# Chemical crosslinking extends and complements UV crosslinking in analysis of RNA/DNA nucleic acid–protein interaction sites by mass spectrometry

**DOI:** 10.1101/2024.08.29.610268

**Authors:** Luisa M. Welp, Alexander Wulf, Aleksandar Chernev, Yehor Horokhovskyi, Sergei Moshkovskii, Olexandr Dybkov, Piotr Neumann, Martin Pašen, Arslan Siraj, Monika Raabe, Henri Göthert, James L. Walshe, Deliana A. Infante, Ana C. de A.P. Schwarzer, Achim Dickmanns, Sven Johannsson, Jana Schmitzova, Ingo Wohlgemuth, Eugen Netz, Yi He, Kai Fritzemeier, Bernard Delanghe, Rosa Viner, Seychelle M. Vos, Elisa Oberbeckmann, Katherine E. Bohnsack, Markus T. Bohnsack, Patrick Cramer, Ralf Ficner, Oliver Kohlbacher, Juliane Liepe, Timo Sachsenberg, Henning Urlaub

## Abstract

UV (ultra-violet) crosslinking with mass spectrometry (XL-MS) has been established for identifying RNA- and DNA-binding proteins along with their domains and amino acids involved. Here, we explore chemical XL-MS for RNA-protein, DNA-protein, and nucleotide-protein complexes *in vitro* and *in vivo*. We introduce a specialized nucleotide-protein-crosslink search engine, NuXL, for robust and fast identification of such crosslinks at amino acid resolution. Chemical XL-MS complements UV XL-MS by generating different crosslink species, increasing crosslinked protein yields *in vivo* almost four-fold, and thus it expands the structural information accessible via XL-MS. Our workflow facilitates integrative structural modelling of nucleic acid–protein complexes and adds spatial information to the described RNA-binding properties of enzymes, for which crosslinking sites are often observed close to their cofactor-binding domains. *In vivo* UV and chemical XL-MS data from *E. coli* cells analysed by NuXL establish a comprehensive nucleic acid–protein crosslink inventory with crosslink sites at amino acid level for more than 1500 proteins. Our new workflow combined with the dedicated NuXL search engine identified RNA crosslinks that cover most RNA-binding proteins, with DNA and RNA crosslinks detected in transcriptional repressors and activators.

## INTRODUCTION

Proteins and RNA (or DNA) form functional complexes termed ribonucleoproteins, RNPs (or deoxyribonucleoproteins, DNPs), which drive and organize key cellular processes. For studying RNA-protein interactions, UV crosslinking combined with mass spectrometry (UV XL-MS) has been established as a standard method, in which the identification of crosslinked RNA-protein heteroconjugates is based either on the inherent reactivity of pyrimidines (mainly uracil) or on UV-reactive derivatives, 4-thiouridine or 6-thioguanine (1–8). In earlier studies, we demonstrated that UV XL-MS can identify the sites of RNA/DNA-protein crosslinking at amino acid resolution and the compositions of the crosslinked (oligo)nucleotide adducts in samples of varying complexity (2, 9–12). Other compelling MS-based studies of the RNA-protein interactome in cells have contributed to this research field by establishing dedicated workflows for UV XL-MS. RBDmap and pCLAP protocols have been used to identify RNA-binding proteins and binding domains by identification of peptides neighbouring crosslinked peptides using sequential proteolytic digestion (13, 14). Similarly, the CAPRI protocol identifies RNA-crosslinked proteins and peptides neighbouring crosslinked peptides (5). XRNAX and OOPS use UV crosslinking with organic phase separation to identify RNA-crosslinked proteins and peptides by MS (7, 8). TRAPP uses silica-based enrichment of UV-crosslinked proteins and for its variant iTRAPP, resolution is increased to the amino acid level (6). Using (p)RBS-ID protocol, Bae *et al.* established chemical digestion of the RNA in UV-crosslinked samples to generate mononucleosides, thus facilitating identification of crosslinked peptides by minimizing the database search space of the MS crosslinking data (3, 4). CLIR-MS identifies UV-crosslinked amino acids and the crosslinked nucleotide in the RNA sequence by using stable isotope-labelled RNA *in vitro* (15).

In DNP analysis, UV XL-MS is much less well-established, owing to the reduced UV reactivity of double-stranded DNA compared to RNA (10, 16). Alternatively, DNP XL-MS studies use chemicals that react with purines in the DNA (17–22). Chemical crosslinking has also been applied successfully in several biochemical studies of RNA-protein interactions (23), but barely combined with larger XL-MS studies although the potential of alkylating reagents for protein-RNA and DNA crosslinking has been systematically investigated by Scalabrin *et al.* in 2018 (24). To increase coverage of interaction sites we set out to complement UV XL-MS by introducing chemical crosslinking with alkylating agents (mechlorethamine/nitrogen mustard, NM; 1,2:3,4-diepoxybutane, DEB; and formaldehyde, FA), identifying the crosslinking sites, and to compare the results with those from UV XL-MS. We developed NuXL, a specialized, open-source cross-link search tool, with integrated false-discovery rate (FDR) control at the crosslink-spectrum match (CSM) level, and extensive reporting of cross-link results tailored for the analysis of protein-nucleic acid complexes. We demonstrate the feasibility of using XL-MS with NuXL for the integrative modelling of RNPs *in vitro,* based on the combination of UV and chemical crosslink sites with high-resolution structural data from cryo-EM and crystallography. Moreover, using NuXL we could identify more than 1,500 proteins with their chemically or UV-crosslinked amino acids to RNA, DNA, or single nucleotides in *E. coli*. The chemically and UV-induced crosslinking sites identified on proteins confirm that chemical crosslinking complements and extends UV crosslinking at the amino acid and nucleotide levels.

## MATERIALS AND METHODS

### Crosslinker safety measures

The chemical crosslinkers used in this study (mechlorethamine/nitrogen mustard, NM; 1,2:3,4-diepoxybutane, DEB; and formaldehyde, FA) are toxic compounds that must always be handled in accordance with the relevant safety regulations. For this study, we kept purchase and use to a minimum.

### UV, DEB, NM and FA *E. coli* ribosome crosslinking and sample processing

70S *E. coli* ribosomes prepared by sucrose-gradient centrifugation as described elsewhere (25) were kindly provided by Prof. Dr. Marina Rodnina (Department of Physical Biochemistry, Max Planck Institute for Multidisciplinary Sciences, Göttingen, Germany) and were stored in 20 mM HEPES pH 7.5, 100 mM KCl, 7 mM MgCl_2_, at –80°C. The protein content of the preparation was 13.3 g/L. 50 µg (3.8 µL) of ribosomes were diluted to 100 µL with the same buffer before crosslinking. For UV XL, samples were pipetted into 96-well polypropylene solid bottom microplates (Greiner Bio-One, 655209) and irradiated for 10 min at a 2 cm distance from four parallel UVC Germicidal T5 lamps (Sylvania; G8W; 0000501; λ=254 nm) as part of an in-house built crosslinking apparatus. DEB crosslinking was performed at a final concentration of 50 mM DEB (Sigma; 202533) for 10 min, 37°C, 300 rpm. NM crosslinking was performed at a final concentration of 10 mM NM (Sigma; 122564) for 10 min, 37°C, 300 rpm. FA crosslinking was performed at 1% [v/v] FA (Thermo Scientific™, 28906) for 10 min, 37°C, 300 rpm. Optimal concentrations of chemical crosslinkers, incubation time and UV irradiation time in order to yield a maximum of crosslinked spectra matches (CSMs) in XL-MS analysis of isolated and crosslinked ribosomes was evaluated in independent experiments (data not shown). All chemical crosslinking reactions were quenched with 50 mM Tris/HCl pH 7.5 and incubation for 5 min, RT, 300 rpm. After crosslinking and quenching, samples were EtOH-precipitated (75% [v/v] EtOH, 0.3 M NaOAc) and pellets were washed once with 80% [v/v] EtOH. Pellets were dissolved in 45 µL 8 M urea in 50 mM Tris/HCl pH 7.5. Urea concentration was diluted to 1 M using 50 mM Tris/HCl pH 7.5. 1mM MgCl_2_, 250 U Pierce™ Universal nuclease for Cell Lysis (Thermo Scientific™, 88700), 100 U nuclease P1 (NEB, M0660S) were added, following incubation for 3 h, 37°C, 300 rpm. Trypsin was added at a 1:20 enzyme-to-ribosome ratio (2.5 µg; Promega, V5111) following incubation ON, 37°C, 300 rpm. Samples were acidified with 2-5 µL 10% [v/v] formic acid to pH 5. 100 U nuclease P1 were added, following incubation for 30 min, 37°C, 300 rpm. 10% [v/v] formic acid and 100% acetonitrile (ACN) were added to reach a final pH of 2 and ACN concentration of 2%. Nucleotide removal was performed using C18 stage tips (Harvard Apparatus; 74-4107) according to manufacturer’s instructions. Briefly, columns were equilibrated with each 400 µL of 100% ACN; 50% [v/v] ACN, 0.1% [v/v] formic acid; and three times 0.1% [v/v] formic acid. Samples were loaded twice and washed two times with 0.1% [v/v] formic acid and once with 5% [v/v] ACN, 0.1% [v/v] formic acid. Elution was performed using twice 150 µL of 50% [v/v] ACN, 0.1% [v/v] formic acid and once 100 µL of 80% [v/v] ACN, 0.1% [v/v] formic acid. All centrifugation steps were performed for 2 min at 600*xg*. Peptides were dried in a speed vac concentrator. Enrichment of crosslinked peptide- (oligo)nucleotides was performed as described in (2) using Titansphere TiO_2_ 5 µm beads (GL Sciences; 5020-75000). Buffer A contained 5% [v/v] glycerol in buffer B; buffer B contained 80% [v/v] ACN, 5% [v/v] trifluoroacetic acid (TFA); buffer B2 contained 60% [v/v] ACN and 0.1% [v/v] TFA; buffer C contained 0.3 M NH_4_OH, pH 10.5. Columns were self-packed by adding approximately 3 mg TiO_2_ beads in 100 % MeOH to p10 pipette tips equipped with a coffee filter insert at the tip (described in more detail by Qamar et al (26)). For each replicate, two columns were used. Columns were equilibrated with 60 µL of each 100% MeOH, buffer B and buffer A. Dried samples were dissolved in 60 µL buffer A and loaded twice and washed three times with buffer A, three times with buffer B and once with buffer B2. Elution was performed in three steps of each adding 40 µL buffer C. Elution tubes contained 20 µL of 10% [v/v] FA for pH neutralization. All centrifugation steps were performed for 2-5 min at 900 × *g*, to prevent columns from running dry. Eluted crosslinked peptide-(oligo)nucleotides were dried in a speed vac concentrator. Once dry, the samples were dissolved first in 10% [v/v] ACN, 0.1% [v/v] formic acid and diluted to 2% [v/v] ACN, 0.1% [v/v] formic acid and subjected to LC-MS/MS measurements.

### Purification of recombinant Hsh49 and Cus1 from *S. cerevisiae*

Recombinant protein complex Cus1-HSH49 was co-expressed from the pETDuet vector in Rosetta II in 2xYT medium ON at 18°C. Purification was performed as described with some modifications (27). Cells were lysed in 50 mM HEPES-NaOH, pH 7.5, 400 mM NaCl, 500 mM urea, 15% glycerol, 5 mM 2-Mercaptoethanol, 20 mM imidazole supplemented with protease inhibitors EDTA free (Roche) using sonication. 5kU Pierce™ Universal nuclease for Cell Lysis (Thermo Scientific™, 88700) was added, following incubation for 15 min on ice. Cell debris was removed by centrifugation 17.000 rpm at 4°C for 1 h (A27 rotor; Thermo Scientific™). The supernatant was loaded on a 5-ml His Trap HP Ni-NTA column (Cytiva) equilibrated with lysis buffer A1. Proteins were eluted with a linear imidazole gradient of 20 to 400 mM in buffer B1 (50 mM HEPES-NaOH, pH 7.5, 200 mM NaCl, 400 mM imidazole, 500 mM urea, 10% glycerol, 5 mM 2-mercaptoethanol) and the main peak fractions containing both proteins were collected and further processed. His-tag was removed by on-column TEV-digestion and sample was loaded on the pre-packed His Trap HP Ni-NTA column (5 mL, Cytiva). The flow-through was collected and loaded on a pre-packed Heparin HP column (3×5 mL, Cytiva). The protein complex was eluted with a linear gradient of 0-100% of 20 mM HEPES-NaOH, pH 7.5, 1M NaCl, 10% glycerol, 5 mM 2-mercaptoethanol. The peak fractions containing purified proteins were pooled and concentrated.

A fragment of *S. cerevisiae* gene encoding nucleotides 1–47 of the U2 snRNA (ggACGAATCTCTTTGCCTTTTGGCTTAGATCAAGTGTAGTATCTGTTCTtctag) was placed under T7 promoter and cloned into EcoRI/XbaI sites of the pUC57 vector. The plasmid was linearized with XbaI and the U2-47Nt snRNA was transcribed *in vitro* with T7 polymerase (NEB). After transcription, DNA was digested with RQ1 DNAse and RNA was purified by Microspin G50 spin columns (Cytiva), and precipitated with ethanol.

Hsh49-Cus1-RNA complexes were reconstituted on ice for 30 min at a protein:RNA molar ratio of 4:1.

### Expression and purification of *Saccharomyces pombe* Dnmt2

*Saccharomyces pombe* Dnmt2 (spDnmt2) was produced as described before (28) with several modifications. The protein was expressed from the pGEX 6P-3 vector as a GST-spDnmt2 fusion protein in *Escherichia coli* BL21(DE3) cells using autoinduction medium (ZYM-5052 medium, 2 mM MgSO_4_ replaced by 1 mM MgCl_2_, supplemented with 100 µM ZnCl_2_) (28). Cells were grown at 37°C for 3 h followed by 60-66 h at 16°C. Cells were disrupted by microfluidization (EmulsiFlex-C3, Avestin) in 50 mM Tris-HCl pH 7.5, 150 mM NaCl, 1 M LiCl, 2 mM DTT and the crude lysate was cleared by ultracentrifugation at 50,000×g for 30 min at 4°C. The cleared supernatant was loaded onto a GSTrap FF column (Cytiva) in 50 mM Tris-HCl pH 7.5, 150 mM NaCl, 2 mM DTT and GST-spDnmt2 was eluted with 30 mM reduced glutathione in the same buffer. The eluted protein was incubated with PreScission Protease (1:80 w/w) to cleave off the GST-tag while dialysing against 50 mM Tris-HCl pH 7.5, 100 mM NaCl, 2 mM DTT for 20 h at 4°C (6-8 kDa MWCO, SpectraPor 1 dialysis membrane) and buffer was exchanged three times. GST was removed via a HiTrap Heparin HP column (Cytiva) in 50 mM Tris-HCl pH 7.5, 100 mM NaCl, 2 mM DTT using a gradient of 0.1-2 M NaCl. The protein was concentrated using an Amicon ultrafiltration device (10 kDa MWCO, Merck) and further purified via a HiLoad Superdex 75 pg in 50 mM Tris-HCl pH 7.5, 150 mM NaCl, 2 mM DTT. Purified spDnmt2 was concentrated to 10 mg/mL and stored at −80°C until further use. All chromatographic steps were performed at 4°C. Purified spDnmt2 was dialysed against 50 mM HEPES pH 7.5, 150 mM NaCl, 2 mM DTT for 16°C at 4°C (6-8 kDa MWCO, SpectraPor 1 dialysis membrane) before complex formation with sp-tRNA-Asp.

### *In vitro* transcription and purification of *Saccharomyces pombe* tRNA-Asp

Synthetic DNA oligonucleotides were used as a template for run-off in vitro transcription (IVT) of *S. pombe*tRNA-Asp (sptRNA-Asp) (Fwd: 5’-TAATACGACTCACTATAGGGTCTCCTTTAGTATAGGGGTAGTACACAAGCCTGTCACGCTTGCAGCCCGGGTTCGAATCCCGGAGGGAGAG-3’; Rev: 5’-CTCTCCCTCCGGGATTCGAACCCGGGCTGCAAGCGTGACAGGCTTGTGTACTACCCCTATACTAAAGGAGACCCTATAGTGAGTCGTATTA-3’). The complementary DNA oligonucleotides were annealed at 100 µM for 5 min at 95°C followed by snap cooling on ice. The annealed template was used in IVT reactions containing 1× Transcription Buffer (Thermo Scientific), 30 mM MgCl_2_, 5 mM DTT, 0.002 U/µL Pyrophosphatase (M0361, NEB), 6 mM rNTPs each, 40 ng/µL annealed template, 5 U/uL T7 RNA Polymerase (Thermo Scientific). Reactions were performed at 37°C for 16 h followed by incubation with 0.1 U/µL TURBO DNase (Invitrogen) at 37°C for 30 min. IVTs were subjected to ethanol precipitation in anhydrous alcohol with 300 mM NaOAc pH 5.2 at −20°C for 16 h. The RNA was pelleted by centrifugation at 20,000×g for 30 min at 4°C, washed with 70% ethanol, air-dried, and resuspended in 20 mM HEPES pH 7.5, 50 mM KCl. IVTs were loaded onto a ResourceQ column (Cytiva), washed with 20% buffer B (20 mM HEPES pH 7.5, 2 M KCl), and eluted in a gradient of 20-35% buffer B. Eluted fractions were precipitated in anhydrous alcohol with 300 mM NaOAc pH 5.2 at −20°C for 16 h. The RNA was pelleted by centrifugation at 20,000×g for 30 min at 4°C, washed with 70% ethanol, air-dried, and resuspended in nuclease-free water. Fractions were analysed in denaturing (7 M urea) 10% polyacrylamide gels and pooled for complex formation with spDnmt2.

### Purification of human NELF complex

The human NELF complex and NELF ΔA, ΔE and ΔAE tentacle mutants were purified as previously described (29). An RNA corresponding to the HIV-1 stem loop (TAR) was purchased from Integrated DNA technologies with the following sequence 5’-CCAGAUCUGAGCCUGGGAGCUCUCUGG-3’. The RNA was folded by incubating of 80 µM of the RNA in a buffer containing 100 mM NaCl, 20 mM HEPES/NaOH pH 7.4, 3 mM MgCl_2_ at 95°C for 3 min, and transferring to ice for 10 min. NELF (∼12 µM final) and RNA (∼80 µM final) were mixed and incubated for 30 min at 4°C. A complex of NELF + TAR RNA was separated from free RNA by size exclusion chromatography (SEC) using a Superose6 Increase 3.2/300 column (Cytiva) in a final buffer containing 20 mM HEPES/NaOH pH 7.4, 150 mM NaCl, 10% glycerol, 1 mM DTT. Peak fractions were analysed by polyacrylamide gel electrophoresis and fractions containing both NELF and TAR RNA pooled for further analysis. Fluorescence anisotropy was performed as previously described (29).

### UV and chemical crosslinking of Hsh49-Cus1-U2 snRNA, Dnmt2-tRNA^Asp^ and NELF-HIV TAR RNA *in vitro* reconstituted complexes and peptide-RNA (oligo)nucleotide enrichment

Samples containing each a maximum of 50 µg reconstituted Hsh49-Cus1-U2 snRNA, Dnmt2-tRNA^Asp^, or NELF:TAR complex were crosslinked and processed as described above for *E. coli* ribosomes with modifications in crosslinking and digestion conditions. Complex concentrations for the crosslinking reaction were approximately 2 mM (Hsh49-Cus1-snRNA), 13.1 µM (Dnmt2-tRNA^Asp^), and 1.6 µM (NELF:TAR), respectively. For Hsh49-Cus1 – U2 snRNA UV crosslinking, samples were irradiated for 10 min as described above, following EtOH precipitation and washing as described for isolated ribosomes. For DEB and NM crosslinking, 1, 5, 10, and 50 mM DEB 1, 5, and 10 mM NM were added, following incubation at 37°C for 10 min and precipitation as described for UV crosslinked samples. Samples were dissolved in 4 M urea, 50 mM Tris/HCl pH 7.5, and diluted to reach 1 M urea, 50 mM Tris/HCl pH 7.5. 2 µg of each Ambion™ RNase A (Invitrogen™, AM2270) and RNase T1 (Invitrogen™, AM2283) were added, following incubation at 37°C for 2 h. Afterwards, 1 mM MgCl_2_, and 750 U of Benzonase® Nuclease (Millipore, 70664) were added, following incubation at 37°C, for another 2 h. Trypsin digestion, nucleotide removal using C18 stage tips, and crosslinked peptide- (oligo)nucleotide enrichment were performed as described above for *E. coli* ribosomes. For Dnmt2-tRNA^Asp^ UV-crosslinking, samples were irradiated for 10 min with consecutive ethanol-precipitation as described for isolated ribosomes. DEB crosslinking was performed using 1, 5, 10, and 50 mM DEB and NM crosslinking of Dnmt2-tRNA^Asp^ was performed using 1 and 10 mM NM following incubation for 10 min at 37°C. DEB and NM crosslinked Dnmt2-tRNA^Asp^ complexes were ethanol precipitated as described above and after ON incubation at −20°C washed with 80% ice-cold ethanol. Subsequently, samples were digested in 1 M urea, 50 mM Tris/HCl pH 7.5, 1 mM MgCl_2_, using 1 µg of each Ambion™ RNase A, 1 µg of RNase T1, and 250 U Benzonase® Nuclease HC (Millipore, 71205) for 2 h, at 37°C (Benzonase was not applied for digestion of RNA in Hsh49-Cus1-U2 snRNA complex). Trypsin or chymotrypsin (Promega, V1061) were added at a 1:20 enzyme-to-protein ratio following incubation ON at 37°C, 300 rpm. The rest of the protocol is identical to what is described for purified *E. coli* ribosomes. The reconstituted NELF-TAR complex was crosslinked by UV irradiation for 10 minutes or by adding 1, 5, 10, and 50 mM DEB, or 1, 5, and 10 mM NM and the TAR-RNA element was digested as described for Hsh49-Cus1 complex. All crosslinked peptide-(oligo)nucleotide samples were dissolved in 2% [v/v] ACN 0.05% [v/v] TFA for LC-MS/MS measurements.

### UV and chemical crosslinking of reconstituted *Saccharomyces cerevisiae* mononucleosomes

Mononucleosomes were reconstituted using the 601 DNA sequence (30) and purified recombinant *S. cerevisiae* histones octamers. Reconstitution and histone octamers purification was done as described previously with minor modifications (31). In brief, DNA and histone octamers were mixed in a 1:1:1 ratio in a high salt buffer (10 mM HEPES-NaOH pH 7.6, 2 M NaCl, 0.5 mM EDTA), transferred to Slide-A-lyzer MINI Dialysis devices (Thermo Scientific) and dialyzed via salt gradient dialysis over 16 hours into a low salt buffer (10 mM HEPES-NaOH pH 7.6, 2 M NaCl, 0.5 mM EDTA). After salt gradient dialysis, the sample was centrifuged at 10,000 g for 1 minute to remove precipitates. The DNA concentration was measured using a spectrophotometer, and nucleosome formation was confirmed by running approximately 100 ng of nucleosomes on a 3-12% NativePAGE (Invitrogen). DNA was visualized by SYBR Gold staining.

UV crosslinking of 30 µg mononucleosomes was performed as described above for *E. coli* ribosomes with minor changes. DEB, NM, and FA crosslinking were performed using 50 mM DEB (we note, that in this case only 50 mM DEB yield crosslinks, while lower concentrations gave no results), or 1 mM NM, or 2% FA, respectively, and incubation for 10 min, 37°C, 300 rpm. For quenching, 50 mM Tris/HCl pH 7.5 was added following incubation for 5 min, RT, 300 rpm. Crosslinked protein-nucleic acid conjugates were EtOH precipitated. DNA was digested in 1 M urea, 50 mM Tris/HCl pH 7.5 with 1mM MgCl_2_, 500 U Pierce™ Universal nuclease for Cell Lysis (Thermo Scientific™, 88700), 200 U nuclease P1 (NEB, M0660S) and incubation for 3 h, 37°C, 300 rpm. Protein digest, nucleotide removal and TiO_2_ based peptide-DNA- (oligo)nucleotide enrichment was performed as described above for *E. coli* ribosomes. Two TiO_2_ columns were used per replicate. Peptides were dried and dissolved like *E. coli* ribosomes for LC-MS/MS measurements.

### *In vivo* UV and chemical crosslinking of *E. coli* cells

*E. coli* BL21 DE3 cells were grown in LB medium to OD_600_ ≈ 0.6. 15 mL of cell culture (corresponding to roughly 7 x 10^9^ cells) were used as starting material per biological replicate for UV, DEB, and NM crosslinking. UV, DEB, and NM crosslinking were performed in medium. Cells were UV crosslinked for 10 min as described above in glass petri dishes (d = 12 cm, vol. cell suspension ≈ 11 mL, ht. cell suspension ≈ 1 mm) on an ice-cold metal block for two replicates or at RT for one replicate. DEB crosslinking was performed using 50 mM DEB for 10 min, 37°C, 300 rpm in 15 mL cell culture. NM crosslinking was performed with 10 mM NM for 10 min, 37°C, 300 rpm in 15 mL cell culture. DEB and NM crosslinking reactions were quenched with final 50 mM Tris/HCl pH 7.5 and incubation for 5 min, RT, 300 rpm. FA crosslinking was performed at 1% [v/v] FA for 10 min, 37°C, 300 rpm in 15 mL cell culture, followed by quenching using 400 mM Tris/HCl pH 7.5 for 10 min, RT, 300 rpm. Cells were pelleted at 3000*×g* for 10 min and the medium was removed. For UV, DEB and NM crosslinked cells, S30/S100 fractionation by ultracentrifugation was performed as follows: Cell pellets were dissolved in 100 µL of 20 mM Tris/HCl pH 7.6, 100 mM NH_4_Cl, 10.5 mM MgCl_2_, 3 mM β-mercaptoethanol, 0.4 mg/mL lysozyme, 10 U/mL RQ1 DNase (Promega, M6101) and cOmplete™ EDTA-free protease inhibitor cocktail (Roche, COEDTAF-RO). Cells were first incubated at 37°C for 30 min and subsequently mechanically disrupted by sonication in a water bath for 15 min, in alternating 30 s ‘ON’ and 30 s ‘OFF’ cycles at the highest intensity setting (Bioruptor, Diagenode). Three volumes of B-PER™ bacterial protein extraction reagent (Thermo Scientific™, 78243) were added and samples were briefly mixed by inverting, following incubation at RT for 5 min and another mixing by inverting. After another 5 min incubation at RT, samples were centrifuged at 40,000*xg* for 30 min for S30 fractions, and for another 30 min at 330,000*xg* for S100 fractions. S30 and S100 fractions were EtOH precipitated as described above, pellets were washed and dissolved in 8 M urea, 50 mM Tris/HCl pH 7.5. LysC (Promega, V1671) was added at a 1:100 enzyme-to-protein ratio, following incubation at 25°C, ON, 300 rpm. Urea concentration was diluted to 1 M using 50 mM Tris/HCl pH 7.5. Trypsin was added at a 1:50 enzyme-to-protein ratio, following incubation ON, 30°C, 300 rpm. Samples were EtOH precipitated as described above and dissolved in 4 M urea, 50 mM Tris/HCl pH 7.5. Urea concentration was diluted to 1 M using 50 mM Tris/HCl pH 7.5 and 1mM MgCl_2_, 500-625 U Pierce™ Universal nuclease, 200-300 U nuclease P1 were added, following incubation for 3 h, 37°C, 300 rpm. Trypsin was added at a 1:50 enzyme-to-protein ratio, following incubation ON, 37°C, 300 rpm. Samples were acidified with formic acid to pH 3-4 and 2% [v/v] ACN was added. Sample clean-up was performed using C18 stage tips as described above. Enrichment of crosslinked peptide-(oligo)nucleotides was performed using 6 to 8 TiO_2_ columns as described above. Eluted crosslinked peptide-(oligo)nucleotides were dried in a speed vac and fractionated by high pH reversed-phase fractionation (see below).

FA crosslinked *E. coli* cell pellets were resuspended in 300 μL PBS containing 9 mg of lysozyme, following incubation for 5 min, RT. 8 M urea, 100 mM HEPES-NaOH pH 7.9, 20 mM EDTA was added, and cells were mechanically lysed by sonication, in alternating 30 s on- and 30 s off-cycles at highest intensity output (Bioruptor, Diagenode). Samples were diluted to 1 M urea and trypsin was added at an estimated 1:20 enzyme-to-protein ratio, following incubation at 37°C, ON, 300 rpm. 5 ml of cell lysate was used for silica enrichment using RNeasy Maxi Kit (Qiagen, 75162) according to manufacturer’s instructions. Each centrifugation step was performed at 3,000*×g*. Briefly, 3.8 volumes of RLT Buffer, supplemented with β-mercaptoethanol and 2.8 volumes of EtOH were added to the cell lysate. Samples were loaded and washed once with RW1 Buffer, and twice with RPE Buffer. Peptides were eluted twice with H_2_O, and 1 M urea, 12.5 mM HEPES-NaOH pH 7.9 were added. Trypsin was added at an estimated 1:20 enzyme-to-protein ratio, following incubation for 2 h at 37°C, 300 rpm, following another round of RNA purification using RNeasy Maxi Kit as described above. Samples were adjusted to 1M urea, 12.5 mM HEPES-NaOH pH 7.9. 1mM MgCl_2_, 1mM ZnCl_2_, 1250 U Pierce™ Universal nuclease, 50 U RNase I, 300 U nuclease P1, 2 U RNase T1, and 200 U RNase A were added, following incubation at 37°C, ON, 300 rpm. Sample clean-up was performed using C18 stage tips as described above. The mixture containing (crosslinked) peptides and RNA oligonucleotides was resuspended in 50 mM NaOAc and 80 U nuclease P1 and 4 U Antarctic phosphatase were added, following incubation at 30°C, 2 h, 300 rpm. Sample clean-up was performed using self-made C18 columns (p200 tips containing three layers of Empore™ C18 SPE Disks (Supelco, 66883-U)). Eluted FA crosslinked peptide-(oligo)nucleotides were dried in a speed vac concentrator, dissolved in 2% [v/v] ACN 0.05% [v/v] TFA, and subjected to LC-MS/MS measurements.

### High pH reversed phase liquid chromatography (LC) sample fractionation

Enriched UV-, DEB-, NM-crosslinked, or whole proteome peptides from *E. coli* were dissolved in 35 µL 10 mM NH_4_OH pH 10, 5% [v/v] ACN. Peptides were loaded onto an Xbridge C18 column (Waters, 186003128) using an Agilent 1100 series chromatography system. A flow rate of 60 µl/min was set up using a buffer system consisting of 10 mM NH_4_OH pH 10 (buffer A) and 10 mM NH_4_OH pH10, 80% [v/v] ACN (buffer B). The column was equilibrated with 5% buffer B and developed over 64 min using the following gradient: 5% buffer B (0-7 min), 8-30% buffer B (8-42 min), 30-50% buffer B (43-50 min), 90-95% buffer B (51-56 min), 5% buffer B (57-64 min). The first 6 min were collected as flow-through fraction, followed by 48 × 1 min fractions, which were reduced to 12 fractions by concatenated pooling for crosslink preparations and to 24 fractions for whole proteome. Fractions were dried in a speed vac concentrator and dissolved like *E. coli* ribosomes for LC-MS/MS measurements.

### Proteome of non-crosslinked *E. coli* cells

For the reference proteome of *E. coli* cells, cells were treated as described above except that no crosslinking and no TiO_2_ purification step was performed. The digested peptides were separated by high pH reverse-phase HPLC as described above and the resultant fractions were measured by LC-MS as described below.

### LC-MS/MS analyses

All crosslinked peptide-(oligo)nucleotide samples were injected onto a C18 PepMap100-trapping column (0.3 x 5 mm, 5 μm, Thermo Scientific™) connected to an in-house packed C18 analytical column (75 μm x 300 mm; Reprosil-Pur 120C18-AQ, 1.9 μm, Dr Maisch GmbH). Columns were equilibrated using 98% buffer A (0.1% [v/v] formic acid), 2% buffer B (80% [v/v] ACN, 0.08% [v/v] formic acid). Liquid chromatography was performed using an UltiMate-3000 RSLC nanosystem (Thermo Scientific™). Purified, crosslinked *E. coli* ribosome samples, yeast nucleosome samples, and high pH reversed phase LC fractions derived from *E. coli* cell UV, DEB and NM crosslinked samples were analysed for 58 min using a linear gradient (10% to 45% buffer B (80% [v/v] ACN, 0.1% [v/v] formic acid) in 44 min). Hsh49-Cus1-snRNA and NELF complex samples were analysed for 58 min using a linear gradient from 12% to 50% buffer B in 43 min. Dnmt1-tRNA^Asp^ samples were analysed for 58 min using a linear gradient from 12% to 50% buffer B in 43 min or from 2% to 48% buffer B in 44 min. FA XL-MS *E. coli* samples were analysed for 118 min using a linear gradient from 8% to 44% buffer B in 104 min. All measurements ended with an approximately 5 min high-percentage buffer B washing step. Eluting peptides were analysed on an Orbitrap Exploris 480 (Thermo Scientific™; *E. coli* ribosomes, UV, DEB and NM XL-MS *E. coli* cells, yeast nucleosomes, *E. coli* proteome, Dnmt2-tRNA^Asp^, NELF complex, Hsh49-Cus1-snRNA), Q-Exactive HF-X (Thermo Scientific™; UV XL-MS *E. coli* cells), or Orbitrap Fusion (Thermo Scientific™; FA XL-MS *E. coli* cells) instrument. The following MS settings were used for *E. coli* ribosome, UV, DEB and NM crosslinked *E. coli* cell, yeast nucleosome, Hsh49-Cus1-snRNA, NELF complex, Dnmt2-tRNA^Asp^ samples: MS1 scan range, 350–1600 *m/z*; MS1 resolution, 120,000 FWHM; AGC target MS1, 1E6; maximum injection time MS1, 60 ms; intensity threshold, 1E4; isolation window, 1.6 Th; normalized collision energy, 30% (DEB, NM, FA) or 28% (UV); charge states, 2+ to 6+; dynamic exclusion, 9 s; Top 20 most abundant precursors were selected for fragmentation; MS2 resolution, 30,000, AGC target MS2, 1E5; maximum injection time MS2, 120 ms. For FA XL-MS *E. coli* samples, the same settings as for *E. coli* ribosomes were used, except for: MS1 scan range, 350–1580 *m/z*; maximum injection time MS1, 50 ms; intensity threshold, 2E5; isolation window, 1.4 Th; normalized collision energy, 36%; dynamic exclusion, 20 s; Cycle time (top speed method), 3 s; AGC target MS2, 1.1E5; maximum injection time MS2, 80 ms.

### XL-MS data analysis using NuXL

*E. coli* ribosome, Hsh49-Cus1, Dnmt2 and NELF *in vitro* complexes, yeast *S. cerevisae* mononucleosomes*, E. coli in vivo* MS raw data were analysed with the NuXL node in the OpenMS open-source C++ library (version OpenMS-3.0.0-pre-HEAD-2022-04-27-Win64) or in Proteome Discoverer using the following databases: i) the reference proteome canonical sequence database for *E. coli* strain downloaded from the UniProt repository on 10.03.2022 (4400 protein sequences) including MaxQuant contaminants (version 2.0.3.0.) for *E. coli* ribosome and *E. coli in vivo* MS data, ii) Hsh49-Cus1 protein sequences for Hsh49-Cus1 data, iii) *S. pombe* Dnmt2 protein sequence plus MaxQuant contaminants (version 2.0.3.0.) for Dnmt2 data, iv) human NELF subunit protein sequences for NELF data, and v) a custom subset of 75 proteins extracted from the reference proteome canonical sequence database for *Saccharomyces cerevisiae* (UP000002311) from the UniProt repository released on 09.01.2025 plus MaxQuant contaminants (version 2.0.3.0.) for yeast nucleosome data. The subset of 75 *S. cerevisiae* protein sequences was generated for NuXL search because the proteome analysis of the starting nucleosome preparation identified (in addition to the core histones) these 75 other *S. cerevisiae* proteins, which could also potentially have been crosslinked. The following NuXL presets were selected: *E. coli* ribosome, RNA-UV (UCGA), RNA-DEB, RNA-NM, RNA-FA; Hsh49-Cus1, RNA-UV (UCGA), RNA-DEB, RNA-NM; Dnmt2, RNA-UV (UCGA), RNA-DEB, RNA-NM; NELF, RNA-UV (UCGA), RNA-DEB, RNA-NM; yeast nucleosome, DNA-UV, DNA-DEB, DNA-NM, DNA-FA; *E. coli in vivo*, RNA-UV (UCGA), RNA-DEB, RNA-NM, RNA-FA, DNA-UV, DNA-DEB, DNA-NM, DNA-FA. Other parameters were set to default or as follows: enzyme, Trypsin/P, or Chymotrypsin/P (only for Dnmt2-tRNA^Asp^ datasets derived from chymotryptic cleavage); max. missed cleavages, 2; min. peptide length, 5; max. peptide length, 30; peptideFDR, 0.01; nucleotide adduct length, 2 (for XL-MS datasets from *E. coli* cells) or 3 (for all other datasets); dynamic modifications, Oxidation (Met). Crosslink MS2 spectra of the *in vitro* reconstituted complexes (Hsh49-Cus1, Dnmt2, and NELF data, yeast mononucleosomes) were manually curated using OpenMS TOPPView. Further details on XL-MS data processing in NuXL can be found in the Supplementary Methods section. Exemplary run times of NuXL and effects of simple and extended presets on the numbers of identified crosslinks are provided in Supplementary Results.

### Label-free quantitative MS data analysis using MaxQuant

Proteomics data from *E. coli* cells were analysed with MaxQuant (version 2.0.3.0) (32) using the reference proteome canonical sequence database for *E. coli* K12 strain downloaded from the UniProt repository on 10.03.2022 (4400 protein sequences) including MaxQuant contaminants (version 2.0.3.0.). Search parameters were set to default or as follows: enzyme, Trypsin/P; max. missed cleavages, 2; dynamic modifications, Oxidation (Met), Acetylation (protein N-term); static modification, Carbamidomethylation (Cys).

### Downstream data analysis

Barplots were generated using R Studio (version 2024.04.2 Build 764), violin plots were generated using Python (version 3.10.3). Protein 3D structures were plotted in PyMOL Molecular Graphics System (version 3.1.0) using Python (version 3.10.3) scripts for crosslink mapping and distance calculations. Figure 1 was created with BioRender.com.

**Figure 1.**
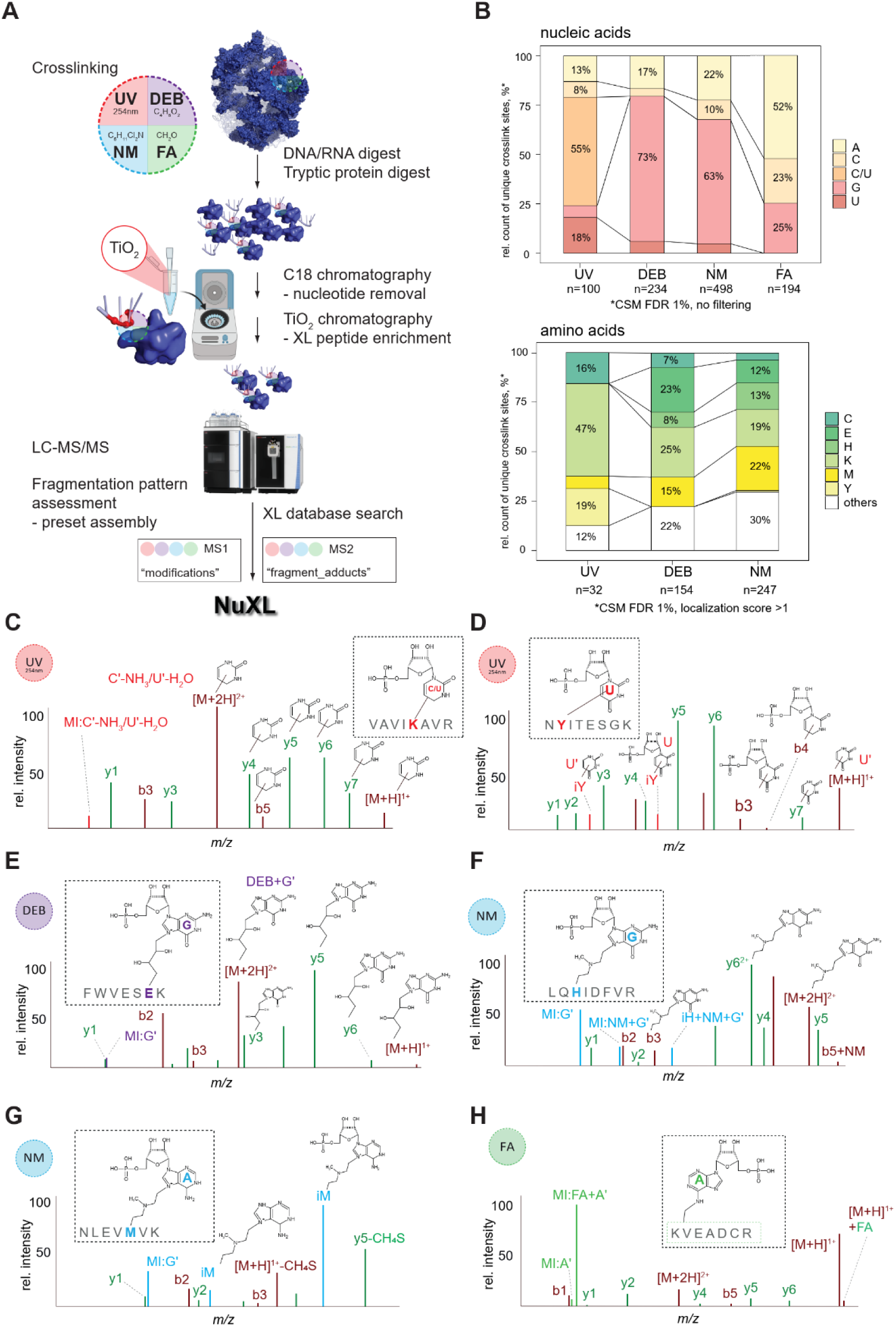
UV and chemical XL-MS workflow for identification of UV and chemical (oligo)nucleotide peptide crosslinks. **(A)** XL-MS processing and analysis workflow. Purified *E. coli* ribosomes were crosslinked with UV, DEB, NM, or FA. DNA, RNA, and proteins were digested enzymatically, and free nucleotides were removed by C18 chromatography. Crosslinked peptides were enriched by TiO_2_ chromatography and analysed by LC-MS/MS. Spectra were evaluated manually and crosslinker-specific adducts were assembled into presets. MS data was then analysed using NuXL. **(B)** Upper panel, stacked bar plot (normalised to 100%) of relative counts of observed crosslinked nucleobases in unique crosslinked peptide-(oligo)nucleotide pairs resulting from UV, DEB, NM and FA XL-MS of *E. coli* ribosomes. CSMs were filtered for 1% FDR. Ambiguous, isobaric C–NH_3_/U–H_2_O crosslinks were counted as C/U. Labels are displayed only for values of 7% and above. Lower panel, 100% stacked bar plot of relative counts of observed crosslinked amino acids in unique crosslinked peptide-(oligo)nucleotide pairs resulting from UV, DEB, NM XL-MS of *E. coli* ribosomes. CSMs were filtered for 1% FDR and a localisation score ≥1. Labels are displayed only for values of 7% and above. FA data are excluded from the visualisation due to low numbers of properly assigned crosslinked amino acids. Absolute CSM counts for each bar are depicted below the plots. **(C-H)** Schematic crosslink MS2 spectra for UV-, DEB-, NM- and FA-crosslinked peptide–RNA(oligo)nucleotides. Peptide fragment b- and y-type ions are shown in wine red and green, respectively. **(C)** UV crosslink MS2 spectrum of VAVIKAVR-CMP-NH_3_/UMP-H_2_O. Marker ion (C*’*–NH_3_ or U*’*–H_2_O), precursor ion, and fragment ions modified with C*’*–NH_3_ or U*’*–H_2_O, as well as unmodified fragment ions indicate a crosslink located at Lys-5. **(D)** UV crosslink MS2 spectrum of NYITESGK-UMP. Immonium ion of tyrosine (iY) modified with U’, precursor ion, and fragment ions modified with UMP or U*’,* as well as unmodified fragment ions indicate a crosslink located at Tyr-2. **(E)** DEB crosslink MS2 spectrum of peptide–RNA(oligo)nucleotide FWVESEK-DEB-GMP. Precursor ions, and fragment ions modified with DEB+G*’* as well as unmodified fragment ions indicate crosslink localisation at glutamic acid-6. **(F)** NM crosslink MS2 spectrum of crosslinked peptide-RNA(oligo)nucleotide LQHIDFVR-NM-GMP. Marker ion (NM+G’), immonium ion of histidine (iH) modified with NM+G’, precursor ion, and fragment ions modified with NM+G’, as well as unmodified fragment ions indicate crosslink localisation at histidine-3. **(G)** NM crosslink MS2 spectrum of crosslinked peptide-RNA(oligo)nucleotide NLEVMVK-NM-AMP. Immonium ions of methionine modified with NM+G’ and NM+GMP, methionine specific loss (-CH_4_S) on the precursor ion and fragment ion y5, as well as unmodified fragment ions indicate crosslink localisation at methionine-5. **(H)** FA crosslink MS2 spectrum of crosslinked peptide-RNA(oligo)nucleotide KVEADCR-FA-AMP. Marker ions (FA plus adenine, FA+A*’*; A*’*), and precursor ion modified with FA identify AMP crosslink. Unmodified fragment ions cover the peptide sequence. Unprocessed mass spectra as visualized from NuXL outputs by TOPPView spectra viewer (70) are represented in Supplementary Figure S2.

### *sp*Dnmt2-tRNA^Asp^ docking-based model generation

The previously published assembly of *sp*Dnmt2-tRNA^Asp^ was generated as described in Johannsson et al. (33). It combined *de novo* RNA modelling protocol (34) with molecular *in silico* docking experiments yielding 3D models of *S. pombe* tRNA^Asp^-Dnmt2 (34, 35).

### Generation of *sp*Dnmt2-tRNA^Asp^ models with AlphaFold3

Complexes of *sp*Dnmt2-tRNA^Asp^ were predicted by AlphaFold3 (AF3, https://alphafoldserver.com/ (36)). To enhance structural diversity in predicted models, 60 individual runs with differing random seeds were executed, yielding 300 predictions.

### Verification of *sp*Dnmt2-tRNA^Asp^ models using XL-MS data

All 300 Alpha Fold 3-based predictions, along the previously published assembly (“OLD”), were individually assessed for compatibility with cross-linking mass spectrometry (XL-MS) data using an adapted protocol (37). The cross-linking dataset was filtered to retain only cross-links with a cross-link spectral match (CSM) count of >2 per site. Each cross-link was evaluated for compatibility with every tRNA^Asp^-Dnmt2 model by computing Euclidean distances between the protein residue (known position) and the potentially linked nucleotide(s) (unknown position), selected based on distance-based criteria. Two sets of cross-link occurrences were identified for each model: (i) SINGLE, comprising the shortest-distance cross-link pairs between the known protein residues and the closest nucleotides within the defined threshold; and (ii) ALL, including all potential cross-link occurrences within the threshold. Distance thresholds were set according to the cross-linker type: UV (10 Å), DEB, and NM (15 Å). Distances were measured between the N3 atom (for U or C nucleotides) or N7 atom (for other nucleotides) of tRNA and the Cα atom of the cross-linked protein residue. The number of cross-link occurrences within these thresholds was termed “occurrences_d0” (where d0 represents no additional slack beyond the threshold distance). To account for potential conformational flexibility of both protein and tRNA, that was unpredictable by AF3 or *in silico* modelling/docking, additional protein residue-nucleotide pairs were identified by incrementally increasing the threshold distance by 2, 3, and 5 Å, designated as d2, d3, and d5, respectively. For all predicted tRNA^Asp^–Dnmt2 complexes, both single (d0, d2, d3, d5) and multiple (ALLd0, ALLd2, ALLd3, ALLd5) cross-link occurrences were determined and subjected to further analysis.

To identify models supporting the highest number of cross-links, we compiled a dataset integrating both ideal (d0) and distance-slacked (d2, d3, d5) single and multiple cross-link occurrences. A custom Bash/Awk script was developed to systematically rank models based on numerical sorting of multiple specified columns, ensuring an unbiased weighting. The script operates in two key steps. During the first step (termed normalization), the maximum value for each specified column was identified to prevent dominance by a single variable. The second step concentrated on scoring and ranking. Here, each row (tRNA^Asp^–Dnmt2 model) was assigned a score reflecting how many of its column values (d0, d2, d3, d5, ALLd0, ALLd2, ALLd3, ALLd5) match the maxima. Rows were then sorted in descending order, with models exhibiting the highest number of maximal values ranked highest. This approach ensured progressive and independent contributions from all criteria.

Our analysis demonstrated that AF3-based predictions exhibit the highest compatibility with crosslinking data, providing compelling evidence of the unprecedented accuracy achieved by AI-driven AF3 models.

### Protein annotation sources

The protein sequences were retrieved from UniProt on 2022-03-10 (38) and UniProt identifiers were used to transfer the information from additional sources. Domain annotation was downloaded from InterPro (39). In addition to that, single-residue annotation of conserved and functionally important residues was obtained using the InterProScan software version 5.56-89.0 on the above mentioned FASTA protein sequences, with residue annotation mode enabled and minimal feature size set to 1 (40). Manual review identified 202 relevant InterPro signatures that were manually grouped into 20 categories, corresponding to the molecule class that the protein feature is interacting with (i.e., ATP, FAD/NAD/NADP). Secondary structure was predicted from protein sequences using the S4PRED (41) using the default settings. Residue-level features were transferred from the DESCRIBEPROT (42) and the continuous scores were binarized into feature and non-feature using the default database thresholds.

Protein abundance in ppm for *E.coli* K12 strain and integrated abundance across strains were downloaded from the PaxDb (43). In addition to the pre-computed reference abundance in parts per million (ppm), the MS1 intensities derived from label-free quantification of the non-crosslinked control *E. coli* dataset were converted into ppm using the PaxDb web-interface. The annotation of RNA-related proteins was derived based on RBP2GO database of putative RNA-binding proteins (44).

### Sampling and statistical testing of feature enrichment

Statistical testing was performed to quantify the enrichment of amino-acid (AA) composition, secondary structure and other residue features. The foreground was defined as a set of crosslinked peptide residues, either exactly at the crosslink site or belonging to the whole peptide. On the site level, only those crosslink-spectrum matches (CSMs) with the localization threshold ≥ 1 were kept. The number of peptides in the foreground set corresponds to the number of the CSMs per identified sequence. The background was derived by an *in silico* tryptic cleavage of the proteome. When comparing the crosslinked residue distributions, one random residue per background peptide was chosen.

Since not all the possible tryptic peptides are observable in the MS experiment and 85% of identified experimental sequences were previously reported in PeptideAtlas (45) and PaxDb (43), a mass-spectrometry-informed background was defined as a union of all the experimental sequences and all the tryptic *E. coli* peptides from PeptideAtlas (45) and PaxDb (43), and was used to sample the background distribution, omitting the non-detectable peptides. A global background consisting of all the MS-detectable peptides was used for the 3D distance enrichment analysis, whilst for feature count enrichment analysis, samples were drawn with replacement from the background distribution to match the experimental distribution of peptide lengths and missed cleavages. This propensity score-based covariate matching was done via matchRanges (46) using the “nearest” matching option.

### Quantification and correction of the protein abundance bias

In order to assess whether the crosslink detection is favouring more abundant proteins we correlated the available protein abundance metrics (below) to the RBP2GO score and the number of observed CSMs in the crosslinking data. When comparing the abundance distributions of the proteins annotated in RBP2GO, the RNA-related proteins appeared to be on average more abundant that other proteins. From this, we concluded that the differences in intracellular protein abundance could affect the identification of crosslinked spectra and the CSM counts of proteins.

We have modified the sampling scheme to compensate for the abundance bias, by associating every peptide with 4 metrics of the source protein abundance: PaxDB integrated (ppm) and E. coli K12 reference abundance (ppm), log10-transformed label-free MS1 intensity of our non-crosslinked *E. coli* proteome control and quantile-normalised ppm values derived from the label-free MS1 intensity of our non-crosslinked *E. coli* proteome control. A combination of multiple features was chosen in order to obtain a balanced abundance estimate informed by both in-house (own proteome data, see above) and public data and cover the most proteins. The abundance-corrected background resulted in comparable amino acid enrichment values, but the frequency of putative RNA-binding residues was reduced, supporting the hypothesis of RNA-related proteins being naturally high-abundant in *E. coli*.

### Statistical testing for residue feature enrichment

The difference of the distribution mean feature counts per residue between the foreground and background was assessed via a t-test as implemented in R language (47). In addition, since most of the protein features are distributed not normally, non-parametric goodness-of-fit tests (48) are used to validate the statistical significance. Furthermore, since many features are discrete, the discrete adaptations of nonparametric tests are used (49, 50). The significance threshold for the feature enrichment was set to the adjusted p-value of 0.01 (Benjamini-Hochberg method (51)) to be surpassed by both the t-test and non-parametric goodness-of-fit test. Sampling was repeated 10 times and the resulting statistics were aggregated by mean. The log2FoldChanges between the foreground and background distribution mean feature counts per residue were formatted into a matrix and plotted via the “pheatmap” R library (52).

### 3D distance analysis

The protein structures were downloaded from the AlphaFold Protein Structure Database (53, 54) and 3D Euclidean distances were retrieved between the pairs of Cα atoms using the Xwalk (55) version 0.6 with maximal distance 100 Å. All the amino acids with min. localisation score ≥ 1 were checked for the presence of annotated protein features in the 10 Å radius. Since some of the features of interest occur infrequently, the background was not sampled from the proteome, but a full set of all annotated residues was used as a background distribution. A combination of a t-test and nonparametric goodness-of-fit-test was applied to test for enrichment, same as for the feature counts enrichment.

### Over-representation analysis

The STRING database was used to define the gene sets for *E. coli*. All the identified proteins with at least 1 unique peptide were grouped into sets corresponding to different crosslinking methods each (56). The hypergeometric test for term over-representation was done via the “fgsea” R package (57). A union set of every protein identified in any of RNA-XL datasets or in the non-crosslinked control dataset was used as an enrichment background. The minimal gene set size was set to 4 and maximal to 1000. After testing and p-value adjustment with the BH method, a set of non-redundant terms was derived (51). All the terms that pass the significance threshold of 0.05 p.adj BH were tested to not enrich on the background of each other, using the “fgsea” package implementation.

### Gene set over-representation heatmaps

The over-represented STRING terms were selected for heatmap display as long as they passed the p.adj BH significance threshold of 0.01 except the Domains set, where it was set to 0.05. In addition, a query and background set overlap size of at least 5 proteins was required. Upon the filtering, the manually selected top representative terms were visualised using the “pheatmap” R library (52).

### Protein feature and coverage plots

We have compiled a residue-level annotation of E. coli proteome and visualised the co-localization of protein features and RNA-XL coverage. Tryptic cleavage sites correspond to Lys and Arg residues in the proteins. Protein annotation was converted into per-residue GenomicRanges format using the protein coordinates and used for further plotting (58).

All the CSMs with the localization score ≥ 1 and their source peptides were plotted for all RNA-XL datasets except FA, where no localization information is available. The positions of RNA-XL sites and peptide coverage were plotted along the protein sequence together with the domain and residue annotation (58–63).

### Logo plots

Crosslink site motifs were derived by centering all CSMs with the localization threshold ≥ 1 around the XL-sites, compared to the background of all MS-detectable peptides in *E.coli* and plotted via “Logolas” R package with the default settings (64).

### HTH motif analysis

Proteins with HTH-motif entries were taken from the InterPro (39) domain annotation. For every HTH motif type, the average count of crosslinkable amino acids (C, E, F, H, K, M, Y based on observed frequencies) in the motif was calculated, and the fraction of crosslinked amino acids was computed, together with the CSM counts per amino acid.

## RESULTS

### Assessment of MS2 spectra and crosslinked species from UV and chemical XL-MS

Our first goal was to identify DEB-, NM-, and FA-induced crosslink sites at the amino acid and nucleotide level in MS analysis. To this end, purified *E. coli* ribosomes were subjected to *in vitro* UV, DEB, NM, and FA crosslinking, nuclease, and trypsin digestion, followed by C18 clean-up for removal of non-crosslinked (oligo)nucleotides and TiO_2_ bead-based peptide-RNA (oligo)nucleotide enrichment followed LC-MS analysis (Figure 1A). We manually annotated MS1 and MS2 data, applying known mass-adduct settings for UV and calculating the mass of hypothetical crosslinker-RNA (oligo)nucleotide adducts of various compositions, based on the detected mass adducts and on previous studies suggesting possible reaction mechanisms (Figure 1A, Supplementary Figures S1 and S2, Supplementary Table S1)(2,63–66). Individual mass adducts on the precursor and fragment ions and several marker ions (crosslinked immonium, nucleobase, and nucleotide ions) were assembled and guided the development of a fast, reliable FDR-controlled computational analysis tool for identifying crosslinking sites, which we named NuXL.

Using NuXL, we determined the occurrence of predominant crosslink types by comparing of counts of crosslink-spectrum matches (CSMs) (Figure 1B). For UV, the majority of peptide–RNA (oligo)nucleotides, in absolute CSM numbers, were crosslinks to Lys, Tyr, and Cys. When normalized by the frequencies of the occurrence of the corresponding amino acids in ribosomal proteins (Supplementary Figure S3A), Cys and Tyr were found to be the amino acids most frequently crosslinked, in agreement with earlier studies (4, 65, 66) followed by Lys and Met (Supplementary Figure S3A). Other aromatic amino acids, such as Phe and Trp, as well as His (all recognized before as preferred UV crosslinking sites (4, 65) were barely found in isolated ribosomes. Lys crosslinks have been previously identified by Kramer et al. (2) and Bae et al. (4). Most UV crosslinks through Lys involved uracil or cytosine: either uracil with neutral loss of H_2_O, or cytosine with loss of NH_3_ (these products are isobaric; Figure 1C; Supplementary Table S2). Cytosine and uracil crosslinks are only distinguishable from one another when no neutral loss of NH_3_ or H_2_O is identified. Significantly fewer CSMs of UV crosslinks matched with peptides crosslinked to adenosine or guanosine (Figure 1B; Supplementary Table S2). Crosslinks of uracil to Tyr did not reveal loss of H_2_O (Figure 1D, Supplementary Table S2).

Unlike UV XL-MS, CSMs from DEB and NM XL-MS ribosome data identified mainly peptides crosslinked to purines. According to unique CSM numbers, 73% of DEB- and 63% of NM-induced crosslinks involved guanine. Adenine was involved in 17% DEB and 22% NM crosslinks. Only a few of the chemical crosslinks were observed to pyrimidines. Both DEB and NM crosslinks, in absolute CSM numbers, were mainly formed to Lys, Met, Glu, His, and Cys, with preferential crosslinking to Glu or Met (Figure 1B; Supplementary Table S2). Normalizing the absolute CSM numbers by the frequencies of the respective amino acids in the ribosomal proteins revealed Cys, Met, and His as preferred residues for NM and DEB crosslinking (Supplementary Figure S3A). MS2 spectra of peptides crosslinked with DEB unambiguously revealed Glu as crosslinked amino acid (Figure 1E), which differs notably from the reactivity of mainly aromatic or basic residues in other types of protein–RNA crosslinking. MS2 spectra of His crosslinks show prominent immonium ions of His with mass adducts of the crosslinker remnant and/or crosslinker remnant plus the crosslinked nucleobase/nucleotide of adenine and guanine (Figure 1F). For peptides with Met crosslinked by DEB/NM, we frequently observed specific loss of C_2_H_2_S from Met on precursors and fragment ions (Figure 1G). As a result, the MS2 spectra contained high-intensity marker ions comprising the linker remnant of DEB/NM, the crosslinked nucleobase/nucleotide plus C_2_H_2_S. Furthermore, marker ions of only DEB/NM plus C_2_H_2_S were observed.

In MS2, spectra of FA crosslinks differed fundamentally from those of DEB and NM. FA MS2 spectra revealed specific and highly abundant marker ions (FA plus nucleobase/nucleotide) and comprehensive peptide sequence coverage by unmodified fragment ions (Figure 1H). The absence of crosslinked peptide fragment ions, presumably due to the unstable peptide-FA bond, simplified the MS2 spectra and improved database search efficiency, but prevents assignment of the crosslinked amino acid (67). Therefore, FA presets contained only a small number of fragment adducts (Supplementary Table S1). 52% of unique FA crosslinks involved adenine, followed by 25% guanine, and 23% cytosine (Figure 1B; Supplementary Table S2).

In conclusion, chemical XL-MS using DEB, NM, or FA adds complementary information by crosslinking amino acids mainly to purines, while UV crosslinks amino acids (mainly) to pyrimidines. The incorporation of assembled crosslink mass adducts within dedicated presets in NuXL facilitates rapid, FDR-controlled assignment of crosslinking sites.

### NuXL – a nucleic acid–protein XL-MS search engine for automated and sensitive crosslink site identification

Our novel software tool NuXL identifies crosslinked peptide-RNA/DNA (oligo)nucleotides and crosslinked amino acids rapidly and sensitively (for more details, see Supplementary Methods). NuXL is an open-source software and has been developed using the OpenMS software framework (68). NuXL is available as standalone version as part of the upcoming OpenMS version and also available as a free node for the Thermo Scientific Proteome Discoverer (PD) software platform (for online tutorials, see https://openms.de/applications/nuxl/) (69). NuXL generates extensive reports at the CSM, peptide, and protein levels, including output for visualization of fragment annotations in the separate viewer application TOPPView for the standalone version of NuXL and in PD (70). NuXL allows the user to choose between crosslinker-specific presets (conventional UV, DEB, NM, FA, 4SU, and 6SG; Figure 2A and Supplementary Table S1) that describe the relation between fragment adducts and cross-linked precursor adducts. In the standalone version of NuXL, these presets can be changed by the user to support novel protocols. The NuXL algorithm processes XL-MS data by MS2 spectra pre-processing, theoretical spectrum generation and spectrum-matching, crosslinked amino acid localization, scoring, and subsequent protein inference (Figure 2B; see Supplementary Methods for extended description). For spectrum matching and scoring, we designed subscores that consider XL-MS-specific spectrum features: precursor ions with crosslinker-specific (oligo)nucleotide mass adducts and their associated neutral losses; precursor adduct and cross-linked nucleotide-dependent MS2 fragment ions with (oligo)nucleotide mass adducts (including losses); and singly charged marker ions (i.e., unmodified or chemically crosslinked nucleobase, nucleoside, nucleotide and immonium ions; Figure 2C; Supplementary Table S3). To increase CSM identification rates, subscores are combined and rescored using the Percolator algorithm (71). Lastly, the CSM-FDR is calculated; protein groups are inferred from all peptides. NuXL localizes crosslink sites from the pattern of unmodified and crosslinked fragment ions and from the presence of crosslinked immonium ions. Spectra with peak annotations can be visualized in TOPPView and in PD. Comprehensive tutorials are provided online for both the standalone version and the PD version of NuXL (https://openms.de/applications/nuxl/).

**Figure 2.**
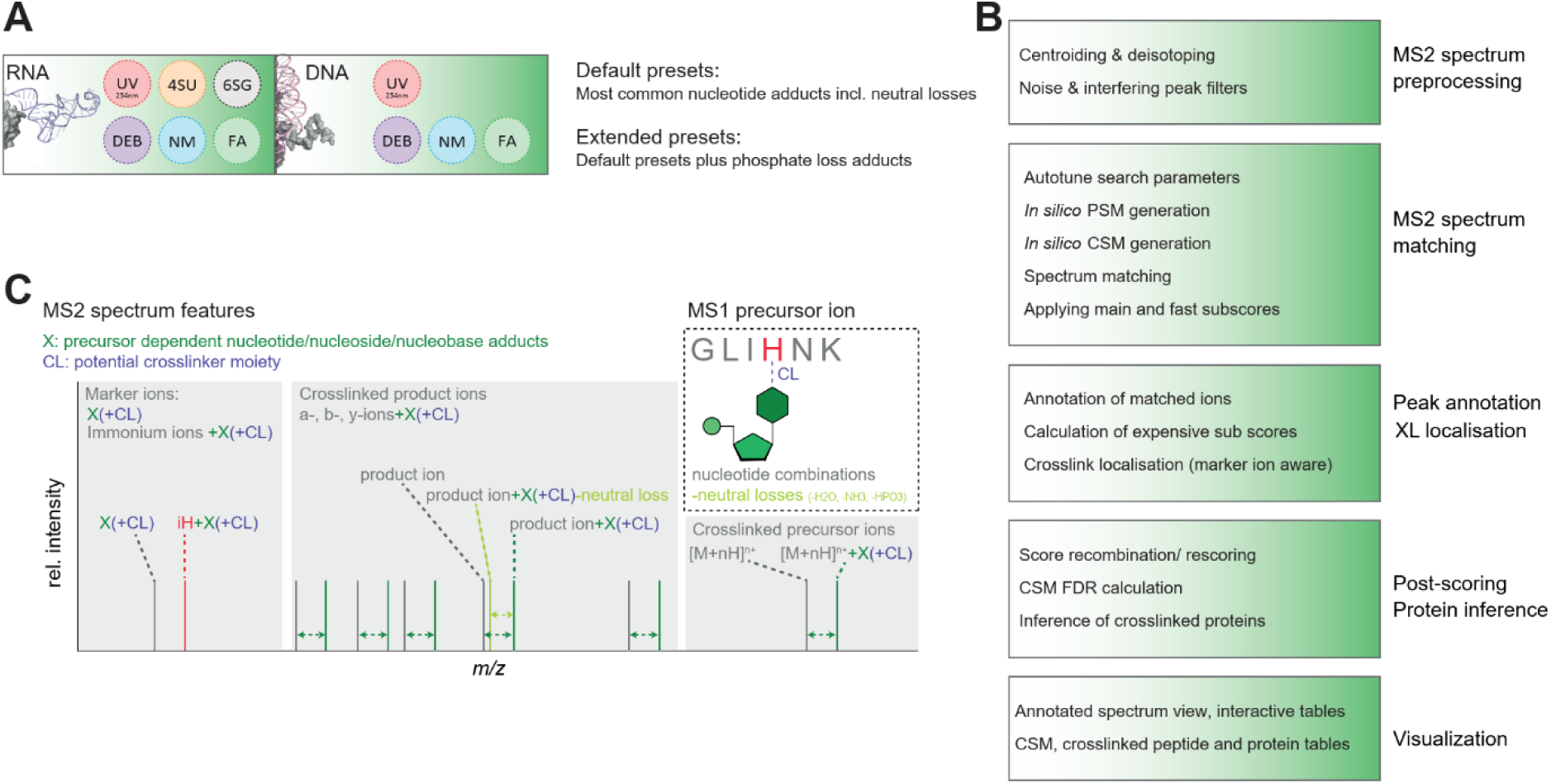
NuXL software tool for the analysis of XL-MS data. **(A)** Crosslinking reagents supported by NuXL database search tool. Available RNA-, DNA- and crosslinker-specific default and extended NuXL presets. **(B)** NuXL pipeline. XL-MS data-processing includes MS2 spectra pre-processing and spectrum-matching; peak annotation and localisation of crosslinked amino acid; post-scoring using the Percolator algorithm and protein inference; visualization of the annotated spectra and generation of tables of results. **(C)** MS2 spectrum features considered in (sub)score calculation. The main score and subscores used by the Percolator algorithm consider the presence and intensities of MS1 and MS2 precursor ions with crosslinker-specific (oligo)nucleotide mass adducts (X(+CL)) including various oligonucleotide combinations and related neutral losses, precursor adduct-dependent MS2 fragment ions with unmodified or chemically crosslinked (oligo)nucleotide mass adducts (a/b/y-ions+X(+CL)), and singly charged marker ions (nucleobase/nucleoside/nucleotide, X(+CL); immonium ions, iX+X(+CL)).

We evaluated NuXL performance using datasets generated for this study including *in vitro* and *in vivo* UV, DEB, NM and FA XL-MS data (Supplementary Tables S2, S4-6). Compared with our previous approach in RNP^xl^, the XL-MS data analysis time with NuXL is decreased by several orders of magnitude (for working times see Supplementary Results); this is achieved by implementing NuXL as a specialized search engine with full control over the internal search process, fragment-ion generation, scoring, optimized spectrum comparison, parallelization and caching of MS2 spectra generated *in silico* (2, 72). Score combination of XL-specific subscores (for details, see Supplementary Methods) and rescoring using Percolator (71) is fundamental to NuXL’s sensitivity and improved crosslink identification rates at the CSM level for all protocols (1.2- to 8.1-fold higher identification rates). We compared NuXL to the established MSFragger custom offset workflow (XRNAX MassOffset (8) plus Percolator rescoring, mass shifts from NuXL presets) using our *E. coli* crosslink data. Our findings revealed a high degree of overlap in identified crosslinks (mean 81% shared CSMs; 88% shared proteins, across all protocols). NuXL showed a higher yield in CSMs. Averaged among different crosslinkers for *E. coli* dataset, 104% CSMs were uniquely identified by NuXL vs. 19% CSMs uniquely identified by MSFragger; 74% distinct proteins were unique to NuXL vs. 12% unique to MSFragger, given that MSFragger output is taken as 100%. When we compared NuXL with MSFragger-Labile search (73), NuXL achieved comparable or higher gains (28% CSMs unique to NuXL vs. 23% CSMs unique to MSFragger; 36% distinct proteins unique to NuXL vs. 15% unique to MSFragger). Percentages are relative to the total number of CSMs or proteins reported by the MSFragger protocols. For further details on the search engine comparison, see Supplementary Results and Supplementary Figure S4.

### UV combined with chemical crosslinking provides spatial constraints in structural modelling of protein-RNA complexes

UV XL-MS resolved at the amino-acid level provides high-resolution spatial information on RNA interaction interfaces of proteins, which proved to be valuable for understanding the three-dimensional arrangement of biomolecular complexes (2, 9–11, 65, 74). Our initial crosslinking experiments with isolated *E. coli* ribosomes demonstrated that chemical crosslinkers connect different amino acids and nucleotides from those connected by UV crosslinking. We therefore applied UV, DEB and NM XL-MS and NuXL for the analysis of different protein–RNA complexes reconstituted *in vitro*, Hsh49/Cus1-U2 snRNA (27), Dnmt2-tRNA^Asp^ (33), and NELF complex HIV TAR RNA (75), highlighting XL MS’s role in structural analysis, particularly for RNPs in which the 3D protein structure, but not the 3D RNA structure, was resolved. FA XL-MS was excluded in this set of experiments because of its limited ability to provide crosslink site information.

We analysed yeast spliceosomal protein Hsh49 in complex with Cus1 and U2 47nt-snRNA (26). Hsh49 contains two RNA recognition motifs (RRM), and thus is a suitable example of a protein with a well-studied RNA-binding domain (27, 76). We UV-irradiated the reconstituted Hsh49/Cus1-U2 RNA complex, or titrated it with chemical crosslinkers at various concentrations, and analysed the XL-MS data. For all crosslinking conditions, we identified crosslink sites with CSM numbers increasing gradually with increasing concentrations of the chemical crosslinker. UV and chemical crosslinking sites are located in both RRM domains of Hsh49 as well as in its regions not resolved, and in Cus1 protein (Figure 3A, Supplementary Figure S5, Supplementary Table S4). The N-terminal RRM domain (RRM1) and the loop structure between two domains of Hsh49 contained more crosslinked amino acids shared between at least two crosslinkers (Figure 3A), while only DEB crosslinking revealed interaction of the RRM2 with U2 47nt snRNA. This result may be consistent with the observation that the N-terminal RRM of Hsh49 was shown to be more important for U2 snRNA binding than the C-terminal one (27). The loop region connecting RRM1 and RRM2 of Hsh49 was crosslinked more with chemicals, than with UV (in terms of CSMs). The importance of loop region connecting two RRMs was recently shown by Keil et al (11) for another yeast RNA-binding protein, Npl3, where mutation of a crosslinked amino acid in such a loop region had severe impact on the function of the protein and, eventually, on the growth of the yeast cells.

**Figure 3.**
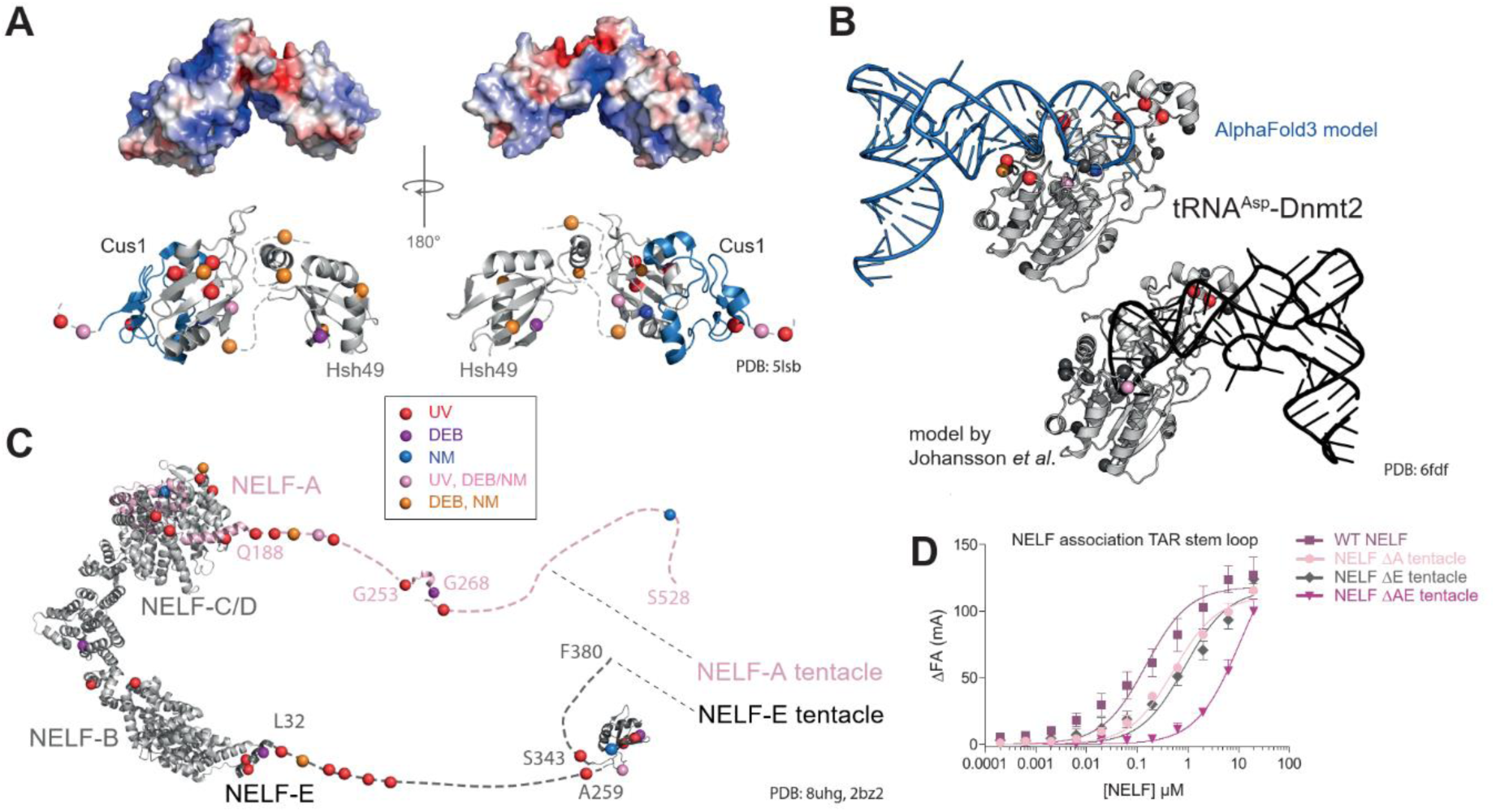
UV, DEB, NM XL-MS in integrated structural modelling. **(A)** 3D structure of yeast spliceosomal protein Hsh49 in complex with Cus1 (PDB: 5lsb (27)). Surface representation of electrostatic potentials depicted as a colour gradient of ±5.0 k_B_T/e. Hsh49-Cus1 RNA- (oligo)nucleotide-crosslinked amino acids (Cɑ) colour-coded depending on crosslinkers are shown based on CSMs filtered for 1% FDR, localization score > 0, and CSM counts of crosslinked peptides >2. In lower panels, Hsh49 and Cus1 proteins are shown in grey and blue, respectively. The parts of chains not resolved structurally are shown schematically by dashed line. Crosslinking was done at 10 min UV irradiation, 10 mM DEB, and 1 mM NM, respectively. **(B)** 3D model of *S. pombe* methyltransferase Dnmt2 (PDB: 6fdf (33)) in complex with tRNA^Asp^. Black cartoon tRNA representation: the published model based on UV XL-MS data. Blue cartoon representation: a new model built using AlphaFold3 (36) based on UV and DEB/NM XL-MS data based on CSMs filtered for 1% FDR, localisation score ≥0, CSM score >2. RNA-(oligo)nucleotide-crosslinked amino acids that fit to each model are highlighted as in panel (A), while crosslinked amino acids that do not fit are shown in black. Crosslinking was done at 10 min UV irradiation, 50 mM DEB, and 10 mM NM, respectively. **(C)** 3D structure of human negative elongation factor (NELF) complex (PDB: 8uhg (78), NELF complex as a part of paused transcription complex Pol II-DSIF-NELF; 2bz2 (111), RRM domain of NELF-E) including RNA-(oligo)nucleotide-crosslinked amino acids color-coded and filtered as in panel (A). Intrinsically disordered regions (tentacles) of NELF-A and NELF-E are indicated by pink and grey dashed lines including relative RNA-(oligo)nucleotide-crosslinked amino acid positions. NELF-A and NELF-E are shown in pink and dark-grey, respectively, Both NELF-B and NELF-C/D are shown in light grey. The NELF-A tentacle part with a determined structure is spatially located in correspondence with the whole structure of the NELF complex; in contrast, the spatial position of the RRM domain in the NELF-E tentacle is unknown and shown arbitrarily. Crosslinking was done at 10 min UV irradiation, 50 mM DEB, and 10 mM NM, respectively. **(D)** Binding of WT NELF, NELF ΔA tentacle, NELF ΔE tentacle, and ΔAE tentacle to the TAR stem-loop measured by fluorescence anisotropy.

We aimed for determining how the *in vitro* crosslinking results of the Hsh49/Cus1-U2 RNA complex correspond to the existing structures of this complex in spliceosomes. However, in the only available 3D structure of yeast spliceosome containing Hsh49-Cus1 complex (77), the proteins are located in poorly resolved regions with multiple RNA regions not modelled at all. Crosslinking of reconstituted spliceosomal complexes containing Hsh49-Cus1 may further improve understanding of their structural interaction with RNAs in spliceosomes.

We also applied UV, DEB and NM XL-MS to obtain additional spatial restraints for the support of *in silico* models of *S. pombe* methyltransferase Dnmt2 in complex with its cognate tRNA^Asp^ (33). We titrated the reconstituted complex with various chemical crosslinker concentrations and analysed the XL-MS data (Supplementary Figure S6). Recent advances in AI-driven structure prediction, particularly AlphaFold3 (36), have enabled unprecedented accuracy in protein-RNA complexes modeling. Motivated by these improvements, we revisited the previously published atomic model of *S. pombe* methyltransferase Dnmt2 in complex with its cognate tRNA^Asp^ (27). To explore potential structural diversity, we generated 300 spDnmt2– tRNA^Asp^ complexes using AlphaFold3, varying the initial random seed. Remarkably, all predictions converged on a consistent binding mode, with only minor variations in tRNA loop conformations (Supplementary Figure S7). This AlphaFold3-derived model reveals a markedly different tRNA^Asp^-Dnmt2 interaction compared to the previously proposed model, which was based solely on UV crosslinking data (Figure 3B, Supplementary Figure S7). In the revised model, Adenine 37 is deeply positioned within the cavity containing the methyl-donating S-adenosylmethionine (SAH), while Cytosine 38, the target nucleotide for modification, is placed above the ribose moiety of SAH. The biological relevance of this prediction is further reinforced by the placement of Guanine 34 directly above Cytosine 38, structurally elucidating the enhanced catalytic efficiency of Dnmt2 in the presence of the hypermodified nucleoside queuosine (Q) at position 34 of tRNA^Asp^. Remarkably, despite the fact that AlphaFold 3 prediction was generated without spatial constraints or prior definition of the protein binding site and its function, the model exhibits a markedly improved fit to cross-linking data compared to previous model. In details, the AF3-based model provided single occurrences of crosslinks fit to the model (occurrences within threshold distances to the tRNA: d0=6, d2=10, d3=11, d5=12, see Materials and methods section for more detail) and multiple (ALLd0=6, ALLd2=10, ALLd3=16, ALLd5=20). In the previously published model, there were fewer single occurrences (d0=4, d2=6, d3=8, d5=9) and multiple occurrences (ALLd0=6, ALLd2=6, ALLd3=9, ALLd5=11). Consequently, chemical cross-linking provides additional spatial constraints that complement UV cross-linking and further enhance experimental mass spectrometry (MS)-based validation of structural models, establishing it as an unprecedented tool for modelling ribonucleoprotein (RNP) complexes.

Finally, we analysed the negative elongation factor (NELF) complex with its four subunits NELF- A/B/D/E, in complex with HIV TAR RNA (29). For this, we titrated the reconstituted complex with the chemical crosslinkers in various concentrations and performed XL-MS analysis (Figure 3C, Supplementary Figure S8, Supplementary Table S4). We identified numerous crosslink sites in the so-called tentacle regions located at the C-termini of NELF-A and NELF-E, most of which are disordered. The tentacle of NELF-E contains an RRM domain and crosslinks by all types of crosslinker to the RNA (Figure 3C). A recent study modelled the structure of the residues 253– 266 in the NELF-A tentacle using CryoEM data (78) and highlighted its role in interaction with the conservative protrusion of the RNA polymerase II subunit, RPB2. Residues in within this particular NELF-A region are believed to interact with the DNA–RNA duplex in the complex (78). Accordingly, three crosslinks generated by UV and DEB are clustered around this region in the NELF-TAR interaction (Figure 3C). In addition, fluorescence anisotropy experiments with mutants of the NELF complex showed significantly reduced binding of TAR to RNA upon deletion of intrinsically disordered regions of either NELF-A or NELF-E, and an even greater loss in binding affinity when both regions were deleted (Figure 3D).

With regards to database search space of crosslinks for these and other samples of low complexity (i.e., mononucleosomes, see below) we used mainly a database comprising the proteins within the sample, if preparations of protein expressed *in vitro* revealed sufficient purity, which can be monitored by proteome analysis of the preparation of the reconstituted protein–RNA/DNA complex). We recommend that in these cases the MS2 spectra, and hence the crosslinks need to be validated by applying additional filters or manual inspection (for example, with TOPPView output or PD output), as relying only on the FDR is not possible in these cases. According to our observations, the search against extended databases, e.g., MaxQuant contaminants, in absence of these contaminants in the samples of *in vitro* reconstituted complexes, could decrease the sensitivity of the searches, such that crosslinks with a small peptide moiety can be wrongly attributed to irrelevant peptides, despite the high-quality spectra. Overall, our data demonstrate that UV and chemical crosslinking cover different sites within RNPs reconstituted *in vitro*, and hence are valuable for mapping extended RNA-binding interfaces in proteins and for providing suitable restraints for in silico docking of RNA to proteins.

### NuXL facilitates identification of UV and chemical *crosslinking sites* in *E. coli*

To assess chemical XL-MS and NuXL *in vivo*, we applied our method to *E. coli* cells. After crosslinking with UV, DEB, NM, or FA, cells were lysed, enriched for crosslinked peptide- (oligo)nucleotides, and analysed with LC-MS. We note that for FA XL-MS we used a different workflow based on silica enrichment subsequent to TRIZOL extraction of intact (crosslinked) RNA (see Materials and Methods and Discussion). NuXL identified 4,924 unique crosslinked peptides across 1,502 proteins, with varying numbers of peptides and proteins identified by each XL-MS method (1,007, 2,715, 2,805 and 670 crosslinked peptides, and 393, 952, 1,186 and 302 crosslinked proteins, were identified by UV, DEB, NM and FA XL-MS, respectively) (Figure 4A, Supplementary Table S5). 162 proteins were found crosslinked under all four crosslinking conditions. In accordance with the results for crosslinking of isolated *E. coli* ribosomes (Supplementary Table S2), UV crosslinked mostly pyrimidines (51% cytosine/uracil, 39% uracil, 2% cytosine) and the chemical crosslinkers most frequently gave purine crosslinks (63%, 70%, 51% guanine, and 32%, 26%, 38% adenine for DEB, NM, and FA, respectively; Figure 4B). The most frequently identified crosslinked amino acids in unique crosslinks in UV XL-MS *E. coli* data were Lys (41%), Tyr (14%), and Cys (7%). For DEB XL-MS, we identified Met (42%), His (14%), and Lys (11%); for NM XL-MS, Met (64%), His (9%), and Cys (5%) (Figure 4B). After normalizing the absolute CSM numbers by the amino acid frequencies of their occurrence in the sequences of all crosslinked *E. coli* proteins, Cys, His, and all aromatic amino acids were found to be the most frequently UV-crosslinked amino acids, which is consistent with results of previous studies (4, 65) (Supplementary Figure S3B). Since we identified crosslinking sites of Phe and Trp in the *E. coli* XL-MS data, but not in isolated ribosomes (Supplementary Table S2), we conclude that their absence in *E. coli* ribosomes may be a specific characteristic of ribosomal RNA-binding proteins. This observation can be partially attributed to the reduced frequencies of Phe and Trp in the sequences of 58 *E. coli* ribosomal proteins when compared to the overall background of the whole proteome (3.1% vs. 3.8% for Phe, and 0.6% vs. 1.5% for Trp, Fisher exact test p-value for both < 0.01). Crosslinking by UV, DEB, and NM in *E. coli* dataset at specific positions was independent on the surrounding sequence context as shown in Figure 4C.

**Figure 4.**
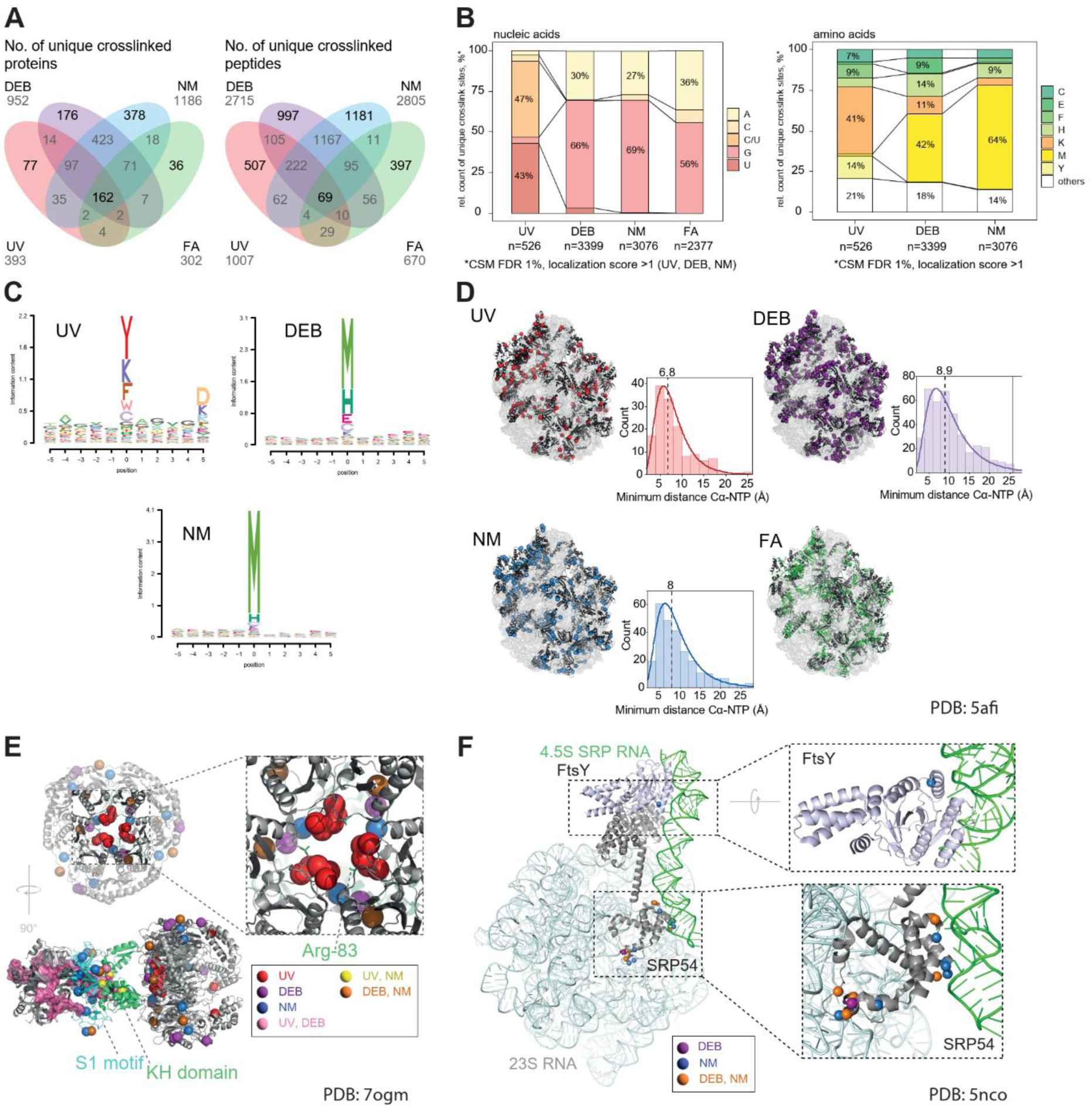
*In vivo* UV and chemical XL-MS of *Escherichia coli* cells. **(A)** Venn diagrams of unique crosslinked proteins (left) and peptides (right) for UV, DEB, NM and FA XL-MS of *E. coli* cells. Identifications are filtered for 1% FDR at CSM level. **(B)** Left panel, stacked bar plot (normalised to 100%) of relative counts of observed crosslinked nucleobases in unique crosslinked peptide-(oligo)nucleotide pairs resulting from UV, DEB, NM and FA XL-MS of *E. coli* ribosomes. CSMs were filtered for 1% FDR. Ambiguous, isobaric C–NH_3_/U–H_2_O crosslinks were counted as C/U. Labels are displayed only for values of 7% and above. Right panel, 100% stacked bar plot of relative counts of observed crosslinked amino acids in unique crosslinked peptide-(oligo)nucleotide pairs resulting from UV, DEB, NM XL-MS of *E. coli* ribosomes. CSMs were filtered for 1% FDR and a localisation score ≥1. Labels are displayed only for values of 7% and above. FA data are excluded from the visualisation due to low numbers of properly assigned crosslinked amino acids. Absolute CSM counts for each bar are depicted below the plots. **(C)** Logo plots of amino acid positional frequency enrichment around the RNA-XL sites for UV, DEB and NM. The localisation score threshold of ≥1 was used to compare the detected CSM counts per position against the amino acid frequency in the MS-detectable *E. coli* tryptic peptides. **(D)** Cartoon representation of cryo-EM 3D structure of *E. coli* 70S ribosome-EF-TU complex (PDB: 5afi (79)) and histograms of crosslink distances. In the structures, RNA-(oligo)nucleotide-crosslinked amino acids are highlighted as spheres (Cα atoms) in red (UV), purple (DEB), or blue (NM); or as green cartoon (FA). Crosslink sites are based on CSMs filtered for 1% FDR for all conditions, and localisation score ≥1 for UV, DEB and NM. Histograms display distances (in Å) between UV (left), DEB (centre left) and NM (centre right) crosslinked amino acids (Cɑ) and closest atom of the closest nucleotide in the *E. coli* 70S ribosome-EF-TU complex structure. **(E)** Cartoon representation of the cryo-electron microscopy structure of a trimer of *E. coli* polynucleotide phosphorylase (PNPase; grey cartoon) in complex with six copies of *E. coli* RNA-binding protein Hfq (grey cartoon) and the 3’external transcribed spacer RNA of leuZ (pink cartoon/surface; PDB: 7ogm (81)). PNPase KH domain and S1 motif are highlighted in limegreen and cyan, respectively. The Arg-83 residue involved in RNA binding is highlighted by green sticks (80). Alpha-carbon atoms (Cɑ) of RNA-(oligo)nucleotide crosslinked amino acids are highlighted as red (UV), purple (DEB), blue (NM), pink (UV and DEB), yellow (UV and NM), or orange (DEB and NM) spheres. Crosslinks are filtered for a localisation score ≥1. Crosslinks mapped on the structure of another member of *E. coli* degradasome, enolase, are shown in Supplementary Figure S9. **(F)** Cartoon representation of the cryoEM structure of *E. coli* ribosome with signal recognition particle bound; for simplicity, ribosomal proteins are not shown (PDB 5nco (84)). Color scheme of chemical crosslinks is as in panel E. Mapped crosslink positions are based on CSMs filtered for 1% FDR, and localisation score >0.

We mapped *in vivo* UV, DEB, and NM crosslink sites and FA crosslinked peptides of ribosomal proteins onto the 70S *E. coli* ribosome structure (Figure 4D) (79). For this, we filtered crosslink sites for localisation score ≥1, as judging from manual spectrum validation, this threshold secures reliable amino acid localisation. We also calculated distances between Cα atoms of crosslinked amino acids and the closest atom of the closest nucleotide for UV, DEB and NM. The median distances found for UV, DEB, and NM were 6.8, 8.9, and 8 Å, respectively, which agrees well with calculated spacer lengths and consideration of structural flexibility (we accept maximum distances of 10, 15 and 15 Å for UV, DEB and NM, respectively) and underscores the specificity of chemical crosslinking for *in vivo* crosslink analysis (Figure 4D).

We also located the *in vivo* XL-MS sites of the polynucleotide phosphorylase (PNPase) on available 3D structures of the protein trimer in complex with six copies of HfQ and RNA (Figure 4E) (80, 81). PNPase UV-crosslinking sites are positioned in the cavity of the PNPase tetramer core, and their locations align with the amino acids responsible for the RNA degradation activity of the PNPase (81). Despite the fact there are amino acids, such as lysines in the cavity that are potentially chemically crosslinkable, we did not identify these as crosslinked to RNA. This might indicate that spatial constraints in this cavity/channel, when occupied with RNA, exist that prevent penetration of the chemical crosslinker NM and DEB inside the cavity/channel. Chemical crosslinking sites are located mainly in, or very close to the S1/KH domain of the PNPase that binds to (small) RNAs and guides them towards degradation. PNPase forms the *E. coli* degradasome with proteins RNaseE, RhlB, and enolase and we also identified crosslinking sites in these proteins (Supplementary Table S5, Supplementary File S1) (82). Enolase also plays an essential part in the glycolysis pathway; the crosslink sites are scattered across the enzyme, with no visible tendency to cluster, for example, at positively charged surface patches. This finding seems to contradict the specific binding of nucleotides between the enolase dimer, even though there we found several crosslinks located in the cleft between two enolase monomers in close vicinity to the active site of the enzyme (Supplementary Figure S9). In addition, it was shown that, in mammalian cells, enolase can bind hundreds of mRNAs which regulate its enzymatic activity, the effect which could be conservative through different kingdoms of life (83). Hence, the different crosslink sites might reflect the various contacts of the different RNAs.

To illustrate the capacity of the chemical crosslinker to link double-stranded RNA to a protein, we mapped chemical crosslinks on a structure of signal recognition particle (SRP) in contact with 23S ribosomal RNA and 4.5S SRP RNA (Figure 4F) (84). Crosslinked amino acids are in alpha-helices of SRP54 and FtsY proteins in immediate proximity either to 4.5S RNA of SPR or to 23S rRNA.

### Properties of crosslinked *E. coli* proteins

Because of the large number of detected crosslinking sites, we aimed to assess whether crosslinking is associated i) with the abundance of crosslinked proteins in the *E. coli* proteome (Supplementary Table S6), thus possibly reflecting a “crowding effect” (i.e., less specific crosslinking of only highly abundant proteins to nucleic acid in the crowded cellular environment), in particular when chemical crosslinkers are used, and ii) with the molecular features of the crosslinked proteins. For the latter, we integrated residue properties and 3D structure annotations (see below).

To investigate the abundance of crosslinked proteins detected across XL-MS methods, we applied label-free MS to *E. coli* cells grown under the same conditions as for our crosslinking experiments and also used abundance values available for *E. coli* K12 in the PaxDb (43) and PaxDb_integrated across strains (Supplementary Figure S10A, see also Materials and Methods). In general, we observed a moderate 0.42-0.53 Pearson correlation values between the number of CSMs and protein abundance and 0.48-0.54 when comparing the number of CSMs to the RBP2GO score (Supplementary Figure S10B). According to STRING (56) and RBP2GO (44) annotation, putative RNA-binding proteins are naturally highly abundant in *E. coli,* while most proteins without RNA-related functions have lower abundance. Nevertheless, there also exists a group of high-abundant proteins without annotated RNA-related functions. Our own proteome analysis is mainly in agreement with this annotation; however, we detected a small subset of abundant non-RNA-binders as well, possibly reflecting molecular crowding. With regard to the crosslinked proteins, we observed that DEB and NM crosslinking covered most annotated RNA-binders, whilst UV and especially FA identified predominantly top-abundant proteins. Crosslinks were mainly identified in high-abundance RNA-binder proteins, but also (upon DEB and NM treatment) in the low-abundant putative RNA-binders and the above-mentioned subset of non-RNA-binder abundant proteins. We compared the iBAQ values of identified proteins in our *E. coli* proteome study and the number of potentially crosslinkable peptides (which is higher in abundant RNA-binders). DEB and NM crosslinking managed to identify less abundant and crosslink-reactive proteins (Supplementary Figure S10C). We observed crosslinking only for a small fraction of other high-abundant proteins and no stochastic crosslinking of lower-abundant proteins without RNA-related functions.

When comparing the difference between the RNA-binding and non-binding protein abundance (Supplementary Figure S10D), we observe that the correlation of abundance and RNA-binding in *E.coli* proteome is reflected in all datasets, biasing the FA the strongest, followed by UV, and with NM and DEB being the least affected by molecular crowding.

We statistically tested the enrichment of amino acid residue features of crosslinked amino acids and peptides against an expected background distribution of *E. coli* peptides, considering peptide length and tryptic missed cleavages and protein abundance (see Methods for details). The DESCRIBEPROT database protein annotations were transferred for all identified crosslinked peptides (for FA XL-MS) and sites (for UV, DEB and NM XL-MS) at the residue level. We found that secondary structure elements were only mildly associated with crosslinking events (Figure 5A, Supplementary Table S7). The analysis of residue properties revealed the under-representation of putative disordered residues at the crosslink sites, but an overall enrichment of disordered RNA and DNA binders on the crosslinked peptides (Figure 5A). The strongest enrichment was observed for specific amino acids at crosslinked sites (Supplementary Figure S11), confirming the amino acid preferences displayed in Figure 1B and Figure 4B. We investigated the correlation of abundance and RNA-binding by sampling the background peptide distribution with regard to the protein abundance, resulting in a more abundant background compared against the detected peptide set. We observed that residue properties, such as solvent exposure and evolutionary conservation of the crosslinked amino acids did not change much, whilst the enrichment of RNA-binding decreased, yet retaining the moderate enrichment upon almost full abundance bias compensation (Supplementary Figure S11).

**Figure 5.**
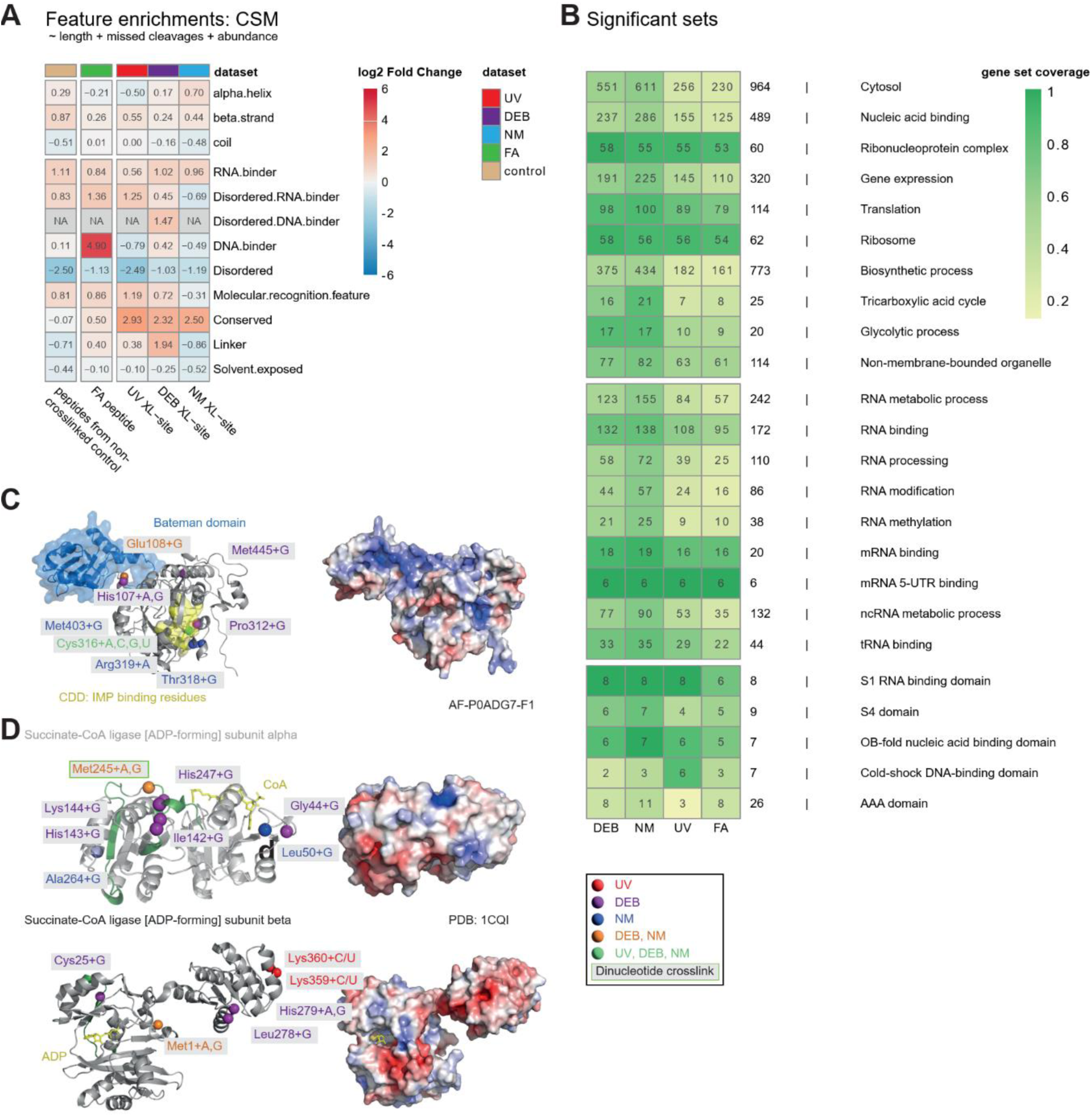
Crosslink-assisted functional and structural protein characterization. **(A)** A heatmap of log2 Fold Changes between the experimental and an *in-silico* background distribution mean feature counts per residue. The background sampling was performed with regards to the experimentally observed peptide length and missed cleavage distribution and source protein quantity to compensate for the abundance bias. The non-crosslinked control and FA datasets were analysed on the peptide level, whilst the UV, NM and DEB enrichment was analysed on crosslinked residue level. The NA values correspond to the cases where the sampling or statistical testing could not be performed due to the feature distribution violating the method assumptions. **(B)** A gene set over-representation by the crosslinked protein sets. Reported are the terms significant in at least one crosslinking method (p.adjusted BH <= 0.01 for all sets and 0.05 for protein domain sets). The gene set coverage is colour coded, with the number of identified proteins noted in the heatmap cells and the full gene set size next to the corresponding rows **(C)** Cartoon/surface representation and electrostatic surface of AlphaFold-predicted 3D structure of *E. coli* inosine-5’-monophosphate dehydrogenase (AF-P0ADG7-F1) with RNA-(oligo)nucleotide-crosslinked amino acids (Cɑ) highlighted as purple (DEB), blue (NM), orange (DEB and NM) or lime-green (UV, DEB and NM) spheres. Mapped crosslink positions are based on CSMs filtered for 1% FDR, and localisation score ≥1. The Bateman domain is highlighted as a transparent blue surface. Conserved residues (derived from conserved domain database CDD) involved in IMP binding are highlighted by a yellow transparent surface. **(D)** Cartoon representation and electrostatic surface of the crystal structure of succinate-CoA ligase subunit alpha (upper panel) with bound coenzyme A (yellow sticks) and subunit beta (lower panel) with bound ADP (yellow sticks) (PDB: 1cqi (112)). Crosslinks are mapped as in **C** with FA-crosslinked peptide displayed as green cartoon and UV-crosslinked amino acids (Cɑ) highlighted as red spheres. Met-245 crosslink found to a dinucleotide is marked by a green-outlined box.

Taken together, the preferential specificity of RNA-binding protein detection, the lack of crosslinkable residue saturation and the enrichment of RNA-binding residues beyond abundance bias all strongly support the specificity of our (chemical) crosslinking methods (Supplementary Figures S3B, S10A,C, S11) and confirm the importance of the observation that RNA-binding proteins are naturally highly abundant in *E. coli*.

It is important to note that the proteome of *E. coli* exhibits significant plasticity depending on the growth conditions (85). In this work, we characterized RNA-binding proteins in nutrient-rich media where cells strongly allocate their resources to translation. This may result in higher concentrations of RNA-binding proteins. Under starving conditions that are closer to natural situations, the distribution of different functional groups of proteins may differ from our observations.

Further, we analysed crosslinked protein sets under different MS-XL conditions for gene ontology (GO) term enrichment using STRING database annotations (Figure 5B, Supplementary Table S8). The most enriched GO terms are related to the cytosol, nucleic acid binding, and translation and those for biosynthetic processes. Regarding the latter, we found almost all proteins involved in glycolysis and the tricarboxylic acid cycle (TCA) and their crosslinked amino acids linked to mono- and dinucleotides. Other studies of the RNA interactome in *E. coli* and eukaryotes also identified metabolic enzymes crosslinked to RNA, but barely with their crosslinked amino acid (Supplementary Figure S12A) (6, 83, 86, 87). In glycolysis, 27% of enzymes were crosslinked across all XL-MS conditions, and 80% under at least one (Supplementary Figure S13A). Over half of these enzymes were linked to dinucleotides or cytosine/uracil, indicating RNA-binding properties. Similar trends were observed in the TCA pathway (Supplementary Figure S13B). Our analysis also revealed an overrepresentation of coding and non-coding RNA-related and metabolic terms and domains, including RNA-binding domains, as well as S1 and OB fold domains, and cofactor-binding domains such as the AAA domain (Figure 5B).

To globally determine if (RNA) crosslinking in cofactor-binding metabolic proteins occurs at consistent or varying sites, we mapped UV, DEB, and NM crosslink sites onto the sequences and structures of proteins (Supplementary File S1, Supplementary Figure S14). We assessed co-localization of crosslinked residues and functionally important amino acids from DESCRIBEPROT and InterPro databases. We compared the 3D distances between C_α_ of the crosslinked amino acids and the C_α_ of residues with annotated features against a background distribution (see Materials and Methods for details) and found that RNA-binding residues, disordered regions, conserved residues and molecular recognition features (i.e., protein interactions) were enriched within a 4 Å radius of crosslink sites (Supplementary Figure S14A-G, Supplementary Table S7, Supplementary Table S9). Crosslinked sites were significantly closer to RNA-binding regions and further away from DNA-binding areas (see below). Additionally, crosslink sites were enriched adjacent to coenzyme A and FAD/NAD/NADP binding sites. Crosslink sites across all methods were close to ATP-binding residues (Supplementary Figure S14H-L).

To further provide structural insights, we show protein structures with their bound mononucleotides/co-enzyme A in Figure 5C and D. For inosine 5’-monophosphate dehydrogenase (IMDH; AF-P0ADG7-F1), an enzyme involved in guanosine-nucleotide metabolism, amino acids crosslinked chemically to adenine and guanine are in close proximity to the Bateman domain of the predicted 3D structure (Figure 5C). The Bateman domain is known to be involved in RNA and ssDNA binding (88, 89). In addition, several crosslinks close to conserved residues in the active site, which binds IMP, were identified. The crosslink data support the suggestion of regulatory roles for RNA associated with ATP and GTP binding in the Bateman domain (90). We further mapped crosslinks to the structure of succinyl-CoA synthetase subunits alpha and beta (Figure 5D). In SUCD, crosslinks were identified close to bound coenzyme A, including a dinucleotide crosslink of unambiguous RNA origin. In SUCC, crosslinks are rather scattered across the whole structure, without any apparent tendency to cluster at the ADP-binding site. This may provide a hint toward the possible involvement of SUCC in RNA-related processes, which could bring RNA molecules close to the complex and lead to more dispersed crosslinks across this relatively small protein.

### Chemical *XL-*MS identification of protein-DNA crosslinks – DNA-specific NuXL settings

We adapted our chemical NuXL presets for the analysis of protein-DNA crosslink sites by comparing chemical and UV XL-MS of 601 DNA sequence (30) to yeast histone octamers as part of an *in vitro* reconstituted nucleosome (Figure 6, Supplementary Table S10). Sample processing, including crosslink enrichment, was performed as described above for the analysis of *E. coli* ribosomes. In addition to peptides crosslinked to the nucleobase moiety of DNA, we observed precursor-ion mass adducts of 178.00311 Da (C_5_H_7_O_5_P) or 196.01368 Da (C_5_H_9_O_6_P), matching deoxyribose-phosphate adducts with and without loss of H_2_O, respectively. Most likely, these are derived from abasic sites in the DNA generated by the crosslinker reaction and lead to the generation of a reactive aldehyde that crosslinks to the proteins (Supplementary Figure S15A-C) (91, 92). We did not observe a similar species in protein-RNA crosslinking.

**Figure 6.**
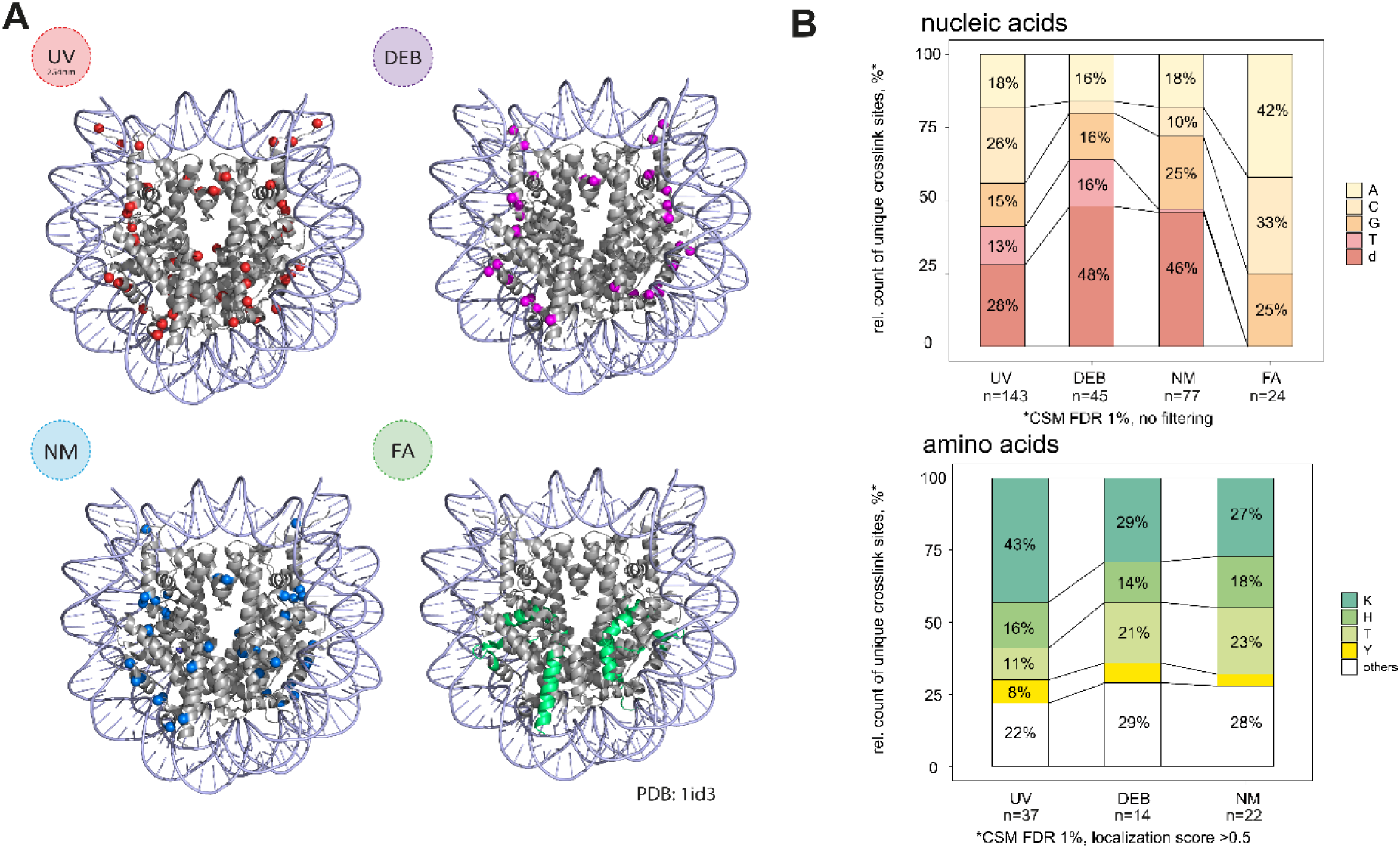
Protein–DNA UV and chemical XL-MS of *S. cerevisiae* nucleosomes. (**A)** *S. cerevisiae* nucleosome core-particle structure (PDB: 1id3 (113)) with DNA-(oligo)nucleotide-crosslinked amino acids (Cɑ) highlighted as red (UV), purple (DEB), or blue (NM) spheres based on CSMs filtered for 1% FDR, localization scores >0, and CSM counts of crosslinked peptides >2. For FA, whole crosslinked peptides filtered for 1% FDR, and CSM counts >2 are shown in green. **(B)** Upper panel, stacked bar plot (normalised to 100%) of relative counts of observed crosslinked nucleobases in unique crosslinked peptide-(oligo)nucleotide pairs resulting from UV, DEB, NM and FA XL-MS of *S. cerevisiae* nucleosomes. CSMs were filtered for 1% FDR. Labels are displayed only for values of 7% and above. Lower panel, 100% stacked bar plot of relative counts of observed crosslinked amino acids in unique crosslinked peptide- (oligo)nucleotide pairs resulting from UV, DEB, NM XL-MS of *E. coli* ribosomes. CSMs were filtered for 1% FDR and a localisation score ≥0.5. Labels are displayed only for values of 7% and above. FA data are excluded from the visualisation due to low numbers of properly assigned crosslinked amino acids. Absolute CSM counts for each bar are depicted below the plots.

UV, DEB, NM, and FA XL-MS sites were mapped to structures of the *S. cerevisiae* nucleosome core particle (Figure 6A). For UV, DEB, and NM, most crosslinked amino acids were found to be clustered in close proximity to the DNA. UV XL-MS mostly identified unique crosslinks to deoxyribose (28%) and cytosine (26%) with lysine as the most frequently crosslinked amino acid (43%; Figure 6B). DEB XL-MS identifies mostly crosslinks to deoxyribose (48% of unique crosslinks), followed by all bases except cytosine (all 16%) involving mainly lysine (28%) and threonine (21%) as crosslinked amino acids (Figure 6B). For NM XL-MS, deoxyribose crosslinks were also the most strongly represented (46%), and guanine was a next preferential moiety for crosslinking (25%). Amino-acid preferences of NM were similar to those of DEB. In the UV-irradiated sample, in addition to lysine and threonine, histidine and tyrosine were frequently crosslinked in histone sequences. The discrepancy between crosslinked amino acid frequencies in nucleosomes (Figure 6B) and in RNA crosslinking samples (Figures 1B,4B) may be explained by the sequences of the four yeast histones, which contain no cysteines and only one methionine (except for the starting residues). We measured distances between Cα atoms of crosslinked amino acid residues that were resolved in the yeast nucleosome structure (Figure 6A) and the closest atom of the closest nucleotide. The median of amino-acid-dNTP distances was 8.5, 7.6 and 8.1 Å for UV, DEB and NM, respectively. No significant difference between crosslinkers was observed in contrast to *E. coli* RNA crosslinking data, most likely, due to much fewer numbers of nucleosomal crosslinks. As in RNA–protein XL-MS experiments, in DNA– protein FA XL-MS the crosslinked amino acid could barely be identified in most cases. The yield (CSM and unique crosslinked peptides) of FA crosslinking was lower than that of the other crosslinking methods; one unique crosslinked peptide derived from H2A, three from H2B, one from H3, and one from H4. FA XL-MS identified no deoxyribose as crosslinked residue in contrast to the other chemical crosslinkers, which were dominant in NM and DEB, and hence more crosslinked peptides were identified with these chemicals (Supplementary Table S10).

### Crosslinking of DNA-binding proteins in *E. coli*

We searched our *in vivo* crosslink *E. coli* dataset with NuXL and DNA presets (Supplementary Table S11). We identified the cold shock domain containing Csp protein family members crosslinked to RNA and DNA according to chaperone properties in binding single-stranded RNA and DNA (93, 94). The crosslinking sites to RNA and DNA are at the same amino acids in these proteins. A similar pattern is present in the ssDNA binding protein SSB, in which the same residues crosslink to DNA and RNA. We also identified RNA and DNA crosslinks in the nucleoid double-stranded (ds)DNA-binding proteins H-NS, DBHA/DBHB, FIS, and IHF (Figure 7A). We mapped IHFA and IHFB crosslinks to the EM 3D structure of the Cas1-Cas2-IHF-DNA holo-complex (Figure 7A). The UV and chemical RNA crosslinking sites are located in close proximity to the dsDNA in the structure. Potential RNA-binding was previously suggested in acid-resistance studies (95). All other nucleoid proteins show RNA and DNA crosslinking at the same positions in the 3D structures, which indicates RNA-binding properties of nucleoid proteins in general (Figure 7A).

**Figure 7.**
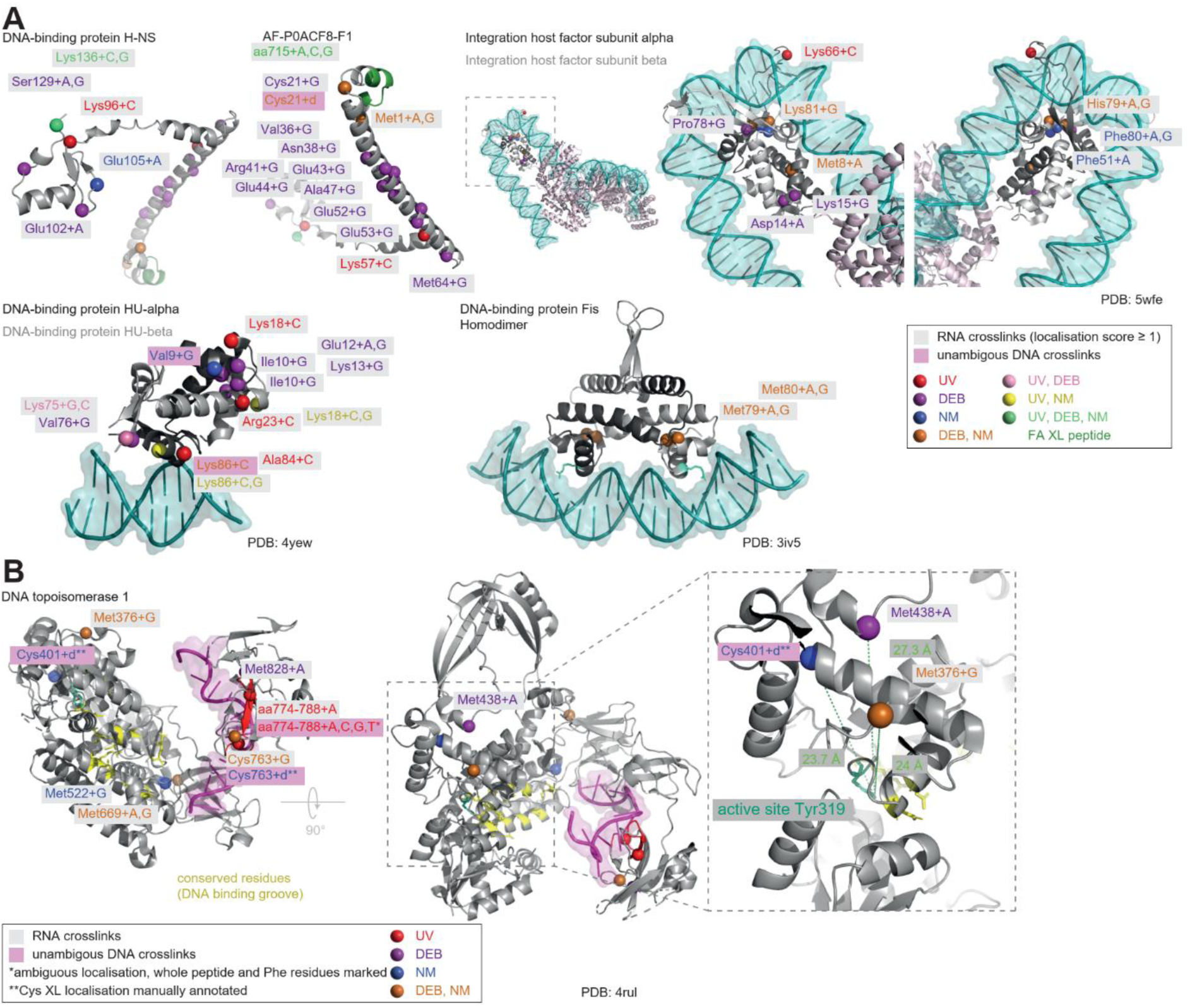
Examples of crosslinked DNA-binding proteins in E. coli. **(A)** Cartoon representations of nucleoid proteins HN-S (AF-P0ACF8-F1), integration host factor (subunit alpha and beta; *PDB:* 5wfe (114)), HU (alpha and beta; PDB: 4yew (115)), and FIS (as homodimer; PDB: 3iv5 (116)) crosslinked to RNA (and DNA) in *E. coli*. DNA is displayed as dark-cyan cartoon/surface. RNA- and unambiguous DNA-(oligo)nucleotide-crosslinked amino acids (Cɑ) are highlighted as red (UV), purple (DEB), blue (NM), light-pink (UV and DEB), yellow (UV and NM), orange (DEB and NM) or lime-green (UV, DEB and NM) spheres. FA-crosslinked peptides are displayed as green cartoon. RNA- and unambiguous DNA-crosslink labels are coloured grey and purple, respectively. Labels indicate crosslinked amino acids and directly crosslinked nucleic acid moiety. Crosslink sites are based on CSMs filtered for 1% FDR for all conditions, and localisation score ≥ 1 for UV, DEB and NM peptide-RNA(oligonucleotide)-crosslinks. **(B)** Cartoon representation of the crystal structure of topoisomerase I (PDB: 4rul (117)) with crosslinks and labels as in A. In the case of UV-crosslinked peptide spanning amino acids 774 to 788, cartoon is coloured in red and Cɑ of putatively crosslinked phenylalanines Phe-780 and Phe-786 are highlighted by spheres. For Cys-401, and Cys-763, the crosslink sites were manually annotated based on the reactivity of cysteine residues. Crosslinks are not filtered based on localisation score. Conserved residues of the DNA binding groove (CDD1: cd00186) and the active site Tyr-319 are highlighted by yellow and cyan sticks, respectively. The distances between Cɑ of crosslinked residues Cys-401 (23.7 Å) and Met-376 (24 Å) and the hydroxyl group of the active site Tyr-319 are indicated. A, adenine; G, guanine; C, cytosine; T, thyrosine; d, deoxyribose-phosphate.

UV XL-MS generally unambiguously distinguishes between RNA and DNA in crosslinks, because thymidine and deoxyribose crosslinks generate DNA-specific mass adducts. In contrast, many chemical crosslinks, involving purines, are ambiguous as deoxyguanosine and adenosine are isobaric (Supplementary Table S12). The detection of adducts of two or more nucleotides harbouring cytidine, uridine, or thymidine reduces such ambiguity. Also, deoxyribose crosslinks originated unambiguously from DNA (see above). However, their presence, in particular in a cellular context, must be still considered with caution. It is not entirely clear whether crosslinking of these reactive deoxyribose derivatives might be (entirely) reversible and could lead to crosslinking to proteins not necessarily involved in direct DNA binding. In the protein TOP1, which releases negative supercoiling in DNA, we identified unambiguous UV crosslinks to deoxydinucleotides, within a peptide encompassing amino acid positions 774-788 (Figure 7B) (96). The peptide contains Phe780 and Phe786, which are very close to the ssDNA in the 3D structure. Ambiguous crosslinks derived from chemical crosslinking (Met376, Met438, Met522, and Met669), were all located near conserved residues involved in DNA binding (96). Deoxyribose crosslinks were found to Cys401 and Cys763 in the DNA- and ssDNA-binding region, respectively.

Overall, we identified 70 crosslinked DNA-binding transcriptional regulators (according to the Regulon database (97)) in *E. coli*, exclusively after chemical crosslinking. Crosslinking sites were found within or close to the described DNA-binding domains of these regulators, such as the helix-turn-helix (HTH) motif, OmpR/PhoB-type domain, or the LytTr DNA-binding domain. Among 29 transcriptional regulators with an HTH motif, in 18 proteins the crosslinking sites are not located in the HTH motif, but in N- or C-termini of the protein. These potentially novel functional regions have not yet been described as nucleic-acid binding before. Importantly, among the all (chemically) crosslinkable amino acids in transcription regulators containing HTH motifs, only a minority were crosslinked, but they were identified by multiple CSMs in XL-MS, supporting the biological/structural relevance of these identifications (Supplementary Table S13). Owing to the ambiguity in DNA/RNA chemical crosslinking, we cannot conclude finally whether the crosslinks in the HTH domain or in other, as yet undefined, domains originated from RNA or from DNA.

Recently, the RNA binding properties of HTH transcription regulators have been confirmed in *Staphylococcus aureus* (98). According to this, we found that the MaR-type HTH domain containing transcriptional repressor MprA (Supplementary File S1) chemically crosslinked within its conserved site of the HTH domain (residues 87–121) to RNA (or DNA) and with residues at its partly disordered C- and N-termini outside the HTH domain (Supplementary Figure S16). Since MprA is not expressed in high abundance in our *E. coli* samples, it appears unlikely that the chemical crosslinking is due to a “crowding effect”, i.e., non-specific crosslinking to RNA or DNA owing to a high abundance of the protein. Predictions of pathogenicity due to missense mutations affecting the protein structure, conducted with AlphaMissense (99), suggest a mild pathogenicity upon mutation in the protein termini, which points towards a possible functional interaction of these particular protein regions with RNA.

## DISCUSSION

In a protein-centric analysis, we regard the detection/identification of crosslinked amino acids by XL-MS as the most direct approach to localizing the interactions of proteins with RNA and/or DNA. Several excellent proteome-wide crosslinking studies applied UV XL-MS and obtained remarkable inventories of proteins bound to (m)RNAs (5–7, 13, 14).

Compared to previous work, our study makes three major contributions: (i) establishing chemical XL-MS as a method to increase the number of crosslinks, (ii) introduce a dedicated search engine for crosslink identification and localization, and (iii) reporting the to-date largest data set of crosslinking sites for *E. coli* proteins.

UV crosslinking is the most frequently applied method for studying RNPs by XL-MS, while chemical XL-MS has so far been favored for application to DNPs (17, 100). UV XL-MS requires, for example, 6-thio-G or purine derivatives incorporated into cellular RNAs in order to capture RNP interaction with bases other than uracil (3, 101). Very recently it was demonstrated that 4-thio thymidine can be incorporated into cellular DNA, and subsequent UV irradiation followed by XL-MS revealed a large number of proteins crosslinked to DNA and allowed identification of their crosslinking sites (102). We show that the reaction with alkylating chemicals, nitrogen mustard, diepoxybutane, and formaldehyde, offers an alternative, as guanine and adenosine are linked to proteins. We anticipate a broad application of chemical XL-MS in global interaction mapping and also in the modelling of DNA/RNA to its binding proteins in RNPs. We show that chemical crosslinking is efficient *in vivo* and *in vitro*. Compared with UV irradiation, it is relatively sensitive to the composition of the sample. For example, primary amino groups containing components in the sample, such as Tris buffer, may inhibit chemical crosslinking; other components in the cell media can also react with chemical crosslinker. In general, it is recommended to choose the optimum concentration of these crosslinkers in each *in vitro* and *in vivo* sample by titration with various crosslinker concentrations.

Recently, chemical, i.e. FA-based crosslinking in XL-MS studies (103) and also in CLIP studies, termed fCLIP (104) have been applied as well and shown that it extends UV crosslinking in terms of numbers of crosslinked proteins but is also highly suitable to map RNA sequences that are in contact to dsRNA-binding proteins (105). Yet, it is not entirely solved whether chemical crosslinks occur mainly to primary RNA-binding proteins. It might be well the case that more indirect (“associated”) binders, which might interact with a primary RNA-binding protein chemically crosslinks to RNA in close vicinity to the primary RNA binder as well. Our own XL-MS *E. coli* data show that 501 crosslinked peptide sequences are common between UV and chemical crosslinks, revealing that chemical crosslinking also identifies primary RNA binders, if those proteins that UV crosslink to RNA are considered as primary RNA-binders based on the “zero-length” nature of UV crosslinking. Moreover, when structures of protein-dsRNA and protein-DNA complexes are available, we show here that chemicals can crosslink amino acid in contact to dsRNA and dsDNA (Figure 3B, 6A, 7A).

The first search engine for nucleotide-protein crosslinks, RNPxl, was restricted to UV XL-MS and required additional manual evaluation/annotation of XL-MS data (2, 72). Such shortcomings have been overcome in our tool NuXL. While general-purpose peptide search engines such as MSFragger (106) have also been applied in several UV-XL MS studies, NuXL has been developed specifically for protein-nucleotide crosslink assignment.

NuXL also allows the incorporation of other protein-nucleotide crosslinkers, for example, SDA, whose NHS ester group reacts with lysine residues in RNPs and upon photoactivation forms a carbene nucleotide in close vicinity to the derivatized lysine residue (Supplementary Figure S17A-C for reaction mechanism, *adduct sum formulae*, and MS2).

We compared our results with results from three other UV XL-MS studies (TRAPP (6), OOPS (7), and RIC (107)) of protein-RNA interactions in *E. coli* (Supplementary Figure S12B). Our results overlap substantially with these studies (Supplementary Figure S12B), but our dataset additionally reveals the sites of crosslinking for all identified proteins. In addition, we detected 548 crosslinked proteins and their crosslinking sites that were not identified in previous studies. 74% of these proteins were detected under only one of the four crosslinking conditions (12%, 20%, 39%, and 3% for UV, DEB, NM, and FA XL-MS, respectively; Supplementary Figure S12C), emphasizing the advantage of the combination of UV and chemical MS-XL. For example, crosslinking sites were identified in 9 of 10 hitherto poorly characterized RBPs described in the RIC (107) study and in 12 of 17 putative RBPs investigated in a recent OOPS study (108); in nine of these exclusively upon chemical crosslinking.

Unlike in the above-mentioned studies, we did not apply a dedicated purification of RNA after crosslinking in all experiments, except for formaldehyde crosslinking of *E. coli* cells (see below). Instead, we chose a comprehensive approach aimed to enrich any (deoxy)ribonucleotide crosslink species to show the feasibility of chemical crosslinking and NuXL in RBP and DBP analysis. Based on this, particularly chemical crosslinks show ambiguities, i.e., crosslinks that could be derived from RNA, DNA, mononucleotides, or enzymatic co-factors. Also, one cannot exclude the possibility that crosslinks derived from shorter RNA oligonucleotides originated from degradation of intact RNA. Nonetheless, our data largely overlap with RNA-crosslinked proteins in other *E. coli* studies and we show that crosslinks of RNA origin coincide with nucleotide/co-factor-binding sites. Our workflow is advantageous, for comparing crosslink sites originating from RNA or DNA with assigned nucleotide/co-factor-binding sites in a single experiment. The identification of nearly all proteins of glycolysis and the TCA cycle and their crosslinking sites (with the key enzymes crosslinked by UV, DEB, NM, and FA) strengthen the view that metabolic enzymes are moonlighting RNA-binding proteins, as previously reported in pro- and eukaryotes (86, 87). For formaldehyde XL-MS of *E. coli* cells we applied a silica-based enrichment strategy of intact (crosslinked) RNA (see Materials and Methods). The reason for this was that upon application of our general XL-MS workflow the yield of FA-crosslinked peptide-(oligo)nucleotide conjugates was surprisingly low (for which we do not yet have a compelling explanation). Importantly, this particular dataset, which shows crosslinked metabolic enzymes, strongly indicates that these must have derived from intact RNA rather than from single nucleotides or co-factors due to the presence of crosslinked dinucleotide.

Alkylating reagents such as those used in this study form intra- and interstrand crosslinks in DNA and also connect bases in RNA (109). We can only speculate that such crosslinks theoretically influence chemical protein–RNA crosslinking if it occurs within sites where the proteins bind and hence would crosslink. Such a scenario would also occur in protein–DNA crosslinking.

In this study, we could not (yet) systematically compare DNA and RNA crosslinking (owing to the ambiguous nature of possible A and dG crosslinking), including nucleotide crosslink reactivity. Our statistical analysis shows that RNA-binding proteins are highly abundant in *E. coli* cells in accordance with the abundance distribution of RNA and DNA (20% and 3% of the cell’s dry mass, respectively) (110). Accordingly, we demonstrated a statistically significant enrichment towards the putative RNA-binding sites on primary and tertiary protein structure levels, but not for DNA-binding sites (Figure 5A; Supplementary Figure S11). Hence, we infer that, at least under the growth conditions applied in this study, i.e., growing and crosslinking *E. coli* in nutrient-rich media where cells strongly allocate their resources to transcription and translation proteins are more prone to crosslinking with RNA than with DNA.

## Supporting information

Supplementary Methods, Results and Figures

## DATA AVAILABILITY

The datasets (.raw, .mzML, .idxML, and .txt files) generated and analysed for this study are deposited in the MassIVE repository and submitted to ProteomeXchange Consortium (118) and are available using the following identifiers MSV000094879 (MassIVE), PXD052614 (ProteomeXchange). Dnmt2-tRNA^Asp^ models and Supplementary Table S5 and S11 were deposited via figshare and are available under the following link: https://figshare.com/s/6b2091a24a327ed546e4.

## SUPPLEMENTARY DATA

Supplementary Data are available at NAR online.

## ACKNOWLEDGEMENTS

We thank Tamara Dehne, Sabine König, Annika Reinelt, and Ralf Pflanz for excellent technical assistance and support in LC-MS/MS analyses. We are grateful for Fanni Bazsó’s excellent contribution to FA XL-MS *E. coli* experiments and corresponding method development.

## AUTHORS CONTRIBUTIONS

Conceptualized of the study: L.M.W., A.C., O.K., J.L., T.S., H.U..; NuXL software tool development: A.C., A.S., E.N., Y. Hi, K.F., B.D., R.V., O.K., T.S.; *in vitro* complex reconstitution, crosslinking, XL-MS, NuXL data analysis: L.M.W., A.W., S.M., O.D., P.N., H.G., J.W., D.A.I., A.C.deA.P.S., A.D., S.J., J.S., I.W., S.M.V, E.O., K.E.B., M.B., P.C., R.F., O.K., T.S., H.U.; molecular modelling: P.N., A.D., R.F.; *in vivo* crosslinking, XL-MS, NuXL data analysis: L.M.W., A.W., A.C., S.M., O.D., A.S., M.R., Y.He, R.V., O.K., T.S., H.U.; *E. coli* proteome and LC-MS data analysis: L.M.W., Y.Hor., M.P., M.R., J.L.; computational XL-MS, LC-MS data analysis and annotation: Y.Hor., M.P., J.L.; manuscript writing: L.M.W., Y.Hor., S.M., J.L., T.S., H.U., with input from all other authors.

