## Supplementary Methods, Results and Figures for "Chemical crosslinking extends and complements UV crosslinking in analysis of RNA/DNA nucleic acid–protein interaction sites by mass spectrometry"

† Joint Authors

### SUPPLEMENTARY TEXT

### SUPPLEMENTARY METHODS

#### XL-MS data processing in NuXL

##### Raw data conversion

Thermo raw data was converted to the mzML format using freely available software (ThermoRawFileParser, Thermo Scientific™; and msconvert/pwiz (1)).

##### Spectra preprocessing

Removal of peak interference: Precursor purities were calculated as the intensity ratio between all isotopic peaks corresponding to the precursor target *m/z* and the total intensity of peaks in the precursor isolation window. All interfering peaks also found in the product MS/MS were removed so as to reduce random hits to these peaks.

##### Intensity processing

Zero-intensity peaks were removed; intensities were normalised to a maximum intensity of one.

##### Deisotoping

Mass-to-charge differences between fragment ions were compared with the expected isotope distance for charges 1 to 3, but not higher than the precursor charge. Wherever at least two consecutive peaks matched the expected distance within a mass tolerance window of 10 ppm, a hypothetical isotope pattern is found. Because low-intensity (e.g., noise) peaks can lead to random matches, we additionally required that fragment intensities approximately matched the theoretical isotope pattern; otherwise, we did not consider the hypothetical isotope pattern for deisotoping. A simple approximation of the averaging model that disallows isotopic intensities to increase after the second isotopic peak proved sufficient. If a pattern passed the test, all fragments corresponding to higher-isotopes were removed and their intensities were added to the monoisotopic peak that is annotated with the charge. Fragments not part of any isotopic pattern were retained and annotated with charge one.

Depending on the protocol and instrument settings, we often observed one or several high-intensity peptide precursor-derived ions in an MS/MS that might have carried nucleotide fragment adducts. As these are highly indicative of a cross-link spectrum, we excluded all high-intensity peaks that could start an isotope pattern according to the criteria above from removal. For the same reasons, we excluded peaks in the typical marker-ion range (<150 *m/z*) from removal.

#### Noise Filter

The top 20 highest-intensity fragments within each window of 75 *m/z* were retained, to remove local noise clusters. We then limited the spectrum complexity by retaining the 400 highest-intensity ones.

#### Identification

The protein database (in FASTA format) is digested in silico by using one of the supported enzyme settings. Decoy peptides were generated deterministically by shuffling peptide sequences randomly between cutting sites (excluding the C-terminal residue) up to 100 times. If the target and shuffled sequence (except for the terminal residue) did not share any prefixes or suffixes it was selected as decoy. Otherwise, after 100 attempts, the sequence that minimised the similarity to its target peptide was selected. Peptides were processed in parallel by using OpenMP as parallelisation backend. For each peptide all features/peptidoforms were generated by applying fixed, variable and RNA/DNA precursor adducts as modifications. The resulting set of masses was then queried against the experimental precursor masses. If the mass of a feature/peptidoform matched an experimental precursor mass (within the specified mass tolerance window), it was a candidate considered for MS/MS comparison.

For every candidate, theoretical fragment spectra were generated. For both peptides and cross-linked peptides, we generated a-, b- and y-ion ladders without additional fragment adduct mass. This spectrum was cached and re-used for all oligonucleotide variants, speeding up the process of spectrum generation. For cross-links, we additionally added all shifted ladder ions such that fragment adducts were compatible with the observed precursor adduct and cross-linked nucleotide. In addition, singly charged marker ions, cross-linked immonium ions and cross-linked precursor ions were generated. If more than one nucleotide had been configured as cross-linkable, fragment spectra were generated for all of them.

#### Autotune of search tolerances and identification-based filtering of peptides

Choosing precursor and fragment mass tolerances that reflect the quality of instrument calibration can greatly improve search results, as it can reduce the number of incorrectly assigned peaks or prevent missed assignments due to overstrict mass tolerance. NuXL implements an autotuning step that estimates precursor and fragment mass tolerances using high-scoring peptides from an initial calibration search. Optionally, NuXL can exclude confidently assigned peptide spectra from the search against cross-link candidates, similar to our ID filter approach described previously (2).

**Spectrum comparison and scoring**

The generated spectrum of a candidate is aligned to the measured tandem mass spectrum to determine matching fragment ions within the specified fragment-mass tolerance. The NuXL score is the primary scoring metric used for identifying cross-linked peptides. It is computed as the sum of match-odds scores, which quantify the likelihood of observed fragment matches occurring by chance. Specifically, NuXL calculates two separate match-odds scores: one for fragment ions without fragment adducts (e.g., non-cross-linked peptide fragments or fragments that completely lost the precursor adduct upon fragmentation) and one for fragments that retain the adduct. The summed value of the log-transformed scores represents the overall NuXL score. In addition, NuXL refines scoring using several subscores capturing cross-link specific fragmentation patterns, such as marker ions or shifted immonium ions, are calculated and used in semi-supervised score calibration to improve the discriminative power of the Percolator (3) algorithm (see Supplementary Table S3 for a list of calculated scores).

**Localisation**

After the main identification loop, cross-links were localised by using ions that still carried nucleotide adducts after fragmentation. Similarly to classical PTM localisation, consecutive prefix (or suffix) ions without and with mass shift constitute site-determining ions that allow one to pinpoint the cross-linking site. In cross-linking we may also observe several ions of the same prefix (or suffix) with and without fragment adducts. Not uncommonly, we observed a particular prefix ion with several fragment adducts that differed in composition. These mixture spectra of complete loss and multiple partial losses were attributed to the lability of the cross-linked nucleotide. While basic and aromatic residues are more susceptible to cross-linking, nearly all residues can cross-link upon UV irradiation (4). NuXL does not restrict the subset of residues during localisation but considers all sites with site determining ions as potentially cross-linked. When potential sites are next to each other, we discarded the site corresponding to the longer prefix/suffix ion if it had a lower intensity. As site score, we assigned the summed intensity of the site determining prefix/suffix ions that carry nucleotide mass shifts. Analogously, we considered all sites corresponding to shifted immonium ions to be site-determining and report their summed intensity over one or over several different observed fragment adducts. In cases of ambiguous localisation (e.g., immonium ion with fragment adducts that match to multiple positions in the peptide; several sites with site-determining prefix/suffix ions) all sites are reported. If no site-determining ions were found, no localisation is reported for that particular CSM. Owing to the lower specificity for individual residues as compared with classical post-translational modifications (PTMs), developing a probability-based scoring function is challenging. Consequently, we adopted a simple intensity-based scoring scheme. We determined that localisations are typically reliable when their intensities exceed the base-peak intensity, corresponding to a localisation score greater than 1.

**Score combination and FDR-estimation of cross-link spectrum matches**

NuXL integrates main and subscores by using the semi-supervised learning approach of the Percolator algorithm to optimise scoring and confidence estimation. Standard false discovery rate (FDR) estimation by target-decoy competition, applied to all spectrum matches, can lead to over- or underestimation of q-values for modified peptides (5, 6). Similarly, the FDR in the group of cross-links (CSM-FDR) typically deviates from the FDR at the spectrum-match level. To account for these differences, NuXL calculates separate q-values (group FDRs) for cross-linked spectrum matches (CSMs) and regular peptide-spectrum matches (PSMs) using the Percolator SVM score. Across a wide range of experiments, this approach has consistently improved the identification rates of cross-linked peptides (data not shown). After processing, q-values are reported as the main score in the final results.

**Identification of nucleic acid–protein crosslinks by MSFragger search engine**

We employed MSFragger v4.0 using FragPipe v20.0 (5) for crosslink identifications, configuring variable modifications to 15.9949 (M) and permitting a maximum of two variable modifications per peptide. Two relevant workflows were used, XRNAX (7) and MSFragger-Labile (8). In both workflows, we set MSFragger mass offsets for crosslink modifications to the ones defined in the NuXL presets. Mining for diagnostic and fragment ions was enabled for the MSFragger-Labile workflow.

### SUPPLEMENTARY RESULTS

#### NuXL run times with simple and extended presets

The NuXL software version used in this work (version OpenMS-3.0.0-pre-HEAD-2022-04-27-Win64) was tested on representative *E. coli* crosslinking data: a merged mzML file, including 13 raw files from fractions of *E. coli* UV crosslinks (2.38 Gb) created by the FileMerger function of OpenMS, was searched against *E. coli* genomic database version as used in this work with RNA-UV(UCGA) and RNA-UV(UCGA-Extended) presets. In addition, a similar mzML file from *E. coli* NM crosslinks (2.87 Gb) was searched against the same database with RNA-NM and RNA-NM presets. Searches were run on Dell Optiplex PC with Intel(R) Core(TM) i7-14700K 3.40 GHz and 32.0 Gb of installed RAM. UV data searches took 54 min 47 s (54:47) with UCGA presets and 01 hour 06 min 15 s (01:06:15) with extended presets, respectively. NM data searches took 01:14:14 with NM presets and 02:03:57 with extended presets, respectively.

Alternatively, for 3 mzML files made of raw files of the UV-crosslinked Hsh49Cus1 preparation (all 868 Mb), NuXL ran for about 7 min against the same database with UCGA presets by the same PC. Similar NM files were searched for 6 min with NM presets.

To assess further the influence of variable modifications, we performed a search on a human sample using the RNA-FA protocol preset and human protein database, allowing for a maximum of two oxidation modifications. When running single-threaded on an AMD Ryzen 7 PRO 4750U processor, the search is completed in 24 minutes. Utilising multi-threading with 20 threads reduced the runtime to approximately 7 minutes, with peak memory usage reaching around 850 Mb. In comparison, performing the search without variable modifications was only approximately 20% faster and was nearly identical in respect of to memory requirement.

The extension of UV presets led to minimal addition in running time, in contrast to NM presets, for which the time doubled owing to the greater number of combinations. The running times on regular PCs were sufficiently short to be practical for typical research applications.

#### Effects of simple and extended presets to crosslink identification yields

To estimate the effects of simplified or extended presets on the number of identifications, we compared outputs from the searches described above. Based on unique crosslinked peptide sequences, UV(UCGA) vs. UV (UCGA-Extended), 78.4% of identifications (497 crosslinked peptides) were shared with 61 (9.6%) and 76 (12.0%) peptides identified only with UV(UCGA) and with UV (UCGA-Extended), respectively. Margins of the diagram contained mostly crosslinks represented by single mass spectra (mean CSM count for UV(UCGA) 1.15 vs. 6.68 for shared crosslinked peptides, Wilcoxon-test *p* value 3.7e-09; mean CSM for UV (UCGA-Extended) 1.24 vs. 7.02 for shared crosslinked peptides, Wilcoxon-test *p* value 3.5e-08). It means that changes in outputs of different presets corresponded to low score hits which did not influence much on the interpretation of results.

When NM and NM-Extended presets were compared, 90.4% (2326) of unique crosslinked peptide identifications were shared with 125 (4.9%) and 122 (4.7%) of them identified only with NM and with NM-Extended, respectively. As for UV crosslink searches, margins of the diagram contained mostly crosslinks represented by single mass spectra (mean CSM count for NM 1.09 vs. 5.24 for shared crosslinked peptides, Wilcoxon-test *p* value < 2e-16; mean CSM for NM- Extended 1.3 vs. 5.46 for shared crosslinked peptides, Wilcoxon-test *p* value 9.9e-15). Extended presets did not give much advantage when searching crosslinks proteome-wide, but they represent an option for advanced users to identify some specific crosslinks in less complex mixtures.

**Comparison of NuXL and MSFragger outputs for crosslinked samples**

Comparison of outputs of MSFragger's XRNAX (7) and Labile (8) workflows with corresponding NuXL outputs made on *in vivo E. coli* crosslinking datasets, the MSFragger-Labile workflow provided higher overlaps with NuXL results in terms of CSMs, unique peptides and proteins (see Results section in the main text). For the further representation, MSFragger-Labile results were considered at the level of unique crosslinked peptides and compared with the same results of NuXL as reported here. For all crosslinkers except FA, the numbers of crosslinked peptides identified only by NuXL exceeded the corresponding numbers of crosslinks identified by MSFragger (Supplementary Figure S4). For NM and FA crosslinkers, the majorities of identifications were shared between two search engines. In contrast, for UV and DEB, significant additions were provided by NuXL in comparison with MSFragger-Labile workflow. In absence of synthetic mixtures of nucleotide–peptide crosslinks that are currently not available for the field, it is not possible to benchmark outputs of any search engine to estimate real shares of false discoveries. However, we curated identifications of UV-generated crosslinks specific for NuXL in the dataset of S30 fraction of *E. coli* cells, replicate 2, manually. Sequences of many crosslinks were covered by specific fragment ions which confirmed the presence and localisation of nucleotide adducts (a few examples are provided as Supplementary File S2).

### SUPPLEMENTARY FIGURES

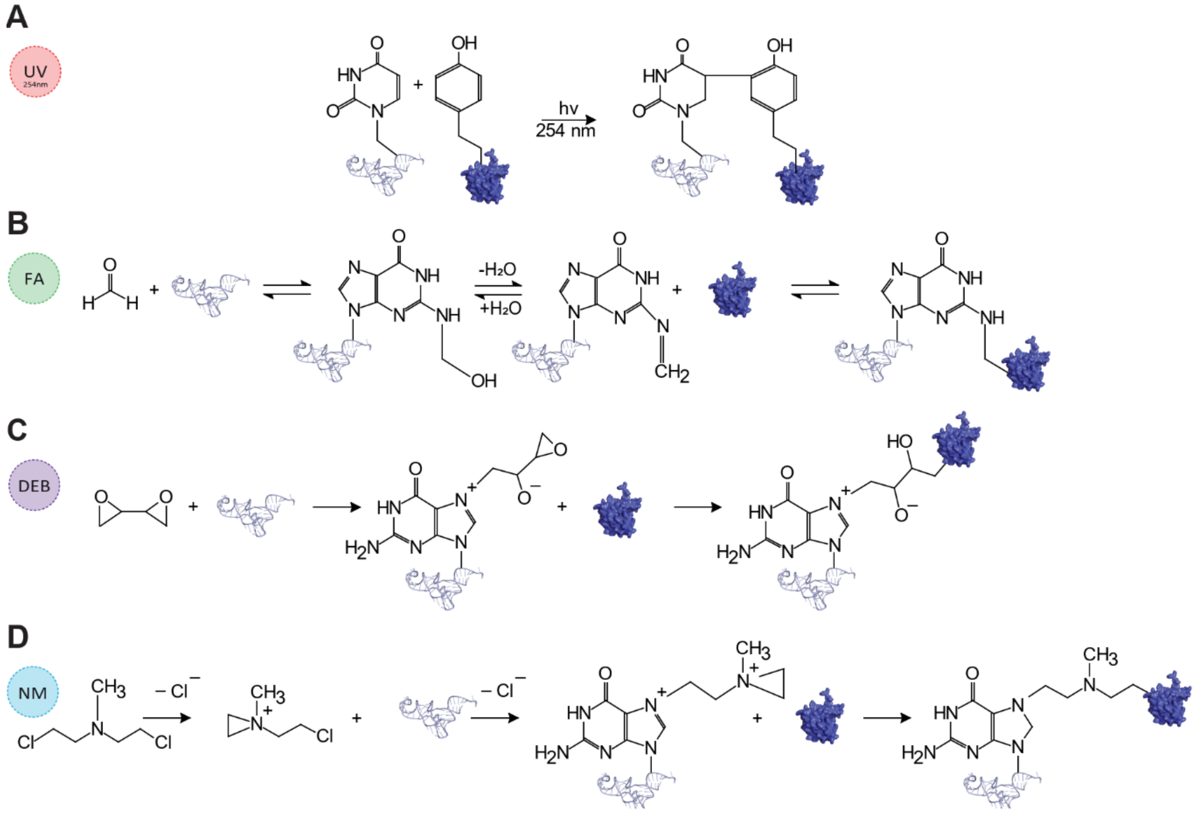


**Supplementary Figure S1.** Proposed chemical protein–RNA crosslinking reaction mechanisms: **(A)** UV light at λ=254 nm (adapted from Kramer *et al.* (9)), **(B)** formaldehyde (FA) (adapted from Hoffman *et al.* (10)), **(C)** 1,2:3,4-diepoxybutane (DEB) (adapted from Tretyakova *et al.* (11)), **(D)** mechlorethamine (nitrogen mustard, NM) (adapted from Tretyakova *et al.* (11)).

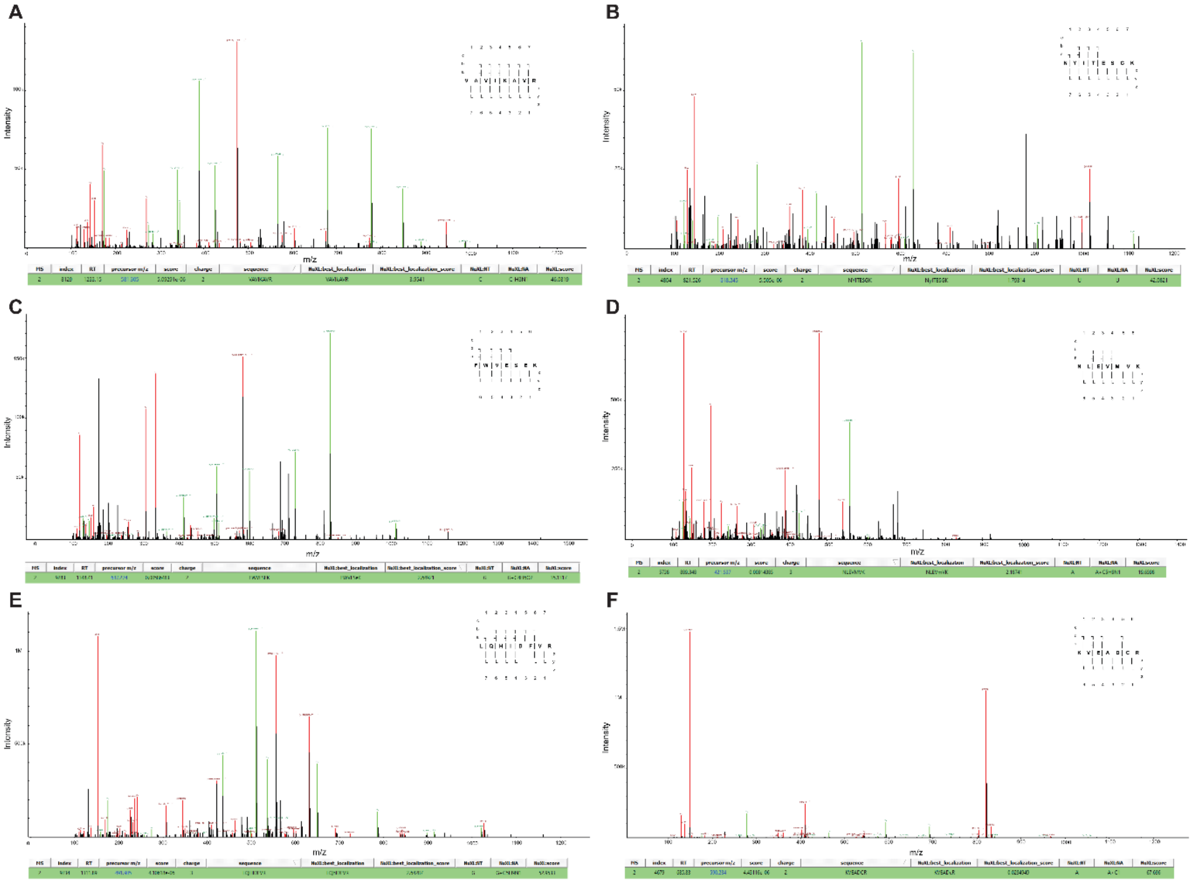

**Supplementary Figure S2.** Examples of MS2 spectra for UV-, DEB-, NM- and FA-crosslinked peptide–RNA(oligo)nucleotides visualised by TOPPView spectra viewer from NuXL outputs. The simplified schematic representation of the same mass spectra is shown in Figure 1C-H. **(A)** UV crosslink MS2 spectrum of VAVIKAVR-CMP-$NH_3$/UMP-$H_2O$. **(B)** UV crosslink MS2 spectrum of NYITESGK-UMP. **(C)** DEB crosslink MS2 spectrum of peptide–RNA(oligo)nucleotide FWVESEK-DEB-GMP. **(D)** NM crosslink MS2 spectrum of crosslinked peptide–RNA(oligo)nucleotide LQHIDFVR-NM-GMP. **(E)** NM crosslink MS2 spectrum of crosslinked peptide–RNA(oligo)nucleotide NLEVMVK-NM-AMP. **(F)** FA crosslink MS2 spectrum of crosslinked peptide–RNA(oligo)nucleotide KVEADCR-FA-AMP.

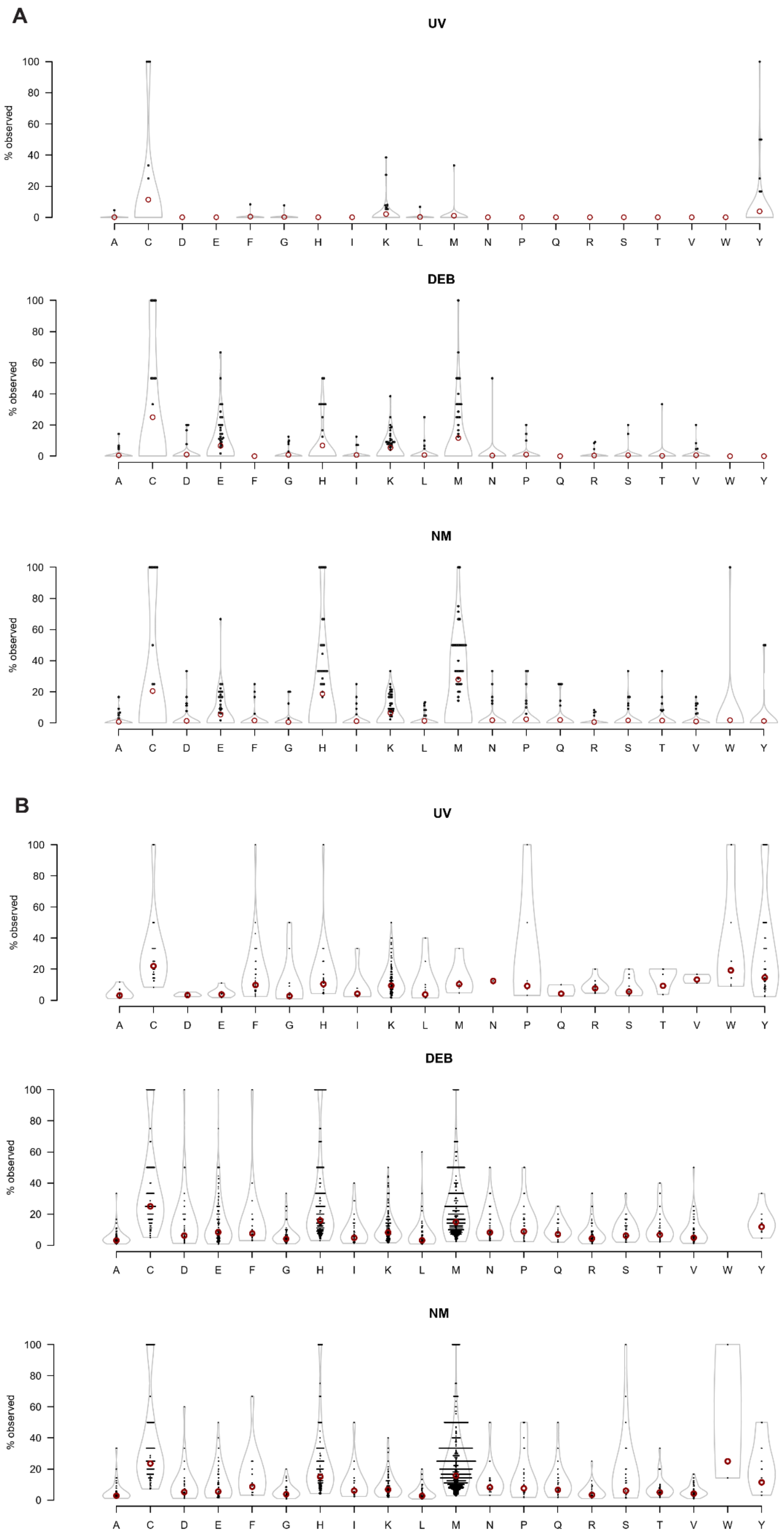
A
UV
DEB
NM
B
UV
DEB
NM
% observed
A C D E F G H I K L M N P Q R S T V W Y

**Supplementary Figure S3.** Occurrence of RNA crosslinking for individual amino-acid residues normalised by the background frequencies of the corresponding amino acids in crosslinked proteins. Only crosslinks with a localisation score ≥1 are included. Data are presented for UV, DEB and NM crosslinkers. FA data are not included, as for these a localisation score is assigned to only a minority of crosslinks for FA. Black dots represent percentages for each protein, and red circles represent all proteins combined. **(A)** Amino-acid crosslinking frequencies in isolated *E. coli* ribosomes normalised by amino-acid frequencies in 57 *E. coli* ribosomal proteins. **(B)** Amino-acid crosslinking frequencies in *E. coli* cells normalised by amino-acid frequencies in all crosslinked proteins.

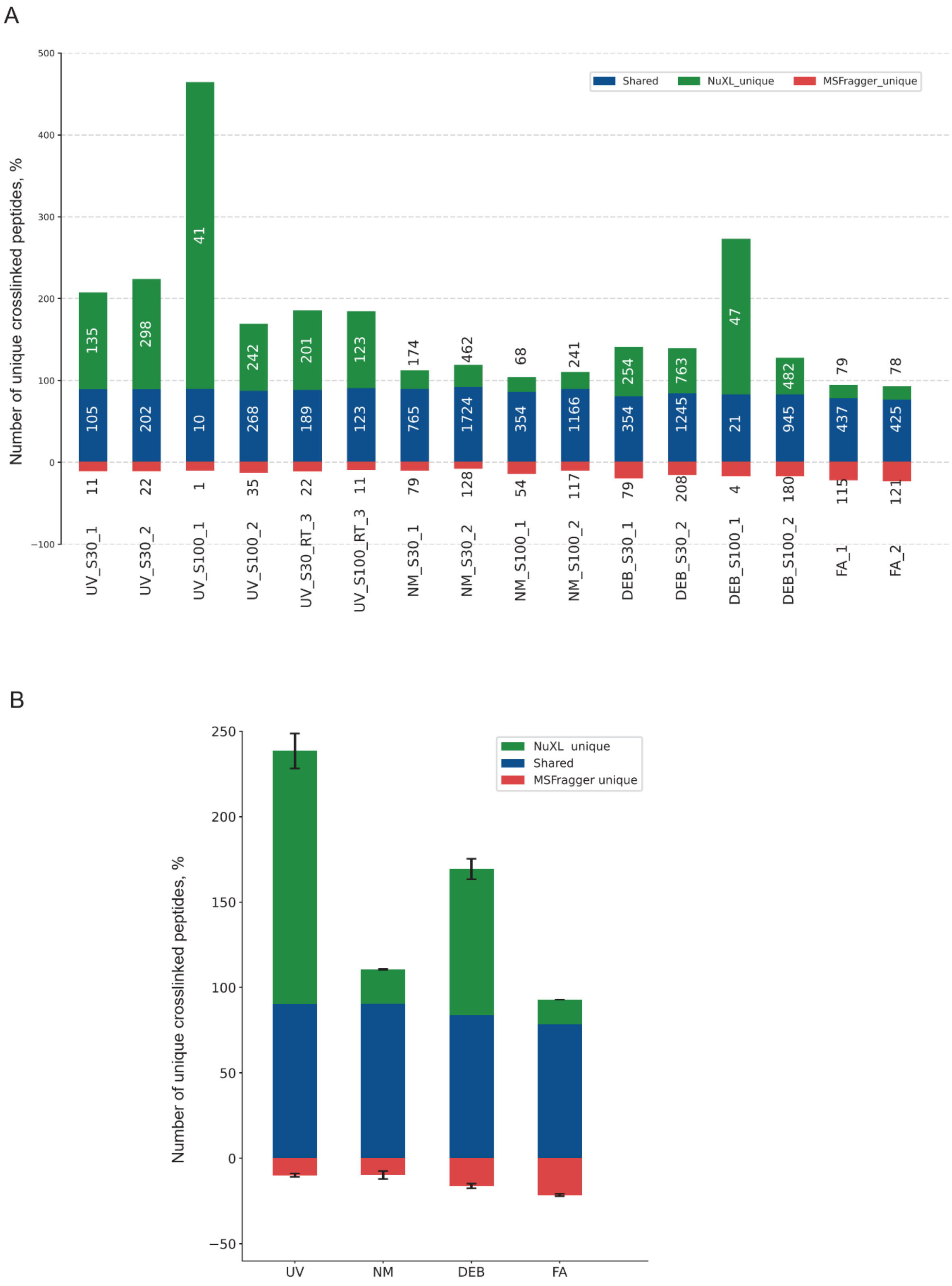


**Supplementary Figure S4.** Comparison of NuXL and MSFragger-Labile (8) workflow outputs for crosslinked *E. coli* cell datasets (Supplementary Table S5) in terms of identification of unique crosslinked peptides (1% FDR calculated at the peptide sequence-level). Percentages represent the fraction of crosslinked peptides exclusive to NuXL, exclusive to MSFragger, or found by both search engines. Percentages are calculated relative to the total number of MSFragger-identified crosslink peptides for the different crosslinking protocols (UV, NM, DEB, FA) and replicates. Only crosslink peptides that map to a single protein are considered. **(A)** Percentages of crosslinked peptide numbers identified in datasets of different crosslinking methods and replicates shown separately. The absolute numbers of identifications are provided. **(B)** The same percentages averaged across crosslinking methods and replicates.

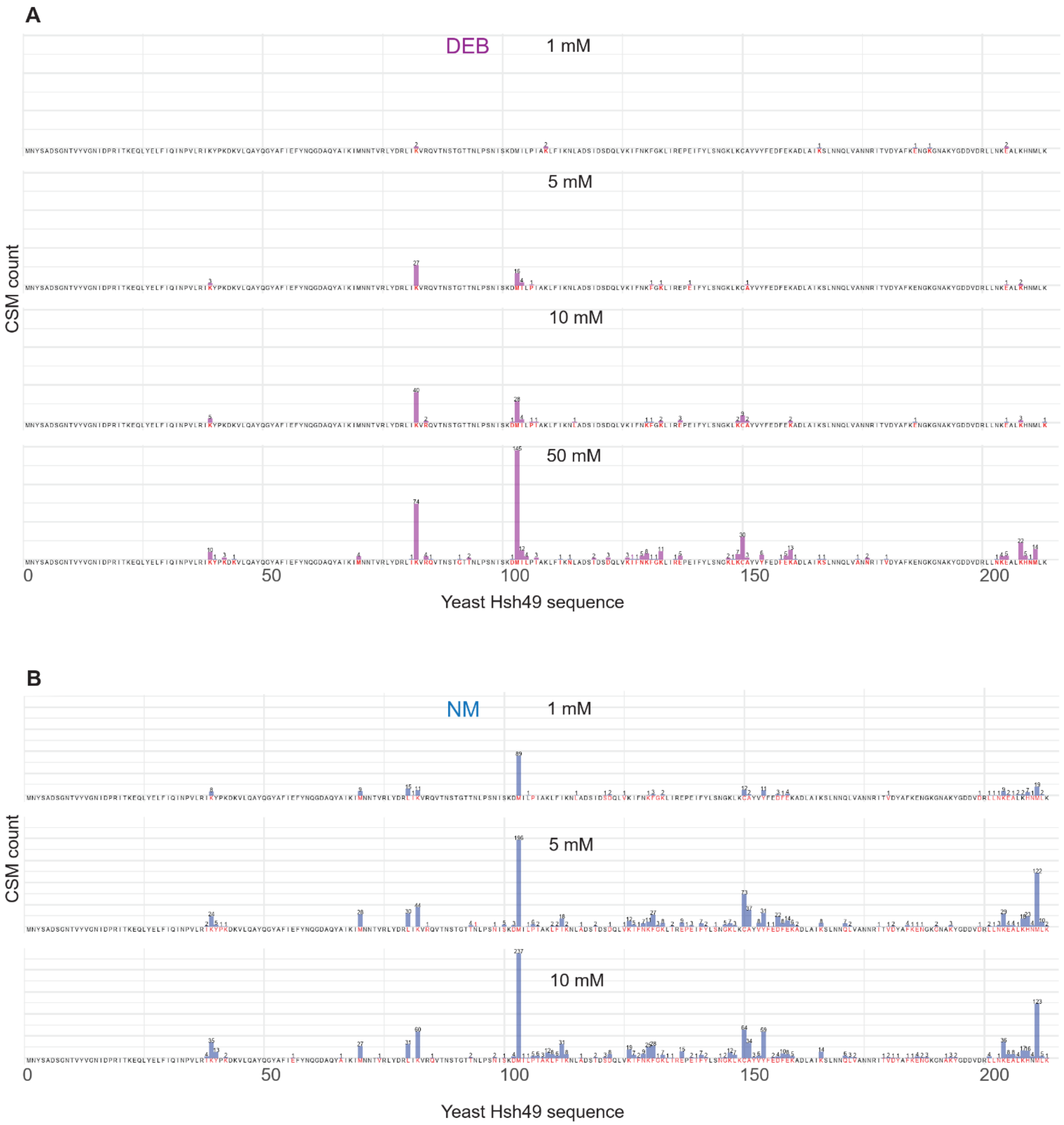


**Supplementary Figure S5.** Peptide–RNA crosslinking of Hsh49 protein in Hsh49-Cus1-U2 47Nt-snRNA complex at different concentrations of DEB **(A)** and NM **(B)**. Data were obtained according to the corresponding Materials and Methods section, with crosslinker concentrations of 1 mM, 5 mM, 10 mM, and 50 mM DEB, and 1 mM, 5 mM, and 10 mM NM. Results are shown with CSM counts based on best amino-acid localisation assigned by NuXL (localisation score > 0).

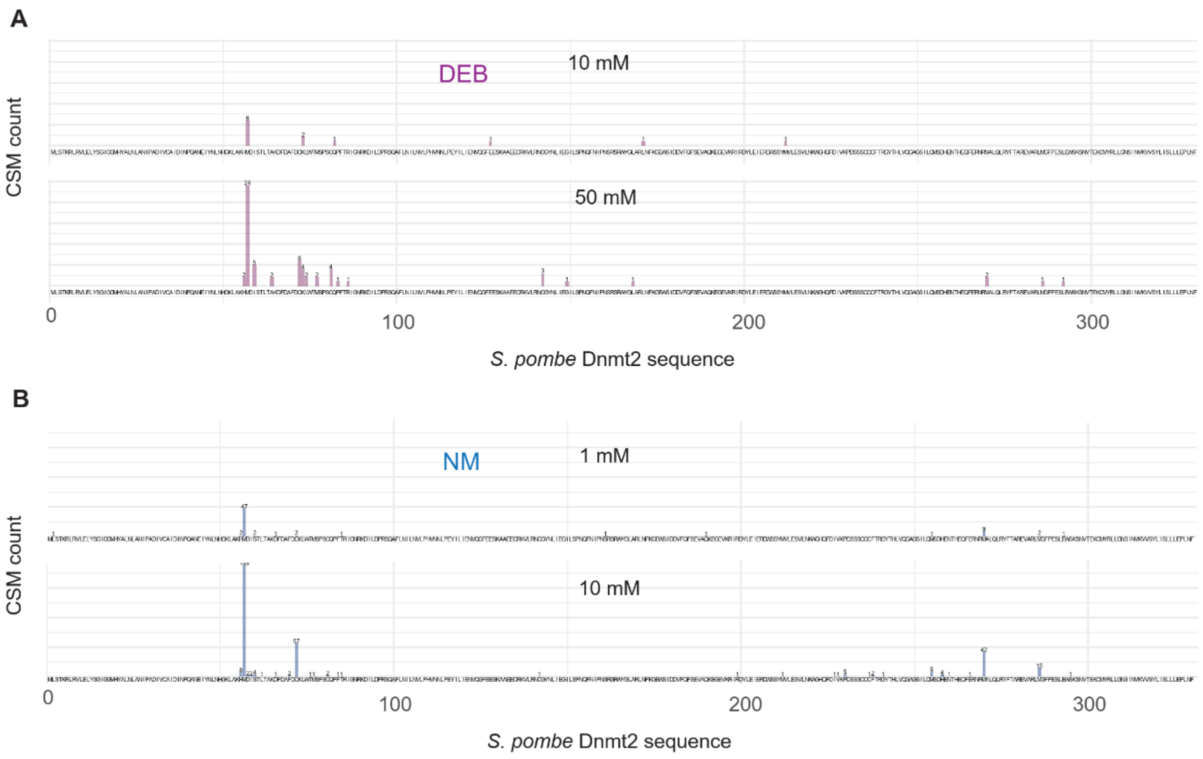


**Supplementary Figure S6.** Nucleotide-peptide crosslinking of *S. pombe* Dnmt2 protein in Dnmt2-tRNA$^{Asp}$ complex by different concentrations of DEB **(A)** and NM **(B)** chemicals (titration by crosslinkers). Data were obtained according to the corresponding Materials and methods section with crosslinker concentrations of 1 mM, 5 mM, 10 mM, and 50 mM DEB, and 1 mM and 10 mM NM. Results are shown as CSM counts based on best localization positions assigned by NuXL (localization score > 0).

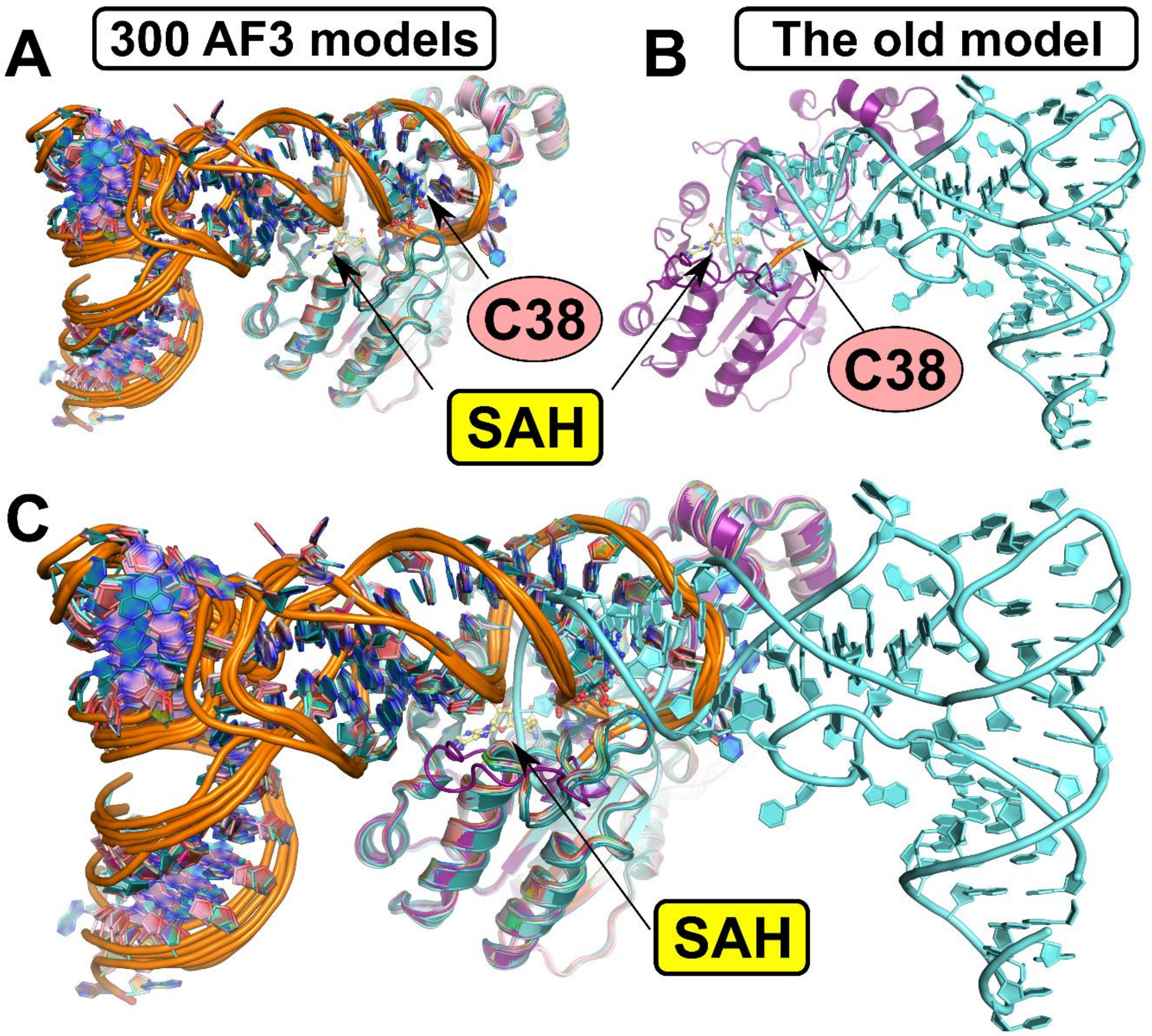


**Supplementary Figure S7.** Comparison of AlphaFold3 (AF3)-based (12) Dnmt2-tRNA$^{Asp}$ models with the previously published U-crosslink-based ("old") model (13). The S-adenosylhomocysteine (SAH) molecule is depicted in a ball-and-stick representation, with carbon atoms coloured yellow. The position of cytidine 38 (C38) is indicated.

**(A)** Superposition of 300 AF3-based predictions. **(B)** Previous UV crosslink-based model (13).

**(C)** Overlay of the models presented in (A) and (B).

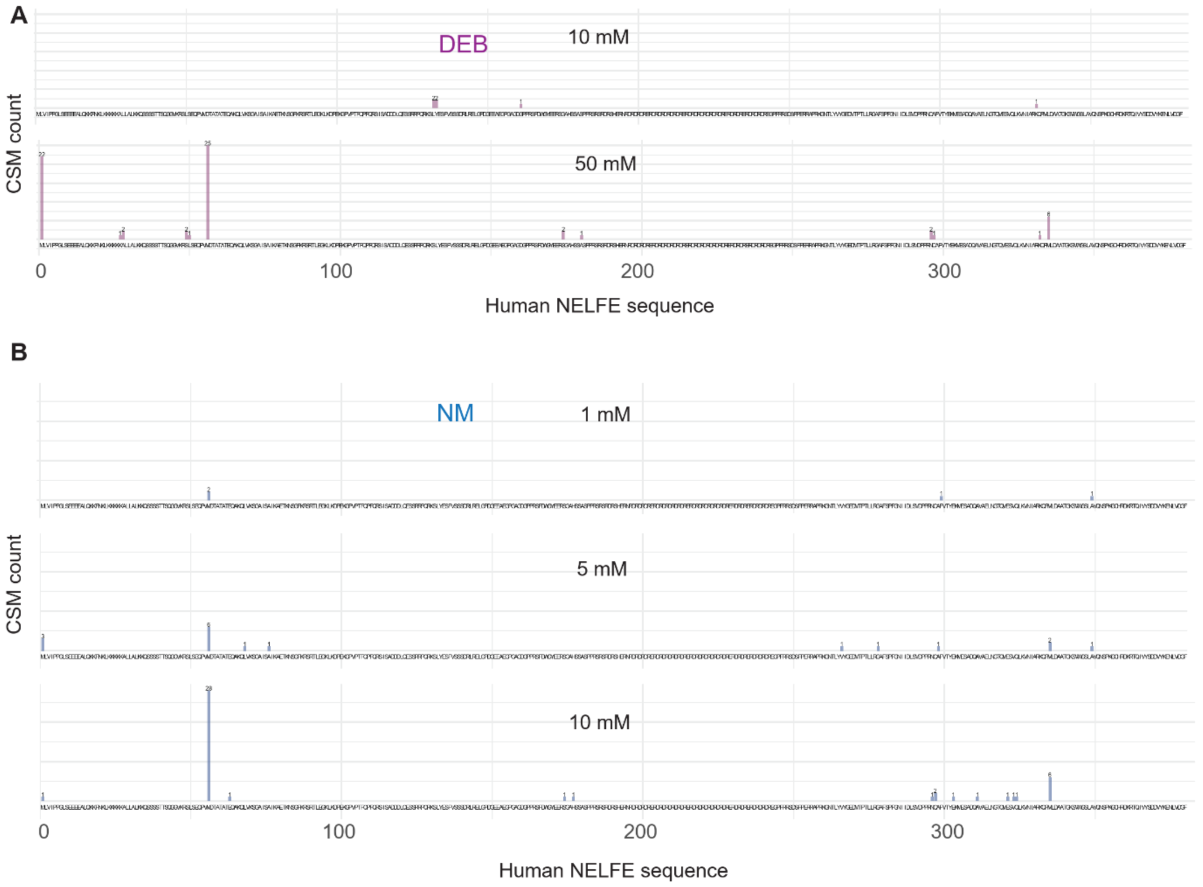


**Supplementary Figure S8.** Nucleotide-peptide crosslinking of human NELF-E protein in NELF-TAR complex by different concentrations of DEB **(A)** and NM **(B)** chemicals (titration by crosslinkers). Data were obtained according to the corresponding Materials and methods section with crosslinker concentrations of 10 mM and 50 mM DEB (1 mM and 5 mM DEB did not crosslink NELF-E protein), and 1 mM, 5 mM, and 10 mM NM. Results are shown as CSM counts based on best localization positions assigned by NuXL (localization score > 0).

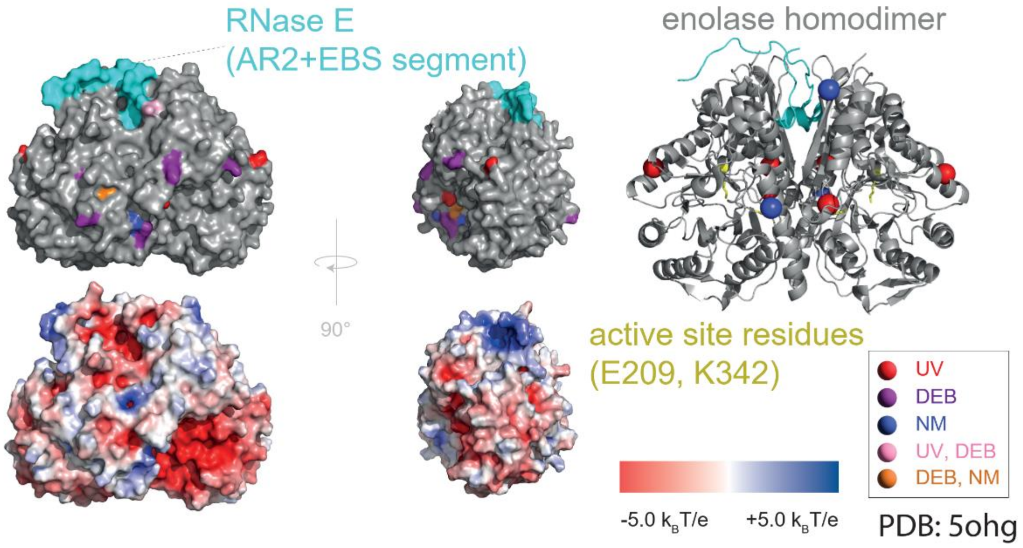


**Supplementary Figure S9.** Surface/cartoon representation and electrostatic surface of the crystal structure of enolase homodimer (grey surface/cartoon) in complex with RNaseE fragment (cyan surface/cartoon; PDB: 5ogh (14)). RNA-(oligo)nucleotide-crosslinked amino acids (Cα) are highlighted as purple (DEB), blue (NM), orange (DEB and NM) or light pink (UV, DEB and NM) surface or spheres. Mapped crosslink positions are based on CSMs filtered for 1% FDR, and localisation score ≥1.

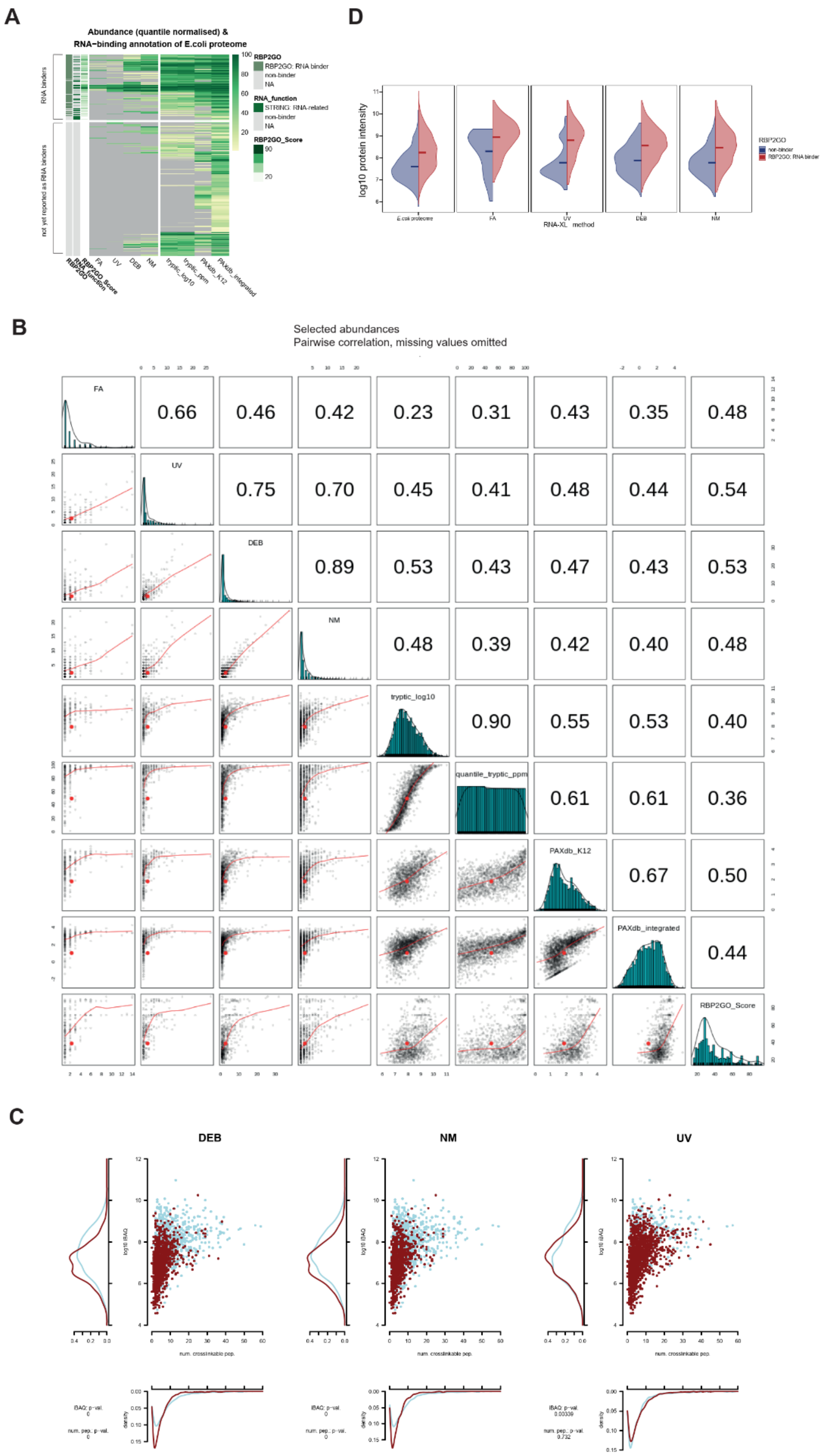


**Supplementary Figure S10.** Crosslinked protein abundances. **(A)** Heatmap of protein abundance. Plotted are the column-wise quantile-normalised CSM counts for RNA crosslinks, tryptic protein abundance and PaxDB (15) protein quantities for the *E.coli* proteome. The annotation on the left shows whether a protein is a putative RNA-binder in the RBP2GO database (16), whether it has RNA-related functions in STRING ontology (17) and the corresponding RBP2GO score. Missing values are plotted in grey. **(B)** Correlation matrix of protein abundance metrics. Pearson correlations were computed among the non-missing value pairs. **(C)** A scatterplot of protein-wise number of crosslinkable peptides plotted against the tryptic protein iBAQ abundance (18) by RNA-crosslinking method (red, detected; blue, not detected). **(D)** Distributions of the *E. coli* protein abundance measured by label-free MS1 intensity and stratified by the annotation given in Caudron-Herger *et al.* (16), and detection across the crosslinking methods. The figure highlights the distribution means.

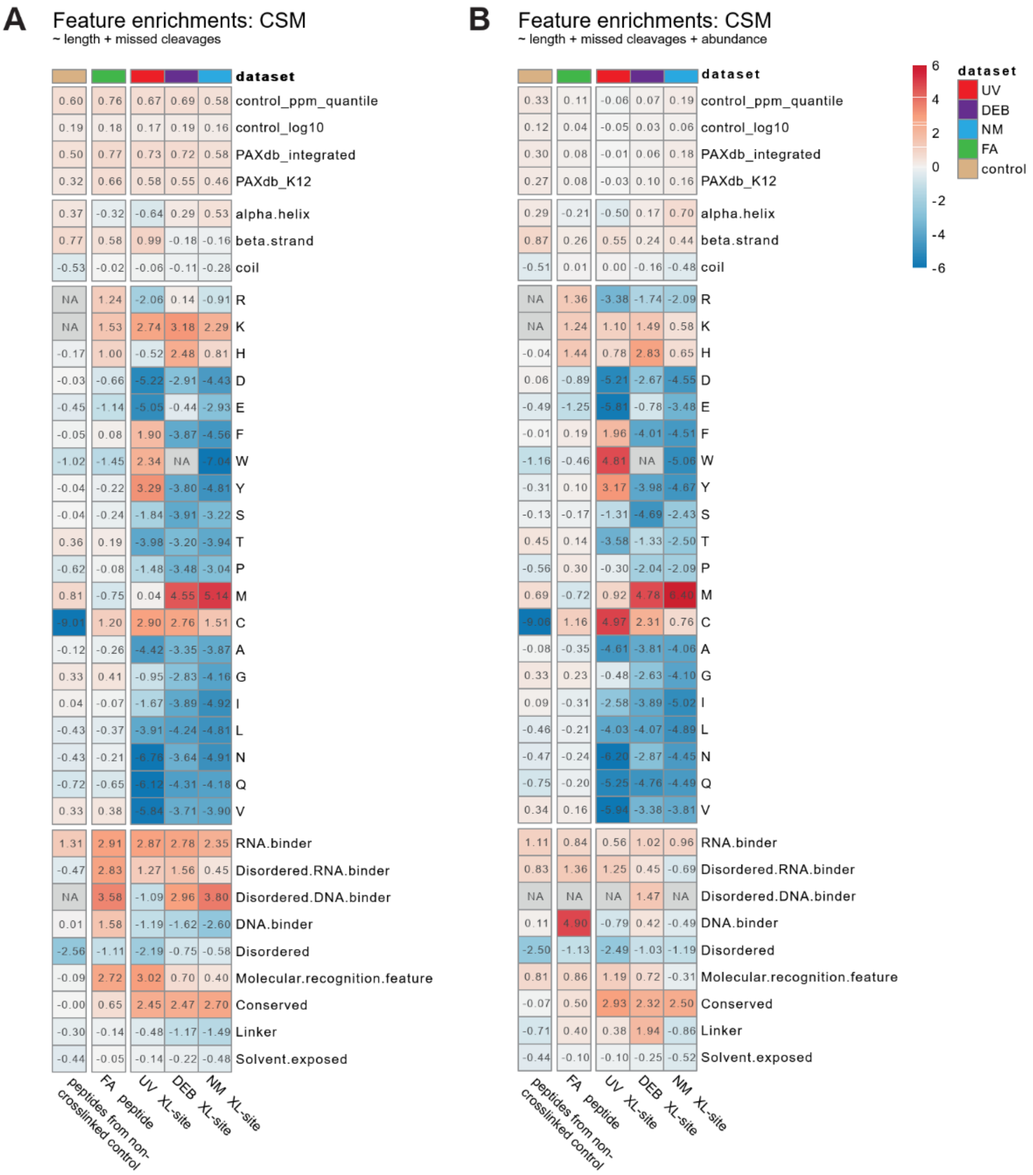


**Supplementary Figure S11.** The log2FoldChanges between the experimental and an *in-silico* background distribution mean feature counts per residue. The background sampling was performed with regard to: **(A)** the experimentally observed peptide length and missed cleavage distribution and **(B)** the experimentally observed peptide length, missed cleavage distribution and source protein abundance. The tryptic and FA datasets were enriched at the peptide level, whilst the UV, NM and DEB enrichment was done at the crosslinked-residue level. The NA values correspond to the cases where the sampling or statistical testing could not be performed owing to the feature distribution violating the method's assumptions. Crosslinker-specific feature enrichment. Control, non-crosslinked *E. coli* proteome.

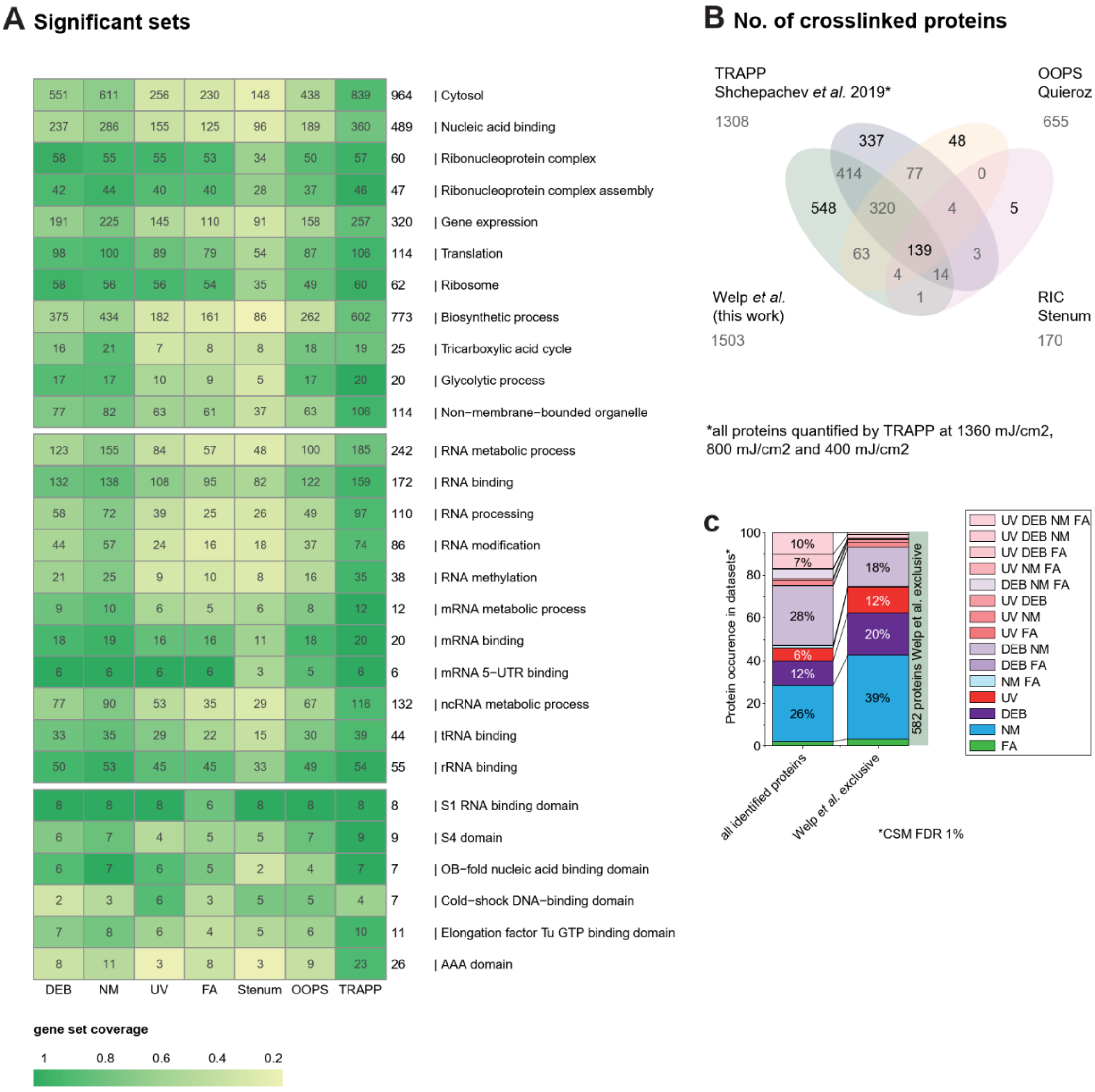


**Supplementary Figure S12.** Protein–RNA XL-MS dataset comparison. **(A)** GO term and domain gene set coverage by *E. coli* UV, DEB, NM and FA XL-MS datasets presented in this study and *E. coli* XL-MS datasets from (6-8). The terms are selected from those significantly enriched in at least one of the UV, DEB, NM or FA XL-MS datasets (padj, ≤ 0.01 for GO terms; padj, ≤ 0.05 for domains). **(B)** Venn diagram of *E. coli* RNA-crosslinked proteins identified in this study compared with proteins identified as RNA-binding in Stenum *et al.* (19) (RIC, protocol used), Quieroz *et al.* (20) (OOPS) and/or Shchepachev *et al.* (21) (TRAPP). **(C)** 100% stacked bar plot of all RNA-crosslinked proteins identified in this study and proteins found crosslinked to RNA but not identified as interaction partners in OOPS or TRAPP (excluding data from the present manuscript). Relative numbers of proteins found to be crosslinked in UV, DEB, NM and FA are indicated.

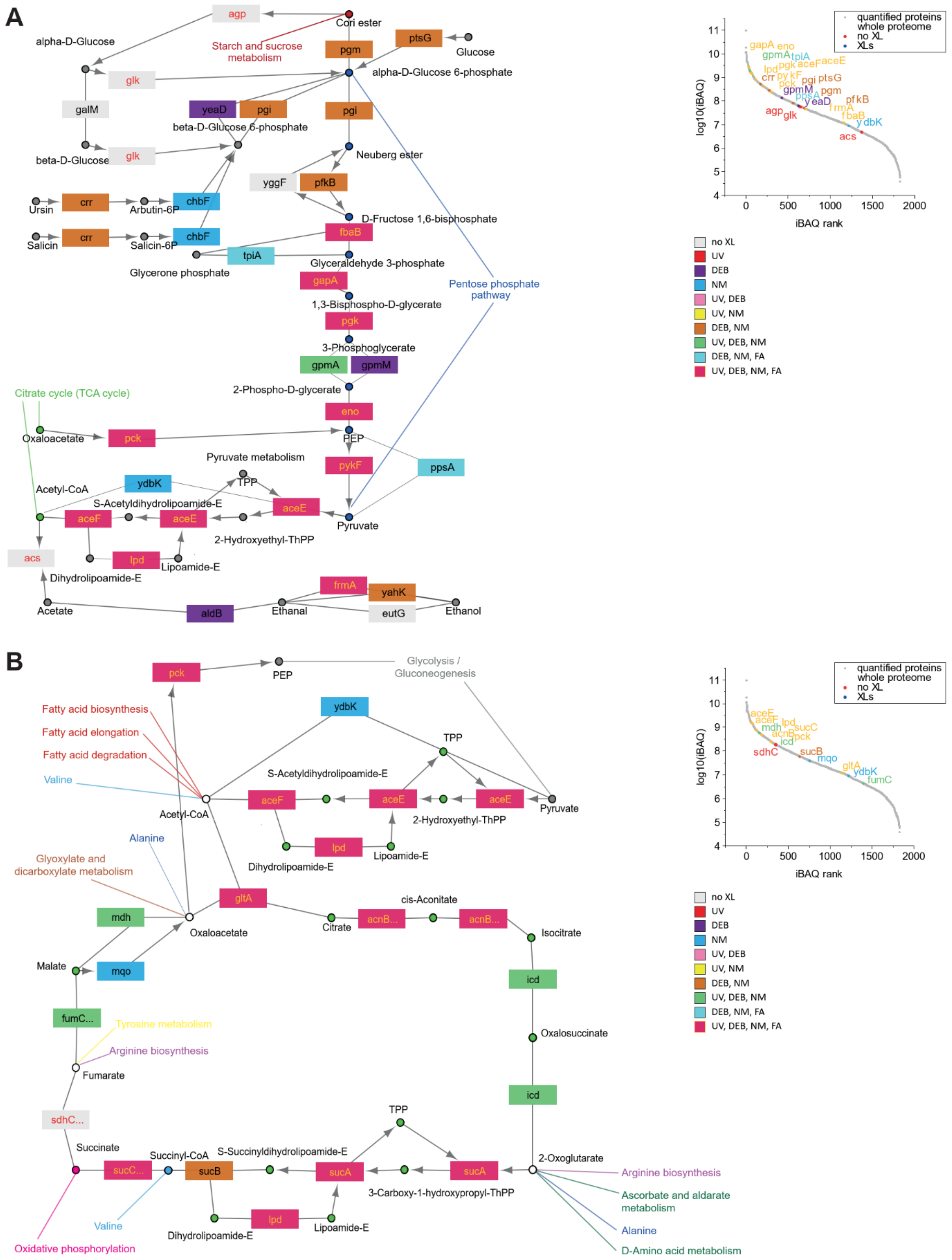


**Supplementary Figure S13.** Coverage of glycolysis- and TCA-cycle-involved enzymes by protein–RNA XL-MS. **(A)** KEGG pathway of glycolysis and gluconeogenesis. Genes are indicated by rectangles. Colours denote respective protein occurrence in UV, DEB, NM and/or

FA dataset. Grey boxes indicate proteins not found crosslinked. Genes highlighted by red font colour were not identified as crosslinked, but were quantified in the proteomics dataset. On the right: Ranked plot of proteins quantified in the *E. coli* proteomics control experiment. Font colour code matches the rectangle colour code in KEGG pathway. Glycolysis-involved proteins not found crosslinked are indicated by red font colour. **(B)** KEGG pathway of the tricarboxylic acid cycle (TCA-cycle) as in (A). On the right: Ranked plot as in (A) with TCA-cycle-involved proteins highlighted as in (A).

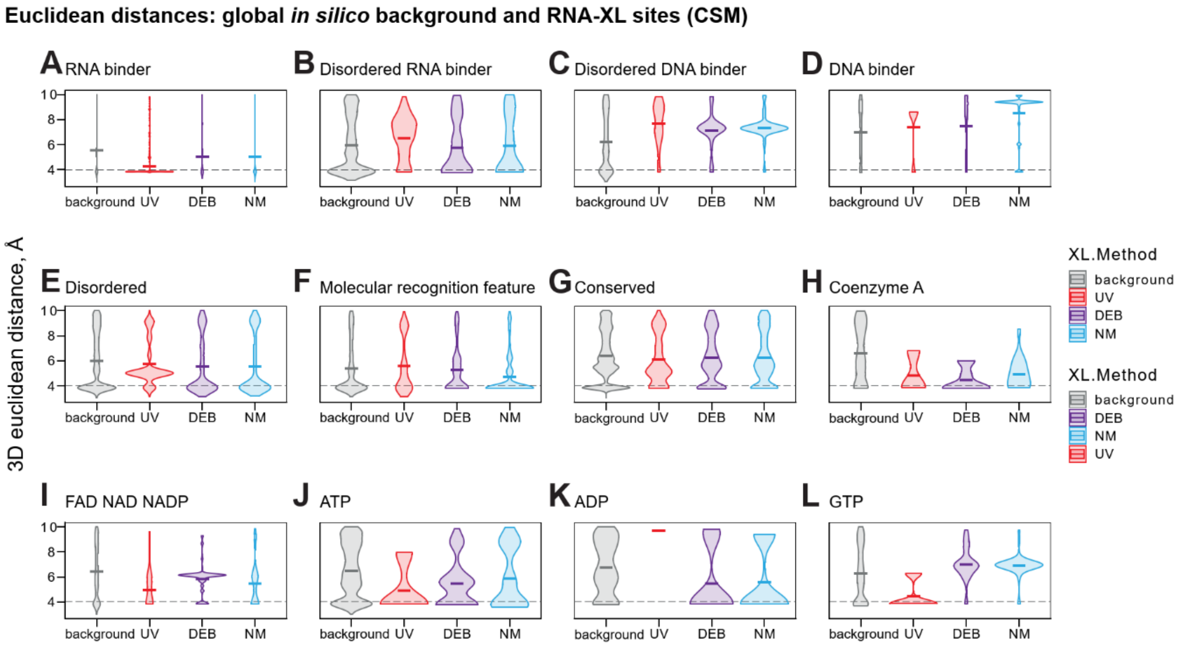


**Supplementary Figure S14.** 3D distances from crosslinked amino acids to protein sequence features. The distributions of 3D Euclidean distances between the Cα atom pairs of crosslinked residues and nearest feature residue Cα annotated in DESCRIBEPROT (22) **(A–G)** and InterPro (23) (**H–L**). The figure highlights the group means. The 4 Å range is marked, corresponding to the typical distance to the nearest residue across the proteome.

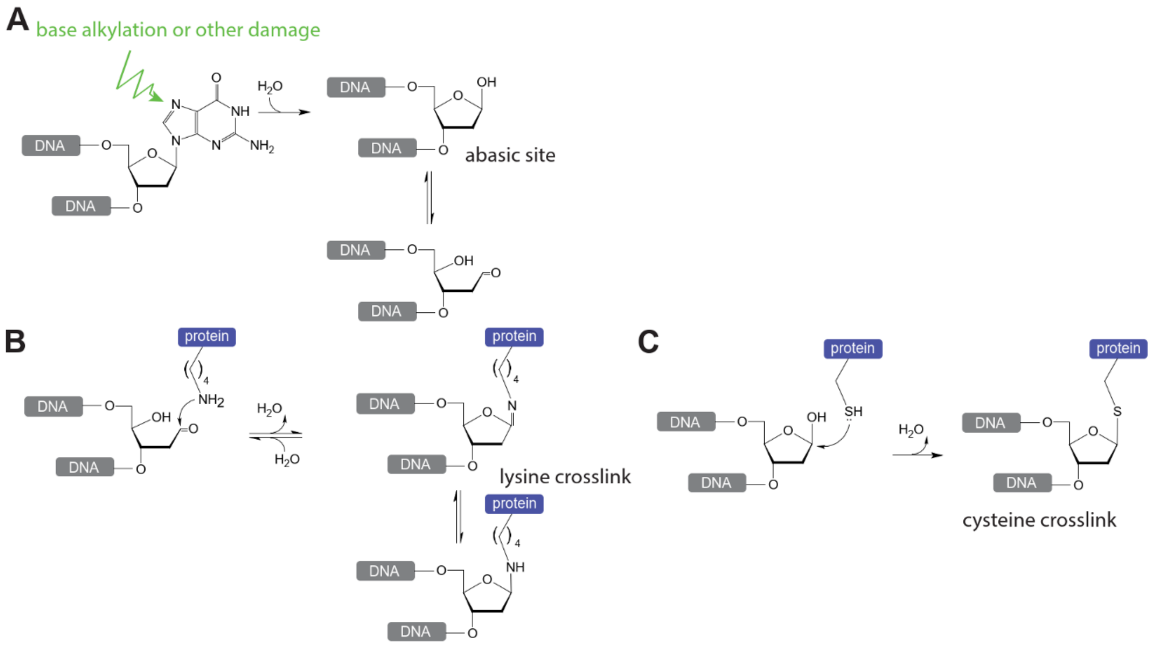


**Supplementary Figure S15.** Depurination and abasic site reactions in DNA. **(A)** Guanosine base alkylation or other forms of base damage weakening N-glyosidic bond, leading to base hydrolysis. Abasic sites exist in ring-opened aldehyde and the closed hemi-acetal form. **(B)** The acetal form of an abasic site can be attacked at the C1' position by the nucleophilic ε-amino group of lysines to create a crosslink via a reversible Schiff-base intermediate (24). **(C)** The acetal form of an abasic site can be attacked at the C1' position by the nucleophilic thiol group in cysteines to form a stable covalent linkage (25).

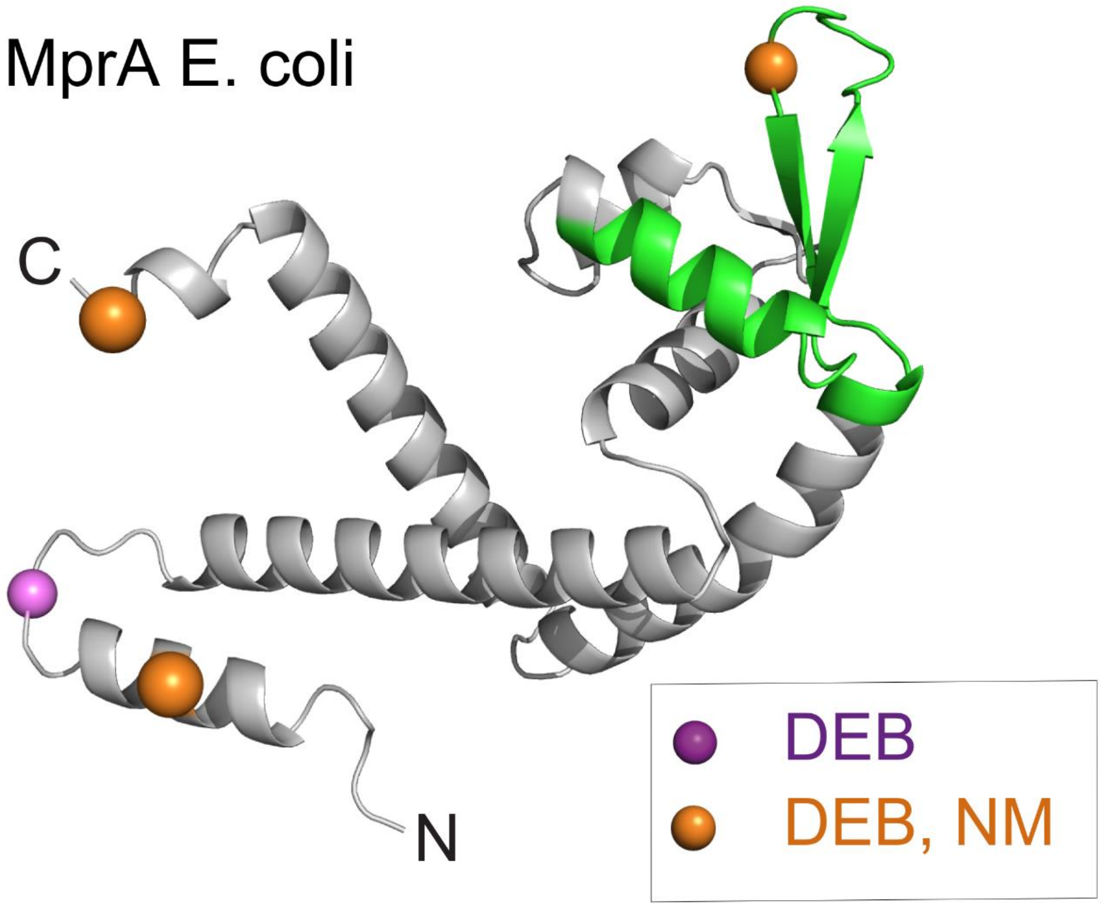


**Supplementary Figure S16.** Cartoon representation and electrostatic surface of the AlphaFold (12) predicted structure of E. coli MprA transcription regulator (Uniprot ID P0ACR9|MPRA_ECOLI) (Oligo)nucleotide-crosslinked amino acids (Cα) are highlighted as purple (DEB), and orange (DEB and NM) spheres. HTH domain is shown in green. Mapped crosslink positions are based on CSMs filtered for 1% FDR, and localisation score ≥1. All four crosslinked amino acid positions were characterized by CSM counts >3 with RNA NuXL presets. Two of them shown by bigger spheres (Met-11, Met-175) were also identified using DNA NuXL presets and also met the requirements of localisation score ≥1, and CSM counts >3.

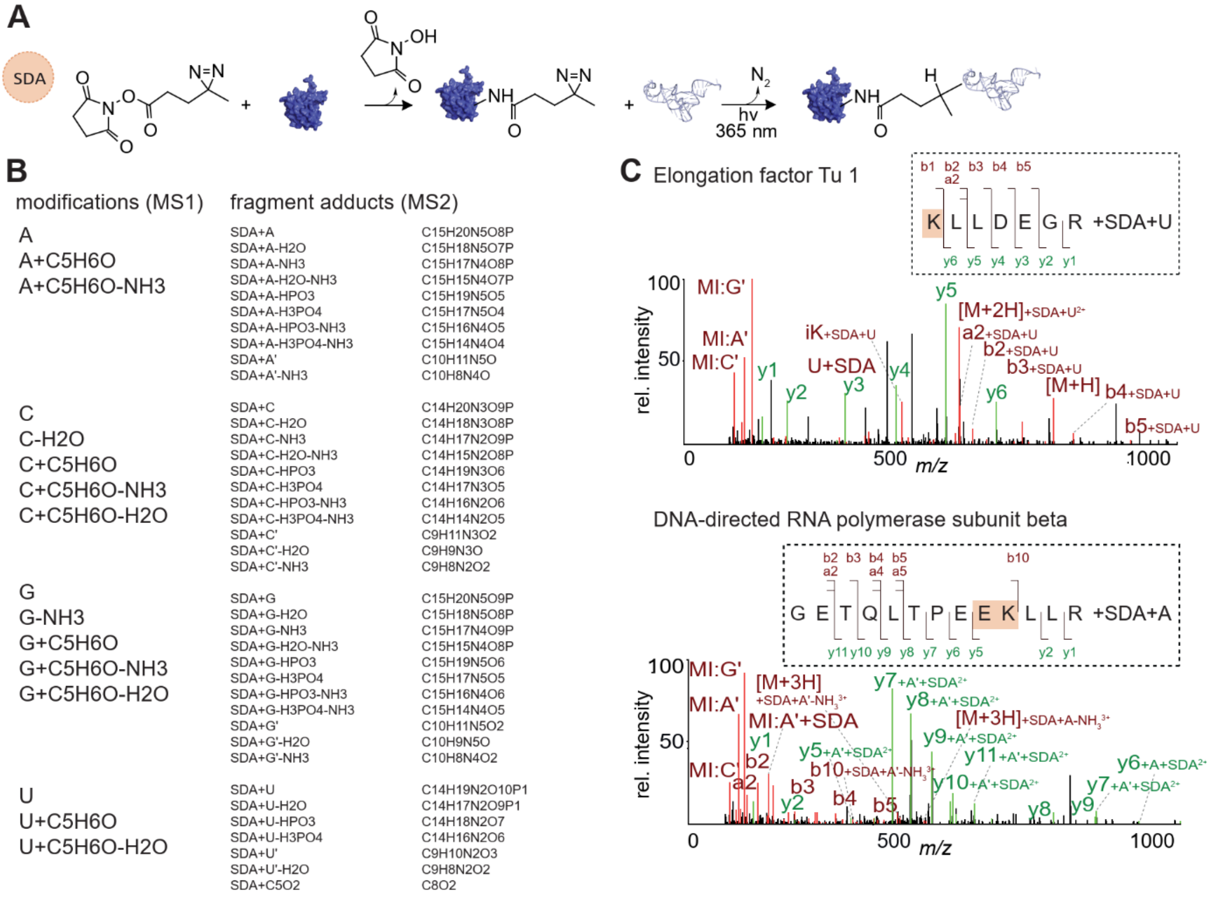


**Supplementary Figure S17.** Protein–nucleic acid XL-MS using SDA. **(A)** Proposed chemical protein–RNA crosslinking reaction mechanism for SDA. **(B)** Sum formula of SDA RNA-crosslink adducts for NuXL analysis. Expected modifications on MS1 level and fragment adducts in MS2 scans. **(C)** MS2 crosslink spectra for SDA-crosslinked peptide–RNA-(oligo)nucleotides from *E. coli* ribosomes resulting from SDA XL-MS and NuXL analysis using settings listed in B. Upper panel: MS2 spectrum of elongation factor Tu 1 peptide KLLDEGR crosslinked to uridine-monophosphate by SDA. Lower panel: MS2 spectrum of DNA-directed RNA polymerase subunit beta peptide GETQLTPEEKLLR crosslinked to adenosine-monophosphate by SDA. Annotated peaks are highlighted in green (y-ions) or red (marker ions and a-/b-ions). Crosslinked amino acids are highlighted in orange.

### SUPPLEMENTARY TABLES

**Supplementary Table S1**. Crosslinker-specific presets in NuXL. Table including all sum formulae for RNA/DNA adducts defined as presets in NuXL. Sheets contain different presets. Columns include: target_nucleotides, definitions (sum formulae) of monophosphate-nucleotides/deoxyribose; modifications, definitions (sum formulae) of RNA/DNA adducts on peptide precursor (MS1); fragment_adducts, definitions (sum formulae) of RNA/DNA adducts on peptide fragments (MS2).

**Supplementary Table S2**. Protein–RNA crosslinks in *E. coli* ribosomes. Table including RNA crosslink-spectrum matches (CSMs) from UV, DEB, NM and FA XL-MS data from purified *E. coli* ribosomes. CSMs are filtered for 1% FDR and rescored by Percolator. Among other values, the table includes: idxml, source file; index, spectrum index number; RT, retention time; precursor *m/z*; score, main score calibrated by percolator algorithm using subscores; PeakAnnotations, MS2 peaks *m/z* values, annotations and intensities; NuXL:NA, RNA adduct on peptide precursor (MS1); NuXL:NT, crosslinked nucleobase.

**Supplementary Table S3**. List of all subscores used in Percolator-based (3) rescoring. List of all subscores used in Percolator-based rescoring.

**Supplementary Table S4.** Protein–RNA crosslinks in *in vitro* protein–RNA complexes. Table including RNA crosslink-spectrum matches (CSMs) from UV, DEB, NM and FA XL-MS data from *in vitro* analysed protein–RNA complexes. Sheets include results from: Hsh49-Cus1-U2RNA; Dnmt2-tRNA$^{Asp}$, NELF-TAR complexes separately. CSMs are filtered for 1% FDR and rescored by Percolator. Among other values, the table includes: idxml, source file; index, spectrum index number; RT, retention time; precursor *m/z*; score, main score calibrated by percolator algorithm using subscores; PeakAnnotations, MS2 peaks *m/z* values, annotations and intensities; NuXL:NA, RNA adduct on peptide precursor (MS1); NuXL:NT, crosslinked nucleobase. The amino-acid positions are indicated according to the numbering in the applied protein sequence files (see uploaded fasta files in ProteomeXchange).

**Supplementary Table S5**. Protein–RNA crosslinks from *E. coli*. Table including RNA crosslink-spectrum matches (CSMs) from UV, DEB, NM and FA XL-MS data from *E. coli* cells. CSMs are filtered for 1% FDR and rescored by Percolator (3). Among other values, the table includes:

fraction, S30/S100 fraction of lysate; idxml, source file; index, spectrum index number; RT, retention time; precursor *m/z*; score, main score calibrated by Percolator algorithm using subscores; PeakAnnotations, MS2 peaks *m/z* values, annotations and intensities; NuXL:NA, RNA adduct on peptide precursor (MS1); NuXL:NT, crosslinked nucleobase. File available via figshare under the following link: https://figshare.com/s/6b2091a24a327ed546e4.

**Supplementary Table S6**. Label-free quantitative *E. coli* proteomics results. Table including modified proteingroups.txt output from MaxQuant search (26) of *E. coli* label-free quantitative proteomics dataset. *Inter alia*, columns include iBAQ values (18) for identified proteins.

**Supplementary Table S7**. Amino acid / sequence feature tests. Table with enrichment analysis of RNA-crosslinks against sequence features of *E. coli* proteome. Secondary structure counts from S4PRED, amino-acid counts, feature counts from DESCRIBEPROT (22) and InterPro (23) are included together with the distances to the next feature in 10 Å radius (1D, 3D-euclidean and 3D-SAS distances). The analysis is performed on the detected peptide and crosslinked residue levels, with and without consideration of crosslink-spectrum match counts in a variety of background assumptions. Residue-level enrichment only considers the CSMs with the NuXL best localisation threshold ≥ 1. Provided are the average values of the detected crosslinks, expected background values, *t*-test outputs, nonparametric goodness-of-fit test outputs, adjusted *p* values for both tests (BH procedure) and sample sizes.

**Supplementary Table S8**. Gene set overrepresentation analysis. Table with a hypergeometric test overrepresentation analysis of identified *E. coli* proteins across crosslinking methods. The proteins were filtered for the detection of at least 1 unique peptide sequence. The table shows the gene set names together with the overlap size, significance of the overlap and the estimates of the gene set redundancy.

**Supplementary Table S9**. Proximal feature enrichment. Table with significance of co-localisation of crosslinked residues and annotated protein sequence features. The table shows the outcomes of binomial tests with feature frequency weights: set sizes of foreground and background within and outside the 4 Å radius, the *p* value of enrichment, adjusted *p* value (BH method) and 0.95 confidence interval.

**Supplementary Table S10**. Protein–DNA crosslinks in *Saccharomyces cerevisiae* nucleosomes. Table including DNA crosslink-spectrum matches (CSMs) from UV, DEB, NM and FA XL-MS data from purified yeast *S. cerevisiae* nucleosomes. CSMs are filtered for 1% FDR and rescored by Percolator. Among other values, the table includes: idxml, source file; index, spectrum index number; RT, retention time; precursor *m/z*; score, main score calibrated by percolator algorithm using subscores; PeakAnnotations, MS2 peaks *m/z* values, annotations and intensities; NuXL:NA, DNA adduct on peptide precursor (MS1); NuXL:NT, crosslinked nucleobase/deoxyribose.

**Supplementary Table S11**. Protein–DNA crosslinks from *E. coli*. Table including DNA crosslink-spectrum matches (CSMs) from UV, DEB, NM and FA XL-MS data from *E. coli* cells. CSMs are filtered for 1% FDR and rescored by Percolator (3). Among other values, the table includes: fraction, S30/S100 fraction of lysate; idxml, source file; index, spectrum index number; RT, retention time; precursor *m/z*; score, main score calibrated by percolator algorithm using subscores; PeakAnnotations, MS2 peaks *m/z* values, annotations and intensities; NuXL:NA, DNA adduct on peptide precursor (MS1); NuXL:NT, crosslinked nucleobase. File available via figshare through the following link: https://figshare.com/s/6b2091a24a327ed546e4.

**Supplementary Table S12**. Ambiguous RNA/DNA mass adducts in XL-MS. Table containing ambiguous RNA/DNA mass adducts in XL-MS. Sum formulae corresponding to ambiguous mass adducts are listed with respective monoisotopic masses and possible RNA/DNA adducts.

**Supplementary Table S13**. A list of crosslinkable HTH-motifs in the *E. coli* proteome.

**SUPPLEMENTARY FILES**

**Supplementary File S1**. Visualisation of RNA-crosslinks in *E. coli*. PDF file displaying 1D sequence representations of all crosslinked proteins from UV, DEB, NM and FA XL-MS *E. coli in vivo* datasets containing at least one crosslink site with a localisation score ≥ 1. The header provides information on: protein accession, protein name, protein abundance ($log_{10}$iBAQ values) from label-free quantitative proteomics dataset presented in this study ("tryptic"), PAXdb (15) K12 strain protein abundance in ppm, PAXdb *E. coli* protein abundance in ppm. For all abundance values, the respective abundance quantile to which the values correspond is given. The plot includes the protein sequence plotted schematically in 1D, including positions of UV, DEB and NM crosslink sites (localisation score ≥1) and indicated CSM counts (y axis), UV, DEB, NM and FA crosslinked peptide positions, and potential tryptic cleavages.

**Supplementary File S2**. Example of MS2 spectra of UV-induced *E. coli* peptide–RNA crosslinks (S30 fraction, 2 replicate) identified by NuXL, but not by MSFragger-Labile workflow, visualised by OpenMS ToppView from NuXL output. C-$NH_3$ additions to the peptide may also represent U-$H_2O$. NuXL outputs are provided in Supplementary Table S5.
