## Supplementary File S1 for "Chemical crosslinking extends and complements UV crosslinking in analysis of RNA/DNA nucleic acid–protein interaction sites by mass spectrometry"

P60422
RL2_ECOLI 50S ribosomal protein L2

− Abundance:
tryptic [log10 Intensity]: 9.26 (Q 93)
PAXdb K12 strain [ppm]: 3.89 (Q 100)
PAXdb E.coli [ppm]: 3.33 (Q 98)

− RNA functions:
Ribosomal Proteins L2, RNA binding domain; RNA binding; rRNA binding

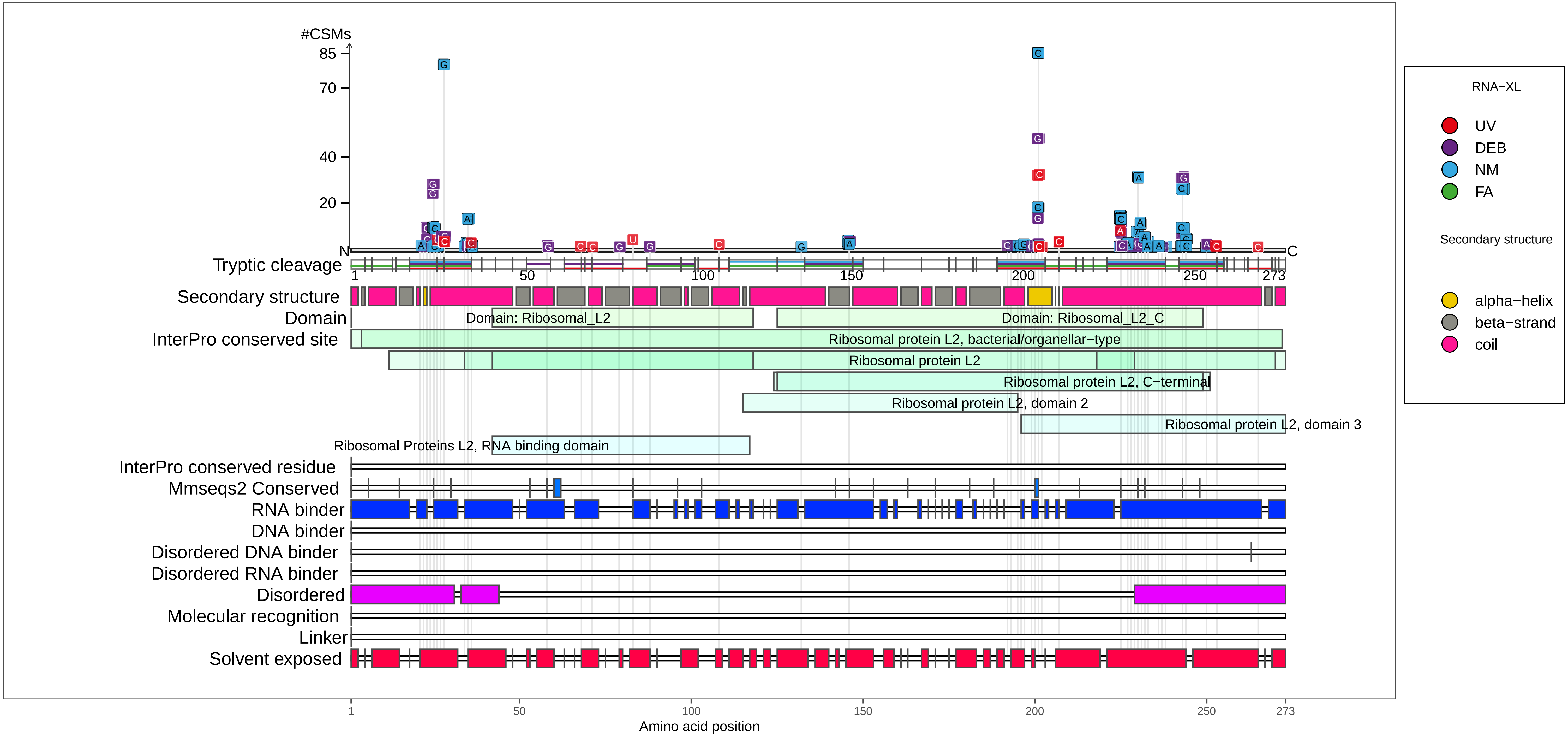

P0A7V3
RS3_ECOLI 30S ribosomal protein S3

− Abundance:
tryptic [log10 Intensity]: 9.66 (Q 98)
PAXdb K12 strain [ppm]: 3.87 (Q 100)
PAXdb E.coli [ppm]: 3.45 (Q 98)

− RNA functions:
mRNA binding; RNA binding; rRNA binding

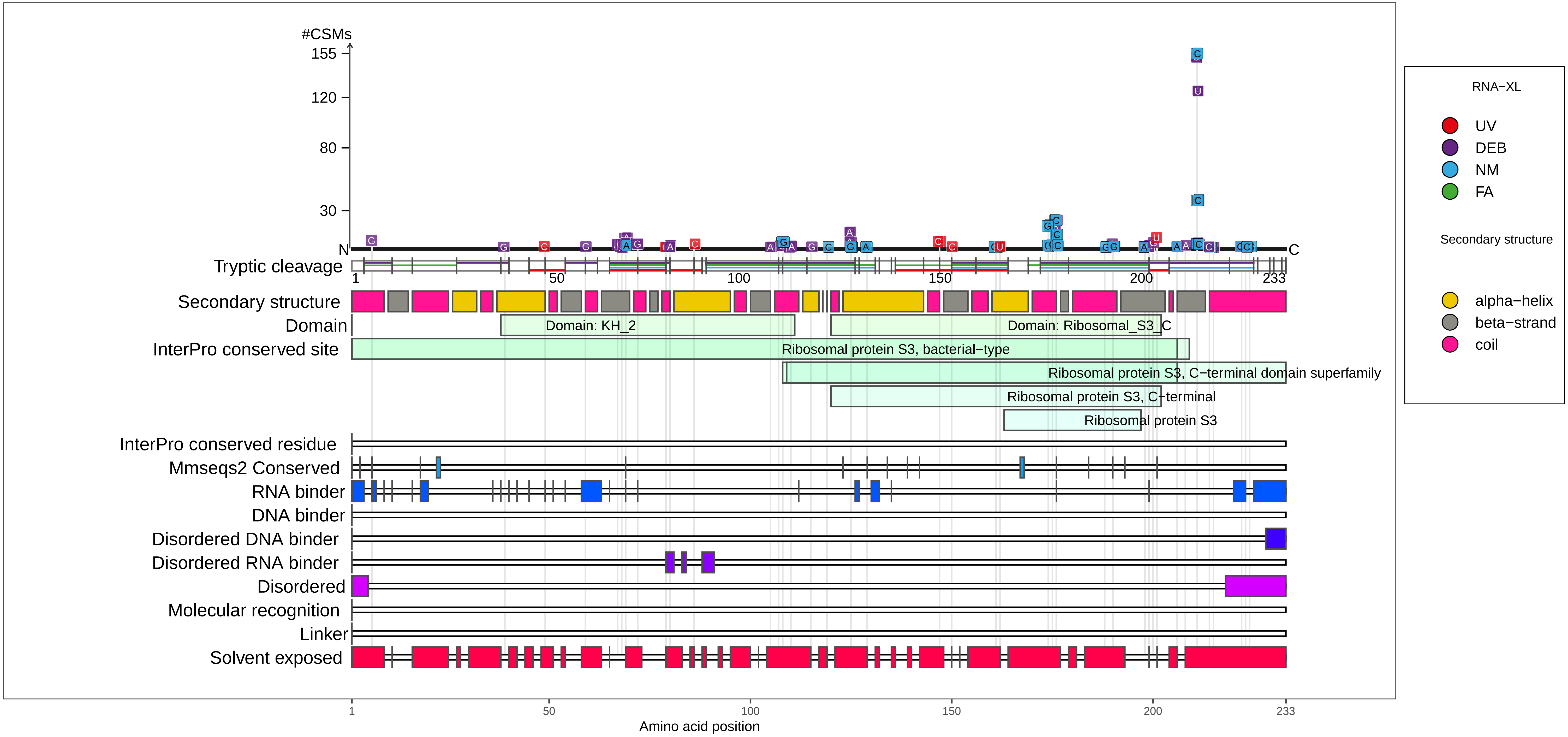

P0A6Y8
DNAK_ECOLI Chaperone protein DnaK

− Abundance:
tryptic [log10 Intensity]: 9.7 (Q 98)
PAXdb K12 strain [ppm]: 3.78 (Q 99)
PAXdb E.coli [ppm]: 3.89 (Q 100)

− RNA functions: not annotated

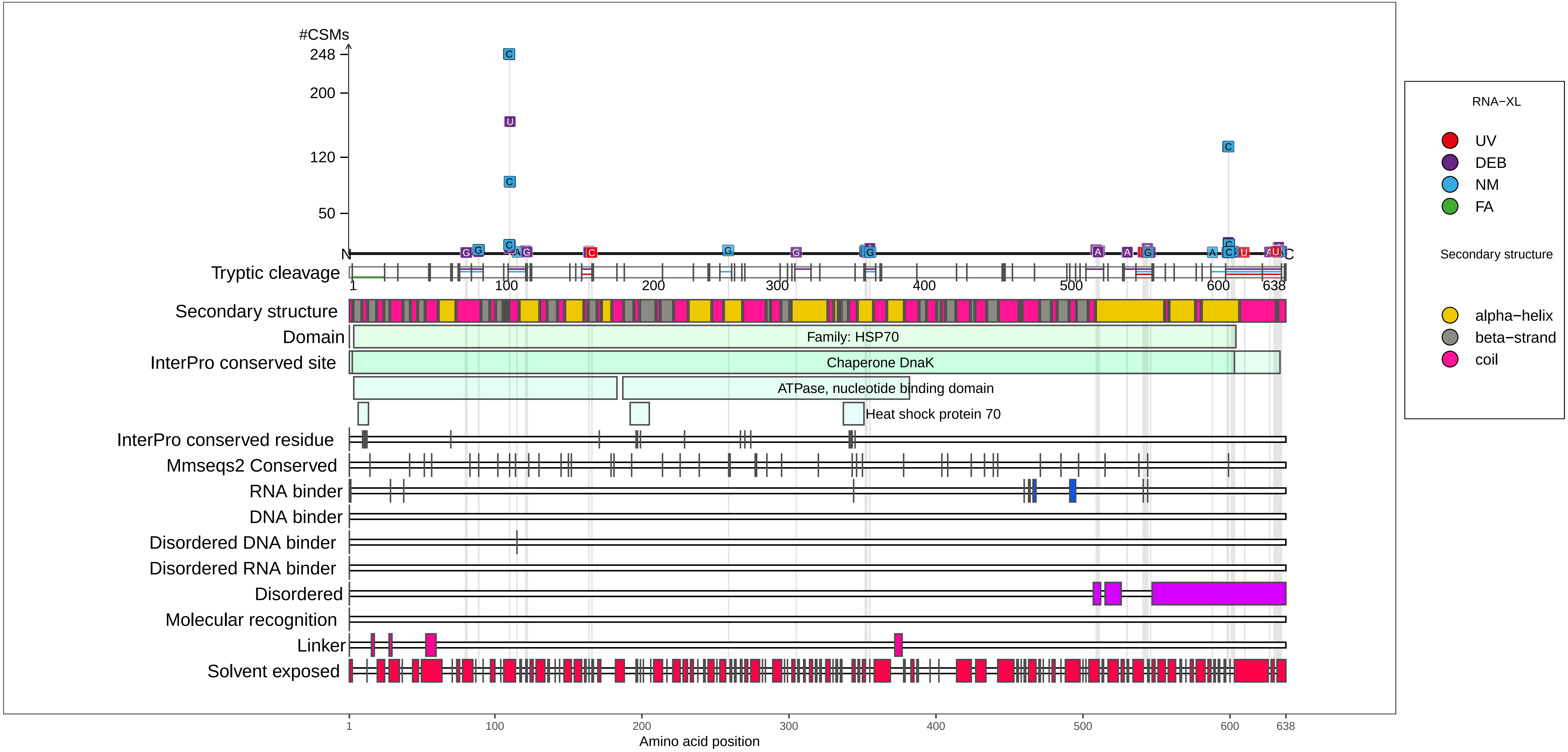

P0CE47
EFTU1_ECOLI Elongation factor Tu 1

− Abundance:
tryptic [log10 Intensity]: NA (Q NA)
PAXdb K12 strain [ppm]: 3.7 (Q 99)
PAXdb E.coli [ppm]: 4.27 (Q 100)

− RNA functions:
RNA binding

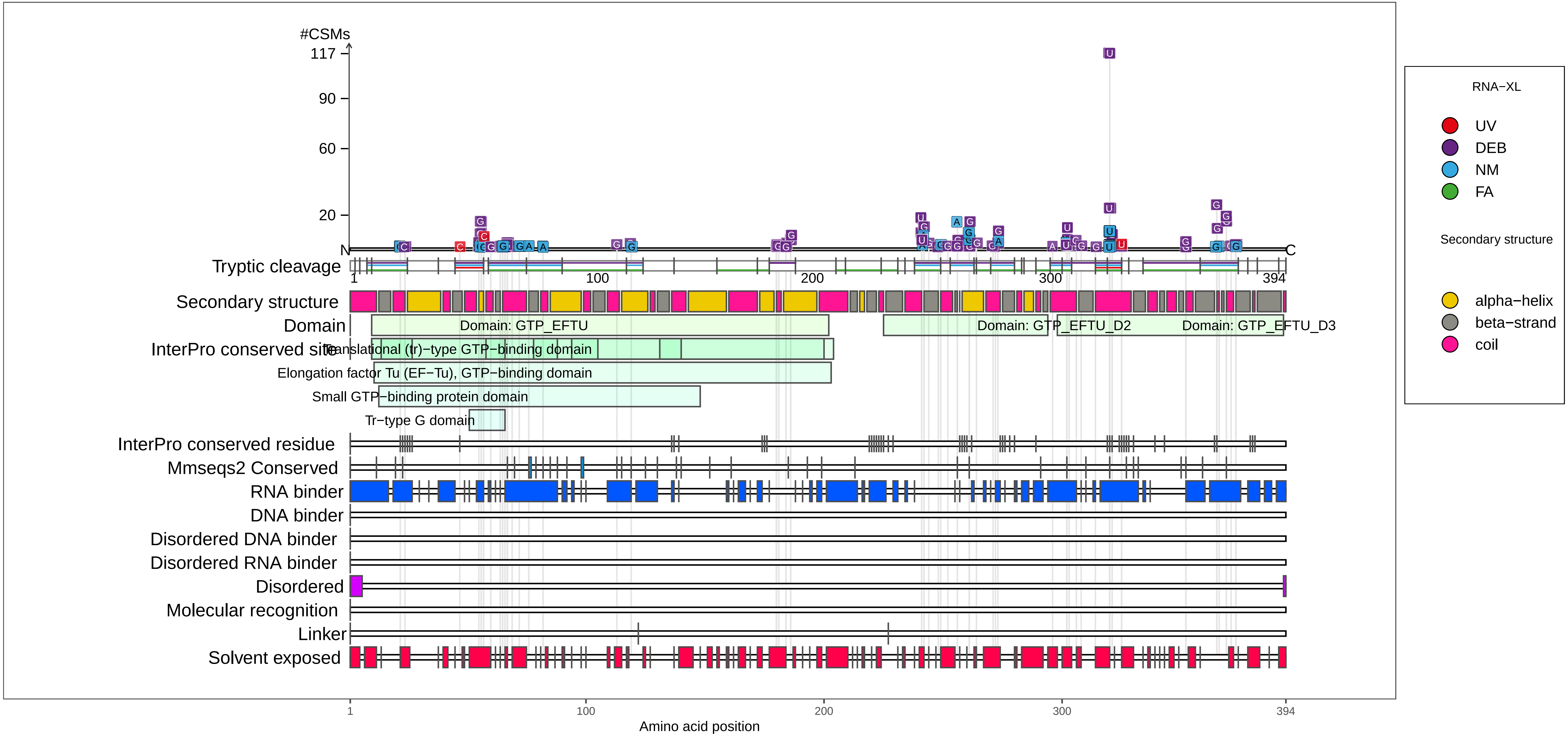

P0A6M8
EFG_ECOLI Elongation factor G

− Abundance:
tryptic [log10 Intensity]: 10.41 (Q 100)
PAXdb K12 strain [ppm]: 3.24 (Q 94)
PAXdb E.coli [ppm]: 3.48 (Q 99)

− RNA functions:
RNA binding

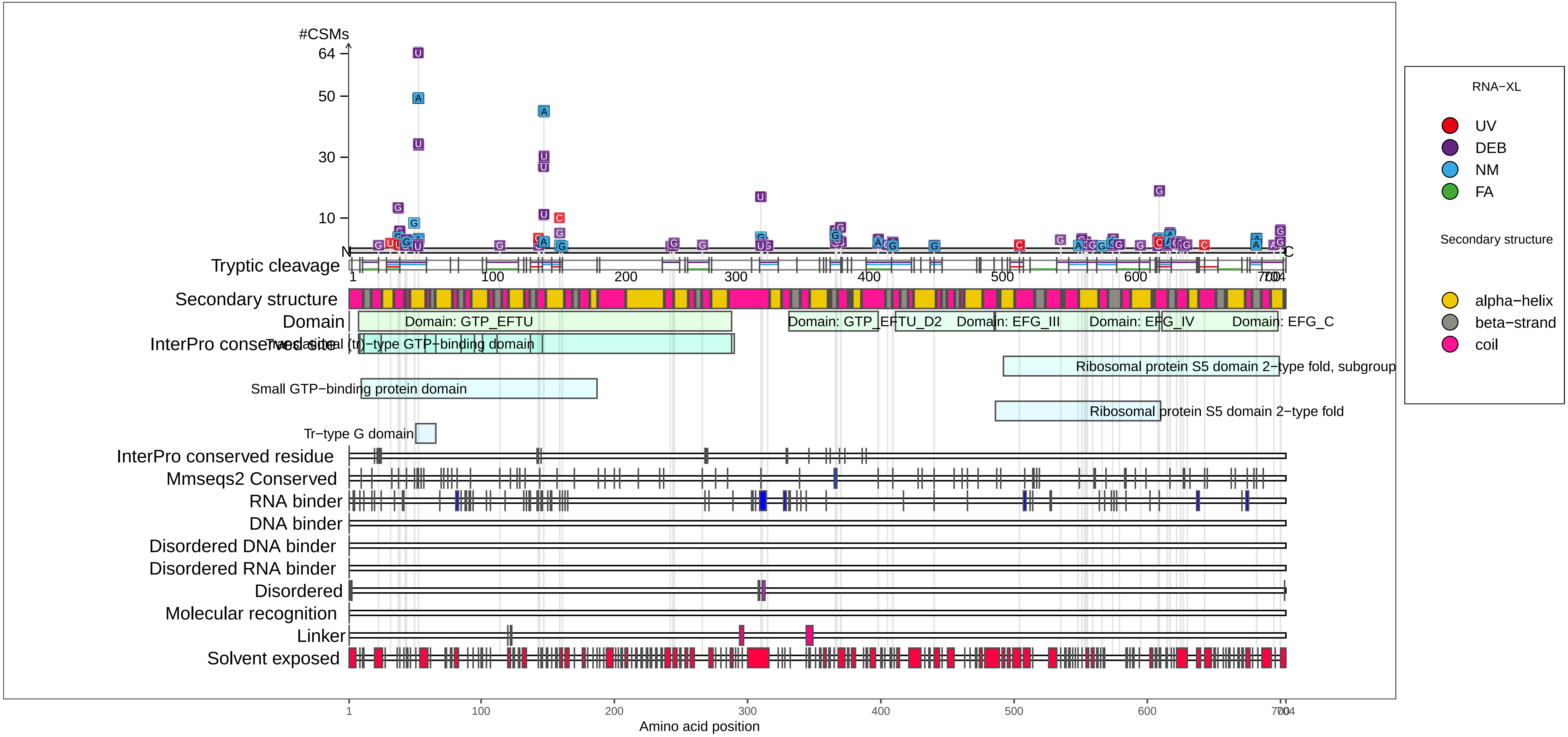

P0AG67
RS1_ECOLI 30S ribosomal protein S1

− Abundance:
tryptic [log10 Intensity]: 10.07 (Q 100)
PAXdb K12 strain [ppm]: 3.53 (Q 98)
PAXdb E.coli [ppm]: 3.38 (Q 98)

− RNA functions:
mRNA binding; RNA binding; S1 RNA binding domain

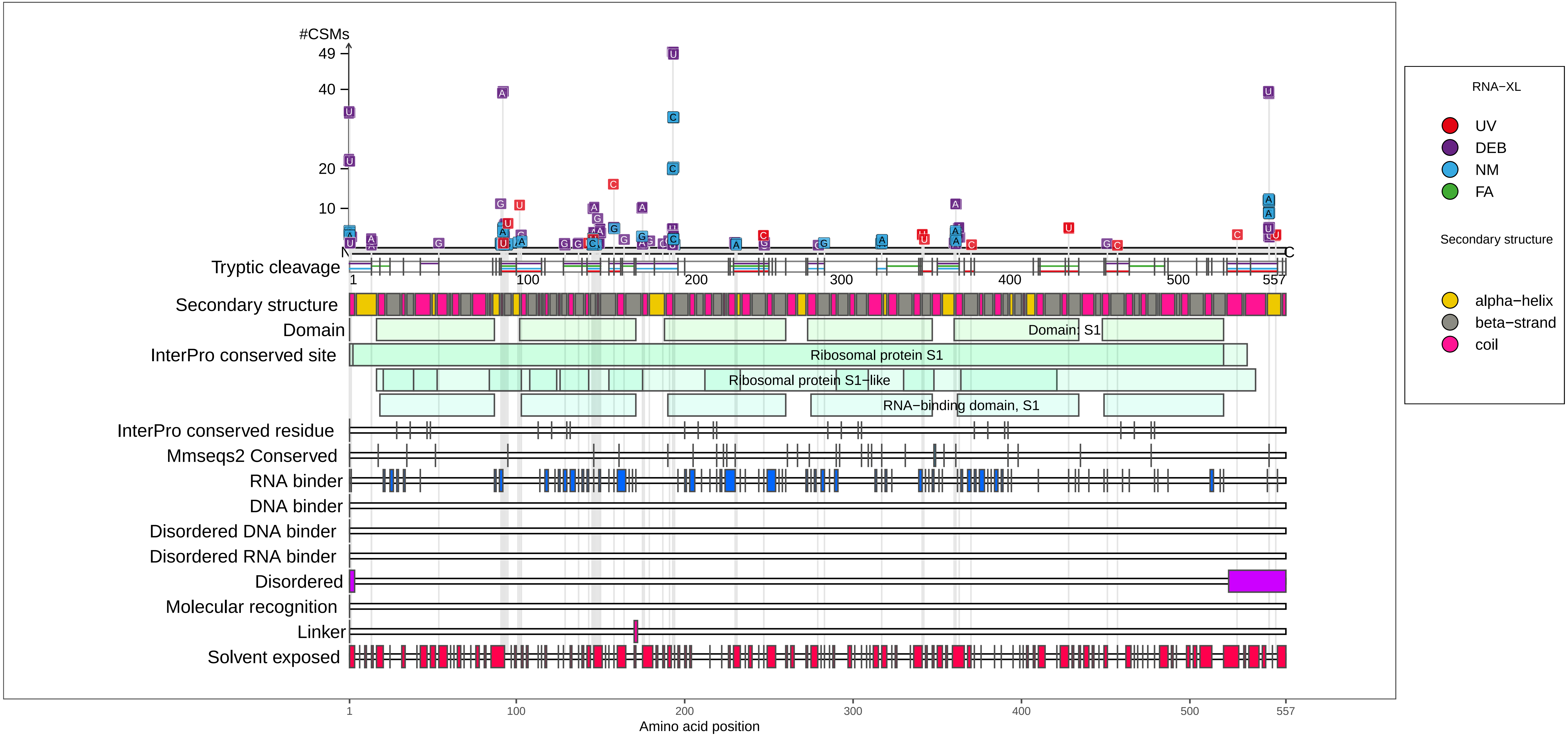

P61175
RL22_ECOLI 50S ribosomal protein L22

− Abundance:
tryptic [log10 Intensity]: 9.75 (Q 98)
PAXdb K12 strain [ppm]: 4.03 (Q 100)
PAXdb E.coli [ppm]: 3.62 (Q 100)

− RNA functions:
RNA binding; rRNA binding

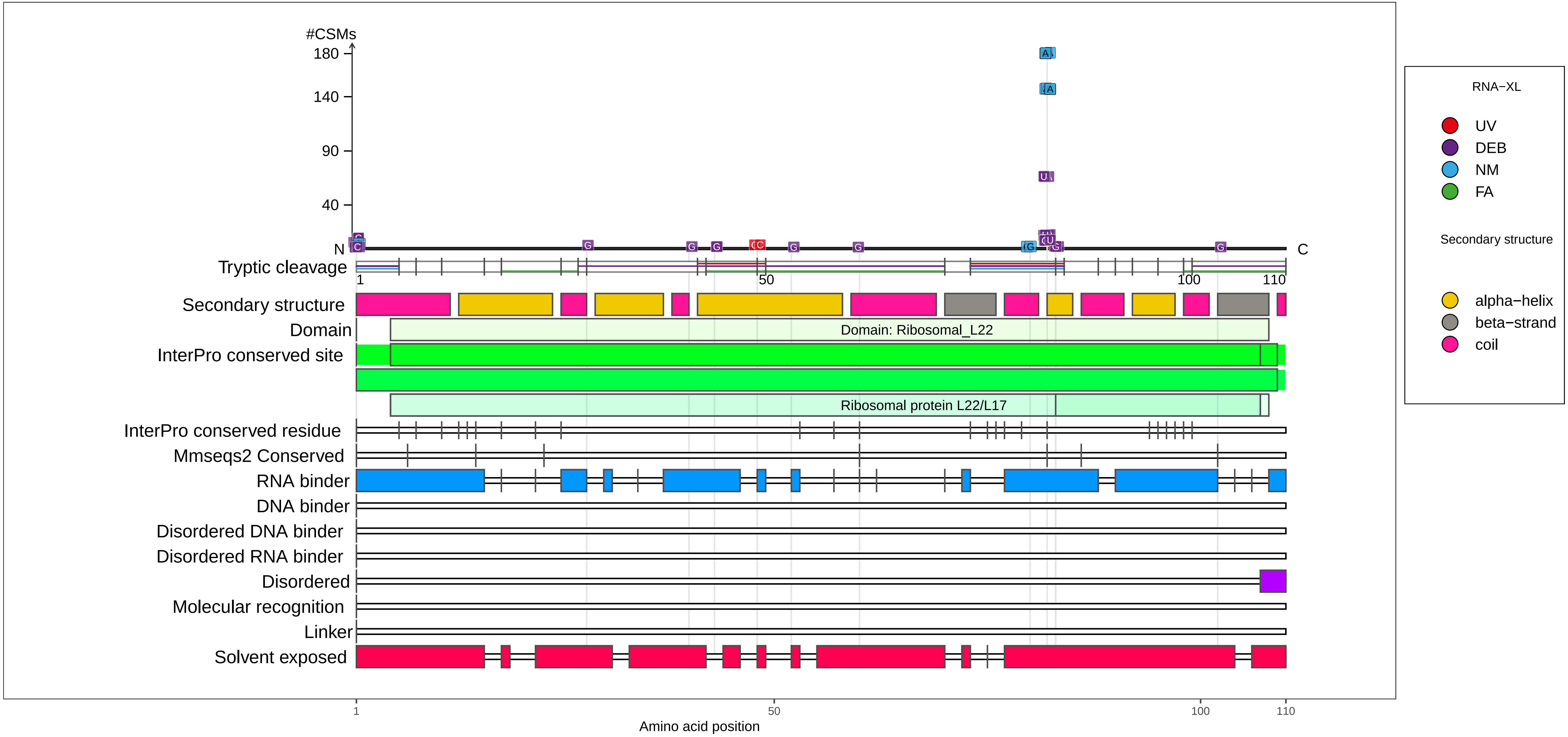

P0ADY3
RL14_ECOLI 50S ribosomal protein L14

− Abundance:
tryptic [log10 Intensity]: 8.73 (Q 82)
PAXdb K12 strain [ppm]: 3.68 (Q 99)
PAXdb E.coli [ppm]: 3.57 (Q 99)

− RNA functions:
RNA binding; rRNA binding

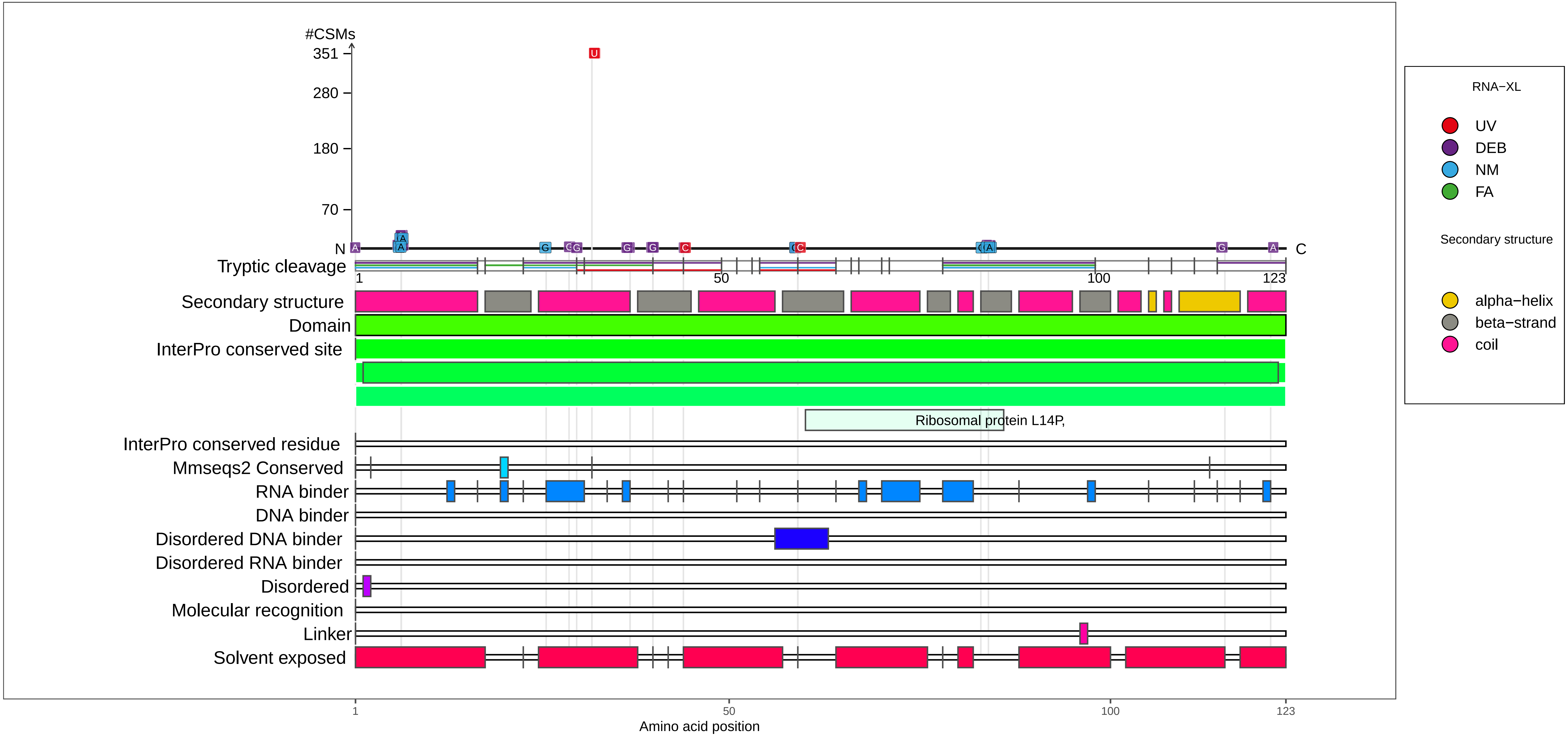

P0A7L0
RL1_ECOLI 50S ribosomal protein L1

− Abundance:
tryptic [log10 Intensity]: 9.33 (Q 94)
PAXdb K12 strain [ppm]: 3.48 (Q 97)
PAXdb E.coli [ppm]: 3.37 (Q 98)

− RNA functions:
ncRNA metabolic process; ncRNA processing; RNA binding; RNA metabolic process
RNA processing; rRNA binding; rRNA metabolic process; rRNA processing; tRNA binding

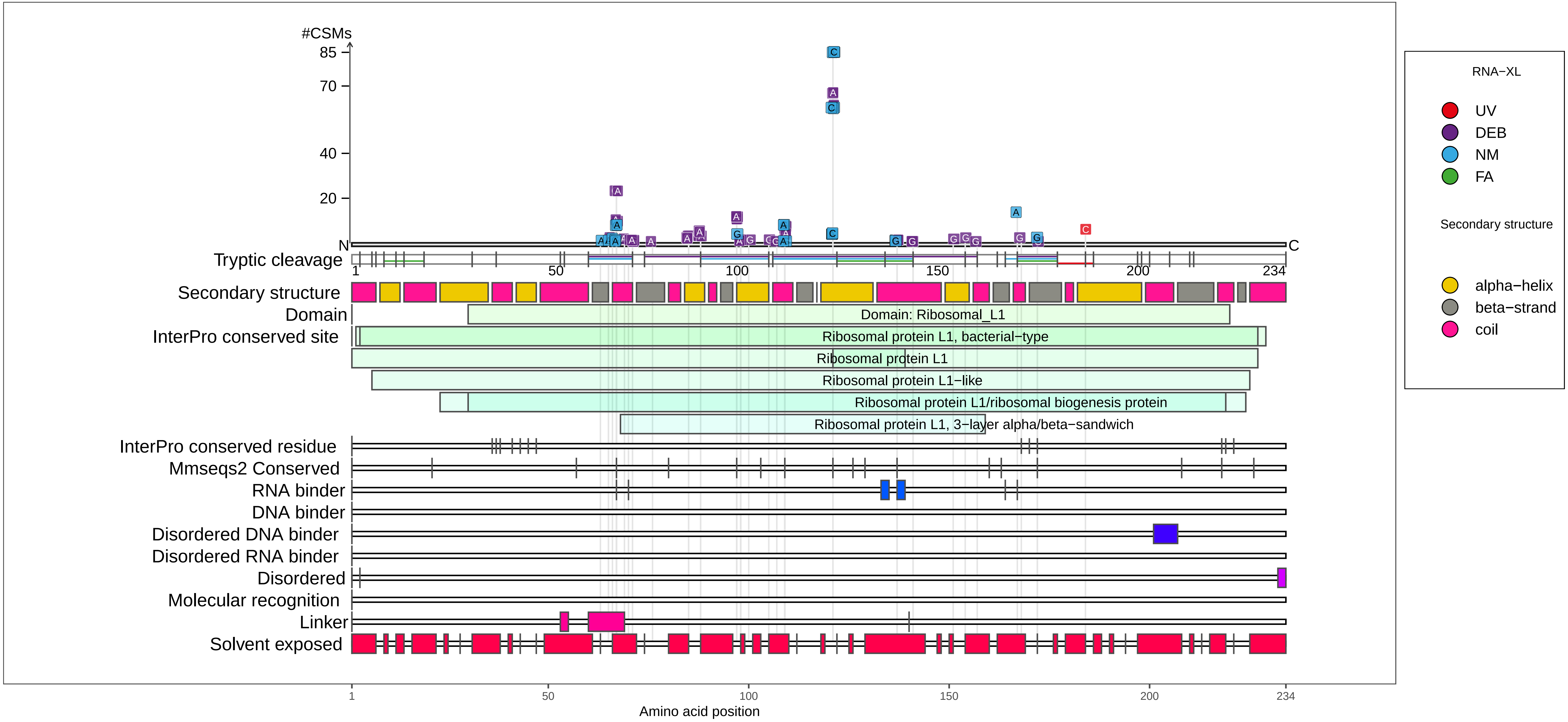

P0A850
TIG_ECOLI Trigger factor

− Abundance:
tryptic [log10 Intensity]: 10.15 (Q 100)
PAXdb K12 strain [ppm]: 3.53 (Q 98)
PAXdb E.coli [ppm]: 3.52 (Q 99)

− RNA functions: not annotated

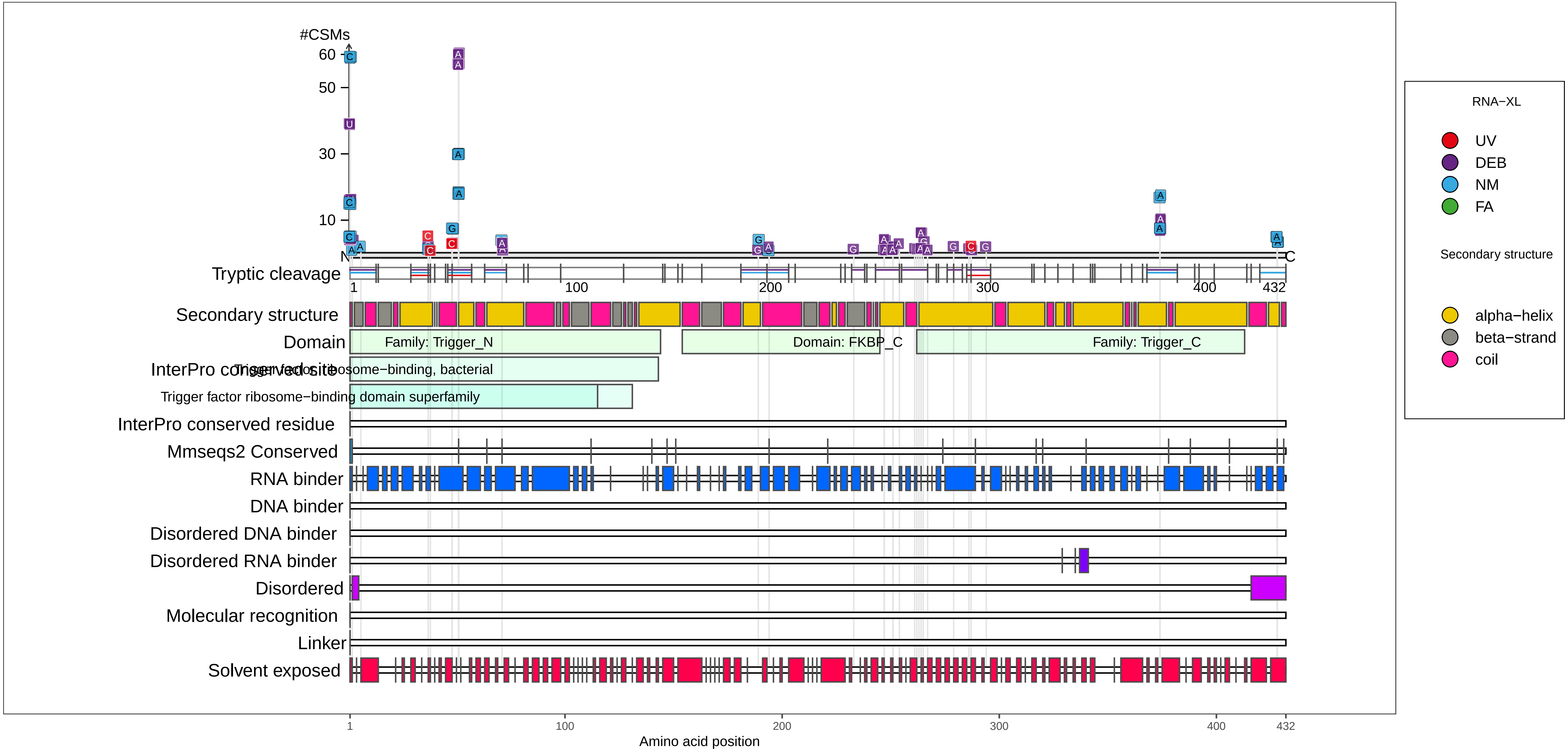

P0A7R9
RS11_ECOLI 30S ribosomal protein S11

− Abundance:
tryptic [log10 Intensity]: 8.89 (Q 86)
PAXdb K12 strain [ppm]: 3.47 (Q 97)
PAXdb E.coli [ppm]: 3.04 (Q 95)

− RNA functions:
mRNA 5−UTR binding; mRNA binding; ncRNA metabolic process; ncRNA processing; RNA binding
RNA metabolic process; RNA processing; rRNA binding; rRNA metabolic process
rRNA processing

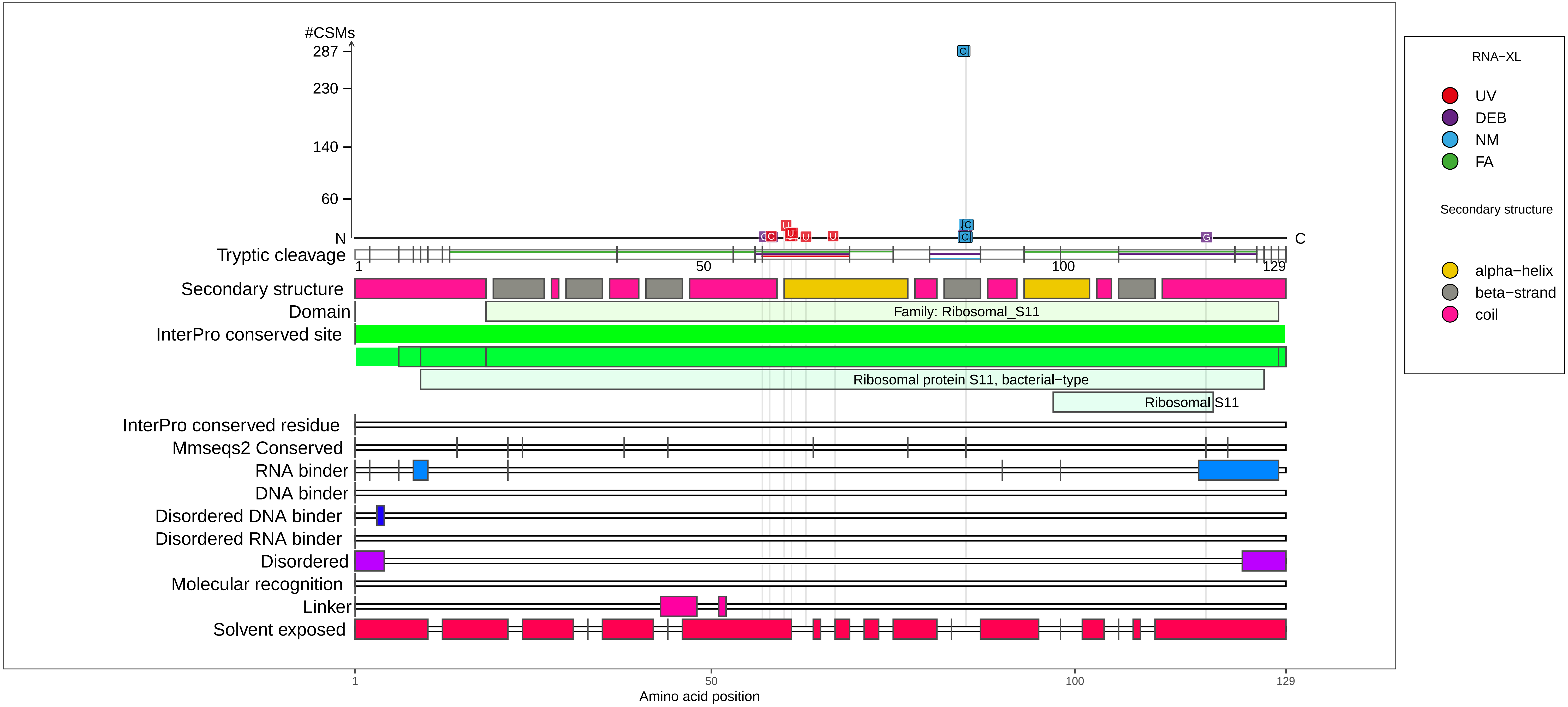

P0A6F5
CH60_ECOLI Chaperonin GroEL

− Abundance:
tryptic [log10 Intensity]: 10.15 (Q 100)
PAXdb K12 strain [ppm]: 3.48 (Q 97)
PAXdb E.coli [ppm]: 3.53 (Q 99)

− RNA functions: not annotated

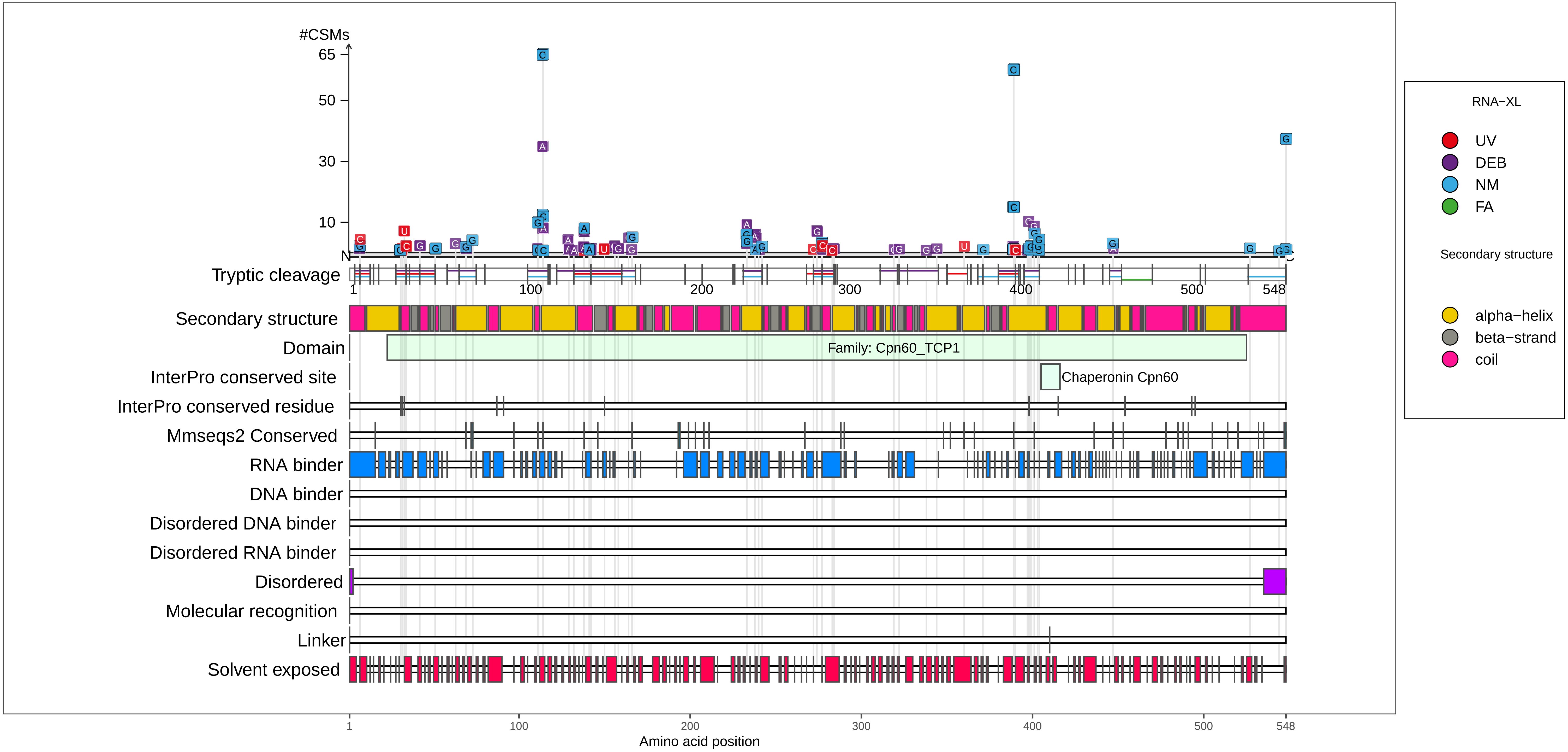

P0A7M9
RL31_ECOLI 50S ribosomal protein L31

− Abundance:
tryptic [log10 Intensity]: 8.83 (Q 84)
PAXdb K12 strain [ppm]: 3.53 (Q 98)
PAXdb E.coli [ppm]: 2.89 (Q 93)

− RNA functions:
RNA binding; rRNA binding

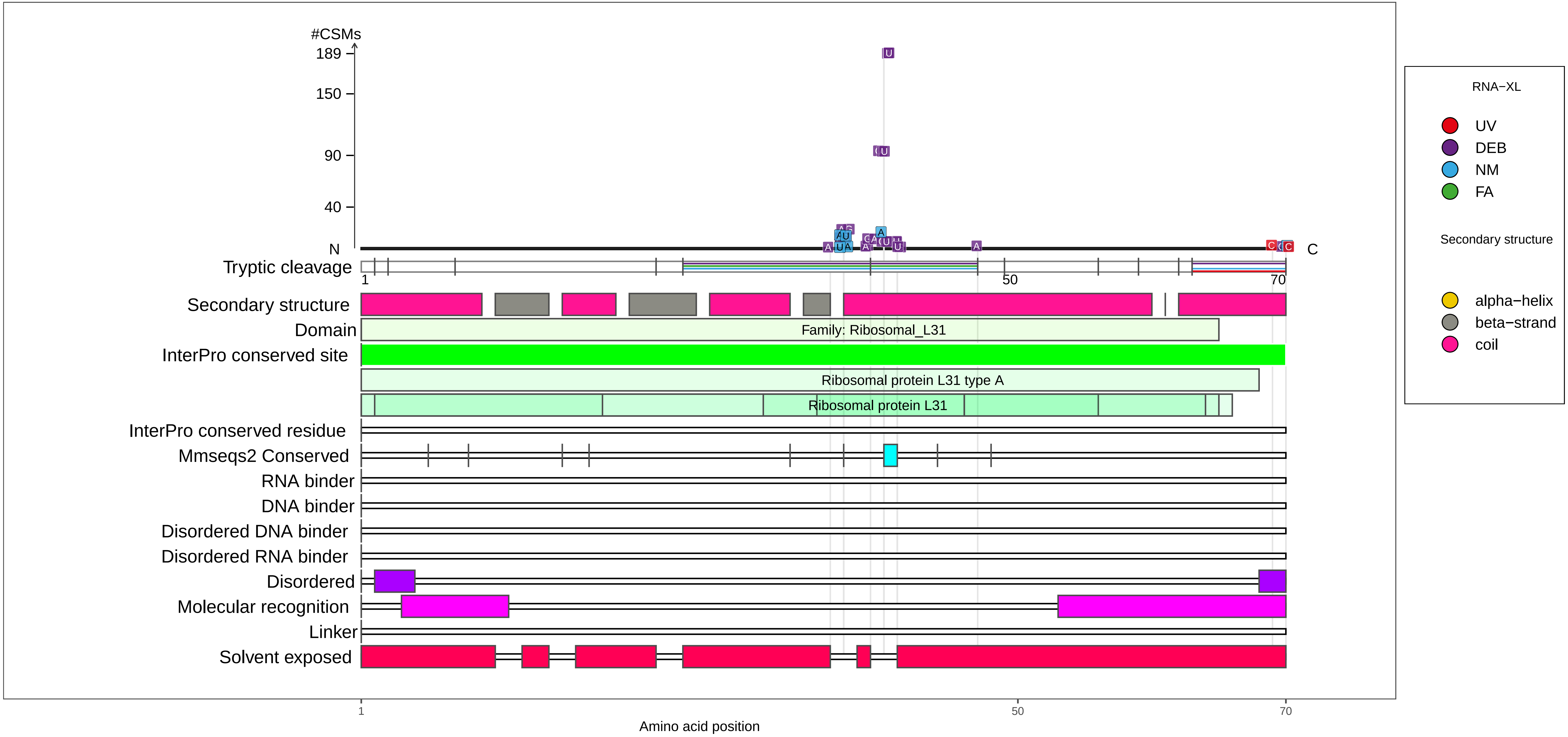

P0A8T7
RPOC_ECOLI DNA−directed RNA polymerase subunit beta'

− Abundance:
tryptic [log10 Intensity]: 9.82 (Q 99)
PAXdb K12 strain [ppm]: 3.18 (Q 94)
PAXdb E.coli [ppm]: 2.9 (Q 93)

− RNA functions:
5−3 RNA polymerase activity; DNA−directed 5−3 RNA polymerase activity
RNA biosynthetic process; RNA metabolic process; RNA polymerase activity
RNA polymerase Rpb1, domain 1; RNA polymerase Rpb1, domain 2; RNA polymerase Rpb1, domain 3
RNA polymerase Rpb1, domain 4; RNA polymerase Rpb1, domain 5

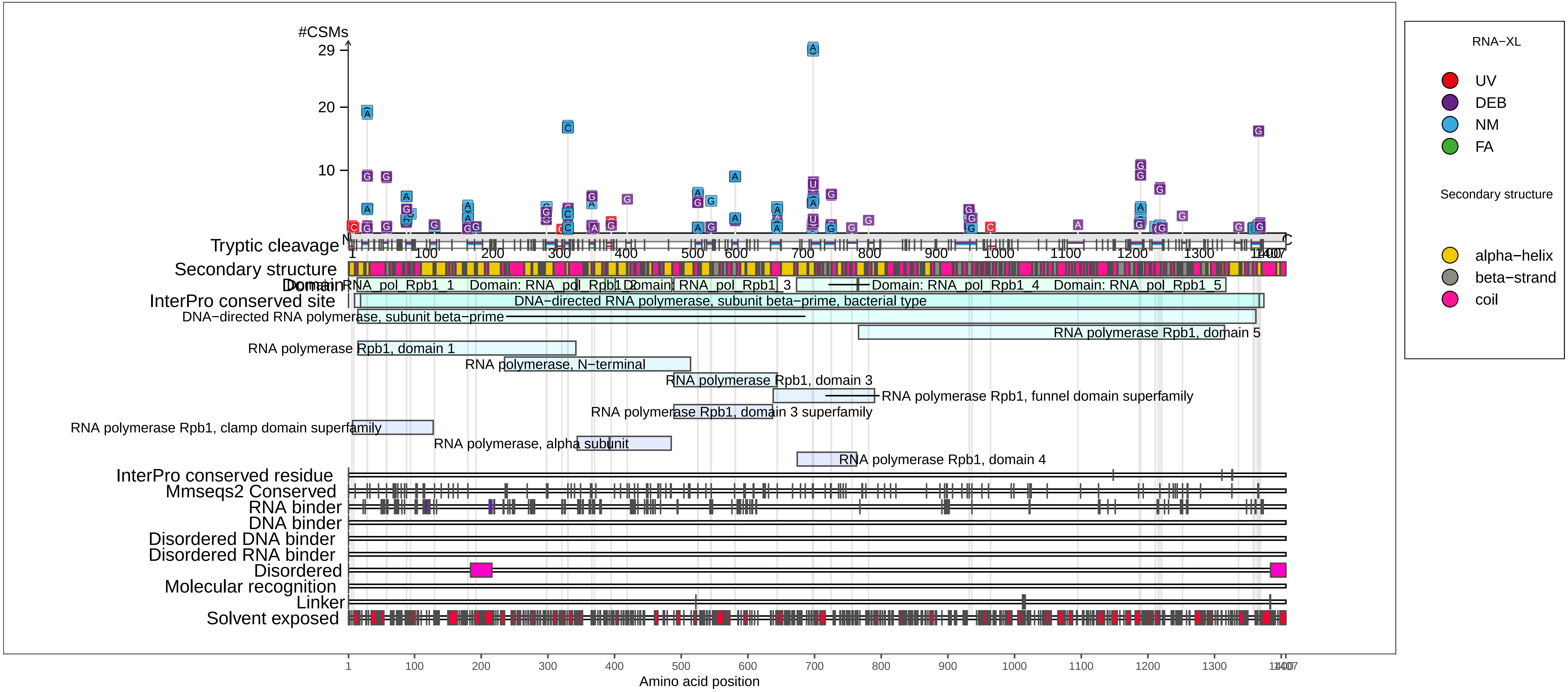

P0AA10
RL13_ECOLI 50S ribosomal protein L13
− Abundance:
tryptic [log10 Intensity]: 9.11 (Q 91)
PAXdb K12 strain [ppm]: 3.67 (Q 99)
PAXdb E.coli [ppm]: 3.44 (Q 98)

− RNA functions:
mRNA binding; RNA binding; rRNA binding

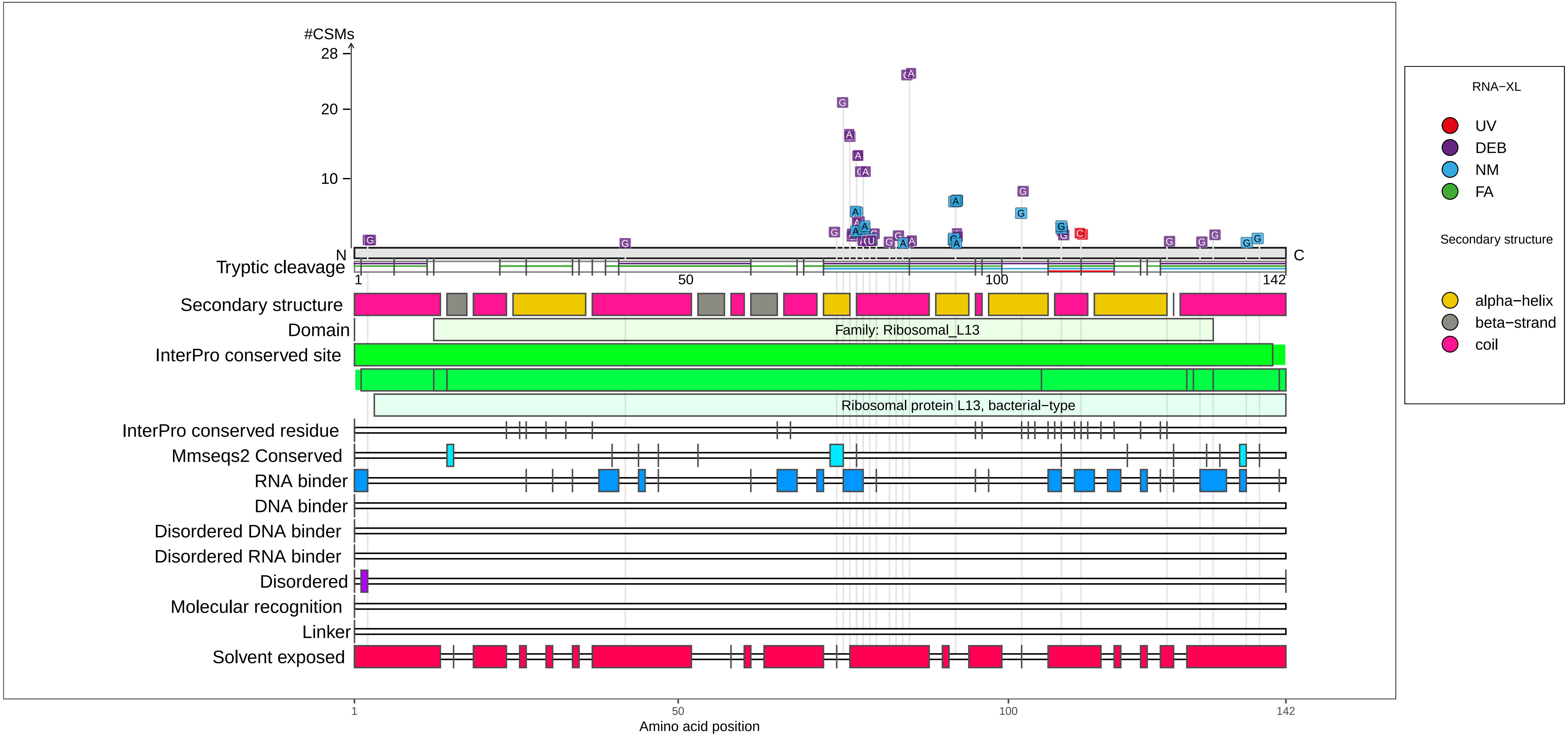

P0A7S9
RS13_ECOLI 30S ribosomal protein S13

− Abundance:
tryptic [log10 Intensity]: 9.56 (Q 97)
PAXdb K12 strain [ppm]: 3.69 (Q 99)
PAXdb E.coli [ppm]: 3.59 (Q 100)

− RNA functions:
RNA binding; rRNA binding; tRNA binding

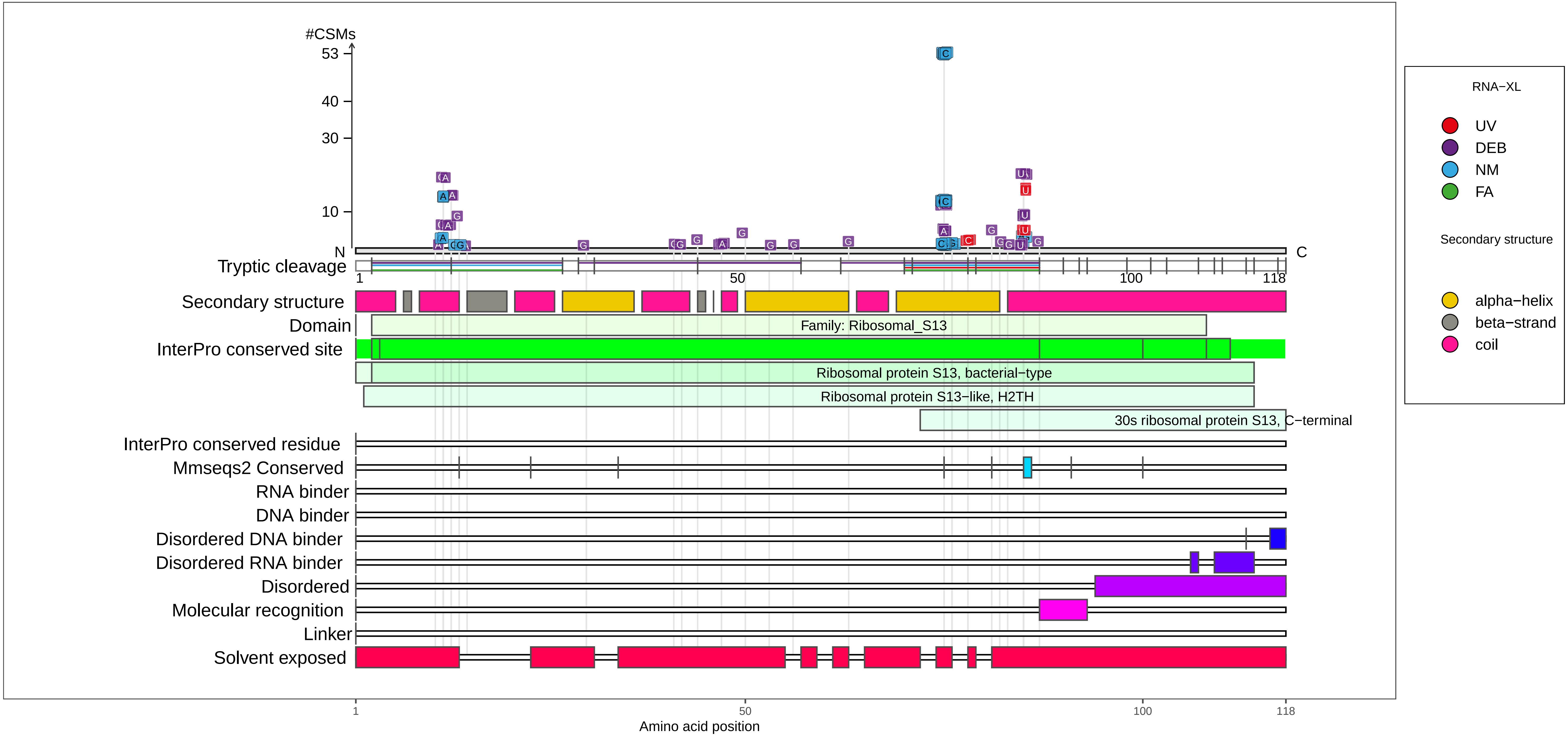

P05055
PNP_ECOLI Polyribonucleotide nucleotidyltransferase

− Abundance:
tryptic [log10 Intensity]: 9.59 (Q 97)
PAXdb K12 strain [ppm]: 2.55 (Q 79)
PAXdb E.coli [ppm]: 3.43 (Q 98)

− RNA functions:
mRNA catabolic process; mRNA metabolic process
Polyribonucleotide nucleotidyltransferase, RNA binding domain; RNA binding; RNA catabolic process; RNA metabolic process
RNA phosphodiester bond hydrolysis
RNA phosphodiester bond hydrolysis, exonucleolytic; RNA processing; S1 RNA binding domain

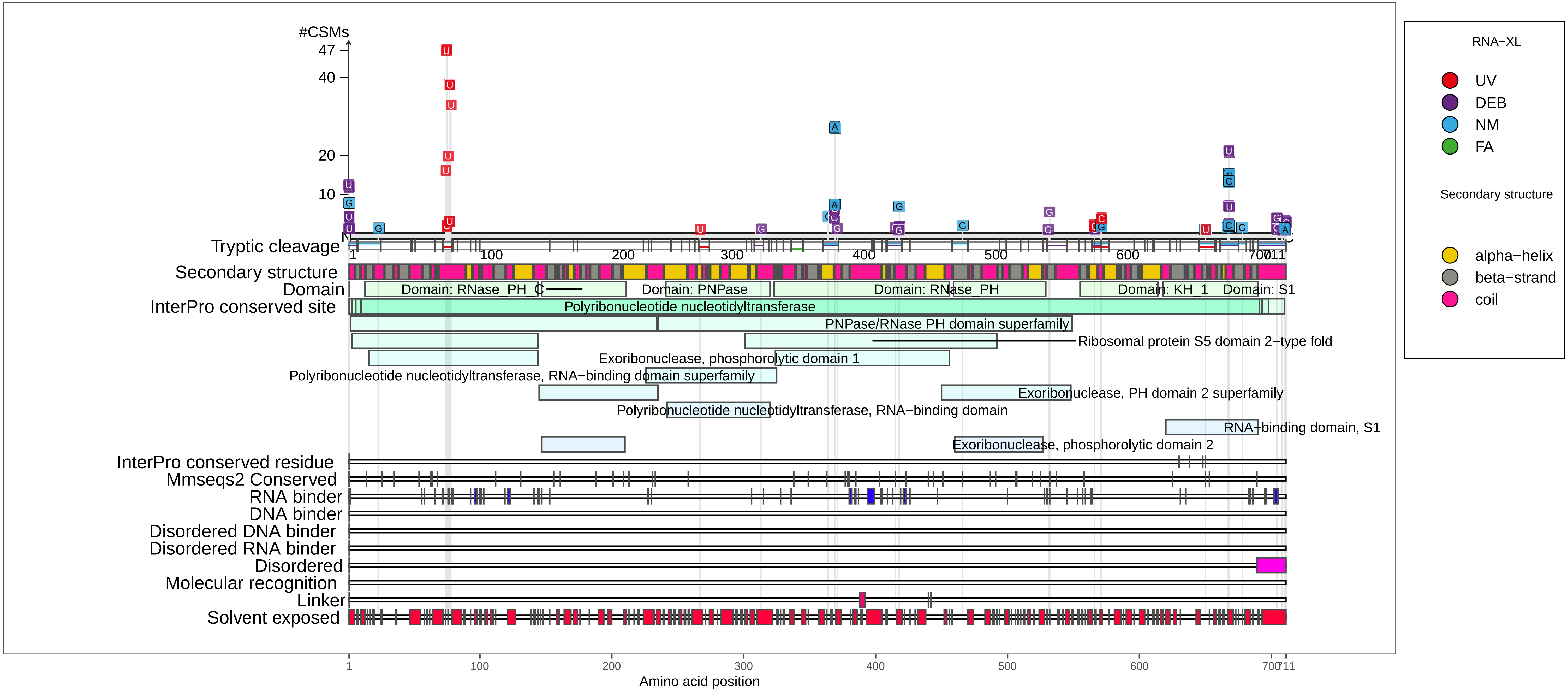

P0A9P0
DLDH_ECOLI Dihydrolipoyl dehydrogenase

− Abundance:
tryptic [log10 Intensity]: 9.67 (Q 98)
PAXdb K12 strain [ppm]: 3.46 (Q 97)
PAXdb E.coli [ppm]: 3.4 (Q 98)

− RNA functions: not annotated

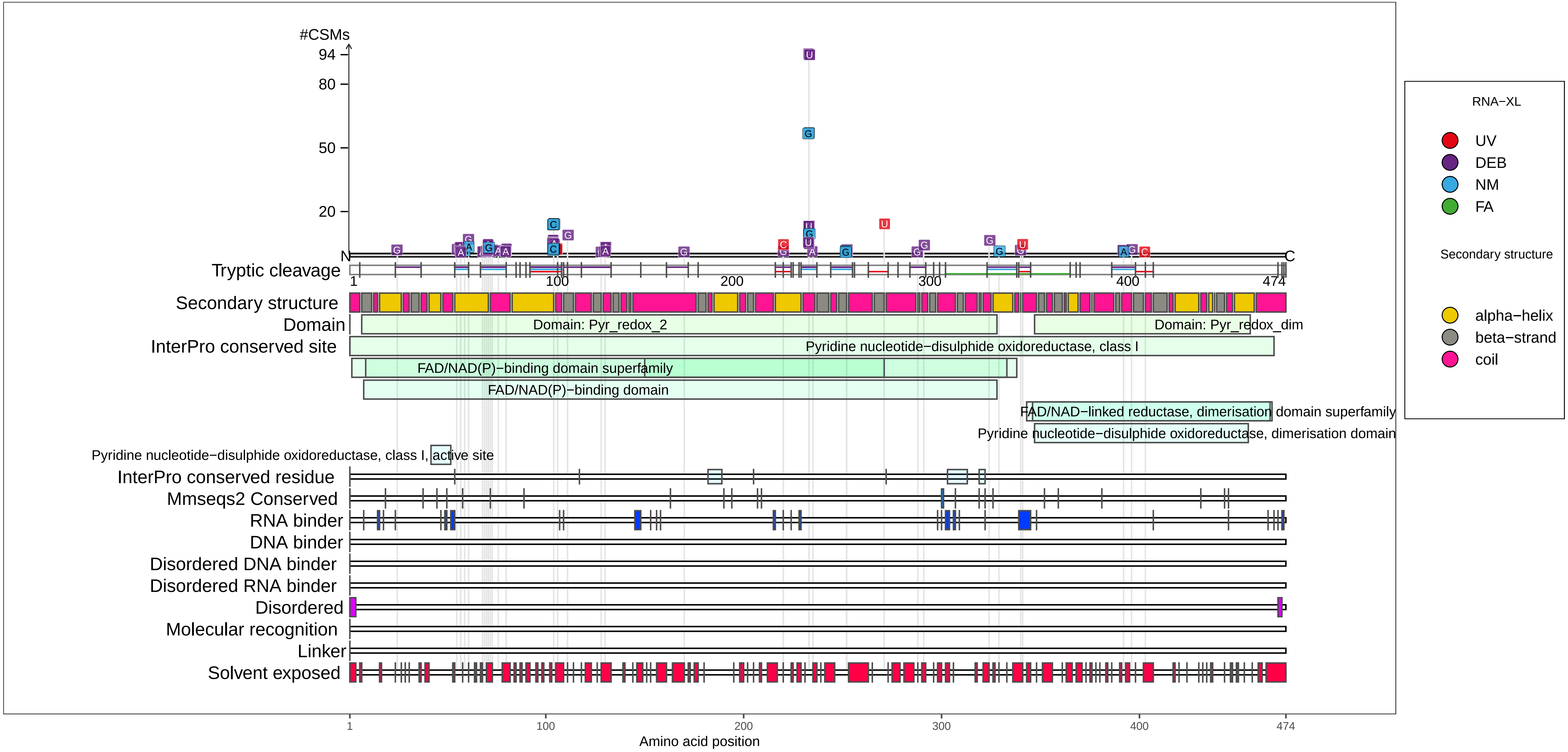

P63284
CLPB_ECOLI Chaperone protein ClpB

− Abundance:
tryptic [log10 Intensity]: 9.37 (Q 95)
PAXdb K12 strain [ppm]: 3.48 (Q 97)
PAXdb E.coli [ppm]: 2.97 (Q 94)

− RNA functions: not annotated

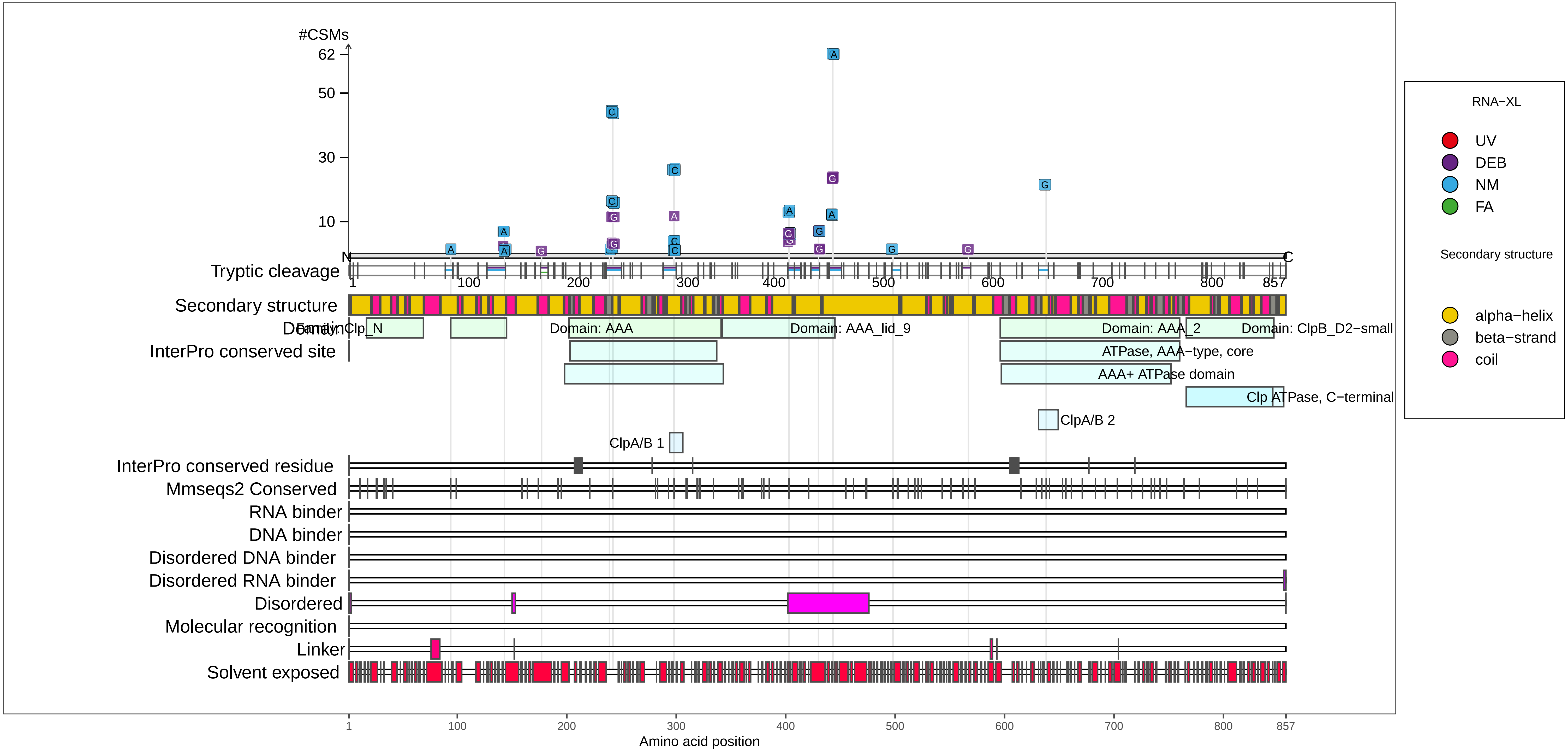

P0A7W1
RS5_ECOLI 30S ribosomal protein S5

− Abundance:
tryptic [log10 Intensity]: 9.19 (Q 92)
PAXdb K12 strain [ppm]: 3.8 (Q 100)
PAXdb E.coli [ppm]: 3.89 (Q 100)

− RNA functions:
RNA binding; rRNA binding

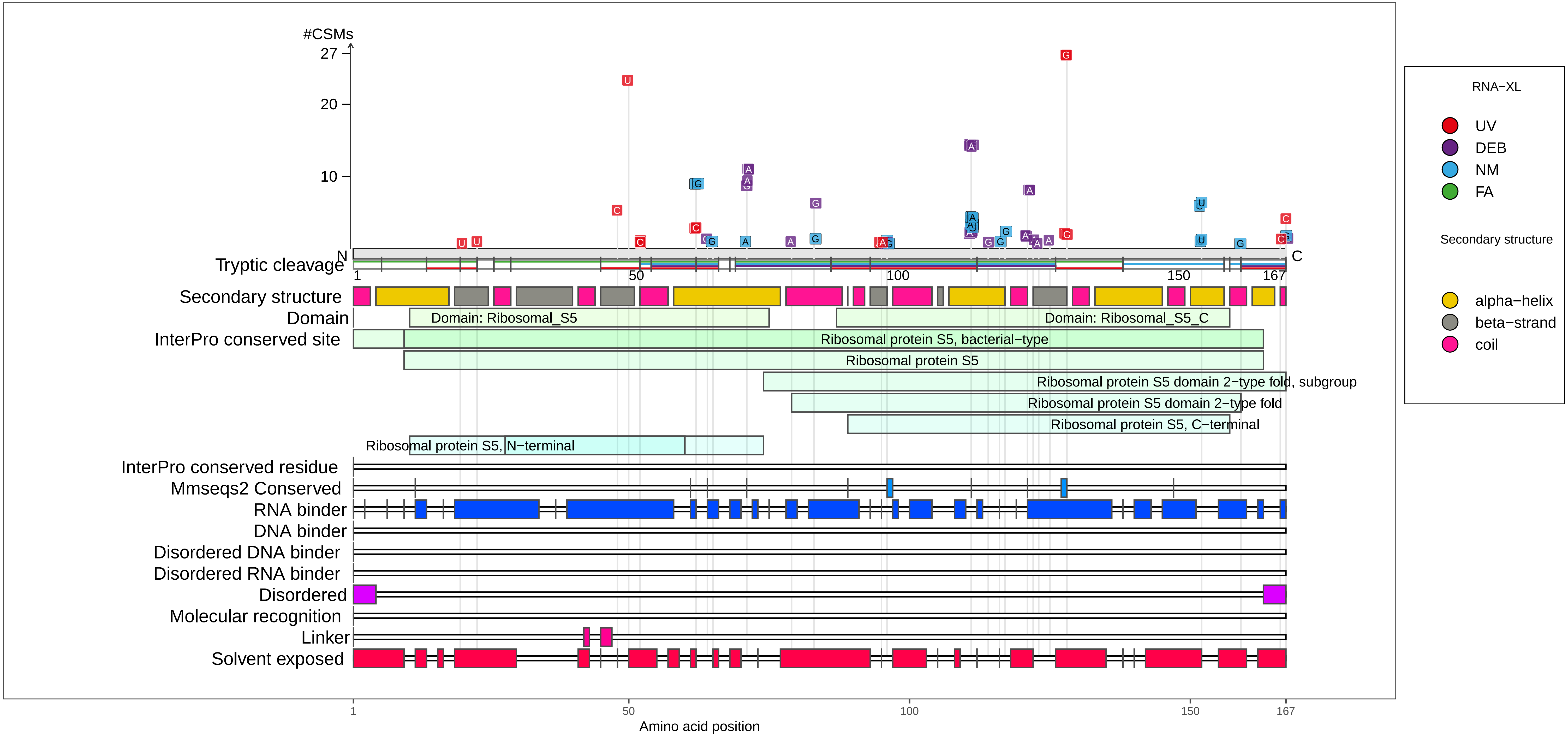

P0A8V2
RPOB_ECOLI DNA−directed RNA polymerase subunit beta

− Abundance:
tryptic [log10 Intensity]: 9.69 (Q 98)
PAXdb K12 strain [ppm]: 2.92 (Q 89)
PAXdb E.coli [ppm]: 3.25 (Q 97)

− RNA functions:
5−3 RNA polymerase activity; DNA−directed 5−3 RNA polymerase activity
RNA biosynthetic process; RNA metabolic process; RNA polymerase activity
RNA polymerase beta subunit; RNA polymerase beta subunit external 1 domain; RNA polymerase Rpb2, domain 2
RNA polymerase Rpb2, domain 3; RNA polymerase Rpb2, domain 6
RNA polymerase Rpb2, domain 7

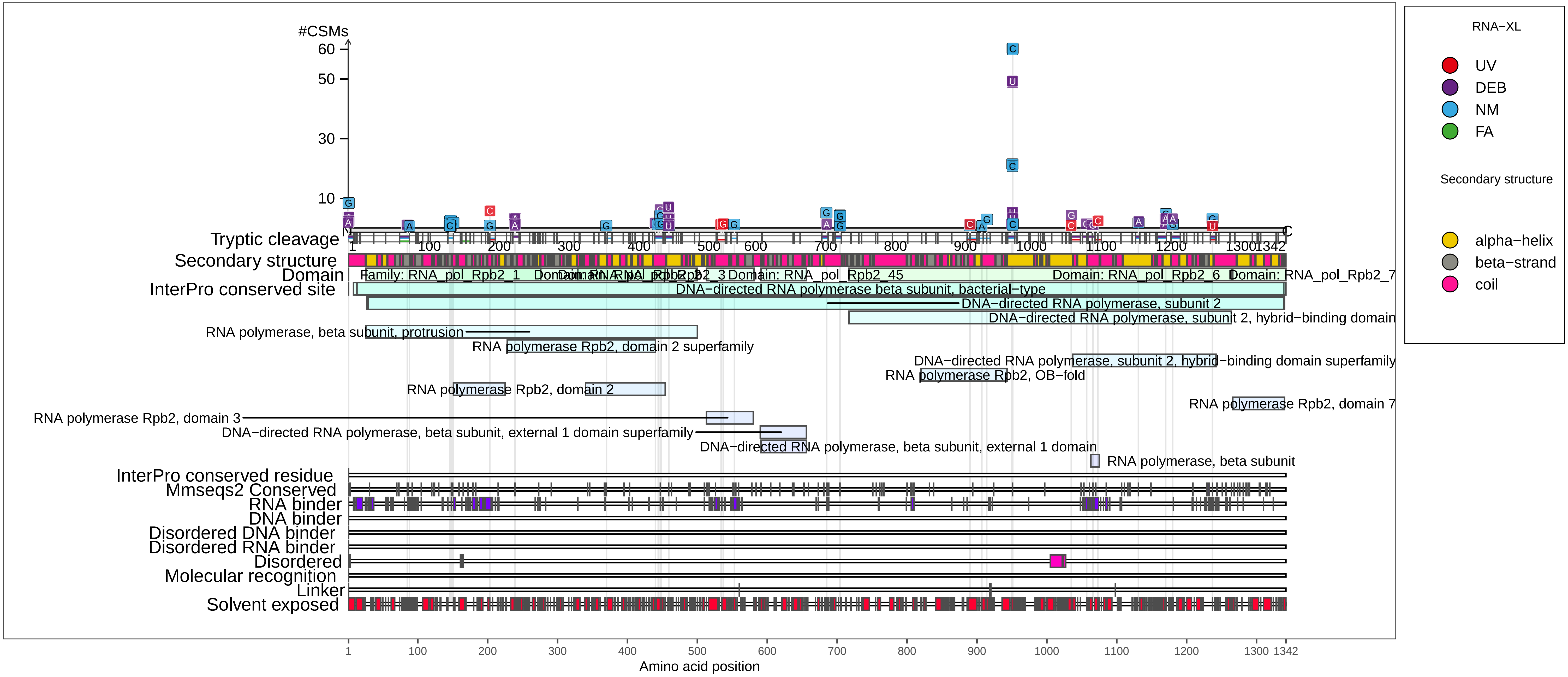

P02359
RS7_ECOLI 30S ribosomal protein S7

− Abundance:
tryptic [log10 Intensity]: 9.12 (Q 91)
PAXdb K12 strain [ppm]: 3.86 (Q 100)
PAXdb E.coli [ppm]: 3.72 (Q 100)

− RNA functions:
mRNA binding; RNA binding; rRNA binding; tRNA binding

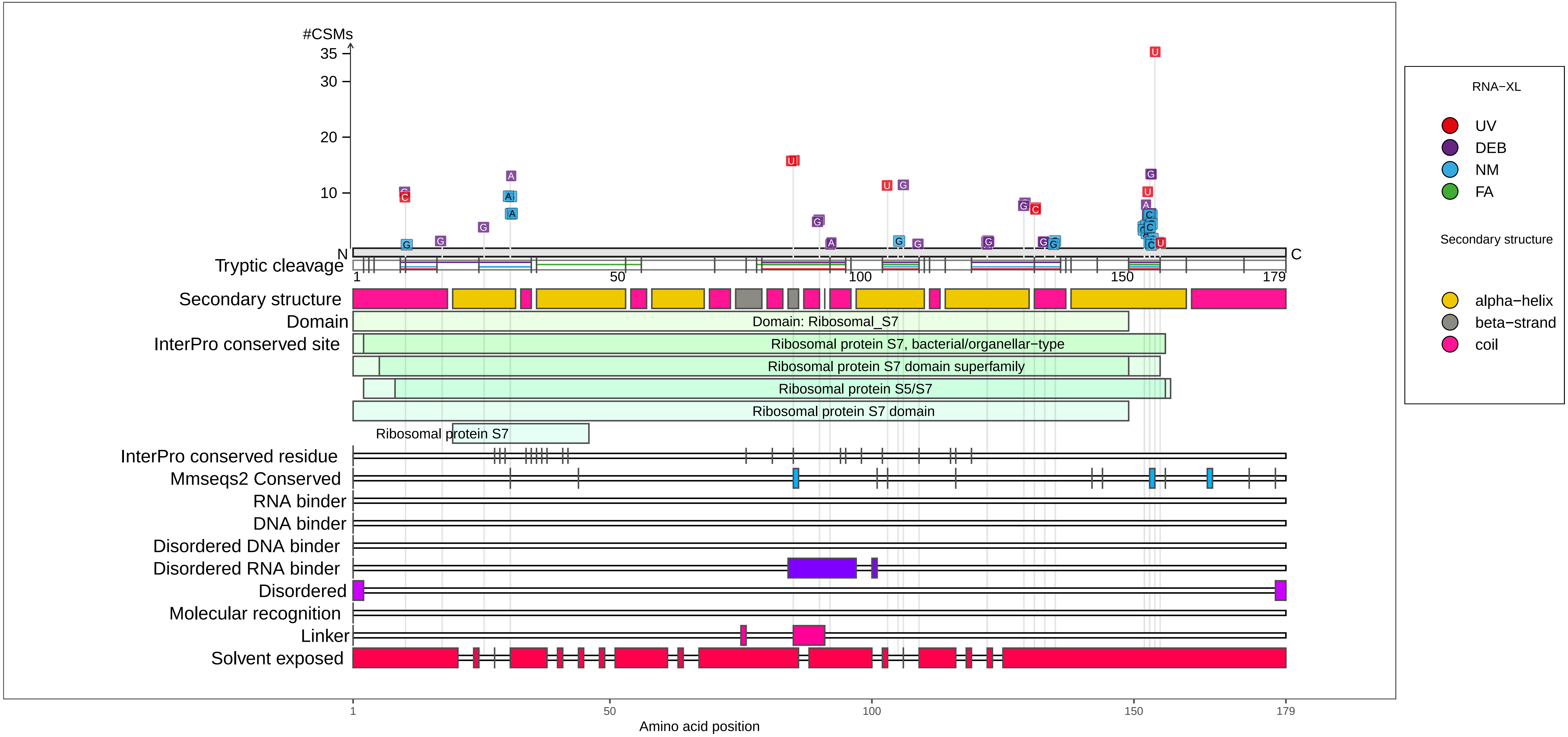

P0A6P1
EFTS_ECOLI Elongation factor Ts

− Abundance:
tryptic [log10 Intensity]: 9.7 (Q 98)
PAXdb K12 strain [ppm]: 3.4 (Q 96)
PAXdb E.coli [ppm]: 3.65 (Q 100)

− RNA functions:
RNA binding

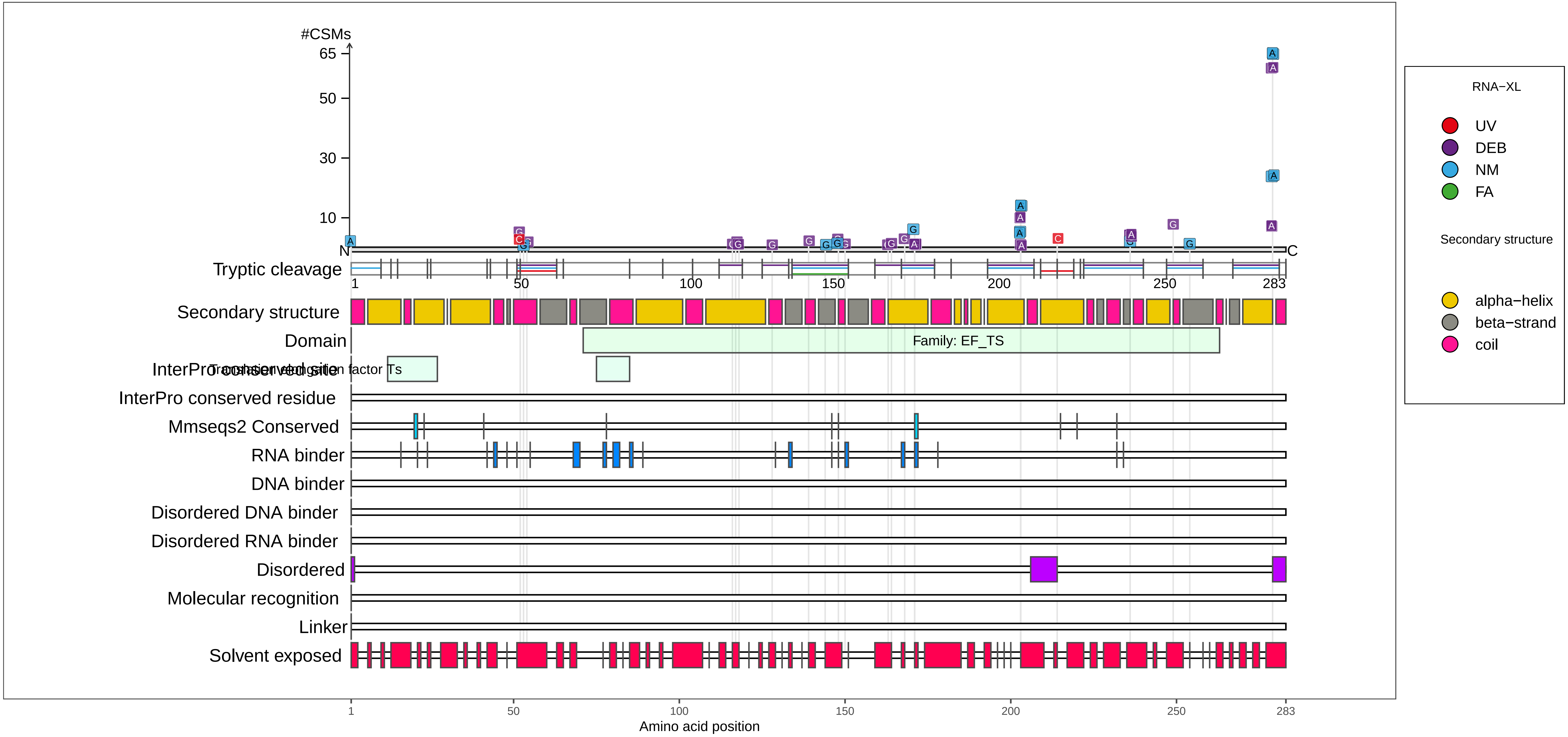

P0A7S3
RS12_ECOLI 30S ribosomal protein S12

− Abundance:
tryptic [log10 Intensity]: 8.53 (Q 76)
PAXdb K12 strain [ppm]: 2.85 (Q 87)
PAXdb E.coli [ppm]: 3.06 (Q 95)

− RNA functions:
RNA binding; RNA folding; RNA metabolic process; RNA processing; RNA splicing
RNA splicing, via transesterification reactions
RNA splicing, via transesterification reactions with guanosine as nucleophile; rRNA binding; tRNA binding

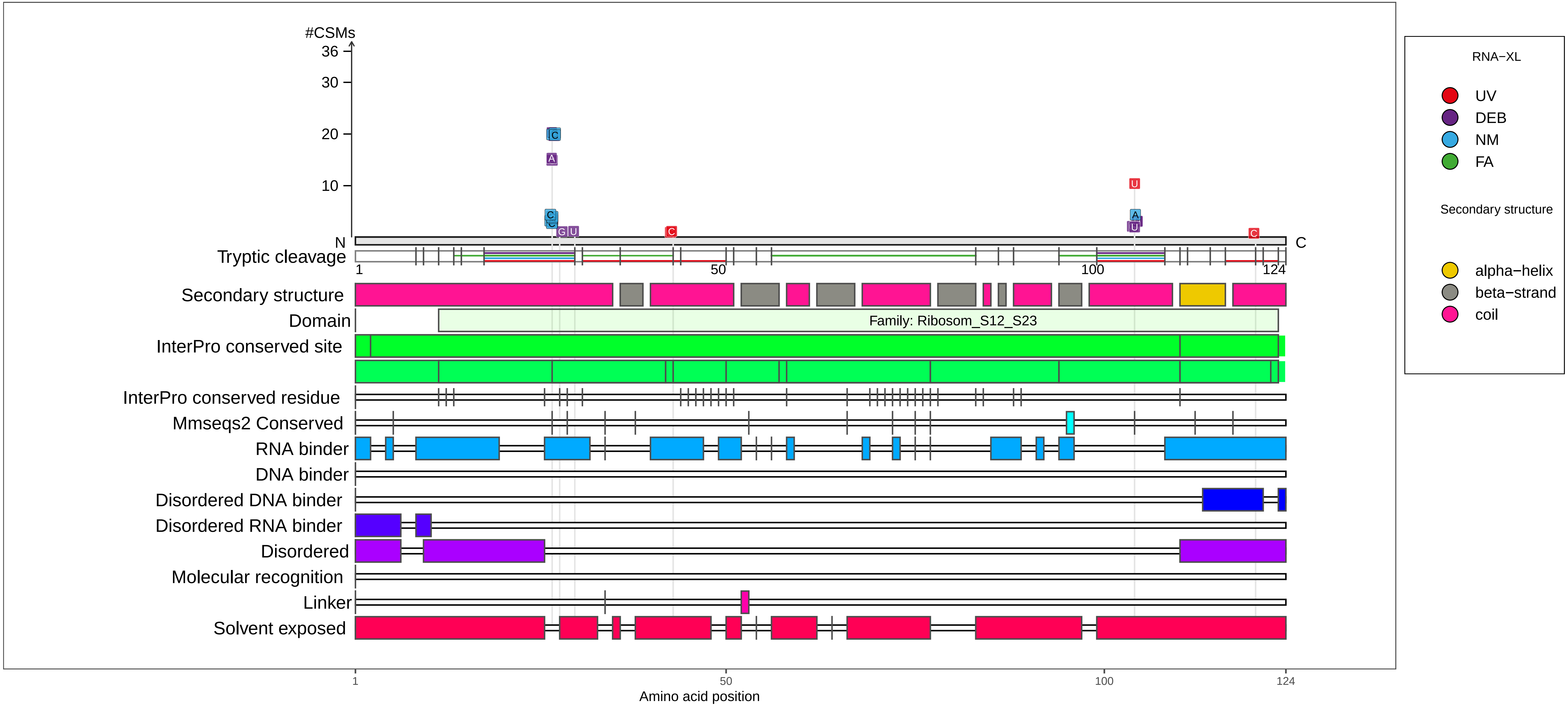

P60624
RL24_ECOLI 50S ribosomal protein L24

− Abundance:
tryptic [log10 Intensity]: 9.65 (Q 97)
PAXdb K12 strain [ppm]: 3.74 (Q 99)
PAXdb E.coli [ppm]: 3.57 (Q 99)

− RNA functions:
RNA binding; rRNA binding

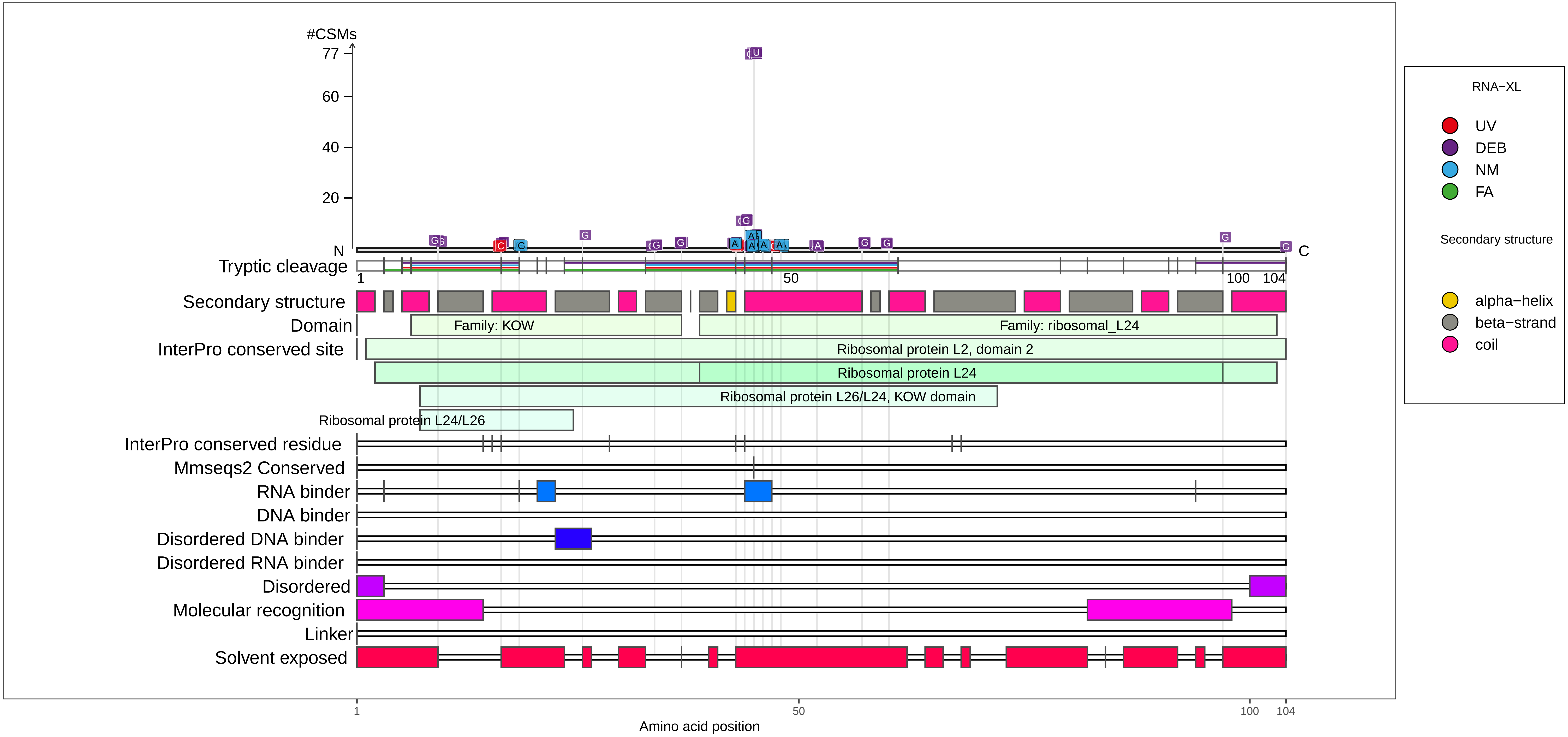

P0A7V0
RS2_ECOLI 30S ribosomal protein S2

− Abundance:
tryptic [log10 Intensity]: 9.45 (Q 96)
PAXdb K12 strain [ppm]: 3.71 (Q 99)
PAXdb E.coli [ppm]: 3.45 (Q 98)

− RNA functions: not annotated

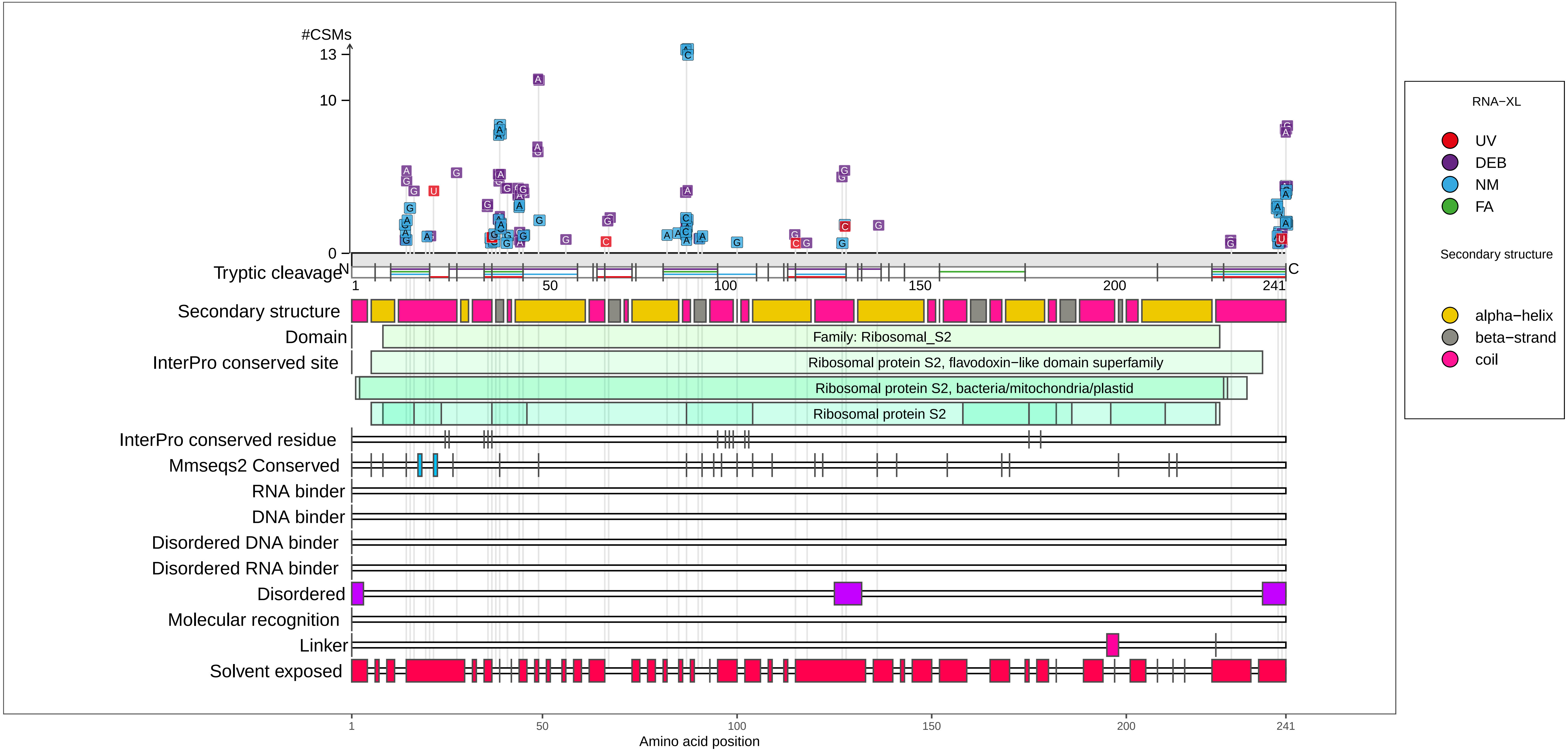

P68919
RL25_ECOLI 50S ribosomal protein L25

− Abundance:
tryptic [log10 Intensity]: 8.93 (Q 87)
PAXdb K12 strain [ppm]: 3.79 (Q 99)
PAXdb E.coli [ppm]: 3.84 (Q 100)

− RNA functions:
5S rRNA binding; RNA binding; rRNA binding

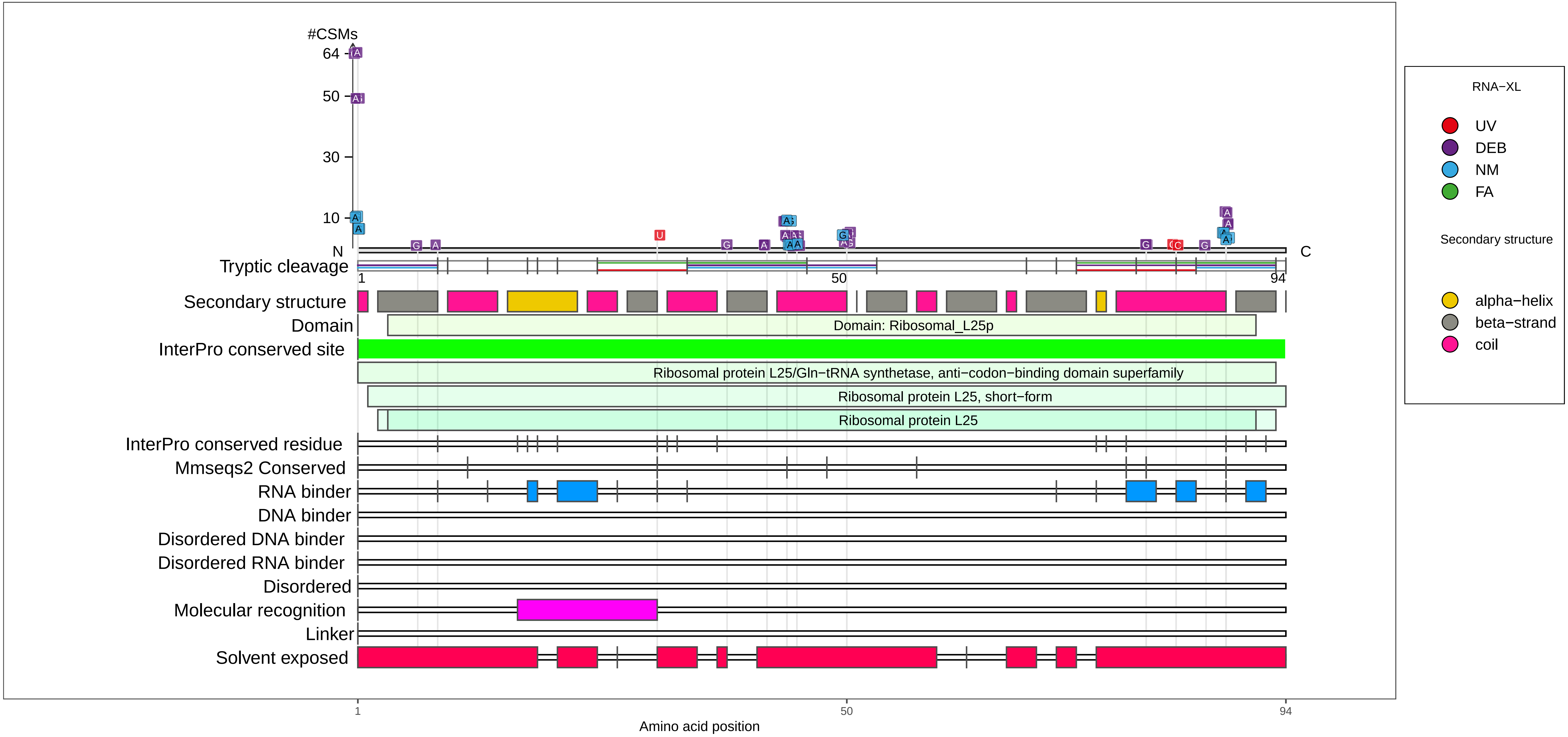

P0AG59
RS14_ECOLI 30S ribosomal protein S14

− Abundance:
tryptic [log10 Intensity]: 8.34 (Q 71)
PAXdb K12 strain [ppm]: 3.8 (Q 100)
PAXdb E.coli [ppm]: 3.35 (Q 98)

− RNA functions:
RNA binding; rRNA binding

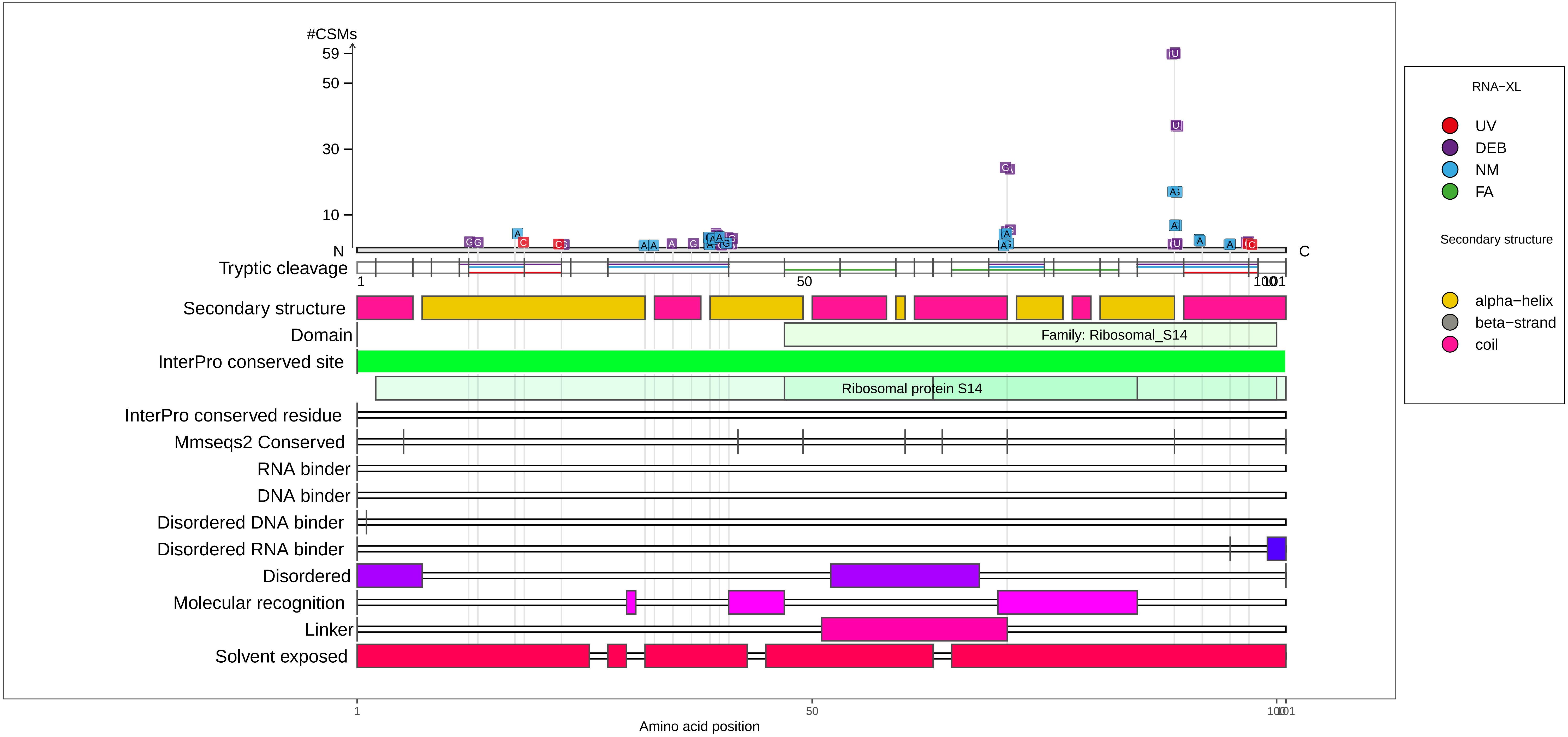

P0A7N4
RL32_ECOLI 50S ribosomal protein L32

− Abundance:
tryptic [log10 Intensity]: 8.26 (Q 68)
PAXdb K12 strain [ppm]: 3.87 (Q 100)
PAXdb E.coli [ppm]: 3.47 (Q 99)

− RNA functions: not annotated

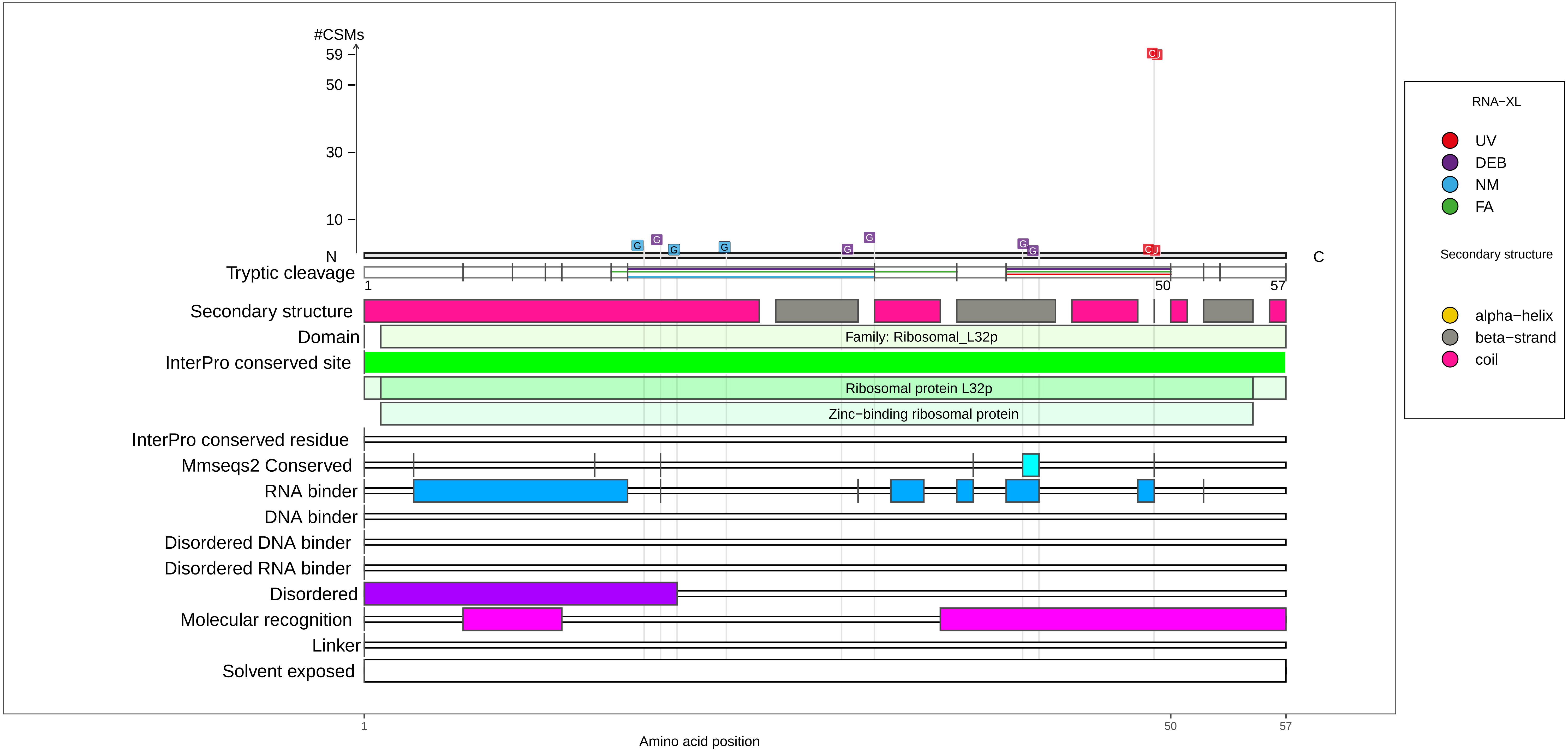

P76177
YDGH_ECOLI Protein YdgH

− Abundance:
tryptic [log10 Intensity]: 8.35 (Q 71)
PAXdb K12 strain [ppm]: 2.94 (Q 89)
PAXdb E.coli [ppm]: 2.84 (Q 92)

− RNA functions: not annotated

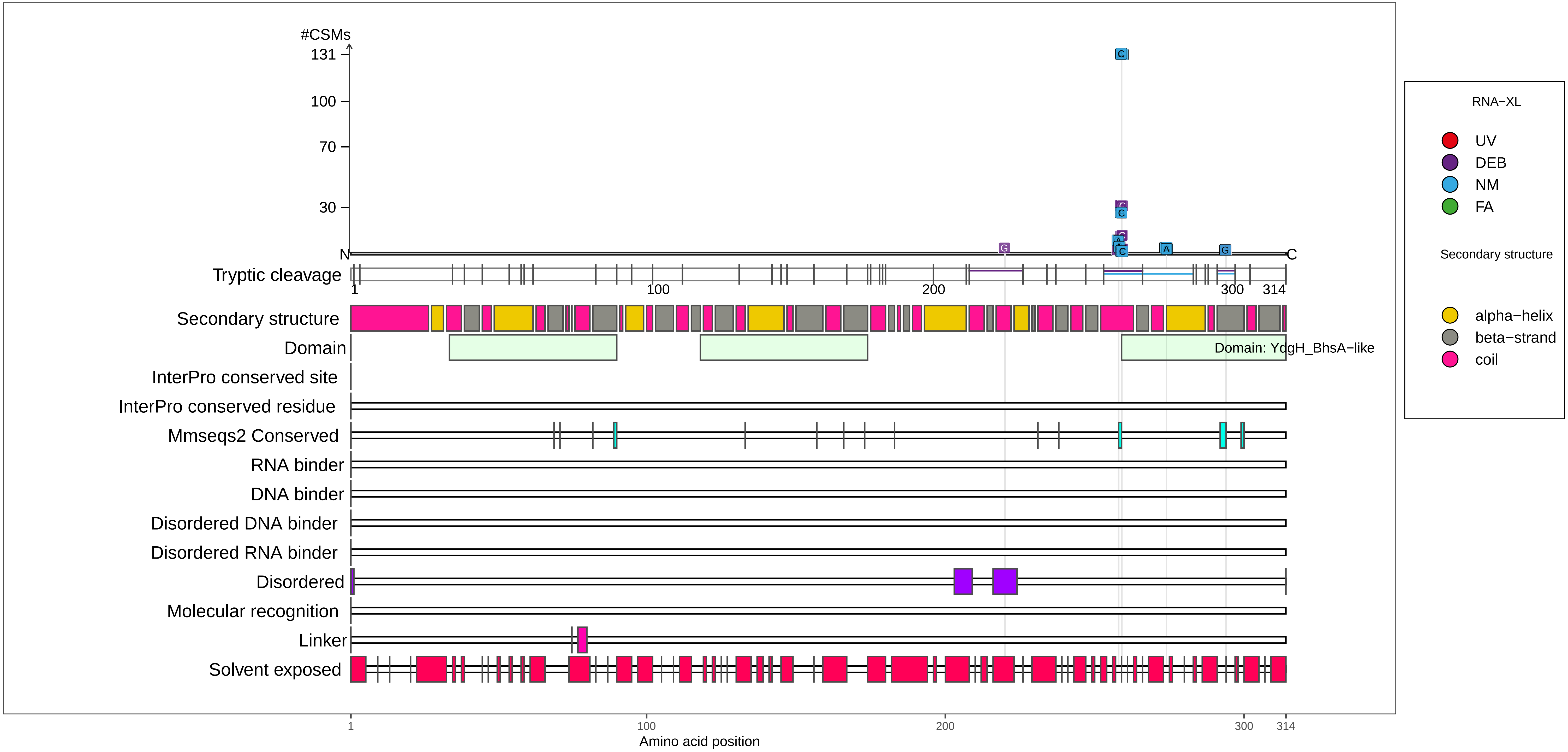

P0A7V8
RS4_ECOLI 30S ribosomal protein S4

− Abundance:
tryptic [log10 Intensity]: 9.07 (Q 90)
PAXdb K12 strain [ppm]: 3.57 (Q 98)
PAXdb E.coli [ppm]: 3.5 (Q 99)

− RNA functions:
mRNA 5−UTR binding; mRNA binding; RNA binding; RNA biosynthetic process
RNA metabolic process; rRNA binding

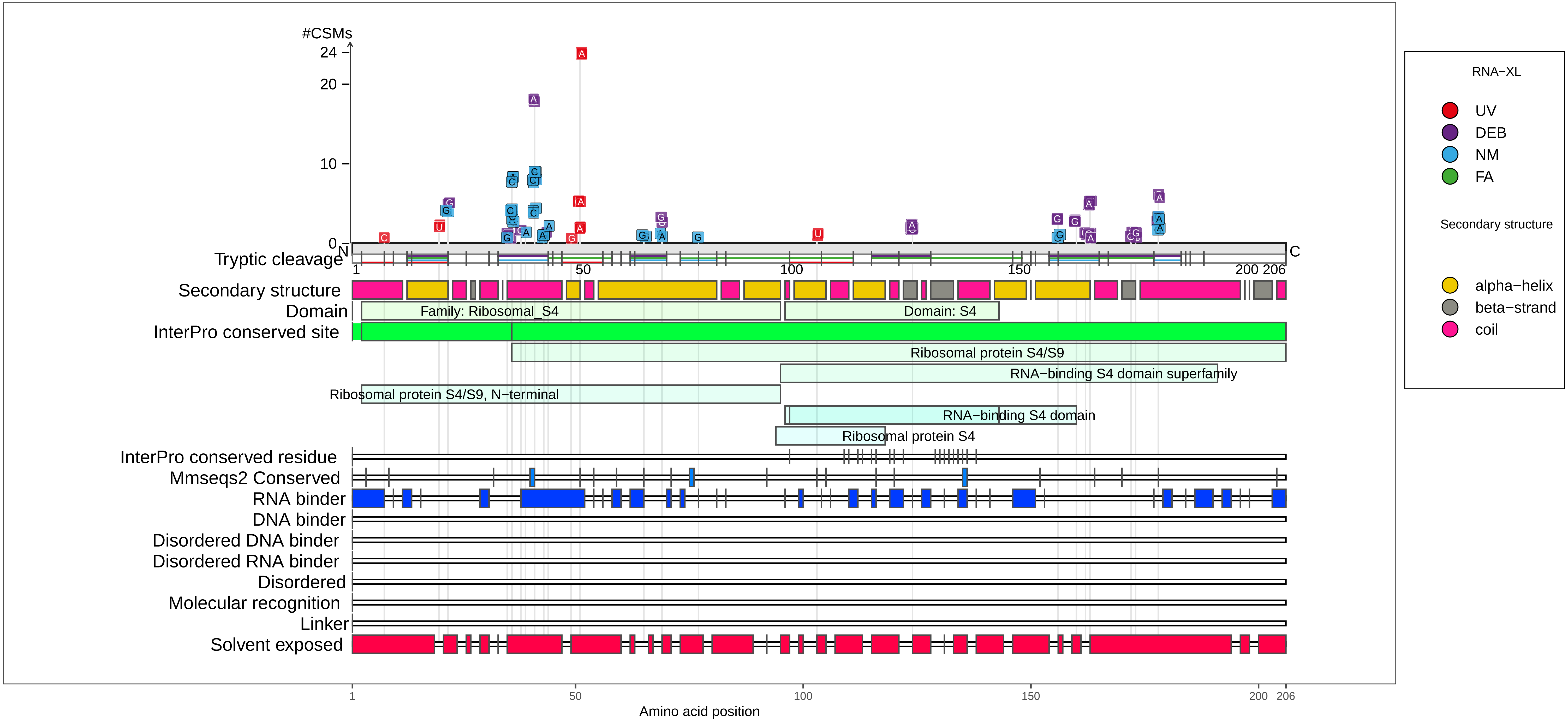

P0A707
IF3_ECOLI Translation initiation factor IF−3
− Abundance:
tryptic [log10 Intensity]: 8.76 (Q 82)
PAXdb K12 strain [ppm]: 3.48 (Q 97)
PAXdb E.coli [ppm]: 2.97 (Q 94)

− RNA functions:
RNA binding

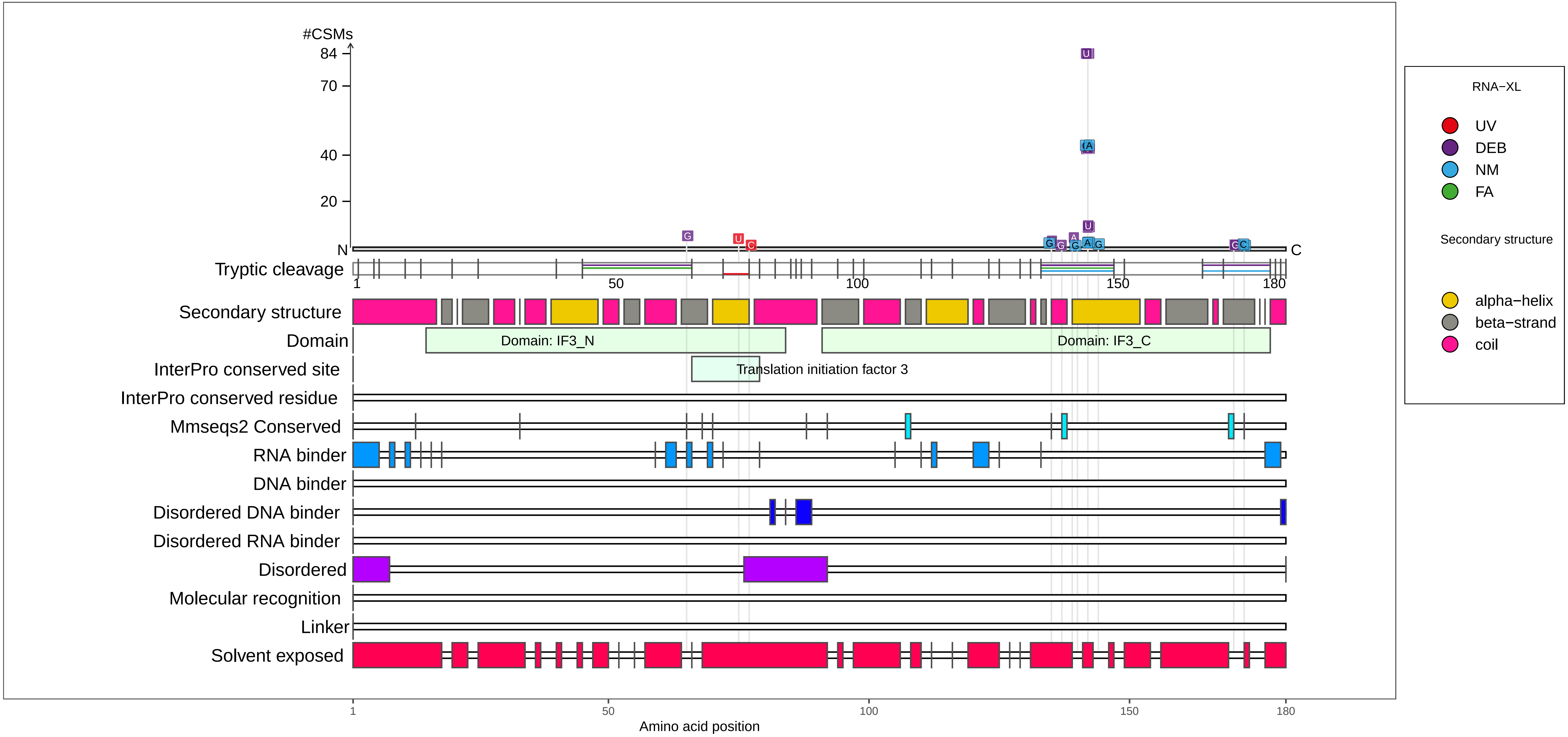

P0AFG8
ODP1_ECOLI Pyruvate dehydrogenase E1 component

− Abundance:
tryptic [log10 Intensity]: 10.3 (Q 100)
PAXdb K12 strain [ppm]: 3.47 (Q 97)
PAXdb E.coli [ppm]: 3.5 (Q 99)

− RNA functions: not annotated

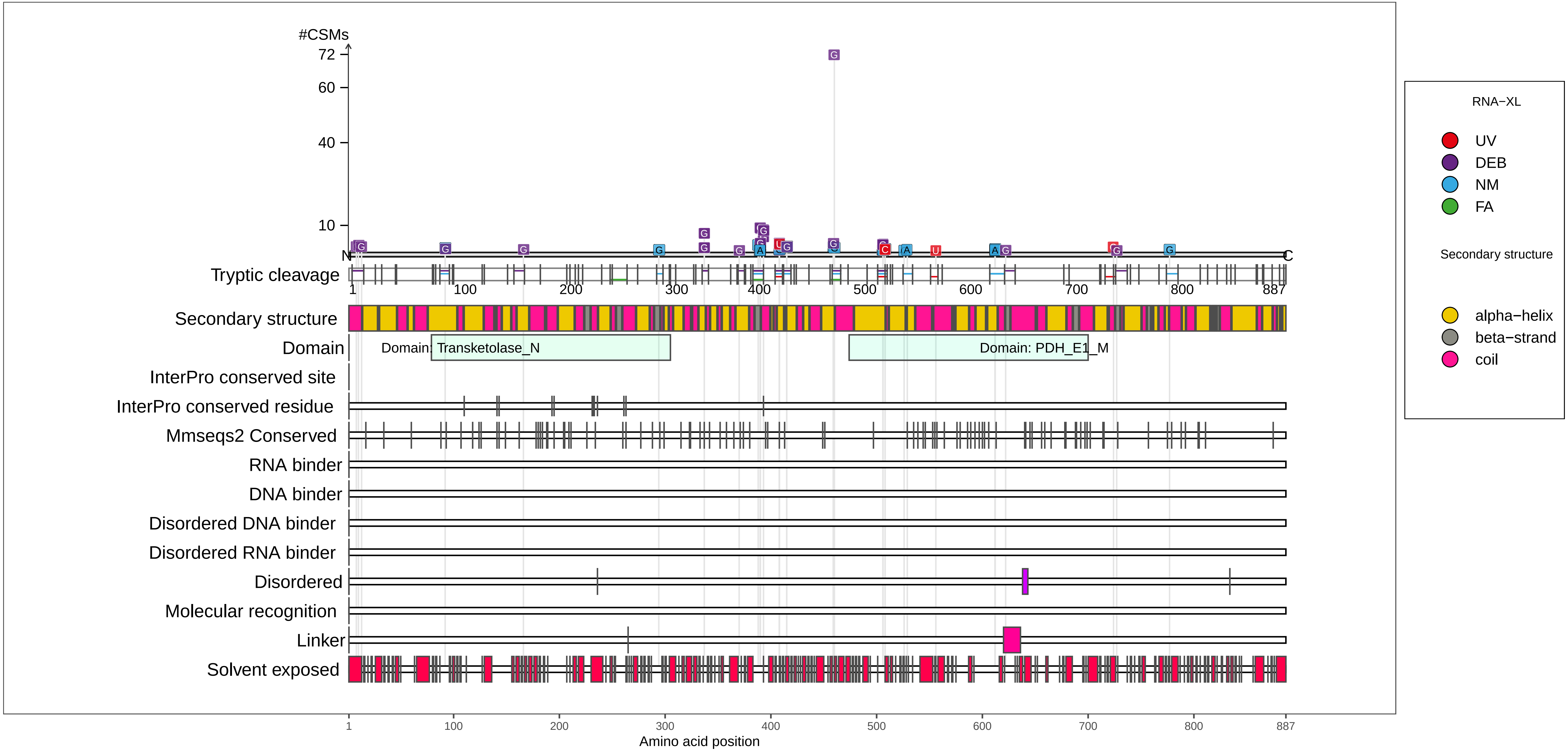

P00957
SYA_ECOLI Alanine−−tRNA ligase

− Abundance:
tryptic [log10 Intensity]: 9.68 (Q 98)
PAXdb K12 strain [ppm]: 3.01 (Q 91)
PAXdb E.coli [ppm]: 2.78 (Q 91)

− RNA functions:
alanine−tRNA ligase activity; alanyl−tRNA aminoacylation
aminoacyl−tRNA editing activity; aminoacyl−tRNA ligase activity
aminoacyl−tRNA metabolism involved in translational fidelity; ncRNA metabolic process; ncRNA processing; RNA binding
RNA metabolic process; RNA modification; RNA processing; Ser−tRNA(Ala) hydrolase activity
Threonyl and Alanyl tRNA synthetase second additional domain; tRNA aminoacylation
tRNA aminoacylation for protein translation; tRNA binding; tRNA metabolic process
tRNA modification; tRNA processing; tRNA synthetases class II (A)

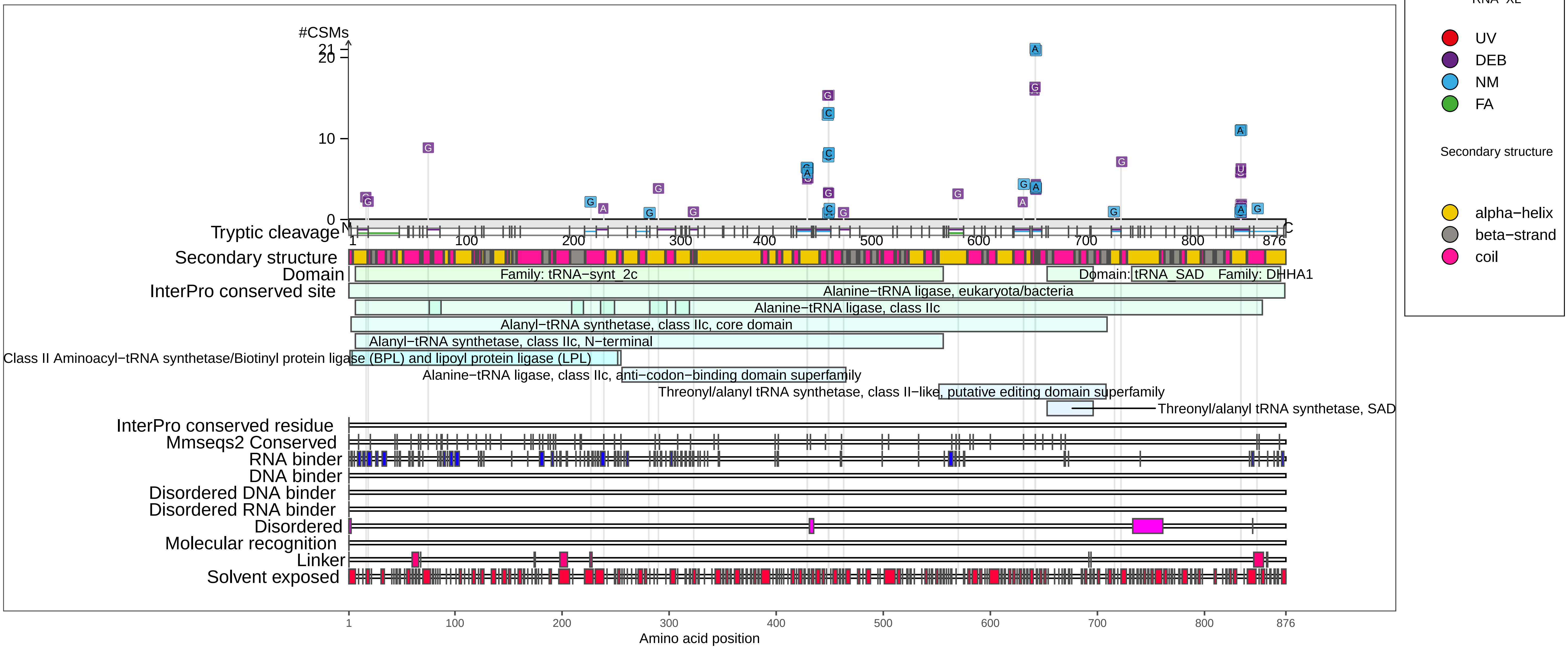

P0A705
IF2_ECOLI Translation initiation factor IF−2

− Abundance:
tryptic [log10 Intensity]: 9.78 (Q 99)
PAXdb K12 strain [ppm]: 3.25 (Q 95)
PAXdb E.coli [ppm]: 2.67 (Q 89)

− RNA functions:
RNA binding

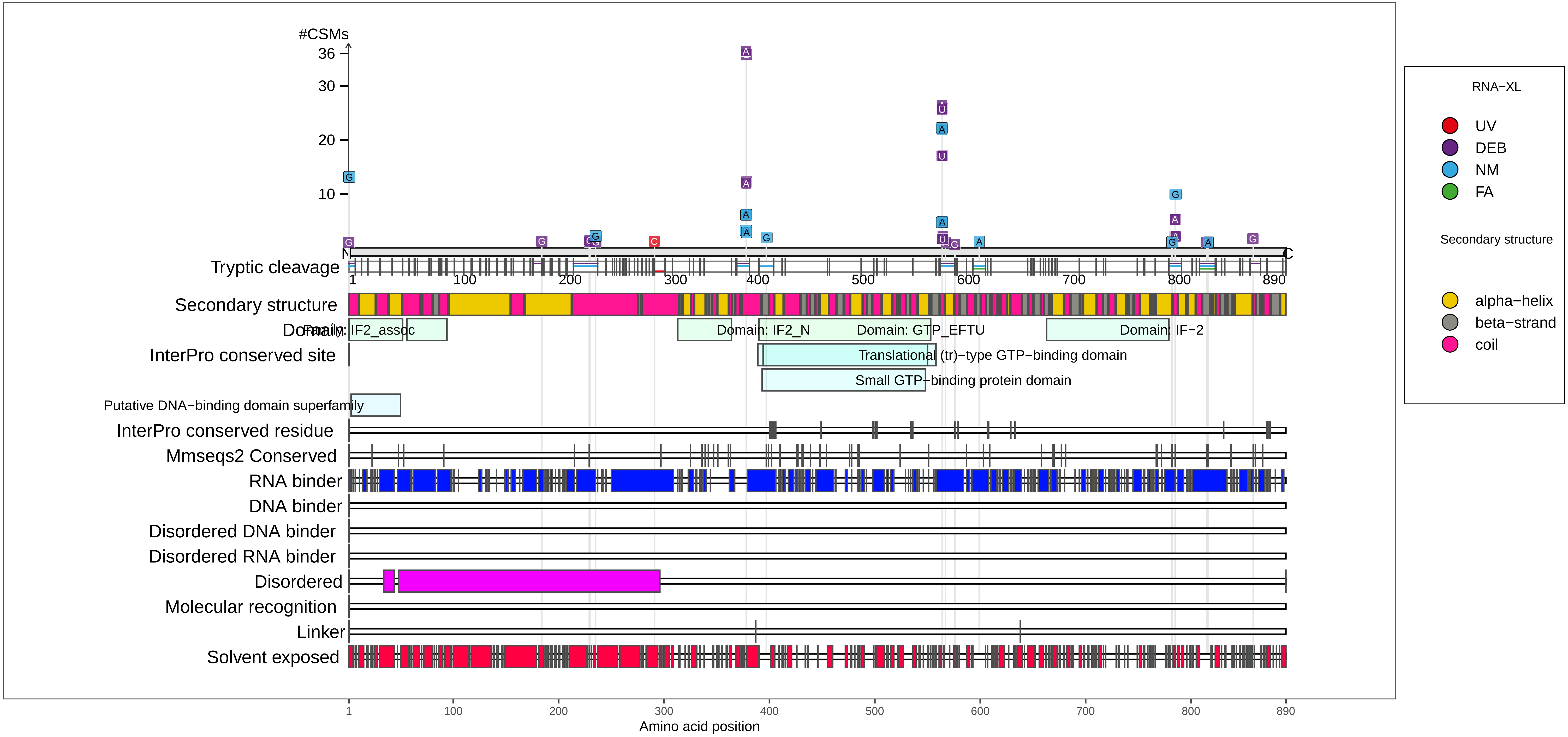

P0A9Y6
CSPC_ECOLI Cold shock−like protein CspC

− Abundance:
tryptic [log10 Intensity]: 9.87 (Q 99)
PAXdb K12 strain [ppm]: 3.46 (Q 97)
PAXdb E.coli [ppm]: 4.32 (Q 100)

− RNA functions:
RNA binding

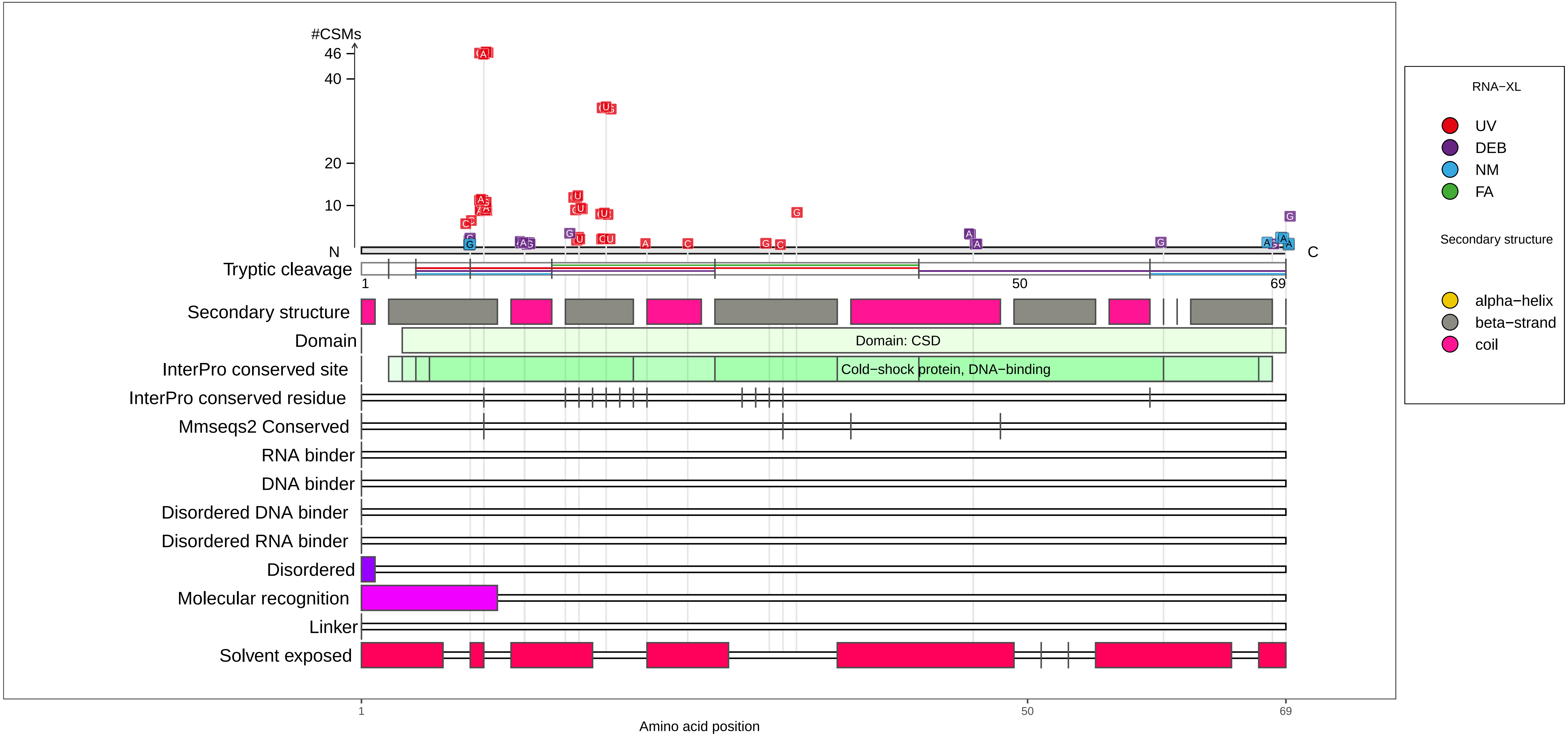

P0A9B2
G3P1_ECOLI Glyceraldehyde−3−phosphate dehydrogenase A

− Abundance:
tryptic [log10 Intensity]: 10.17 (Q 100)
PAXdb K12 strain [ppm]: 3.25 (Q 95)
PAXdb E.coli [ppm]: 4.03 (Q 100)

− RNA functions: not annotated

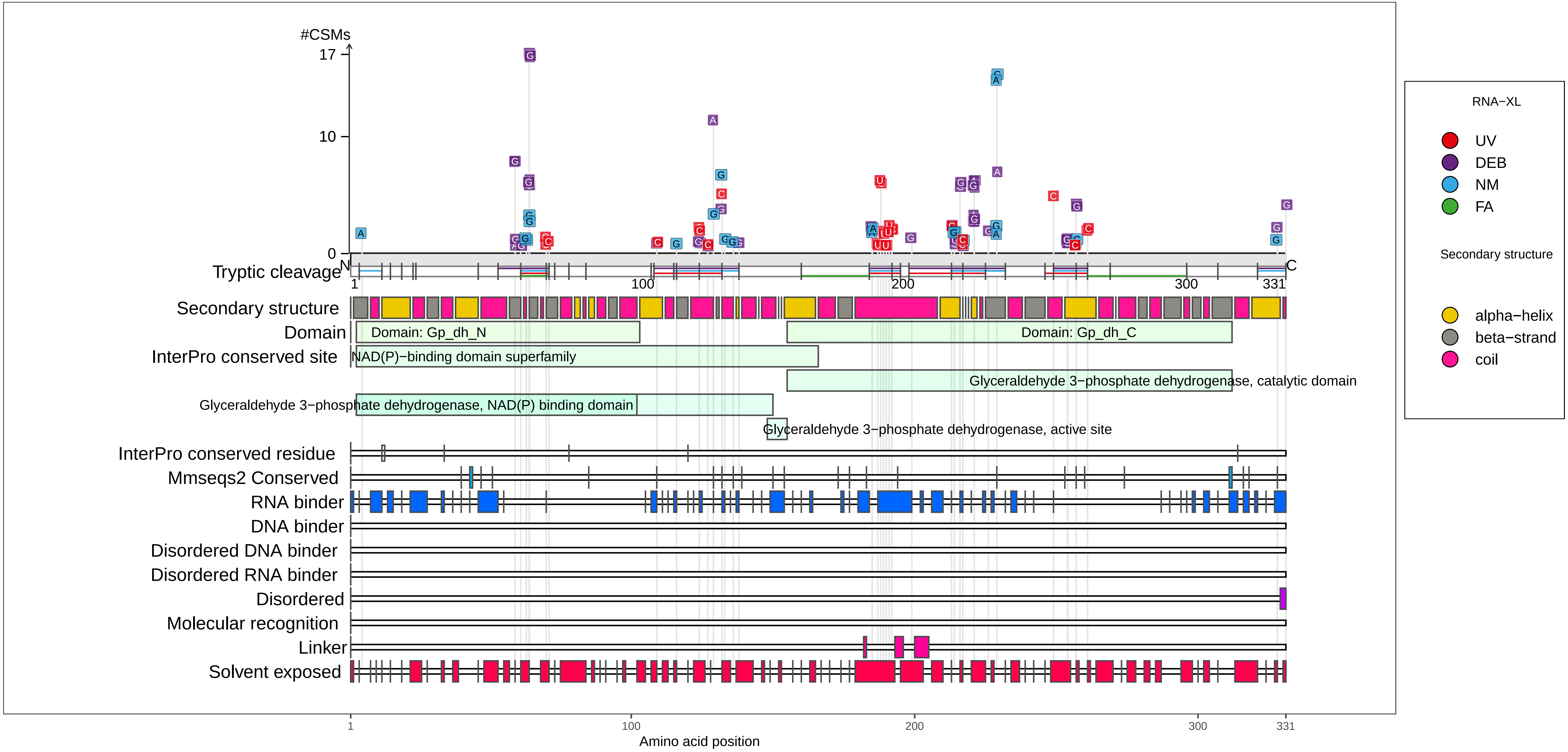

P62399
RL5_ECOLI 50S ribosomal protein L5

− Abundance:
tryptic [log10 Intensity]: 9.56 (Q 97)
PAXdb K12 strain [ppm]: 3.79 (Q 99)
PAXdb E.coli [ppm]: 3.56 (Q 99)

− RNA functions:
5S rRNA binding; RNA binding; rRNA binding; tRNA binding

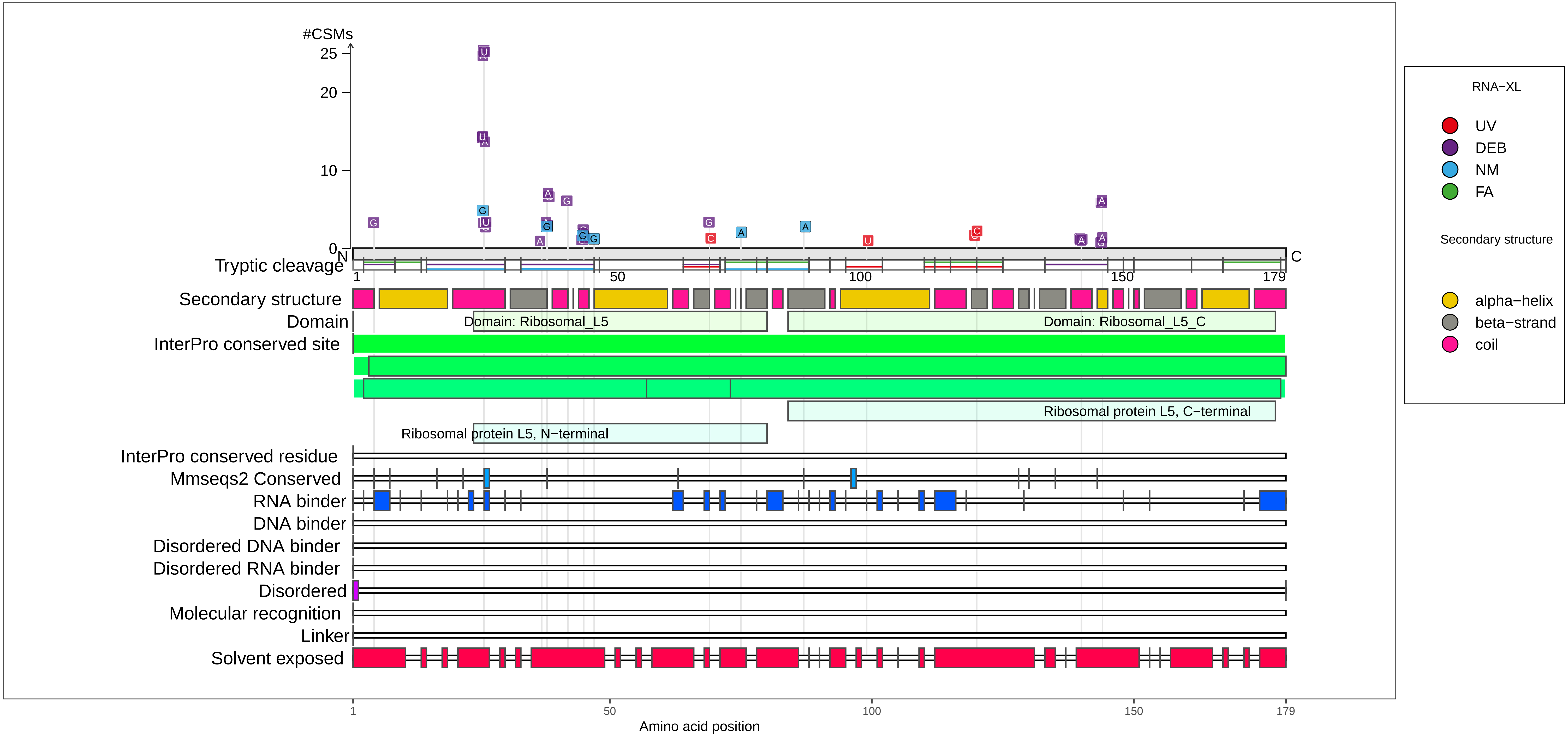

P02413
RL15_ECOLI 50S ribosomal protein L15

− Abundance:
tryptic [log10 Intensity]: 9.45 (Q 96)
PAXdb K12 strain [ppm]: 3.41 (Q 96)
PAXdb E.coli [ppm]: 3.45 (Q 99)

− RNA functions:
RNA binding; rRNA binding

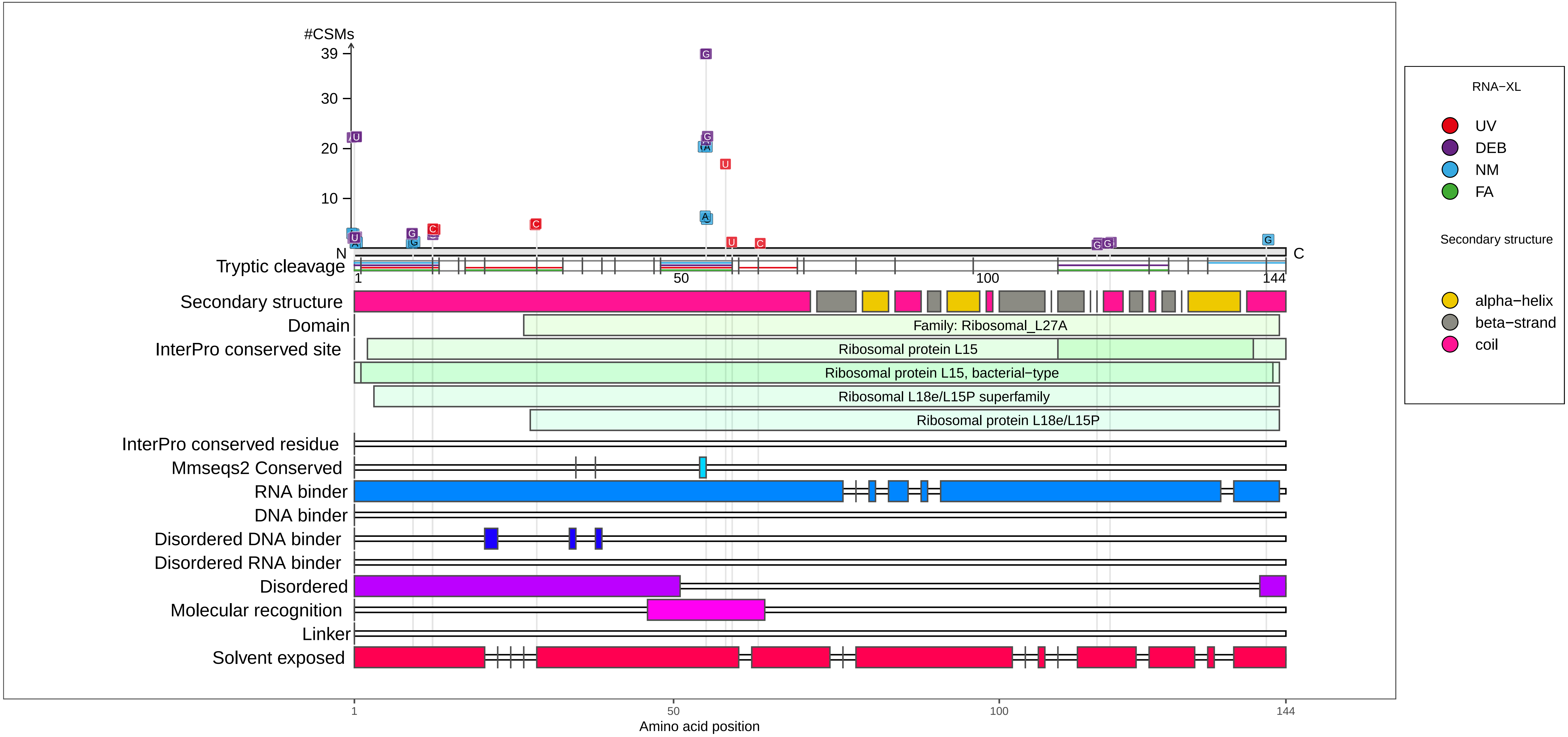

P0AE08
AHPC_ECOLI Alkyl hydroperoxide reductase C

− Abundance:
tryptic [log10 Intensity]: 9.82 (Q 99)
PAXdb K12 strain [ppm]: 4.01 (Q 100)
PAXdb E.coli [ppm]: 3.6 (Q 100)

− RNA functions: not annotated

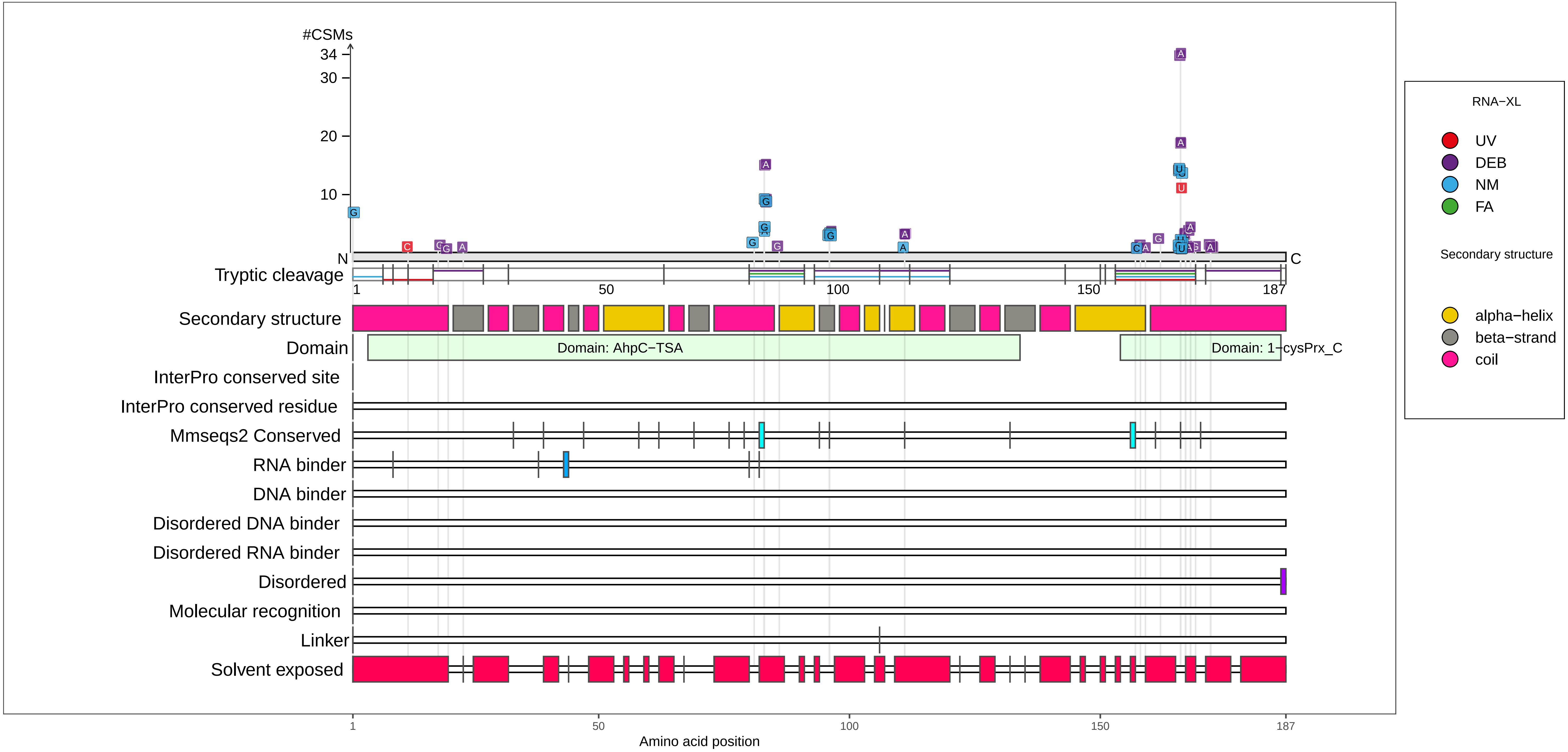

P00954
SYW_ECOLI Tryptophan−−tRNA ligase

− Abundance:
tryptic [log10 Intensity]: 8.6 (Q 79)
PAXdb K12 strain [ppm]: 3.33 (Q 96)
PAXdb E.coli [ppm]: 2.83 (Q 92)

− RNA functions:
aminoacyl−tRNA ligase activity; ncRNA metabolic process; RNA metabolic process
tRNA aminoacylation; tRNA aminoacylation for protein translation; tRNA metabolic process
tRNA synthetases class I (W and Y); tryptophan−tRNA ligase activity
tryptophanyl−tRNA aminoacylation

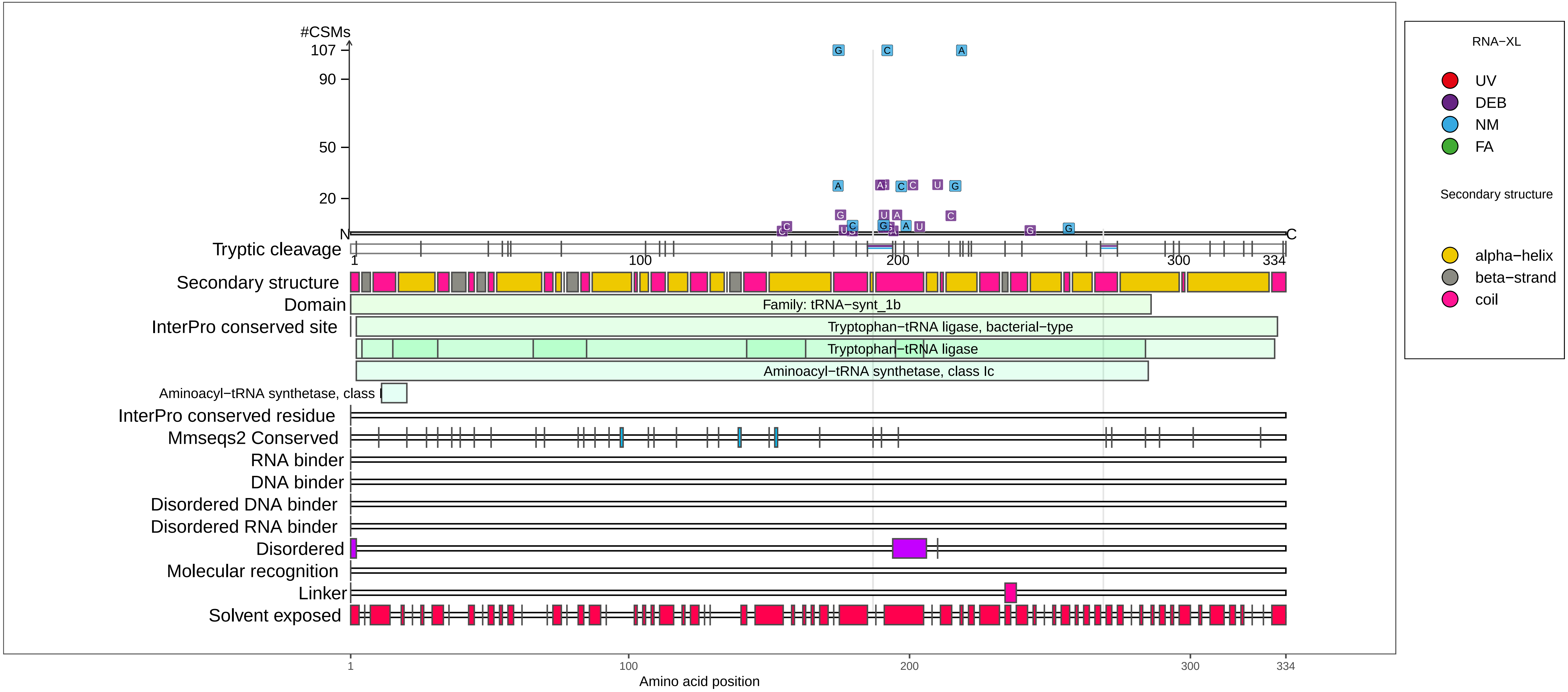

P0A7L8
RL27_ECOLI 50S ribosomal protein L27

− Abundance:
tryptic [log10 Intensity]: 8.89 (Q 86)
PAXdb K12 strain [ppm]: 3.89 (Q 100)
PAXdb E.coli [ppm]: 3.24 (Q 97)

− RNA functions:
RNA binding; rRNA binding; tRNA binding

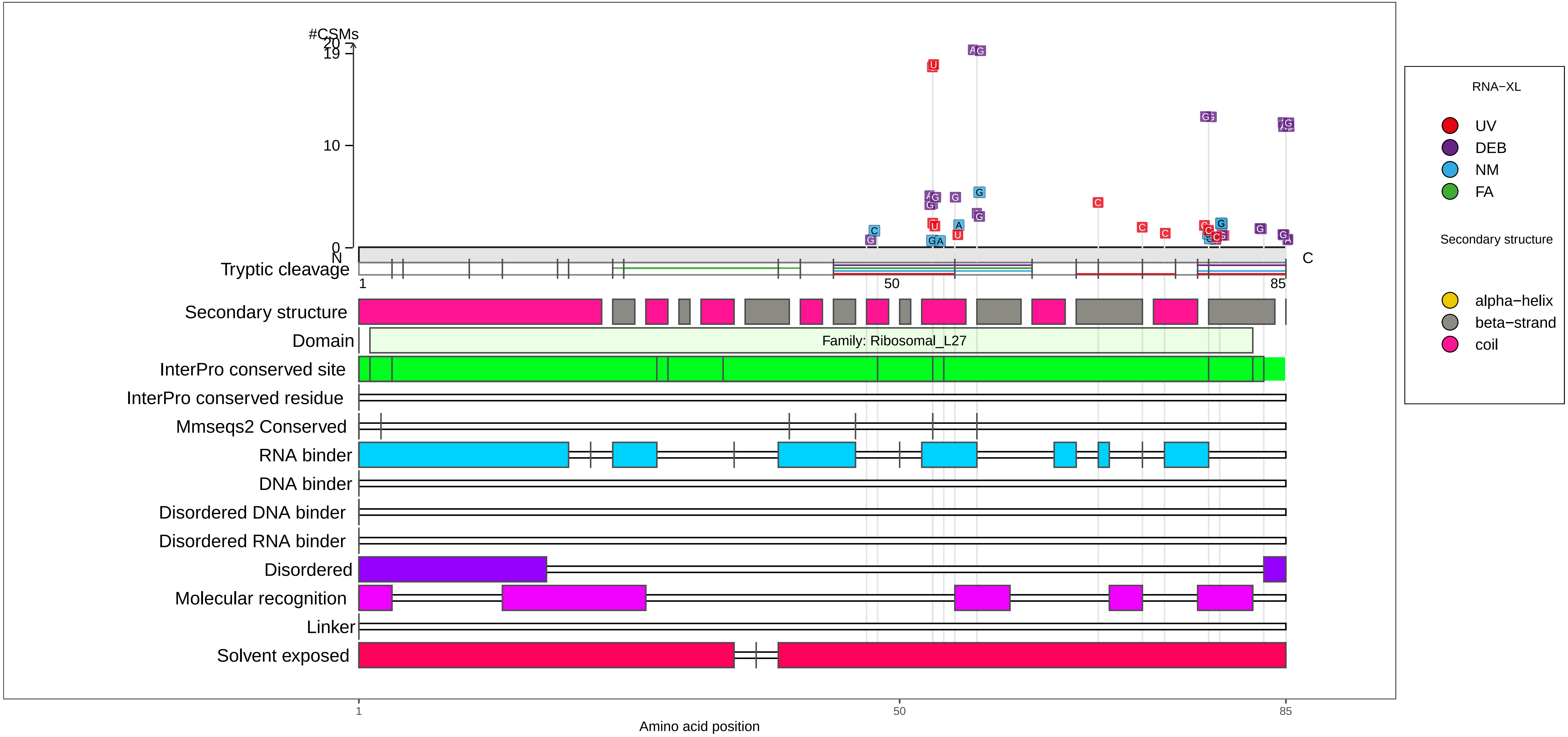

P0ABB4
ATPB_ECOLI ATP synthase subunit beta

− Abundance:
tryptic [log10 Intensity]: 10.24 (Q 100)
PAXdb K12 strain [ppm]: 2.94 (Q 89)
PAXdb E.coli [ppm]: 3.62 (Q 100)

− RNA functions: not annotated

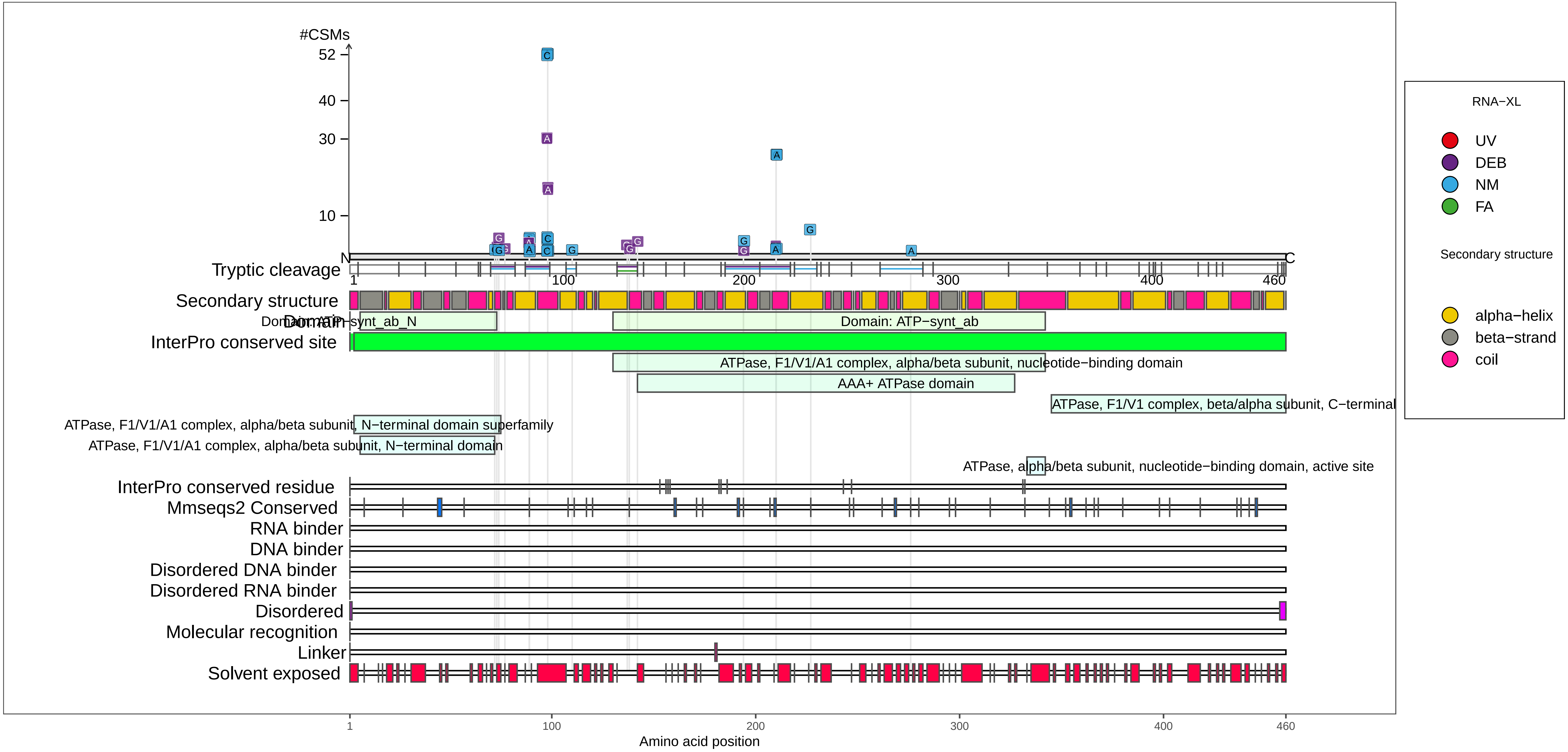

P0A8B5
YBAB_ECOLI Nucleoid−associated protein YbaB
− Abundance:
tryptic [log10 Intensity]: 8.79 (Q 83)
PAXdb K12 strain [ppm]: 2.8 (Q 86)
PAXdb E.coli [ppm]: 3.28 (Q 97)

− RNA functions: not annotated

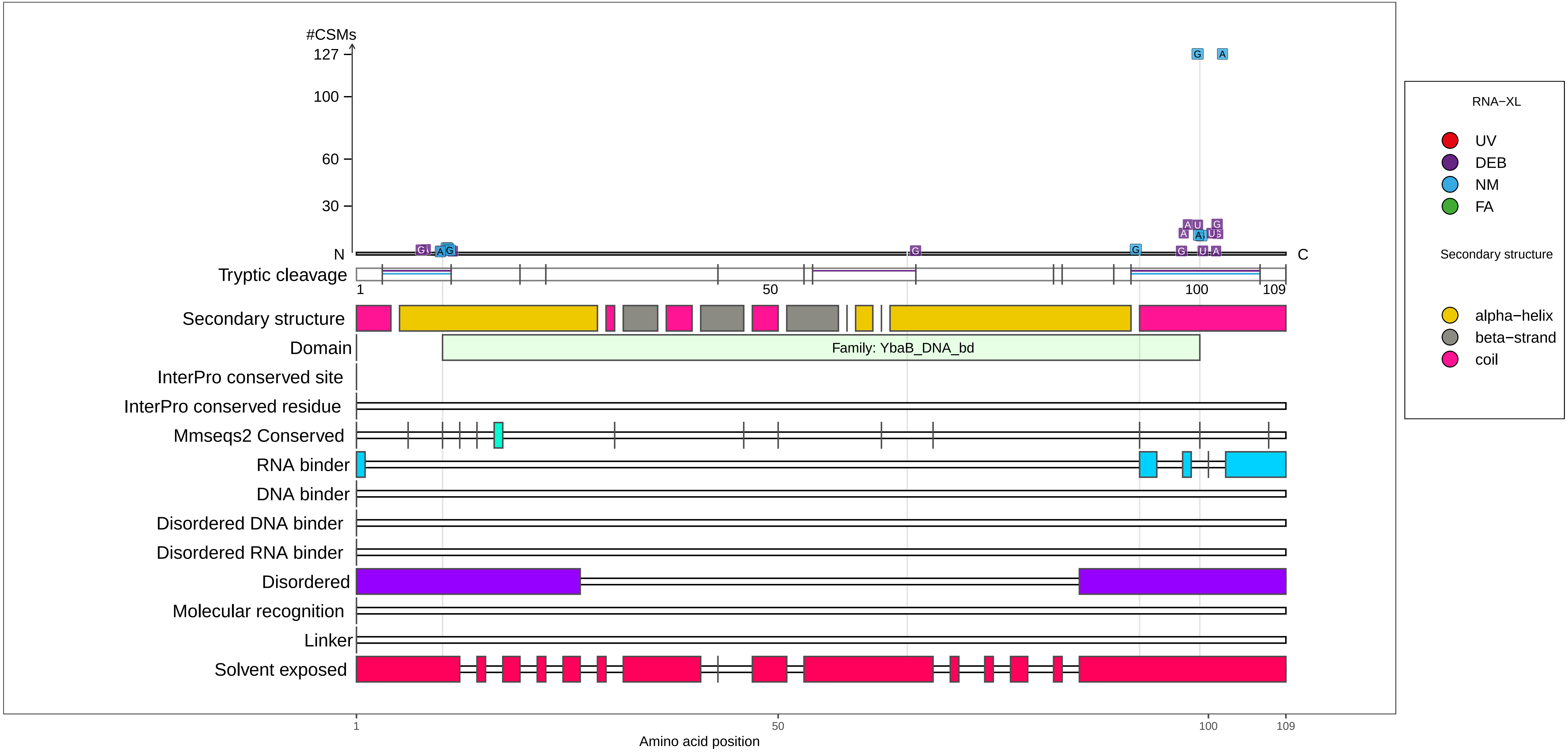

P0A7X3
RS9_ECOLI 30S ribosomal protein S9

− Abundance:
tryptic [log10 Intensity]: 9.19 (Q 92)
PAXdb K12 strain [ppm]: 3.71 (Q 99)
PAXdb E.coli [ppm]: 3.25 (Q 97)

− RNA functions:
RNA binding; tRNA binding

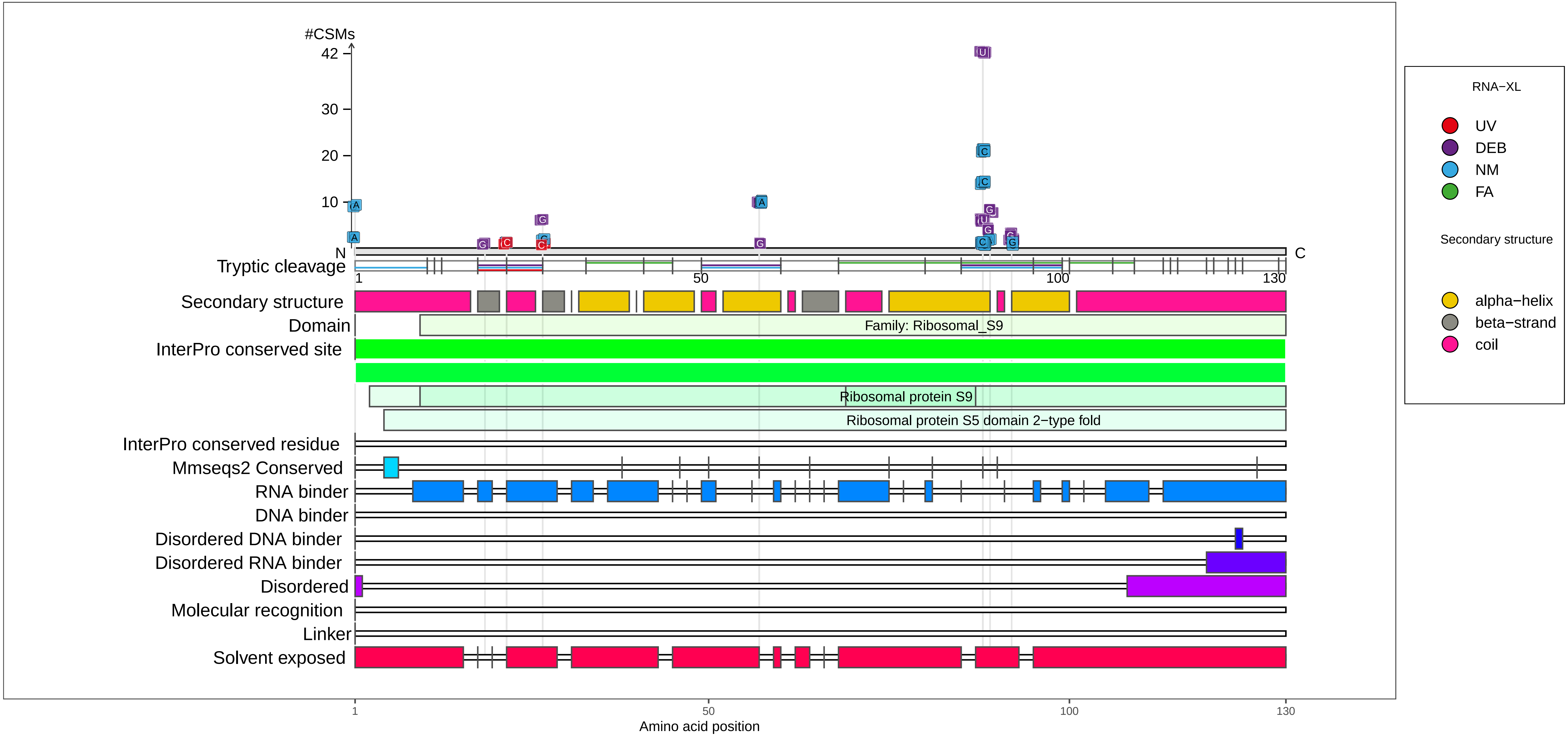

P0A6D7
AROK_ECOLI Shikimate kinase 1

− Abundance:
tryptic [log10 Intensity]: 9 (Q 88)
PAXdb K12 strain [ppm]: 2.81 (Q 86)
PAXdb E.coli [ppm]: 2.32 (Q 81)

− RNA functions: not annotated

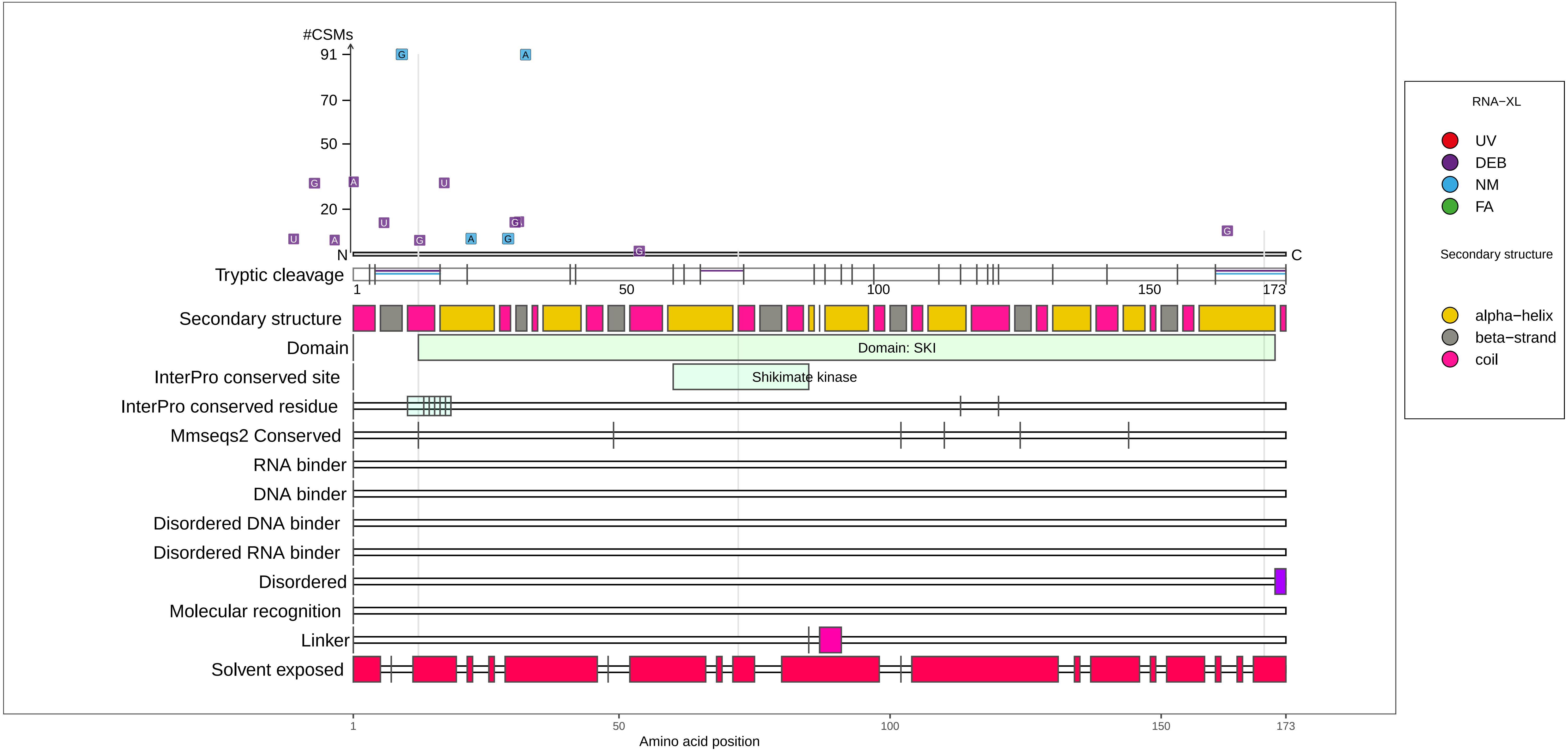

P0A7Z4
RPOA_ECOLI DNA−directed RNA polymerase subunit alpha

− Abundance:
tryptic [log10 Intensity]: 9.79 (Q 99)
PAXdb K12 strain [ppm]: 3.34 (Q 96)
PAXdb E.coli [ppm]: 3.16 (Q 96)

− RNA functions:
5−3 RNA polymerase activity
Bacterial RNA polymerase, alpha chain C terminal domain; DNA−directed 5−3 RNA polymerase activity; RNA biosynthetic process
RNA metabolic process; RNA polymerase activity; RNA polymerase Rpb3/Rpb11 dimerisation domain
RNA polymerase Rpb3/RpoA insert domain

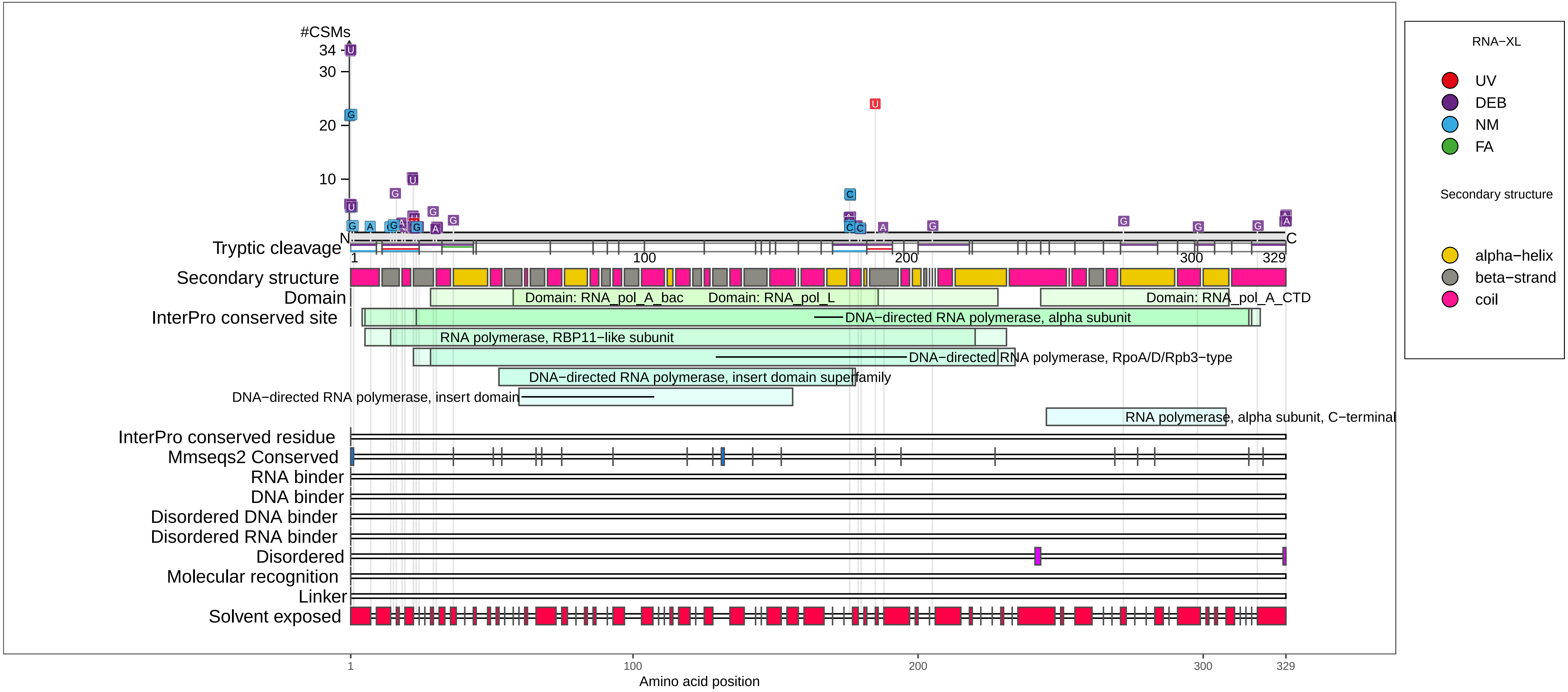

P45577
PROQ_ECOLI RNA chaperone ProQ

− Abundance:
tryptic [log10 Intensity]: 8.59 (Q 78)
PAXdb K12 strain [ppm]: 3.45 (Q 97)
PAXdb E.coli [ppm]: 2.78 (Q 91)

− RNA functions:
mRNA 3−UTR binding; mRNA binding; RNA binding; RNA strand annealing activity
RNA strand−exchange activity

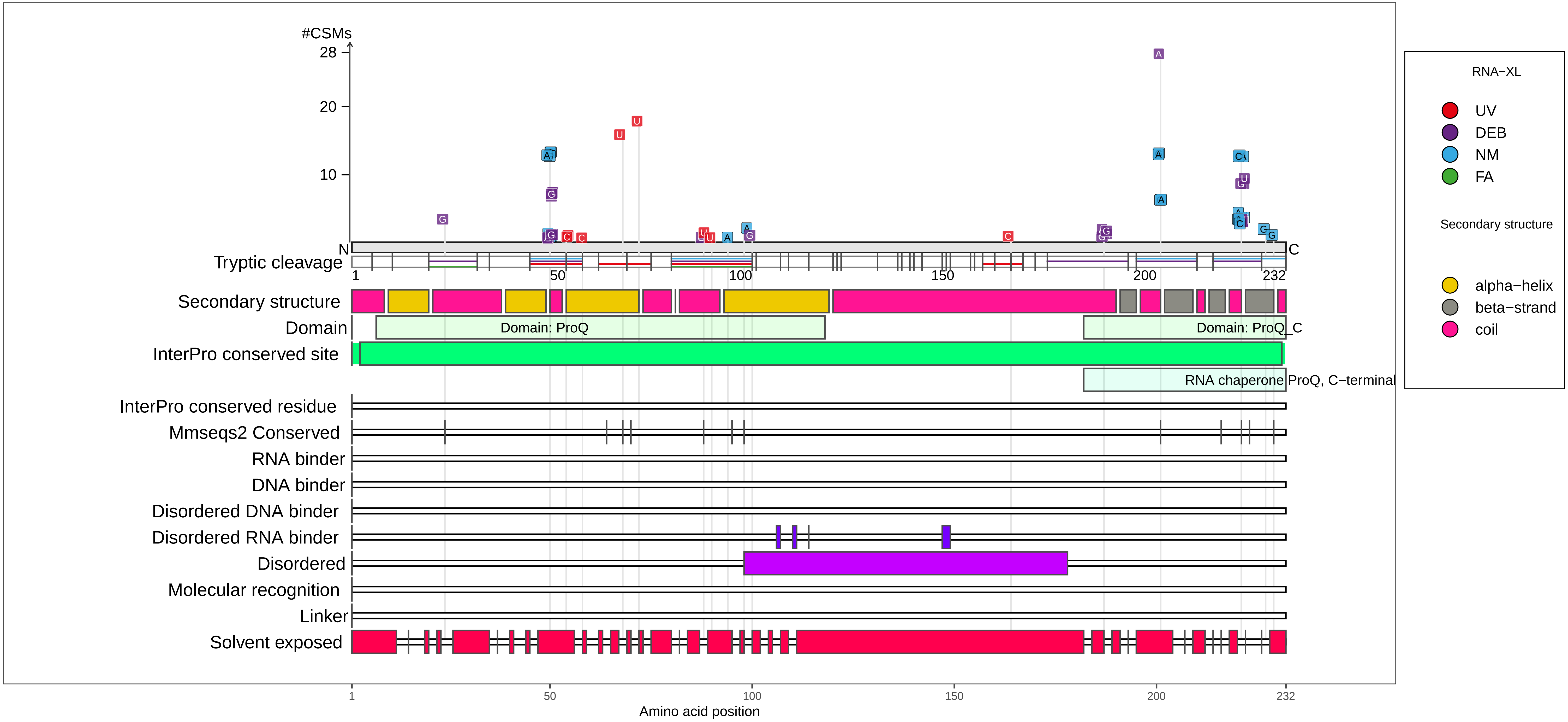

P0A7R5
RS10_ECOLI 30S ribosomal protein S10

− Abundance:
tryptic [log10 Intensity]: 9.4 (Q 95)
PAXdb K12 strain [ppm]: 3.82 (Q 100)
PAXdb E.coli [ppm]: 3.45 (Q 98)

− RNA functions:
RNA binding; tRNA binding

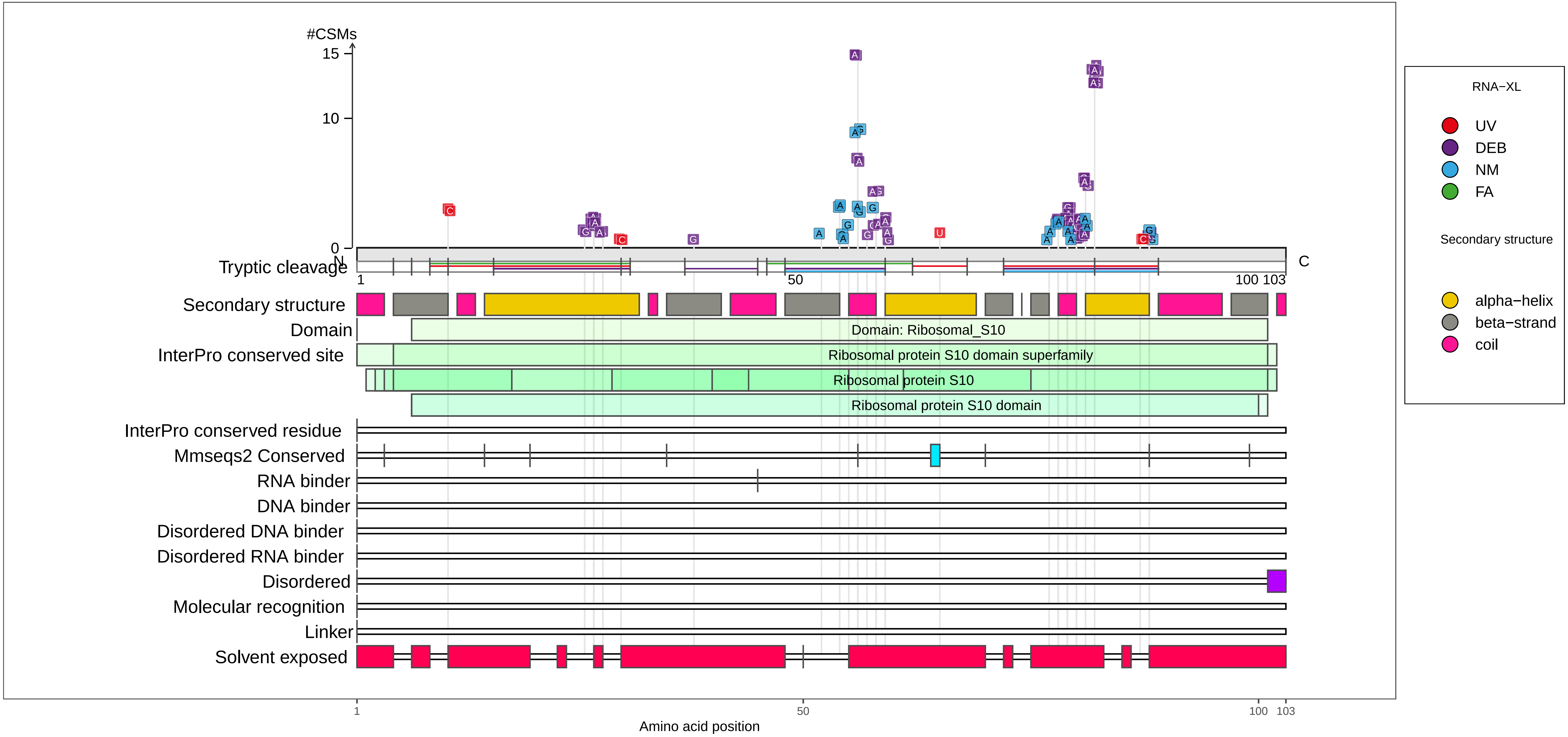

P0A8M3
SYT_ECOLI Threonine−−tRNA ligase

− Abundance:
tryptic [log10 Intensity]: 9.3 (Q 94)
PAXdb K12 strain [ppm]: 2.06 (Q 62)
PAXdb E.coli [ppm]: 3.29 (Q 97)

− RNA functions:
aminoacyl−tRNA editing activity; aminoacyl−tRNA ligase activity
aminoacyl−tRNA metabolism involved in translational fidelity; mRNA 5−UTR binding; mRNA binding
ncRNA metabolic process; RNA binding; RNA metabolic process; threonine−tRNA ligase activity
Threonyl and Alanyl tRNA synthetase second additional domain
threonyl−tRNA aminoacylation; tRNA aminoacylation; tRNA aminoacylation for protein translation; tRNA binding
tRNA metabolic process; tRNA synthetase class II core domain (G, H, P, S and T)

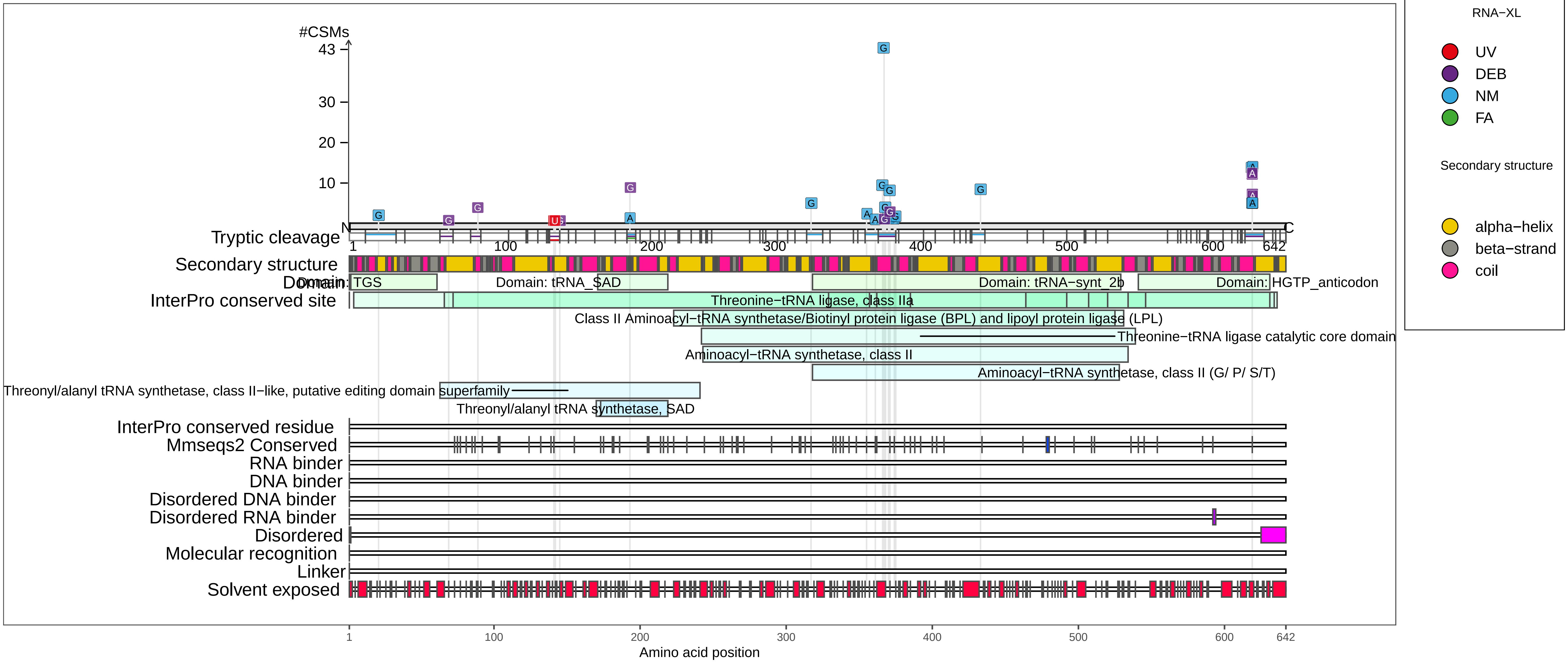

P0AFF6
NUSA_ECOLI Transcription termination/antitermination protein NusA

− Abundance:
tryptic [log10 Intensity]: 9.04 (Q 89)
PAXdb K12 strain [ppm]: 3.17 (Q 94)
PAXdb E.coli [ppm]: 3.21 (Q 97)

− RNA functions:
RNA binding; RNA biosynthetic process; RNA metabolic process; RNA polymerase binding
RNA polymerase core enzyme binding; S1 RNA binding domain

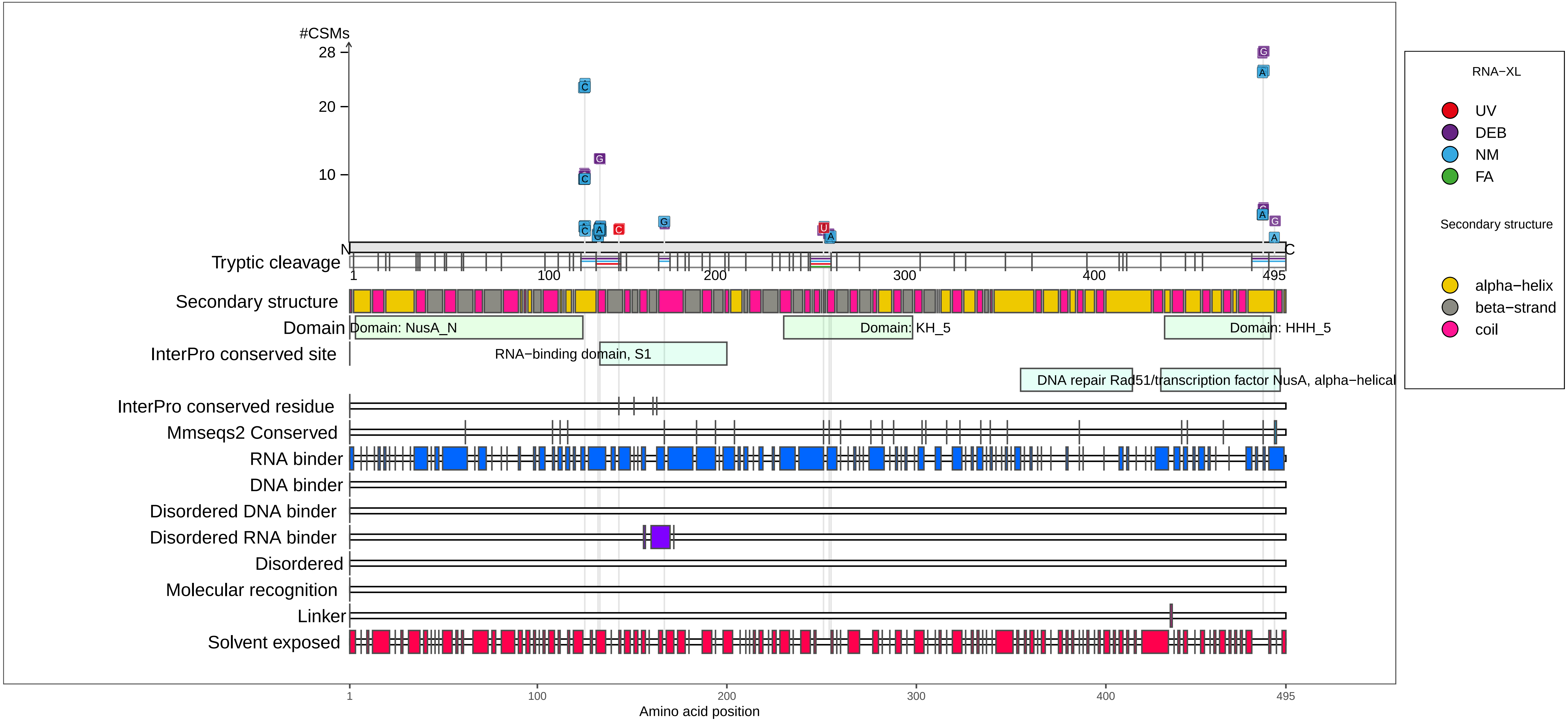

P60438
RL3_ECOLI 50S ribosomal protein L3

− Abundance:
tryptic [log10 Intensity]: 9.42 (Q 96)
PAXdb K12 strain [ppm]: 3.38 (Q 96)
PAXdb E.coli [ppm]: 3.49 (Q 99)

− RNA functions:
RNA binding; rRNA binding

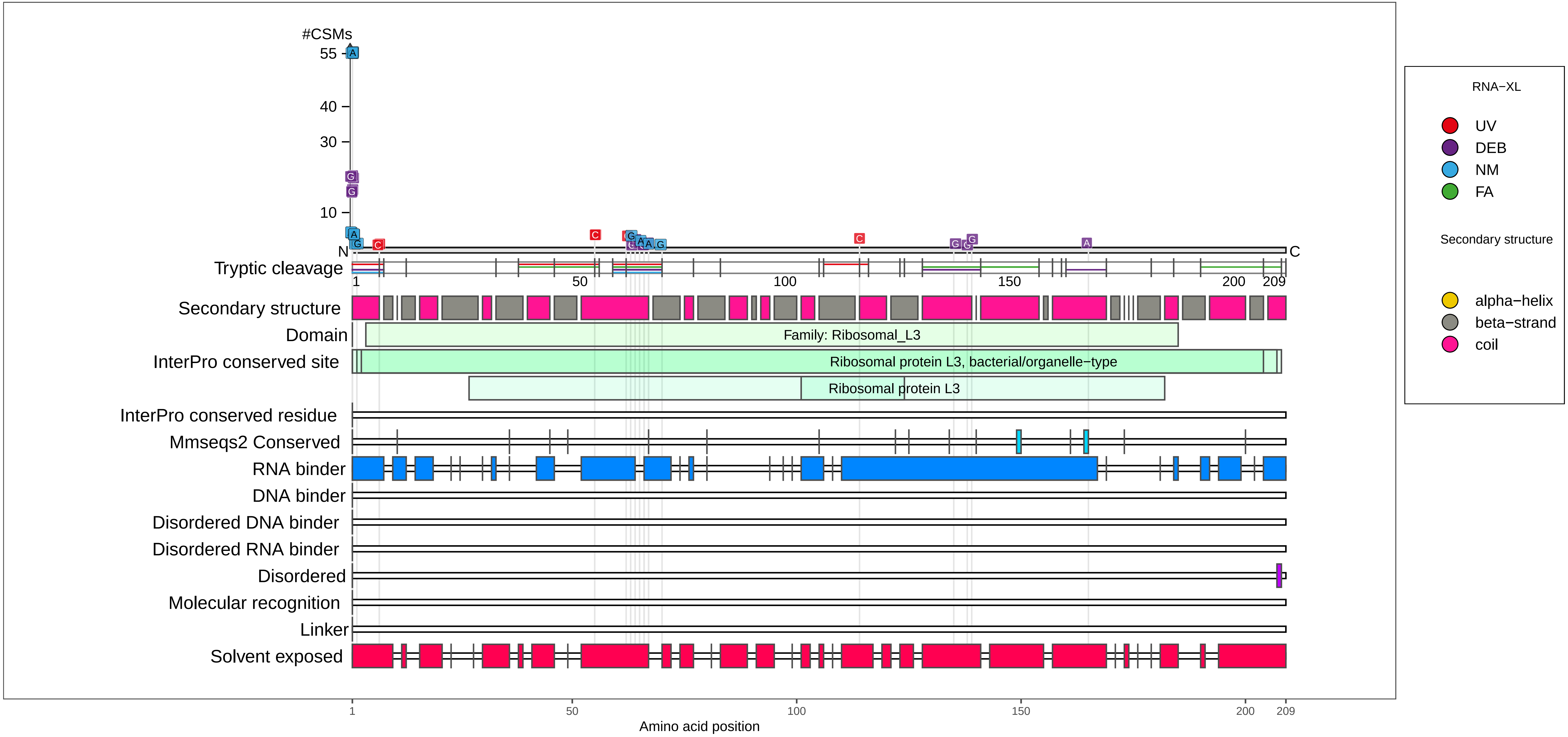

P09373
PFLB_ECOLI Formate acetyltransferase 1

− Abundance:
tryptic [log10 Intensity]: 9.93 (Q 99)
PAXdb K12 strain [ppm]: 3.15 (Q 93)
PAXdb E.coli [ppm]: 3.52 (Q 99)

− RNA functions: not annotated

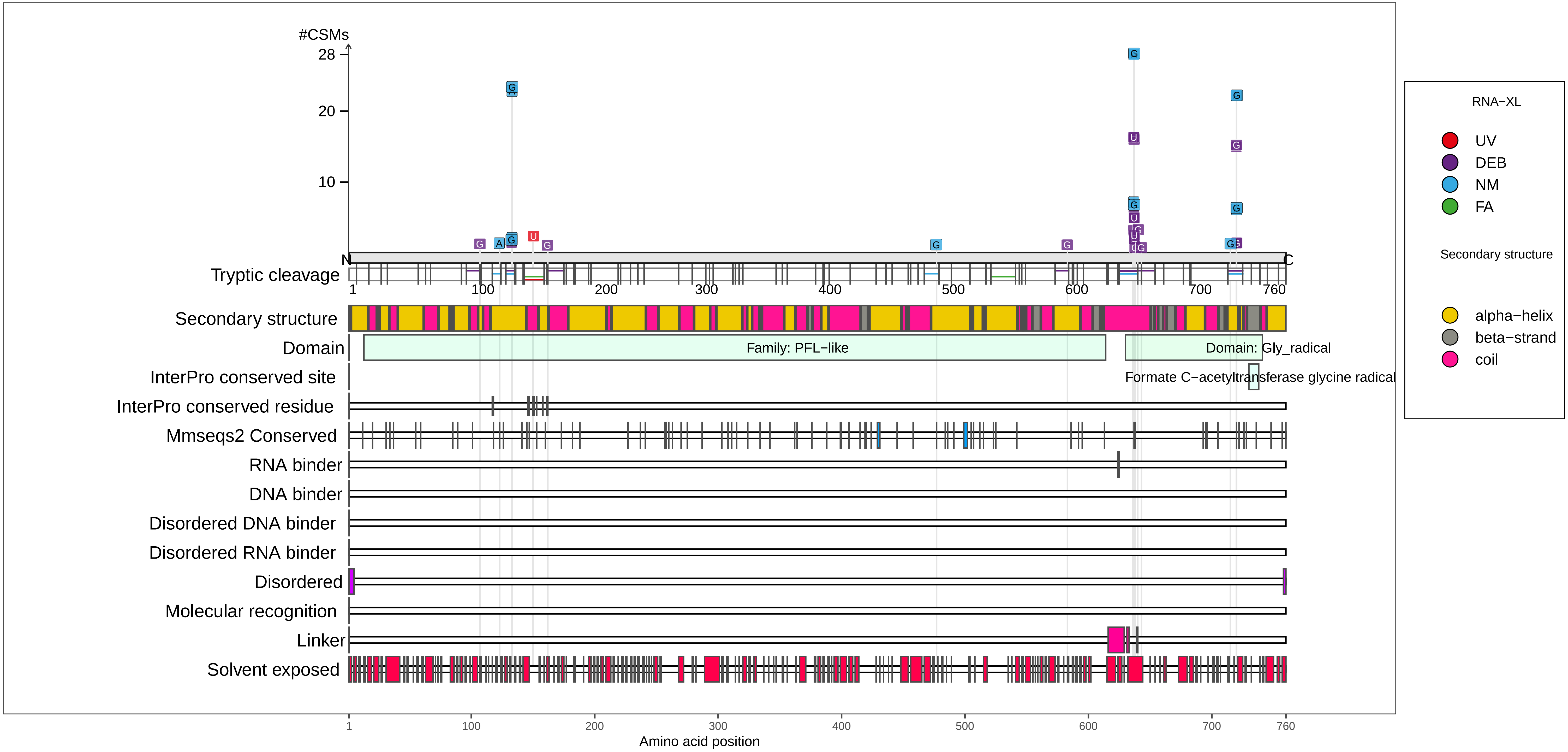

P0A978
CSPG_ECOLI Cold shock−like protein CspG

− Abundance:
tryptic [log10 Intensity]: 9.15 (Q 92)
PAXdb K12 strain [ppm]: 2.85 (Q 87)
PAXdb E.coli [ppm]: 1.73 (Q 65)

− RNA functions:
RNA binding

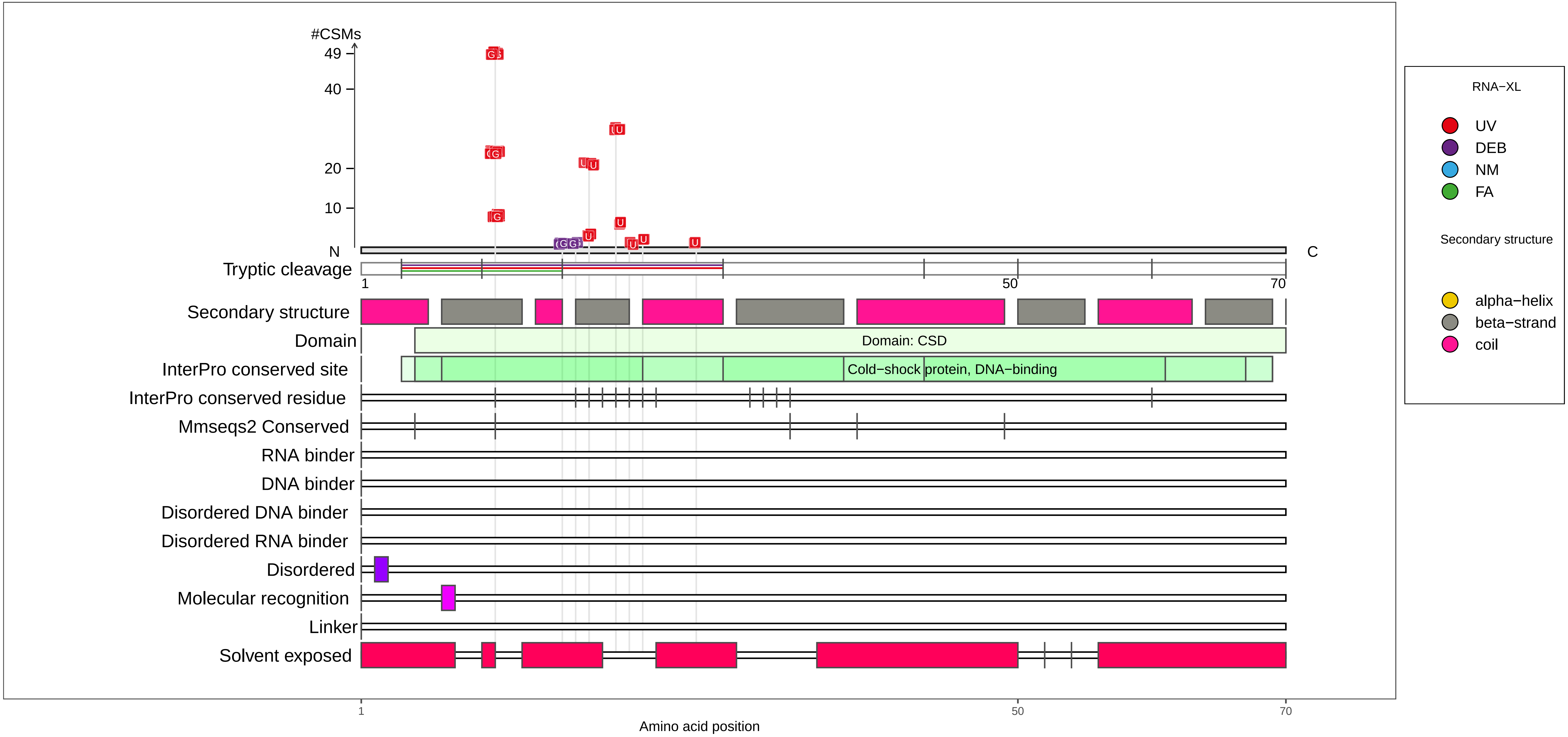

P0A6F3
GLPK_ECOLI Glycerol kinase

− Abundance:
tryptic [log10 Intensity]: 9.91 (Q 99)
PAXdb K12 strain [ppm]: 3.28 (Q 95)
PAXdb E.coli [ppm]: 2.75 (Q 91)

− RNA functions: not annotated

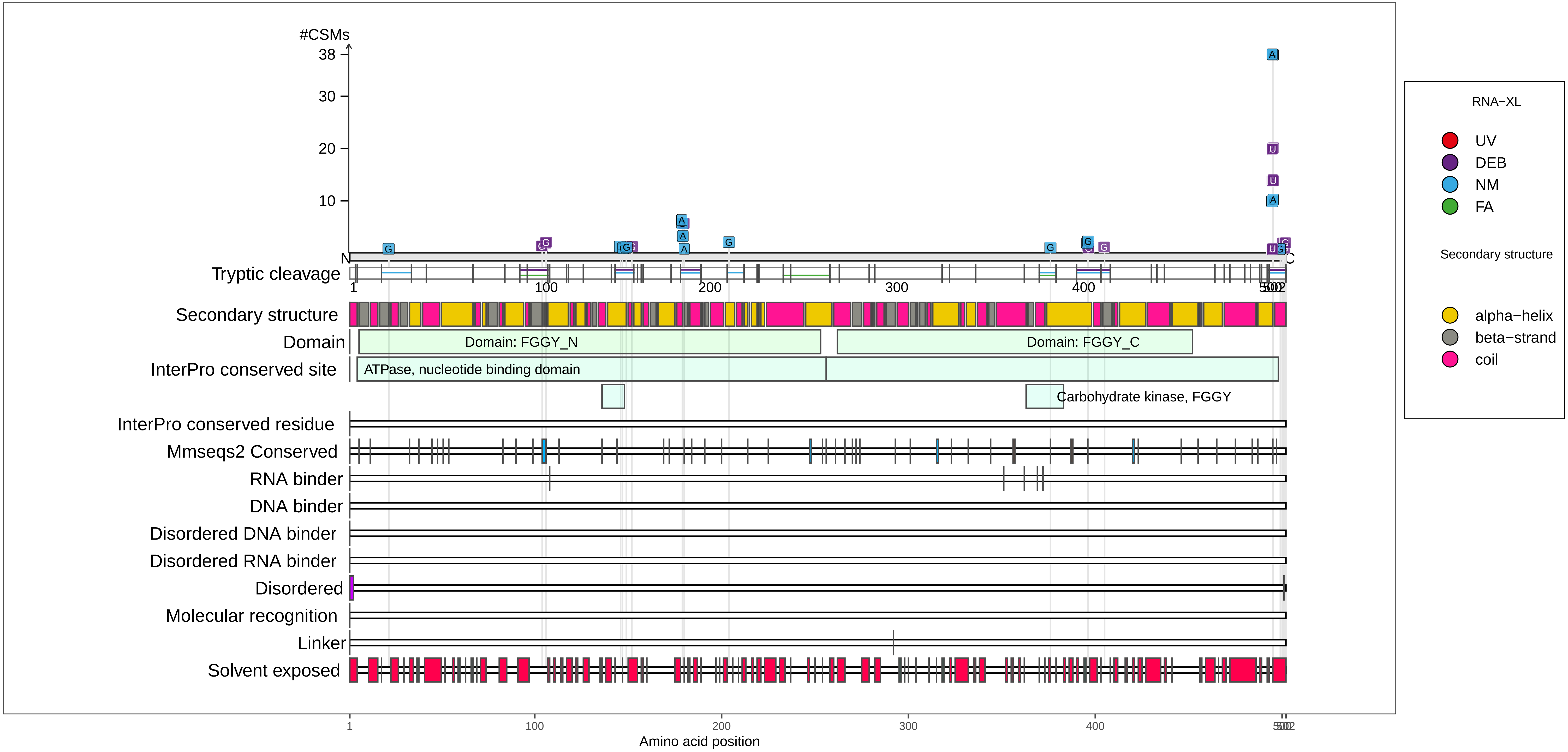

P0A805
RRF_ECOLI Ribosome−recycling factor

− Abundance:
tryptic [log10 Intensity]: 8.51 (Q 76)
PAXdb K12 strain [ppm]: 3.53 (Q 98)
PAXdb E.coli [ppm]: 3.08 (Q 96)

− RNA functions: not annotated

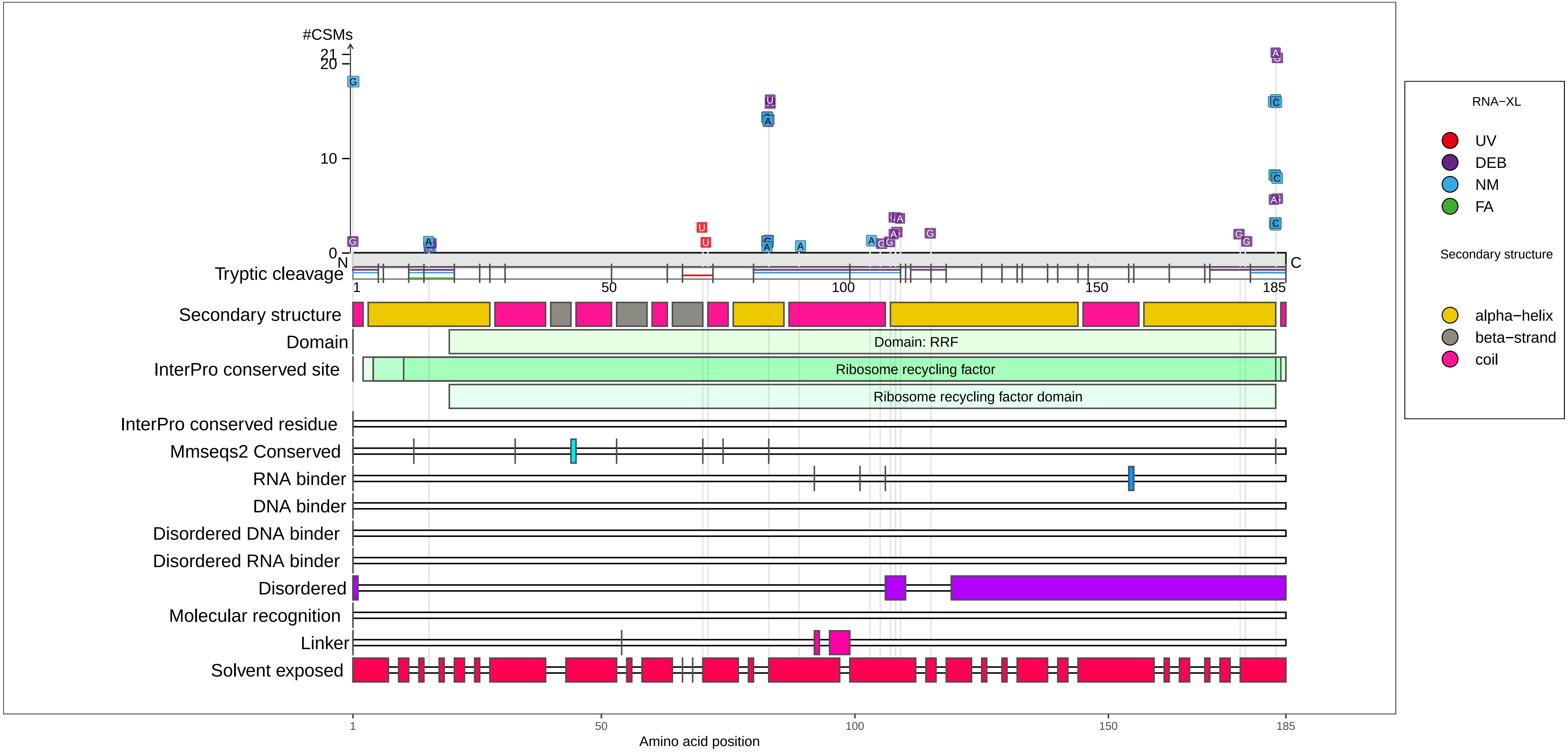

P06959
ODP2_ECOLI Dihydrolipoyllysine−residue acetyltransferase component of pyruvate dehydrogenase complex

− Abundance:
tryptic [log10 Intensity]: 10.12 (Q 100)
PAXdb K12 strain [ppm]: 3.12 (Q 93)
PAXdb E.coli [ppm]: 3.45 (Q 99)

− RNA functions: not annotated

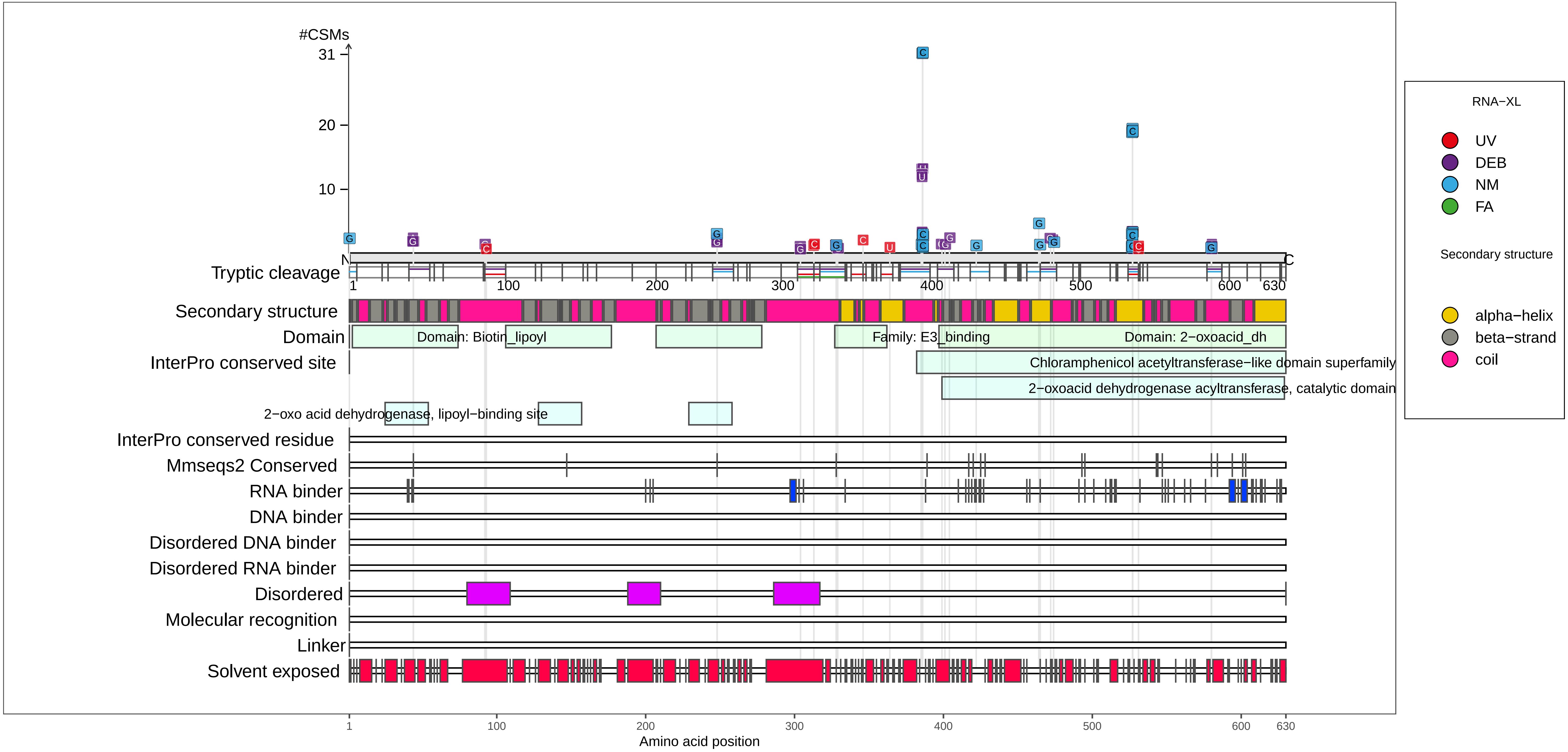

P0ACR9
MPRA_ECOLI Transcriptional repressor MprA

− Abundance:
tryptic [log10 Intensity]: 7.76 (Q 47)
PAXdb K12 strain [ppm]: 2.79 (Q 86)
PAXdb E.coli [ppm]: 2.16 (Q 76)

− RNA functions:
RNA biosynthetic process; RNA metabolic process

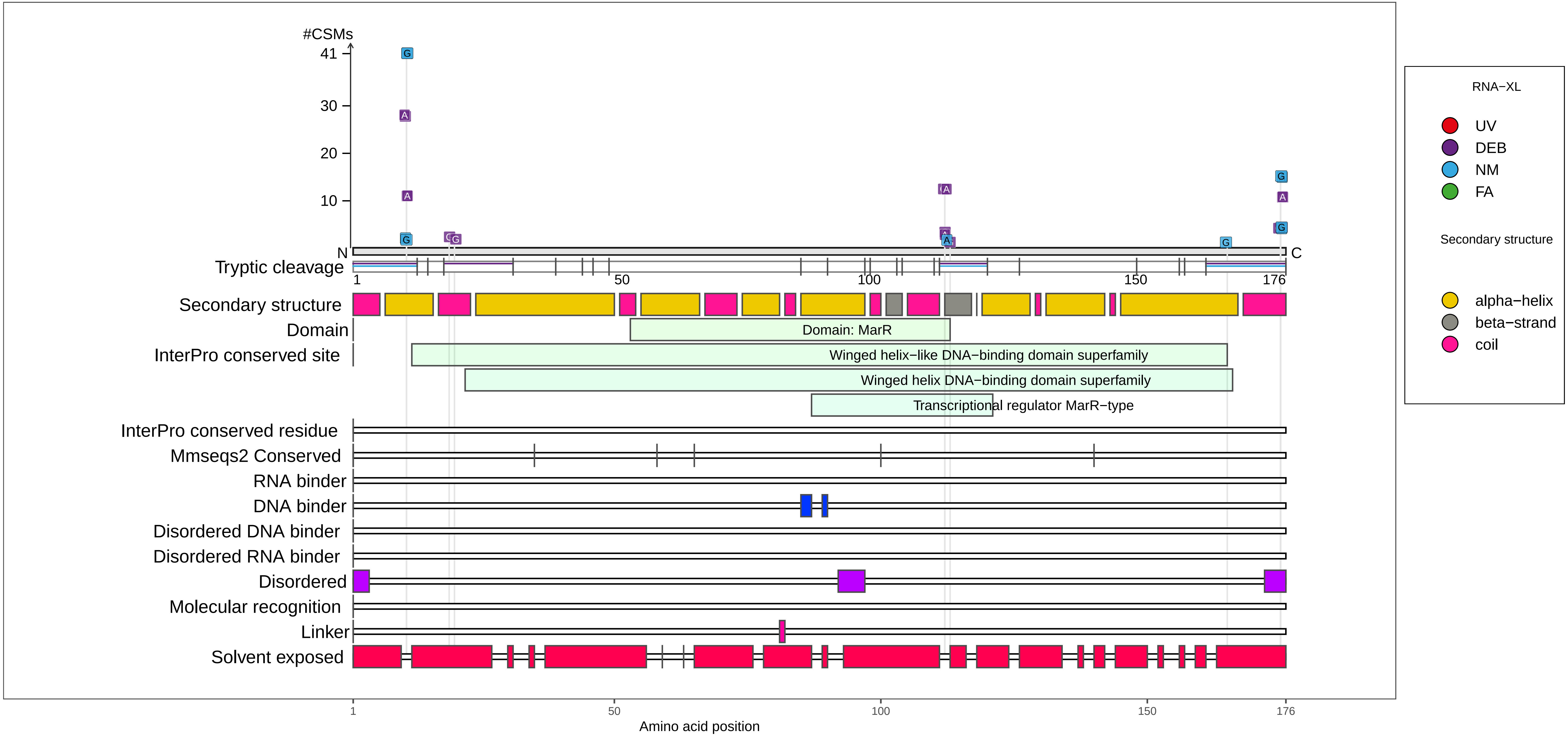

P24182
ACCC_ECOLI Biotin carboxylase

− Abundance:
tryptic [log10 Intensity]: 9.14 (Q 91)
PAXdb K12 strain [ppm]: 3.47 (Q 97)
PAXdb E.coli [ppm]: 2.4 (Q 83)

− RNA functions: not annotated

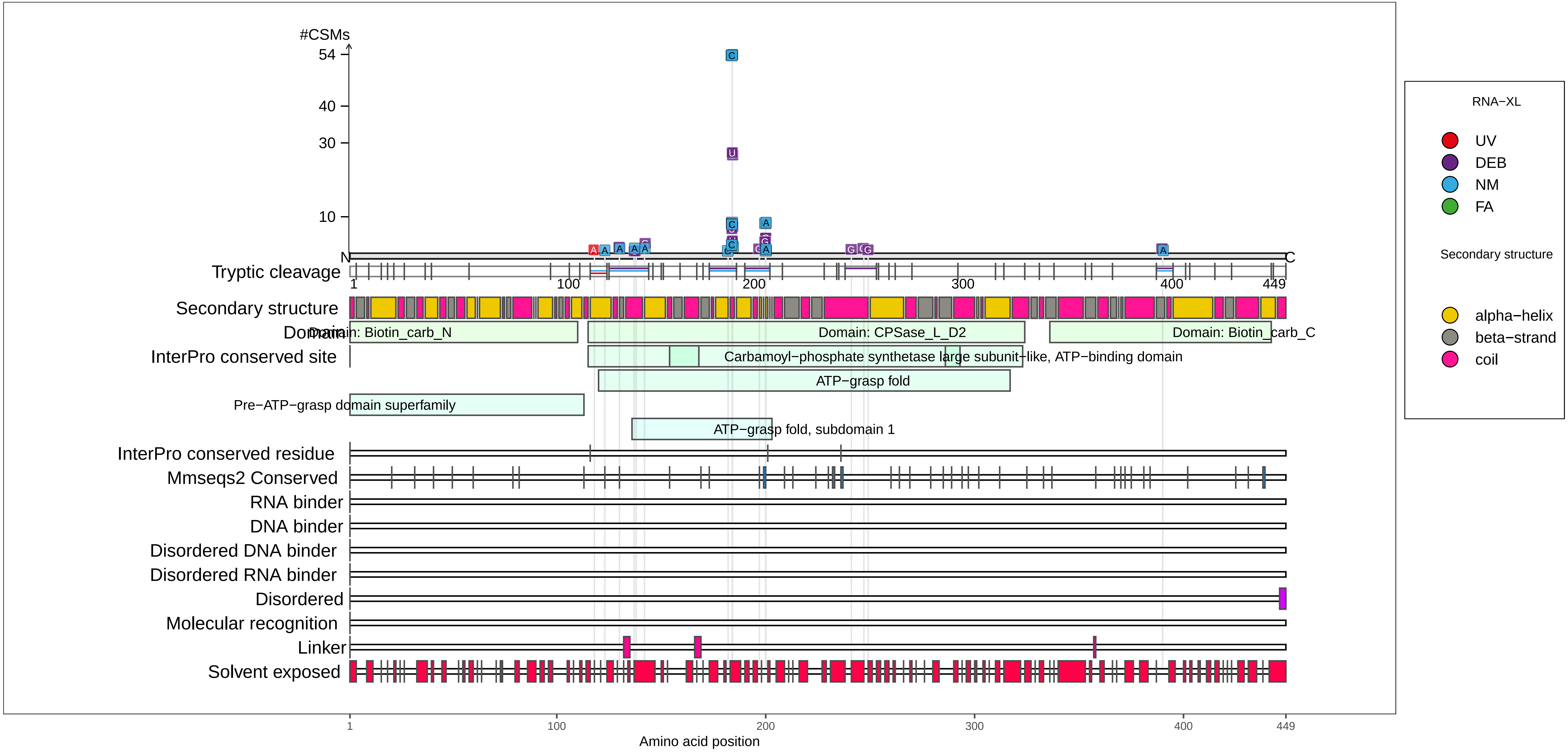

P0A8M0
SYN_ECOLI Asparagine−−tRNA ligase

− Abundance:
tryptic [log10 Intensity]: 9.34 (Q 95)
PAXdb K12 strain [ppm]: 3.26 (Q 95)
PAXdb E.coli [ppm]: 2.78 (Q 91)

− RNA functions:
aminoacyl−tRNA ligase activity; asparagine−tRNA ligase activity
asparaginyl−tRNA aminoacylation; ncRNA metabolic process; RNA metabolic process; tRNA aminoacylation
tRNA aminoacylation for protein translation; tRNA metabolic process
tRNA synthetases class II (D, K and N)

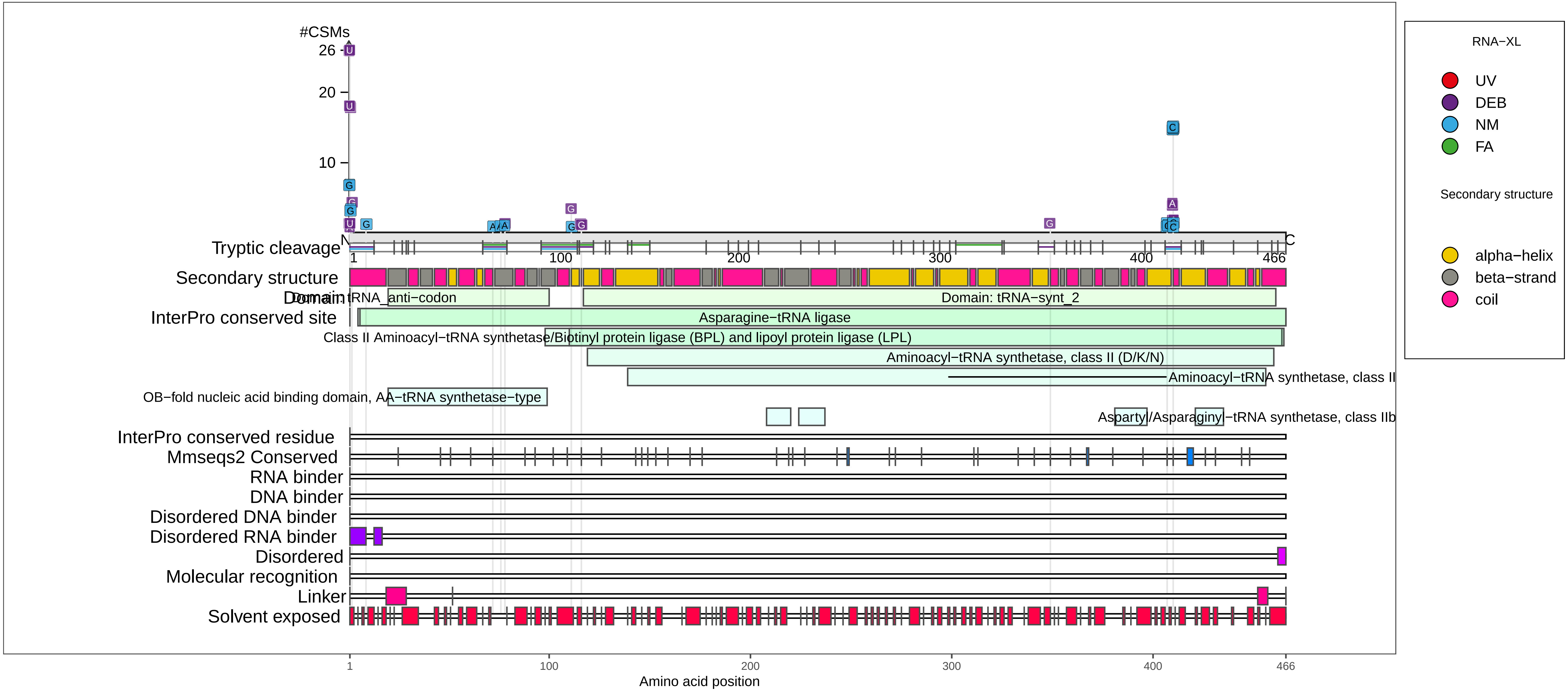

P0A982
CSPH_ECOLI Cold shock−like protein CspH

− Abundance:
tryptic [log10 Intensity]: NA (Q NA)
PAXdb K12 strain [ppm]: NA (Q NA)
PAXdb E.coli [ppm]: −1.48 (Q 4)

− RNA functions:
RNA binding

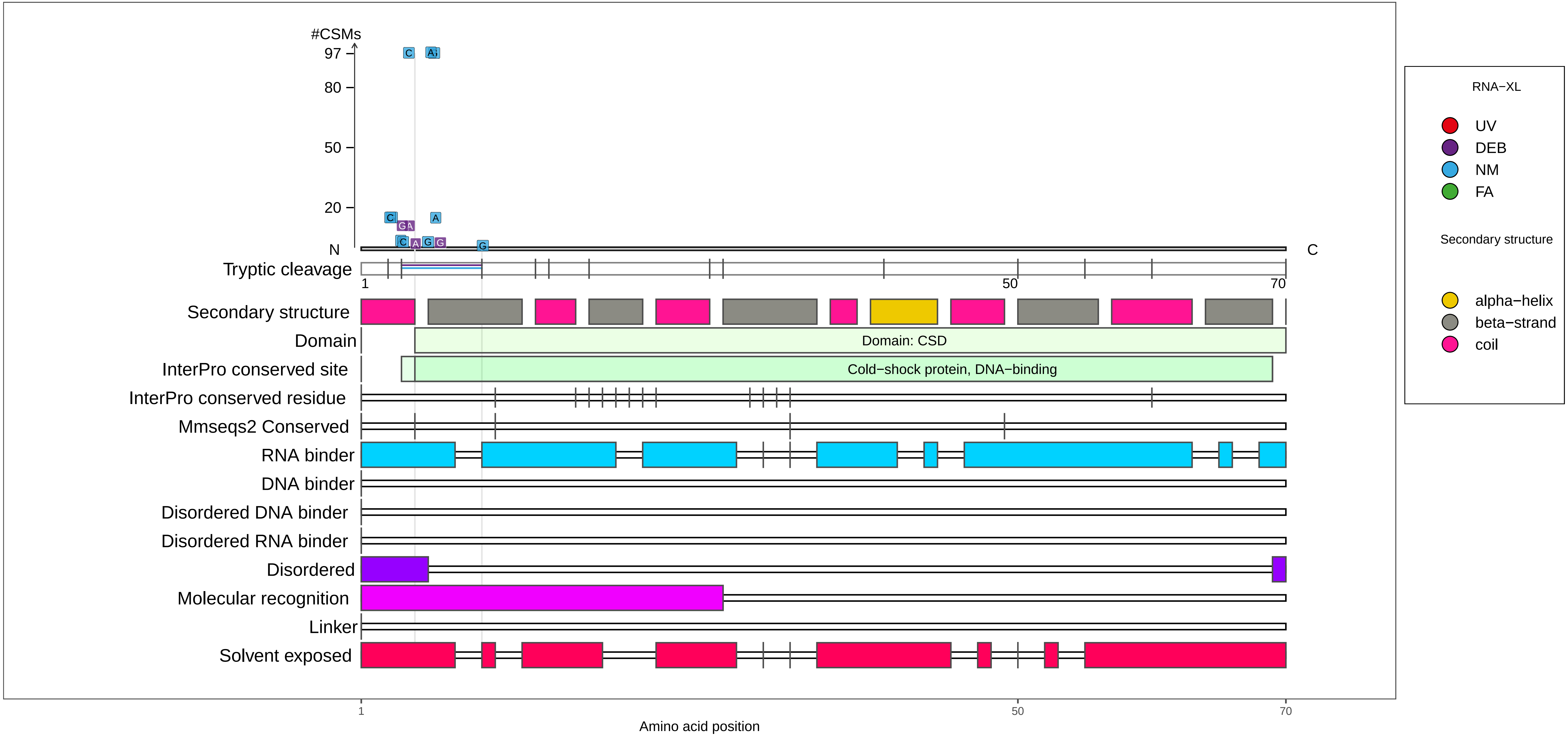

P0A7K6
RL19_ECOLI 50S ribosomal protein L19

− Abundance:
tryptic [log10 Intensity]: 9.12 (Q 91)
PAXdb K12 strain [ppm]: 3.73 (Q 99)
PAXdb E.coli [ppm]: 3.34 (Q 98)

− RNA functions:
RNA binding; rRNA binding

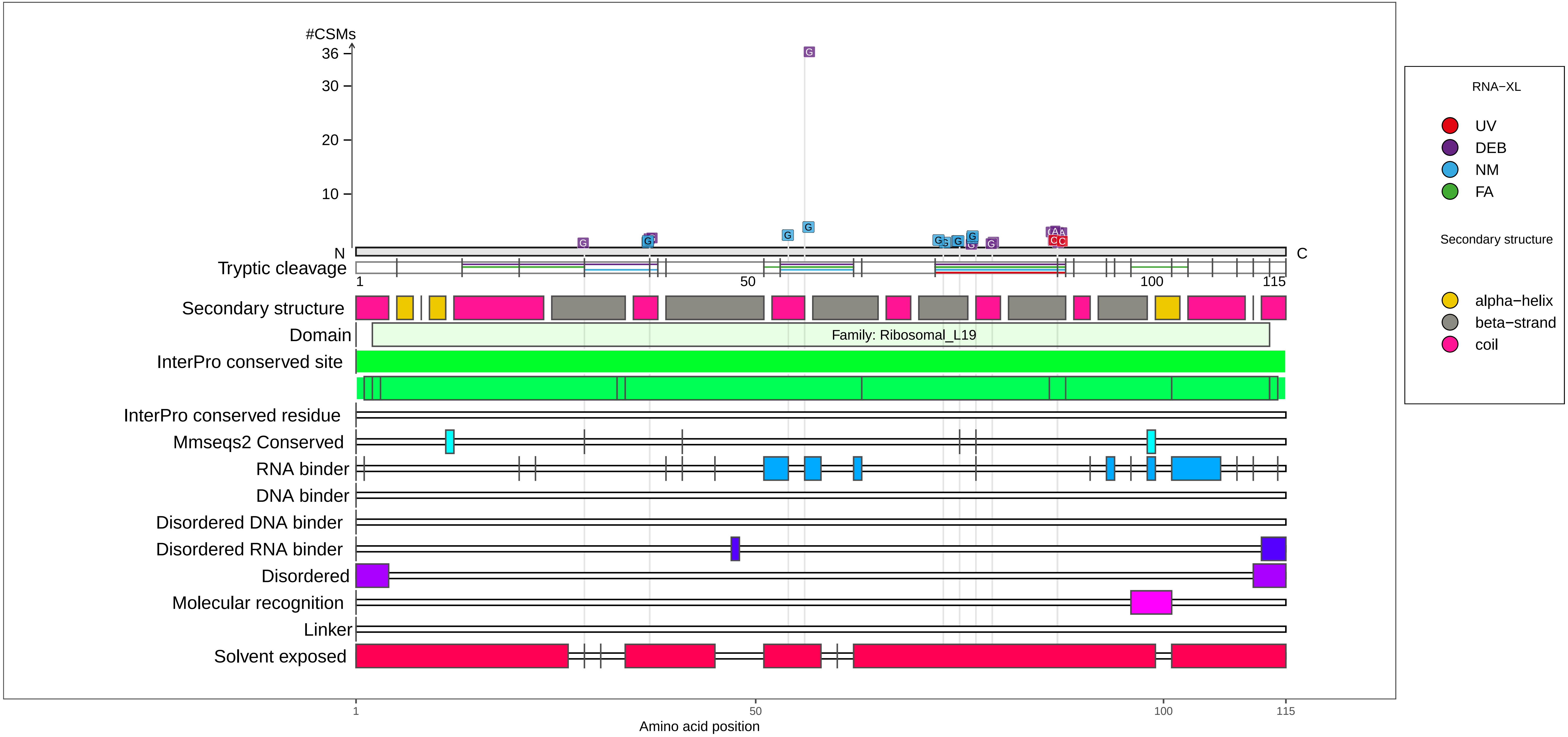

P0A9A6
FTSZ_ECOLI Cell division protein FtsZ

− Abundance:
tryptic [log10 Intensity]: 9.05 (Q 90)
PAXdb K12 strain [ppm]: 2.98 (Q 90)
PAXdb E.coli [ppm]: 2.95 (Q 94)

− RNA functions: not annotated

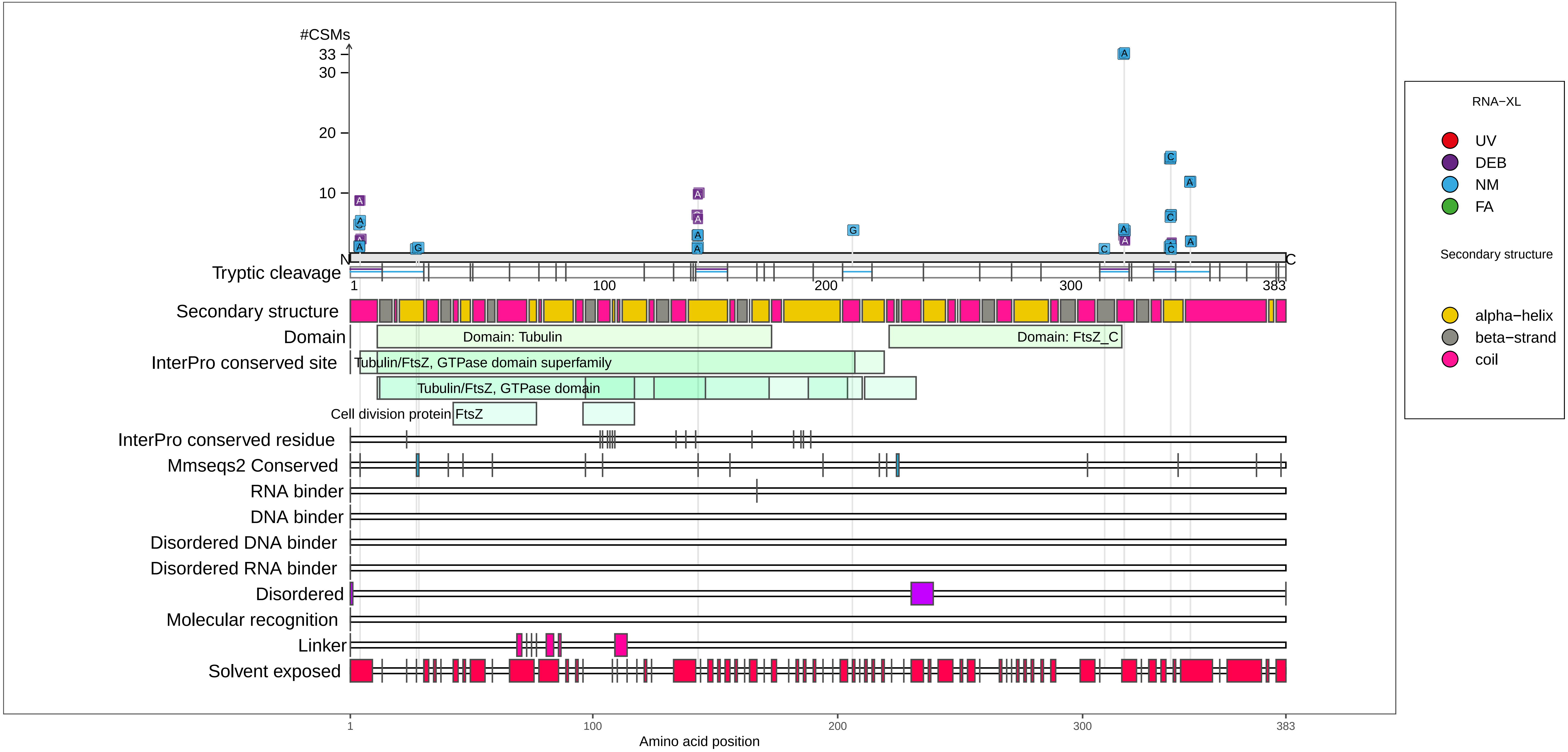

P0A825
GLYA_ECOLI Serine hydroxymethyltransferase

− Abundance:
tryptic [log10 Intensity]: 9.34 (Q 95)
PAXdb K12 strain [ppm]: 3.54 (Q 98)
PAXdb E.coli [ppm]: 3.7 (Q 100)

− RNA functions: not annotated

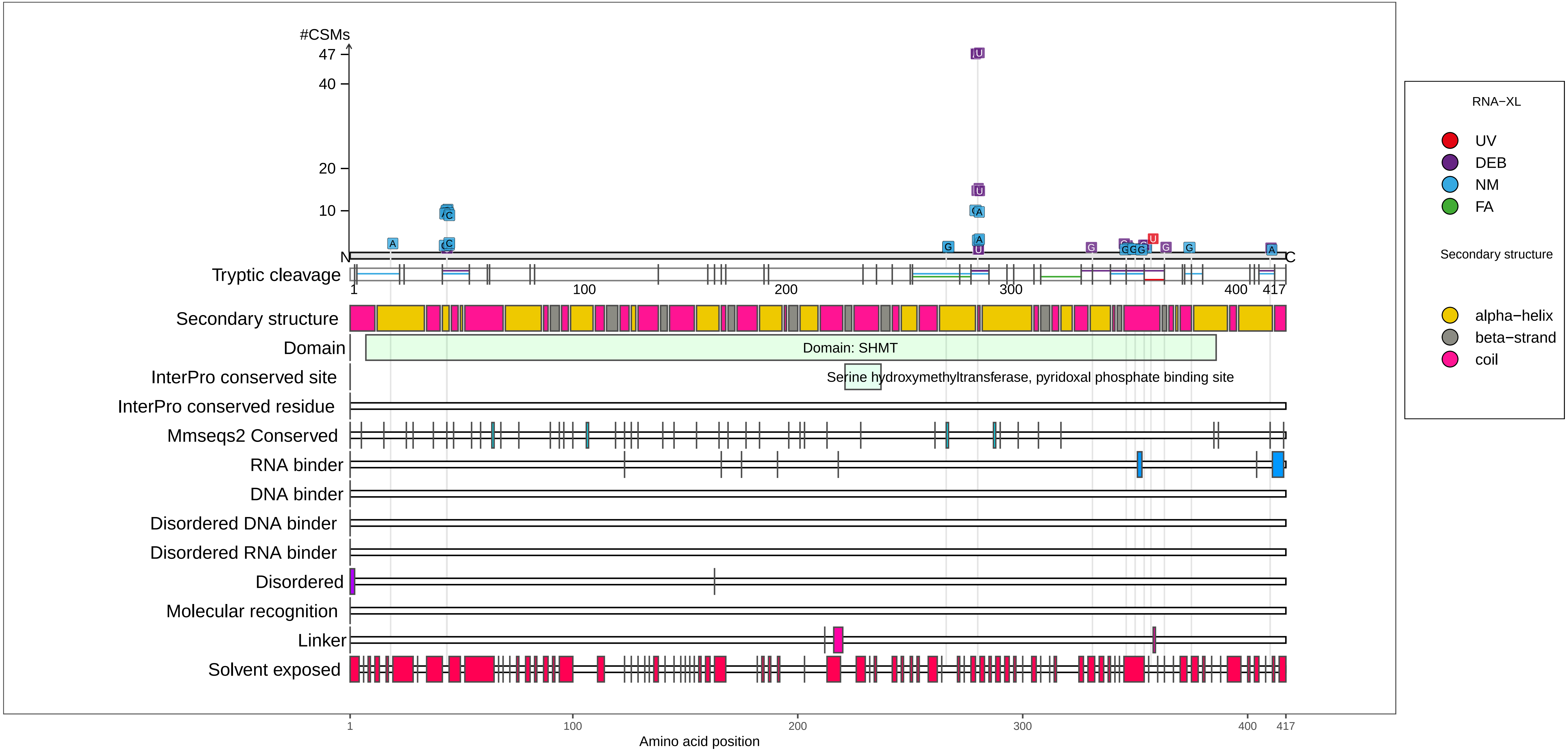

P0ADY7
RL16_ECOLI 50S ribosomal protein L16

− Abundance:
tryptic [log10 Intensity]: 8.89 (Q 86)
PAXdb K12 strain [ppm]: 3.85 (Q 100)
PAXdb E.coli [ppm]: 3.16 (Q 96)

− RNA functions:
RNA binding; rRNA binding; tRNA binding

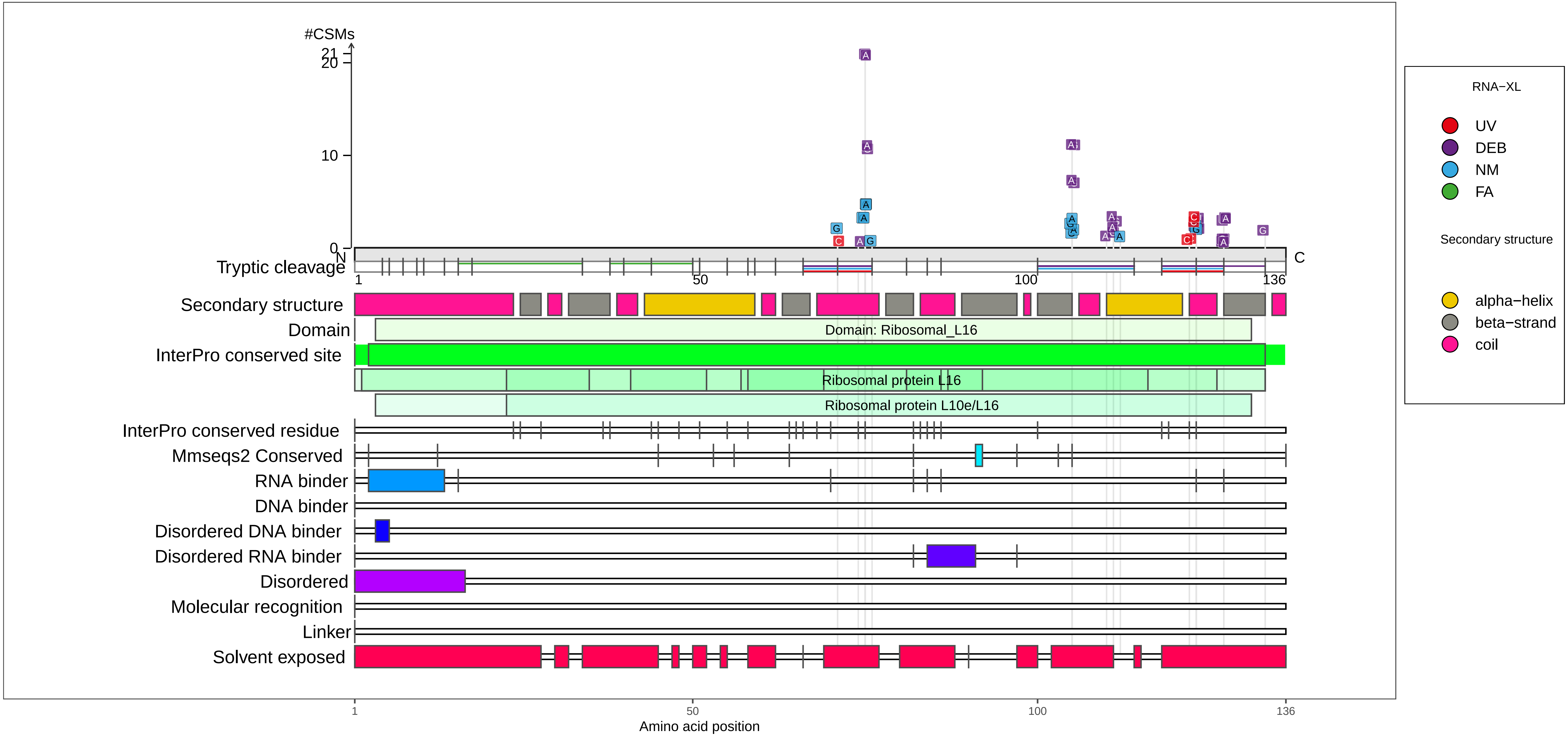

P0A9L3
FKBB_ECOLI FKBP−type 22 kDa peptidyl−prolyl cis−trans isomerase

− Abundance:
tryptic [log10 Intensity]: 8.32 (Q 70)
PAXdb K12 strain [ppm]: 2.71 (Q 83)
PAXdb E.coli [ppm]: 2.97 (Q 94)

− RNA functions: not annotated

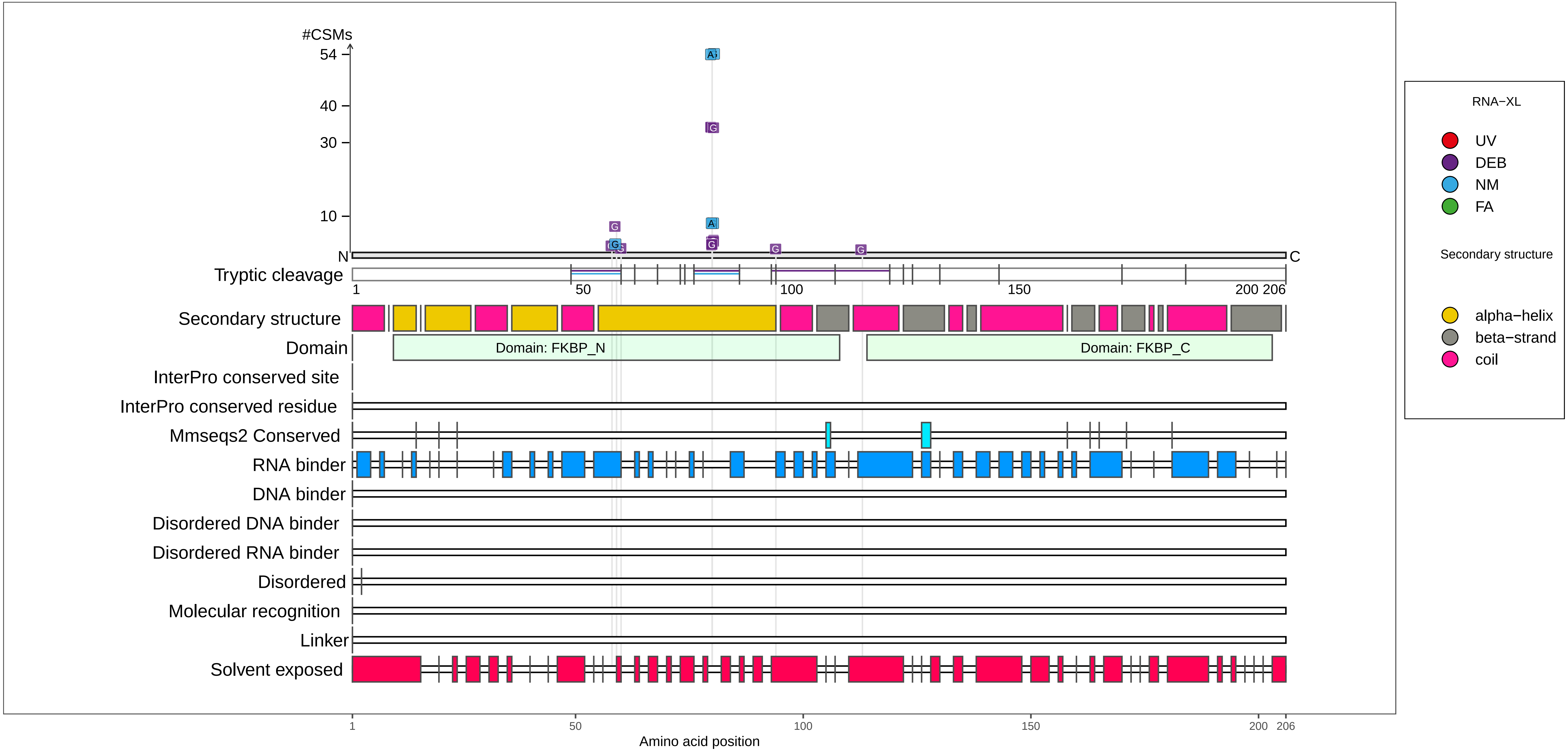

P0ABB0
ATPA_ECOLI ATP synthase subunit alpha

− Abundance:
tryptic [log10 Intensity]: 9.73 (Q 98)
PAXdb K12 strain [ppm]: 2.94 (Q 89)
PAXdb E.coli [ppm]: 3.39 (Q 98)

− RNA functions: not annotated

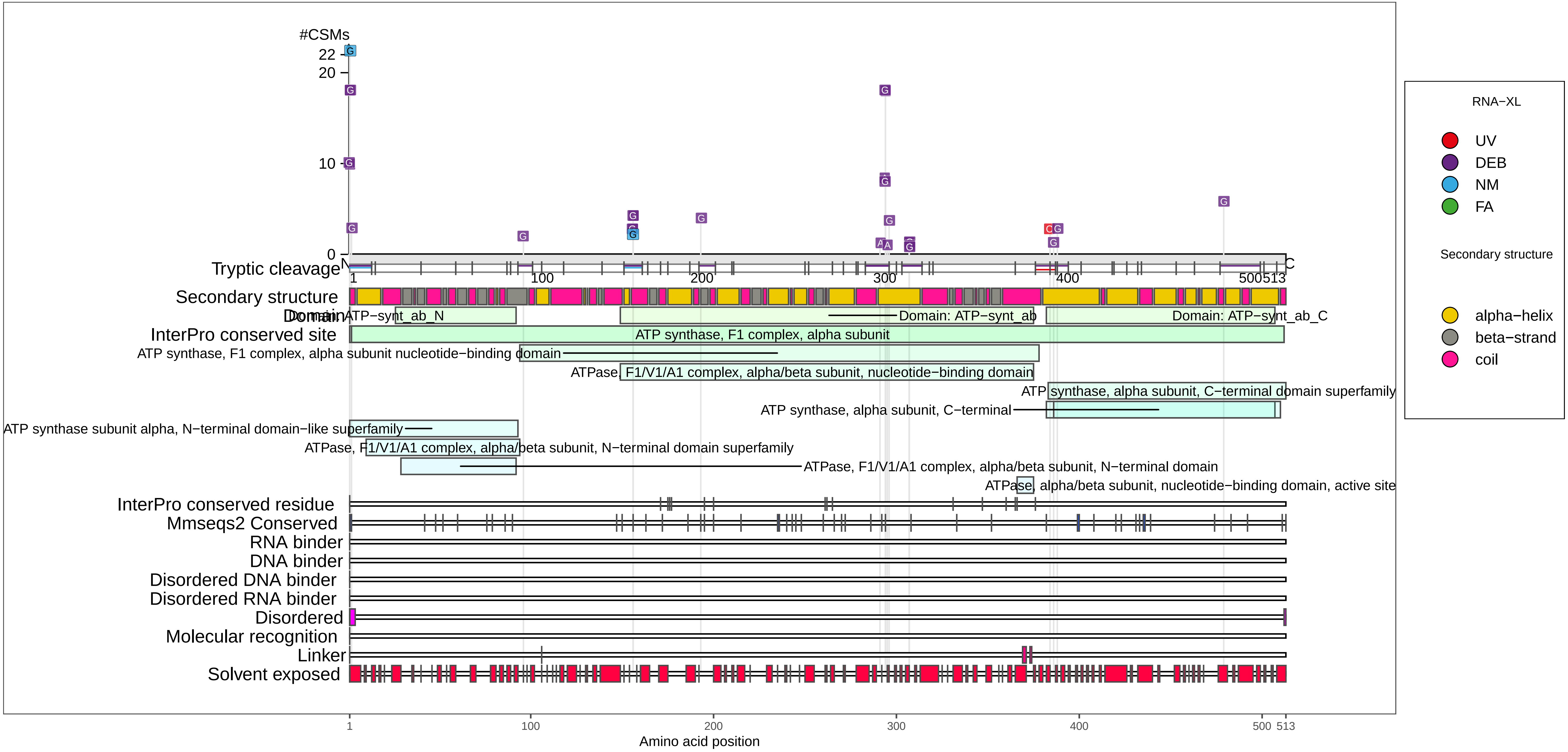

P0A7T3
RS16_ECOLI 30S ribosomal protein S16

− Abundance:
tryptic [log10 Intensity]: 9.14 (Q 91)
PAXdb K12 strain [ppm]: 3.66 (Q 98)
PAXdb E.coli [ppm]: 3.45 (Q 99)

− RNA functions: not annotated

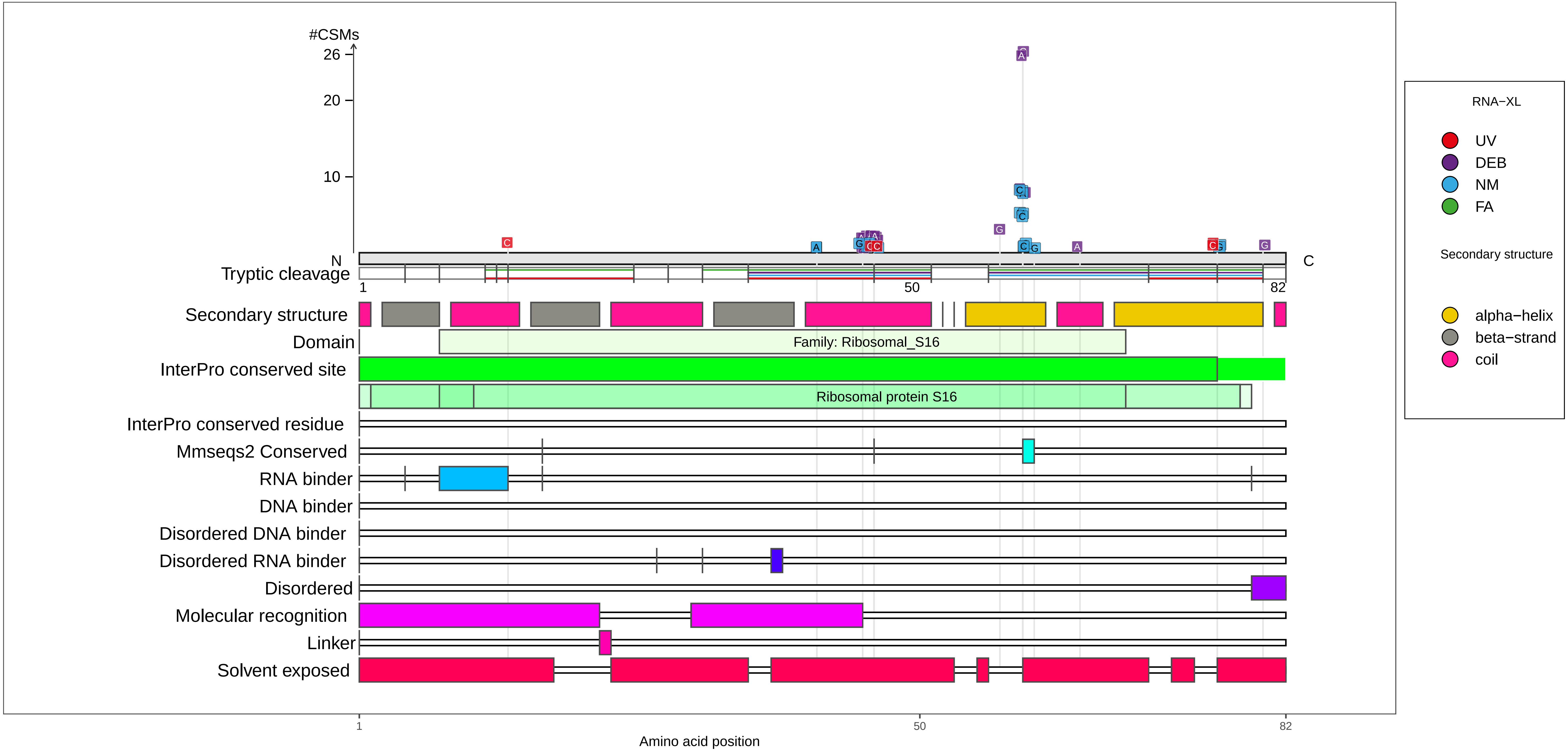

P00579
RPOD_ECOLI RNA polymerase sigma factor RpoD

− Abundance:
tryptic [log10 Intensity]: 8.98 (Q 88)
PAXdb K12 strain [ppm]: 2.99 (Q 91)
PAXdb E.coli [ppm]: 2.57 (Q 87)

− RNA functions:
RNA biosynthetic process; RNA metabolic process

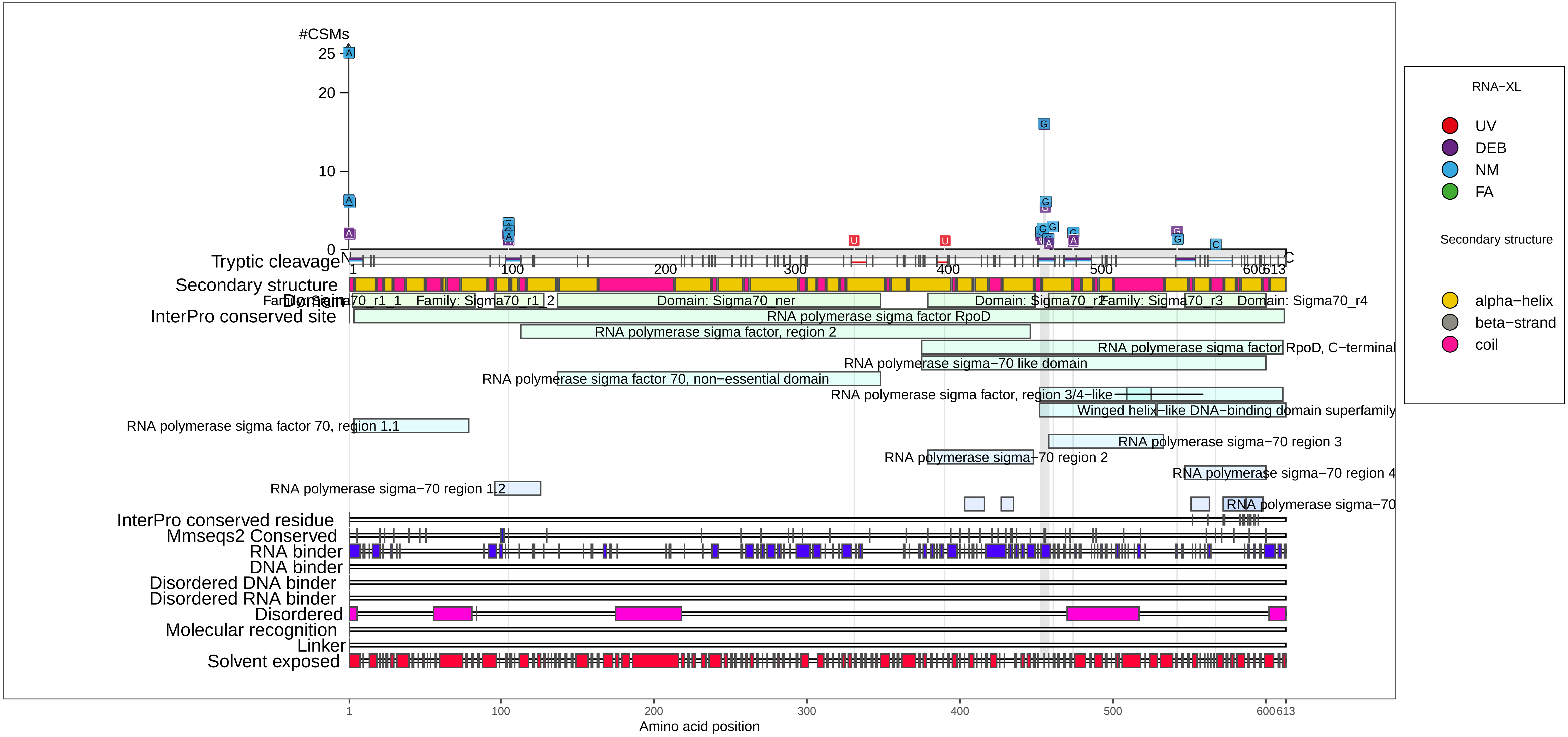

P02358
RS6_ECOLI 30S ribosomal protein S6

− Abundance:
tryptic [log10 Intensity]: 8.99 (Q 88)
PAXdb K12 strain [ppm]: 3.01 (Q 91)
PAXdb E.coli [ppm]: 3.48 (Q 99)

− RNA functions:
mRNA 5−UTR binding; mRNA binding; RNA binding; rRNA binding

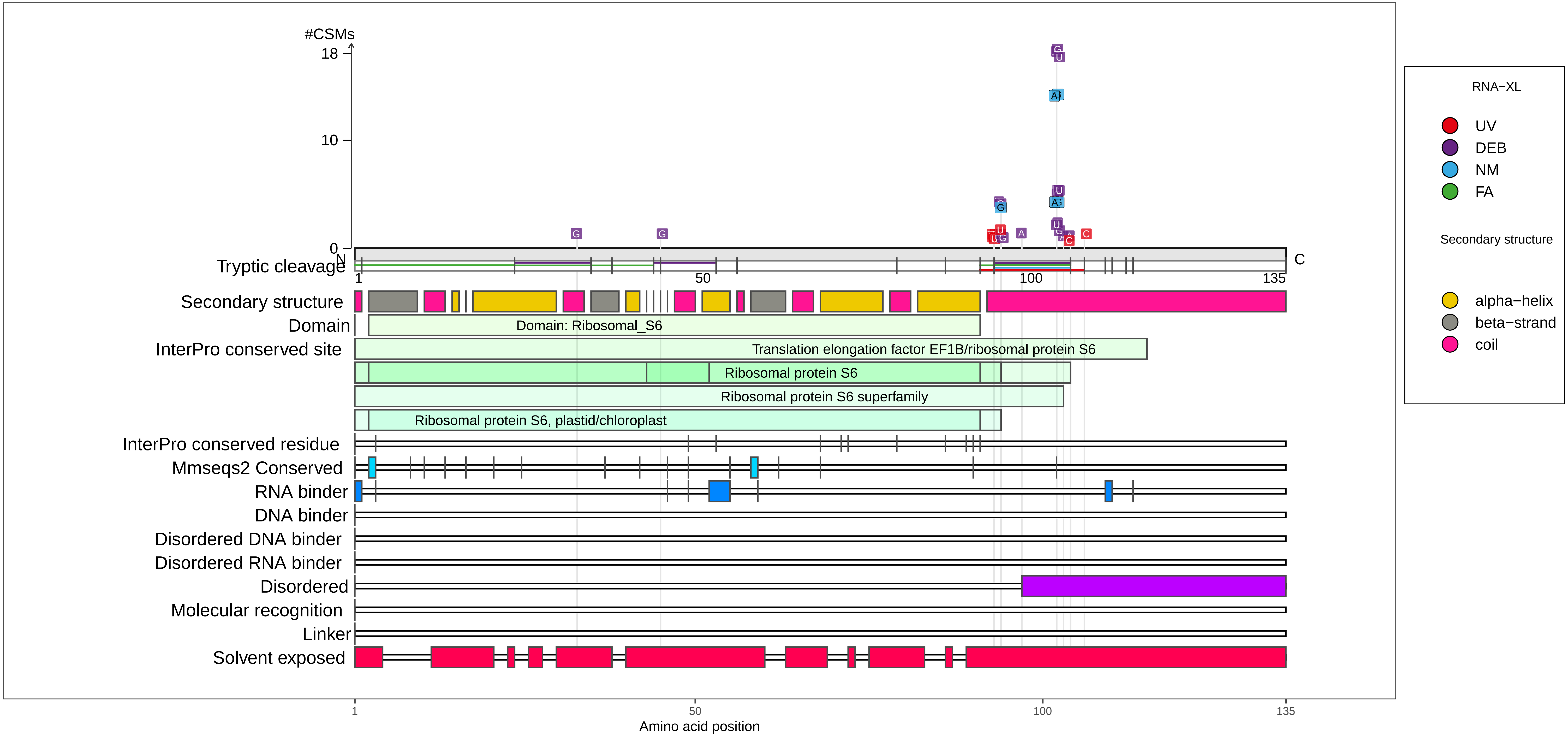

P68679
RS21_ECOLI 30S ribosomal protein S21
− Abundance:
tryptic [log10 Intensity]: 8.65 (Q 80)
PAXdb K12 strain [ppm]: 4.2 (Q 100)
PAXdb E.coli [ppm]: 3.54 (Q 99)

− RNA functions:
RNA binding; rRNA binding

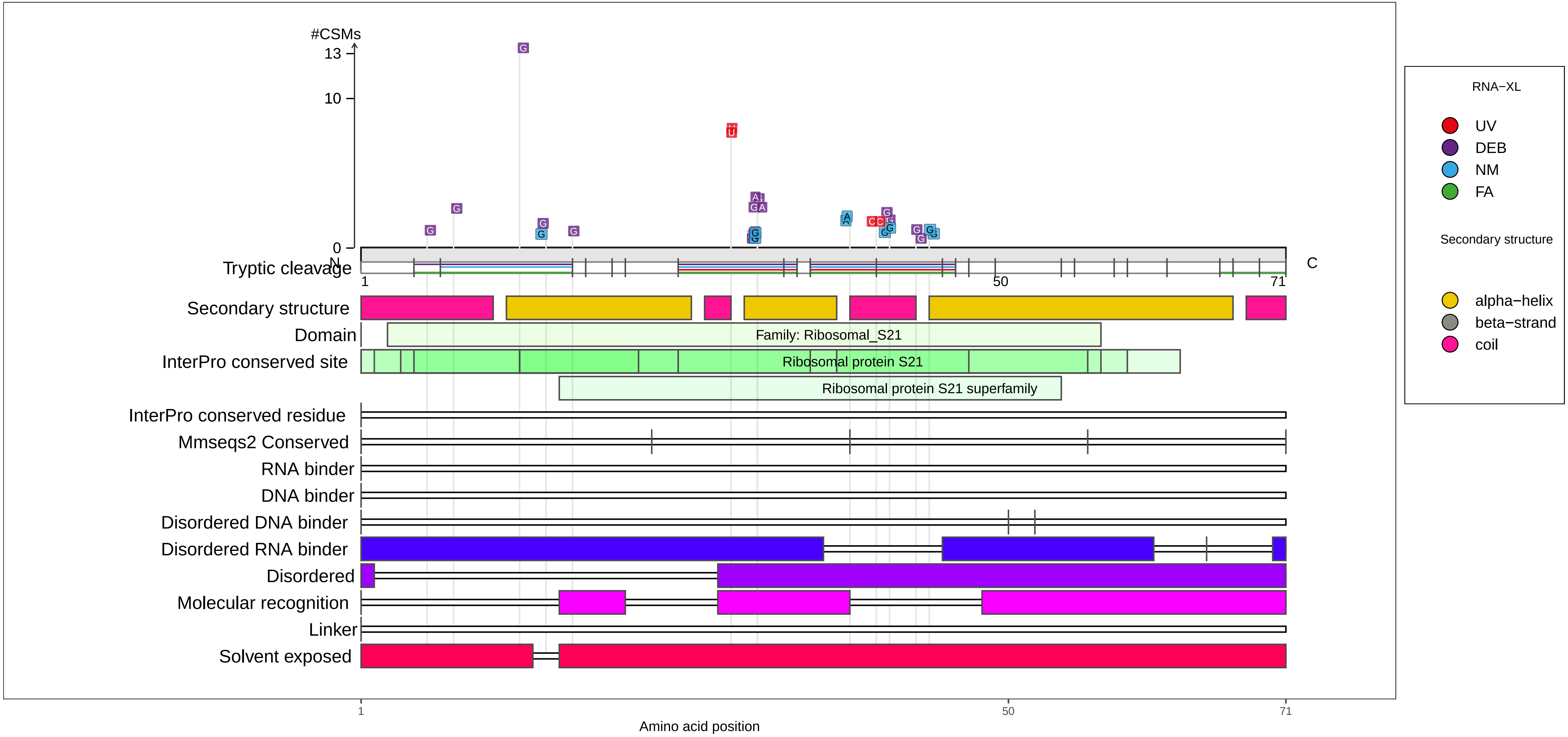

P00961
SYGB_ECOLI Glycine−−tRNA ligase beta subunit

− Abundance:
tryptic [log10 Intensity]: 8.74 (Q 82)
PAXdb K12 strain [ppm]: 3 (Q 91)
PAXdb E.coli [ppm]: 2.71 (Q 90)

− RNA functions:
aminoacyl−tRNA ligase activity; arginine−tRNA ligase activity
arginyl−tRNA aminoacylation; glycine−tRNA ligase activity; glycyl−tRNA aminoacylation
Glycyl−tRNA synthetase beta subunit; ncRNA metabolic process; RNA metabolic process; tRNA aminoacylation
tRNA aminoacylation for protein translation; tRNA metabolic process

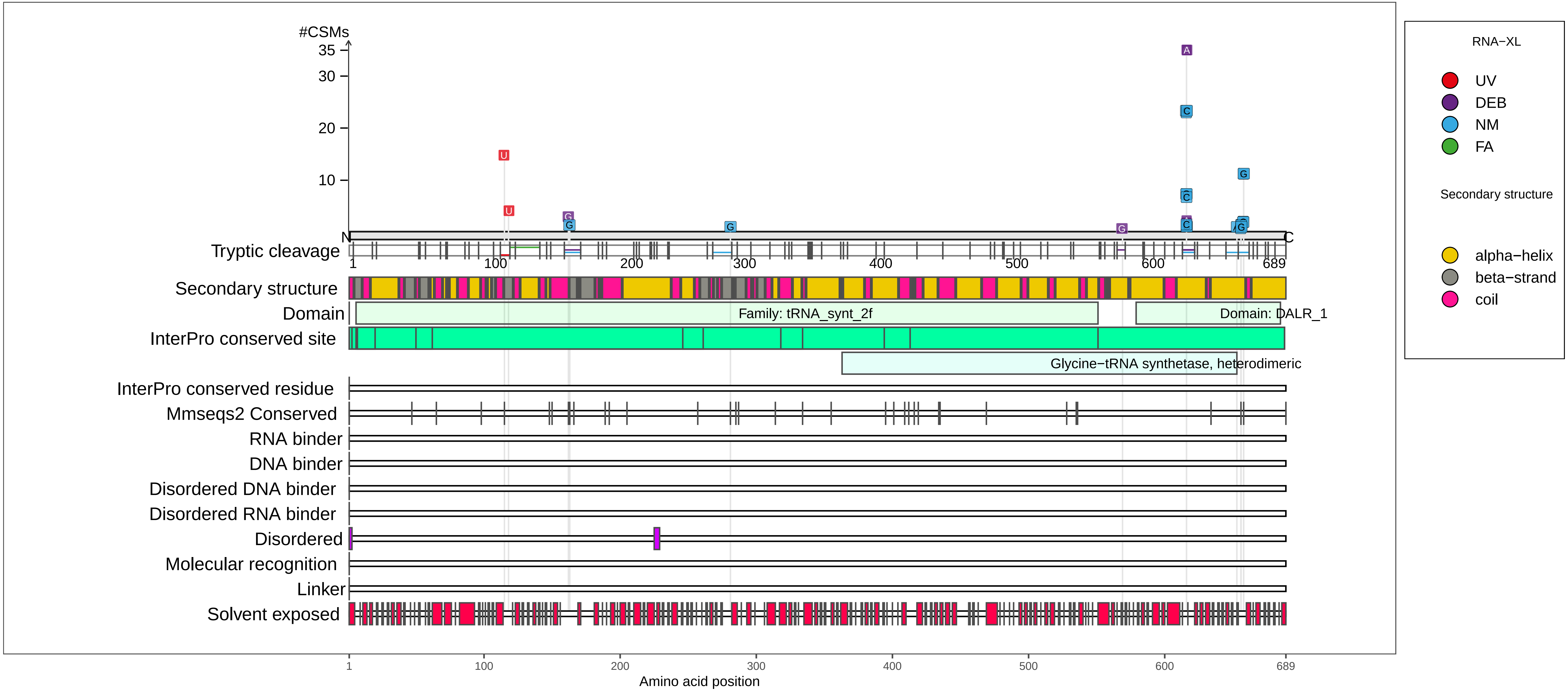

P0A7U3
RS19_ECOLI 30S ribosomal protein S19

− Abundance:
tryptic [log10 Intensity]: 8.56 (Q 78)
PAXdb K12 strain [ppm]: 3.73 (Q 99)
PAXdb E.coli [ppm]: 3.6 (Q 100)

− RNA functions:
RNA binding; rRNA binding; tRNA binding

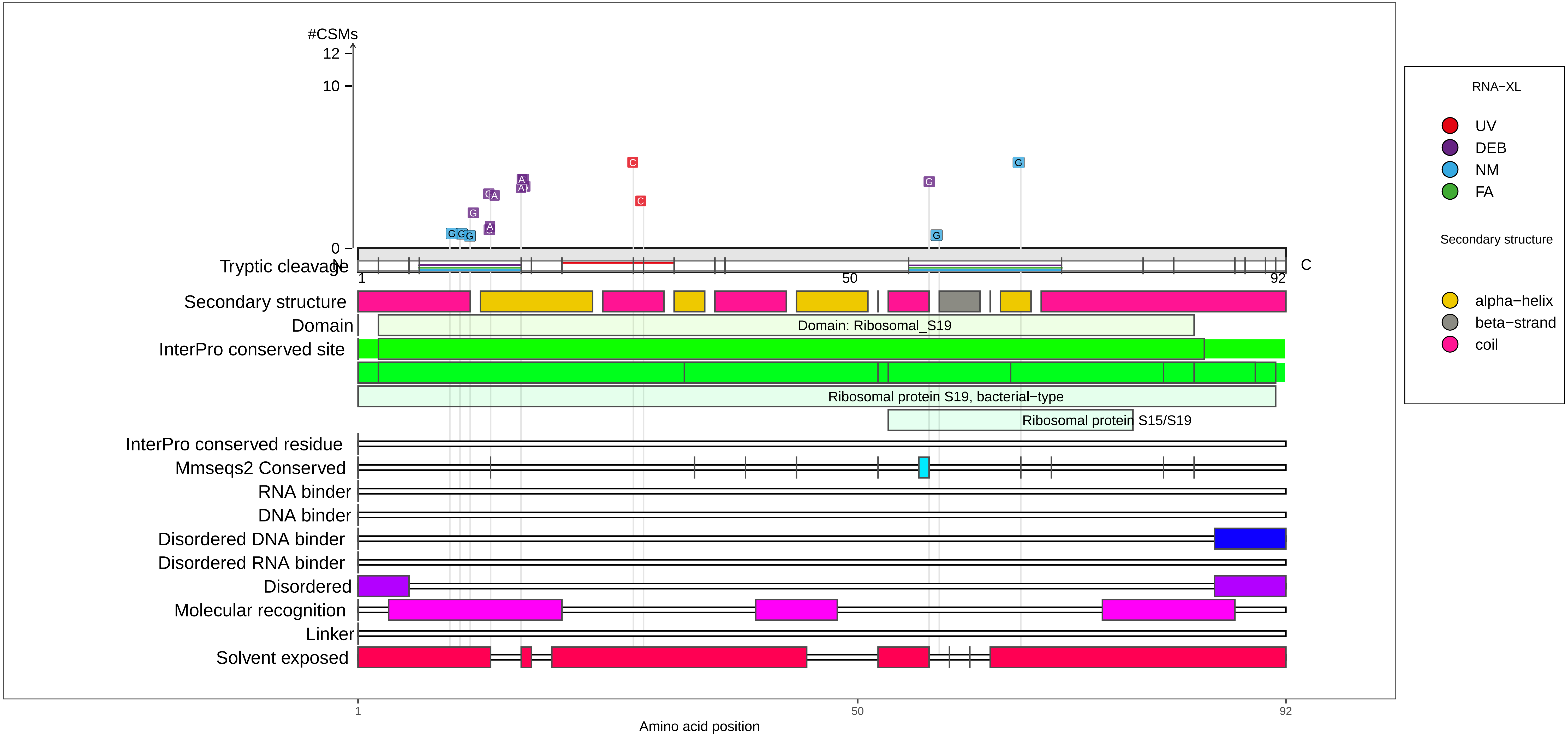

P0A6A8
ACP_ECOLI Acyl carrier protein

− Abundance:
tryptic [log10 Intensity]: 9.19 (Q 92)
PAXdb K12 strain [ppm]: 3.67 (Q 99)
PAXdb E.coli [ppm]: 3.53 (Q 99)

− RNA functions: not annotated

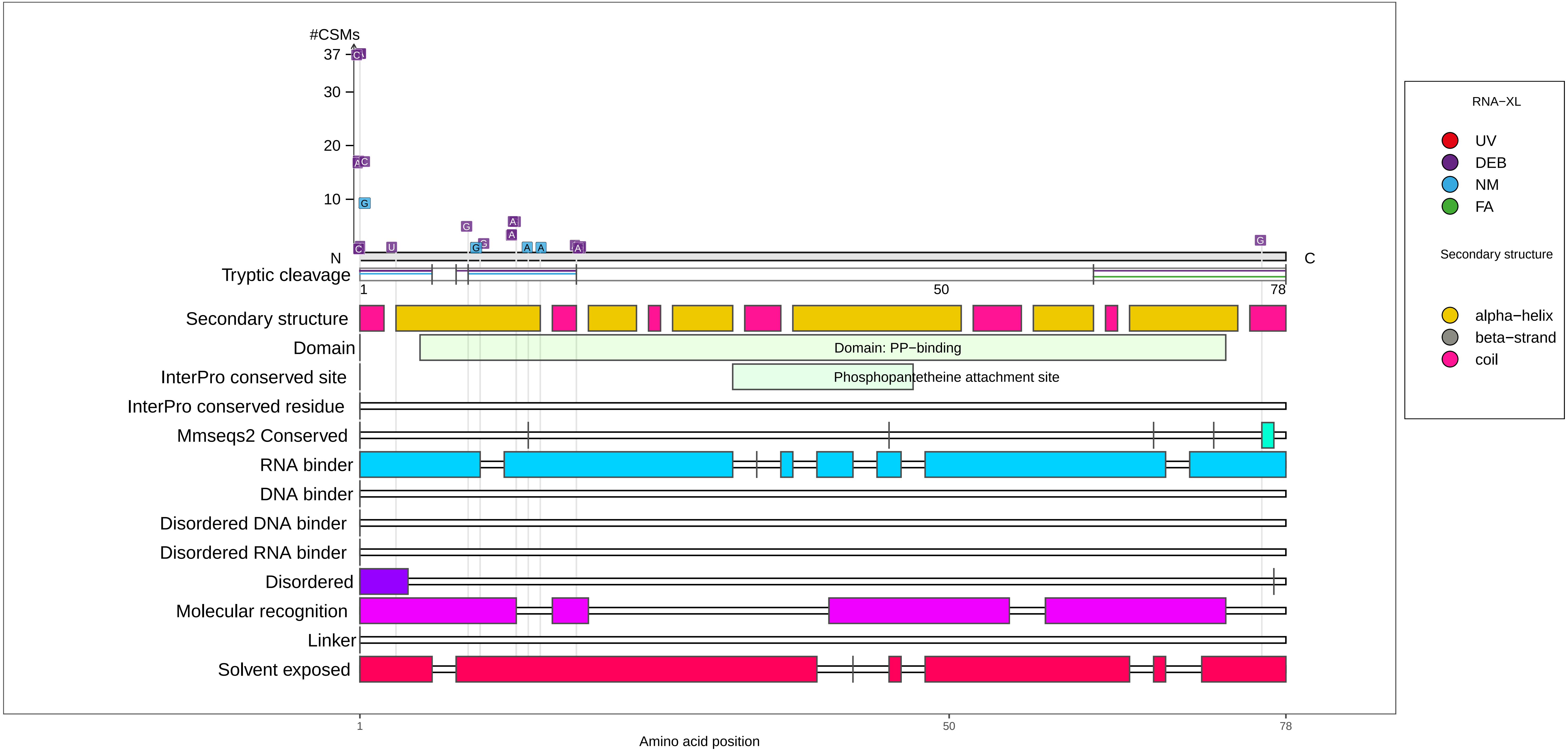

P25665
METE_ECOLI 5−methyltetrahydropteroyltriglutamate−−homocysteine methyltransferase

− Abundance:
tryptic [log10 Intensity]: 9.26 (Q 93)
PAXdb K12 strain [ppm]: 2.96 (Q 90)
PAXdb E.coli [ppm]: 3.18 (Q 96)

− RNA functions: not annotated

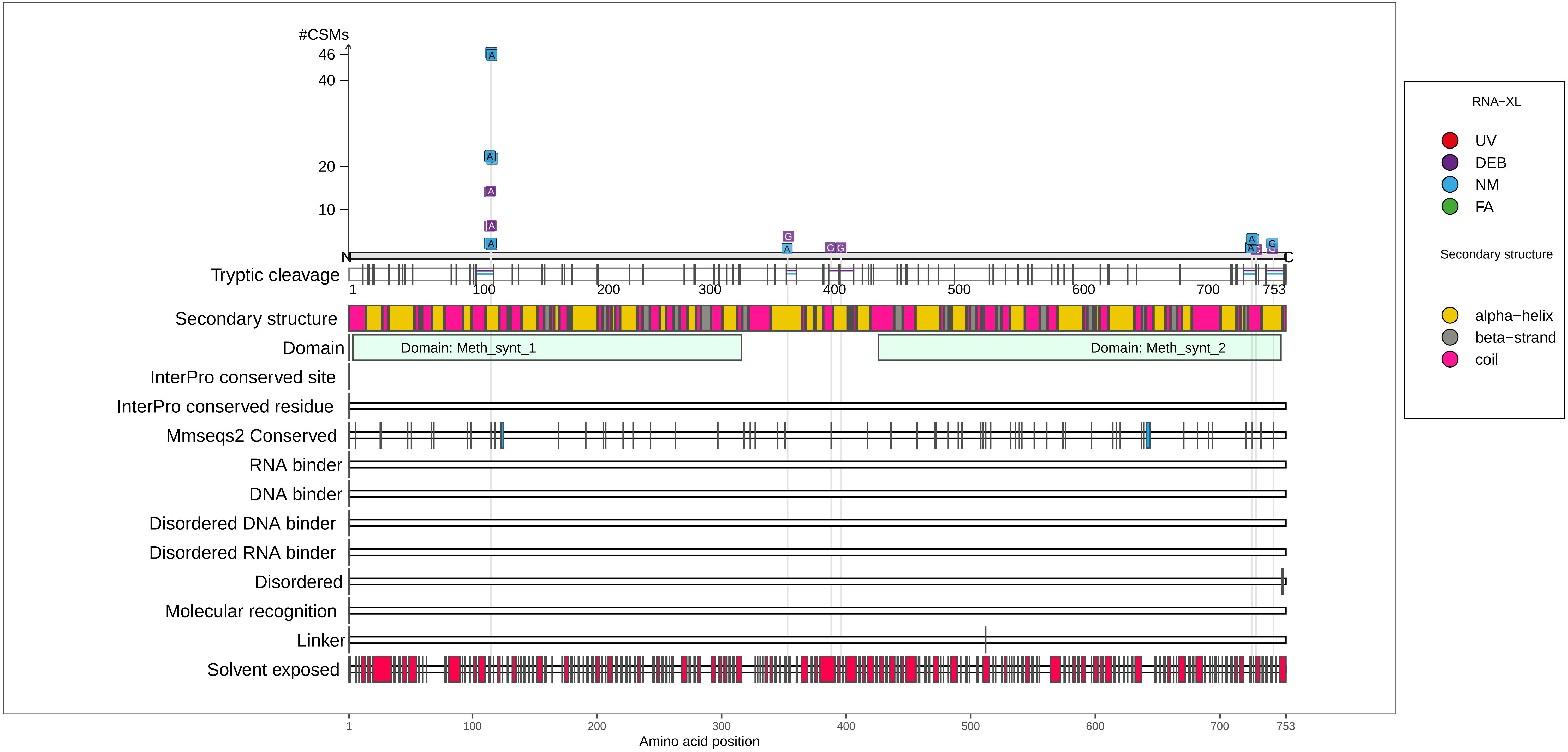

P0A6K6
DEOB_ECOLI Phosphopentomutase

− Abundance:
tryptic [log10 Intensity]: 9.44 (Q 96)
PAXdb K12 strain [ppm]: 3.28 (Q 95)
PAXdb E.coli [ppm]: 2.95 (Q 94)

− RNA functions: not annotated

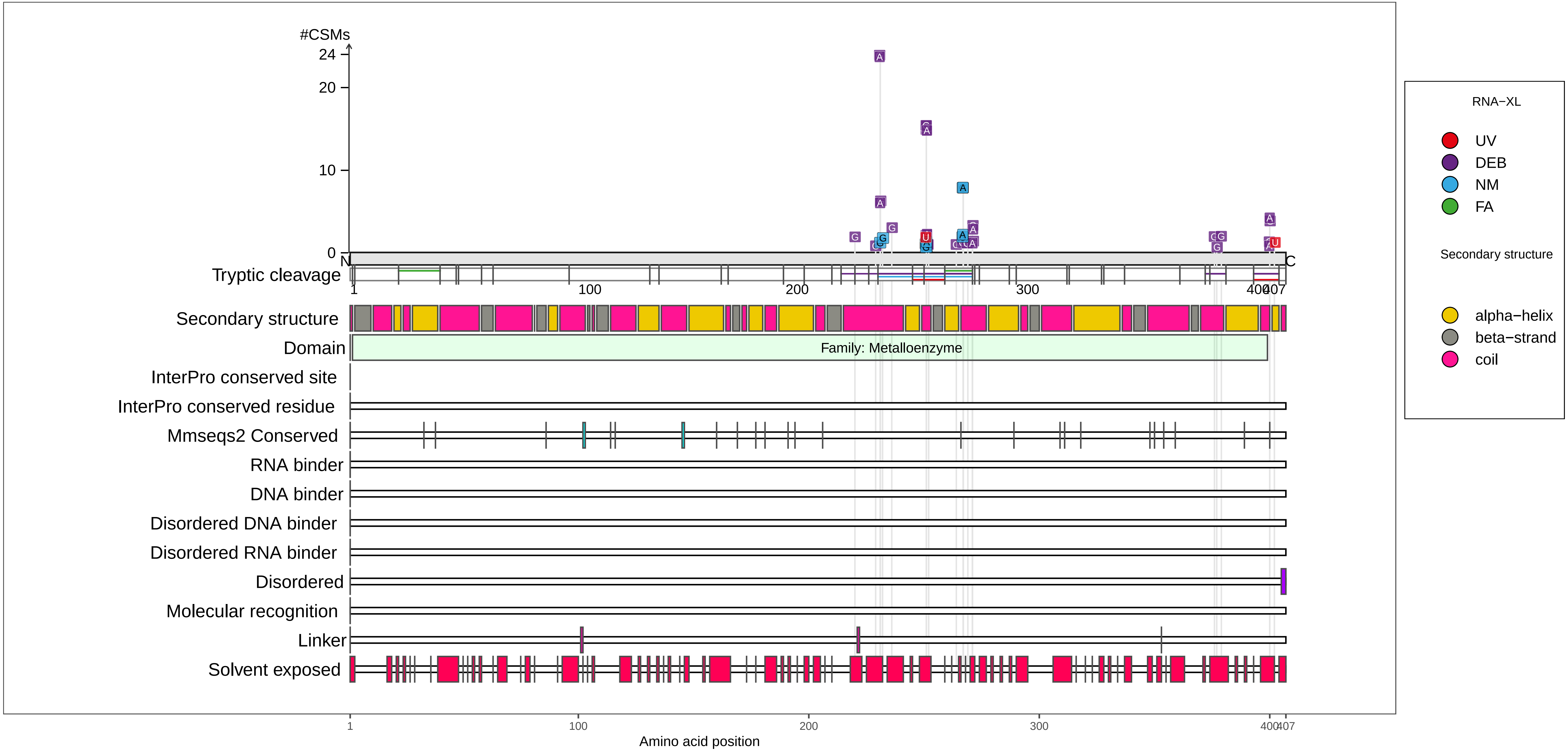

P0A7N9
RL33_ECOLI 50S ribosomal protein L33

− Abundance:
tryptic [log10 Intensity]: 10.61 (Q 100)
PAXdb K12 strain [ppm]: 3.75 (Q 99)
PAXdb E.coli [ppm]: 3.2 (Q 96)

− RNA functions:
RNA binding; tRNA binding

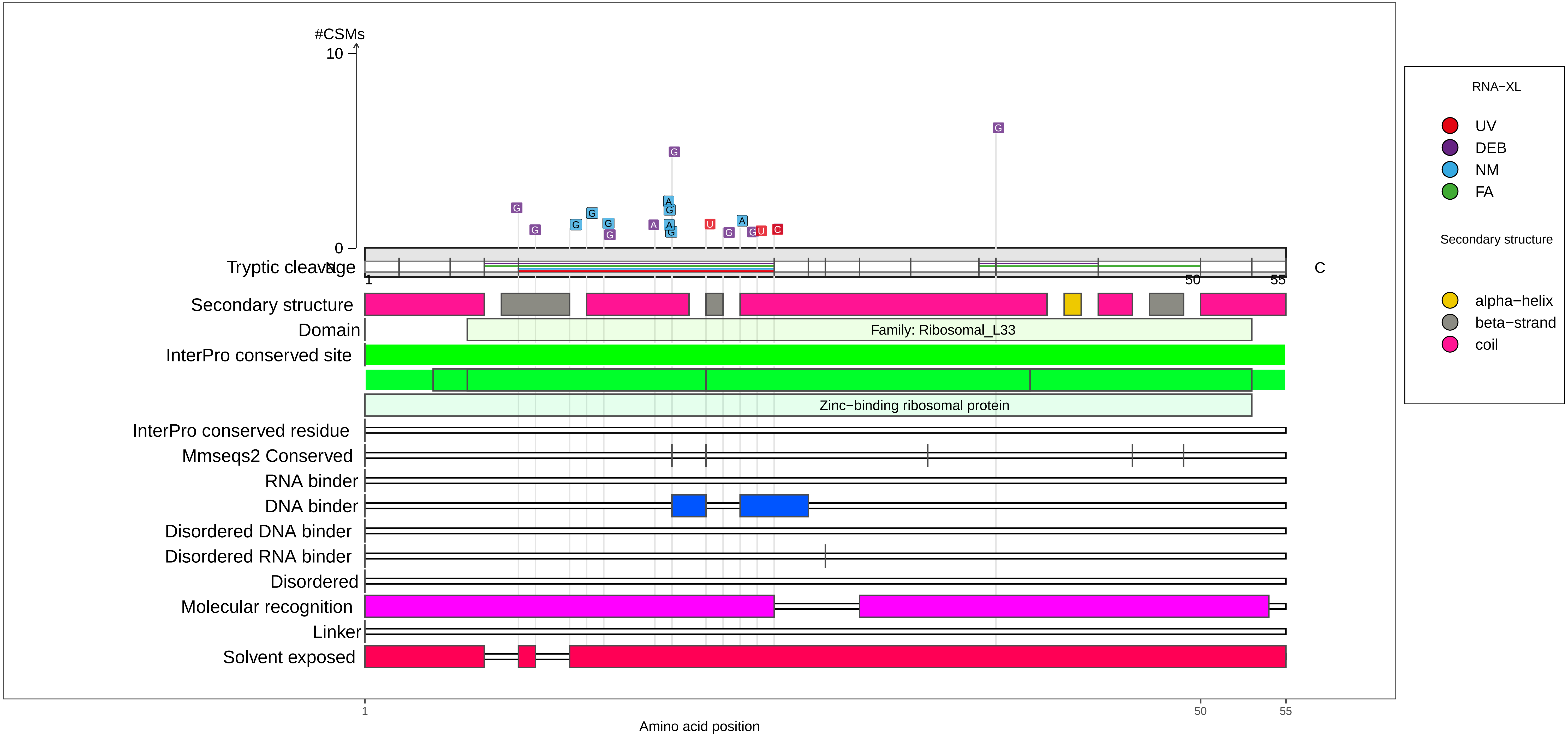

P0A7R1
RL9_ECOLI 50S ribosomal protein L9

− Abundance:
tryptic [log10 Intensity]: 9.46 (Q 96)
PAXdb K12 strain [ppm]: 3.61 (Q 98)
PAXdb E.coli [ppm]: 3.51 (Q 99)

− RNA functions:
RNA binding; rRNA binding

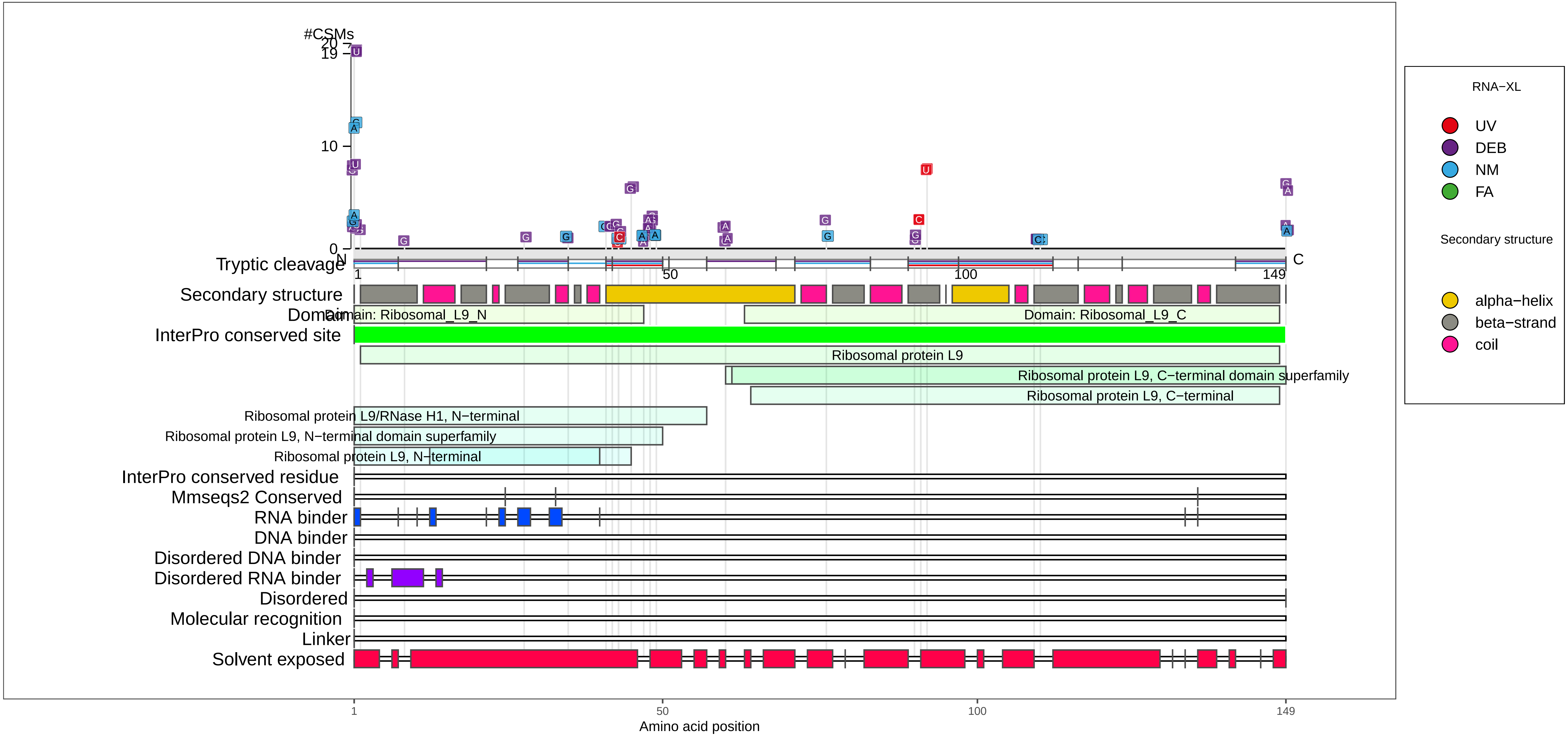

P0A7J3
RL10_ECOLI 50S ribosomal protein L10

− Abundance:
tryptic [log10 Intensity]: 9.09 (Q 90)
PAXdb K12 strain [ppm]: 3.58 (Q 98)
PAXdb E.coli [ppm]: 3.44 (Q 98)

− RNA functions:
RNA binding; rRNA binding

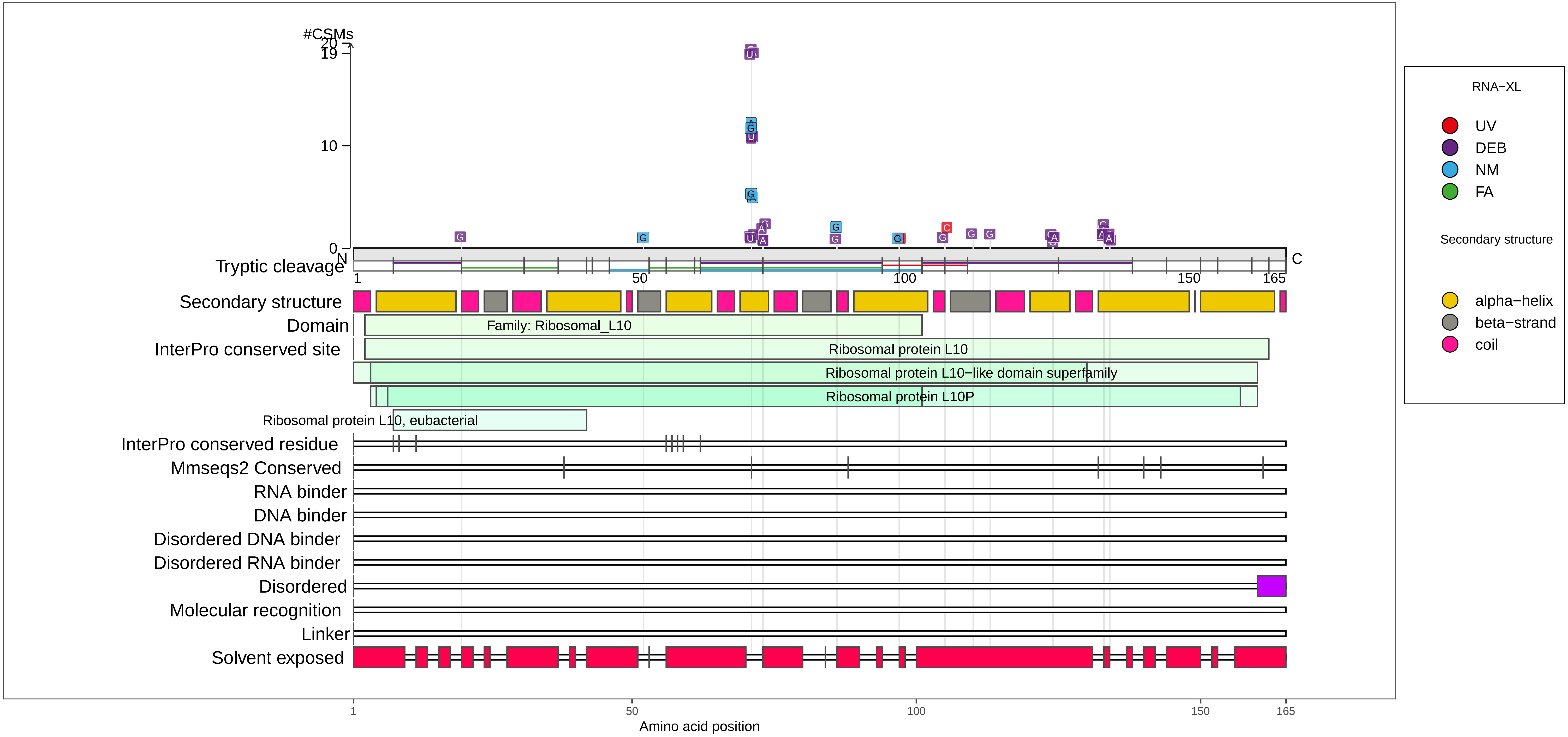

P60723
RL4_ECOLI 50S ribosomal protein L4
− Abundance:
tryptic [log10 Intensity]: 9.3 (Q 94)
PAXdb K12 strain [ppm]: 3.44 (Q 97)
PAXdb E.coli [ppm]: 3.32 (Q 98)

− RNA functions:
RNA binding; RNA biosynthetic process; RNA metabolic process; rRNA binding

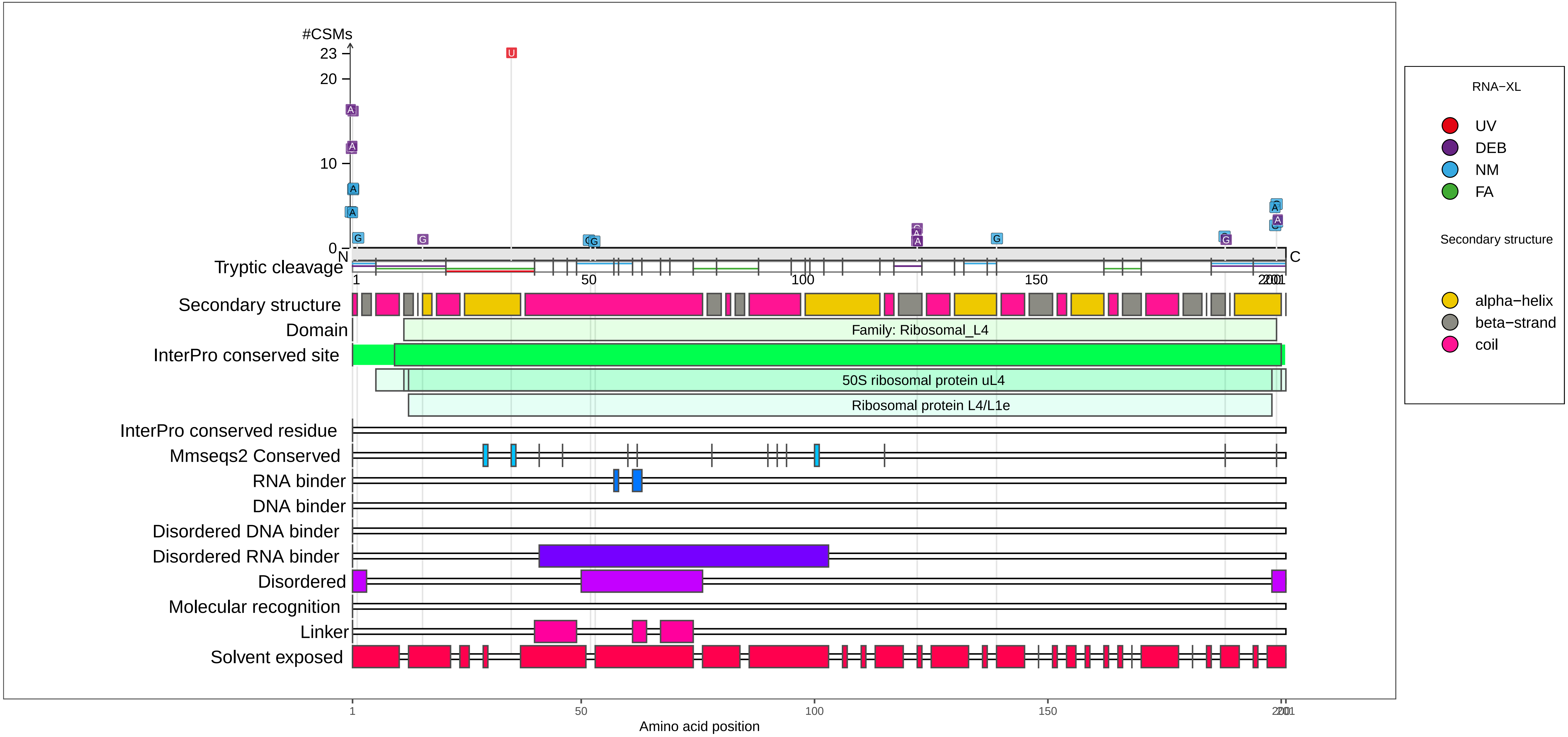

P69441
KAD_ECOLI Adenylate kinase

− Abundance:
tryptic [log10 Intensity]: 9.05 (Q 90)
PAXdb K12 strain [ppm]: 3.7 (Q 99)
PAXdb E.coli [ppm]: 3.41 (Q 98)

− RNA functions: not annotated

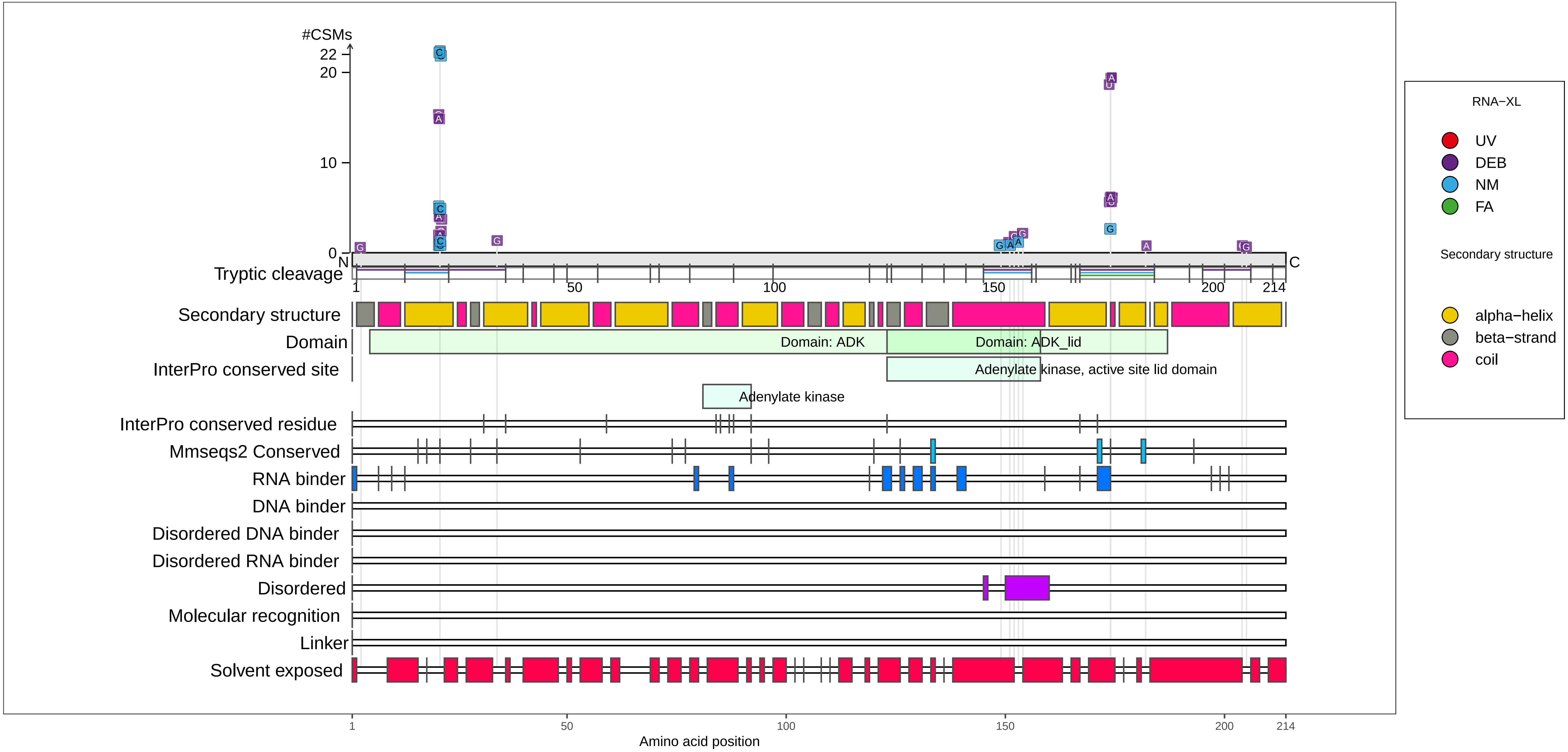

P0A7M2
RL28_ECOLI 50S ribosomal protein L28

− Abundance:
tryptic [log10 Intensity]: 8.99 (Q 88)
PAXdb K12 strain [ppm]: 3.71 (Q 99)
PAXdb E.coli [ppm]: 3.5 (Q 99)

− RNA functions:
RNA binding; rRNA binding

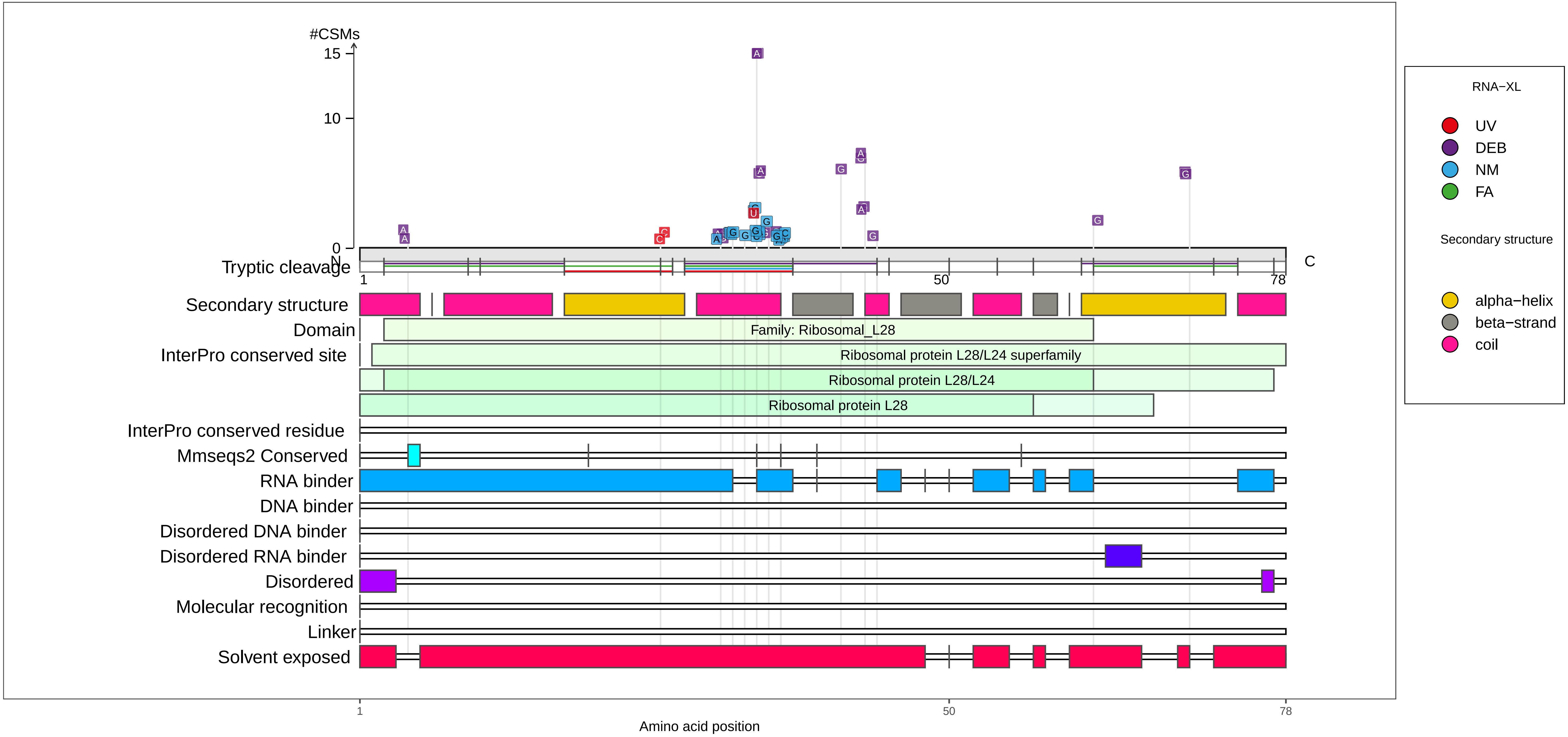

P0A7W7
RS8_ECOLI 30S ribosomal protein S8

− Abundance:
tryptic [log10 Intensity]: 9.33 (Q 94)
PAXdb K12 strain [ppm]: 3.34 (Q 96)
PAXdb E.coli [ppm]: 3.31 (Q 97)

− RNA functions:
RNA binding; rRNA binding

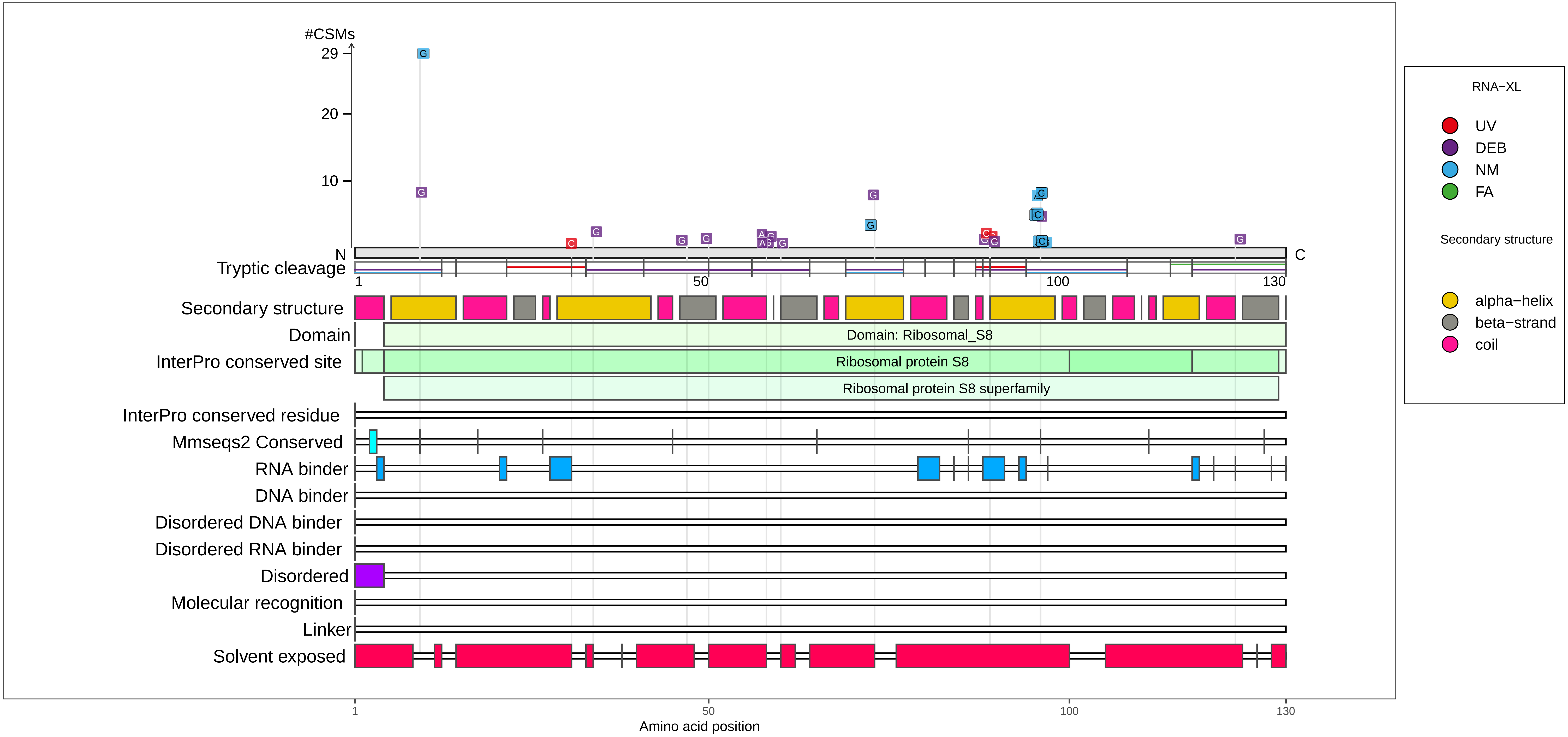

P07813
SYL_ECOLI Leucine−−tRNA ligase

− Abundance:
tryptic [log10 Intensity]: 9.26 (Q 93)
PAXdb K12 strain [ppm]: 2.91 (Q 89)
PAXdb E.coli [ppm]: 2.79 (Q 92)

− RNA functions:
aminoacyl−tRNA editing activity; aminoacyl−tRNA ligase activity
aminoacyl−tRNA metabolism involved in translational fidelity; Anticodon−binding domain of tRNA
leucine−tRNA ligase activity; leucyl−tRNA aminoacylation; Leucyl−tRNA synthetase, Domain 2
ncRNA metabolic process; RNA metabolic process; tRNA aminoacylation
tRNA aminoacylation for protein translation; tRNA metabolic process
tRNA synthetases class I (I, L, M and V); tRNA synthetases class I (M)

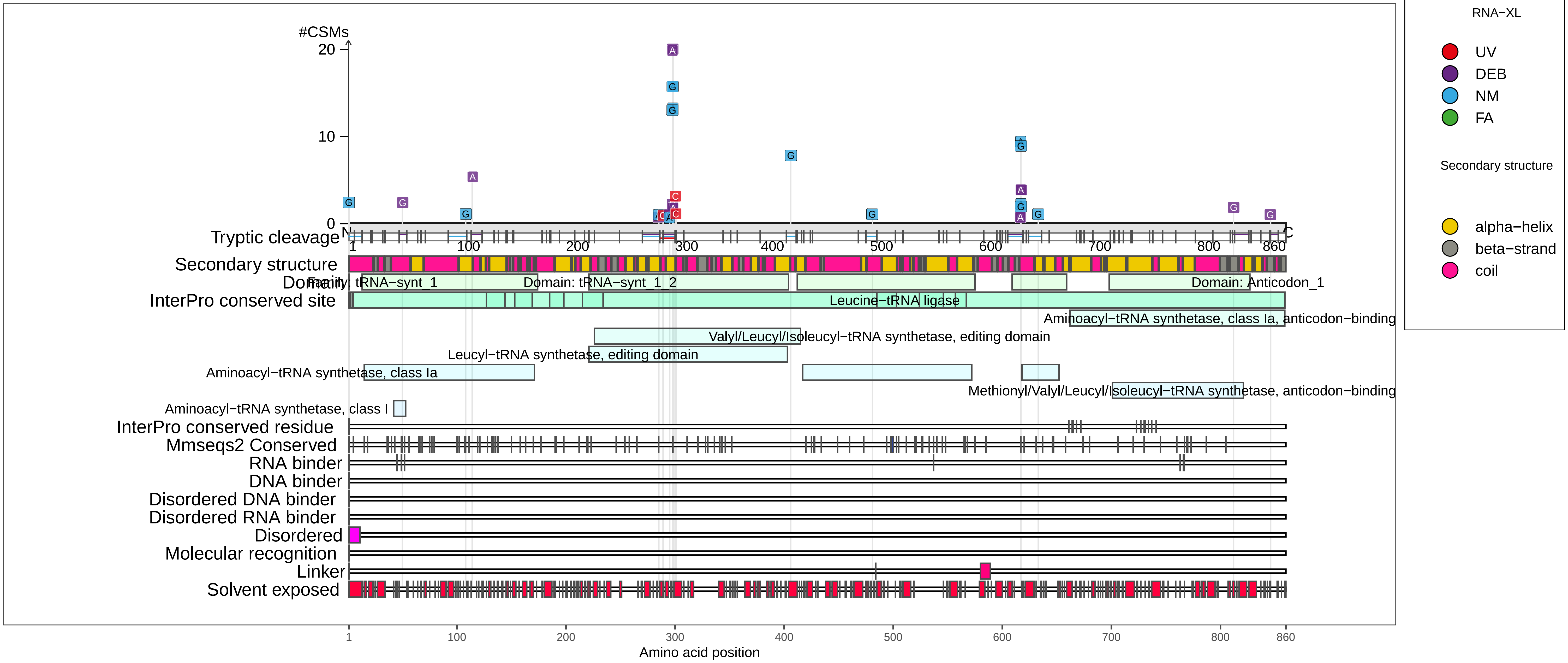

P0A9P4
TRXB_ECOLI Thioredoxin reductase

− Abundance:
tryptic [log10 Intensity]: 9.18 (Q 92)
PAXdb K12 strain [ppm]: 3.24 (Q 94)
PAXdb E.coli [ppm]: 2.62 (Q 88)

− RNA functions: not annotated

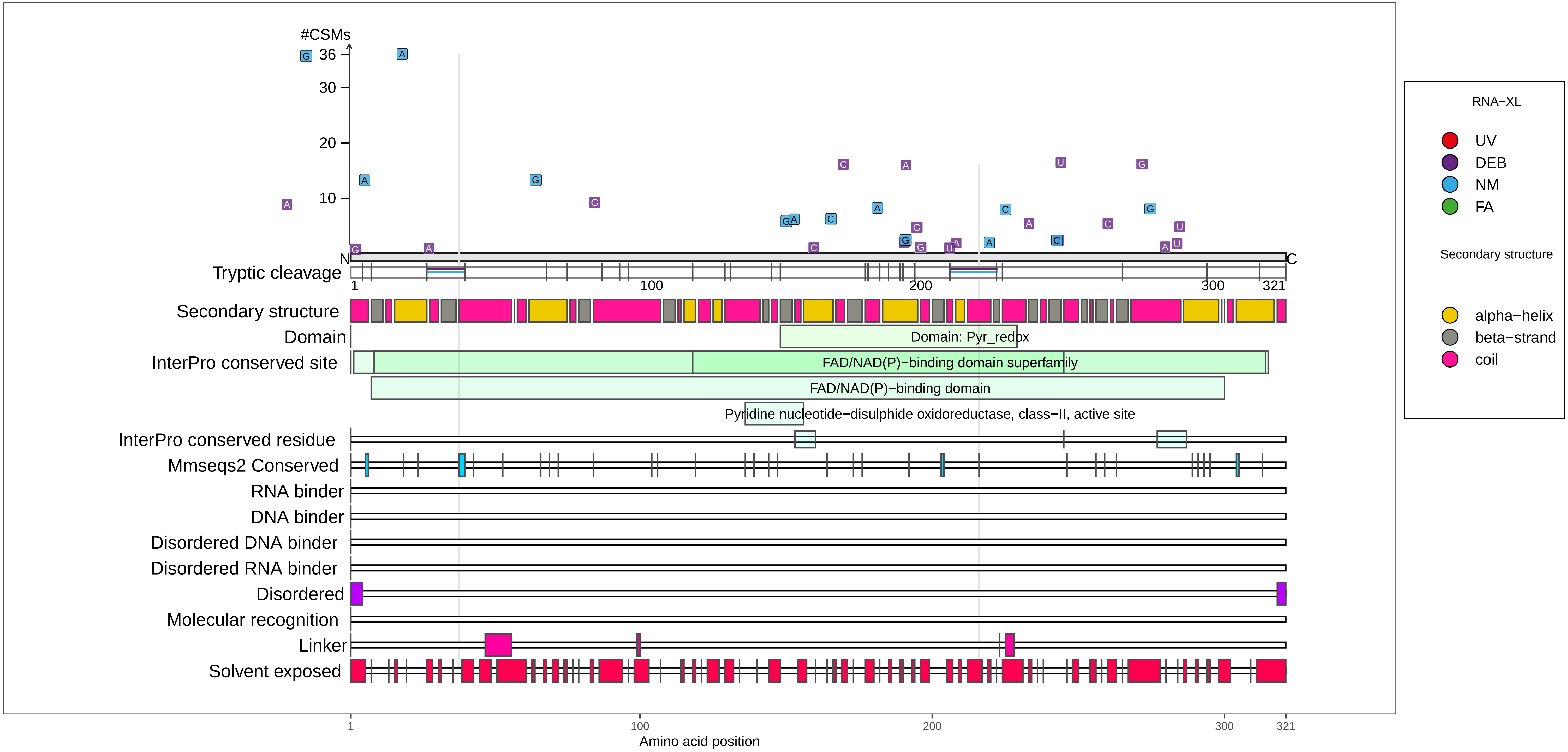

P0AFG0
NUSG_ECOLI Transcription termination/antitermination protein NusG

− Abundance:
tryptic [log10 Intensity]: 9.04 (Q 89)
PAXdb K12 strain [ppm]: 3.03 (Q 91)
PAXdb E.coli [ppm]: 3.03 (Q 95)

− RNA functions:
RNA biosynthetic process; RNA metabolic process

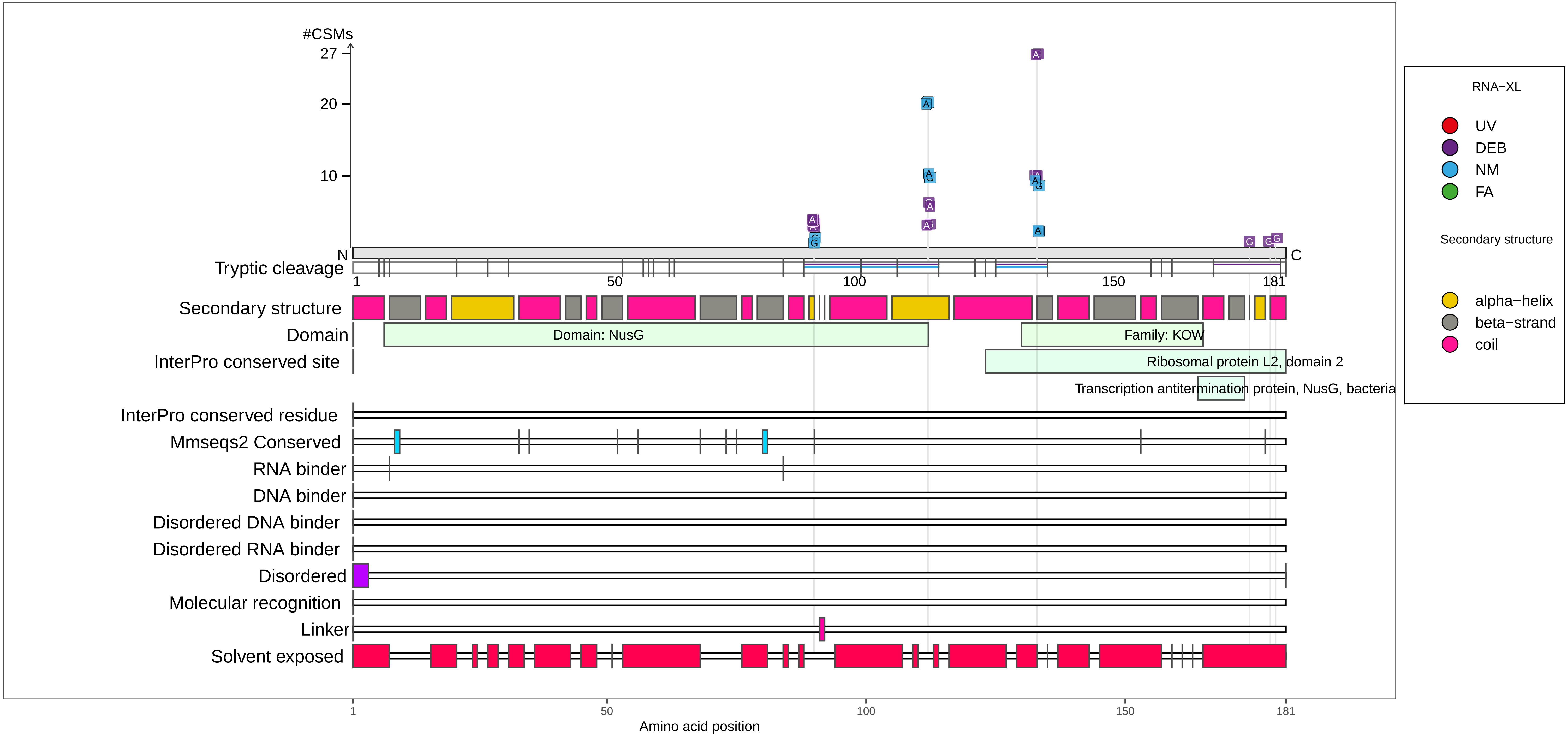

P0C0V0
DEGP_ECOLI Periplasmic serine endoprotease DegP

− Abundance:
tryptic [log10 Intensity]: 8.81 (Q 84)
PAXdb K12 strain [ppm]: 2.86 (Q 88)
PAXdb E.coli [ppm]: 2.77 (Q 91)

− RNA functions: not annotated

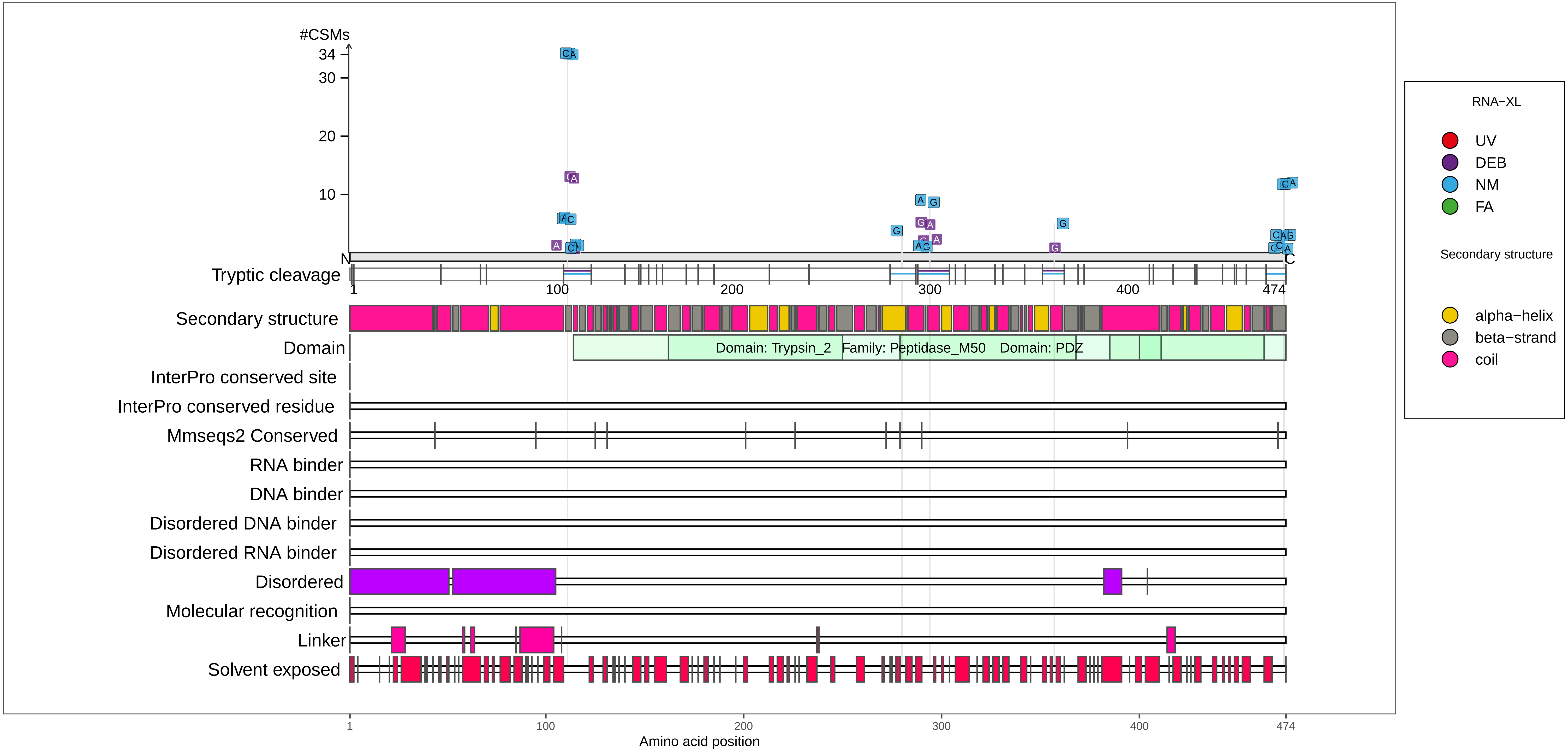

P07012
RF2_ECOLI Peptide chain release factor RF2

− Abundance:
tryptic [log10 Intensity]: 8.93 (Q 86)
PAXdb K12 strain [ppm]: 2.98 (Q 90)
PAXdb E.coli [ppm]: 2.51 (Q 86)

− RNA functions:
RNA binding

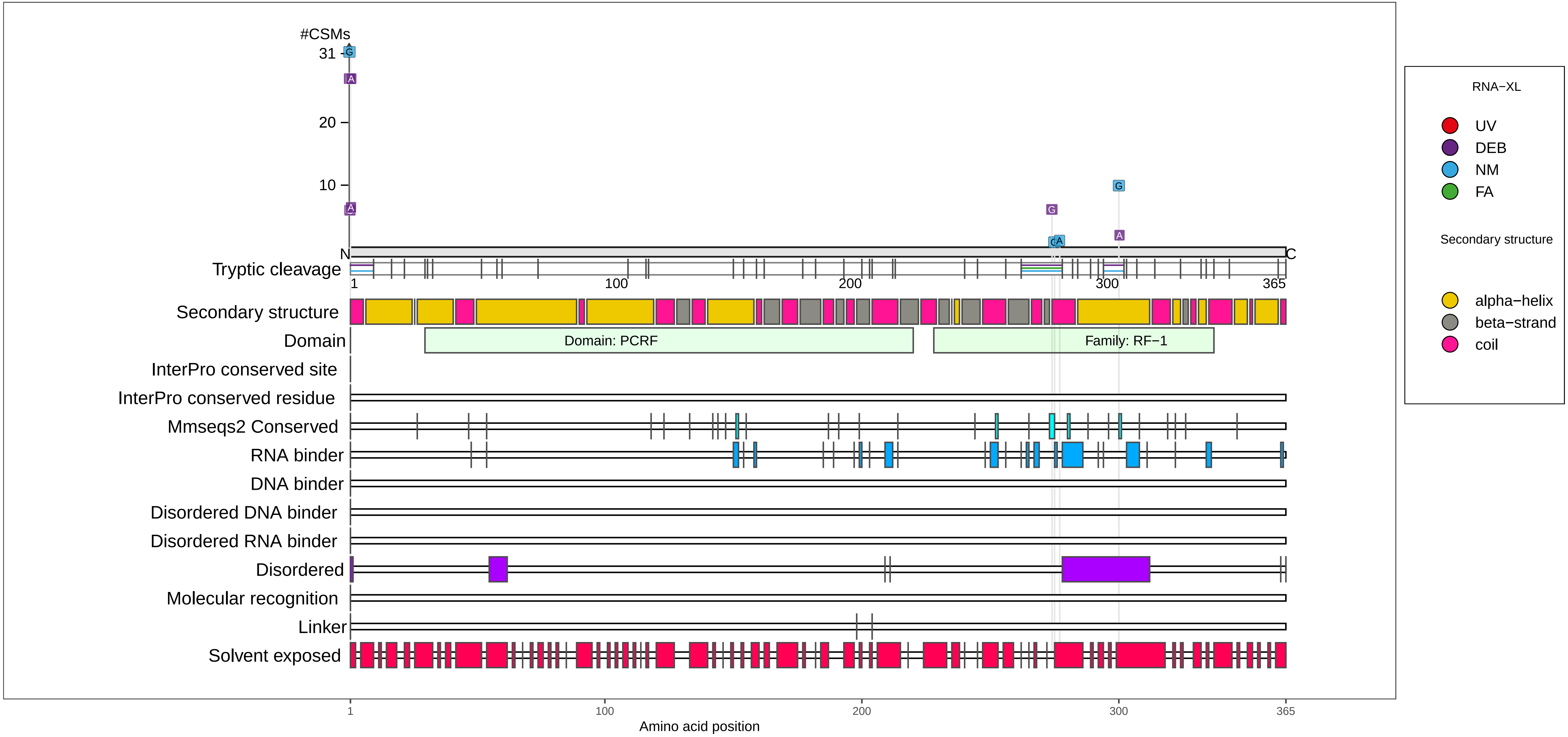

P0A9Q7
ADHE_ECOLI Aldehyde−alcohol dehydrogenase

− Abundance:
tryptic [log10 Intensity]: 8.73 (Q 82)
PAXdb K12 strain [ppm]: 3.05 (Q 91)
PAXdb E.coli [ppm]: 3.22 (Q 97)

− RNA functions: not annotated

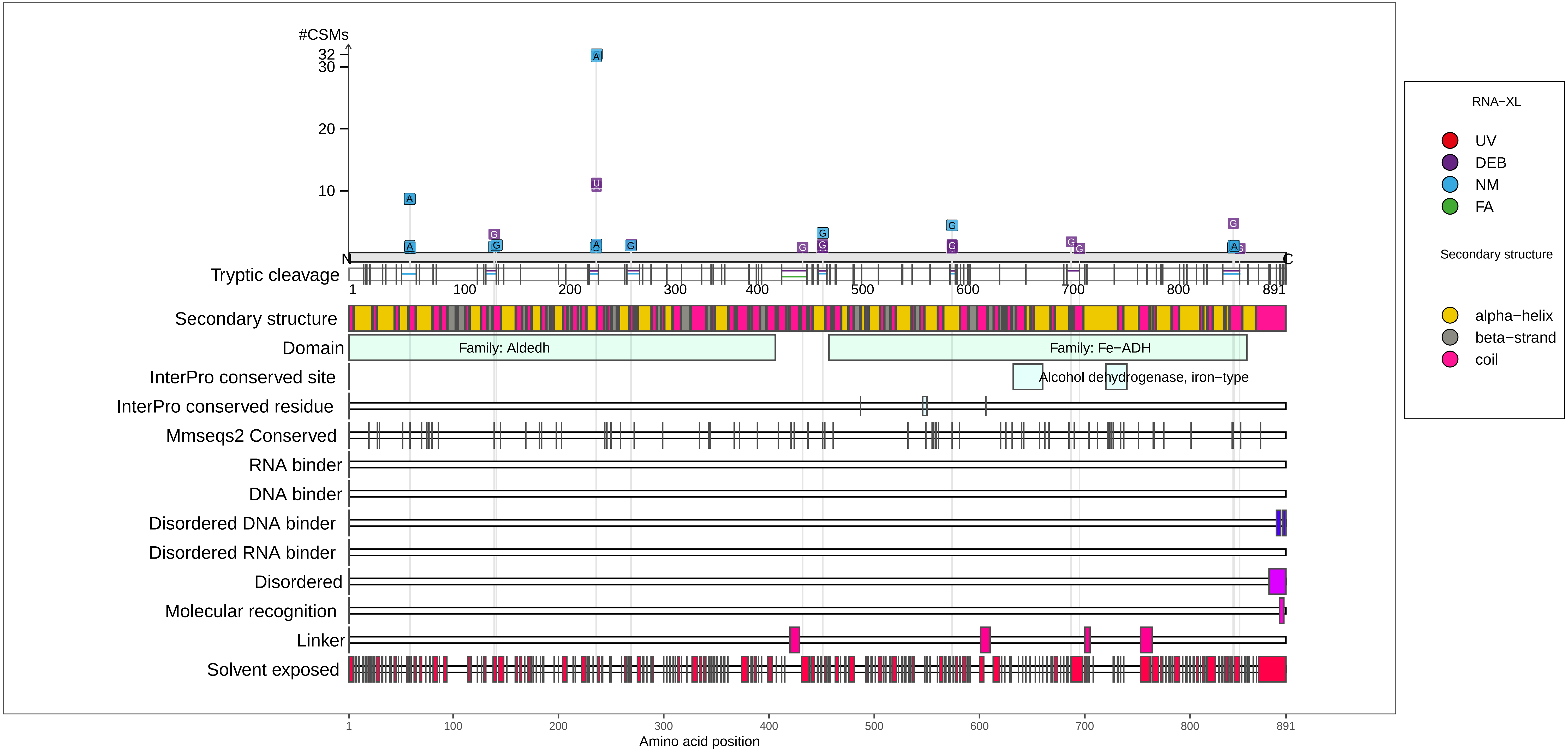

P0A910
OMPA_ECOLI Outer membrane protein A

− Abundance:
tryptic [log10 Intensity]: 10.15 (Q 100)
PAXdb K12 strain [ppm]: 3.17 (Q 94)
PAXdb E.coli [ppm]: 3.55 (Q 99)

− RNA functions: not annotated

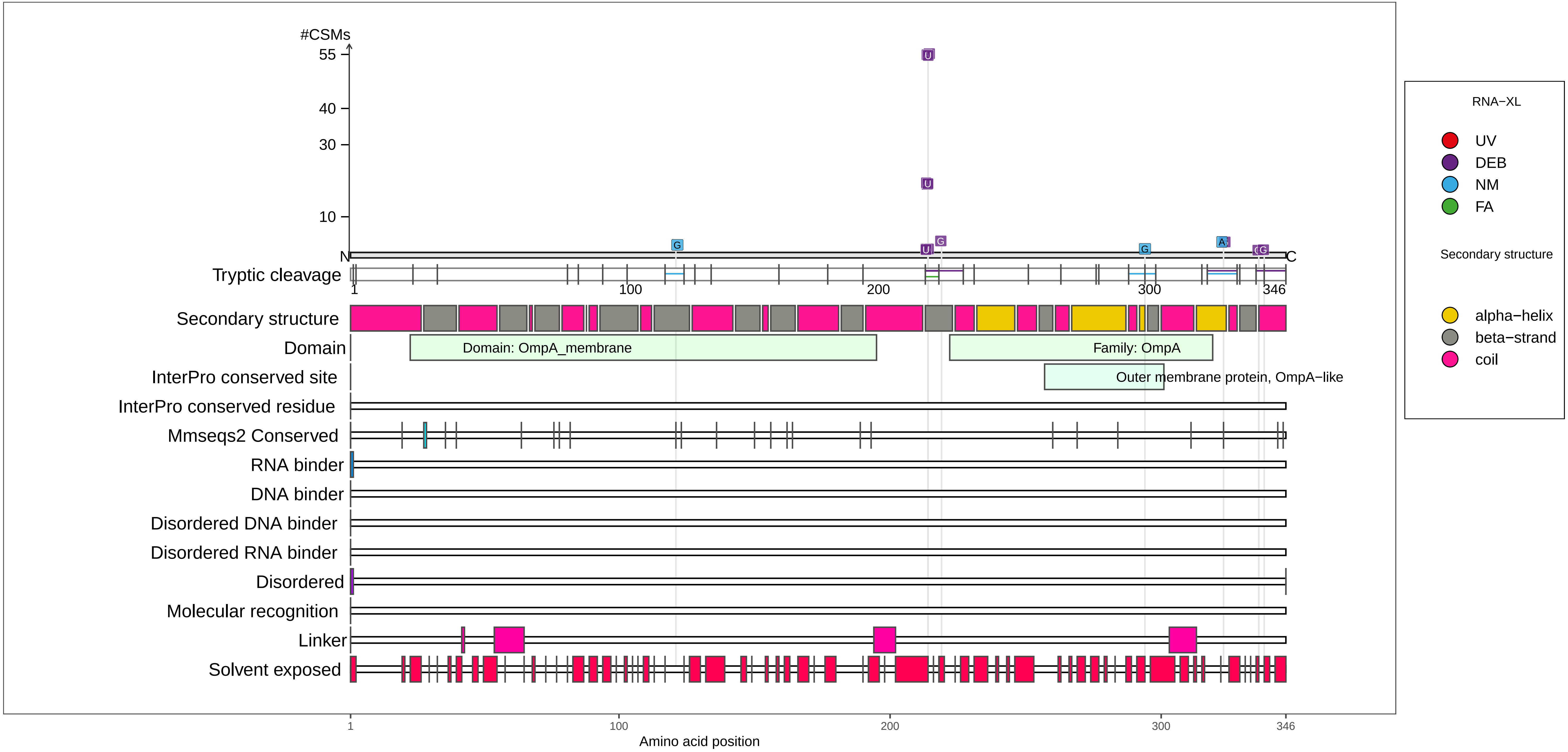

P0ABH7
CISY_ECOLI Citrate synthase

− Abundance:
tryptic [log10 Intensity]: 9.52 (Q 97)
PAXdb K12 strain [ppm]: 2.98 (Q 90)
PAXdb E.coli [ppm]: 3.03 (Q 95)

− RNA functions: not annotated

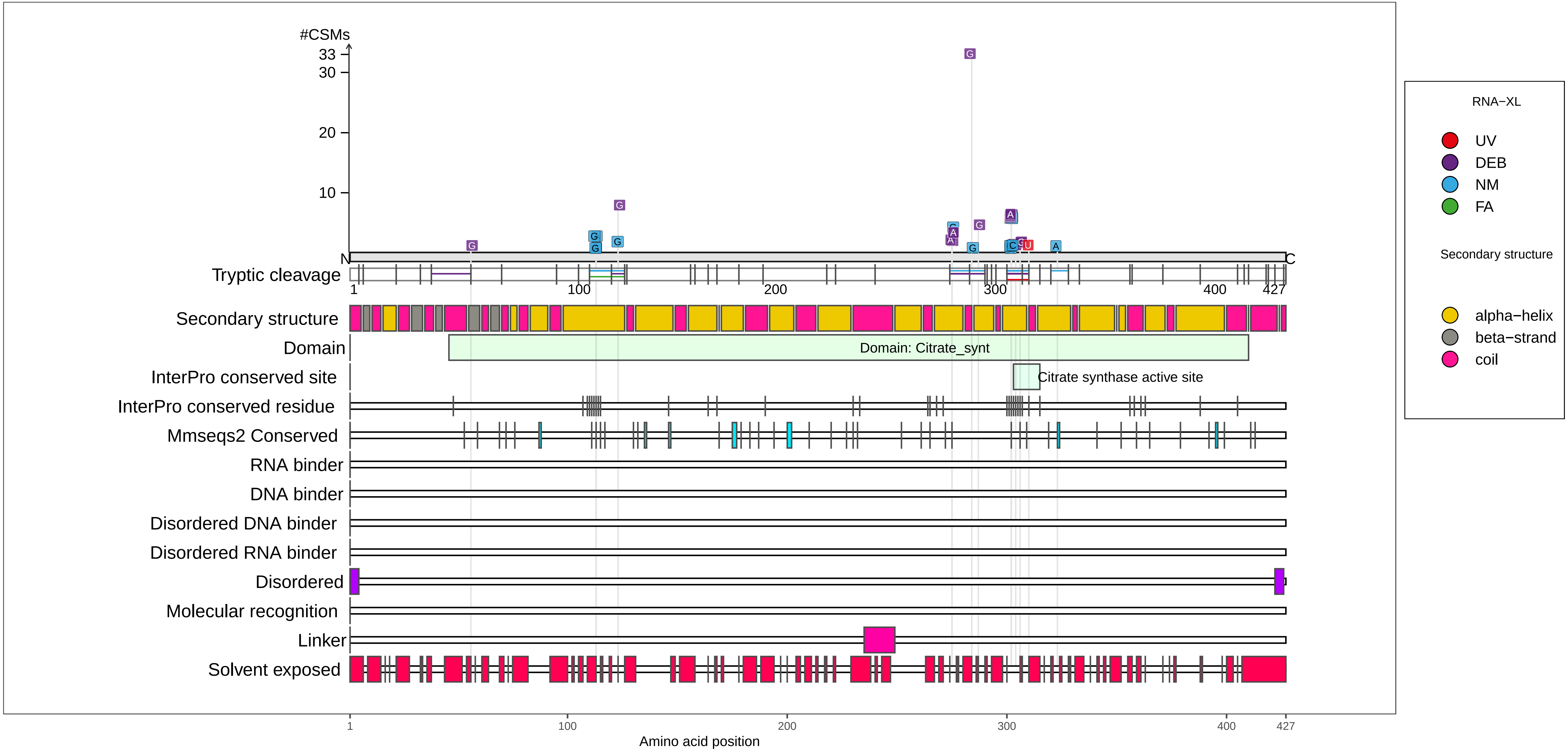

P0AG55
RL6_ECOLI 50S ribosomal protein L6

− Abundance:
tryptic [log10 Intensity]: 9.05 (Q 89)
PAXdb K12 strain [ppm]: 3.67 (Q 99)
PAXdb E.coli [ppm]: 3.43 (Q 98)

− RNA functions:
RNA binding; rRNA binding

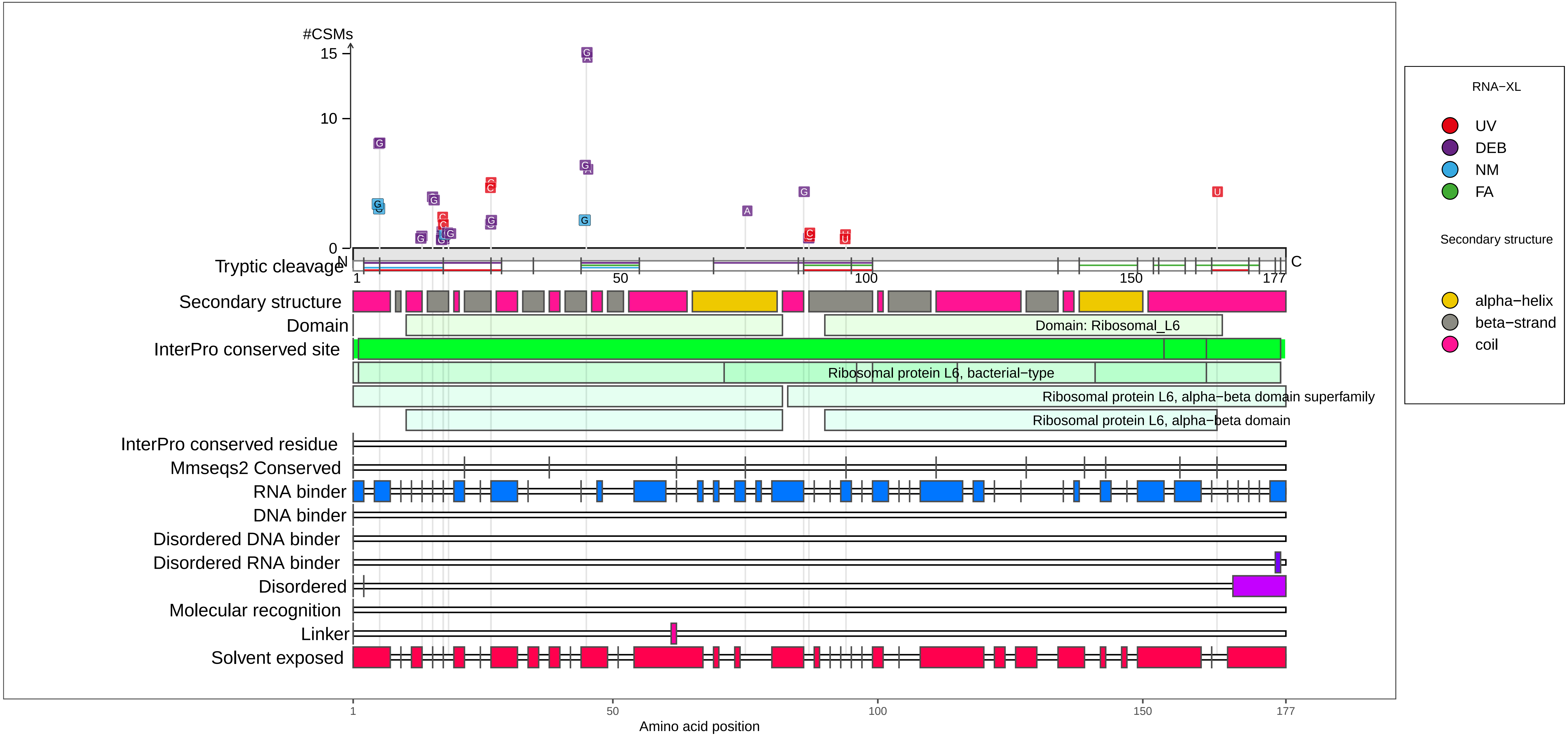

P0A7K2
RL7_ECOLI 50S ribosomal protein L7/L12

− Abundance:
tryptic [log10 Intensity]: 10.08 (Q 100)
PAXdb K12 strain [ppm]: 3.8 (Q 99)
PAXdb E.coli [ppm]: 3.86 (Q 100)

− RNA functions: not annotated

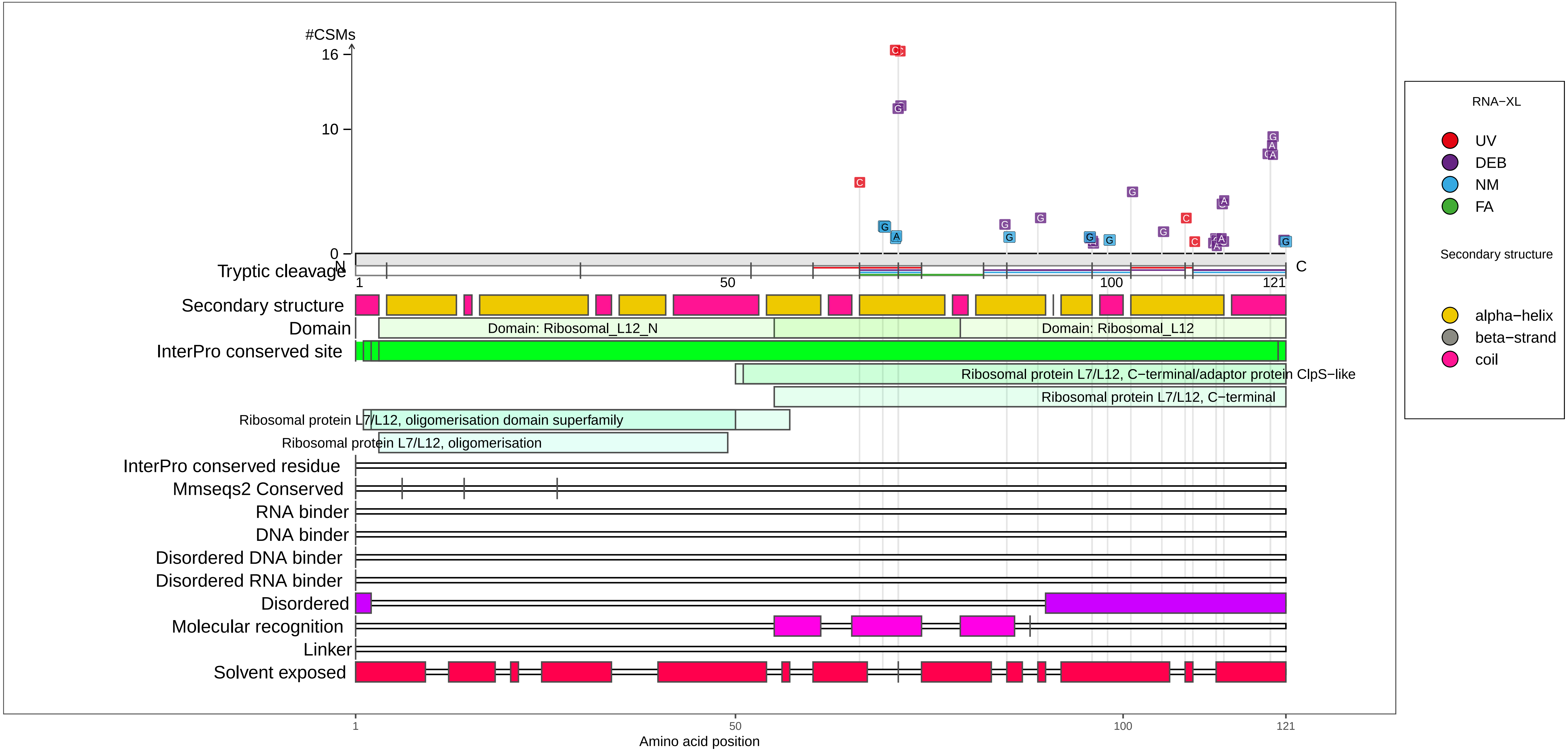

P0A858
TPIS_ECOLI Triosephosphate isomerase

− Abundance:
tryptic [log10 Intensity]: 10.02 (Q 99)
PAXdb K12 strain [ppm]: 2.58 (Q 80)
PAXdb E.coli [ppm]: 3.54 (Q 99)

− RNA functions: not annotated

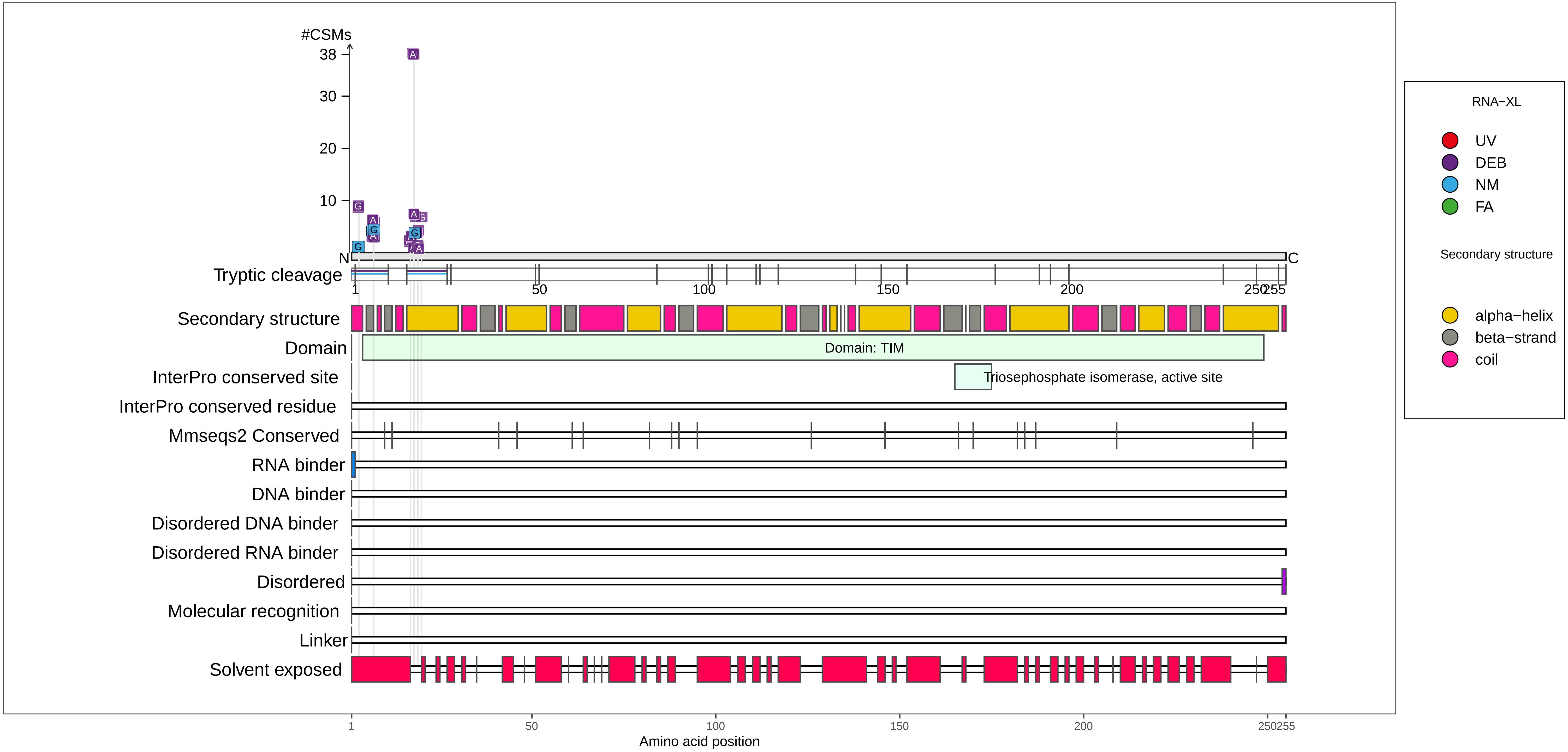

P0A8N3
SYK1_ECOLI Lysine−−tRNA ligase

− Abundance:
tryptic [log10 Intensity]: 9.64 (Q 97)
PAXdb K12 strain [ppm]: 3.15 (Q 93)
PAXdb E.coli [ppm]: 2.92 (Q 93)

− RNA functions:
aminoacyl−tRNA ligase activity; lysine−tRNA ligase activity
lysyl−tRNA aminoacylation; ncRNA metabolic process; RNA binding; RNA metabolic process; tRNA aminoacylation
tRNA aminoacylation for protein translation; tRNA binding; tRNA metabolic process
tRNA synthetases class II (D, K and N)

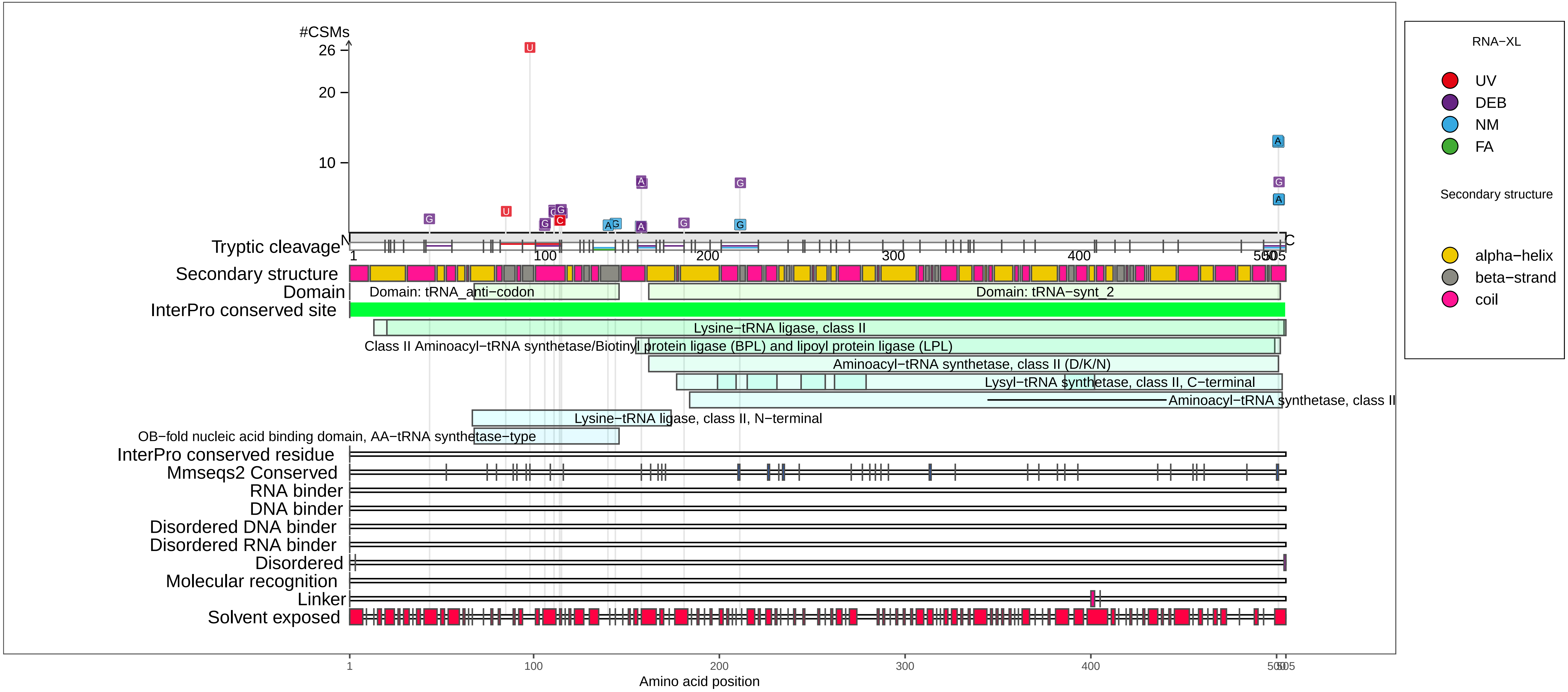

P0A998
FTNA_ECOLI Bacterial non−heme ferritin

− Abundance:
tryptic [log10 Intensity]: 9.36 (Q 95)
PAXdb K12 strain [ppm]: 2.57 (Q 79)
PAXdb E.coli [ppm]: 2.83 (Q 92)

− RNA functions: not annotated

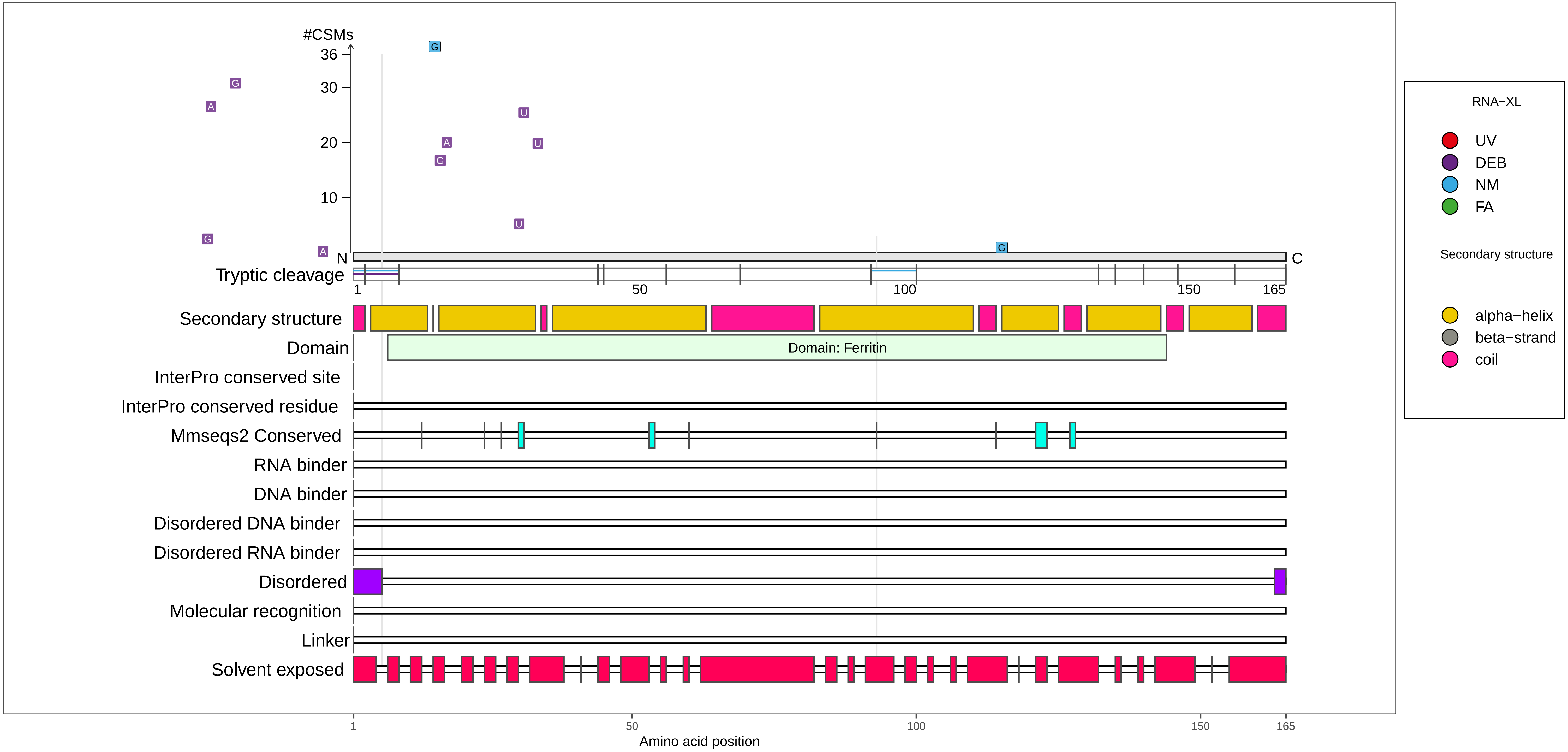

P0AAT6
IOJAP_ECOLI Ribosomal silencing factor RsfS

− Abundance:
tryptic [log10 Intensity]: 8.17 (Q 64)
PAXdb K12 strain [ppm]: 2.53 (Q 78)
PAXdb E.coli [ppm]: 2.36 (Q 82)

− RNA functions: not annotated

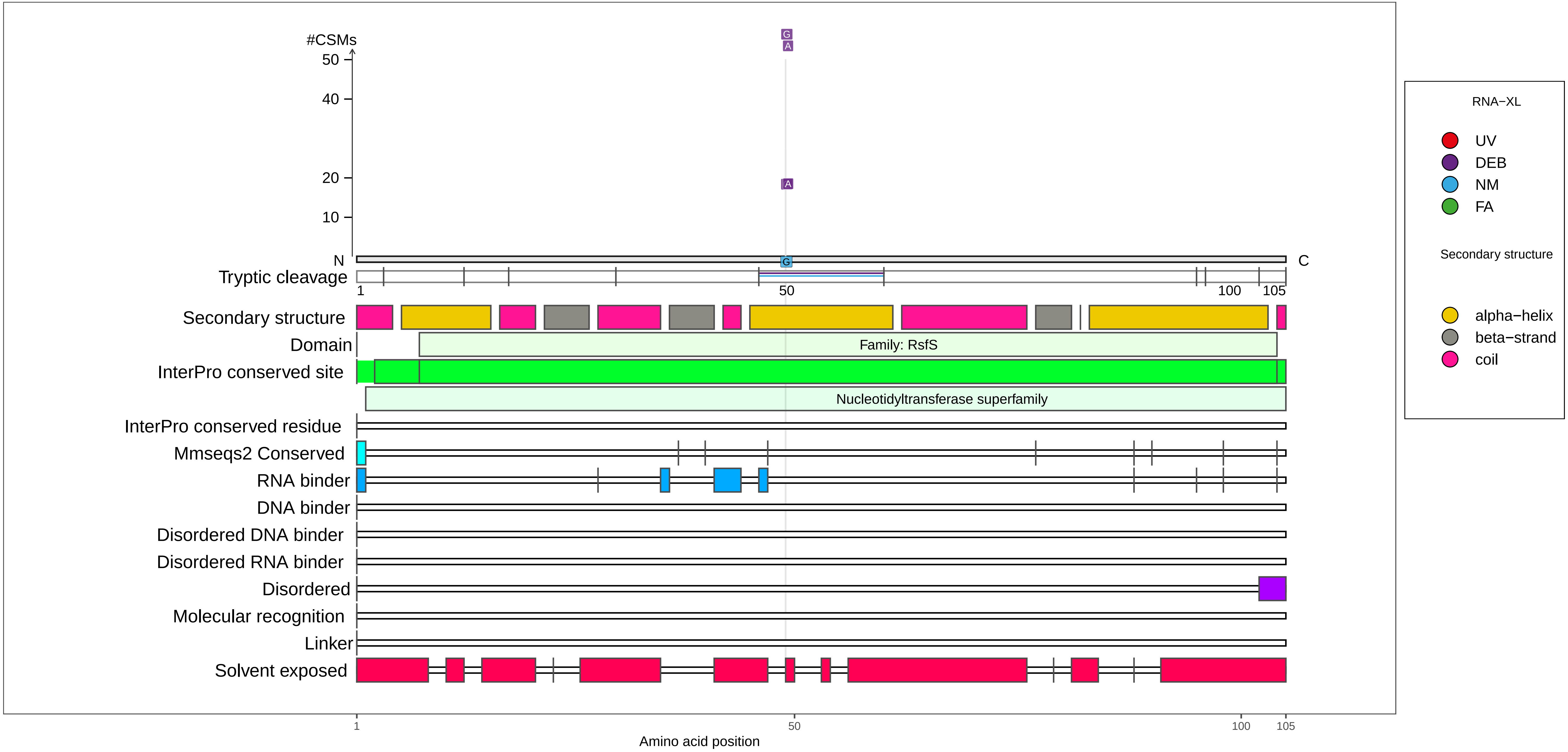

P0AG30
RHO_ECOLI Transcription termination factor Rho

− Abundance:
tryptic [log10 Intensity]: 8.54 (Q 77)
PAXdb K12 strain [ppm]: 3.4 (Q 96)
PAXdb E.coli [ppm]: 3.73 (Q 100)

− RNA functions:
Rho termination factor, RNA−binding domain; RNA binding; RNA biosynthetic process
RNA metabolic process; RNA−dependent ATPase activity

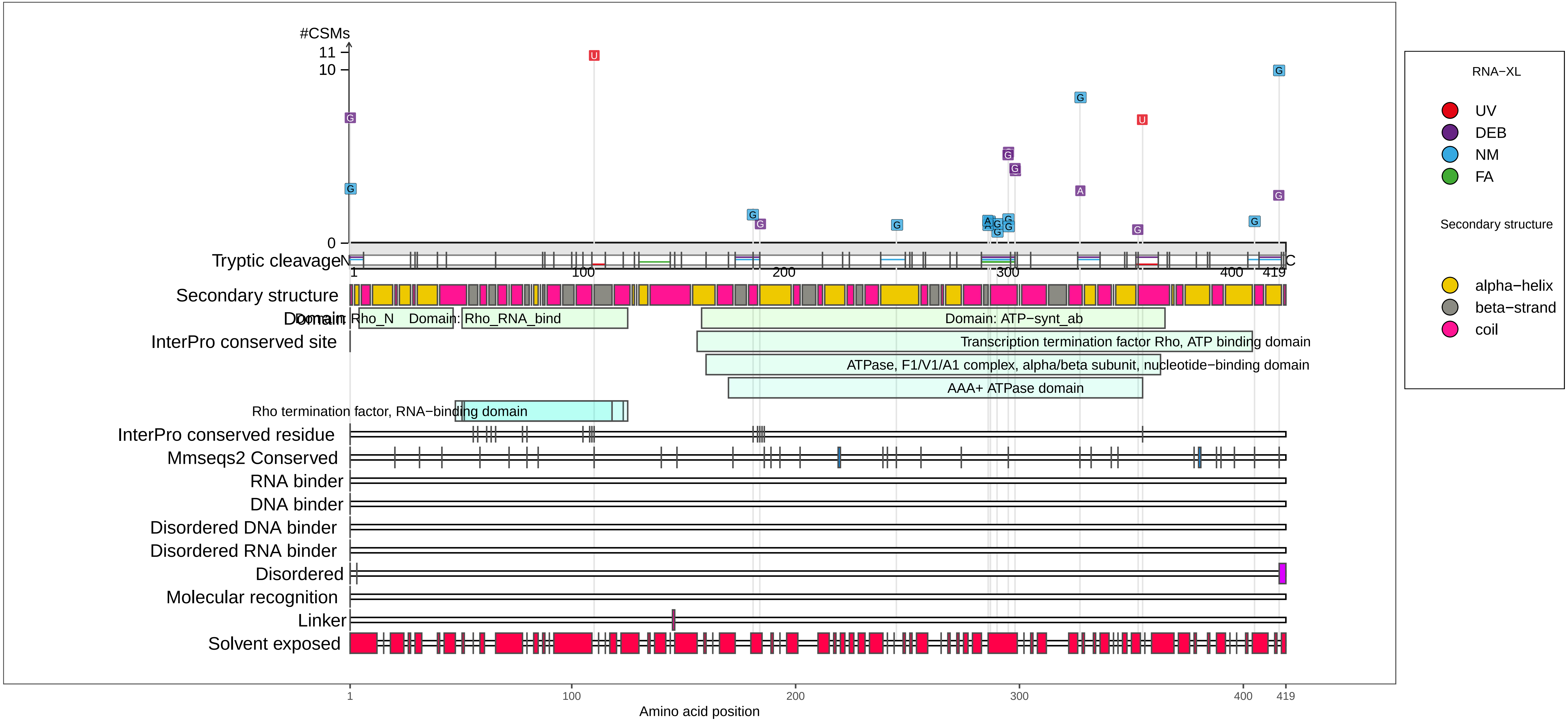

P0AG44
RL17_ECOLI 50S ribosomal protein L17

− Abundance:
tryptic [log10 Intensity]: 8.55 (Q 77)
PAXdb K12 strain [ppm]: 3.85 (Q 100)
PAXdb E.coli [ppm]: 3.33 (Q 98)

− RNA functions: not annotated

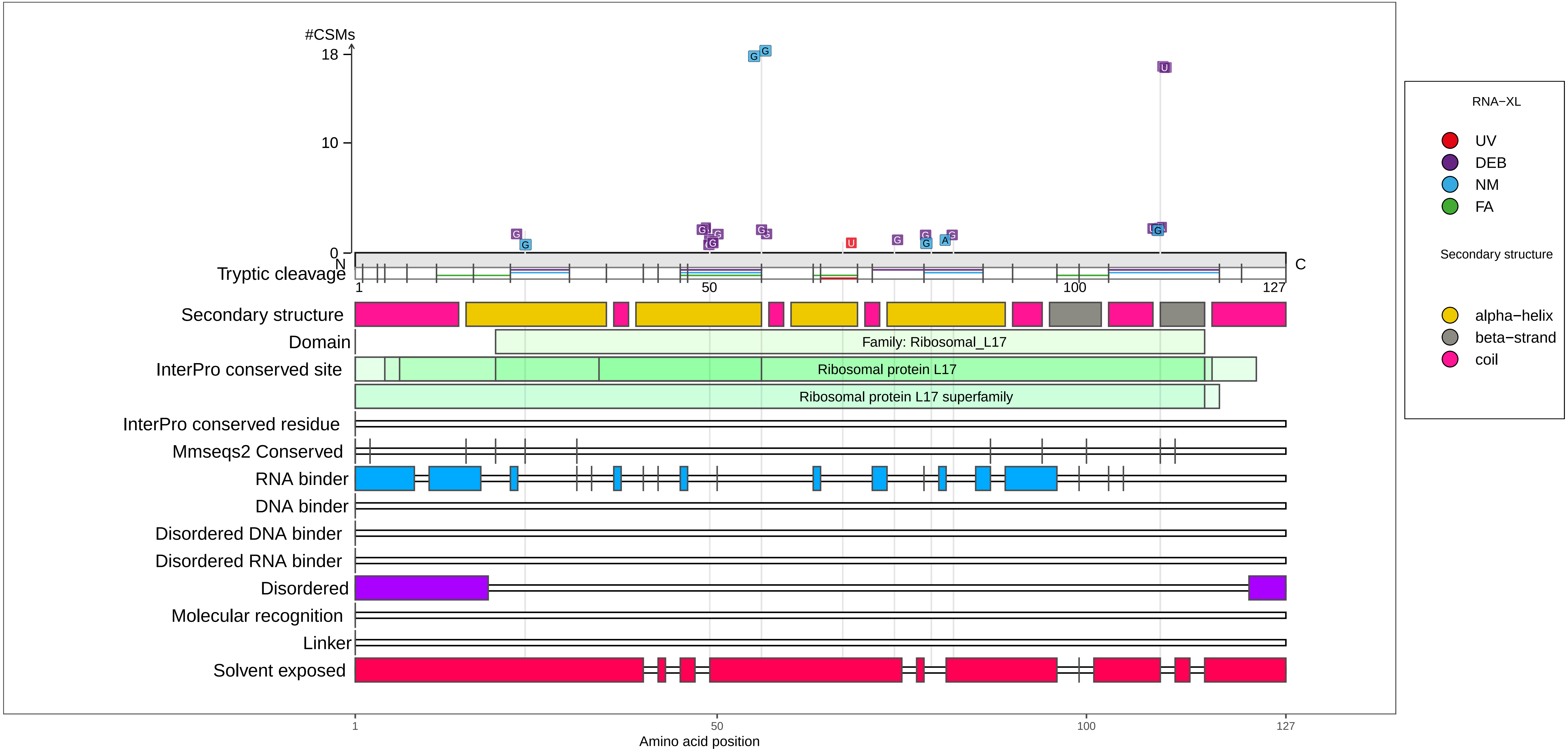

P05793
ILVC_ECOLI Ketol−acid reductoisomerase (NADP(+))

− Abundance:
tryptic [log10 Intensity]: 9.75 (Q 98)
PAXdb K12 strain [ppm]: 2.99 (Q 91)
PAXdb E.coli [ppm]: 3.28 (Q 97)

− RNA functions: not annotated

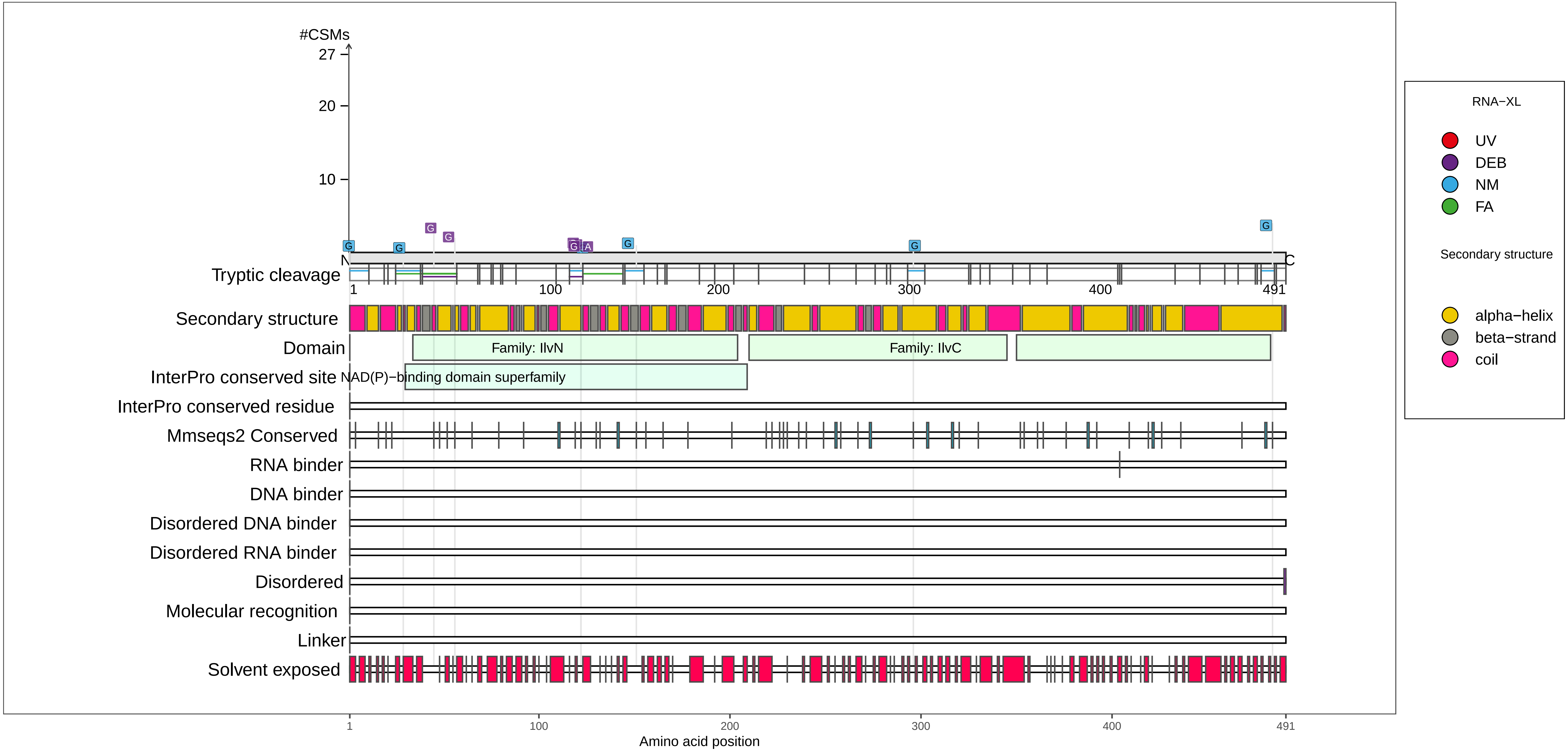

P61517
CAN_ECOLI Carbonic anhydrase 2

− Abundance:
tryptic [log10 Intensity]: 9.61 (Q 97)
PAXdb K12 strain [ppm]: 2.5 (Q 78)
PAXdb E.coli [ppm]: 2.73 (Q 90)

− RNA functions: not annotated

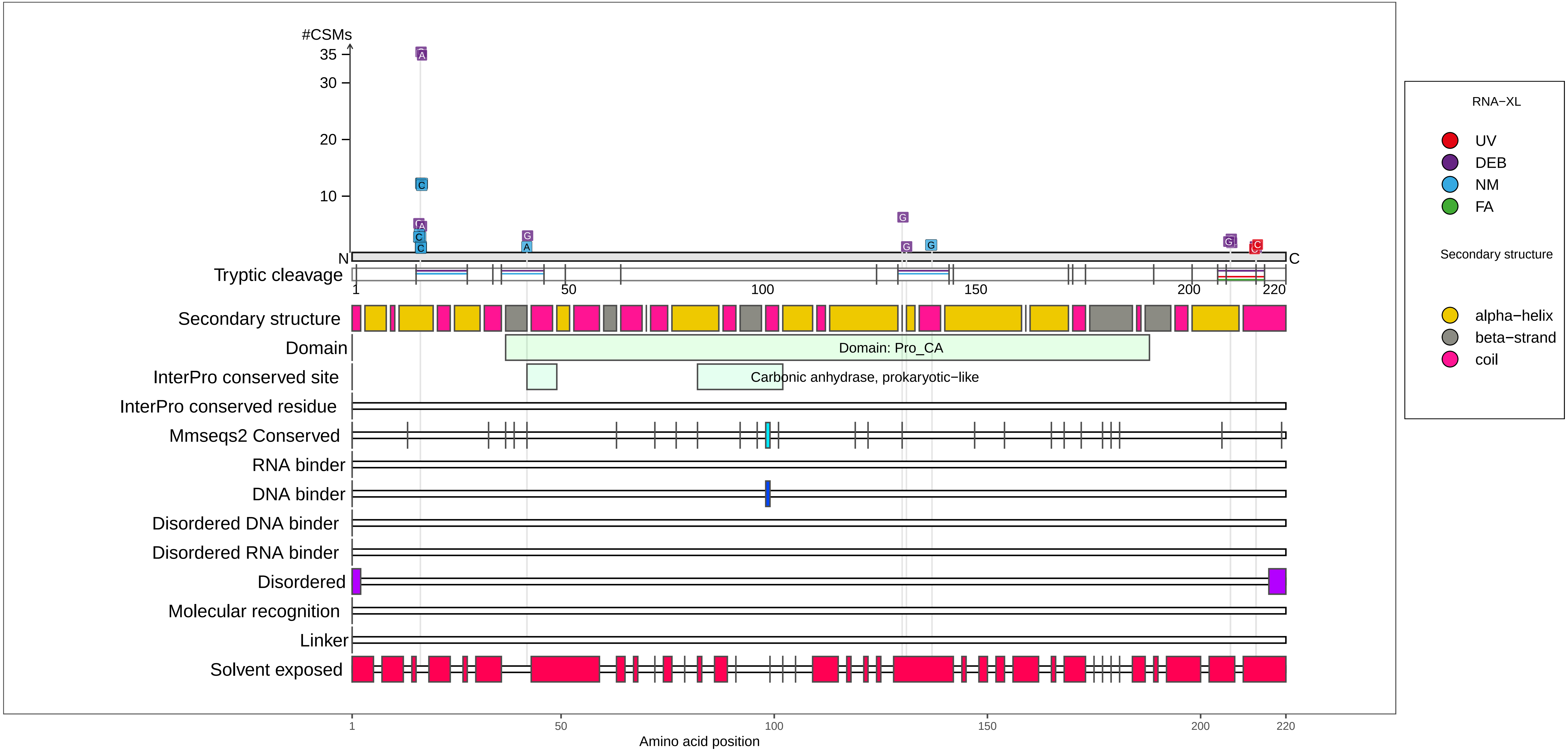

P75937
FLGE_ECOLI Flagellar hook protein FlgE

− Abundance:
tryptic [log10 Intensity]: 8.02 (Q 58)
PAXdb K12 strain [ppm]: 2.62 (Q 81)
PAXdb E.coli [ppm]: 1.17 (Q 52)

− RNA functions: not annotated

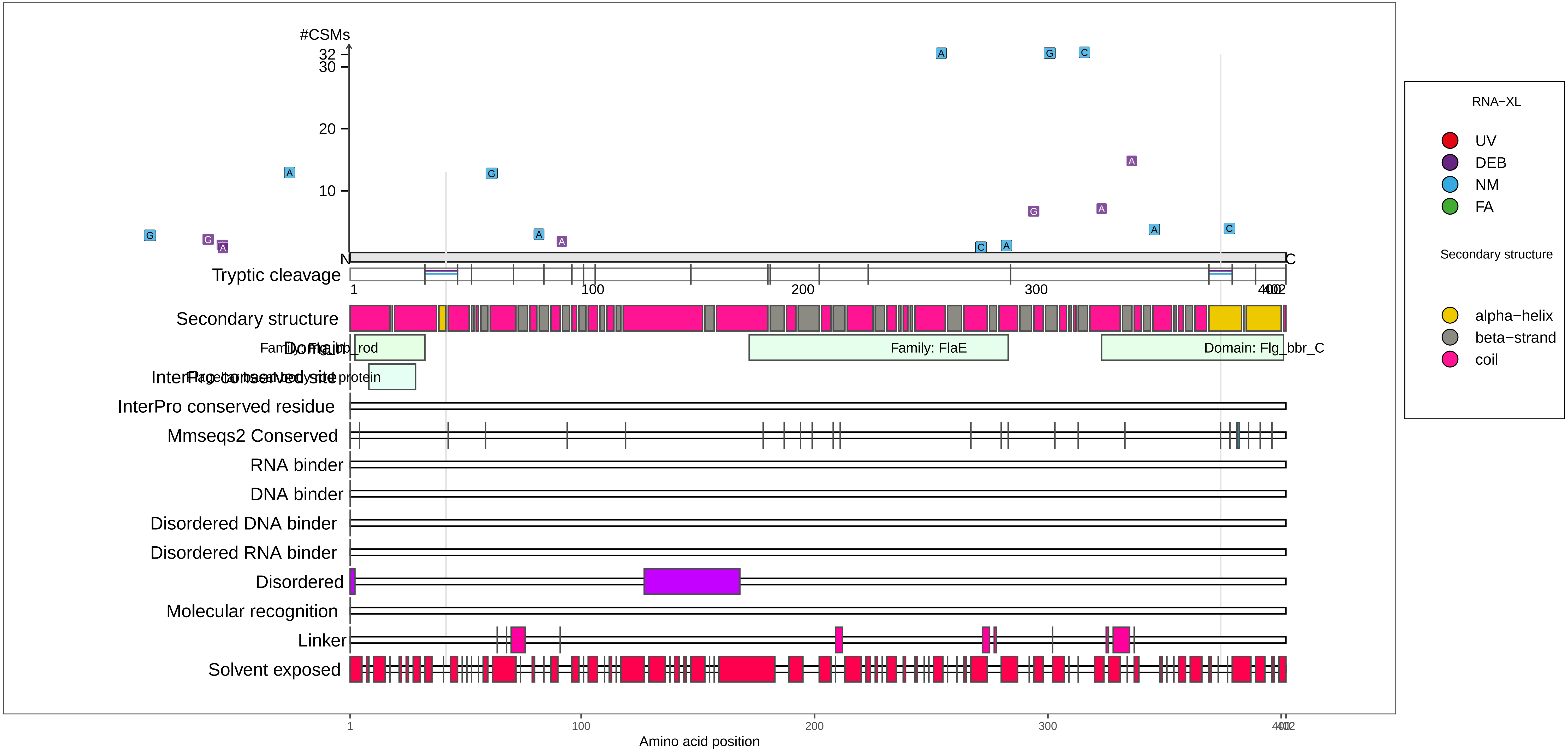

P0A940
BAMA_ECOLI Outer membrane protein assembly factor BamA

− Abundance:
tryptic [log10 Intensity]: 9.14 (Q 91)
PAXdb K12 strain [ppm]: 2 (Q 60)
PAXdb E.coli [ppm]: 2.28 (Q 80)

− RNA functions: not annotated

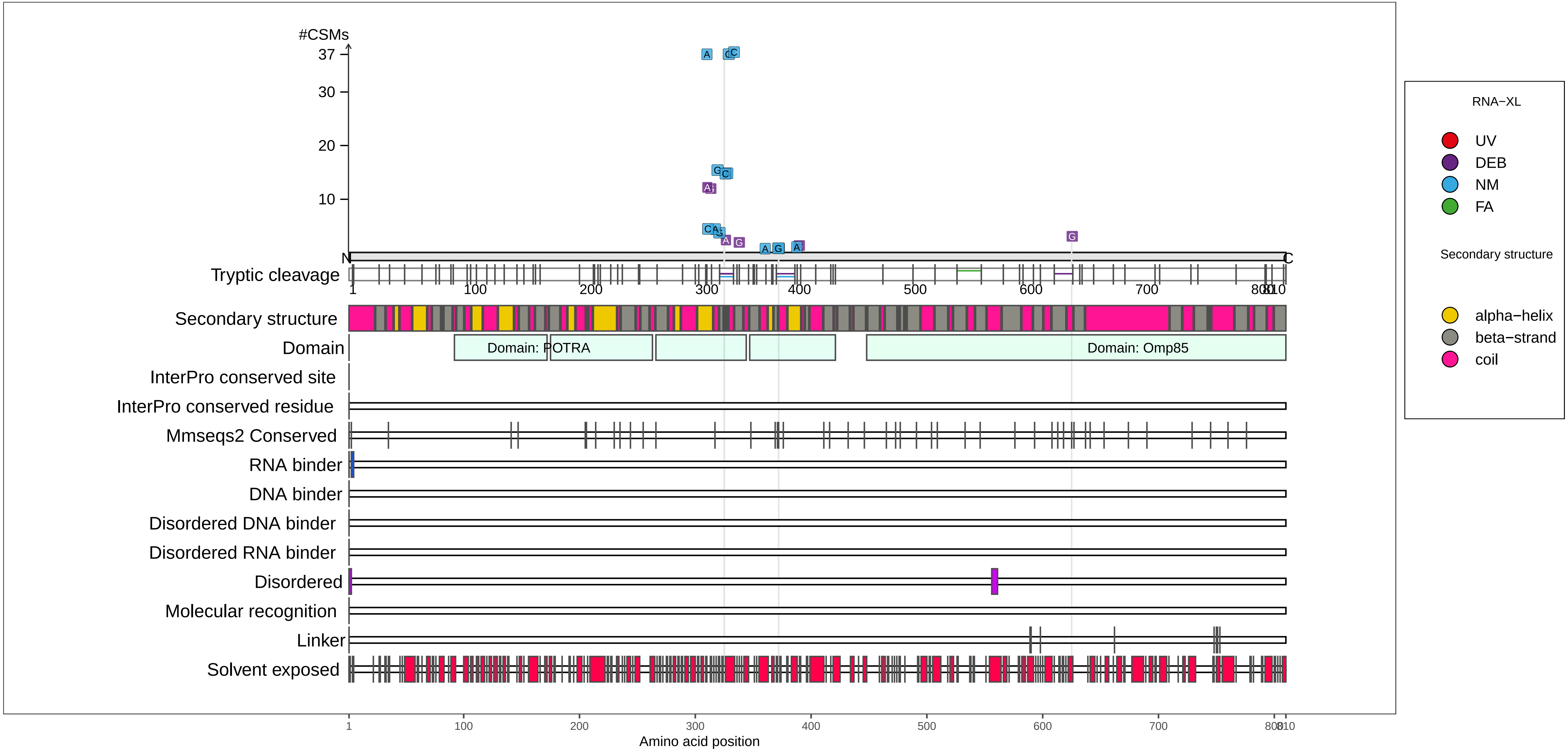

P09030
EX3_ECOLI Exodeoxyribonuclease III

− Abundance:
tryptic [log10 Intensity]: 8.21 (Q 66)
PAXdb K12 strain [ppm]: 2.77 (Q 85)
PAXdb E.coli [ppm]: 2.42 (Q 84)

− RNA functions: not annotated

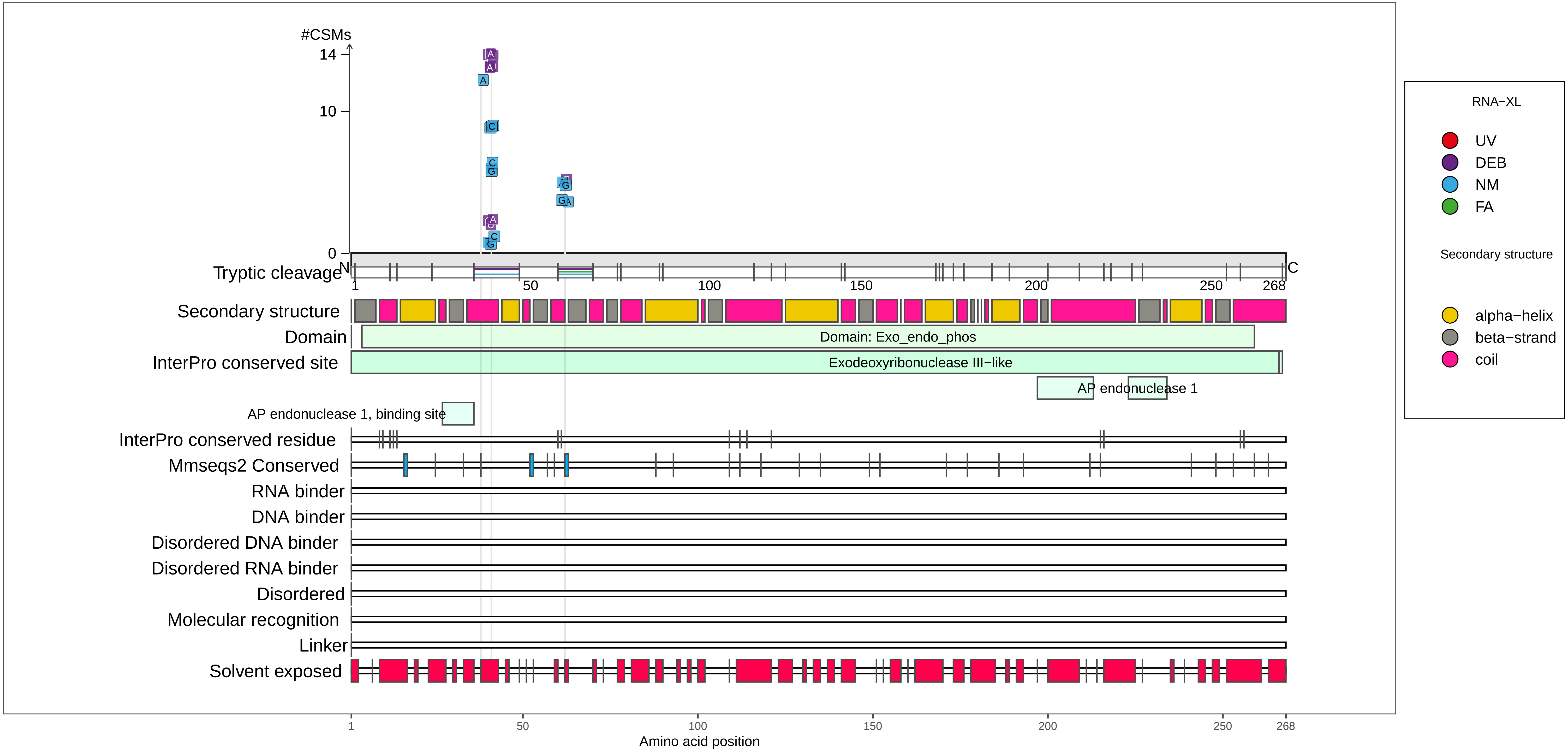

P36683
ACNB_ECOLI Aconitate hydratase B

− Abundance:
tryptic [log10 Intensity]: 9.47 (Q 96)
PAXdb K12 strain [ppm]: 3.11 (Q 93)
PAXdb E.coli [ppm]: 3.36 (Q 98)

− RNA functions:
mRNA 3−UTR binding; mRNA binding; RNA binding

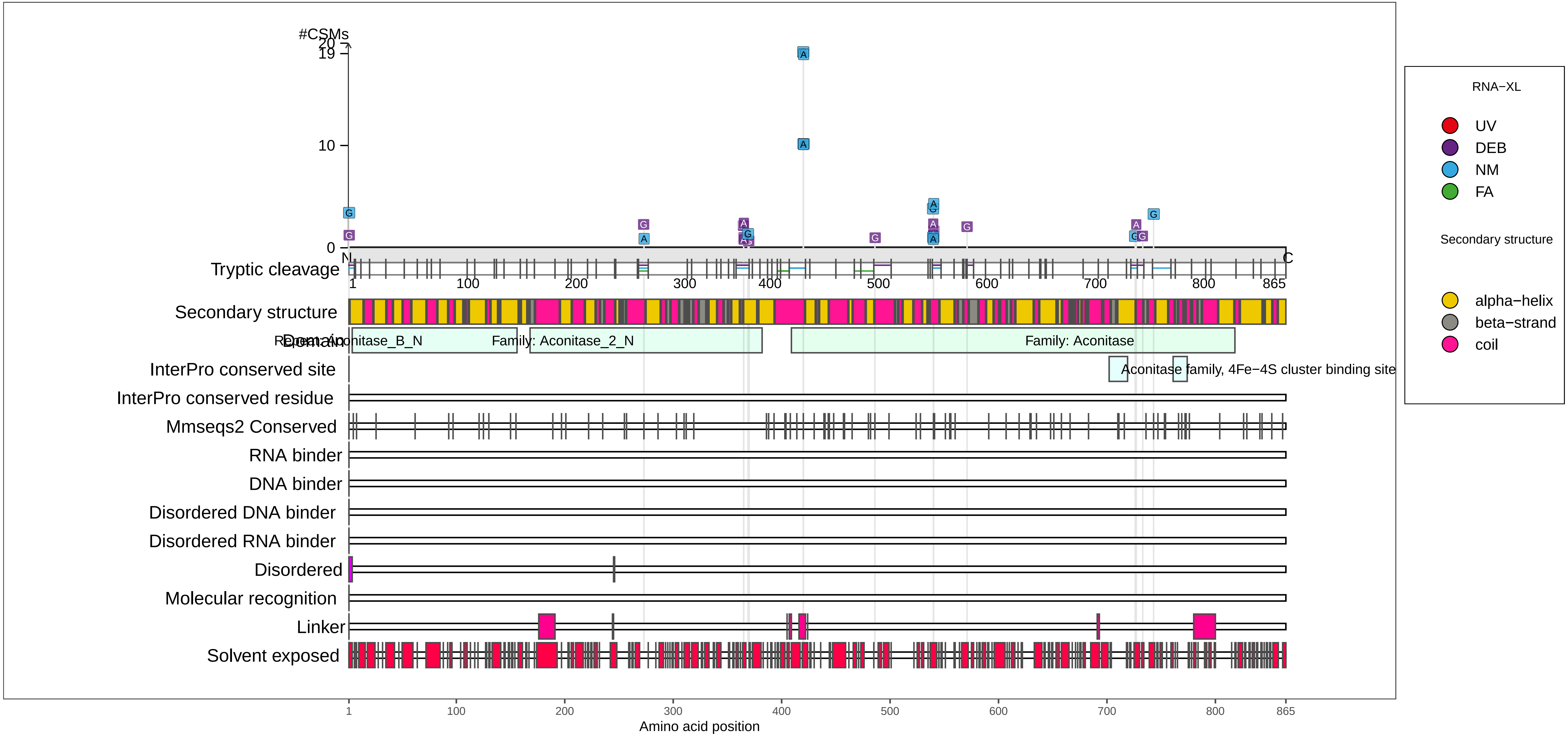

P0A7G2
RBFA_ECOLI 30S ribosome−binding factor

− Abundance:
tryptic [log10 Intensity]: 8.03 (Q 59)
PAXdb K12 strain [ppm]: 3.08 (Q 92)
PAXdb E.coli [ppm]: 2.81 (Q 92)

− RNA functions:
ncRNA metabolic process; ncRNA processing; RNA metabolic process; RNA processing
rRNA metabolic process; rRNA processing

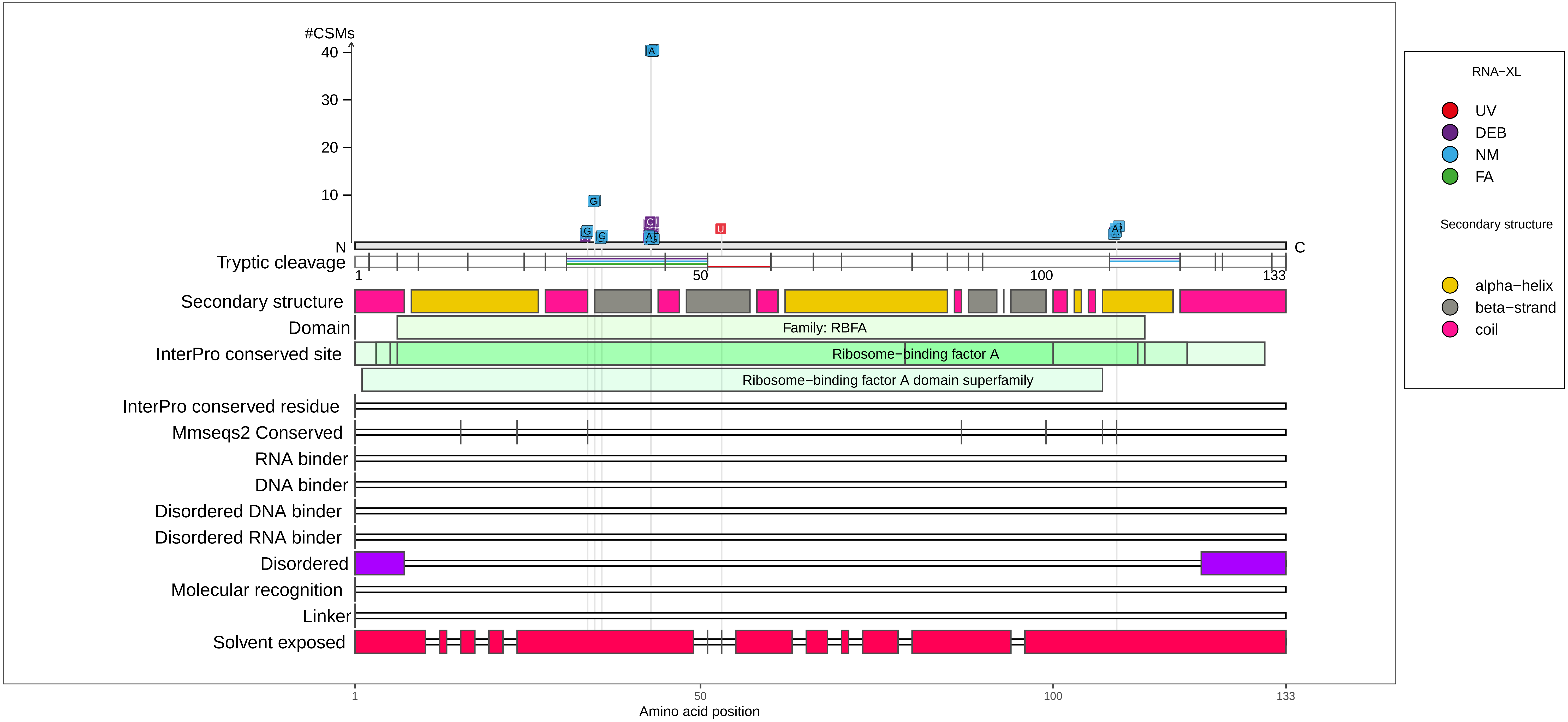

P21513
RNE_ECOLI Ribonuclease E

− Abundance:
tryptic [log10 Intensity]: 8.93 (Q 87)
PAXdb K12 strain [ppm]: 2.48 (Q 77)
PAXdb E.coli [ppm]: 2.43 (Q 84)

− RNA functions:
7S RNA binding; DEAD/H−box RNA helicase binding; mRNA catabolic process
mRNA metabolic process; ncRNA 5−end processing; ncRNA metabolic process; ncRNA processing
Ribosomal protein S1−like RNA−binding domain; RNA 5−end processing; RNA binding
RNA catabolic process; RNA metabolic process; RNA phosphodiester bond hydrolysis
RNA phosphodiester bond hydrolysis, endonucleolytic; RNA processing; rRNA 5−end processing
rRNA metabolic process; rRNA processing; S1 RNA binding domain; tRNA metabolic process
tRNA processing

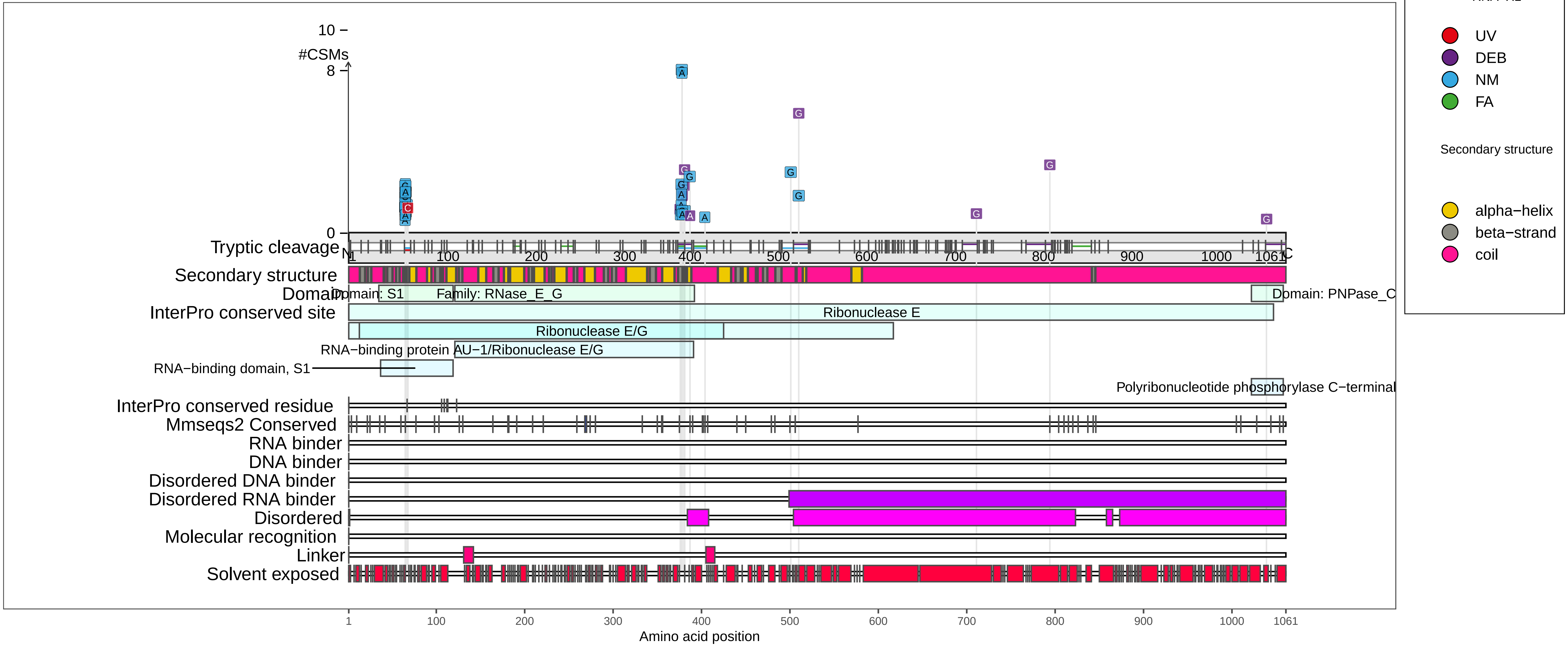

P0AAI3
FTSH_ECOLI ATP−dependent zinc metalloprotease FtsH

− Abundance:
tryptic [log10 Intensity]: 8.55 (Q 77)
PAXdb K12 strain [ppm]: 2.56 (Q 79)
PAXdb E.coli [ppm]: 2.66 (Q 89)

− RNA functions: not annotated

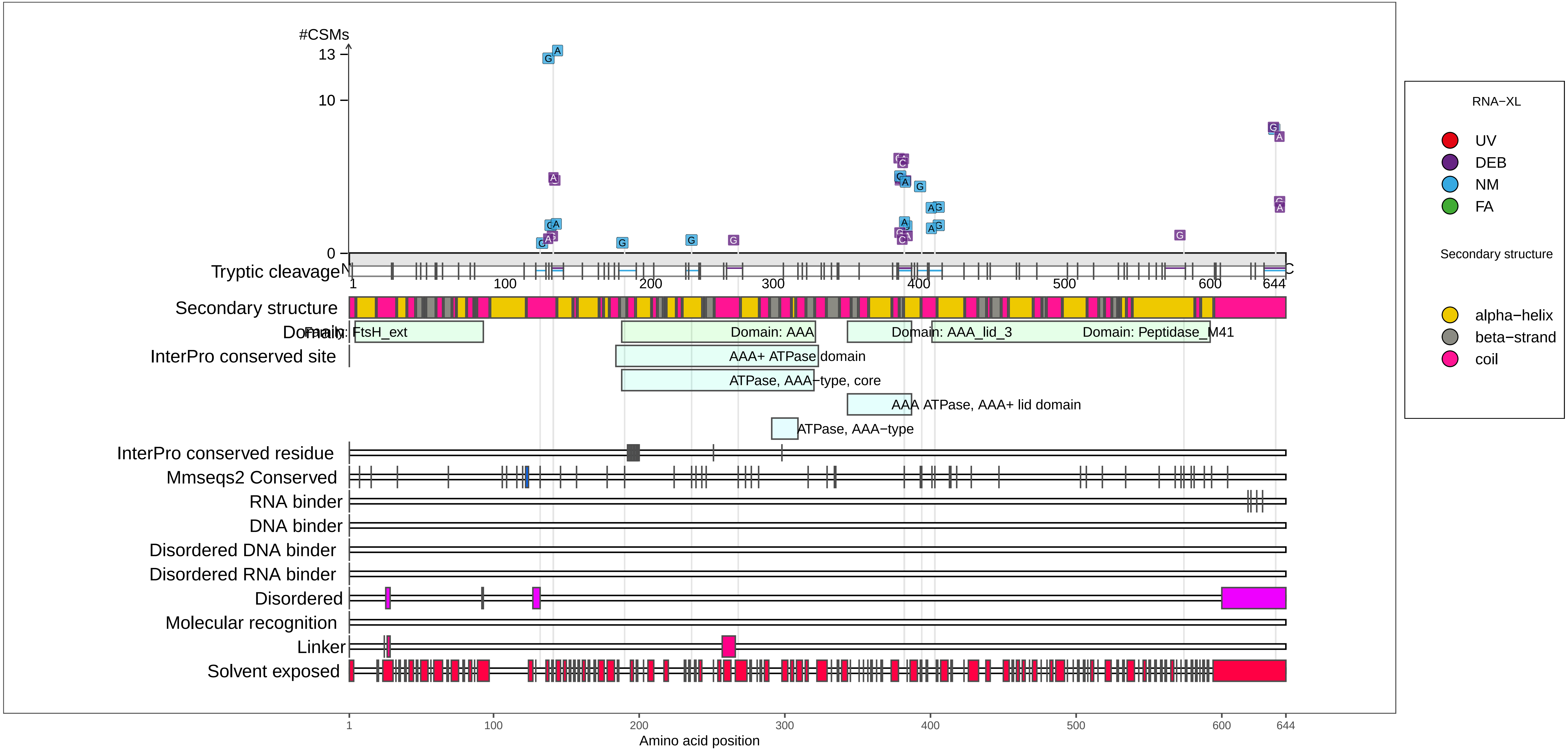

P12758
UDP_ECOLI Uridine phosphorylase

− Abundance:
tryptic [log10 Intensity]: 9.14 (Q 91)
PAXdb K12 strain [ppm]: 3.03 (Q 91)
PAXdb E.coli [ppm]: 3.18 (Q 96)

− RNA functions: not annotated

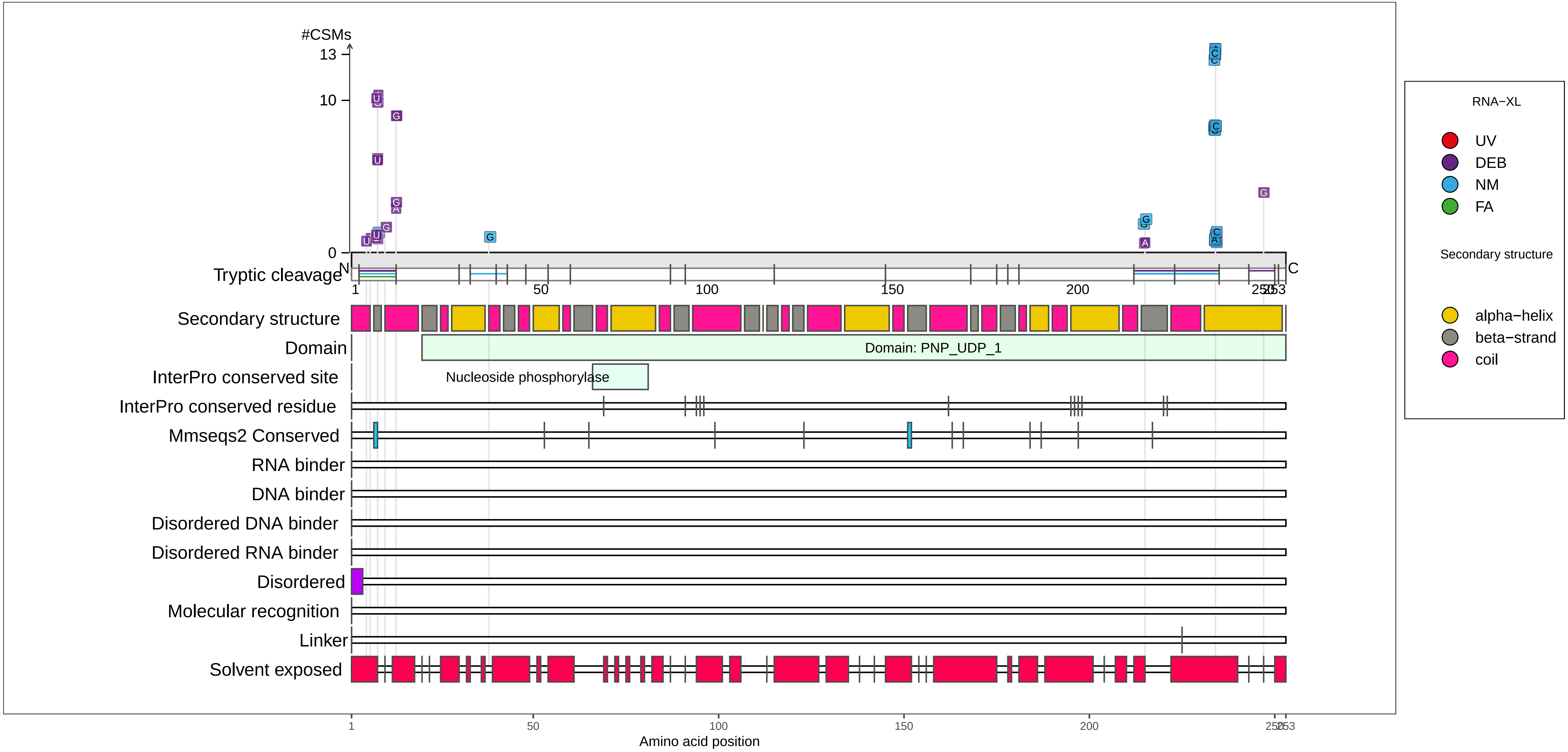

P23538
PPSA_ECOLI Phosphoenolpyruvate synthase

− Abundance:
tryptic [log10 Intensity]: 8.59 (Q 78)
PAXdb K12 strain [ppm]: 2.35 (Q 72)
PAXdb E.coli [ppm]: 2.89 (Q 93)

− RNA functions: not annotated

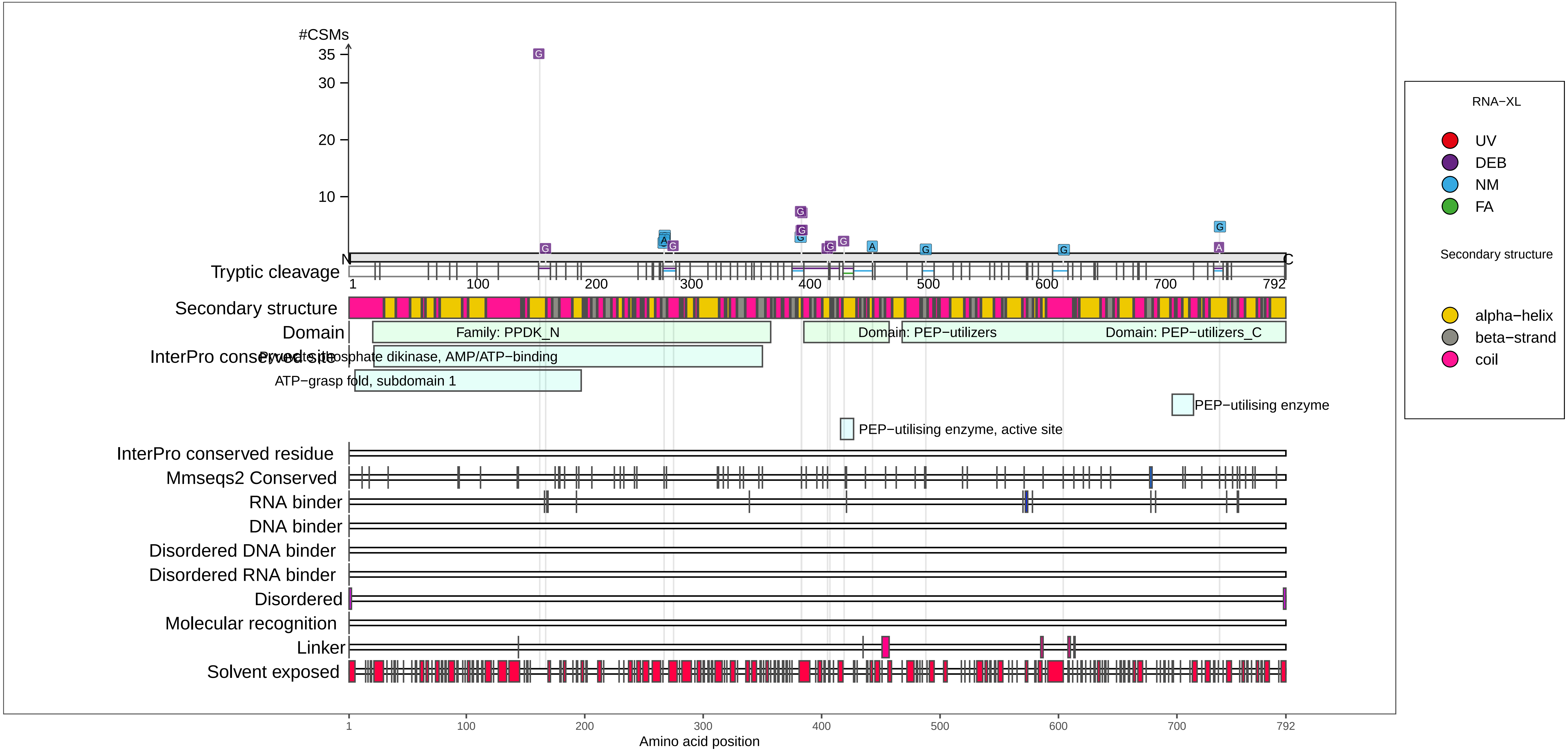

P0ACJ8
CRP_ECOLI cAMP−activated global transcriptional regulator CRP

− Abundance:
tryptic [log10 Intensity]: 8.67 (Q 80)
PAXdb K12 strain [ppm]: 2.95 (Q 90)
PAXdb E.coli [ppm]: 2.92 (Q 93)

− RNA functions:
RNA biosynthetic process; RNA metabolic process

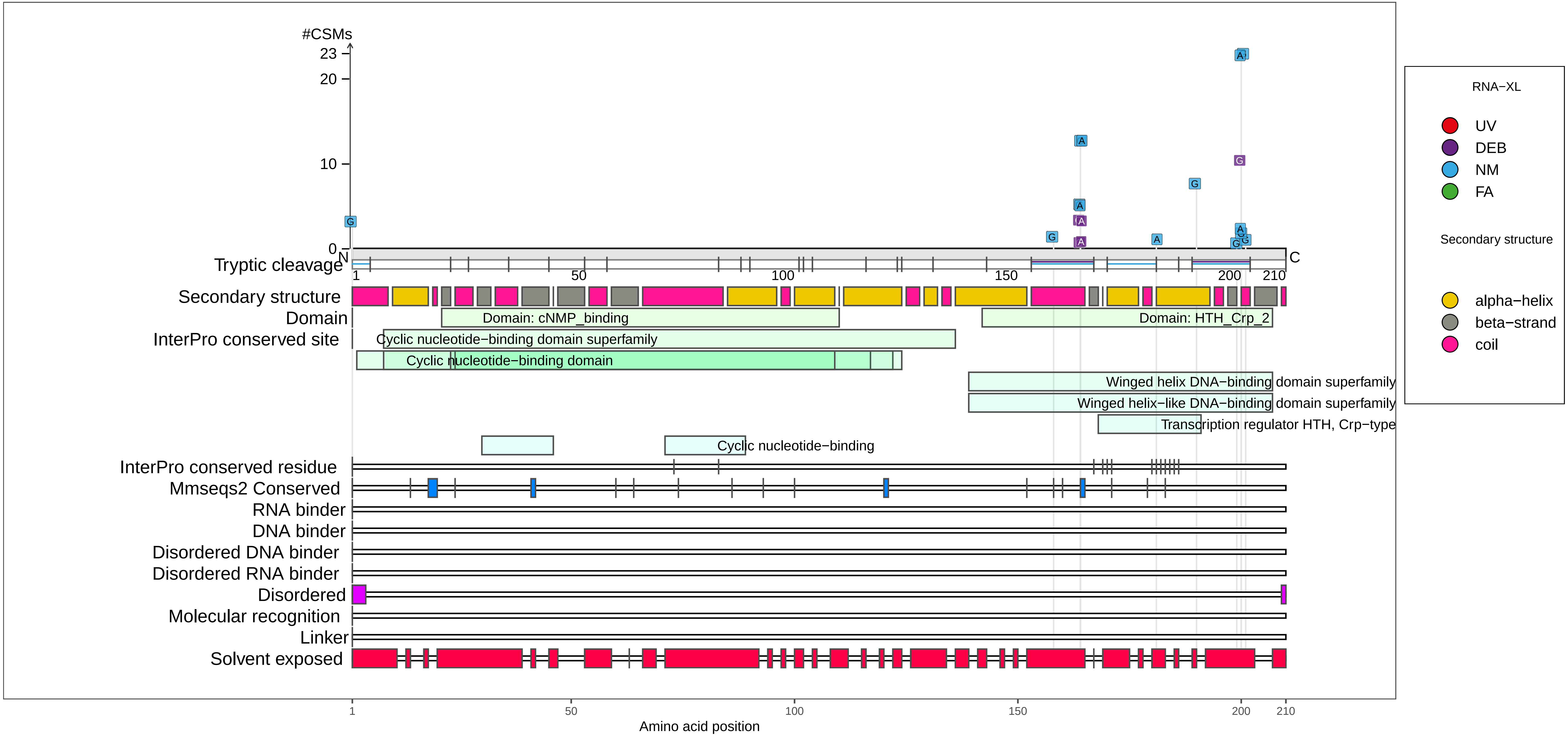

P0AGE9

SUCD_ECOLI Succinate−−CoA ligase [ADP−forming] subunit alpha

− Abundance:
tryptic [log10 Intensity]: 9.07 (Q 90)
PAXdb K12 strain [ppm]: 3.32 (Q 95)
PAXdb E.coli [ppm]: 3.31 (Q 97)

− RNA functions: not annotated

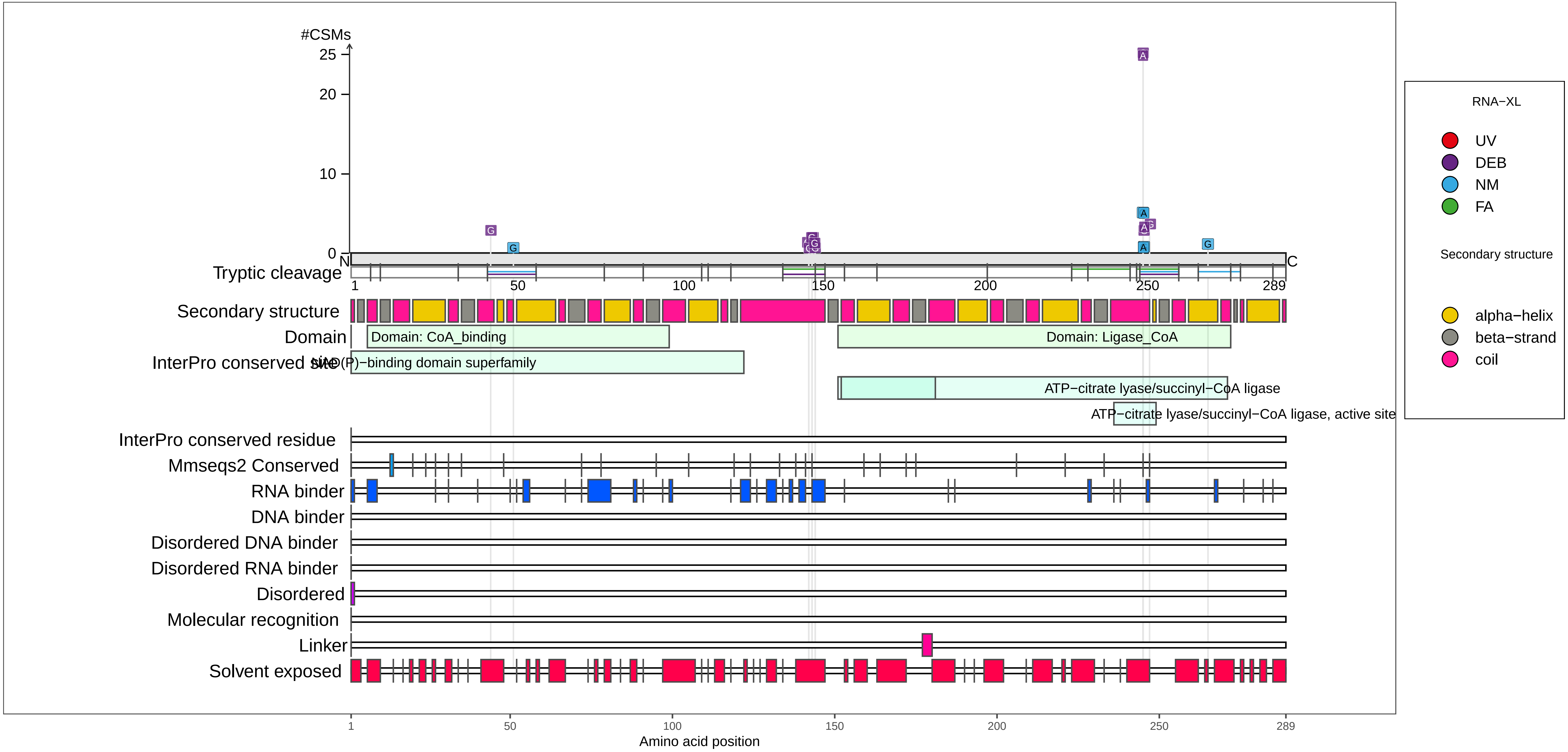

P0A7J7
RL11_ECOLI 50S ribosomal protein L11

− Abundance:
tryptic [log10 Intensity]: 9.33 (Q 94)
PAXdb K12 strain [ppm]: 3.51 (Q 98)
PAXdb E.coli [ppm]: 3.28 (Q 97)

− RNA functions:
Ribosomal protein L11, RNA binding domain; RNA binding; rRNA binding

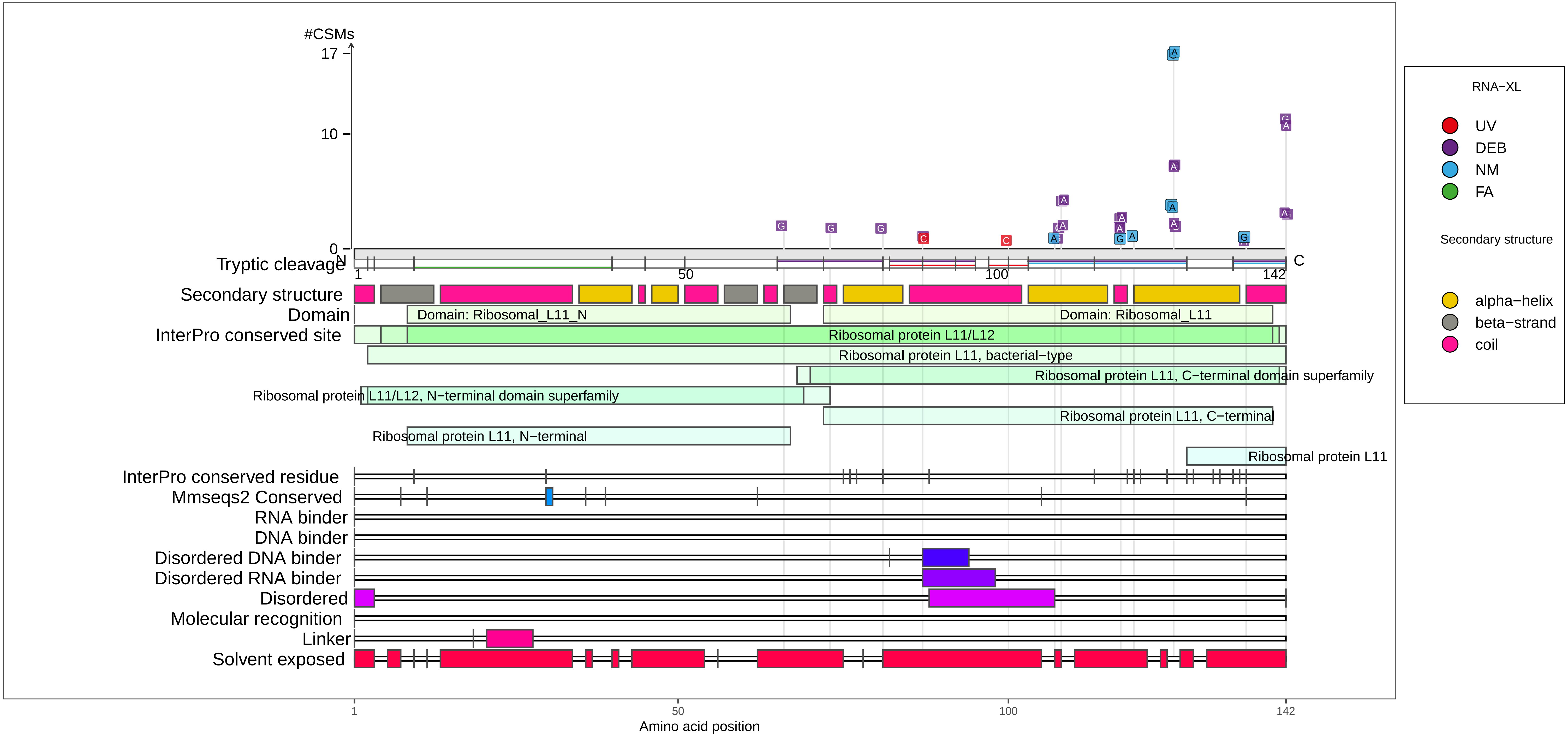

P0A7U7
RS20_ECOLI 30S ribosomal protein S20

− Abundance:
tryptic [log10 Intensity]: 8.9 (Q 86)
PAXdb K12 strain [ppm]: 3.85 (Q 100)
PAXdb E.coli [ppm]: 3.49 (Q 99)

− RNA functions:
RNA binding; rRNA binding

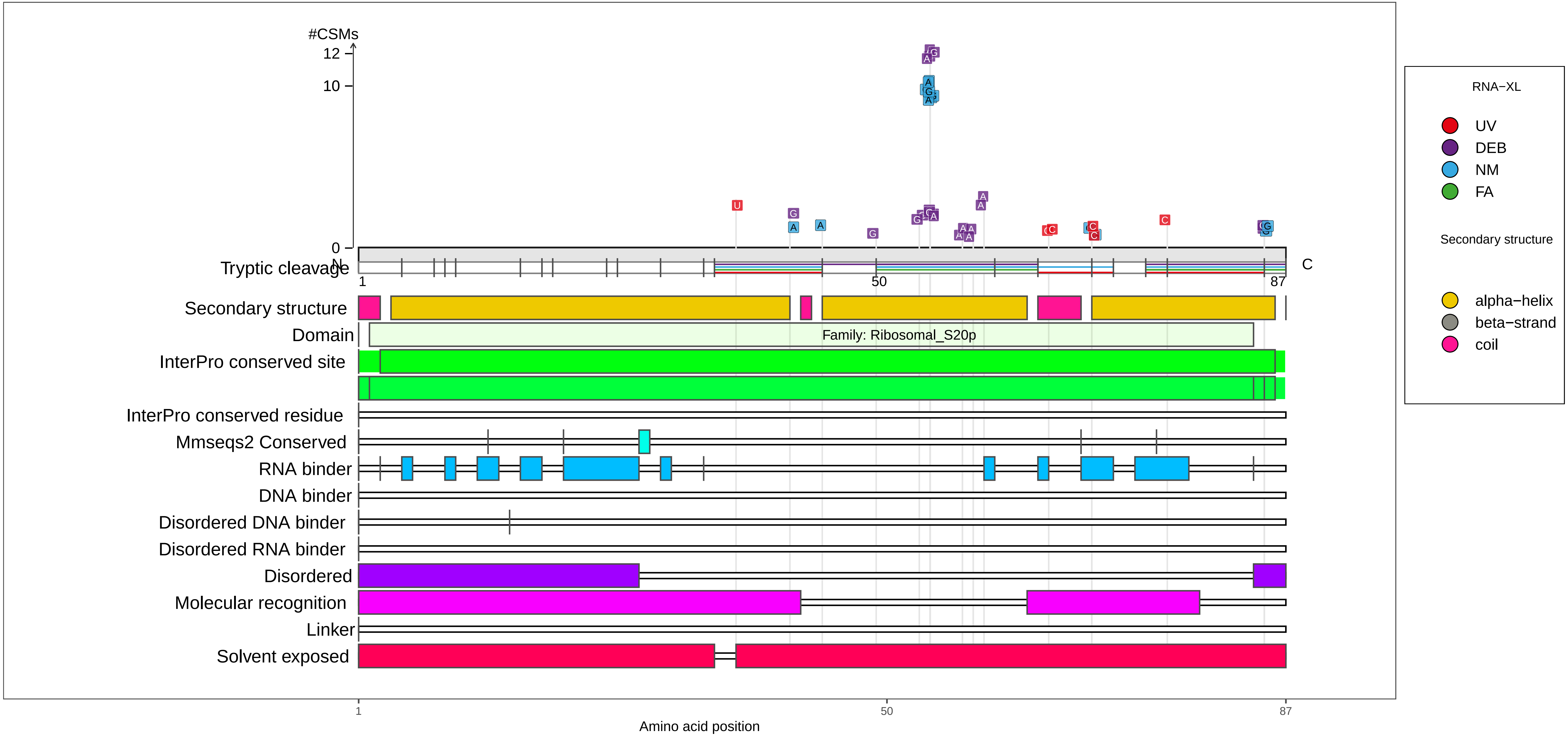

P26646
ACUI_ECOLI Probable acrylyl−CoA reductase AcuI

− Abundance:
tryptic [log10 Intensity]: 8.64 (Q 80)
PAXdb K12 strain [ppm]: 2.75 (Q 84)
PAXdb E.coli [ppm]: 2.63 (Q 89)

− RNA functions: not annotated

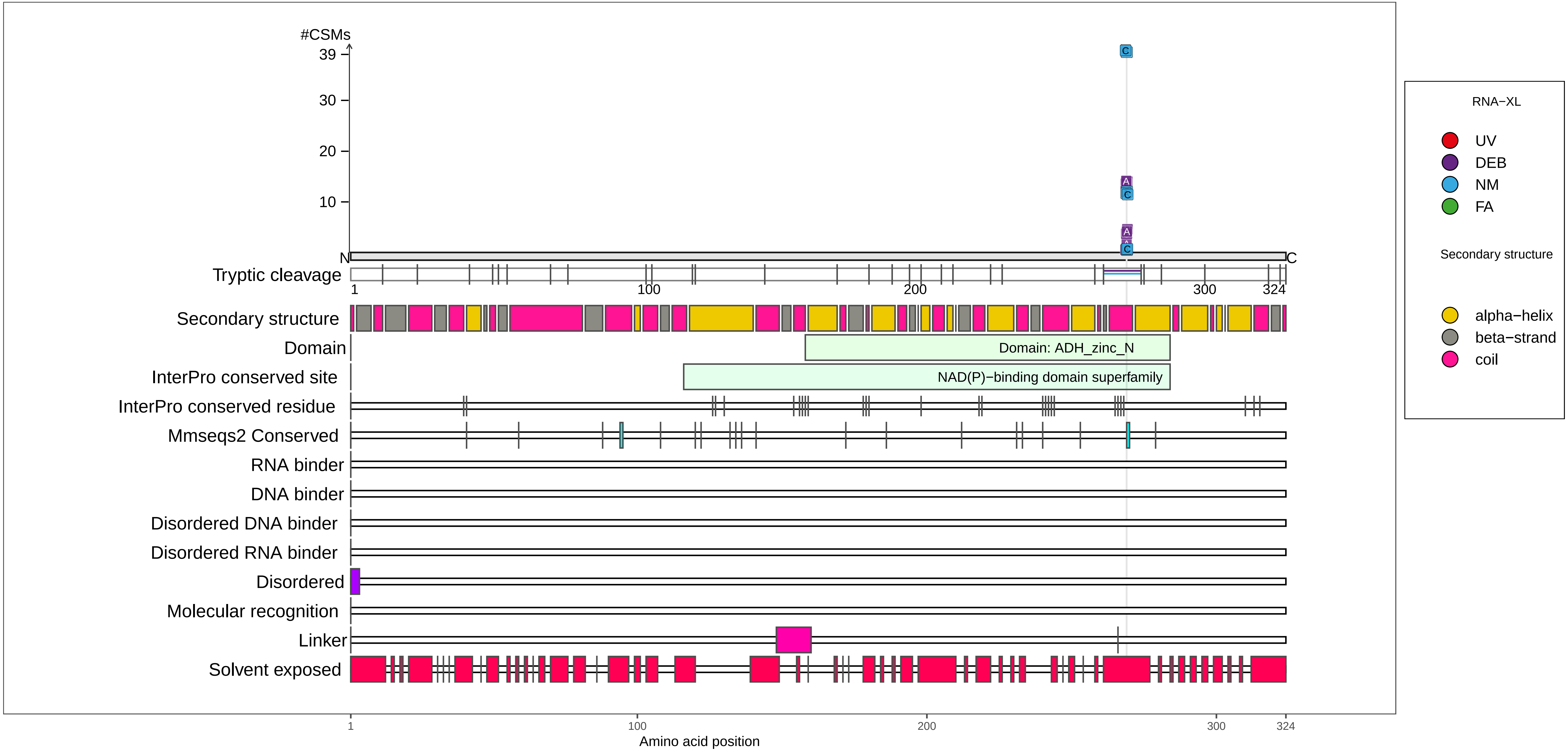

P07118
SYV_ECOLI Valine−−tRNA ligase

− Abundance:
tryptic [log10 Intensity]: 9.77 (Q 99)
PAXdb K12 strain [ppm]: 2.76 (Q 85)
PAXdb E.coli [ppm]: 2.67 (Q 89)

− RNA functions:
aminoacyl−tRNA editing activity; aminoacyl−tRNA ligase activity
aminoacyl−tRNA metabolism involved in translational fidelity; Anticodon−binding domain of tRNA
Leucyl−tRNA synthetase, Domain 2; ncRNA metabolic process; RNA metabolic process
tRNA aminoacylation; tRNA aminoacylation for protein translation; tRNA metabolic process
tRNA synthetases class I (I, L, M and V); tRNA synthetases class I (M)
valine−tRNA ligase activity; Valyl tRNA synthetase tRNA binding arm; valyl−tRNA aminoacylation

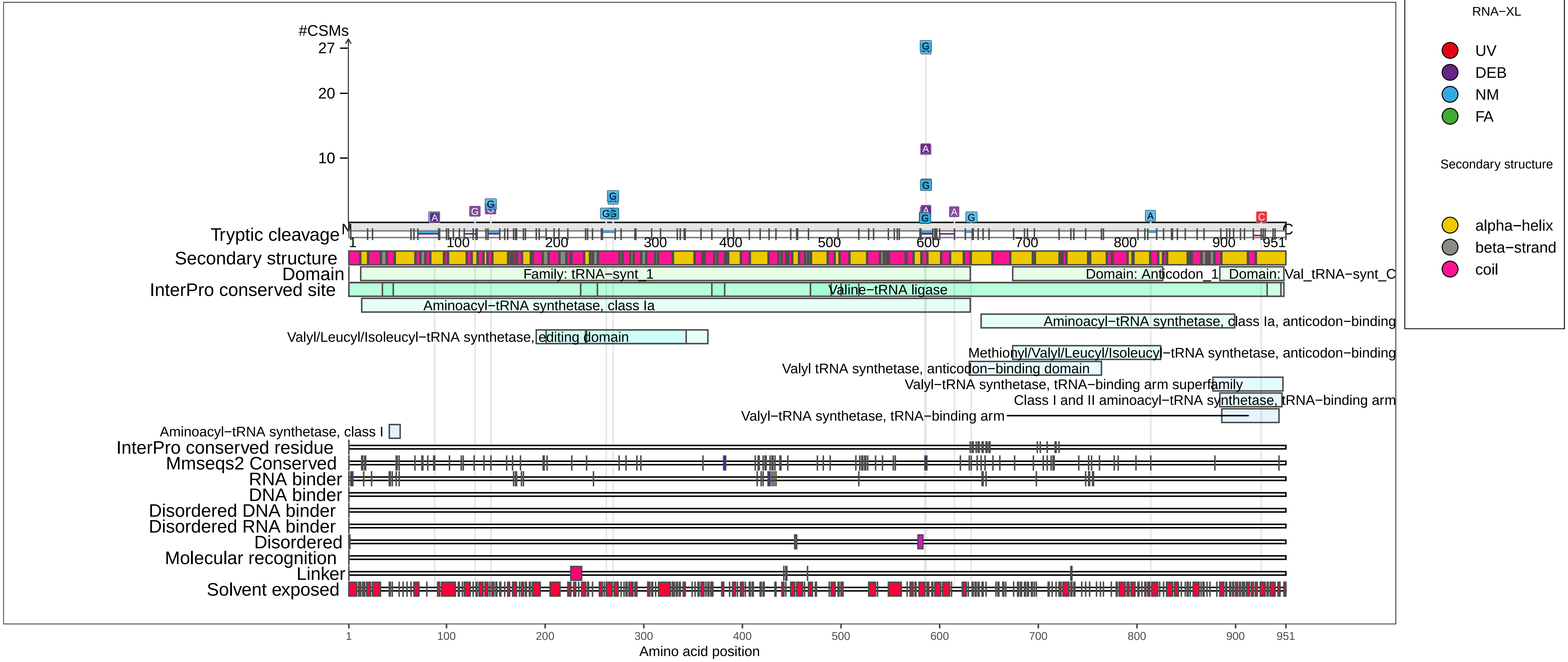

P0AG63
RS17_ECOLI 30S ribosomal protein S17

− Abundance:
tryptic [log10 Intensity]: 9.84 (Q 99)
PAXdb K12 strain [ppm]: 3.83 (Q 100)
PAXdb E.coli [ppm]: 2.92 (Q 94)

− RNA functions:
RNA binding; rRNA binding

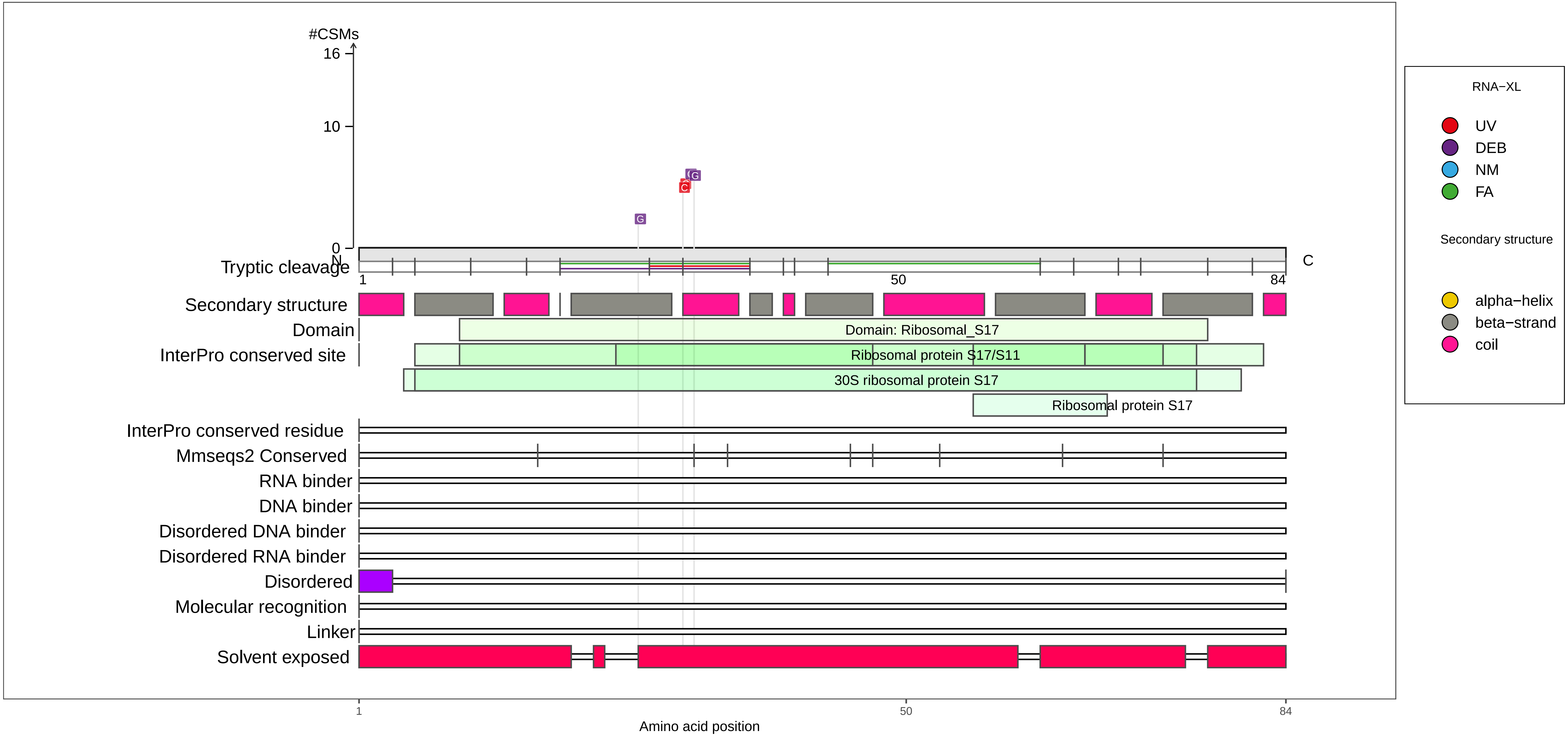

P35340
AHPF_ECOLI Alkyl hydroperoxide reductase subunit F

− Abundance:
tryptic [log10 Intensity]: 9.34 (Q 95)
PAXdb K12 strain [ppm]: 2.69 (Q 83)
PAXdb E.coli [ppm]: 2.45 (Q 84)

− RNA functions: not annotated

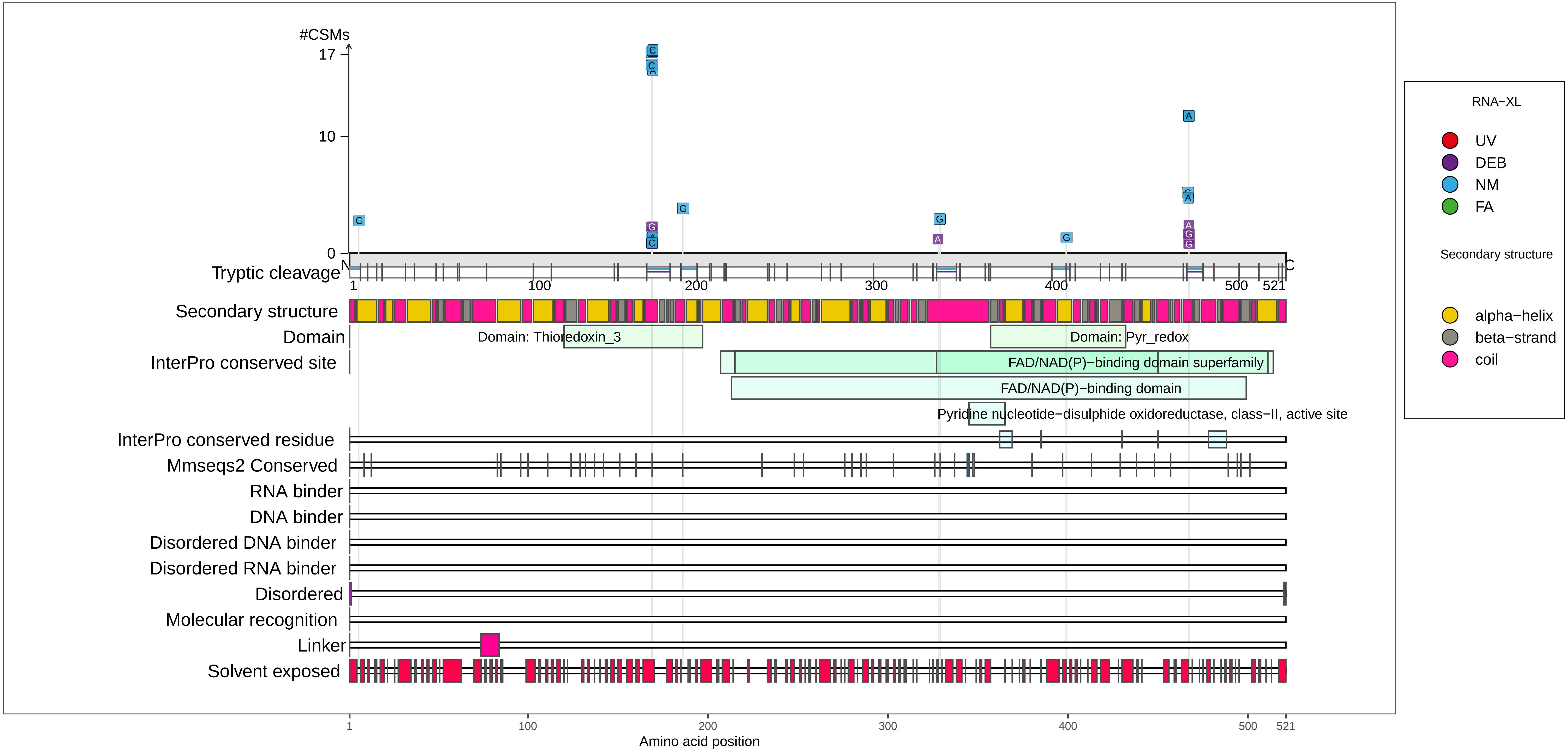

P0AGD7
SRP54_ECOLI Signal recognition particle protein

− Abundance:
tryptic [log10 Intensity]: 6.56 (Q 3)
PAXdb K12 strain [ppm]: 2.57 (Q 79)
PAXdb E.coli [ppm]: 2.31 (Q 81)

− RNA functions:
7S RNA binding; RNA binding

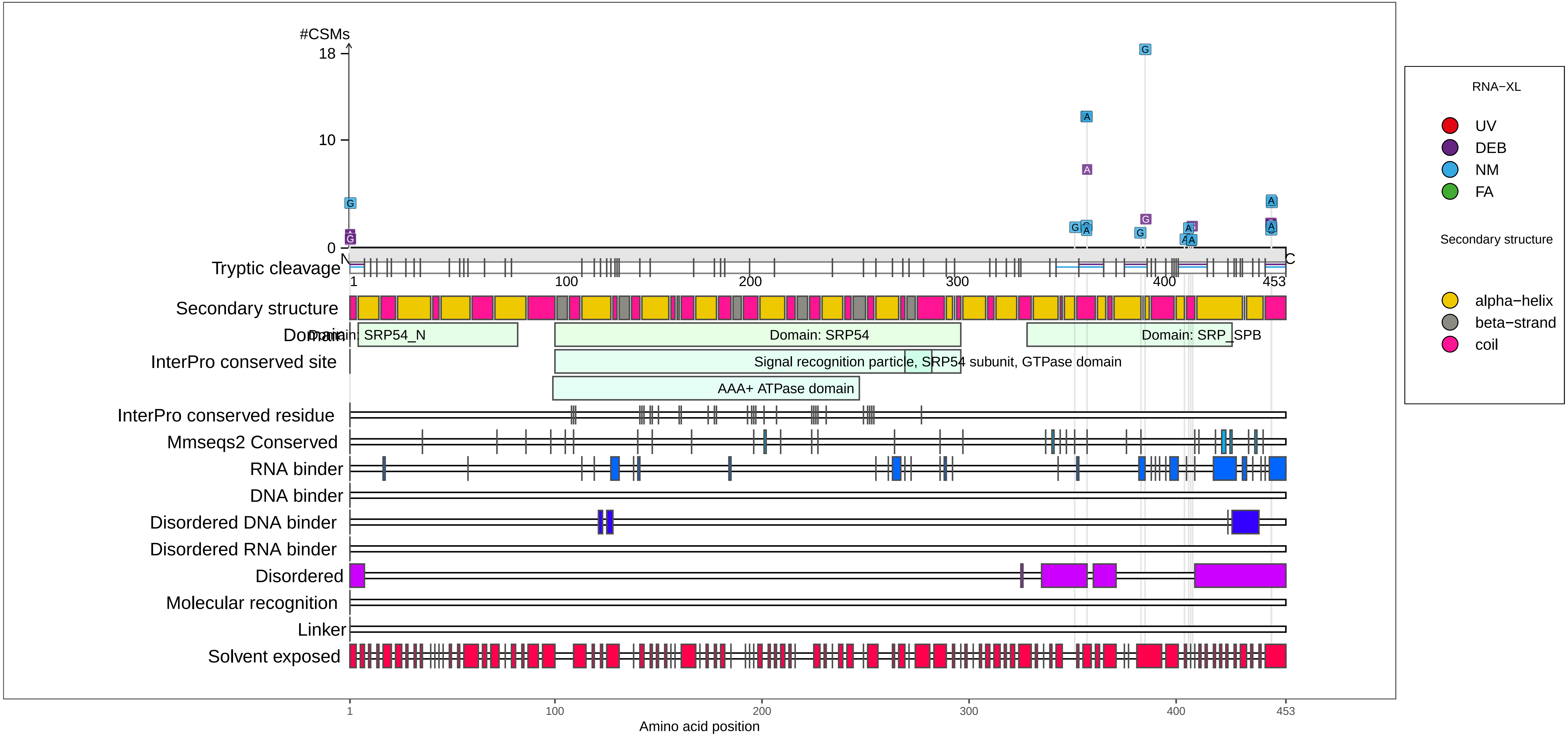

P13035
GLPD_ECOLI Aerobic glycerol−3−phosphate dehydrogenase

− Abundance:
tryptic [log10 Intensity]: 9.67 (Q 98)
PAXdb K12 strain [ppm]: 2.25 (Q 68)
PAXdb E.coli [ppm]: 2.55 (Q 87)

− RNA functions: not annotated

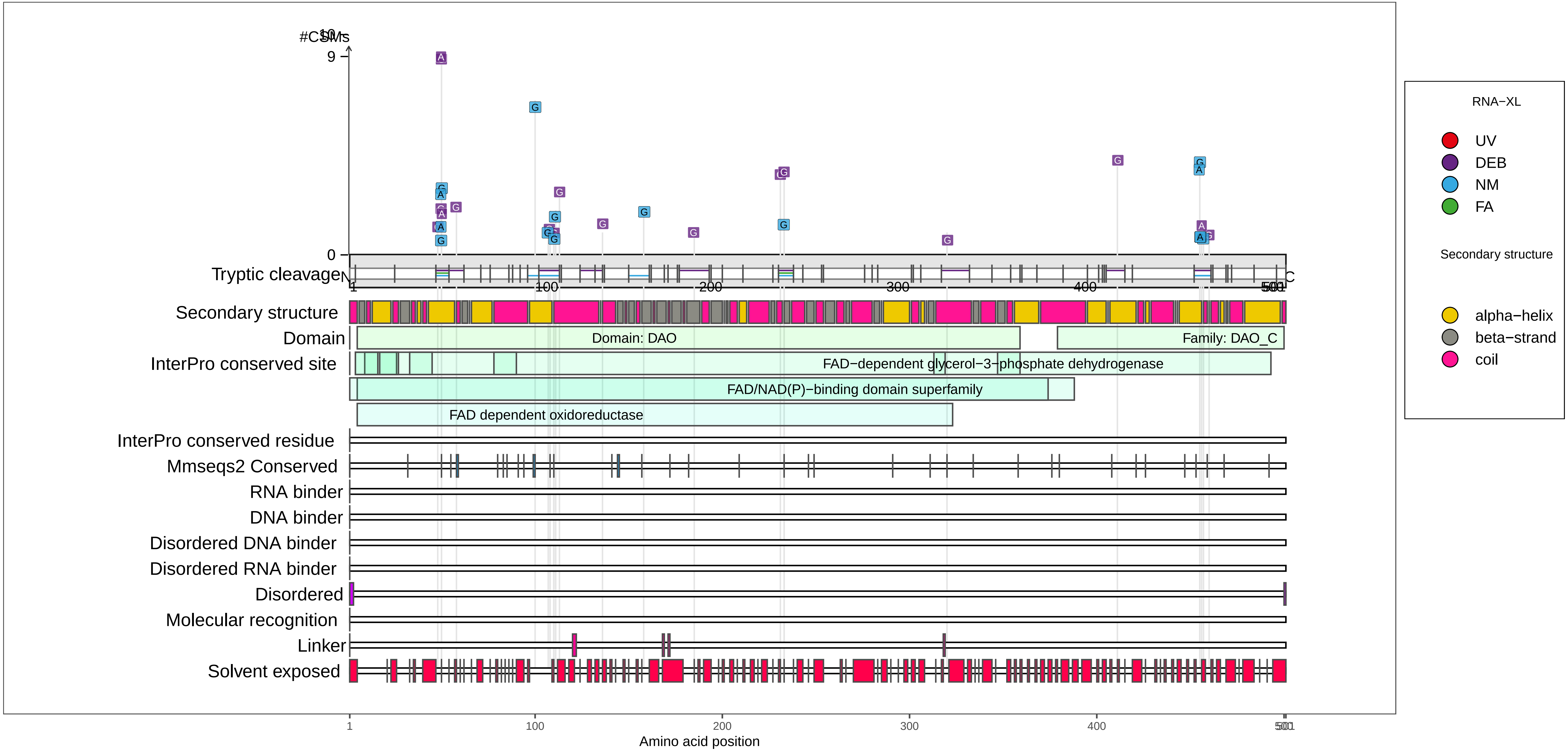

P0A799
PGK_ECOLI Phosphoglycerate kinase

− Abundance:
tryptic [log10 Intensity]: 9.45 (Q 96)
PAXdb K12 strain [ppm]: 3.17 (Q 94)
PAXdb E.coli [ppm]: 3.74 (Q 100)

− RNA functions: not annotated

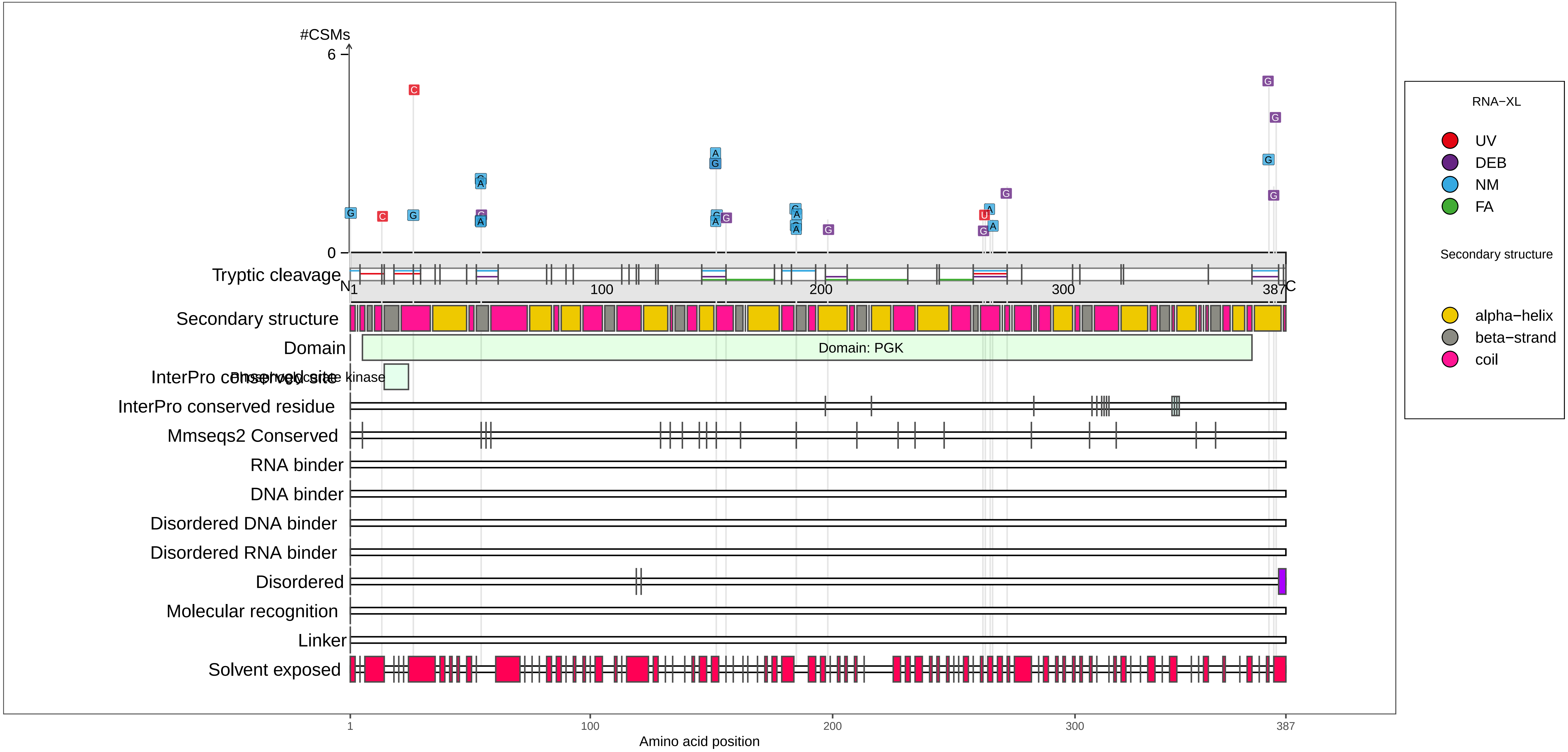

P69222
IF1_ECOLI Translation initiation factor IF−1

− Abundance:
tryptic [log10 Intensity]: 8.73 (Q 82)
PAXdb K12 strain [ppm]: 3.5 (Q 97)
PAXdb E.coli [ppm]: 3.3 (Q 97)

− RNA functions:
RNA binding; rRNA binding

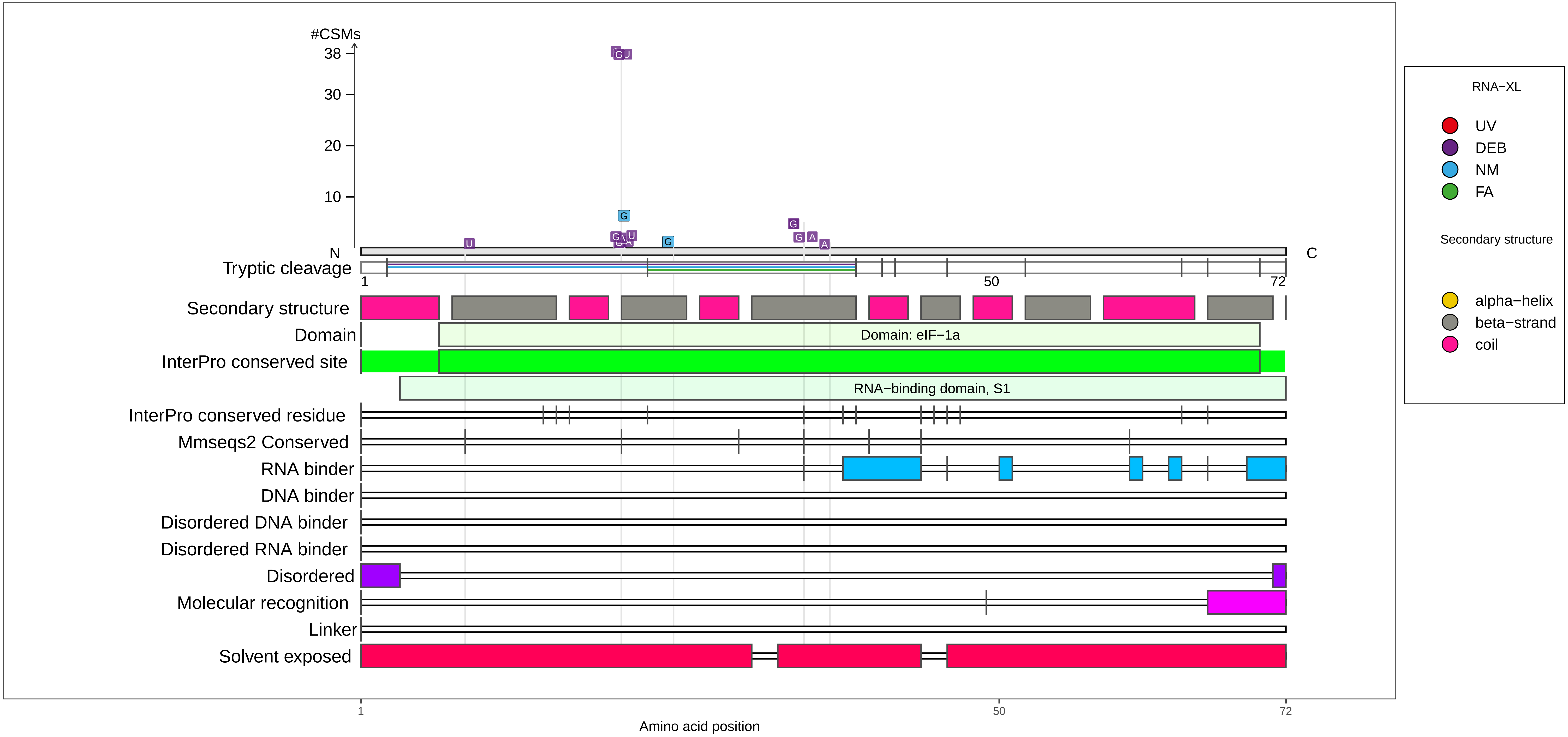

P69776
LPP_ECOLI Major outer membrane lipoprotein Lpp

− Abundance:
tryptic [log10 Intensity]: 10.6 (Q 100)
PAXdb K12 strain [ppm]: 2.77 (Q 85)
PAXdb E.coli [ppm]: 3.58 (Q 100)

− RNA functions: not annotated

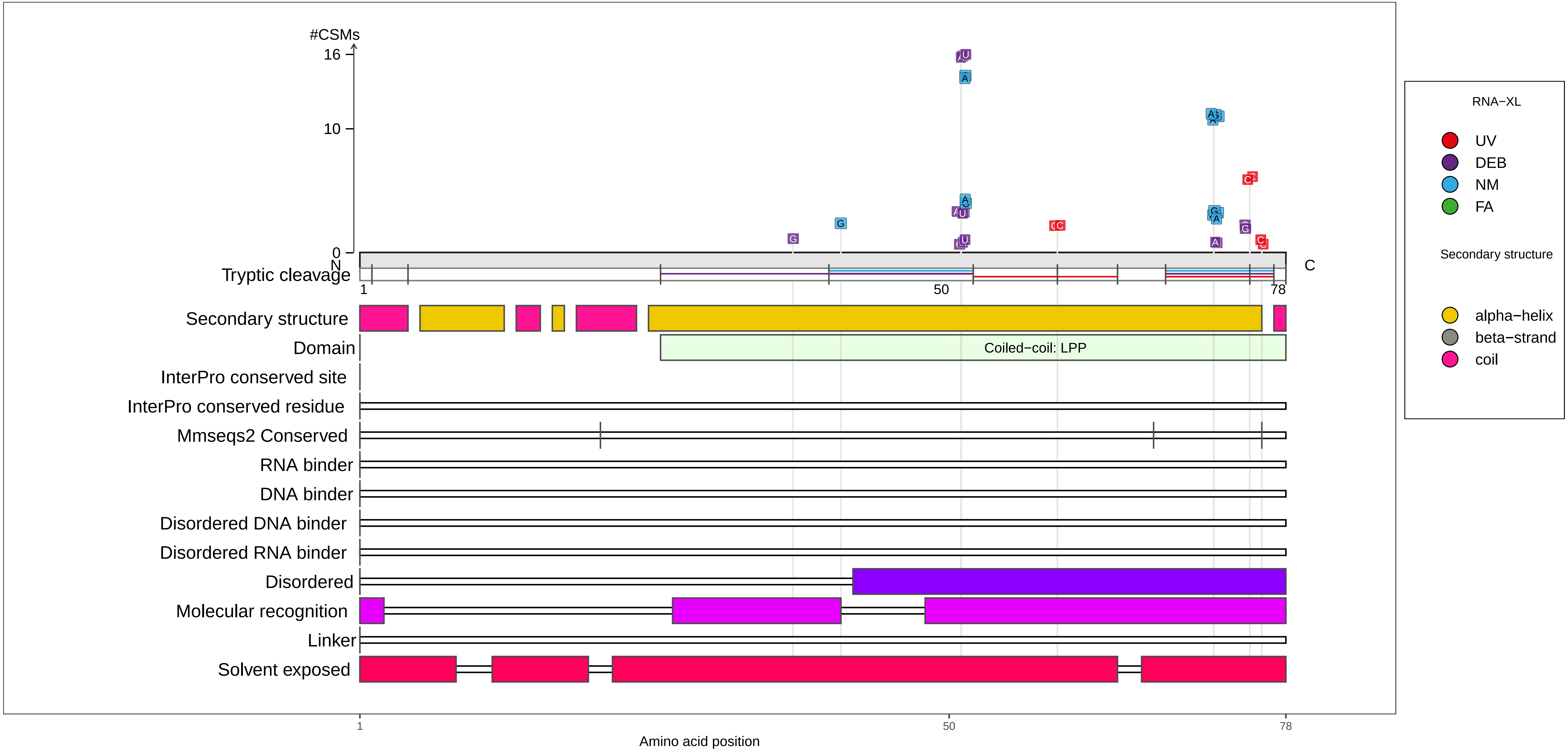

P00956
SYI_ECOLI Isoleucine−−tRNA ligase

− Abundance:
tryptic [log10 Intensity]: 9.76 (Q 99)
PAXdb K12 strain [ppm]: 3 (Q 91)
PAXdb E.coli [ppm]: 2.74 (Q 90)

− RNA functions:
aminoacyl−tRNA editing activity; aminoacyl−tRNA ligase activity
aminoacyl−tRNA metabolism involved in translational fidelity; Anticodon−binding domain of tRNA
isoleucine−tRNA ligase activity; isoleucyl−tRNA aminoacylation
Leucyl−tRNA synthetase, Domain 2; ncRNA metabolic process; RNA binding; RNA metabolic process; tRNA aminoacylation
tRNA aminoacylation for protein translation; tRNA binding; tRNA metabolic process
tRNA synthetases class I (C) catalytic domain
tRNA synthetases class I (I, L, M and V); tRNA synthetases class I (K); tRNA synthetases class I (M)

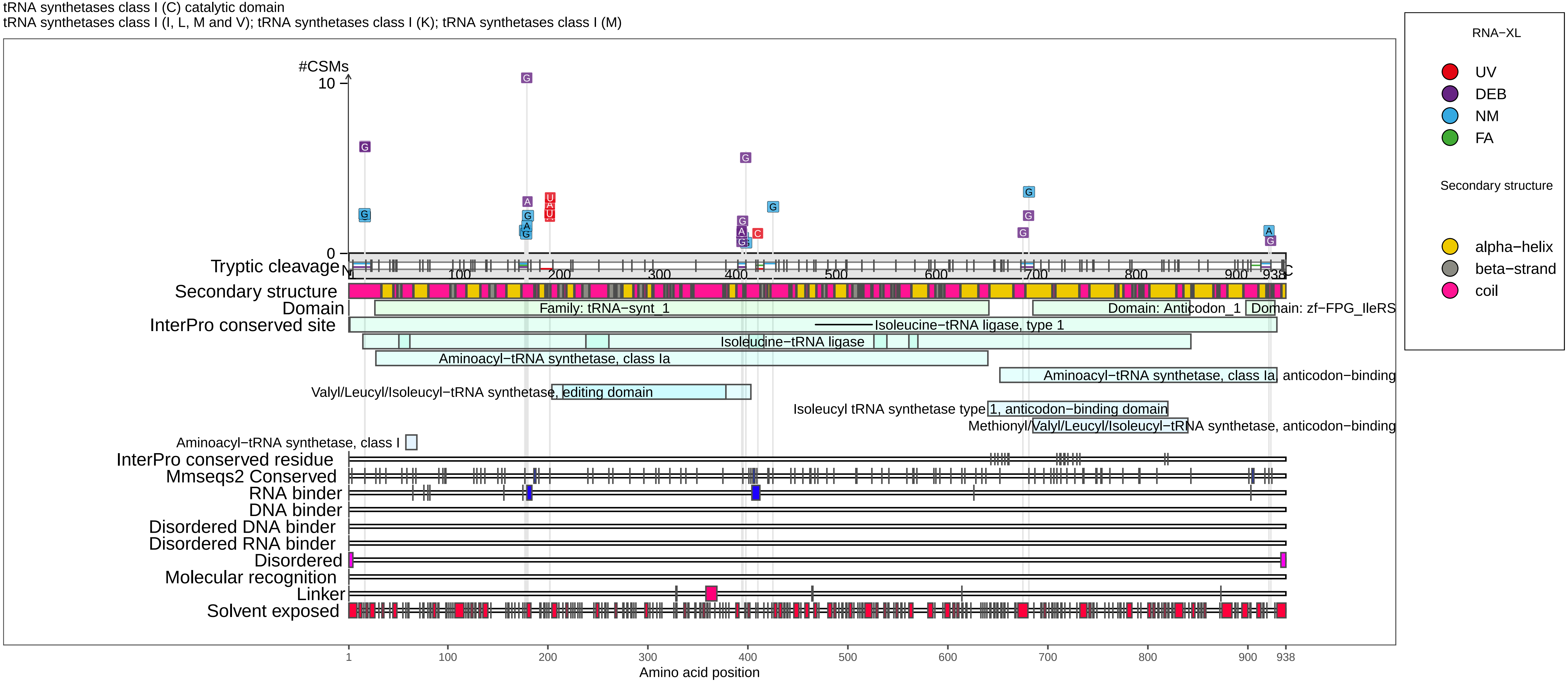

P63020
NFUA_ECOLI Fe/S biogenesis protein NfuA

− Abundance:
tryptic [log10 Intensity]: 8.33 (Q 70)
PAXdb K12 strain [ppm]: 2.66 (Q 82)
PAXdb E.coli [ppm]: 3.09 (Q 96)

− RNA functions: not annotated

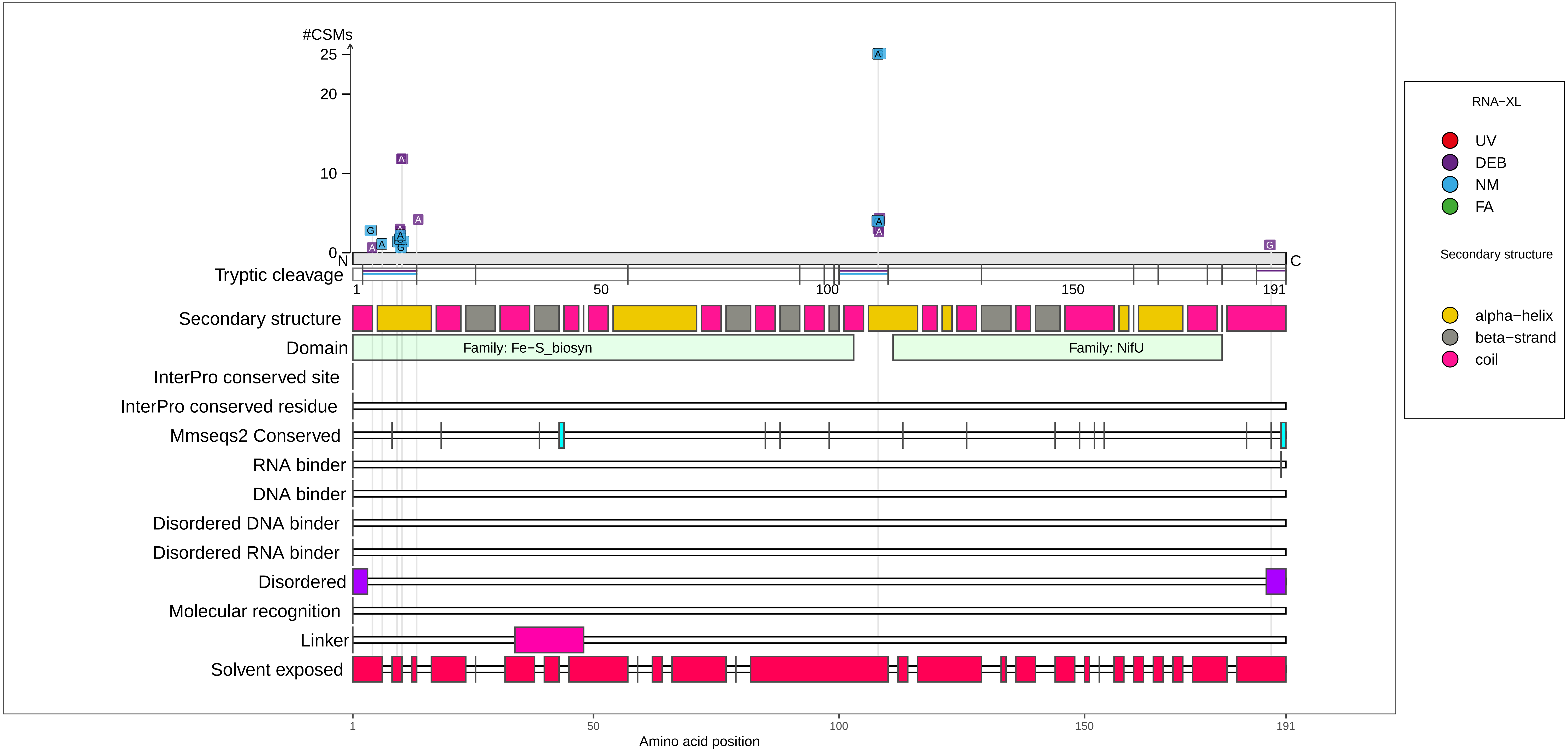

P0A6H5
HSLU_ECOLI ATP−dependent protease ATPase subunit HslU

− Abundance:
tryptic [log10 Intensity]: 9.37 (Q 95)
PAXdb K12 strain [ppm]: 3.26 (Q 95)
PAXdb E.coli [ppm]: 2.8 (Q 92)

− RNA functions: not annotated

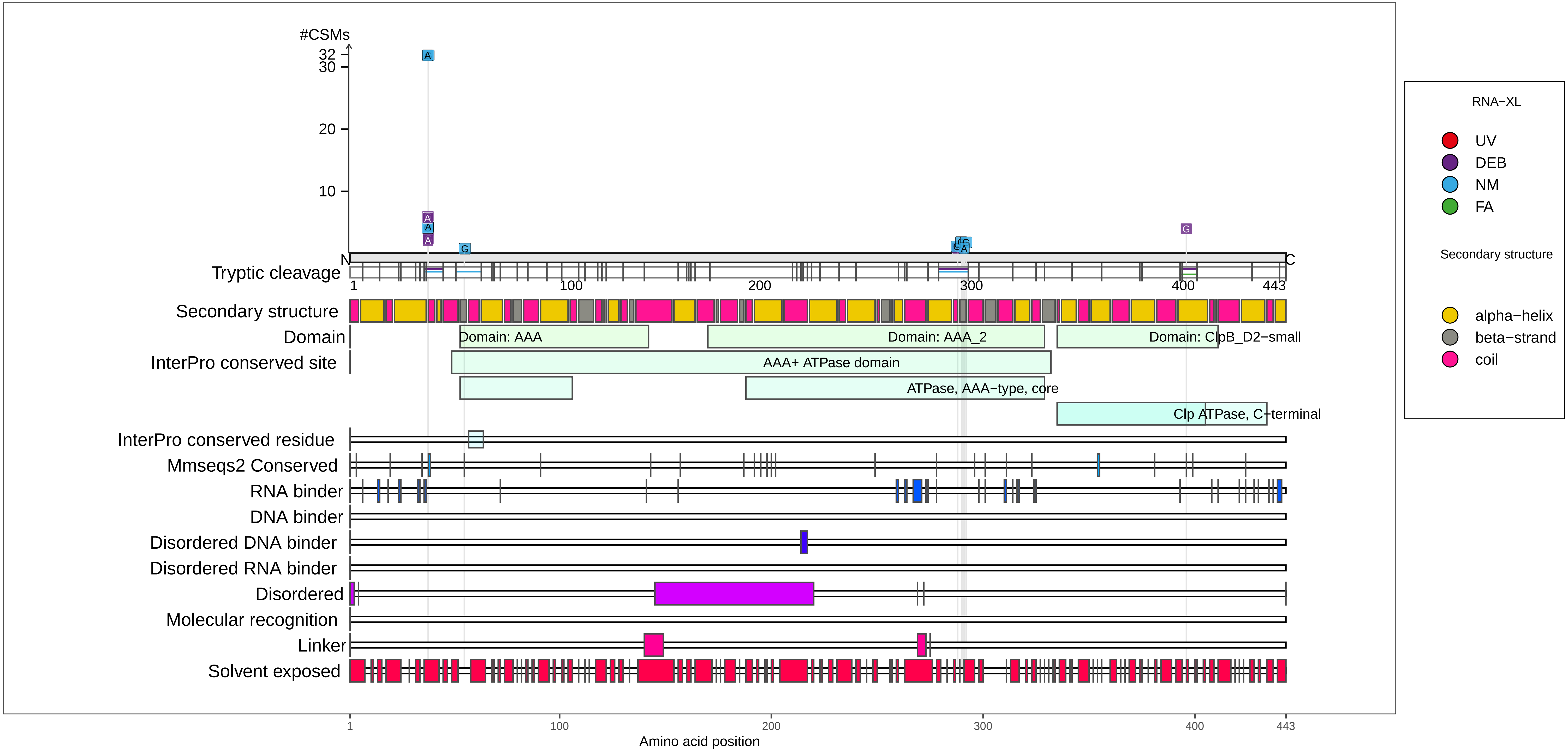

P0A7T7
RS18_ECOLI 30S ribosomal protein S18

− Abundance:
tryptic [log10 Intensity]: 8.82 (Q 84)
PAXdb K12 strain [ppm]: 3.88 (Q 100)
PAXdb E.coli [ppm]: 3.48 (Q 99)

− RNA functions:
mRNA 5−UTR binding; mRNA binding; RNA binding; rRNA binding

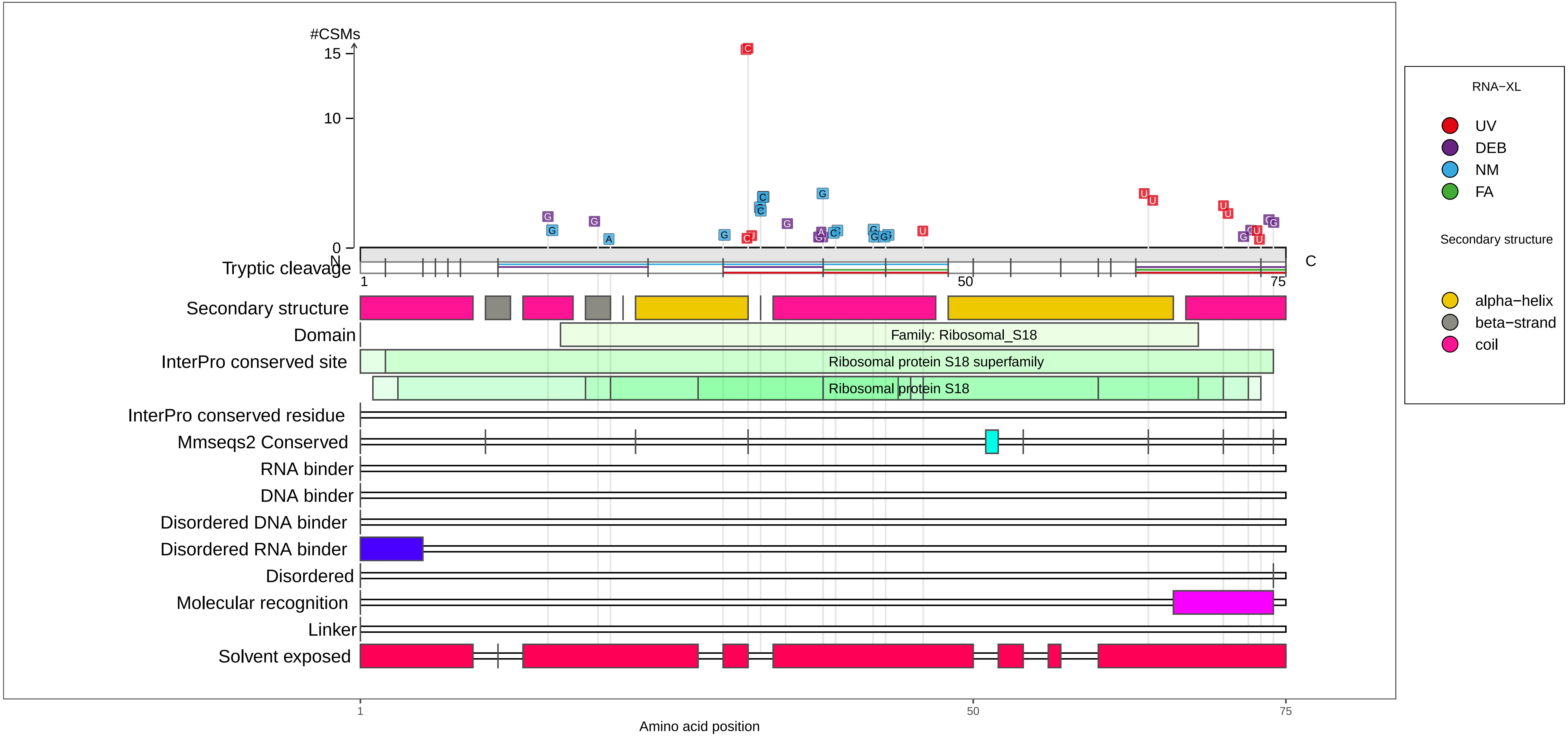

P0AD61
KPYK1_ECOLI Pyruvate kinase I

− Abundance:
tryptic [log10 Intensity]: 9.41 (Q 95)
PAXdb K12 strain [ppm]: 2.78 (Q 86)
PAXdb E.coli [ppm]: 3.18 (Q 96)

− RNA functions: not annotated

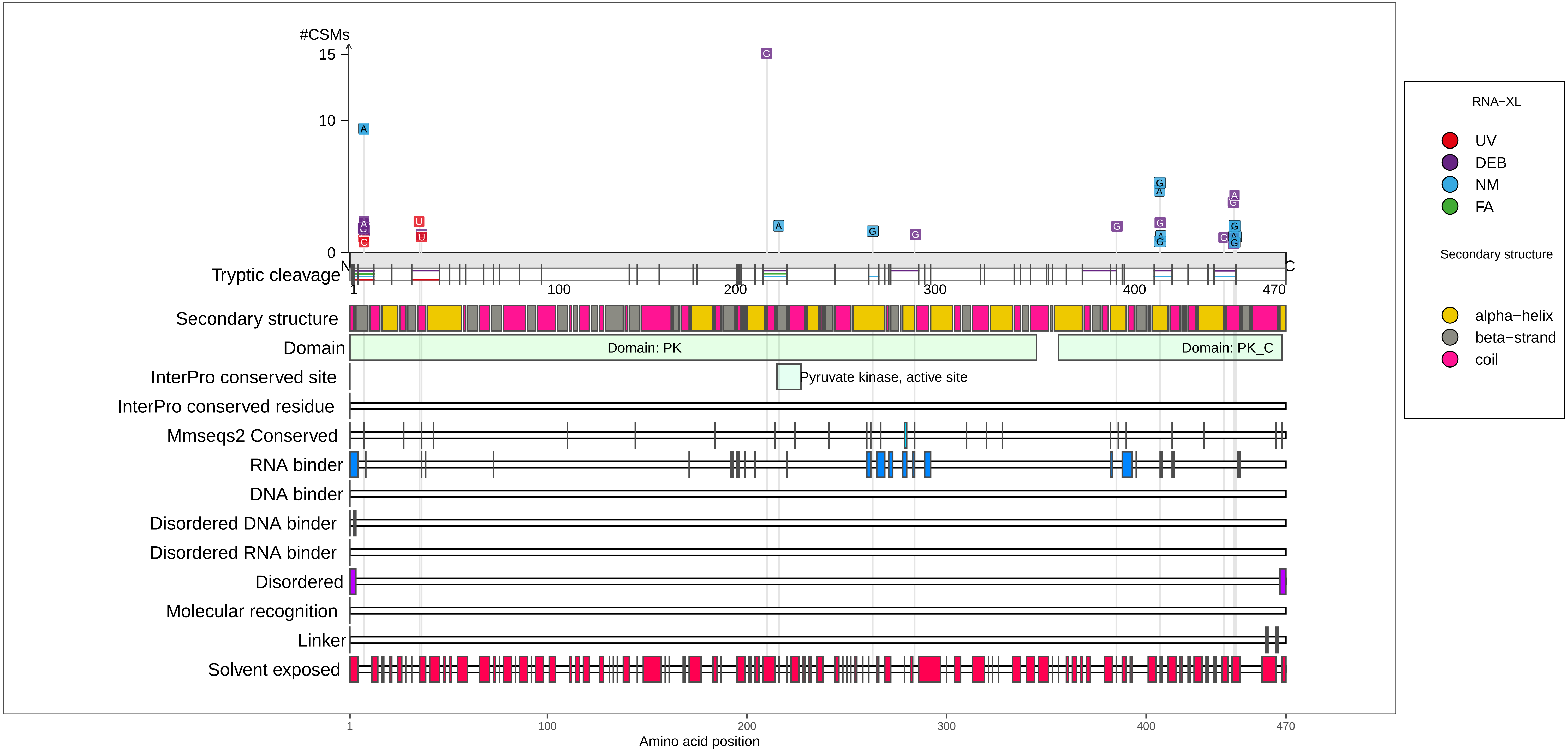

P37744
RMLA1_ECOLI Glucose−1−phosphate thymidylyltransferase 1

− Abundance:
tryptic [log10 Intensity]: 8.4 (Q 72)
PAXdb K12 strain [ppm]: 2.39 (Q 74)
PAXdb E.coli [ppm]: 2.39 (Q 83)

− RNA functions: not annotated

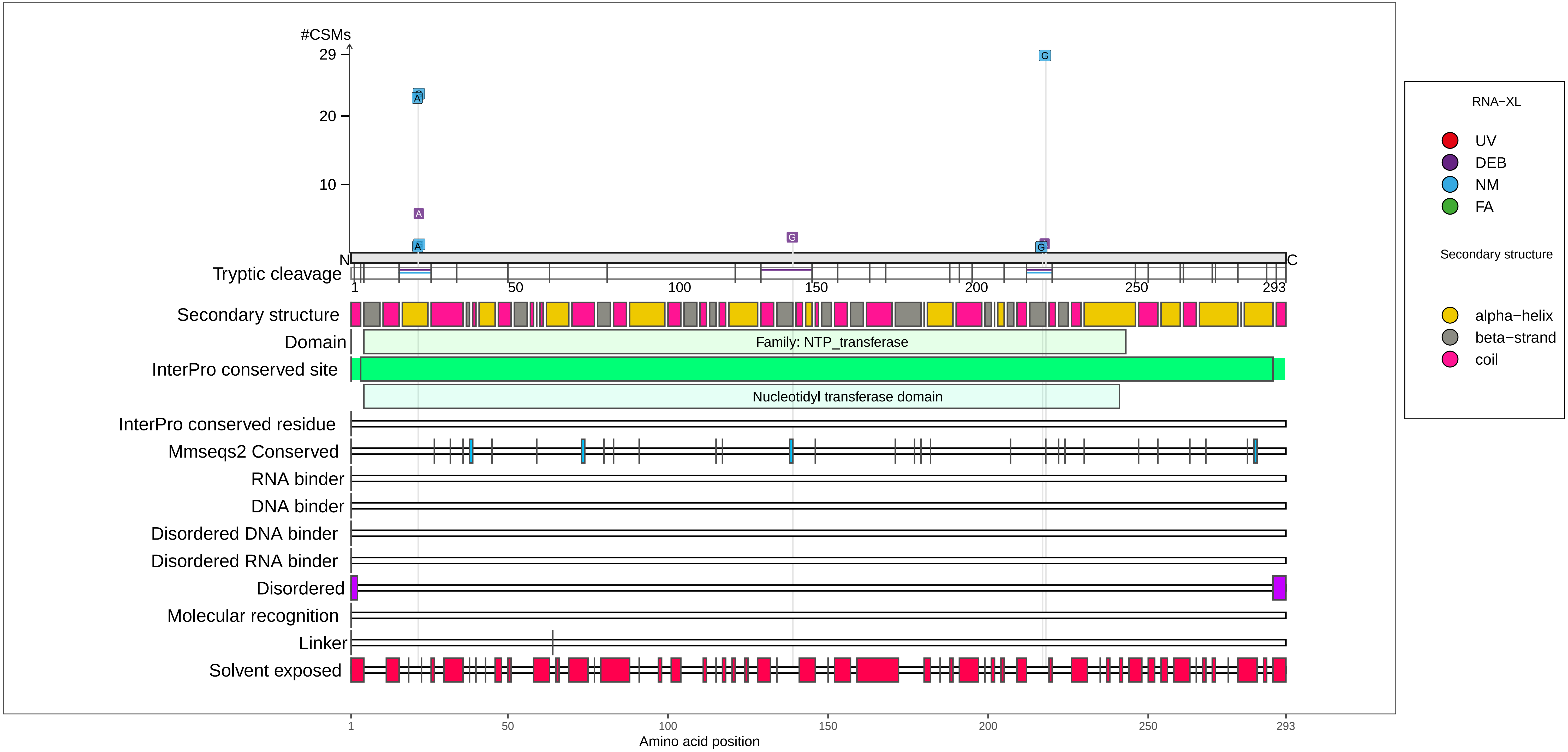

P07001
PNTA_ECOLI NAD(P) transhydrogenase subunit alpha

− Abundance:
tryptic [log10 Intensity]: 9 (Q 88)
PAXdb K12 strain [ppm]: 1.3 (Q 26)
PAXdb E.coli [ppm]: 2.4 (Q 83)

− RNA functions: not annotated

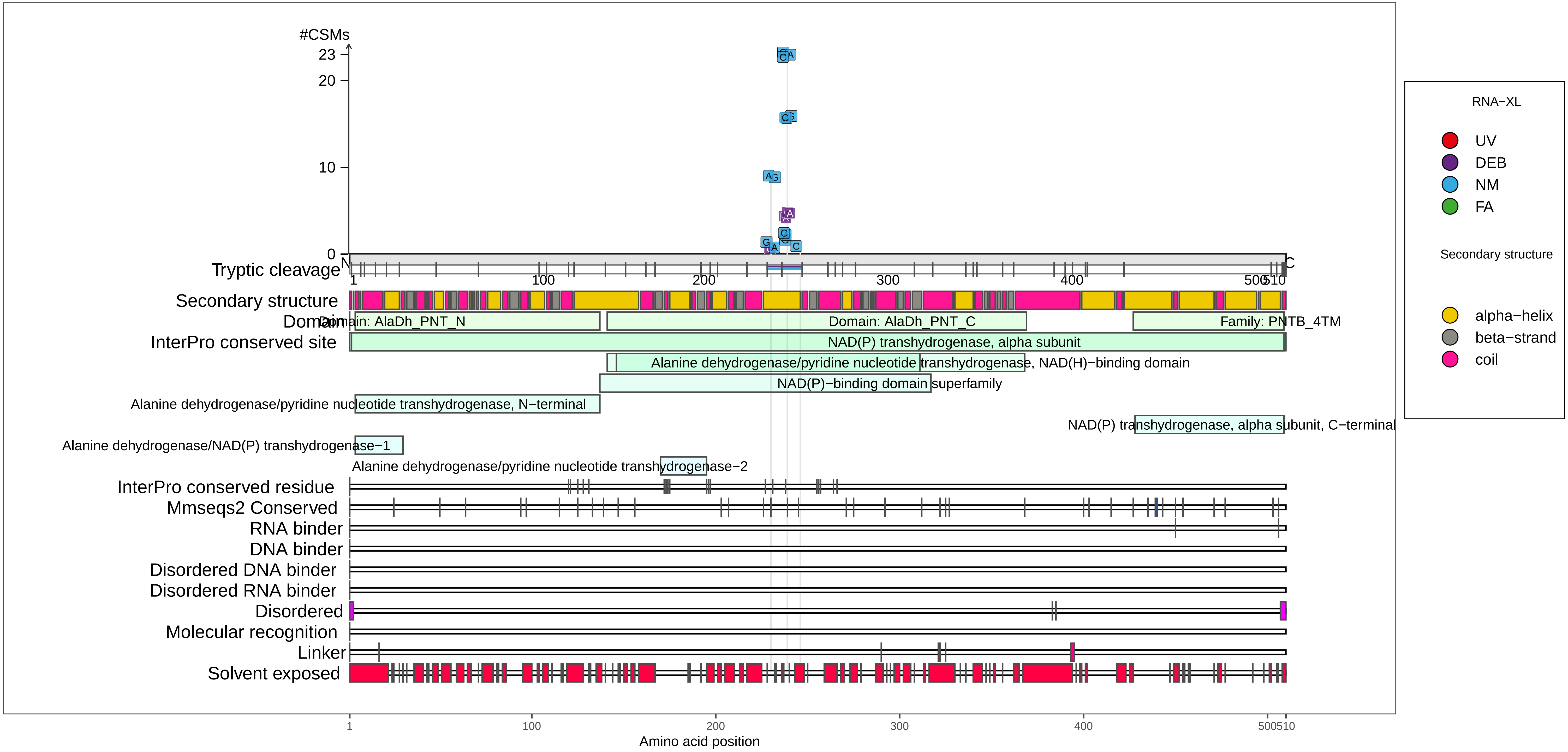

P07650
TYPH_ECOLI Thymidine phosphorylase

− Abundance:
tryptic [log10 Intensity]: 8.93 (Q 86)
PAXdb K12 strain [ppm]: 2.61 (Q 81)
PAXdb E.coli [ppm]: 2.19 (Q 77)

− RNA functions: not annotated

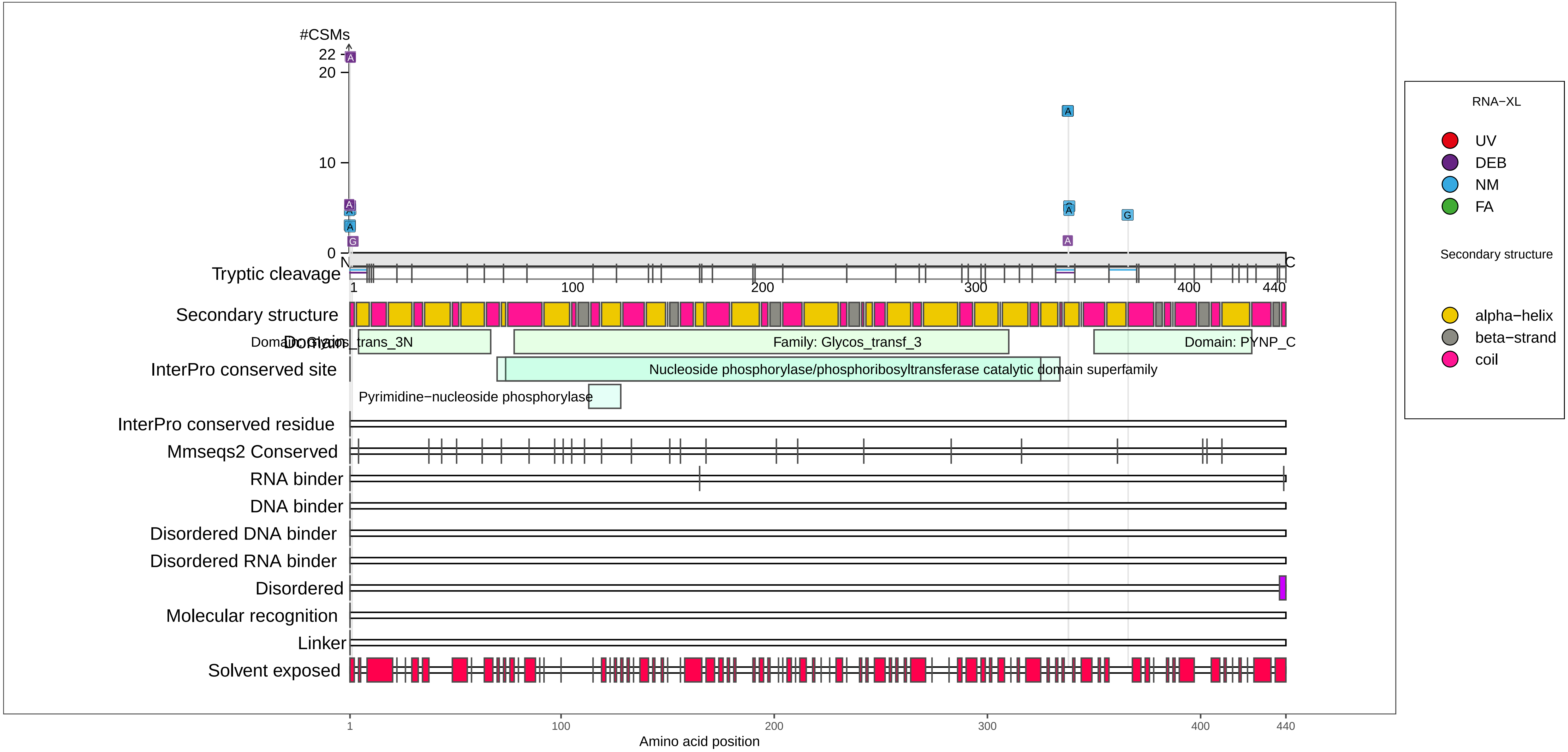

P21889
SYD_ECOLI Aspartate−−tRNA ligase

− Abundance:
tryptic [log10 Intensity]: 9.25 (Q 93)
PAXdb K12 strain [ppm]: 2.37 (Q 73)
PAXdb E.coli [ppm]: 2.9 (Q 93)

− RNA functions:
aminoacyl−tRNA ligase activity; aspartate−tRNA ligase activity
aspartyl−tRNA aminoacylation; ncRNA metabolic process; RNA metabolic process; tRNA aminoacylation
tRNA aminoacylation for protein translation; tRNA metabolic process
tRNA synthetases class II (D, K and N)

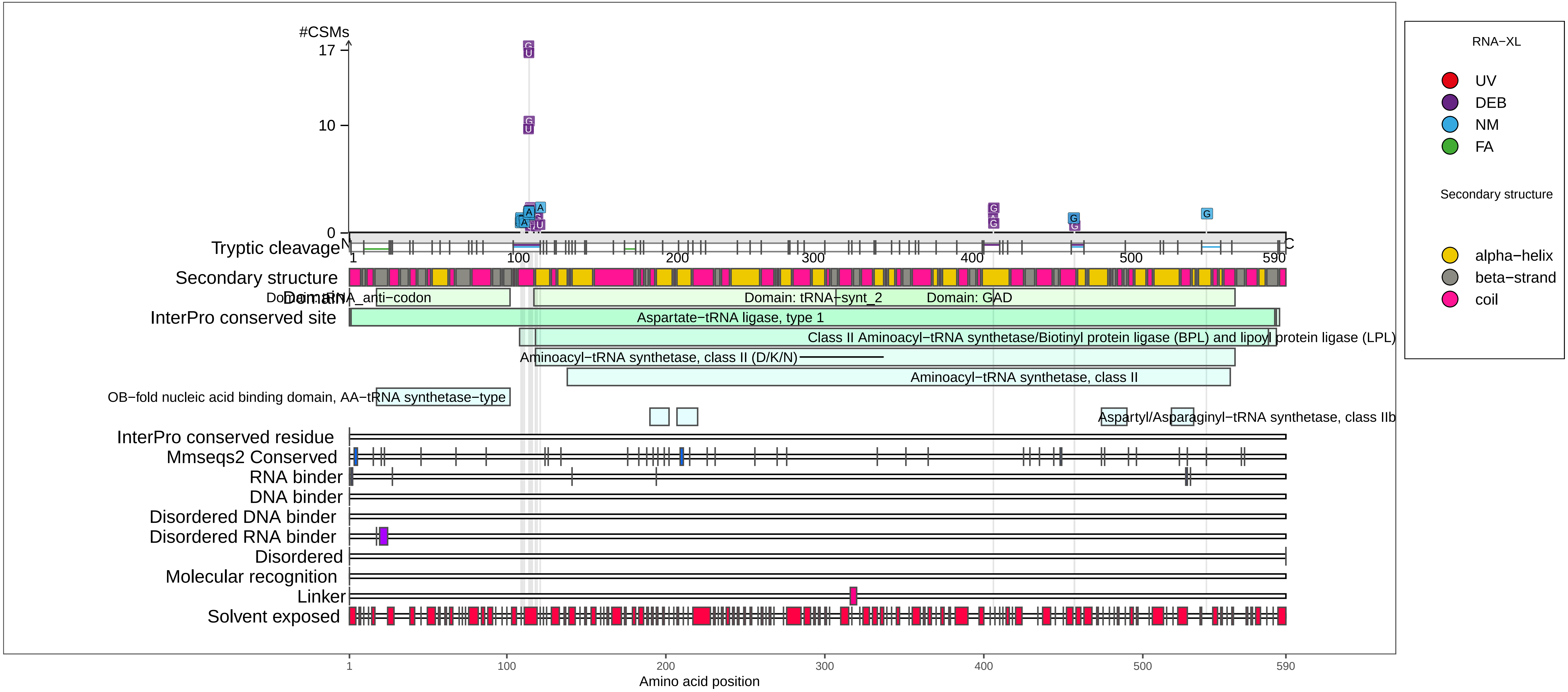

P0A8N5
SYK2_ECOLI Lysine−−tRNA ligase, heat inducible

− Abundance:
tryptic [log10 Intensity]: 9.36 (Q 95)
PAXdb K12 strain [ppm]: 3.3 (Q 95)
PAXdb E.coli [ppm]: 2.34 (Q 82)

− RNA functions:
aminoacyl−tRNA ligase activity; lysine−tRNA ligase activity
lysyl−tRNA aminoacylation; ncRNA metabolic process; RNA binding; RNA metabolic process; tRNA aminoacylation
tRNA aminoacylation for protein translation; tRNA binding; tRNA metabolic process
tRNA synthetases class II (D, K and N)

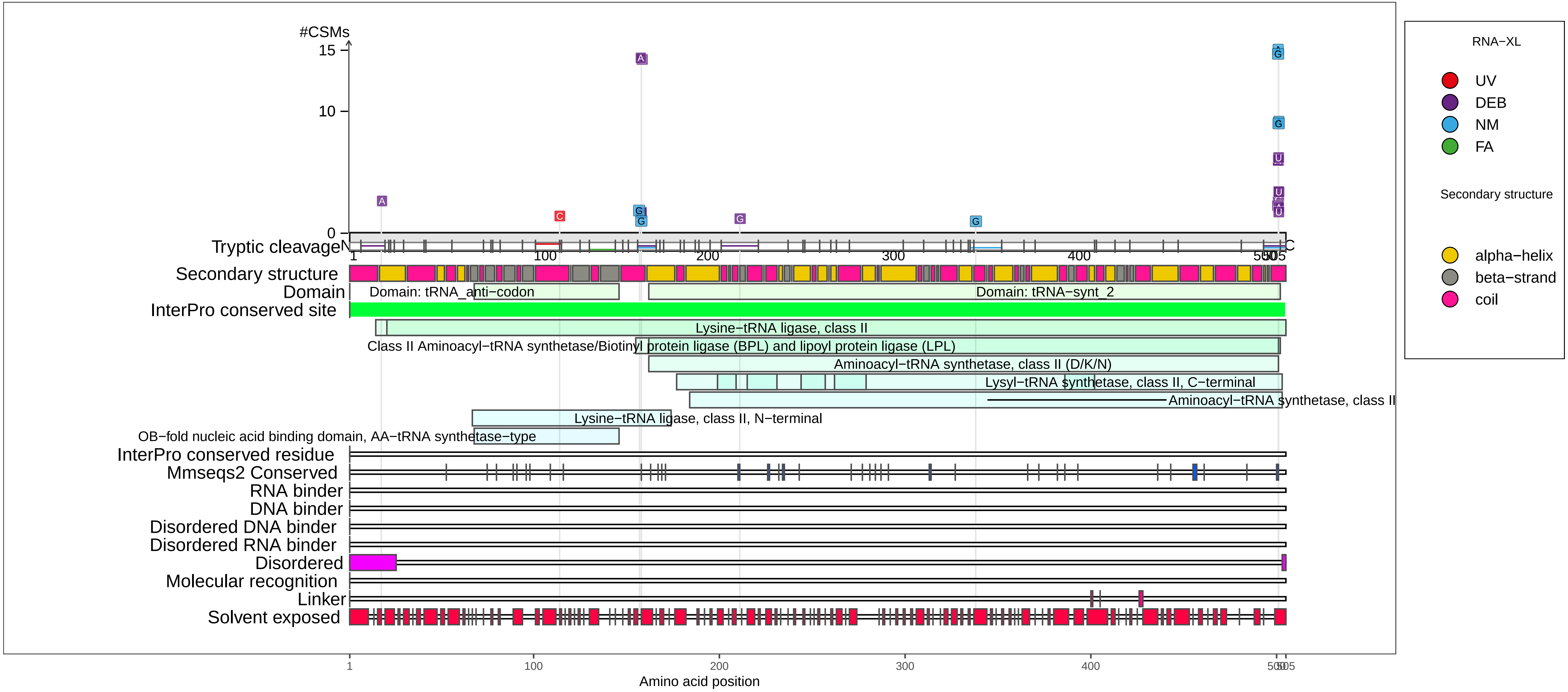

P17117
NFSA_ECOLI Oxygen−insensitive NADPH nitroreductase

− Abundance:
tryptic [log10 Intensity]: 8.93 (Q 87)
PAXdb K12 strain [ppm]: 2.86 (Q 88)
PAXdb E.coli [ppm]: 2 (Q 72)

− RNA functions: not annotated

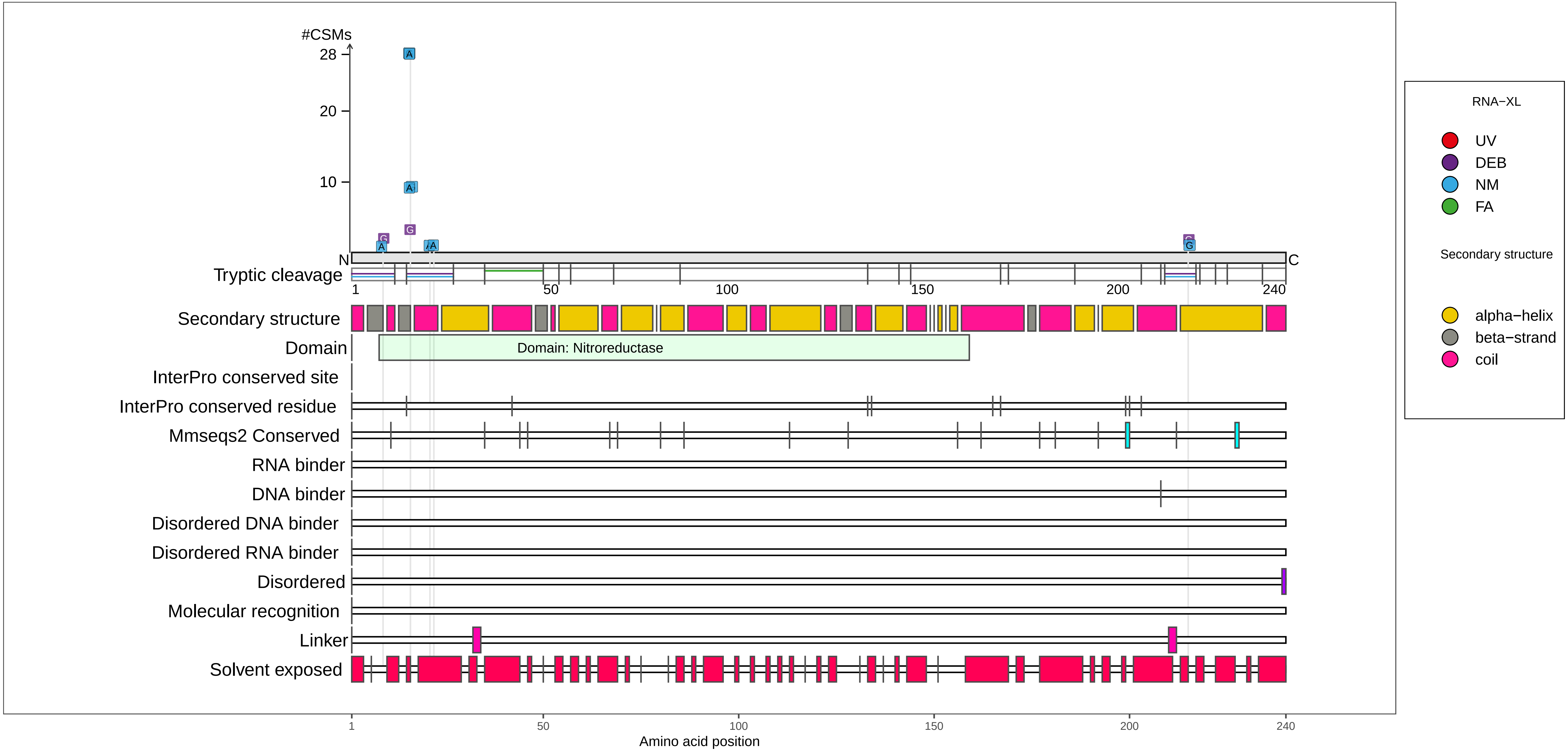

P38489
NFSB_ECOLI Oxygen−insensitive NAD(P)H nitroreductase

− Abundance:
tryptic [log10 Intensity]: 8.98 (Q 88)
PAXdb K12 strain [ppm]: 3.52 (Q 98)
PAXdb E.coli [ppm]: 2.84 (Q 92)

− RNA functions: not annotated

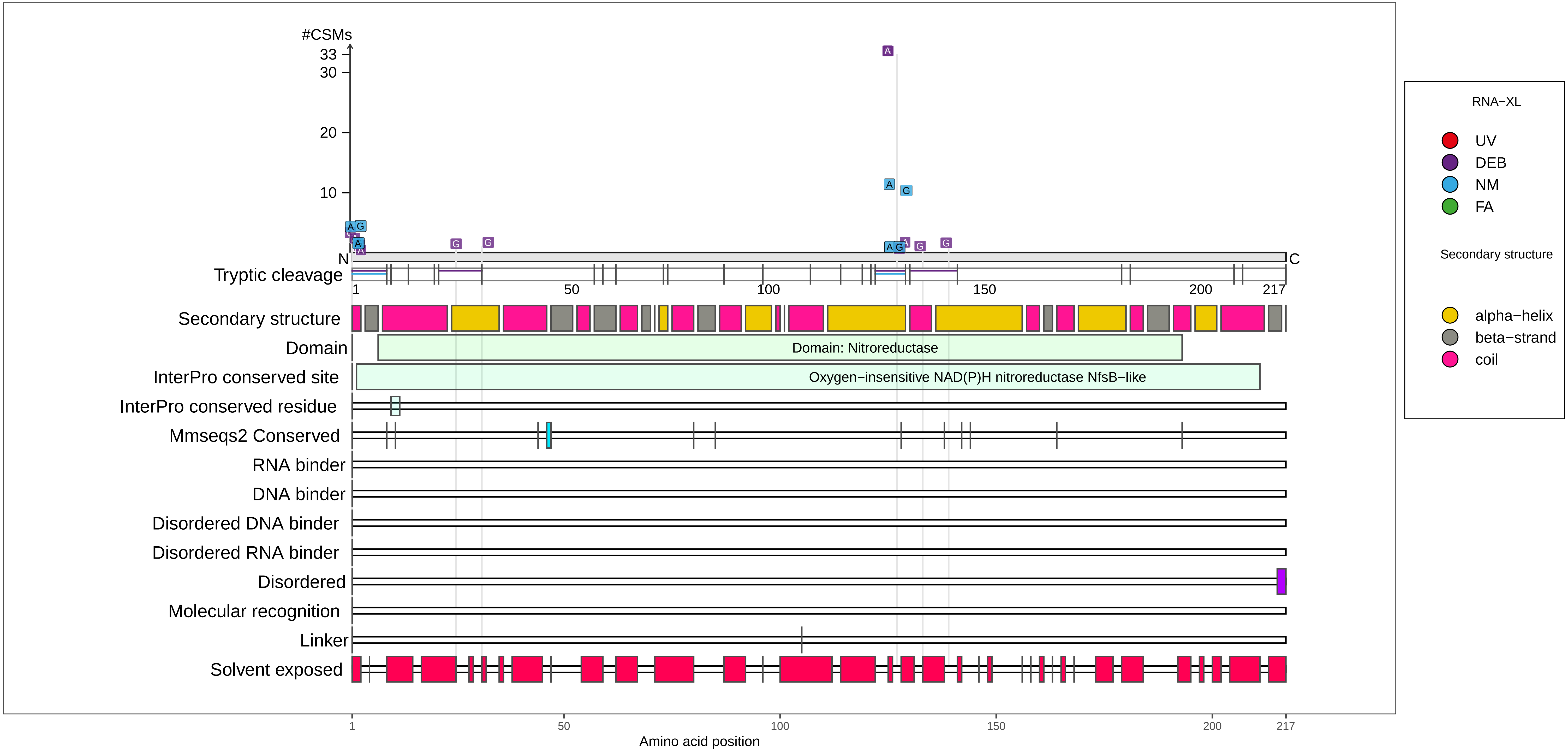

P0A6P9
ENO_ECOLI Enolase

− Abundance:
tryptic [log10 Intensity]: 9.84 (Q 99)
PAXdb K12 strain [ppm]: 3.79 (Q 99)
PAXdb E.coli [ppm]: 3.69 (Q 100)

− RNA functions: not annotated

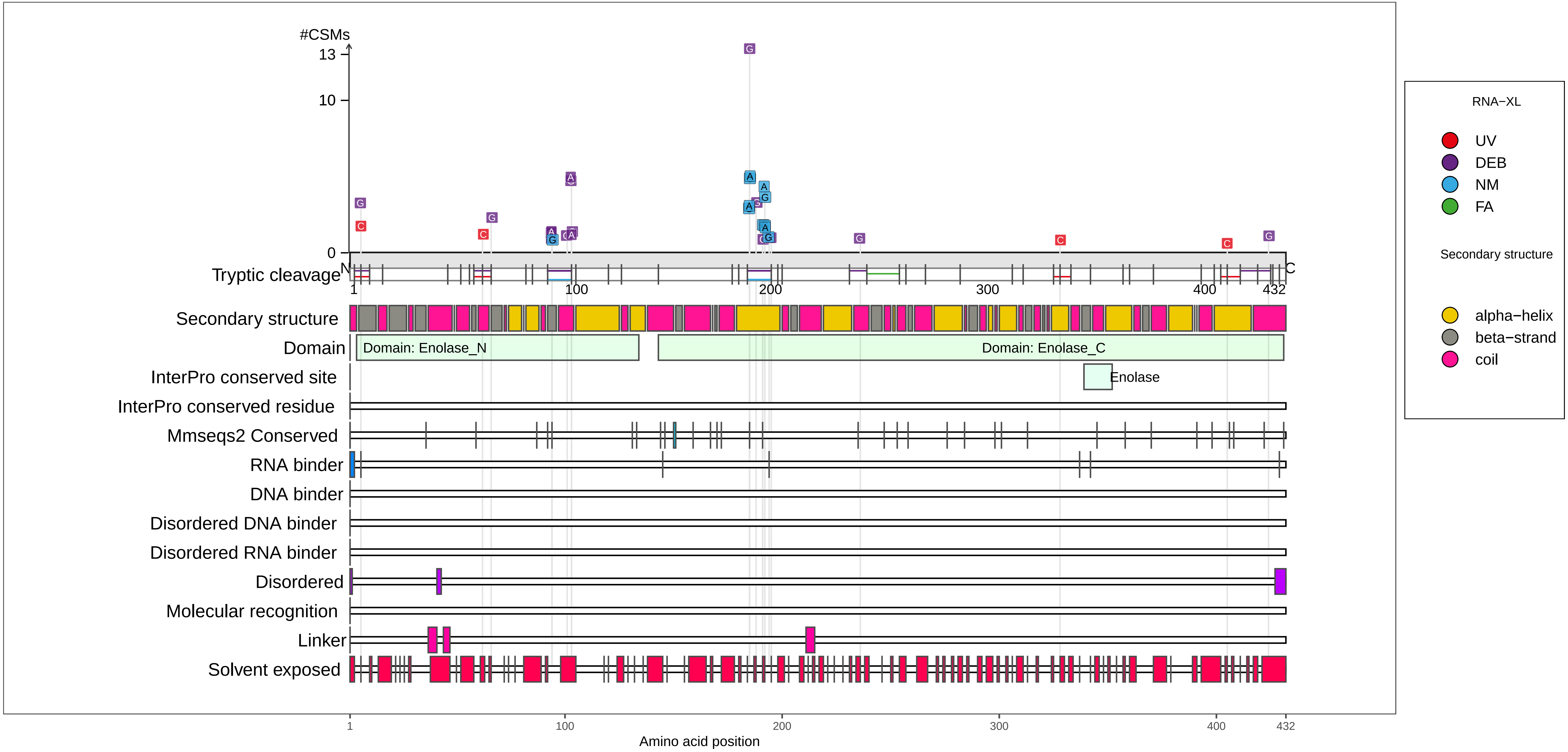

P0A817
METK_ECOLI S−adenosylmethionine synthase
− Abundance:
tryptic [log10 Intensity]: 9.37 (Q 95)
PAXdb K12 strain [ppm]: 2.85 (Q 87)
PAXdb E.coli [ppm]: 3.21 (Q 97)

− RNA functions: not annotated

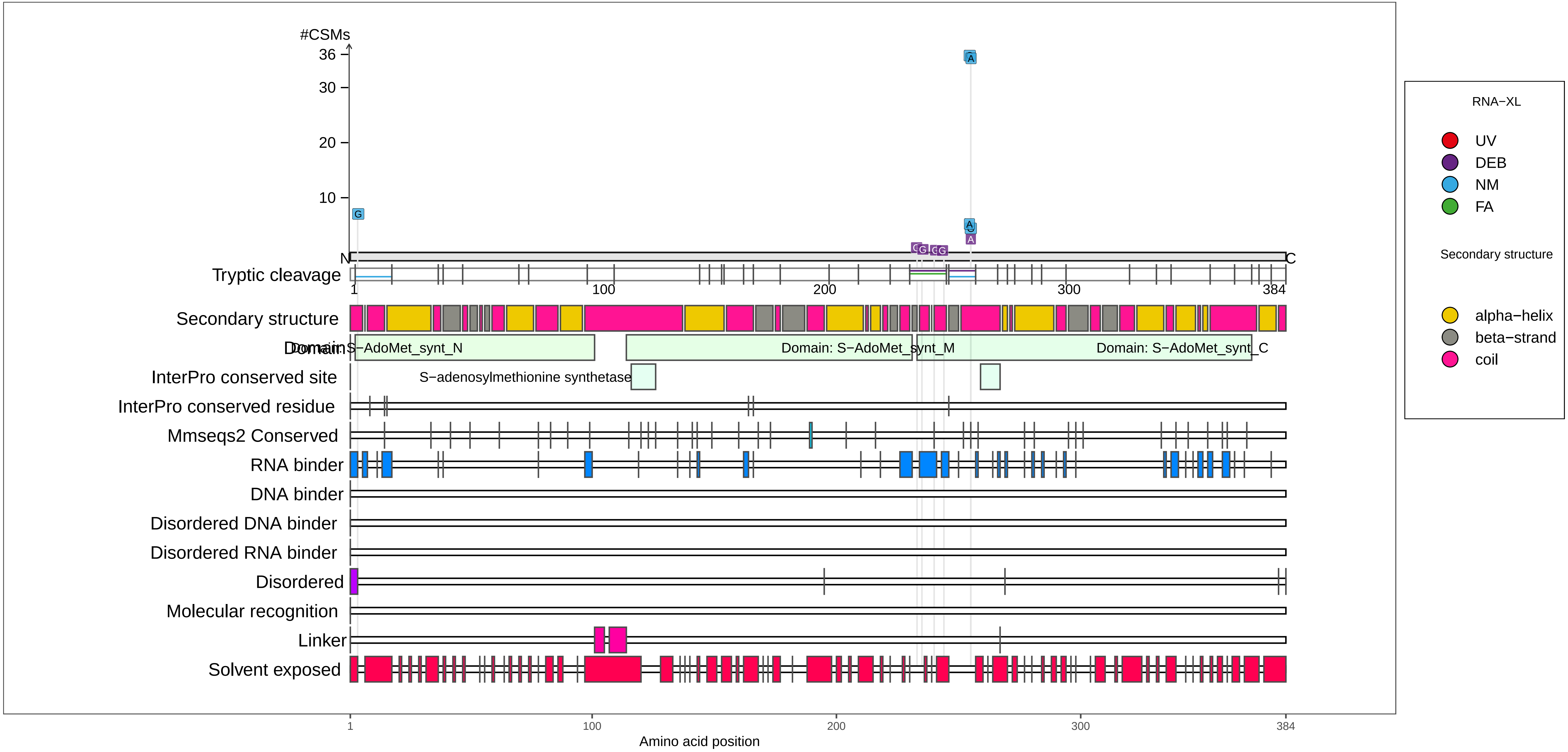

P0AGJ9
SYY_ECOLI Tyrosine−−tRNA ligase

− Abundance:
tryptic [log10 Intensity]: 7.94 (Q 56)
PAXdb K12 strain [ppm]: 3.05 (Q 92)
PAXdb E.coli [ppm]: 2.46 (Q 85)

− RNA functions:
aminoacyl−tRNA ligase activity; ncRNA metabolic process; RNA binding
RNA metabolic process; S4 RNA−binding domain; tRNA aminoacylation
tRNA aminoacylation for protein translation; tRNA metabolic process; tRNA synthetases class I (W and Y)
tyrosine−tRNA ligase activity; tyrosyl−tRNA aminoacylation

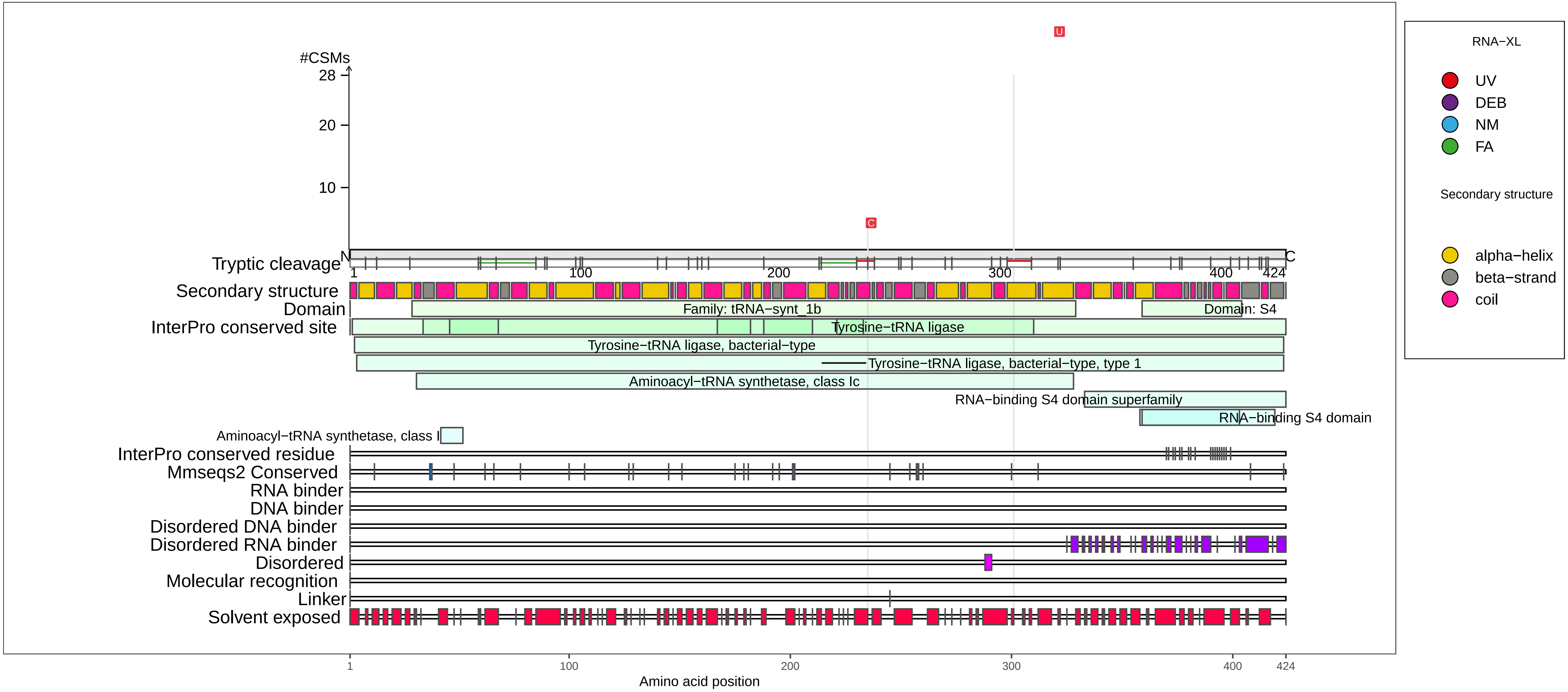

P08400
PHOR_ECOLI Phosphate regulon sensor protein PhoR

− Abundance:
tryptic [log10 Intensity]: NA (Q NA)
PAXdb K12 strain [ppm]: NA (Q NA)
PAXdb E.coli [ppm]: 1.45 (Q 58)

− RNA functions: not annotated

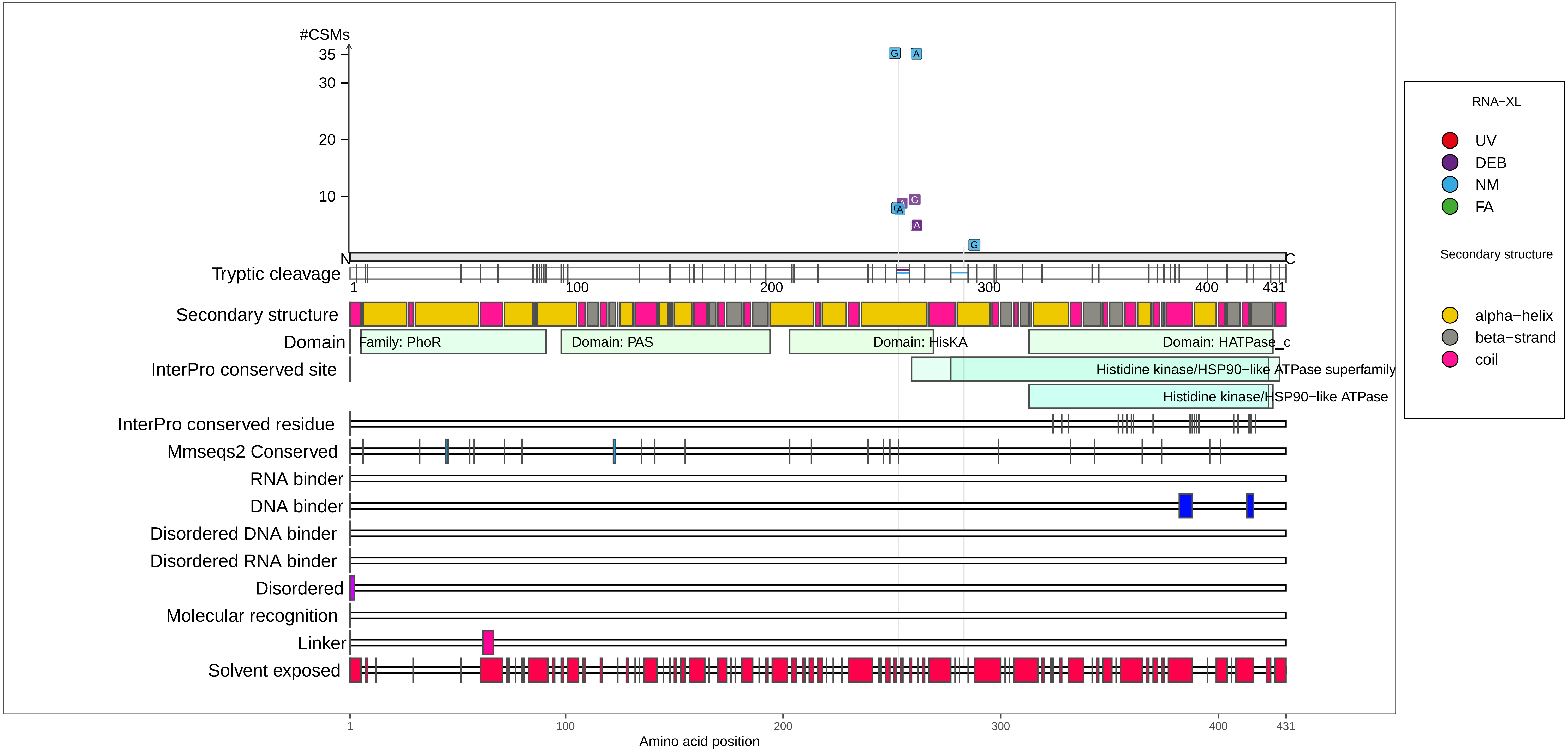

P0A7E5
PYRG_ECOLI CTP synthase

− Abundance:
tryptic [log10 Intensity]: 9.42 (Q 96)
PAXdb K12 strain [ppm]: 3.23 (Q 94)
PAXdb E.coli [ppm]: 2.32 (Q 81)

− RNA functions: not annotated

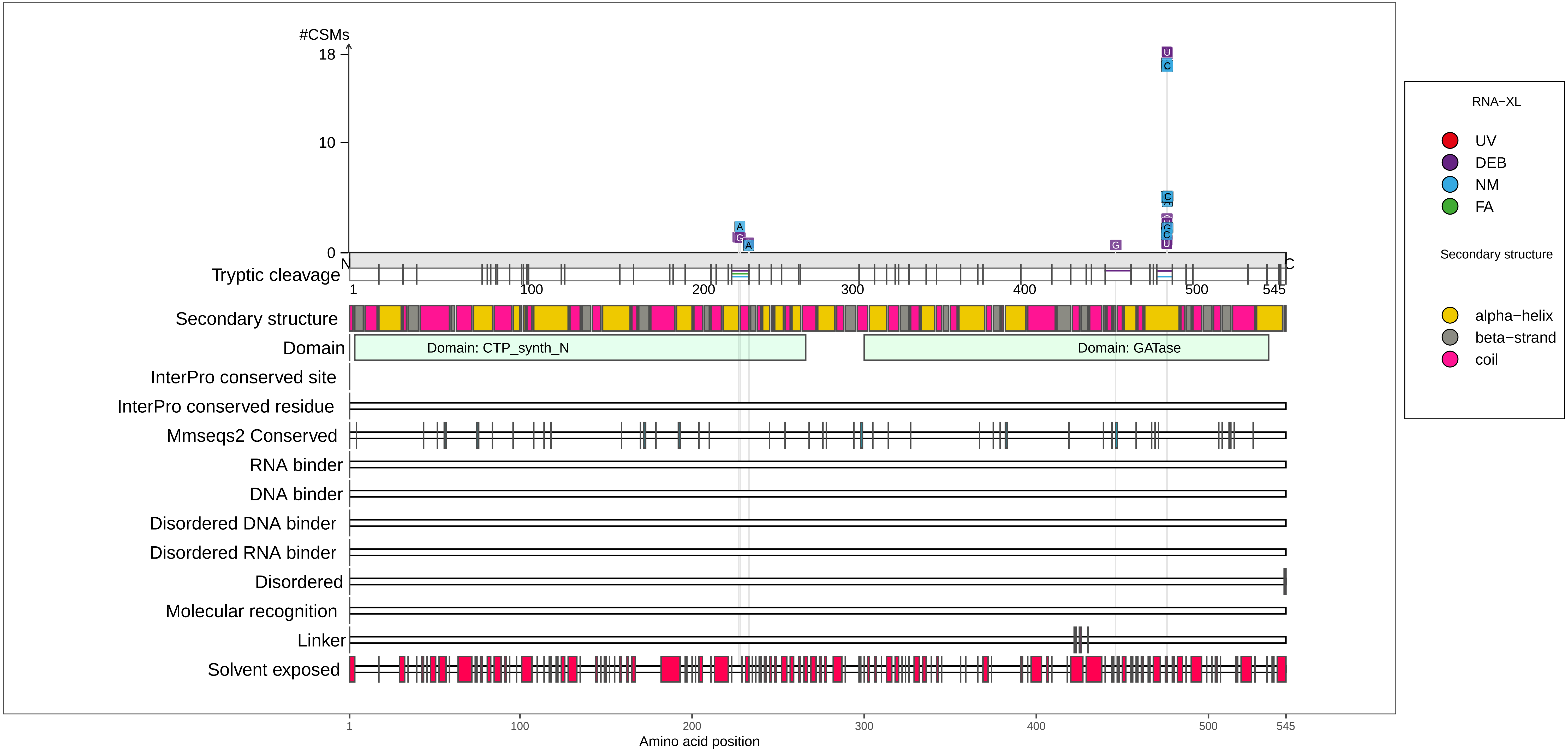

P0A8P3
FETP_ECOLI Probable Fe(2+)−trafficking protein

− Abundance:
tryptic [log10 Intensity]: 8.29 (Q 69)
PAXdb K12 strain [ppm]: 2.7 (Q 83)
PAXdb E.coli [ppm]: 3.16 (Q 96)

− RNA functions: not annotated

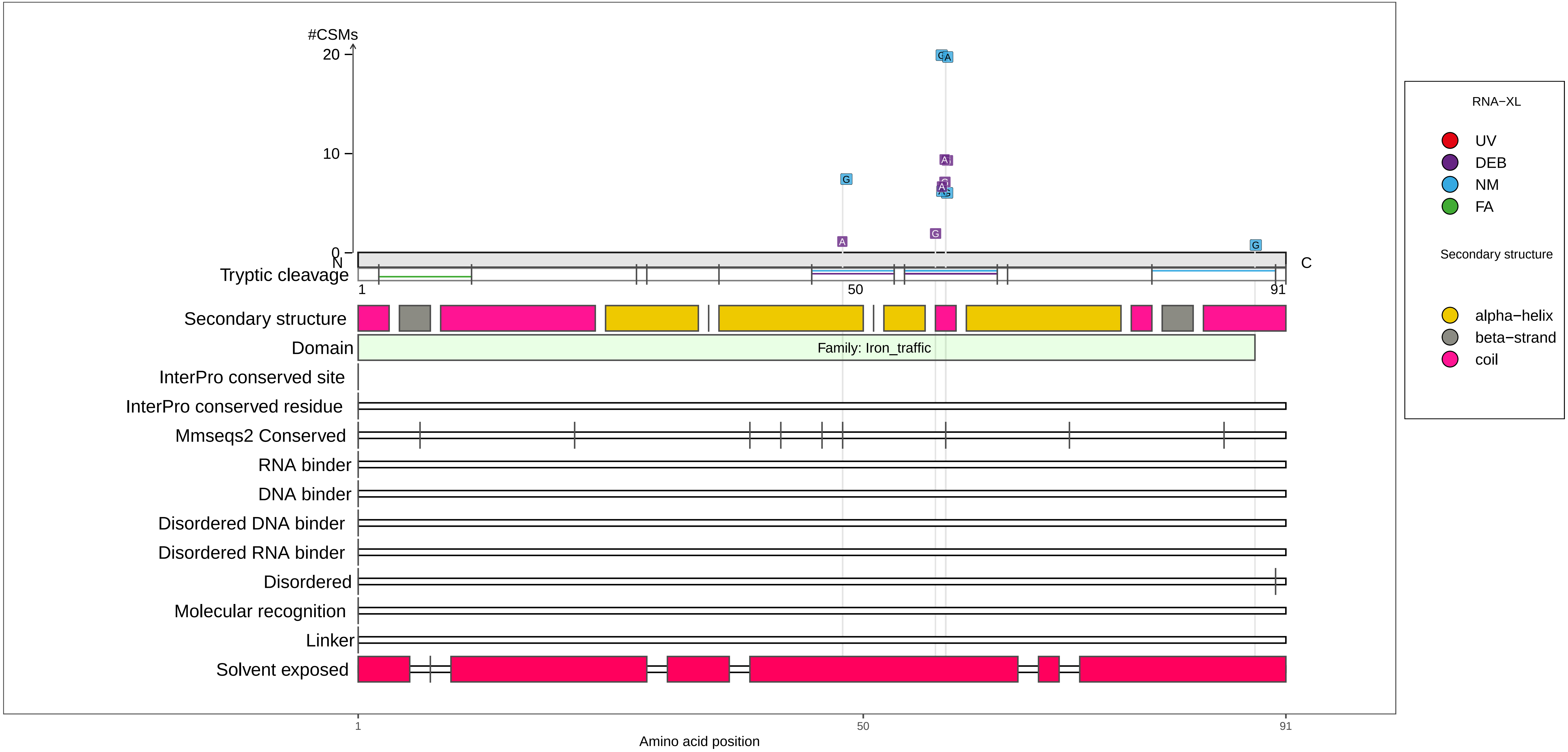

P09372
GRPE_ECOLI Protein GrpE

− Abundance:
tryptic [log10 Intensity]: 8.79 (Q 83)
PAXdb K12 strain [ppm]: 3.06 (Q 92)
PAXdb E.coli [ppm]: 3.01 (Q 95)

− RNA functions: not annotated

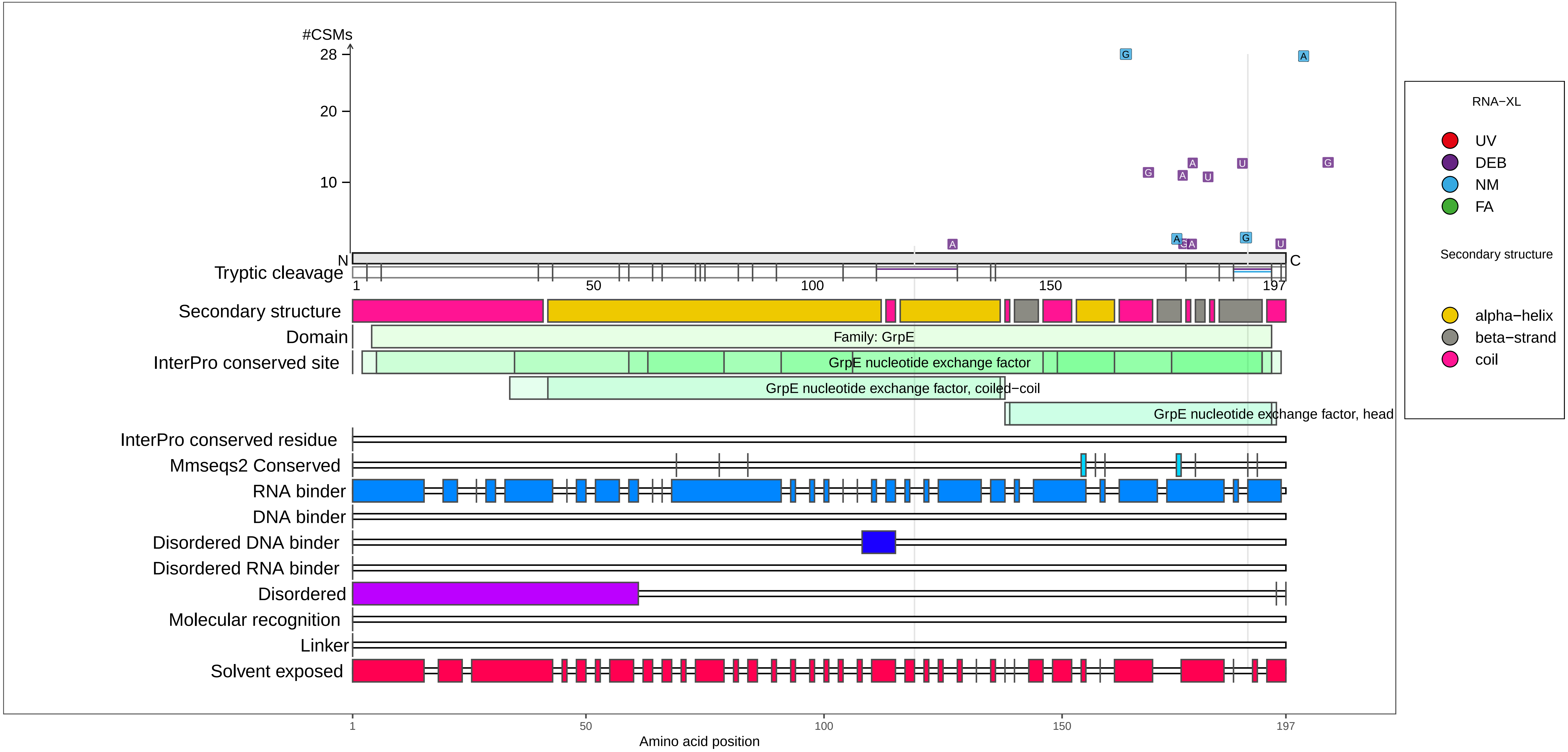

P0A9Z1
GLNB_ECOLI Nitrogen regulatory protein P−II 1

− Abundance:
tryptic [log10 Intensity]: 7.76 (Q 47)
PAXdb K12 strain [ppm]: 2.32 (Q 71)
PAXdb E.coli [ppm]: 2.66 (Q 89)

− RNA functions: not annotated

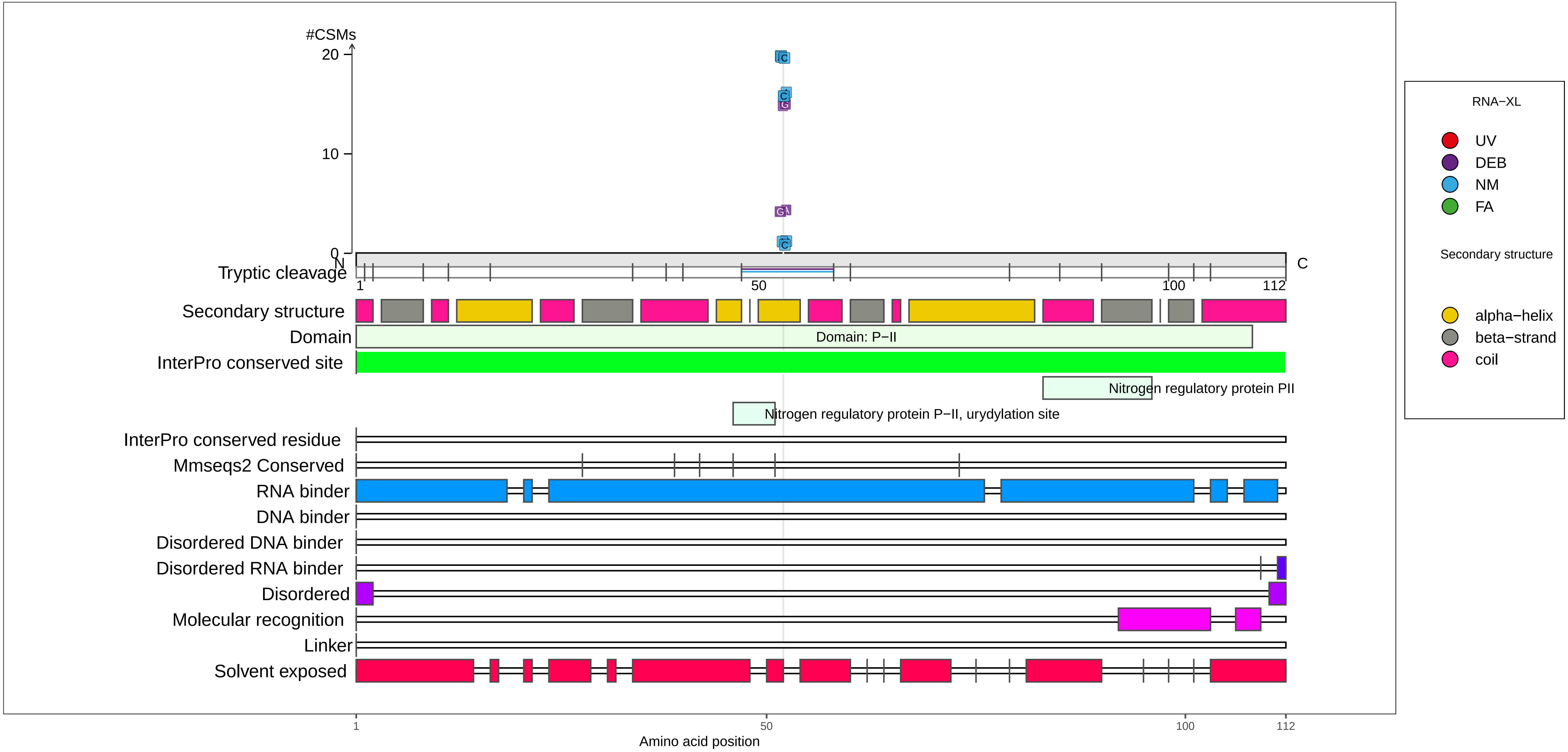

P0A836
SUCC_ECOLI Succinate−−CoA ligase [ADP−forming] subunit beta

− Abundance:
tryptic [log10 Intensity]: 9.18 (Q 92)
PAXdb K12 strain [ppm]: 3.25 (Q 95)
PAXdb E.coli [ppm]: 3.27 (Q 97)

− RNA functions: not annotated

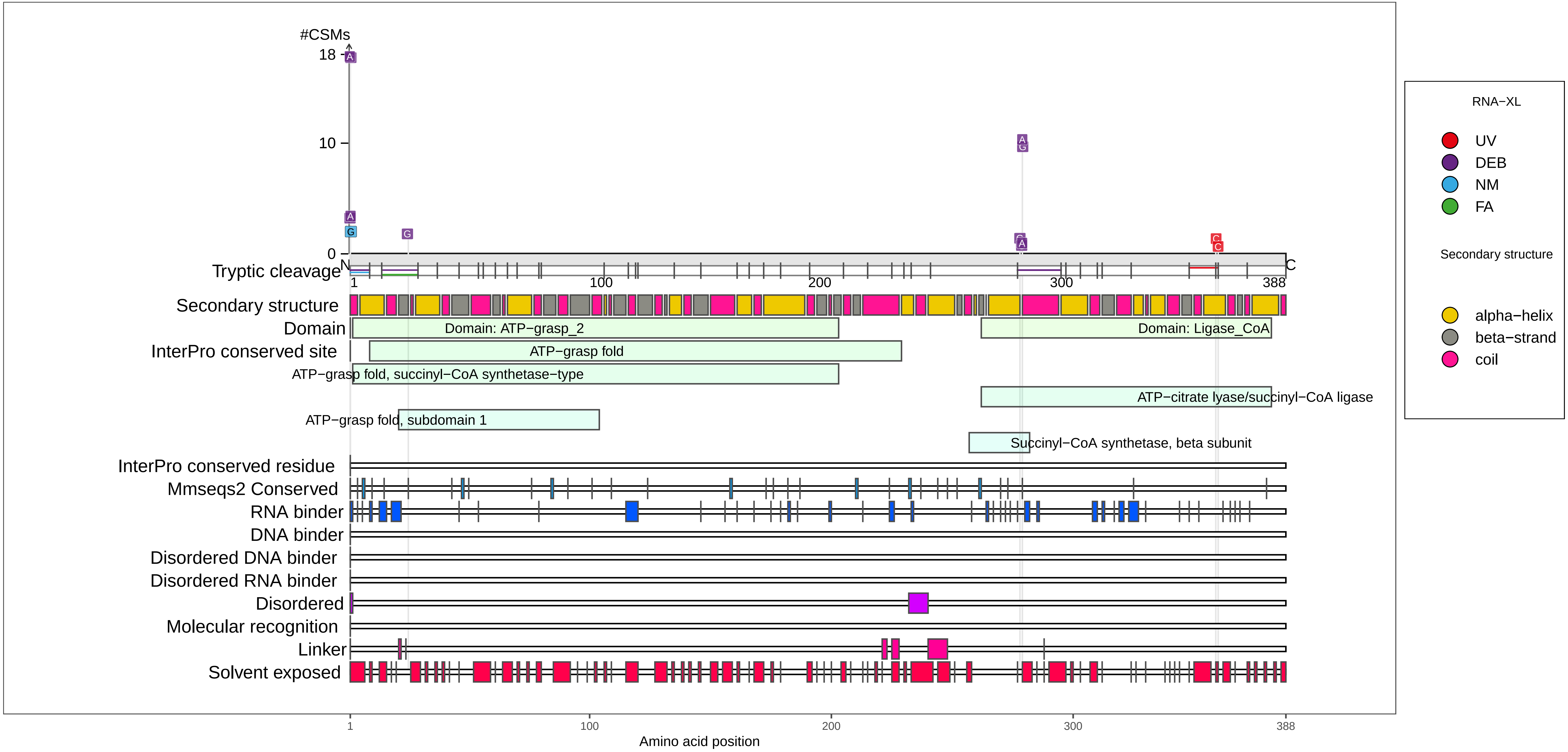

P21599
KPYK2_ECOLI Pyruvate kinase II

− Abundance:
tryptic [log10 Intensity]: 9.19 (Q 93)
PAXdb K12 strain [ppm]: 3.17 (Q 93)
PAXdb E.coli [ppm]: 2.91 (Q 93)

− RNA functions: not annotated

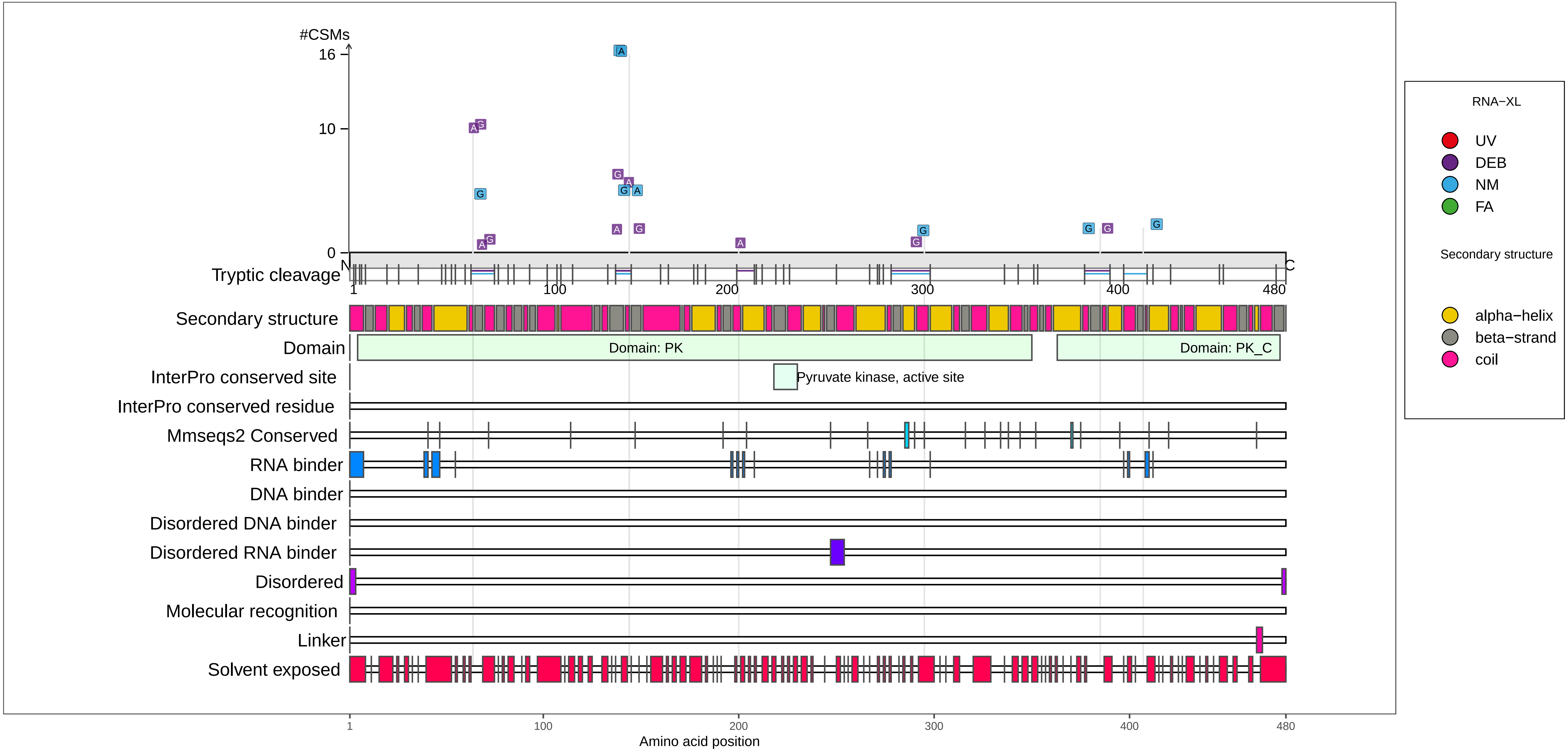

P25745
MNMA_ECOLI tRNA−specific 2−thiouridylase MnmA

− Abundance:
tryptic [log10 Intensity]: 8.49 (Q 75)
PAXdb K12 strain [ppm]: 2.21 (Q 67)
PAXdb E.coli [ppm]: 2.31 (Q 80)

− RNA functions:
ncRNA metabolic process; ncRNA processing; RNA binding; RNA metabolic process
RNA modification; RNA processing; tRNA binding; tRNA metabolic process; tRNA methyl transferase
tRNA modification; tRNA processing; tRNA thio−modification
tRNA wobble base modification; tRNA wobble position uridine thiolation; tRNA wobble uridine modification

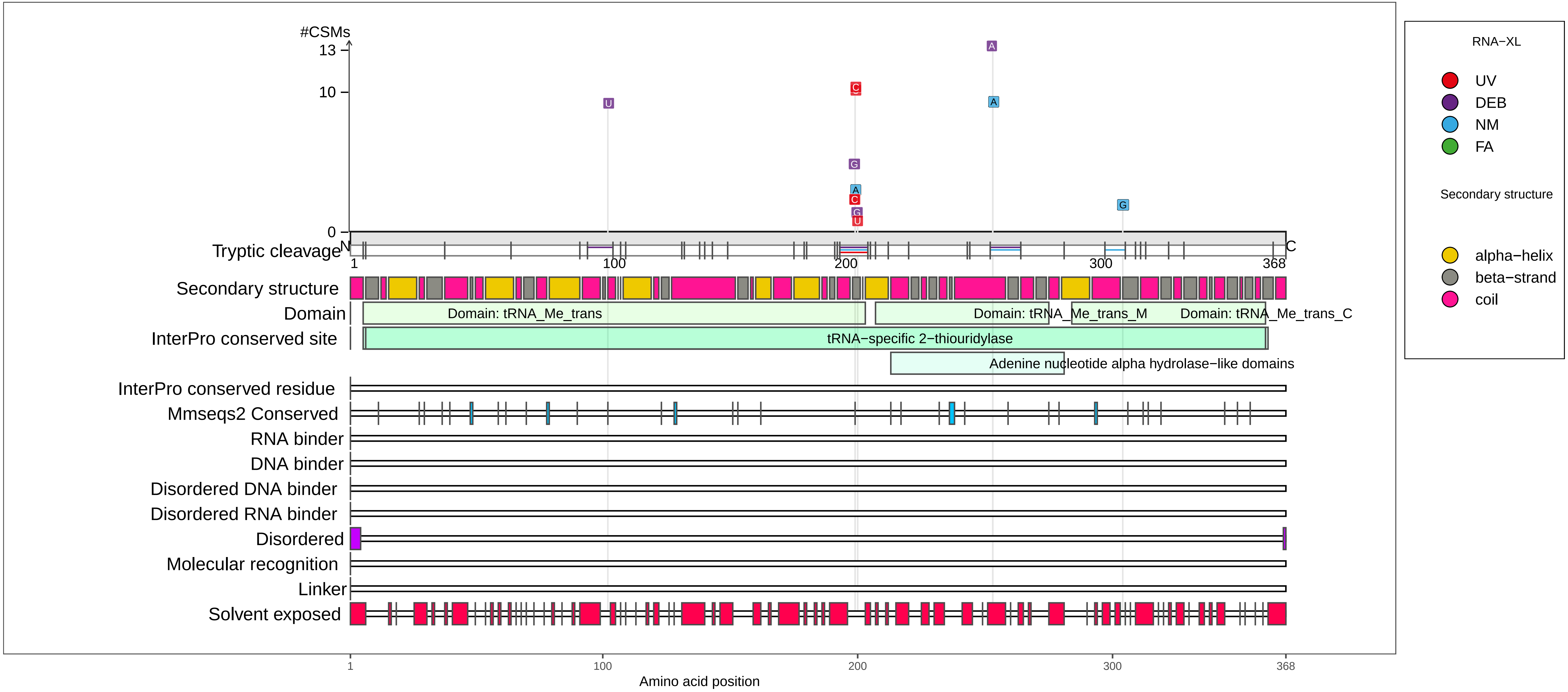

P0C018
RL18_ECOLI 50S ribosomal protein L18

− Abundance:
tryptic [log10 Intensity]: 8.7 (Q 81)
PAXdb K12 strain [ppm]: 3.89 (Q 100)
PAXdb E.coli [ppm]: 3.39 (Q 98)

− RNA functions:
5S rRNA binding; RNA binding; rRNA binding

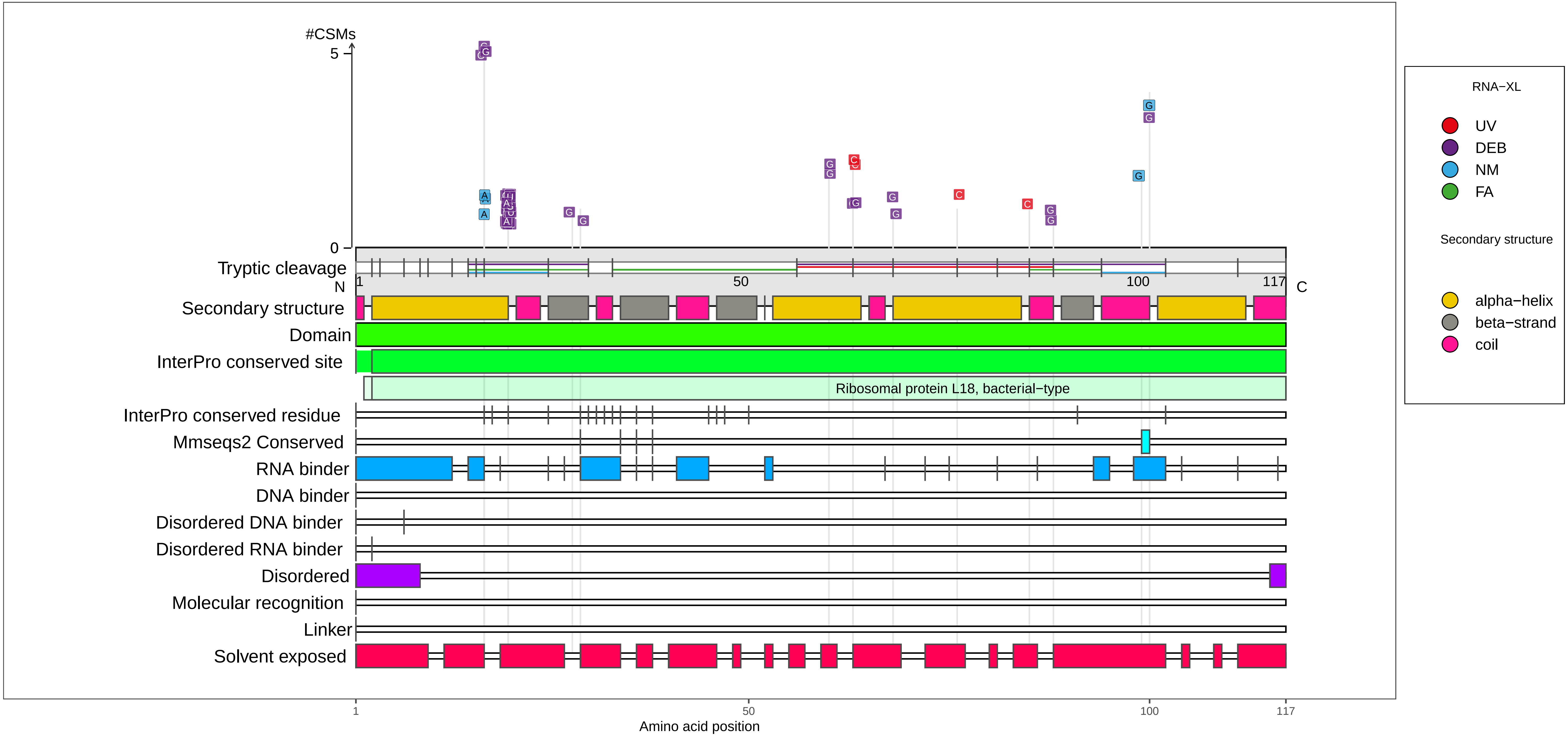

P0A7B8
HSLV_ECOLI ATP−dependent protease subunit HslV

− Abundance:
tryptic [log10 Intensity]: 8.52 (Q 76)
PAXdb K12 strain [ppm]: 3.03 (Q 91)
PAXdb E.coli [ppm]: 2.16 (Q 76)

− RNA functions: not annotated

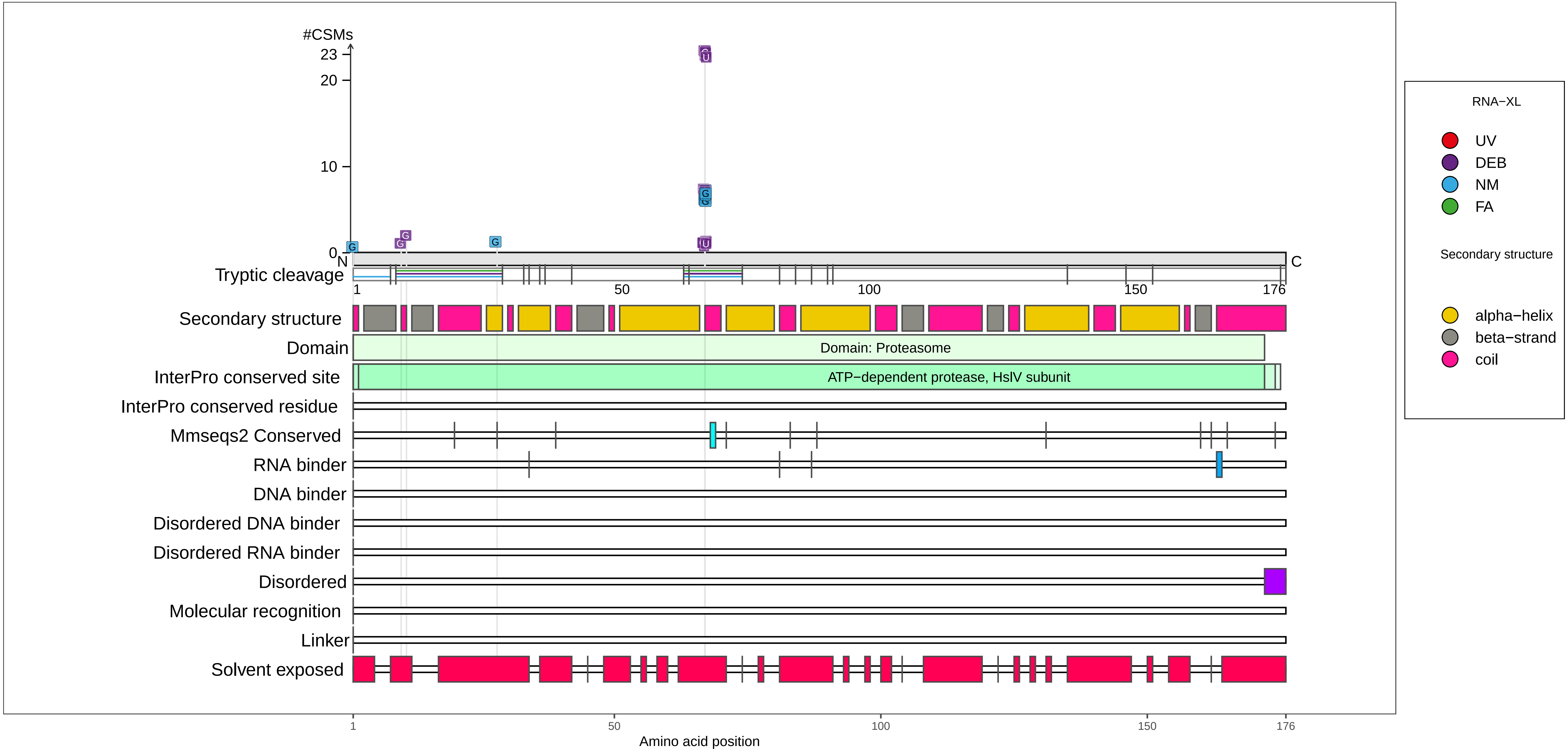

P16095
SDHL_ECOLI L−serine dehydratase 1

− Abundance:
tryptic [log10 Intensity]: 8.53 (Q 77)
PAXdb K12 strain [ppm]: 2.39 (Q 74)
PAXdb E.coli [ppm]: 1.44 (Q 58)

− RNA functions: not annotated

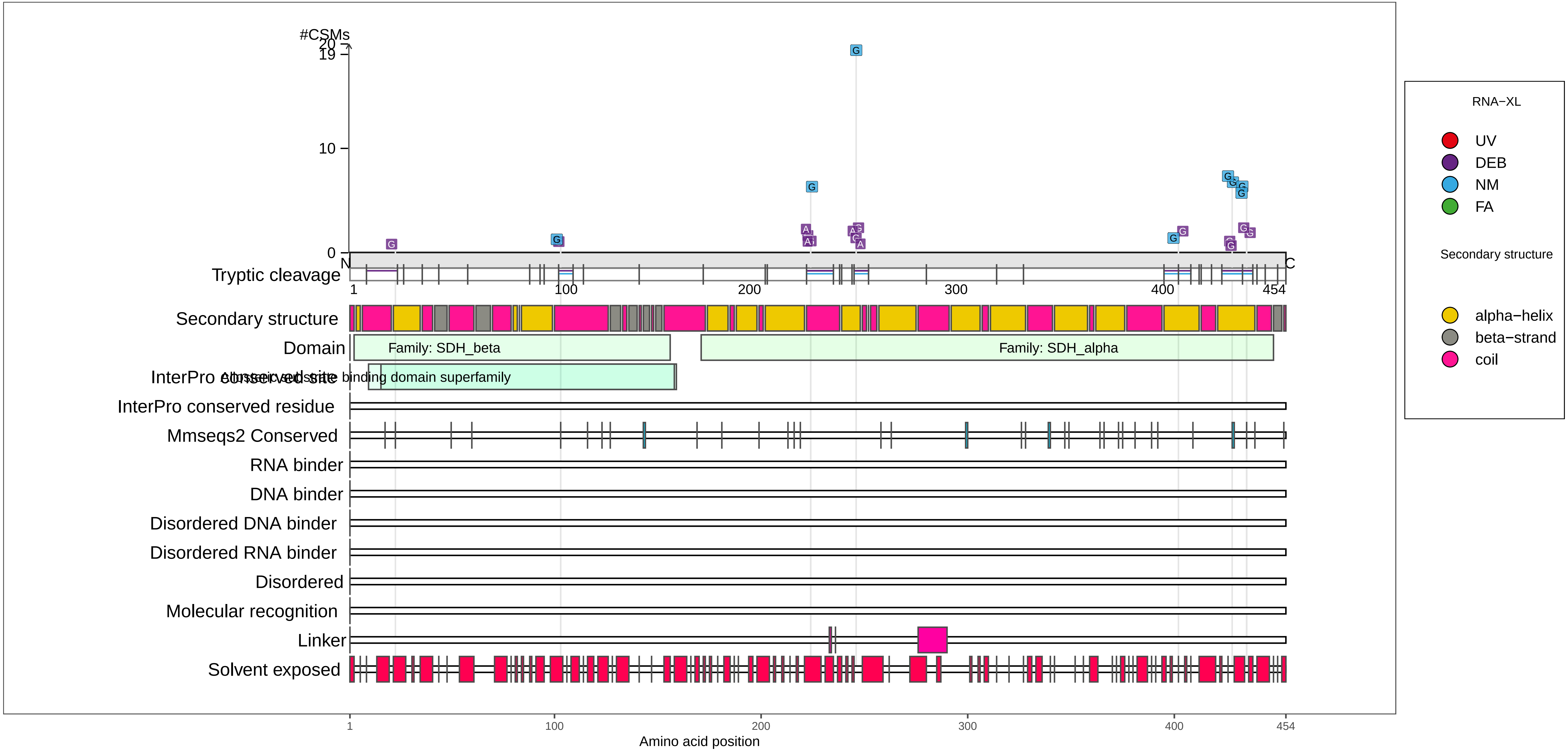

P0AB71
ALF_ECOLI Fructose−bisphosphate aldolase class 2
− Abundance:
tryptic [log10 Intensity]: 10.21 (Q 100)
PAXdb K12 strain [ppm]: 3.52 (Q 98)
PAXdb E.coli [ppm]: 3.5 (Q 99)

− RNA functions: not annotated

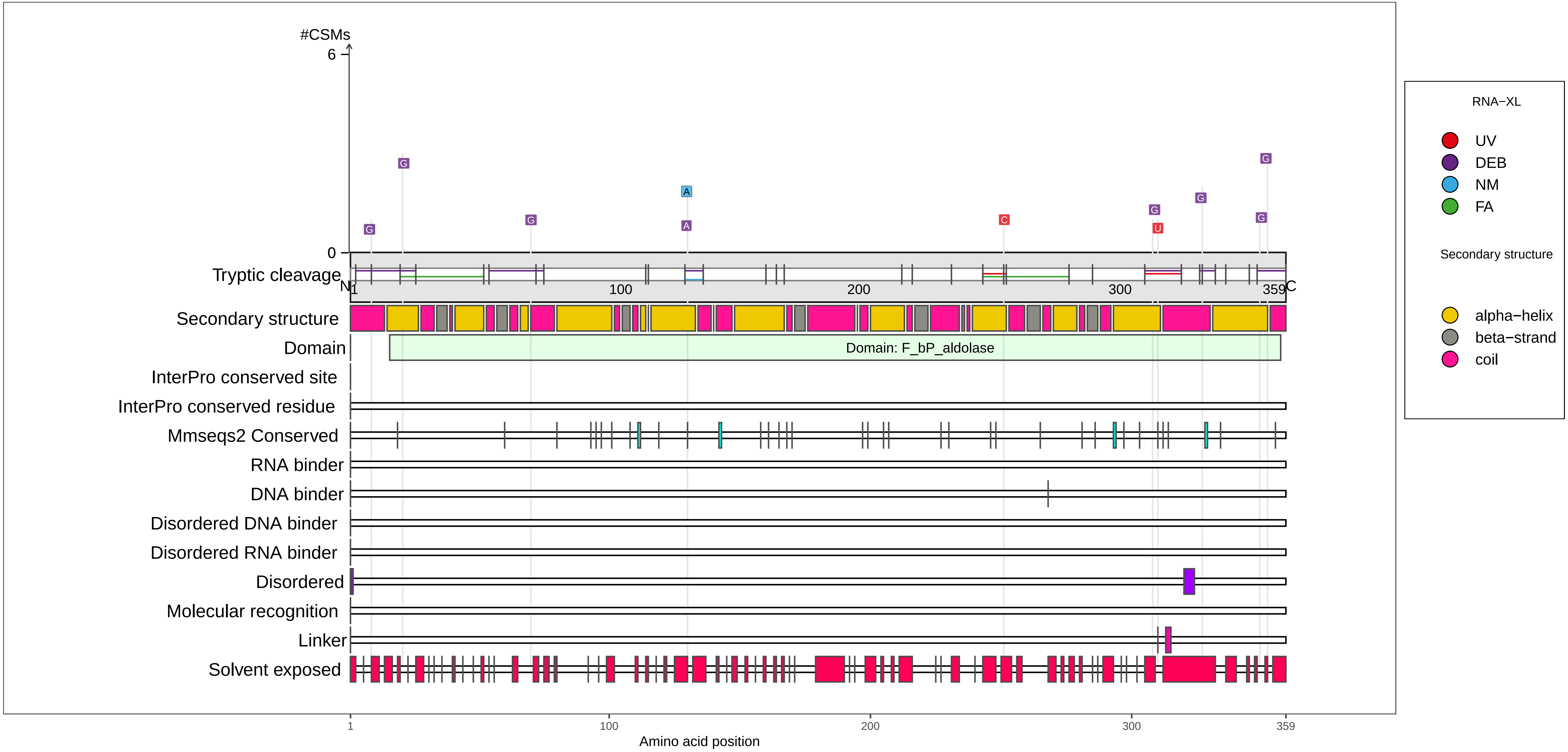

P0A8L1
SYS_ECOLI Serine−−tRNA ligase

− Abundance:
tryptic [log10 Intensity]: 9.34 (Q 95)
PAXdb K12 strain [ppm]: 3.25 (Q 95)
PAXdb E.coli [ppm]: 2.93 (Q 94)

− RNA functions:
aminoacyl−tRNA ligase activity; ncRNA metabolic process; RNA biosynthetic process
RNA metabolic process; selenocysteinyl−tRNA(Sec) biosynthetic process
serine−tRNA ligase activity; seryl−tRNA aminoacylation; Seryl−tRNA synthetase N−terminal domain
tRNA aminoacylation; tRNA aminoacylation for protein translation; tRNA metabolic process
tRNA synthetase class II core domain (G, H, P, S and T)

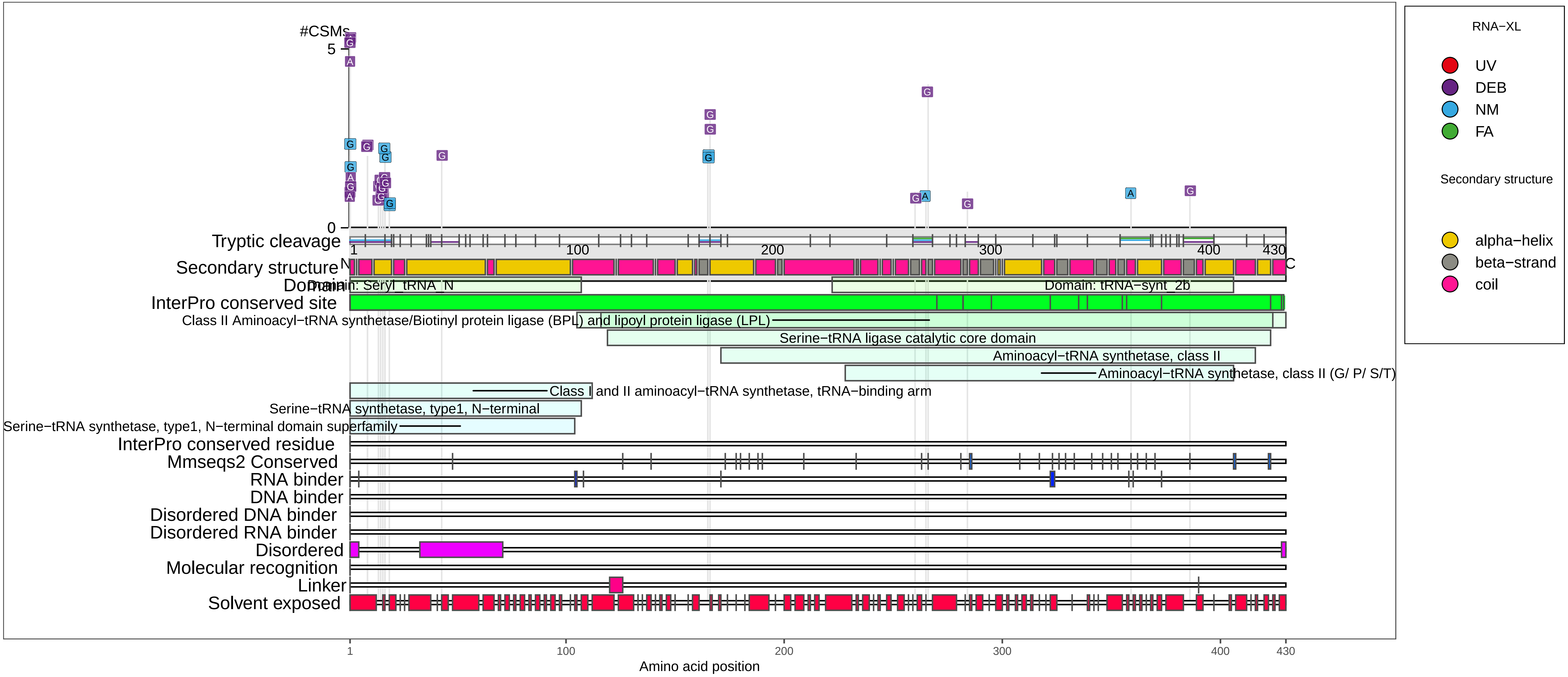

P0AEI1
MIAB_ECOLI tRNA−2−methylthio−N(6)−dimethylallyladenosine synthase

− Abundance:
tryptic [log10 Intensity]: 8.78 (Q 83)
PAXdb K12 strain [ppm]: 1.45 (Q 35)
PAXdb E.coli [ppm]: 1.5 (Q 60)

− RNA functions:
ncRNA metabolic process; ncRNA processing; RNA metabolic process; RNA methylation
RNA modification; RNA processing; tRNA metabolic process; tRNA methylation
tRNA methylthiolation; tRNA modification; tRNA processing

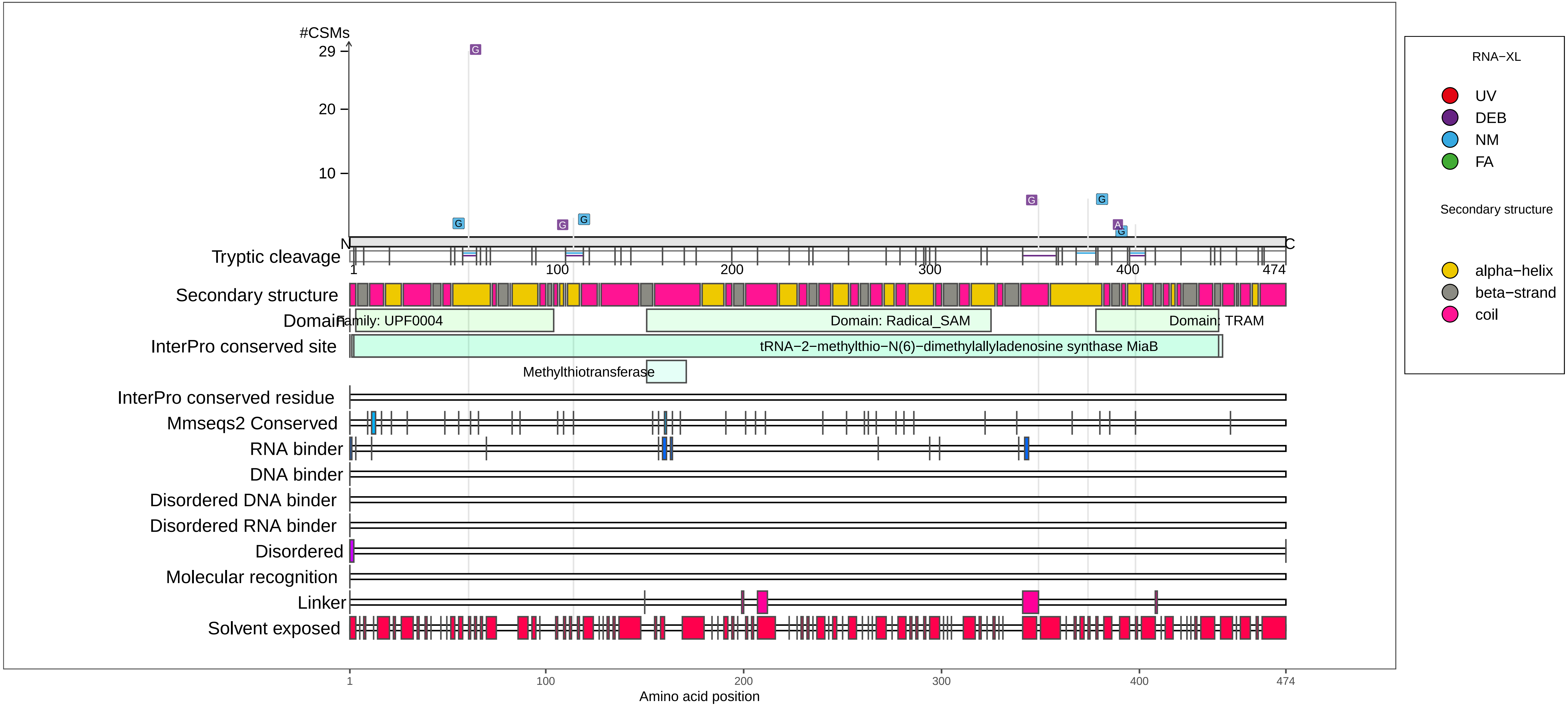

P77718
THII_ECOLI tRNA sulfurtransferase

− Abundance:
tryptic [log10 Intensity]: 7.62 (Q 41)
PAXdb K12 strain [ppm]: 1.31 (Q 27)
PAXdb E.coli [ppm]: 1.51 (Q 60)

− RNA functions:
ncRNA metabolic process; ncRNA processing; RNA adenylyltransferase activity
RNA binding; RNA metabolic process; RNA modification; RNA processing
tRNA 4−thiouridine biosynthesis; tRNA adenylyltransferase activity; tRNA binding; tRNA metabolic process
tRNA modification; tRNA processing; tRNA thio−modification

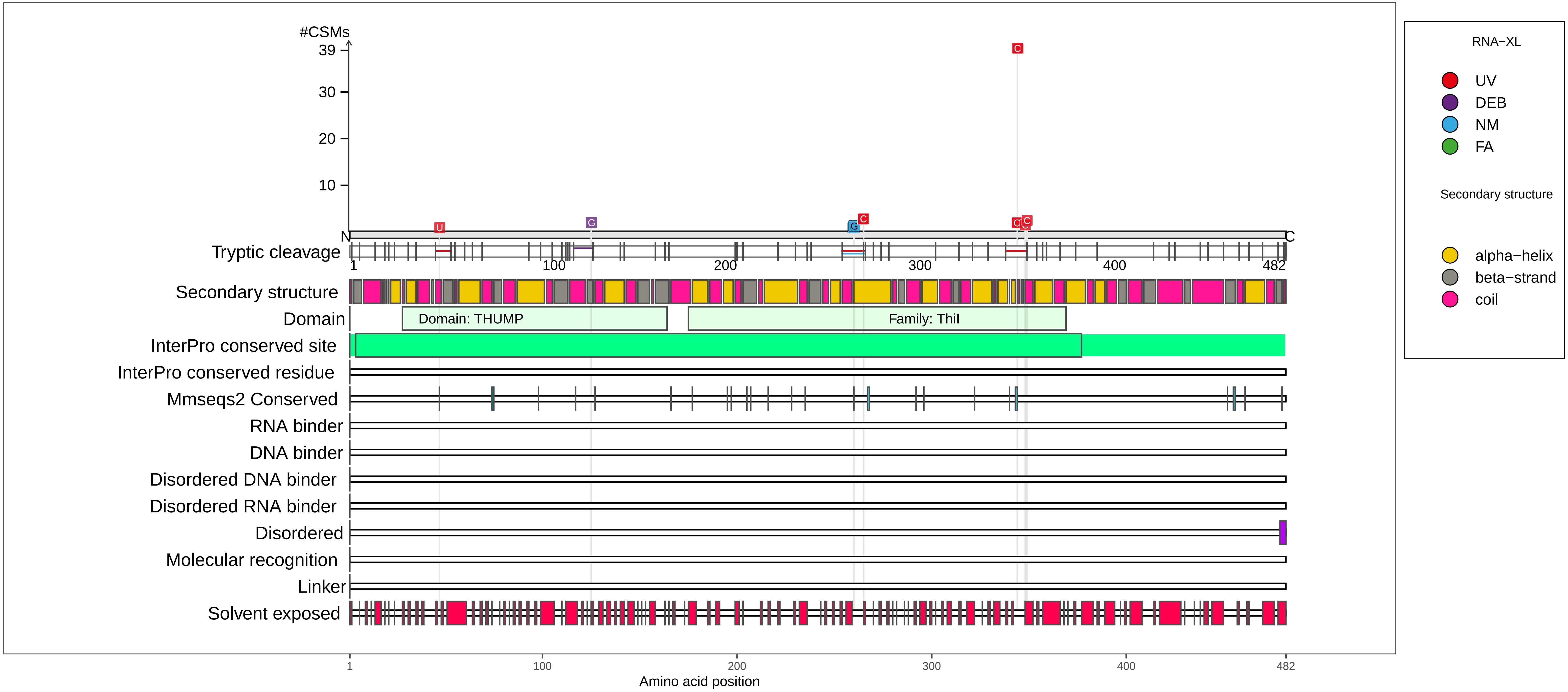

P08312
SYFA_ECOLI Phenylalanine−−tRNA ligase alpha subunit

− Abundance:
tryptic [log10 Intensity]: 9.12 (Q 91)
PAXdb K12 strain [ppm]: 2.97 (Q 90)
PAXdb E.coli [ppm]: 2.61 (Q 88)

− RNA functions:
Aminoacyl tRNA synthetase class II, N−terminal domain
aminoacyl−tRNA ligase activity; ncRNA metabolic process; phenylalanine−tRNA ligase activity
phenylalanine−tRNA ligase complex; phenylalanyl−tRNA aminoacylation; RNA binding; RNA metabolic process
tRNA aminoacylation; tRNA aminoacylation for protein translation; tRNA binding
tRNA metabolic process; tRNA synthetases class II core domain (F)

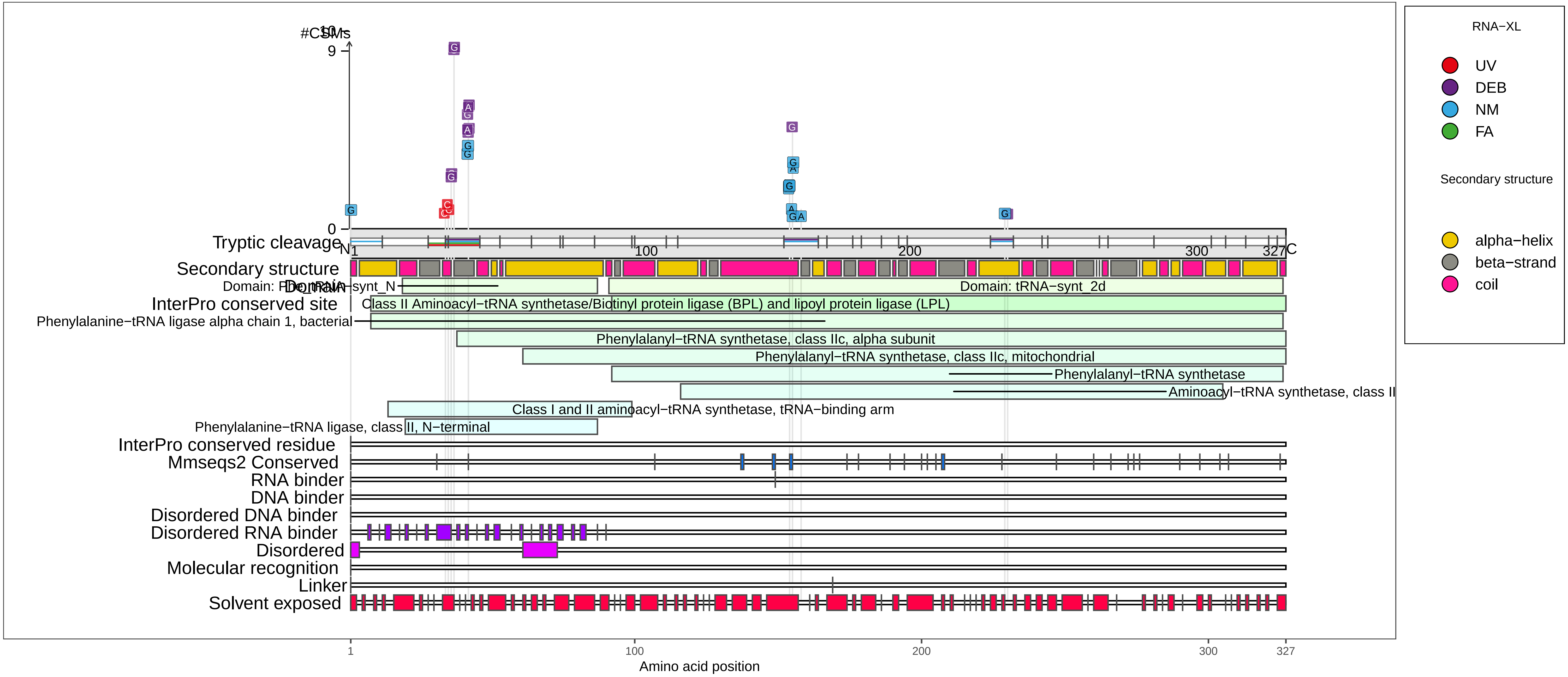

P0A9K9
SLYD_ECOLI FKBP−type peptidyl−prolyl cis−trans isomerase SlyD

− Abundance:
tryptic [log10 Intensity]: 9.87 (Q 99)
PAXdb K12 strain [ppm]: 3.39 (Q 96)
PAXdb E.coli [ppm]: 3.2 (Q 96)

− RNA functions: not annotated

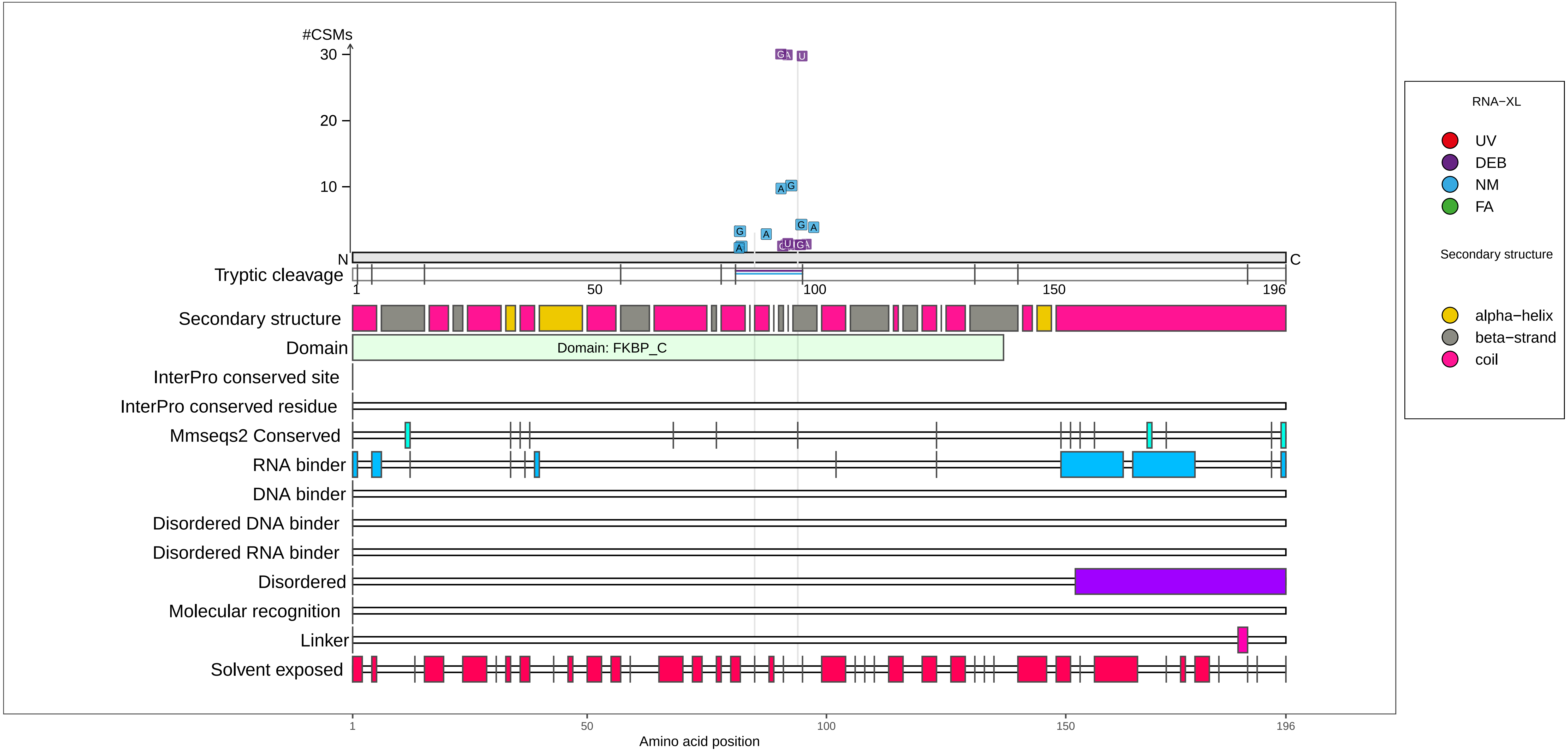

P04805
SYE_ECOLI Glutamate−−tRNA ligase

− Abundance:
tryptic [log10 Intensity]: 9.05 (Q 89)
PAXdb K12 strain [ppm]: 2.92 (Q 89)
PAXdb E.coli [ppm]: 2.56 (Q 87)

− RNA functions:
aminoacyl−tRNA ligase activity; glutamate−tRNA ligase activity
glutamyl−tRNA aminoacylation; ncRNA metabolic process; RNA binding; RNA metabolic process
tRNA aminoacylation; tRNA aminoacylation for protein translation; tRNA binding; tRNA metabolic process
tRNA synthetases class I (E and Q), catalytic domain

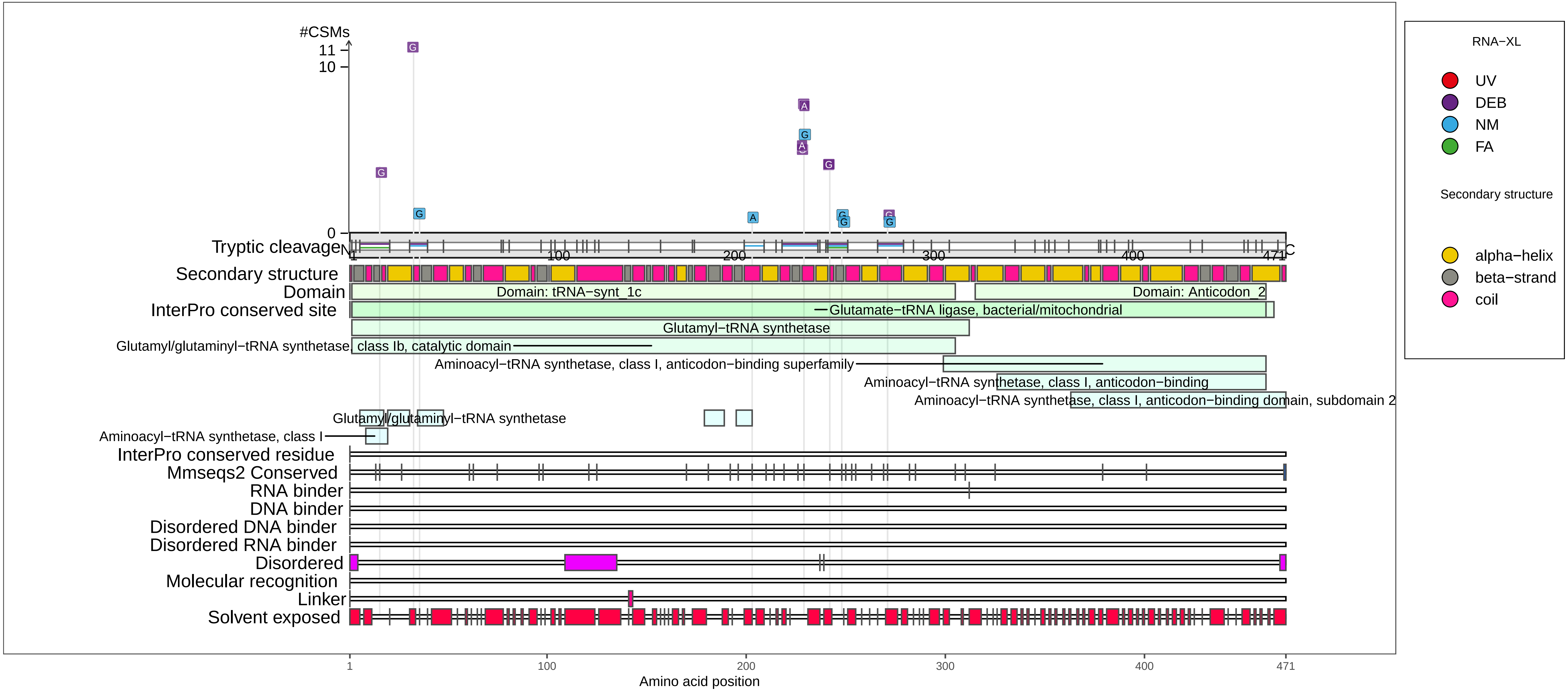

P00490
PHSM_ECOLI Maltodextrin phosphorylase

− Abundance:
tryptic [log10 Intensity]: 8.8 (Q 83)
PAXdb K12 strain [ppm]: 1.85 (Q 55)
PAXdb E.coli [ppm]: 2.17 (Q 77)

− RNA functions: not annotated

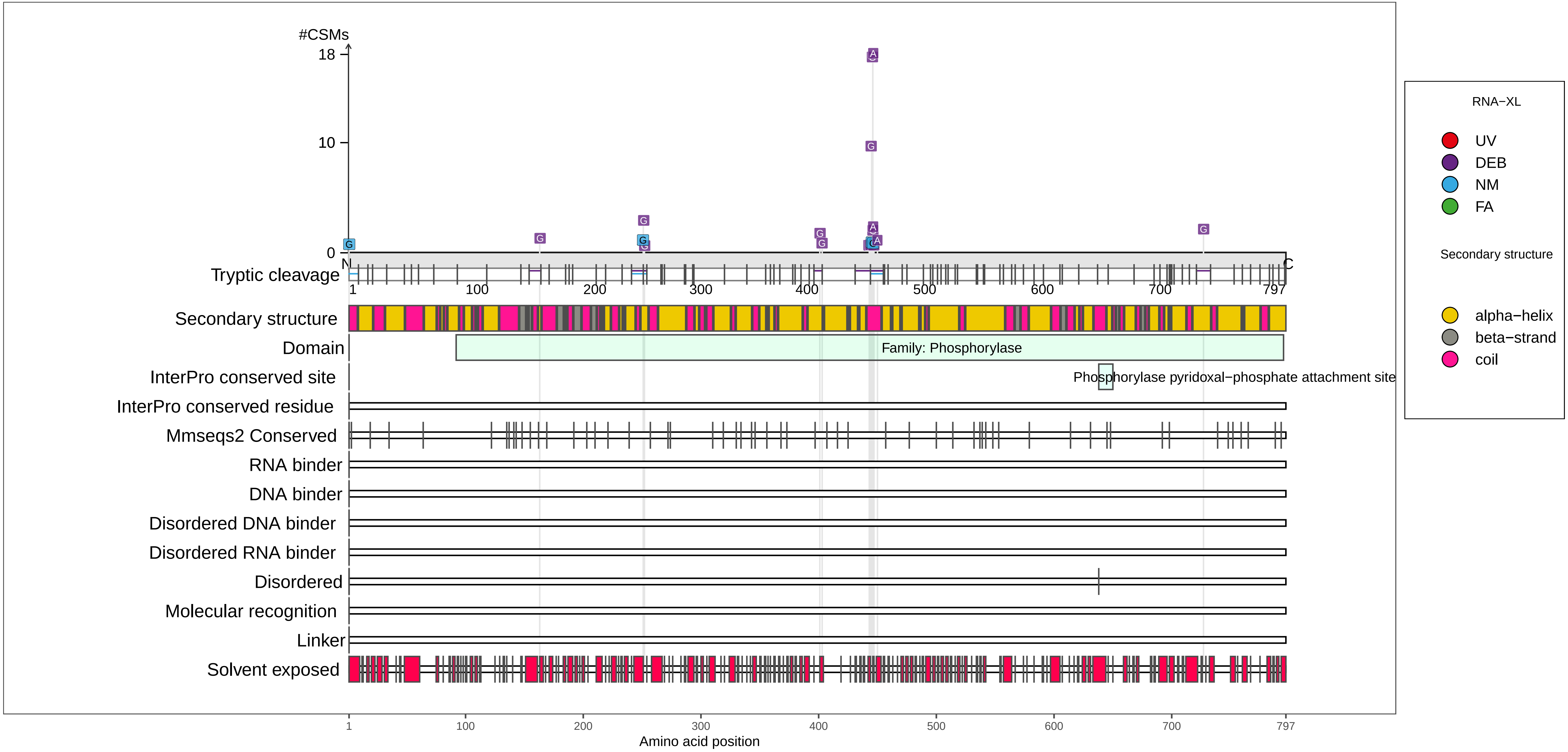

P00960
SYGA_ECOLI Glycine−−tRNA ligase alpha subunit

− Abundance:
tryptic [log10 Intensity]: 9.38 (Q 95)
PAXdb K12 strain [ppm]: 2.2 (Q 67)
PAXdb E.coli [ppm]: 2.57 (Q 87)

− RNA functions:
aminoacyl−tRNA ligase activity; glycine−tRNA ligase activity
glycyl−tRNA aminoacylation; Glycyl−tRNA synthetase alpha subunit; ncRNA metabolic process
RNA metabolic process; tRNA aminoacylation; tRNA aminoacylation for protein translation
tRNA metabolic process

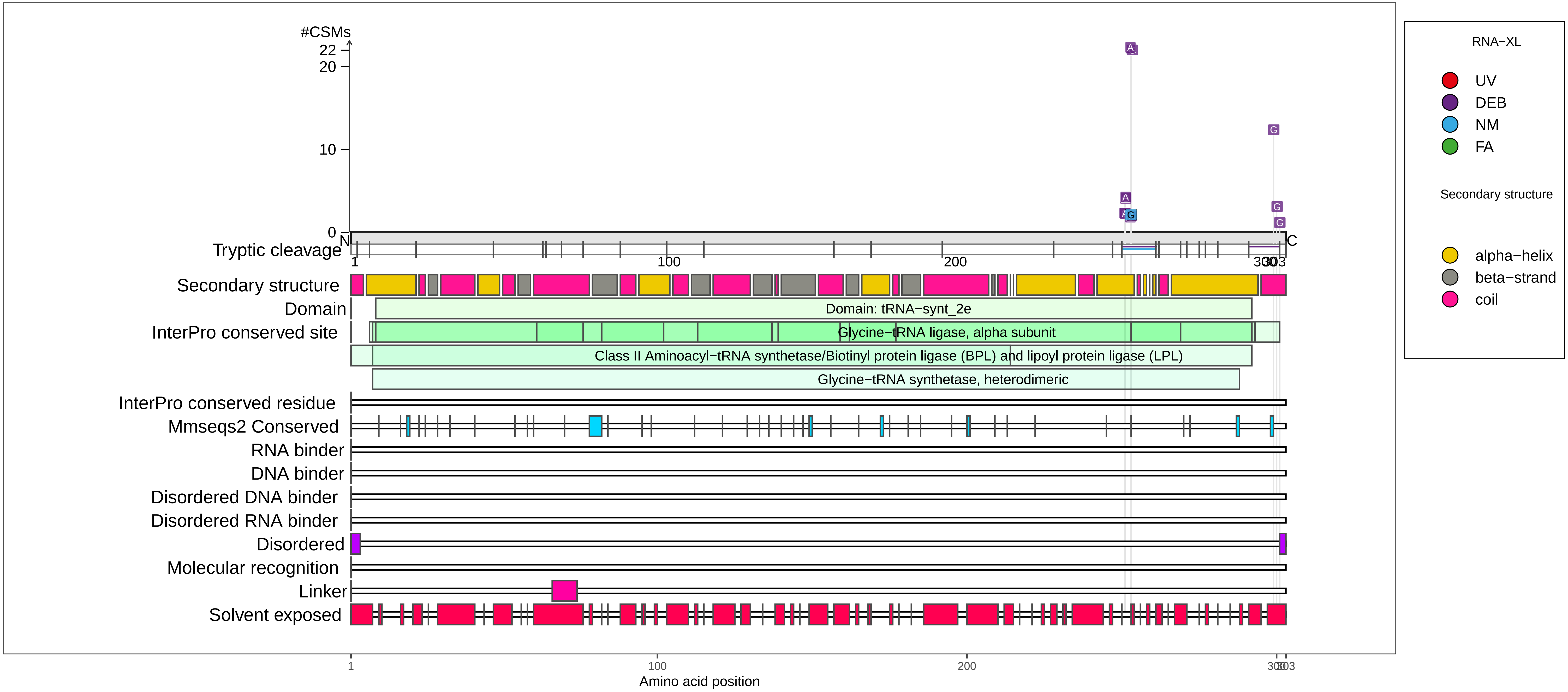

P0A6F9
CH10_ECOLI Co−chaperonin GroES

− Abundance:
tryptic [log10 Intensity]: 9.49 (Q 96)
PAXdb K12 strain [ppm]: 3.64 (Q 98)
PAXdb E.coli [ppm]: 3.85 (Q 100)

− RNA functions: not annotated

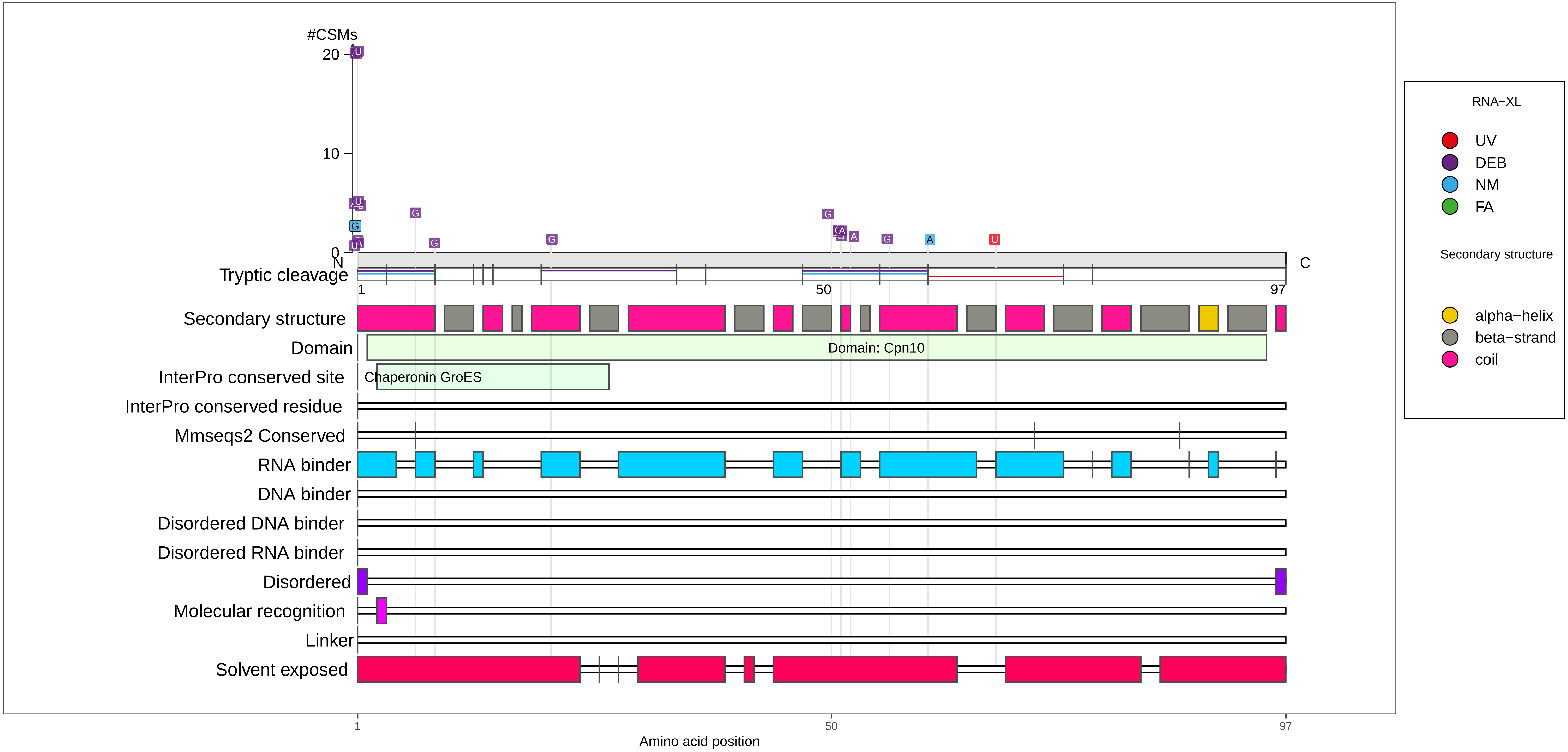

P0ABP8
DEOD_ECOLI Purine nucleoside phosphorylase DeoD−type

− Abundance:
tryptic [log10 Intensity]: 9.22 (Q 93)
PAXdb K12 strain [ppm]: 2.76 (Q 85)
PAXdb E.coli [ppm]: 3.2 (Q 96)

− RNA functions: not annotated

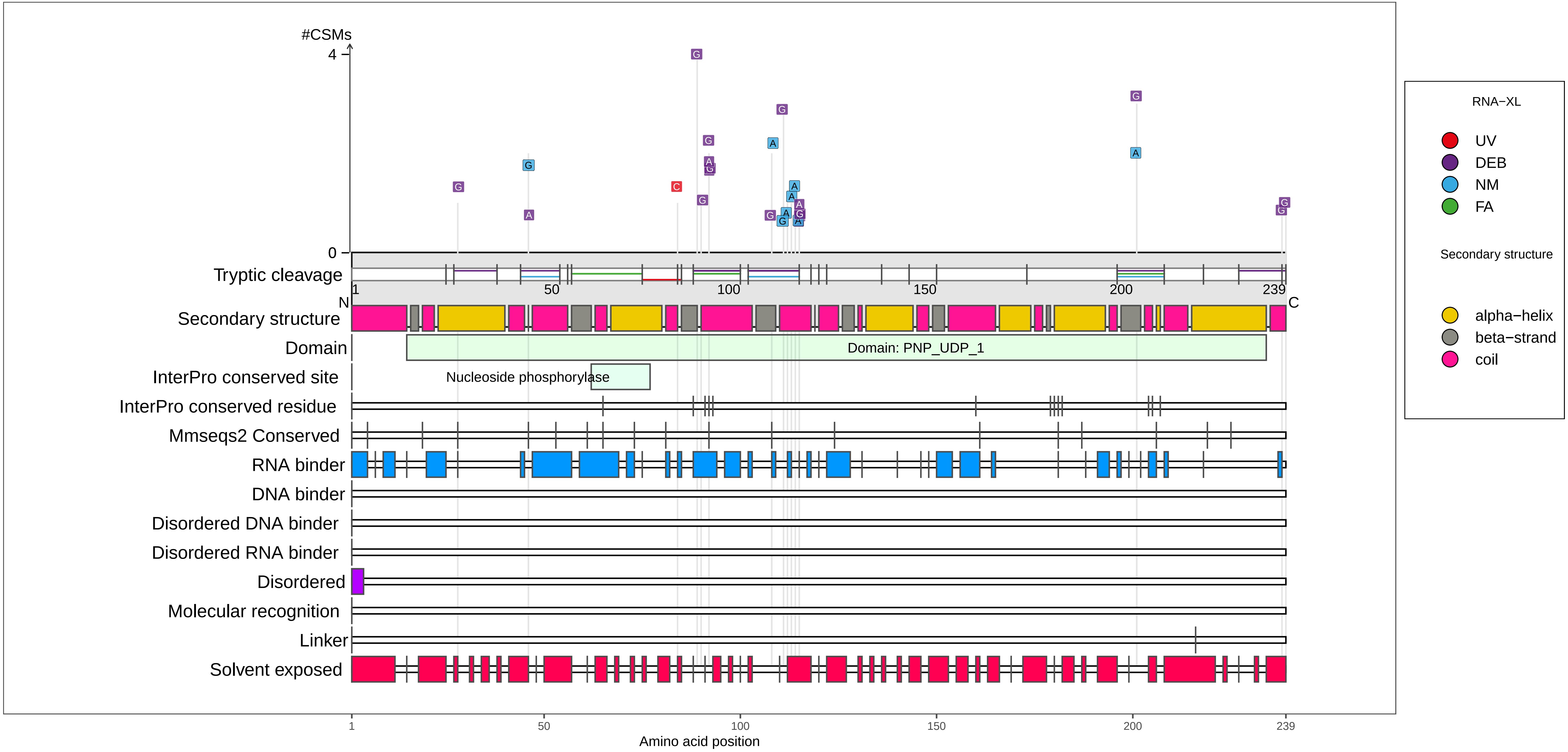

P0ADZ0
RL23_ECOLI 50S ribosomal protein L23

− Abundance:
tryptic [log10 Intensity]: 8.78 (Q 83)
PAXdb K12 strain [ppm]: 2.89 (Q 88)
PAXdb E.coli [ppm]: 2.81 (Q 92)

− RNA functions:
RNA binding; rRNA binding

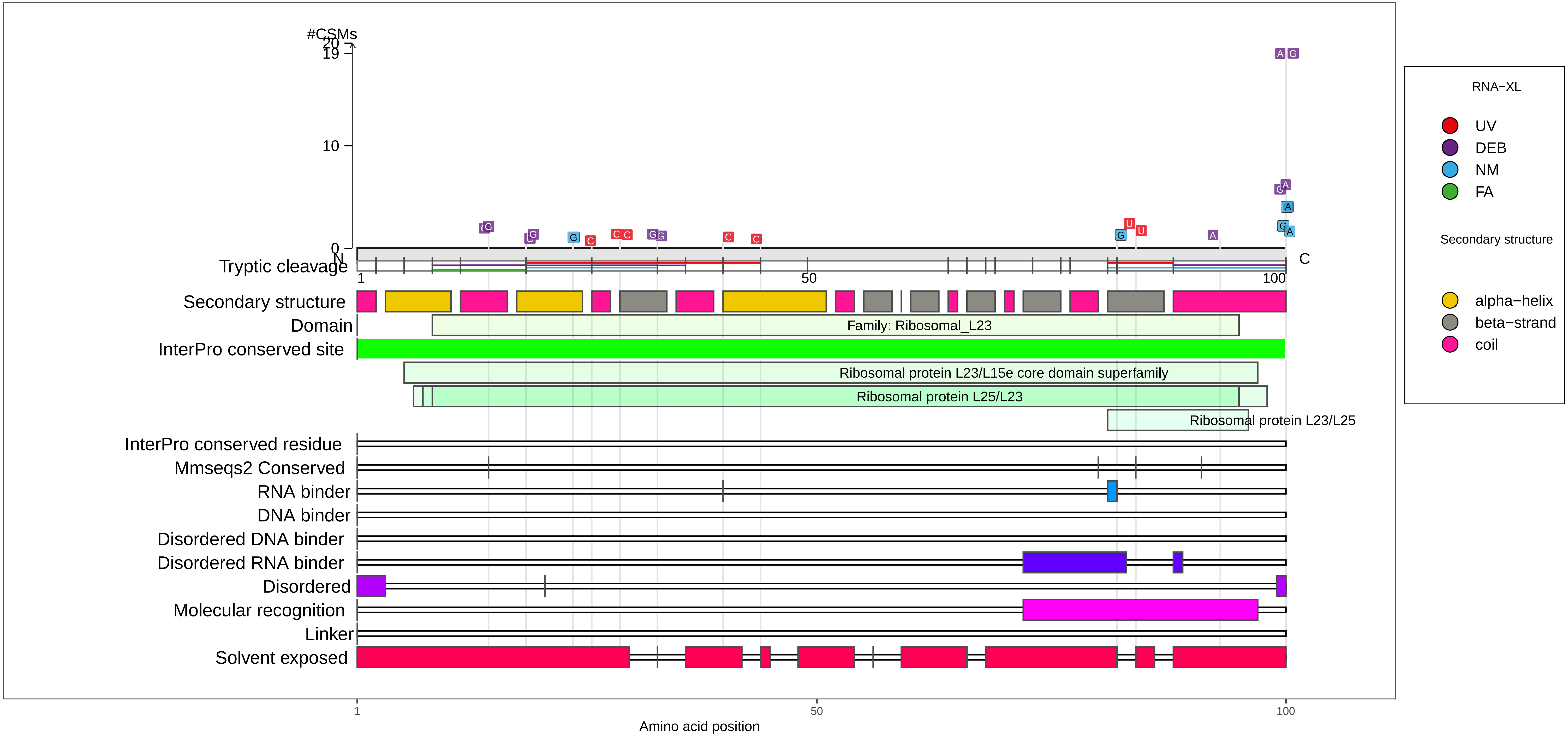

P11875
SYR_ECOLI Arginine−−tRNA ligase

− Abundance:
tryptic [log10 Intensity]: 8.36 (Q 71)
PAXdb K12 strain [ppm]: 2.16 (Q 66)
PAXdb E.coli [ppm]: 2.23 (Q 78)

− RNA functions:
aminoacyl−tRNA ligase activity; arginine−tRNA ligase activity
Arginyl tRNA synthetase N terminal domain; arginyl−tRNA aminoacylation; ncRNA metabolic process
RNA metabolic process; tRNA aminoacylation; tRNA aminoacylation for protein translation
tRNA metabolic process; tRNA synthetases class I (R)

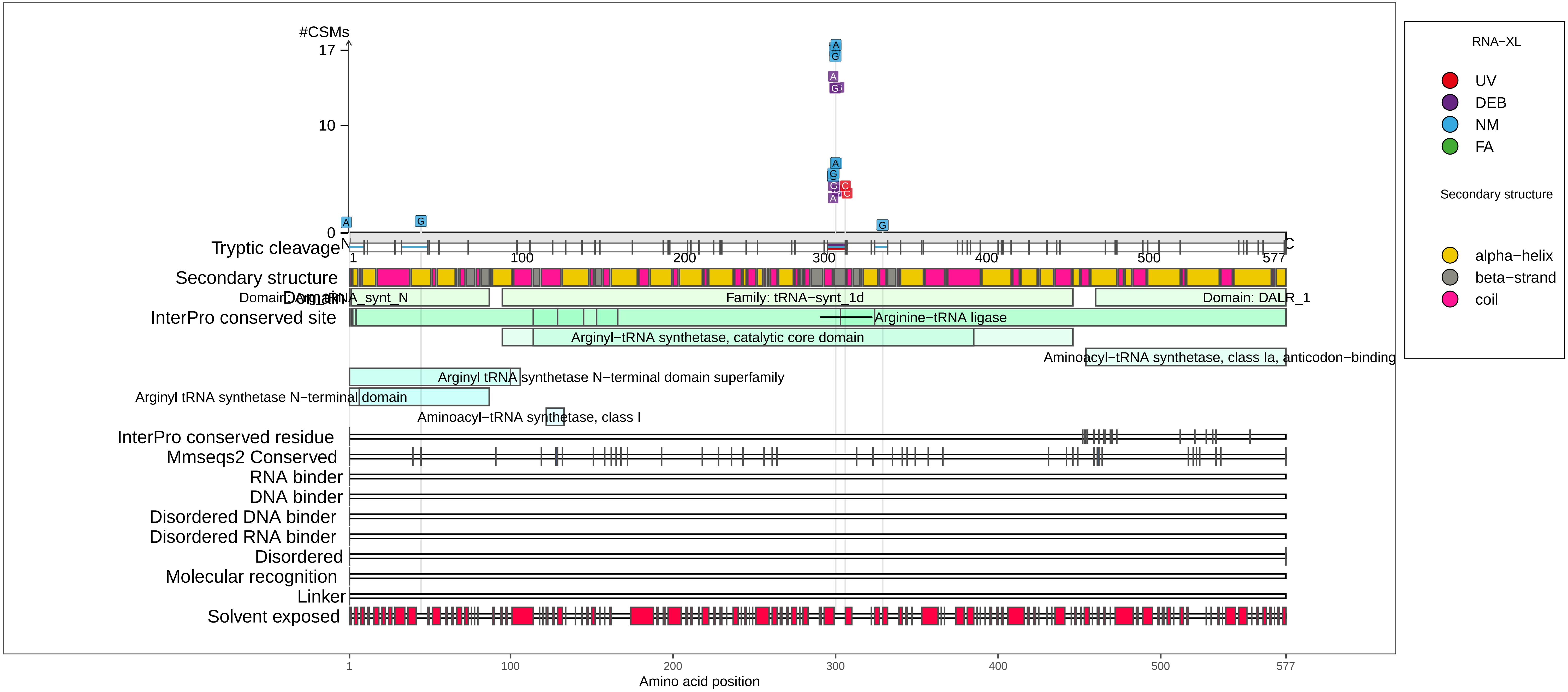

P0A6Y1
IHFB_ECOLI Integration host factor subunit beta

− Abundance:
tryptic [log10 Intensity]: 7.85 (Q 51)
PAXdb K12 strain [ppm]: 3.42 (Q 96)
PAXdb E.coli [ppm]: 3.18 (Q 96)

− RNA functions:
RNA biosynthetic process; RNA metabolic process

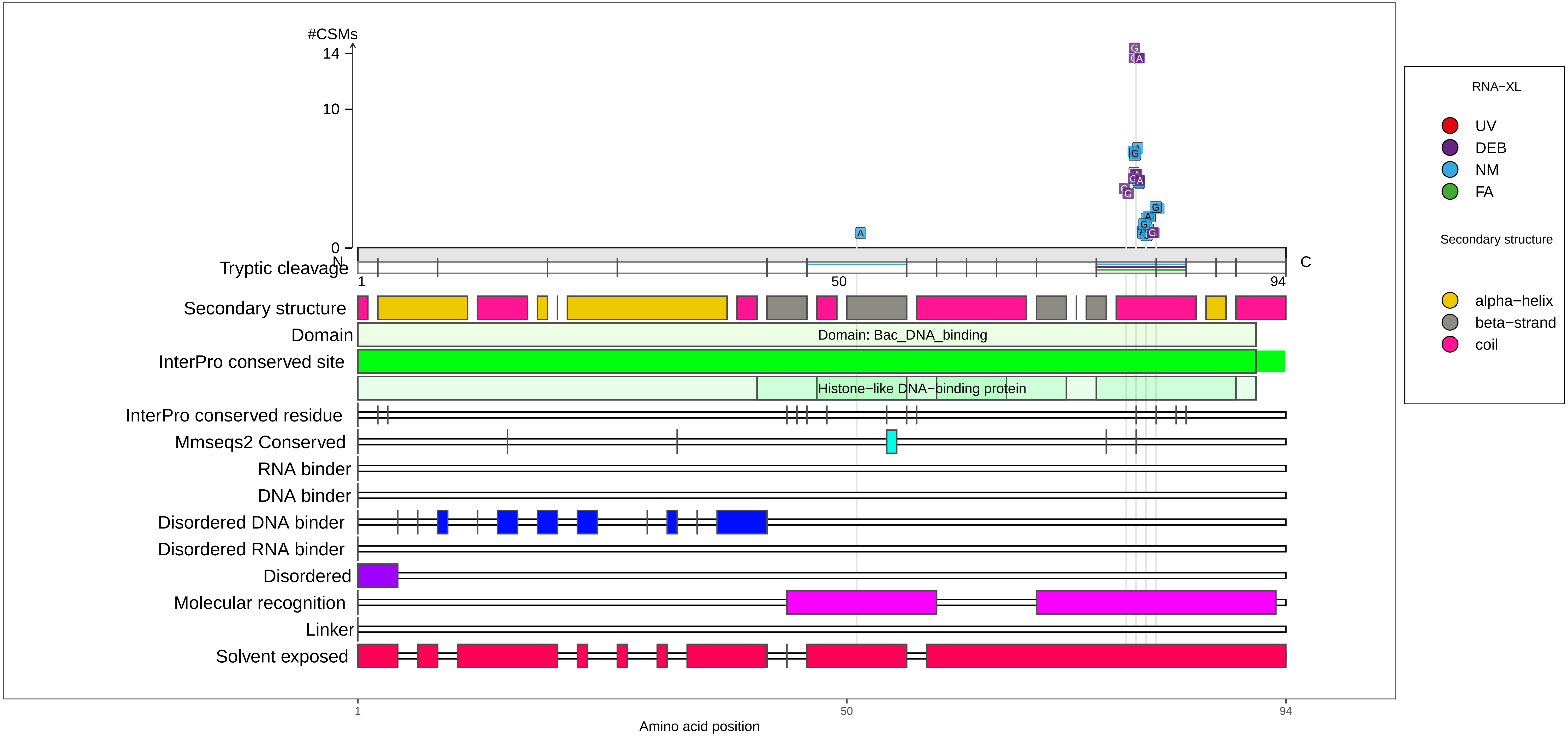

P0A7D4
PURA_ECOLI Adenylosuccinate synthetase

− Abundance:
tryptic [log10 Intensity]: 9.69 (Q 98)
PAXdb K12 strain [ppm]: 3.38 (Q 96)
PAXdb E.coli [ppm]: 3.32 (Q 98)

− RNA functions: not annotated

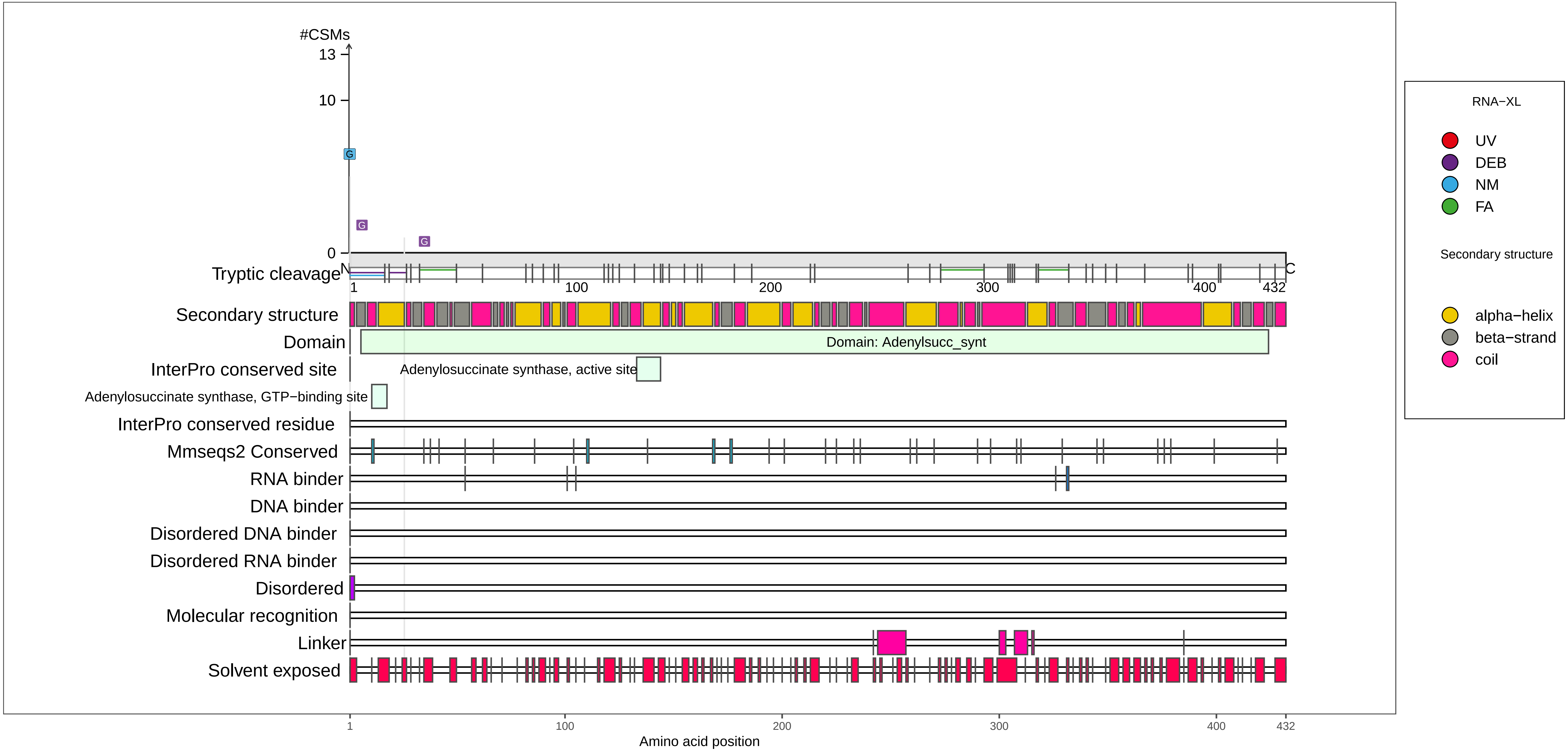

P0ABA4
ATPD_ECOLI ATP synthase subunit delta

− Abundance:
tryptic [log10 Intensity]: 9 (Q 88)
PAXdb K12 strain [ppm]: 2.87 (Q 88)
PAXdb E.coli [ppm]: 3.01 (Q 95)

− RNA functions: not annotated

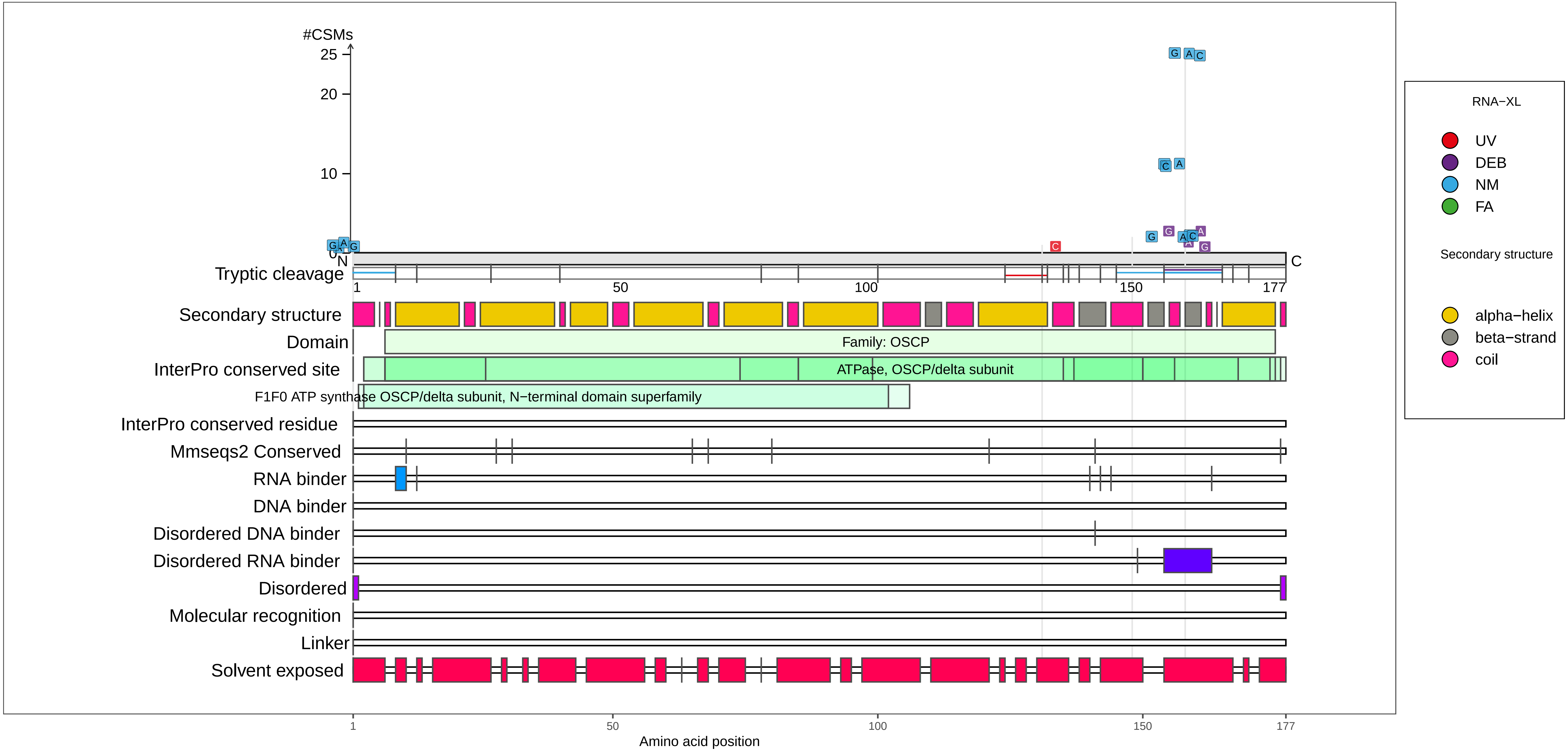

P76268
KDGR_ECOLI Transcriptional regulator KdgR

− Abundance:
tryptic [log10 Intensity]: 9.03 (Q 89)
PAXdb K12 strain [ppm]: 2.36 (Q 72)
PAXdb E.coli [ppm]: 2.48 (Q 85)

− RNA functions: not annotated

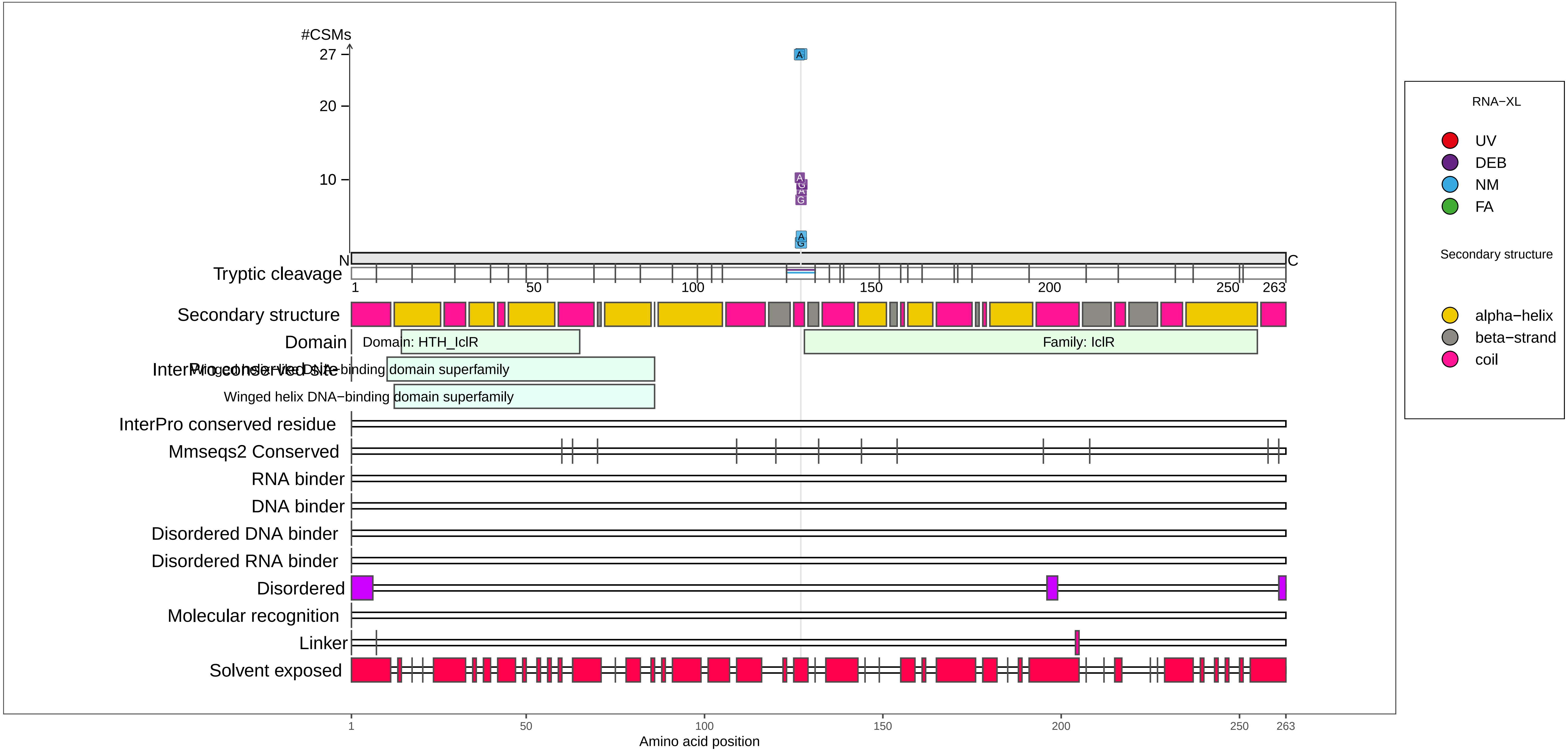

P0ADG7
IMDH_ECOLI Inosine−5'−monophosphate dehydrogenase

− Abundance:
tryptic [log10 Intensity]: 9.54 (Q 97)
PAXdb K12 strain [ppm]: 2.83 (Q 87)
PAXdb E.coli [ppm]: 2.94 (Q 94)

− RNA functions: not annotated

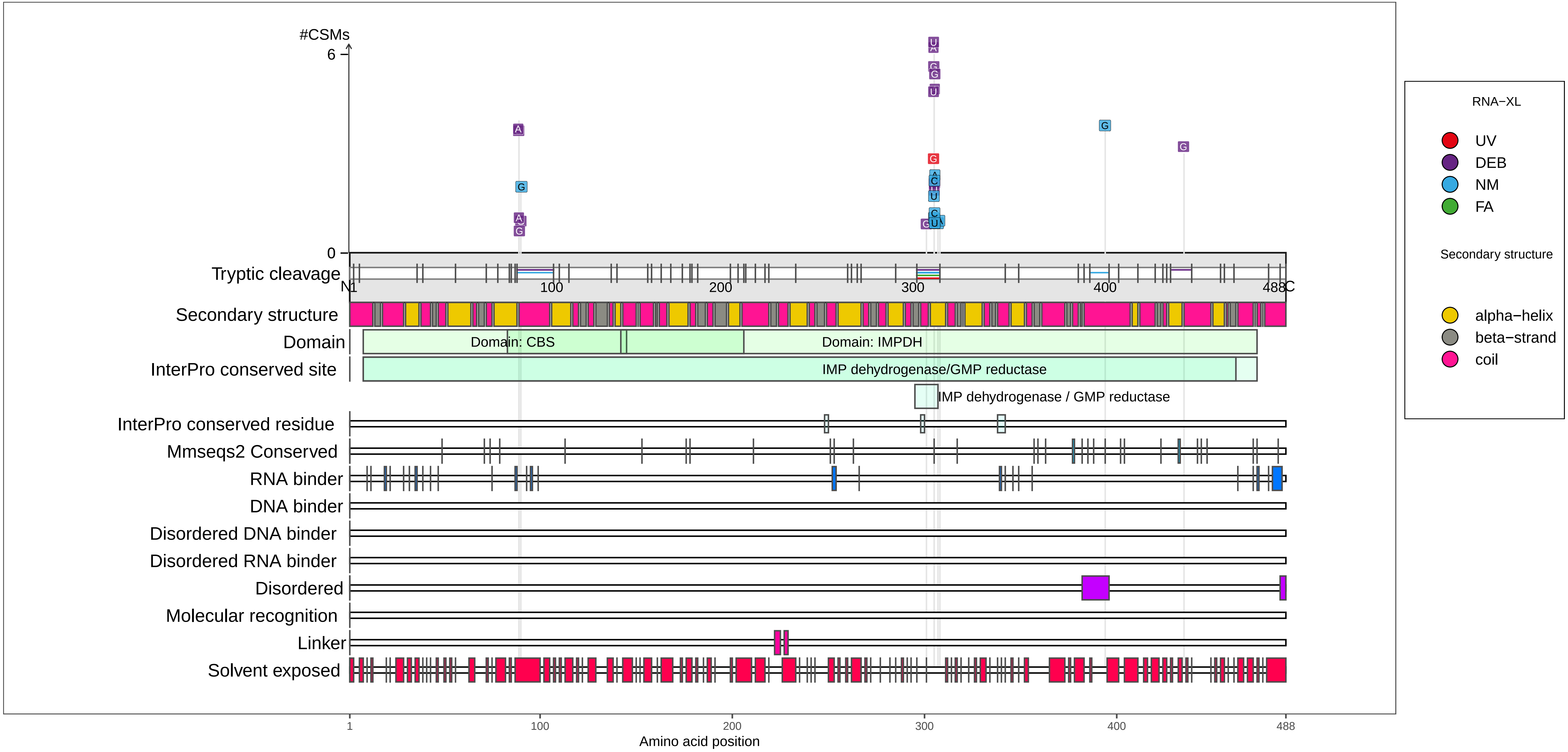

P0ACF8
HNS_ECOLI DNA−binding protein H−NS

− Abundance:
tryptic [log10 Intensity]: 9.73 (Q 98)
PAXdb K12 strain [ppm]: 3.95 (Q 100)
PAXdb E.coli [ppm]: 3.82 (Q 100)

− RNA functions:
RNA binding

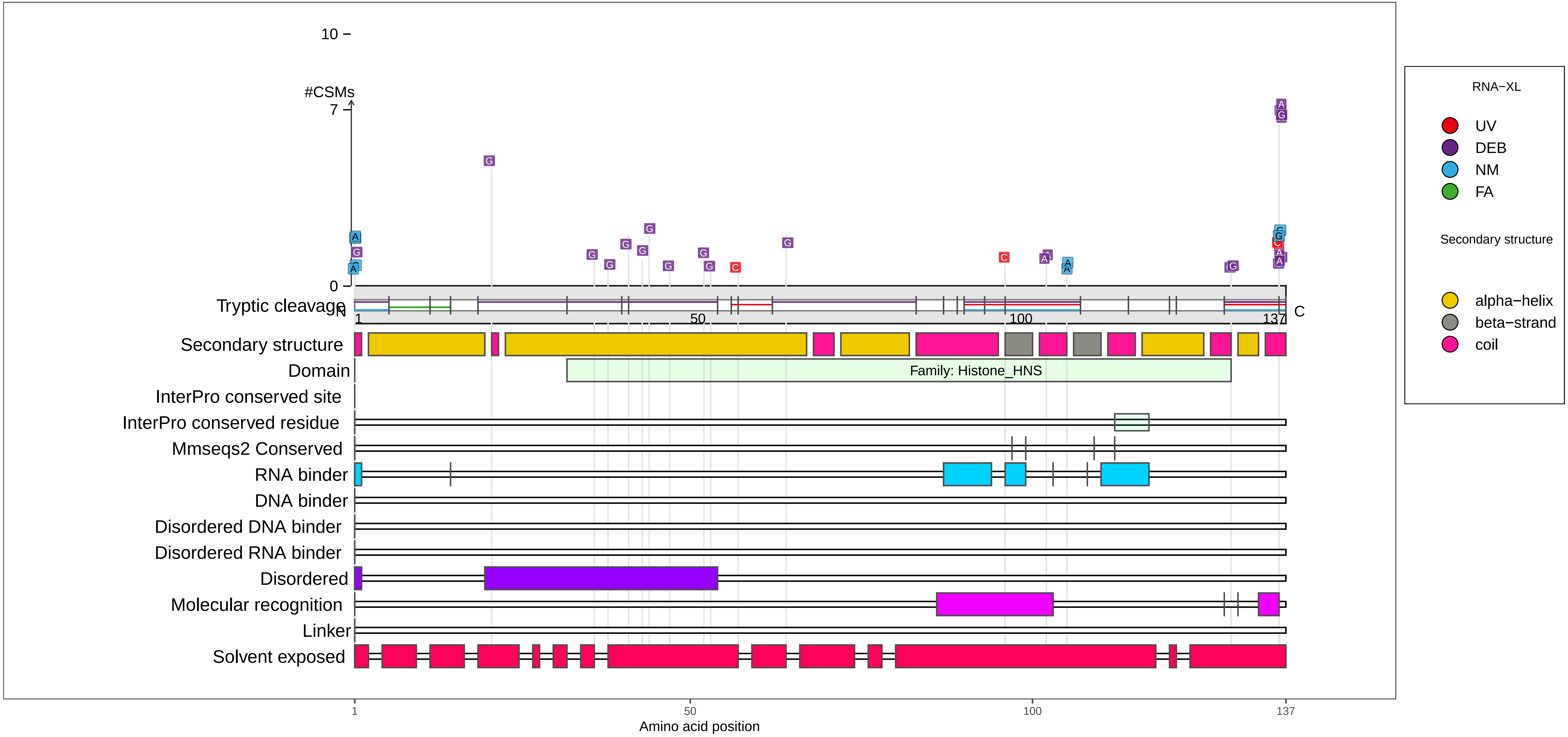

P0ADN2
YIFE_ECOLI UPF0438 protein YifE

− Abundance:
tryptic [log10 Intensity]: 8.72 (Q 81)
PAXdb K12 strain [ppm]: 3.47 (Q 97)
PAXdb E.coli [ppm]: 2.98 (Q 94)

− RNA functions: not annotated

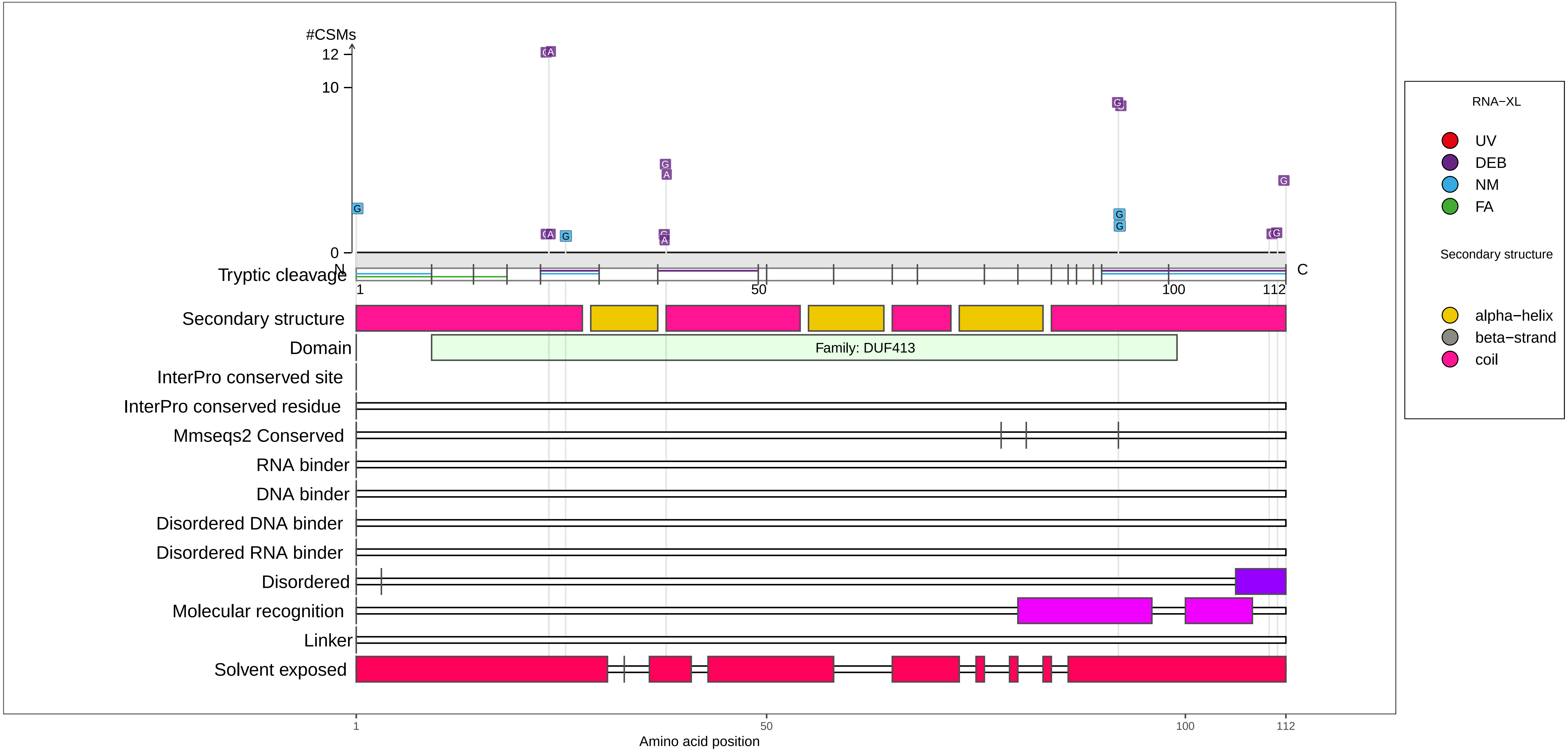

P00509
AAT_ECOLI Aspartate aminotransferase

− Abundance:
tryptic [log10 Intensity]: 9.75 (Q 98)
PAXdb K12 strain [ppm]: 3.43 (Q 96)
PAXdb E.coli [ppm]: 3.24 (Q 97)

− RNA functions: not annotated

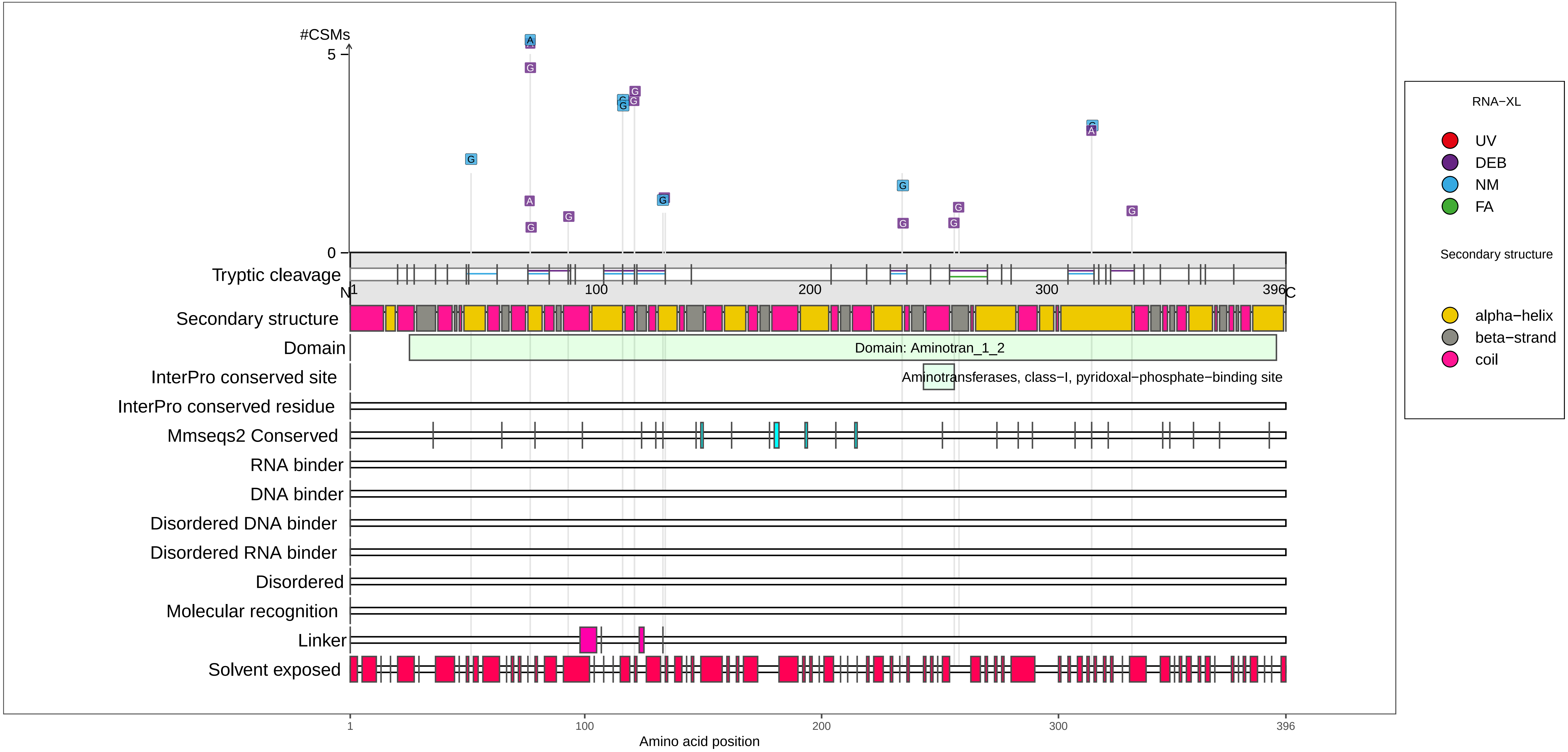

P07649
TRUA_ECOLI tRNA pseudouridine synthase A

− Abundance:
tryptic [log10 Intensity]: 8.89 (Q 86)
PAXdb K12 strain [ppm]: 2.11 (Q 64)
PAXdb E.coli [ppm]: 1.18 (Q 52)

− RNA functions:
ncRNA metabolic process; ncRNA processing; RNA binding; RNA metabolic process
RNA modification; RNA processing; tRNA binding; tRNA metabolic process; tRNA modification
tRNA processing; tRNA pseudouridine synthase; tRNA pseudouridine synthase activity
tRNA pseudouridine synthesis

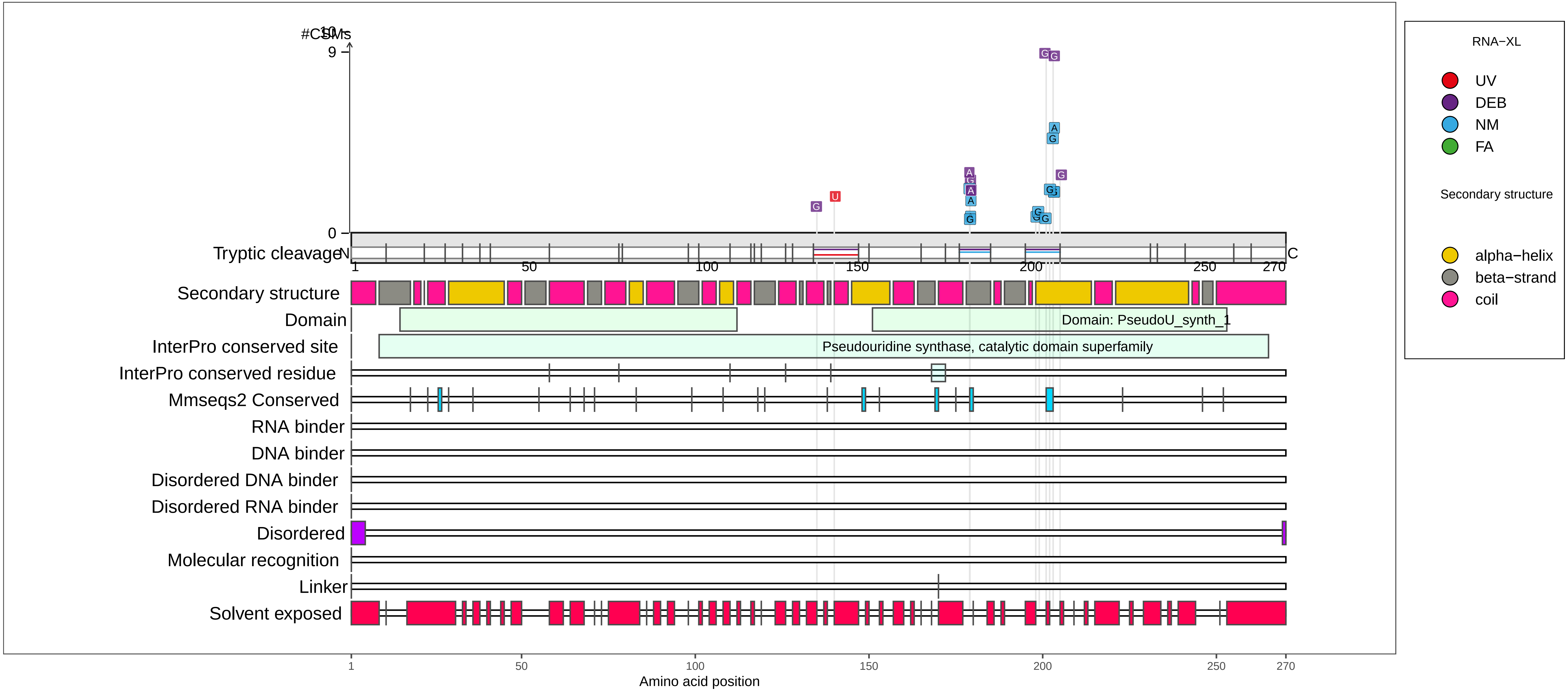

P0A7M6
RL29_ECOLI 50S ribosomal protein L29

− Abundance:
tryptic [log10 Intensity]: 9.05 (Q 89)
PAXdb K12 strain [ppm]: 3.37 (Q 96)
PAXdb E.coli [ppm]: 3.37 (Q 98)

− RNA functions:
RNA binding; rRNA binding

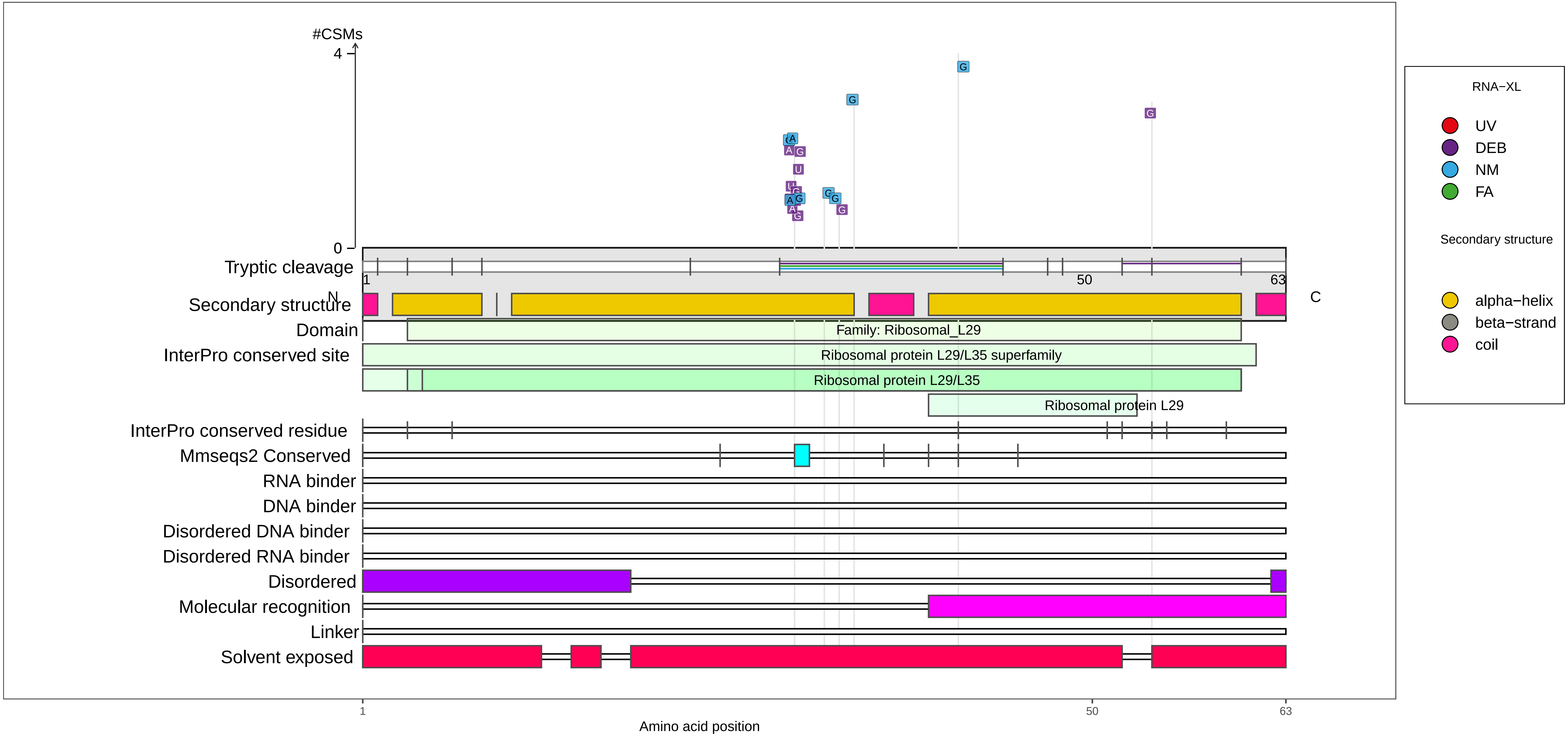

P0ABJ9
CYDA_ECOLI Cytochrome bd−I ubiquinol oxidase subunit 1

− Abundance:
tryptic [log10 Intensity]: 10.13 (Q 100)
PAXdb K12 strain [ppm]: 2.32 (Q 70)
PAXdb E.coli [ppm]: 3.12 (Q 96)

− RNA functions: not annotated

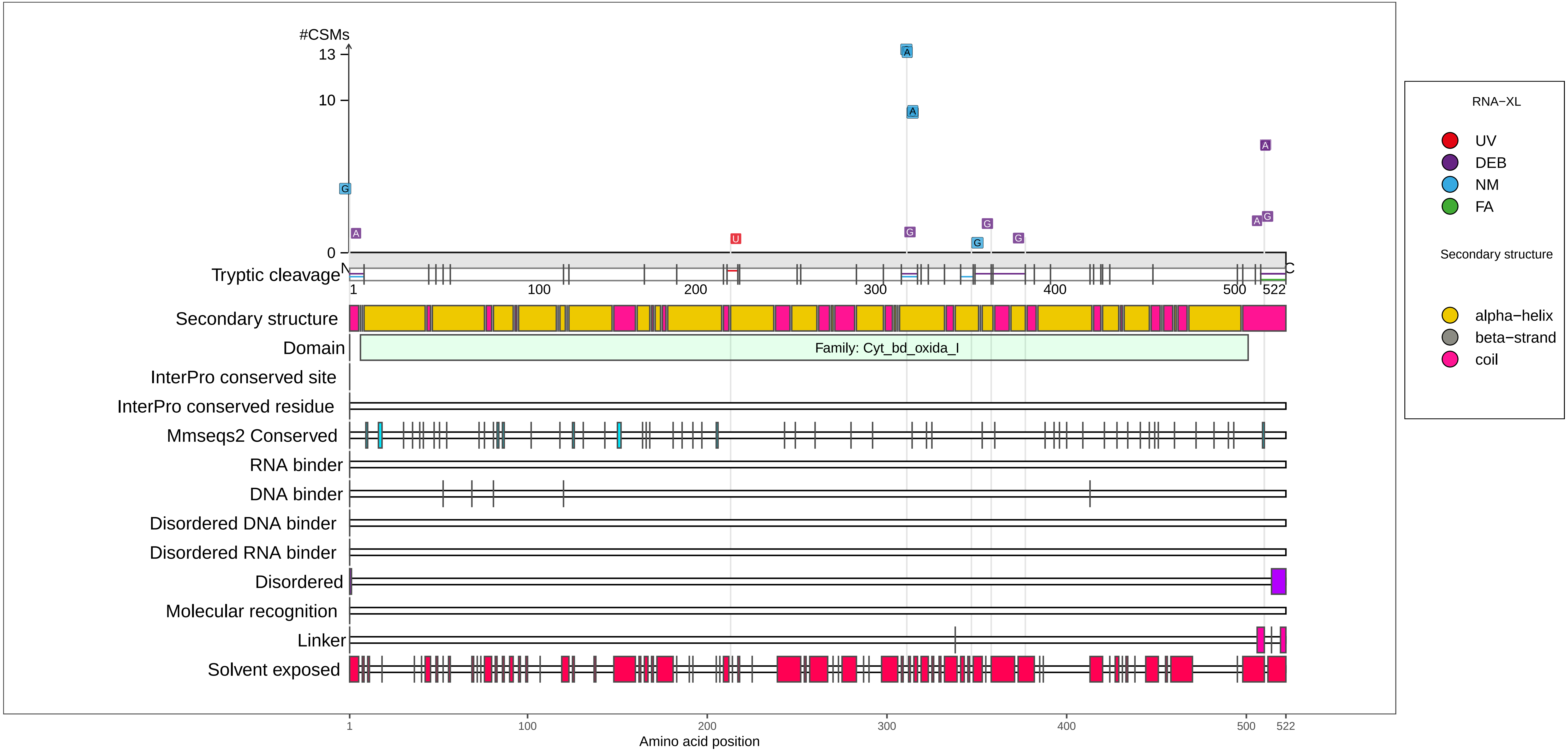

P33570
TKT2_ECOLI Transketolase 2

− Abundance:
tryptic [log10 Intensity]: 7.69 (Q 44)
PAXdb K12 strain [ppm]: 2.61 (Q 81)
PAXdb E.coli [ppm]: 2.82 (Q 92)

− RNA functions: not annotated

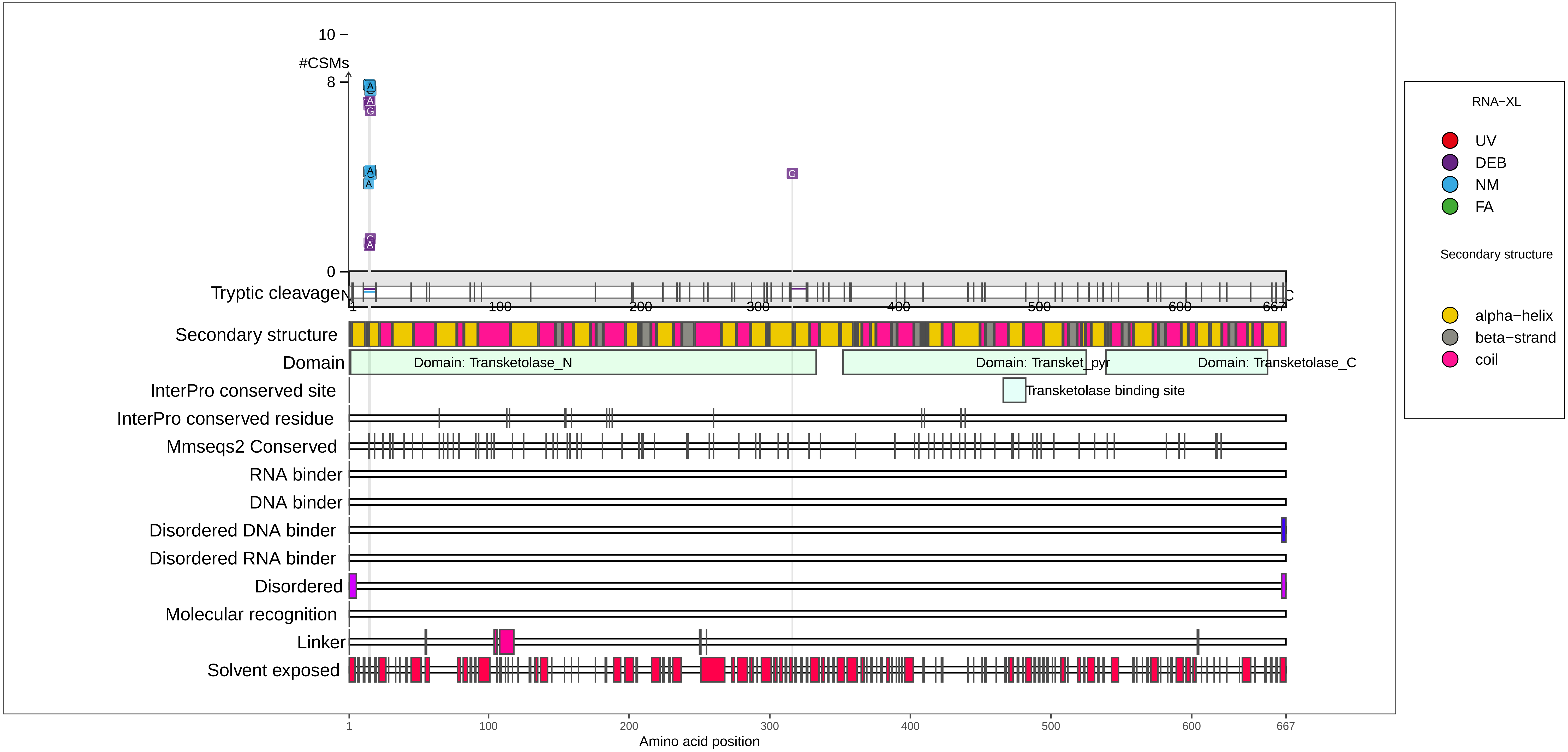

P0AB91
AROG_ECOLI Phospho−2−dehydro−3−deoxyheptonate aldolase, Phe−sensitive

− Abundance:
tryptic [log10 Intensity]: 8.52 (Q 76)
PAXdb K12 strain [ppm]: 3.14 (Q 93)
PAXdb E.coli [ppm]: 2.7 (Q 90)

− RNA functions: not annotated

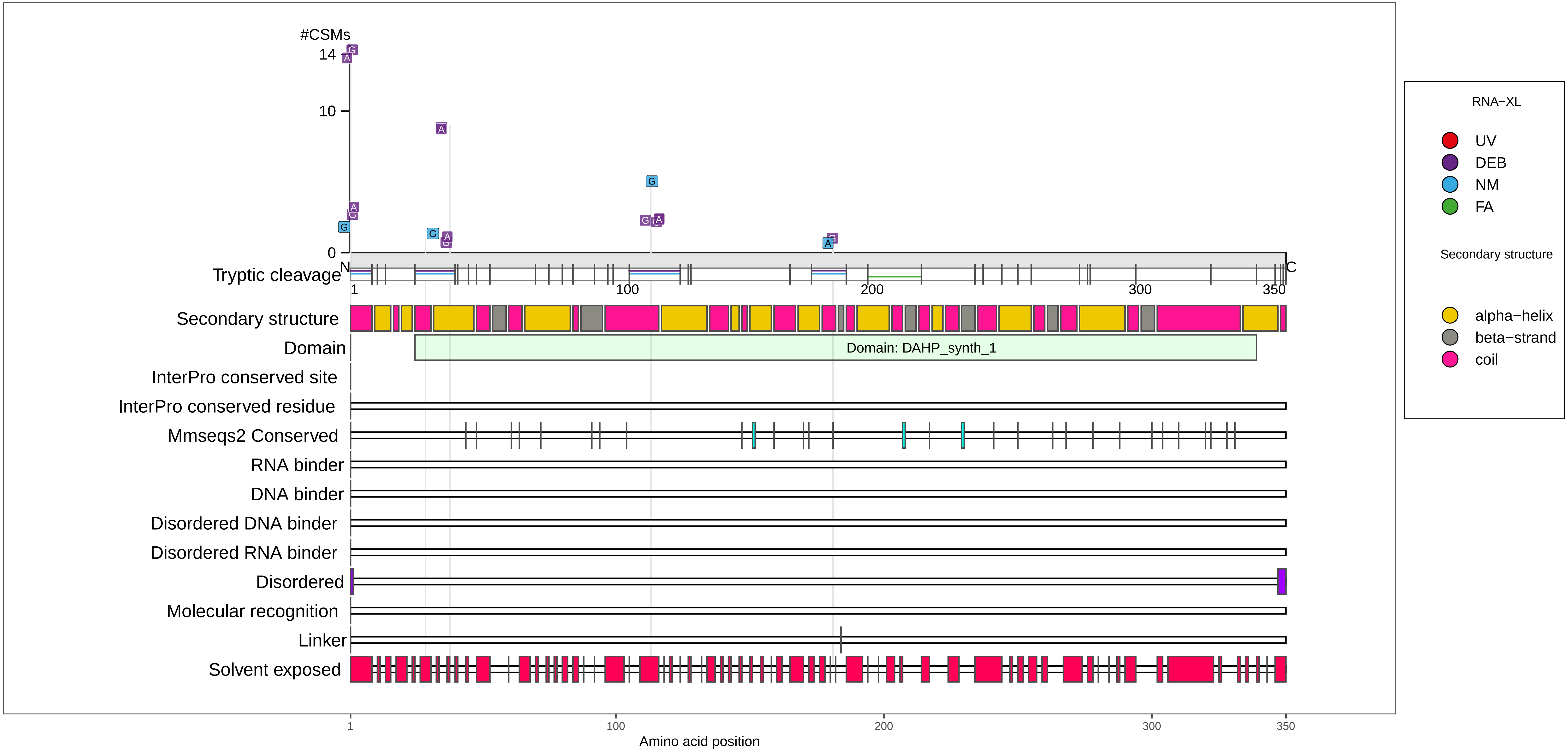

P0ABZ6
SURA_ECOLI Chaperone SurA

− Abundance:
tryptic [log10 Intensity]: 8.37 (Q 71)
PAXdb K12 strain [ppm]: 3.07 (Q 92)
PAXdb E.coli [ppm]: 2.76 (Q 91)

− RNA functions: not annotated

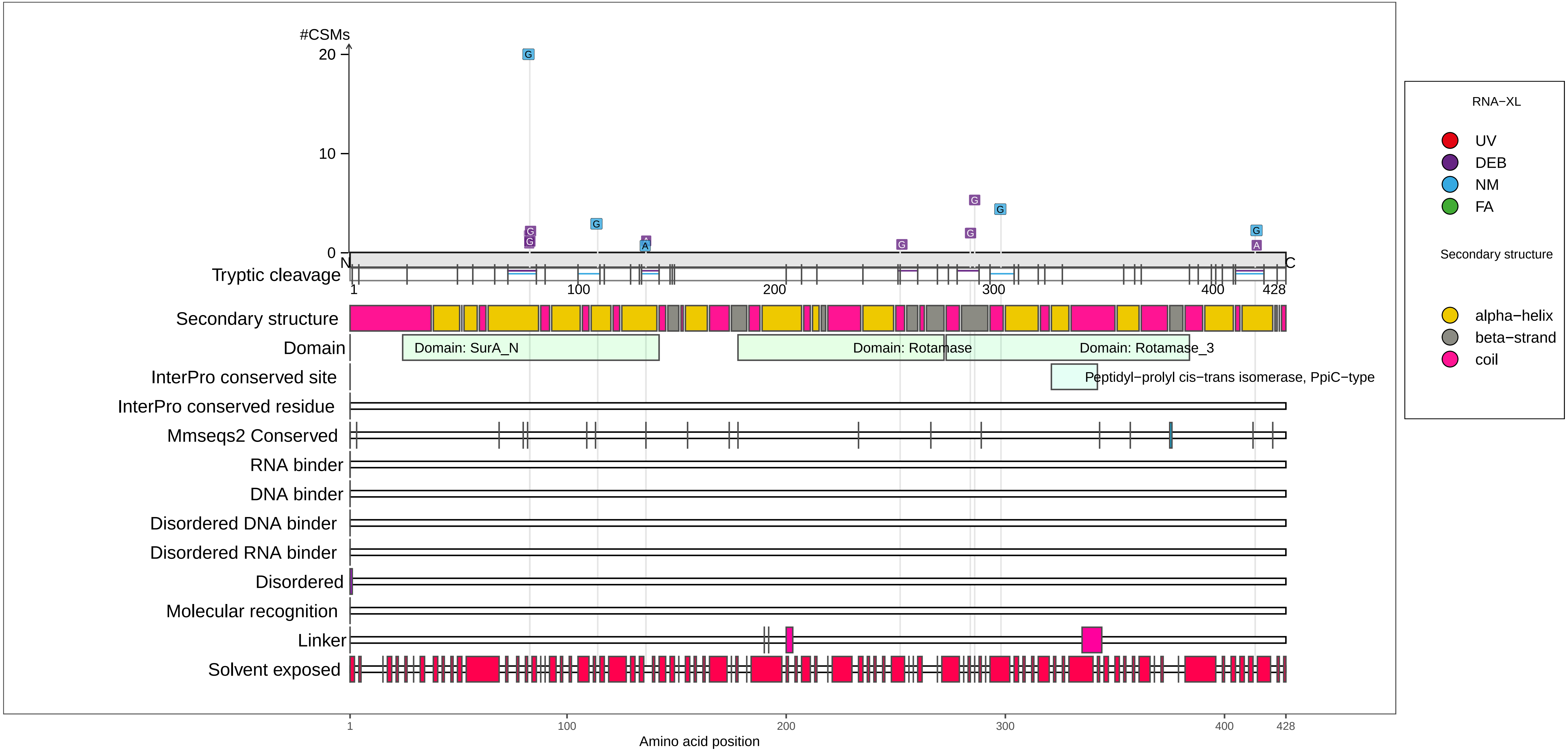

P0AEK4
FABI_ECOLI Enoyl−[acyl−carrier−protein] reductase [NADH] FabI

− Abundance:
tryptic [log10 Intensity]: 9.75 (Q 98)
PAXdb K12 strain [ppm]: 3.29 (Q 95)
PAXdb E.coli [ppm]: 3.31 (Q 97)

− RNA functions: not annotated

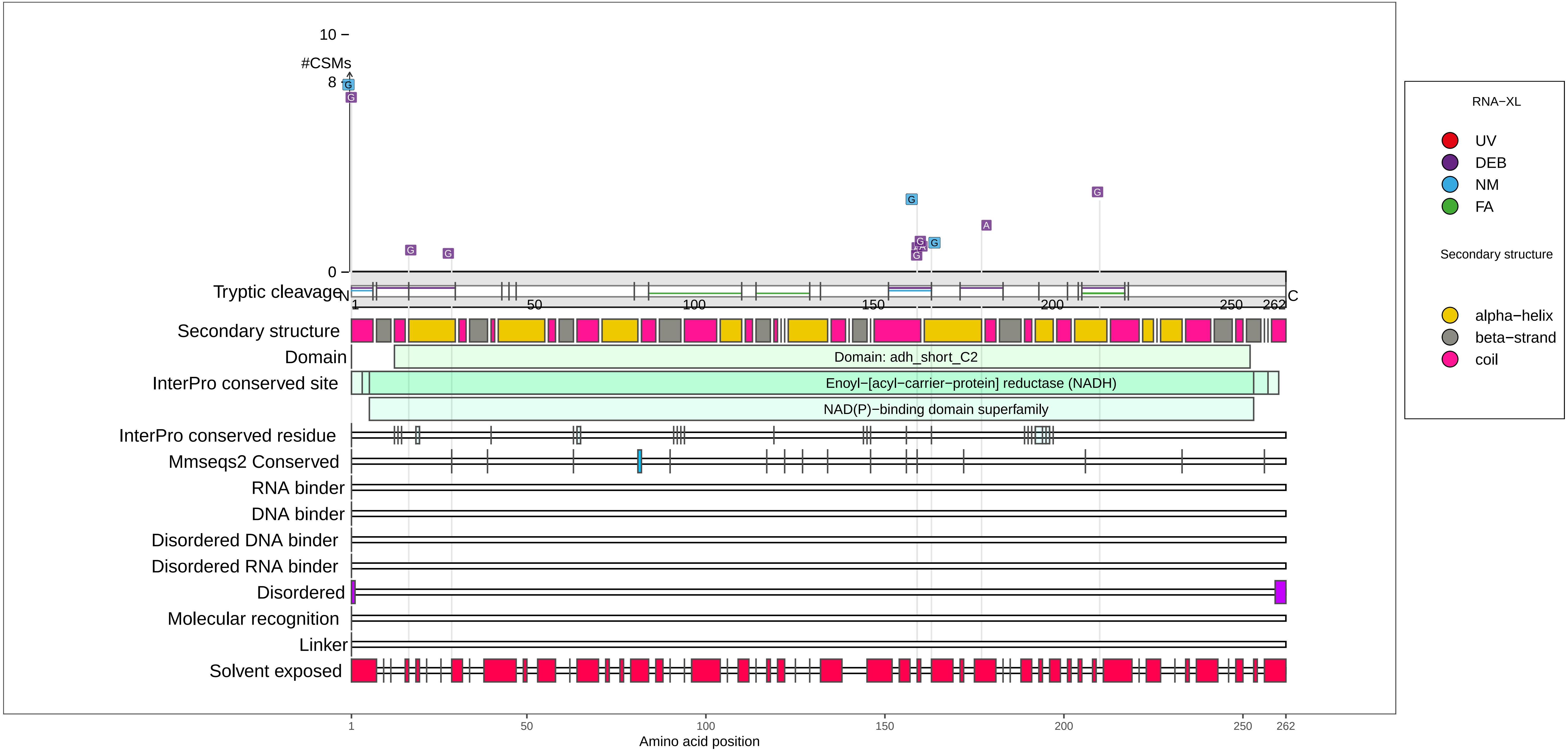

P0A6B7
ISCS_ECOLI Cysteine desulfurase IscS

− Abundance:
tryptic [log10 Intensity]: 9.17 (Q 92)
PAXdb K12 strain [ppm]: 3.19 (Q 94)
PAXdb E.coli [ppm]: 3.4 (Q 98)

− RNA functions:
ncRNA metabolic process; ncRNA processing; RNA metabolic process; RNA modification
RNA processing; tRNA metabolic process; tRNA modification; tRNA processing
tRNA pseudouridine synthesis

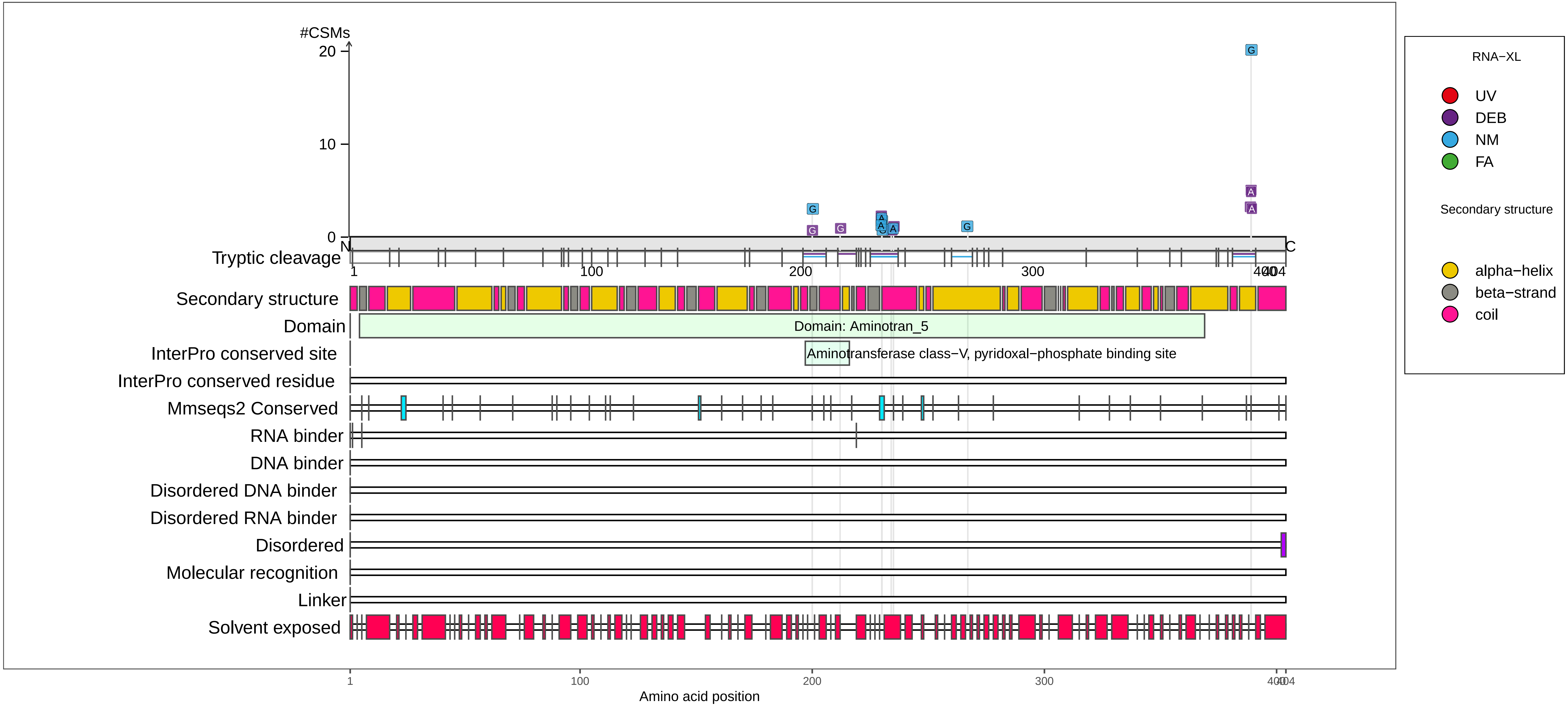

P0A6P5
DER_ECOLI GTPase Der

− Abundance:
tryptic [log10 Intensity]: 7.51 (Q 35)
PAXdb K12 strain [ppm]: 2.46 (Q 76)
PAXdb E.coli [ppm]: 2.4 (Q 83)

− RNA functions: not annotated

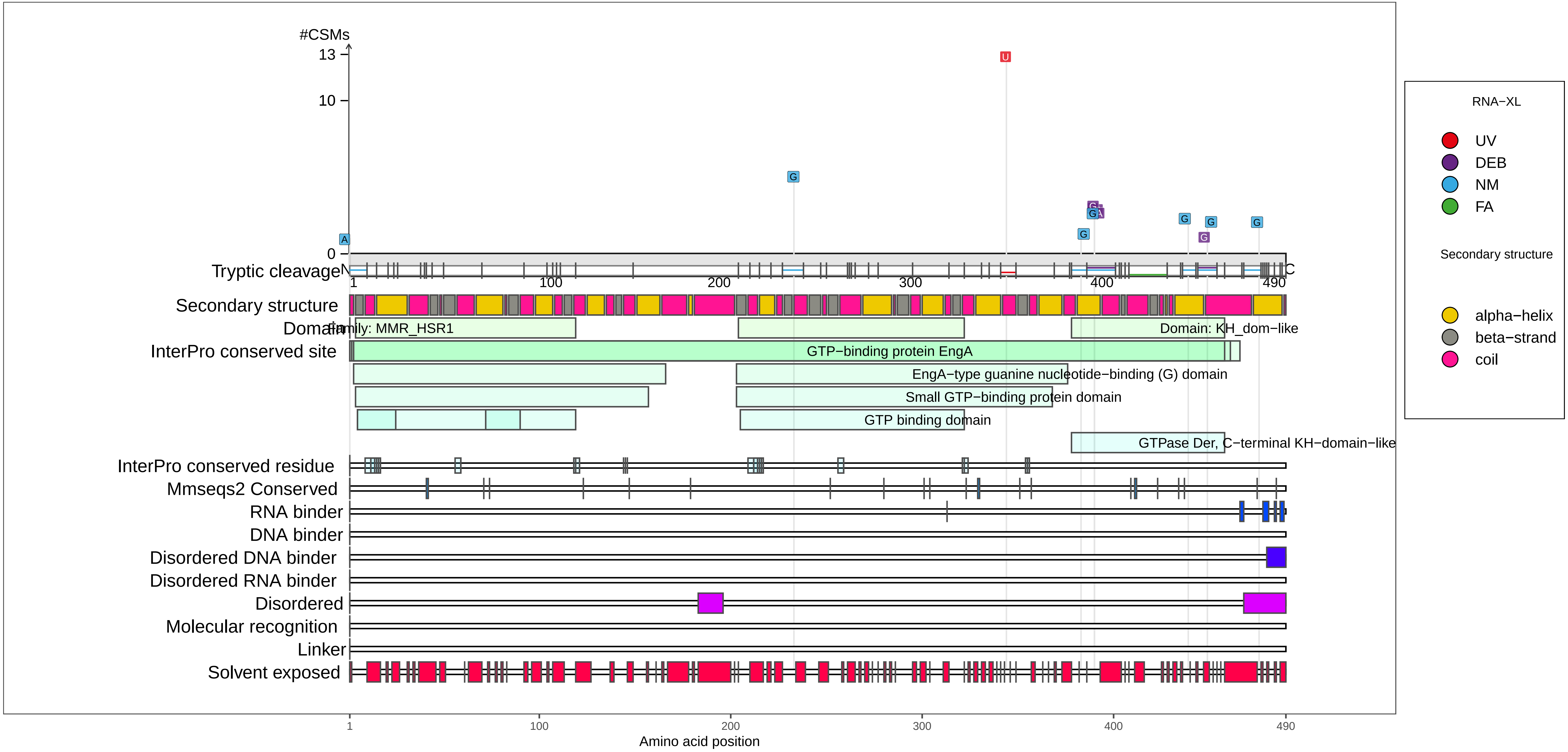

P0AGJ5
YFIF_ECOLI Uncharacterized tRNA/rRNA methyltransferase YfiF

− Abundance:
tryptic [log10 Intensity]: 8.46 (Q 74)
PAXdb K12 strain [ppm]: 1.61 (Q 45)
PAXdb E.coli [ppm]: 2.39 (Q 83)

− RNA functions:
RNA 2−O ribose methyltransferase substrate binding; RNA binding
RNA metabolic process; RNA methylation; RNA methyltransferase activity; RNA modification; RNA processing
SpoU rRNA Methylase family

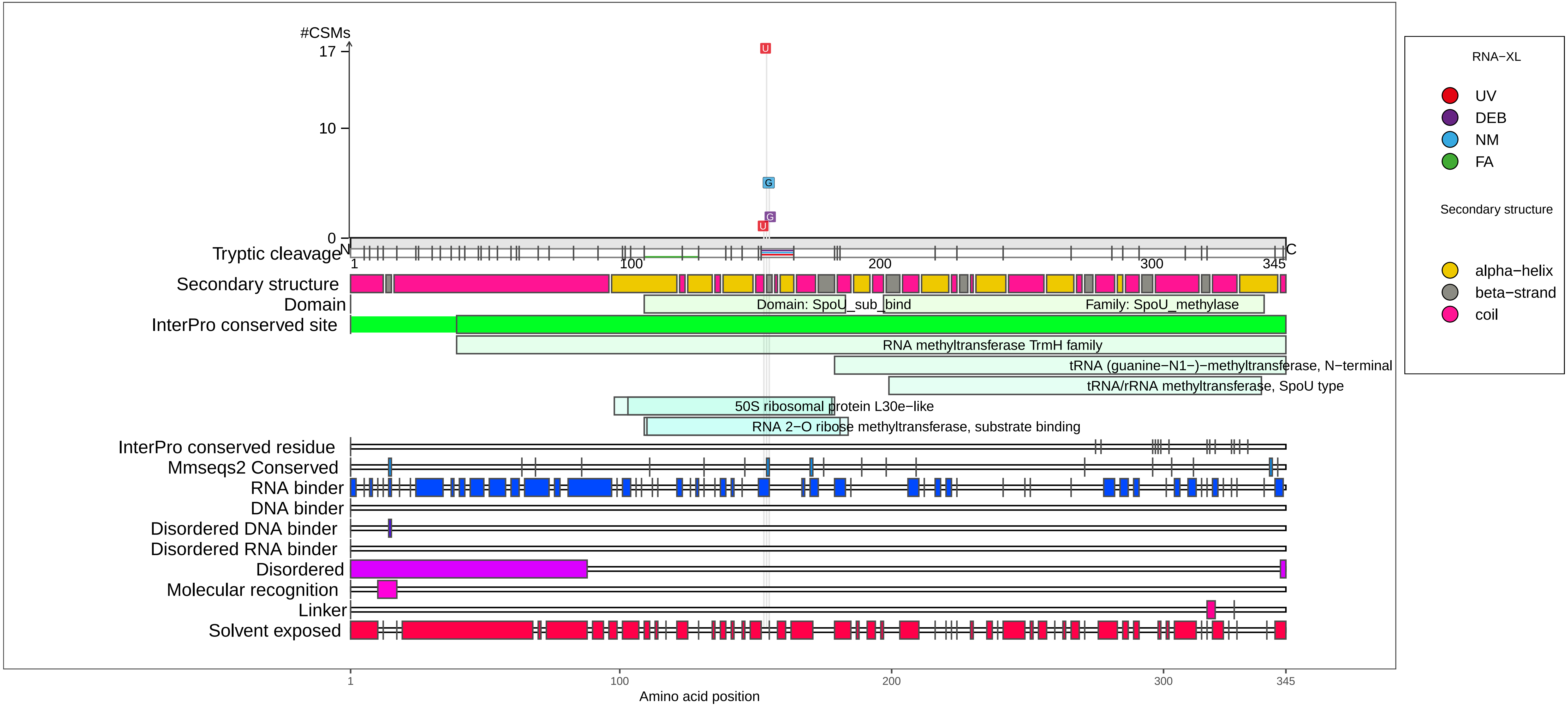

P37634
RLMJ_ECOLI Ribosomal RNA large subunit methyltransferase J

− Abundance:
tryptic [log10 Intensity]: 7.76 (Q 47)
PAXdb K12 strain [ppm]: NA (Q NA)
PAXdb E.coli [ppm]: 1.55 (Q 61)

− RNA functions:
23S rRNA (adenine(2030)−N(6))−methyltransferase activity; ncRNA metabolic process
ncRNA processing; Ribosomal RNA large subunit methyltransferase D, RlmJ; RNA binding
RNA metabolic process; RNA methylation; RNA methyltransferase activity
RNA modification; RNA processing; rRNA (adenine−N6−)−methyltransferase activity
rRNA (adenine) methyltransferase activity; rRNA base methylation; rRNA metabolic process; rRNA methylation
rRNA methyltransferase activity; rRNA modification; rRNA processing

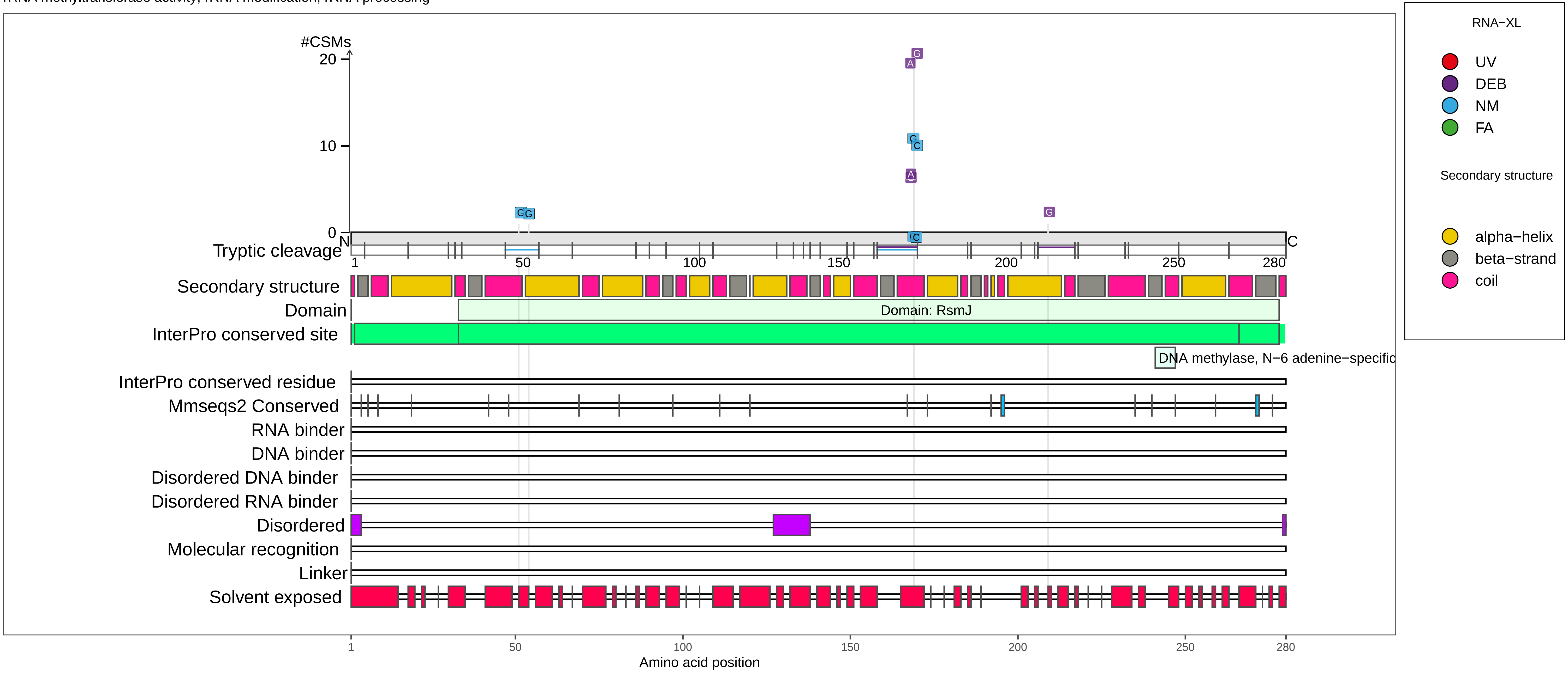

P00968
CARB_ECOLI Carbamoyl−phosphate synthase large chain

− Abundance:
tryptic [log10 Intensity]: 9.04 (Q 89)
PAXdb K12 strain [ppm]: 3.08 (Q 92)
PAXdb E.coli [ppm]: 2.57 (Q 87)

− RNA functions: not annotated

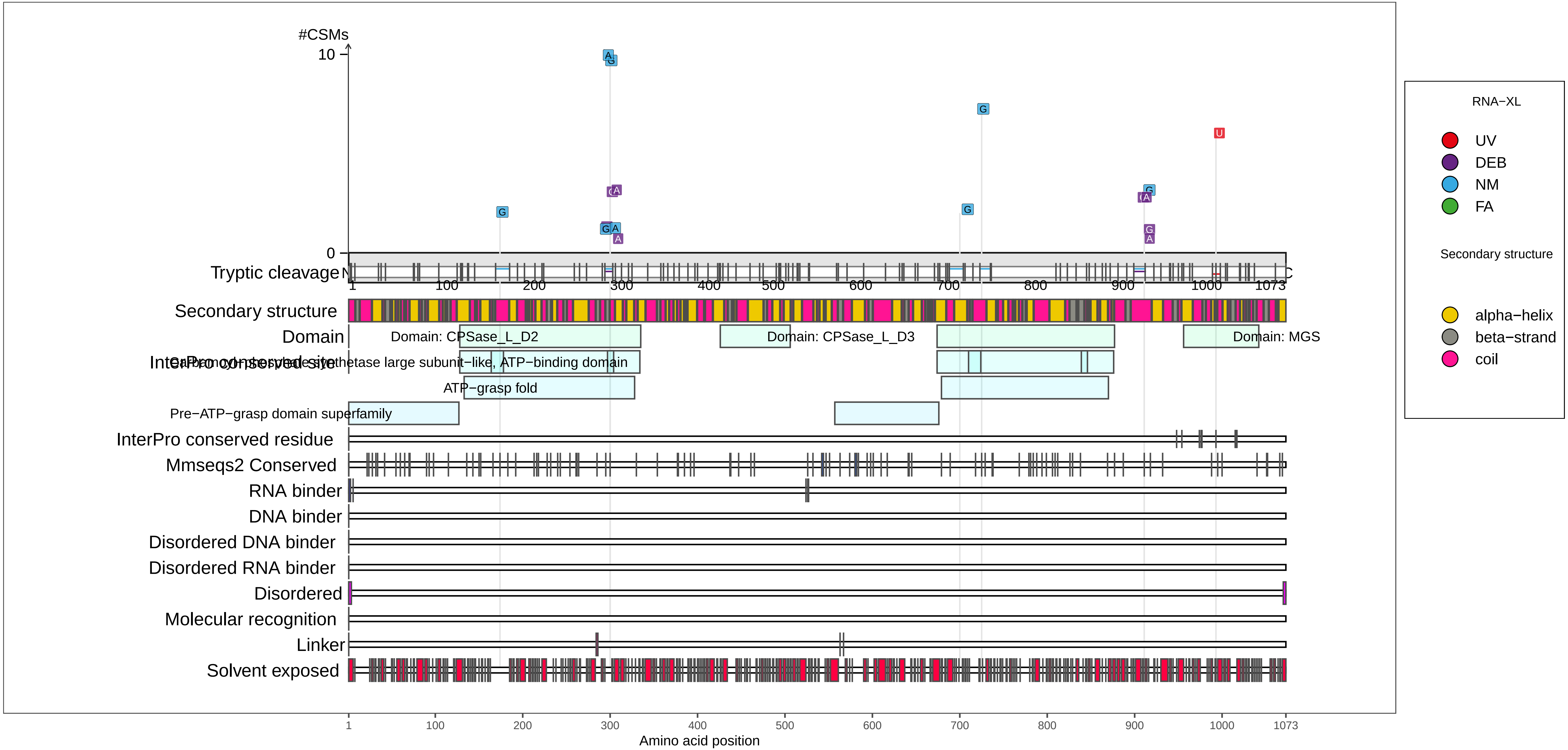

P0A870
TALB_ECOLI Transaldolase B

− Abundance:
tryptic [log10 Intensity]: 9.67 (Q 98)
PAXdb K12 strain [ppm]: 3.09 (Q 92)
PAXdb E.coli [ppm]: 3.33 (Q 98)

− RNA functions: not annotated

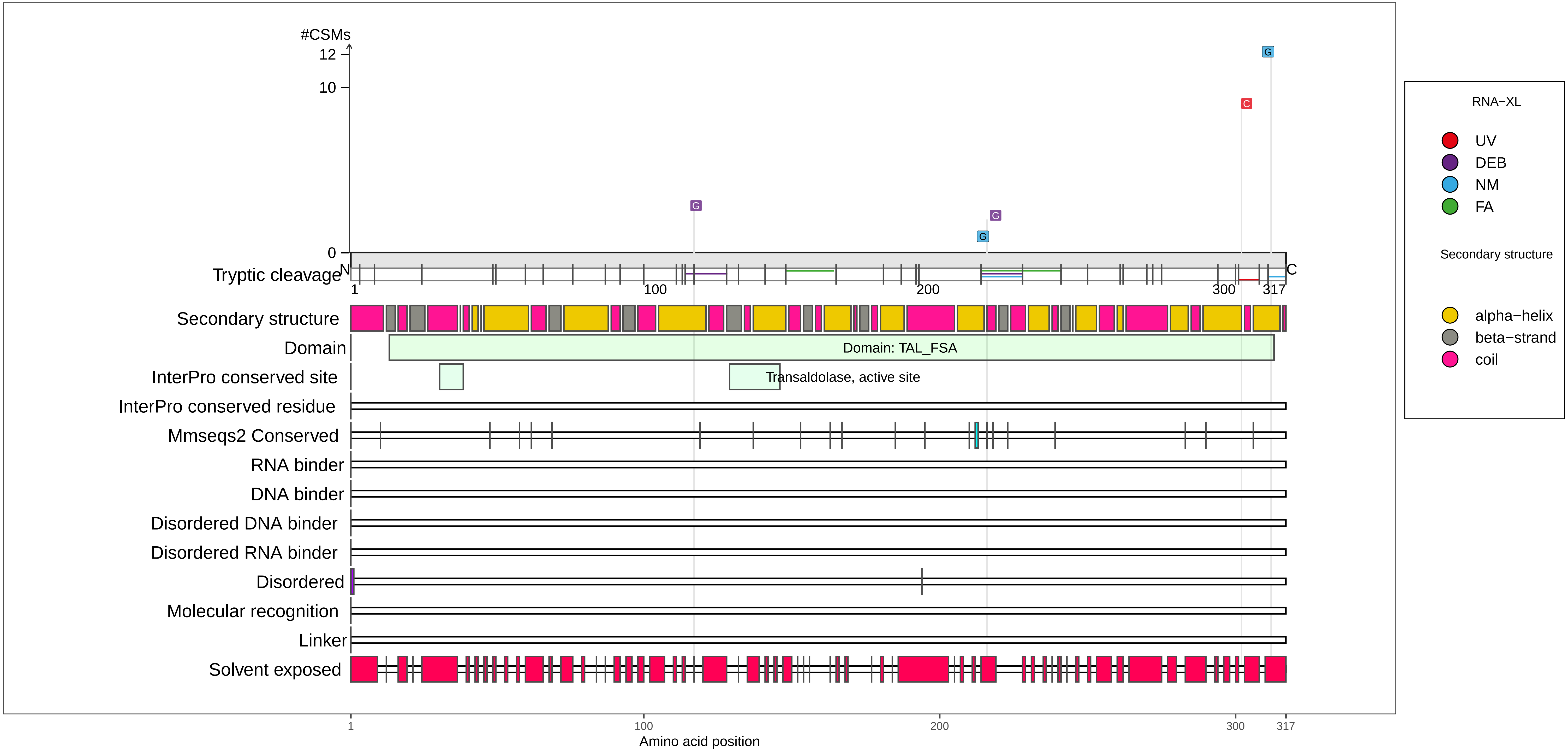

P0DTT0

BIPA_ECOLI 50S ribosomal subunit assembly factor BipA

− Abundance:
tryptic [log10 Intensity]: 9.85 (Q 99)
PAXdb K12 strain [ppm]: 3.13 (Q 93)
PAXdb E.coli [ppm]: 2.73 (Q 90)

− RNA functions: not annotated

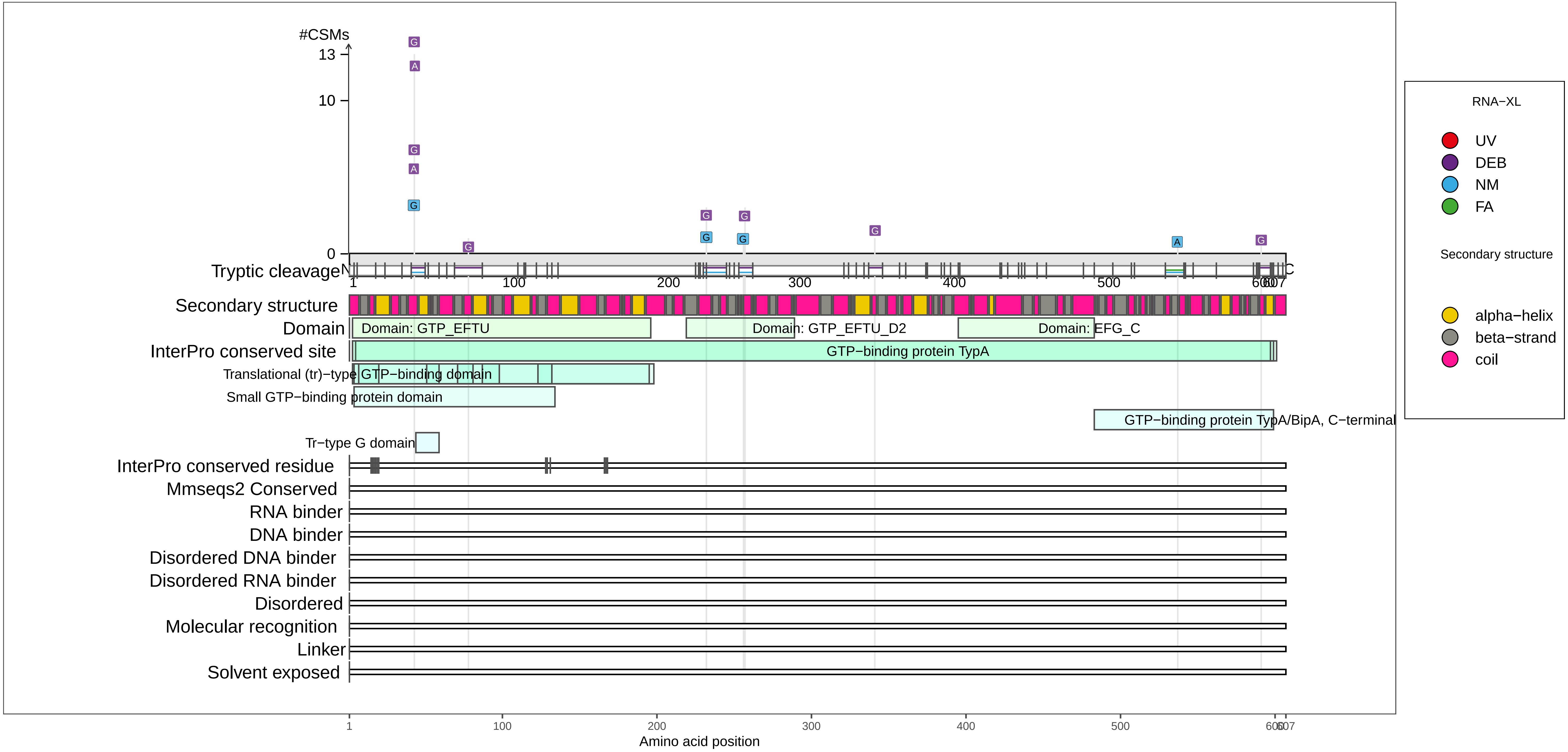

P0ACW6
YDCH_ECOLI Uncharacterized protein YdcH
− Abundance:
tryptic [log10 Intensity]: 6.94 (Q 11)
PAXdb K12 strain [ppm]: 2.77 (Q 85)
PAXdb E.coli [ppm]: 2.69 (Q 90)

− RNA functions: not annotated

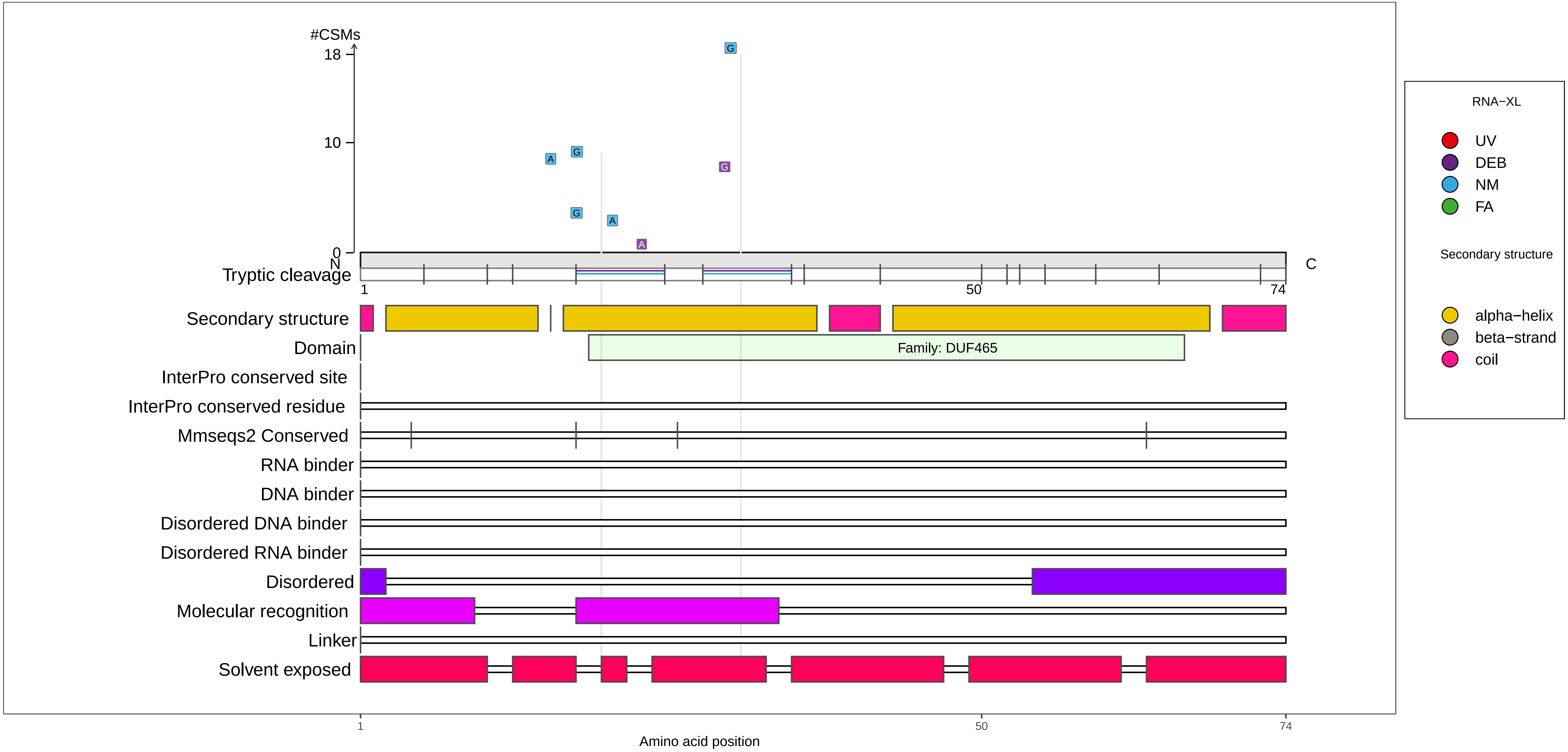

P0ADW3
YHCB_ECOLI Inner membrane protein YhcB

− Abundance:
tryptic [log10 Intensity]: 8.54 (Q 77)
PAXdb K12 strain [ppm]: 2.41 (Q 74)
PAXdb E.coli [ppm]: 2.75 (Q 91)

− RNA functions: not annotated

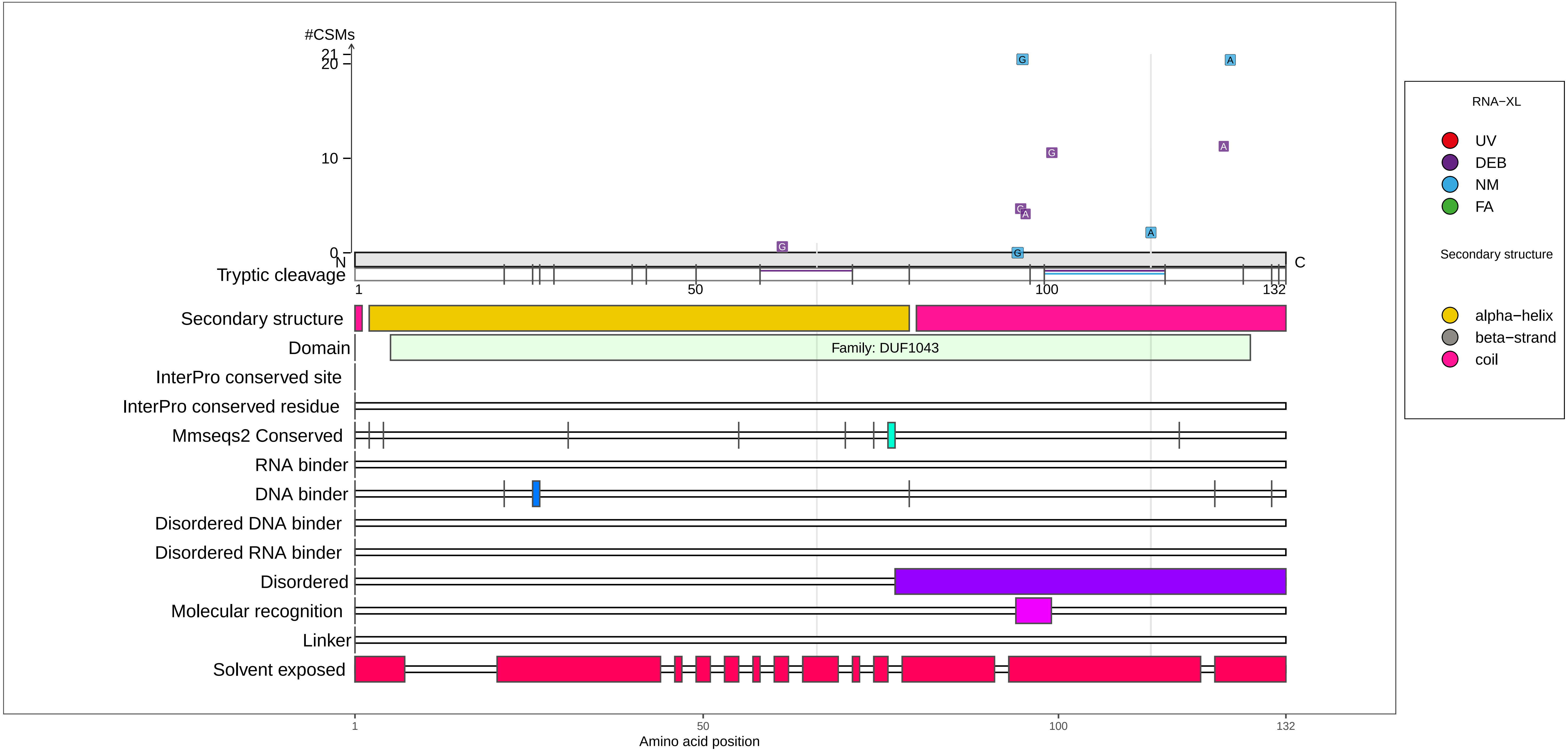

P0A953
FABB_ECOLI 3−oxoacyl−[acyl−carrier−protein] synthase 1

− Abundance:
tryptic [log10 Intensity]: 9.48 (Q 96)
PAXdb K12 strain [ppm]: 3.29 (Q 95)
PAXdb E.coli [ppm]: 3.19 (Q 96)

− RNA functions: not annotated

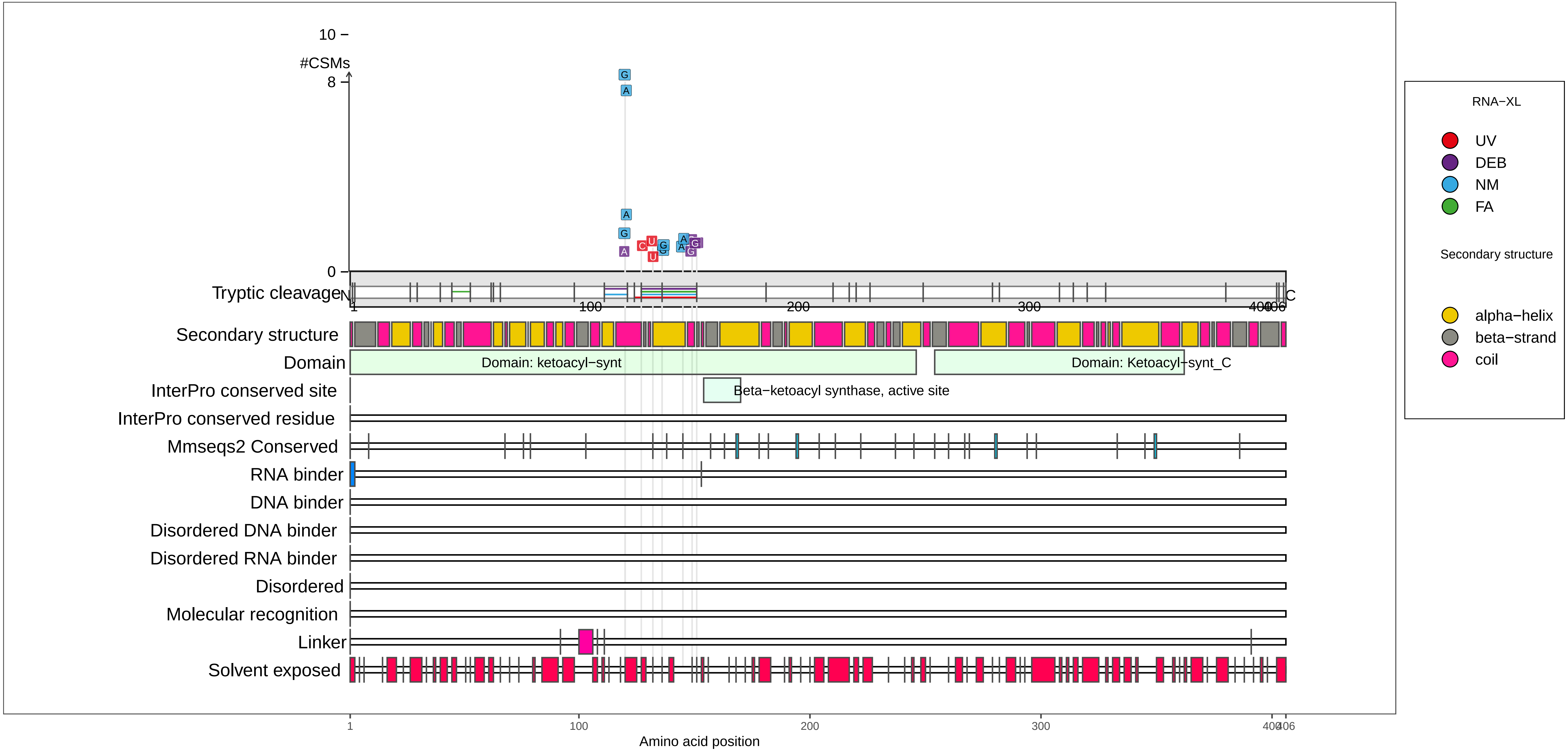

P0A9J8
CMPDT_ECOLI Bifunctional chorismate mutase/prephenate dehydratase

− Abundance:
tryptic [log10 Intensity]: 8.08 (Q 61)
PAXdb K12 strain [ppm]: 2.2 (Q 67)
PAXdb E.coli [ppm]: 2.38 (Q 83)

− RNA functions: not annotated

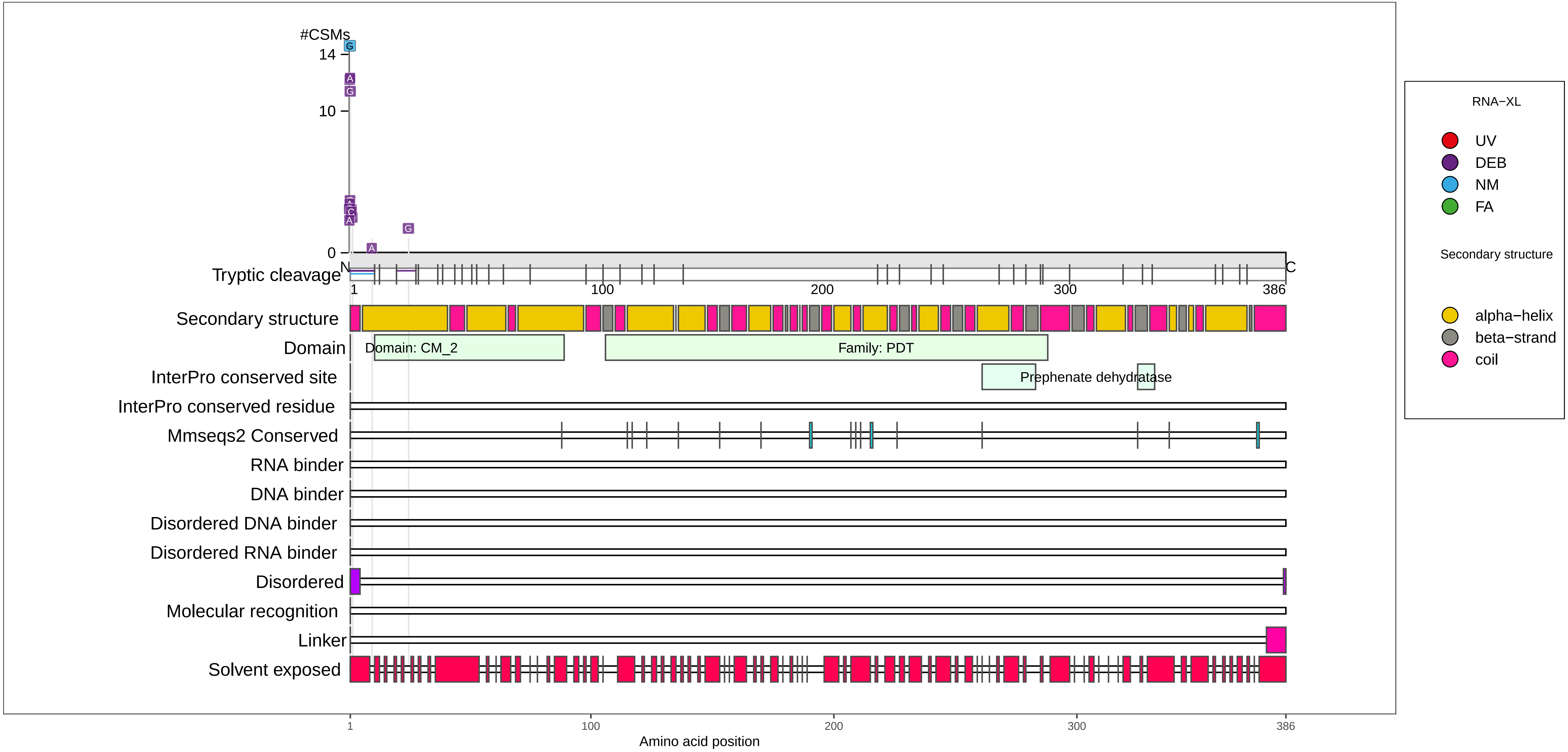

P0A9P6
DEAD_ECOLI ATP−dependent RNA helicase DeaD

− Abundance:
tryptic [log10 Intensity]: 7.69 (Q 44)
PAXdb K12 strain [ppm]: 2.66 (Q 82)
PAXdb E.coli [ppm]: 2.29 (Q 80)

− RNA functions:
DbpA RNA binding domain; mRNA stabilization; RNA binding; RNA catabolic process
RNA helicase activity; RNA metabolic process; RNA stabilization
RNA strand annealing activity

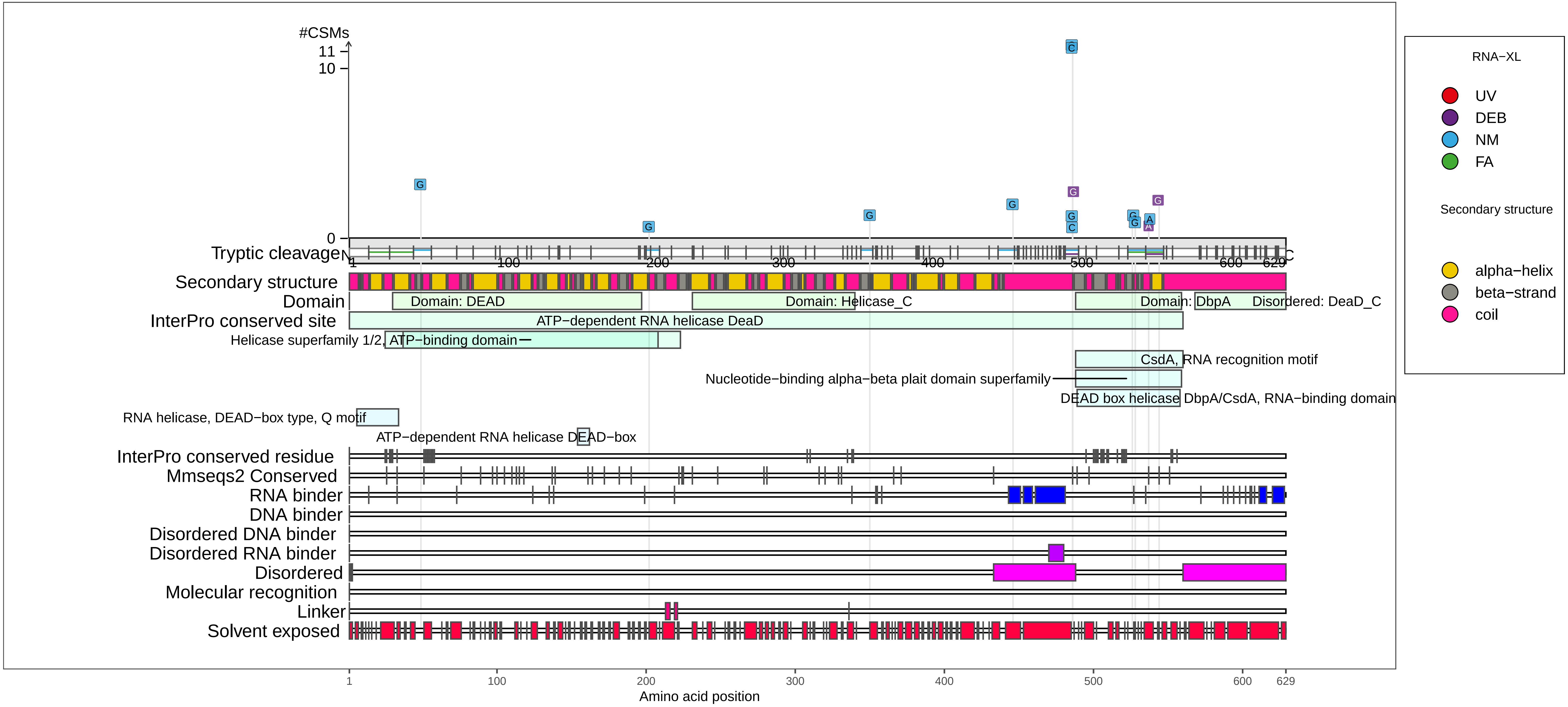

P0A9X9
CSPA_ECOLI Cold shock protein CspA

− Abundance:
tryptic [log10 Intensity]: 9.33 (Q 95)
PAXdb K12 strain [ppm]: 3.49 (Q 97)
PAXdb E.coli [ppm]: 3.24 (Q 97)

− RNA functions:
RNA binding; RNA biosynthetic process; RNA metabolic process

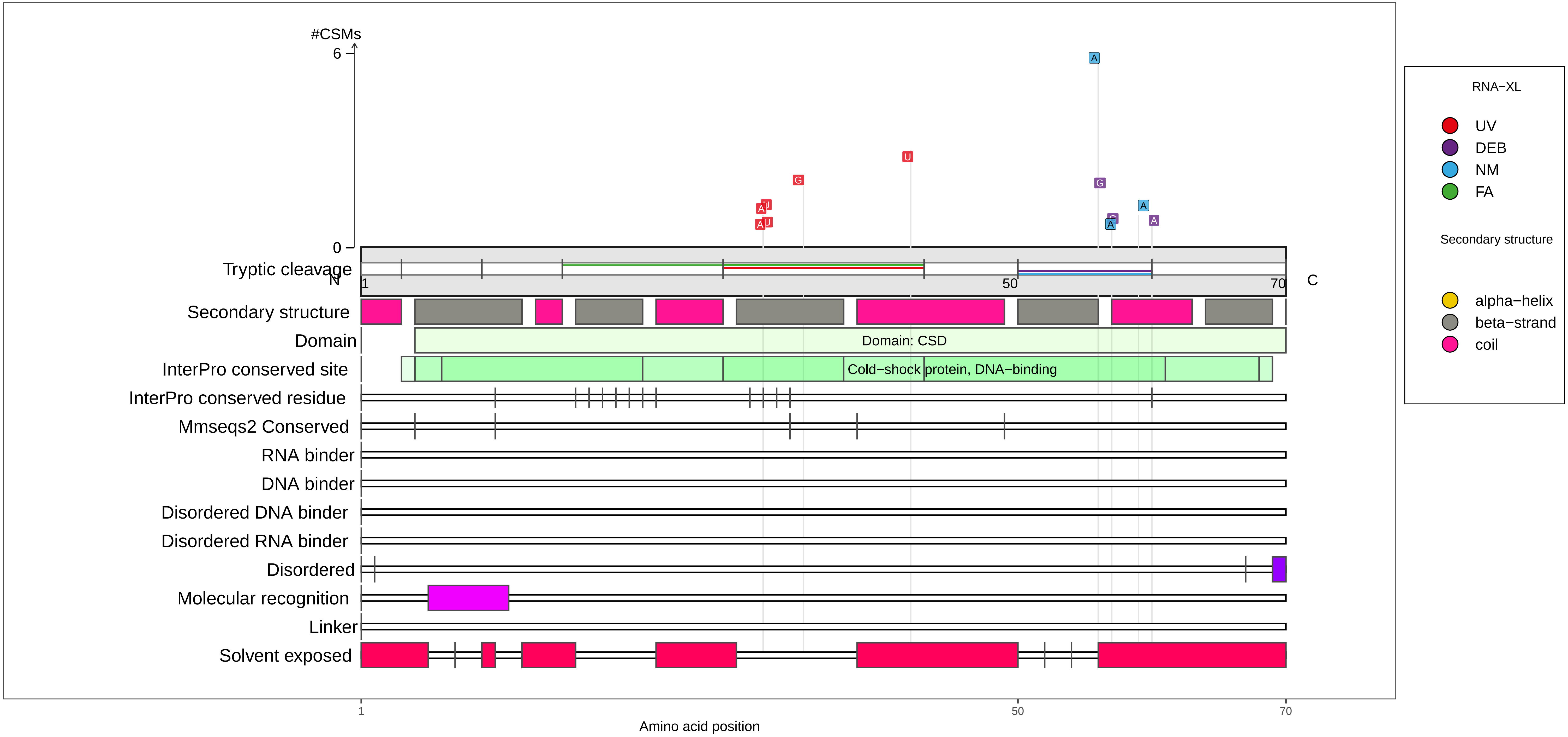

P0ADZ4
RS15_ECOLI 30S ribosomal protein S15

− Abundance:
tryptic [log10 Intensity]: 8.83 (Q 84)
PAXdb K12 strain [ppm]: 2.69 (Q 83)
PAXdb E.coli [ppm]: 3.22 (Q 97)

− RNA functions:
mRNA binding; mRNA binding involved in posttranscriptional gene silencing
RNA binding; RNA binding involved in posttranscriptional gene silencing; rRNA binding

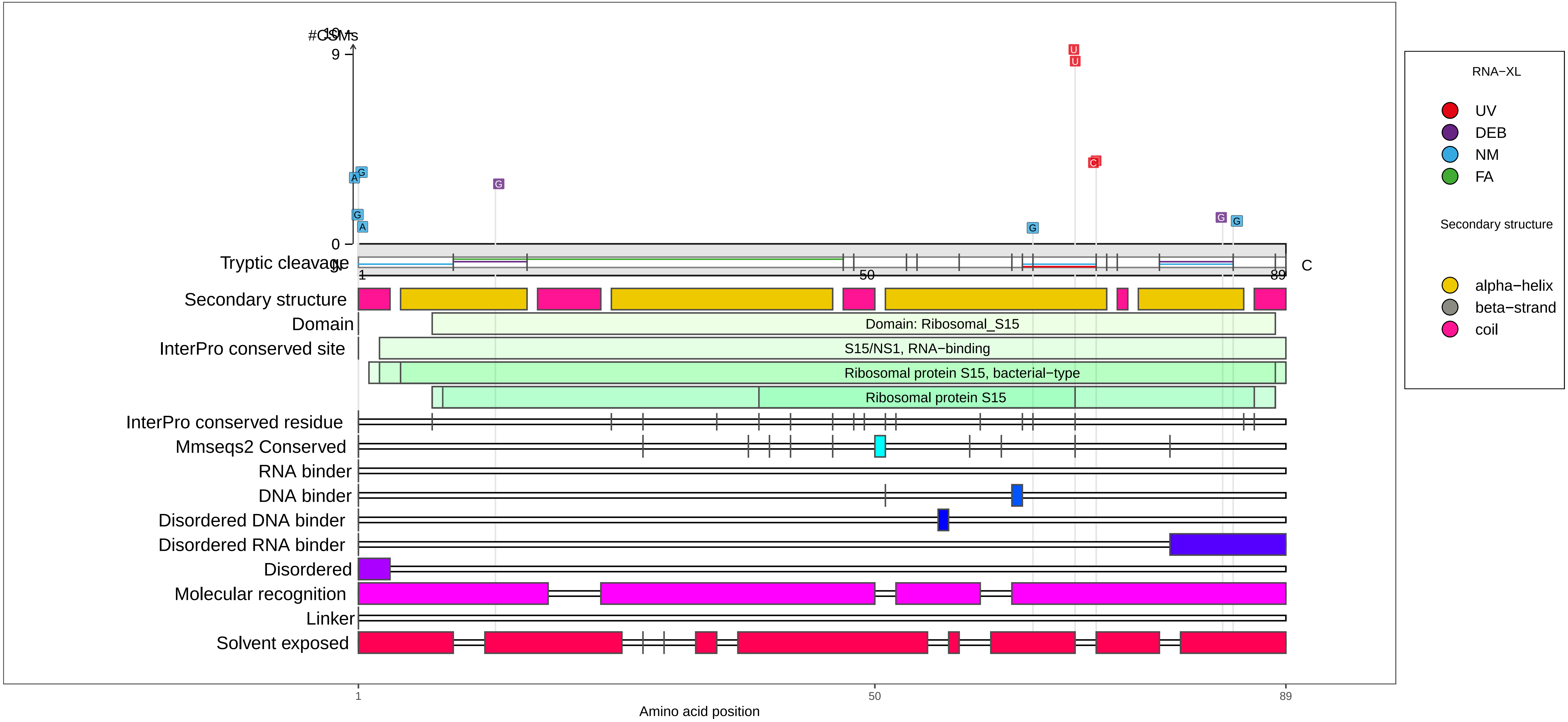

P13029
KATG_ECOLI Catalase−peroxidase

− Abundance:
tryptic [log10 Intensity]: 8.91 (Q 86)
PAXdb K12 strain [ppm]: 2.82 (Q 87)
PAXdb E.coli [ppm]: 3.04 (Q 95)

− RNA functions: not annotated

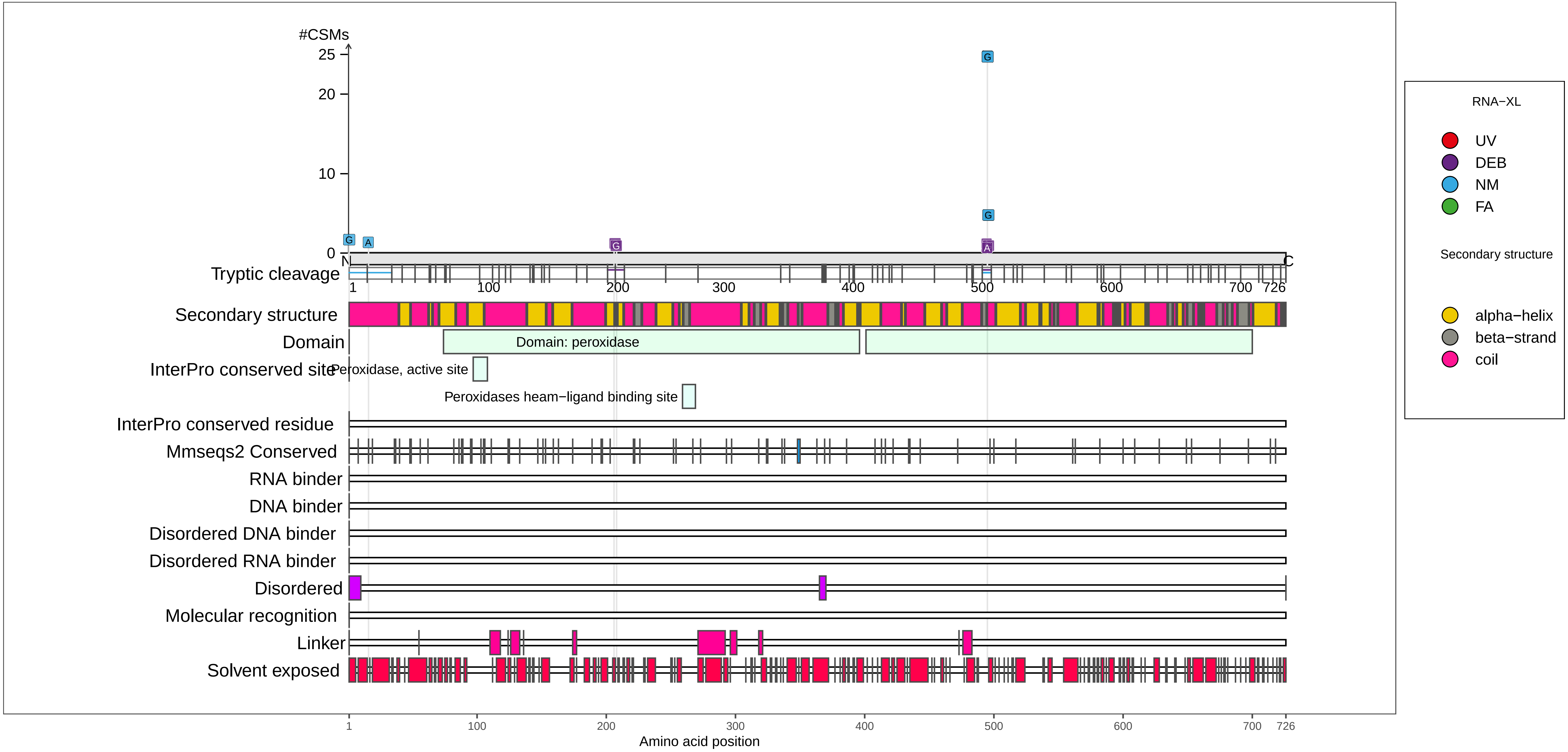

P27302
TKT1_ECOLI Transketolase 1

− Abundance:
tryptic [log10 Intensity]: 10.15 (Q 100)
PAXdb K12 strain [ppm]: 3.1 (Q 92)
PAXdb E.coli [ppm]: 3.04 (Q 95)

− RNA functions: not annotated

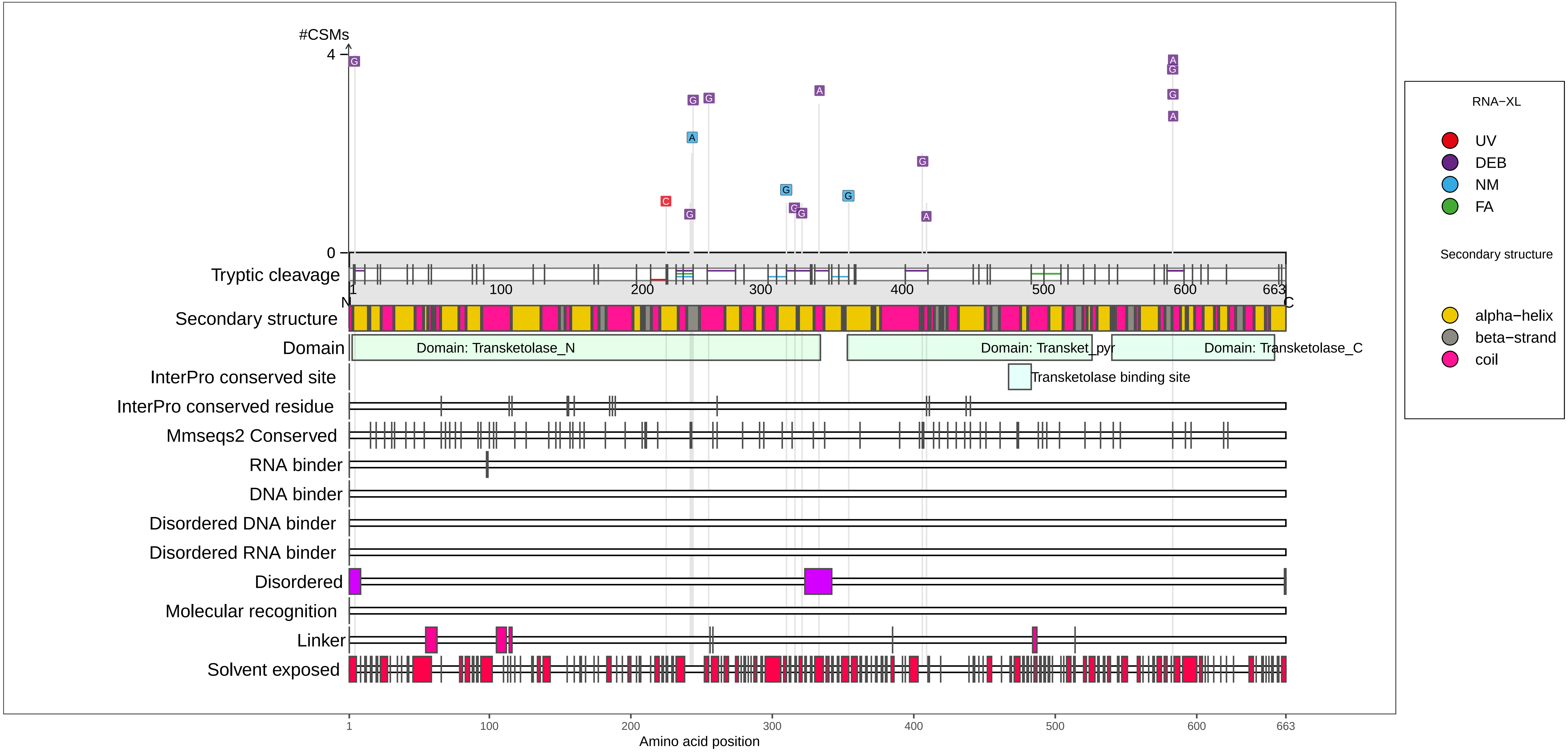

P60390
RSMH_ECOLI Ribosomal RNA small subunit methyltransferase H

− Abundance:
tryptic [log10 Intensity]: 7.04 (Q 15)
PAXdb K12 strain [ppm]: 2.55 (Q 79)
PAXdb E.coli [ppm]: 2.06 (Q 74)

− RNA functions:
ncRNA metabolic process; ncRNA processing; RNA metabolic process; RNA methylation
RNA methyltransferase activity; RNA modification; RNA processing
rRNA (cytosine−N4−)−methyltransferase activity; rRNA (cytosine) methyltransferase activity
rRNA base methylation; rRNA metabolic process; rRNA methylation; rRNA methyltransferase activity
rRNA modification; rRNA processing

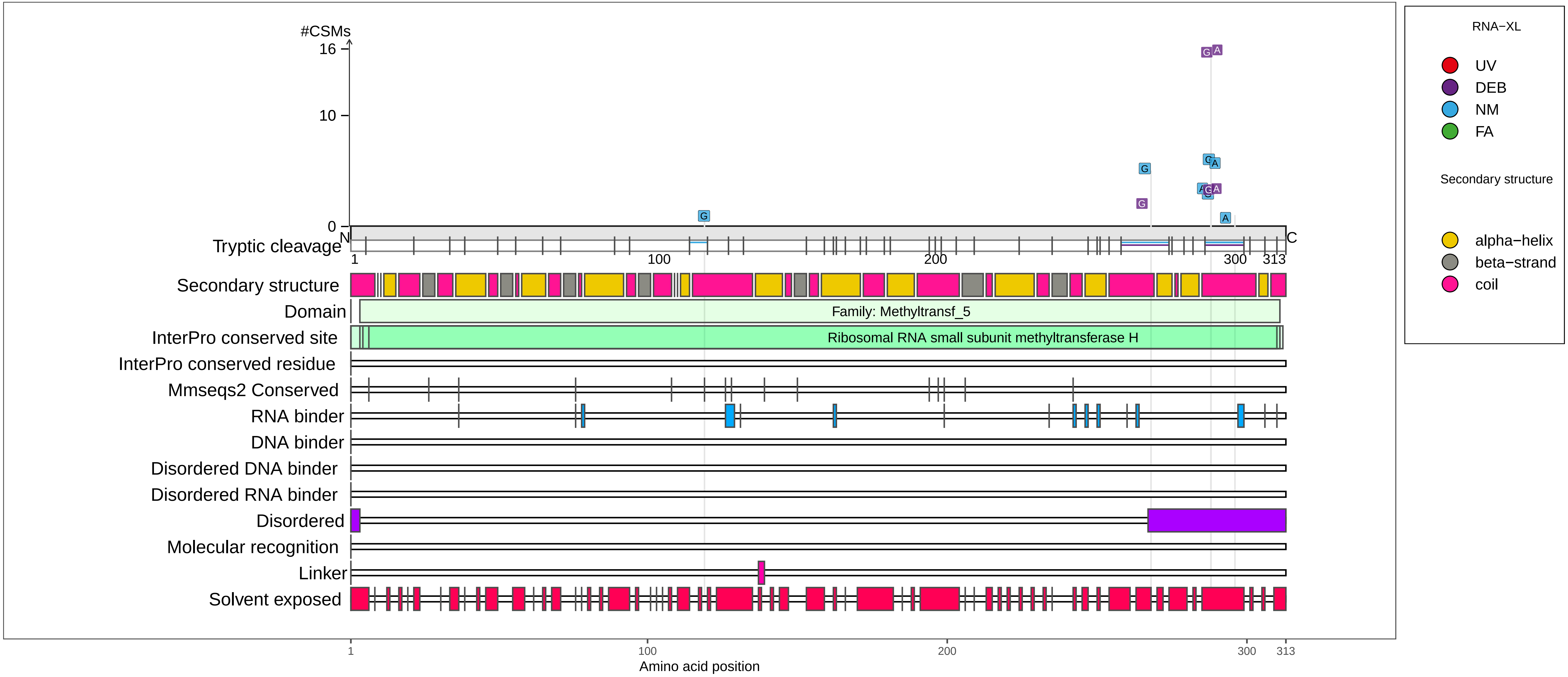

P60785
LEPA_ECOLI Elongation factor 4

− Abundance:
tryptic [log10 Intensity]: 8.87 (Q 85)
PAXdb K12 strain [ppm]: 2.06 (Q 63)
PAXdb E.coli [ppm]: 2.49 (Q 85)

− RNA functions:
RNA binding

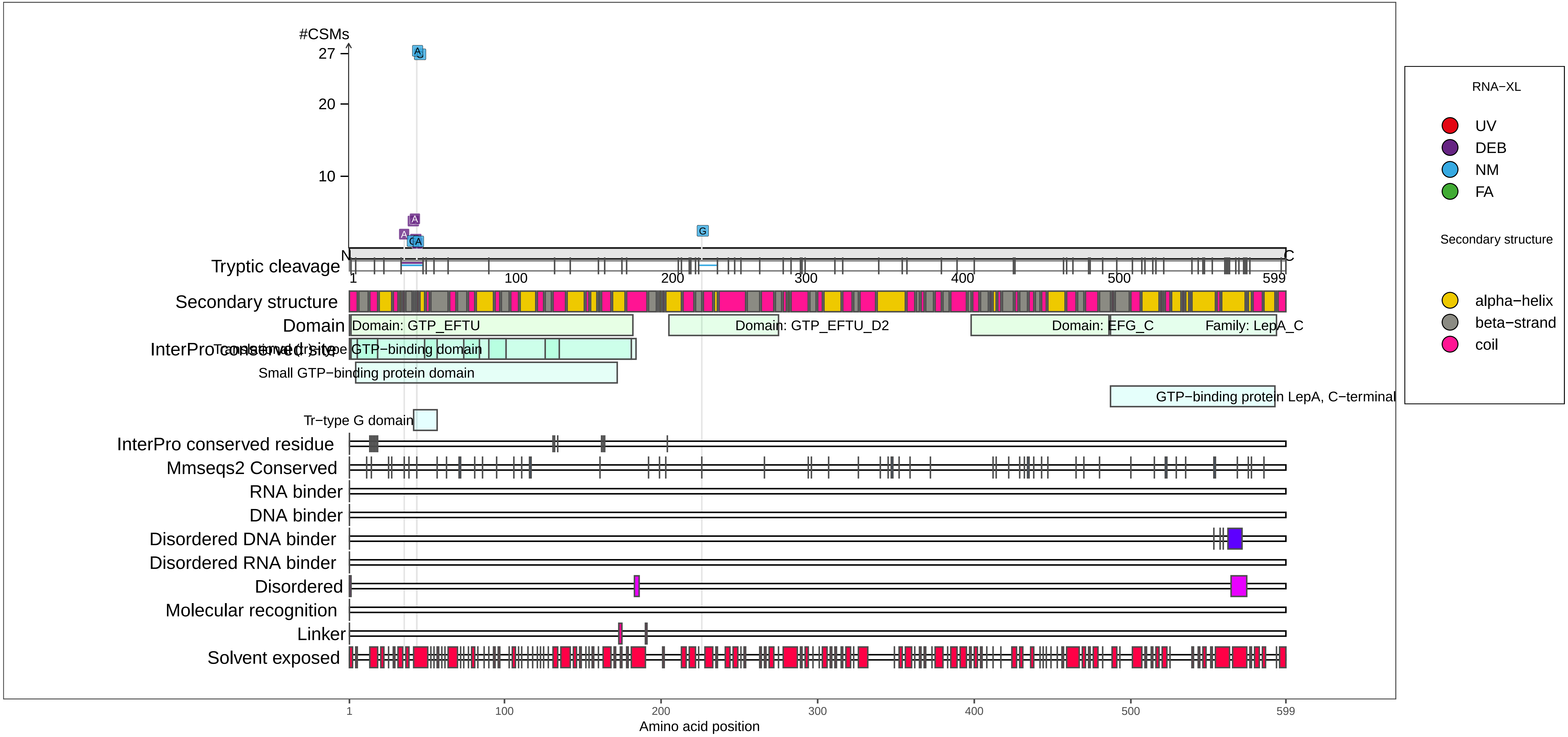

P0A7A9
IPYR_ECOLI Inorganic pyrophosphatase

− Abundance:
tryptic [log10 Intensity]: 9.14 (Q 91)
PAXdb K12 strain [ppm]: 2.82 (Q 86)
PAXdb E.coli [ppm]: 3.42 (Q 98)

− RNA functions: not annotated

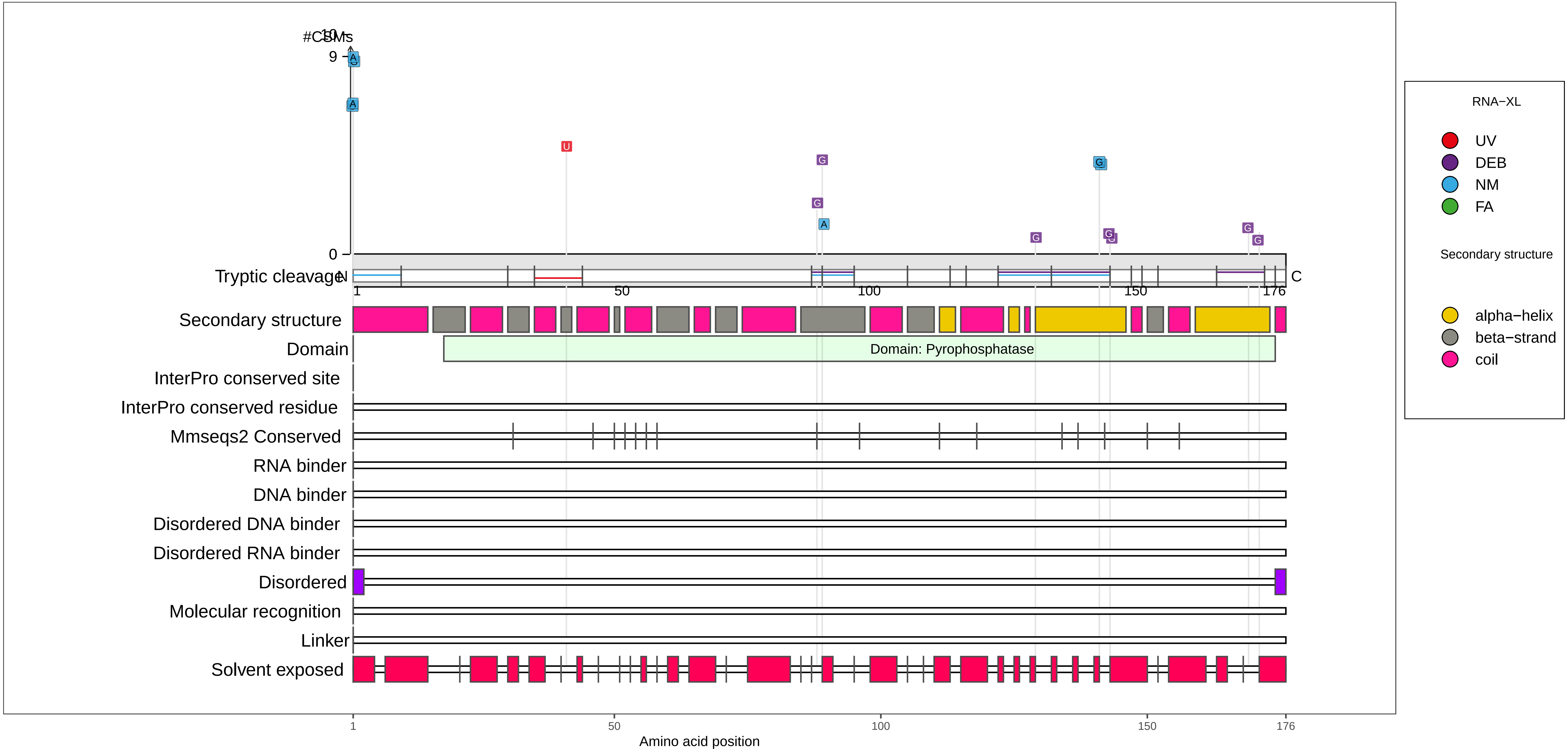

P08839
PT1_ECOLI Phosphoenolpyruvate−protein phosphotransferase

− Abundance:
tryptic [log10 Intensity]: 9.02 (Q 89)
PAXdb K12 strain [ppm]: 3.27 (Q 95)
PAXdb E.coli [ppm]: 3.04 (Q 95)

− RNA functions: not annotated

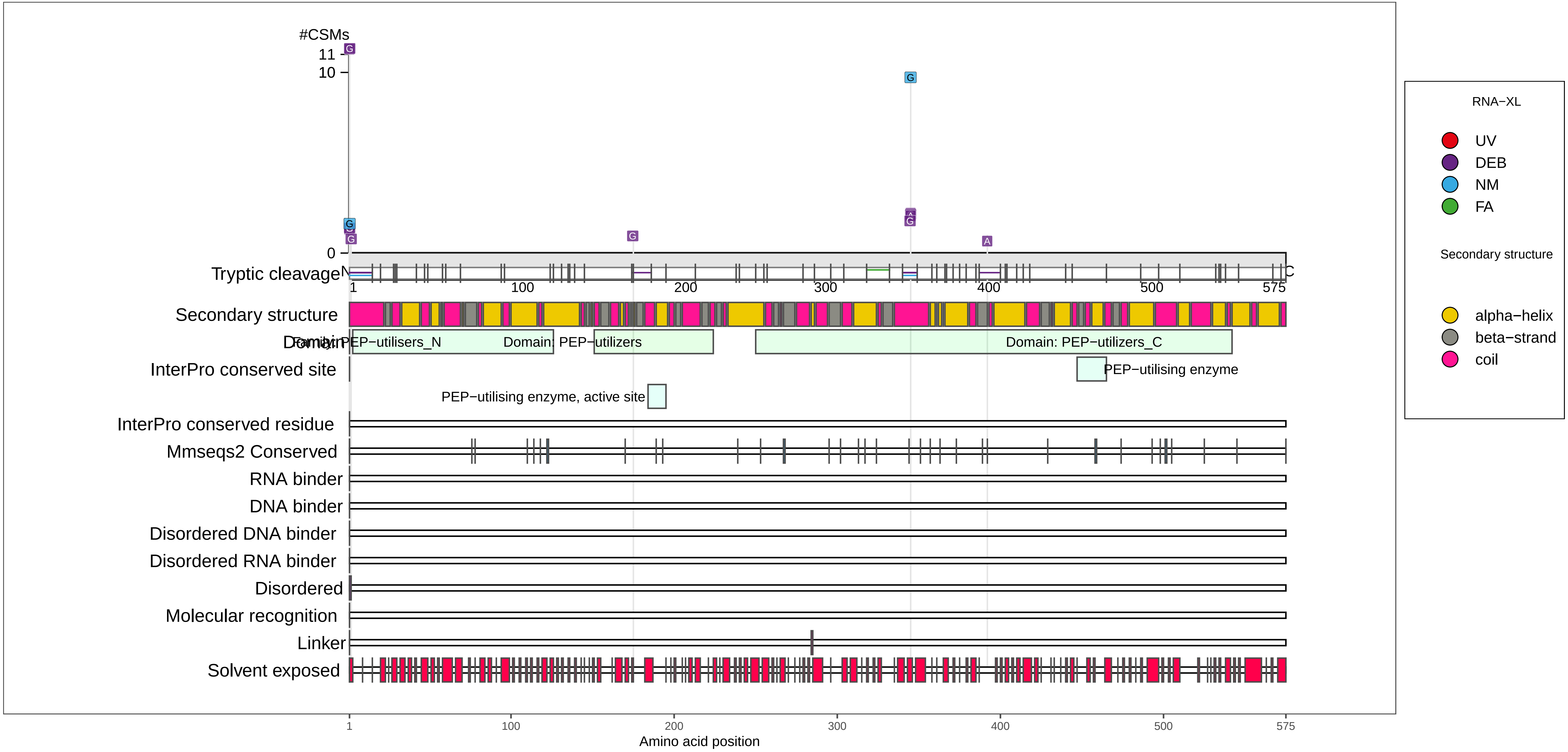

P0A749
MURA_ECOLI UDP−N−acetylglucosamine 1−carboxyvinyltransferase

− Abundance:
tryptic [log10 Intensity]: 7.31 (Q 25)
PAXdb K12 strain [ppm]: 2.57 (Q 79)
PAXdb E.coli [ppm]: 2.27 (Q 79)

− RNA functions: not annotated

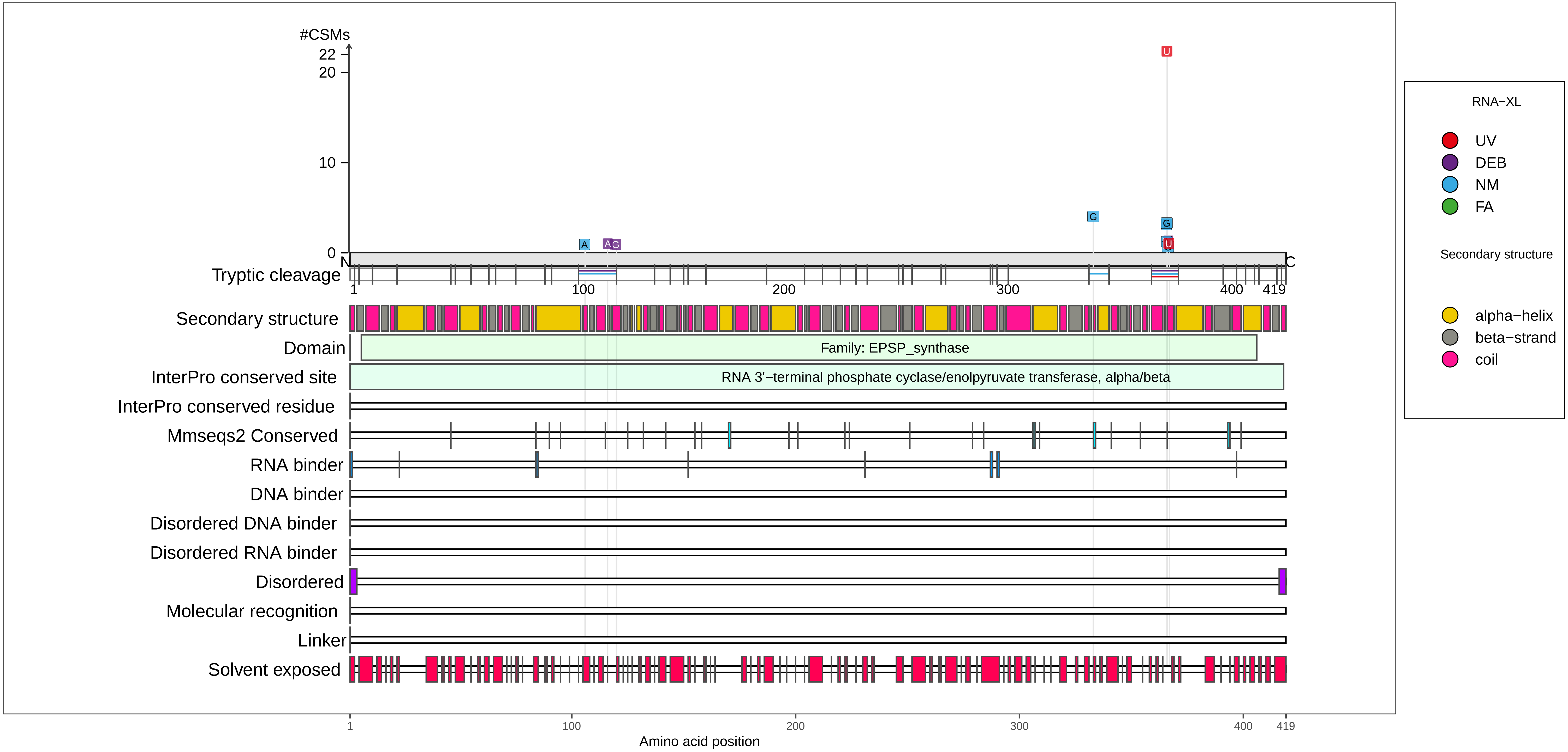

P0C037
PPNP_ECOLI Pyrimidine/purine nucleoside phosphorylase

− Abundance:
tryptic [log10 Intensity]: 8.23 (Q 67)
PAXdb K12 strain [ppm]: 2.42 (Q 75)
PAXdb E.coli [ppm]: 2.85 (Q 92)

− RNA functions: not annotated

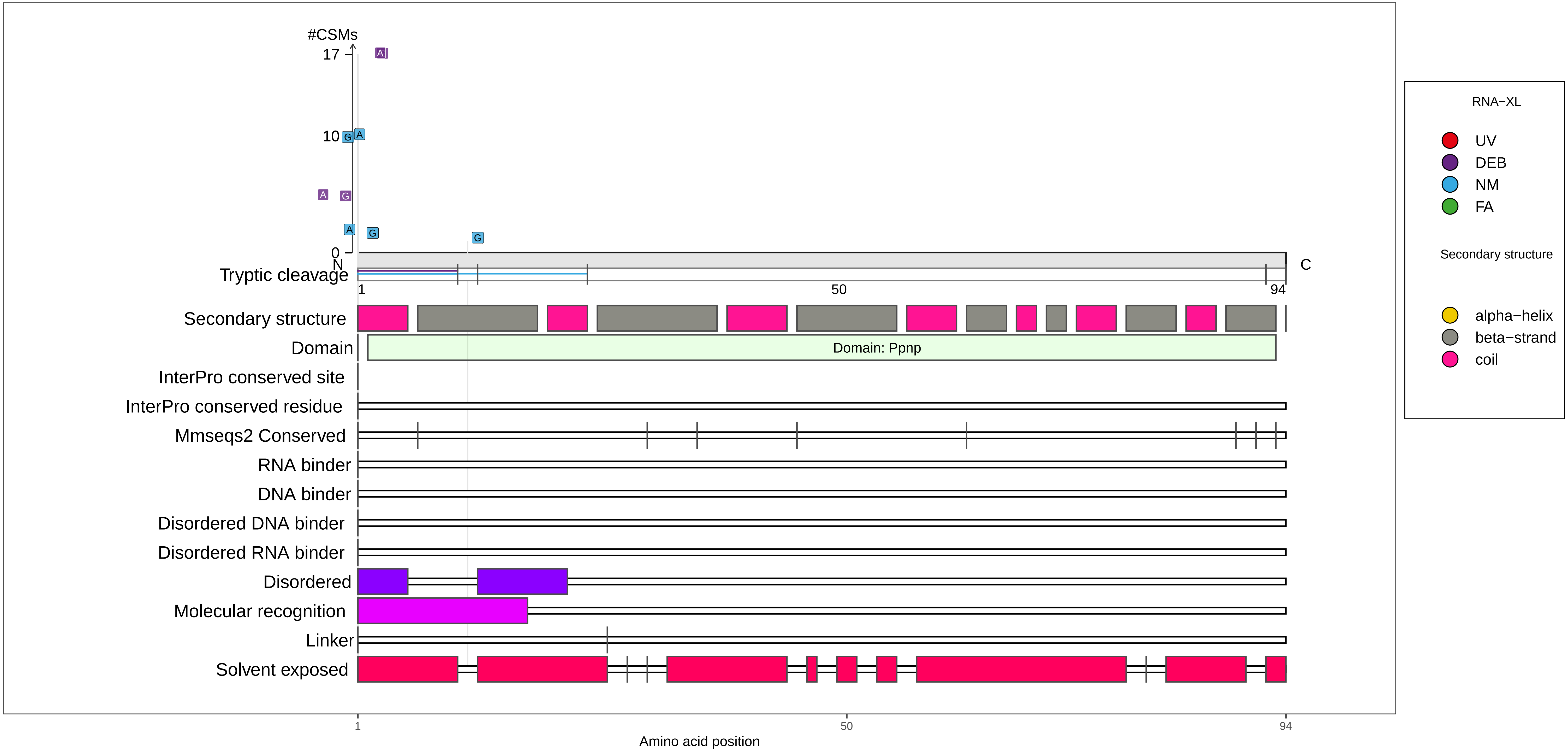

P45565
AIS_ECOLI Lipopolysaccharide core heptose(II)−phosphate phosphatase

− Abundance:
tryptic [log10 Intensity]: 8.98 (Q 88)
PAXdb K12 strain [ppm]: 3.22 (Q 94)
PAXdb E.coli [ppm]: 1.69 (Q 64)

− RNA functions: not annotated

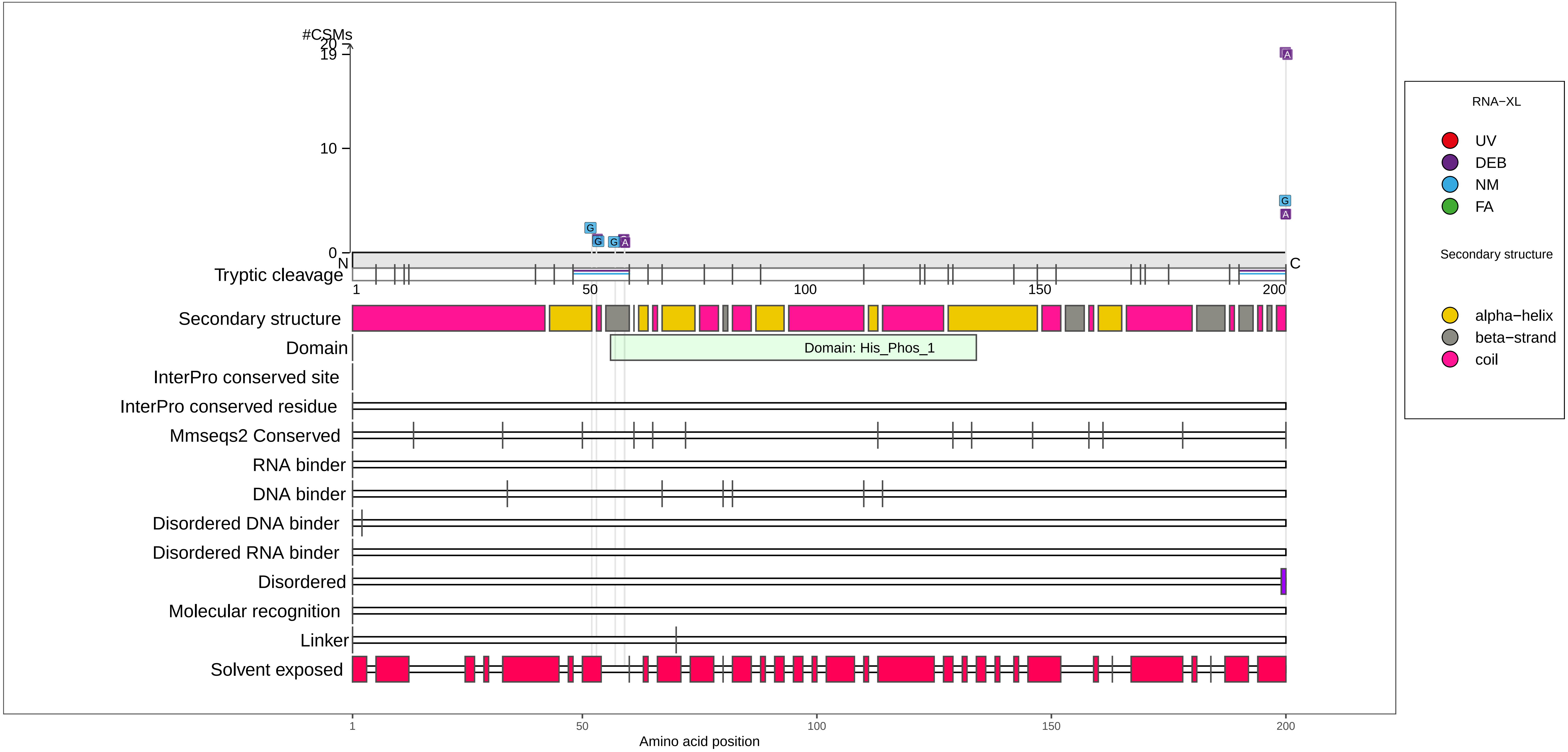

P77398
ARNA_ECOLI Bifunctional polymyxin resistance protein ArnA

− Abundance:
tryptic [log10 Intensity]: 9.03 (Q 89)
PAXdb K12 strain [ppm]: 2.86 (Q 88)
PAXdb E.coli [ppm]: 1.73 (Q 65)

− RNA functions: not annotated

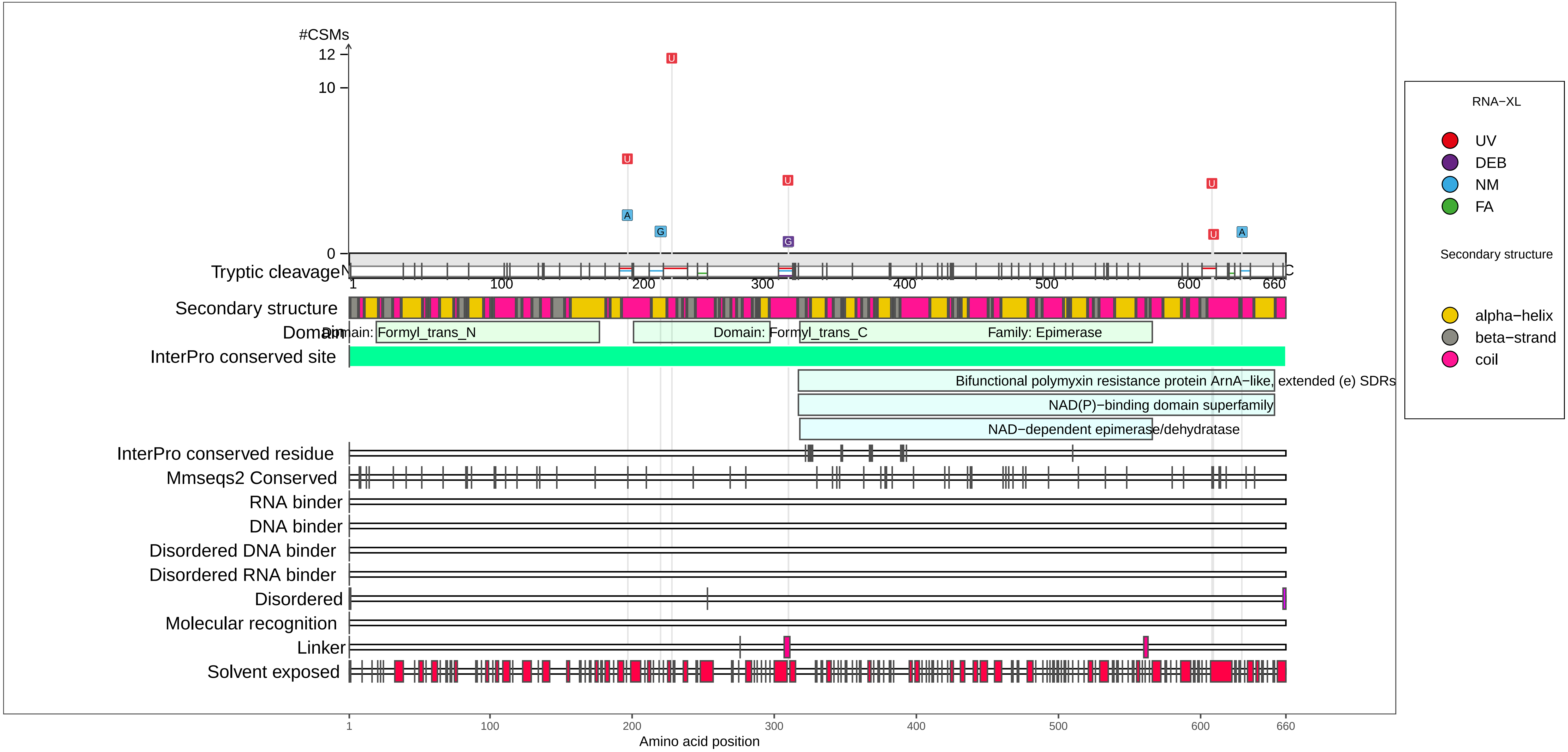

P0AE88
CPXR_ECOLI Transcriptional regulatory protein CpxR

− Abundance:
tryptic [log10 Intensity]: 7.71 (Q 44)
PAXdb K12 strain [ppm]: NA (Q NA)
PAXdb E.coli [ppm]: 2.77 (Q 91)

− RNA functions: not annotated

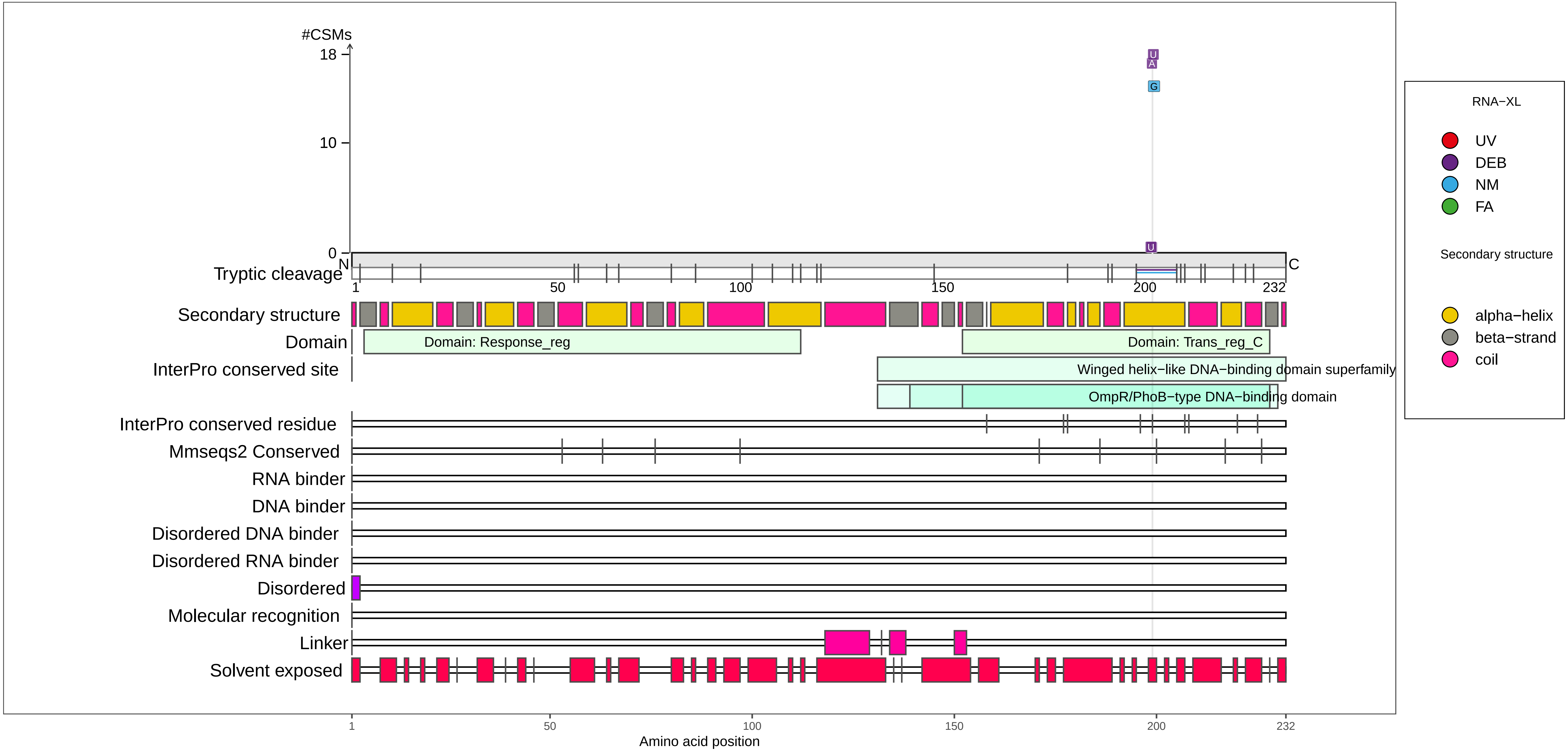

P0AFD1
NUOE_ECOLI NADH−quinone oxidoreductase subunit E

− Abundance:
tryptic [log10 Intensity]: 8.46 (Q 74)
PAXdb K12 strain [ppm]: 2.07 (Q 63)
PAXdb E.coli [ppm]: 2.46 (Q 84)

− RNA functions: not annotated

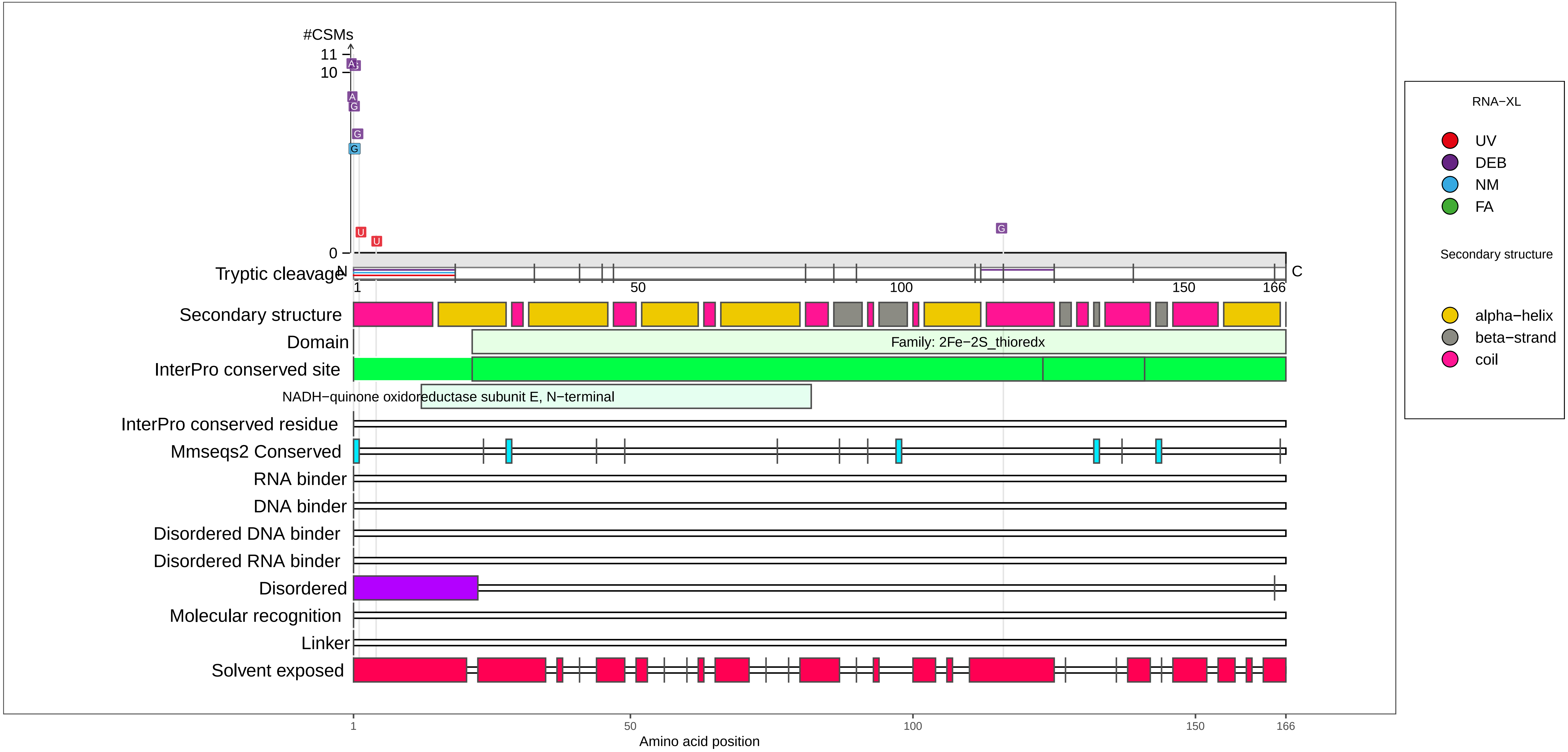

P32662
GPH_ECOLI Phosphoglycolate phosphatase

− Abundance:
tryptic [log10 Intensity]: 7.94 (Q 55)
PAXdb K12 strain [ppm]: 2.05 (Q 62)
PAXdb E.coli [ppm]: 1.97 (Q 71)

− RNA functions: not annotated

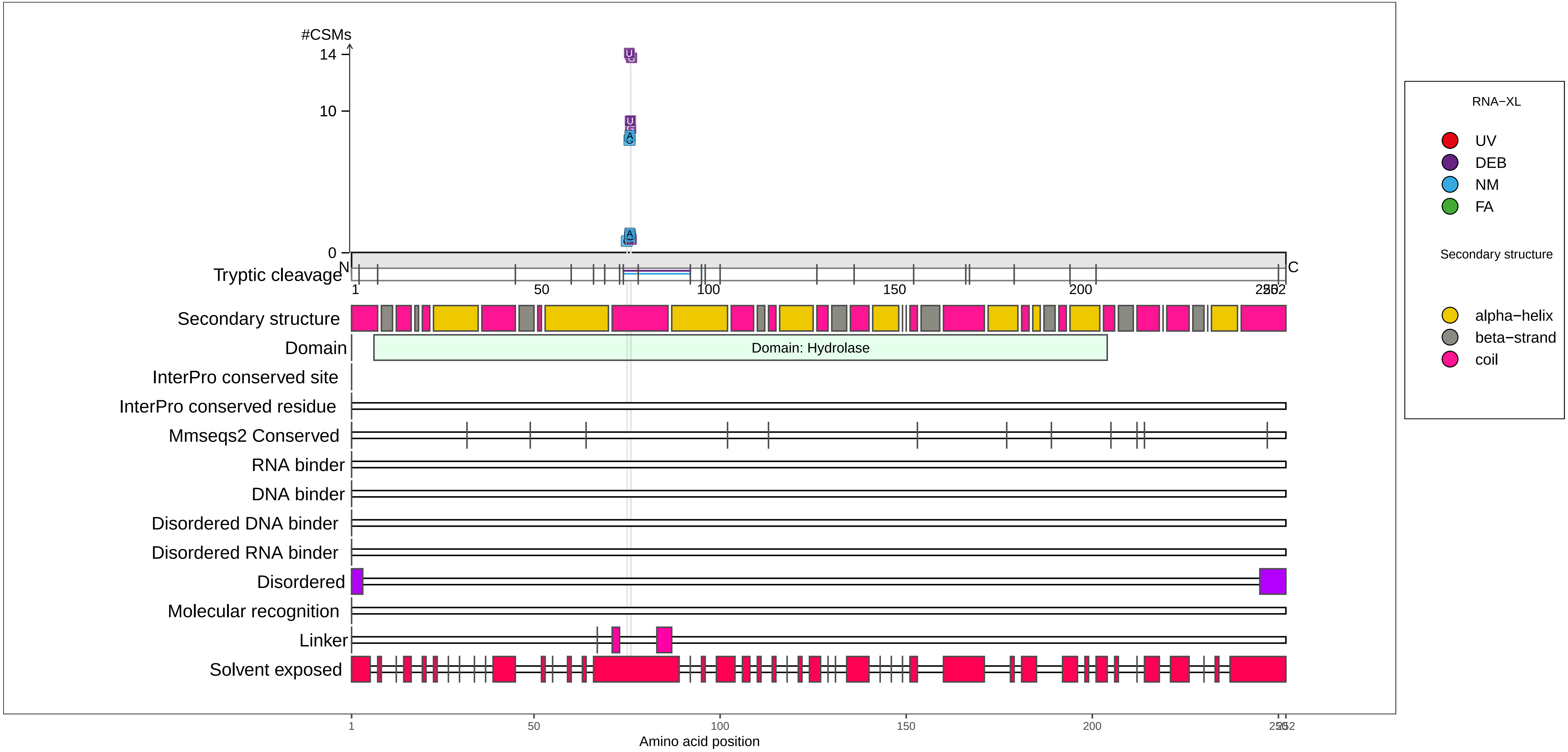

P07004
PROA_ECOLI Gamma−glutamyl phosphate reductase

− Abundance:
tryptic [log10 Intensity]: 8.59 (Q 78)
PAXdb K12 strain [ppm]: 2.39 (Q 74)
PAXdb E.coli [ppm]: 2.58 (Q 87)

− RNA functions: not annotated

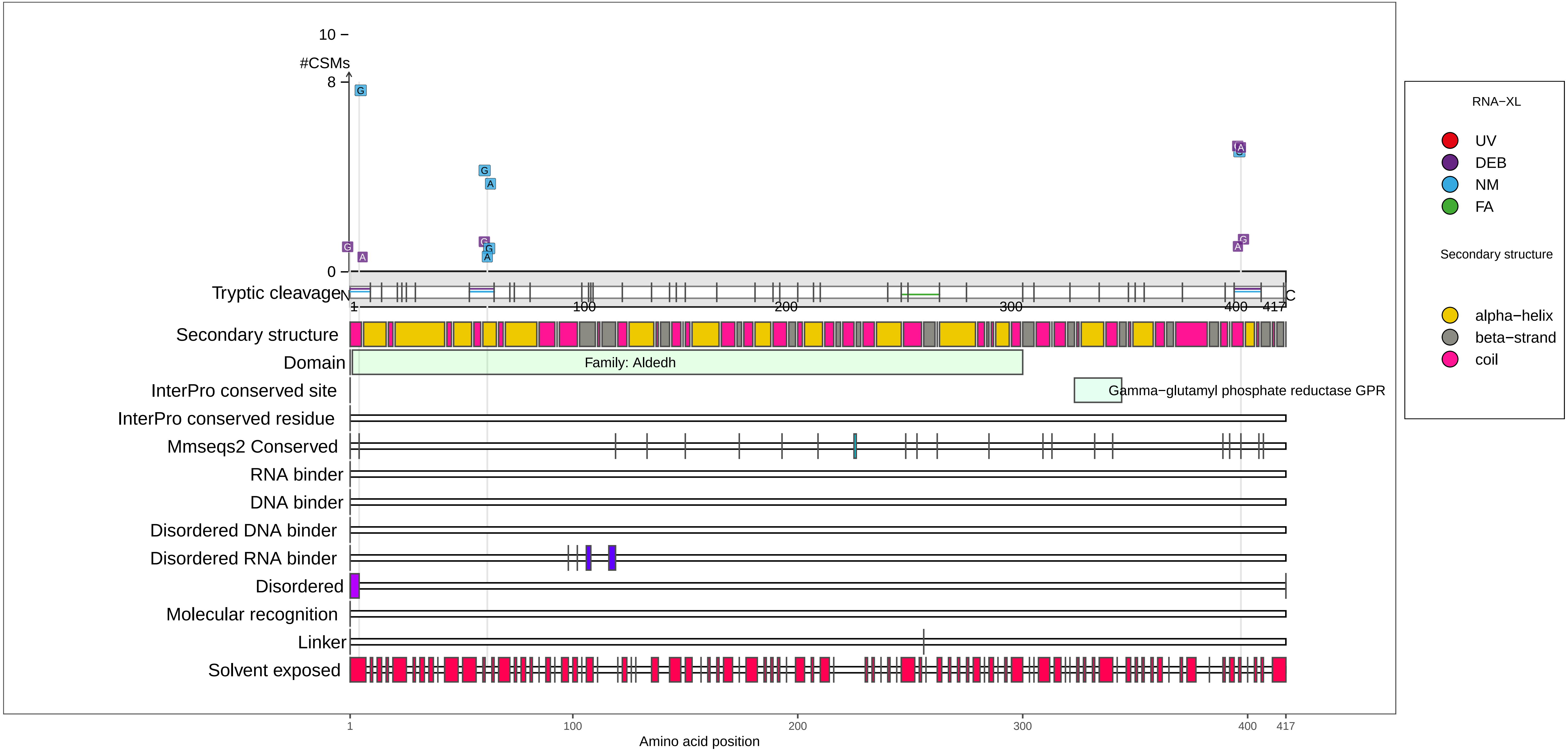

P0A7Q6
RL36_ECOLI 50S ribosomal protein L36

− Abundance:
tryptic [log10 Intensity]: NA (Q NA)
PAXdb K12 strain [ppm]: NA (Q NA)
PAXdb E.coli [ppm]: 2.63 (Q 89)

− RNA functions: not annotated

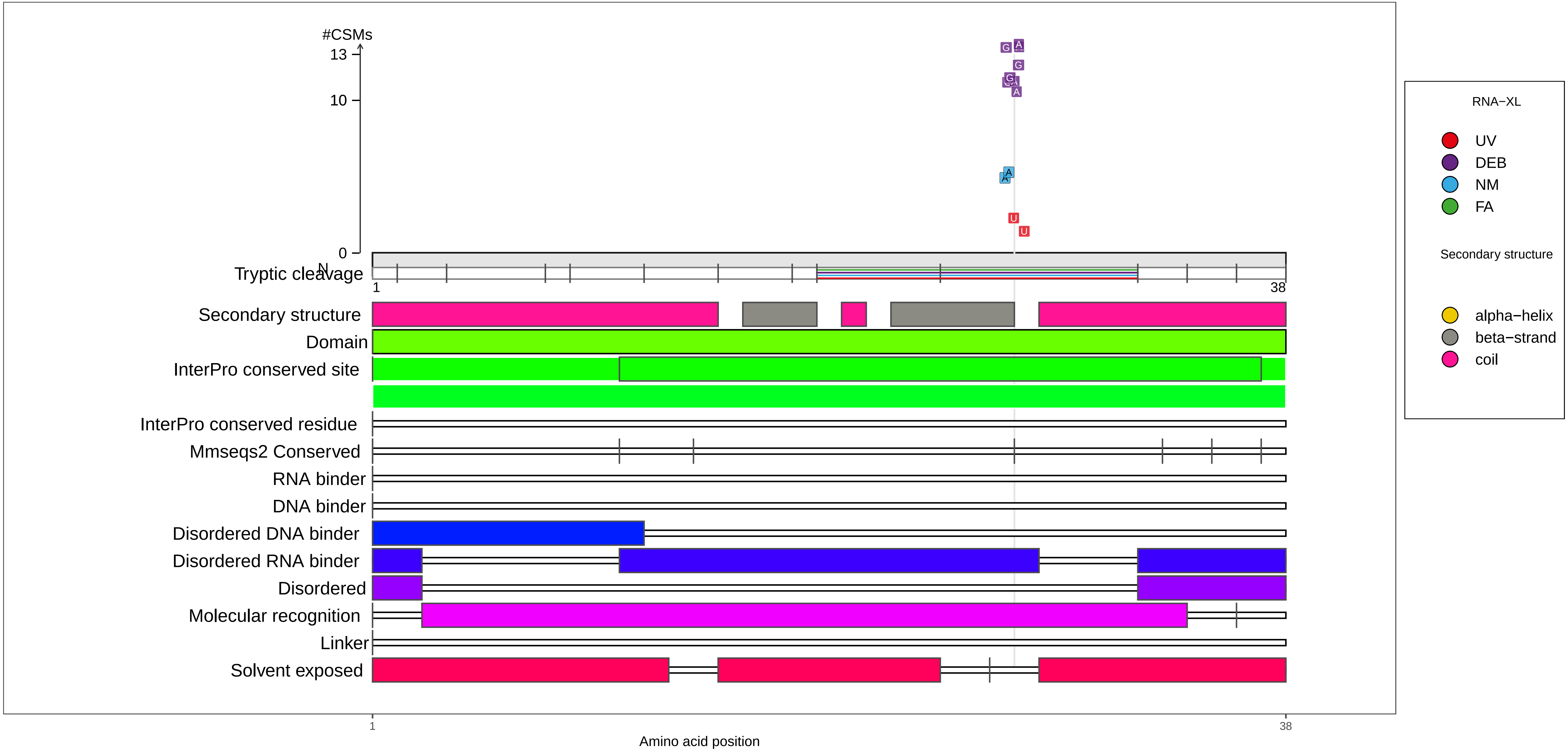

P60906
SYH_ECOLI Histidine−−tRNA ligase

− Abundance:
tryptic [log10 Intensity]: 9.35 (Q 95)
PAXdb K12 strain [ppm]: 2.96 (Q 90)
PAXdb E.coli [ppm]: 2.7 (Q 90)

− RNA functions:
aminoacyl−tRNA ligase activity; histidine−tRNA ligase activity
histidyl−tRNA aminoacylation; Histidyl−tRNA synthetase; ncRNA metabolic process; RNA metabolic process
tRNA aminoacylation; tRNA aminoacylation for protein translation; tRNA metabolic process
tRNA synthetase class II core domain (G, H, P, S and T)

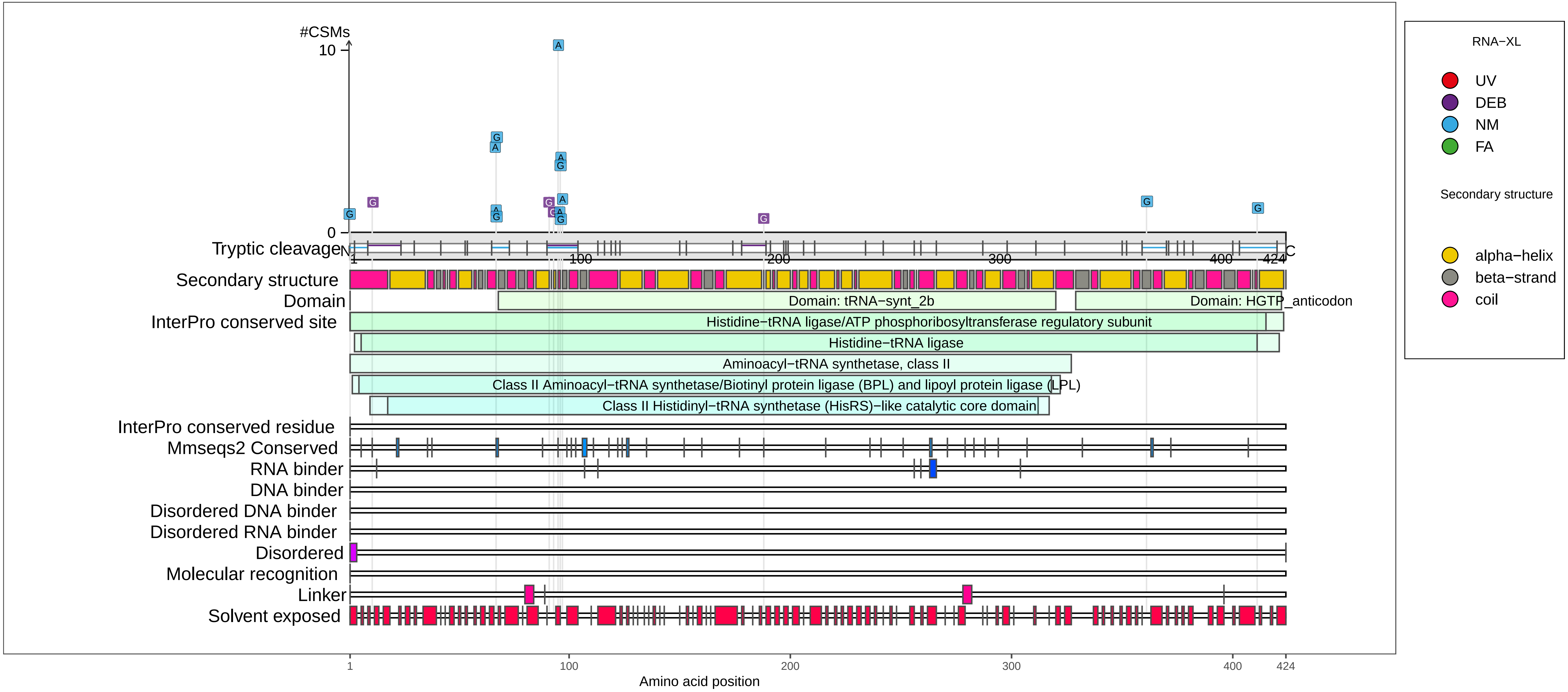

P69913
CSRA_ECOLI Carbon storage regulator

− Abundance:
tryptic [log10 Intensity]: 9.33 (Q 94)
PAXdb K12 strain [ppm]: 3.49 (Q 97)
PAXdb E.coli [ppm]: 3.33 (Q 98)

− RNA functions:
mRNA 5−UTR binding; mRNA binding; mRNA catabolic process; mRNA metabolic process
mRNA stabilization; RNA binding; RNA catabolic process; RNA metabolic process
RNA stabilization

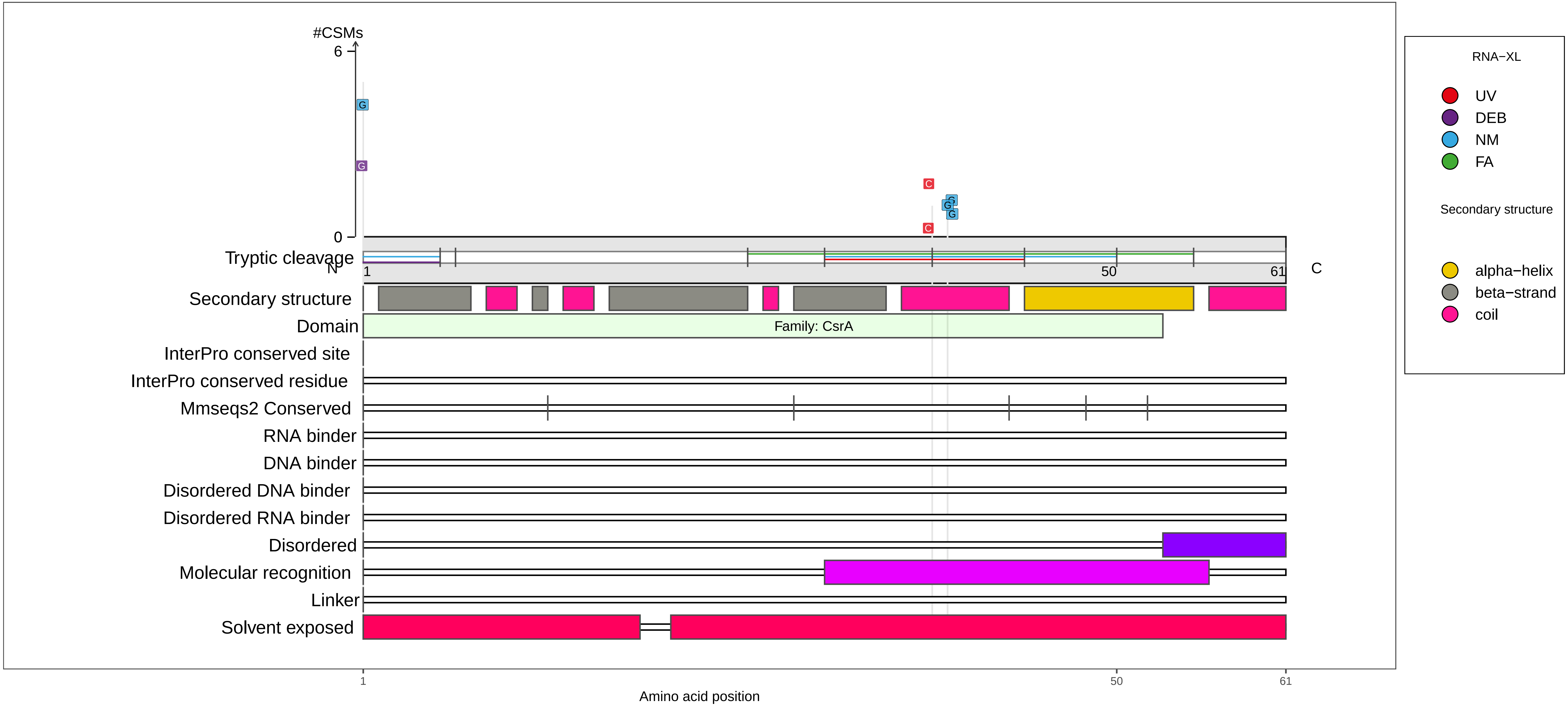

P0A6A3
ACKA_ECOLI Acetate kinase

− Abundance:
tryptic [log10 Intensity]: 9.49 (Q 96)
PAXdb K12 strain [ppm]: 3.24 (Q 94)
PAXdb E.coli [ppm]: 2.99 (Q 94)

− RNA functions: not annotated

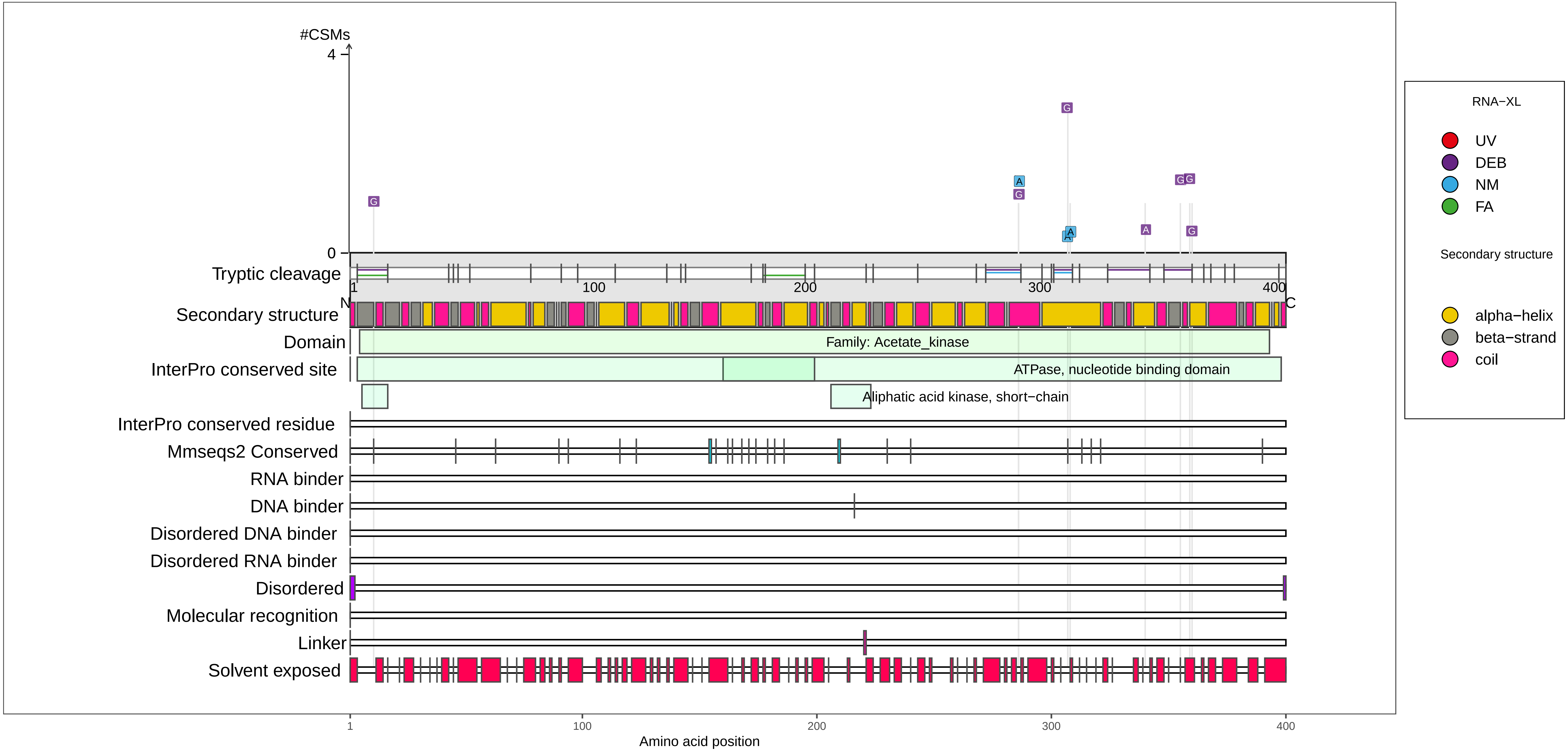

P0A8J8
RHLB_ECOLI ATP−dependent RNA helicase RhlB

− Abundance:
tryptic [log10 Intensity]: 8.47 (Q 74)
PAXdb K12 strain [ppm]: 2.32 (Q 71)
PAXdb E.coli [ppm]: 2.41 (Q 83)

− RNA functions:
RNA binding; RNA catabolic process; RNA helicase activity; RNA metabolic process

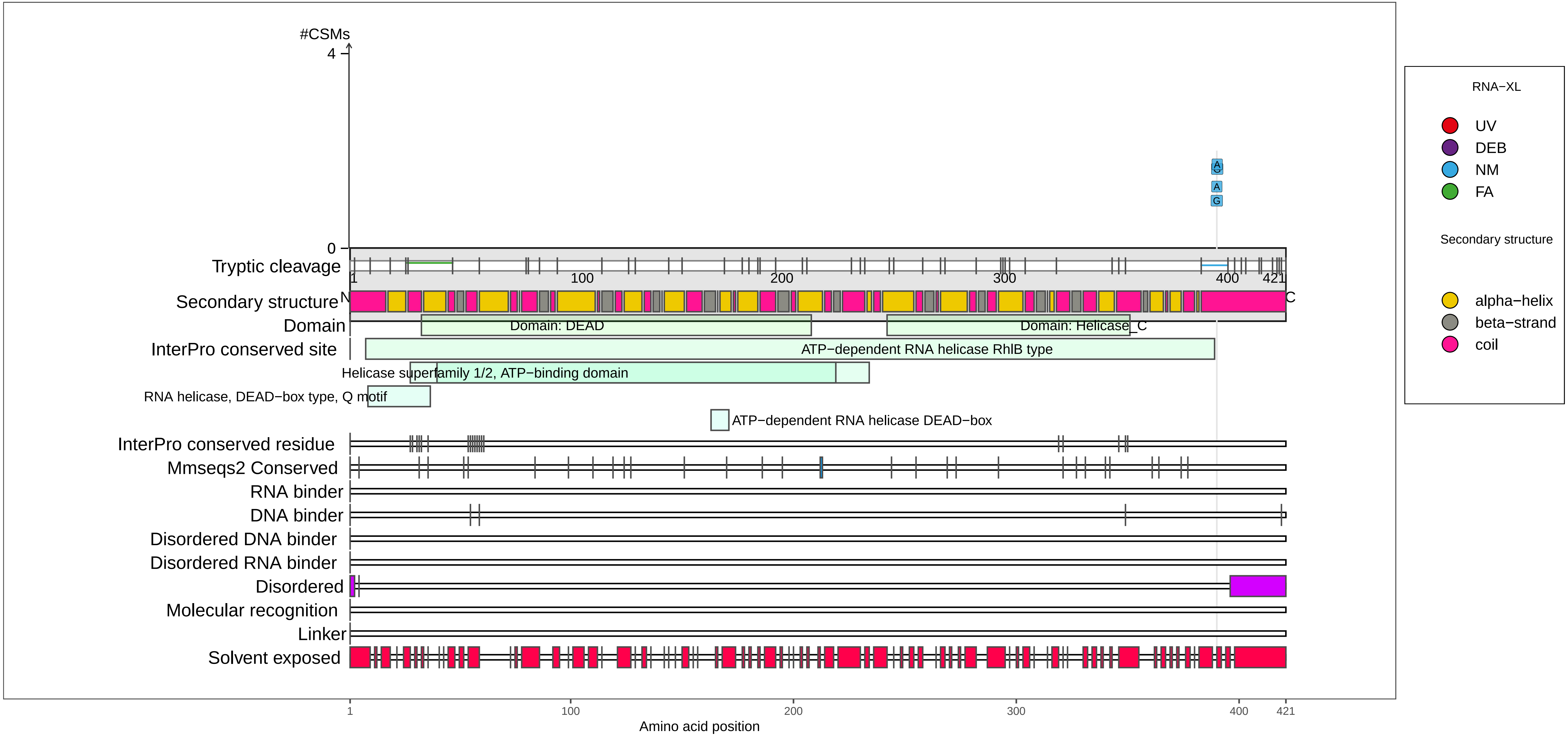

P0A912
PAL_ECOLI Peptidoglycan−associated lipoprotein

− Abundance:
tryptic [log10 Intensity]: 9.39 (Q 95)
PAXdb K12 strain [ppm]: 1.56 (Q 42)
PAXdb E.coli [ppm]: 2.56 (Q 87)

− RNA functions: not annotated

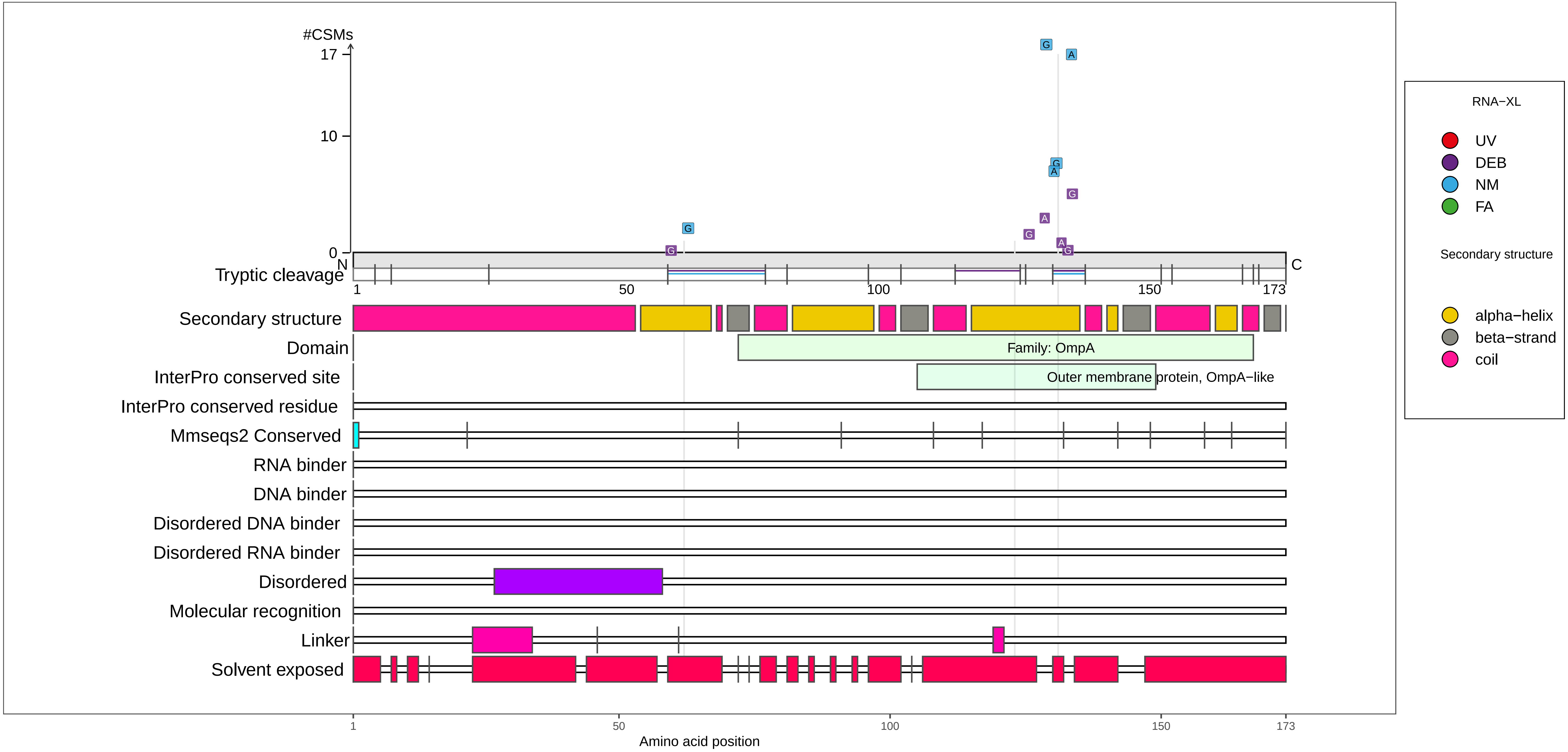

P0AA04
PTHP_ECOLI Phosphocarrier protein HPr

− Abundance:
tryptic [log10 Intensity]: 10.03 (Q 99)
PAXdb K12 strain [ppm]: 3.89 (Q 100)
PAXdb E.coli [ppm]: 3.19 (Q 96)

− RNA functions: not annotated

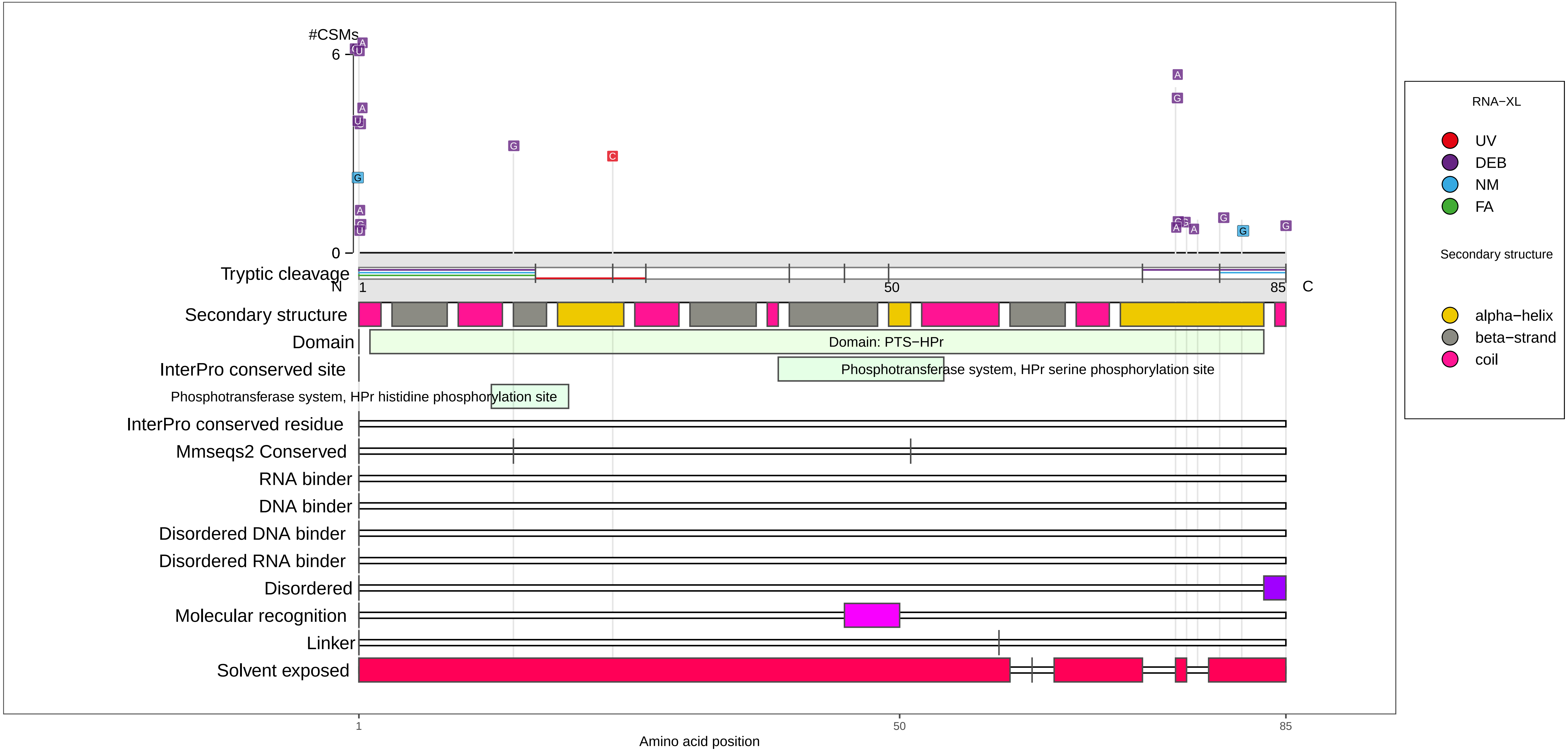

P0AFC3
NUOA_ECOLI NADH−quinone oxidoreductase subunit A

− Abundance:
tryptic [log10 Intensity]: 9.53 (Q 97)
PAXdb K12 strain [ppm]: 2.28 (Q 69)
PAXdb E.coli [ppm]: 2.56 (Q 87)

− RNA functions: not annotated

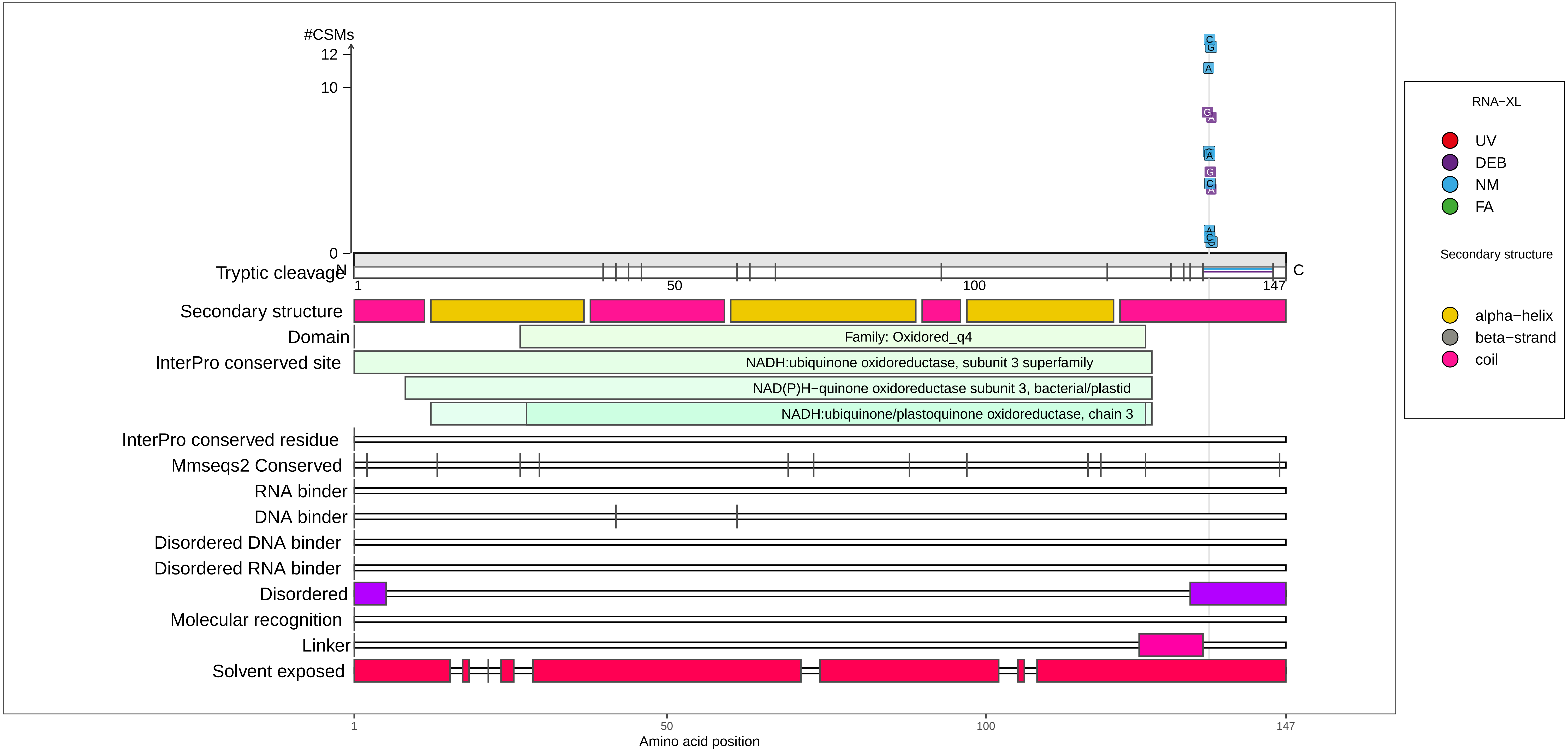

P18843
NADE_ECOLI NH(3)−dependent NAD(+) synthetase

− Abundance:
tryptic [log10 Intensity]: 8.56 (Q 77)
PAXdb K12 strain [ppm]: 2.96 (Q 90)
PAXdb E.coli [ppm]: 2.68 (Q 89)

− RNA functions: not annotated

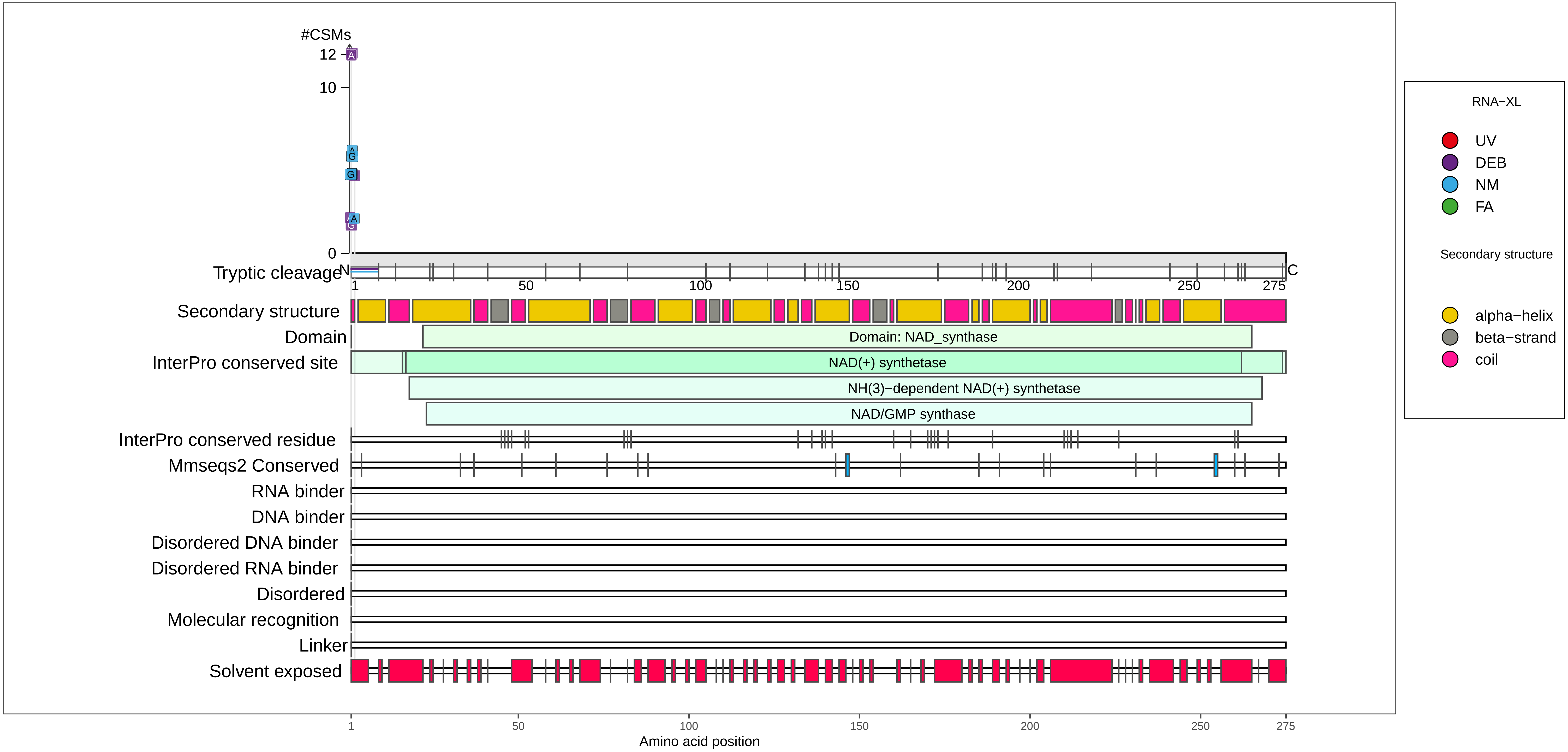

P23839
YICC_ECOLI UPF0701 protein YicC

− Abundance:
tryptic [log10 Intensity]: 8.35 (Q 71)
PAXdb K12 strain [ppm]: 2.43 (Q 75)
PAXdb E.coli [ppm]: 2.27 (Q 79)

− RNA functions:
ncRNA catabolic process; ncRNA metabolic process; ncRNA processing; RNA binding
RNA catabolic process; RNA metabolic process; RNA phosphodiester bond hydrolysis
RNA phosphodiester bond hydrolysis, exonucleolytic; RNA processing; rRNA catabolic process
rRNA metabolic process; rRNA processing; tRNA binding; tRNA metabolic process
tRNA nucleotidyltransferase activity; tRNA processing

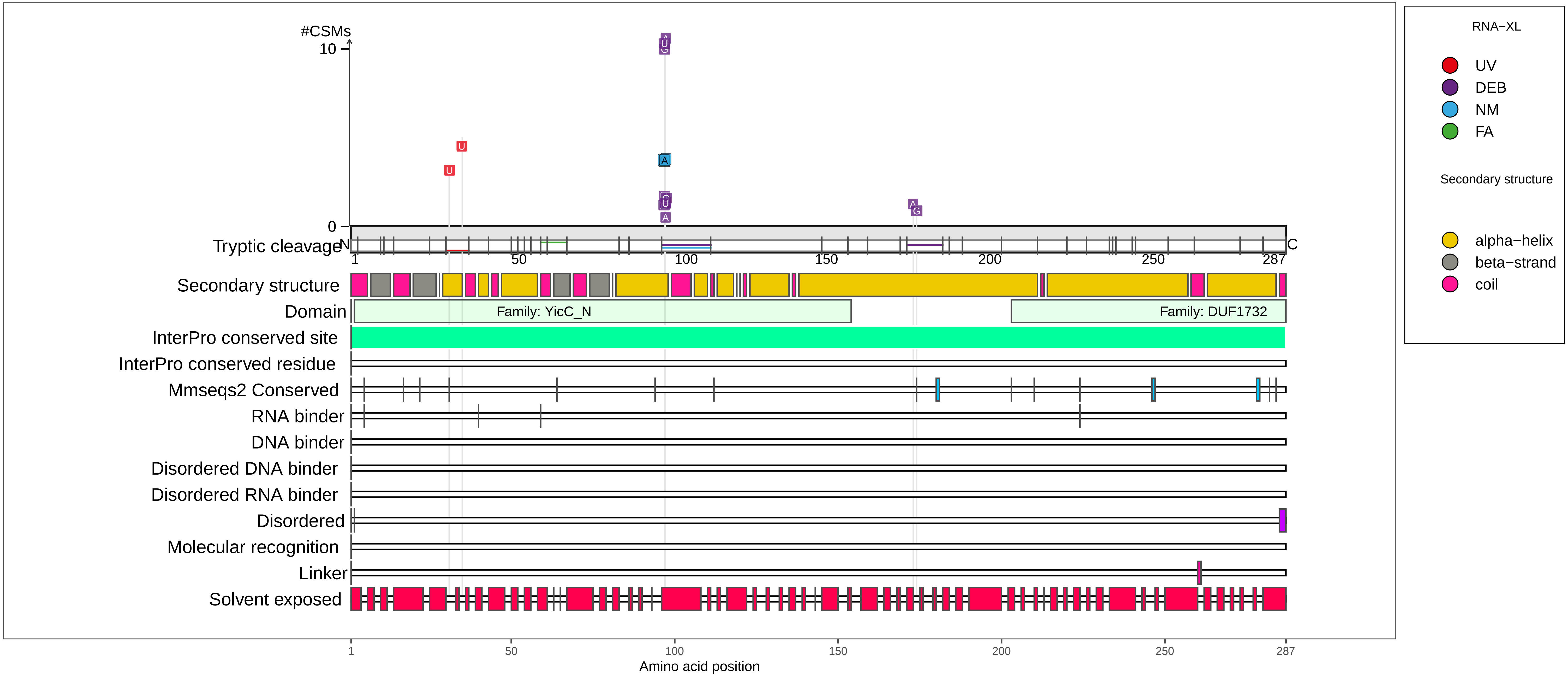

P0A6F1
CARA_ECOLI Carbamoyl−phosphate synthase small chain

− Abundance:
tryptic [log10 Intensity]: 9.06 (Q 90)
PAXdb K12 strain [ppm]: 3.05 (Q 91)
PAXdb E.coli [ppm]: 3.1 (Q 96)

− RNA functions: not annotated

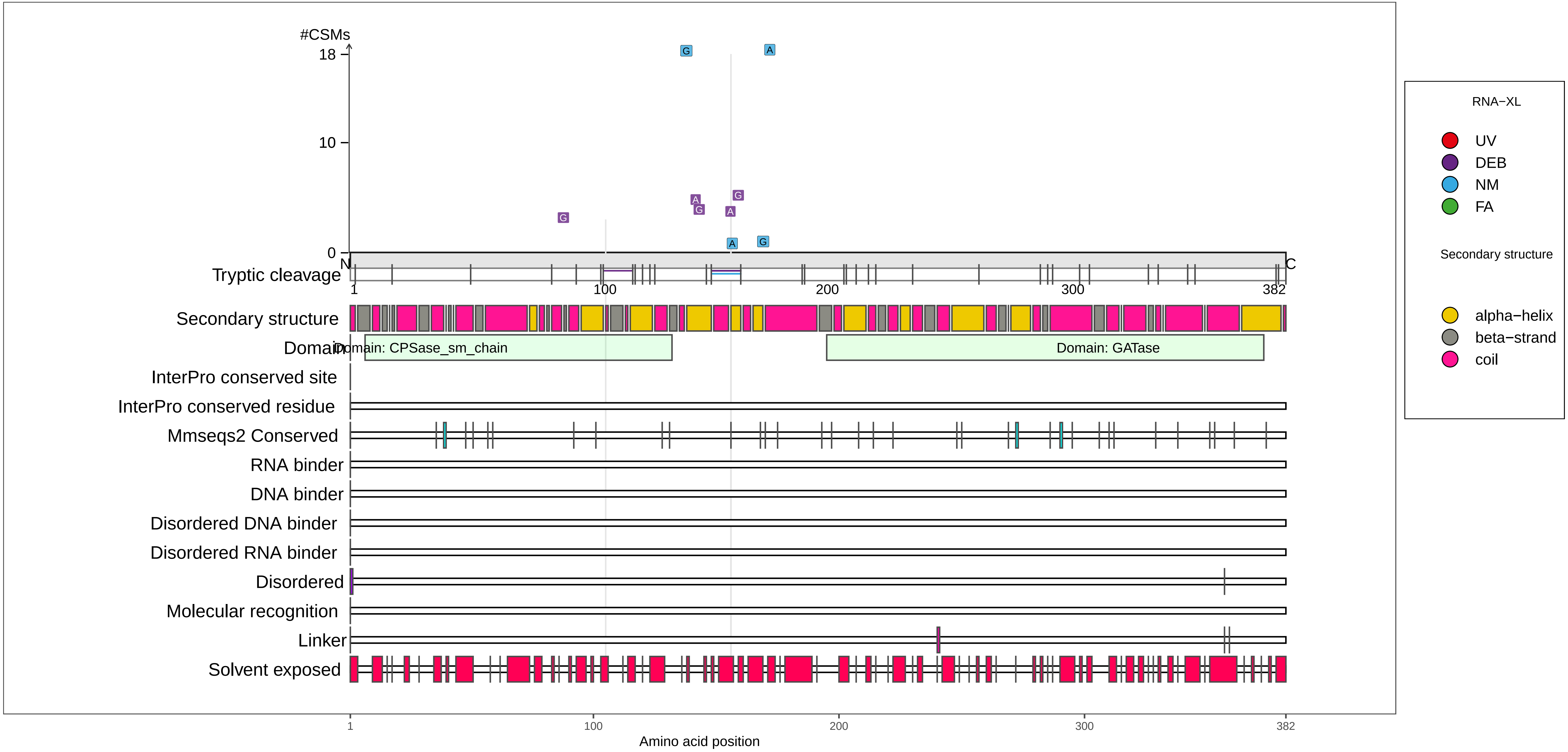

P0A6L4
NANA_ECOLI N−acetylneuraminate lyase

− Abundance:
tryptic [log10 Intensity]: 9.31 (Q 94)
PAXdb K12 strain [ppm]: NA (Q NA)
PAXdb E.coli [ppm]: 1.58 (Q 62)

− RNA functions: not annotated

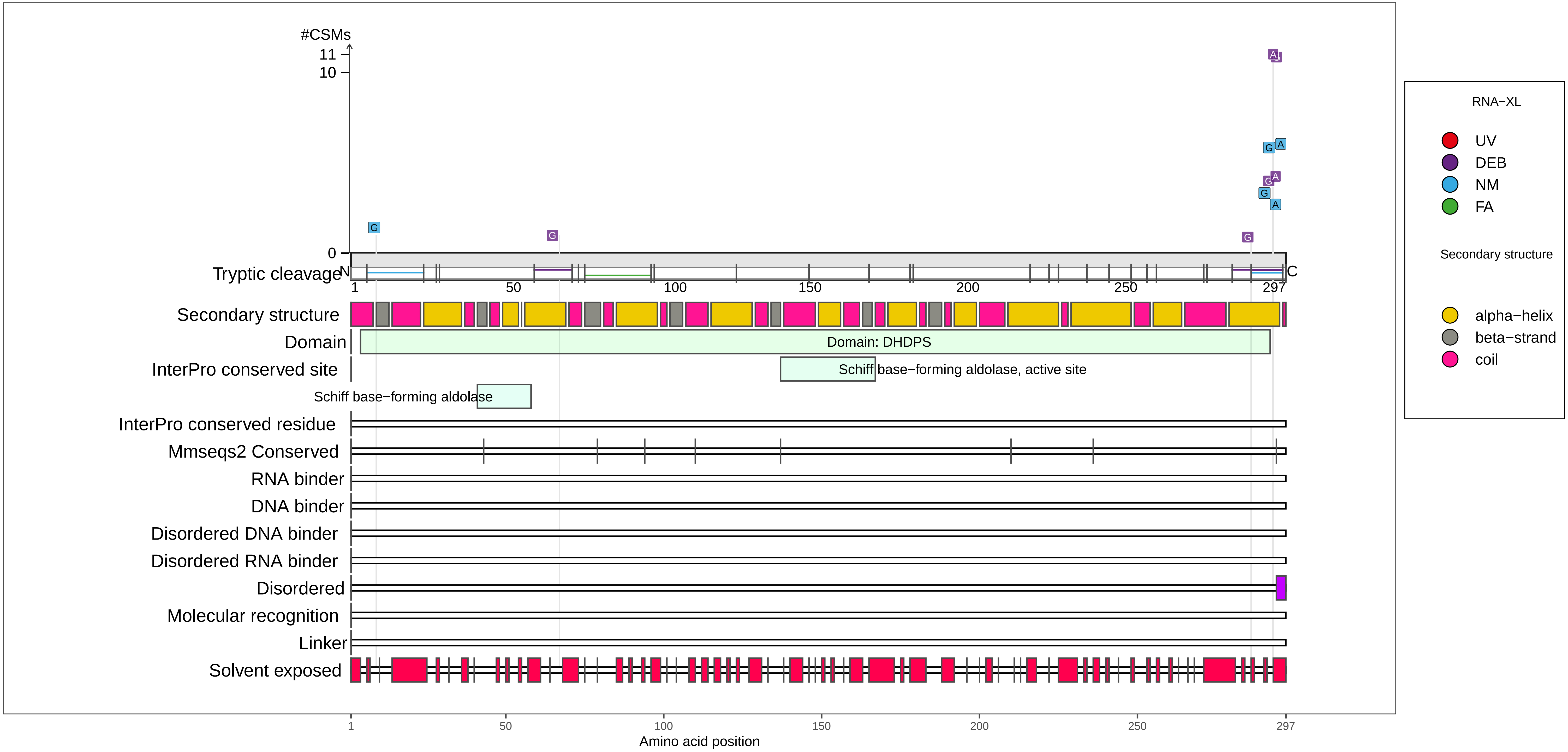

P0A905
SLYB_ECOLI Outer membrane lipoprotein SlyB

− Abundance:
tryptic [log10 Intensity]: 9.87 (Q 99)
PAXdb K12 strain [ppm]: 2.4 (Q 74)
PAXdb E.coli [ppm]: 3.01 (Q 95)

− RNA functions: not annotated

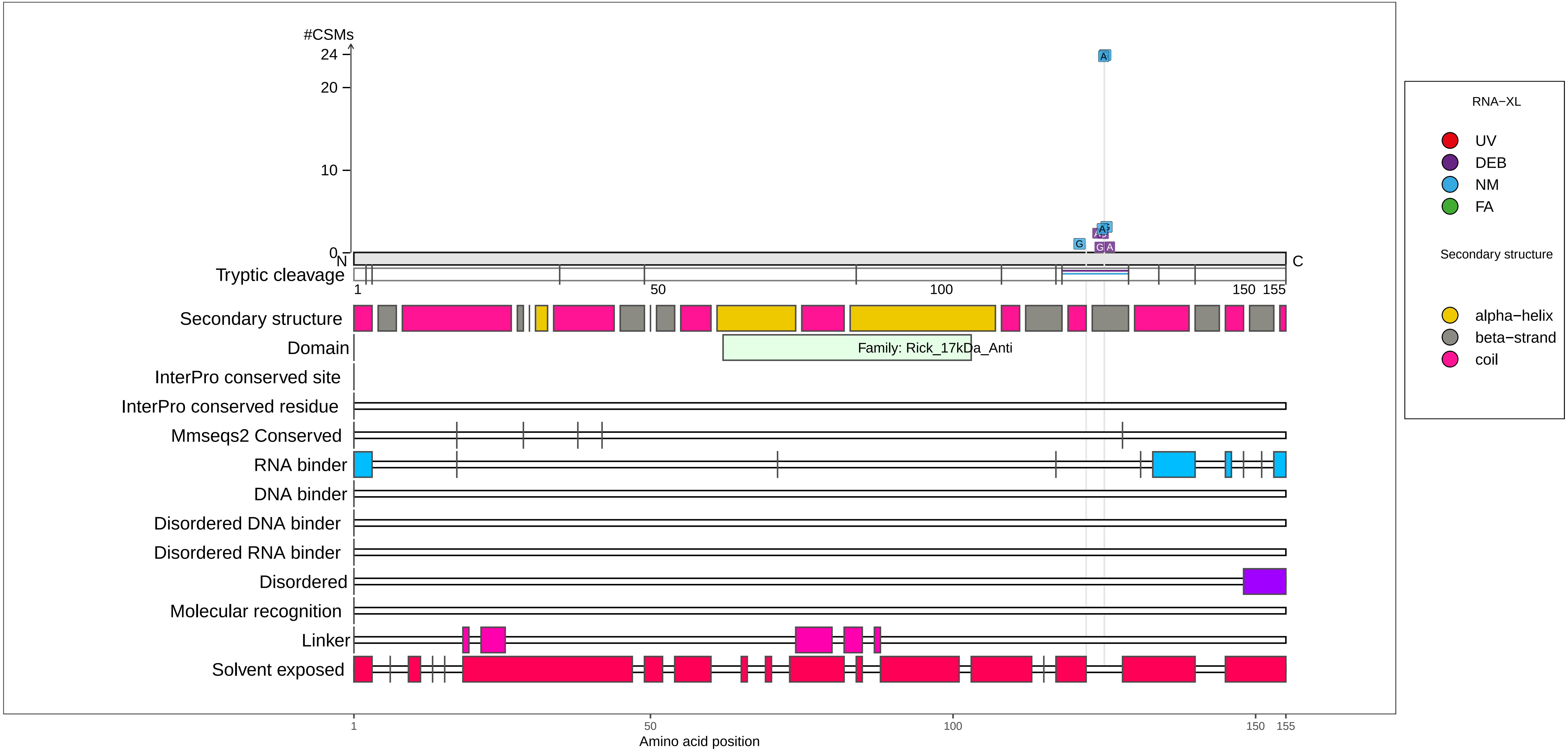

P0AC92
GNSA_ECOLI Protein GnsA

− Abundance:
tryptic [log10 Intensity]: 8.43 (Q 73)
PAXdb K12 strain [ppm]: 3.5 (Q 98)
PAXdb E.coli [ppm]: 2.08 (Q 74)

− RNA functions: not annotated

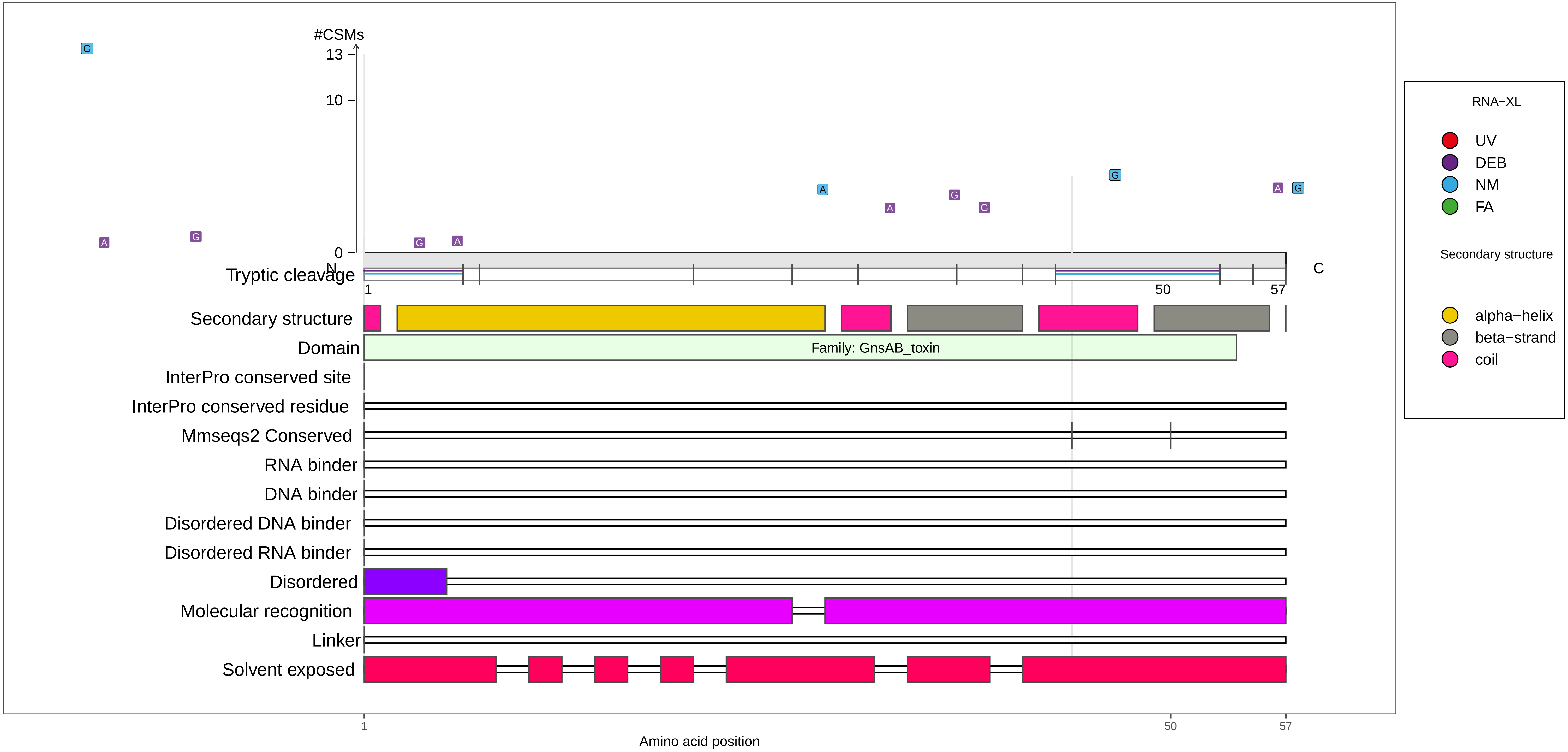

P0ACE0
MBHM_ECOLI Hydrogenase−2 large chain

− Abundance:
tryptic [log10 Intensity]: 8.59 (Q 78)
PAXdb K12 strain [ppm]: 2.03 (Q 61)
PAXdb E.coli [ppm]: 1.74 (Q 65)

− RNA functions: not annotated

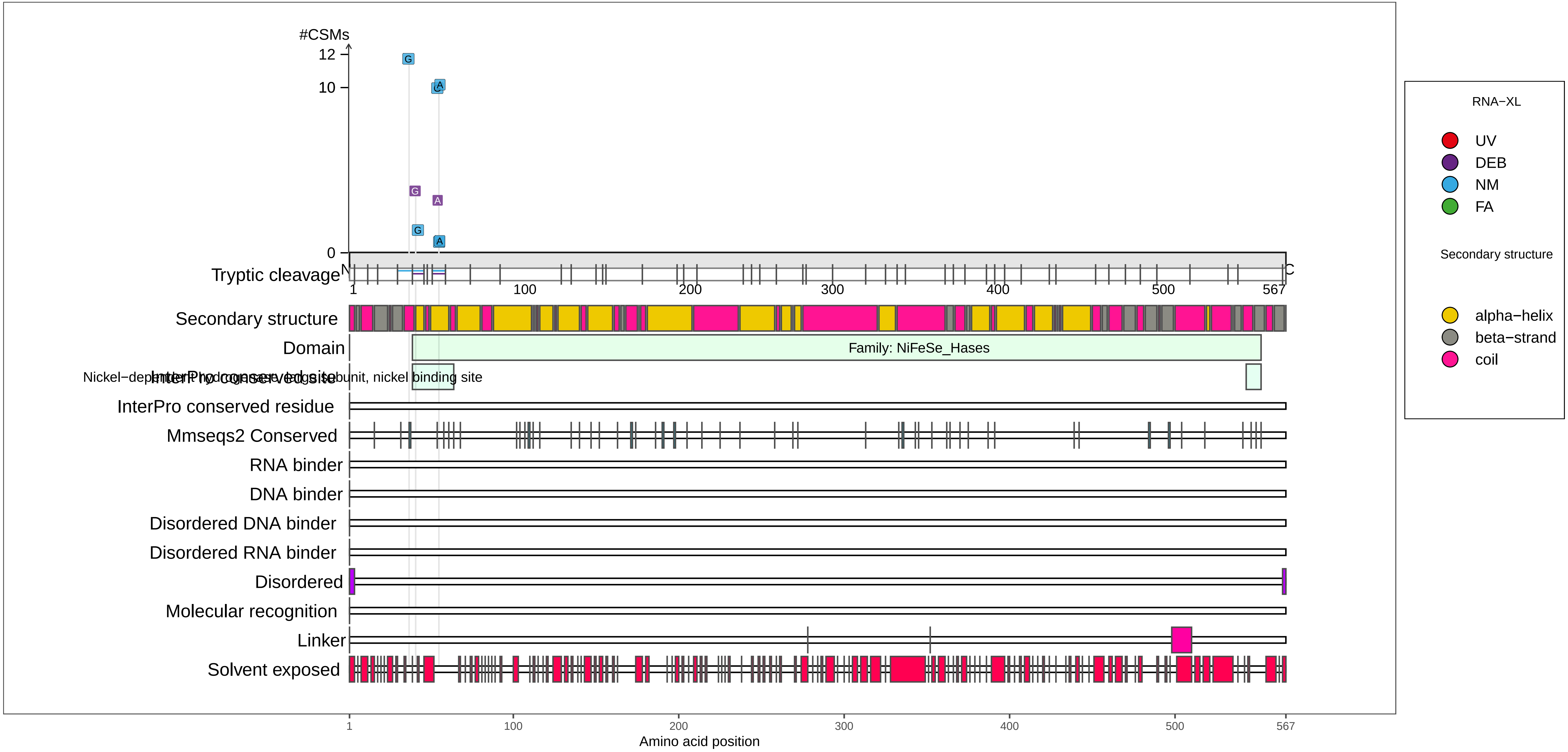

P0ADY1
PPID_ECOLI Periplasmic chaperone PpiD

− Abundance:
tryptic [log10 Intensity]: 8.66 (Q 80)
PAXdb K12 strain [ppm]: 2.53 (Q 78)
PAXdb E.coli [ppm]: 2.47 (Q 85)

− RNA functions: not annotated

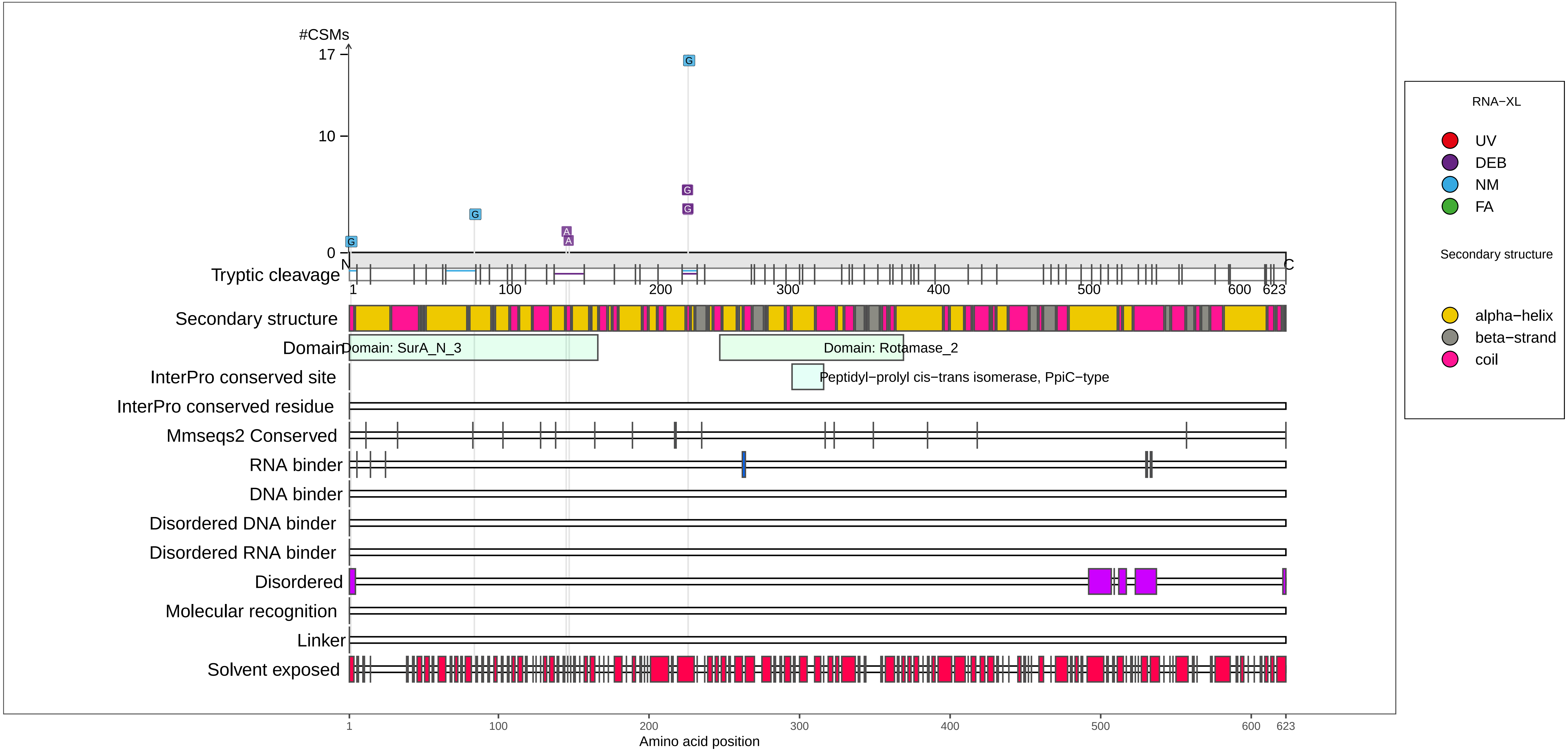

P0AEG4
DSBA_ECOLI Thiol:disulfide interchange protein DsbA

− Abundance:
tryptic [log10 Intensity]: 8.48 (Q 75)
PAXdb K12 strain [ppm]: 2.4 (Q 74)
PAXdb E.coli [ppm]: 2.61 (Q 88)

− RNA functions: not annotated

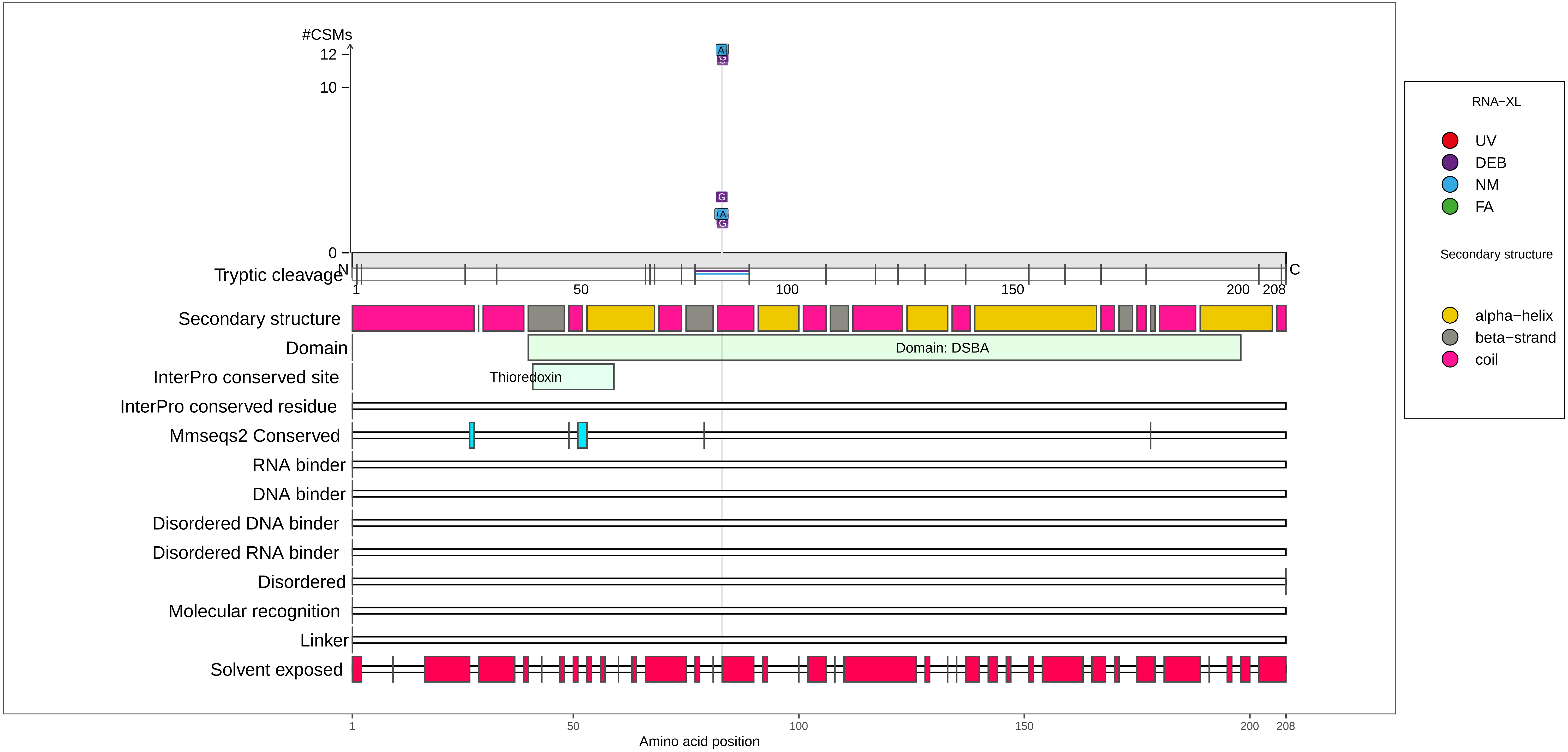

P33602
NUOG_ECOLI NADH−quinone oxidoreductase subunit G

− Abundance:
tryptic [log10 Intensity]: 9.64 (Q 97)
PAXdb K12 strain [ppm]: 2.5 (Q 78)
PAXdb E.coli [ppm]: 2.54 (Q 86)

− RNA functions: not annotated

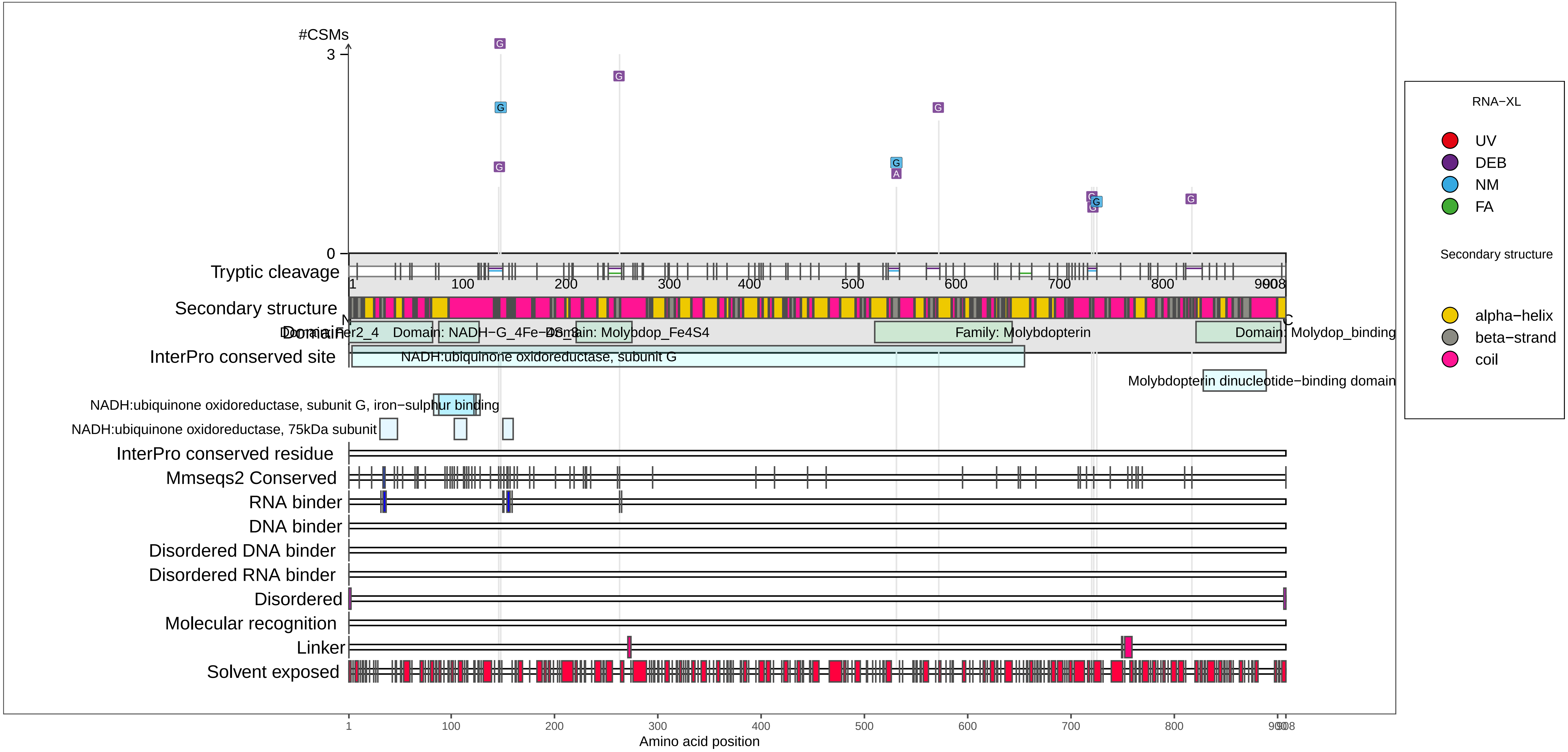

P40710
NLPE_ECOLI Lipoprotein NlpE

− Abundance:
tryptic [log10 Intensity]: 8.38 (Q 72)
PAXdb K12 strain [ppm]: 1.24 (Q 22)
PAXdb E.coli [ppm]: 1.04 (Q 48)

− RNA functions: not annotated

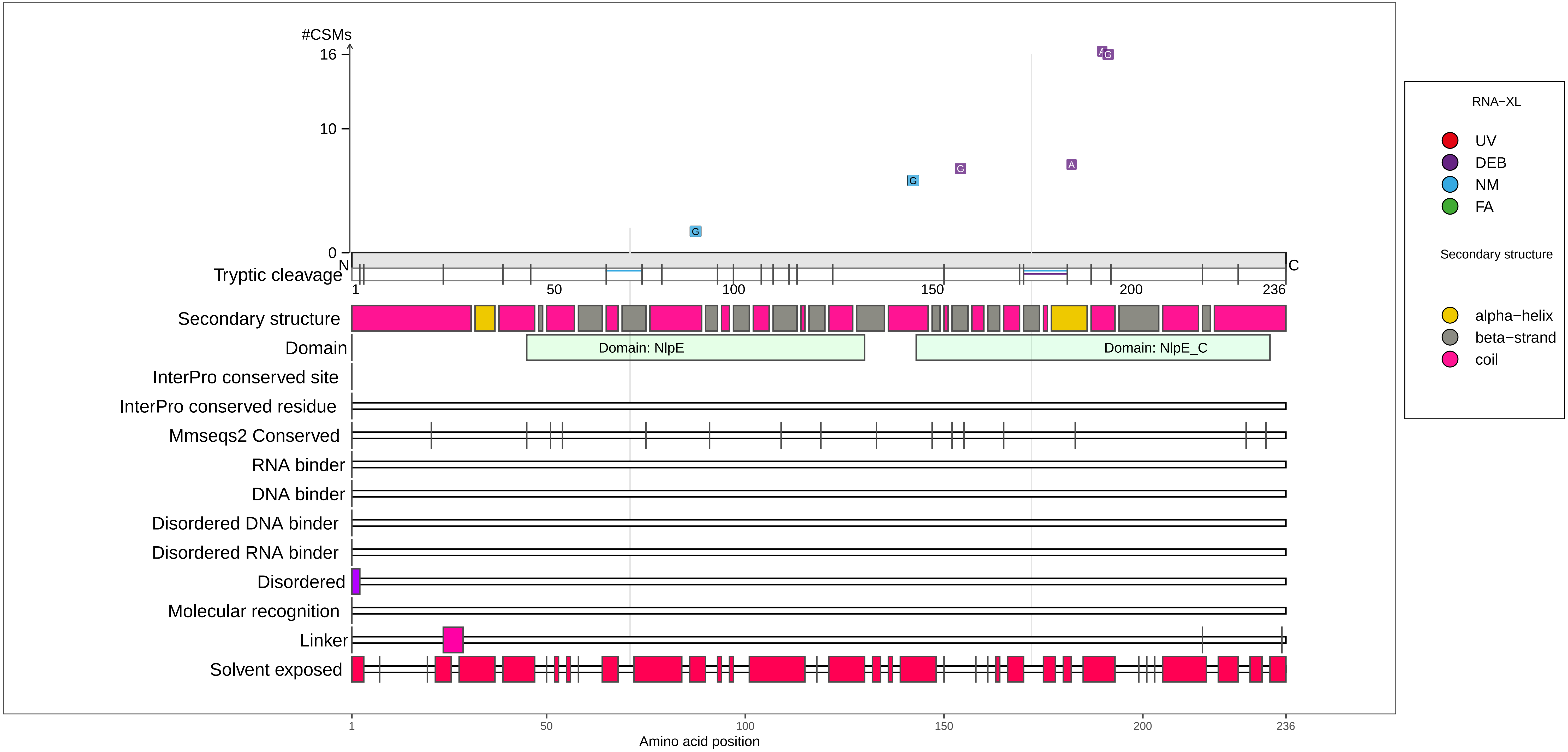

P0AAR3
YBAK_ECOLI Cys−tRNA(Pro)/Cys−tRNA(Cys) deacylase YbaK

− Abundance:
tryptic [log10 Intensity]: 7.76 (Q 47)
PAXdb K12 strain [ppm]: 2.84 (Q 87)
PAXdb E.coli [ppm]: 2.22 (Q 78)

− RNA functions:
aminoacyl−tRNA editing activity; Aminoacyl−tRNA editing domain
aminoacyl−tRNA metabolism involved in translational fidelity; Cys−tRNA(Pro) hydrolase activity
ncRNA metabolic process; RNA metabolic process; tRNA metabolic process

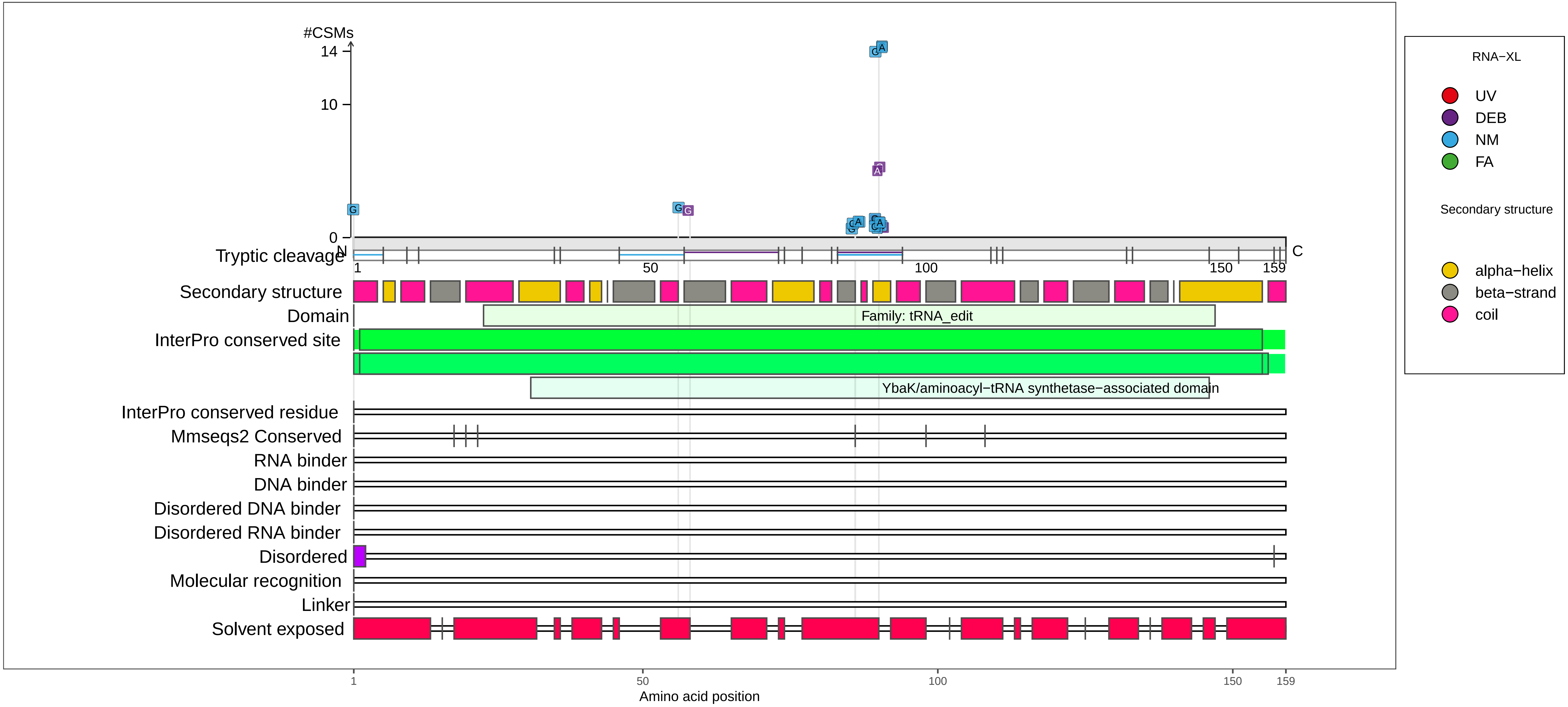

P0ACB2
HEM2_ECOLI Delta−aminolevulinic acid dehydratase
− Abundance:
tryptic [log10 Intensity]: 9.6 (Q 97)
PAXdb K12 strain [ppm]: 2.62 (Q 81)
PAXdb E.coli [ppm]: 2.1 (Q 75)

− RNA functions: not annotated

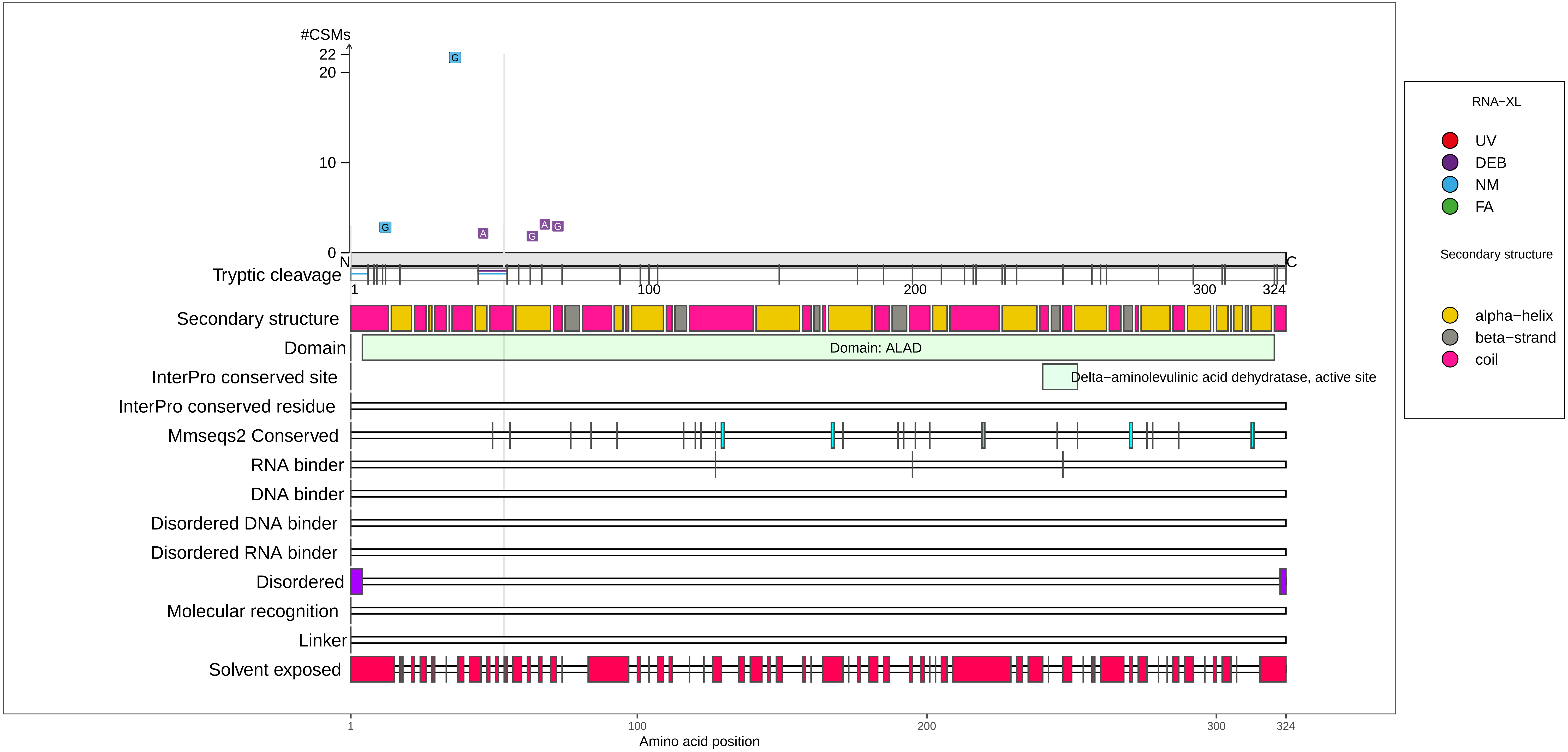

P0AFG6
ODO2_ECOLI Dihydrolipoyllysine−residue succinyltransferase component of 2−oxoglutarate dehydrogenase complex

− Abundance:
tryptic [log10 Intensity]: 9.19 (Q 92)
PAXdb K12 strain [ppm]: 3.71 (Q 99)
PAXdb E.coli [ppm]: 2.92 (Q 93)

− RNA functions: not annotated

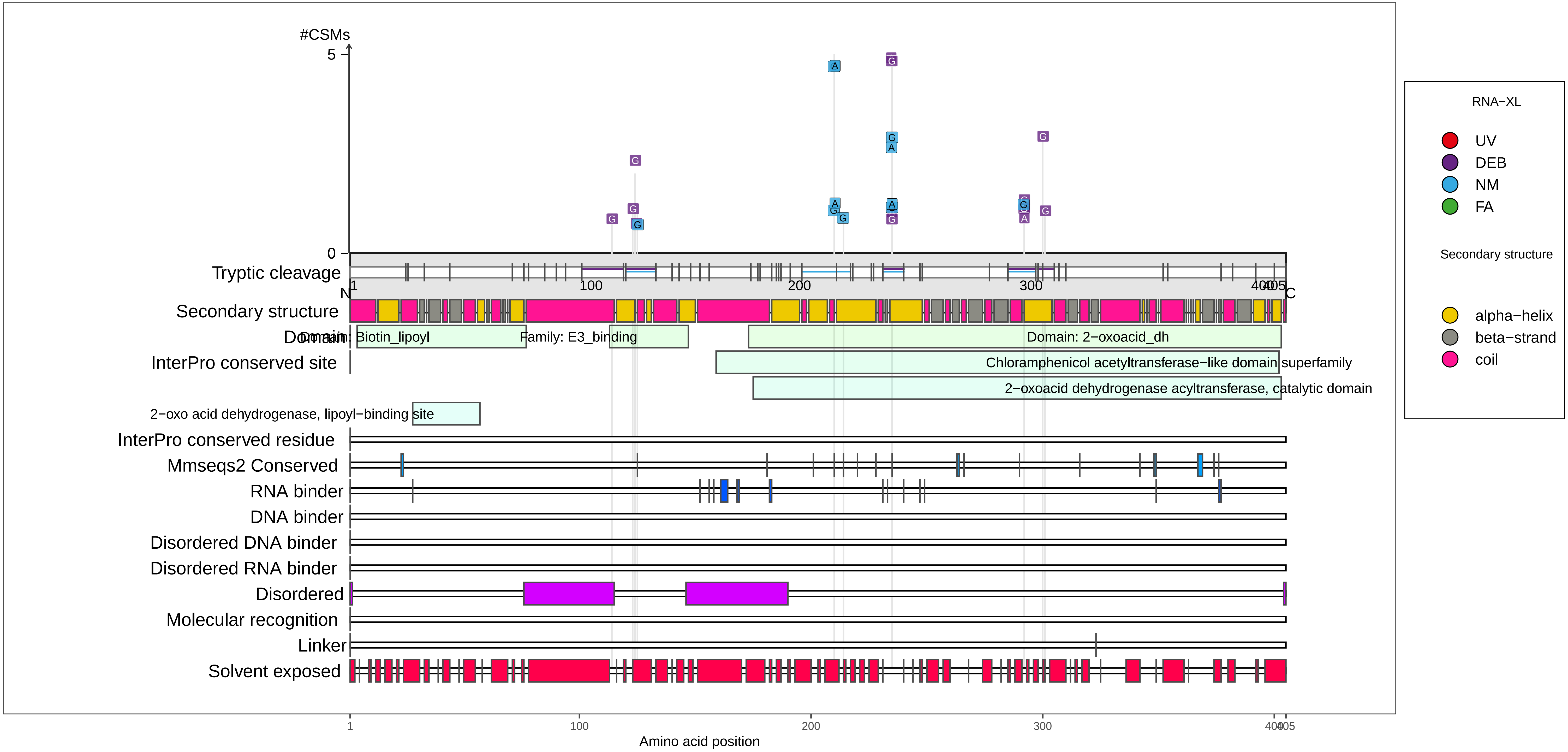

P11557
DAMX_ECOLI Cell division protein DamX

− Abundance:
tryptic [log10 Intensity]: 9.25 (Q 93)
PAXdb K12 strain [ppm]: 1.25 (Q 23)
PAXdb E.coli [ppm]: 2.24 (Q 78)

− RNA functions: not annotated

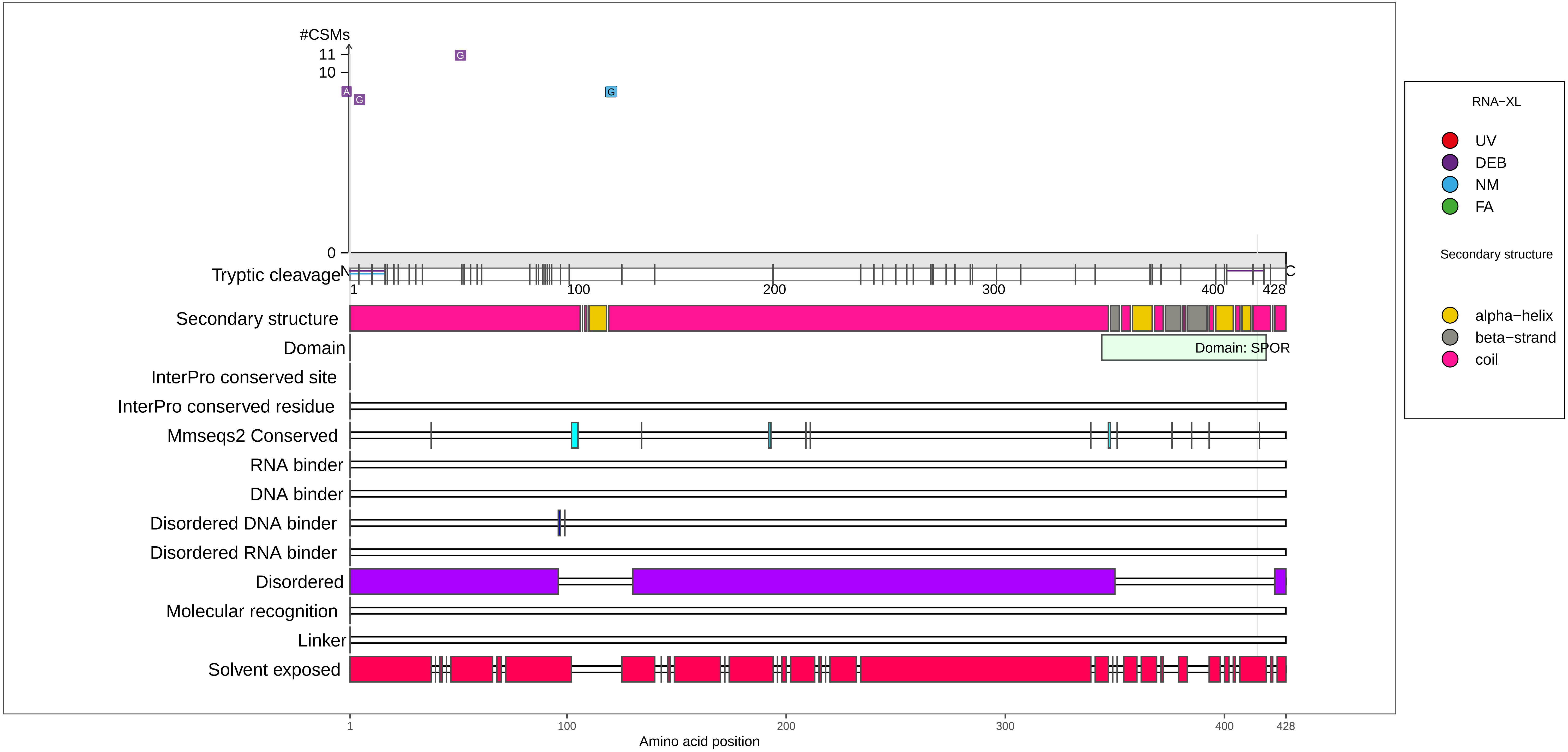

P27434
RODZ_ECOLI Cytoskeleton protein RodZ

− Abundance:
tryptic [log10 Intensity]: 8.31 (Q 70)
PAXdb K12 strain [ppm]: 2.54 (Q 78)
PAXdb E.coli [ppm]: 2.41 (Q 83)

− RNA functions: not annotated

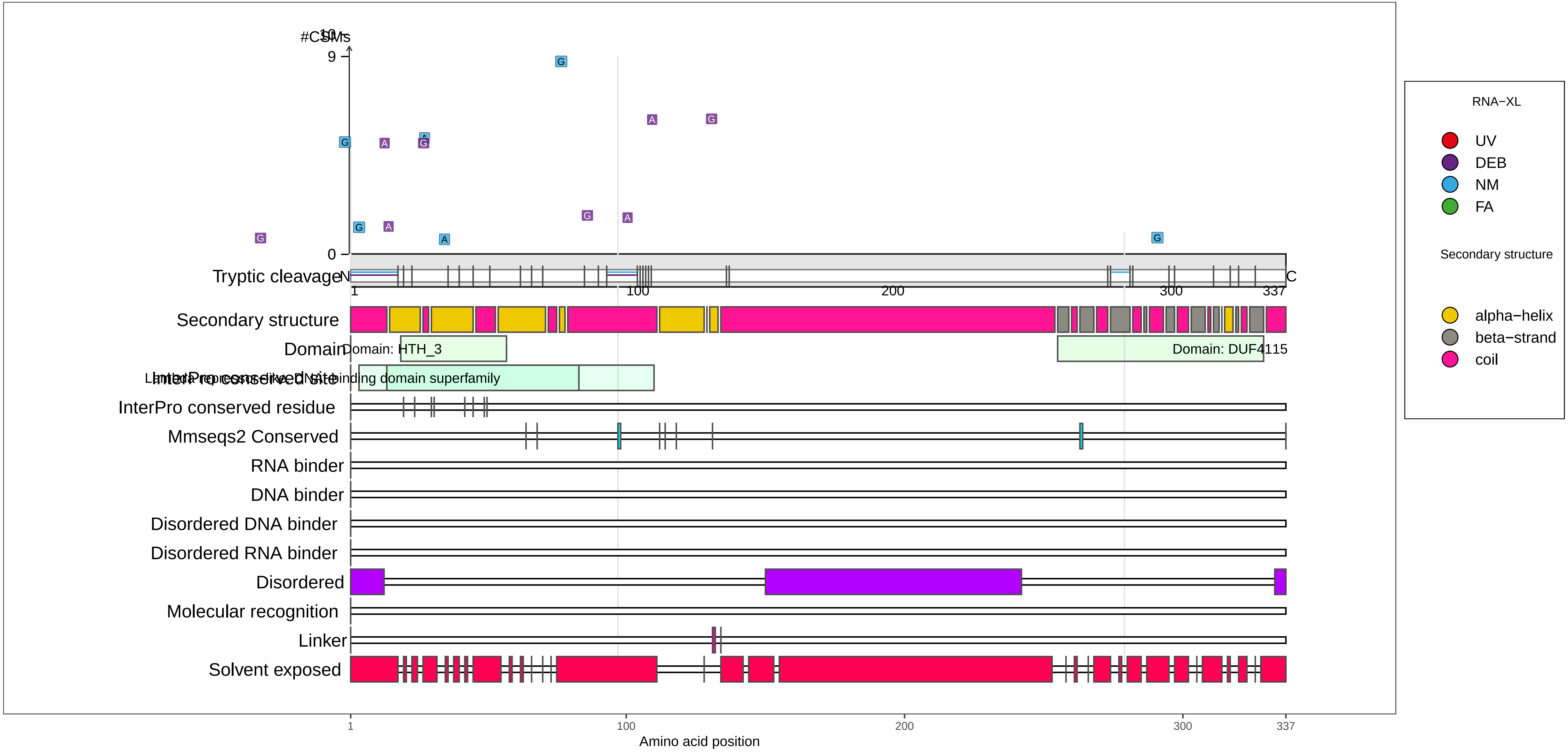

P36938
PGM_ECOLI Phosphoglucomutase

− Abundance:
tryptic    [log10 Intensity]:  8.62 (Q 79)
PAXdb K12 strain   [ppm]:  2.77 (Q 85)
PAXdb E.coli     [ppm]:  2.8 (Q 92)

− RNA functions: not annotated

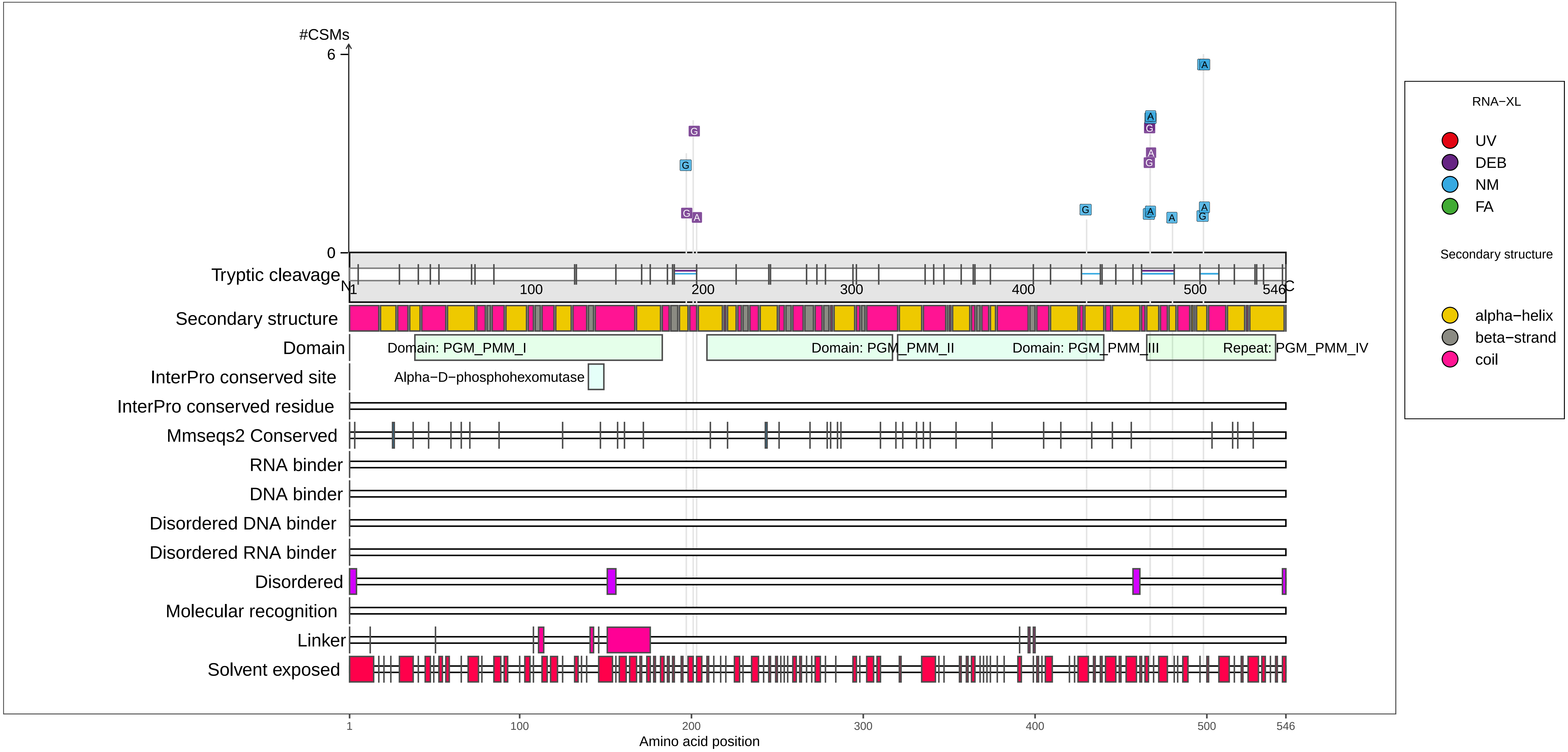

P65556
YFCD_ECOLI Uncharacterized Nudix hydrolase YfcD

− Abundance:
tryptic [log10 Intensity]: 8.39 (Q 72)
PAXdb K12 strain [ppm]: NA (Q NA)
PAXdb E.coli [ppm]: 2.39 (Q 83)

− RNA functions: not annotated

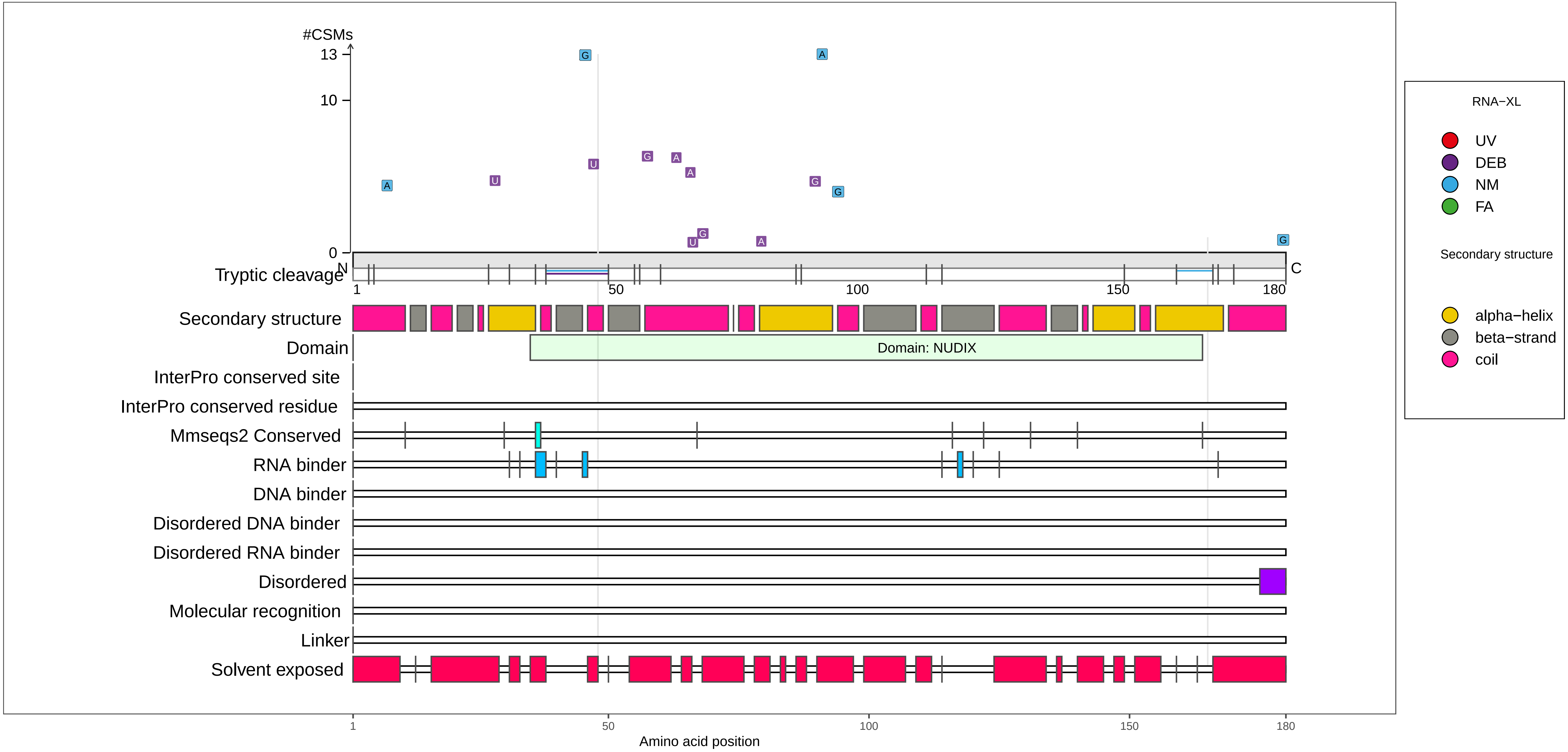

P76046
YCJX_ECOLI Uncharacterized protein YcjX

− Abundance:
tryptic [log10 Intensity]: 7.68 (Q 43)
PAXdb K12 strain [ppm]: 1.18 (Q 19)
PAXdb E.coli [ppm]: 1.26 (Q 54)

− RNA functions: not annotated

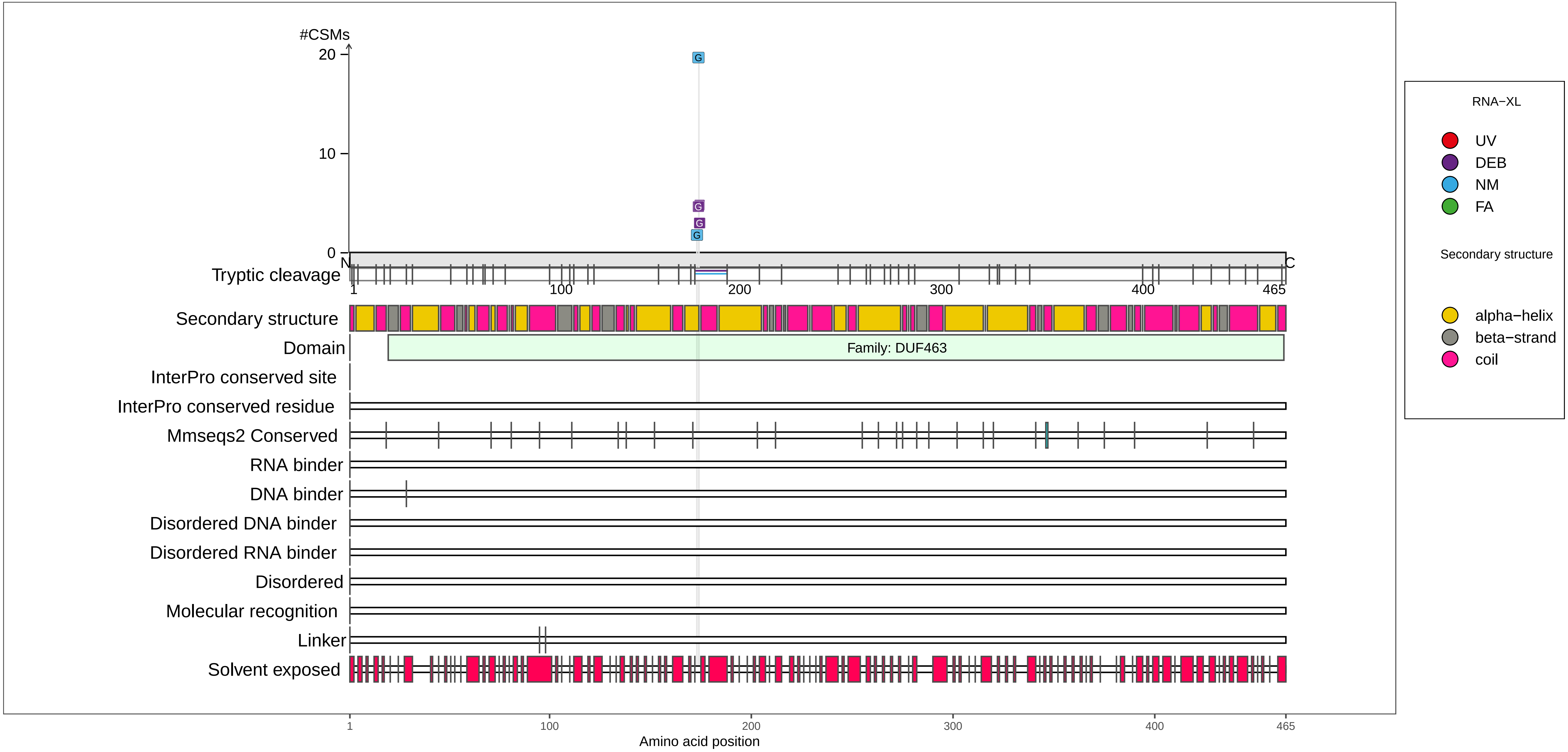

P00959
SYM_ECOLI Methionine−−tRNA ligase

− Abundance:
tryptic [log10 Intensity]: 8.96 (Q 87)
PAXdb K12 strain [ppm]: 2.43 (Q 75)
PAXdb E.coli [ppm]: 2.68 (Q 89)

− RNA functions:
aminoacyl−tRNA ligase activity; methionine−tRNA ligase activity
methionyl−tRNA aminoacylation; ncRNA metabolic process; Putative tRNA binding domain; RNA binding
RNA metabolic process; tRNA aminoacylation; tRNA aminoacylation for protein translation
tRNA binding; tRNA metabolic process; tRNA synthetases class I (C) catalytic domain
tRNA synthetases class I (I, L, M and V); tRNA synthetases class I (M)

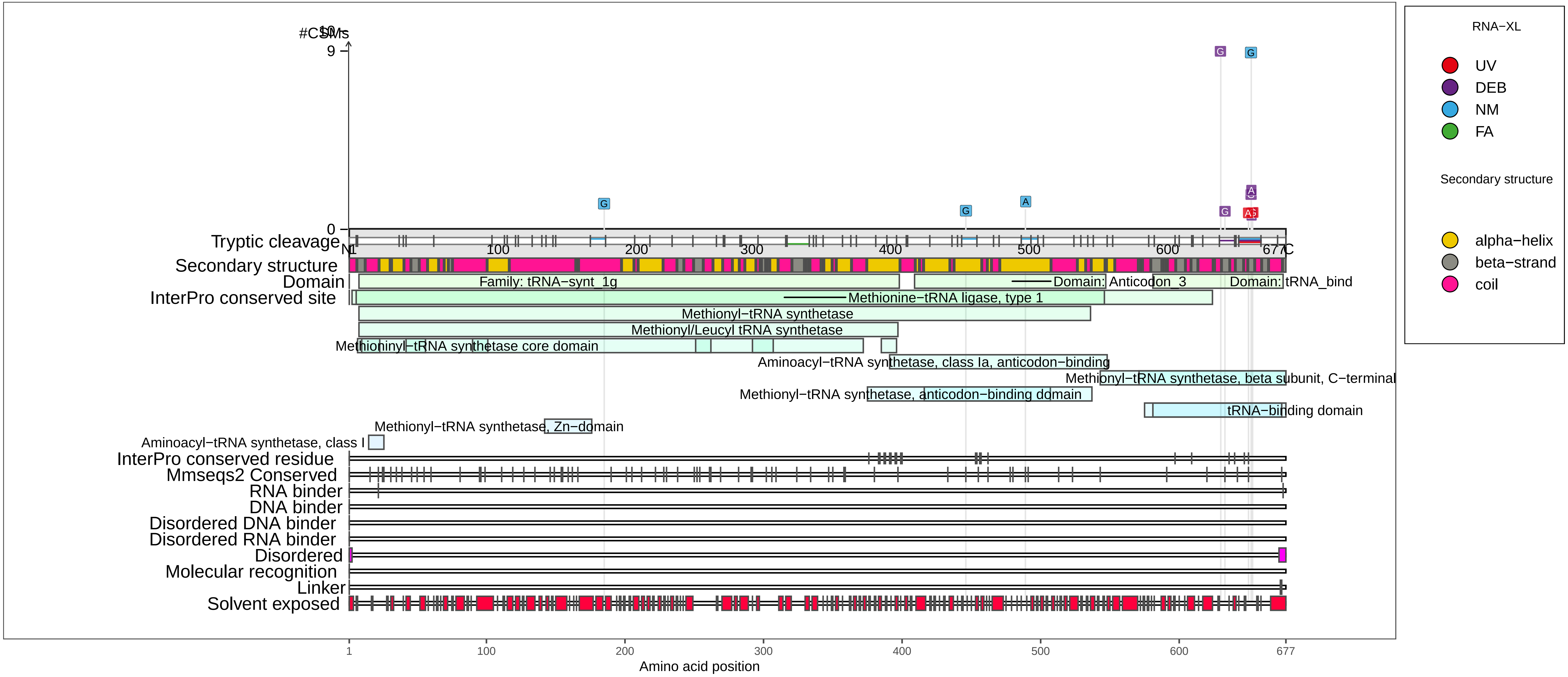

P0AB38
LPOB_ECOLI Penicillin−binding protein activator LpoB

− Abundance:
tryptic [log10 Intensity]: 8.2 (Q 65)
PAXdb K12 strain [ppm]: 2.16 (Q 65)
PAXdb E.coli [ppm]: 2.22 (Q 78)

− RNA functions: not annotated

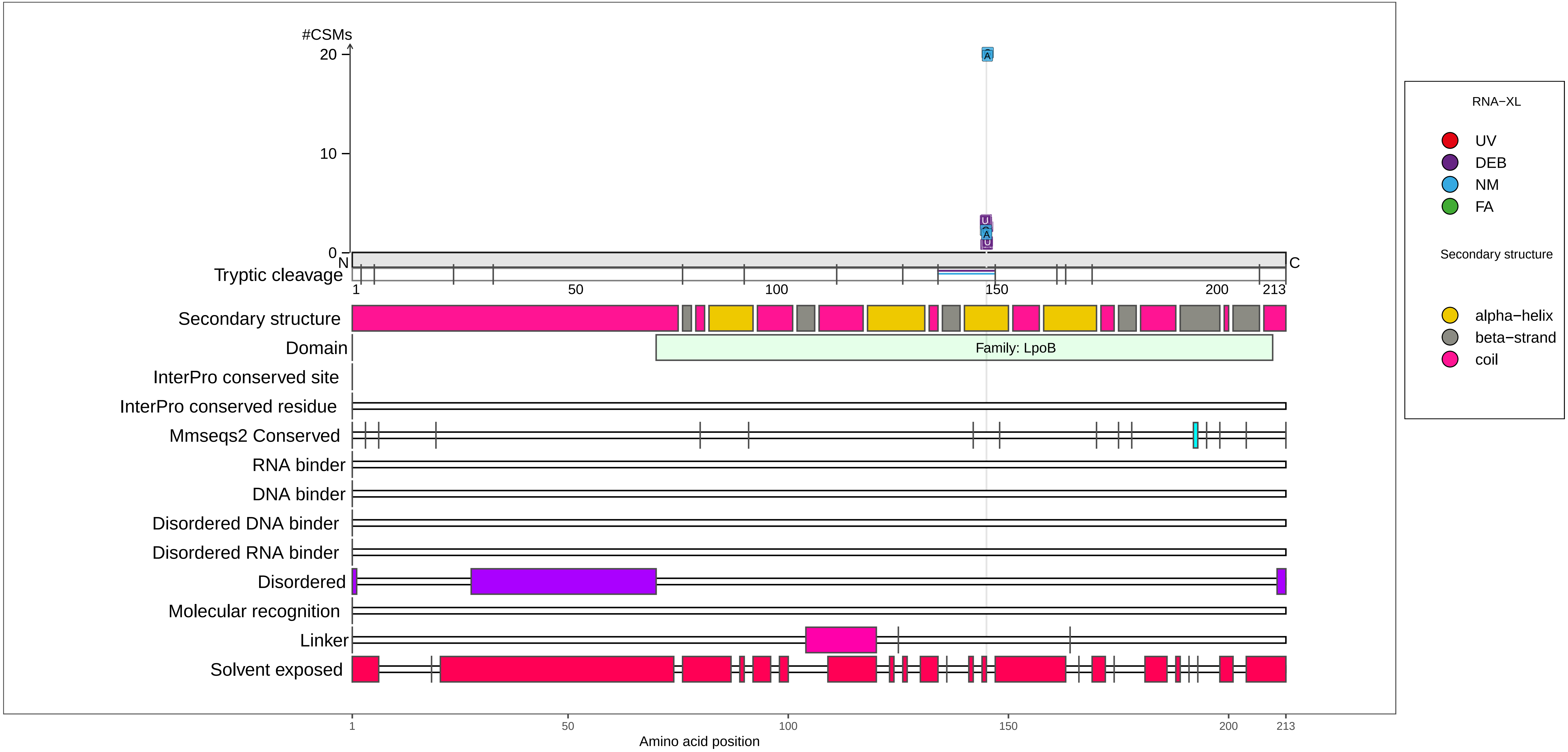

P0A9C5
GLN1B_ECOLI Glutamine synthetase

− Abundance:
tryptic [log10 Intensity]: 9.07 (Q 90)
PAXdb K12 strain [ppm]: 2.97 (Q 90)
PAXdb E.coli [ppm]: 2.89 (Q 93)

− RNA functions: not annotated

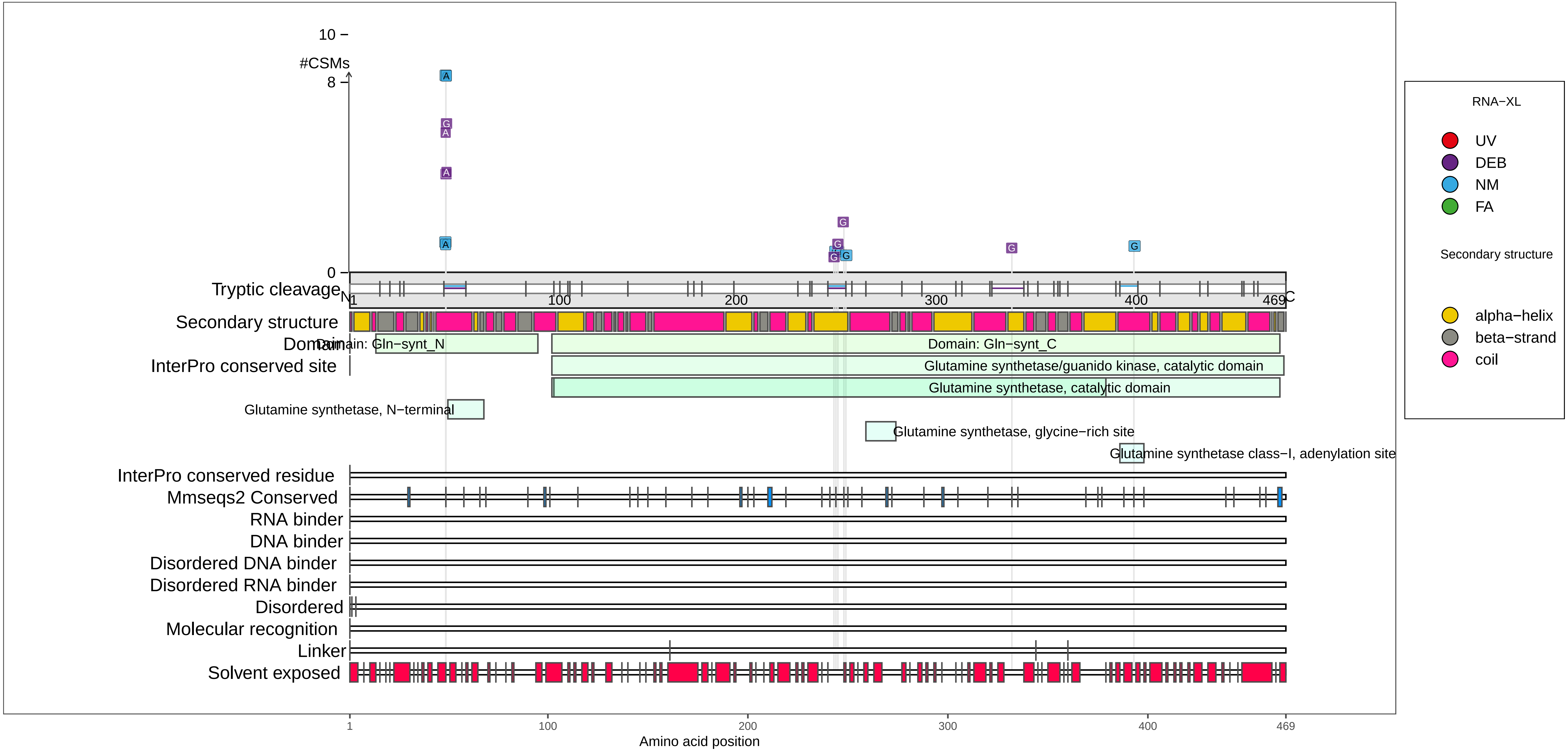

P0AB24
EFEO_ECOLI Iron uptake system component EfeO

− Abundance:
tryptic [log10 Intensity]: 8.44 (Q 73)
PAXdb K12 strain [ppm]: 2.8 (Q 86)
PAXdb E.coli [ppm]: 2.09 (Q 74)

− RNA functions: not annotated

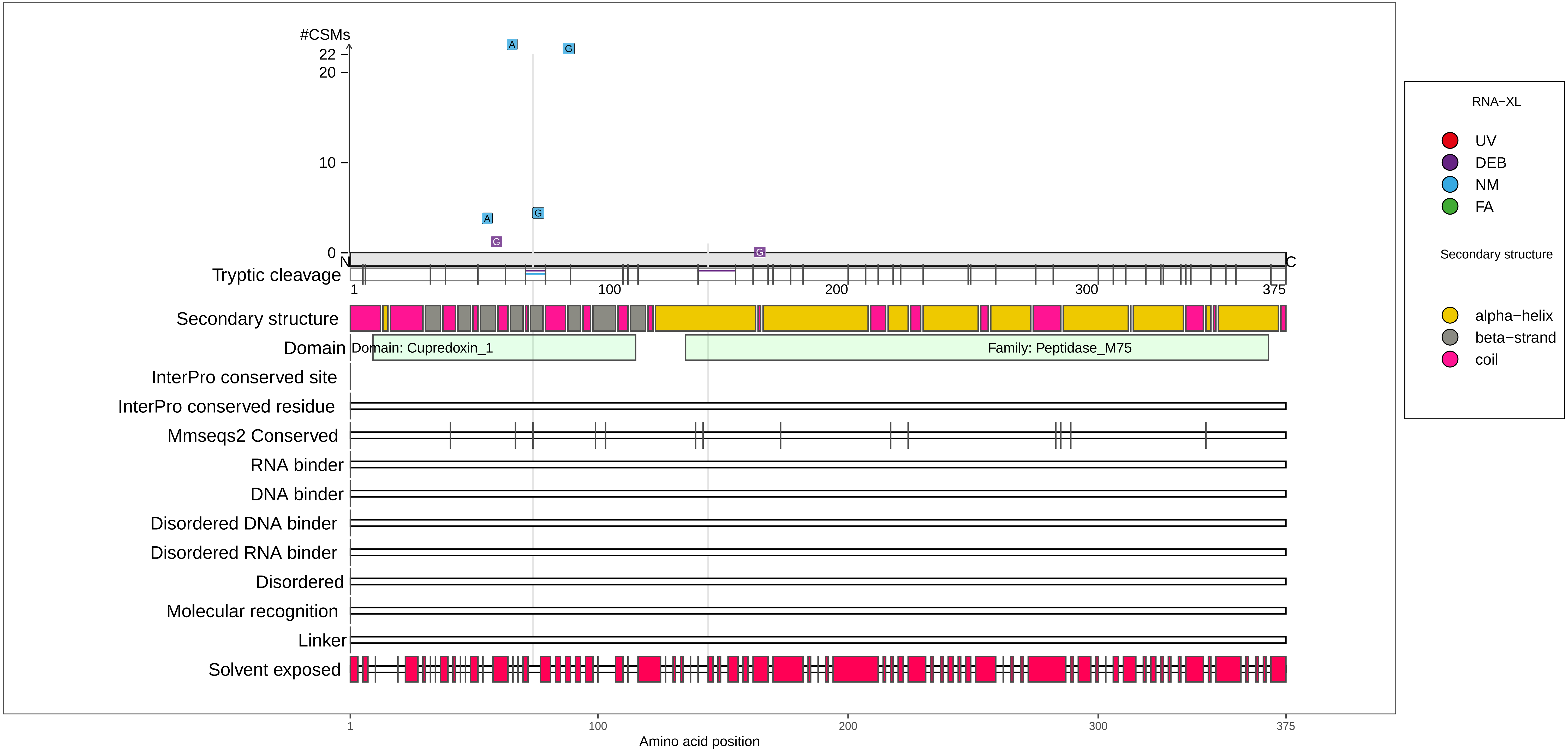

P0ACD4
ISCU_ECOLI Iron−sulfur cluster assembly scaffold protein IscU

− Abundance:
tryptic [log10 Intensity]: 8.15 (Q 63)
PAXdb K12 strain [ppm]: 2.98 (Q 90)
PAXdb E.coli [ppm]: 3.27 (Q 97)

− RNA functions: not annotated

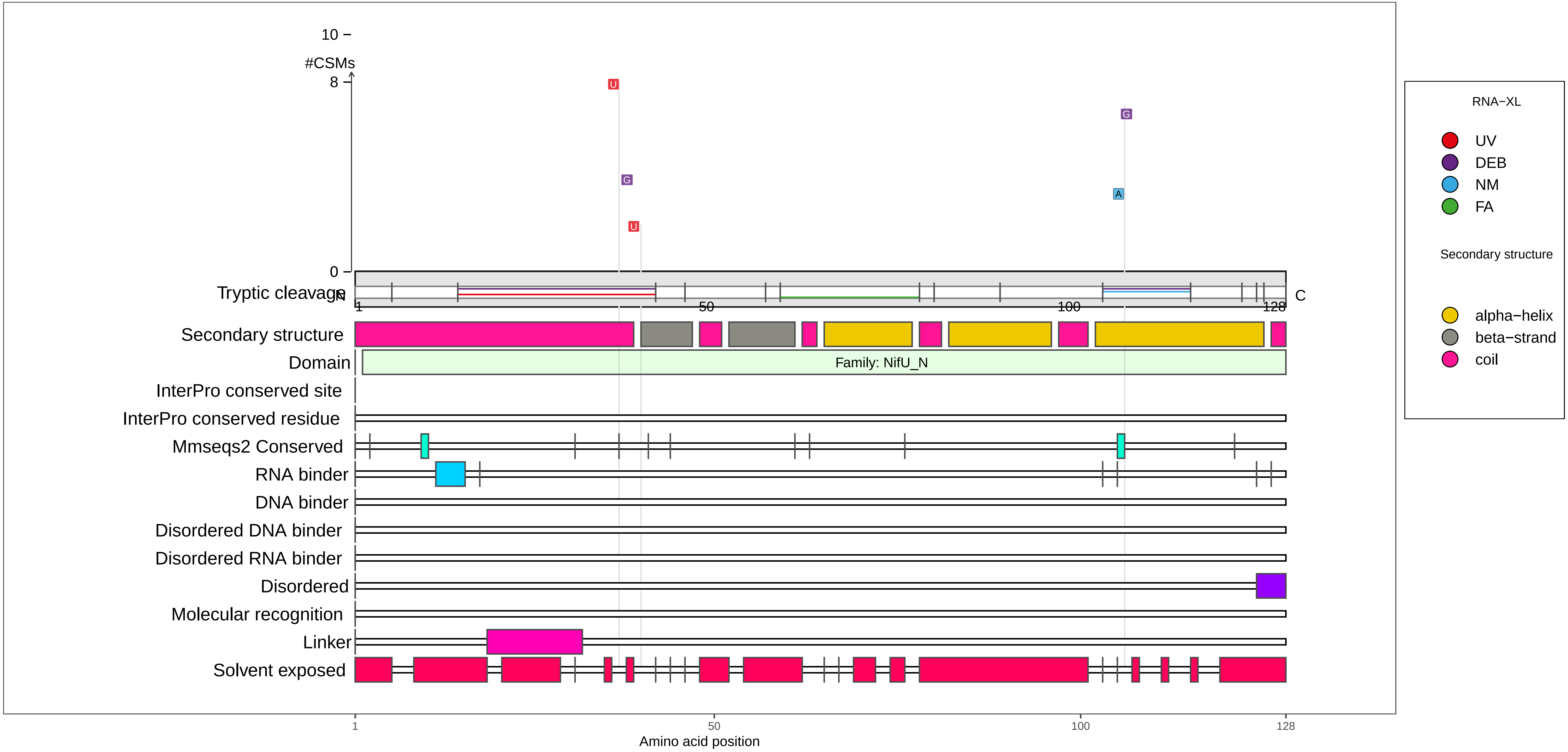

P0ACQ4
OXYR_ECOLI Hydrogen peroxide−inducible genes activator

− Abundance:
tryptic [log10 Intensity]: 8.06 (Q 60)
PAXdb K12 strain [ppm]: 2.36 (Q 72)
PAXdb E.coli [ppm]: 2.38 (Q 83)

− RNA functions: not annotated

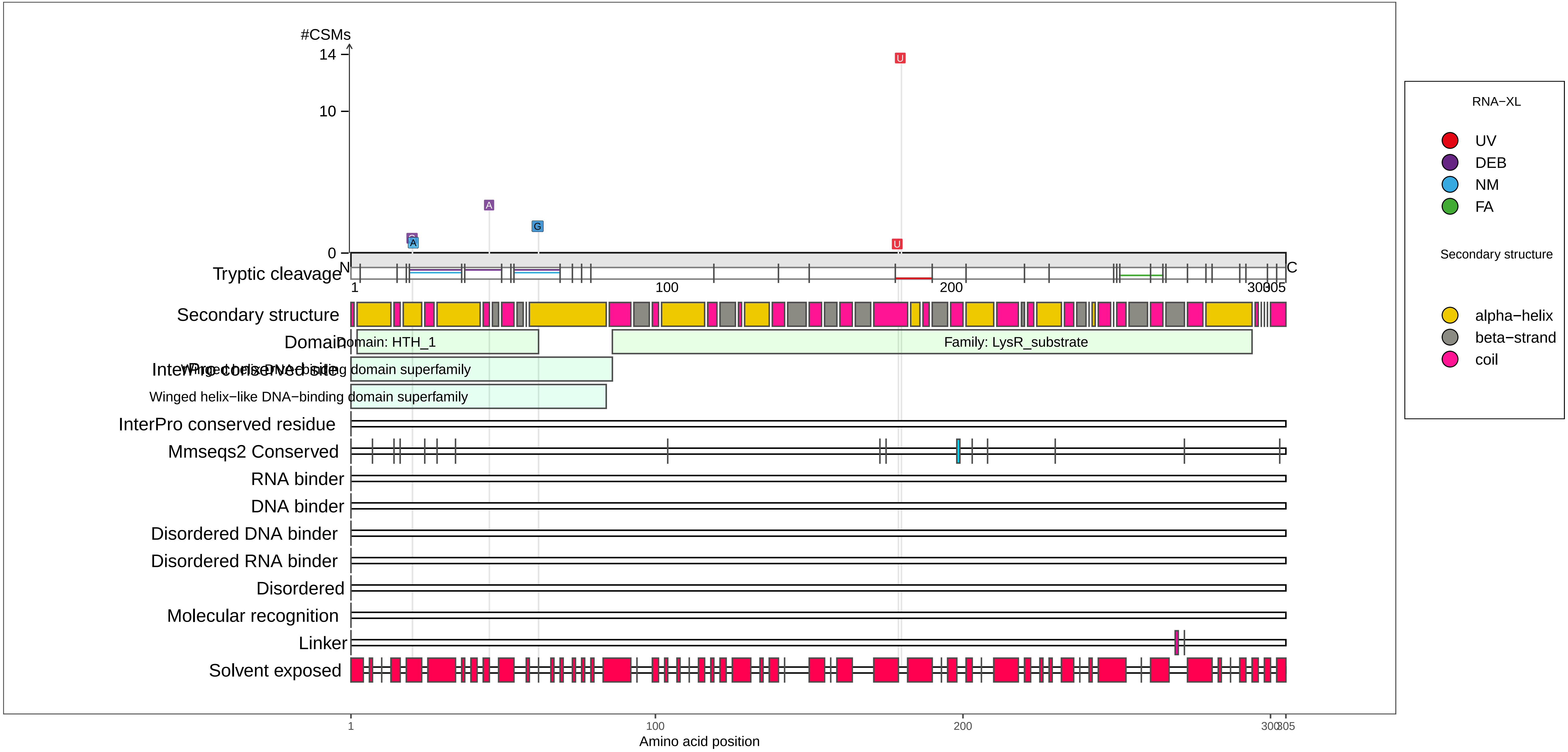

P0AD49
YFIA_ECOLI Ribosome−associated inhibitor A

− Abundance:
tryptic [log10 Intensity]: 8.69 (Q 81)
PAXdb K12 strain [ppm]: 2.9 (Q 88)
PAXdb E.coli [ppm]: 3.35 (Q 98)

− RNA functions:
RNA binding; rRNA binding

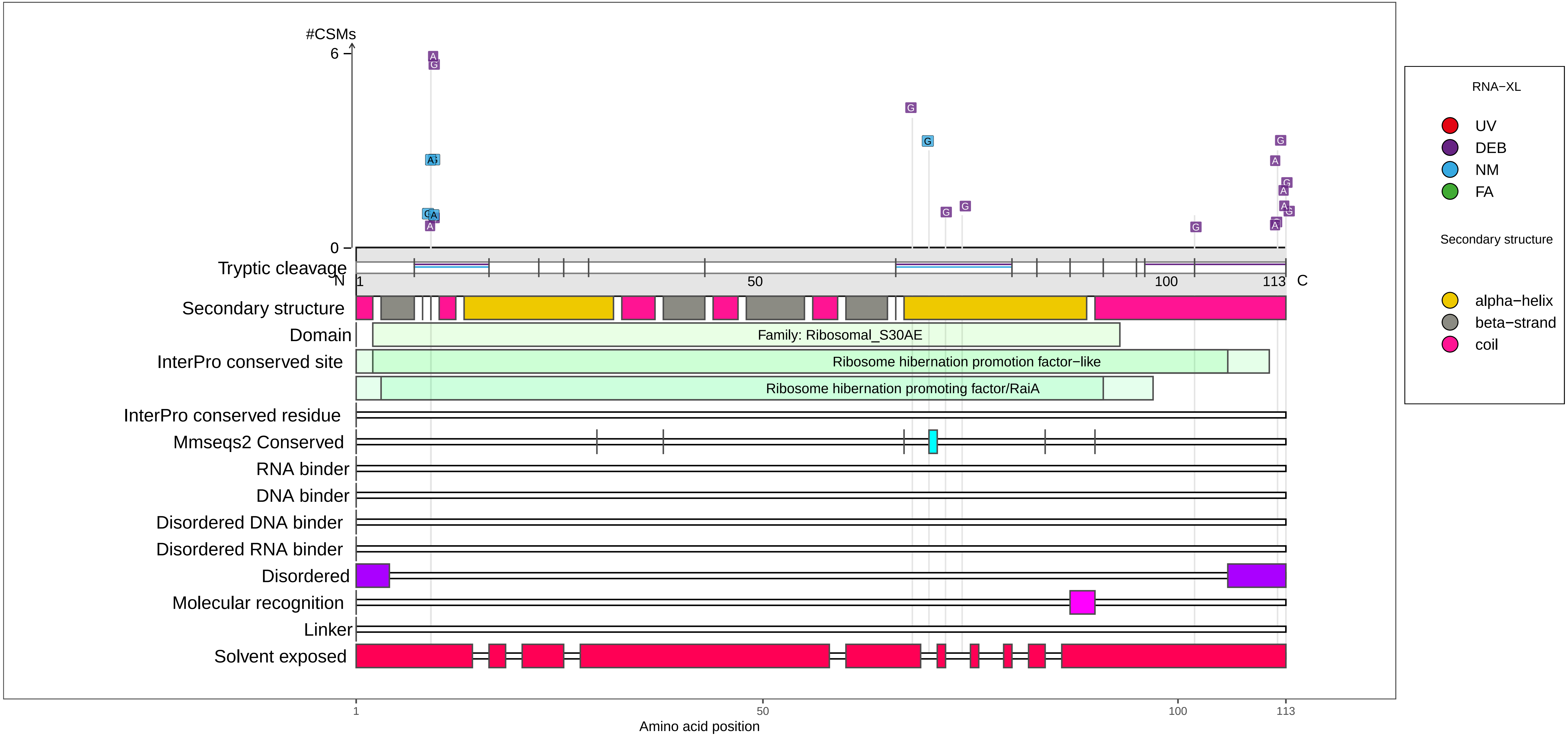

P0AG48
RL21_ECOLI 50S ribosomal protein L21

− Abundance:
tryptic [log10 Intensity]: 9.08 (Q 90)
PAXdb K12 strain [ppm]: 3.52 (Q 98)
PAXdb E.coli [ppm]: 3.13 (Q 96)

− RNA functions:
RNA binding; rRNA binding

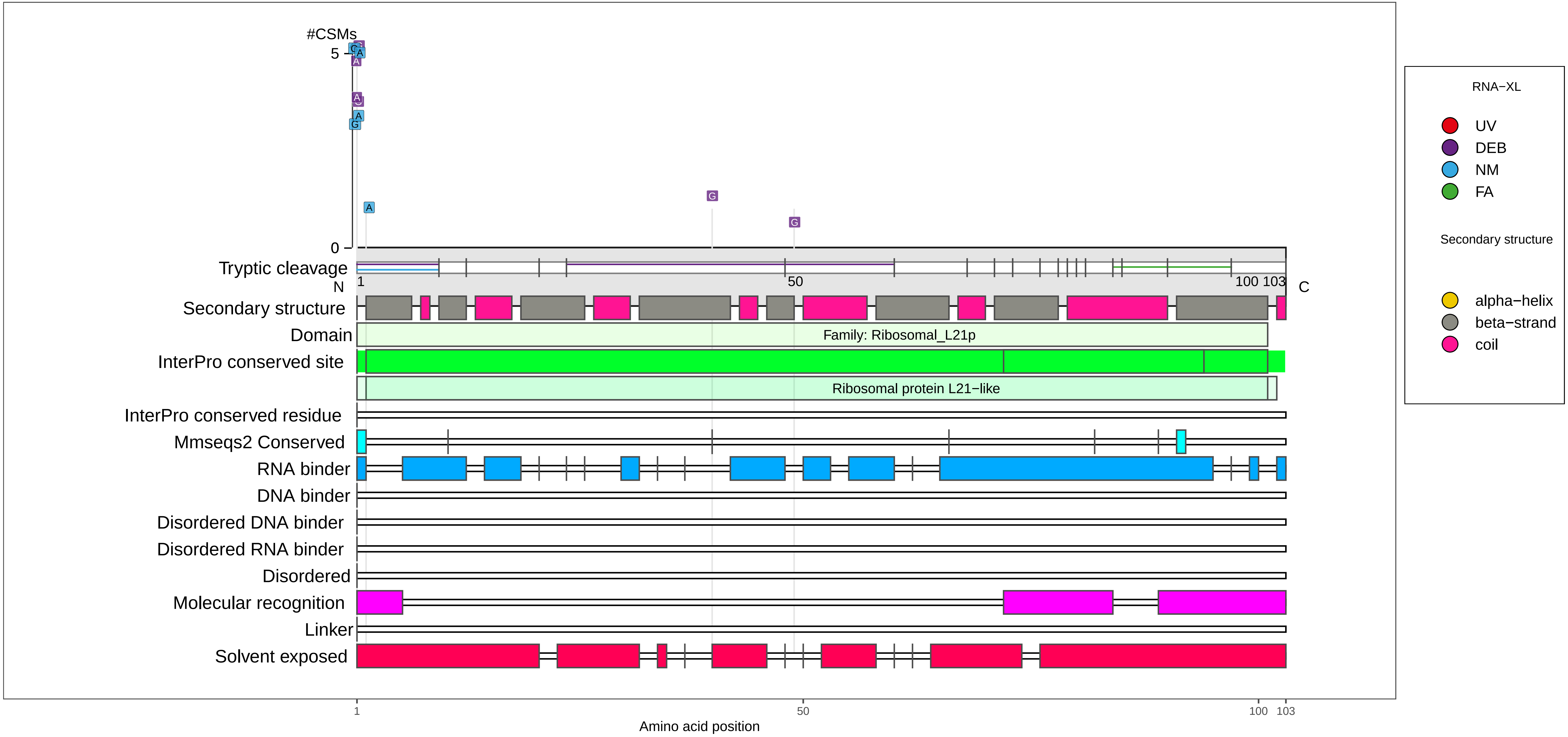

P10408
SECA_ECOLI Protein translocase subunit SecA

− Abundance:
tryptic [log10 Intensity]: 9.11 (Q 91)
PAXdb K12 strain [ppm]: 2.55 (Q 79)
PAXdb E.coli [ppm]: 2.59 (Q 88)

− RNA functions: not annotated

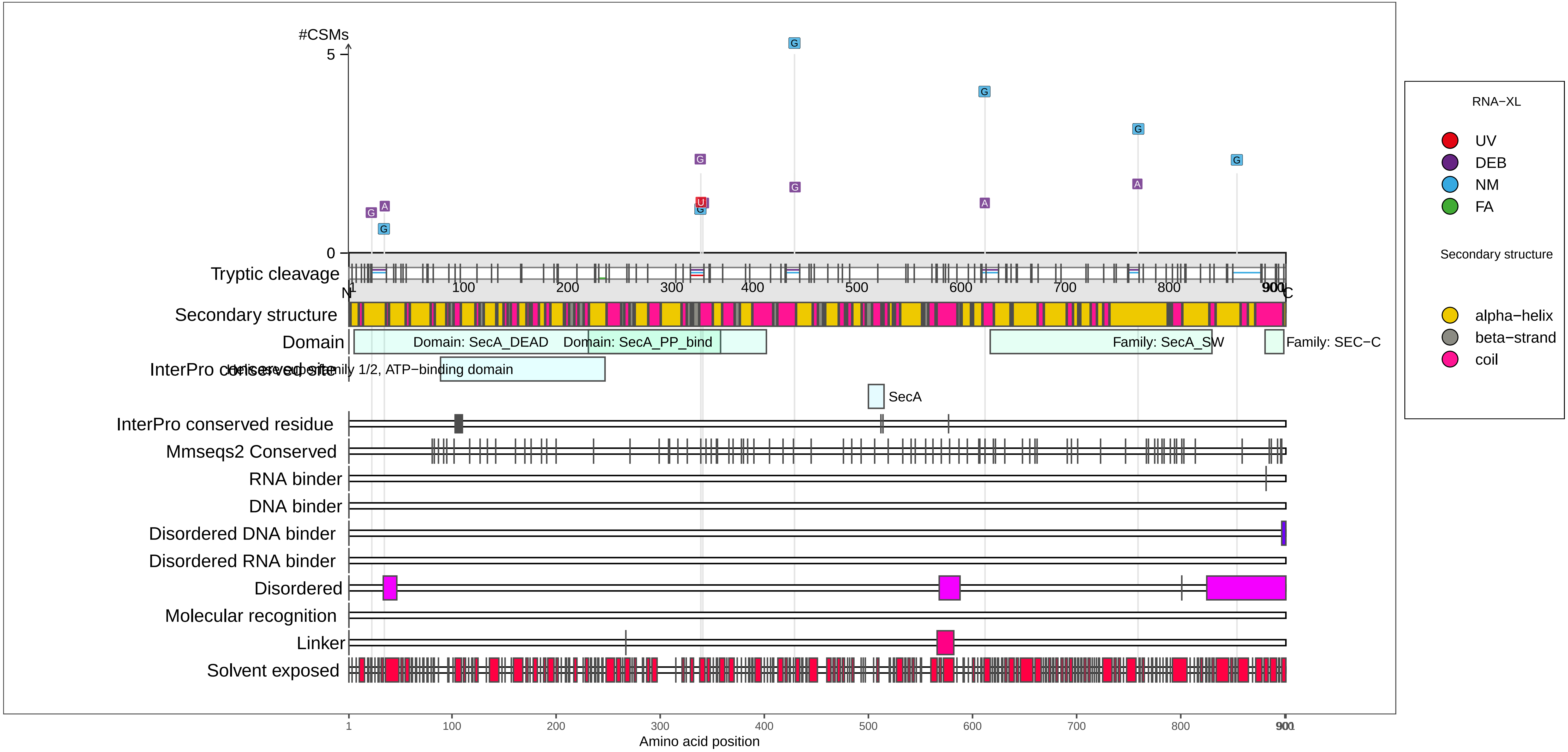

P45750
HOFP_ECOLI DNA utilization protein HofP

− Abundance:
tryptic [log10 Intensity]: NA (Q NA)
PAXdb K12 strain [ppm]: NA (Q NA)
PAXdb E.coli [ppm]: −0.11 (Q 22)

− RNA functions: not annotated

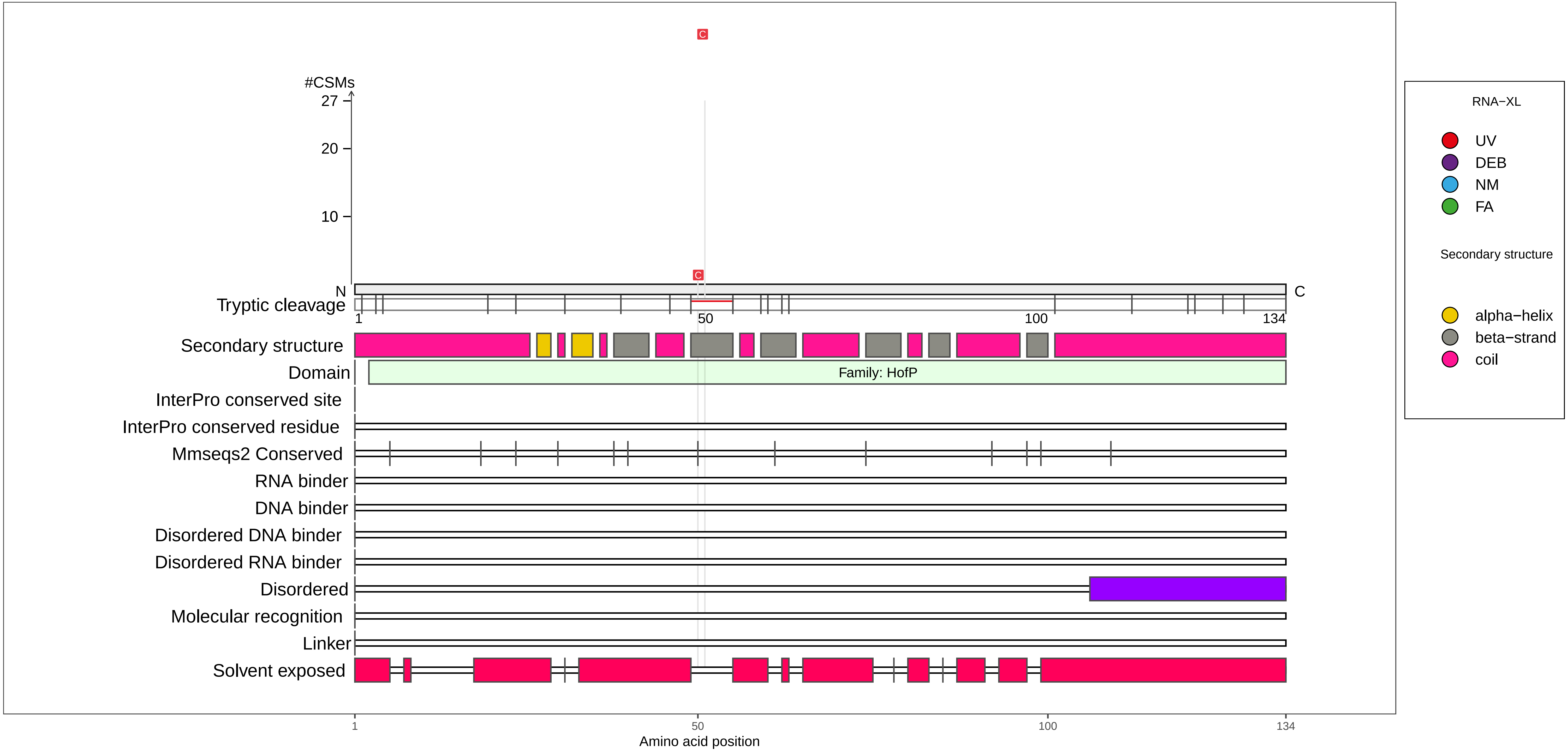

P46837
YHGF_ECOLI Protein YhgF

− Abundance:
tryptic [log10 Intensity]: 7.85 (Q 52)
PAXdb K12 strain [ppm]: 2.64 (Q 82)
PAXdb E.coli [ppm]: 1.89 (Q 69)

− RNA functions:
mRNA binding; Ribosomal protein S1−like RNA−binding domain; RNA binding
S1 RNA binding domain

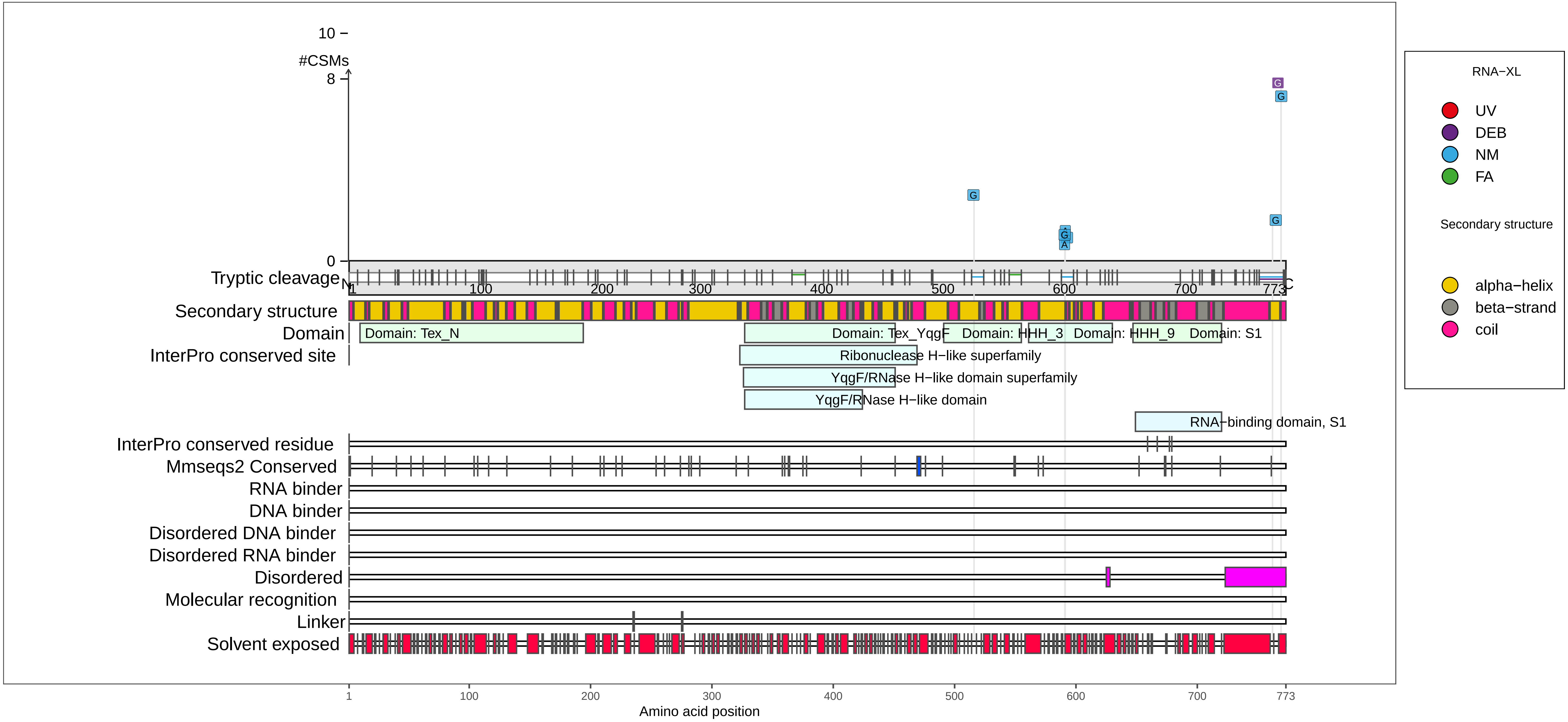

P0ABA6
ATPG_ECOLI ATP synthase gamma chain

− Abundance:
tryptic [log10 Intensity]: 8.53 (Q 77)
PAXdb K12 strain [ppm]: 2.36 (Q 72)
PAXdb E.coli [ppm]: 3.04 (Q 95)

− RNA functions: not annotated

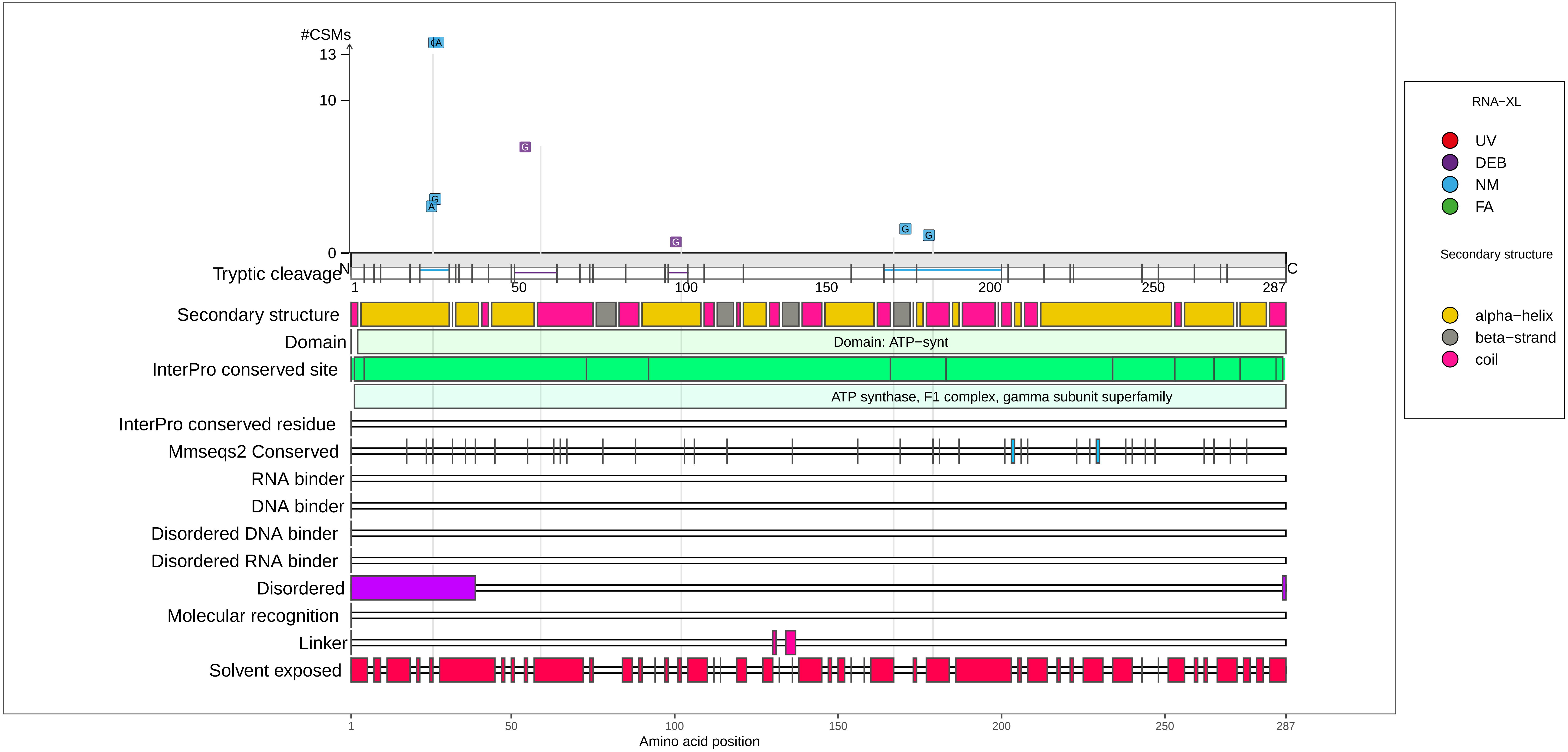

P17169
GLMS_ECOLI Glutamine−−fructose−6−phosphate aminotransferase [isomerizing]

− Abundance:
tryptic [log10 Intensity]: 9.69 (Q 98)
PAXdb K12 strain [ppm]: 3.12 (Q 93)
PAXdb E.coli [ppm]: 2.5 (Q 85)

− RNA functions: not annotated

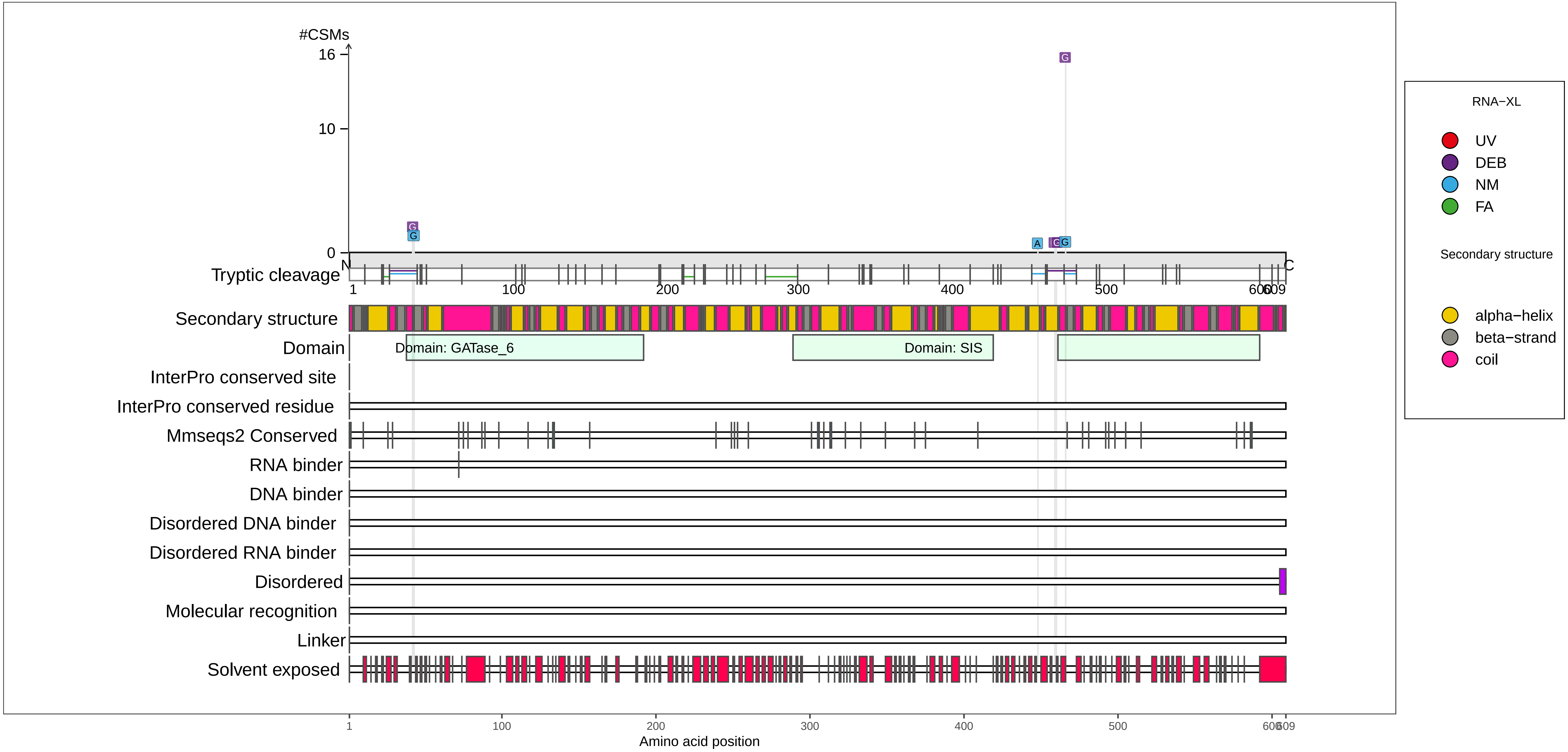

P30850
RNB_ECOLI Exoribonuclease 2

− Abundance:
tryptic [log10 Intensity]: 8.3 (Q 69)
PAXdb K12 strain [ppm]: 2.76 (Q 85)
PAXdb E.coli [ppm]: 2.16 (Q 76)

− RNA functions:
mRNA catabolic process; mRNA metabolic process
Ribosomal protein S1−like RNA−binding domain; RNA binding; RNA catabolic process; RNA metabolic process
RNA phosphodiester bond hydrolysis; RNA phosphodiester bond hydrolysis, exonucleolytic
S1 RNA binding domain

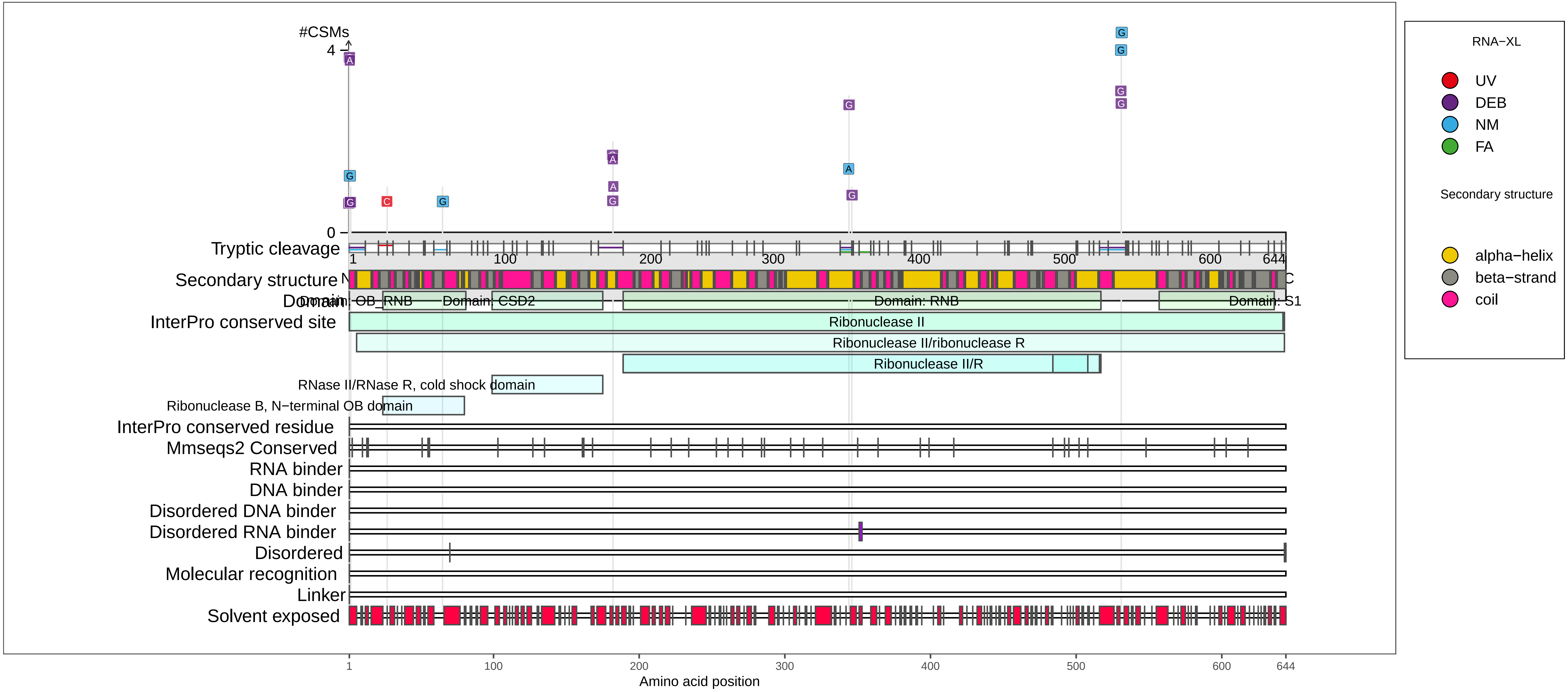

P0A715
KDSA_ECOLI 2−dehydro−3−deoxyphosphooctonate aldolase
− Abundance:
tryptic [log10 Intensity]: 8.73 (Q 82)
PAXdb K12 strain [ppm]: 3.45 (Q 97)
PAXdb E.coli [ppm]: 3.07 (Q 95)

− RNA functions: not annotated

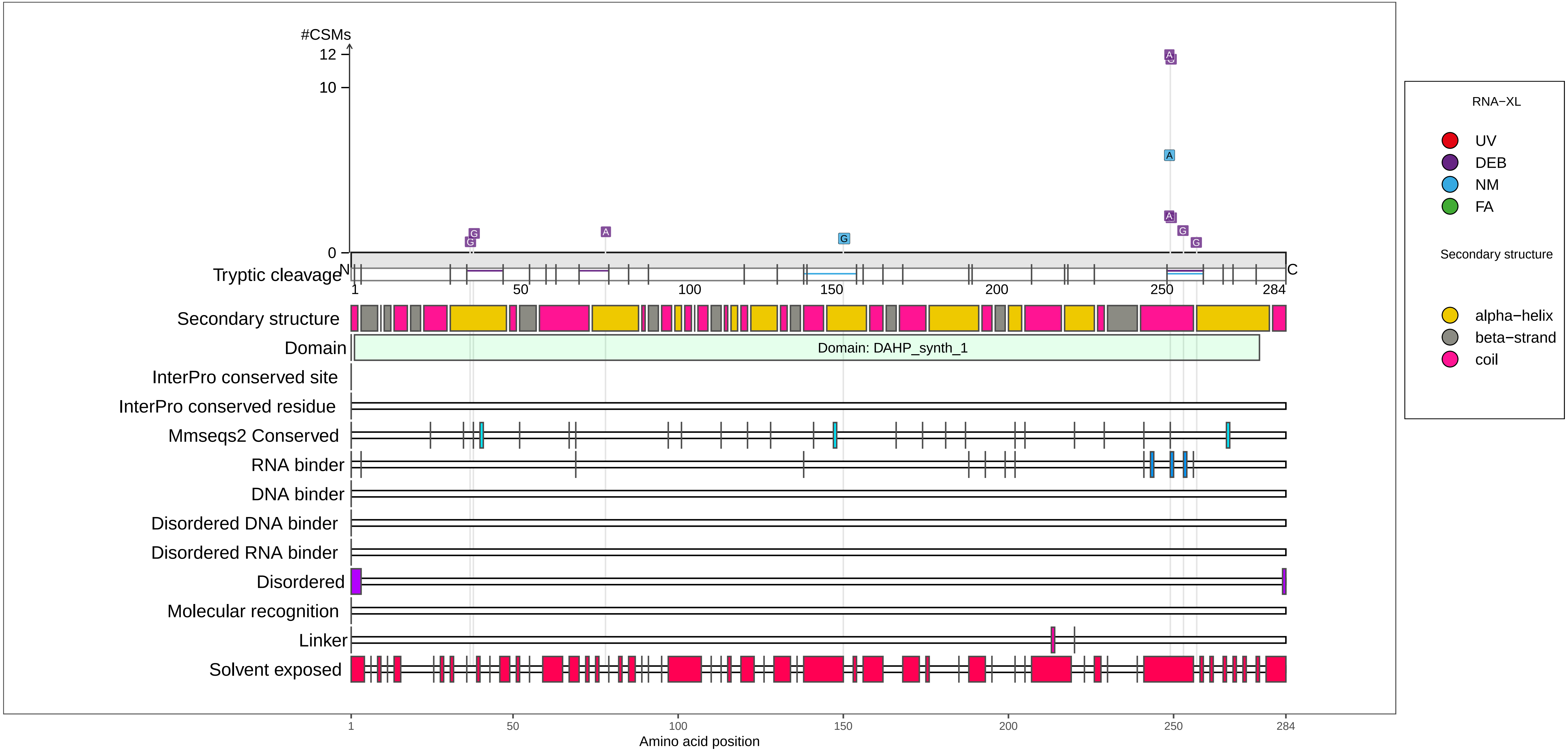

P0A7G6
RECA_ECOLI Protein RecA

− Abundance:
tryptic [log10 Intensity]: 7.58 (Q 39)
PAXdb K12 strain [ppm]: 2.86 (Q 88)
PAXdb E.coli [ppm]: 2.7 (Q 90)

− RNA functions: not annotated

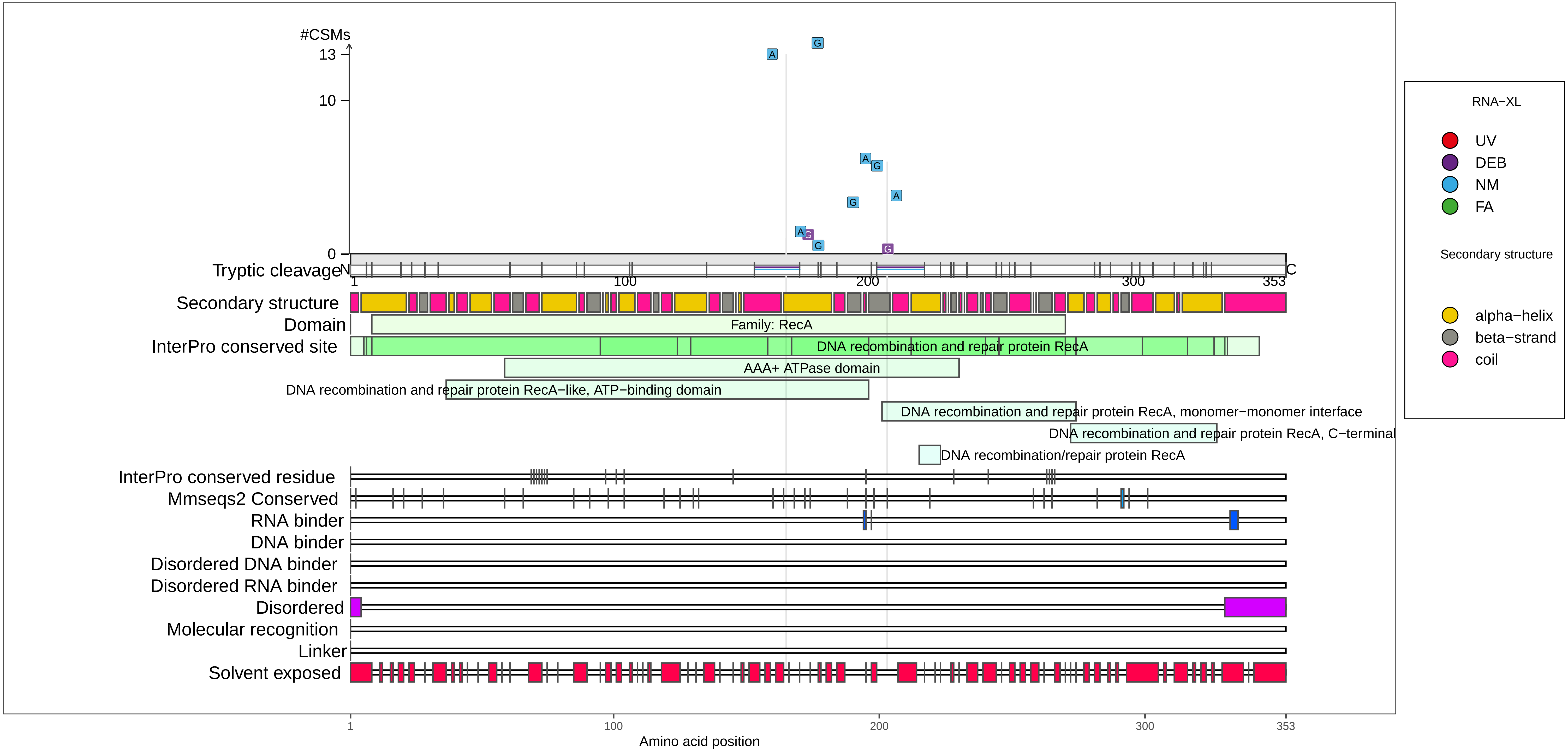

P0A7L3
RL20_ECOLI 50S ribosomal protein L20

− Abundance:
tryptic [log10 Intensity]: 9.04 (Q 89)
PAXdb K12 strain [ppm]: 3.02 (Q 91)
PAXdb E.coli [ppm]: 3.36 (Q 98)

− RNA functions:
mRNA binding; RNA binding; rRNA binding

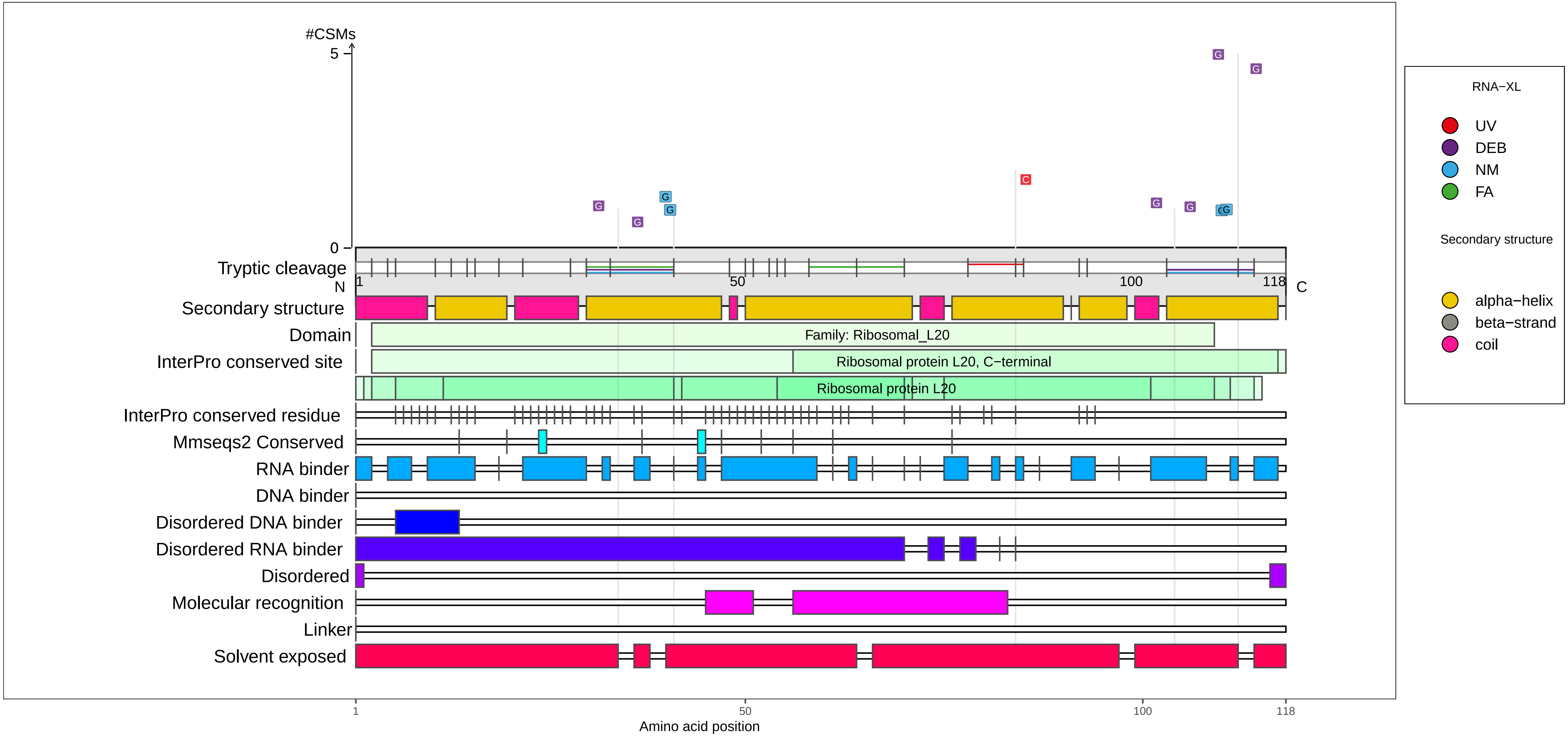

P0A862
TPX_ECOLI Thiol peroxidase

− Abundance:
tryptic [log10 Intensity]: 9.19 (Q 92)
PAXdb K12 strain [ppm]: 3.19 (Q 94)
PAXdb E.coli [ppm]: 3.39 (Q 98)

− RNA functions: not annotated

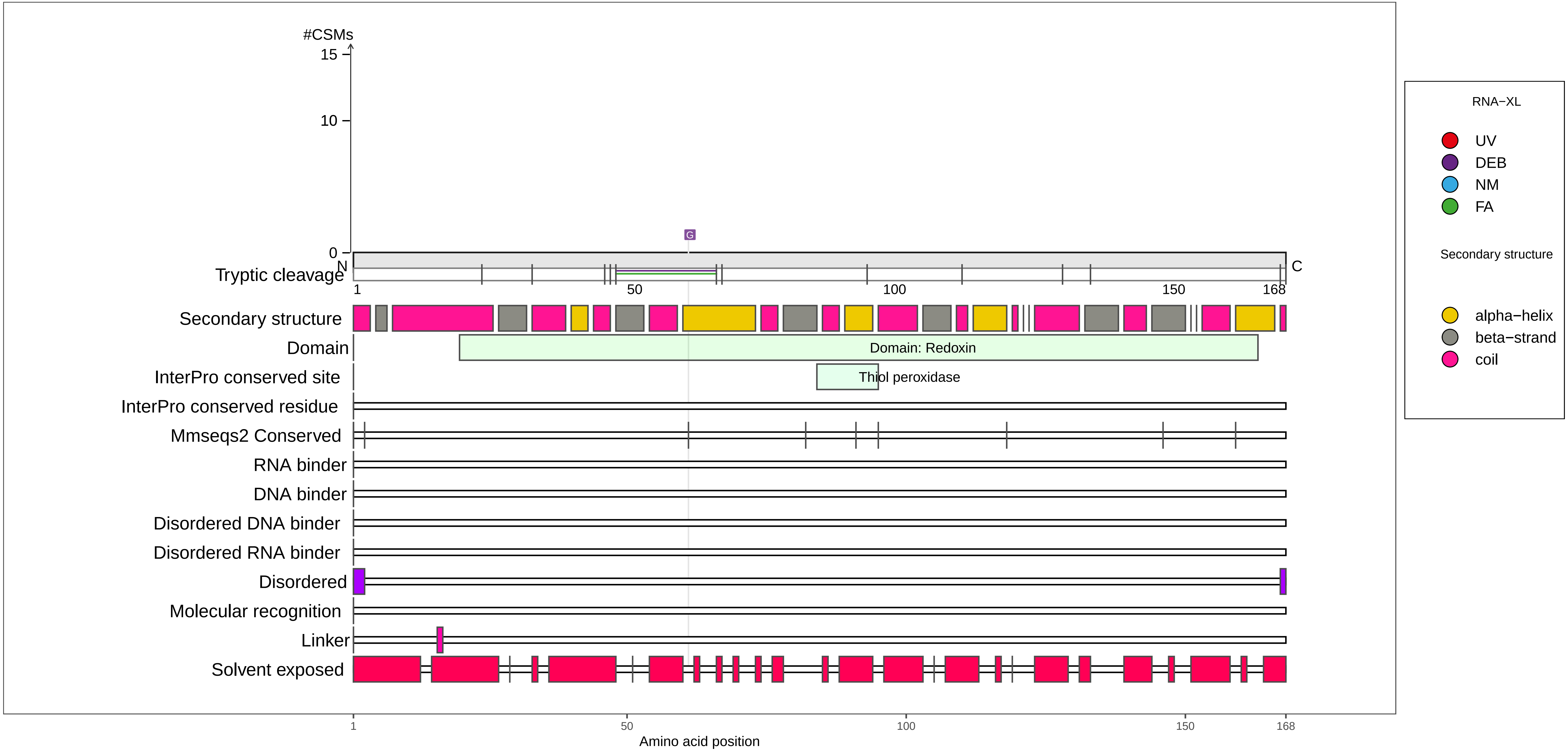

P0A8A0
YEBC_ECOLI Probable transcriptional regulatory protein YebC

− Abundance:
tryptic [log10 Intensity]: 7.94 (Q 56)
PAXdb K12 strain [ppm]: 2.95 (Q 89)
PAXdb E.coli [ppm]: 2.83 (Q 92)

− RNA functions: not annotated

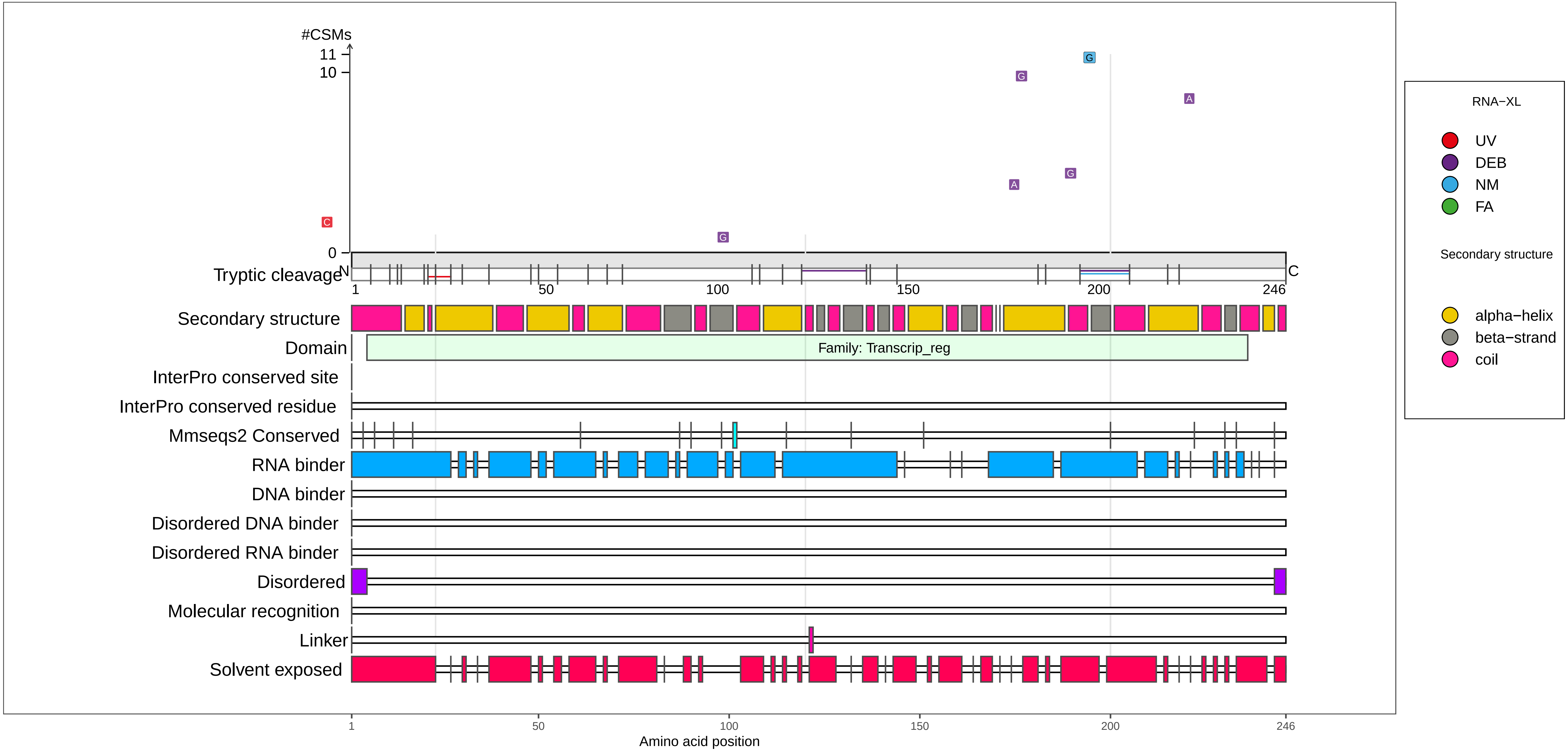

P0AGD3
SODF_ECOLI Superoxide dismutase [Fe]

− Abundance:
tryptic [log10 Intensity]: 9.9 (Q 99)
PAXdb K12 strain [ppm]: 2.98 (Q 90)
PAXdb E.coli [ppm]: 2.92 (Q 94)

− RNA functions: not annotated

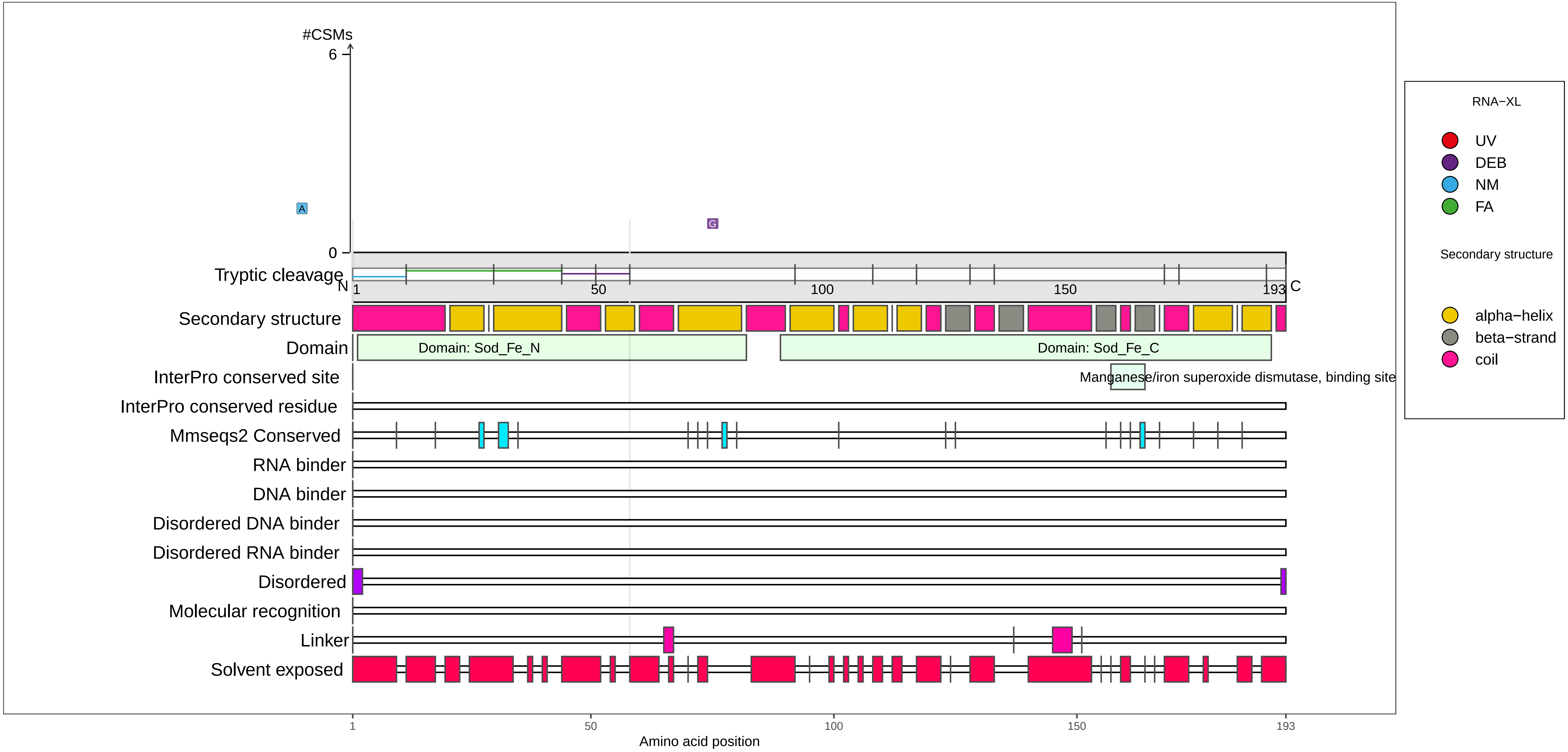

P16659
SYP_ECOLI Proline−−tRNA ligase

− Abundance:
tryptic [log10 Intensity]: 8.73 (Q 82)
PAXdb K12 strain [ppm]: 2.99 (Q 90)
PAXdb E.coli [ppm]: 2.73 (Q 90)

− RNA functions:
Ala−tRNA(Pro) hydrolase activity; aminoacyl−tRNA editing activity
Aminoacyl−tRNA editing domain; aminoacyl−tRNA ligase activity
aminoacyl−tRNA metabolism involved in translational fidelity; ncRNA metabolic process; proline−tRNA ligase activity
prolyl−tRNA aminoacylation; RNA metabolic process; tRNA aminoacylation
tRNA aminoacylation for protein translation; tRNA metabolic process
tRNA synthetase class II core domain (G, H, P, S and T)

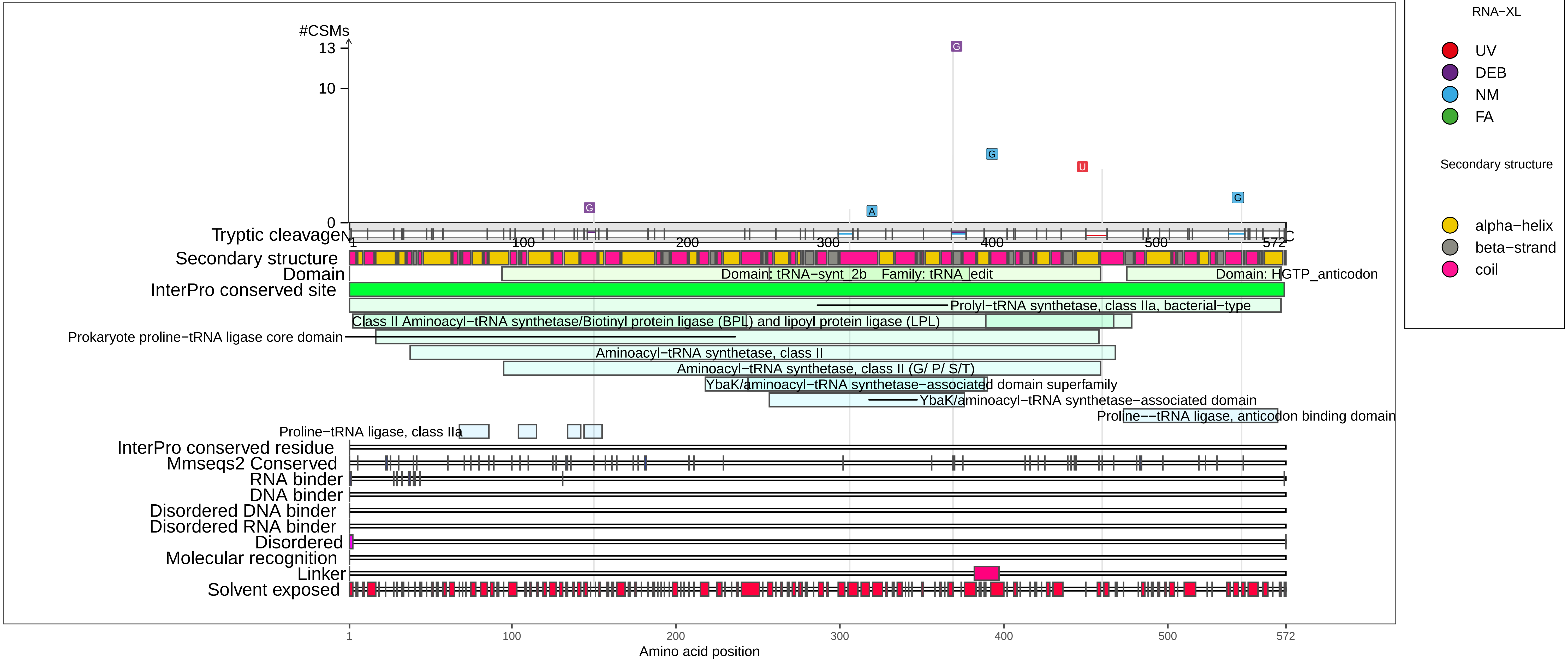

P21165
PEPQ_ECOLI Xaa−Pro dipeptidase

− Abundance:
tryptic [log10 Intensity]: 8.83 (Q 84)
PAXdb K12 strain [ppm]: 2.39 (Q 74)
PAXdb E.coli [ppm]: 2.66 (Q 89)

− RNA functions: not annotated

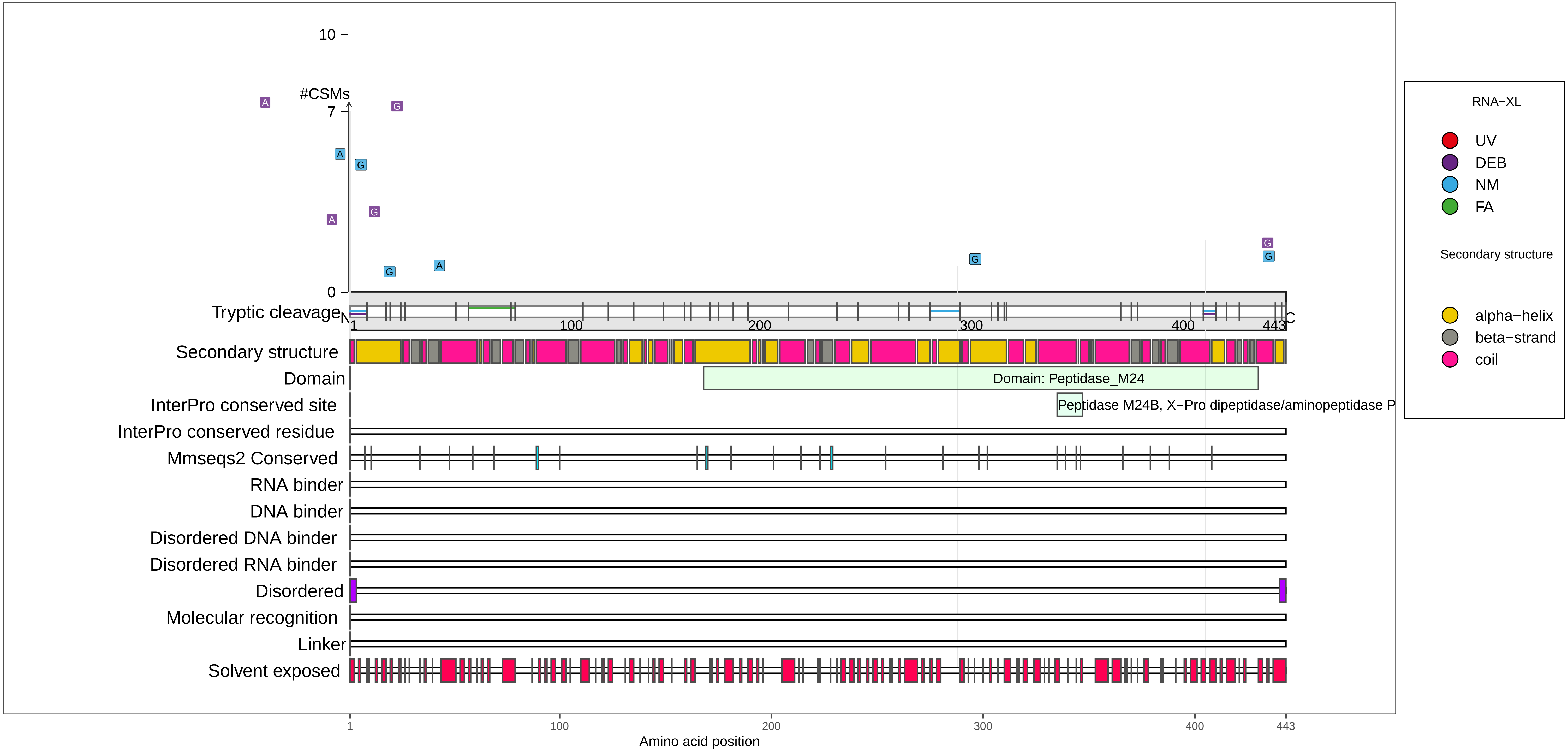

P45464
LPOA_ECOLI Penicillin−binding protein activator LpoA

− Abundance:
tryptic [log10 Intensity]: 7.76 (Q 47)
PAXdb K12 strain [ppm]: 1.24 (Q 23)
PAXdb E.coli [ppm]: 1.84 (Q 68)

− RNA functions: not annotated

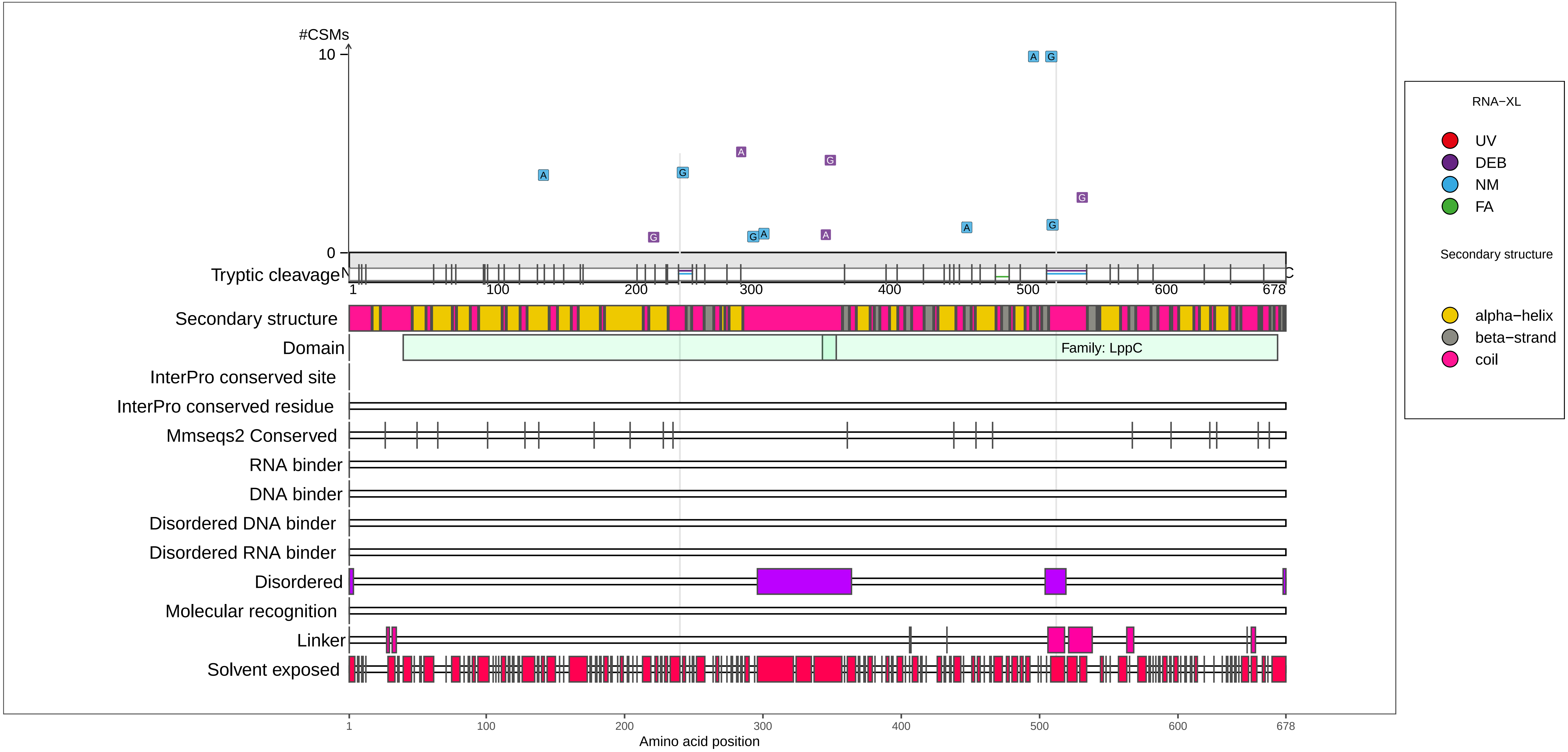

P63386
MLAF_ECOLI Intermembrane phospholipid transport system ATP−binding protein MlaF

− Abundance:
tryptic [log10 Intensity]: 7.29 (Q 25)
PAXdb K12 strain [ppm]: 1.72 (Q 50)
PAXdb E.coli [ppm]: 1.46 (Q 59)

− RNA functions: not annotated

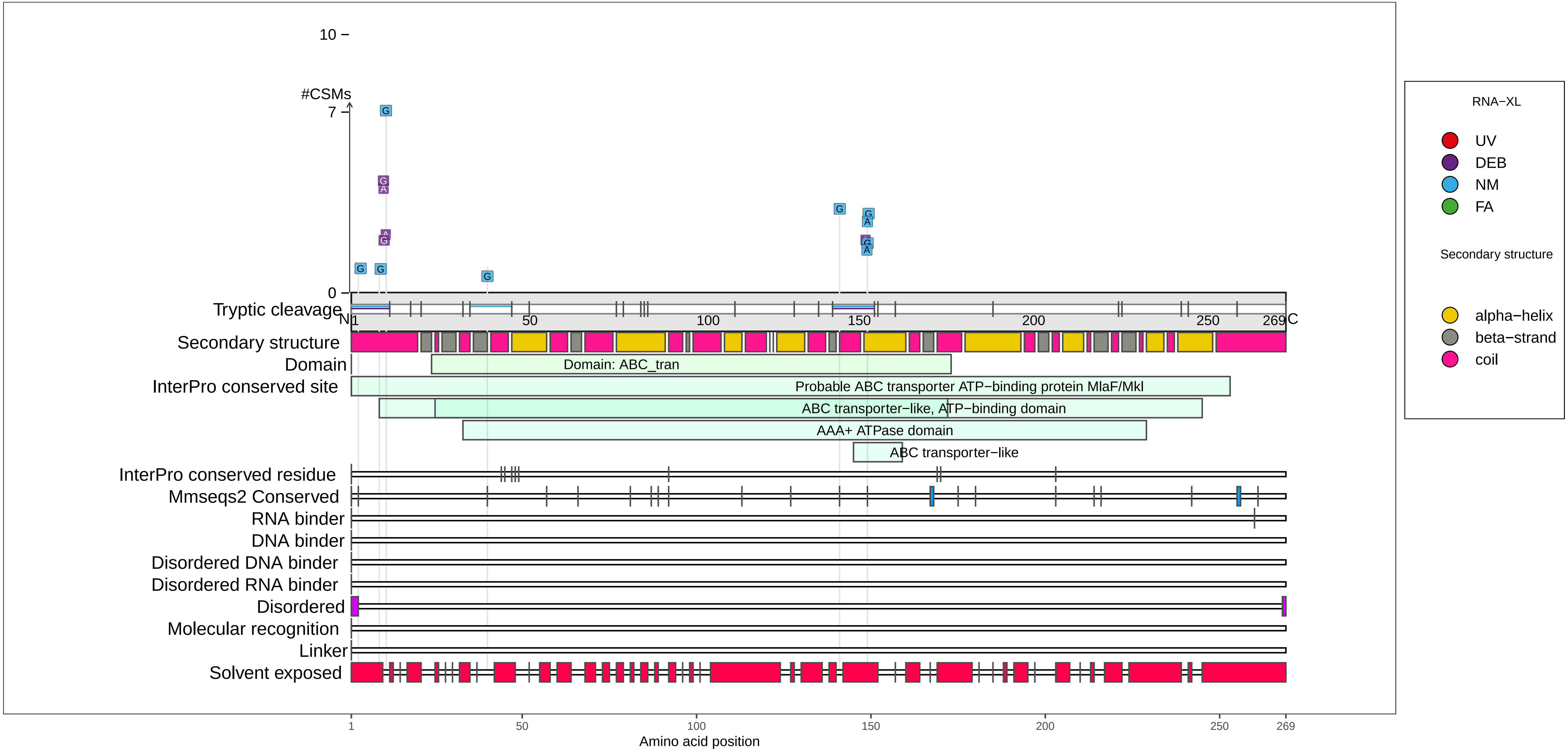

P0A6D3
AROA_ECOLI 3−phosphoshikimate 1−carboxyvinyltransferase

− Abundance:
tryptic [log10 Intensity]: 8.52 (Q 76)
PAXdb K12 strain [ppm]: 2.04 (Q 62)
PAXdb E.coli [ppm]: 2.23 (Q 78)

− RNA functions: not annotated

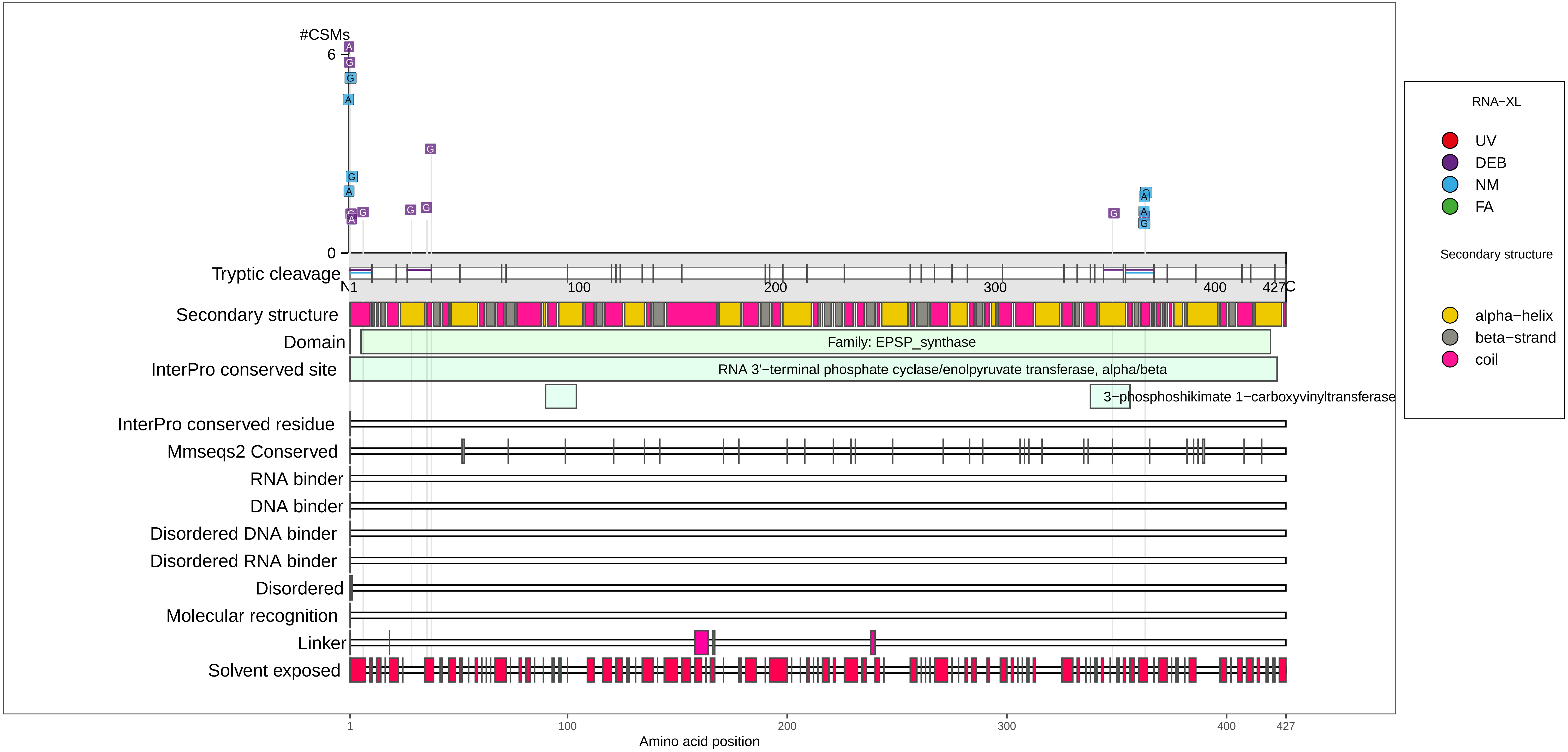

P0A6W5
GREA_ECOLI Transcription elongation factor GreA
− Abundance:
tryptic [log10 Intensity]: 8.46 (Q 74)
PAXdb K12 strain [ppm]: 3.31 (Q 95)
PAXdb E.coli [ppm]: 2.85 (Q 92)

− RNA functions:
RNA biosynthetic process; RNA metabolic process; RNA polymerase binding

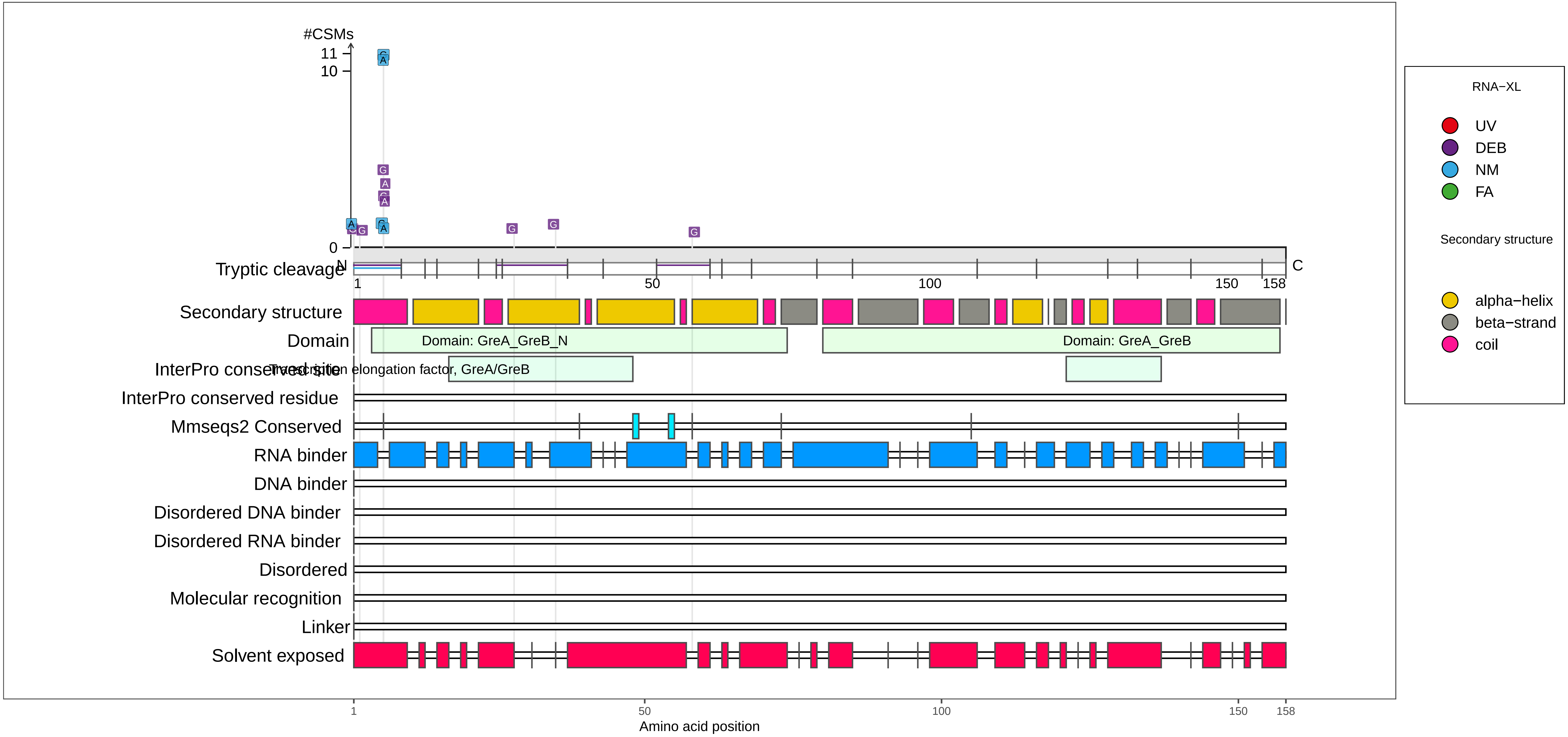

P0A847
TGT_ECOLI Queuine tRNA−ribosyltransferase

− Abundance:
tryptic [log10 Intensity]: 8.01 (Q 58)
PAXdb K12 strain [ppm]: 2.08 (Q 63)
PAXdb E.coli [ppm]: 1.46 (Q 59)

− RNA functions:
ncRNA metabolic process; ncRNA processing; Queuine tRNA−ribosyltransferase
RNA metabolic process; RNA modification; RNA processing; tRNA metabolic process; tRNA modification
tRNA processing; tRNA wobble base modification; tRNA wobble guanine modification
tRNA−guanine transglycosylation

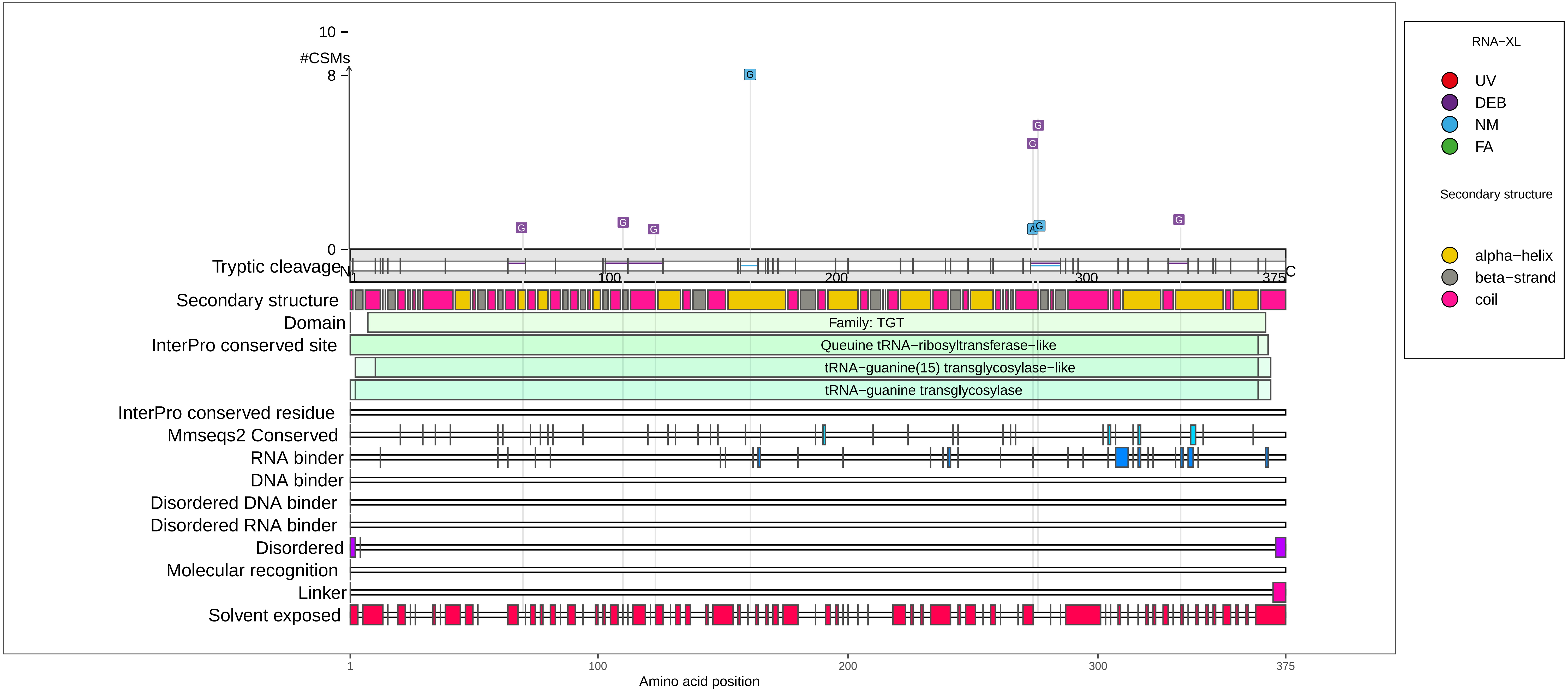

P0AA39
RLUC_ECOLI Ribosomal large subunit pseudouridine synthase C

− Abundance:
tryptic [log10 Intensity]: 7.43 (Q 32)
PAXdb K12 strain [ppm]: 1.31 (Q 27)
PAXdb E.coli [ppm]: 1.75 (Q 66)

− RNA functions:
ncRNA metabolic process; ncRNA processing; RNA binding; RNA metabolic process
RNA modification; RNA processing; RNA pseudouridylate synthase; rRNA metabolic process
rRNA modification; rRNA processing; rRNA pseudouridine synthase activity
rRNA pseudouridine synthesis

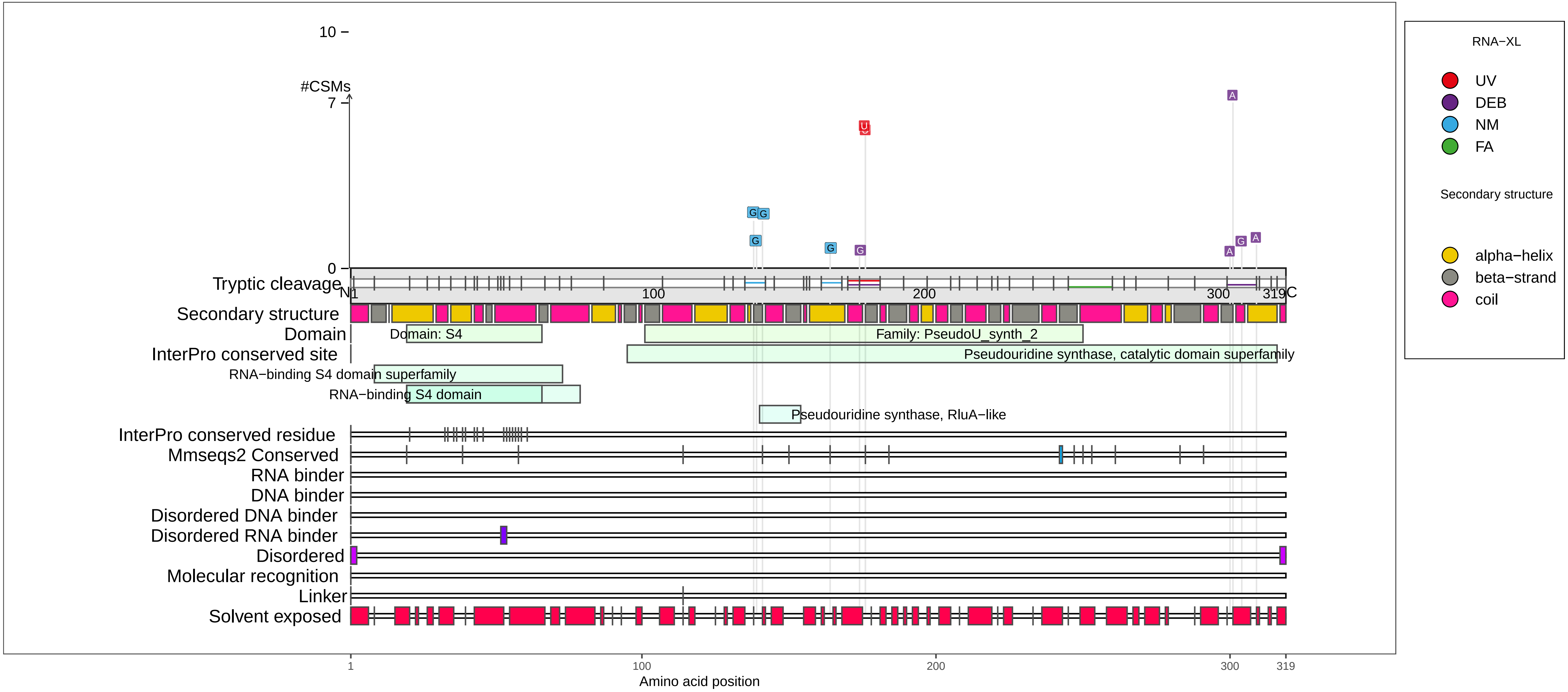

P23882
FMT_ECOLI Methionyl−tRNA formyltransferase

− Abundance:
tryptic [log10 Intensity]: 8.21 (Q 66)
PAXdb K12 strain [ppm]: 2.28 (Q 69)
PAXdb E.coli [ppm]: 2.34 (Q 82)

− RNA functions:
charged−tRNA amino acid modification; methionyl−tRNA formyltransferase activity
ncRNA metabolic process; ncRNA processing; RNA metabolic process; RNA modification
RNA processing; tRNA metabolic process; tRNA modification; tRNA processing

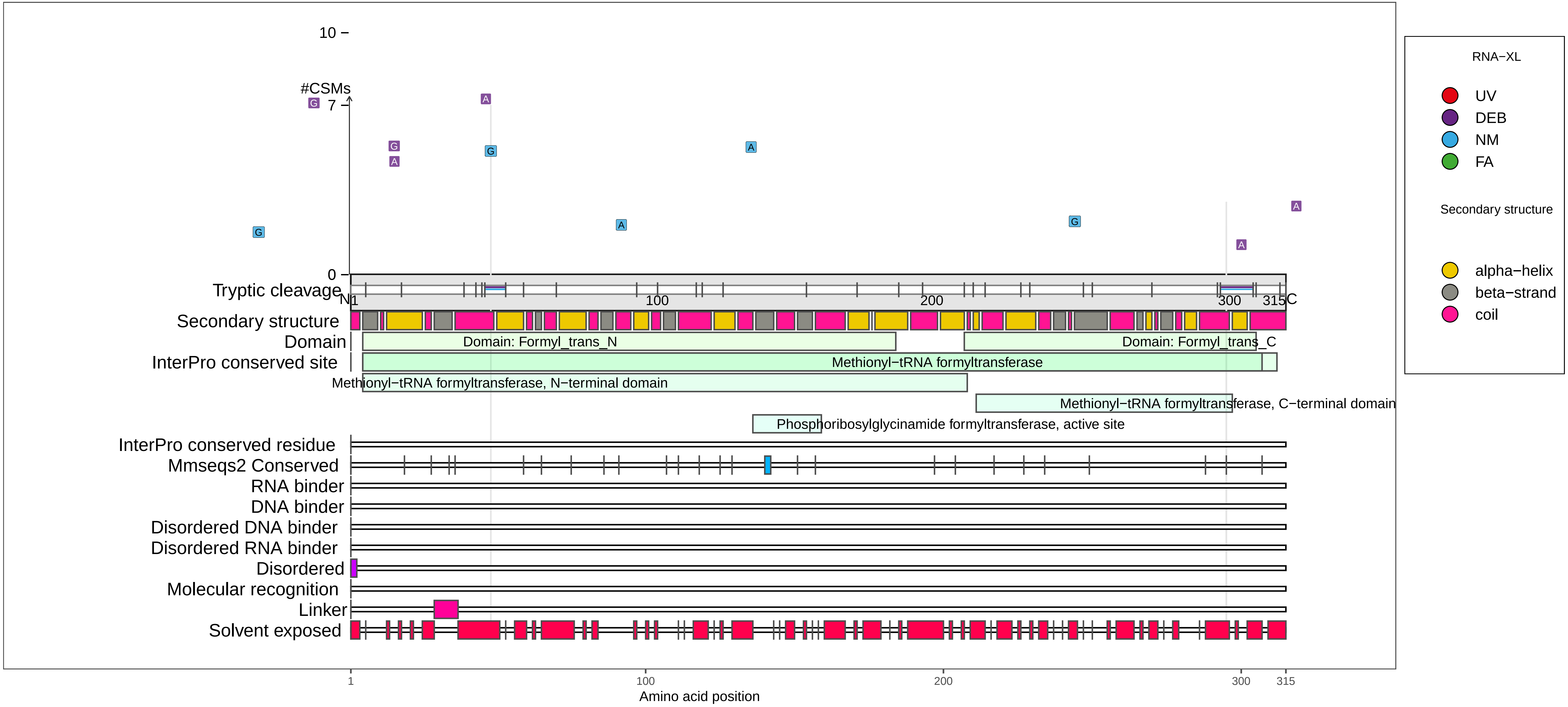

P33232
LLDD_ECOLI L−lactate dehydrogenase

− Abundance:
tryptic [log10 Intensity]: 8.49 (Q 75)
PAXdb K12 strain [ppm]: 1.21 (Q 21)
PAXdb E.coli [ppm]: 1.91 (Q 70)

− RNA functions: not annotated

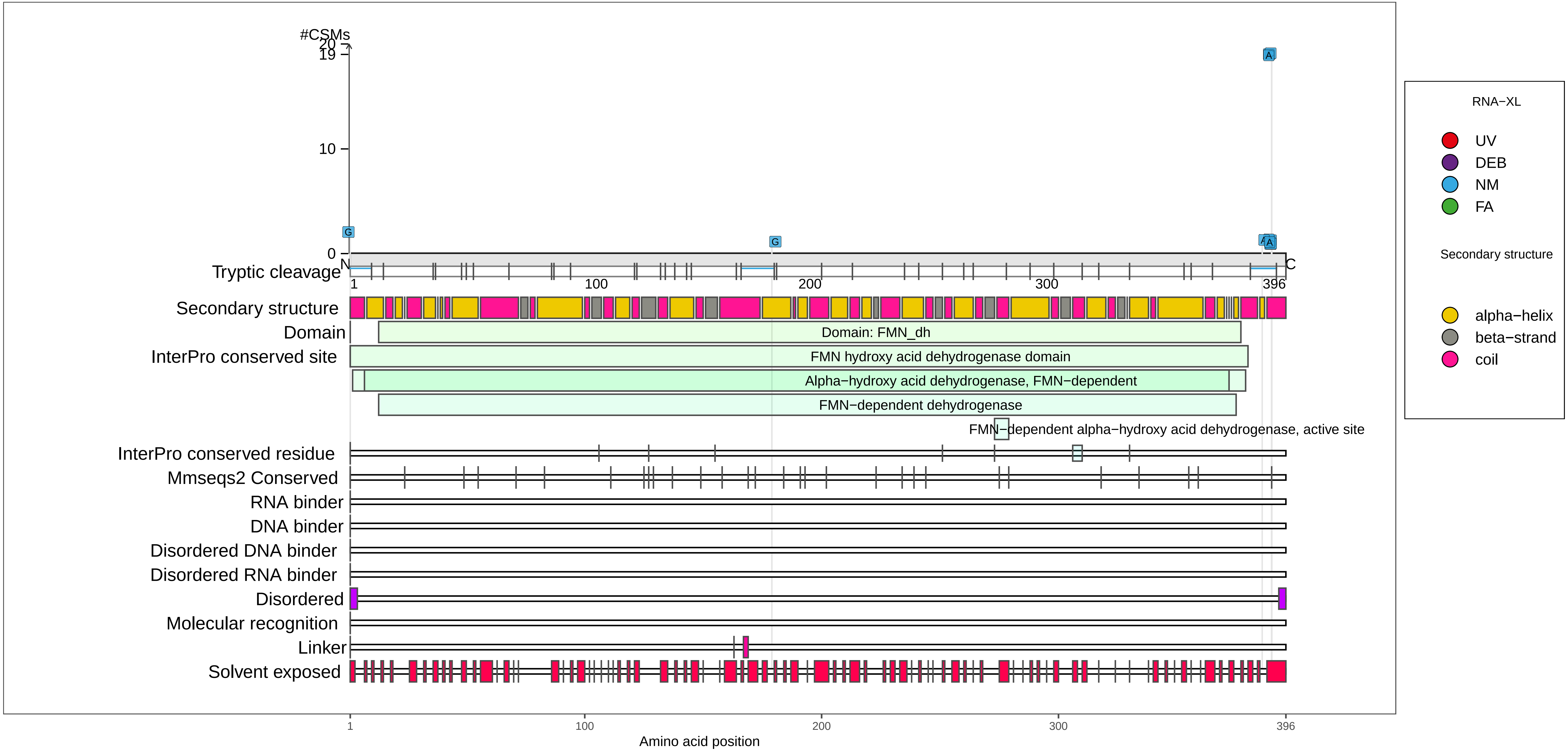

P60560
GUAC_ECOLI GMP reductase

− Abundance:
tryptic [log10 Intensity]: 8.34 (Q 71)
PAXdb K12 strain [ppm]: 1.19 (Q 19)
PAXdb E.coli [ppm]: 2.52 (Q 86)

− RNA functions: not annotated

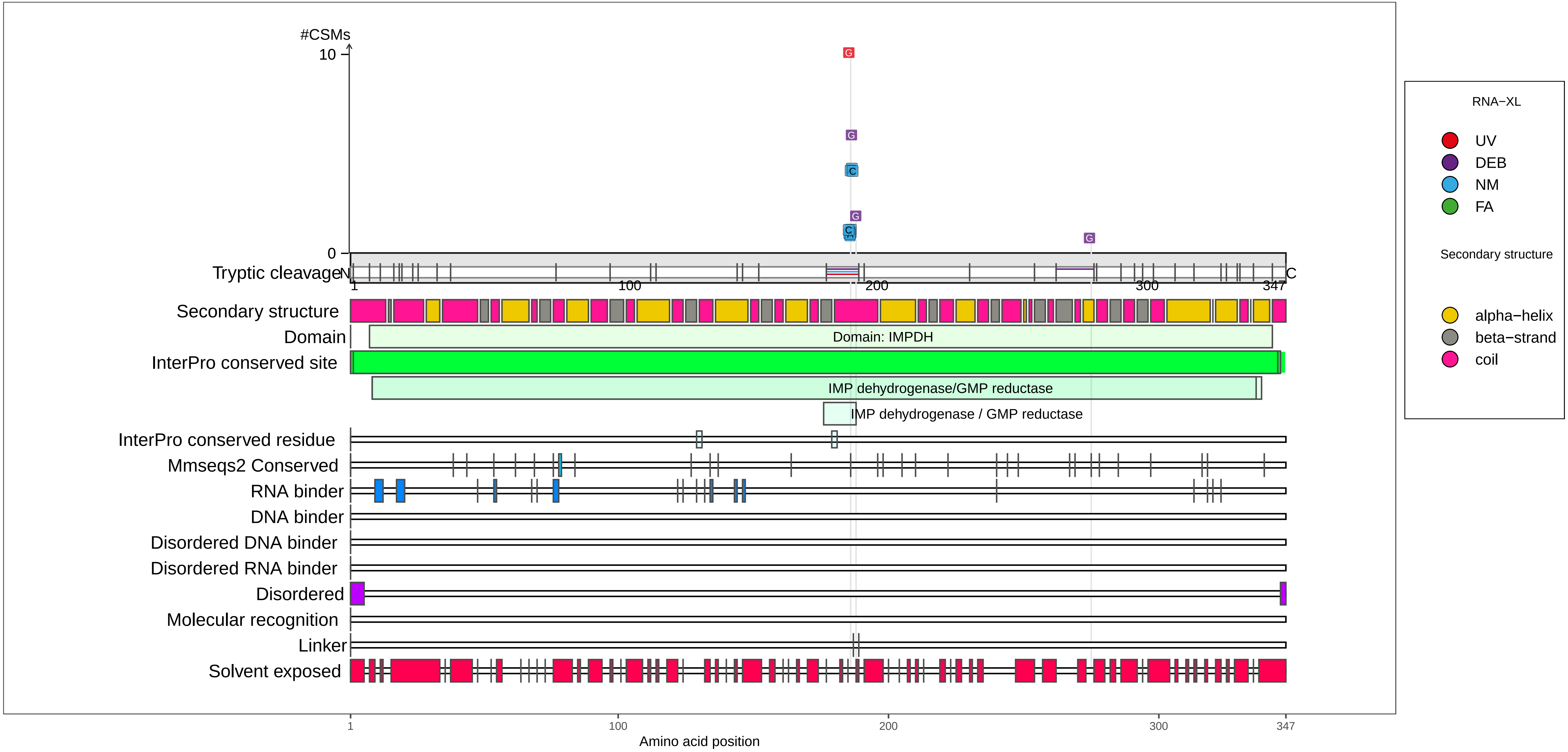

P67087
RSMI_ECOLI Ribosomal RNA small subunit methyltransferase I

− Abundance:
tryptic [log10 Intensity]: 6.89 (Q 9)
PAXdb K12 strain [ppm]: 2.78 (Q 85)
PAXdb E.coli [ppm]: 2.02 (Q 73)

− RNA functions:
ncRNA metabolic process; ncRNA processing; RNA 2−O−methyltransferase activity
RNA metabolic process; RNA methylation; RNA methyltransferase activity; RNA modification
RNA processing; rRNA (cytosine−2−O−)−methyltransferase activity
rRNA (cytosine) methyltransferase activity; rRNA 2−O−methylation; rRNA metabolic process; rRNA methylation
rRNA methyltransferase activity; rRNA modification; rRNA processing

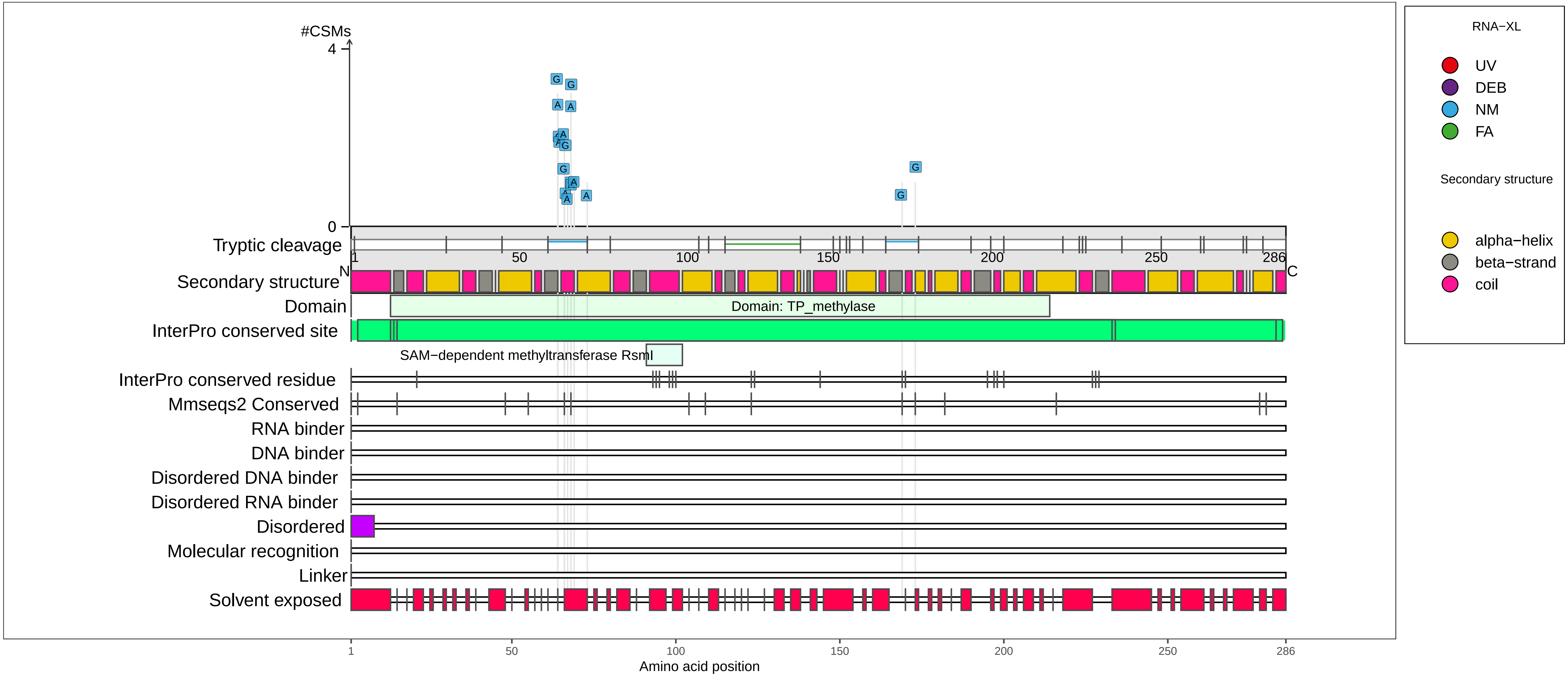

P05042
FUMC_ECOLI Fumarate hydratase class II

− Abundance:
tryptic [log10 Intensity]: 7.91 (Q 54)
PAXdb K12 strain [ppm]: 2.5 (Q 78)
PAXdb E.coli [ppm]: 2.81 (Q 92)

− RNA functions: not annotated

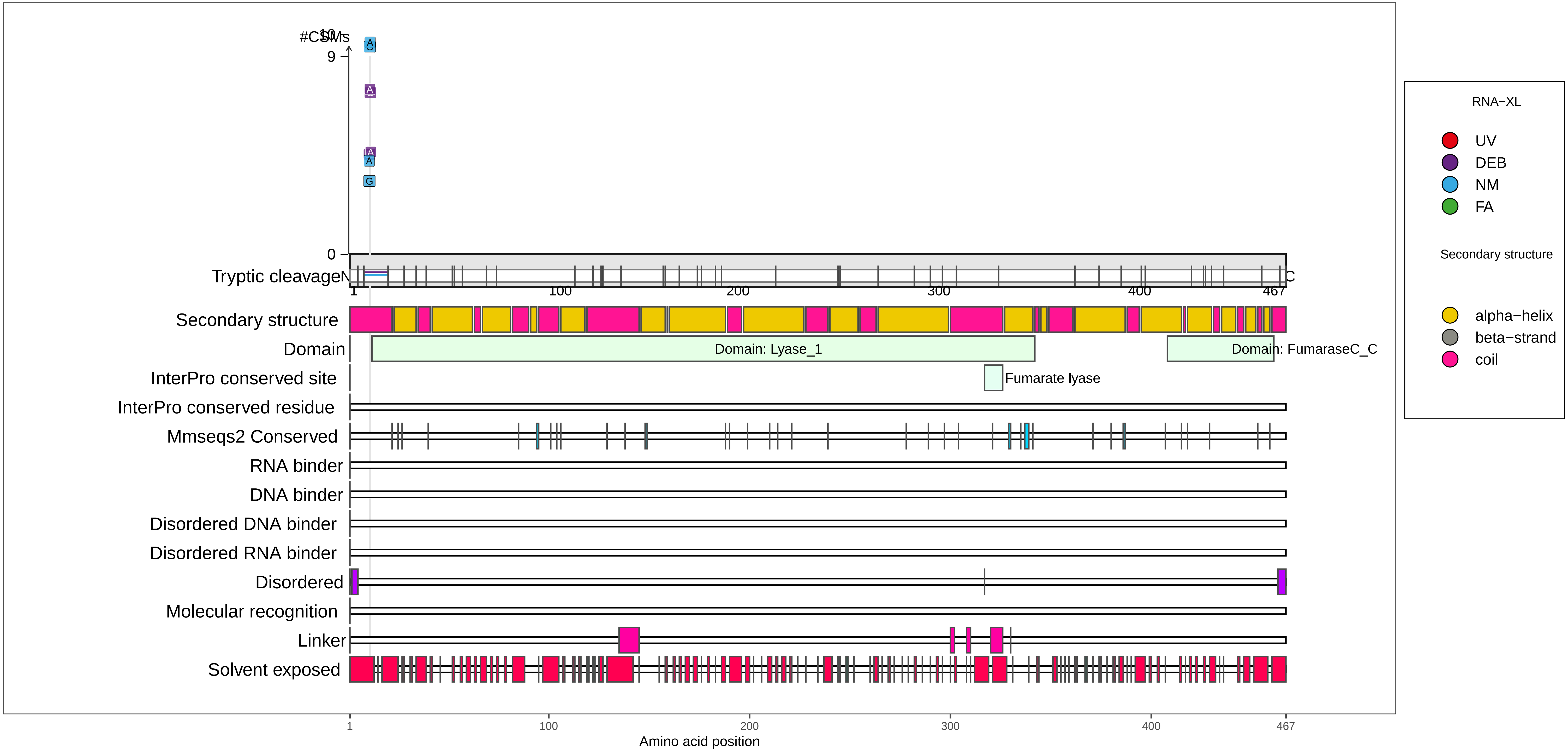

P0A6L2
DAPA_ECOLI 4−hydroxy−tetrahydrodipicolinate synthase

− Abundance:
tryptic [log10 Intensity]: 8.97 (Q 87)
PAXdb K12 strain [ppm]: 1.79 (Q 53)
PAXdb E.coli [ppm]: 2.42 (Q 83)

− RNA functions: not annotated

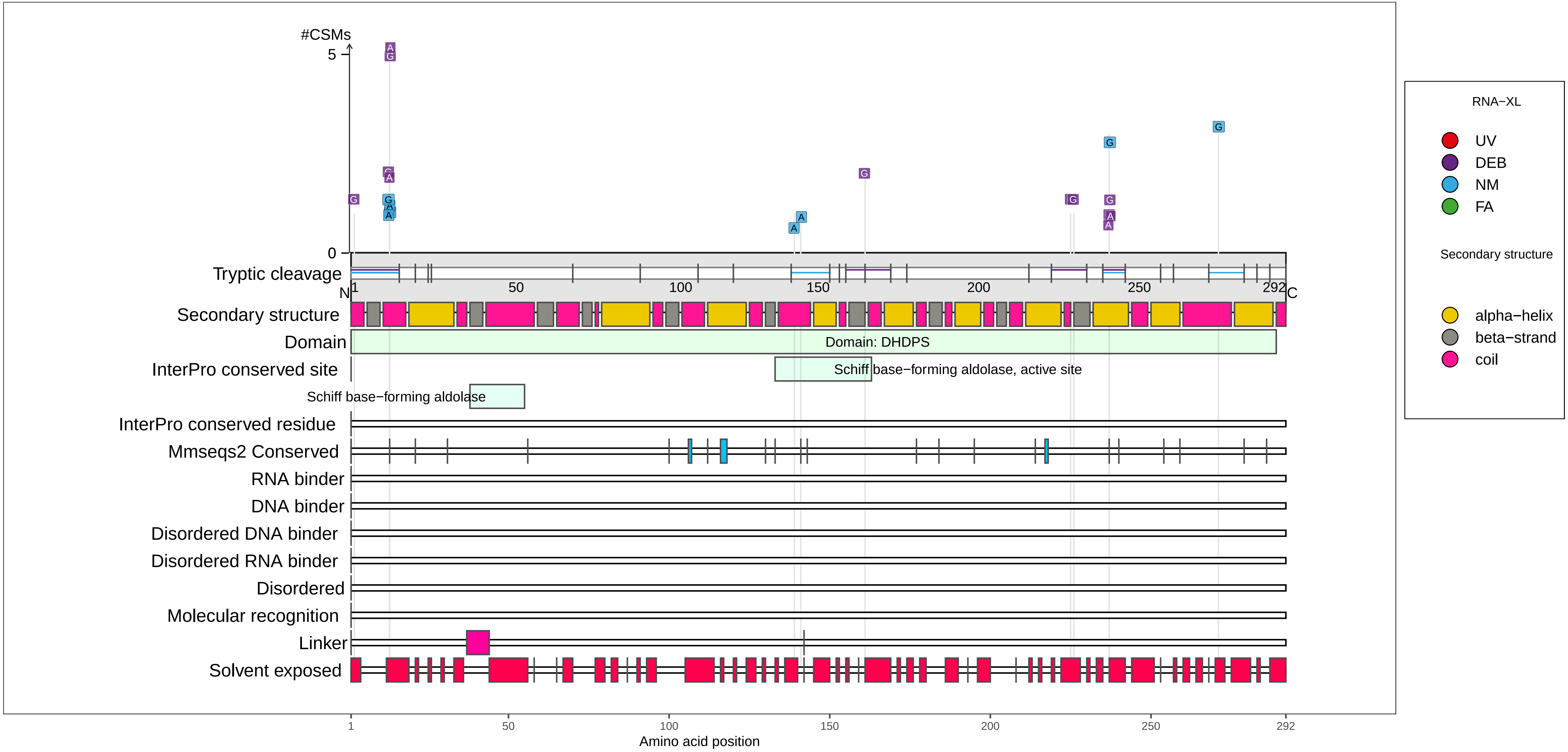

P0A734
MINE_ECOLI Cell division topological specificity factor

− Abundance:
tryptic [log10 Intensity]: 8.09 (Q 61)
PAXdb K12 strain [ppm]: 2.36 (Q 72)
PAXdb E.coli [ppm]: 2.96 (Q 94)

− RNA functions: not annotated

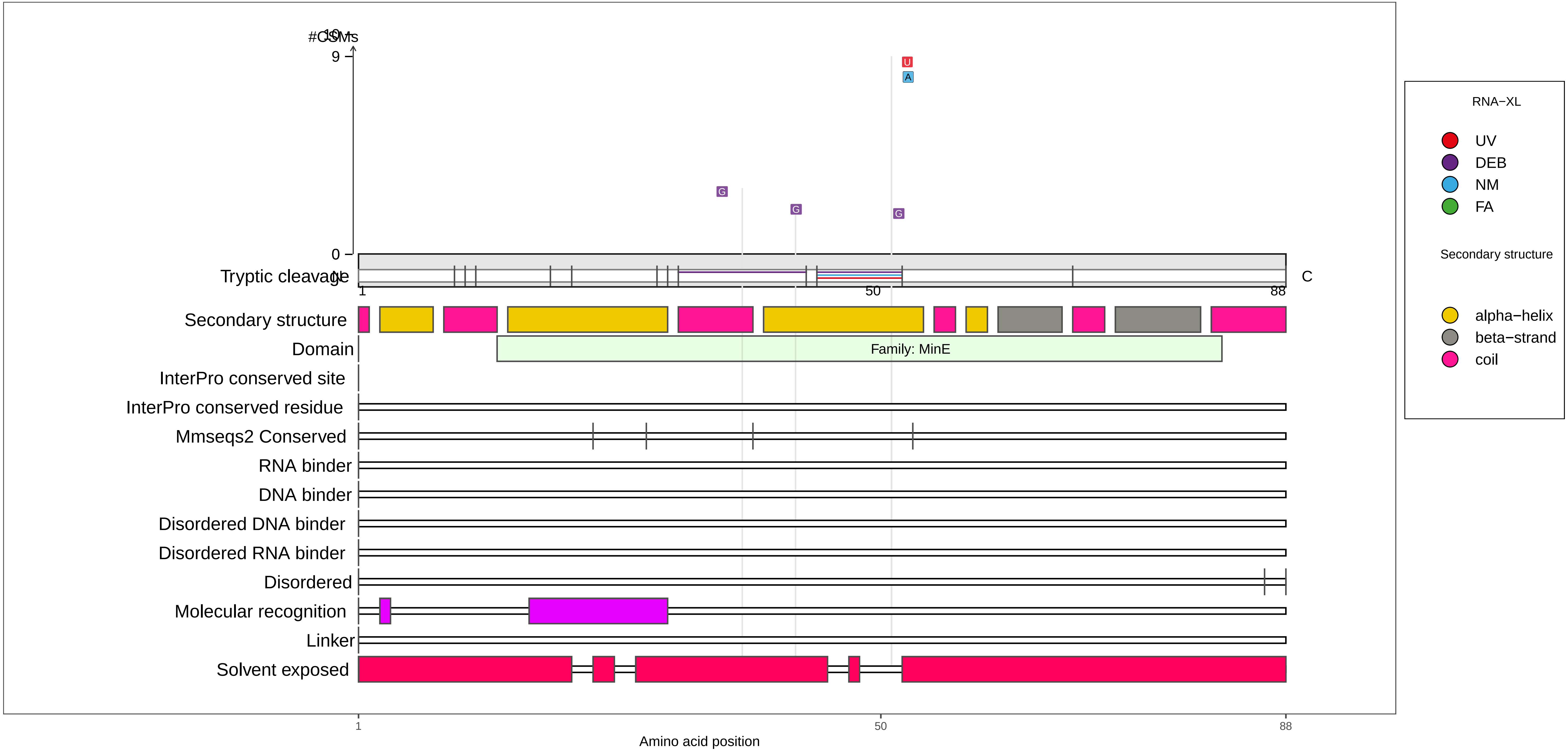

P0A9M5
XGPT_ECOLI Xanthine−guanine phosphoribosyltransferase

− Abundance:
tryptic [log10 Intensity]: 8.89 (Q 86)
PAXdb K12 strain [ppm]: 1.99 (Q 60)
PAXdb E.coli [ppm]: 2.11 (Q 75)

− RNA functions: not annotated

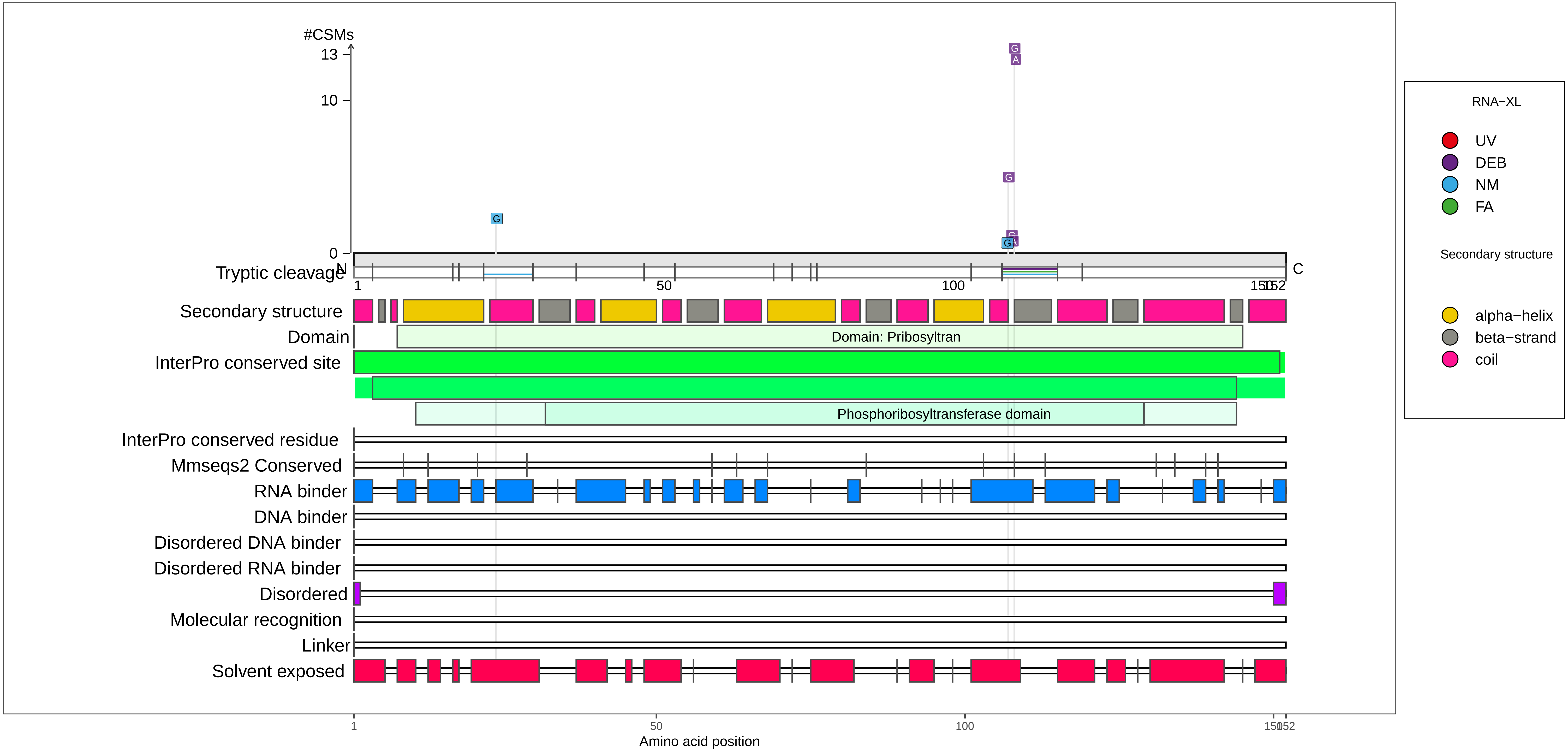

P0A9S3
GATD_ECOLI Galactitol 1−phosphate 5−dehydrogenase

− Abundance:
tryptic [log10 Intensity]: 8.9 (Q 86)
PAXdb K12 strain [ppm]: 1.92 (Q 58)
PAXdb E.coli [ppm]: 2.81 (Q 92)

− RNA functions: not annotated

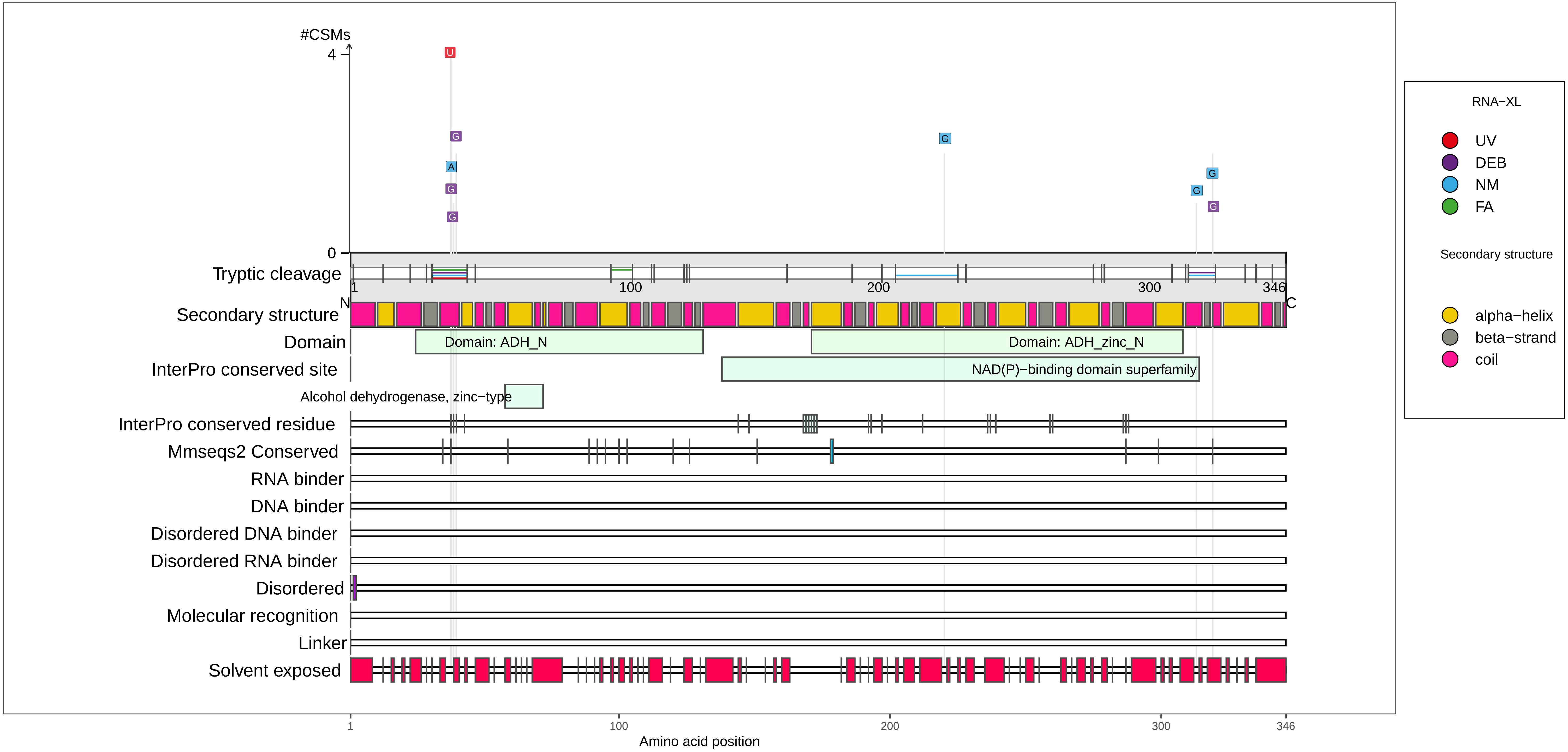

P0AB55
YCII_ECOLI Protein YciI
− Abundance:
tryptic [log10 Intensity]: 7.75 (Q 47)
PAXdb K12 strain [ppm]: 3.91 (Q 100)
PAXdb E.coli [ppm]: 2.97 (Q 94)

− RNA functions: not annotated

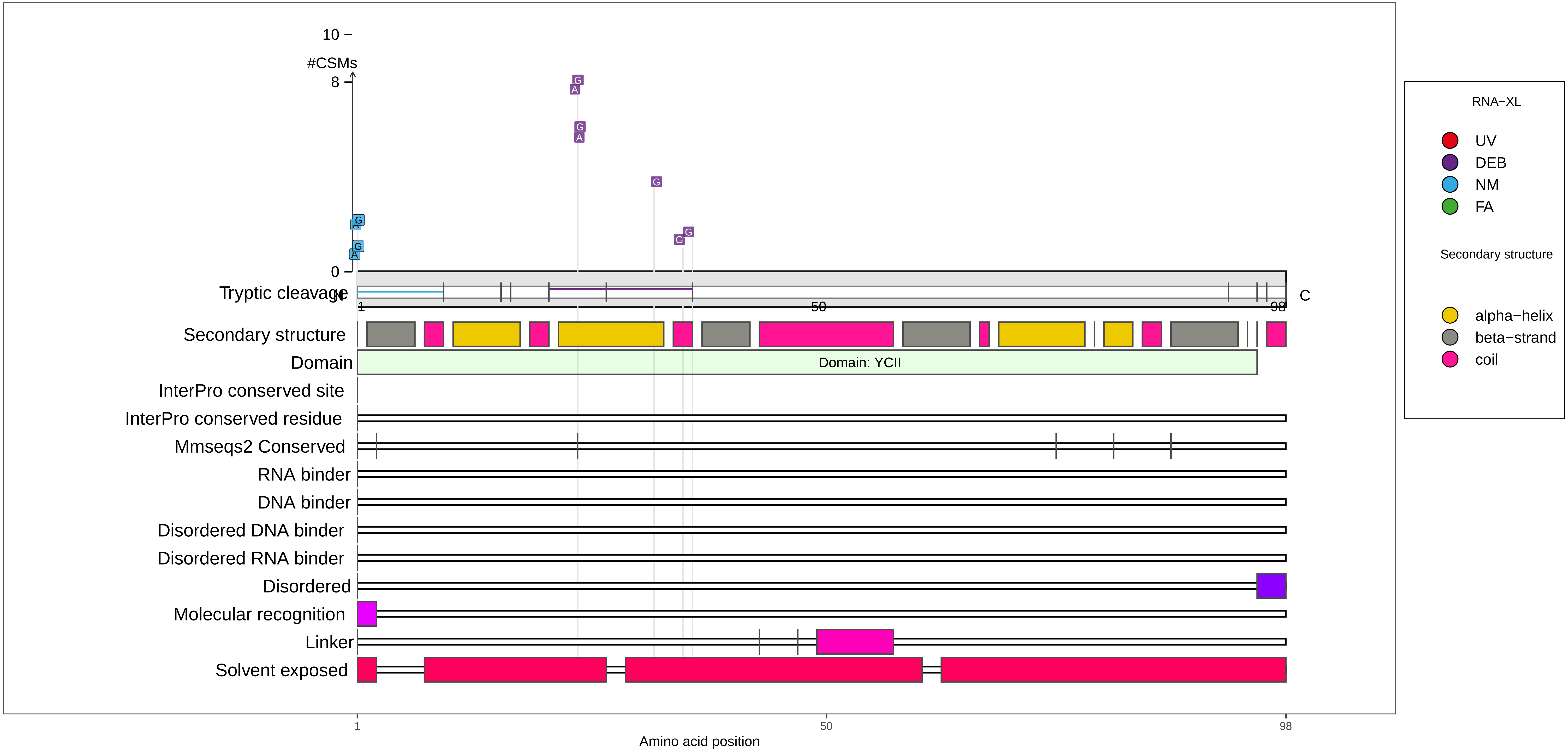

P0ADZ7
YAJC_ECOLI Sec translocon accessory complex subunit YajC

− Abundance:
tryptic [log10 Intensity]: 9.23 (Q 93)
PAXdb K12 strain [ppm]: 2.32 (Q 71)
PAXdb E.coli [ppm]: 3.58 (Q 99)

− RNA functions: not annotated

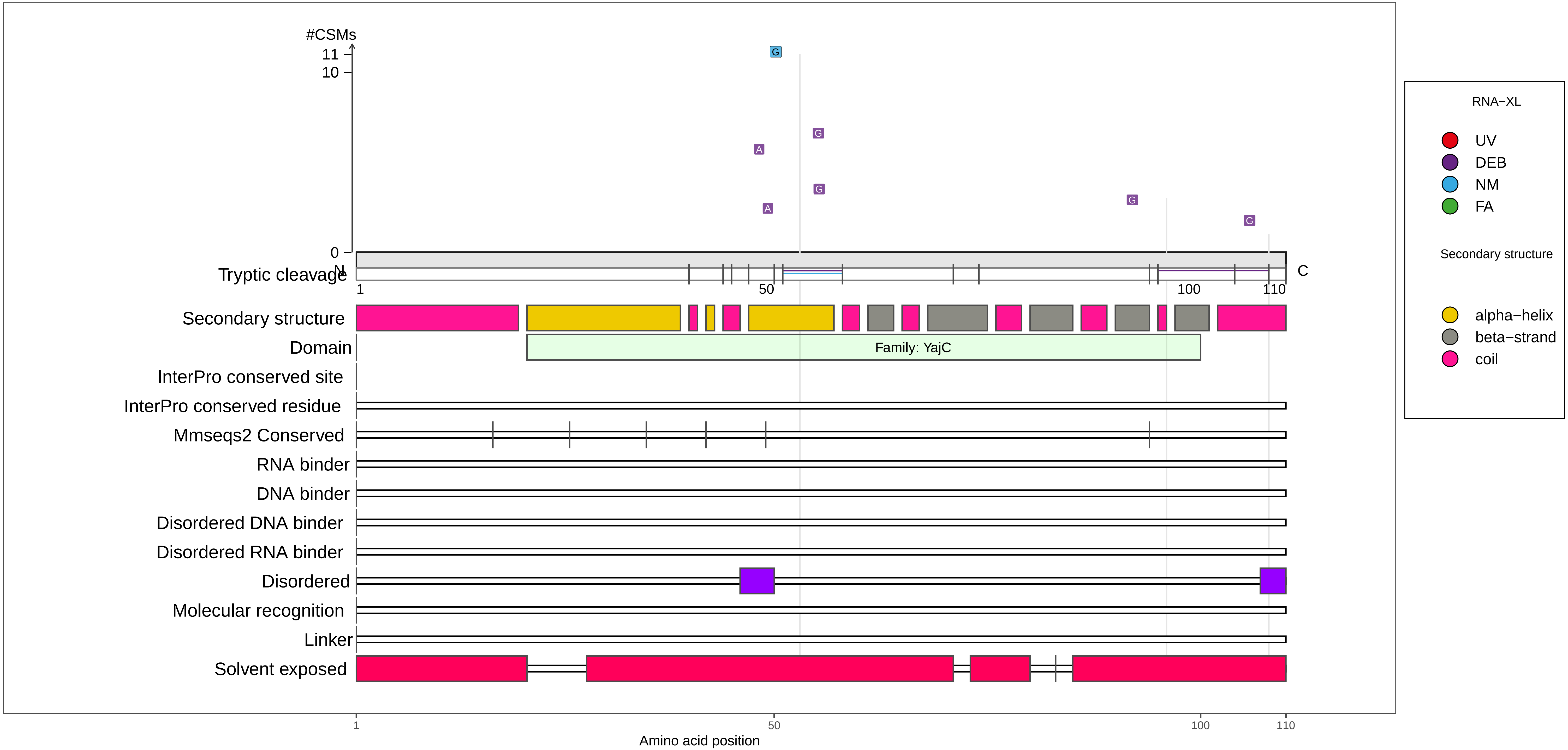

P21170
SPEA_ECOLI Biosynthetic arginine decarboxylase

− Abundance:
tryptic [log10 Intensity]: 8.9 (Q 86)
PAXdb K12 strain [ppm]: 2.87 (Q 88)
PAXdb E.coli [ppm]: 2.37 (Q 82)

− RNA functions: not annotated

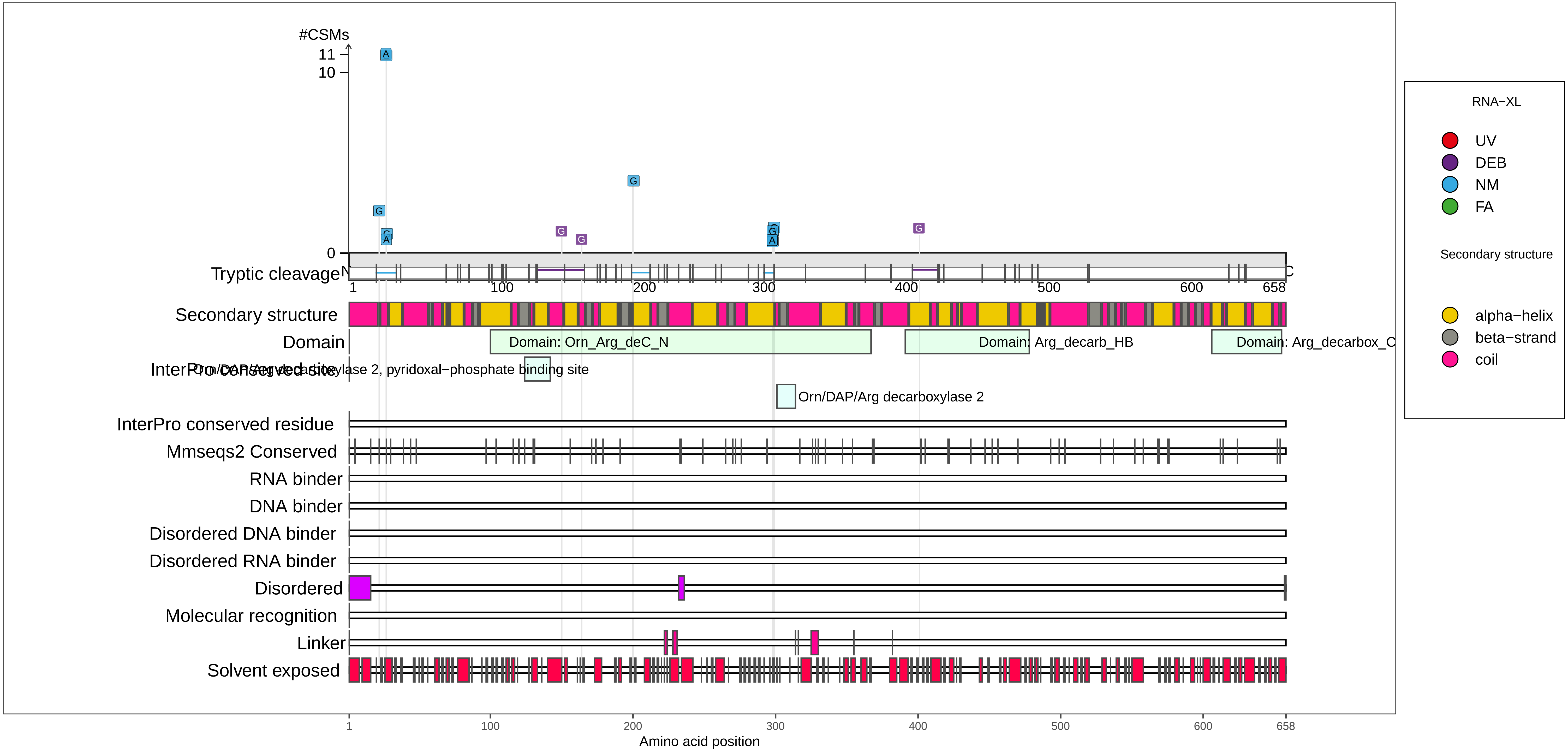

P25516
ACNA_ECOLI Aconitate hydratase A
− Abundance:
tryptic [log10 Intensity]: 8.48 (Q 75)
PAXdb K12 strain [ppm]: 2.9 (Q 88)
PAXdb E.coli [ppm]: 2.61 (Q 88)

− RNA functions:
mRNA 3−UTR binding; mRNA binding; RNA binding

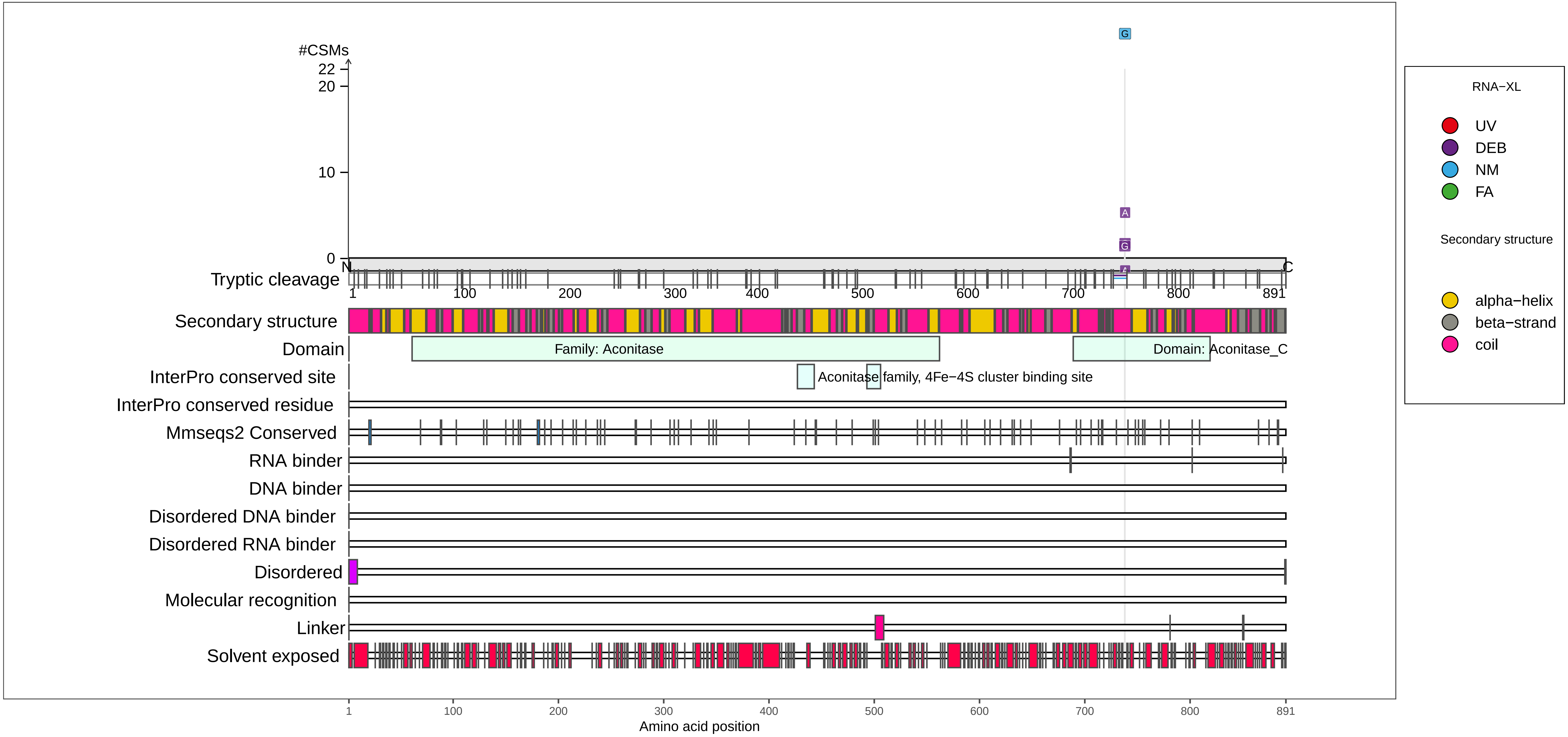

P64559
SDHE_ECOLI FAD assembly factor SdhE

− Abundance:
tryptic [log10 Intensity]: 8.63 (Q 79)
PAXdb K12 strain [ppm]: NA (Q NA)
PAXdb E.coli [ppm]: 1.88 (Q 69)

− RNA functions: not annotated

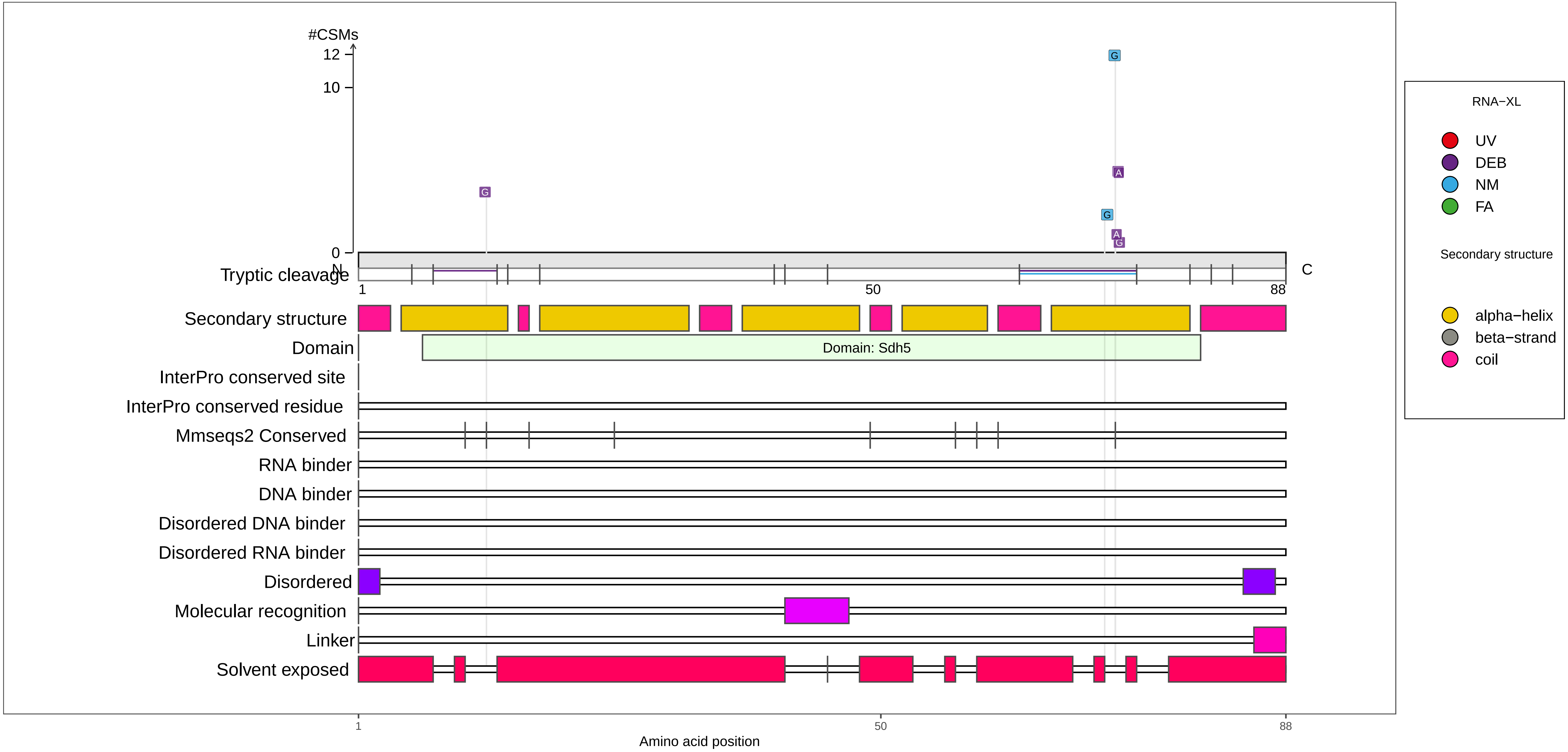

P75864
RLMKL_ECOLI Ribosomal RNA large subunit methyltransferase K/L

− Abundance:
tryptic [log10 Intensity]: 7.88 (Q 52)
PAXdb K12 strain [ppm]: 1.92 (Q 57)
PAXdb E.coli [ppm]: 1.3 (Q 54)

− RNA functions:
23S rRNA (guanine(2445)−N(2))−methyltransferase activity; ncRNA metabolic process
ncRNA processing; Putative RNA methylase family UPF0020; RNA (guanine−N7)−methylation
RNA binding; RNA metabolic process; RNA methylation; RNA methyltransferase activity
RNA modification; RNA processing; rRNA (guanine−N2−)−methyltransferase activity
rRNA (guanine−N7−)−methyltransferase activity; rRNA (guanine−N7)−methylation
rRNA (guanine) methyltransferase activity; rRNA base methylation; rRNA metabolic process
rRNA methylation; rRNA methyltransferase activity; rRNA modification; rRNA processing

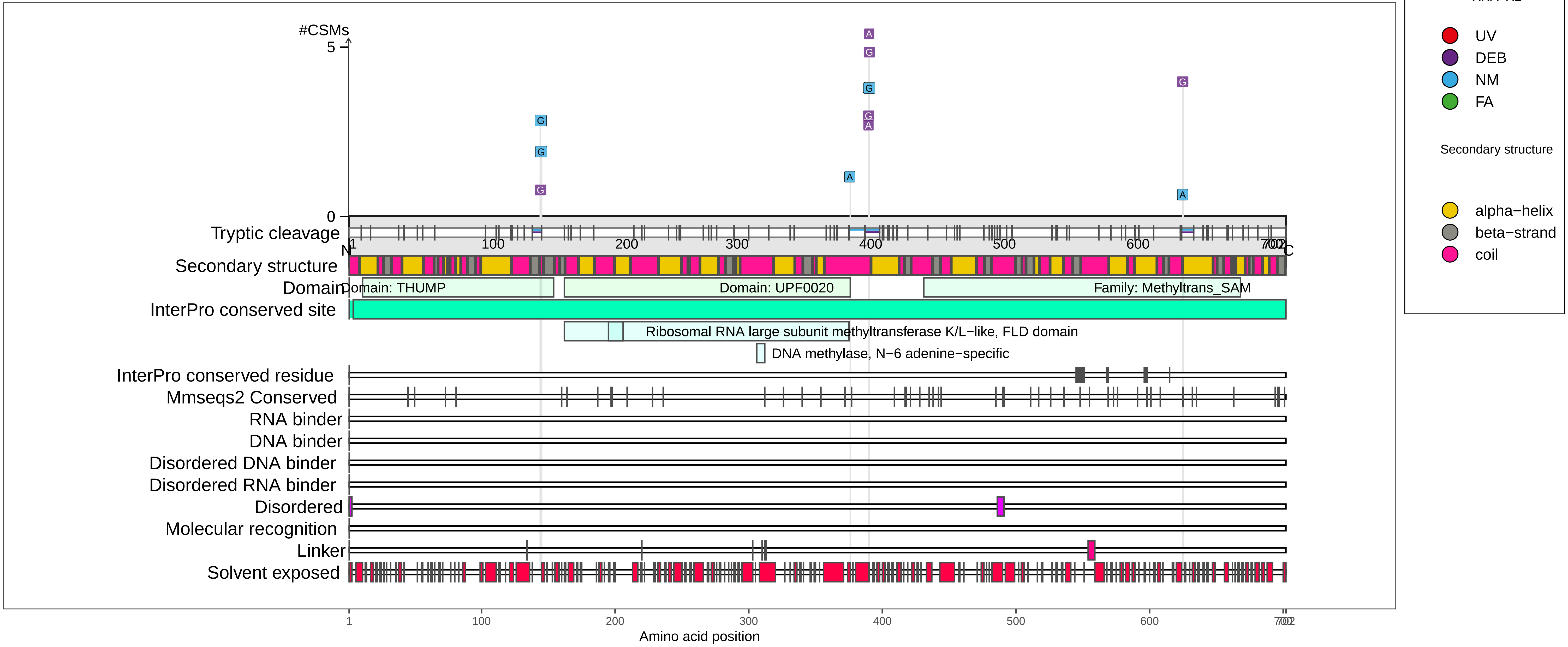

P05791
ILVD_ECOLI Dihydroxy−acid dehydratase

− Abundance:
tryptic [log10 Intensity]: 8.97 (Q 87)
PAXdb K12 strain [ppm]: 2.6 (Q 80)
PAXdb E.coli [ppm]: 1.99 (Q 72)

− RNA functions: not annotated

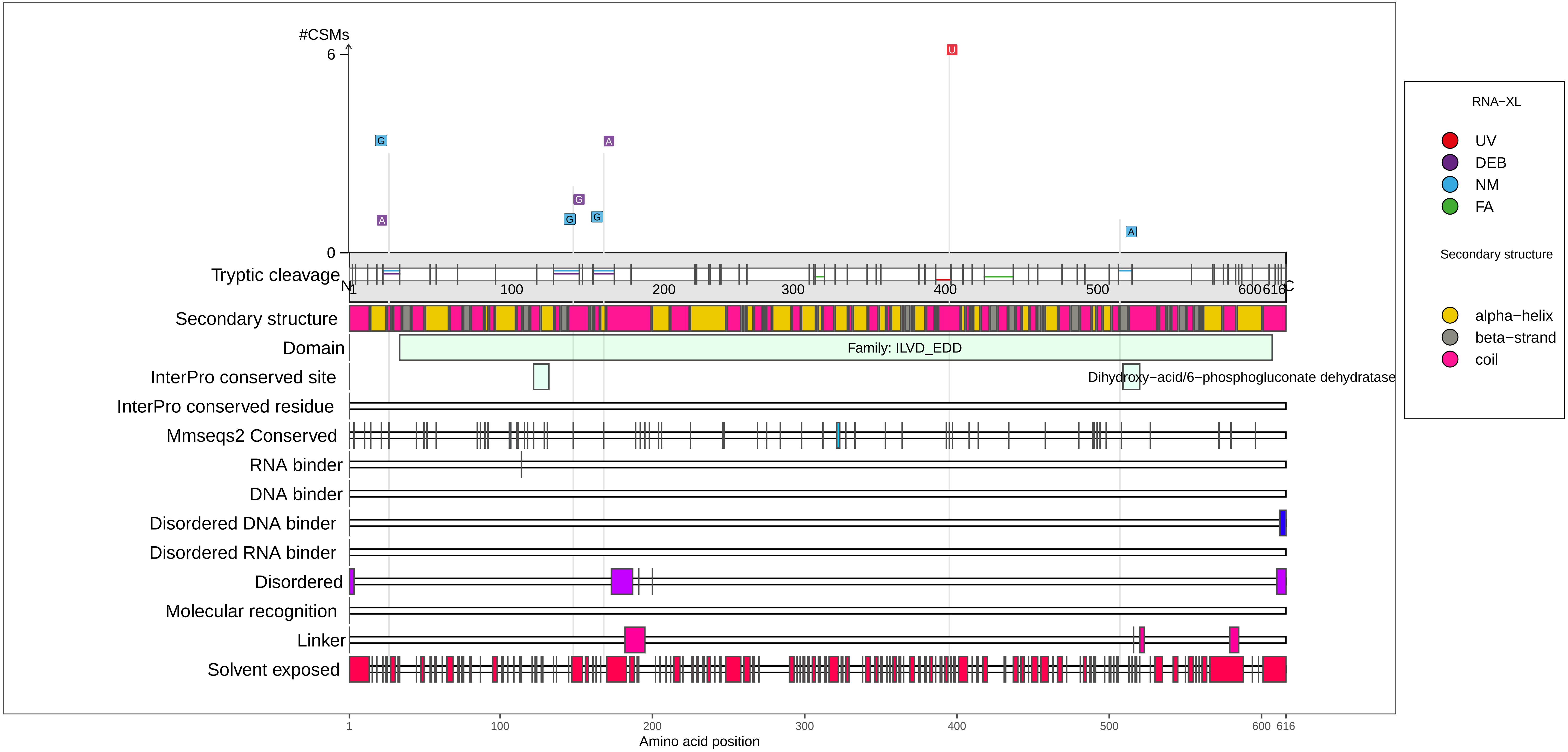

P06149
DLD_ECOLI Quinone−dependent D−lactate dehydrogenase

− Abundance:
tryptic [log10 Intensity]: 9.01 (Q 88)
PAXdb K12 strain [ppm]: 1.33 (Q 28)
PAXdb E.coli [ppm]: 2.46 (Q 84)

− RNA functions: not annotated

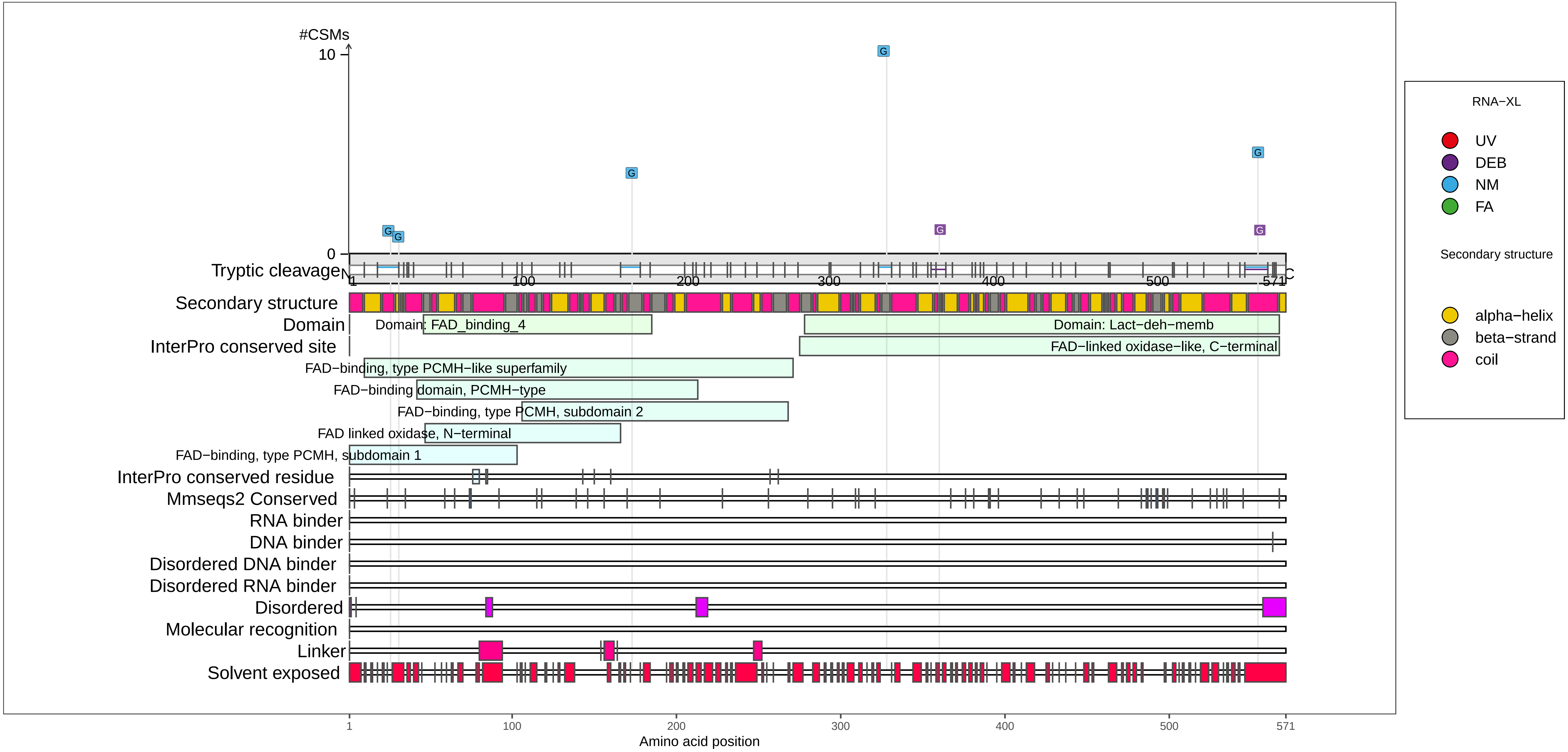

P0A6P7
ENGB_ECOLI Probable GTP−binding protein EngB

− Abundance:
tryptic [log10 Intensity]: 7.39 (Q 29)
PAXdb K12 strain [ppm]: 2.75 (Q 84)
PAXdb E.coli [ppm]: 2.2 (Q 77)

− RNA functions: not annotated

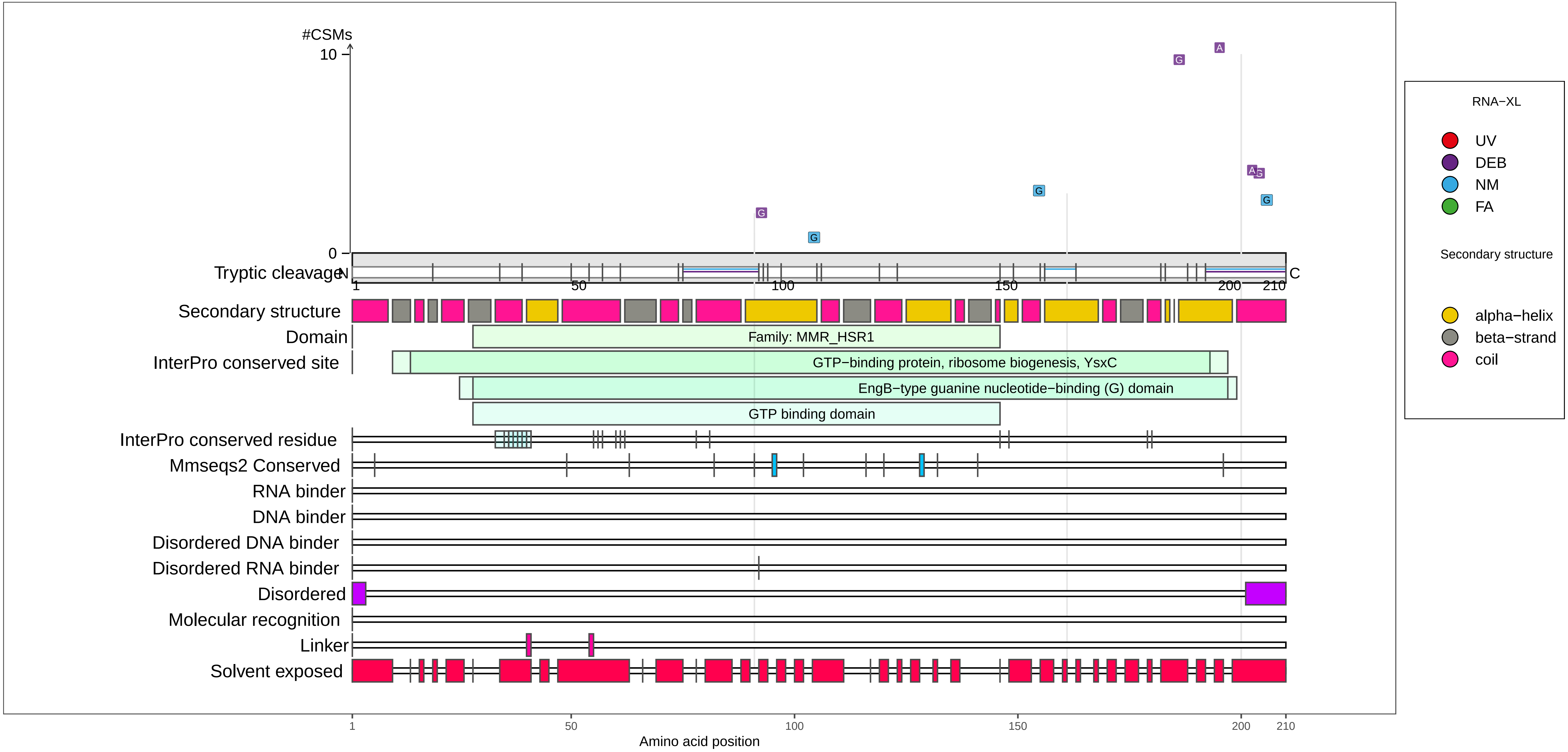

P0A6R0
FABH_ECOLI 3−oxoacyl−[acyl−carrier−protein] synthase 3

− Abundance:
tryptic [log10 Intensity]: 8.68 (Q 81)
PAXdb K12 strain [ppm]: 2.85 (Q 87)
PAXdb E.coli [ppm]: 2.79 (Q 91)

− RNA functions: not annotated

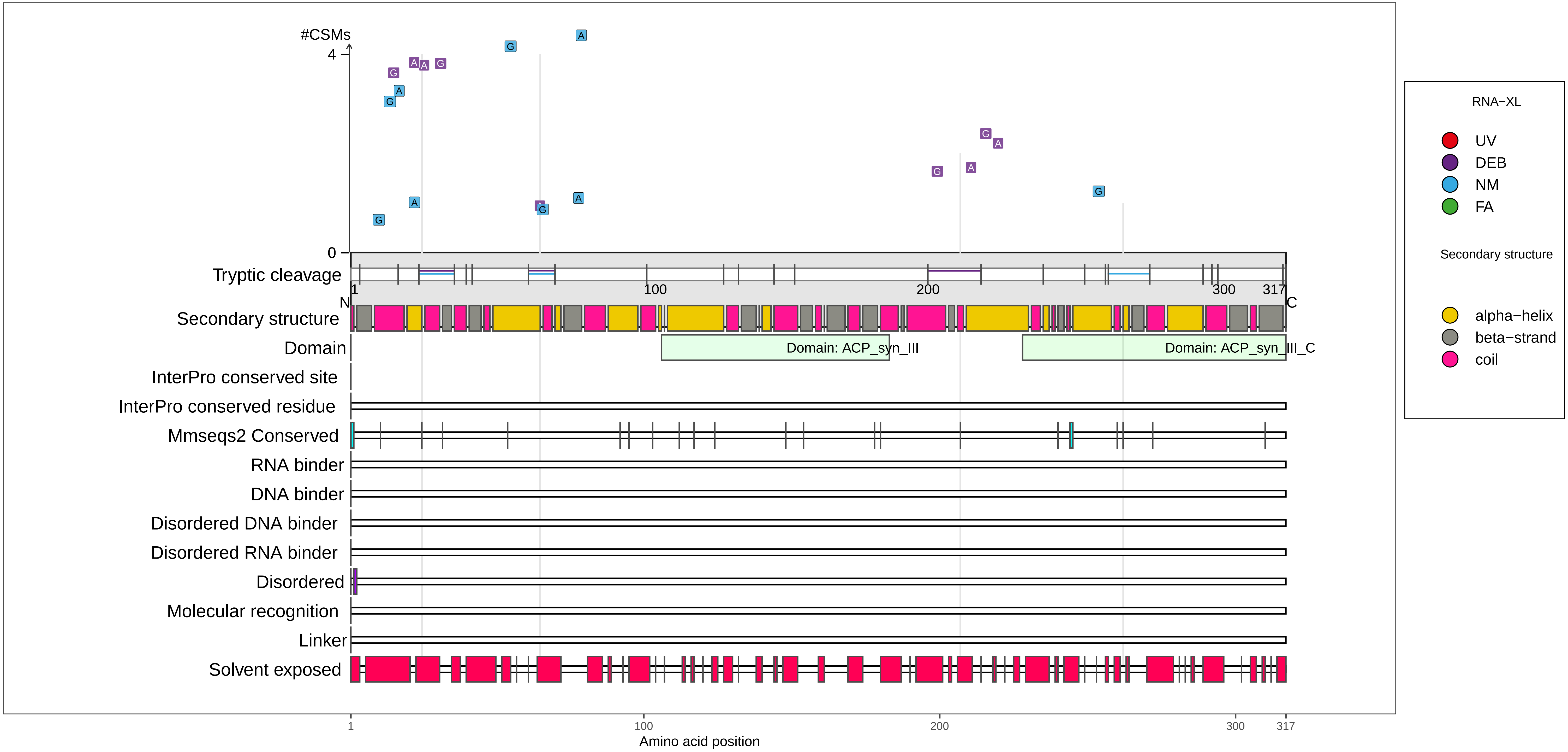

P0A9D8
DAPD_ECOLI 2,3,4,5−tetrahydropyridine−2,6−dicarboxylate N−succinyltransferase

− Abundance:
tryptic [log10 Intensity]: 9.49 (Q 96)
PAXdb K12 strain [ppm]: 3.55 (Q 98)
PAXdb E.coli [ppm]: 2.99 (Q 94)

− RNA functions: not annotated

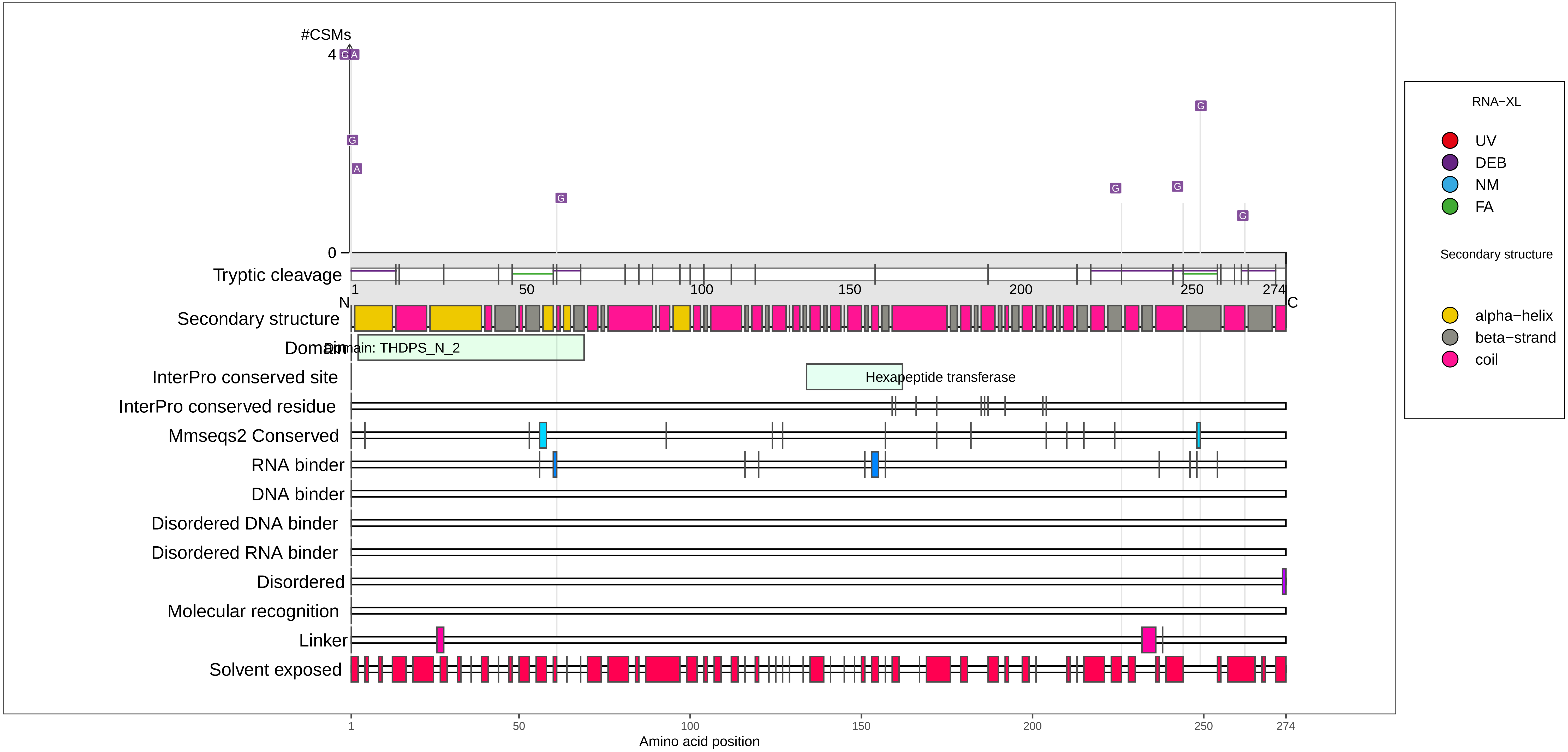

P0A9M2
HPRT_ECOLI Hypoxanthine phosphoribosyltransferase

− Abundance:
tryptic [log10 Intensity]: 9.15 (Q 92)
PAXdb K12 strain [ppm]: 3.34 (Q 96)
PAXdb E.coli [ppm]: 2.37 (Q 82)

− RNA functions: not annotated

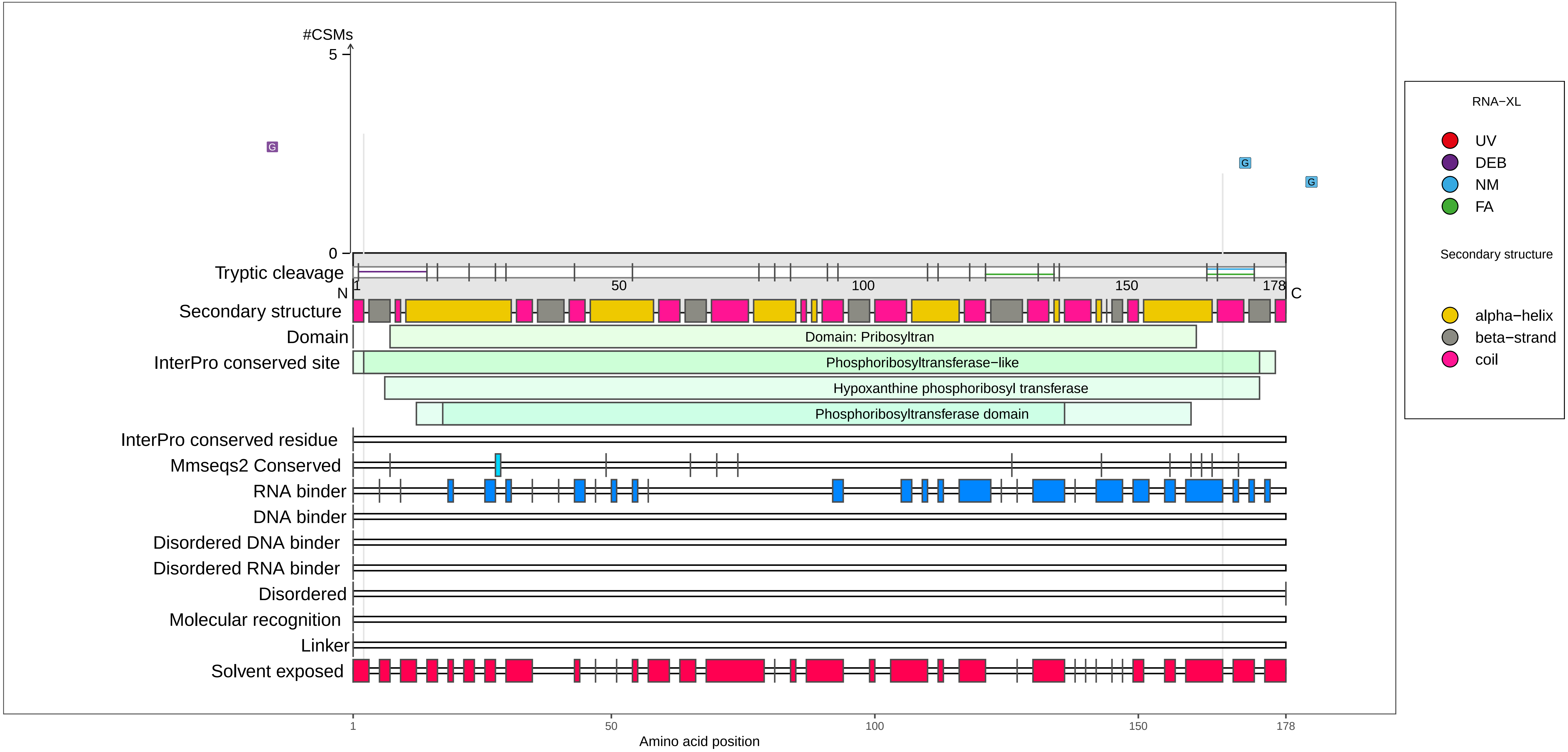

P0A9X4
MREB_ECOLI Cell shape−determining protein MreB

− Abundance:
tryptic [log10 Intensity]: 7.24 (Q 22)
PAXdb K12 strain [ppm]: 3.14 (Q 93)
PAXdb E.coli [ppm]: 2.99 (Q 95)

− RNA functions: not annotated

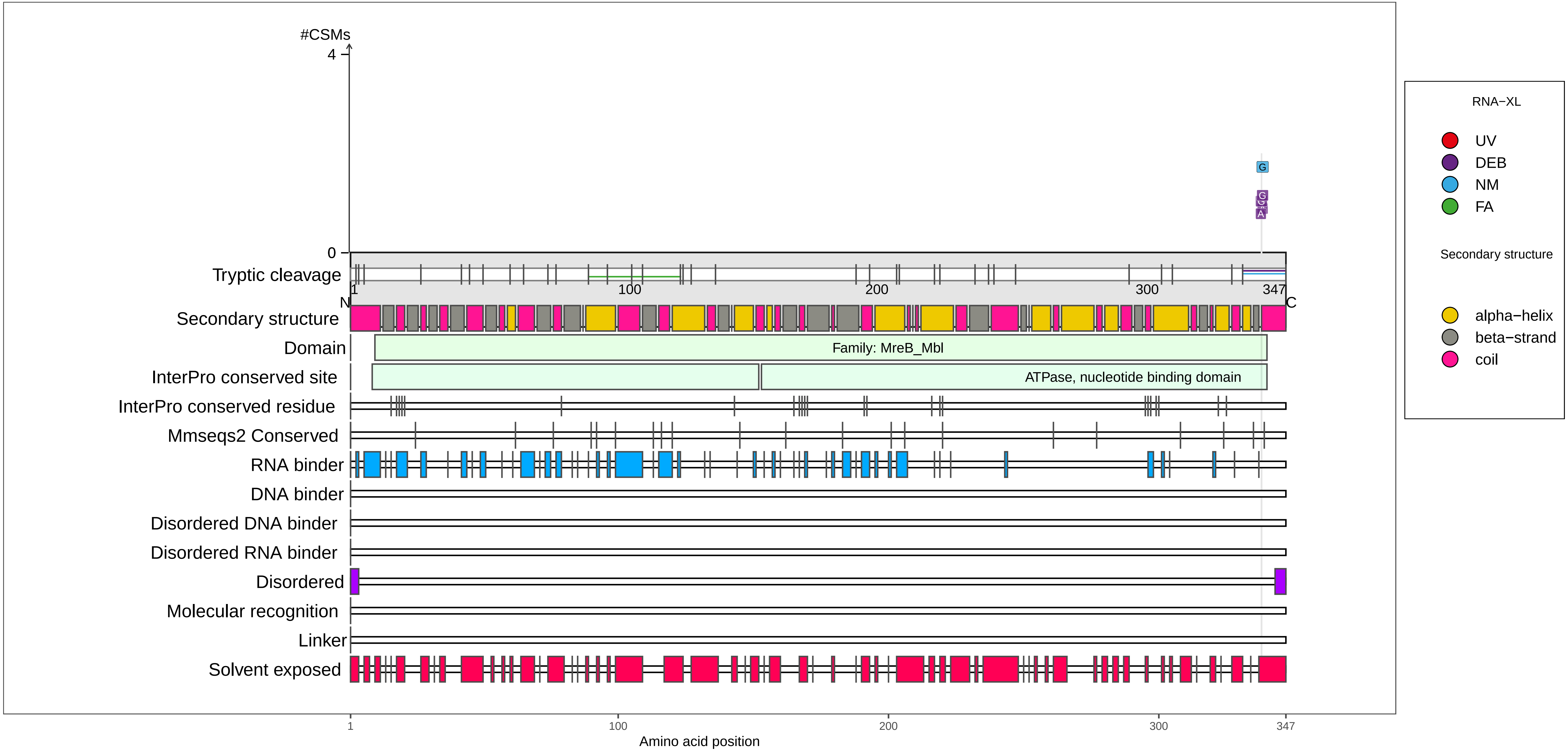

P0AB28
YCED_ECOLI Large ribosomal RNA subunit accumulation protein YceD

− Abundance:
tryptic [log10 Intensity]: 7.88 (Q 53)
PAXdb K12 strain [ppm]: 2.16 (Q 65)
PAXdb E.coli [ppm]: 2.24 (Q 78)

− RNA functions: not annotated

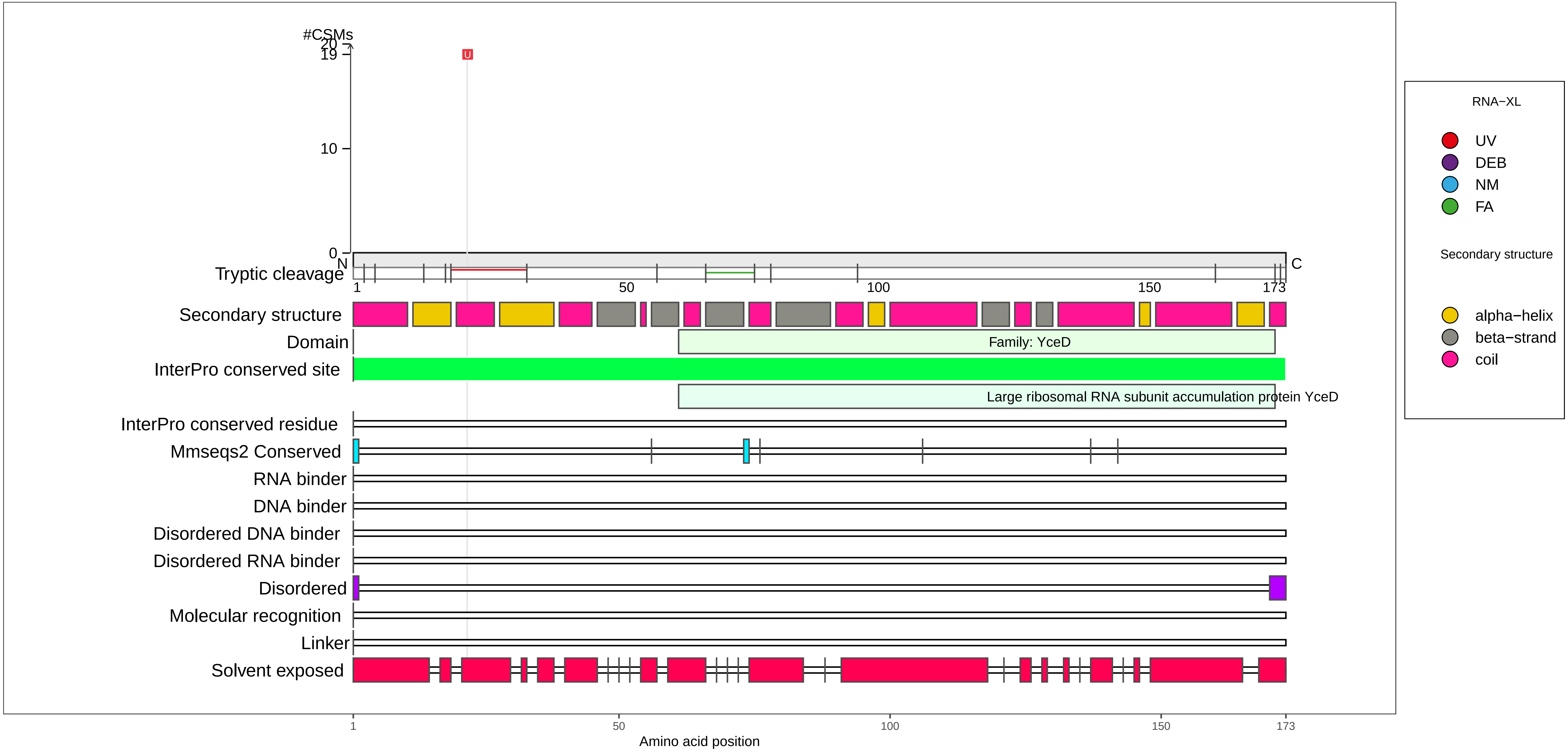

P0AB89
PUR8_ECOLI Adenylosuccinate lyase

− Abundance:
tryptic [log10 Intensity]: 8.75 (Q 82)
PAXdb K12 strain [ppm]: 3 (Q 91)
PAXdb E.coli [ppm]: 2.8 (Q 92)

− RNA functions: not annotated

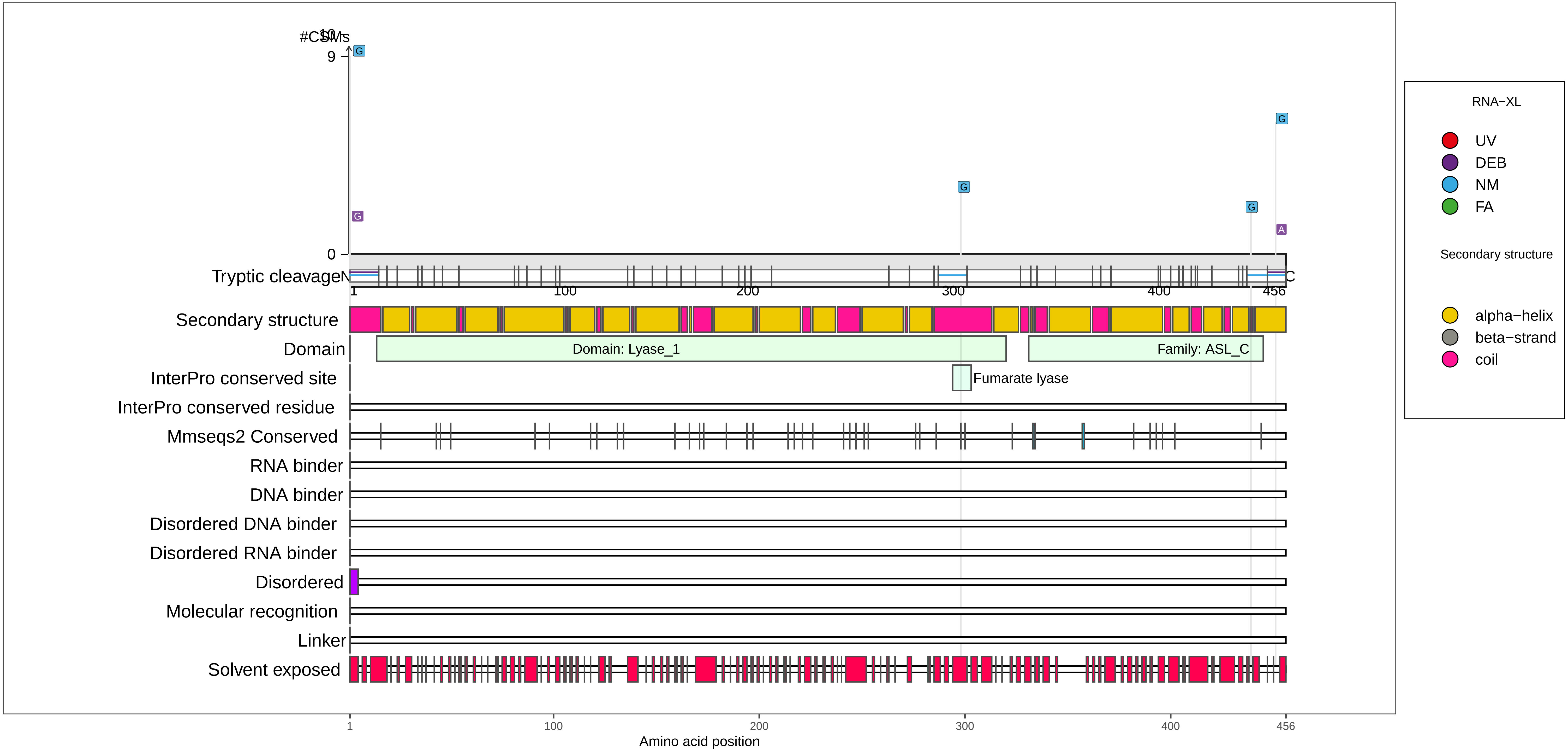

P0ACF0
DBHA_ECOLI DNA−binding protein HU−alpha

− Abundance:
tryptic [log10 Intensity]: 9.62 (Q 97)
PAXdb K12 strain [ppm]: 3.93 (Q 100)
PAXdb E.coli [ppm]: 3.67 (Q 100)

− RNA functions:
RNA biosynthetic process; RNA metabolic process

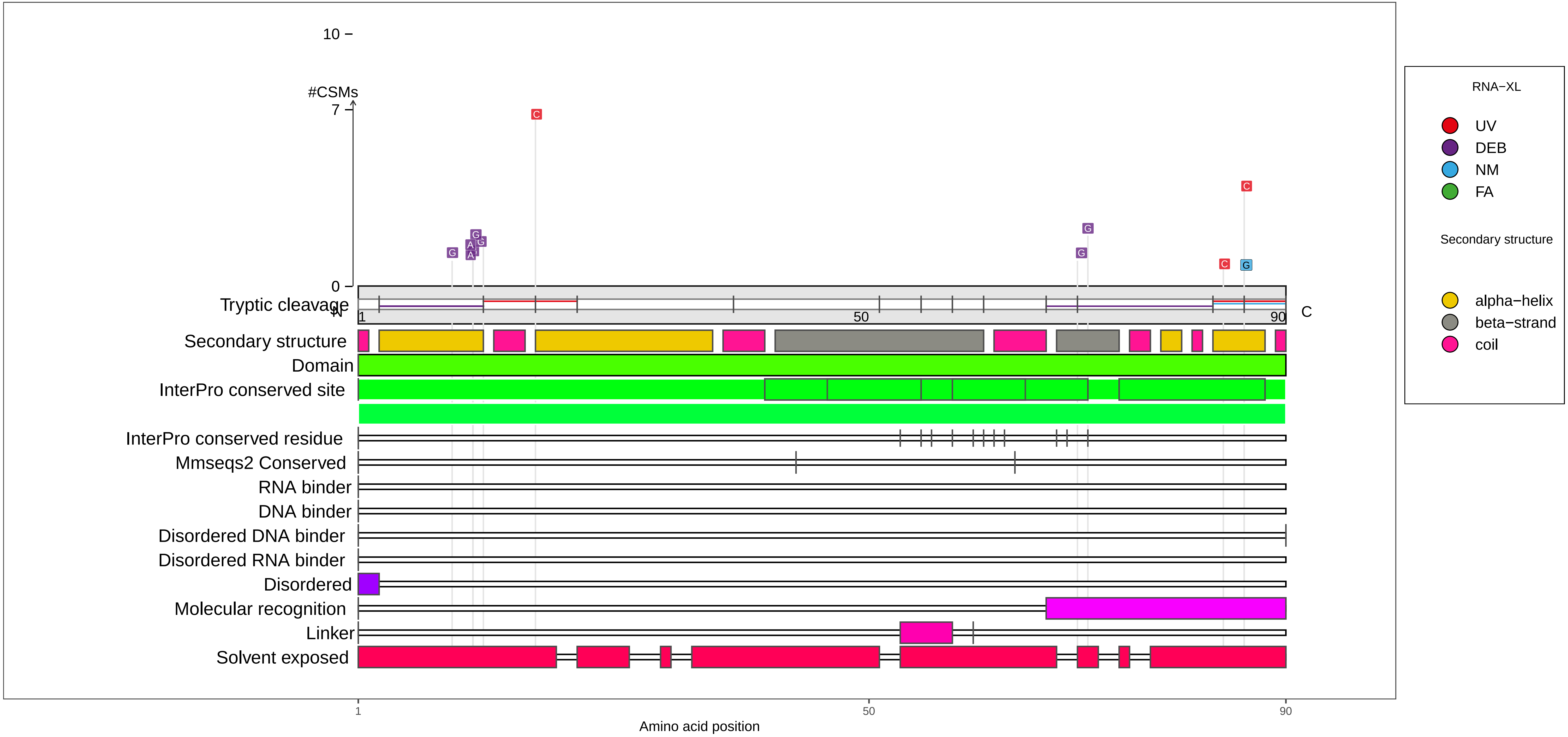

P15288
PEPD_ECOLI Cytosol non−specific dipeptidase
− Abundance:
tryptic [log10 Intensity]: 9.28 (Q 94)
PAXdb K12 strain [ppm]: 3.25 (Q 95)
PAXdb E.coli [ppm]: 2.99 (Q 94)

− RNA functions:
RNA binding; RNA ligase activity; RNA metabolic process; RNA processing

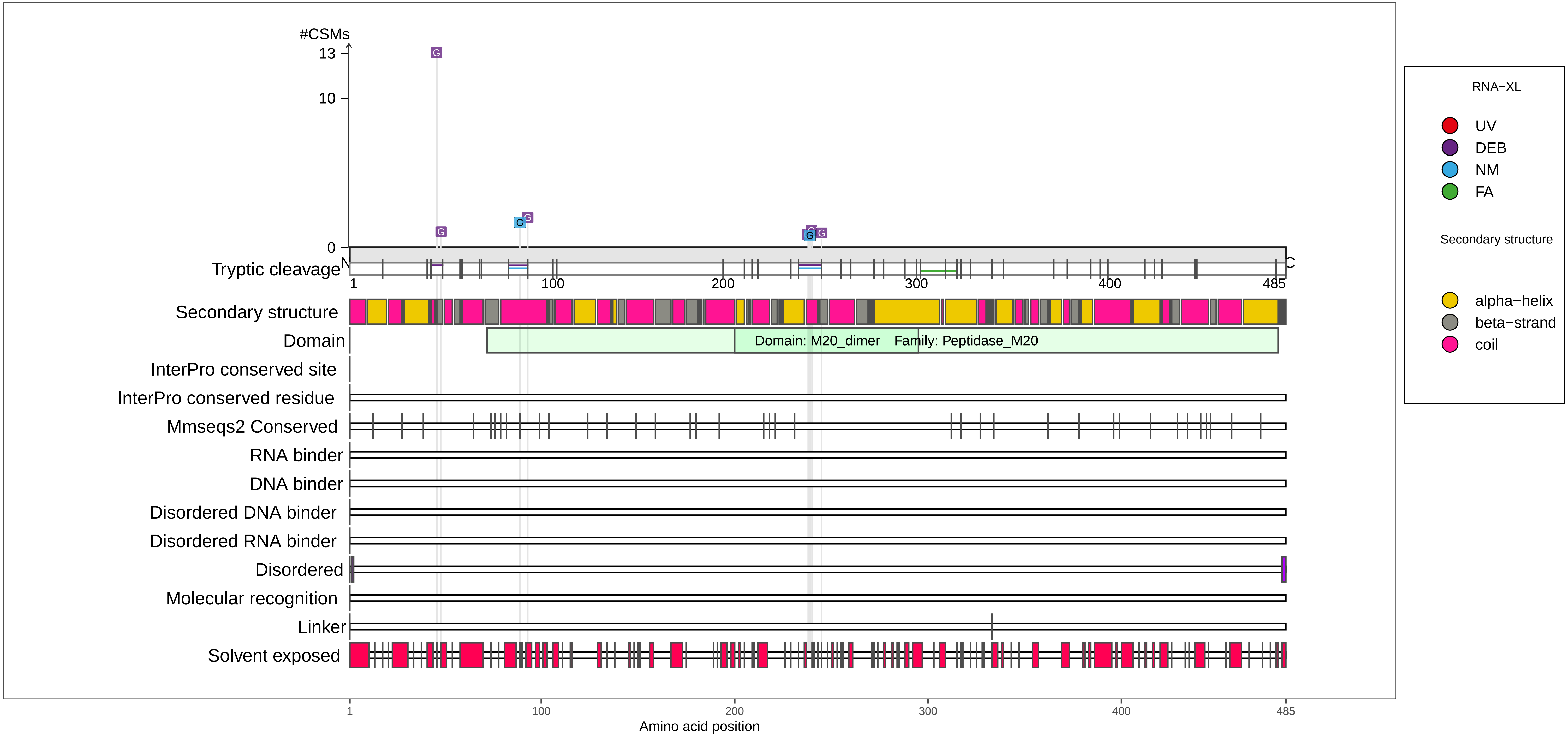

P37765
RLUB_ECOLI Ribosomal large subunit pseudouridine synthase B

− Abundance:
tryptic [log10 Intensity]: 7.48 (Q 34)
PAXdb K12 strain [ppm]: 1.93 (Q 58)
PAXdb E.coli [ppm]: 2.2 (Q 77)

− RNA functions:
ncRNA metabolic process; ncRNA processing; RNA binding; RNA metabolic process
RNA modification; RNA processing; RNA pseudouridylate synthase; rRNA metabolic process
rRNA modification; rRNA processing; rRNA pseudouridine synthase activity
rRNA pseudouridine synthesis

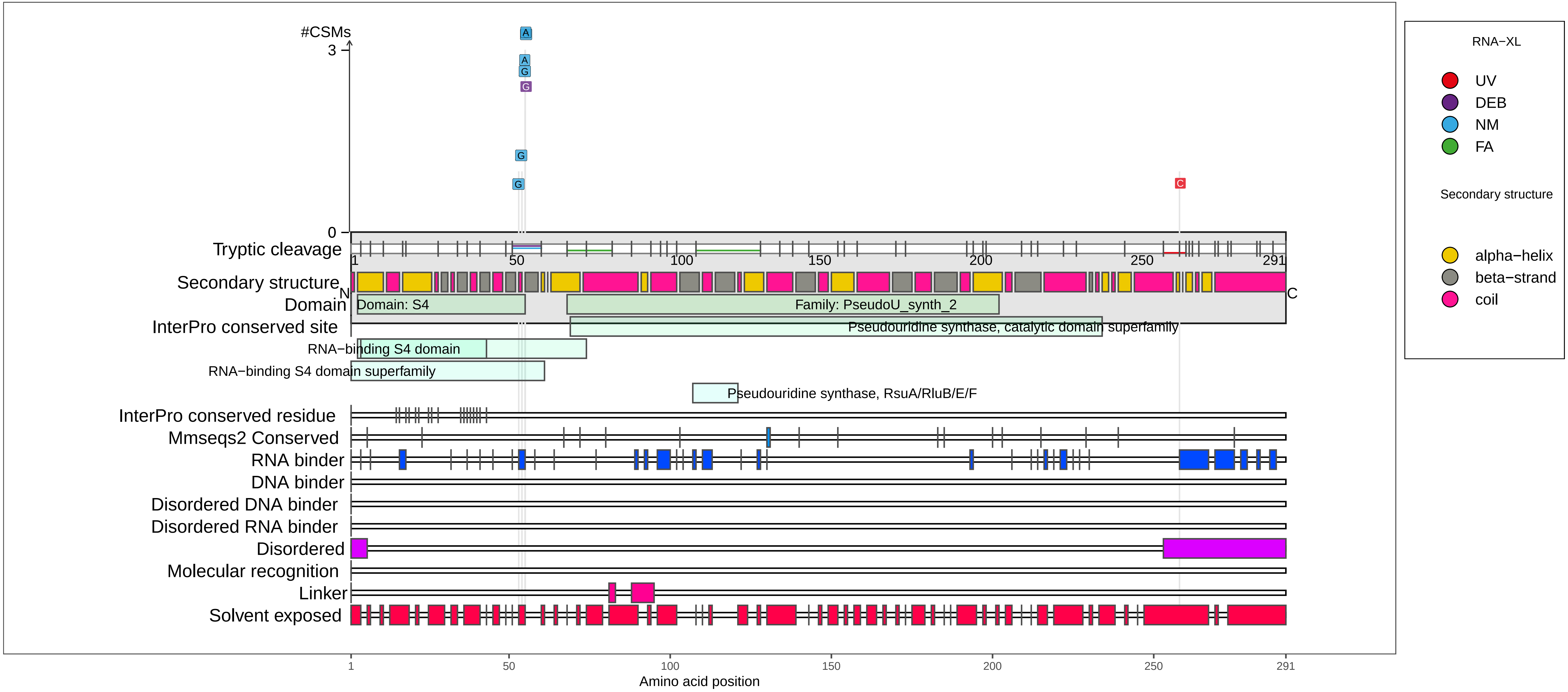

P08506
DACC_ECOLI D−alanyl−D−alanine carboxypeptidase DacC

− Abundance:
tryptic [log10 Intensity]: 8.22 (Q 66)
PAXdb K12 strain [ppm]: 1.39 (Q 32)
PAXdb E.coli [ppm]: 2.47 (Q 85)

− RNA functions: not annotated

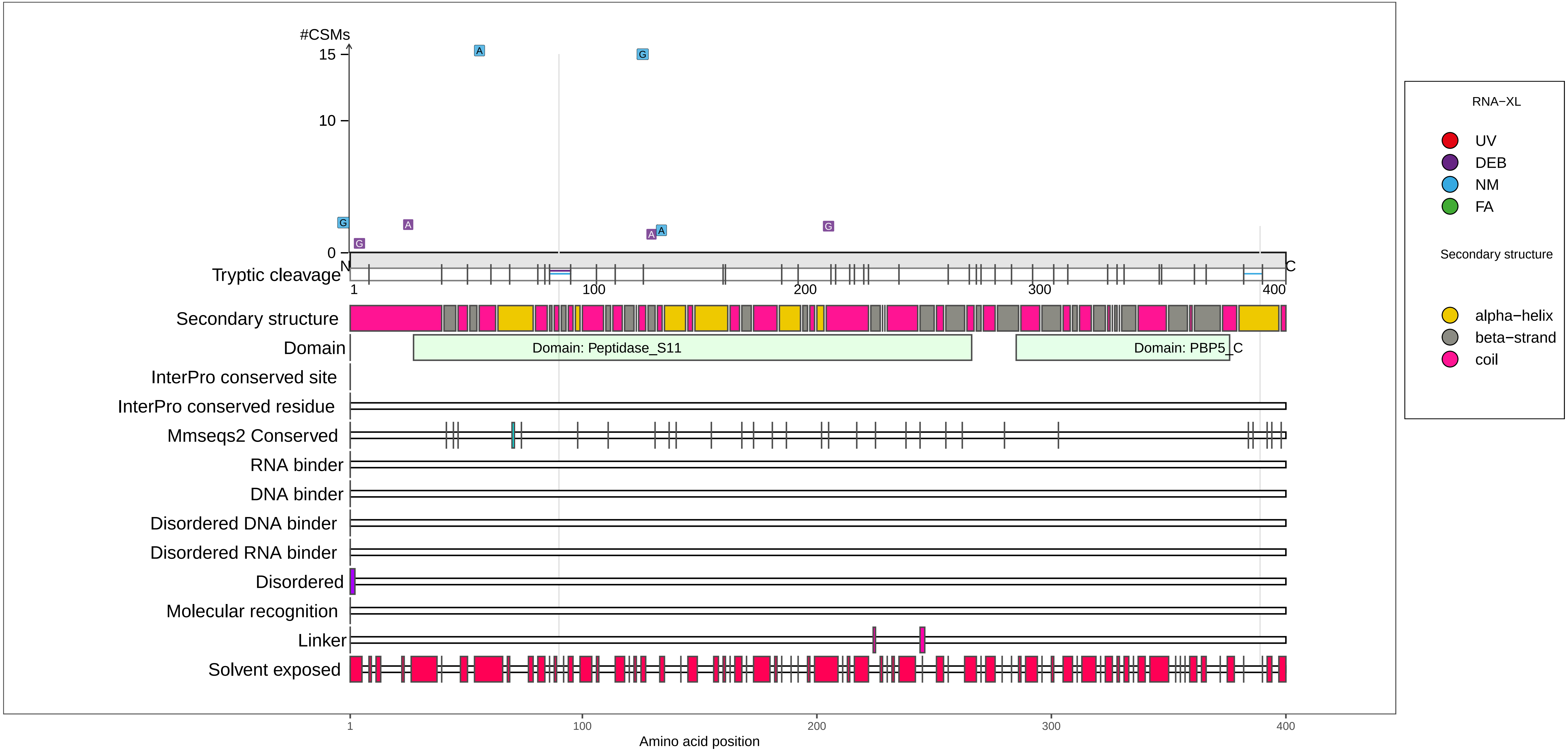

P0A6E4
ASSY_ECOLI Argininosuccinate synthase

− Abundance:
tryptic [log10 Intensity]: 9.02 (Q 89)
PAXdb K12 strain [ppm]: 2.96 (Q 90)
PAXdb E.coli [ppm]: 2.74 (Q 91)

− RNA functions: not annotated

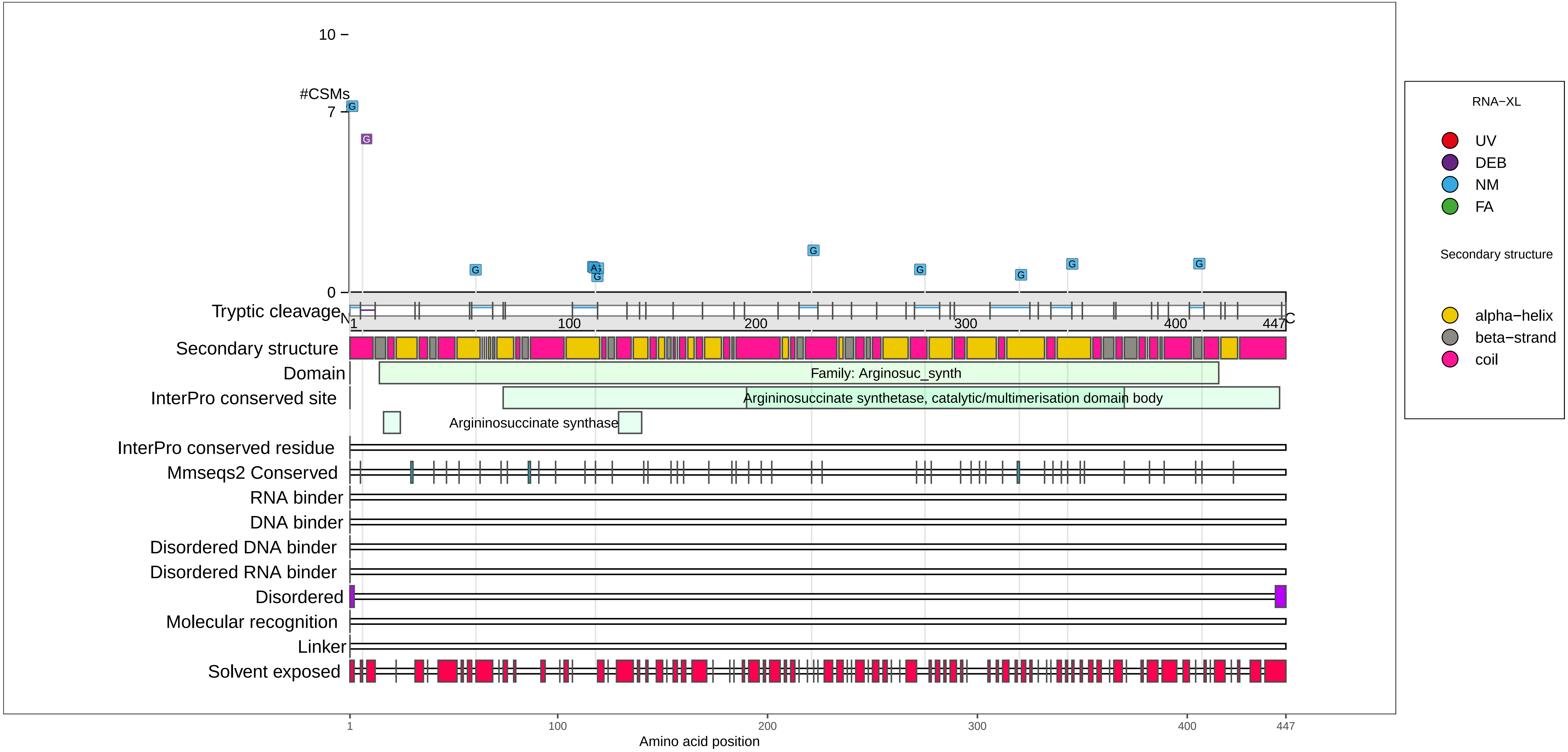

P0A9A9
FUR_ECOLI Ferric uptake regulation protein

− Abundance:
tryptic [log10 Intensity]: 8.52 (Q 76)
PAXdb K12 strain [ppm]: 3.15 (Q 93)
PAXdb E.coli [ppm]: 2.92 (Q 94)

− RNA functions: not annotated

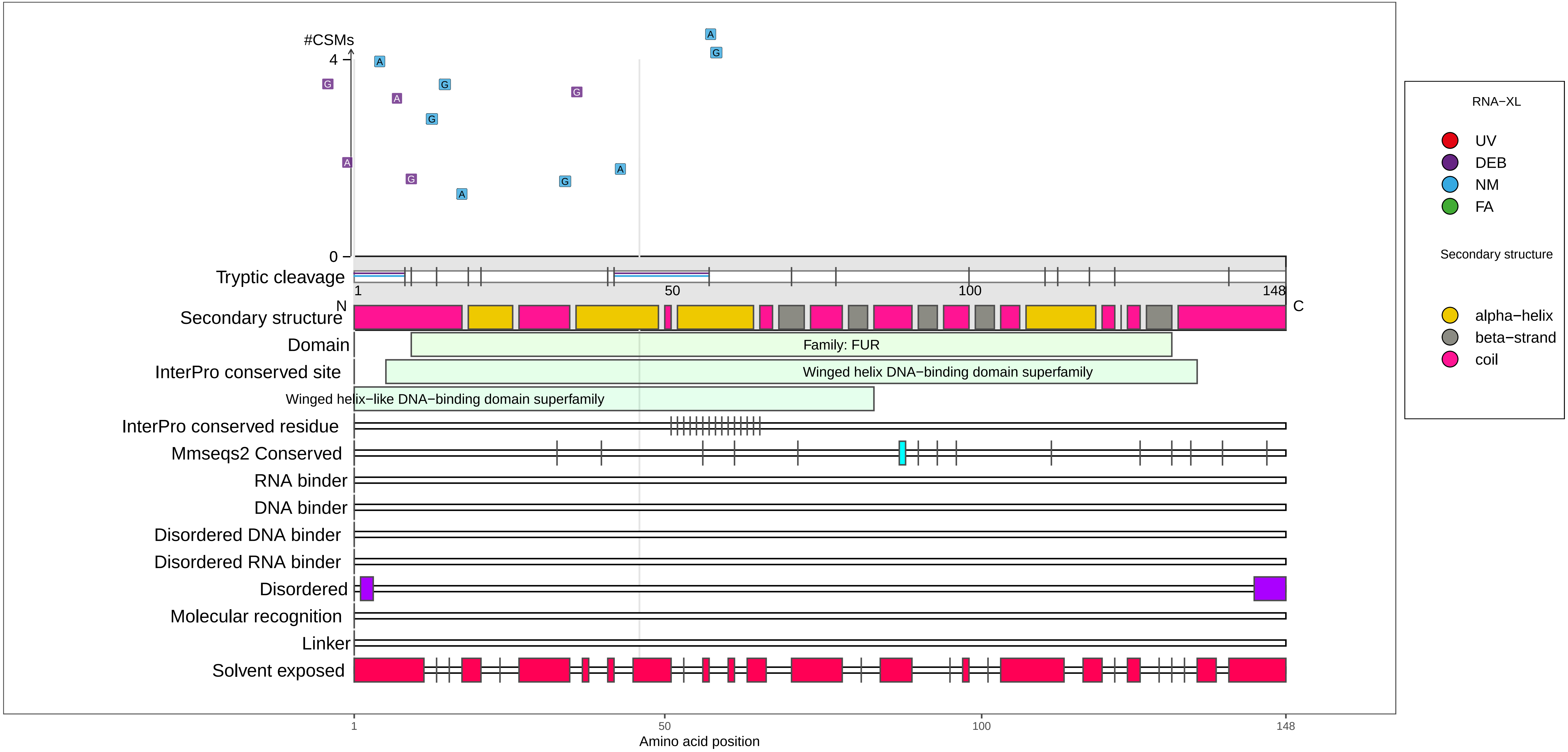

P0A9M8
PTA_ECOLI Phosphate acetyltransferase

− Abundance:
tryptic [log10 Intensity]: 9.19 (Q 92)
PAXdb K12 strain [ppm]: 3.23 (Q 94)
PAXdb E.coli [ppm]: 2.88 (Q 93)

− RNA functions: not annotated

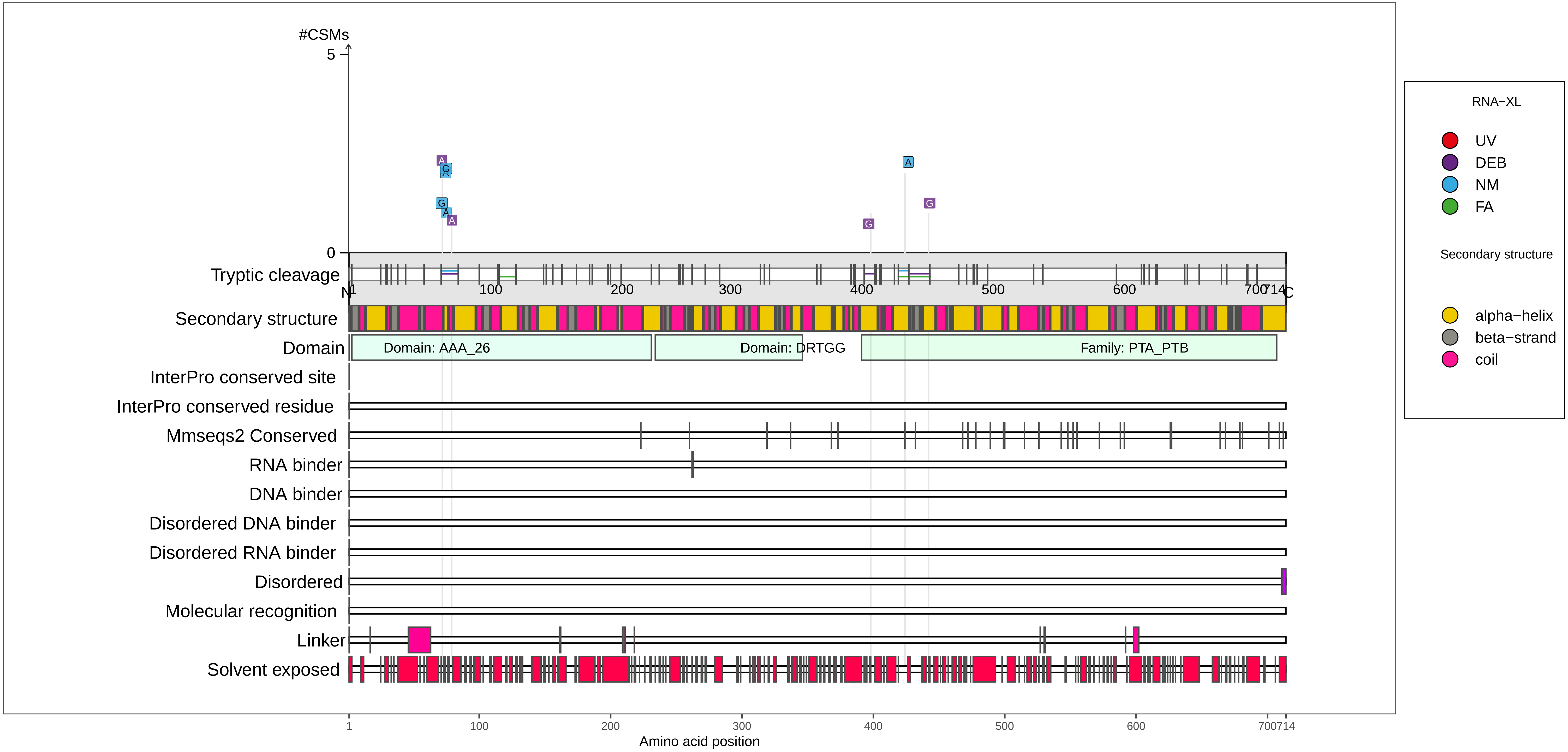

P0A9U3
YBIT_ECOLI Probable ATP−binding protein YbiT

− Abundance:
tryptic [log10 Intensity]: 9.15 (Q 92)
PAXdb K12 strain [ppm]: 2.44 (Q 75)
PAXdb E.coli [ppm]: 2.25 (Q 79)

− RNA functions: not annotated

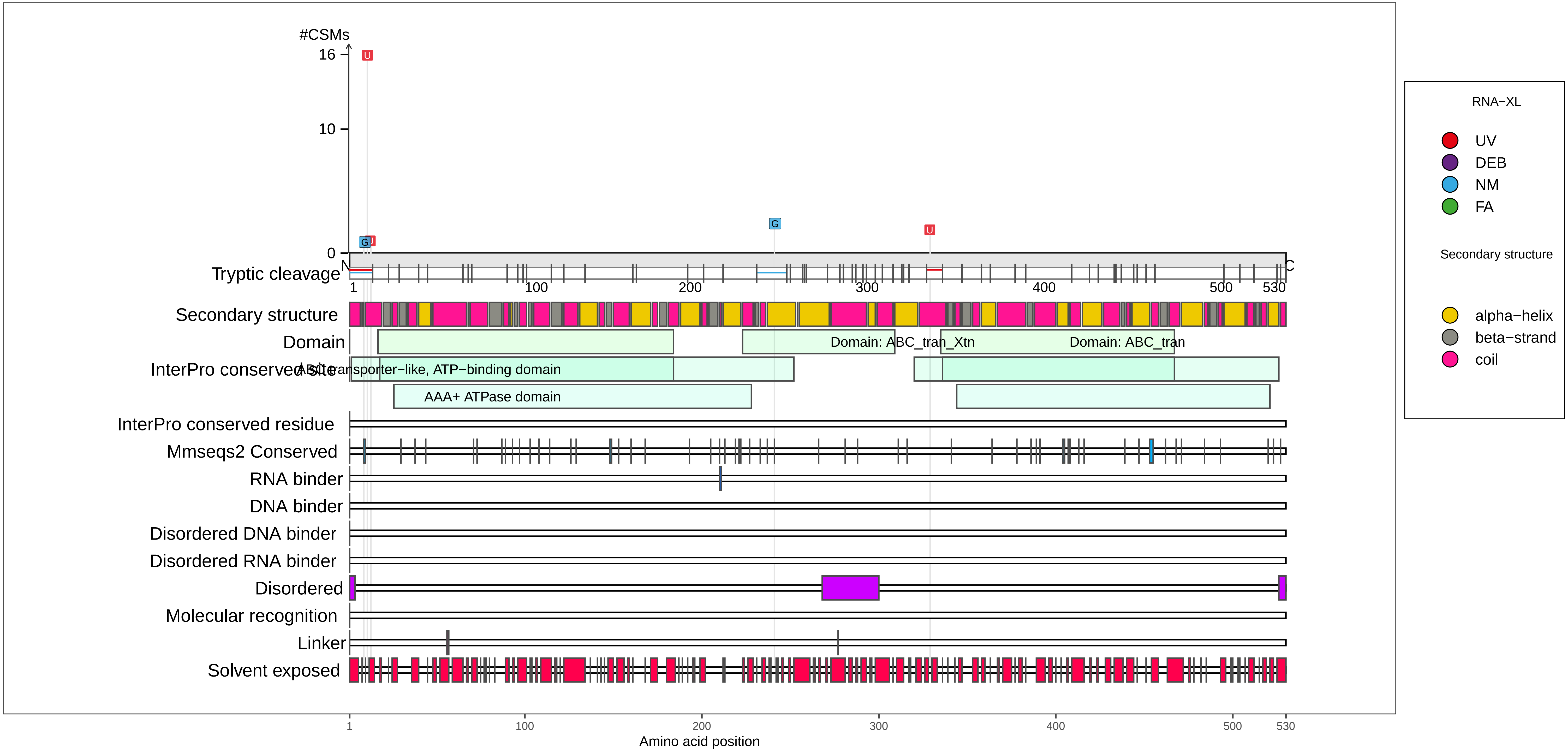

P0ABD5
ACCA_ECOLI Acetyl−coenzyme A carboxylase carboxyl transferase subunit alpha

− Abundance:
tryptic [log10 Intensity]: 8.48 (Q 75)
PAXdb K12 strain [ppm]: 2.78 (Q 85)
PAXdb E.coli [ppm]: 2.88 (Q 93)

− RNA functions: not annotated

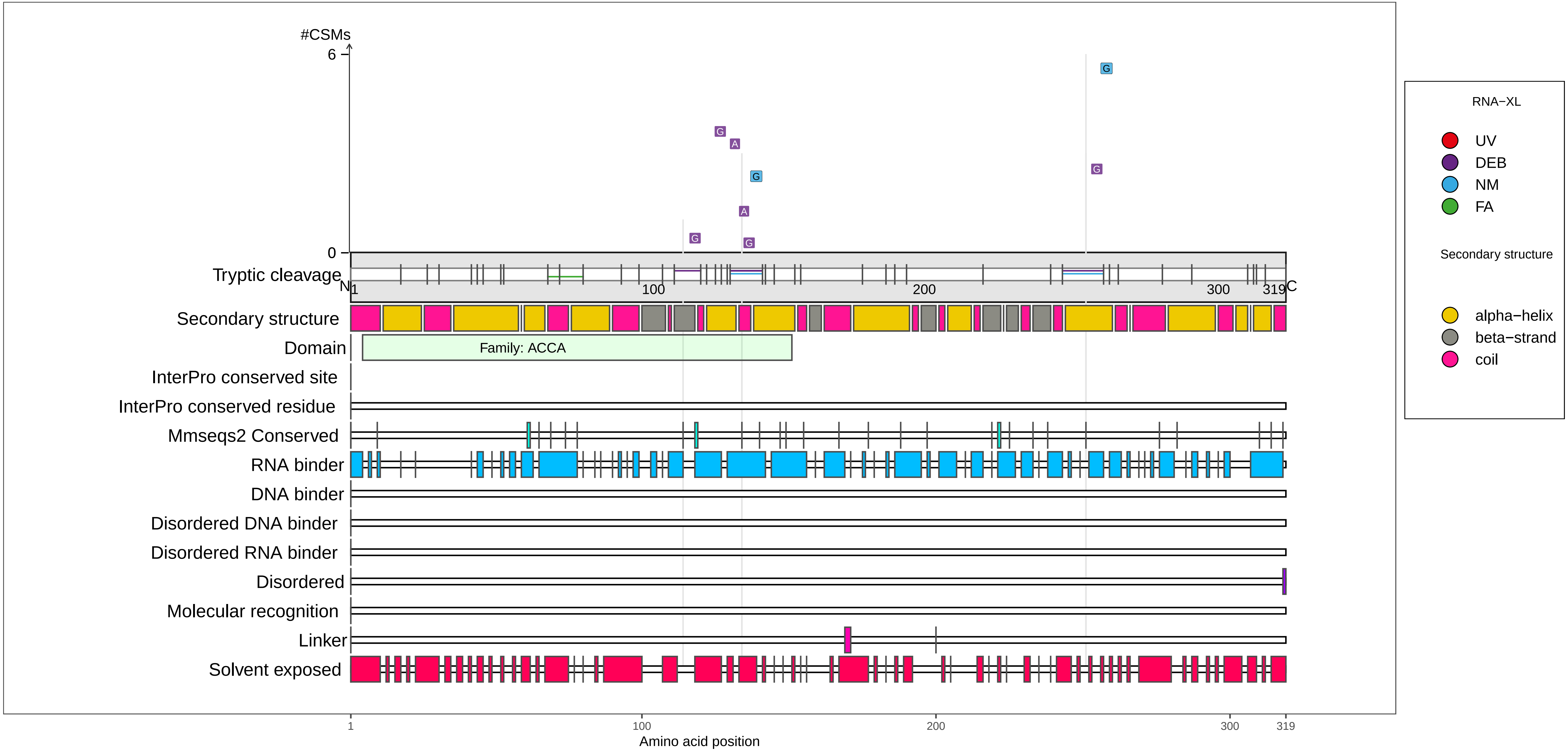

P60757
HIS1_ECOLI ATP phosphoribosyltransferase

− Abundance:
tryptic [log10 Intensity]: 8.14 (Q 63)
PAXdb K12 strain [ppm]: 2.76 (Q 85)
PAXdb E.coli [ppm]: 2.52 (Q 86)

− RNA functions: not annotated

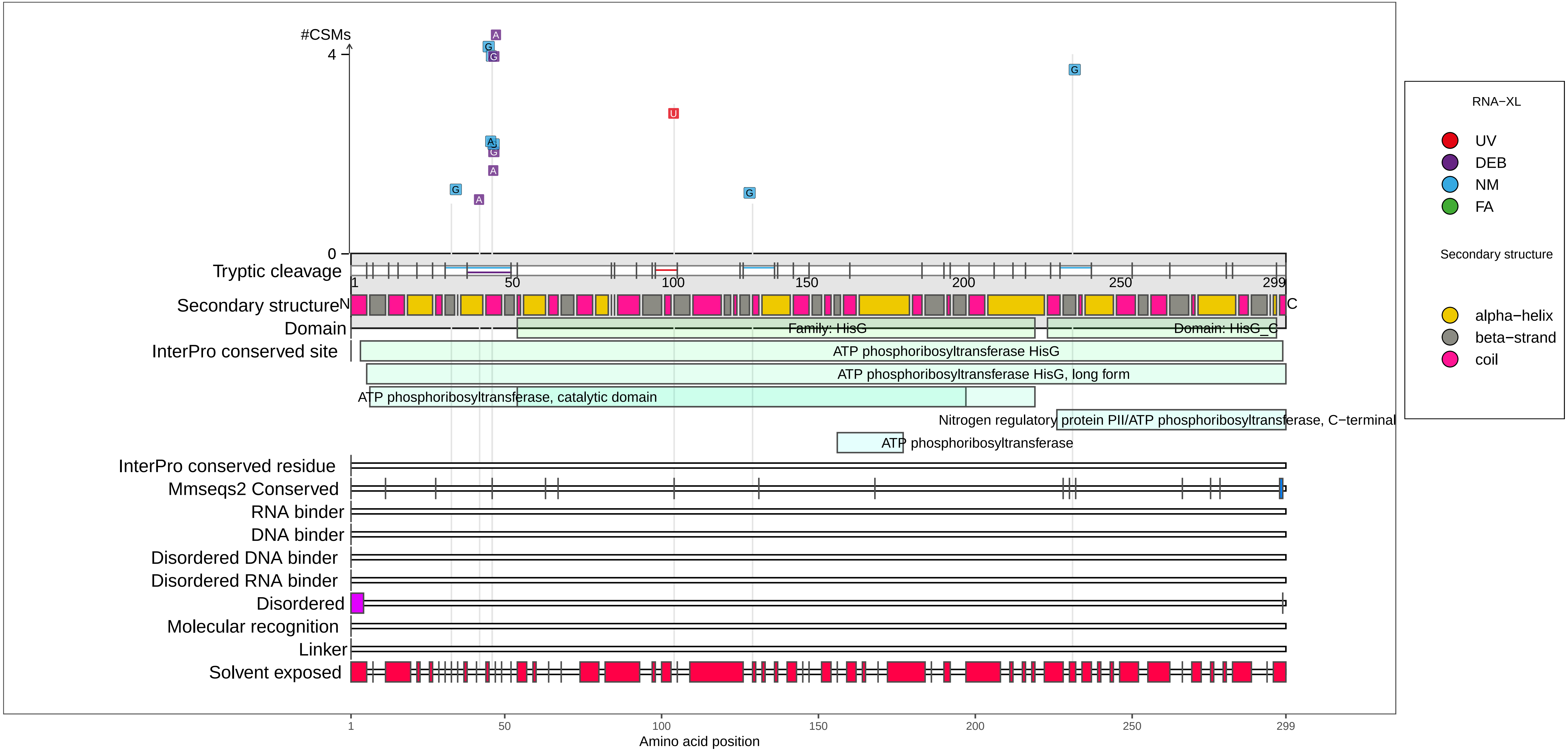

P69228
BAER_ECOLI Transcriptional regulatory protein BaeR

− Abundance:
tryptic [log10 Intensity]: 7.47 (Q 33)
PAXdb K12 strain [ppm]: 1.55 (Q 41)
PAXdb E.coli [ppm]: 1.83 (Q 68)

− RNA functions:
RNA biosynthetic process; RNA metabolic process

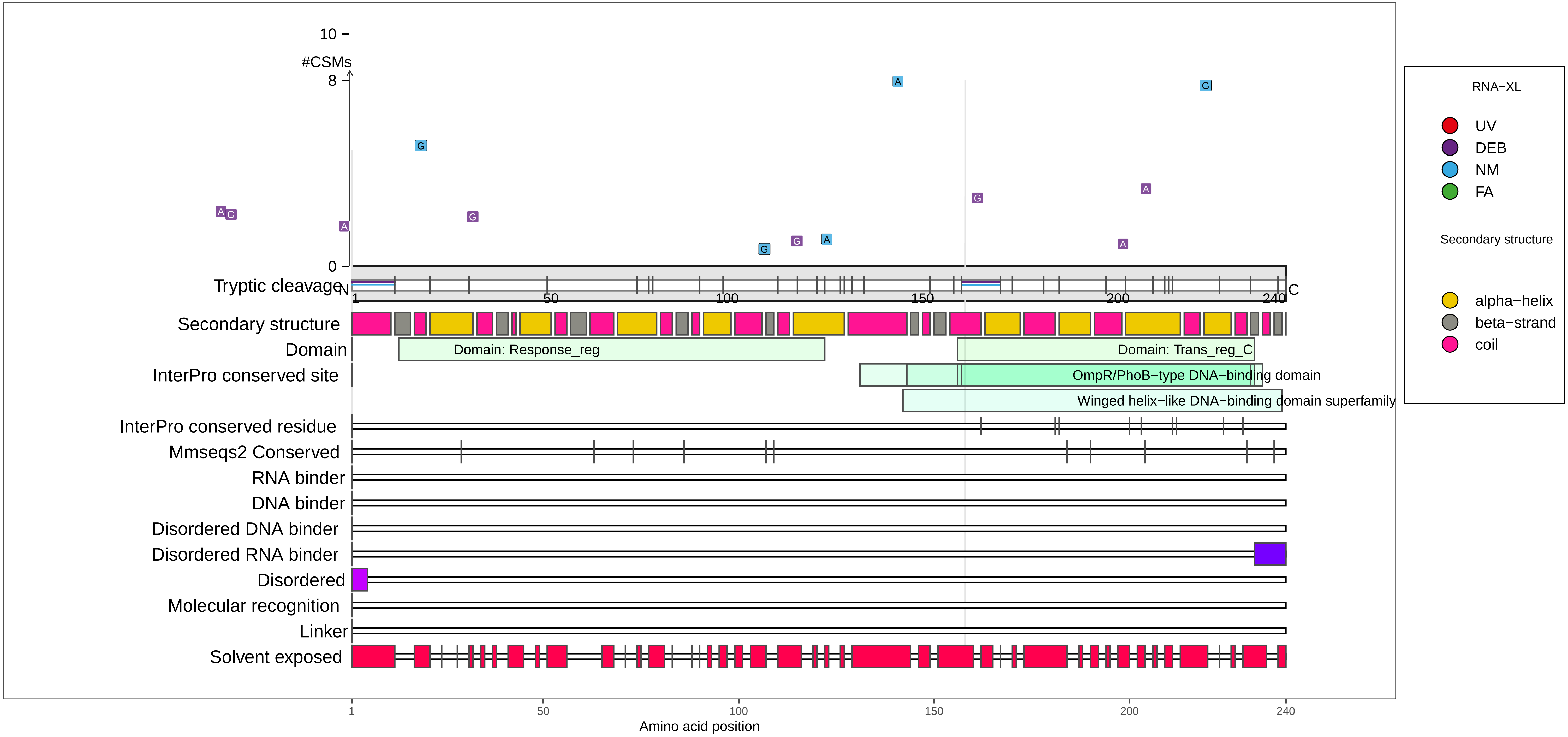

P69797
PTNAB_ECOLI PTS system mannose−specific EIIAB component

− Abundance:
tryptic [log10 Intensity]: 8.81 (Q 84)
PAXdb K12 strain [ppm]: 2.49 (Q 77)
PAXdb E.coli [ppm]: 3.21 (Q 97)

− RNA functions: not annotated

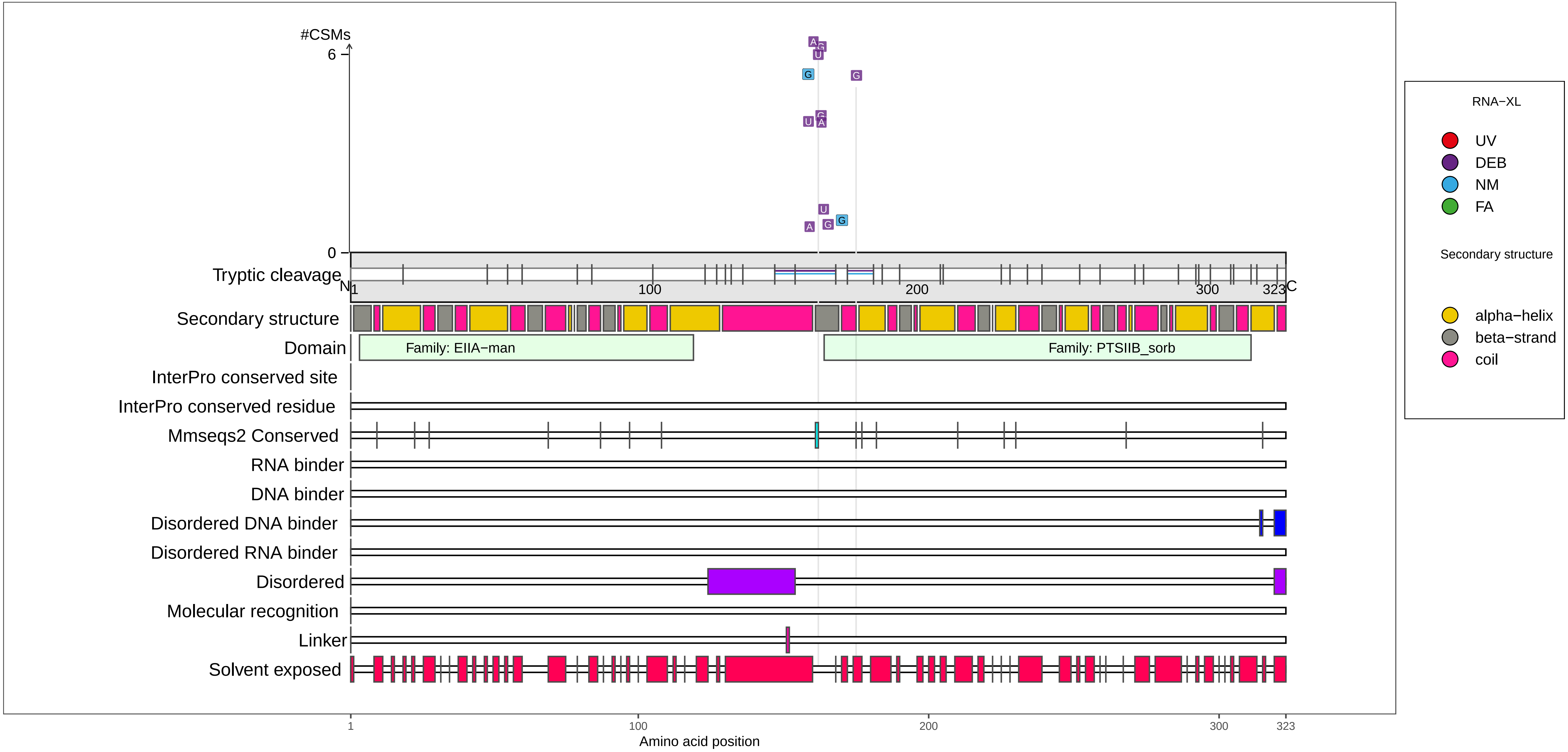

P00393
NDH_ECOLI Type II NADH:quinone oxidoreductase
− RNA functions: not annotated

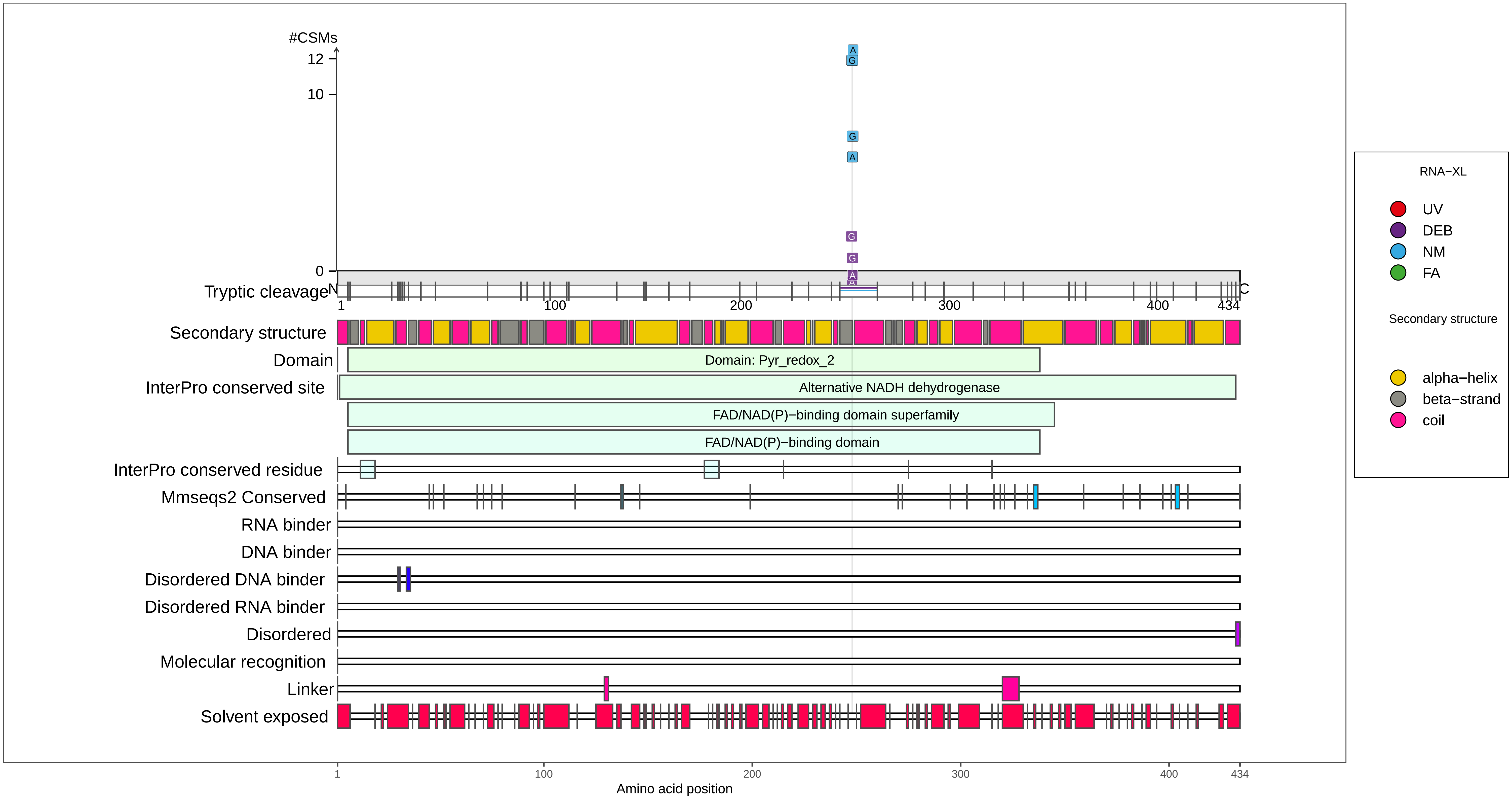

P06992
RSMA_ECOLI Ribosomal RNA small subunit methyltransferase A

− Abundance:
tryptic [log10 Intensity]: 7.51 (Q 36)
PAXdb K12 strain [ppm]: 2.43 (Q 75)
PAXdb E.coli [ppm]: 1.97 (Q 71)

− RNA functions:
16S rRNA (adenine(1518)−N(6)/adenine(1519)−N(6))−dimethyltransferase activity
mRNA binding; ncRNA metabolic process; ncRNA processing
Ribosomal RNA adenine dimethylase; RNA binding; RNA metabolic process; RNA methylation; RNA methyltransferase activity
RNA modification; RNA processing; rRNA (adenine−N6−)−methyltransferase activity
rRNA (adenine−N6,N6−)−dimethyltransferase activity
rRNA (adenine) methyltransferase activity; rRNA base methylation; rRNA binding; rRNA metabolic process; rRNA methylation
rRNA methyltransferase activity; rRNA modification; rRNA processing

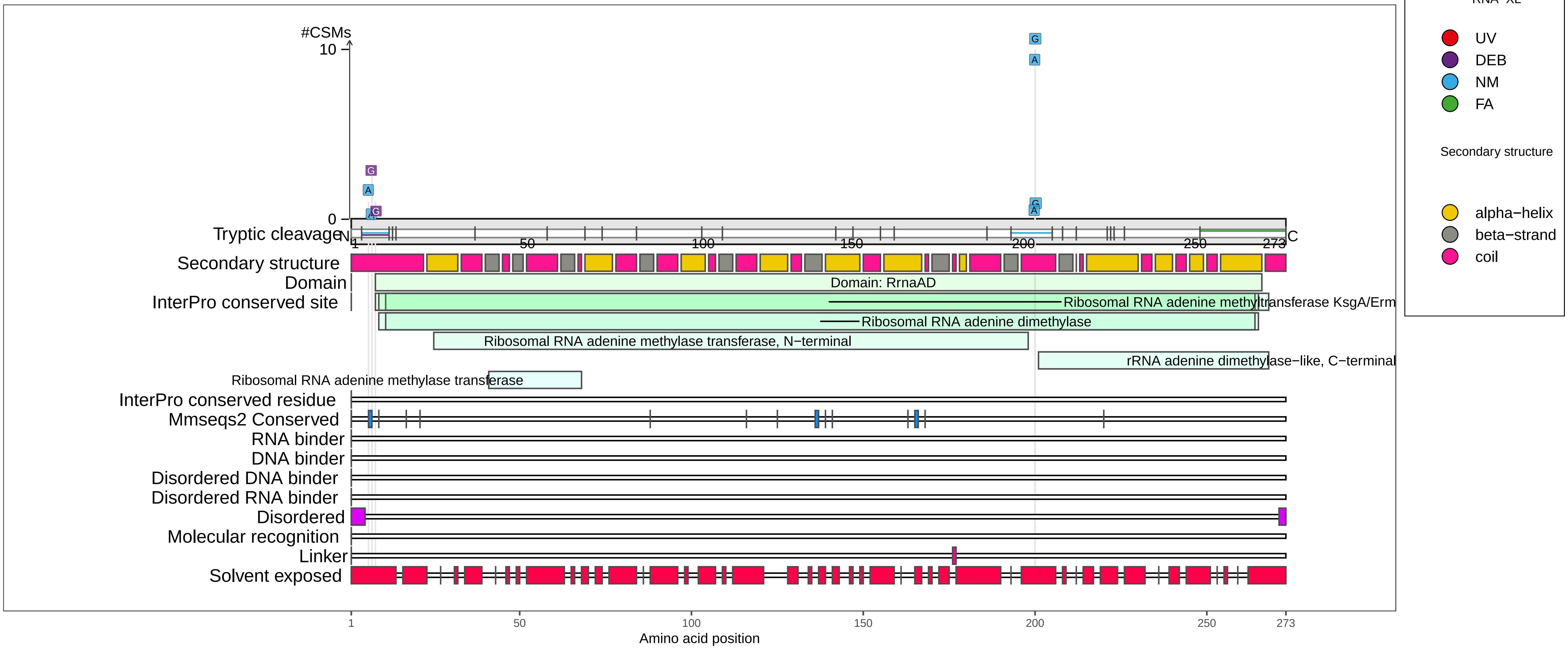

P0A6J5
DADA_ECOLI D−amino acid dehydrogenase

− Abundance:
tryptic [log10 Intensity]: 7.33 (Q 26)
PAXdb K12 strain [ppm]: 1.96 (Q 59)
PAXdb E.coli [ppm]: 1.98 (Q 72)

− RNA functions: not annotated

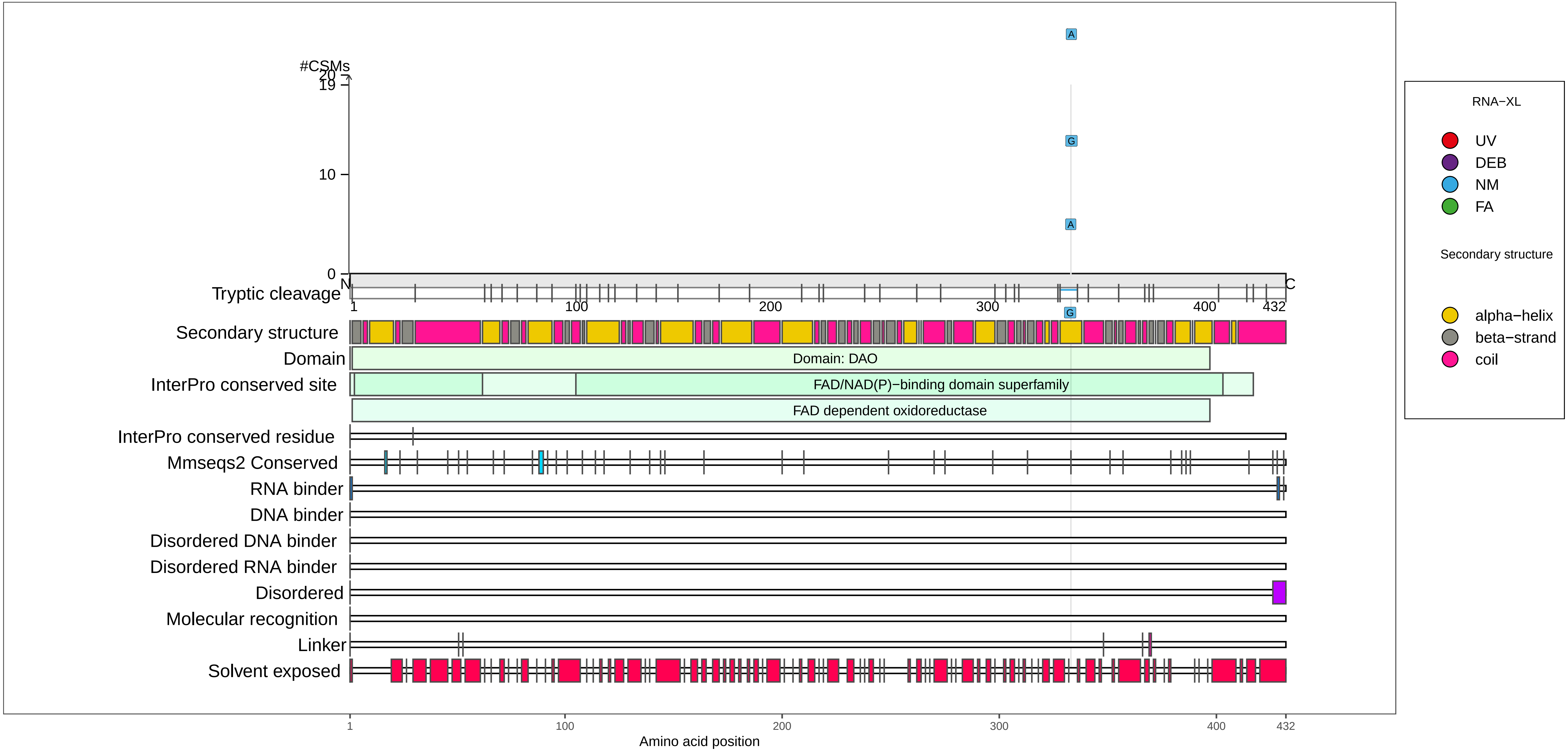

P0A832
SSRP_ECOLI SsrA−binding protein

− Abundance:
tryptic [log10 Intensity]: 7.86 (Q 52)
PAXdb K12 strain [ppm]: 2.32 (Q 71)
PAXdb E.coli [ppm]: 1.9 (Q 70)

− RNA functions:
RNA binding

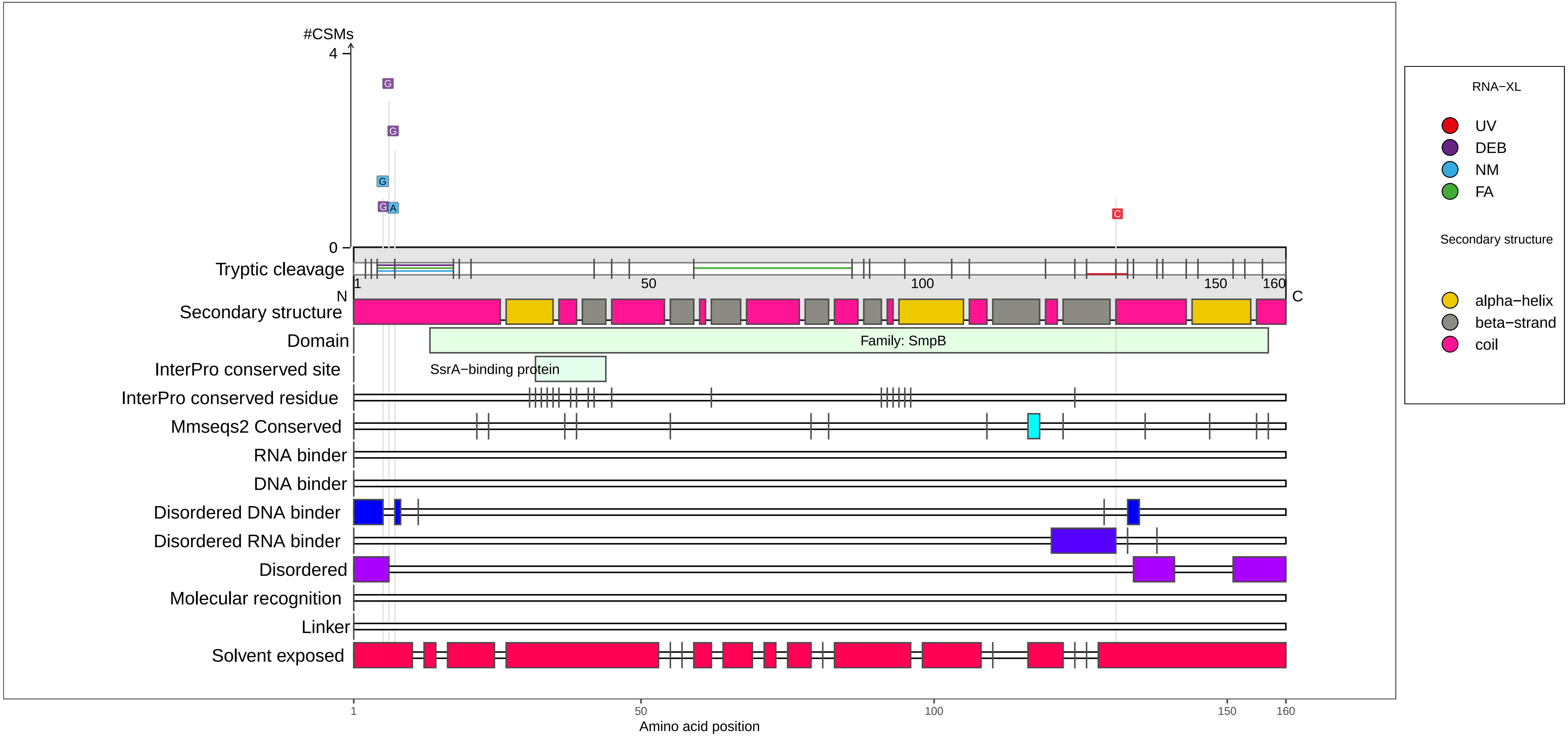

P0ACJ0
LRP_ECOLI Leucine−responsive regulatory protein

− Abundance:
tryptic [log10 Intensity]: 8.19 (Q 65)
PAXdb K12 strain [ppm]: 3.08 (Q 92)
PAXdb E.coli [ppm]: 3.05 (Q 95)

− RNA functions: not annotated

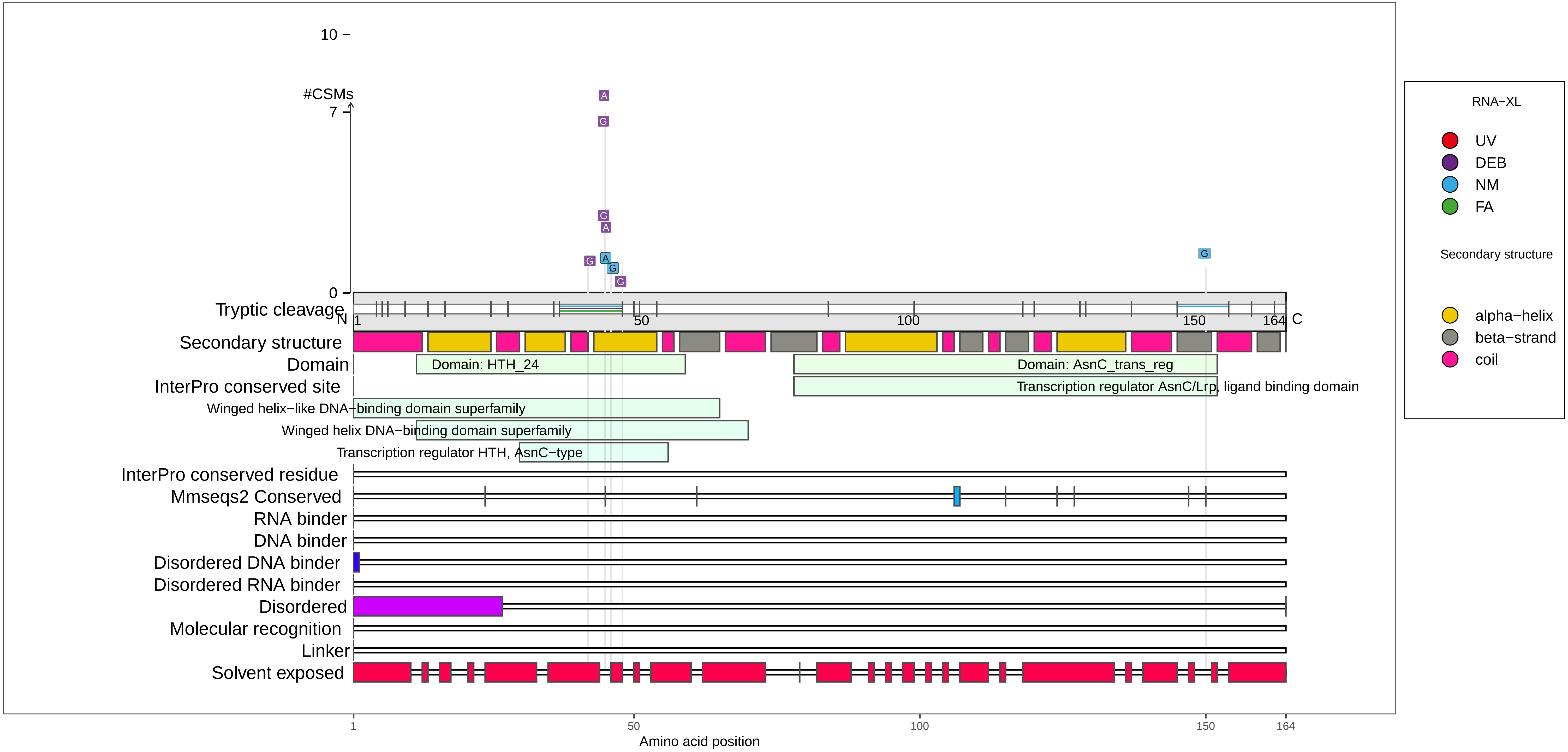

P13024
FDHE_ECOLI Protein FdhE

− Abundance:
tryptic [log10 Intensity]: 8.17 (Q 64)
PAXdb K12 strain [ppm]: 2.32 (Q 71)
PAXdb E.coli [ppm]: 1.72 (Q 65)

− RNA functions: not annotated

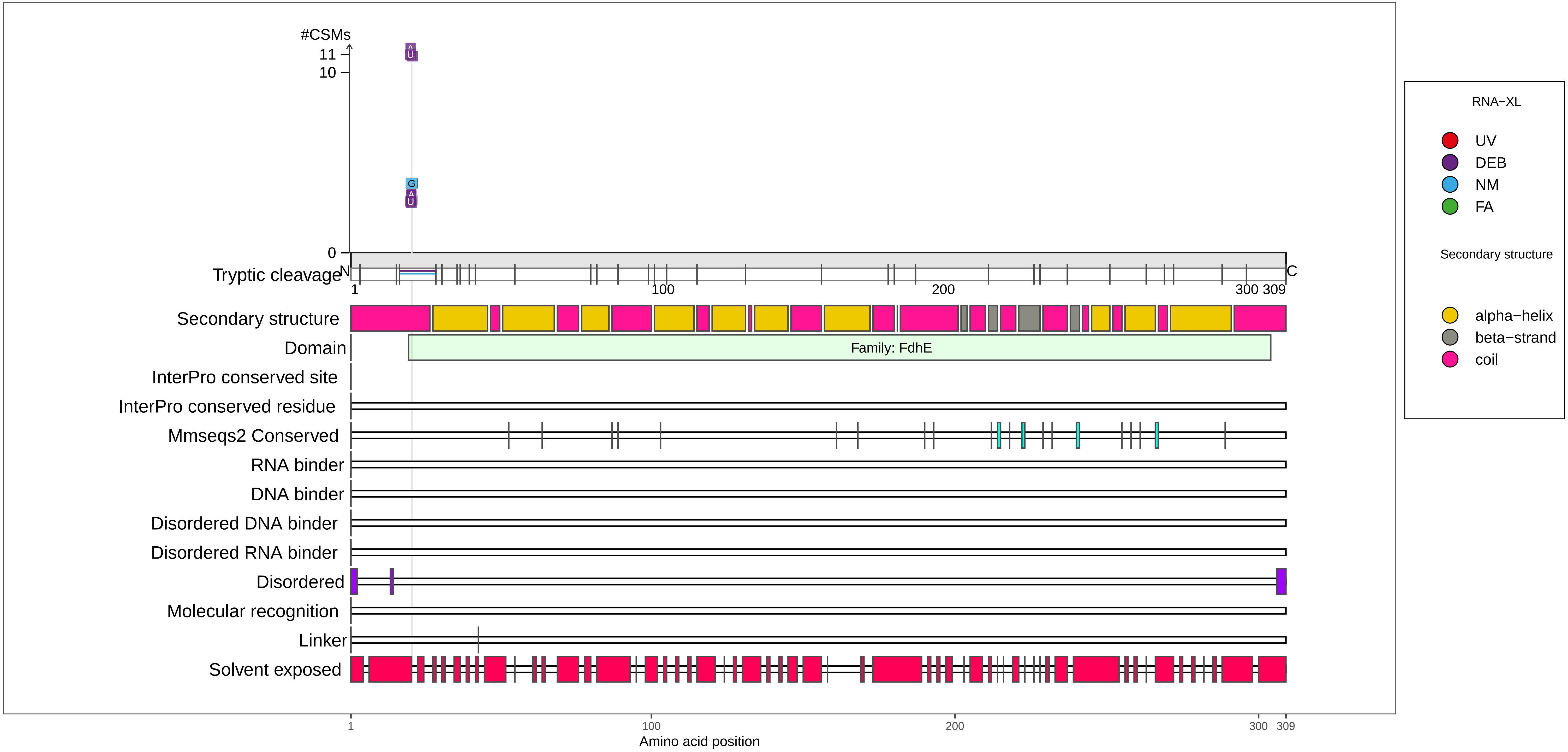

P24232
HMP_ECOLI Flavohemoprotein

− Abundance:
tryptic [log10 Intensity]: 8.14 (Q 63)
PAXdb K12 strain [ppm]: 1.52 (Q 39)
PAXdb E.coli [ppm]: 1.29 (Q 54)

− RNA functions: not annotated

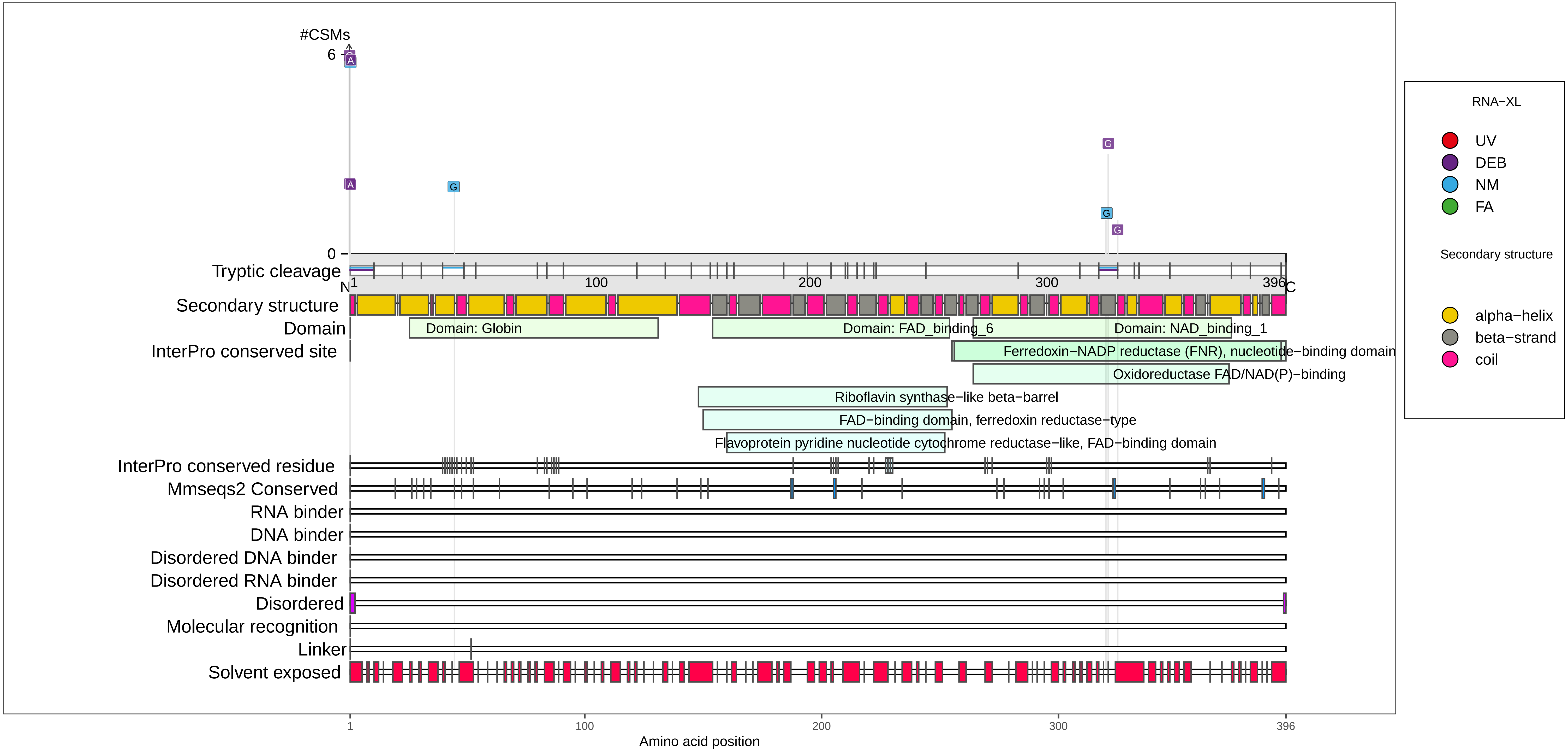

P63224
GMHA_ECOLI Phosphoheptose isomerase

− Abundance:
tryptic [log10 Intensity]: 9.17 (Q 92)
PAXdb K12 strain [ppm]: 2.45 (Q 76)
PAXdb E.coli [ppm]: 2.93 (Q 94)

− RNA functions: not annotated

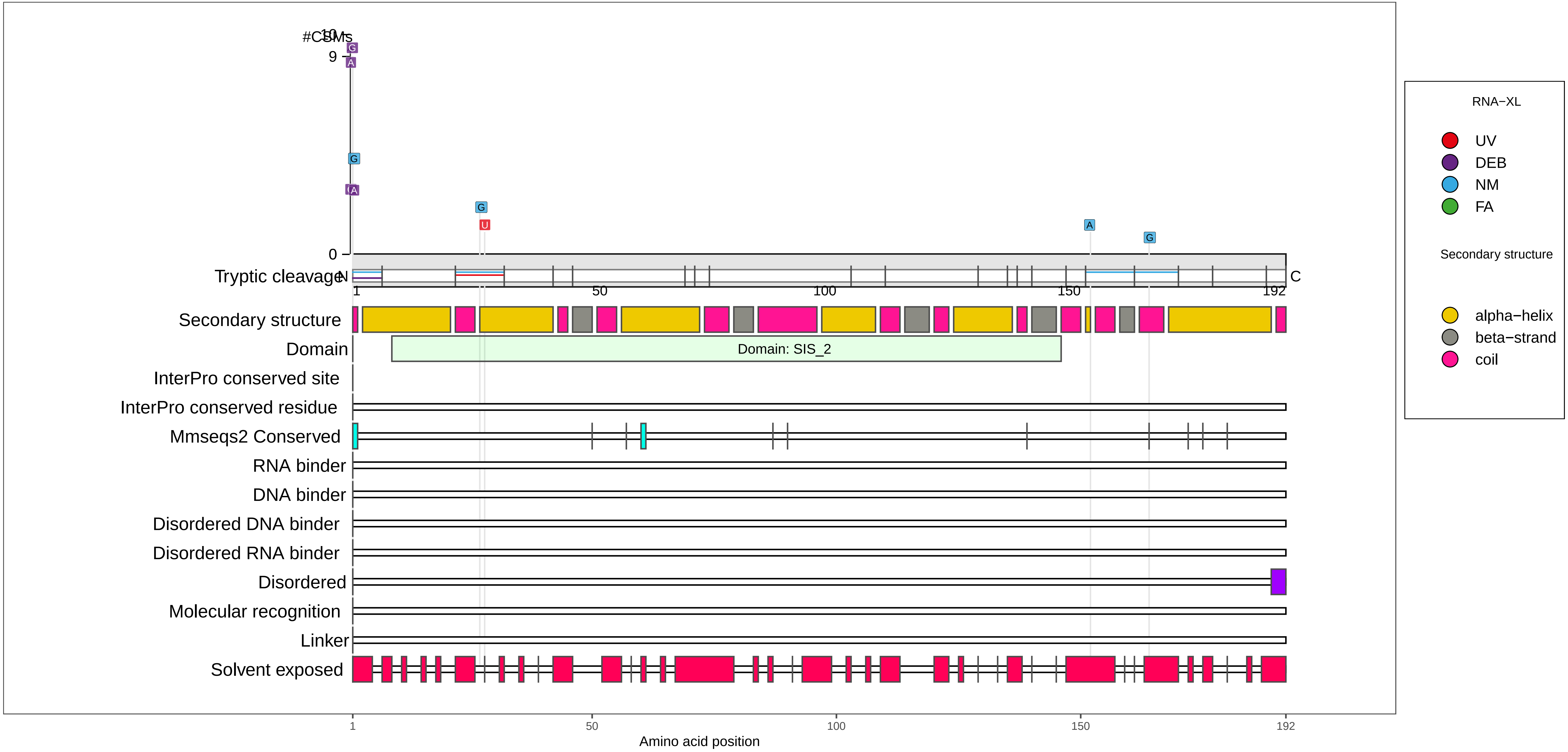

P76576
YFGM_ECOLI Ancillary SecYEG translocon subunit

− Abundance:
tryptic [log10 Intensity]: 8.73 (Q 82)
PAXdb K12 strain [ppm]: 1.9 (Q 57)
PAXdb E.coli [ppm]: 2.49 (Q 85)

− RNA functions: not annotated

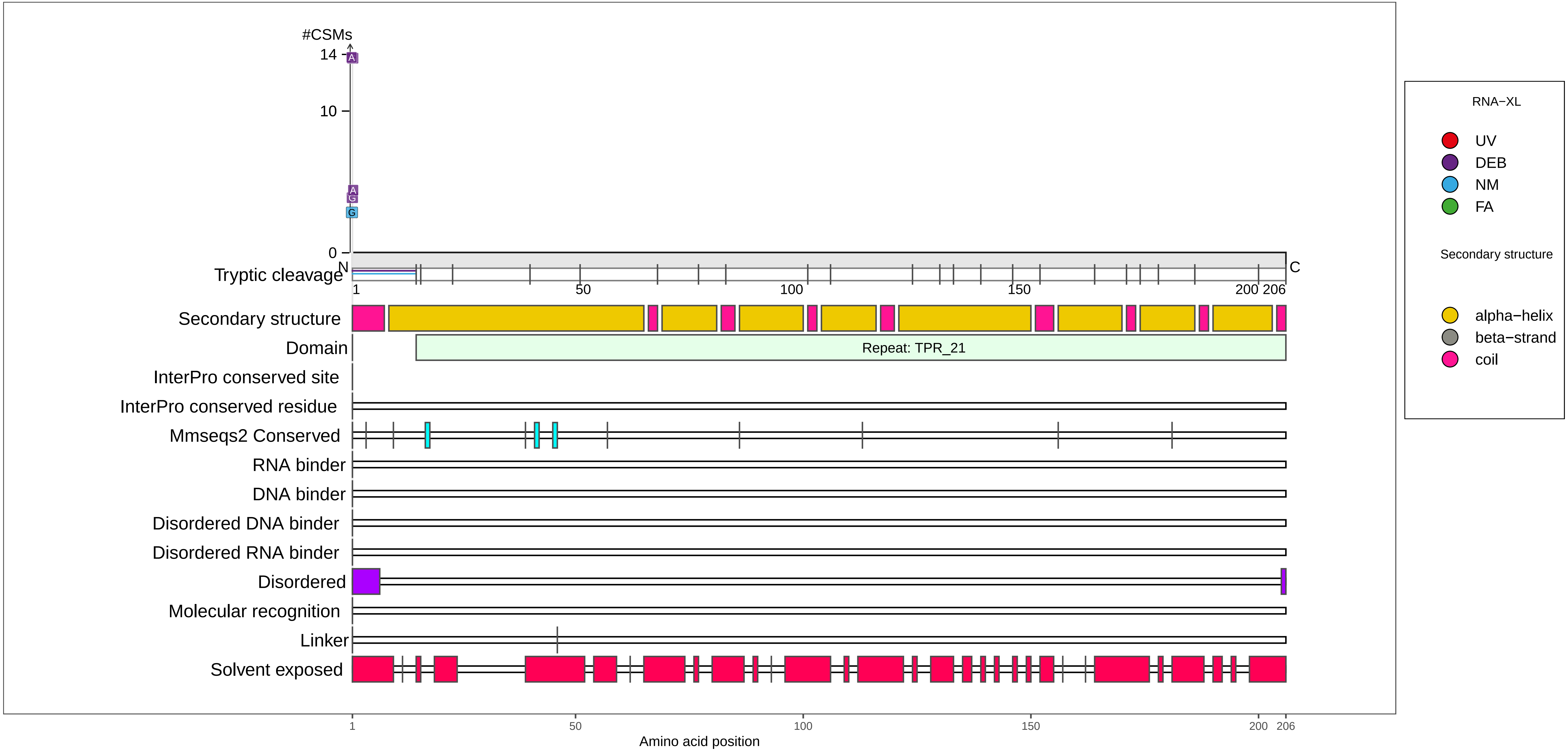

P0A6T9
GCSH_ECOLI Glycine cleavage system H protein

− Abundance:
tryptic [log10 Intensity]: 8.24 (Q 67)
PAXdb K12 strain [ppm]: 3.67 (Q 98)
PAXdb E.coli [ppm]: 2.59 (Q 88)

− RNA functions: not annotated

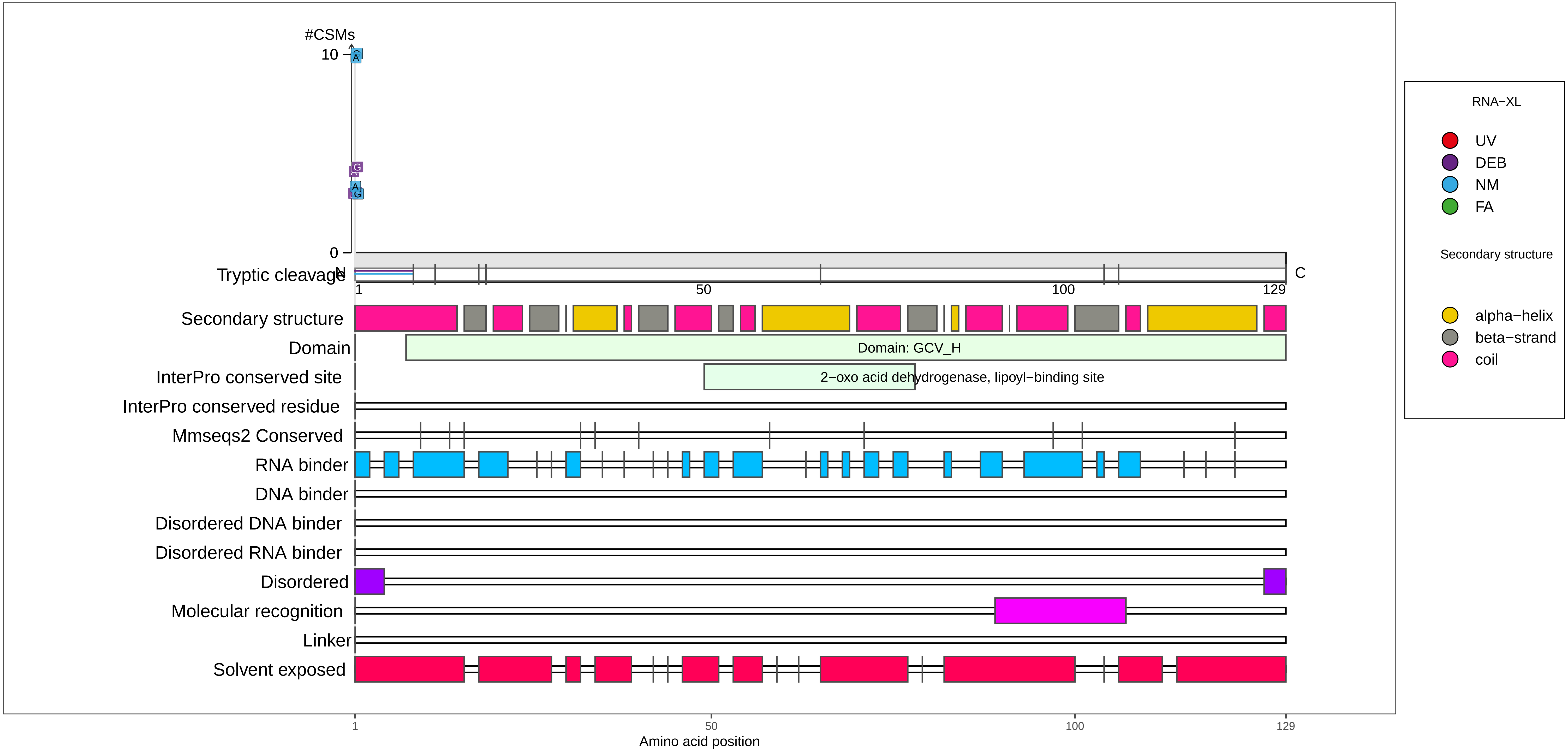

P0A6Z3
HTPG_ECOLI Chaperone protein HtpG

− Abundance:
tryptic [log10 Intensity]: 9.75 (Q 98)
PAXdb K12 strain [ppm]: 3.16 (Q 93)
PAXdb E.coli [ppm]: 2.67 (Q 89)

− RNA functions: not annotated

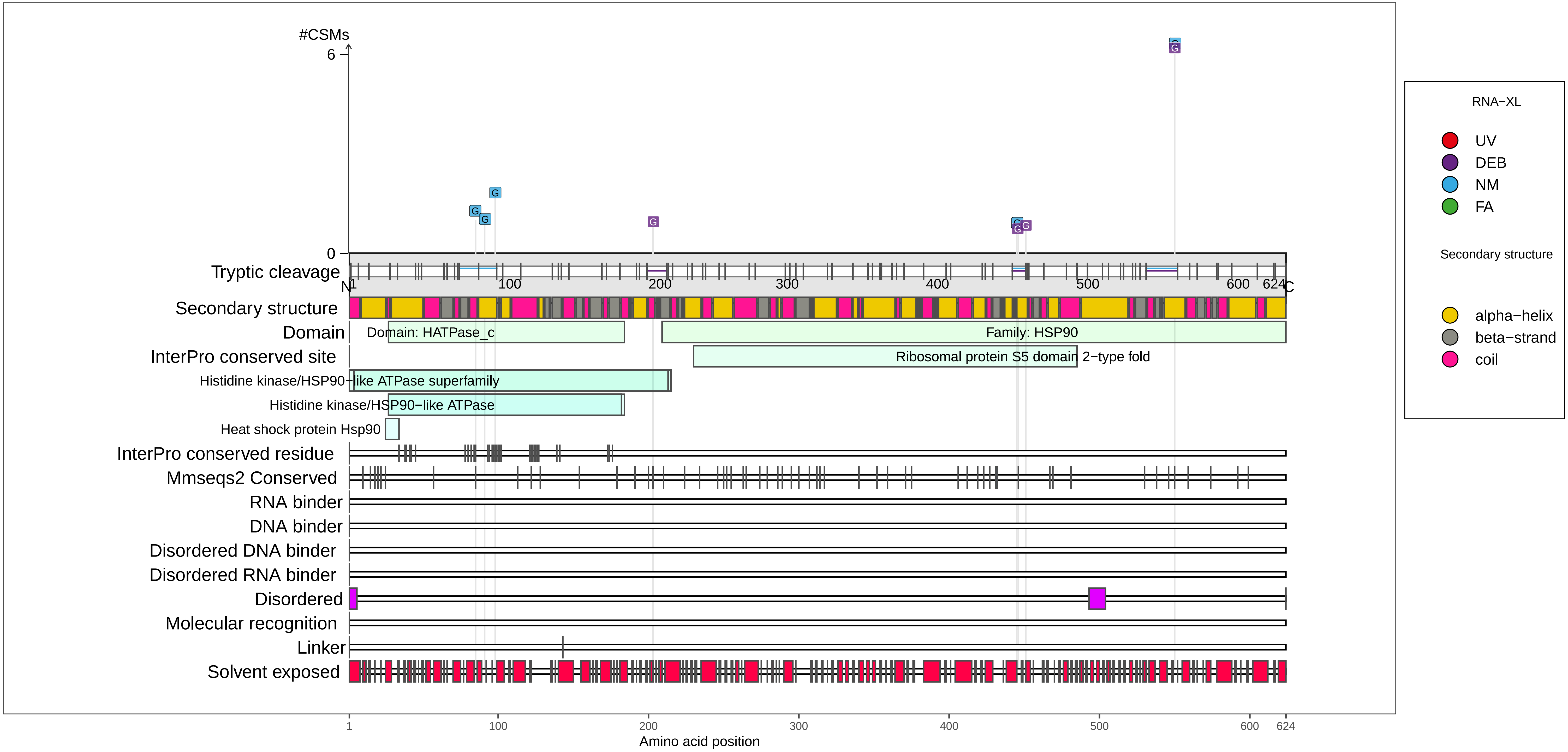

P0A959
ALAA_ECOLI Glutamate−pyruvate aminotransferase AlaA

− Abundance:
tryptic [log10 Intensity]: 7 (Q 13)
PAXdb K12 strain [ppm]: 2.18 (Q 66)
PAXdb E.coli [ppm]: 2.04 (Q 73)

− RNA functions: not annotated

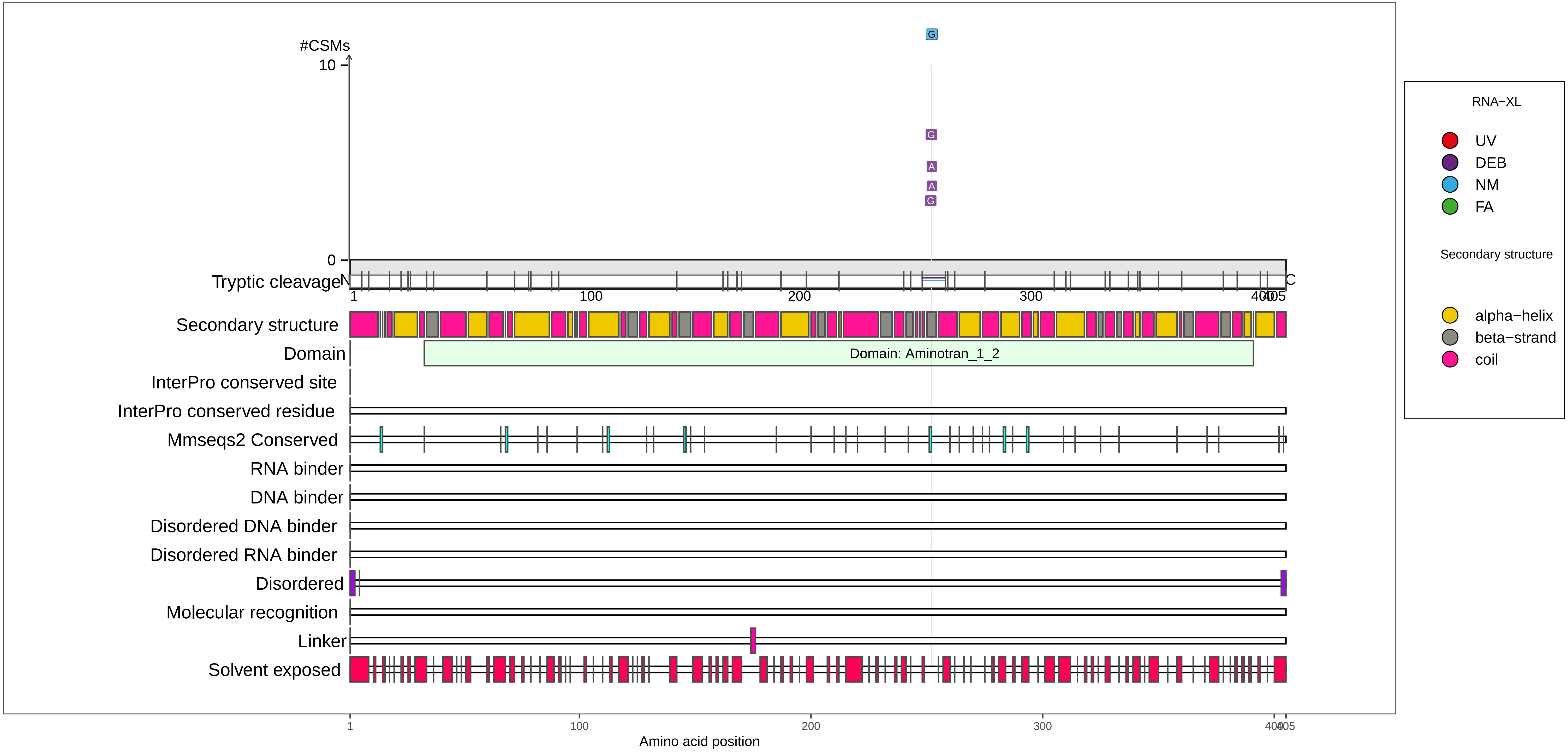

P0ABC7
HFLK_ECOLI Modulator of FtsH protease HflK

− Abundance:
tryptic [log10 Intensity]: 8.64 (Q 80)
PAXdb K12 strain [ppm]: 2.48 (Q 77)
PAXdb E.coli [ppm]: 2.59 (Q 88)

− RNA functions: not annotated

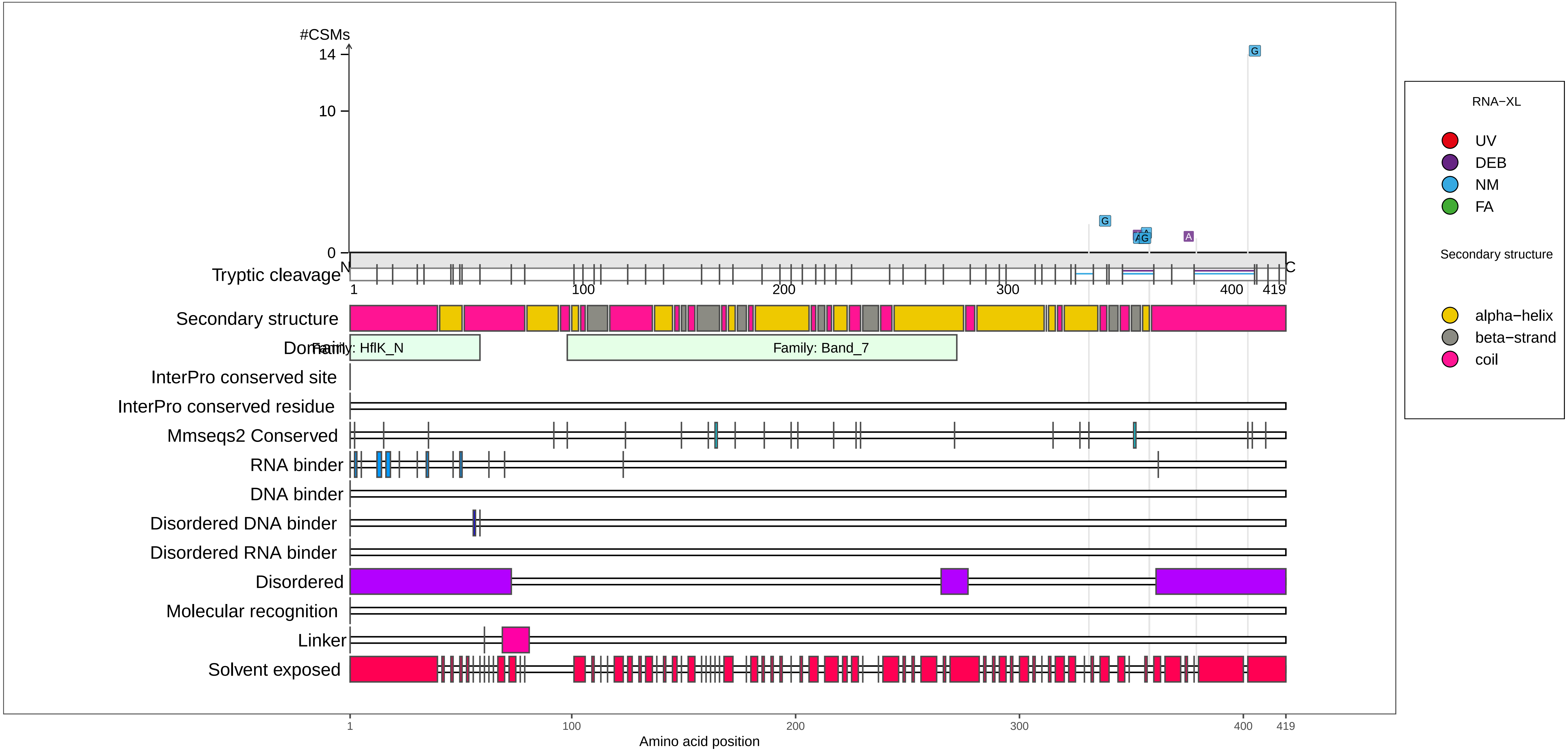

P0ABH9
CLPA_ECOLI ATP−dependent Clp protease ATP−binding subunit ClpA

− Abundance:
tryptic [log10 Intensity]: 7.88 (Q 53)
PAXdb K12 strain [ppm]: 2.45 (Q 76)
PAXdb E.coli [ppm]: 2.72 (Q 90)

− RNA functions: not annotated

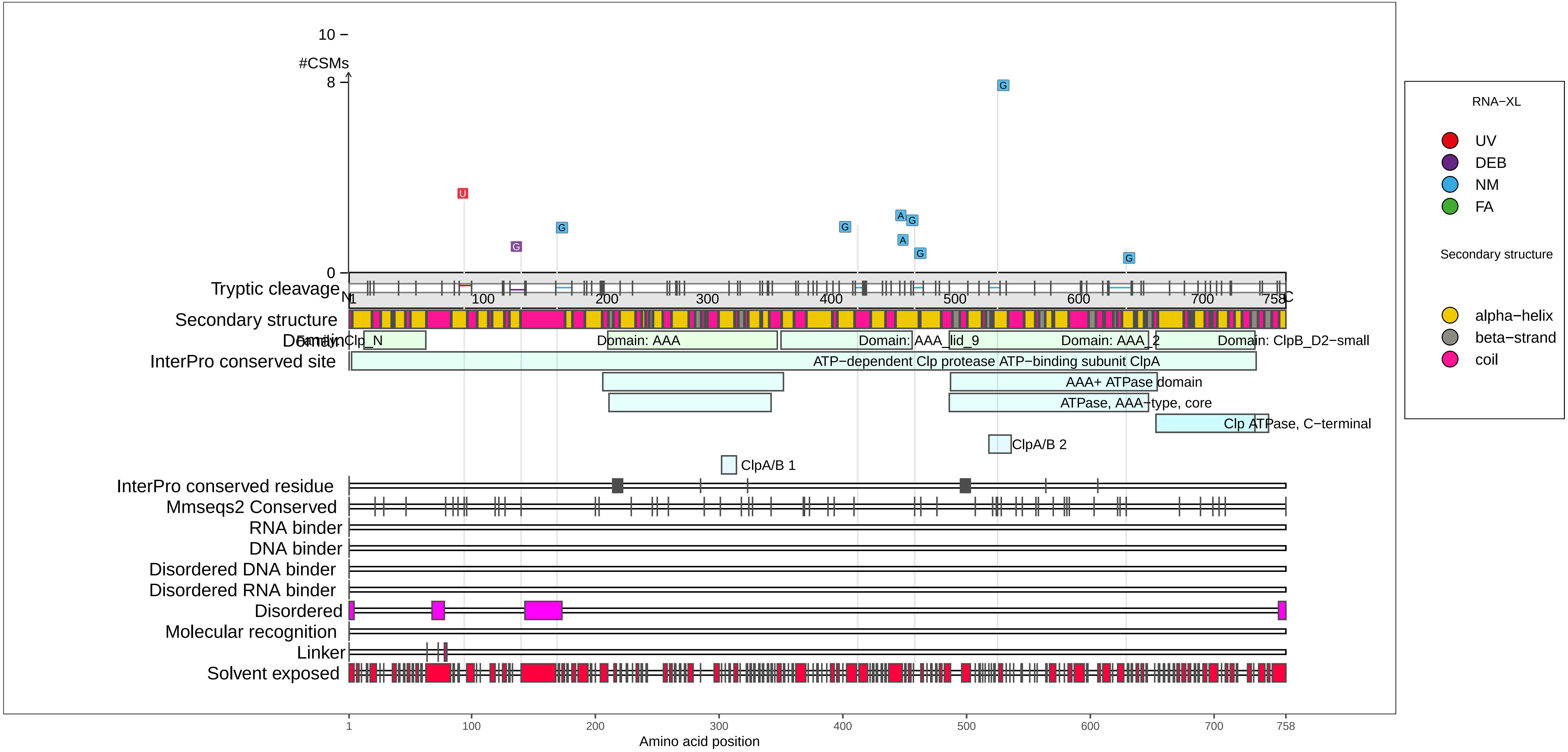

P31224
ACRB_ECOLI Multidrug efflux pump subunit AcrB
− Abundance:
tryptic [log10 Intensity]: 9.03 (Q 89)
PAXdb K12 strain [ppm]: NA (Q NA)
PAXdb E.coli [ppm]: 1.72 (Q 65)

− RNA functions: not annotated

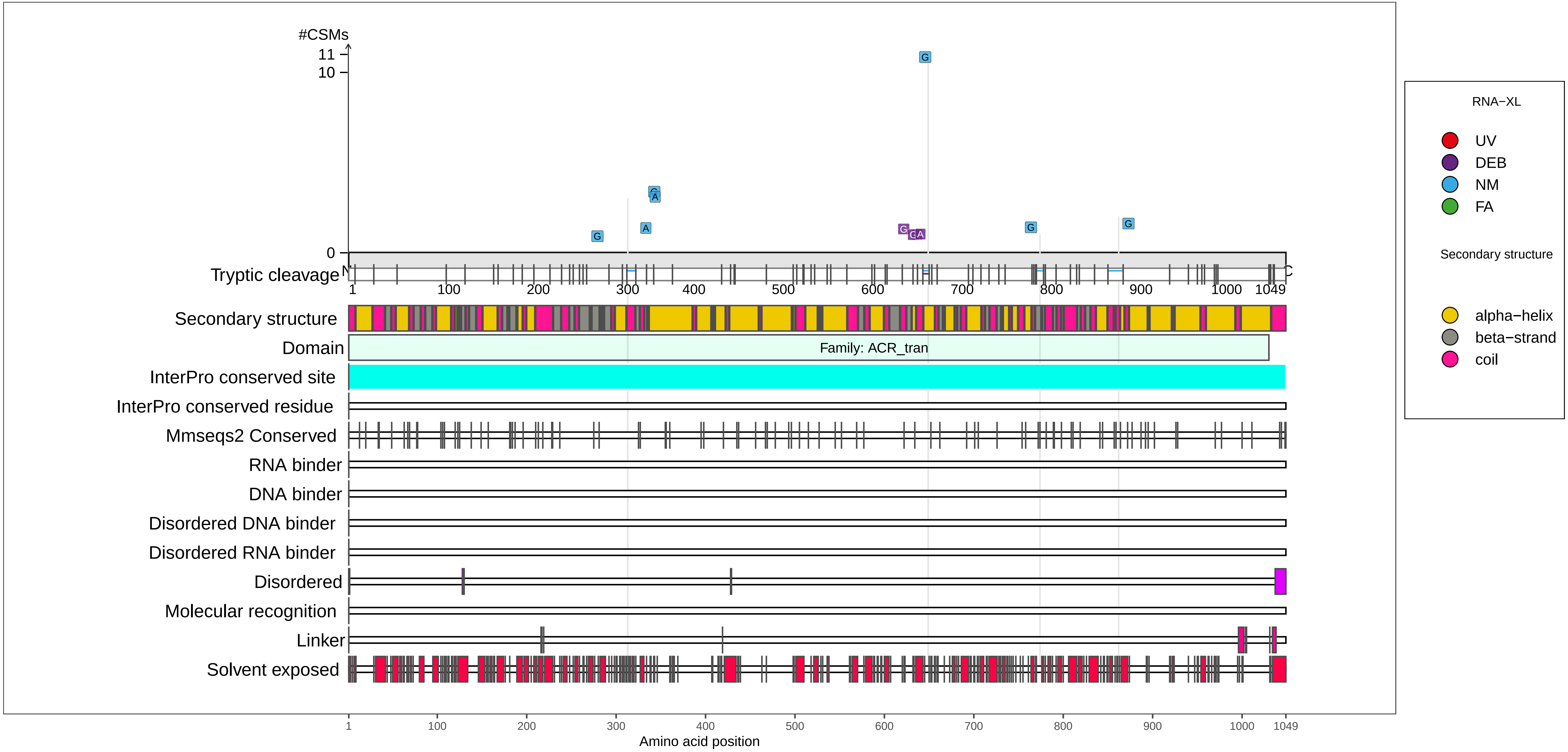

P33136
OPGG_ECOLI Glucans biosynthesis protein G

− Abundance:
tryptic [log10 Intensity]: 8.39 (Q 72)
PAXdb K12 strain [ppm]: 2.29 (Q 69)
PAXdb E.coli [ppm]: 2.43 (Q 84)

− RNA functions: not annotated

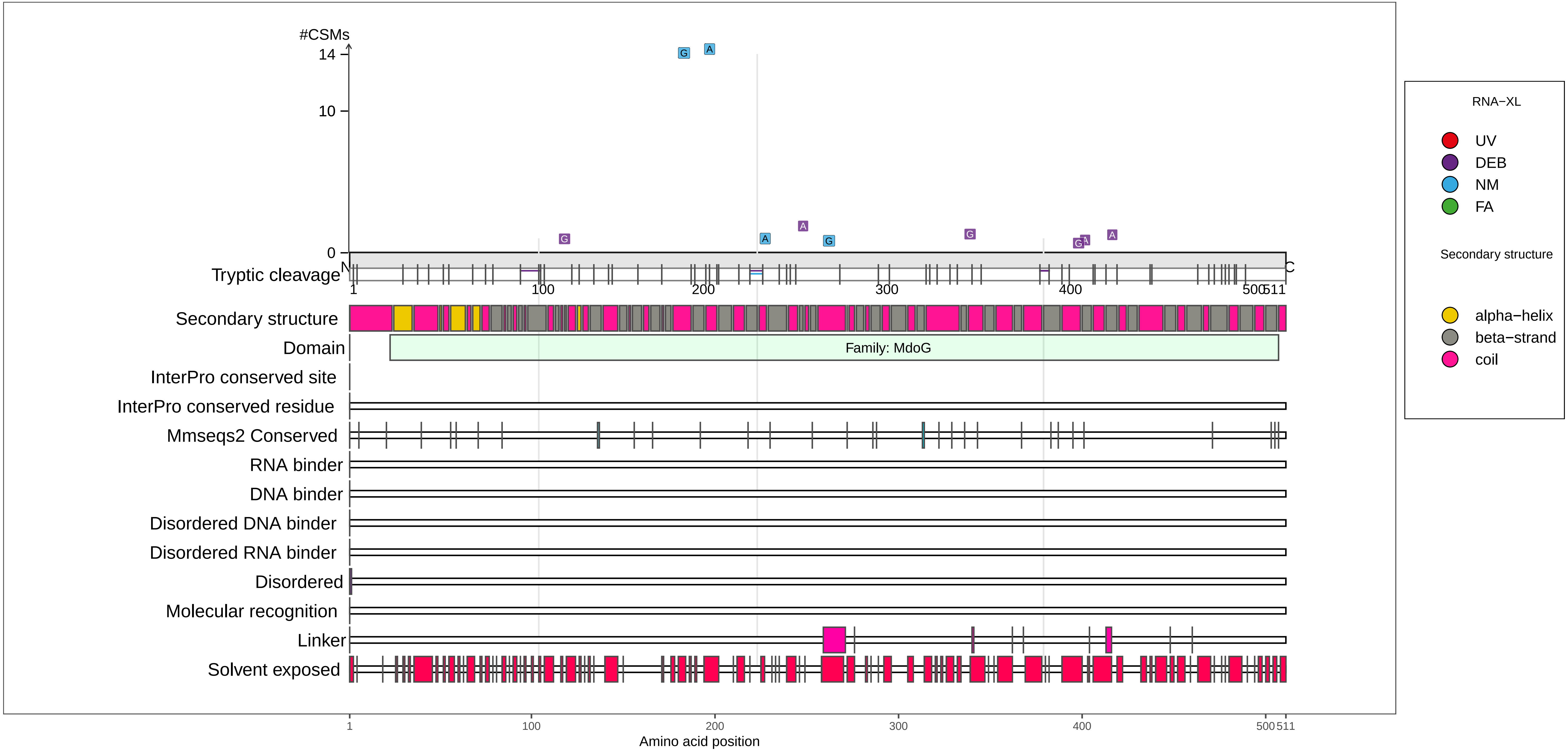

P62768
YAEH_ECOLI UPF0325 protein YaeH

− Abundance:
tryptic [log10 Intensity]: 7.67 (Q 43)
PAXdb K12 strain [ppm]: 3.06 (Q 92)
PAXdb E.coli [ppm]: 3.13 (Q 96)

− RNA functions: not annotated

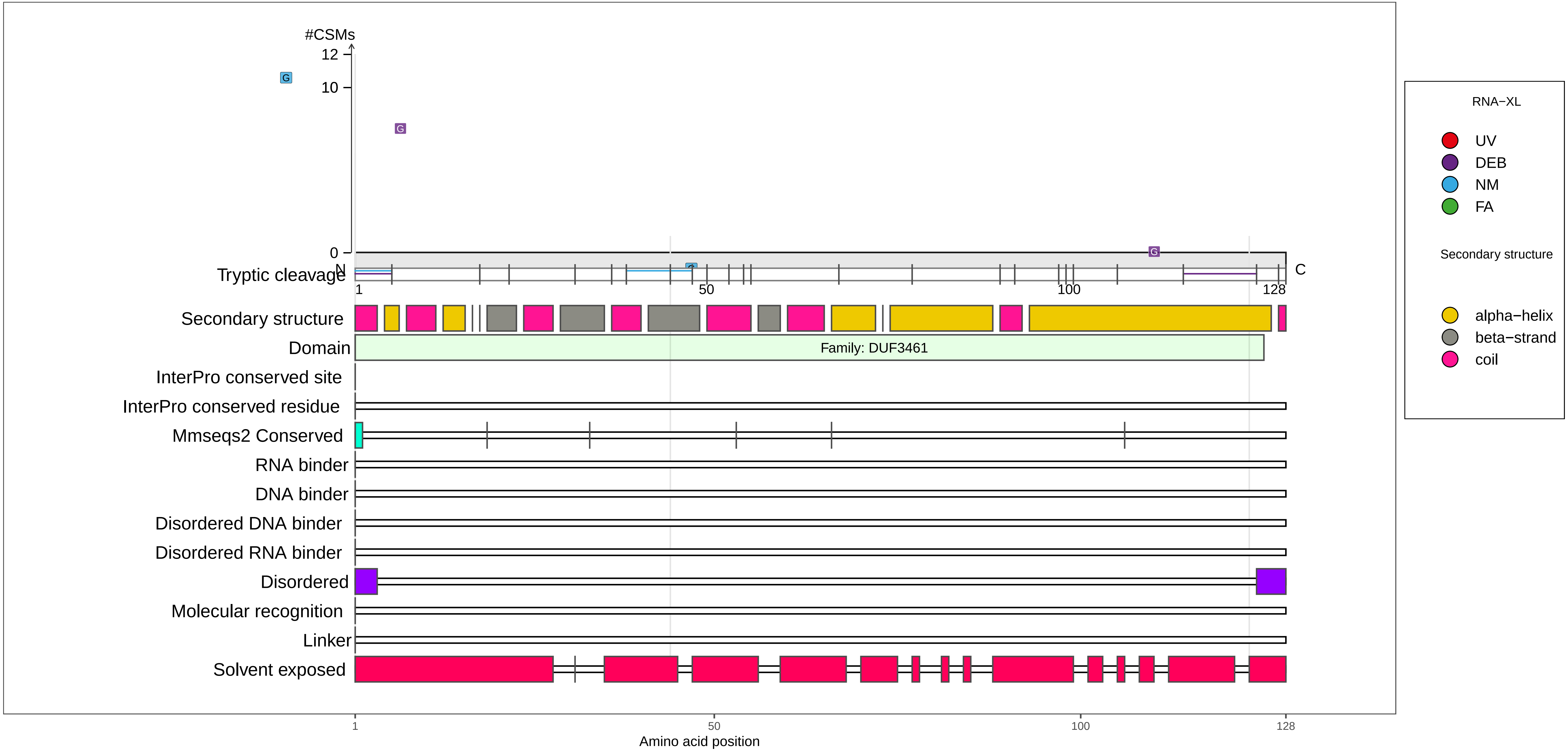

P77488

DXS_ECOLI 1−deoxy−D−xylulose−5−phosphate synthase

− Abundance:
tryptic [log10 Intensity]: 7.48 (Q 34)
PAXdb K12 strain [ppm]: 1.95 (Q 59)
PAXdb E.coli [ppm]: 2.25 (Q 79)

− RNA functions: not annotated

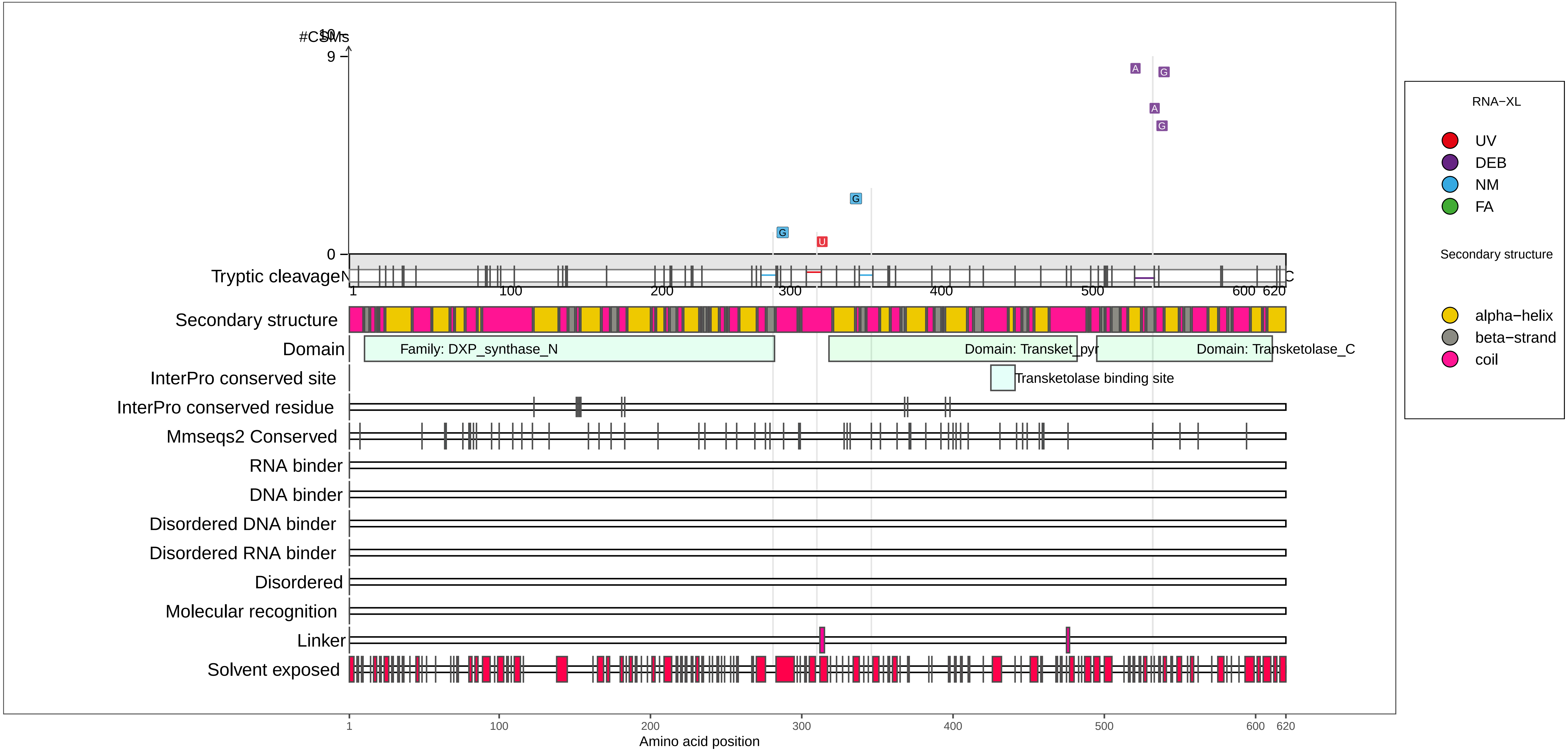

P00448
SODM_ECOLI Superoxide dismutase [Mn]

− Abundance:
tryptic [log10 Intensity]: 9.02 (Q 89)
PAXdb K12 strain [ppm]: 2.41 (Q 74)
PAXdb E.coli [ppm]: 3.73 (Q 100)

− RNA functions: not annotated

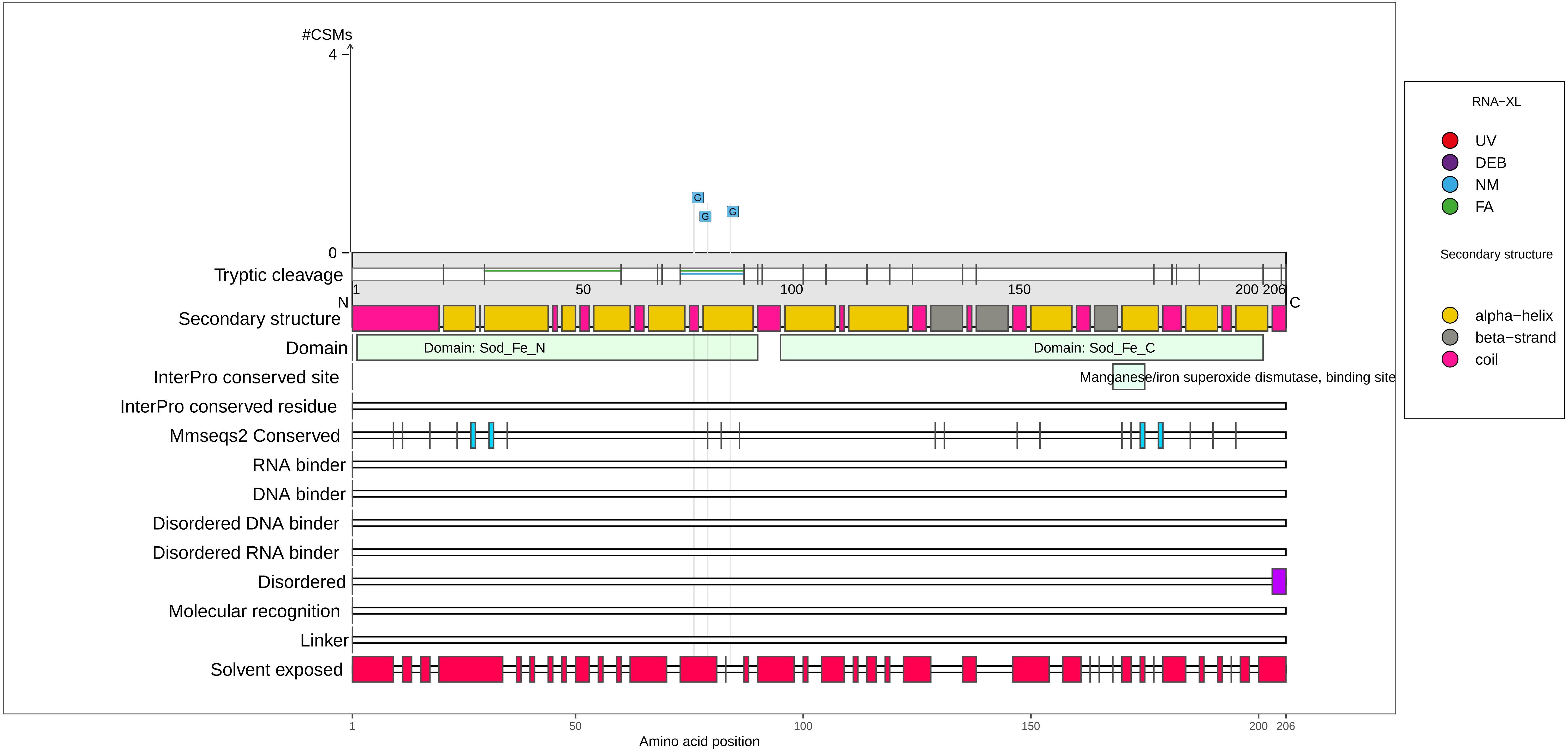

P00946
MANA_ECOLI Mannose−6−phosphate isomerase

− Abundance:
tryptic [log10 Intensity]: 8.17 (Q 64)
PAXdb K12 strain [ppm]: 2.78 (Q 85)
PAXdb E.coli [ppm]: 2.56 (Q 87)

− RNA functions: not annotated

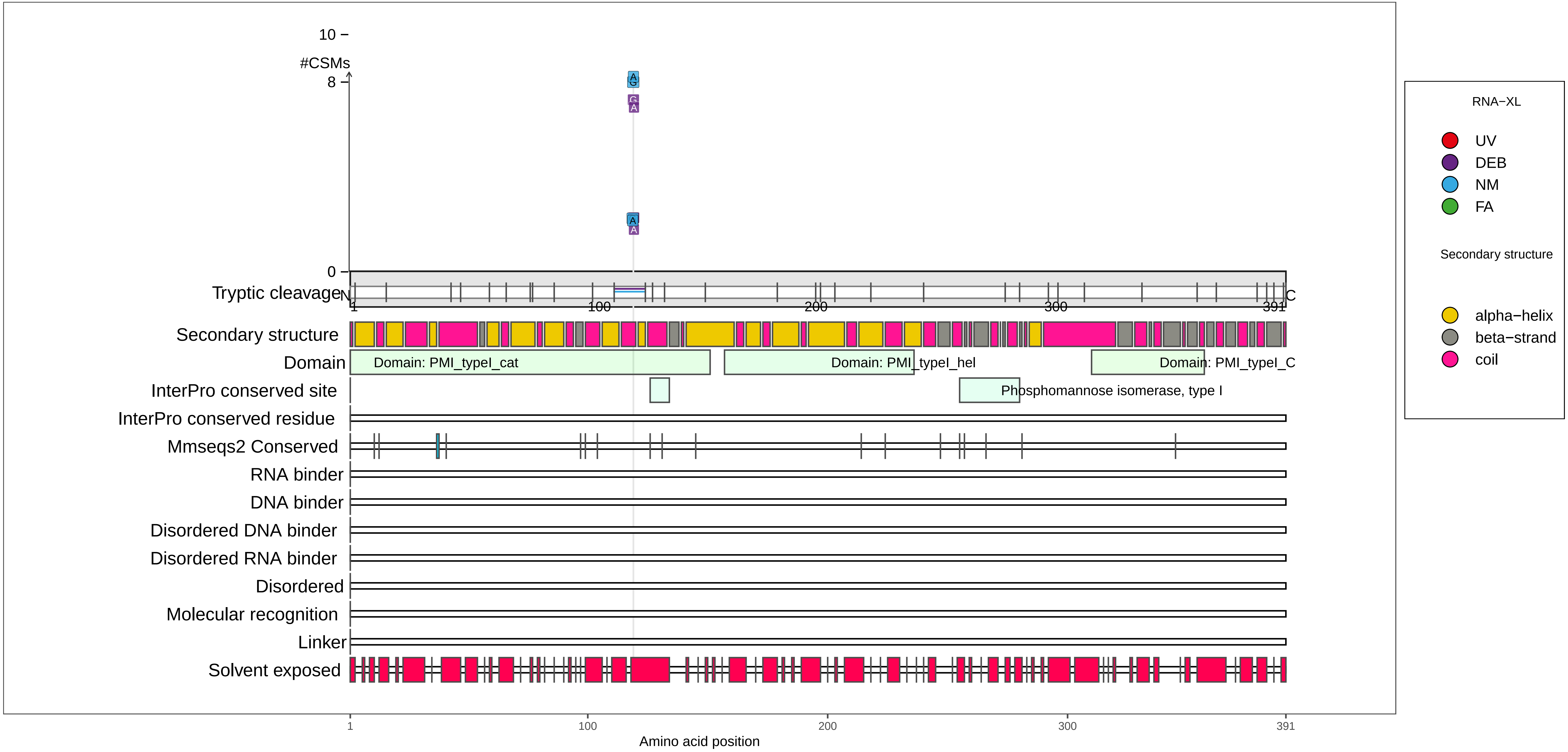

P04079
GUAA_ECOLI GMP synthase [glutamine−hydrolyzing]

− Abundance:
tryptic [log10 Intensity]: 9.46 (Q 96)
PAxdb K12 strain [ppm]: 1.94 (Q 58)
PAxdb E.coli [ppm]: 3.05 (Q 95)

− RNA functions:
tRNA methyl transferase

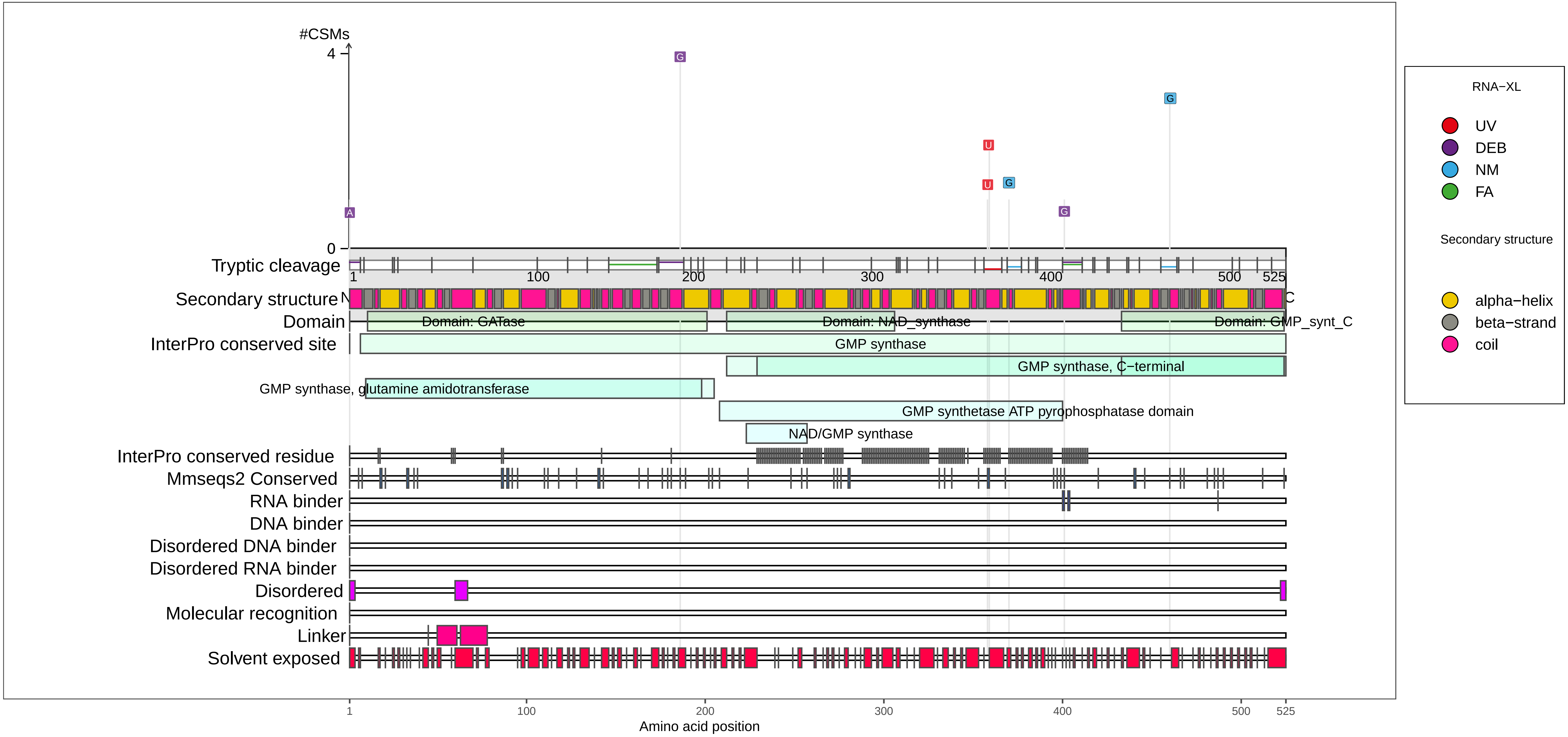

P0A7Z0
RPIA_ECOLI Ribose−5−phosphate isomerase A

− Abundance:
tryptic [log10 Intensity]: 8.6 (Q 79)
PAXdb K12 strain [ppm]: 3.49 (Q 97)
PAXdb E.coli [ppm]: 2.85 (Q 93)

− RNA functions: not annotated

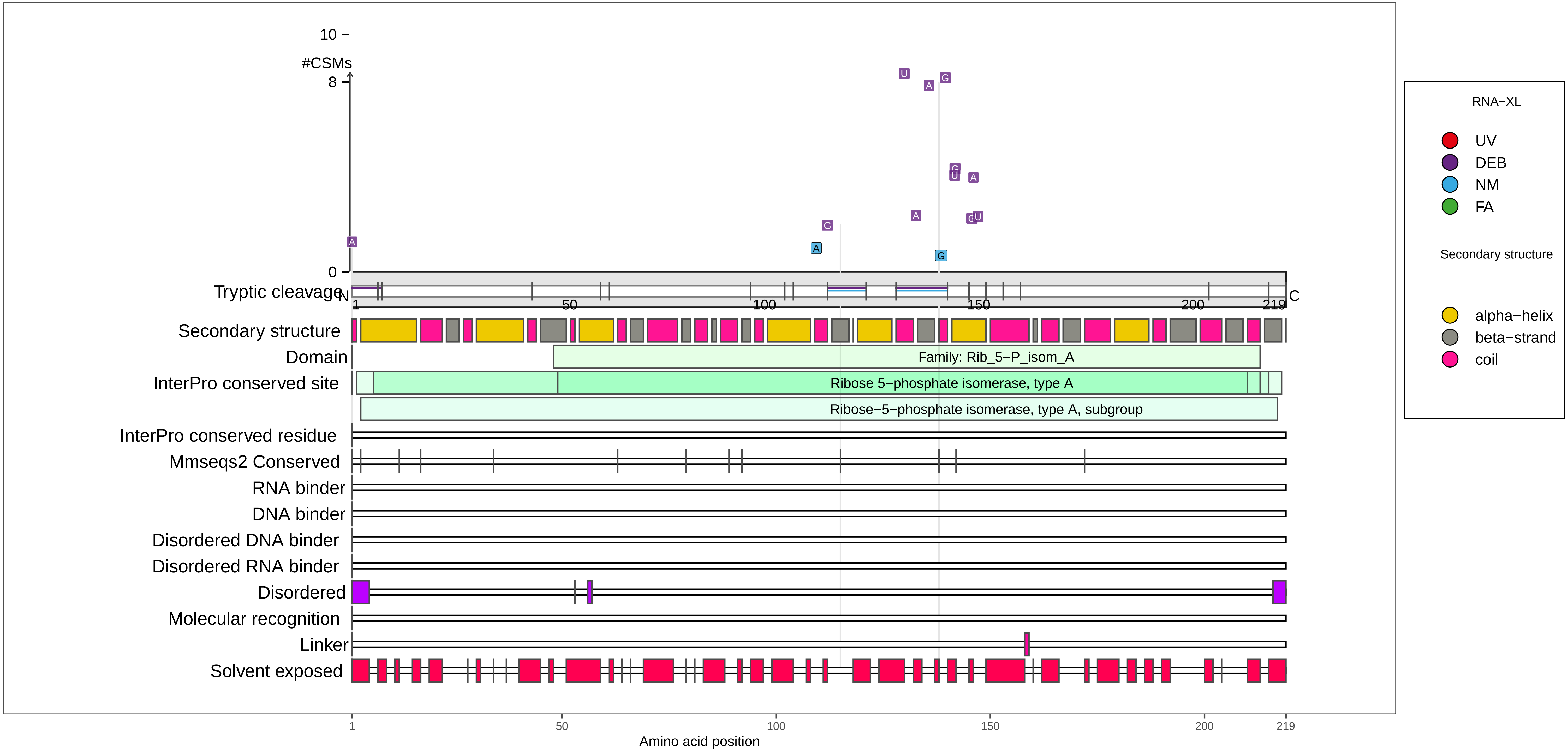

P0ADC1
LPTE_ECOLI LPS−assembly lipoprotein LptE

− Abundance:
tryptic [log10 Intensity]: 9.23 (Q 93)
PAXdb K12 strain [ppm]: 2.44 (Q 76)
PAXdb E.coli [ppm]: 2.05 (Q 73)

− RNA functions: not annotated

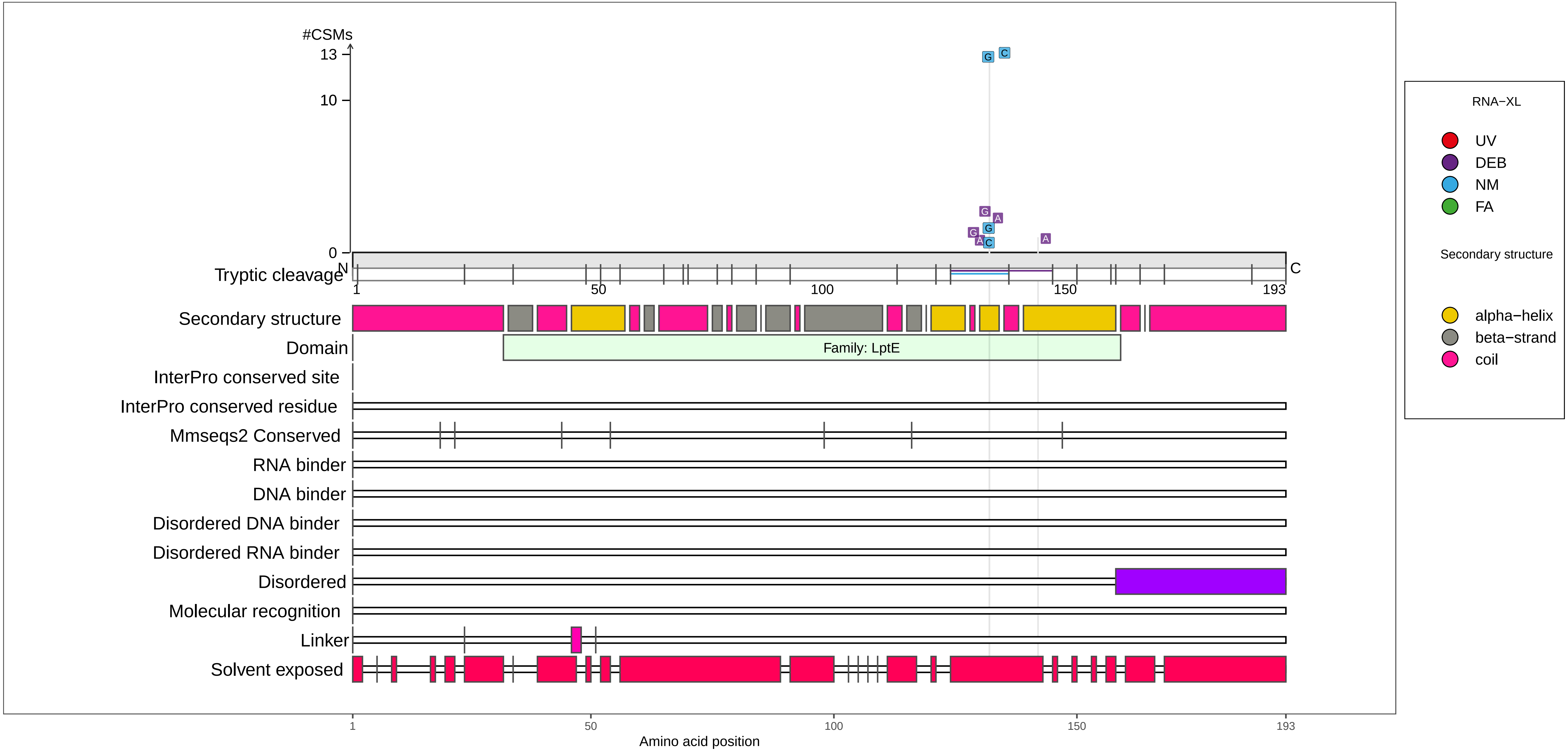

P0AFC7
NUOB_ECOLI NADH−quinone oxidoreductase subunit B

− Abundance:
tryptic [log10 Intensity]: 8.47 (Q 74)
PAXdb K12 strain [ppm]: 1.95 (Q 59)
PAXdb E.coli [ppm]: 2.59 (Q 88)

− RNA functions: not annotated

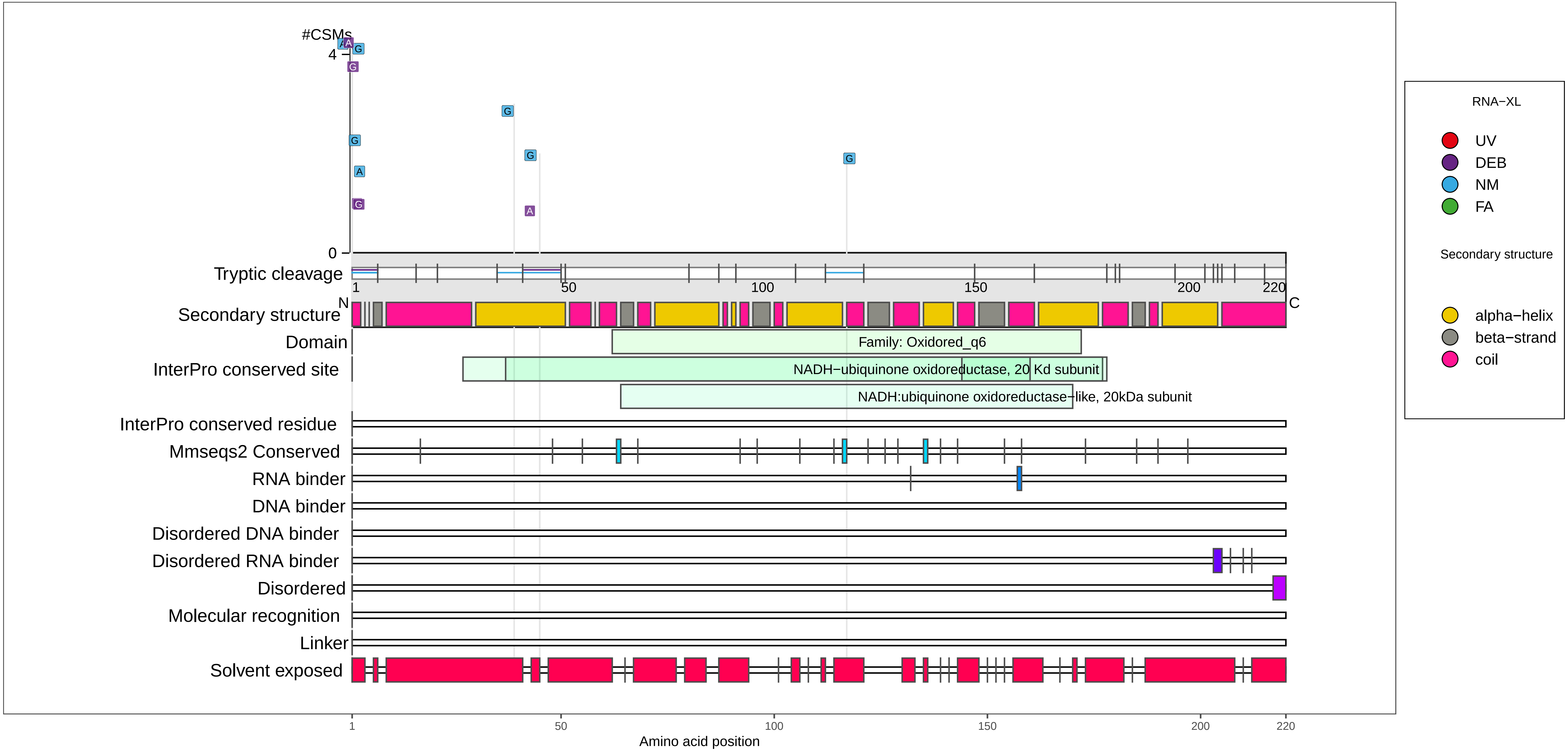

P0AFD6
NUOI_ECOLI NADH−quinone oxidoreductase subunit I

− Abundance:
tryptic [log10 Intensity]: 8.56 (Q 77)
PAXdb K12 strain [ppm]: 2.96 (Q 90)
PAXdb E.coli [ppm]: 2.45 (Q 84)

− RNA functions: not annotated

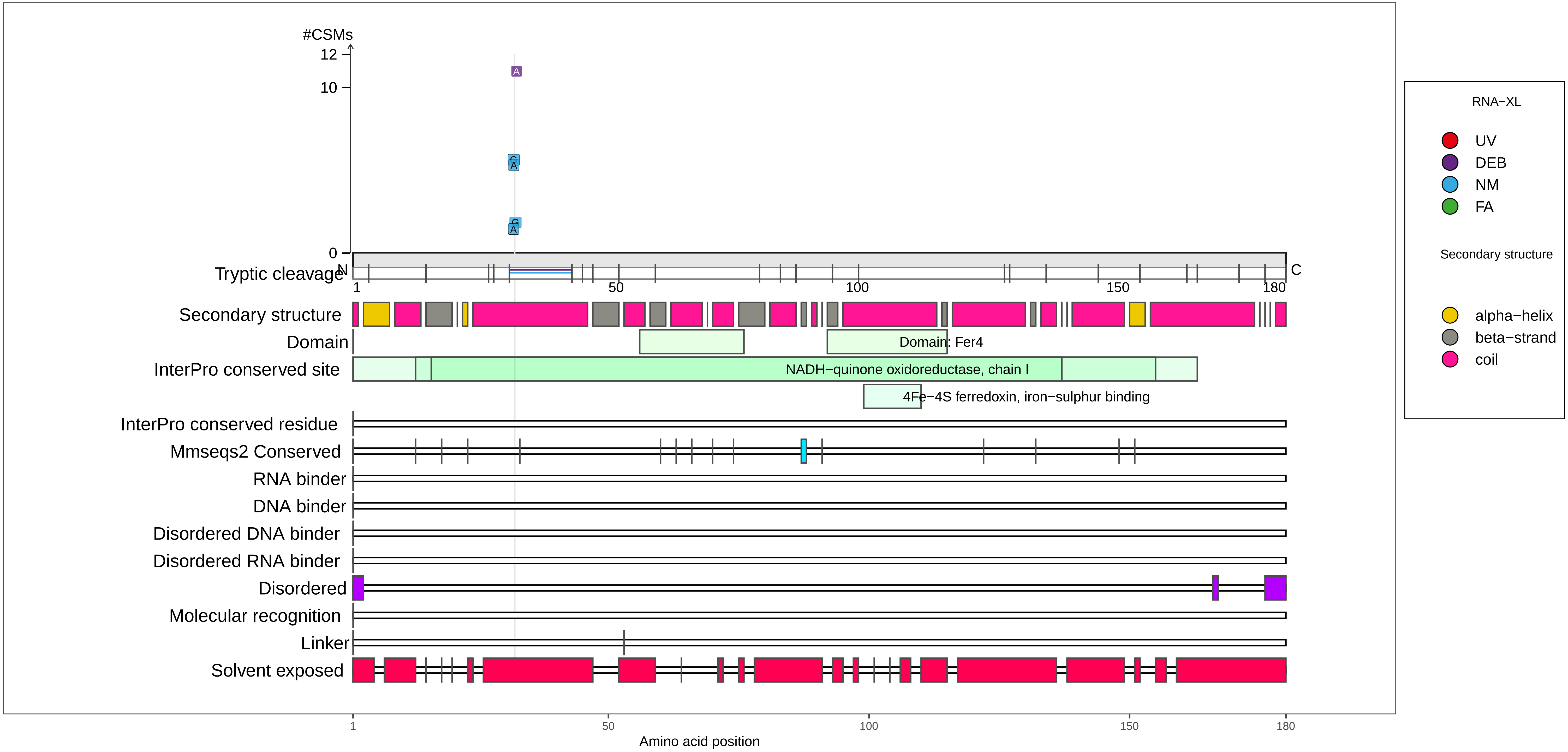

P0C8J8
GATZ_ECOLI D−tagatose−1,6−bisphosphate aldolase subunit GatZ

− Abundance:
tryptic [log10 Intensity]: 9.49 (Q 96)
PAXdb K12 strain [ppm]: 2.66 (Q 82)
PAXdb E.coli [ppm]: 3.48 (Q 99)

− RNA functions: not annotated

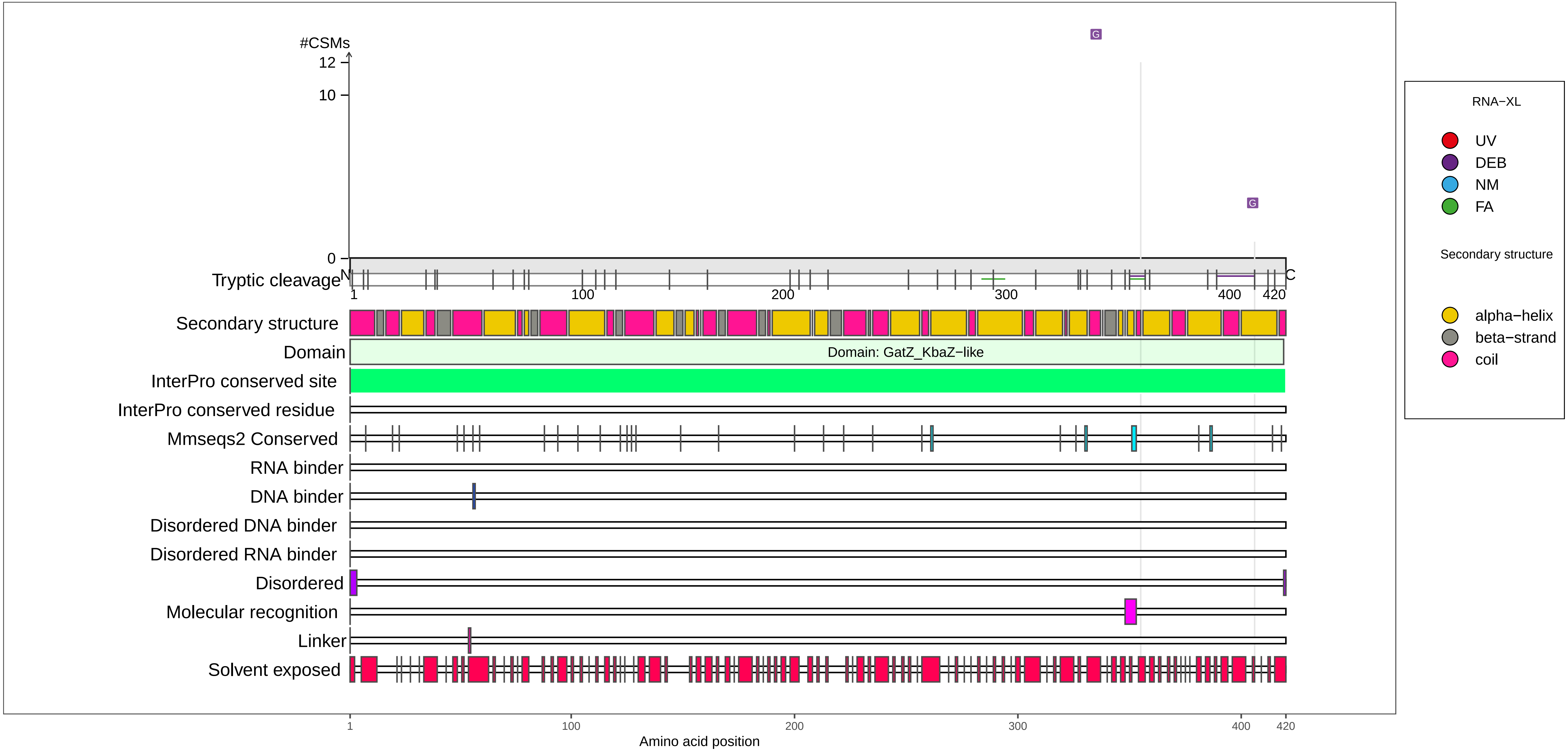

P15639
PUR9_ECOLI Bifunctional purine biosynthesis protein PurH

− Abundance:
tryptic [log10 Intensity]: 7.83 (Q 51)
PAXdb K12 strain [ppm]: 2.79 (Q 86)
PAXdb E.coli [ppm]: 2.55 (Q 87)

− RNA functions: not annotated

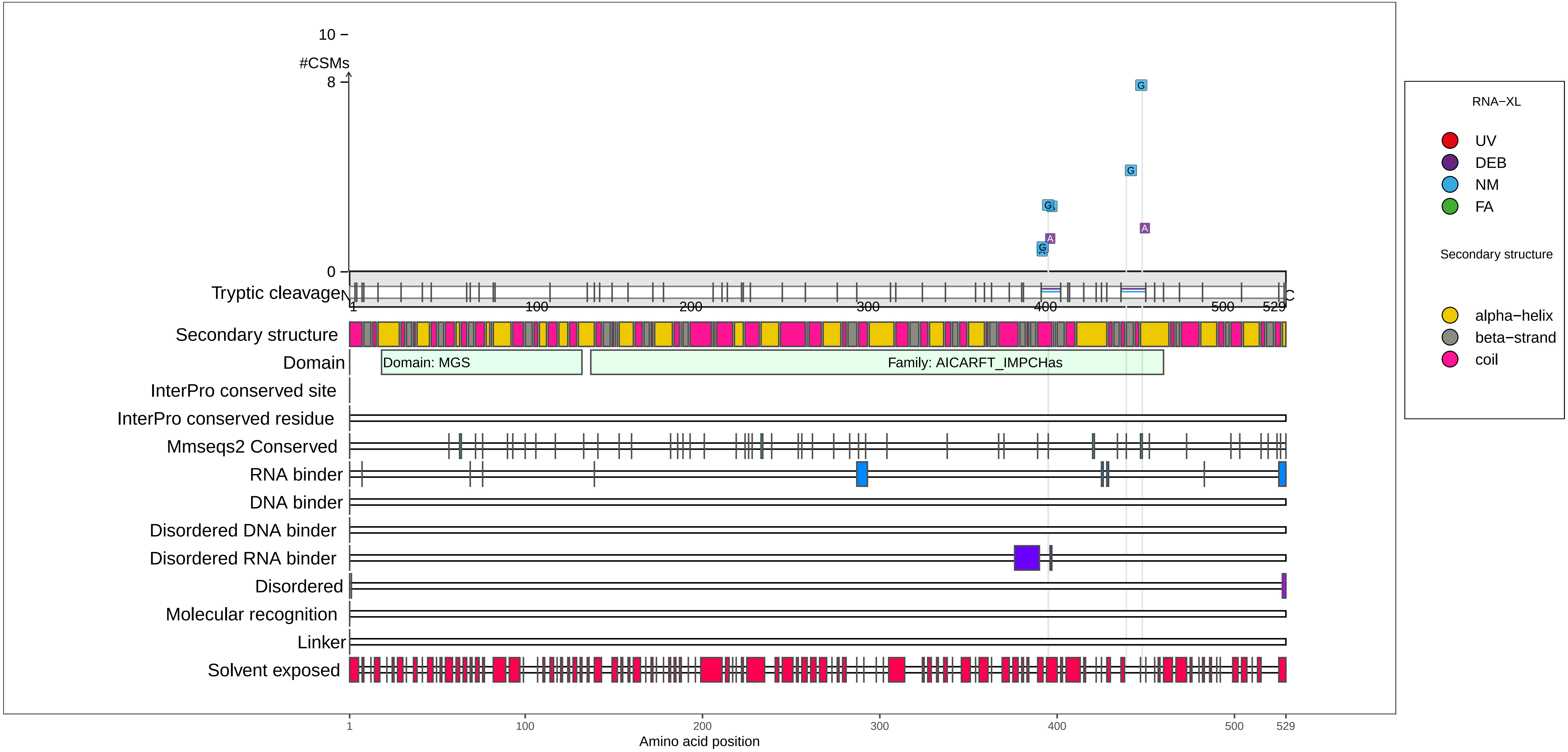

P68206
YJBJ_ECOLI UPF0337 protein YjbJ

− Abundance:
tryptic [log10 Intensity]: NA (Q NA)
PAXdb K12 strain [ppm]: 4.46 (Q 100)
PAXdb E.coli [ppm]: 3.07 (Q 96)

− RNA functions: not annotated

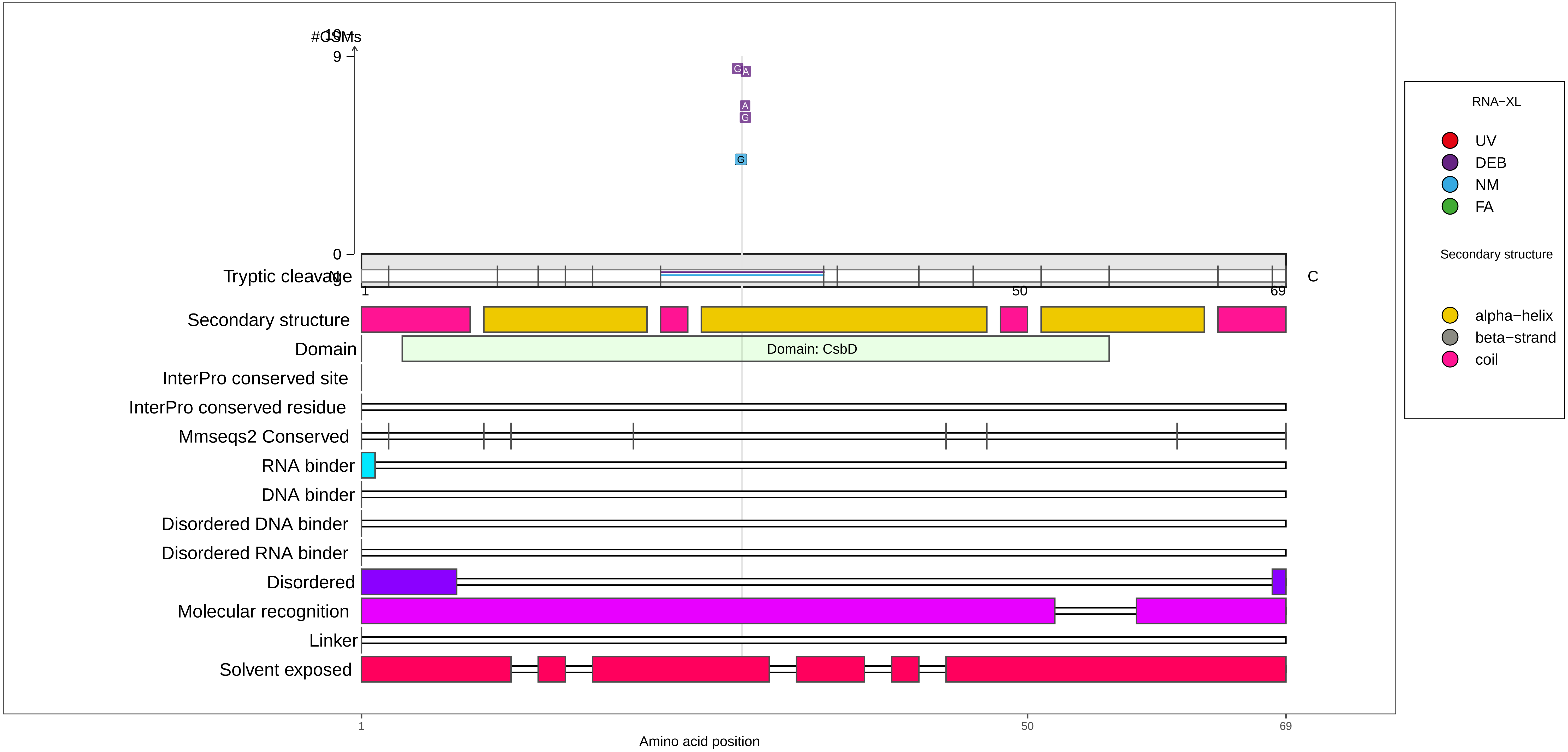

P03018
UVRD_ECOLI DNA helicase II

− Abundance:
tryptic [log10 Intensity]: 7.78 (Q 48)
PAXdb K12 strain [ppm]: 1.83 (Q 54)
PAXdb E.coli [ppm]: 1.89 (Q 69)

− RNA functions: not annotated

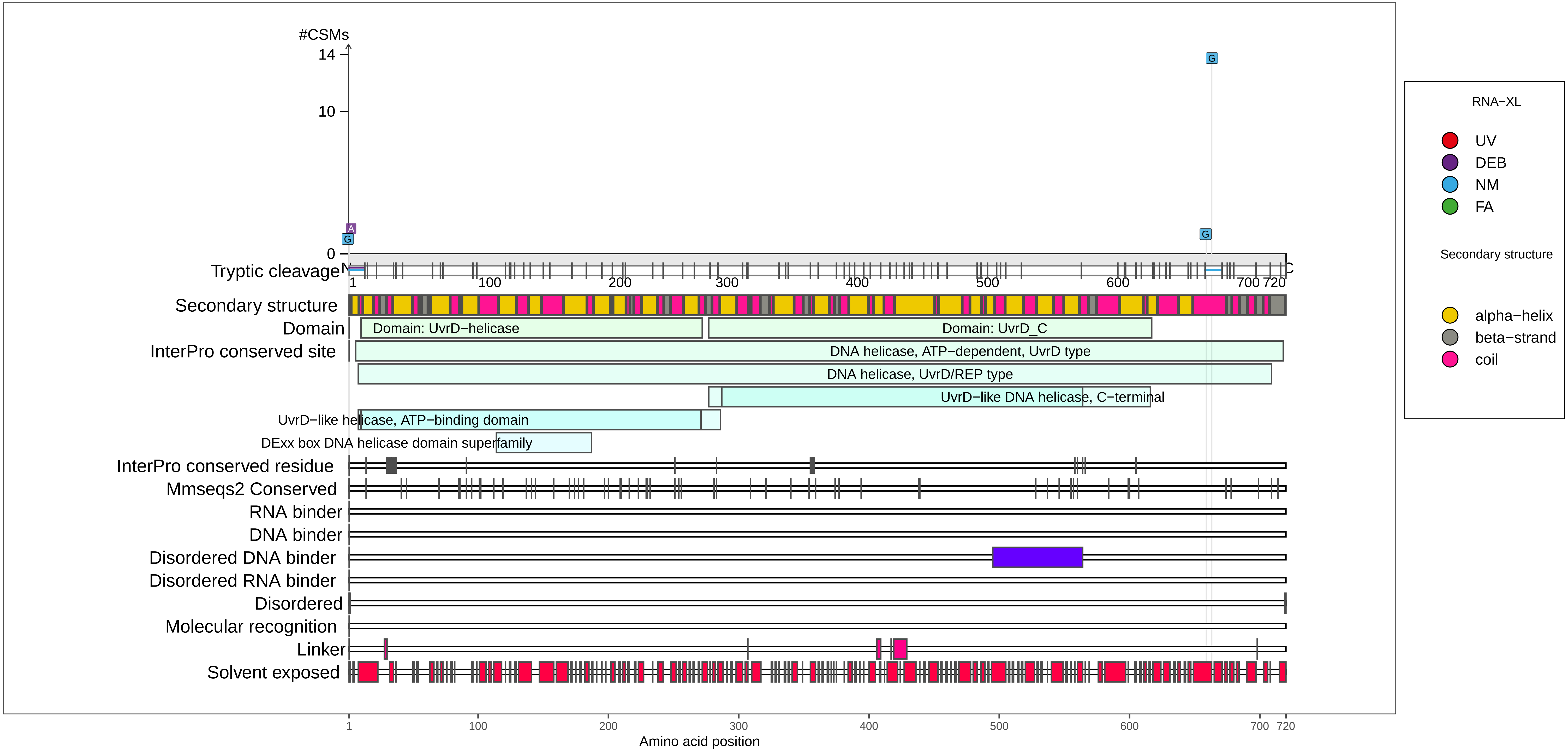

P05459
PDXB_ECOLI Erythronate−4−phosphate dehydrogenase

− Abundance:
tryptic [log10 Intensity]: 8.76 (Q 83)
PAXdb K12 strain [ppm]: 2.64 (Q 82)
PAXdb E.coli [ppm]: 2.33 (Q 81)

− RNA functions: not annotated

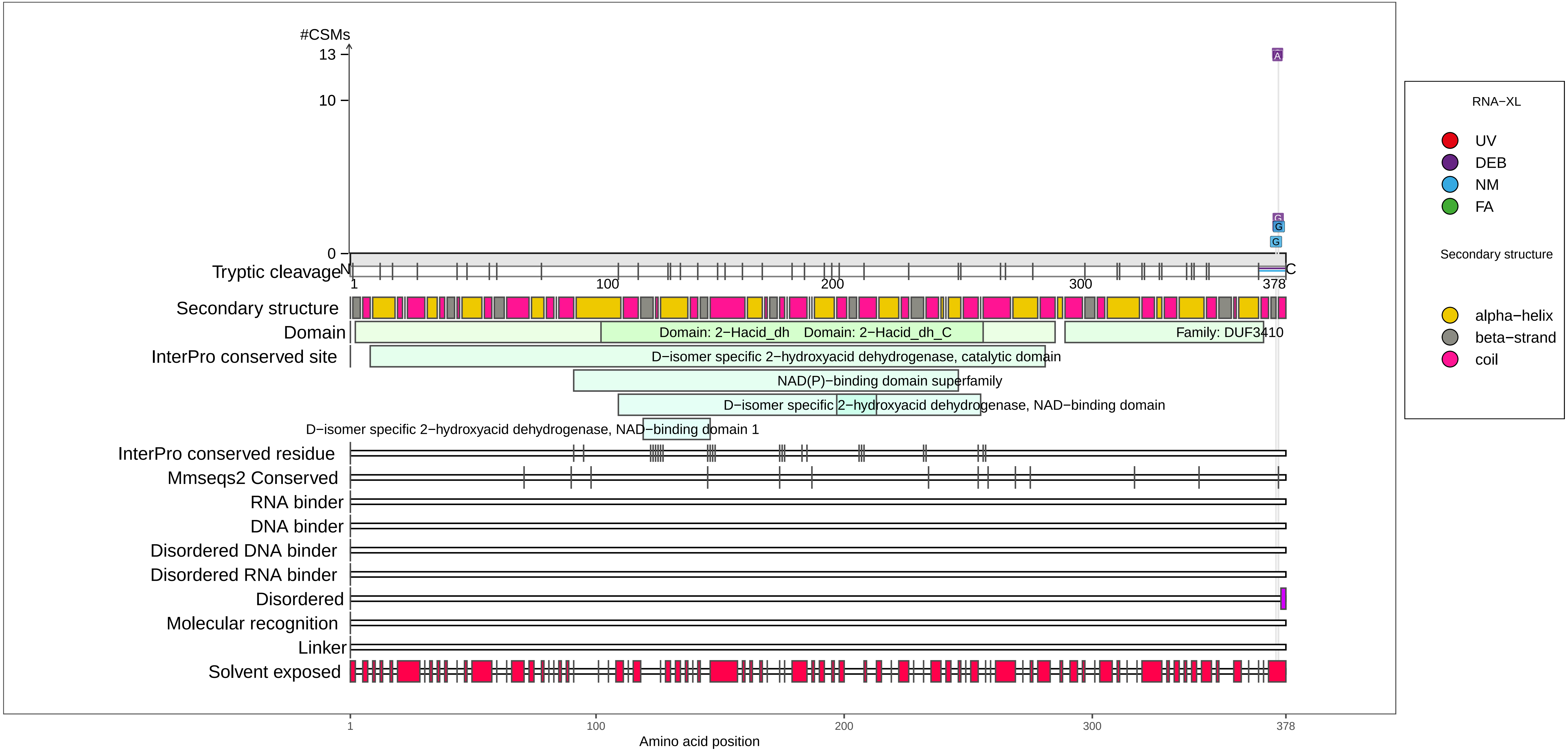

P0A968
CSPD_ECOLI Cold shock−like protein CspD

− Abundance:
tryptic [log10 Intensity]: 6.98 (Q 13)
PAXdb K12 strain [ppm]: 1.63 (Q 46)
PAXdb E.coli [ppm]: 2.87 (Q 93)

− RNA functions:
RNA binding

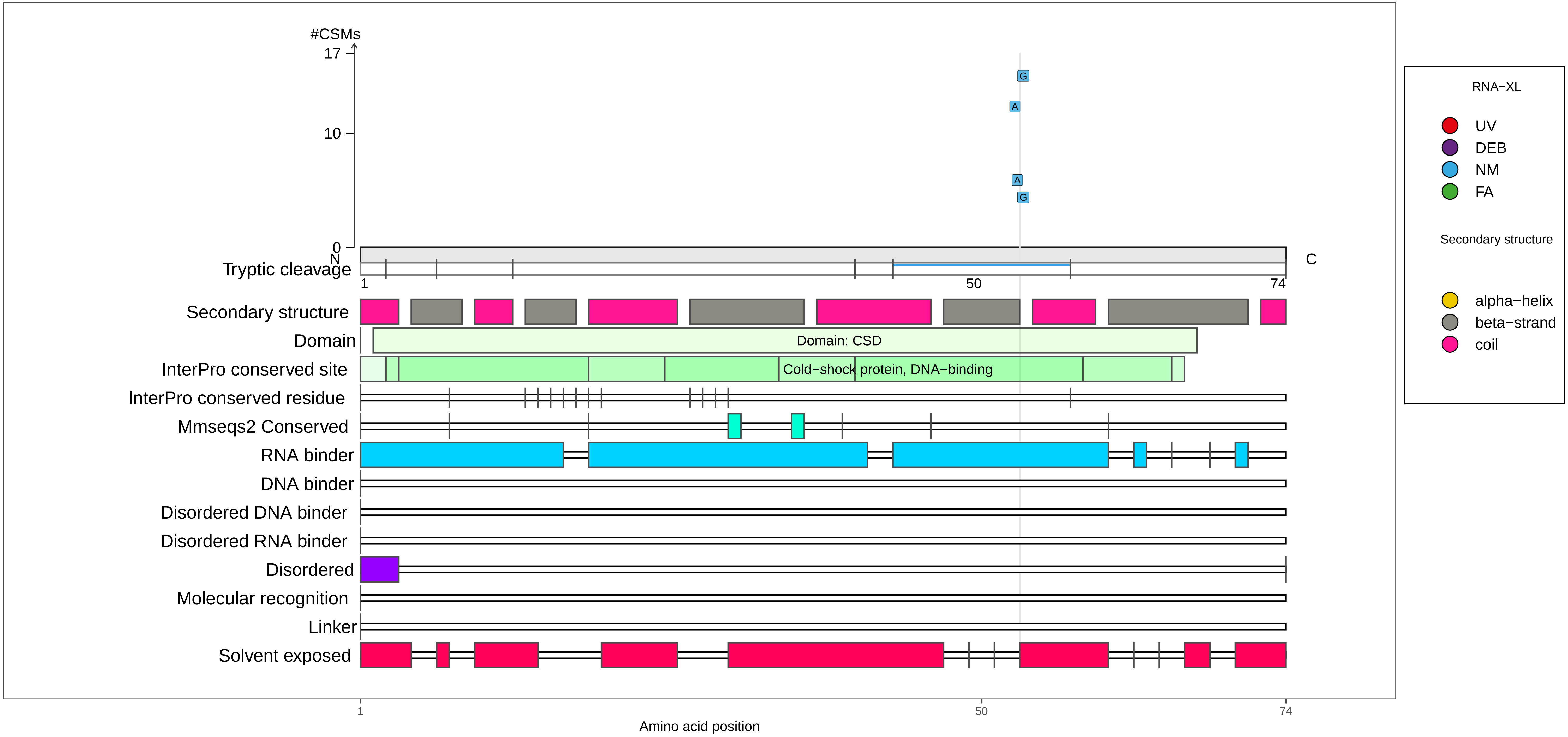

P0ACN7
CYTR_ECOLI HTH−type transcriptional repressor CytR

− Abundance:
tryptic [log10 Intensity]: 7.32 (Q 26)
PAXdb K12 strain [ppm]: 2.13 (Q 65)
PAXdb E.coli [ppm]: 0.42 (Q 34)

− RNA functions: not annotated

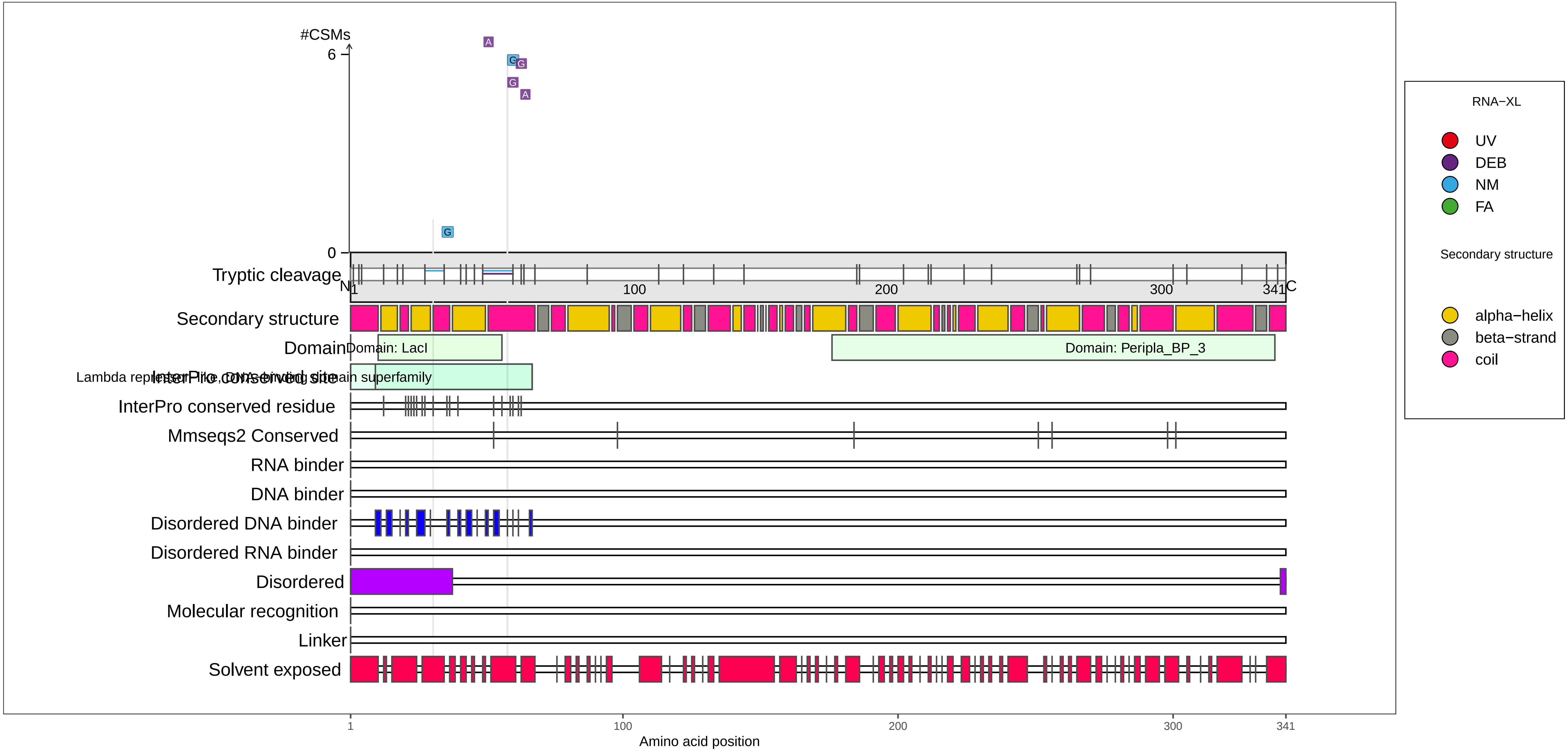

P0ADK8
YIBL_ECOLI Uncharacterized protein YibL

− Abundance:
tryptic [log10 Intensity]: 7.29 (Q 25)
PAXdb K12 strain [ppm]: 3.01 (Q 91)
PAXdb E.coli [ppm]: 2.38 (Q 83)

− RNA functions: not annotated

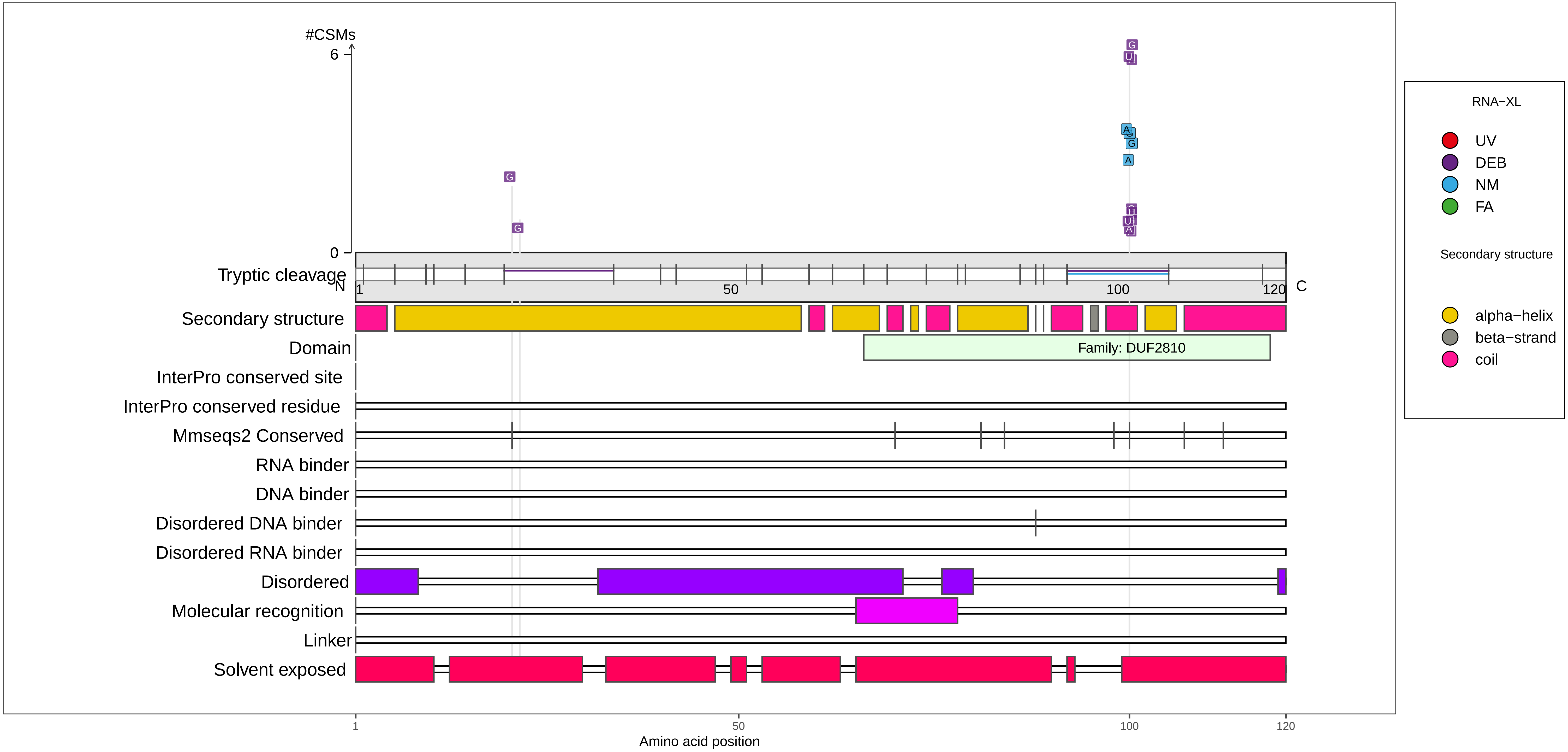

P0AEY5
MDAB_ECOLI NADPH:quinone oxidoreductase MdaB

− Abundance:
tryptic [log10 Intensity]: 8.57 (Q 78)
PAXdb K12 strain [ppm]: NA (Q NA)
PAXdb E.coli [ppm]: 2.12 (Q 75)

− RNA functions: not annotated

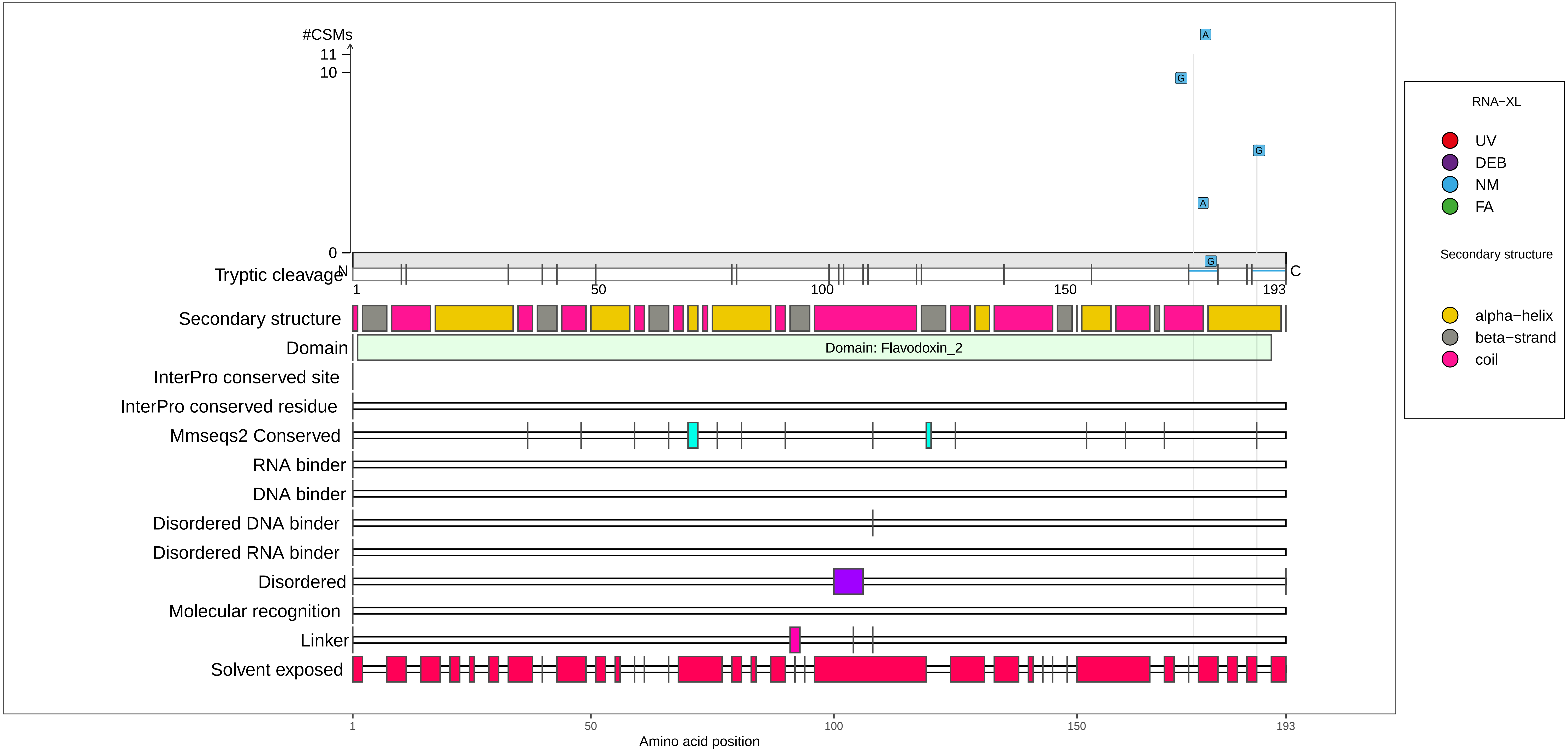

P0AFH8
OSMY_ECOLI Osmotically−inducible protein Y

− Abundance:
tryptic [log10 Intensity]: 7.52 (Q 36)
PAXdb K12 strain [ppm]: 3.77 (Q 99)
PAXdb E.coli [ppm]: 2.85 (Q 92)

− RNA functions: not annotated

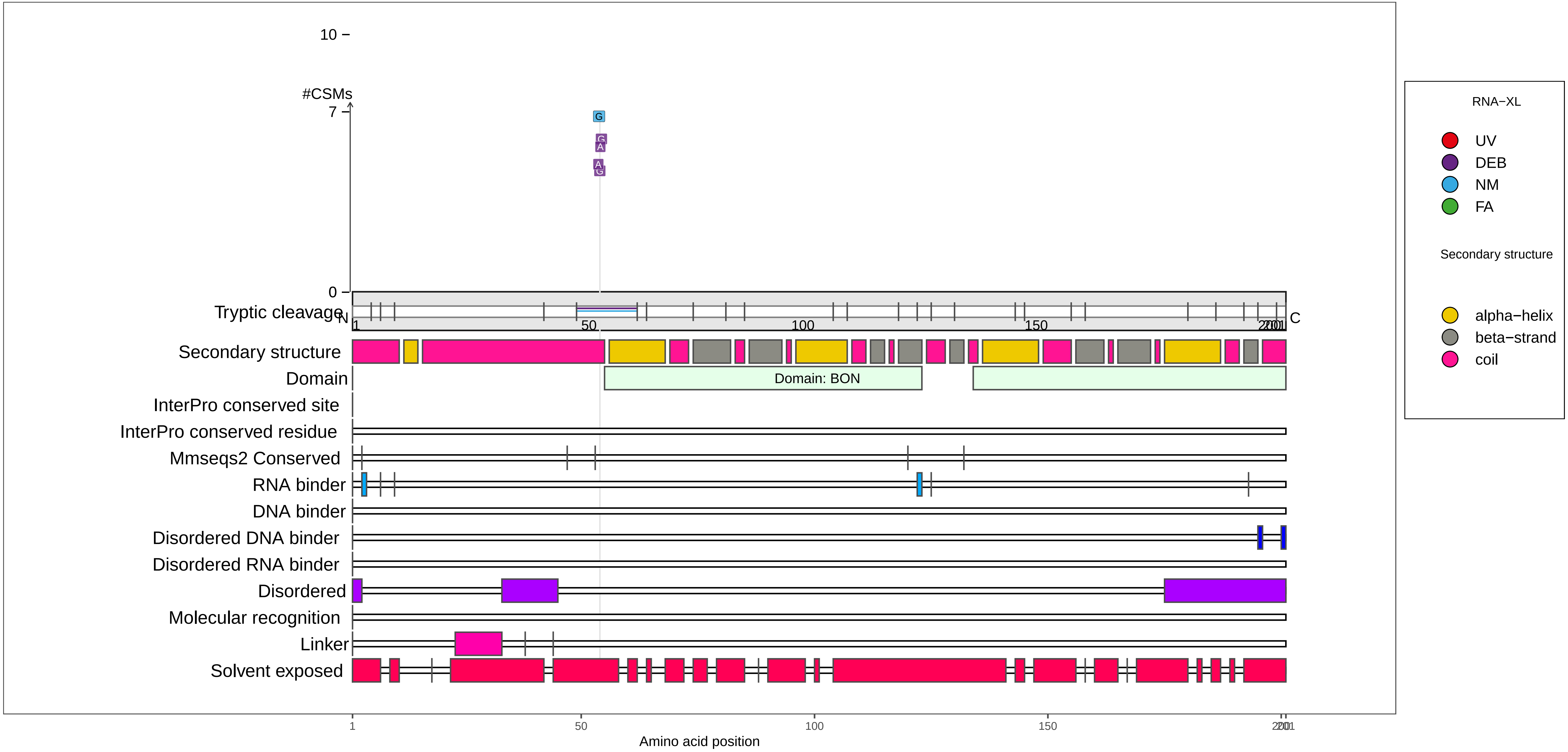

P0AGB0
SERB_ECOLI Phosphoserine phosphatase

− Abundance:
tryptic [log10 Intensity]: 8.79 (Q 83)
PAXdb K12 strain [ppm]: 1.45 (Q 35)
PAXdb E.coli [ppm]: 1.26 (Q 54)

− RNA functions: not annotated

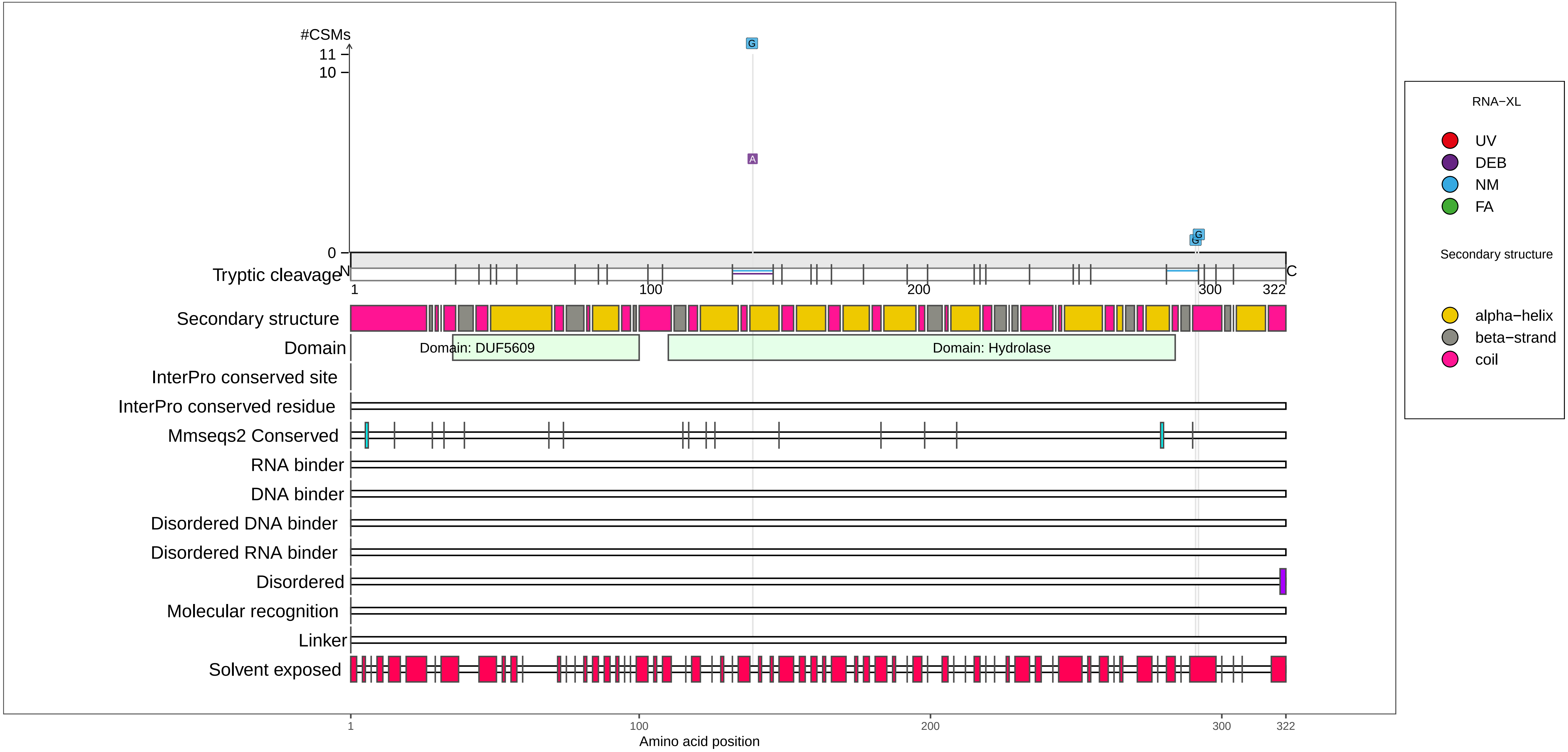

P27828
WECB_ECOLI UDP−N−acetylglucosamine 2−epimerase

− Abundance:
tryptic [log10 Intensity]: 8.07 (Q 60)
PAXdb K12 strain [ppm]: 1.02 (Q 12)
PAXdb E.coli [ppm]: 1.31 (Q 55)

− RNA functions: not annotated

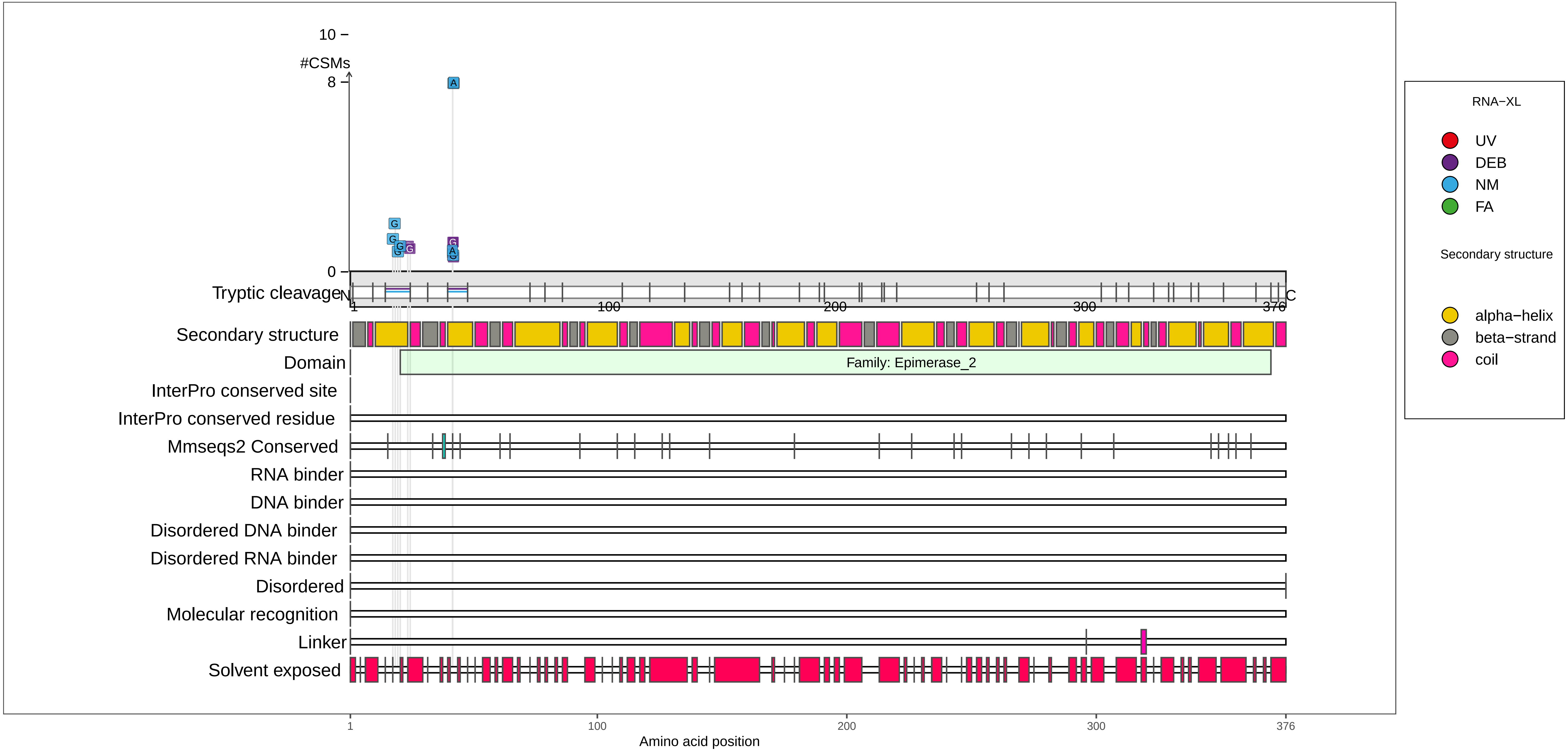

P60716
LIPA_ECOLI Lipoyl synthase

− Abundance:
tryptic [log10 Intensity]: 9.01 (Q 88)
PAXdb K12 strain [ppm]: 2.8 (Q 86)
PAXdb E.coli [ppm]: 2.23 (Q 78)

− RNA functions: not annotated

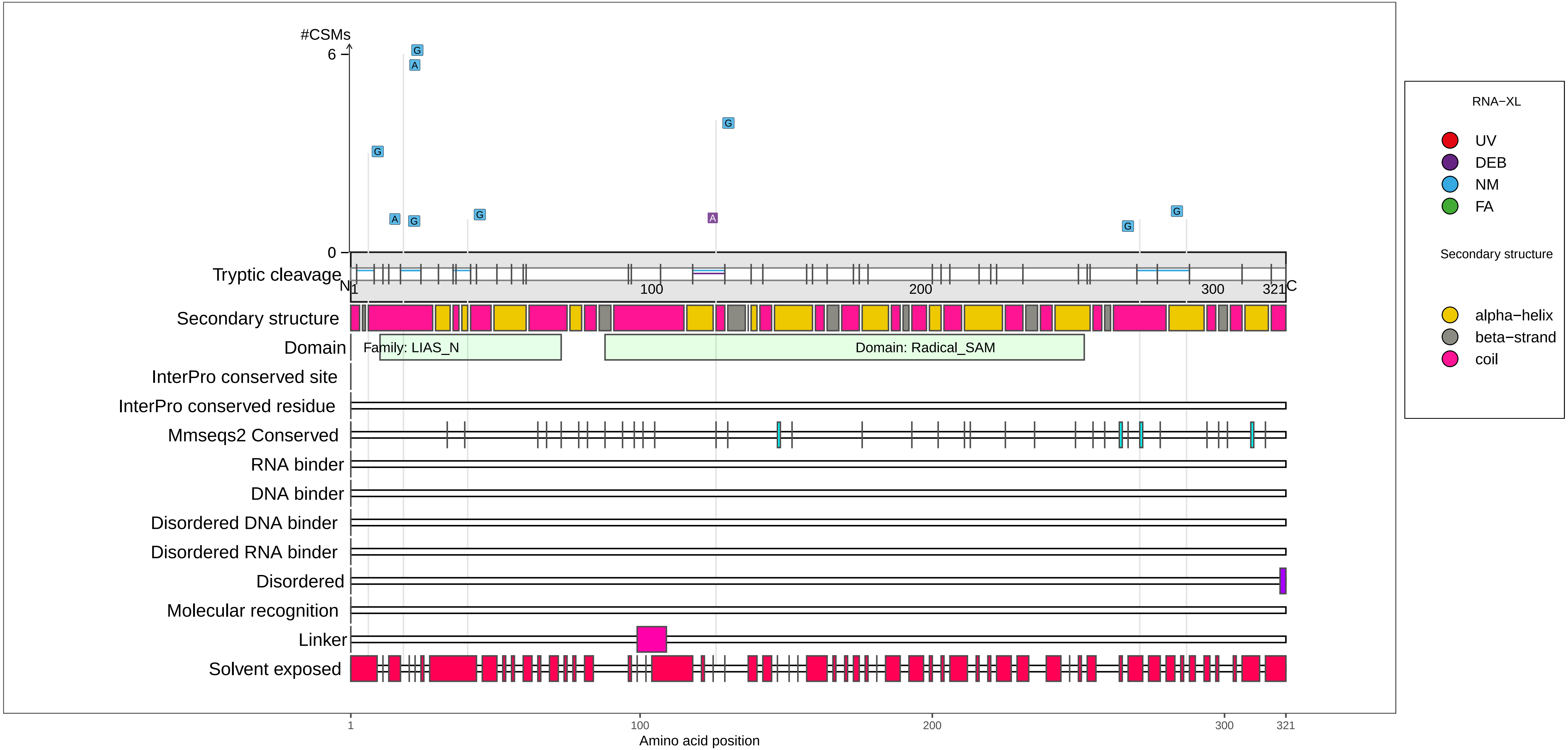

P77754
SPY_ECOLI Periplasmic chaperone Spy

− Abundance:
tryptic [log10 Intensity]: 6.7 (Q 5)
PAXdb K12 strain [ppm]: 2.37 (Q 73)
PAXdb E.coli [ppm]: 2.56 (Q 87)

− RNA functions: not annotated

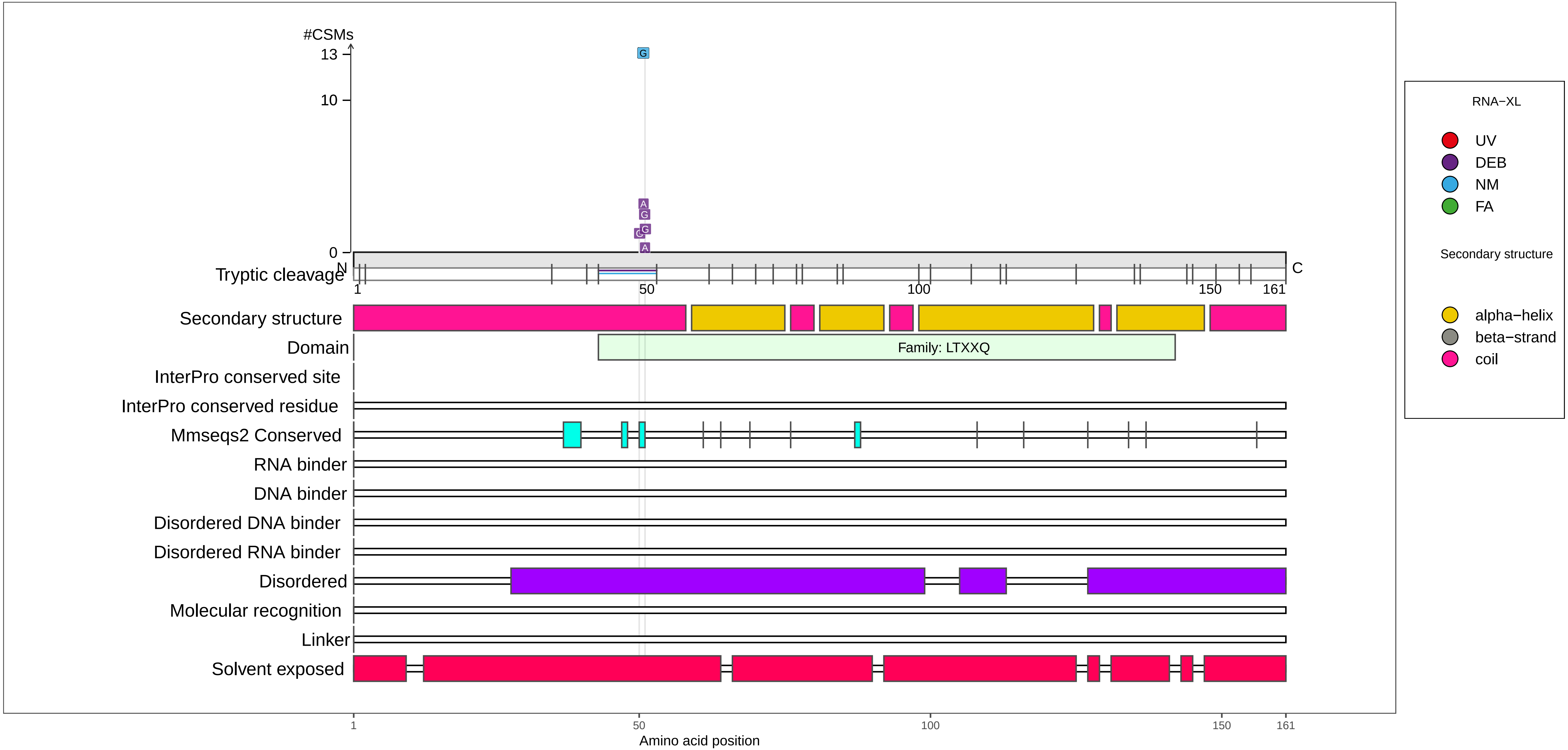

P08957
T1MK_ECOLI Type I restriction enzyme EcoKI methylase subunit

− Abundance:
tryptic [log10 Intensity]: 7.88 (Q 53)
PAXdb K12 strain [ppm]: 2.13 (Q 65)
PAXdb E.coli [ppm]: 1.66 (Q 63)

− RNA functions: not annotated

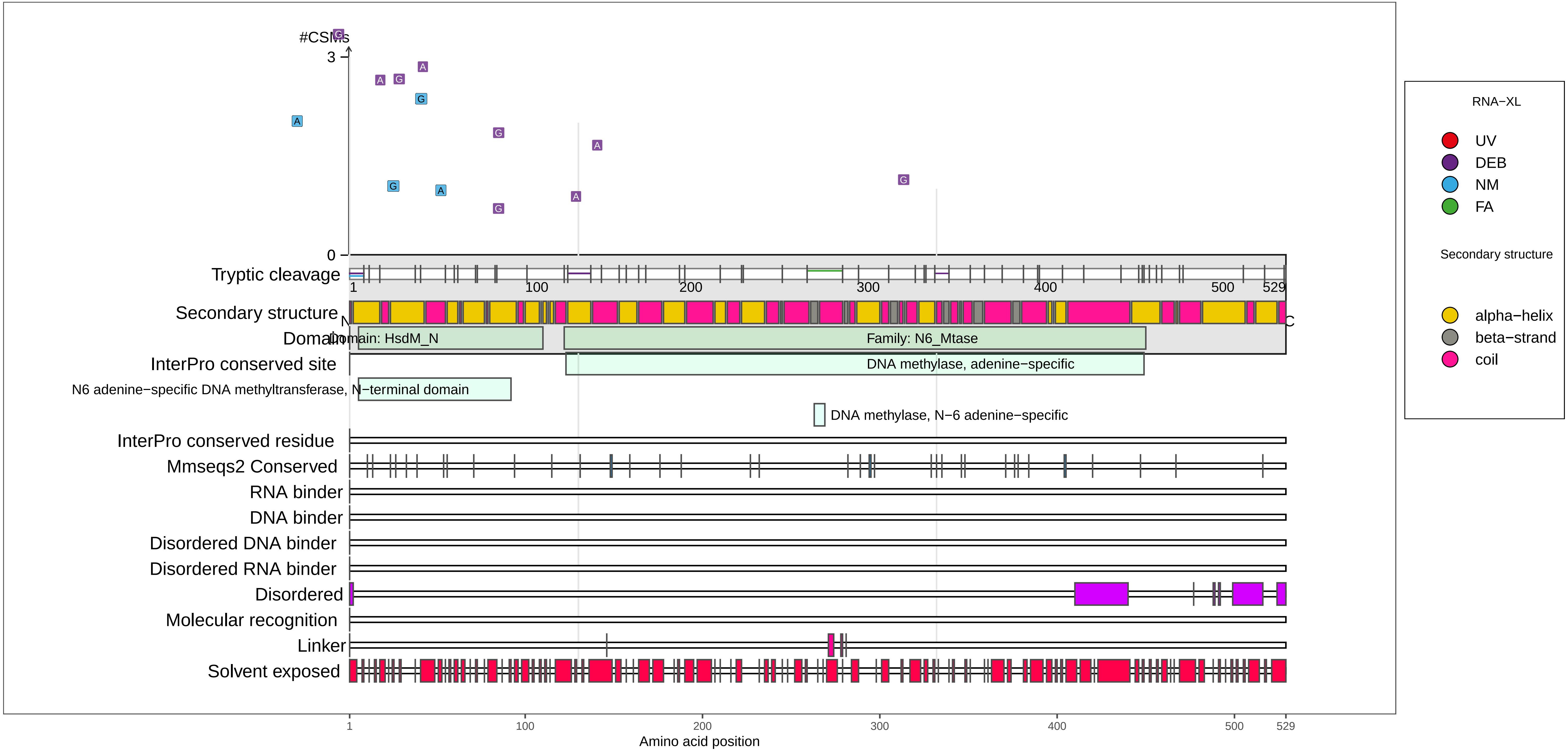

P0A744
MSRA_ECOLI Peptide methionine sulfoxide reductase MsrA

− Abundance:
tryptic [log10 Intensity]: 7.94 (Q 56)
PAXdb K12 strain [ppm]: 2.78 (Q 85)
PAXdb E.coli [ppm]: 2.57 (Q 87)

− RNA functions: not annotated

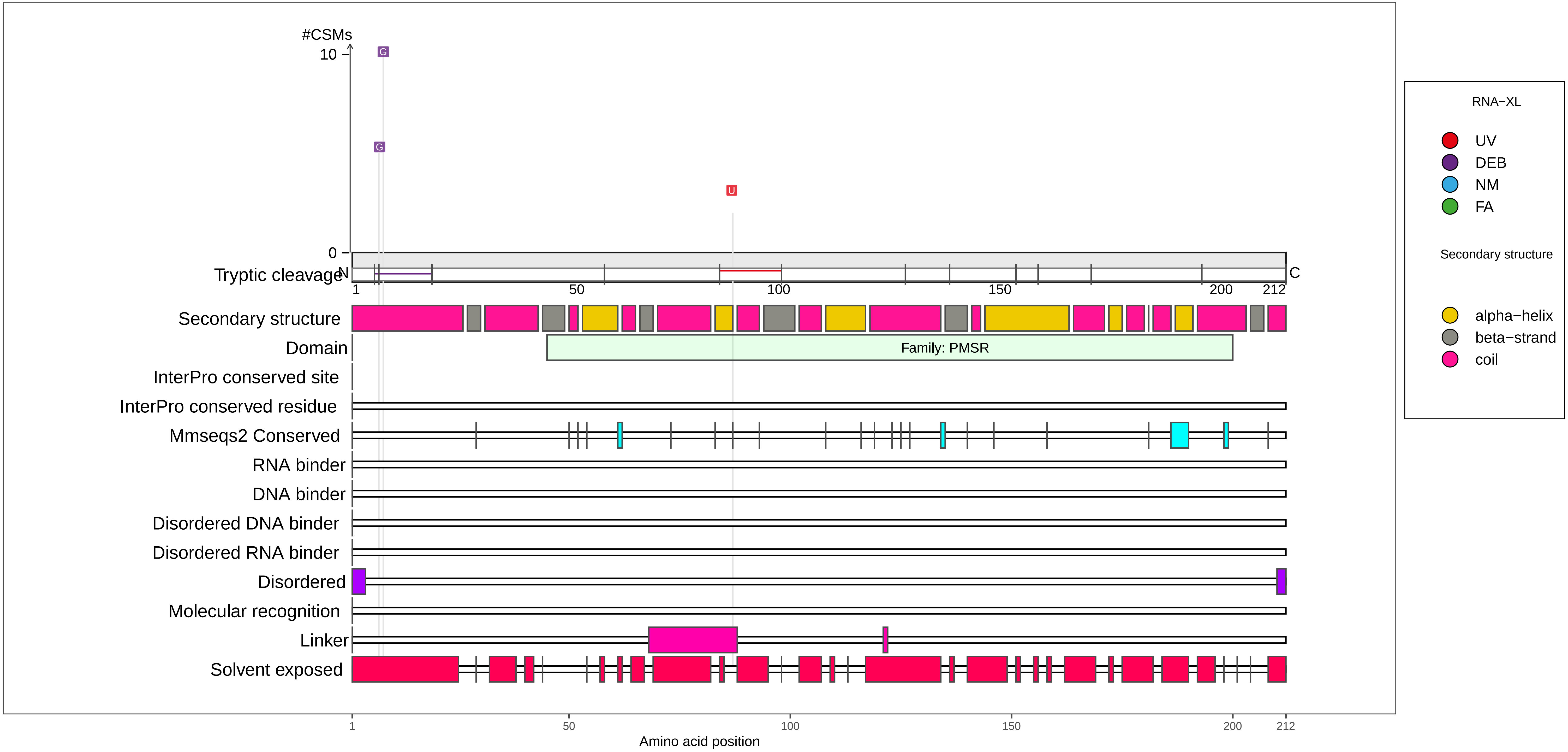

P0A7B5
PROB_ECOLI Glutamate 5−kinase

− Abundance:
tryptic [log10 Intensity]: 8.93 (Q 87)
PAXdb K12 strain [ppm]: 2.36 (Q 72)
PAXdb E.coli [ppm]: 2.13 (Q 76)

− RNA functions:
RNA binding

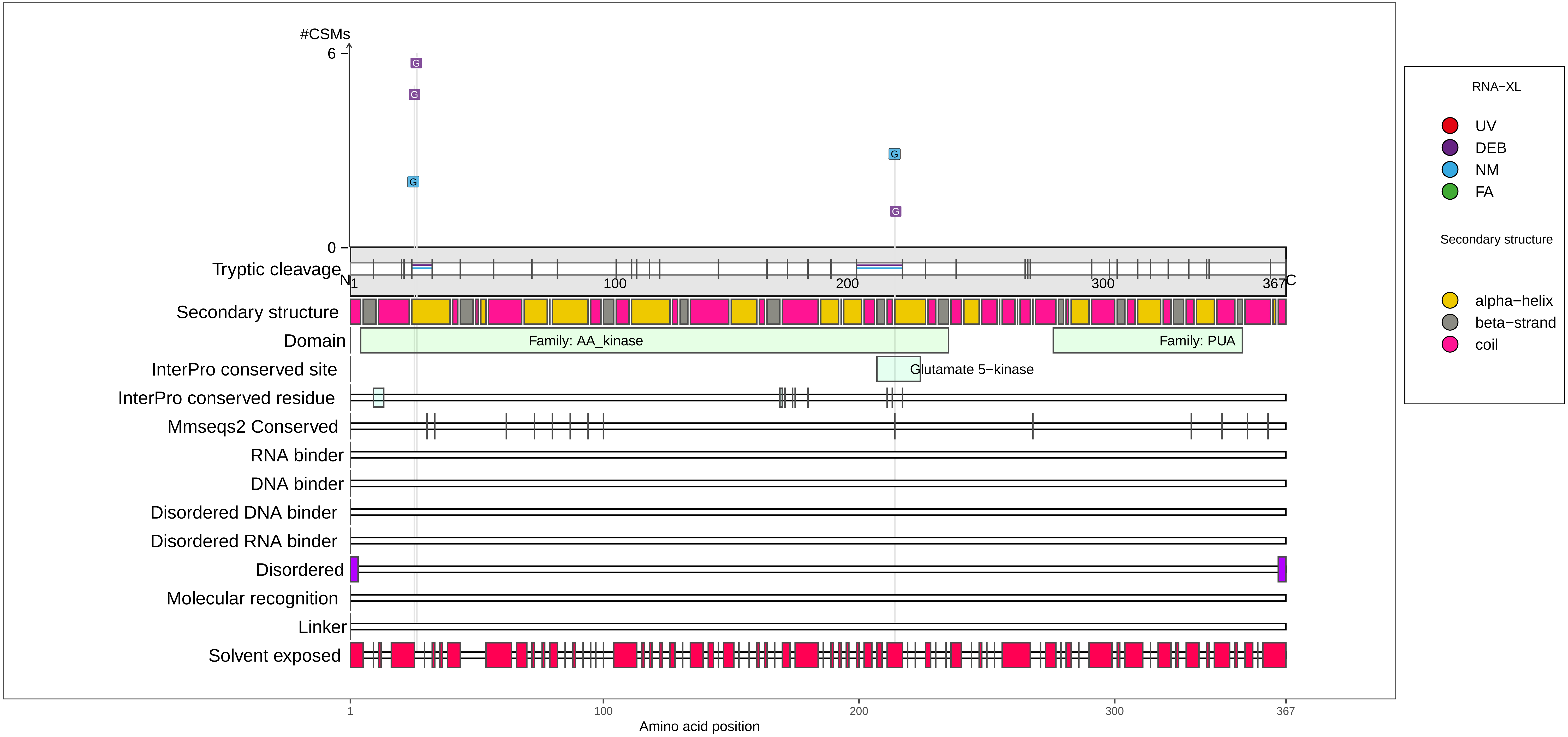

P0A8F0
UPP_ECOLI Uracil phosphoribosyltransferase

− Abundance:
tryptic [log10 Intensity]: 9.08 (Q 90)
PAXdb K12 strain [ppm]: 3.14 (Q 93)
PAXdb E.coli [ppm]: 3.07 (Q 95)

− RNA functions: not annotated

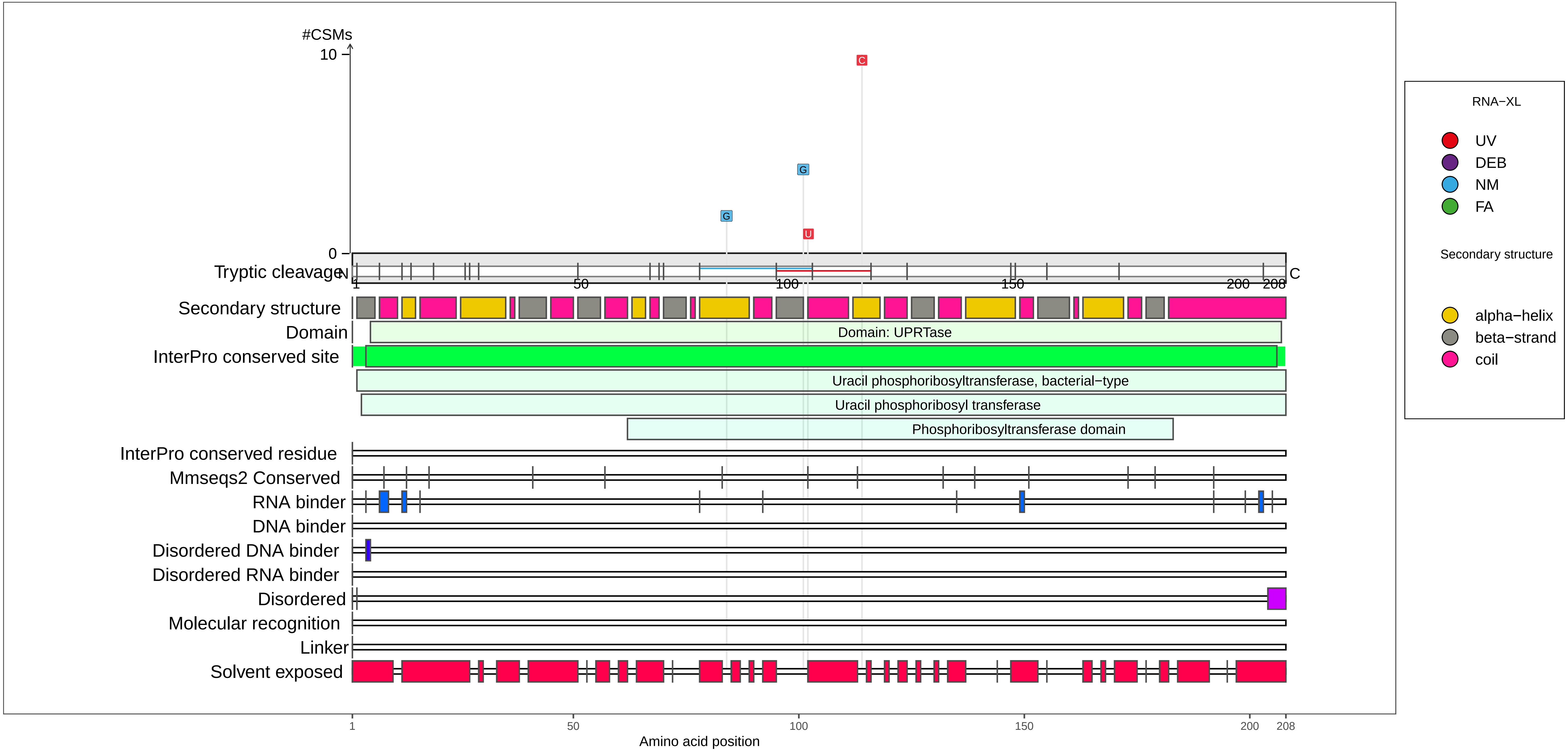

P0A8W2
SLYA_ECOLI Transcriptional regulator SlyA

− Abundance:
tryptic [log10 Intensity]: 7.36 (Q 27)
PAXdb K12 strain [ppm]: 2.24 (Q 68)
PAXdb E.coli [ppm]: 2.61 (Q 88)

− RNA functions: not annotated

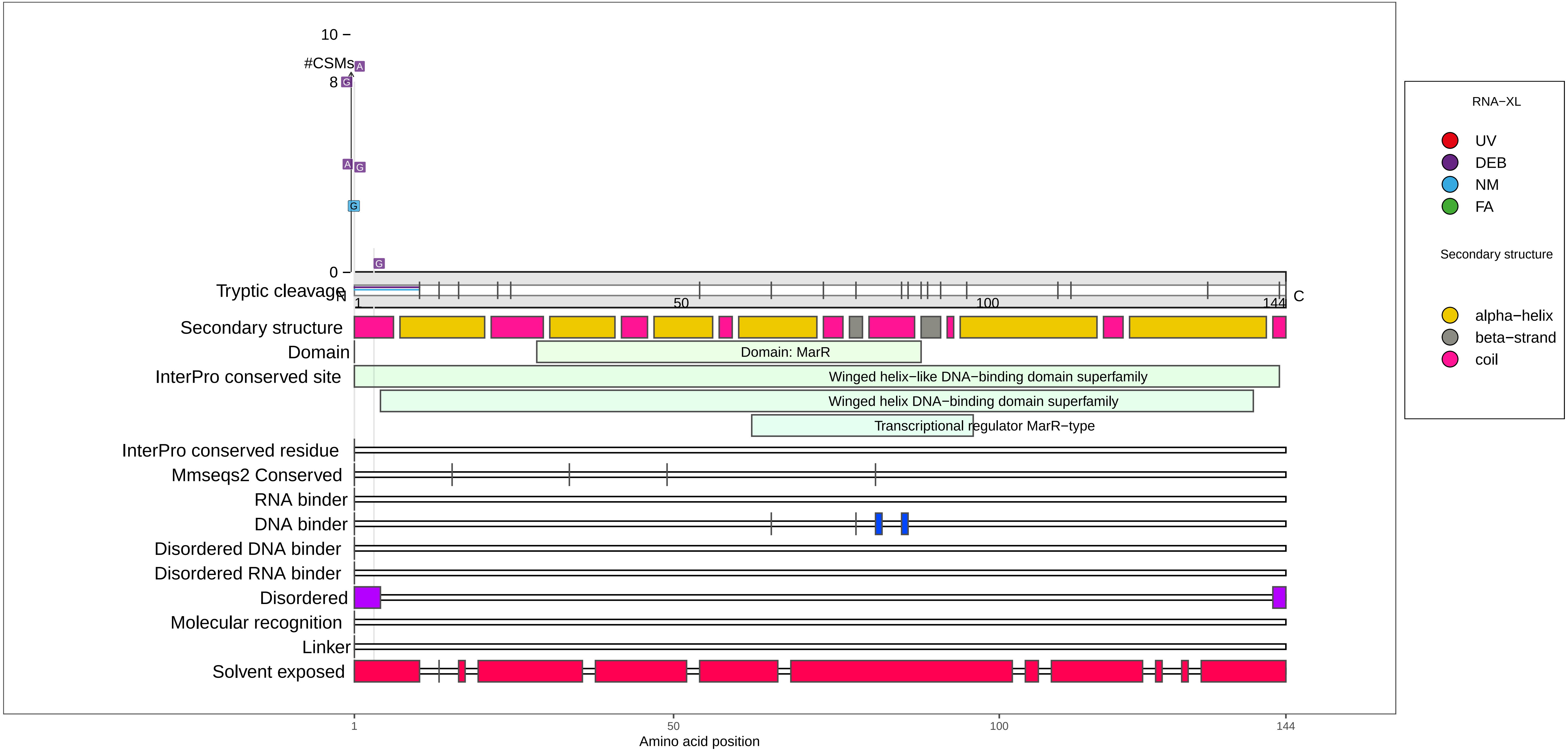

P0ABQ2
GARR_ECOLI 2−hydroxy−3−oxopropionate reductase

− Abundance:
tryptic [log10 Intensity]: 8.37 (Q 71)
PAXdb K12 strain [ppm]: 2.1 (Q 64)
PAXdb E.coli [ppm]: 1.74 (Q 66)

− RNA functions: not annotated

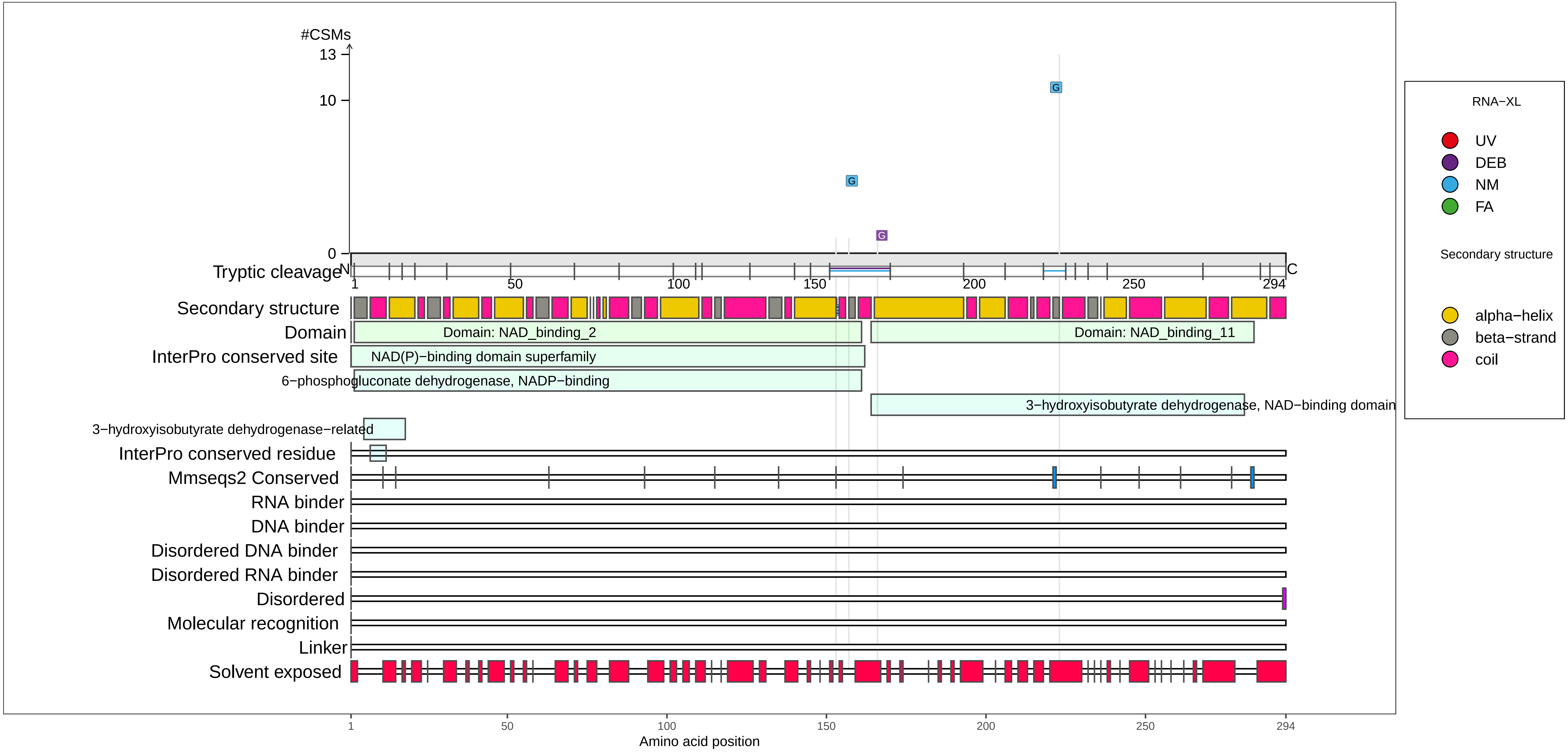

P0AEI4
RIMO_ECOLI Ribosomal protein S12 methylthiotransferase RimO

− Abundance:
tryptic [log10 Intensity]: 8.22 (Q 66)
PAXdb K12 strain [ppm]: 1.97 (Q 59)
PAXdb E.coli [ppm]: 1.3 (Q 55)

− RNA functions:
ncRNA metabolic process; ncRNA processing; RNA metabolic process; RNA modification
RNA processing; tRNA metabolic process; tRNA modification; tRNA processing

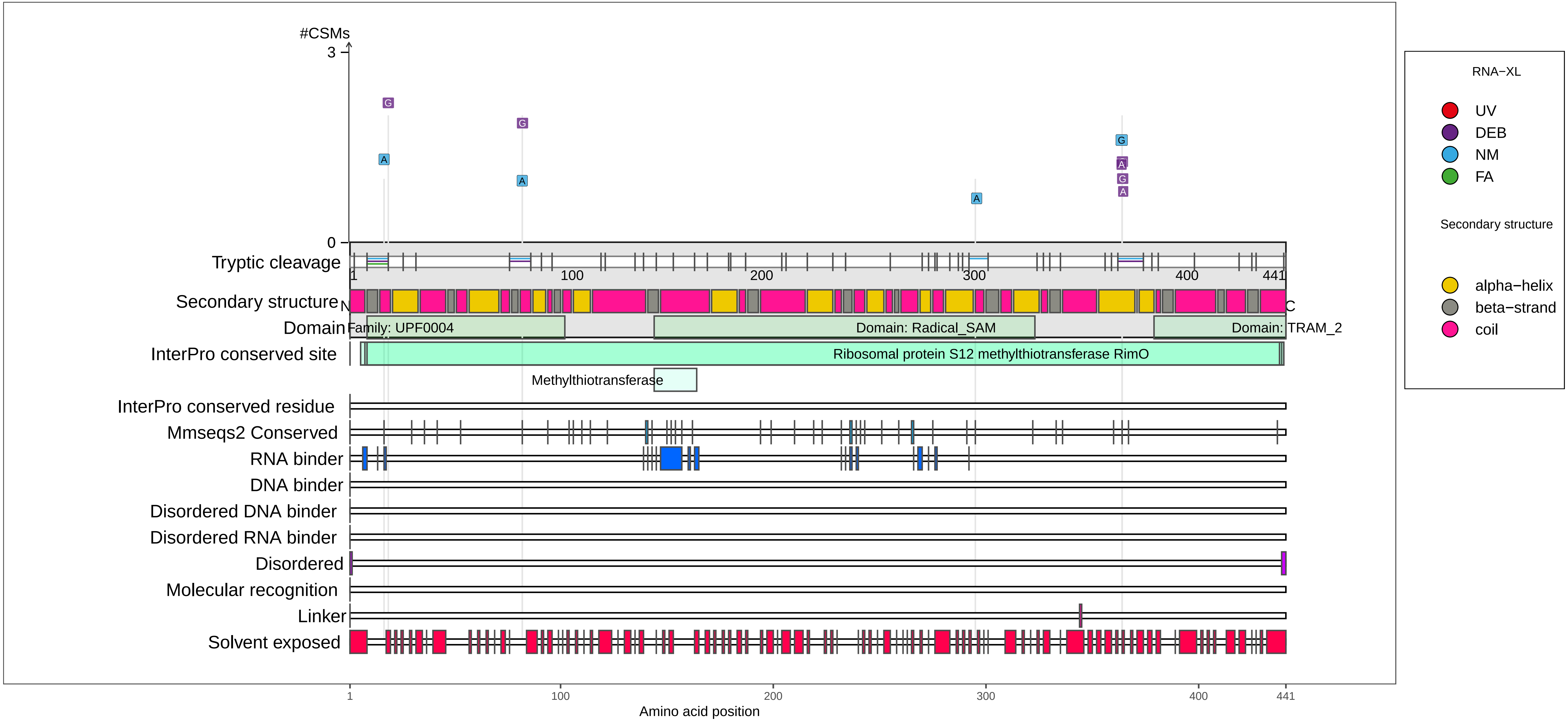

P0AFG3
ODO1_ECOLI 2−oxoglutarate dehydrogenase E1 component

− Abundance:
tryptic [log10 Intensity]: 8.84 (Q 85)
PAXdb K12 strain [ppm]: 3.11 (Q 92)
PAXdb E.coli [ppm]: 2.91 (Q 93)

− RNA functions: not annotated

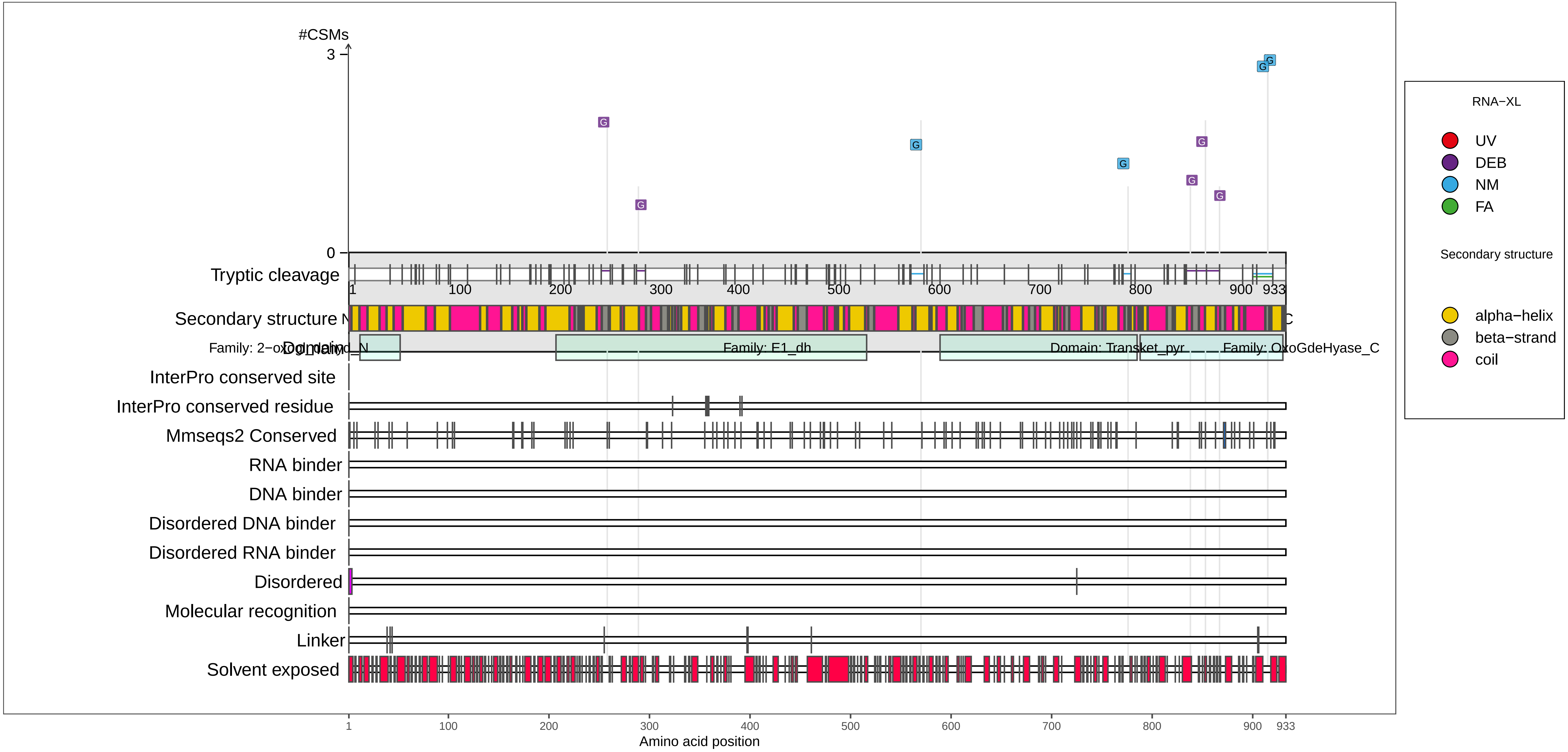

P0AG51
RL30_ECOLI 50S ribosomal protein L30

− Abundance:
tryptic [log10 Intensity]: 8.87 (Q 85)
PAXdb K12 strain [ppm]: 3.42 (Q 96)
PAXdb E.coli [ppm]: 3.46 (Q 99)

− RNA functions: not annotated

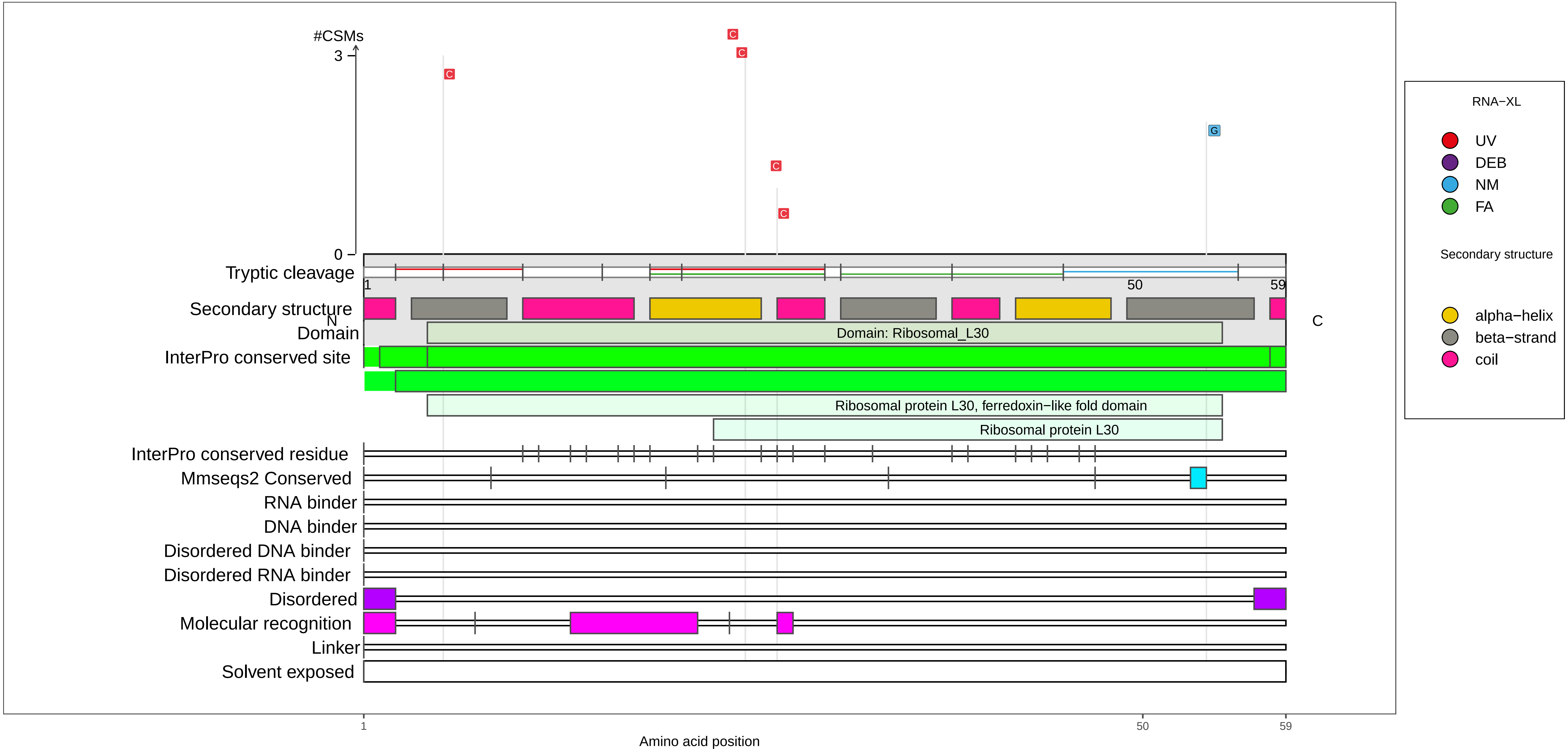

P0AGJ7
TRML_ECOLI tRNA (cytidine(34)−2'−O)−methyltransferase

− Abundance:
tryptic [log10 Intensity]: 7.4 (Q 30)
PAXdb K12 strain [ppm]: 1.23 (Q 22)
PAXdb E.coli [ppm]: 1.16 (Q 51)

− RNA functions:
ncRNA metabolic process; ncRNA processing; RNA 2−O−methyltransferase activity
RNA binding; RNA metabolic process; RNA methylation; RNA methyltransferase activity
RNA modification; RNA processing; SpoU rRNA Methylase family
tRNA (cytosine−2−O−)−methyltransferase activity; tRNA (cytosine) methyltransferase activity
tRNA (uracil−2−O−)−methyltransferase activity; tRNA (uracil) methyltransferase activity
tRNA 2−O−methyltransferase activity; tRNA metabolic process; tRNA methylation
tRNA methyltransferase activity; tRNA modification; tRNA nucleoside ribose methylation; tRNA processing

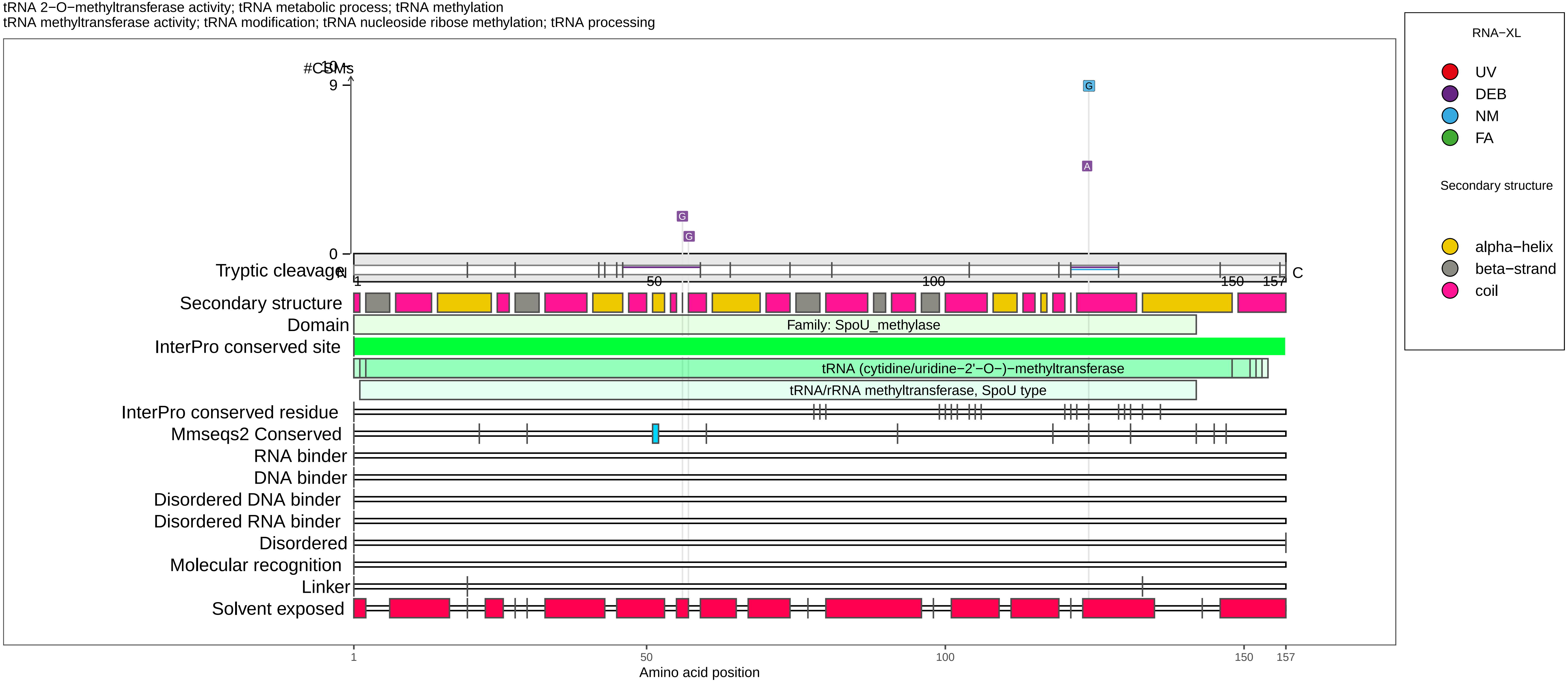

P17445
BETB_ECOLI Betaine aldehyde dehydrogenase

− Abundance:
tryptic [log10 Intensity]: 7.79 (Q 49)
PAXdb K12 strain [ppm]: 1.53 (Q 40)
PAXdb E.coli [ppm]: 1.12 (Q 51)

− RNA functions: not annotated

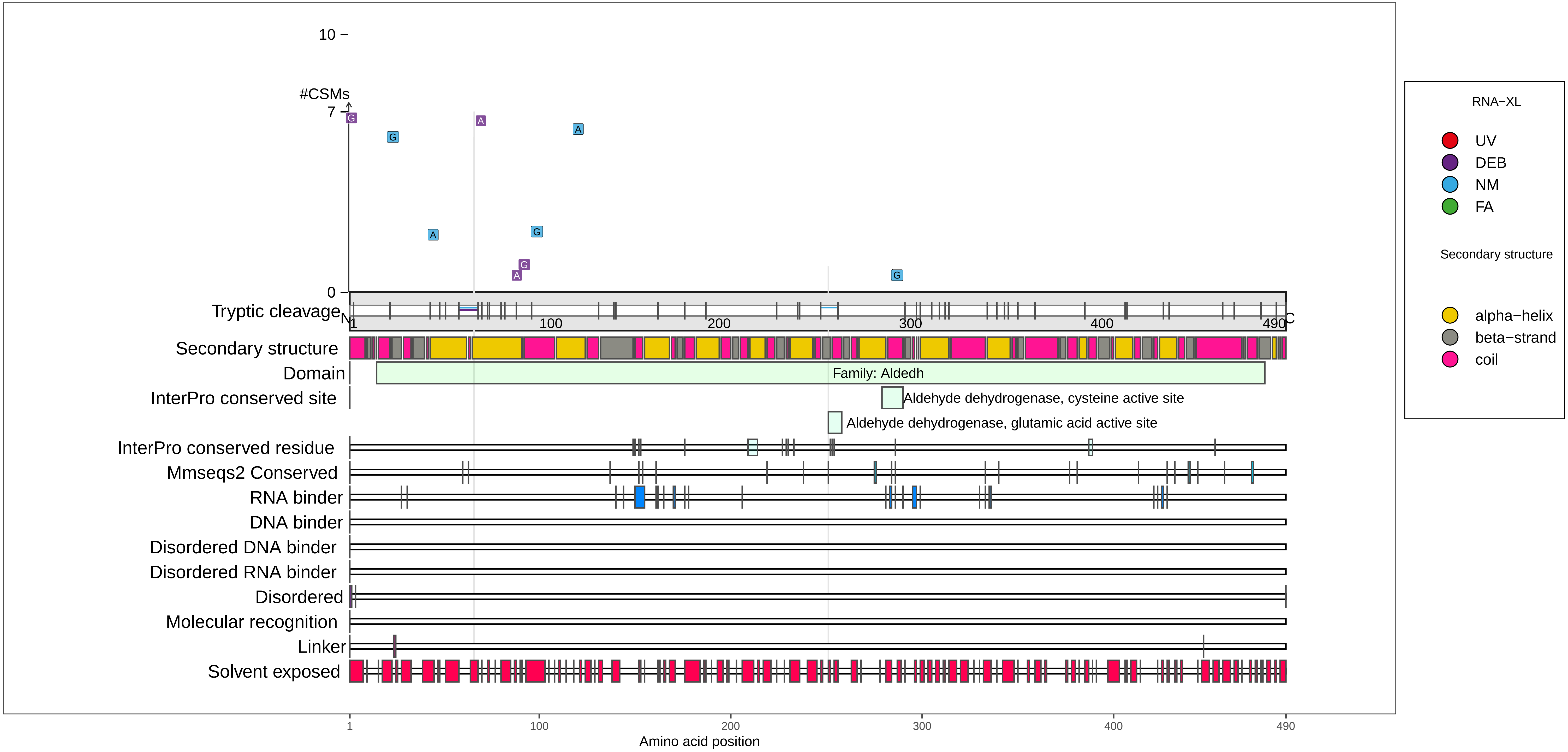

P27248
GCST_ECOLI Aminomethyltransferase

− Abundance:
tryptic [log10 Intensity]: 9.49 (Q 96)
PAXdb K12 strain [ppm]: 3.2 (Q 94)
PAXdb E.coli [ppm]: 2.29 (Q 80)

− RNA functions: not annotated

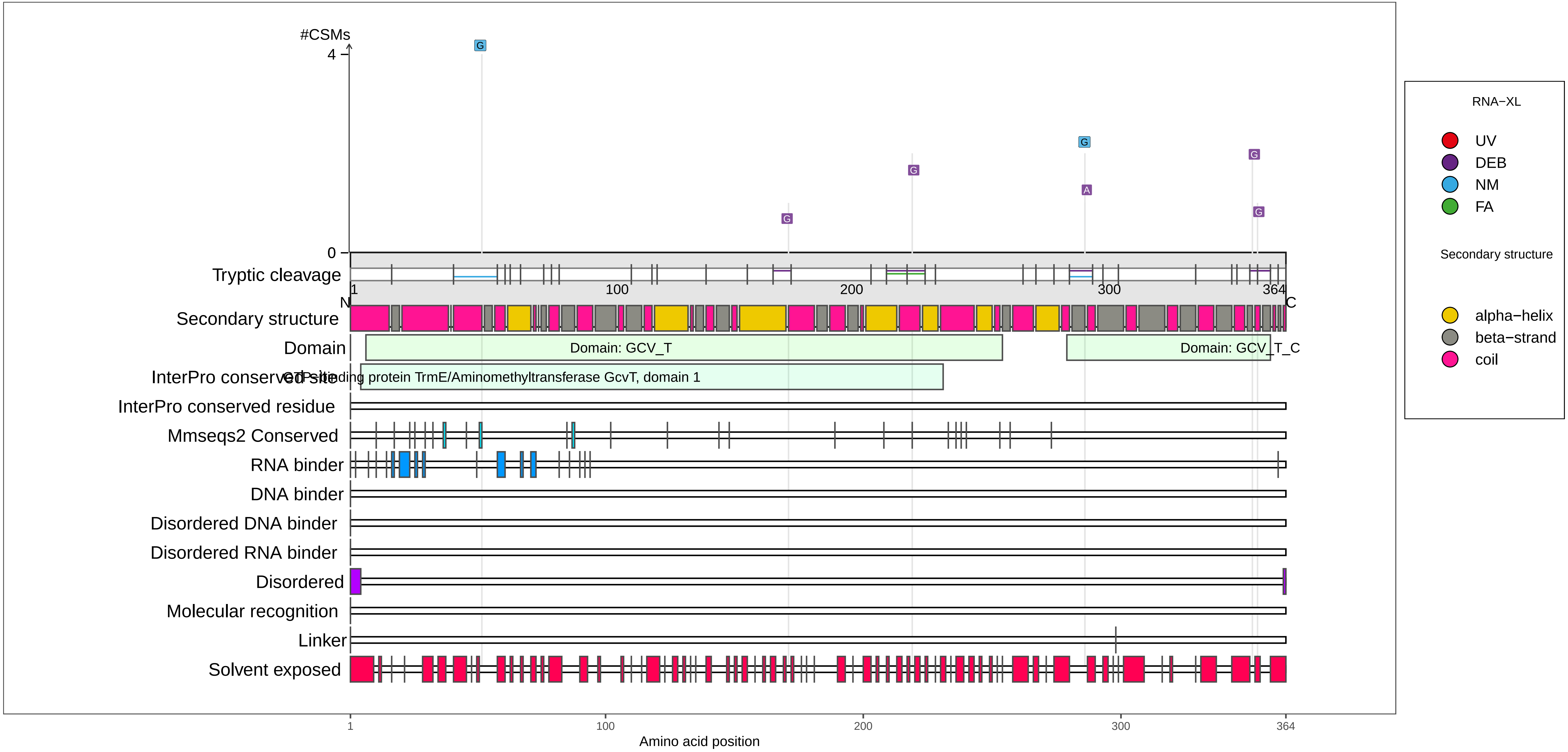

P0AB77
KBL_ECOLI 2−amino−3−ketobutyrate coenzyme A ligase
− Abundance:
tryptic [log10 Intensity]: 8.96 (Q 87)
PAXdb K12 strain [ppm]: 3.06 (Q 92)
PAXdb E.coli [ppm]: 2.22 (Q 78)

− RNA functions: not annotated

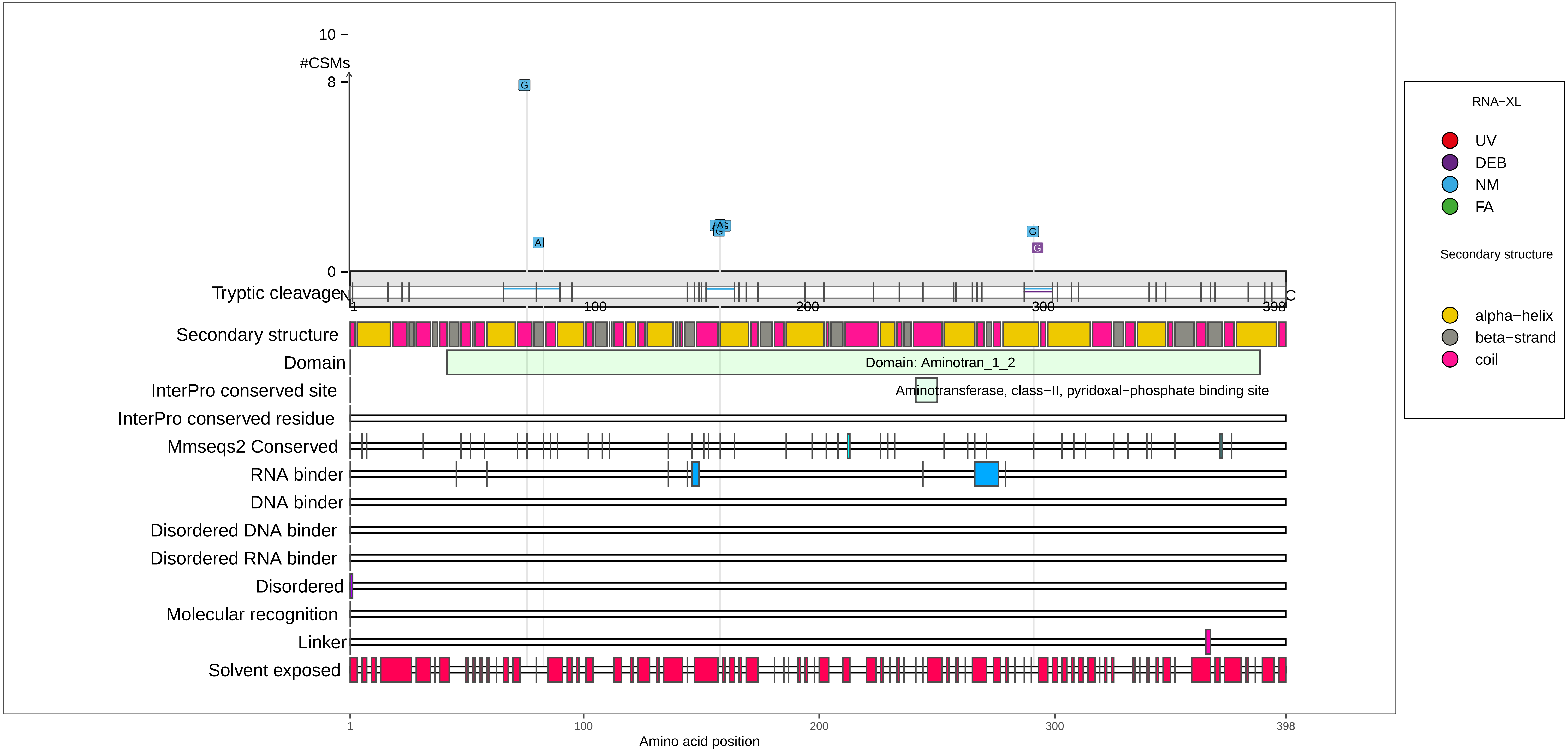

P0ACE7
HINT_ECOLI Purine nucleoside phosphoramidase

− Abundance:
tryptic [log10 Intensity]: 8.2 (Q 65)
PAXdb K12 strain [ppm]: 2.56 (Q 79)
PAXdb E.coli [ppm]: 3 (Q 95)

− RNA functions:
Scavenger mRNA decapping enzyme C−term binding

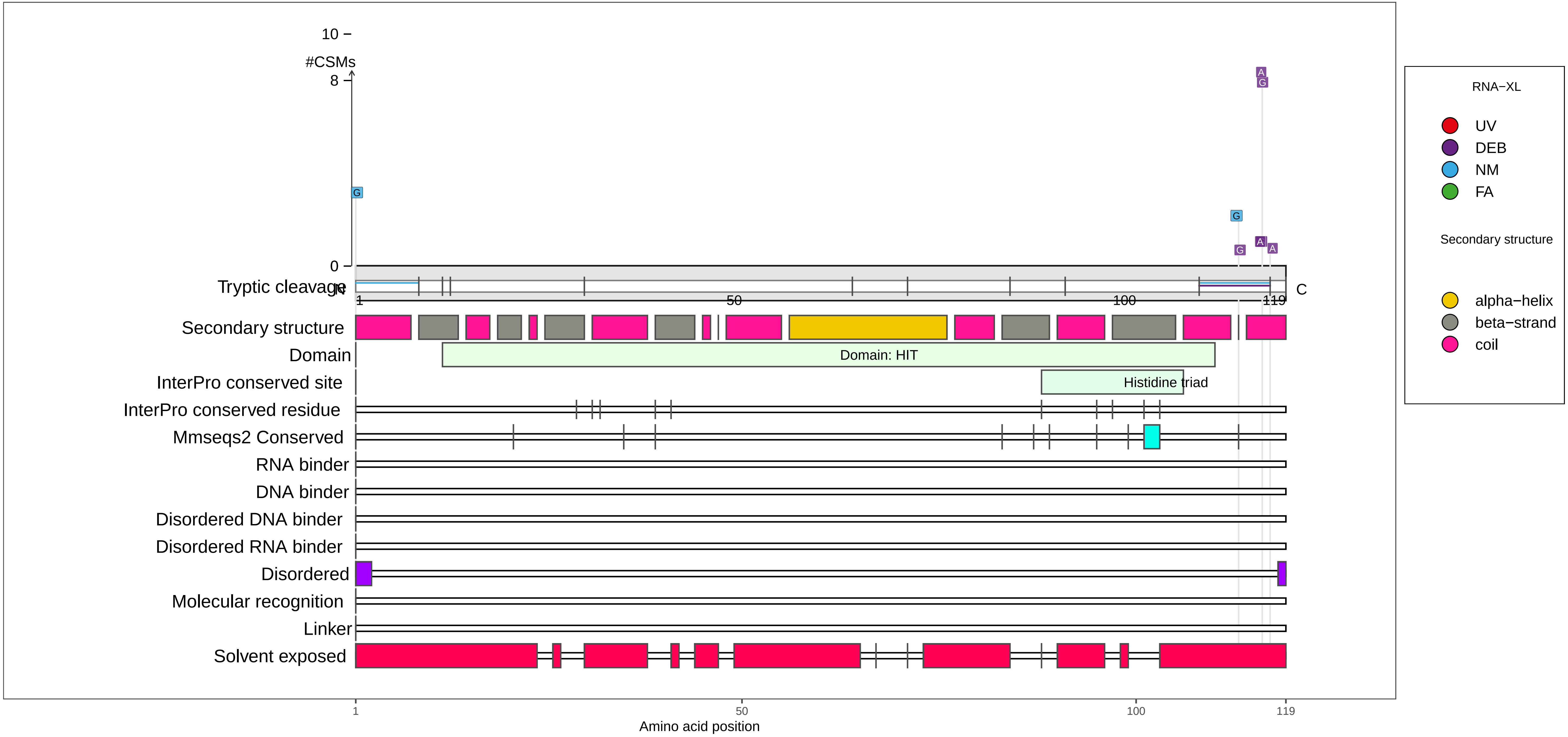

P0AFM6
PSPA_ECOLI Phage shock protein A
− Abundance:
tryptic [log10 Intensity]: 8.46 (Q 74)
PAXdb K12 strain [ppm]: 2.57 (Q 80)
PAXdb E.coli [ppm]: 2.73 (Q 90)

− RNA functions: not annotated

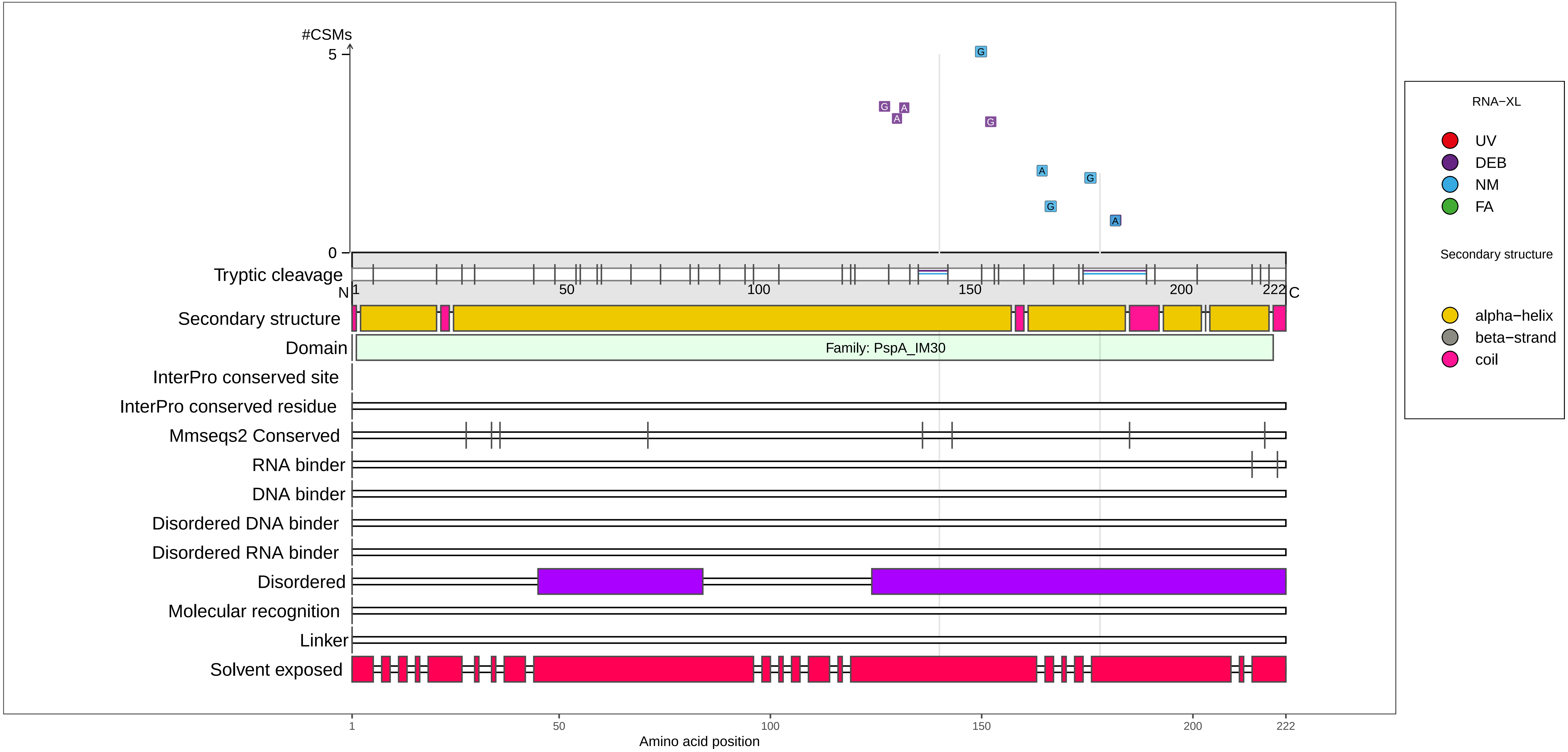

P13001
BIOH_ECOLI Pimeloyl−[acyl−carrier protein] methyl ester esterase

− Abundance:
tryptic [log10 Intensity]: 7.86 (Q 52)
PAXdb K12 strain [ppm]: NA (Q NA)
PAXdb E.coli [ppm]: 1.29 (Q 54)

− RNA functions: not annotated

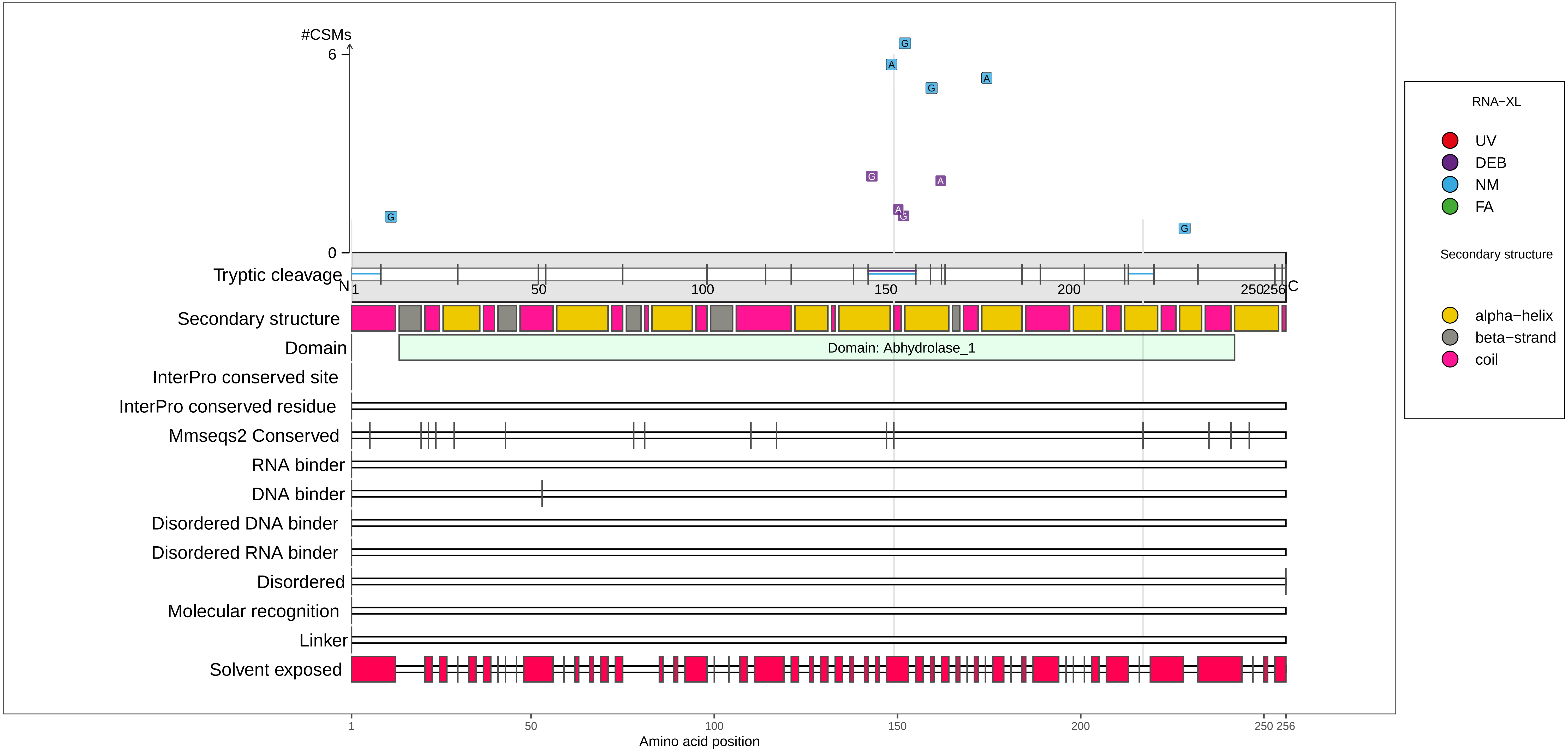

P22333
ADD_ECOLI Adenosine deaminase

− Abundance:
tryptic [log10 Intensity]: 8.3 (Q 69)
PAXdb K12 strain [ppm]: 2.25 (Q 68)
PAXdb E.coli [ppm]: 2.15 (Q 76)

− RNA functions: not annotated

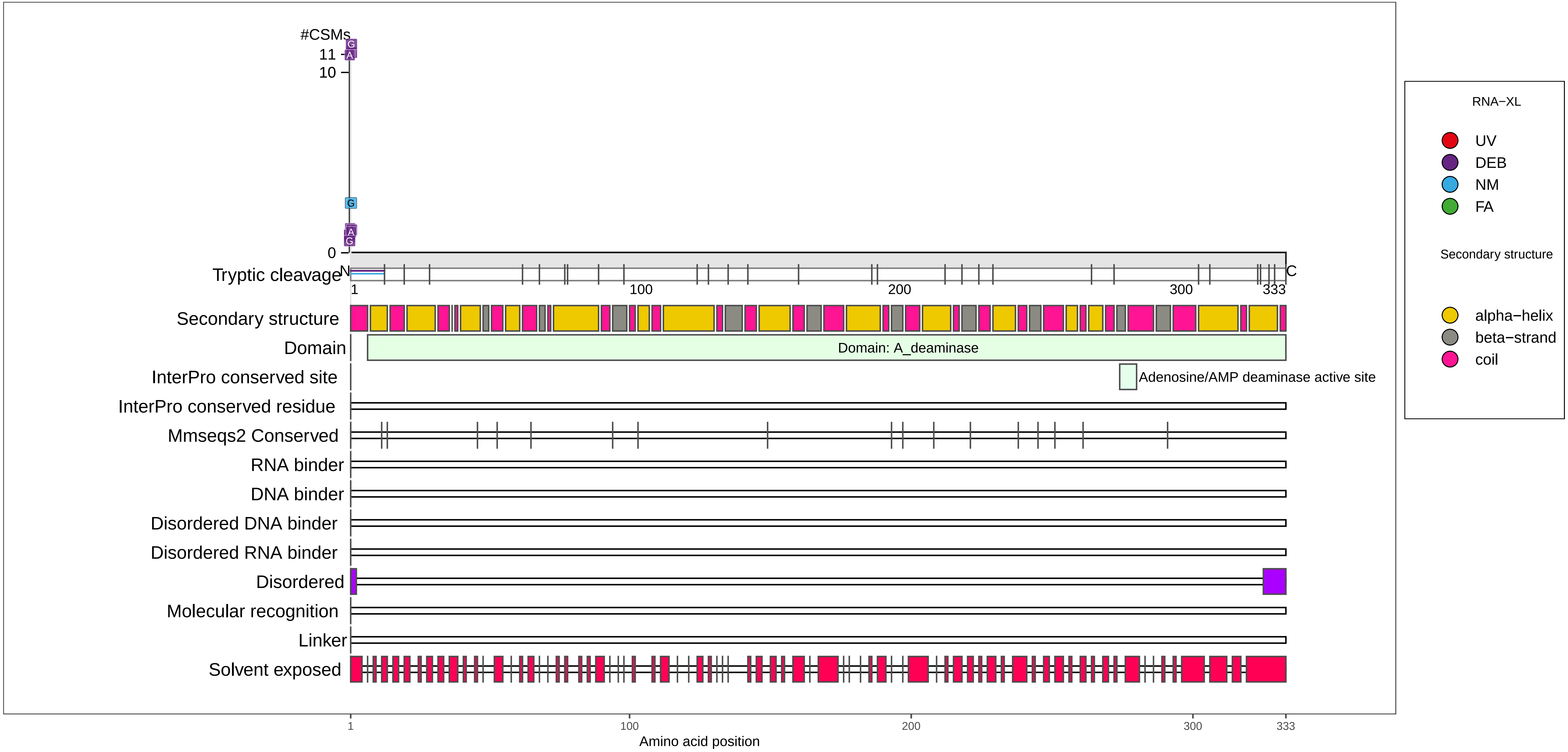

P23843
OPPA_ECOLI Periplasmic oligopeptide−binding protein

− Abundance:
tryptic [log10 Intensity]: 8.53 (Q 76)
PAXdb K12 strain [ppm]: 2.81 (Q 86)
PAXdb E.coli [ppm]: 3.15 (Q 96)

− RNA functions: not annotated

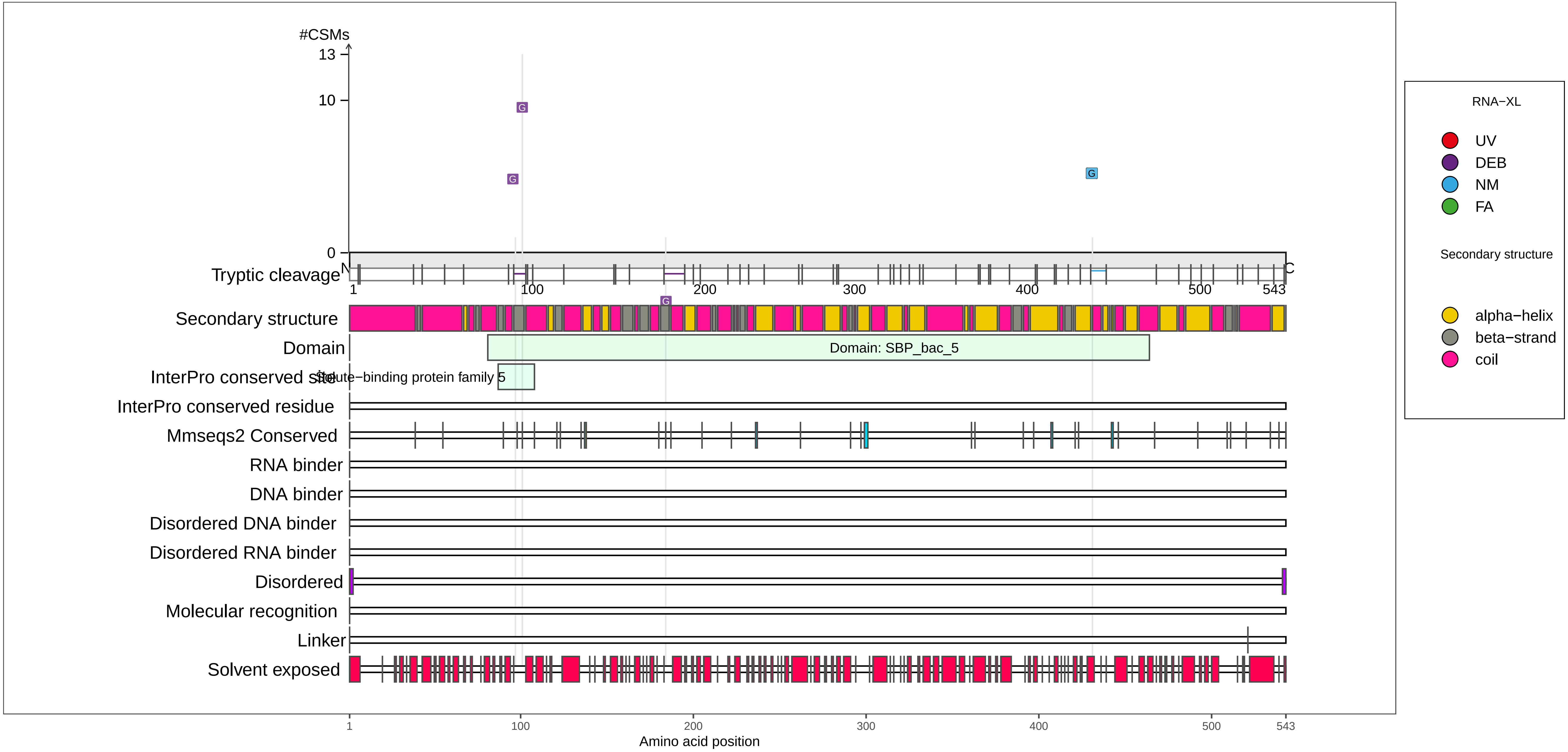

P23865
PRC_ECOLI Tail−specific protease
− Abundance:
tryptic [log10 Intensity]: 8.04 (Q 59)
PAXdb K12 strain [ppm]: 1.59 (Q 44)
PAXdb E.coli [ppm]: 2.29 (Q 80)

− RNA functions: not annotated

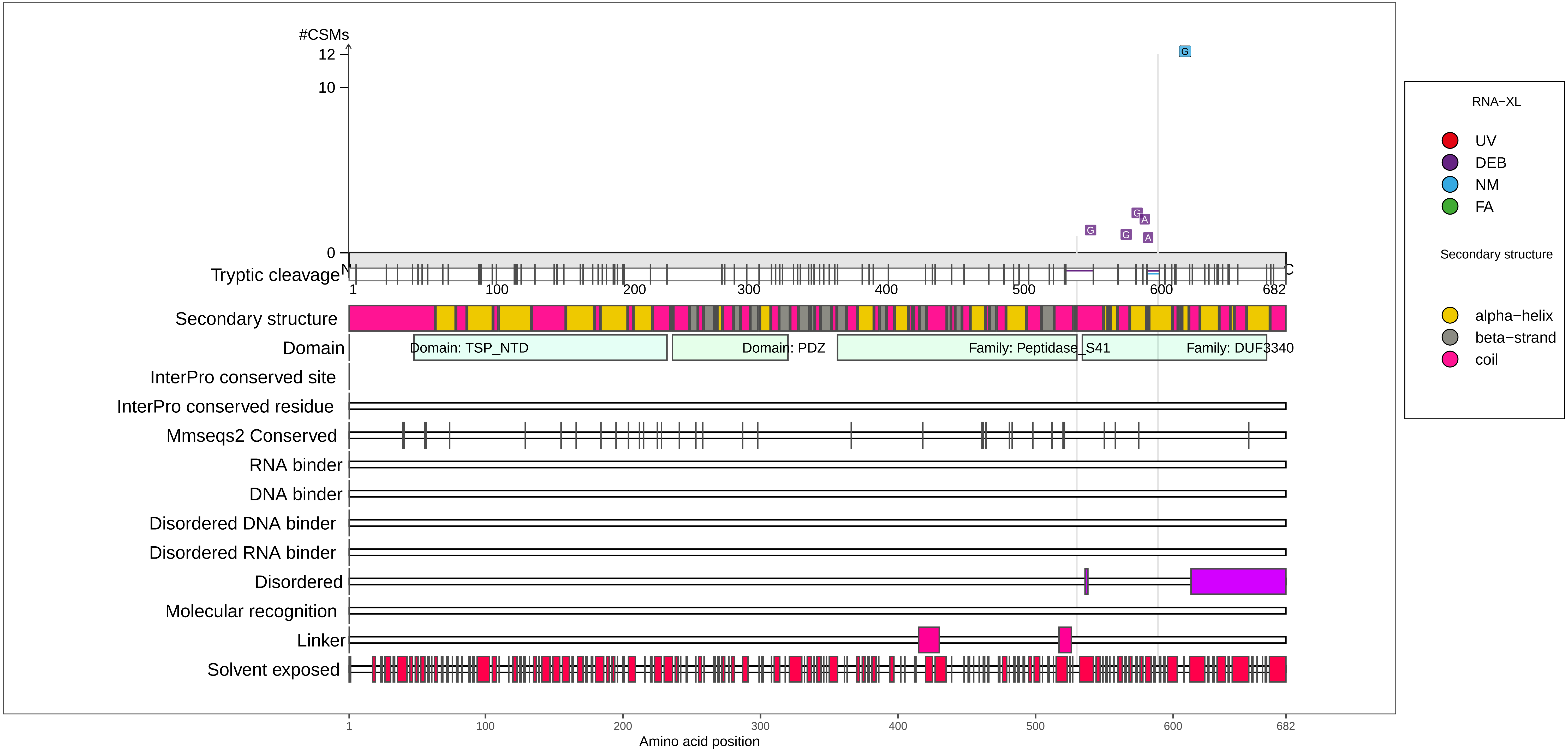

P30859
ARTI_ECOLI Putative ABC transporter arginine−binding protein 2

− Abundance:
tryptic [log10 Intensity]: 8.67 (Q 80)
PAXdb K12 strain [ppm]: 2.71 (Q 84)
PAXdb E.coli [ppm]: 2.94 (Q 94)

− RNA functions: not annotated

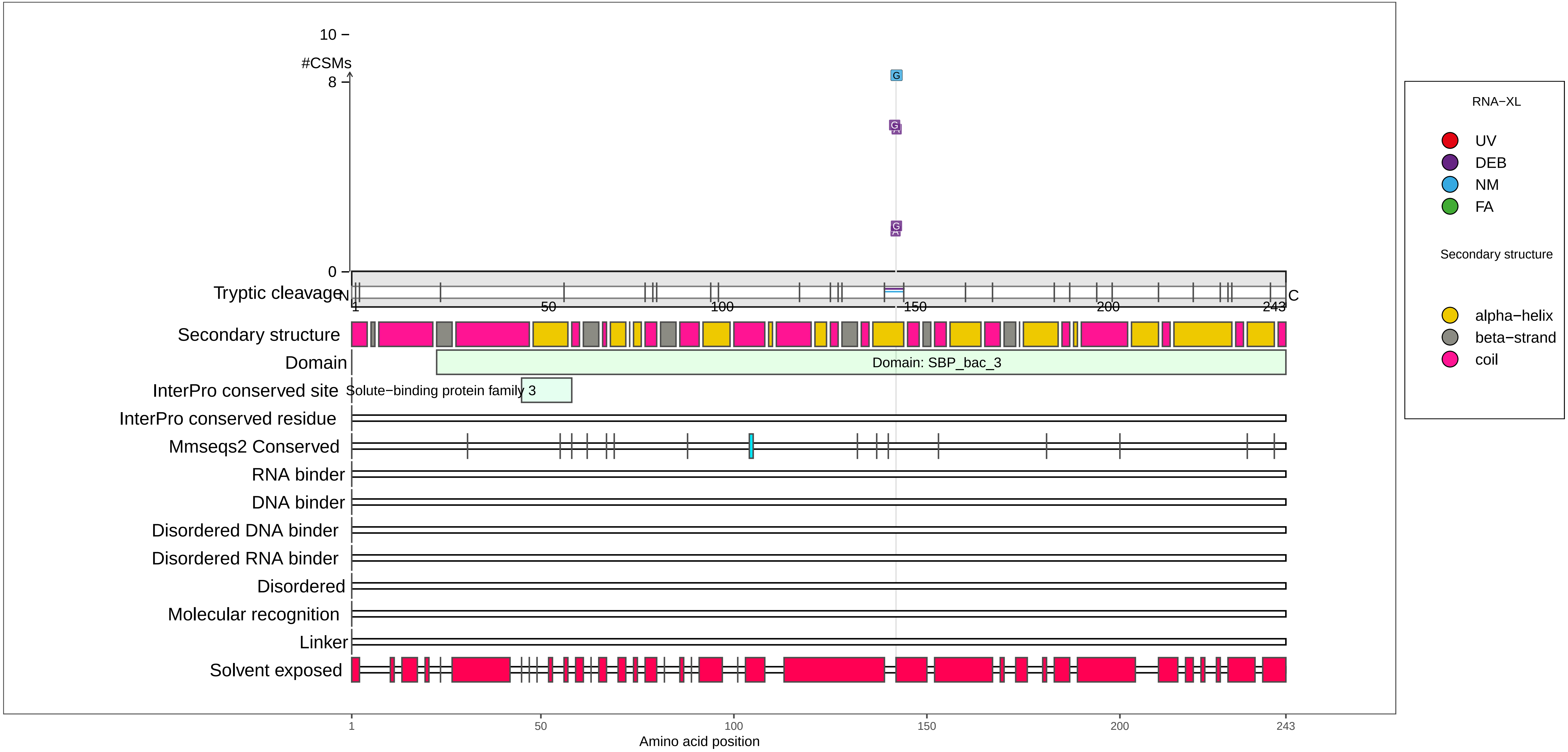

P75694
YAHO_ECOLI Uncharacterized protein YahO

− Abundance:
tryptic [log10 Intensity]: 8.11 (Q 62)
PAXdb K12 strain [ppm]: NA (Q NA)
PAXdb E.coli [ppm]: 3.17 (Q 96)

− RNA functions: not annotated

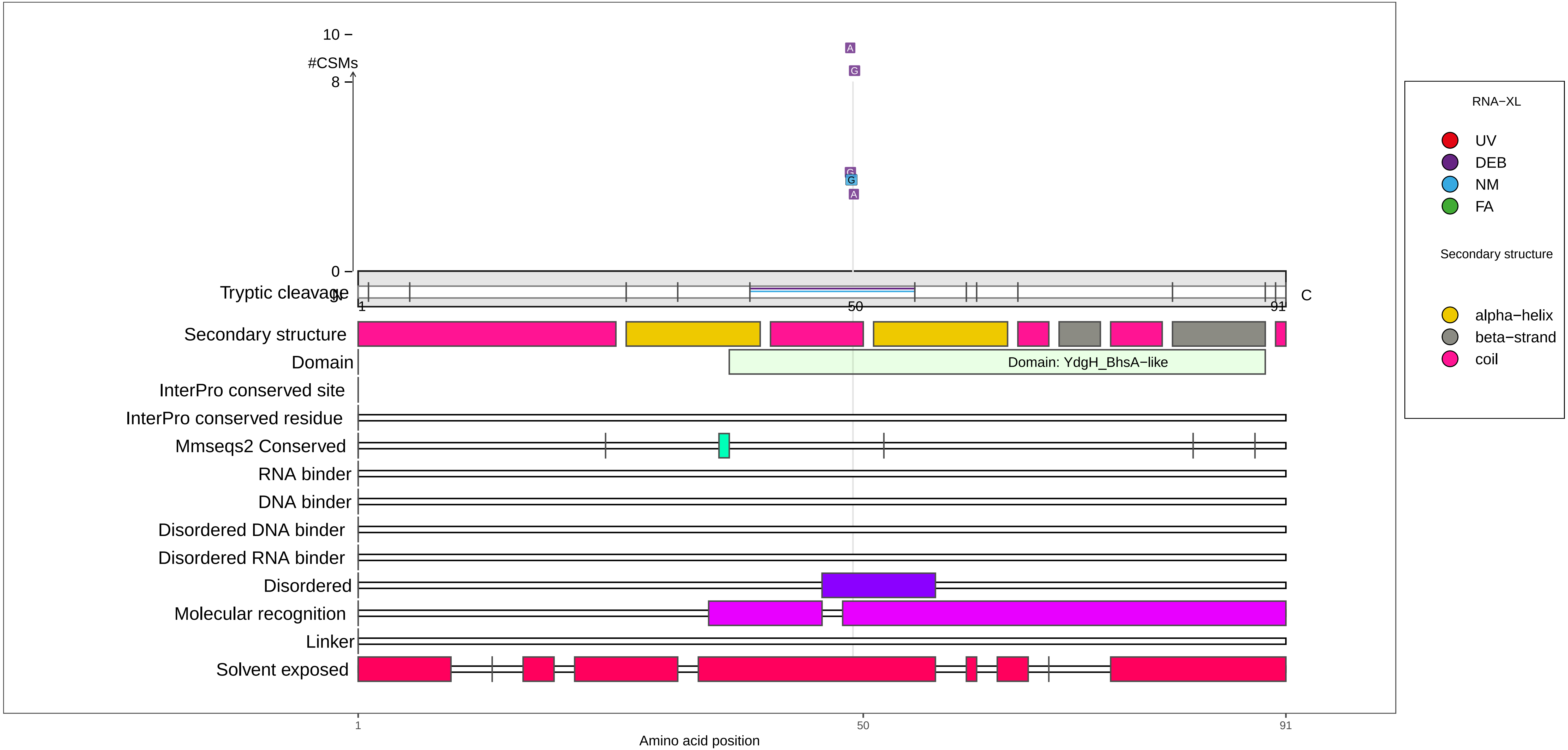

P77552
YDHQ_ECOLI Uncharacterized protein YdhQ

− Abundance:
tryptic [log10 Intensity]: 8.21 (Q 66)
PAXdb K12 strain [ppm]: 1.49 (Q 38)
PAXdb E.coli [ppm]: 0.68 (Q 40)

− RNA functions: not annotated

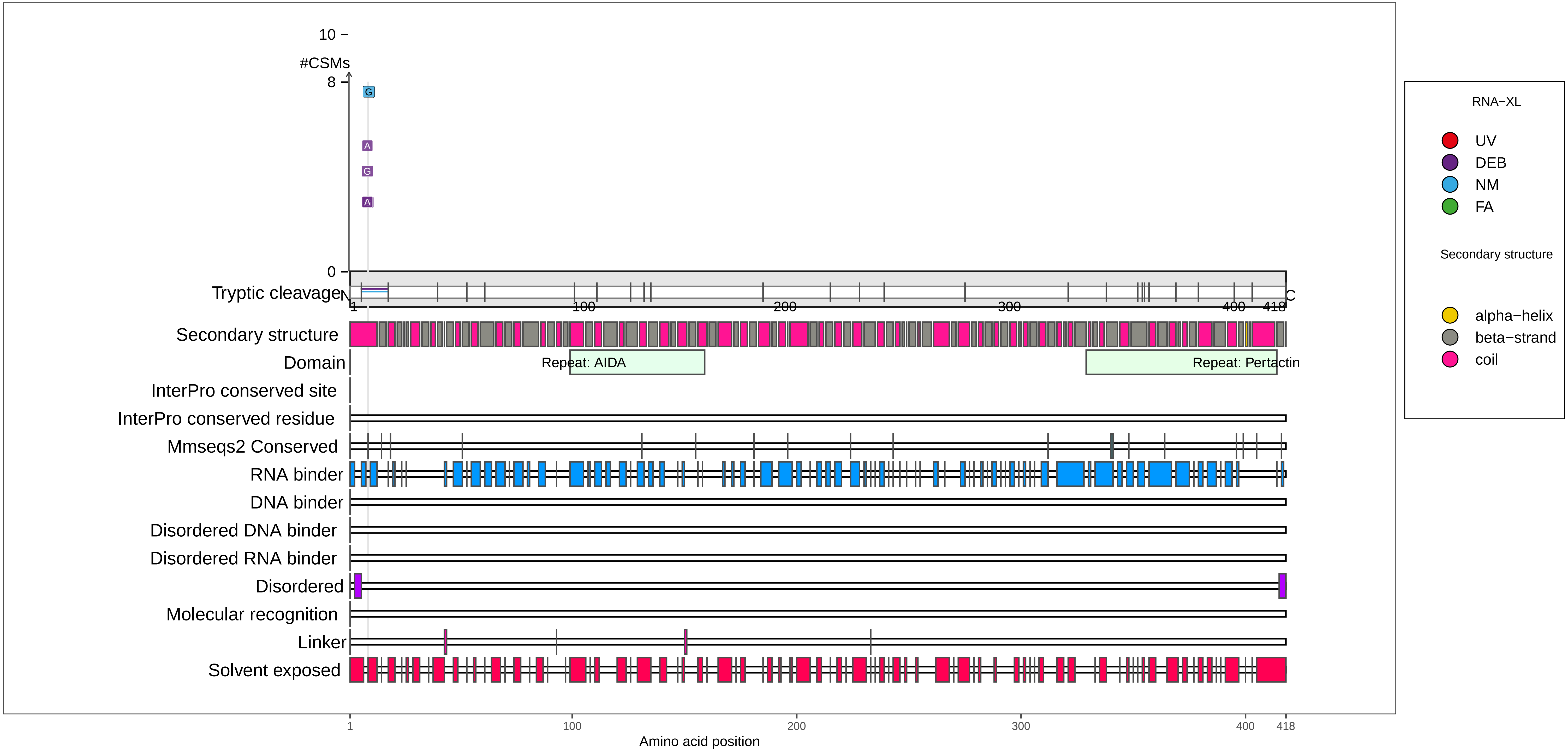

P00350
6PGD_ECOLI 6−phosphogluconate dehydrogenase, decarboxylating

− Abundance:
tryptic [log10 Intensity]: 9.41 (Q 95)
PAXdb K12 strain [ppm]: 3.03 (Q 91)
PAXdb E.coli [ppm]: 3.18 (Q 96)

− RNA functions: not annotated

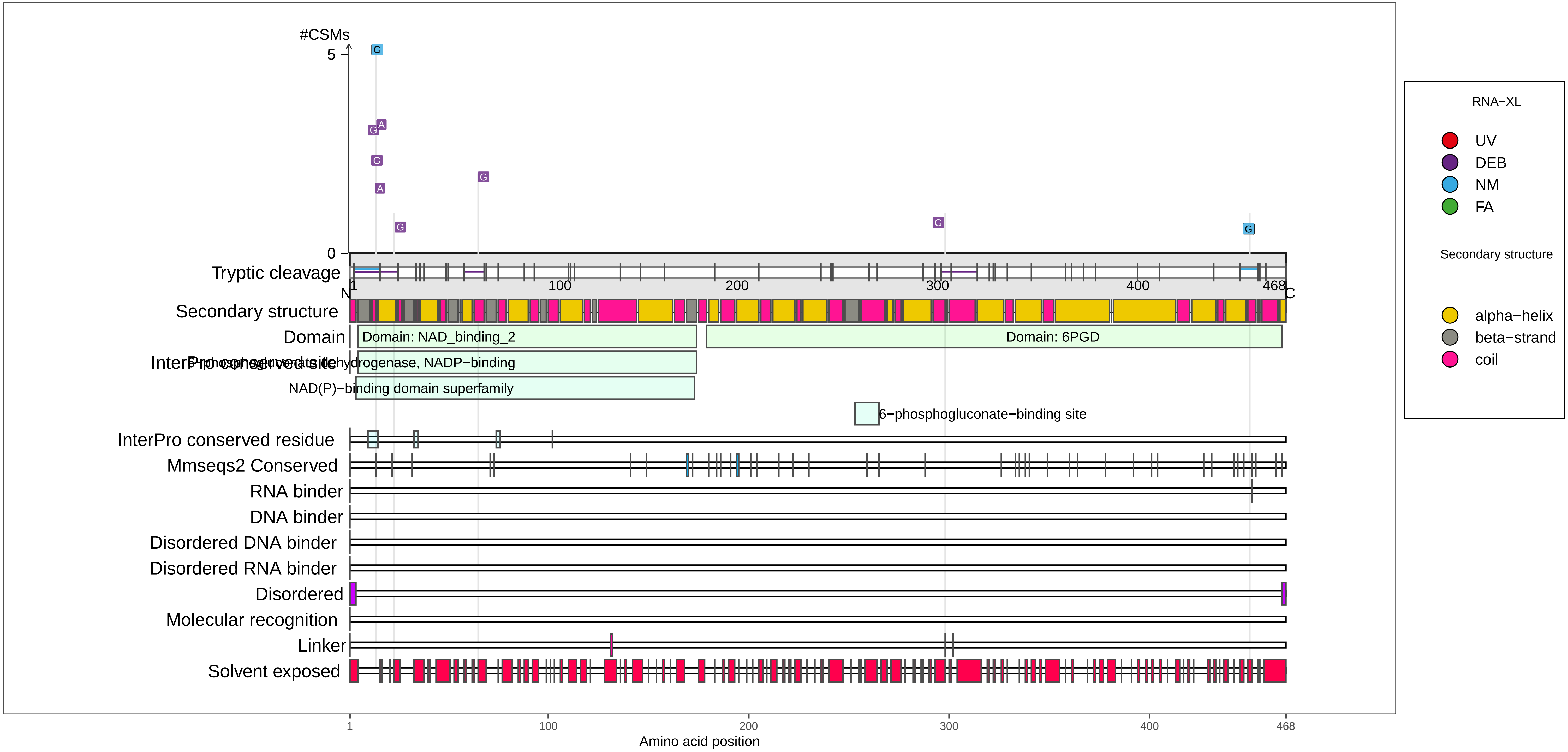

P0ABJ1
CYOA_ECOLI Cytochrome bo(3) ubiquinol oxidase subunit 2

− Abundance:
tryptic [log10 Intensity]: 9.75 (Q 98)
PAXdb K12 strain [ppm]: 2.08 (Q 63)
PAXdb E.coli [ppm]: 2.32 (Q 81)

− RNA functions: not annotated

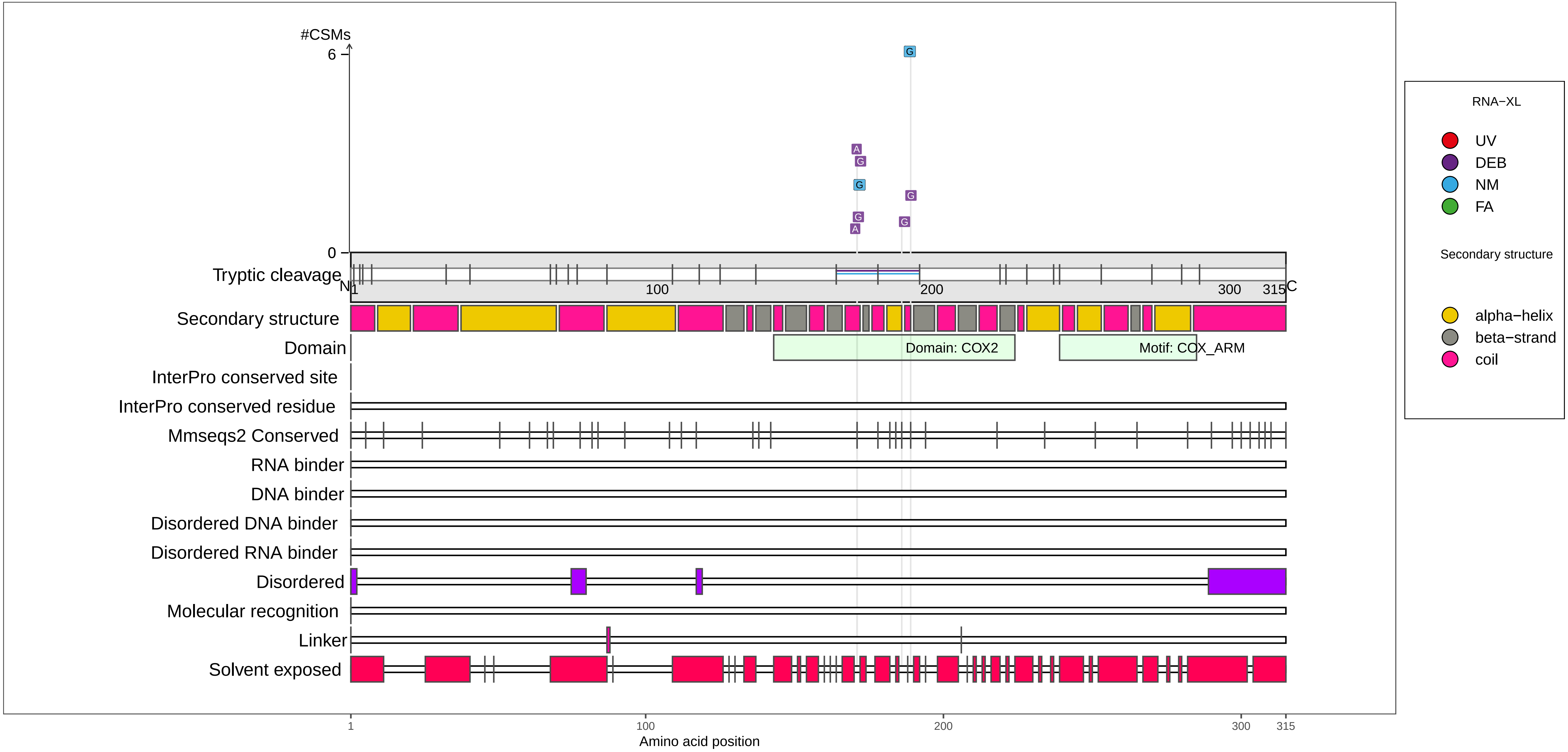

P0AEM4
FLGM_ECOLI Negative regulator of flagellin synthesis

− Abundance:
tryptic [log10 Intensity]: 7.56 (Q 38)
PAXdb K12 strain [ppm]: 3.09 (Q 92)
PAXdb E.coli [ppm]: 2.51 (Q 86)

− RNA functions: not annotated

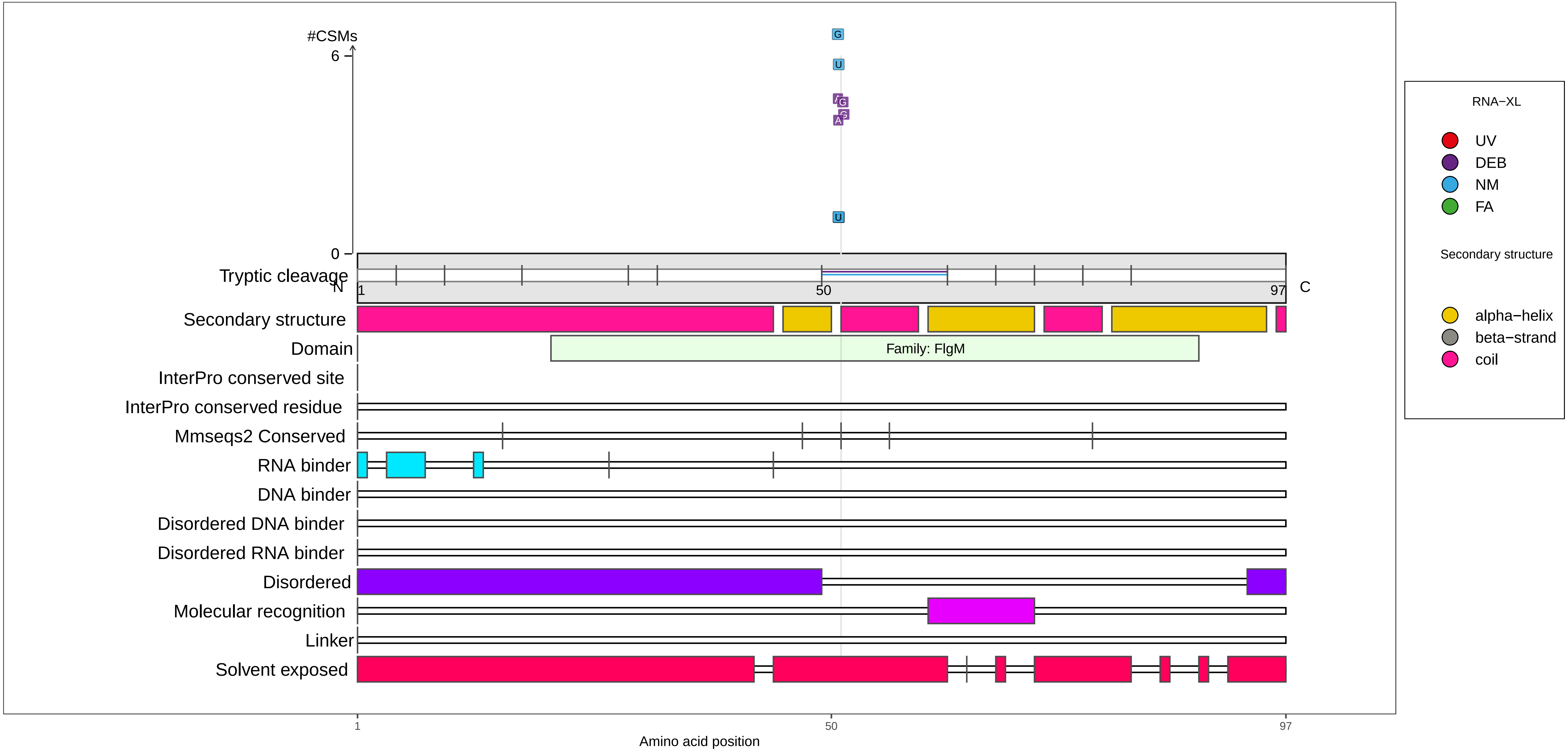

P23522
GARL_ECOLI 5−keto−4−deoxy−D−glucarate aldolase

− Abundance:
tryptic [log10 Intensity]: 7.71 (Q 45)
PAXdb K12 strain [ppm]: 1.38 (Q 31)
PAXdb E.coli [ppm]: 0.92 (Q 46)

− RNA functions: not annotated

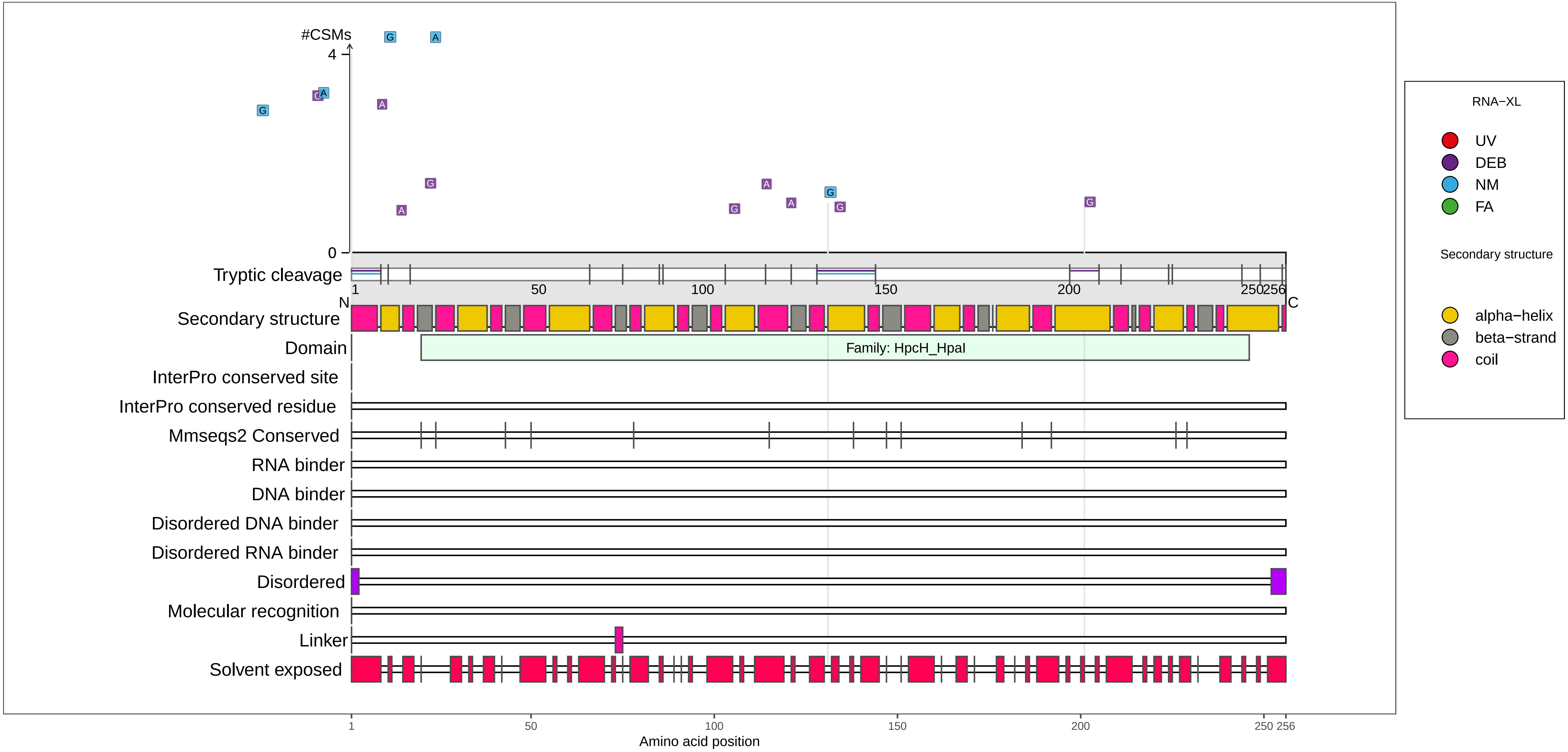

P23721
SERC_ECOLI Phosphoserine aminotransferase

− Abundance:
tryptic [log10 Intensity]: 8.99 (Q 88)
PAXdb K12 strain [ppm]: 2.42 (Q 75)
PAXdb E.coli [ppm]: 3.15 (Q 96)

− RNA functions: not annotated

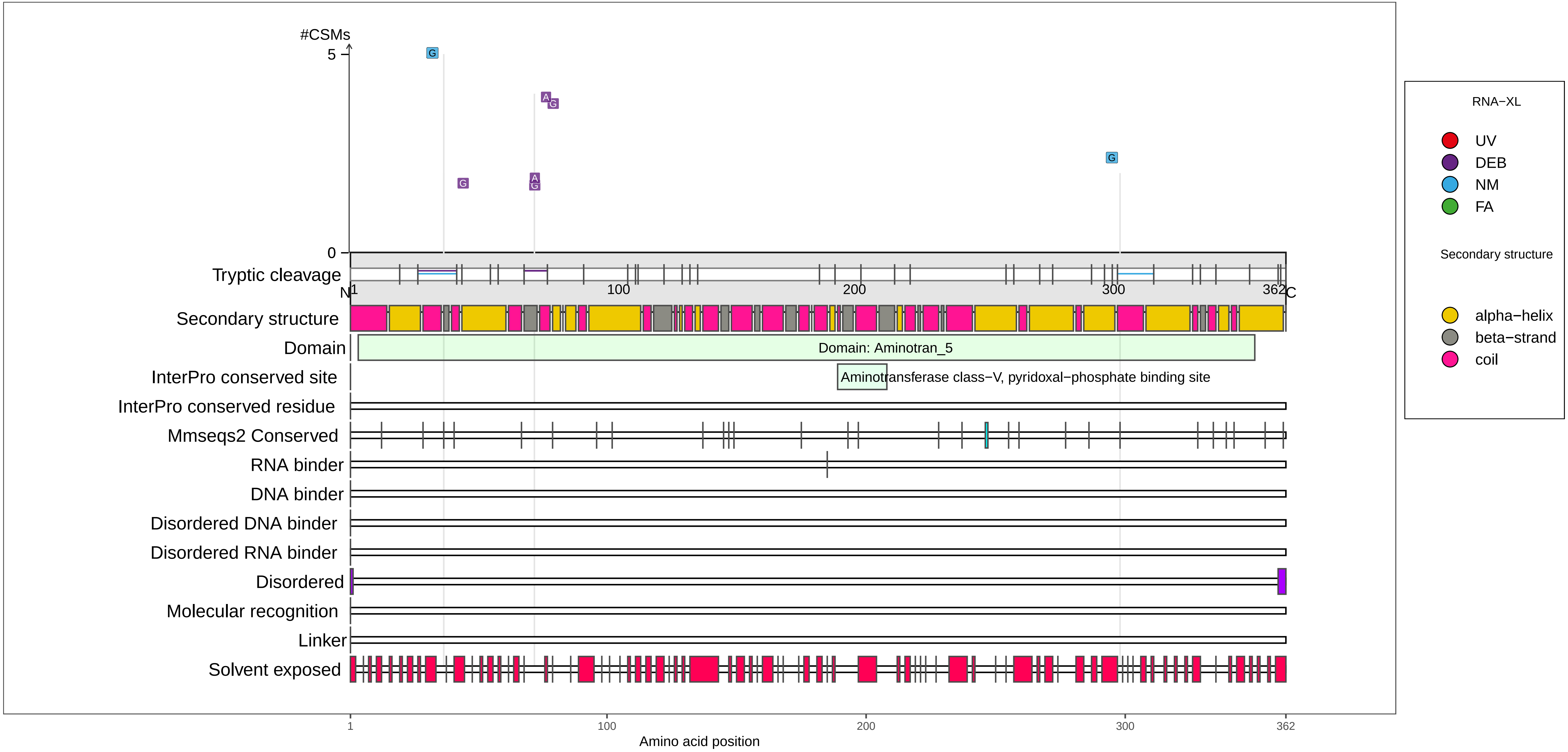

P25740

RFAG_ECOLI Lipopolysaccharide core biosynthesis protein RfaG

− Abundance:
tryptic [log10 Intensity]: 7.08 (Q 16)
PAXdb K12 strain [ppm]: 1.17 (Q 19)
PAXdb E.coli [ppm]: 0.48 (Q 36)

− RNA functions: not annotated

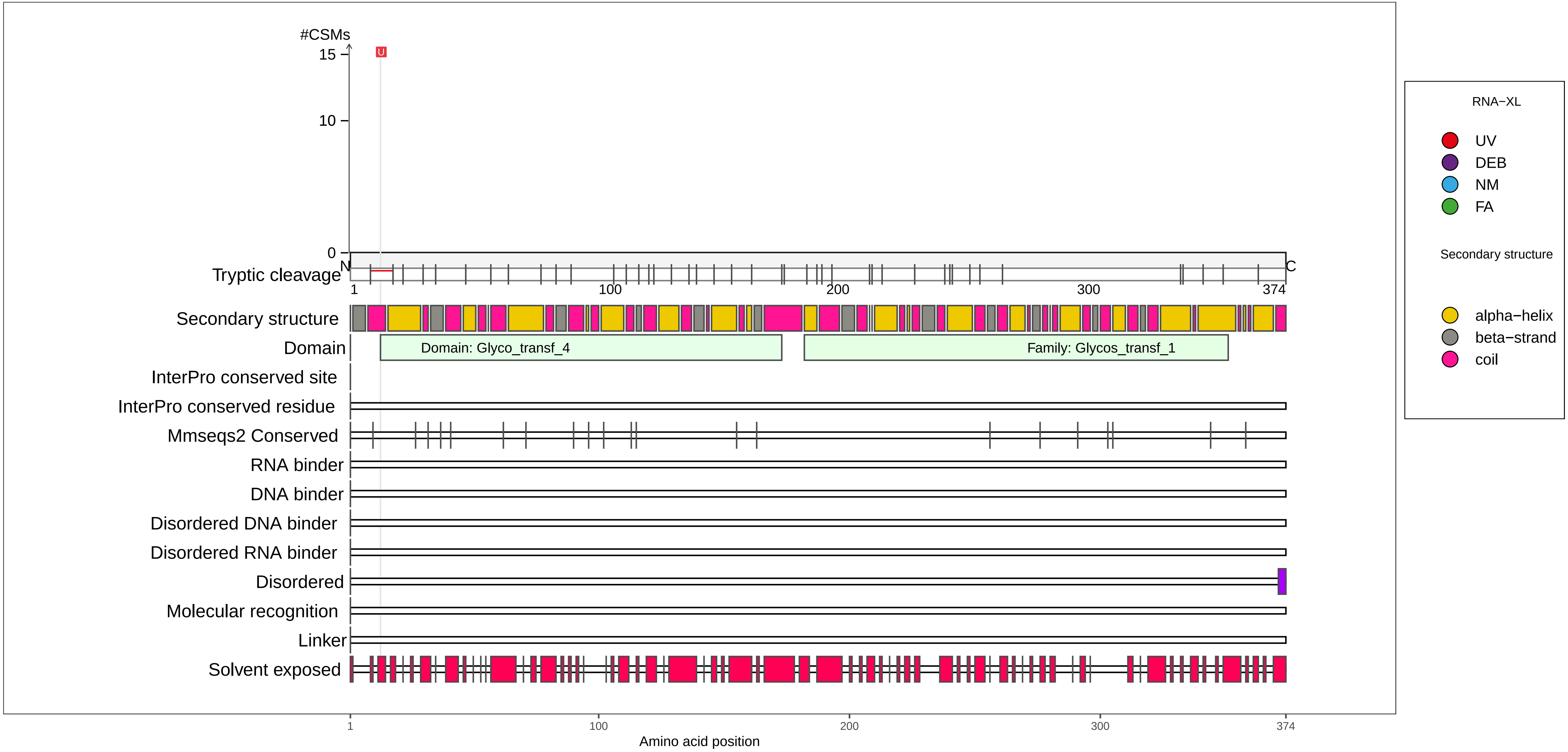

P36979
RLMN_ECOLI Dual−specificity RNA methyltransferase RlmN

− Abundance:
tryptic [log10 Intensity]: 7.66 (Q 43)
PAXdb K12 strain [ppm]: 1.17 (Q 19)
PAXdb E.coli [ppm]: 1.96 (Q 71)

− RNA functions:
ncRNA metabolic process; ncRNA processing; RNA binding; RNA metabolic process
RNA methylation; RNA methyltransferase activity; RNA modification; RNA processing
rRNA (adenine−C2−)−methyltransferase activity; rRNA (adenine) methyltransferase activity
rRNA base methylation; rRNA binding; rRNA metabolic process; rRNA methylation
rRNA methyltransferase activity; rRNA modification; rRNA processing
tRNA (adenine−C2−)−methyltransferase activity; tRNA (adenine) methyltransferase activity; tRNA binding
tRNA metabolic process; tRNA methylation; tRNA methyltransferase activity; tRNA modification
tRNA processing

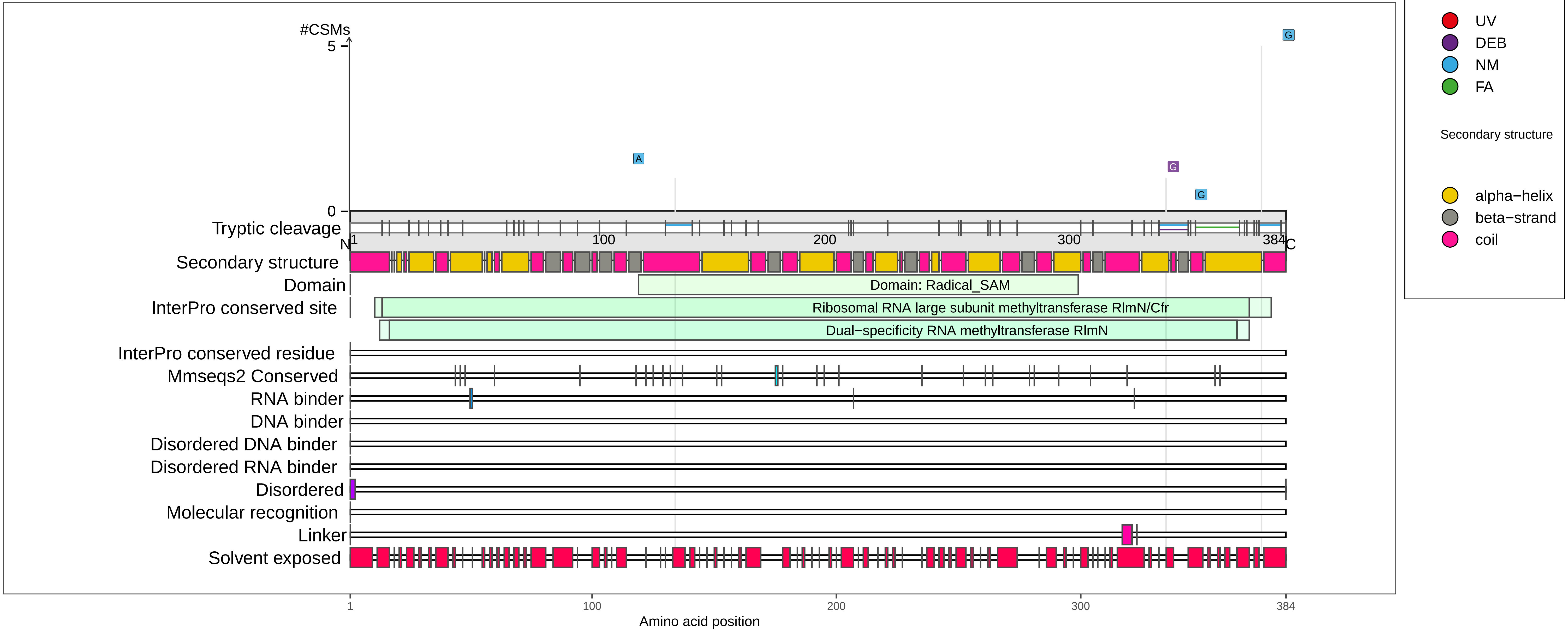

P37666
GHRB_ECOLI Glyoxylate/hydroxypyruvate reductase B

− Abundance:
tryptic [log10 Intensity]: 8.41 (Q 73)
PAXdb K12 strain [ppm]: 2.61 (Q 81)
PAXdb E.coli [ppm]: 2.53 (Q 86)

− RNA functions: not annotated

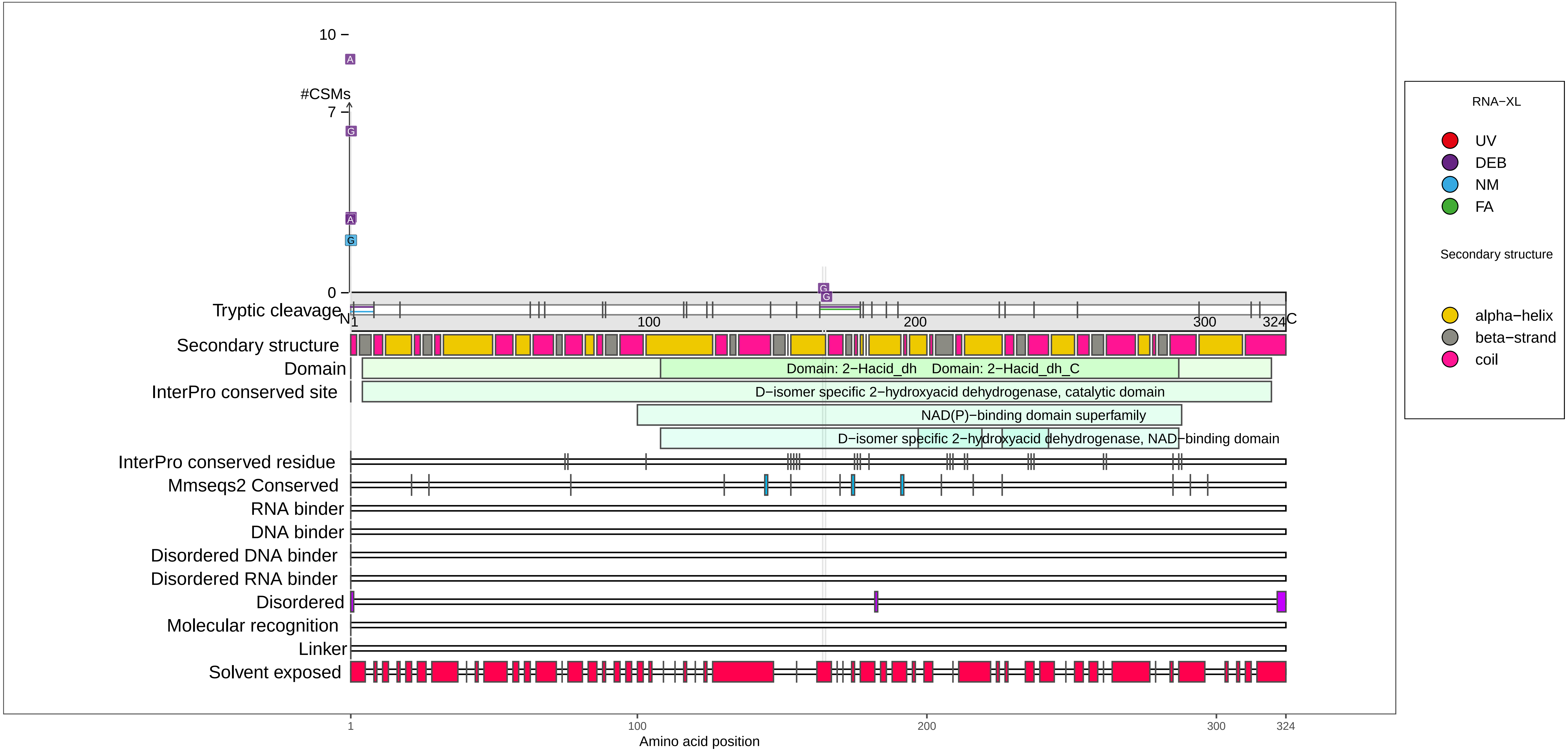

P62623
ISPH_ECOLI 4−hydroxy−3−methylbut−2−enyl diphosphate reductase

− Abundance:
tryptic [log10 Intensity]: 8.84 (Q 85)
PAXdb K12 strain [ppm]: NA (Q NA)
PAXdb E.coli [ppm]: 2.09 (Q 74)

− RNA functions: not annotated

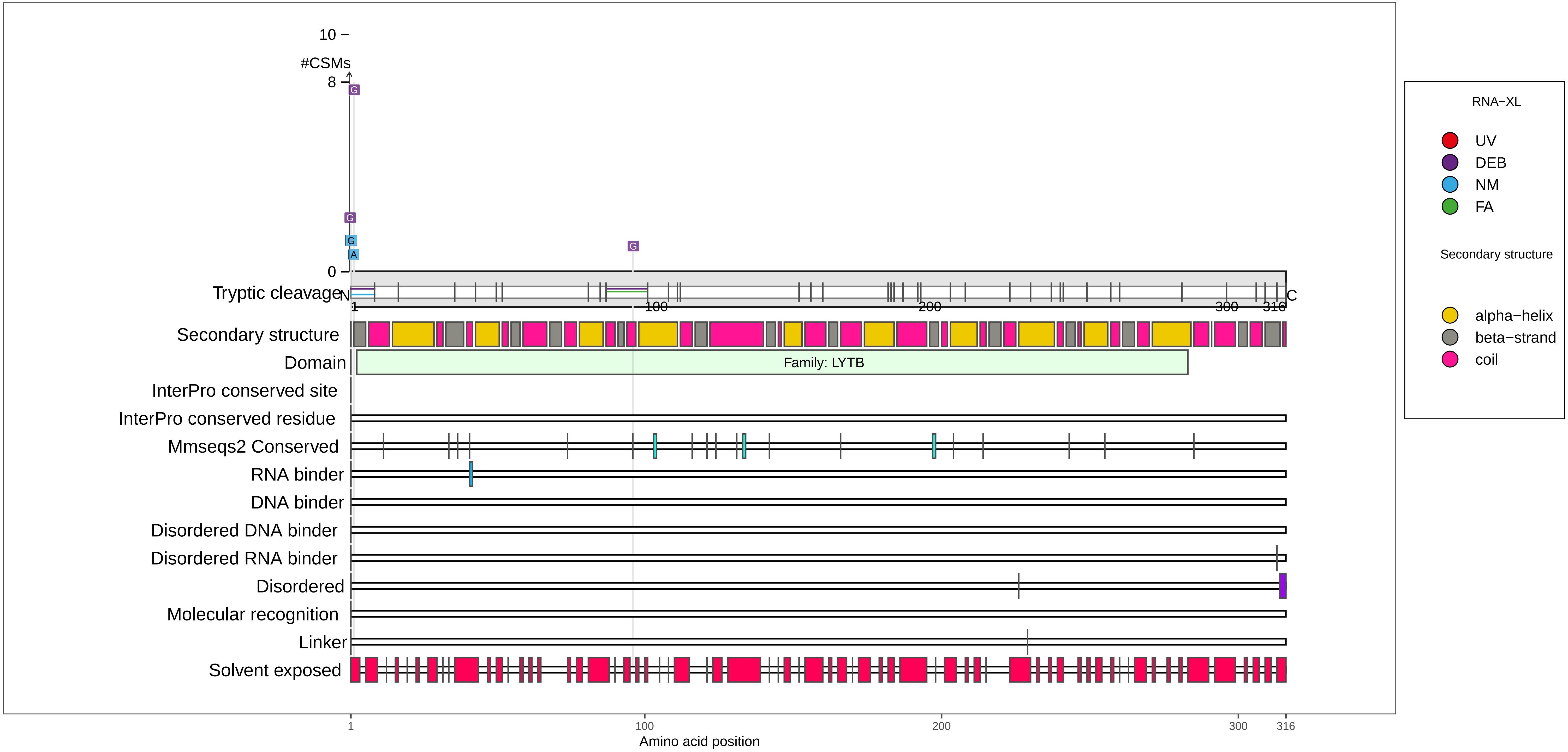

P67603
AC4CH_ECOLI N(4)−acetylcytidine amidohydrolase
− RNA functions: not annotated

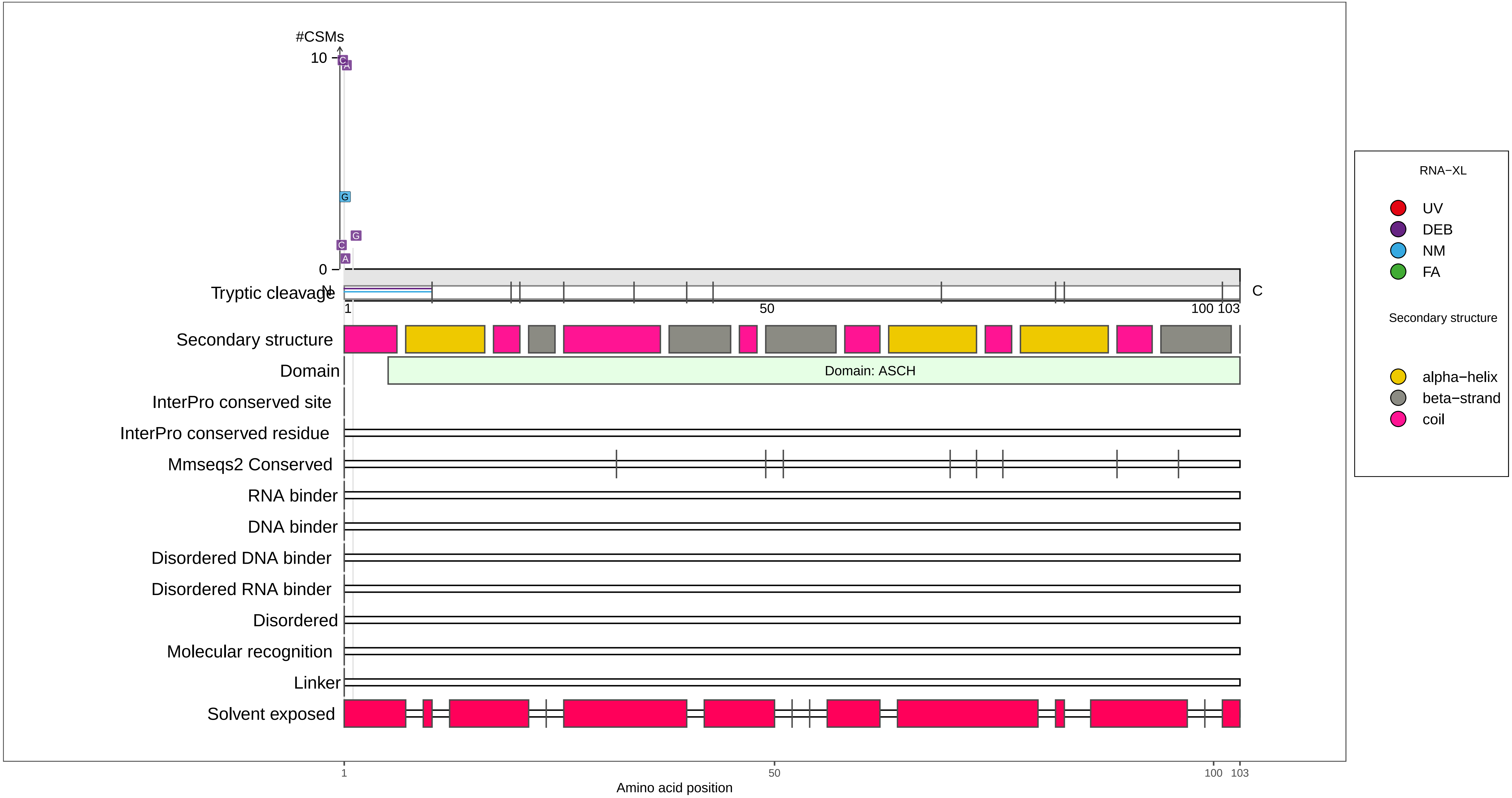

P69924
RIR2_ECOLI Ribonucleoside−diphosphate reductase 1 subunit beta

− Abundance:
tryptic [log10 Intensity]: 8.86 (Q 85)
PAXdb K12 strain [ppm]: 2.32 (Q 71)
PAXdb E.coli [ppm]: 2.22 (Q 78)

− RNA functions: not annotated

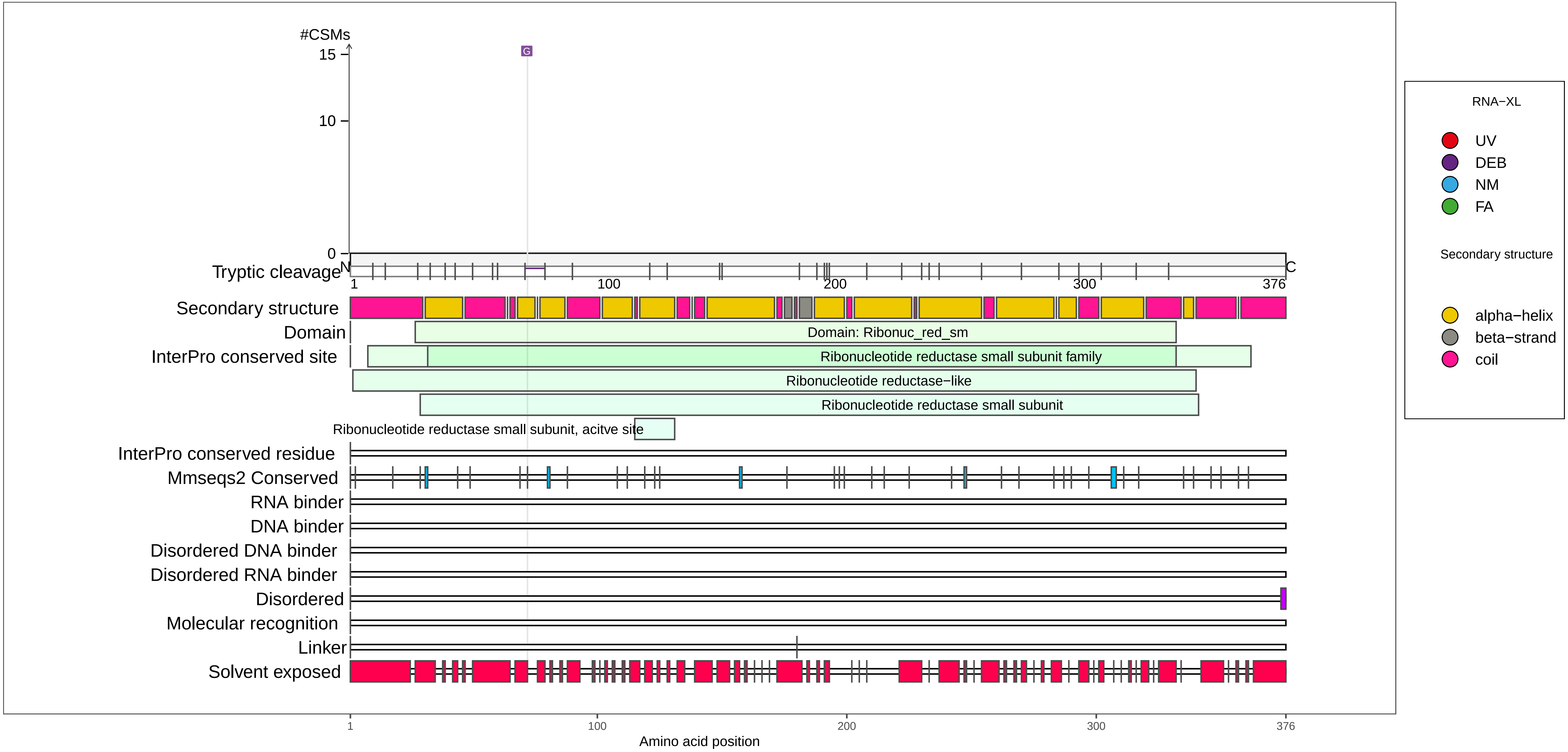

P76373
UDG_ECOLI UDP−glucose 6−dehydrogenase

− Abundance:
tryptic [log10 Intensity]: 6.7 (Q 5)
PAXdb K12 strain [ppm]: 2.27 (Q 69)
PAXdb E.coli [ppm]: 1.15 (Q 51)

− RNA functions: not annotated

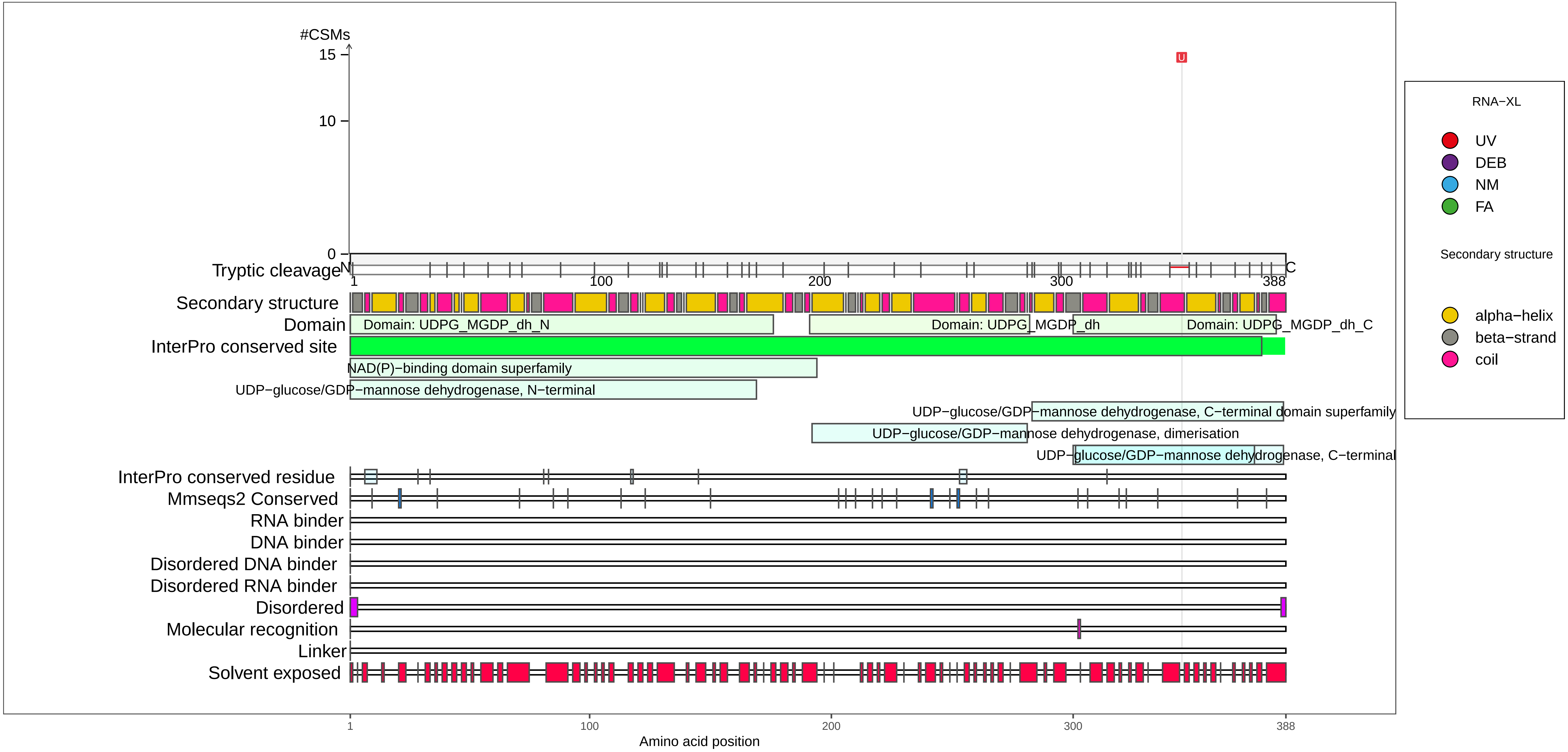

P04995
EX1_ECOLI Exodeoxyribonuclease I

− Abundance:
tryptic [log10 Intensity]: 7.95 (Q 56)
PAXdb K12 strain [ppm]: 1.26 (Q 24)
PAXdb E.coli [ppm]: 1.21 (Q 52)

− RNA functions:
RNA metabolic process; RNA phosphodiester bond hydrolysis
RNA phosphodiester bond hydrolysis, exonucleolytic

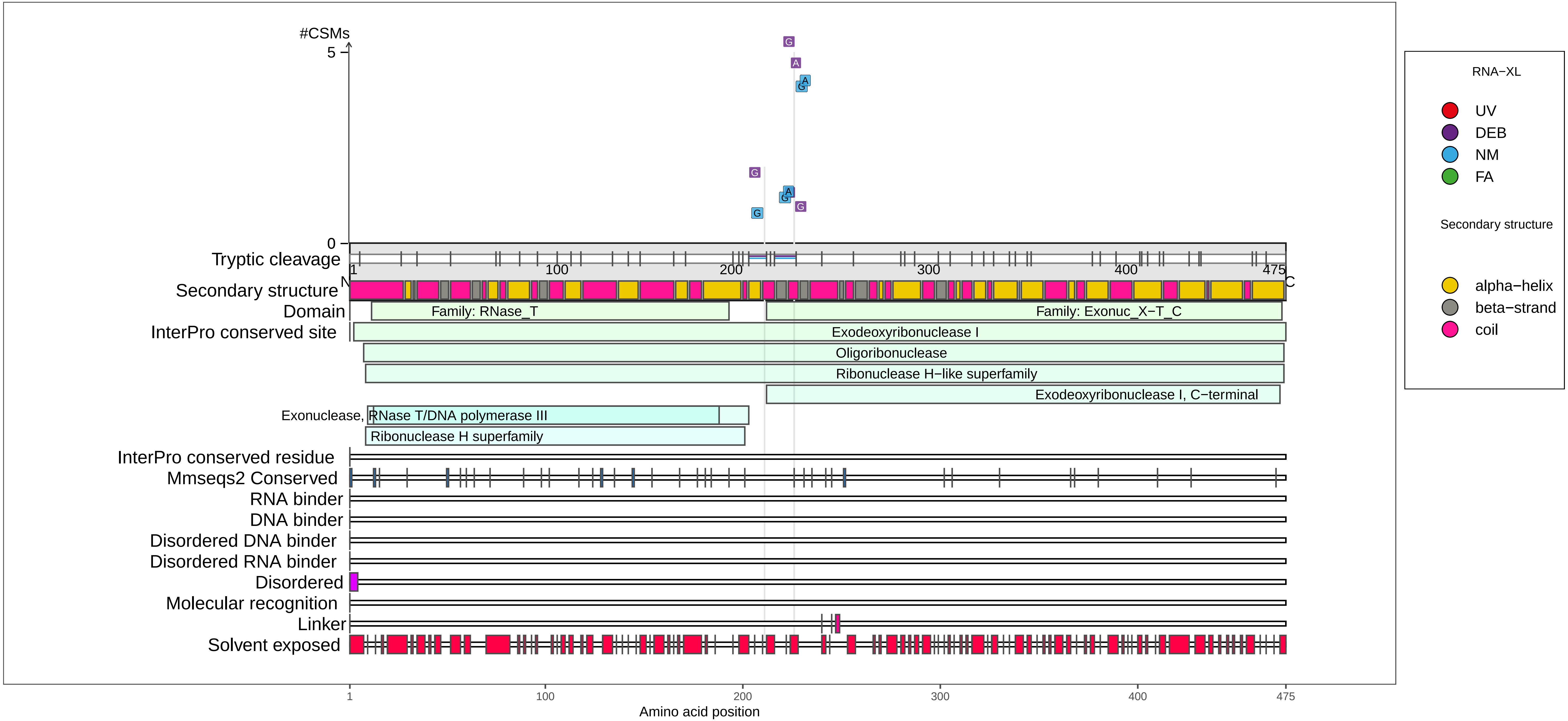

P09394
GLPQ_ECOLI Glycerophosphodiester phosphodiesterase, periplasmic

− Abundance:
tryptic [log10 Intensity]: 9.44 (Q 96)
PAXdb K12 strain [ppm]: 2.86 (Q 87)
PAXdb E.coli [ppm]: 2.29 (Q 80)

− RNA functions: not annotated

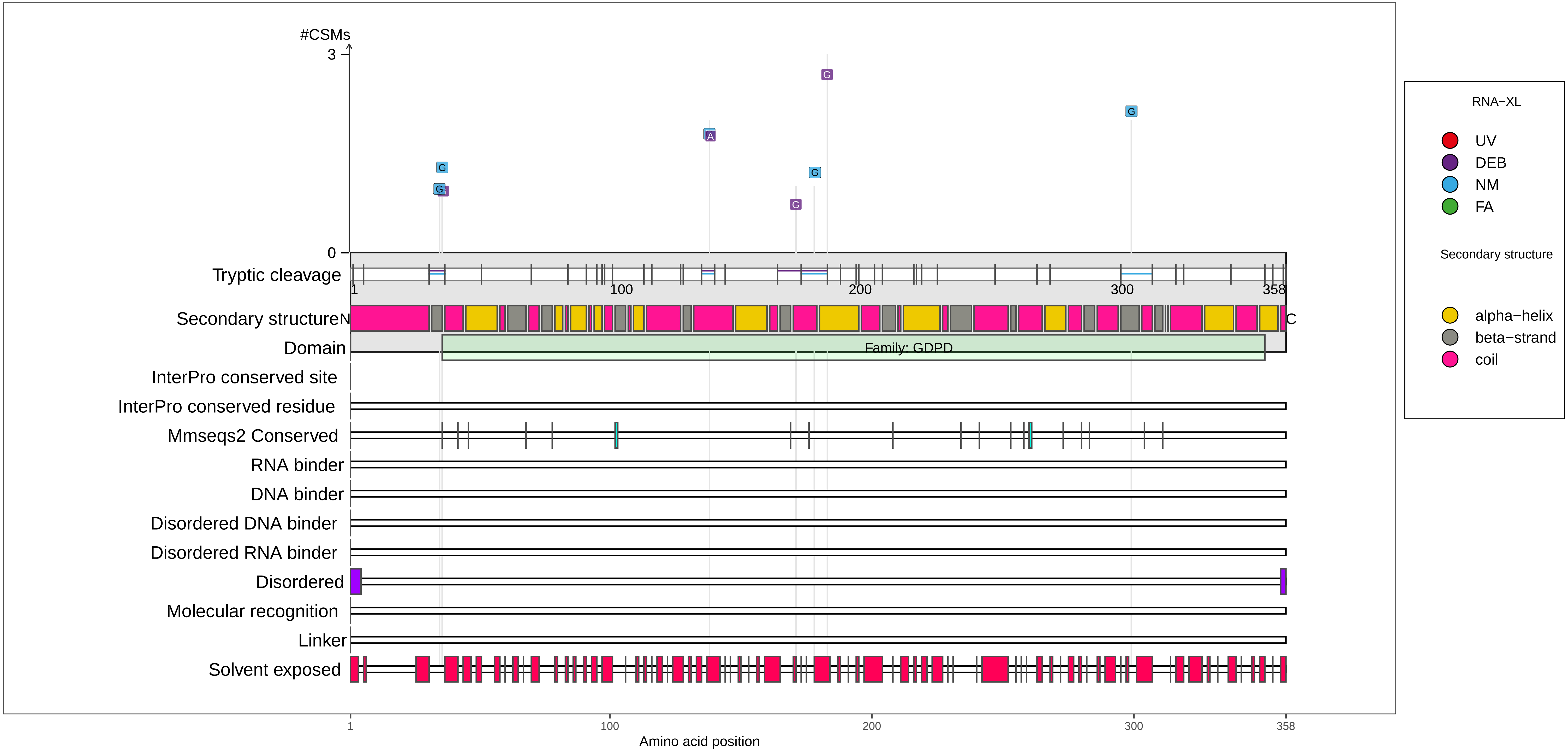

P0A796
PFKA_ECOLI ATP−dependent 6−phosphofructokinase isozyme 1

− Abundance:
tryptic [log10 Intensity]: 8.63 (Q 79)
PAXdb K12 strain [ppm]: 2.29 (Q 69)
PAXdb E.coli [ppm]: 2.9 (Q 93)

− RNA functions: not annotated

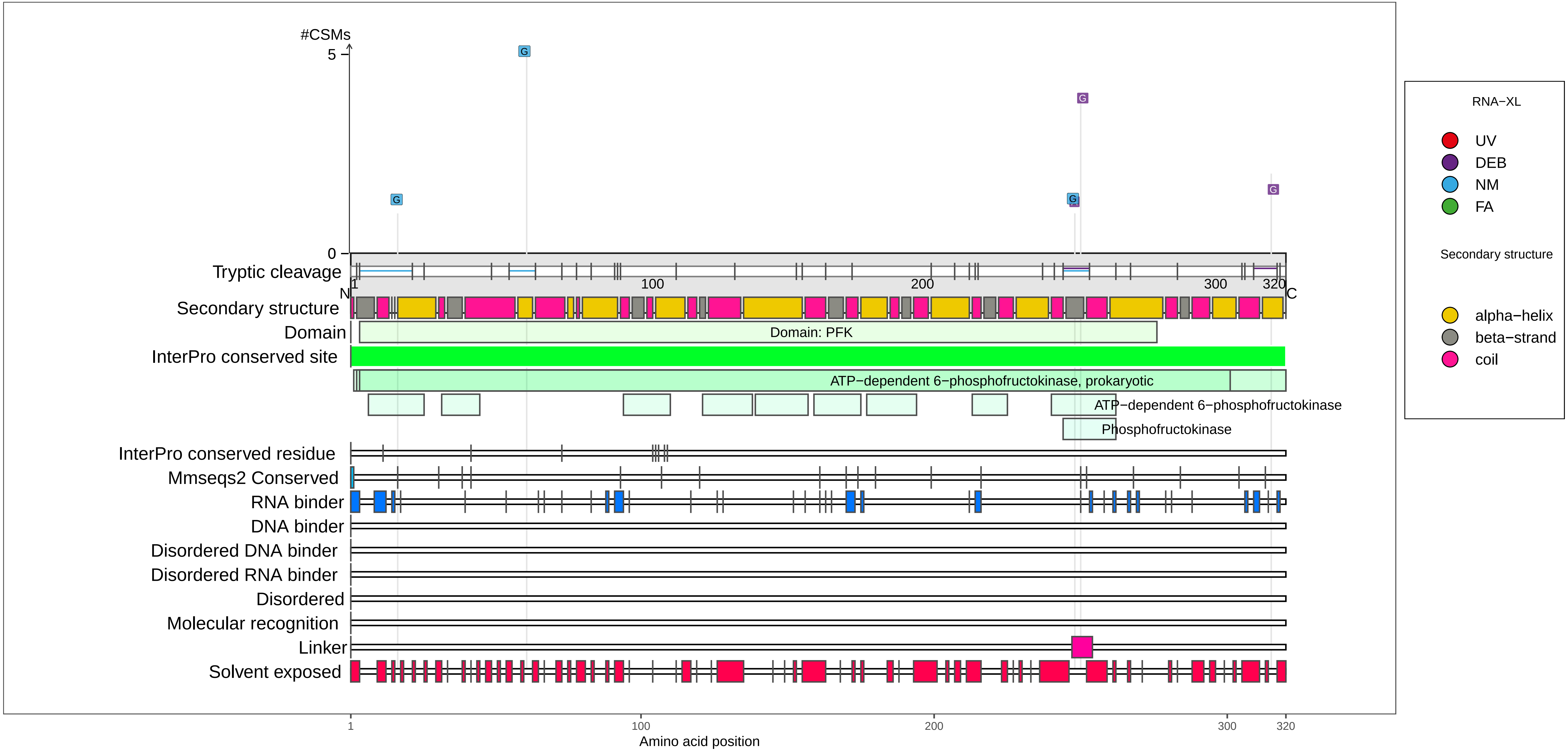

P0A7Y0
RNC_ECOLI Ribonuclease 3

− Abundance:
tryptic [log10 Intensity]: 7.86 (Q 52)
PAXdb K12 strain [ppm]: 2.6 (Q 80)
PAXdb E.coli [ppm]: 2.43 (Q 84)

− RNA functions:
Double−stranded RNA binding motif; mRNA metabolic process; mRNA processing
ncRNA catabolic process; ncRNA metabolic process; ncRNA processing; RNA binding
RNA catabolic process; RNA metabolic process; RNA phosphodiester bond hydrolysis
RNA phosphodiester bond hydrolysis, endonucleolytic; RNA processing; rRNA binding; rRNA catabolic process
rRNA metabolic process; rRNA processing; tRNA metabolic process; tRNA processing

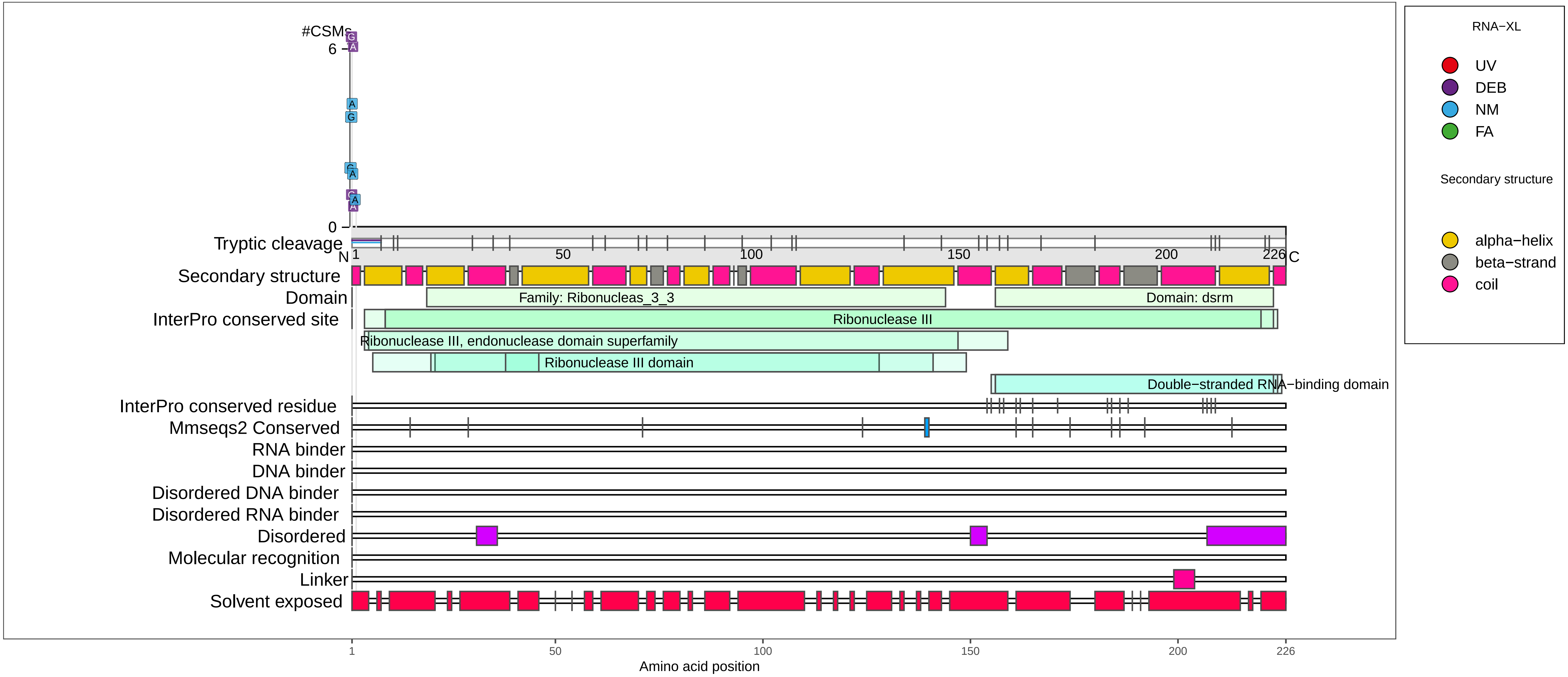

P0A8W8
YFBU_ECOLI UPF0304 protein YfbU

− Abundance:
tryptic [log10 Intensity]: 9.66 (Q 98)
PAXdb K12 strain [ppm]: 2.55 (Q 79)
PAXdb E.coli [ppm]: 2.76 (Q 91)

− RNA functions: not annotated

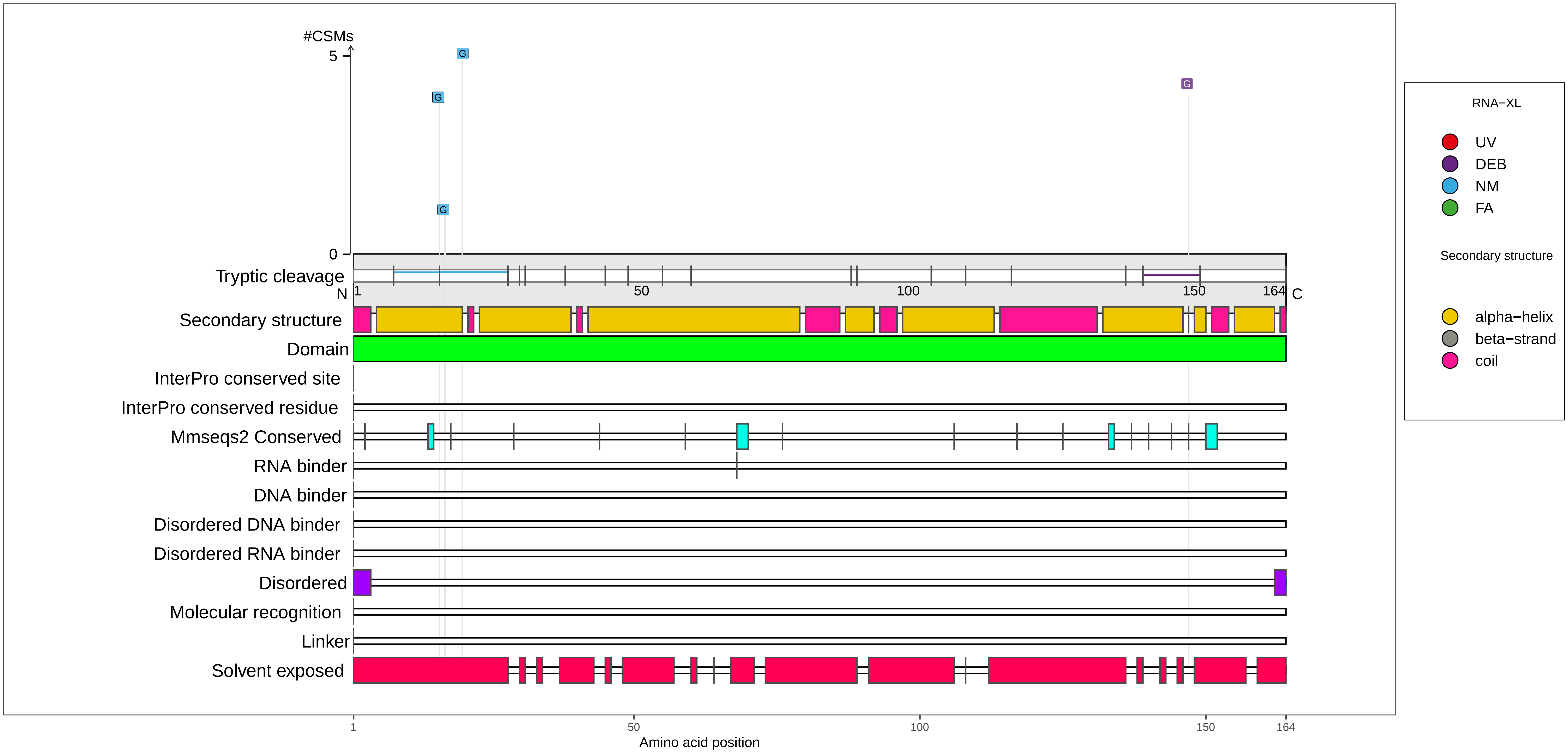

P0A8X0
YJGA_ECOLI UPF0307 protein YjgA

− Abundance:
tryptic [log10 Intensity]: 8.1 (Q 61)
PAXdb K12 strain [ppm]: 2.6 (Q 80)
PAXdb E.coli [ppm]: 2.6 (Q 88)

− RNA functions: not annotated

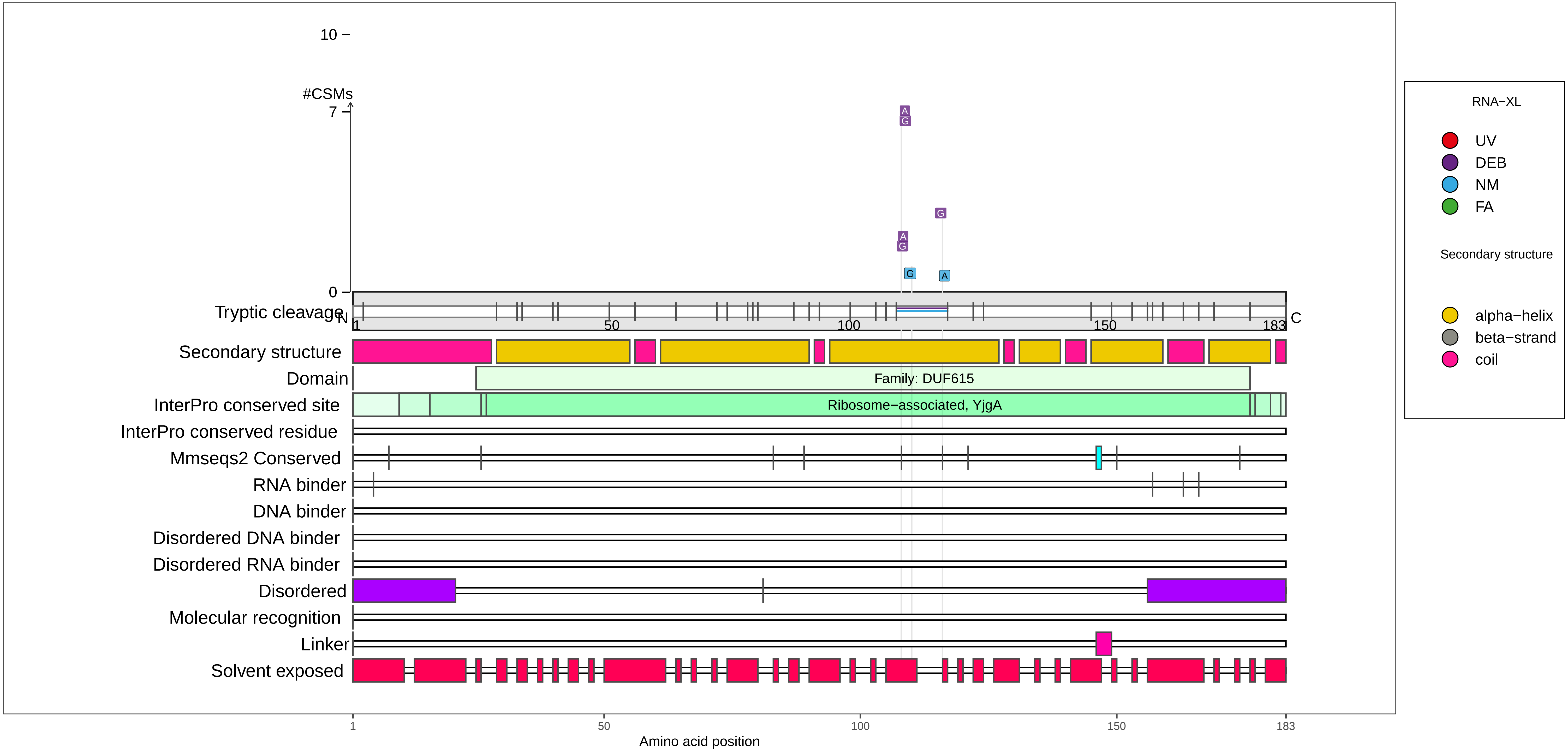

P0ABF1
PCNB_ECOLI Poly(A) polymerase I

− Abundance:
tryptic [log10 Intensity]: 7.56 (Q 38)
PAXdb K12 strain [ppm]: 1.95 (Q 59)
PAXdb E.coli [ppm]: 1.87 (Q 69)

− RNA functions:
mRNA 3−end processing; mRNA metabolic process; mRNA polyadenylation; mRNA processing
Probable RNA and SrmB− binding site of polymerase A; RNA 3−end processing; RNA binding
RNA metabolic process; RNA modification; RNA polyadenylation; RNA processing

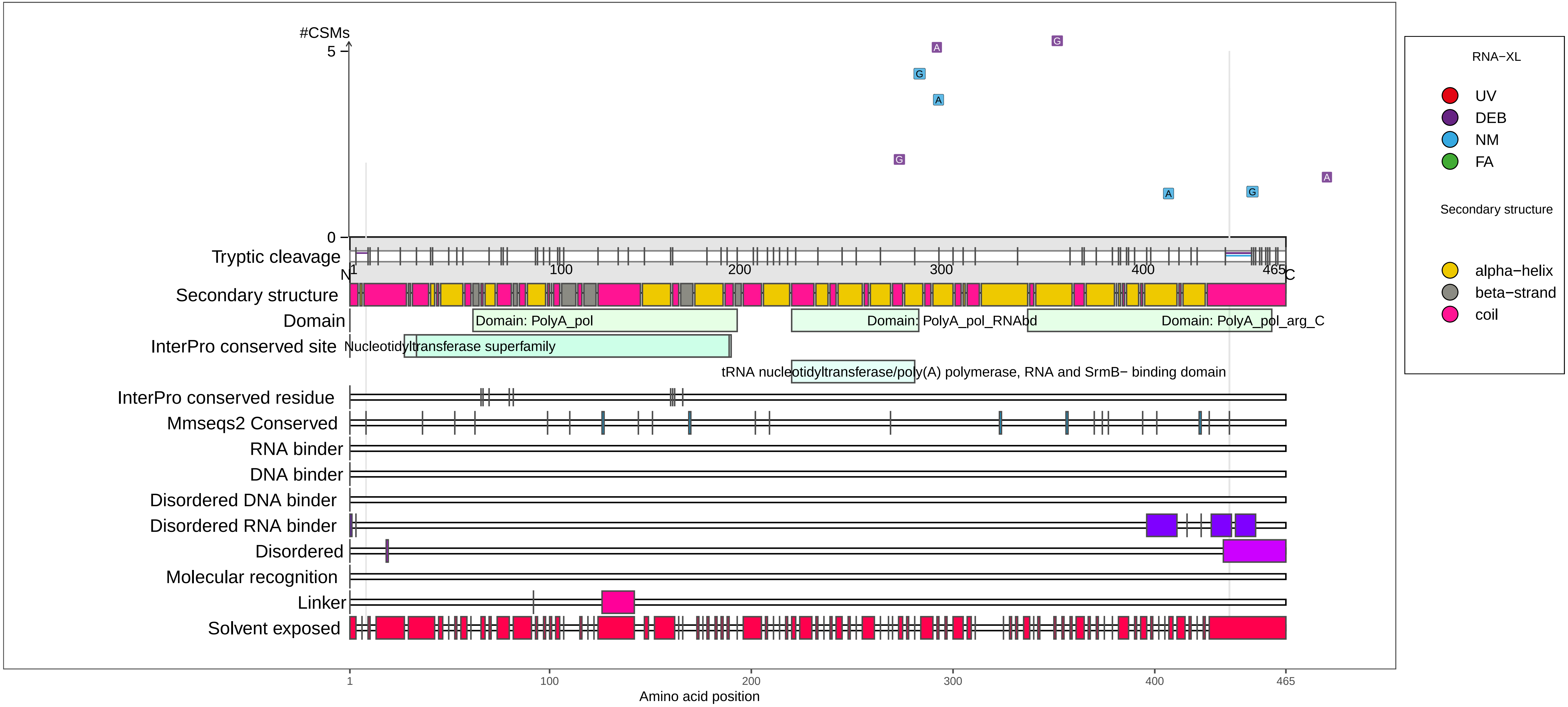

P0AC81
LGUL_ECOLI Lactoylglutathione lyase

− Abundance:
tryptic [log10 Intensity]: 7.98 (Q 57)
PAXdb K12 strain [ppm]: 1.69 (Q 49)
PAXdb E.coli [ppm]: 2.59 (Q 88)

− RNA functions: not annotated

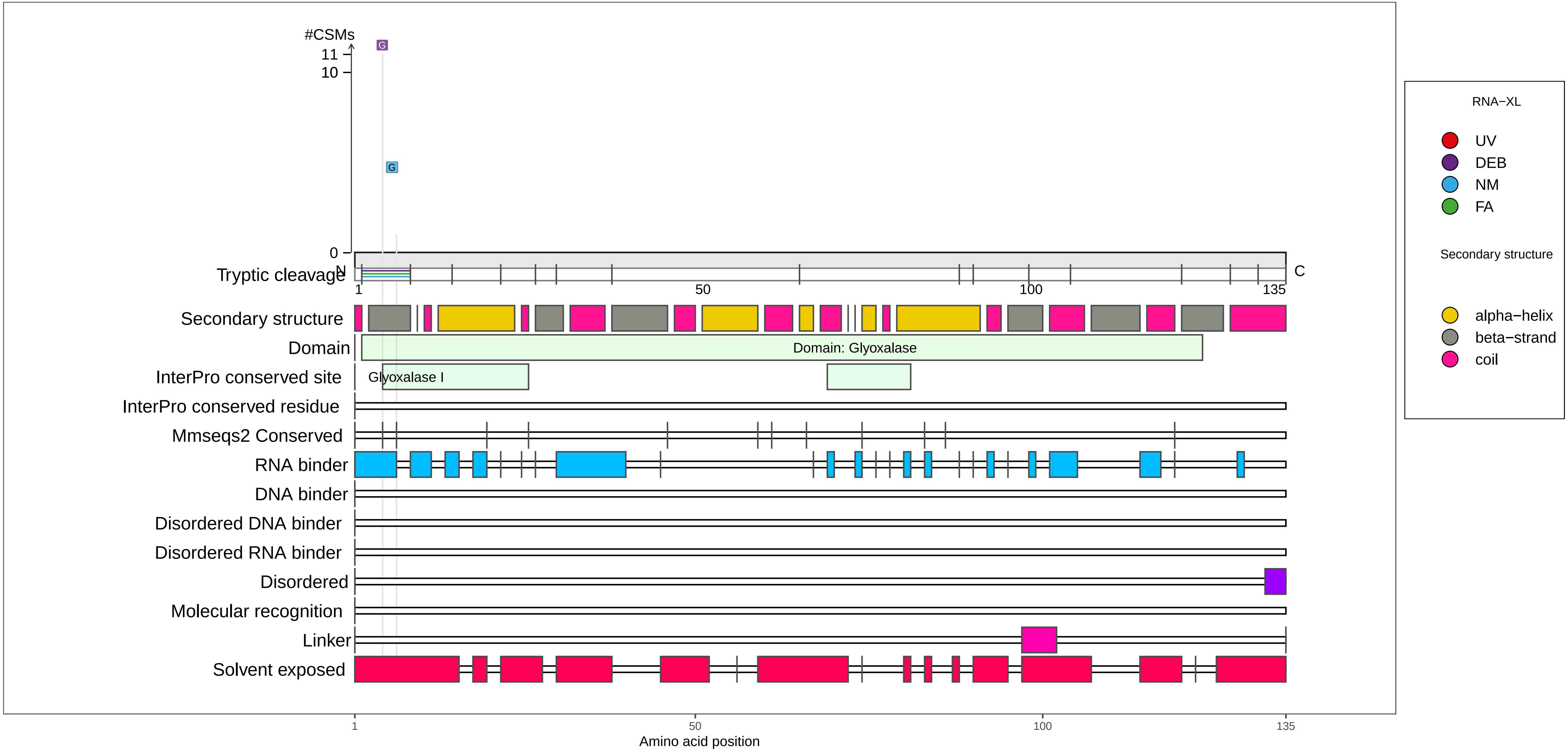

P0ADP7
UBIJ_ECOLI Ubiquinone biosynthesis accessory factor UbiJ

− Abundance:
tryptic [log10 Intensity]: 7.92 (Q 55)
PAXdb K12 strain [ppm]: NA (Q NA)
PAXdb E.coli [ppm]: 1.45 (Q 59)

− RNA functions: not annotated

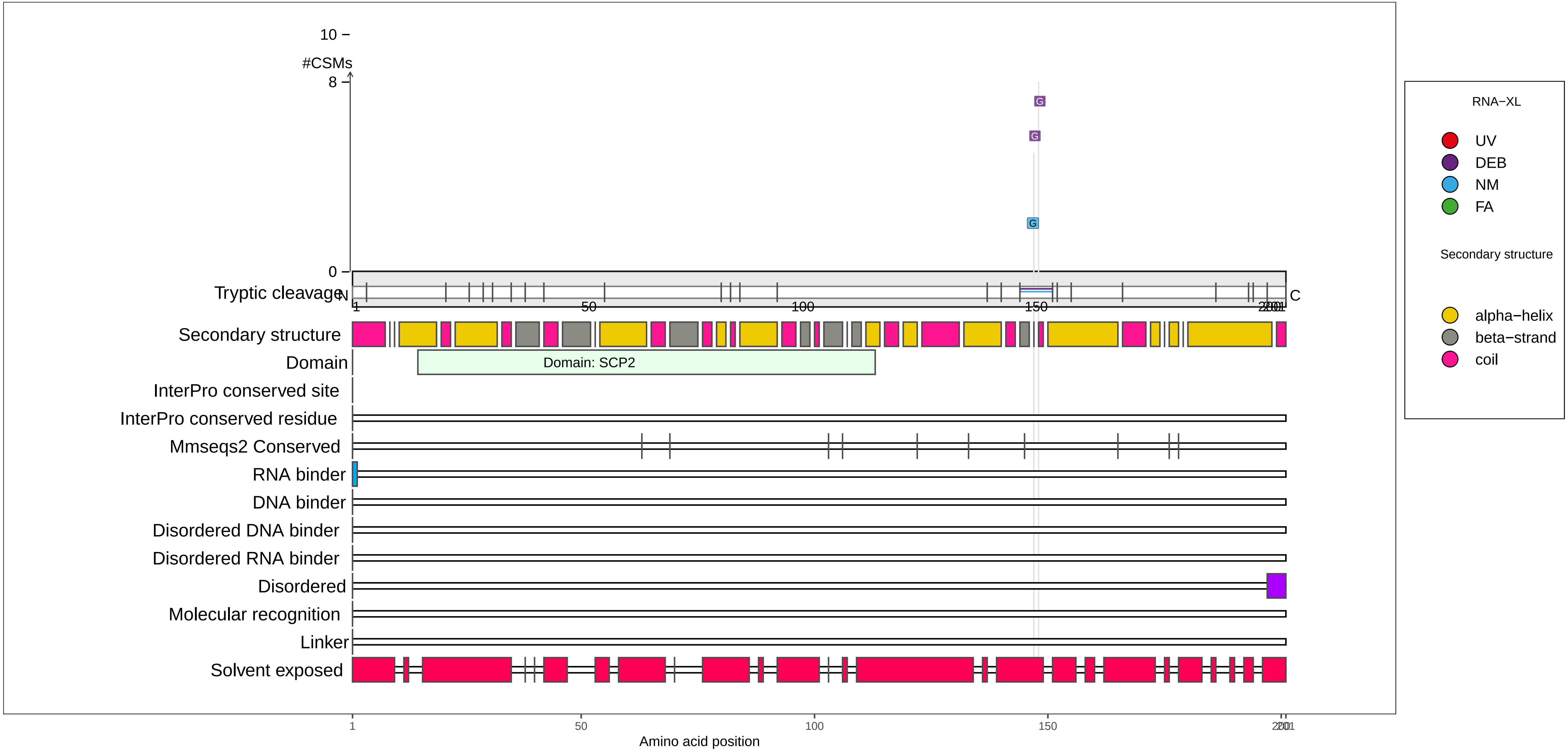

P0AFX9
RSEB_ECOLI Sigma−E factor regulatory protein RseB

− Abundance:
tryptic [log10 Intensity]: 8.26 (Q 68)
PAXdb K12 strain [ppm]: 2.5 (Q 78)
PAXdb E.coli [ppm]: 2.06 (Q 74)

− RNA functions: not annotated

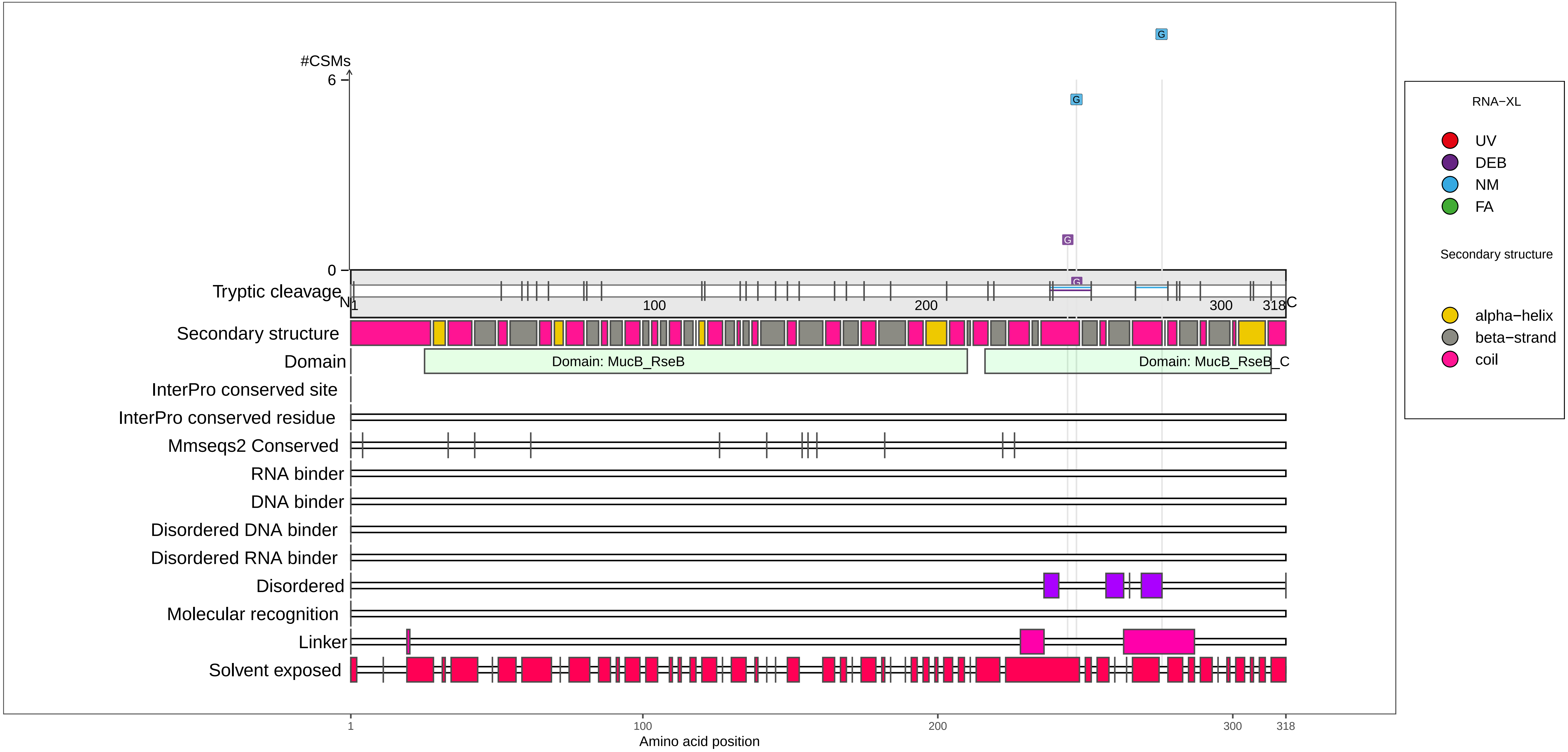

P0AFZ3
SSPB_ECOLI Stringent starvation protein B

− Abundance:
tryptic [log10 Intensity]: 7.83 (Q 51)
PAXdb K12 strain [ppm]: 1.66 (Q 48)
PAXdb E.coli [ppm]: 2.55 (Q 87)

− RNA functions:
RNA binding

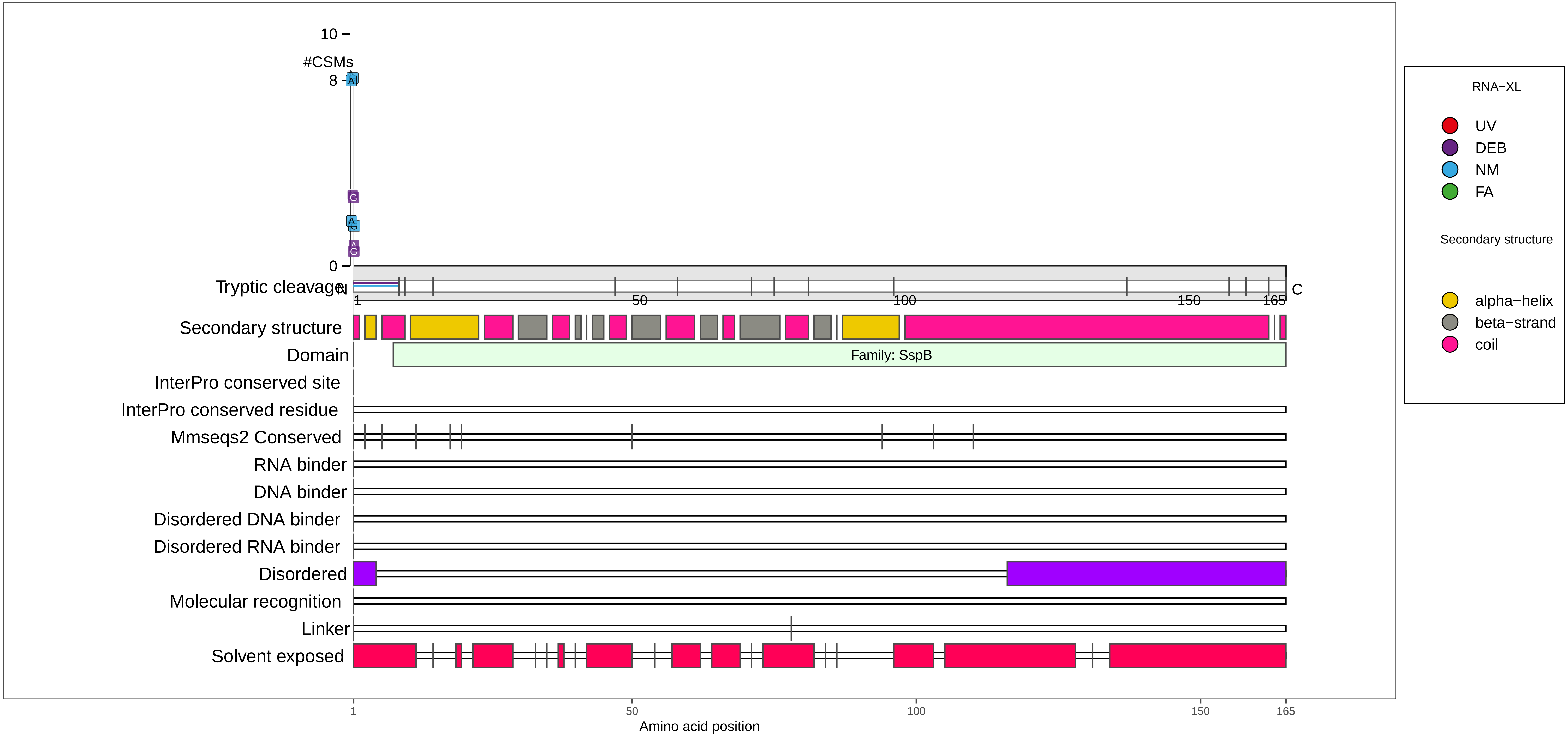

P21888
SYC_ECOLI Cysteine−−tRNA ligase

− Abundance:
tryptic [log10 Intensity]: 9.01 (Q 88)
PAXdb K12 strain [ppm]: 2.85 (Q 87)
PAXdb E.coli [ppm]: 2.54 (Q 86)

− RNA functions:
aminoacyl−tRNA ligase activity; cysteine−tRNA ligase activity
cysteinyl−tRNA aminoacylation; ncRNA metabolic process; RNA metabolic process
This DALR domain is found in cysteinyl−tRNA−synthetases.; tRNA aminoacylation
tRNA aminoacylation for protein translation; tRNA metabolic process; tRNA synthetases class I (C) catalytic domain
tRNA synthetases class I (I, L, M and V); tRNA synthetases class I (K)
tRNA synthetases class I (M)

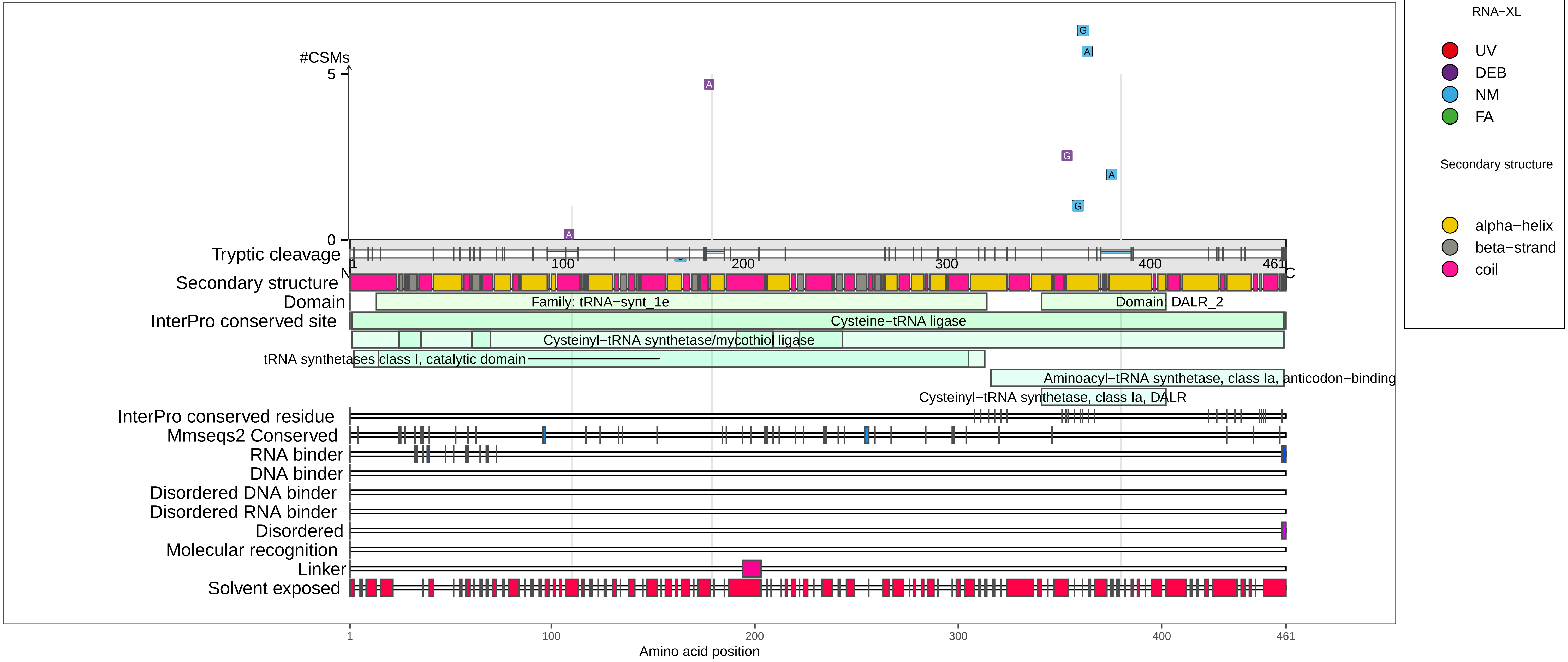

P27306
STHA_ECOLI Soluble pyridine nucleotide transhydrogenase

− Abundance:
tryptic [log10 Intensity]: 7.81 (Q 49)
PAXdb K12 strain [ppm]: 2.57 (Q 79)
PAXdb E.coli [ppm]: 2.39 (Q 83)

− RNA functions: not annotated

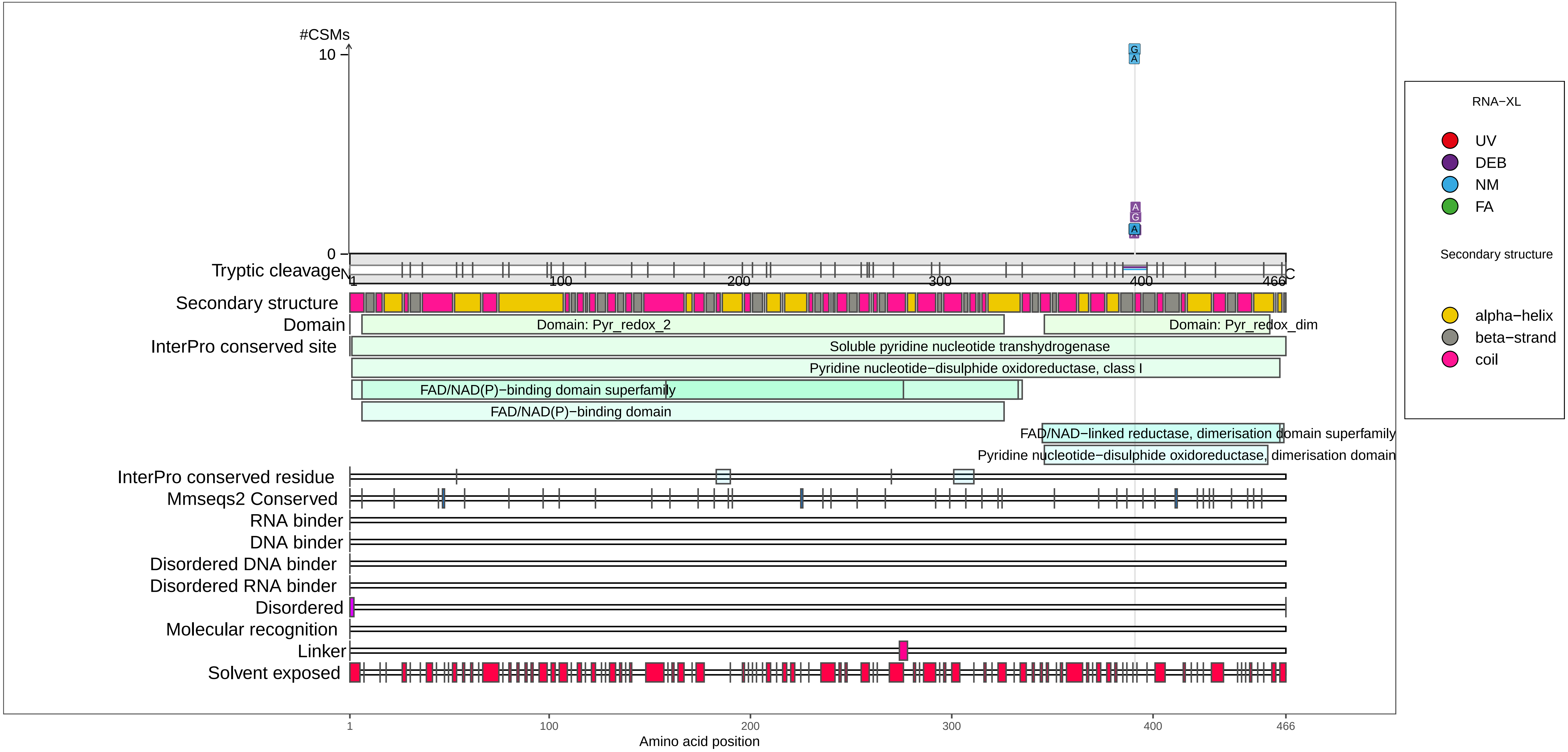

P37648
YHJJ_ECOLI Protein YhjJ

− Abundance:
tryptic [log10 Intensity]: 8.35 (Q 71)
PAXdb K12 strain [ppm]: 2.43 (Q 75)
PAXdb E.coli [ppm]: 1.91 (Q 70)

− RNA functions: not annotated

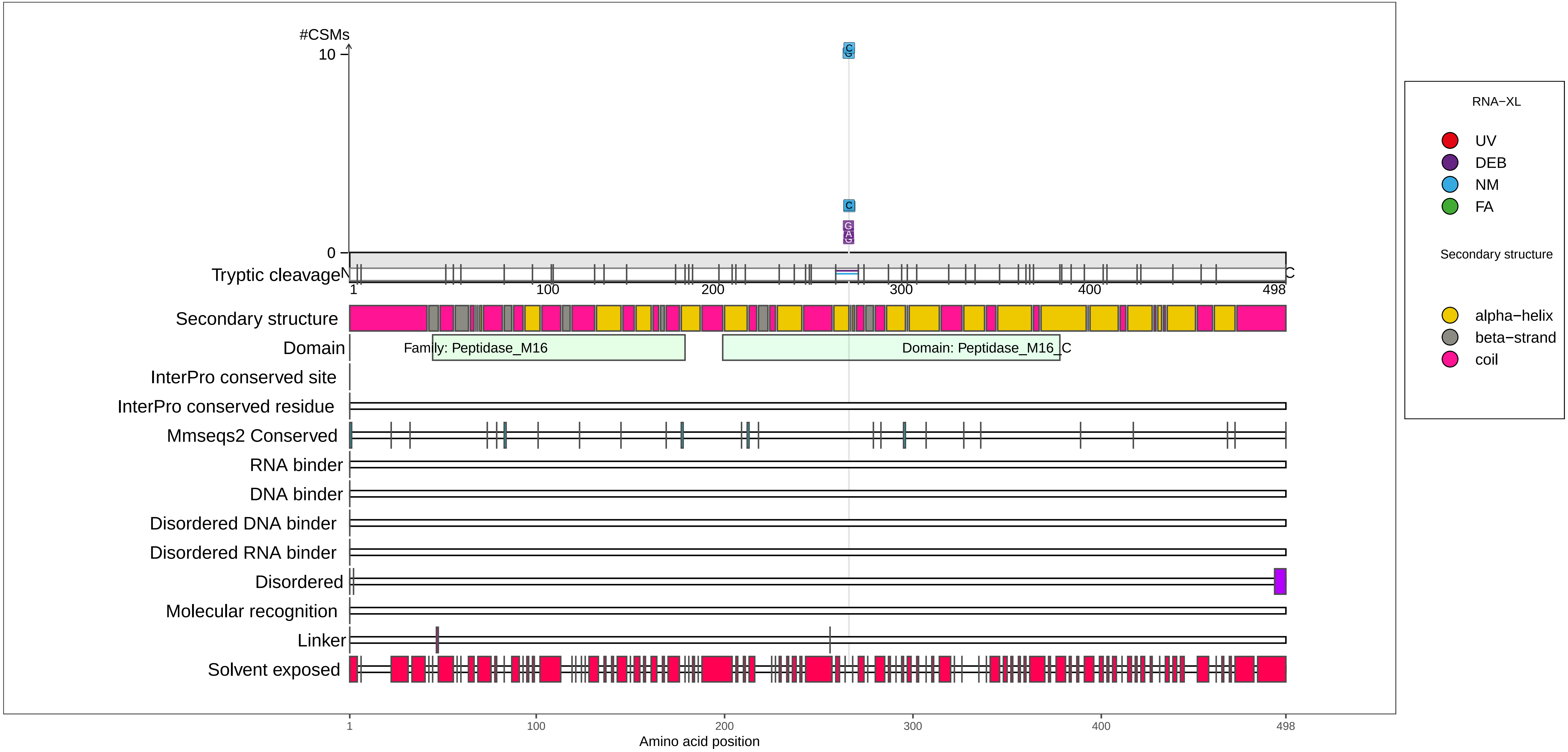

P39831
YDFG_ECOLI NADP−dependent 3−hydroxy acid dehydrogenase YdfG

− Abundance:
tryptic [log10 Intensity]: 8.36 (Q 71)
PAXdb K12 strain [ppm]: 3.02 (Q 91)
PAXdb E.coli [ppm]: 2.9 (Q 93)

− RNA functions: not annotated

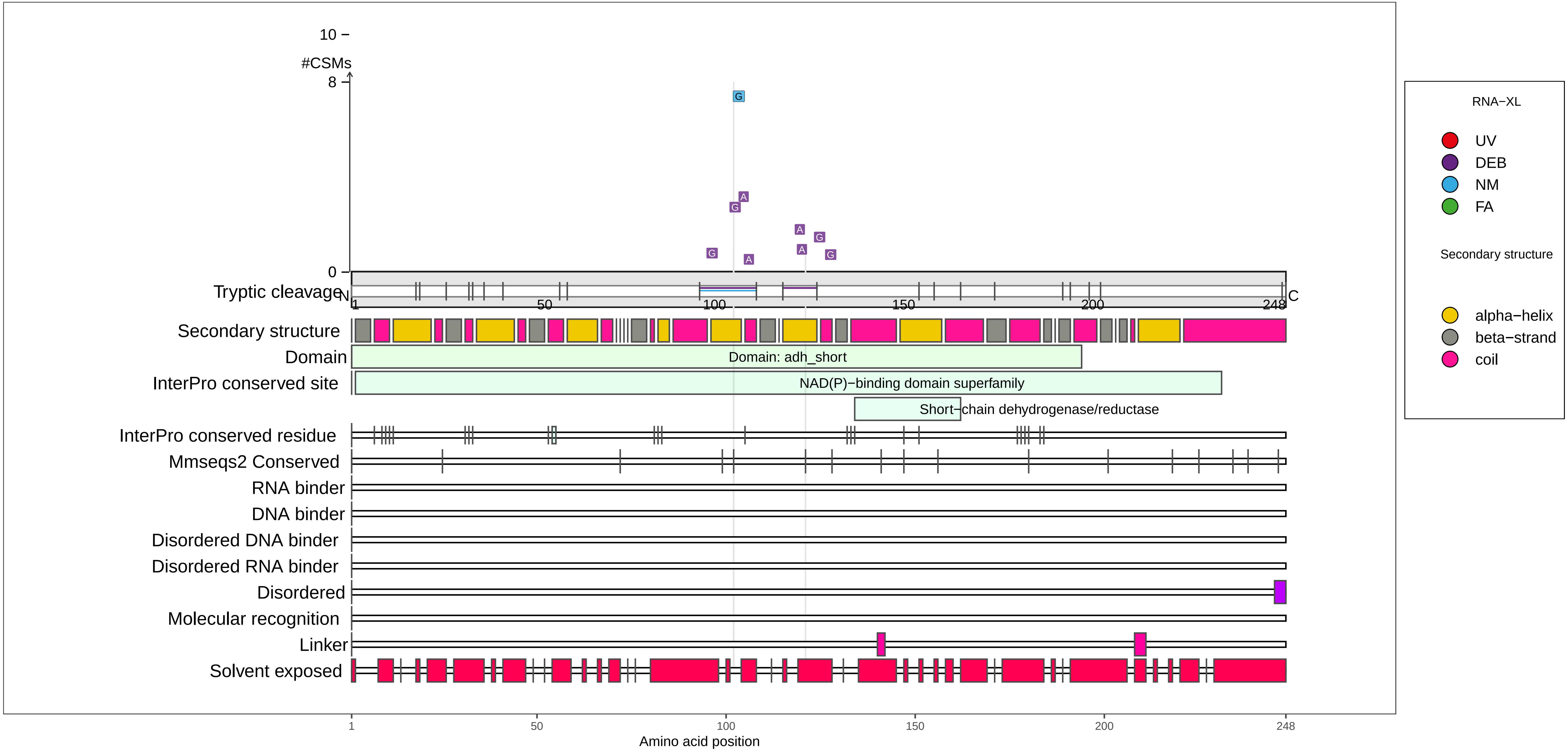

P43672
UUP_ECOLI ATP−binding protein Uup

− Abundance:
tryptic [log10 Intensity]: 8.77 (Q 83)
PAXdb K12 strain [ppm]: 1.32 (Q 28)
PAXdb E.coli [ppm]: 1.68 (Q 64)

− RNA functions: not annotated

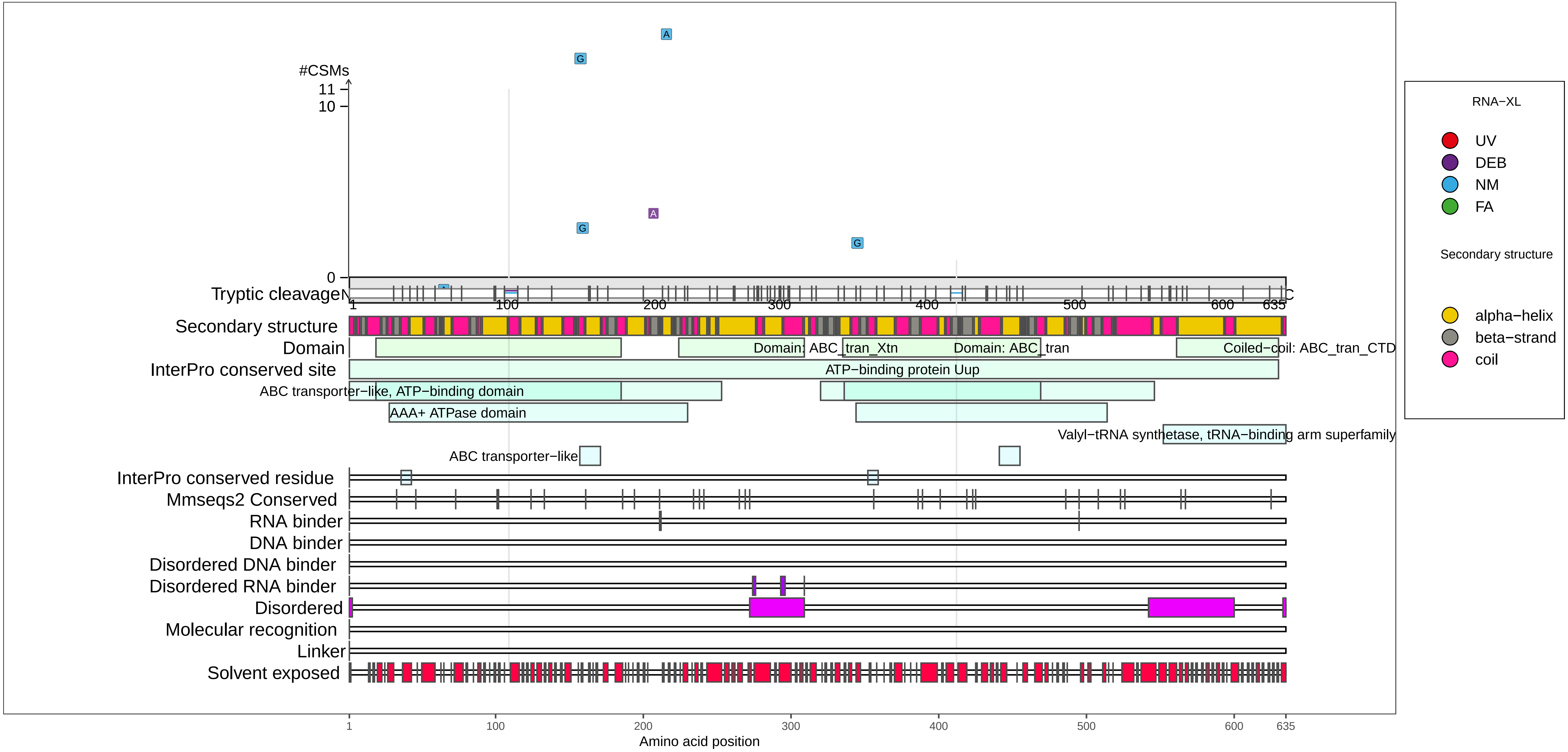

P61316
LOLA_ECOLI Outer−membrane lipoprotein carrier protein

− Abundance:
tryptic [log10 Intensity]: 8.06 (Q 60)
PAXdb K12 strain [ppm]: 2.62 (Q 81)
PAXdb E.coli [ppm]: 2.56 (Q 87)

− RNA functions: not annotated

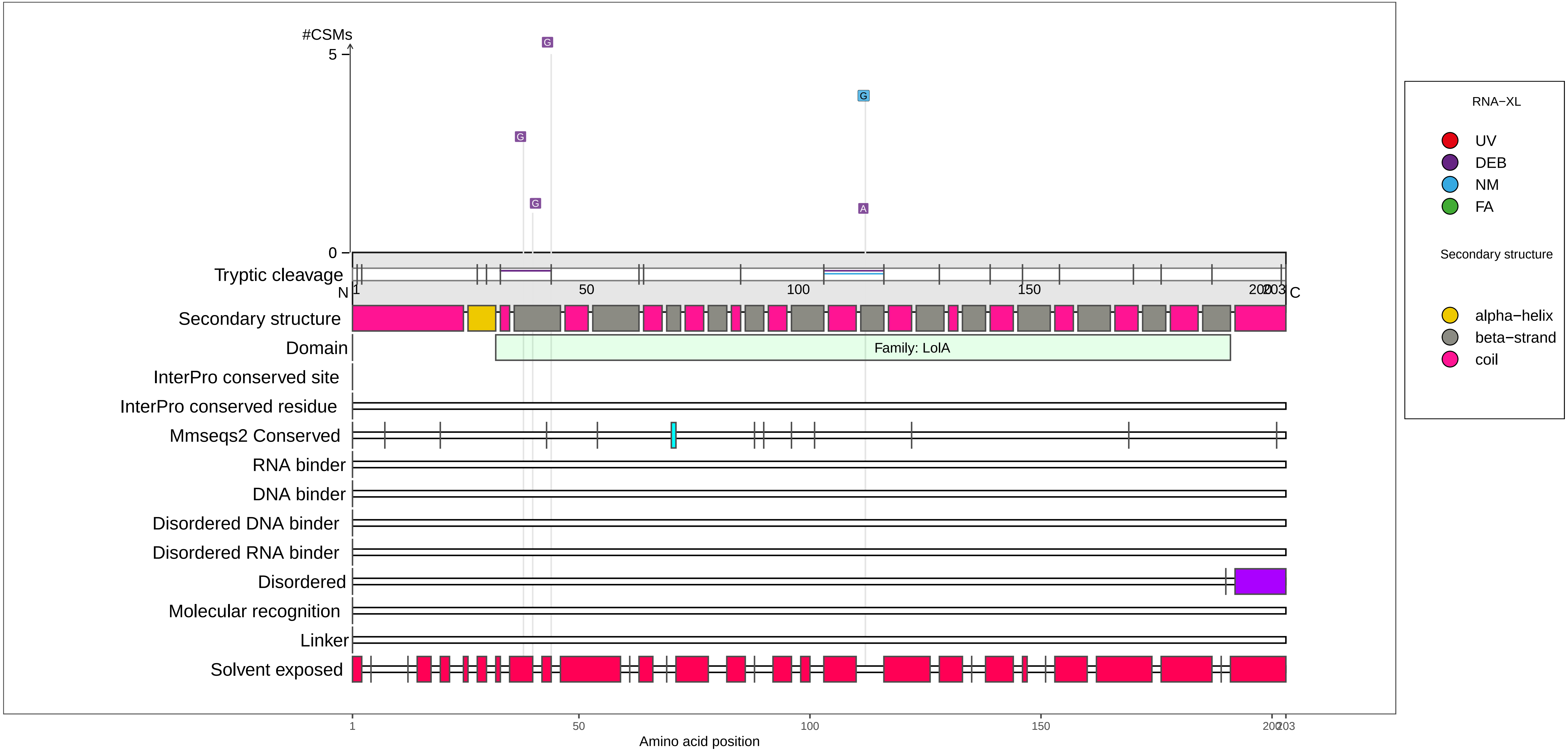

P69829
PTSN_ECOLI Nitrogen regulatory protein

− Abundance:
tryptic [log10 Intensity]: 8.01 (Q 58)
PAXdb K12 strain [ppm]: NA (Q NA)
PAXdb E.coli [ppm]: 2.57 (Q 87)

− RNA functions: not annotated

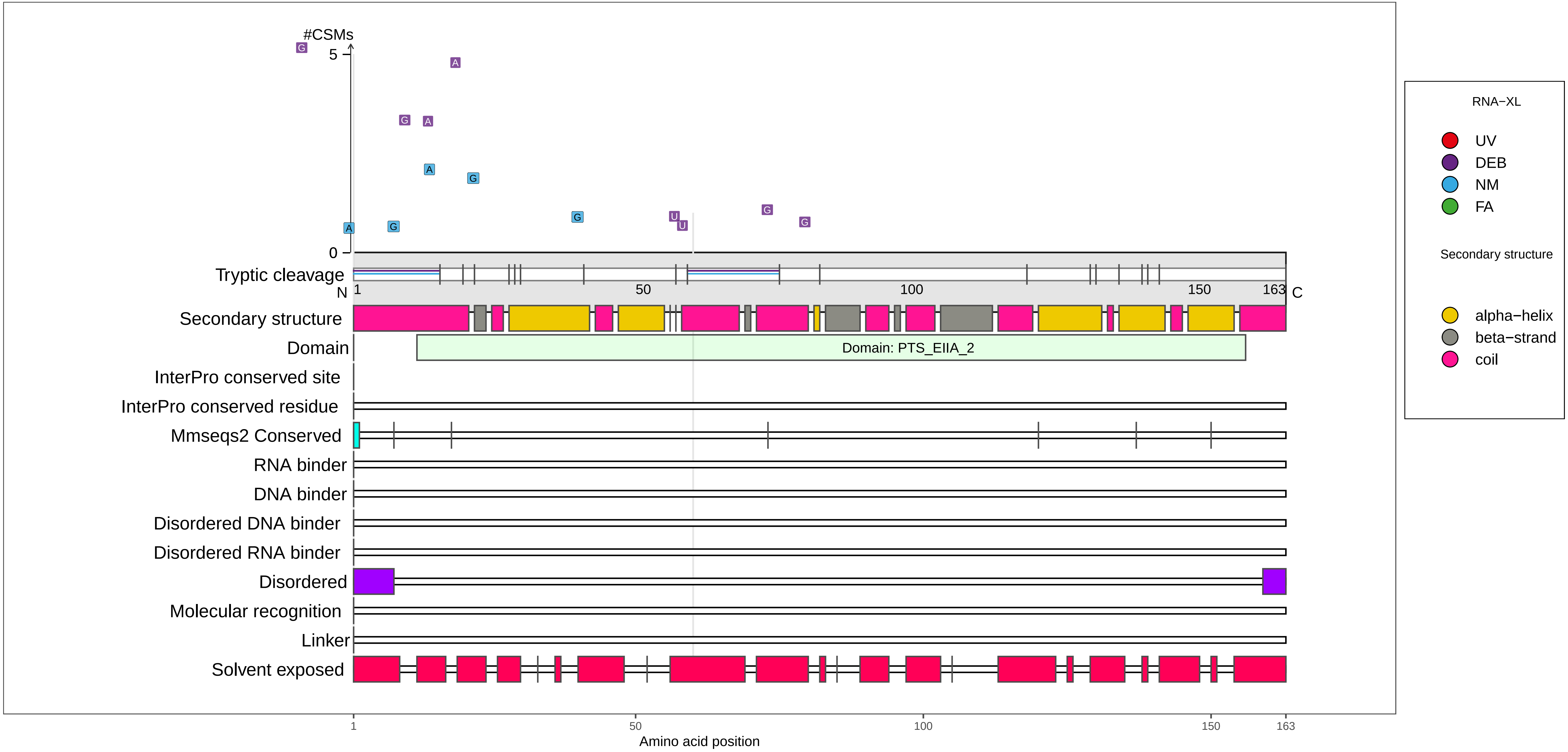

P75829
YBJX_ECOLI Uncharacterized protein YbjX

− Abundance:
tryptic [log10 Intensity]: 7.56 (Q 38)
PAXdb K12 strain [ppm]: 0.88 (Q 7)
PAXdb E.coli [ppm]: 1.35 (Q 56)

− RNA functions: not annotated

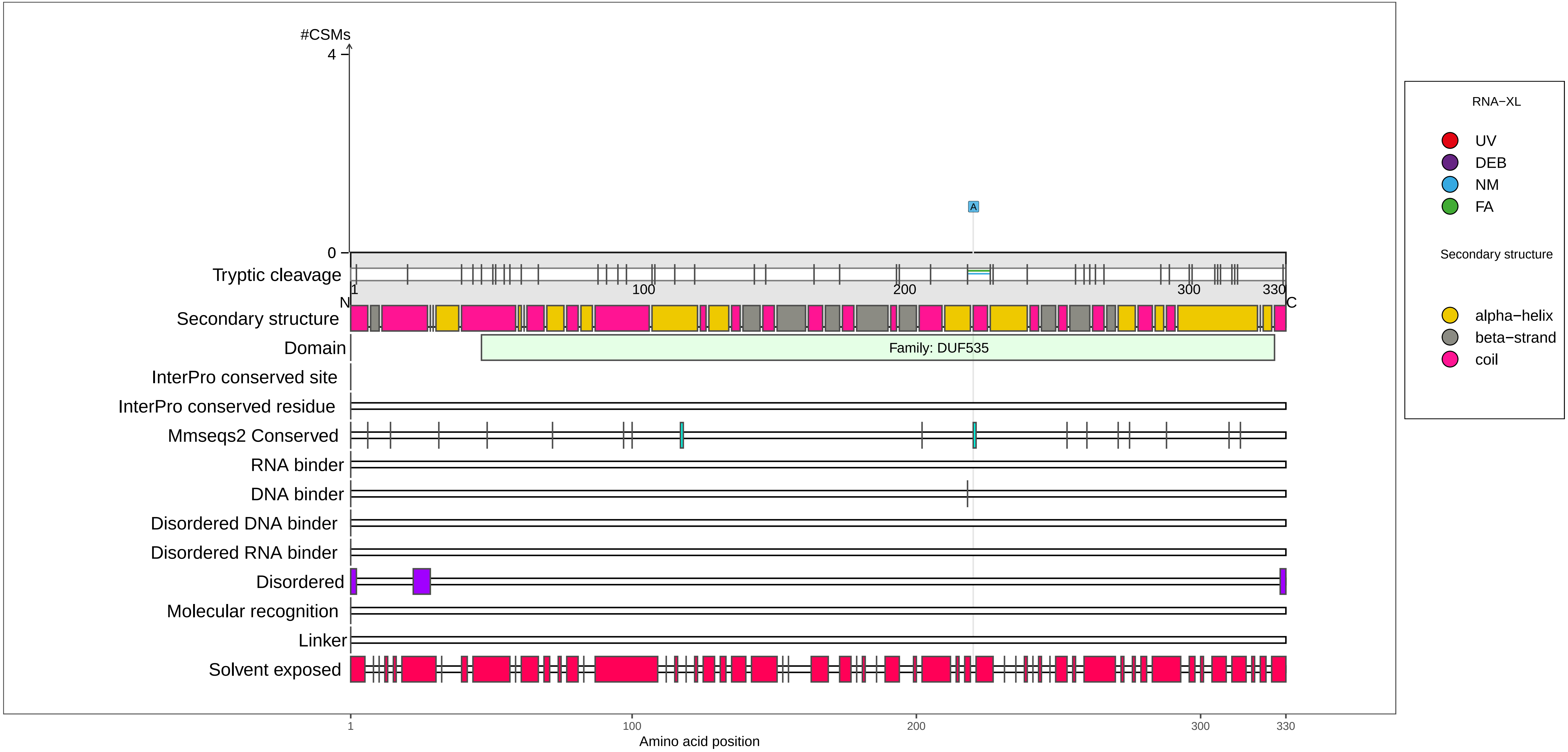

P02931
OMPF_ECOLI Outer membrane porin F

− Abundance:
tryptic [log10 Intensity]: 10.37 (Q 100)
PAXdb K12 strain [ppm]: NA (Q NA)
PAXdb E.coli [ppm]: 2.96 (Q 94)

− RNA functions: not annotated

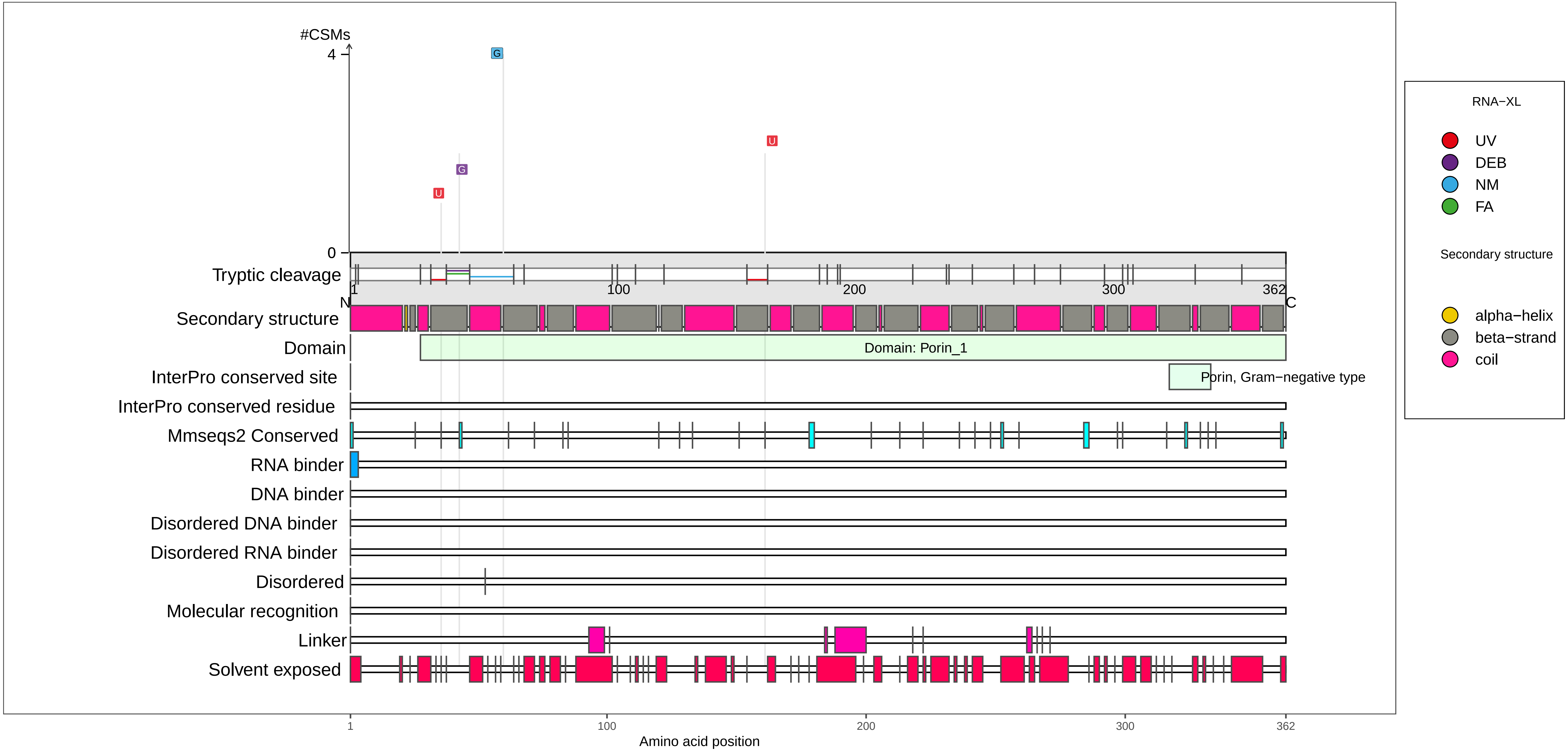

P08200
IDH_ECOLI Isocitrate dehydrogenase [NADP]

− Abundance:
tryptic [log10 Intensity]: 8.86 (Q 85)
PAXdb K12 strain [ppm]: 3.57 (Q 98)
PAXdb E.coli [ppm]: 3.47 (Q 99)

− RNA functions: not annotated

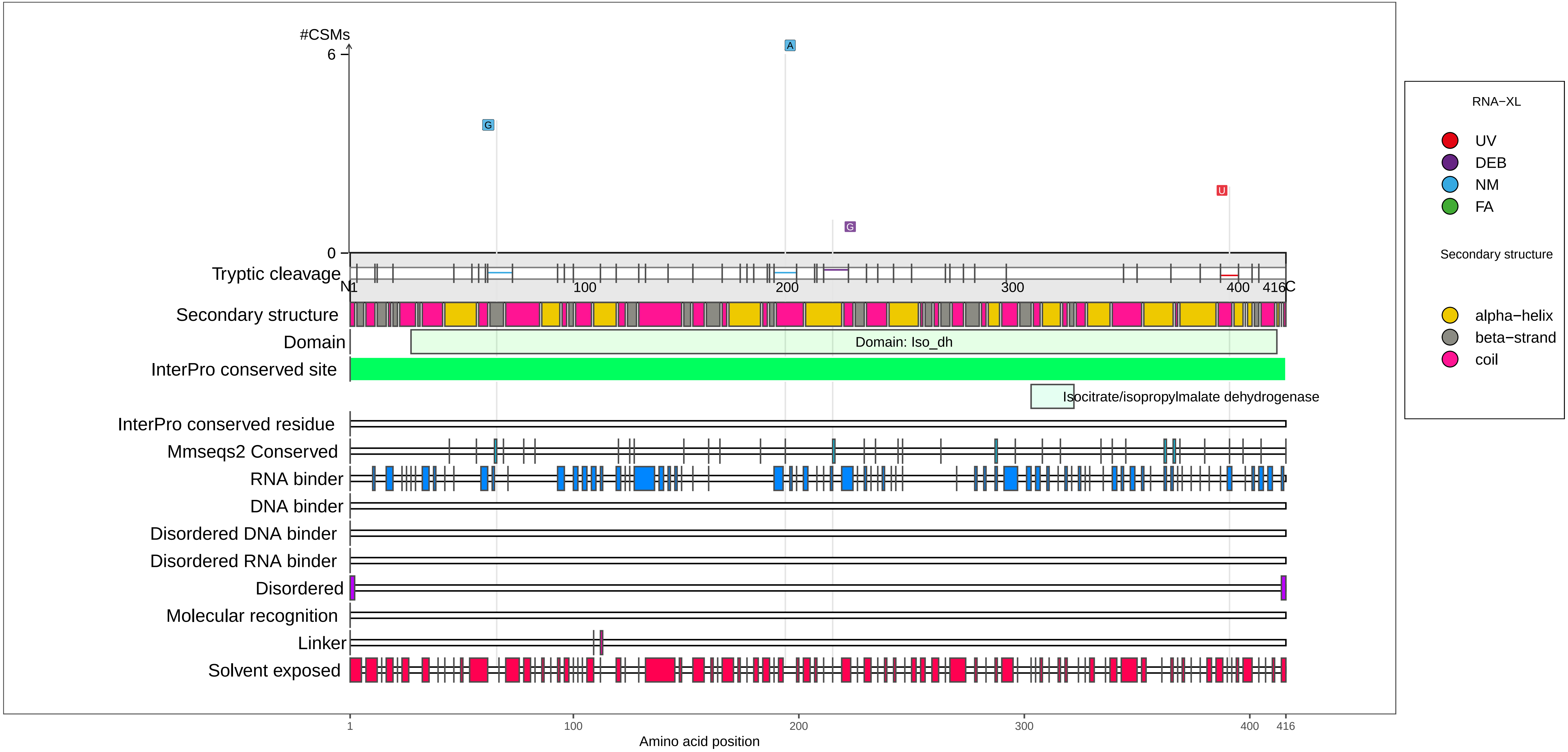

P0A6Q3
FABA_ECOLI 3−hydroxydecanoyl−[acyl−carrier−protein] dehydratase

− Abundance:
tryptic [log10 Intensity]: 9.63 (Q 97)
PAXdb K12 strain [ppm]: 2.76 (Q 85)
PAXdb E.coli [ppm]: 3.31 (Q 98)

− RNA functions: not annotated

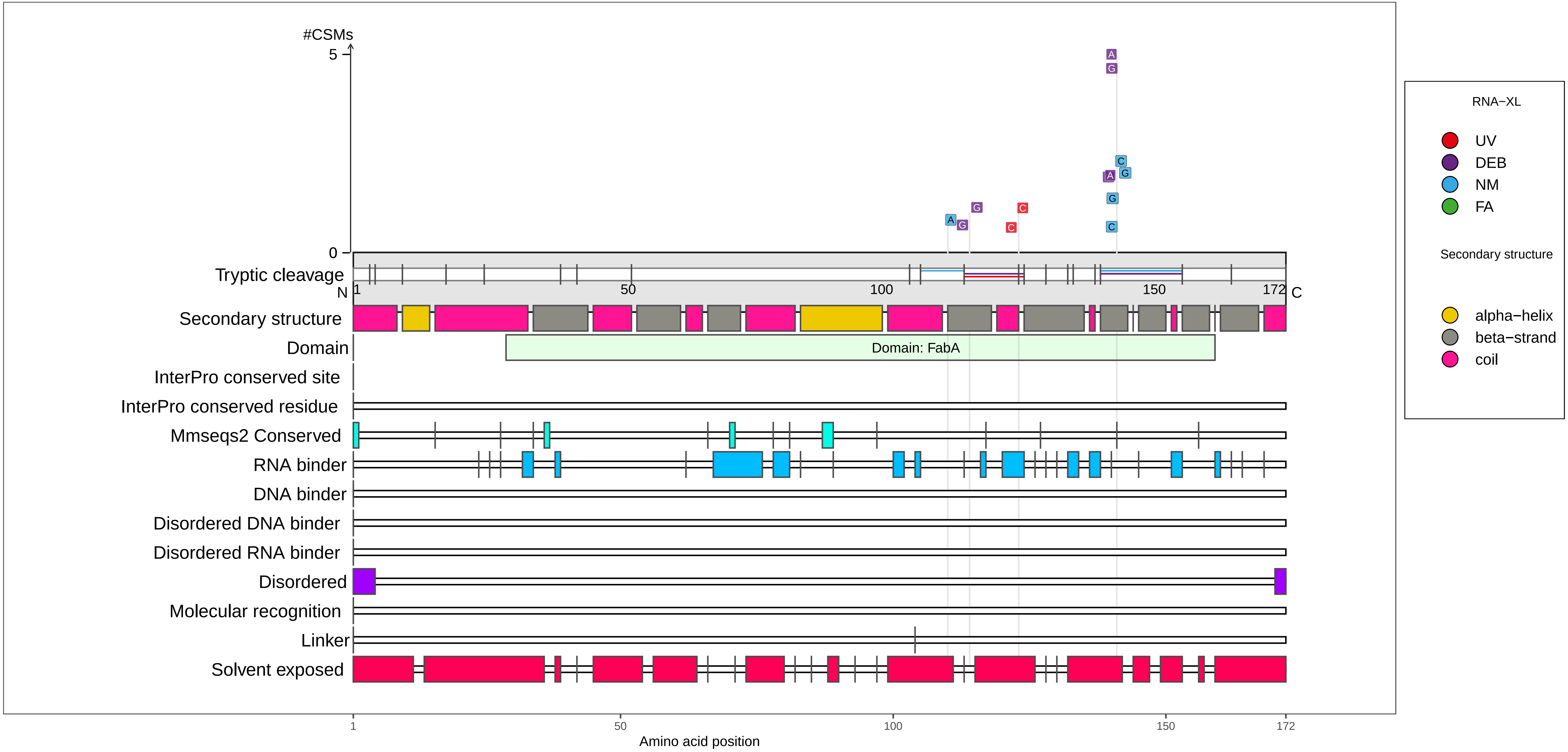

P0A6T5
GCH1_ECOLI GTP cyclohydrolase 1

− Abundance:
tryptic [log10 Intensity]: 9.2 (Q 93)
PAXdb K12 strain [ppm]: 1.34 (Q 29)
PAXdb E.coli [ppm]: 2.84 (Q 92)

− RNA functions: not annotated

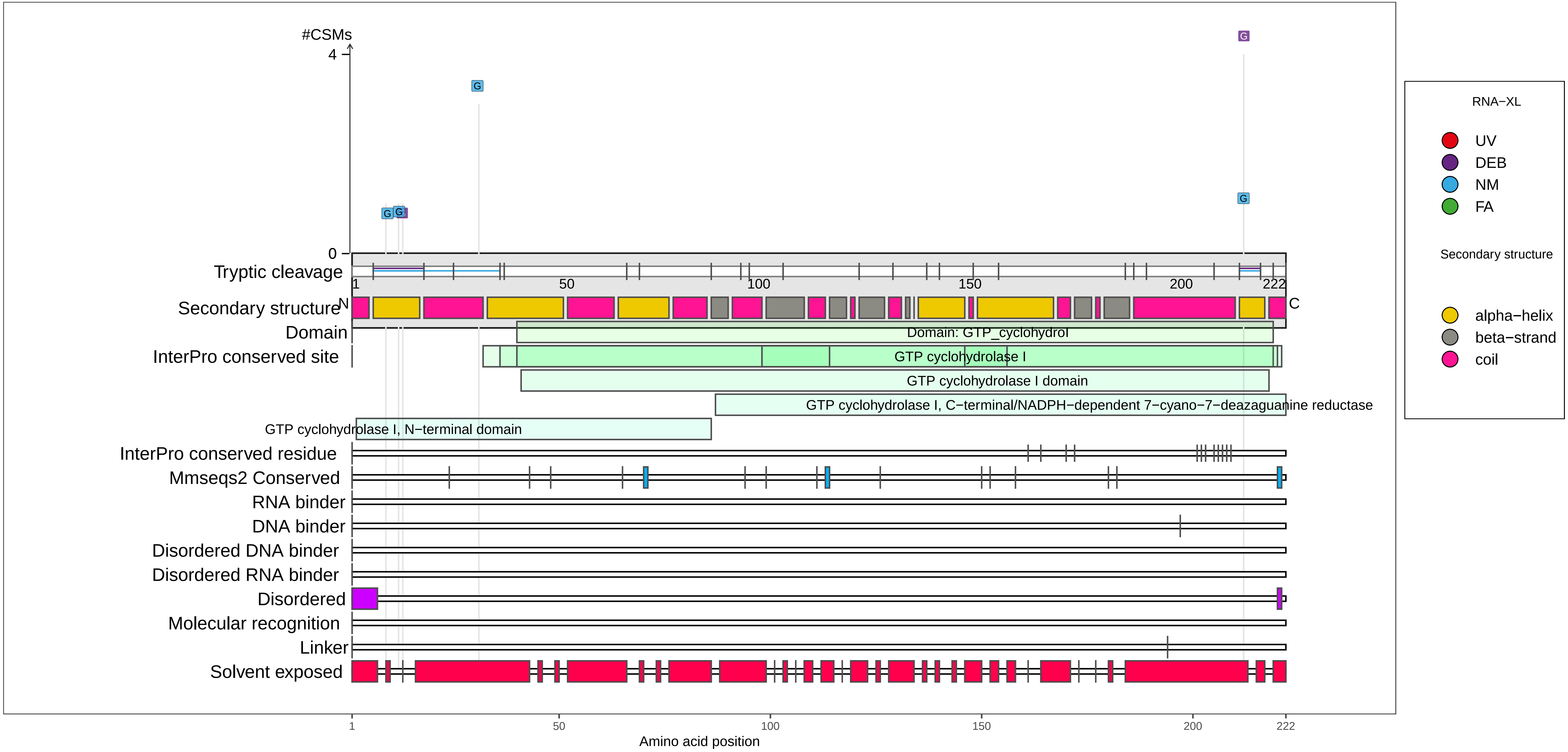

P0A6U3
MNMG_ECOLI tRNA uridine 5−carboxymethylaminomethyl modification enzyme MnmG

− Abundance:
tryptic [log10 Intensity]: 8.12 (Q 62)
PAXdb K12 strain [ppm]: 1.85 (Q 55)
PAXdb E.coli [ppm]: 1.42 (Q 57)

− RNA functions:
ncRNA metabolic process; ncRNA processing; RNA metabolic process; RNA methylation
RNA modification; RNA processing; tRNA metabolic process; tRNA methylation; tRNA modification
tRNA processing; tRNA wobble base modification; tRNA wobble uridine modification

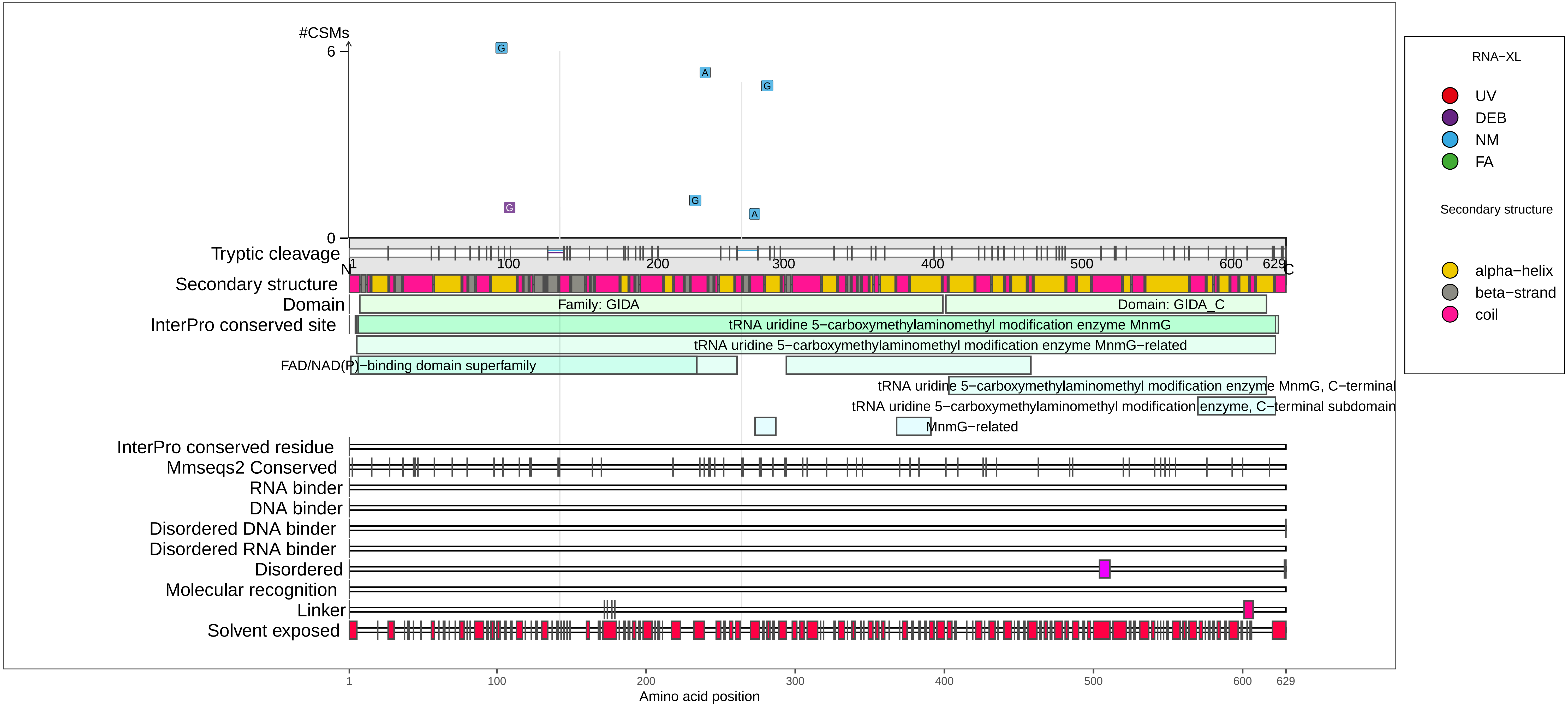

P0A6X7
IHFA_ECOLI Integration host factor subunit alpha
− Abundance:
tryptic [log10 Intensity]: 8.11 (Q 62)
PAXdb K12 strain [ppm]: 3.06 (Q 92)
PAXdb E.coli [ppm]: 3.43 (Q 98)

− RNA functions:
RNA biosynthetic process; RNA metabolic process

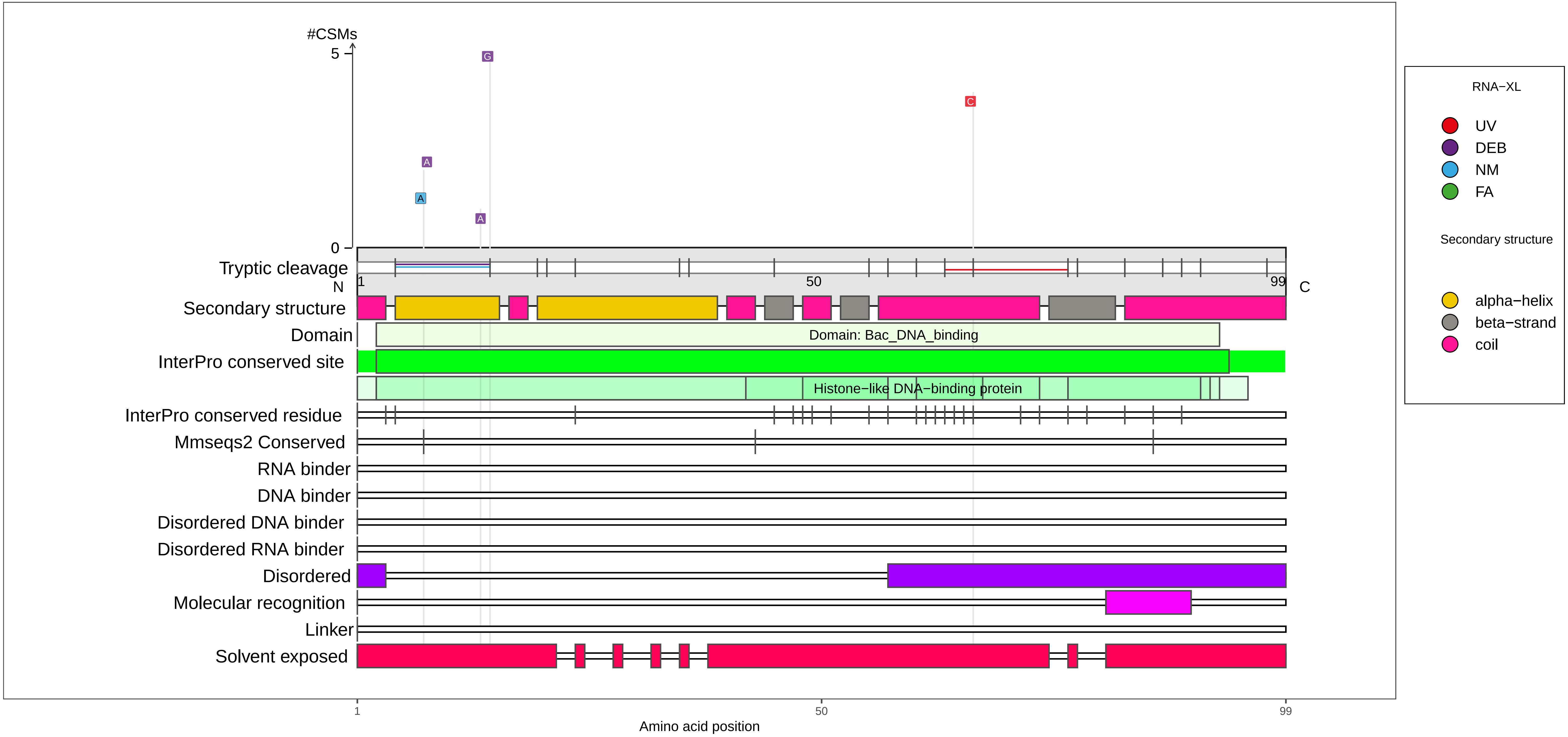

P0A7P5
RL34_ECOLI 50S ribosomal protein L34

− Abundance:
tryptic [log10 Intensity]: 8.46 (Q 74)
PAXdb K12 strain [ppm]: 3.48 (Q 97)
PAXdb E.coli [ppm]: 3.19 (Q 96)

− RNA functions: not annotated

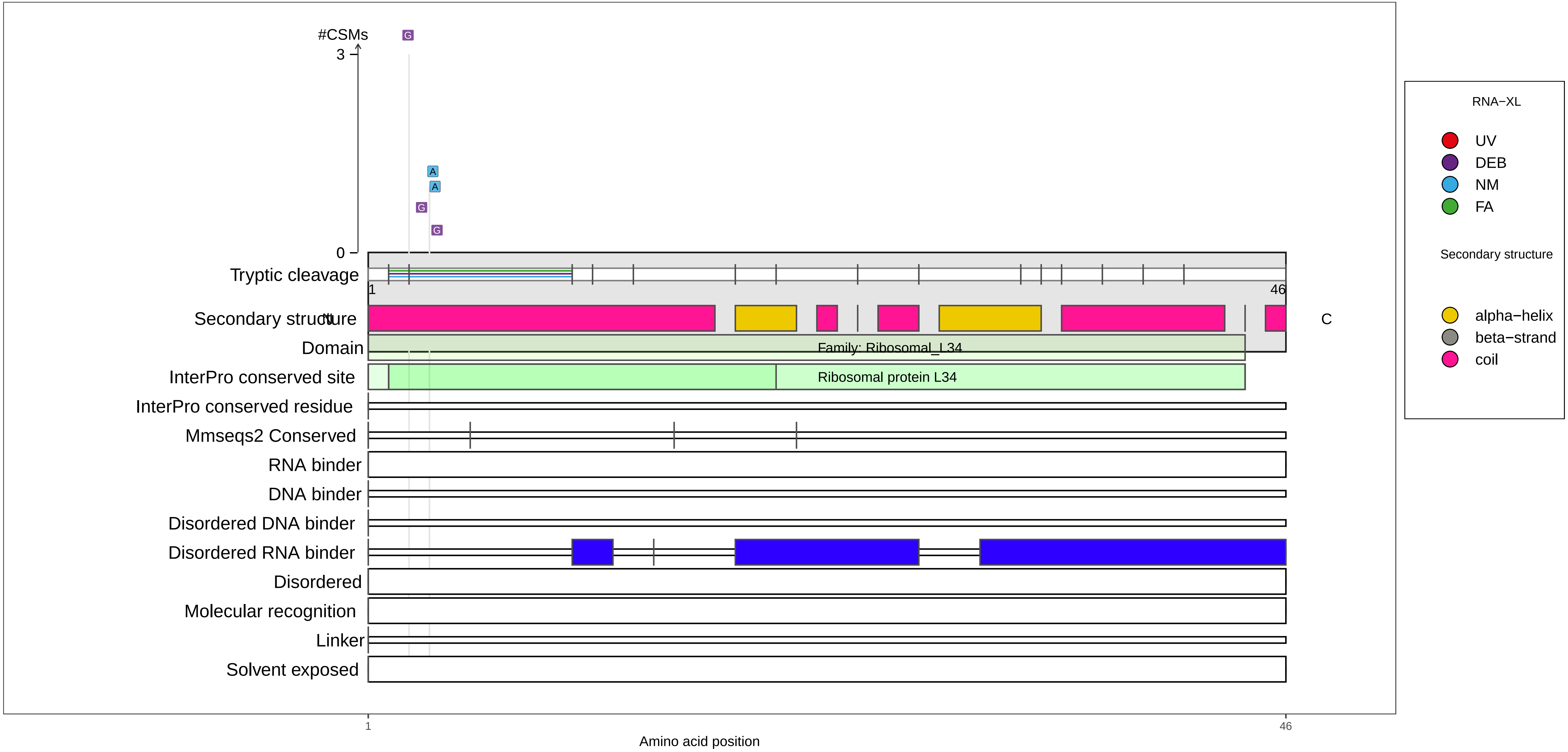

P0AAD6
SDAC_ECOLI Serine transporter SdaC

− Abundance:
tryptic [log10 Intensity]: 6.75 (Q 6)
PAXdb K12 strain [ppm]: 2.14 (Q 65)
PAXdb E.coli [ppm]: 1.99 (Q 72)

− RNA functions: not annotated

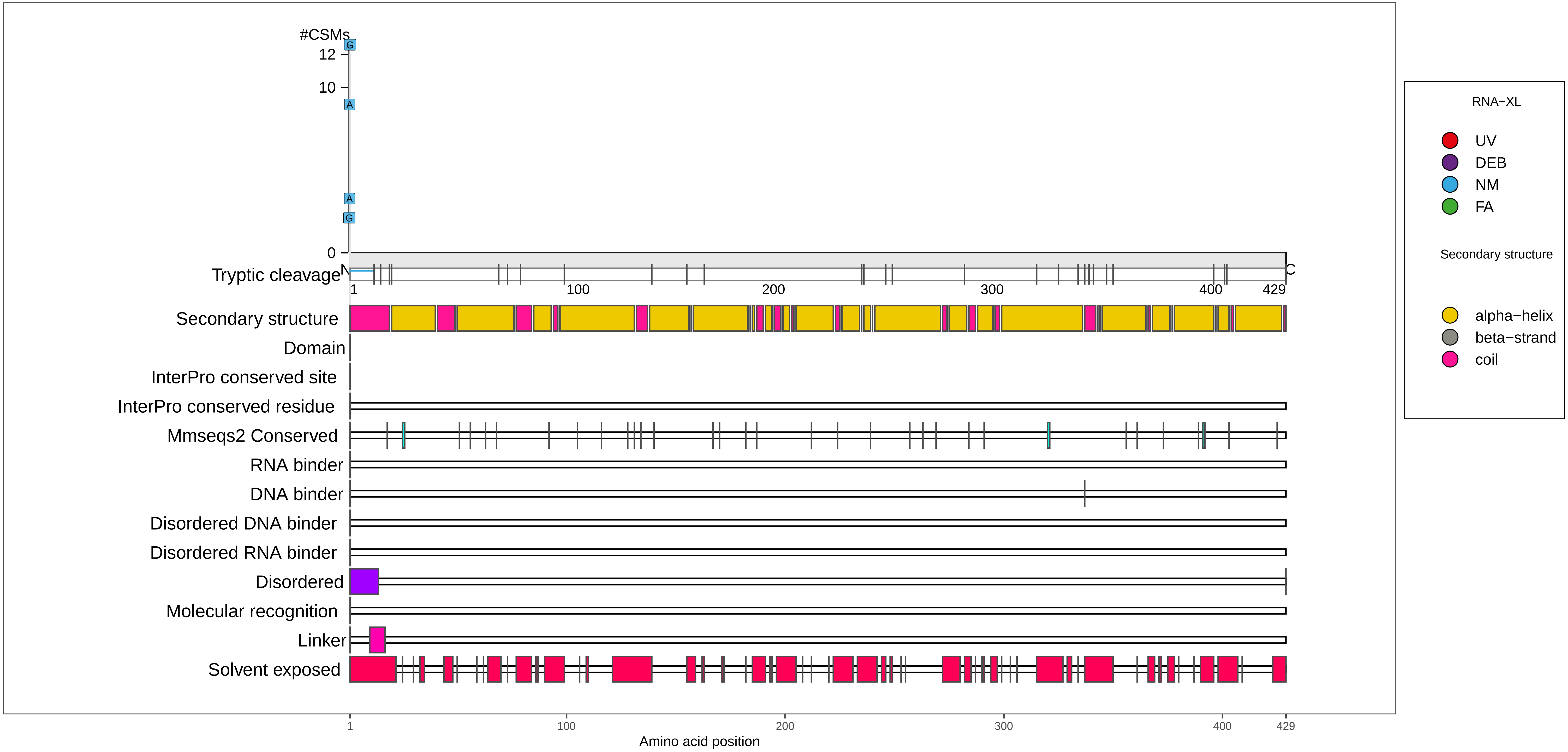

P0ABS1
DKSA_ECOLI RNA polymerase−binding transcription factor DksA

− Abundance:
tryptic [log10 Intensity]: 9.18 (Q 92)
PAXdb K12 strain [ppm]: 2.78 (Q 85)
PAXdb E.coli [ppm]: 2.75 (Q 91)

− RNA functions: not annotated

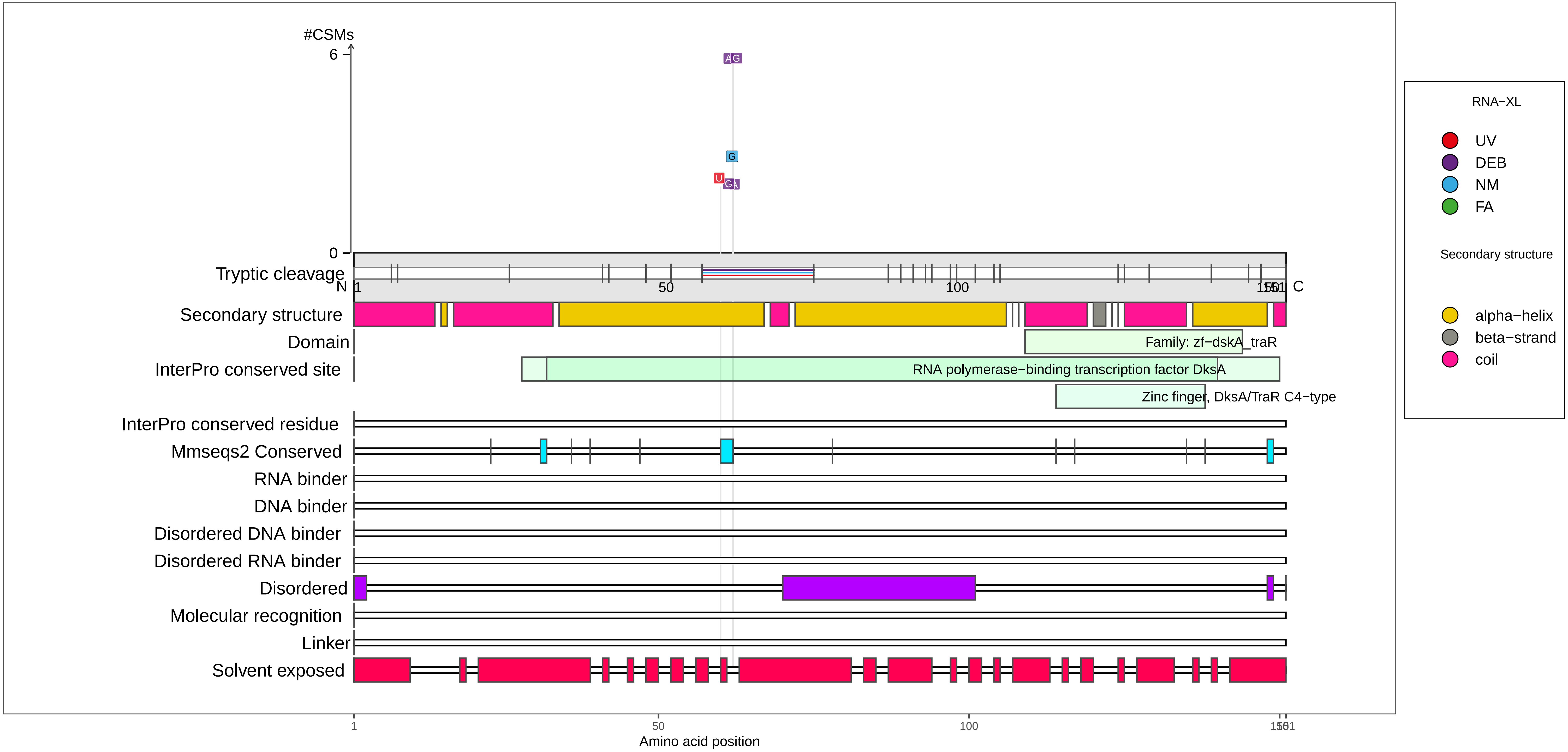

P0ABU2
YCHF_ECOLI Ribosome−binding ATPase YchF

− Abundance:
tryptic [log10 Intensity]: 8.42 (Q 73)
PAXdb K12 strain [ppm]: 2.85 (Q 87)
PAXdb E.coli [ppm]: 2.42 (Q 83)

− RNA functions: not annotated

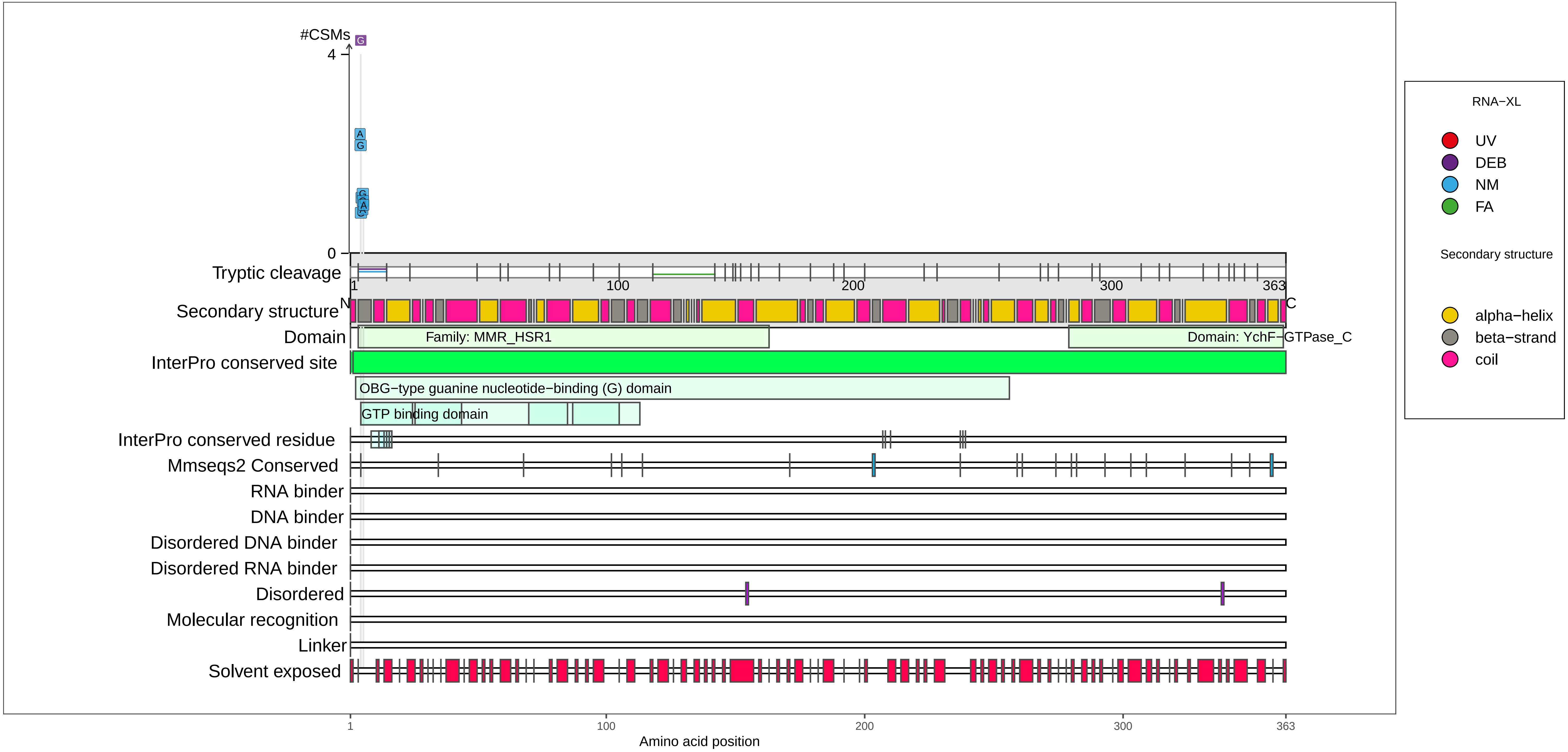

P0ACY1
YDJA_ECOLI Putative NAD(P)H nitroreductase YdjA

− Abundance:
tryptic [log10 Intensity]: 8.26 (Q 68)
PAXdb K12 strain [ppm]: 3.19 (Q 94)
PAXdb E.coli [ppm]: 2.61 (Q 88)

− RNA functions: not annotated

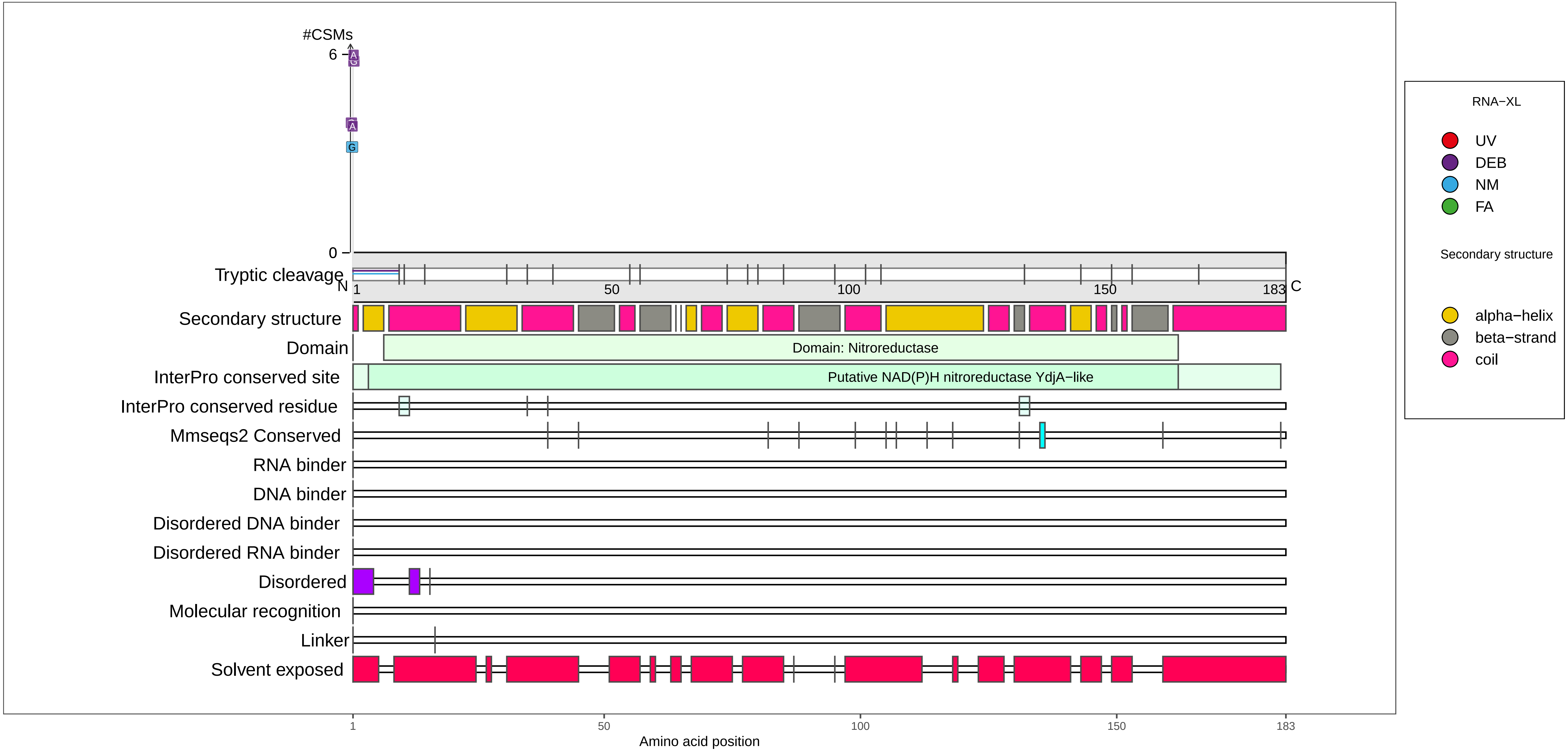

P0ADT5
YGIC_ECOLI Putative acid−−amine ligase YgiC

− Abundance:
tryptic [log10 Intensity]: 8.41 (Q 73)
PAXdb K12 strain [ppm]: 1.56 (Q 42)
PAXdb E.coli [ppm]: 1.3 (Q 55)

− RNA functions: not annotated

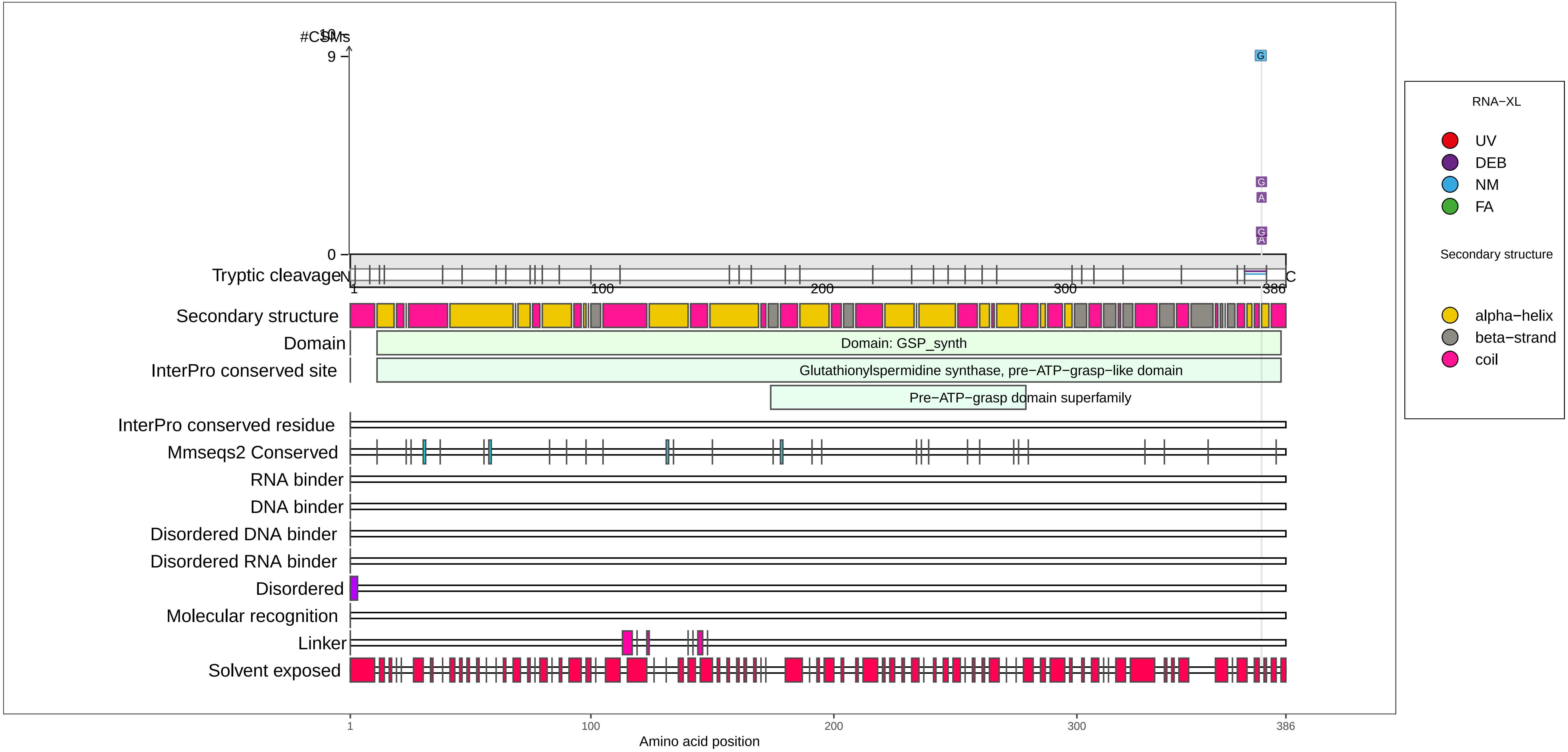

P0ADU2
YGIN_ECOLI Probable quinol monooxygenase YgiN

− Abundance:
tryptic [log10 Intensity]: 8.16 (Q 63)
PAXdb K12 strain [ppm]: 2.96 (Q 90)
PAXdb E.coli [ppm]: 2.71 (Q 90)

− RNA functions: not annotated

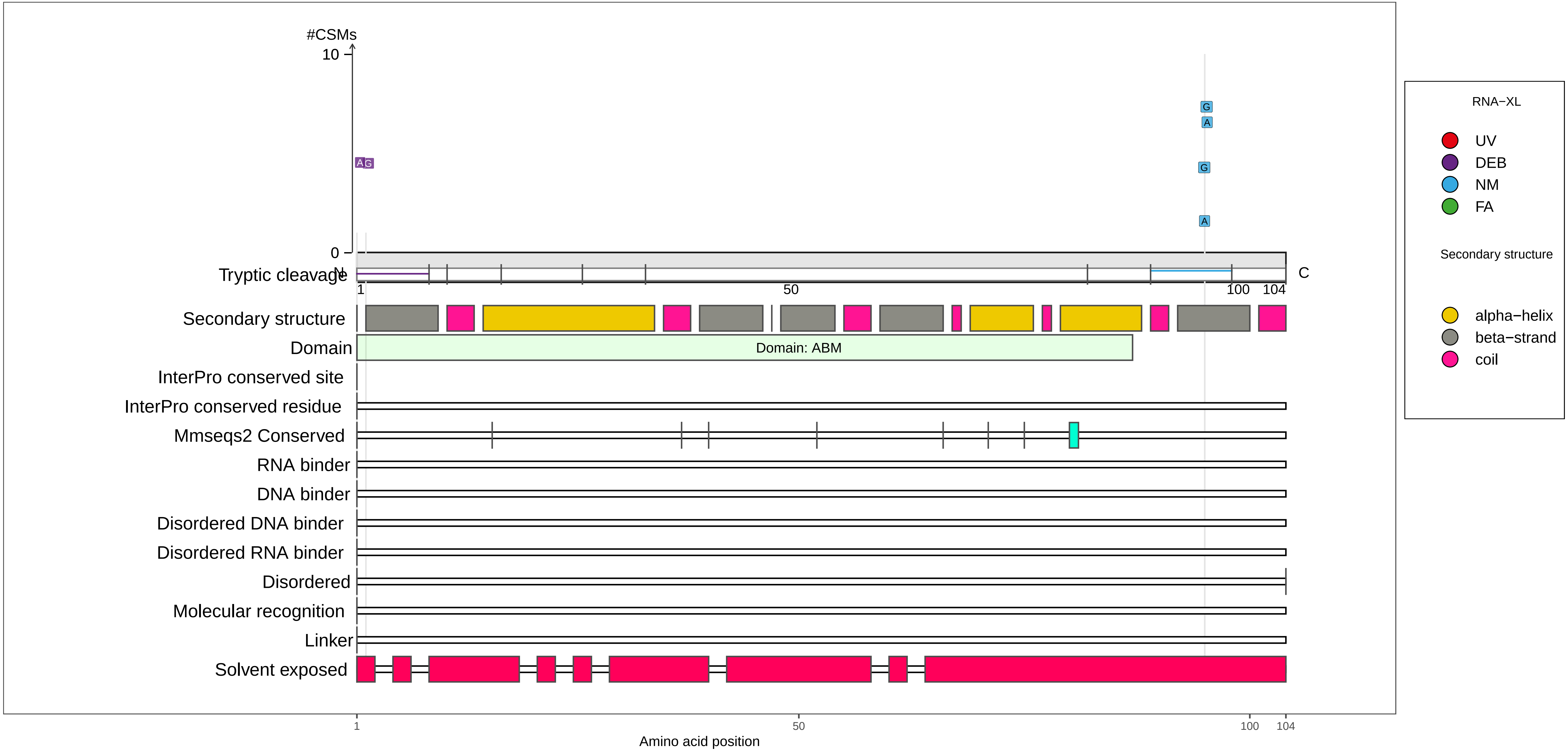

P0AG00
WZZE_ECOLI ECA polysaccharide chain length modulation protein

− Abundance:
tryptic [log10 Intensity]: 8.31 (Q 70)
PAXdb K12 strain [ppm]: NA (Q NA)
PAXdb E.coli [ppm]: 1.67 (Q 64)

− RNA functions: not annotated

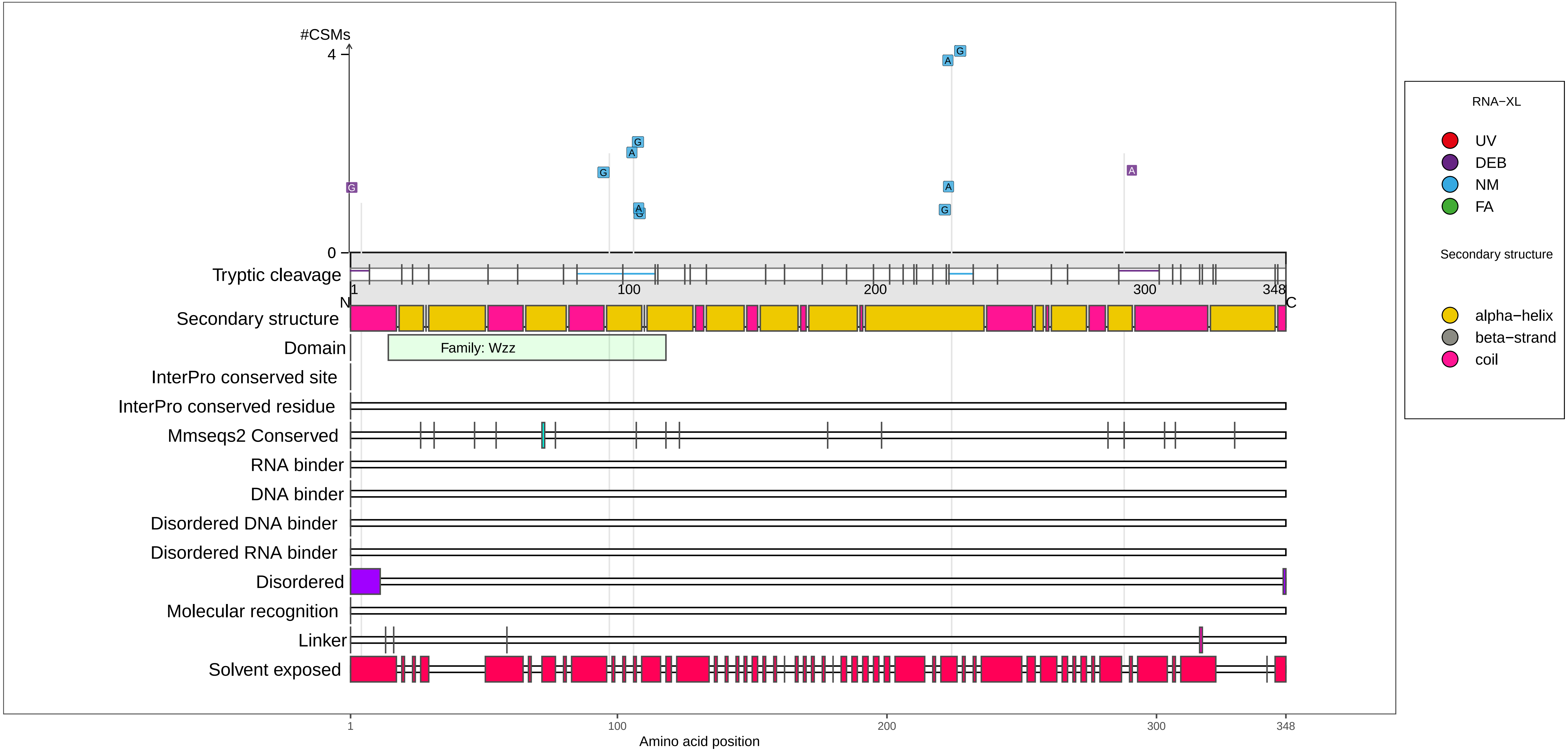

P23909
MUTS_ECOLI DNA mismatch repair protein MutS

− Abundance:
tryptic [log10 Intensity]: 6.63 (Q 4)
PAXdb K12 strain [ppm]: 1.57 (Q 43)
PAXdb E.coli [ppm]: 0.99 (Q 47)

− RNA functions: not annotated

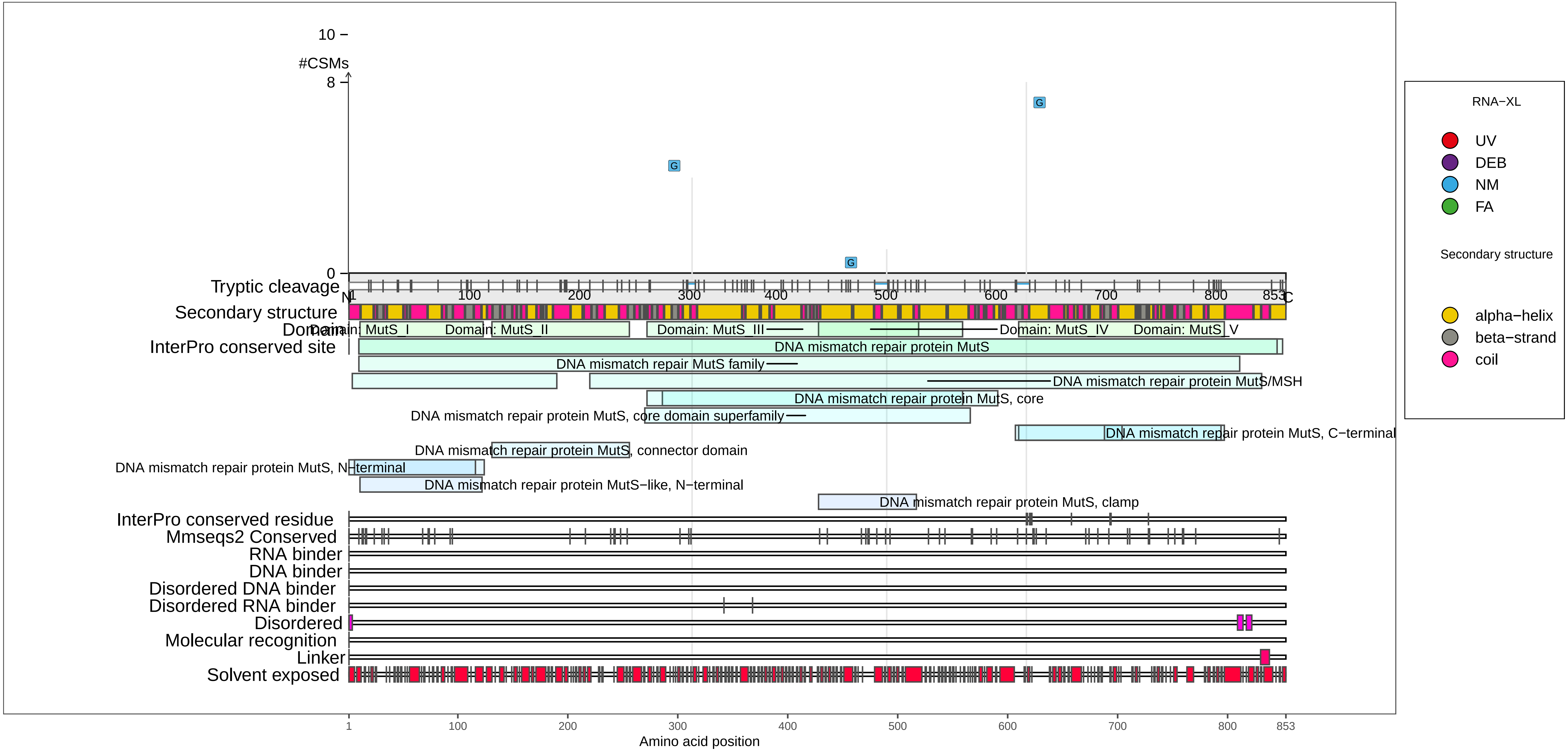

P39406
RSMC_ECOLI Ribosomal RNA small subunit methyltransferase C

− Abundance:
tryptic [log10 Intensity]: 8.04 (Q 59)
PAXdb K12 strain [ppm]: 1.33 (Q 28)
PAXdb E.coli [ppm]: 1.71 (Q 64)

− RNA functions:
16S rRNA (guanine(1207)−N(2))−methyltransferase activity; ncRNA metabolic process
ncRNA processing; RNA metabolic process; RNA methylation
RNA methyltransferase activity; RNA modification; RNA processing; rRNA (guanine−N2−)−methyltransferase activity
rRNA (guanine) methyltransferase activity; rRNA base methylation; rRNA metabolic process
rRNA methylation; rRNA methyltransferase activity; rRNA modification; rRNA processing

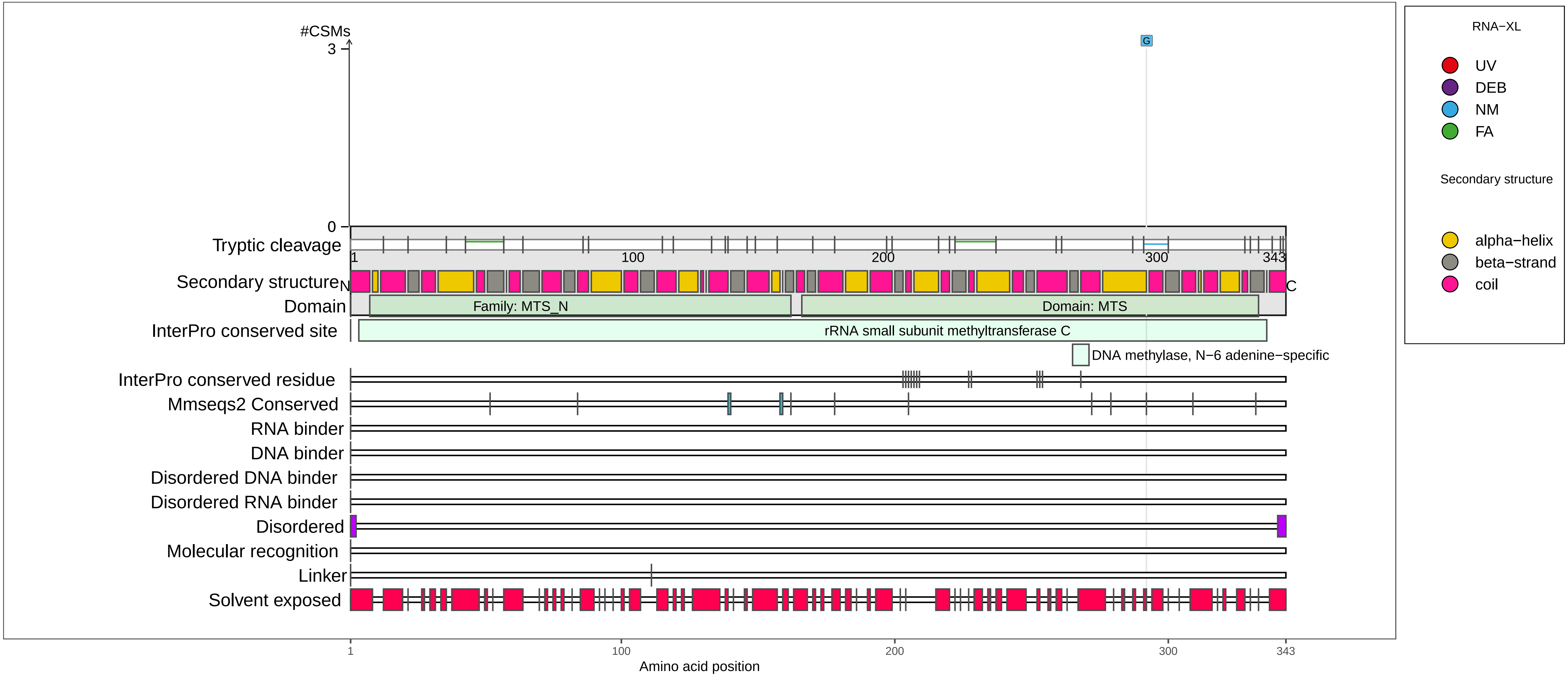

P60472
UPPS_ECOLI Ditrans,polycis−undecaprenyl−diphosphate synthase ((2E,6E)−farnesyl−diphosphate specific)

− Abundance:
tryptic [log10 Intensity]: 6.74 (Q 6)
PAXdb K12 strain [ppm]: 1.87 (Q 56)
PAXdb E.coli [ppm]: 1.78 (Q 66)

− RNA functions: not annotated

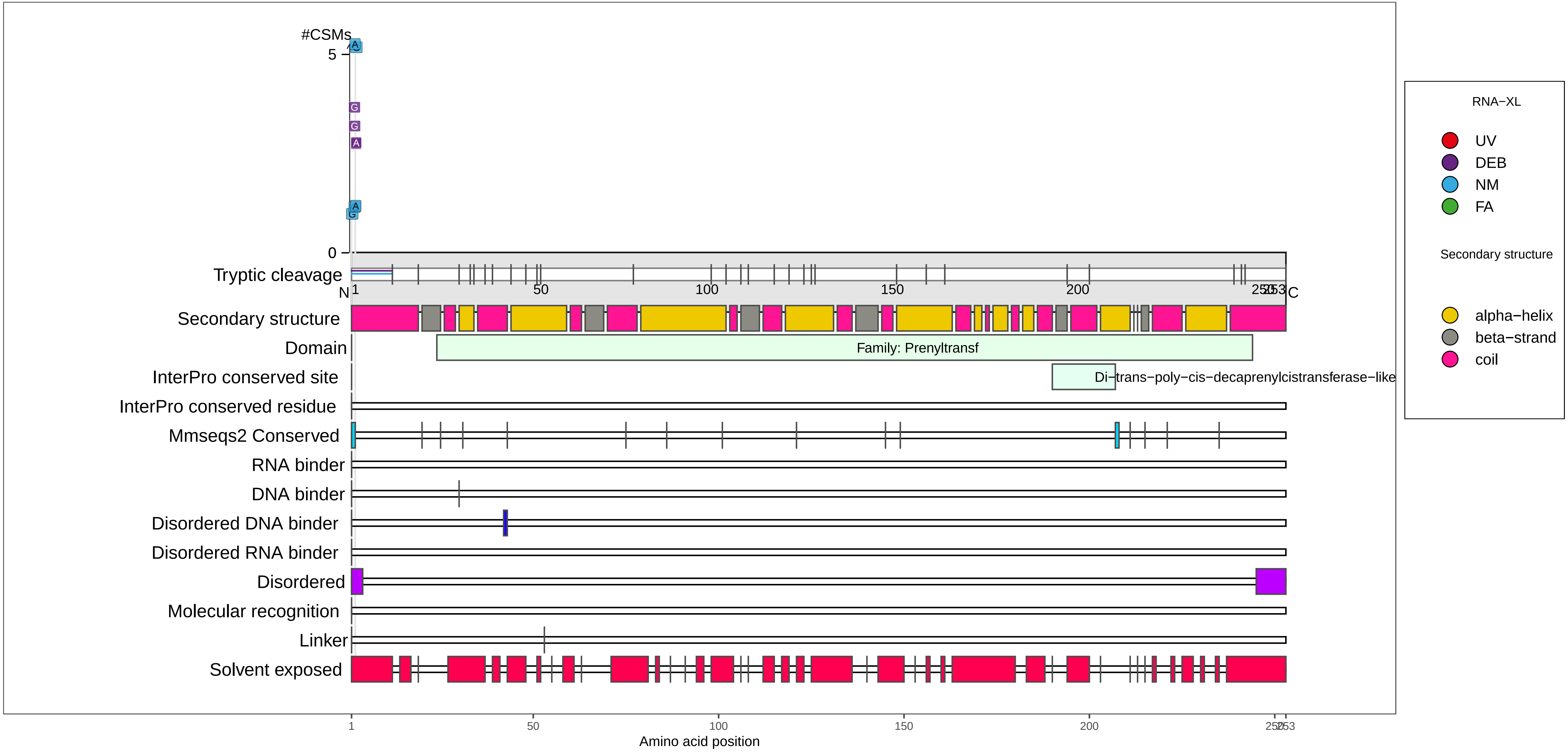

P62707
GPMA_ECOLI 2,3−bisphosphoglycerate−dependent phosphoglycerate mutase

− Abundance:
tryptic [log10 Intensity]: 9.1 (Q 91)
PAXdb K12 strain [ppm]: 3.61 (Q 98)
PAXdb E.coli [ppm]: 3.65 (Q 100)

− RNA functions: not annotated

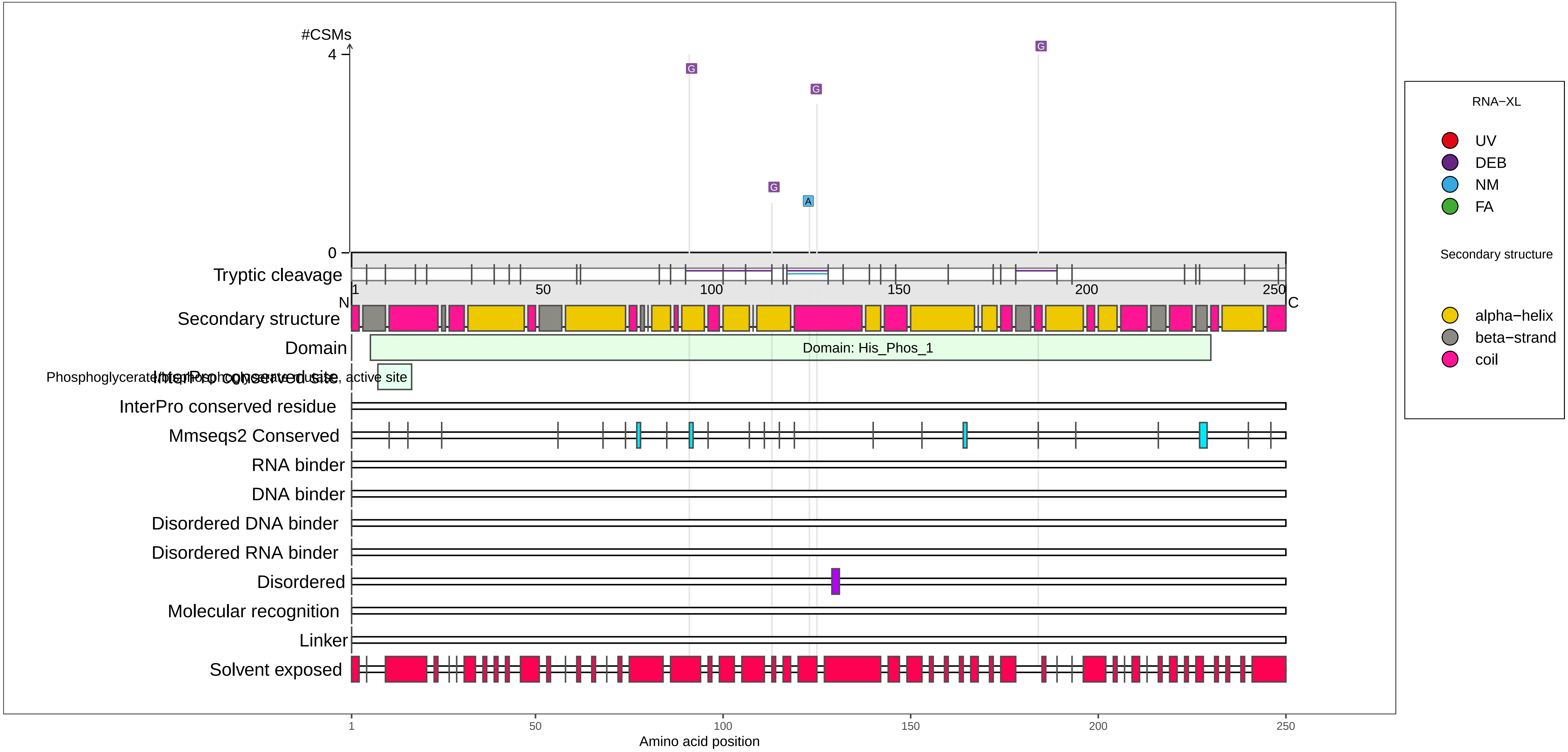

P69503
APT_ECOLI Adenine phosphoribosyltransferase

− Abundance:
tryptic [log10 Intensity]: 9.33 (Q 94)
PAXdb K12 strain [ppm]: 2.44 (Q 75)
PAXdb E.coli [ppm]: 2.42 (Q 84)

− RNA functions: not annotated

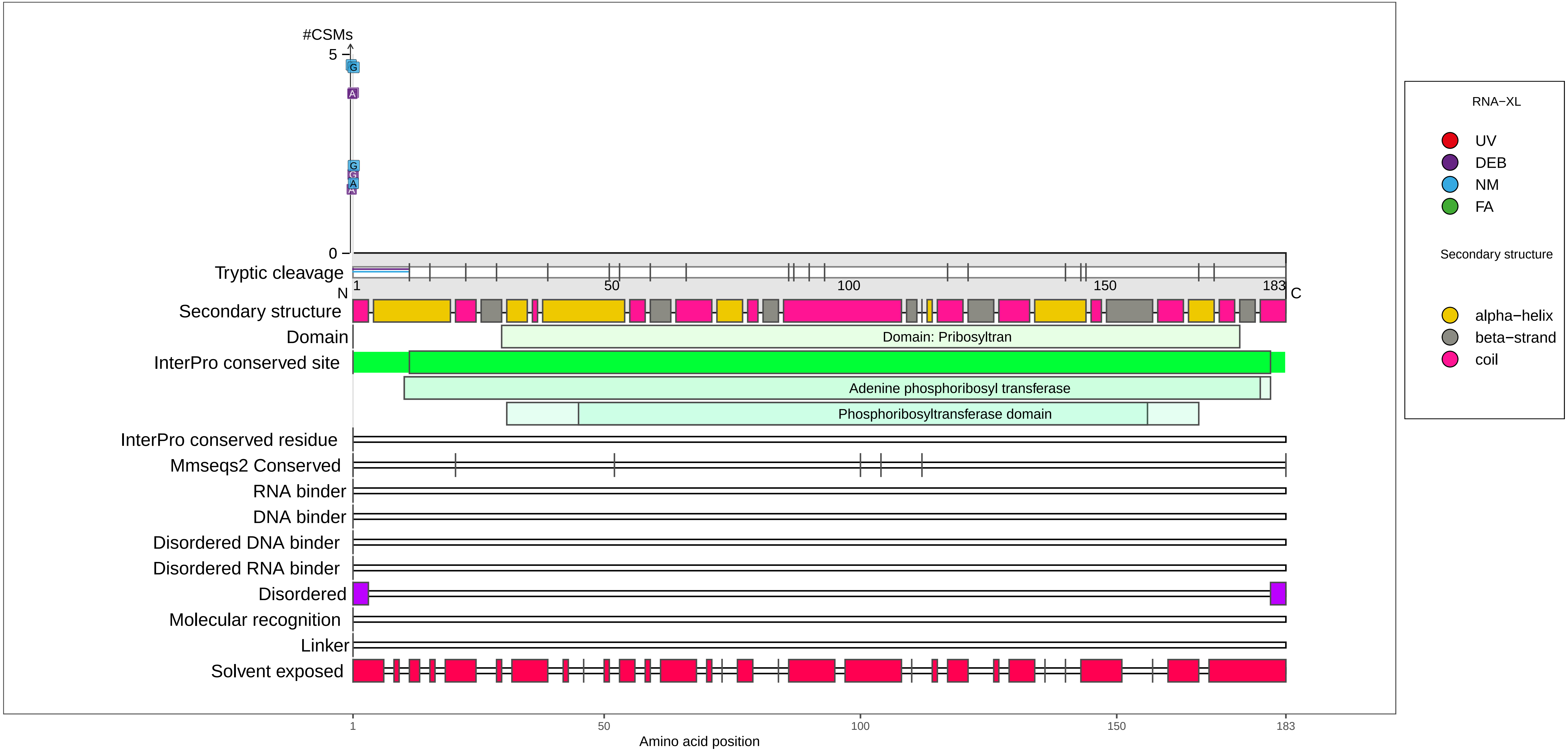

P69786
PTGCB_ECOLI PTS system glucose−specific EIICB component

− Abundance:
tryptic [log10 Intensity]: 9.43 (Q 96)
PAXdb K12 strain [ppm]: 2.05 (Q 62)
PAXdb E.coli [ppm]: 2.54 (Q 86)

− RNA functions: not annotated

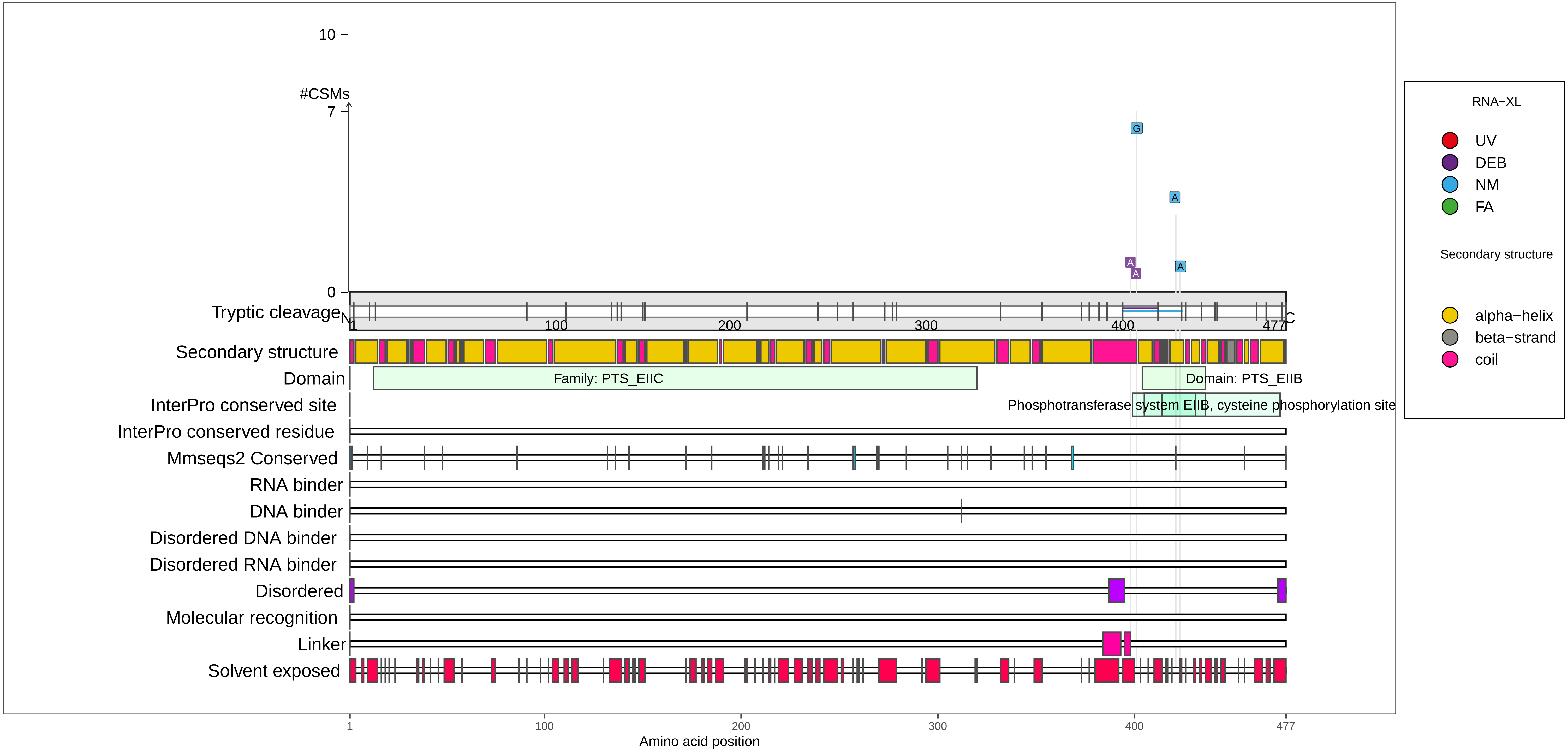

P04968
ILVA_ECOLI L−threonine dehydratase biosynthetic IlvA

− Abundance:
tryptic [log10 Intensity]: 8.67 (Q 81)
PAXdb K12 strain [ppm]: 1.5 (Q 38)
PAXdb E.coli [ppm]: 1.55 (Q 61)

− RNA functions: not annotated

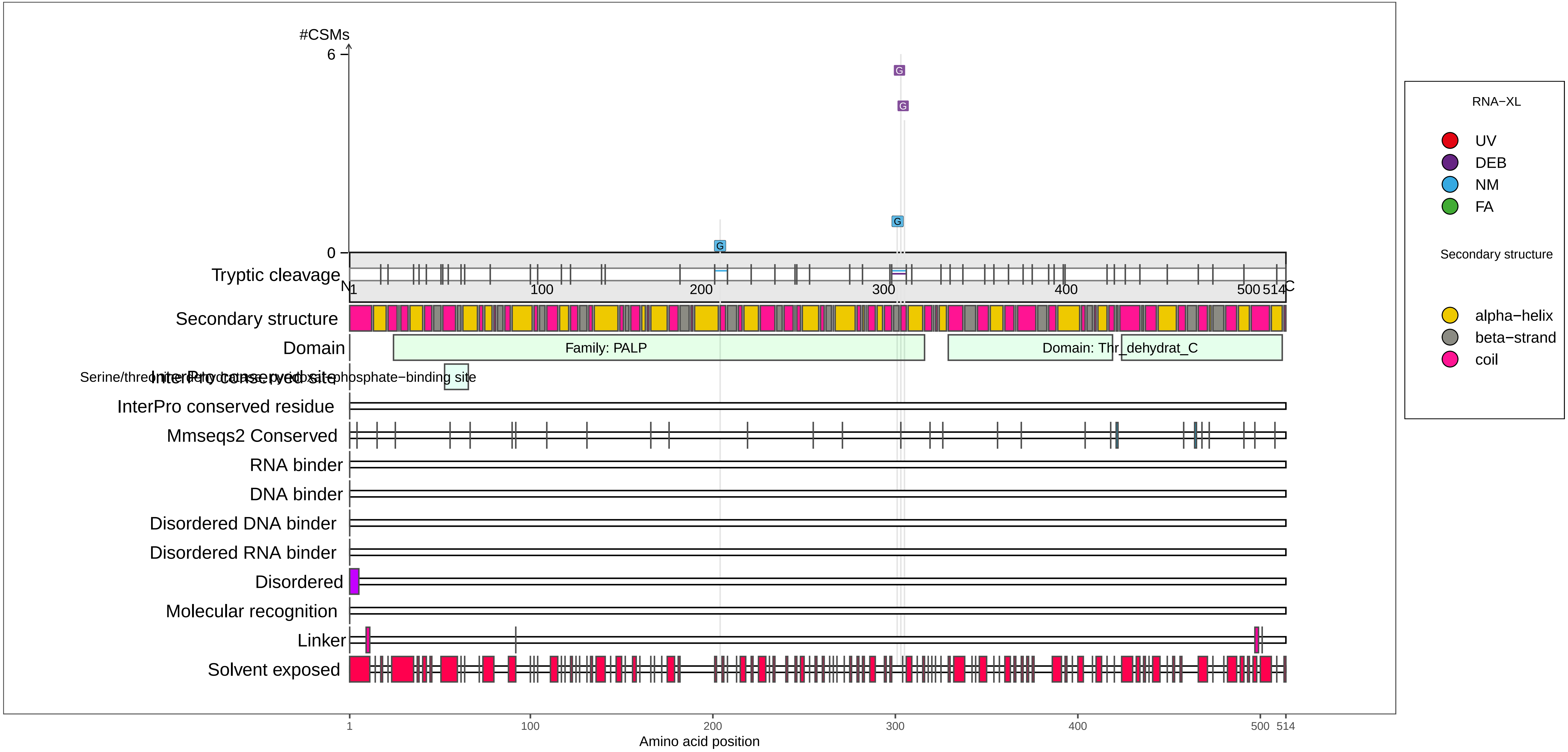

P09323
PTW3C_ECOLI PTS system N−acetylglucosamine−specific EIICBA component

− Abundance:
tryptic [log10 Intensity]: 8.72 (Q 81)
PAXdb K12 strain [ppm]: 1.42 (Q 34)
PAXdb E.coli [ppm]: 2.09 (Q 74)

− RNA functions: not annotated

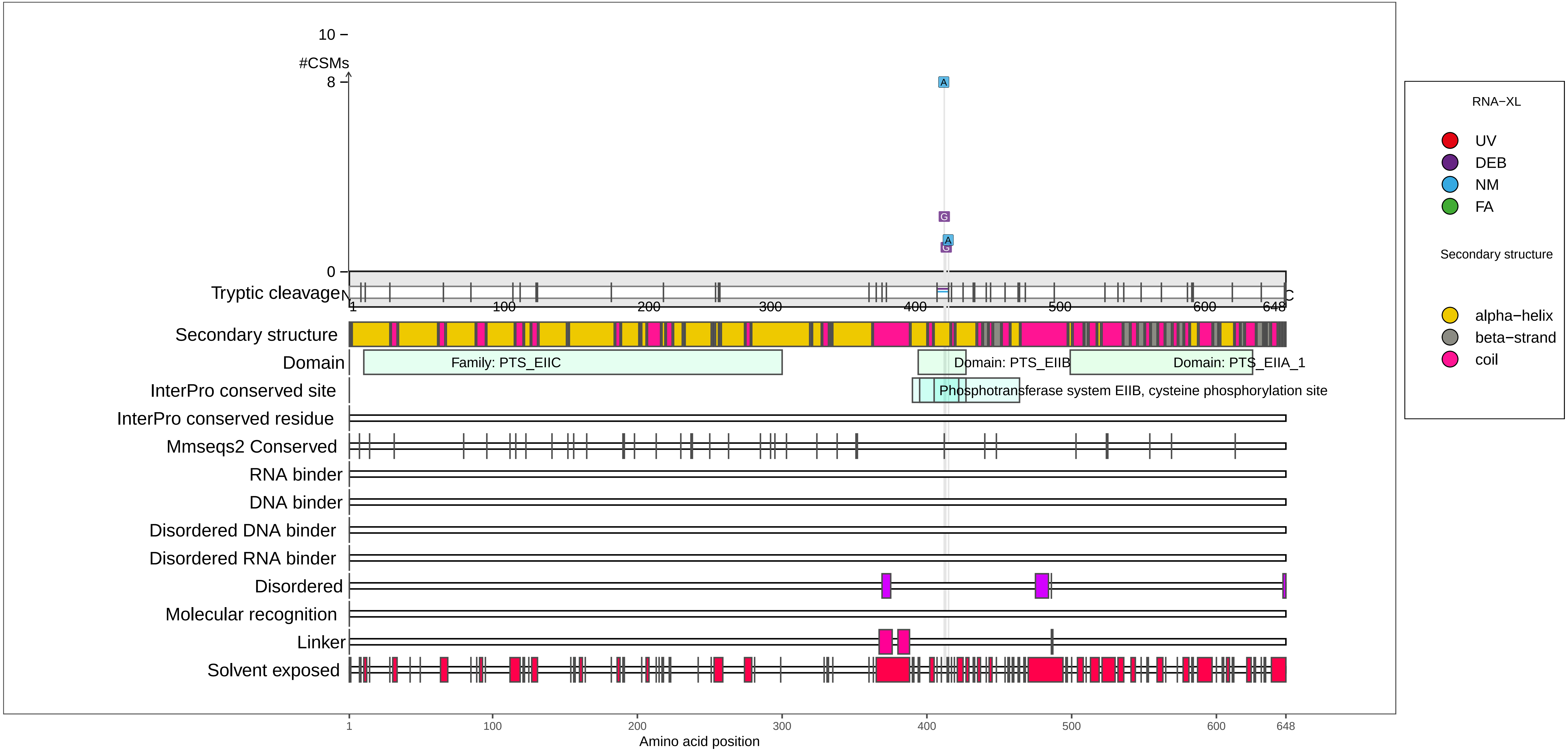

P0A717
KPRS_ECOLI Ribose−phosphate pyrophosphokinase

− Abundance:
tryptic [log10 Intensity]: 8.43 (Q 73)
PAXdb K12 strain [ppm]: 3.16 (Q 93)
PAXdb E.coli [ppm]: 3.07 (Q 96)

− RNA functions: not annotated

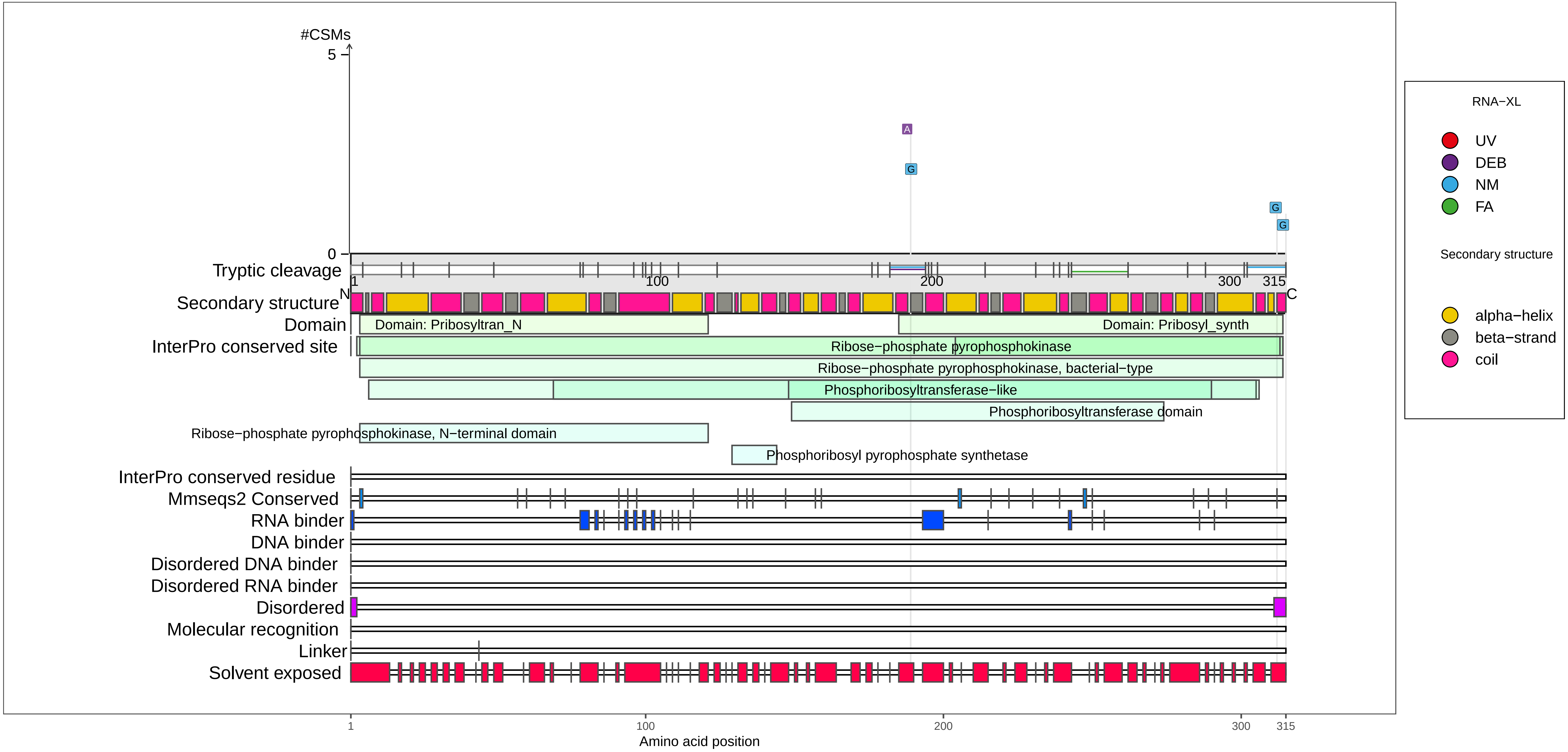

P0A887
UBIE_ECOLI Ubiquinone/menaquinone biosynthesis C−methyltransferase UbiE

− Abundance:
tryptic [log10 Intensity]: 7.21 (Q 21)
PAXdb K12 strain [ppm]: 2.72 (Q 84)
PAXdb E.coli [ppm]: 2.44 (Q 84)

− RNA functions: not annotated

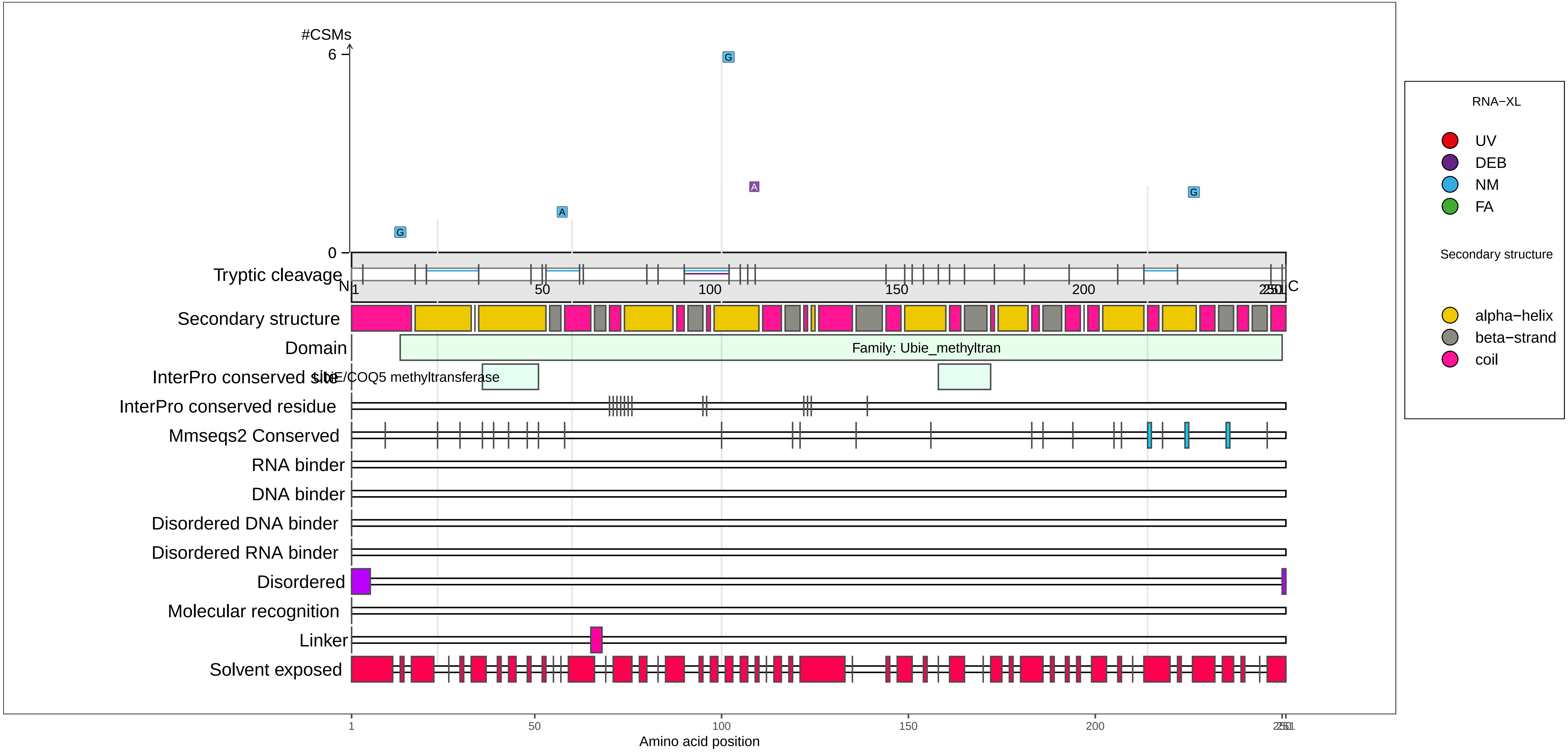

P0AA25
THIO_ECOLI Thioredoxin 1

− Abundance:
tryptic [log10 Intensity]: 9.11 (Q 91)
PAXdb K12 strain [ppm]: 3.17 (Q 94)
PAXdb E.coli [ppm]: 3.3 (Q 97)

− RNA functions: not annotated

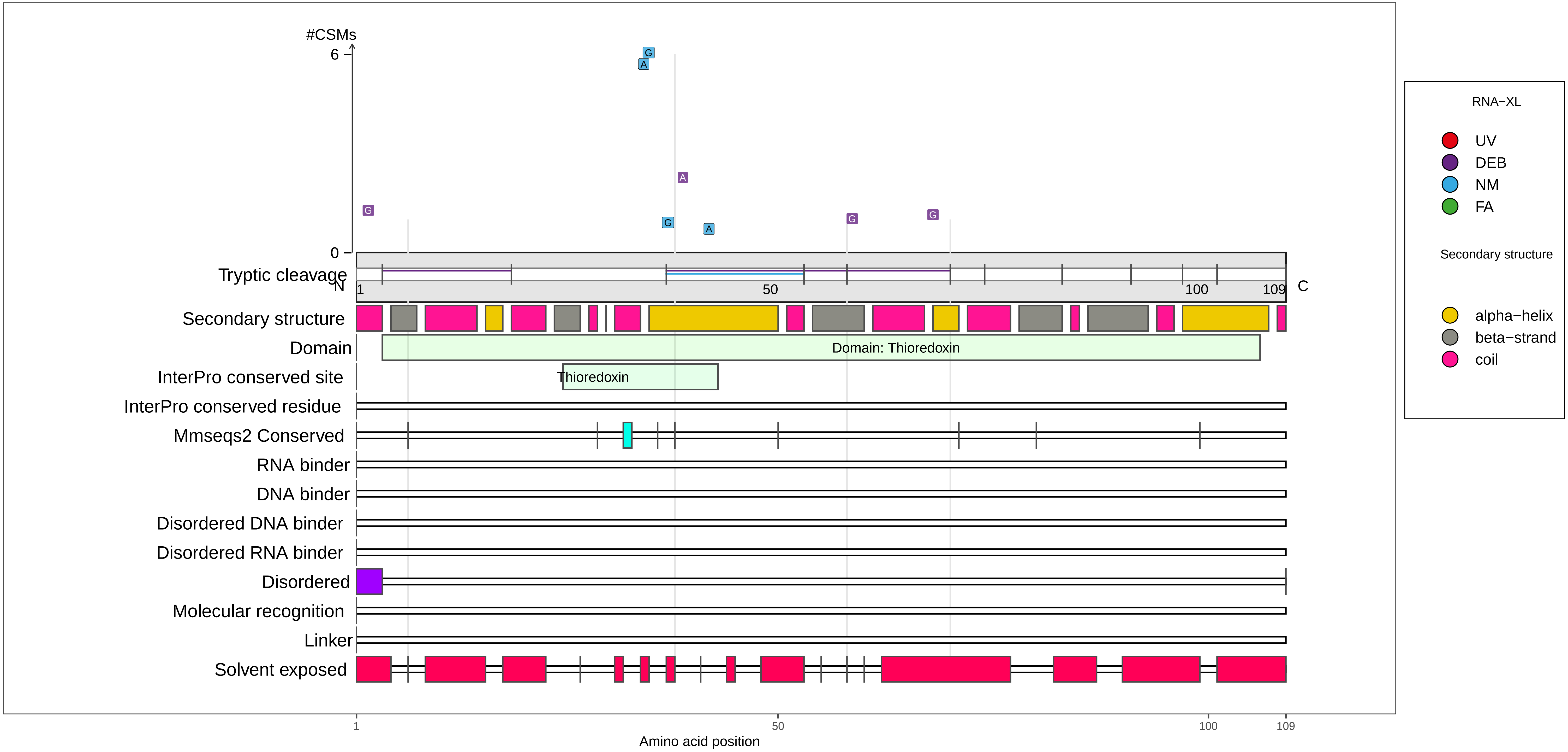

P0AAI5
FABF_ECOLI 3−oxoacyl−[acyl−carrier−protein] synthase 2

− Abundance:
tryptic [log10 Intensity]: 9.58 (Q 97)
PAXdb K12 strain [ppm]: 2.51 (Q 78)
PAXdb E.coli [ppm]: 2.59 (Q 88)

− RNA functions: not annotated

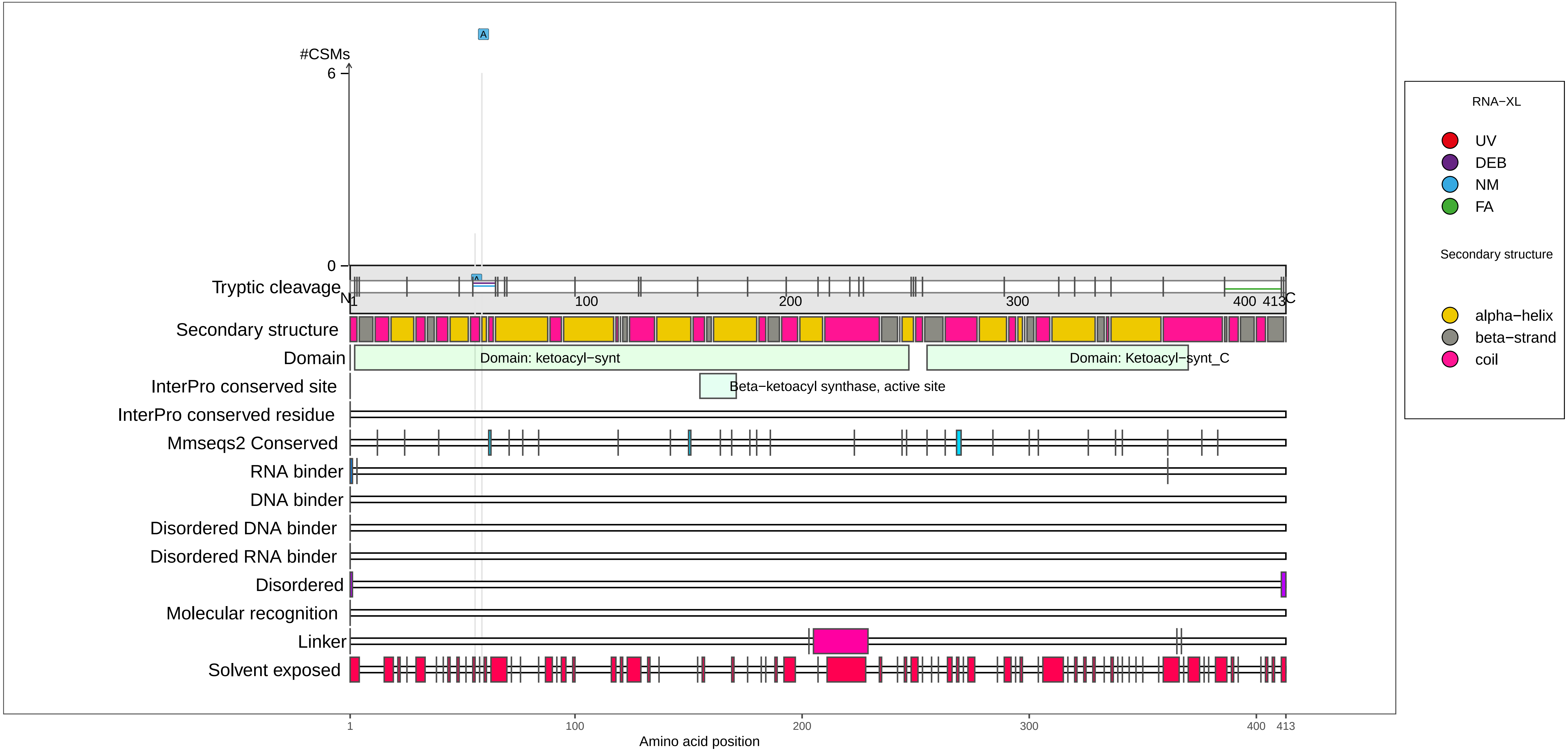

P0ACF4
DBHB_ECOLI DNA−binding protein HU−beta

− Abundance:
tryptic [log10 Intensity]: 9.53 (Q 97)
PAXdb K12 strain [ppm]: 3.83 (Q 100)
PAXdb E.coli [ppm]: 3.91 (Q 100)

− RNA functions:
RNA biosynthetic process; RNA metabolic process

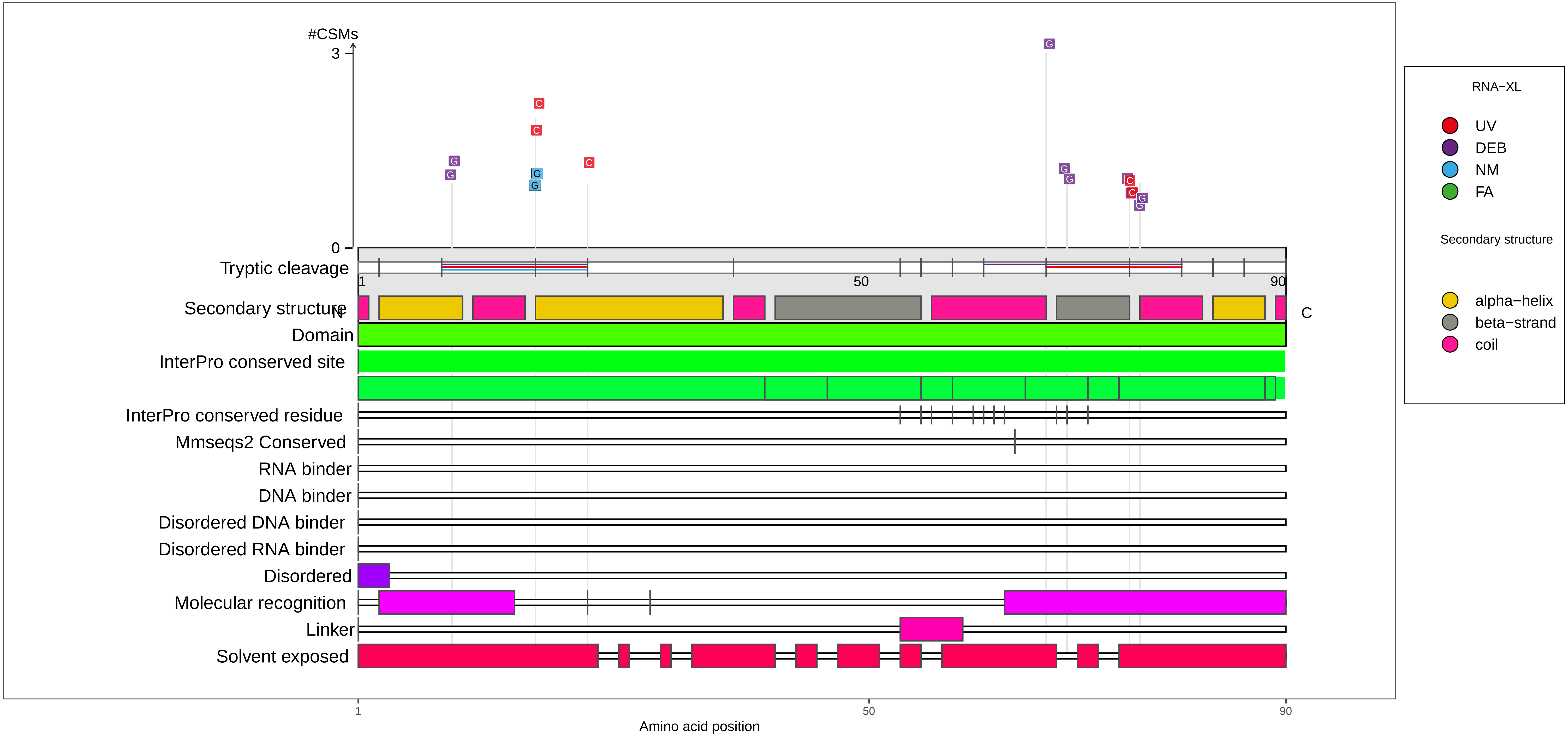

P0AFF0
NUON_ECOLI NADH−quinone oxidoreductase subunit N

− Abundance:
tryptic [log10 Intensity]: 8.81 (Q 84)
PAXdb K12 strain [ppm]: NA (Q NA)
PAXdb E.coli [ppm]: 1.04 (Q 48)

− RNA functions: not annotated

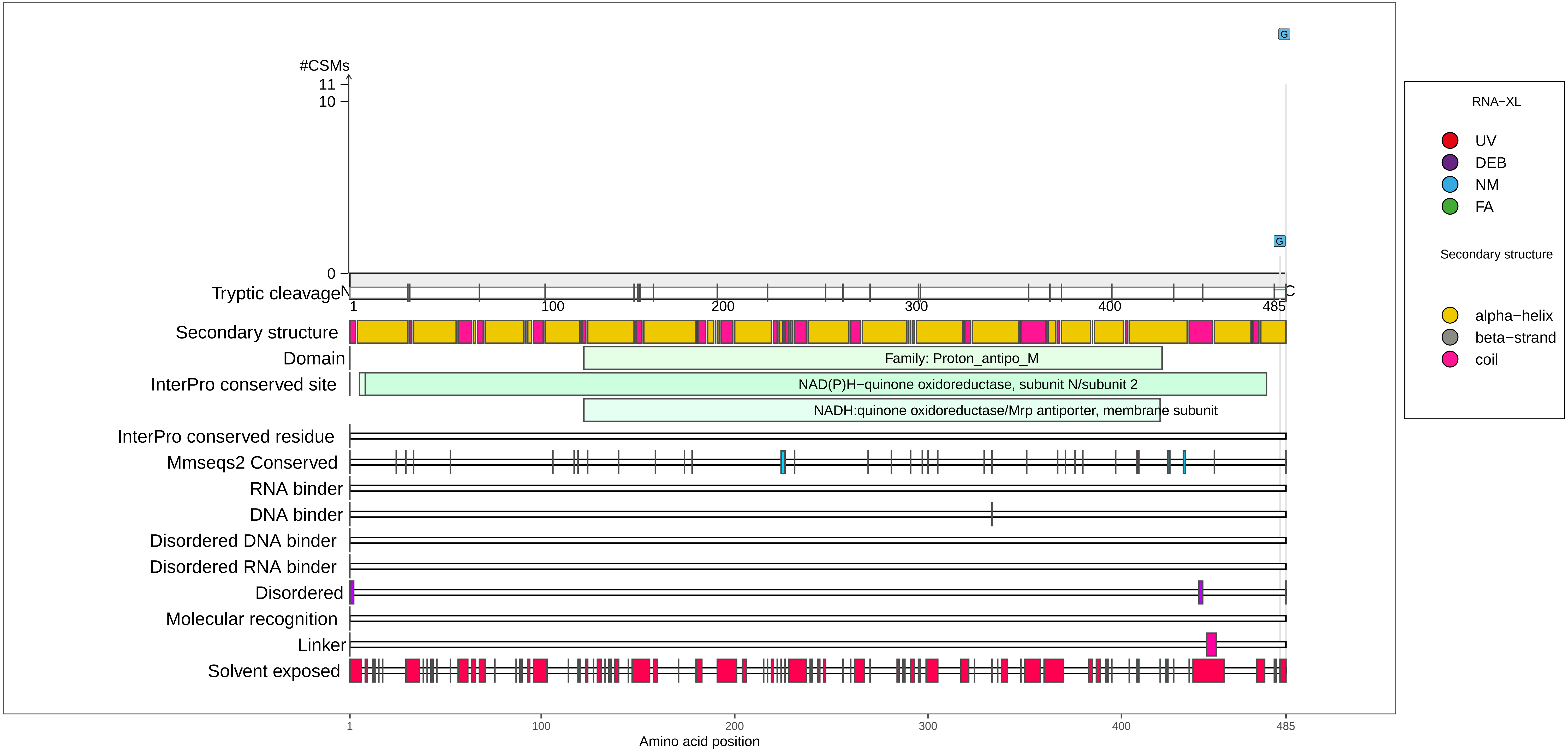

P0AFX4
RSD_ECOLI Regulator of sigma D
− Abundance:
tryptic [log10 Intensity]: 7.48 (Q 34)
PAXdb K12 strain [ppm]: 1.99 (Q 60)
PAXdb E.coli [ppm]: 2.51 (Q 86)

− RNA functions:
Regulator of RNA polymerase sigma(70) subunit, Rsd/AlgQ

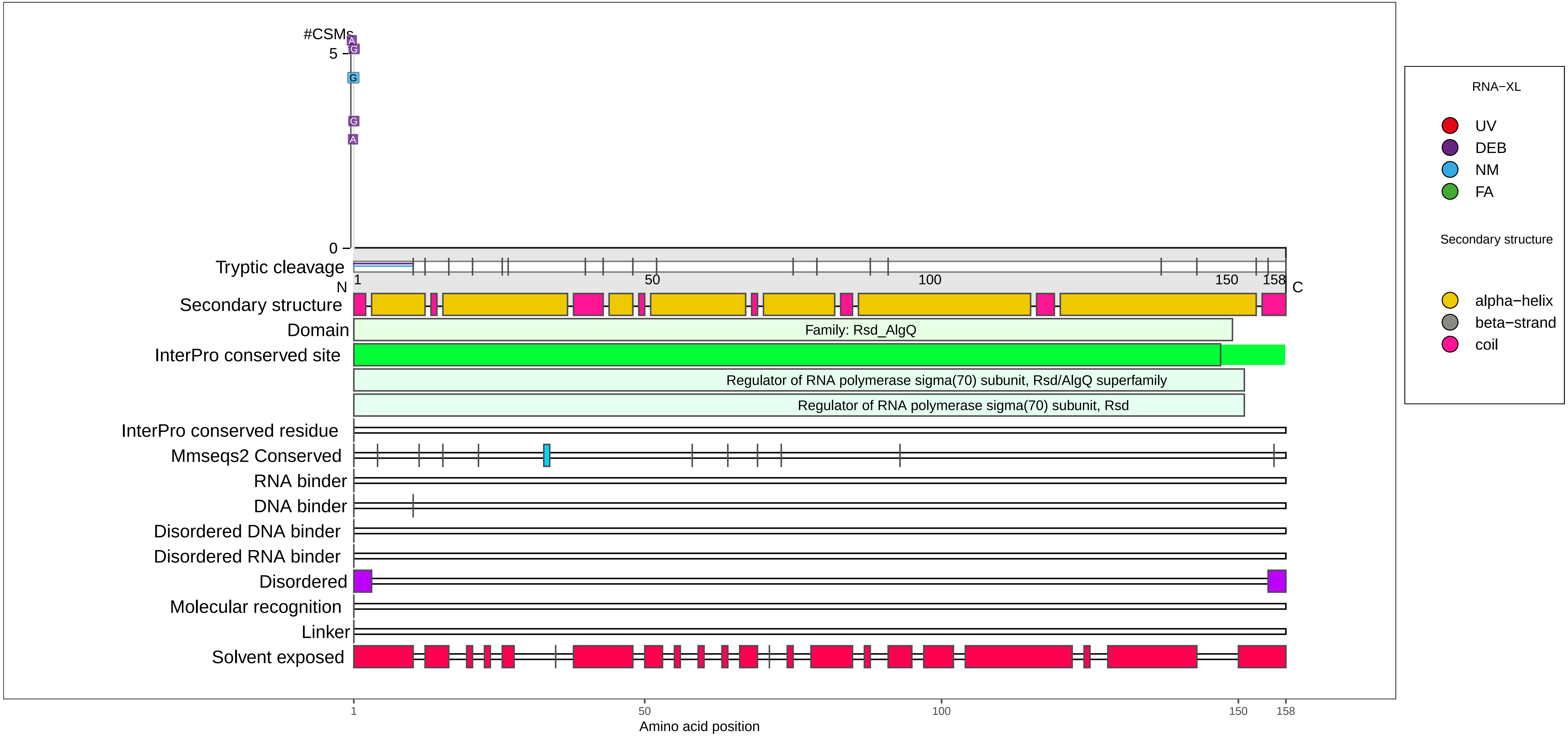

P22259
PCKA_ECOLI Phosphoenolpyruvate carboxykinase (ATP)

− Abundance:
tryptic [log10 Intensity]: 9.21 (Q 93)
PAXdb K12 strain [ppm]: 2.66 (Q 82)
PAXdb E.coli [ppm]: 3.07 (Q 95)

− RNA functions: not annotated

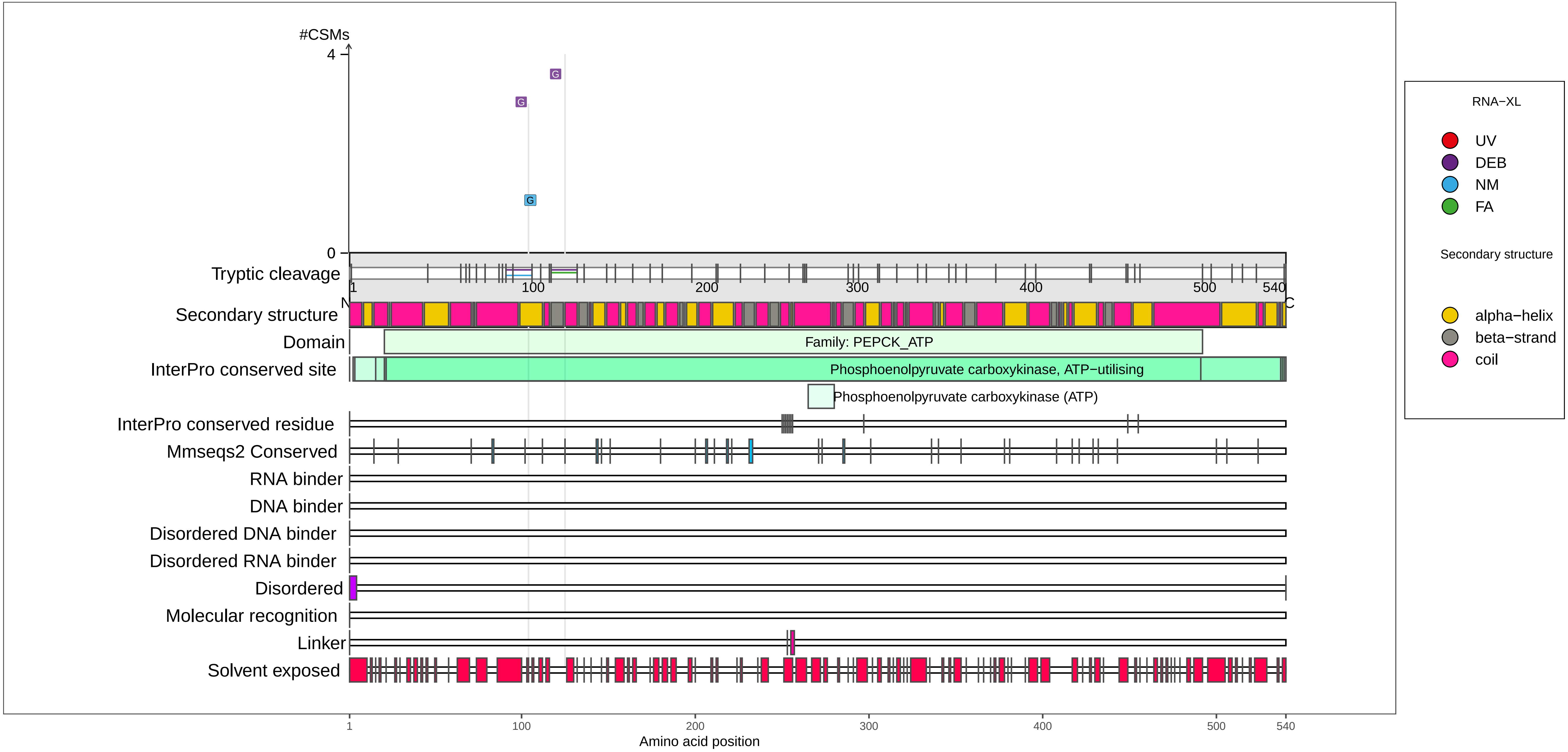

P26648
FTSP_ECOLI Cell division protein FtsP

− Abundance:
tryptic [log10 Intensity]: 7.96 (Q 56)
PAXdb K12 strain [ppm]: 1.66 (Q 48)
PAXdb E.coli [ppm]: 0.65 (Q 40)

− RNA functions: not annotated

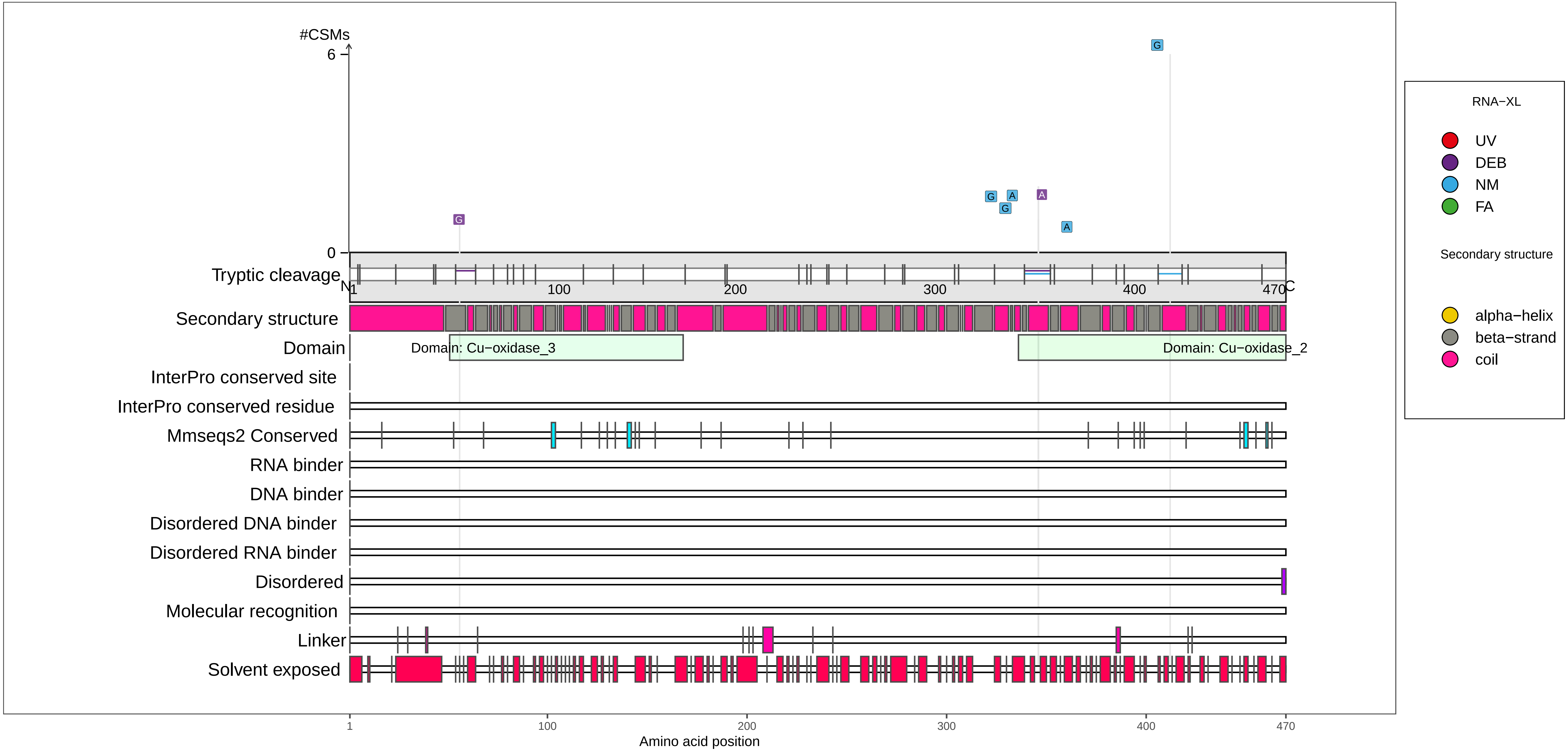

P36929
RSMB_ECOLI Ribosomal RNA small subunit methyltransferase B

− Abundance:
tryptic [log10 Intensity]: 7.98 (Q 57)
PAXdb K12 strain [ppm]: 1.93 (Q 58)
PAXdb E.coli [ppm]: 1.56 (Q 61)

− RNA functions:
16S rRNA methyltransferase RsmB/F; ncRNA metabolic process; ncRNA processing
RNA binding; RNA metabolic process; RNA methylation; RNA methyltransferase activity
RNA modification; RNA processing; rRNA (cytosine−C5−)−methyltransferase activity
rRNA (cytosine) methyltransferase activity; rRNA base methylation; rRNA metabolic process
rRNA methylation; rRNA methyltransferase activity; rRNA modification; rRNA processing

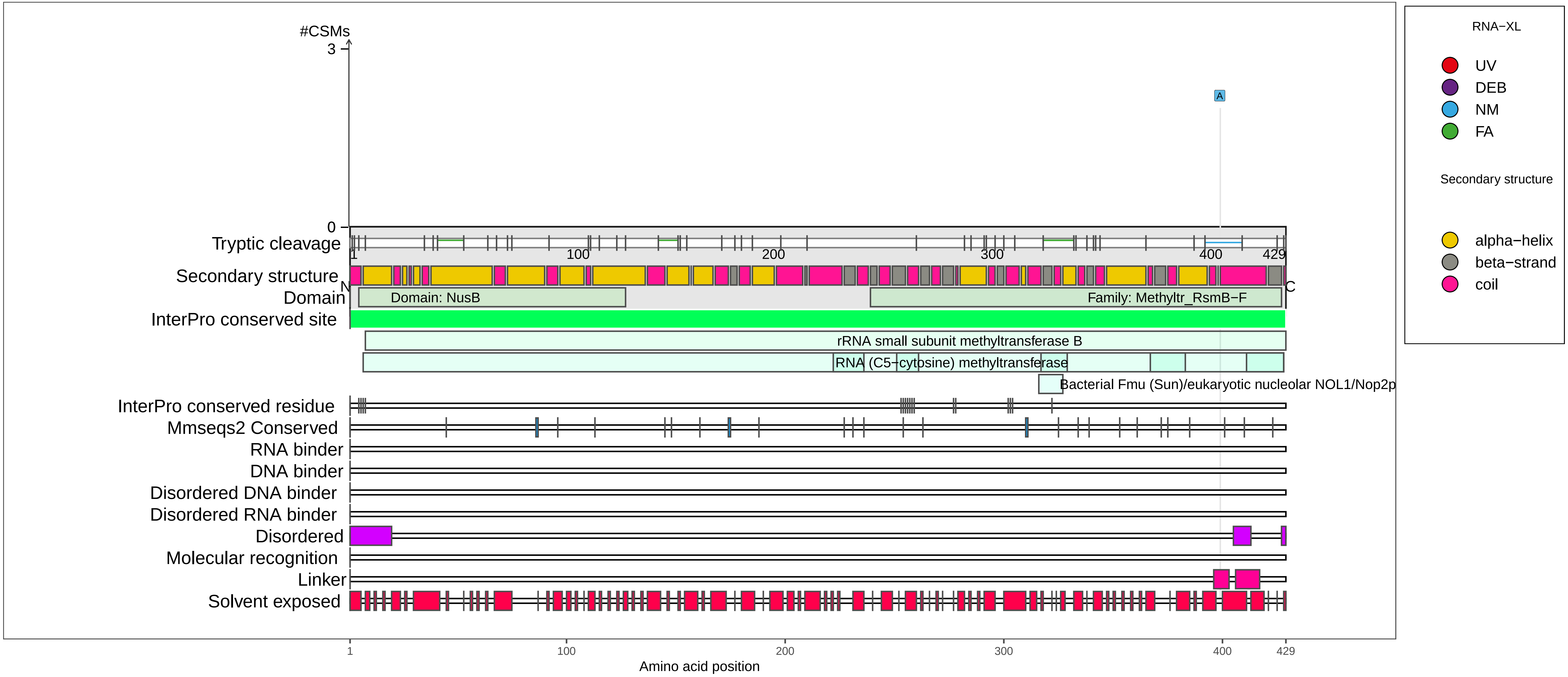

P64588
YQJI_ECOLI Transcriptional regulator YqjI

− Abundance:
tryptic [log10 Intensity]: 7.54 (Q 37)
PAXdb K12 strain [ppm]: NA (Q NA)
PAXdb E.coli [ppm]: 2.29 (Q 80)

− RNA functions: not annotated

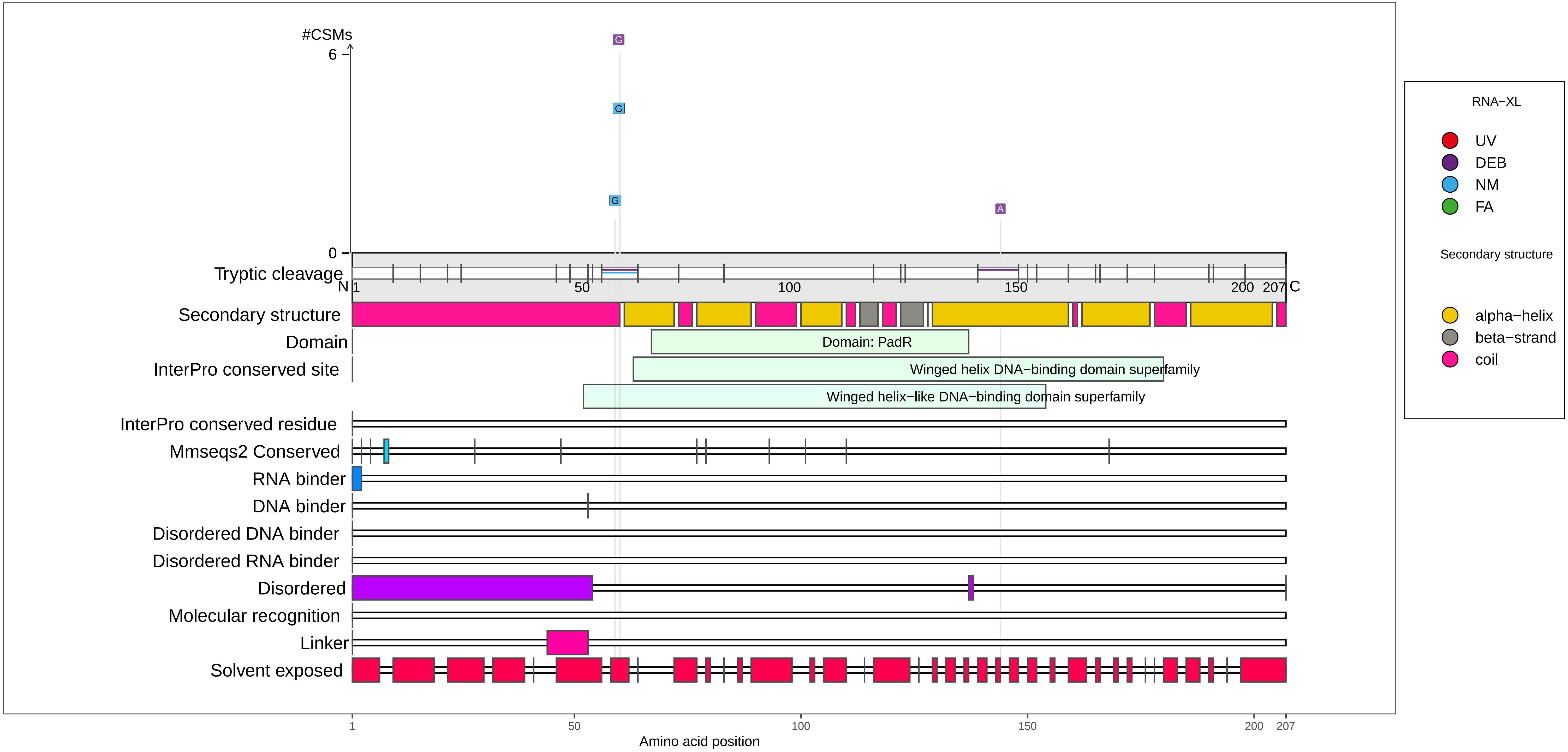

P67080
PLPHP_ECOLI Pyridoxal phosphate homeostasis protein

− Abundance:
tryptic [log10 Intensity]: 7.85 (Q 51)
PAXdb K12 strain [ppm]: NA (Q NA)
PAXdb E.coli [ppm]: 2.31 (Q 81)

− RNA functions: not annotated

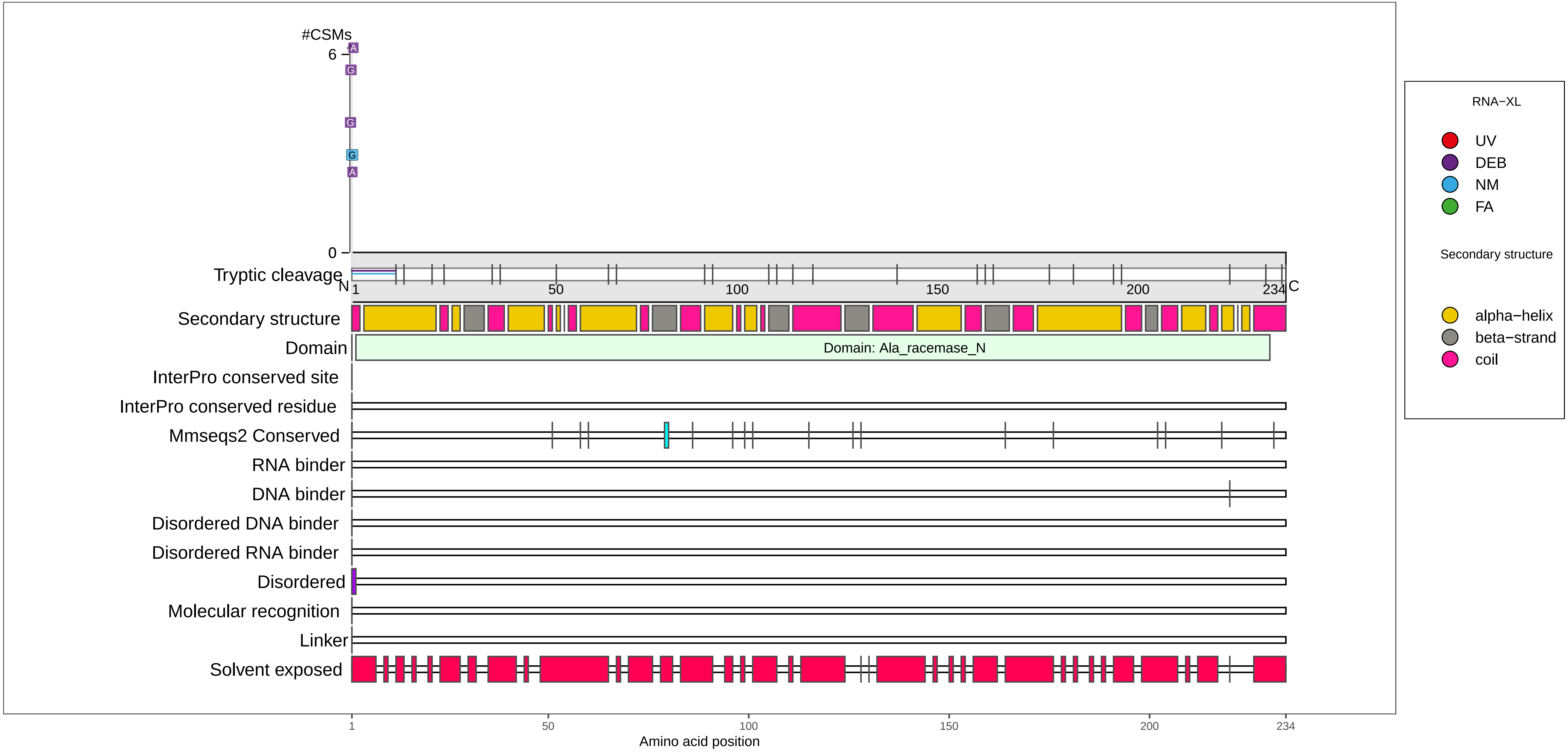

P77150
PDXY_ECOLI Pyridoxal kinase PdxY

− Abundance:
tryptic [log10 Intensity]: 8.55 (Q 77)
PAXdb K12 strain [ppm]: NA (Q NA)
PAXdb E.coli [ppm]: 1.55 (Q 61)

− RNA functions: not annotated

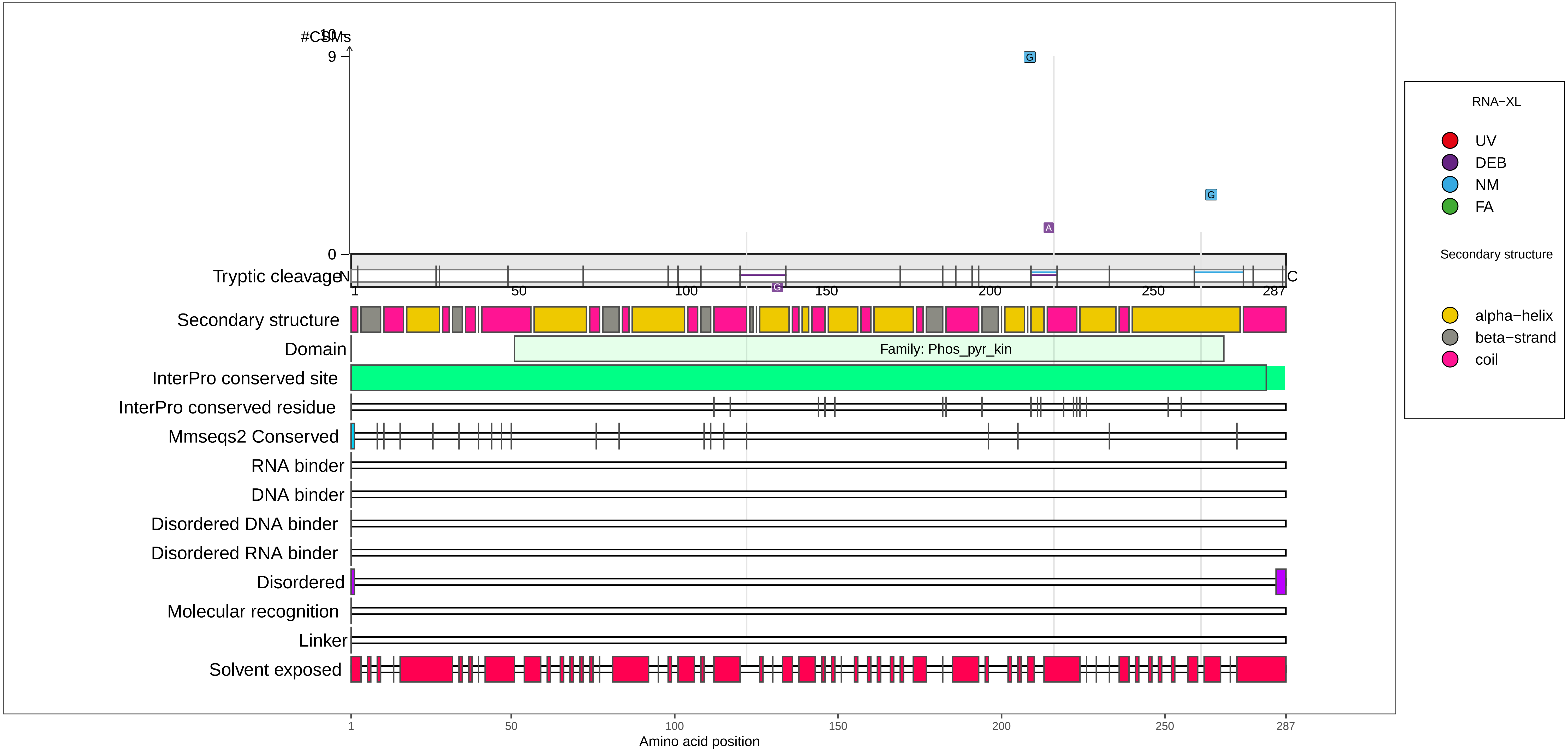

P77173
ZIPA_ECOLI Cell division protein ZipA

− Abundance:
tryptic [log10 Intensity]: 8.41 (Q 72)
PAXdb K12 strain [ppm]: 2.25 (Q 68)
PAXdb E.coli [ppm]: 1.74 (Q 66)

− RNA functions: not annotated

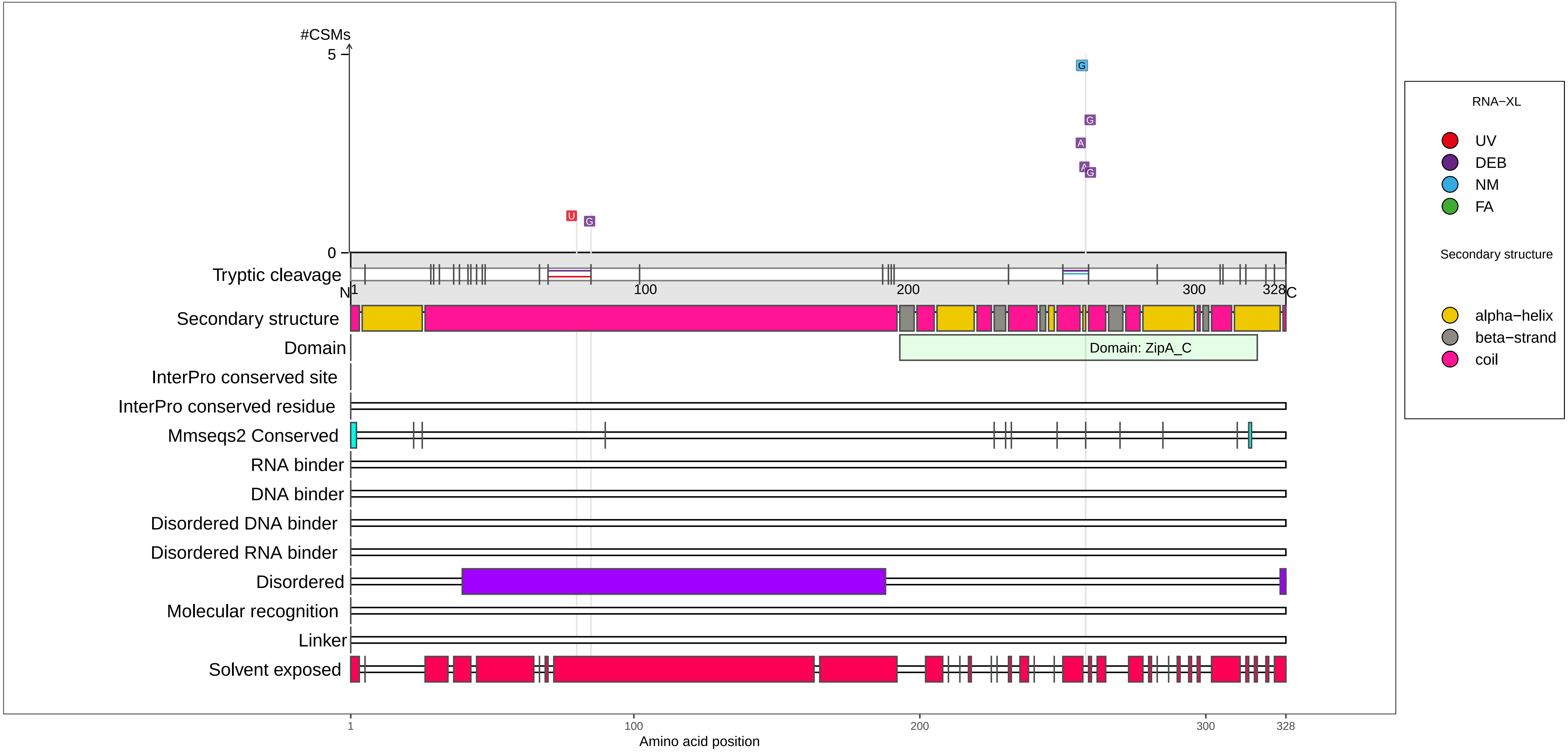

P77258
NEMA_ECOLI N−ethylmaleimide reductase

− Abundance:
tryptic [log10 Intensity]: 8.33 (Q 70)
PAXdb K12 strain [ppm]: 2.63 (Q 81)
PAXdb E.coli [ppm]: 1.27 (Q 54)

− RNA functions: not annotated

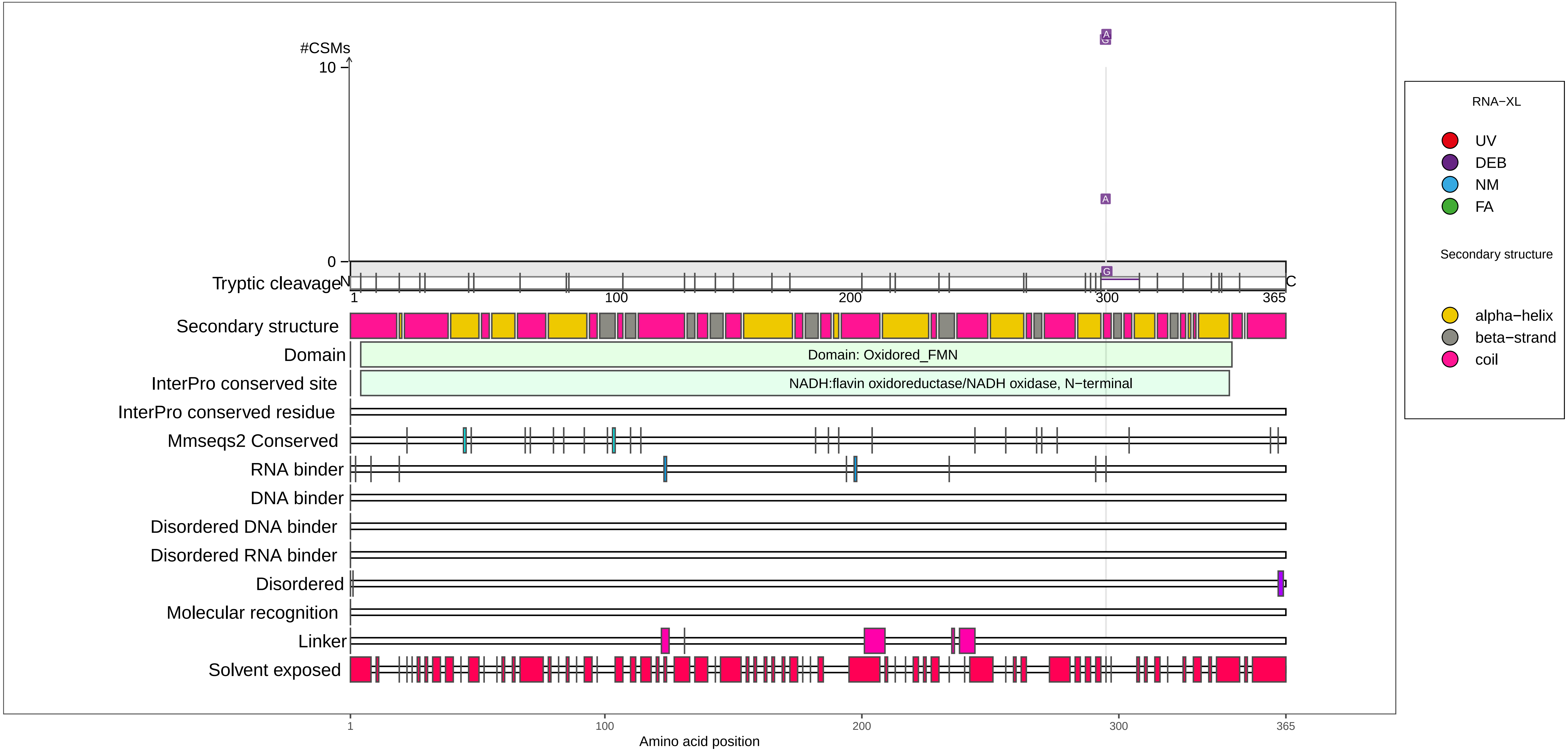

P06610
BTUE_ECOLI Thioredoxin/glutathione peroxidase BtuE

− Abundance:
tryptic [log10 Intensity]: 7.58 (Q 39)
PAXdb K12 strain [ppm]: 2.63 (Q 82)
PAXdb E.coli [ppm]: 2.77 (Q 91)

− RNA functions: not annotated

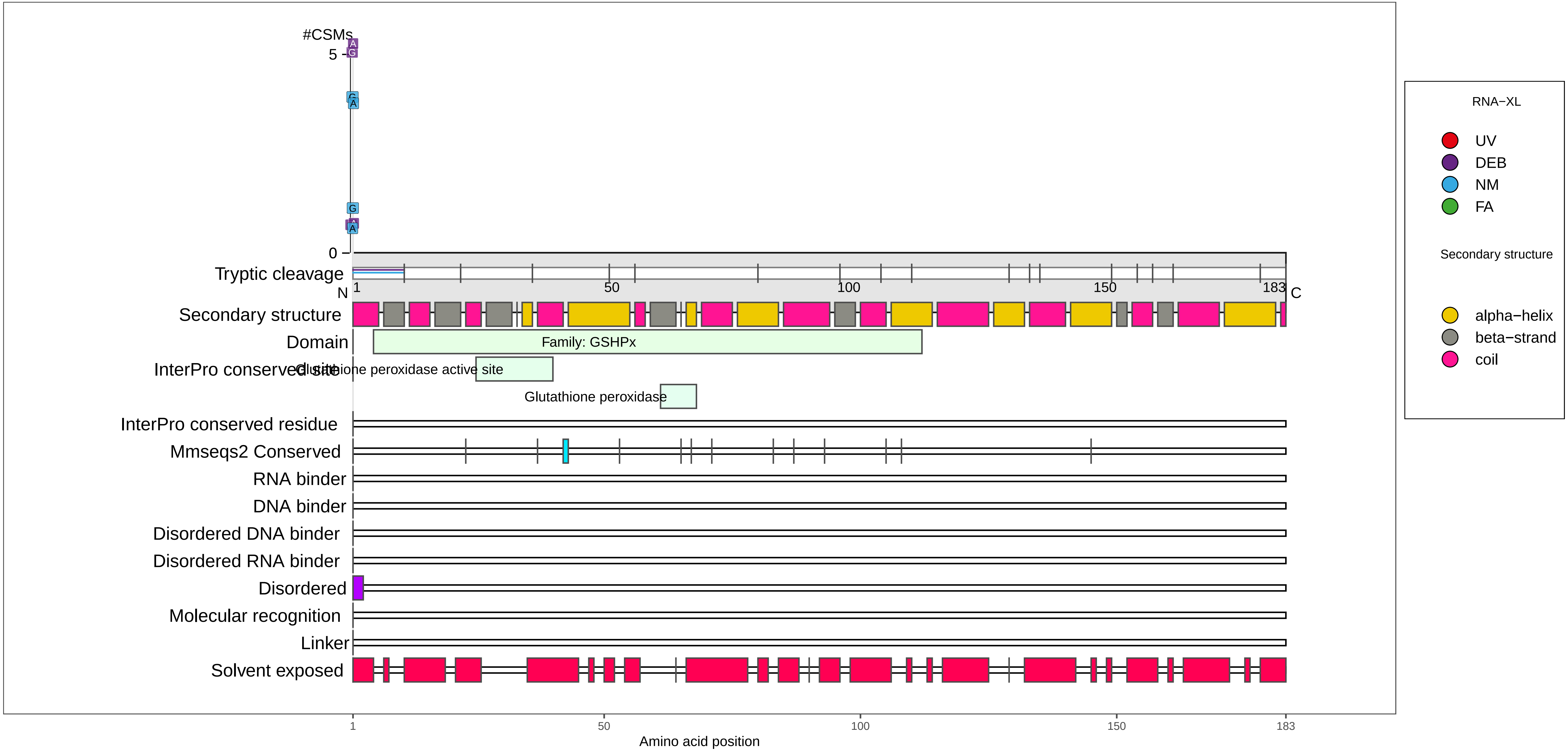

P0A6Z1
HSCA_ECOLI Chaperone protein HscA

− Abundance:
tryptic [log10 Intensity]: 8.64 (Q 80)
PAXdb K12 strain [ppm]: 2.42 (Q 75)
PAXdb E.coli [ppm]: 1.69 (Q 64)

− RNA functions: not annotated

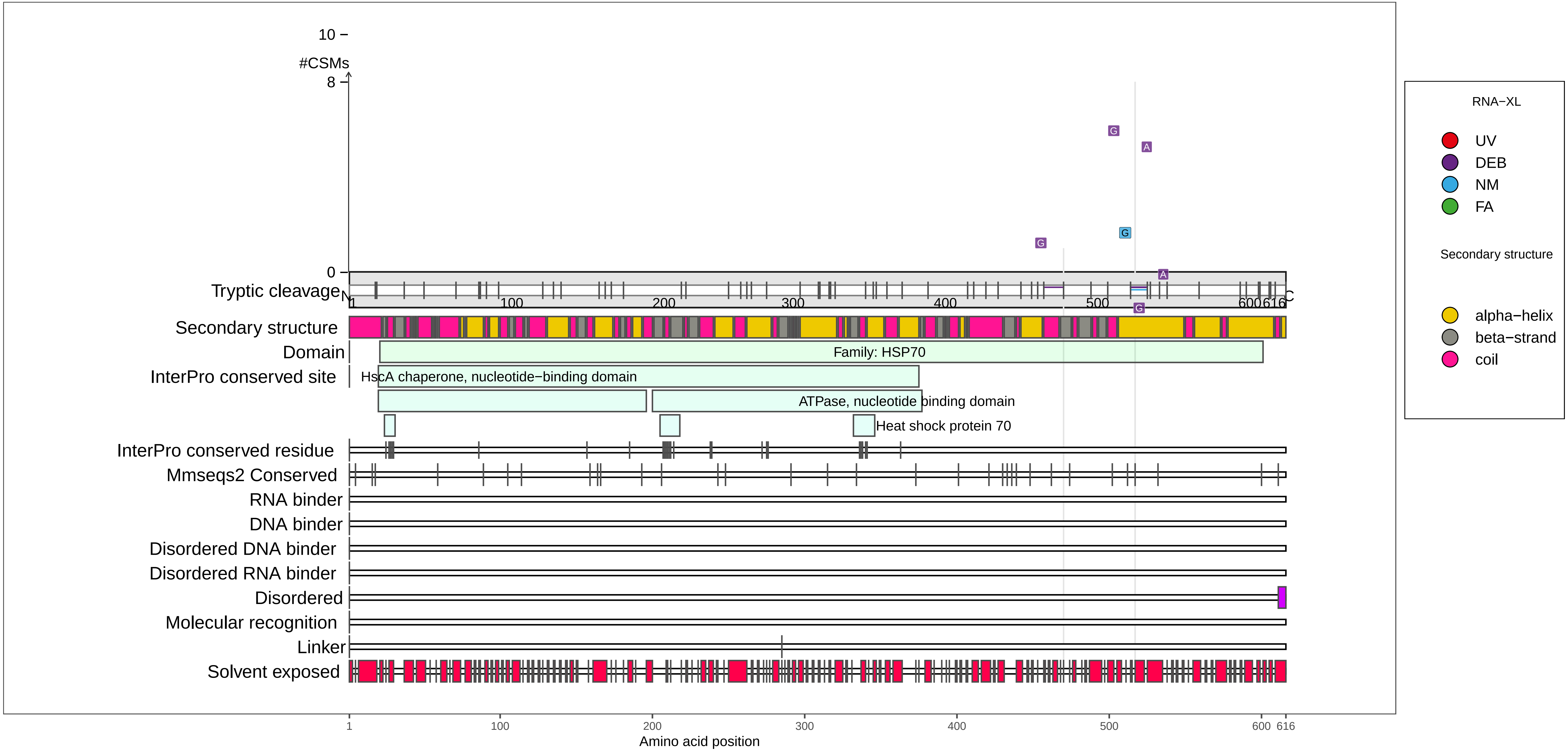

P0A7E9
PYRH_ECOLI Uridylate kinase
− Abundance:
tryptic [log10 Intensity]: 8.76 (Q 83)
PAXdb K12 strain [ppm]: 2.21 (Q 67)
PAXdb E.coli [ppm]: 2.66 (Q 89)

− RNA functions: not annotated

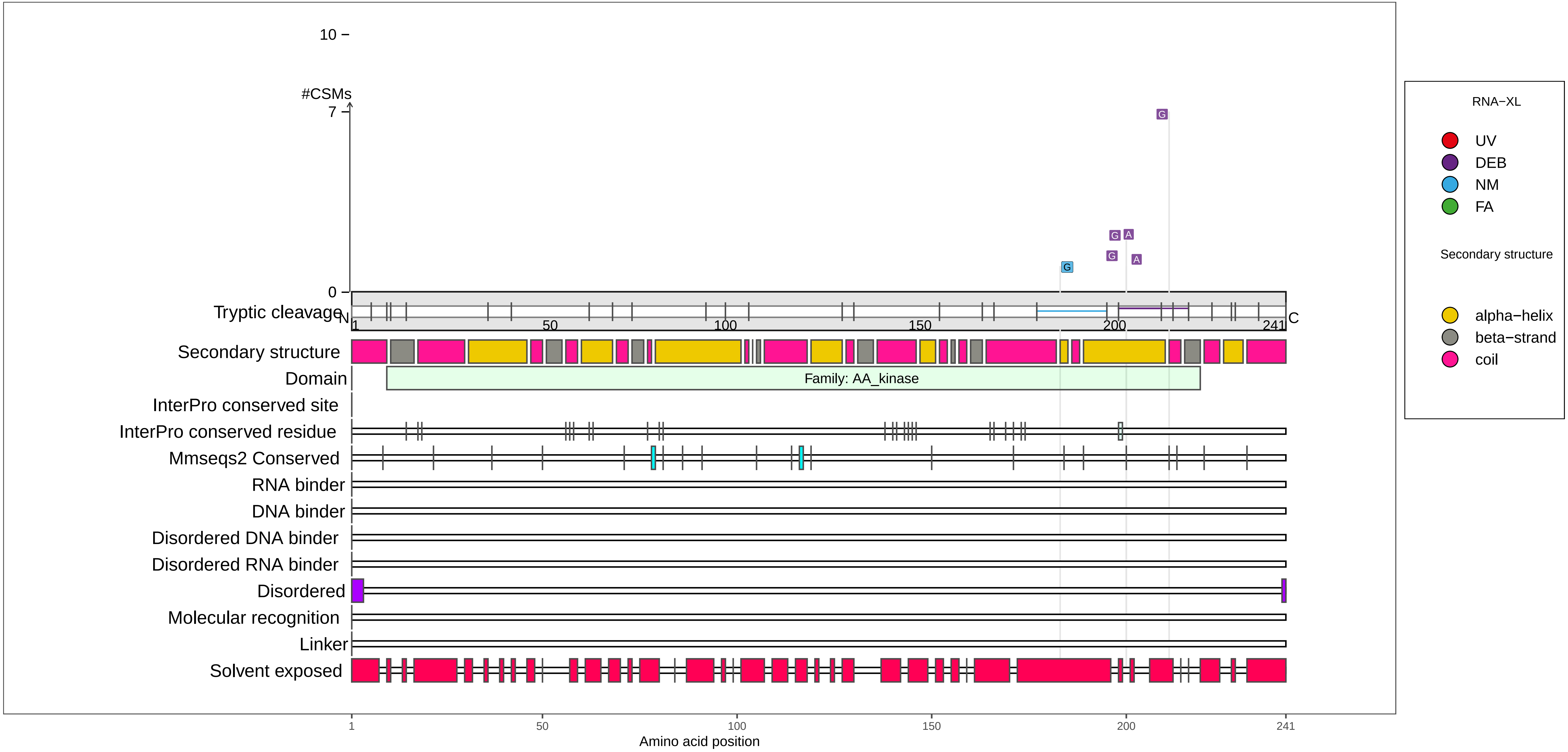

P0A7I4
RF3_ECOLI Peptide chain release factor RF3

− Abundance:
tryptic [log10 Intensity]: 9.08 (Q 90)
PAXdb K12 strain [ppm]: 2.98 (Q 90)
PAXdb E.coli [ppm]: 2.17 (Q 77)

− RNA functions:
RNA binding

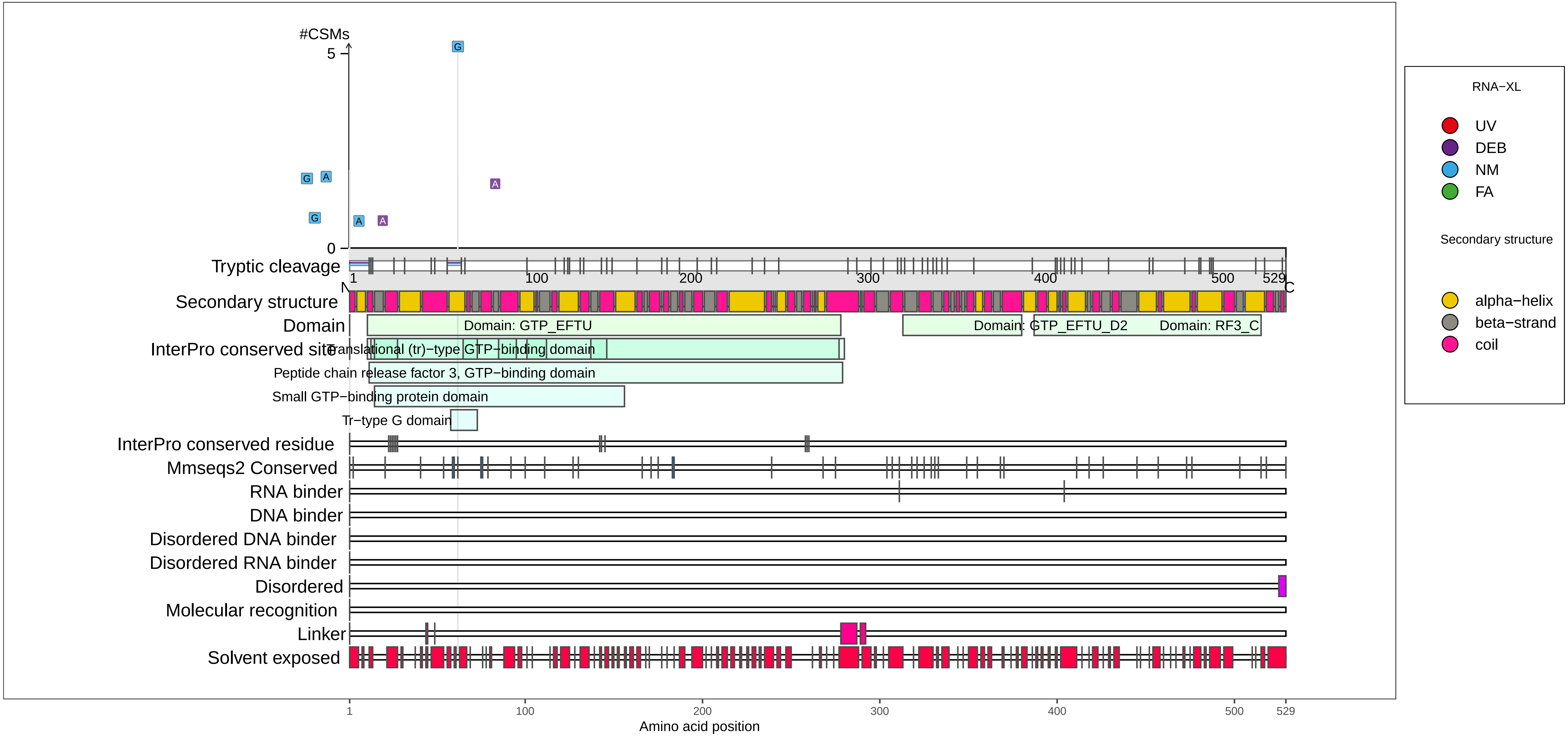

P0A800
RPOZ_ECOLI DNA−directed RNA polymerase subunit omega

− Abundance:
tryptic [log10 Intensity]: 8.84 (Q 84)
PAXdb K12 strain [ppm]: 3.56 (Q 98)
PAXdb E.coli [ppm]: 3.46 (Q 99)

− RNA functions:
5−3 RNA polymerase activity; DNA−directed 5−3 RNA polymerase activity
RNA biosynthetic process; RNA metabolic process; RNA polymerase activity; RNA polymerase complex
RNA polymerase Rpb6

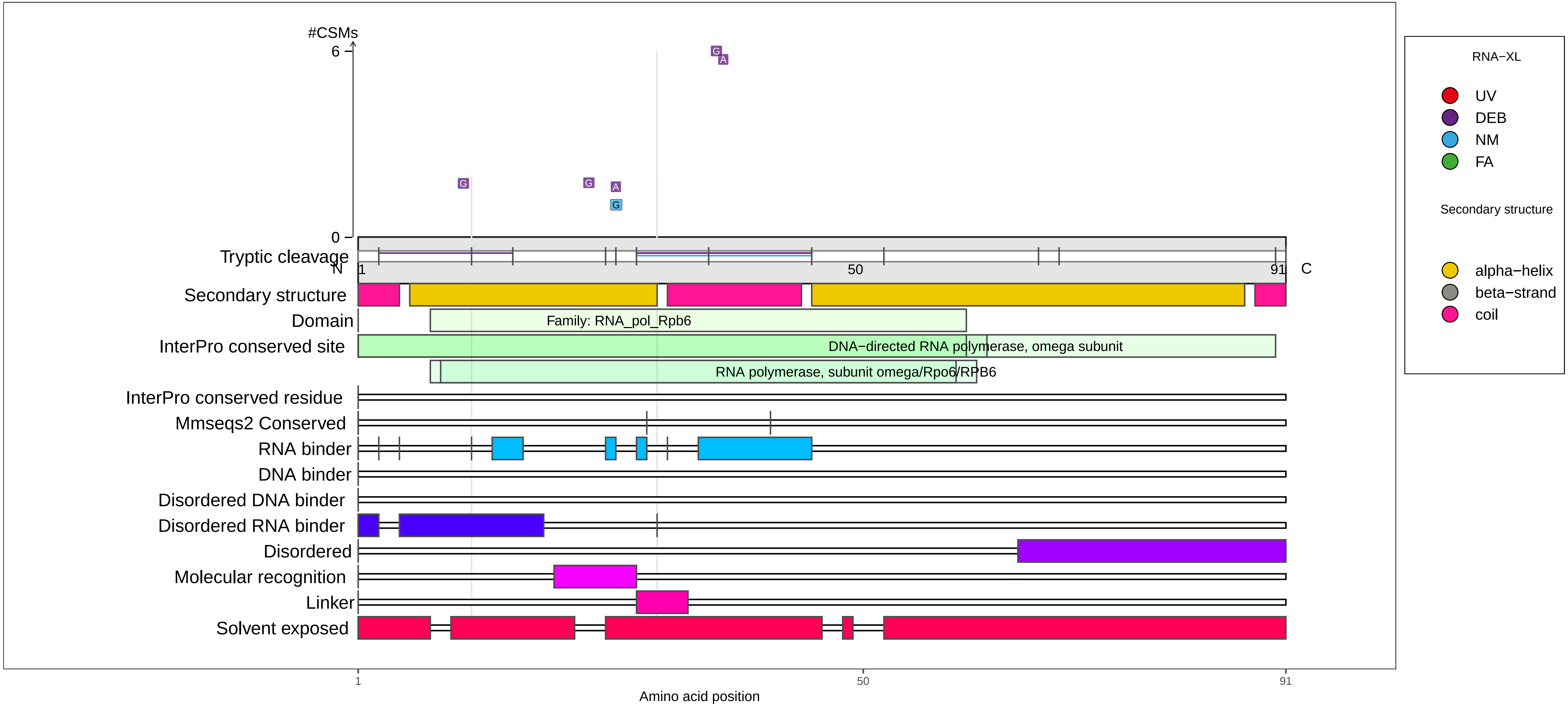

P0A8E7
YAJQ_ECOLI UPF0234 protein YajQ

− Abundance:
tryptic [log10 Intensity]: 8.6 (Q 79)
PAXdb K12 strain [ppm]: 3.31 (Q 95)
PAXdb E.coli [ppm]: 3.07 (Q 95)

− RNA functions:
RNA binding; tRNA binding

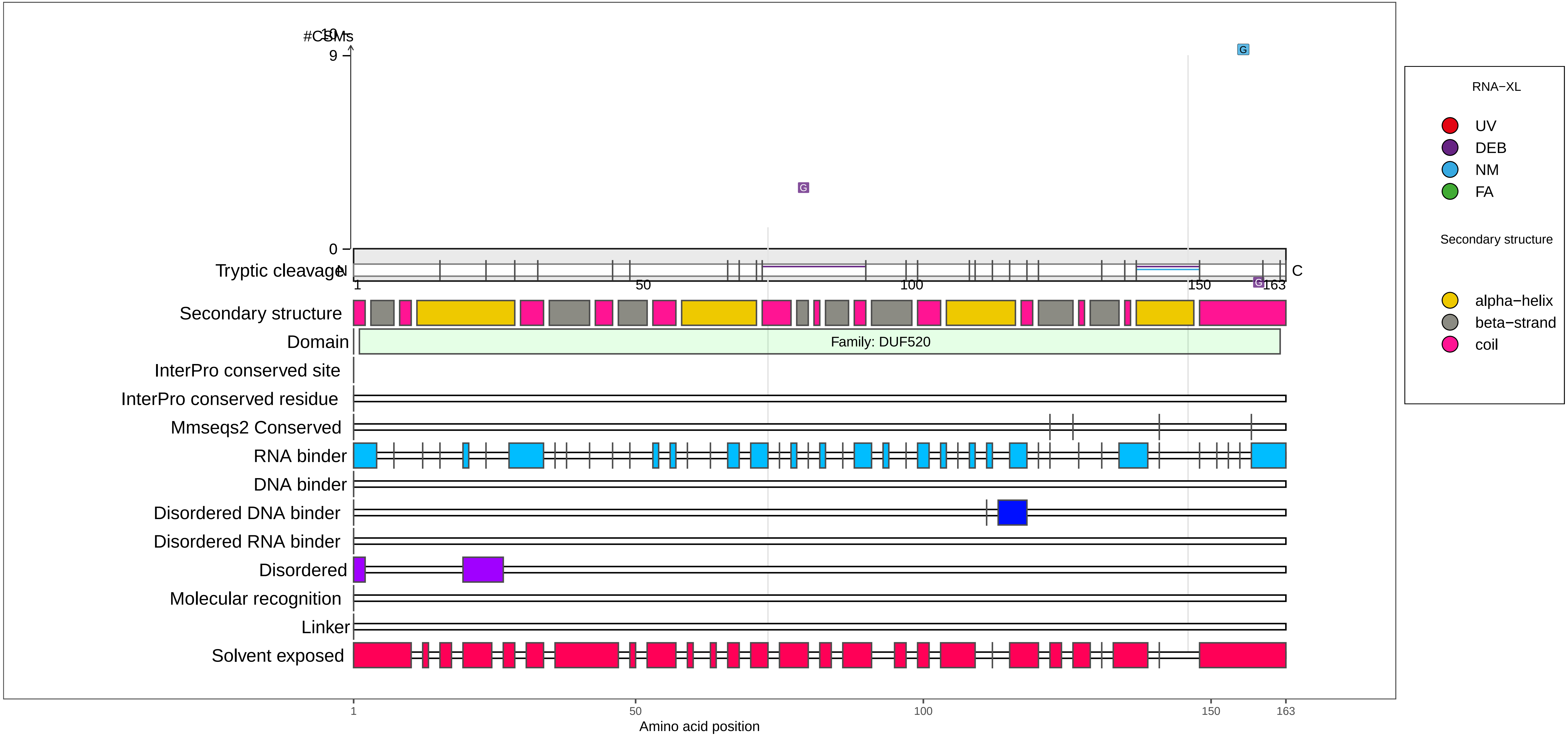

P0A9Q1
ARCA_ECOLI Aerobic respiration control protein ArcA

− Abundance:
tryptic [log10 Intensity]: 8.07 (Q 60)
PAXdb K12 strain [ppm]: 3.13 (Q 93)
PAXdb E.coli [ppm]: 2.97 (Q 94)

− RNA functions: not annotated

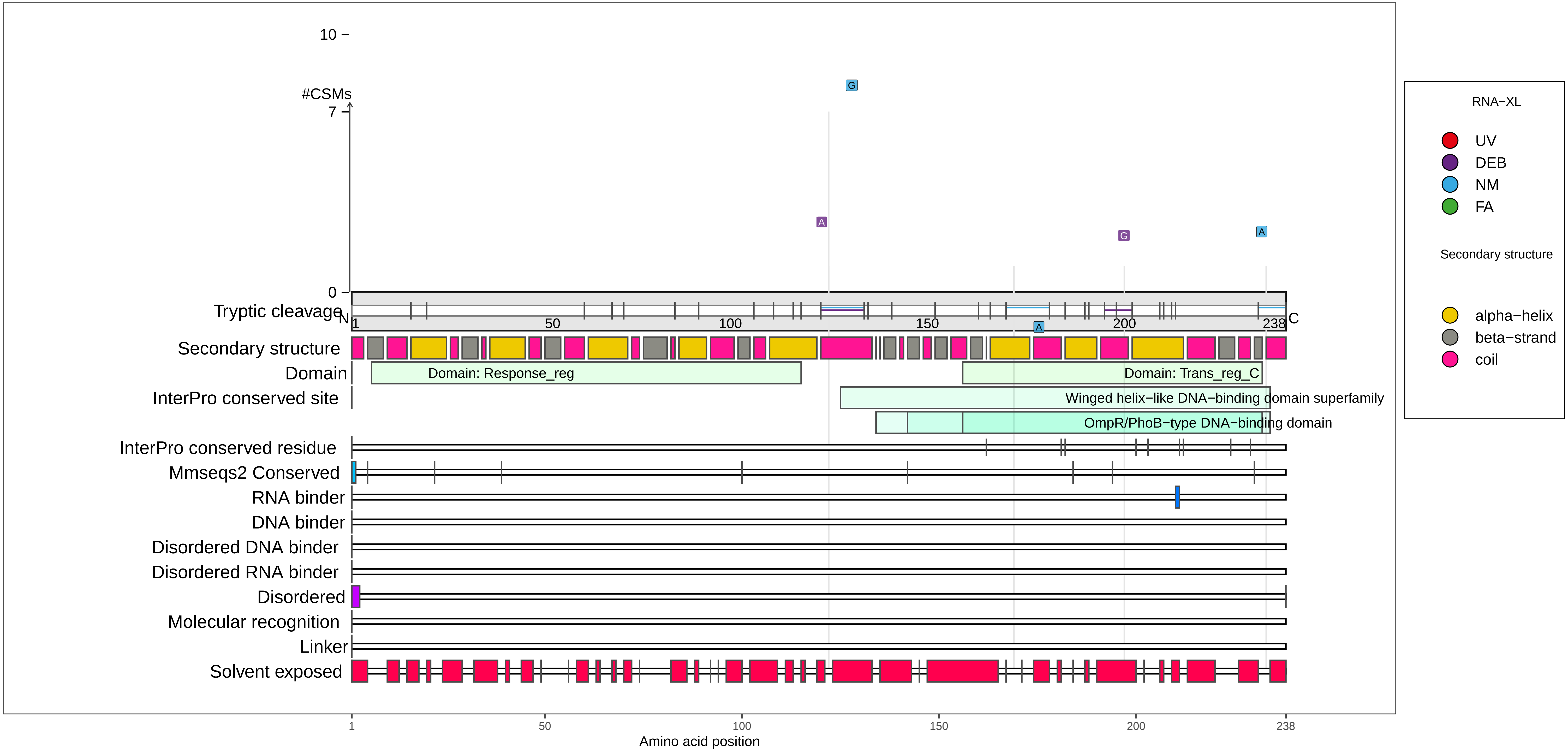

P0ABW9
FLGB_ECOLI Flagellar basal body rod protein FlgB

− Abundance:
tryptic [log10 Intensity]: 6.88 (Q 9)
PAXdb K12 strain [ppm]: NA (Q NA)
PAXdb E.coli [ppm]: −0.24 (Q 20)

− RNA functions: not annotated

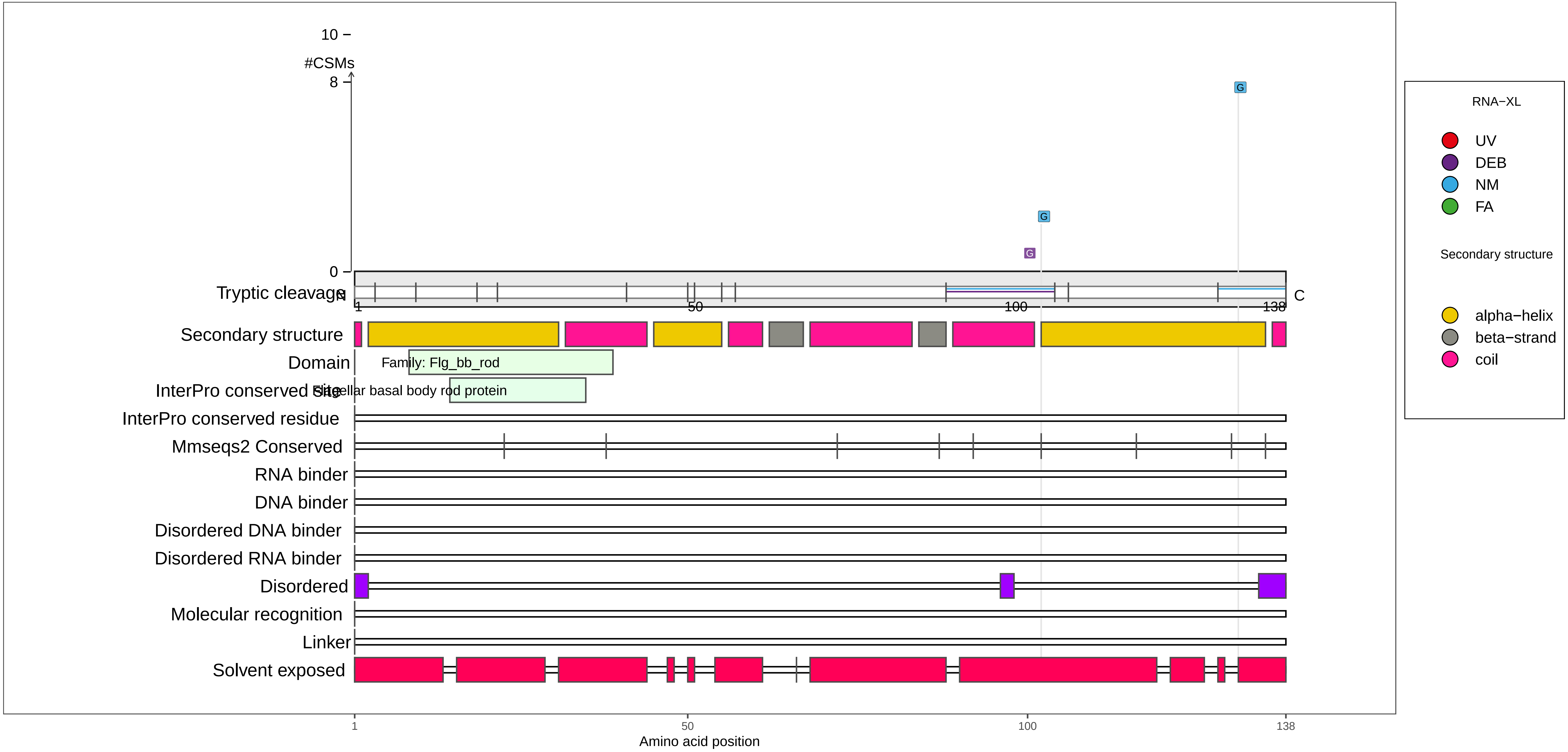

P0AEP3
GALU_ECOLI UTP−−glucose−1−phosphate uridylyltransferase

− Abundance:
tryptic [log10 Intensity]: 8.3 (Q 69)
PAXdb K12 strain [ppm]: 2.72 (Q 84)
PAXdb E.coli [ppm]: 2.6 (Q 88)

− RNA functions: not annotated

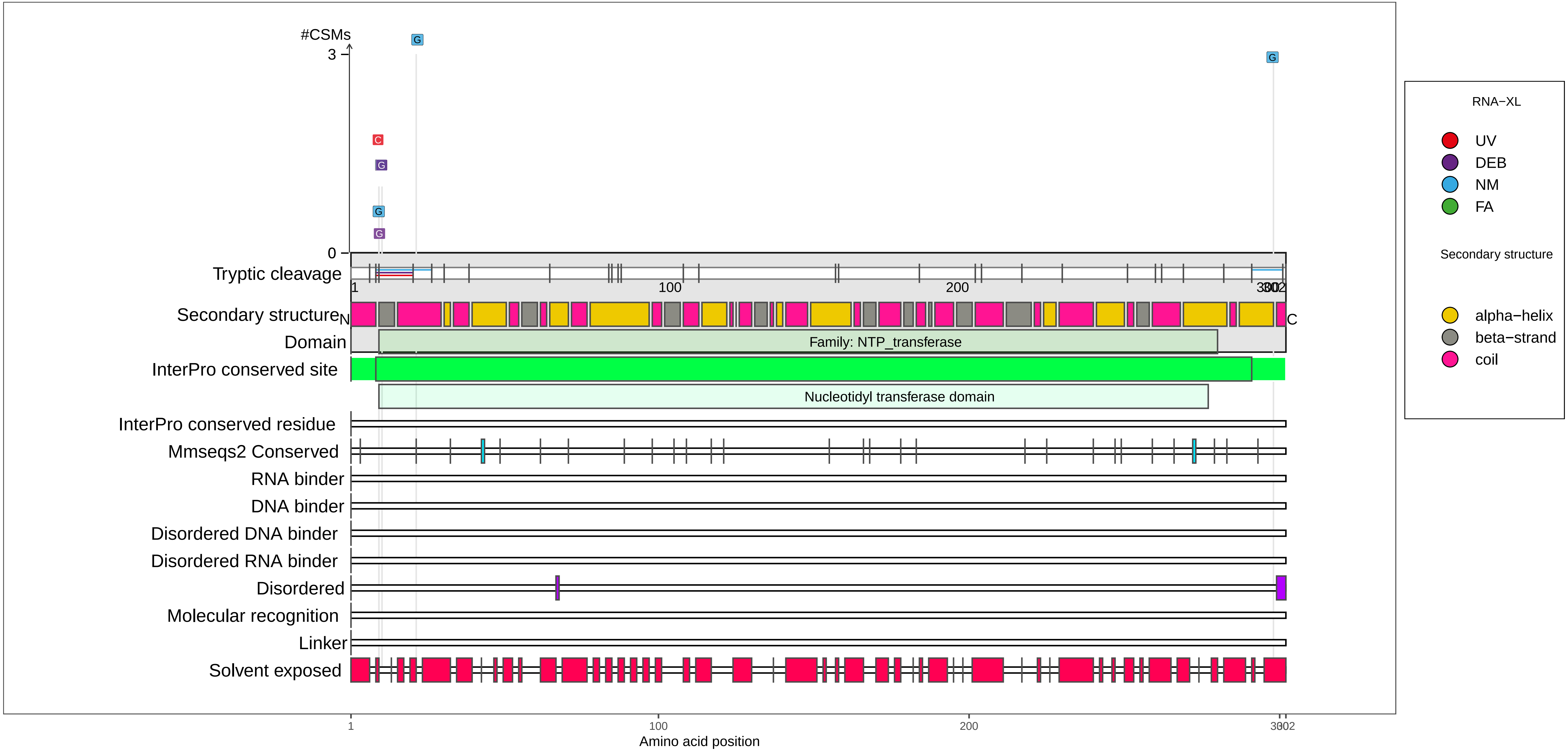

P17952
MURC_ECOLI UDP−N−acetylmuramate−−L−alanine ligase

− Abundance:
tryptic [log10 Intensity]: 8.2 (Q 65)
PAXdb K12 strain [ppm]: 1.91 (Q 57)
PAXdb E.coli [ppm]: 1.69 (Q 64)

− RNA functions: not annotated

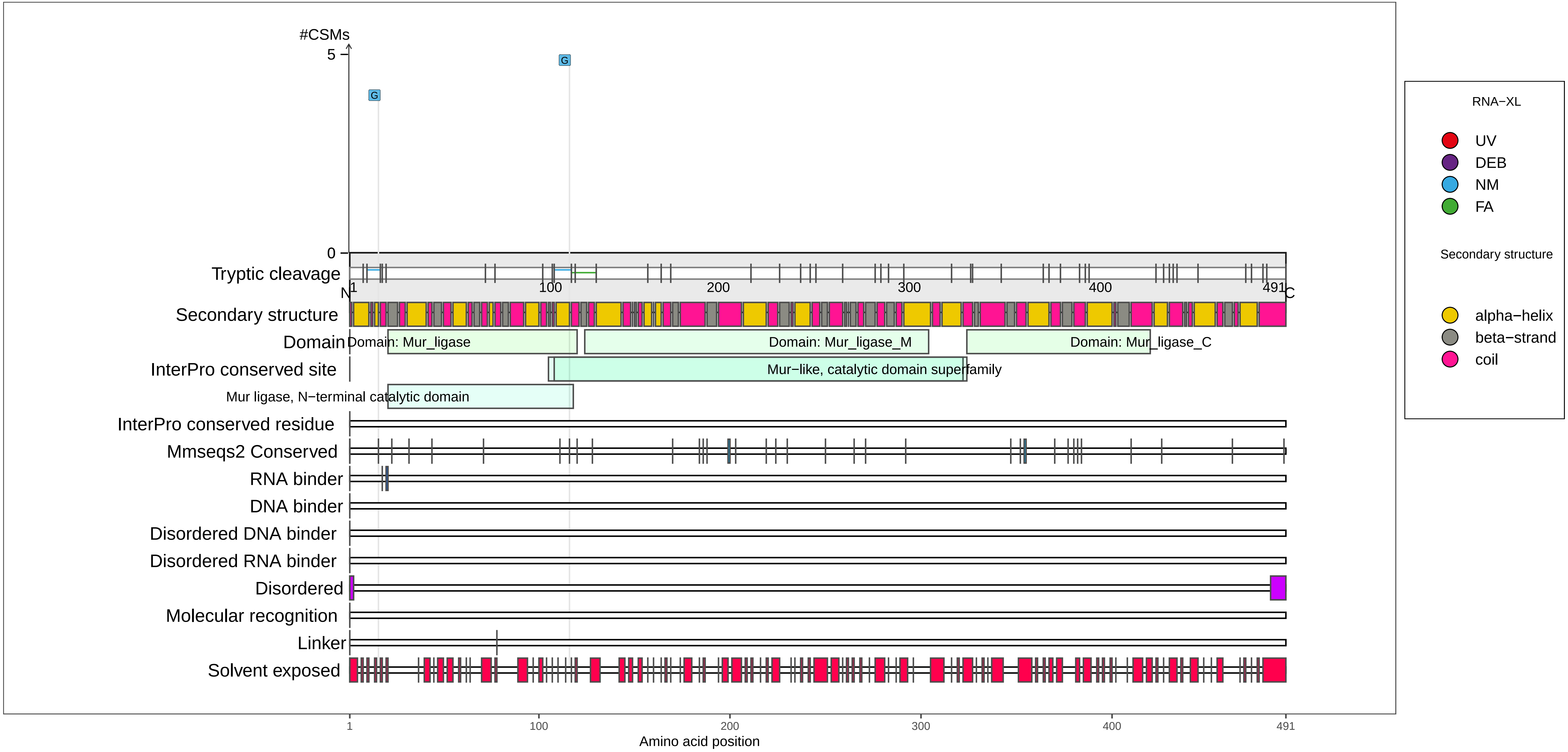

P20083
PARE_ECOLI DNA topoisomerase 4 subunit B

− Abundance:
tryptic [log10 Intensity]: 7.72 (Q 45)
PAXdb K12 strain [ppm]: 1.84 (Q 55)
PAXdb E.coli [ppm]: 1.39 (Q 57)

− RNA functions: not annotated

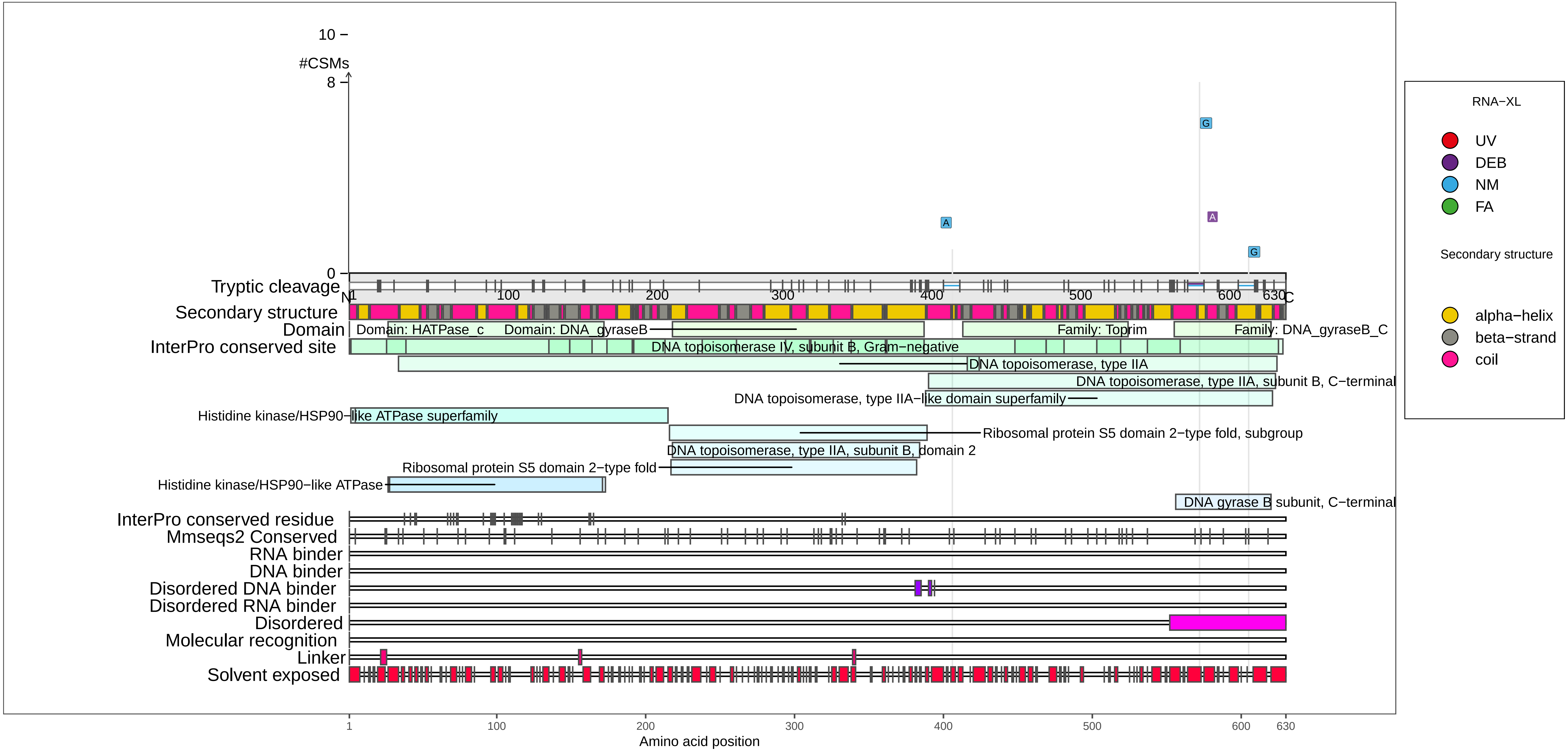

P21499
RNR_ECOLI Ribonuclease R

− Abundance:
tryptic [log10 Intensity]: 8.27 (Q 68)
PAXdb K12 strain [ppm]: 1.94 (Q 58)
PAXdb E.coli [ppm]: 2.43 (Q 84)

− RNA functions:
3−5 RNA helicase activity; mRNA catabolic process; mRNA metabolic process
ncRNA metabolic process; ncRNA processing; RNA binding; RNA catabolic process
RNA helicase activity; RNA metabolic process; RNA phosphodiester bond hydrolysis
RNA phosphodiester bond hydrolysis, exonucleolytic; RNA processing; S1 RNA binding domain

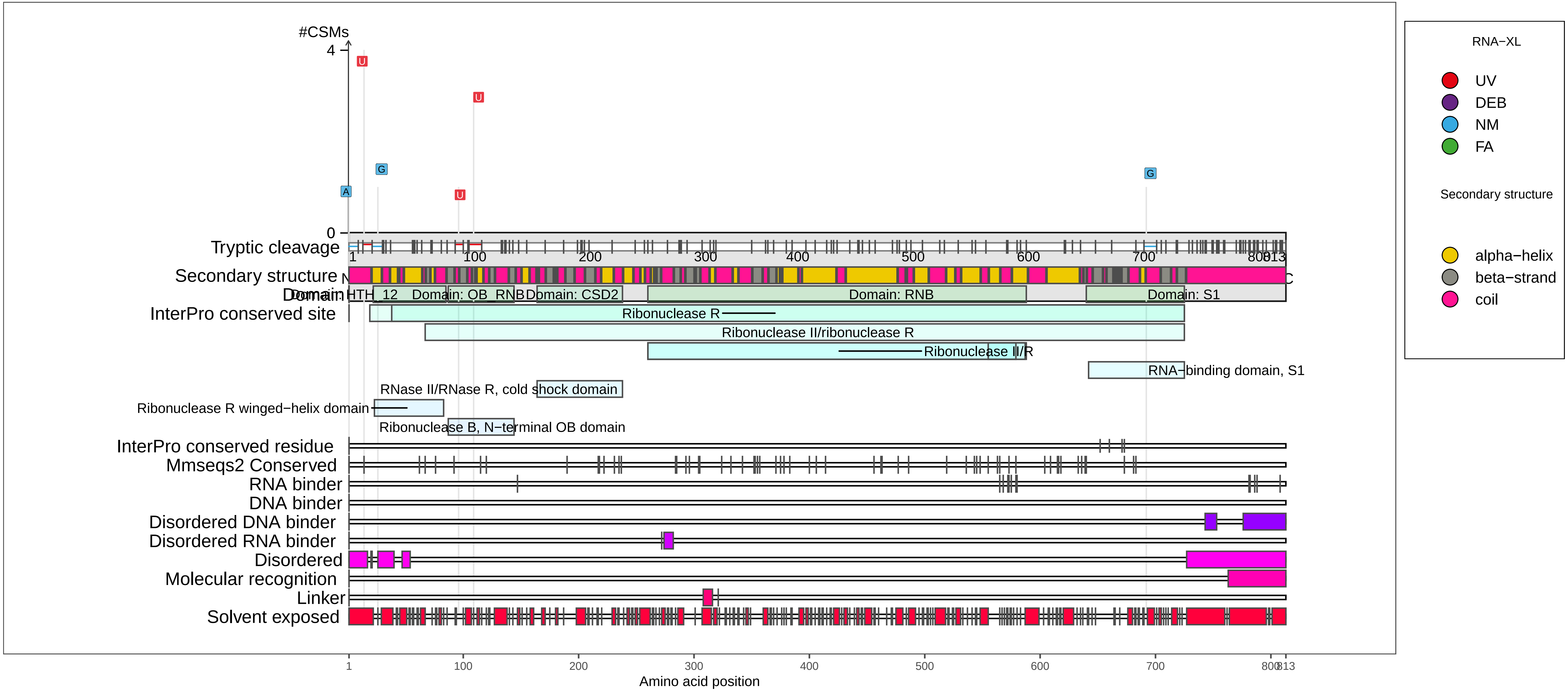

P25397
TEHB_ECOLI Tellurite methyltransferase

− Abundance:
tryptic [log10 Intensity]: 7.76 (Q 47)
PAXdb K12 strain [ppm]: NA (Q NA)
PAXdb E.coli [ppm]: 2.19 (Q 77)

− RNA functions: not annotated

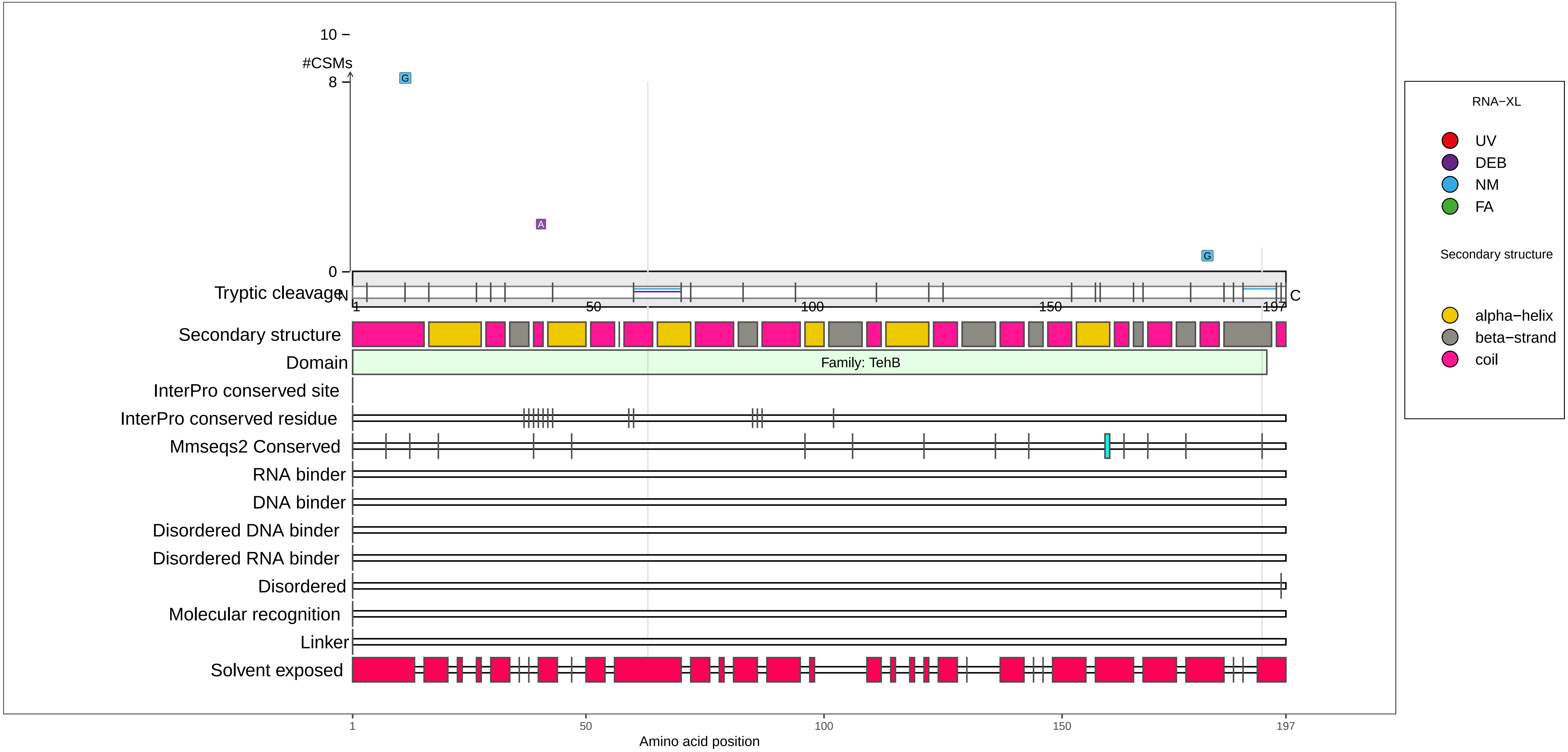

P25714
YIDC_ECOLI Membrane protein insertase YidC

− Abundance:
tryptic [log10 Intensity]: 7.59 (Q 39)
PAXdb K12 strain [ppm]: 2.01 (Q 61)
PAXdb E.coli [ppm]: 1.84 (Q 68)

− RNA functions: not annotated

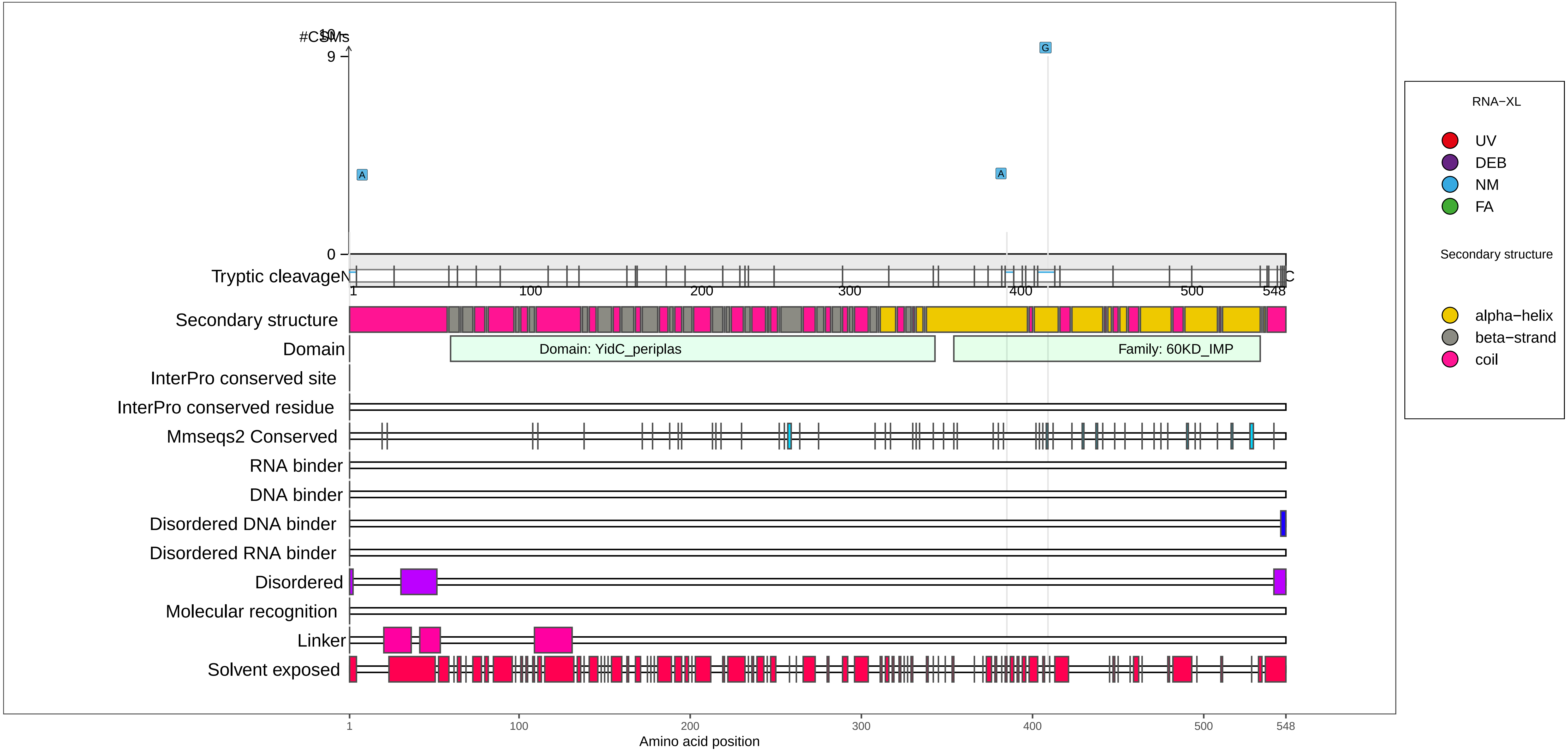

P39337
YJGM_ECOLI Uncharacterized N−acetyltransferase YjgM

− Abundance:
tryptic [log10 Intensity]: 7.14 (Q 18)
PAXdb K12 strain [ppm]: 2.11 (Q 64)
PAXdb E.coli [ppm]: 1.25 (Q 53)

− RNA functions: not annotated

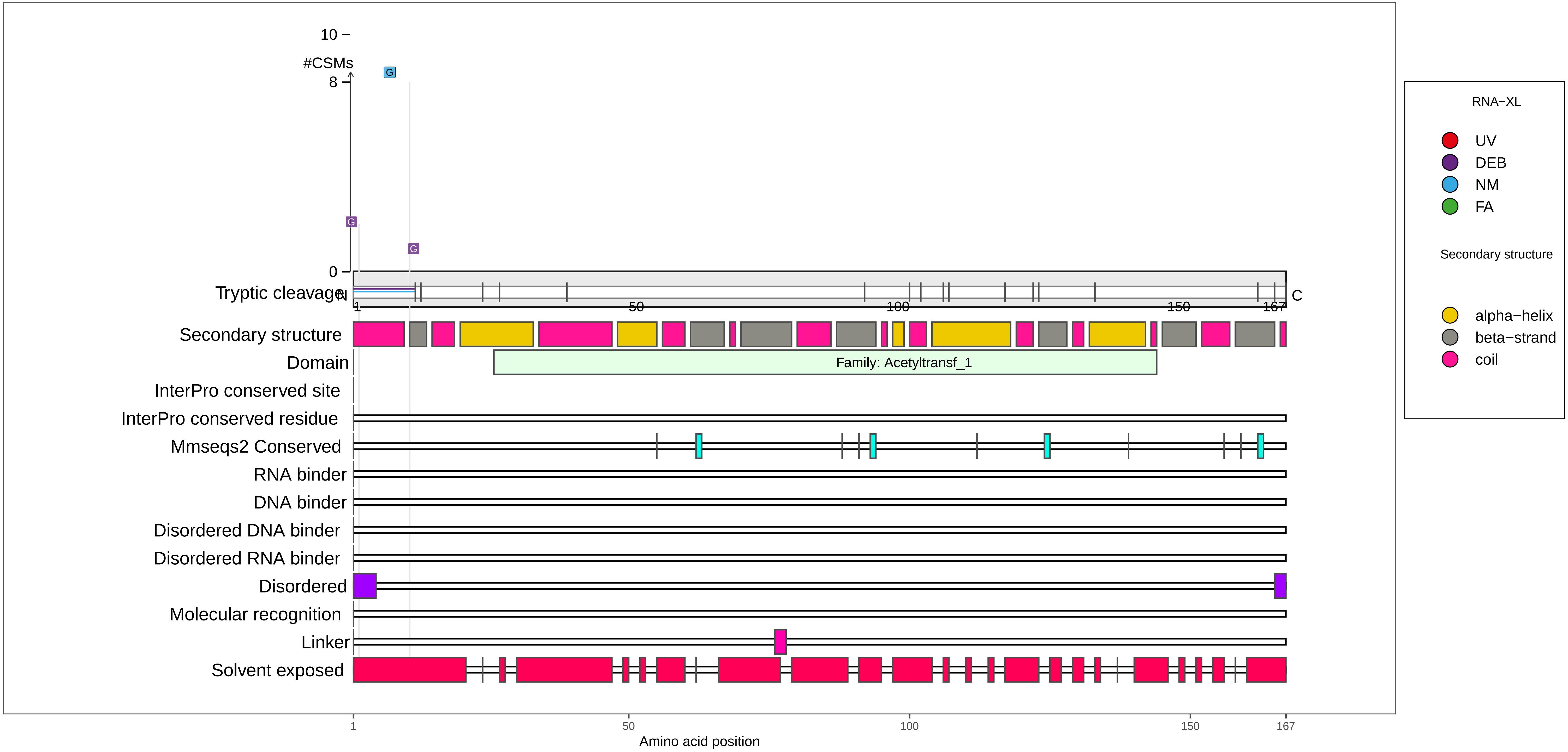

P52643
LDHD_ECOLI D−lactate dehydrogenase

− Abundance:
tryptic [log10 Intensity]: 8.28 (Q 69)
PAXdb K12 strain [ppm]: 2.87 (Q 88)
PAXdb E.coli [ppm]: 2.33 (Q 81)

− RNA functions: not annotated

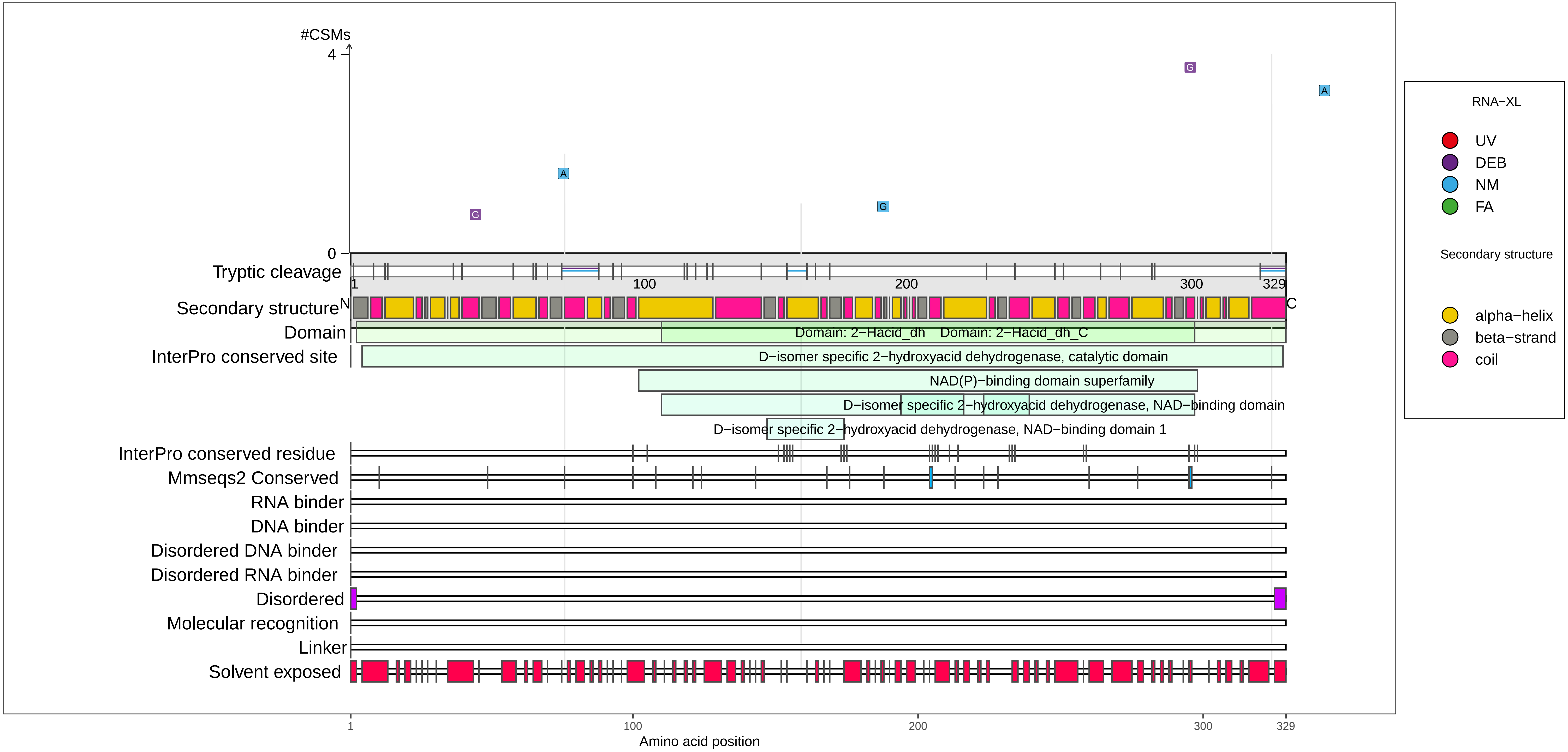

P69811
PTFAH_ECOLI Multiphosphoryl transfer protein

− Abundance:
tryptic [log10 Intensity]: 7.95 (Q 56)
PAXdb K12 strain [ppm]: 2.3 (Q 70)
PAXdb E.coli [ppm]: 2.06 (Q 74)

− RNA functions: not annotated

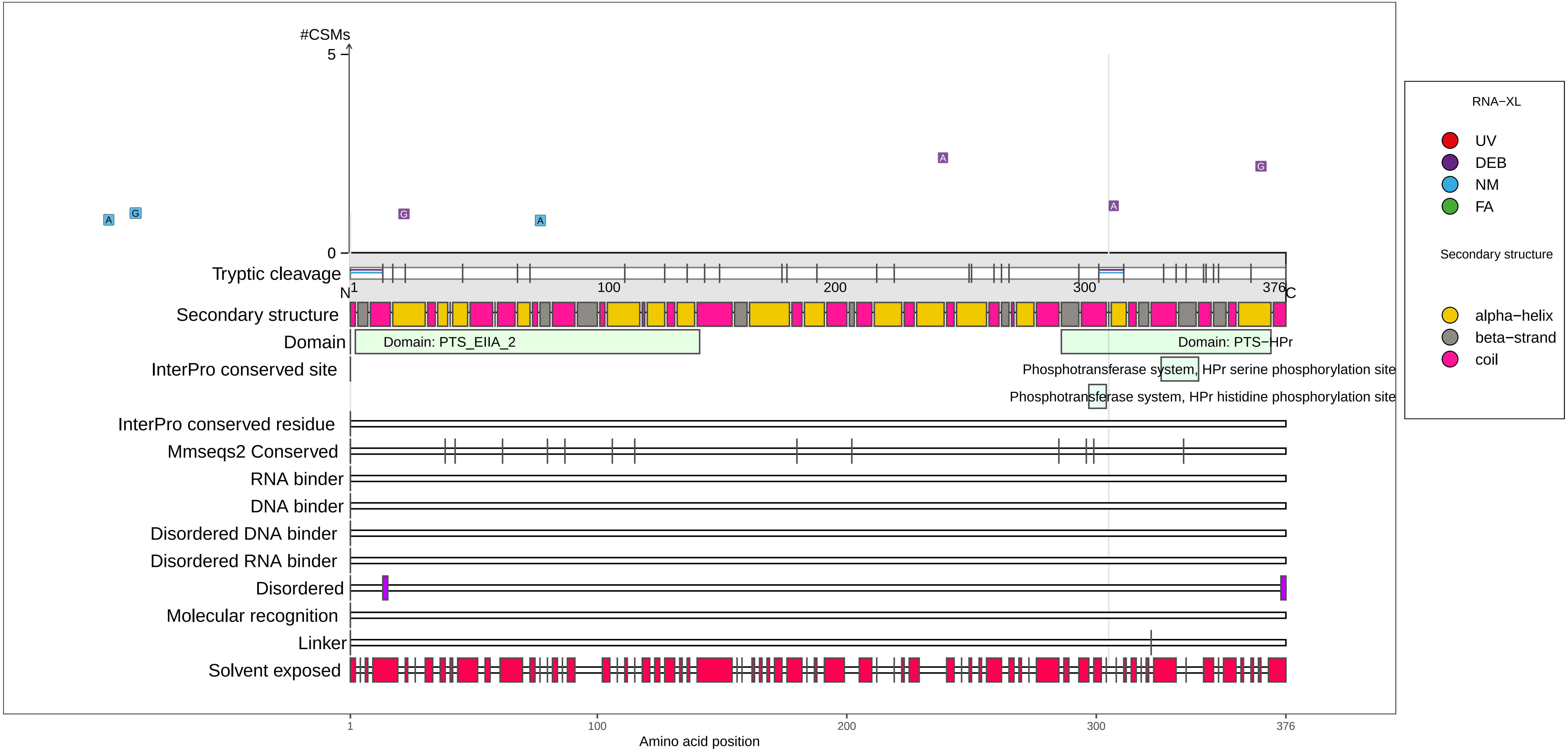

P69831
PTKC_ECOLI PTS system galactitol−specific EIIC component
− RNA functions: not annotated

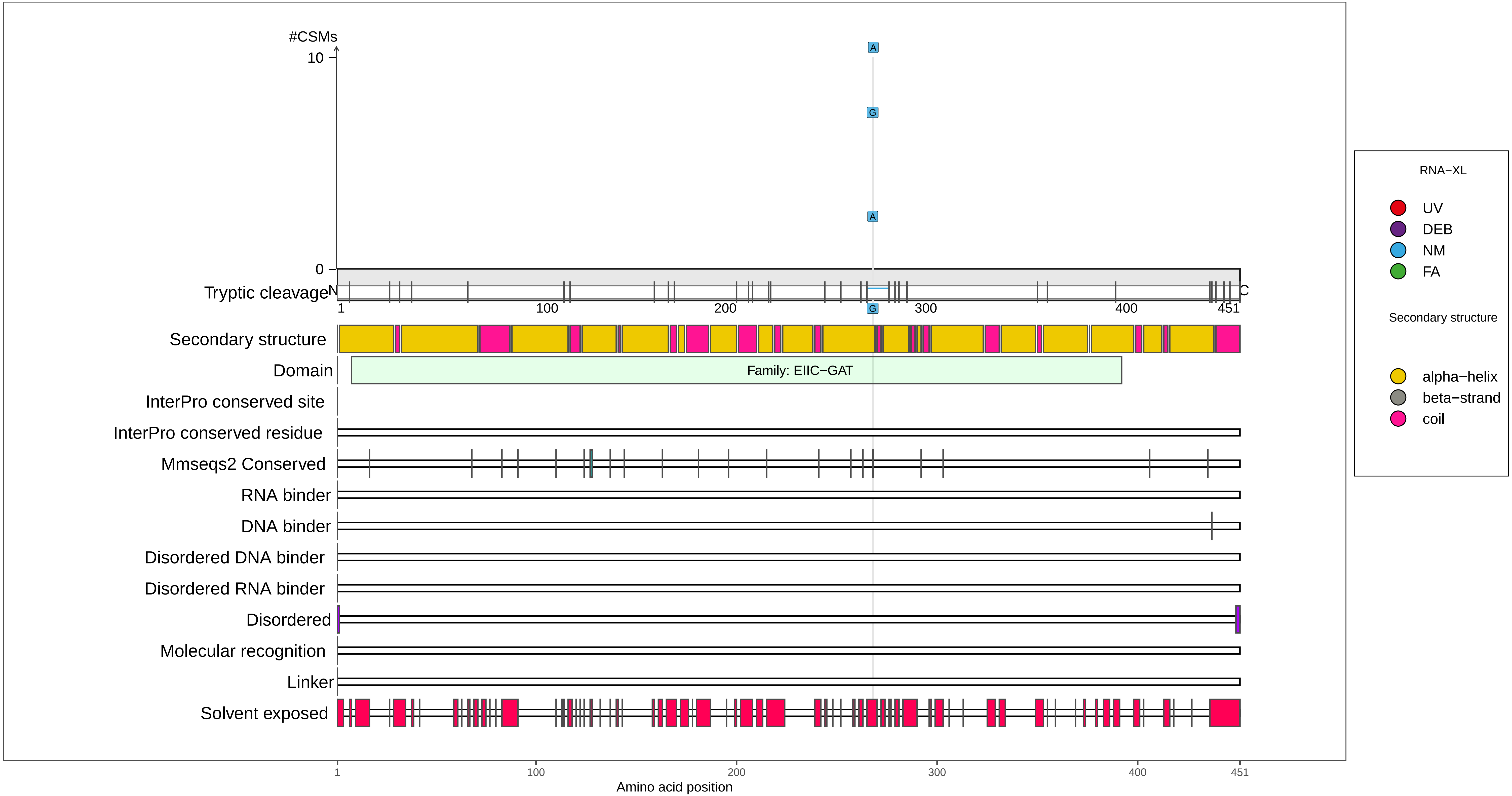

P75745
PXPC_ECOLI 5−oxoprolinase subunit C
− Abundance:
tryptic [log10 Intensity]: 6.93 (Q 10)
PAXdb K12 strain [ppm]: 1.03 (Q 12)
PAXdb E.coli [ppm]: 1.83 (Q 68)

− RNA functions: not annotated

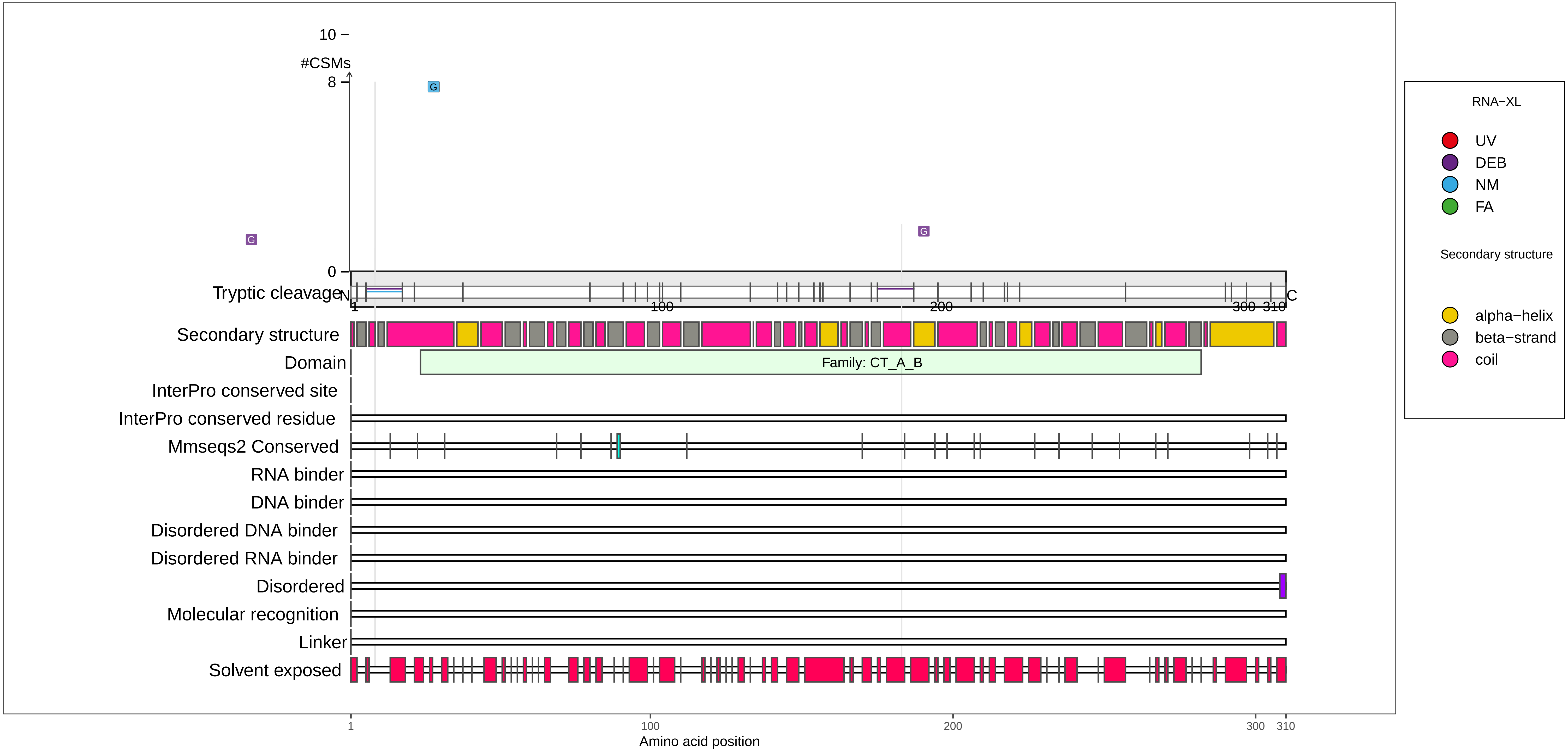

P00582
DPO1_ECOLI DNA polymerase I

− Abundance:
tryptic [log10 Intensity]: 8.85 (Q 85)
PAXdb K12 strain [ppm]: 2.55 (Q 79)
PAXdb E.coli [ppm]: 2.47 (Q 85)

− RNA functions: not annotated

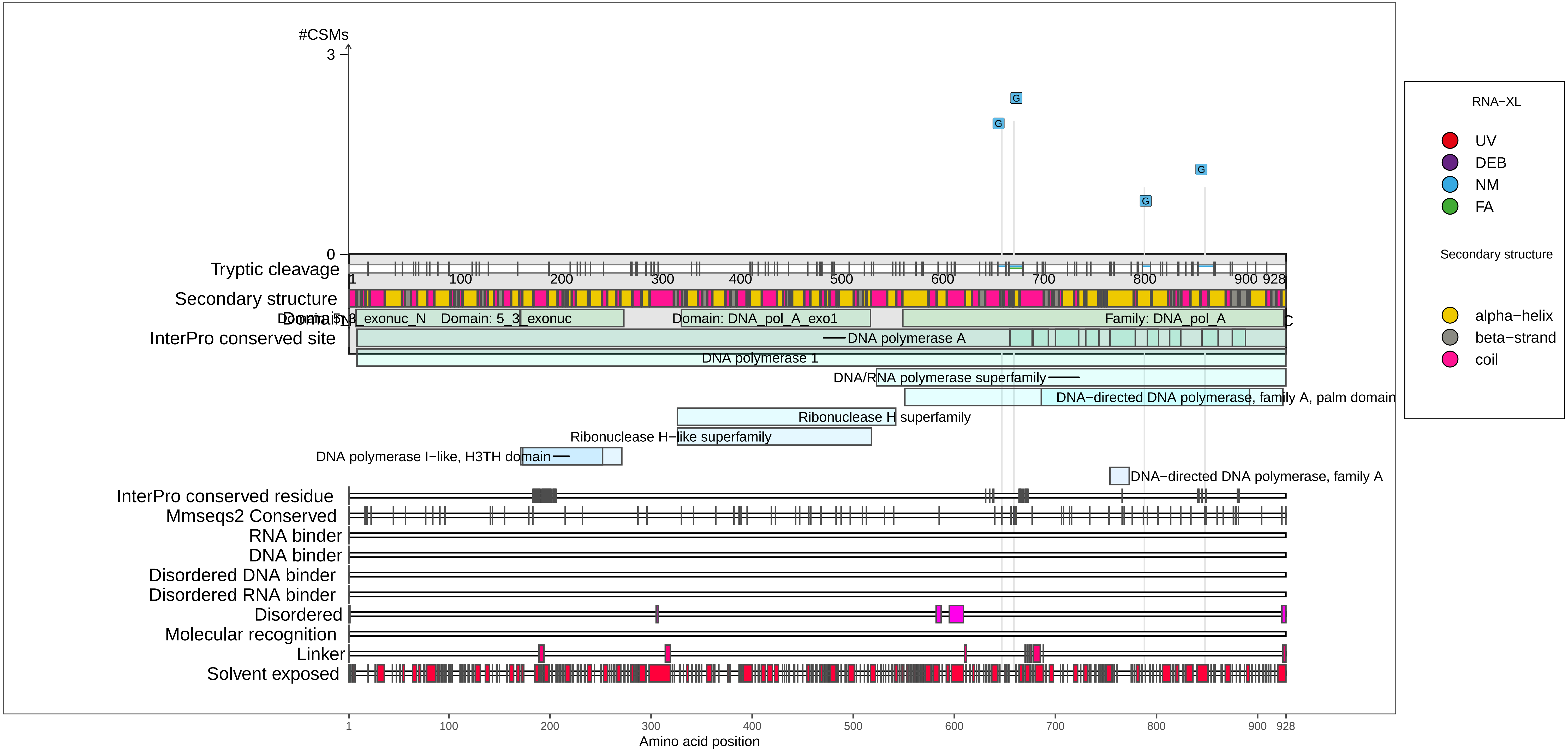

P00926
SDHD_ECOLI D−serine dehydratase

− Abundance:
tryptic [log10 Intensity]: 8.86 (Q 85)
PAXdb K12 strain [ppm]: 1.52 (Q 39)
PAXdb E.coli [ppm]: 1.16 (Q 51)

− RNA functions: not annotated

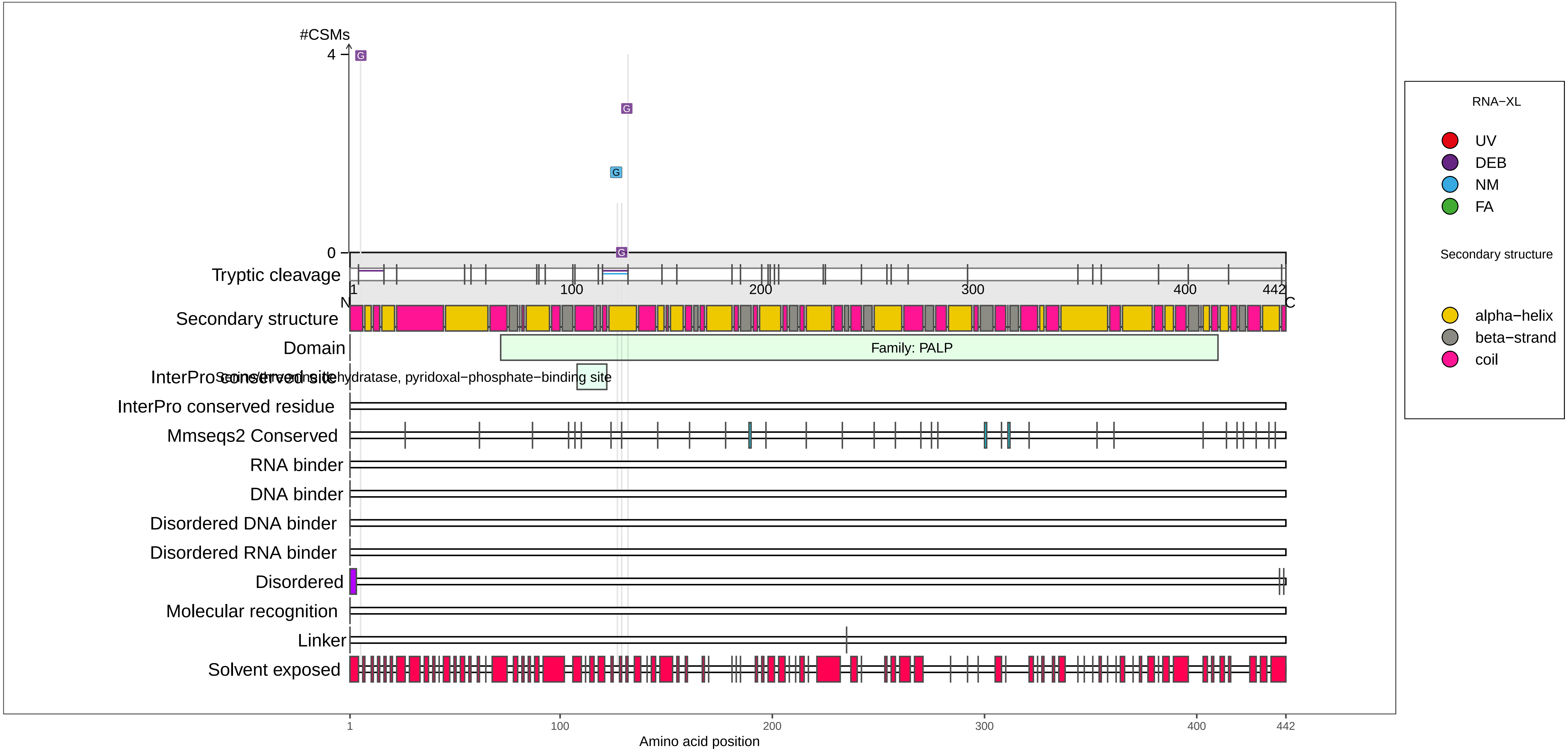

P02918
PBPA_ECOLI Penicillin−binding protein 1A

− Abundance:
tryptic [log10 Intensity]: 8.56 (Q 77)
PAXdb K12 strain [ppm]: 0.91 (Q 8)
PAXdb E.coli [ppm]: 0.94 (Q 46)

− RNA functions: not annotated

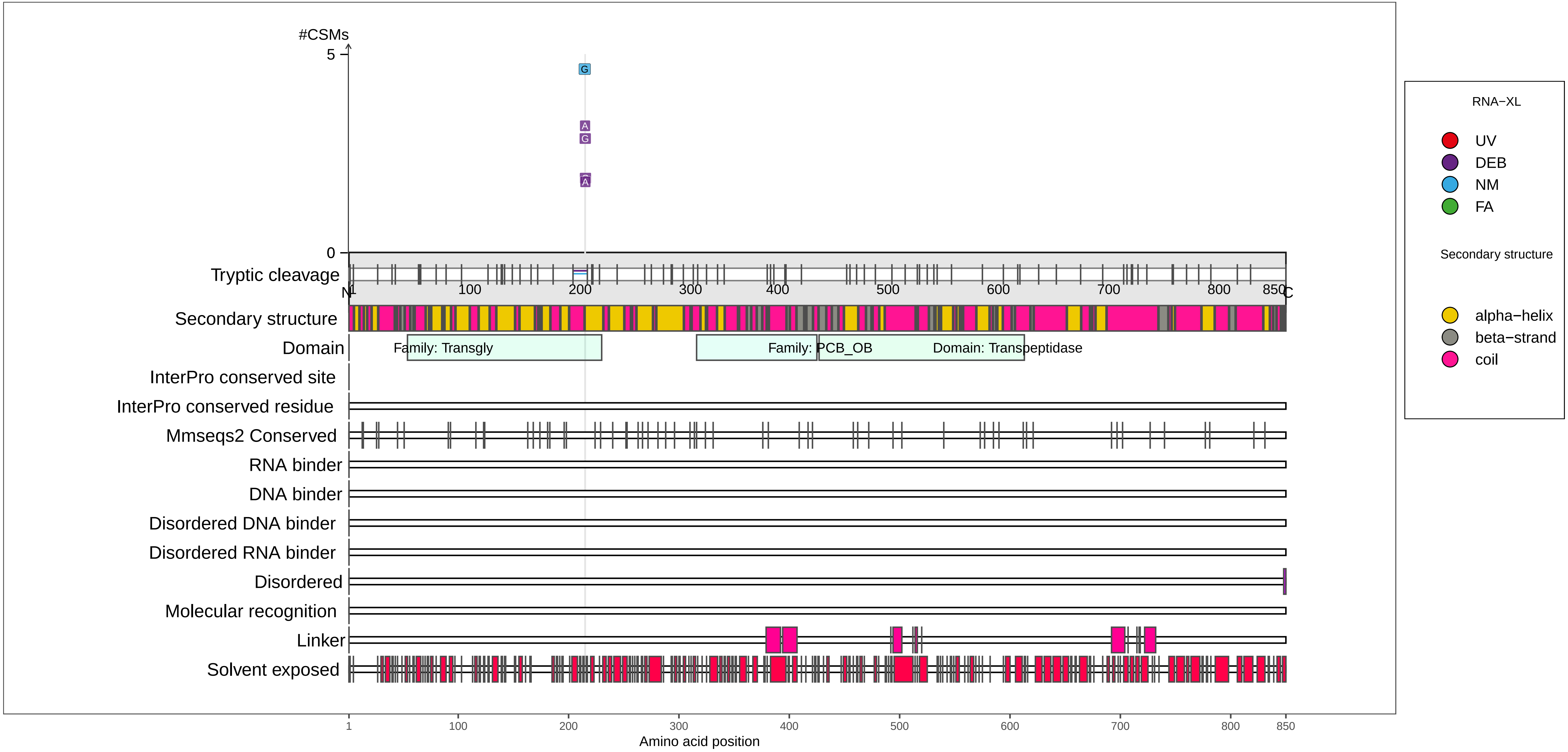

P05020
PYRC_ECOLI Dihydroorotase

− Abundance:
tryptic [log10 Intensity]: 7.37 (Q 28)
PAXdb K12 strain [ppm]: 1.65 (Q 47)
PAXdb E.coli [ppm]: 2.65 (Q 89)

− RNA functions: not annotated

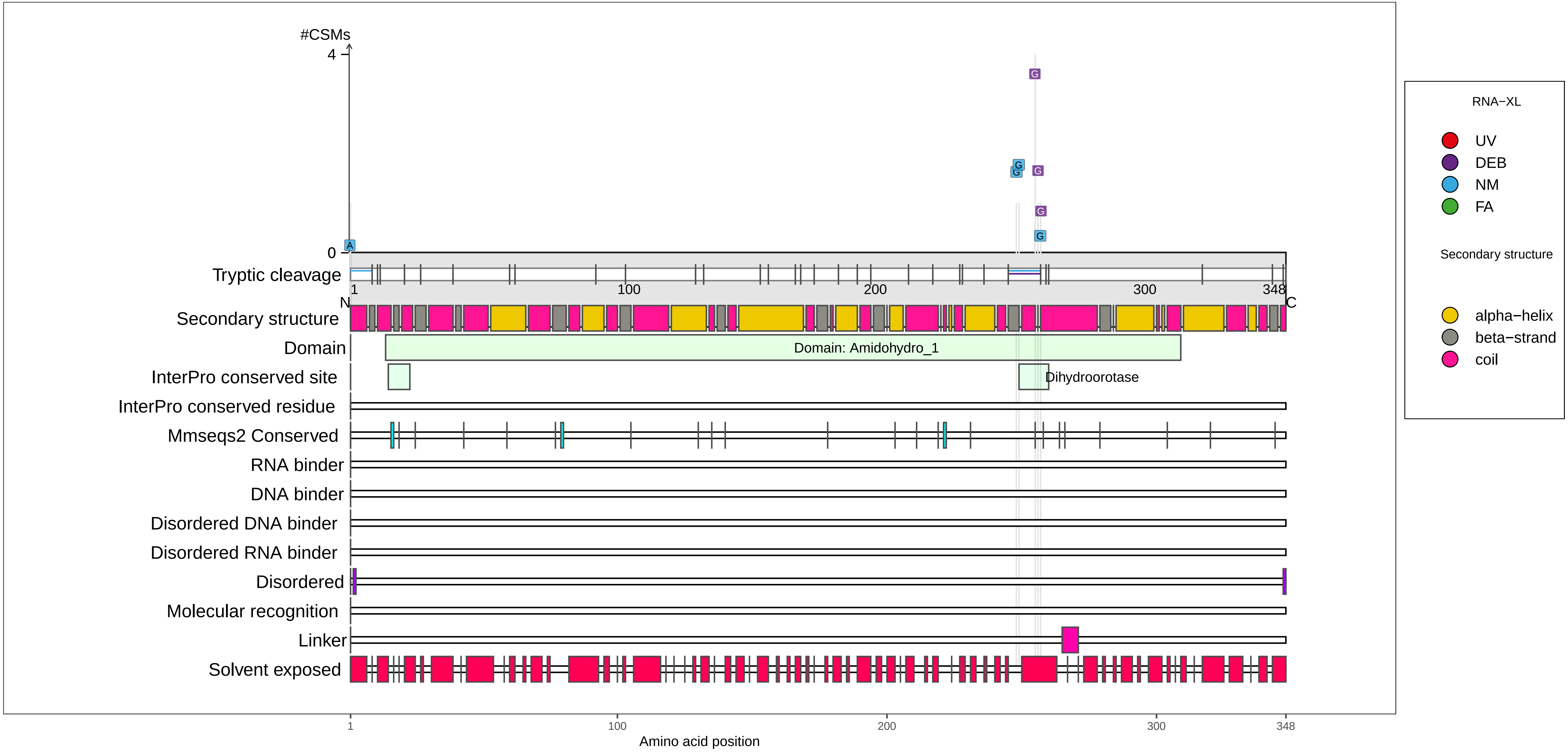

P0A6B4
ALR1_ECOLI Alanine racemase, biosynthetic

− Abundance:
tryptic [log10 Intensity]: 7.81 (Q 49)
PAXdb K12 strain [ppm]: 2.8 (Q 86)
PAXdb E.coli [ppm]: 1.48 (Q 59)

− RNA functions: not annotated

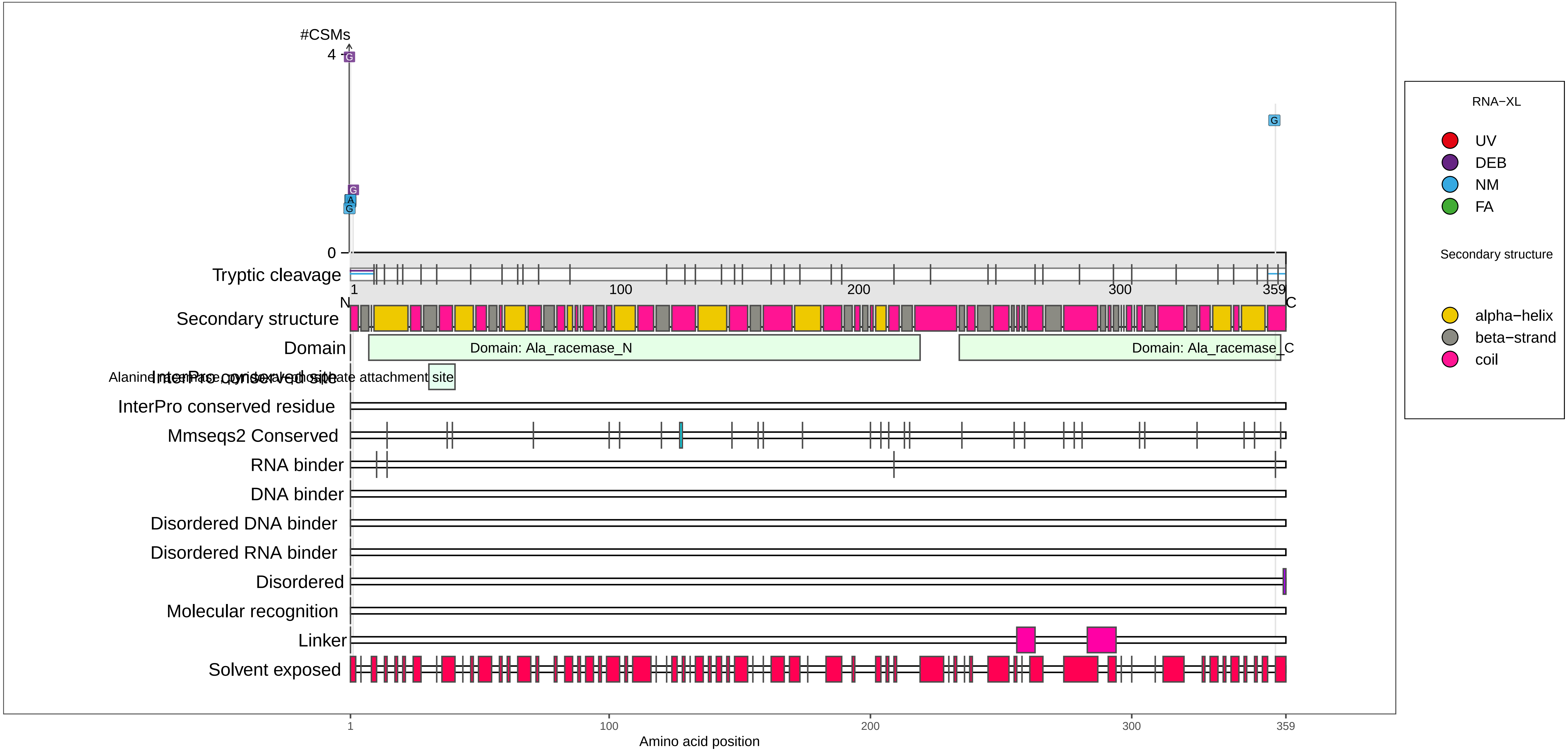

P0A6T1
G6PI_ECOLI Glucose−6−phosphate isomerase

− Abundance:
tryptic [log10 Intensity]: 9.08 (Q 90)
PAXdb K12 strain [ppm]: 2.63 (Q 82)
PAXdb E.coli [ppm]: 3.04 (Q 95)

− RNA functions: not annotated

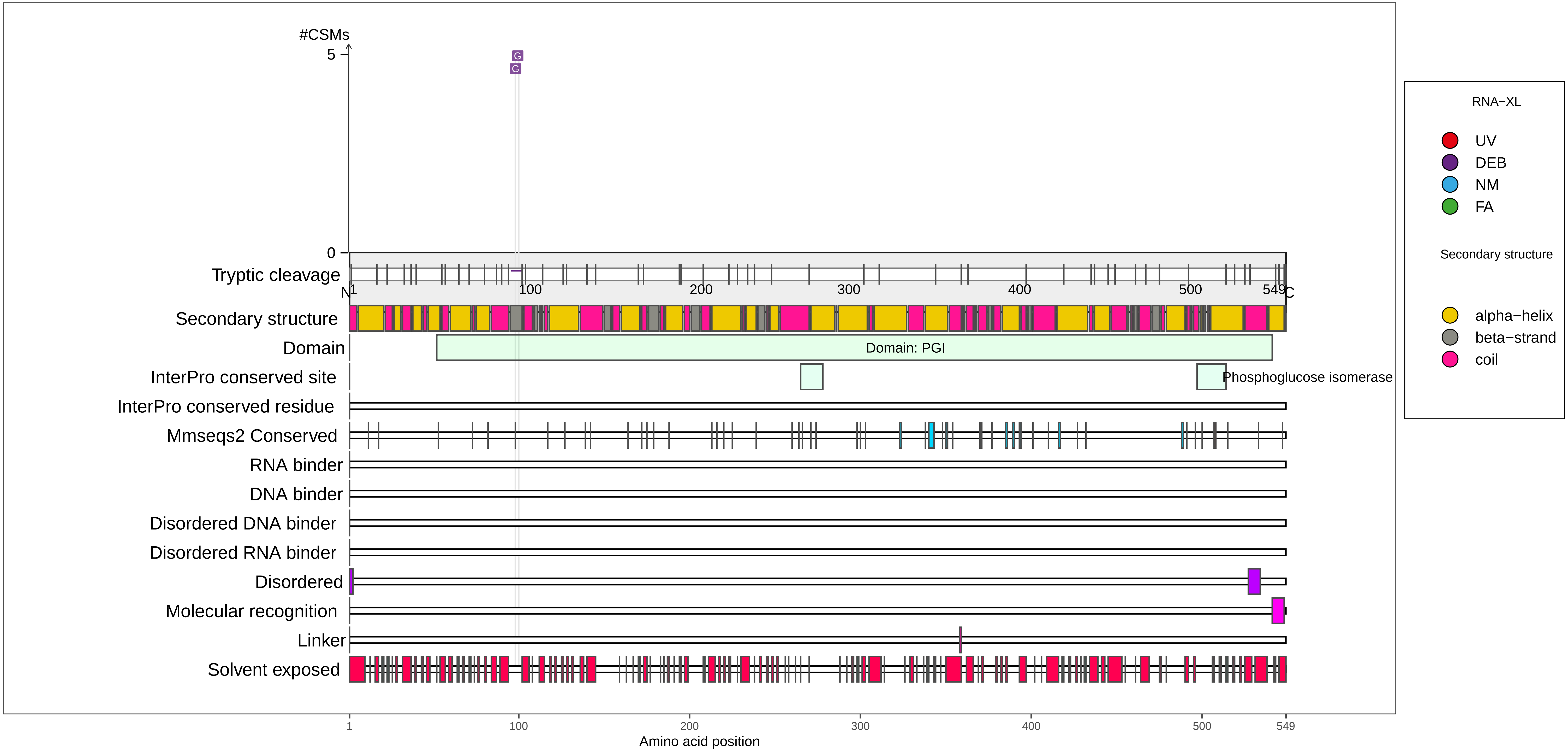

P0A9L8
P5CR_ECOLI Pyrroline−5−carboxylate reductase

− Abundance:
tryptic [log10 Intensity]: 8.62 (Q 79)
PAXdb K12 strain [ppm]: 2.38 (Q 73)
PAXdb E.coli [ppm]: 2.47 (Q 85)

− RNA functions: not annotated

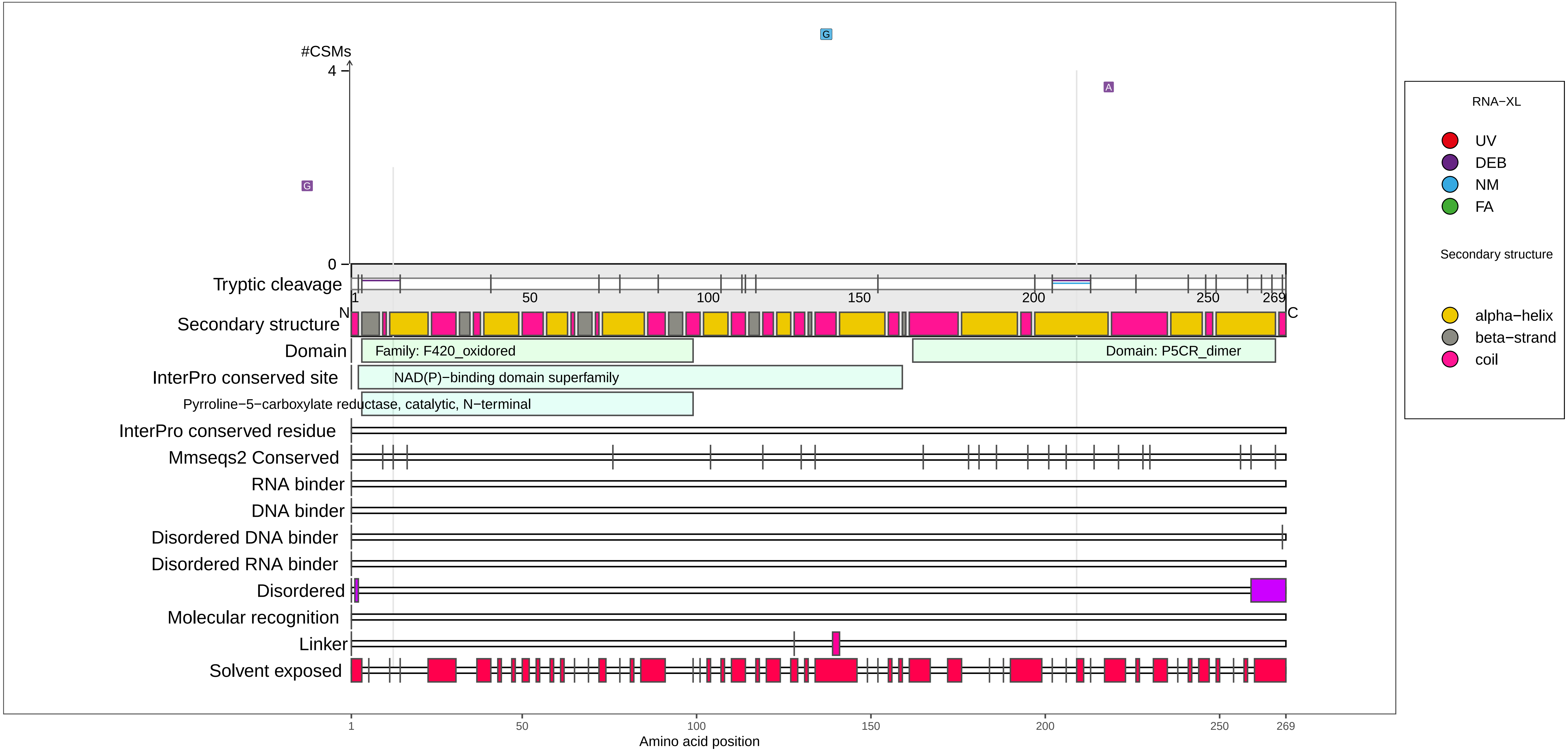

P0AAC0
USPE_ECOLI Universal stress protein E

− Abundance:
tryptic [log10 Intensity]: 8.33 (Q 70)
PAXdb K12 strain [ppm]: 2.08 (Q 63)
PAXdb E.coli [ppm]: 2.89 (Q 93)

− RNA functions: not annotated

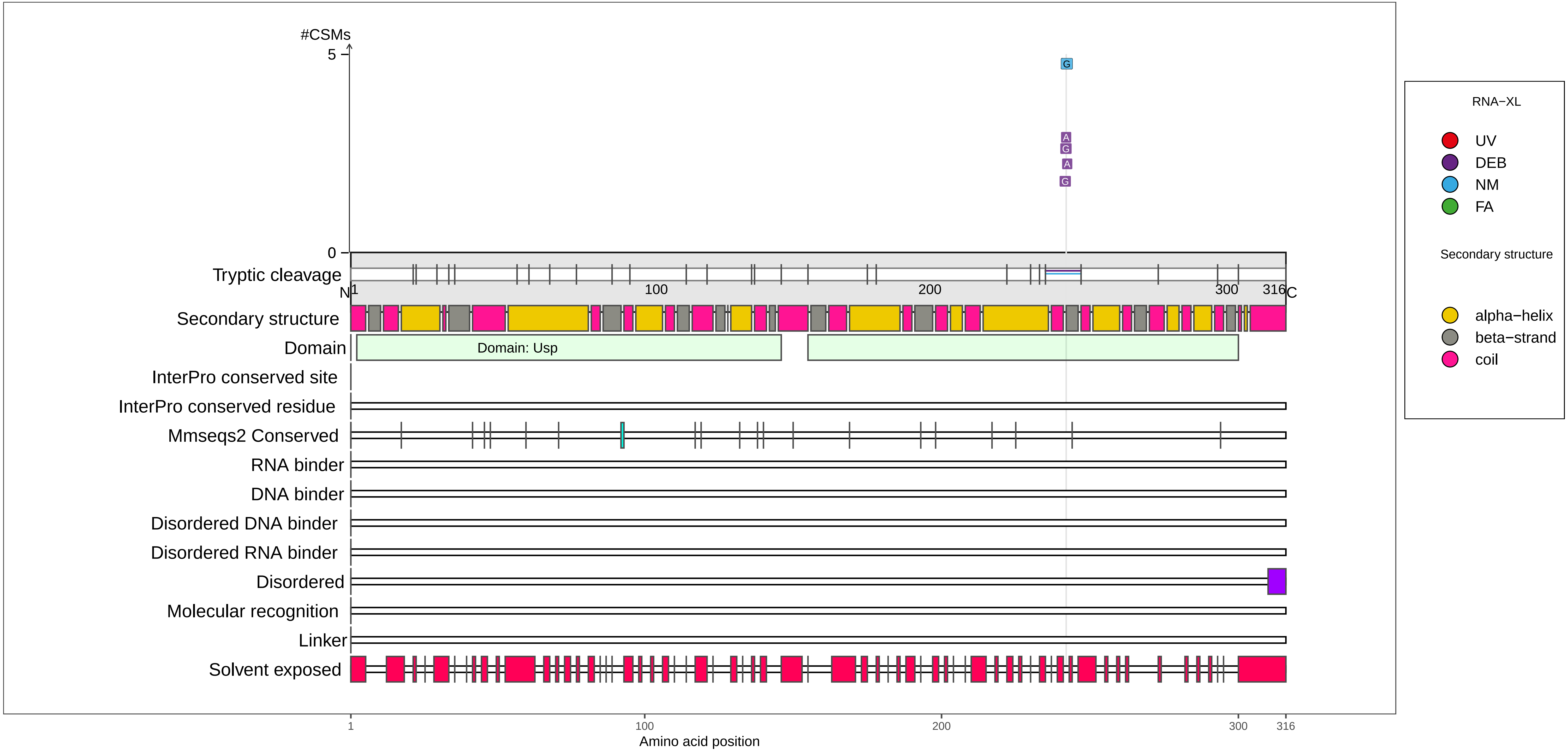

P0AC41
SDHA_ECOLI Succinate dehydrogenase flavoprotein subunit

− Abundance:
tryptic [log10 Intensity]: 8.79 (Q 83)
PAXdb K12 strain [ppm]: 2.28 (Q 69)
PAXdb E.coli [ppm]: 2.8 (Q 92)

− RNA functions: not annotated

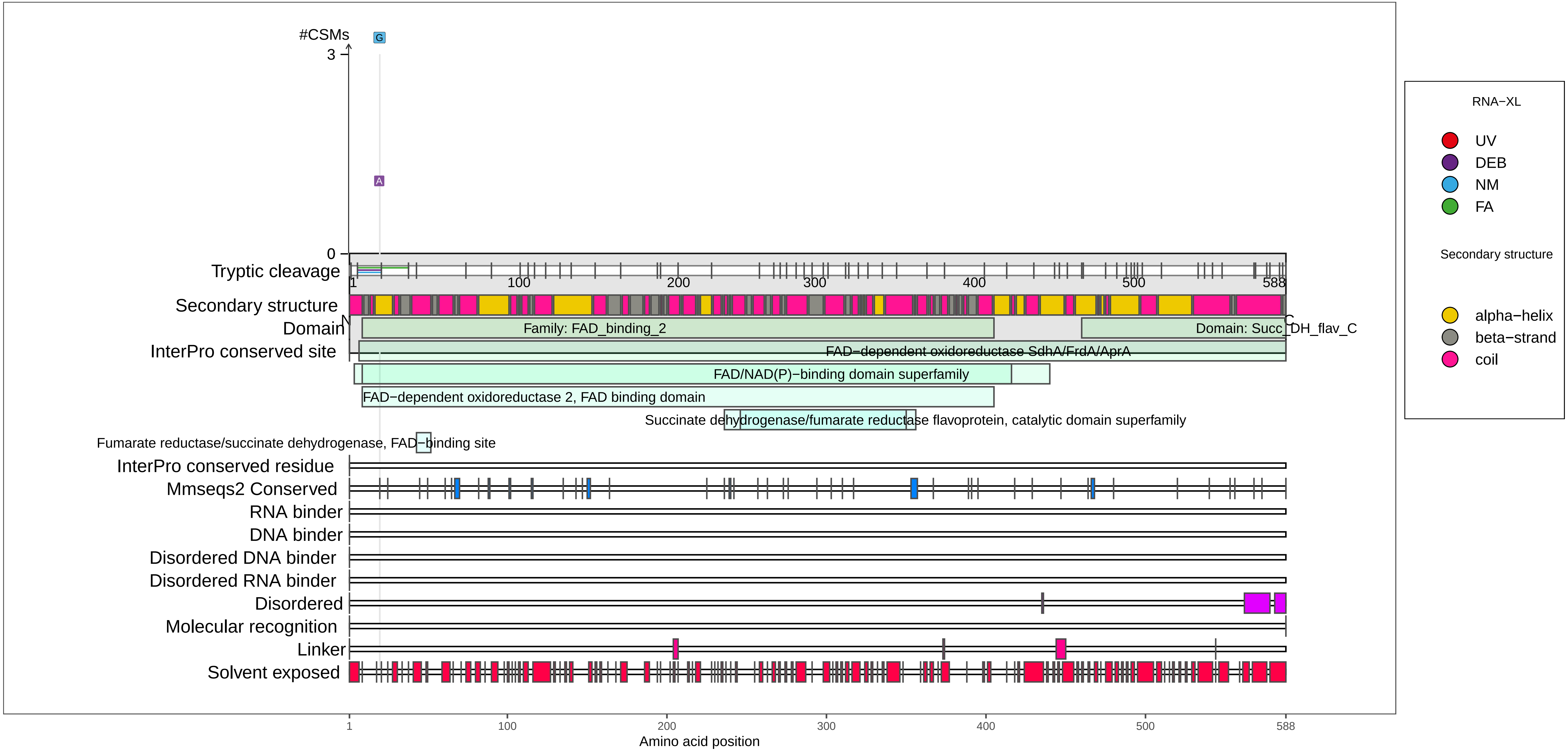

P0AFF2
NUPC_ECOLI Nucleoside permease NupC

− Abundance:
tryptic [log10 Intensity]: 6.85 (Q 8)
PAXdb K12 strain [ppm]: NA (Q NA)
PAXdb E.coli [ppm]: 1.48 (Q 59)

− RNA functions: not annotated

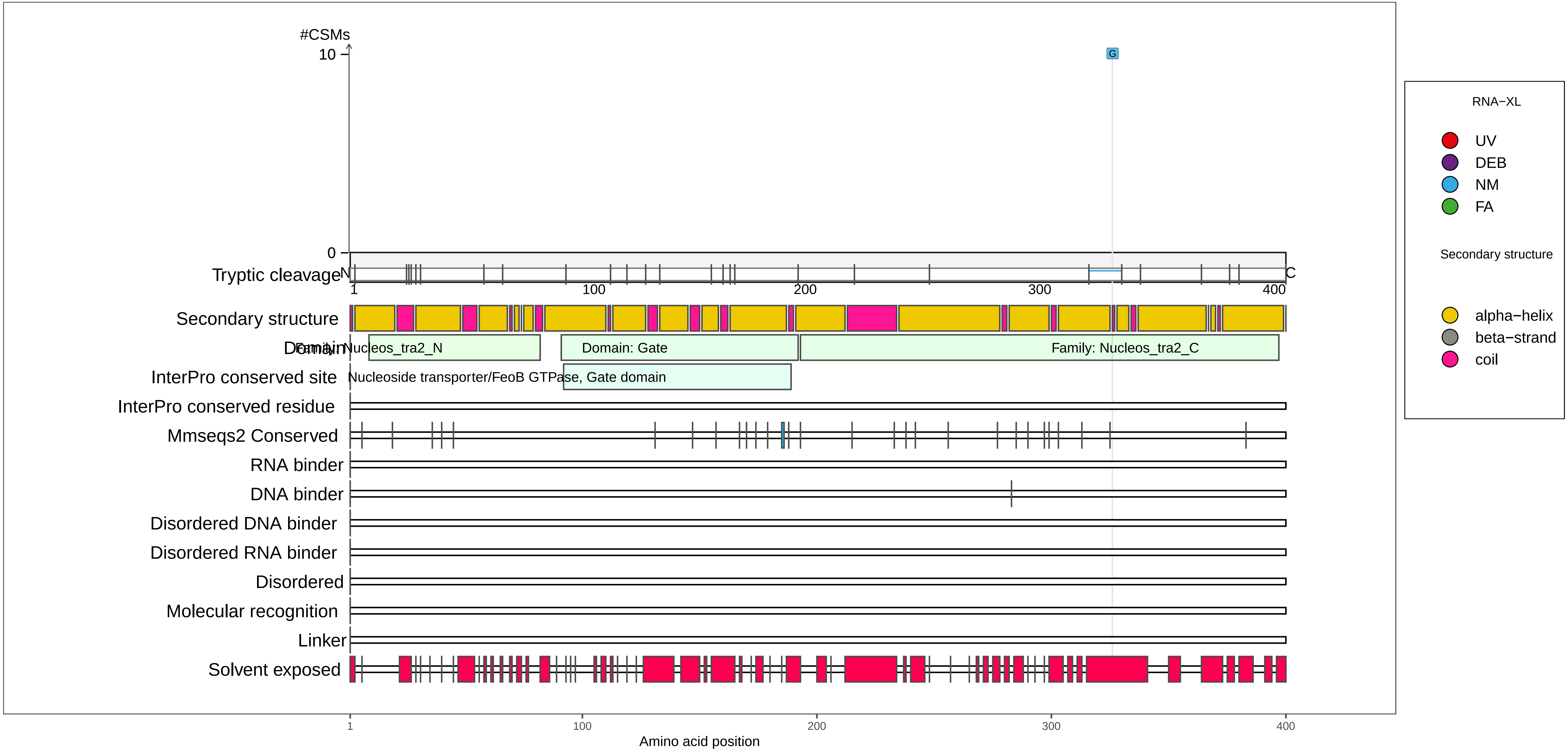

P0AG14
SOHB_ECOLI Probable protease SohB

− Abundance:
tryptic [log10 Intensity]: 7.95 (Q 56)
PAXdb K12 strain [ppm]: 1.31 (Q 27)
PAXdb E.coli [ppm]: 2.24 (Q 79)

− RNA functions: not annotated

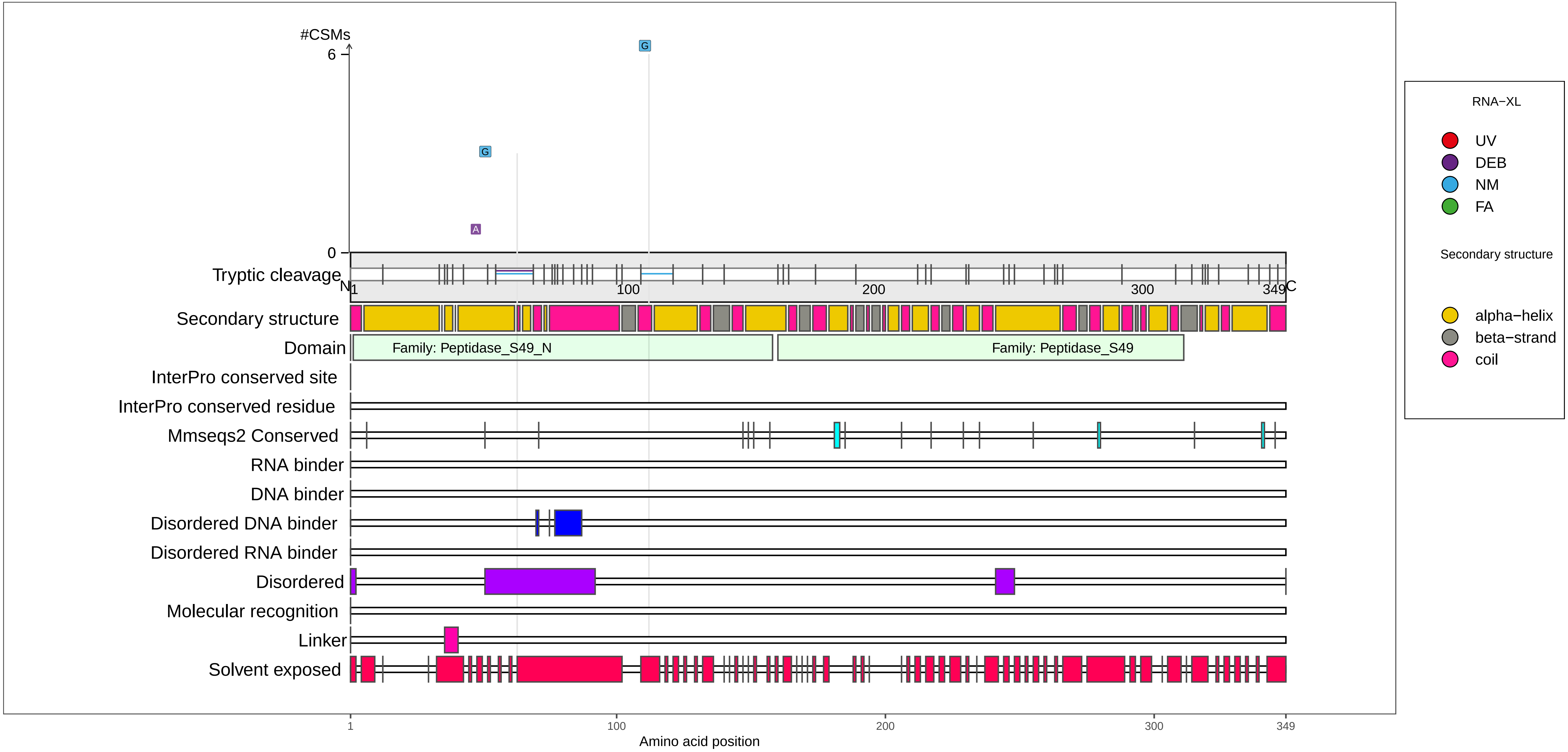

P15977
MALQ_ECOLI 4−alpha−glucanotransferase

− Abundance:
tryptic [log10 Intensity]: 8.28 (Q 68)
PAXdb K12 strain [ppm]: 2.64 (Q 82)
PAXdb E.coli [ppm]: 1.59 (Q 62)

− RNA functions: not annotated

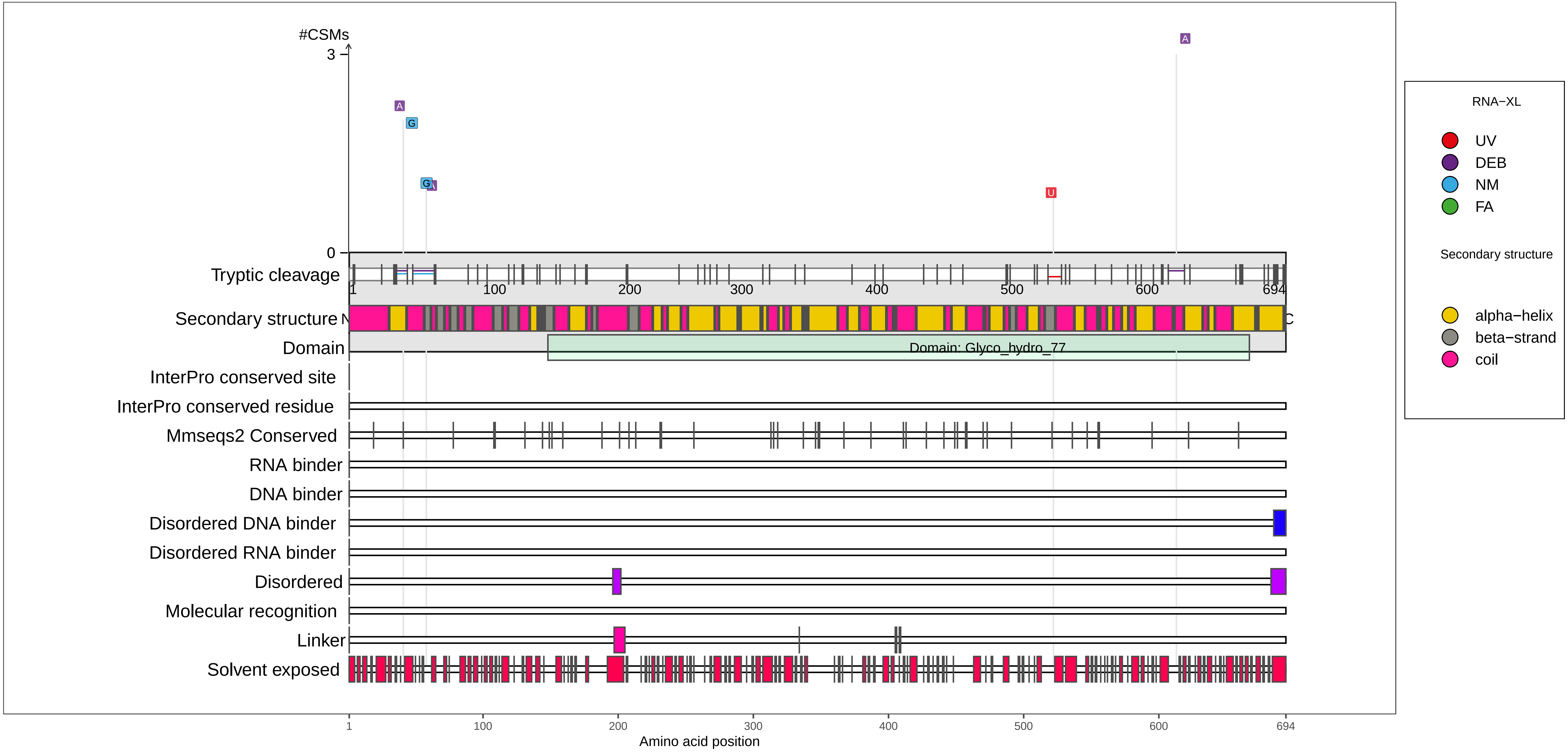

P23869
PPIB_ECOLI Peptidyl−prolyl cis−trans isomerase B

− Abundance:
tryptic [log10 Intensity]: 9.32 (Q 94)
PAXdb K12 strain [ppm]: 3.65 (Q 98)
PAXdb E.coli [ppm]: 3.21 (Q 97)

− RNA functions: not annotated

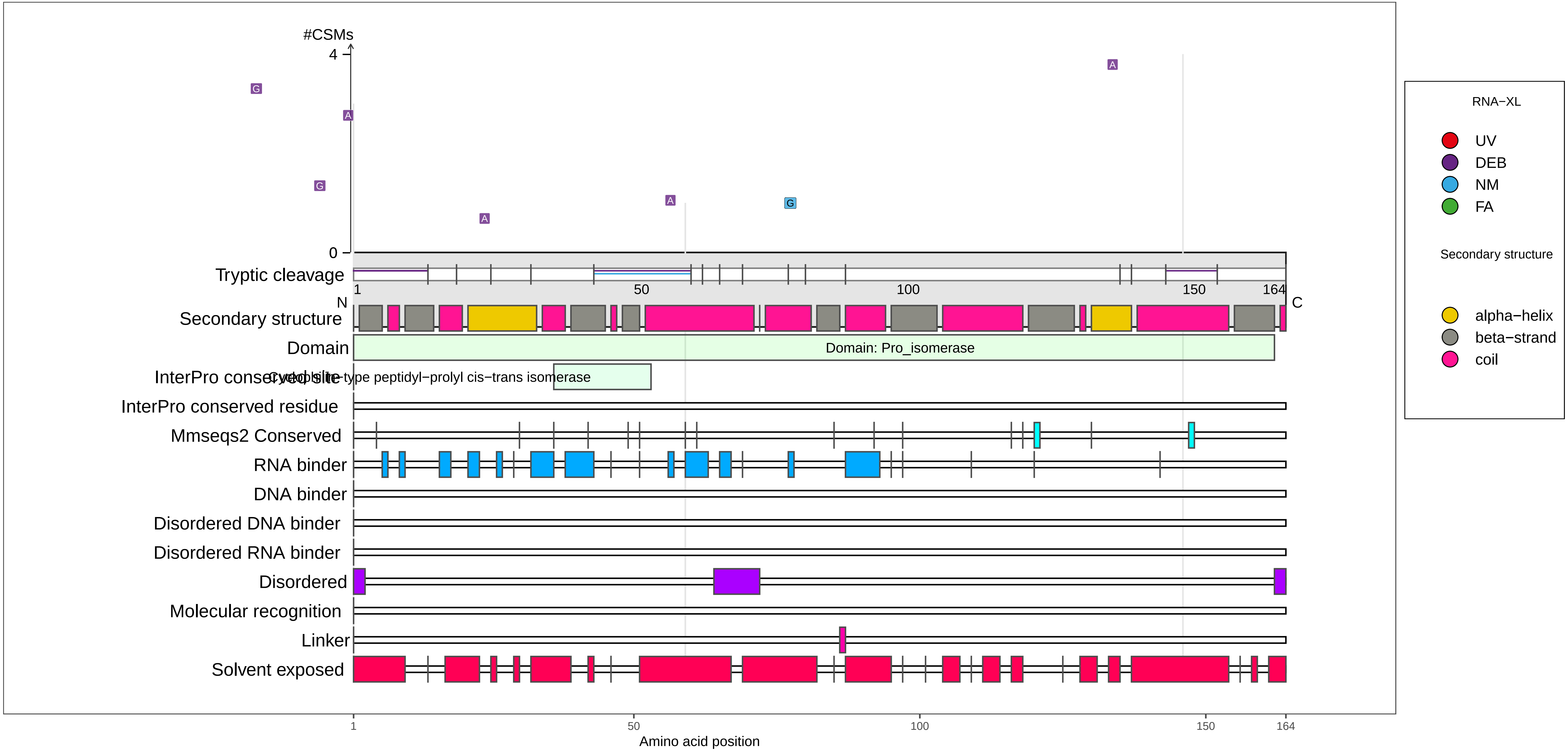

P24178
YFFB_ECOLI Protein YffB

− Abundance:
tryptic [log10 Intensity]: 7.55 (Q 37)
PAXdb K12 strain [ppm]: 2.03 (Q 61)
PAXdb E.coli [ppm]: 2.25 (Q 79)

− RNA functions: not annotated

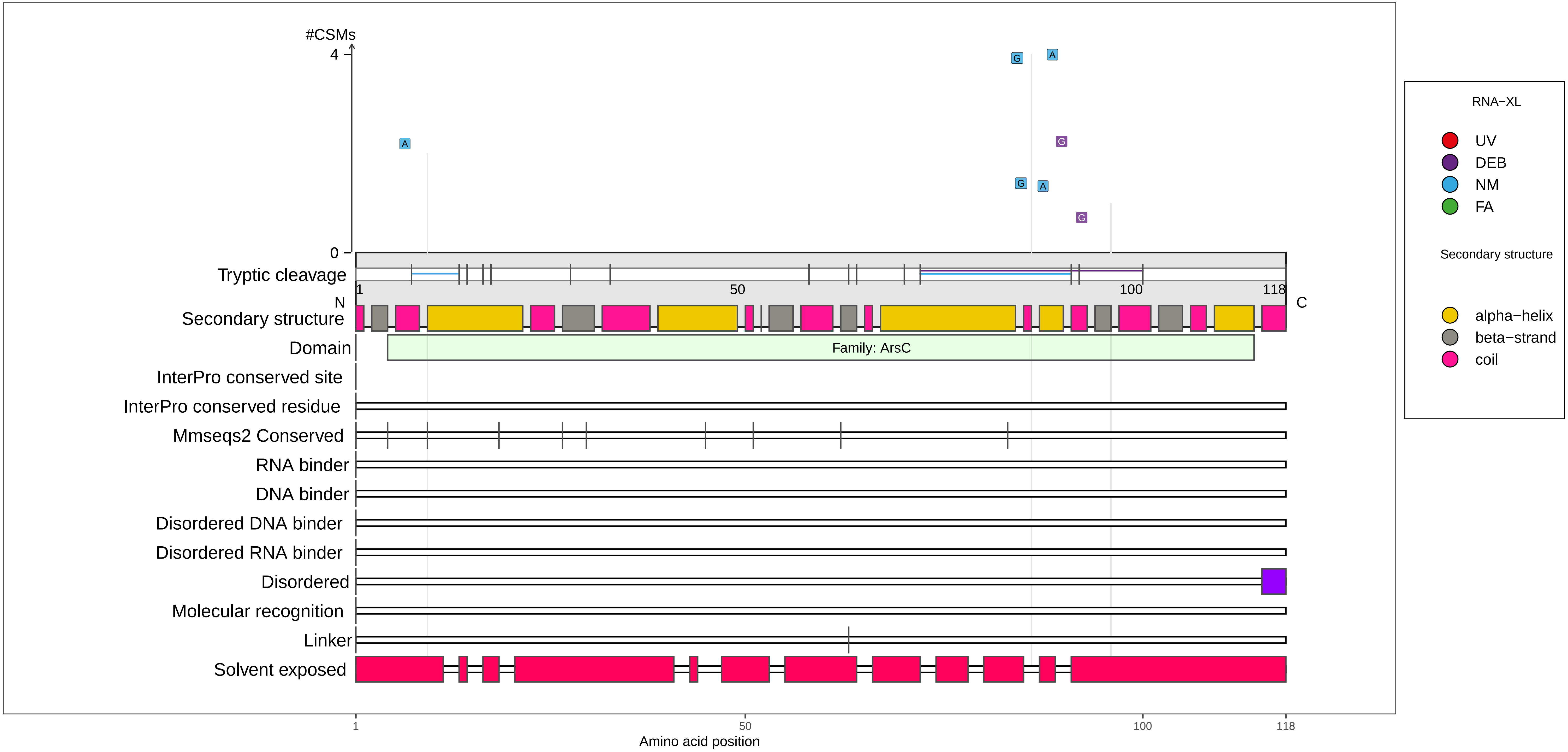

P32712
NRFG_ECOLI Formate−dependent nitrite reductase complex subunit NrfG

− Abundance:
tryptic [log10 Intensity]: NA (Q NA)
PAXdb K12 strain [ppm]: NA (Q NA)
PAXdb E.coli [ppm]: NA (Q NA)

− RNA functions: not annotated

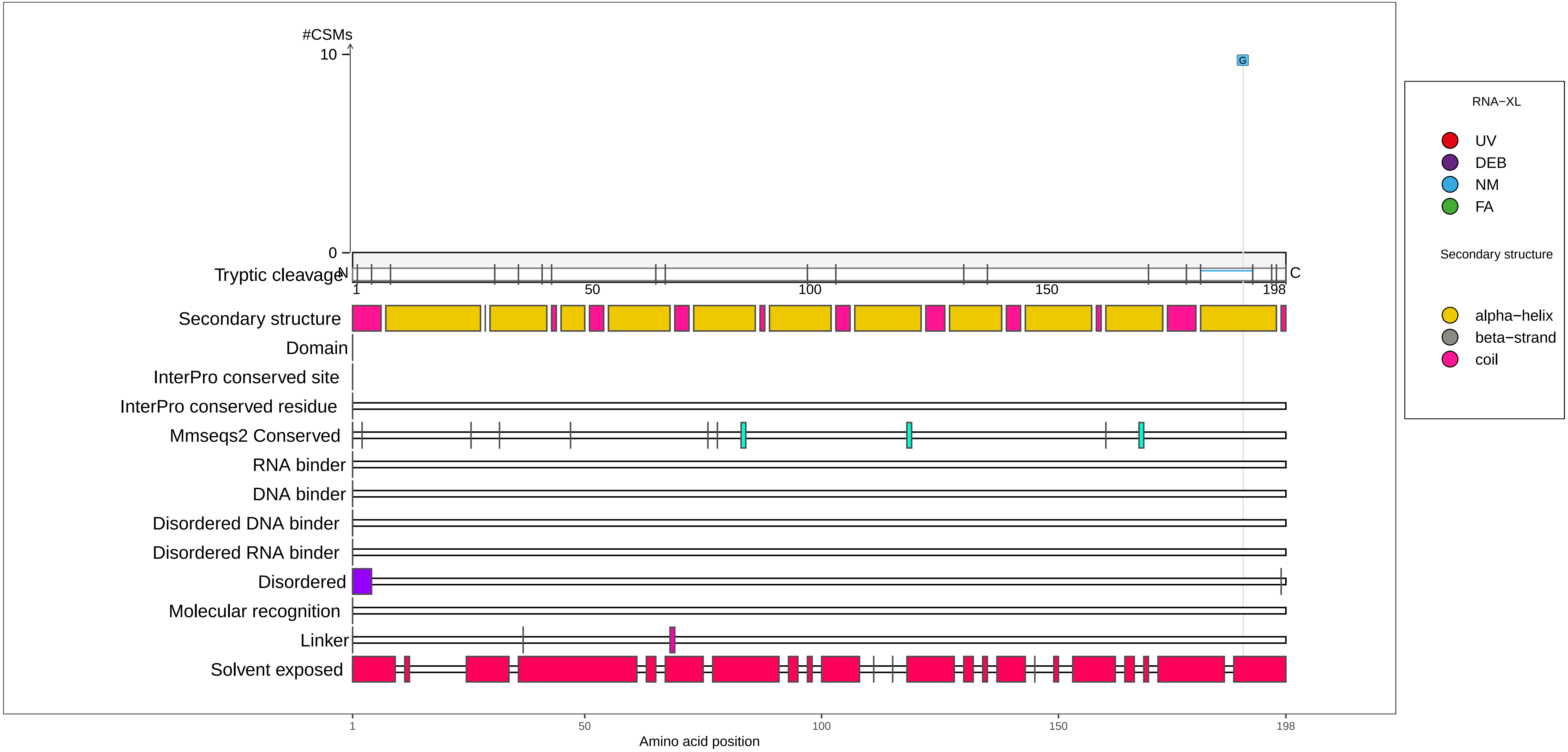

P36930
GATR_ECOLI Putative galactitol utilization operon repressor
− RNA functions: not annotated

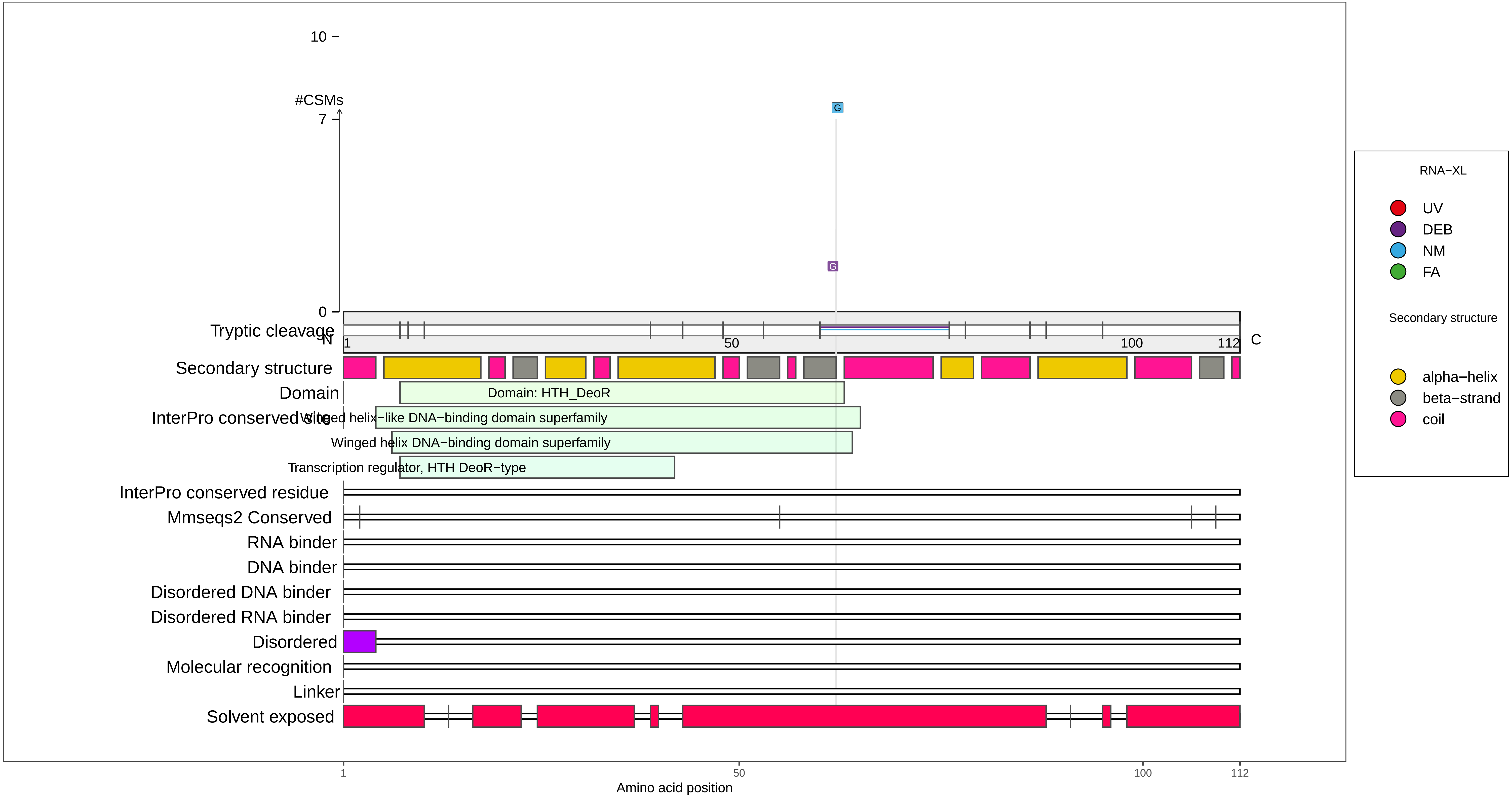

P37671
YIAJ_ECOLI DNA−binding transcriptional repressor YiaJ

− Abundance:
tryptic [log10 Intensity]: 7.6 (Q 40)
PAXdb K12 strain [ppm]: 1.05 (Q 13)
PAXdb E.coli [ppm]: 1.74 (Q 66)

− RNA functions: not annotated

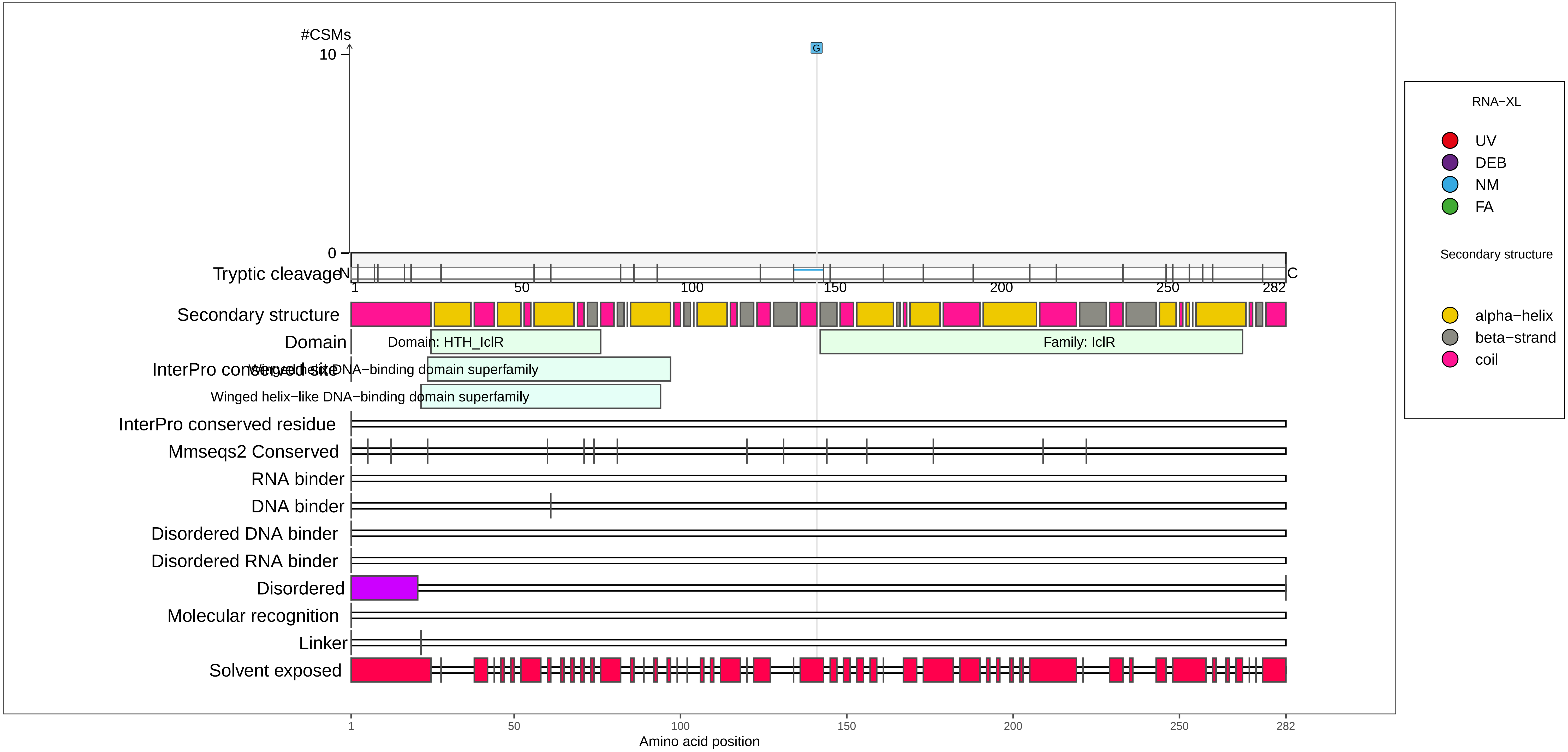

P39172
ZNUA_ECOLI High−affinity zinc uptake system protein ZnuA

− Abundance:
tryptic [log10 Intensity]: 8.23 (Q 67)
PAXdb K12 strain [ppm]: 1.93 (Q 58)
PAXdb E.coli [ppm]: 2.99 (Q 95)

− RNA functions: not annotated

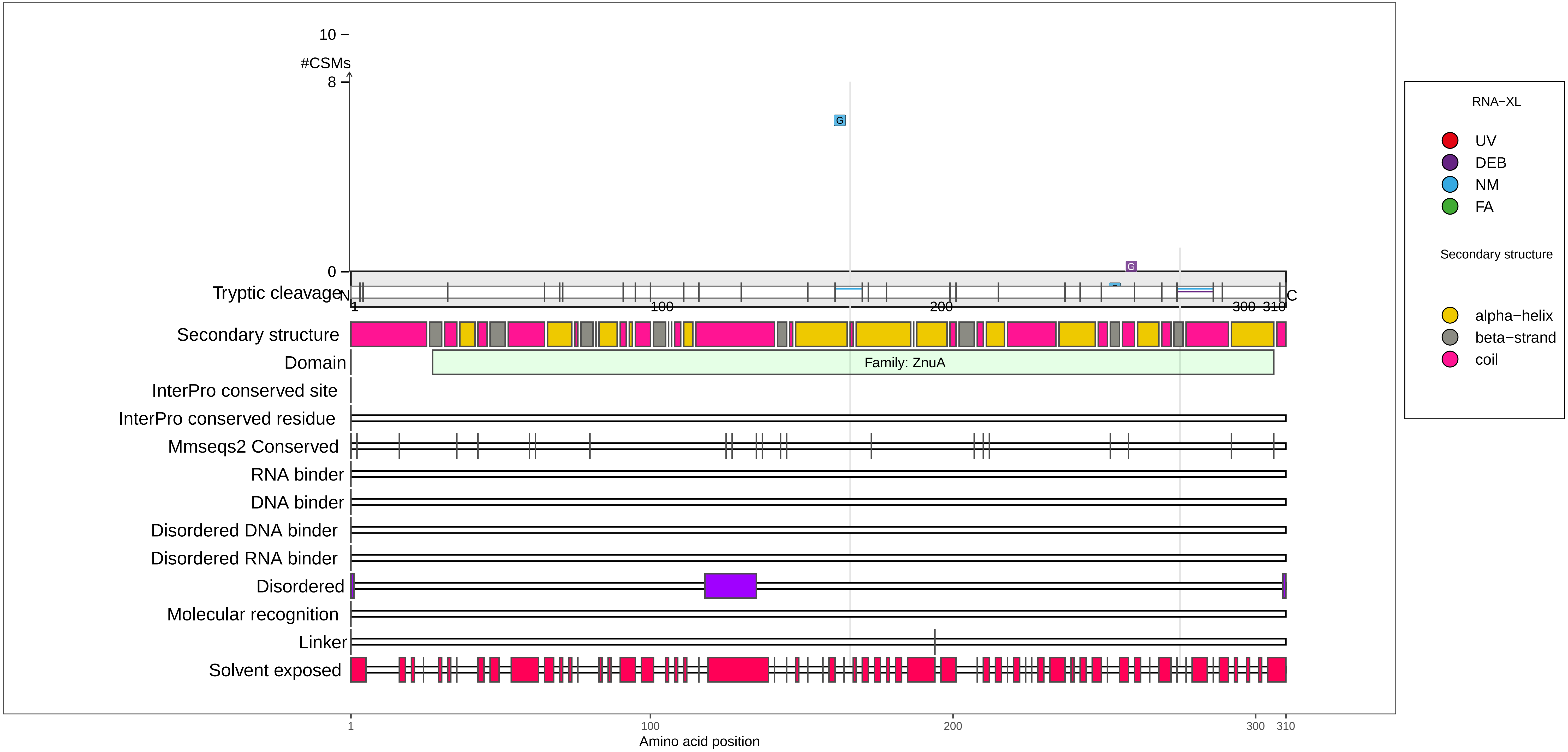

P68767
AMPA_ECOLI Cytosol aminopeptidase

− Abundance:
tryptic [log10 Intensity]: 8.14 (Q 63)
PAXdb K12 strain [ppm]: 1.63 (Q 46)
PAXdb E.coli [ppm]: 2.3 (Q 80)

− RNA functions:
RNA biosynthetic process; RNA metabolic process

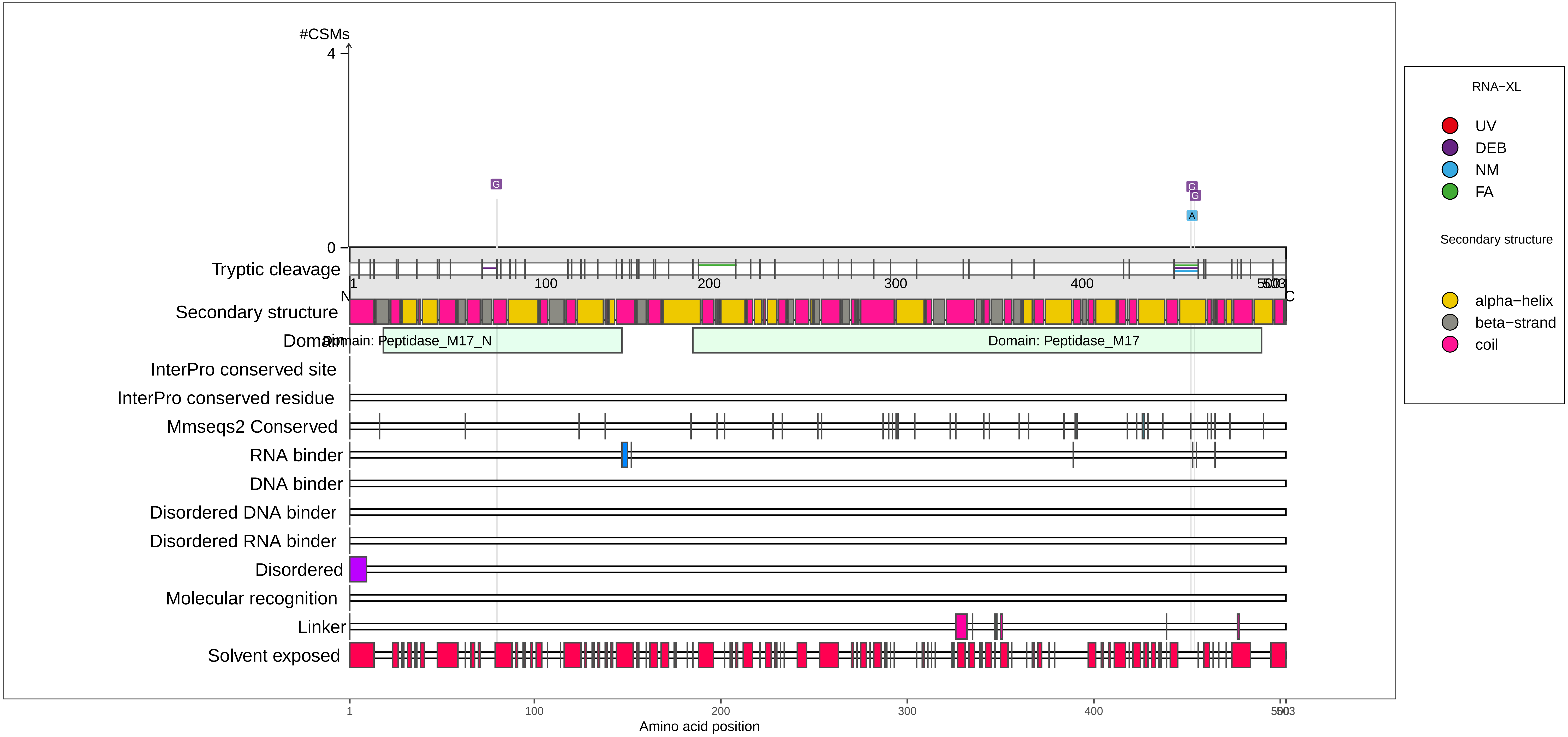

P75876
RLMI_ECOLI Ribosomal RNA large subunit methyltransferase I

− Abundance:
tryptic [log10 Intensity]: 8.04 (Q 59)
PAXdb K12 strain [ppm]: NA (Q NA)
PAXdb E.coli [ppm]: 1.43 (Q 58)

− RNA functions:
ncRNA metabolic process; ncRNA processing; RNA binding; RNA metabolic process
RNA methylation; RNA methyltransferase activity; RNA modification; RNA processing
rRNA (cytosine−C5−)−methyltransferase activity; rRNA (cytosine) methyltransferase activity
rRNA base methylation; rRNA metabolic process; rRNA methylation
rRNA methyltransferase activity; rRNA modification; rRNA processing

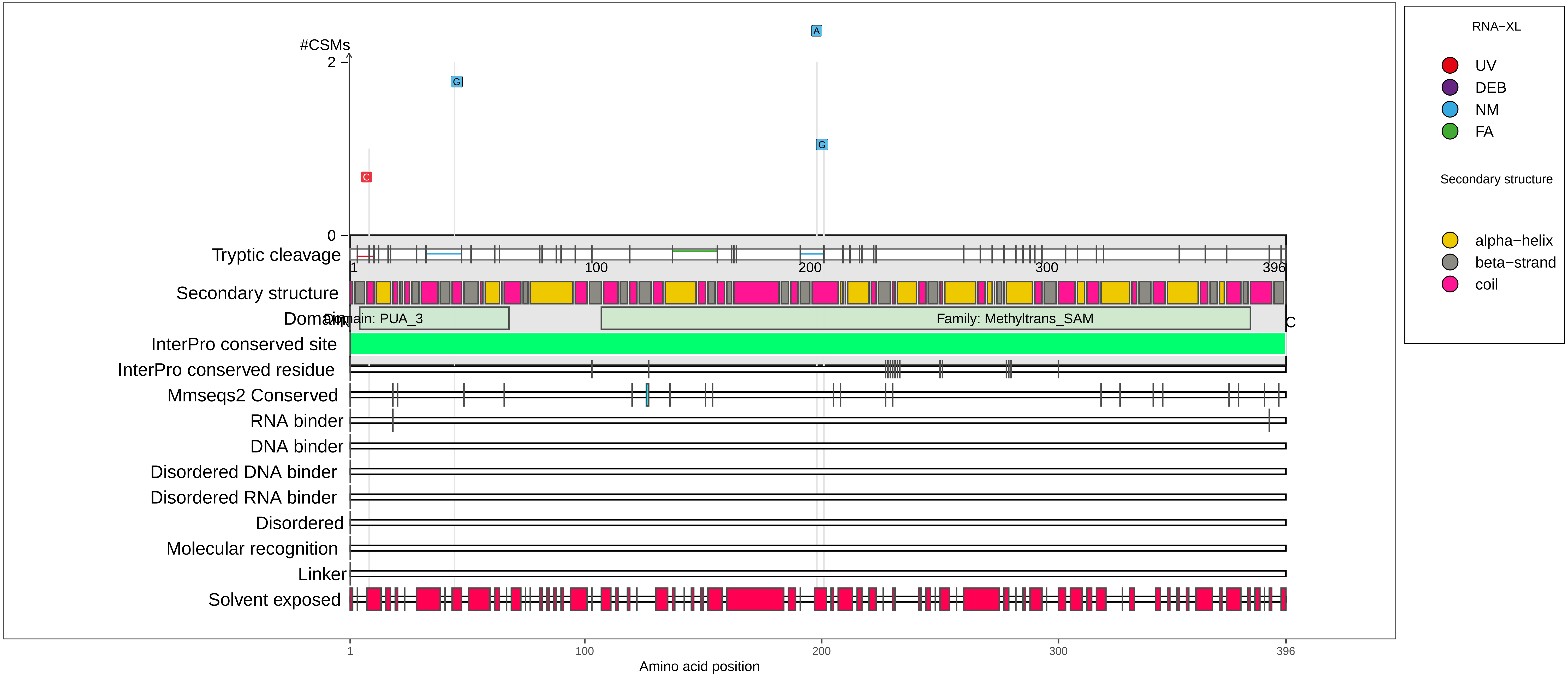

P76423
THIM_ECOLI Hydroxyethylthiazole kinase

− Abundance:
tryptic [log10 Intensity]: 7.76 (Q 47)
PAXdb K12 strain [ppm]: 1.9 (Q 57)
PAXdb E.coli [ppm]: 2.52 (Q 86)

− RNA functions: not annotated

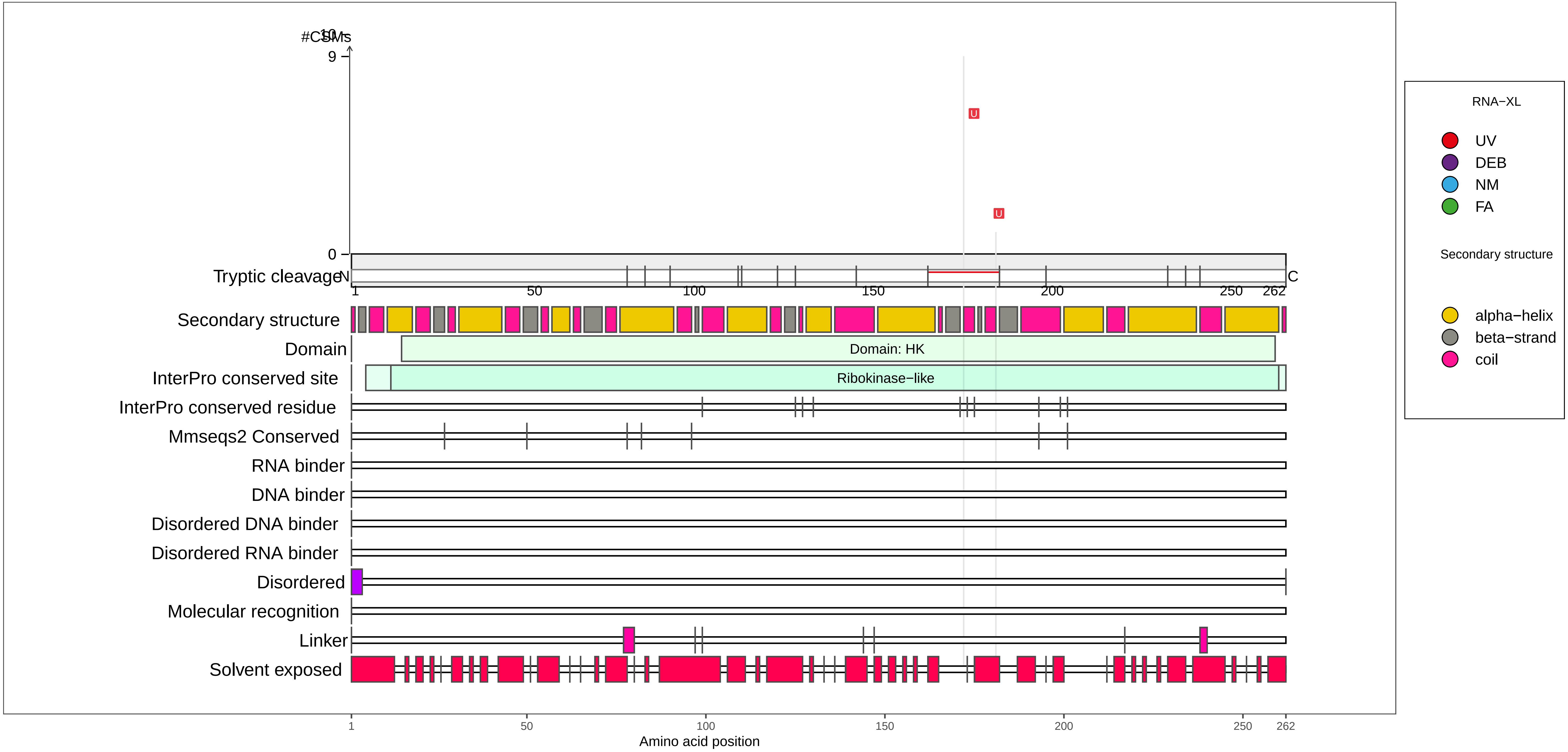

P00370
DHE4_ECOLI NADP−specific glutamate dehydrogenase

− Abundance:
tryptic [log10 Intensity]: 8.61 (Q 79)
PAXdb K12 strain [ppm]: 2.18 (Q 66)
PAXdb E.coli [ppm]: 2.89 (Q 93)

− RNA functions: not annotated

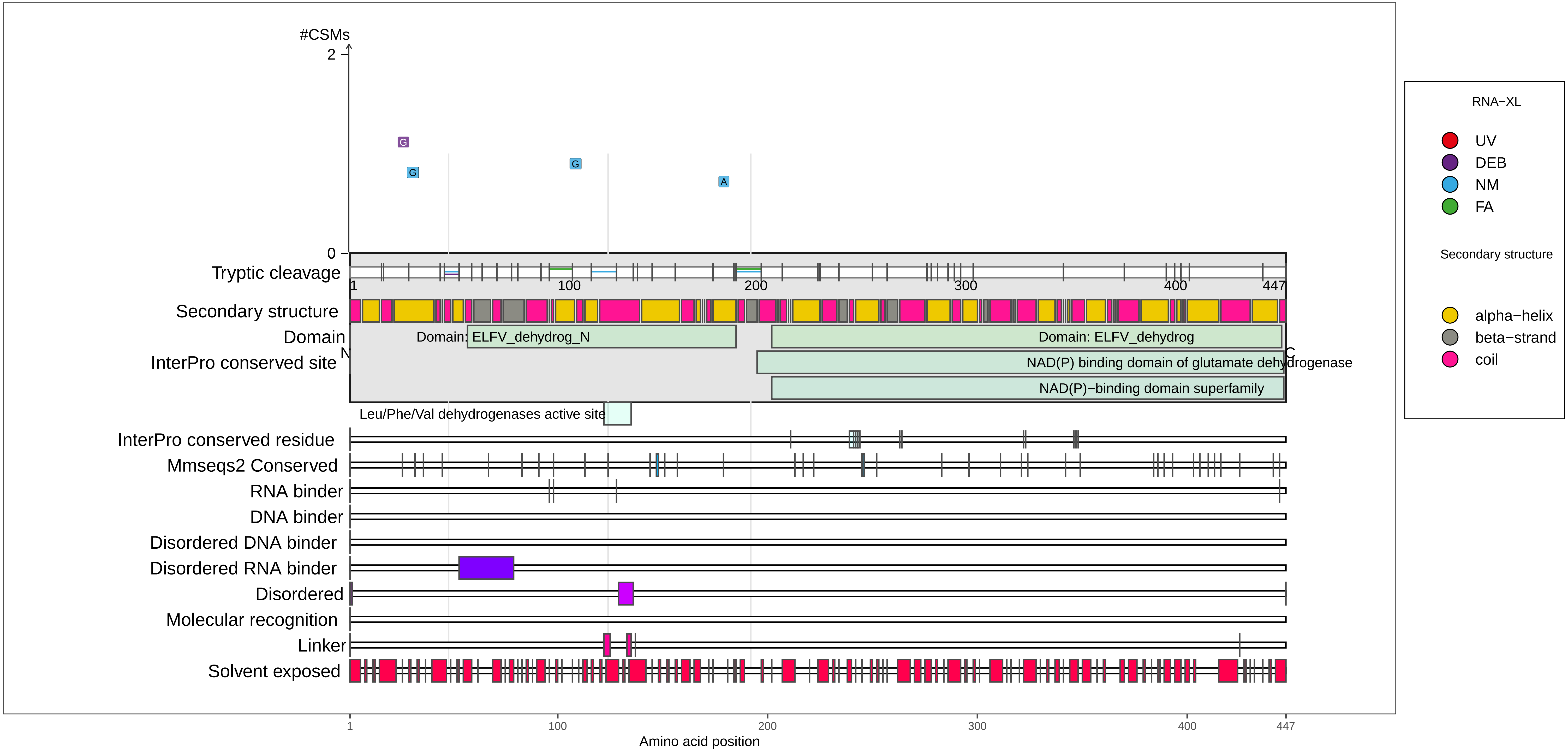

P06612
TOP1_ECOLI DNA topoisomerase 1

− Abundance:
tryptic [log10 Intensity]: 8.57 (Q 78)
PAXdb K12 strain [ppm]: 2.47 (Q 77)
PAXdb E.coli [ppm]: 2.36 (Q 82)

− RNA functions: not annotated

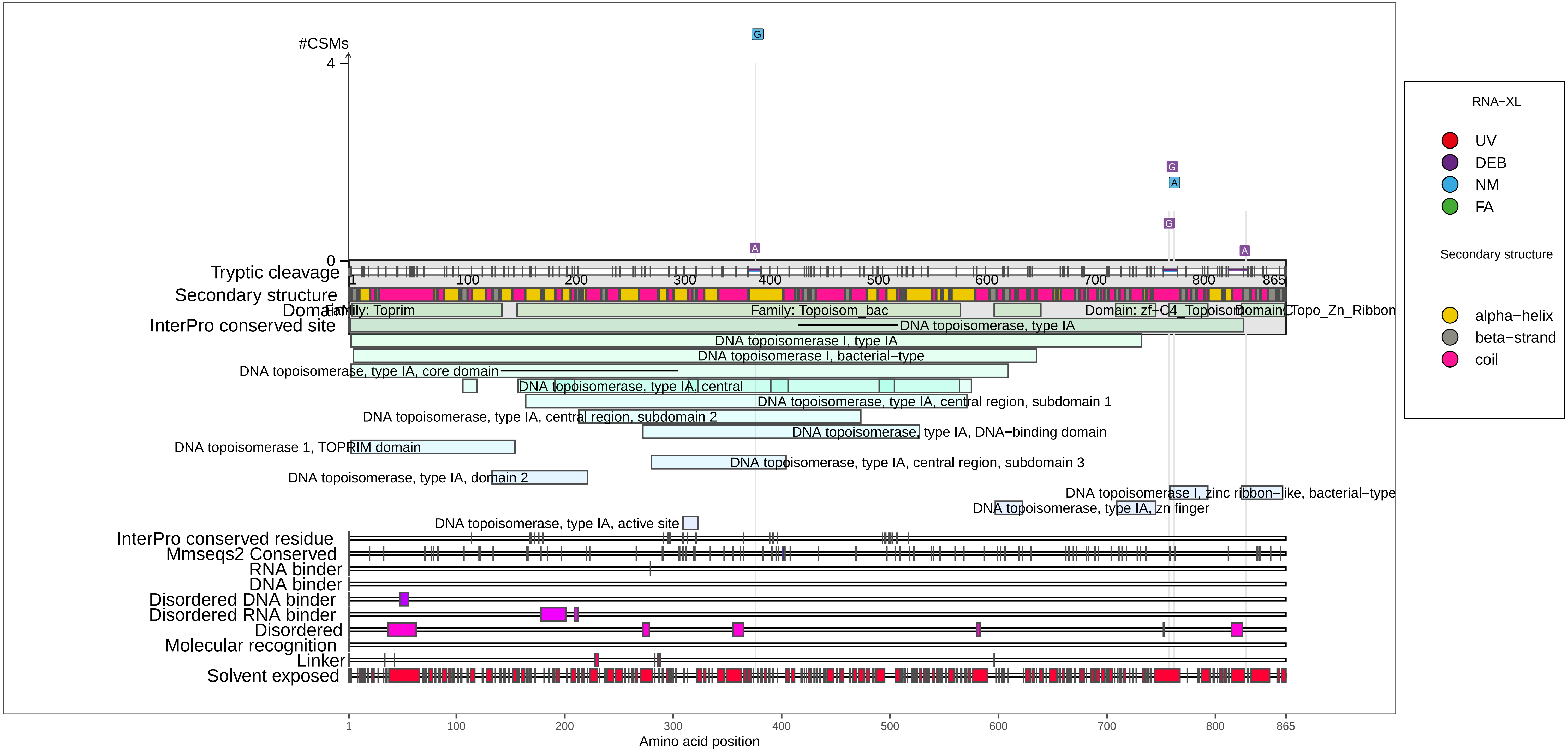

P08331
CPDB_ECOLI 2',3'−cyclic−nucleotide 2'−phosphodiesterase/3'−nucleotidase

− Abundance:
tryptic [log10 Intensity]: 8.56 (Q 77)
PAXdb K12 strain [ppm]: 1.6 (Q 44)
PAXdb E.coli [ppm]: 2.05 (Q 73)

− RNA functions: not annotated

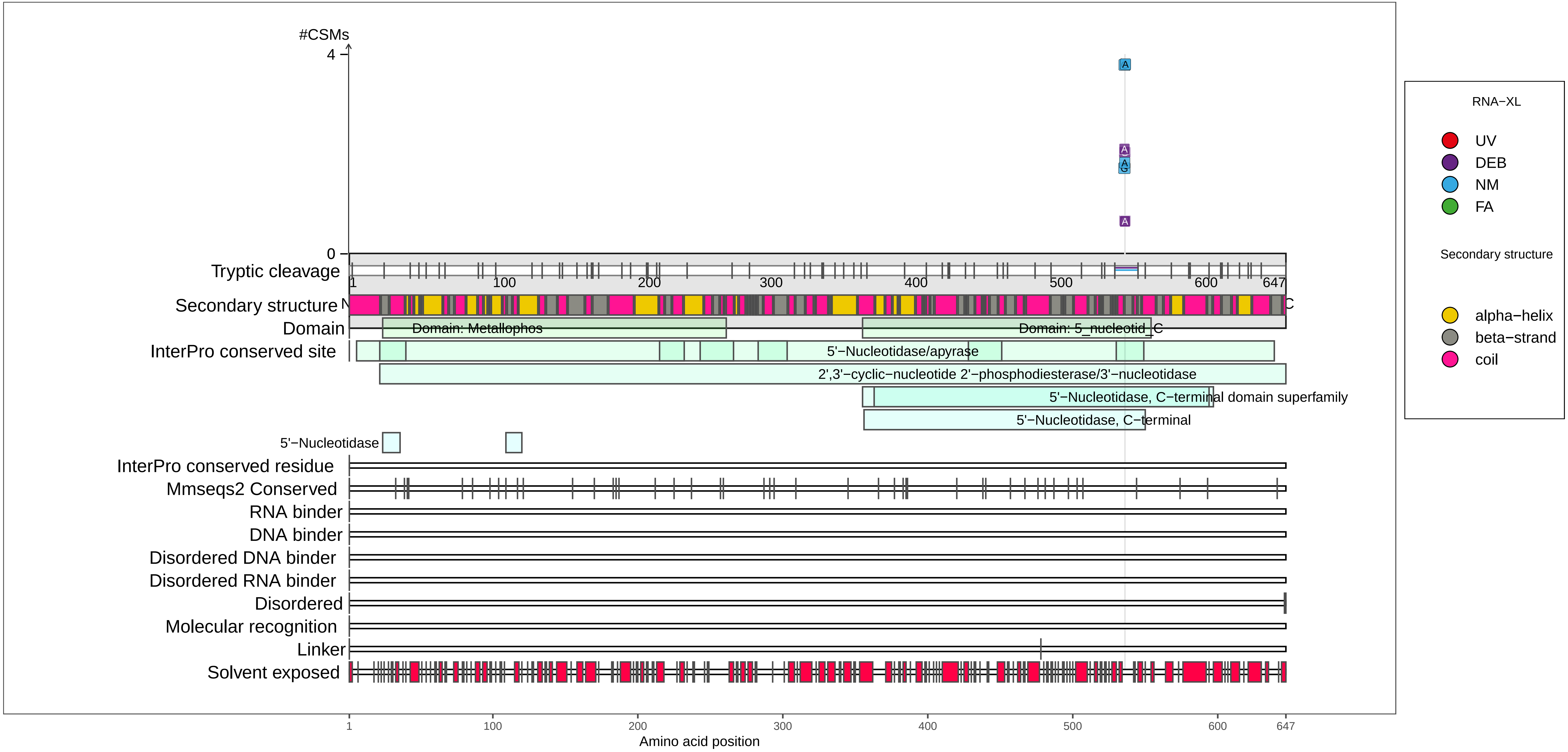

P08337
MUTT_ECOLI 8−oxo−dGTP diphosphatase
− Abundance:
tryptic [log10 Intensity]: 7.4 (Q 30)
PAXdb K12 strain [ppm]: NA (Q NA)
PAXdb E.coli [ppm]: 1.05 (Q 49)

− RNA functions: not annotated

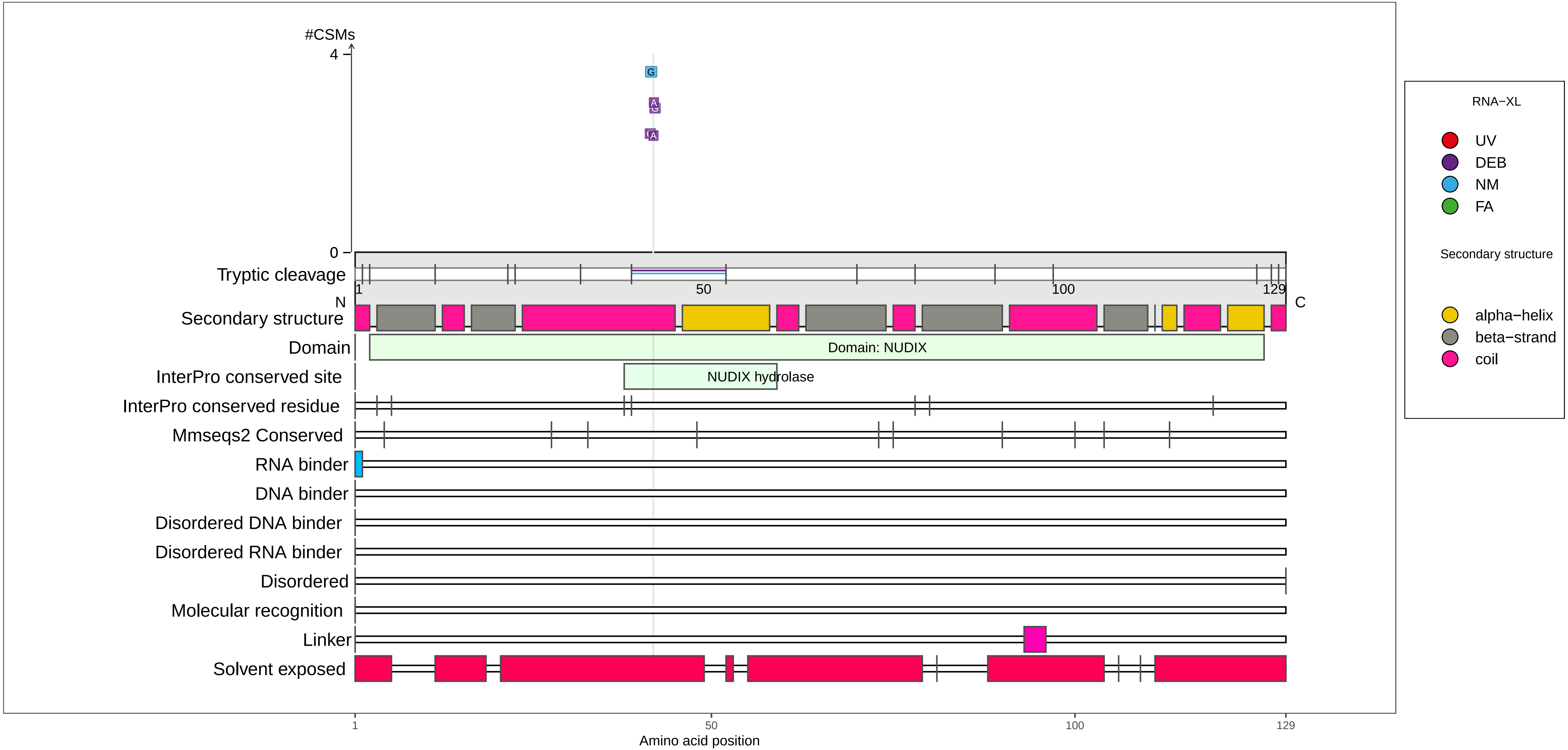

P09831
GLTB_ECOLI Glutamate synthase [NADPH] large chain

− Abundance:
tryptic [log10 Intensity]: 8.83 (Q 84)
PAXdb K12 strain [ppm]: 2.46 (Q 76)
PAXdb E.coli [ppm]: 2.76 (Q 91)

− RNA functions: not annotated

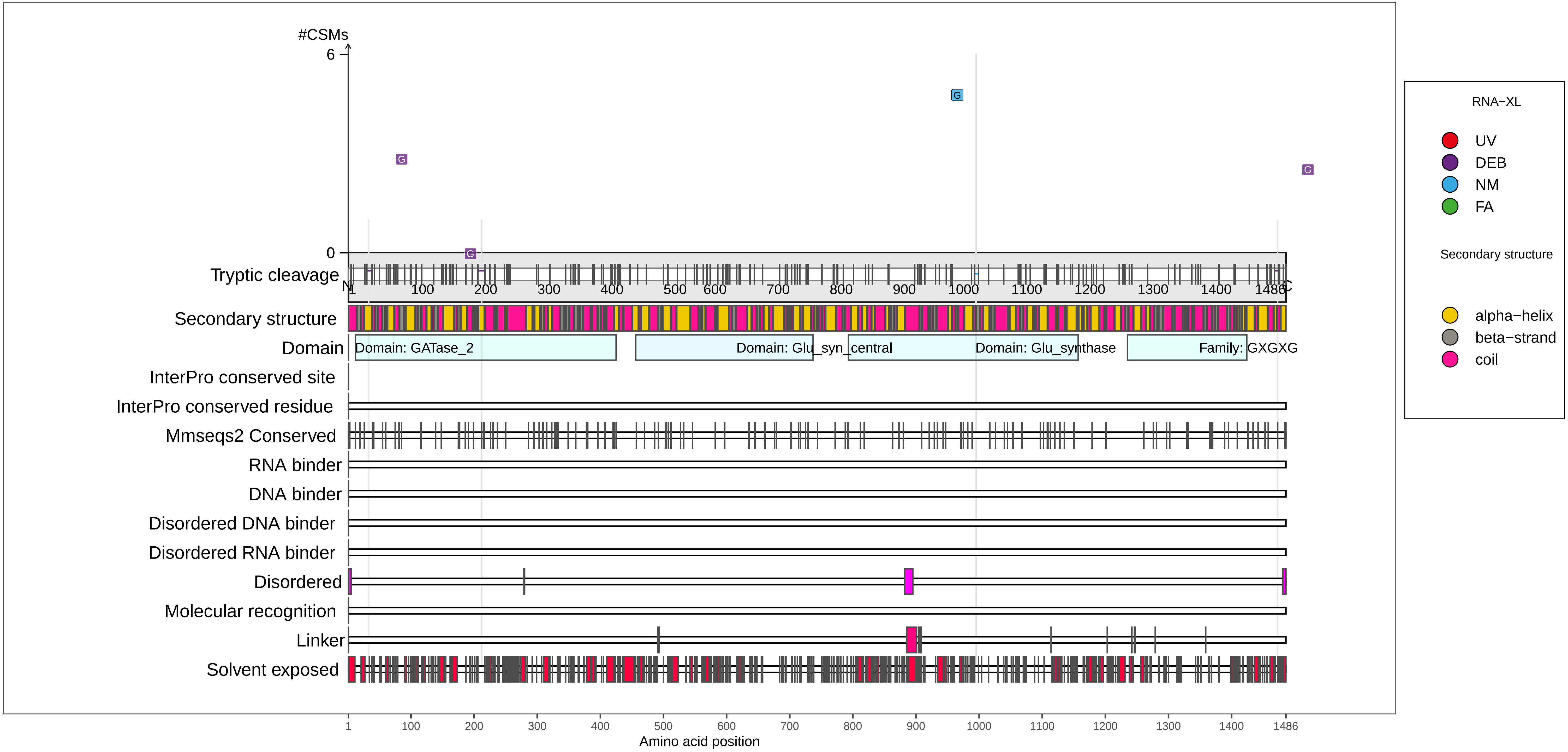

P0A6L0
DEOC_ECOLI Deoxyribose−phosphate aldolase

− Abundance:
tryptic [log10 Intensity]: 8.82 (Q 84)
PAXdb K12 strain [ppm]: 2.31 (Q 70)
PAXdb E.coli [ppm]: 2.67 (Q 89)

− RNA functions: not annotated

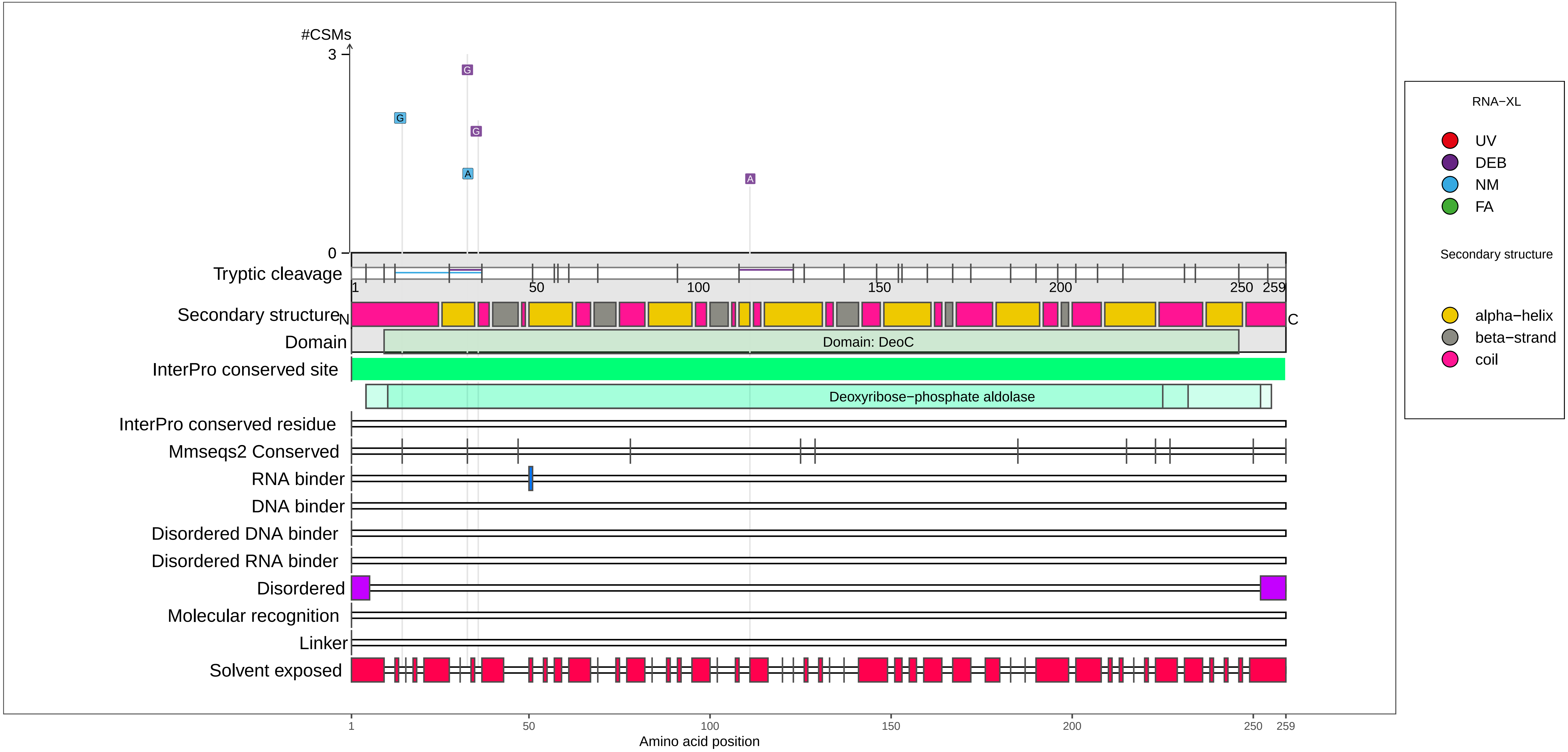

P0A6V1
GLGC_ECOLI Glucose−1−phosphate adenylyltransferase

− Abundance:
tryptic [log10 Intensity]: 8.04 (Q 59)
PAXdb K12 strain [ppm]: 1.84 (Q 55)
PAXdb E.coli [ppm]: 2.34 (Q 82)

− RNA functions: not annotated

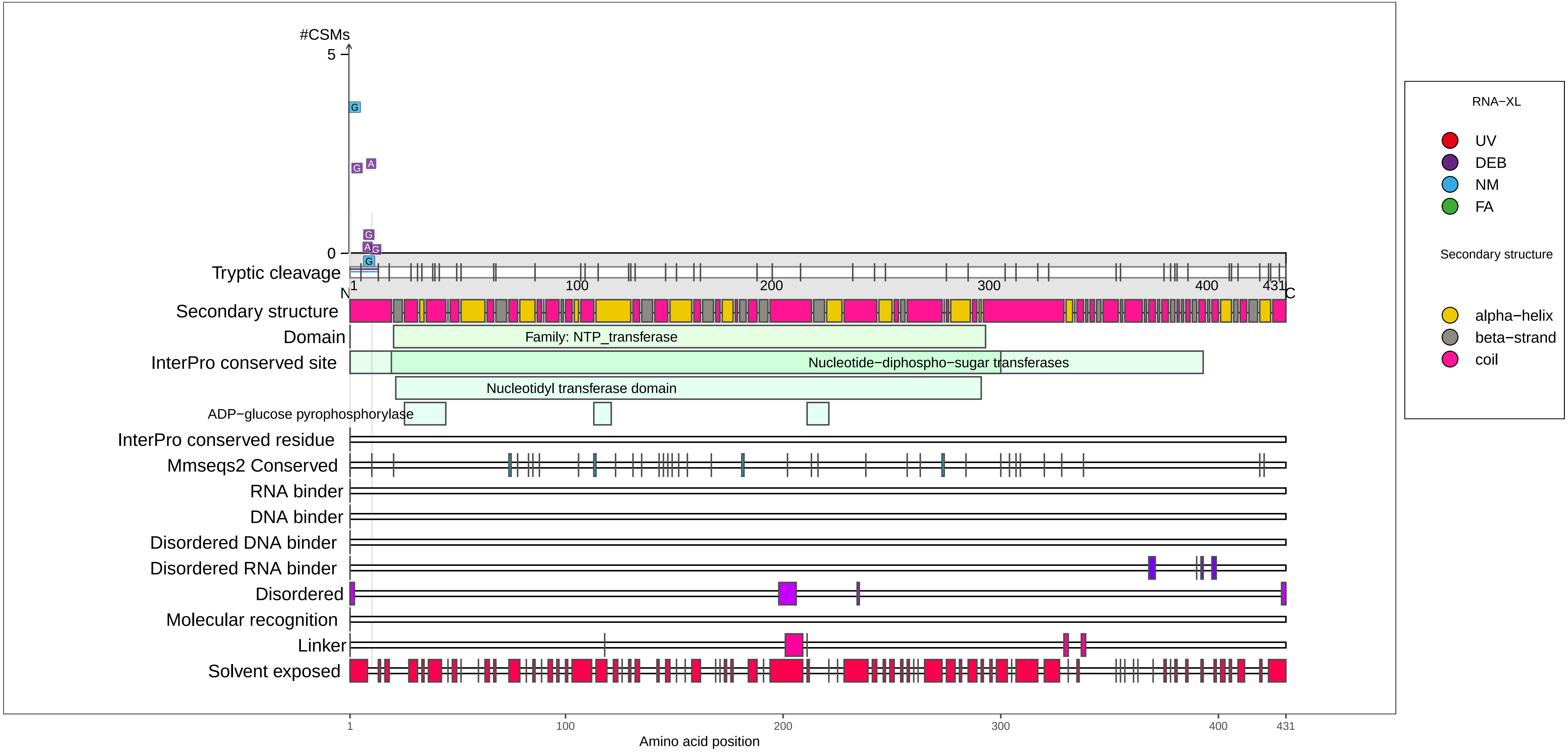

P0A9D4
CYSE_ECOLI Serine acetyltransferase

− Abundance:
tryptic [log10 Intensity]: 8.61 (Q 79)
PAXdb K12 strain [ppm]: 2.45 (Q 76)
PAXdb E.coli [ppm]: 1.96 (Q 71)

− RNA functions: not annotated

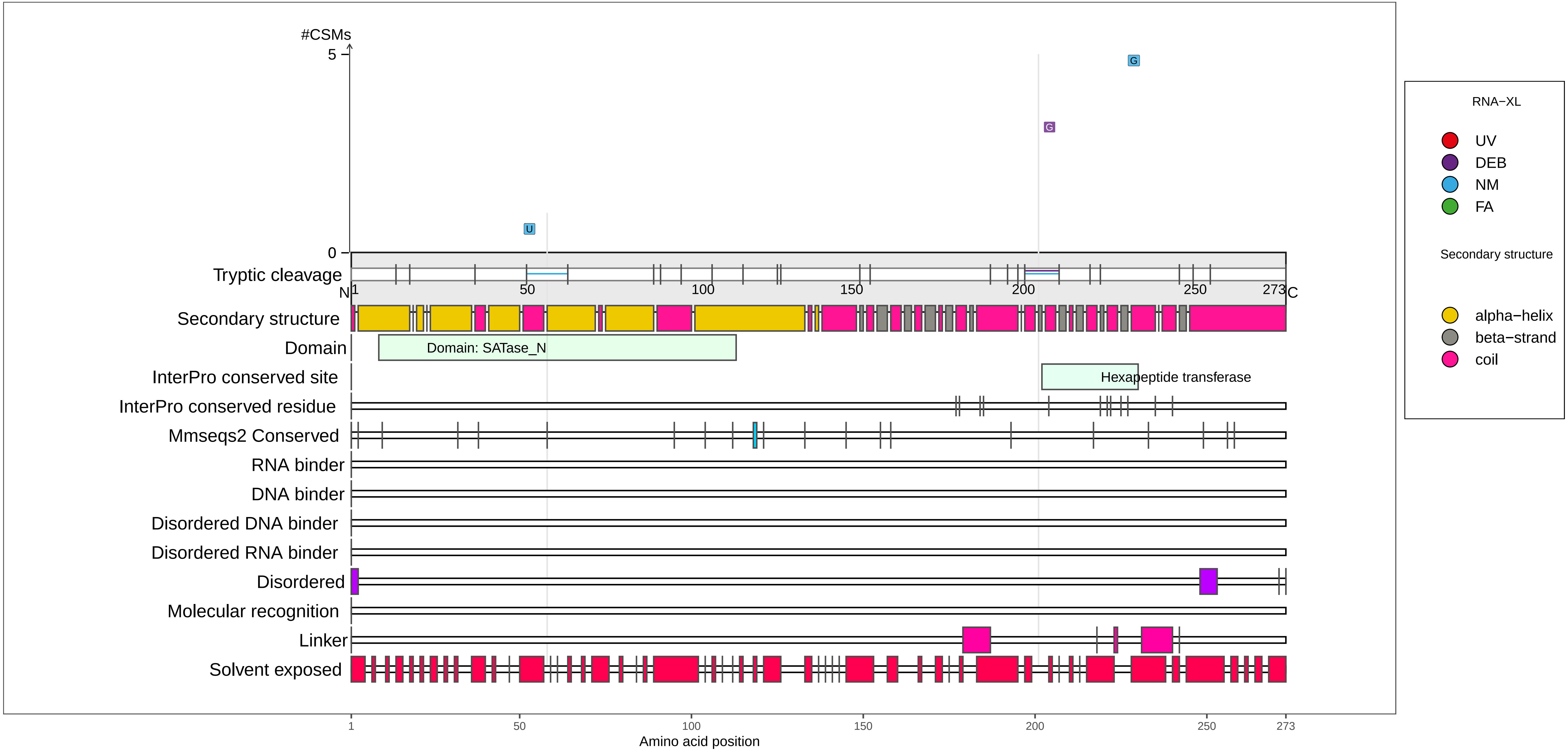

P0A9K3
PHOL_ECOLI PhoH−like protein

− Abundance:
tryptic [log10 Intensity]: 8.57 (Q 78)
PAXdb K12 strain [ppm]: 2.79 (Q 86)
PAXdb E.coli [ppm]: 2.56 (Q 87)

− RNA functions: not annotated

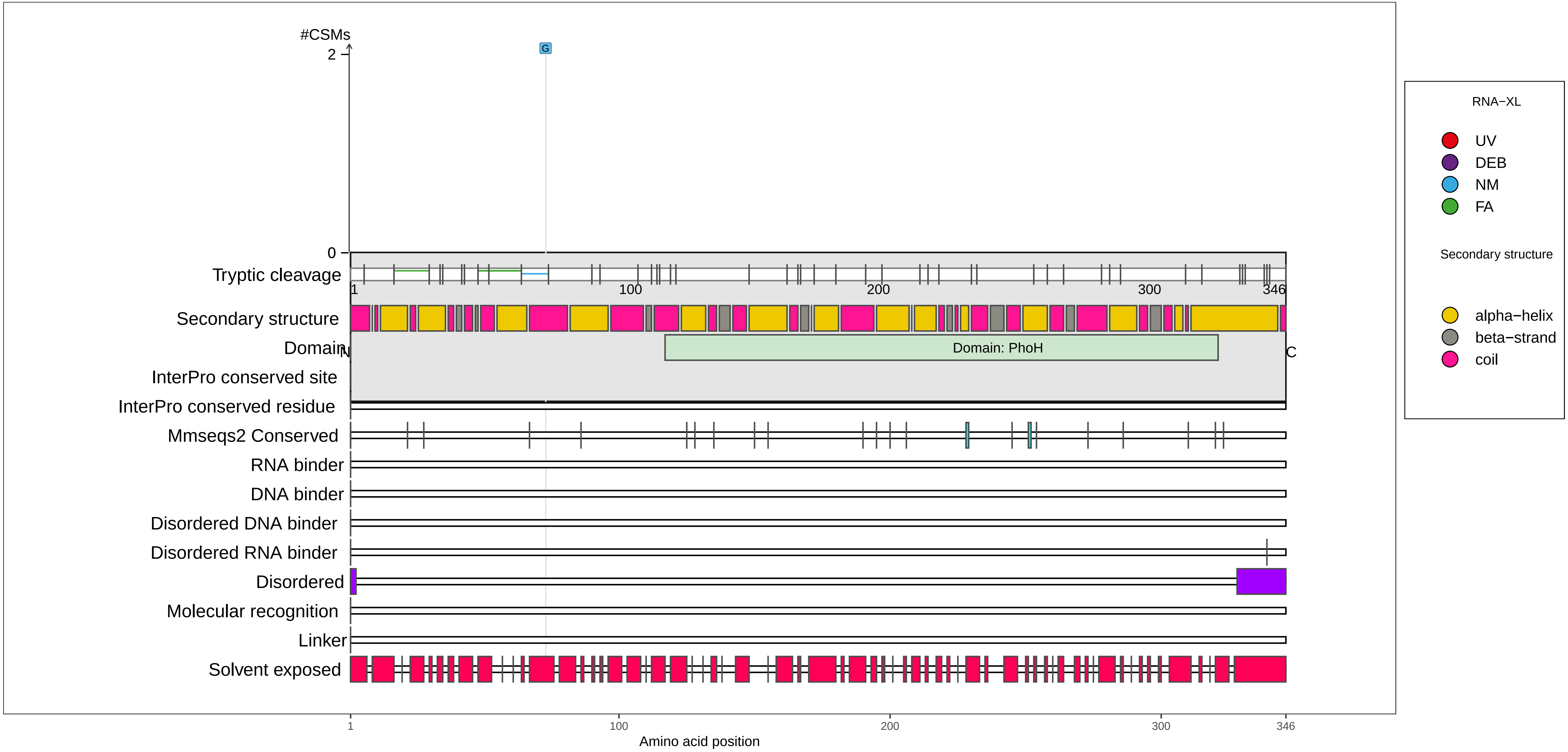

P0AAN1
HYBE_ECOLI Hydrogenase−2 operon protein HybE

− Abundance:
tryptic [log10 Intensity]: 7.45 (Q 32)
PAXdb K12 strain [ppm]: NA (Q NA)
PAXdb E.coli [ppm]: −0.19 (Q 21)

− RNA functions: not annotated

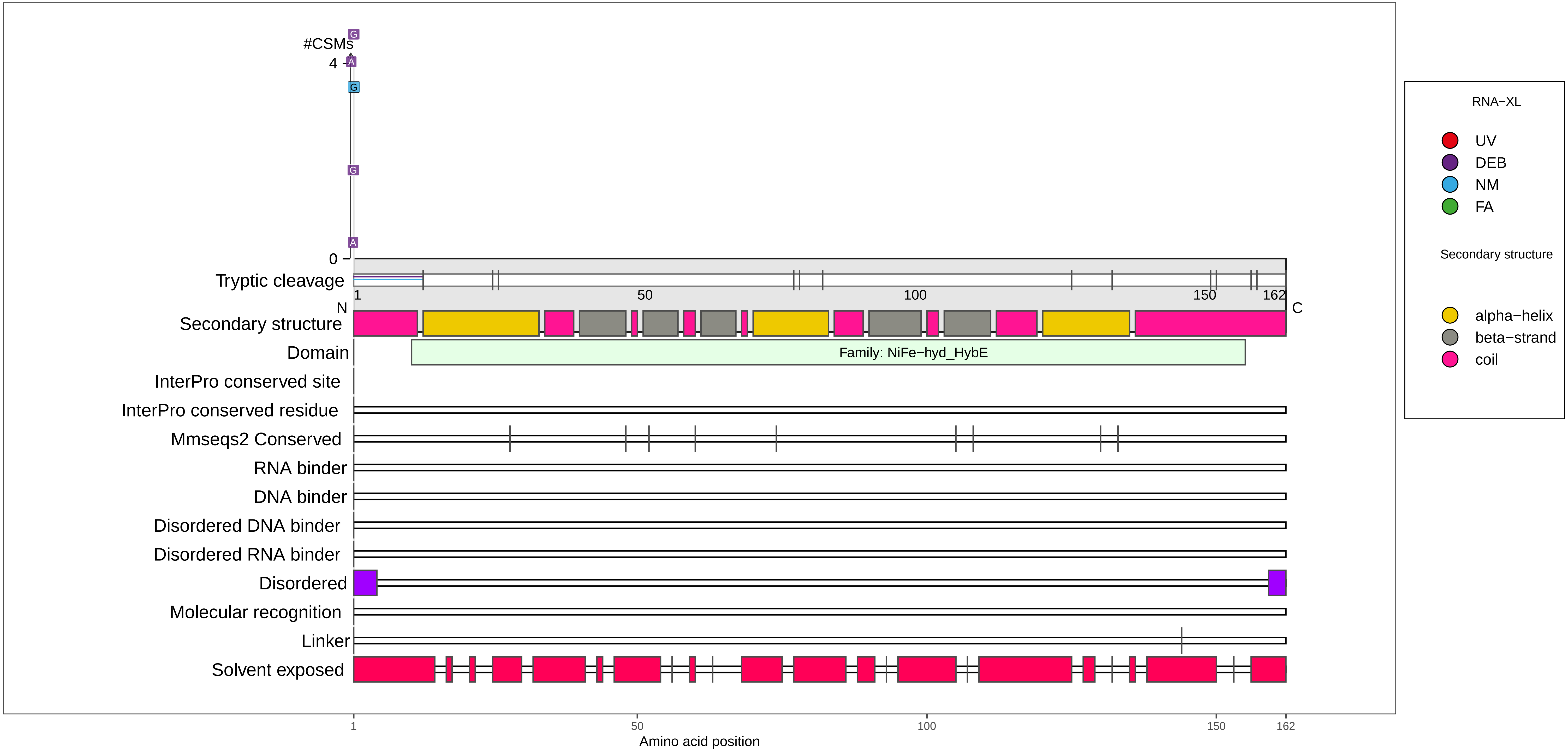

P0ADV5
YHBW_ECOLI Luciferase−like monooxygenase

− Abundance:
tryptic [log10 Intensity]: 8.13 (Q 63)
PAXdb K12 strain [ppm]: 2.82 (Q 87)
PAXdb E.coli [ppm]: 1.14 (Q 51)

− RNA functions: not annotated

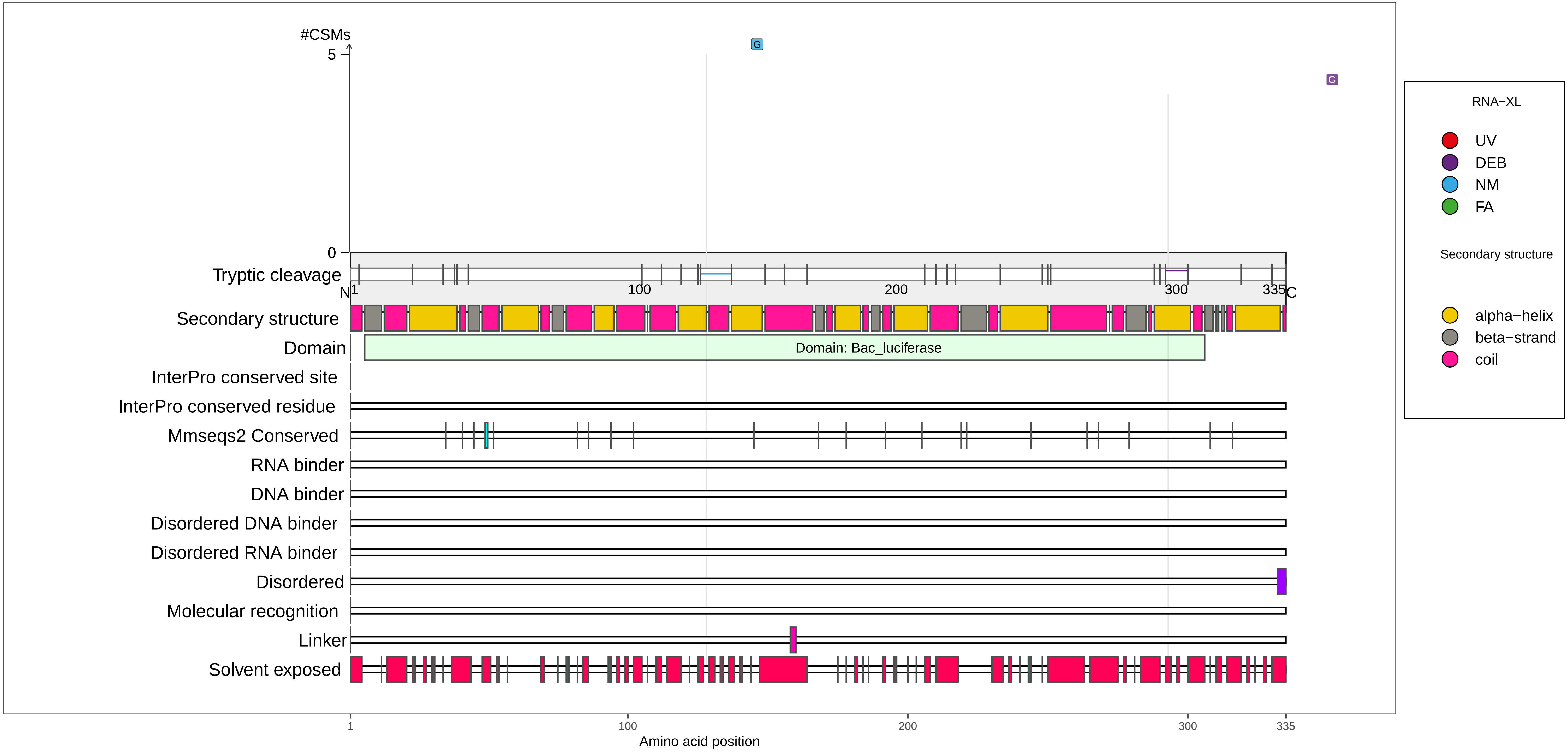

P0AEA8
CYSG_ECOLI Siroheme synthase

− Abundance:
tryptic [log10 Intensity]: 7.62 (Q 41)
PAXdb K12 strain [ppm]: 1.6 (Q 45)
PAXdb E.coli [ppm]: 0.99 (Q 47)

− RNA functions: not annotated

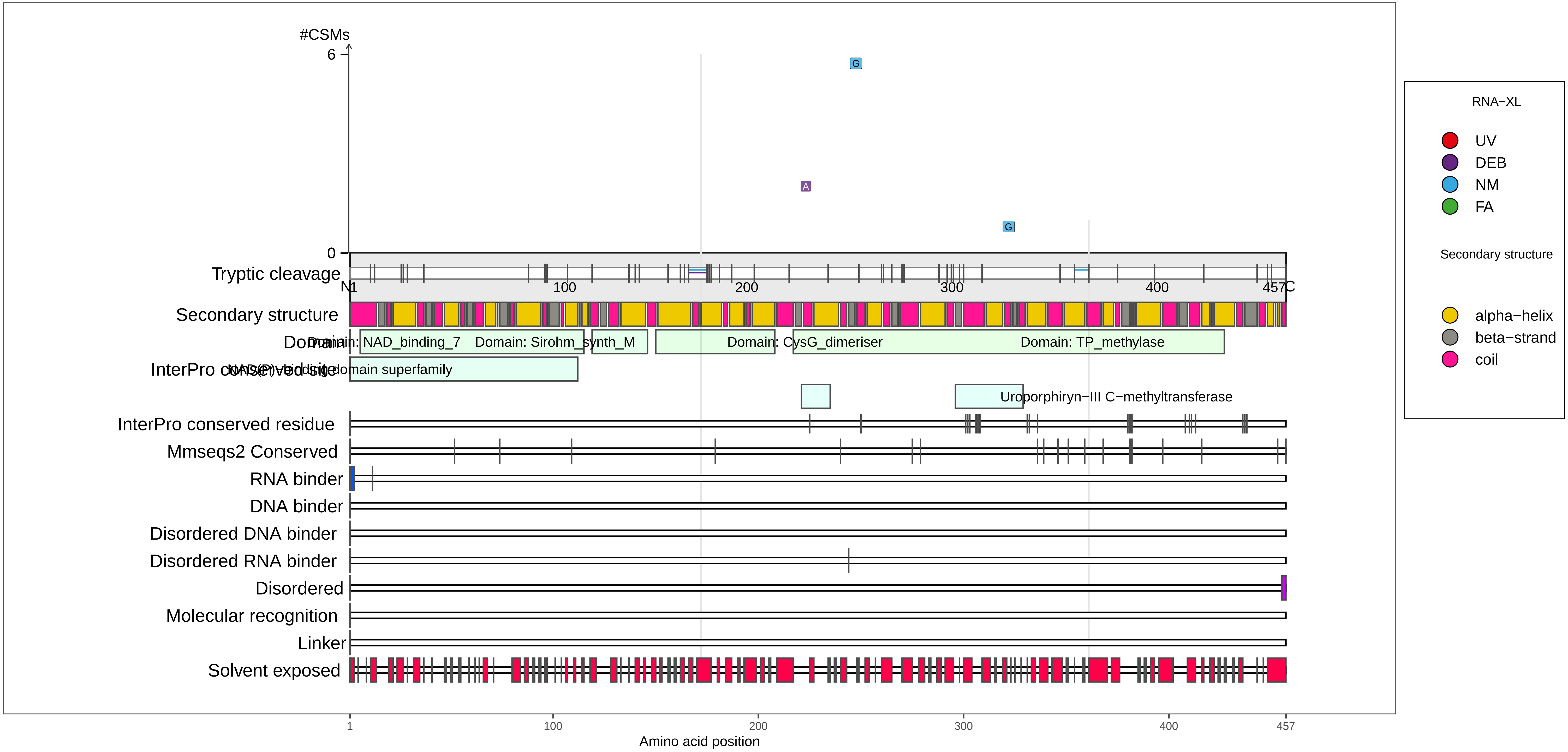

P0AFU8
RISA_ECOLI Riboflavin synthase

− Abundance:
tryptic [log10 Intensity]: 8.13 (Q 63)
PAXdb K12 strain [ppm]: 2.89 (Q 88)
PAXdb E.coli [ppm]: 2.32 (Q 81)

− RNA functions: not annotated

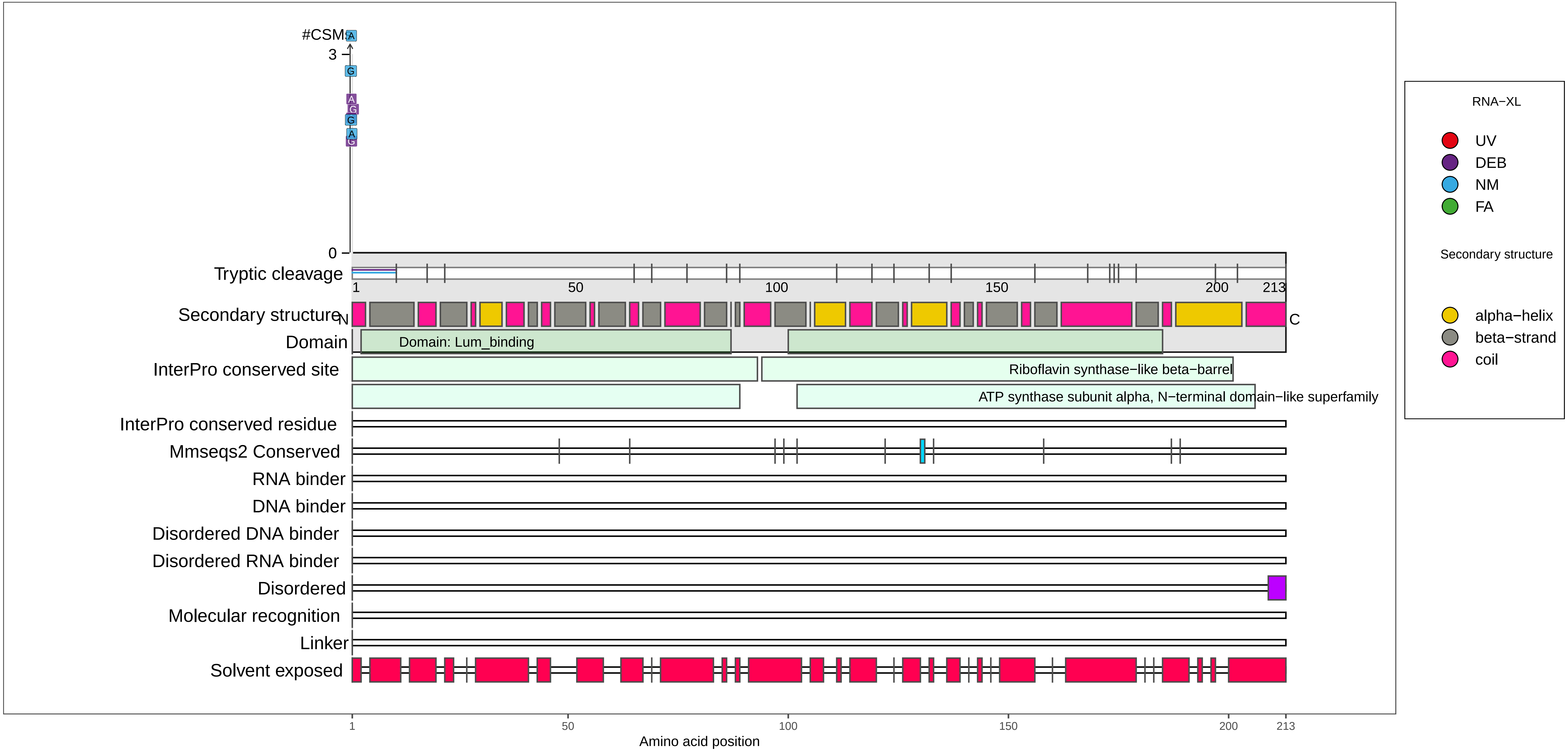

P0AGE0
SSB_ECOLI Single−stranded DNA−binding protein

− Abundance:
tryptic [log10 Intensity]: 9.32 (Q 94)
PAXdb K12 strain [ppm]: NA (Q NA)
PAXdb E.coli [ppm]: 2.7 (Q 90)

− RNA functions: not annotated

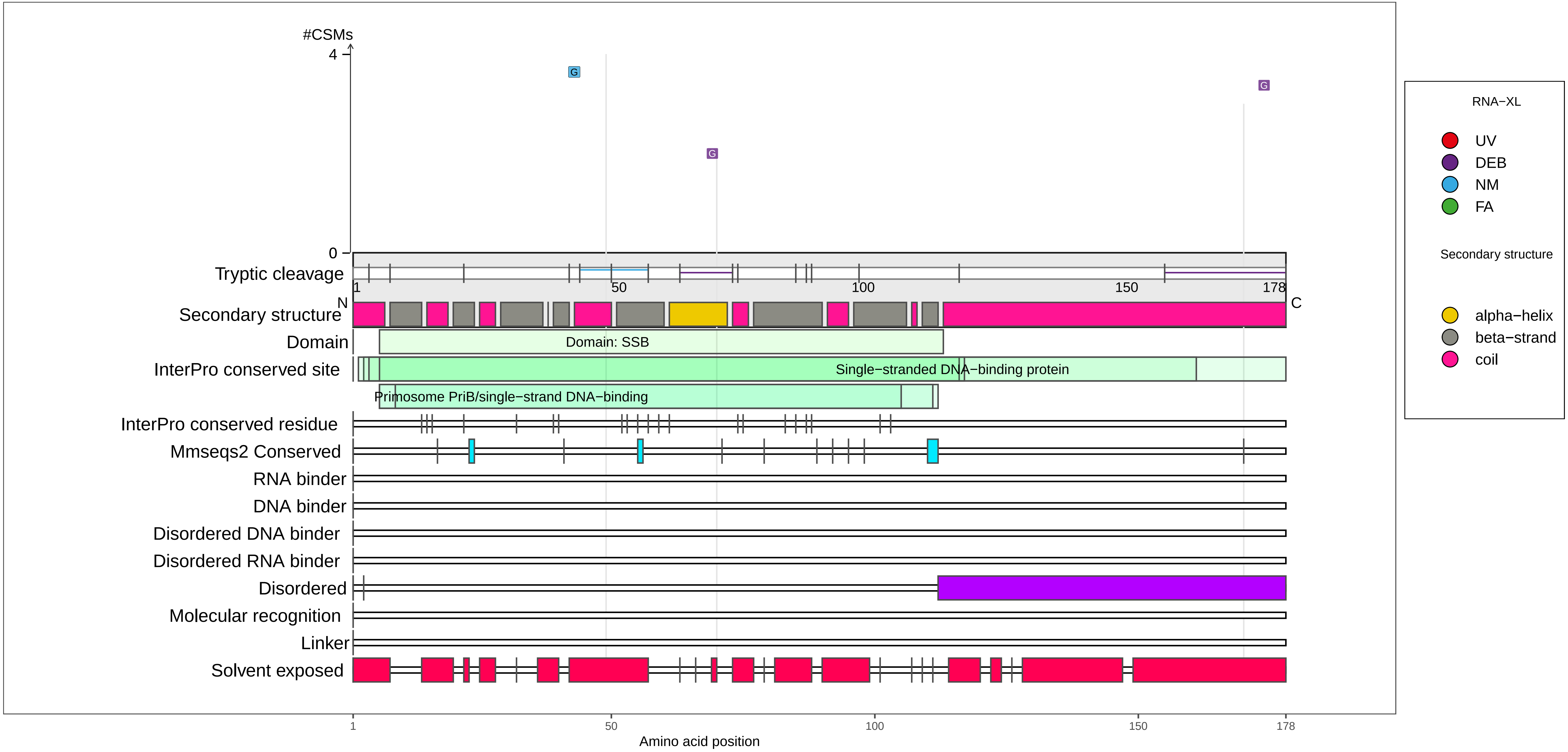

P0AGG8
TLDD_ECOLI Metalloprotease TldD

− Abundance:
tryptic [log10 Intensity]: 8.92 (Q 86)
PAXdb K12 strain [ppm]: 2.32 (Q 70)
PAXdb E.coli [ppm]: 2.44 (Q 84)

− RNA functions: not annotated

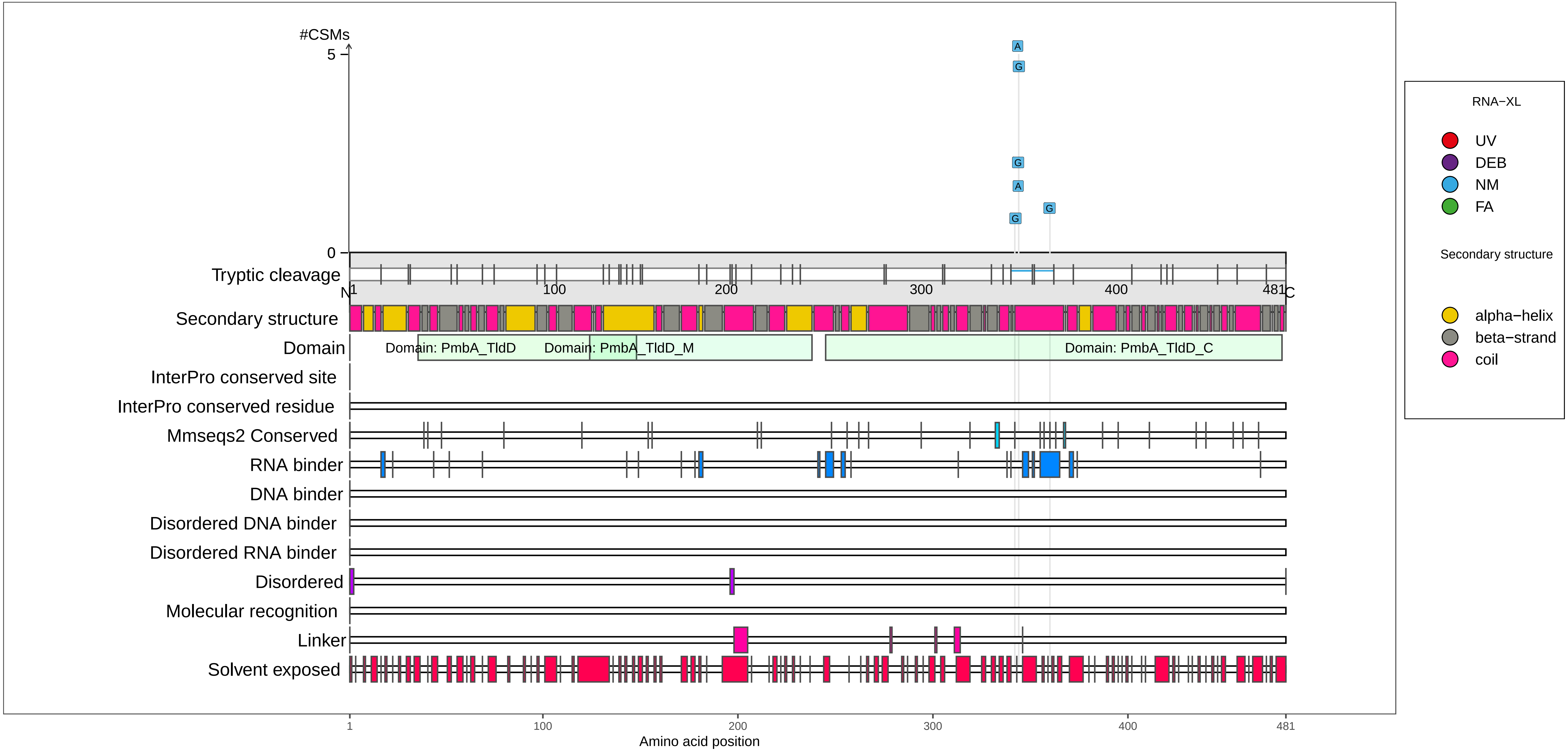

P0CG19
RNPH_ECOLI Truncated inactive ribonuclease PH
− RNA functions: not annotated

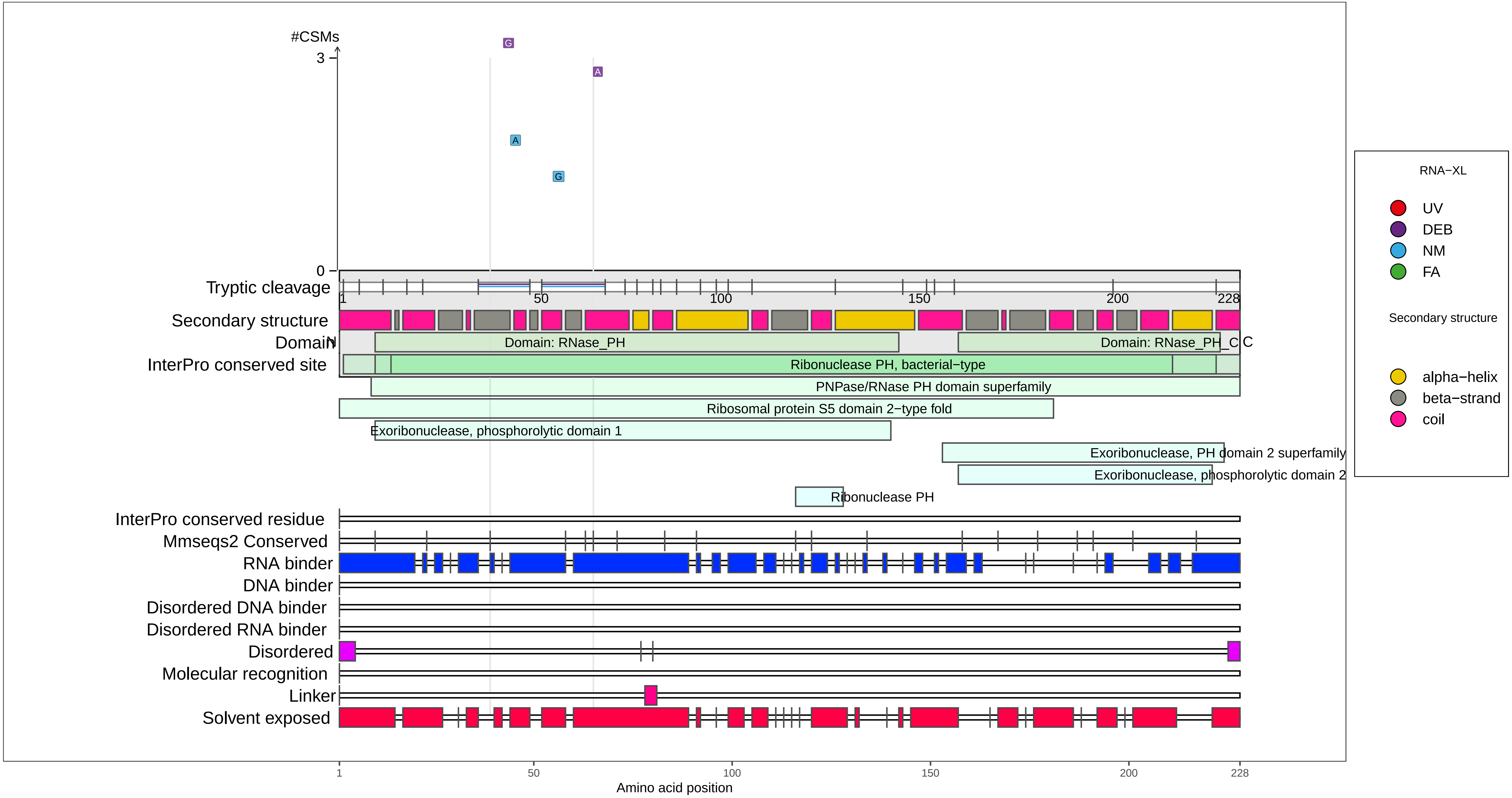

P15877
DHG_ECOLI Quinoprotein glucose dehydrogenase

− Abundance:
tryptic [log10 Intensity]: 8.81 (Q 84)
PAXdb K12 strain [ppm]: 2.5 (Q 77)
PAXdb E.coli [ppm]: 1.74 (Q 65)

− RNA functions: not annotated

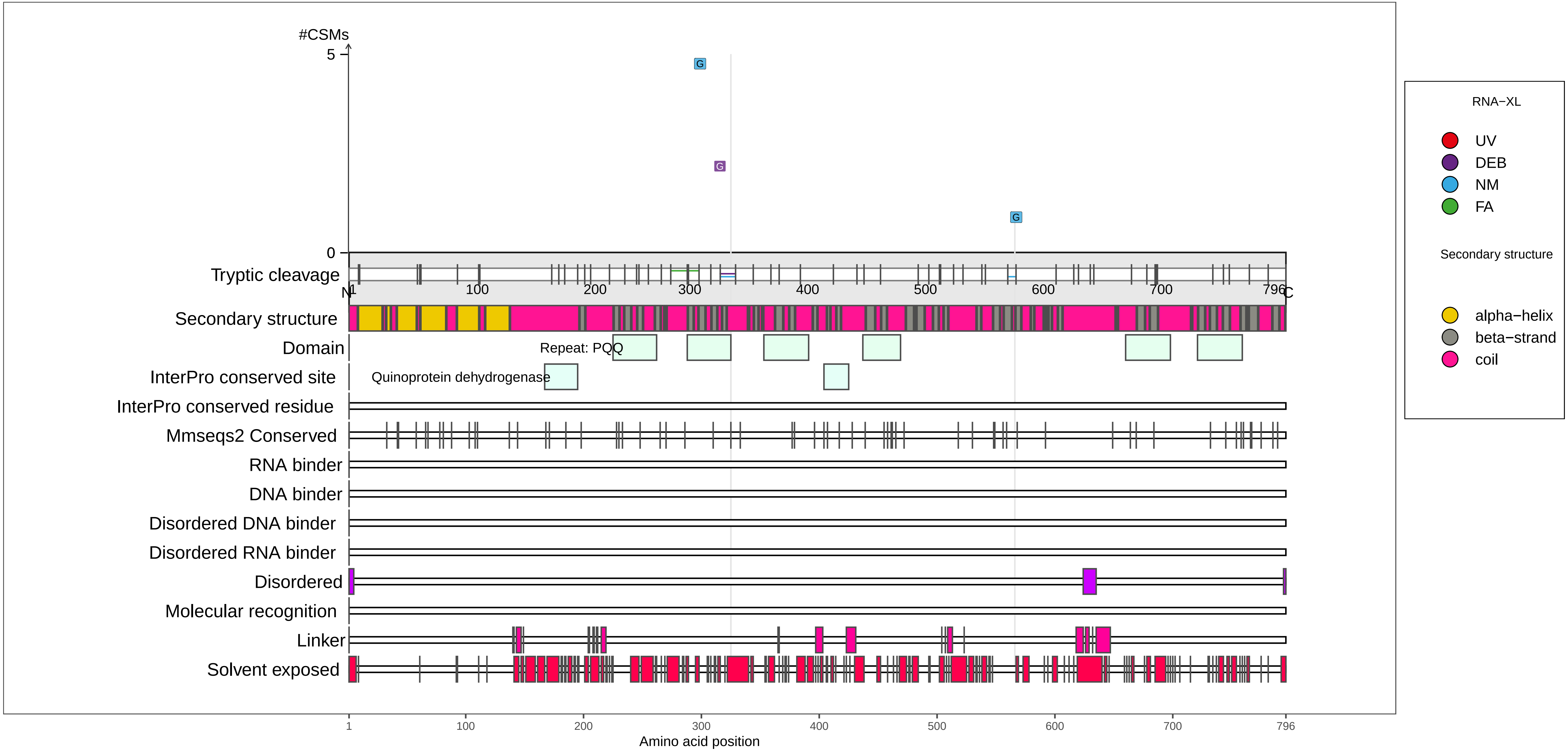

P21189
DPO2_ECOLI DNA polymerase II

− Abundance:
tryptic [log10 Intensity]: NA (Q NA)
PAXdb K12 strain [ppm]: 1.62 (Q 46)
PAXdb E.coli [ppm]: −0.64 (Q 14)

− RNA functions: not annotated

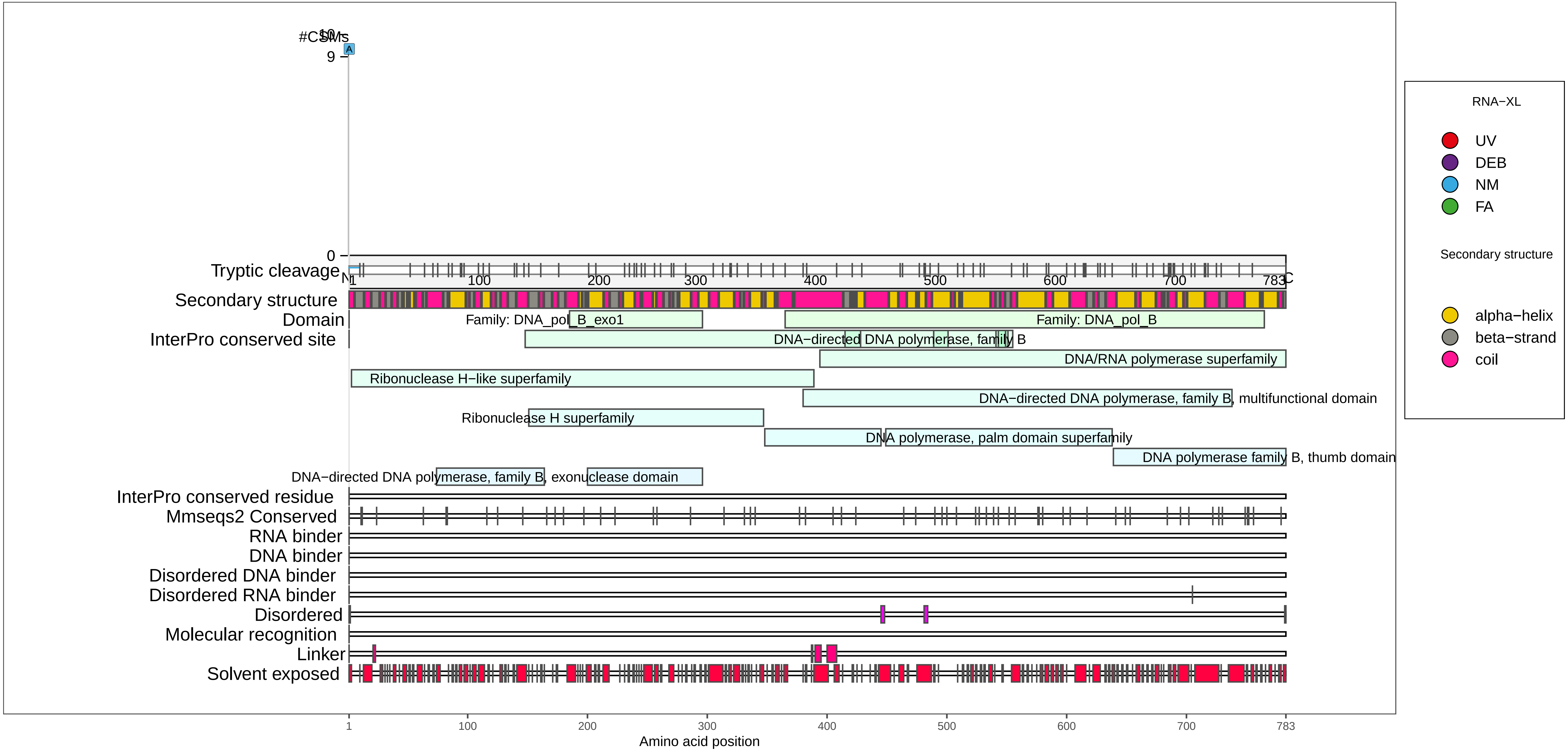

P22523
MUKB_ECOLI Chromosome partition protein MukB

− Abundance:
tryptic [log10 Intensity]: 7.24 (Q 22)
PAXdb K12 strain [ppm]: 2.1 (Q 64)
PAXdb E.coli [ppm]: 2.09 (Q 74)

− RNA functions: not annotated

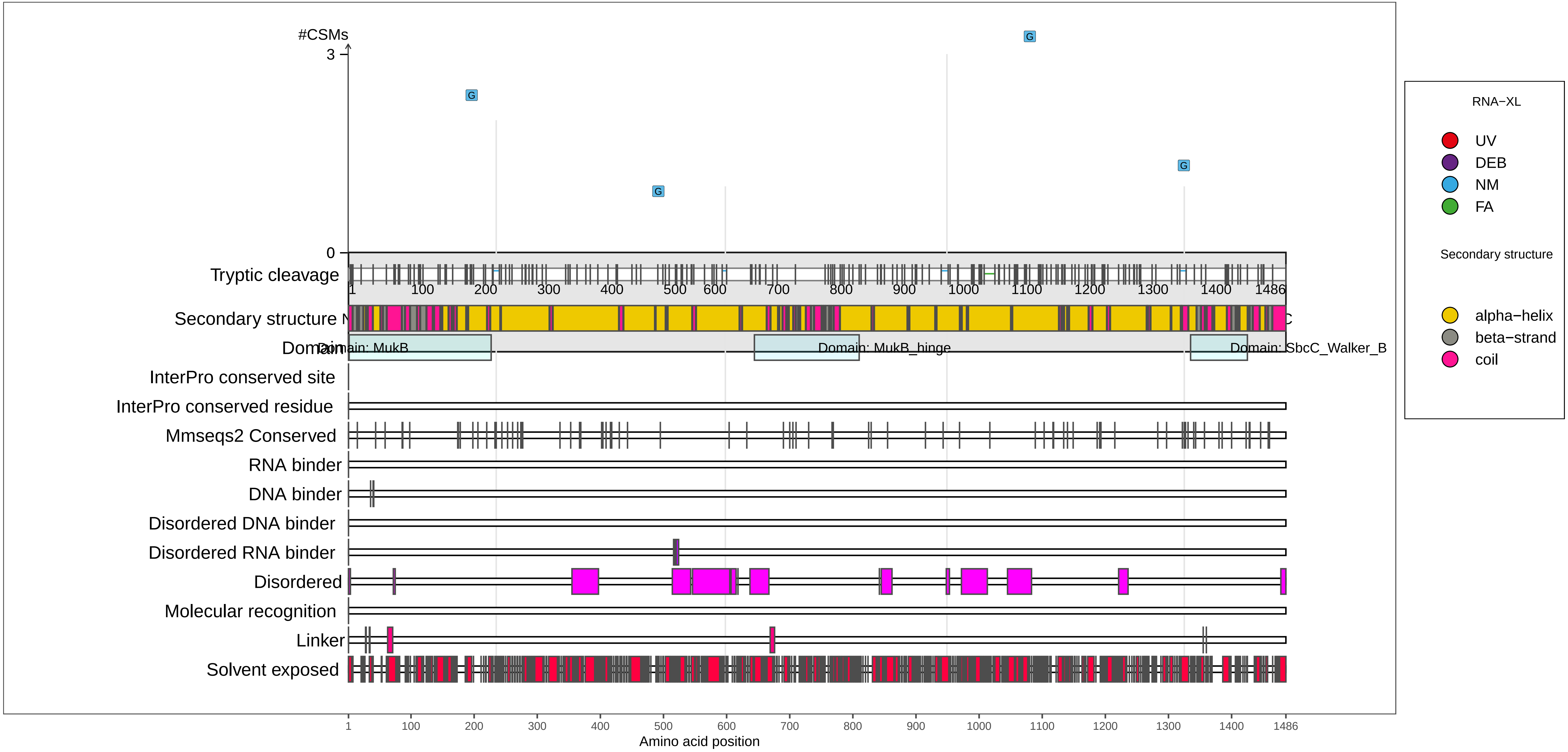

P24175
MANB_ECOLI Phosphomannomutase

− Abundance:
tryptic [log10 Intensity]: 7.26 (Q 23)
PAXdb K12 strain [ppm]: 1.26 (Q 24)
PAXdb E.coli [ppm]: 1.16 (Q 51)

− RNA functions: not annotated

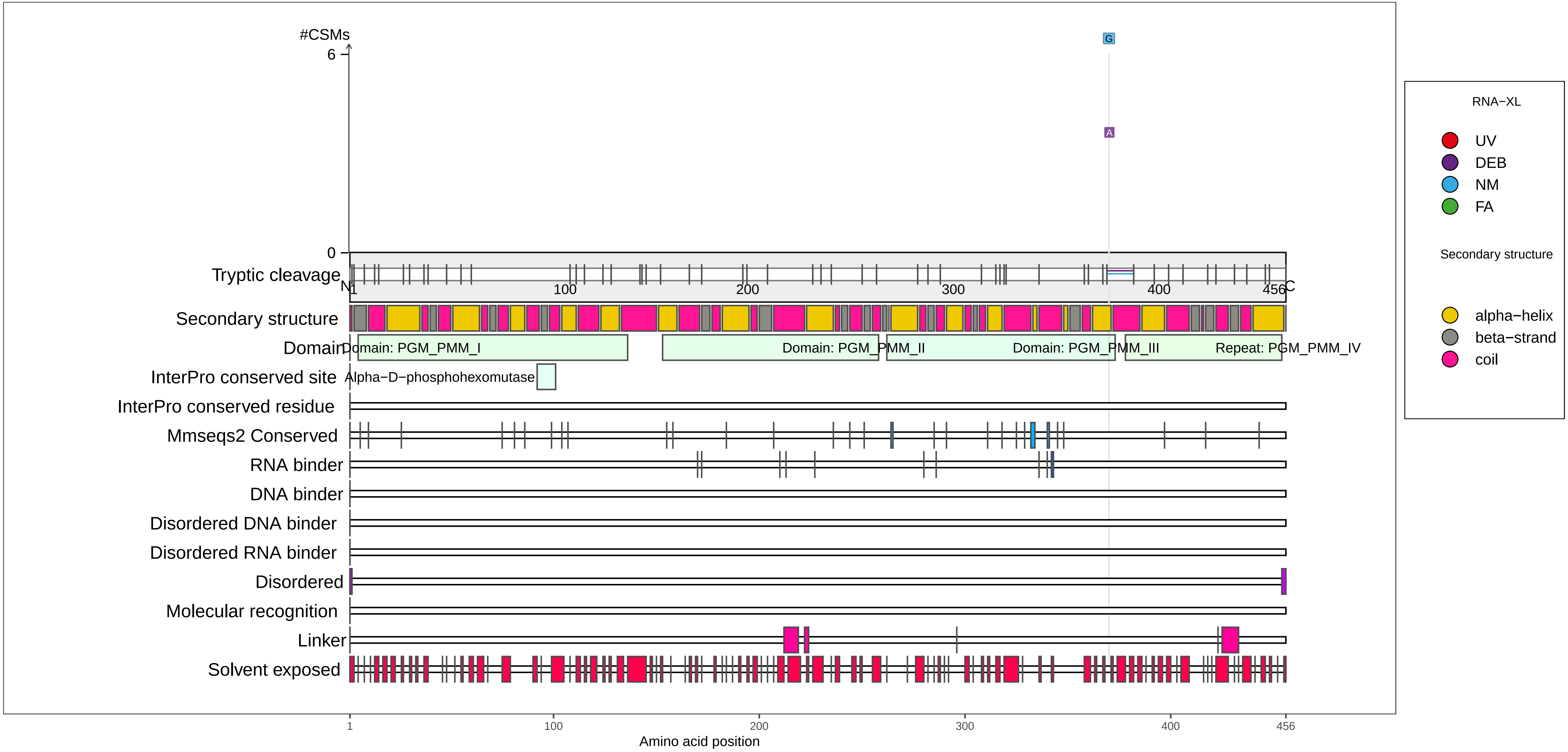

P25539
RIBD_ECOLI Riboflavin biosynthesis protein RibD

− Abundance:
tryptic [log10 Intensity]: 6.99 (Q 13)
PAXdb K12 strain [ppm]: 2.03 (Q 61)
PAXdb E.coli [ppm]: 1.71 (Q 65)

− RNA functions: not annotated

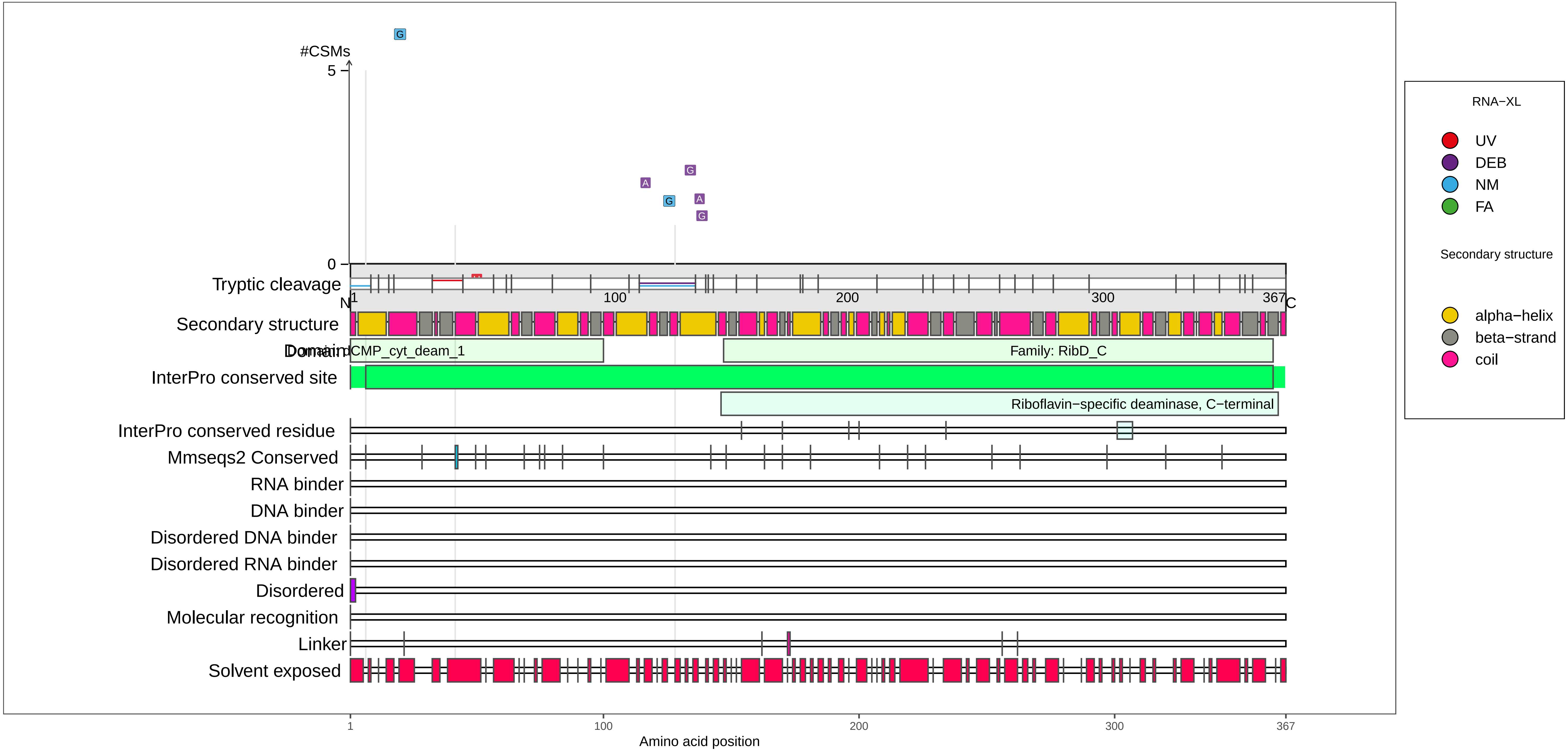

P33195
GCSP_ECOLI Glycine dehydrogenase (decarboxylating)

− Abundance:
tryptic [log10 Intensity]: 9.4 (Q 95)
PAXdb K12 strain [ppm]: 2.63 (Q 81)
PAXdb E.coli [ppm]: 2.51 (Q 86)

− RNA functions: not annotated

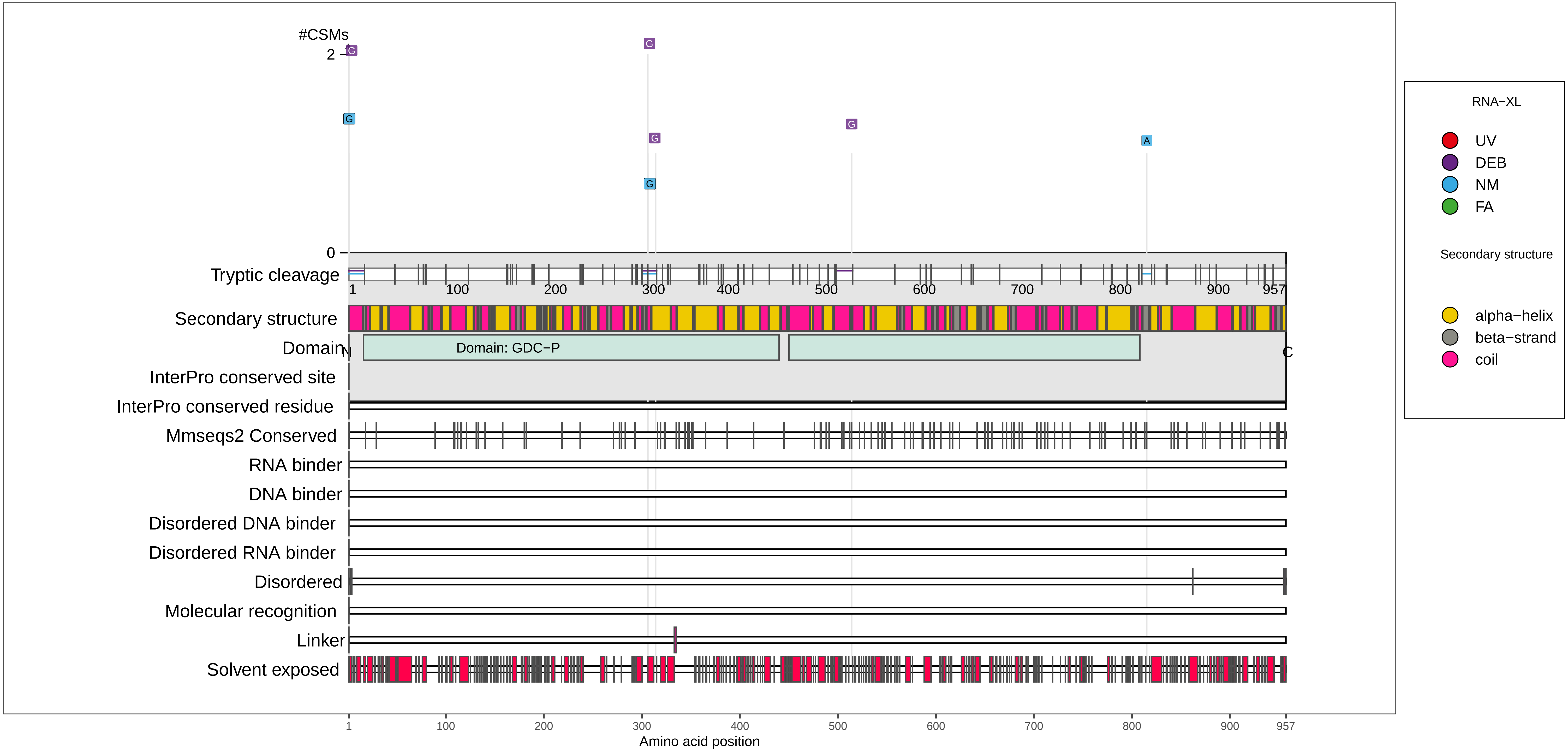

P36767
RDGC_ECOLI Recombination−associated protein RdgC

− Abundance:
tryptic [log10 Intensity]: 8.06 (Q 60)
PAXdb K12 strain [ppm]: 2.92 (Q 89)
PAXdb E.coli [ppm]: 2.04 (Q 73)

− RNA functions: not annotated

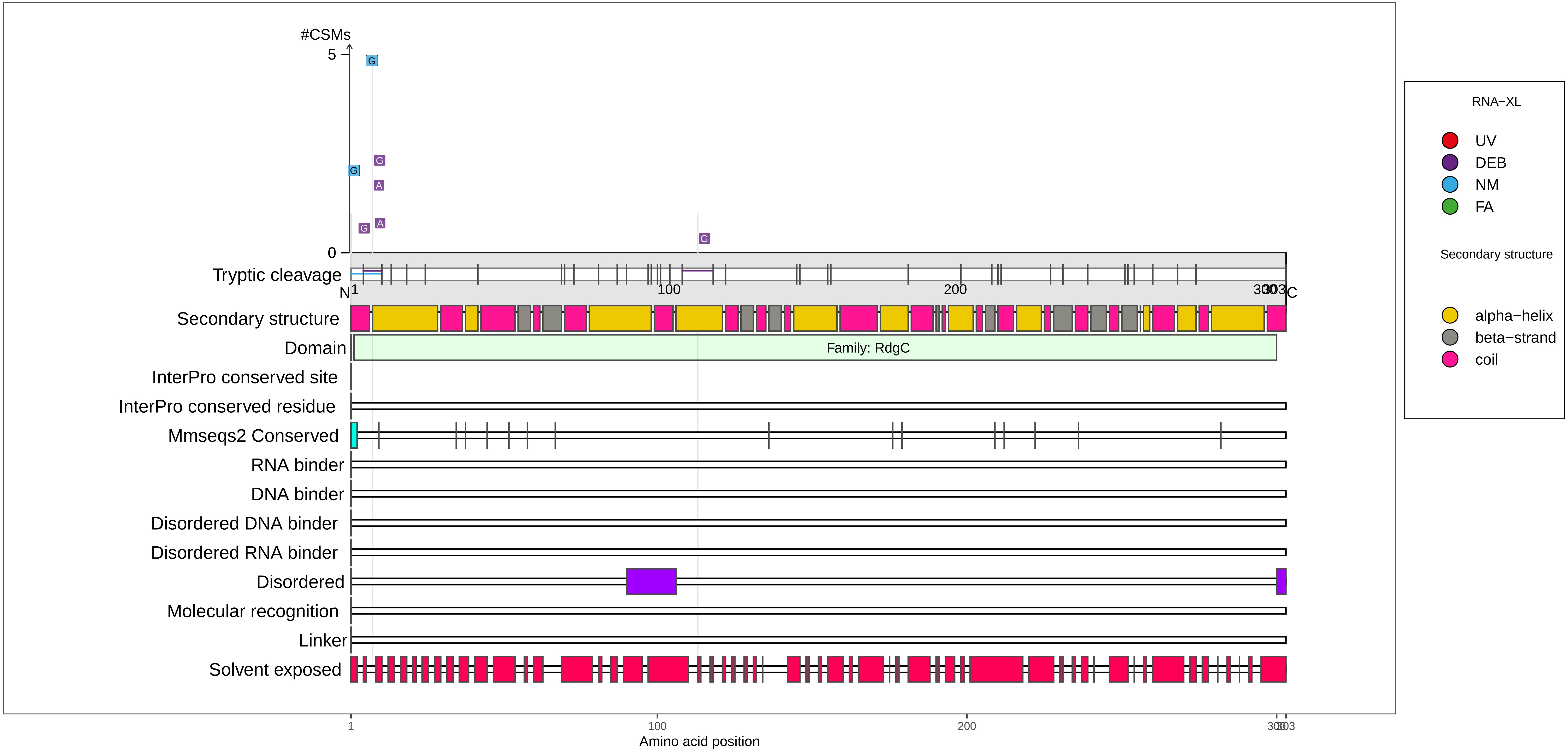

P46859
GNTK_ECOLI Thermoresistant gluconokinase

− Abundance:
tryptic [log10 Intensity]: 7.6 (Q 40)
PAXdb K12 strain [ppm]: 2.39 (Q 74)
PAXdb E.coli [ppm]: 0.91 (Q 45)

− RNA functions: not annotated

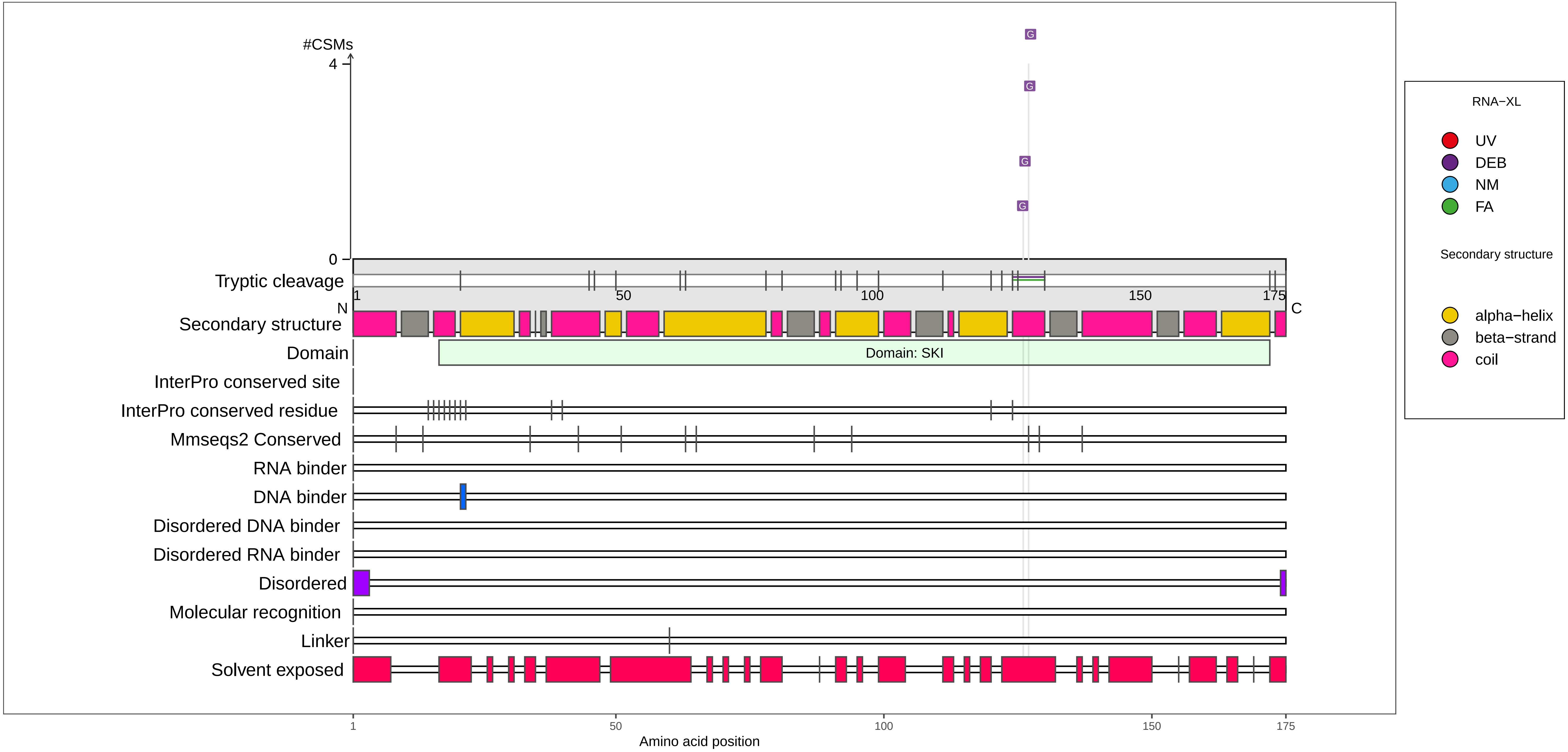

P52647
NIFJ_ECOLI Probable pyruvate−flavodoxin oxidoreductase

− Abundance:
tryptic [log10 Intensity]: 7.78 (Q 48)
PAXdb K12 strain [ppm]: 2.22 (Q 68)
PAXdb E.coli [ppm]: 1.72 (Q 65)

− RNA functions: not annotated

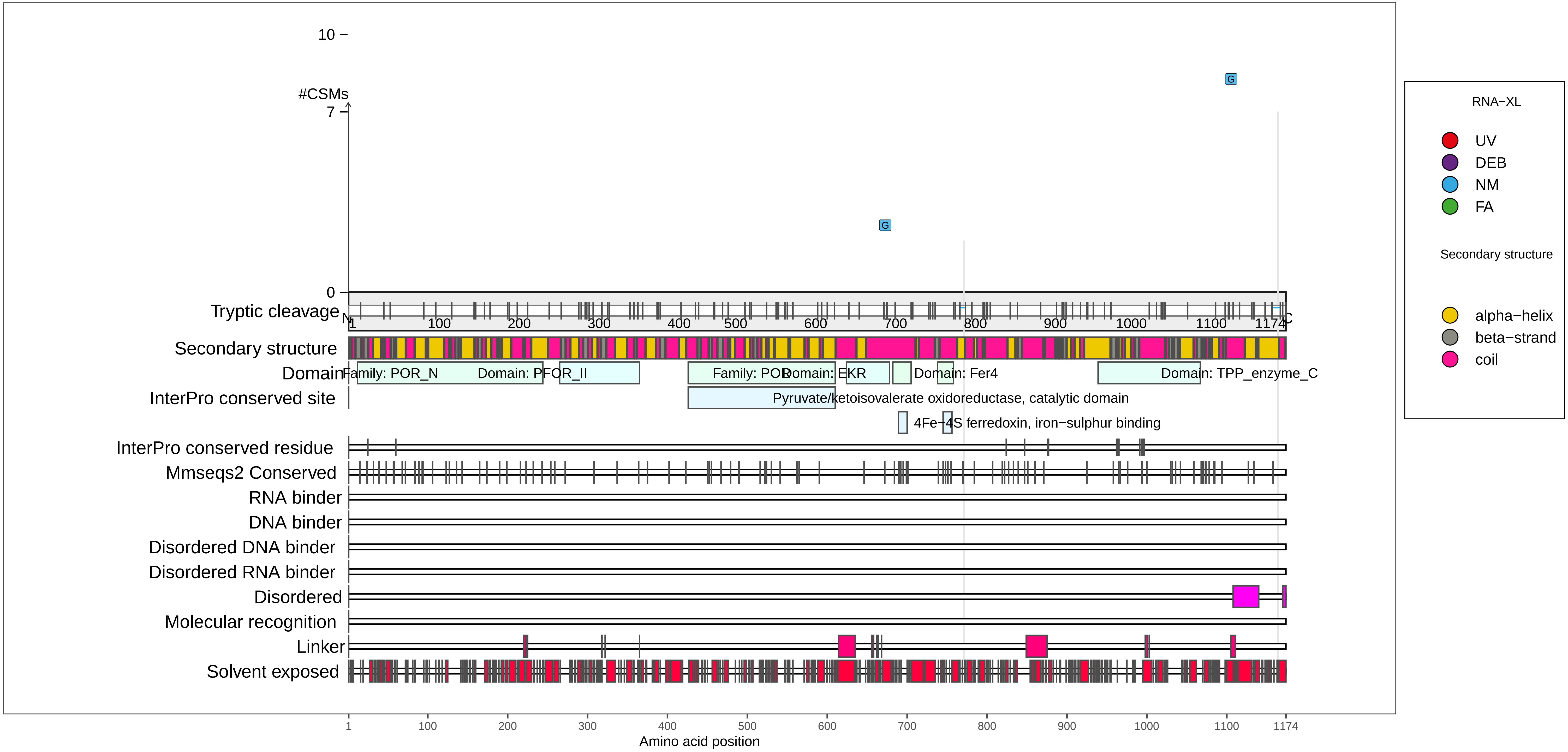

P69490
CCME_ECOLI Cytochrome c−type biogenesis protein CcmE

− Abundance:
tryptic [log10 Intensity]: 7.96 (Q 57)
PAXdb K12 strain [ppm]: 1.26 (Q 23)
PAXdb E.coli [ppm]: 0.18 (Q 29)

− RNA functions: not annotated

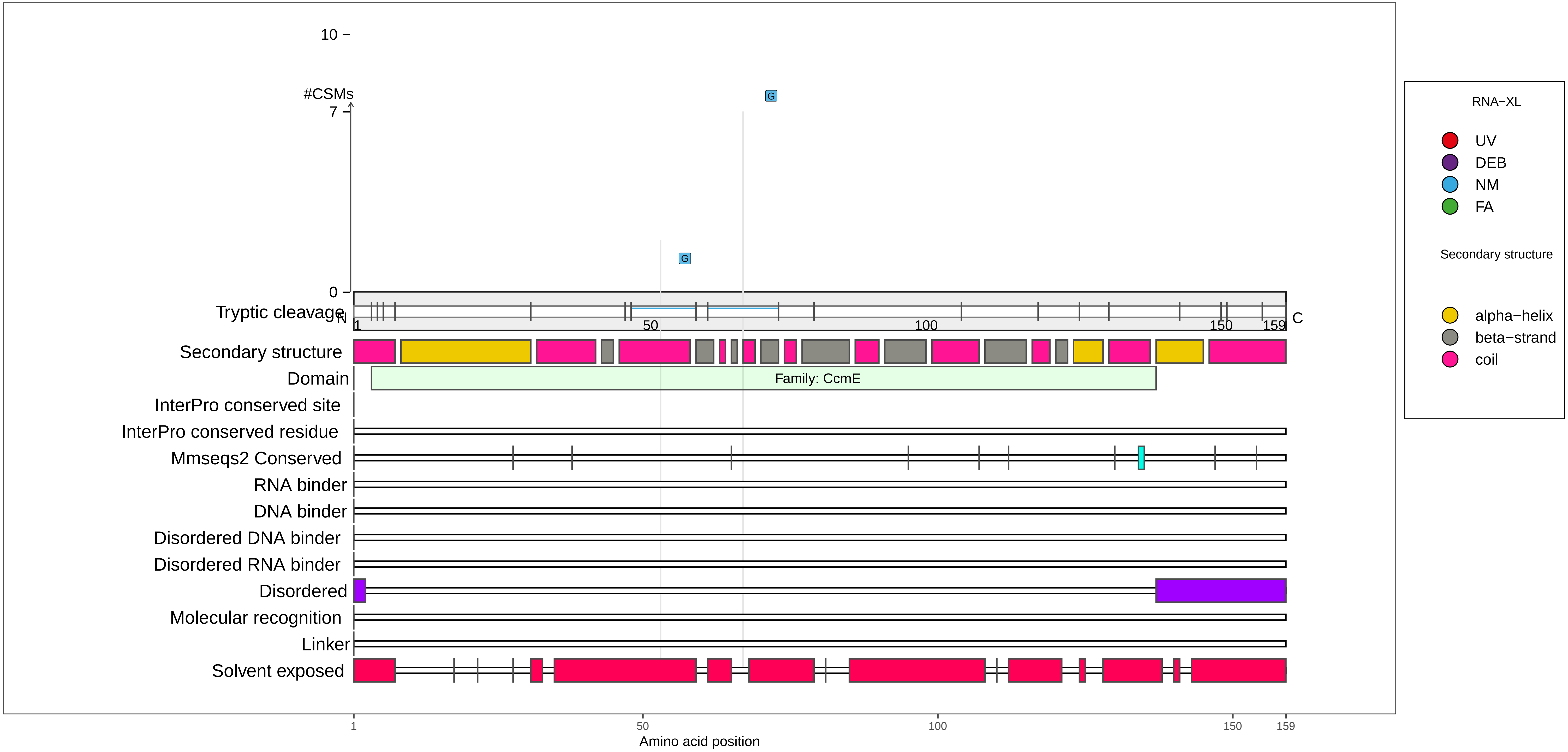

P69783
PTGA_ECOLI PTS system glucose−specific EIIA component

− Abundance:
tryptic [log10 Intensity]: 9.17 (Q 92)
PAXdb K12 strain [ppm]: 3.32 (Q 95)
PAXdb E.coli [ppm]: 3.5 (Q 99)

− RNA functions: not annotated

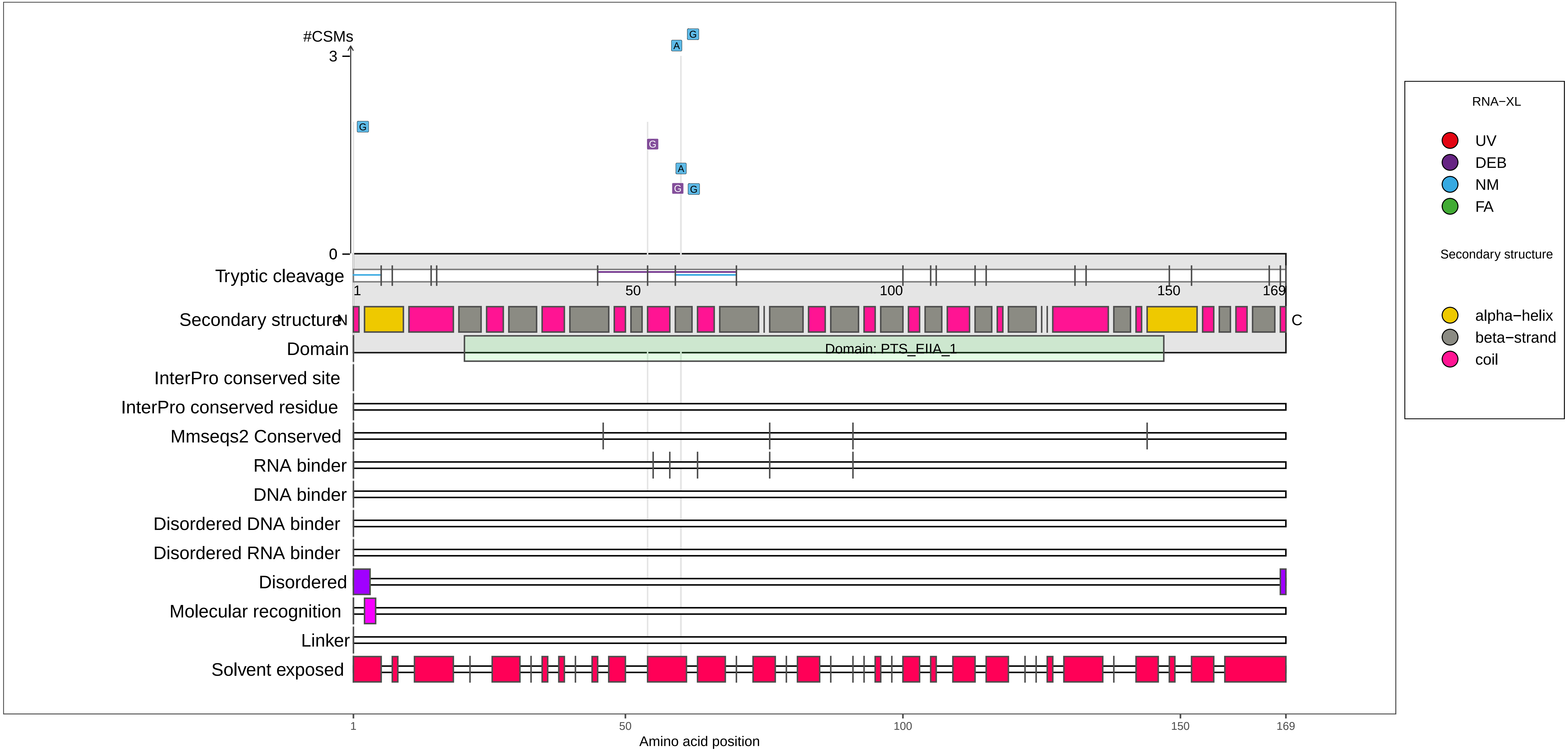

P69805
PTND_ECOLI PTS system mannose−specific EIID component

− Abundance:
tryptic [log10 Intensity]: 8.87 (Q 85)
PAXdb K12 strain [ppm]: 1.51 (Q 39)
PAXdb E.coli [ppm]: 2.58 (Q 88)

− RNA functions: not annotated

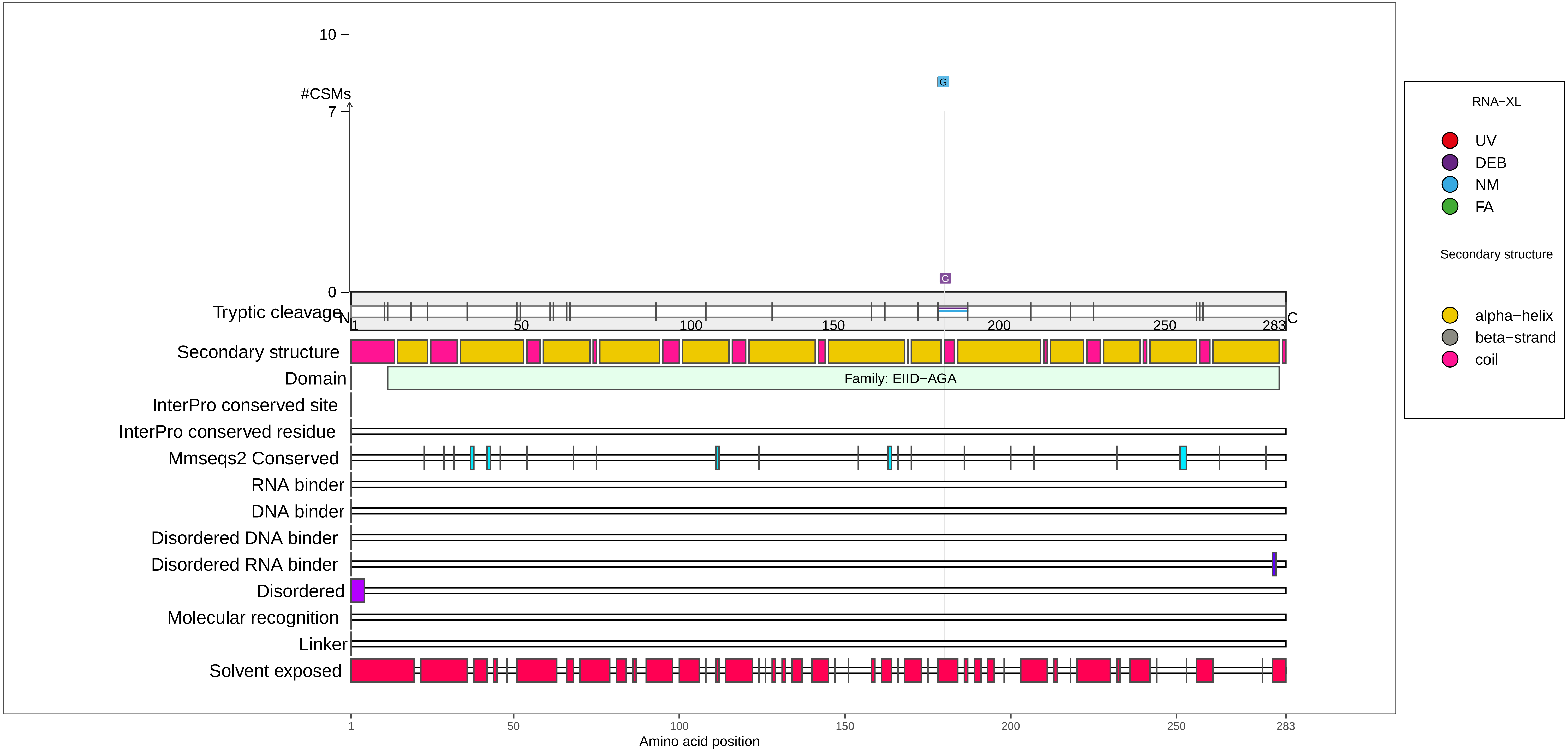

P76341
HIUH_ECOLI 5−hydroxyisourate hydrolase

− Abundance:
tryptic [log10 Intensity]: 7.33 (Q 26)
PAXdb K12 strain [ppm]: NA (Q NA)
PAXdb E.coli [ppm]: 0.95 (Q 47)

− RNA functions: not annotated

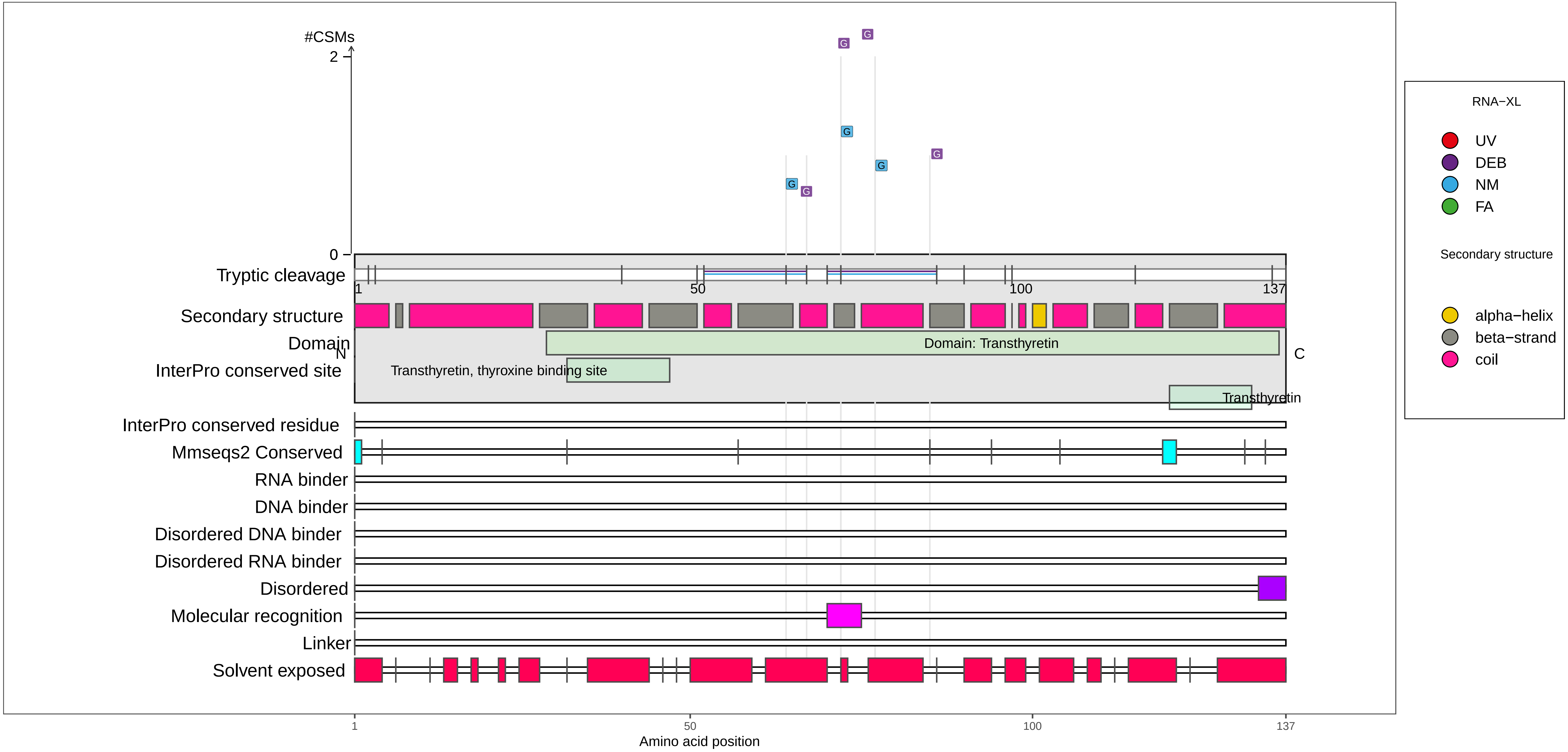

P76558
MAO2_ECOLI NADP−dependent malic enzyme

− Abundance:
tryptic [log10 Intensity]: 8.49 (Q 75)
PAXdb K12 strain [ppm]: 2.55 (Q 79)
PAXdb E.coli [ppm]: 2.83 (Q 92)

− RNA functions: not annotated

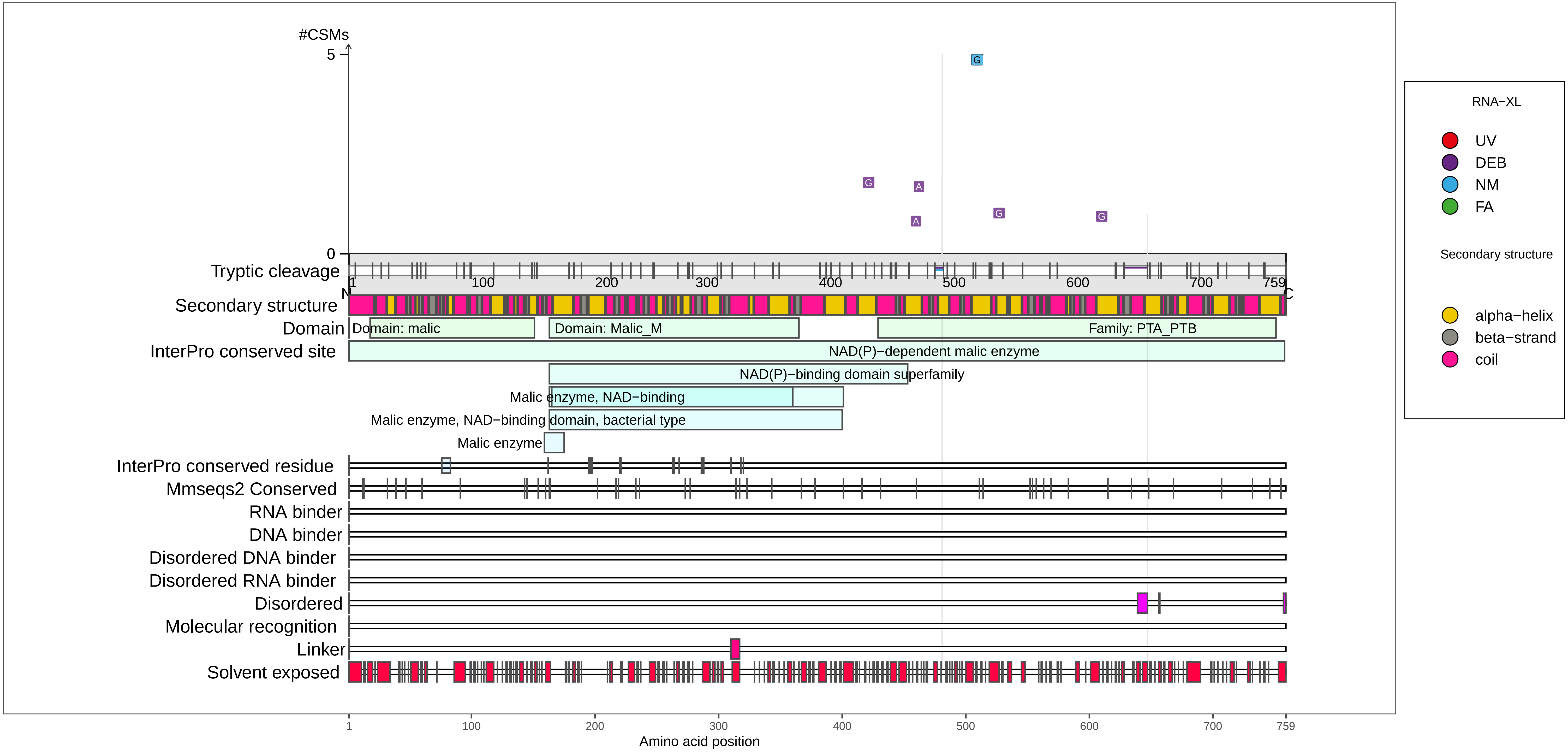

P77781
MENI_ECOLI 1,4−dihydroxy−2−naphthoyl−CoA hydrolase

− Abundance:
tryptic [log10 Intensity]: 7.52 (Q 36)
PAXdb K12 strain [ppm]: 1.75 (Q 51)
PAXdb E.coli [ppm]: 1.51 (Q 60)

− RNA functions: not annotated

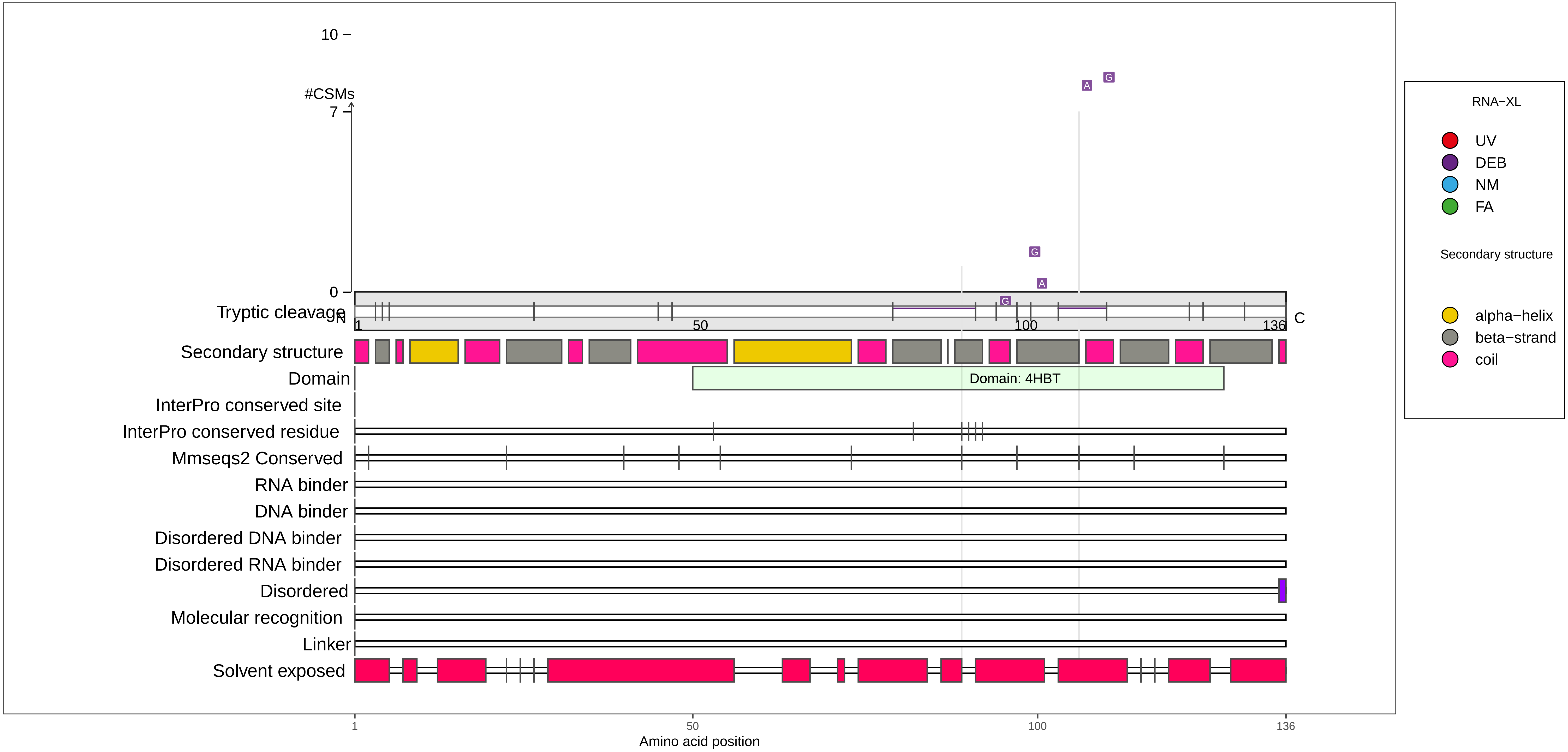

Q46851
GPR_ECOLI L−glyceraldehyde 3−phosphate reductase

− Abundance:
tryptic [log10 Intensity]: 8.07 (Q 60)
PAXdb K12 strain [ppm]: 2.54 (Q 78)
PAXdb E.coli [ppm]: 2.56 (Q 87)

− RNA functions: not annotated

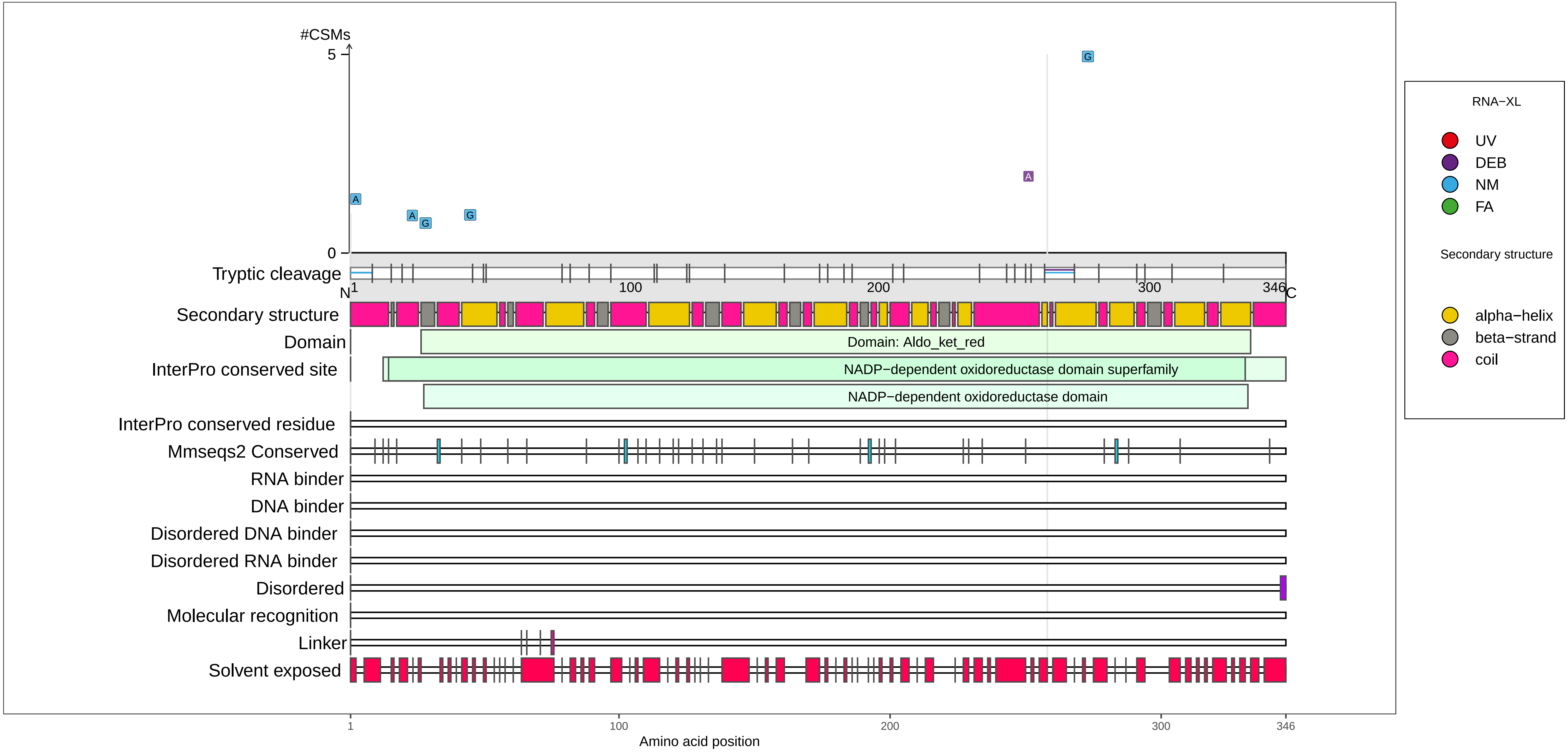

P04036
DAPB_ECOLI 4−hydroxy−tetrahydrodipicolinate reductase

− Abundance:
tryptic [log10 Intensity]: 7.83 (Q 50)
PAXdb K12 strain [ppm]: 1.96 (Q 59)
PAXdb E.coli [ppm]: 2.54 (Q 86)

− RNA functions: not annotated

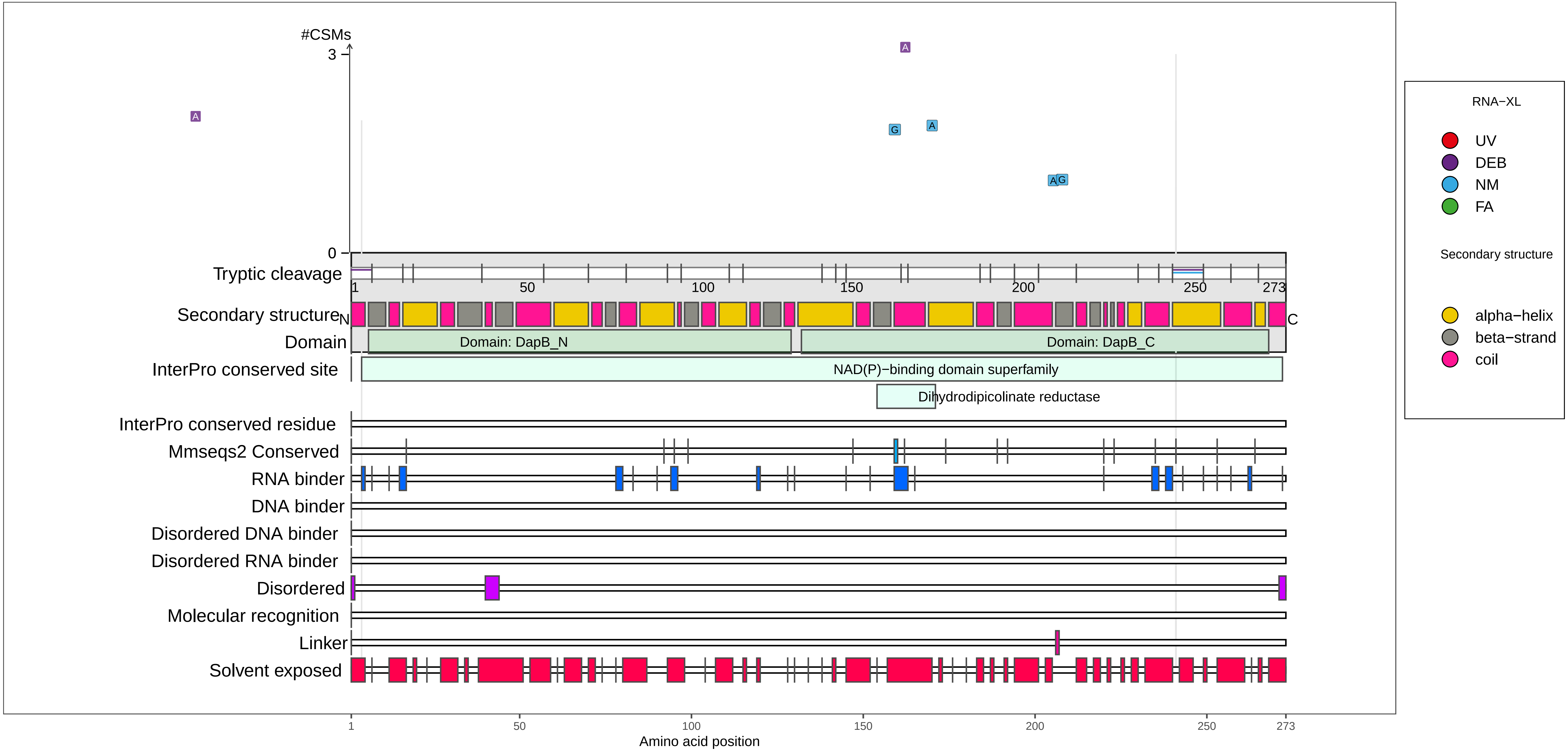

P06616
ERA_ECOLI GTPase Era

− Abundance:
tryptic [log10 Intensity]: 6.93 (Q 11)
PAXdb K12 strain [ppm]: 2.4 (Q 74)
PAXdb E.coli [ppm]: 1.59 (Q 62)

− RNA functions:
RNA binding; rRNA binding

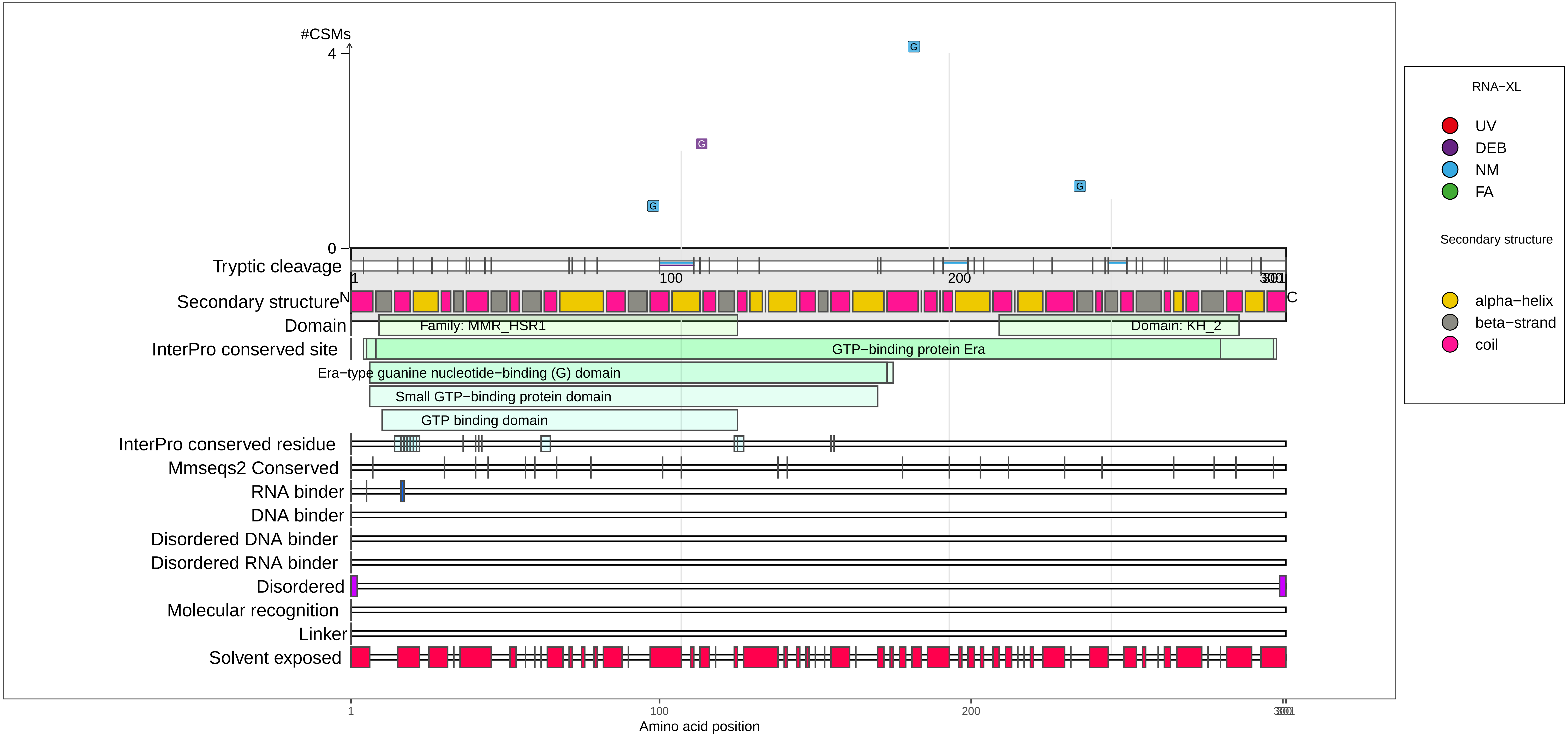

P06983
HEM3_ECOLI Porphobilinogen deaminase

− Abundance:
tryptic [log10 Intensity]: 7.66 (Q 43)
PAXdb K12 strain [ppm]: 1.87 (Q 56)
PAXdb E.coli [ppm]: 1.19 (Q 52)

− RNA functions: not annotated

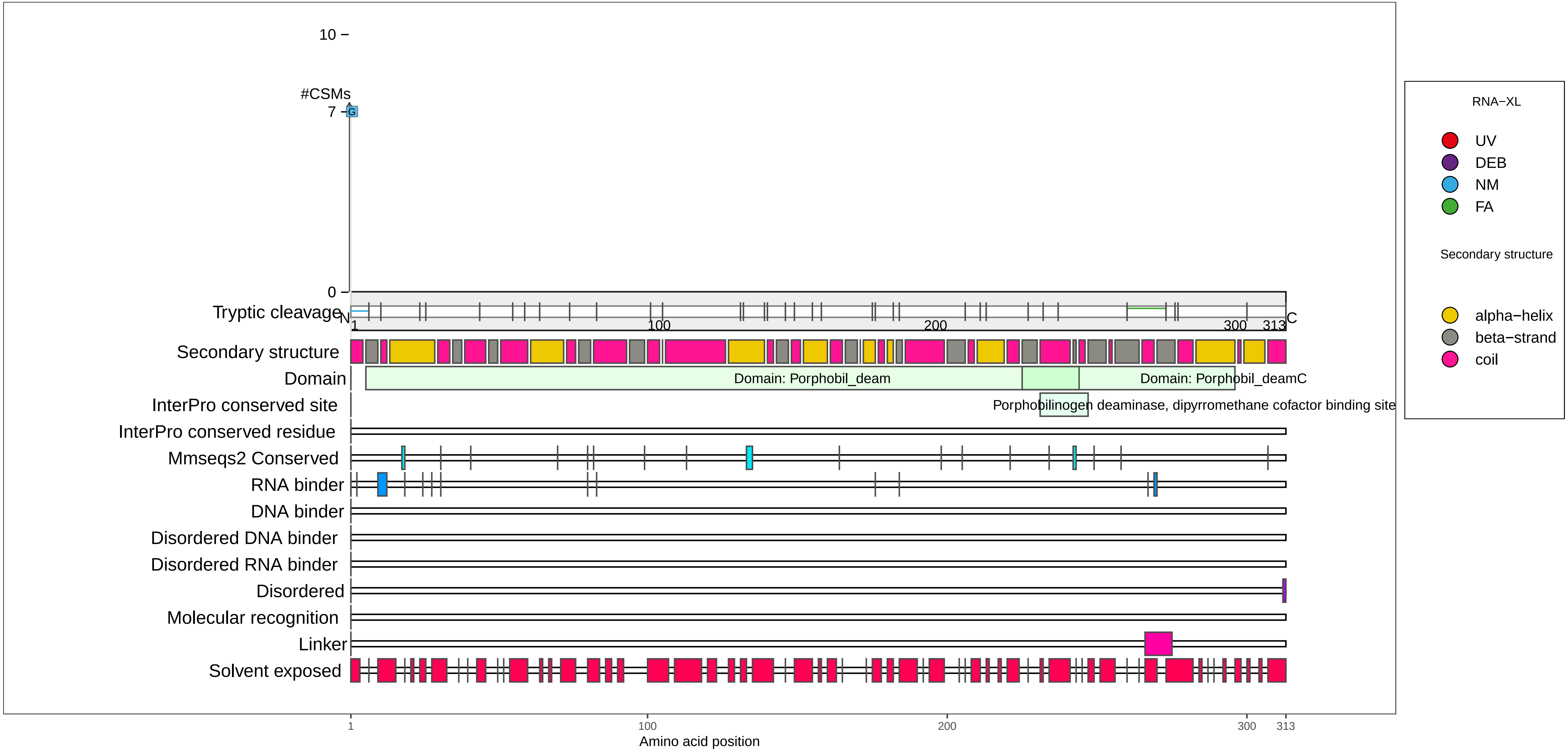

P0A6K1
DAPF_ECOLI Diaminopimelate epimerase

− Abundance:
tryptic [log10 Intensity]: 7.38 (Q 29)
PAXdb K12 strain [ppm]: 2.11 (Q 64)
PAXdb E.coli [ppm]: 2.11 (Q 75)

− RNA functions: not annotated

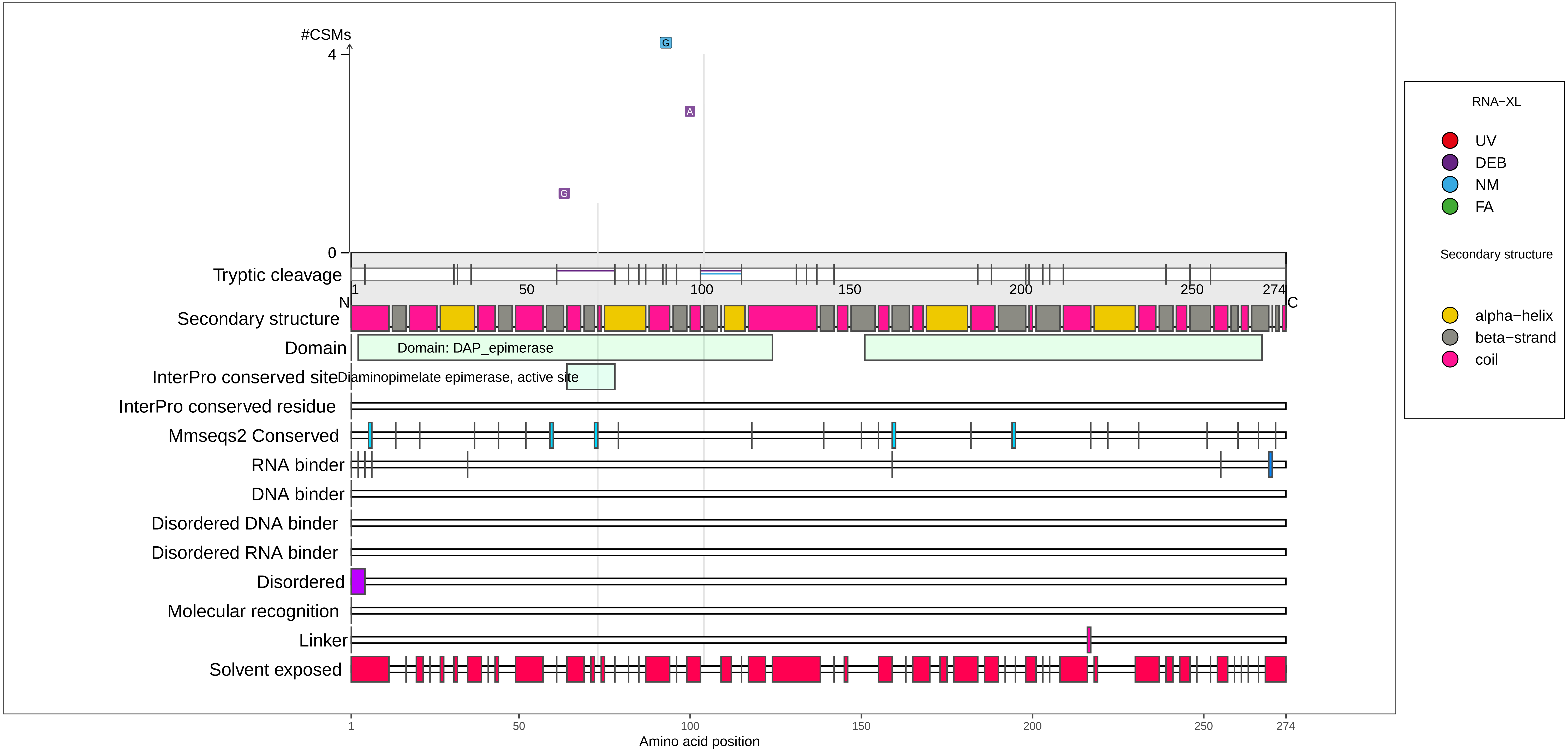

P0A7F6
SPED_ECOLI S−adenosylmethionine decarboxylase proenzyme

− Abundance:
tryptic [log10 Intensity]: 8.66 (Q 80)
PAXdb K12 strain [ppm]: 2.08 (Q 63)
PAXdb E.coli [ppm]: 2.18 (Q 77)

− RNA functions: not annotated

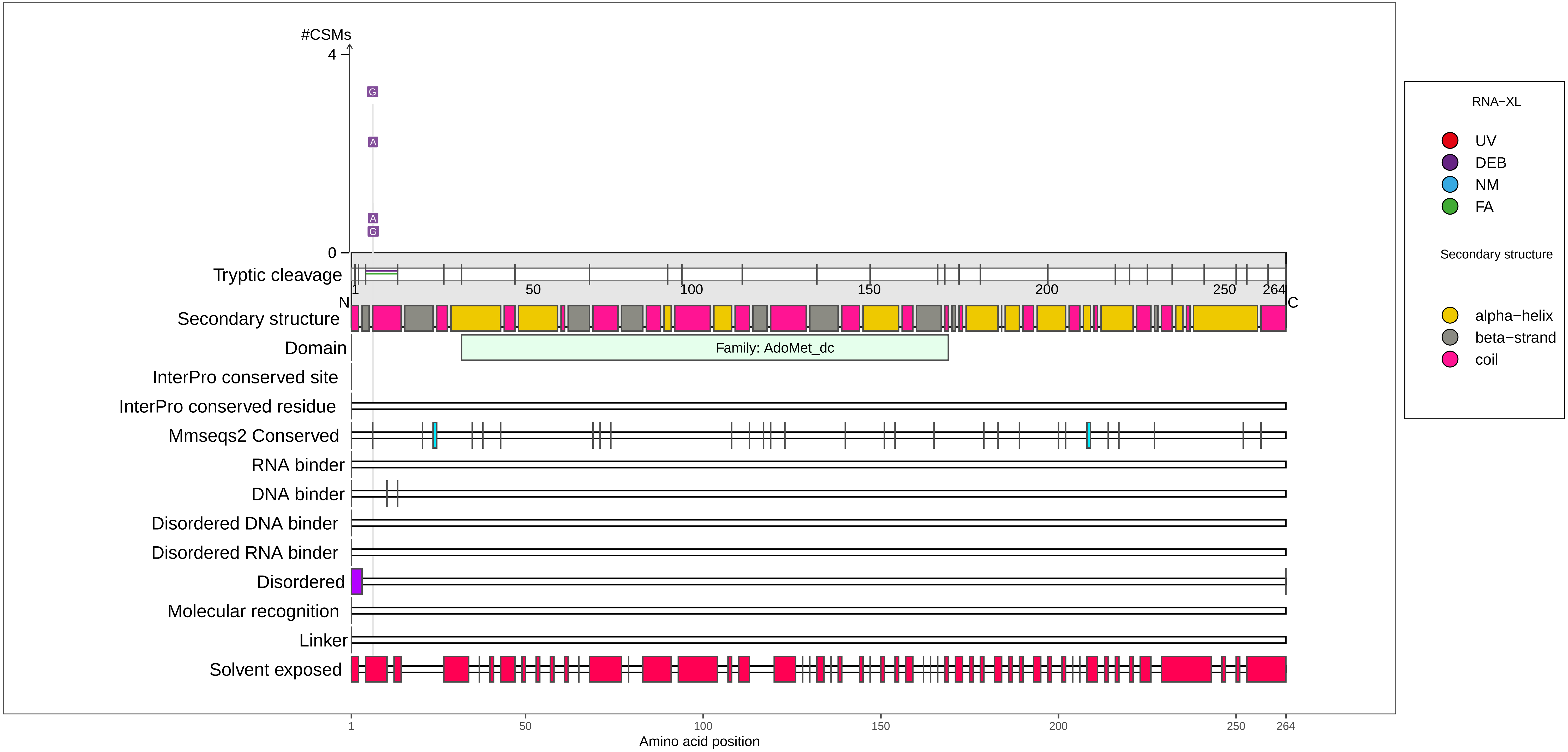

P0A8A8
RIMP_ECOLI Ribosome maturation factor RimP

− Abundance:
tryptic [log10 Intensity]: 9.48 (Q 96)
PAXdb K12 strain [ppm]: 2.21 (Q 67)
PAXdb E.coli [ppm]: 2.09 (Q 75)

− RNA functions: not annotated

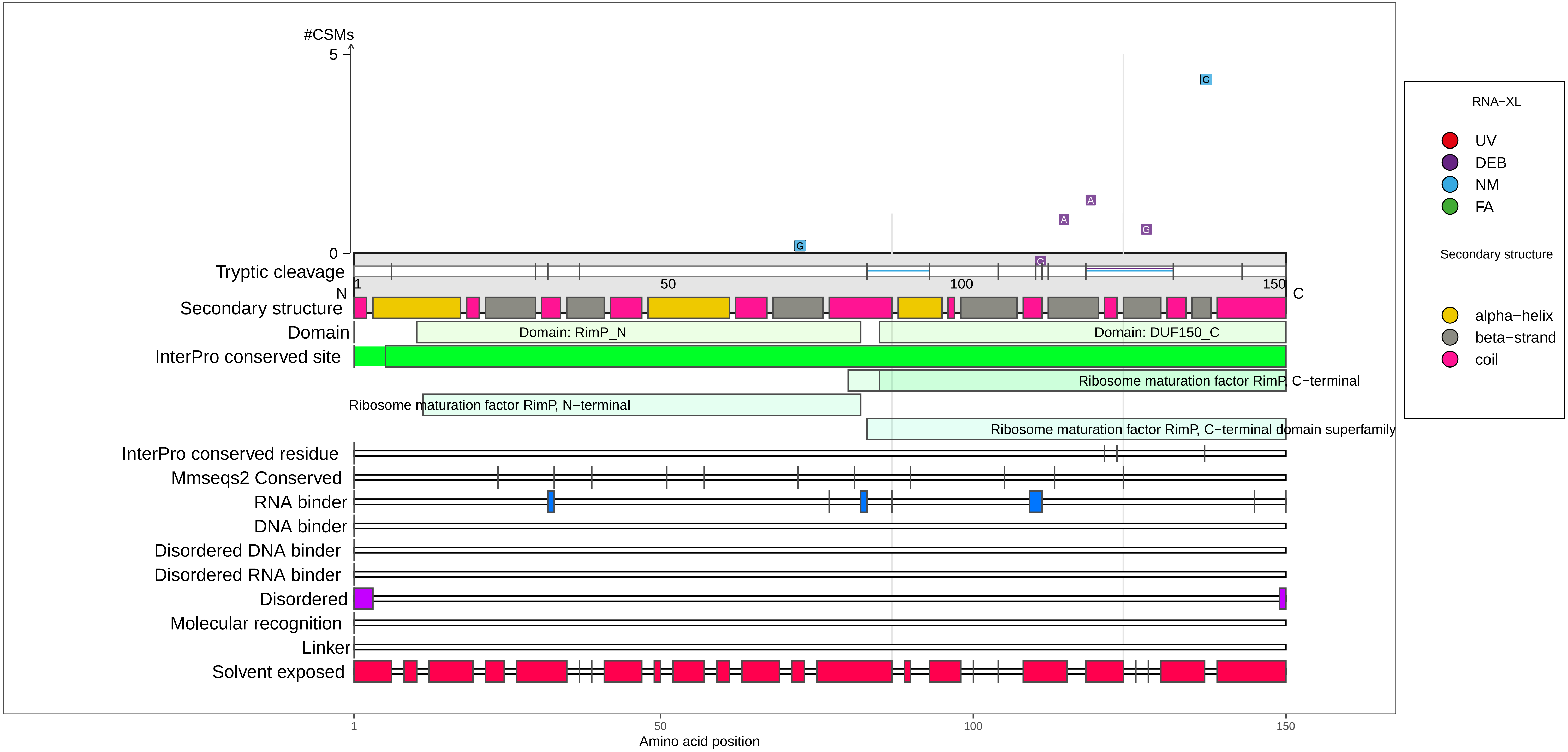

P0A9Q5
ACCD_ECOLI Acetyl−coenzyme A carboxylase carboxyl transferase subunit beta

− Abundance:
tryptic [log10 Intensity]: 8.11 (Q 62)
PAXdb K12 strain [ppm]: 3.09 (Q 92)
PAXdb E.coli [ppm]: 2.58 (Q 87)

− RNA functions:
mRNA binding; RNA binding

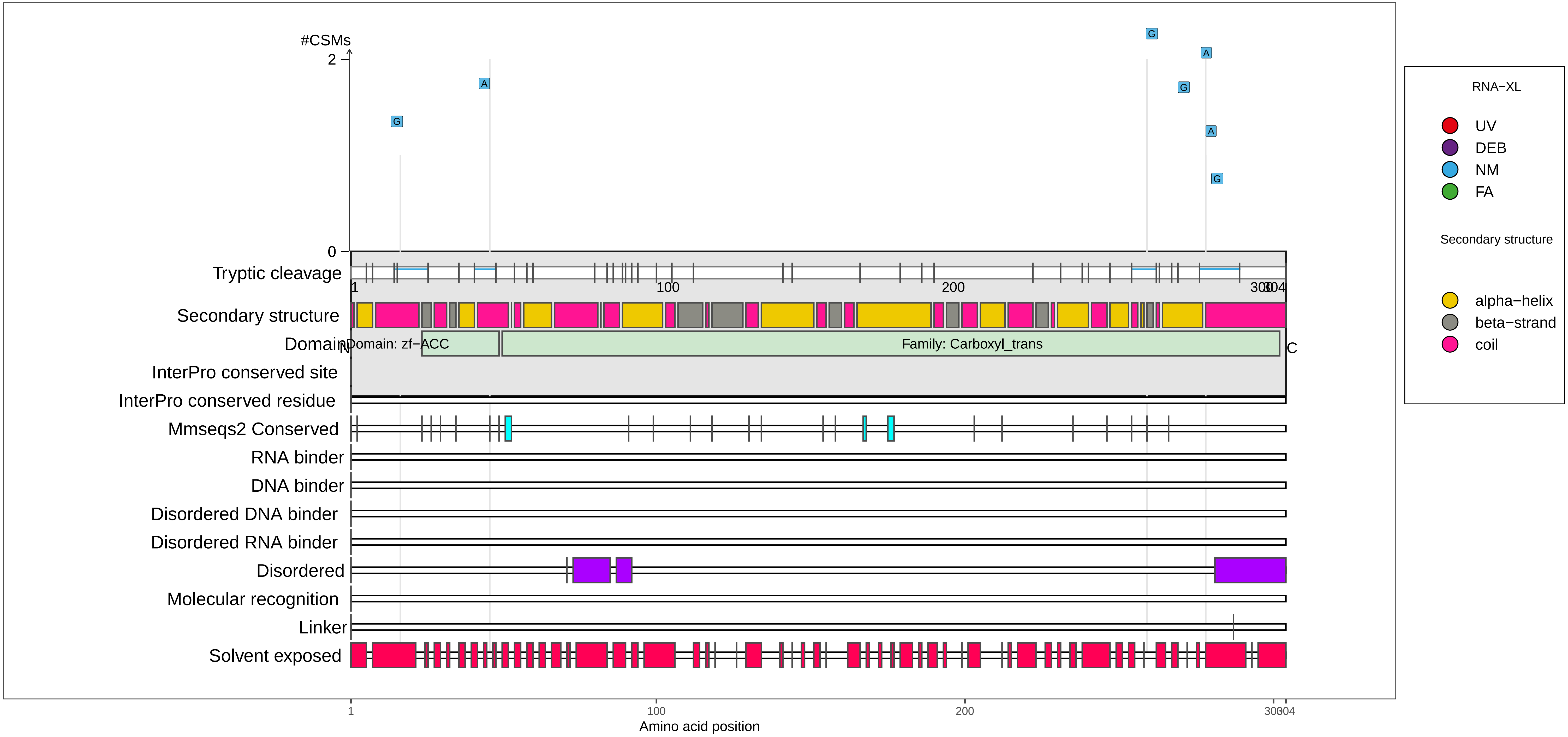

P0AAB6
GALF_ECOLI UTP−−glucose−1−phosphate uridylyltransferase

− Abundance:
tryptic [log10 Intensity]: 7.83 (Q 50)
PAXdb K12 strain [ppm]: 2.78 (Q 85)
PAXdb E.coli [ppm]: 2.97 (Q 94)

− RNA functions: not annotated

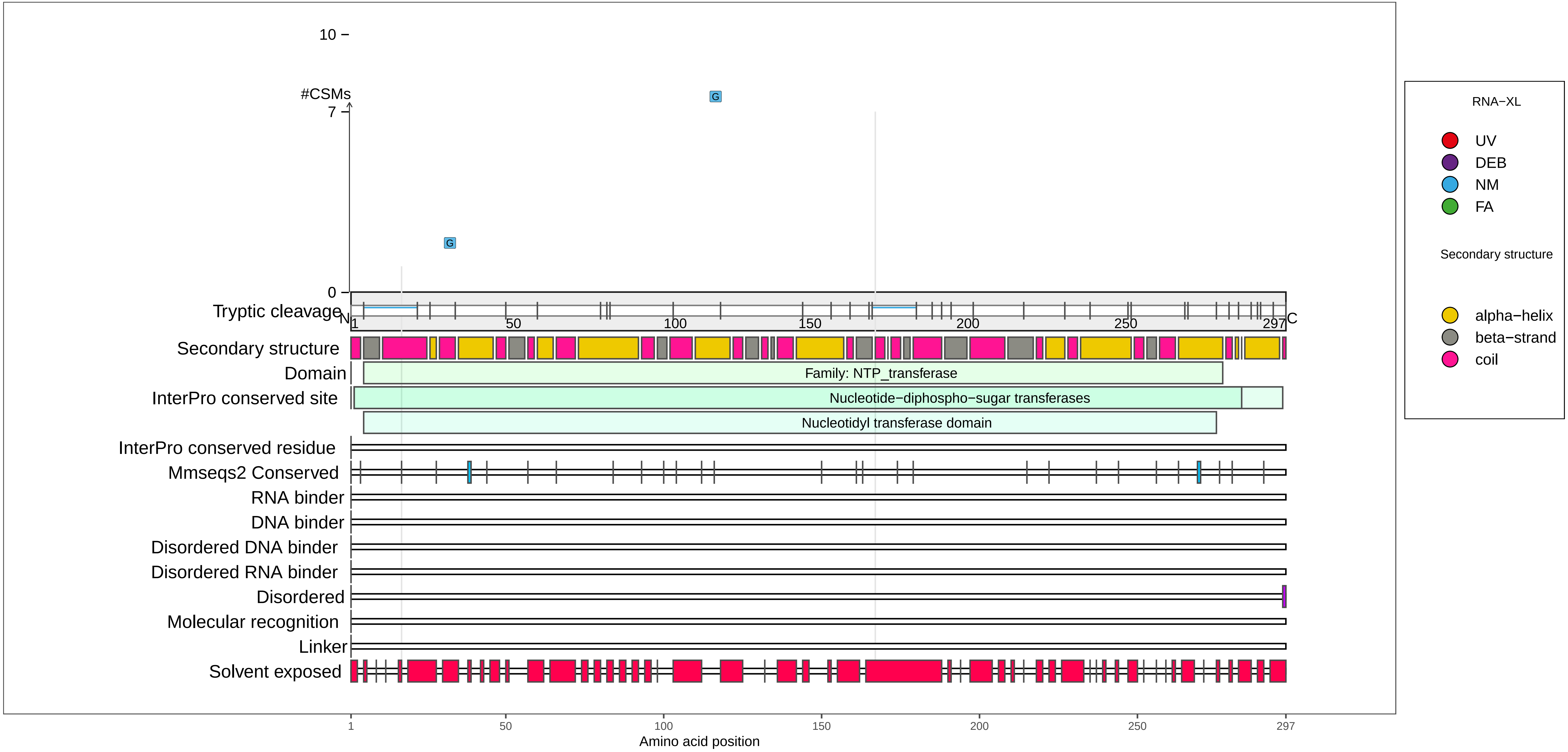

P0AB80
ILVE_ECOLI Branched−chain−amino−acid aminotransferase

− Abundance:
tryptic [log10 Intensity]: 9.05 (Q 89)
PAXdb K12 strain [ppm]: 2.75 (Q 85)
PAXdb E.coli [ppm]: 2.31 (Q 80)

− RNA functions: not annotated

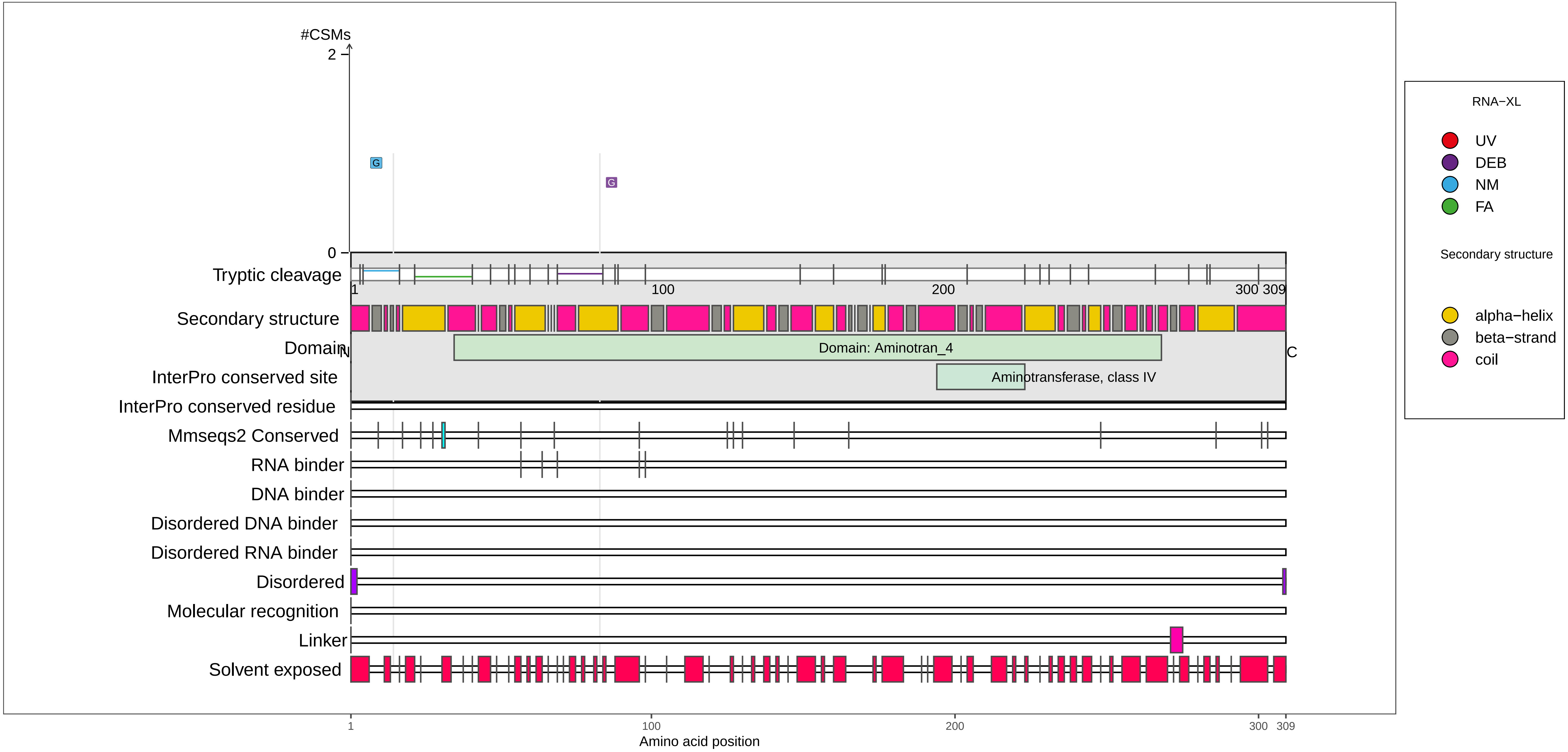

P0AC53
G6PD_ECOLI Glucose−6−phosphate 1−dehydrogenase

− Abundance:
tryptic [log10 Intensity]: 8.95 (Q 87)
PAXdb K12 strain [ppm]: 2.62 (Q 81)
PAXdb E.coli [ppm]: 2.76 (Q 91)

− RNA functions: not annotated

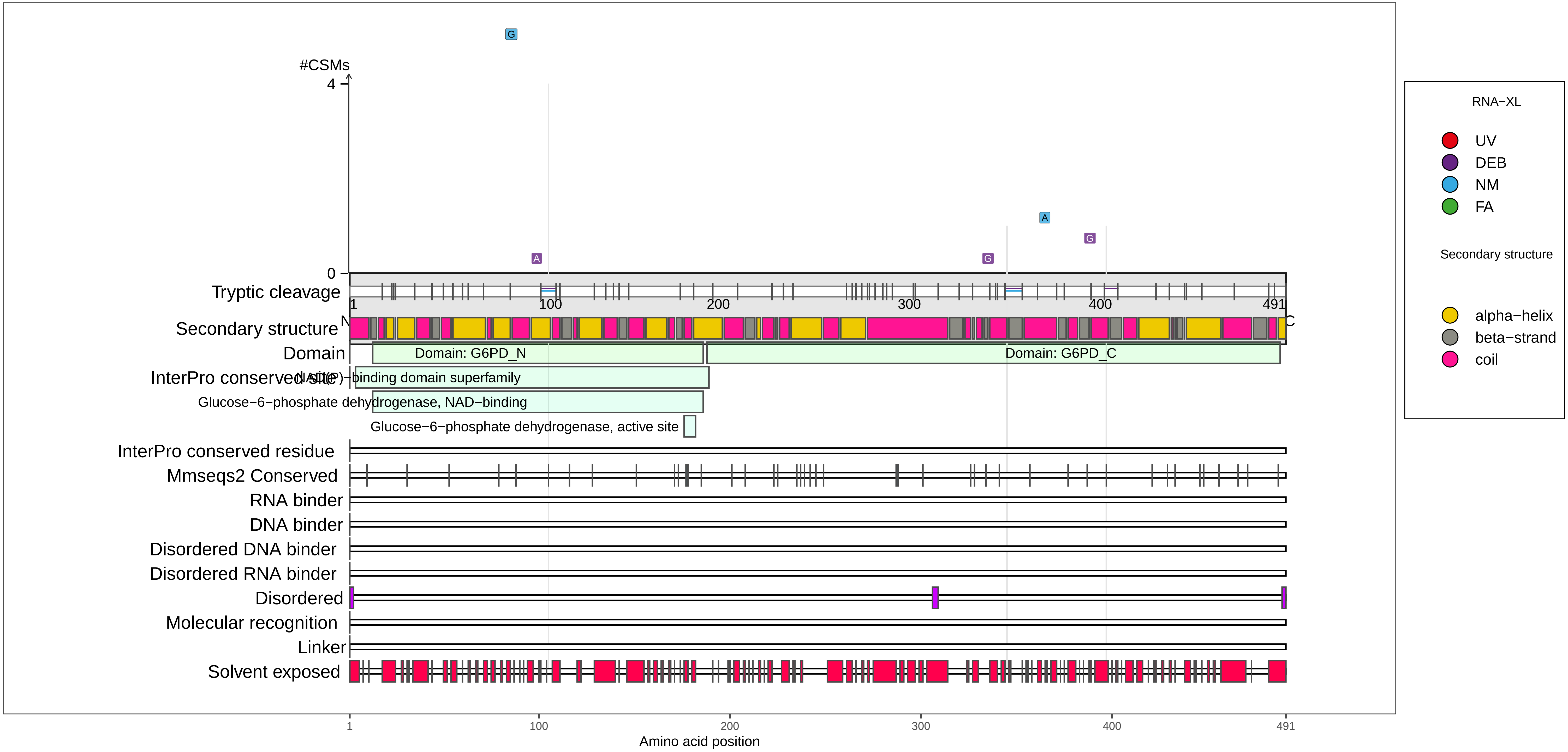

P0ADG1
ILVM_ECOLI Acetolactate synthase isozyme 2 small subunit

− Abundance:
tryptic [log10 Intensity]: 7.47 (Q 33)
PAXdb K12 strain [ppm]: NA (Q NA)
PAXdb E.coli [ppm]: NA (Q NA)

− RNA functions: not annotated

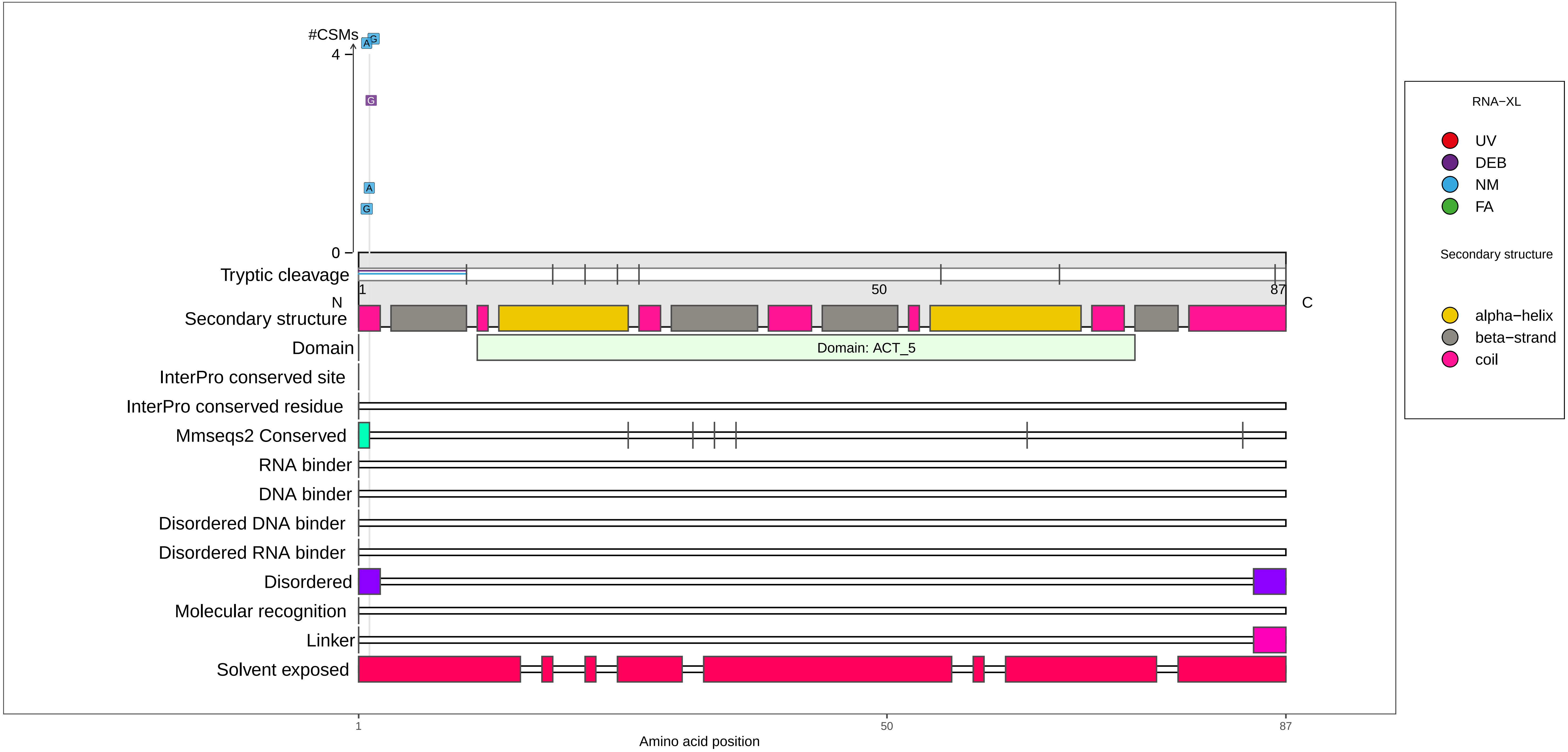

P0ADG4
SUHB_ECOLI Nus factor SuhB

− Abundance:
tryptic [log10 Intensity]: 9.28 (Q 94)
PAXdb K12 strain [ppm]: 2.31 (Q 70)
PAXdb E.coli [ppm]: 2.36 (Q 82)

− RNA functions: not annotated

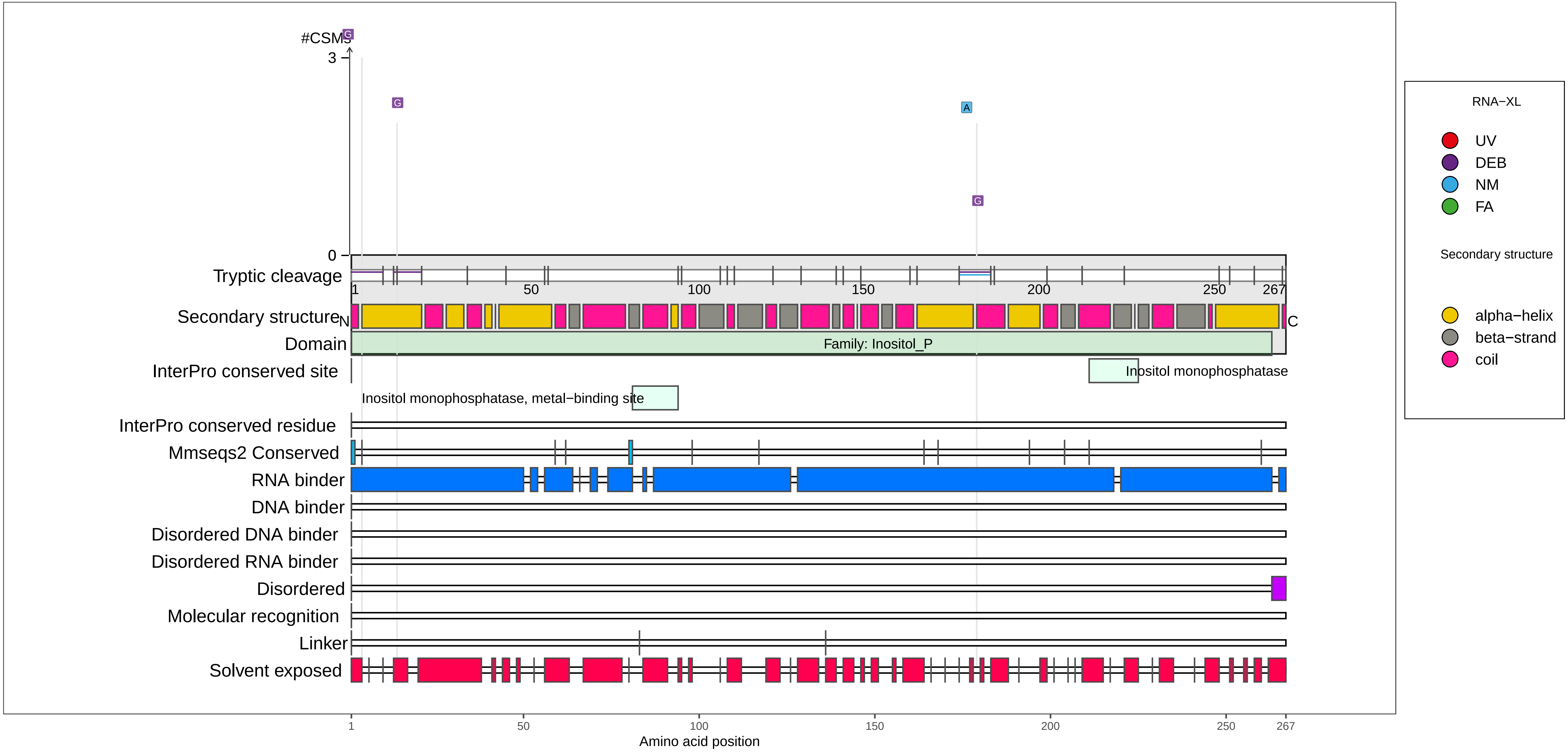

P0ADT2
YGIB_ECOLI UPF0441 protein YgiB

− Abundance:
tryptic [log10 Intensity]: 7.24 (Q 22)
PAXdb K12 strain [ppm]: 1.75 (Q 51)
PAXdb E.coli [ppm]: 1.85 (Q 68)

− RNA functions: not annotated

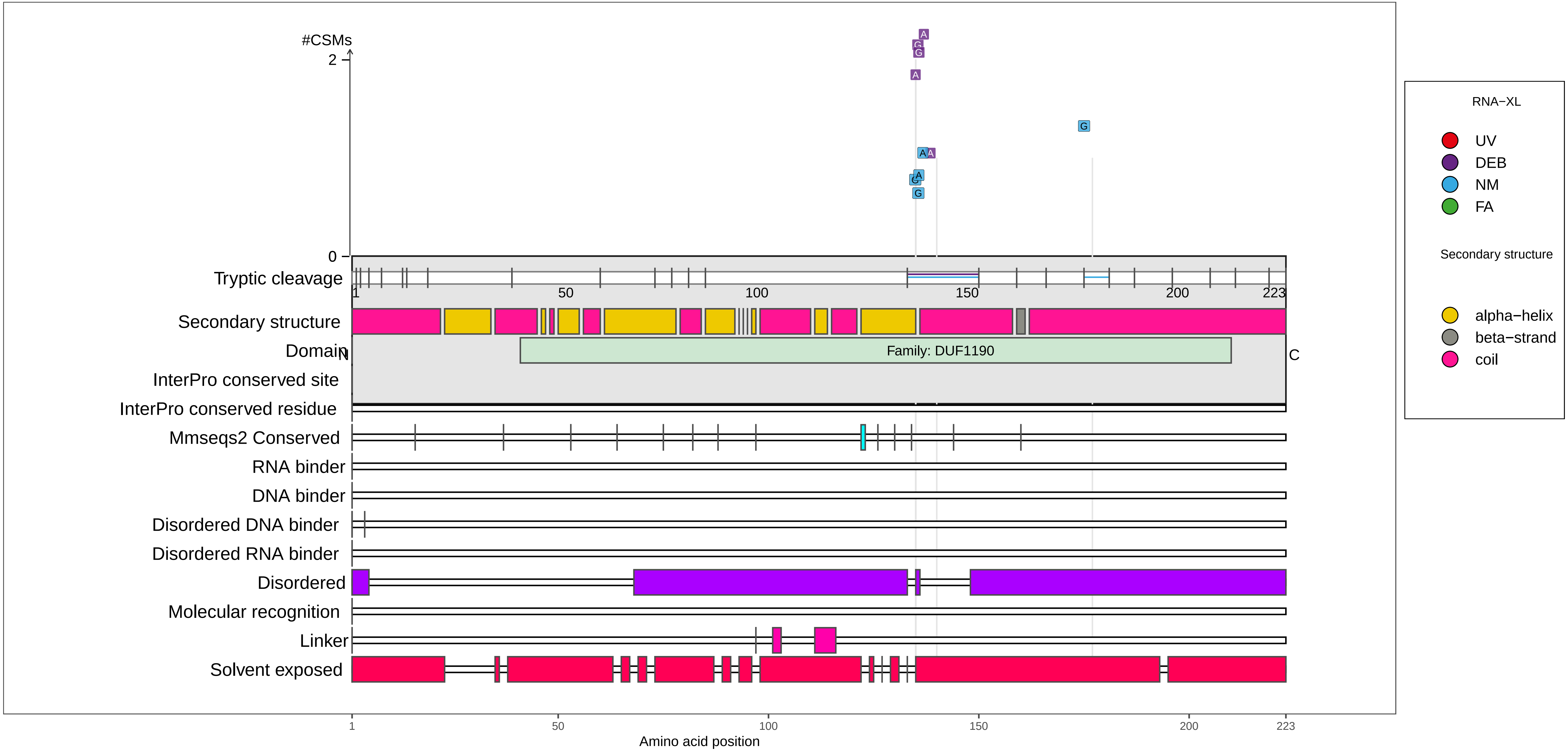

P0AF36
ZAPB_ECOLI Cell division protein ZapB

− Abundance:
tryptic [log10 Intensity]: 8.55 (Q 77)
PAXdb K12 strain [ppm]: 3.37 (Q 96)
PAXdb E.coli [ppm]: 3.6 (Q 100)

− RNA functions: not annotated

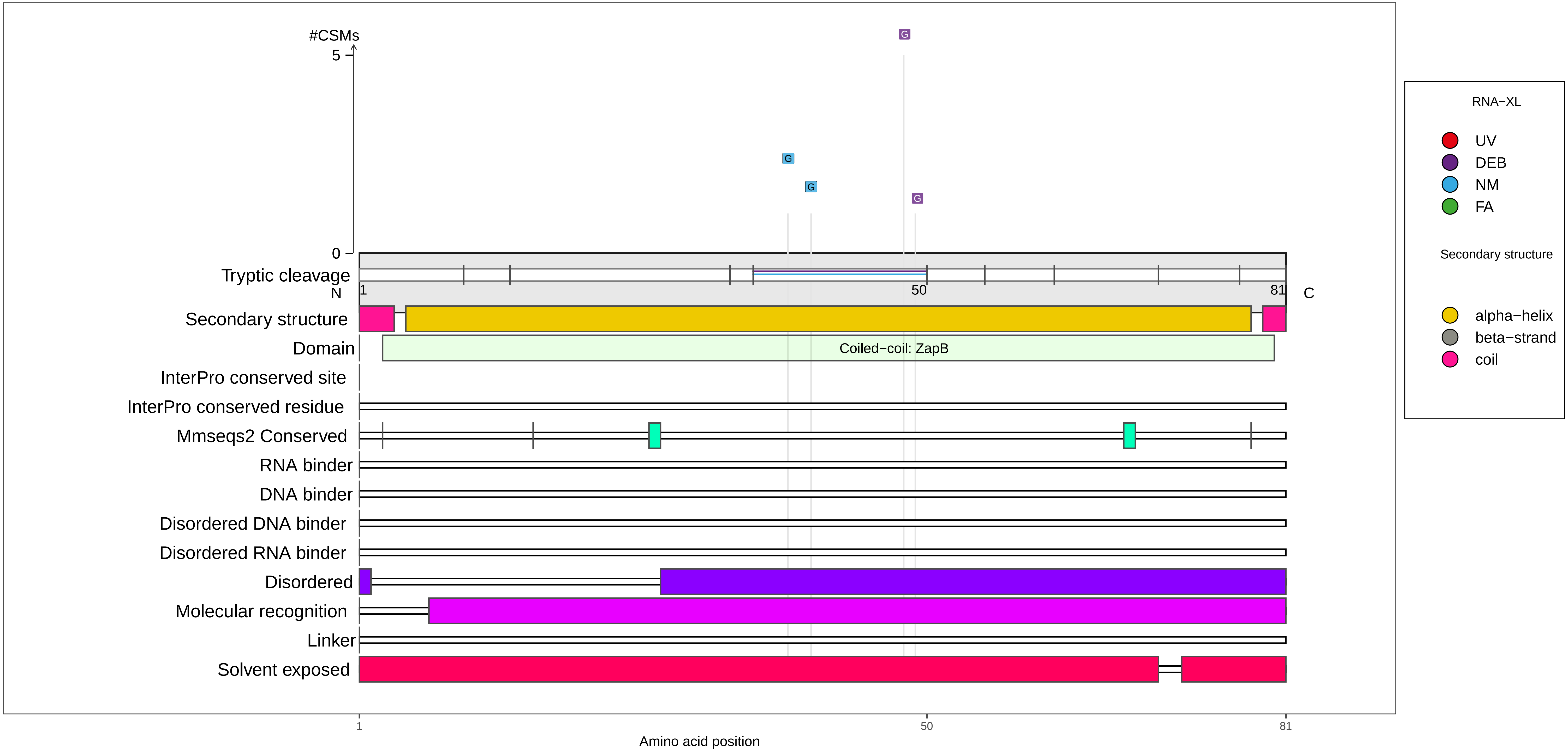

P0AGK4
YHBY_ECOLI RNA−binding protein YhbY

− Abundance:
tryptic [log10 Intensity]: 8.53 (Q 76)
PAXdb K12 strain [ppm]: 2.98 (Q 90)
PAXdb E.coli [ppm]: 3.01 (Q 95)

− RNA functions:
ncRNA 5−end processing; ncRNA metabolic process; ncRNA processing
RNA 5−end processing; RNA binding; RNA metabolic process; RNA processing; rRNA 5−end processing
rRNA metabolic process; rRNA processing

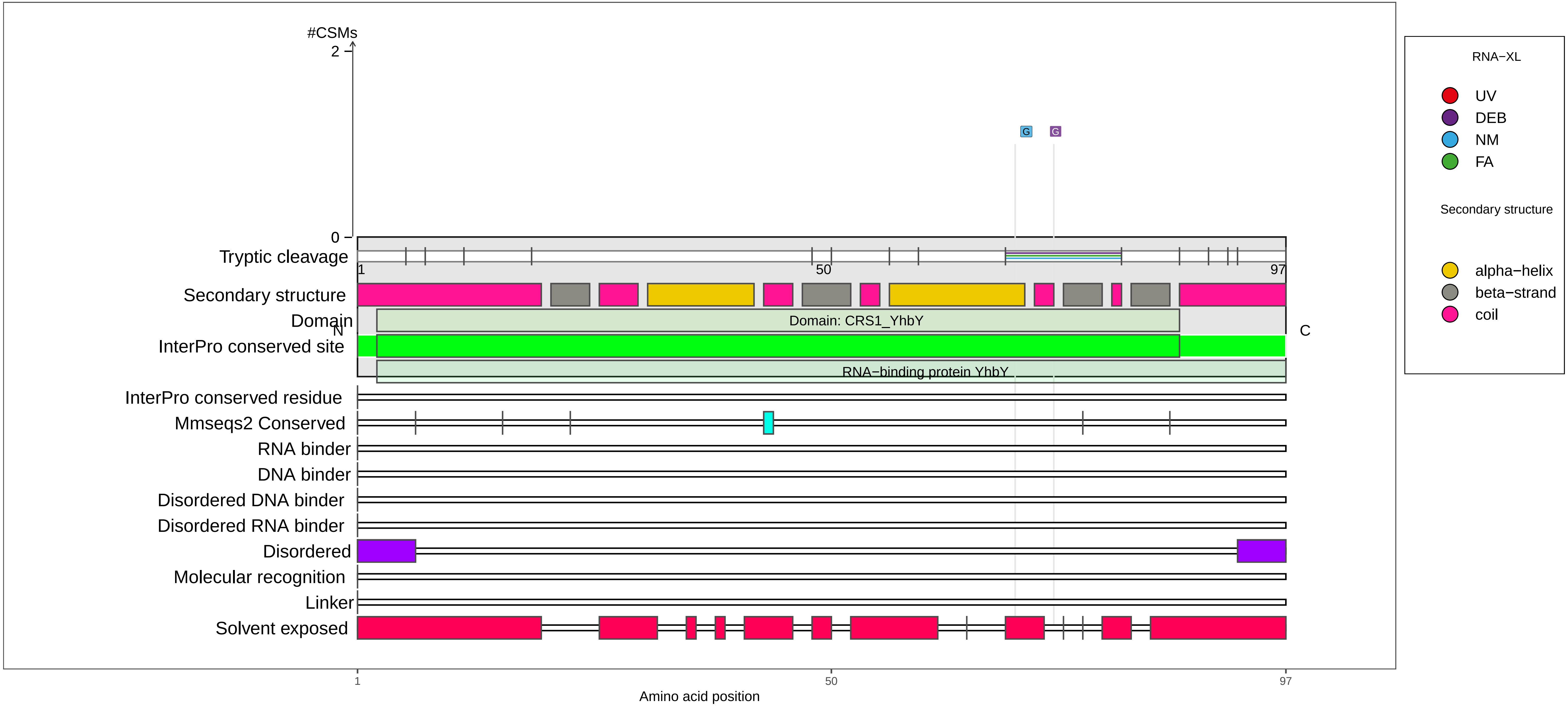

P11447
ARLY_ECOLI Argininosuccinate lyase

− Abundance:
tryptic [log10 Intensity]: 9.49 (Q 96)
PAXdb K12 strain [ppm]: 2.33 (Q 71)
PAXdb E.coli [ppm]: 2.15 (Q 76)

− RNA functions: not annotated

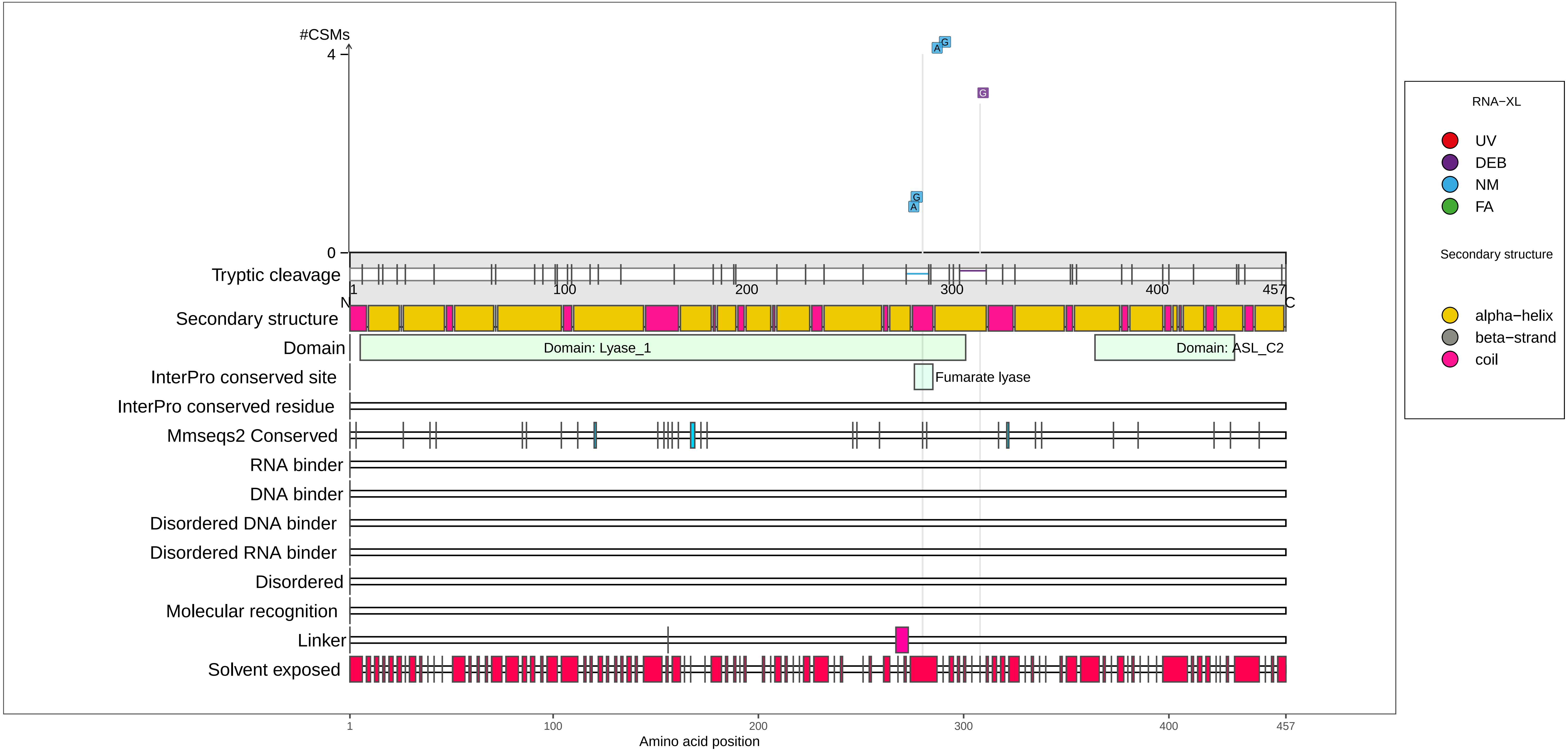

P25748
GALS_ECOLI HTH−type transcriptional regulator GalS

− Abundance:
tryptic [log10 Intensity]: 7.59 (Q 39)
PAXdb K12 strain [ppm]: 1.72 (Q 50)
PAXdb E.coli [ppm]: −0.15 (Q 21)

− RNA functions: not annotated

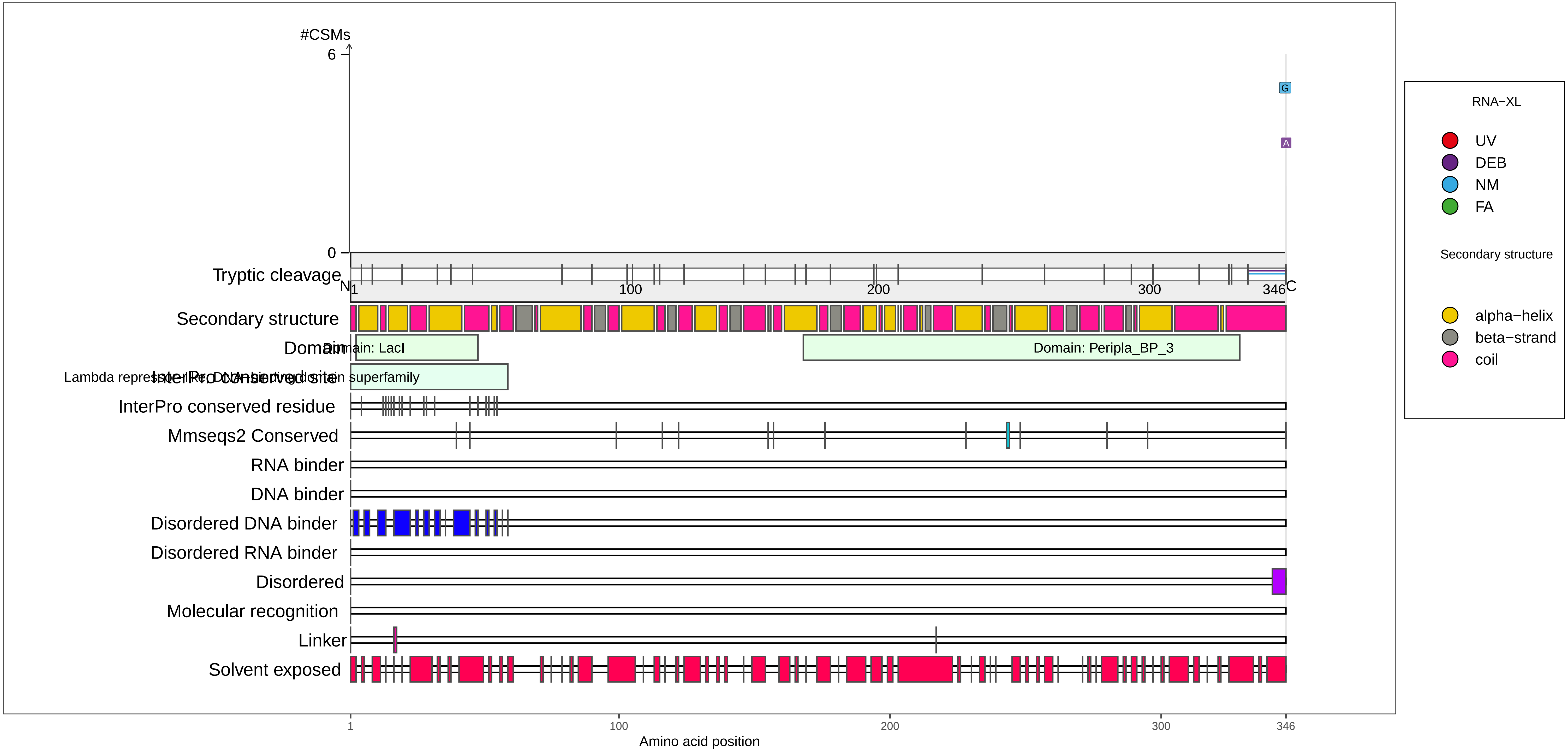

P27848
YIGL_ECOLI Pyridoxal phosphate phosphatase YigL

− Abundance:
tryptic [log10 Intensity]: 7.57 (Q 38)
PAXdb K12 strain [ppm]: 2.62 (Q 81)
PAXdb E.coli [ppm]: 1.42 (Q 58)

− RNA functions: not annotated

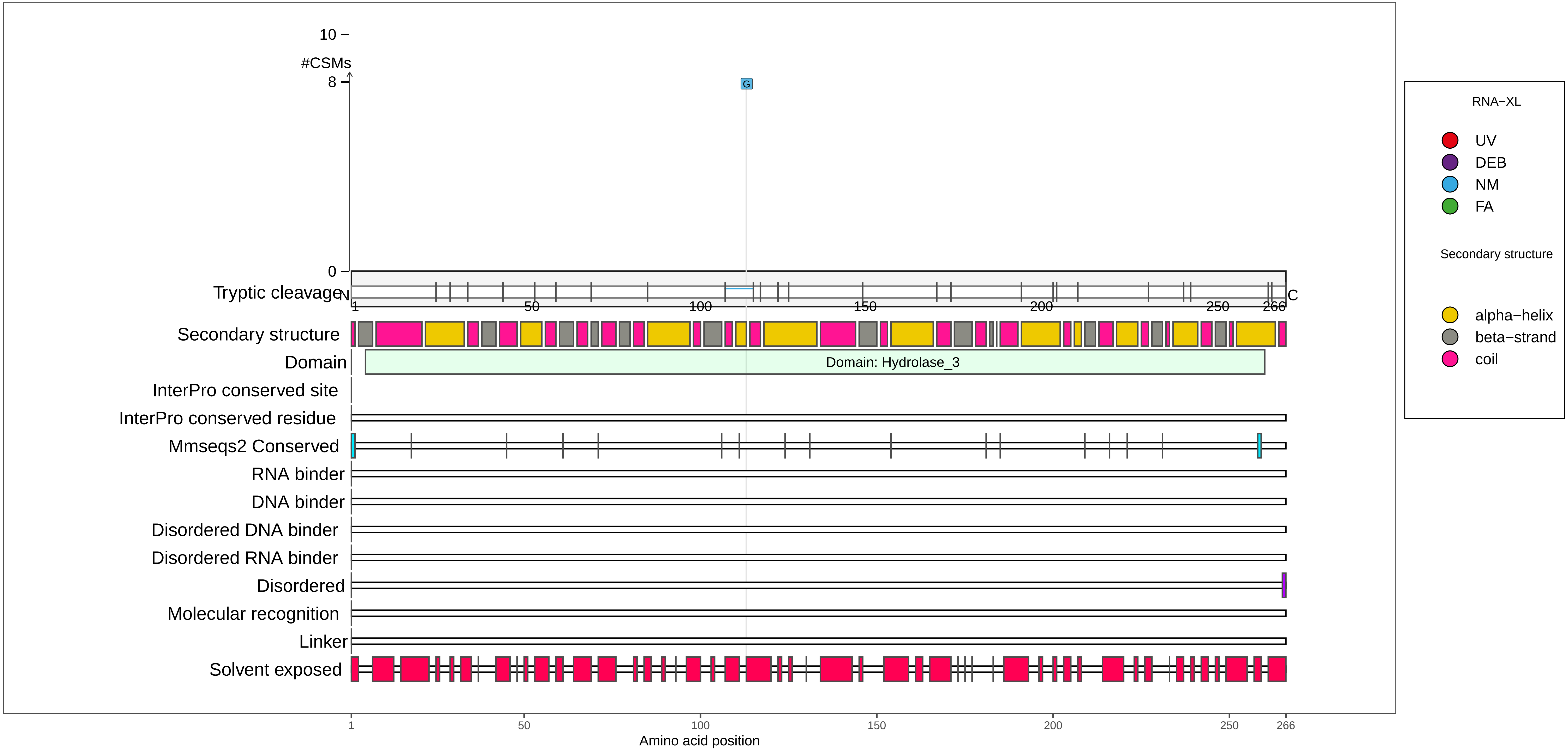

P33599
NUOCD_ECOLI NADH−quinone oxidoreductase subunit C/D

− Abundance:
tryptic [log10 Intensity]: 9.24 (Q 93)
PAXdb K12 strain [ppm]: 2.39 (Q 73)
PAXdb E.coli [ppm]: 2.58 (Q 88)

− RNA functions: not annotated

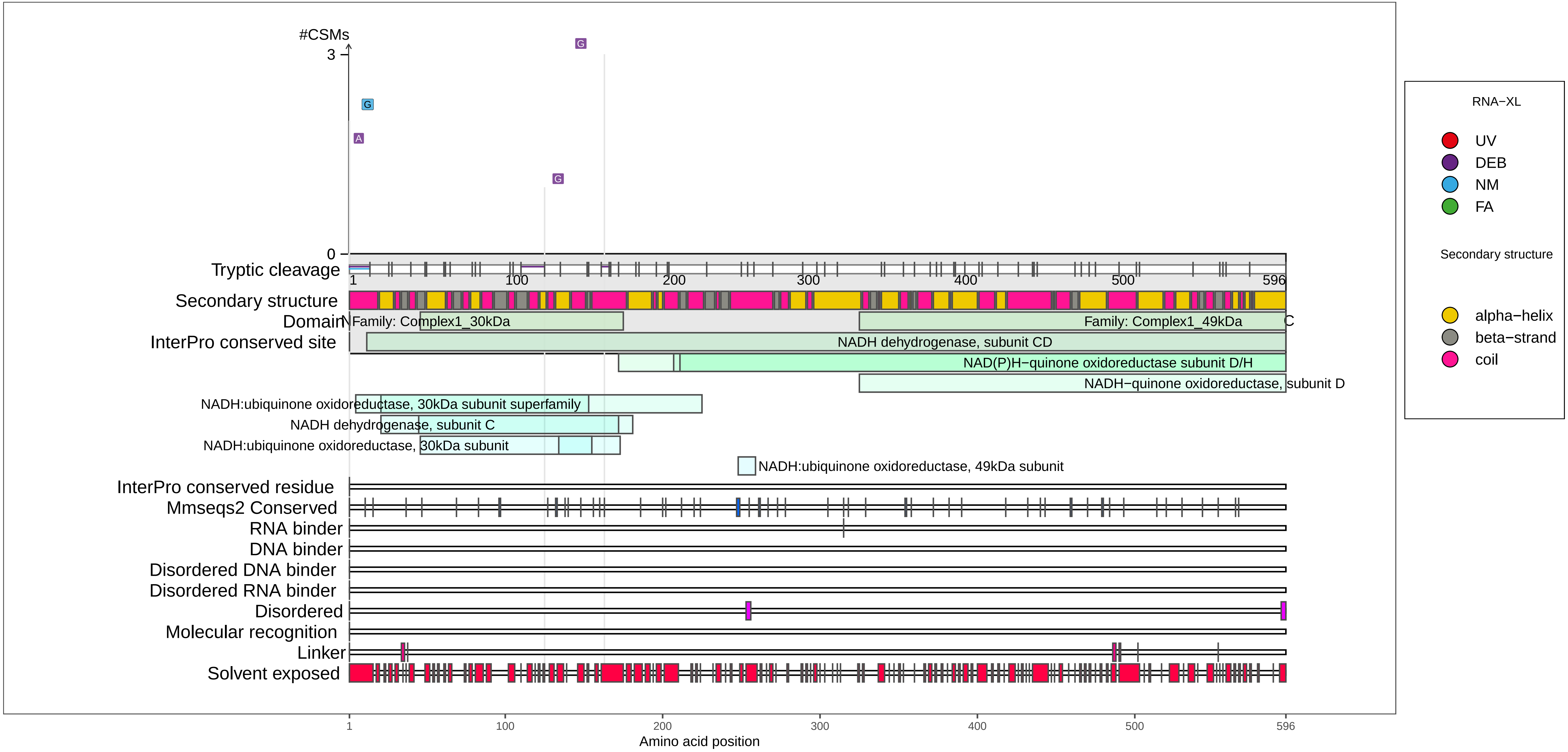

P39212
INSN2_ECOLI Putative transposase InsN for insertion sequence element IS911B
− RNA functions: not annotated

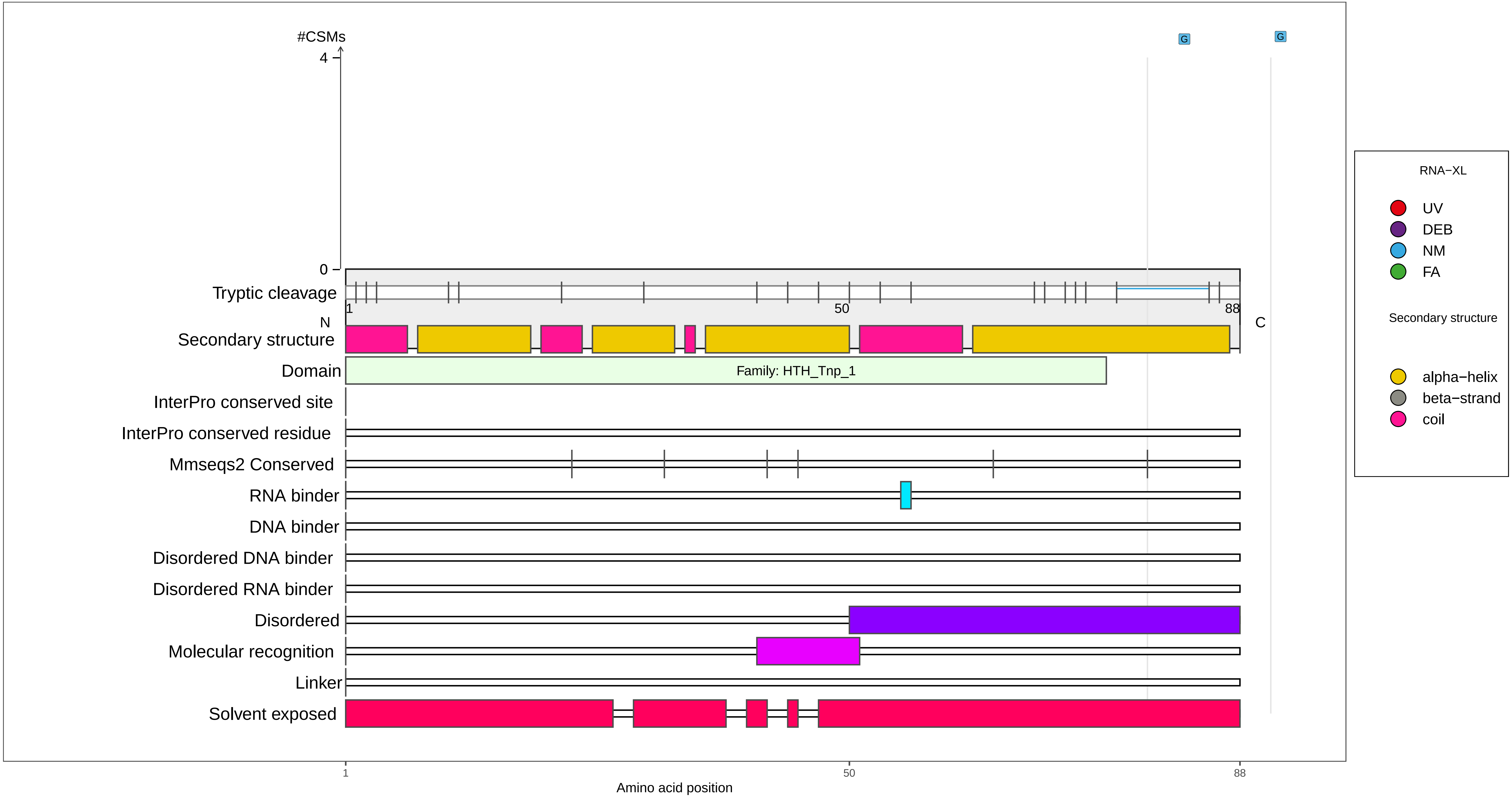

P39286
RSGA_ECOLI Small ribosomal subunit biogenesis GTPase RsgA

− Abundance:
tryptic [log10 Intensity]: 7.65 (Q 43)
PAXdb K12 strain [ppm]: 2.6 (Q 80)
PAXdb E.coli [ppm]: 2.05 (Q 73)

− RNA functions:
RNA binding; rRNA binding

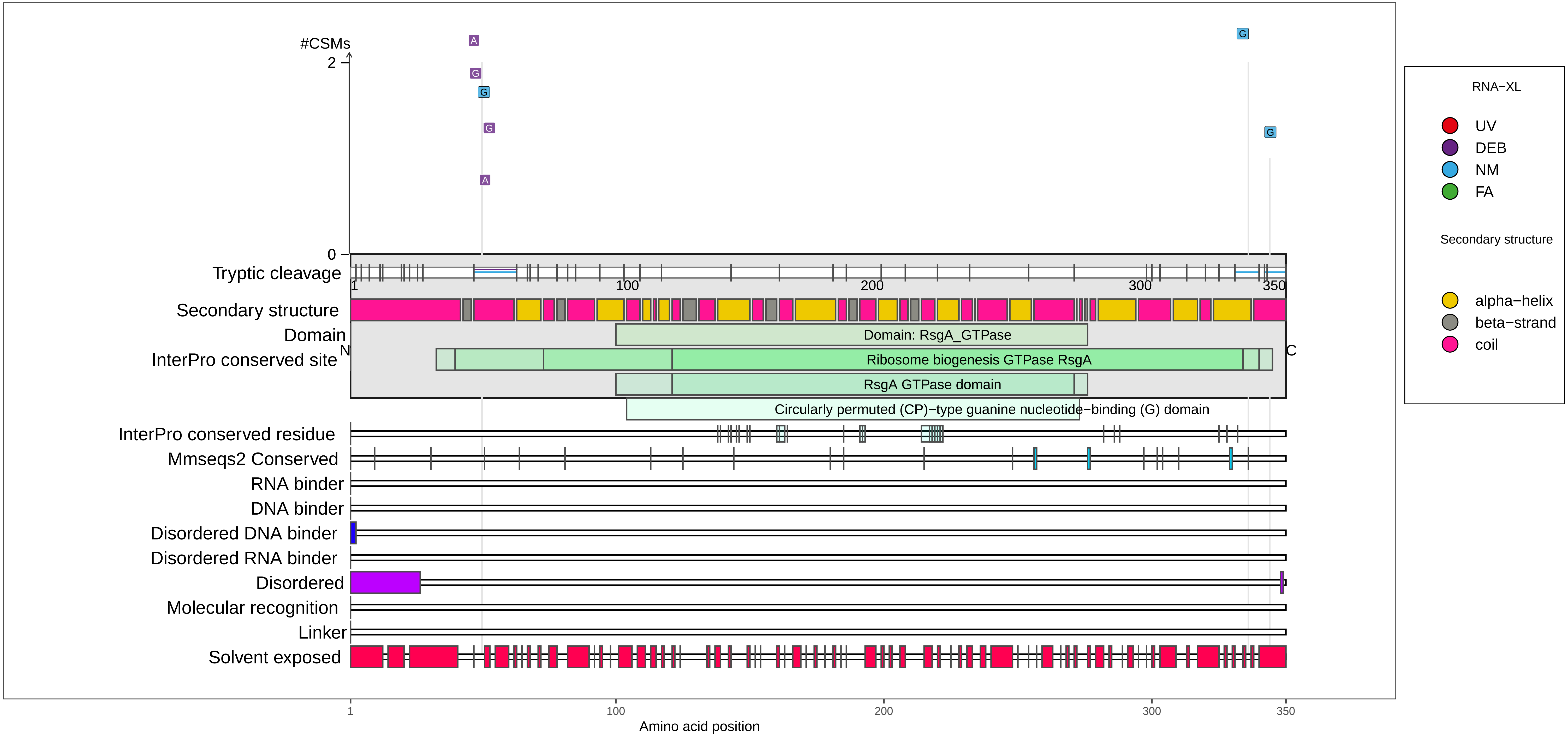

P40874
MTOX_ECOLI N−methyl−L−tryptophan oxidase

− Abundance:
tryptic [log10 Intensity]: 8.31 (Q 70)
PAXdb K12 strain [ppm]: 2.03 (Q 61)
PAXdb E.coli [ppm]: 1.9 (Q 70)

− RNA functions: not annotated

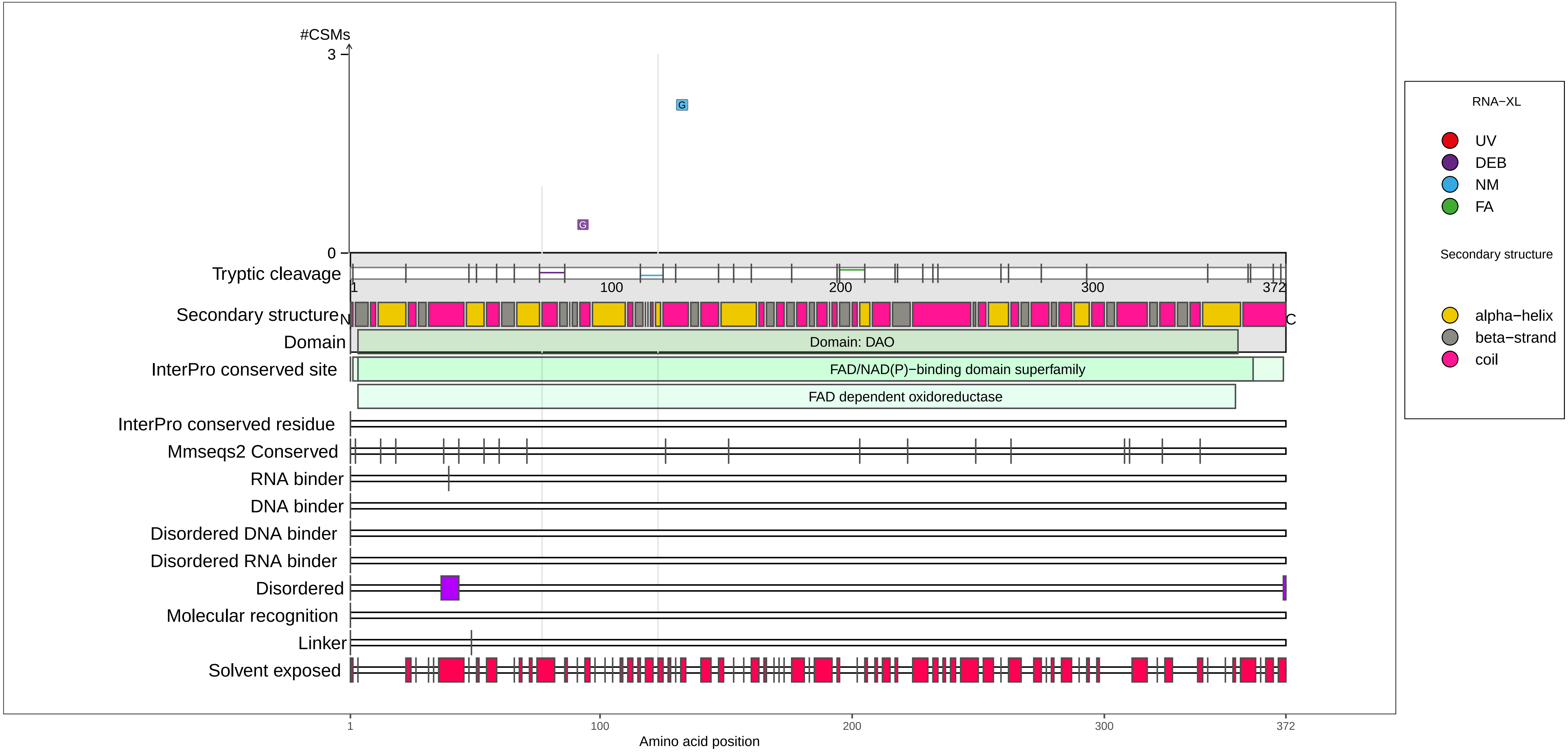

P64506
YEBY_ECOLI Uncharacterized protein YebY

− Abundance:
tryptic [log10 Intensity]: 7.99 (Q 57)
PAXdb K12 strain [ppm]: NA (Q NA)
PAXdb E.coli [ppm]: 1.55 (Q 61)

− RNA functions: not annotated

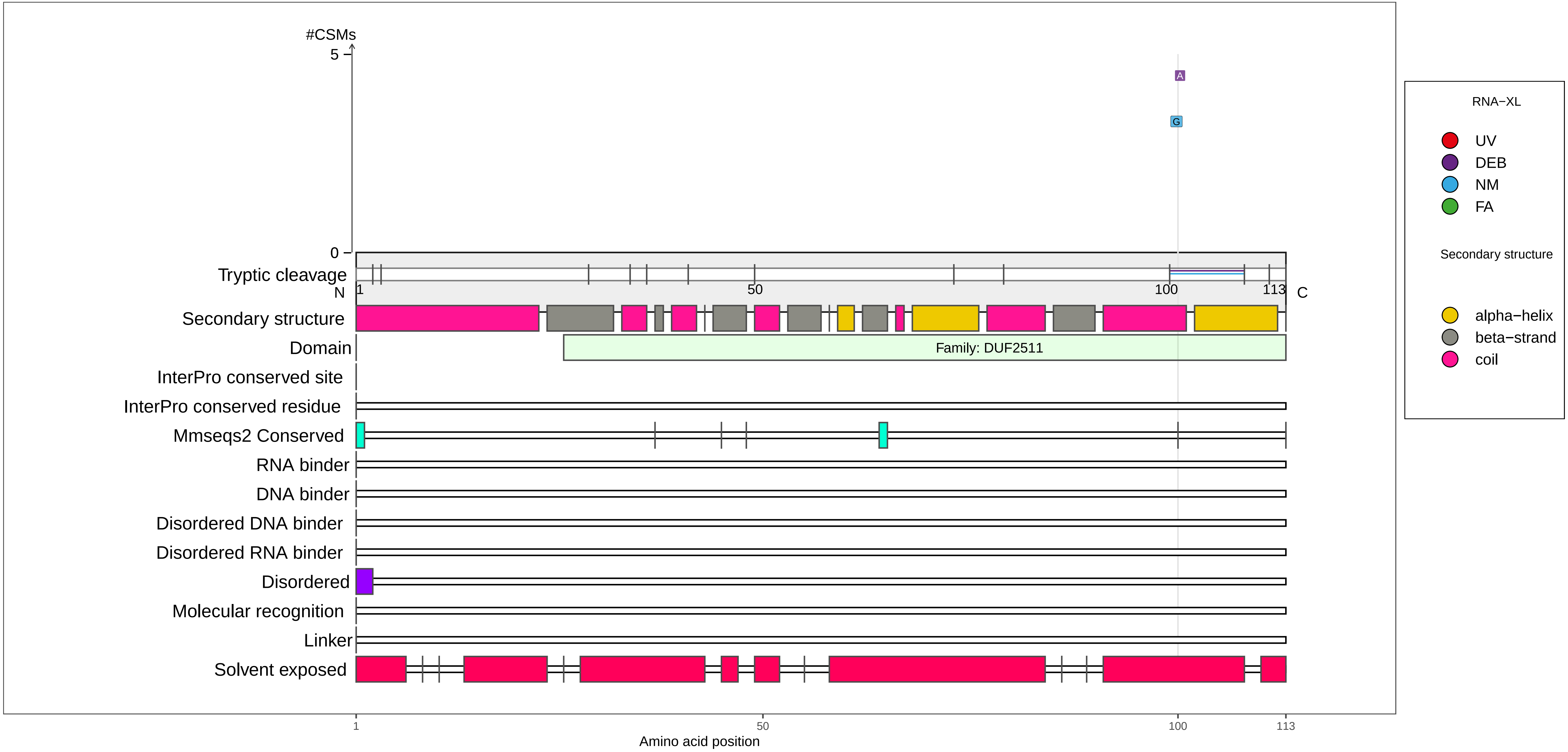

P64540
YFCL_ECOLI Uncharacterized protein YfcL

− Abundance:
tryptic [log10 Intensity]: 8.54 (Q 77)
PAXdb K12 strain [ppm]: 2.05 (Q 62)
PAXdb E.coli [ppm]: 1.79 (Q 67)

− RNA functions: not annotated

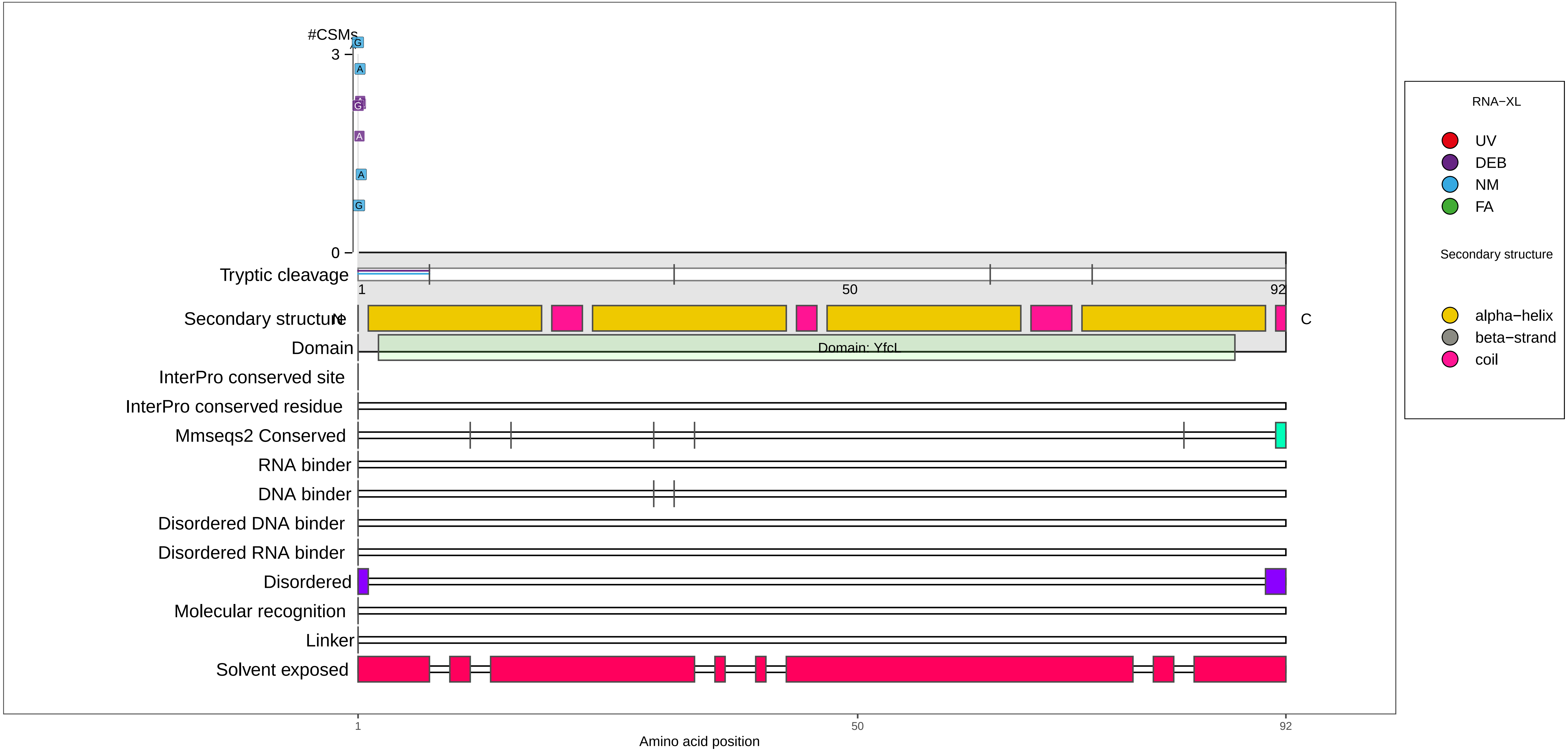

P69828
PTKA_ECOLI PTS system galactitol−specific EIIA component

− Abundance:
tryptic [log10 Intensity]: 7.3 (Q 25)
PAXdb K12 strain [ppm]: 2.06 (Q 62)
PAXdb E.coli [ppm]: 3.36 (Q 98)

− RNA functions: not annotated

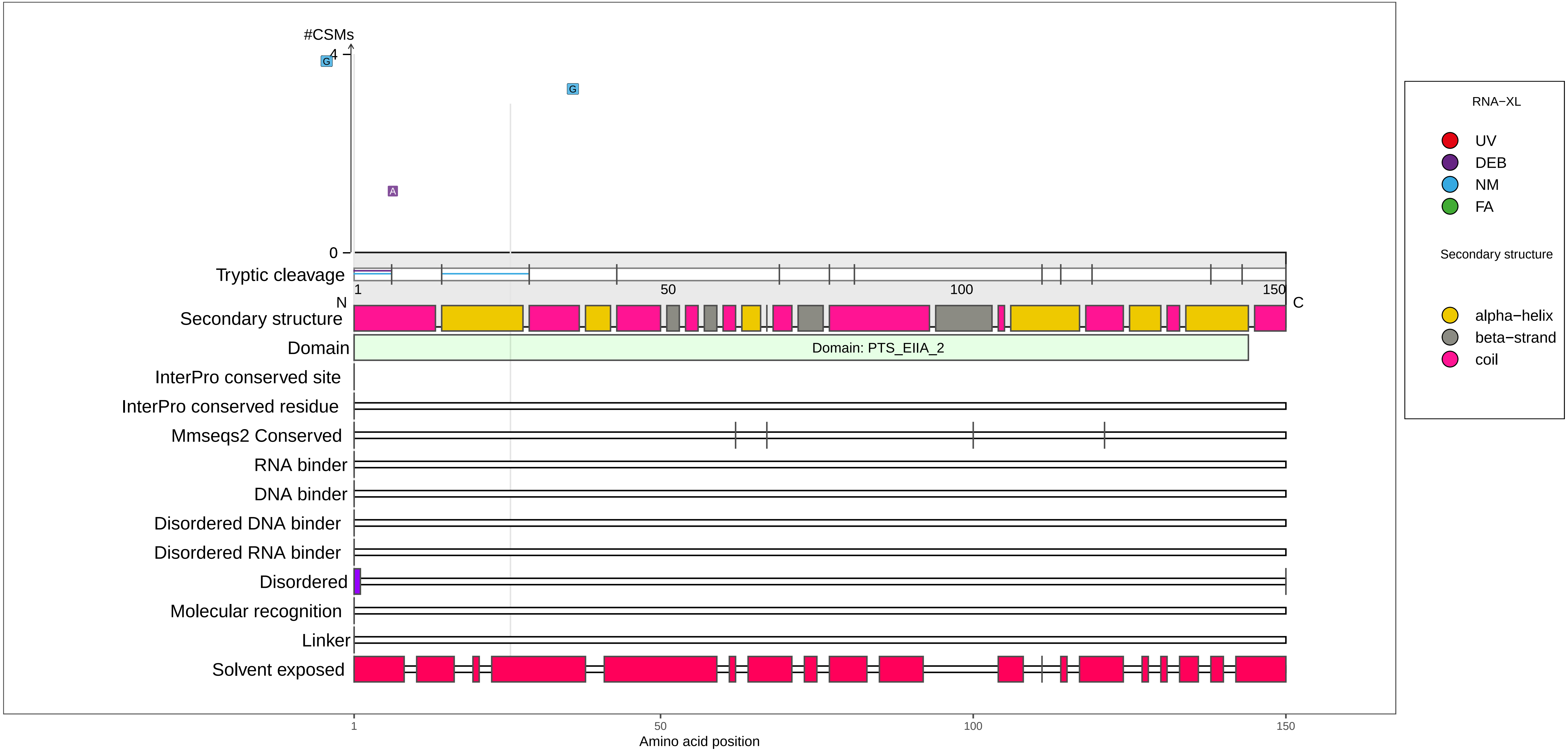

P76034
YCIT_ECOLI Uncharacterized HTH−type transcriptional regulator YciT

− Abundance:
tryptic [log10 Intensity]: 8.06 (Q 60)
PAXdb K12 strain [ppm]: 2.32 (Q 71)
PAXdb E.coli [ppm]: 1.82 (Q 68)

− RNA functions:
RNA biosynthetic process; RNA metabolic process

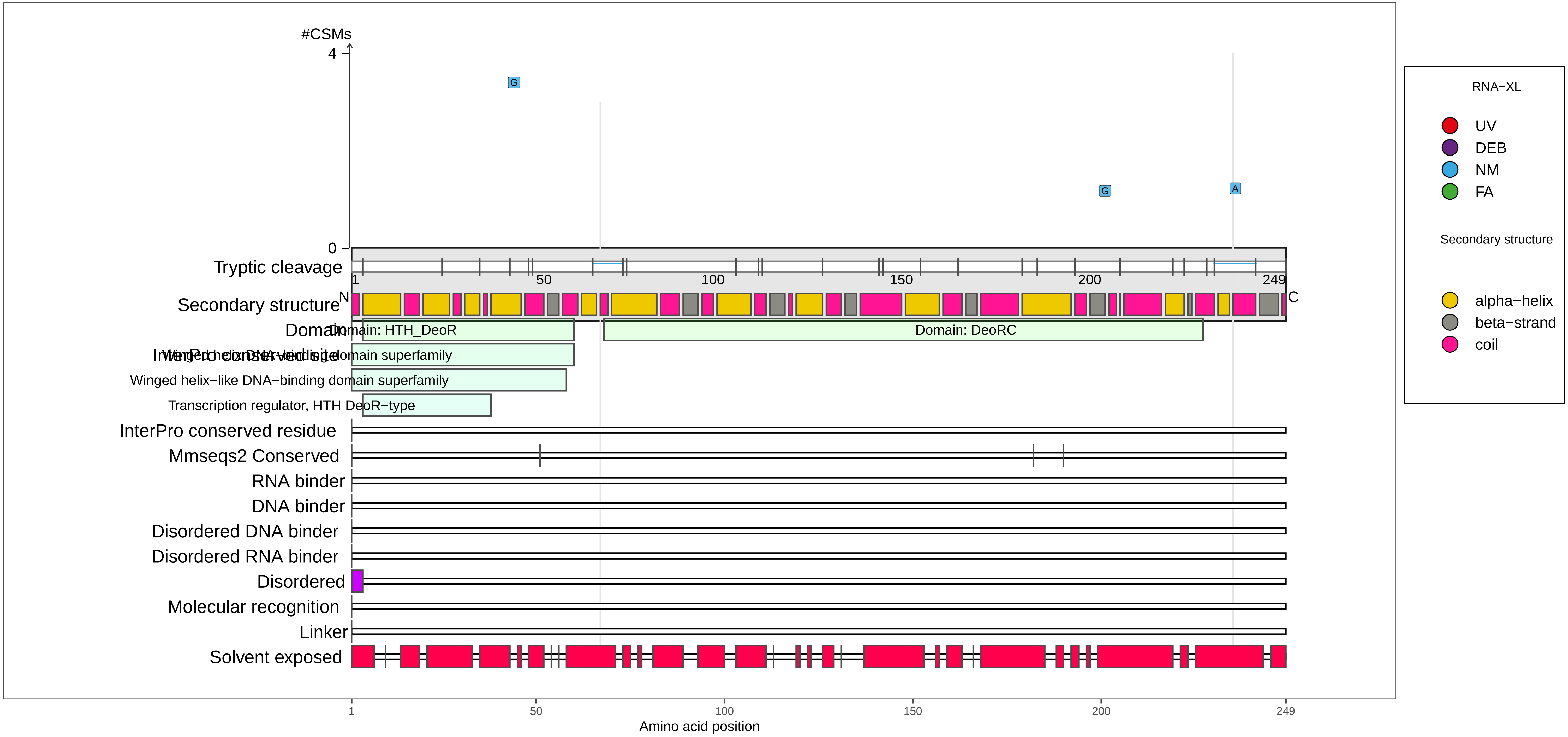

P76342
MSRP_ECOLI Protein−methionine−sulfoxide reductase catalytic subunit MsrP

− Abundance:
tryptic [log10 Intensity]: NA (Q NA)
PAXdb K12 strain [ppm]: 1.16 (Q 18)
PAXdb E.coli [ppm]: 1.18 (Q 52)

− RNA functions: not annotated

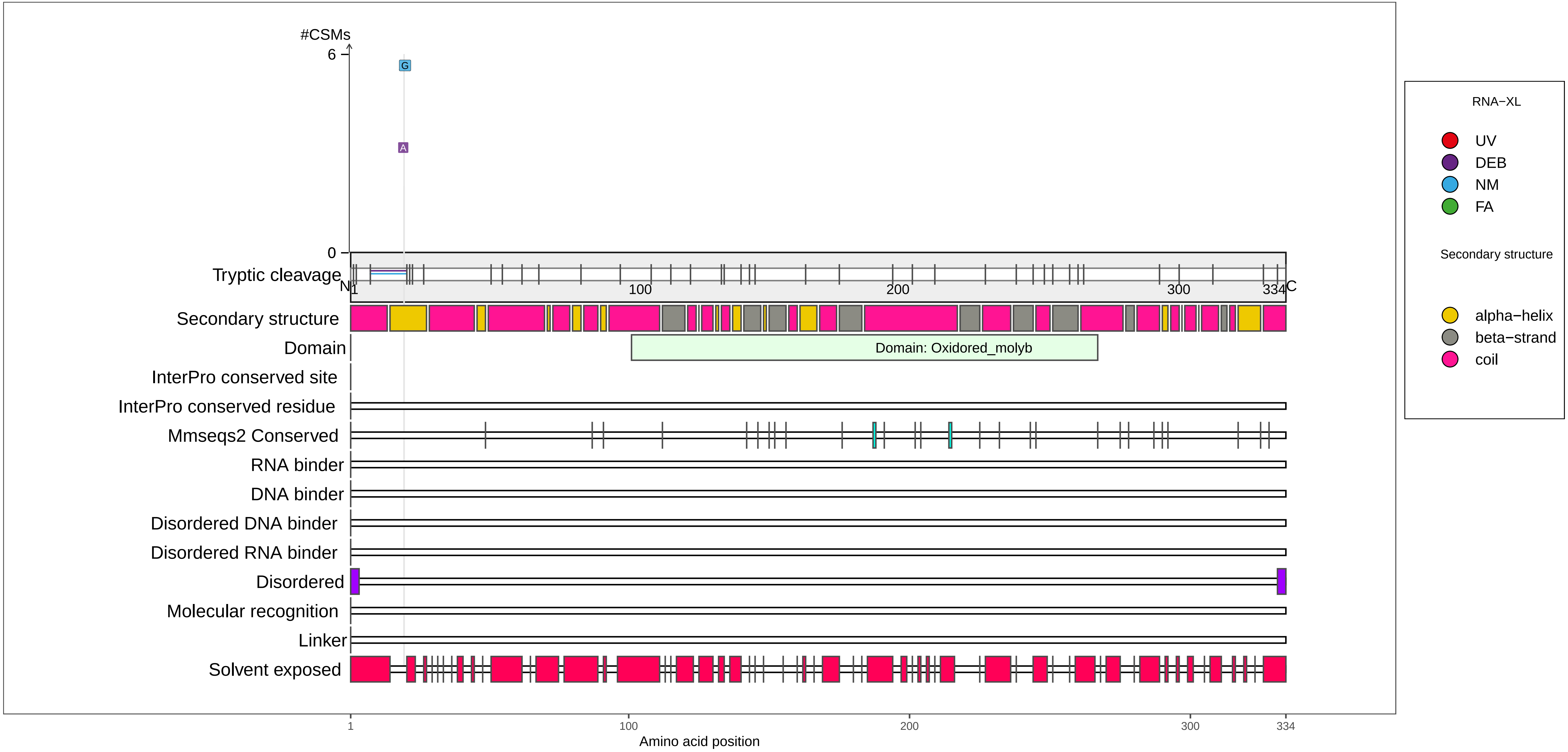

P77804
YDGA_ECOLI Protein YdgA
− Abundance:
tryptic [log10 Intensity]: 8.62 (Q 79)
PAXdb K12 strain [ppm]: 1.99 (Q 60)
PAXdb E.coli [ppm]: 2.66 (Q 89)

− RNA functions: not annotated

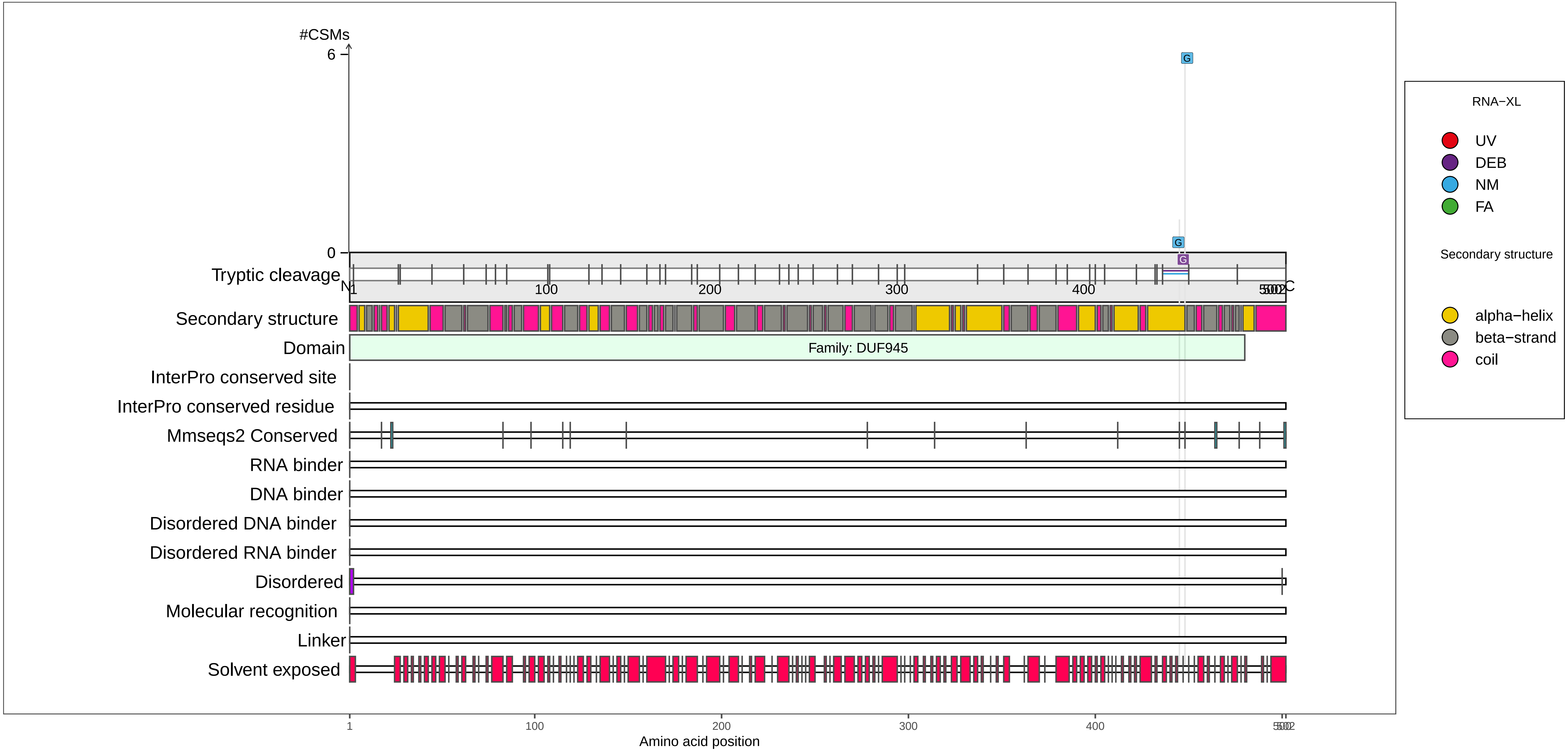

P02919
PBPB_ECOLI Penicillin−binding protein 1B

− Abundance:
tryptic [log10 Intensity]: 7.84 (Q 51)
PAXdb K12 strain [ppm]: 1.57 (Q 42)
PAXdb E.coli [ppm]: 1.31 (Q 55)

− RNA functions: not annotated

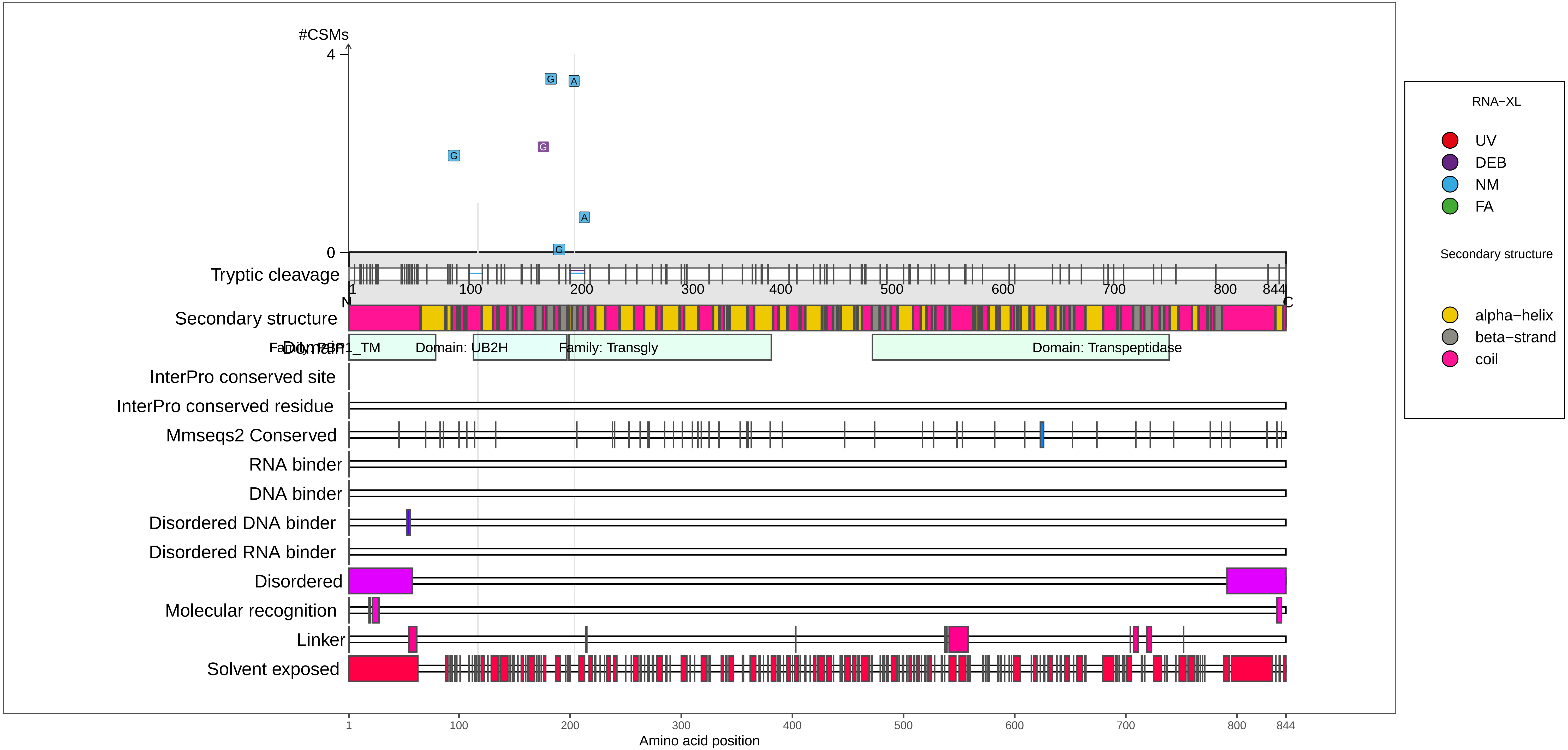

P04425
GSHB_ECOLI Glutathione synthetase

− Abundance:
tryptic [log10 Intensity]: 8.83 (Q 84)
PAXdb K12 strain [ppm]: 2.65 (Q 82)
PAXdb E.coli [ppm]: 2.25 (Q 79)

− RNA functions: not annotated

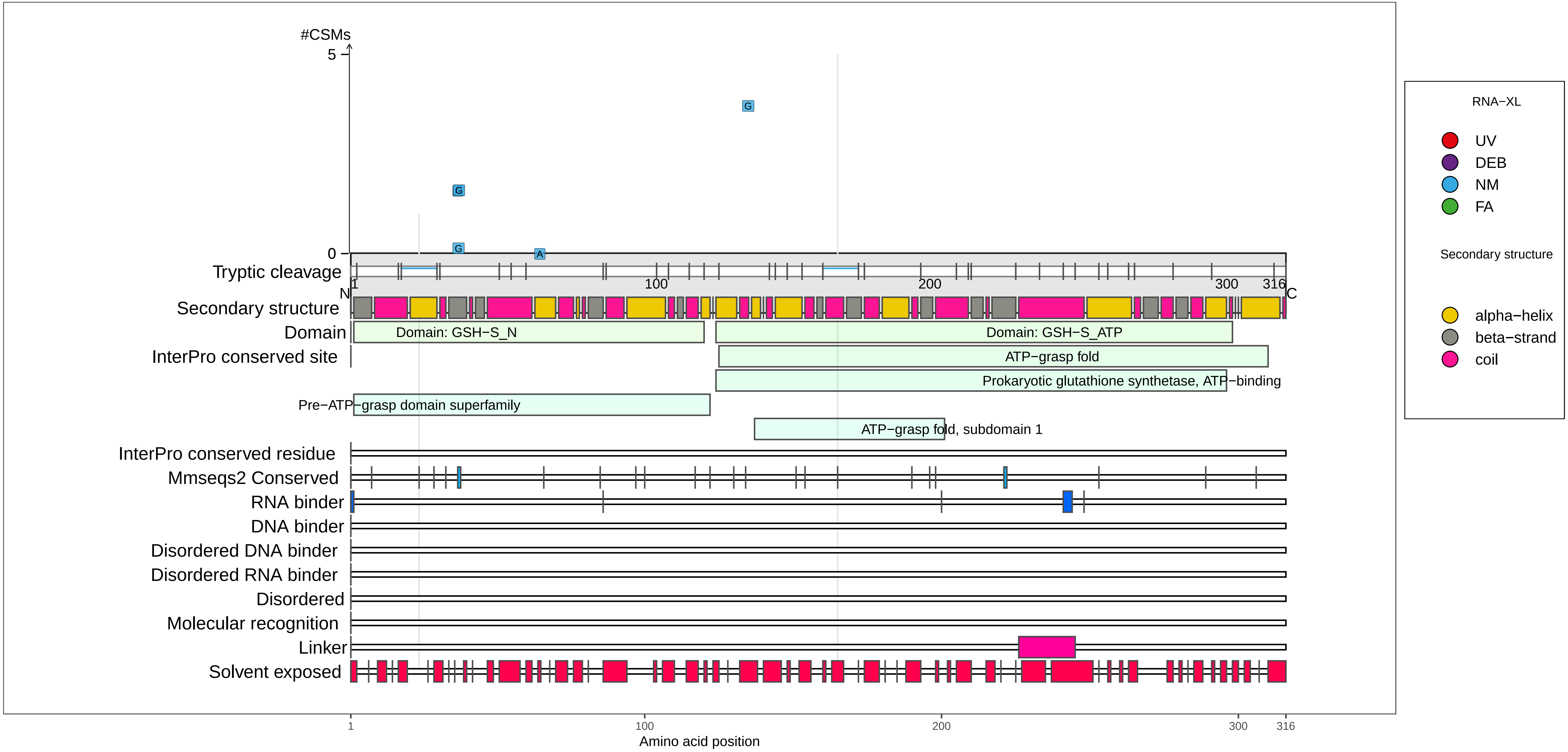

P05458
PTRA_ECOLI Protease 3

− Abundance:
tryptic [log10 Intensity]: 7.77 (Q 48)
PAXdb K12 strain [ppm]: 1.56 (Q 42)
PAXdb E.coli [ppm]: 1.05 (Q 49)

− RNA functions: not annotated

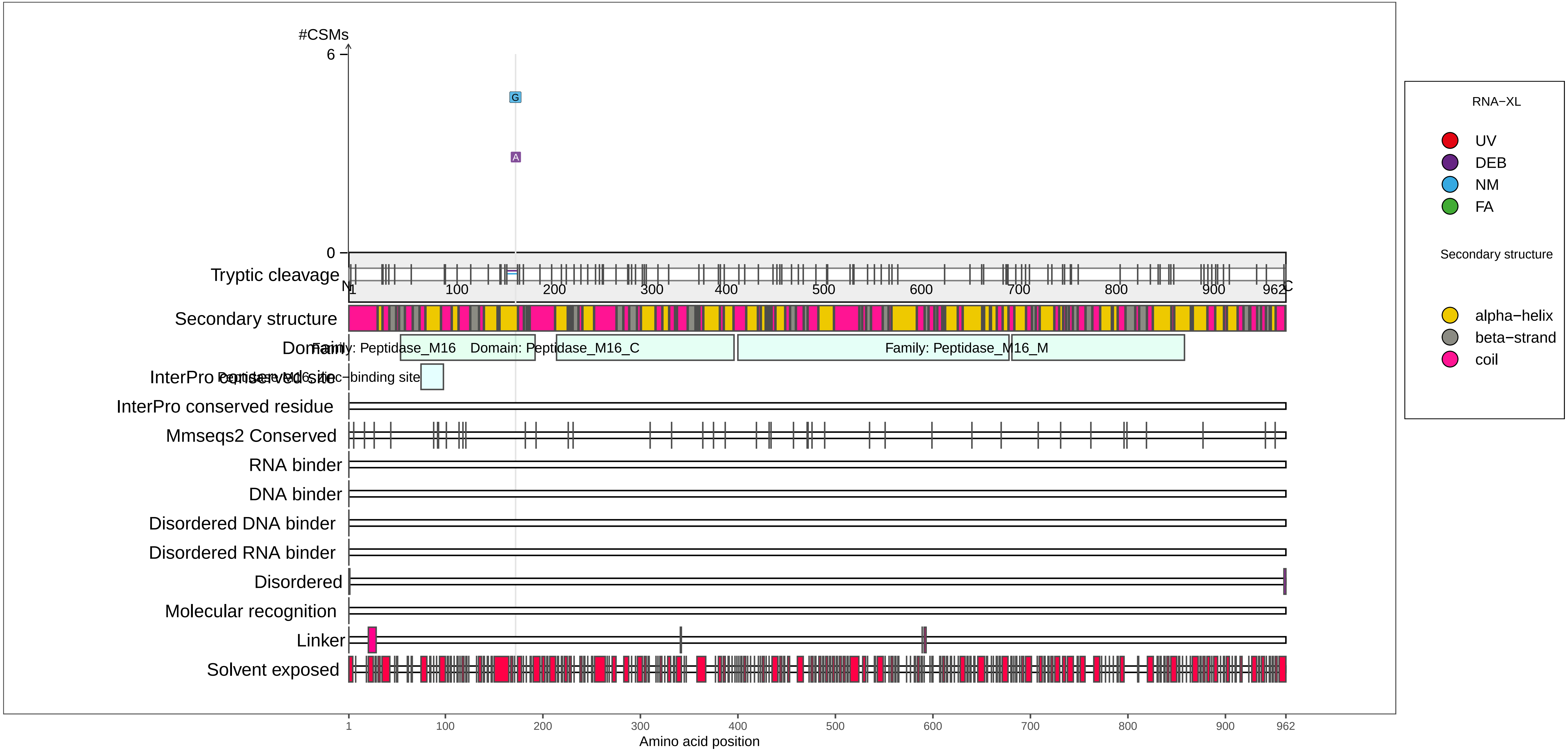

P06715
GSHR_ECOLI Glutathione reductase

− Abundance:
tryptic [log10 Intensity]: 9.13 (Q 91)
PAXdb K12 strain [ppm]: 2.36 (Q 72)
PAXdb E.coli [ppm]: 2.06 (Q 74)

− RNA functions: not annotated

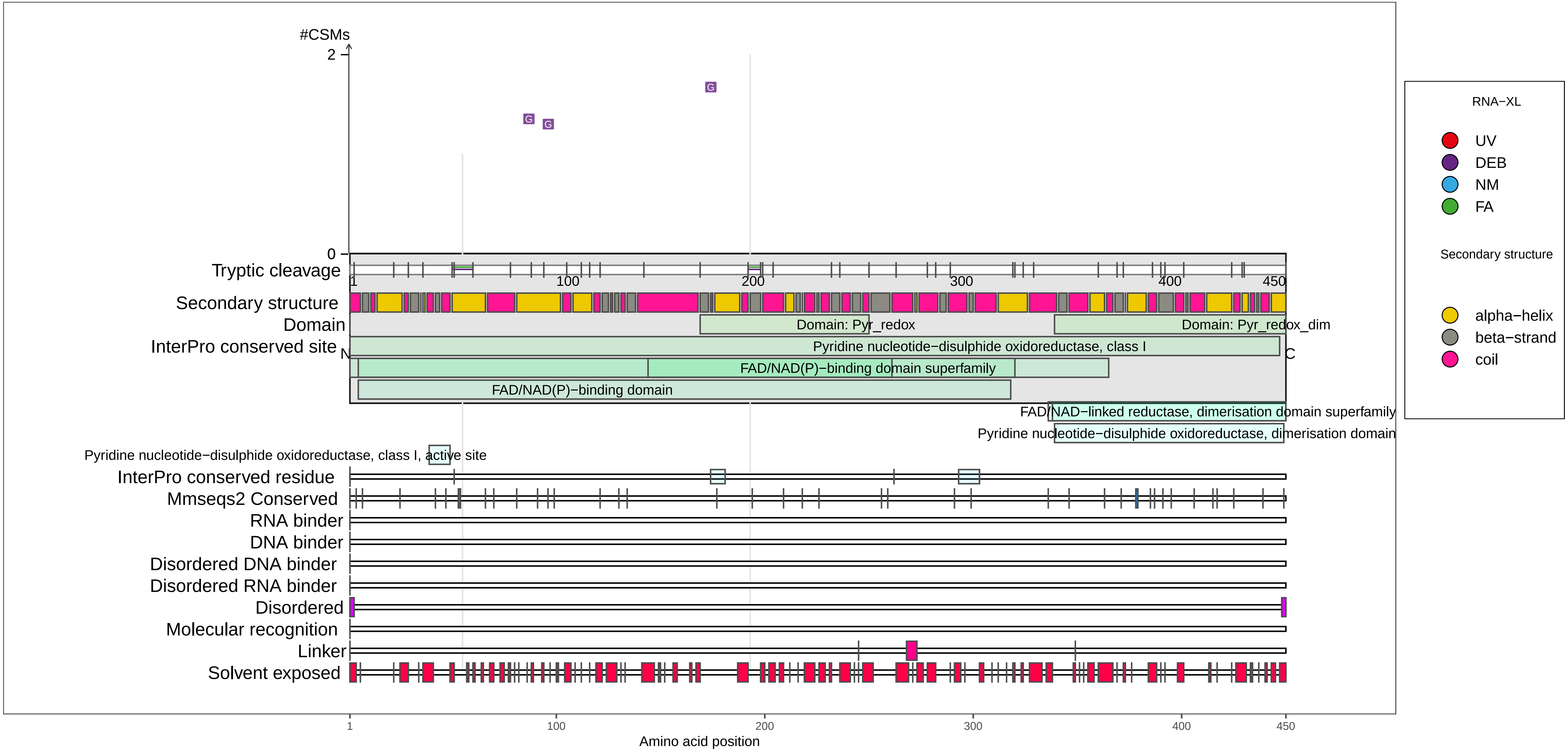

P0A6J8
DDLA_ECOLI D−alanine−−D−alanine ligase A

− Abundance:
tryptic [log10 Intensity]: 8.47 (Q 74)
PAXdb K12 strain [ppm]: 2.5 (Q 77)
PAXdb E.coli [ppm]: 2.11 (Q 75)

− RNA functions: not annotated

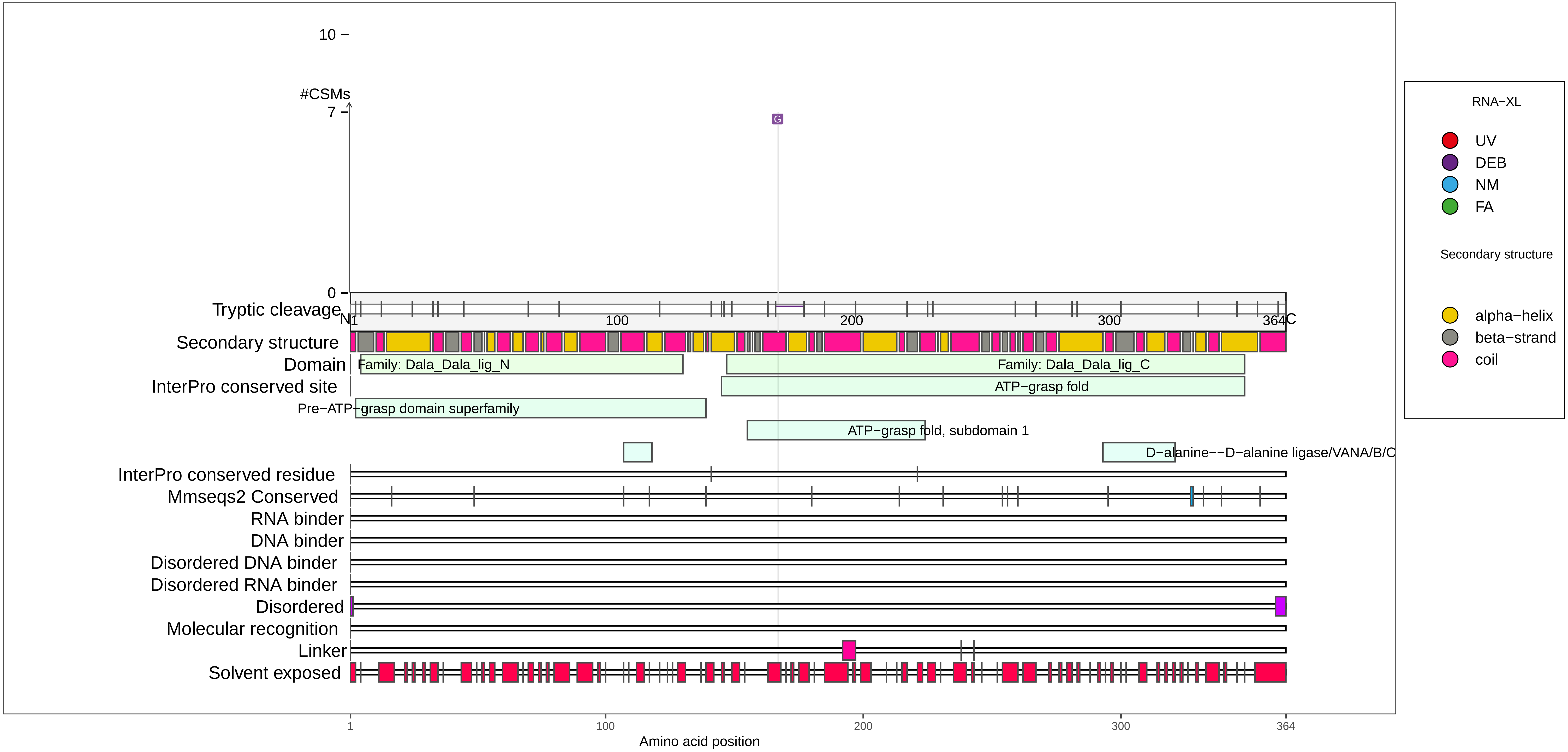

P0A6S0
FLGH_ECOLI Flagellar L−ring protein

− Abundance:
tryptic [log10 Intensity]: 7.88 (Q 53)
PAXdb K12 strain [ppm]: 2.57 (Q 79)
PAXdb E.coli [ppm]: 1.9 (Q 70)

− RNA functions: not annotated

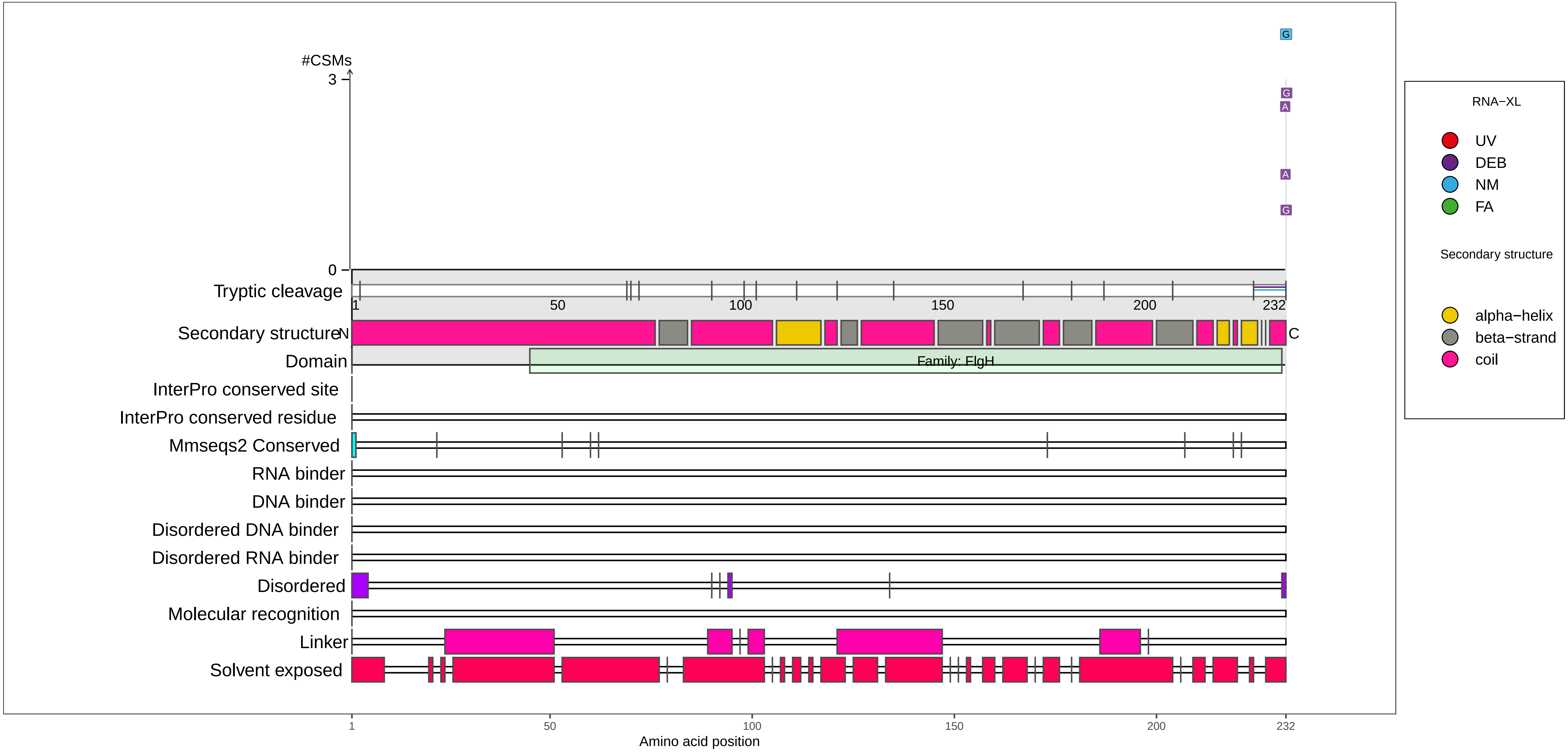

P0A763
NDK_ECOLI Nucleoside diphosphate kinase
− Abundance:
tryptic [log10 Intensity]: 8.83 (Q 84)
PAXdb K12 strain [ppm]: 3.56 (Q 98)
PAXdb E.coli [ppm]: 3.38 (Q 98)

− RNA functions: not annotated

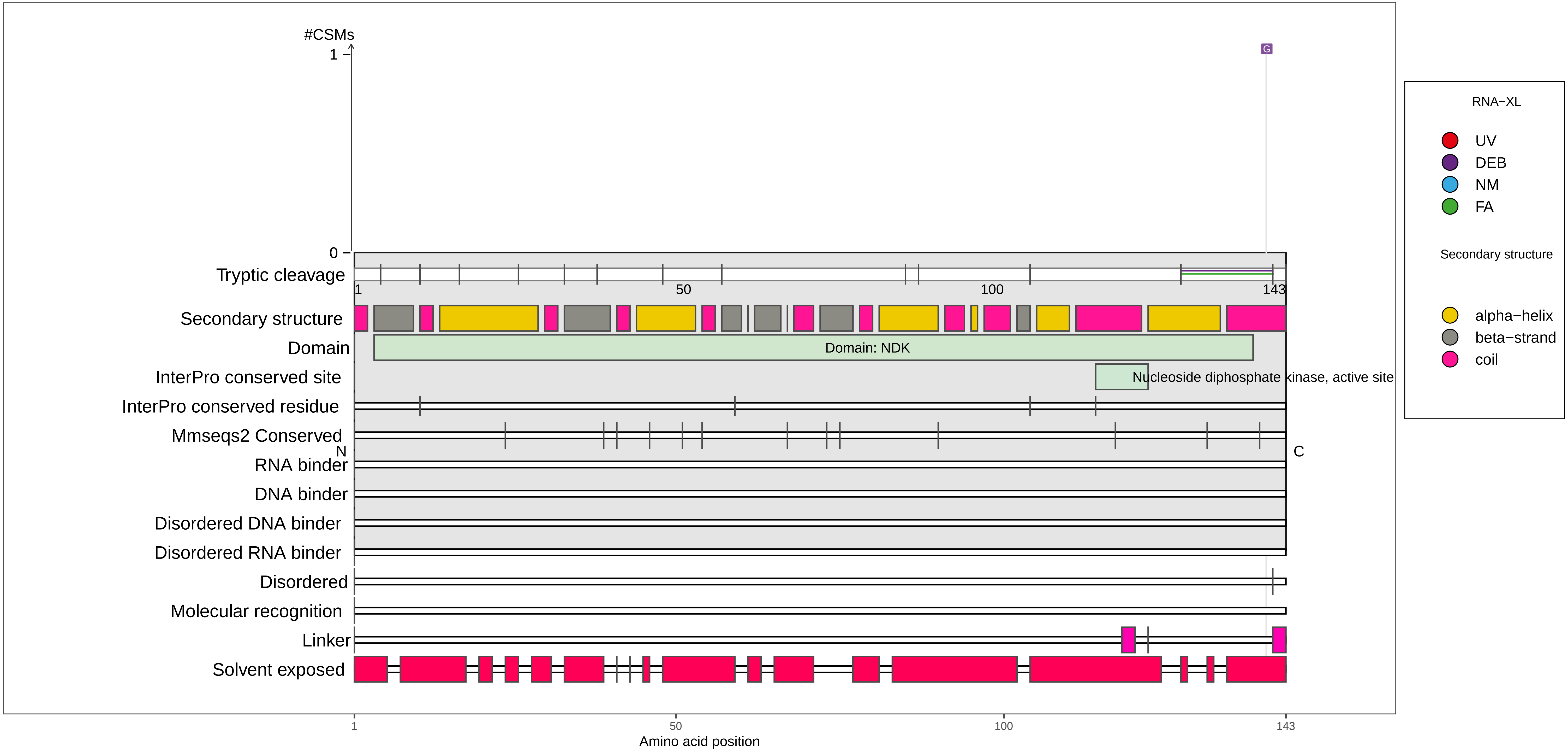

P0A853
TNAA_ECOLI Tryptophanase

− Abundance:
tryptic [log10 Intensity]: 8.36 (Q 71)
PAXdb K12 strain [ppm]: 3.68 (Q 99)
PAXdb E.coli [ppm]: 3.27 (Q 97)

− RNA functions: not annotated

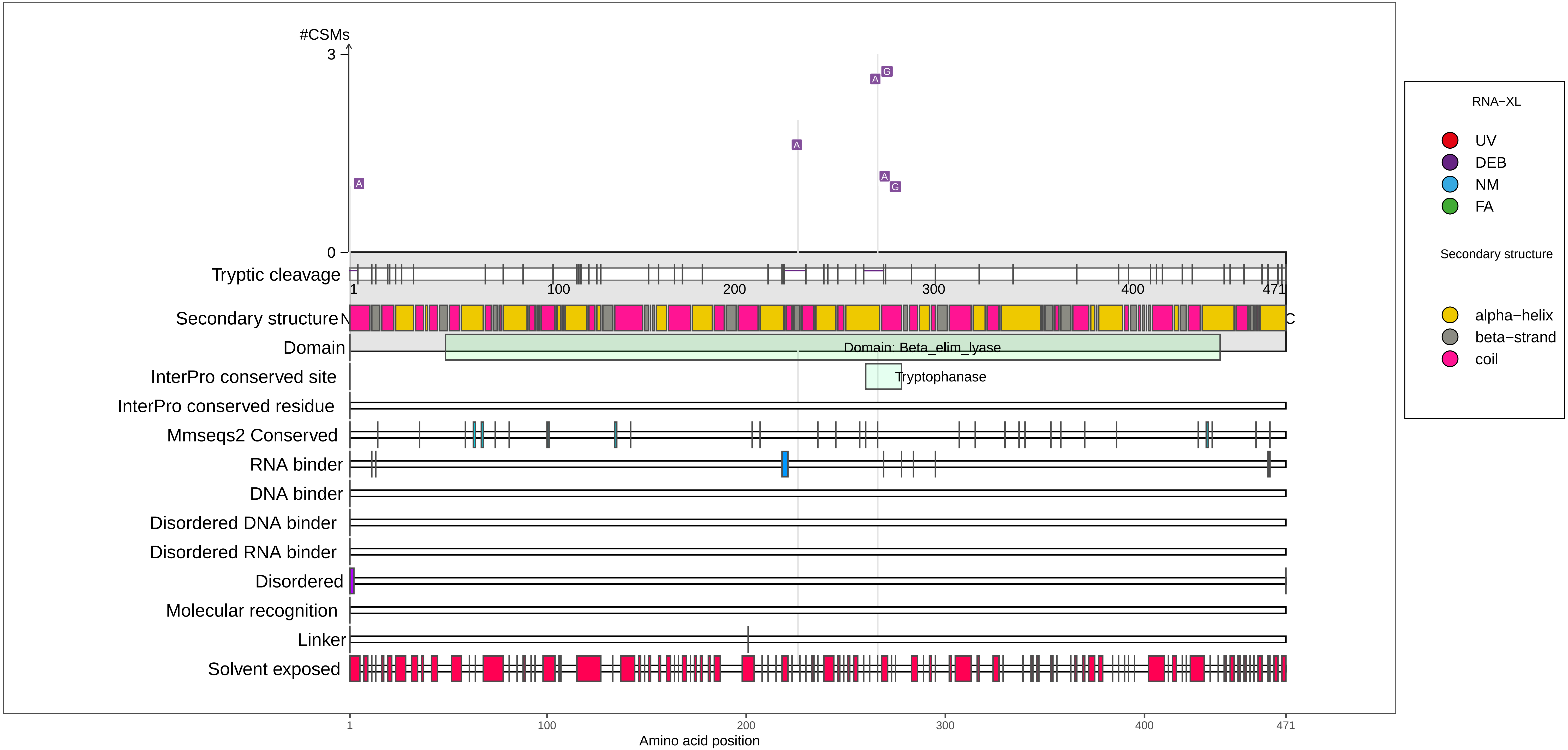

P0A9Q9
DHAS_ECOLI Aspartate−semialdehyde dehydrogenase

− Abundance:
tryptic [log10 Intensity]: 8.85 (Q 85)
PAXdb K12 strain [ppm]: 2.56 (Q 79)
PAXdb E.coli [ppm]: 3.03 (Q 95)

− RNA functions: not annotated

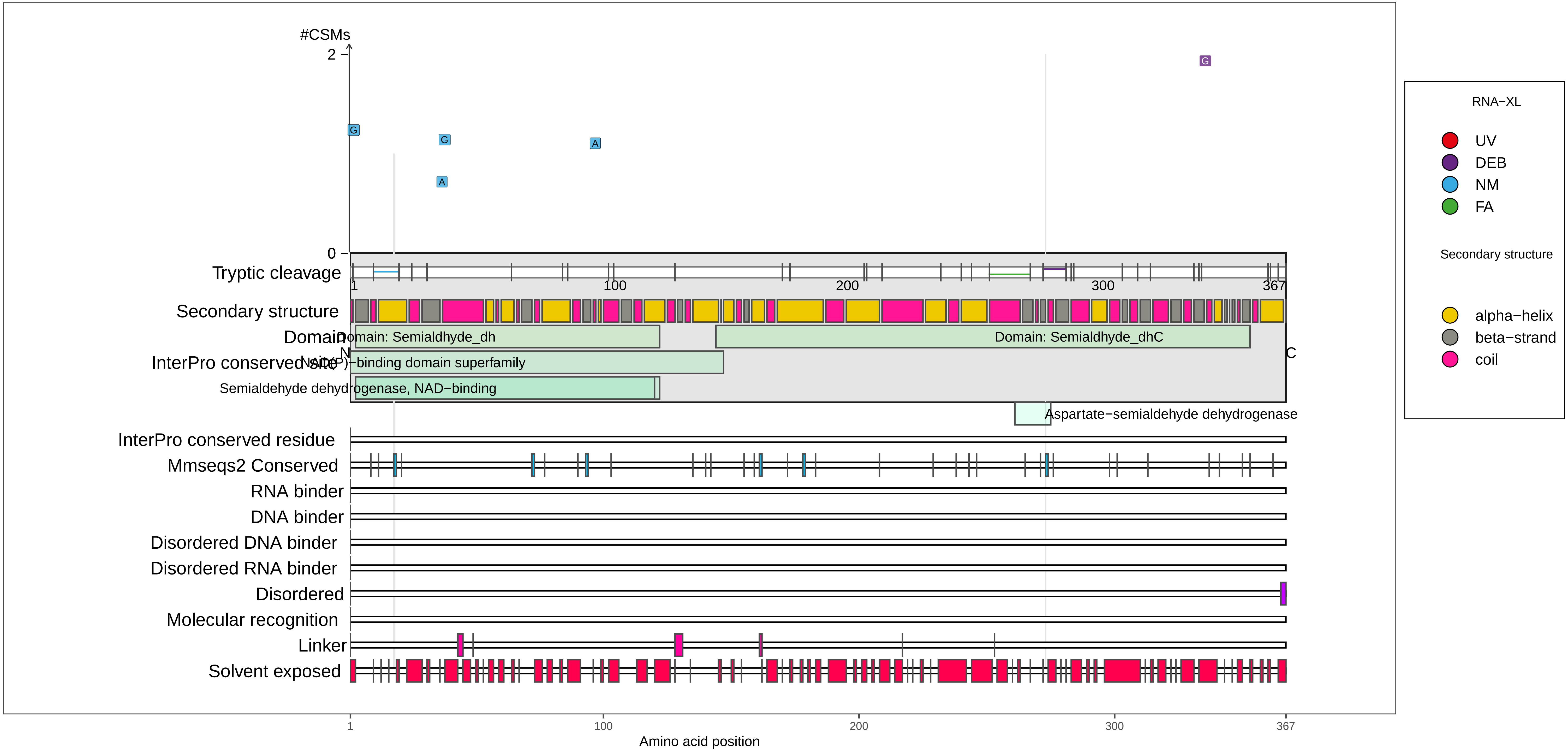

P0AA16
OMPR_ECOLI DNA−binding dual transcriptional regulator OmpR

− Abundance:
tryptic [log10 Intensity]: 6.76 (Q 6)
PAXdb K12 strain [ppm]: 2.65 (Q 82)
PAXdb E.coli [ppm]: 2.8 (Q 92)

− RNA functions: not annotated

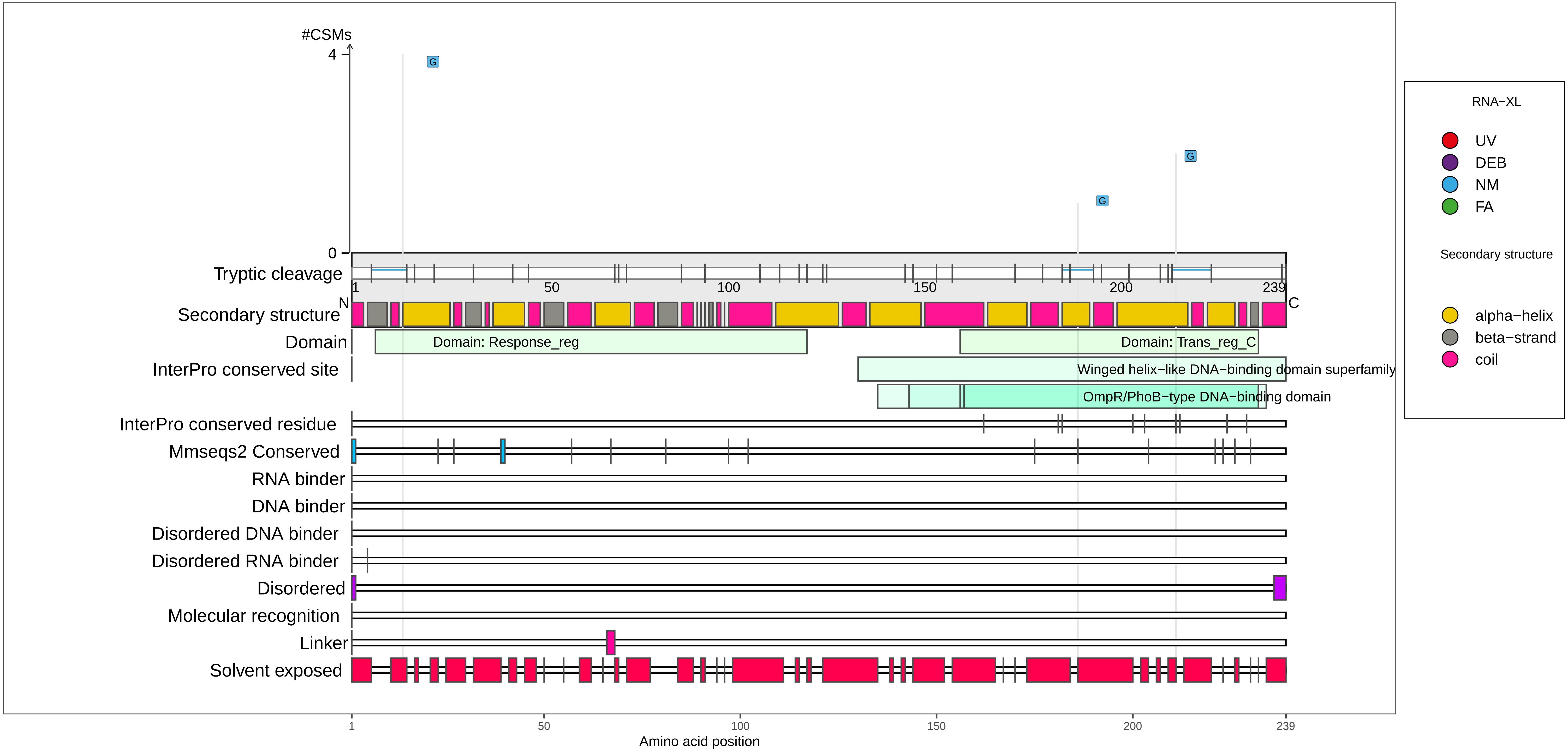

P0ABA0
ATPF_ECOLI ATP synthase subunit b

− Abundance:
tryptic [log10 Intensity]: 9 (Q 88)
PAXdb K12 strain [ppm]: 2.93 (Q 89)
PAXdb E.coli [ppm]: 2.93 (Q 94)

− RNA functions: not annotated

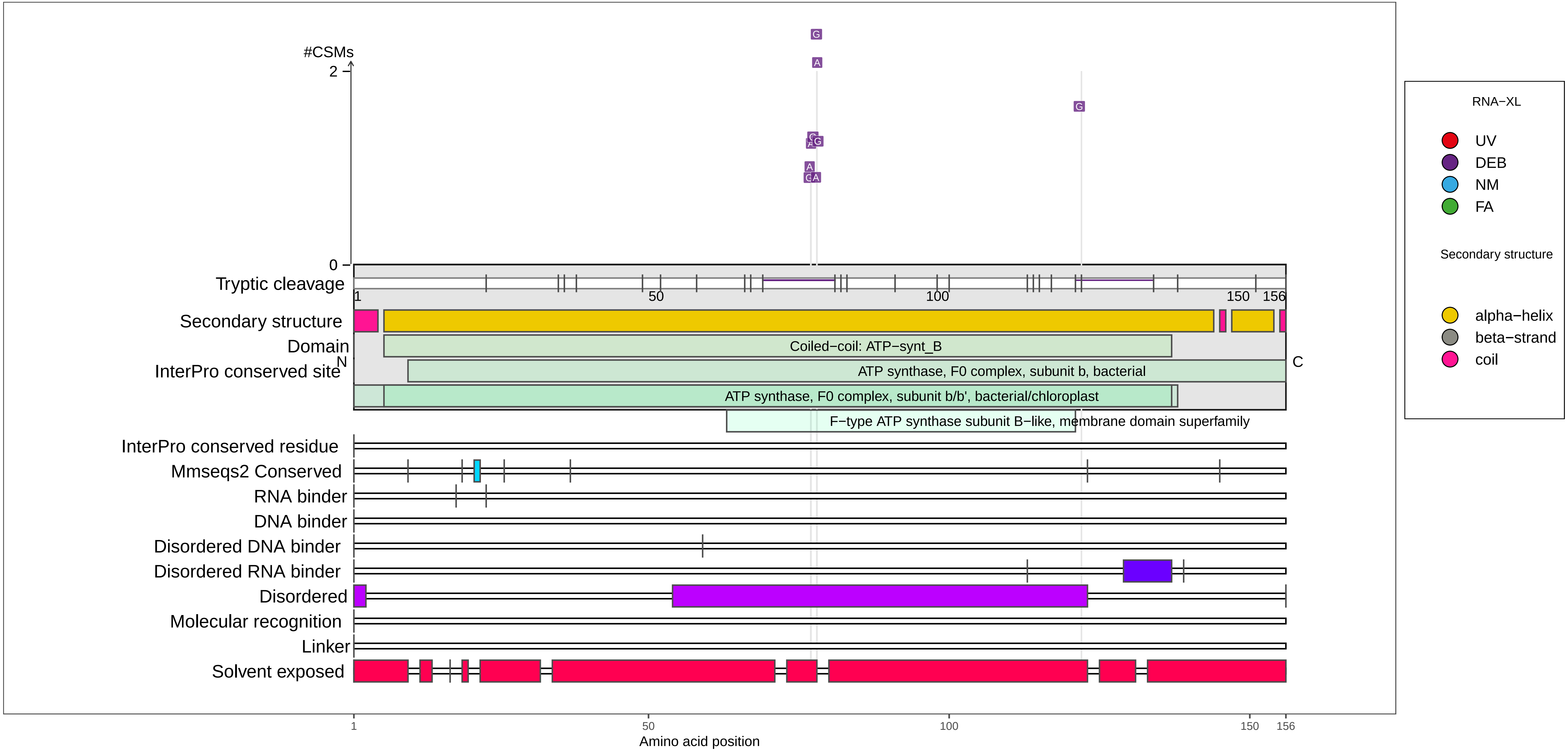

P0ABK5
CYSK_ECOLI Cysteine synthase A

− Abundance:
tryptic [log10 Intensity]: 8.83 (Q 84)
PAXdb K12 strain [ppm]: 2.98 (Q 90)
PAXdb E.coli [ppm]: 3.5 (Q 99)

− RNA functions: not annotated

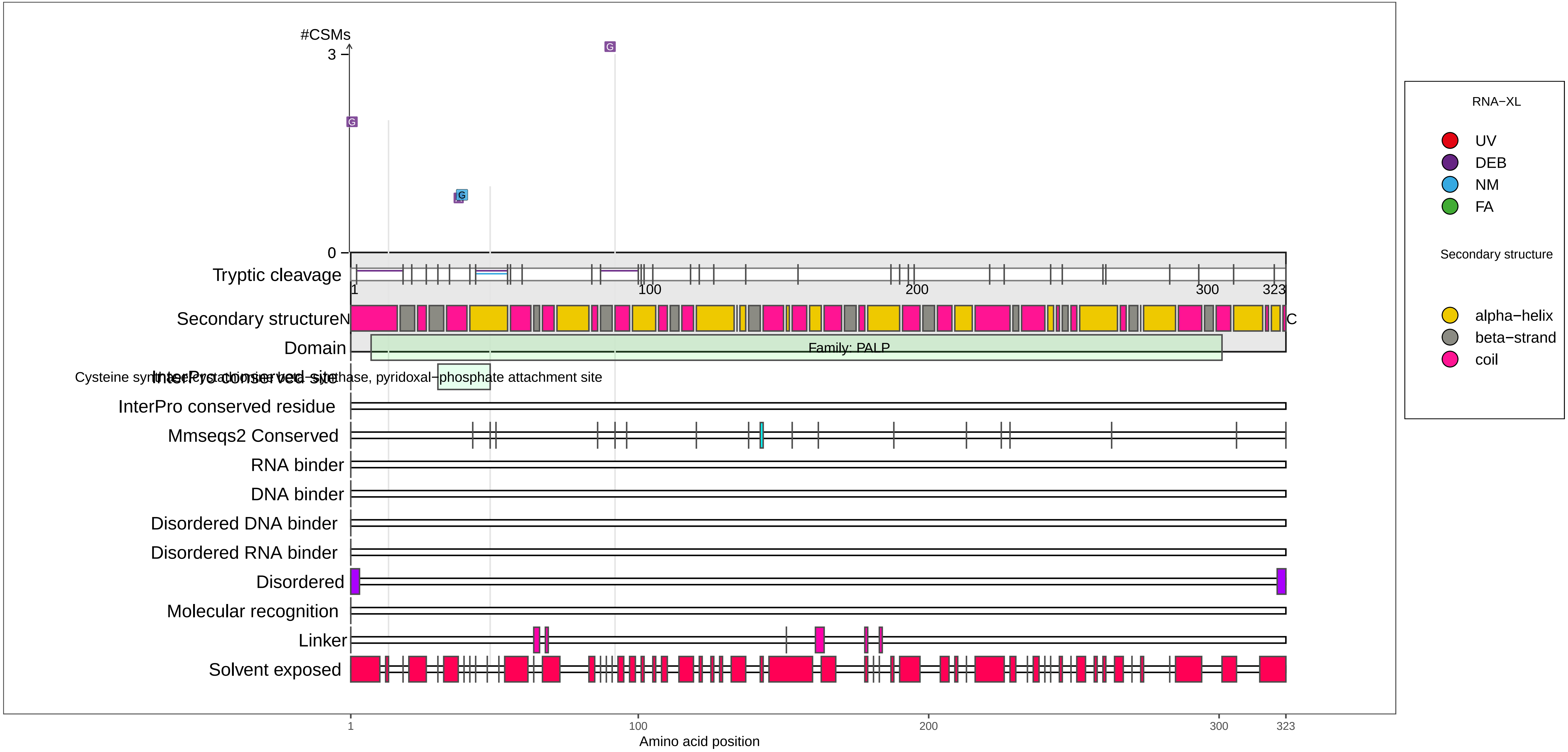

P0ABX5
FLGG_ECOLI Flagellar basal−body rod protein FlgG

− Abundance:
tryptic [log10 Intensity]: 7.48 (Q 34)
PAXdb K12 strain [ppm]: 1.77 (Q 52)
PAXdb E.coli [ppm]: 0.72 (Q 41)

− RNA functions: not annotated

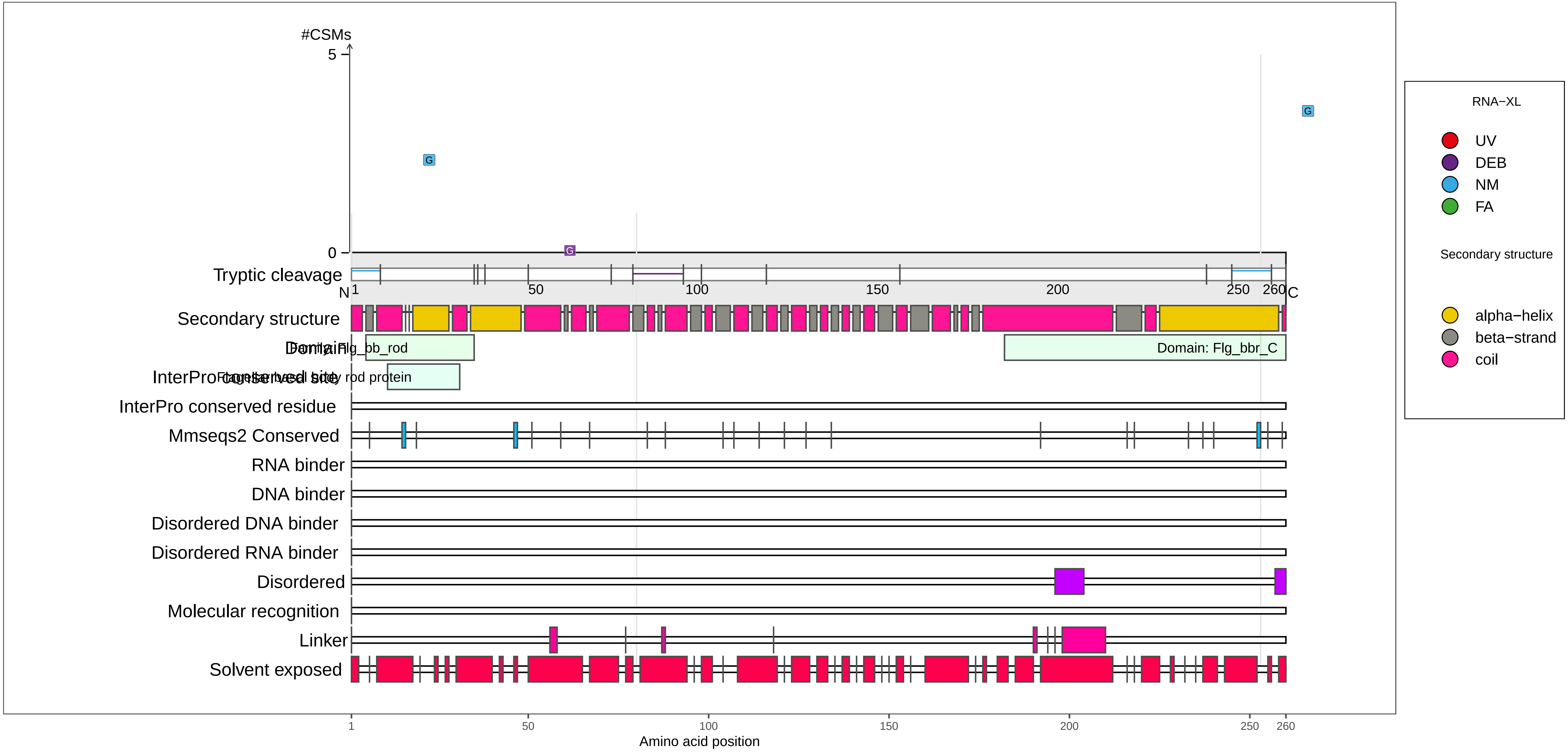

P0AC62
GLRX3_ECOLI Glutaredoxin 3

− Abundance:
tryptic [log10 Intensity]: NA (Q NA)
PAXdb K12 strain [ppm]: 3.11 (Q 92)
PAXdb E.coli [ppm]: 2.8 (Q 92)

− RNA functions: not annotated

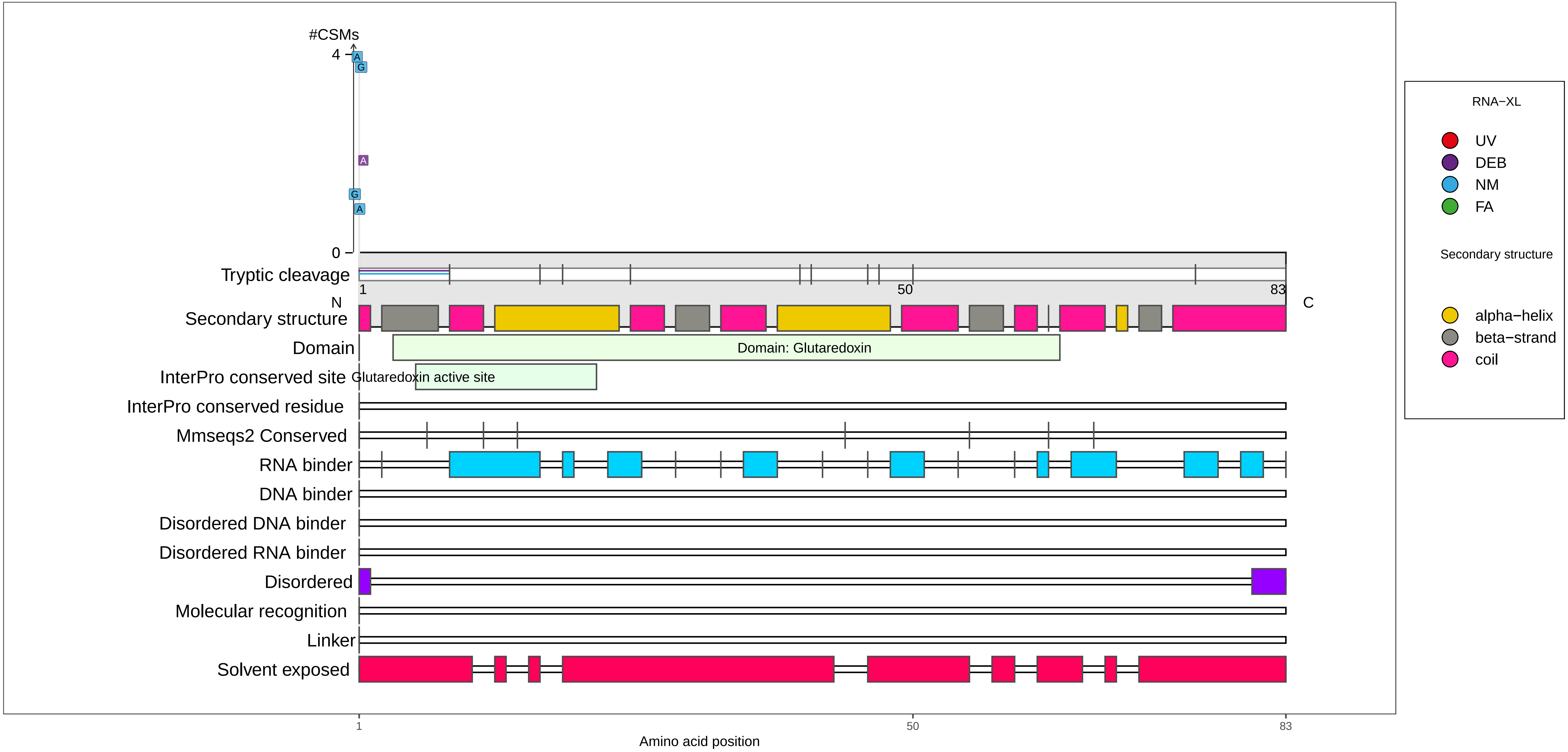

P0AE52
BCP_ECOLI Peroxiredoxin Bcp

− Abundance:
tryptic [log10 Intensity]: 8.78 (Q 83)
PAXdb K12 strain [ppm]: 3.45 (Q 97)
PAXdb E.coli [ppm]: 2.94 (Q 94)

− RNA functions: not annotated

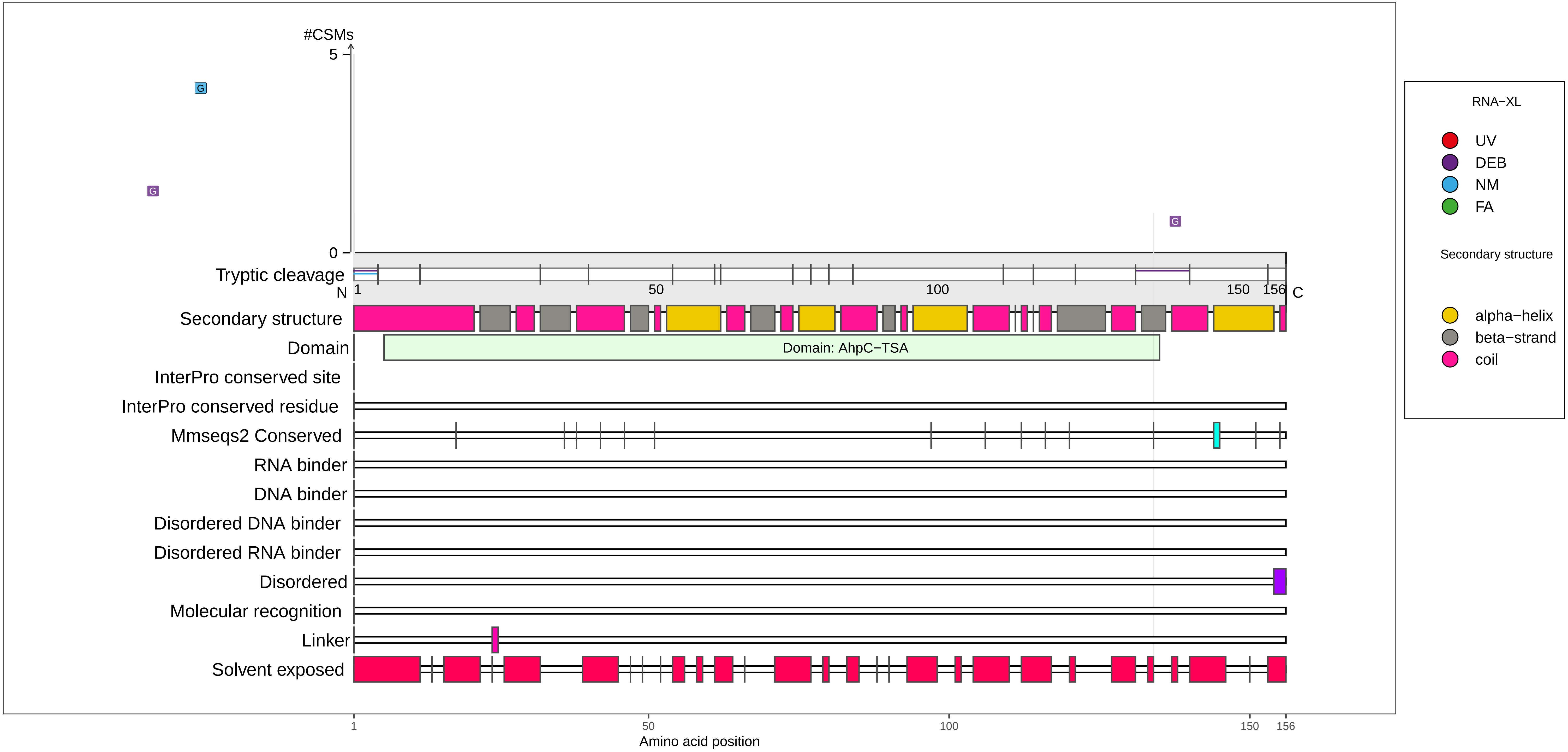

P0AED7
DAPE_ECOLI Succinyl−diaminopimelate desuccinylase

− Abundance:
tryptic [log10 Intensity]: 8.66 (Q 80)
PAXdb K12 strain [ppm]: 0.76 (Q 4)
PAXdb E.coli [ppm]: 1.46 (Q 59)

− RNA functions: not annotated

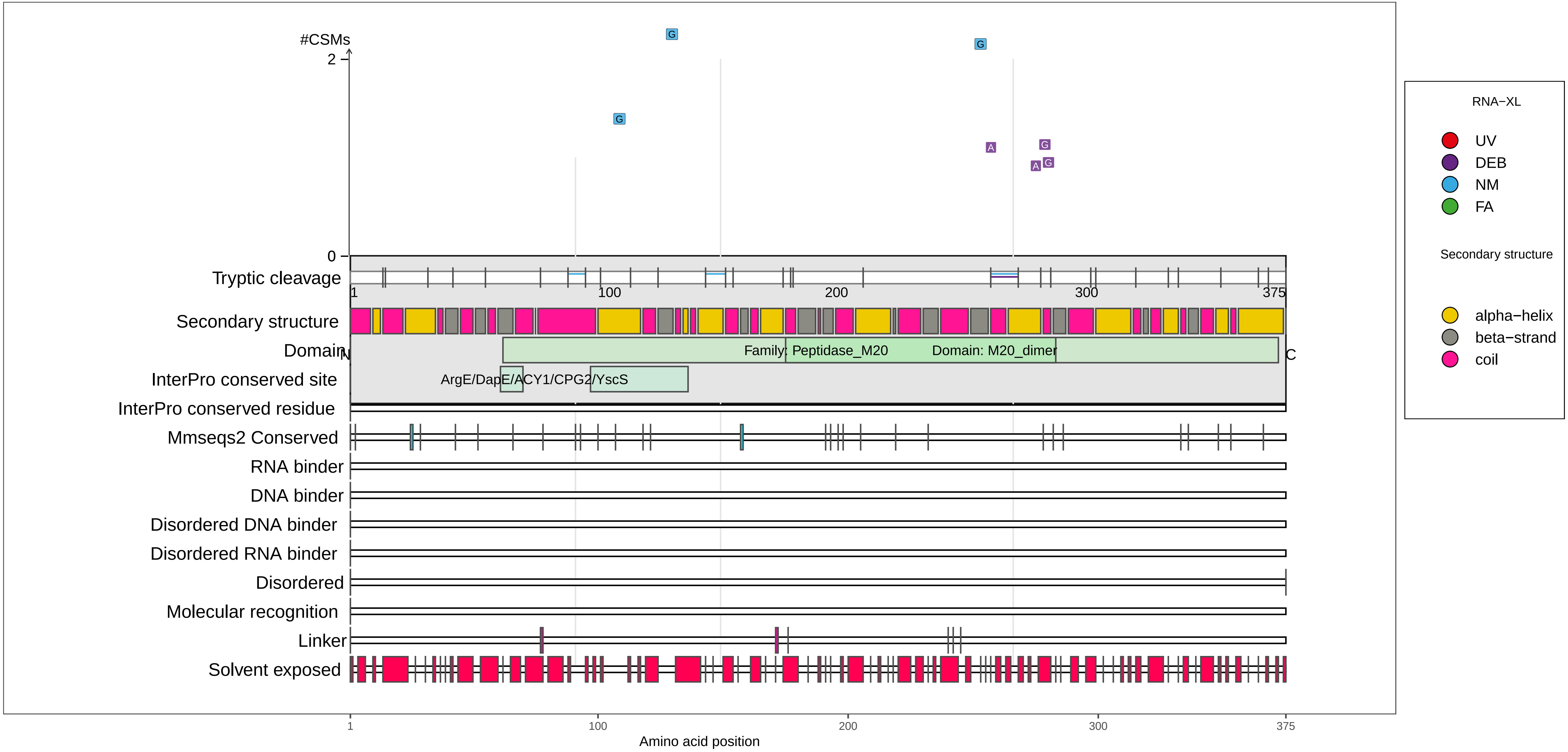

P0AEJ6
EUTB_ECOLI Ethanolamine ammonia−lyase large subunit

− Abundance:
tryptic [log10 Intensity]: 6.98 (Q 12)
PAXdb K12 strain [ppm]: 1.3 (Q 26)
PAXdb E.coli [ppm]: 1.31 (Q 55)

− RNA functions: not annotated

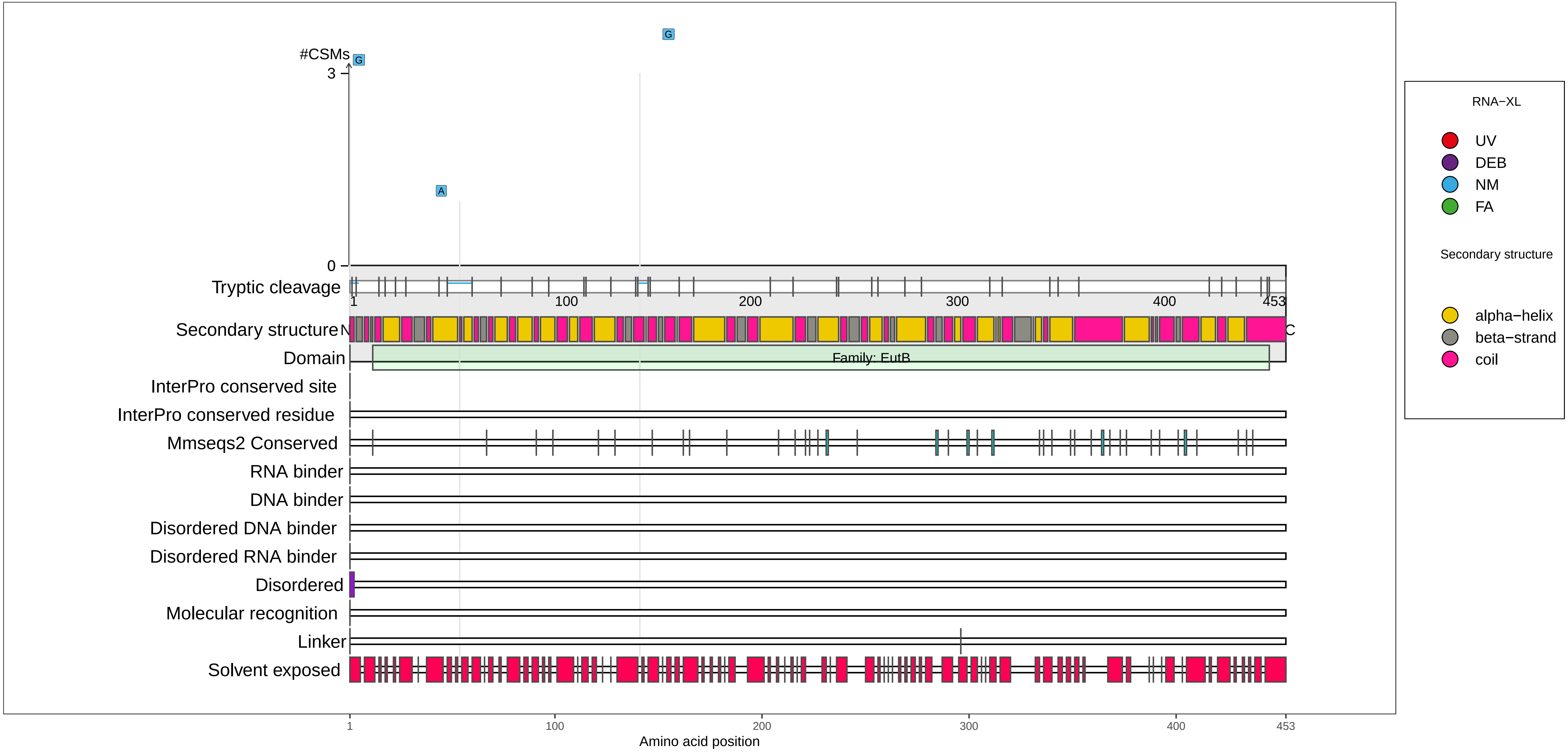

P0AEW4
CPDA_ECOLI 3',5'−cyclic adenosine monophosphate phosphodiesterase CpdA

− Abundance:
tryptic [log10 Intensity]: 7.63 (Q 41)
PAXdb K12 strain [ppm]: 1.99 (Q 60)
PAXdb E.coli [ppm]: 1.79 (Q 67)

− RNA functions: not annotated

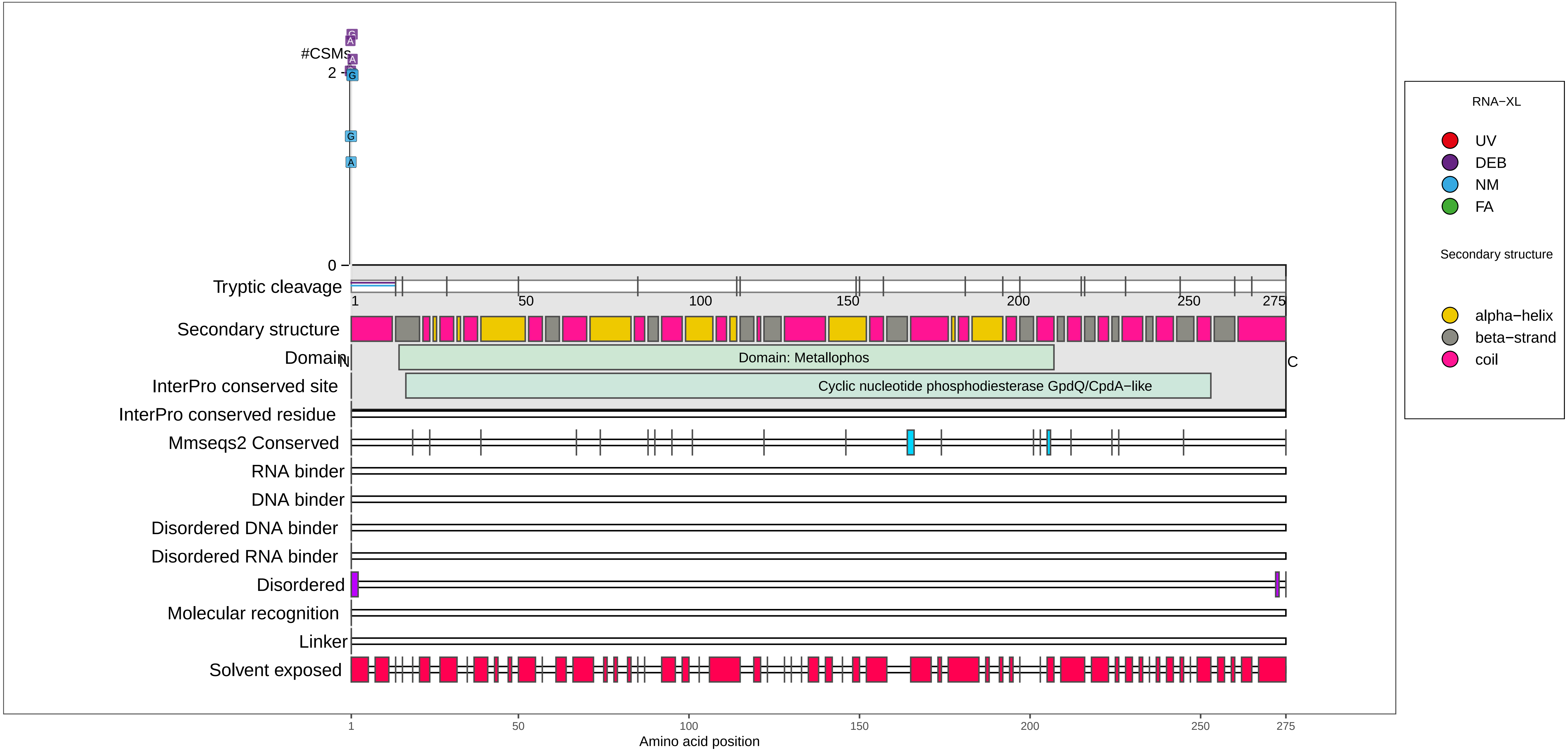

P0AEZ1
METF_ECOLI 5,10−methylenetetrahydrofolate reductase

− Abundance:
tryptic [log10 Intensity]: 7.32 (Q 26)
PAXdb K12 strain [ppm]: 2.18 (Q 66)
PAXdb E.coli [ppm]: 2.47 (Q 85)

− RNA functions: not annotated

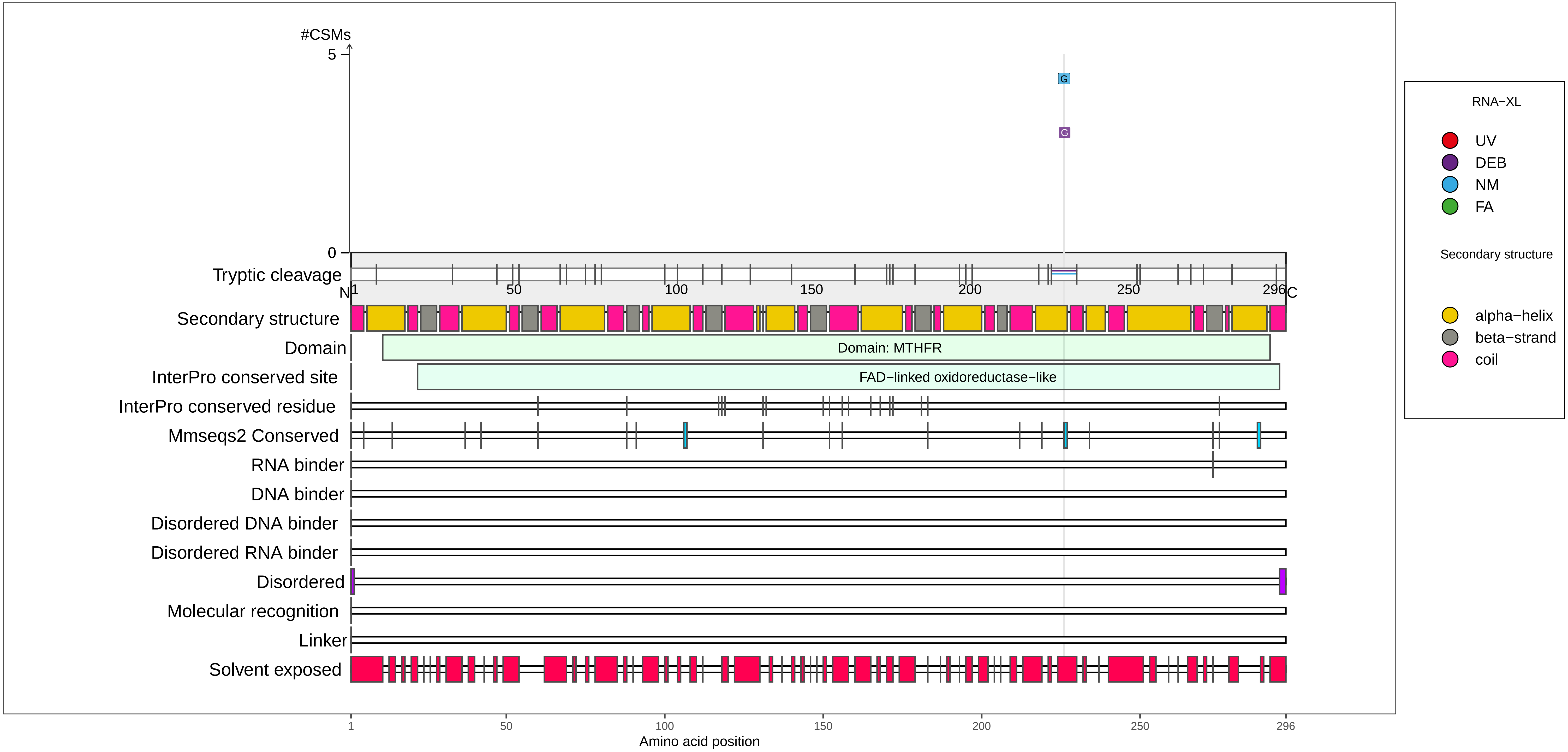

P0AFL3
PPIA_ECOLI Peptidyl−prolyl cis−trans isomerase A

− Abundance:
tryptic [log10 Intensity]: 8.93 (Q 87)
PAXdb K12 strain [ppm]: NA (Q NA)
PAXdb E.coli [ppm]: 2.53 (Q 86)

− RNA functions: not annotated

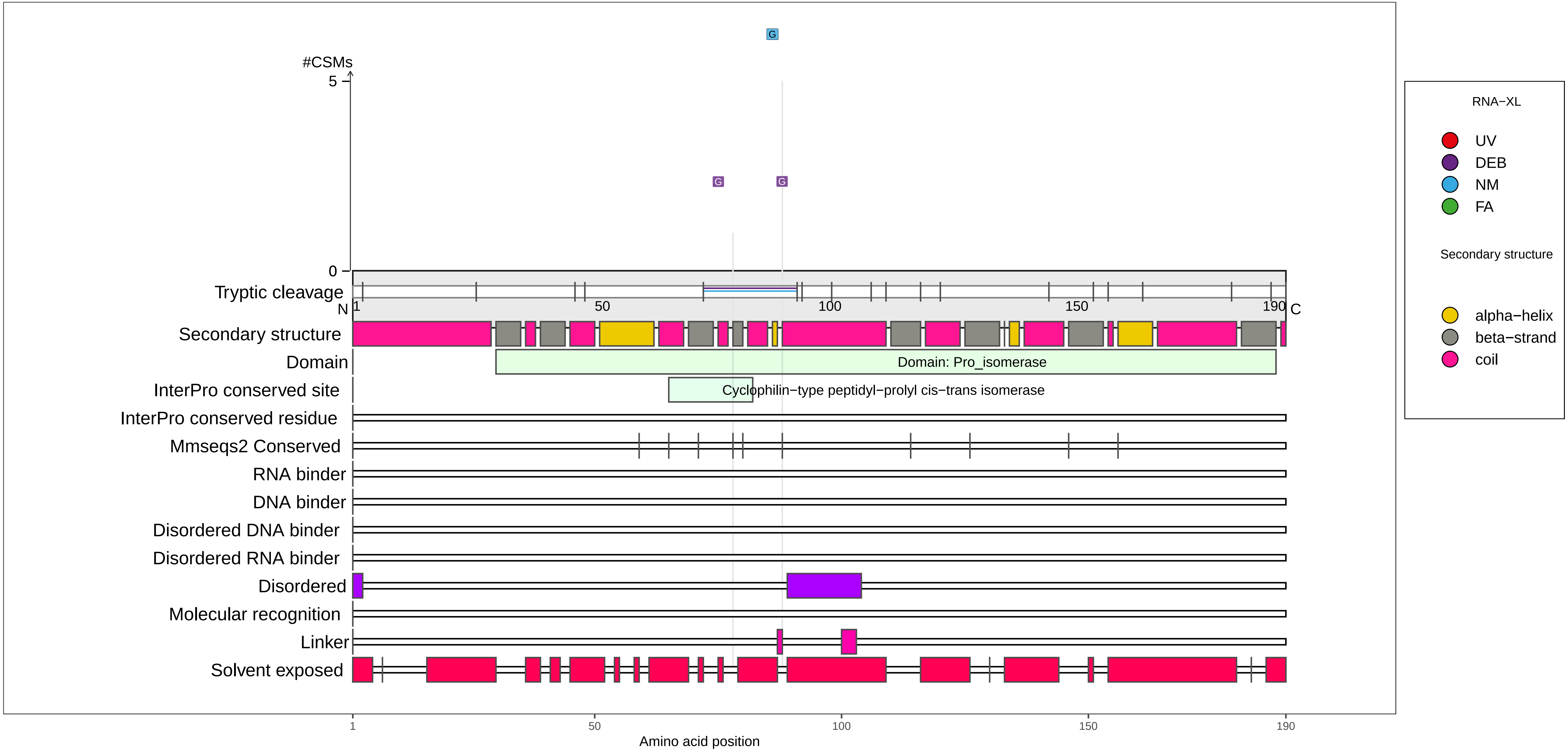

P0AG86
SECB_ECOLI Protein−export protein SecB

− Abundance:
tryptic [log10 Intensity]: 8.86 (Q 85)
PAXdb K12 strain [ppm]: 3.19 (Q 94)
PAXdb E.coli [ppm]: 3.42 (Q 98)

− RNA functions: not annotated

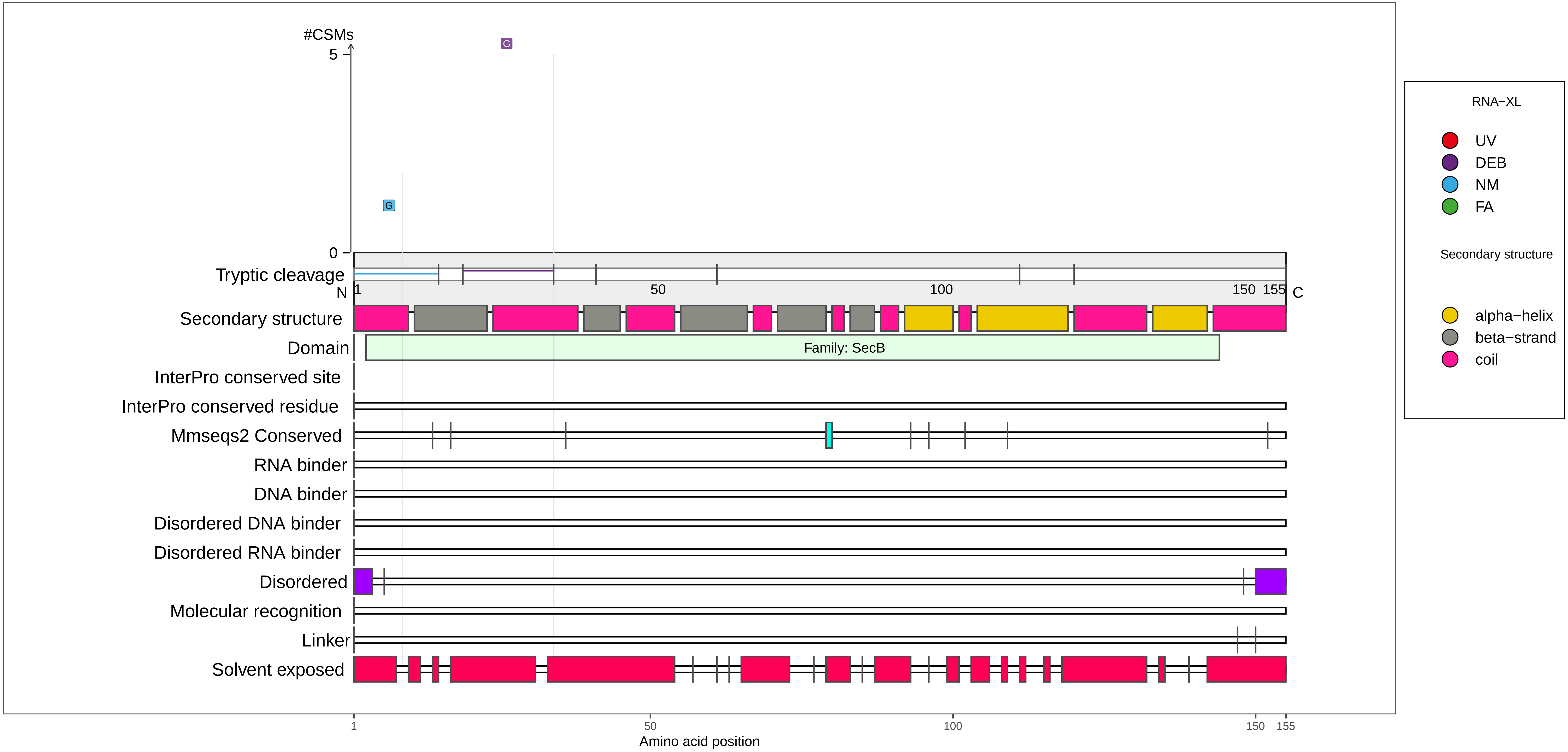

P15042
DNLJ_ECOLI DNA ligase

− Abundance:
tryptic [log10 Intensity]: 8.57 (Q 78)
PAXdb K12 strain [ppm]: 1.54 (Q 40)
PAXdb E.coli [ppm]: 1.91 (Q 70)

− RNA functions: not annotated

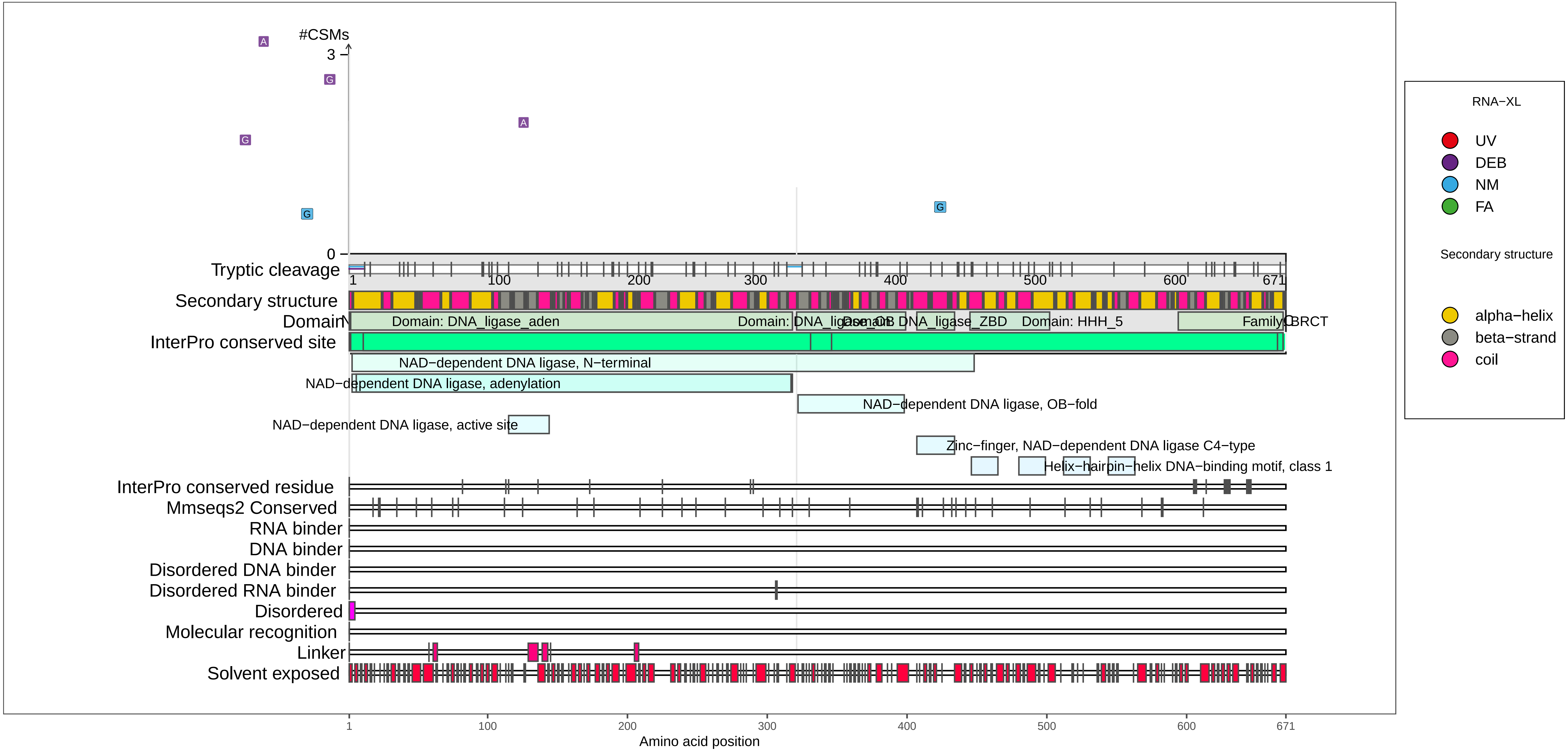

P24171
DCP_ECOLI Dipeptidyl carboxypeptidase

− Abundance:
tryptic [log10 Intensity]: 8.75 (Q 82)
PAXdb K12 strain [ppm]: 1.86 (Q 56)
PAXdb E.coli [ppm]: 2.41 (Q 83)

− RNA functions: not annotated

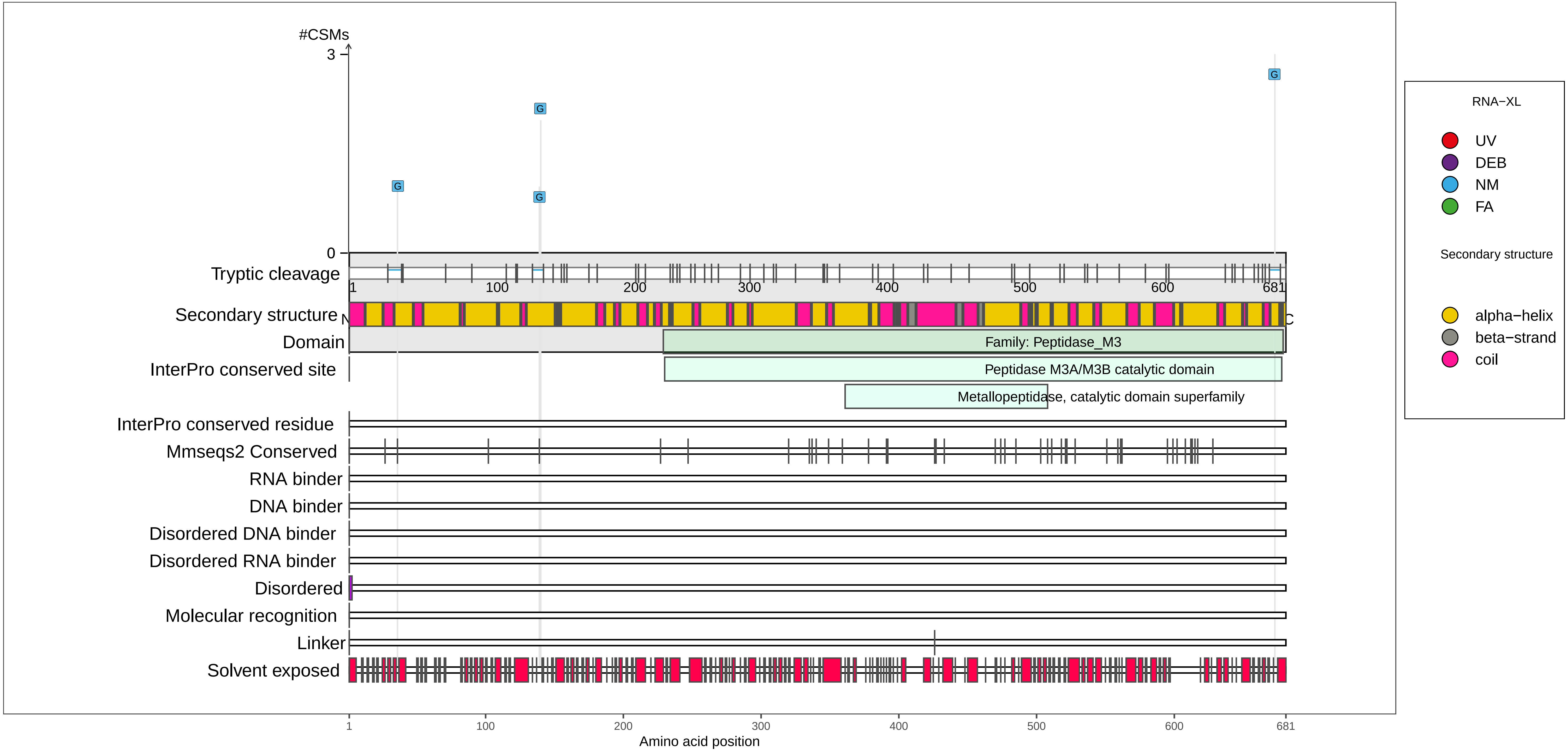

P30958
MFD_ECOLI Transcription−repair−coupling factor

− Abundance:
tryptic [log10 Intensity]: 7.92 (Q 54)
PAXdb K12 strain [ppm]: 2.18 (Q 66)
PAXdb E.coli [ppm]: 1.74 (Q 66)

− RNA functions:
RNA polymerase binding; RNA polymerase core enzyme binding
This domain is found in proteins necessary for strand−specific repair in DNA such as TRCF in Escherichia coli. A lesion in the template strand blocks the RNA polymerase complex (RNAP). The RNAP−DNA−RNA complex is spec

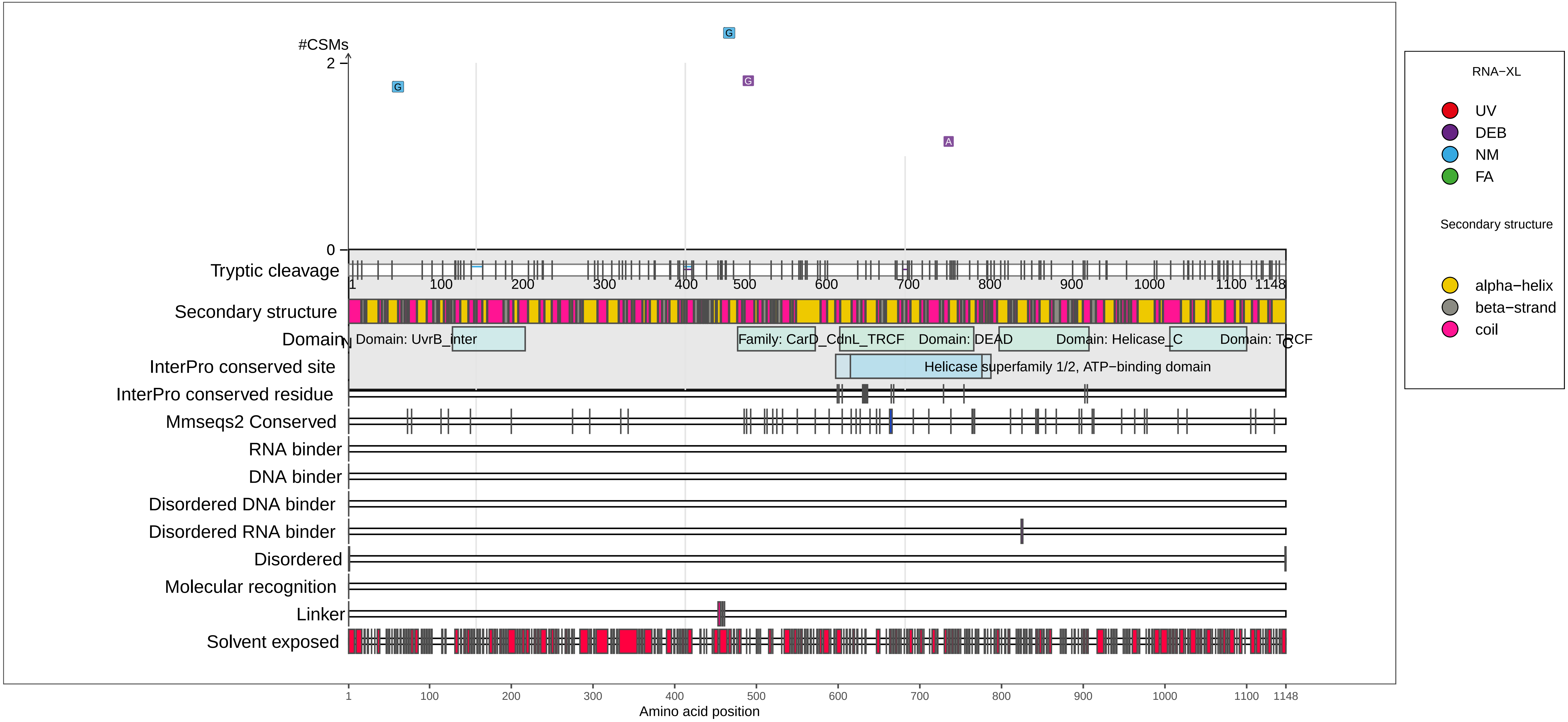

P37051
PURU_ECOLI Formyltetrahydrofolate deformylase

− Abundance:
tryptic [log10 Intensity]: 9.19 (Q 92)
PAXdb K12 strain [ppm]: 2.24 (Q 68)
PAXdb E.coli [ppm]: 2.25 (Q 79)

− RNA functions: not annotated

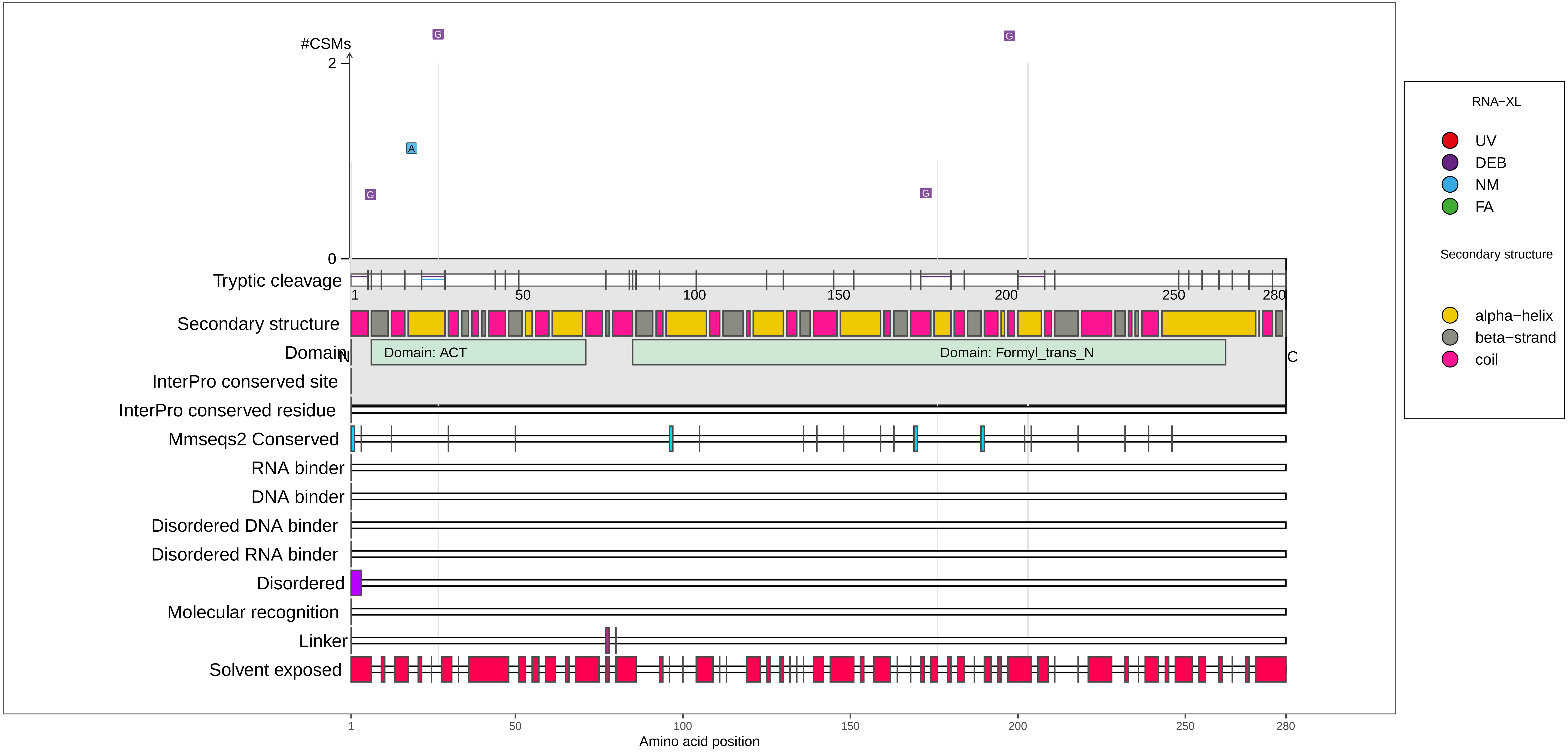

P37692
RFAF_ECOLI ADP−heptose−−LPS heptosyltransferase 2

− Abundance:
tryptic [log10 Intensity]: 7.69 (Q 44)
PAXdb K12 strain [ppm]: 1.83 (Q 54)
PAXdb E.coli [ppm]: 0.71 (Q 41)

− RNA functions: not annotated

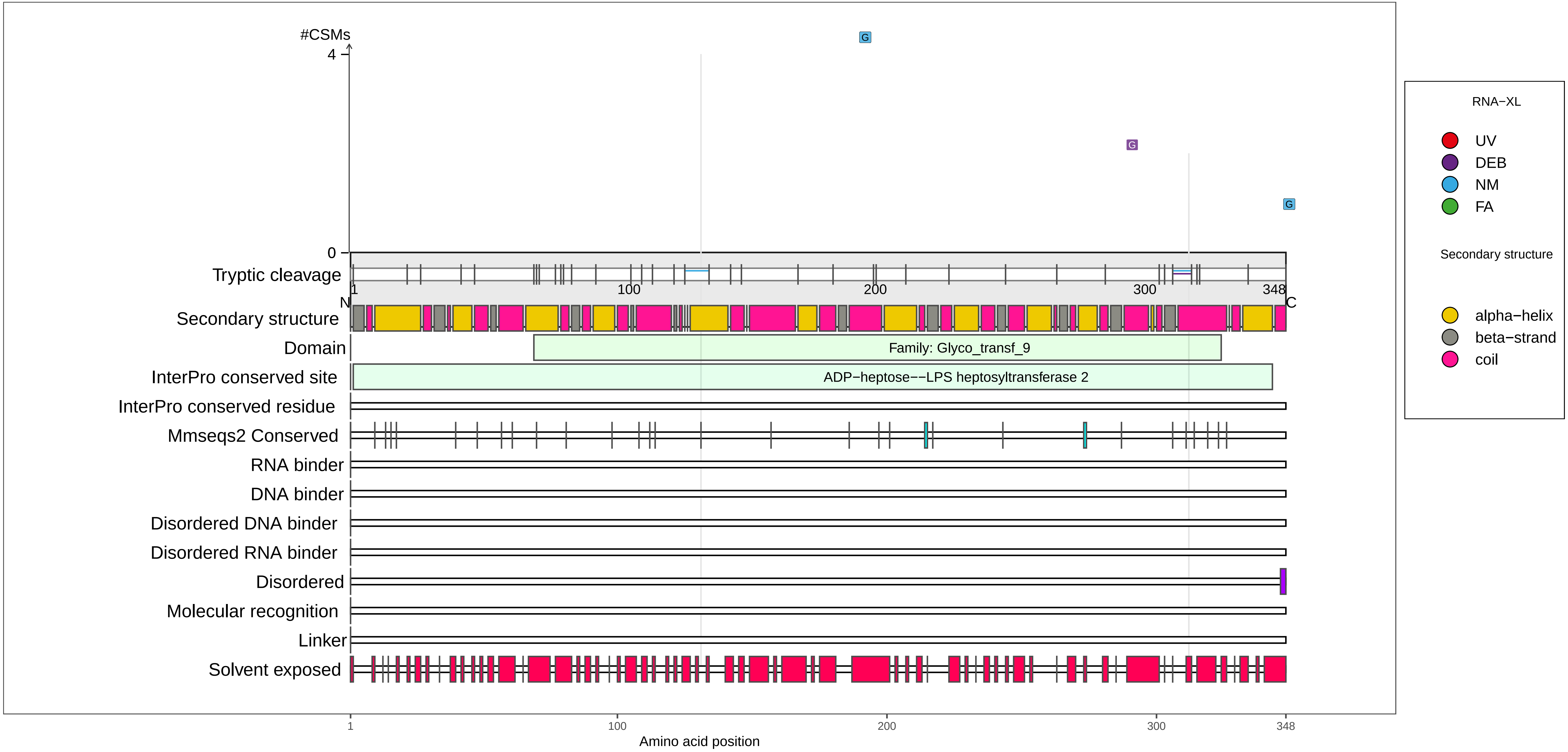

P38054
CUSA_ECOLI Cation efflux system protein CusA

− Abundance:
tryptic [log10 Intensity]: 7.11 (Q 17)
PAXdb K12 strain [ppm]: 1 (Q 11)
PAXdb E.coli [ppm]: 0.77 (Q 42)

− RNA functions: not annotated

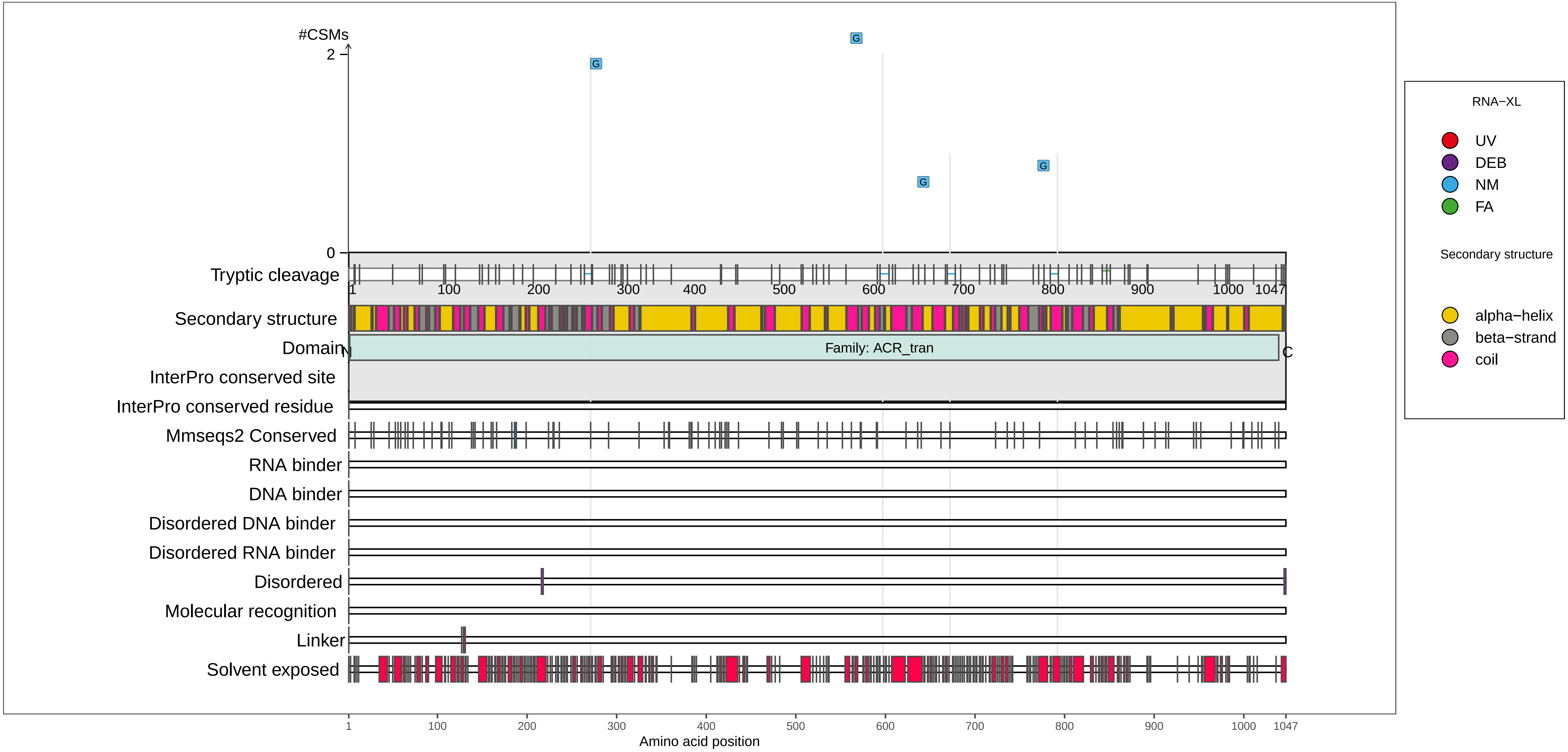

P67660
YHAJ_ECOLI Probable HTH−type transcriptional regulator YhaJ
− Abundance:
tryptic [log10 Intensity]: 7.08 (Q 16)
PAXdb K12 strain [ppm]: NA (Q NA)
PAXdb E.coli [ppm]: 1.87 (Q 69)

− RNA functions:
RNA biosynthetic process; RNA metabolic process

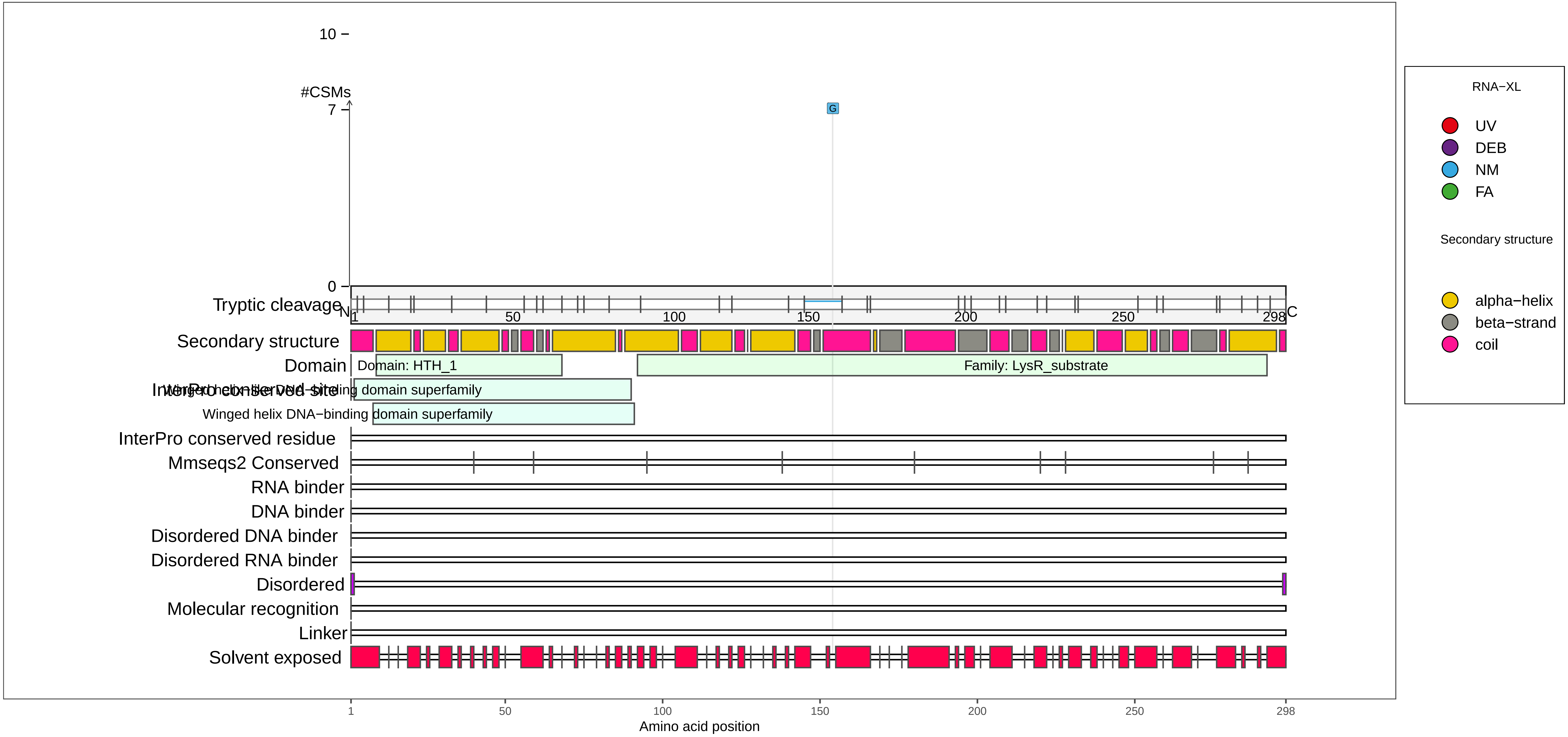

P67699
YDDM_ECOLI Uncharacterized HTH−type transcriptional regulator YddM

− Abundance:
tryptic [log10 Intensity]: NA (Q NA)
PAXdb K12 strain [ppm]: NA (Q NA)
PAXdb E.coli [ppm]: −0.38 (Q 17)

− RNA functions: not annotated

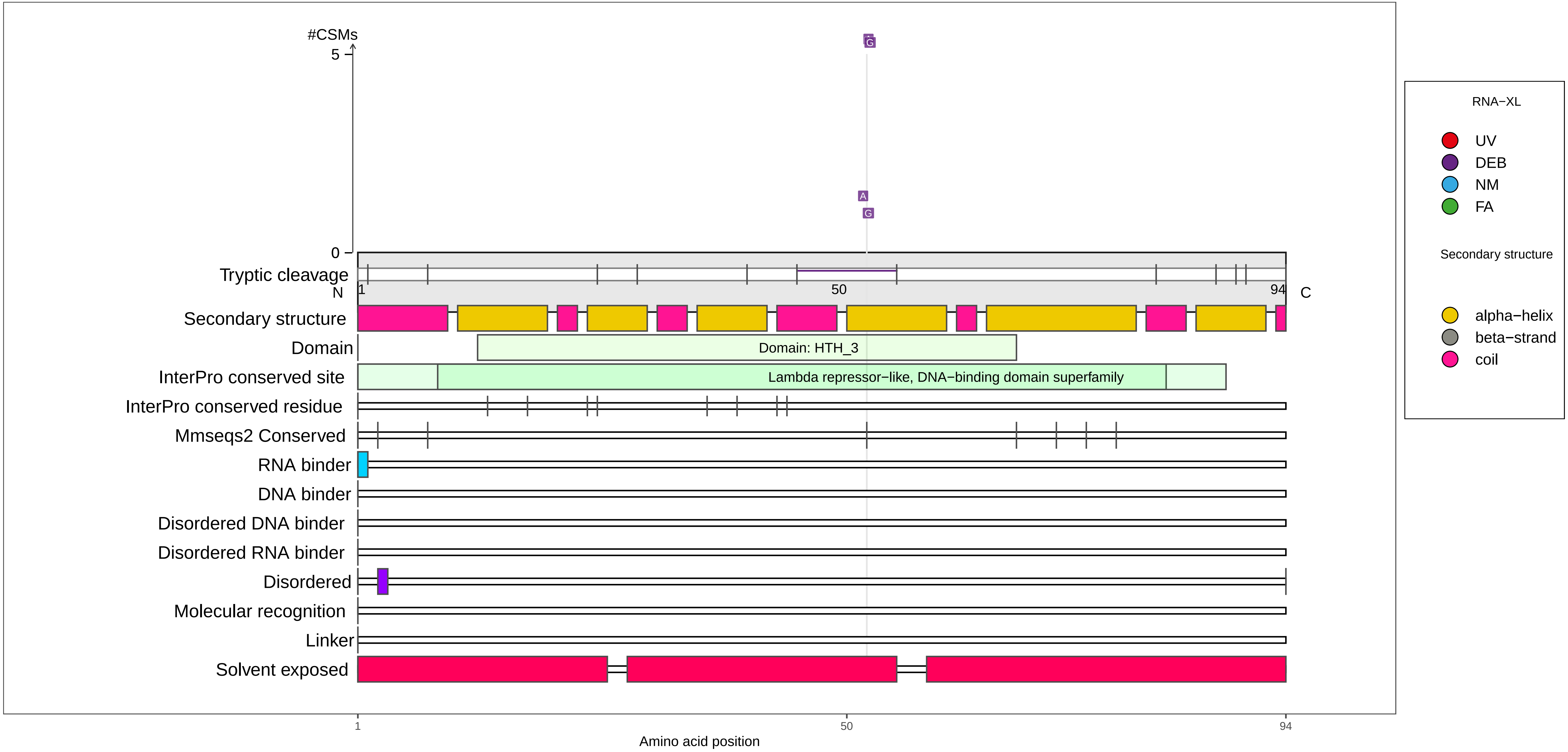

P68688
GLRX1_ECOLI Glutaredoxin 1

− Abundance:
tryptic [log10 Intensity]: 8.47 (Q 74)
PAXdb K12 strain [ppm]: 2.1 (Q 64)
PAXdb E.coli [ppm]: 1.97 (Q 71)

− RNA functions: not annotated

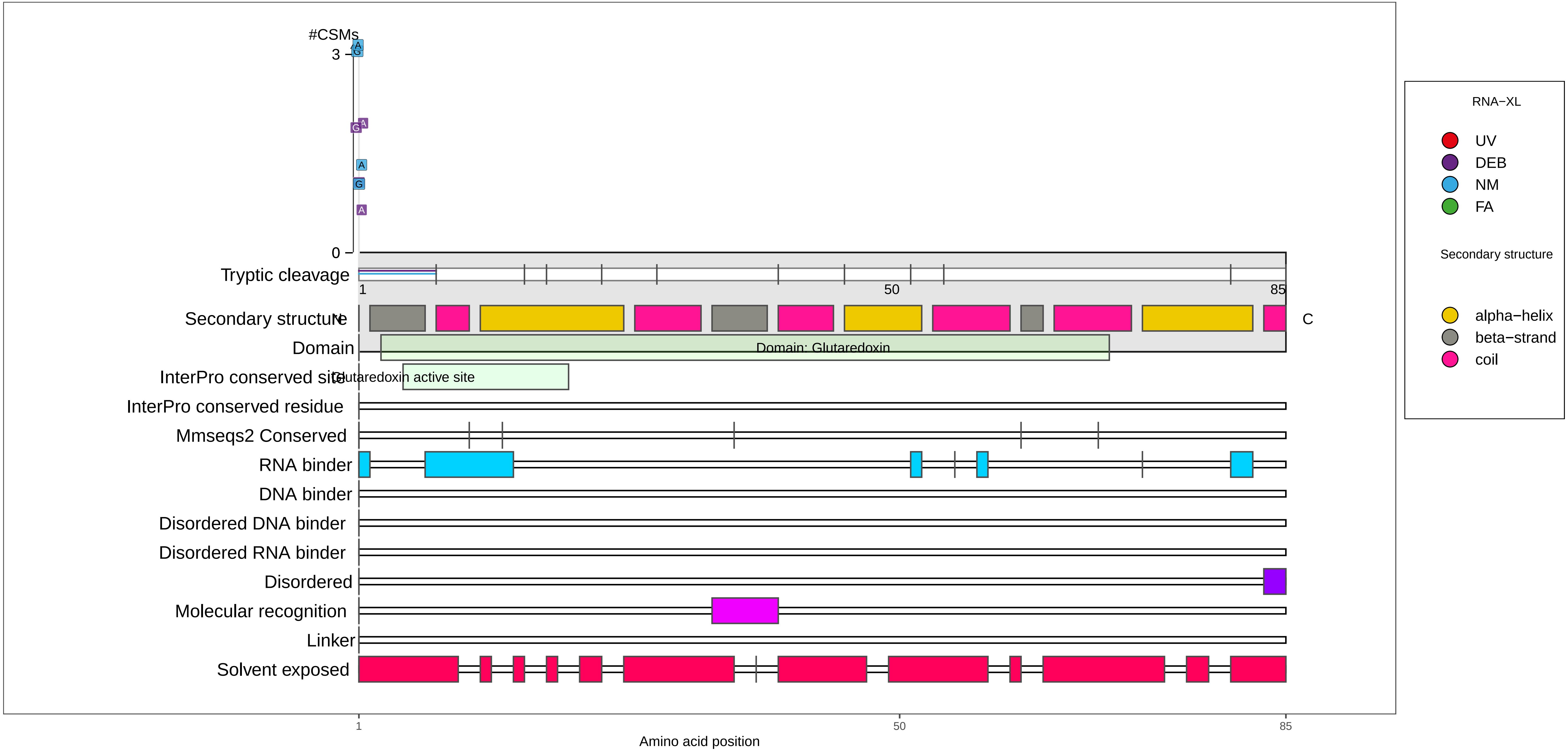

P75863
YCBX_ECOLI Uncharacterized protein YcbX

− Abundance:
tryptic [log10 Intensity]: 8.1 (Q 61)
PAXdb K12 strain [ppm]: 2.1 (Q 64)
PAXdb E.coli [ppm]: 1.82 (Q 68)

− RNA functions: not annotated

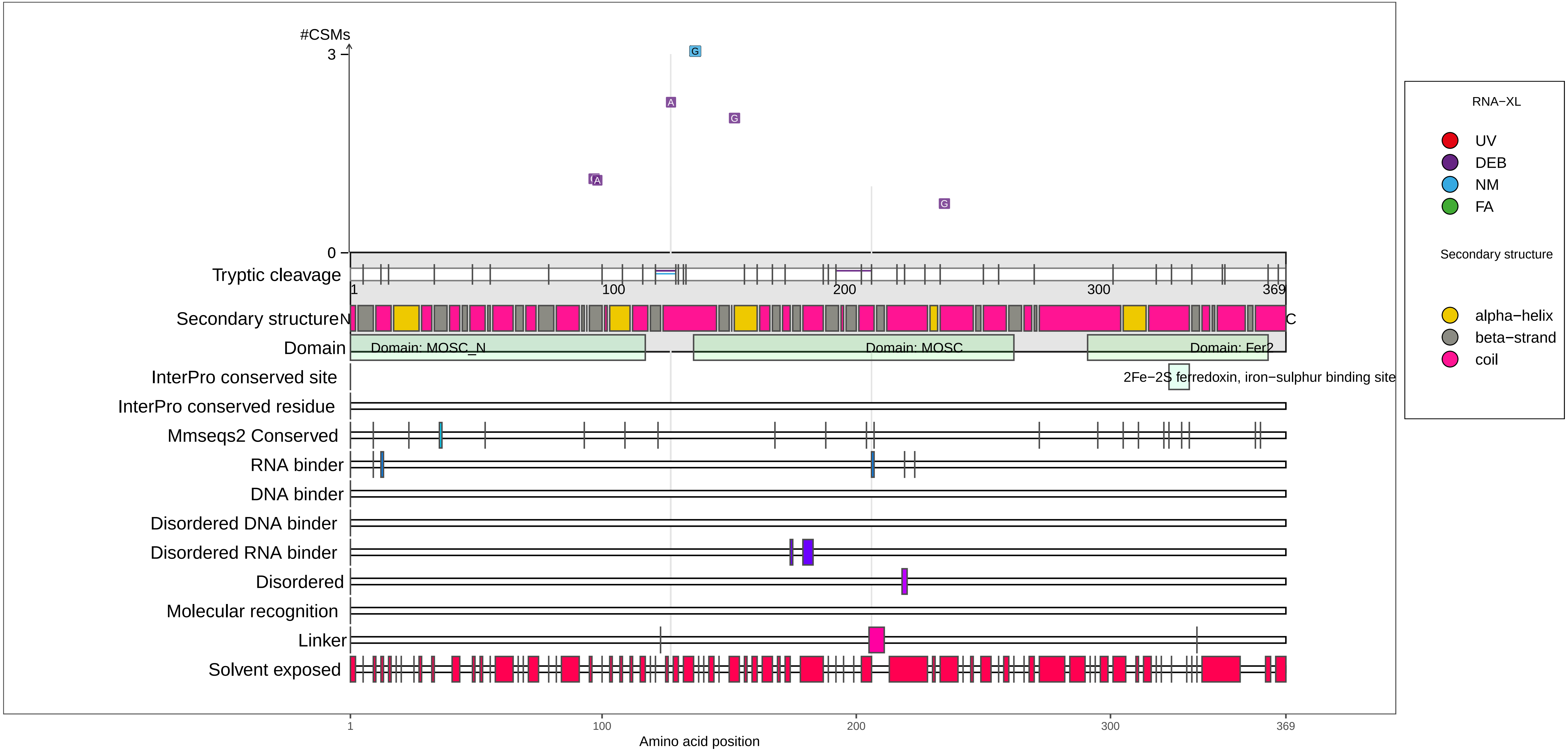

Q46856
YQHD_ECOLI Alcohol dehydrogenase YqhD

− Abundance:
tryptic [log10 Intensity]: 8.39 (Q 72)
PAXdb K12 strain [ppm]: 2.1 (Q 64)
PAXdb E.coli [ppm]: 2.33 (Q 81)

− RNA functions: not annotated

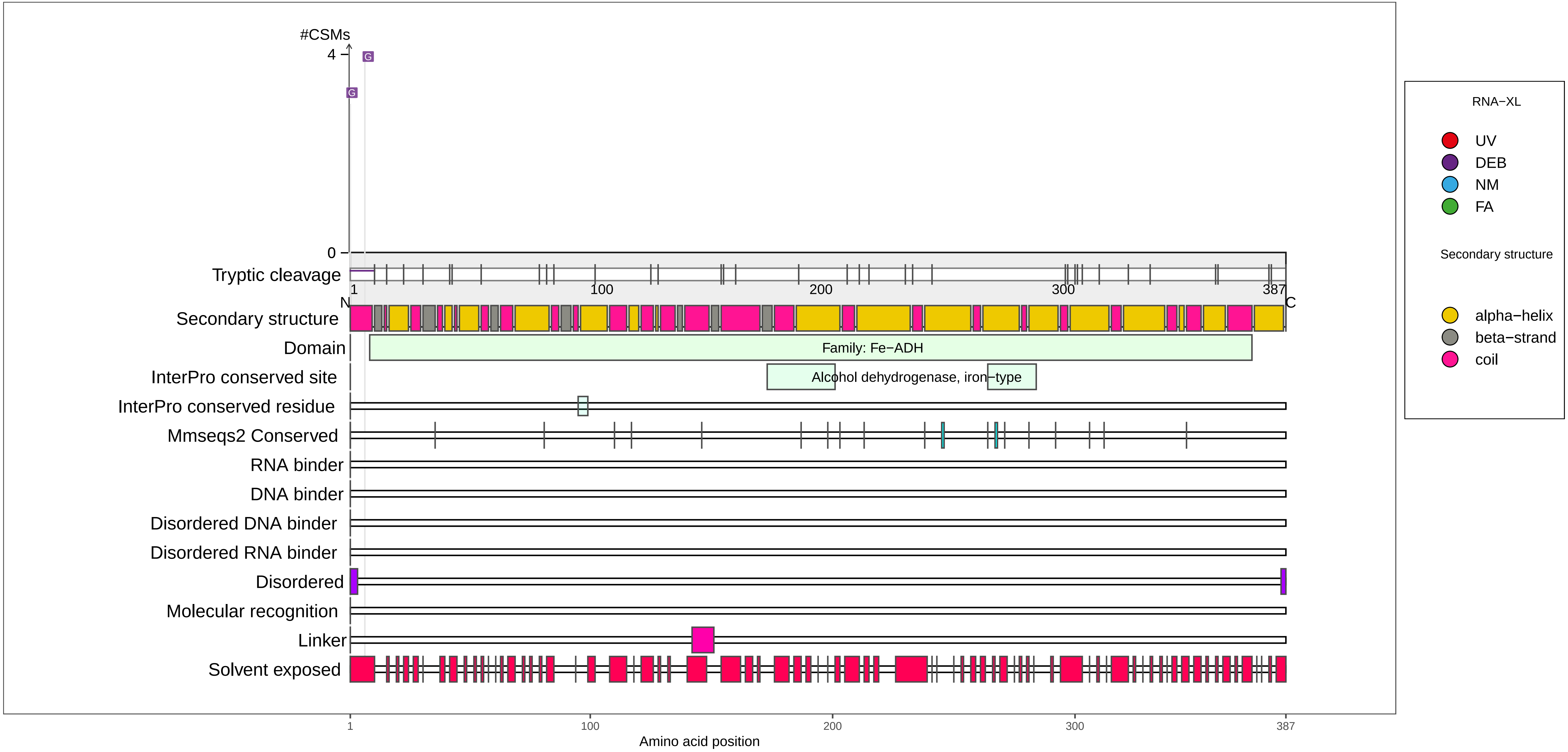

P00561

AK1H_ECOLI Bifunctional aspartokinase/homoserine dehydrogenase 1

− Abundance:
tryptic [log10 Intensity]: 7.13 (Q 18)
PAXdb K12 strain [ppm]: 2.32 (Q 70)
PAXdb E.coli [ppm]: 2.66 (Q 89)

− RNA functions: not annotated

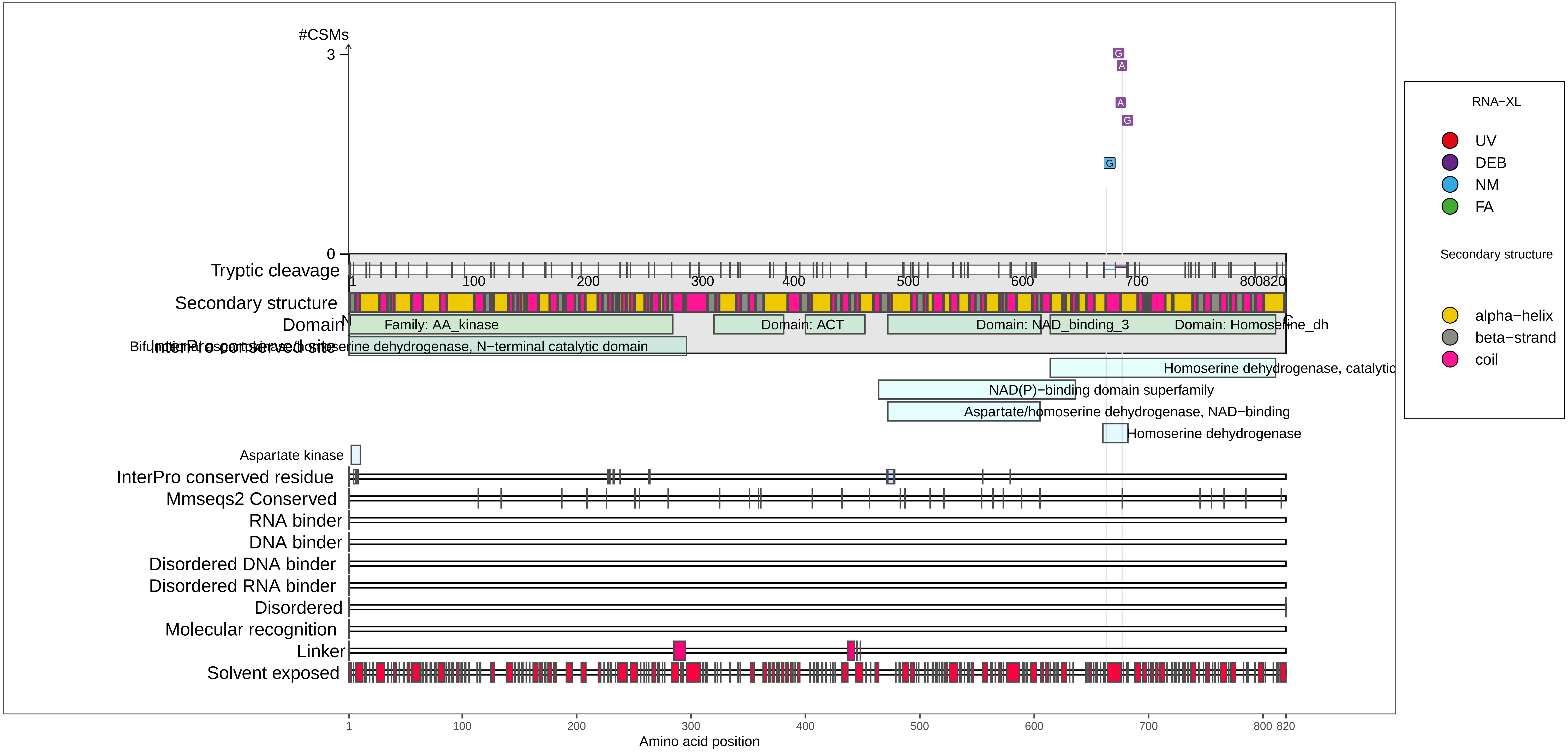

P00962
SYQ_ECOLI Glutamine−−tRNA ligase

− Abundance:
tryptic [log10 Intensity]: 9.03 (Q 89)
PAXdb K12 strain [ppm]: 2.77 (Q 85)
PAXdb E.coli [ppm]: 2.68 (Q 89)

− RNA functions:
aminoacyl−tRNA ligase activity; glutamine−tRNA ligase activity
glutaminyl−tRNA aminoacylation; glutamyl−tRNA aminoacylation; ncRNA metabolic process
RNA metabolic process; tRNA aminoacylation; tRNA aminoacylation for protein translation
tRNA metabolic process; tRNA synthetases class I (E and Q), anti−codon binding domain
tRNA synthetases class I (E and Q), catalytic domain

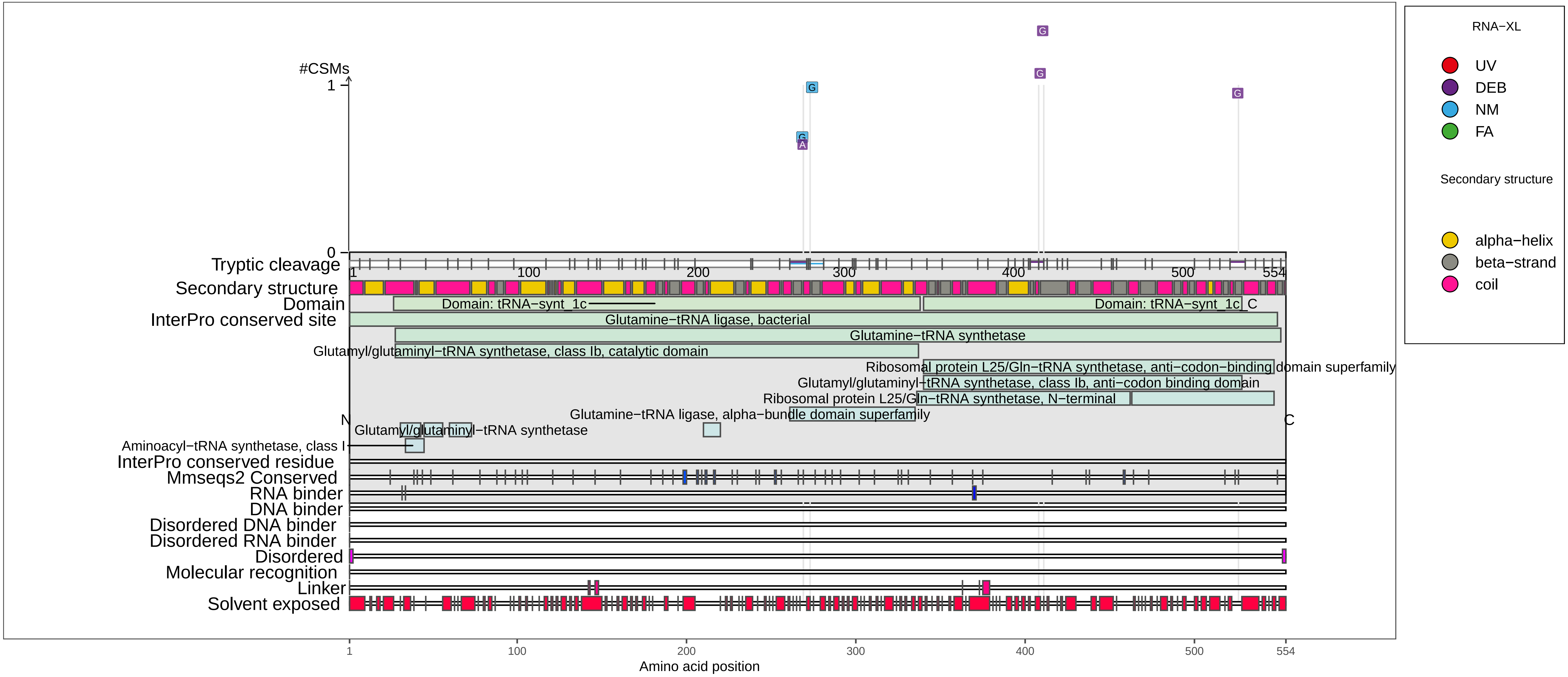

P07014
SDHB_ECOLI Succinate dehydrogenase iron−sulfur subunit

− Abundance:
tryptic [log10 Intensity]: 8.66 (Q 80)
PAXdb K12 strain [ppm]: 2.85 (Q 87)
PAXdb E.coli [ppm]: 2.82 (Q 92)

− RNA functions: not annotated

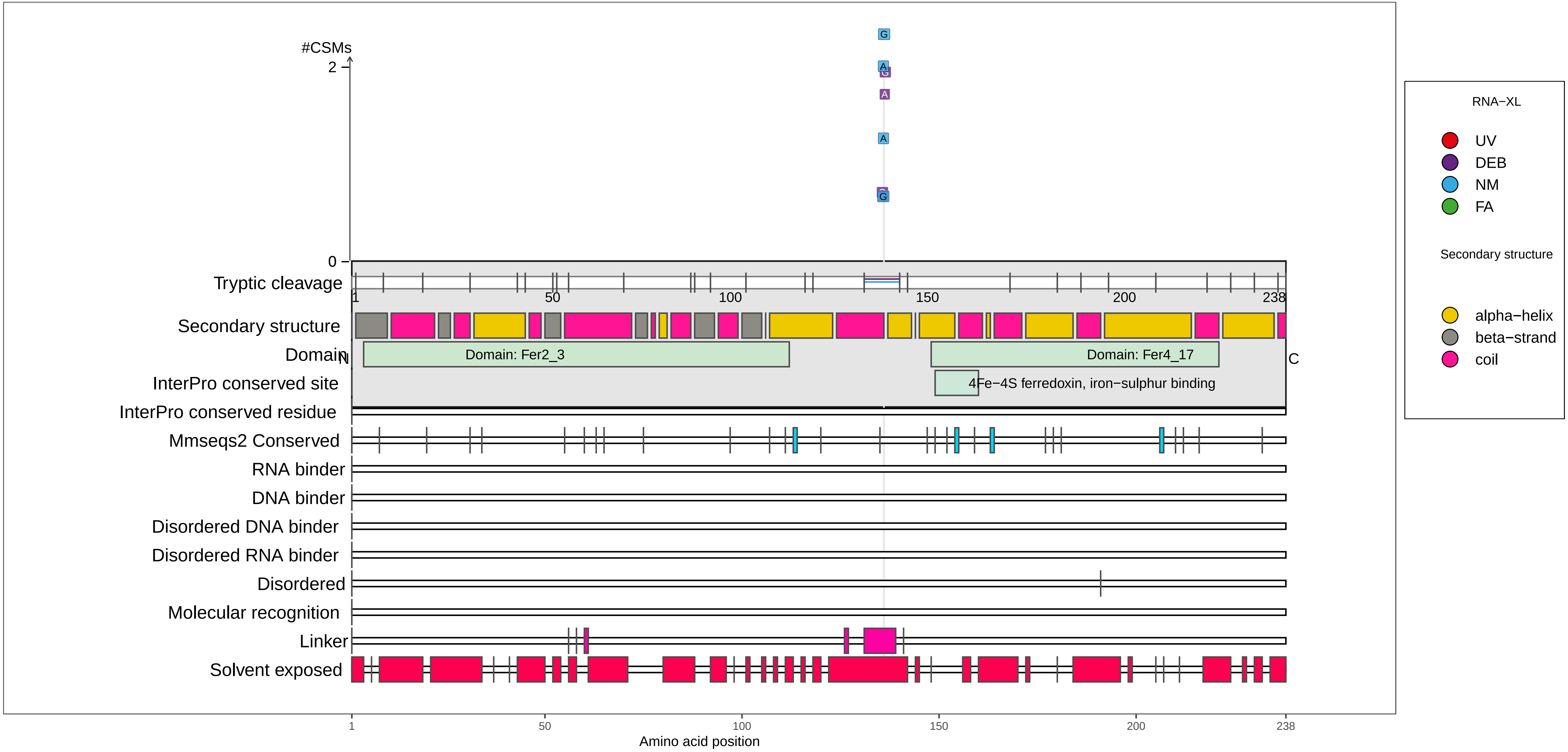

P09158
SPEE_ECOLI Polyamine aminopropyltransferase

− Abundance:
tryptic [log10 Intensity]: 8.61 (Q 79)
PAXdb K12 strain [ppm]: 1.32 (Q 27)
PAXdb E.coli [ppm]: 2 (Q 72)

− RNA functions: not annotated

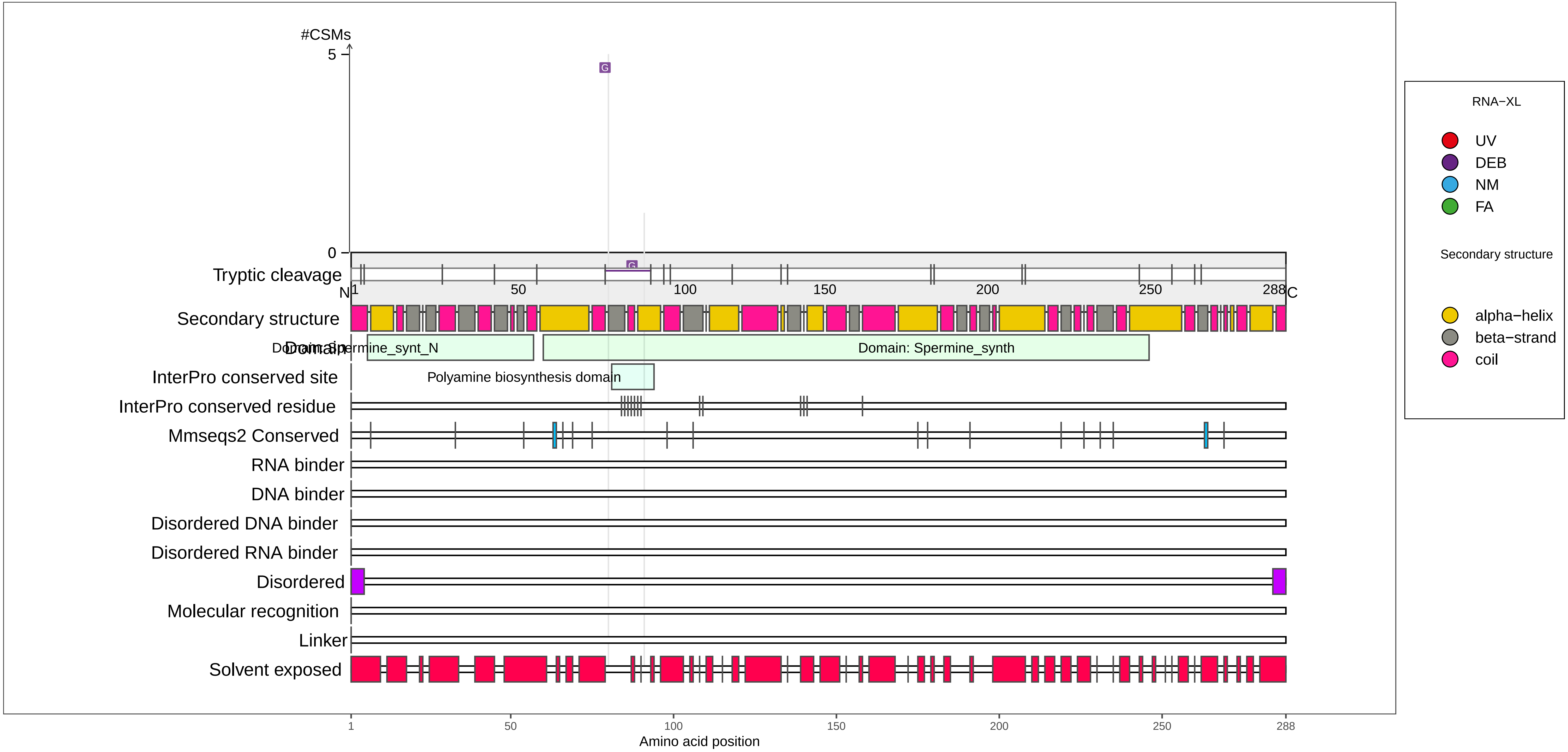

P0A6R3
FIS_ECOLI DNA−binding protein Fis

− Abundance:
tryptic [log10 Intensity]: 9.03 (Q 89)
PAXdb K12 strain [ppm]: 3.33 (Q 96)
PAXdb E.coli [ppm]: 2.39 (Q 83)

− RNA functions:
RNA biosynthetic process; RNA metabolic process

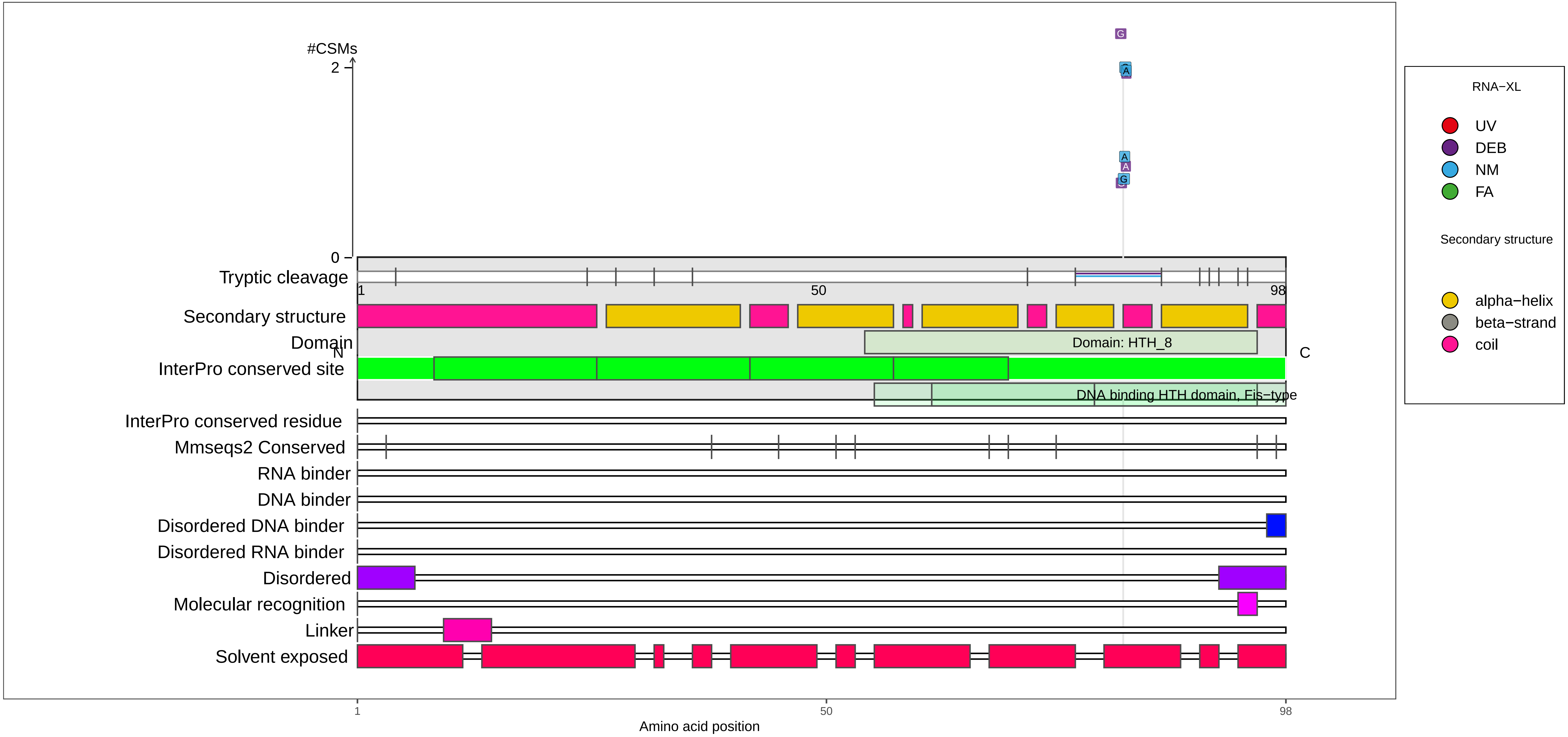

P0A7A2
GPMB_ECOLI Probable phosphoglycerate mutase GpmB

− Abundance:
tryptic [log10 Intensity]: 7.82 (Q 49)
PAXdb K12 strain [ppm]: 2.63 (Q 81)
PAXdb E.coli [ppm]: 1.92 (Q 70)

− RNA functions: not annotated

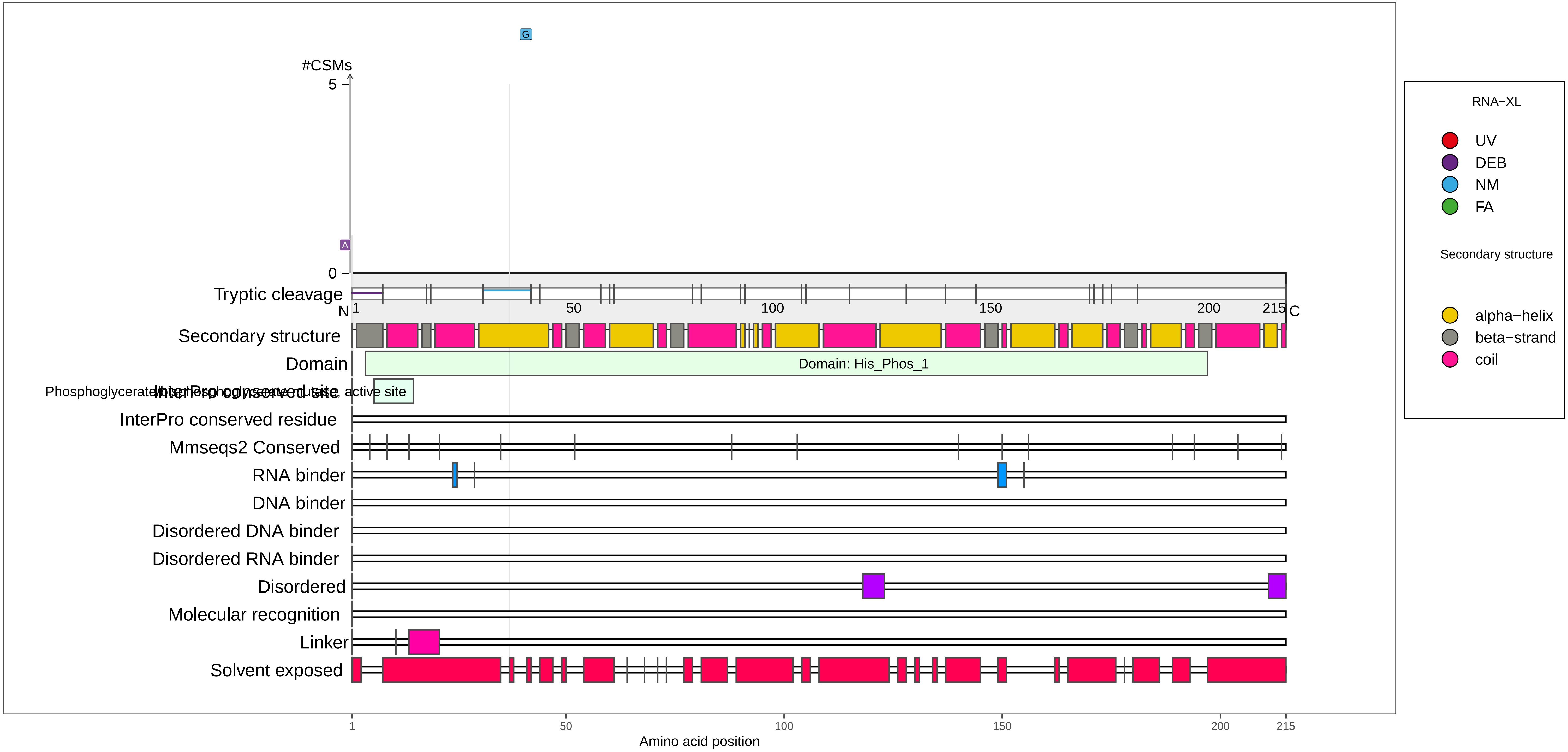

P0A855
TOLB_ECOLI Tol−Pal system protein TolB

− Abundance:
tryptic [log10 Intensity]: 8.72 (Q 82)
PAXdb K12 strain [ppm]: 2.94 (Q 89)
PAXdb E.coli [ppm]: 2.75 (Q 91)

− RNA functions: not annotated

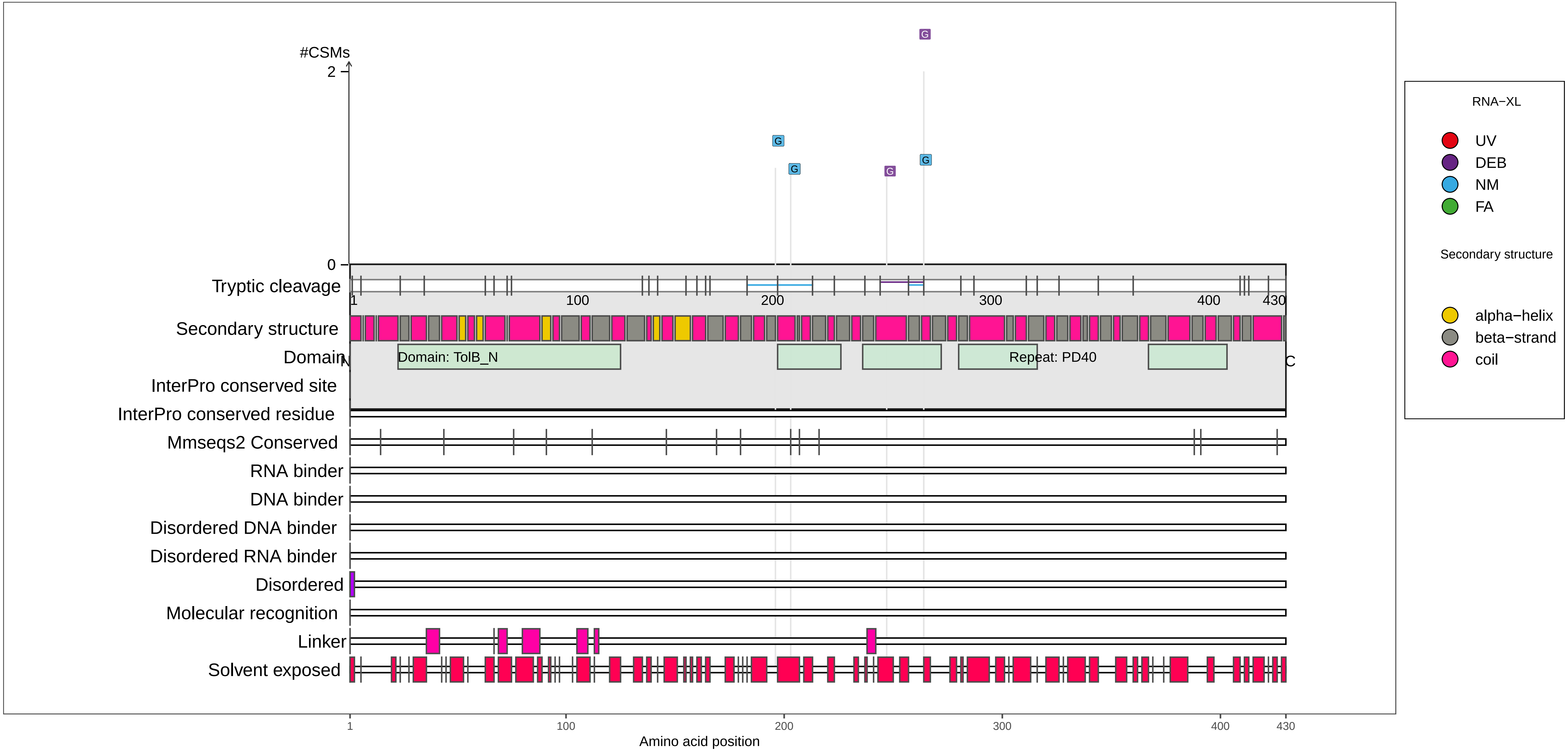

P0A8Y5
YIDA_ECOLI Sugar phosphatase YidA

− Abundance:
tryptic [log10 Intensity]: 8.47 (Q 74)
PAXdb K12 strain [ppm]: 3.15 (Q 93)
PAXdb E.coli [ppm]: 1.7 (Q 64)

− RNA functions: not annotated

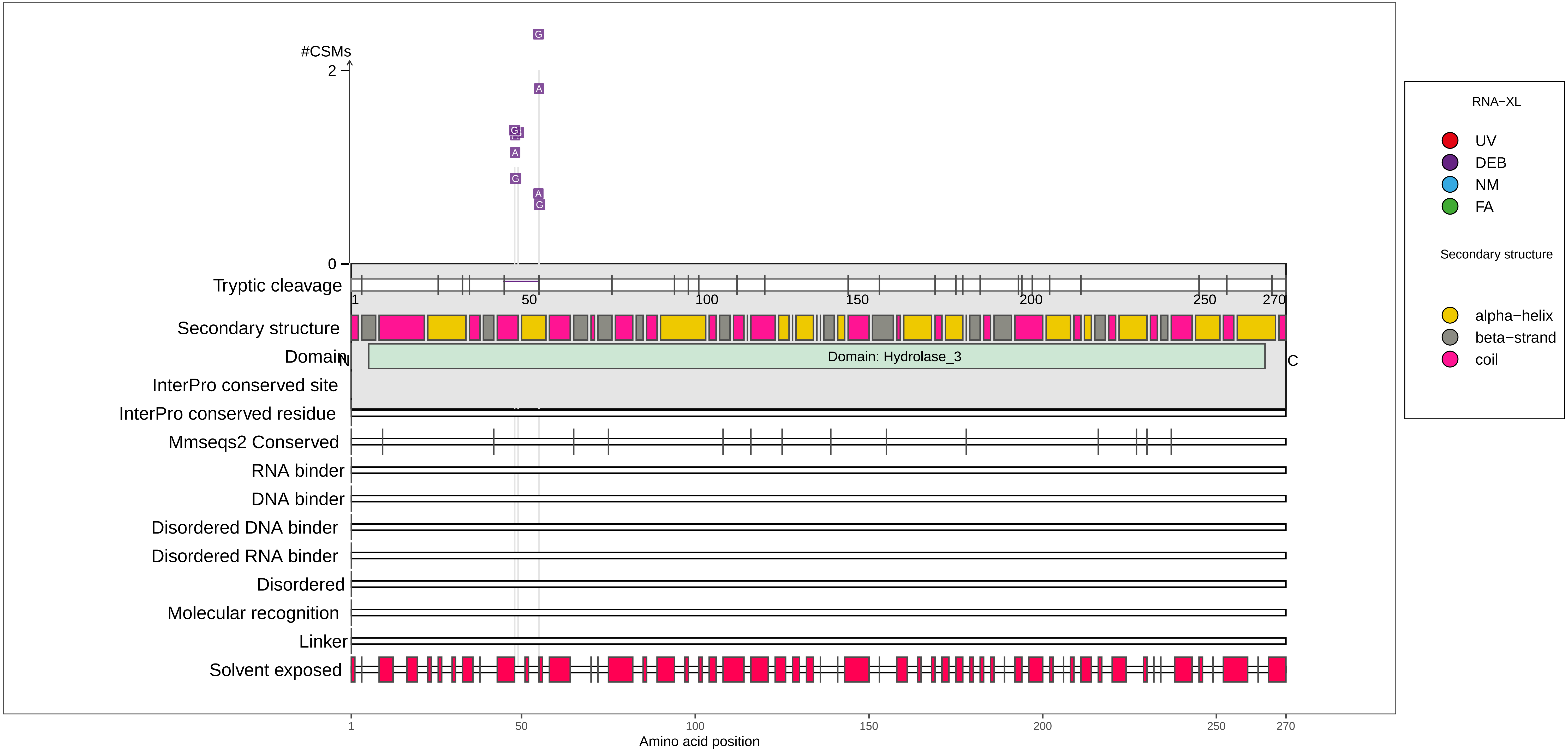

P0A944
RIMI_ECOLI [Ribosomal protein S18]−alanine N−acetyltransferase

− Abundance:
tryptic [log10 Intensity]: 6.67 (Q 5)
PAXdb K12 strain [ppm]: 1.21 (Q 21)
PAXdb E.coli [ppm]: 0.66 (Q 40)

− RNA functions: not annotated

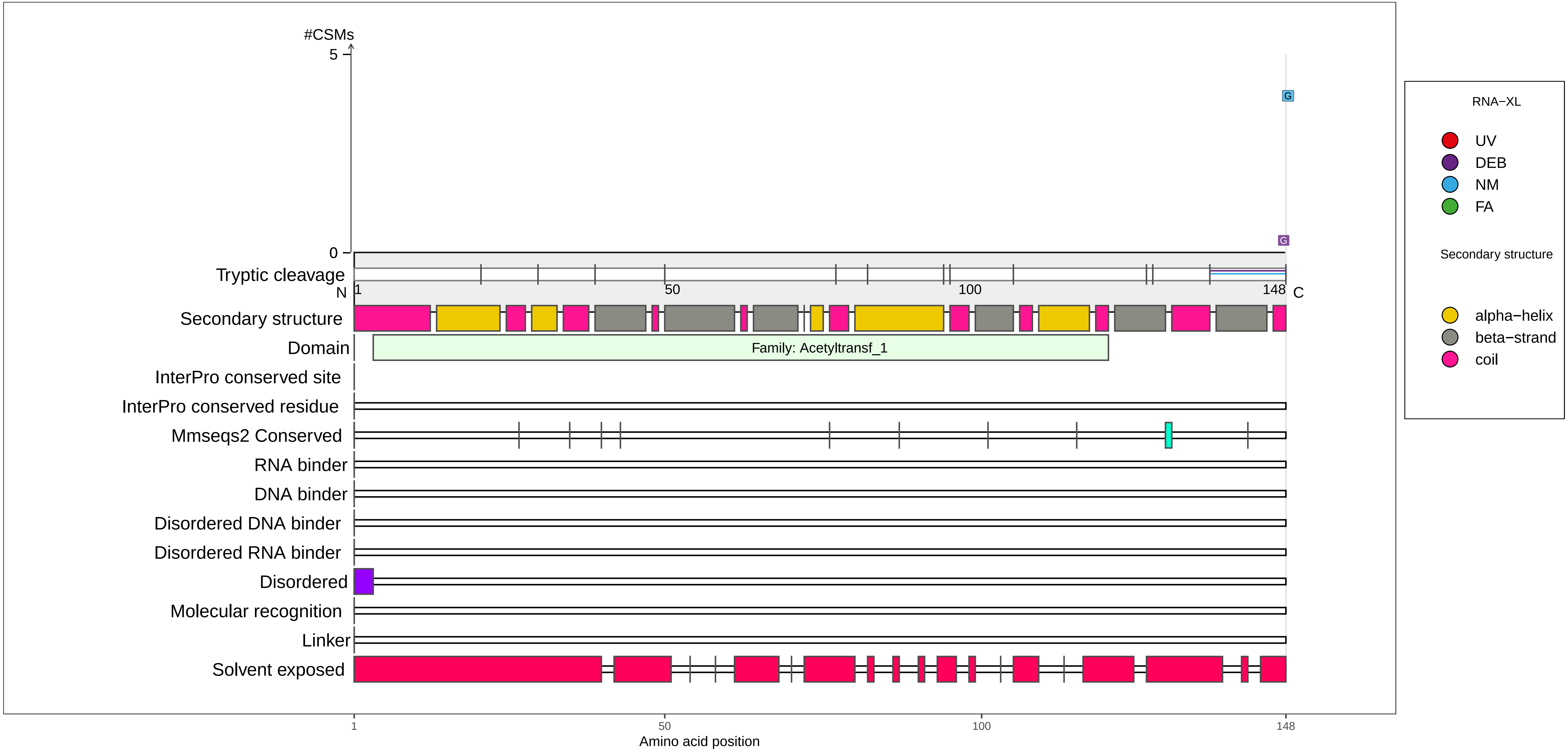

P0A9R4
FER_ECOLI 2Fe−2S ferredoxin

− Abundance:
tryptic [log10 Intensity]: 8 (Q 58)
PAXdb K12 strain [ppm]: 2.69 (Q 83)
PAXdb E.coli [ppm]: 2.59 (Q 88)

− RNA functions: not annotated

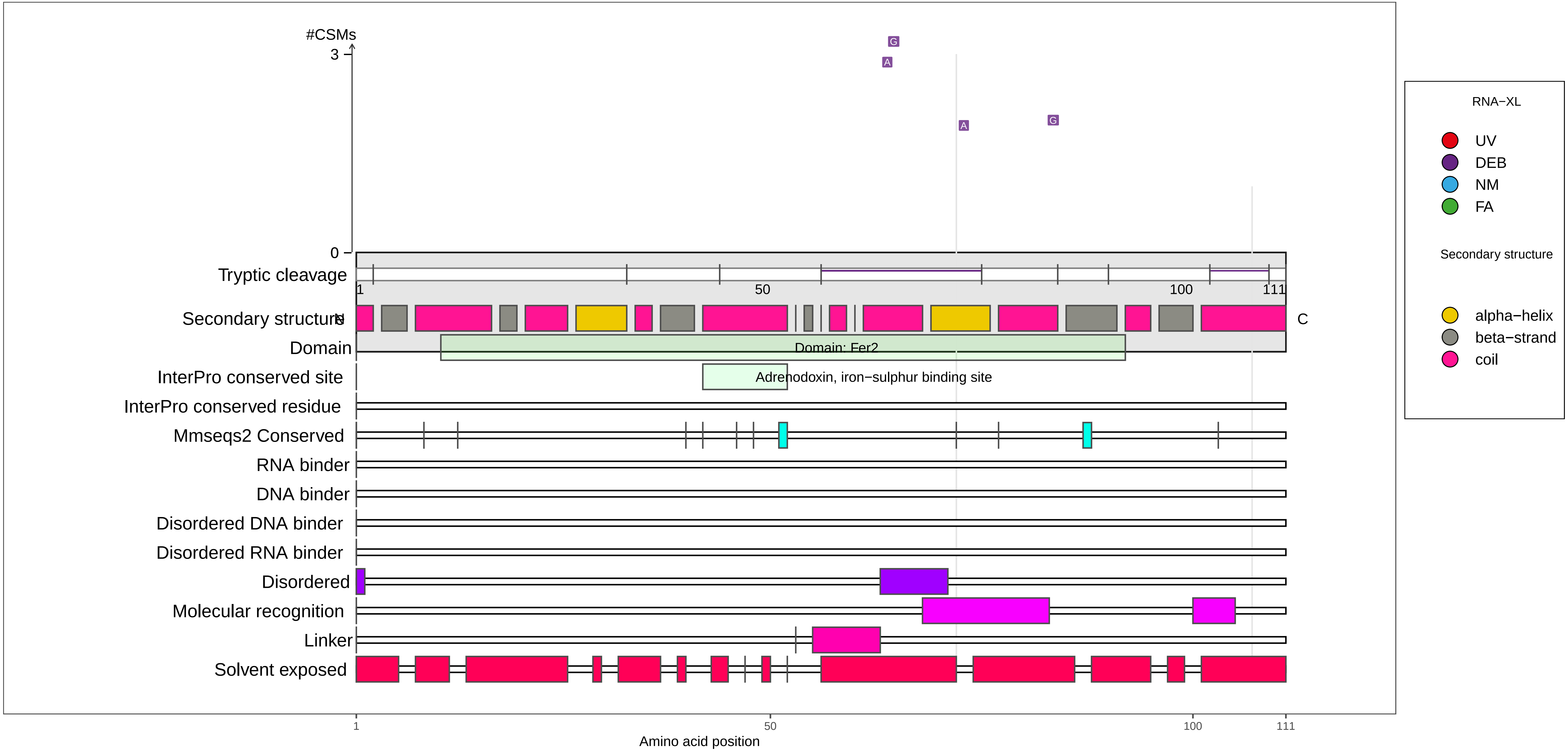

P0ABI8
CYOB_ECOLI Cytochrome bo(3) ubiquinol oxidase subunit 1

− Abundance:
tryptic [log10 Intensity]: 9.41 (Q 95)
PAXdb K12 strain [ppm]: 1.75 (Q 51)
PAXdb E.coli [ppm]: 1.8 (Q 67)

− RNA functions: not annotated

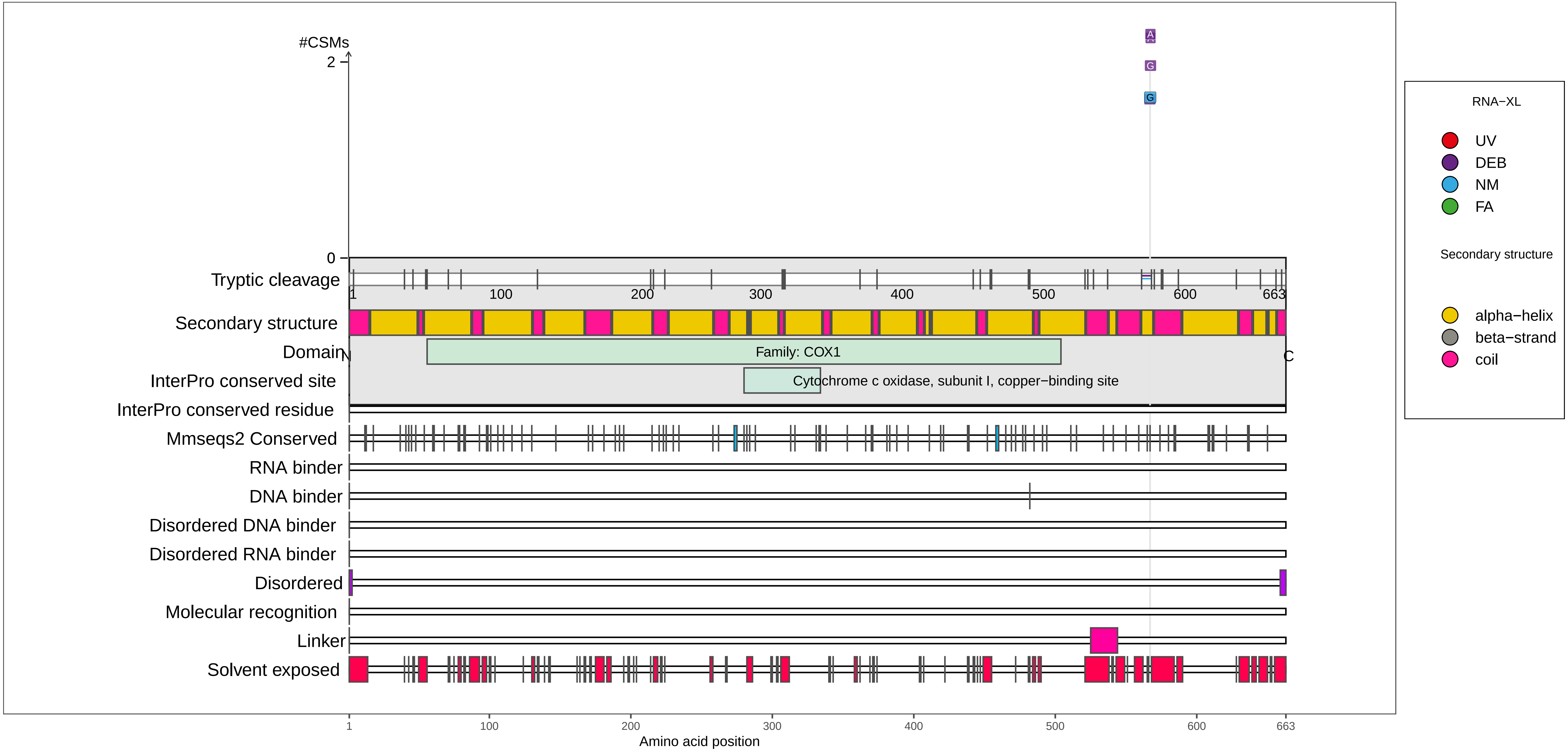

P0ABQ0
COABC_ECOLI Coenzyme A biosynthesis bifunctional protein CoaBC

− Abundance:
tryptic [log10 Intensity]: 7.81 (Q 49)
PAXdb K12 strain [ppm]: 1.78 (Q 53)
PAXdb E.coli [ppm]: 2.15 (Q 76)

− RNA functions: not annotated

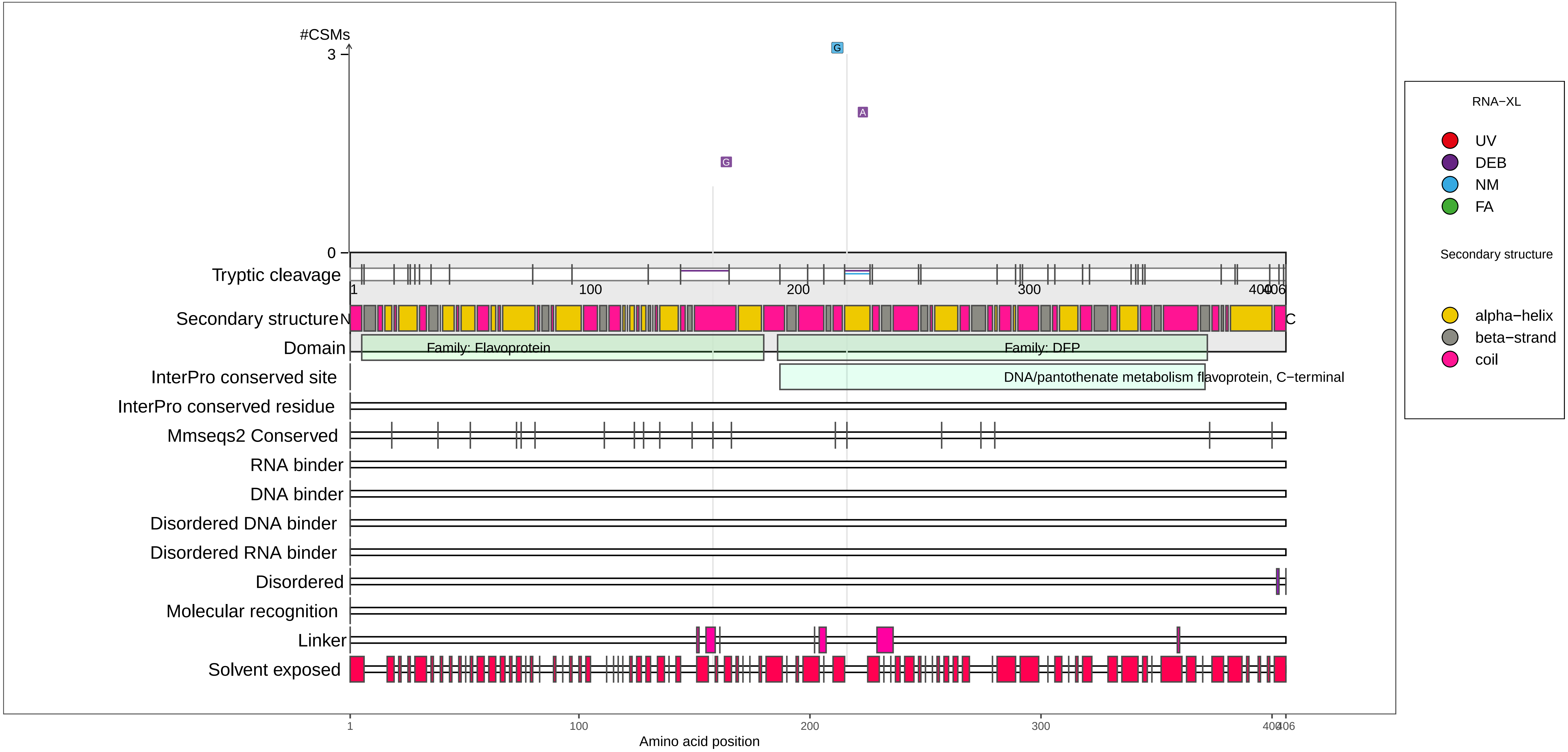

P0ABU0
MENB_ECOLI 1,4−dihydroxy−2−naphthoyl−CoA synthase

− Abundance:
tryptic [log10 Intensity]: 8.22 (Q 66)
PAXdb K12 strain [ppm]: 2.74 (Q 84)
PAXdb E.coli [ppm]: 2.45 (Q 84)

− RNA functions: not annotated

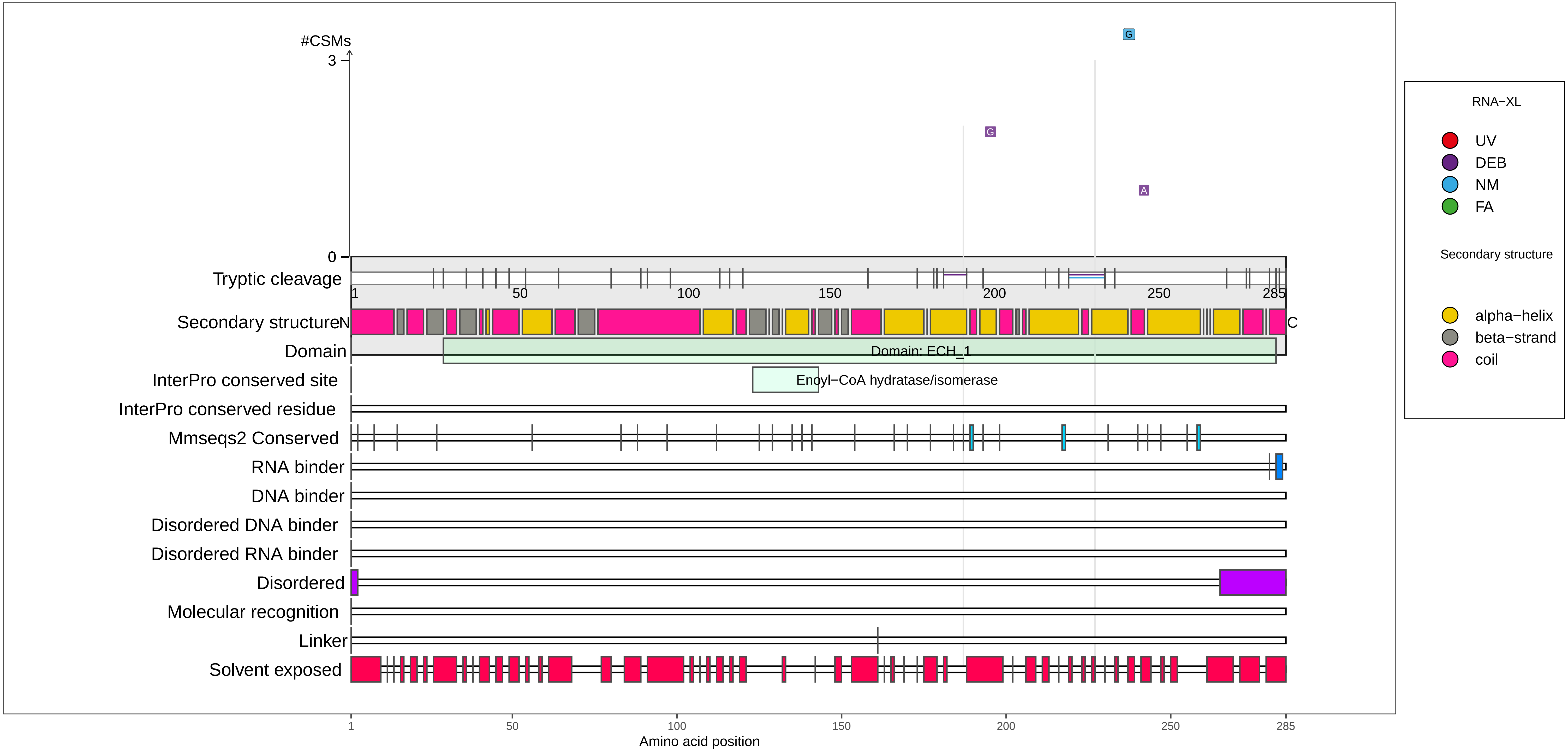

P0AC33
FUMA_ECOLI Fumarate hydratase class I, aerobic

− Abundance:
tryptic [log10 Intensity]: 8.54 (Q 77)
PAXdb K12 strain [ppm]: 2.58 (Q 80)
PAXdb E.coli [ppm]: 2.53 (Q 86)

− RNA functions: not annotated

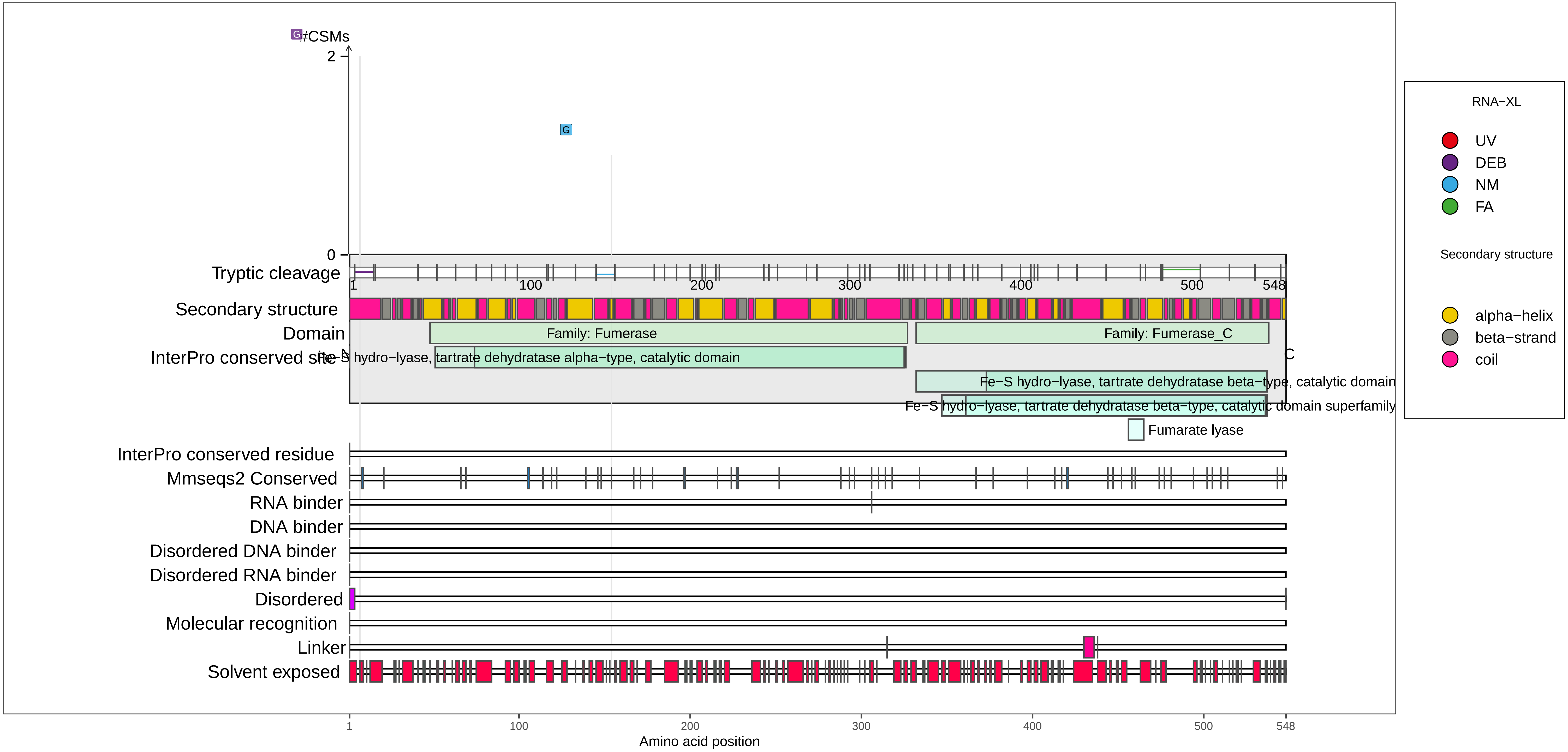

P0AC44
DHSD_ECOLI Succinate dehydrogenase hydrophobic membrane anchor subunit

− Abundance:
tryptic [log10 Intensity]: 8.34 (Q 71)
PAXdb K12 strain [ppm]: 2.14 (Q 65)
PAXdb E.coli [ppm]: 2.45 (Q 84)

− RNA functions: not annotated

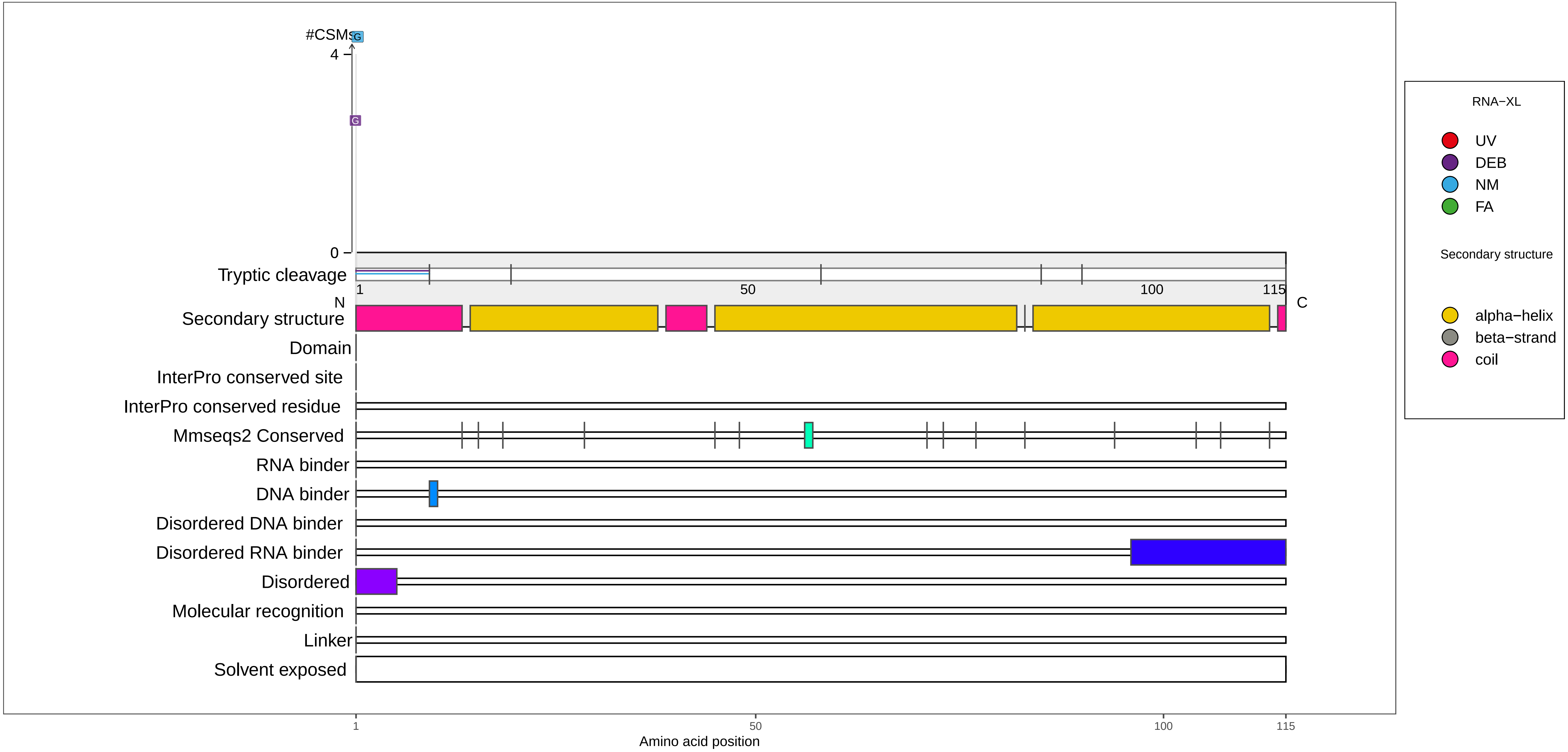

P0ACB7
HEMY_ECOLI Protein HemY

− Abundance:
tryptic [log10 Intensity]: 7.2 (Q 21)
PAXdb K12 strain [ppm]: 1.44 (Q 35)
PAXdb E.coli [ppm]: 1.95 (Q 71)

− RNA functions: not annotated

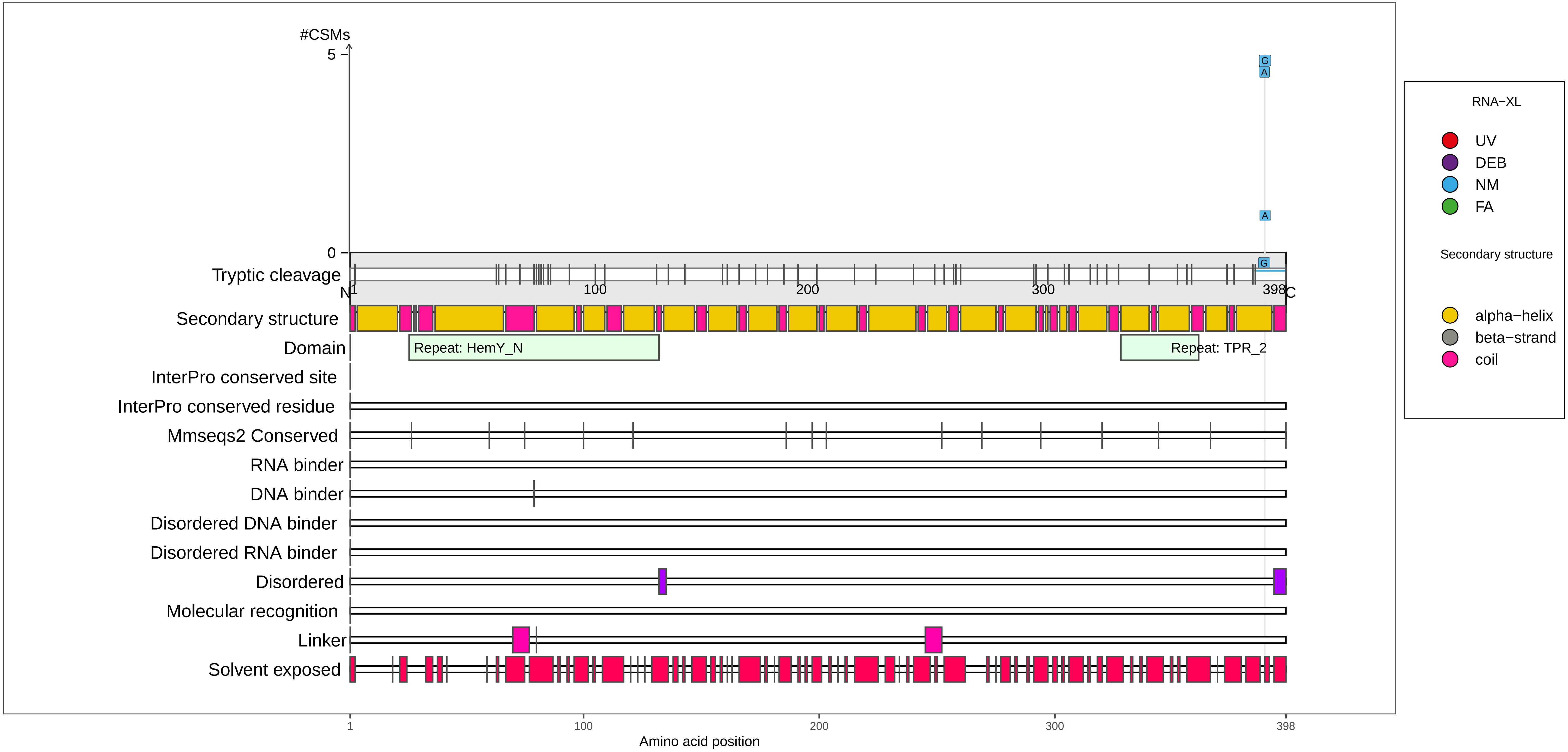

P0ACI0
ROB_ECOLI Right origin−binding protein

− Abundance:
tryptic [log10 Intensity]: 8 (Q 58)
PAXdb K12 strain [ppm]: 2.24 (Q 68)
PAXdb E.coli [ppm]: 2.17 (Q 77)

− RNA functions: not annotated

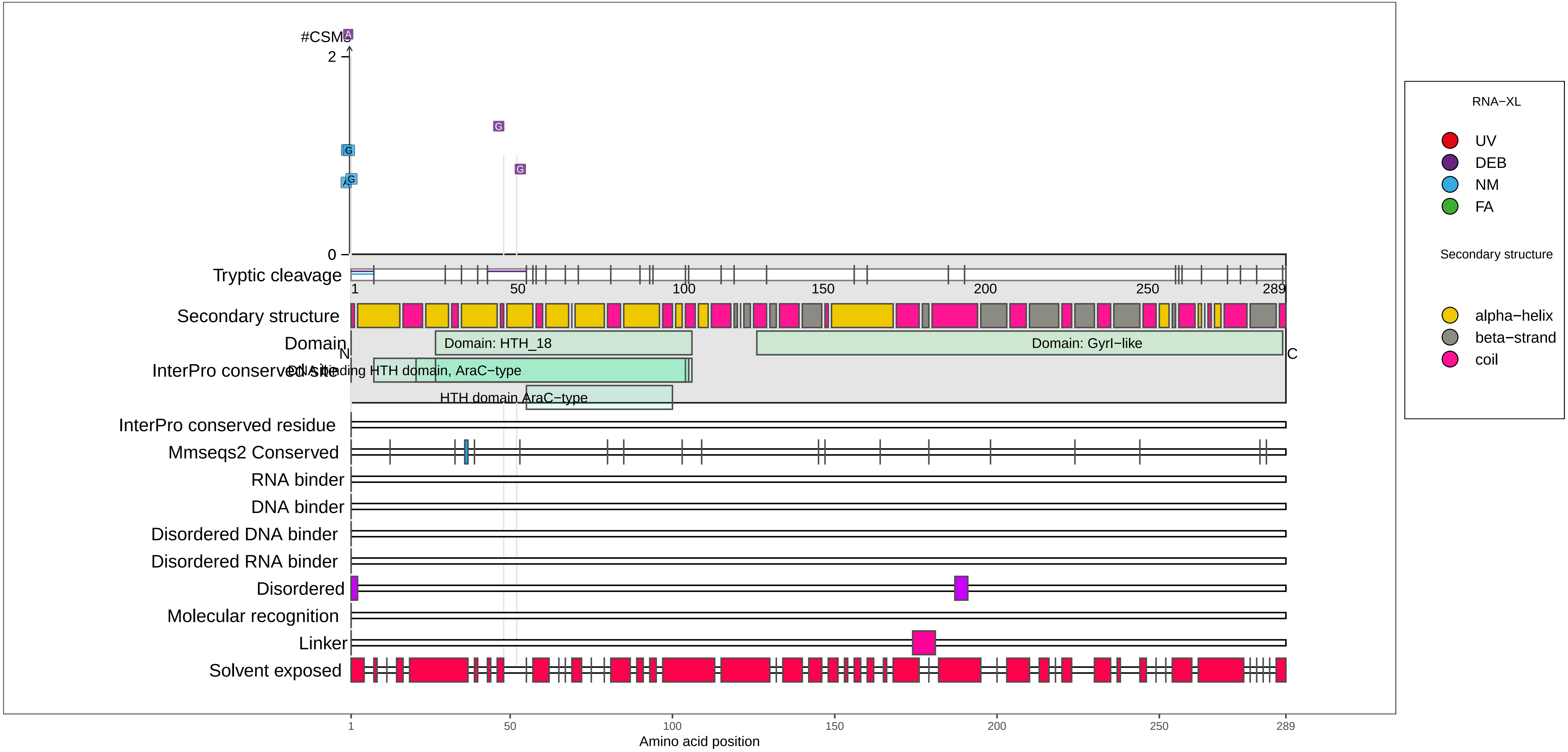

P0ACS5
ZNTR_ECOLI HTH−type transcriptional regulator ZntR

− Abundance:
tryptic [log10 Intensity]: 7.5 (Q 35)
PAXdb K12 strain [ppm]: 1.96 (Q 59)
PAXdb E.coli [ppm]: 0.92 (Q 46)

− RNA functions:
RNA biosynthetic process; RNA metabolic process

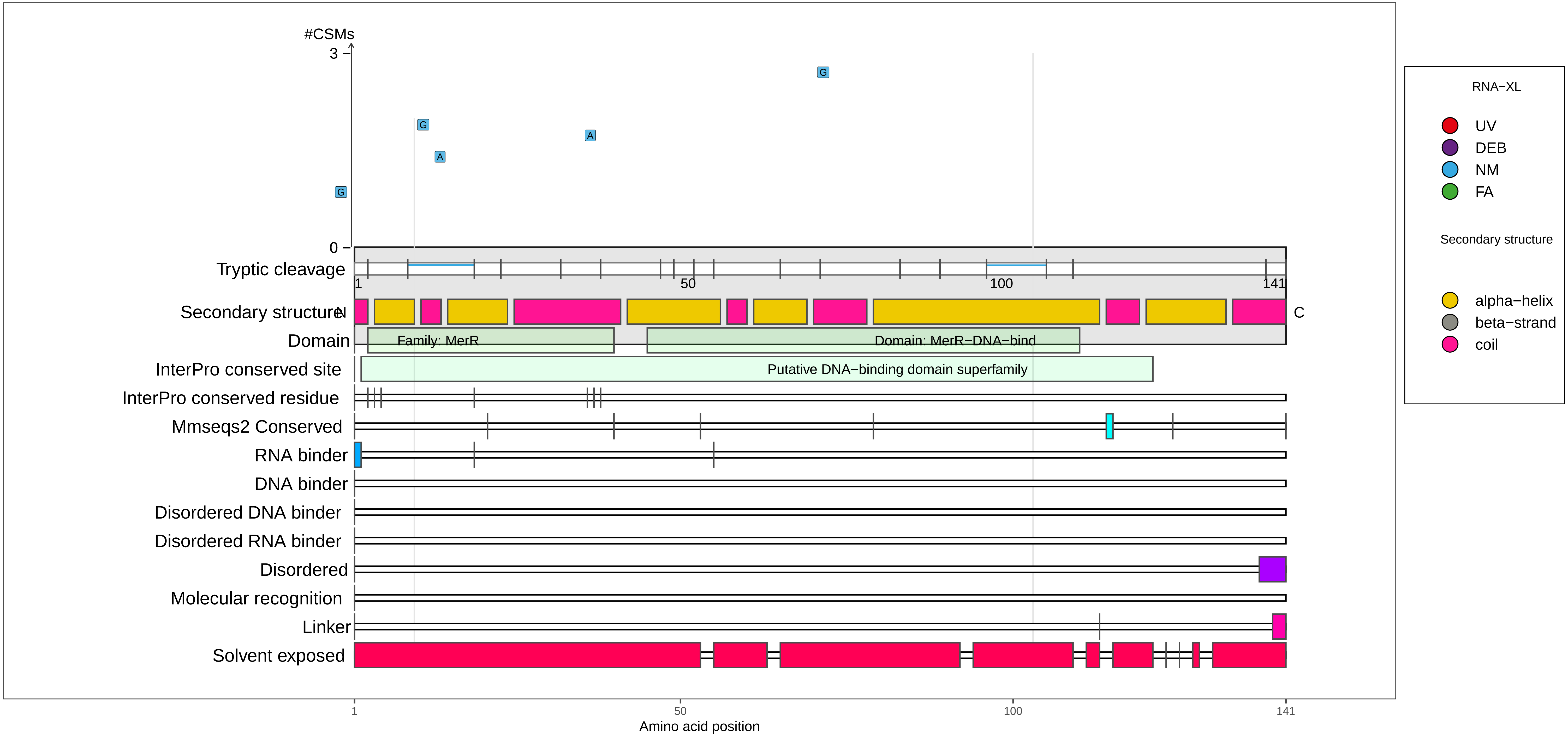

P0ACY9
YEBG_ECOLI Uncharacterized protein YebG

− Abundance:
tryptic [log10 Intensity]: 7.74 (Q 46)
PAXdb K12 strain [ppm]: 2.41 (Q 75)
PAXdb E.coli [ppm]: 2.28 (Q 80)

− RNA functions: not annotated

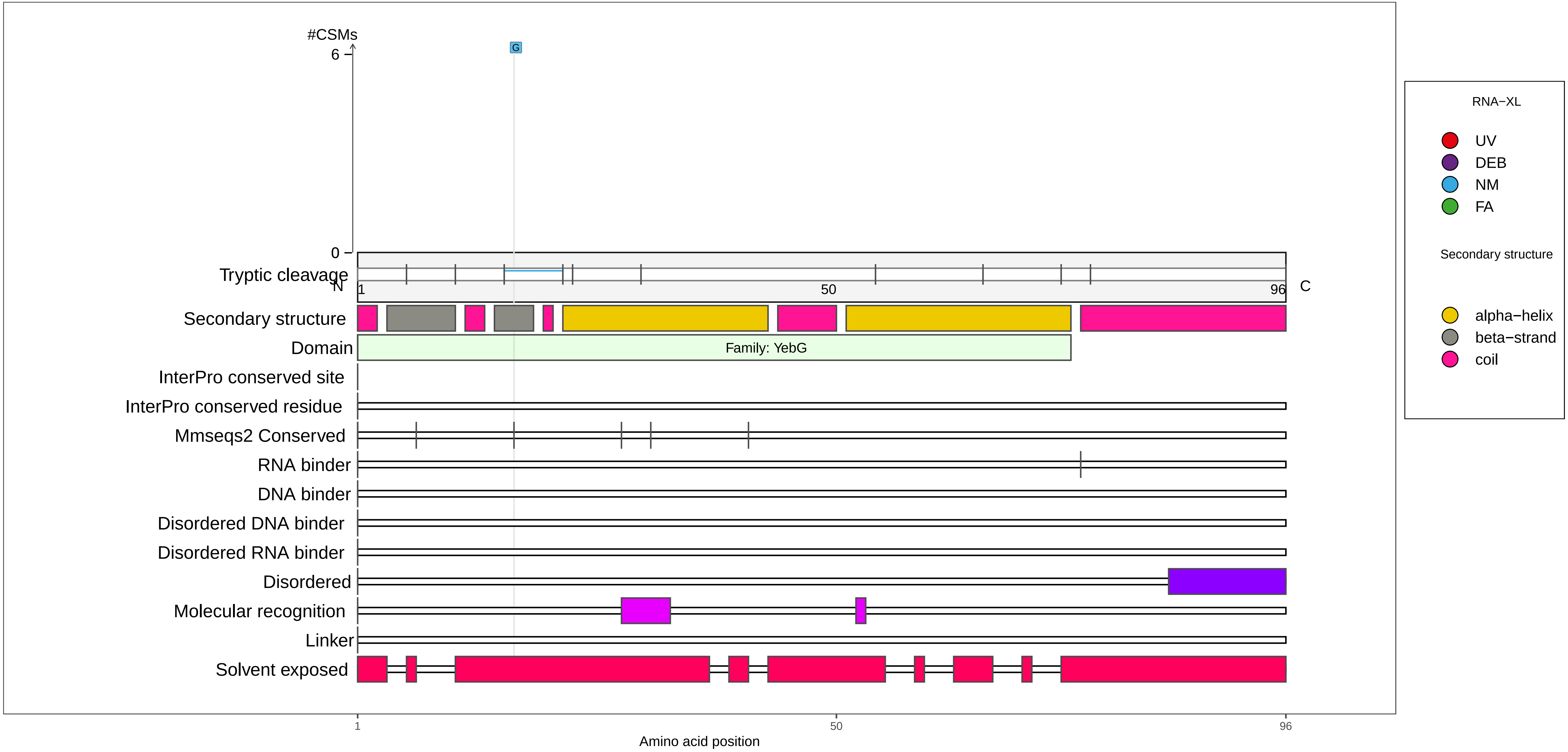

P0AEB2
DACA_ECOLI D−alanyl−D−alanine carboxypeptidase DacA

− Abundance:
tryptic [log10 Intensity]: 8.93 (Q 86)
PAXdb K12 strain [ppm]: 2.3 (Q 70)
PAXdb E.coli [ppm]: 2.36 (Q 82)

− RNA functions: not annotated

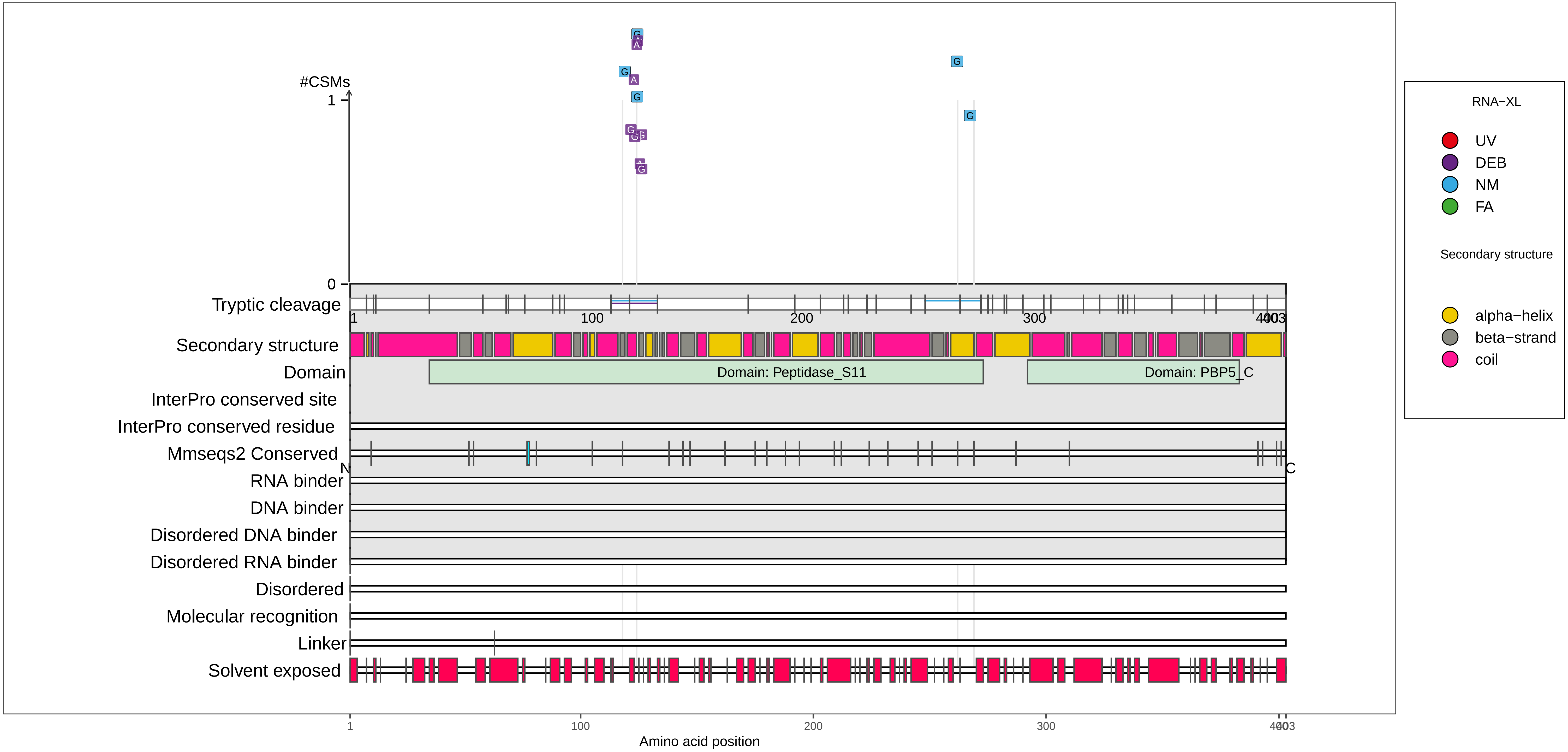

P0AFI5
PBP7_ECOLI D−alanyl−D−alanine endopeptidase

− Abundance:
tryptic [log10 Intensity]: 6.94 (Q 11)
PAXdb K12 strain [ppm]: 1.73 (Q 50)
PAXdb E.coli [ppm]: 0.76 (Q 42)

− RNA functions: not annotated

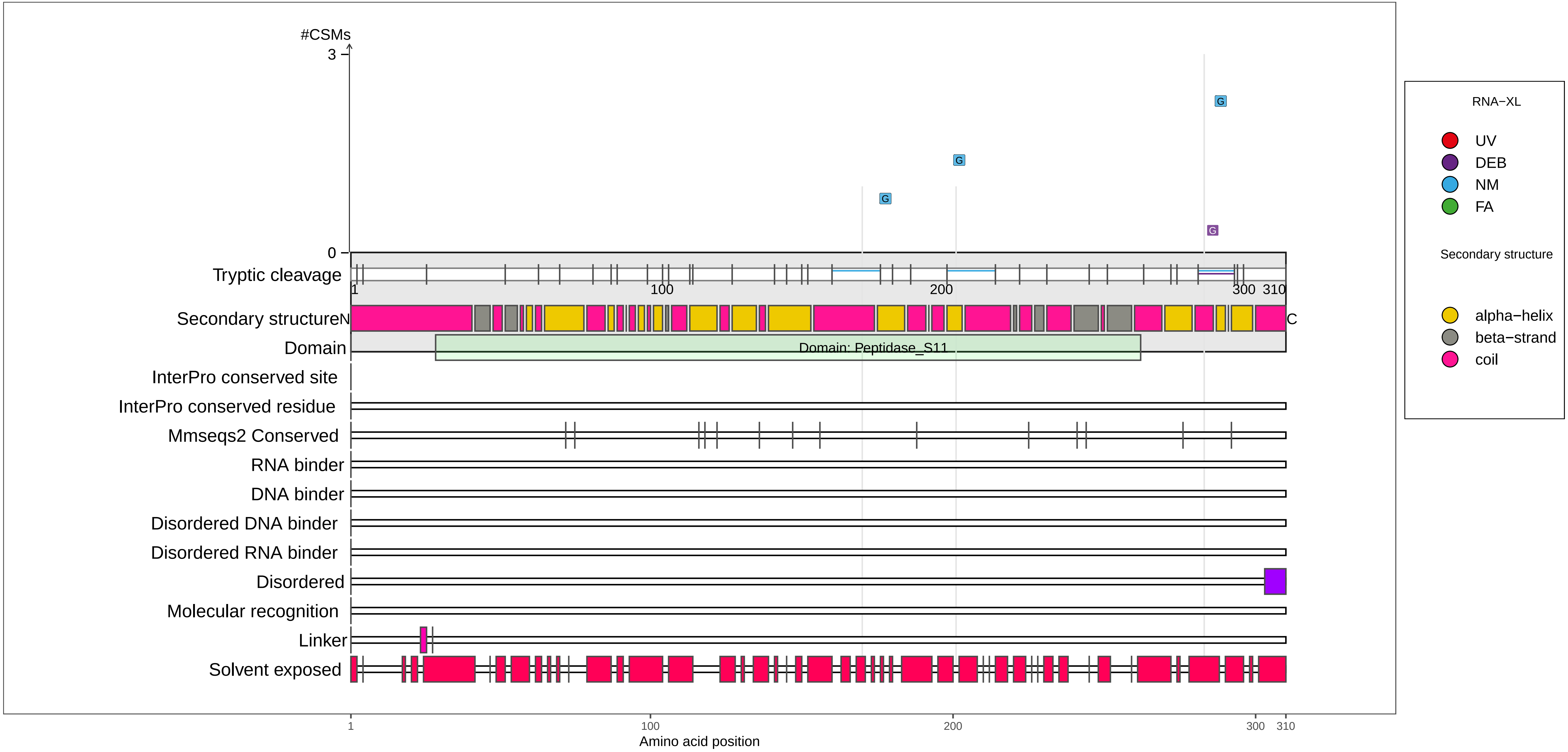

P0AFX7
RSEA_ECOLI Anti−sigma−E factor RseA

− Abundance:
tryptic [log10 Intensity]: 7.45 (Q 32)
PAXdb K12 strain [ppm]: NA (Q NA)
PAXdb E.coli [ppm]: 0.98 (Q 47)

− RNA functions: not annotated

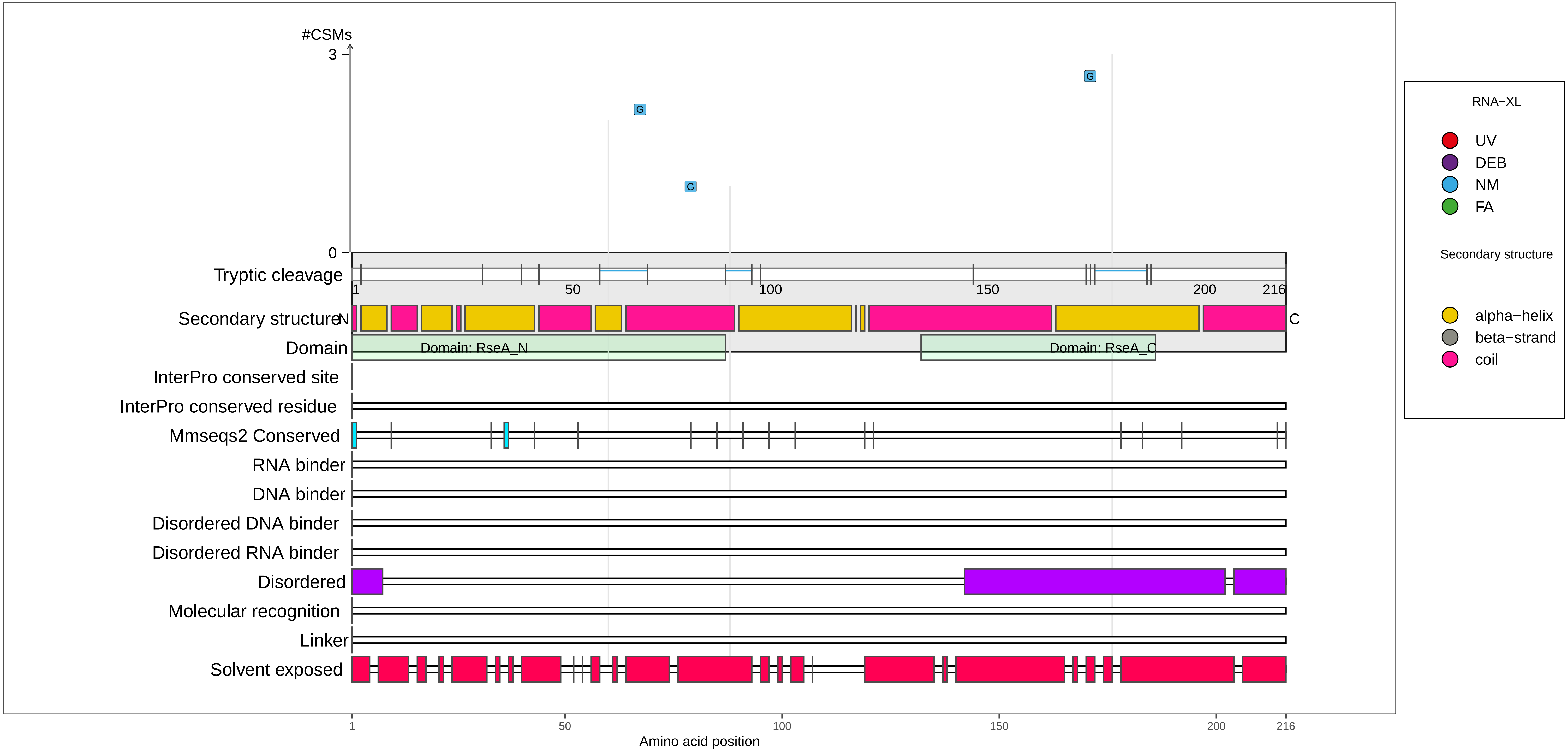

P14407
FUMB_ECOLI Fumarate hydratase class I, anaerobic

− Abundance:
tryptic [log10 Intensity]: 7.16 (Q 19)
PAXdb K12 strain [ppm]: 2.15 (Q 65)
PAXdb E.coli [ppm]: 1.72 (Q 65)

− RNA functions: not annotated

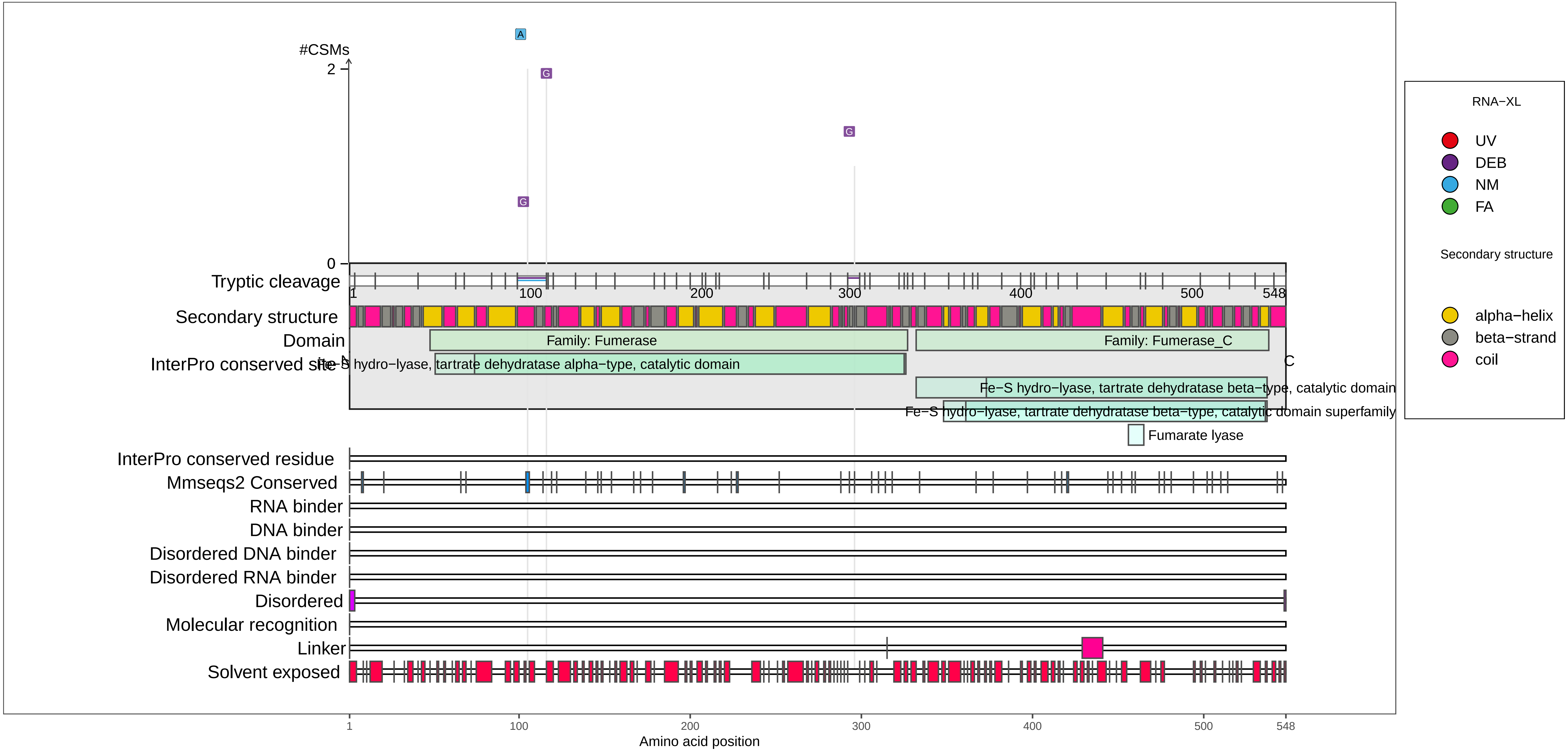

P18196
MINC_ECOLI Septum site−determining protein MinC

− Abundance:
tryptic [log10 Intensity]: 7.43 (Q 32)
PAXdb K12 strain [ppm]: 2.68 (Q 83)
PAXdb E.coli [ppm]: 1.55 (Q 61)

− RNA functions: not annotated

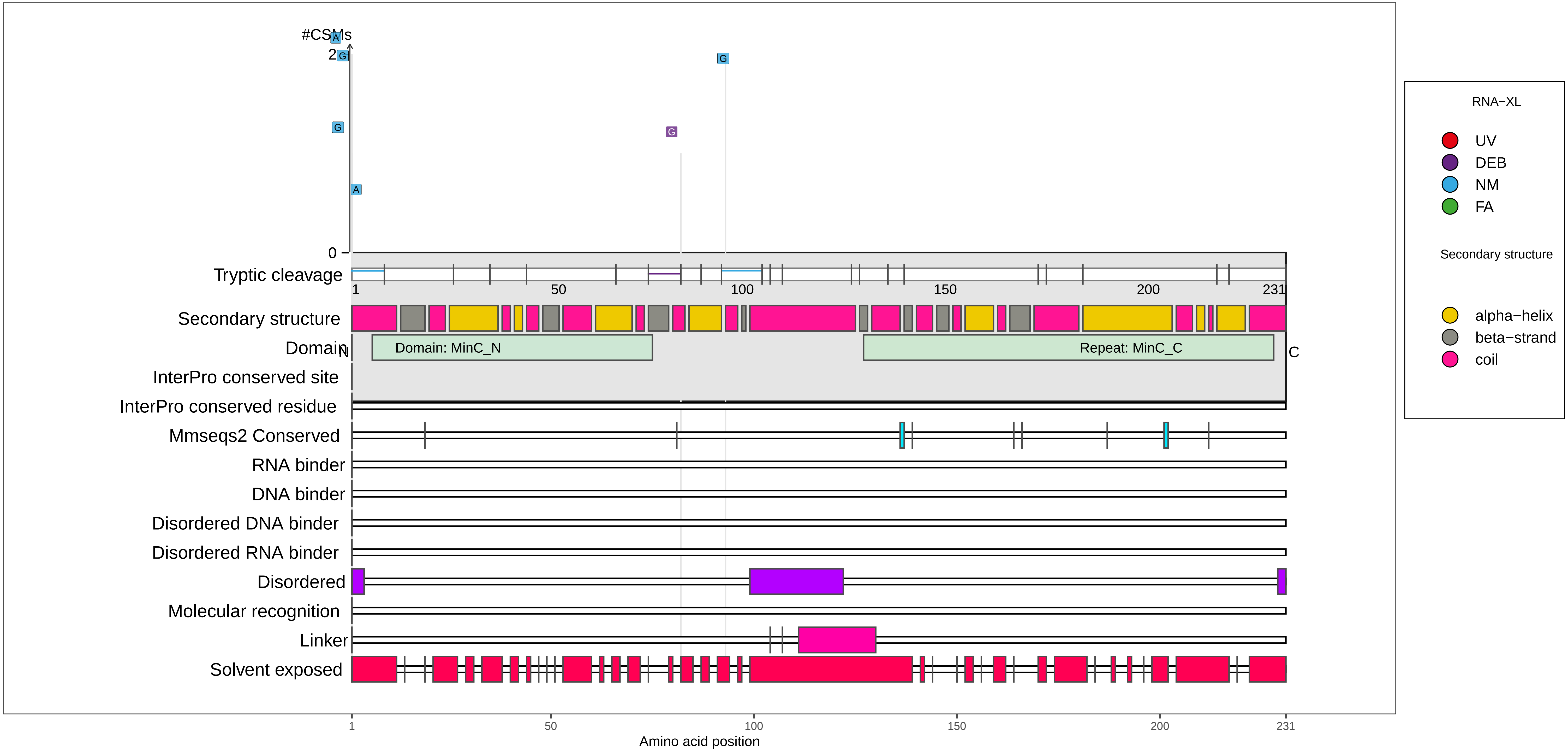

P21645
LPXD_ECOLI UDP−3−O−(3−hydroxymyristoyl)glucosamine N−acyltransferase

− Abundance:
tryptic [log10 Intensity]: 7.31 (Q 25)
PAXdb K12 strain [ppm]: 2.09 (Q 63)
PAXdb E.coli [ppm]: 2.54 (Q 86)

− RNA functions: not annotated

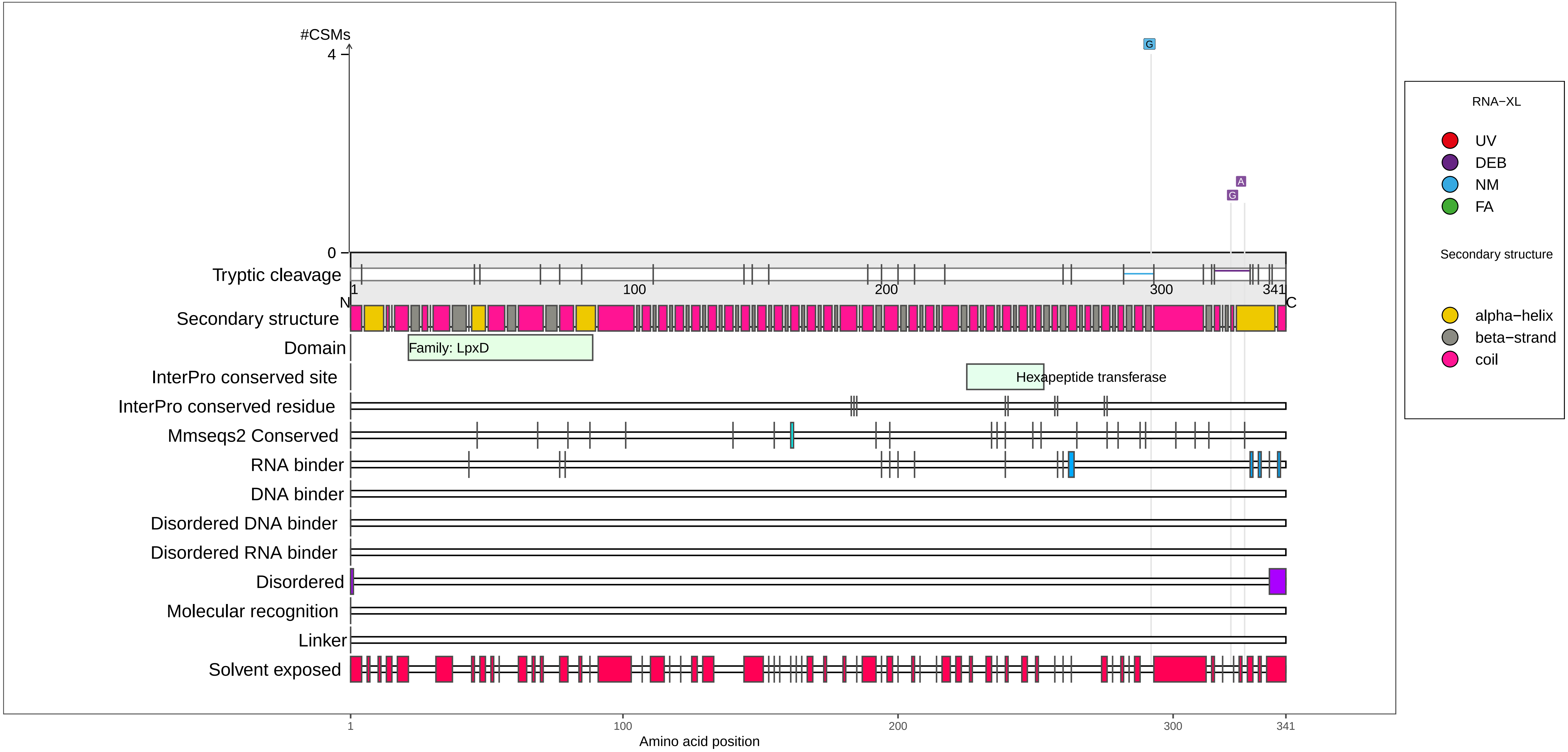

P25519
HFLX_ECOLI GTPase HflX

− Abundance:
tryptic [log10 Intensity]: 7.92 (Q 54)
PAXdb K12 strain [ppm]: 2.33 (Q 71)
PAXdb E.coli [ppm]: 1.9 (Q 69)

− RNA functions:
RNA binding; rRNA binding

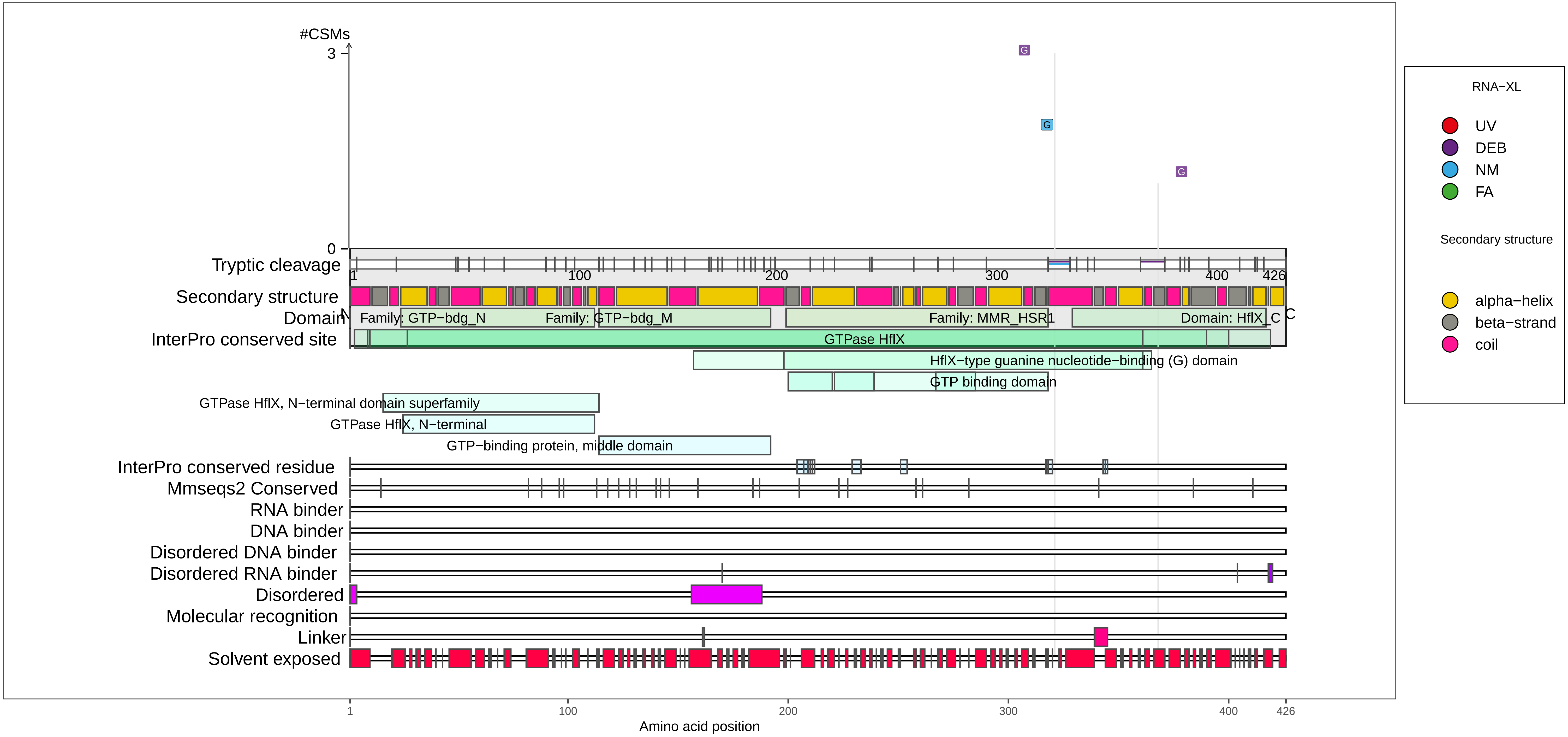

P25552
GPPA_ECOLI Guanosine−5'−triphosphate,3'−diphosphate pyrophosphatase
− Abundance:
tryptic [log10 Intensity]: 8.49 (Q 75)
PAXdb K12 strain [ppm]: 0.78 (Q 4)
PAXdb E.coli [ppm]: 1.81 (Q 67)

− RNA functions: not annotated

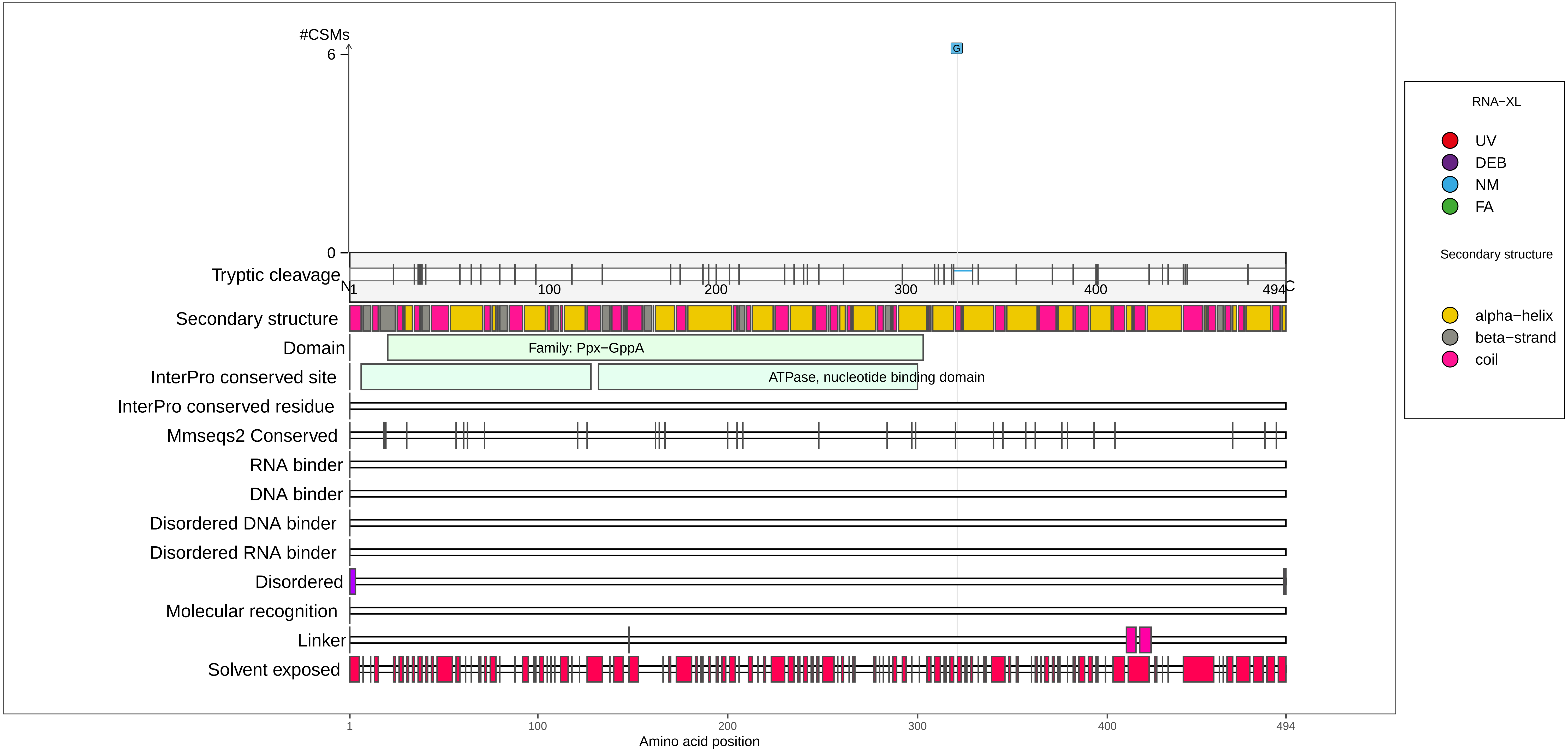

P26616
MAO1_ECOLI NAD−dependent malic enzyme

− Abundance:
tryptic [log10 Intensity]: 8.77 (Q 83)
PAXdb K12 strain [ppm]: 2.59 (Q 80)
PAXdb E.coli [ppm]: 2.47 (Q 85)

− RNA functions: not annotated

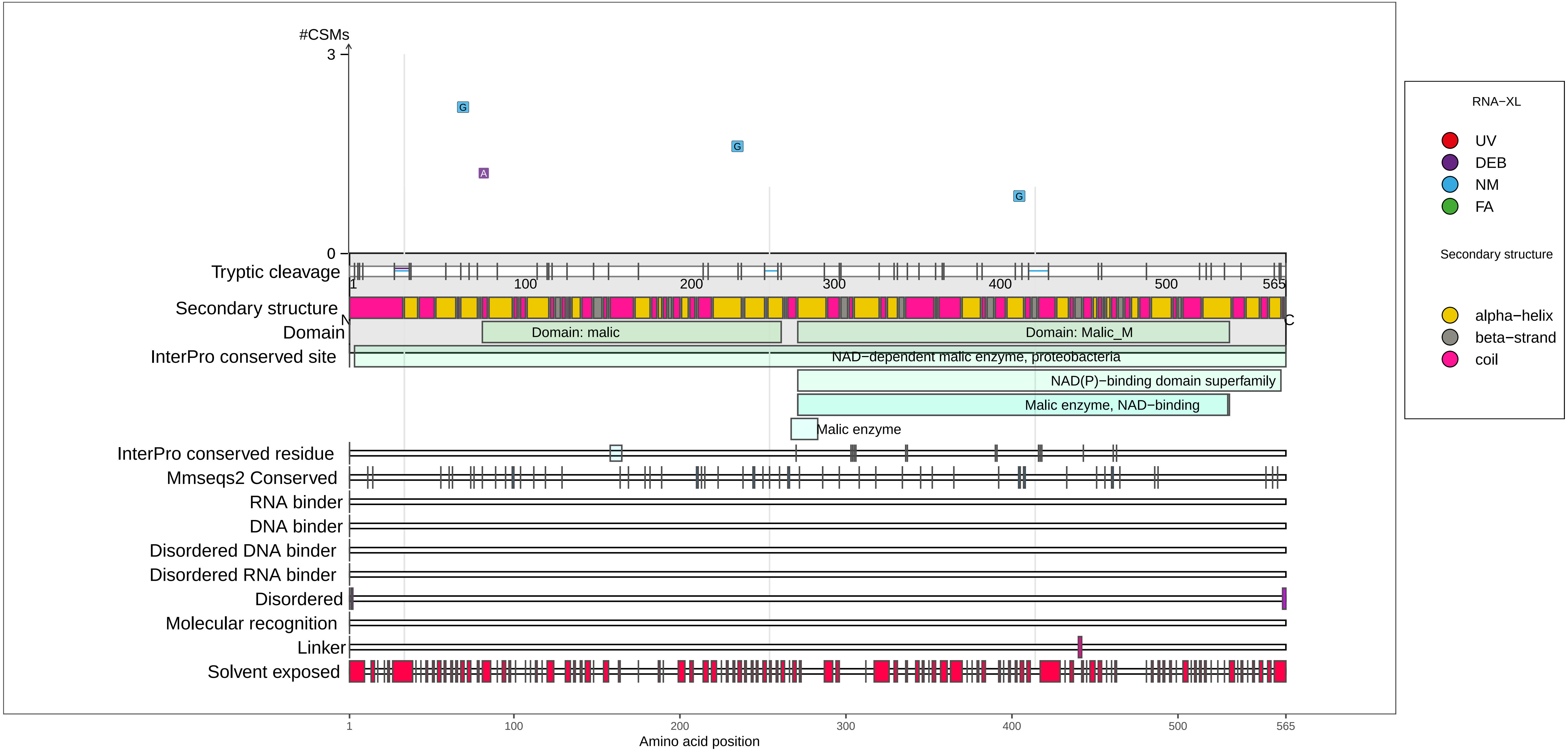

P31455
YIDR_ECOLI Uncharacterized protein YidR

− Abundance:
tryptic [log10 Intensity]: 8.52 (Q 76)
PAXdb K12 strain [ppm]: 1.61 (Q 45)
PAXdb E.coli [ppm]: 1.49 (Q 59)

− RNA functions: not annotated

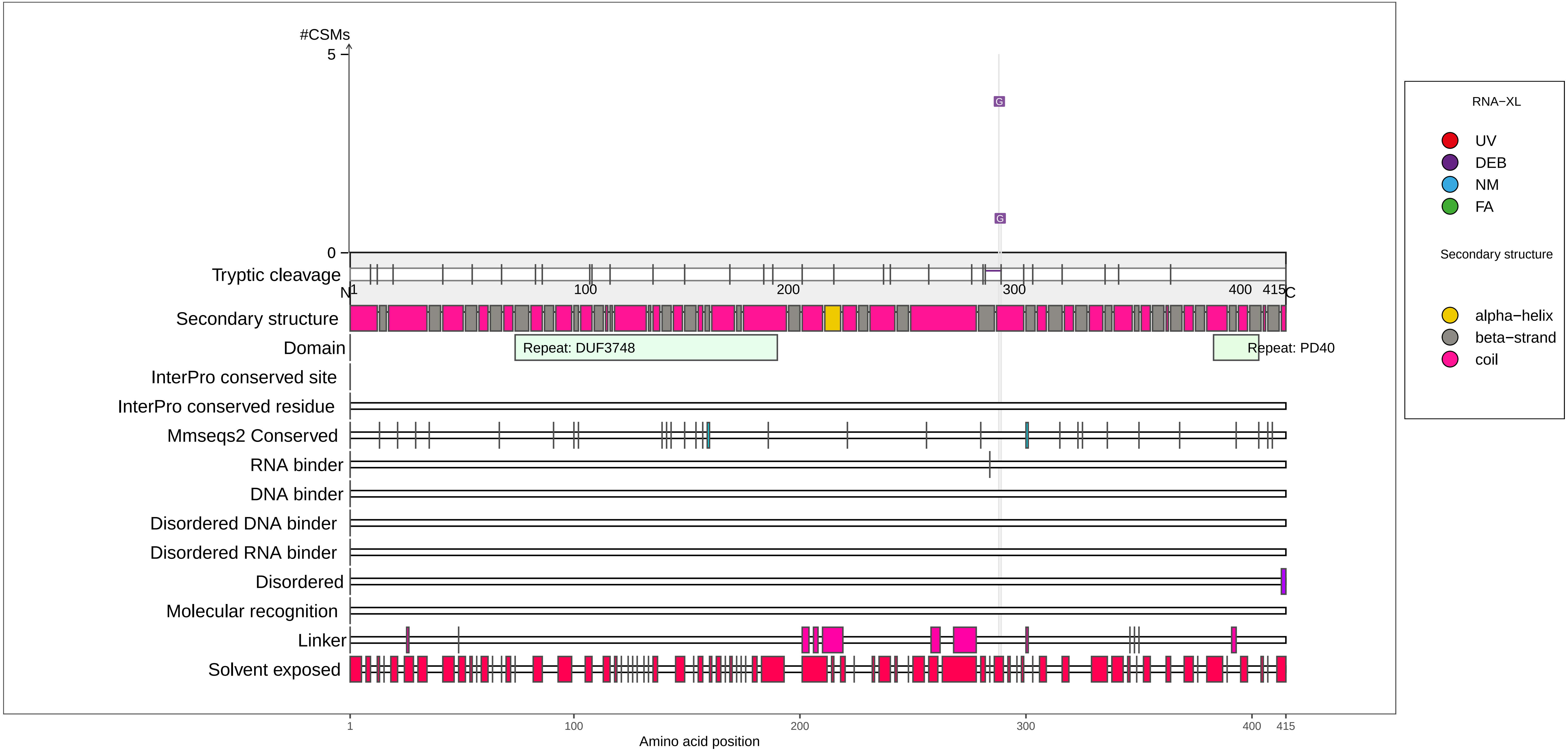

P36649
CUEO_ECOLI Blue copper oxidase CueO

− Abundance:
tryptic [log10 Intensity]: 7.93 (Q 55)
PAXdb K12 strain [ppm]: 1.01 (Q 11)
PAXdb E.coli [ppm]: 2 (Q 72)

− RNA functions: not annotated

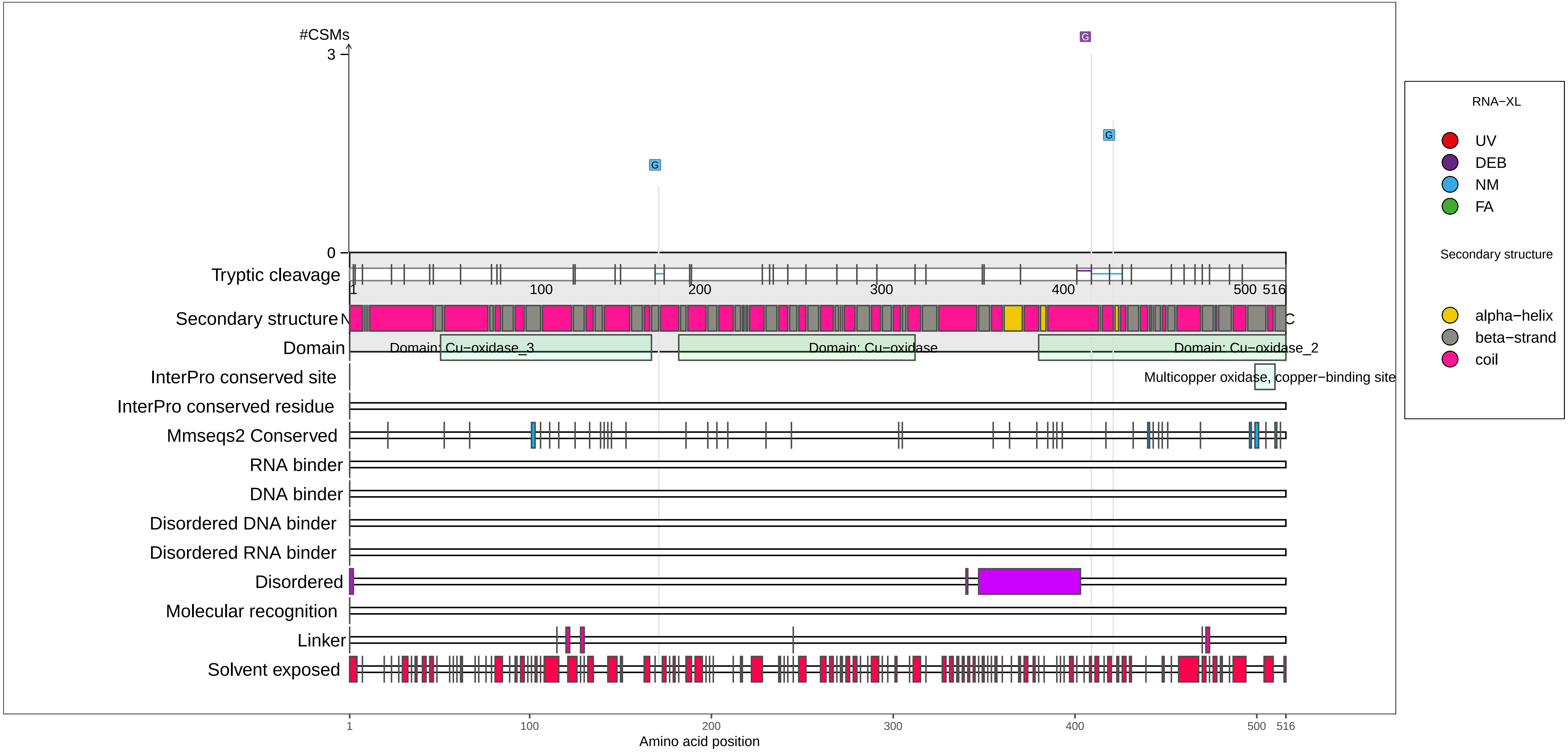

P37147
FXSA_ECOLI UPF0716 protein FxsA

− Abundance:
tryptic [log10 Intensity]: 8.02 (Q 58)
PAXdb K12 strain [ppm]: 2.05 (Q 62)
PAXdb E.coli [ppm]: 0.44 (Q 35)

− RNA functions: not annotated

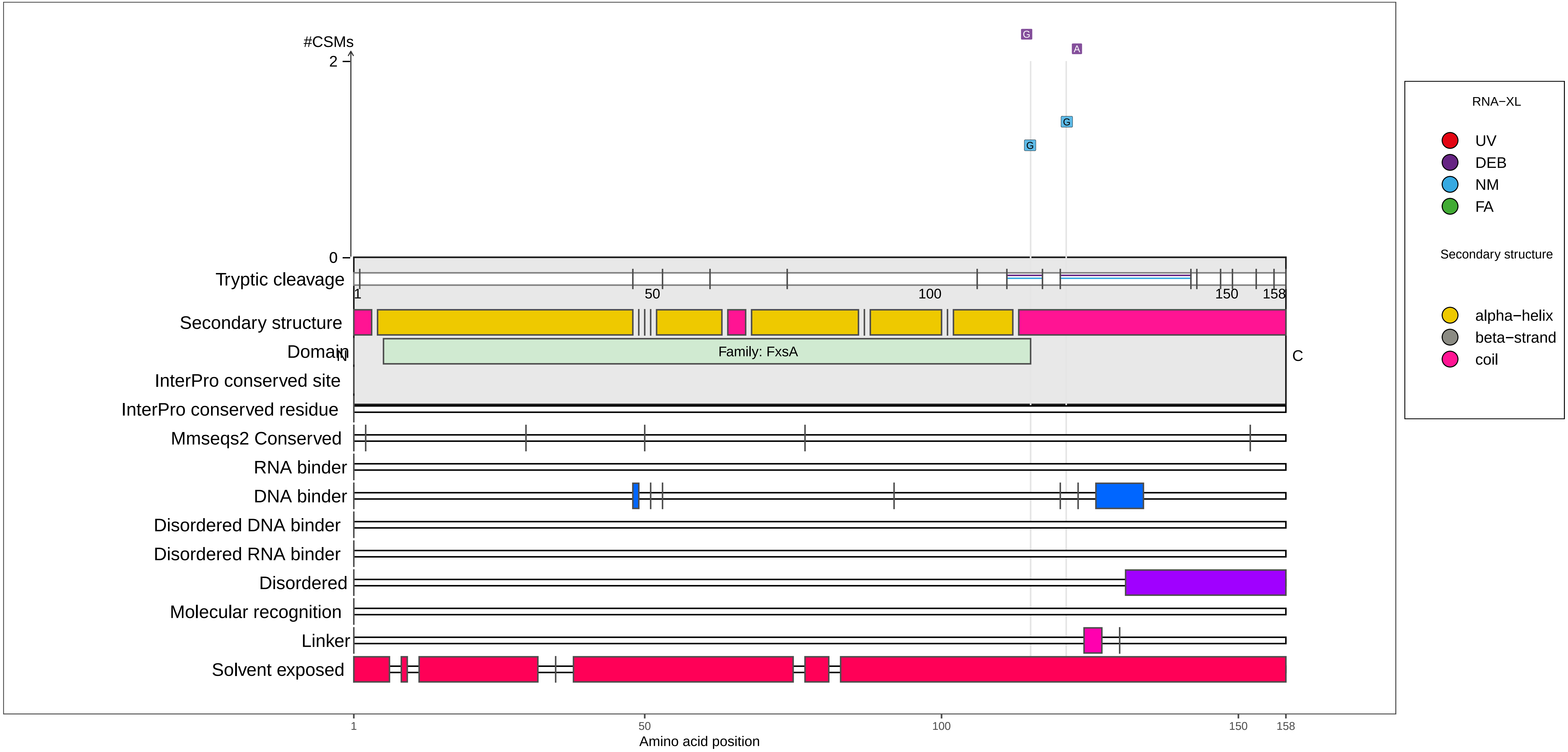

P45395
KDSD_ECOLI Arabinose 5−phosphate isomerase KdsD

− Abundance:
tryptic [log10 Intensity]: 7.78 (Q 48)
PAXdb K12 strain [ppm]: 1.84 (Q 55)
PAXdb E.coli [ppm]: 2.1 (Q 75)

− RNA functions: not annotated

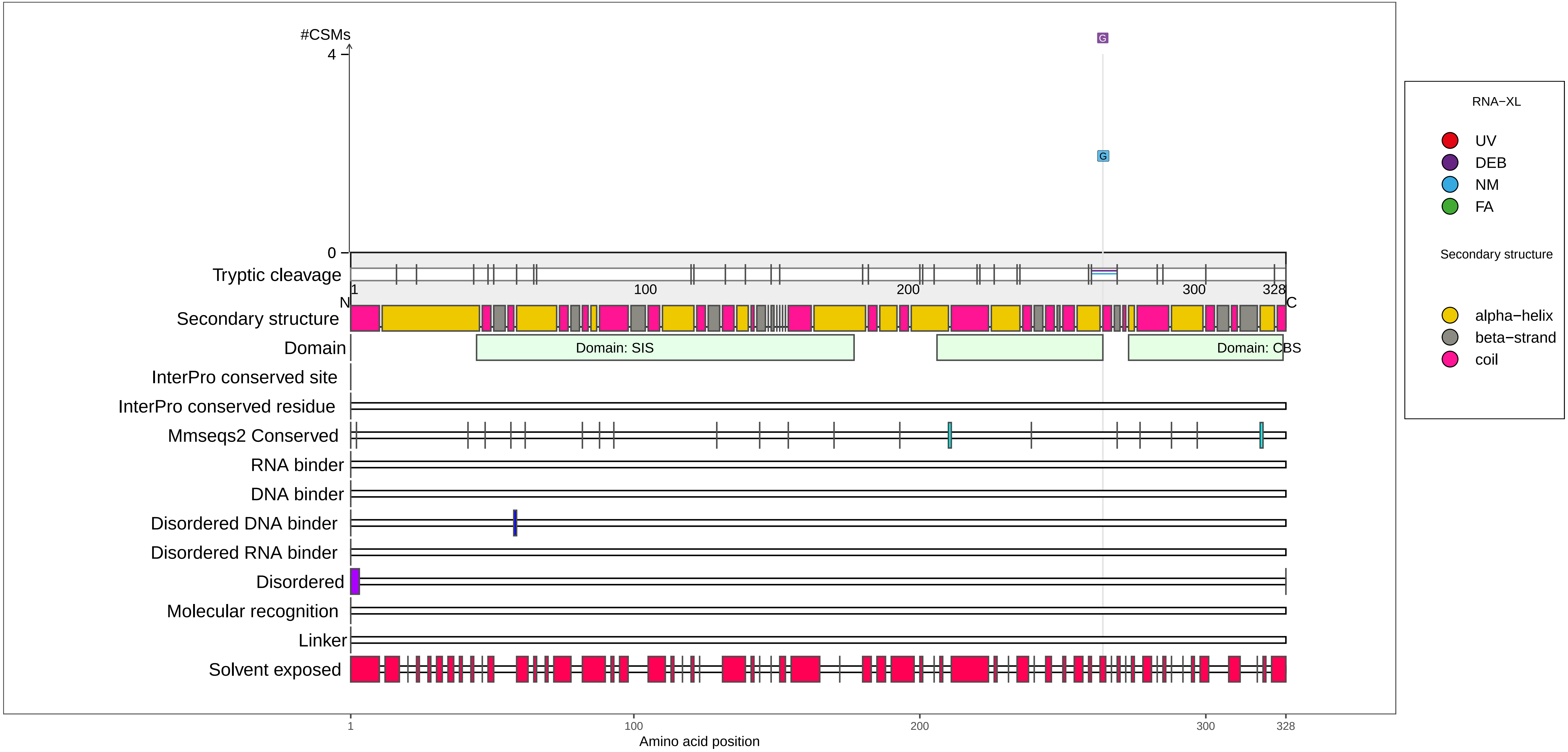

P61320
LOLB_ECOLI Outer−membrane lipoprotein LolB

− Abundance:
tryptic [log10 Intensity]: 8.33 (Q 70)
PAXdb K12 strain [ppm]: 1.41 (Q 33)
PAXdb E.coli [ppm]: 1.89 (Q 69)

− RNA functions: not annotated

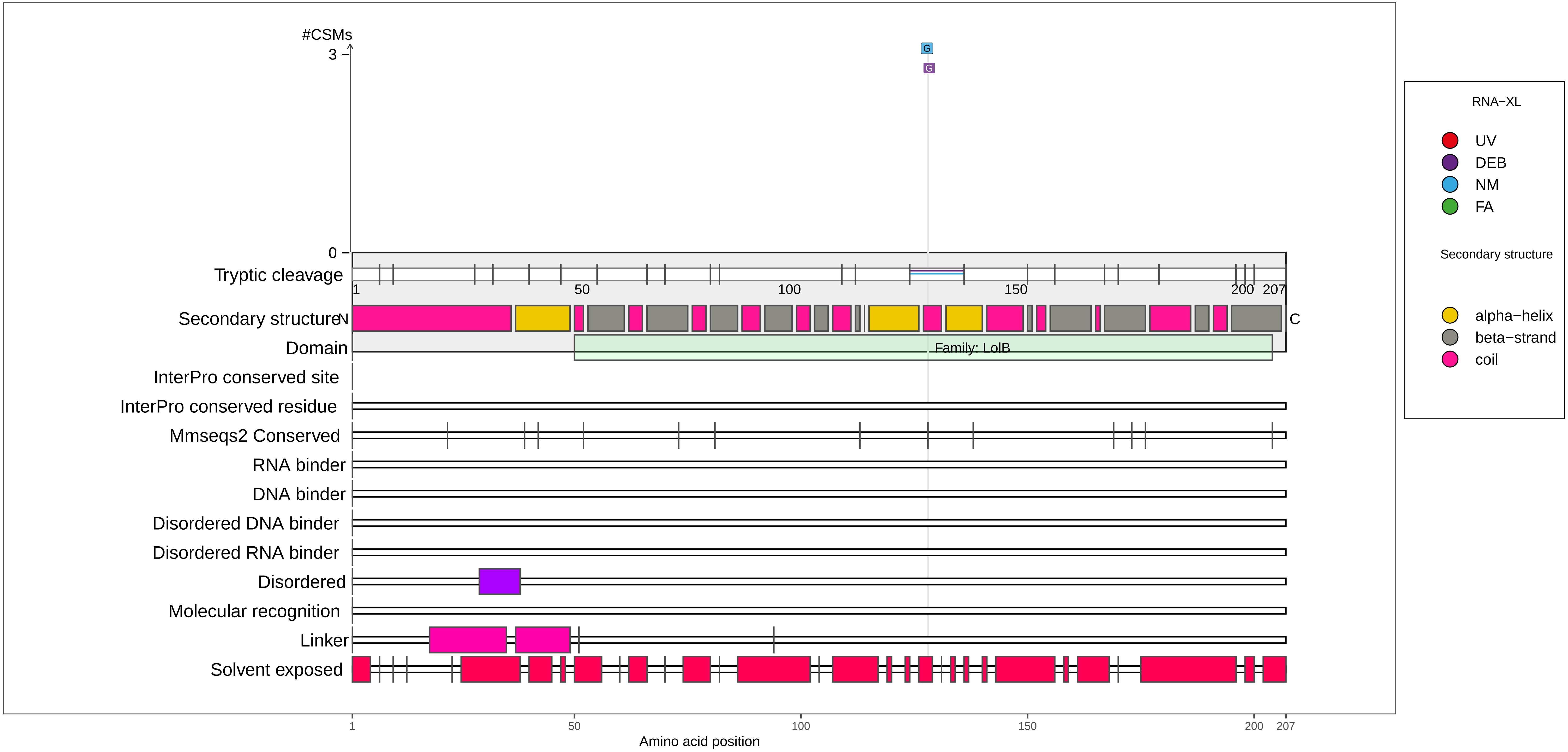

P61714
RISB_ECOLI 6,7−dimethyl−8−ribityllumazine synthase

− Abundance:
tryptic [log10 Intensity]: 9.19 (Q 93)
PAXdb K12 strain [ppm]: 3.13 (Q 93)
PAXdb E.coli [ppm]: 3.19 (Q 96)

− RNA functions: not annotated

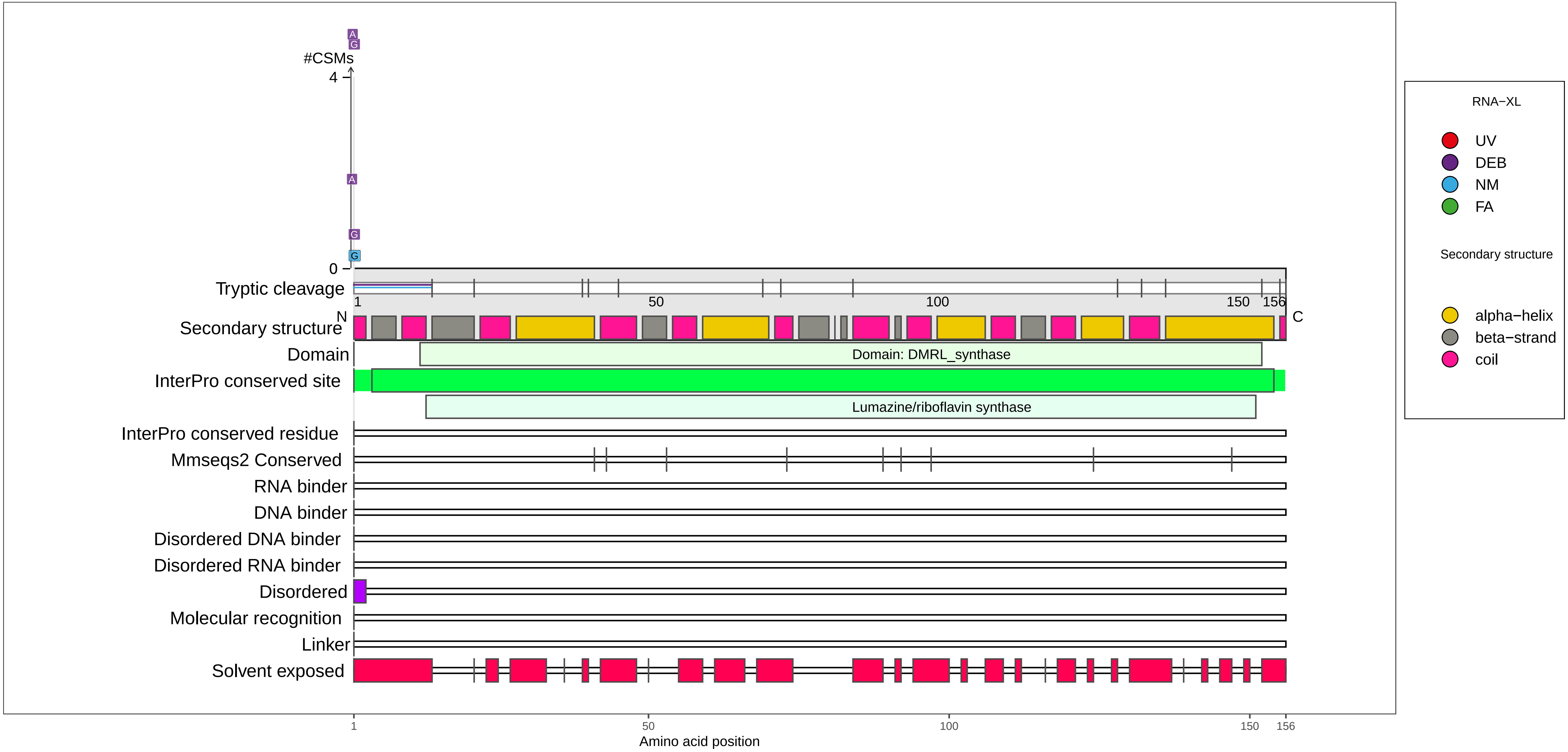

P62620
ISPG_ECOLI 4−hydroxy−3−methylbut−2−en−1−yl diphosphate synthase (flavodoxin)

− Abundance:
tryptic [log10 Intensity]: 8.62 (Q 79)
PAXdb K12 strain [ppm]: 2.72 (Q 84)
PAXdb E.coli [ppm]: 2.44 (Q 84)

− RNA functions: not annotated

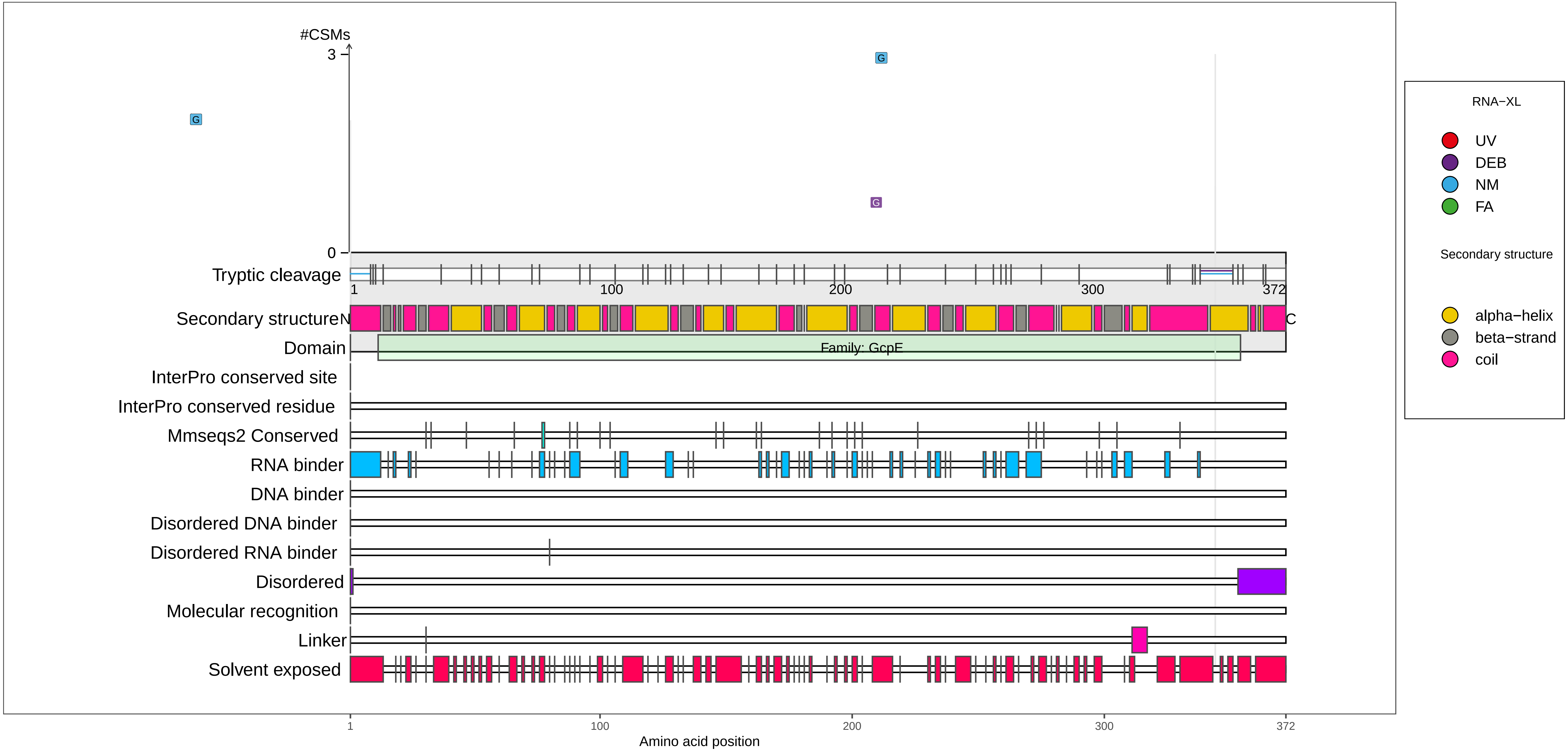

P66817
DIAA_ECOLI DnaA initiator−associating protein DiaA

− Abundance:
tryptic [log10 Intensity]: 7.27 (Q 24)
PAXdb K12 strain [ppm]: 1.43 (Q 34)
PAXdb E.coli [ppm]: 1.47 (Q 59)

− RNA functions: not annotated

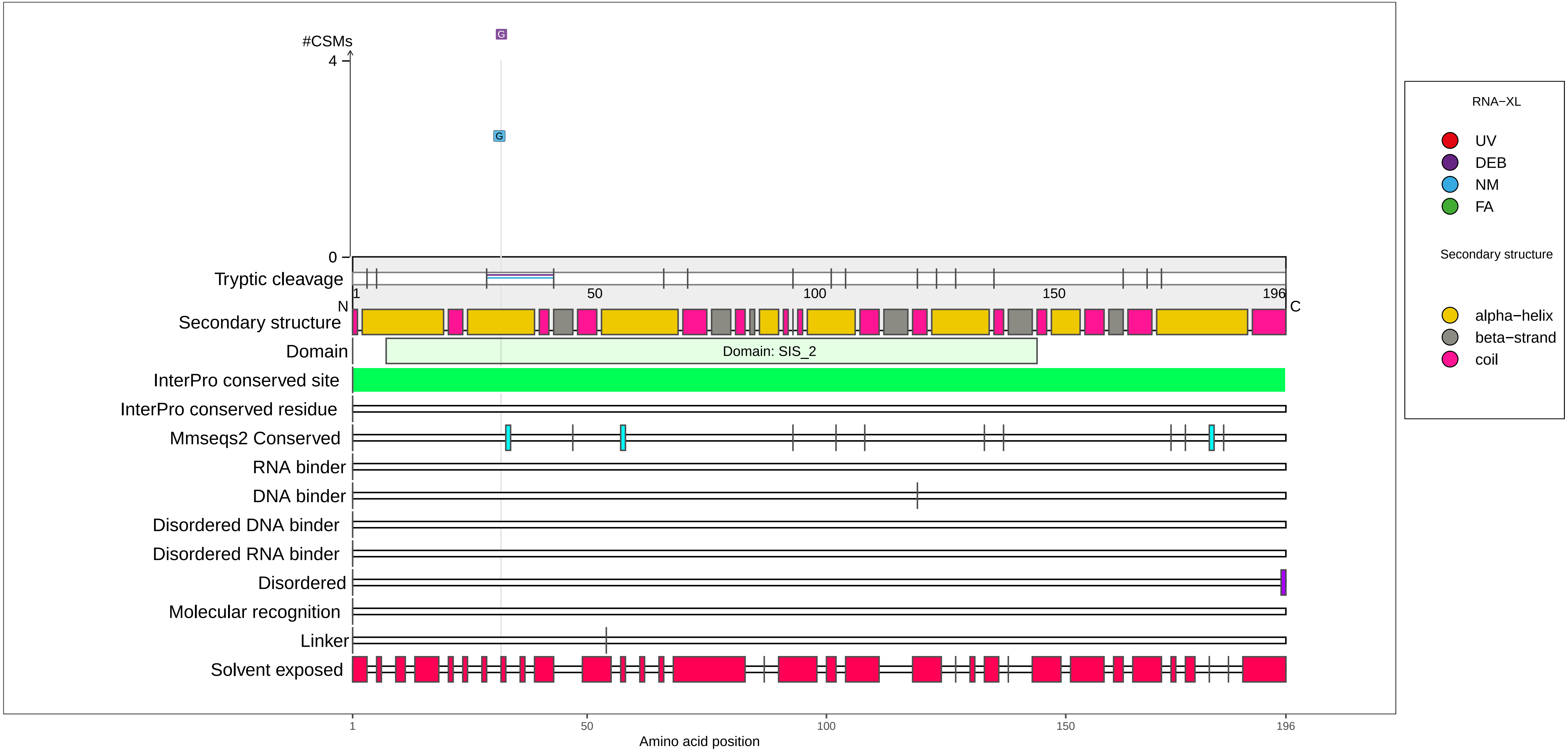

P67762
YHBP_ECOLI UPF0306 protein YhbP

− Abundance:
tryptic [log10 Intensity]: 6.76 (Q 6)
PAXdb K12 strain [ppm]: 2.12 (Q 65)
PAXdb E.coli [ppm]: 0.41 (Q 34)

− RNA functions: not annotated

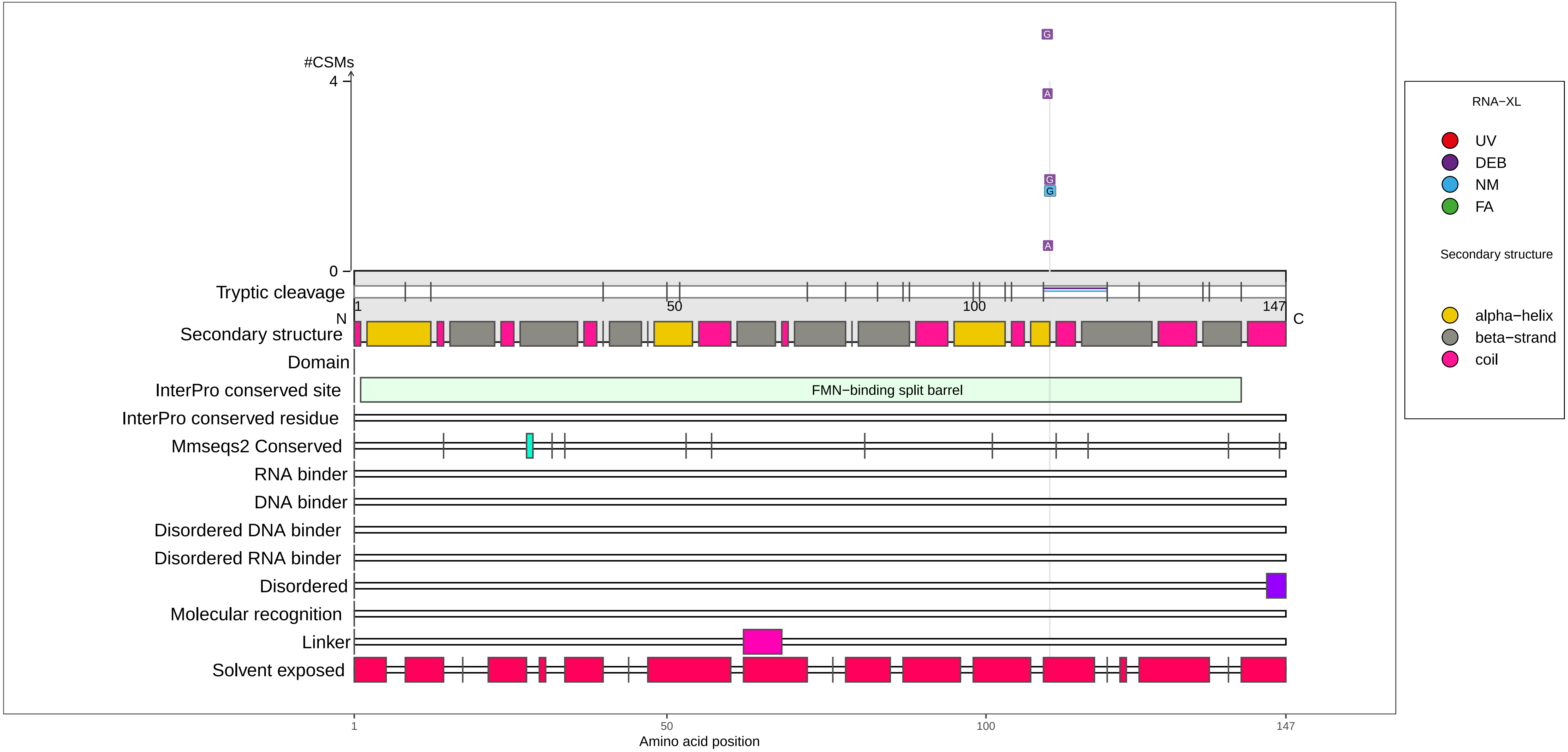

P68739
NFI_ECOLI Endonuclease V

− Abundance:
tryptic [log10 Intensity]: 7.82 (Q 50)
PAXdb K12 strain [ppm]: 1.21 (Q 21)
PAXdb E.coli [ppm]: 1.4 (Q 57)

− RNA functions:
RNA binding; RNA metabolic process; RNA phosphodiester bond hydrolysis
RNA phosphodiester bond hydrolysis, endonucleolytic

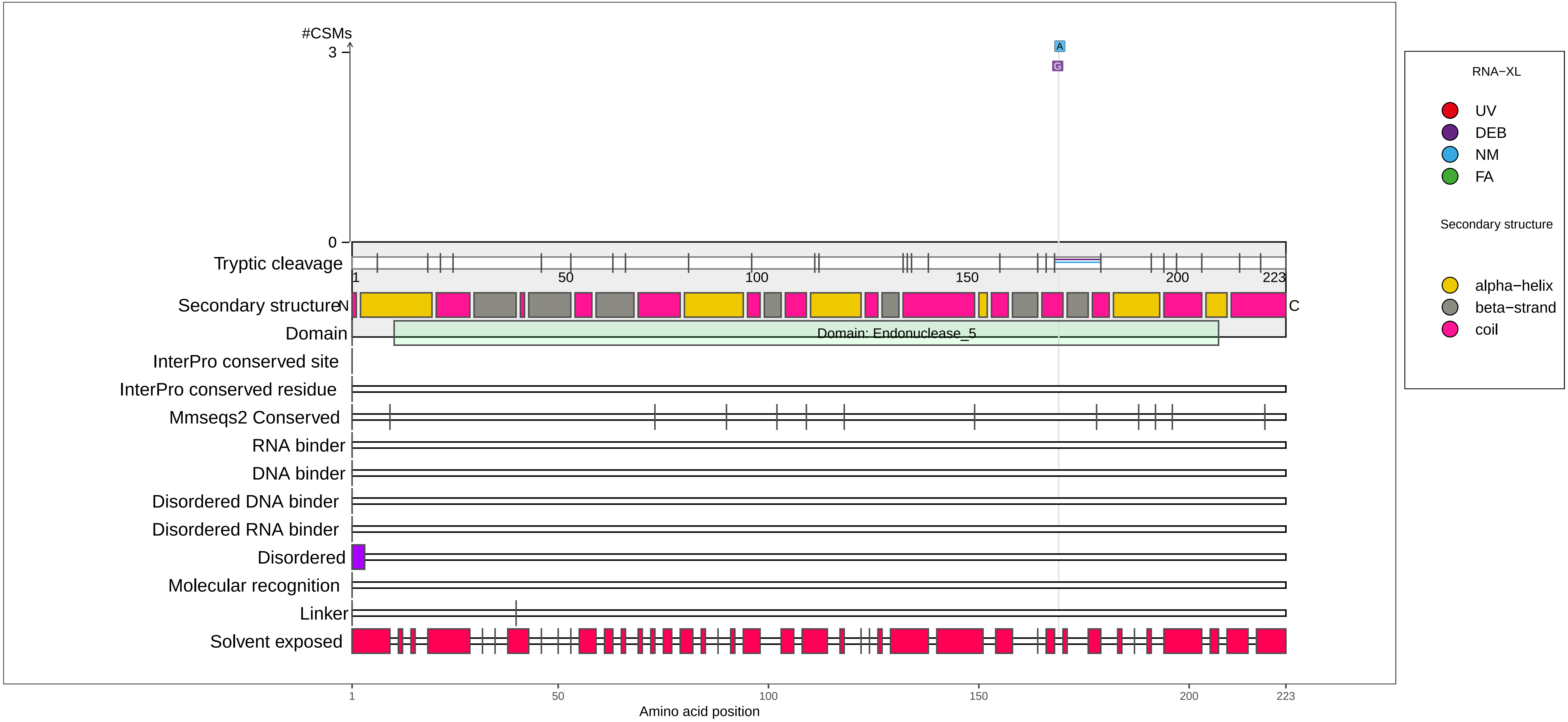

P75728
UBIF_ECOLI 3−demethoxyubiquinol 3−hydroxylase

− Abundance:
tryptic [log10 Intensity]: 7.73 (Q 46)
PAXdb K12 strain [ppm]: 1.39 (Q 32)
PAXdb E.coli [ppm]: 1.96 (Q 71)

− RNA functions: not annotated

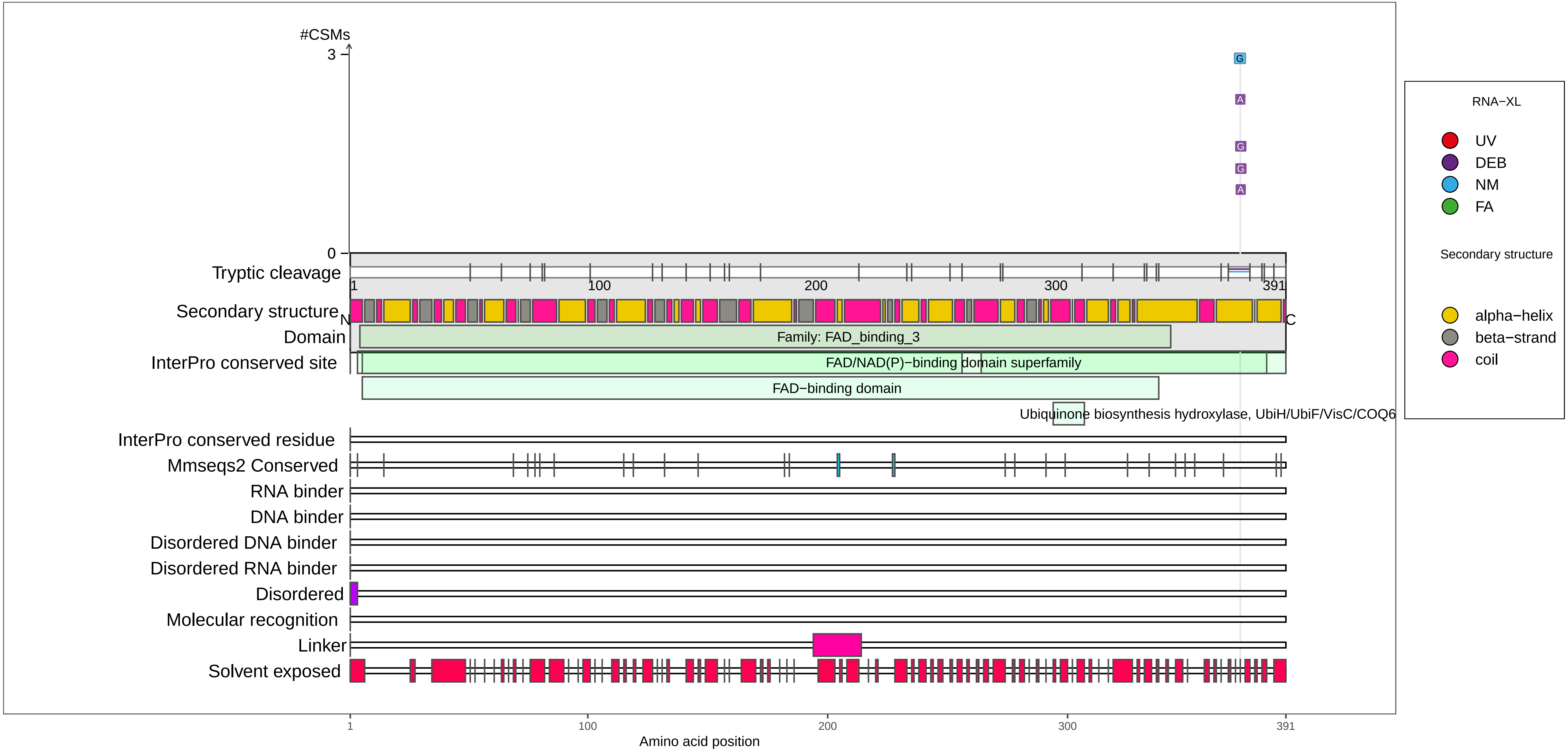

P75830
MACA_ECOLI Macrolide export protein MacA
− Abundance:
tryptic [log10 Intensity]: 7.53 (Q 36)
PAXdb K12 strain [ppm]: 1.24 (Q 22)
PAXdb E.coli [ppm]: 1.2 (Q 52)

− RNA functions: not annotated

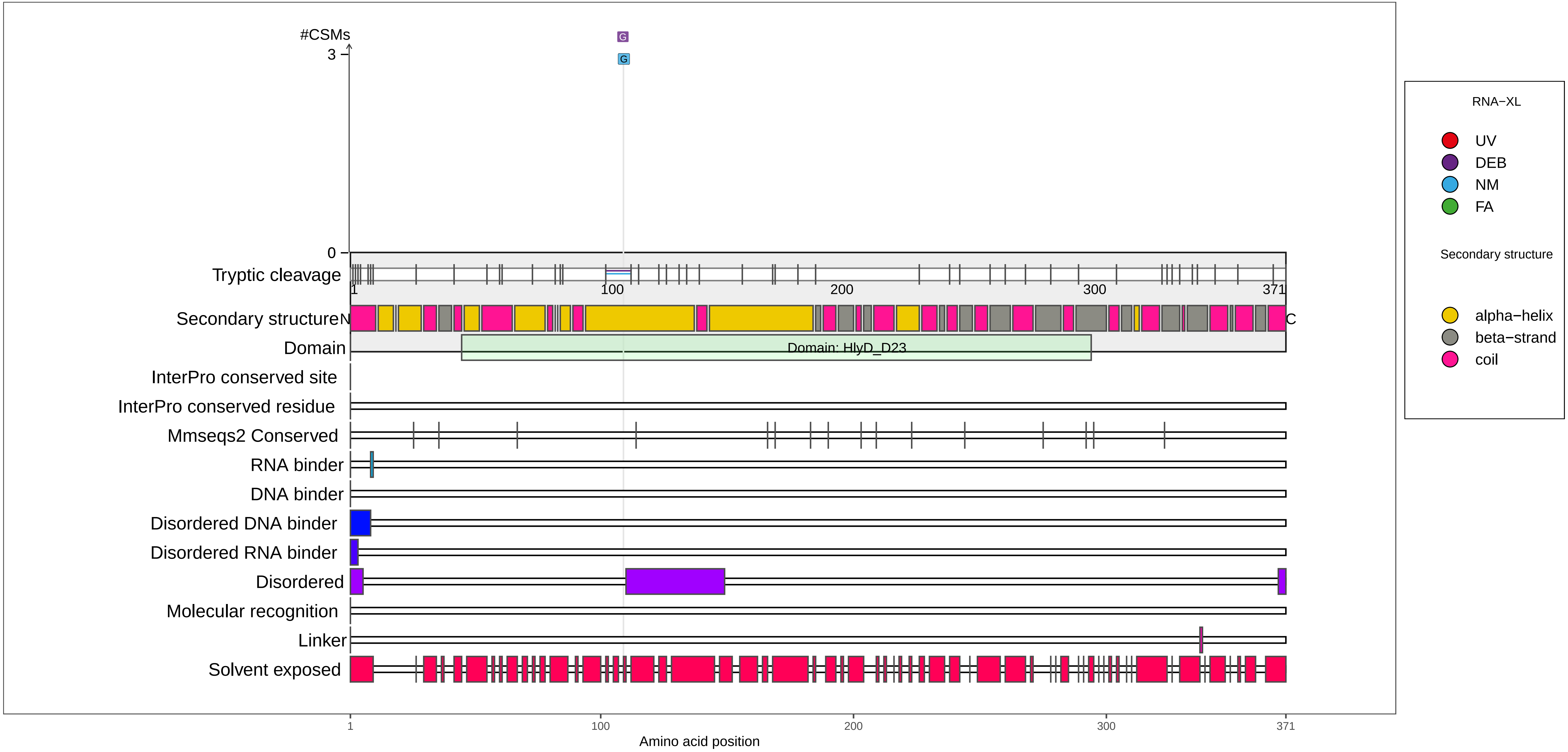

Q57261
TRUD_ECOLI tRNA pseudouridine synthase D

− Abundance:
tryptic [log10 Intensity]: 8.39 (Q 72)
PAXdb K12 strain [ppm]: 2.3 (Q 70)
PAXdb E.coli [ppm]: 1.7 (Q 64)

− RNA functions:
ncRNA metabolic process; ncRNA processing; RNA binding; RNA metabolic process
RNA modification; RNA processing; tRNA metabolic process; tRNA modification; tRNA processing
tRNA pseudouridine synthase activity; tRNA pseudouridine synthase D (TruD)
tRNA pseudouridine synthesis

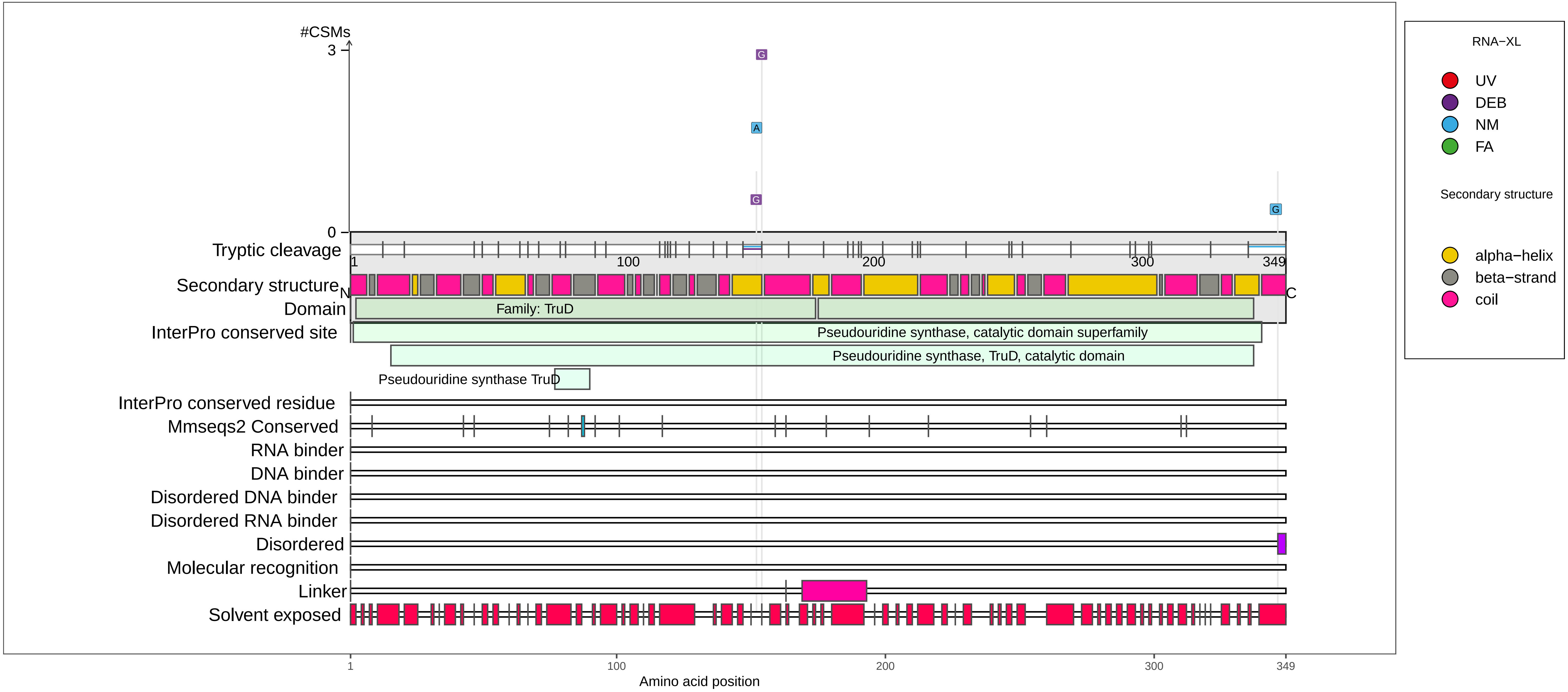

Q93K97
ADPP_ECOLI ADP−ribose pyrophosphatase

− Abundance:
tryptic [log10 Intensity]: 8.38 (Q 72)
PAXdb K12 strain [ppm]: NA (Q NA)
PAXdb E.coli [ppm]: 1.83 (Q 68)

− RNA functions: not annotated

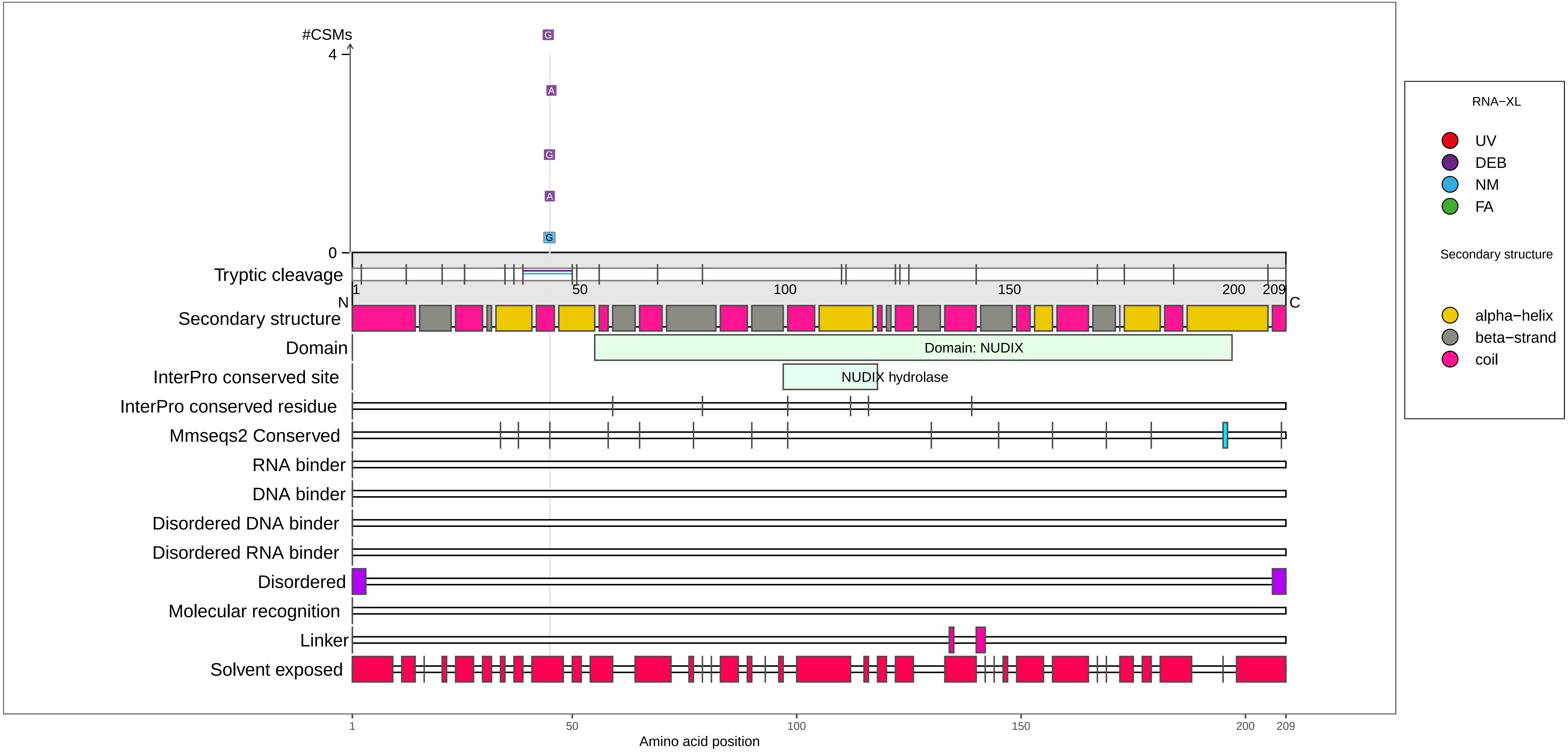

P00452
RIR1_ECOLI Ribonucleoside−diphosphate reductase 1 subunit alpha

− Abundance:
tryptic [log10 Intensity]: 9.14 (Q 91)
PAXdb K12 strain [ppm]: 2.8 (Q 86)
PAXdb E.coli [ppm]: 2.89 (Q 93)

− RNA functions: not annotated

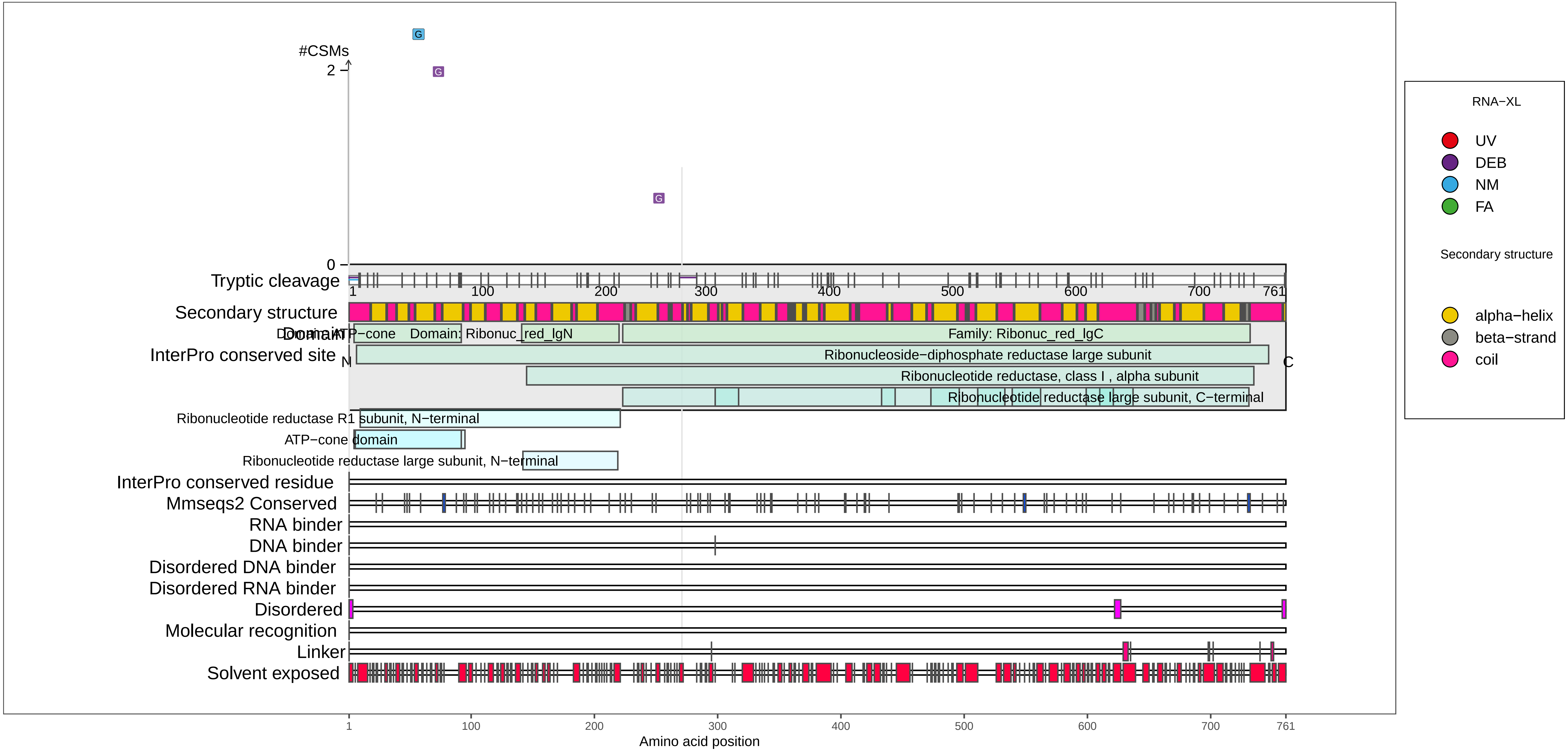

P00887
AROH_ECOLI Phospho−2−dehydro−3−deoxyheptonate aldolase, Trp−sensitive

− Abundance:
tryptic [log10 Intensity]: 7.62 (Q 41)
PAXdb K12 strain [ppm]: 1.45 (Q 35)
PAXdb E.coli [ppm]: 1.42 (Q 58)

− RNA functions: not annotated

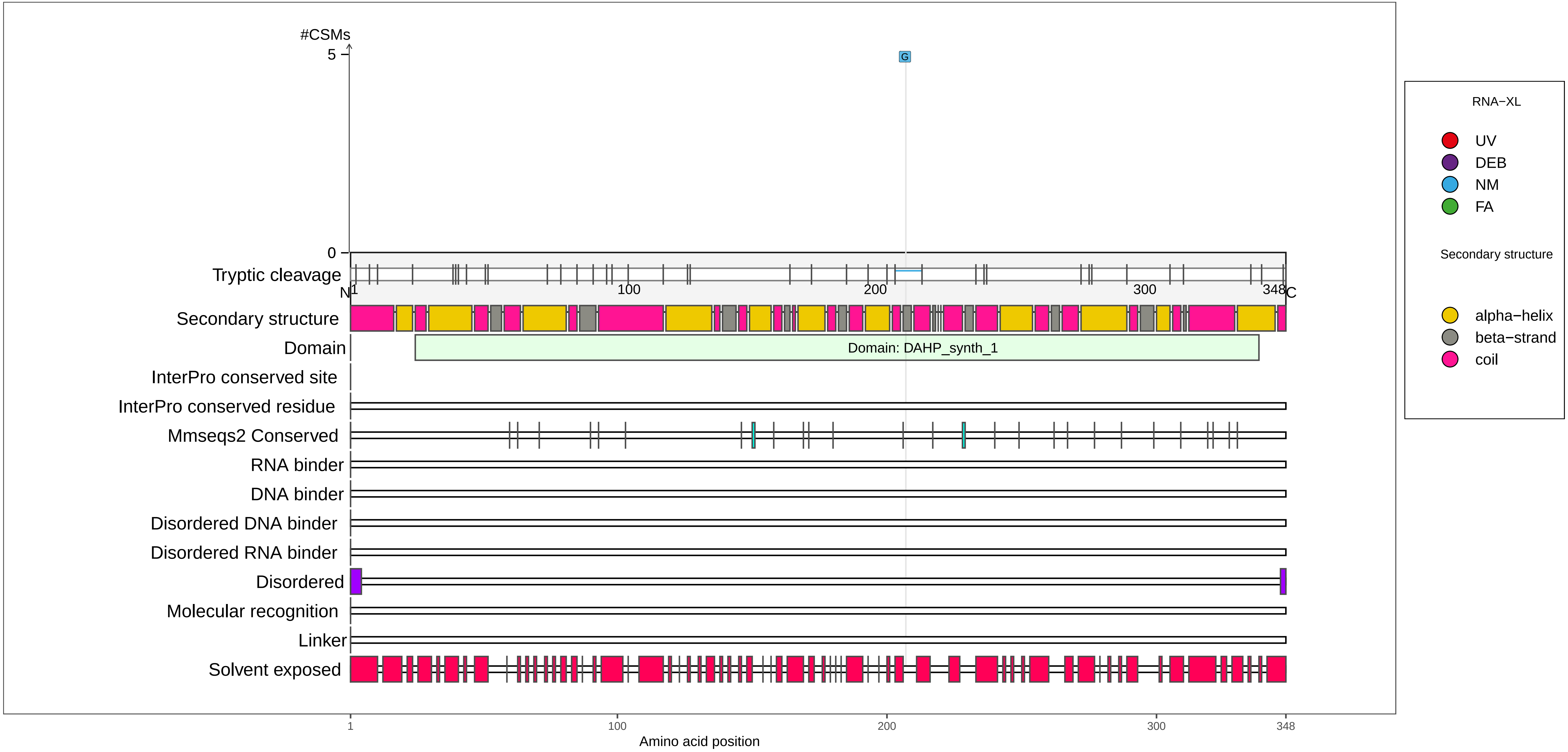

P06710
DPO3X_ECOLI DNA polymerase III subunit tau

− Abundance:
tryptic [log10 Intensity]: 7.06 (Q 15)
PAXdb K12 strain [ppm]: 1.77 (Q 52)
PAXdb E.coli [ppm]: 1.59 (Q 62)

− RNA functions: not annotated

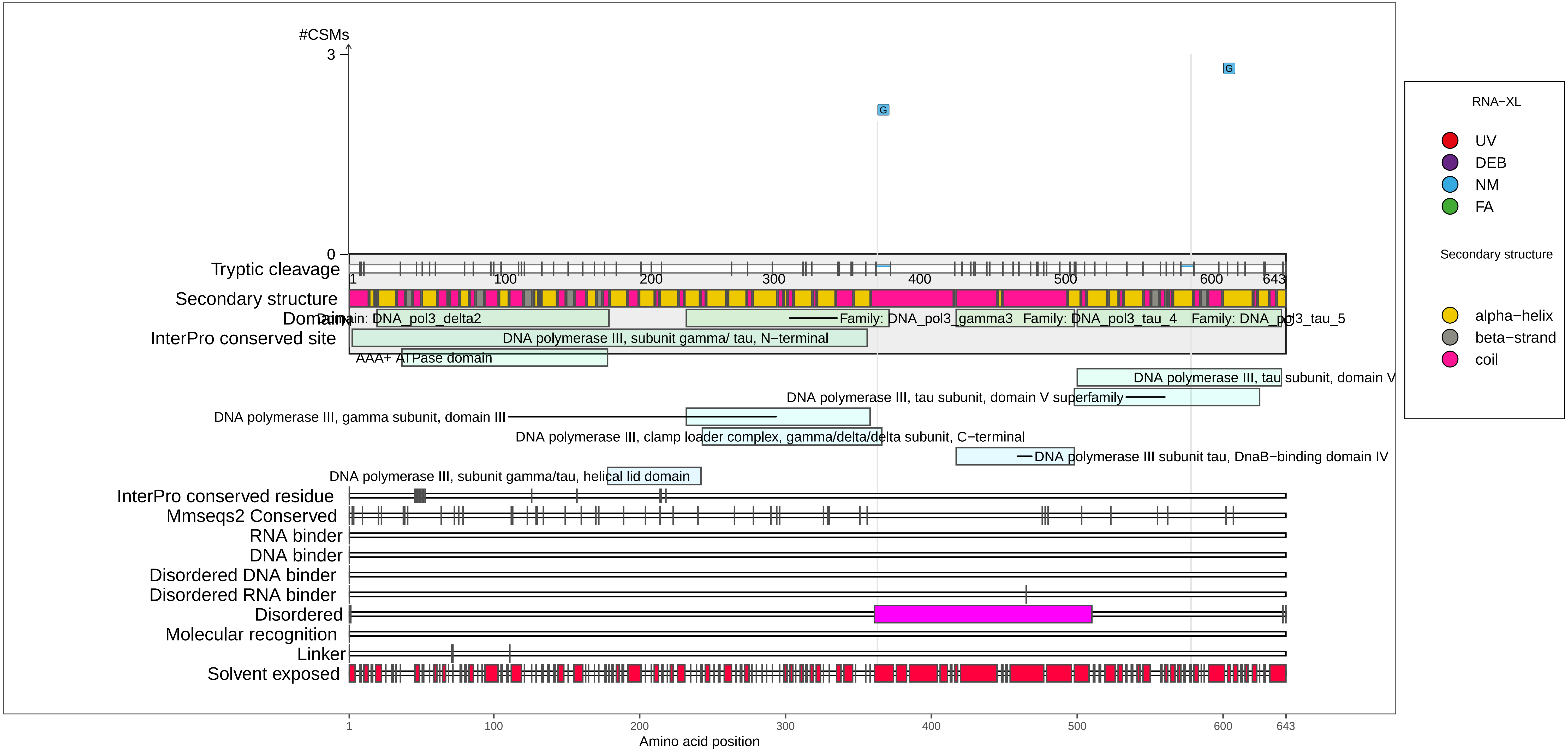

P08142
ILVB_ECOLI Acetolactate synthase isozyme 1 large subunit

− Abundance:
tryptic [log10 Intensity]: 6.44 (Q 2)
PAXdb K12 strain [ppm]: NA (Q NA)
PAXdb E.coli [ppm]: 2.59 (Q 88)

− RNA functions: not annotated

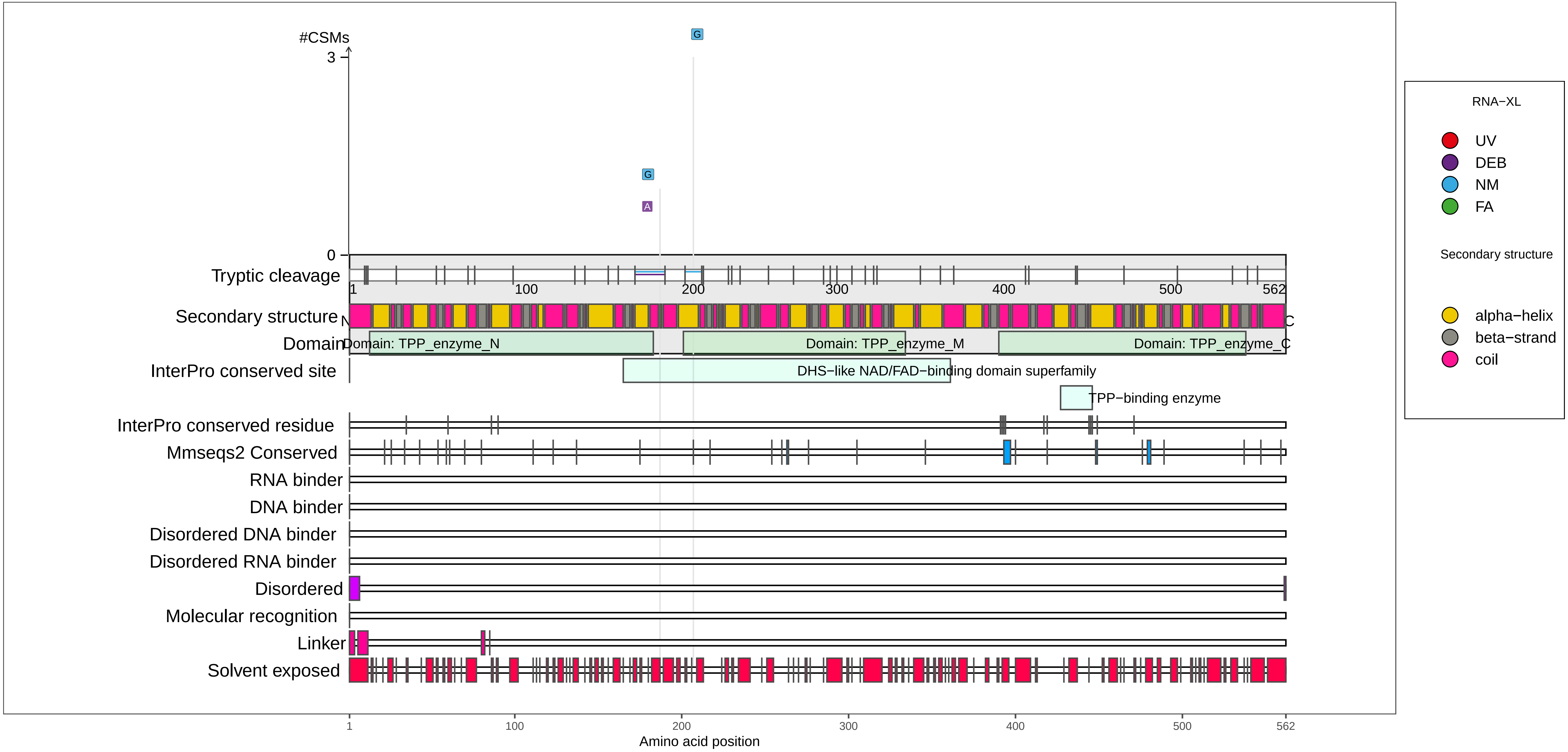

P09980
REP_ECOLI ATP−dependent DNA helicase Rep

− Abundance:
tryptic [log10 Intensity]: NA (Q NA)
PAXdb K12 strain [ppm]: 0.8 (Q 5)
PAXdb E.coli [ppm]: 1 (Q 48)

− RNA functions: not annotated

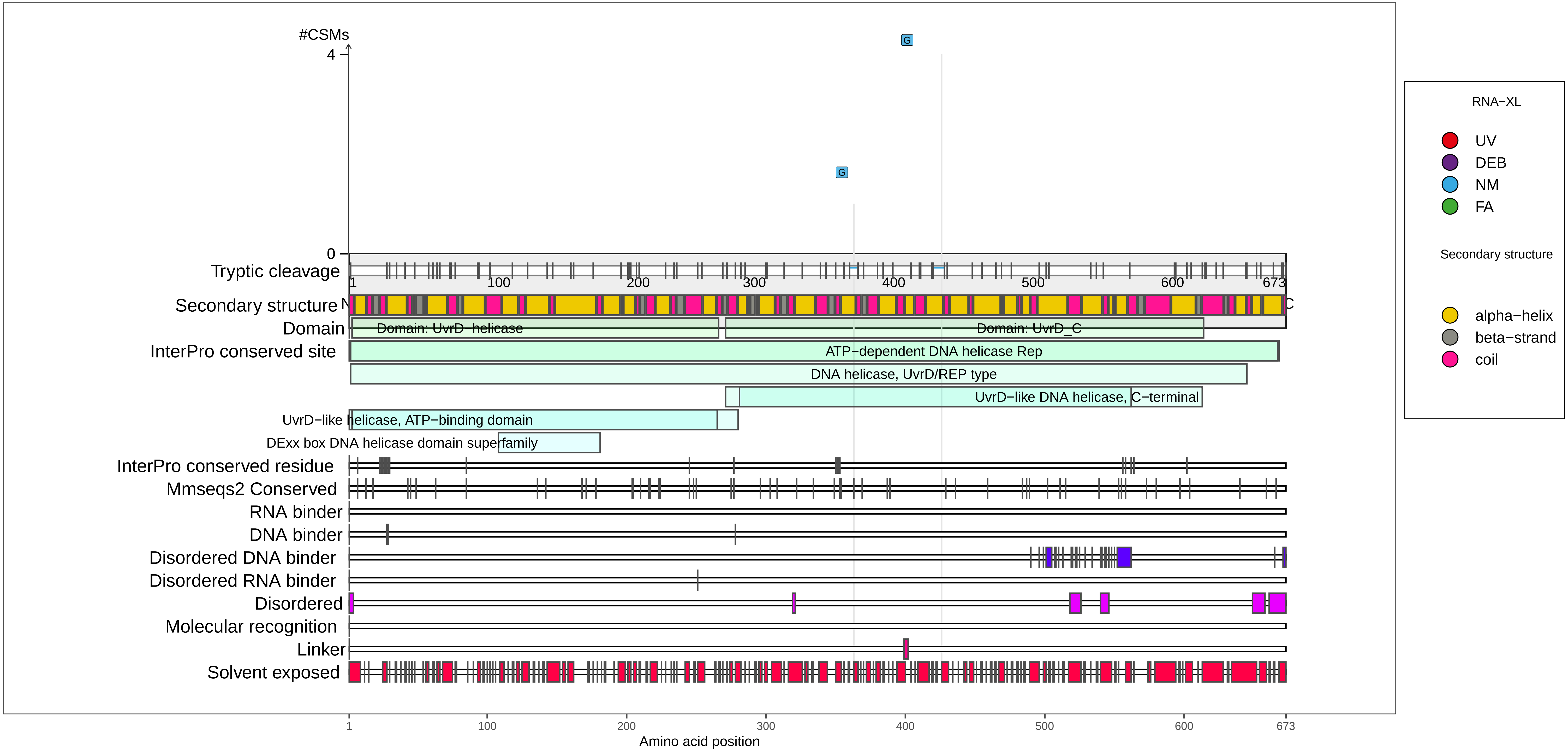

P0A698
UVRA_ECOLI UvrABC system protein A

− Abundance:
tryptic [log10 Intensity]: 8.97 (Q 87)
PAXdb K12 strain [ppm]: 2.35 (Q 72)
PAXdb E.coli [ppm]: 1.92 (Q 70)

− RNA functions: not annotated

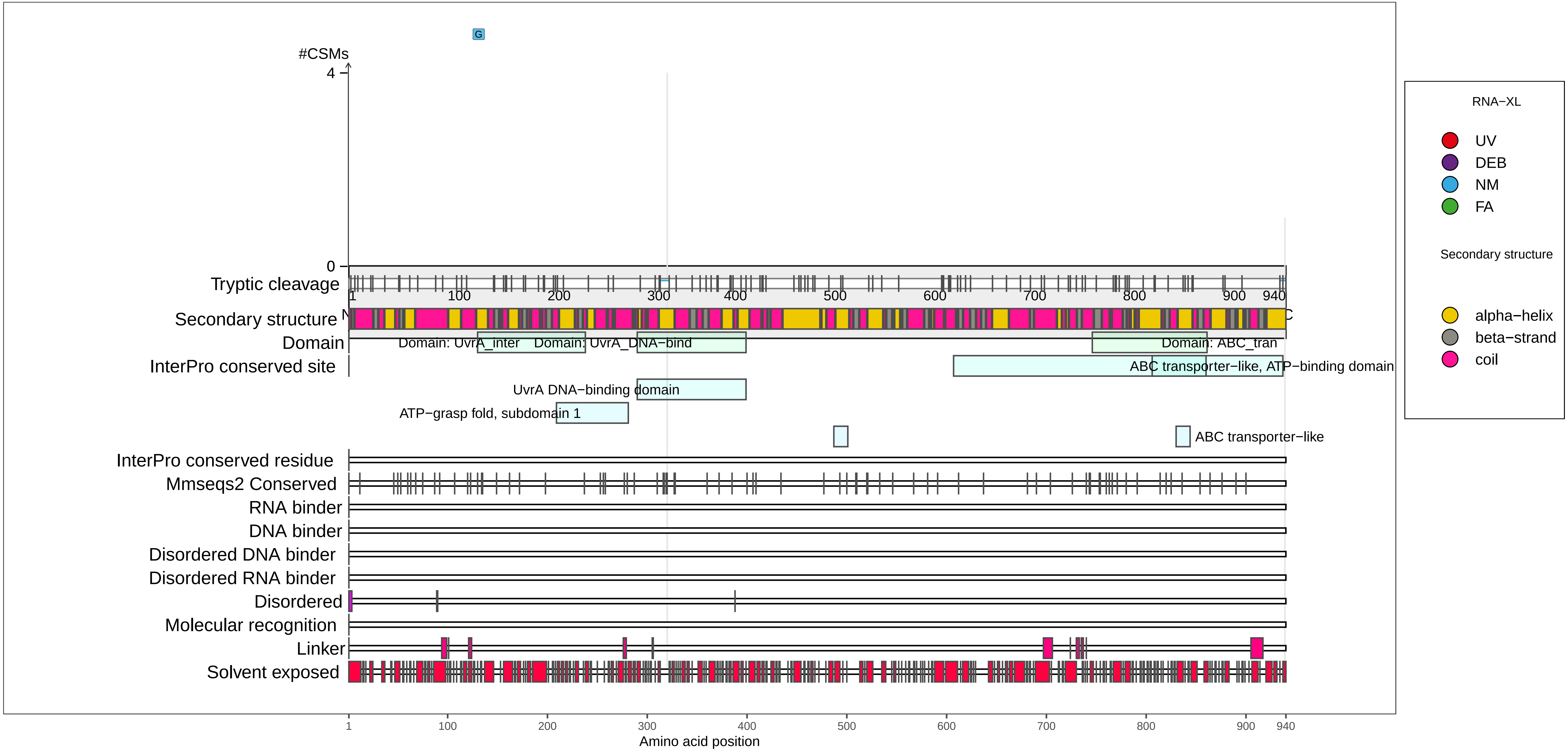

P0A6E1
AROL_ECOLI Shikimate kinase 2

− Abundance:
tryptic [log10 Intensity]: 7.7 (Q 44)
PAXdb K12 strain [ppm]: 1.47 (Q 36)
PAXdb E.coli [ppm]: 1.8 (Q 67)

− RNA functions: not annotated

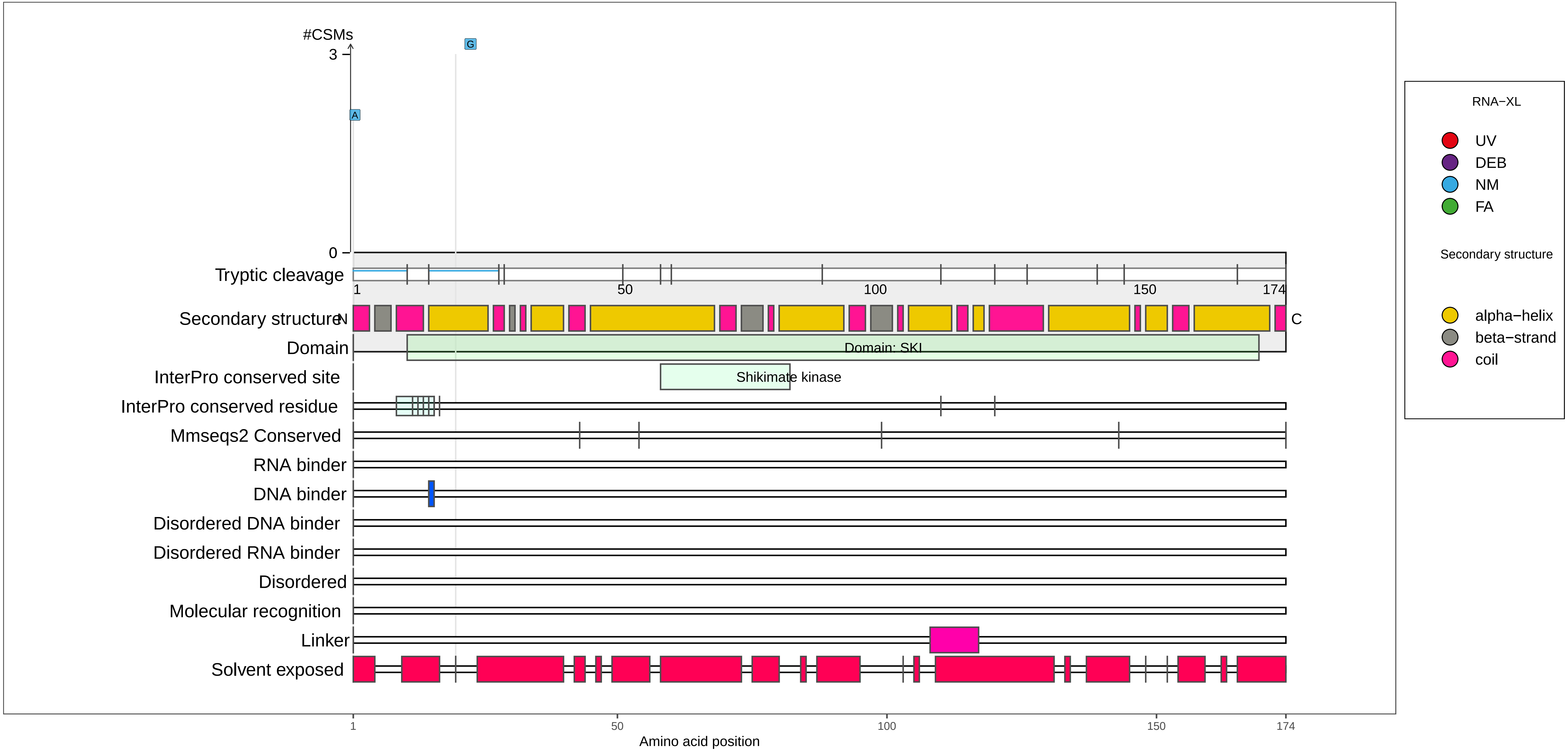

P0A6I0
KCY_ECOLI Cytidylate kinase

− Abundance:
tryptic [log10 Intensity]: 8.09 (Q 61)
PAXdb K12 strain [ppm]: 1.98 (Q 60)
PAXdb E.coli [ppm]: 2.31 (Q 81)

− RNA functions: not annotated

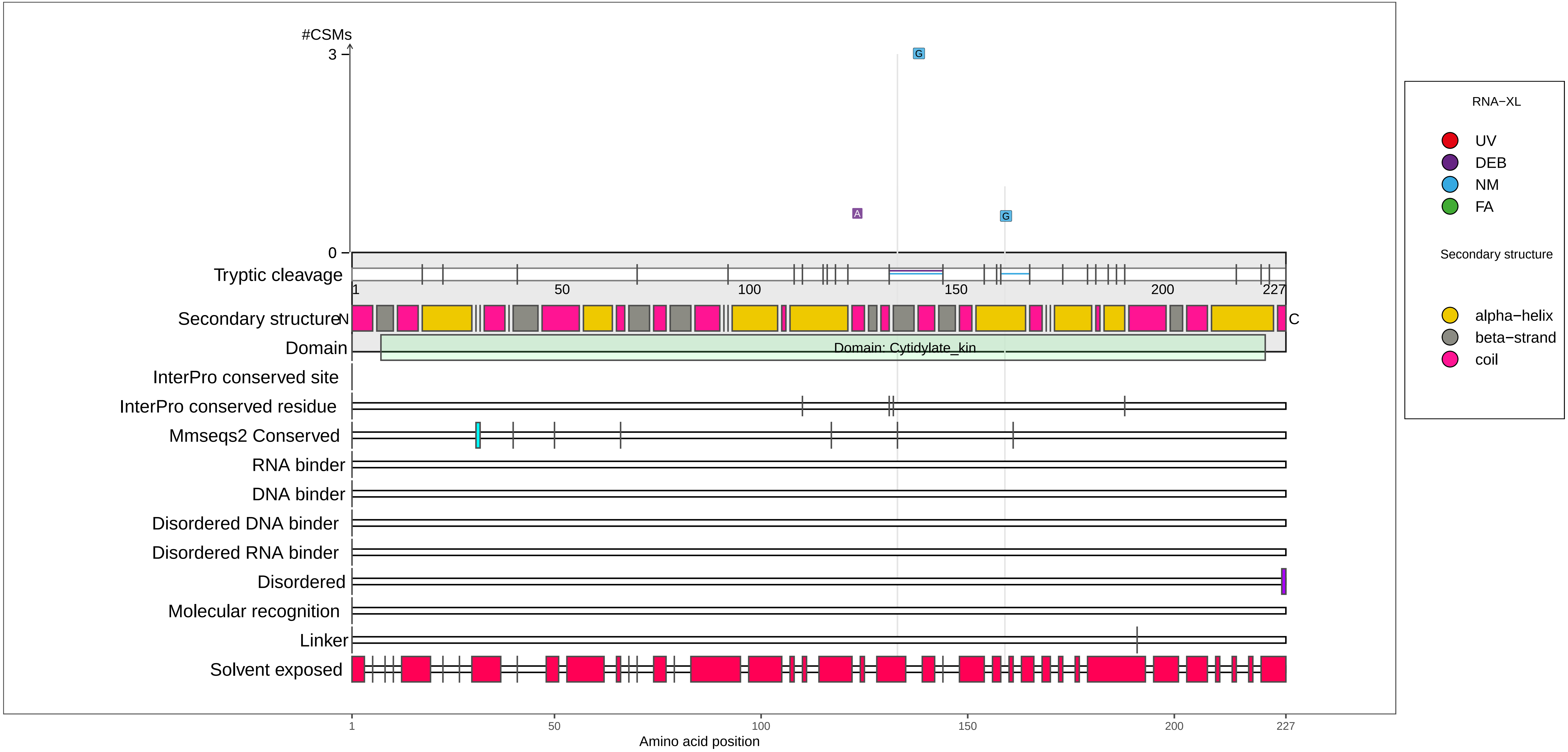

P0A6S3
FLGI_ECOLI Flagellar P−ring protein

− Abundance:
tryptic [log10 Intensity]: 6.7 (Q 5)
PAXdb K12 strain [ppm]: 1.81 (Q 54)
PAXdb E.coli [ppm]: −0.09 (Q 23)

− RNA functions: not annotated

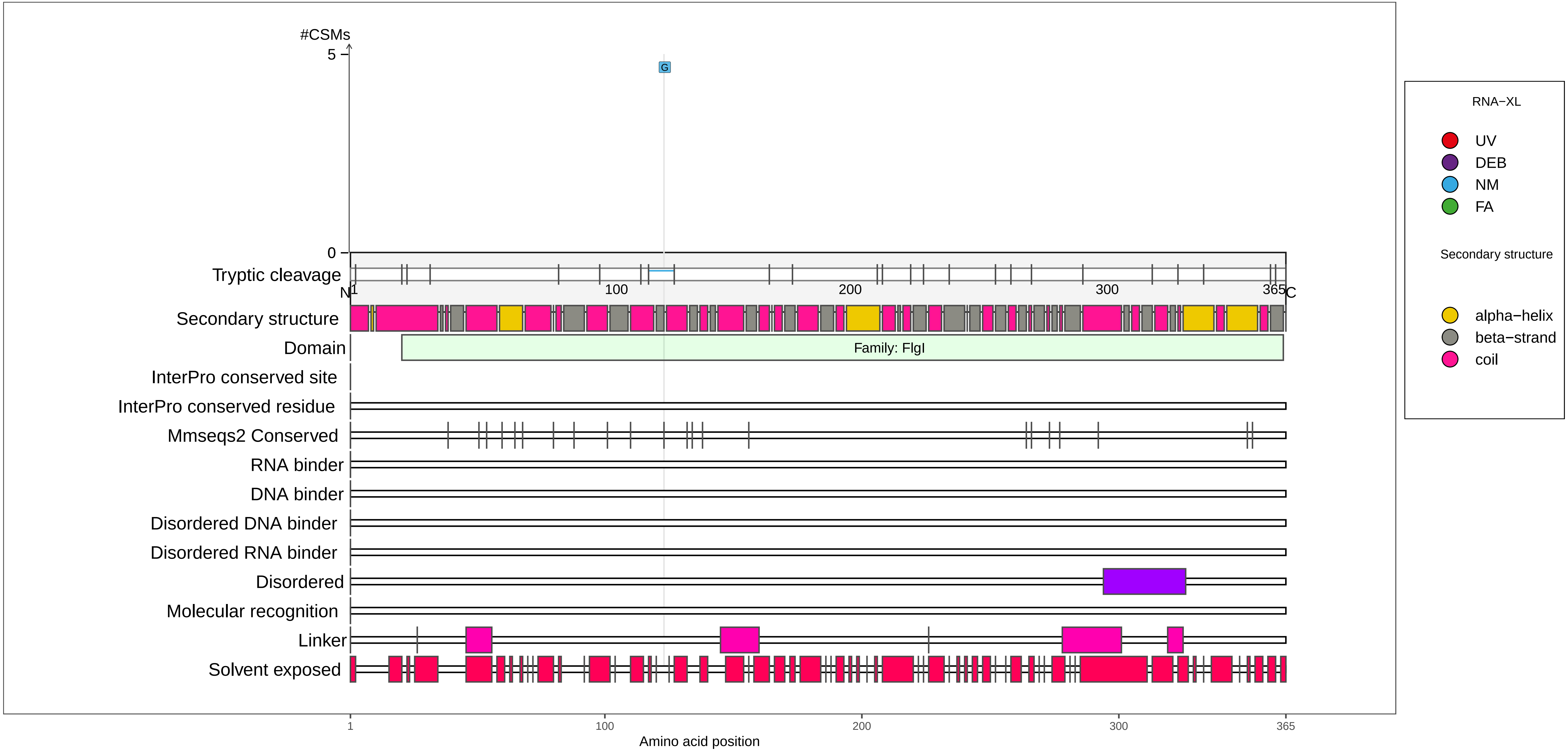

P0A7I7
RIBA_ECOLI GTP cyclohydrolase−2

− Abundance:
tryptic [log10 Intensity]: 9.05 (Q 89)
PAXdb K12 strain [ppm]: 1.35 (Q 29)
PAXdb E.coli [ppm]: 1.74 (Q 66)

− RNA functions: not annotated

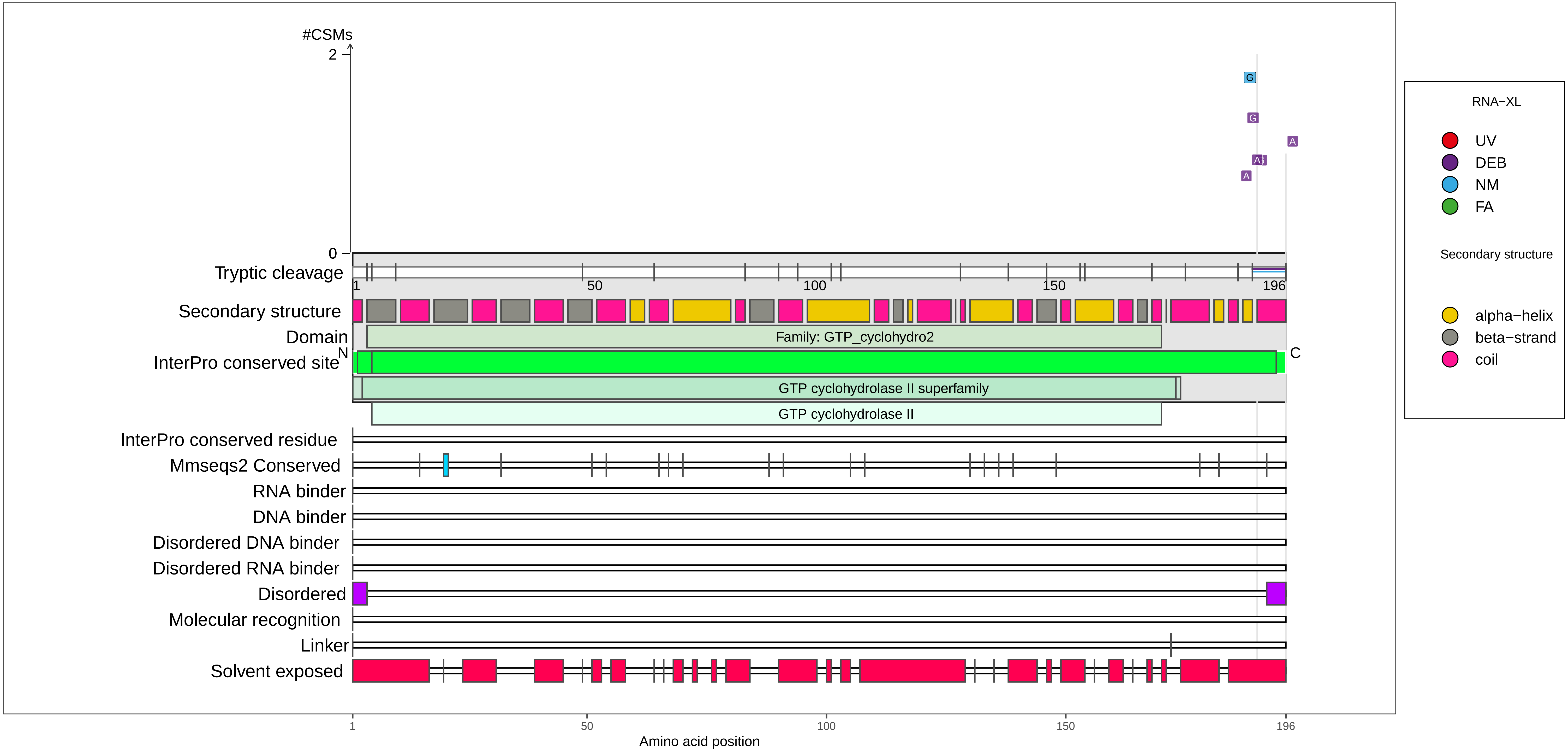

P0A7J0
RIBB_ECOLI 3,4−dihydroxy−2−butanone 4−phosphate synthase

− Abundance:
tryptic [log10 Intensity]: 8.4 (Q 72)
PAXdb K12 strain [ppm]: 2.81 (Q 86)
PAXdb E.coli [ppm]: 2.62 (Q 89)

− RNA functions: not annotated

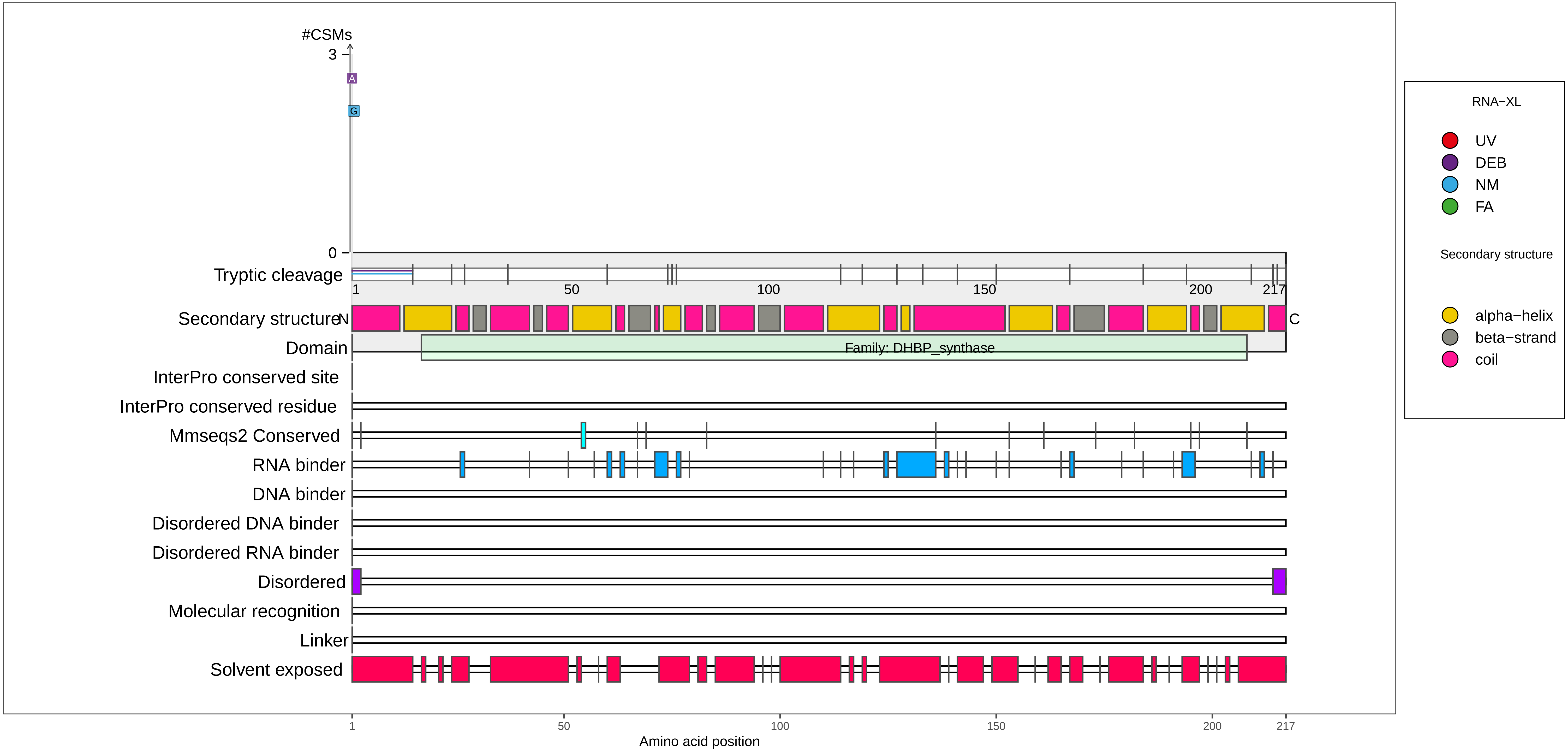

P0A821
SELA_ECOLI L−seryl−tRNA(Sec) selenium transferase

− Abundance:
tryptic [log10 Intensity]: 7.63 (Q 41)
PAXdb K12 strain [ppm]: 1.05 (Q 13)
PAXdb E.coli [ppm]: 0.98 (Q 47)

− RNA functions:
L−seryl−tRNA selenium transferase; L−seryl−tRNASec selenium transferase activity
ncRNA metabolic process; RNA biosynthetic process; RNA metabolic process
selenocysteinyl−tRNA(Sec) biosynthetic process; tRNA metabolic process

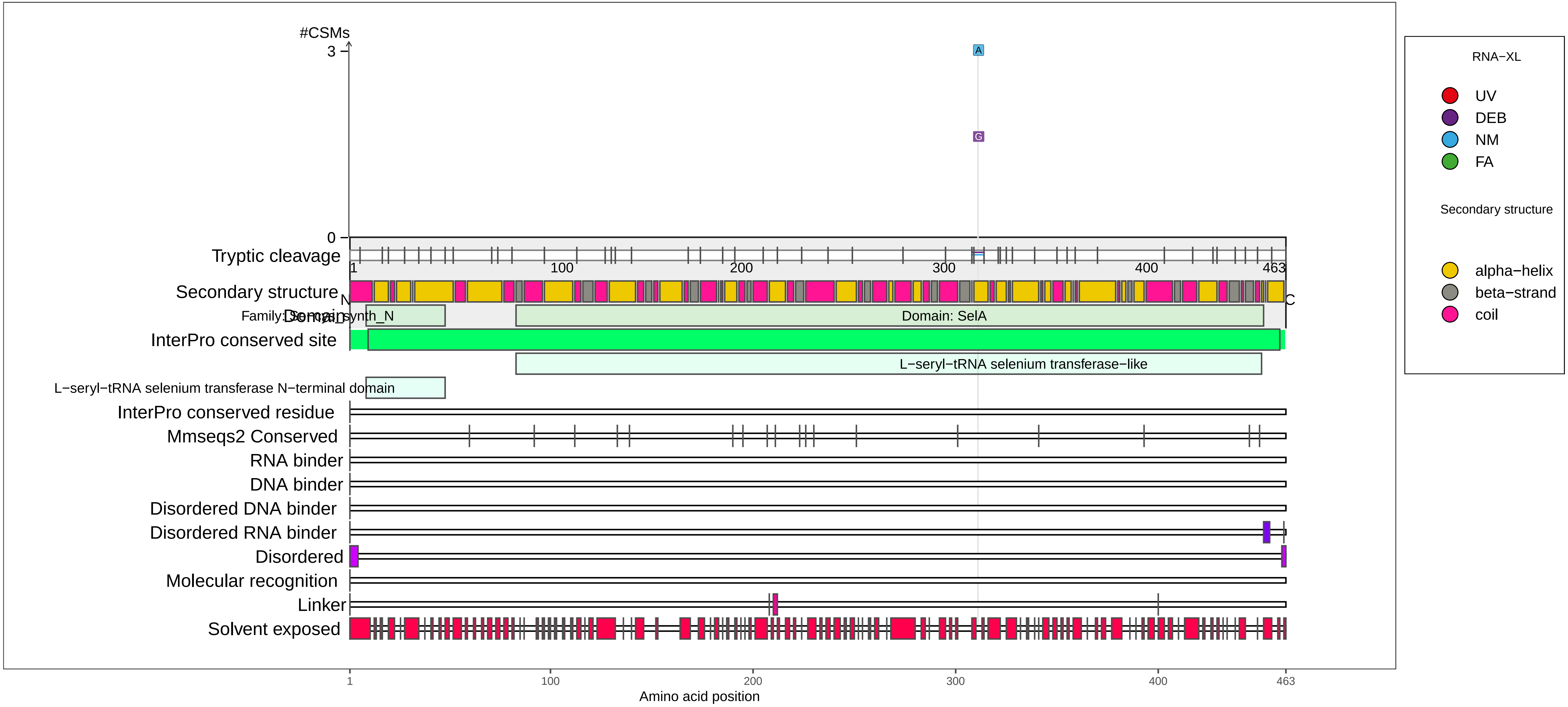

P0A8D3
YAII_ECOLI UPF0178 protein YaiI

− Abundance:
tryptic [log10 Intensity]: 7.64 (Q 42)
PAXdb K12 strain [ppm]: 1.93 (Q 58)
PAXdb E.coli [ppm]: 1.3 (Q 55)

− RNA functions: not annotated

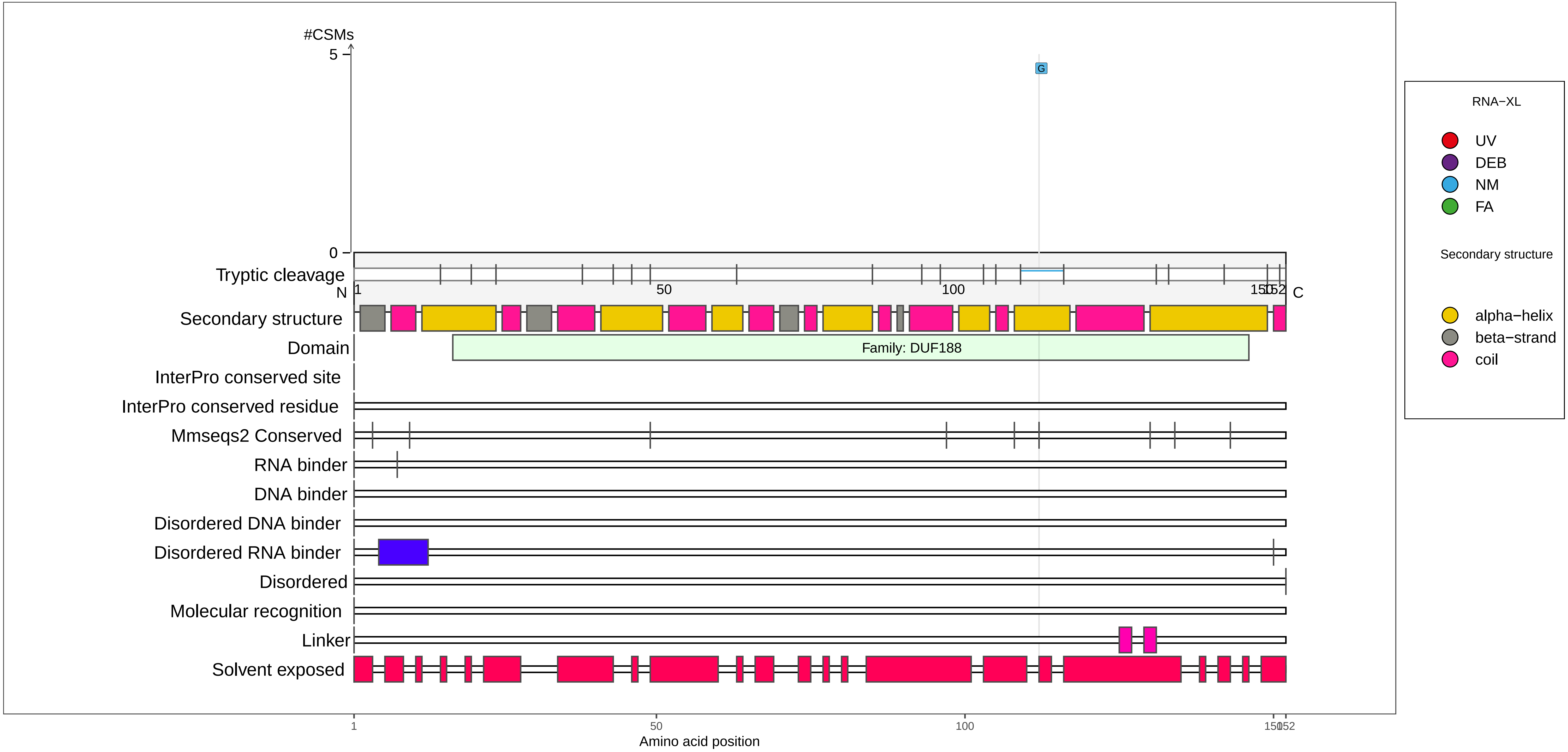

P0A8M6
YEEX_ECOLI UPF0265 protein YeeX

− Abundance:
tryptic [log10 Intensity]: 8.28 (Q 69)
PAXdb K12 strain [ppm]: 3.03 (Q 91)
PAXdb E.coli [ppm]: 3.29 (Q 97)

− RNA functions: not annotated

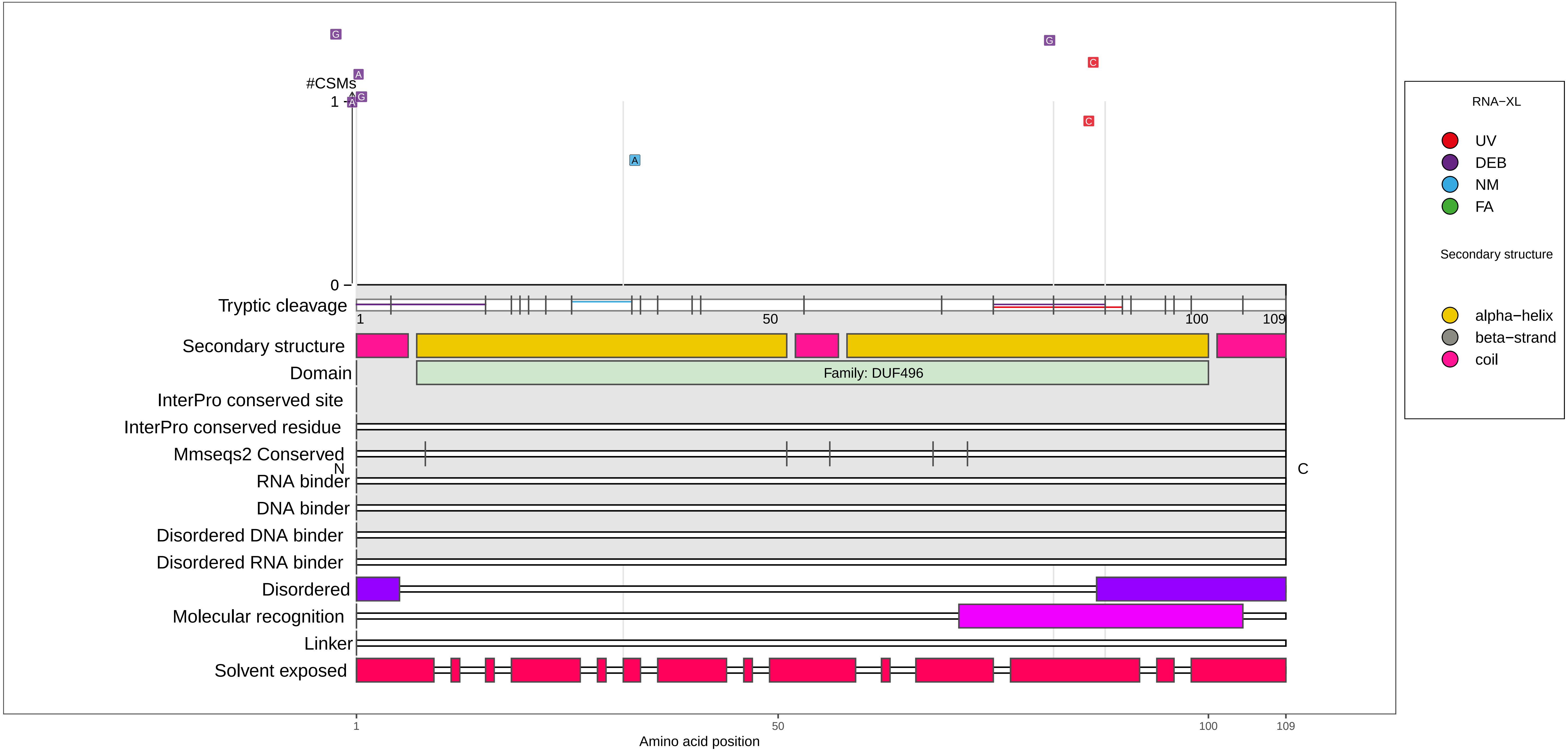

P0A8R4
SLYX_ECOLI Protein SlyX

− Abundance:
tryptic [log10 Intensity]: NA (Q NA)
PAXdb K12 strain [ppm]: 2.36 (Q 73)
PAXdb E.coli [ppm]: 1.88 (Q 69)

− RNA functions: not annotated

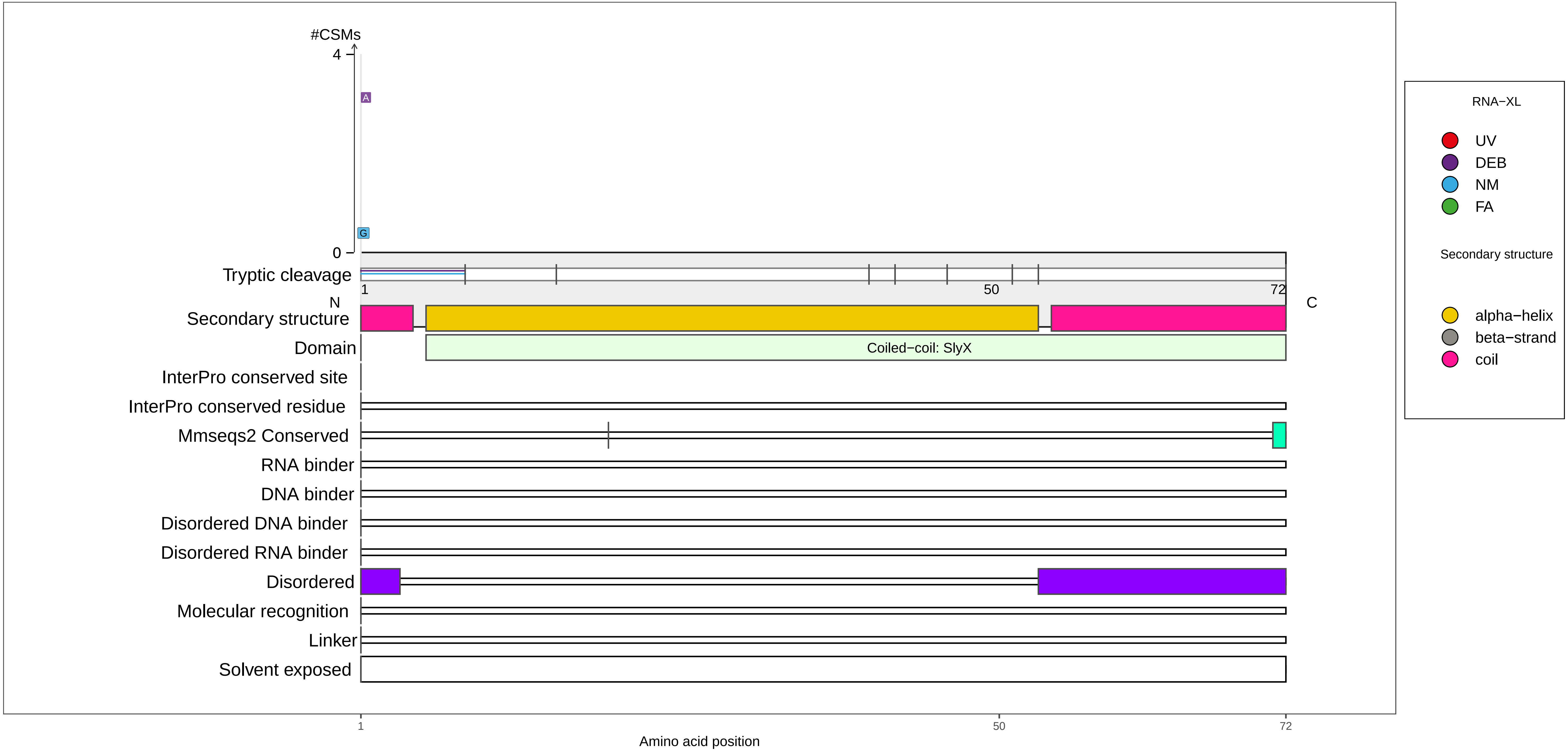

P0A8S1
ARGP_ECOLI HTH−type transcriptional regulator ArgP

− Abundance:
tryptic [log10 Intensity]: NA (Q NA)
PAXdb K12 strain [ppm]: 2.08 (Q 63)
PAXdb E.coli [ppm]: 2.34 (Q 82)

− RNA functions: not annotated

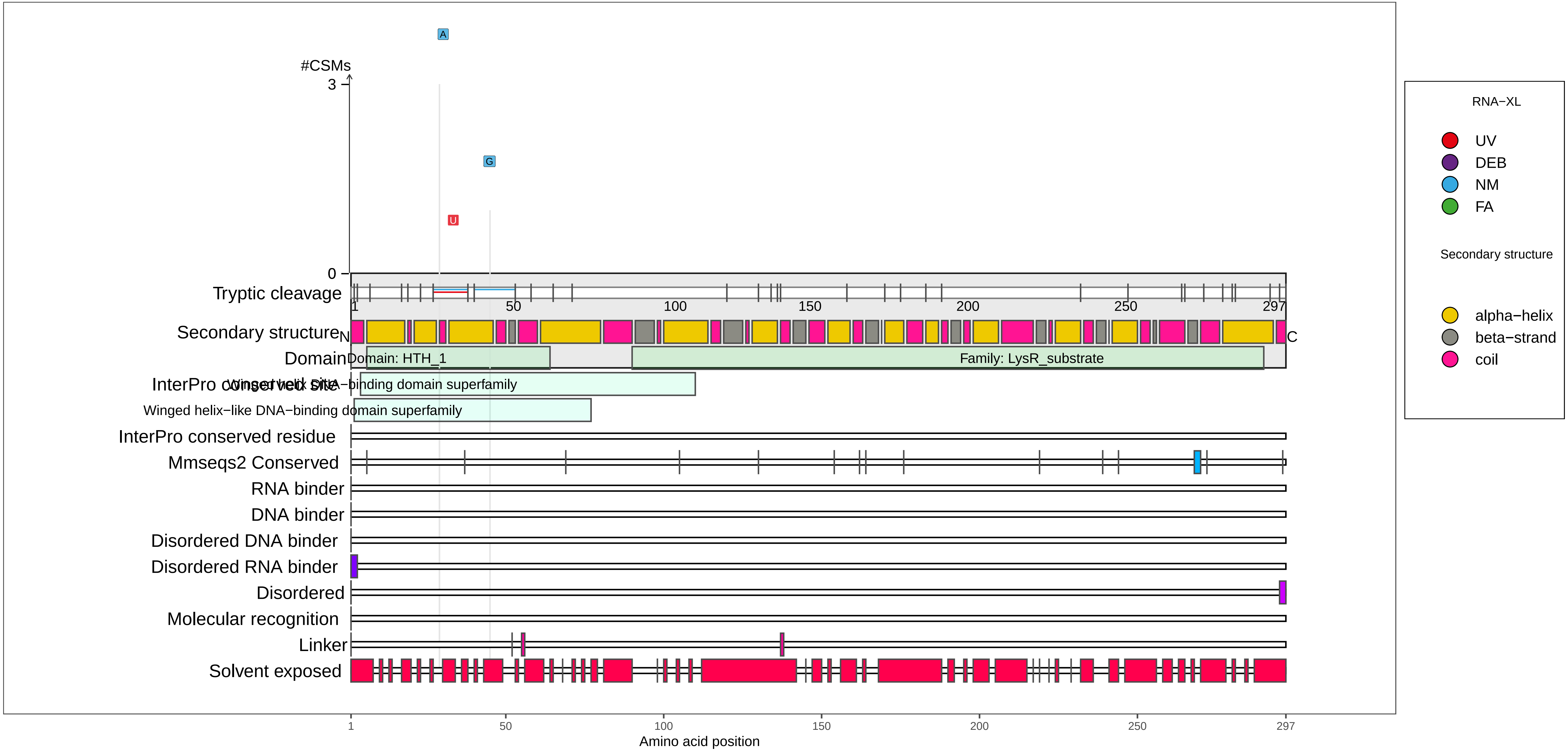

P0A955
ALKH_ECOLI KHG/KDPG aldolase

− Abundance:
tryptic [log10 Intensity]: 8.95 (Q 87)
PAXdb K12 strain [ppm]: 3.15 (Q 93)
PAXdb E.coli [ppm]: 2.93 (Q 94)

− RNA functions: not annotated

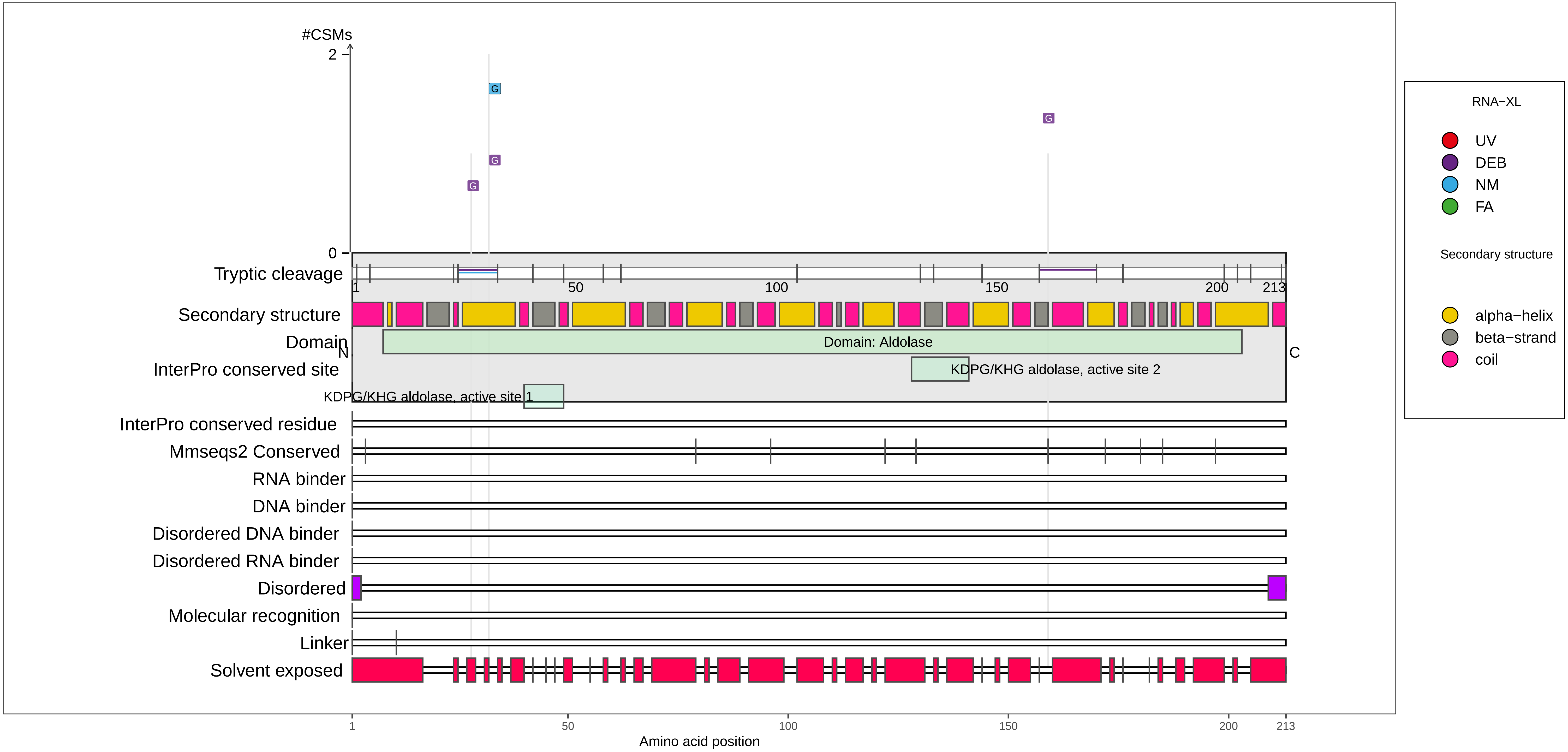

P0A996
GLPC_ECOLI Anaerobic glycerol−3−phosphate dehydrogenase subunit C

− Abundance:
tryptic [log10 Intensity]: 8.11 (Q 62)
PAXdb K12 strain [ppm]: 0.74 (Q 3)
PAXdb E.coli [ppm]: 1.56 (Q 61)

− RNA functions: not annotated

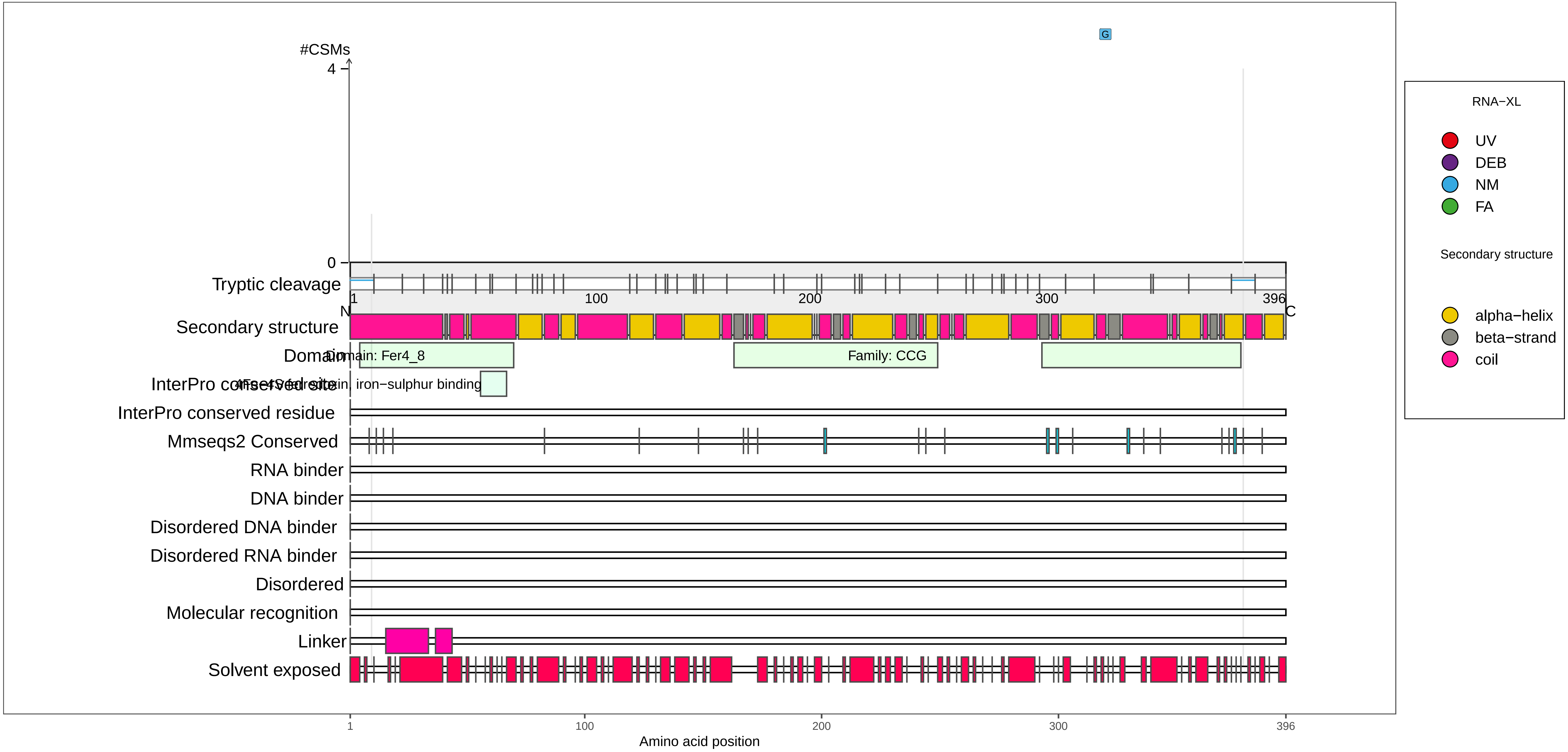

P0A9L5
PPIC_ECOLI Peptidyl−prolyl cis−trans isomerase C

− Abundance:
tryptic [log10 Intensity]: 7.48 (Q 34)
PAXdb K12 strain [ppm]: NA (Q NA)
PAXdb E.coli [ppm]: 2.59 (Q 88)

− RNA functions: not annotated

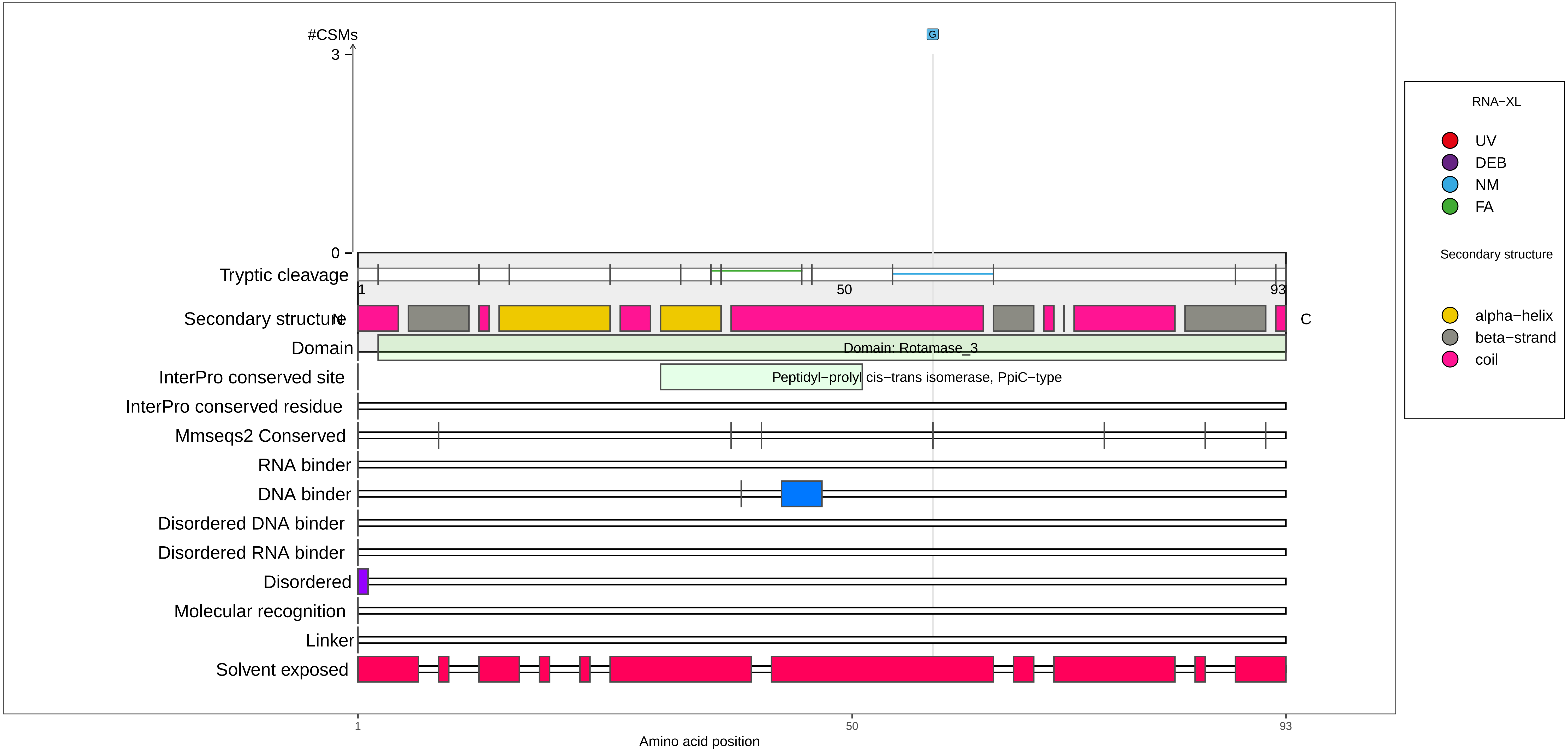

P0A9W6
IBAG_ECOLI Acid stress protein IbaG

− Abundance:
tryptic [log10 Intensity]: 8.09 (Q 61)
PAXdb K12 strain [ppm]: 2.43 (Q 75)
PAXdb E.coli [ppm]: 2.62 (Q 88)

− RNA functions: not annotated

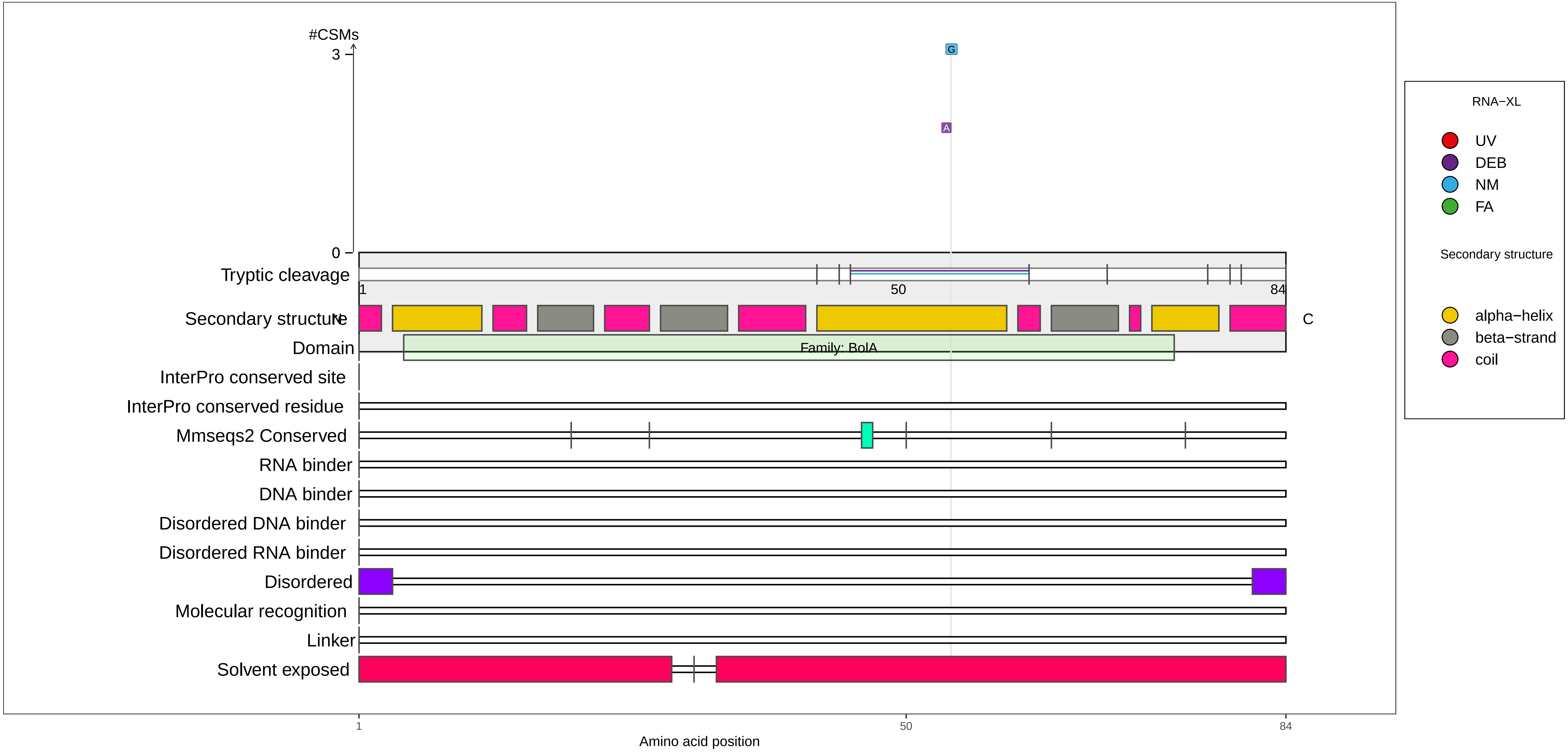

P0AAT9
YBEL_ECOLI Uncharacterized protein YbeL

− Abundance:
tryptic [log10 Intensity]: 7.7 (Q 44)
PAXdb K12 strain [ppm]: 2.37 (Q 73)
PAXdb E.coli [ppm]: 2.86 (Q 93)

− RNA functions: not annotated

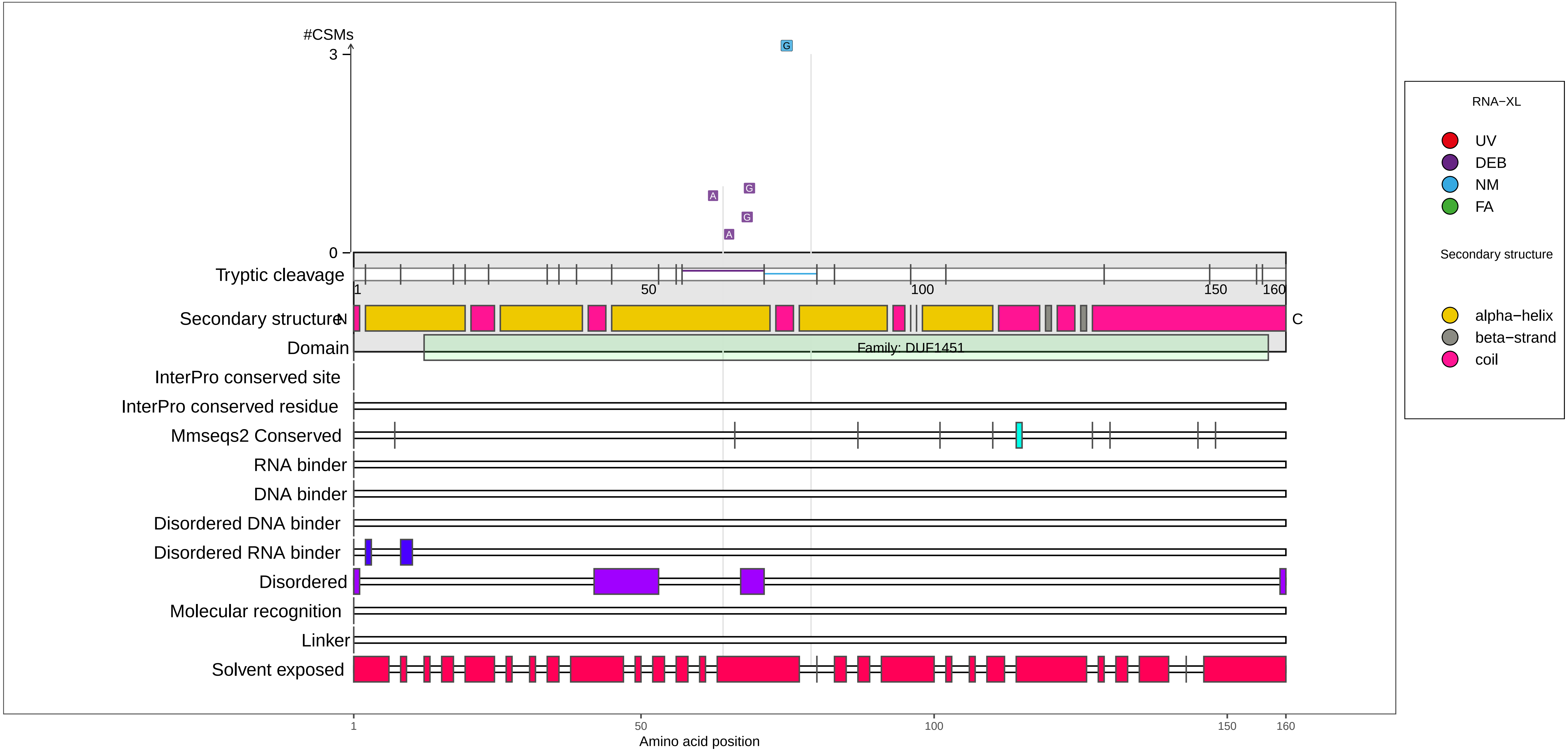

P0ABI4
CORA_ECOLI Magnesium transport protein CorA

− Abundance:
tryptic [log10 Intensity]: 7.13 (Q 18)
PAXdb K12 strain [ppm]: 2.21 (Q 67)
PAXdb E.coli [ppm]: 2.2 (Q 78)

− RNA functions: not annotated

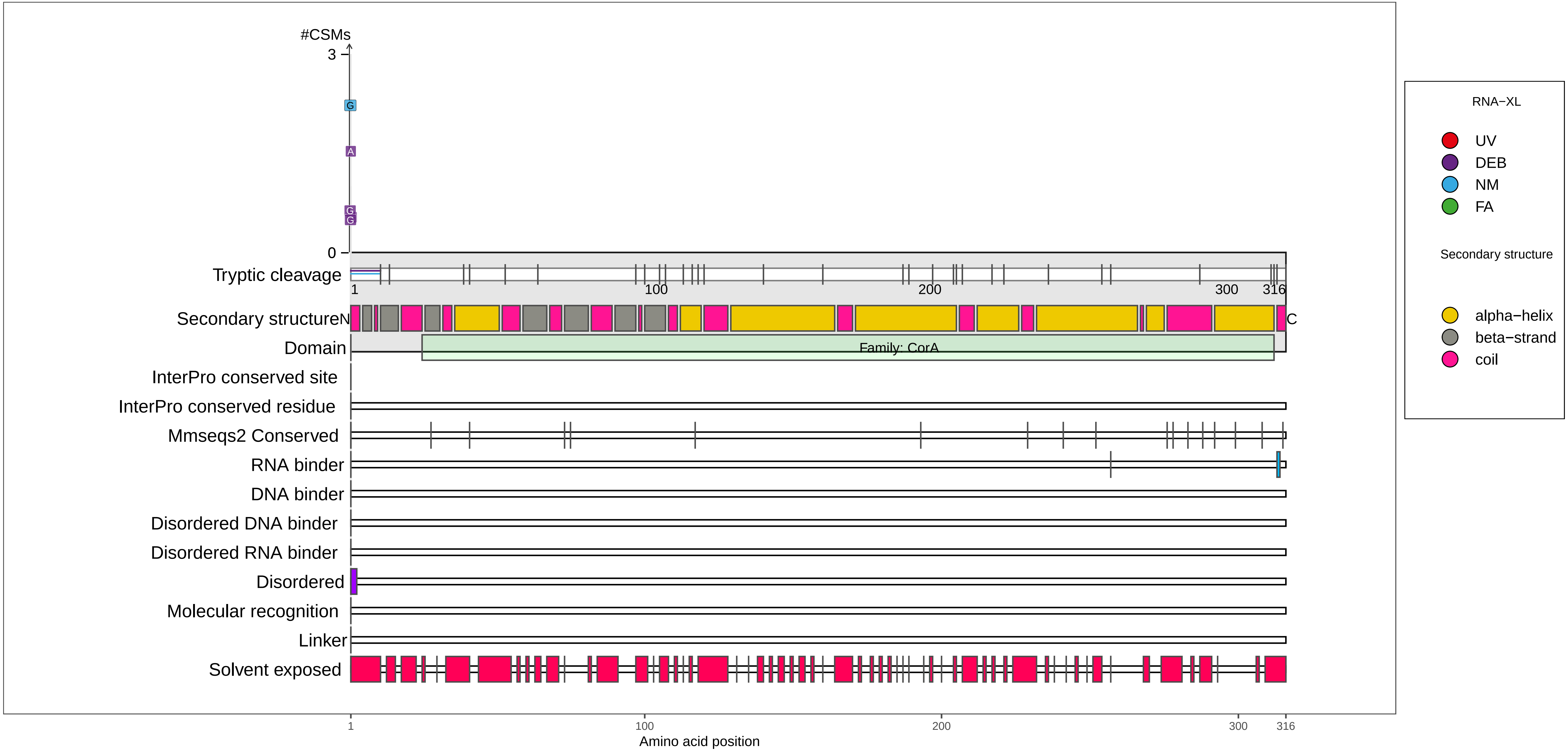

P0ACC7
GLMU_ECOLI Bifunctional protein GlmU

− Abundance:
tryptic [log10 Intensity]: 8.64 (Q 80)
PAXdb K12 strain [ppm]: 1.91 (Q 57)
PAXdb E.coli [ppm]: 2.25 (Q 79)

− RNA functions: not annotated

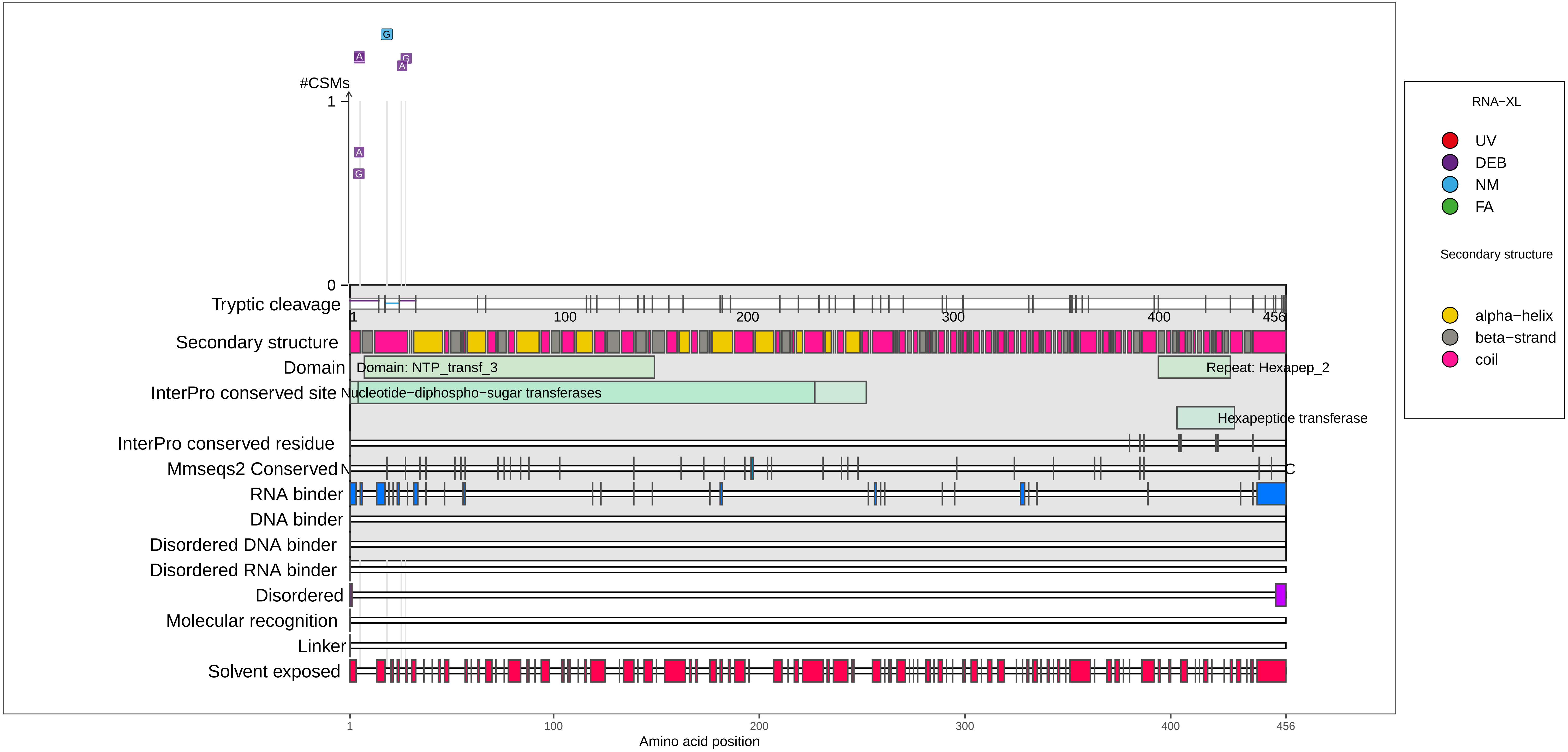

P0ACG8
HSLR_ECOLI Heat shock protein 15

− Abundance:
tryptic [log10 Intensity]: 7.3 (Q 25)
PAXdb K12 strain [ppm]: NA (Q NA)
PAXdb E.coli [ppm]: 1.82 (Q 68)

− RNA functions:
RNA binding

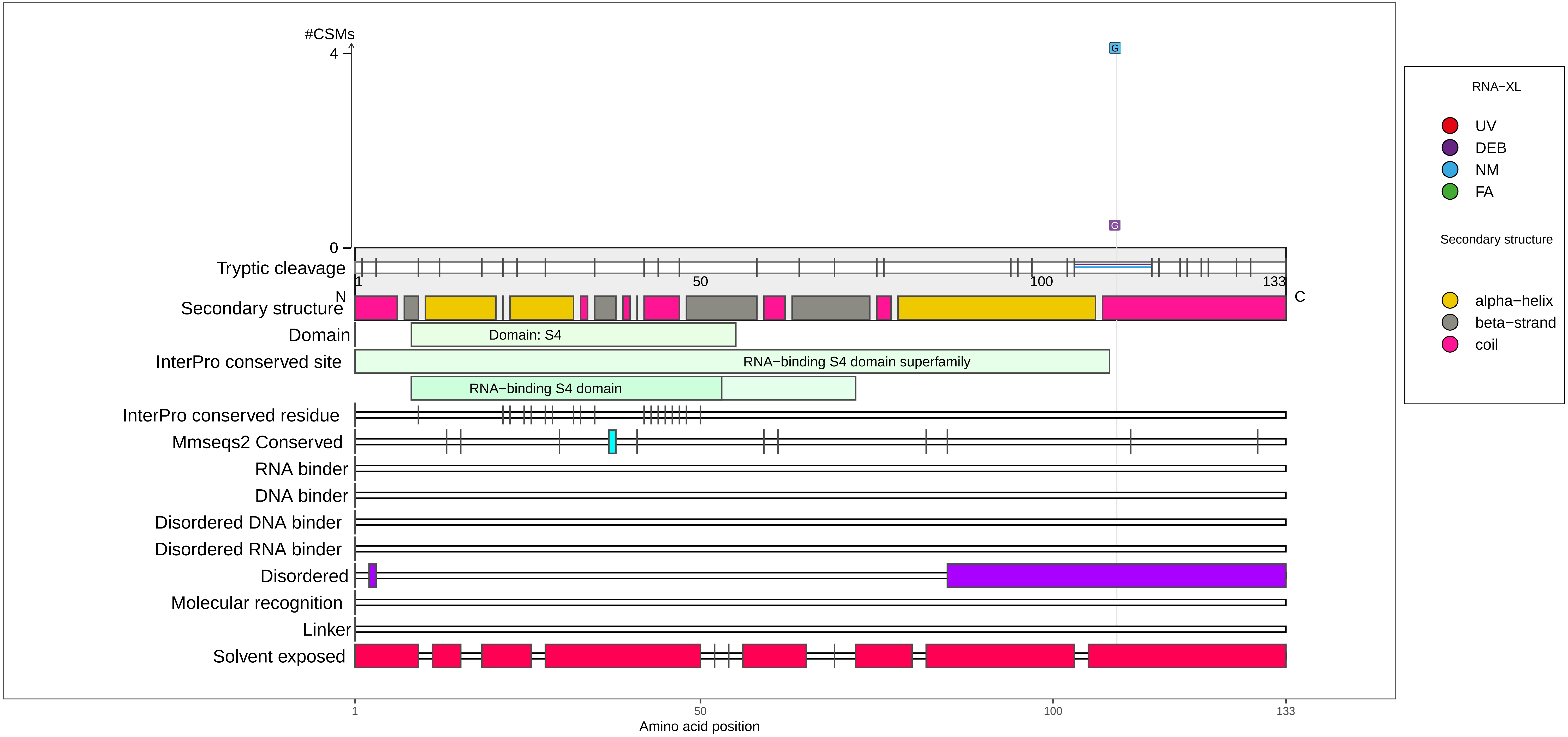

P0ACP7
PURR_ECOLI HTH−type transcriptional repressor PurR

− Abundance:
tryptic [log10 Intensity]: 8.03 (Q 59)
PAXdb K12 strain [ppm]: 1.63 (Q 46)
PAXdb E.coli [ppm]: 2.42 (Q 83)

− RNA functions: not annotated

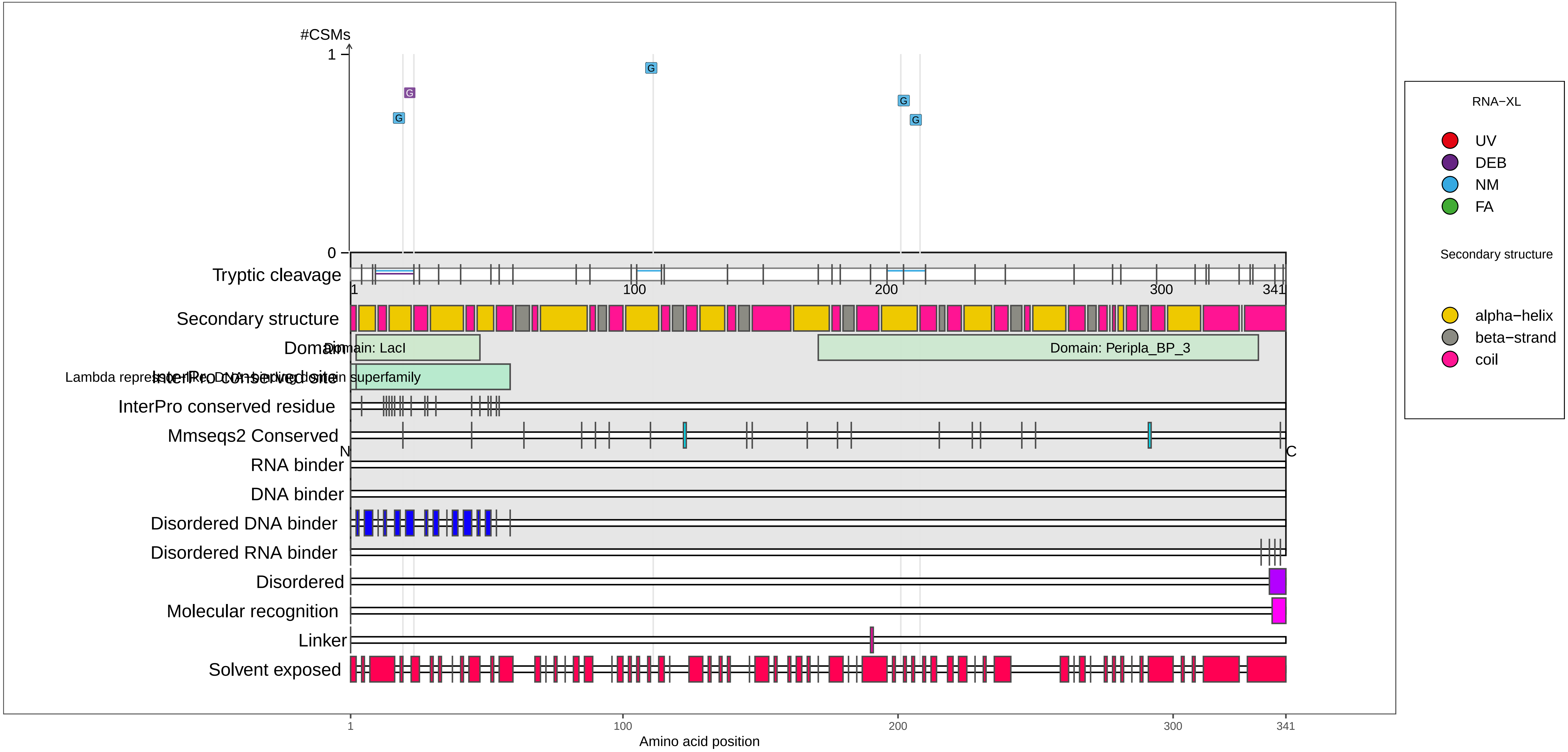

P0AD12
YEEZ_ECOLI Protein YeeZ

− Abundance:
tryptic [log10 Intensity]: 7.32 (Q 26)
PAXdb K12 strain [ppm]: 2.33 (Q 71)
PAXdb E.coli [ppm]: 2.27 (Q 79)

− RNA functions: not annotated

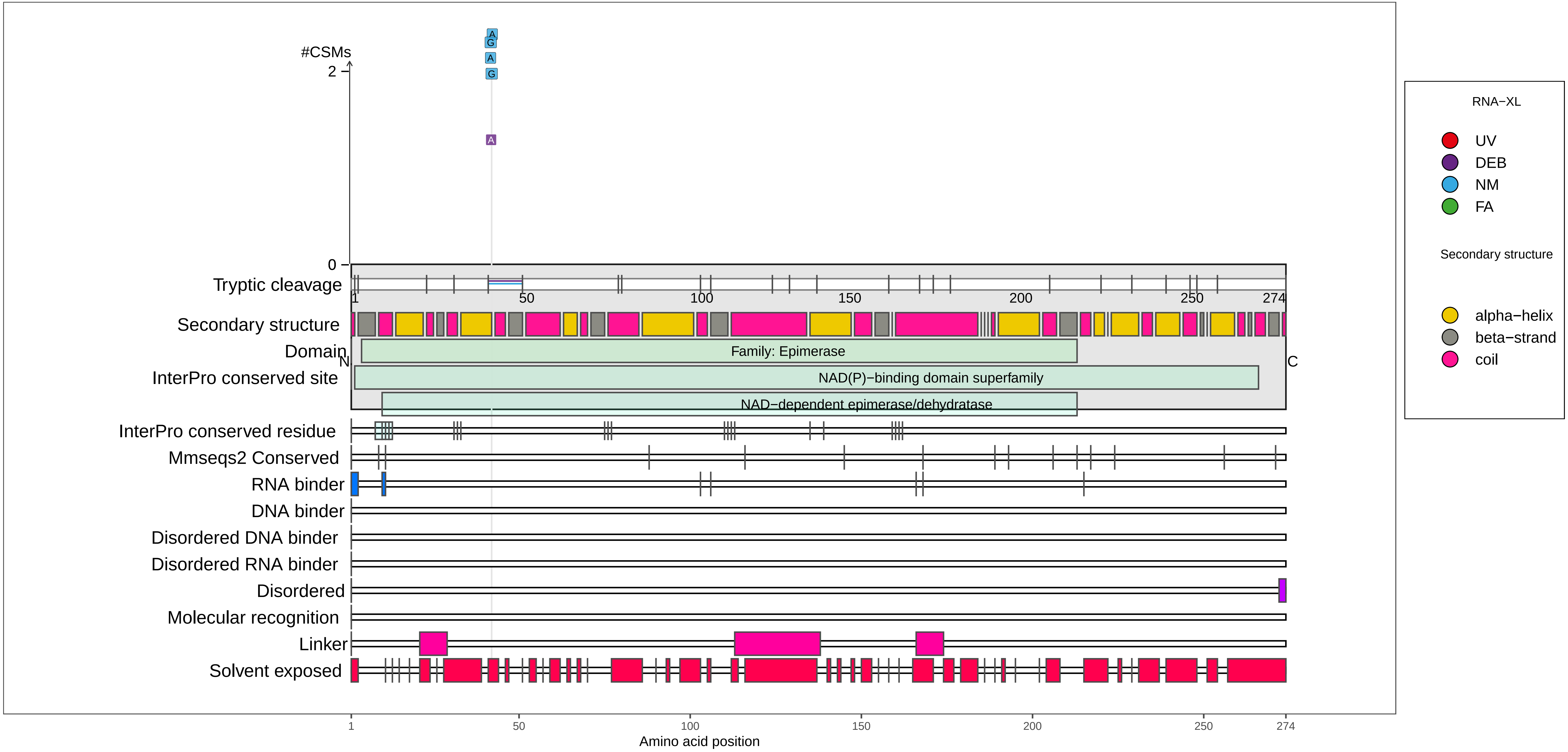

P0ADE8
YGFZ_ECOLI tRNA−modifying protein YgfZ

− Abundance:
tryptic [log10 Intensity]: 8.78 (Q 83)
PAXdb K12 strain [ppm]: 3.19 (Q 94)
PAXdb E.coli [ppm]: 2.64 (Q 89)

− RNA functions:
ncRNA metabolic process; ncRNA processing; RNA metabolic process; RNA modification
RNA processing; tRNA metabolic process; tRNA processing

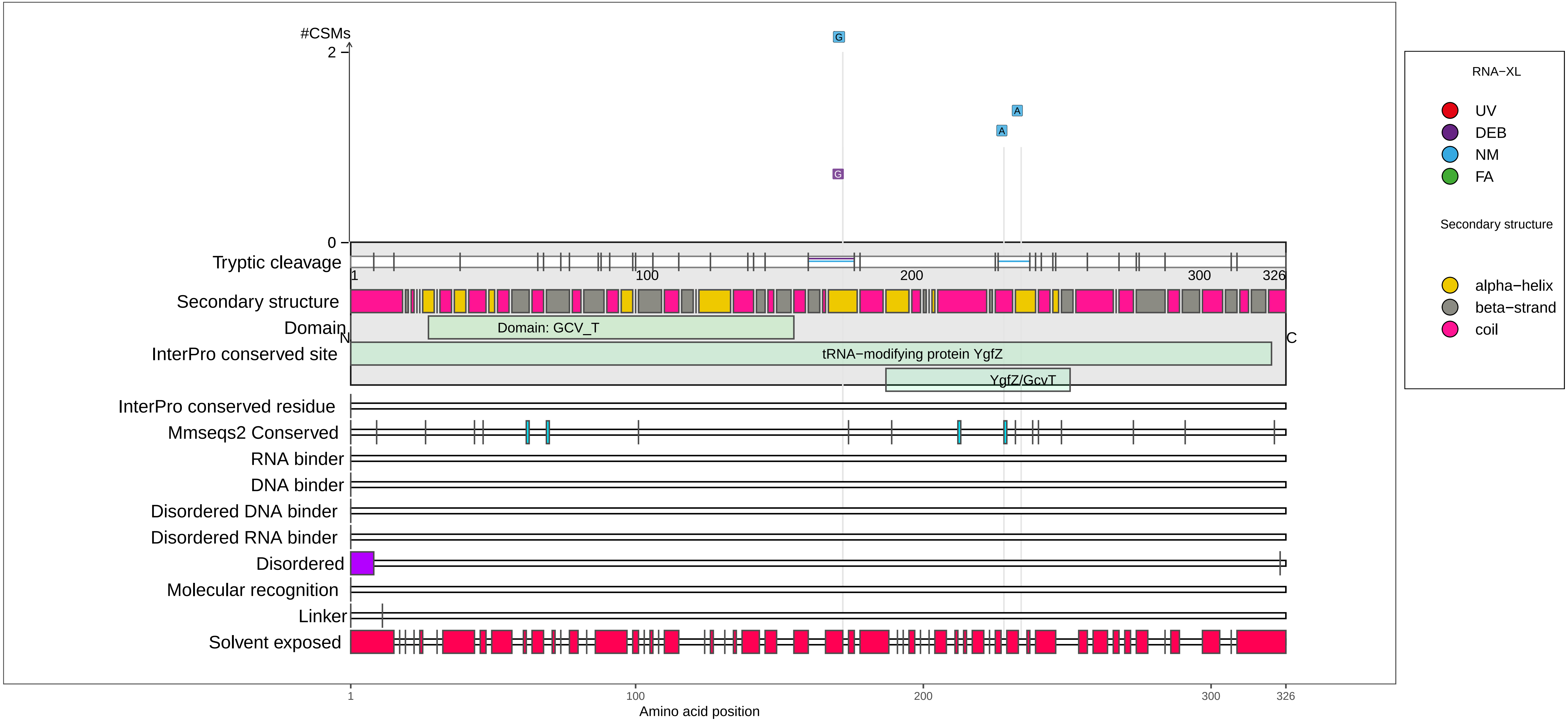

P0ADI7
YECD_ECOLI Isochorismatase family protein YecD

− Abundance:
tryptic [log10 Intensity]: 7.85 (Q 51)
PAXdb K12 strain [ppm]: 2.23 (Q 68)
PAXdb E.coli [ppm]: 2.09 (Q 74)

− RNA functions: not annotated

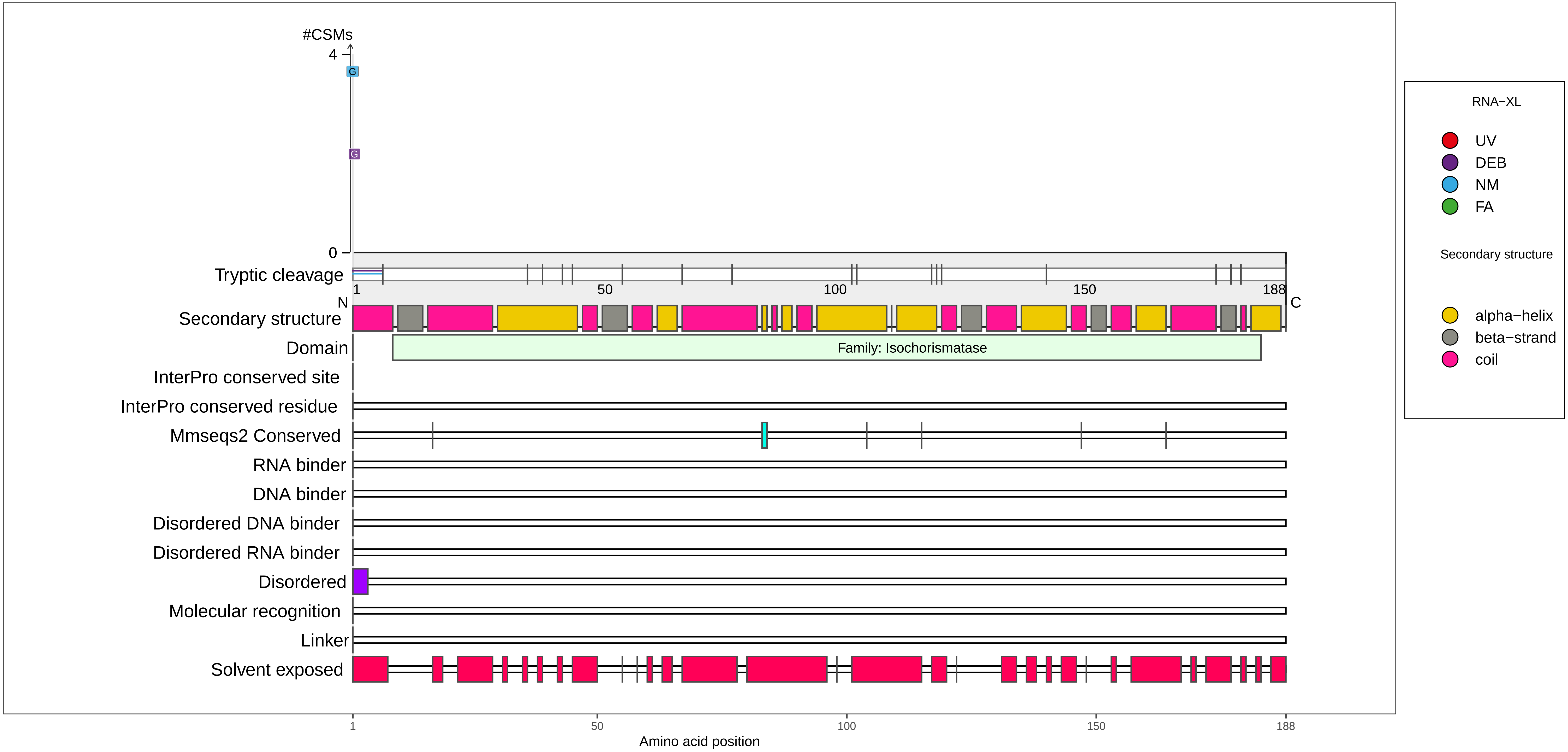

P0ADU5
YGIW_ECOLI Protein YgiW

− Abundance:
tryptic [log10 Intensity]: 8.64 (Q 80)
PAXdb K12 strain [ppm]: 3.41 (Q 96)
PAXdb E.coli [ppm]: 2.83 (Q 92)

− RNA functions: not annotated

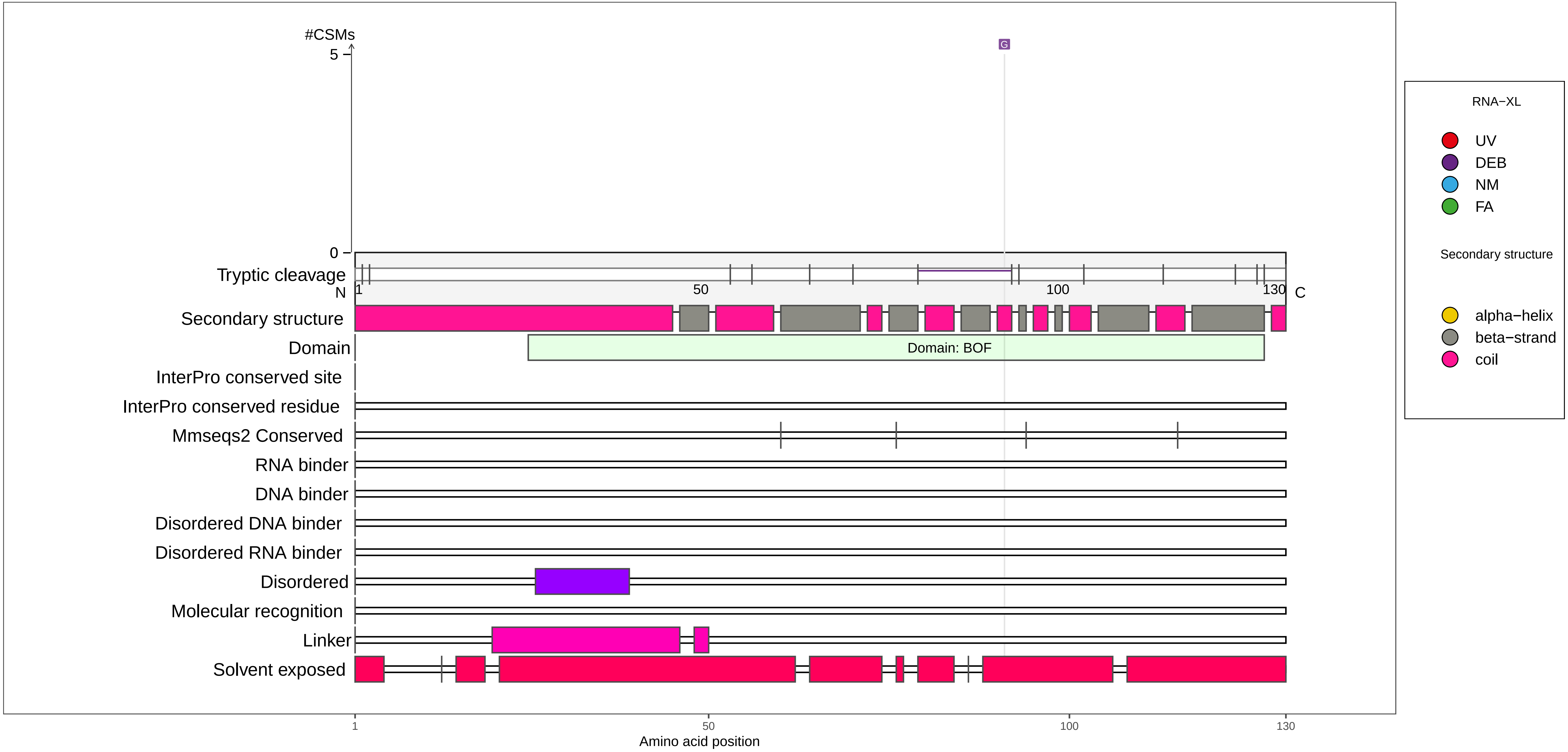

P0AE18
MAP1_ECOLI Methionine aminopeptidase

− Abundance:
tryptic [log10 Intensity]: 9.13 (Q 91)
PAXdb K12 strain [ppm]: 3.18 (Q 94)
PAXdb E.coli [ppm]: 2.71 (Q 90)

− RNA functions: not annotated

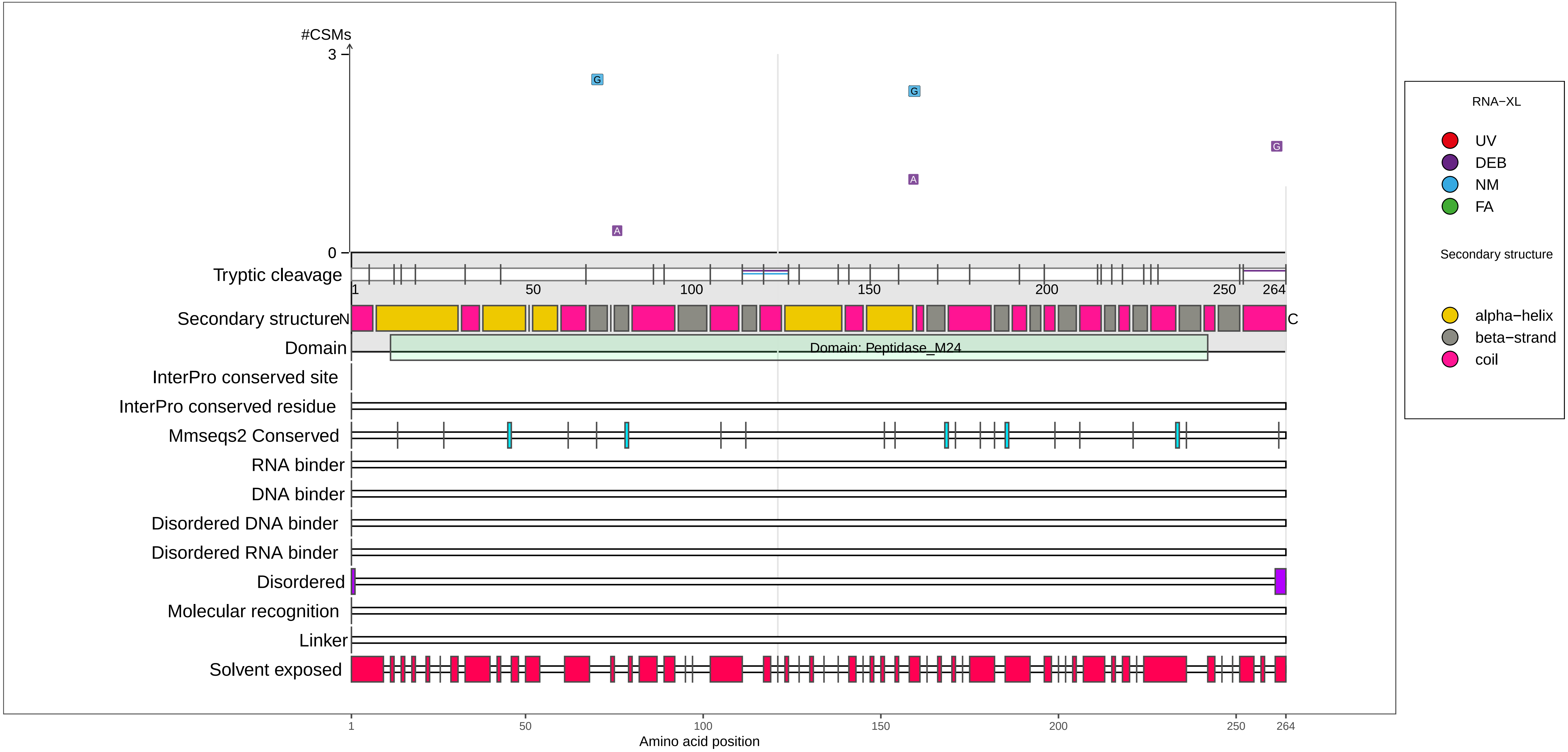

P0AEE1
DCRB_ECOLI Inner membrane lipoprotein DcrB

− Abundance:
tryptic [log10 Intensity]: 8.45 (Q 74)
PAXdb K12 strain [ppm]: 2.5 (Q 78)
PAXdb E.coli [ppm]: 2.44 (Q 84)

− RNA functions: not annotated

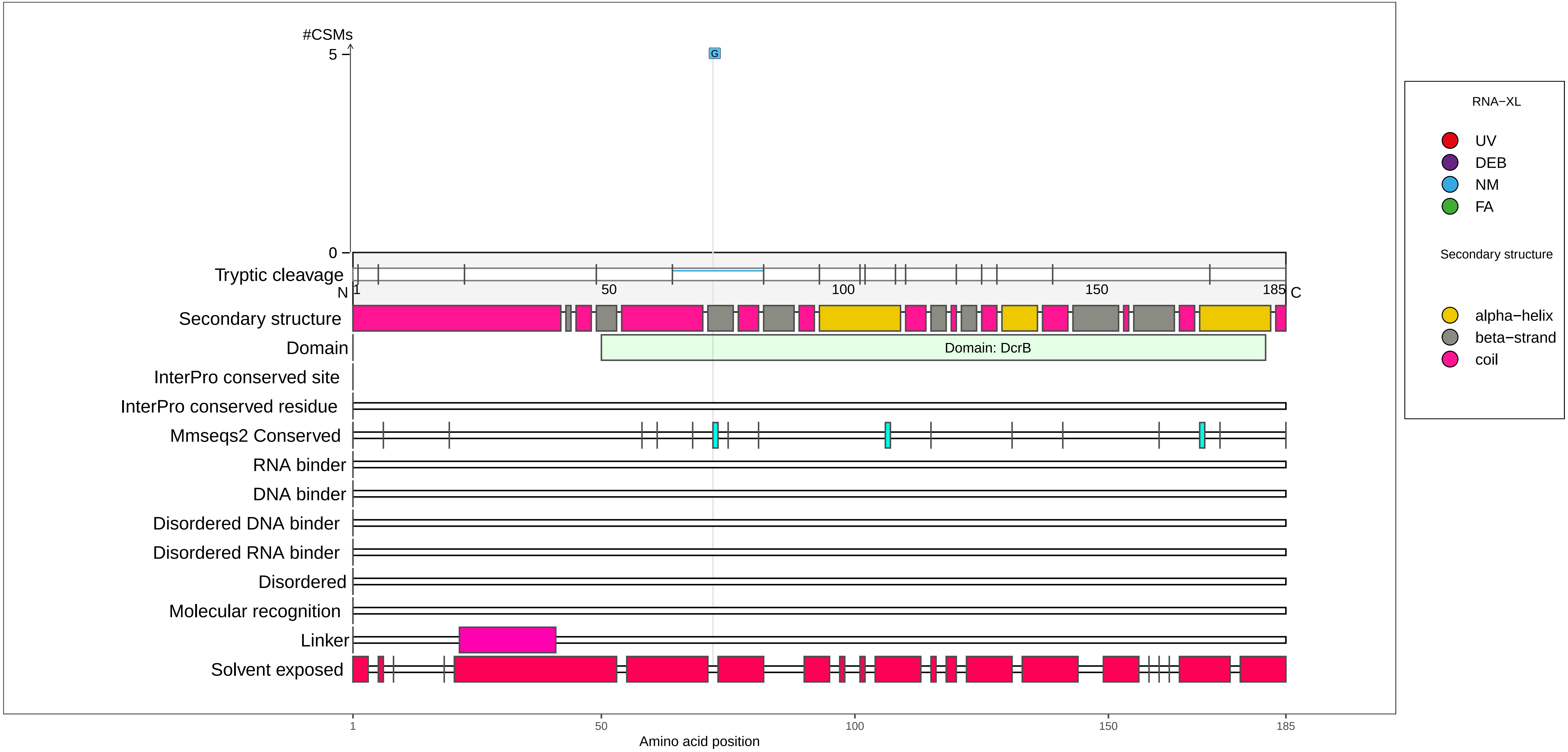

P0AEK2
FABG_ECOLI 3−oxoacyl−[acyl−carrier−protein] reductase FabG

− Abundance:
tryptic [log10 Intensity]: 8.67 (Q 81)
PAXdb K12 strain [ppm]: 3.12 (Q 93)
PAXdb E.coli [ppm]: 3.07 (Q 95)

− RNA functions: not annotated

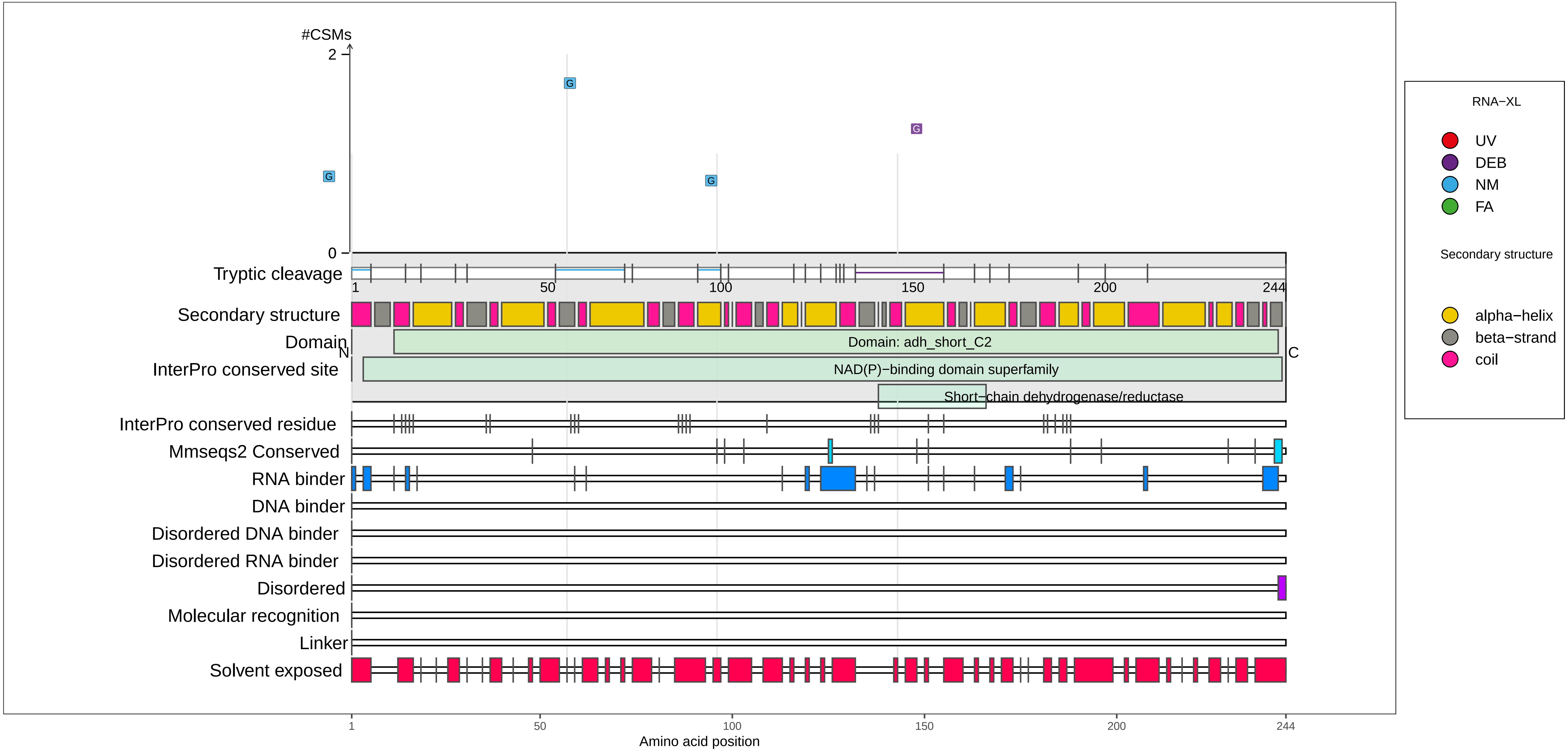

P0AET8
HDHA_ECOLI 7alpha−hydroxysteroid dehydrogenase

− Abundance:
tryptic [log10 Intensity]: 7.35 (Q 27)
PAXdb K12 strain [ppm]: 2.72 (Q 84)
PAXdb E.coli [ppm]: 2.55 (Q 87)

− RNA functions: not annotated

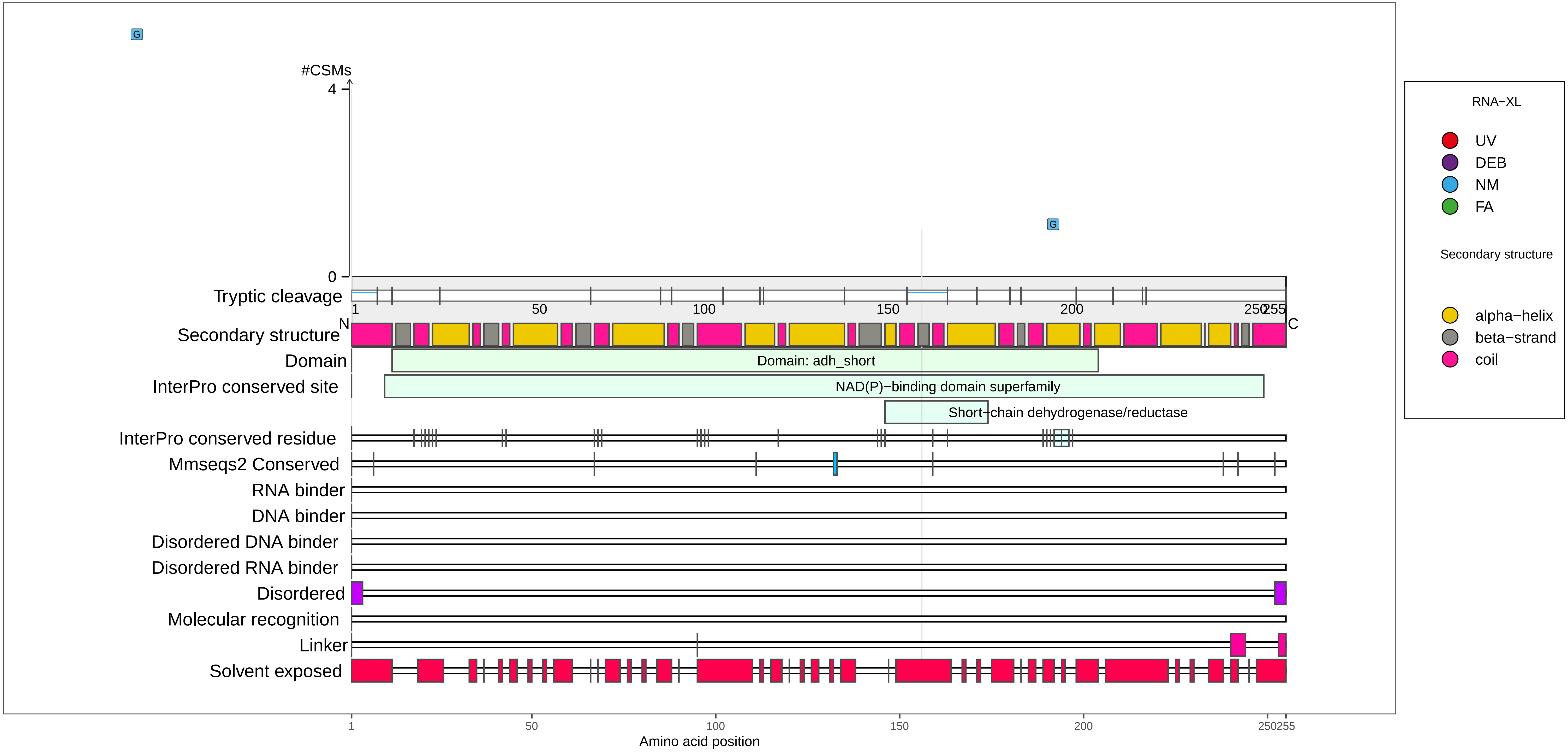

P0AEU7
SKP_ECOLI Chaperone protein Skp

− Abundance:
tryptic [log10 Intensity]: 8.67 (Q 81)
PAXdb K12 strain [ppm]: 3.35 (Q 96)
PAXdb E.coli [ppm]: 3.07 (Q 95)

− RNA functions: not annotated

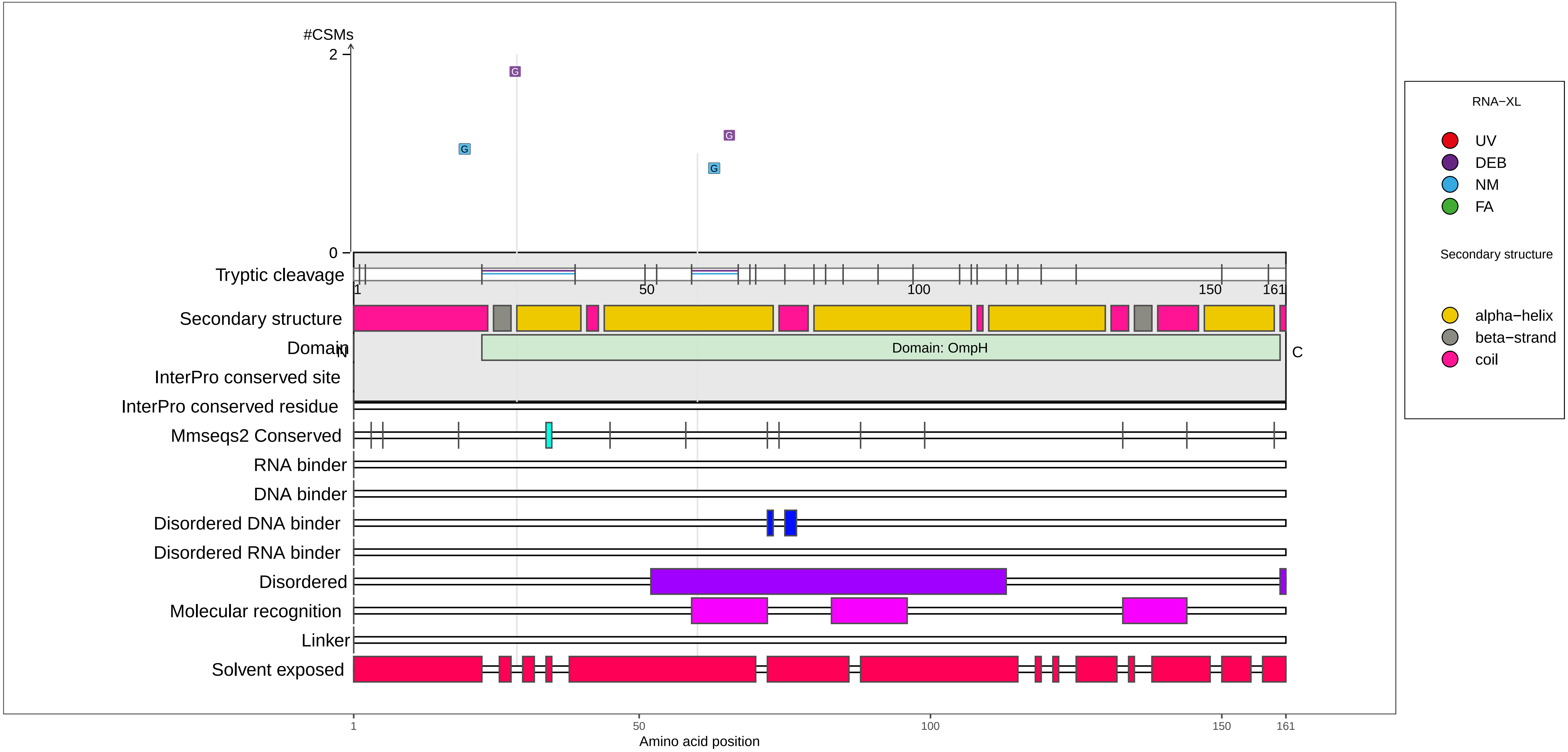

P0AEW6
INGK_ECOLI Guanosine−inosine kinase

− Abundance:
tryptic [log10 Intensity]: 8.3 (Q 69)
PAXdb K12 strain [ppm]: 2.22 (Q 67)
PAXdb E.coli [ppm]: 0.99 (Q 47)

− RNA functions: not annotated

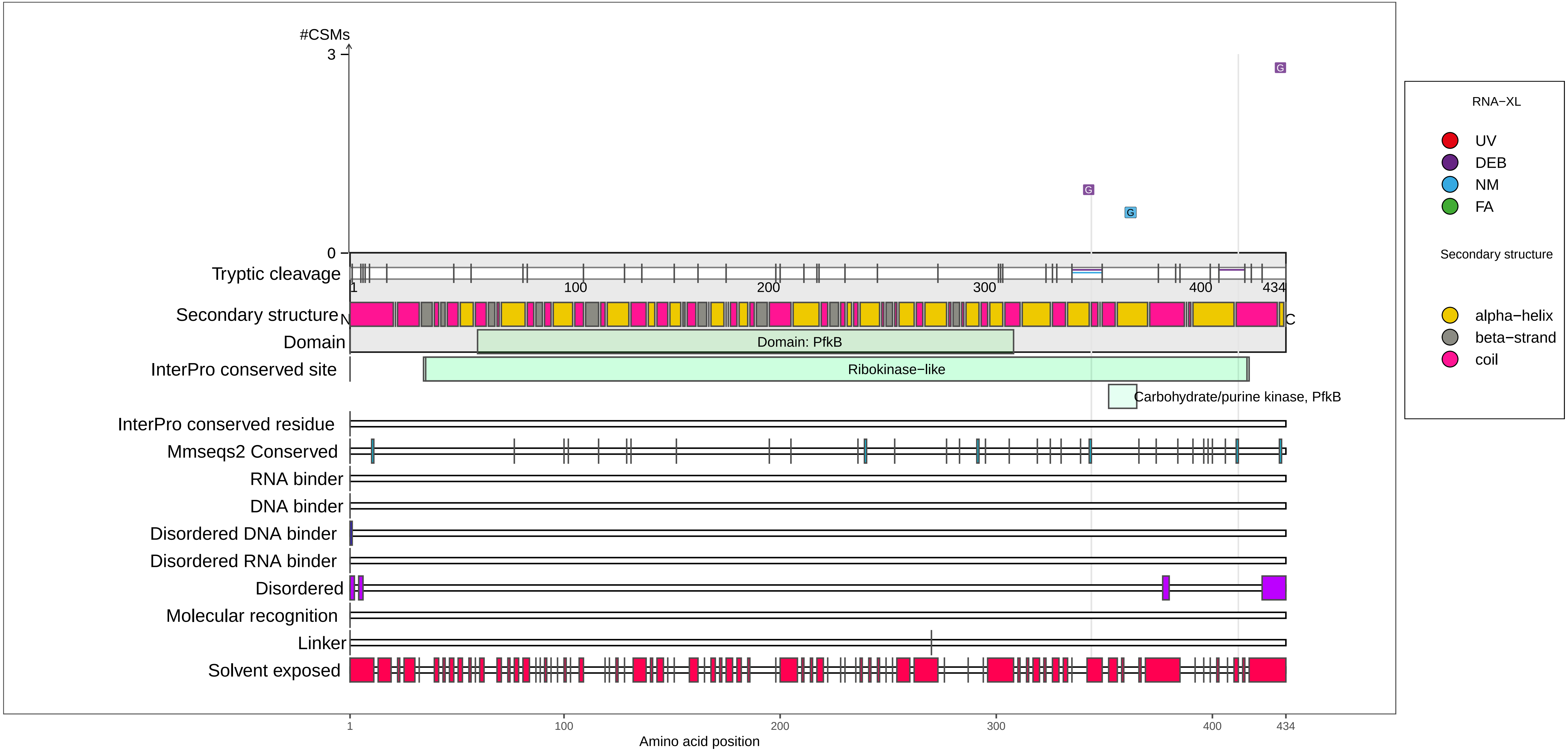

P0AF20
NAGC_ECOLI N−acetylglucosamine repressor

− Abundance:
tryptic [log10 Intensity]: 7.82 (Q 50)
PAXdb K12 strain [ppm]: 1.94 (Q 58)
PAXdb E.coli [ppm]: 2.08 (Q 74)

− RNA functions: not annotated

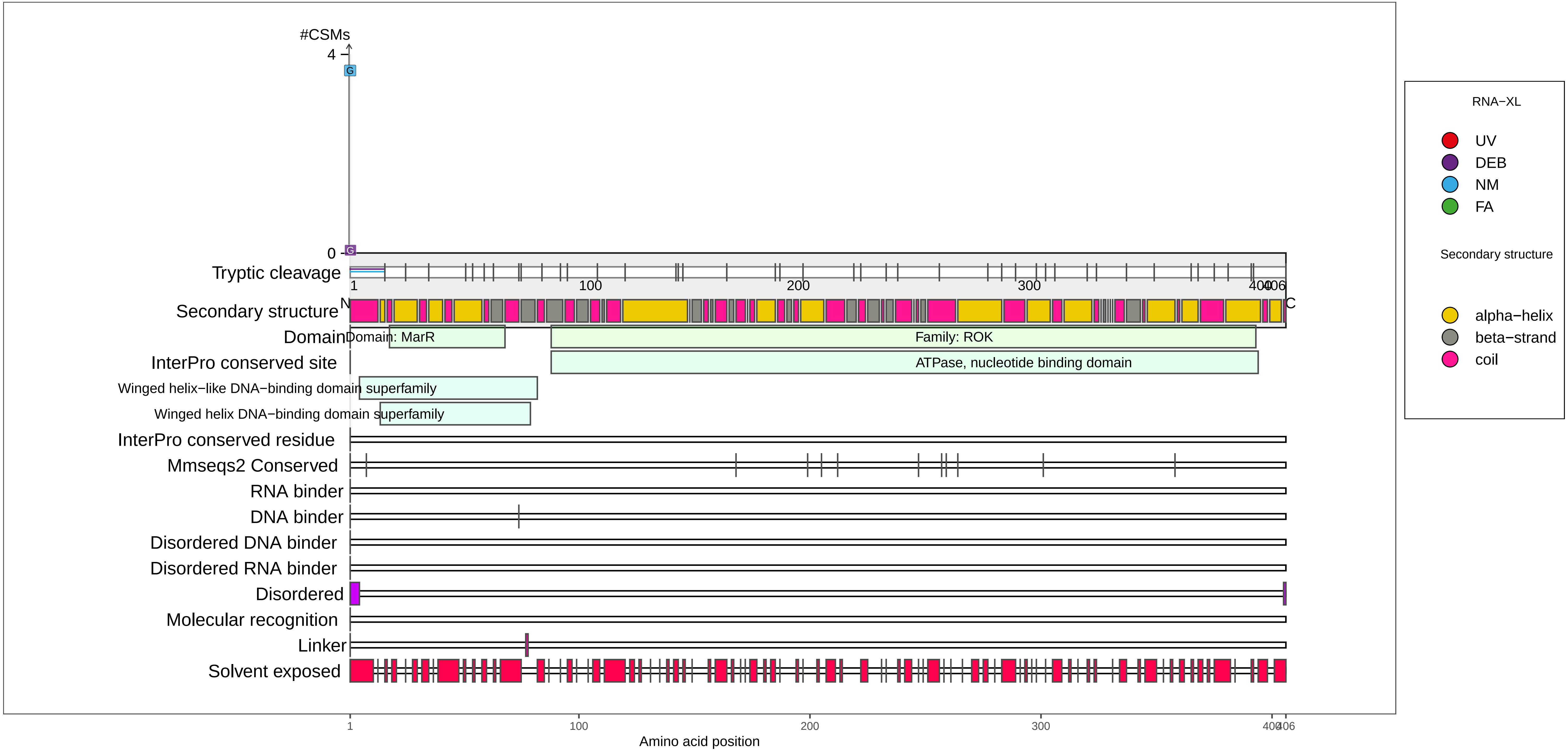

P0AFE8
NUOM_ECOLI NADH−quinone oxidoreductase subunit M

− Abundance:
tryptic [log10 Intensity]: 7.52 (Q 36)
PAXdb K12 strain [ppm]: NA (Q NA)
PAXdb E.coli [ppm]: 0.46 (Q 35)

− RNA functions: not annotated

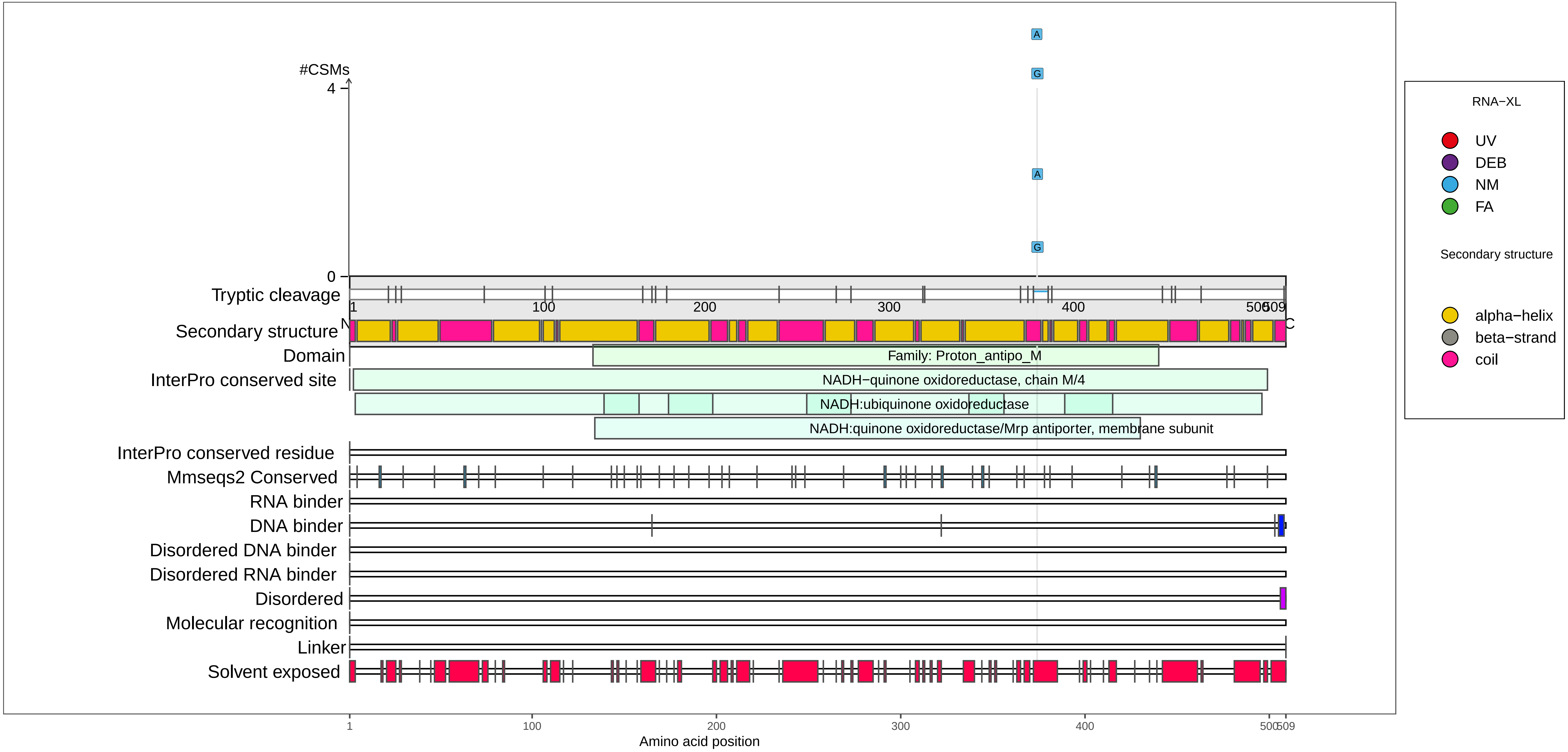

P0AG90
SECD_ECOLI Protein translocase subunit SecD
− Abundance:
tryptic [log10 Intensity]: 7.42 (Q 31)
PAXdb K12 strain [ppm]: 2.03 (Q 61)
PAXdb E.coli [ppm]: 2 (Q 72)

− RNA functions: not annotated

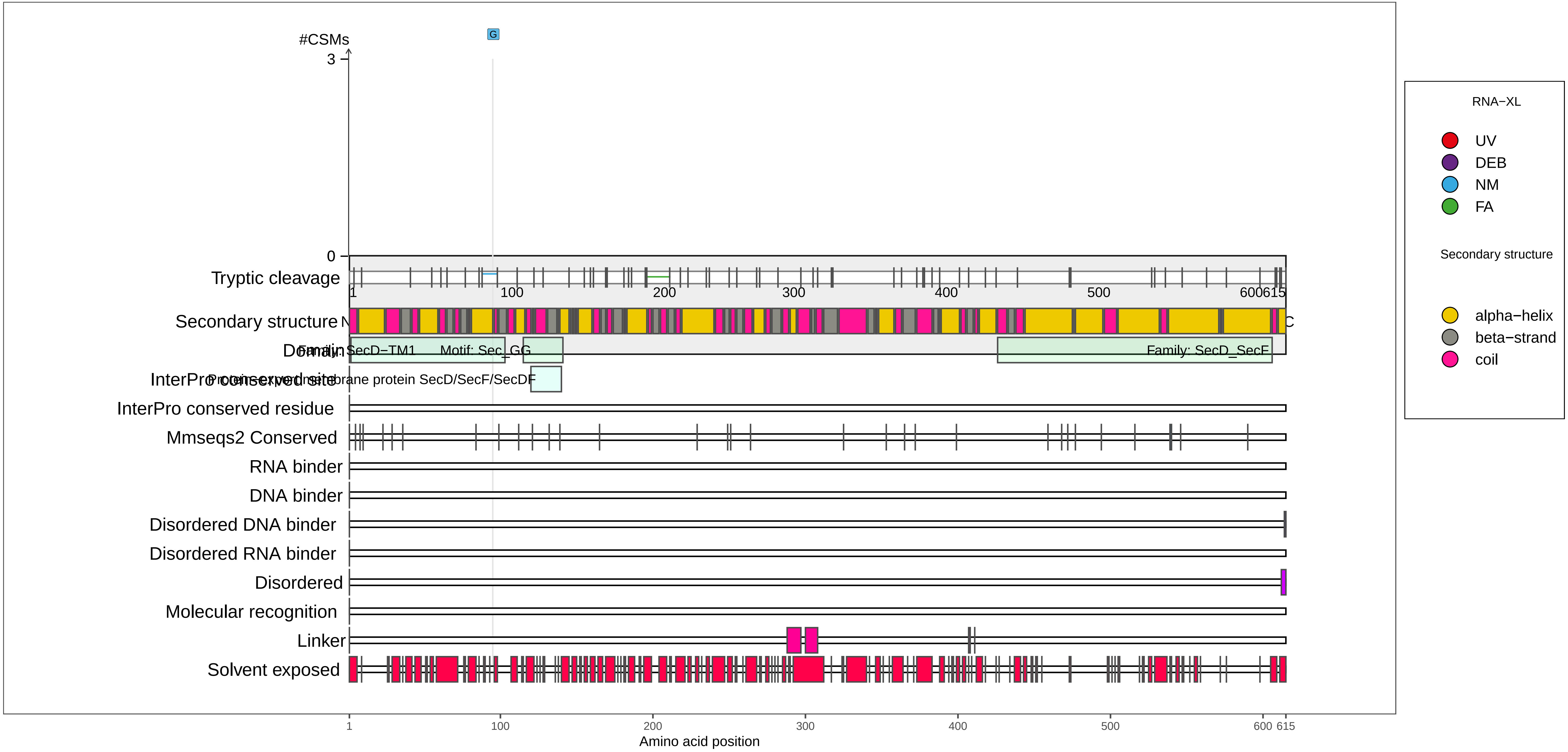

P11446
ARGC_ECOLI N−acetyl−gamma−glutamyl−phosphate reductase

− Abundance:
tryptic [log10 Intensity]: 8.32 (Q 70)
PAXdb K12 strain [ppm]: NA (Q NA)
PAXdb E.coli [ppm]: 1.27 (Q 54)

− RNA functions: not annotated

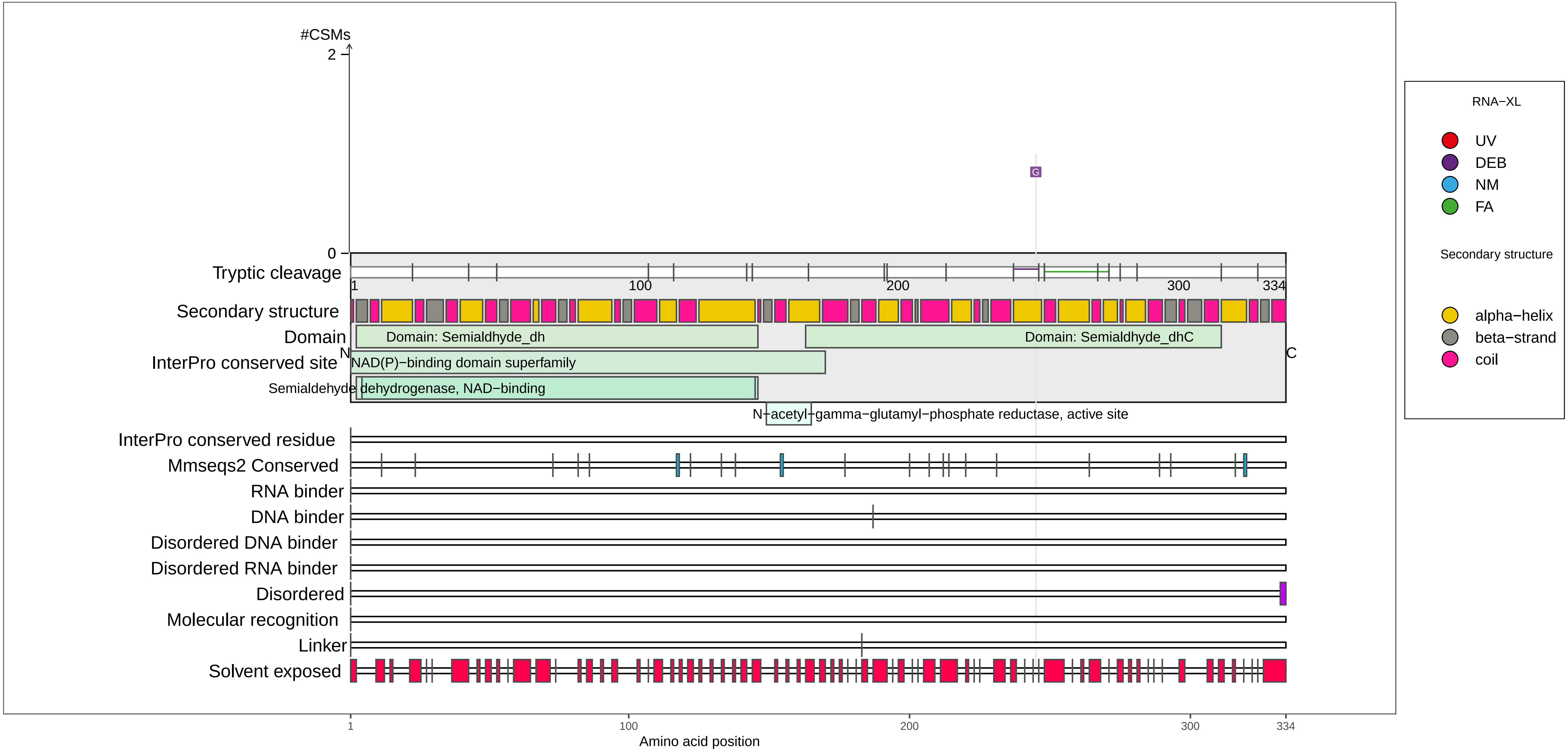

P12008
AROC_ECOLI Chorismate synthase

− Abundance:
tryptic [log10 Intensity]: 7.78 (Q 48)
PAXdb K12 strain [ppm]: 2.17 (Q 66)
PAXdb E.coli [ppm]: 1.98 (Q 72)

− RNA functions: not annotated

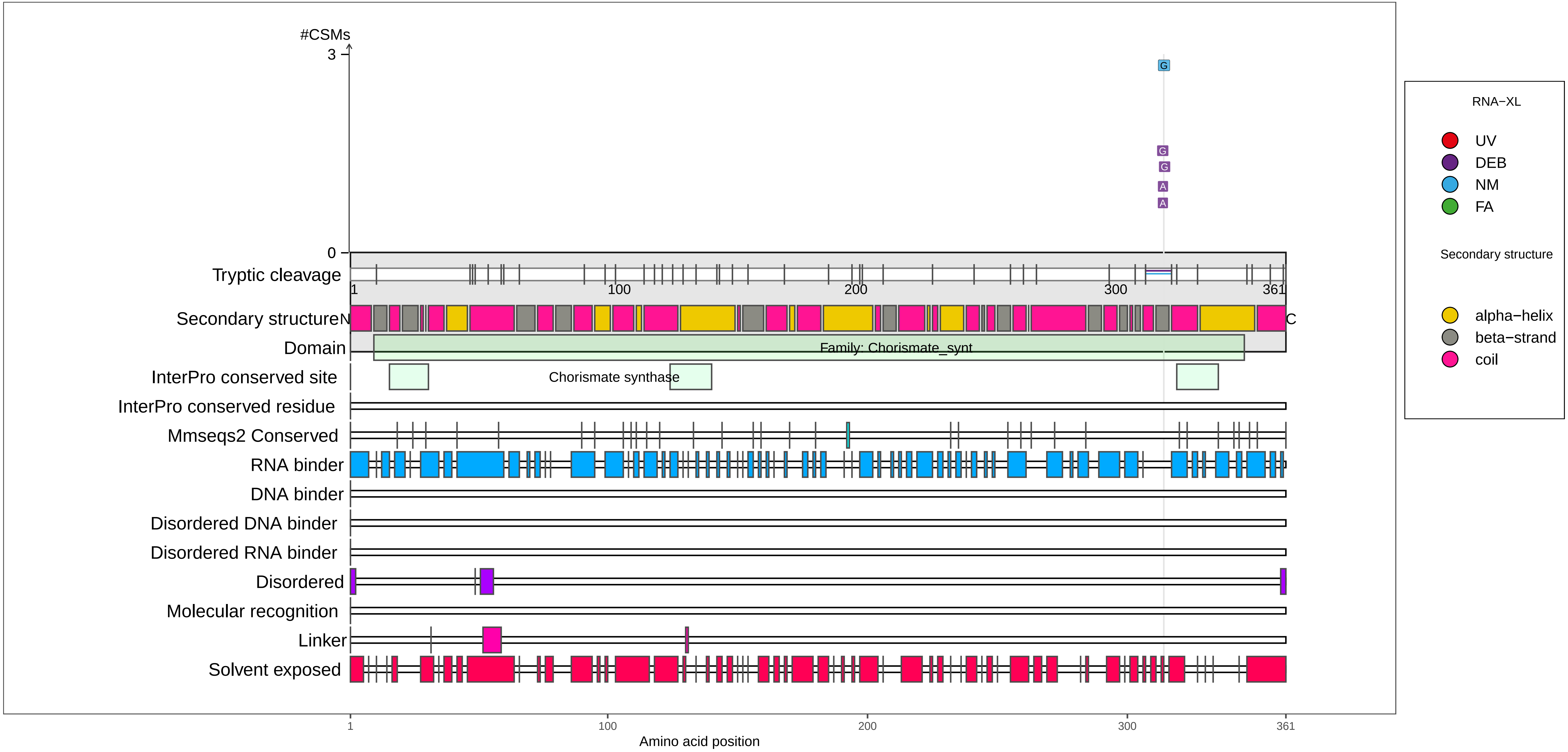

P16528
ICLR_ECOLI Transcriptional repressor IclR

− Abundance:
tryptic [log10 Intensity]: 7.45 (Q 33)
PAXdb K12 strain [ppm]: NA (Q NA)
PAXdb E.coli [ppm]: 0.33 (Q 32)

− RNA functions: not annotated

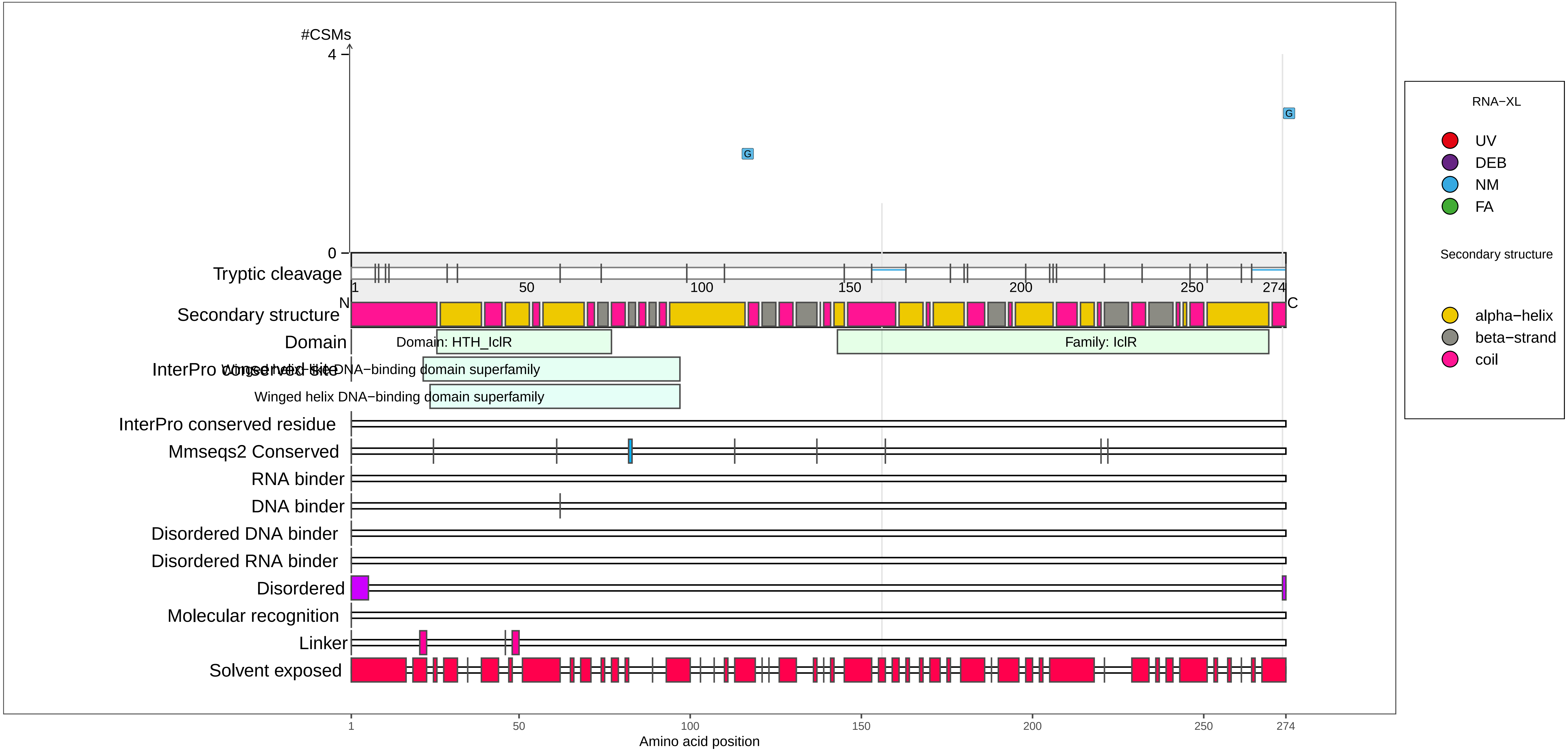

P21507
SRMB_ECOLI ATP−dependent RNA helicase SrmB

− Abundance:
tryptic [log10 Intensity]: 7.75 (Q 46)
PAXdb K12 strain [ppm]: 2.34 (Q 71)
PAXdb E.coli [ppm]: 2.02 (Q 73)

− RNA functions:
RNA binding; RNA helicase activity; RNA strand annealing activity
RNA−dependent ATPase activity

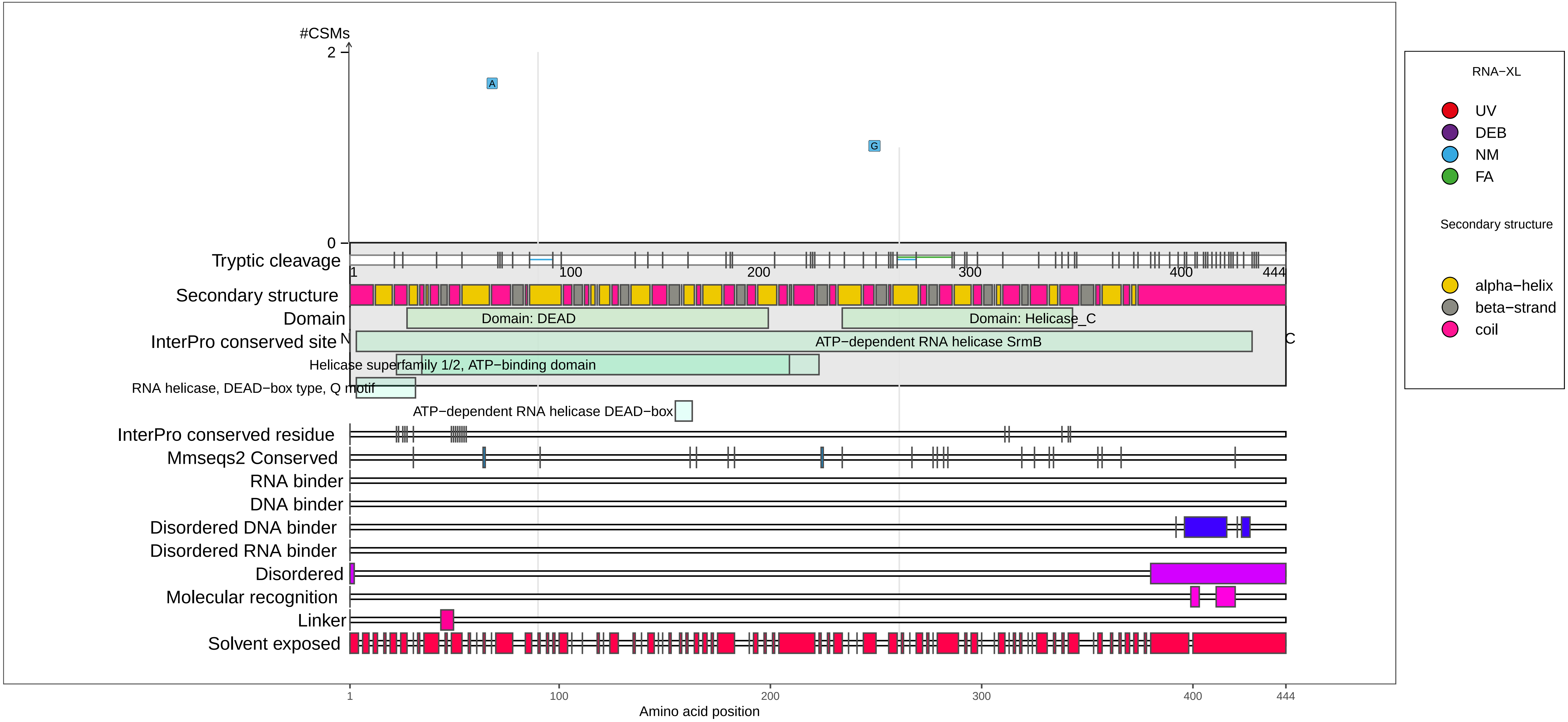

P23827
ECOT_ECOLI Ecotin

− Abundance:
tryptic [log10 Intensity]: 8.19 (Q 65)
PAXdb K12 strain [ppm]: 2.58 (Q 80)
PAXdb E.coli [ppm]: 1.57 (Q 61)

− RNA functions: not annotated

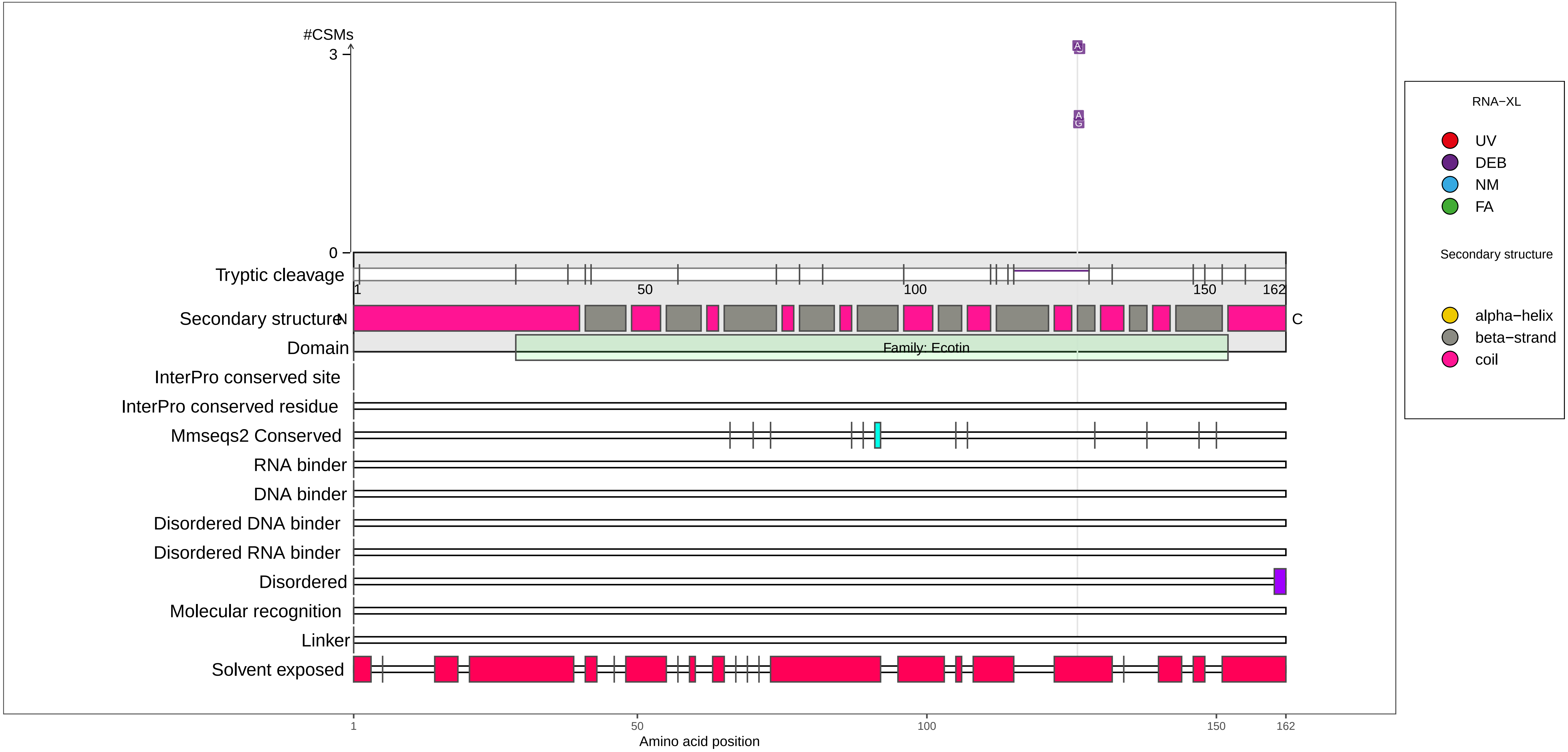

P23830
PSS_ECOLI CDP−diacylglycerol−−serine O−phosphatidyltransferase

− Abundance:
tryptic [log10 Intensity]: 8.25 (Q 67)
PAXdb K12 strain [ppm]: 1.86 (Q 56)
PAXdb E.coli [ppm]: 2.68 (Q 90)

− RNA functions: not annotated

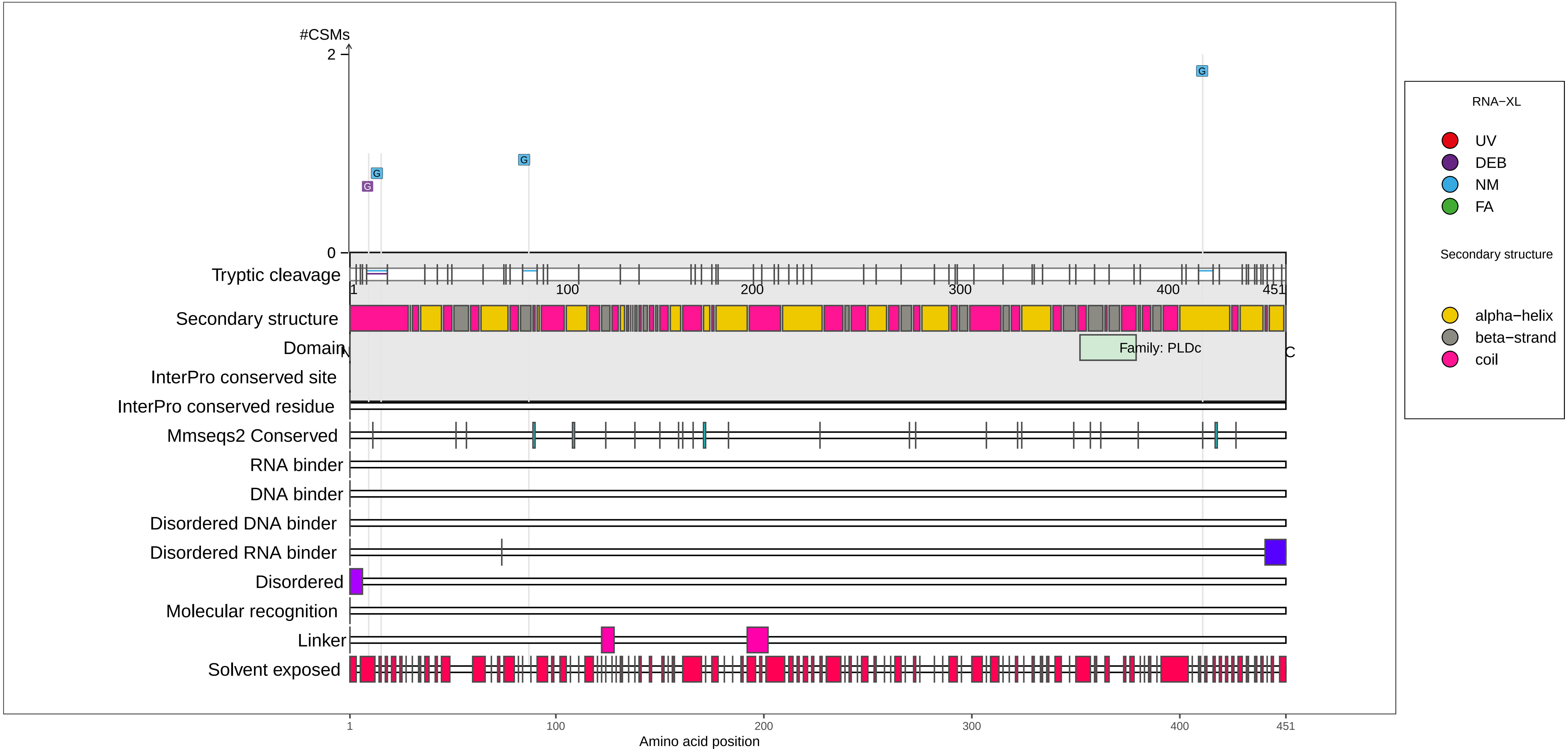

P23893
GSA_ECOLI Glutamate−1−semialdehyde 2,1−aminomutase

− Abundance:
tryptic [log10 Intensity]: 8.3 (Q 69)
PAXdb K12 strain [ppm]: 3 (Q 91)
PAXdb E.coli [ppm]: 2.46 (Q 85)

− RNA functions: not annotated

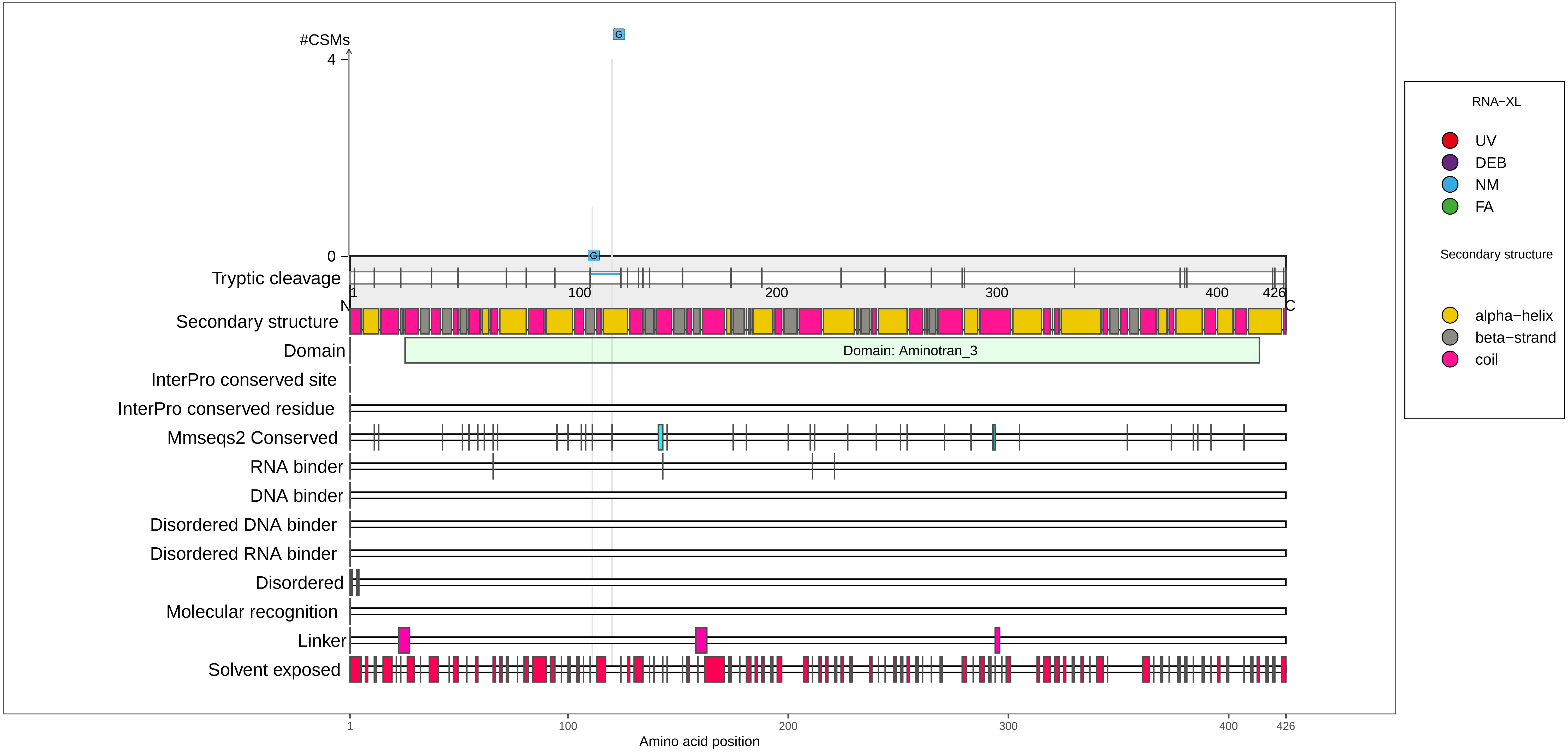

P24186
FOLD_ECOLI Bifunctional protein FolD

− Abundance:
tryptic [log10 Intensity]: 9.05 (Q 89)
PAXdb K12 strain [ppm]: 2.53 (Q 78)
PAXdb E.coli [ppm]: 2.01 (Q 72)

− RNA functions: not annotated

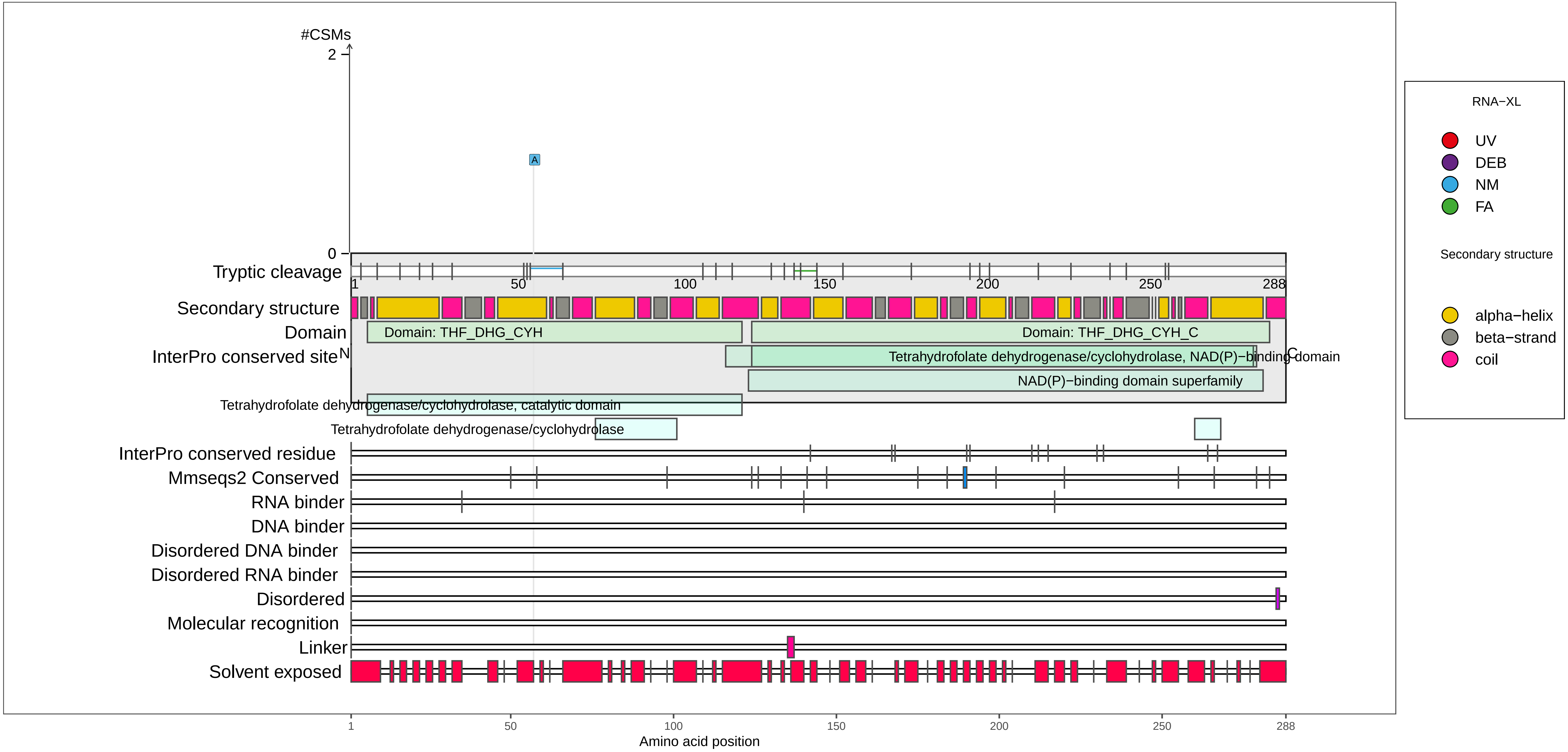

P25522
MNME_ECOLI tRNA modification GTPase MnmE

− Abundance:
tryptic [log10 Intensity]: NA (Q NA)
PAXdb K12 strain [ppm]: 2.15 (Q 65)
PAXdb E.coli [ppm]: 2.6 (Q 88)

− RNA functions:
ncRNA metabolic process; ncRNA processing; RNA metabolic process; RNA methylation
RNA modification; RNA processing; tRNA metabolic process; tRNA methylation; tRNA modification
tRNA processing; tRNA wobble base modification; tRNA wobble uridine modification

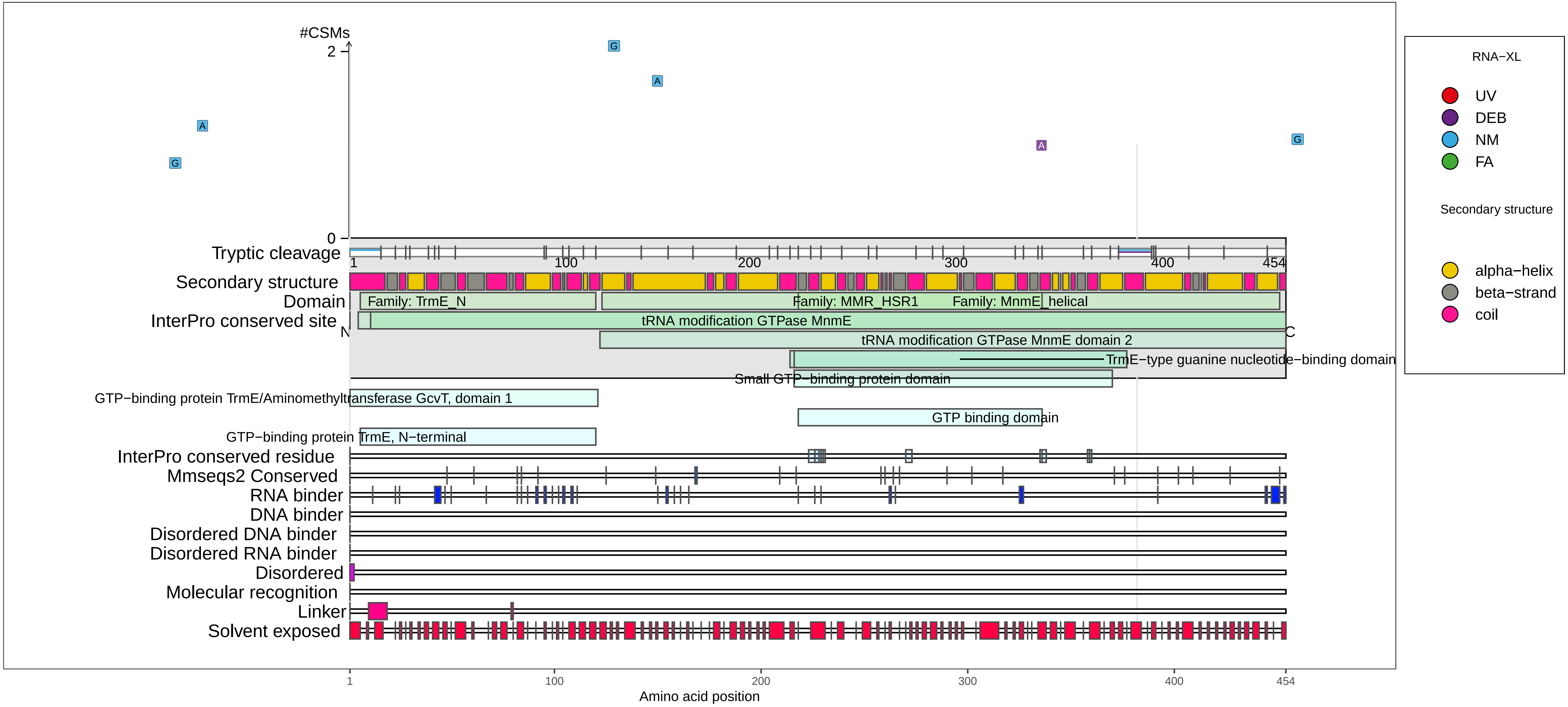

P25888
RHLE_ECOLI ATP−dependent RNA helicase RhlE

− Abundance:
tryptic [log10 Intensity]: 7.58 (Q 39)
PAXdb K12 strain [ppm]: 2.41 (Q 74)
PAXdb E.coli [ppm]: 1.01 (Q 48)

− RNA functions:
RNA binding; RNA helicase activity

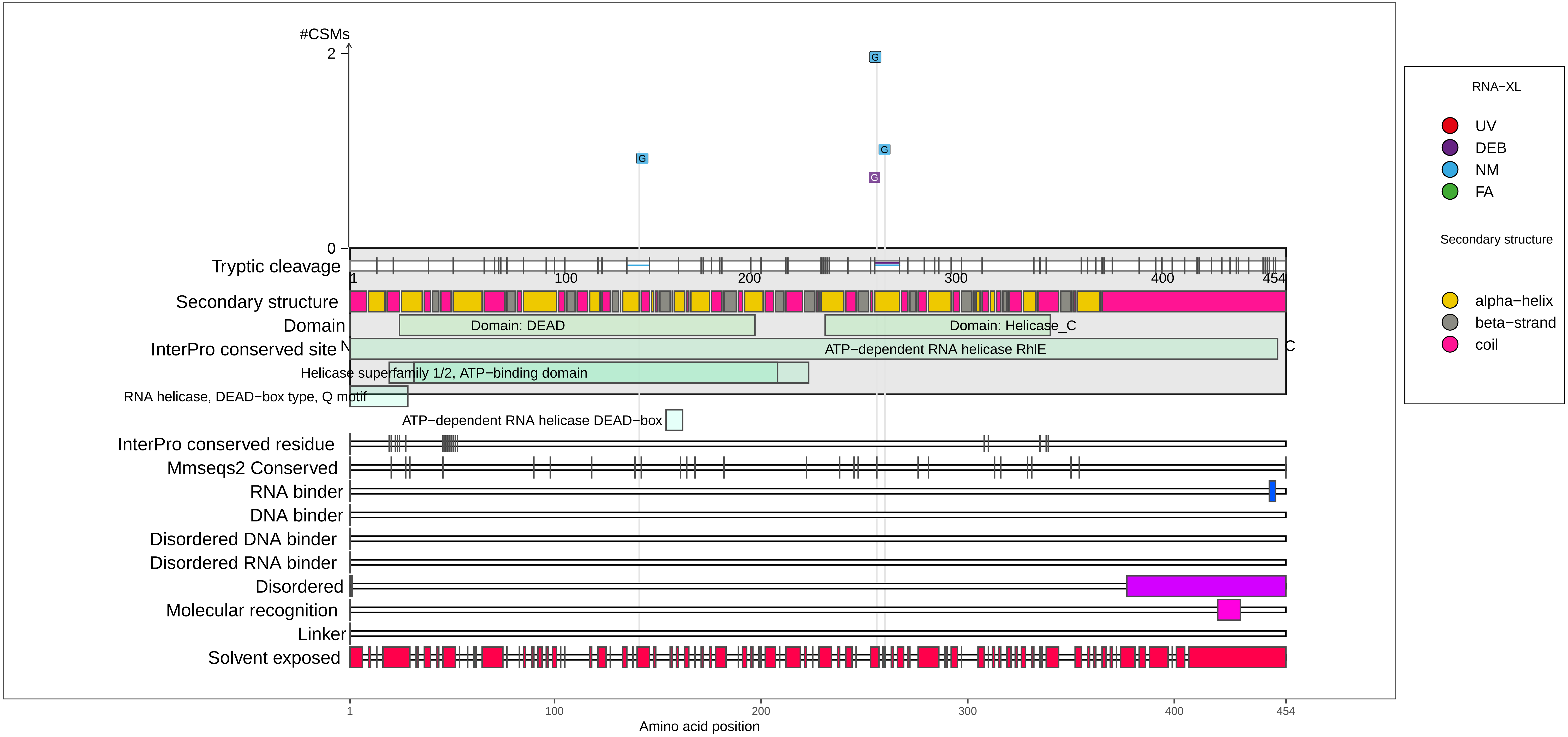

P30871
3PASE_ECOLI Inorganic triphosphatase

− Abundance:
tryptic [log10 Intensity]: 6.83 (Q 8)
PAXdb K12 strain [ppm]: 2.41 (Q 75)
PAXdb E.coli [ppm]: 1.94 (Q 70)

− RNA functions: not annotated

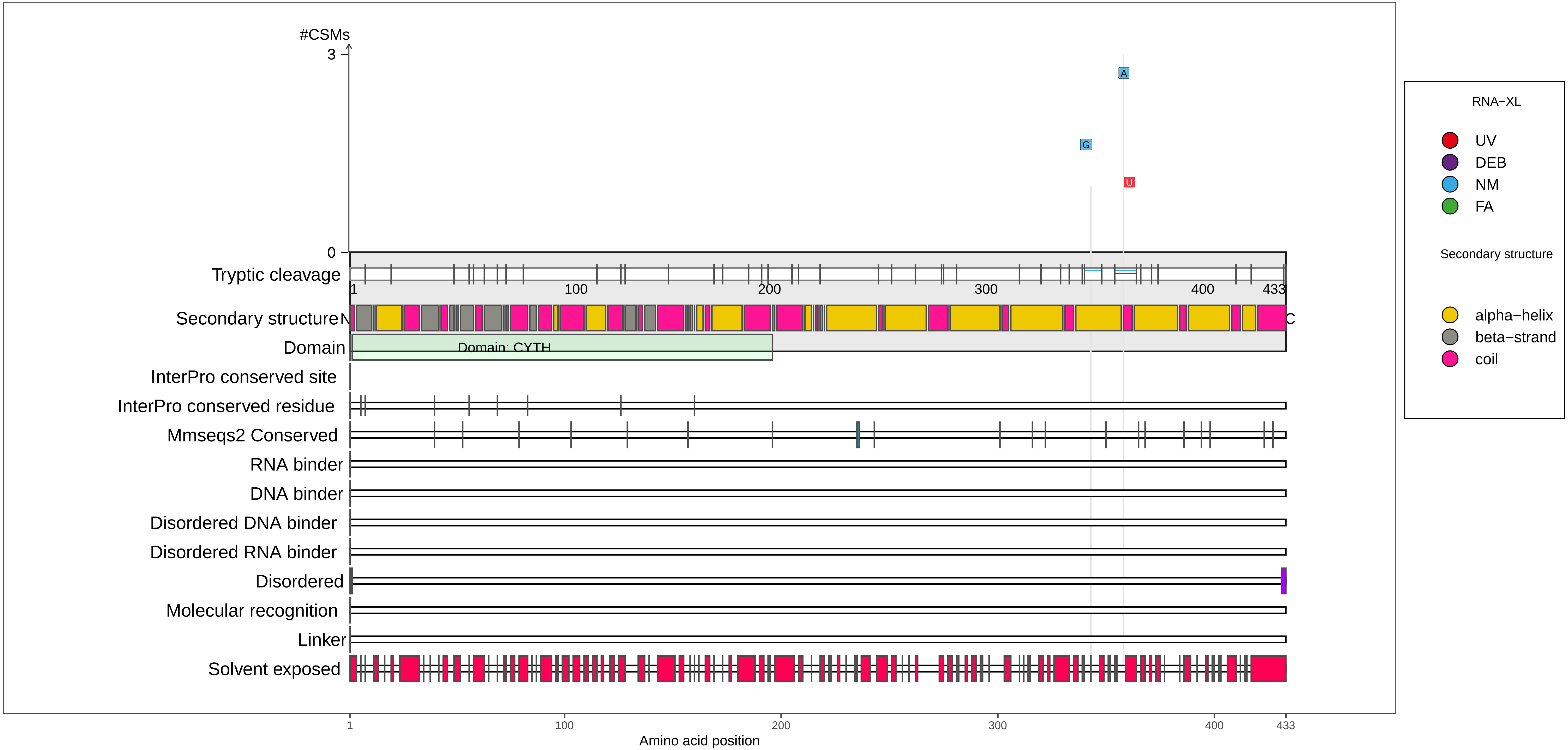

P31545
EFEB_ECOLI Deferrochelatase/peroxidase EfeB

− Abundance:
tryptic [log10 Intensity]: 7.54 (Q 37)
PAXdb K12 strain [ppm]: 1.69 (Q 49)
PAXdb E.coli [ppm]: −0.26 (Q 19)

− RNA functions: not annotated

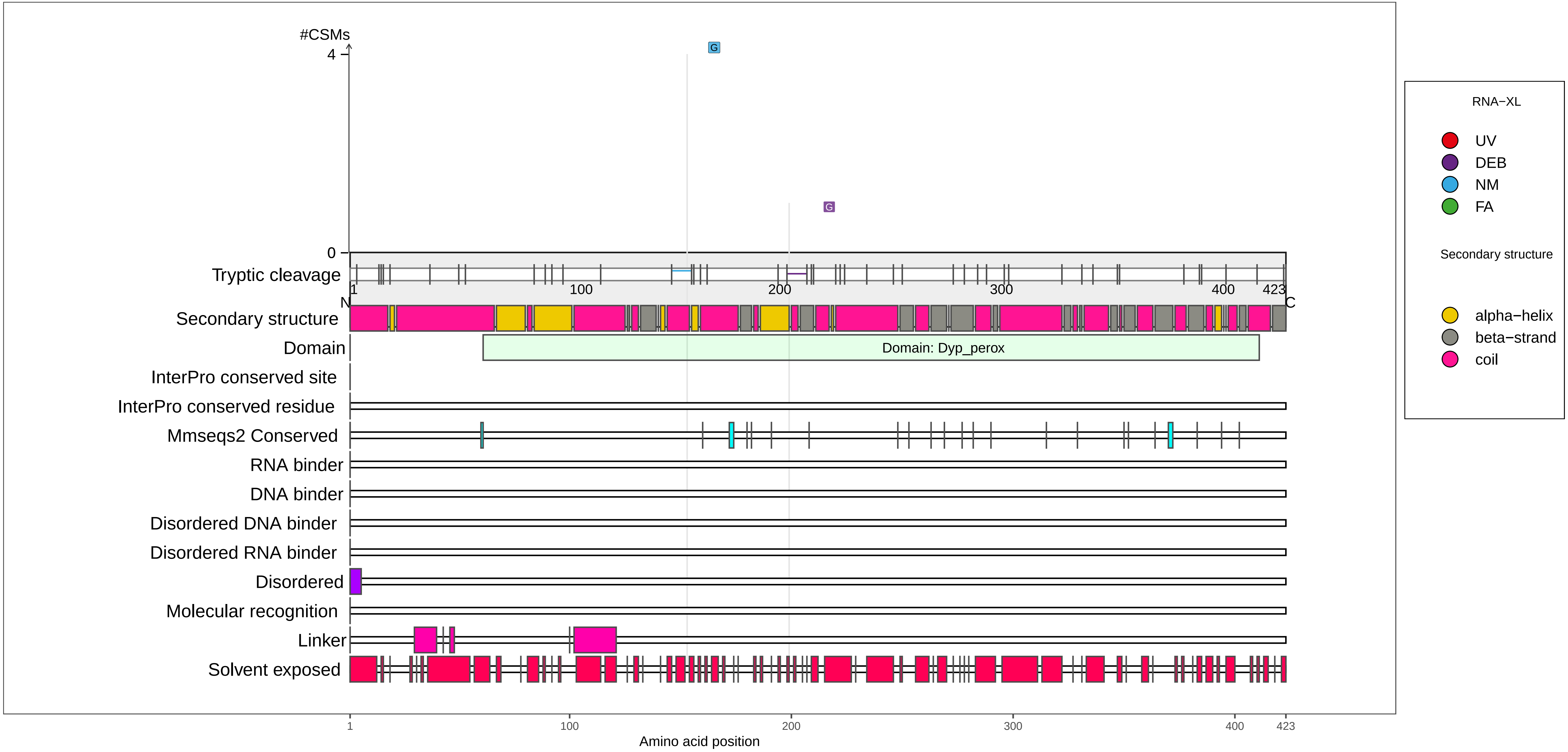

P31979
NUOF_ECOLI NADH−quinone oxidoreductase subunit F

− Abundance:
tryptic [log10 Intensity]: 9.23 (Q 93)
PAXdb K12 strain [ppm]: 2.85 (Q 87)
PAXdb E.coli [ppm]: 2.27 (Q 80)

− RNA functions: not annotated

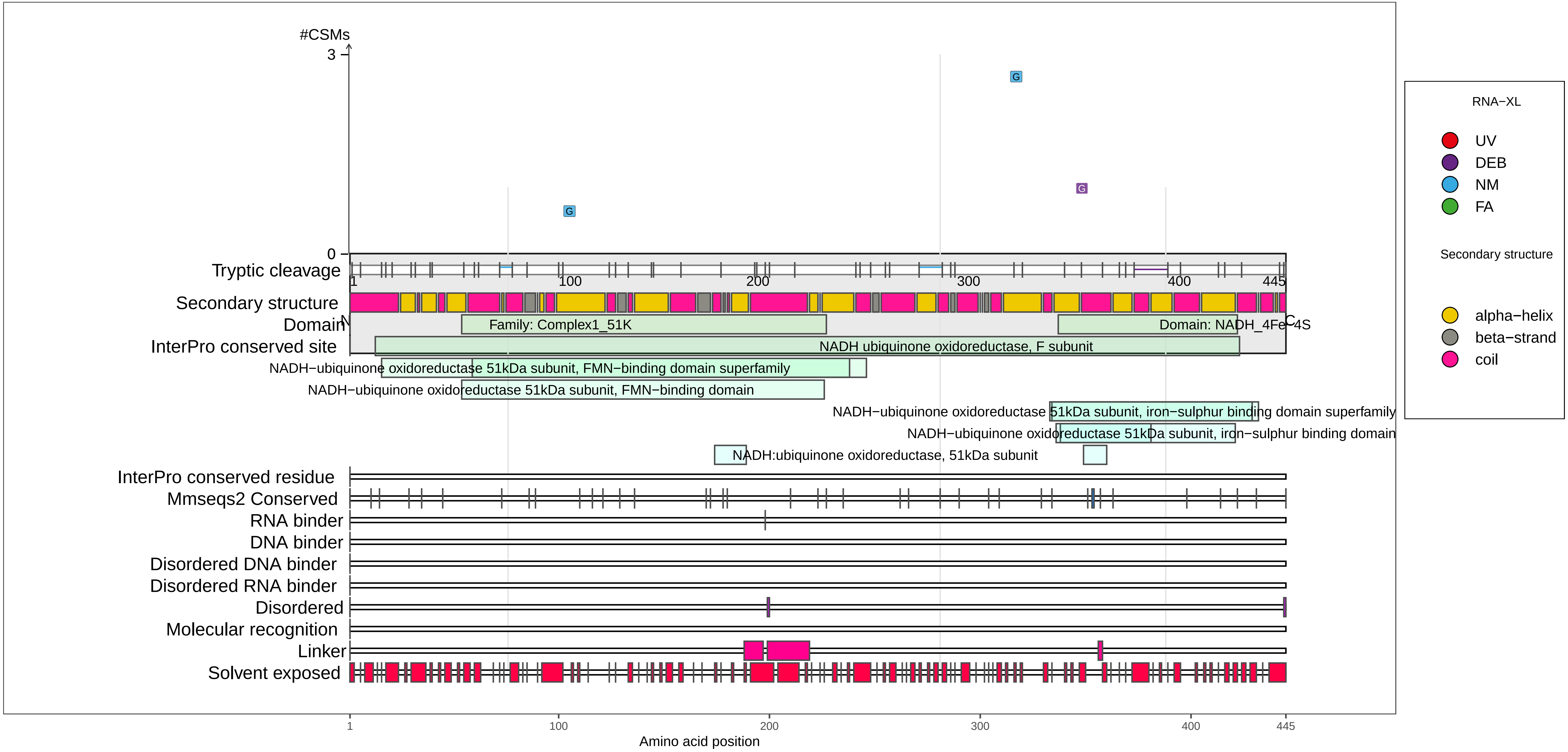

P37095
PEPB_ECOLI Peptidase B
− Abundance:
tryptic [log10 Intensity]: 8.41 (Q 73)
PAXdb K12 strain [ppm]: 2.48 (Q 77)
PAXdb E.coli [ppm]: 2.82 (Q 92)

− RNA functions: not annotated

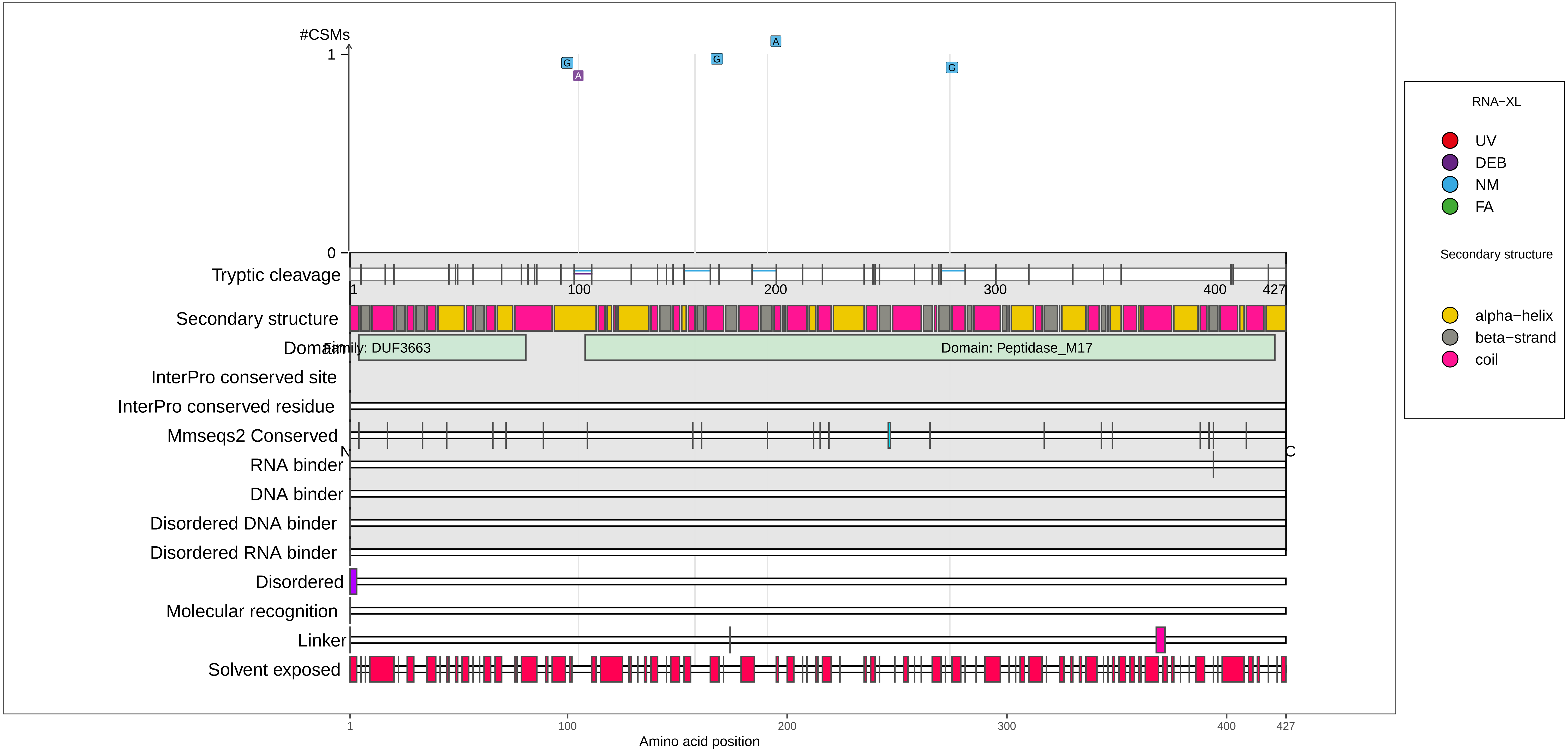

P37188
PTKB_ECOLI PTS system galactitol−specific EIIB component

− Abundance:
tryptic [log10 Intensity]: 8.48 (Q 75)
PAXdb K12 strain [ppm]: NA (Q NA)
PAXdb E.coli [ppm]: 3.79 (Q 100)

− RNA functions: not annotated

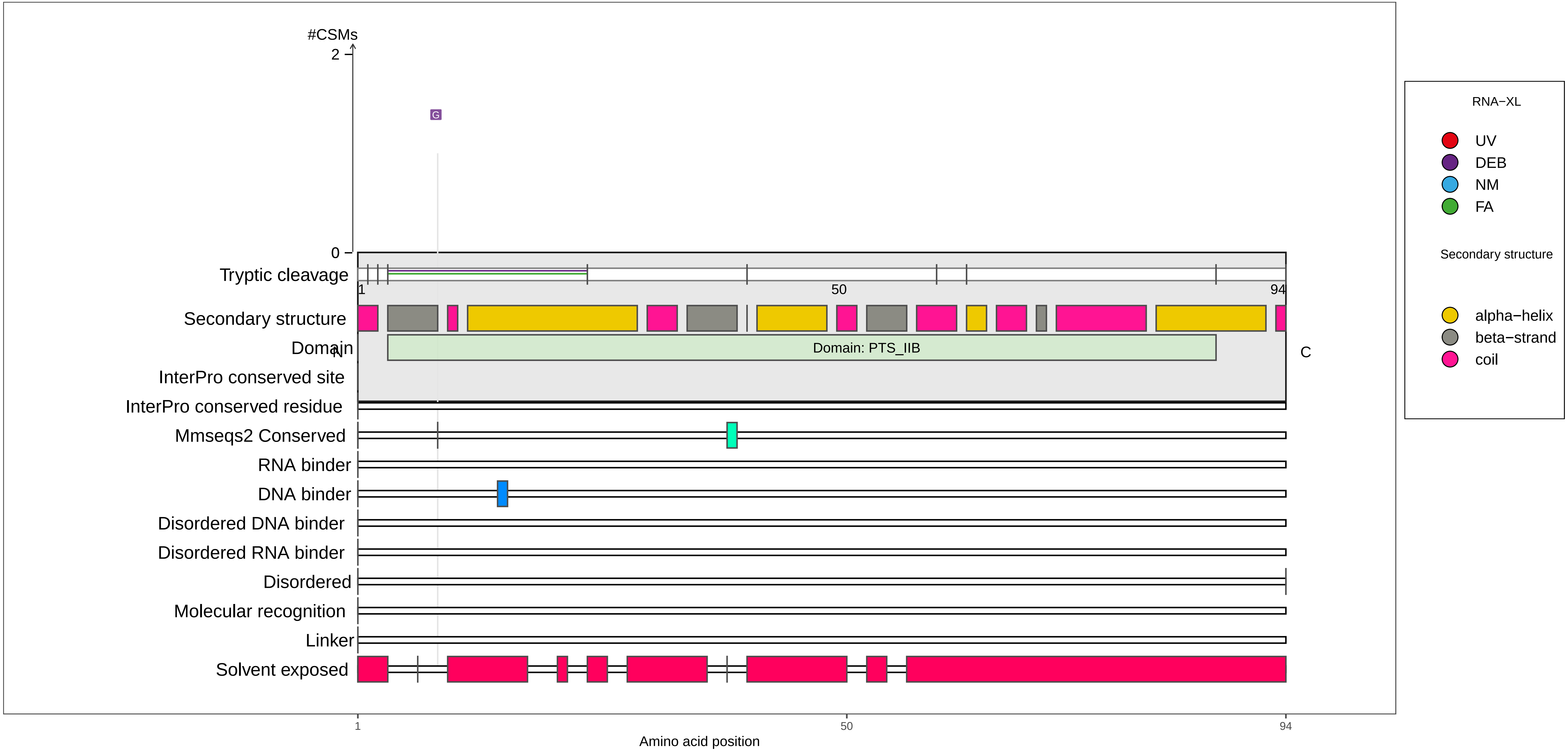

P42596
RLMG_ECOLI Ribosomal RNA large subunit methyltransferase G

− Abundance:
tryptic [log10 Intensity]: 7.86 (Q 52)
PAXdb K12 strain [ppm]: 1.42 (Q 34)
PAXdb E.coli [ppm]: 0.2 (Q 29)

− RNA functions:
23S rRNA (guanine(1835)−N(2))−methyltransferase activity; ncRNA metabolic process
ncRNA processing; RNA metabolic process; RNA methylation
RNA methyltransferase activity; RNA modification; RNA processing; rRNA (guanine−N2−)−methyltransferase activity
rRNA (guanine) methyltransferase activity; rRNA base methylation; rRNA metabolic process
rRNA methylation; rRNA methyltransferase activity; rRNA modification; rRNA processing

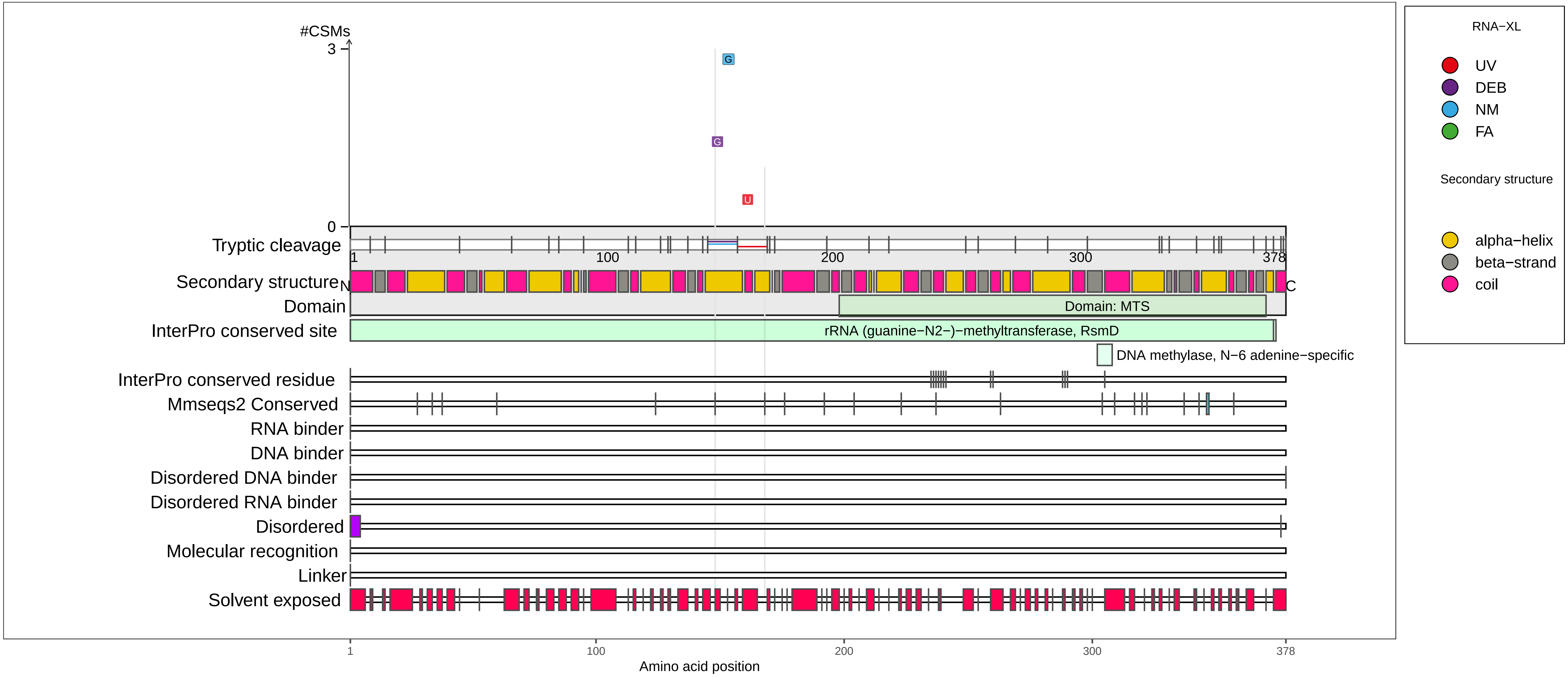

P45568
DXR_ECOLI 1−deoxy−D−xylulose 5−phosphate reductoisomerase

− Abundance:
tryptic [log10 Intensity]: 7.82 (Q 50)
PAXdb K12 strain [ppm]: 1.43 (Q 34)
PAXdb E.coli [ppm]: 0.31 (Q 32)

− RNA functions: not annotated

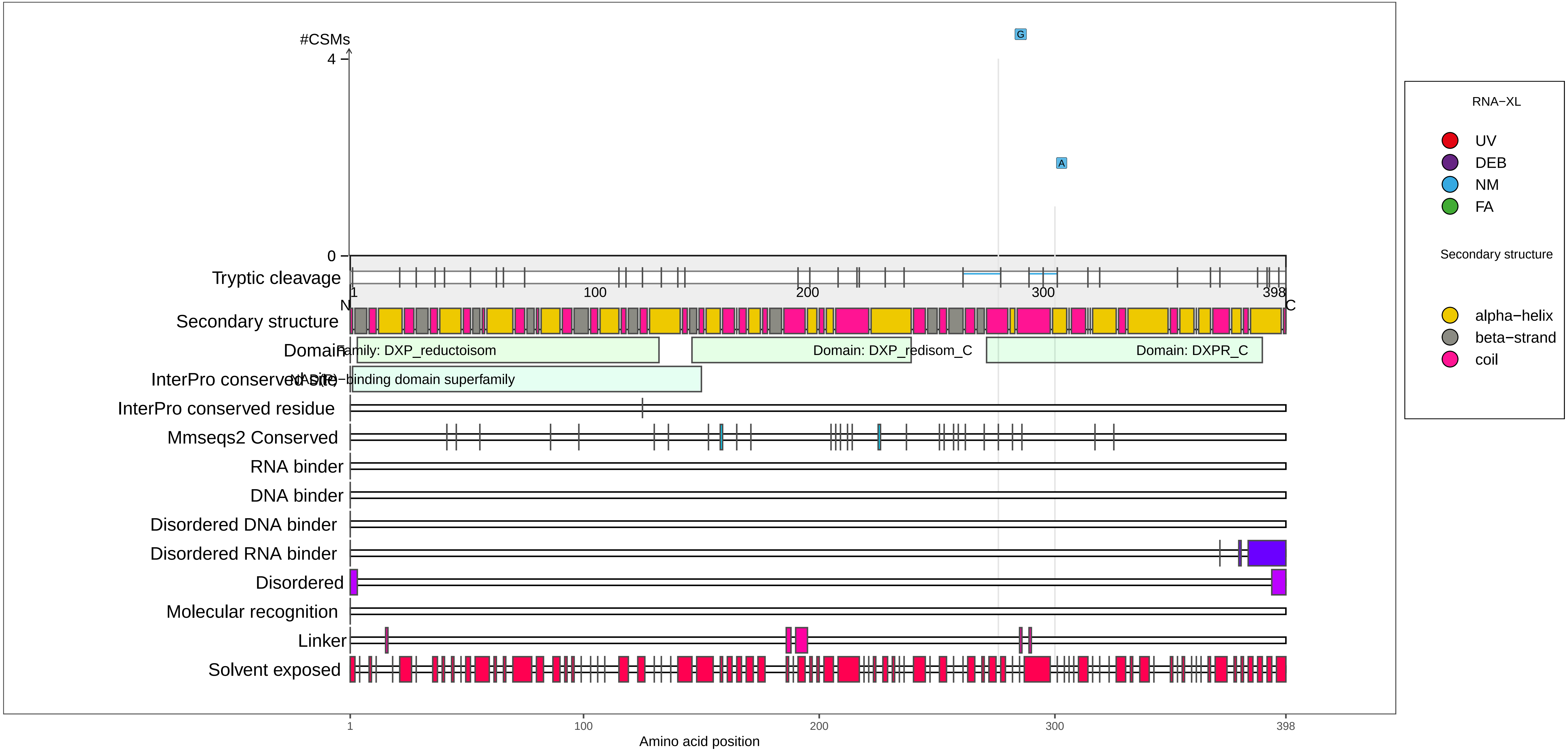

P52060
YGGU_ECOLI UPF0235 protein YggU

− Abundance:
tryptic [log10 Intensity]: 7.09 (Q 16)
PAXdb K12 strain [ppm]: NA (Q NA)
PAXdb E.coli [ppm]: 0.95 (Q 47)

− RNA functions: not annotated

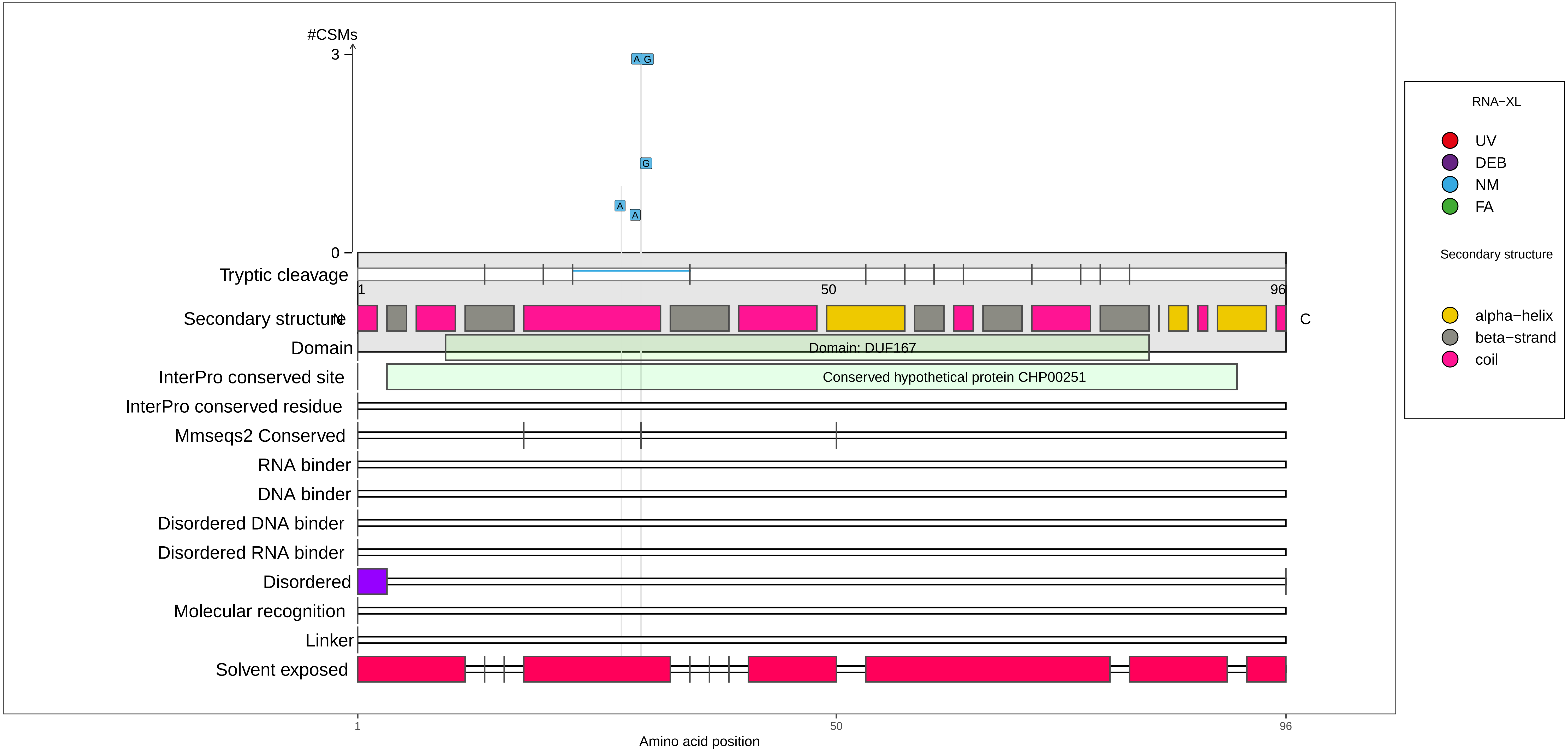

P56580
PTHB_ECOLI PTS system glucitol/sorbitol−specific EIIB component

− Abundance:
tryptic [log10 Intensity]: 7.96 (Q 56)
PAXdb K12 strain [ppm]: NA (Q NA)
PAXdb E.coli [ppm]: 0.15 (Q 28)

− RNA functions: not annotated

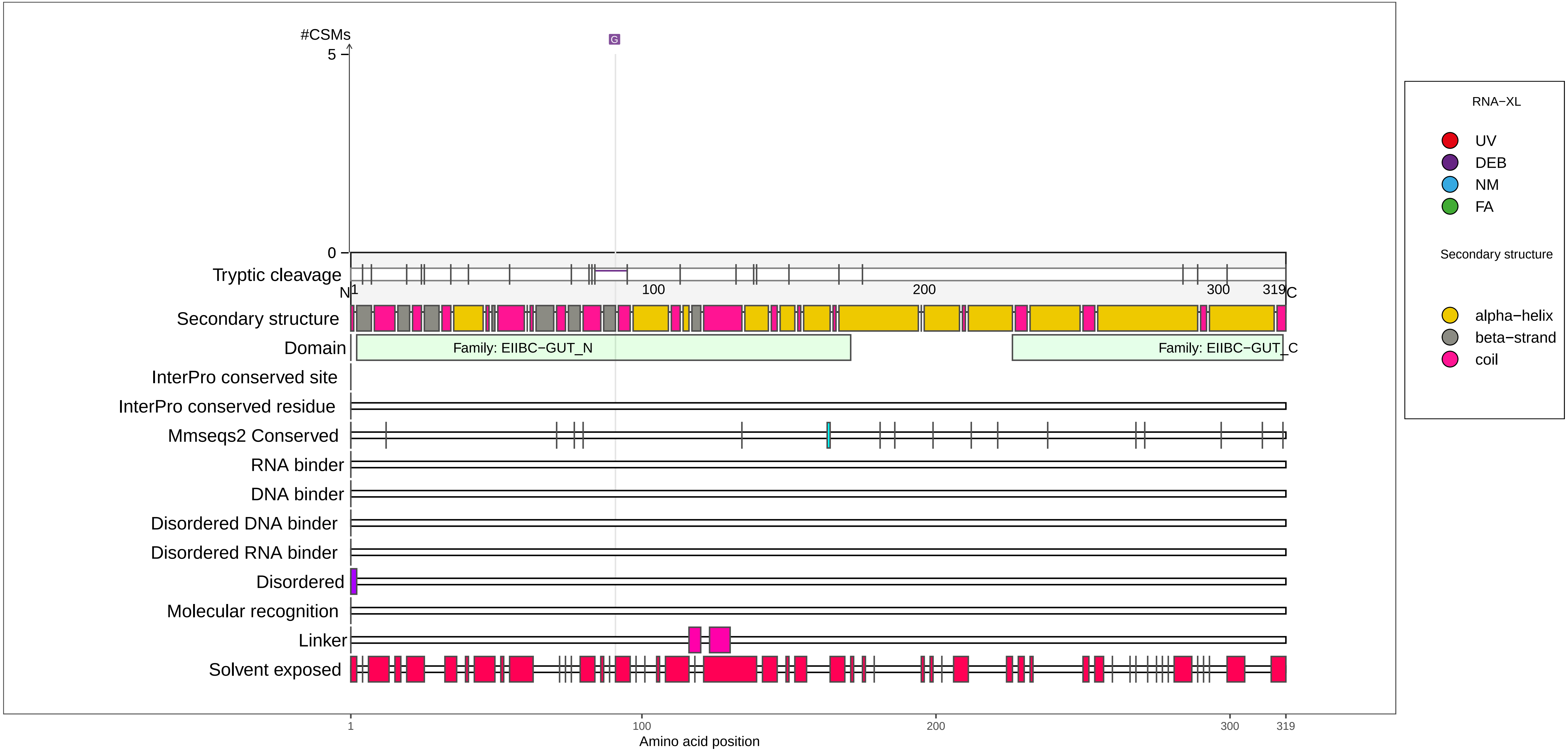

P60340
TRUB_ECOLI tRNA pseudouridine synthase B

− Abundance:
tryptic [log10 Intensity]: 7.48 (Q 34)
PAXdb K12 strain [ppm]: 2.05 (Q 62)
PAXdb E.coli [ppm]: 0.93 (Q 46)

− RNA functions:
mRNA metabolic process; mRNA modification; mRNA pseudouridine synthesis
ncRNA metabolic process; ncRNA processing; RNA binding; RNA folding; RNA metabolic process
RNA modification; RNA processing; tRNA binding; tRNA folding; tRNA metabolic process
tRNA modification; tRNA processing; tRNA pseudouridine synthase activity
tRNA pseudouridine synthesis; tRNA pseudouridylate synthase B C−terminal domain

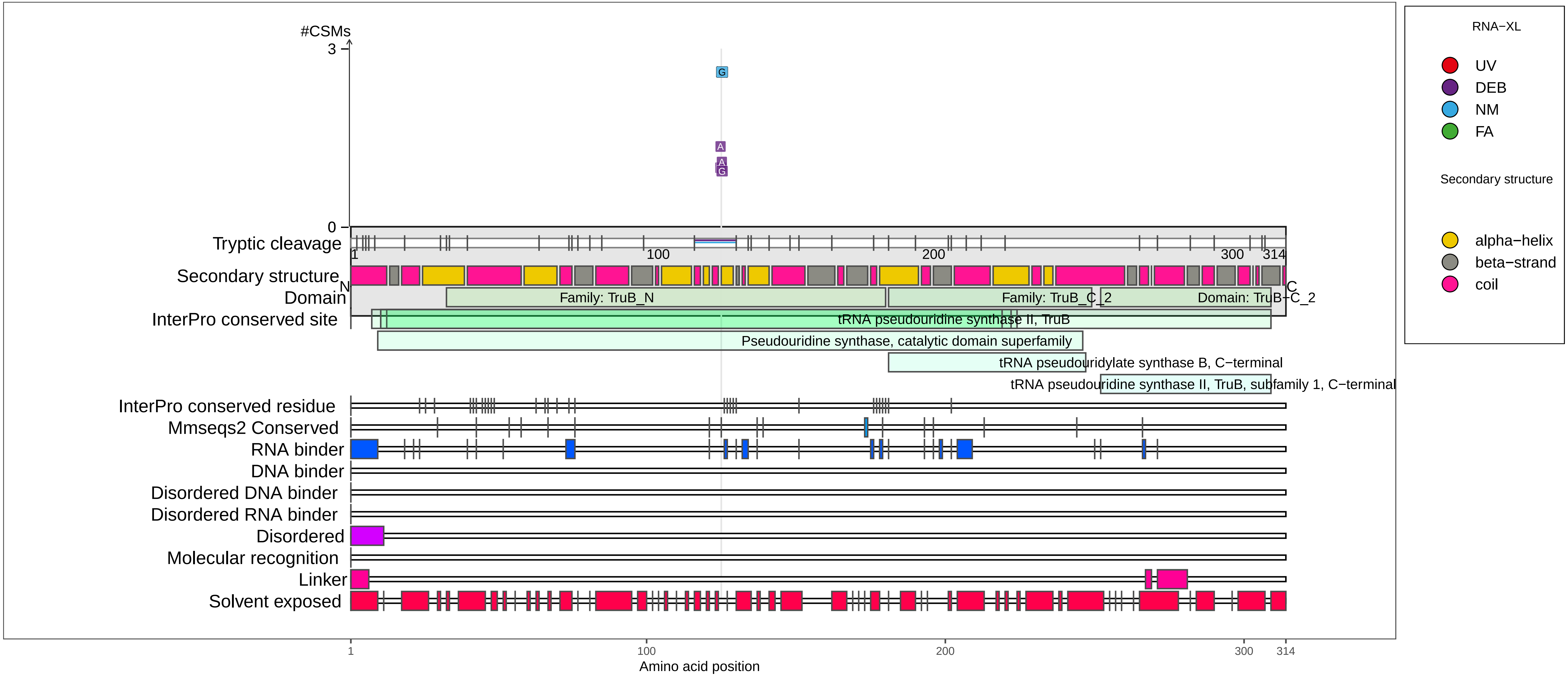

P60651
SPEB_ECOLI Agmatinase

− Abundance:
tryptic [log10 Intensity]: 8.21 (Q 66)
PAXdb K12 strain [ppm]: NA (Q NA)
PAXdb E.coli [ppm]: 2.42 (Q 84)

− RNA functions: not annotated

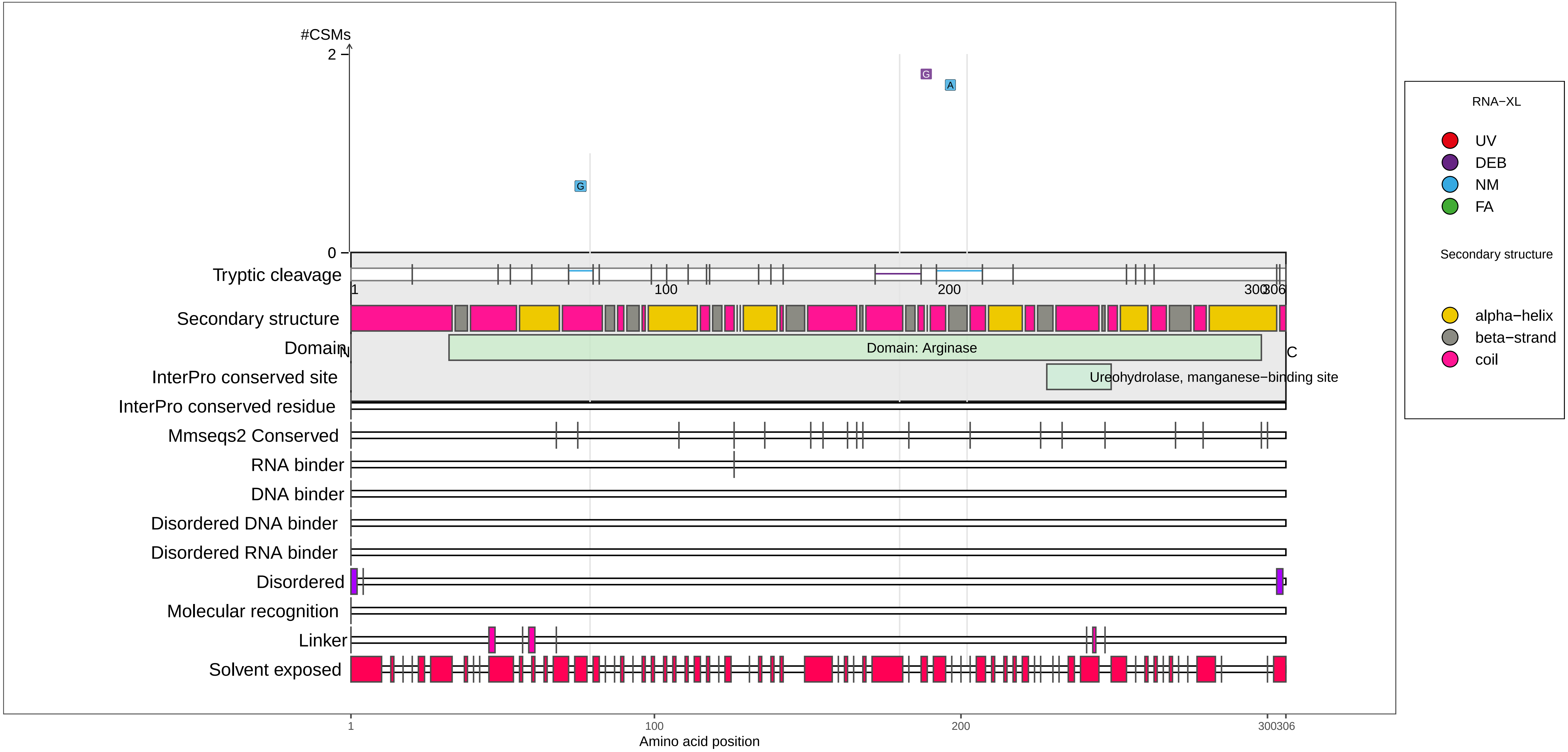

P61889
MDH_ECOLI Malate dehydrogenase

− Abundance:
tryptic [log10 Intensity]: 9.08 (Q 90)
PAXdb K12 strain [ppm]: 3.25 (Q 95)
PAXdb E.coli [ppm]: 3.77 (Q 100)

− RNA functions: not annotated

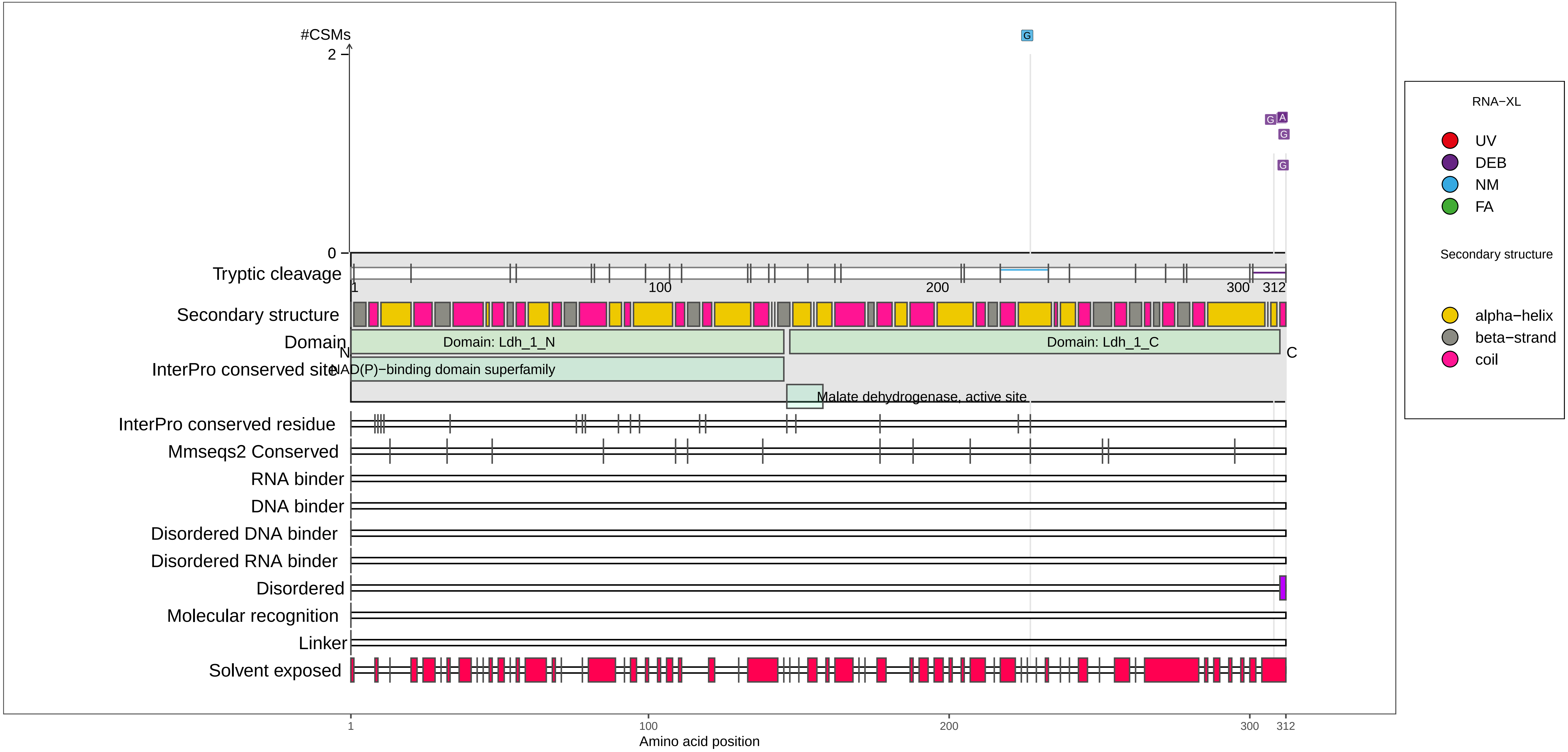

P61949
FLAV_ECOLI Flavodoxin 1

− Abundance:
tryptic [log10 Intensity]: 10.45 (Q 100)
PAXdb K12 strain [ppm]: 2.52 (Q 78)
PAXdb E.coli [ppm]: 3.03 (Q 95)

− RNA functions: not annotated

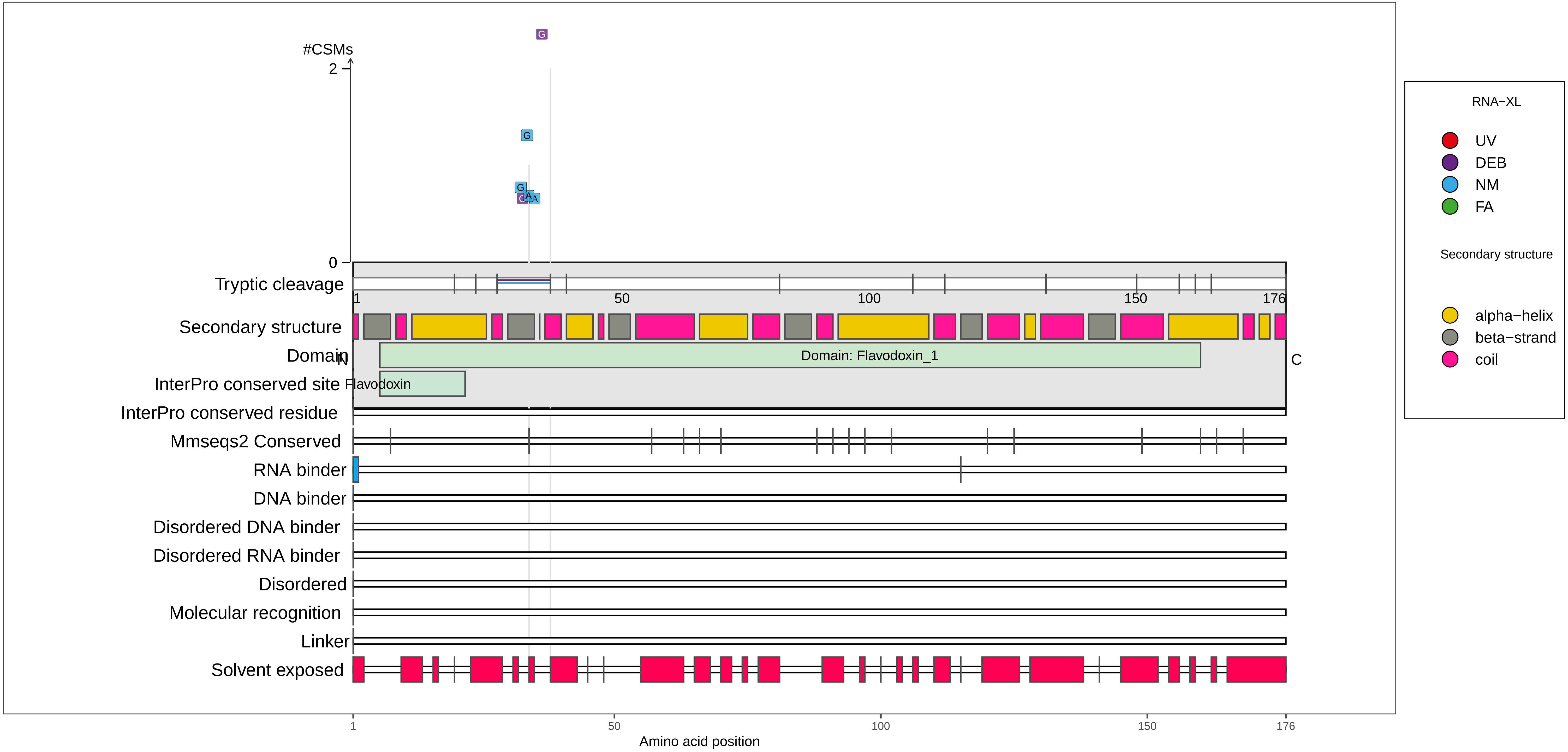

P63389
YHES_ECOLI Probable ATP−binding protein YheS

− Abundance:
tryptic [log10 Intensity]: 8.5 (Q 76)
PAXdb K12 strain [ppm]: 2.12 (Q 65)
PAXdb E.coli [ppm]: 1.17 (Q 52)

− RNA functions: not annotated

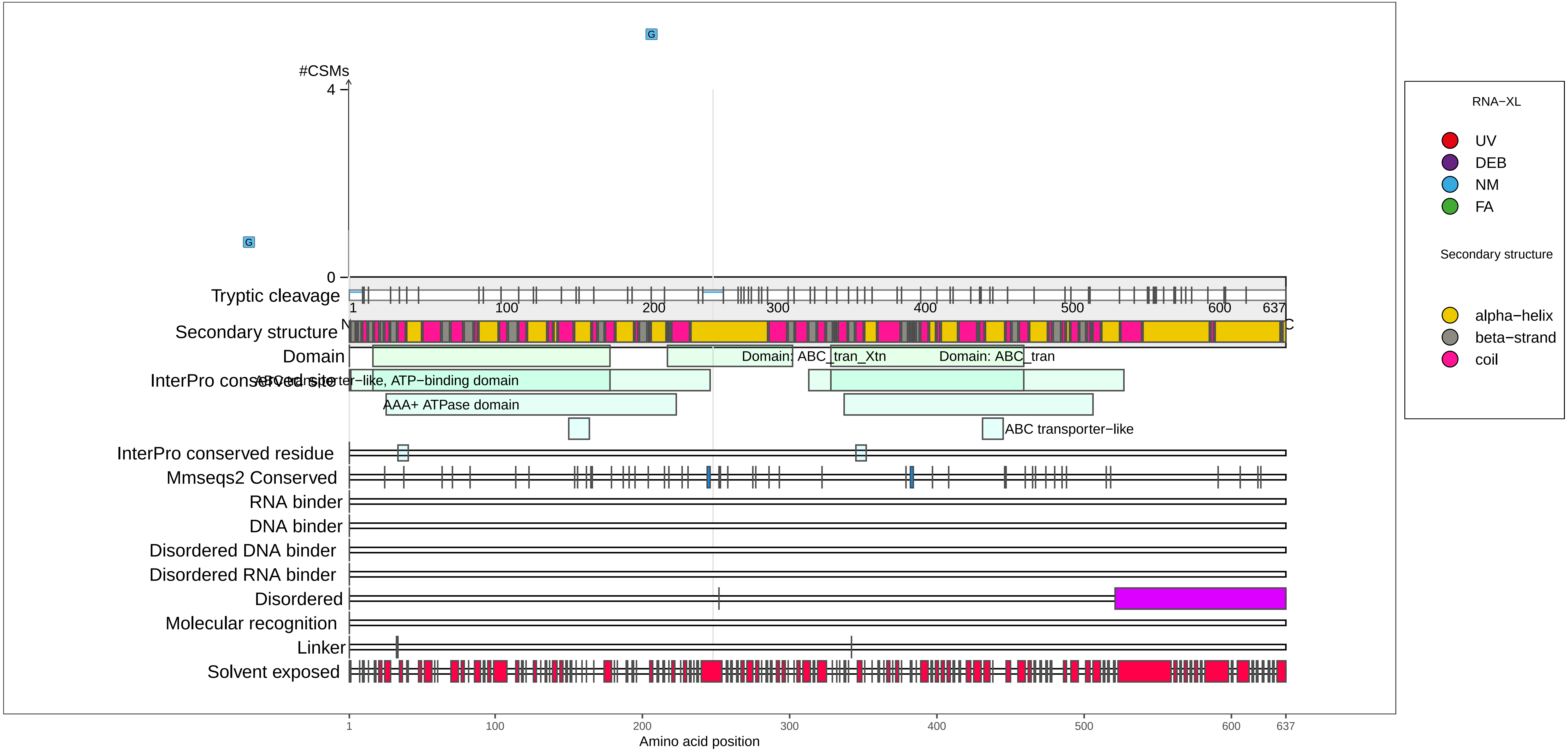

P64451
YDCL_ECOLI Uncharacterized lipoprotein YdcL

− Abundance:
tryptic [log10 Intensity]: 7.54 (Q 37)
PAXdb K12 strain [ppm]: 1.49 (Q 38)
PAXdb E.coli [ppm]: 2.86 (Q 93)

− RNA functions: not annotated

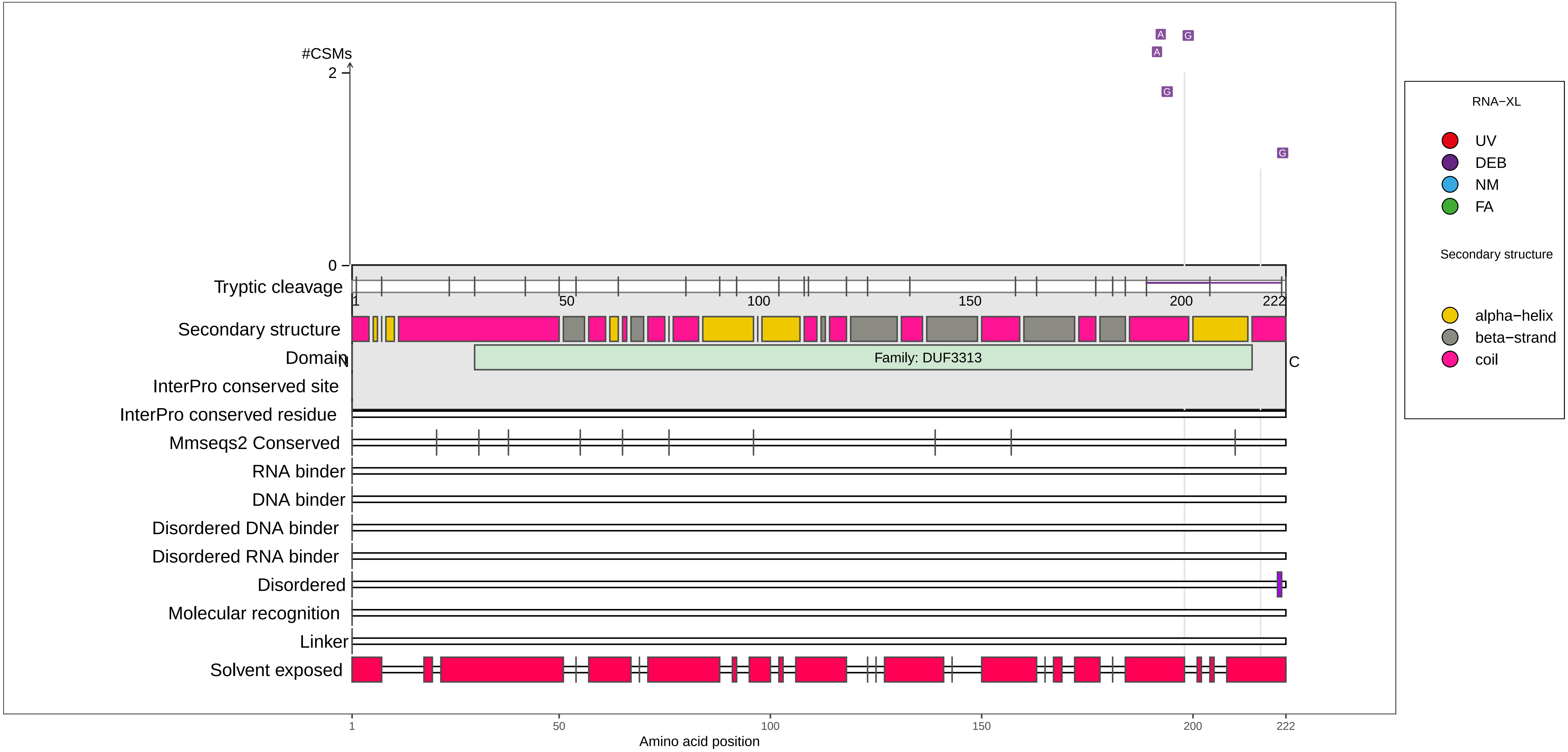

P64548
YFIR_ECOLI Protein YfiR

− Abundance:
tryptic [log10 Intensity]: NA (Q NA)
PAXdb K12 strain [ppm]: 2.03 (Q 62)
PAXdb E.coli [ppm]: −0.08 (Q 23)

− RNA functions: not annotated

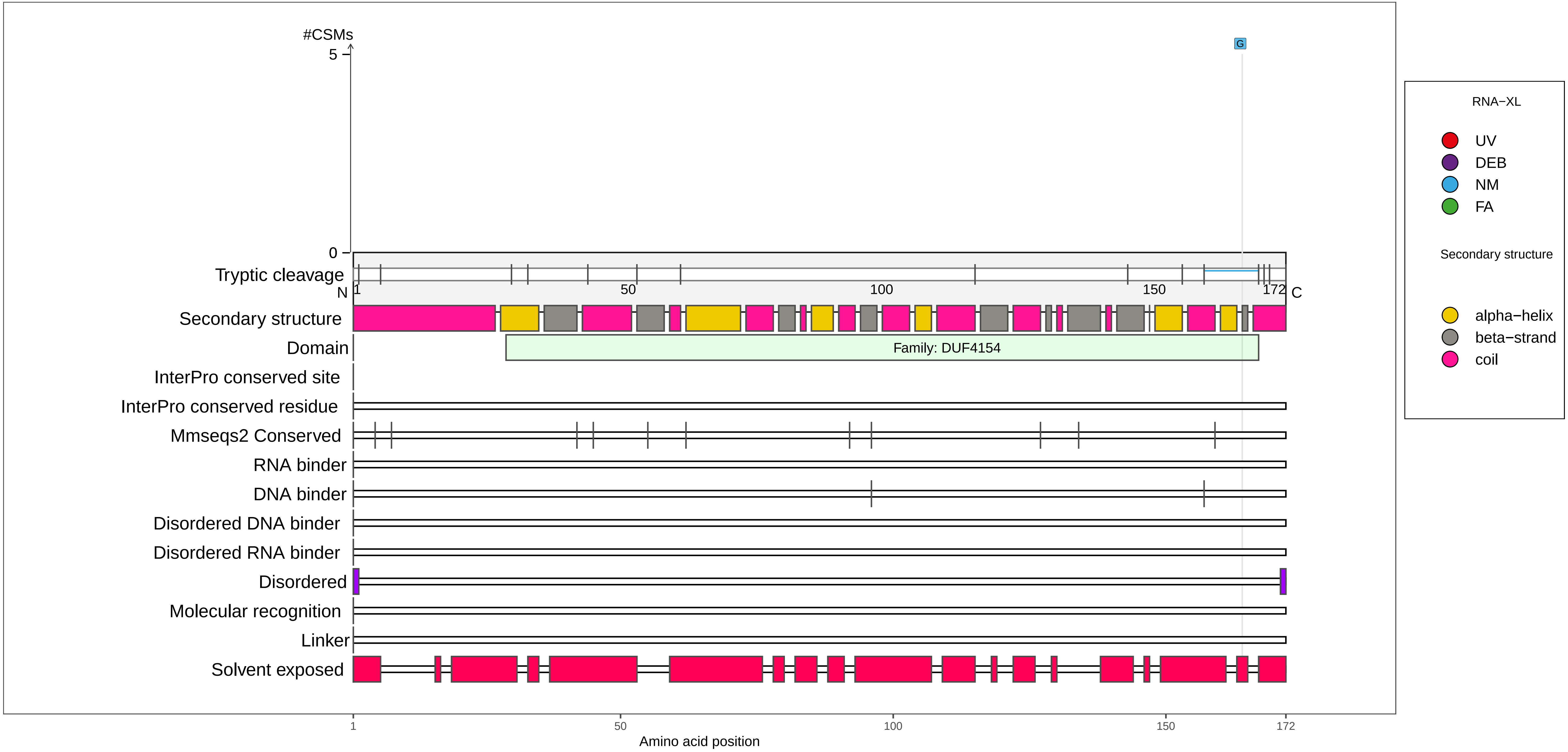

P66948
BEPA_ECOLI Beta−barrel assembly−enhancing protease

− Abundance:
tryptic [log10 Intensity]: 8.42 (Q 73)
PAXdb K12 strain [ppm]: 1.42 (Q 34)
PAXdb E.coli [ppm]: 1.88 (Q 69)

− RNA functions: not annotated

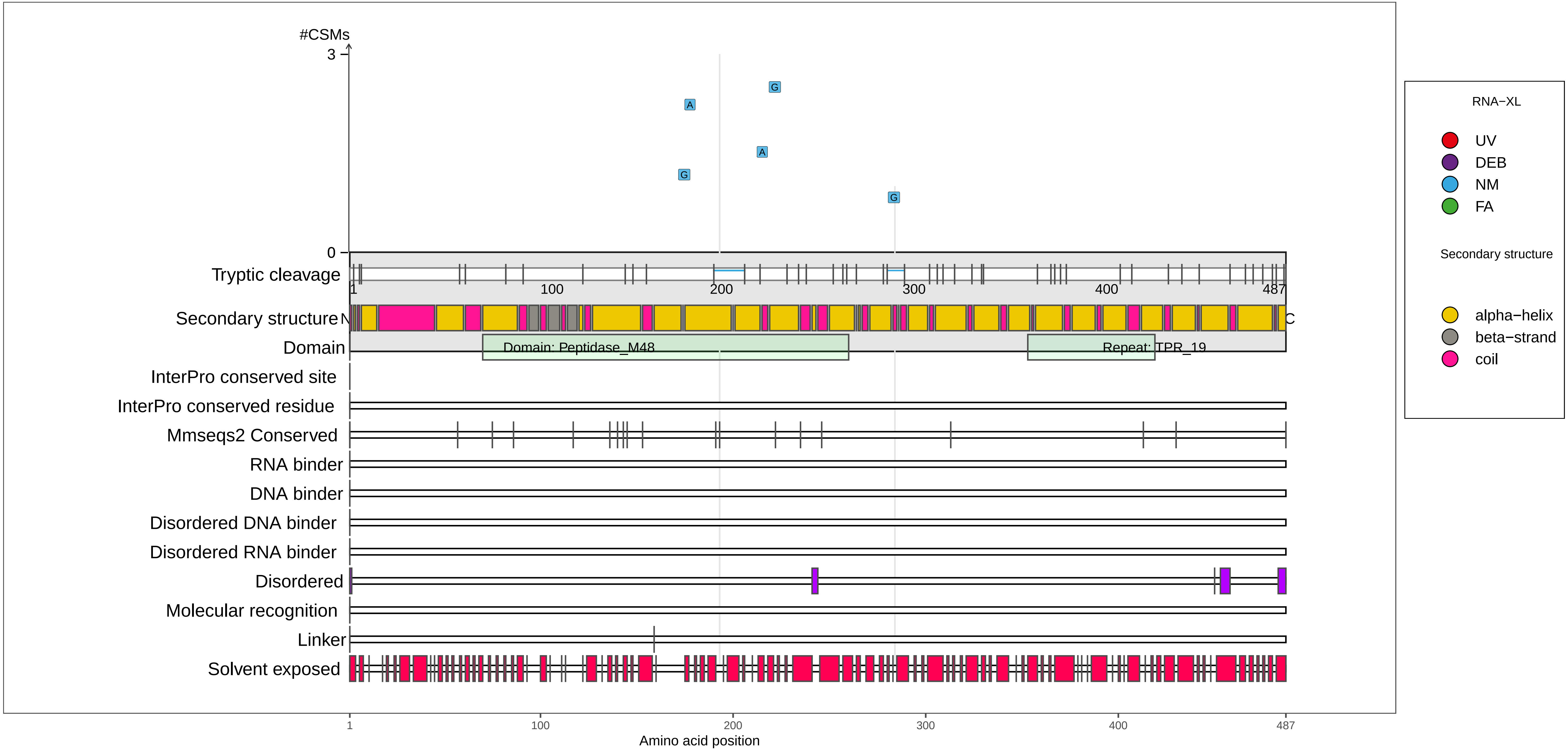

P75691
YAHK_ECOLI Aldehyde reductase YahK

− Abundance:
tryptic [log10 Intensity]: 7.2 (Q 21)
PAXdb K12 strain [ppm]: 0.83 (Q 6)
PAXdb E.coli [ppm]: 3.07 (Q 95)

− RNA functions: not annotated

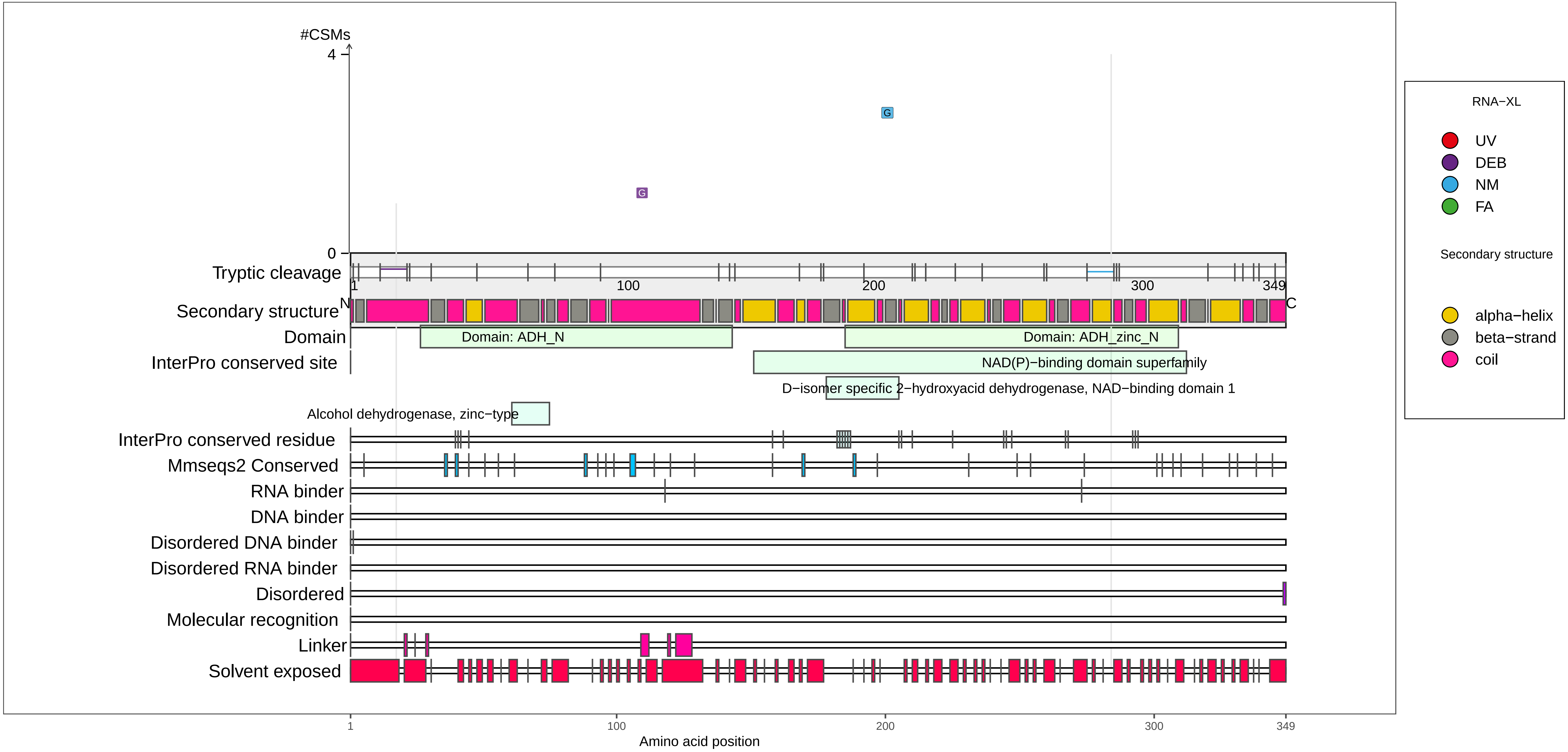

P75797
GSIB_ECOLI Glutathione−binding protein GsiB

− Abundance:
tryptic [log10 Intensity]: 7.86 (Q 52)
PAXdb K12 strain [ppm]: 1.33 (Q 28)
PAXdb E.coli [ppm]: 2.4 (Q 83)

− RNA functions: not annotated

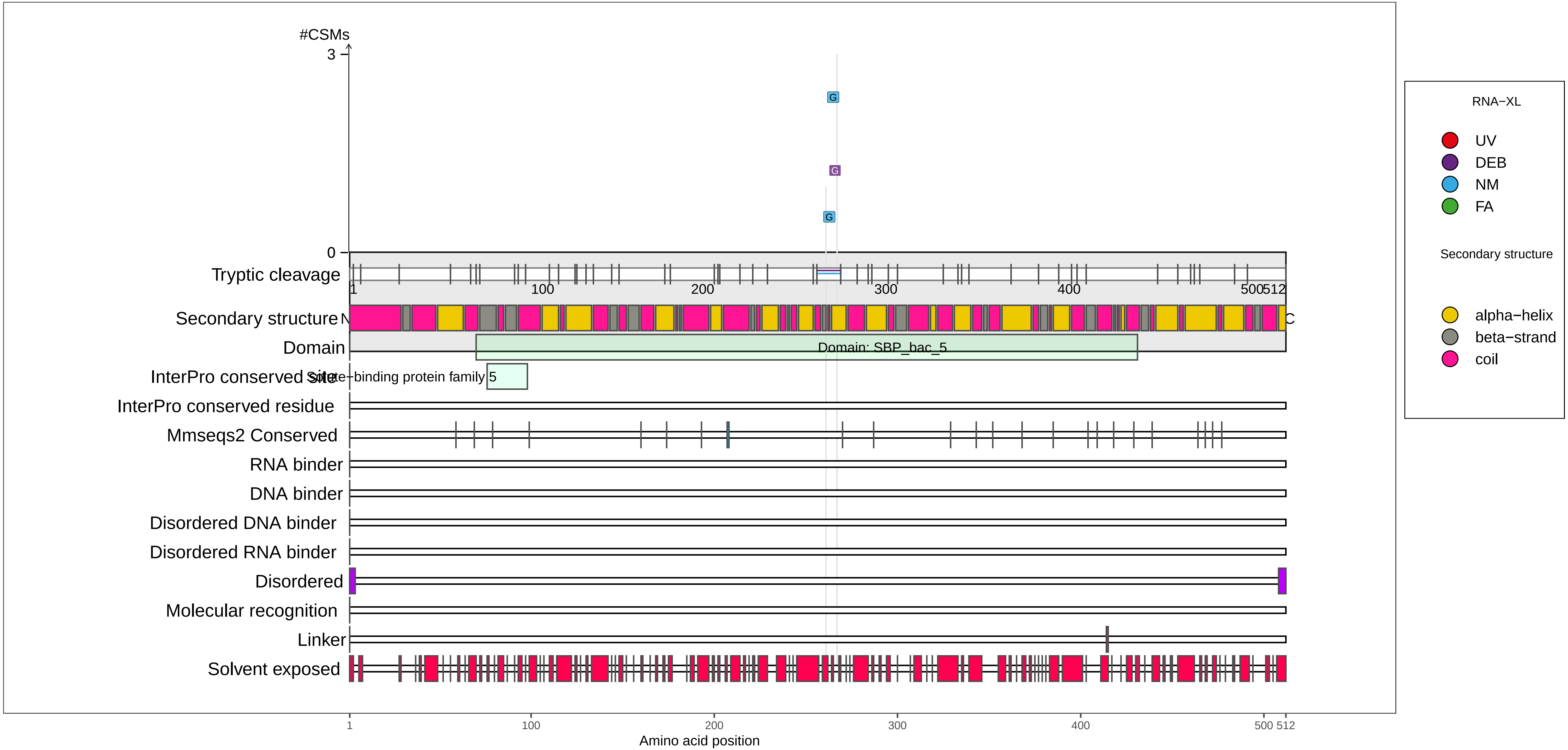

P75938
FLGF_ECOLI Flagellar basal−body rod protein FlgF

− Abundance:
tryptic [log10 Intensity]: 7.14 (Q 18)
PAXdb K12 strain [ppm]: NA (Q NA)
PAXdb E.coli [ppm]: 0.35 (Q 32)

− RNA functions: not annotated

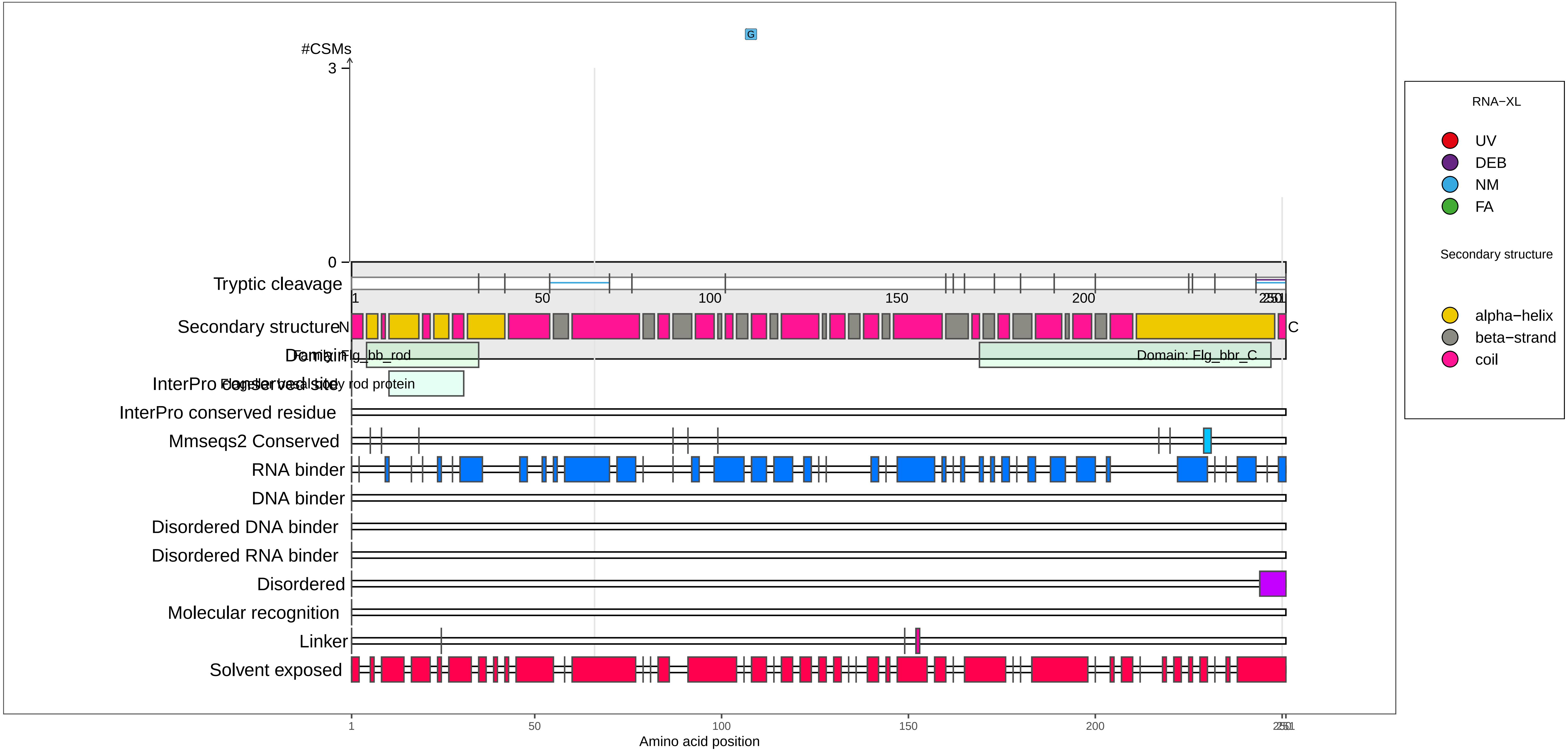

P76104
RLHA_ECOLI 23S rRNA 5−hydroxycytidine C2501 synthase

− Abundance:
tryptic [log10 Intensity]: 6.91 (Q 10)
PAXdb K12 strain [ppm]: 0.69 (Q 3)
PAXdb E.coli [ppm]: 1.41 (Q 57)

− RNA functions:
ncRNA metabolic process; ncRNA processing; RNA metabolic process; RNA modification
RNA processing; rRNA metabolic process; rRNA modification; rRNA processing

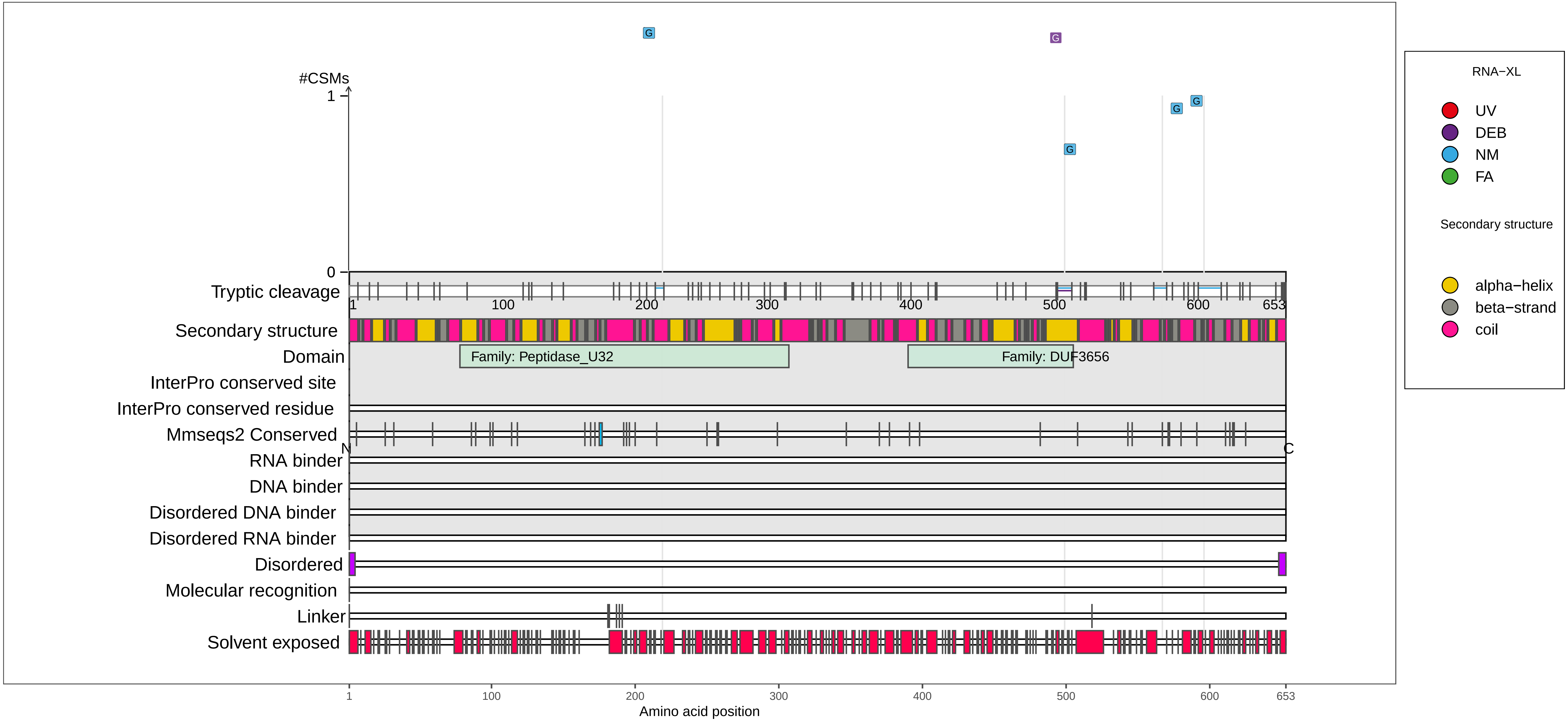

P76297
FLHE_ECOLI Flagellar protein FlhE

− Abundance:
tryptic [log10 Intensity]: 7.9 (Q 54)
PAXdb K12 strain [ppm]: 1.71 (Q 50)
PAXdb E.coli [ppm]: 0.24 (Q 30)

− RNA functions: not annotated

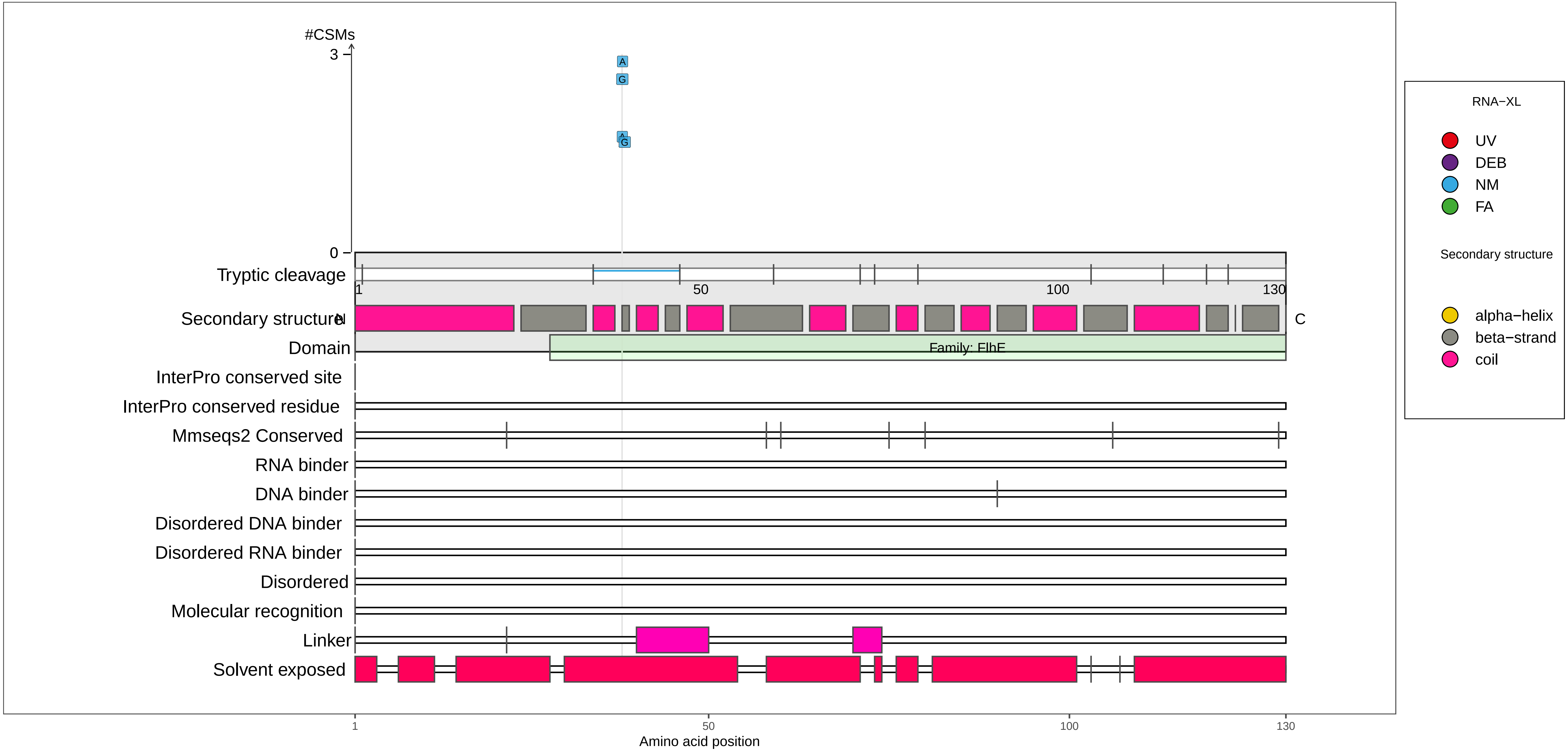

P76403
TRHP_ECOLI tRNA hydroxylation protein P
− RNA functions: not annotated

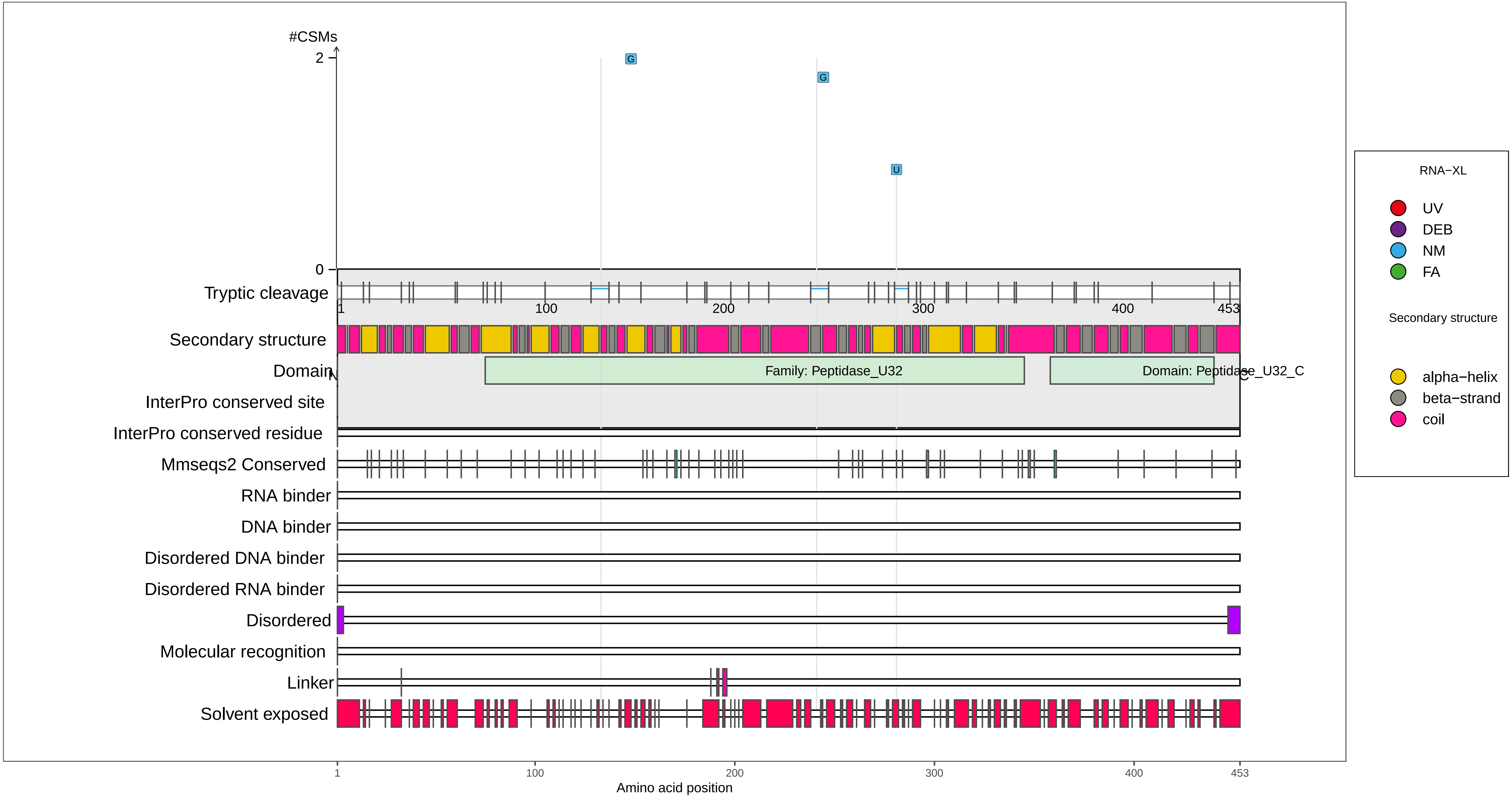

P03023
LACI_ECOLI Lactose operon repressor

− Abundance:
tryptic [log10 Intensity]: 7.7 (Q 44)
PAXdb K12 strain [ppm]: 2.67 (Q 82)
PAXdb E.coli [ppm]: 1.28 (Q 54)

− RNA functions: not annotated

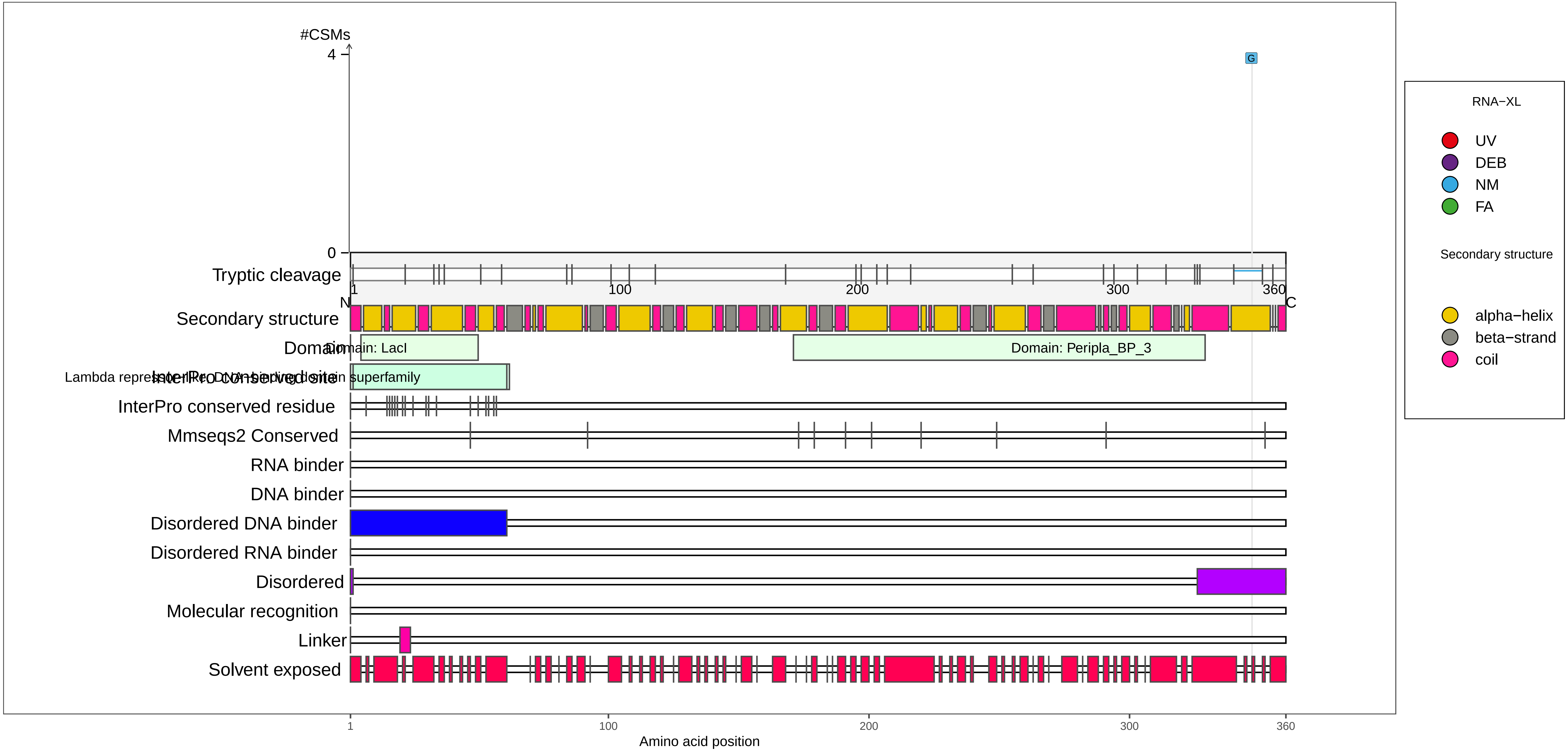

P04391
OTC1_ECOLI Ornithine carbamoyltransferase subunit I

− Abundance:
tryptic [log10 Intensity]: 8.49 (Q 75)
PAXdb K12 strain [ppm]: 0.43 (Q 1)
PAXdb E.coli [ppm]: 2.36 (Q 82)

− RNA functions: not annotated

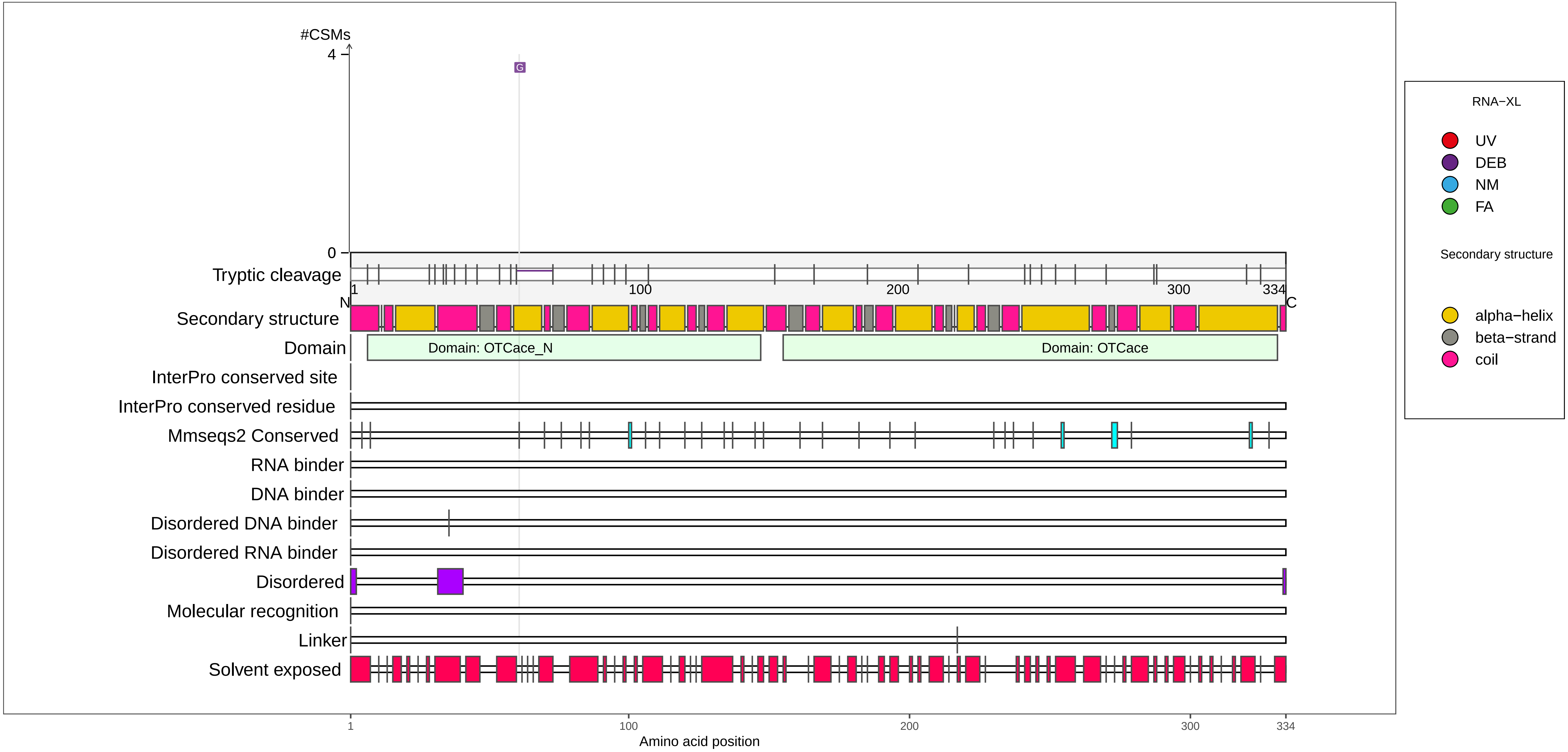

P04825
AMPN_ECOLI Aminopeptidase N

− Abundance:
tryptic [log10 Intensity]: 9.08 (Q 90)
PAXdb K12 strain [ppm]: 2.82 (Q 86)
PAXdb E.coli [ppm]: 2.72 (Q 90)

− RNA functions: not annotated

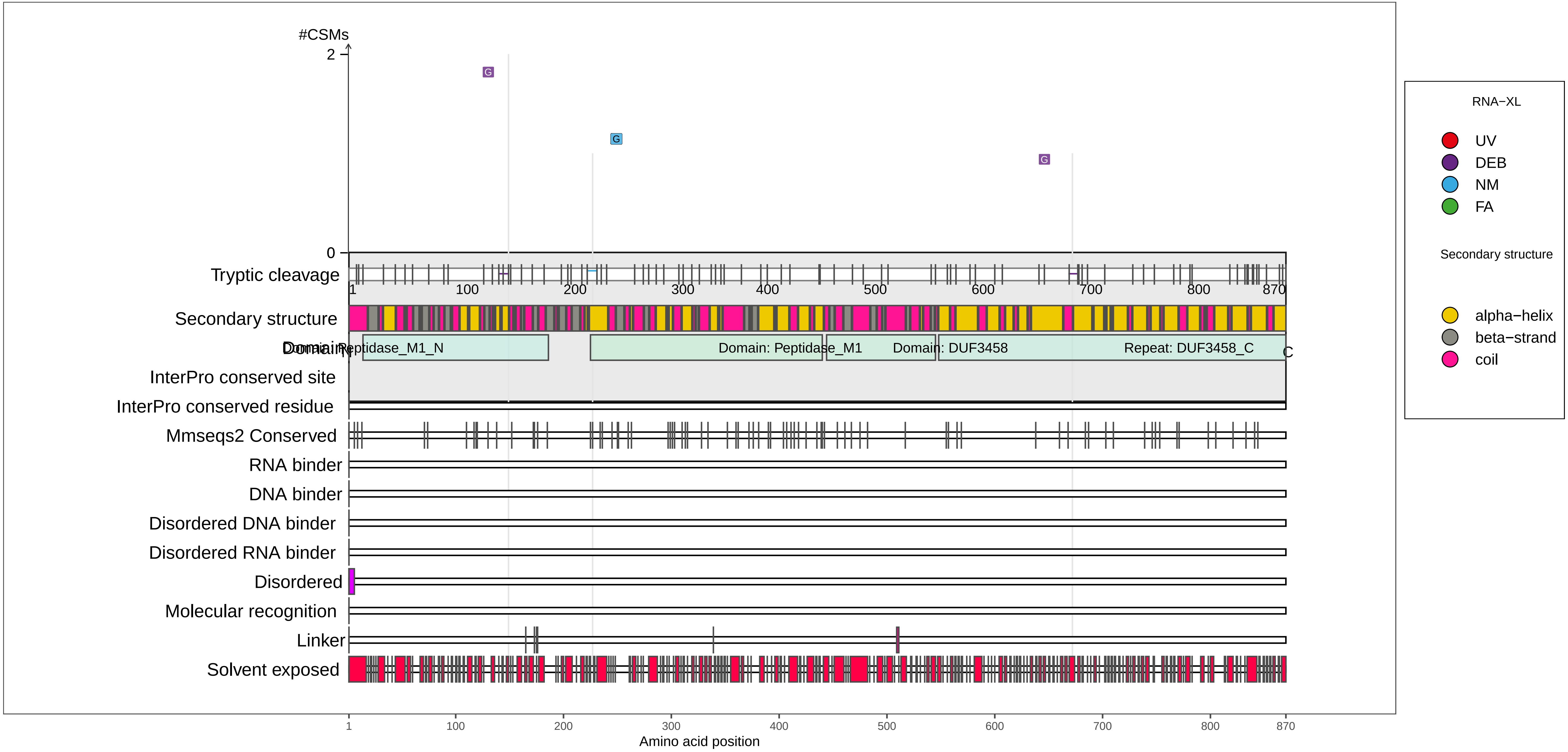

P07017
MCP2_ECOLI Methyl−accepting chemotaxis protein II

− Abundance:
tryptic [log10 Intensity]: NA (Q NA)
PAXdb K12 strain [ppm]: 1.23 (Q 22)
PAXdb E.coli [ppm]: 2.57 (Q 87)

− RNA functions: not annotated

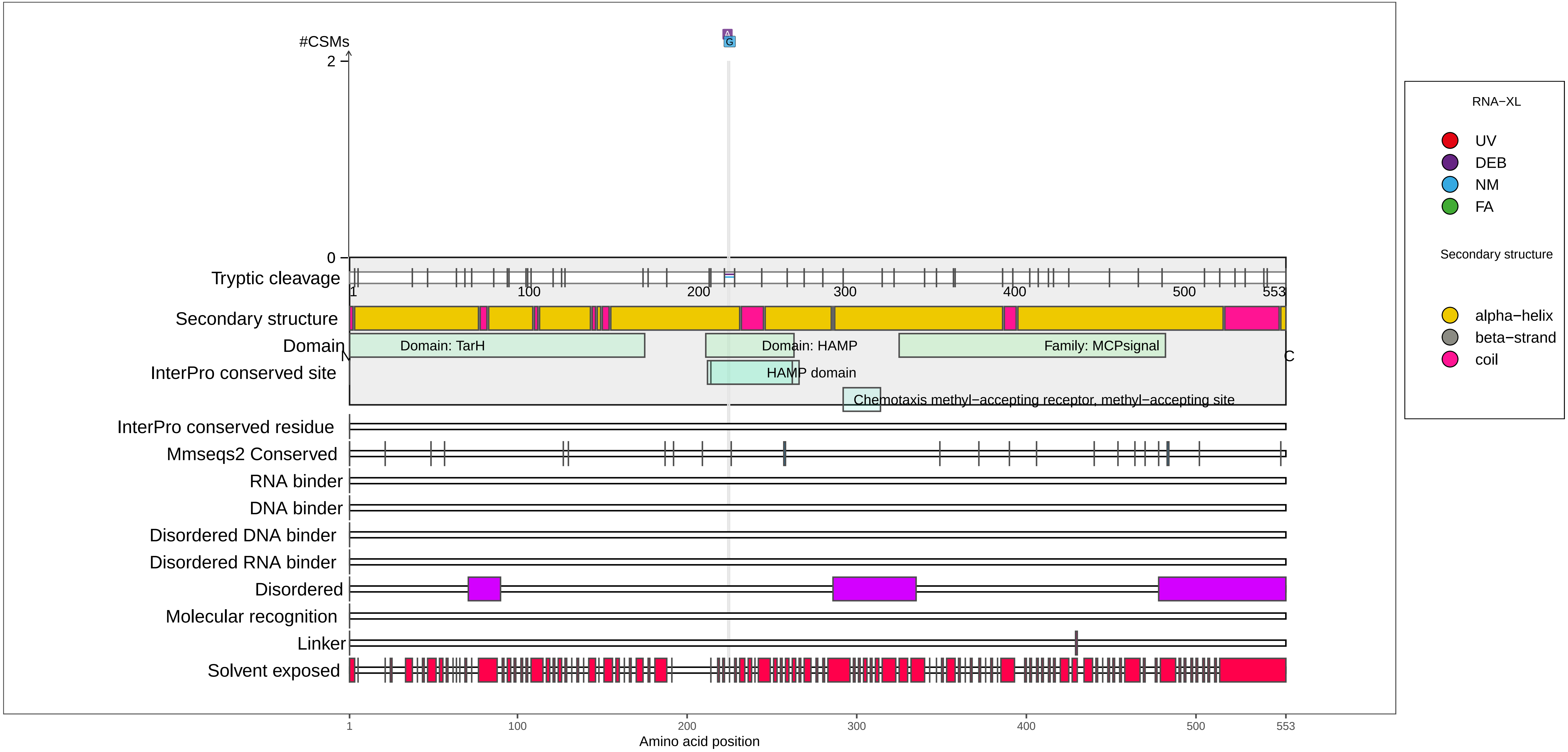

P07395
SYFB_ECOLI Phenylalanine−−tRNA ligase beta subunit

− Abundance:
tryptic [log10 Intensity]: 9.33 (Q 94)
PAXdb K12 strain [ppm]: 2.87 (Q 88)
PAXdb E.coli [ppm]: 2.42 (Q 83)

− RNA functions:
aminoacyl−tRNA ligase activity; ncRNA metabolic process
phenylalanine−tRNA ligase activity; phenylalanine−tRNA ligase complex; phenylalanyl−tRNA aminoacylation
Putative tRNA binding domain; RNA binding; RNA metabolic process; tRNA aminoacylation
tRNA aminoacylation for protein translation; tRNA binding; tRNA metabolic process
tRNA synthetase B5 domain

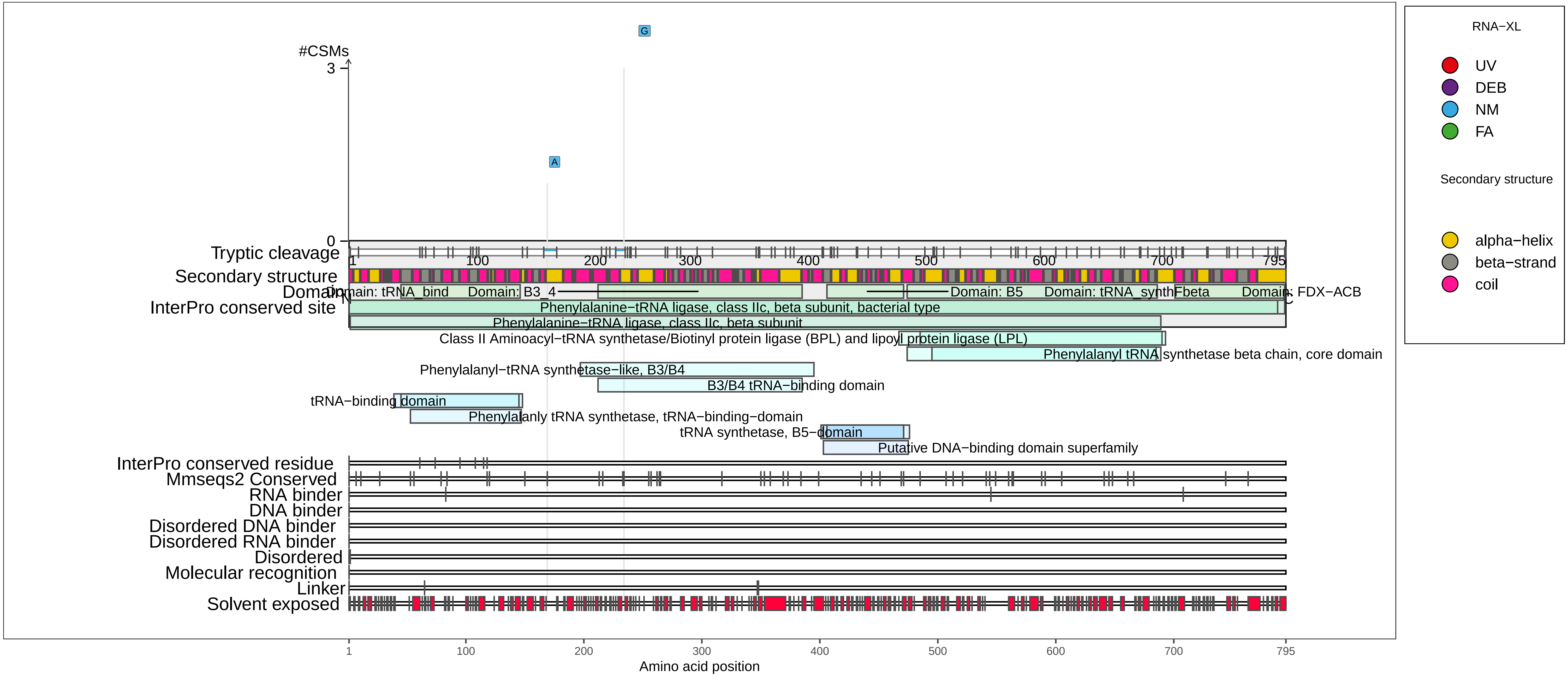

P0A738
MOAC_ECOLI Cyclic pyranopterin monophosphate synthase

− Abundance:
tryptic [log10 Intensity]: 7.68 (Q 44)
PAXdb K12 strain [ppm]: NA (Q NA)
PAXdb E.coli [ppm]: 2.12 (Q 75)

− RNA functions: not annotated

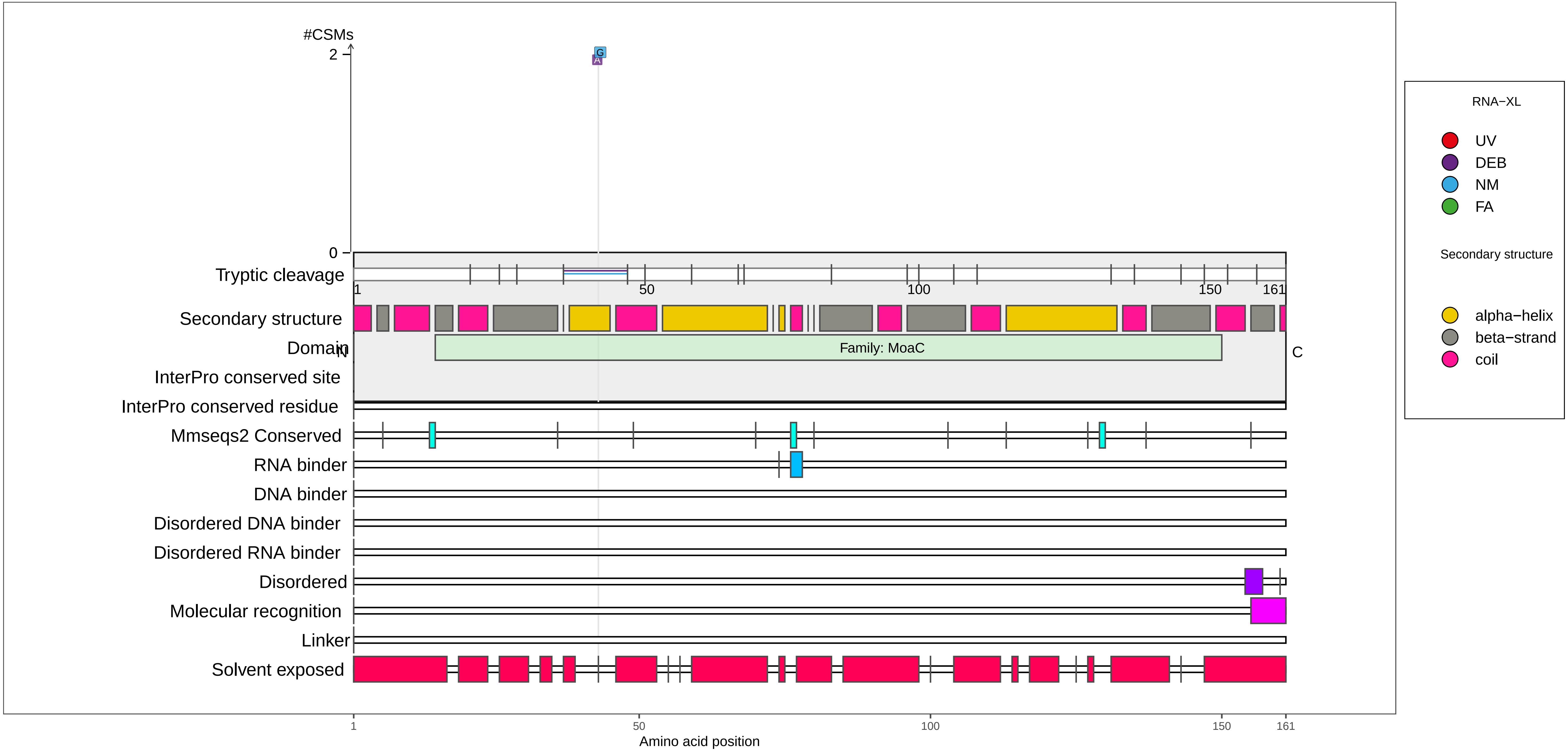

P0A780
NUSB_ECOLI Transcription antitermination protein NusB

− Abundance:
tryptic [log10 Intensity]: 8.94 (Q 87)
PAXdb K12 strain [ppm]: 2.45 (Q 76)
PAXdb E.coli [ppm]: 2.28 (Q 80)

− RNA functions:
RNA binding; RNA biosynthetic process; RNA metabolic process

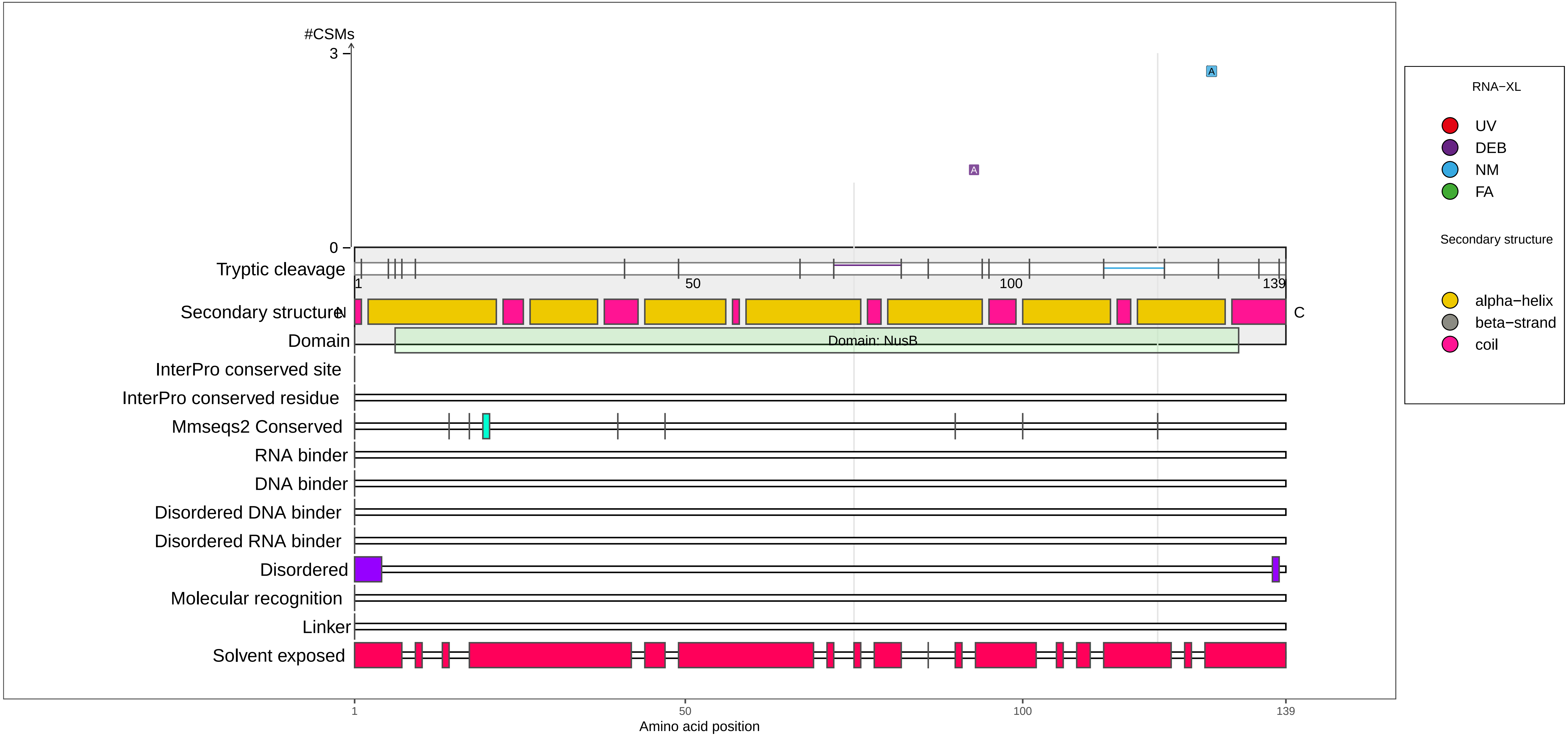

P0A7D1
PTH_ECOLI Peptidyl−tRNA hydrolase
− Abundance:
tryptic [log10 Intensity]: 7.65 (Q 43)
PAXdb K12 strain [ppm]: 1.88 (Q 56)
PAXdb E.coli [ppm]: 1.95 (Q 71)

− RNA functions:
aminoacyl−tRNA hydrolase activity; Peptidyl−tRNA hydrolase

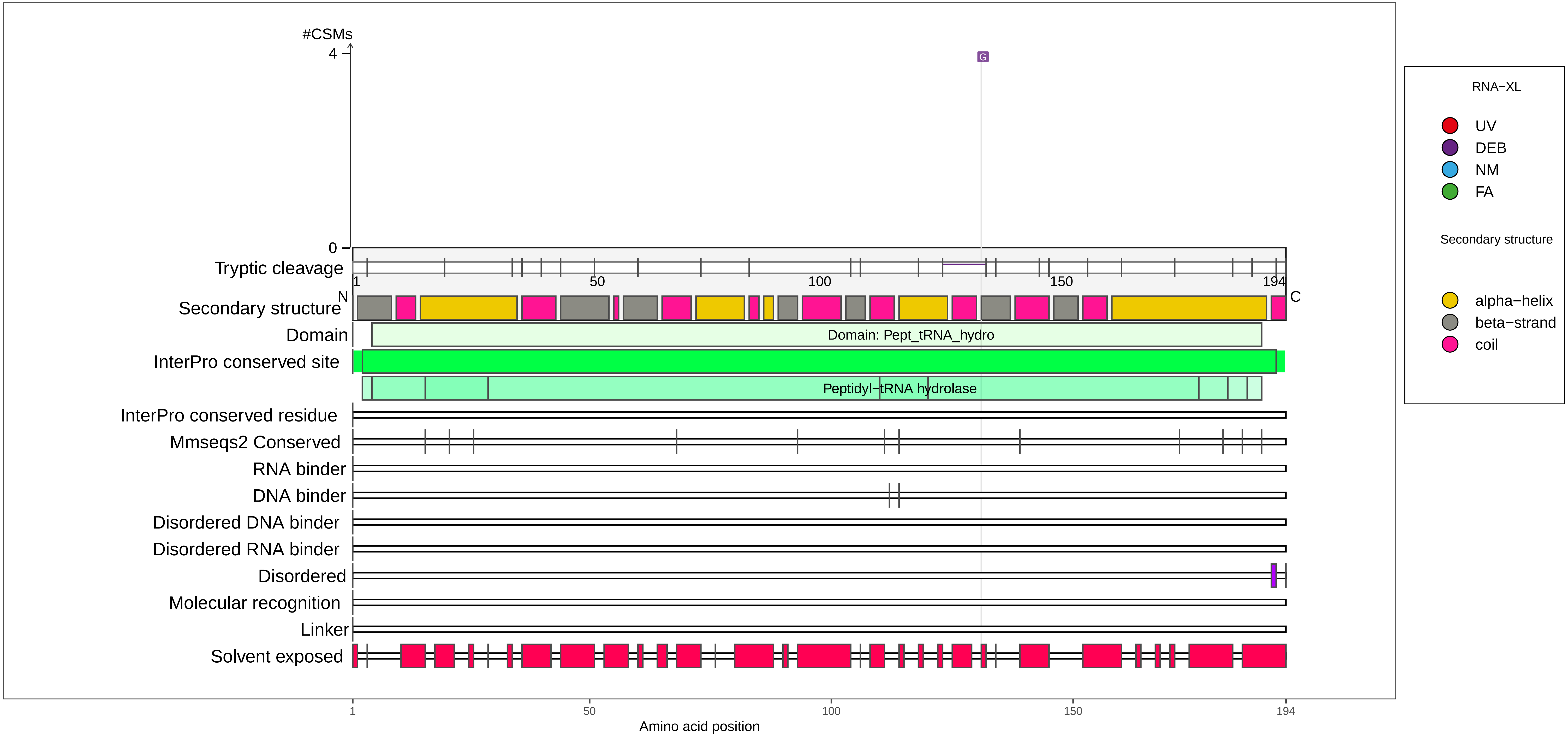

P0A7I0
RF1_ECOLI Peptide chain release factor RF1

− Abundance:
tryptic [log10 Intensity]: 7.99 (Q 57)
PAXdb K12 strain [ppm]: 2.29 (Q 70)
PAXdb E.coli [ppm]: 1.84 (Q 68)

− RNA functions:
RNA binding

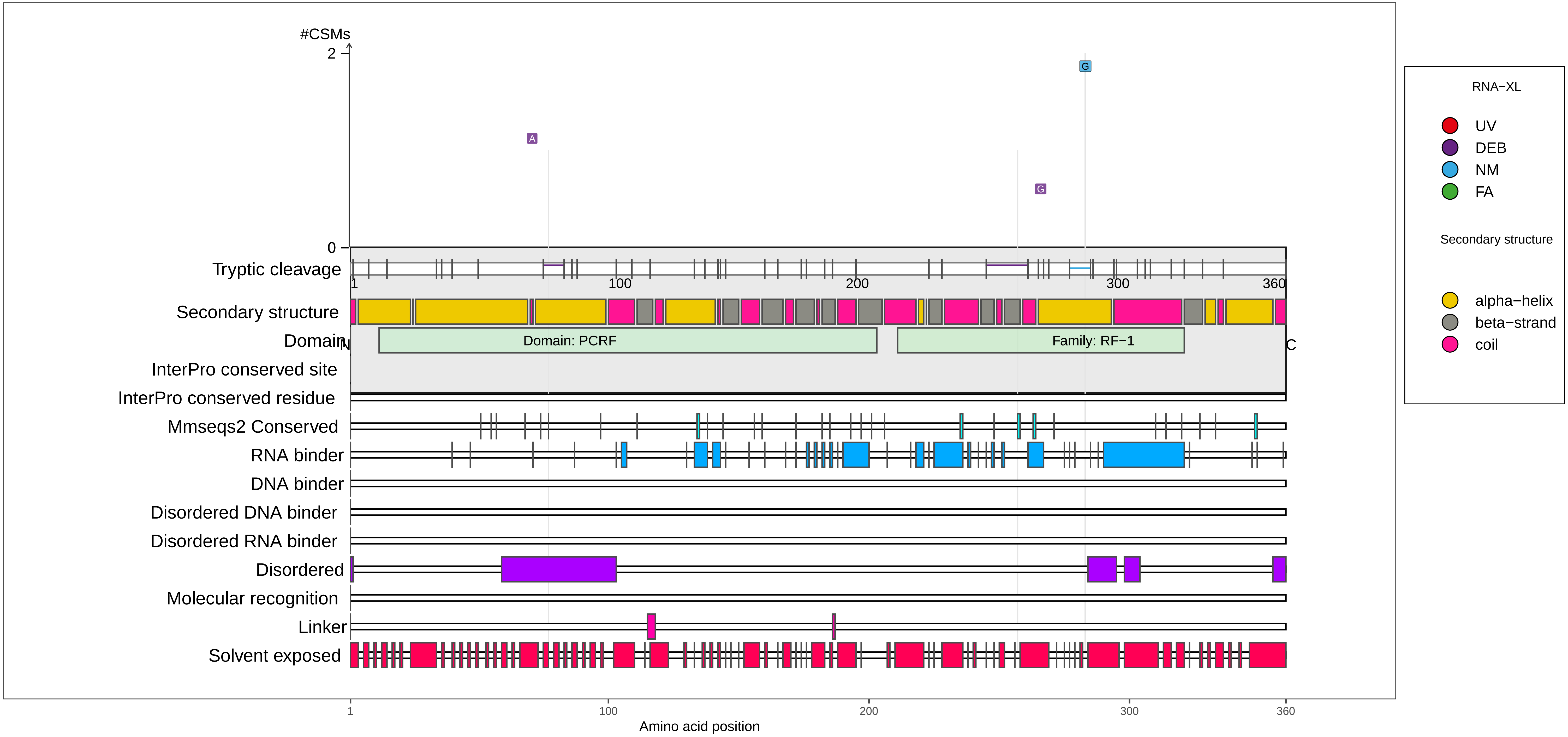

P0A8Q3
FRDD_ECOLI Fumarate reductase subunit D

− Abundance:
tryptic [log10 Intensity]: 7.2 (Q 21)
PAXdb K12 strain [ppm]: NA (Q NA)
PAXdb E.coli [ppm]: 0.37 (Q 33)

− RNA functions: not annotated

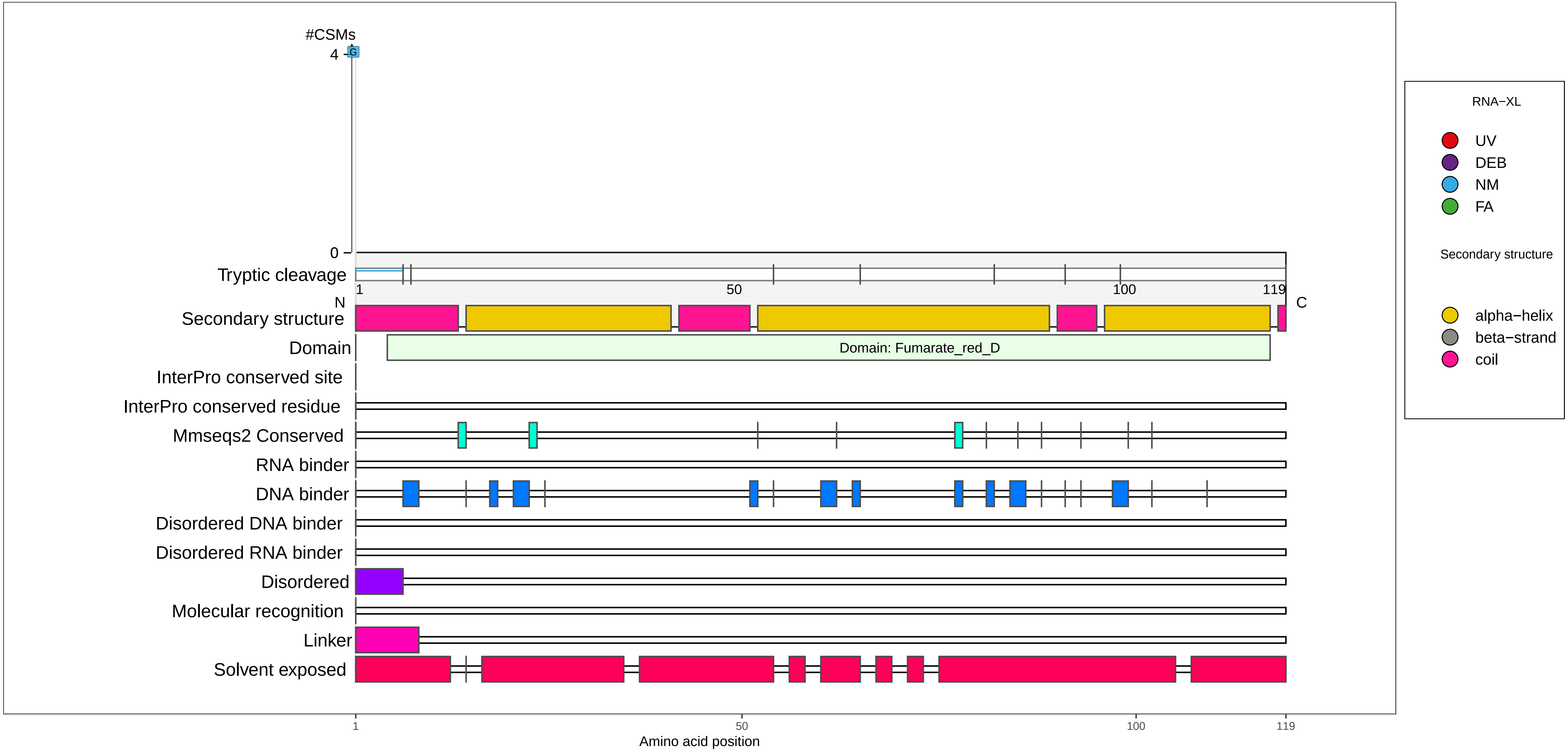

P0A8T1
PRMA_ECOLI Ribosomal protein L11 methyltransferase

− Abundance:
tryptic [log10 Intensity]: 7.82 (Q 50)
PAXdb K12 strain [ppm]: 2.26 (Q 69)
PAXdb E.coli [ppm]: 2.15 (Q 76)

− RNA functions: not annotated

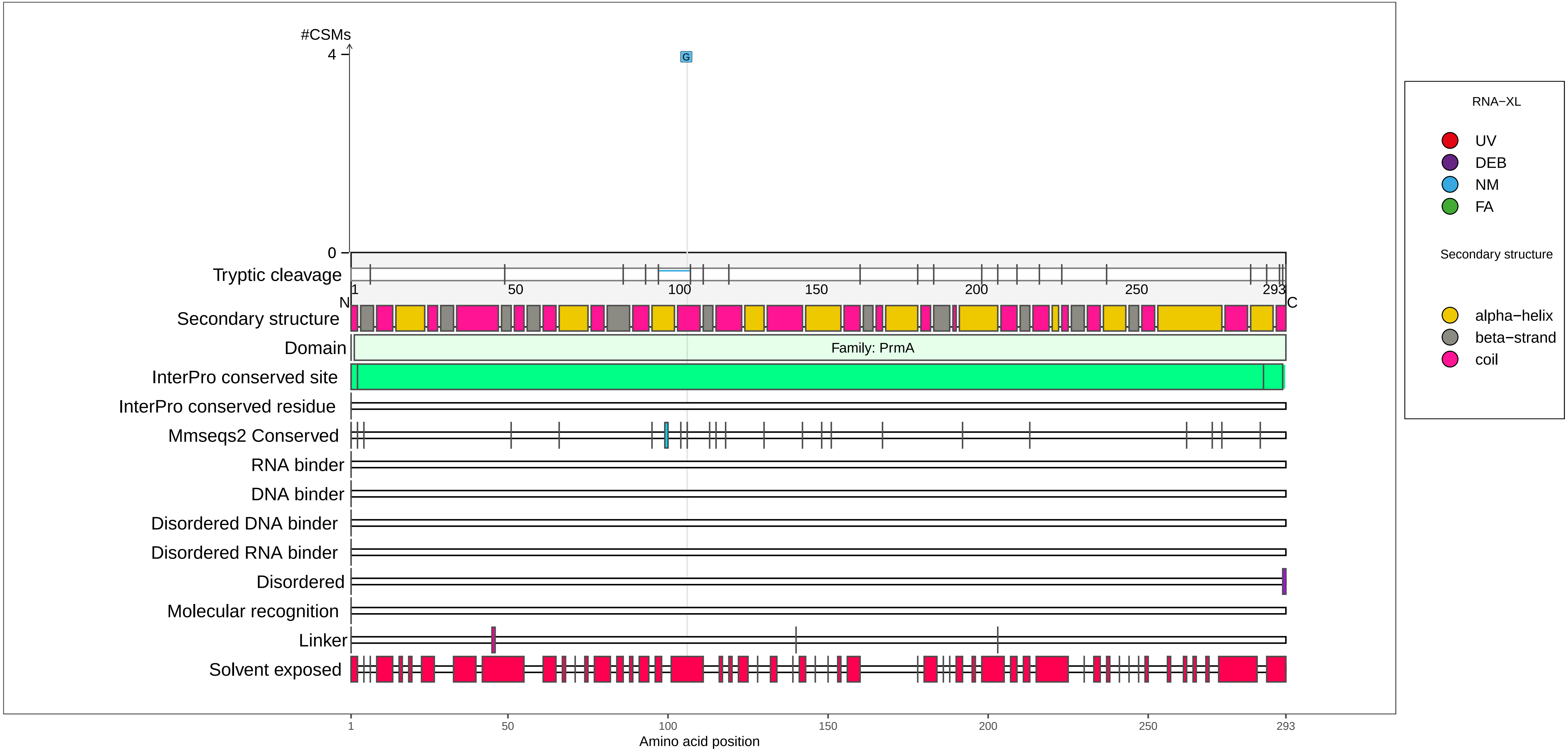

P0A948
RIMJ_ECOLI [Ribosomal protein S5]−alanine N−acetyltransferase

− Abundance:
tryptic [log10 Intensity]: 6.95 (Q 12)
PAXdb K12 strain [ppm]: 1.37 (Q 30)
PAXdb E.coli [ppm]: 1.28 (Q 54)

− RNA functions:
ncRNA metabolic process; ncRNA processing; RNA metabolic process; RNA processing
rRNA metabolic process; rRNA processing

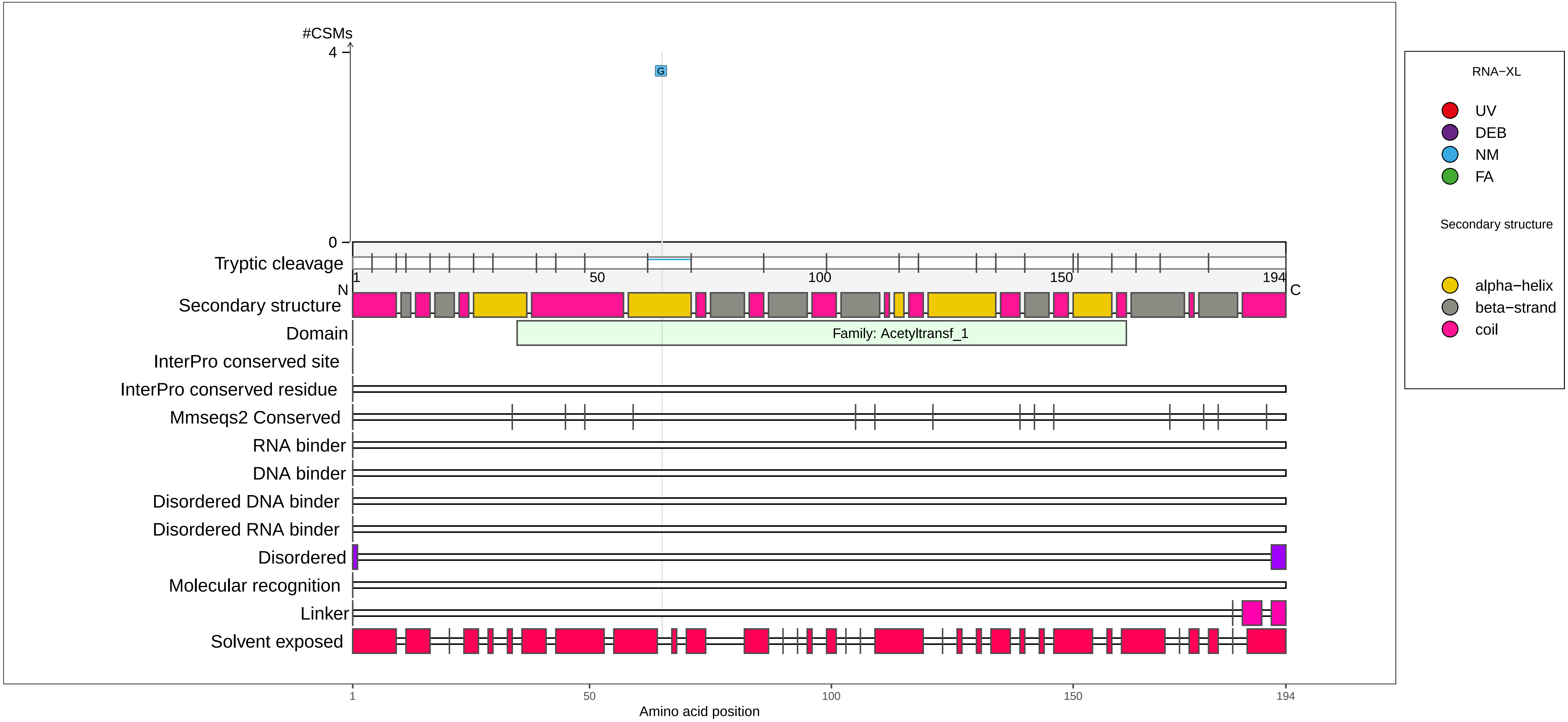

P0A988
DPO3B_ECOLI Beta sliding clamp

− Abundance:
tryptic [log10 Intensity]: 8.46 (Q 74)
PAXdb K12 strain [ppm]: 2.27 (Q 69)
PAXdb E.coli [ppm]: 2.18 (Q 77)

− RNA functions: not annotated

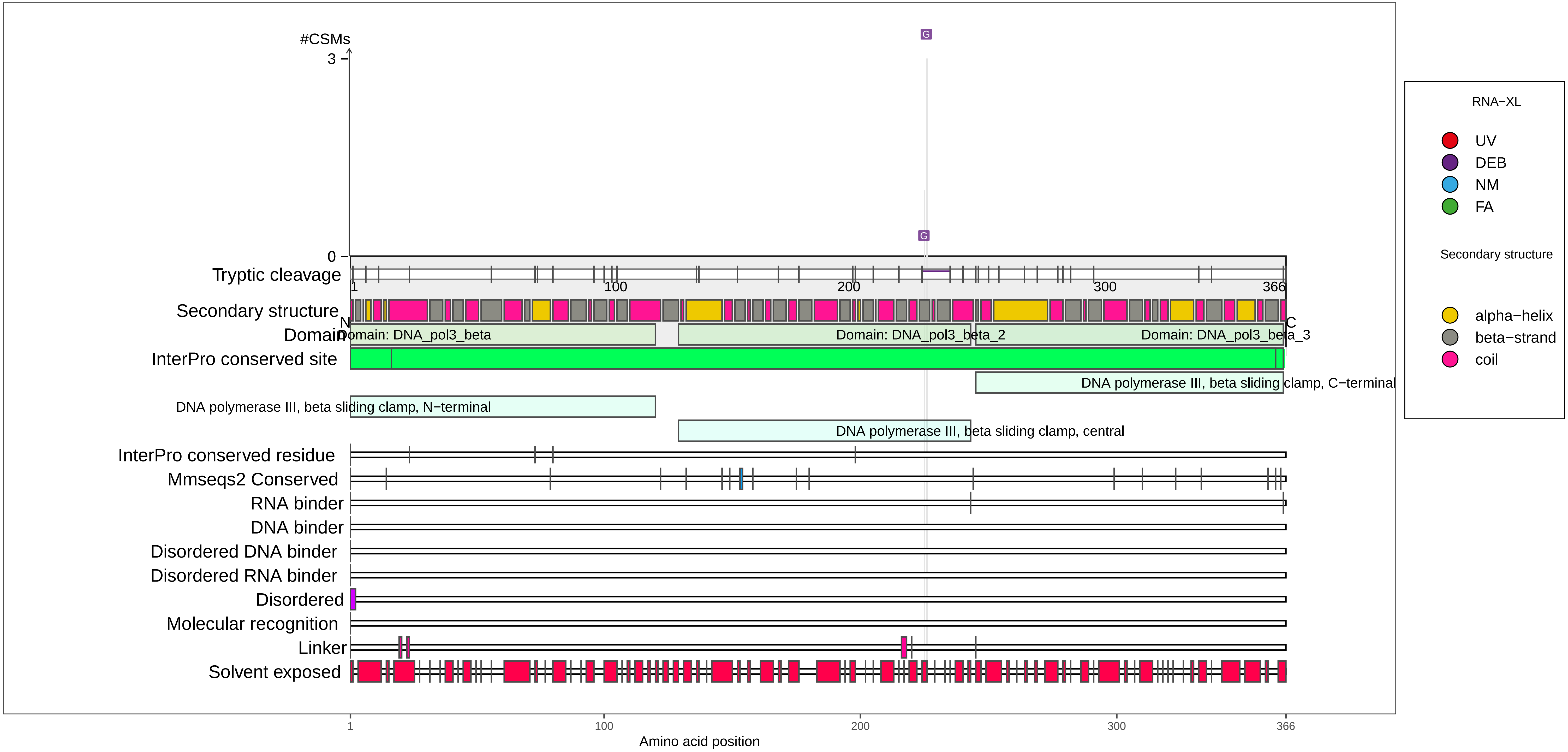

P0A993
F16PA_ECOLI Fructose−1,6−bisphosphatase class 1

− Abundance:
tryptic [log10 Intensity]: 8.38 (Q 72)
PAXdb K12 strain [ppm]: 2.74 (Q 84)
PAXdb E.coli [ppm]: 2.31 (Q 80)

− RNA functions: not annotated

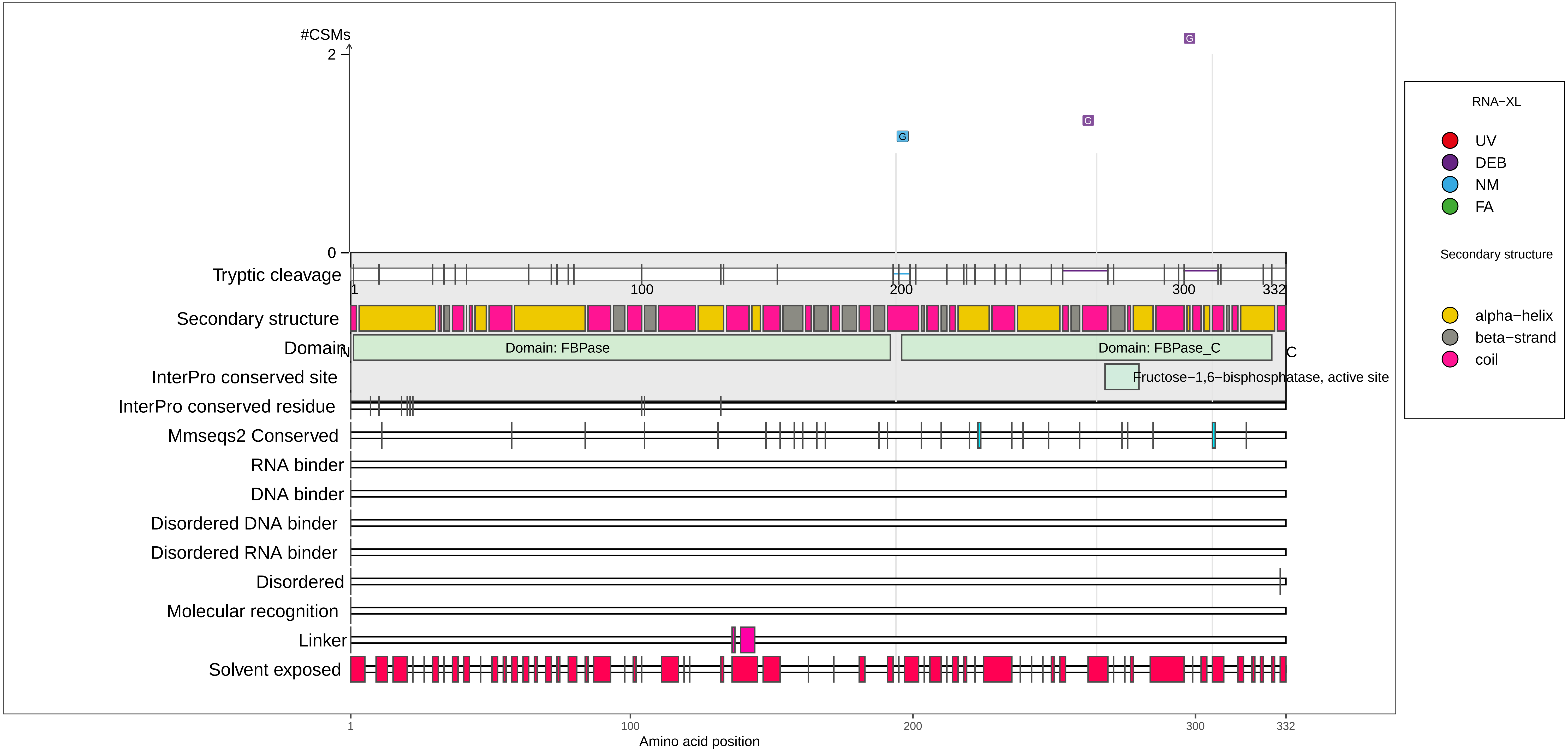

P0AC19
FOLX_ECOLI Dihydroneopterin triphosphate 2'−epimerase

− Abundance:
tryptic [log10 Intensity]: 8.82 (Q 84)
PAXdb K12 strain [ppm]: 2.2 (Q 67)
PAXdb E.coli [ppm]: 2.24 (Q 78)

− RNA functions: not annotated

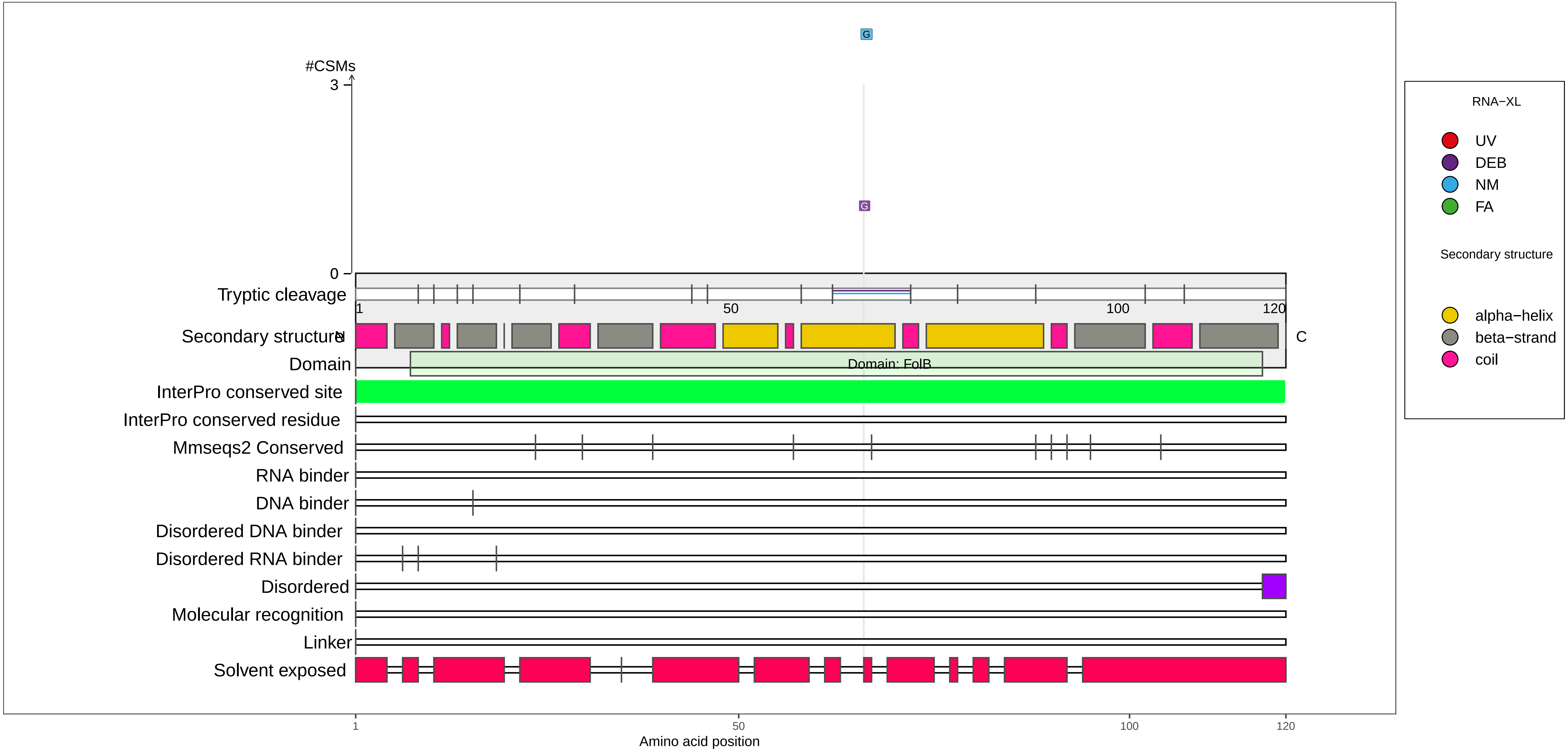

P0AD10
YECJ_ECOLI Uncharacterized protein YecJ

− Abundance:
tryptic [log10 Intensity]: 7.43 (Q 32)
PAXdb K12 strain [ppm]: NA (Q NA)
PAXdb E.coli [ppm]: 2.11 (Q 75)

− RNA functions: not annotated

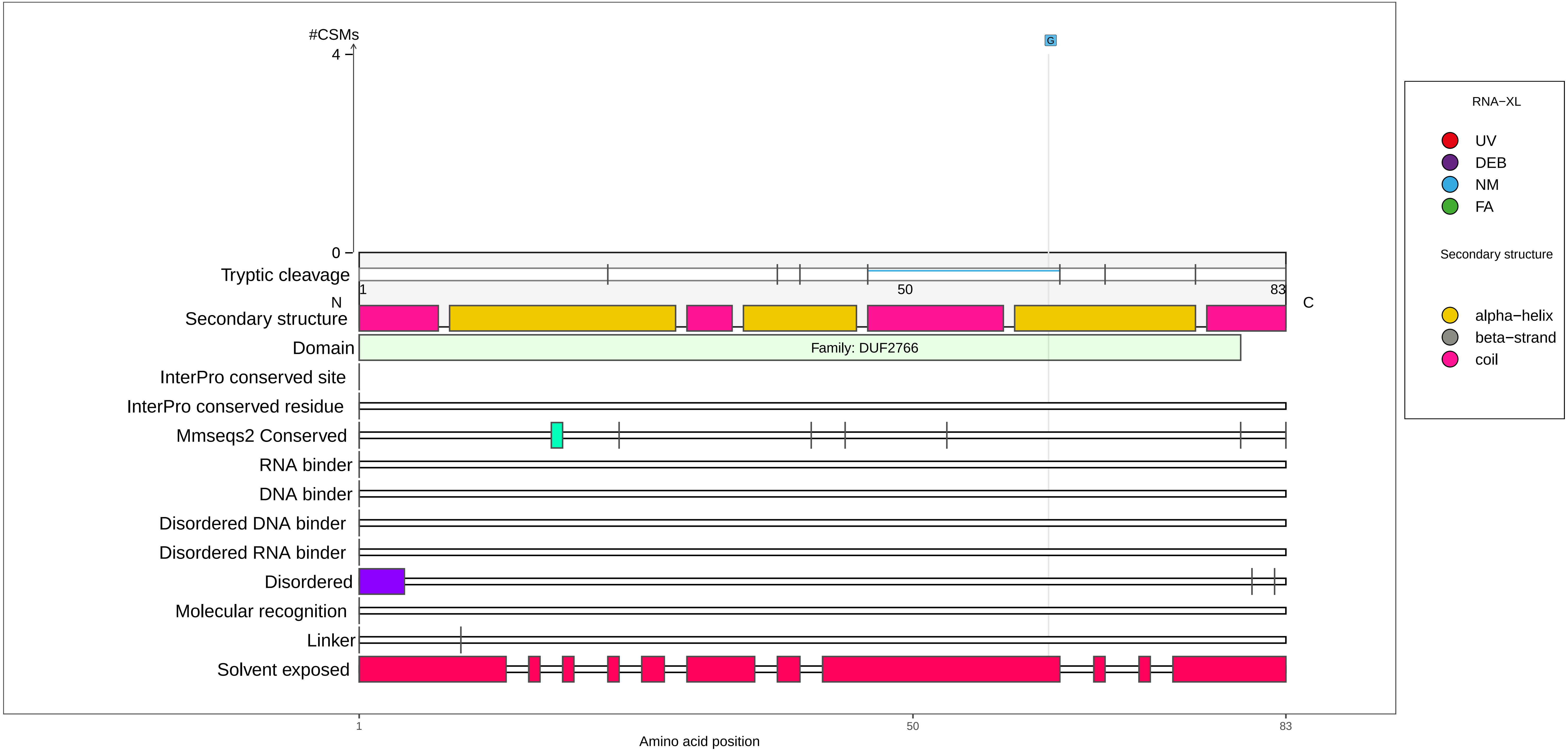

P0ADF6
EDD_ECOLI Phosphogluconate dehydratase

− Abundance:
tryptic [log10 Intensity]: 8.09 (Q 61)
PAXdb K12 strain [ppm]: 2.06 (Q 62)
PAXdb E.coli [ppm]: 1.61 (Q 63)

− RNA functions: not annotated

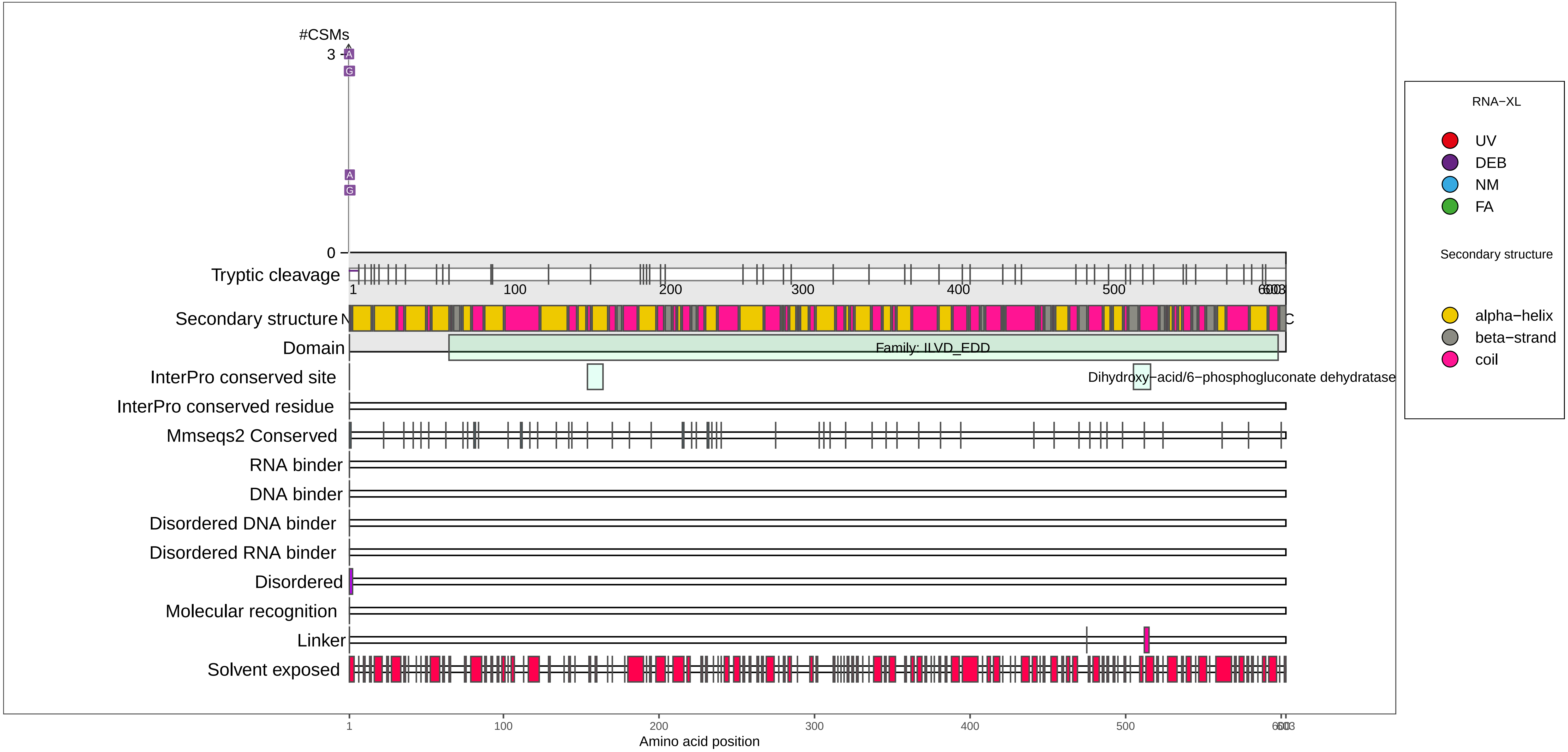

P0ADS6
YGGE_ECOLI Uncharacterized protein YggE

− Abundance:
tryptic [log10 Intensity]: 8.44 (Q 73)
PAXdb K12 strain [ppm]: 1.62 (Q 45)
PAXdb E.coli [ppm]: 2.67 (Q 89)

− RNA functions: not annotated

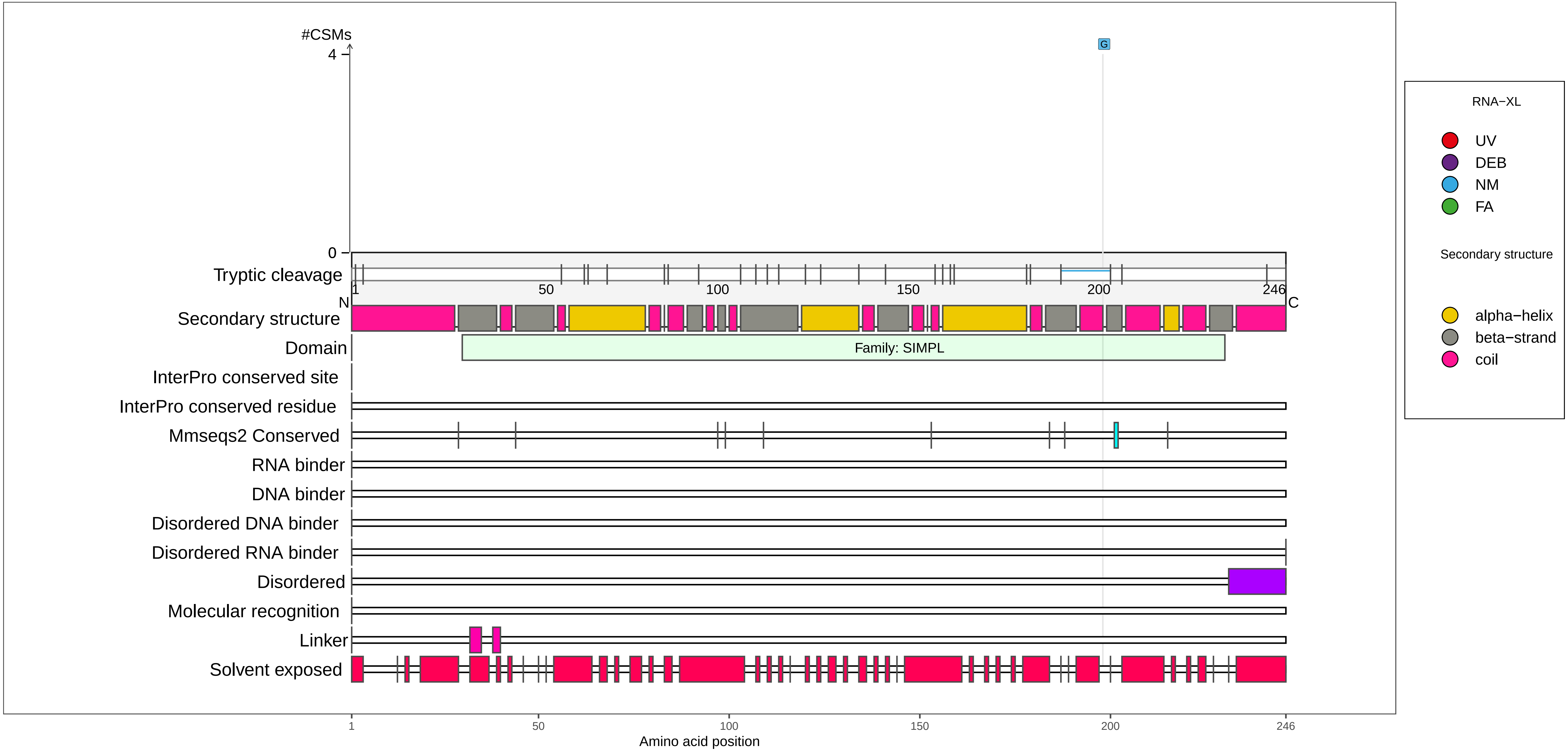

P0ADX9
RSMD_ECOLI Ribosomal RNA small subunit methyltransferase D

− Abundance:
tryptic [log10 Intensity]: NA (Q NA)
PAXdb K12 strain [ppm]: 1.4 (Q 33)
PAXdb E.coli [ppm]: 1.55 (Q 61)

− RNA functions:
16S rRNA (guanine(966)−N(2))−methyltransferase activity; ncRNA metabolic process
ncRNA processing; RNA metabolic process; RNA methylation; RNA methyltransferase activity
RNA modification; RNA processing; rRNA (guanine−N2−)−methyltransferase activity
rRNA (guanine) methyltransferase activity; rRNA base methylation; rRNA metabolic process
rRNA methylation; rRNA methyltransferase activity; rRNA modification; rRNA processing

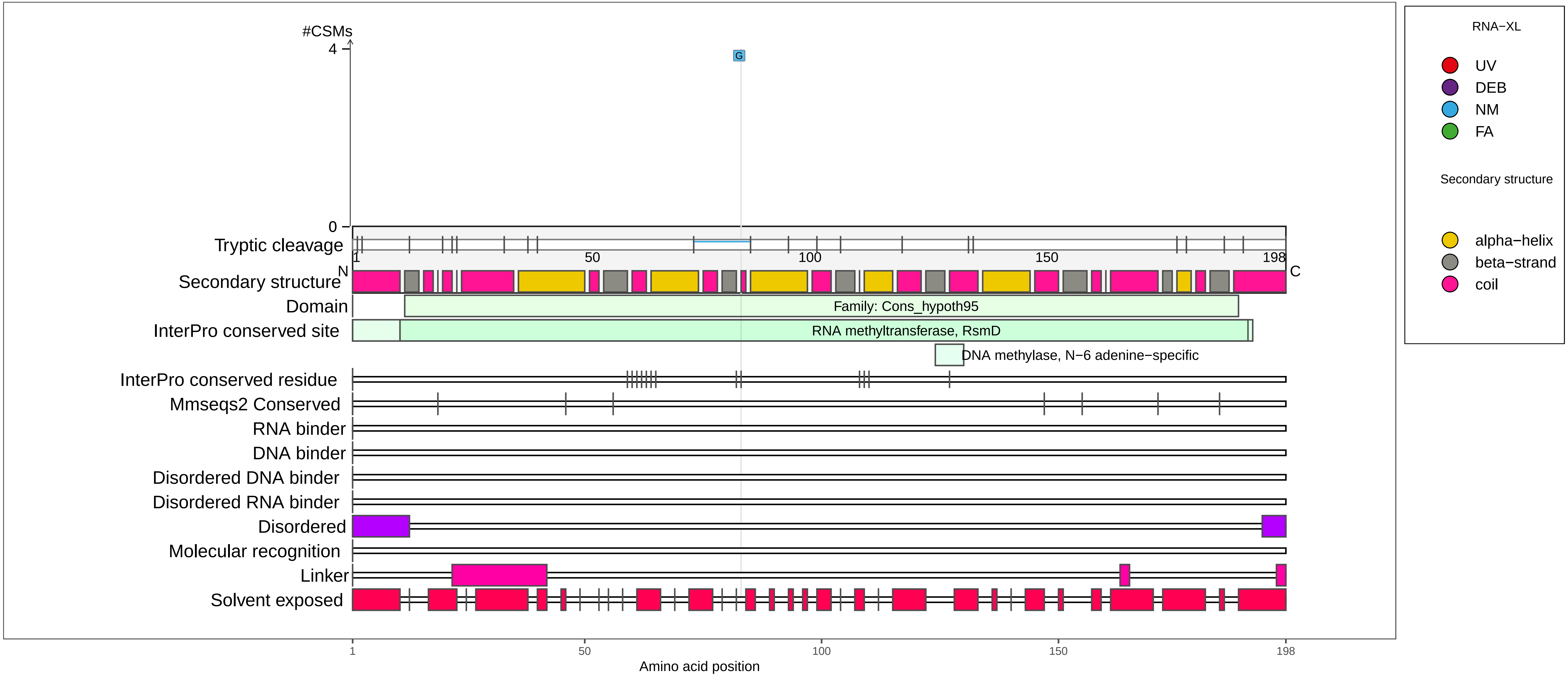

P0AE12
AMN_ECOLI AMP nucleosidase

− Abundance:
tryptic [log10 Intensity]: 8.4 (Q 72)
PAXdb K12 strain [ppm]: 2.1 (Q 64)
PAXdb E.coli [ppm]: 1.7 (Q 64)

− RNA functions: not annotated

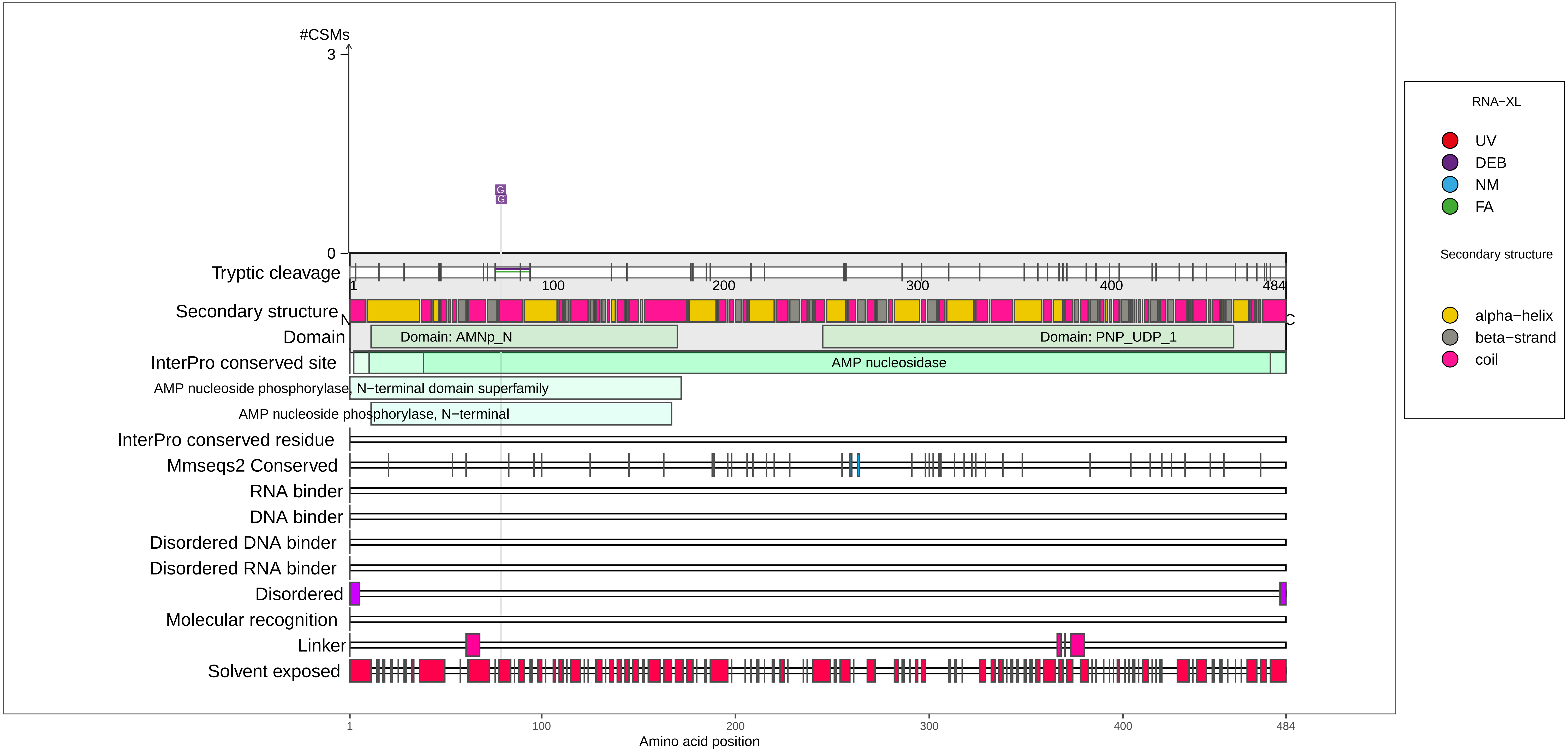

P0AE91
CREA_ECOLI Protein CreA

− Abundance:
tryptic [log10 Intensity]: 7.98 (Q 57)
PAXdb K12 strain [ppm]: 2.46 (Q 76)
PAXdb E.coli [ppm]: 1.77 (Q 66)

− RNA functions: not annotated

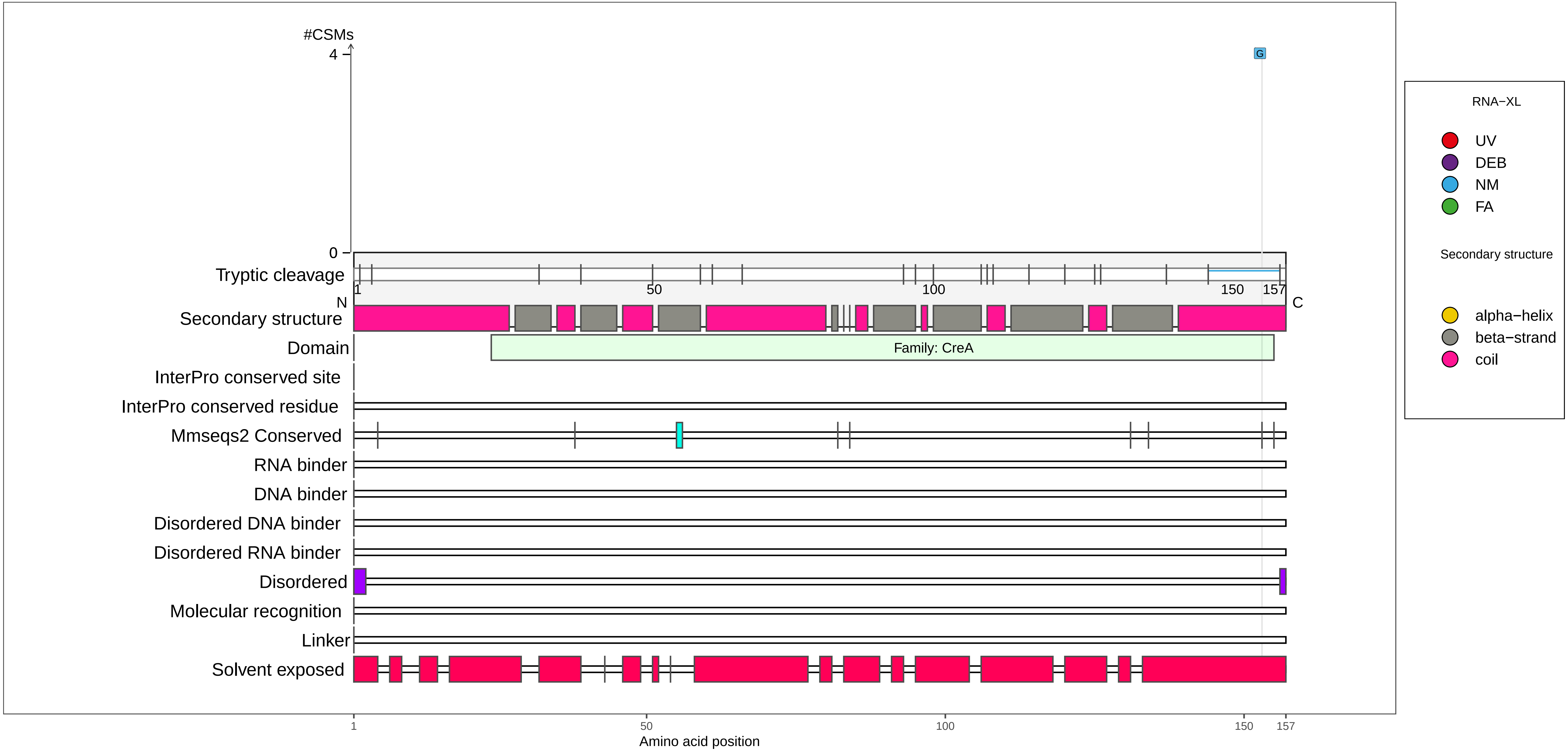

P0AES4
GYRA_ECOLI DNA gyrase subunit A
− Abundance:
tryptic [log10 Intensity]: 8.85 (Q 85)
PAXdb K12 strain [ppm]: 2.71 (Q 84)
PAXdb E.coli [ppm]: 2.67 (Q 89)

− RNA functions:
RNA biosynthetic process; RNA metabolic process

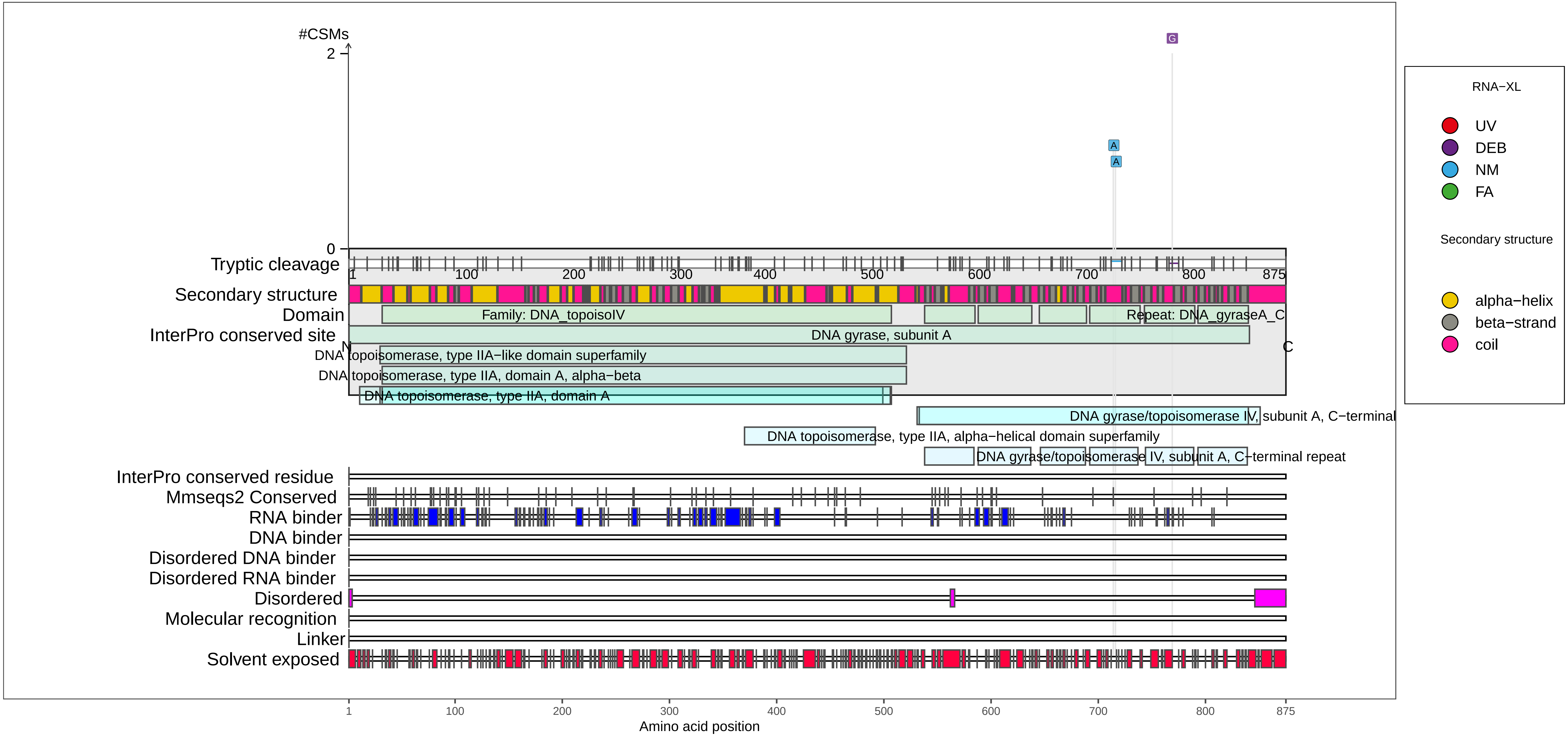

P0AFJ1
YJDM_ECOLI Protein YjdM

− Abundance:
tryptic [log10 Intensity]: 7.53 (Q 37)
PAXdb K12 strain [ppm]: 3.5 (Q 97)
PAXdb E.coli [ppm]: 2.3 (Q 80)

− RNA functions: not annotated

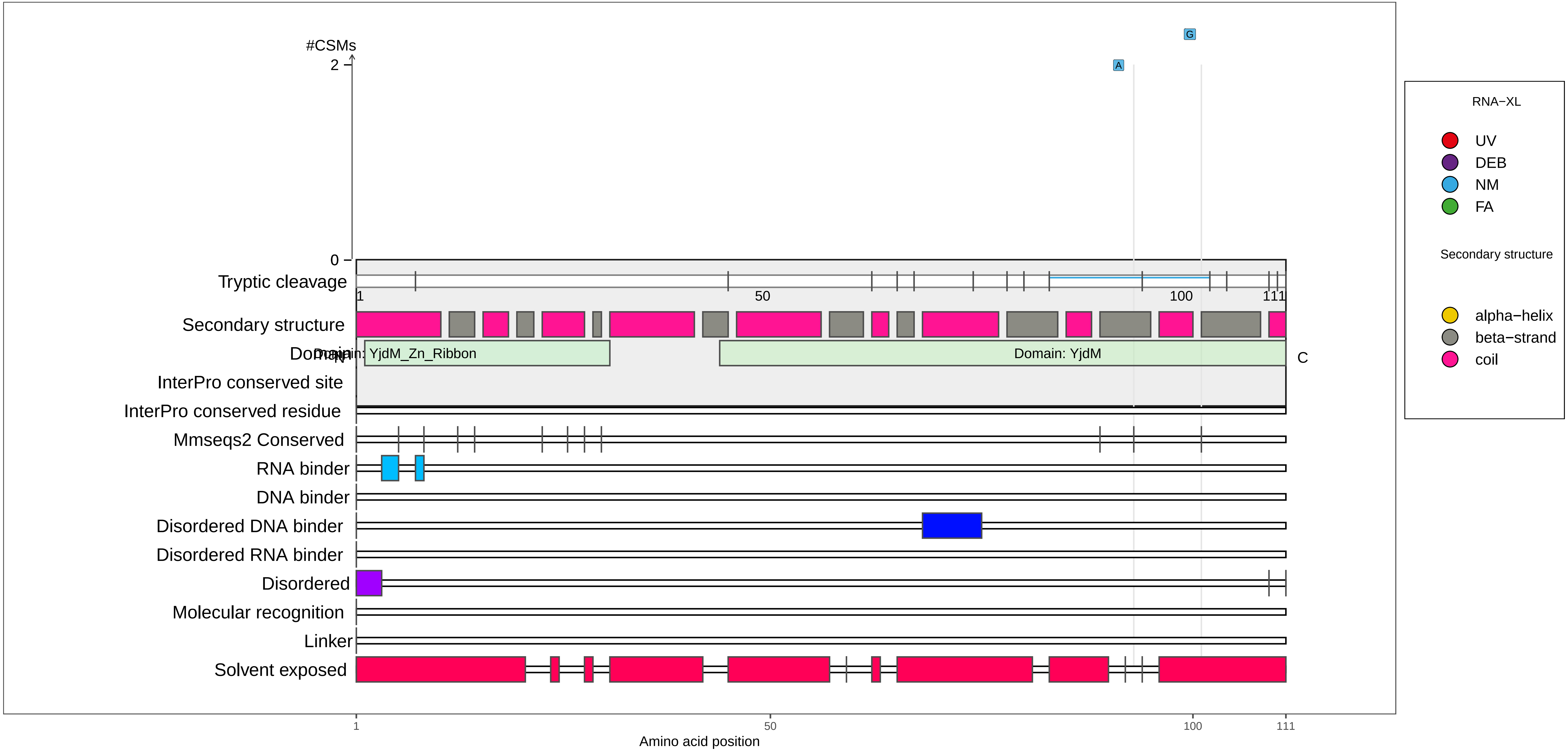

P0AGJ2
TRMH_ECOLI tRNA (guanosine(18)−2'−O)−methyltransferase

− Abundance:
tryptic [log10 Intensity]: 8.17 (Q 64)
PAXdb K12 strain [ppm]: 2.33 (Q 71)
PAXdb E.coli [ppm]: 1.02 (Q 48)

− RNA functions:
ncRNA metabolic process; ncRNA processing; RNA 2−O−methyltransferase activity
RNA binding; RNA metabolic process; RNA methylation; RNA methyltransferase activity
RNA modification; RNA processing; SpoU rRNA Methylase family; SpoU, rRNA methylase, C−terminal
tRNA (guanine) methyltransferase activity
tRNA (guanosine−2−O−)−methyltransferase activity; tRNA 2−O−methyltransferase activity; tRNA binding
tRNA guanine ribose methylation; tRNA metabolic process; tRNA methylation; tRNA methyltransferase activity
tRNA modification; tRNA nucleoside ribose methylation; tRNA processing

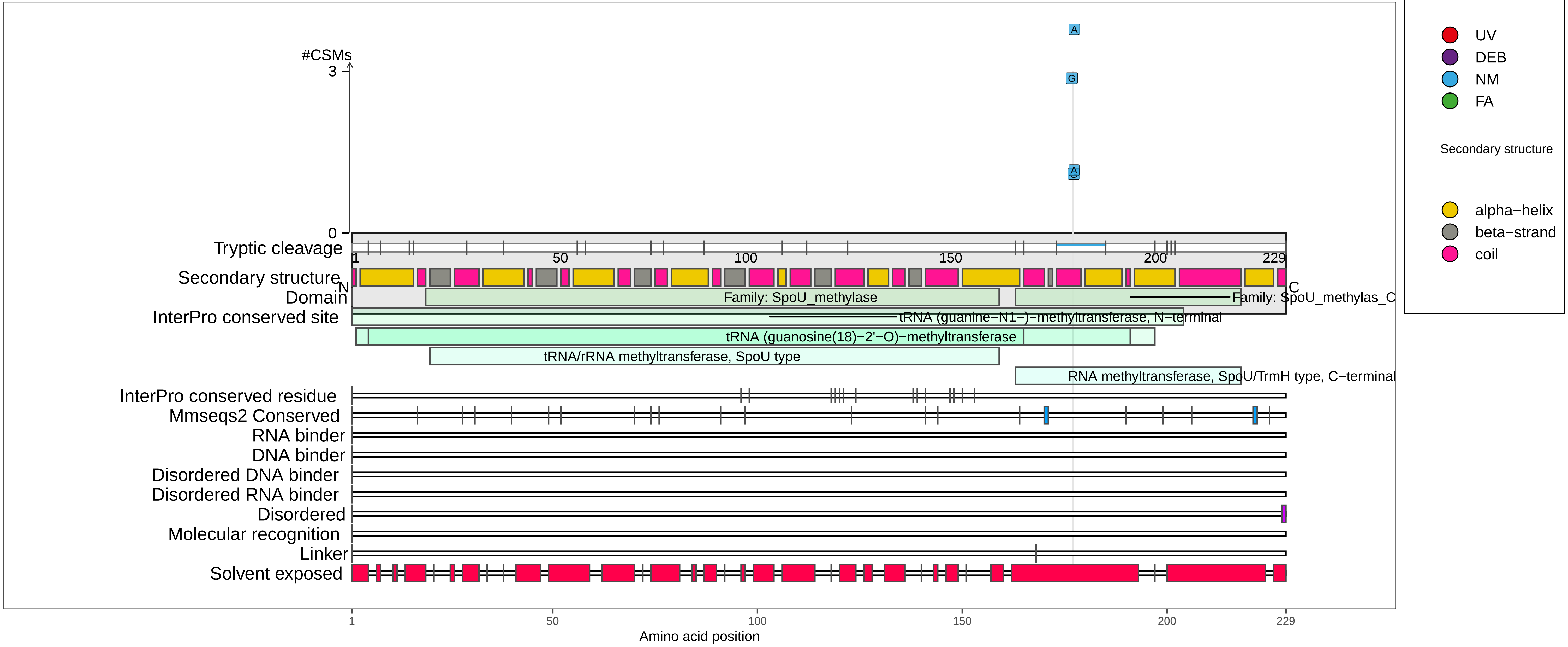

P0C058
IBPB_ECOLI Small heat shock protein IbpB

− Abundance:
tryptic [log10 Intensity]: 7.92 (Q 55)
PAXdb K12 strain [ppm]: NA (Q NA)
PAXdb E.coli [ppm]: 1.99 (Q 72)

− RNA functions: not annotated

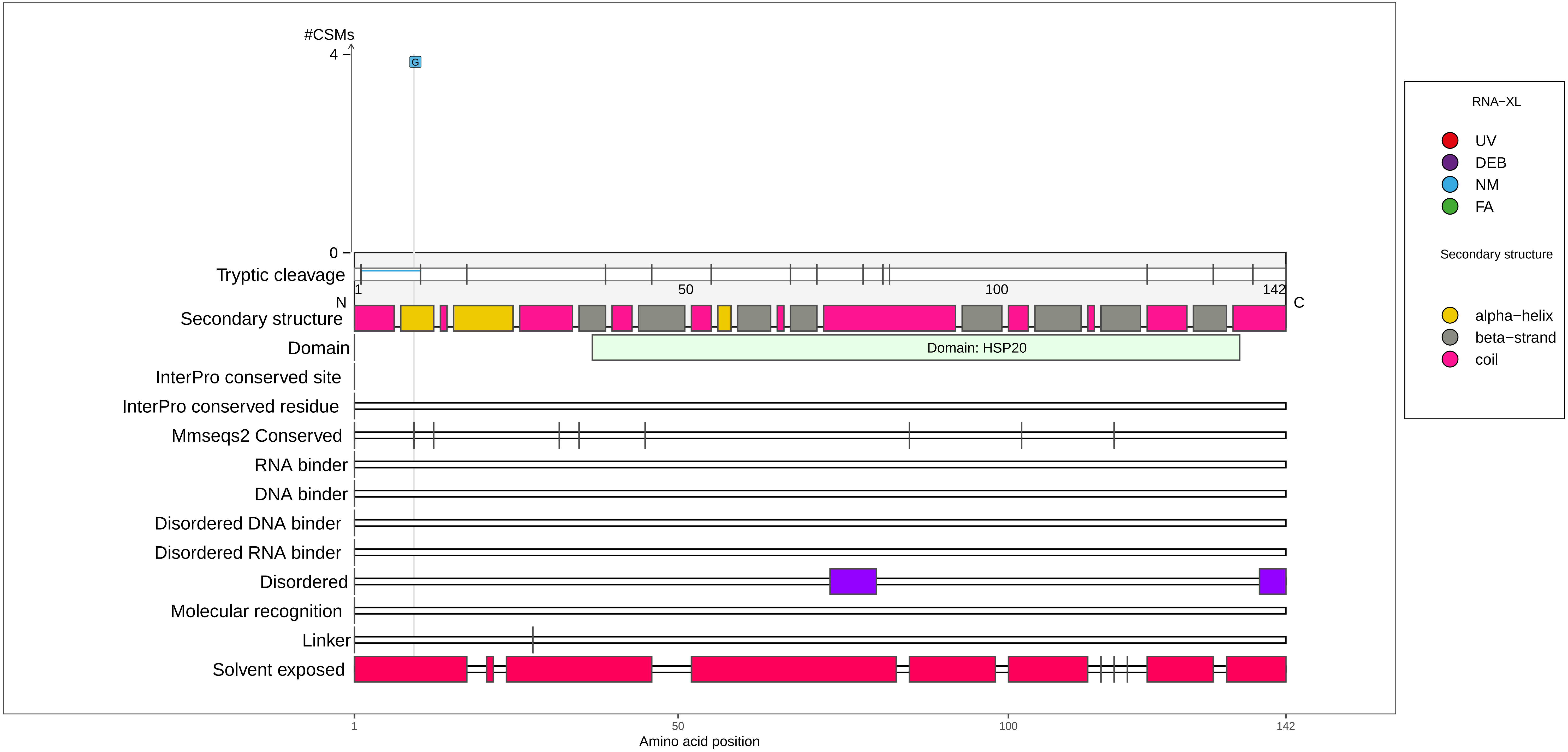

P0C0K3
SRKA_ECOLI Stress response kinase A

− Abundance:
tryptic [log10 Intensity]: 7.13 (Q 18)
PAXdb K12 strain [ppm]: NA (Q NA)
PAXdb E.coli [ppm]: 0.24 (Q 30)

− RNA functions: not annotated

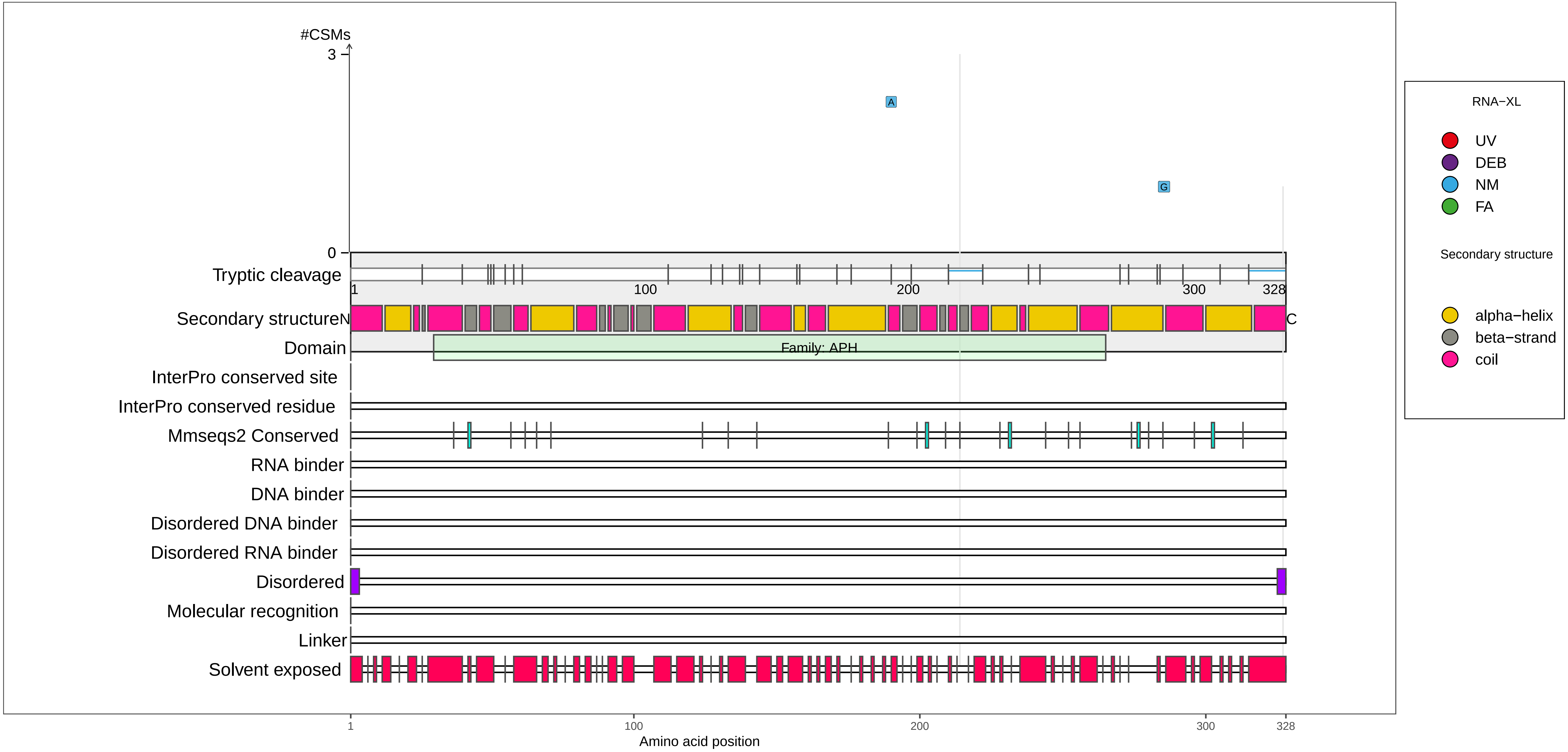

P15254
PUR4_ECOLI Phosphoribosylformylglycinamidine synthase

− Abundance:
tryptic [log10 Intensity]: 8.63 (Q 80)
PAXdb K12 strain [ppm]: 2.7 (Q 83)
PAXdb E.coli [ppm]: 2.57 (Q 87)

− RNA functions: not annotated

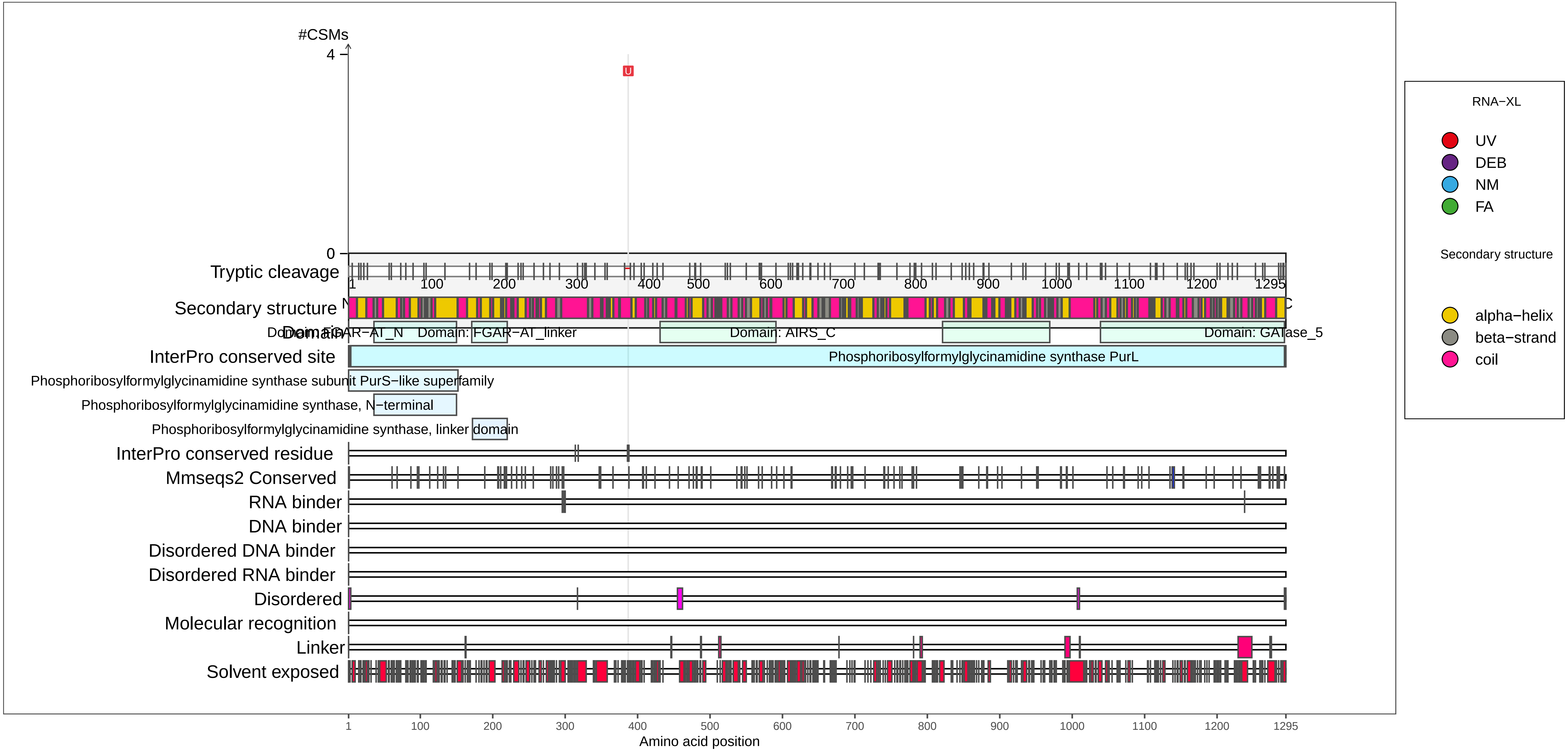

P16926
MREC_ECOLI Cell shape−determining protein MreC

− Abundance:
tryptic [log10 Intensity]: 7.82 (Q 50)
PAXdb K12 strain [ppm]: 1.31 (Q 27)
PAXdb E.coli [ppm]: 1.1 (Q 50)

− RNA functions: not annotated

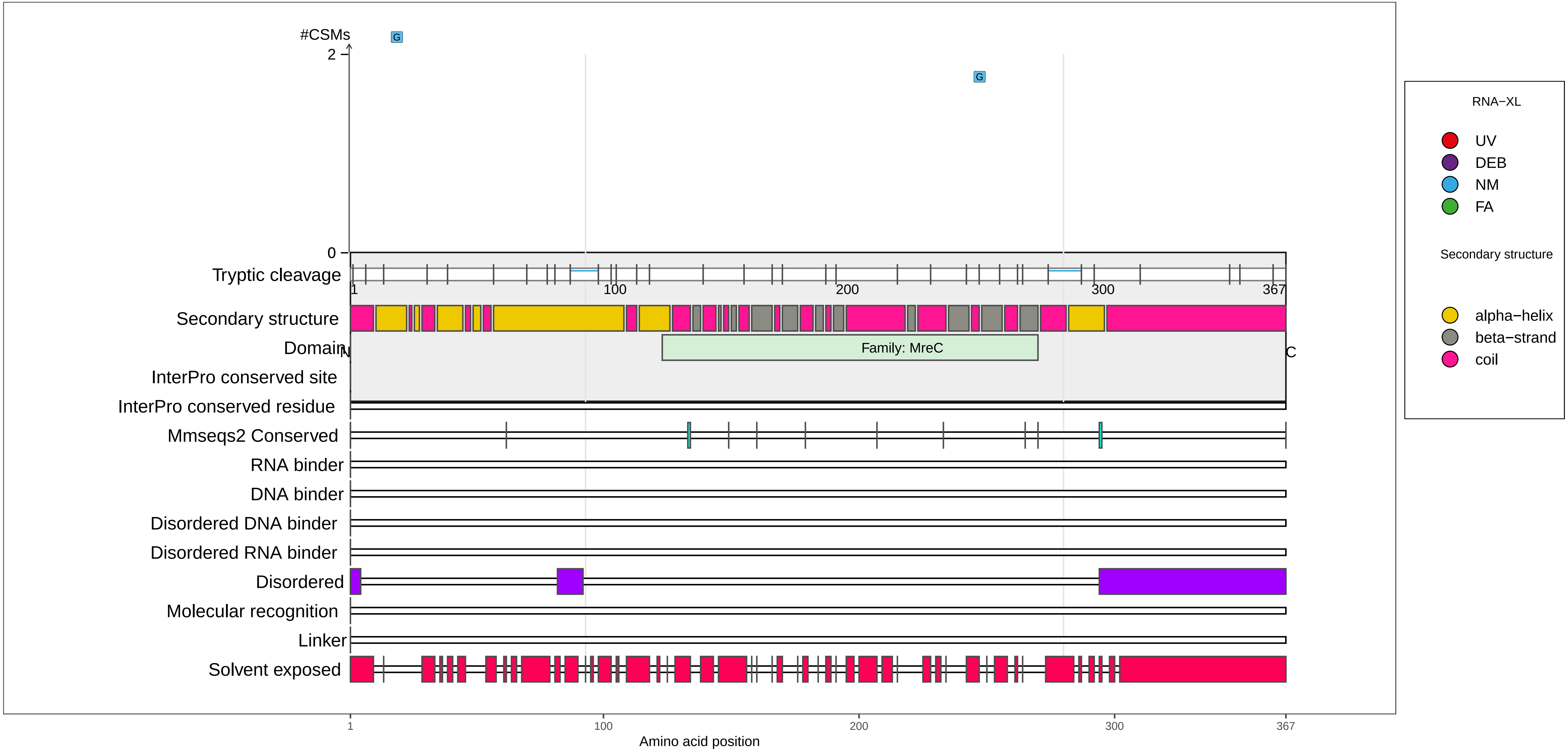

P23003
TRMA_ECOLI tRNA/tmRNA (uracil−C(5))−methyltransferase

− Abundance:
tryptic [log10 Intensity]: 7.83 (Q 51)
PAXdb K12 strain [ppm]: 0.69 (Q 3)
PAXdb E.coli [ppm]: 1.89 (Q 69)

− RNA functions:
ncRNA metabolic process; ncRNA processing; RNA binding; RNA metabolic process
RNA methylation; RNA methyltransferase activity; RNA modification; RNA processing; rRNA binding
S−adenosylmethionine−dependent tRNA (m5U54) methyltransferase activity
tRNA (m5U54) methyltransferase activity; tRNA (Uracil−5−)−methyltransferase
tRNA (uracil) methyltransferase activity; tRNA binding; tRNA metabolic process; tRNA methylation
tRNA methyltransferase activity; tRNA modification; tRNA processing

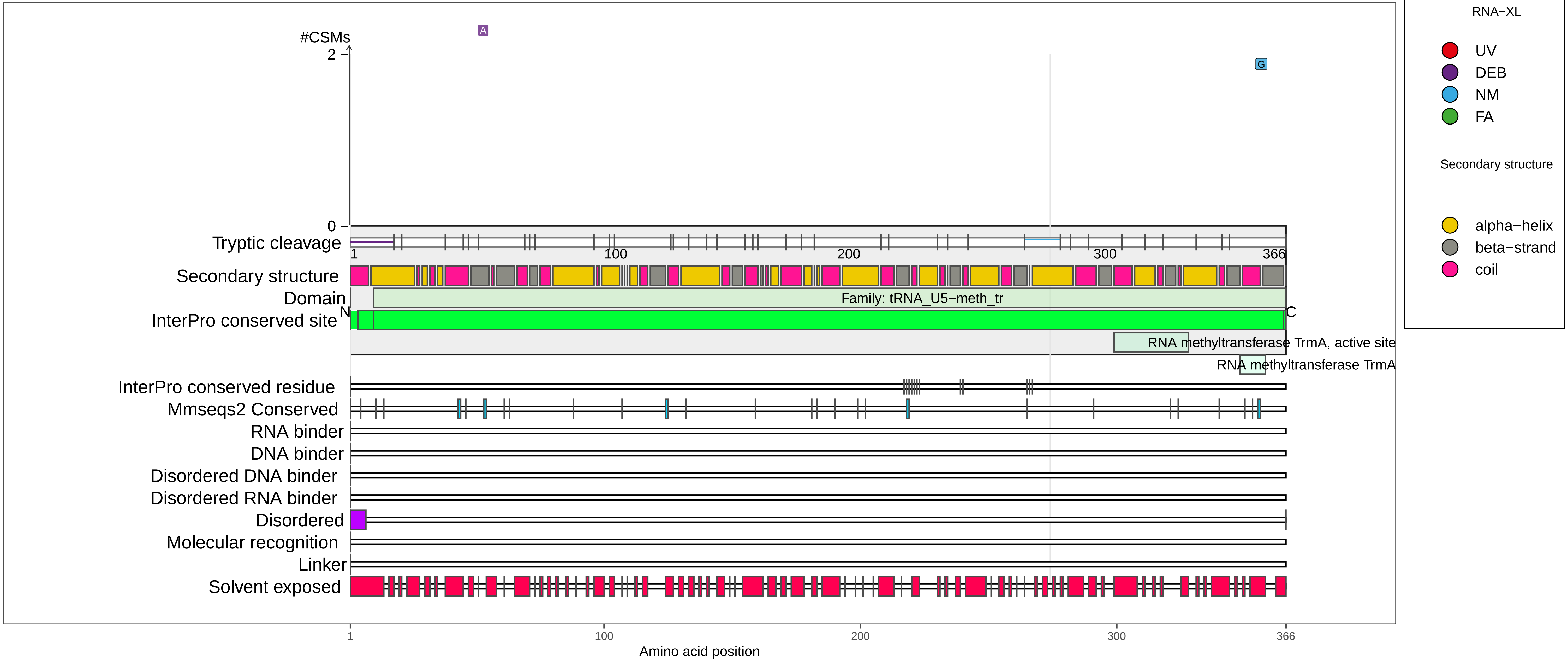

P26365
AMIB_ECOLI N−acetylmuramoyl−L−alanine amidase AmiB

− Abundance:
tryptic [log10 Intensity]: 7.88 (Q 53)
PAXdb K12 strain [ppm]: NA (Q NA)
PAXdb E.coli [ppm]: 0.44 (Q 35)

− RNA functions: not annotated

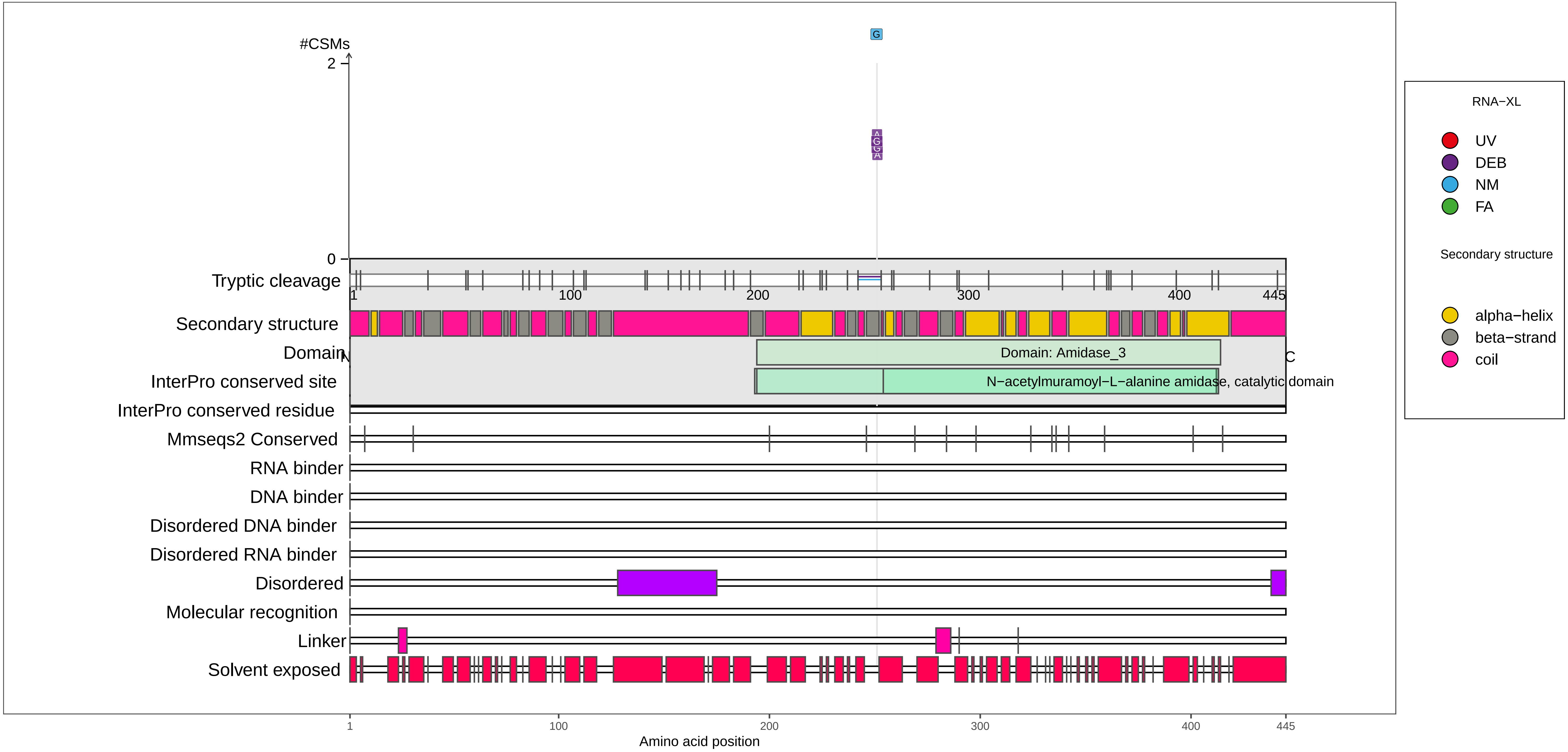

P28303
DINF_ECOLI DNA damage−inducible protein F

− Abundance:
tryptic [log10 Intensity]: 7.28 (Q 24)
PAXdb K12 strain [ppm]: NA (Q NA)
PAXdb E.coli [ppm]: 0.06 (Q 26)

− RNA functions: not annotated

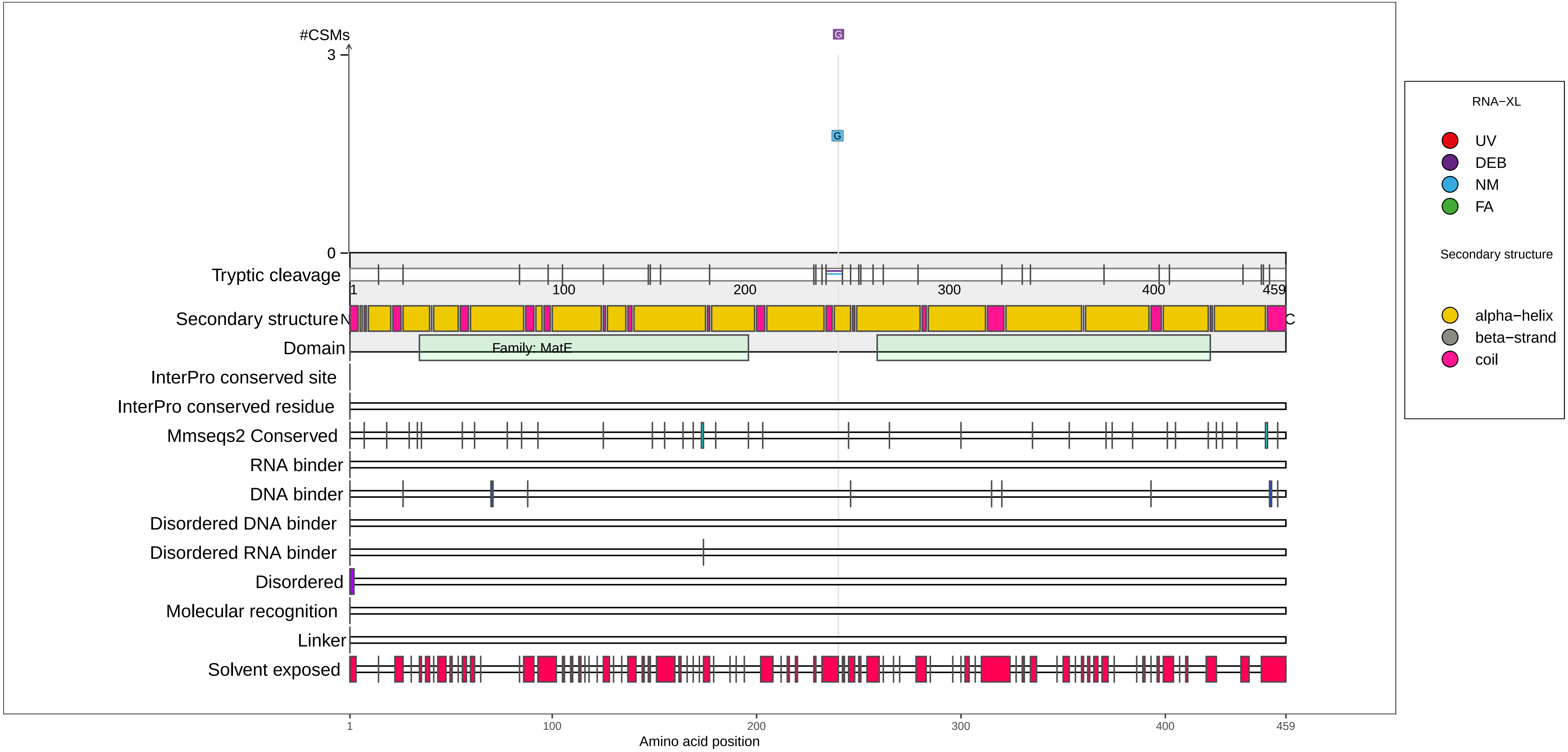

P29680
DCUP_ECOLI Uroporphyrinogen decarboxylase

− Abundance:
tryptic [log10 Intensity]: 8.57 (Q 78)
PAXdb K12 strain [ppm]: 1.9 (Q 57)
PAXdb E.coli [ppm]: 2.18 (Q 77)

− RNA functions: not annotated

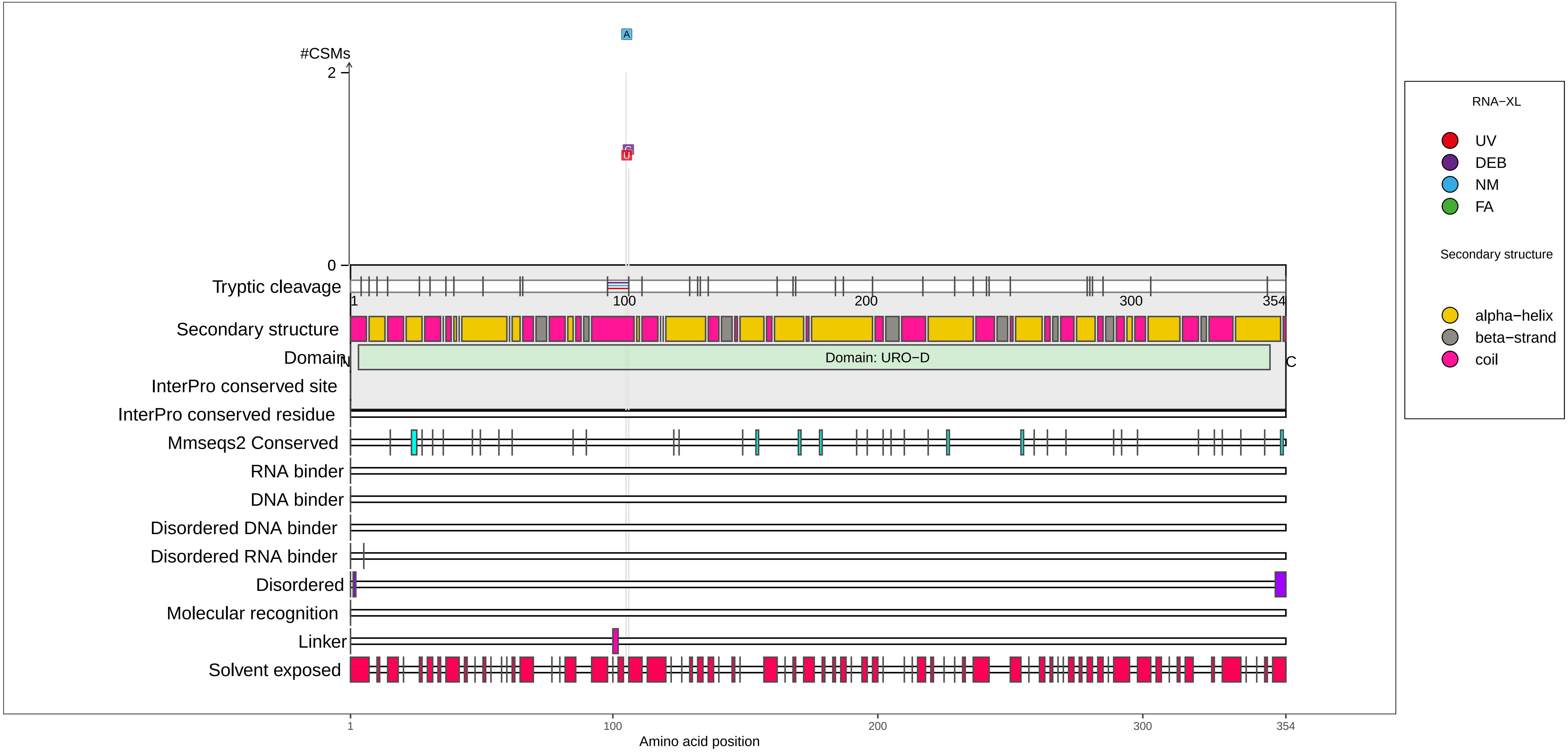

P30744
SDHM_ECOLI L−serine dehydratase 2

− Abundance:
tryptic [log10 Intensity]: 7.59 (Q 40)
PAXdb K12 strain [ppm]: 1.45 (Q 36)
PAXdb E.coli [ppm]: 0.78 (Q 43)

− RNA functions: not annotated

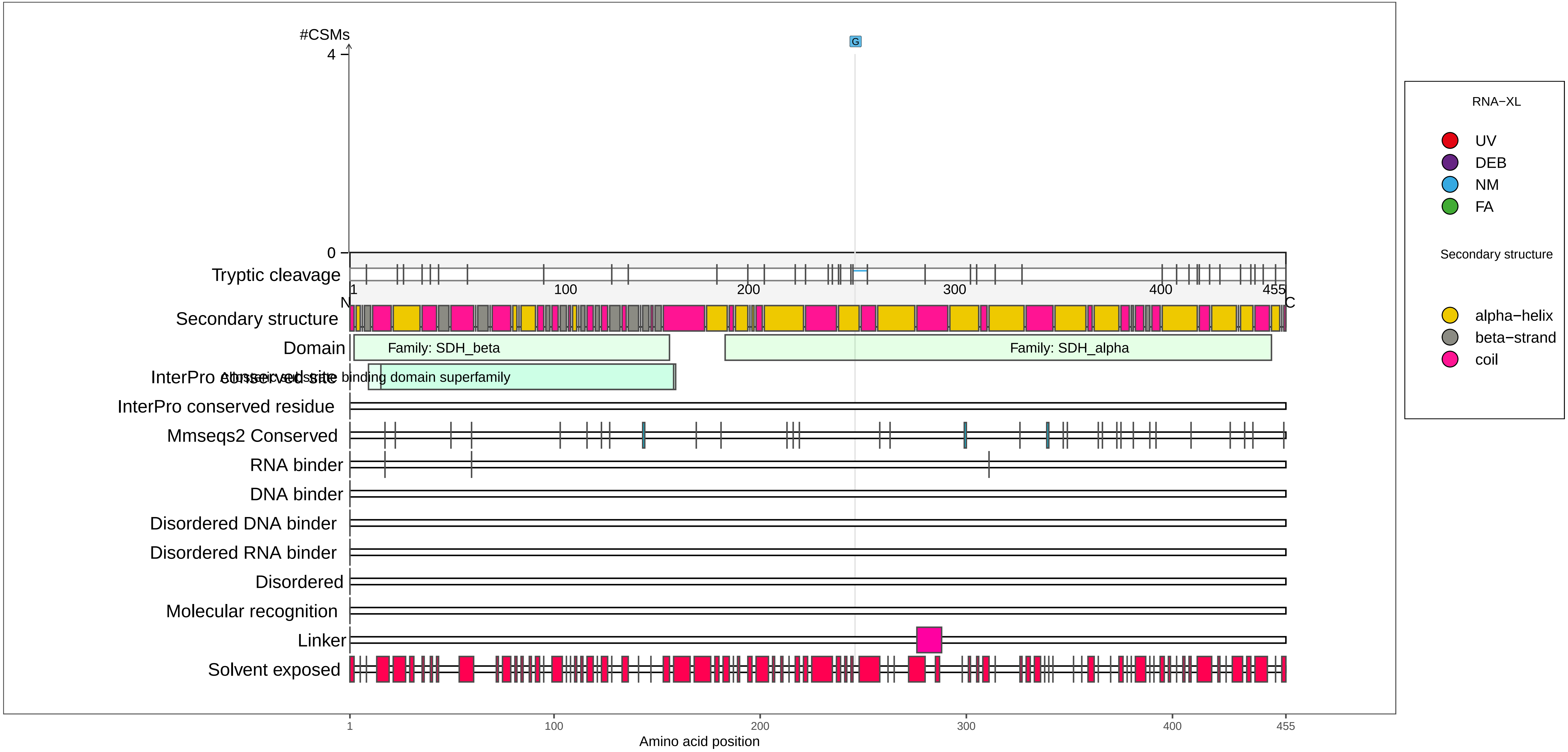

P31808
YCIK_ECOLI Uncharacterized oxidoreductase YciK
− Abundance:
tryptic [log10 Intensity]: 7.09 (Q 17)
PAXdb K12 strain [ppm]: 2.44 (Q 76)
PAXdb E.coli [ppm]: 1.42 (Q 57)

− RNA functions: not annotated

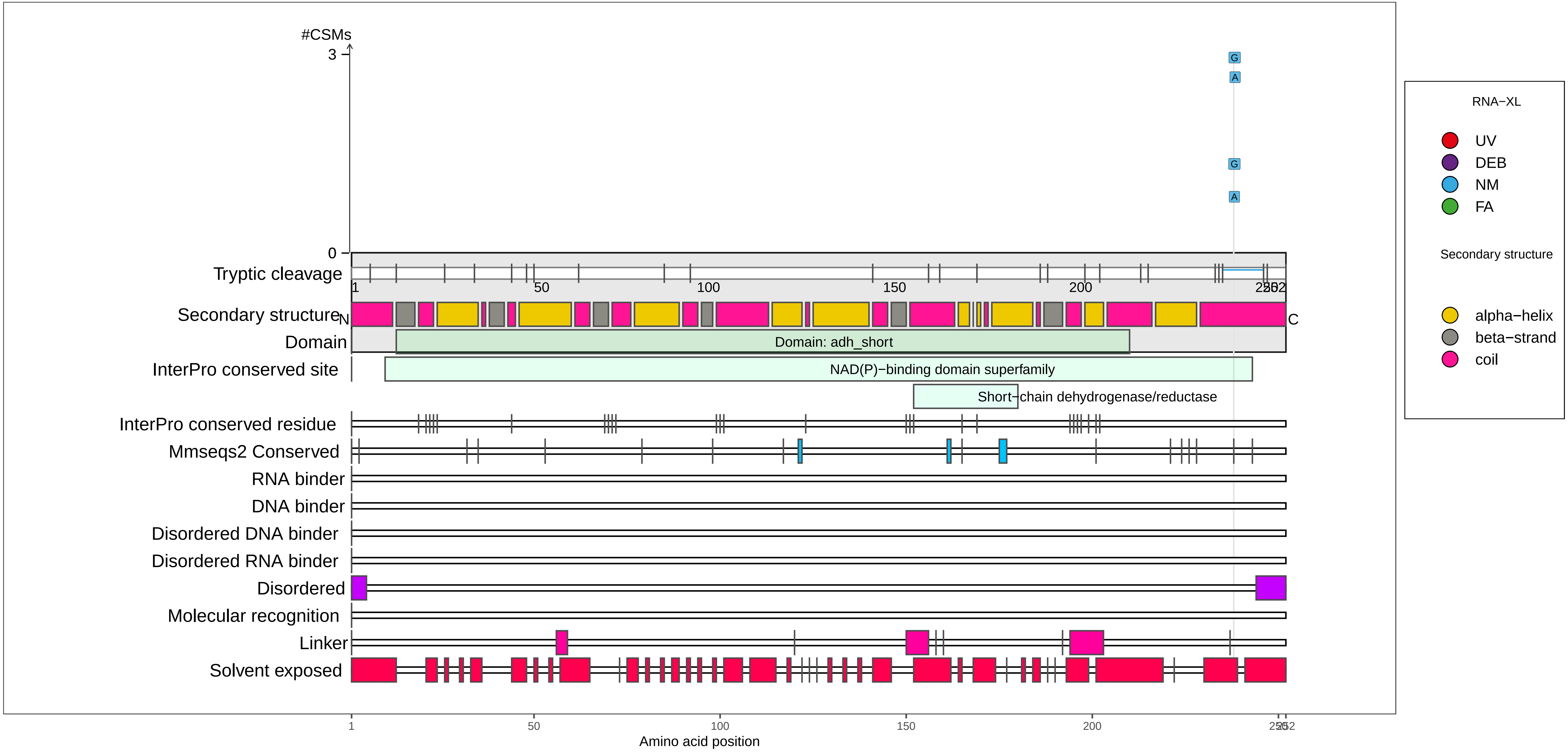

P32157
YIIM_ECOLI Protein YiiM

− Abundance:
tryptic [log10 Intensity]: 7.3 (Q 25)
PAXdb K12 strain [ppm]: 2.31 (Q 70)
PAXdb E.coli [ppm]: 2.11 (Q 75)

− RNA functions: not annotated

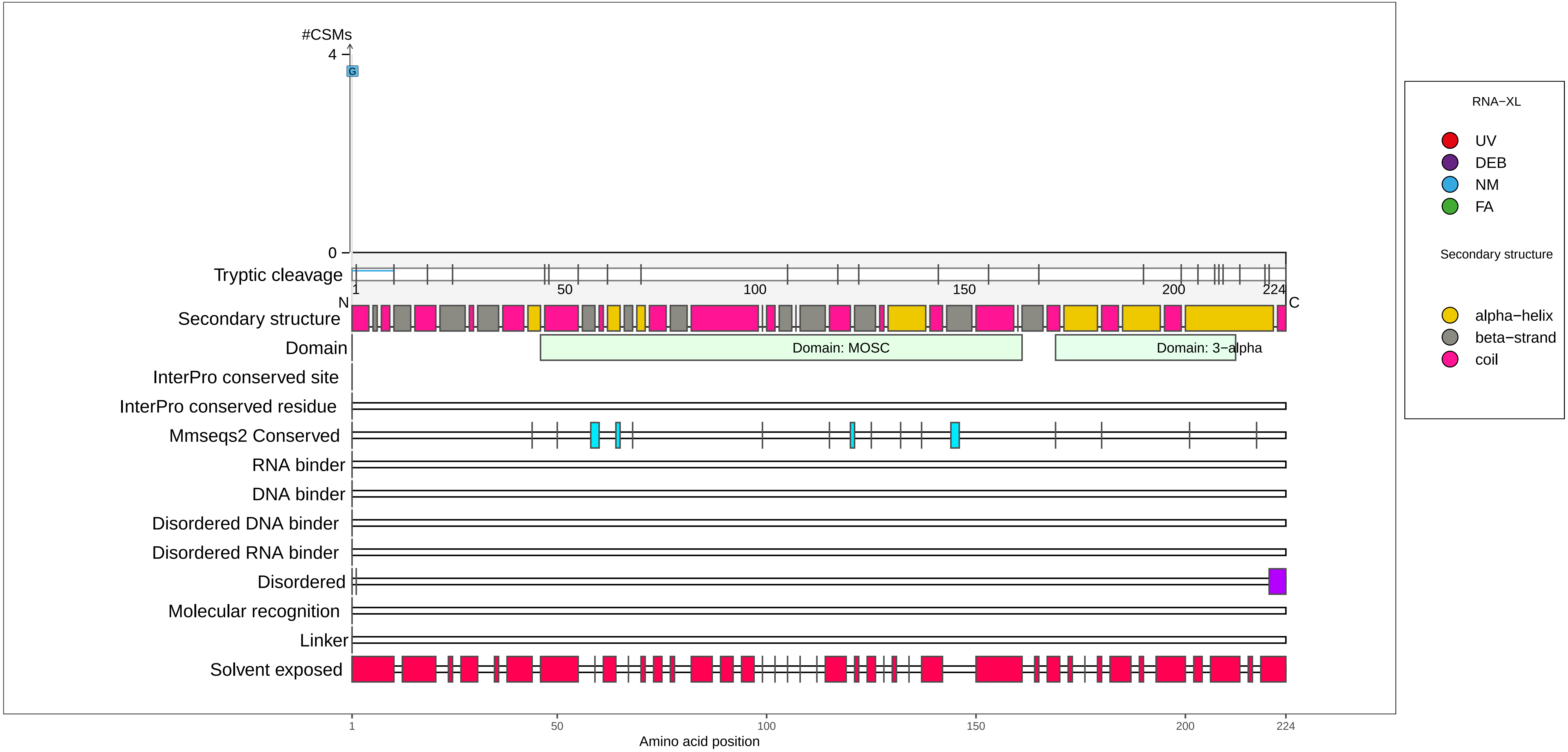

P36672
PTTBC_ECOLI PTS system trehalose−specific EIIBC component

− Abundance:
tryptic [log10 Intensity]: 8.46 (Q 74)
PAXdb K12 strain [ppm]: 1.52 (Q 39)
PAXdb E.coli [ppm]: 1.07 (Q 49)

− RNA functions: not annotated

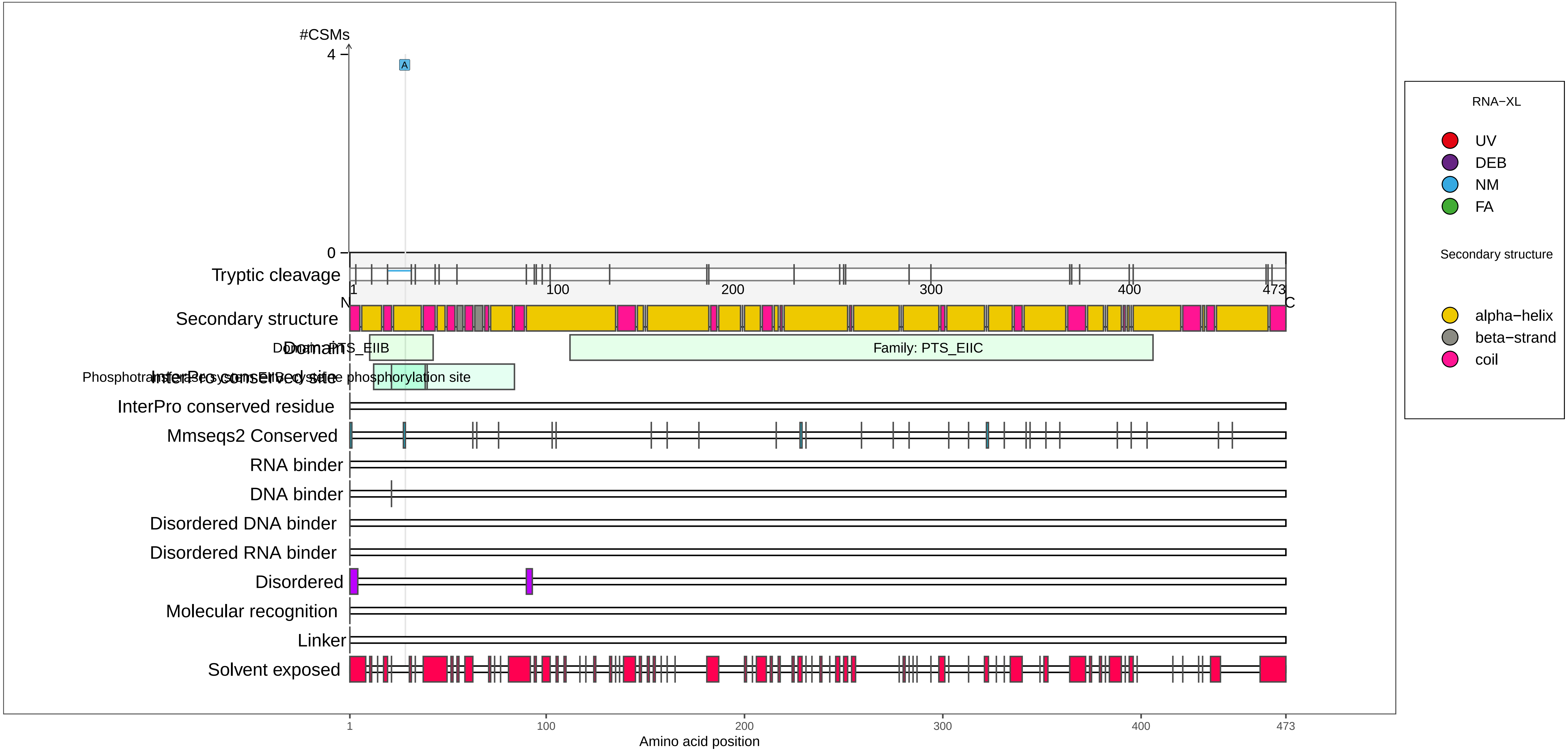

P36680
ZAPD_ECOLI Cell division protein ZapD

− Abundance:
tryptic [log10 Intensity]: 8.68 (Q 81)
PAXdb K12 strain [ppm]: 1.74 (Q 51)
PAXdb E.coli [ppm]: 1.82 (Q 67)

− RNA functions: not annotated

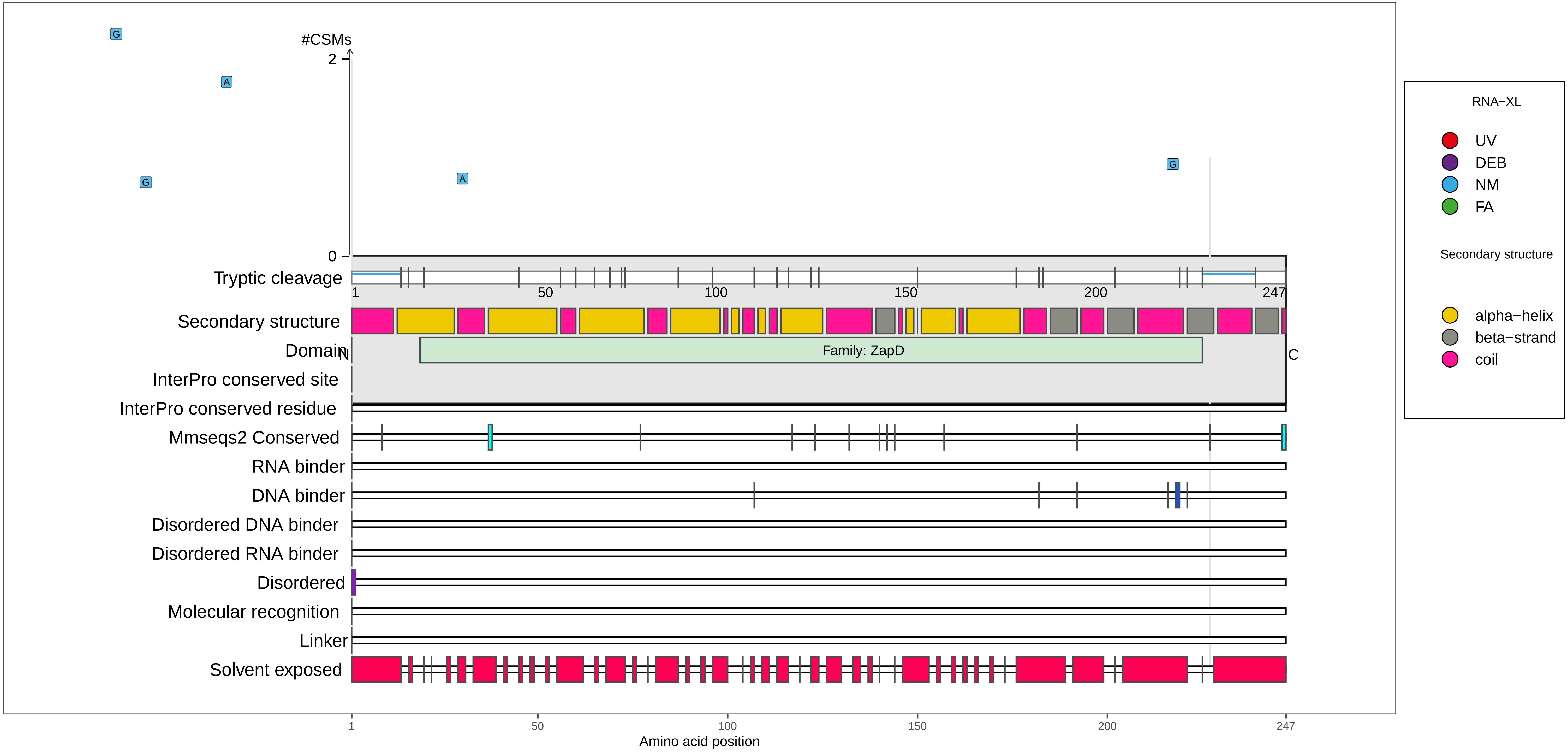

P37024
HRPB_ECOLI ATP−dependent RNA helicase HrpB

− Abundance:
tryptic [log10 Intensity]: 7.47 (Q 33)
PAXdb K12 strain [ppm]: 1.18 (Q 19)
PAXdb E.coli [ppm]: −0.04 (Q 24)

− RNA functions:
RNA helicase activity

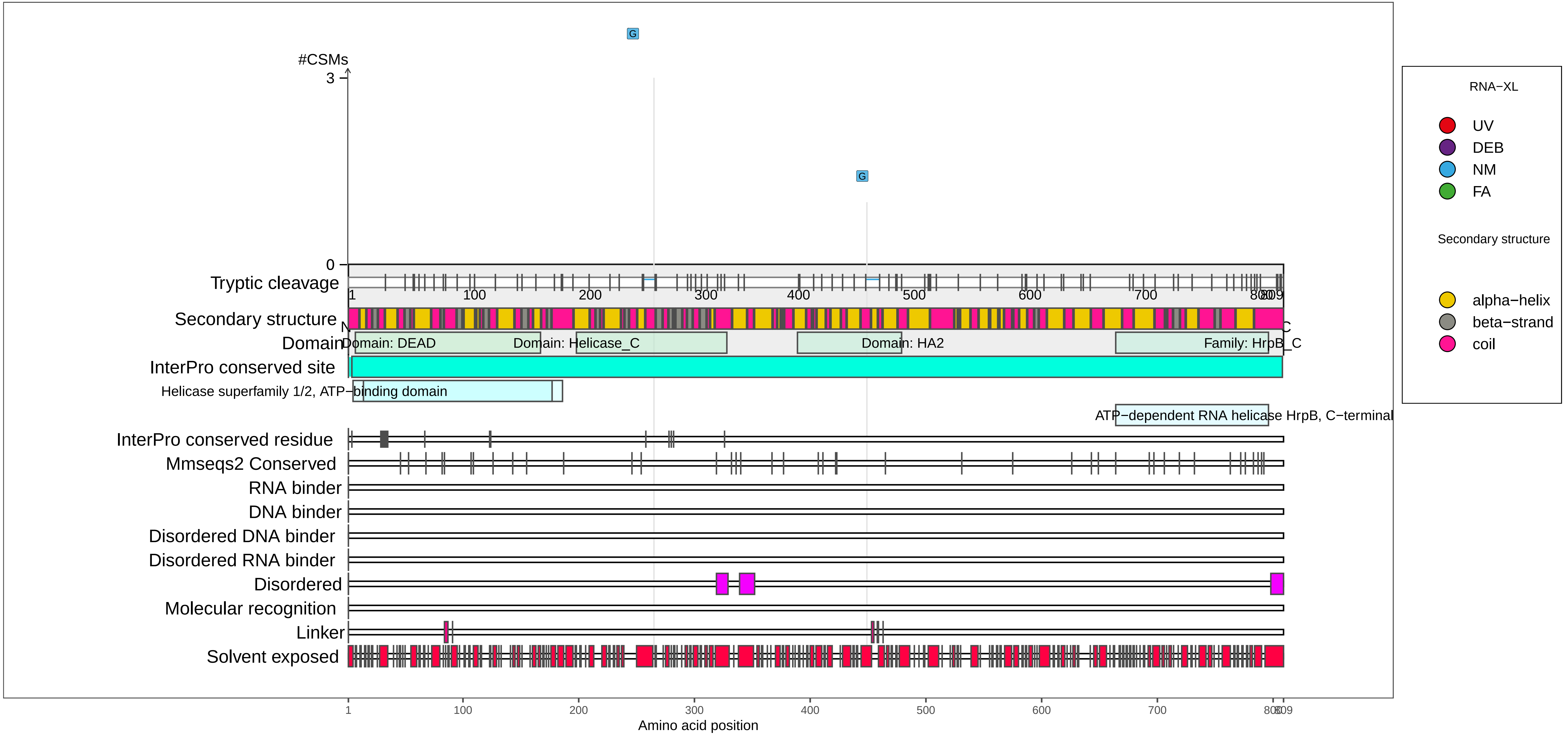

P37128
NUDK_ECOLI GDP−mannose pyrophosphatase

− Abundance:
tryptic [log10 Intensity]: 7.97 (Q 57)
PAXdb K12 strain [ppm]: 2.43 (Q 75)
PAXdb E.coli [ppm]: 1.63 (Q 63)

− RNA functions: not annotated

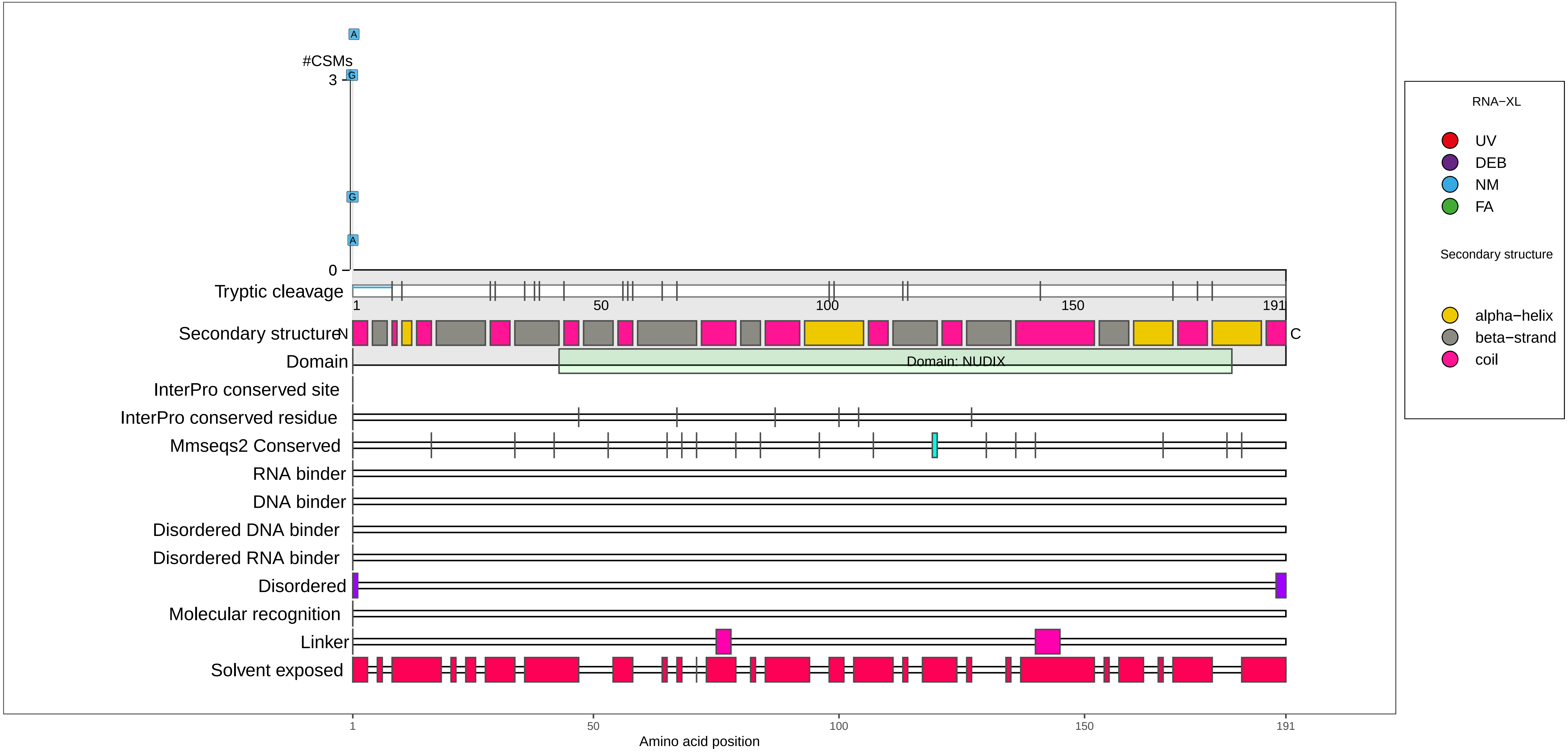

P37590
PMRD_ECOLI Signal transduction protein PmrD

− Abundance:
tryptic [log10 Intensity]: 7.44 (Q 32)
PAXdb K12 strain [ppm]: 2.01 (Q 61)
PAXdb E.coli [ppm]: 1.75 (Q 66)

− RNA functions: not annotated

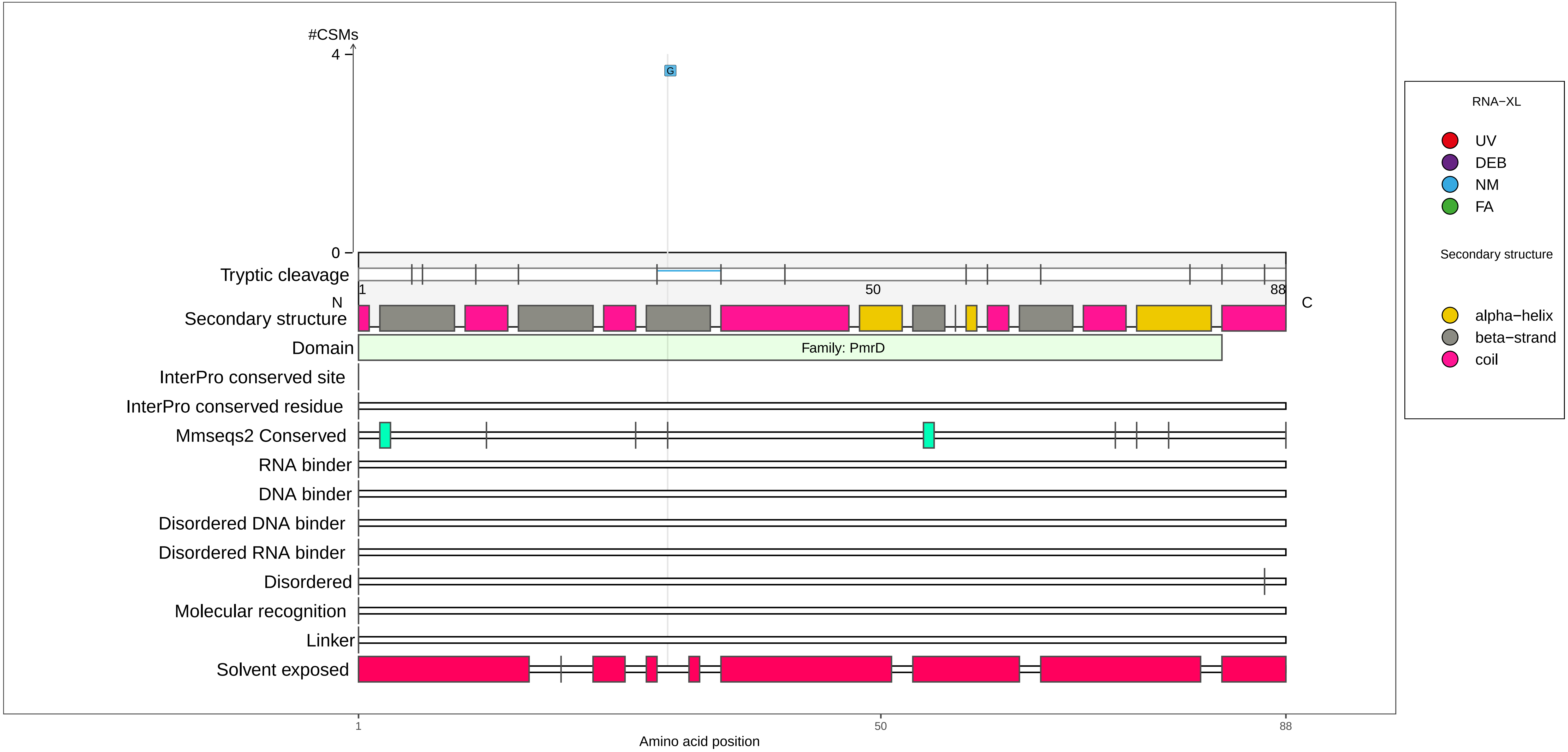

P42641
OBG_ECOLI GTPase ObgE/CgtA

− Abundance:
tryptic [log10 Intensity]: 7.74 (Q 46)
PAXdb K12 strain [ppm]: 1.25 (Q 23)
PAXdb E.coli [ppm]: 2.5 (Q 85)

− RNA functions:
RNA binding; rRNA binding

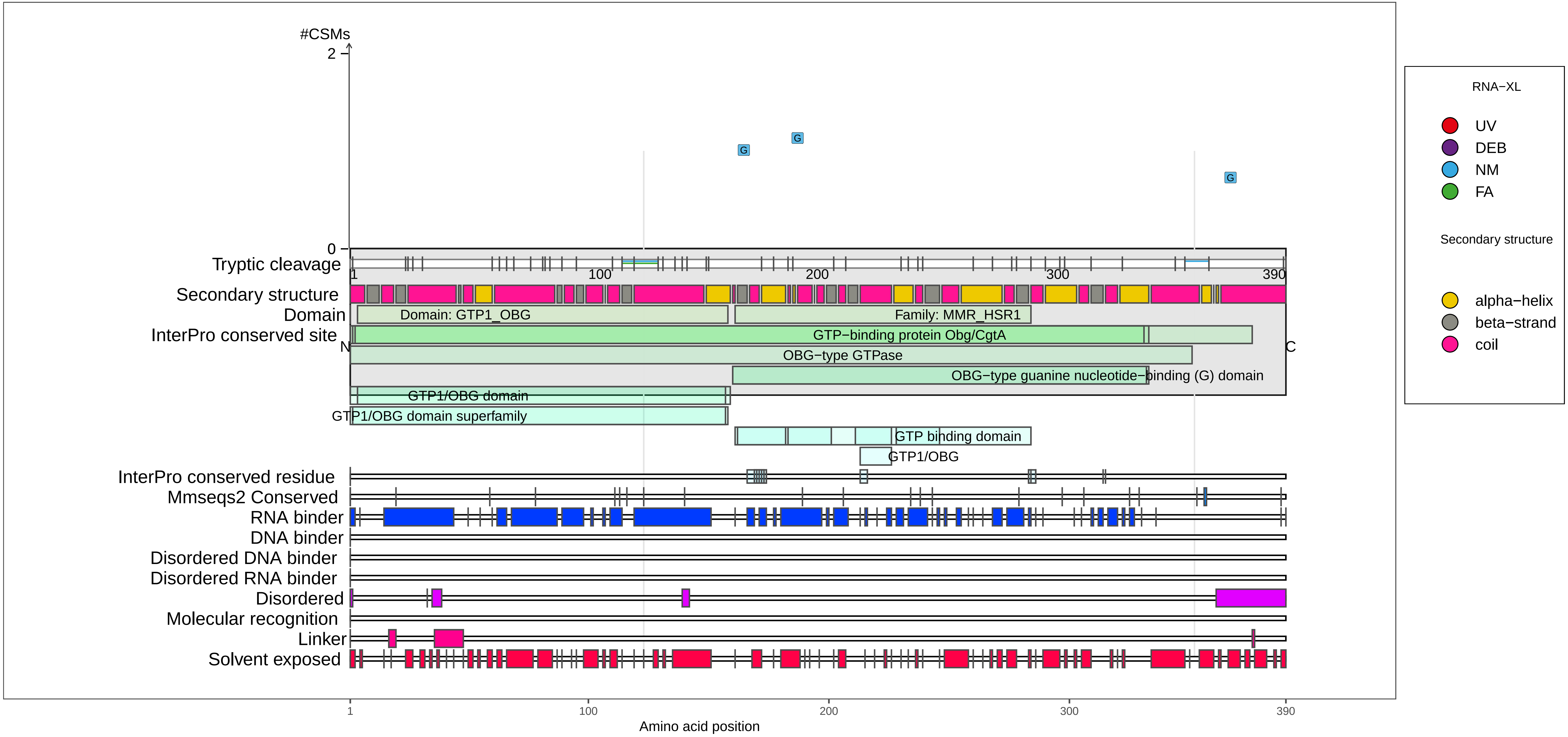

P43671
PQIB_ECOLI Intermembrane transport protein PqiB
− Abundance:
tryptic [log10 Intensity]: 7.72 (Q 45)
PAXdb K12 strain [ppm]: 1.23 (Q 22)
PAXdb E.coli [ppm]: 1.25 (Q 53)

− RNA functions: not annotated

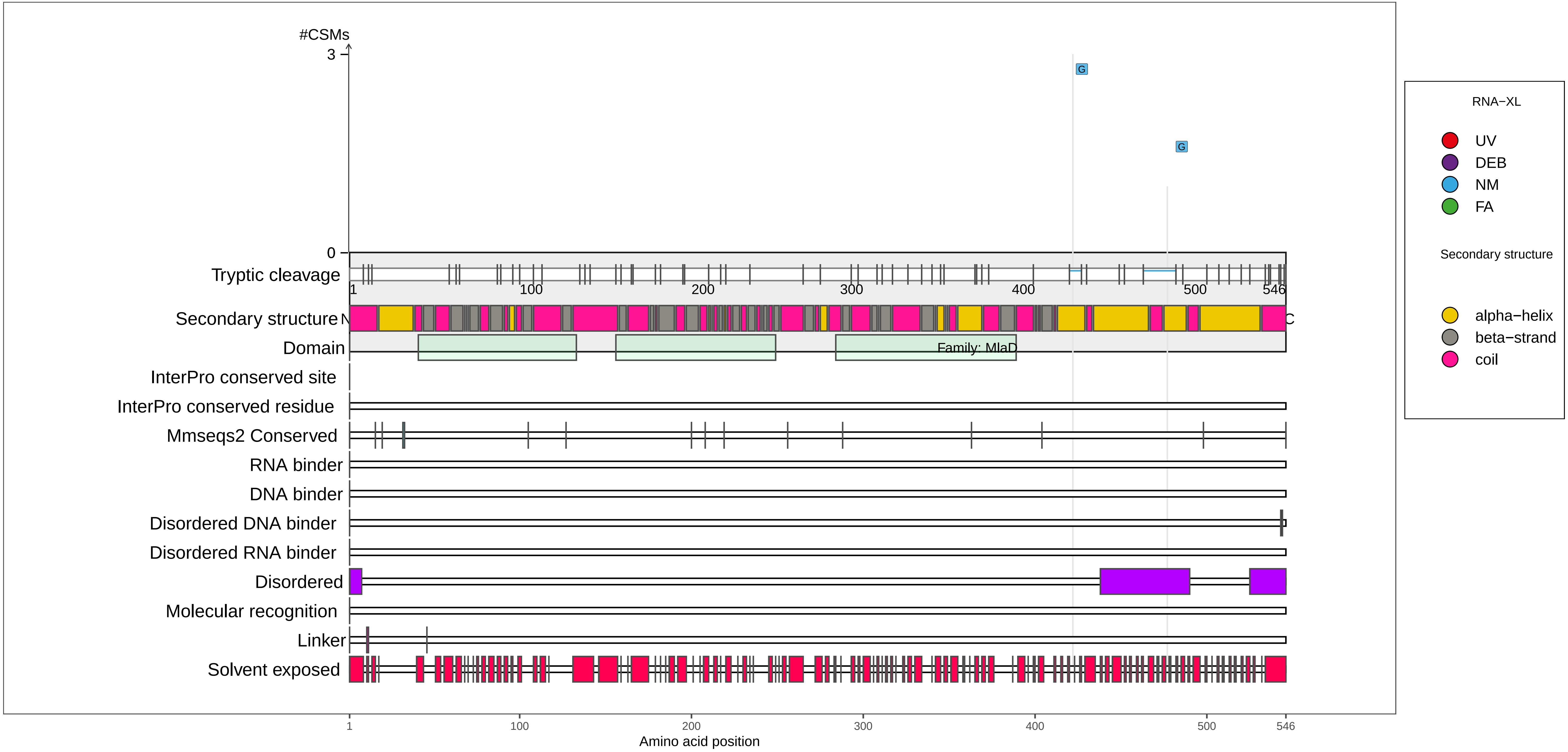

P45578
LUXS_ECOLI S−ribosylhomocysteine lyase

− Abundance:
tryptic [log10 Intensity]: 8.44 (Q 73)
PAXdb K12 strain [ppm]: 3.38 (Q 96)
PAXdb E.coli [ppm]: 3.11 (Q 96)

− RNA functions: not annotated

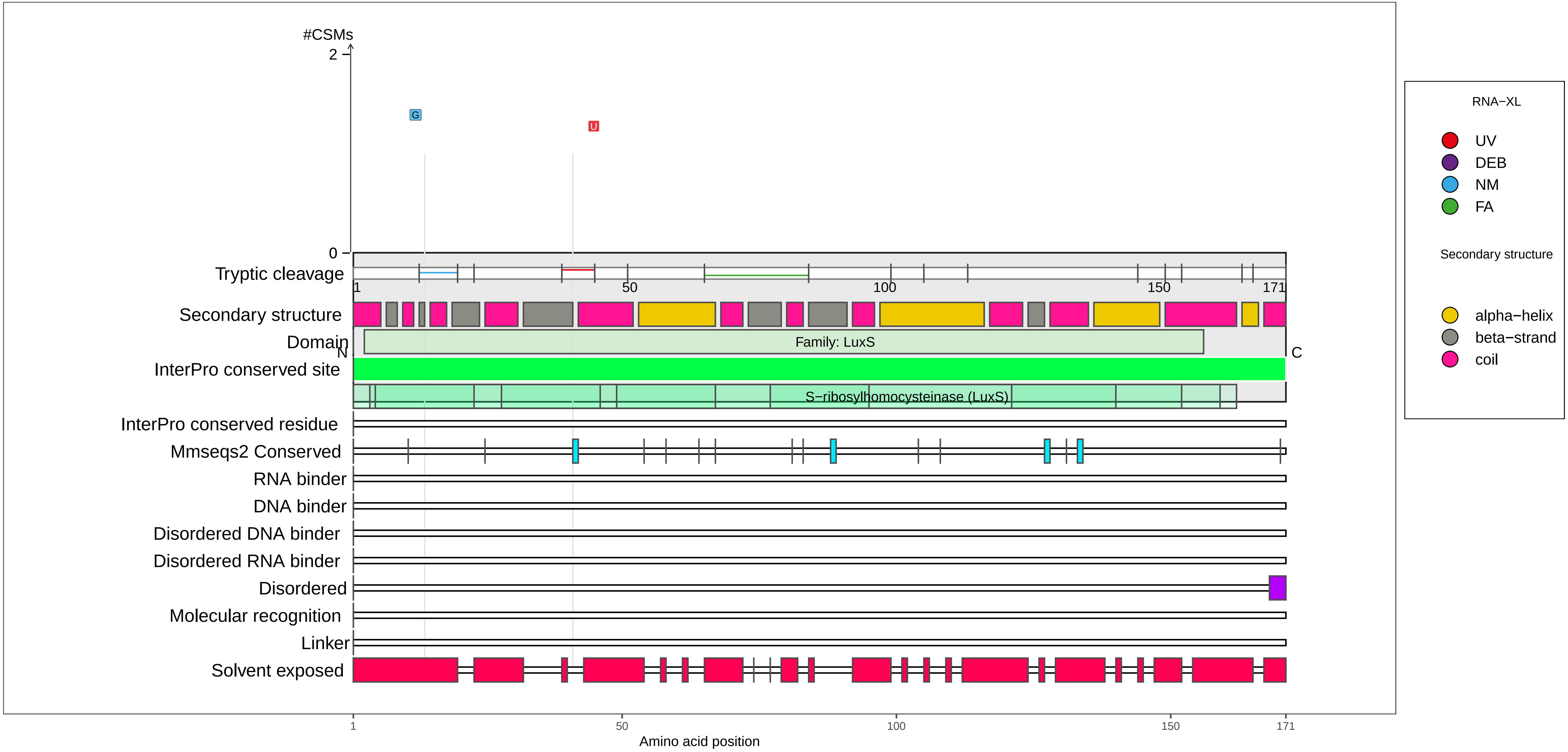

P45955
CPOB_ECOLI Cell division coordinator CpoB

− Abundance:
tryptic [log10 Intensity]: 8.29 (Q 69)
PAXdb K12 strain [ppm]: 2.36 (Q 72)
PAXdb E.coli [ppm]: 2.25 (Q 79)

− RNA functions: not annotated

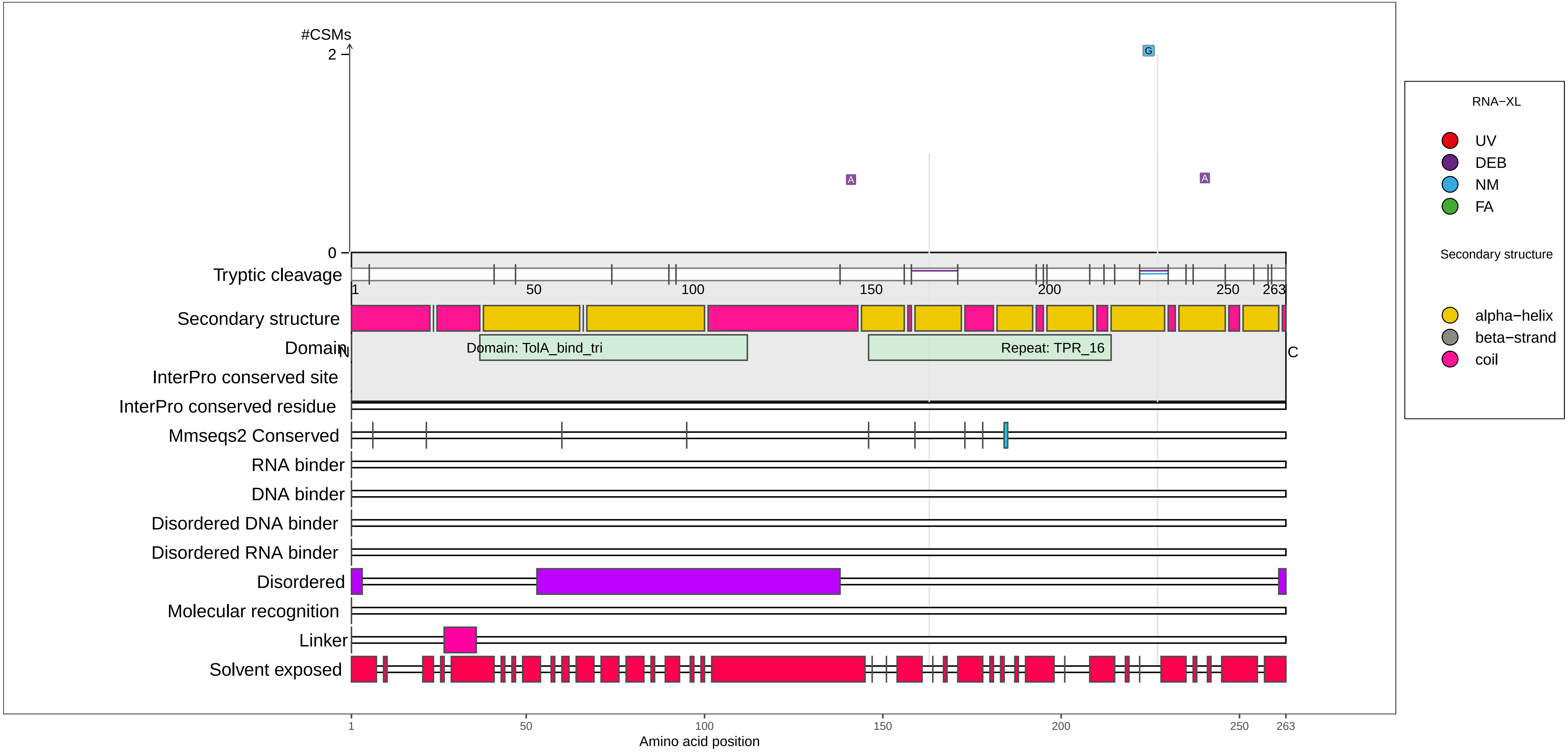

P46889
FTSK_ECOLI DNA translocase FtsK

− Abundance:
tryptic [log10 Intensity]: 7.86 (Q 52)
PAXdb K12 strain [ppm]: 1.6 (Q 45)
PAXdb E.coli [ppm]: 1.28 (Q 54)

− RNA functions: not annotated

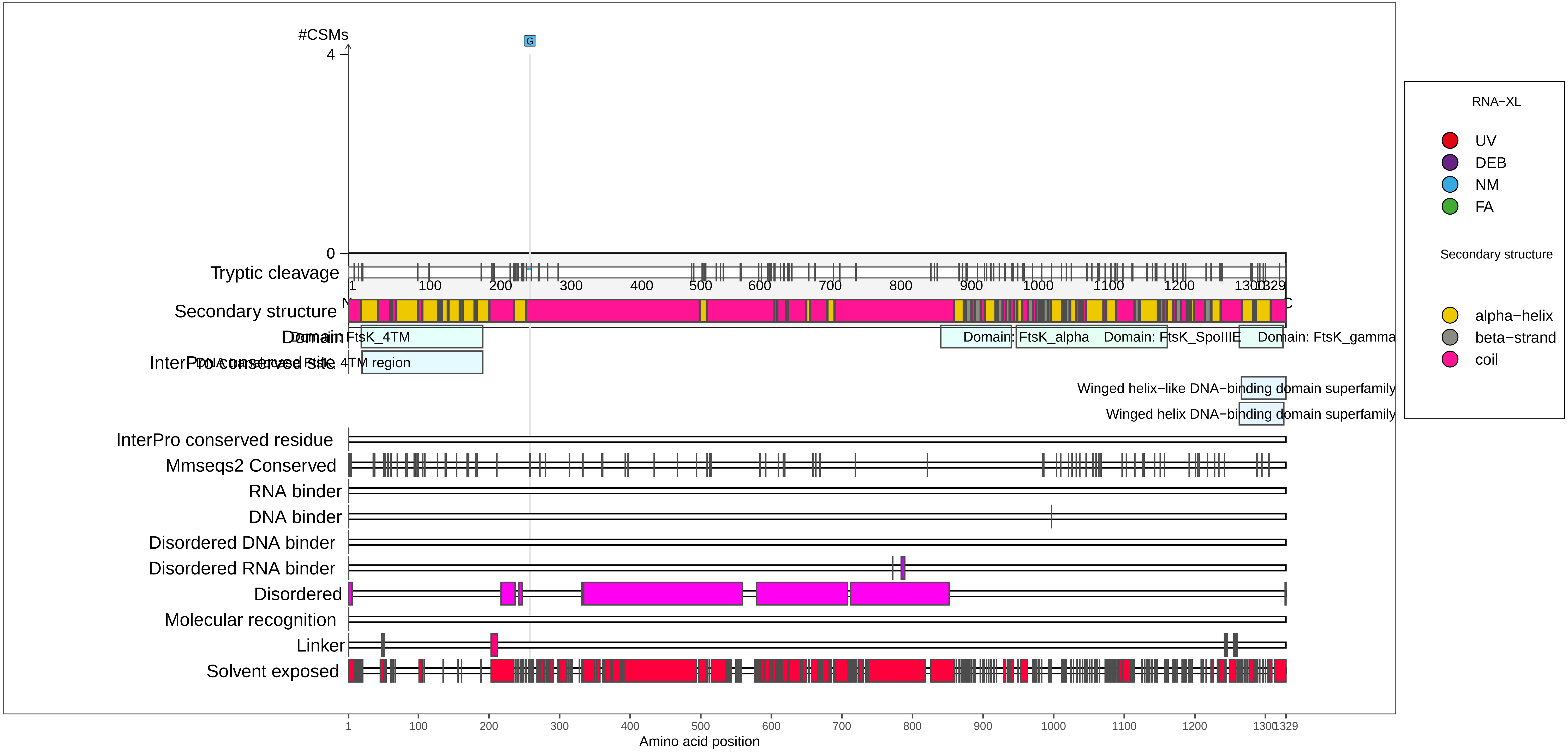

P68066
GRCA_ECOLI Autonomous glycyl radical cofactor

− Abundance:
tryptic [log10 Intensity]: 7.09 (Q 16)
PAXdb K12 strain [ppm]: 2.89 (Q 88)
PAXdb E.coli [ppm]: 3.24 (Q 97)

− RNA functions: not annotated

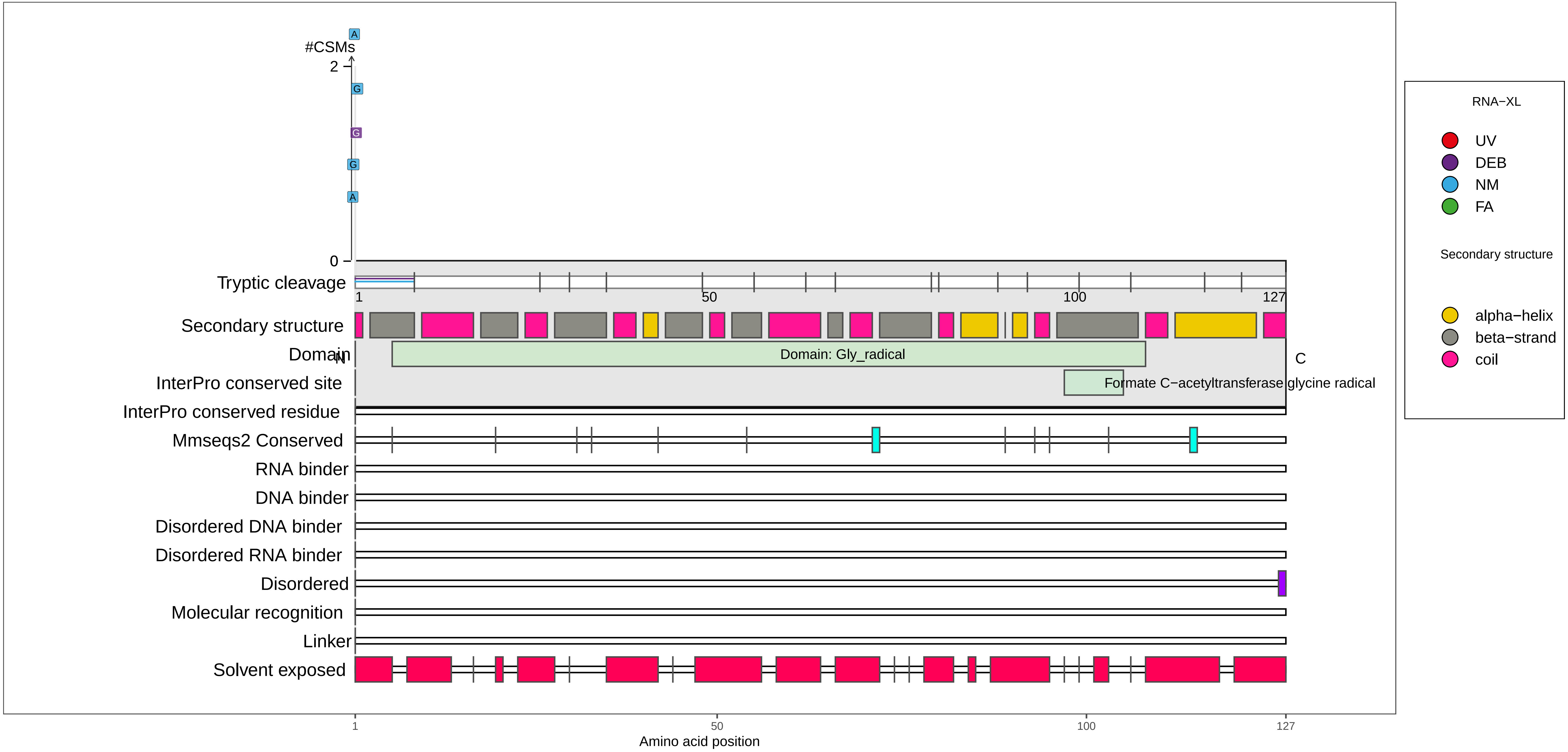

P68191
SRA_ECOLI Stationary−phase−induced ribosome−associated protein

− Abundance:
tryptic [log10 Intensity]: 7.17 (Q 20)
PAXdb K12 strain [ppm]: 2.91 (Q 88)
PAXdb E.coli [ppm]: 3.9 (Q 100)

− RNA functions: not annotated

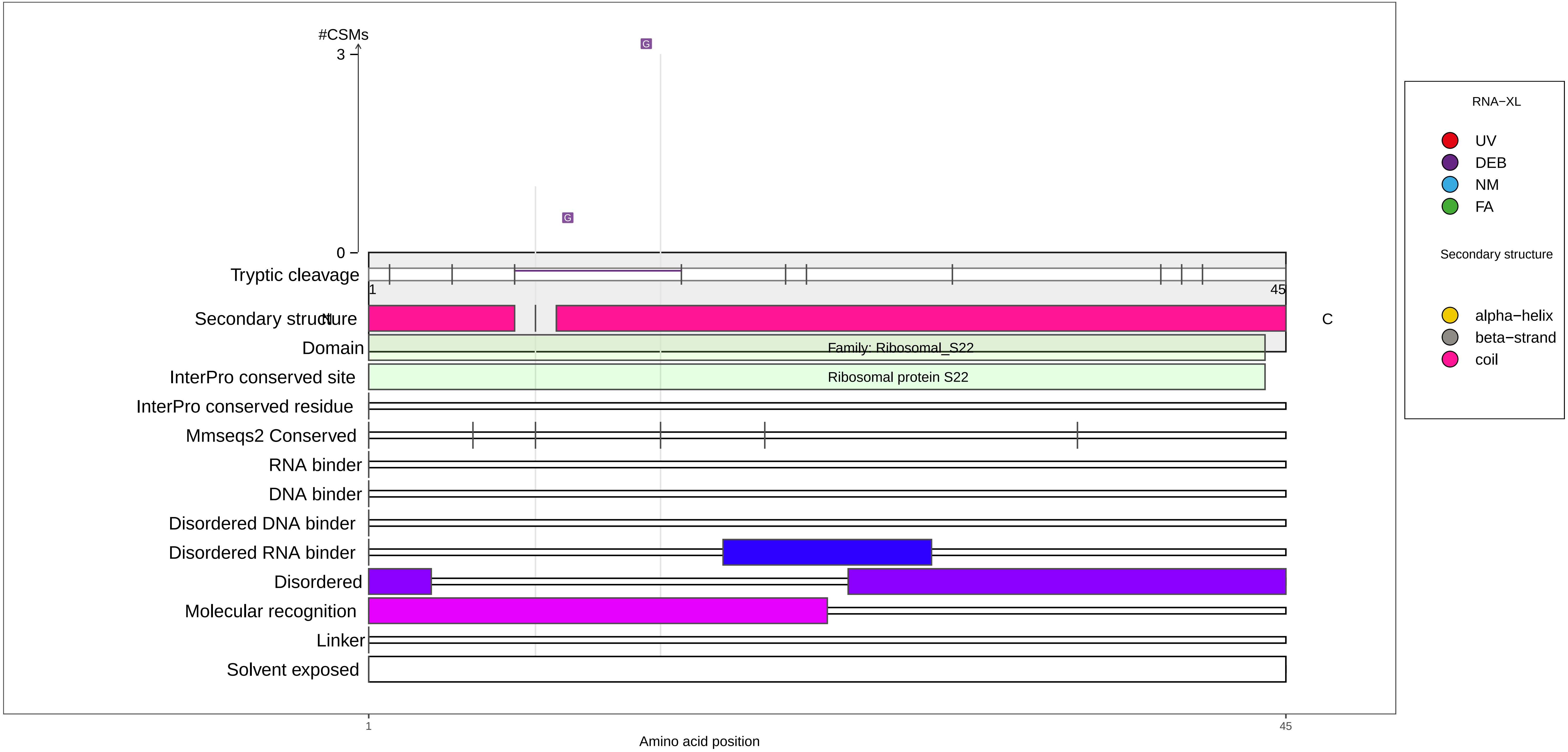

P69874
POTA_ECOLI Spermidine/putrescine import ATP−binding protein PotA

− Abundance:
tryptic [log10 Intensity]: 7.81 (Q 49)
PAXdb K12 strain [ppm]: 1.65 (Q 47)
PAXdb E.coli [ppm]: 1.82 (Q 68)

− RNA functions: not annotated

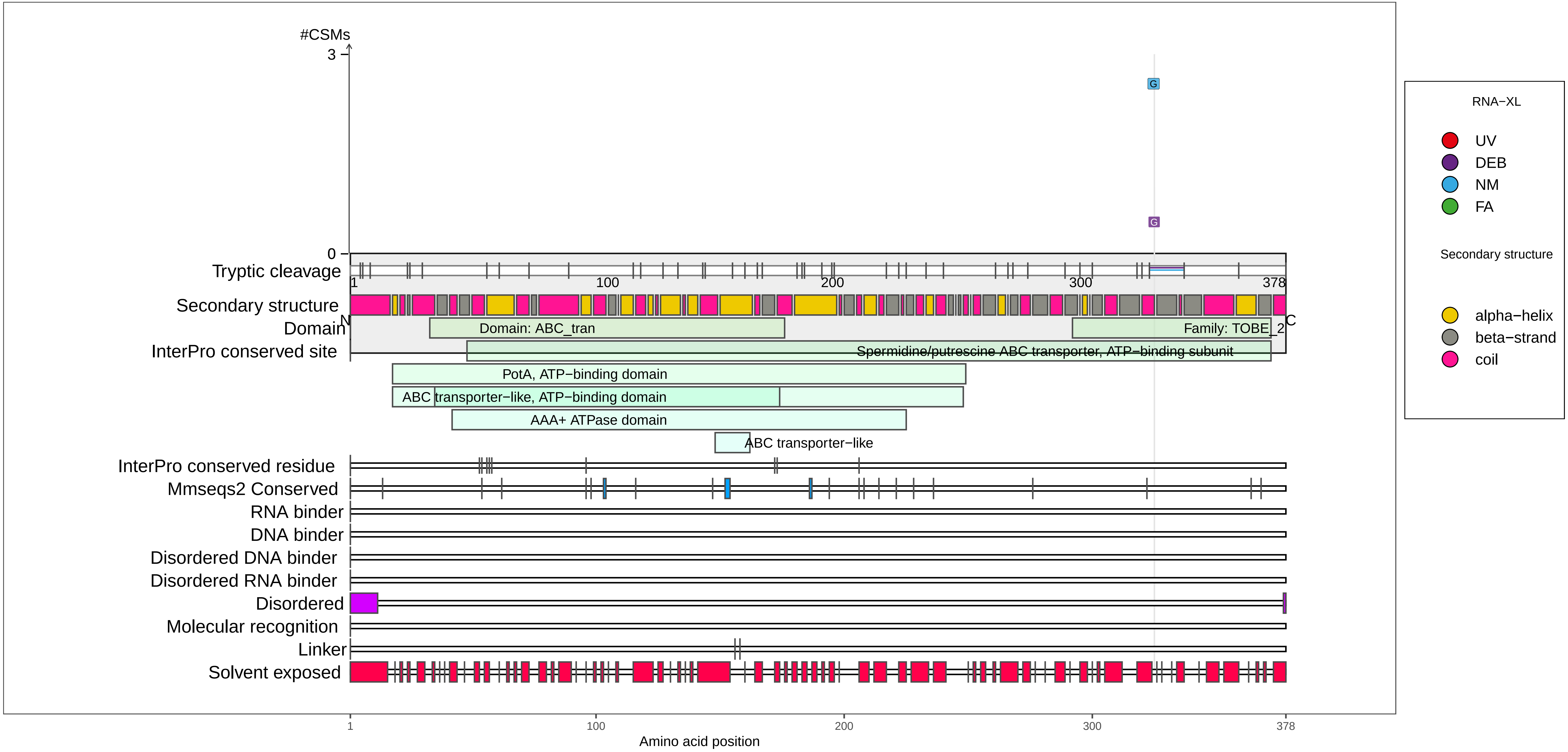

P76256
TSAB_ECOLI tRNA threonylcarbamoyladenosine biosynthesis protein TsaB

− Abundance:
tryptic [log10 Intensity]: 7.82 (Q 50)
PAXdb K12 strain [ppm]: 2.17 (Q 66)
PAXdb E.coli [ppm]: 1.31 (Q 55)

− RNA functions:
ncRNA metabolic process; ncRNA processing; RNA metabolic process; RNA modification
RNA processing; tRNA metabolic process; tRNA modification; tRNA processing
tRNA threonylcarbamoyladenosine metabolic process; tRNA threonylcarbamoyladenosine modification

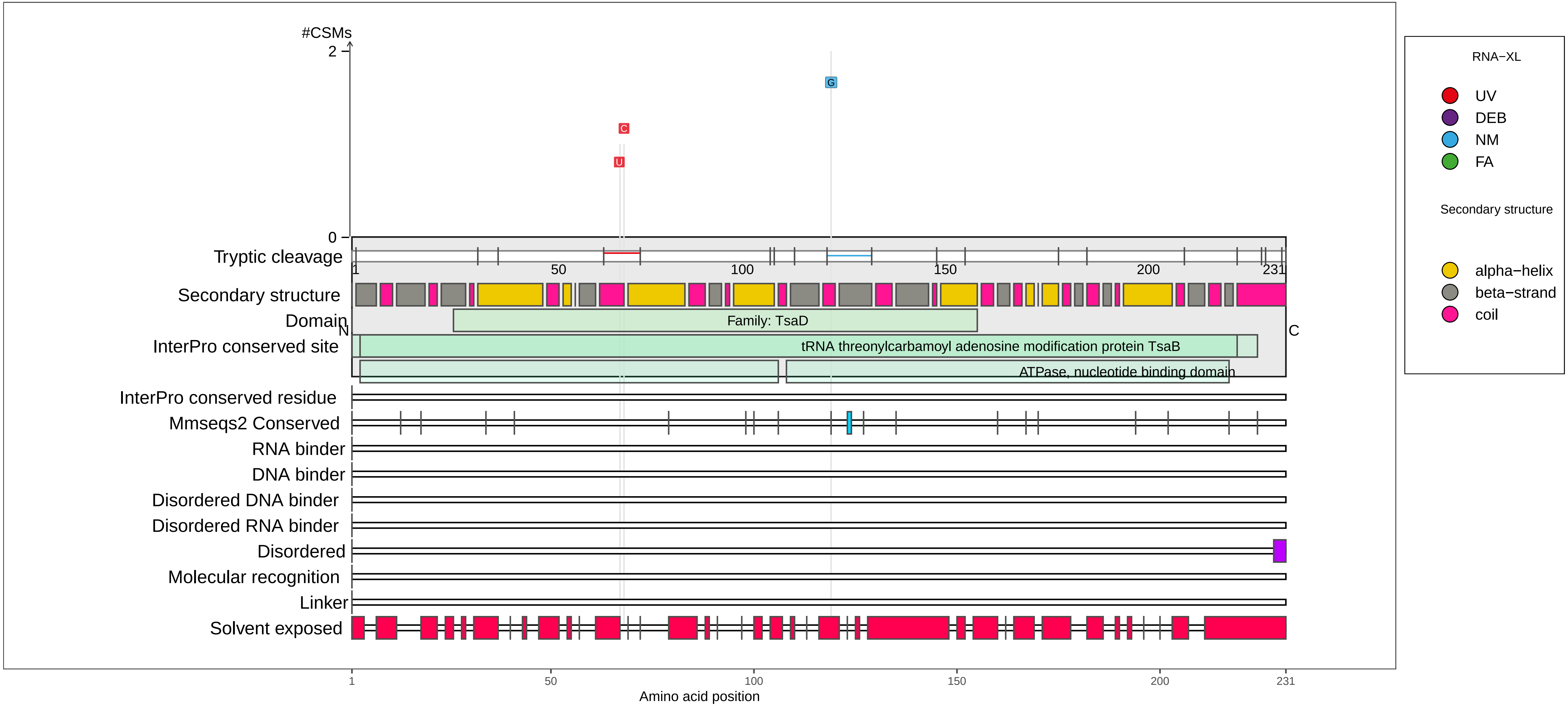

P77202
DSBG_ECOLI Thiol:disulfide interchange protein DsbG

− Abundance:
tryptic [log10 Intensity]: 7.26 (Q 23)
PAXdb K12 strain [ppm]: 0.98 (Q 10)
PAXdb E.coli [ppm]: 1.73 (Q 65)

− RNA functions: not annotated

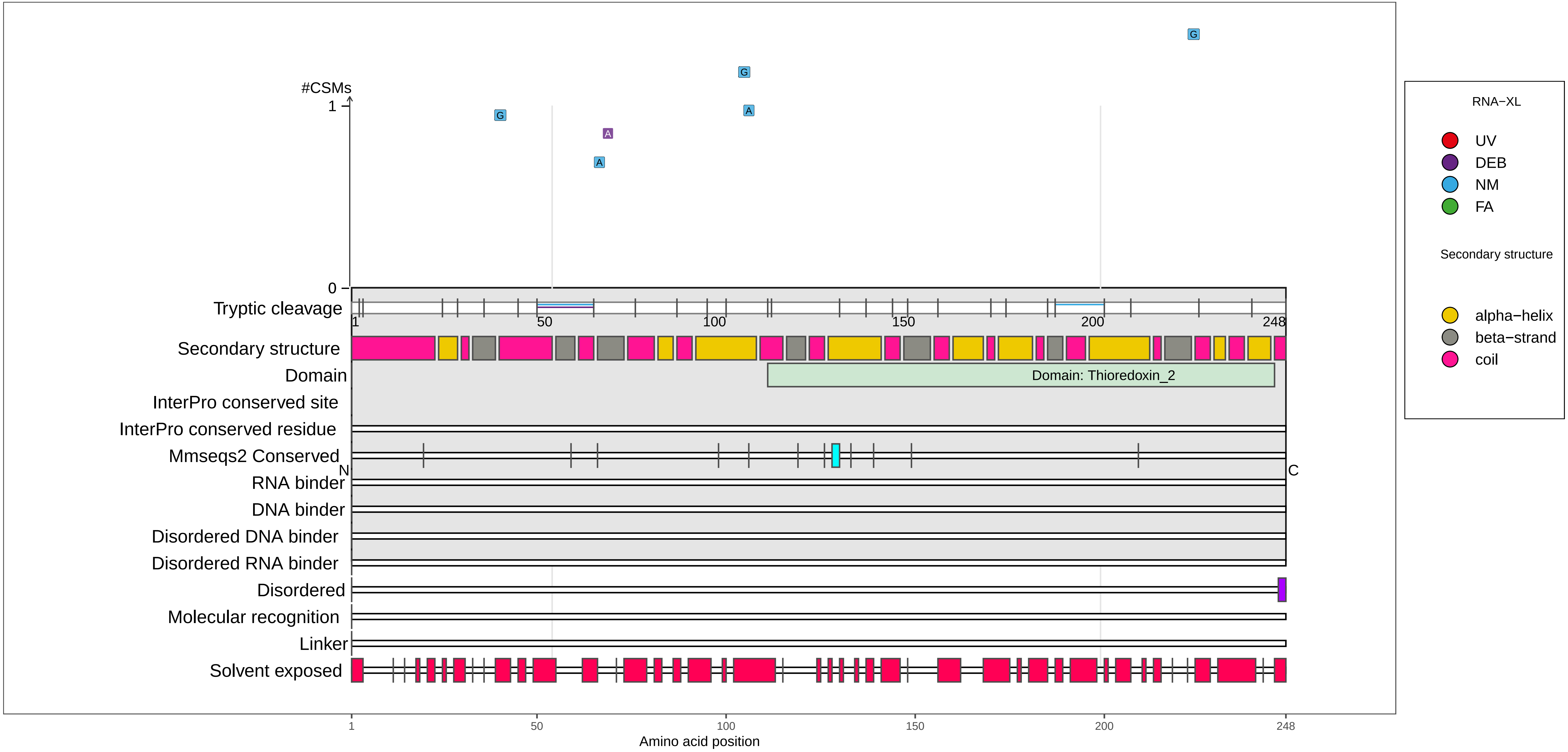

P77615
YCJW_ECOLI Uncharacterized HTH−type transcriptional regulator YcjW

− Abundance:
tryptic [log10 Intensity]: 7.71 (Q 45)
PAXdb K12 strain [ppm]: 1.33 (Q 28)
PAXdb E.coli [ppm]: 0.54 (Q 37)

− RNA functions:
RNA biosynthetic process; RNA metabolic process

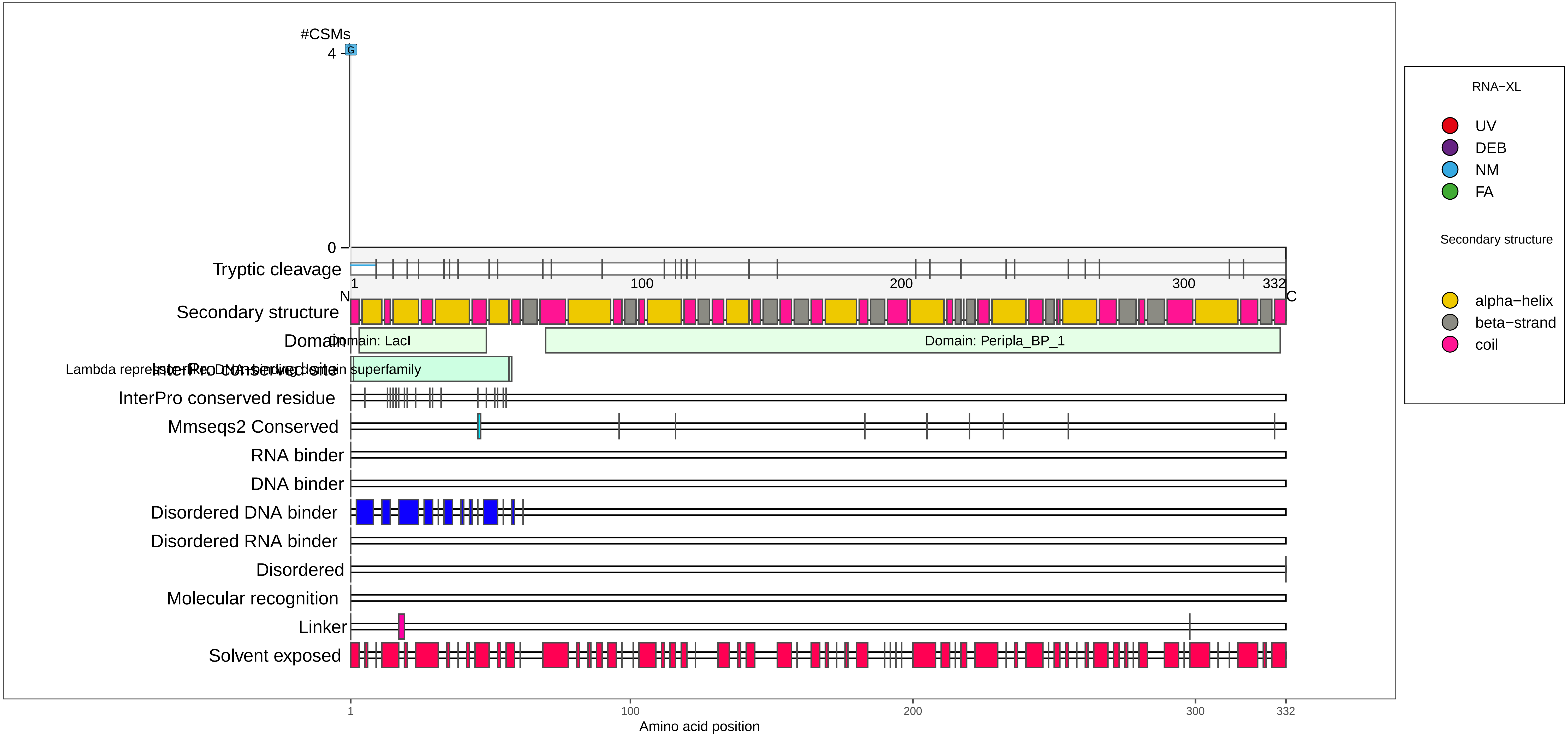

Q46864
MQSA_ECOLI Antitoxin MqsA

− Abundance:
tryptic [log10 Intensity]: 6.51 (Q 3)
PAXdb K12 strain [ppm]: 1.54 (Q 40)
PAXdb E.coli [ppm]: 0.65 (Q 40)

− RNA functions: not annotated

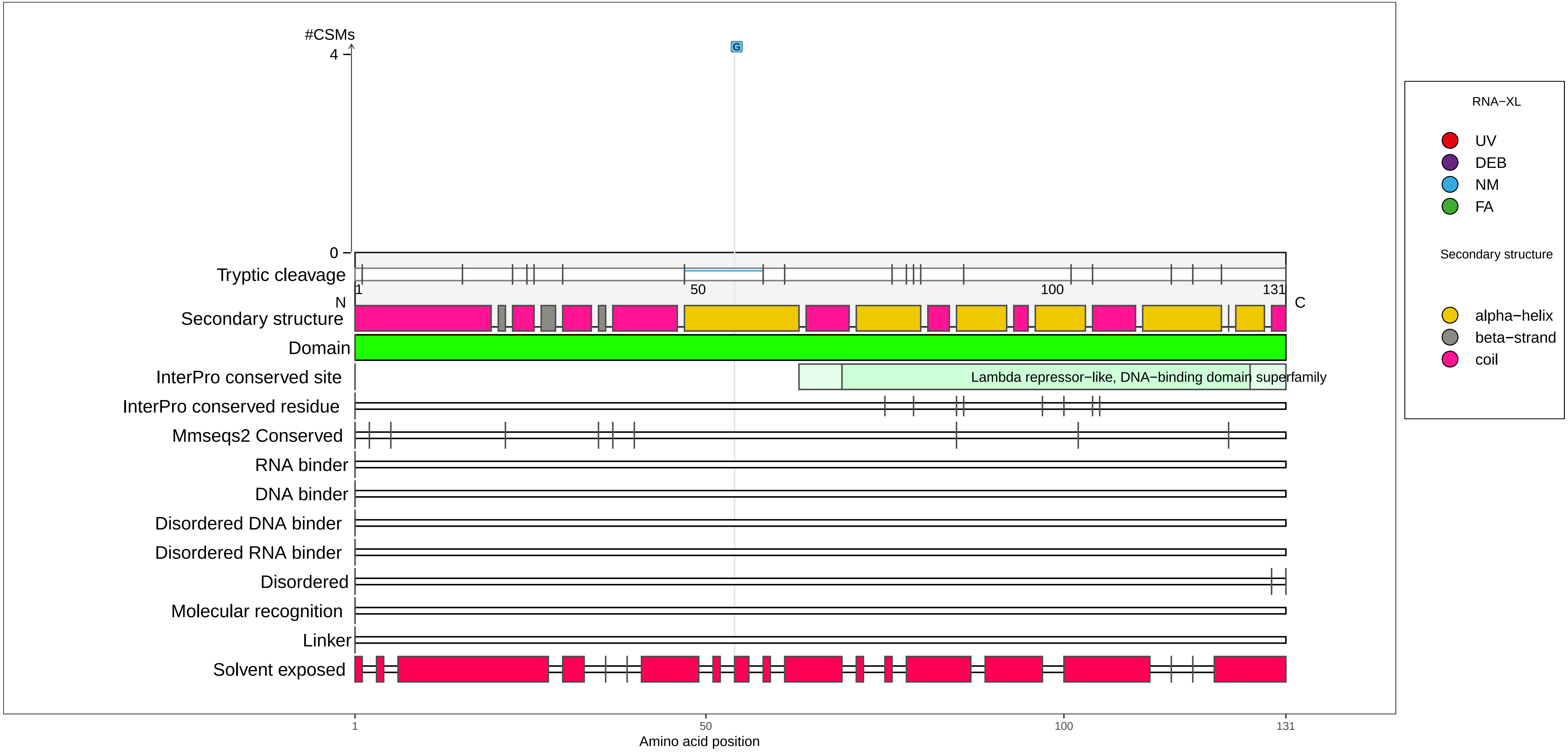

Q47147
YAFJ_ECOLI Putative glutamine amidotransferase YafJ

− Abundance:
tryptic [log10 Intensity]: 8.23 (Q 67)
PAXdb K12 strain [ppm]: 1.77 (Q 52)
PAXdb E.coli [ppm]: 1.05 (Q 49)

− RNA functions: not annotated

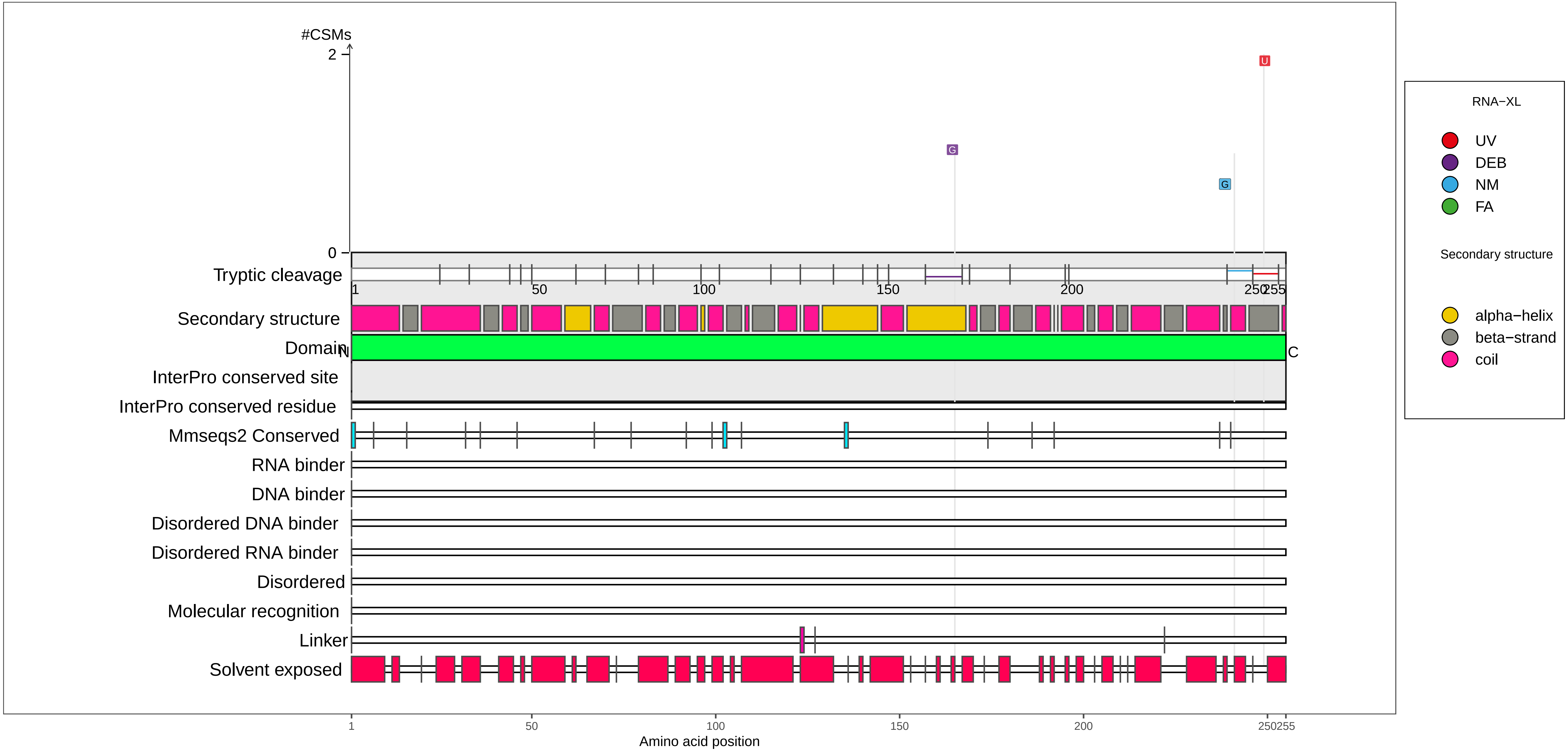

Q47679
YAFV_ECOLI Omega−amidase YafV

− Abundance:
tryptic [log10 Intensity]: 7.29 (Q 24)
PAXdb K12 strain [ppm]: NA (Q NA)
PAXdb E.coli [ppm]: 1.76 (Q 66)

− RNA functions: not annotated

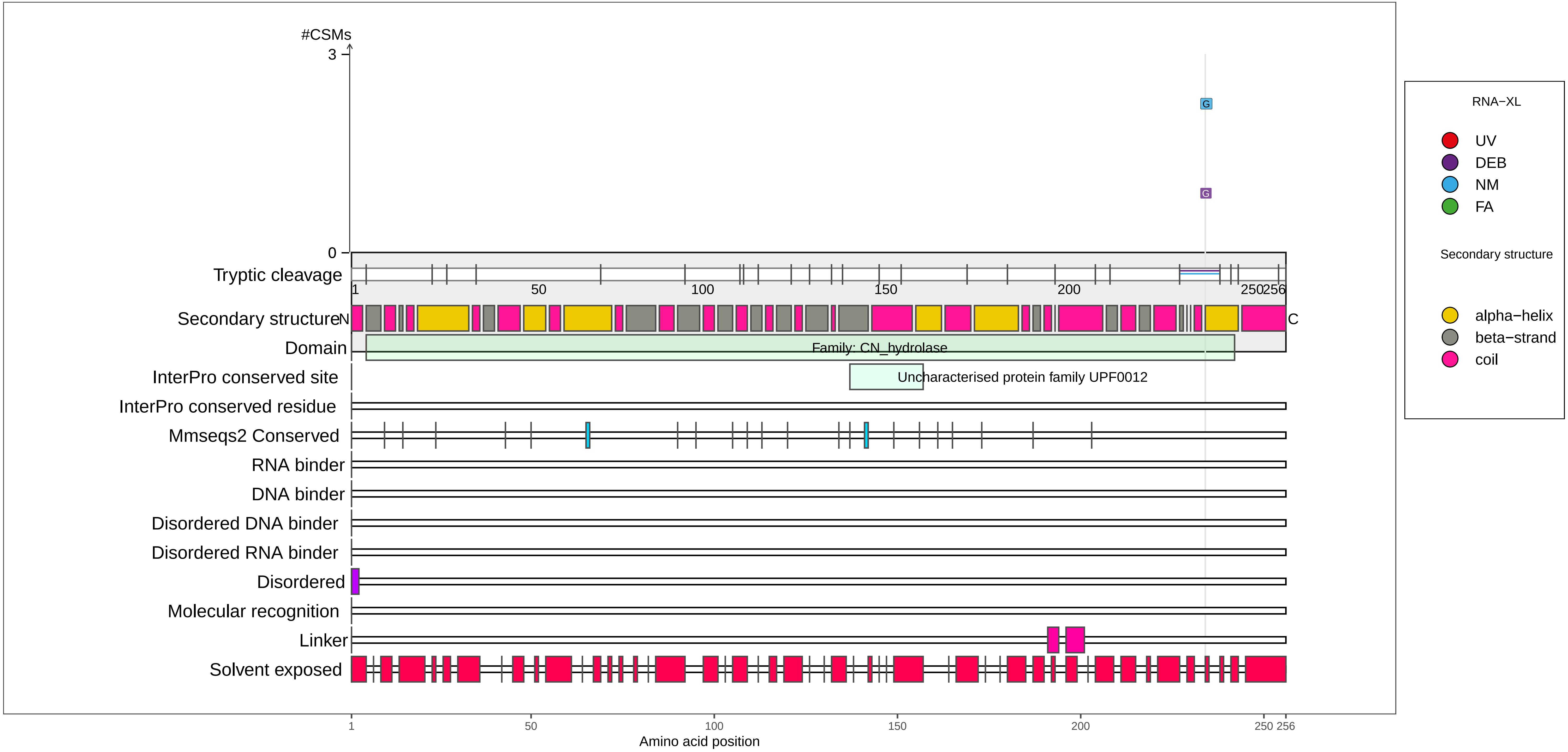

P02924
ARAF_ECOLI L−arabinose−binding periplasmic protein

− Abundance:
tryptic [log10 Intensity]: 7.15 (Q 19)
PAXdb K12 strain [ppm]: 1.67 (Q 48)
PAXdb E.coli [ppm]: 2.54 (Q 86)

− RNA functions: not annotated

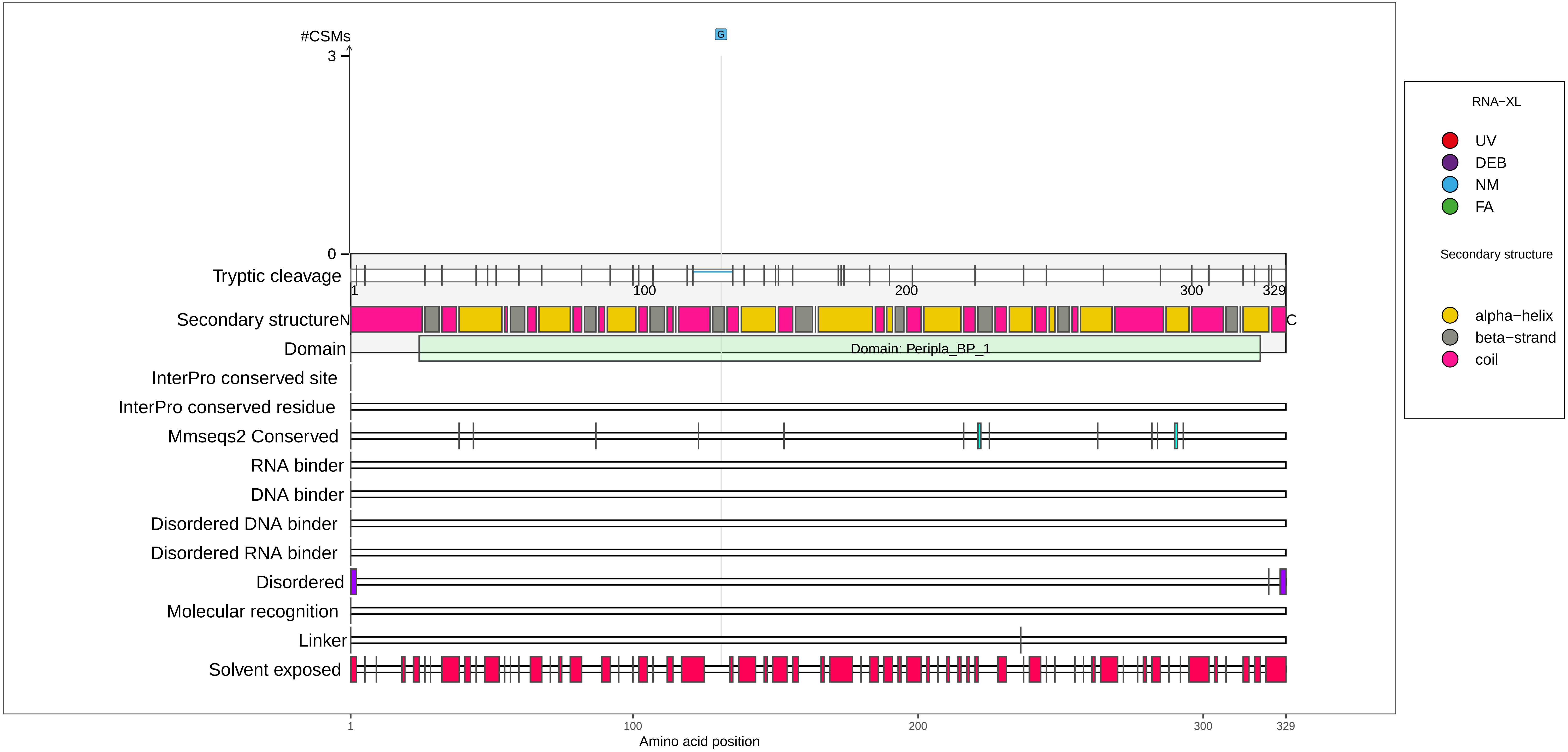

P04693
TYRB_ECOLI Aromatic−amino−acid aminotransferase

− Abundance:
tryptic [log10 Intensity]: 8.34 (Q 71)
PAXdb K12 strain [ppm]: 2.82 (Q 87)
PAXdb E.coli [ppm]: 2.39 (Q 83)

− RNA functions: not annotated

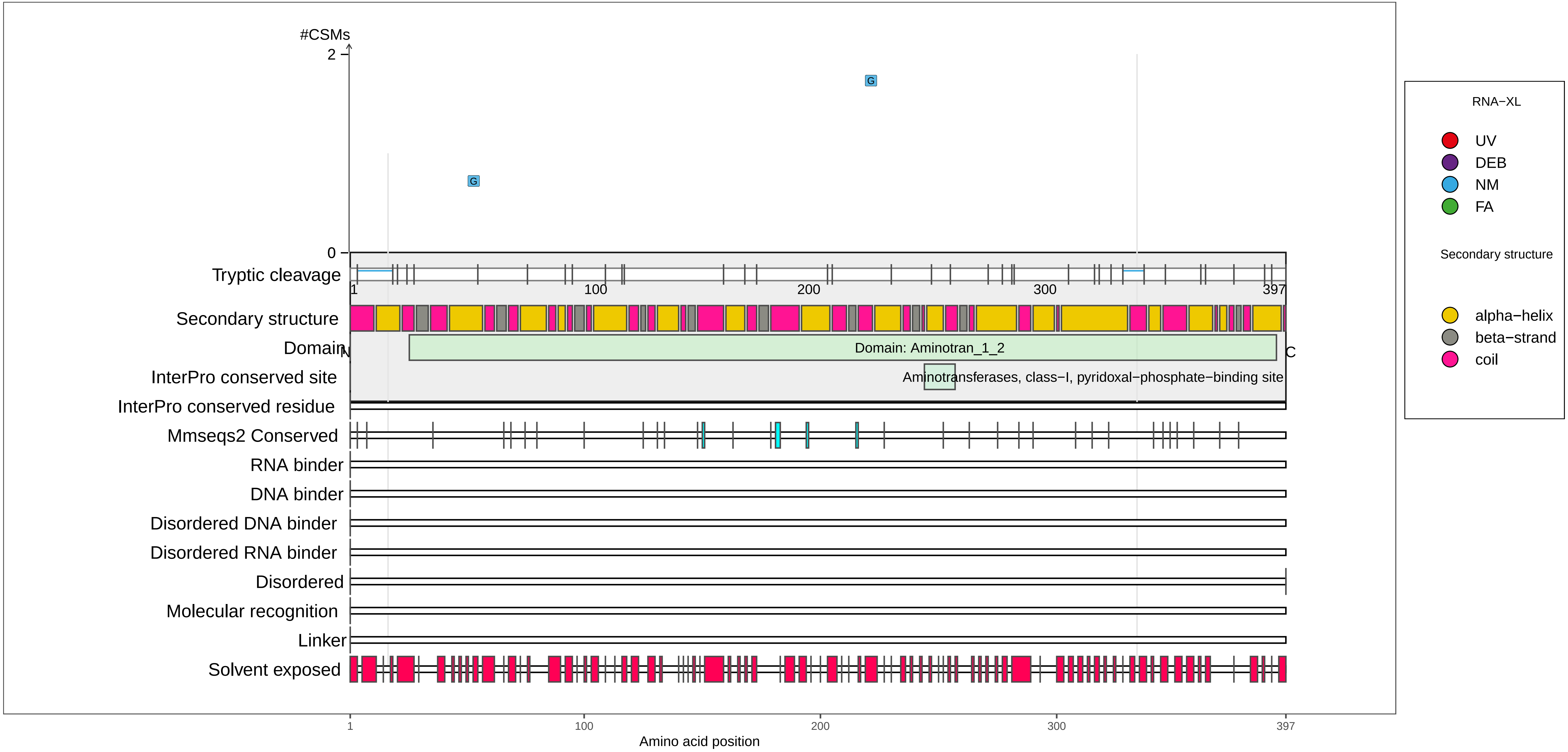

P04994
EX7L_ECOLI Exodeoxyribonuclease 7 large subunit

− Abundance:
tryptic [log10 Intensity]: 8.16 (Q 64)
PAXdb K12 strain [ppm]: 1.28 (Q 25)
PAXdb E.coli [ppm]: 0.33 (Q 32)

− RNA functions: not annotated

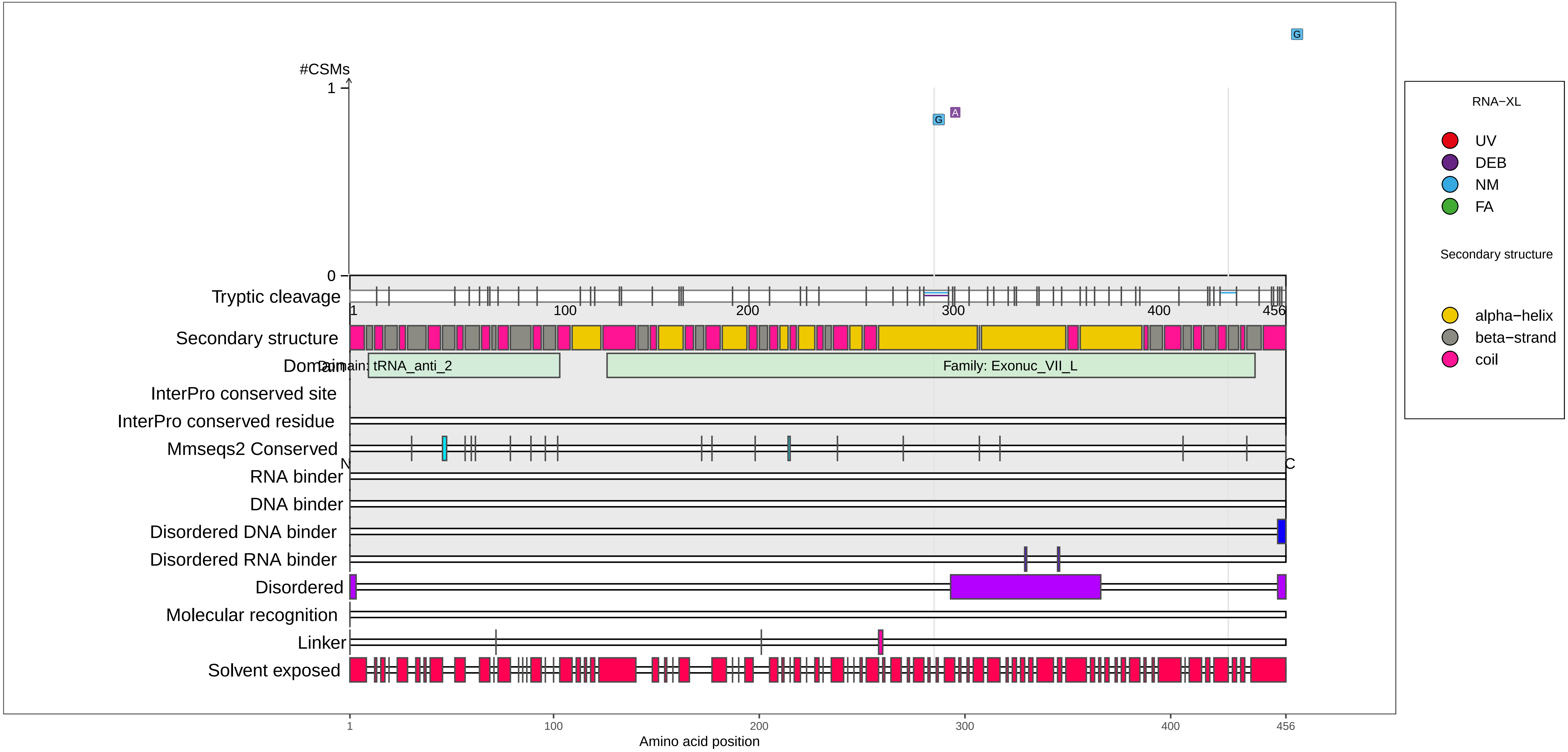

P05804
BGLR_ECOLI Beta−glucuronidase

− Abundance:
tryptic [log10 Intensity]: 7.1 (Q 17)
PAXdb K12 strain [ppm]: 1.76 (Q 52)
PAXdb E.coli [ppm]: 0.55 (Q 37)

− RNA functions: not annotated

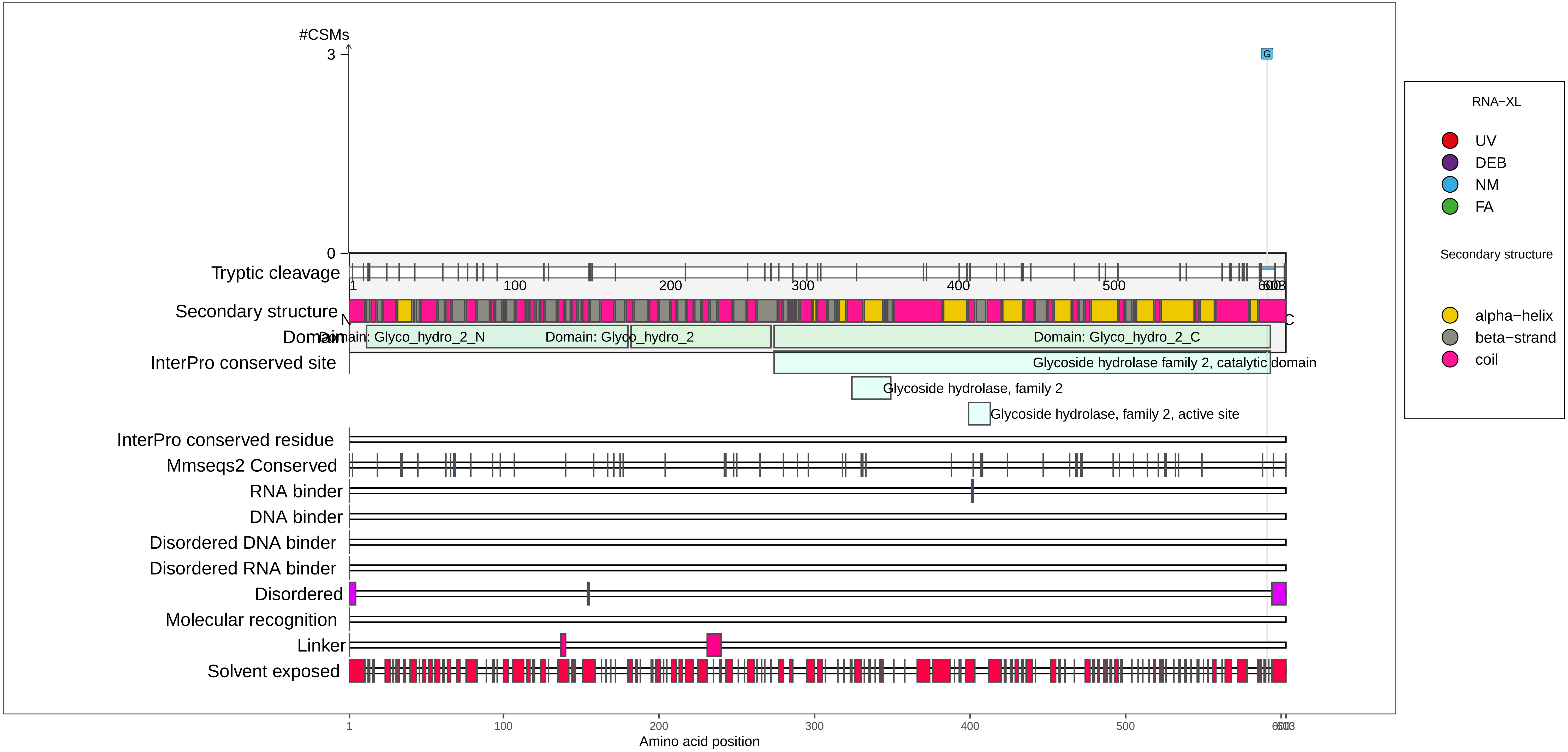

P06966
DICA_ECOLI HTH−type transcriptional regulator DicA
− Abundance:
tryptic [log10 Intensity]: NA (Q NA)
PAXdb K12 strain [ppm]: 1.37 (Q 30)
PAXdb E.coli [ppm]: 1.59 (Q 62)

− RNA functions:
RNA biosynthetic process; RNA metabolic process

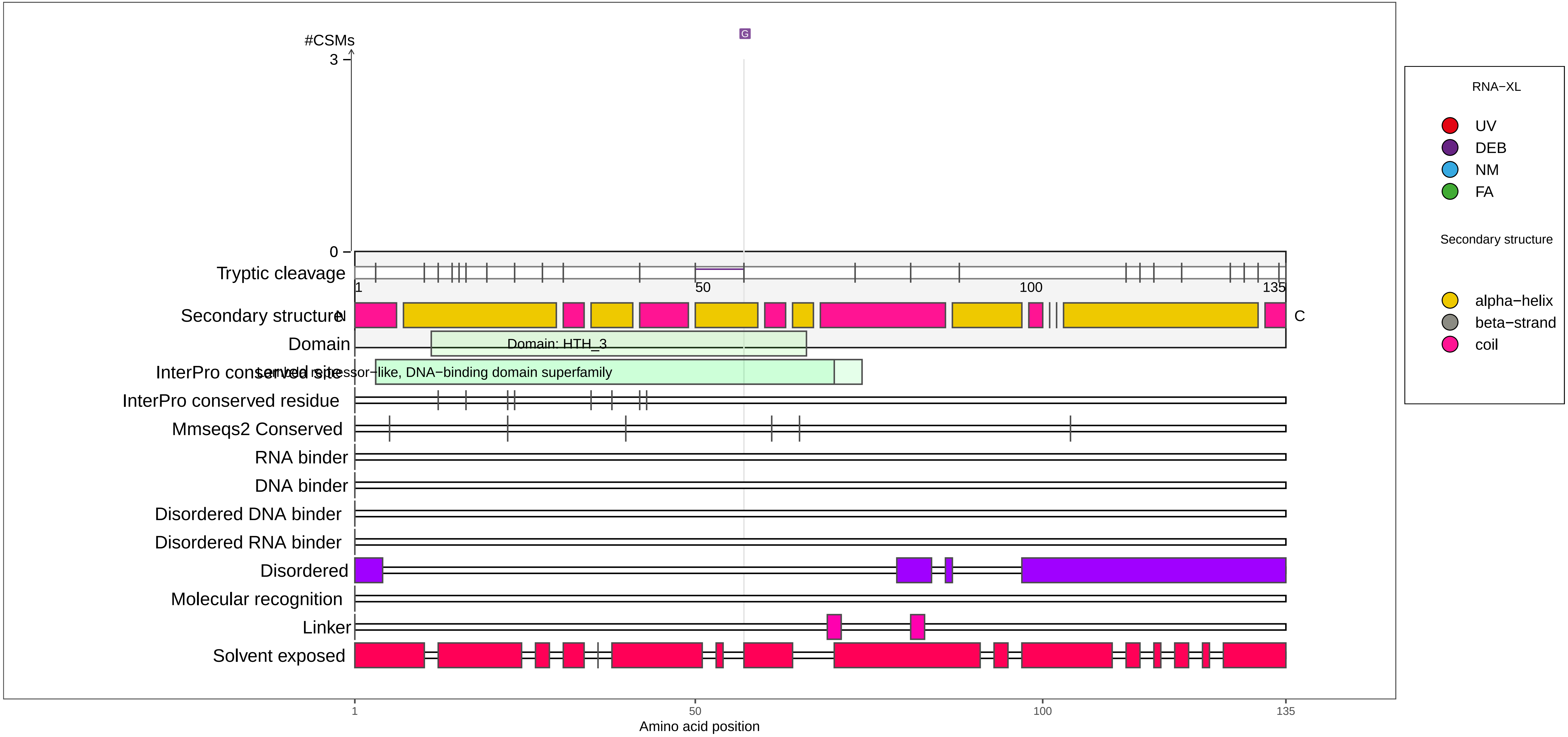

P06999
PFKB_ECOLI ATP−dependent 6−phosphofructokinase isozyme 2

− Abundance:
tryptic [log10 Intensity]: 8.57 (Q 78)
PAXdb K12 strain [ppm]: 2.99 (Q 91)
PAXdb E.coli [ppm]: 2.47 (Q 85)

− RNA functions: not annotated

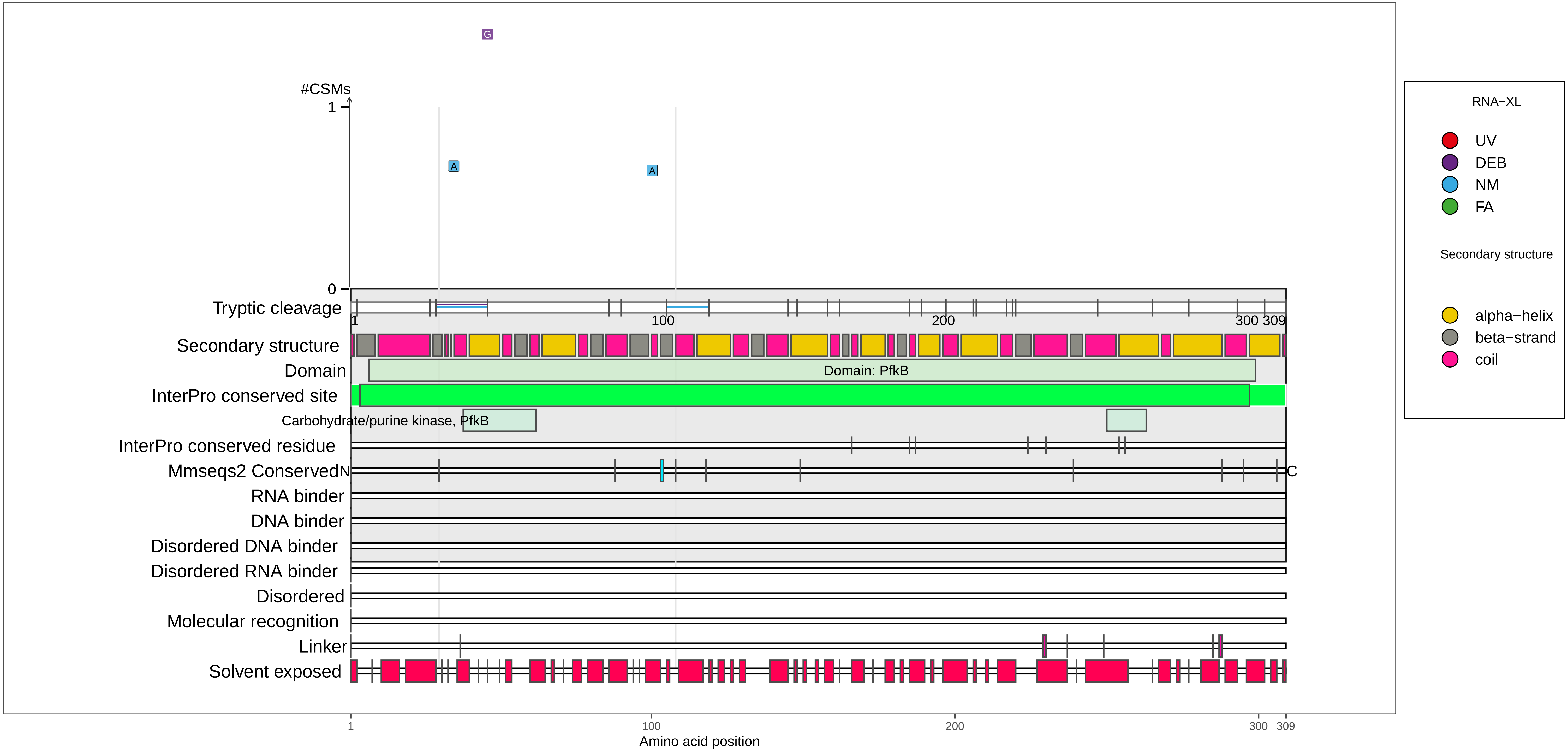

P07913
TDH_ECOLI L−threonine 3−dehydrogenase

− Abundance:
tryptic [log10 Intensity]: 9.3 (Q 94)
PAXdb K12 strain [ppm]: 2.69 (Q 83)
PAXdb E.coli [ppm]: 1.95 (Q 71)

− RNA functions: not annotated

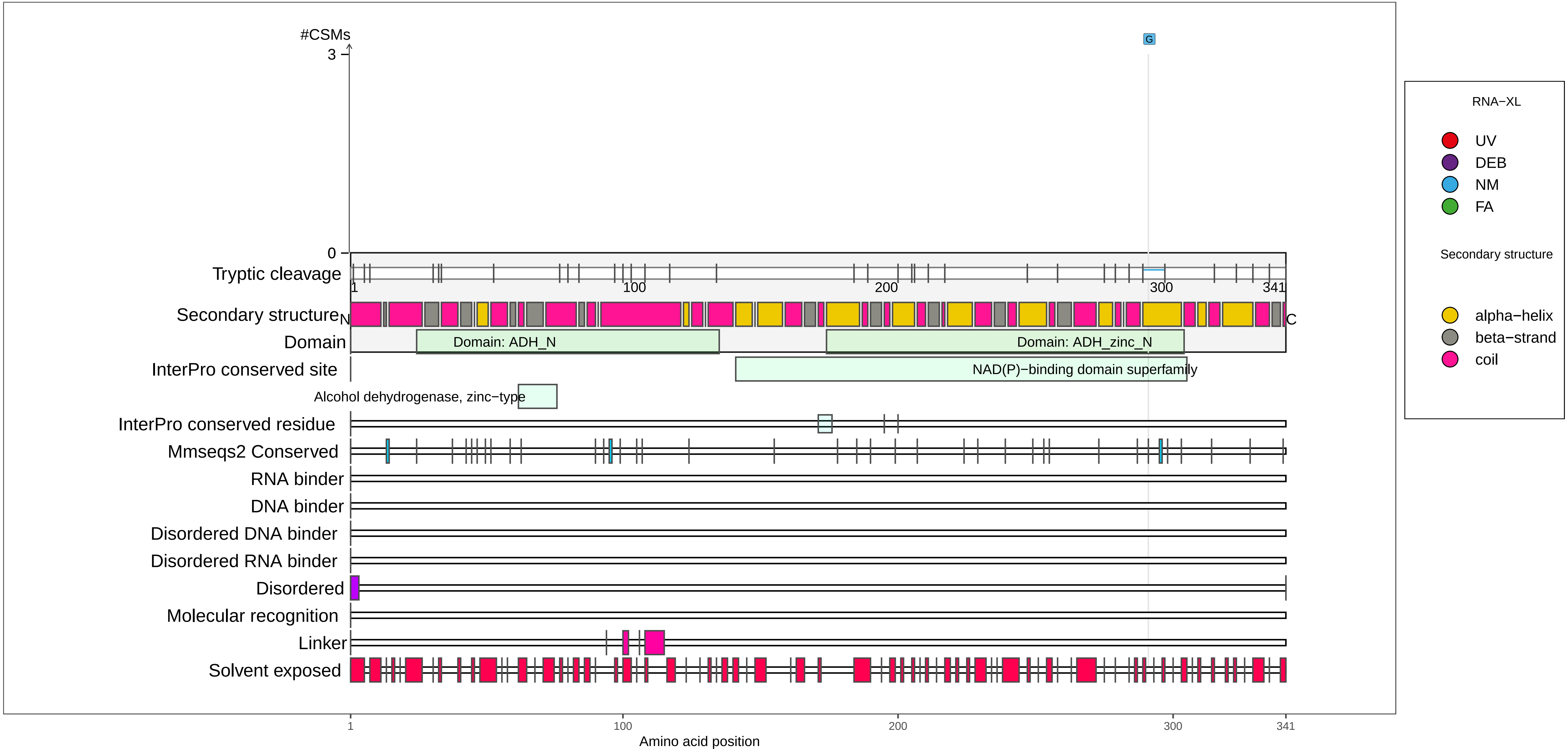

P08194
GLPT_ECOLI Glycerol−3−phosphate transporter

− Abundance:
tryptic [log10 Intensity]: 9.34 (Q 95)
PAXdb K12 strain [ppm]: 2.47 (Q 76)
PAXdb E.coli [ppm]: 1.61 (Q 63)

− RNA functions: not annotated

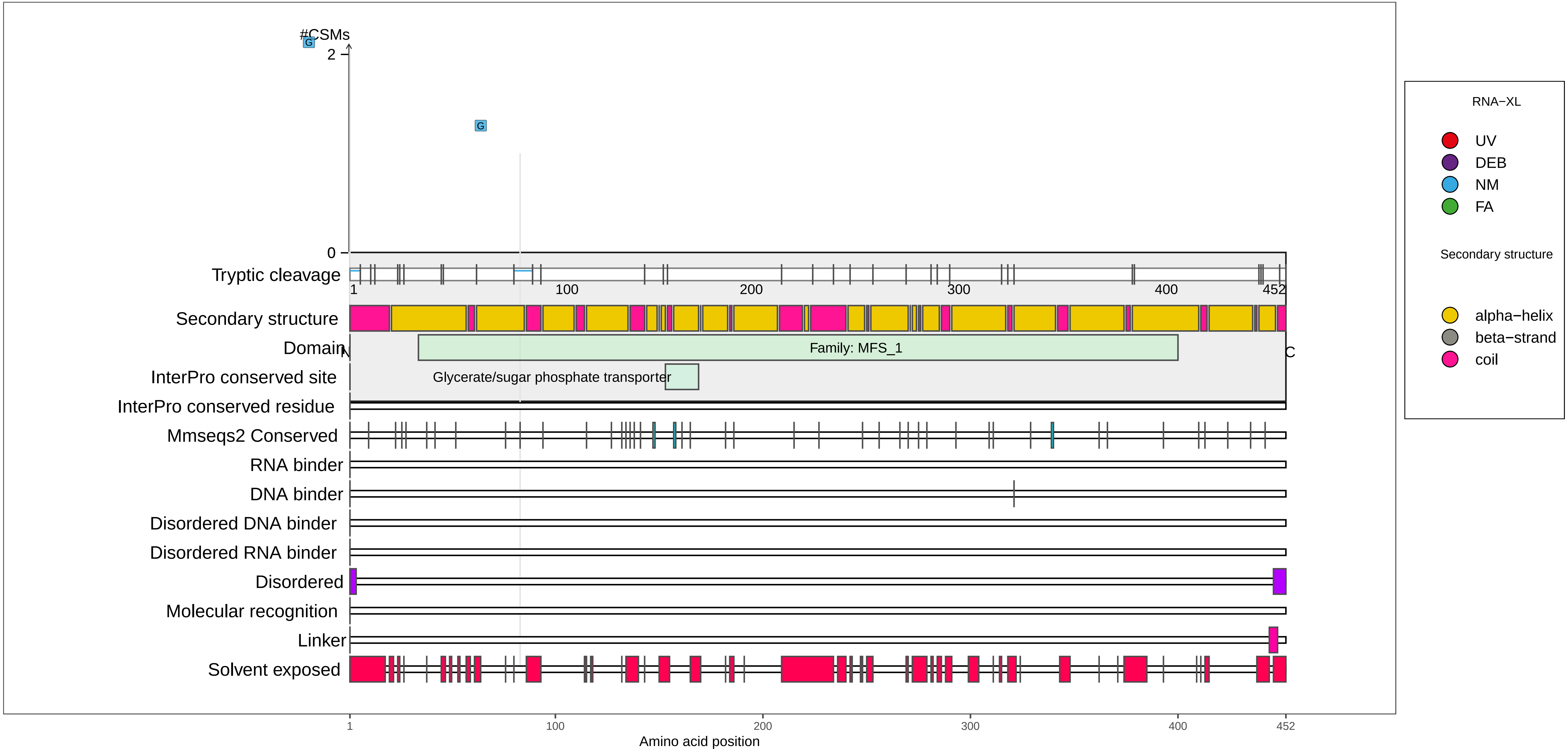

P09832
GLTD_ECOLI Glutamate synthase [NADPH] small chain

− Abundance:
tryptic [log10 Intensity]: 7.65 (Q 43)
PAXdb K12 strain [ppm]: 2.66 (Q 82)
PAXdb E.coli [ppm]: 2.92 (Q 94)

− RNA functions: not annotated

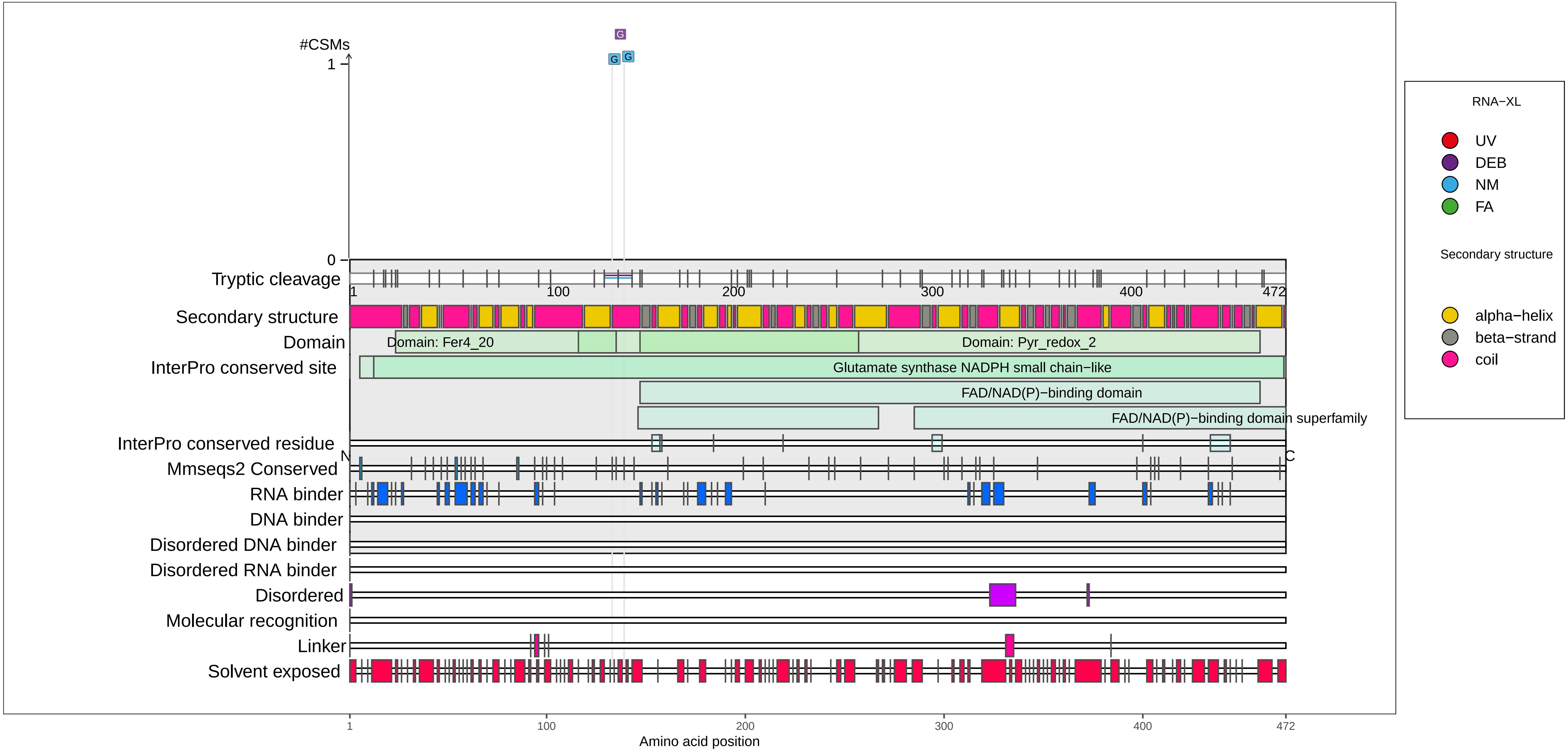

P0A6N8
EFPL_ECOLI Elongation factor P−like protein

− Abundance:
tryptic [log10 Intensity]: 8.57 (Q 78)
PAXdb K12 strain [ppm]: 3.03 (Q 91)
PAXdb E.coli [ppm]: 2.72 (Q 90)

− RNA functions:
RNA binding

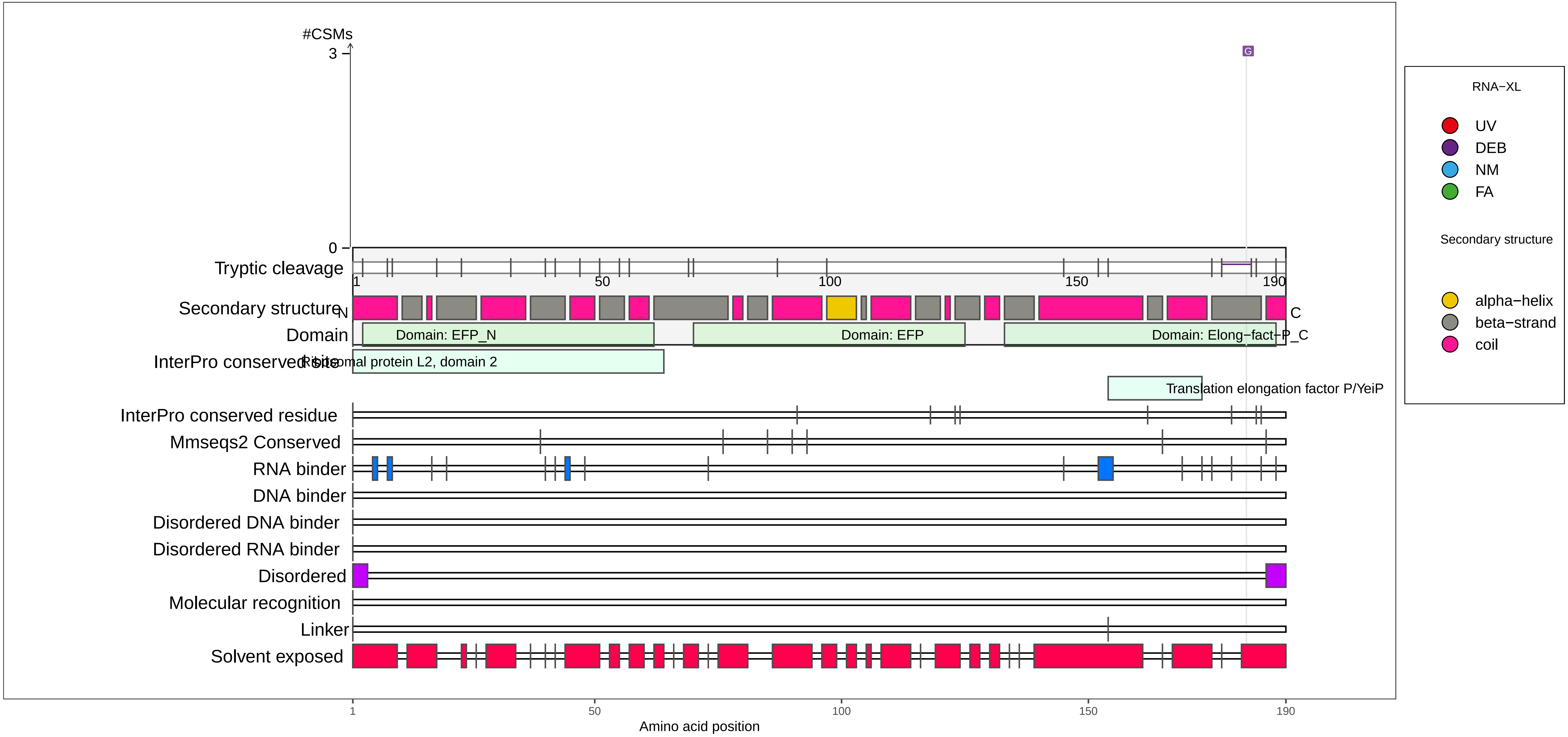

P0A731
MGSA_ECOLI Methylglyoxal synthase

− Abundance:
tryptic [log10 Intensity]: 8.04 (Q 59)
PAXdb K12 strain [ppm]: NA (Q NA)
PAXdb E.coli [ppm]: 2.39 (Q 83)

− RNA functions: not annotated

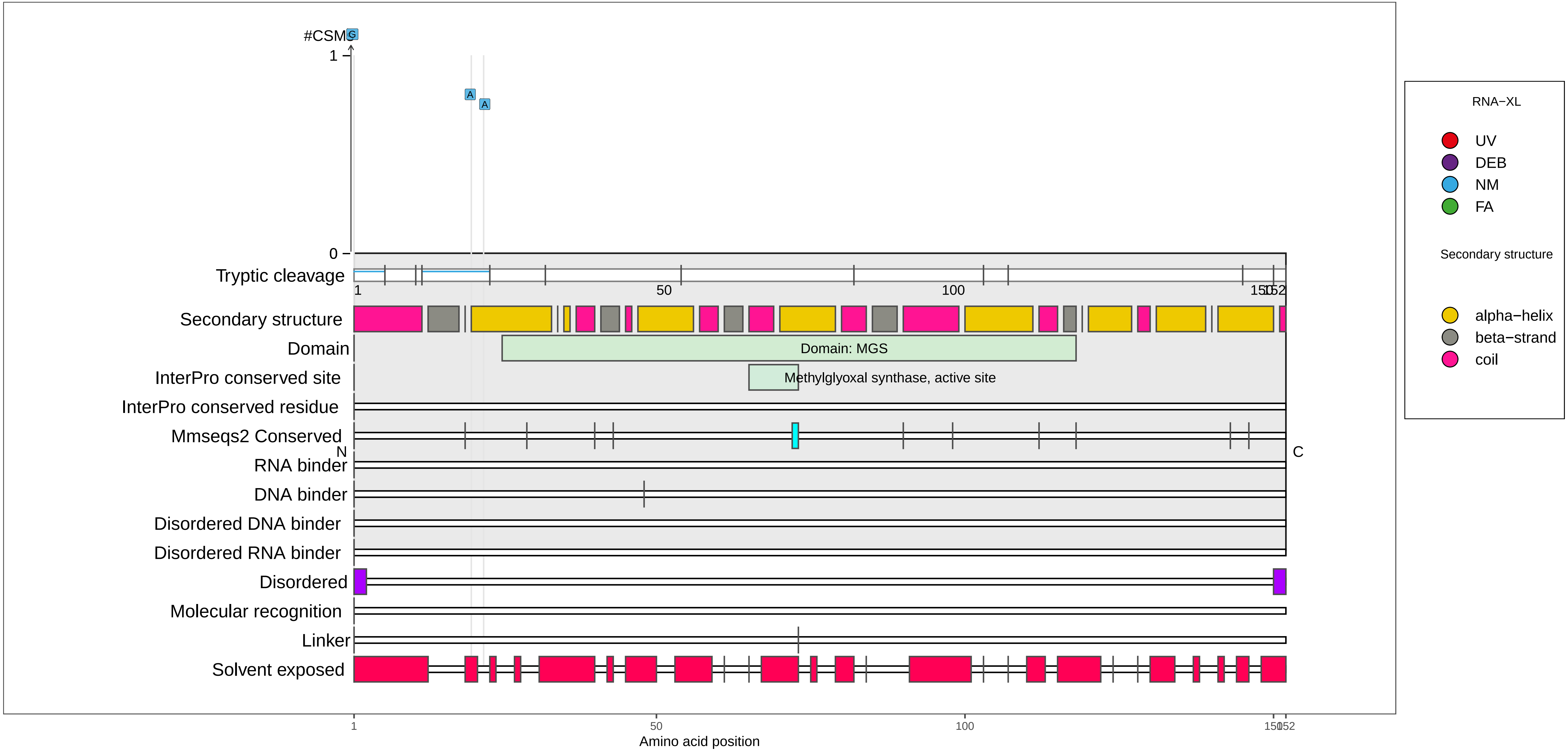

P0A746
MSRB_ECOLI Peptide methionine sulfoxide reductase MsrB
− Abundance:
tryptic [log10 Intensity]: 7.57 (Q 38)
PAXdb K12 strain [ppm]: 2.46 (Q 76)
PAXdb E.coli [ppm]: 2.7 (Q 90)

− RNA functions: not annotated

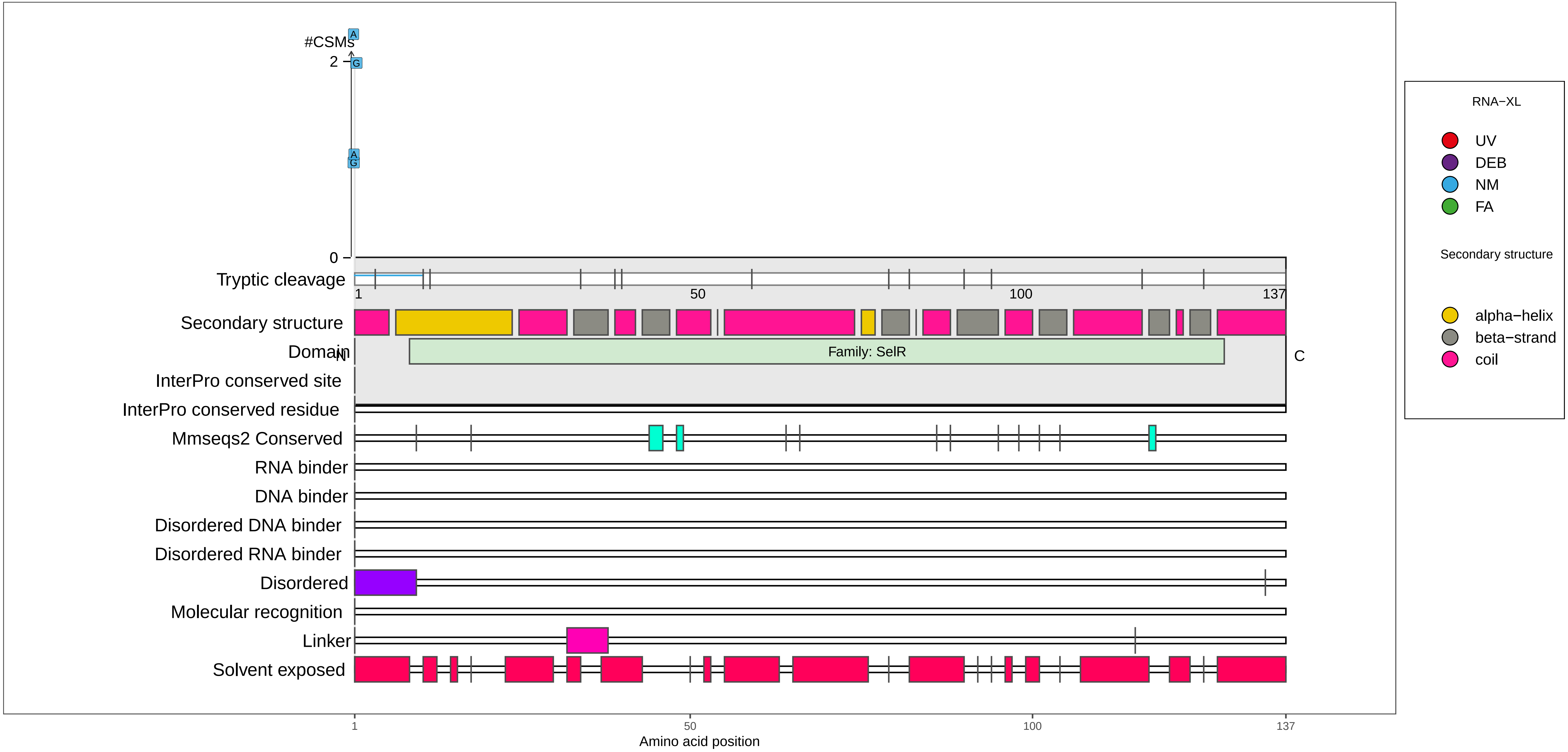

P0A7B3
NADK_ECOLI NAD kinase

− Abundance:
tryptic [log10 Intensity]: 7.58 (Q 39)
PAXdb K12 strain [ppm]: 1.04 (Q 13)
PAXdb E.coli [ppm]: 1.62 (Q 63)

− RNA functions: not annotated

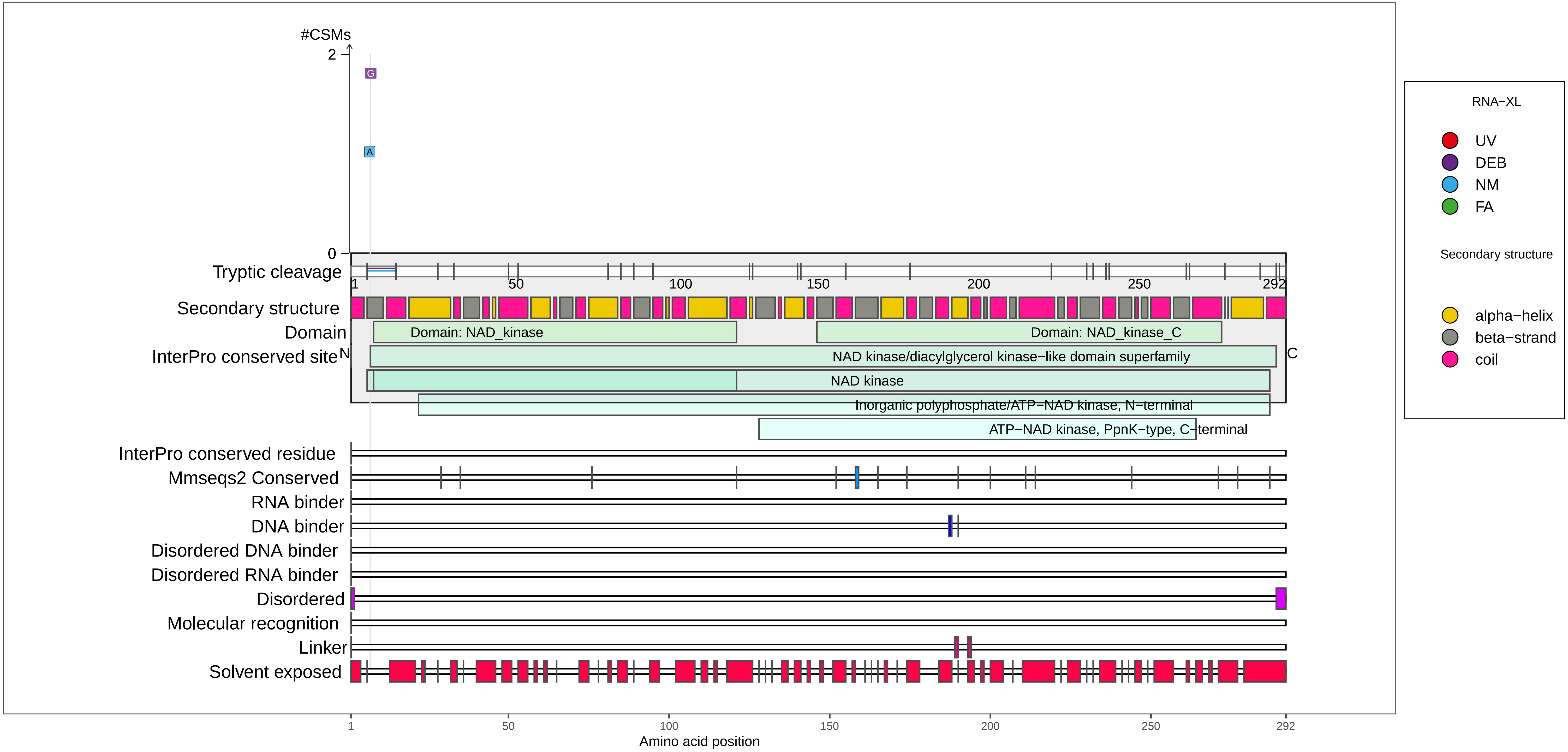

P0A7C2
LEXA_ECOLI LexA repressor
− Abundance:
tryptic [log10 Intensity]: 8.04 (Q 59)
PAXdb K12 strain [ppm]: 2.32 (Q 70)
PAXdb E.coli [ppm]: 1.82 (Q 67)

− RNA functions:
RNA biosynthetic process; RNA metabolic process

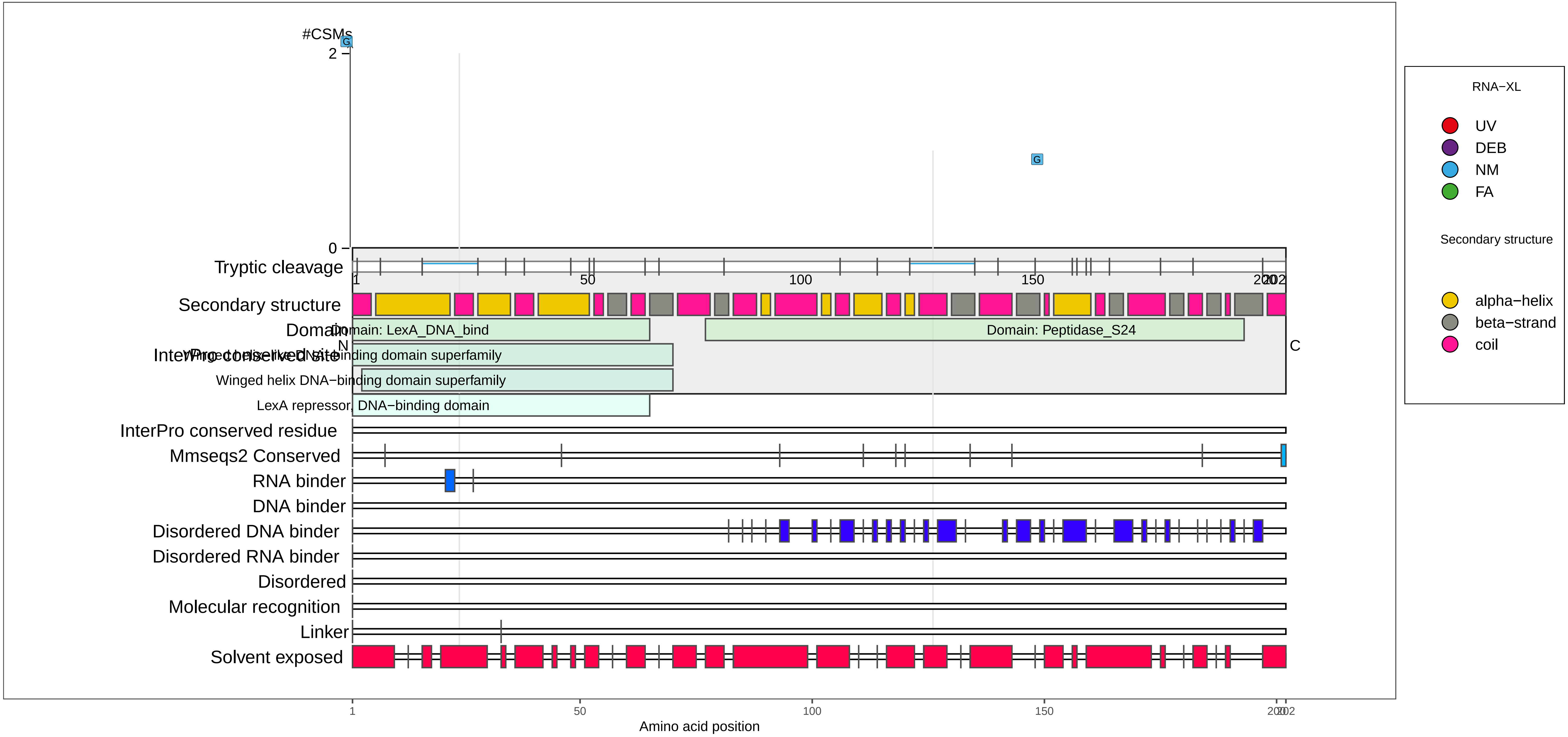

P0A867
TALA_ECOLI Transaldolase A

− Abundance:
tryptic [log10 Intensity]: 7.51 (Q 35)
PAXdb K12 strain [ppm]: 2.73 (Q 84)
PAXdb E.coli [ppm]: 3.1 (Q 96)

− RNA functions: not annotated

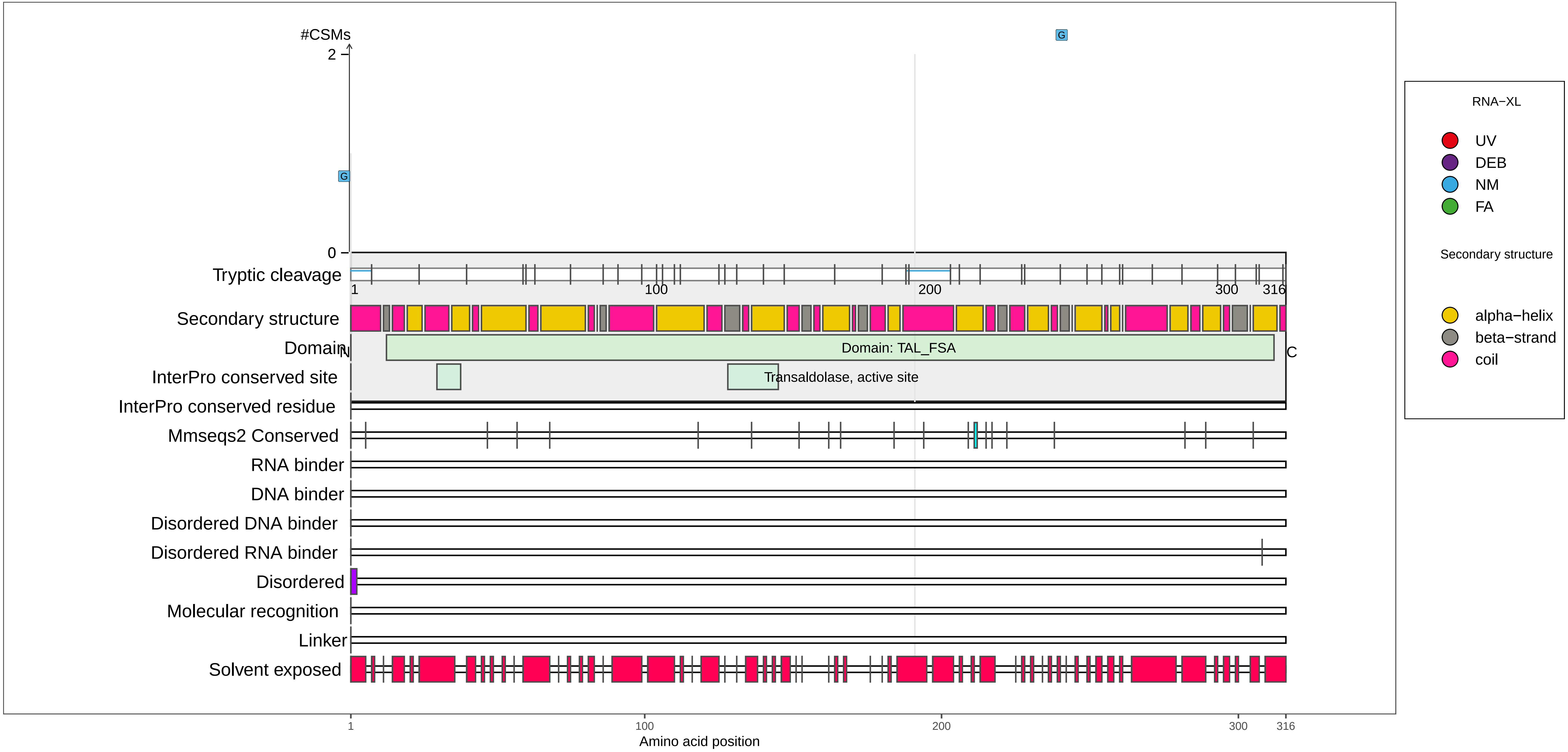

P0A8F8
UVRB_ECOLI UvrABC system protein B

− Abundance:
tryptic [log10 Intensity]: 8.22 (Q 66)
PAXdb K12 strain [ppm]: 2.26 (Q 68)
PAXdb E.coli [ppm]: 1.77 (Q 66)

− RNA functions: not annotated

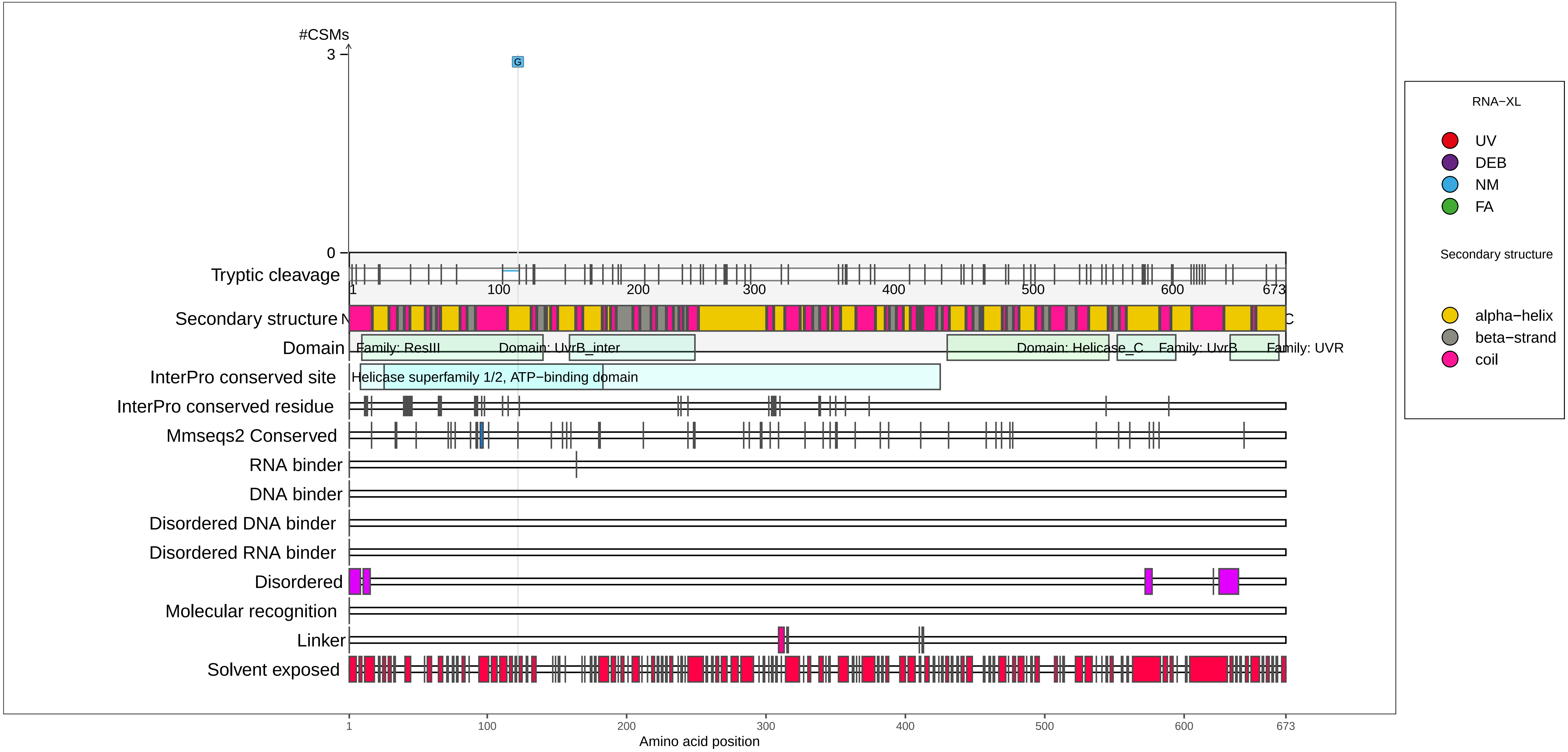

P0A8P6
XERC_ECOLI Tyrosine recombinase XerC

− Abundance:
tryptic [log10 Intensity]: NA (Q NA)
PAXdb K12 strain [ppm]: 1.47 (Q 37)
PAXdb E.coli [ppm]: 0.46 (Q 35)

− RNA functions: not annotated

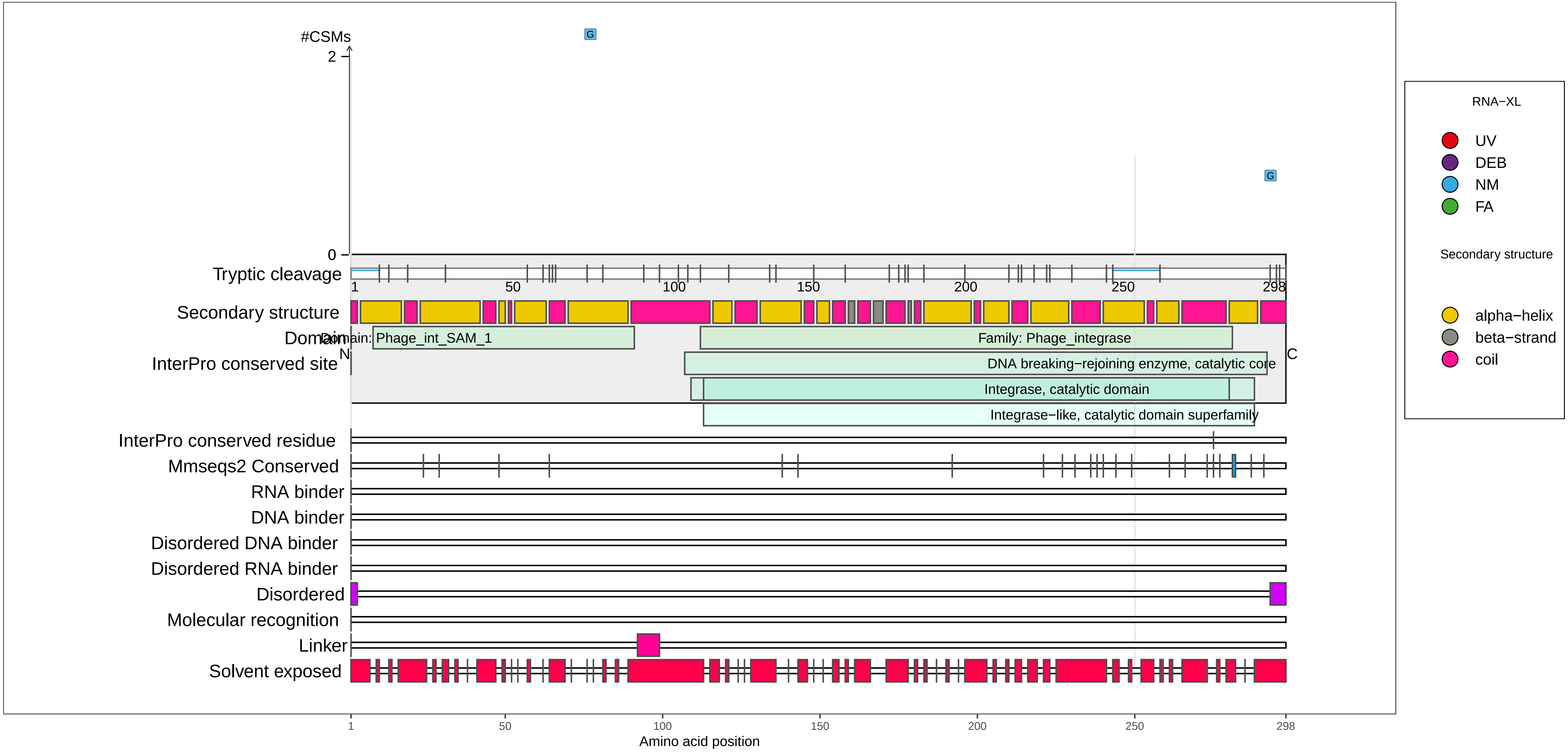

P0A8P8
XERD_ECOLI Tyrosine recombinase XerD

− Abundance:
tryptic [log10 Intensity]: NA (Q NA)
PAXdb K12 strain [ppm]: 0.92 (Q 8)
PAXdb E.coli [ppm]: 0.26 (Q 31)

− RNA functions: not annotated

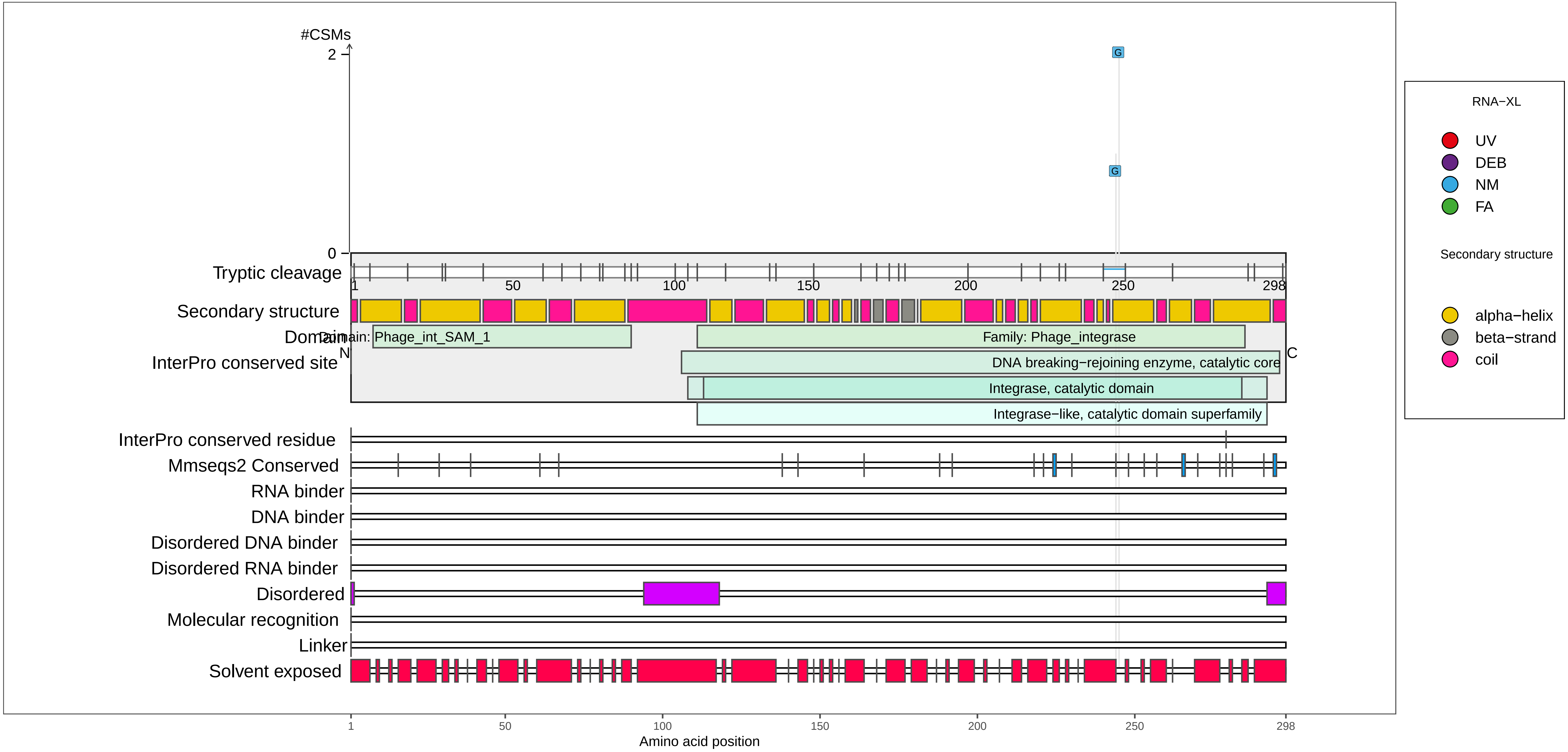

P0A8U0
SYDP_ECOLI Protein Syd

− Abundance:
tryptic [log10 Intensity]: 7.91 (Q 54)
PAXdb K12 strain [ppm]: 2.31 (Q 70)
PAXdb E.coli [ppm]: 1.55 (Q 61)

− RNA functions: not annotated

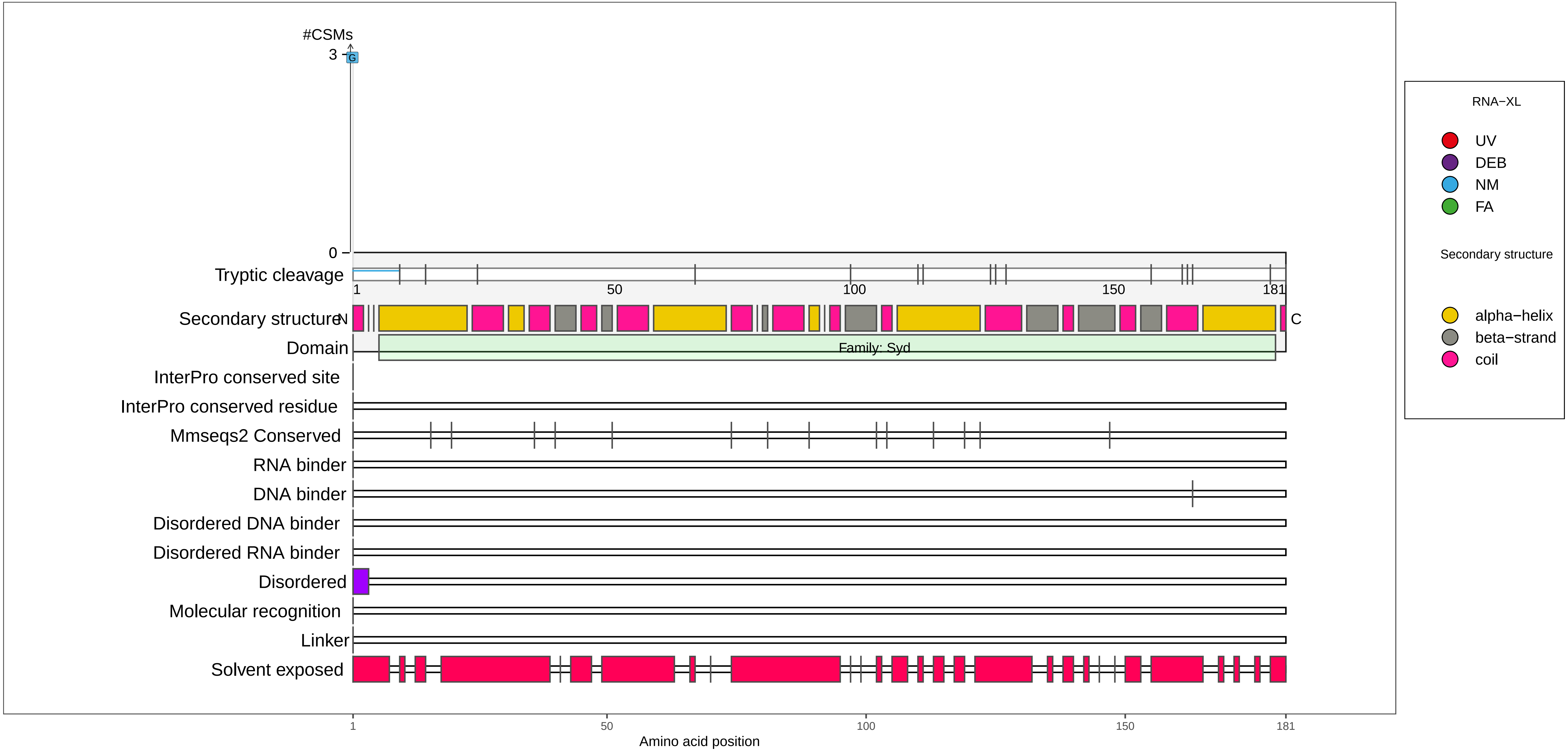

P0A8W0
NANR_ECOLI HTH−type transcriptional repressor NanR
− Abundance:
tryptic [log10 Intensity]: 8.2 (Q 65)
PAXdb K12 strain [ppm]: 1.78 (Q 53)
PAXdb E.coli [ppm]: 1.17 (Q 52)

− RNA functions:
RNA biosynthetic process; RNA metabolic process

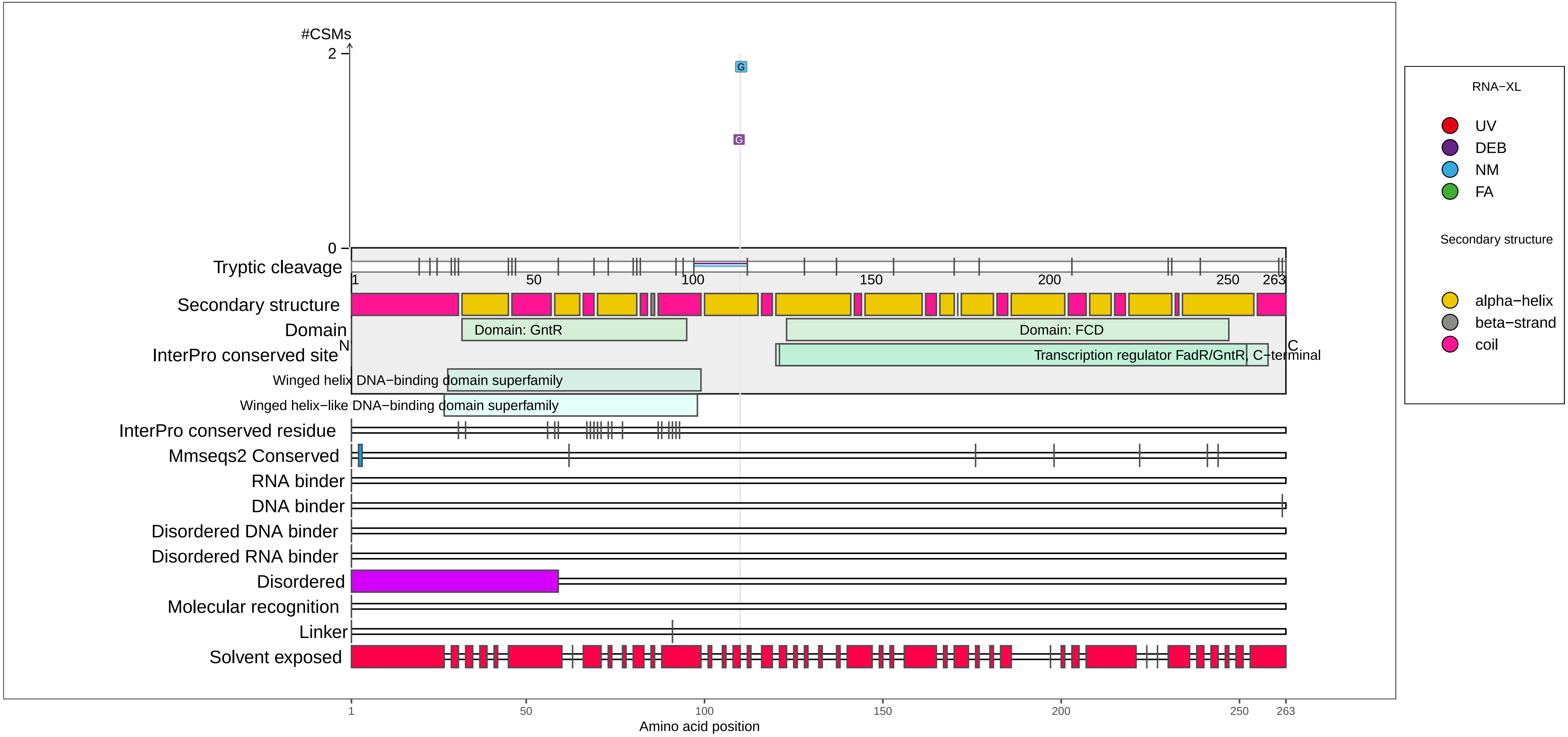

P0A9G4
CUER_ECOLI HTH−type transcriptional regulator CueR

− Abundance:
tryptic [log10 Intensity]: 7.02 (Q 14)
PAXdb K12 strain [ppm]: 1.51 (Q 39)
PAXdb E.coli [ppm]: 0.84 (Q 44)

− RNA functions:
RNA biosynthetic process; RNA metabolic process

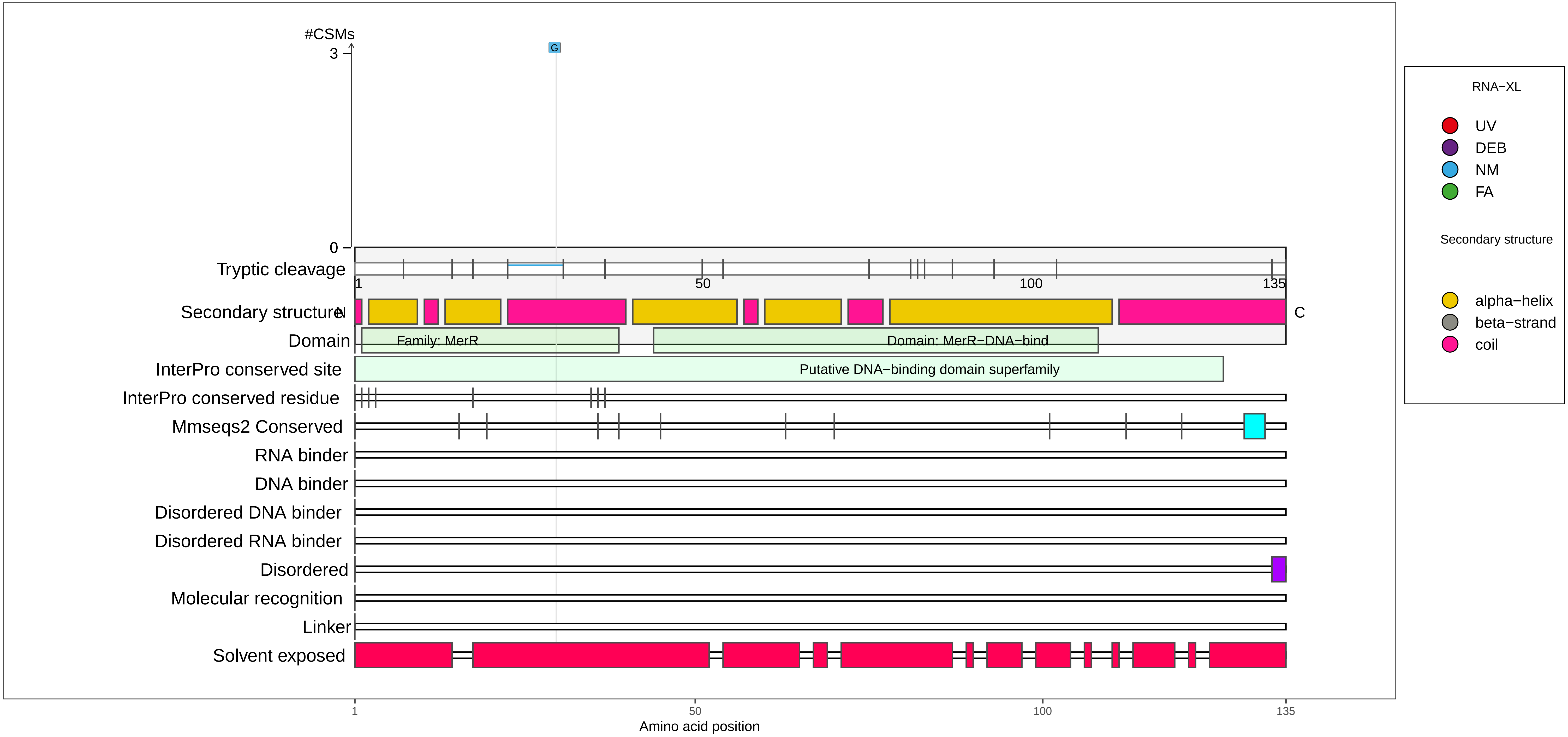

P0A9H3
LDCI_ECOLI Inducible lysine decarboxylase
− Abundance:
tryptic [log10 Intensity]: 8.12 (Q 62)
PAXdb K12 strain [ppm]: NA (Q NA)
PAXdb E.coli [ppm]: 1.25 (Q 53)

− RNA functions: not annotated

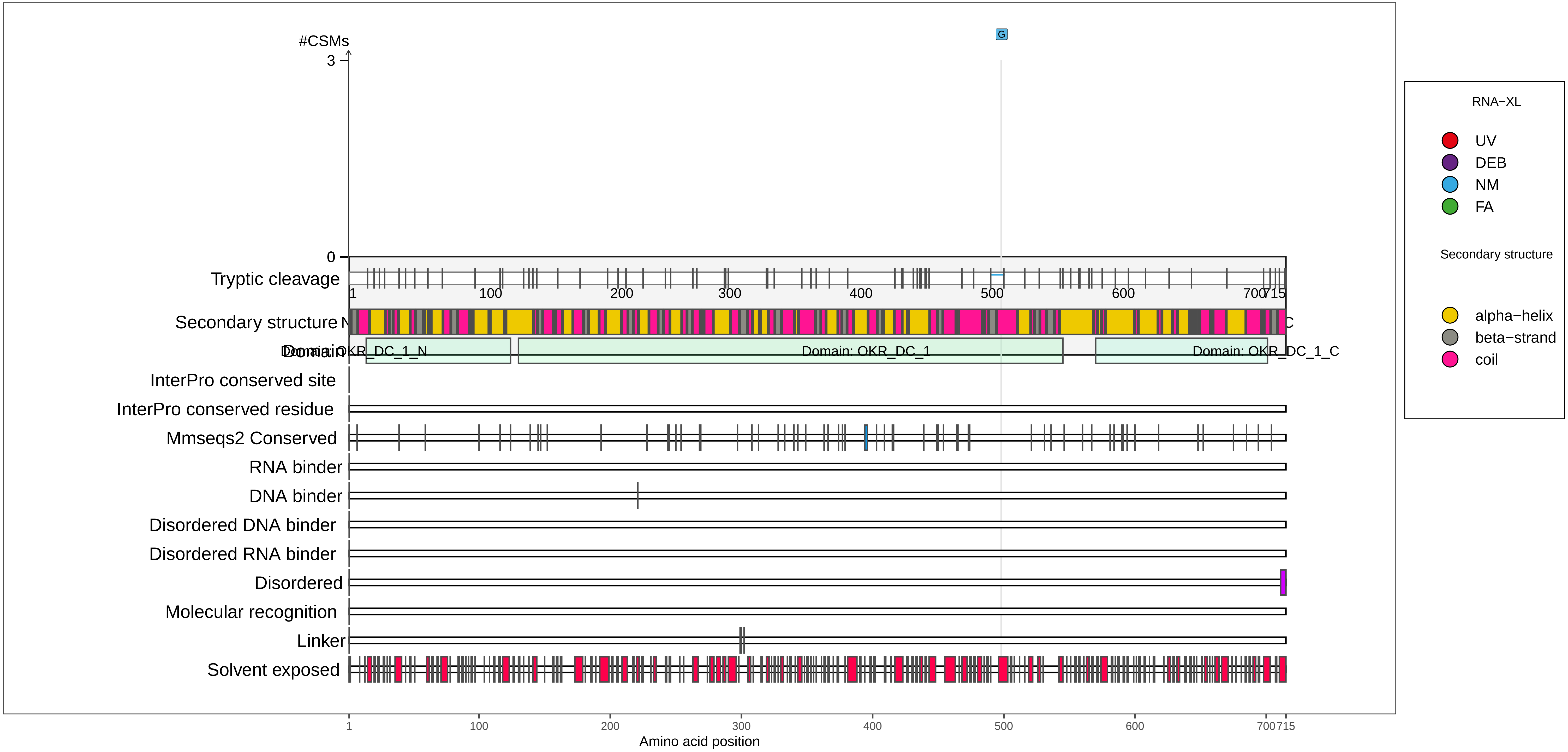

P0A9K7
PHOU_ECOLI Phosphate−specific transport system accessory protein PhoU

− Abundance:
tryptic [log10 Intensity]: 8.45 (Q 73)
PAXdb K12 strain [ppm]: 2.06 (Q 62)
PAXdb E.coli [ppm]: 1.89 (Q 69)

− RNA functions: not annotated

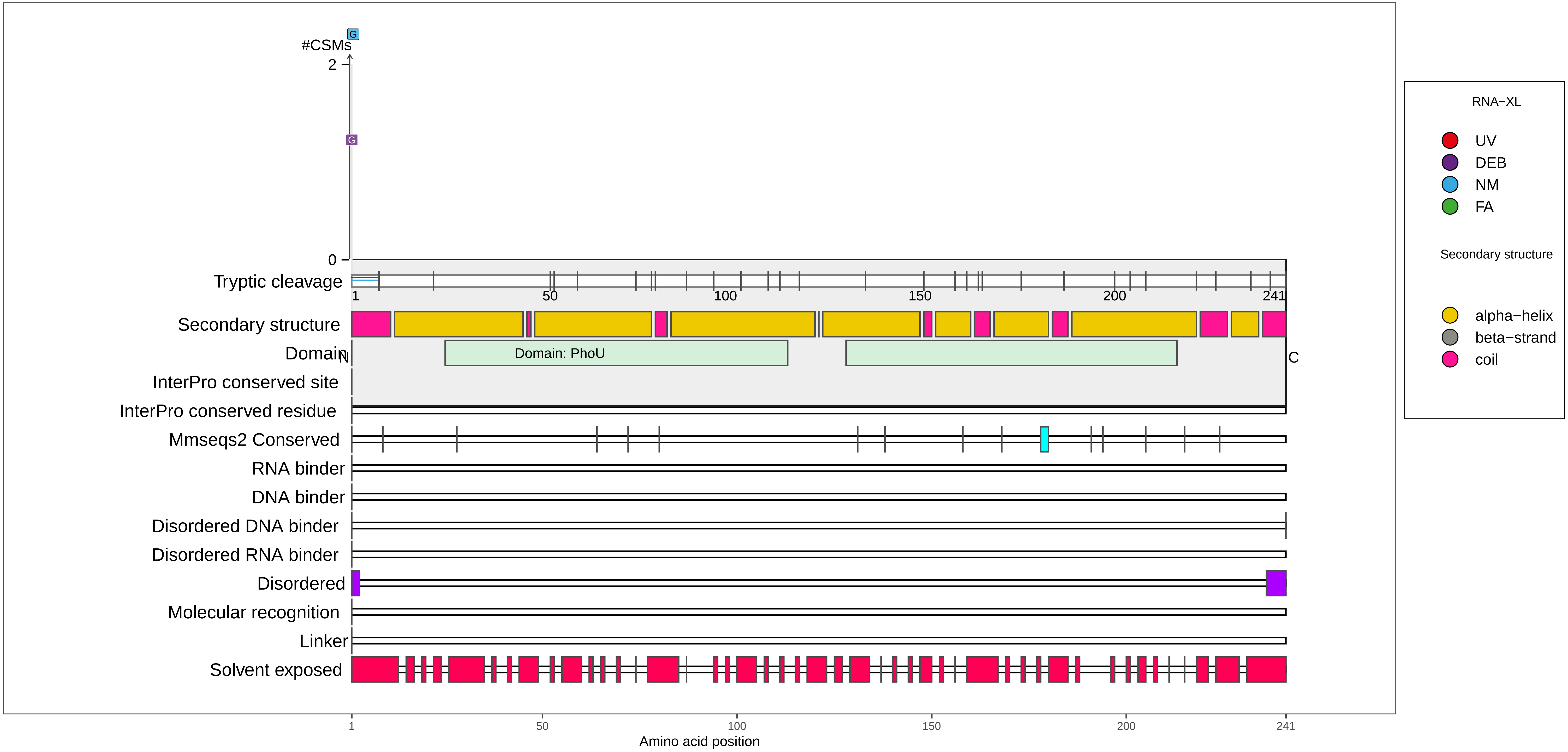

P0A9W9
YRDA_ECOLI Protein YrdA

− Abundance:
tryptic [log10 Intensity]: 8.59 (Q 78)
PAXdb K12 strain [ppm]: 2.94 (Q 89)
PAXdb E.coli [ppm]: 2.53 (Q 86)

− RNA functions: not annotated

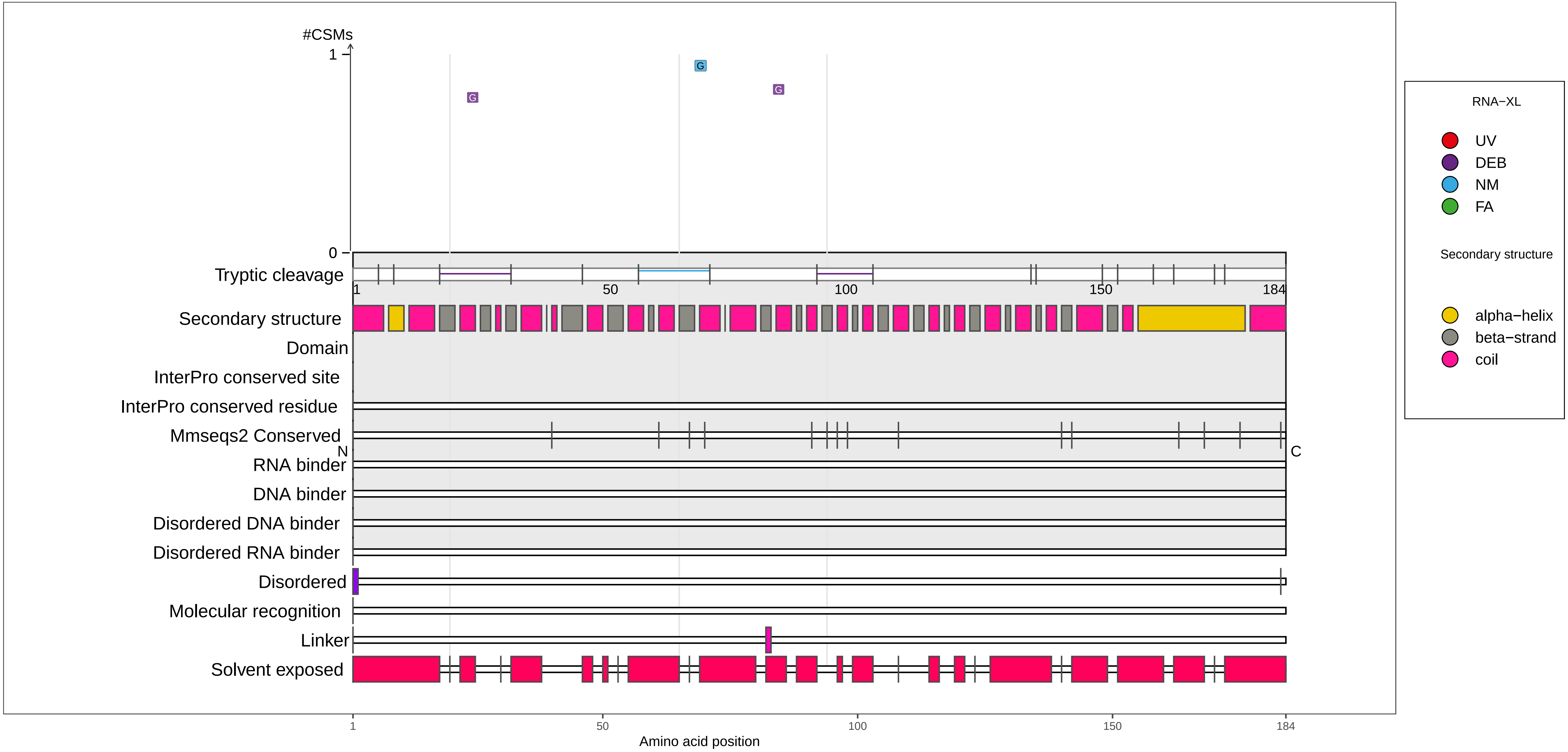

P0AAB8
USPD_ECOLI Universal stress protein D

− Abundance:
tryptic [log10 Intensity]: 8 (Q 58)
PAXdb K12 strain [ppm]: 2.68 (Q 83)
PAXdb E.coli [ppm]: 2.23 (Q 78)

− RNA functions: not annotated

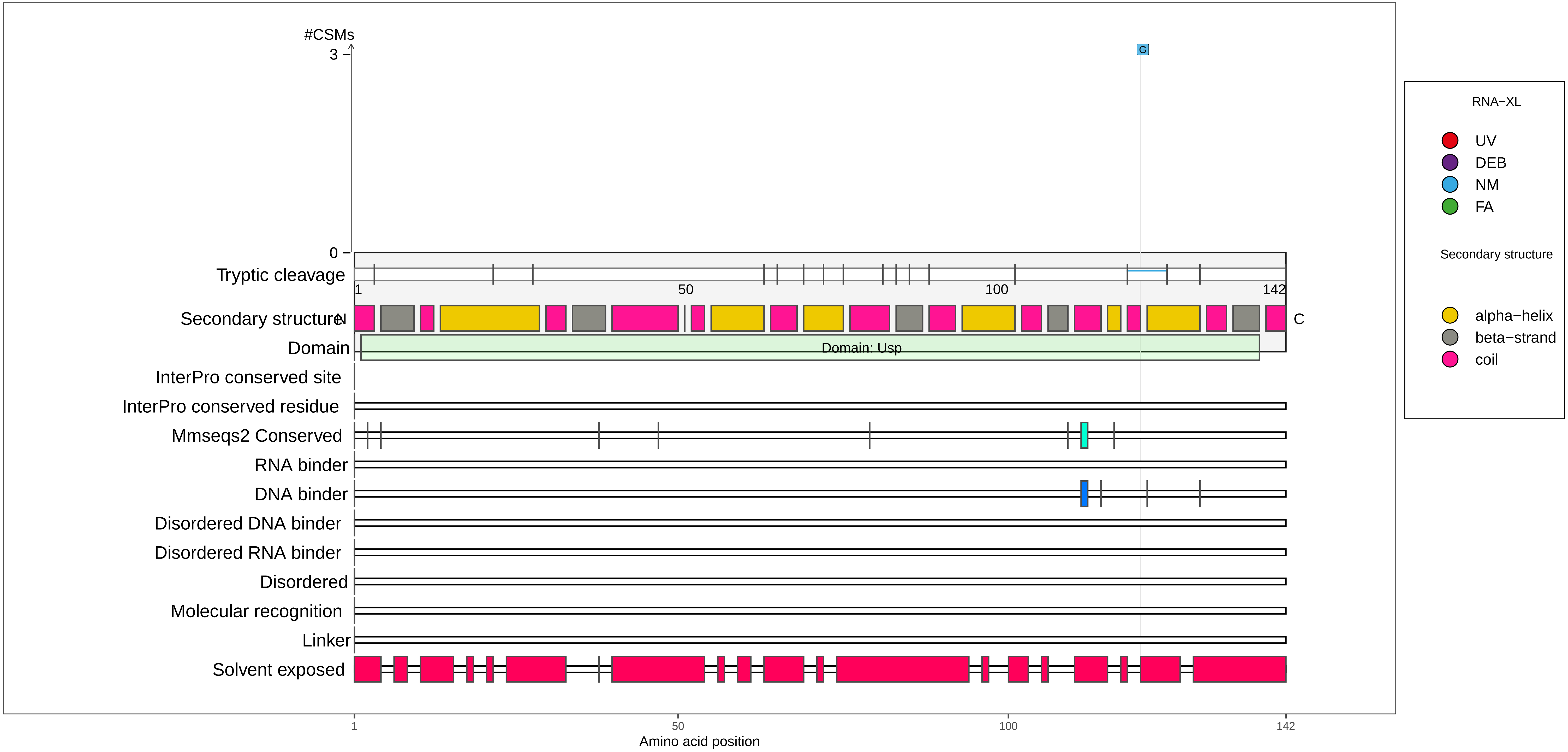

P0AAC8
ISCA_ECOLI Iron−binding protein IscA

− Abundance:
tryptic [log10 Intensity]: 7.88 (Q 53)
PAXdb K12 strain [ppm]: 2.24 (Q 68)
PAXdb E.coli [ppm]: 1.81 (Q 67)

− RNA functions: not annotated

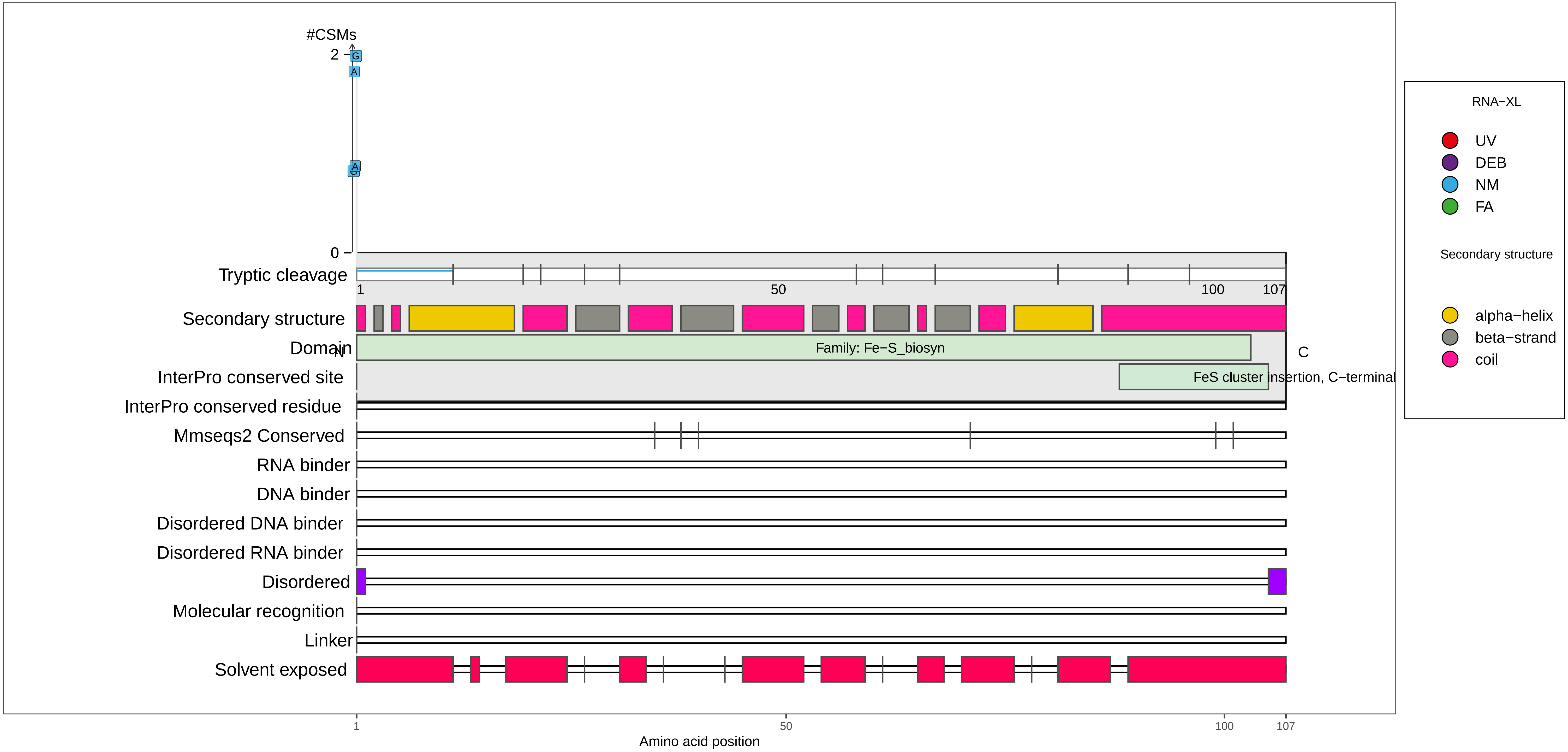

P0AB43
YCGL_ECOLI Protein YcgL

− Abundance:
tryptic [log10 Intensity]: 8.05 (Q 59)
PAXdb K12 strain [ppm]: 2.95 (Q 90)
PAXdb E.coli [ppm]: 2 (Q 72)

− RNA functions: not annotated

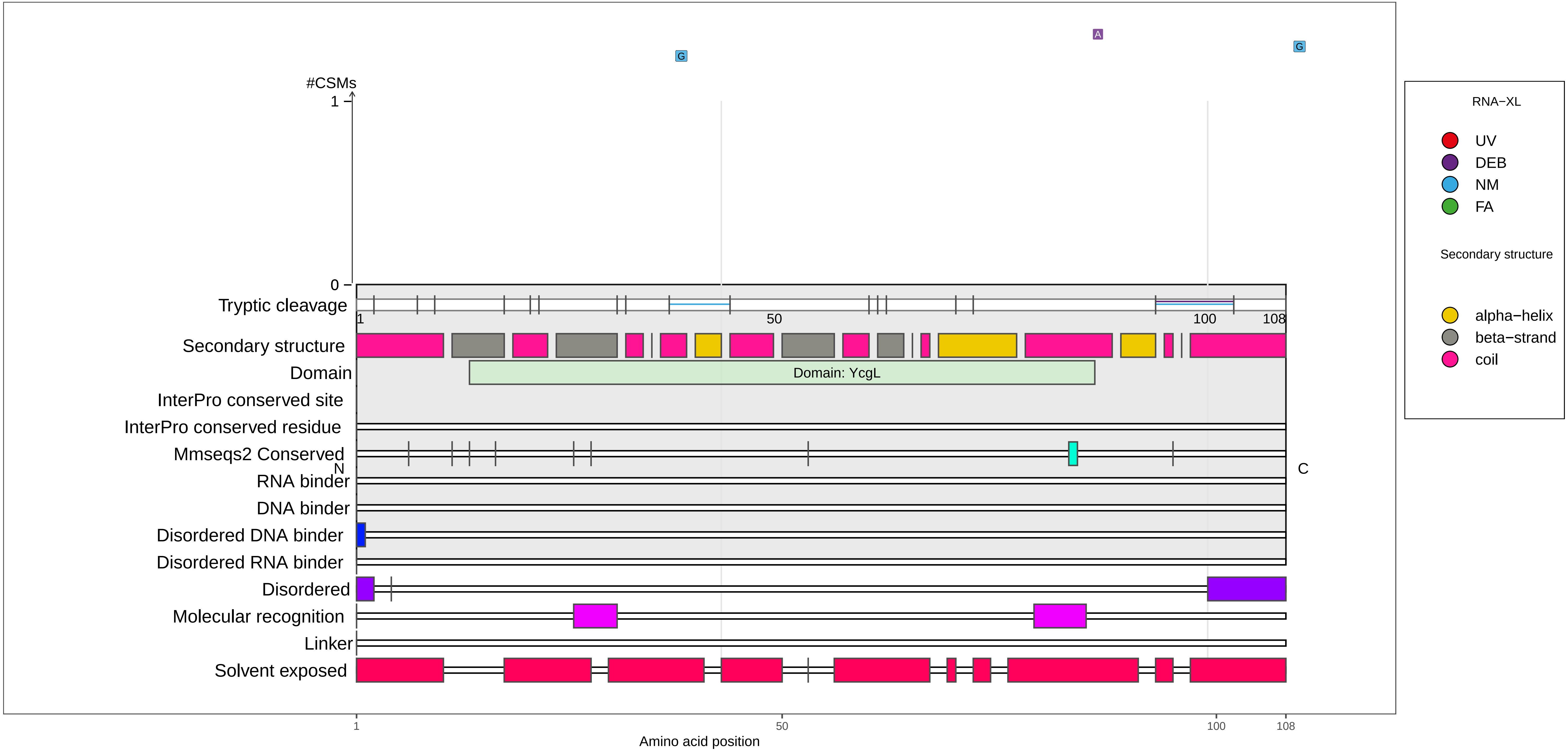

P0AC00
FRLB_ECOLI Fructoselysine 6−phosphate deglycase

− Abundance:
tryptic [log10 Intensity]: 7.6 (Q 40)
PAXdb K12 strain [ppm]: NA (Q NA)
PAXdb E.coli [ppm]: −2.1 (Q 1)

− RNA functions: not annotated

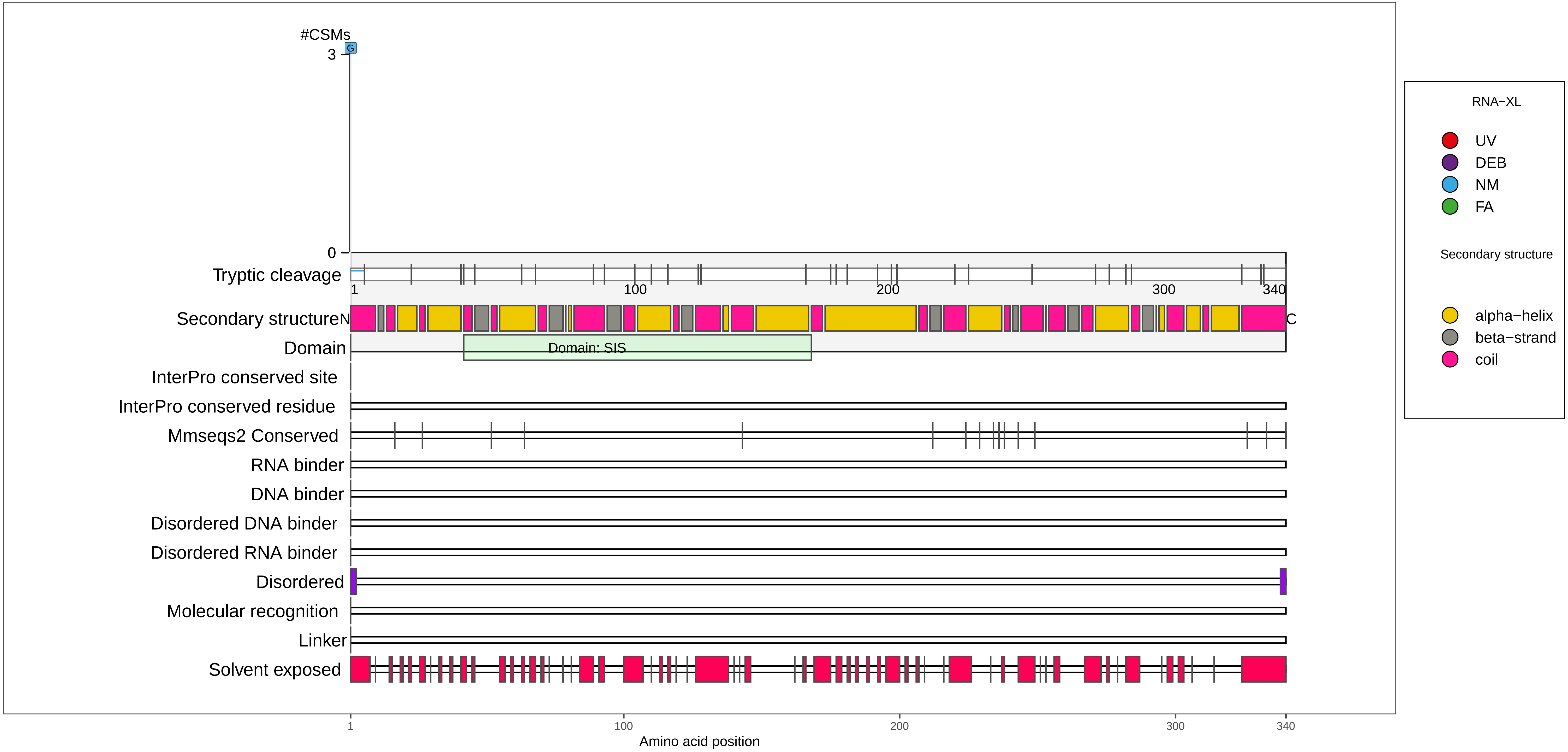

P0AC38
ASPA_ECOLI Aspartate ammonia−lyase

− Abundance:
tryptic [log10 Intensity]: 9.32 (Q 94)
PAXdb K12 strain [ppm]: 2.97 (Q 90)
PAXdb E.coli [ppm]: 2.9 (Q 93)

− RNA functions: not annotated

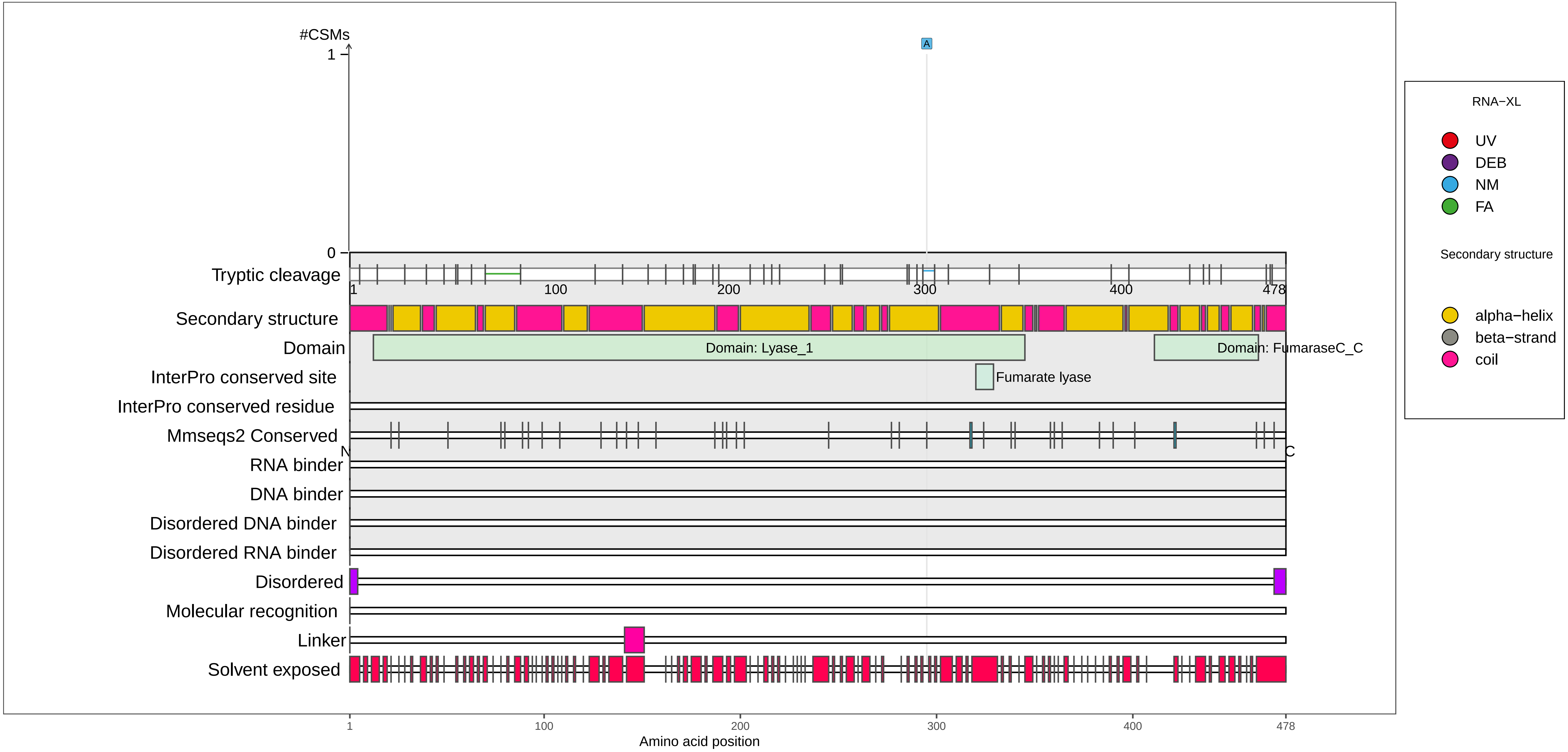

P0ACA7
GSTB_ECOLI Glutathione S−transferase GstB

− Abundance:
tryptic [log10 Intensity]: 8.08 (Q 61)
PAXdb K12 strain [ppm]: 2.27 (Q 69)
PAXdb E.coli [ppm]: 2.92 (Q 93)

− RNA functions: not annotated

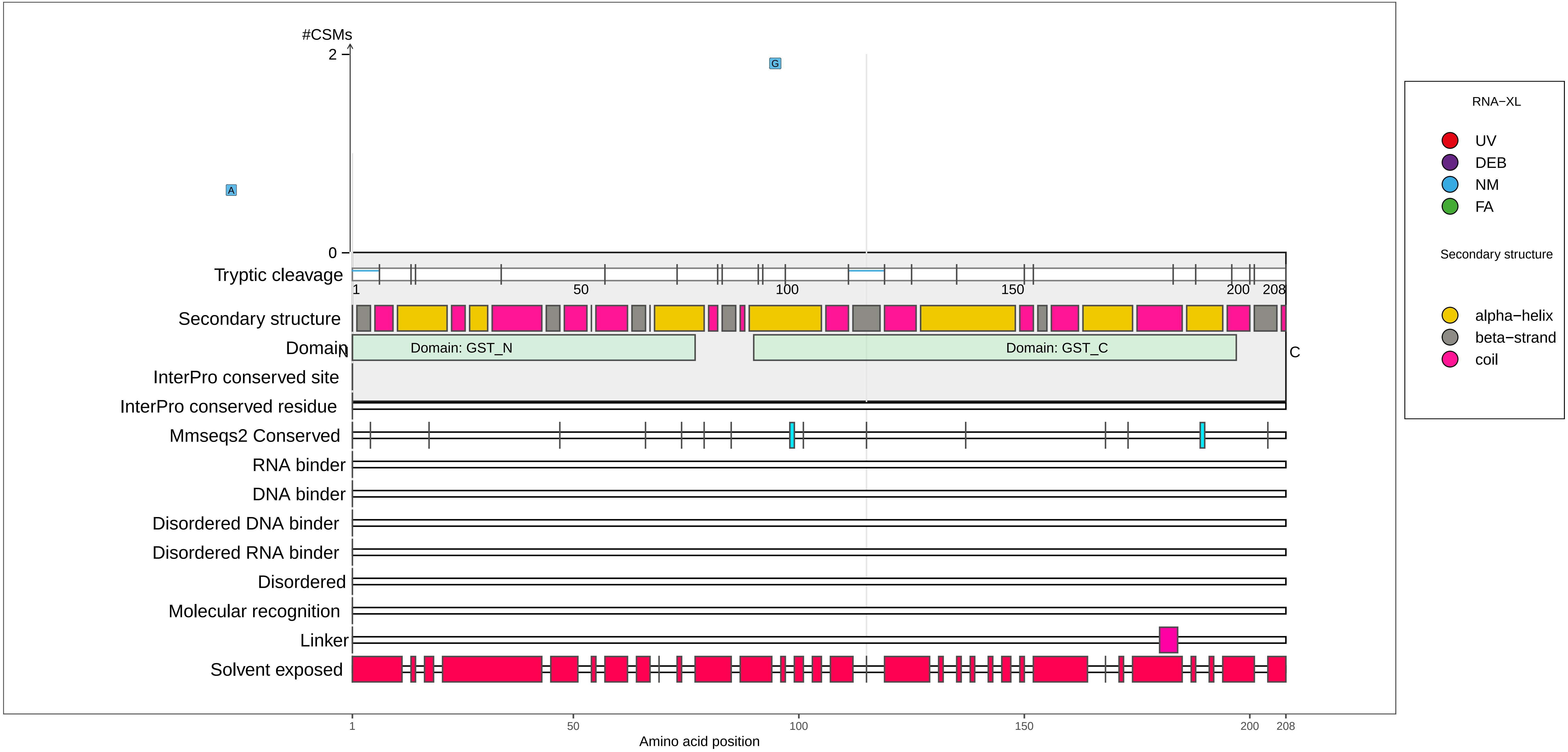

P0ACI6
ASNC_ECOLI Regulatory protein AsnC

− Abundance:
tryptic [log10 Intensity]: 6.81 (Q 7)
PAXdb K12 strain [ppm]: 2.76 (Q 85)
PAXdb E.coli [ppm]: 2.26 (Q 79)

− RNA functions: not annotated

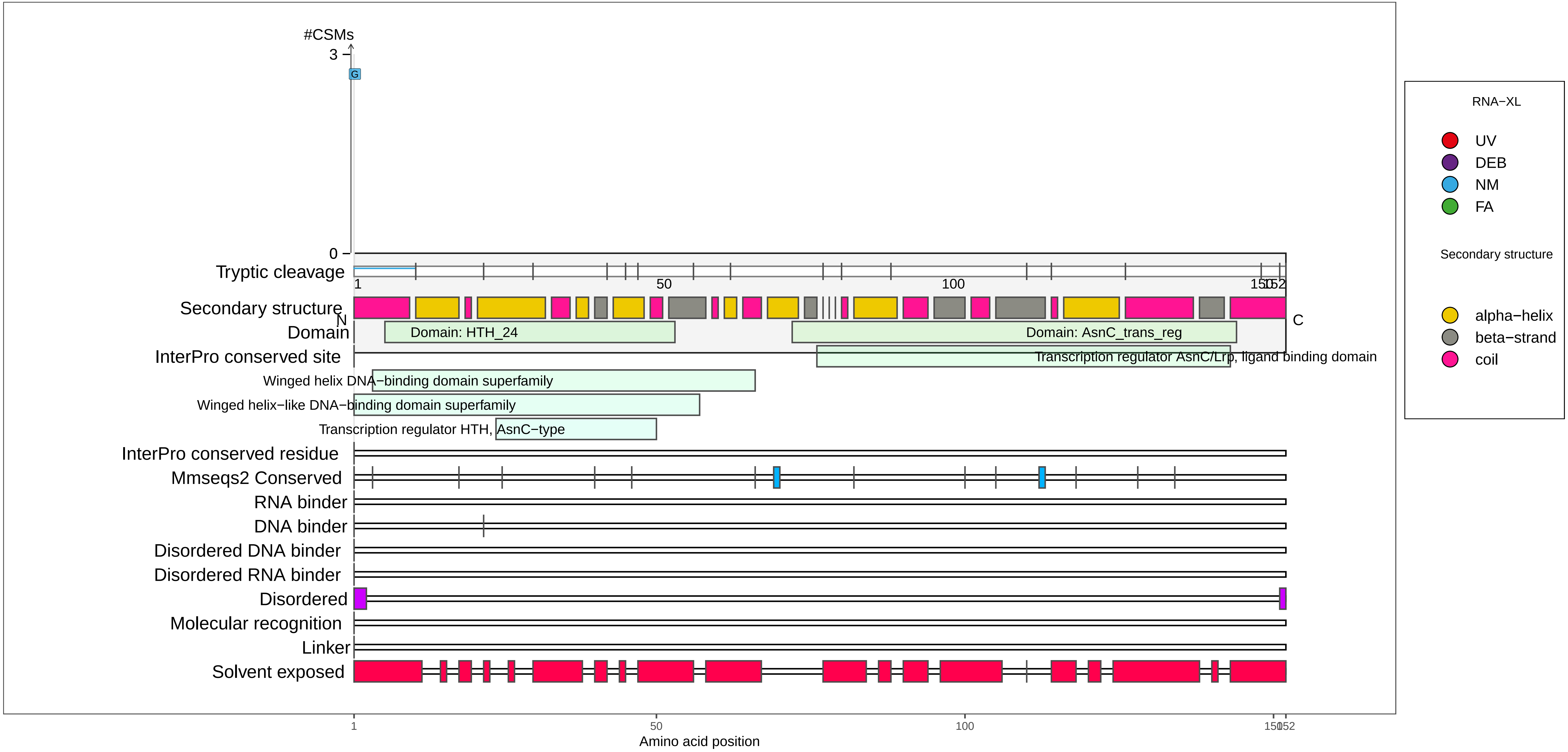

P0ACK5
DEOR_ECOLI Deoxyribose operon repressor

− Abundance:
tryptic [log10 Intensity]: NA (Q NA)
PAXdb K12 strain [ppm]: 1.33 (Q 28)
PAXdb E.coli [ppm]: 0.45 (Q 35)

− RNA functions: not annotated

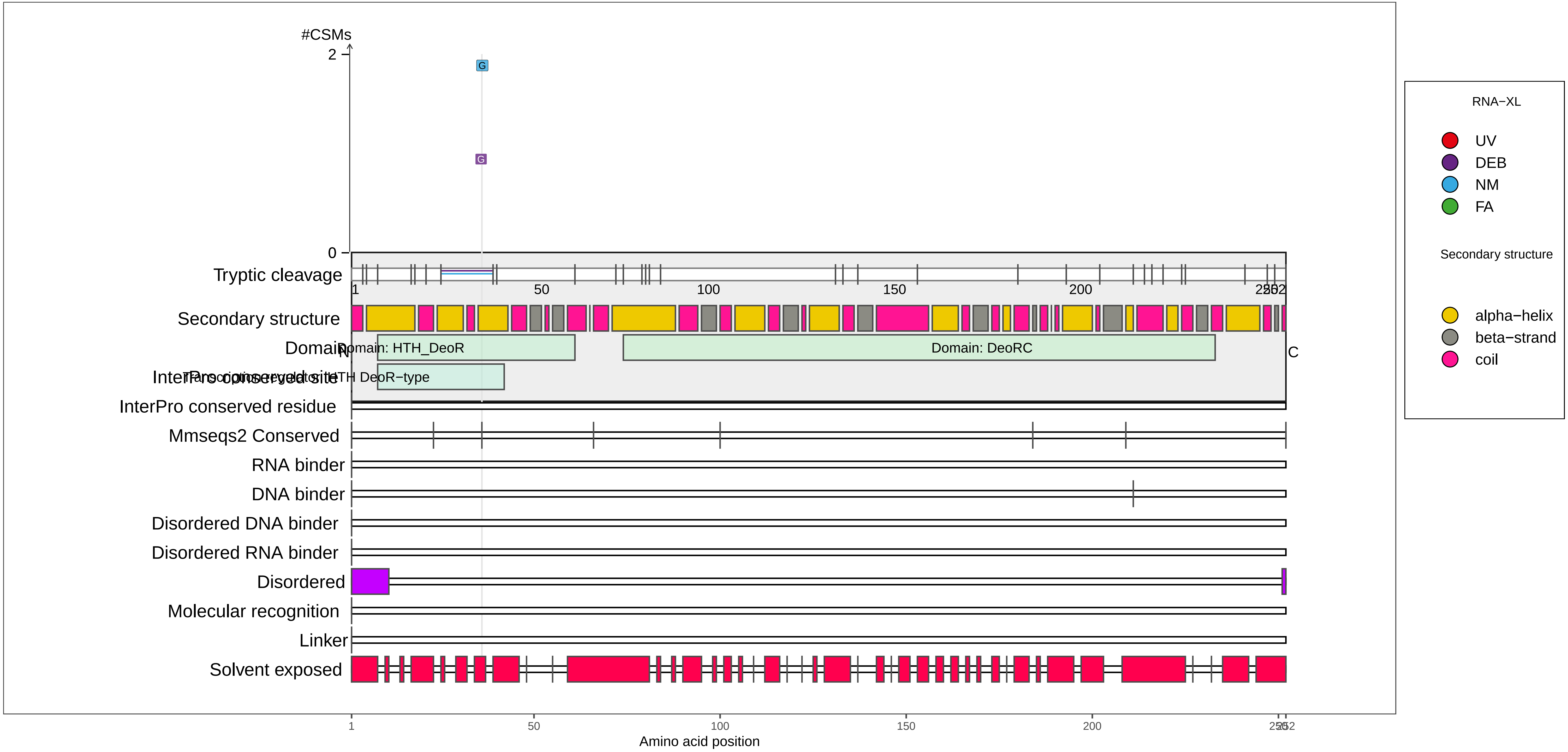

P0ACL7
LLDR_ECOLI Putative L−lactate dehydrogenase operon regulatory protein

− Abundance:
tryptic [log10 Intensity]: 7.16 (Q 19)
PAXdb K12 strain [ppm]: 1.33 (Q 28)
PAXdb E.coli [ppm]: −0.98 (Q 9)

− RNA functions: not annotated

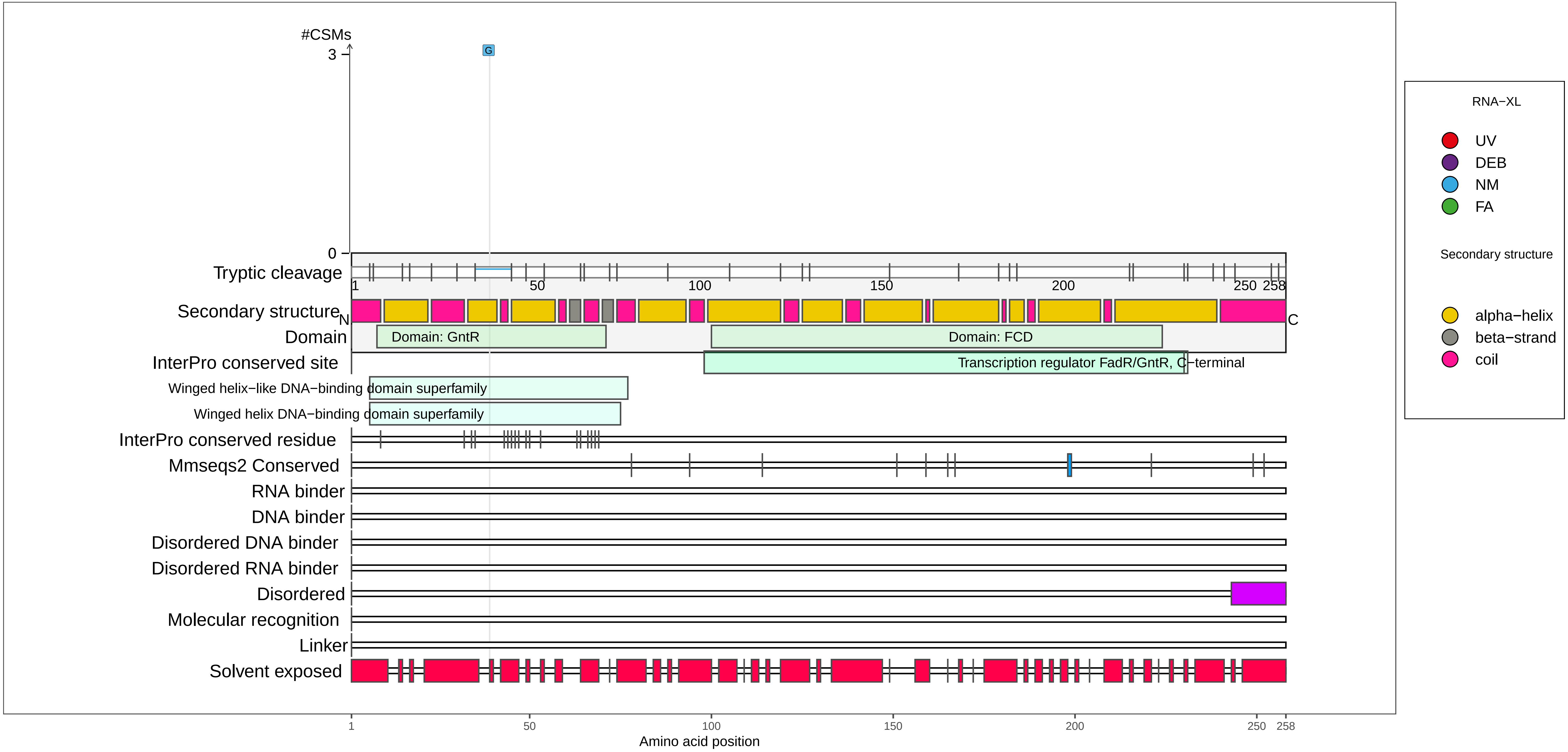

P0ACT6
UIDR_ECOLI HTH−type transcriptional regulator UidR

− Abundance:
tryptic [log10 Intensity]: 6.94 (Q 11)
PAXdb K12 strain [ppm]: 0.93 (Q 8)
PAXdb E.coli [ppm]: 0.74 (Q 42)

− RNA functions:
RNA biosynthetic process; RNA metabolic process

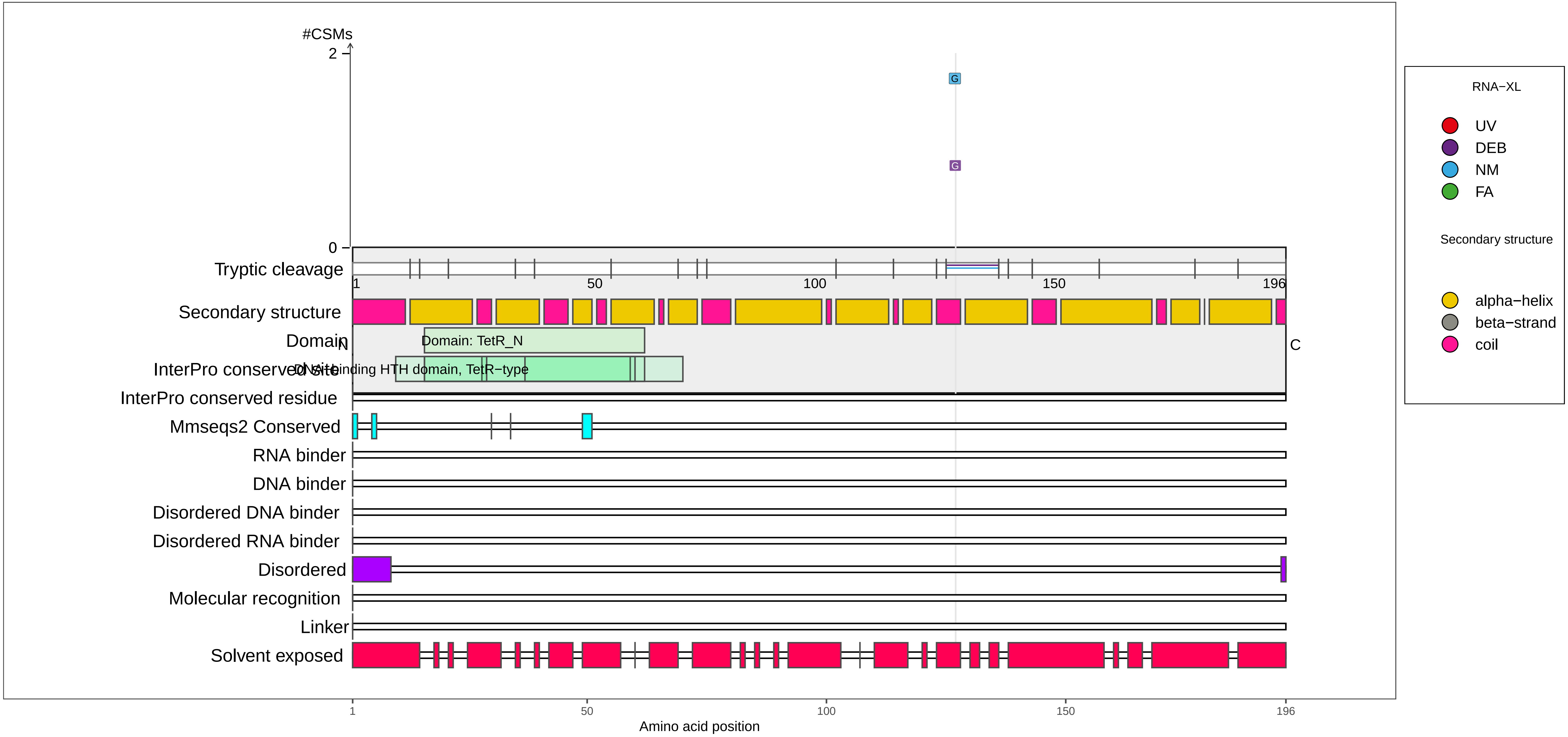

P0ADE6
KBP_ECOLI Potassium binding protein Kbp

− Abundance:
tryptic [log10 Intensity]: 7.13 (Q 18)
PAXdb K12 strain [ppm]: 2.2 (Q 67)
PAXdb E.coli [ppm]: 3.64 (Q 100)

− RNA functions: not annotated

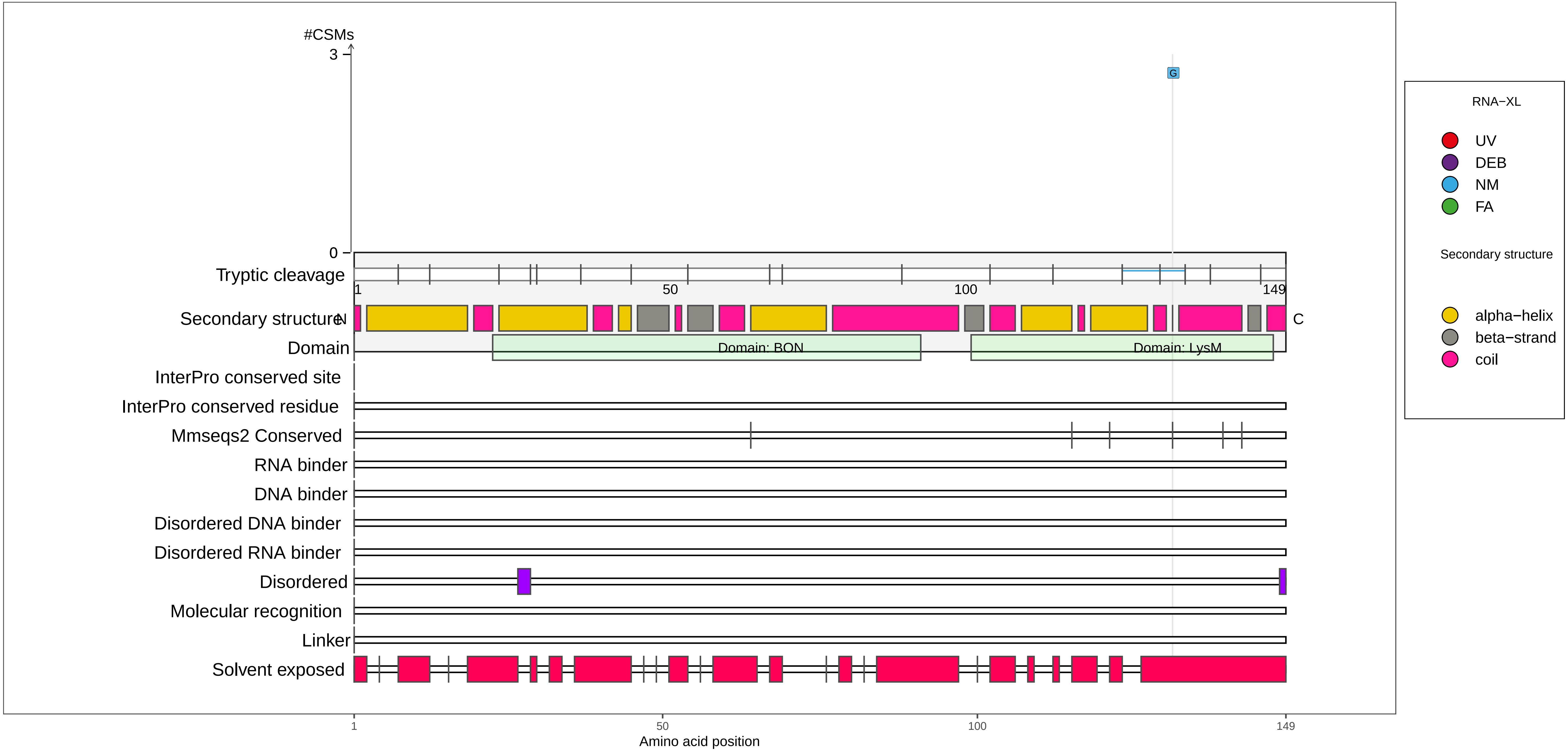

P0ADW8
YHEV_ECOLI Uncharacterized protein YheV

− Abundance:
tryptic [log10 Intensity]: 8.02 (Q 58)
PAXdb K12 strain [ppm]: 2.94 (Q 89)
PAXdb E.coli [ppm]: 1.31 (Q 55)

− RNA functions: not annotated

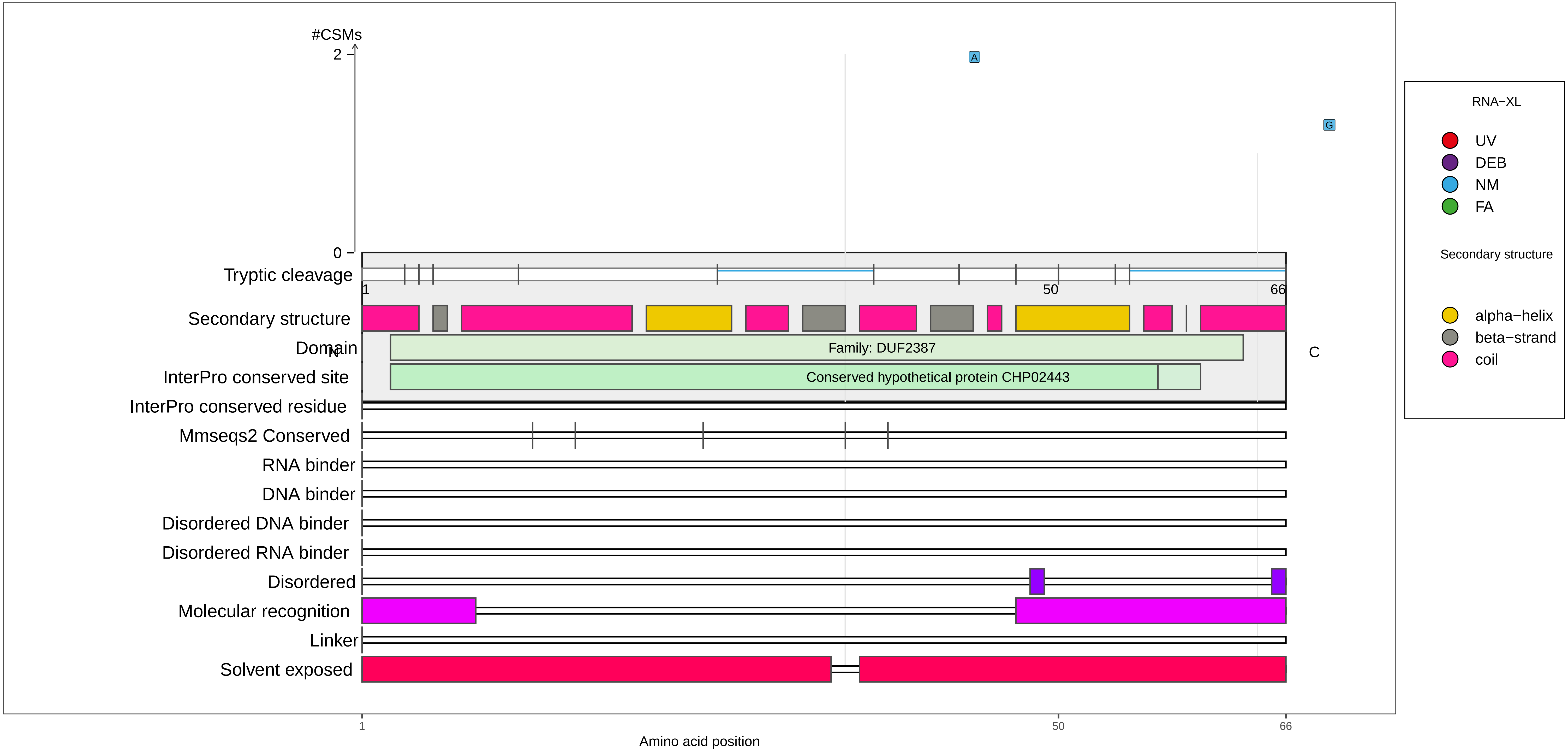

P0AEX9
MALE_ECOLI Maltose/maltodextrin−binding periplasmic protein

− Abundance:
tryptic [log10 Intensity]: 8.94 (Q 87)
PAXdb K12 strain [ppm]: 2.7 (Q 83)
PAXdb E.coli [ppm]: 3.25 (Q 97)

− RNA functions: not annotated

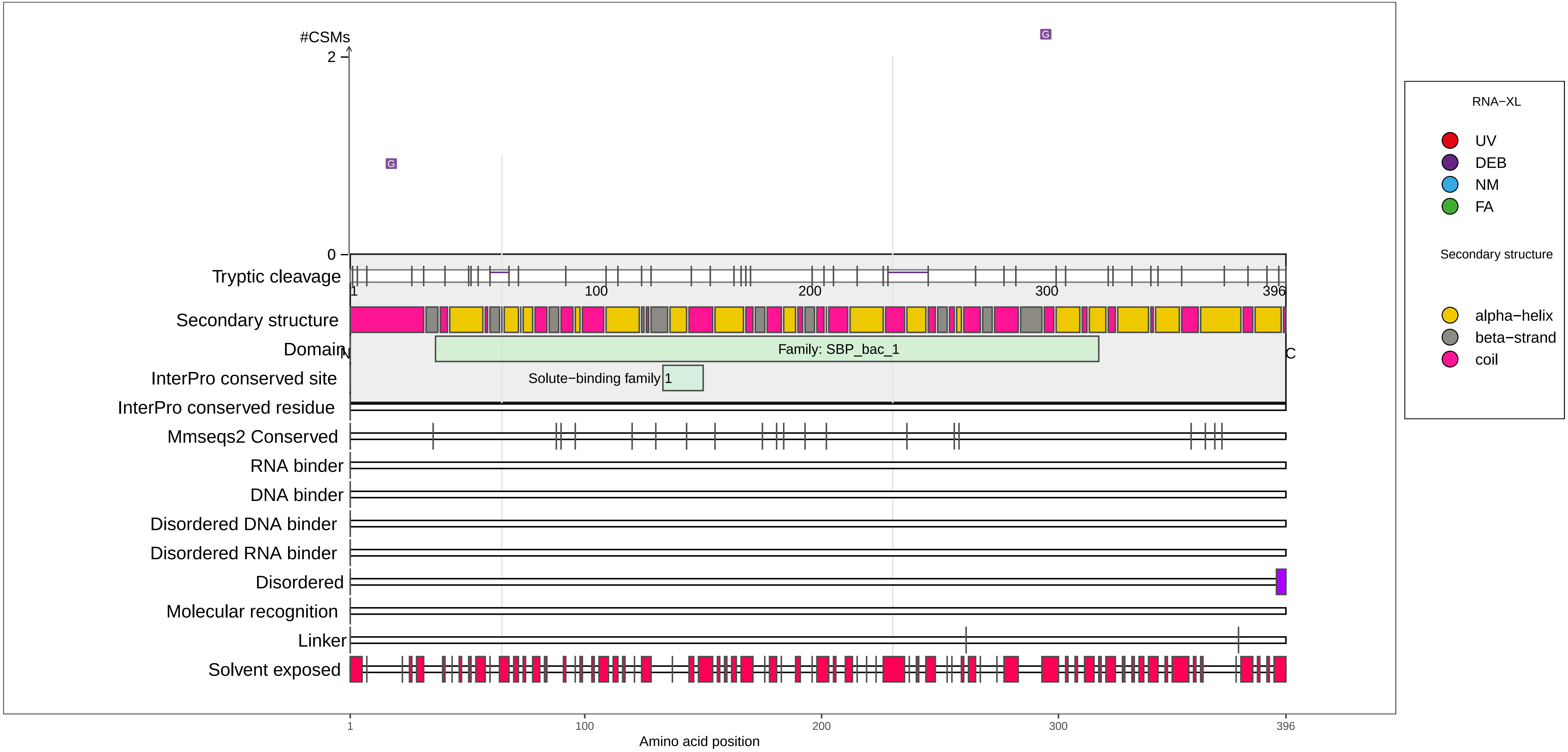

P0AFI7
PDXH_ECOLI Pyridoxine/pyridoxamine 5'−phosphate oxidase

− Abundance:
tryptic [log10 Intensity]: 8.08 (Q 61)
PAXdb K12 strain [ppm]: 2.37 (Q 73)
PAXdb E.coli [ppm]: 2.48 (Q 85)

− RNA functions: not annotated

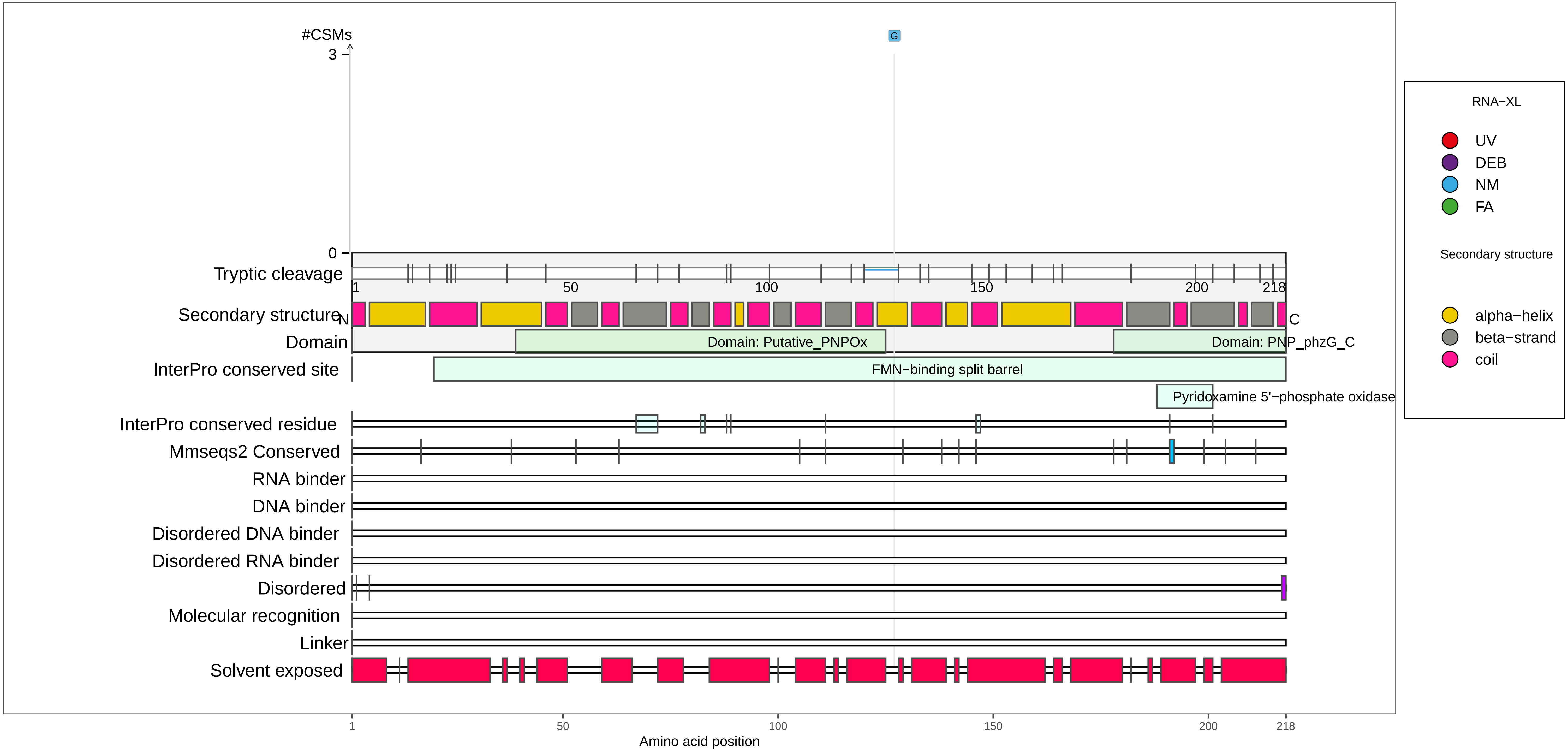

P0AFY8
SEQA_ECOLI Negative modulator of initiation of replication

− Abundance:
tryptic [log10 Intensity]: 7.47 (Q 33)
PAXdb K12 strain [ppm]: 2.74 (Q 84)
PAXdb E.coli [ppm]: 2.34 (Q 82)

− RNA functions: not annotated

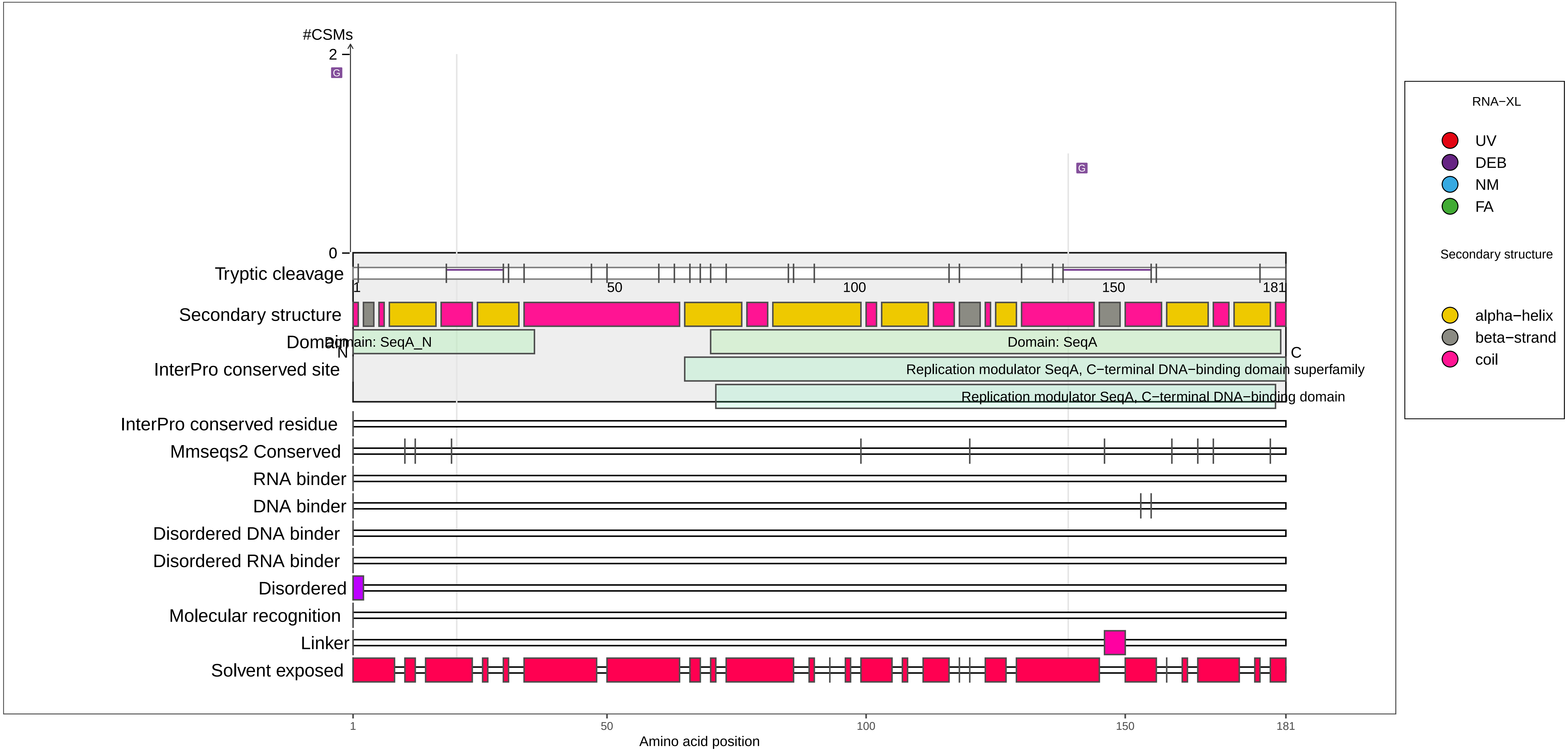

P0AG96
SECE_ECOLI Protein translocase subunit SecE

− Abundance:
tryptic [log10 Intensity]: 8.6 (Q 79)
PAXdb K12 strain [ppm]: 1.62 (Q 45)
PAXdb E.coli [ppm]: 1.56 (Q 61)

− RNA functions: not annotated

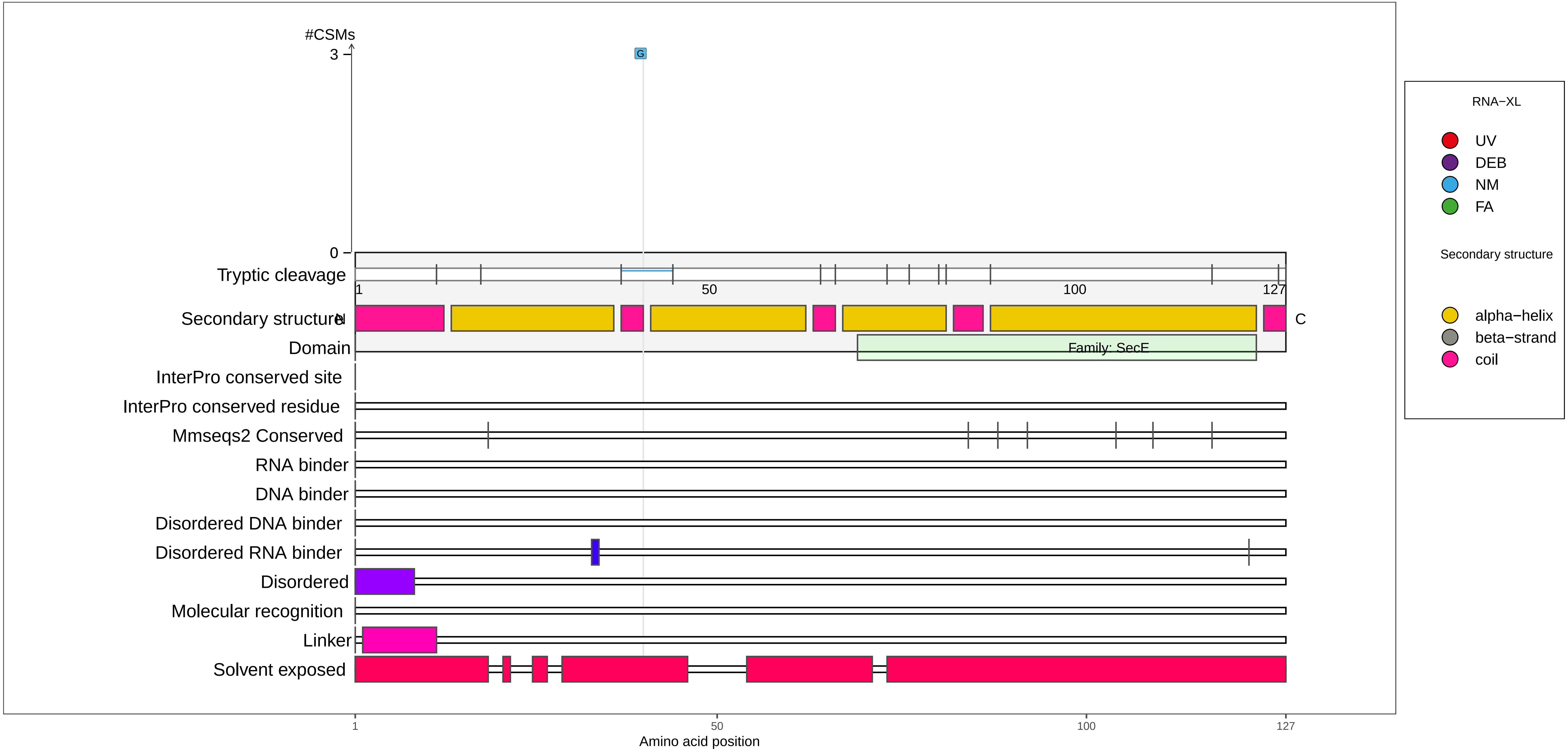

P0AGA2
SECY_ECOLI Protein translocase subunit SecY
− Abundance:
tryptic [log10 Intensity]: 7.41 (Q 30)
PAXdb K12 strain [ppm]: 2.11 (Q 64)
PAXdb E.coli [ppm]: 1.96 (Q 71)

− RNA functions: not annotated

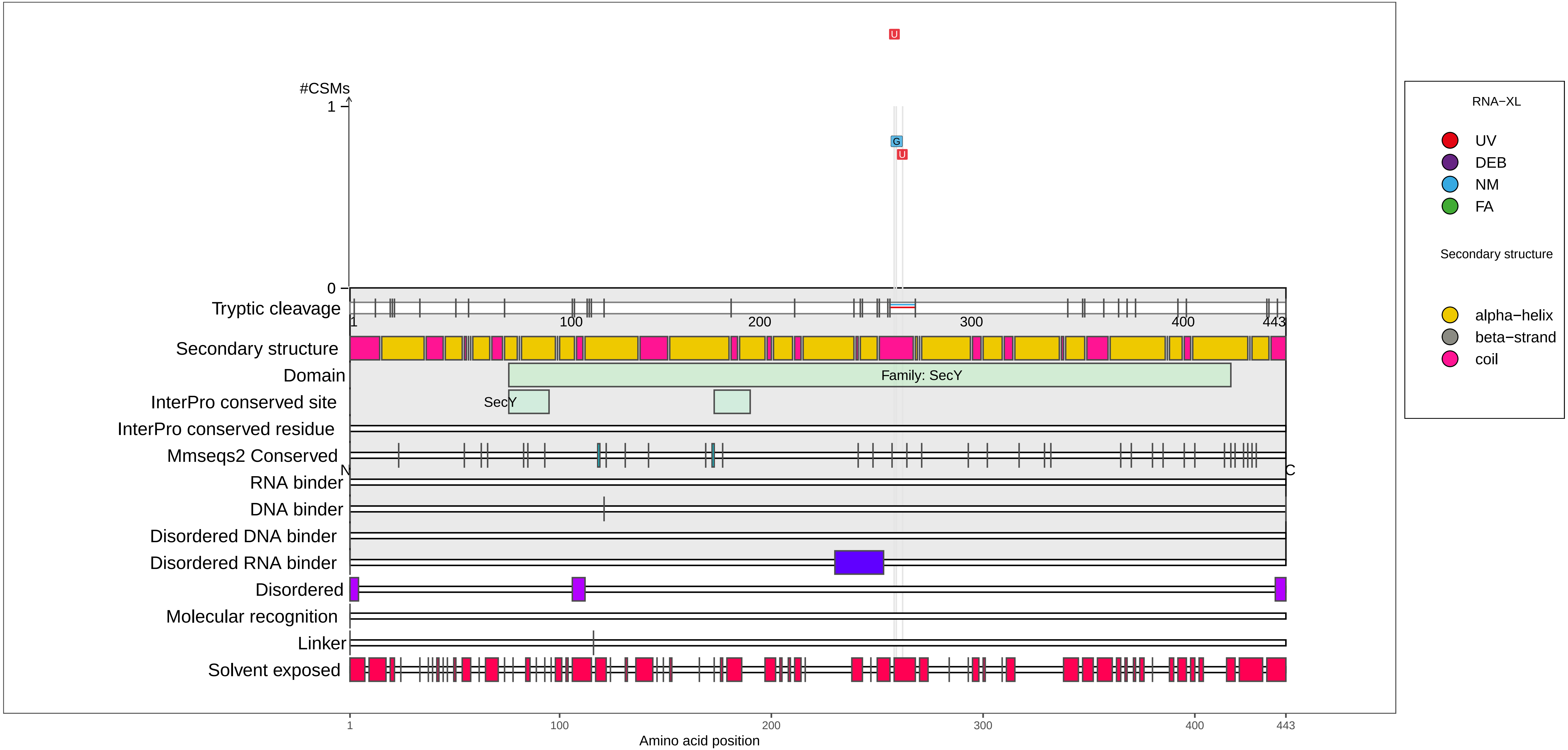

P0C0R7
RLME_ECOLI Ribosomal RNA large subunit methyltransferase E

− Abundance:
tryptic [log10 Intensity]: 7.09 (Q 17)
PAXdb K12 strain [ppm]: 2.49 (Q 77)
PAXdb E.coli [ppm]: 1.73 (Q 65)

− RNA functions:
ncRNA metabolic process; ncRNA processing; RNA 2−O−methyltransferase activity
RNA metabolic process; RNA methylation; RNA methyltransferase activity; RNA modification
RNA processing; rRNA (uridine−2−O−)−methyltransferase activity
rRNA (uridine) methyltransferase activity; rRNA 2−O−methylation; rRNA metabolic process; rRNA methylation
rRNA methyltransferase activity; rRNA modification; rRNA processing

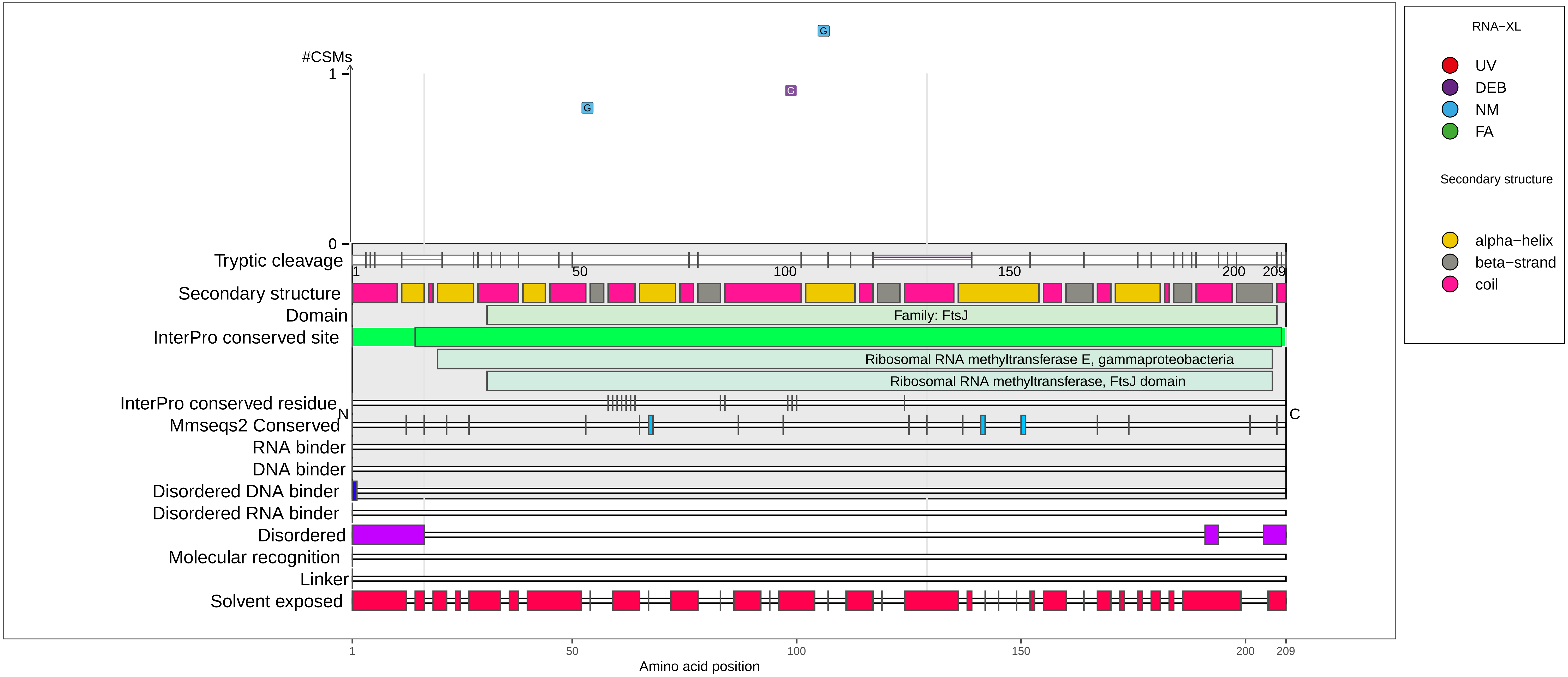

P14900
MURD_ECOLI UDP−N−acetylmuramoylalanine−−D−glutamate ligase

− Abundance:
tryptic [log10 Intensity]: 8.27 (Q 68)
PAXdb K12 strain [ppm]: 1.71 (Q 49)
PAXdb E.coli [ppm]: 1.74 (Q 65)

− RNA functions: not annotated

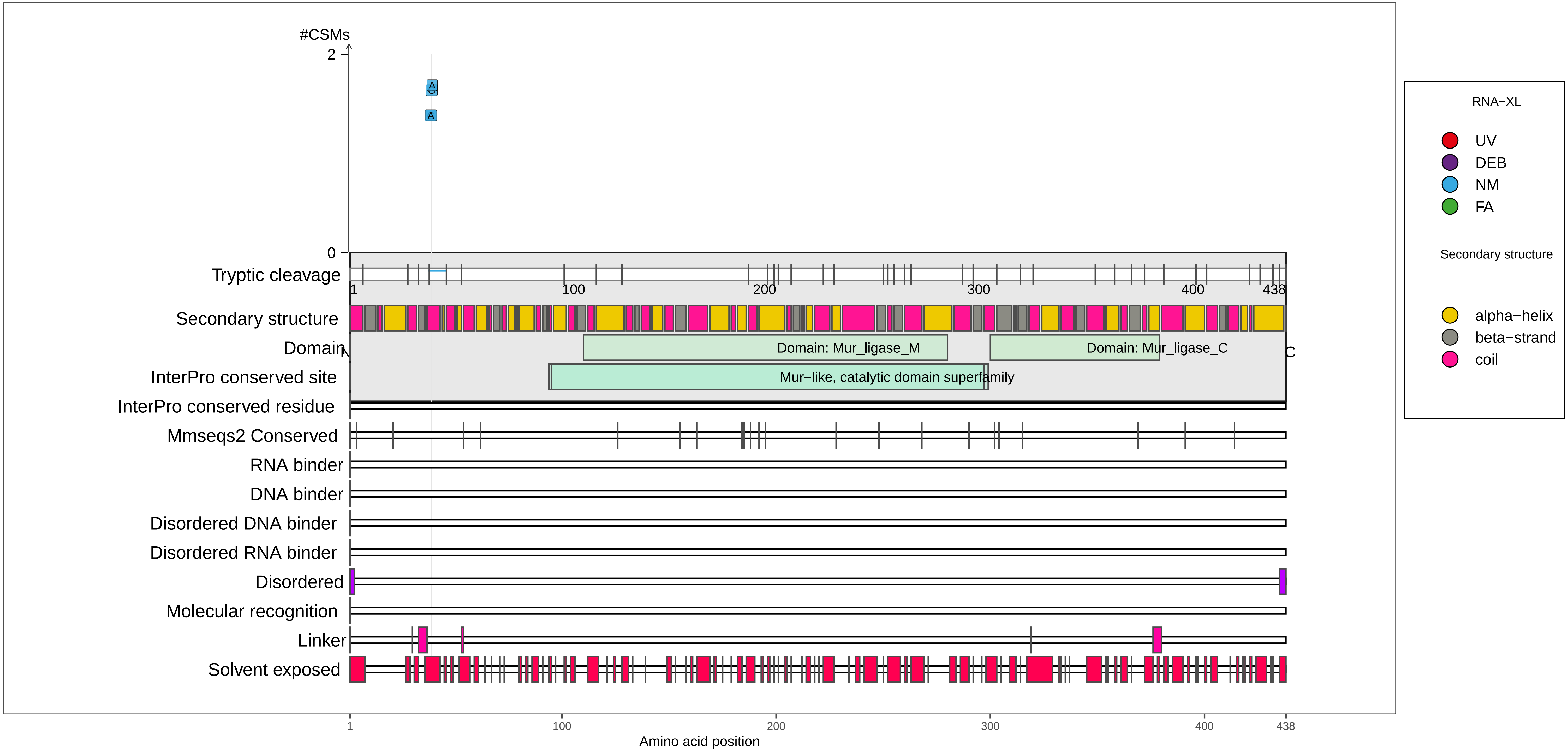

P15034
AMPP_ECOLI Xaa−Pro aminopeptidase

− Abundance:
tryptic [log10 Intensity]: 9.08 (Q 90)
PAXdb K12 strain [ppm]: 2.51 (Q 78)
PAXdb E.coli [ppm]: 2.74 (Q 91)

− RNA functions: not annotated

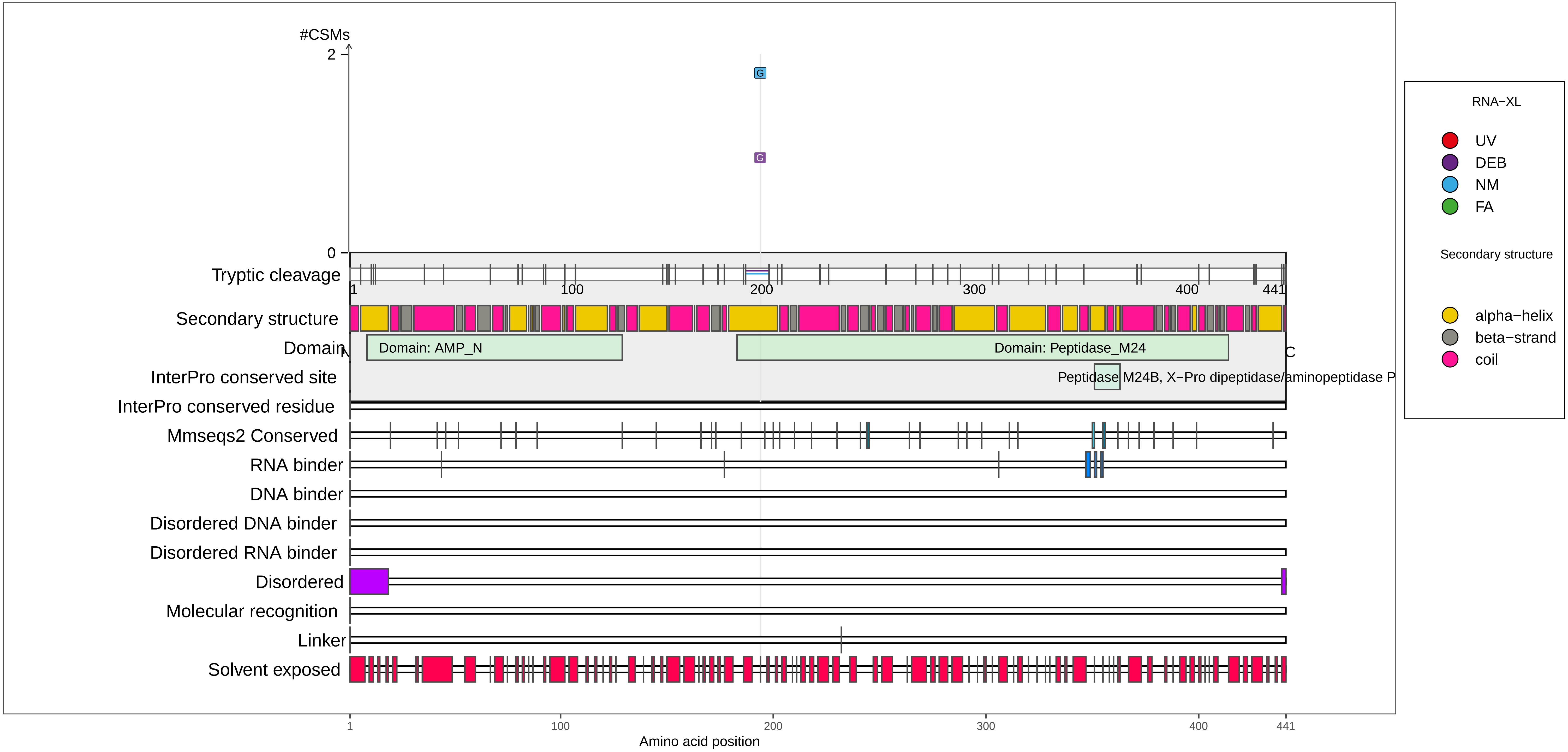

P15723
DGTP_ECOLI Deoxyguanosinetriphosphate triphosphohydrolase

− Abundance:
tryptic [log10 Intensity]: 7.34 (Q 27)
PAXdb K12 strain [ppm]: 0.9 (Q 8)
PAXdb E.coli [ppm]: 0.59 (Q 38)

− RNA functions: not annotated

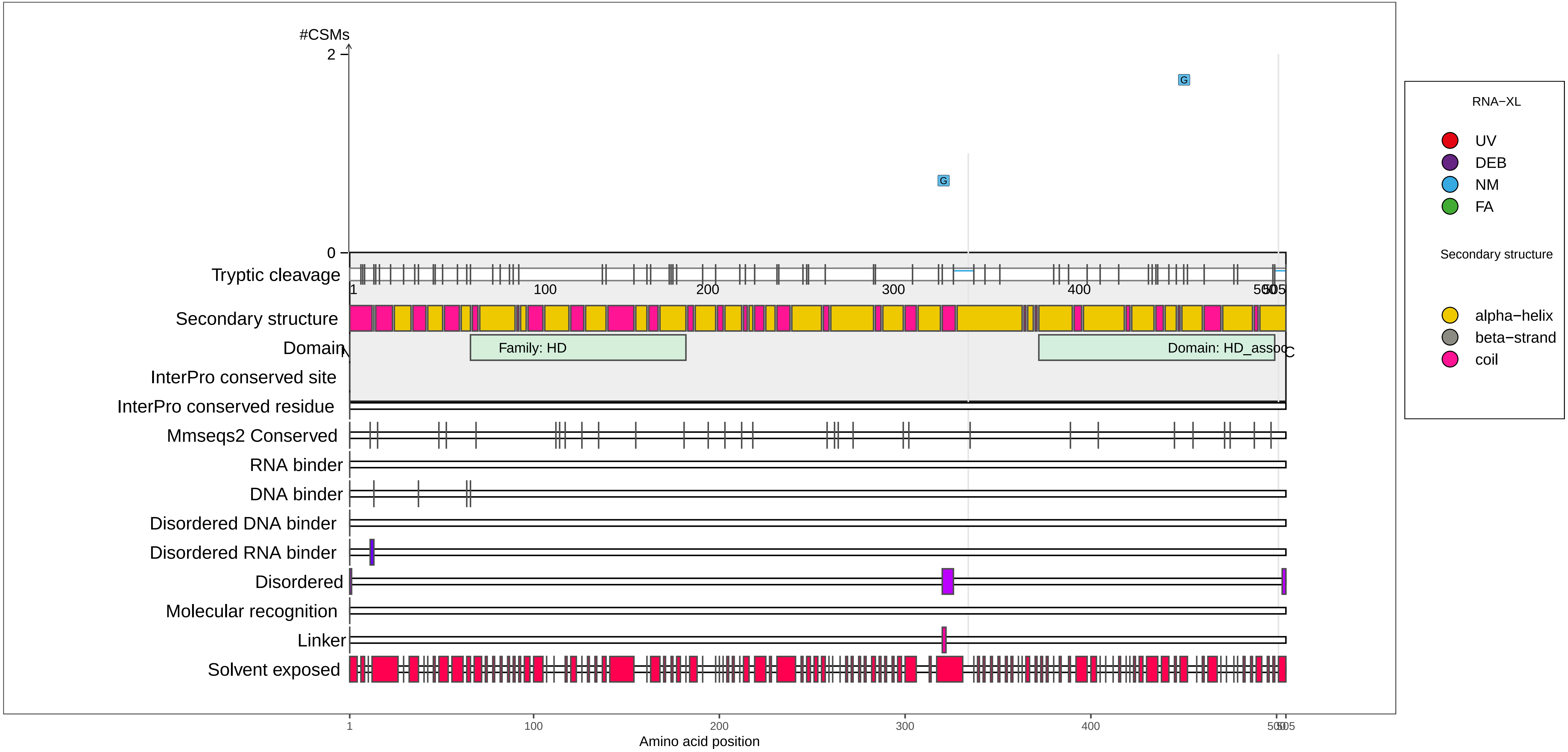

P17846
CYSI_ECOLI Sulfite reductase [NADPH] hemoprotein beta−component

− Abundance:
tryptic [log10 Intensity]: 7 (Q 13)
PAXdb K12 strain [ppm]: 2.81 (Q 86)
PAXdb E.coli [ppm]: 2.56 (Q 87)

− RNA functions: not annotated

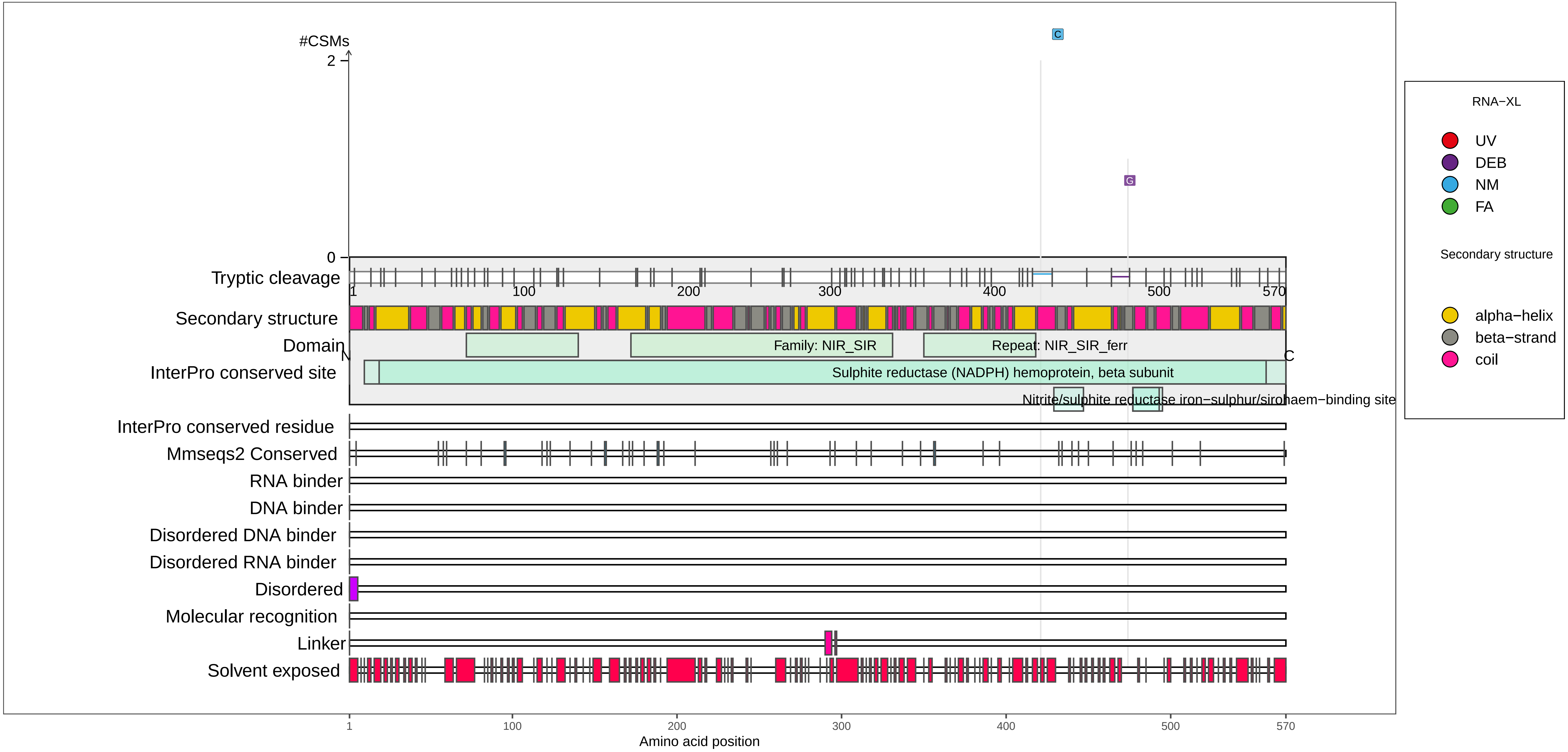

P19768
INSJ_ECOLI Insertion element IS150 protein InsJ

− Abundance:
tryptic [log10 Intensity]: NA (Q NA)
PAXdb K12 strain [ppm]: NA (Q NA)
PAXdb E.coli [ppm]: NA (Q NA)

− RNA functions: not annotated

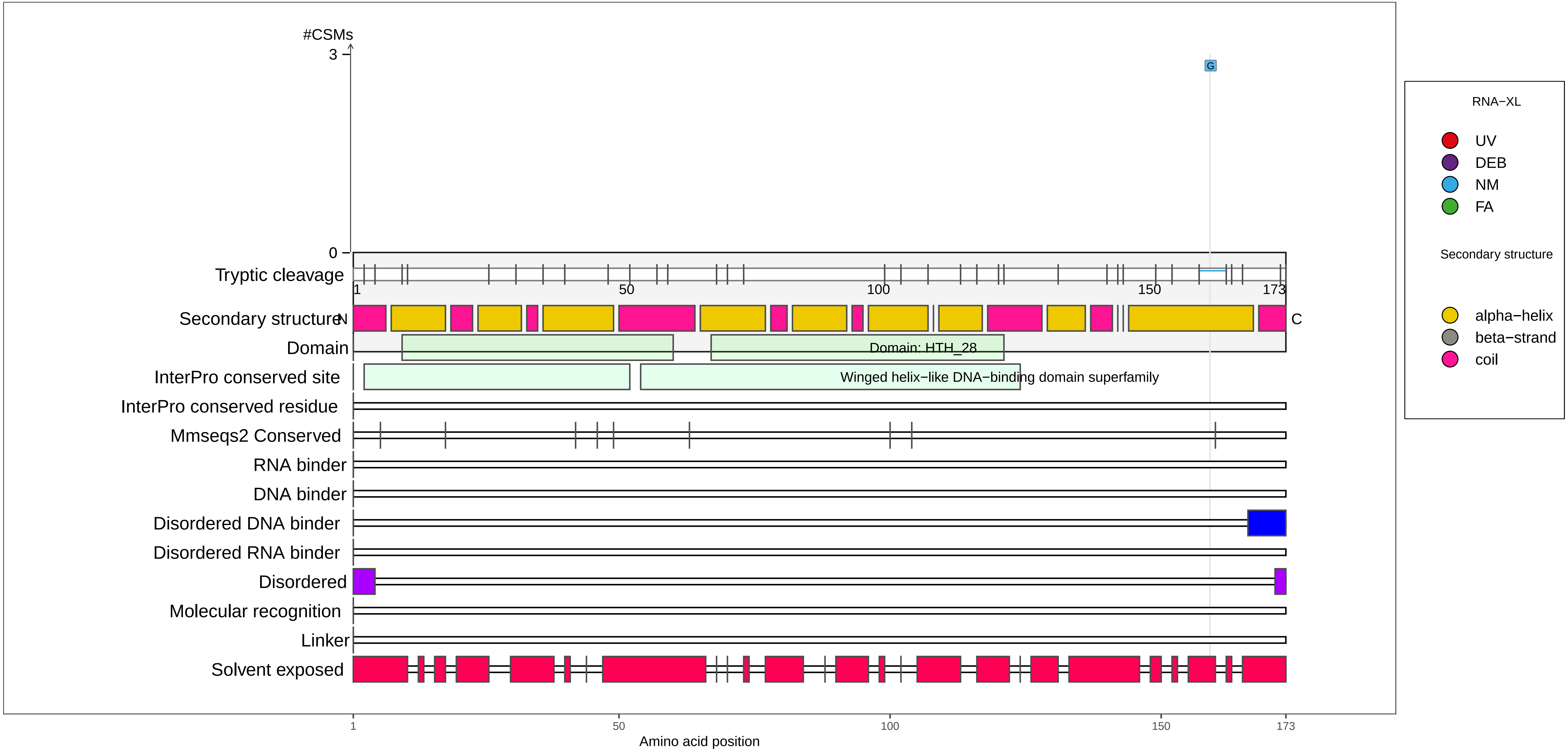

P19934
TOLA_ECOLI Tol−Pal system protein TolA
− Abundance:
tryptic [log10 Intensity]: 7.45 (Q 33)
PAXdb K12 strain [ppm]: 1.93 (Q 58)
PAXdb E.coli [ppm]: 1.25 (Q 53)

− RNA functions: not annotated

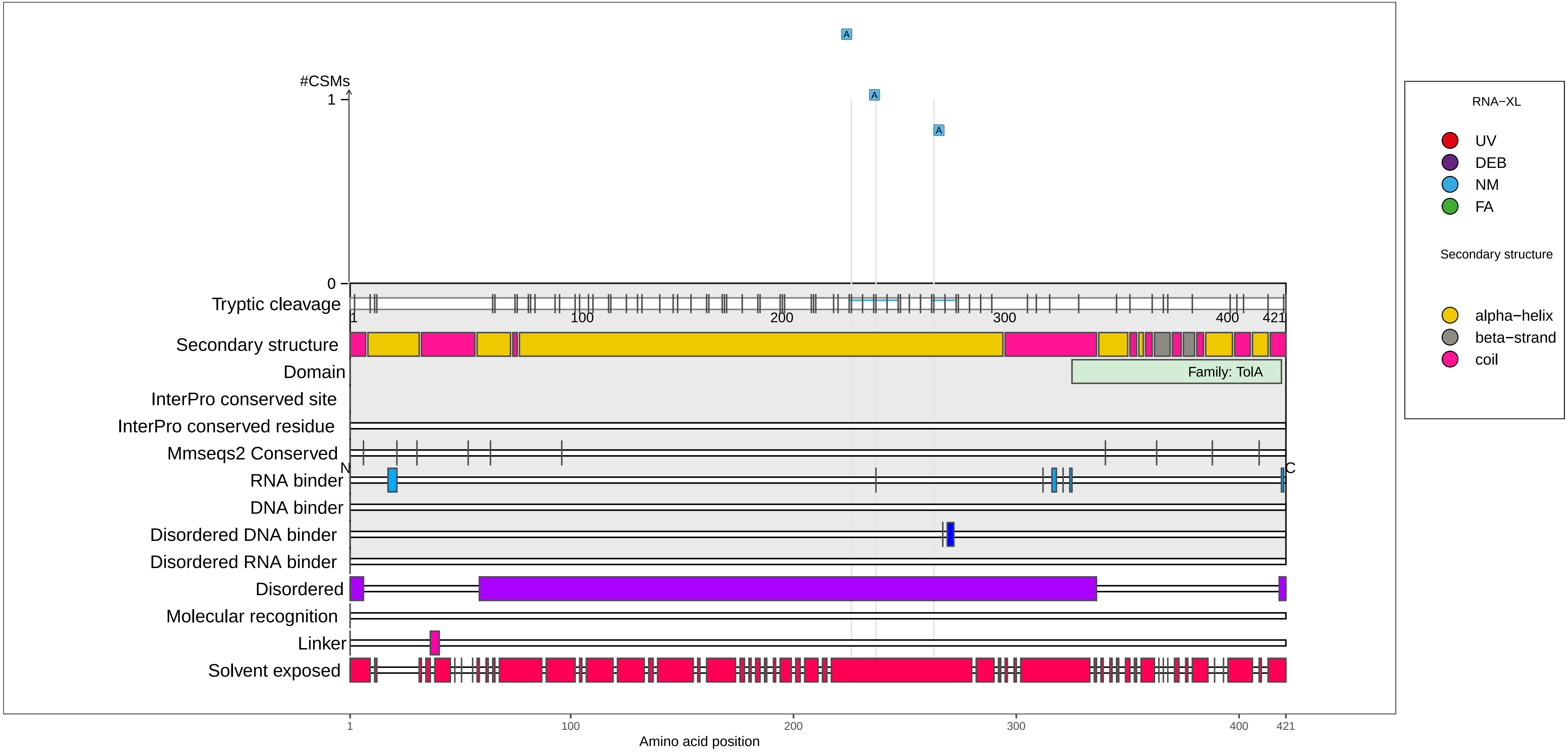

P22939
ISPA_ECOLI Farnesyl diphosphate synthase

− Abundance:
tryptic [log10 Intensity]: 7.17 (Q 20)
PAXdb K12 strain [ppm]: 2.4 (Q 74)
PAXdb E.coli [ppm]: 1.95 (Q 71)

− RNA functions: not annotated

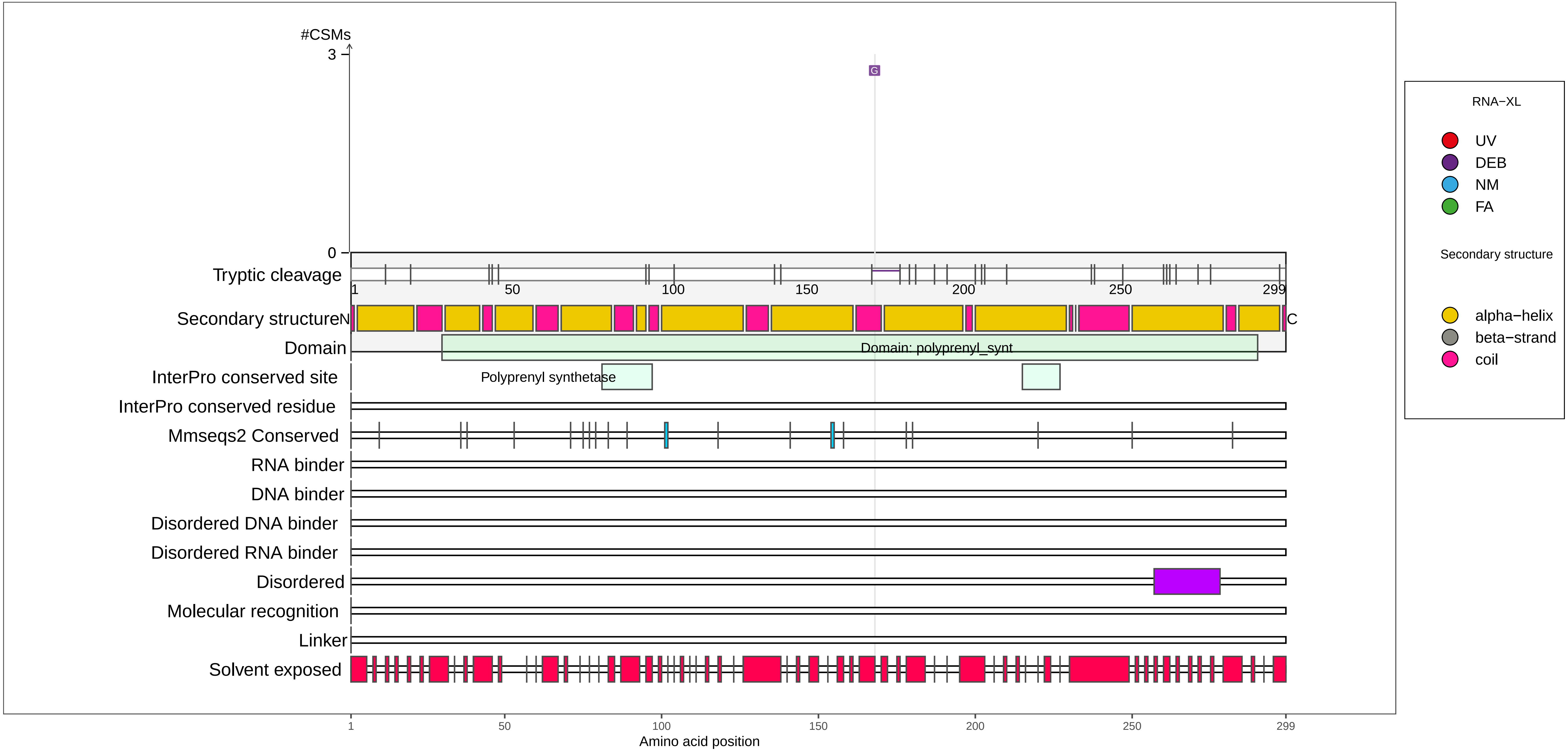

P23908
ARGE_ECOLI Acetylornithine deacetylase

− Abundance:
tryptic [log10 Intensity]: 8.55 (Q 77)
PAXdb K12 strain [ppm]: 1.03 (Q 12)
PAXdb E.coli [ppm]: 2.13 (Q 76)

− RNA functions: not annotated

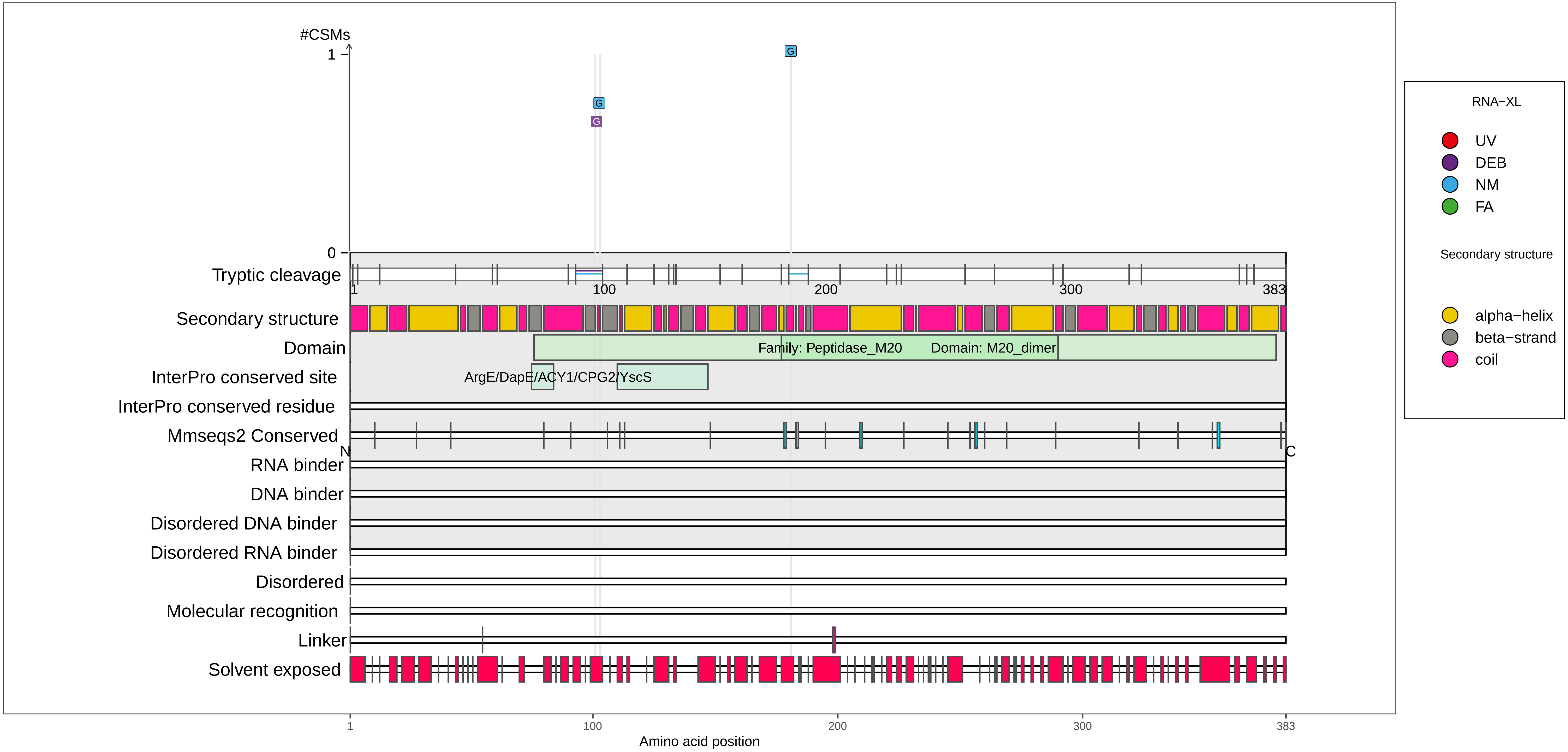

P24197
YGID_ECOLI 4,5−DOPA dioxygenase extradiol

− Abundance:
tryptic [log10 Intensity]: NA (Q NA)
PAXdb K12 strain [ppm]: NA (Q NA)
PAXdb E.coli [ppm]: 0.1 (Q 27)

− RNA functions: not annotated

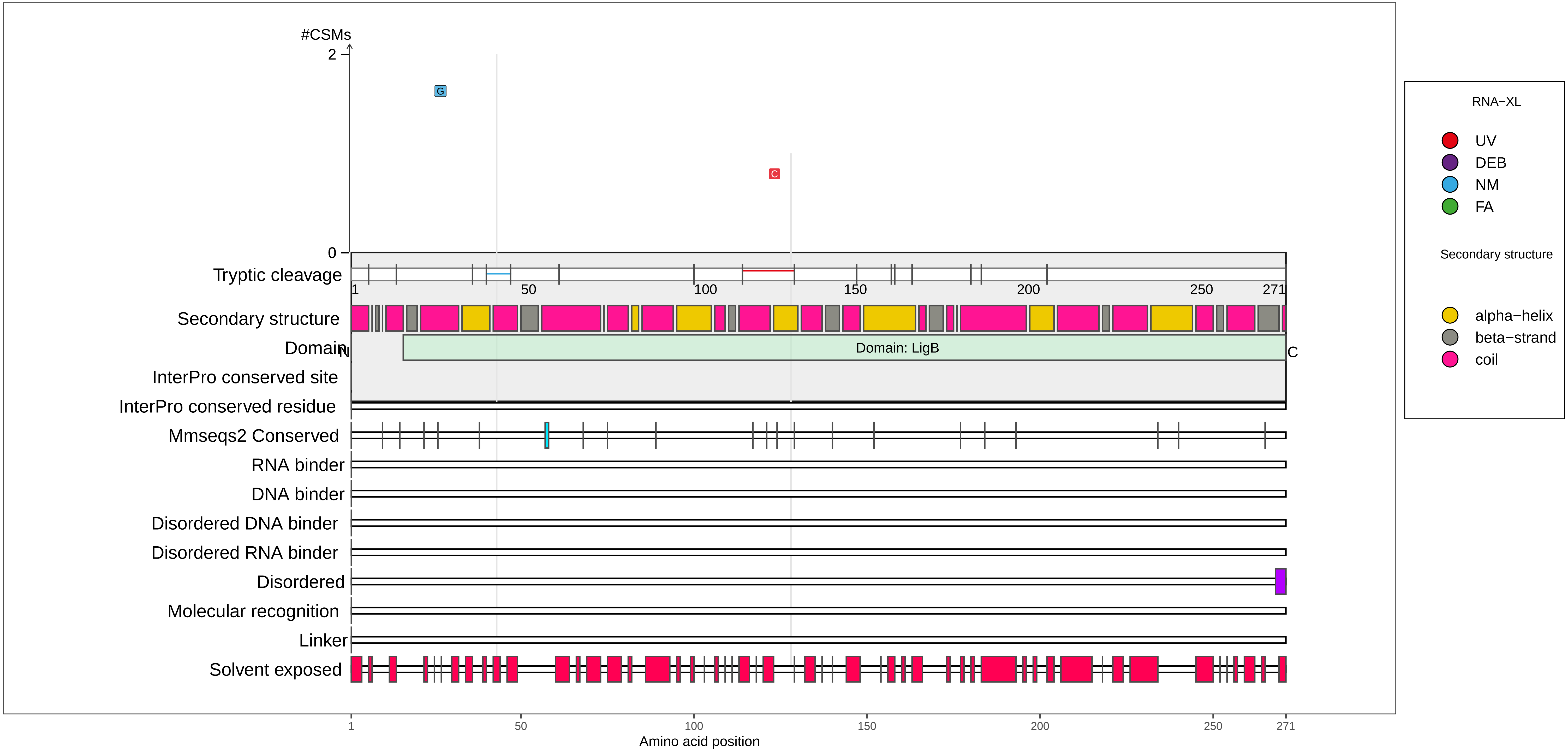

P27838
CYAY_ECOLI Iron−sulfur cluster assembly protein CyaY

− Abundance:
tryptic [log10 Intensity]: 7.92 (Q 55)
PAXdb K12 strain [ppm]: NA (Q NA)
PAXdb E.coli [ppm]: 2.33 (Q 81)

− RNA functions: not annotated

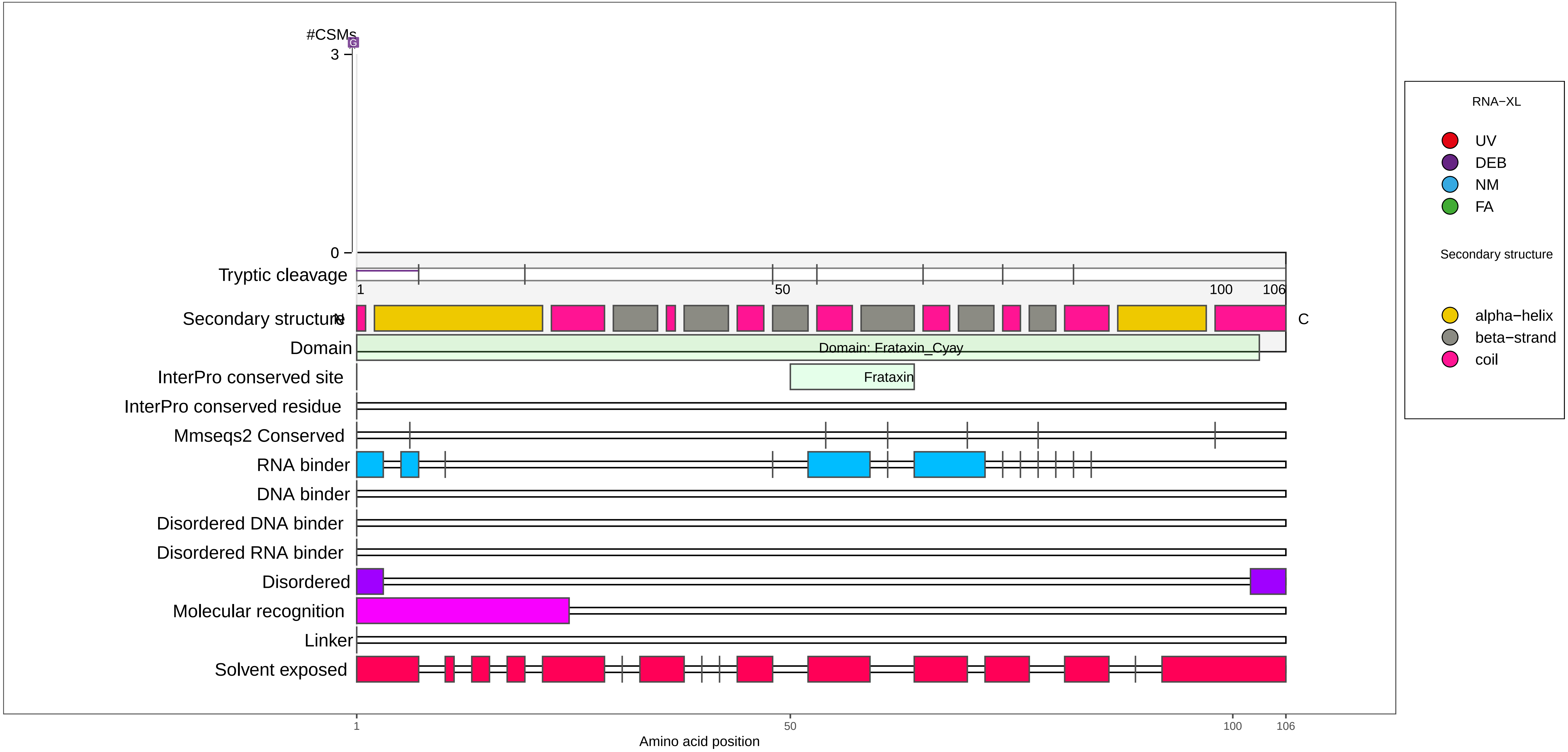

P29744
FLGL_ECOLI Flagellar hook−associated protein 3

− Abundance:
tryptic [log10 Intensity]: 7.2 (Q 21)
PAXdb K12 strain [ppm]: 1.74 (Q 51)
PAXdb E.coli [ppm]: 1.92 (Q 70)

− RNA functions: not annotated

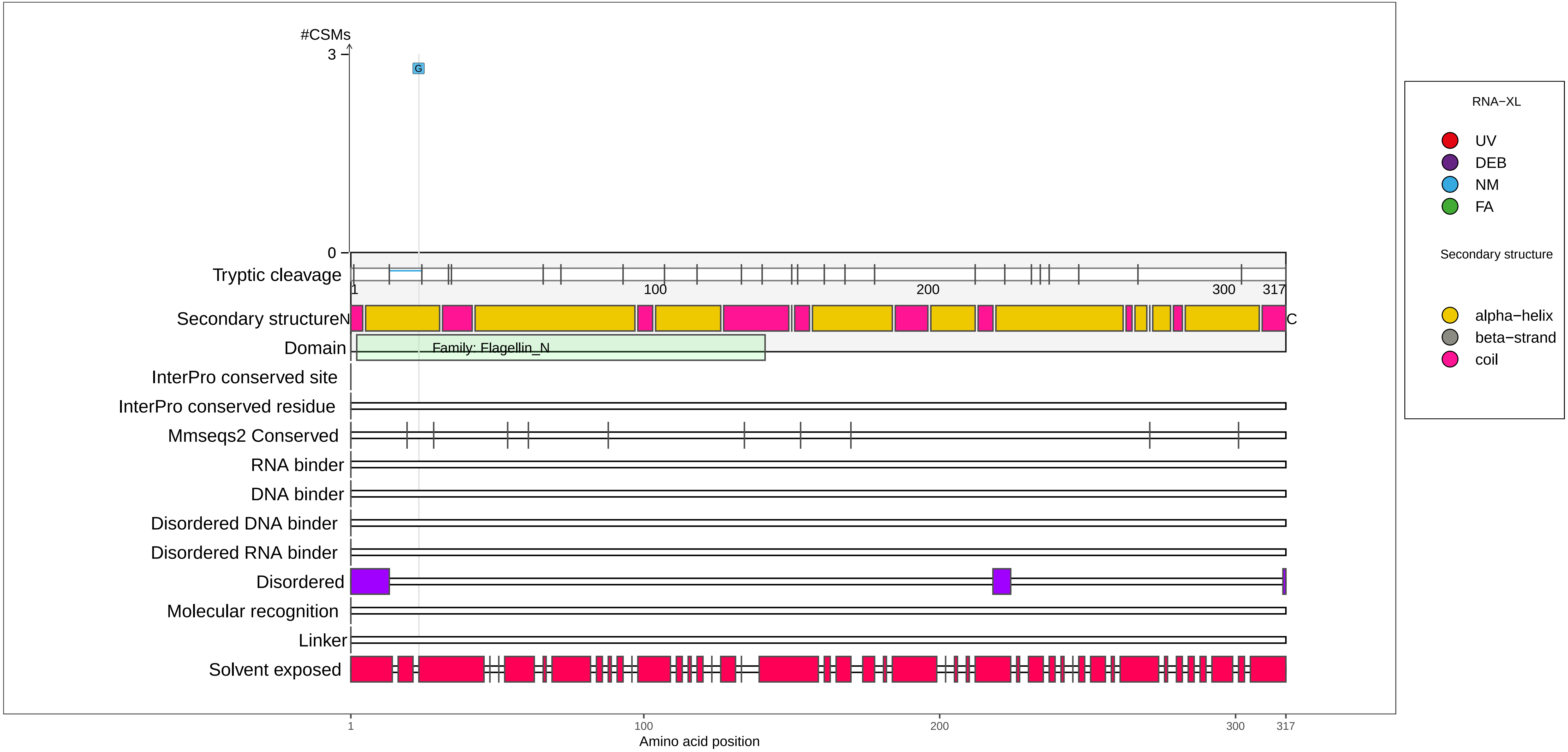

P30136
THIC_ECOLI Phosphomethylpyrimidine synthase

− Abundance:
tryptic [log10 Intensity]: 7.17 (Q 20)
PAXdb K12 strain [ppm]: 0.58 (Q 1)
PAXdb E.coli [ppm]: 1.98 (Q 72)

− RNA functions: not annotated

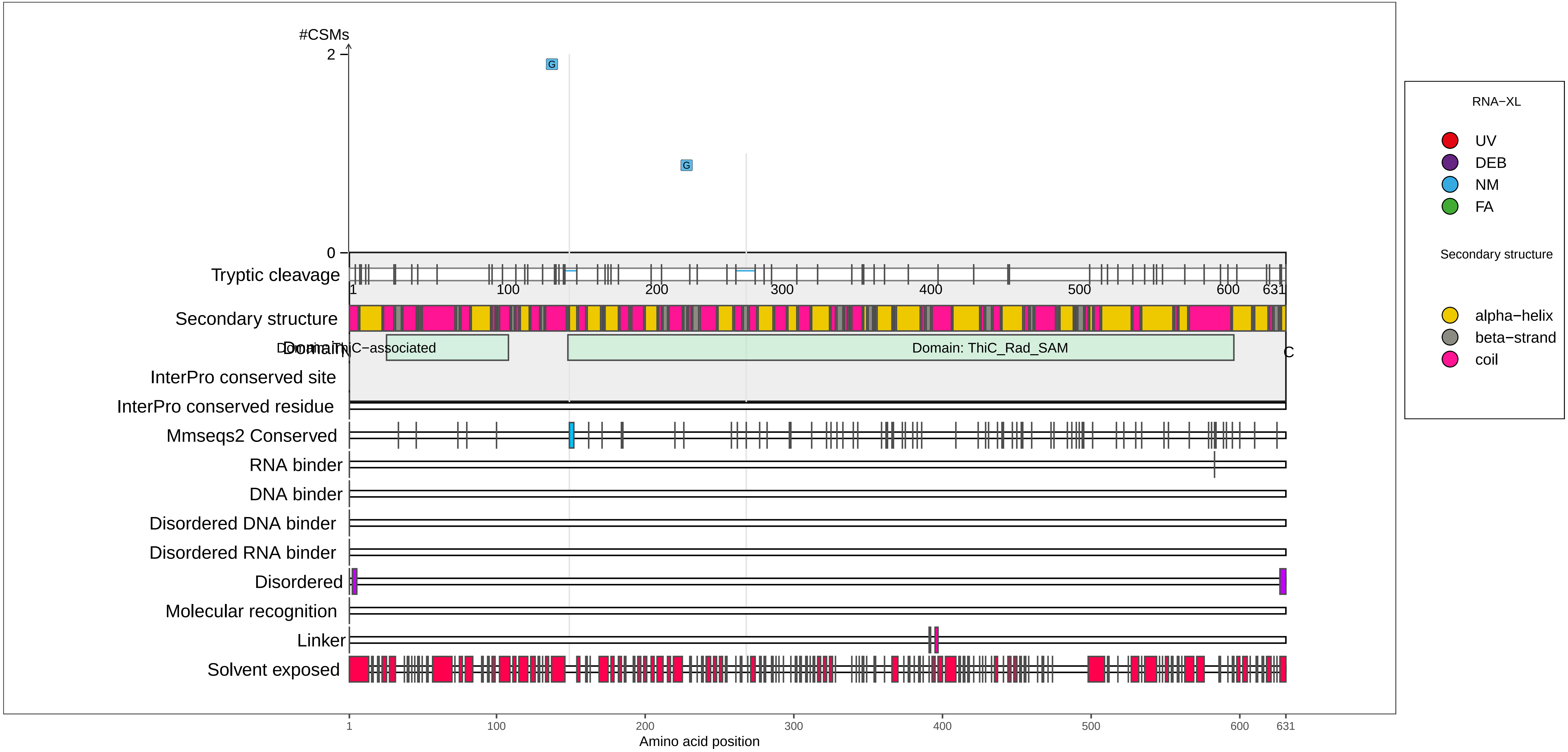

P31554
LPTD_ECOLI LPS−assembly protein LptD
− Abundance:
tryptic [log10 Intensity]: 9.51 (Q 97)
PAXdb K12 strain [ppm]: 0.75 (Q 4)
PAXdb E.coli [ppm]: 1.78 (Q 66)

− RNA functions: not annotated

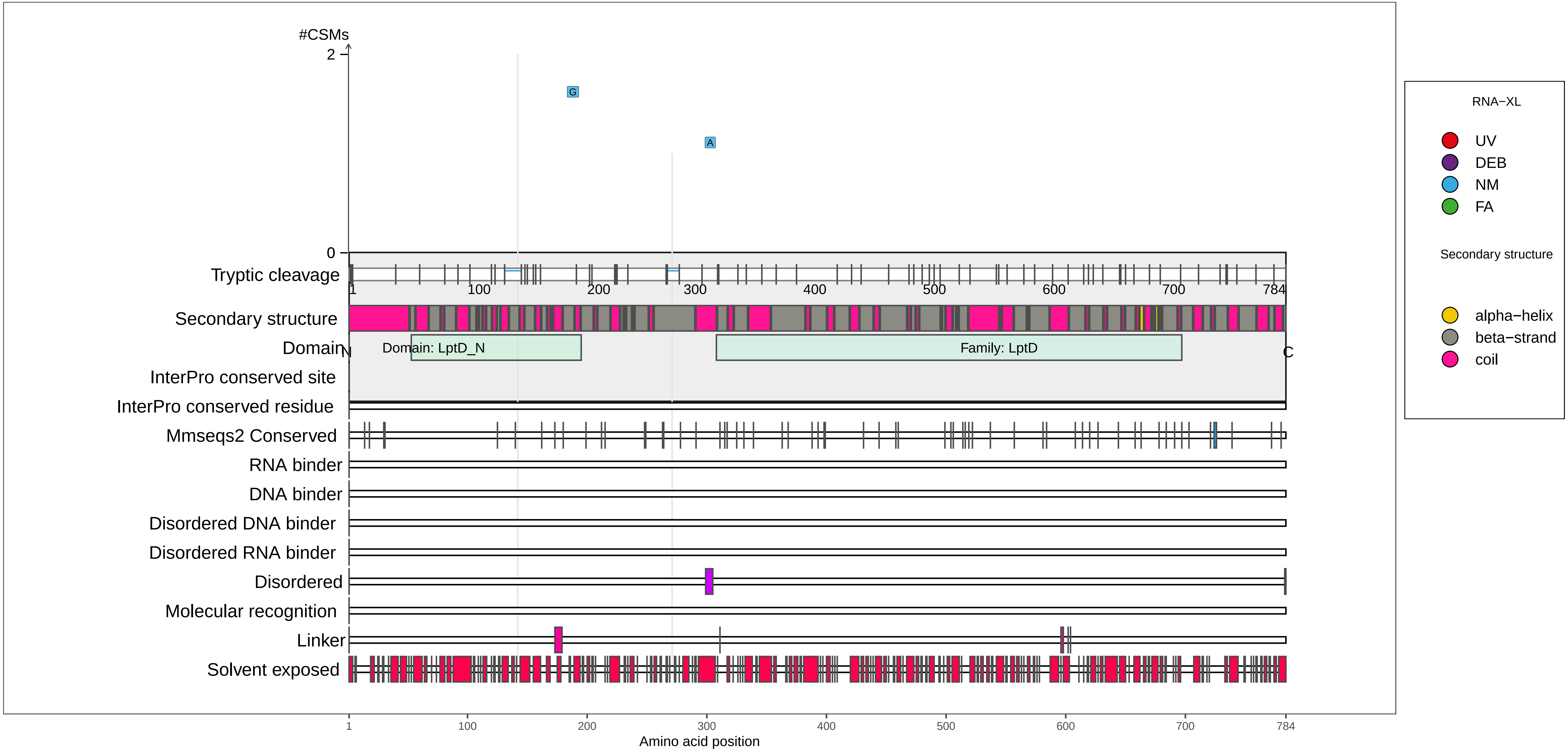

P32151

YIIG_ECOLI Uncharacterized protein YiiG

− Abundance:
tryptic [log10 Intensity]: NA (Q NA)
PAXdb K12 strain [ppm]: 1.21 (Q 21)
PAXdb E.coli [ppm]: −0.31 (Q 19)

− RNA functions: not annotated

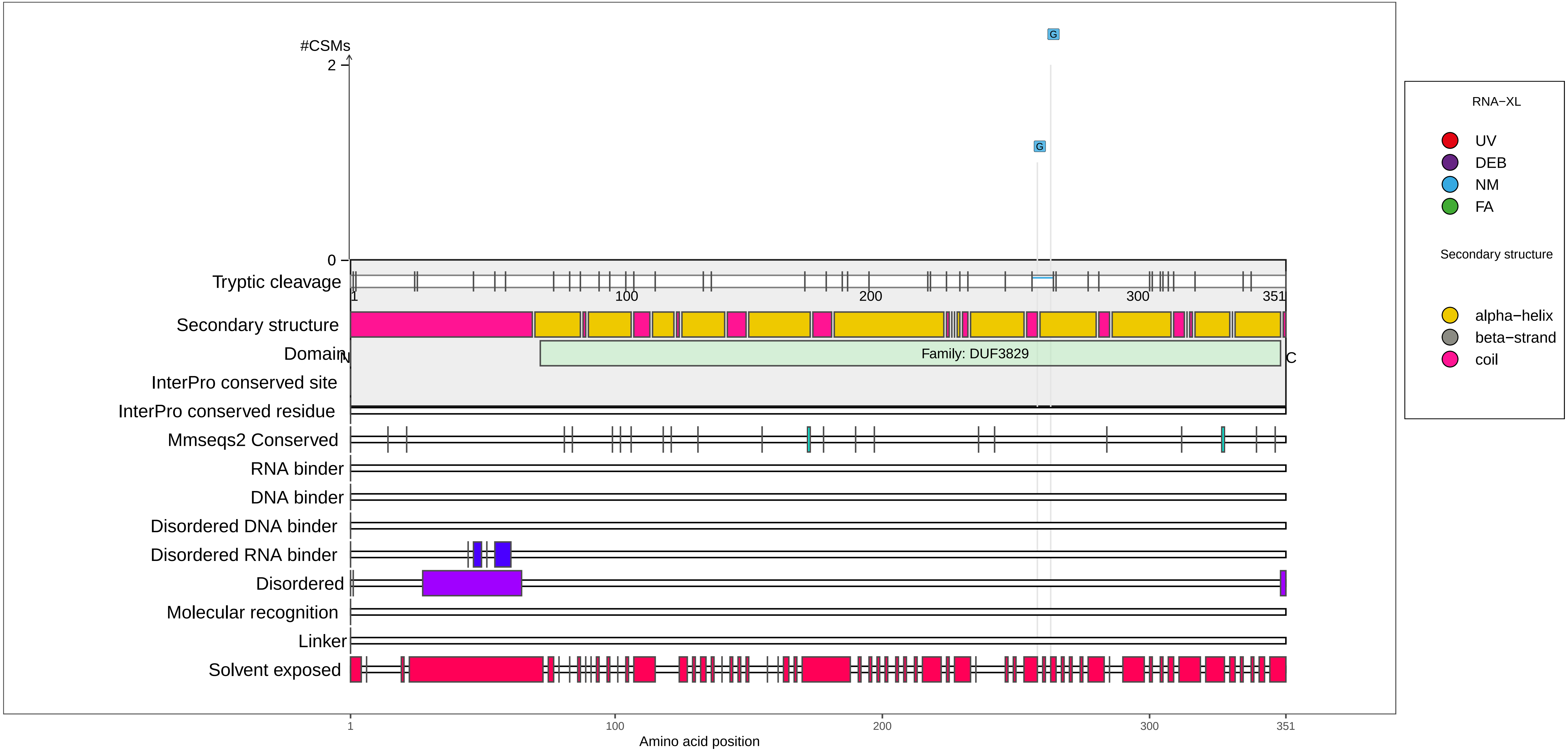

P32695
DUSA_ECOLI tRNA−dihydrouridine(20/20a) synthase

− Abundance:
tryptic [log10 Intensity]: 7.89 (Q 53)
PAXdb K12 strain [ppm]: 0.83 (Q 6)
PAXdb E.coli [ppm]: 1.6 (Q 62)

− RNA functions:
ncRNA metabolic process; ncRNA processing; RNA binding; RNA metabolic process
RNA modification; RNA processing; tRNA binding; tRNA dihydrouridine synthase activity
tRNA dihydrouridine synthesis; tRNA metabolic process; tRNA modification; tRNA processing
tRNA−dihydrouridine20 synthase activity

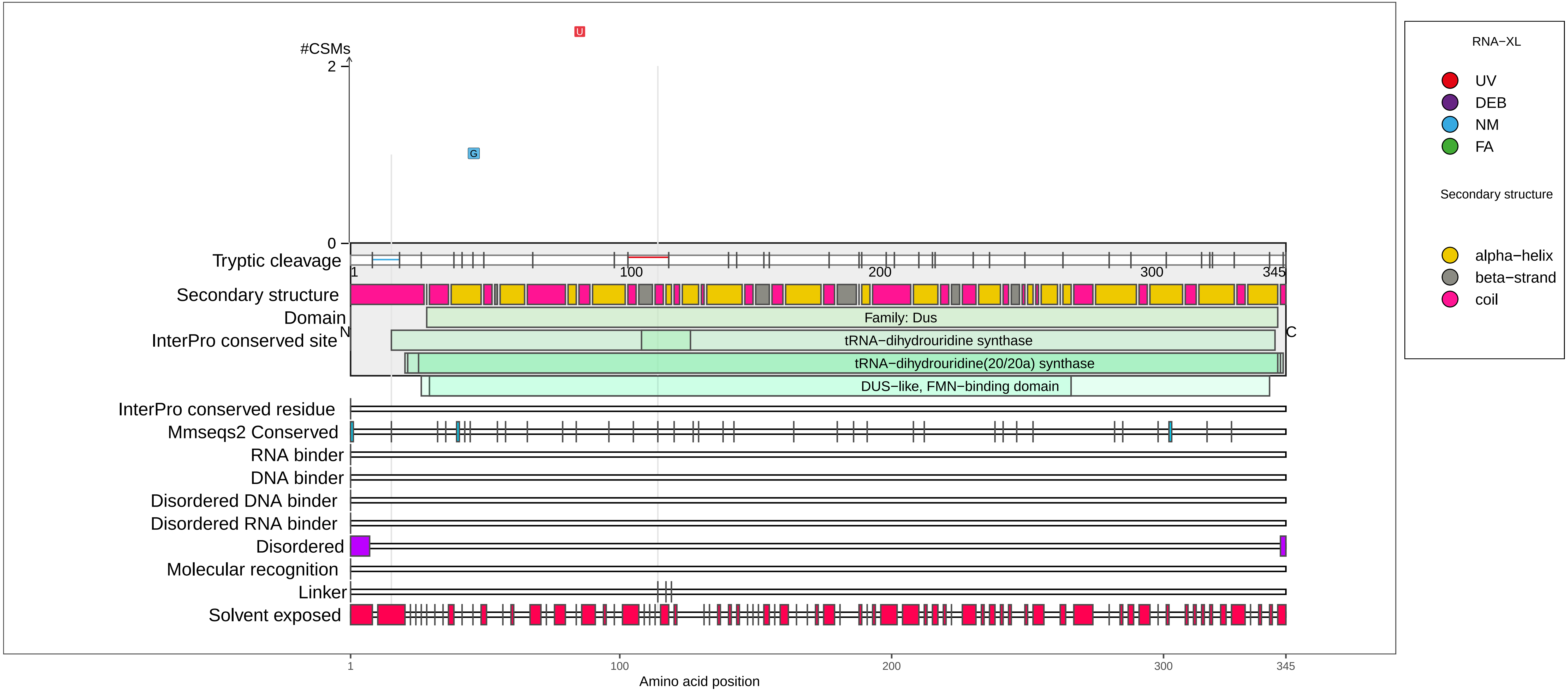

P33030
YEIR_ECOLI Zinc−binding GTPase YeiR

− Abundance:
tryptic [log10 Intensity]: 7.83 (Q 50)
PAXdb K12 strain [ppm]: 2.46 (Q 76)
PAXdb E.coli [ppm]: 0.59 (Q 38)

− RNA functions: not annotated

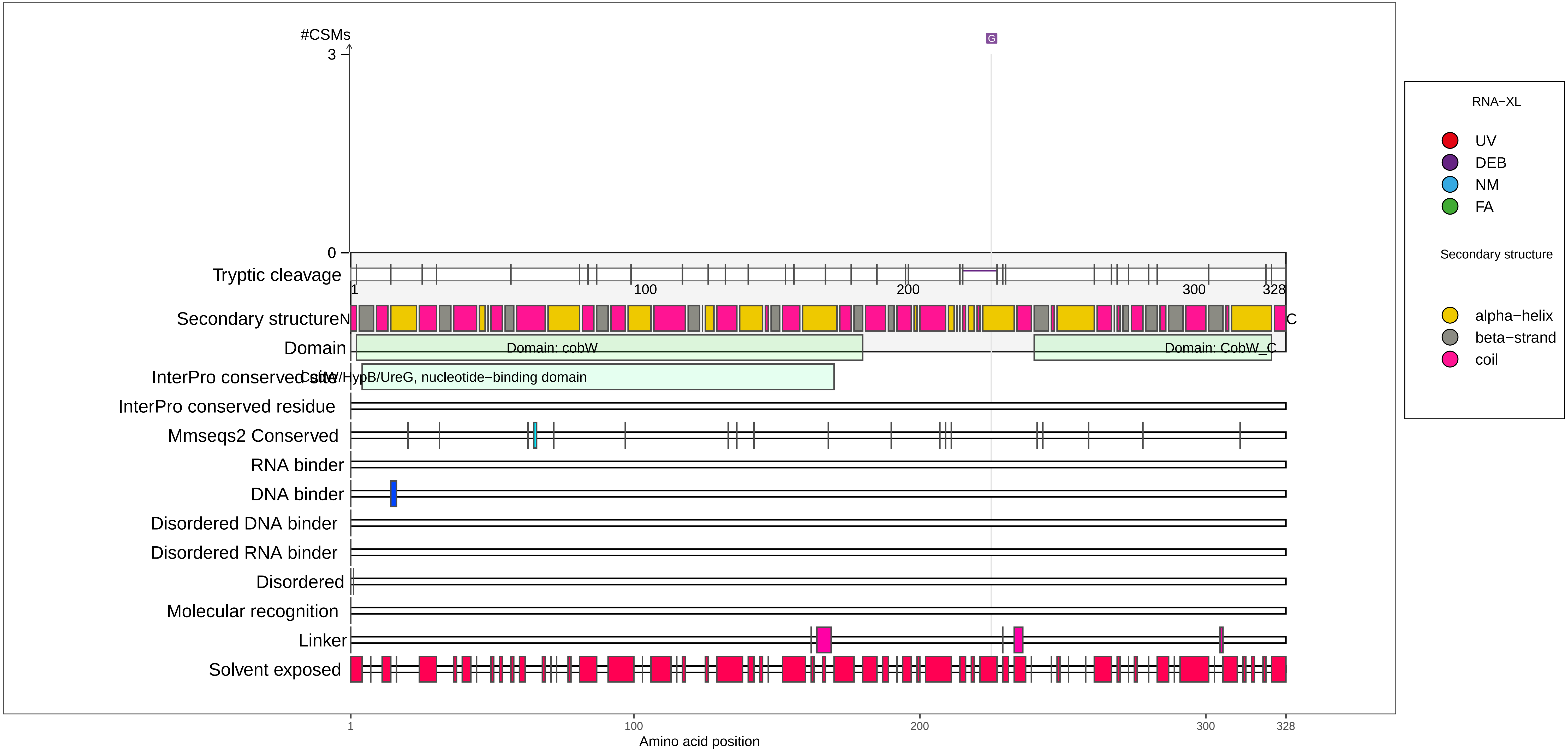

P37440
UCPA_ECOLI Oxidoreductase UcpA

− Abundance:
tryptic [log10 Intensity]: 7.95 (Q 56)
PAXdb K12 strain [ppm]: 2.47 (Q 76)
PAXdb E.coli [ppm]: 2.32 (Q 81)

− RNA functions: not annotated

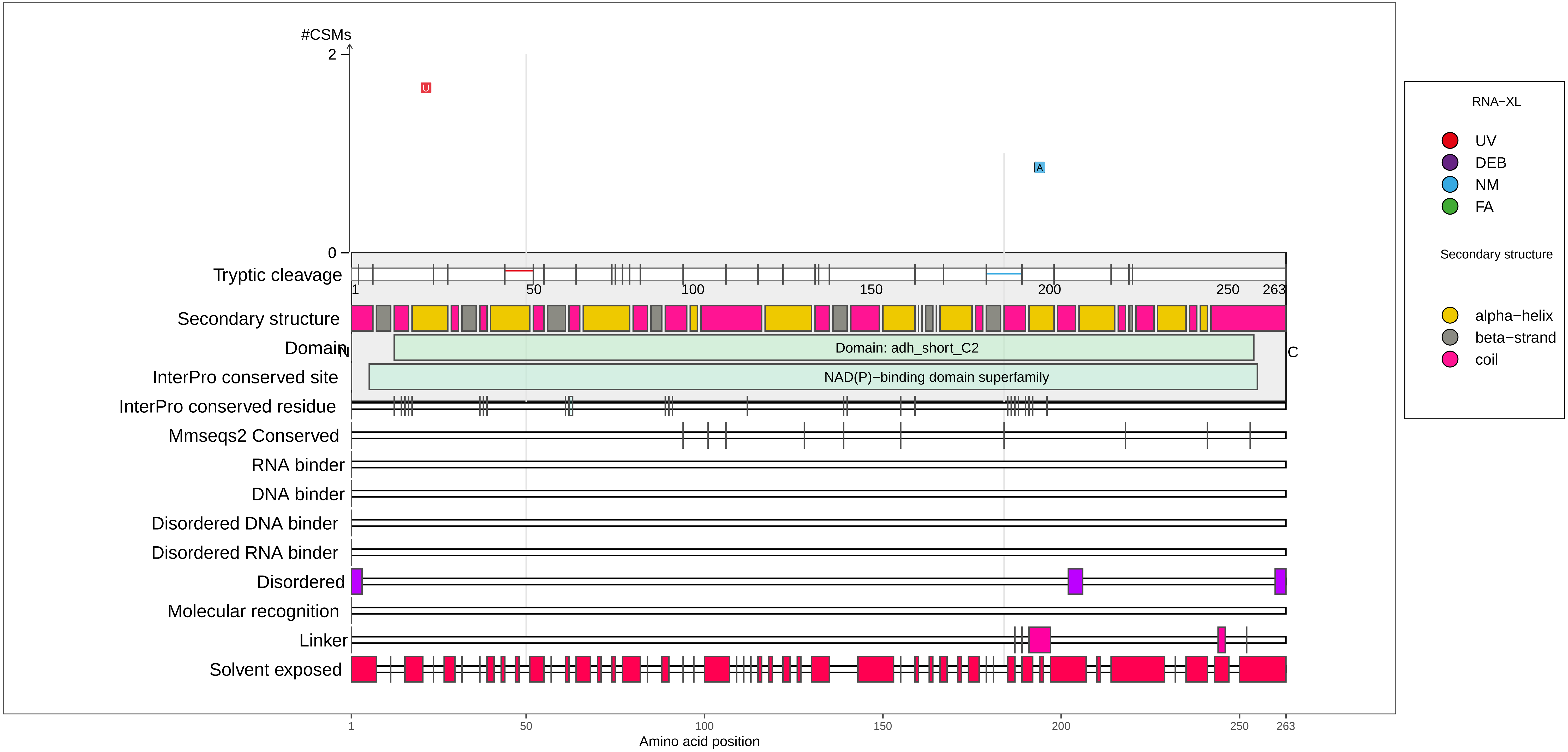

P37650
BCSC_ECOLI Cellulose synthase operon protein C

− Abundance:
tryptic [log10 Intensity]: 7.59 (Q 39)
PAXdb K12 strain [ppm]: 1.01 (Q 11)
PAXdb E.coli [ppm]: 0.81 (Q 43)

− RNA functions: not annotated

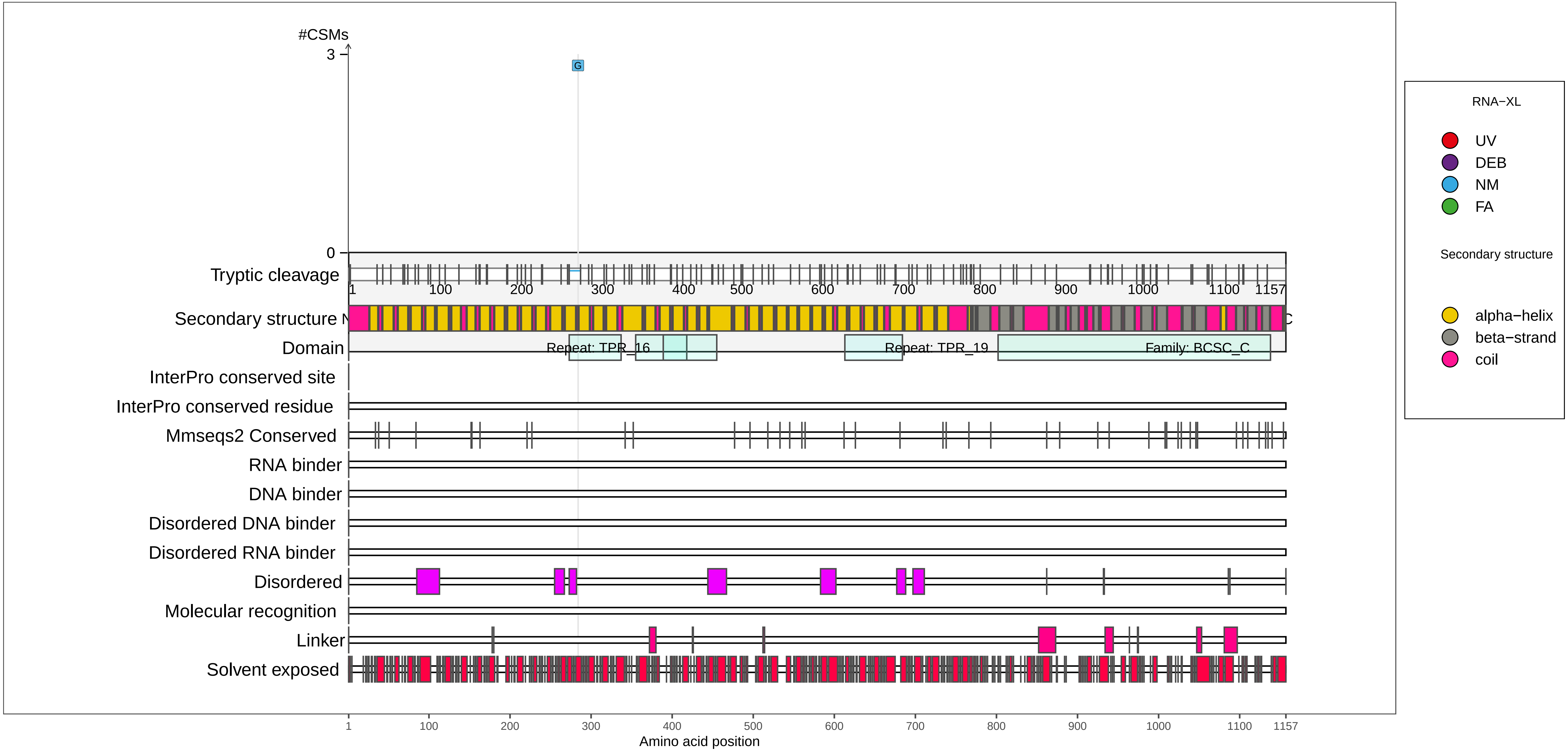

P37665
YIAD_ECOLI Probable lipoprotein YiaD

− Abundance:
tryptic [log10 Intensity]: 9.4 (Q 95)
PAXdb K12 strain [ppm]: 1.97 (Q 59)
PAXdb E.coli [ppm]: 1.6 (Q 62)

− RNA functions: not annotated

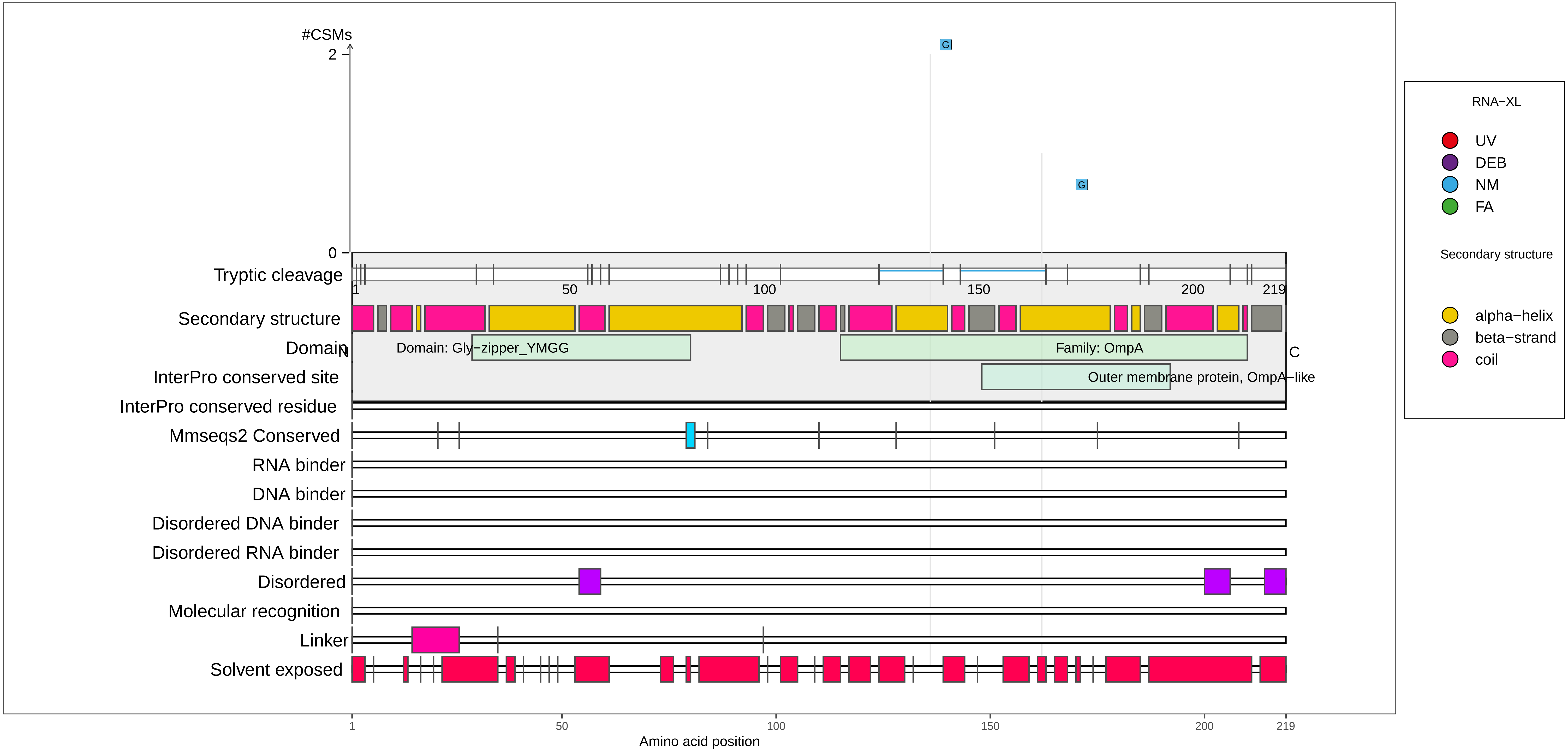

P39177
USPG_ECOLI Universal stress protein UP12

− Abundance:
tryptic [log10 Intensity]: 8.16 (Q 64)
PAXdb K12 strain [ppm]: 2 (Q 60)
PAXdb E.coli [ppm]: 3.16 (Q 96)

− RNA functions: not annotated

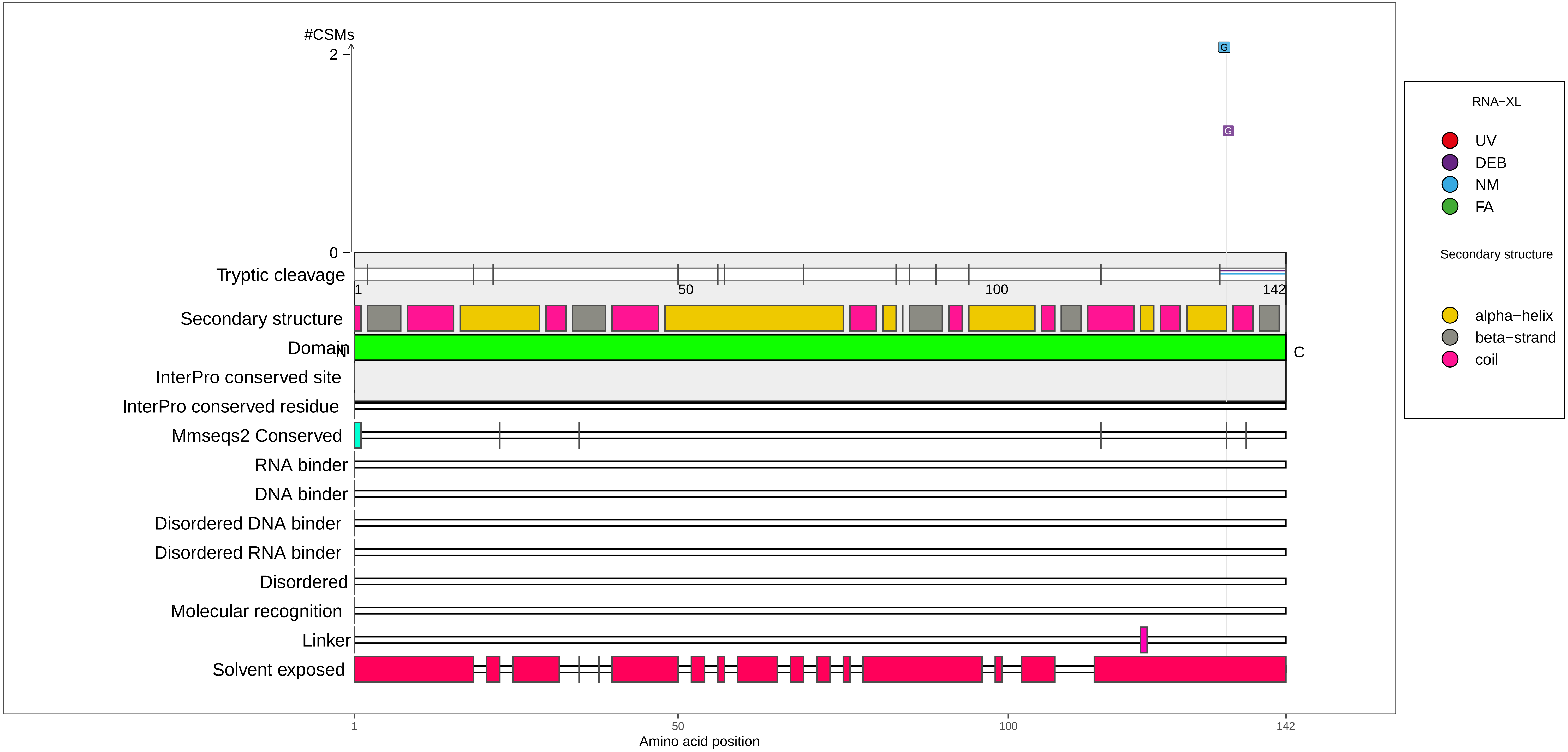

P39300
ULAG_ECOLI Probable L−ascorbate−6−phosphate lactonase UlaG

− Abundance:
tryptic [log10 Intensity]: 7.38 (Q 29)
PAXdb K12 strain [ppm]: 2.6 (Q 80)
PAXdb E.coli [ppm]: 1.3 (Q 54)

− RNA functions: not annotated

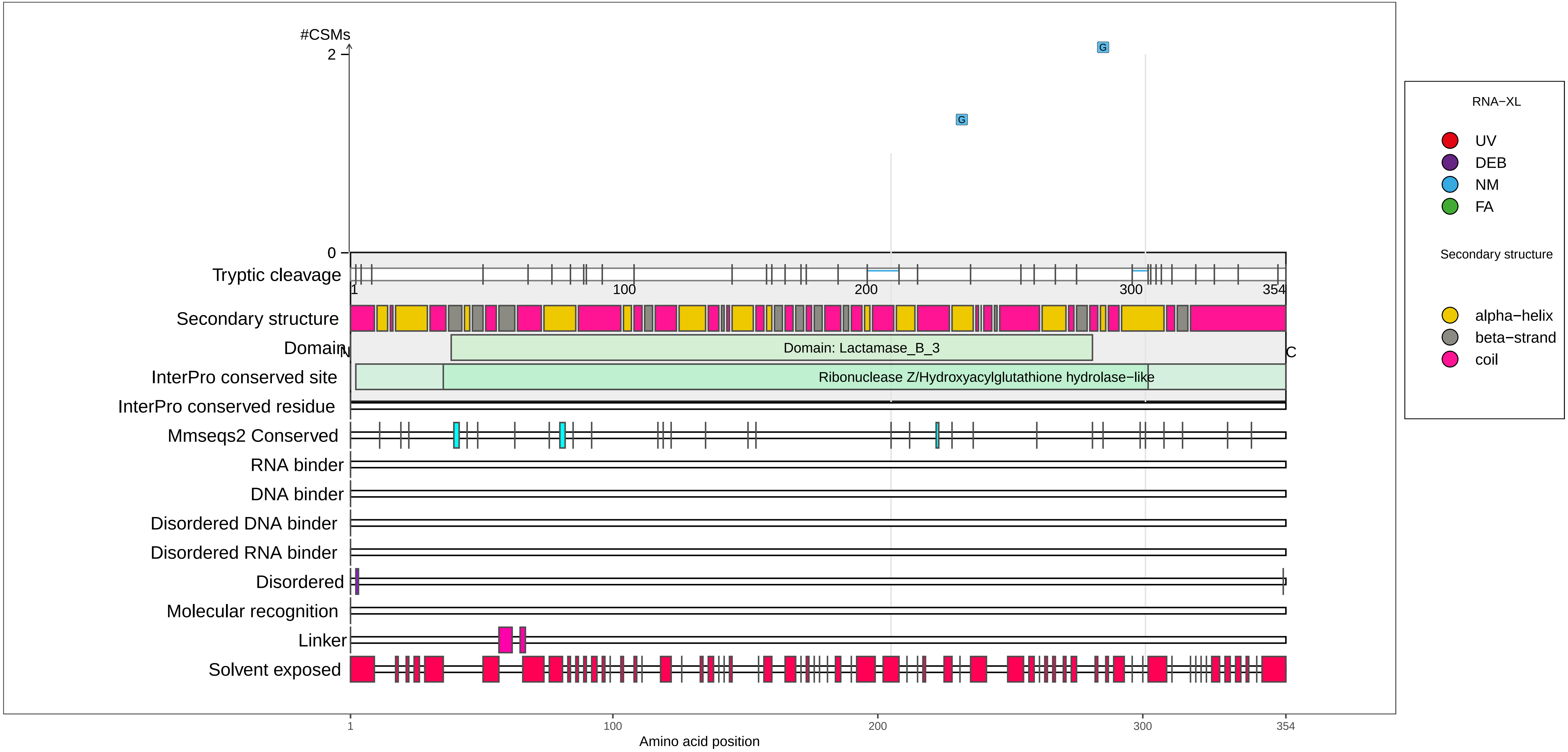

P39346
IDND_ECOLI L−idonate 5−dehydrogenase (NAD(P)(+))

− Abundance:
tryptic [log10 Intensity]: 8.57 (Q 78)
PAXdb K12 strain [ppm]: NA (Q NA)
PAXdb E.coli [ppm]: −1.4 (Q 5)

− RNA functions: not annotated

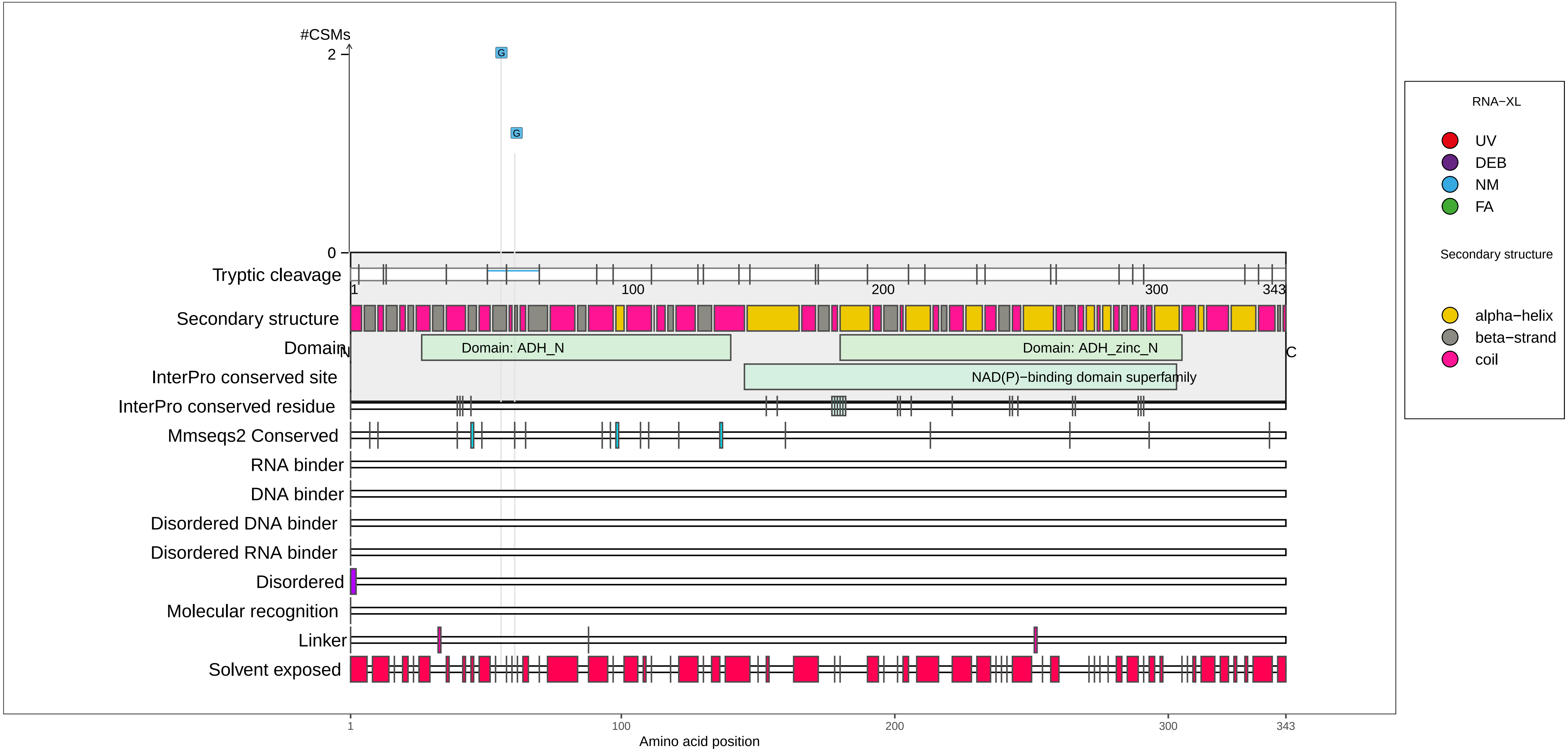

P39353
YJHC_ECOLI Uncharacterized oxidoreductase YjhC

− Abundance:
tryptic [log10 Intensity]: 8.05 (Q 59)
PAXdb K12 strain [ppm]: 1.08 (Q 15)
PAXdb E.coli [ppm]: 0.81 (Q 43)

− RNA functions: not annotated

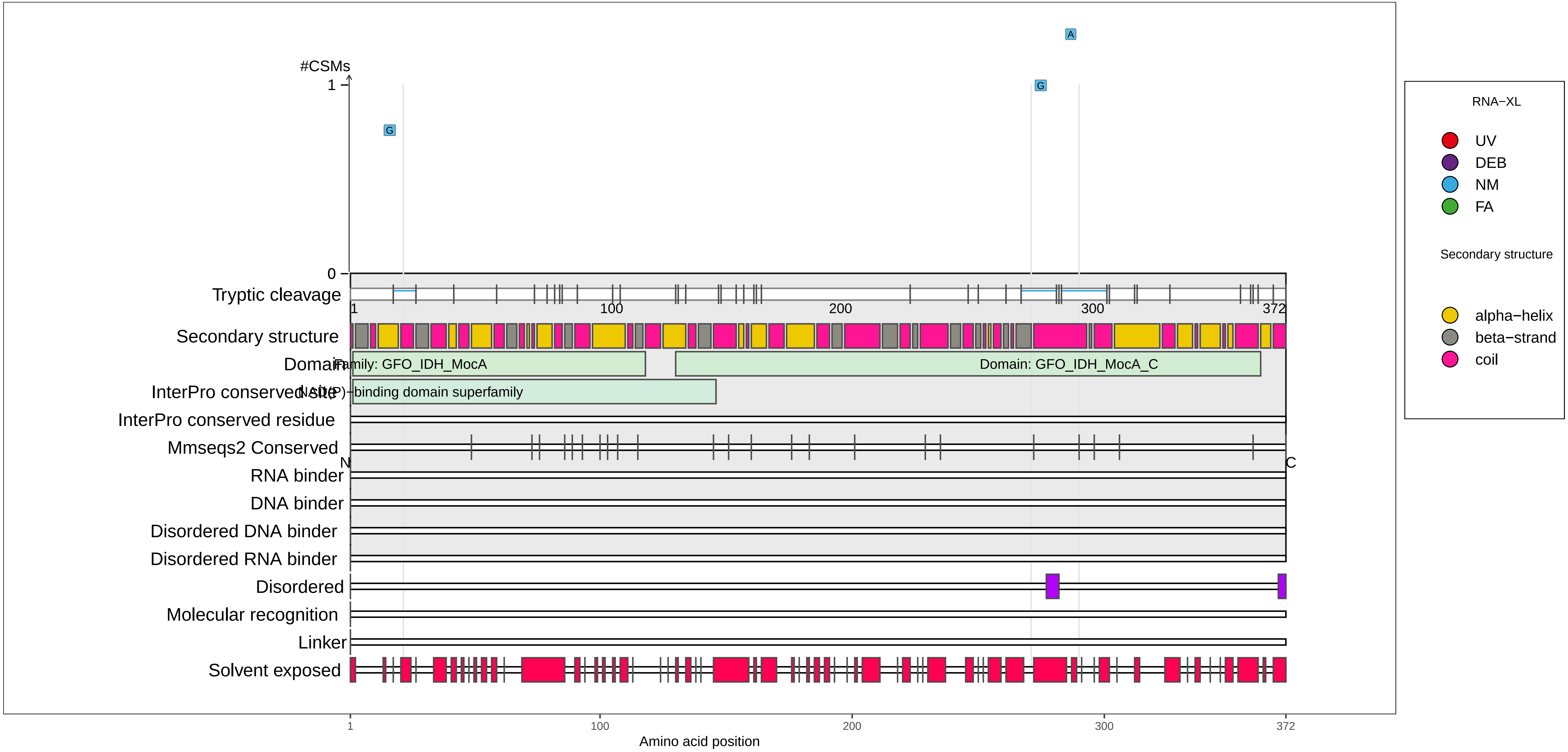

P39401
OPGB_ECOLI Phosphoglycerol transferase I

− Abundance:
tryptic [log10 Intensity]: 8.16 (Q 64)
PAXdb K12 strain [ppm]: 1.55 (Q 41)
PAXdb E.coli [ppm]: 0.3 (Q 32)

− RNA functions: not annotated

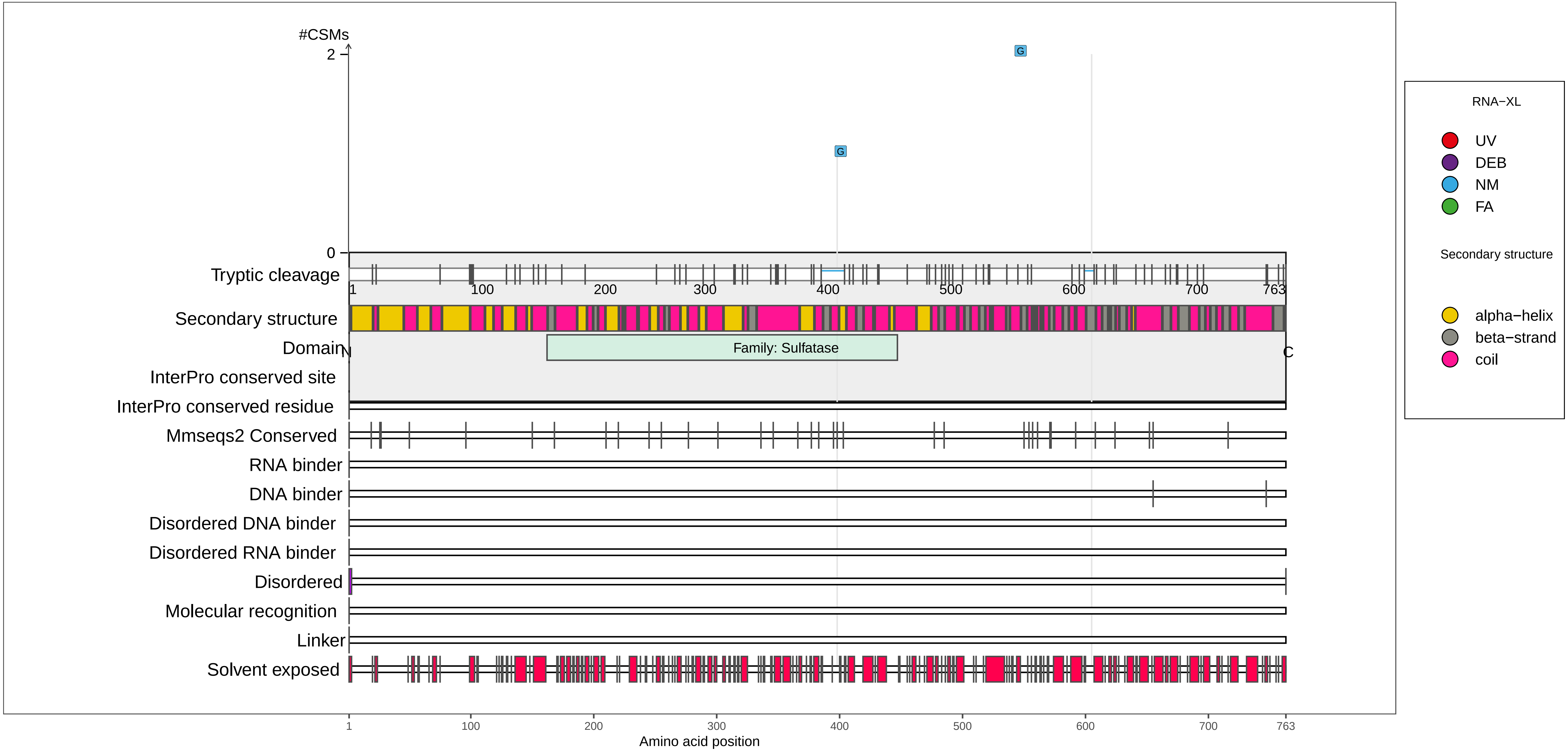

P45523
FKBA_ECOLI FKBP−type peptidyl−prolyl cis−trans isomerase FkpA

− Abundance:
tryptic [log10 Intensity]: 9.51 (Q 97)
PAXdb K12 strain [ppm]: 3.49 (Q 97)
PAXdb E.coli [ppm]: 2.71 (Q 90)

− RNA functions: not annotated

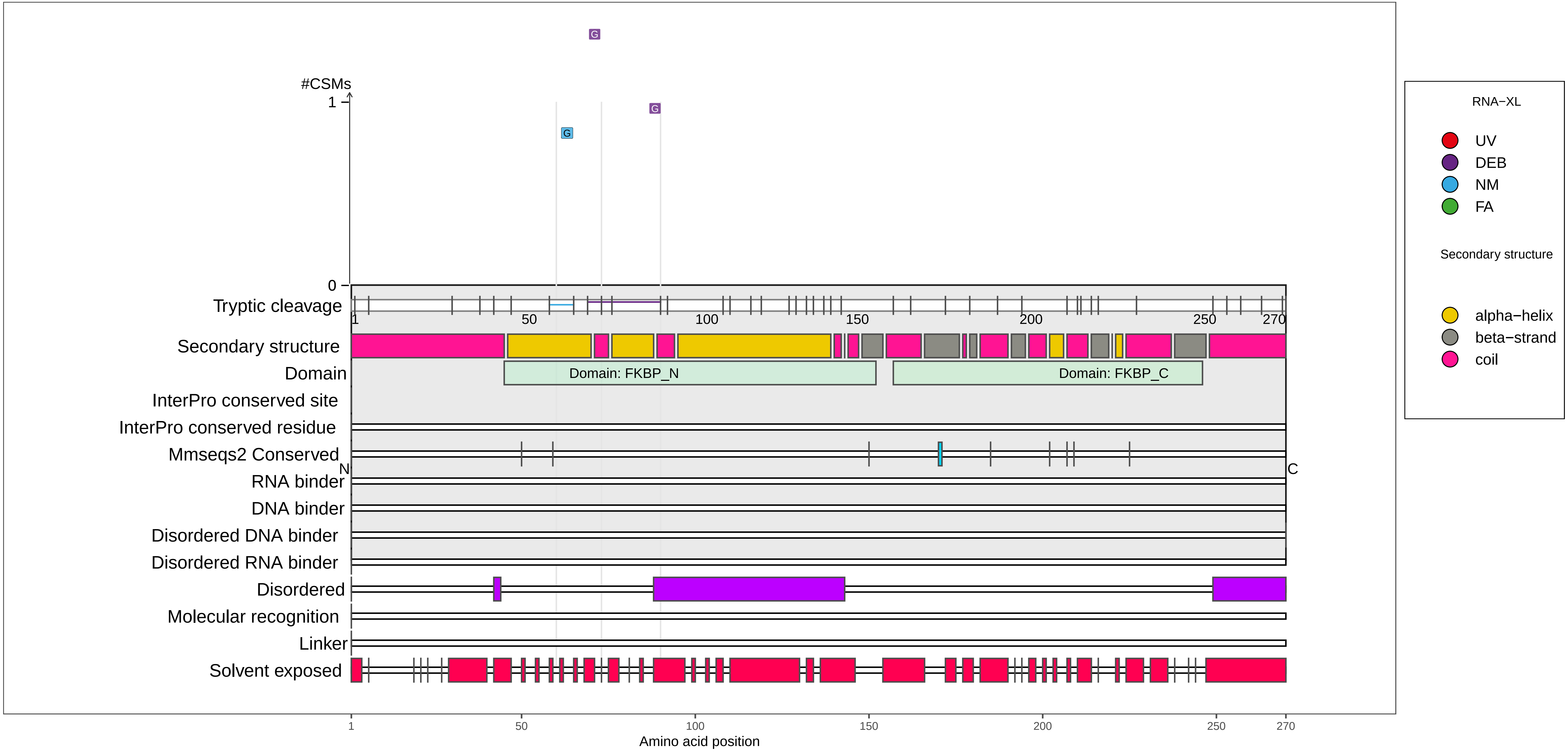

P46068
DSDC_ECOLI HTH−type transcriptional regulator DsdC
− Abundance:
tryptic [log10 Intensity]: NA (Q NA)
PAXdb K12 strain [ppm]: NA (Q NA)
PAXdb E.coli [ppm]: −0.94 (Q 10)

− RNA functions:
RNA biosynthetic process; RNA metabolic process

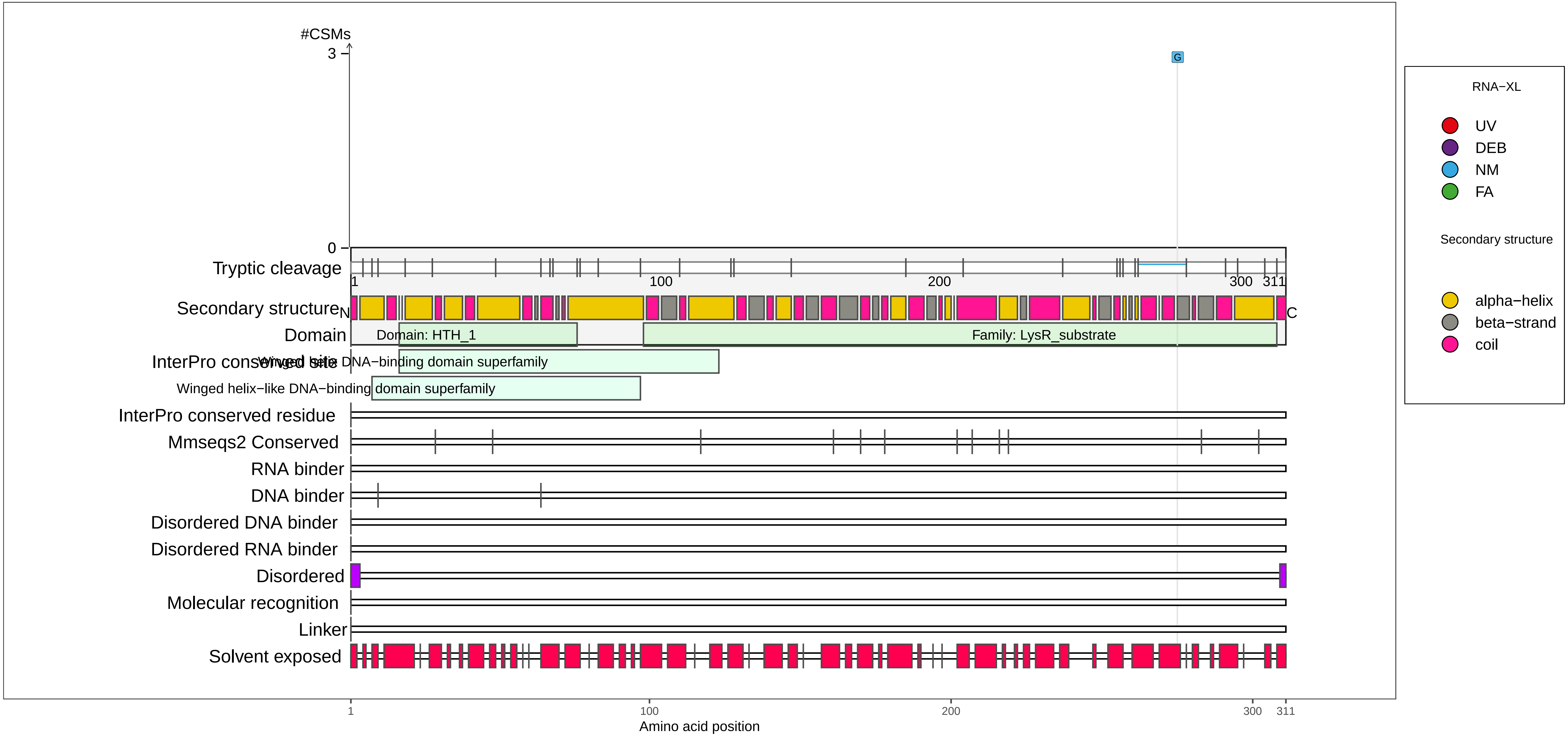

P60546
KGUA_ECOLI Guanylate kinase

− Abundance:
tryptic [log10 Intensity]: 9.1 (Q 91)
PAXdb K12 strain [ppm]: 2.9 (Q 88)
PAXdb E.coli [ppm]: 2.18 (Q 77)

− RNA functions: not annotated

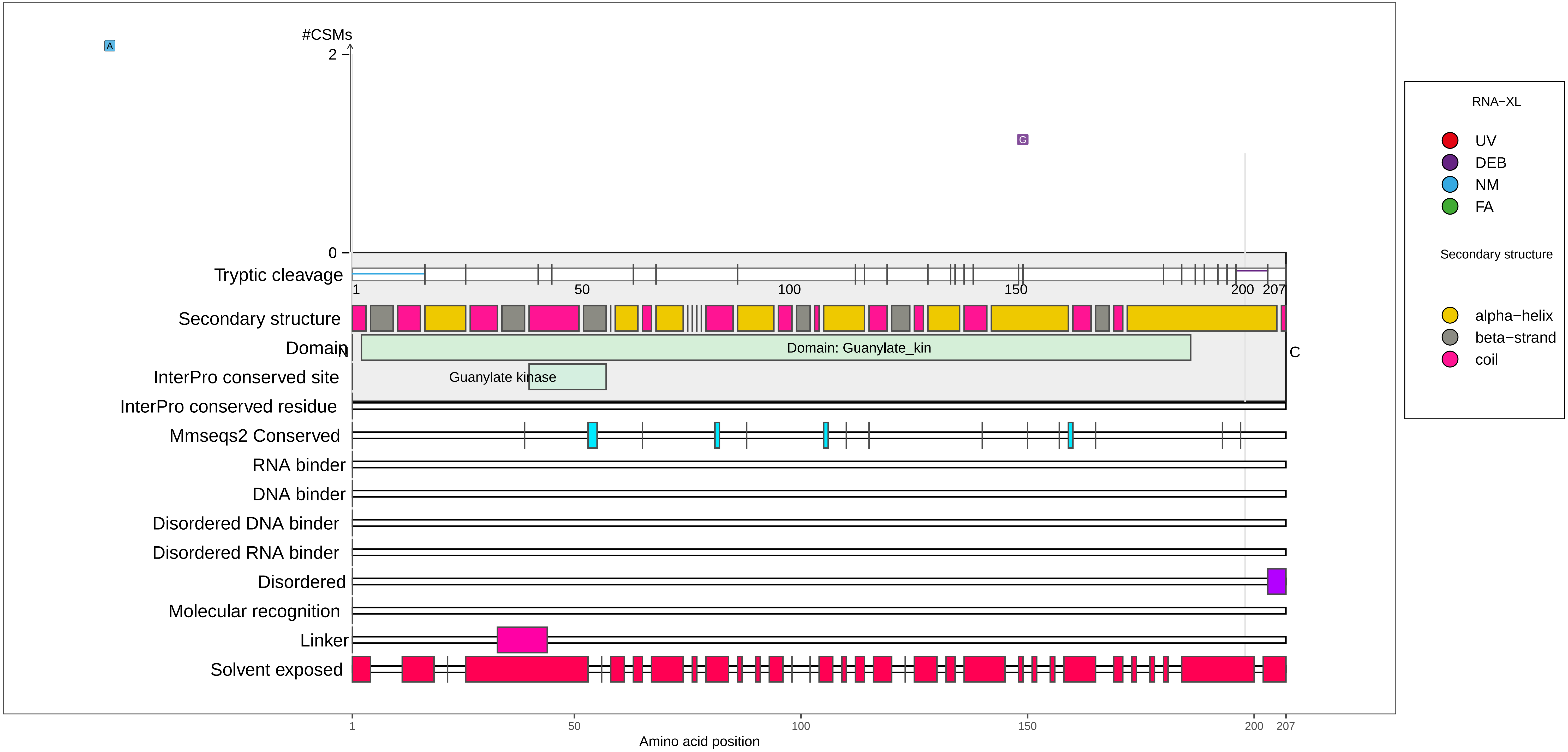

P62066
YCEQ_ECOLI Uncharacterized protein YceQ

− Abundance:
tryptic [log10 Intensity]: NA (Q NA)
PAXdb K12 strain [ppm]: 1.56 (Q 42)
PAXdb E.coli [ppm]: −1.2 (Q 7)

− RNA functions: not annotated

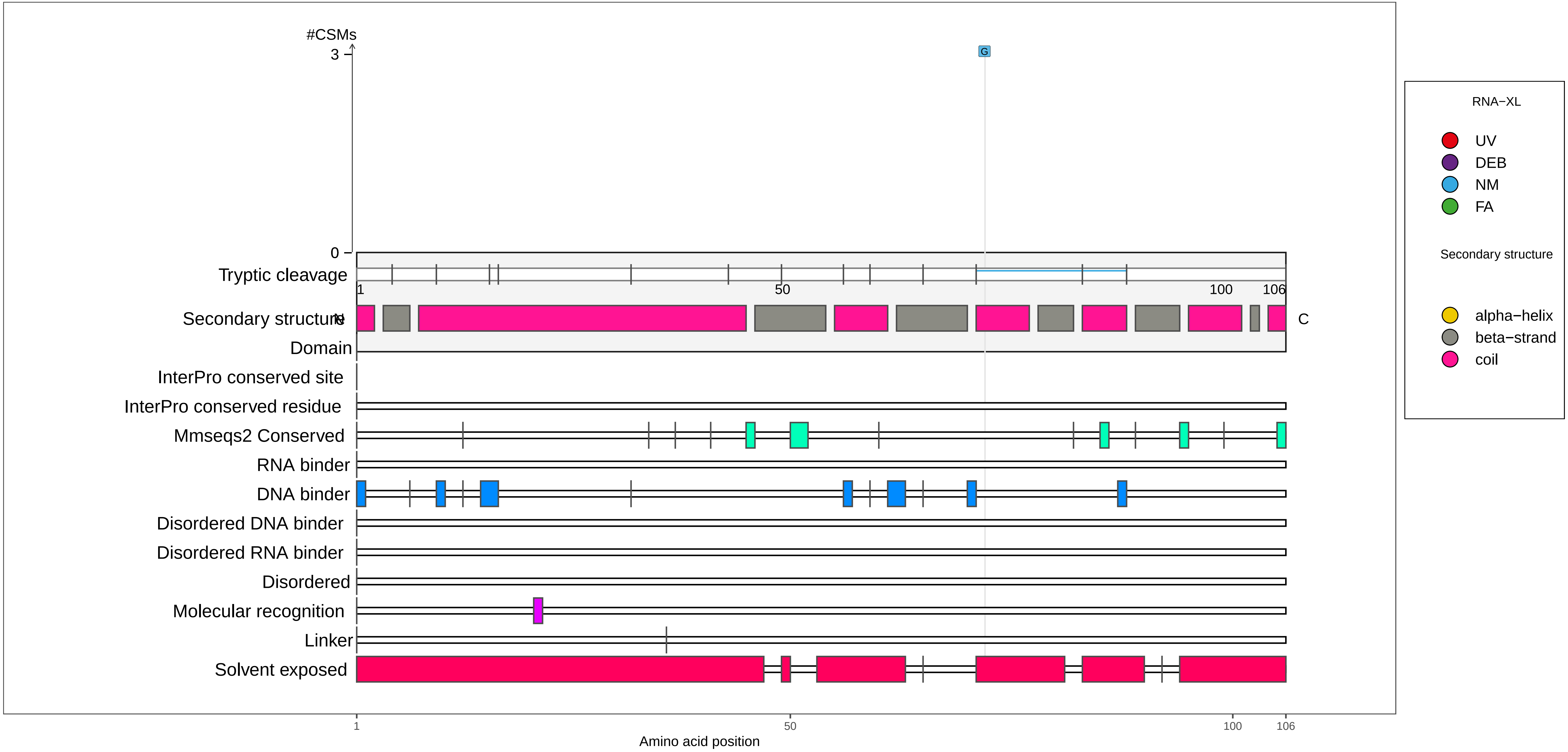

P63177
RLMB_ECOLI 23S rRNA (guanosine−2'−O−)−methyltransferase RlmB

− Abundance:
tryptic [log10 Intensity]: 7.57 (Q 38)
PAXdb K12 strain [ppm]: 2.69 (Q 83)
PAXdb E.coli [ppm]: 1.94 (Q 71)

− RNA functions:
ncRNA metabolic process; ncRNA processing
RNA 2−O ribose methyltransferase substrate binding; RNA 2−O−methyltransferase activity; RNA binding; RNA metabolic process
RNA methylation; RNA methyltransferase activity; RNA modification; RNA processing
rRNA (guanine) methyltransferase activity; rRNA (guanosine−2−O−)−methyltransferase activity
rRNA 2−O−methylation; rRNA metabolic process; rRNA methylation
rRNA methyltransferase activity; rRNA modification; rRNA processing; SpoU rRNA Methylase family

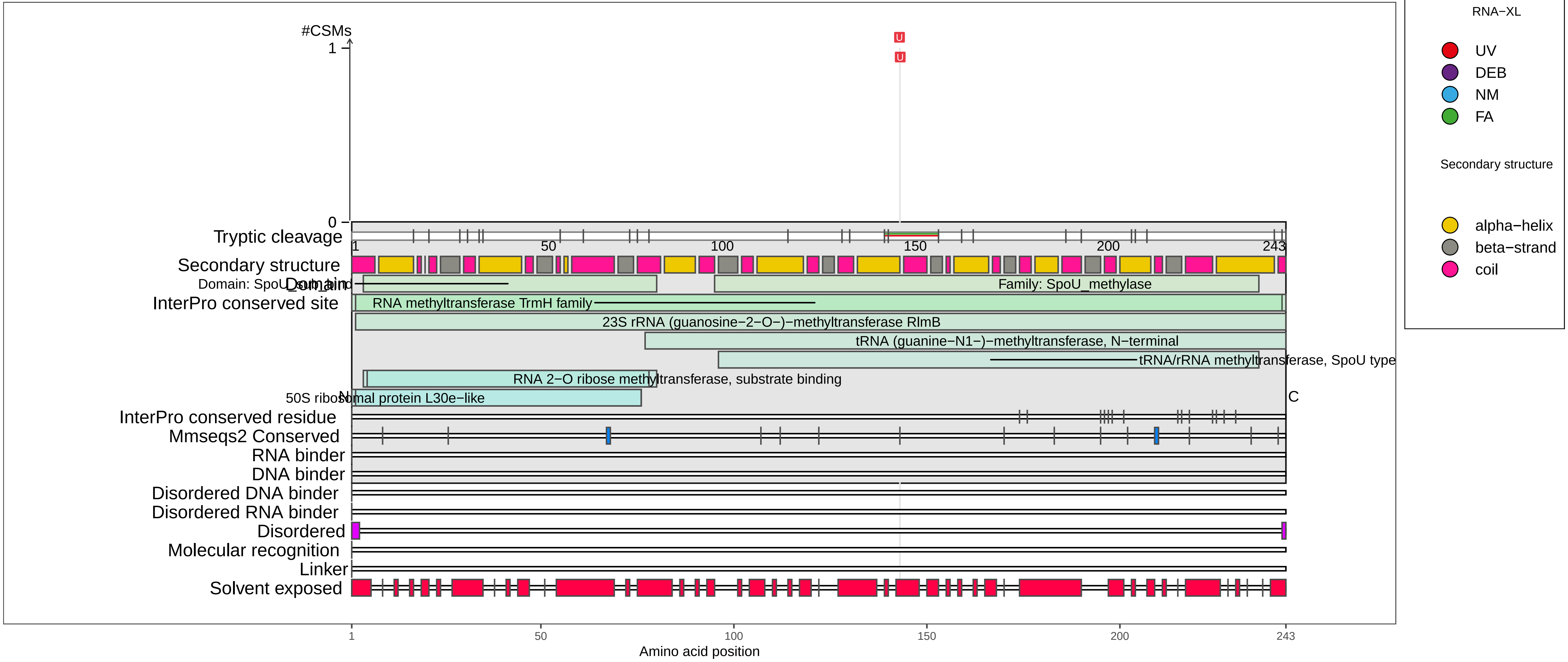

P64574
YGHW_ECOLI Uncharacterized protein YghW

− Abundance:
tryptic [log10 Intensity]: 6.93 (Q 11)
PAXdb K12 strain [ppm]: 1.95 (Q 59)
PAXdb E.coli [ppm]: 0.9 (Q 45)

− RNA functions: not annotated

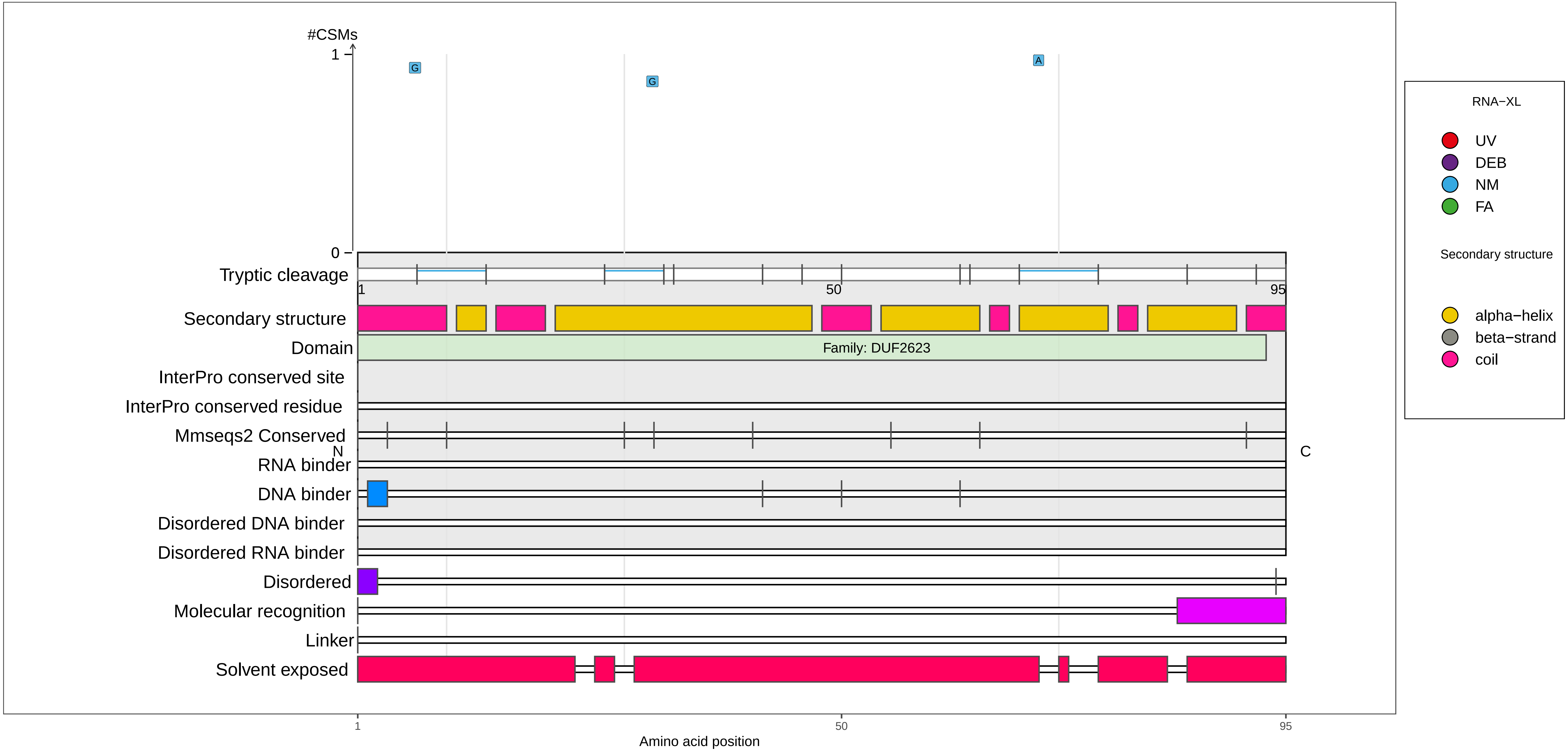

P64581
YQJD_ECOLI Uncharacterized protein YqjD

− Abundance:
tryptic [log10 Intensity]: 8.91 (Q 86)
PAXdb K12 strain [ppm]: 3.06 (Q 92)
PAXdb E.coli [ppm]: 3.37 (Q 98)

− RNA functions: not annotated

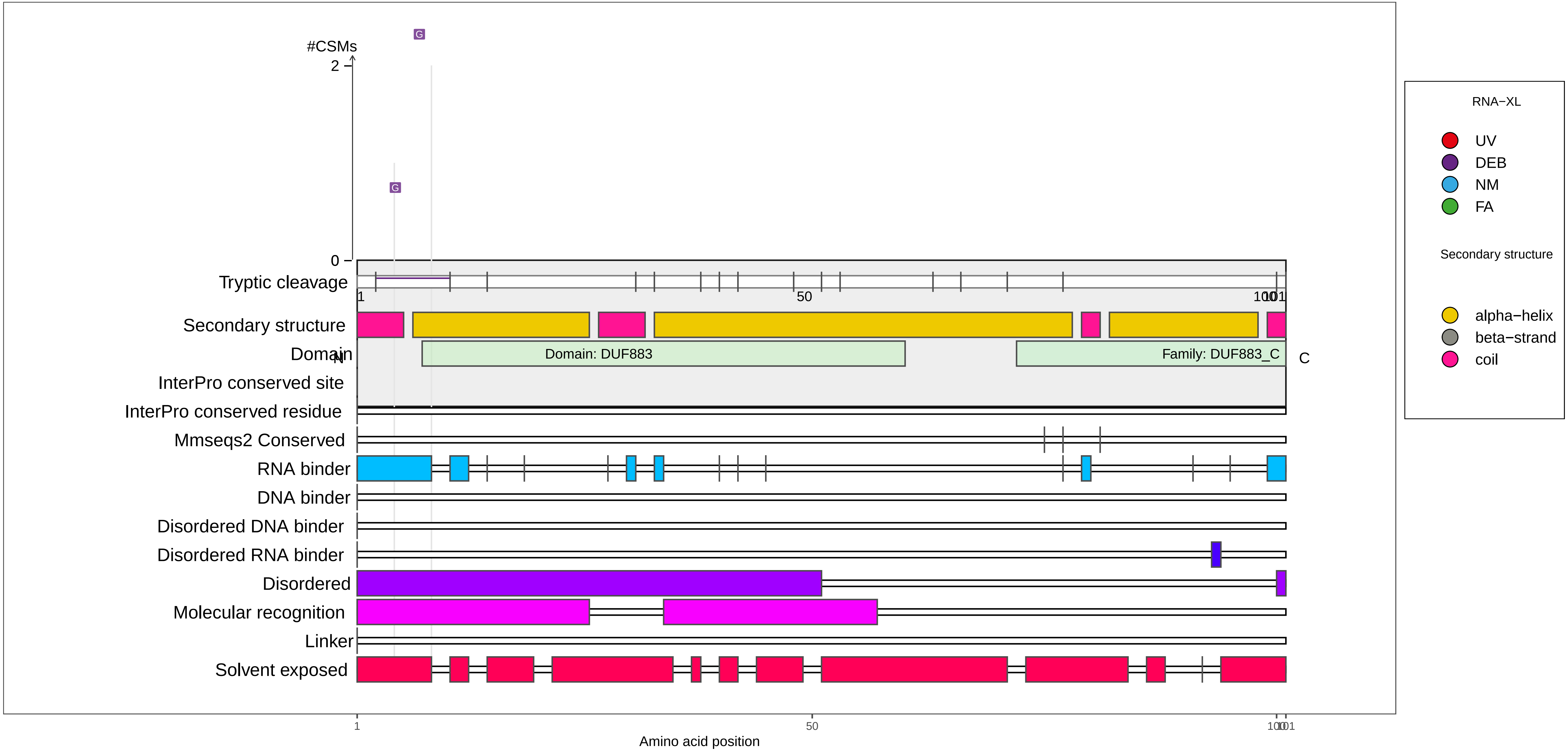

P75792
SUPH_ECOLI Sugar phosphatase YbiV

− Abundance:
tryptic [log10 Intensity]: 7.41 (Q 30)
PAXdb K12 strain [ppm]: 2.03 (Q 61)
PAXdb E.coli [ppm]: 1.65 (Q 63)

− RNA functions: not annotated

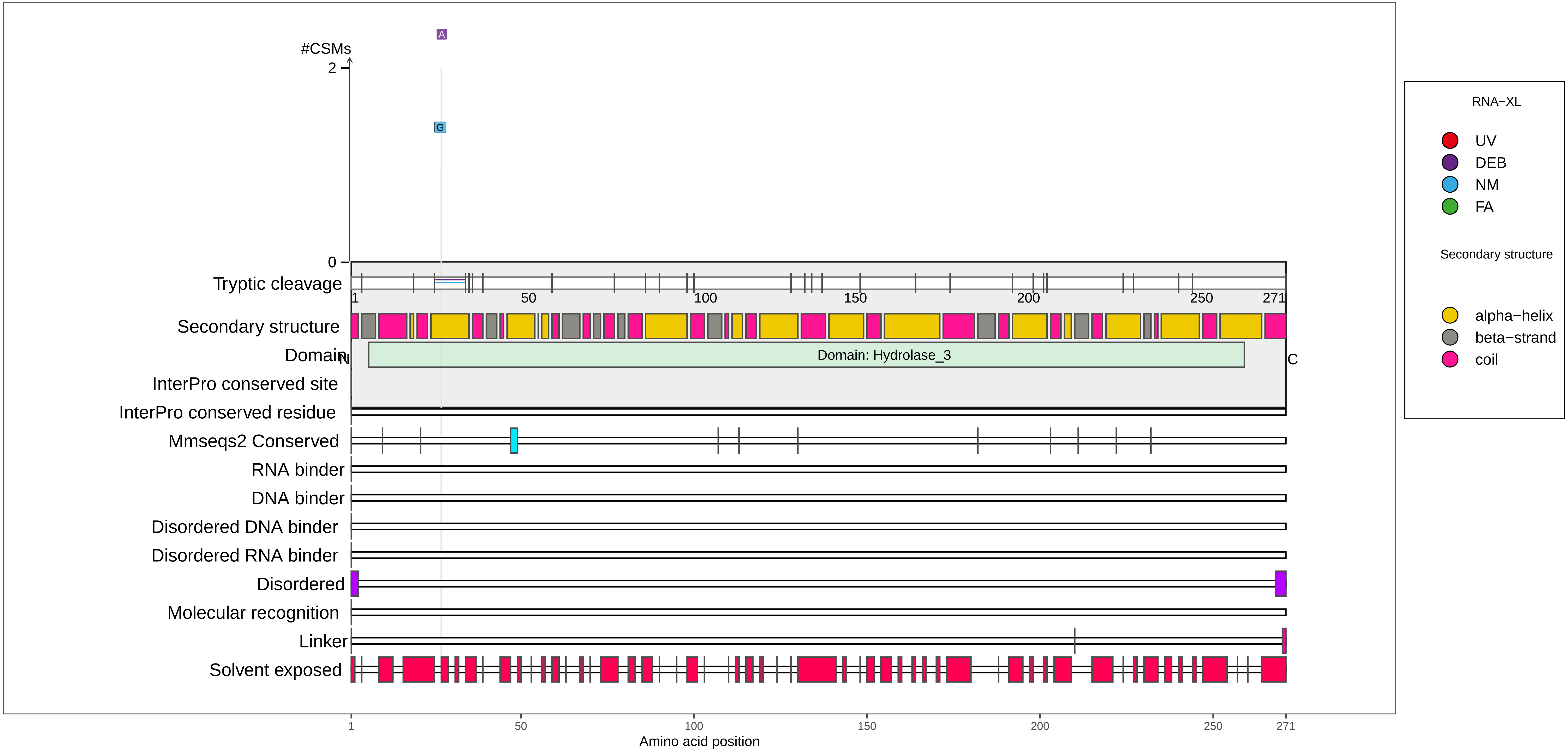

P75821
YBJS_ECOLI Uncharacterized protein YbjS

− Abundance:
tryptic [log10 Intensity]: 6.89 (Q 9)
PAXdb K12 strain [ppm]: 1.35 (Q 29)
PAXdb E.coli [ppm]: 0.62 (Q 39)

− RNA functions: not annotated

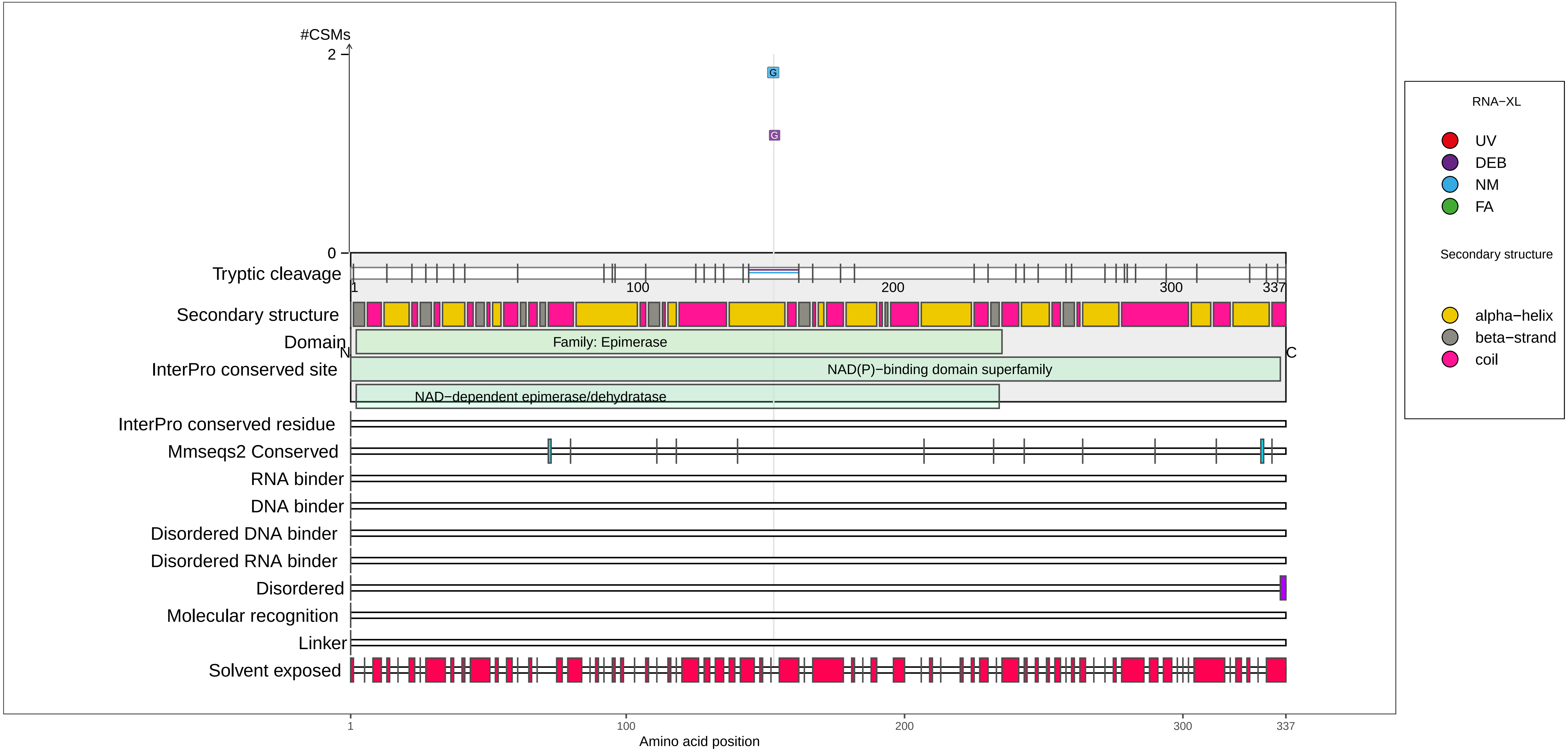

P75874
YCCU_ECOLI Uncharacterized protein YccU

− Abundance:
tryptic [log10 Intensity]: 7.8 (Q 49)
PAXdb K12 strain [ppm]: 2.88 (Q 88)
PAXdb E.coli [ppm]: 2.75 (Q 91)

− RNA functions: not annotated

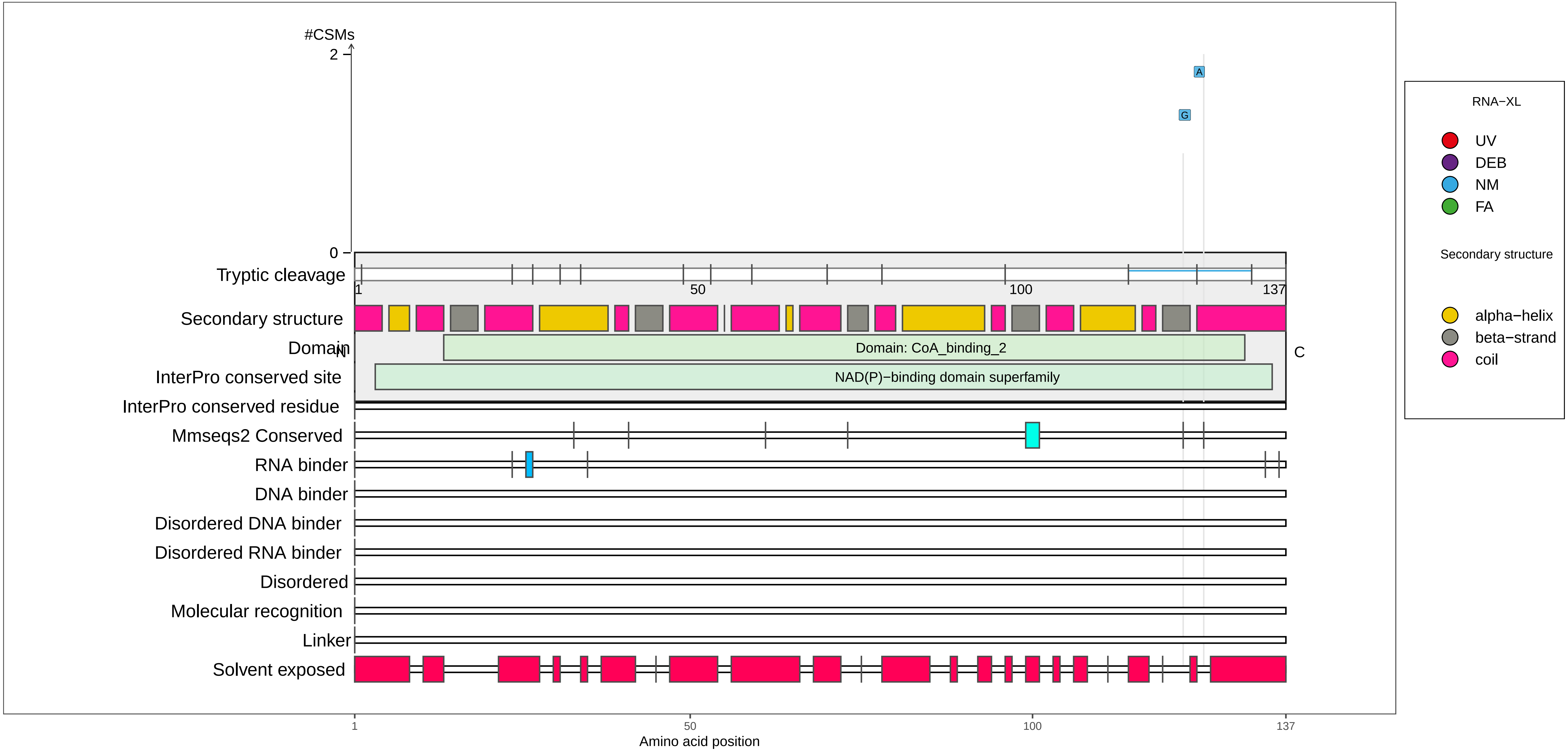

P76231
YEAC_ECOLI Uncharacterized protein YeaC

− Abundance:
tryptic [log10 Intensity]: 7.53 (Q 36)
PAXdb K12 strain [ppm]: NA (Q NA)
PAXdb E.coli [ppm]: 1.68 (Q 64)

− RNA functions: not annotated

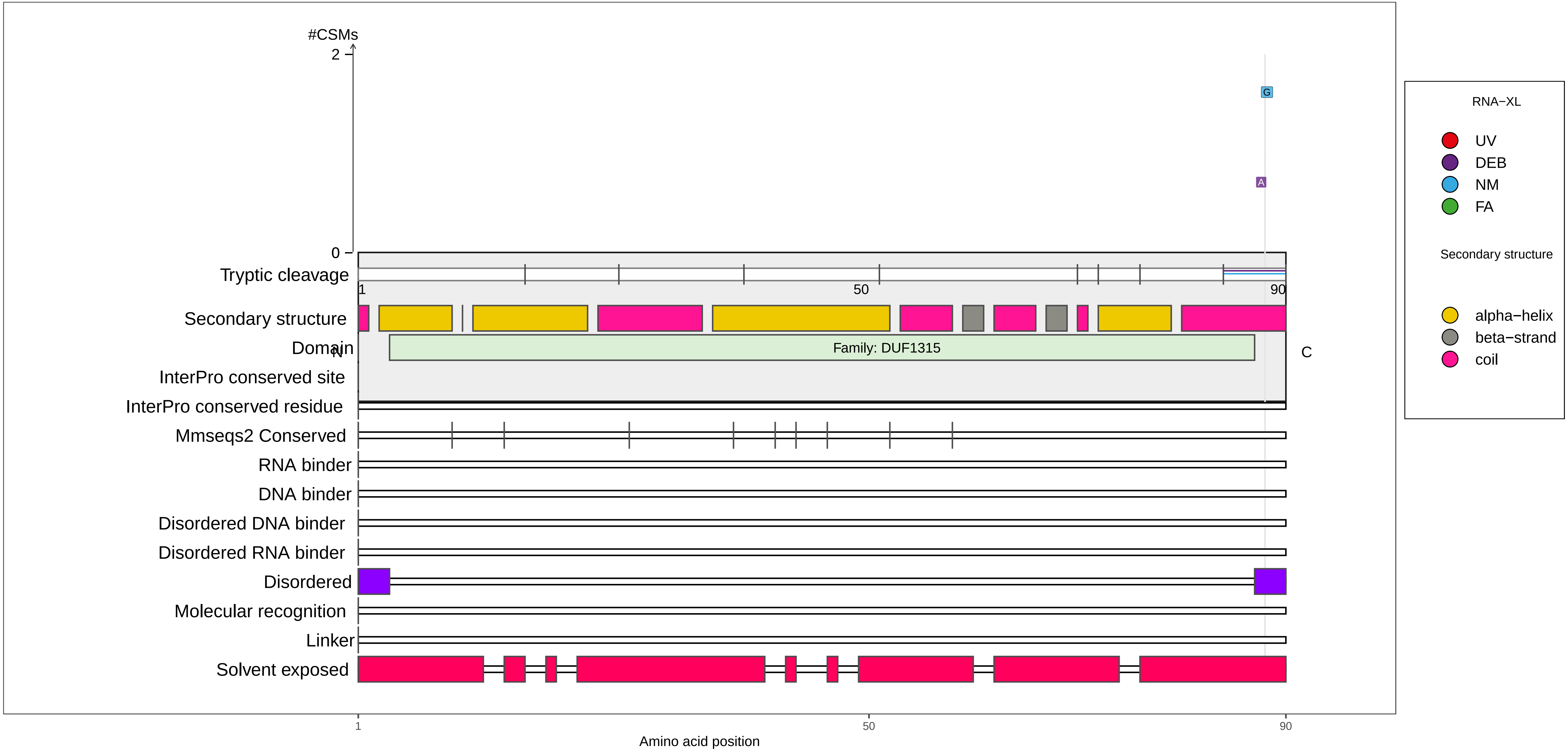

P76469
RHMA_ECOLI 2−keto−3−deoxy−L−rhamnonate aldolase

− Abundance:
tryptic [log10 Intensity]: NA (Q NA)
PAXdb K12 strain [ppm]: NA (Q NA)
PAXdb E.coli [ppm]: 0.1 (Q 27)

− RNA functions: not annotated

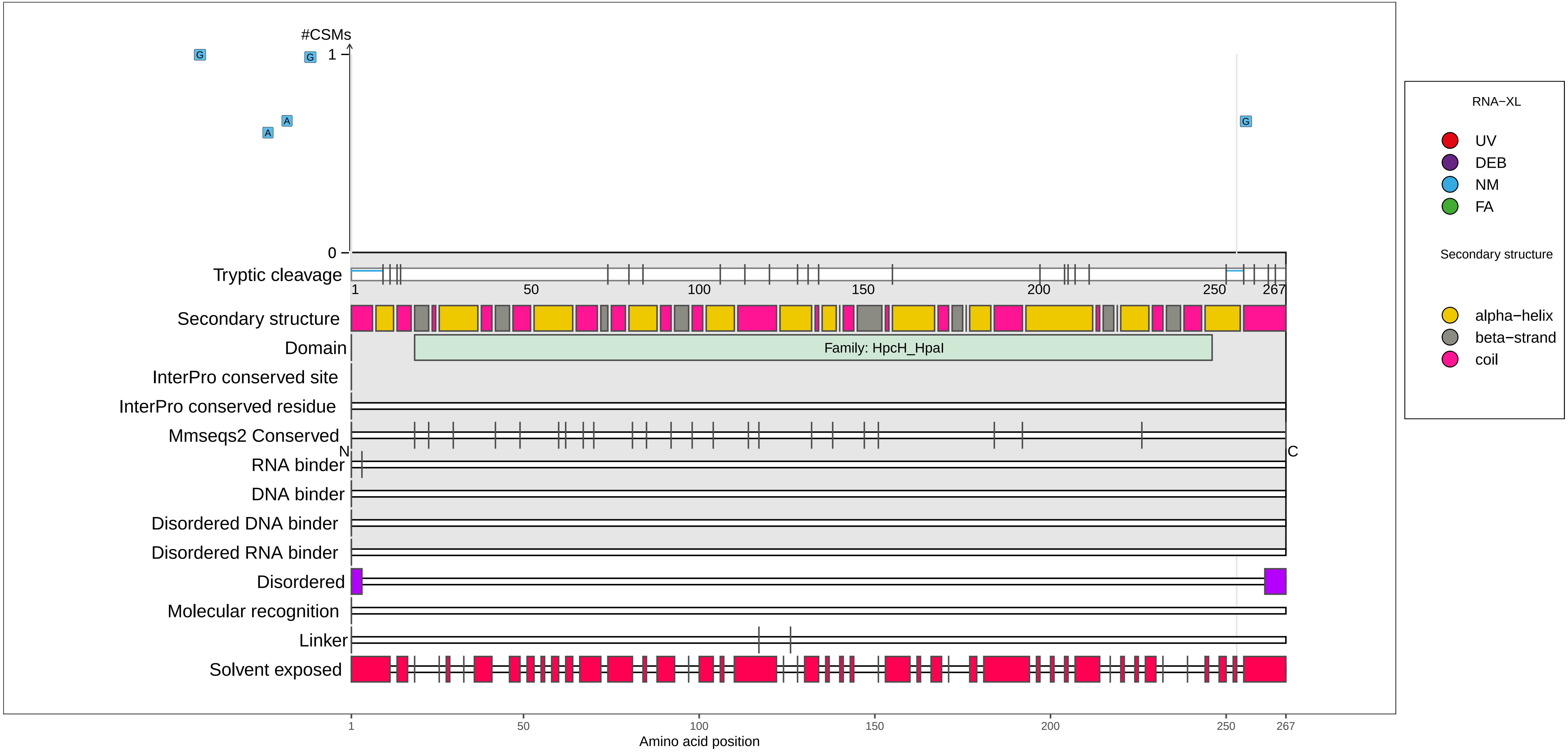

P76562
TMCA_ECOLI tRNA(Met) cytidine acetyltransferase TmcA

− Abundance:
tryptic [log10 Intensity]: 7.09 (Q 16)
PAXdb K12 strain [ppm]: 1.76 (Q 52)
PAXdb E.coli [ppm]: 0.76 (Q 42)

− RNA functions:
ncRNA metabolic process; ncRNA processing; RNA acetylation; RNA binding
RNA metabolic process; RNA modification; RNA processing; tRNA acetylation; tRNA binding
tRNA metabolic process; tRNA modification; tRNA N−acetyltransferase activity; tRNA processing
tRNA wobble base modification; tRNA wobble cytosine modification; tRNA−binding domain

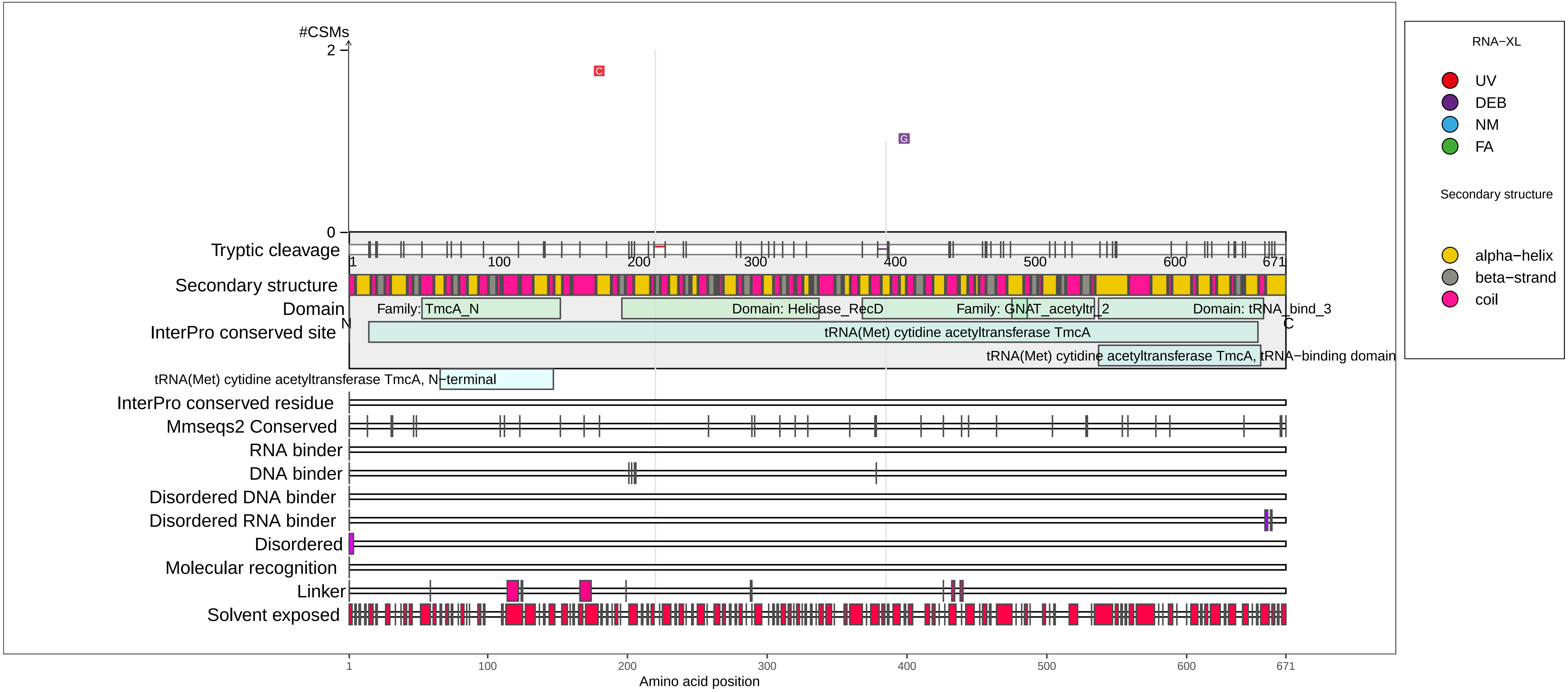

P77526
YFCG_ECOLI Disulfide−bond oxidoreductase YfcG

− Abundance:
tryptic [log10 Intensity]: 6.6 (Q 3)
PAXdb K12 strain [ppm]: 1.39 (Q 32)
PAXdb E.coli [ppm]: 1.81 (Q 67)

− RNA functions: not annotated

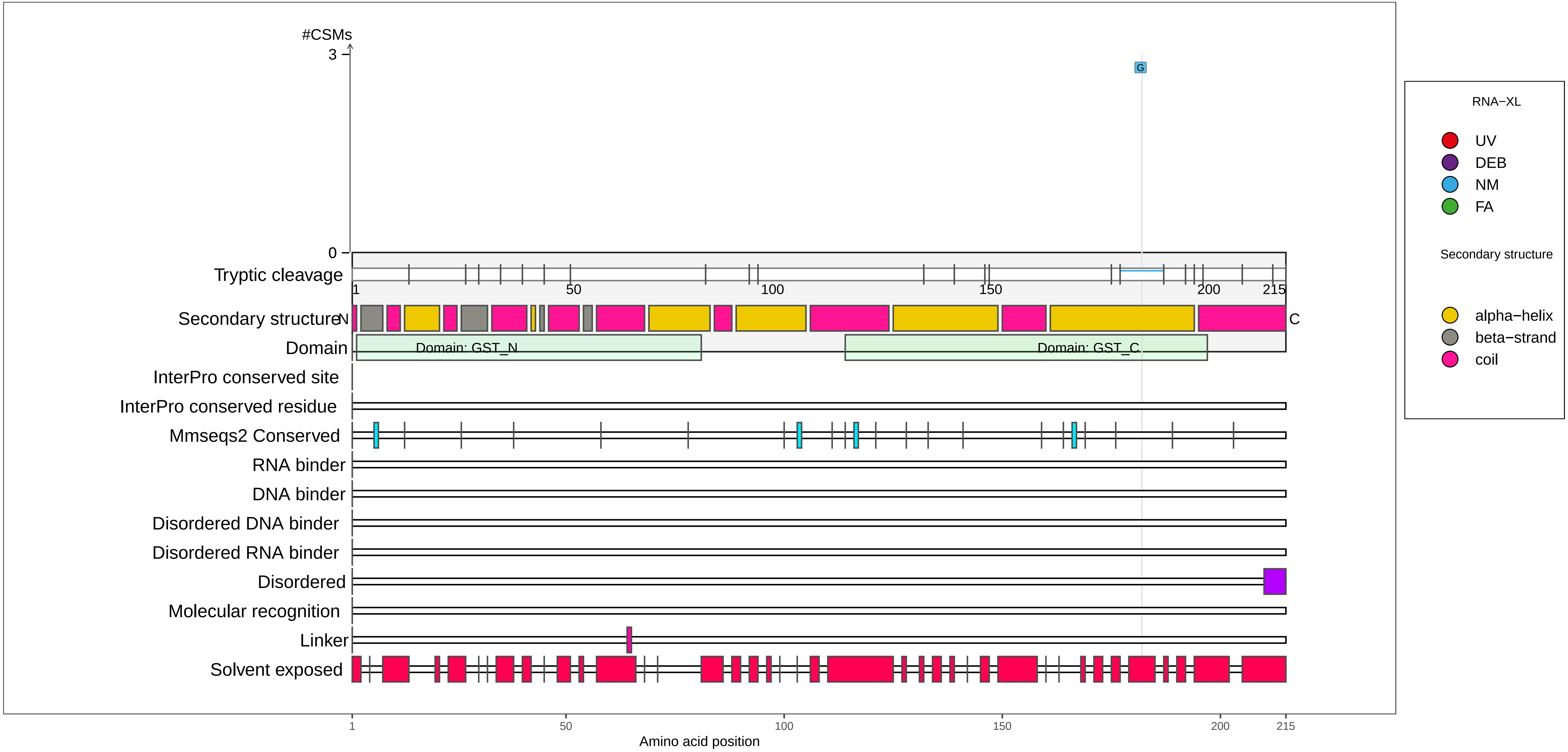

P77756
QUEC_ECOLI 7−cyano−7−deazaguanine synthase

− Abundance:
tryptic [log10 Intensity]: 7.92 (Q 55)
PAXdb K12 strain [ppm]: 2.11 (Q 64)
PAXdb E.coli [ppm]: 1.93 (Q 70)

− RNA functions: not annotated

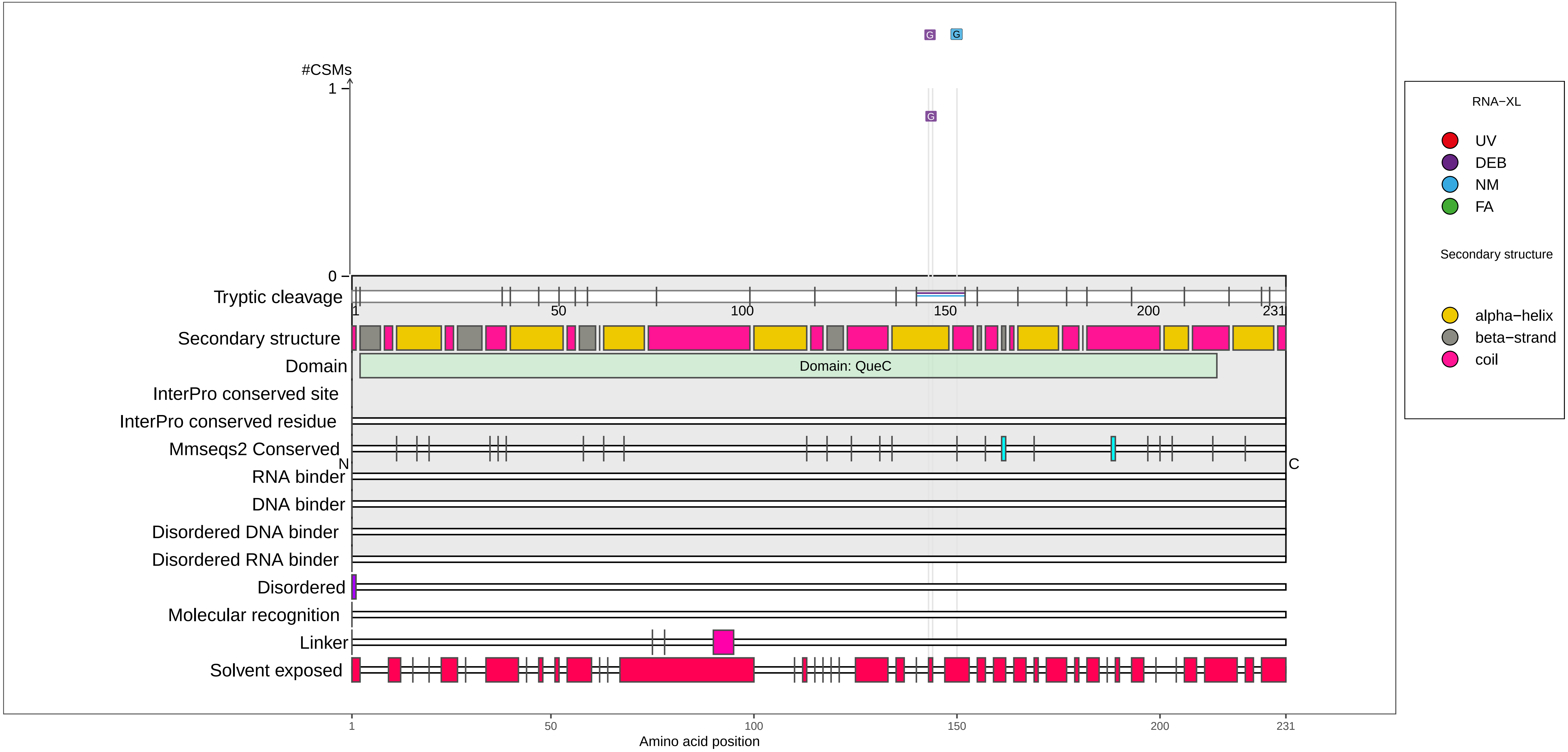

P77783
YNFF_ECOLI Probable dimethyl sulfoxide reductase chain YnfF

− Abundance:
tryptic [log10 Intensity]: 7.79 (Q 48)
PAXdb K12 strain [ppm]: 0.68 (Q 2)
PAXdb E.coli [ppm]: −0.01 (Q 25)

− RNA functions: not annotated

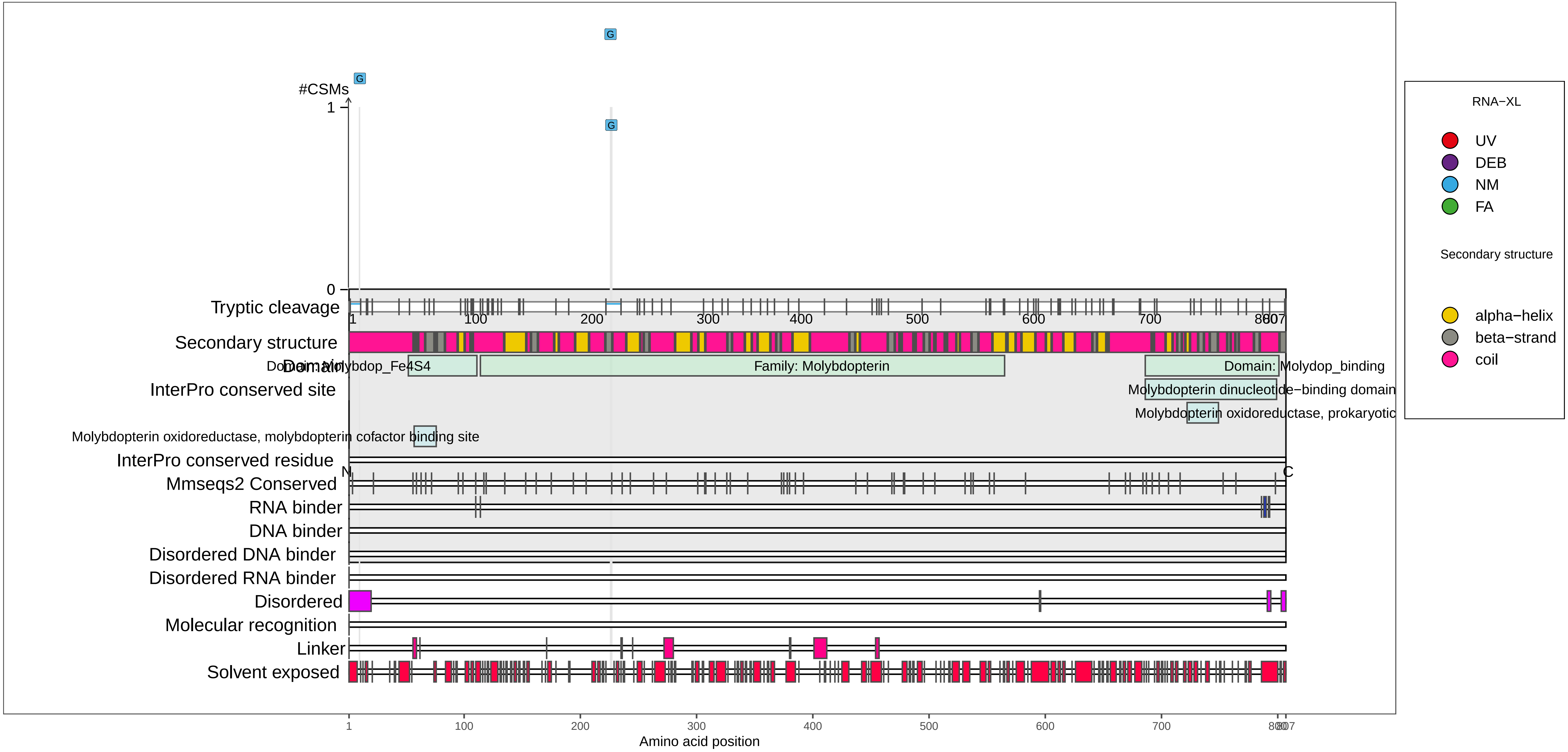

Q46829
BGLA_ECOLI 6−phospho−beta−glucosidase BglA
− Abundance:
tryptic [log10 Intensity]: 9.04 (Q 89)
PAXdb K12 strain [ppm]: 1.68 (Q 48)
PAXdb E.coli [ppm]: 2.3 (Q 80)

− RNA functions: not annotated

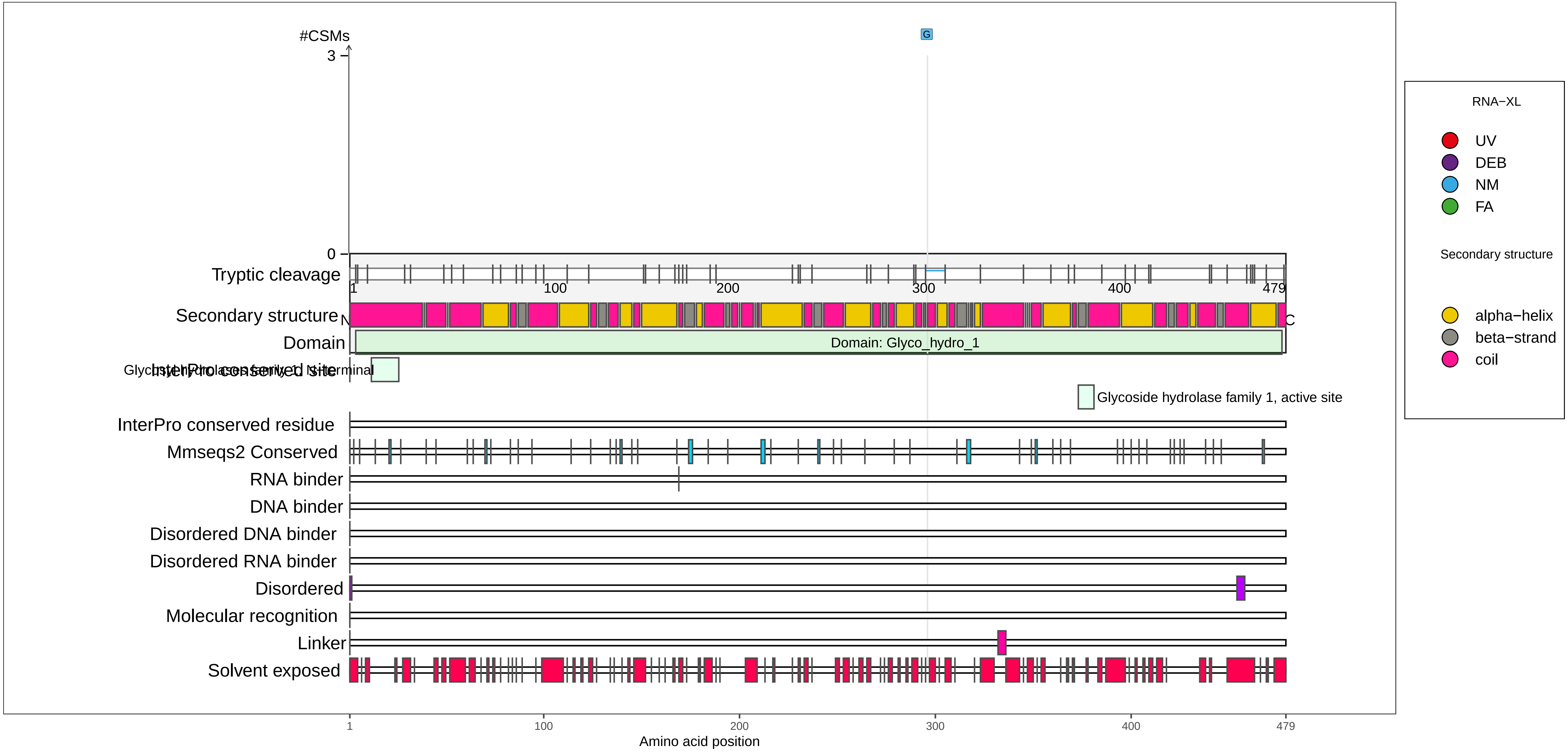

Q46861
YGIQ_ECOLI UPF0313 protein YgiQ
− Abundance:
tryptic [log10 Intensity]: 6.61 (Q 4)
PAXdb K12 strain [ppm]: 0.83 (Q 6)
PAXdb E.coli [ppm]: 0.16 (Q 28)

− RNA functions: not annotated

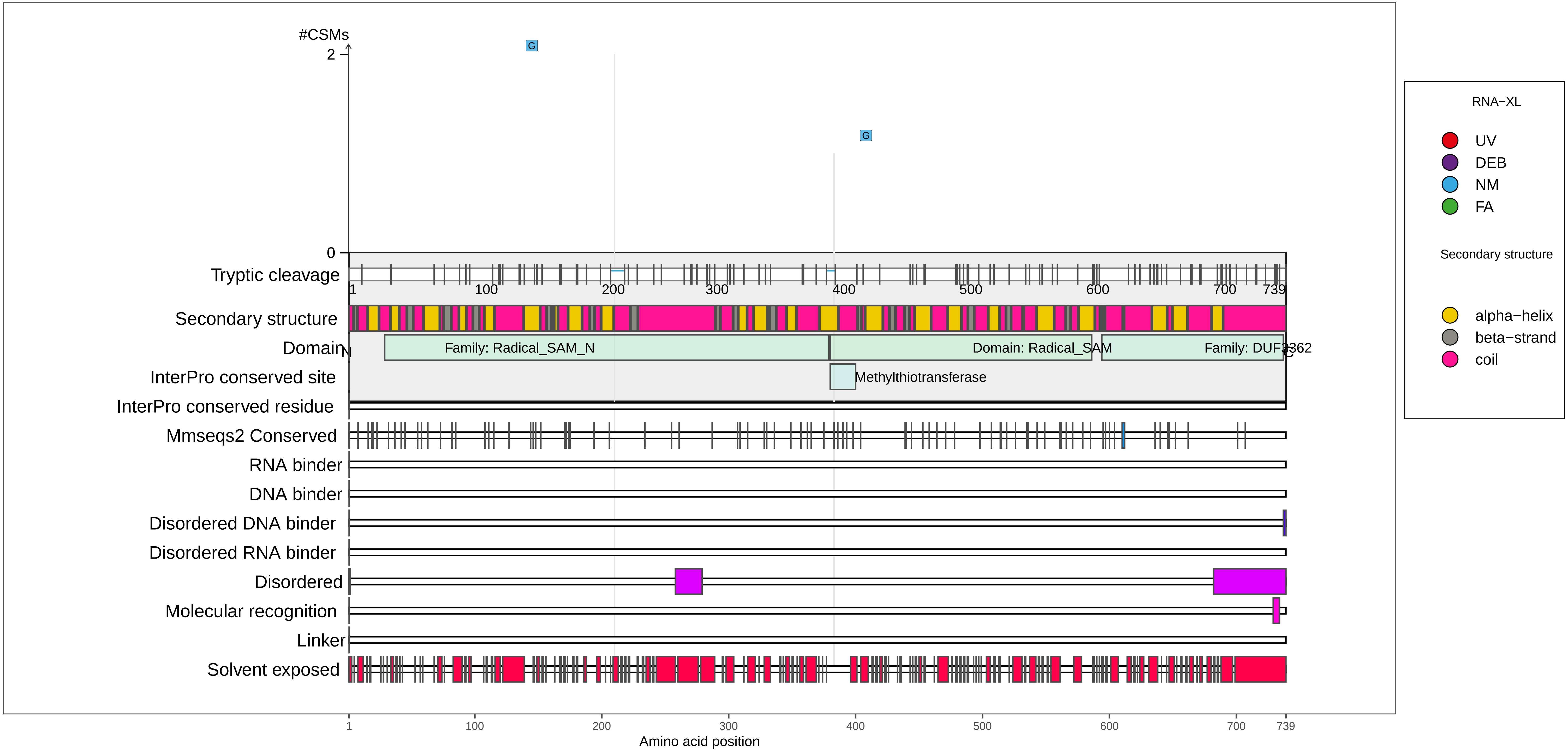

P00803
LEP_ECOLI Signal peptidase I

− Abundance:
tryptic [log10 Intensity]: 8.27 (Q 68)
PAXdb K12 strain [ppm]: NA (Q NA)
PAXdb E.coli [ppm]: 1.85 (Q 68)

− RNA functions: not annotated

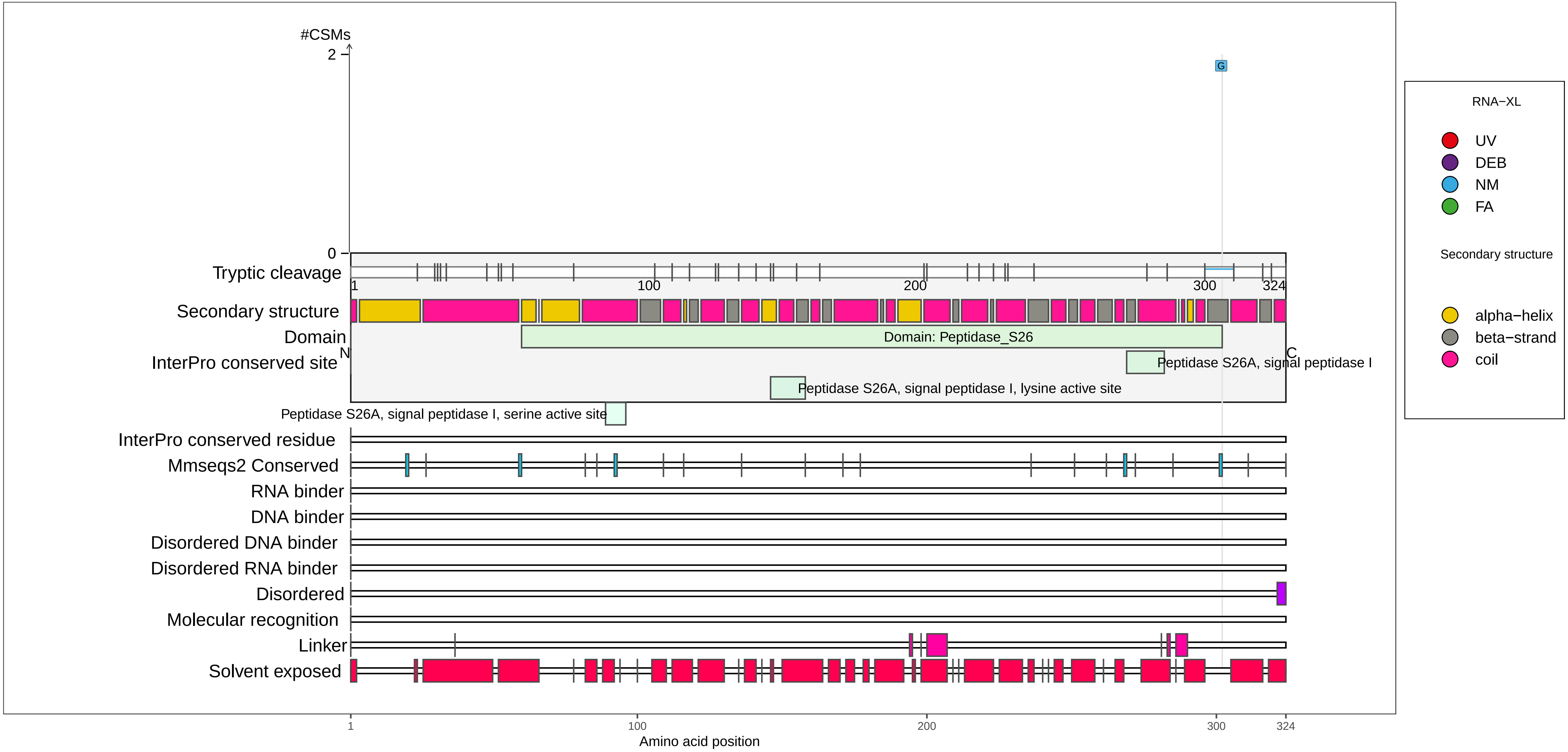

P00934
THRC_ECOLI Threonine synthase

− Abundance:
tryptic [log10 Intensity]: 7.99 (Q 57)
PAXdb K12 strain [ppm]: 2.56 (Q 79)
PAXdb E.coli [ppm]: 2.9 (Q 93)

− RNA functions: not annotated

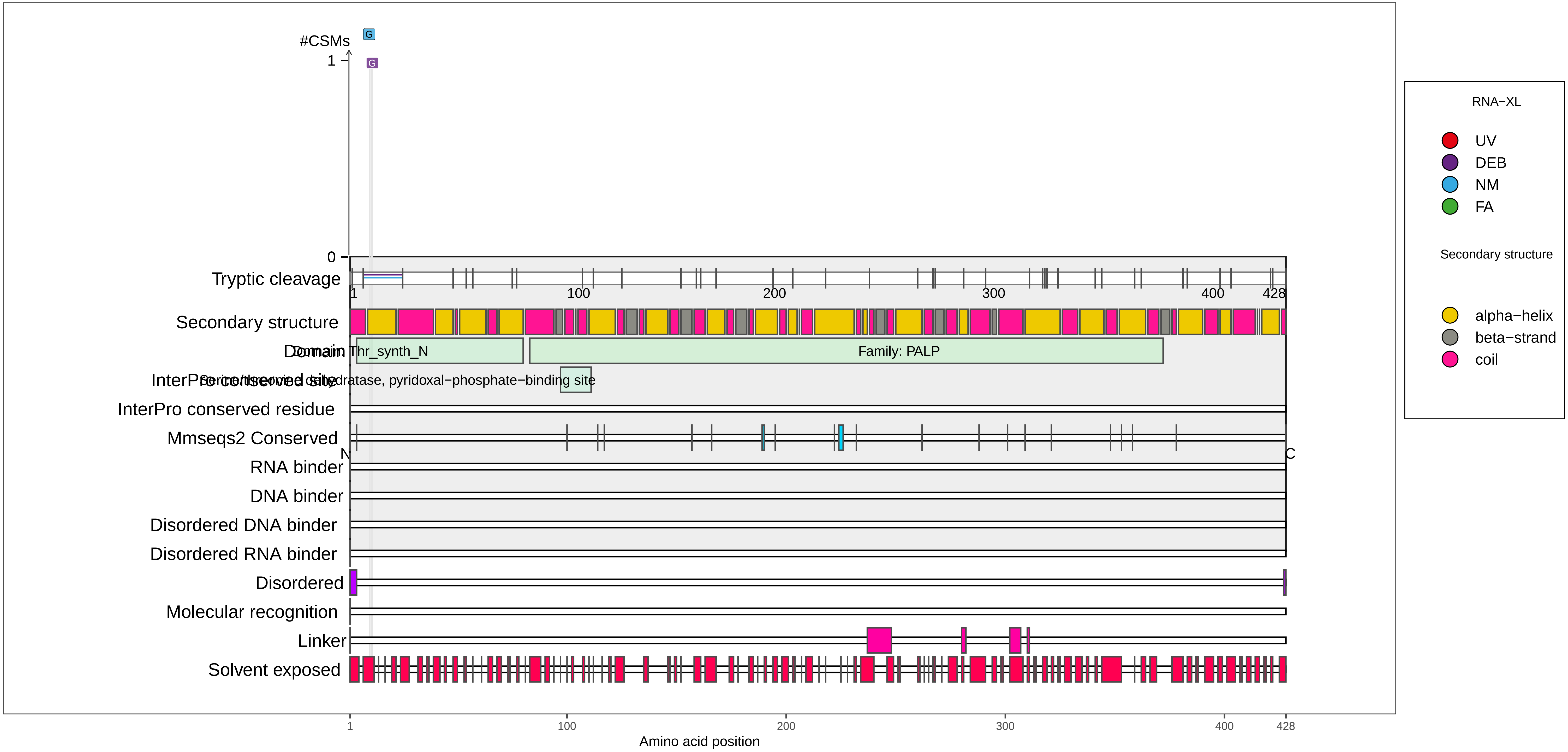

P02925
RBSB_ECOLI Ribose import binding protein RbsB

− Abundance:
tryptic [log10 Intensity]: 8.33 (Q 71)
PAXdb K12 strain [ppm]: 3.16 (Q 93)
PAXdb E.coli [ppm]: 3.26 (Q 97)

− RNA functions: not annotated

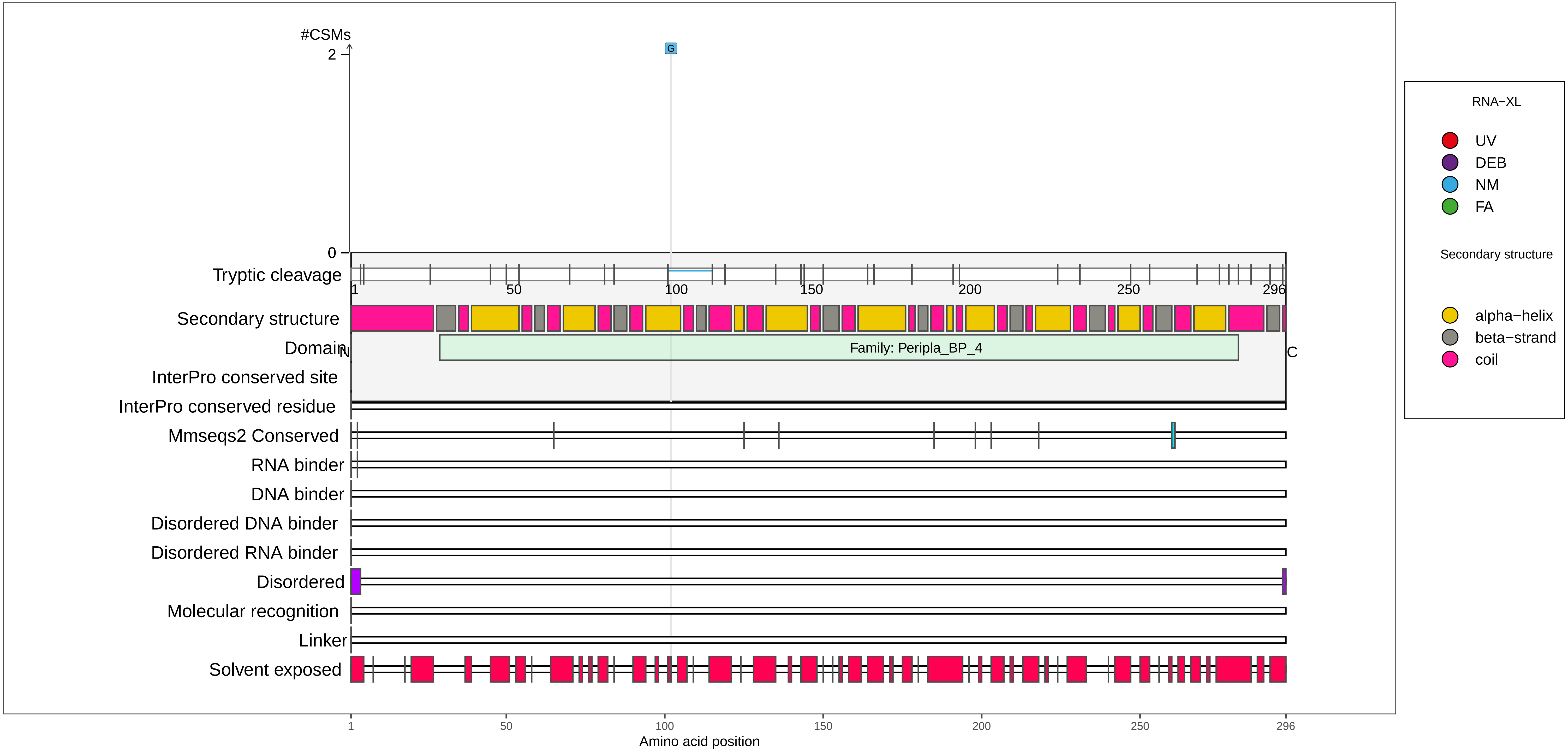

P04335
FRSA_ECOLI Esterase FrsA

− Abundance:
tryptic [log10 Intensity]: 8.73 (Q 82)
PAXdb K12 strain [ppm]: 2.24 (Q 68)
PAXdb E.coli [ppm]: 1.63 (Q 63)

− RNA functions: not annotated

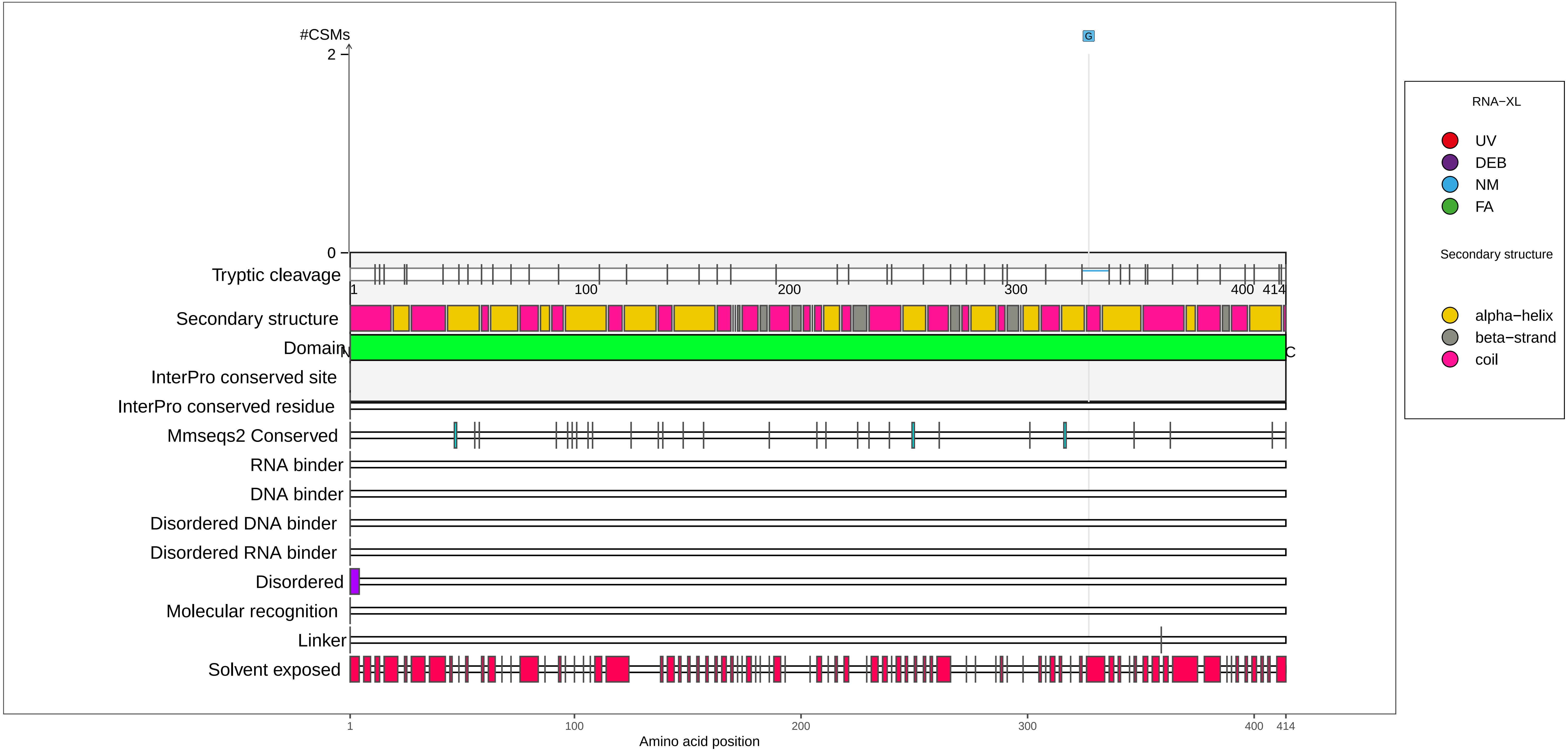

P05707
SRLD_ECOLI Sorbitol−6−phosphate 2−dehydrogenase

− Abundance:
tryptic [log10 Intensity]: 9.06 (Q 90)
PAXdb K12 strain [ppm]: NA (Q NA)
PAXdb E.coli [ppm]: 0.05 (Q 26)

− RNA functions: not annotated

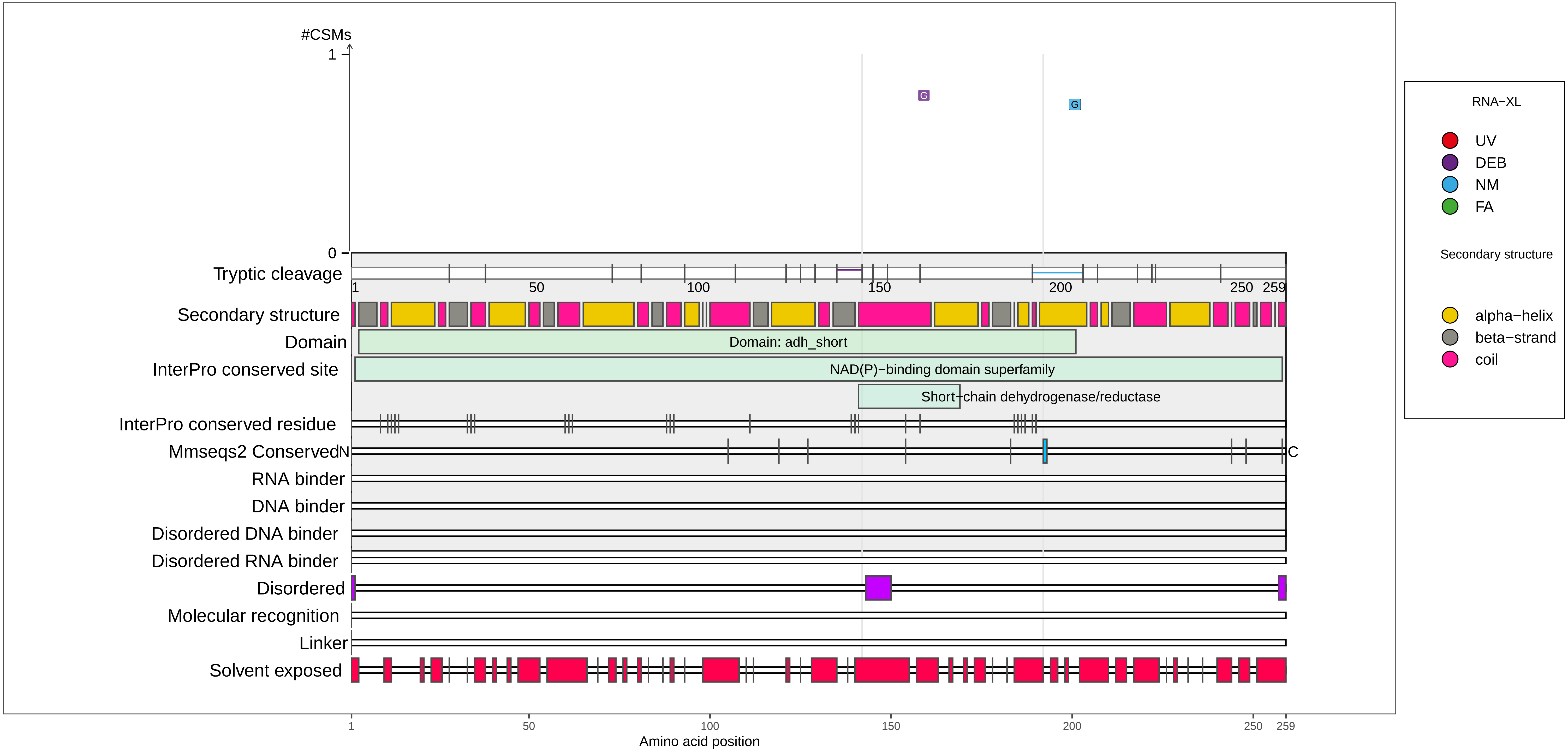

P06960
OTC2_ECOLI Ornithine carbamoyltransferase subunit F

− Abundance:
tryptic [log10 Intensity]: NA (Q NA)
PAXdb K12 strain [ppm]: 0.43 (Q 1)
PAXdb E.coli [ppm]: 1.41 (Q 57)

− RNA functions: not annotated

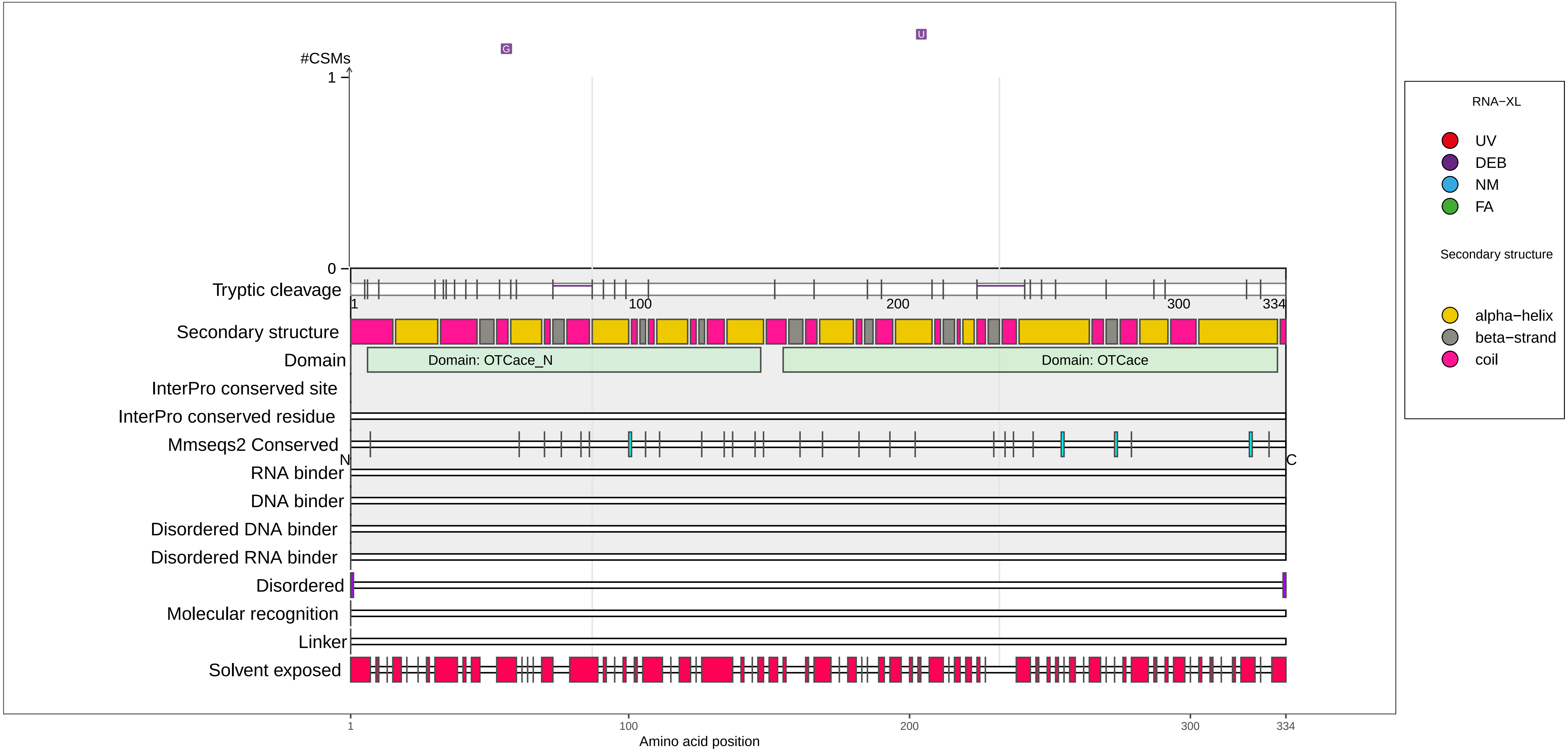

P07623
METAS_ECOLI Homoserine O−succinyltransferase

− Abundance:
tryptic [log10 Intensity]: 7.6 (Q 40)
PAXdb K12 strain [ppm]: 2.12 (Q 65)
PAXdb E.coli [ppm]: 2.19 (Q 77)

− RNA functions: not annotated

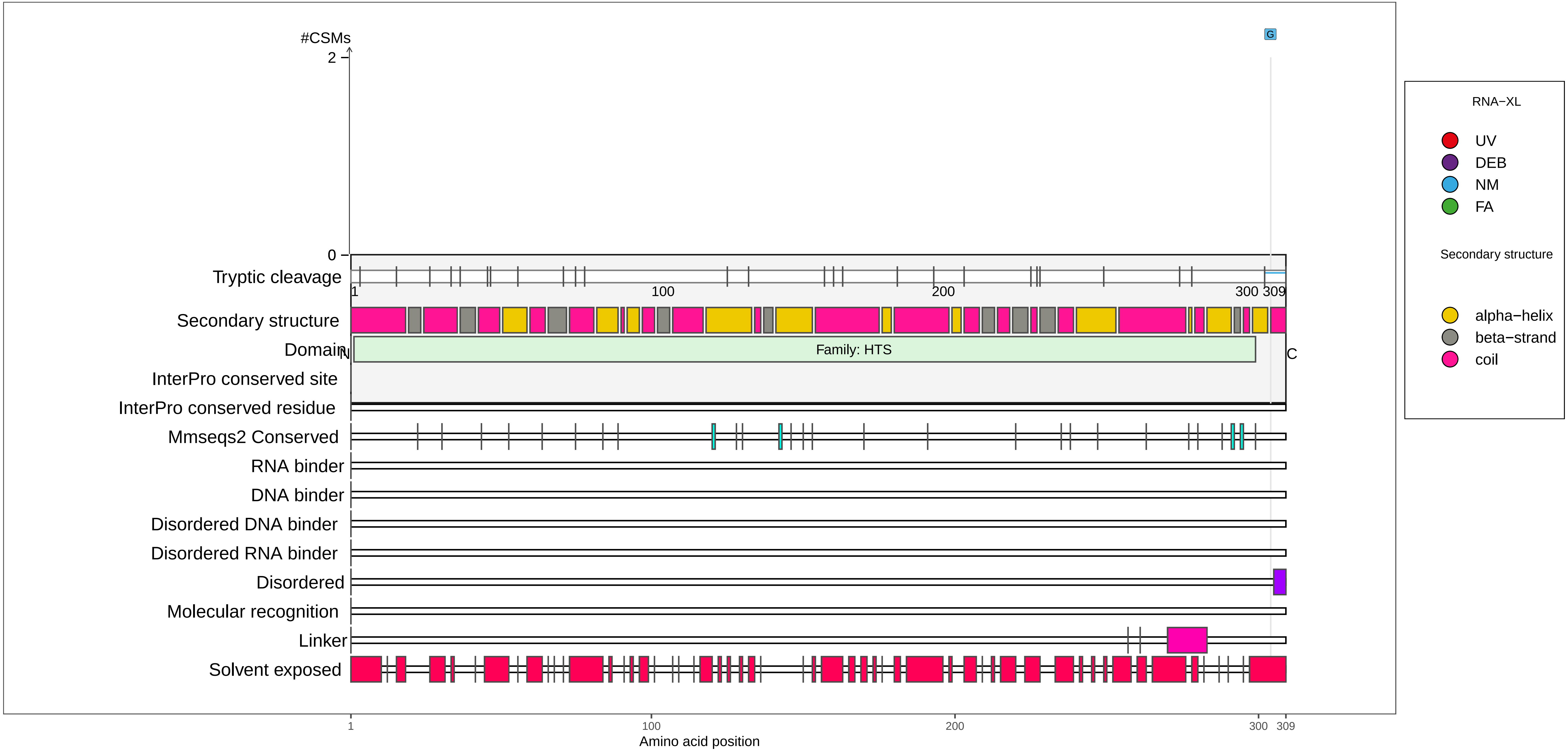

P09155
RND_ECOLI Ribonuclease D

− Abundance:
tryptic [log10 Intensity]: NA (Q NA)
PAXdb K12 strain [ppm]: 1.92 (Q 58)
PAXdb E.coli [ppm]: 1.09 (Q 50)

− RNA functions:
ncRNA 3−end processing; ncRNA metabolic process; ncRNA processing
RNA 3−end processing; RNA metabolic process; RNA phosphodiester bond hydrolysis
RNA phosphodiester bond hydrolysis, exonucleolytic; RNA processing; tRNA 3−end processing
tRNA metabolic process; tRNA processing

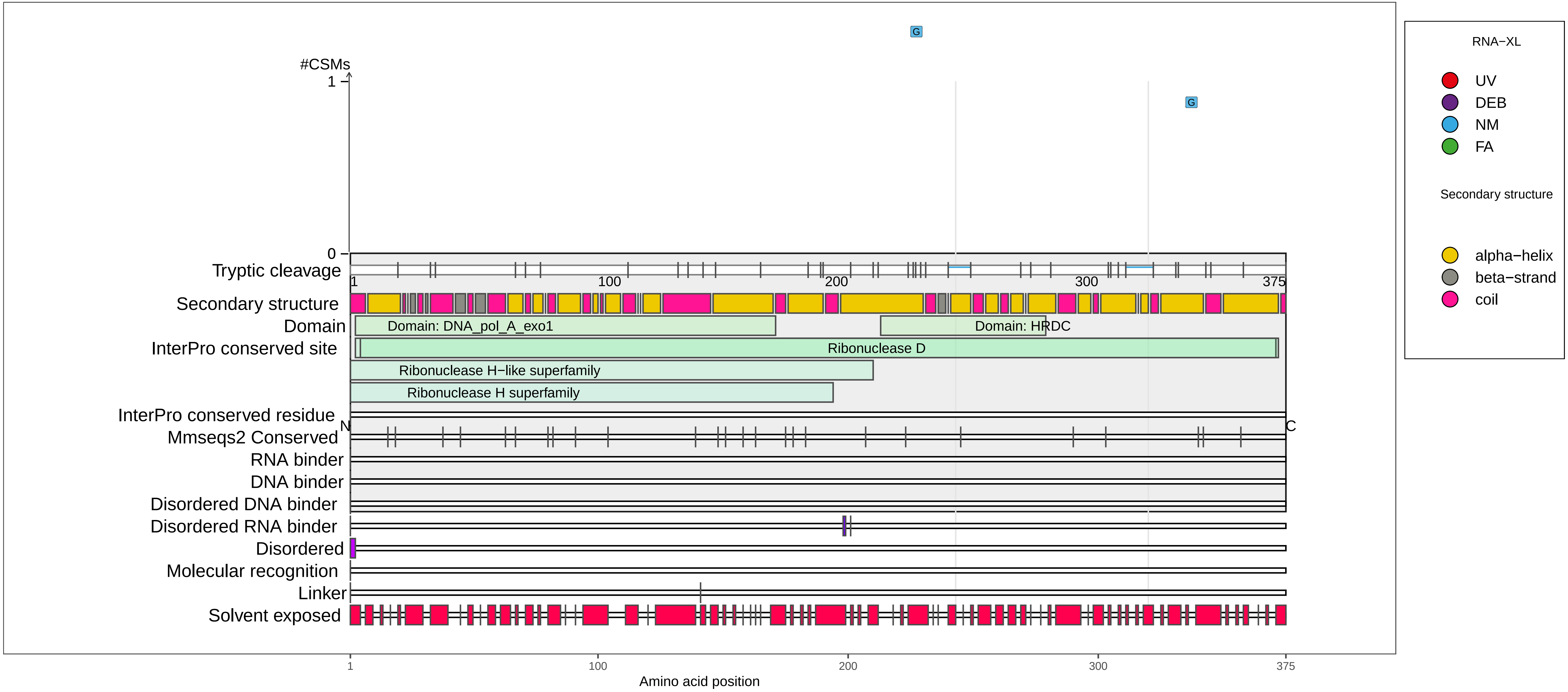

P09549
DEDD_ECOLI Cell division protein DedD

− Abundance:
tryptic [log10 Intensity]: 7.42 (Q 31)
PAXdb K12 strain [ppm]: 2.61 (Q 81)
PAXdb E.coli [ppm]: 1.35 (Q 56)

− RNA functions: not annotated

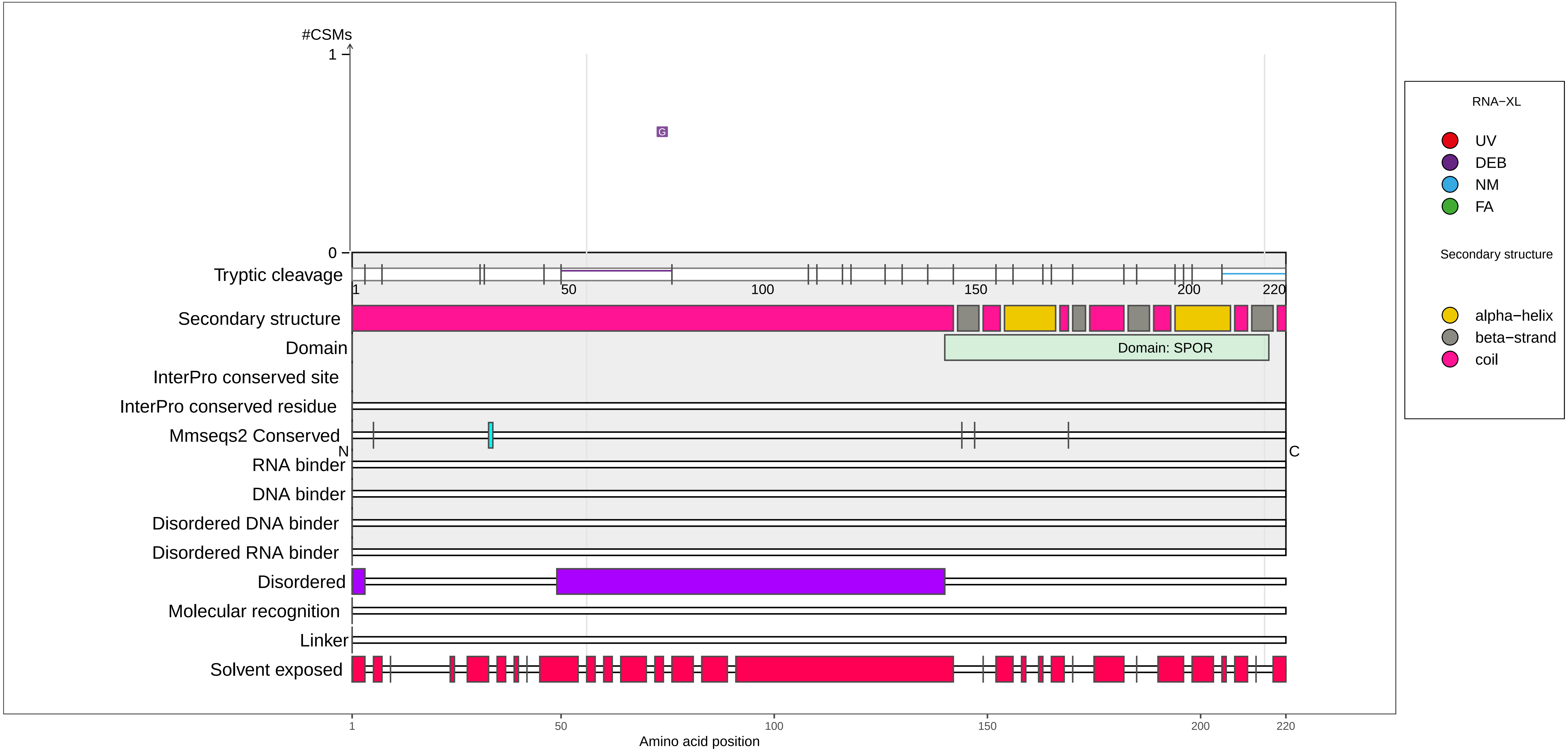

P0A6K3
DEF_ECOLI Peptide deformylase

− Abundance:
tryptic [log10 Intensity]: 8.67 (Q 81)
PAXdb K12 strain [ppm]: 3.14 (Q 93)
PAXdb E.coli [ppm]: 2.55 (Q 87)

− RNA functions: not annotated

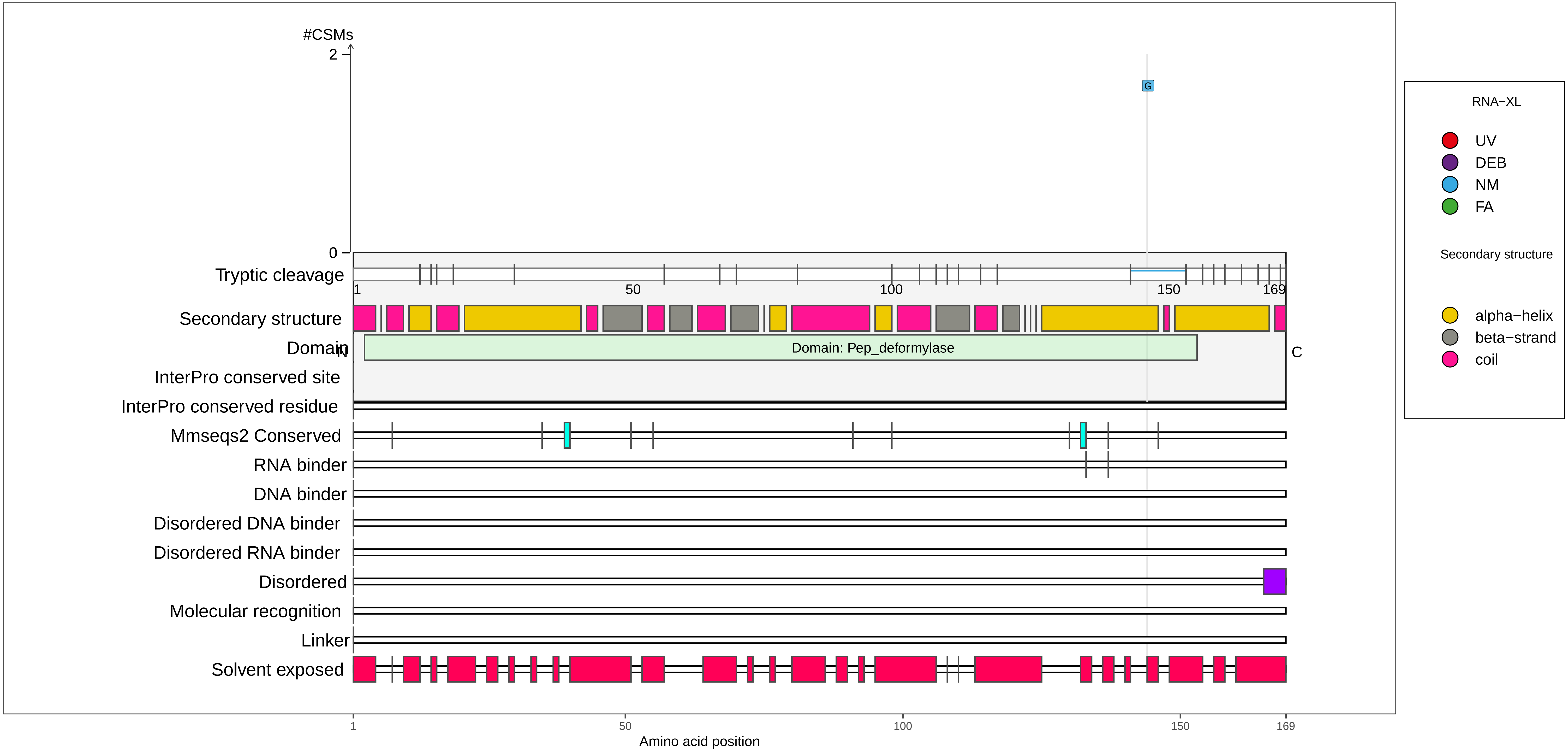

P0A6U5
RSMG_ECOLI Ribosomal RNA small subunit methyltransferase G

− Abundance:
tryptic [log10 Intensity]: 7.73 (Q 46)
PAXdb K12 strain [ppm]: NA (Q NA)
PAXdb E.coli [ppm]: 1.63 (Q 63)

− RNA functions:
ncRNA metabolic process; ncRNA processing; RNA (guanine−N7)−methylation
RNA metabolic process; RNA methylation; RNA methyltransferase activity; RNA modification
RNA processing; rRNA (guanine−N7−)−methyltransferase activity; rRNA (guanine−N7)−methylation
rRNA (guanine) methyltransferase activity; rRNA base methylation
rRNA metabolic process; rRNA methylation; rRNA methyltransferase activity; rRNA modification; rRNA processing
rRNA small subunit methyltransferase G

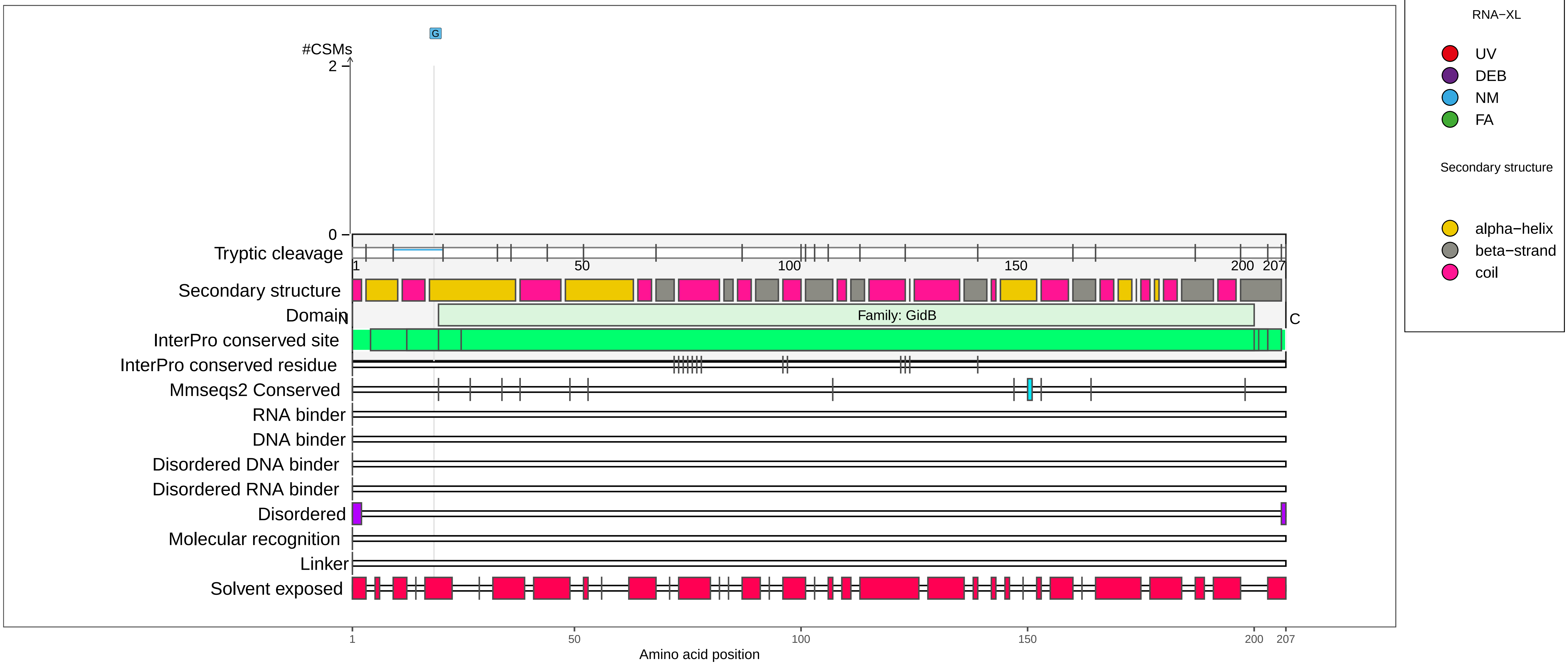

P0A761
NANE_ECOLI Putative N−acetylmannosamine−6−phosphate 2−epimerase

− Abundance:
tryptic [log10 Intensity]: 8.42 (Q 73)
PAXdb K12 strain [ppm]: 1.94 (Q 58)
PAXdb E.coli [ppm]: 1.43 (Q 58)

− RNA functions: not annotated

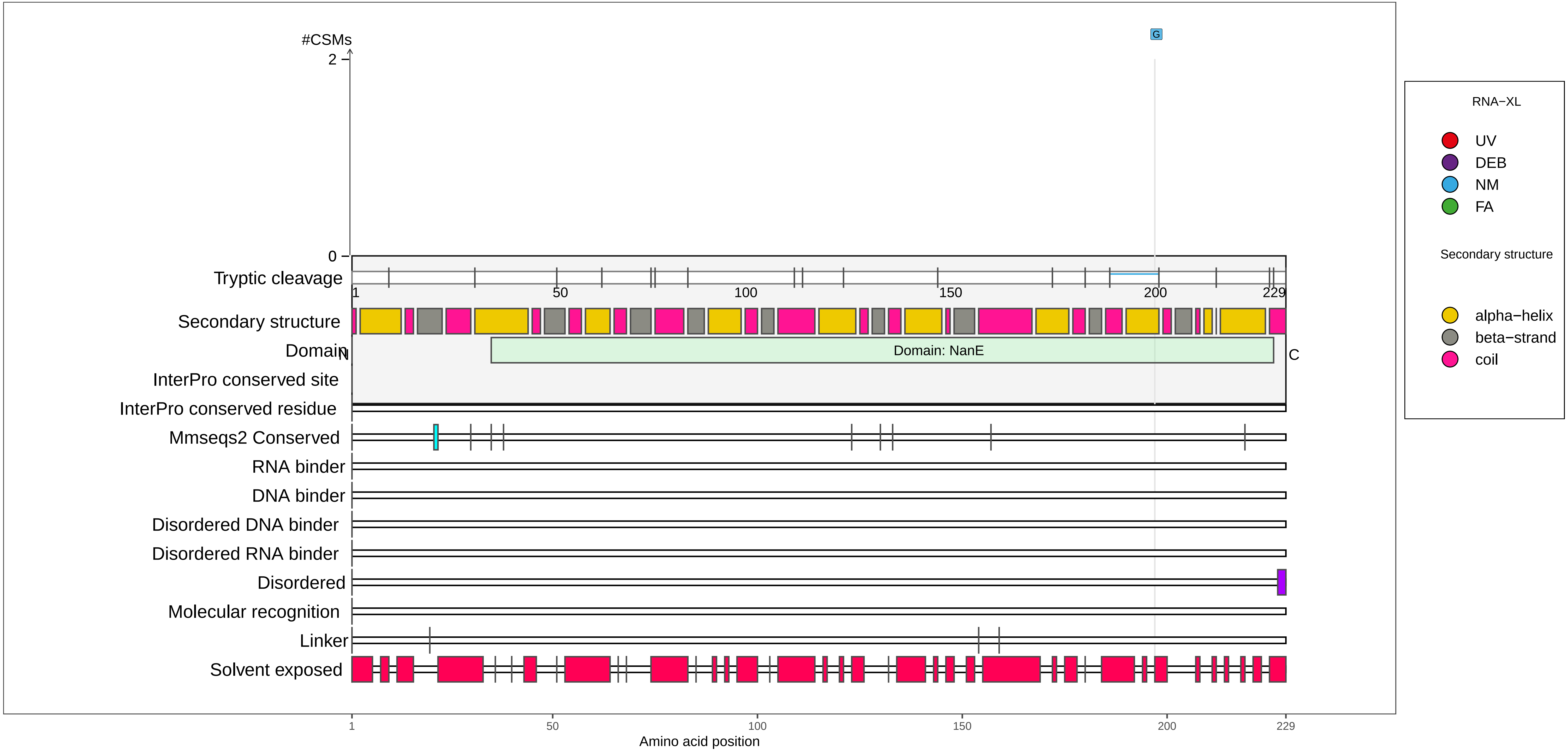

P0A7A5
PIMT_ECOLI Protein−L−isoaspartate O−methyltransferase

− Abundance:
tryptic [log10 Intensity]: 7.38 (Q 28)
PAXdb K12 strain [ppm]: 2.37 (Q 73)
PAXdb E.coli [ppm]: 1.38 (Q 56)

− RNA functions:
Ribosomal RNA adenine dimethylase

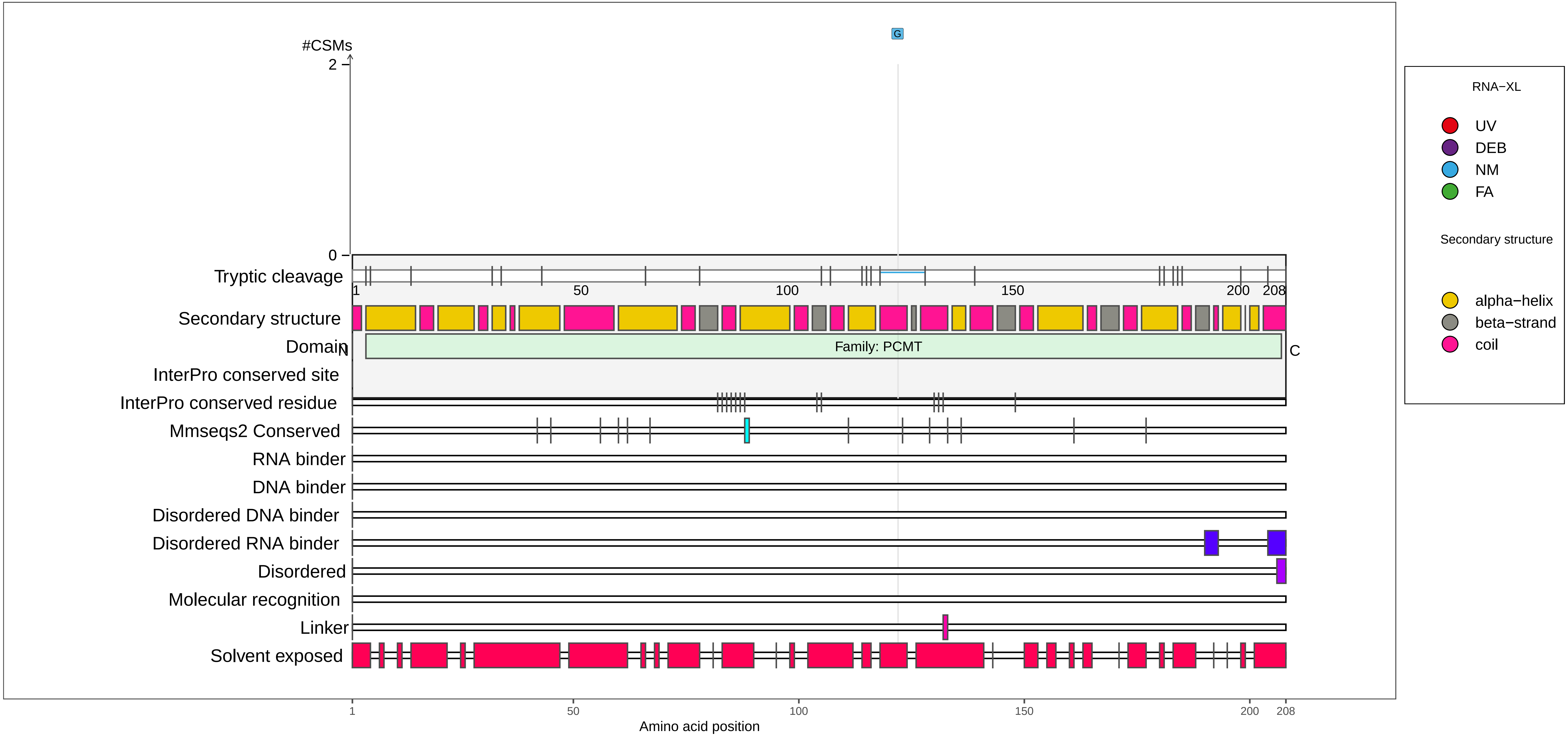

P0A7C6
PEPE_ECOLI Peptidase E

− Abundance:
tryptic [log10 Intensity]: 6.99 (Q 13)
PAXdb K12 strain [ppm]: 1.83 (Q 54)
PAXdb E.coli [ppm]: 2.12 (Q 75)

− RNA functions: not annotated

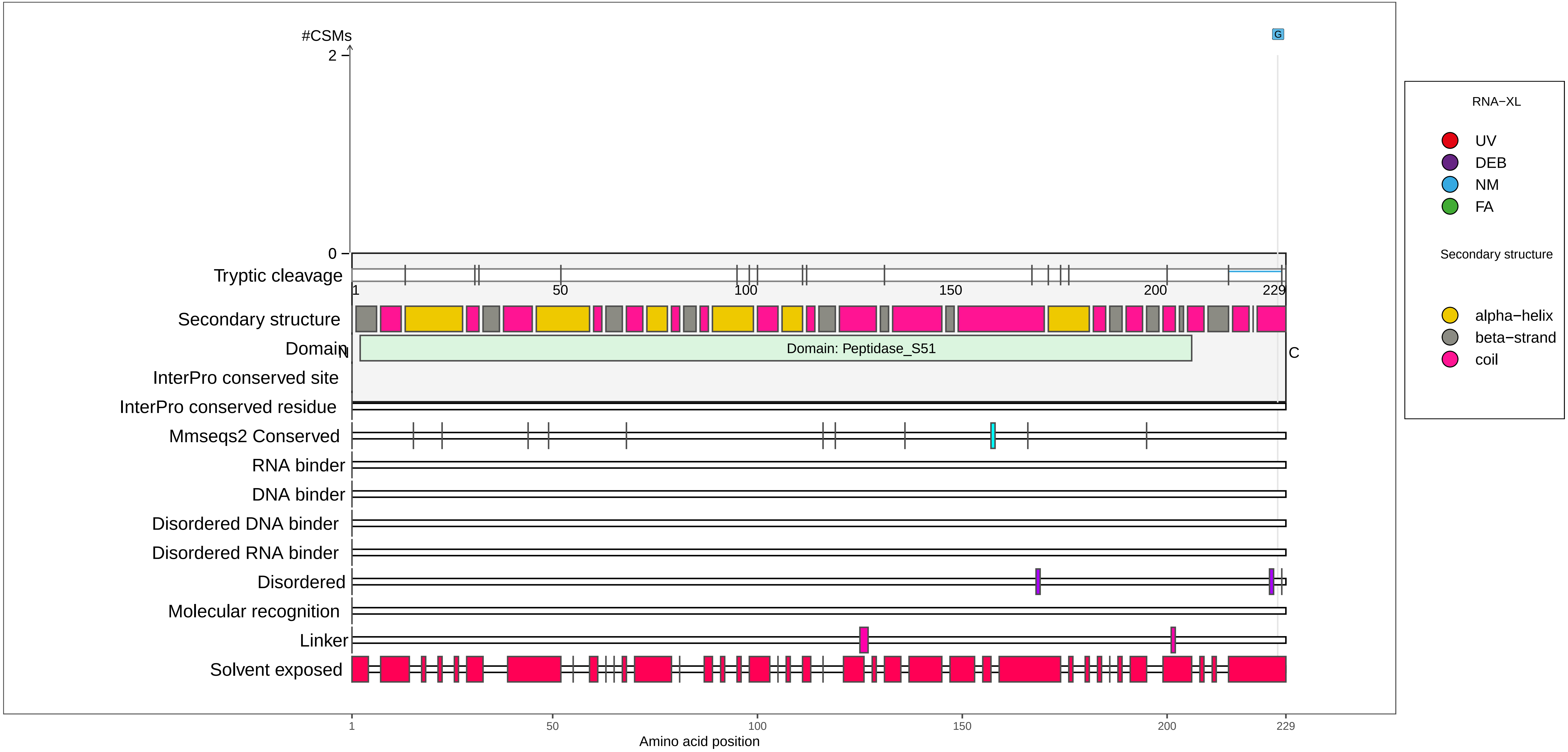

P0A890
TUSA_ECOLI Sulfur carrier protein TusA

− Abundance:
tryptic [log10 Intensity]: 7.48 (Q 34)
PAXdb K12 strain [ppm]: 2.64 (Q 82)
PAXdb E.coli [ppm]: 1.8 (Q 67)

− RNA functions:
ncRNA metabolic process; ncRNA processing; RNA metabolic process; RNA modification
RNA processing; tRNA metabolic process; tRNA modification; tRNA processing
tRNA thio−modification; tRNA wobble base modification; tRNA wobble position uridine thiolation
tRNA wobble uridine modification

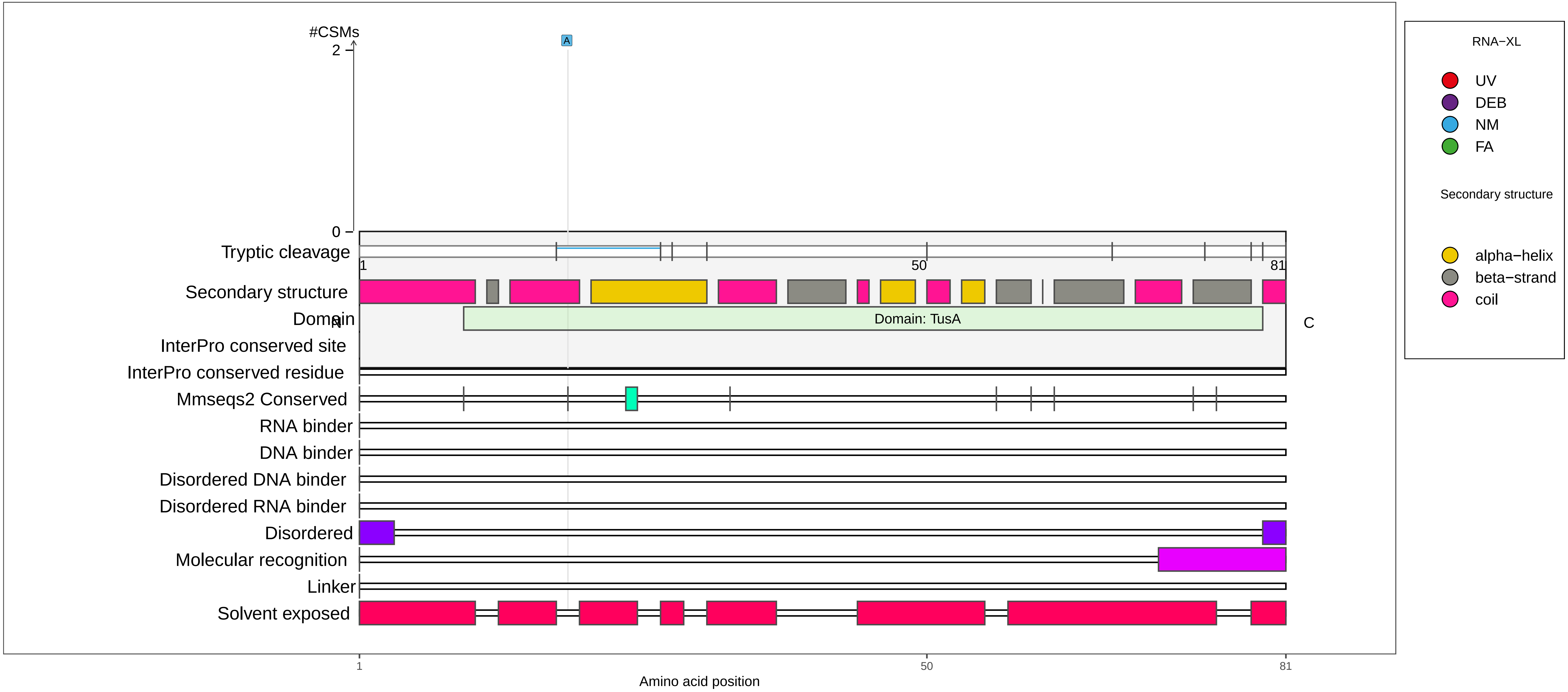

P0A894
RAPZ_ECOLI RNase adapter protein RapZ
− Abundance:
tryptic [log10 Intensity]: 8.15 (Q 63)
PAXdb K12 strain [ppm]: 1.92 (Q 57)
PAXdb E.coli [ppm]: 2.14 (Q 76)

− RNA functions:
RNA binding; RNA destabilization

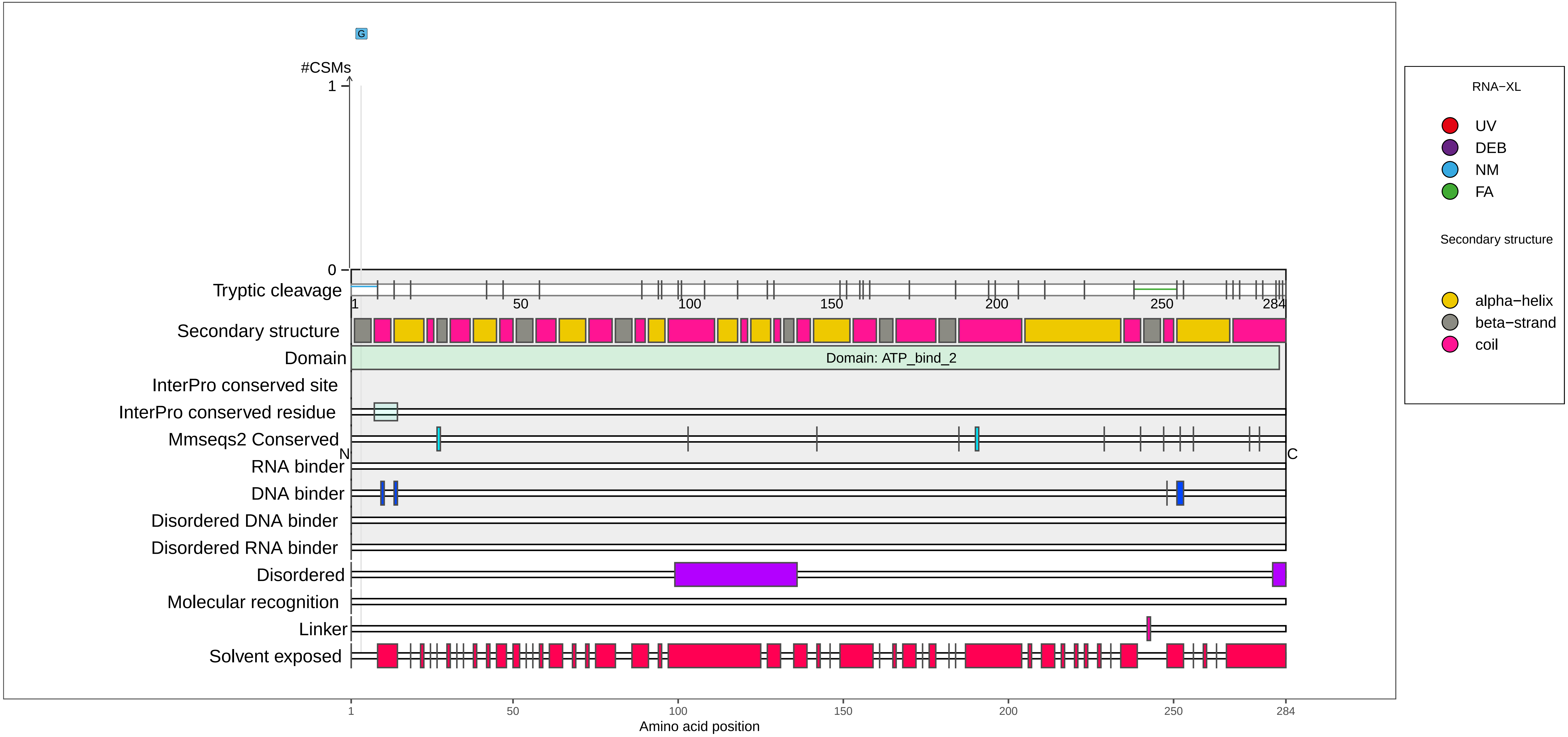

P0A8A2
YEEN_ECOLI Probable transcriptional regulatory protein YeeN

− Abundance:
tryptic [log10 Intensity]: 8.06 (Q 60)
PAXdb K12 strain [ppm]: 2.63 (Q 81)
PAXdb E.coli [ppm]: 2.33 (Q 81)

− RNA functions: not annotated

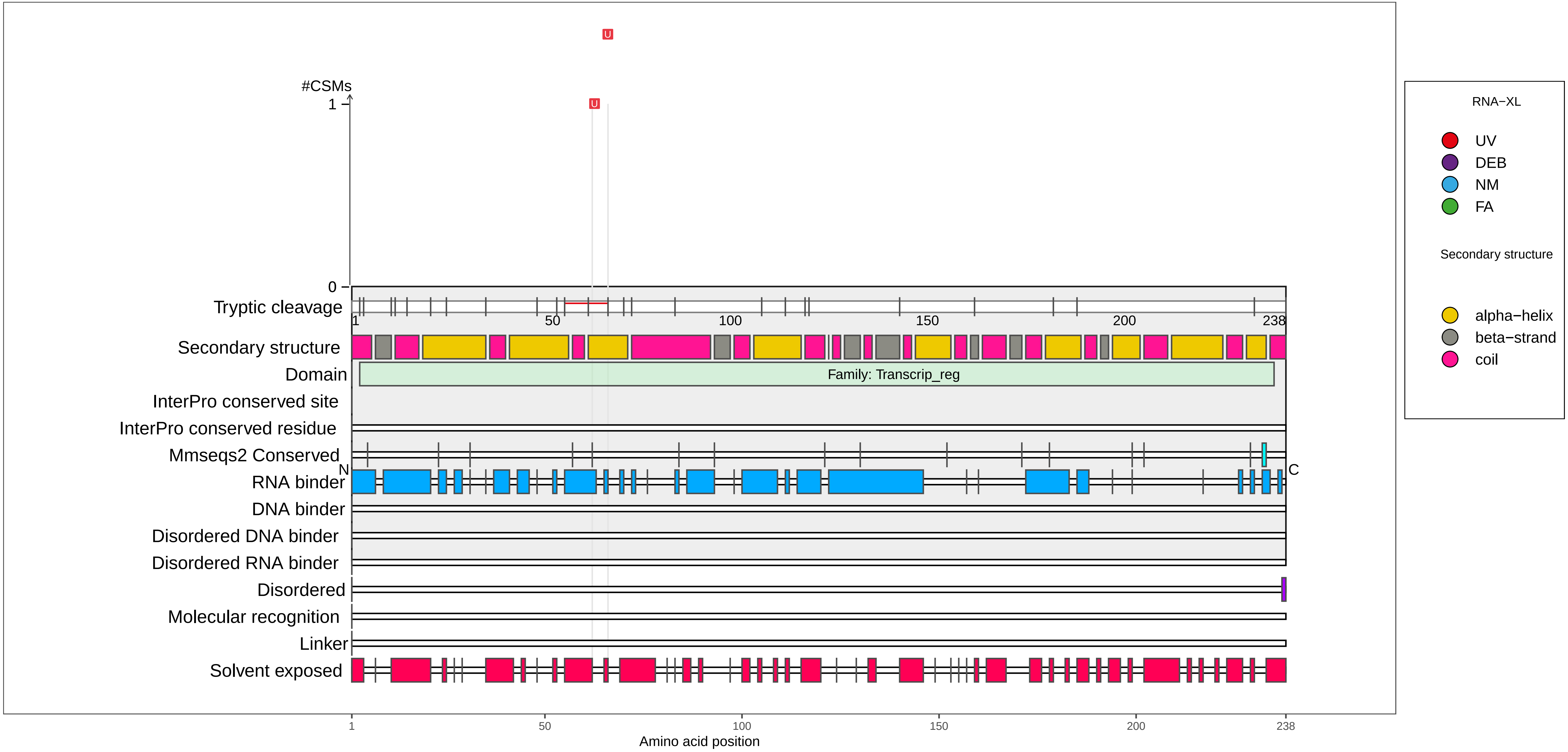

P0A8I5
TRMB_ECOLI tRNA (guanine−N(7)−)−methyltransferase

− Abundance:
tryptic [log10 Intensity]: 7.62 (Q 41)
PAXdb K12 strain [ppm]: 2.94 (Q 89)
PAXdb E.coli [ppm]: 2.28 (Q 80)

− RNA functions:
ncRNA metabolic process; ncRNA processing; RNA (guanine−N7)−methylation
RNA metabolic process; RNA methylation; RNA methyltransferase activity; RNA modification
RNA processing; tRNA (guanine−N7−)−methyltransferase activity; tRNA (guanine−N7)−methylation
tRNA (guanine) methyltransferase activity; tRNA metabolic process; tRNA methylation
tRNA methyltransferase activity; tRNA methyltransferase complex; tRNA modification
tRNA processing

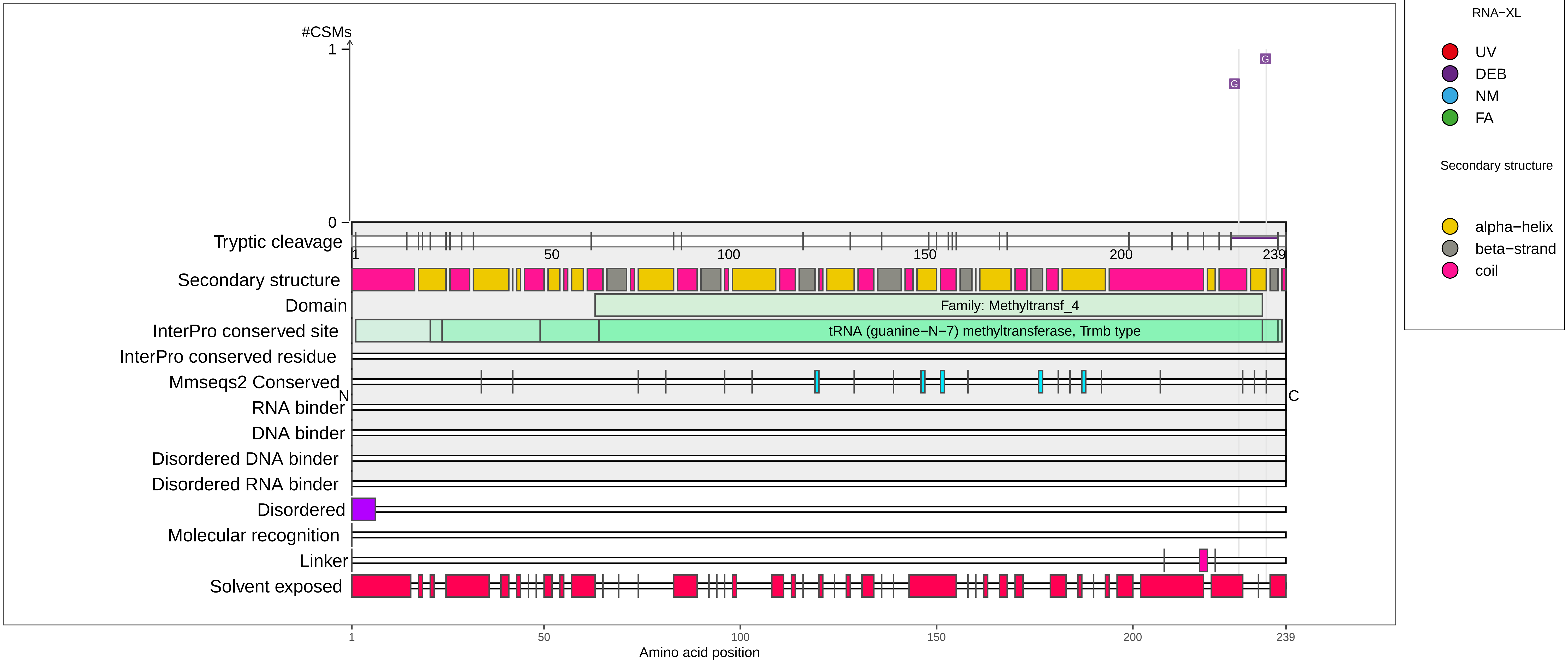

P0A8V6
FADR_ECOLI Fatty acid metabolism regulator protein

− Abundance:
tryptic [log10 Intensity]: 8.02 (Q 58)
PAXdb K12 strain [ppm]: 1.51 (Q 39)
PAXdb E.coli [ppm]: 2.32 (Q 81)

− RNA functions: not annotated

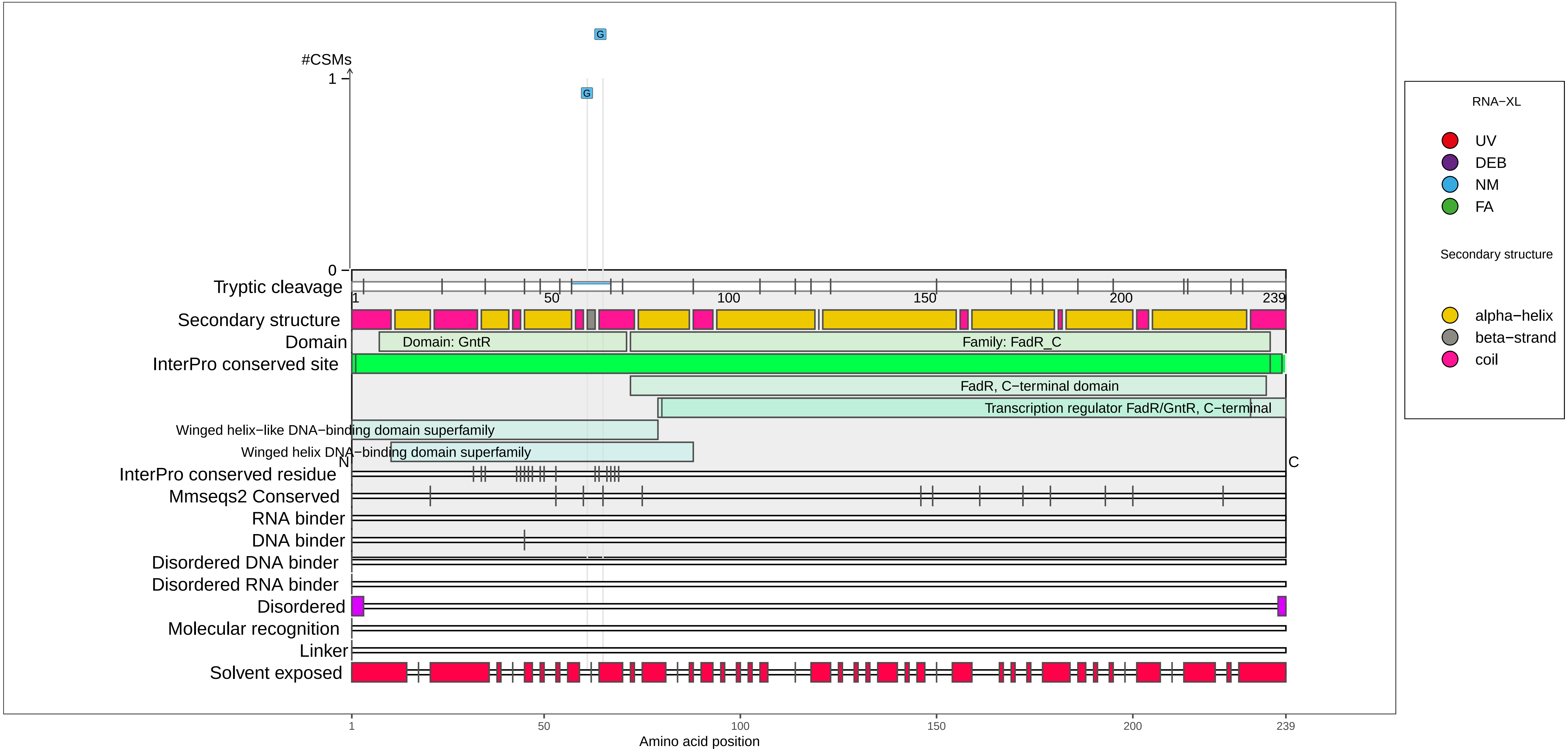

P0A8Y3
YIHX_ECOLI Alpha−D−glucose 1−phosphate phosphatase YihX

− Abundance:
tryptic [log10 Intensity]: 7.93 (Q 55)
PAXdb K12 strain [ppm]: 1.1 (Q 16)
PAXdb E.coli [ppm]: 2.23 (Q 78)

− RNA functions: not annotated

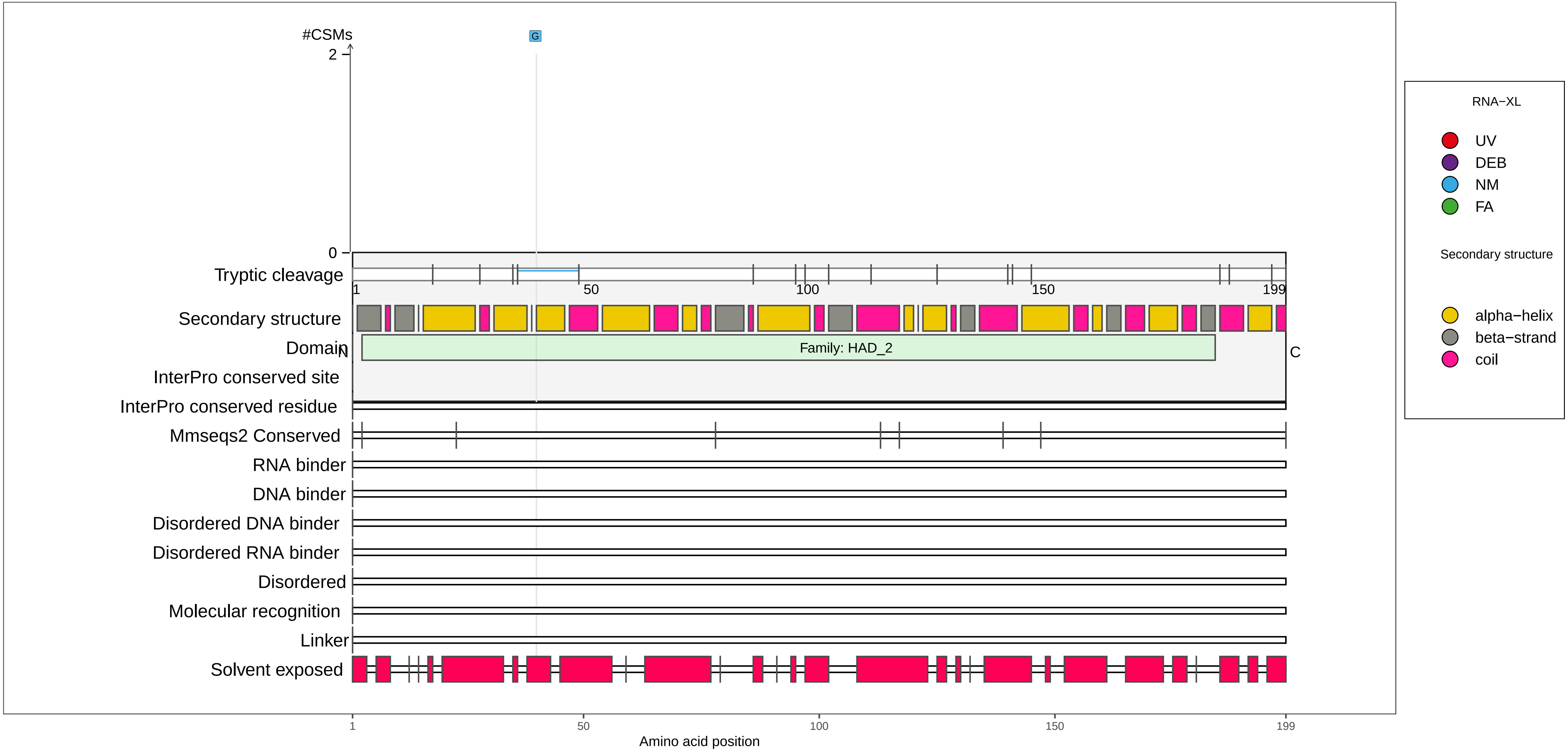

P0A9G6
ACEA_ECOLI Isocitrate lyase

− Abundance:
tryptic [log10 Intensity]: 8.27 (Q 68)
PAXdb K12 strain [ppm]: 2.72 (Q 84)
PAXdb E.coli [ppm]: 3.73 (Q 100)

− RNA functions: not annotated

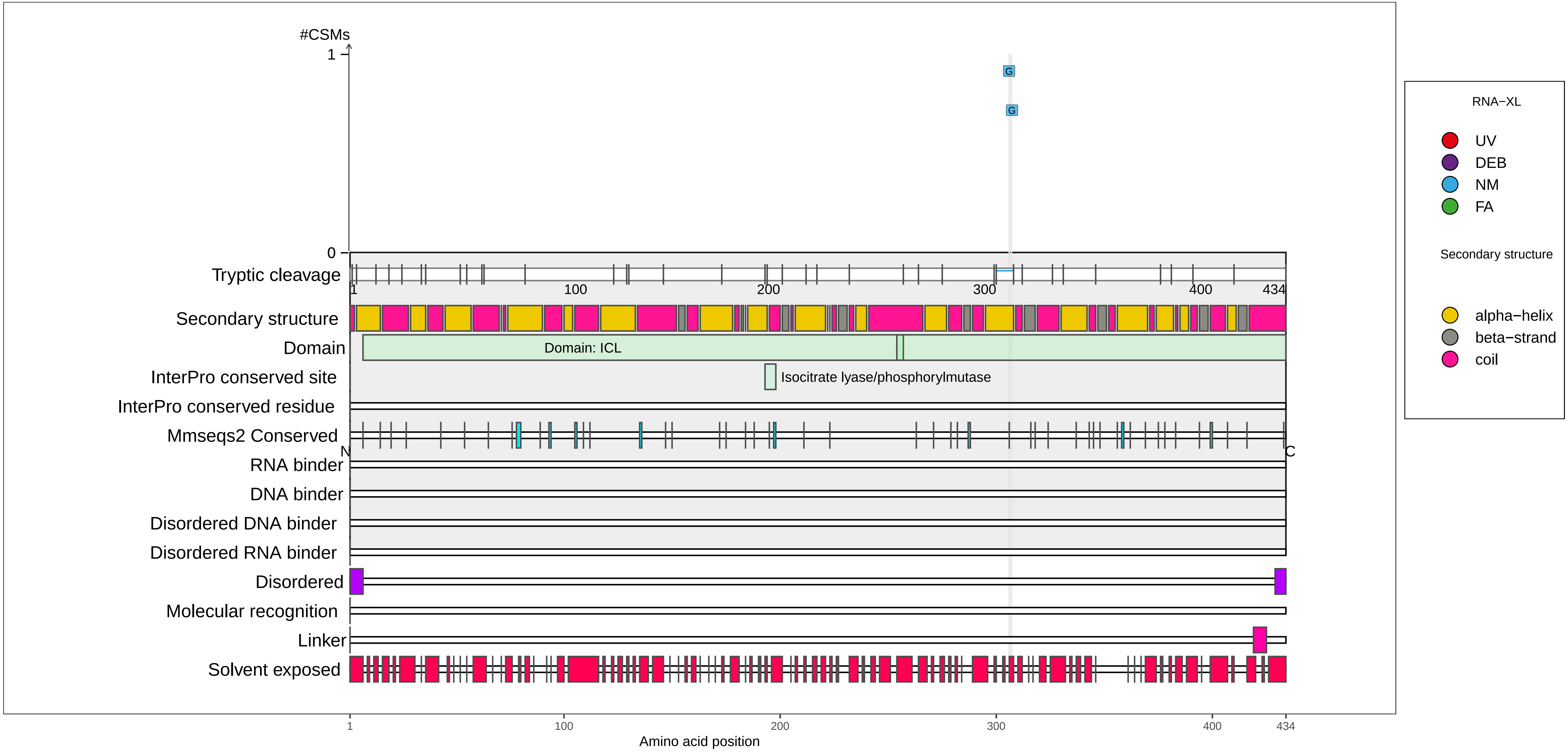

P0A9J0
RNG_ECOLI Ribonuclease G

− Abundance:
tryptic [log10 Intensity]: 8.04 (Q 59)
PAXdb K12 strain [ppm]: 1.61 (Q 45)
PAXdb E.coli [ppm]: 1.19 (Q 52)

− RNA functions:
ncRNA metabolic process; ncRNA processing; RNA binding; RNA metabolic process
RNA phosphodiester bond hydrolysis; RNA processing; rRNA metabolic process; rRNA processing
S1 RNA binding domain

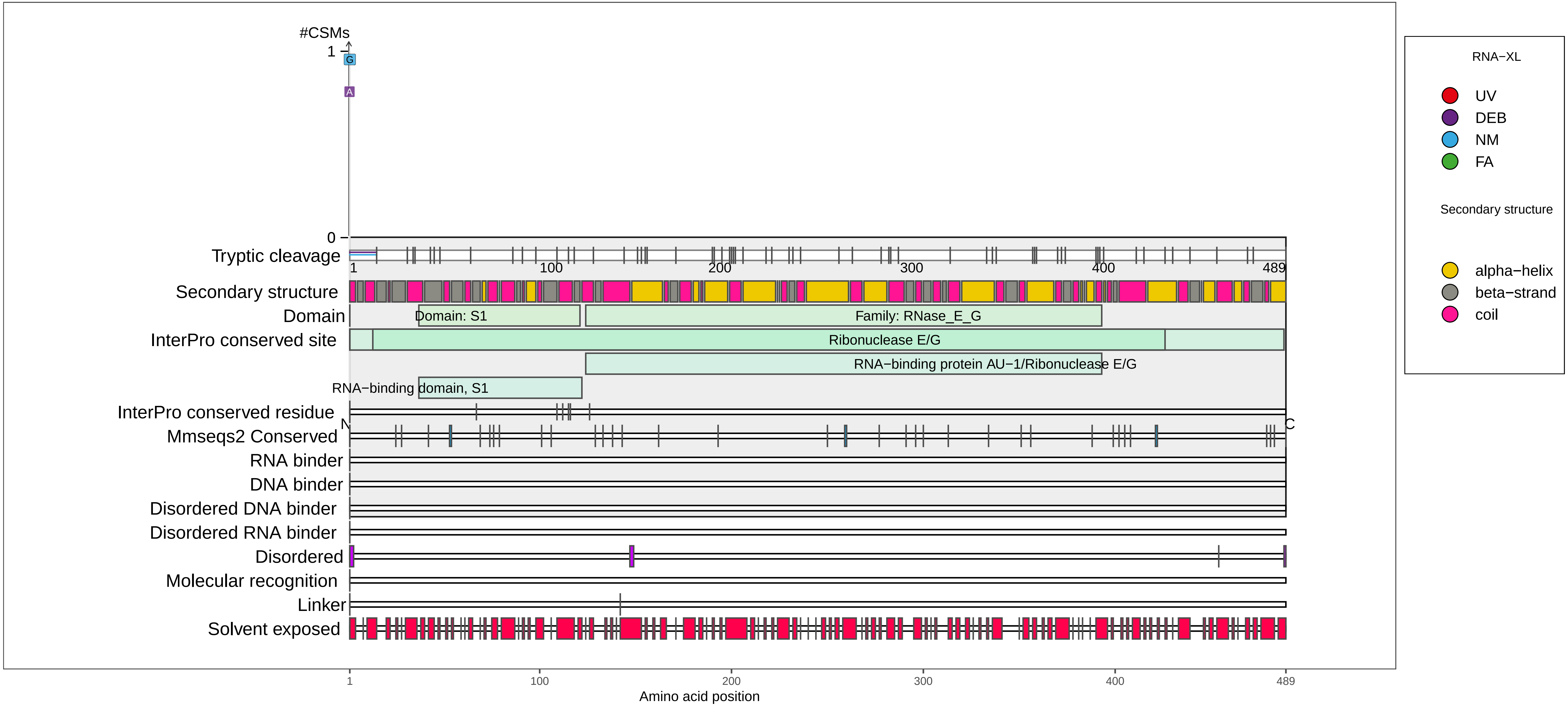

P0A9N4
PFLA_ECOLI Pyruvate formate−lyase 1−activating enzyme
− Abundance:
tryptic [log10 Intensity]: 8.28 (Q 68)
PAXdb K12 strain [ppm]: 2.68 (Q 83)
PAXdb E.coli [ppm]: 1.91 (Q 70)

− RNA functions: not annotated

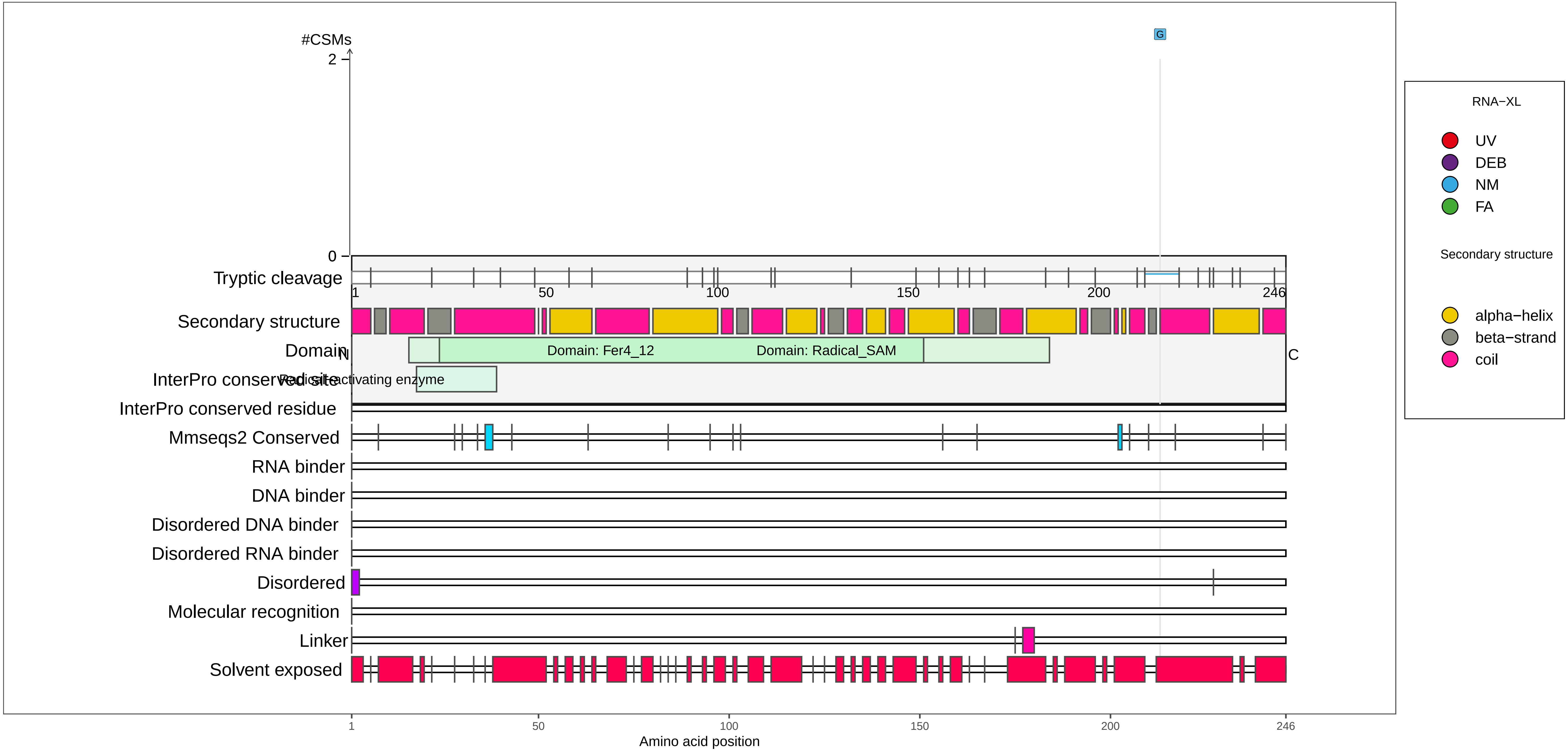

P0A9T0
SERA_ECOLI D−3−phosphoglycerate dehydrogenase

− Abundance:
tryptic [log10 Intensity]: 8.67 (Q 81)
PAXdb K12 strain [ppm]: 3.34 (Q 96)
PAXdb E.coli [ppm]: 3.04 (Q 95)

− RNA functions: not annotated

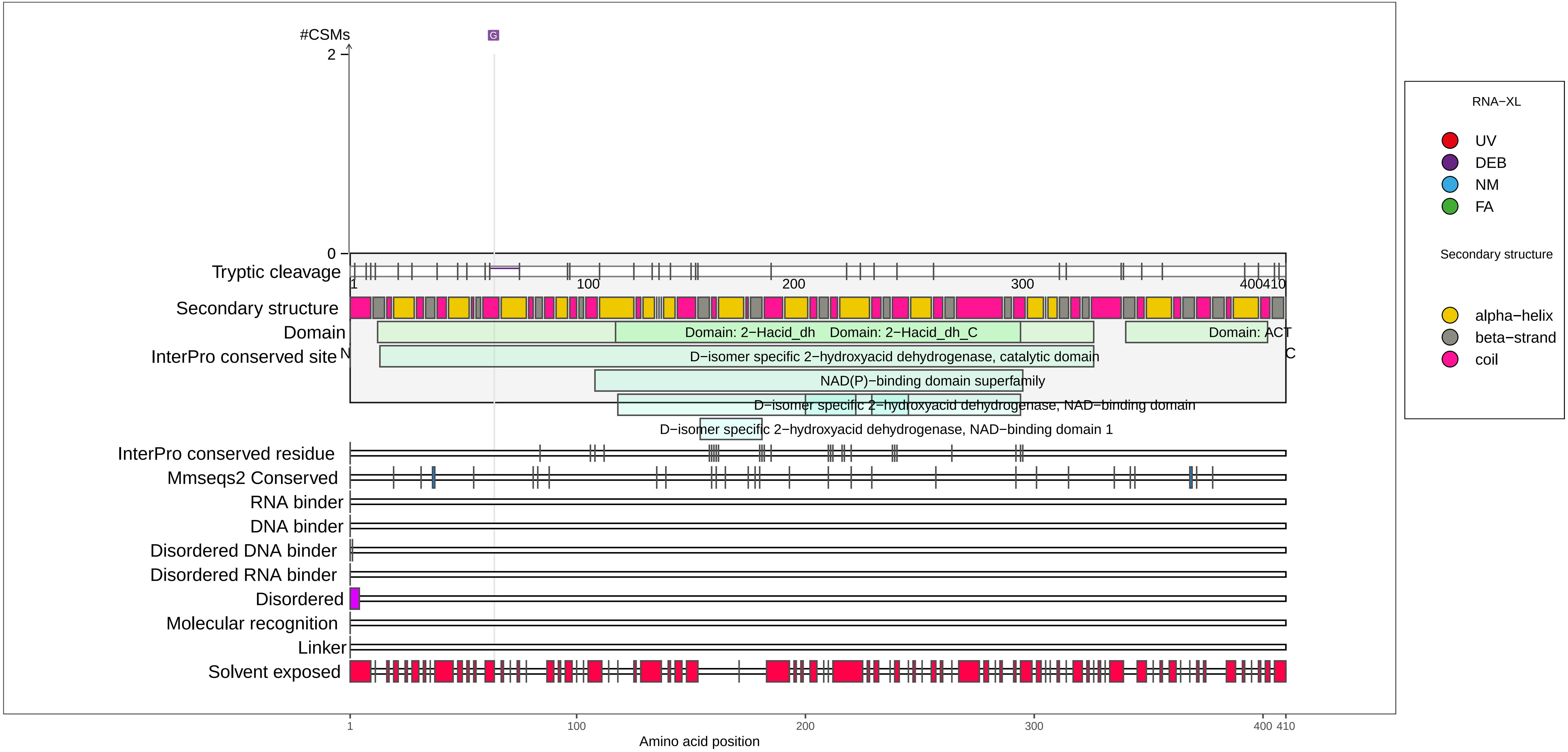

P0A9V1
LPTB_ECOLI Lipopolysaccharide export system ATP−binding protein LptB
− Abundance:
tryptic [log10 Intensity]: 7.87 (Q 52)
PAXdb K12 strain [ppm]: NA (Q NA)
PAXdb E.coli [ppm]: 2.35 (Q 82)

− RNA functions: not annotated

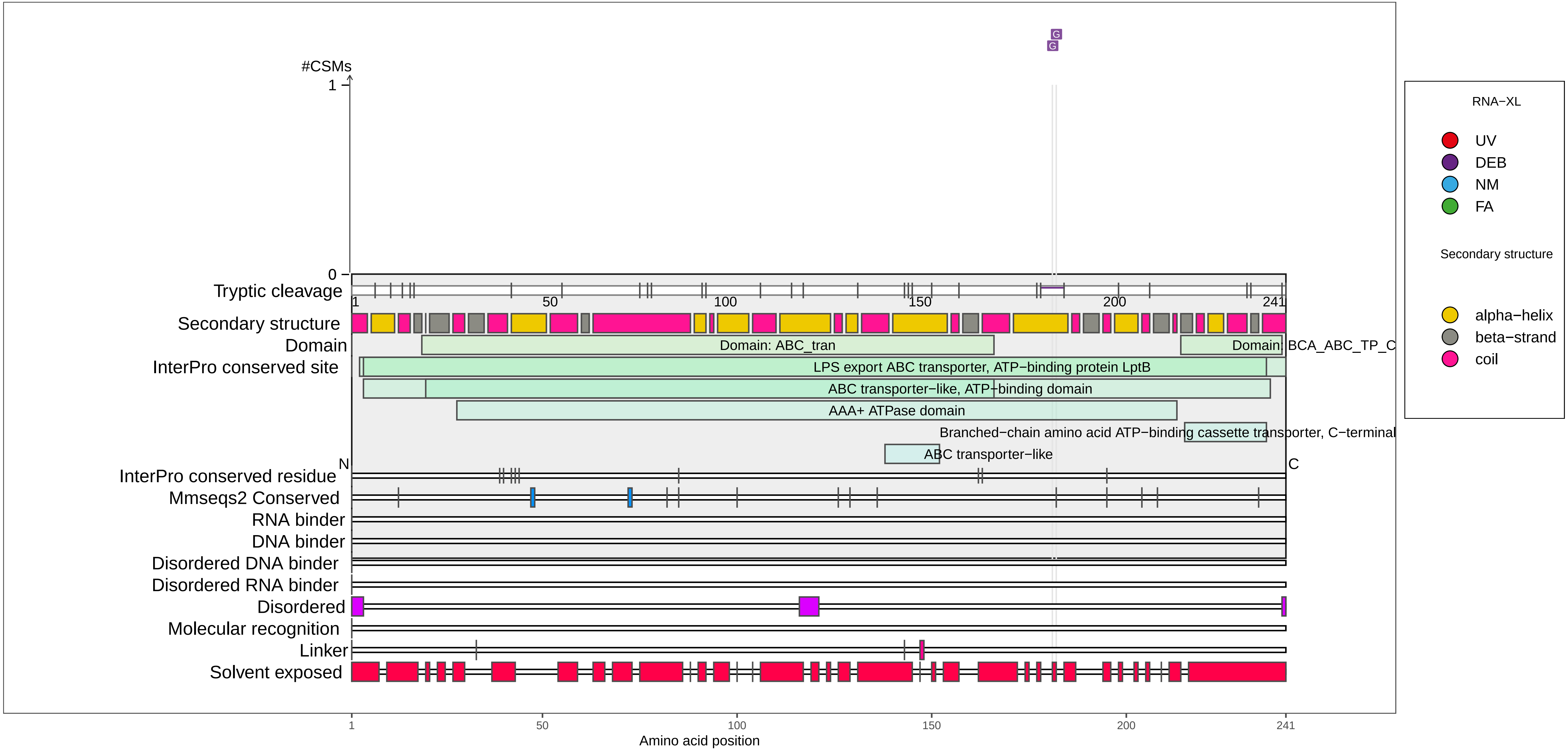

P0AA41
TRUC_ECOLI tRNA pseudouridine synthase C

− Abundance:
tryptic [log10 Intensity]: 6.95 (Q 11)
PAXdb K12 strain [ppm]: NA (Q NA)
PAXdb E.coli [ppm]: 0.1 (Q 27)

− RNA functions:
ncRNA metabolic process; ncRNA processing; RNA binding; RNA metabolic process
RNA modification; RNA processing; RNA pseudouridylate synthase; tRNA metabolic process
tRNA modification; tRNA processing; tRNA pseudouridine synthase activity
tRNA pseudouridine synthesis

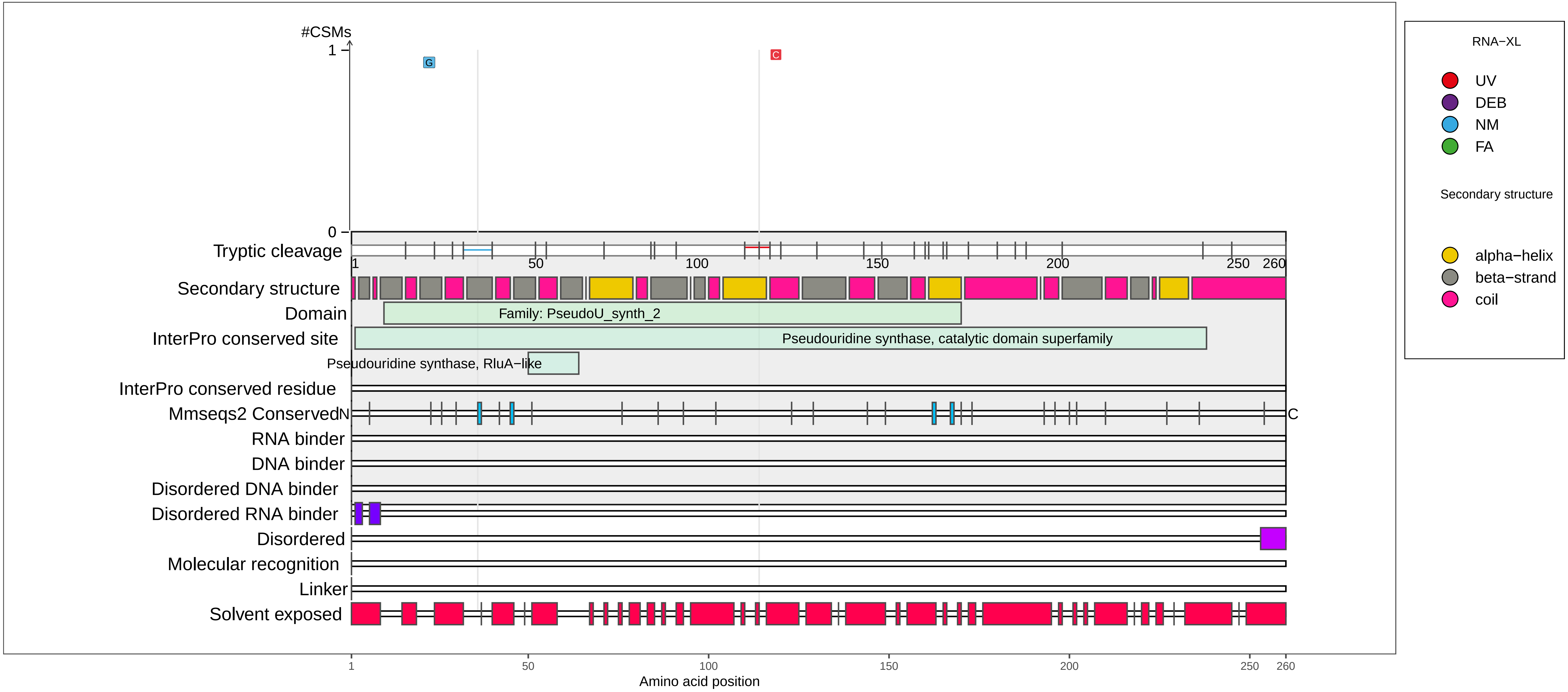

P0AAF6
ARTP_ECOLI Arginine transport ATP−binding protein ArtP

− Abundance:
tryptic [log10 Intensity]: 7.71 (Q 45)
PAXdb K12 strain [ppm]: 2.37 (Q 73)
PAXdb E.coli [ppm]: 2.3 (Q 80)

− RNA functions: not annotated

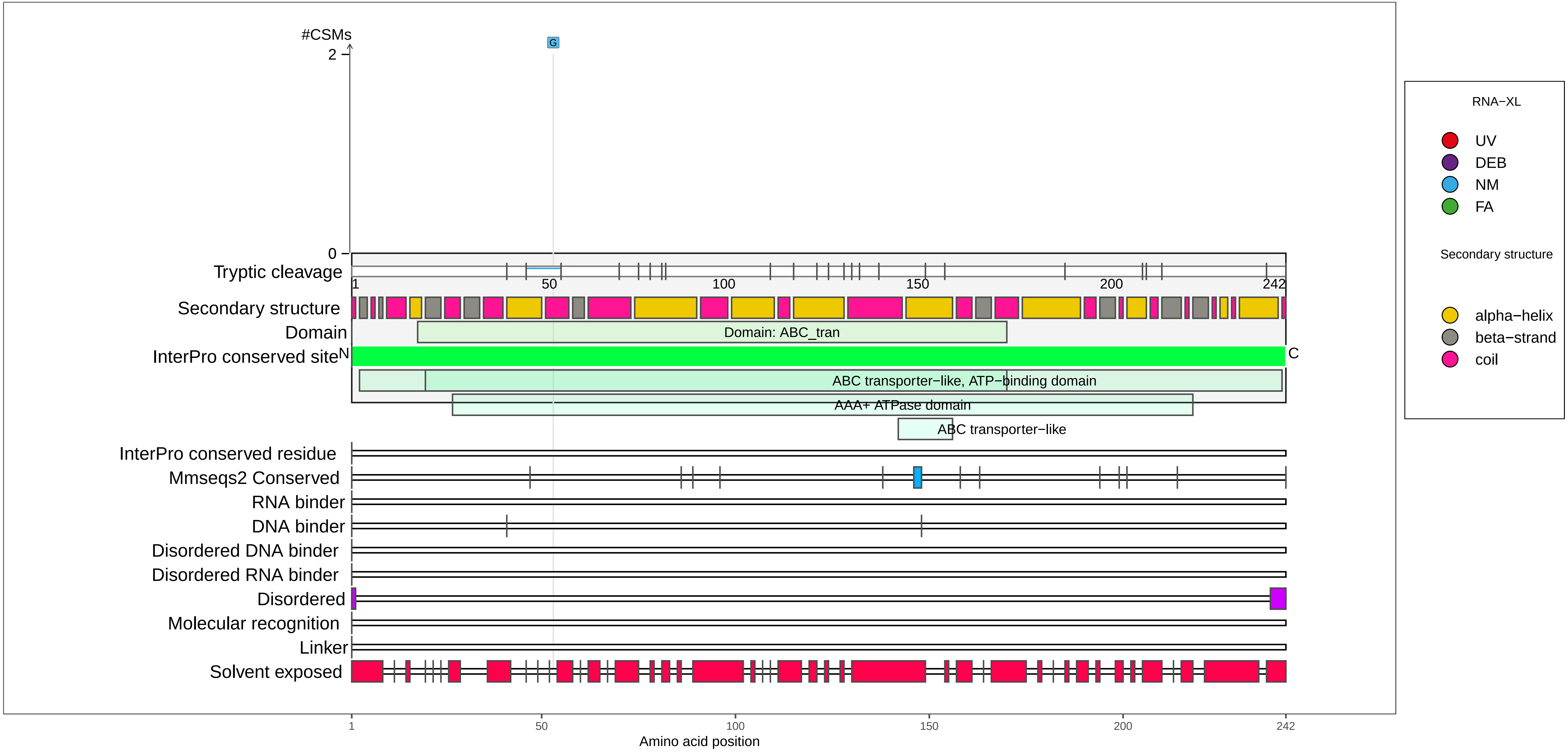

P0AAI9
FABD_ECOLI Malonyl CoA−acyl carrier protein transacylase

− Abundance:
tryptic [log10 Intensity]: 8.55 (Q 77)
PAXdb K12 strain [ppm]: 2.61 (Q 80)
PAXdb E.coli [ppm]: 2.94 (Q 94)

− RNA functions: not annotated

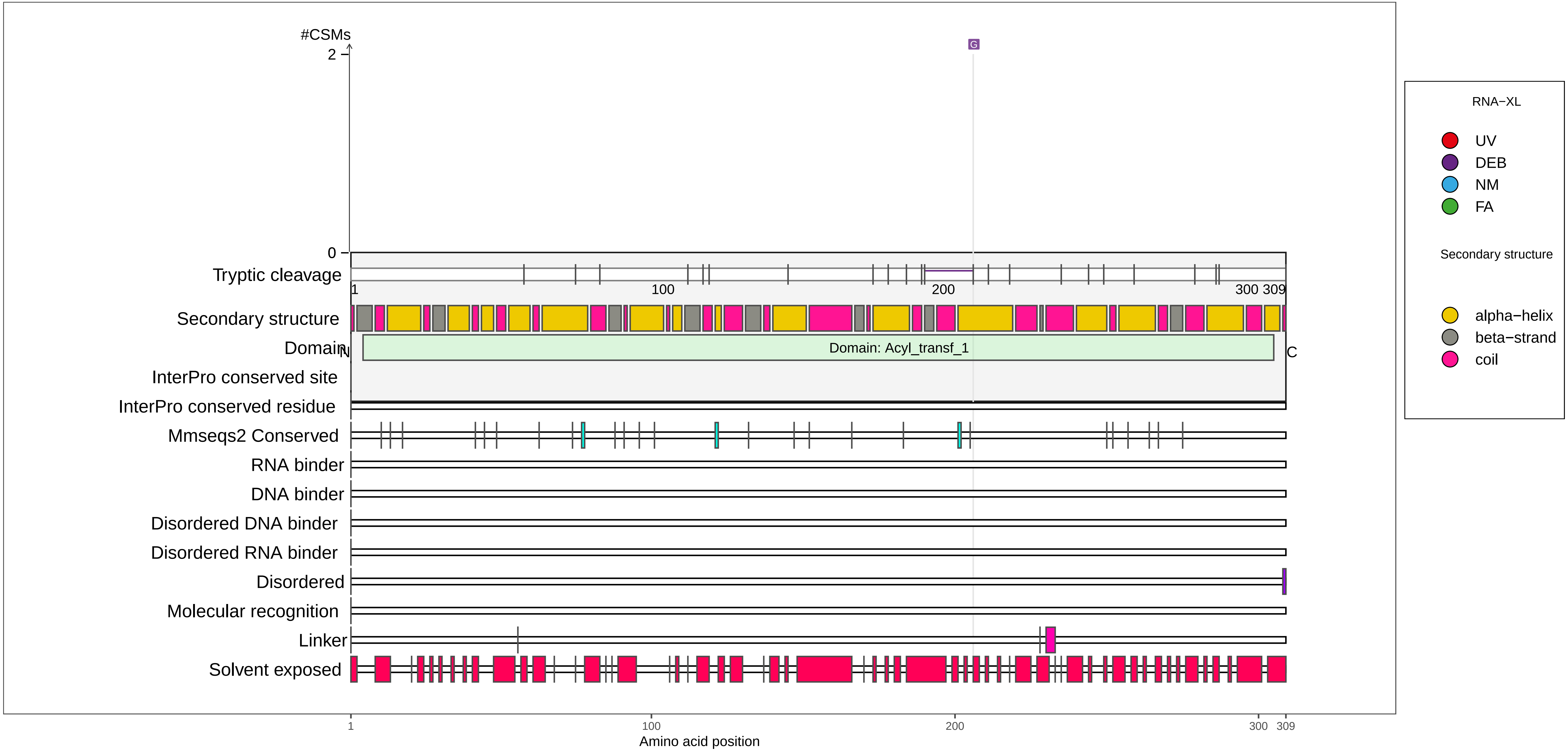

P0AB18
TUSE_ECOLI Sulfurtransferase TusE

− Abundance:
tryptic [log10 Intensity]: 8.08 (Q 61)
PAXdb K12 strain [ppm]: 1.81 (Q 54)
PAXdb E.coli [ppm]: 1.51 (Q 60)

− RNA functions:
ncRNA metabolic process; ncRNA processing; RNA metabolic process; RNA modification
RNA processing; tRNA metabolic process; tRNA modification; tRNA processing
tRNA thio−modification; tRNA wobble base modification; tRNA wobble position uridine thiolation
tRNA wobble uridine modification

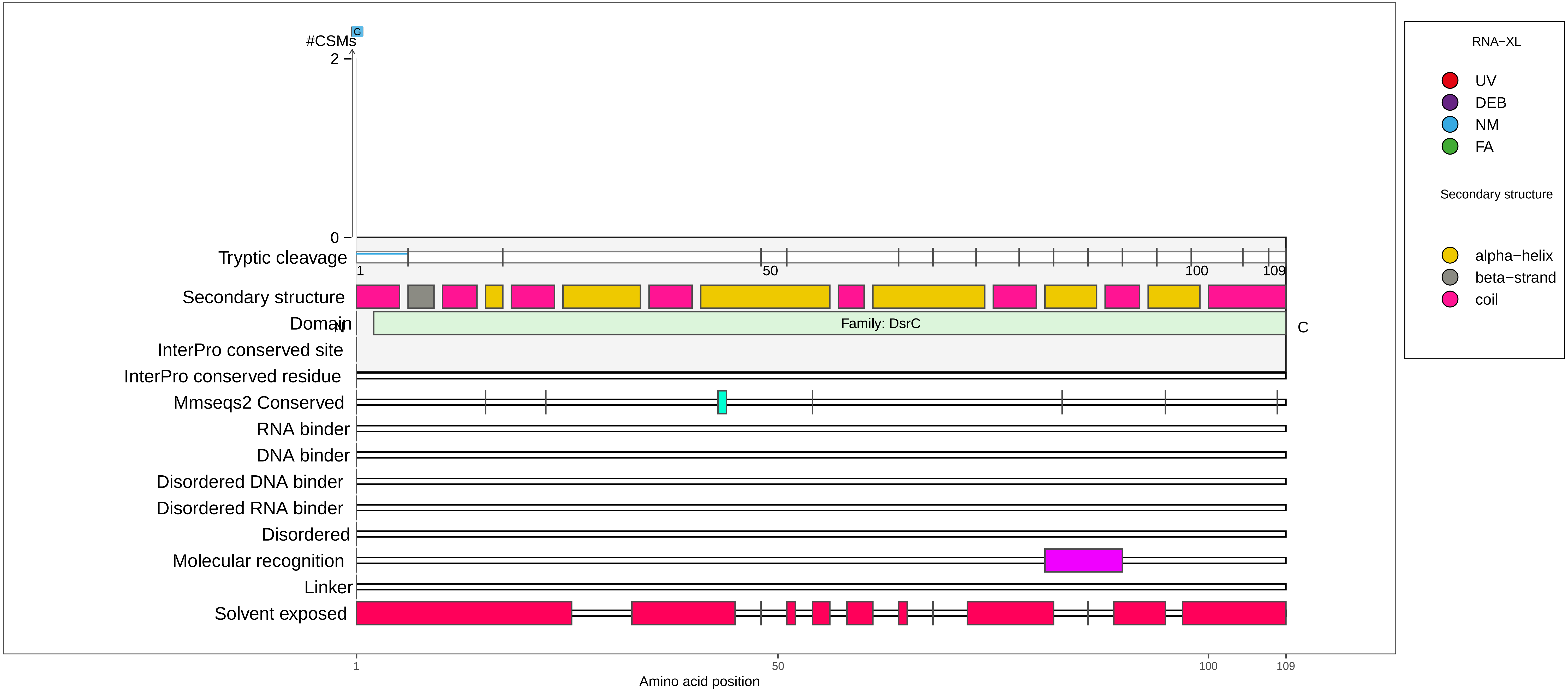

P0ABL3
NAPB_ECOLI Periplasmic nitrate reductase, electron transfer subunit

− Abundance:
tryptic [log10 Intensity]: NA (Q NA)
PAXdb K12 strain [ppm]: NA (Q NA)
PAXdb E.coli [ppm]: −0.36 (Q 18)

− RNA functions: not annotated

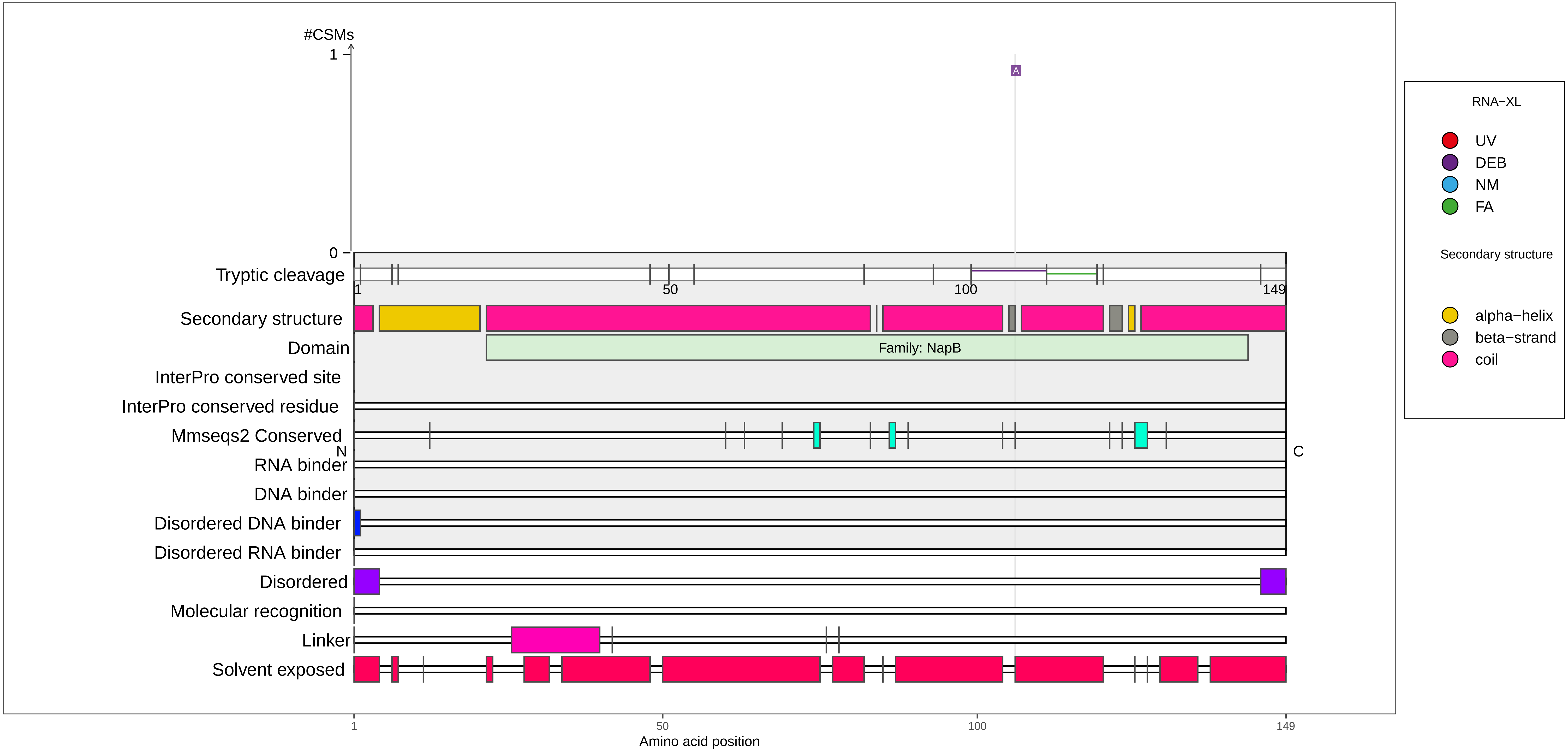

P0ABQ4
DYR_ECOLI Dihydrofolate reductase

− Abundance:
tryptic [log10 Intensity]: 7.09 (Q 17)
PAXdb K12 strain [ppm]: 2.94 (Q 89)
PAXdb E.coli [ppm]: 2.13 (Q 76)

− RNA functions: not annotated

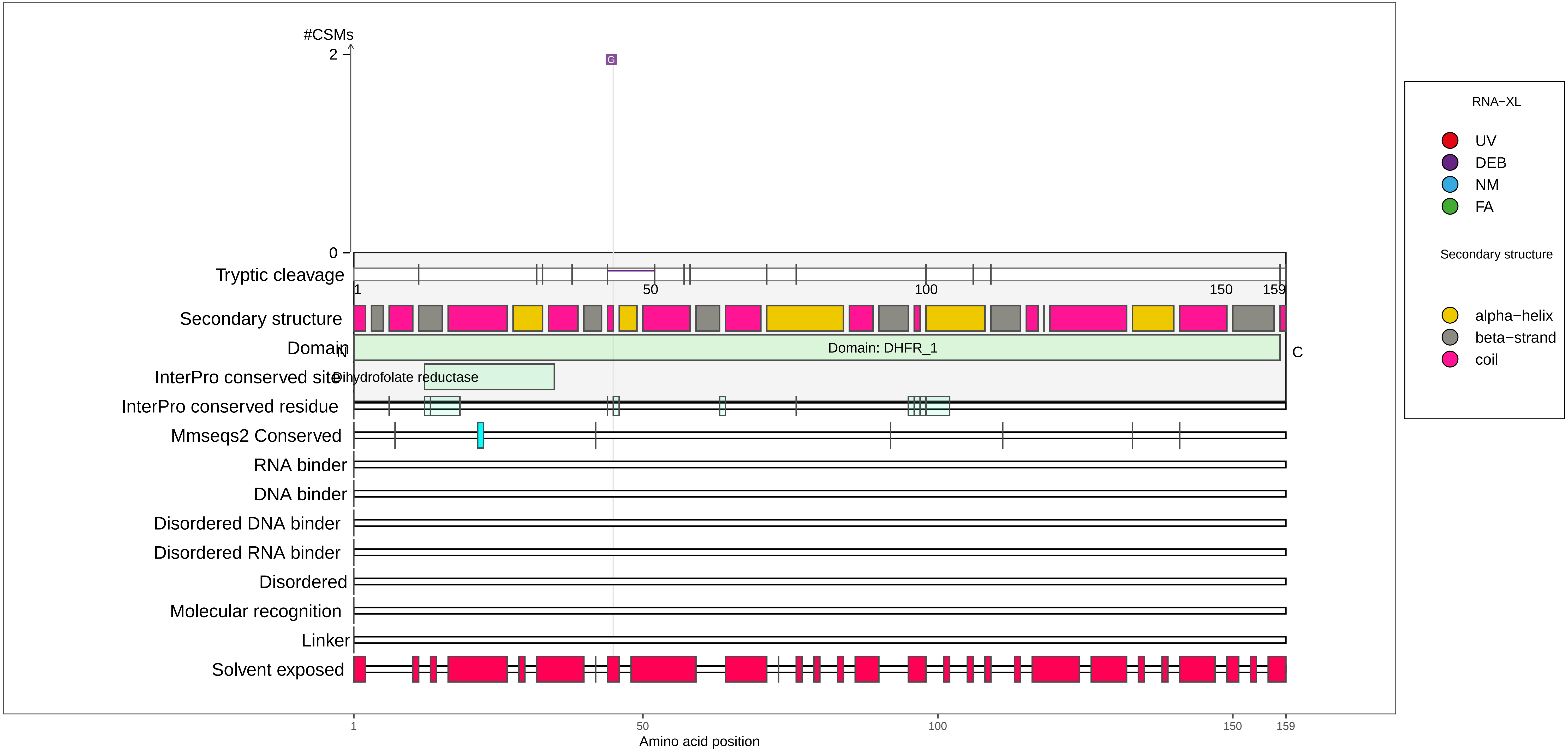

P0ABT5
DUSB_ECOLI tRNA−dihydrouridine synthase B

− Abundance:
tryptic [log10 Intensity]: NA (Q NA)
PAXdb K12 strain [ppm]: NA (Q NA)
PAXdb E.coli [ppm]: −0.11 (Q 22)

− RNA functions:
ncRNA metabolic process; ncRNA processing; RNA binding; RNA metabolic process
RNA modification; RNA processing; tRNA binding; tRNA dihydrouridine synthase activity
tRNA dihydrouridine synthesis; tRNA metabolic process; tRNA modification; tRNA processing

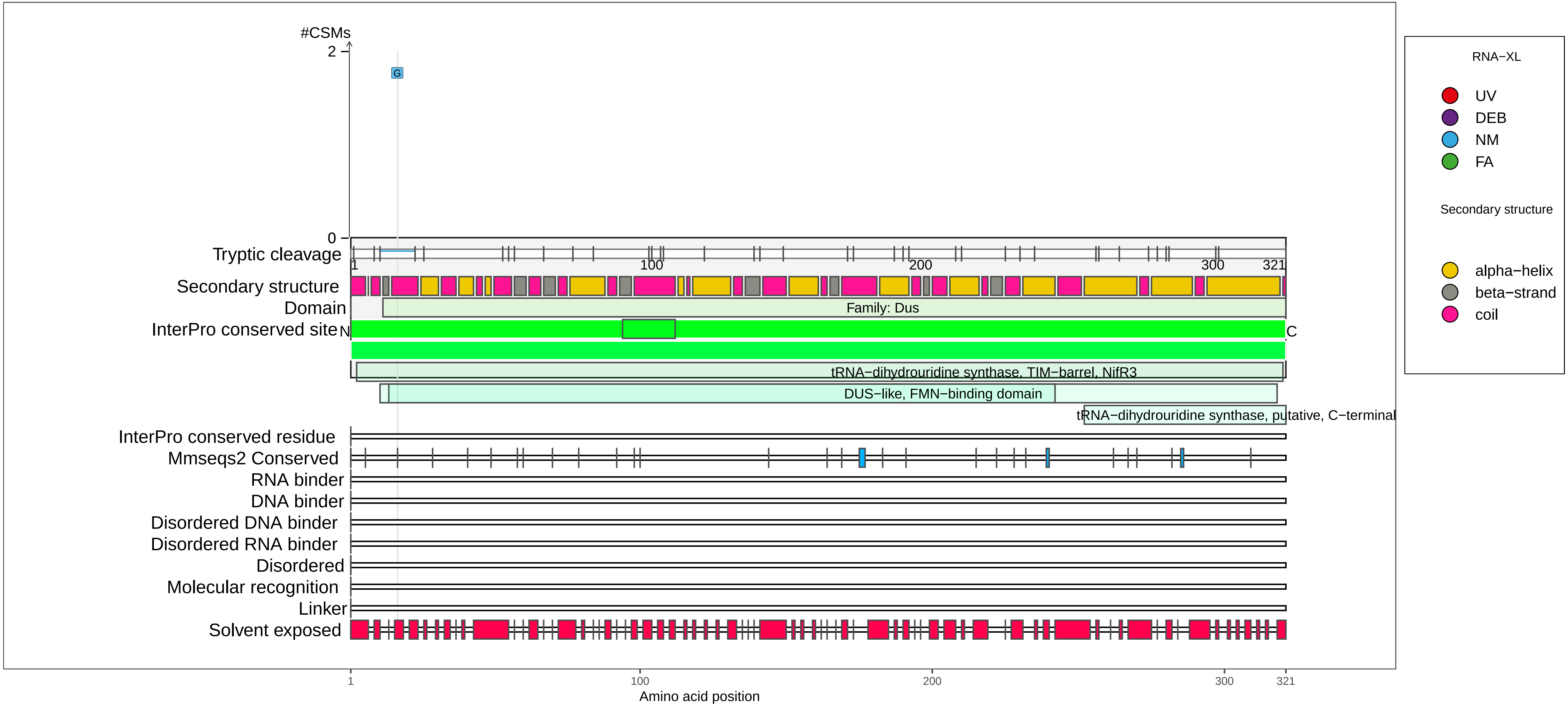

P0AC78
WECA_ECOLI Undecaprenyl−phosphate alpha−N−acetylglucosaminyl 1−phosphate transferase

− Abundance:
tryptic [log10 Intensity]: 6.96 (Q 12)
PAXdb K12 strain [ppm]: NA (Q NA)
PAXdb E.coli [ppm]: −0.52 (Q 15)

− RNA functions: not annotated

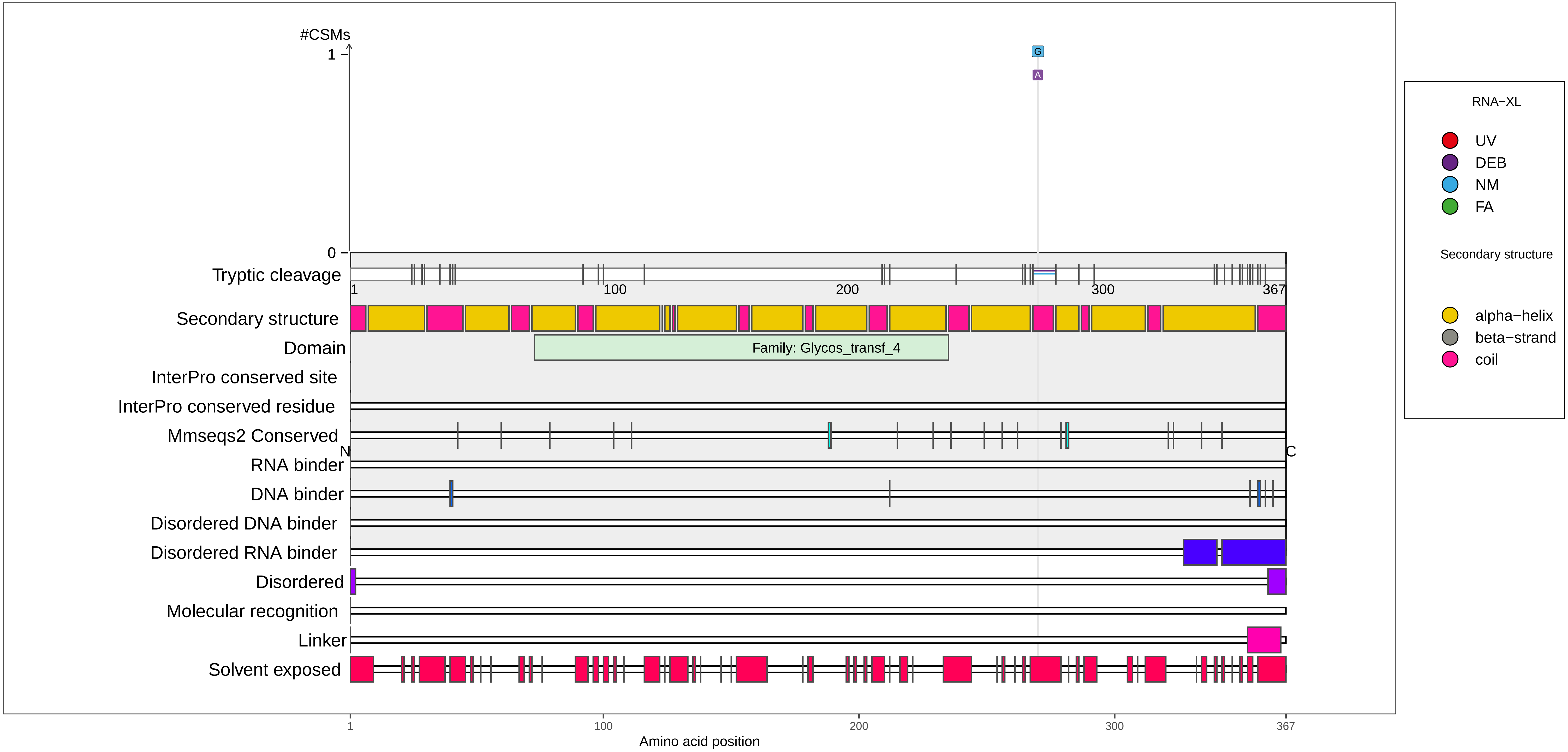

P0AD01
DCUR_ECOLI Transcriptional regulatory protein DcuR

− Abundance:
tryptic [log10 Intensity]: NA (Q NA)
PAXdb K12 strain [ppm]: NA (Q NA)
PAXdb E.coli [ppm]: 0.86 (Q 44)

− RNA functions:
RNA biosynthetic process; RNA metabolic process

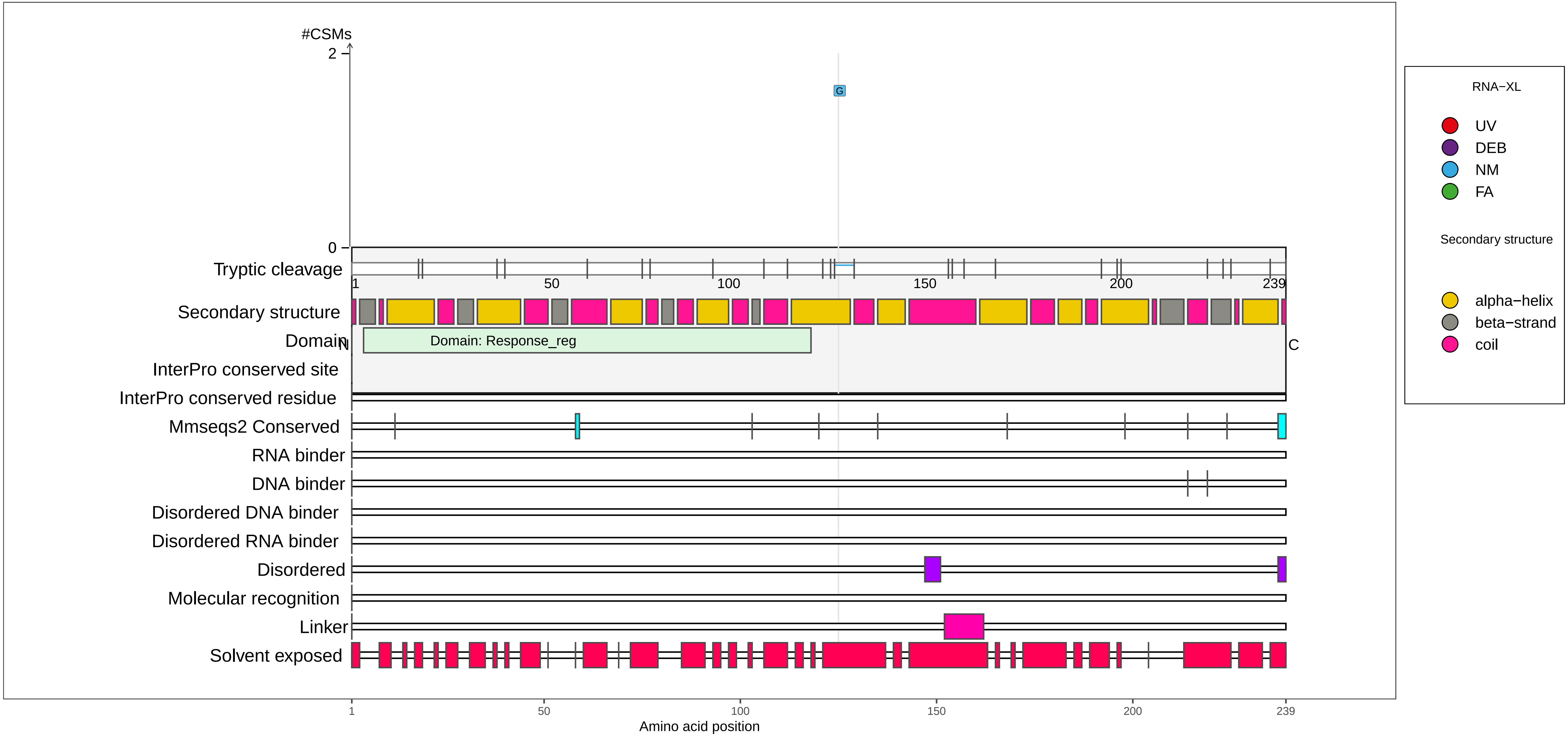

P0AD27
YEJM_ECOLI Inner membrane protein YejM

− Abundance:
tryptic [log10 Intensity]: 6.94 (Q 11)
PAXdb K12 strain [ppm]: NA (Q NA)
PAXdb E.coli [ppm]: 0.52 (Q 36)

− RNA functions: not annotated

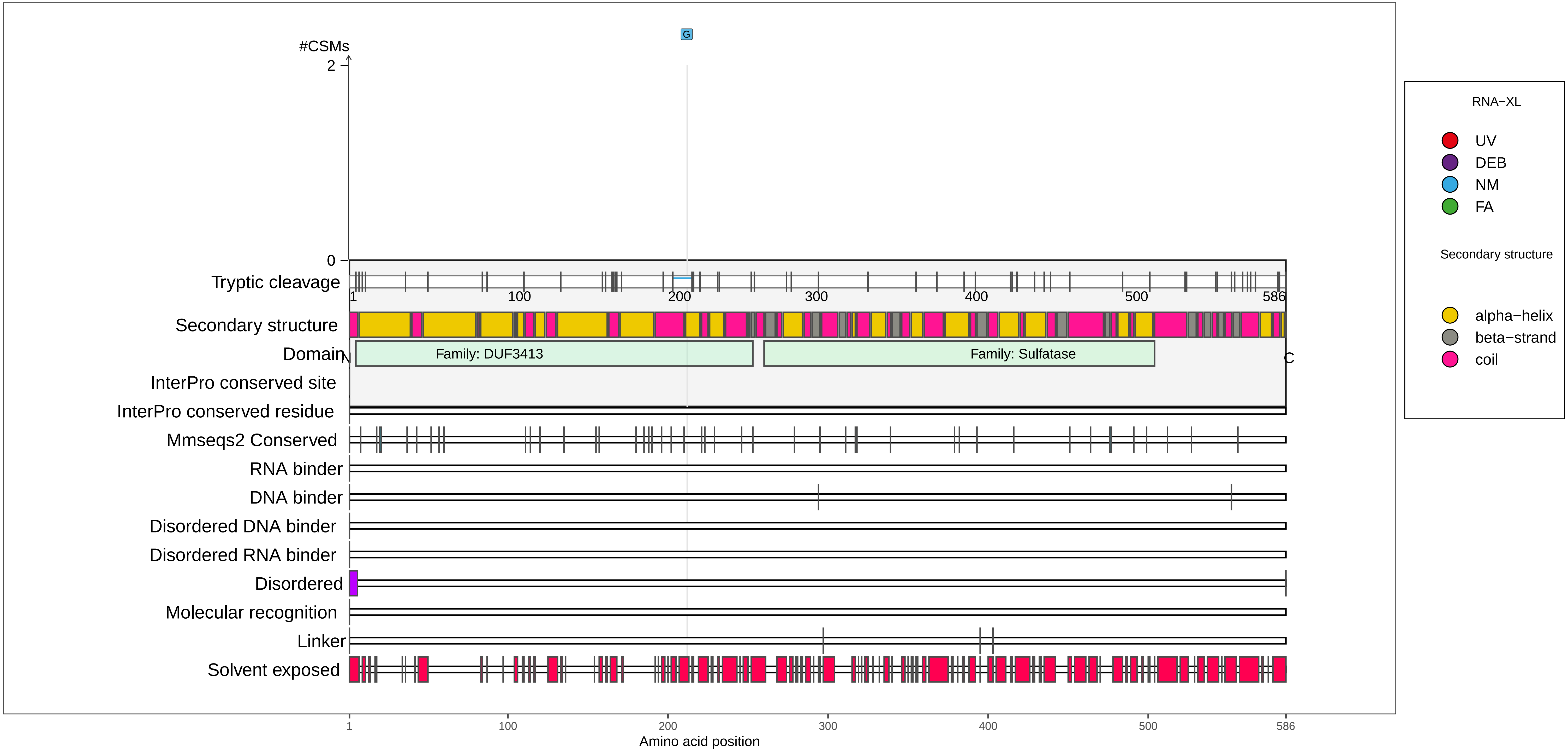

P0ADS2
ZAPA_ECOLI Cell division protein ZapA

− Abundance:
tryptic [log10 Intensity]: 7.52 (Q 36)
PAXdb K12 strain [ppm]: 2.48 (Q 77)
PAXdb E.coli [ppm]: 2.24 (Q 78)

− RNA functions: not annotated

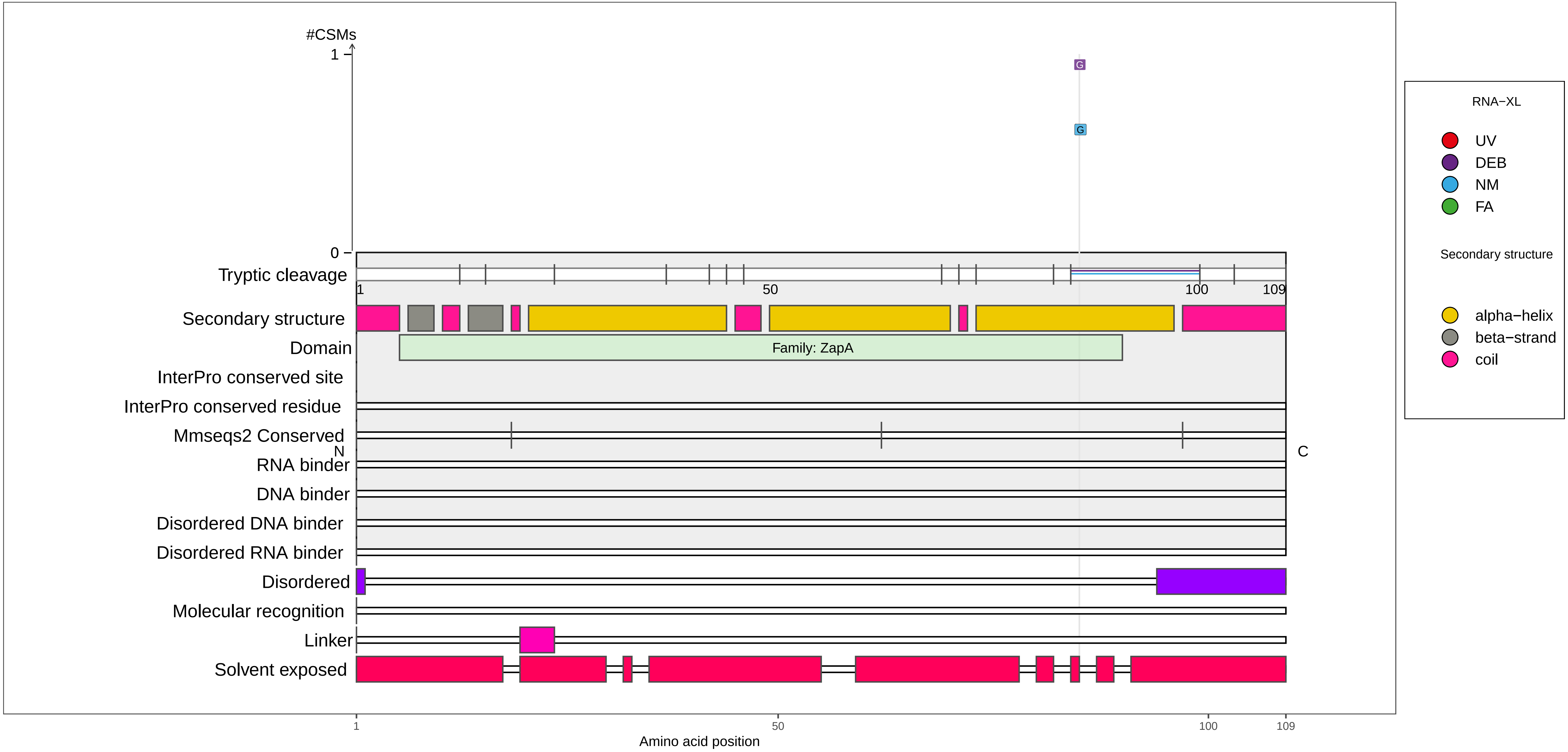

P0ADS9
YGGN_ECOLI Uncharacterized protein YggN

− Abundance:
tryptic [log10 Intensity]: 6.81 (Q 7)
PAXdb K12 strain [ppm]: 1.88 (Q 56)
PAXdb E.coli [ppm]: 2.23 (Q 78)

− RNA functions: not annotated

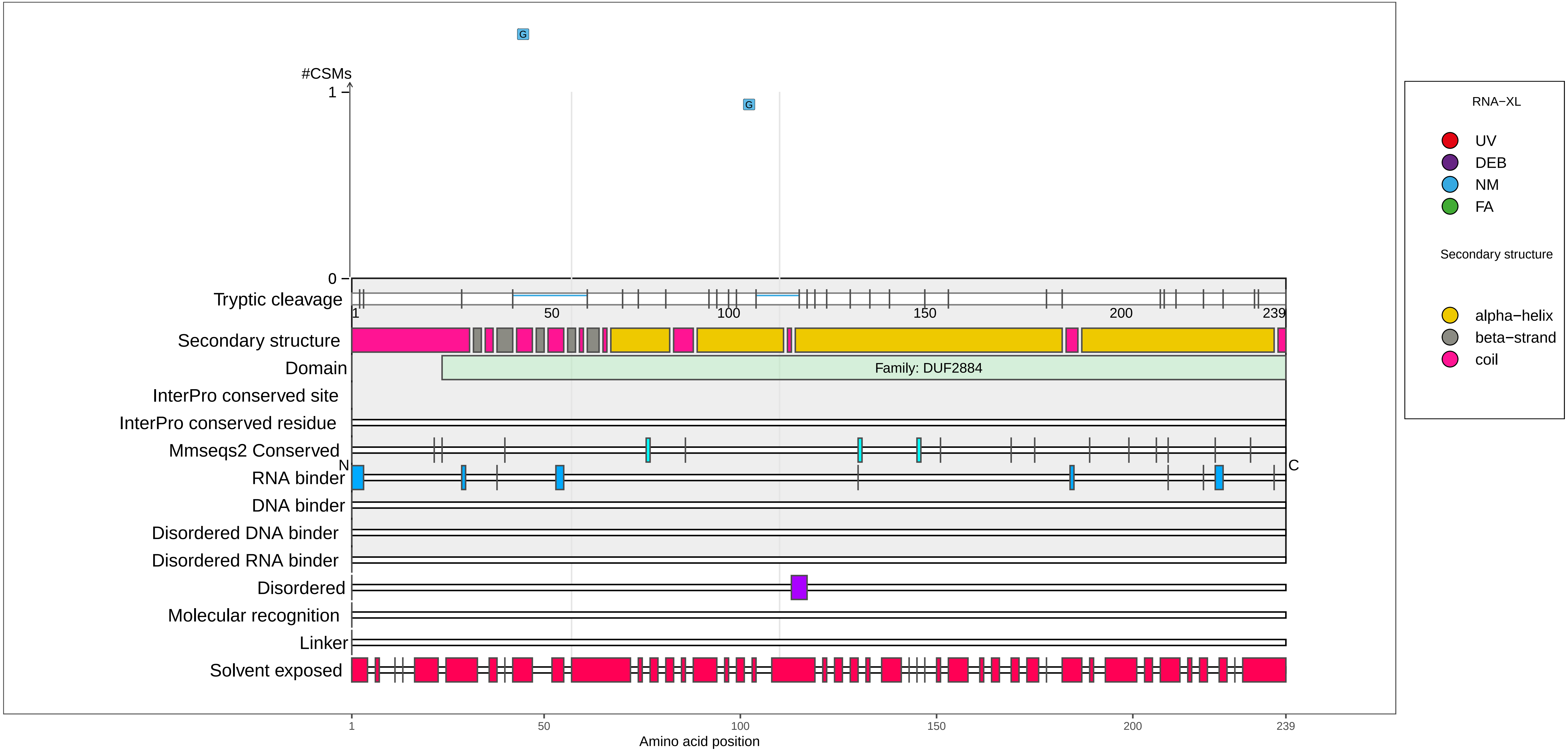

P0AE01
TRMJ_ECOLI tRNA (cytidine/uridine−2'−O−)−methyltransferase TrmJ

− Abundance:
tryptic [log10 Intensity]: 7.95 (Q 56)
PAXdb K12 strain [ppm]: 3.06 (Q 92)
PAXdb E.coli [ppm]: 1.91 (Q 70)

− RNA functions:
ncRNA metabolic process; ncRNA processing; RNA 2−O−methyltransferase activity
RNA binding; RNA metabolic process; RNA methylation; RNA methyltransferase activity
RNA modification; RNA processing; SpoU rRNA Methylase family
tRNA (cytosine−2−O−)−methyltransferase activity; tRNA (cytosine) methyltransferase activity
tRNA (uracil−2−O−)−methyltransferase activity; tRNA (uracil) methyltransferase activity
tRNA 2−O−methyltransferase activity; tRNA metabolic process; tRNA methylation
tRNA methyltransferase activity; tRNA modification; tRNA nucleoside ribose methylation; tRNA processing

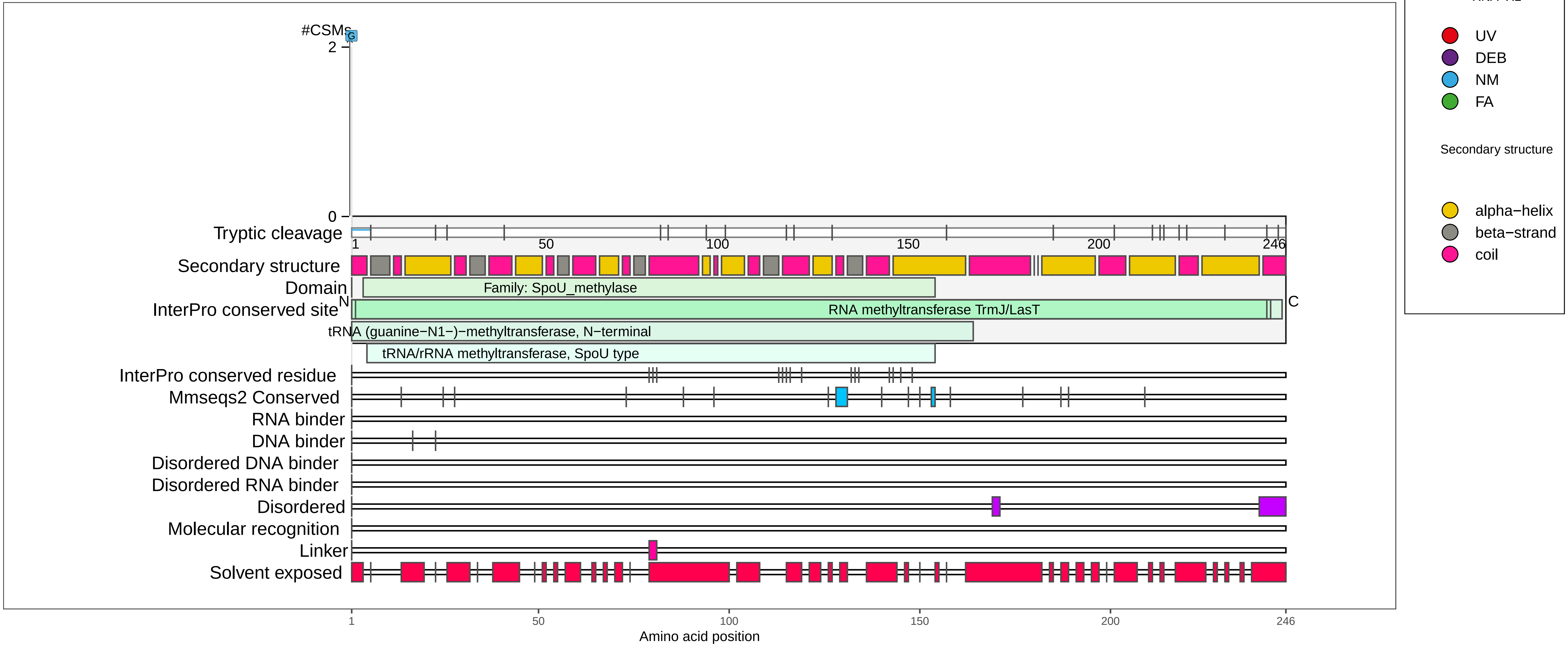

P0AE06
ACRA_ECOLI Multidrug efflux pump subunit AcrA
− Abundance:
tryptic [log10 Intensity]: 8.84 (Q 84)
PAXdb K12 strain [ppm]: 2.26 (Q 69)
PAXdb E.coli [ppm]: 2.44 (Q 84)

− RNA functions: not annotated

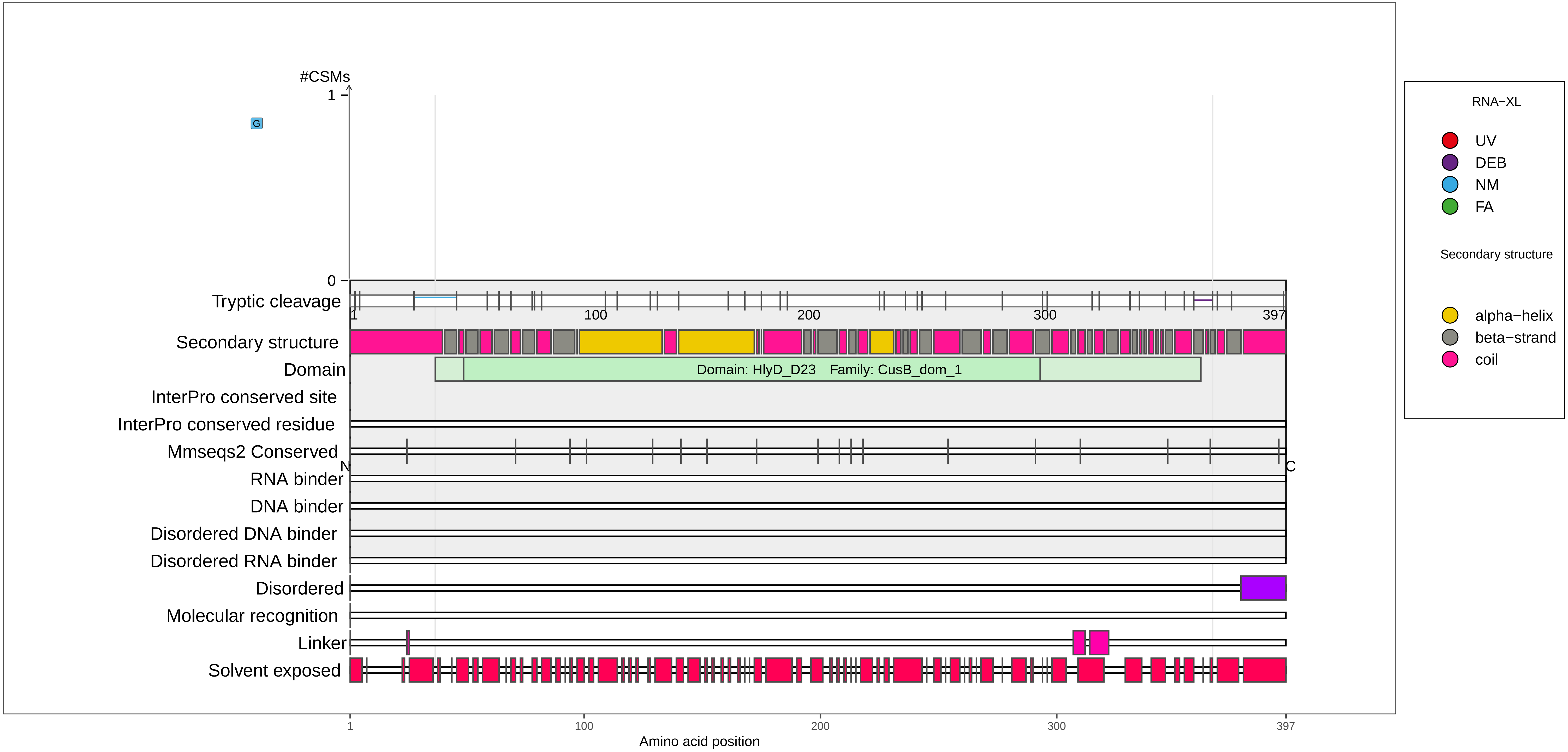

P0AF18
NAGA_ECOLI N−acetylglucosamine−6−phosphate deacetylase

− Abundance:
tryptic [log10 Intensity]: 8.49 (Q 75)
PAXdb K12 strain [ppm]: 2.73 (Q 84)
PAXdb E.coli [ppm]: 2.13 (Q 76)

− RNA functions: not annotated

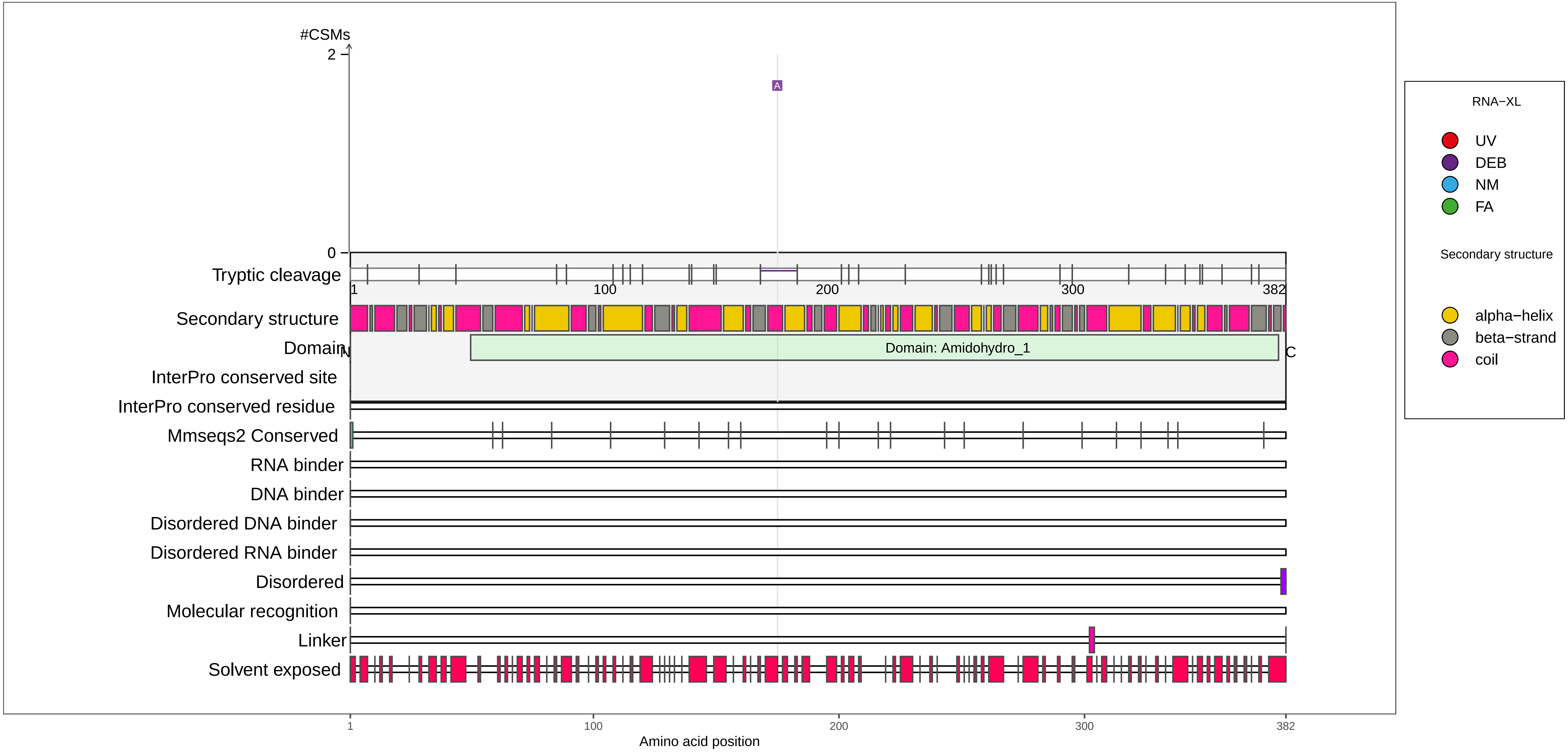

P0AFI2
PARC_ECOLI DNA topoisomerase 4 subunit A

− Abundance:
tryptic [log10 Intensity]: 7.69 (Q 44)
PAXdb K12 strain [ppm]: 1.81 (Q 54)
PAXdb E.coli [ppm]: 1.95 (Q 71)

− RNA functions: not annotated

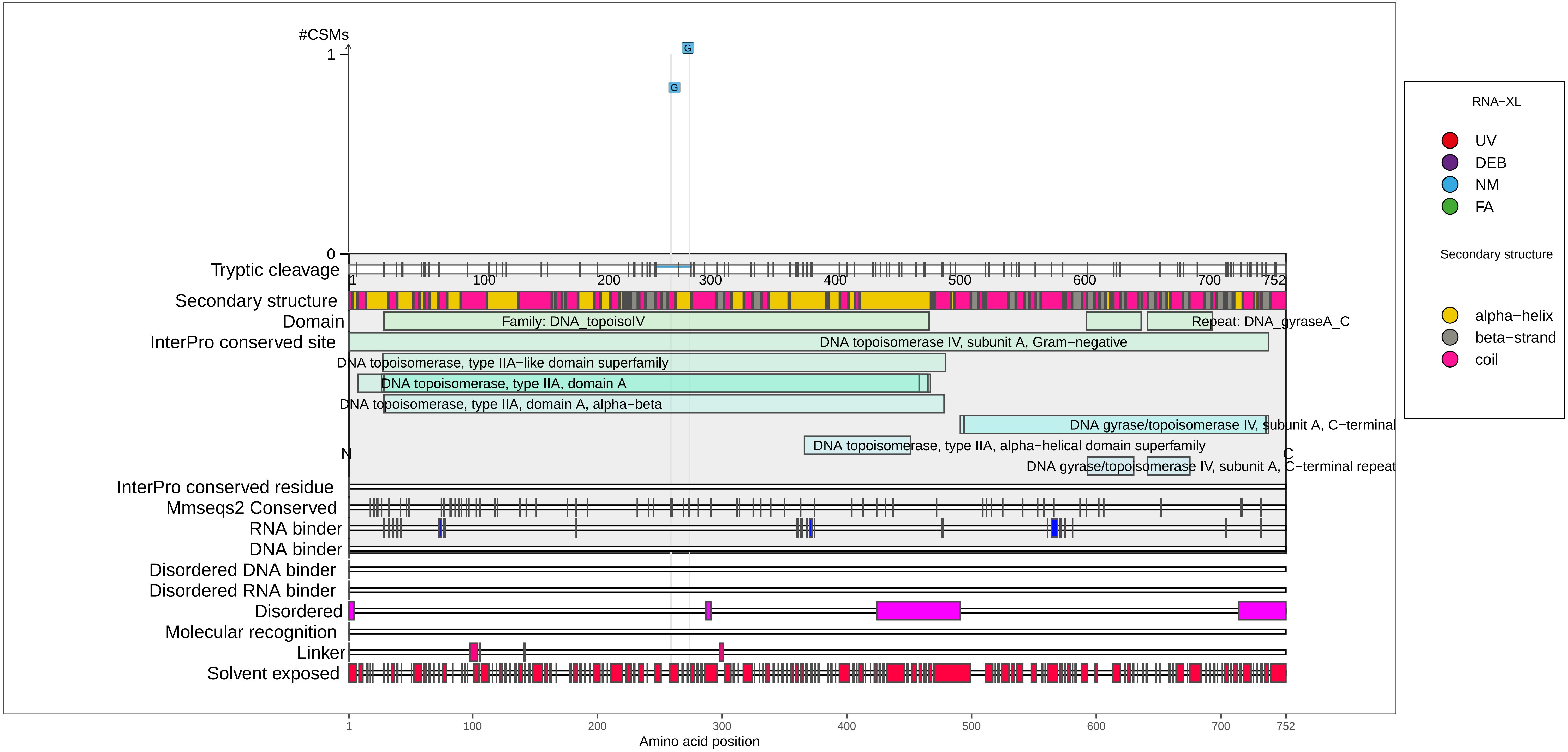

P0AFW0
RFAH_ECOLI Transcription antitermination protein RfaH

− Abundance:
tryptic [log10 Intensity]: 6.22 (Q 1)
PAXdb K12 strain [ppm]: 1.76 (Q 52)
PAXdb E.coli [ppm]: 1.12 (Q 51)

− RNA functions:
RNA biosynthetic process; RNA metabolic process; RNA polymerase binding
RNA polymerase core enzyme binding

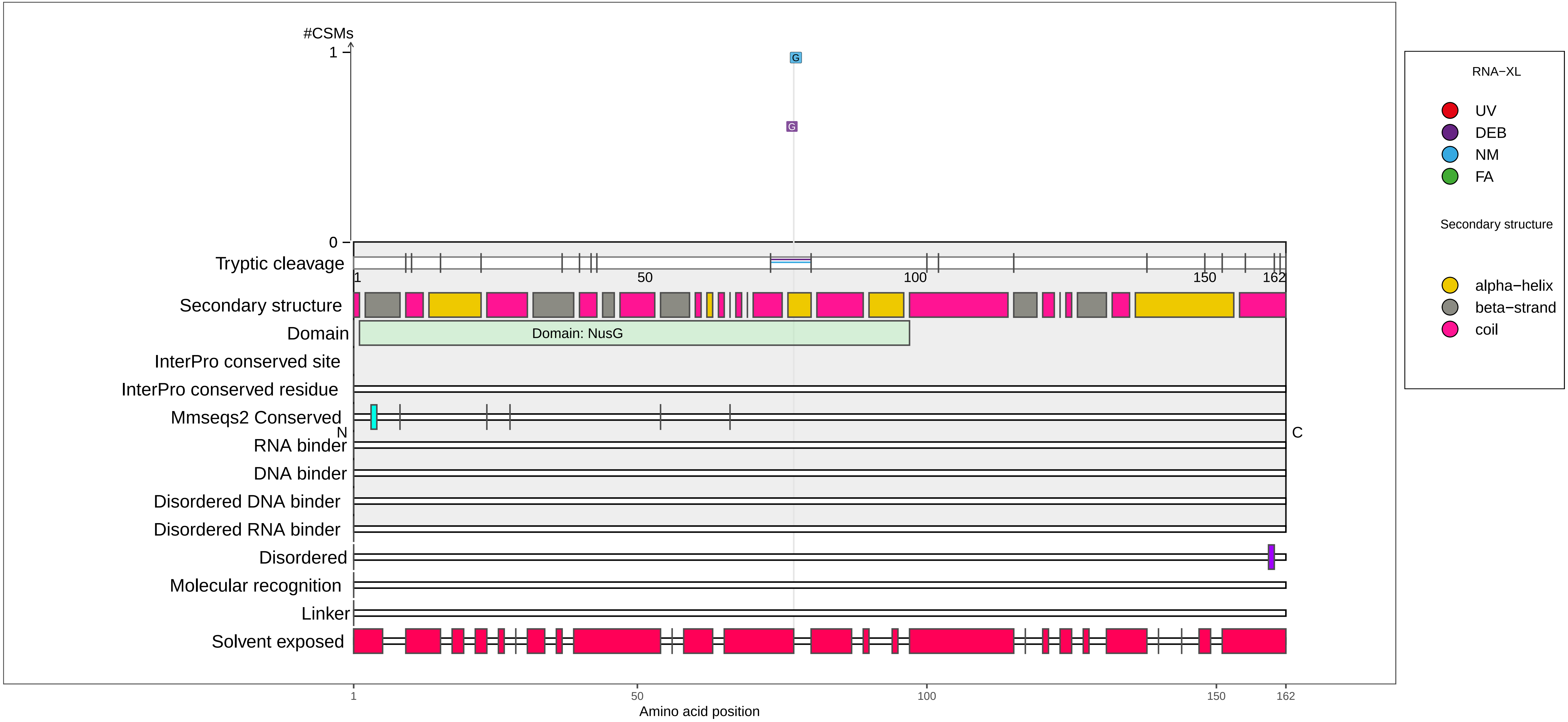

P0AFX0
HPF_ECOLI Ribosome hibernation promoting factor

− Abundance:
tryptic [log10 Intensity]: 8.88 (Q 85)
PAXdb K12 strain [ppm]: 2.36 (Q 72)
PAXdb E.coli [ppm]: 3.29 (Q 97)

− RNA functions: not annotated

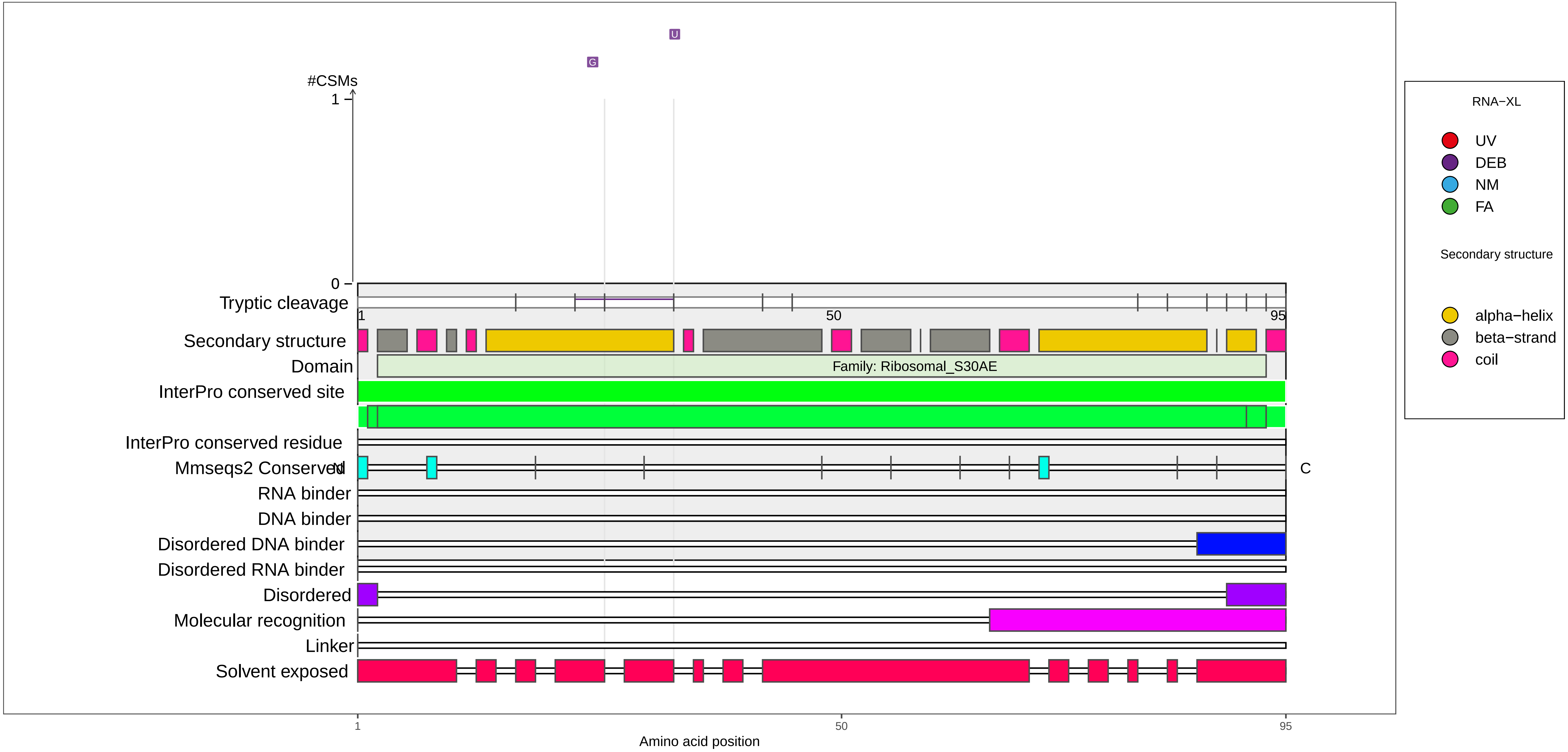

P0AG76
SBCD_ECOLI Nuclease SbcCD subunit D

− Abundance:
tryptic [log10 Intensity]: 7.24 (Q 23)
PAXdb K12 strain [ppm]: NA (Q NA)
PAXdb E.coli [ppm]: −1.01 (Q 9)

− RNA functions: not annotated

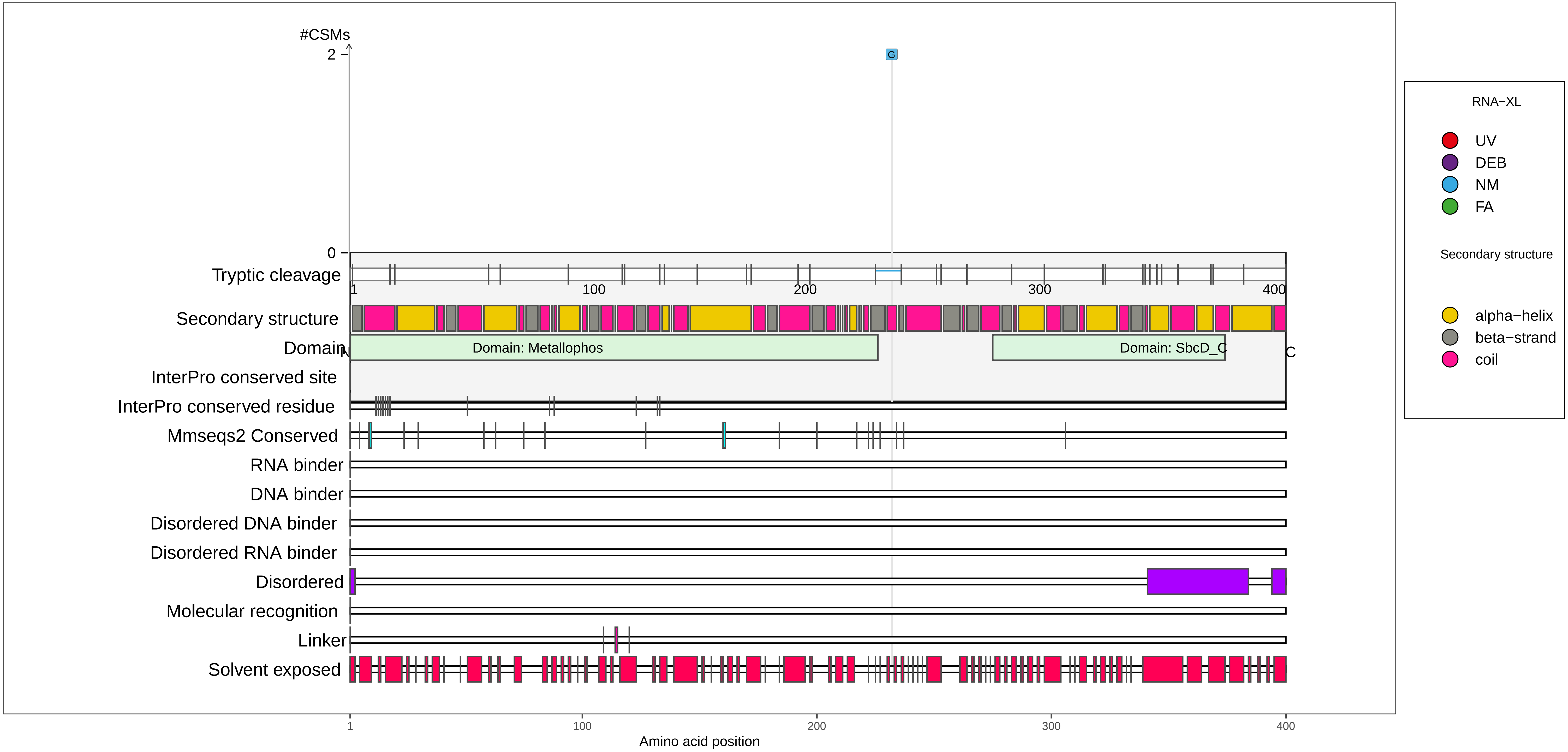

P0AGF2
CSDE_ECOLI Sulfur acceptor protein CsdE

− Abundance:
tryptic [log10 Intensity]: 7.7 (Q 44)
PAXdb K12 strain [ppm]: 1.61 (Q 45)
PAXdb E.coli [ppm]: 1.73 (Q 65)

− RNA functions:
ncRNA metabolic process; ncRNA processing; RNA metabolic process; RNA modification
RNA processing; tRNA metabolic process; tRNA modification; tRNA processing

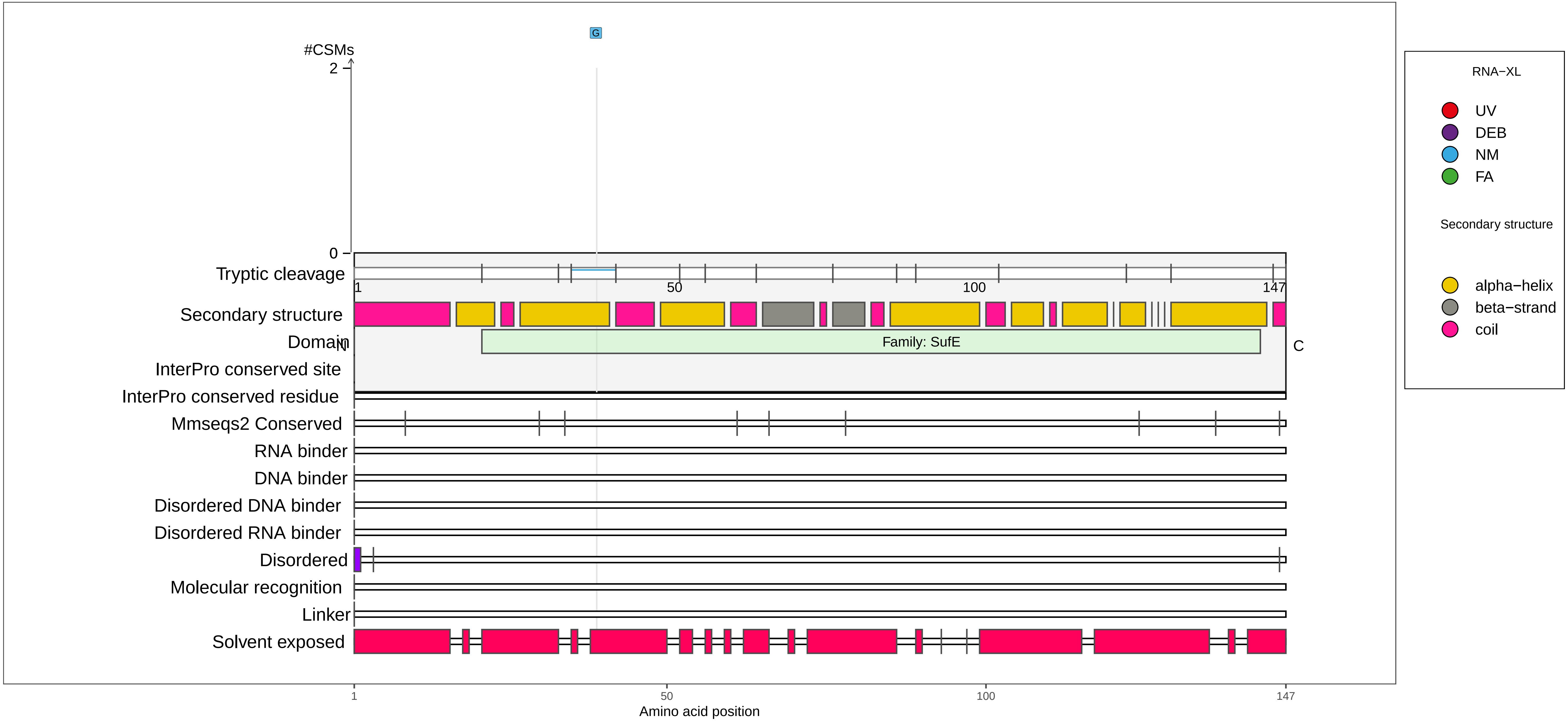

P10442
RNH2_ECOLI Ribonuclease HII

− Abundance:
tryptic [log10 Intensity]: NA (Q NA)
PAXdb K12 strain [ppm]: NA (Q NA)
PAXdb E.coli [ppm]: 1.36 (Q 56)

− RNA functions:
DNA replication, removal of RNA primer; RNA binding; RNA catabolic process
RNA metabolic process; RNA phosphodiester bond hydrolysis
RNA phosphodiester bond hydrolysis, endonucleolytic; RNA−DNA hybrid ribonuclease activity

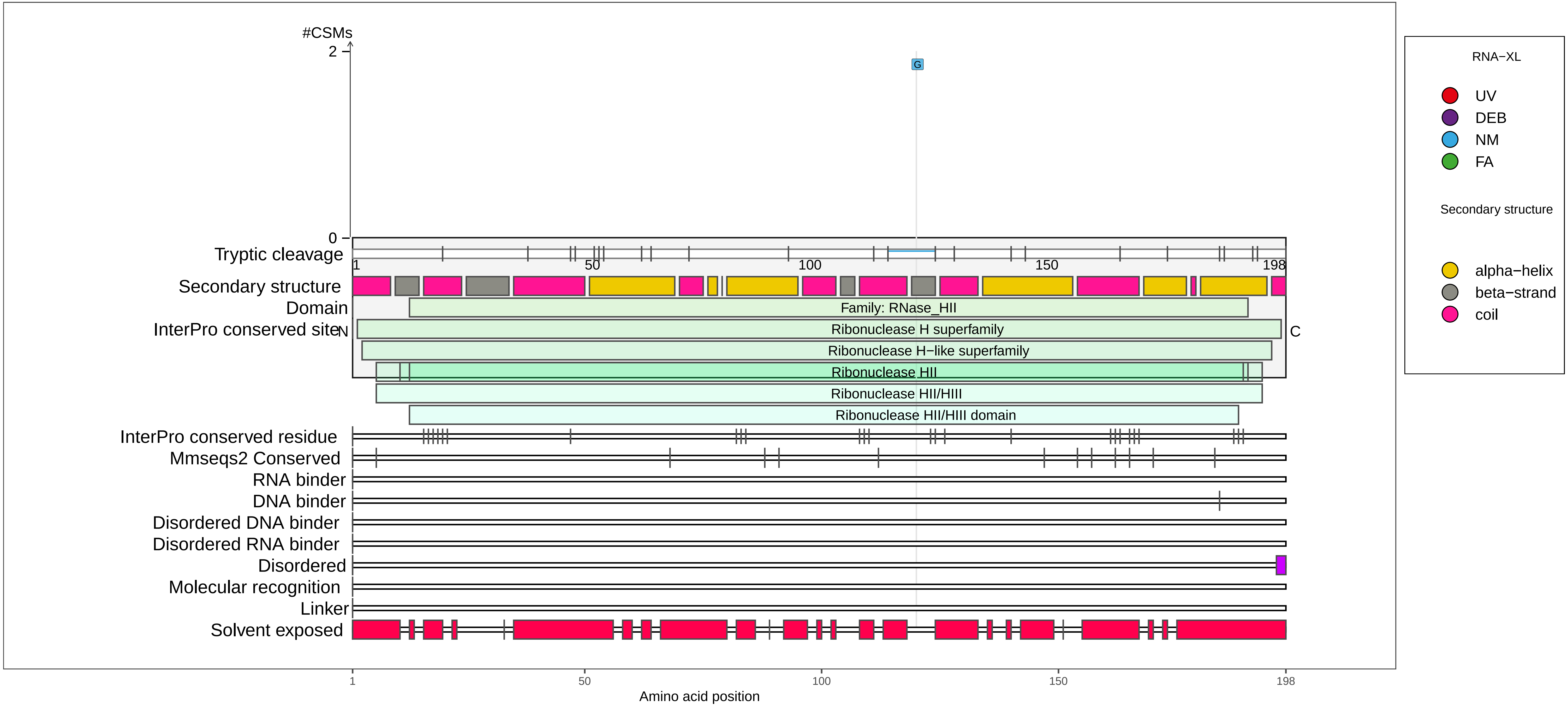

P10443
DPO3A_ECOLI DNA polymerase III subunit alpha

− Abundance:
tryptic [log10 Intensity]: NA (Q NA)
PAXdb K12 strain [ppm]: 1 (Q 11)
PAXdb E.coli [ppm]: 1.26 (Q 54)

− RNA functions: not annotated

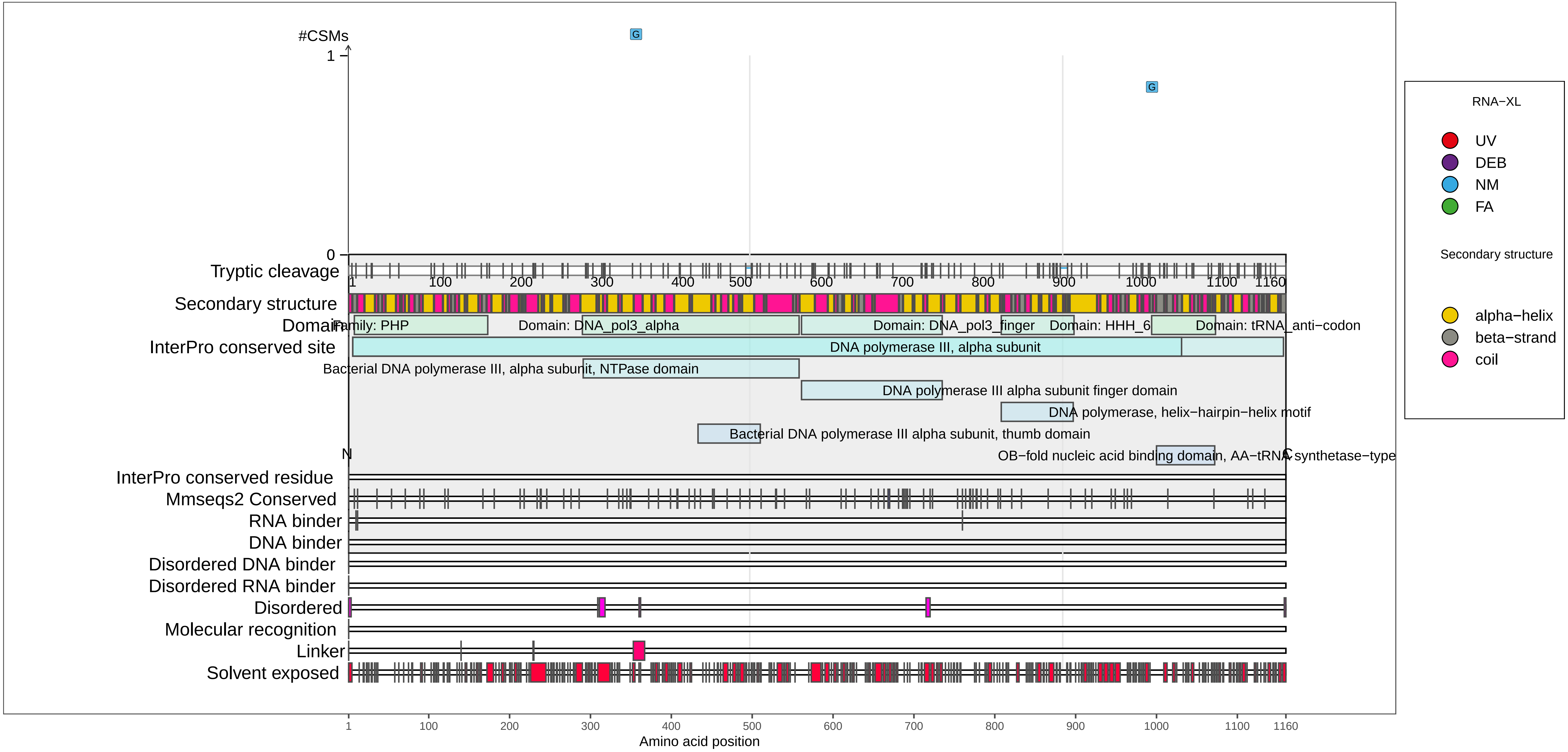

P13009
METH_ECOLI Methionine synthase

− Abundance:
tryptic [log10 Intensity]: 8.34 (Q 71)
PAXdb K12 strain [ppm]: 2.33 (Q 71)
PAXdb E.coli [ppm]: 1.81 (Q 67)

− RNA functions: not annotated

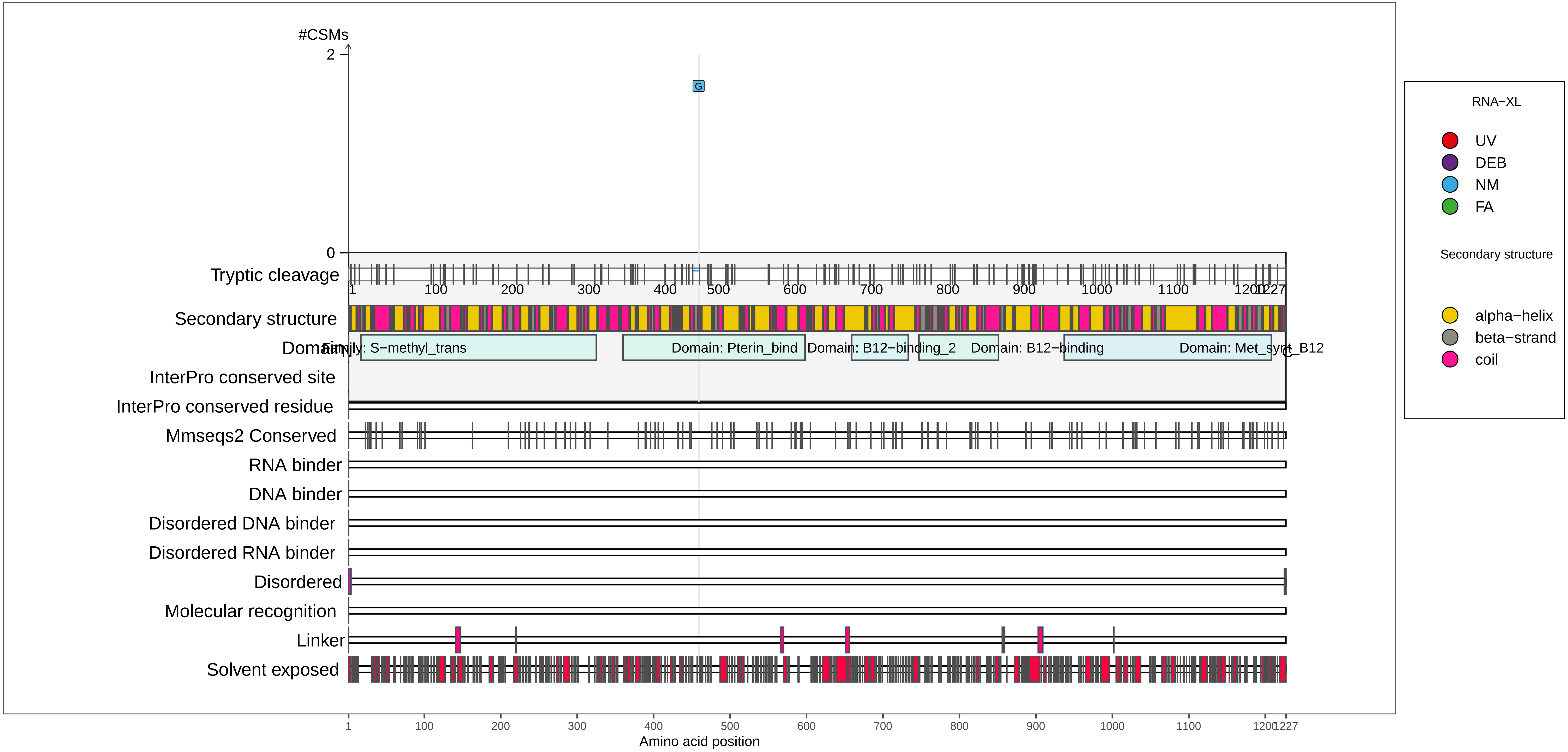

P15770
AROE_ECOLI Shikimate dehydrogenase (NADP(+))

− Abundance:
tryptic [log10 Intensity]: 7.58 (Q 39)
PAXdb K12 strain [ppm]: NA (Q NA)
PAXdb E.coli [ppm]: 1.86 (Q 69)

− RNA functions: not annotated

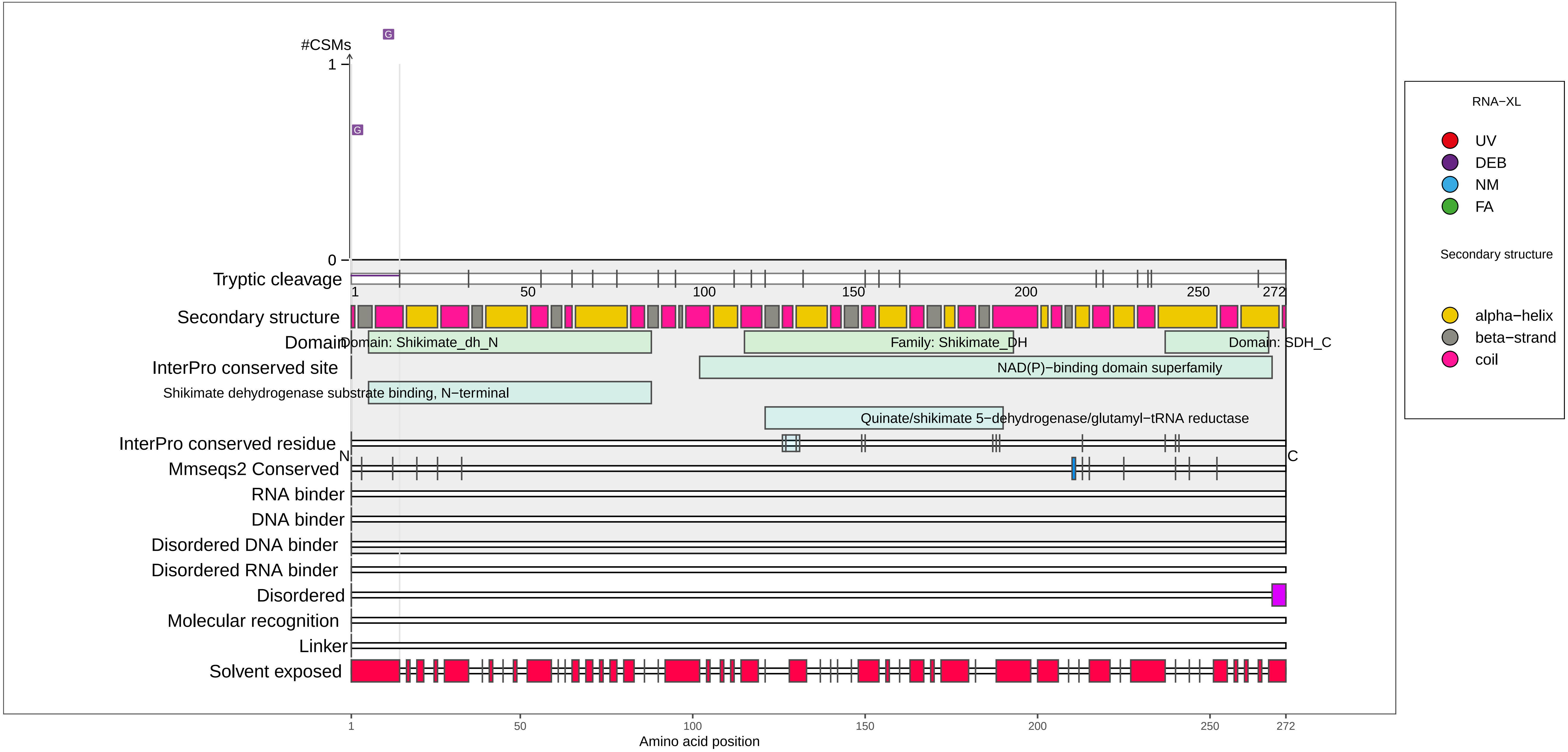

P16703
CYSM_ECOLI Cysteine synthase B

− Abundance:
tryptic [log10 Intensity]: 7.86 (Q 52)
PAXdb K12 strain [ppm]: NA (Q NA)
PAXdb E.coli [ppm]: 2.1 (Q 75)

− RNA functions: not annotated

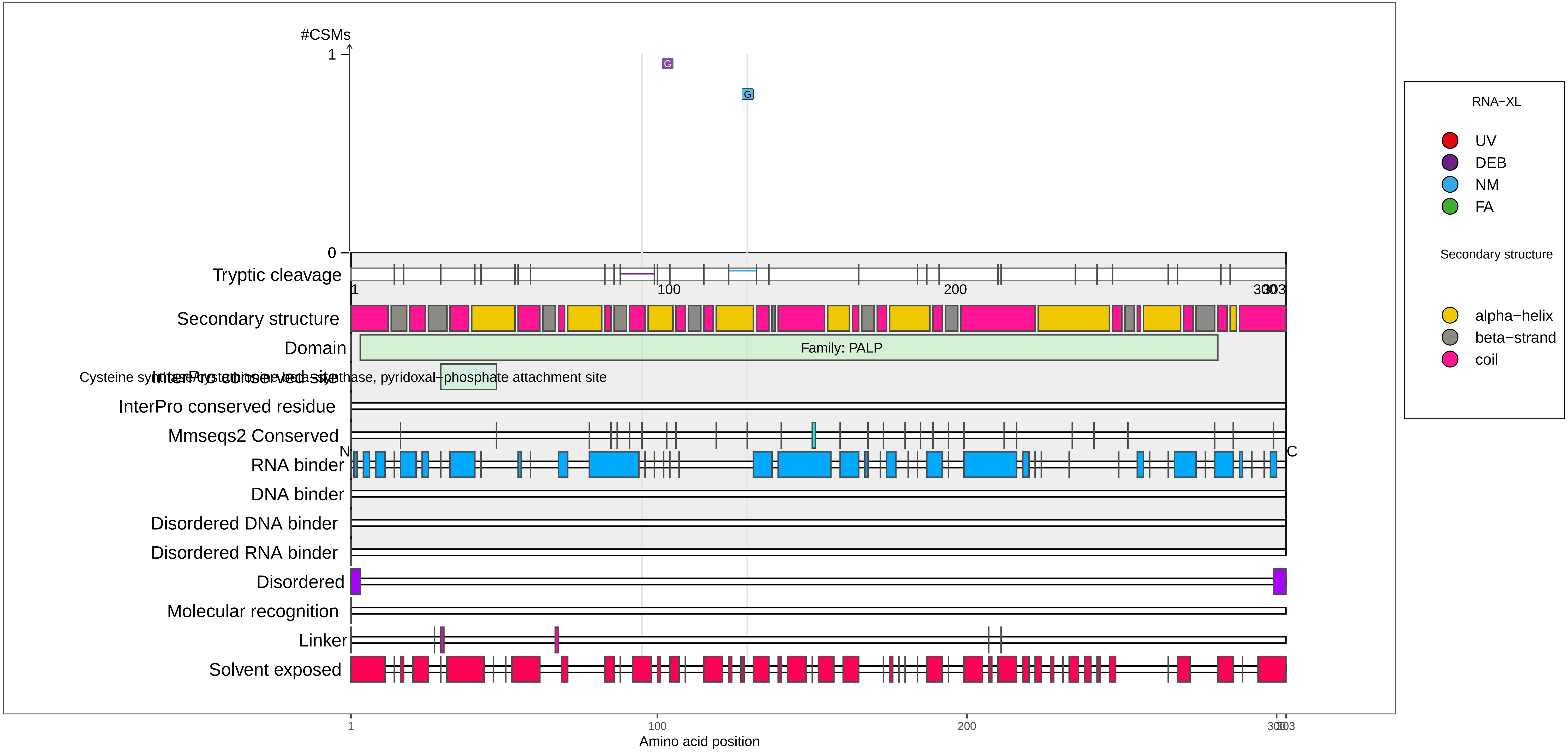

P20099
BISC_ECOLI Biotin sulfoxide reductase

− Abundance:
tryptic [log10 Intensity]: 6.8 (Q 7)
PAXdb K12 strain [ppm]: 1 (Q 11)
PAXdb E.coli [ppm]: 1.2 (Q 52)

− RNA functions: not annotated

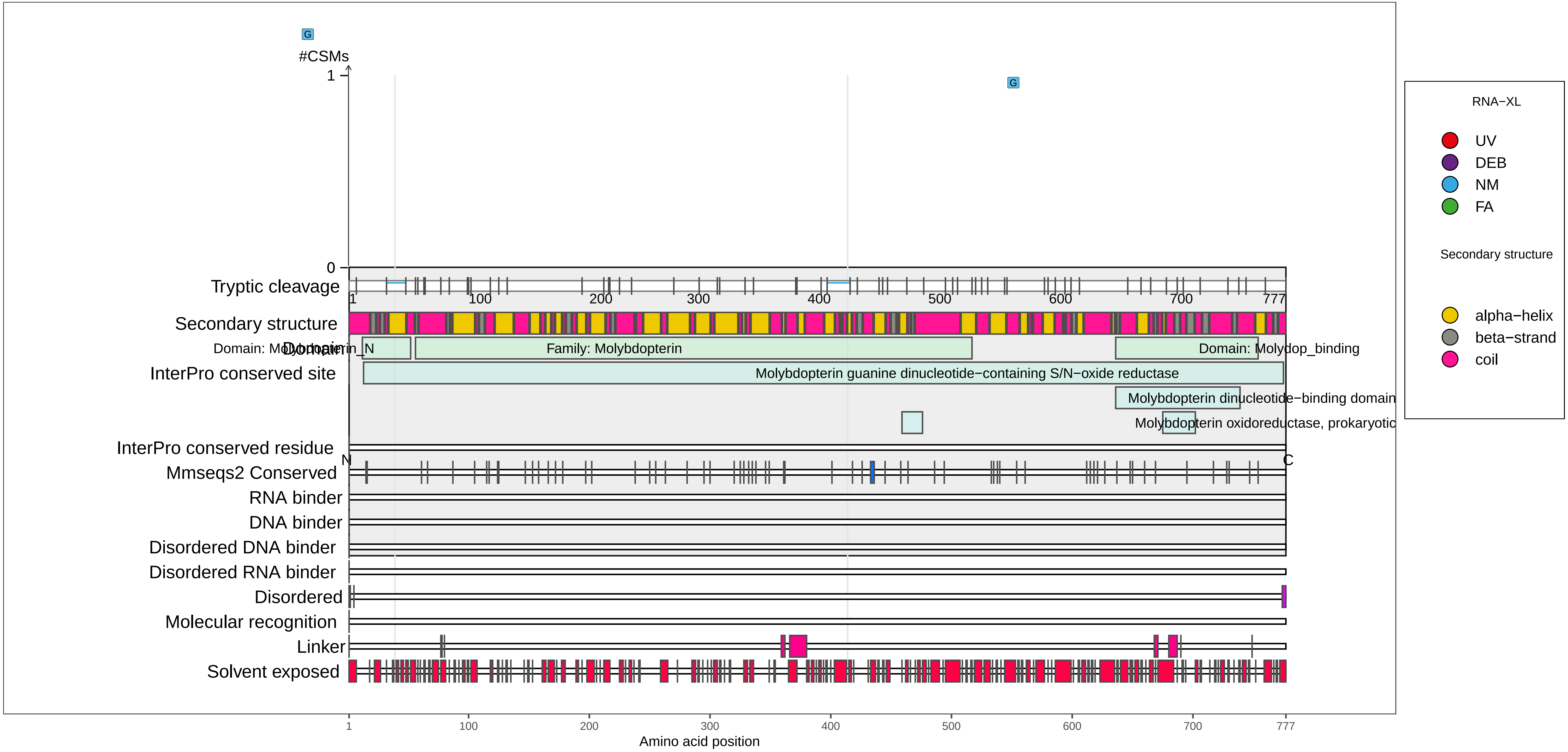

P22525
YCBB_ECOLI Probable L,D−transpeptidase YcbB

− Abundance:
tryptic [log10 Intensity]: 7.12 (Q 17)
PAXdb K12 strain [ppm]: 0.58 (Q 1)
PAXdb E.coli [ppm]: 1.09 (Q 50)

− RNA functions: not annotated

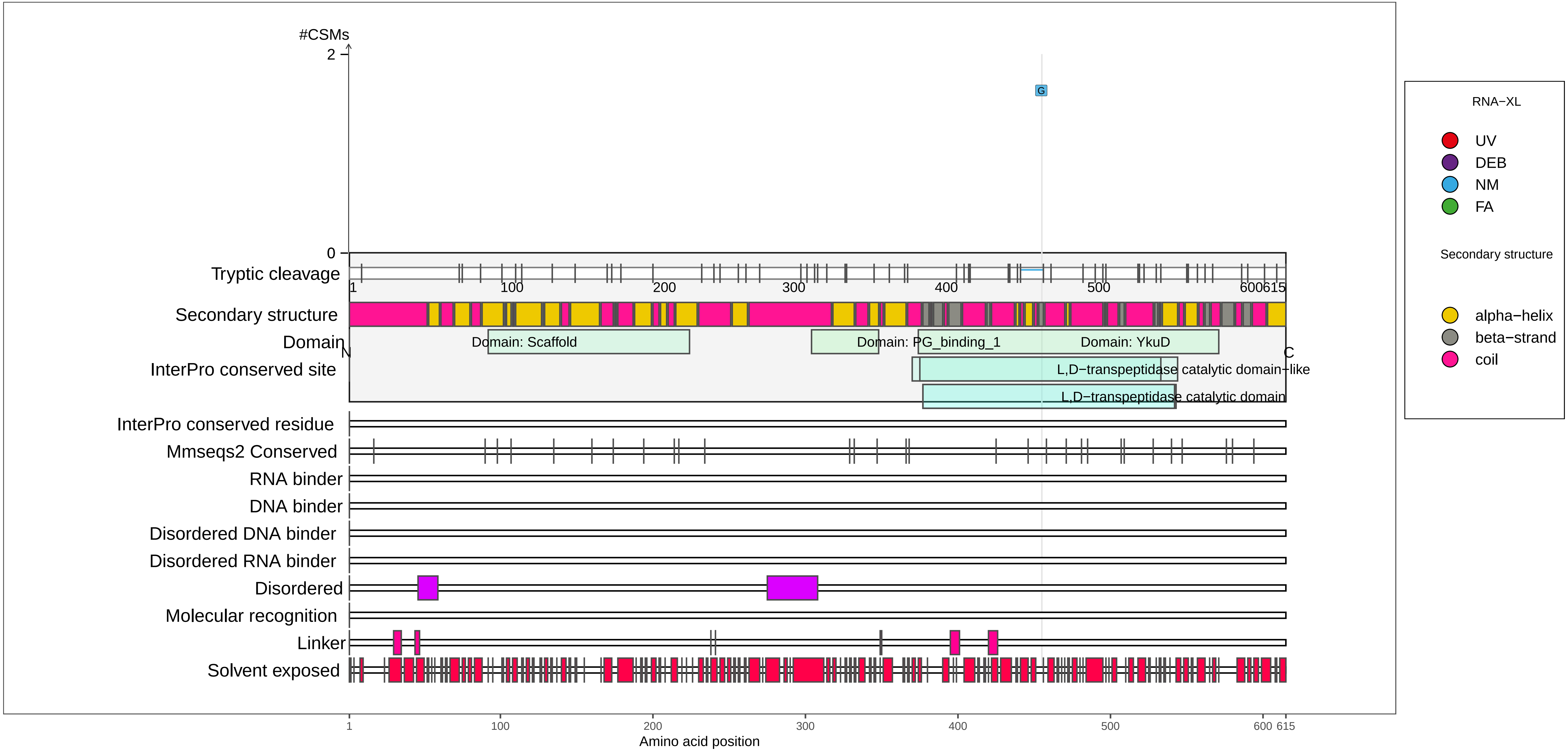

P23930
LNT_ECOLI Apolipoprotein N−acyltransferase
− Abundance:
tryptic [log10 Intensity]: 8.12 (Q 62)
PAXdb K12 strain [ppm]: NA (Q NA)
PAXdb E.coli [ppm]: 0.6 (Q 38)

− RNA functions: not annotated

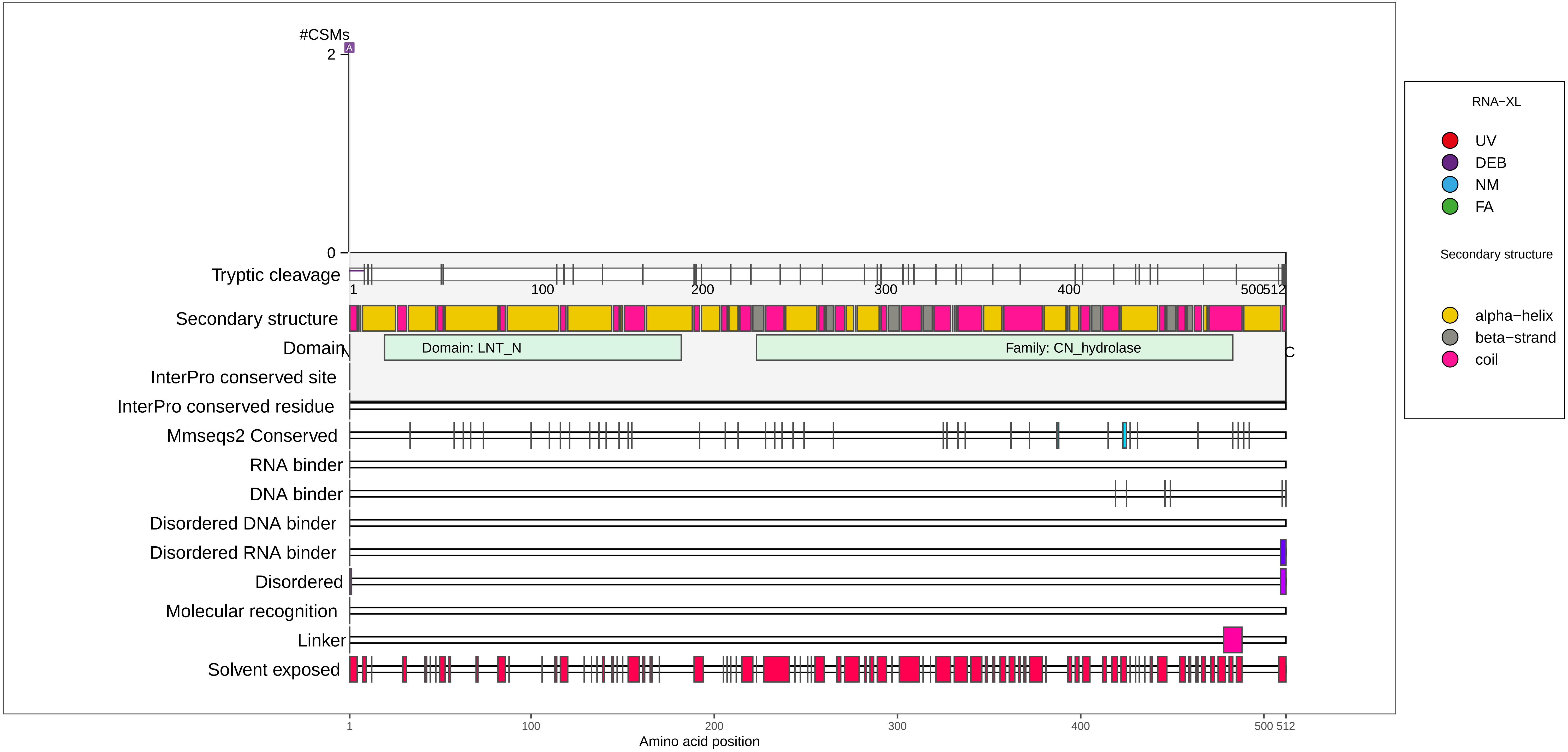

P25746
HFLD_ECOLI High frequency lysogenization protein HflD

− Abundance:
tryptic [log10 Intensity]: 6.84 (Q 8)
PAXdb K12 strain [ppm]: NA (Q NA)
PAXdb E.coli [ppm]: 1.63 (Q 63)

− RNA functions: not annotated

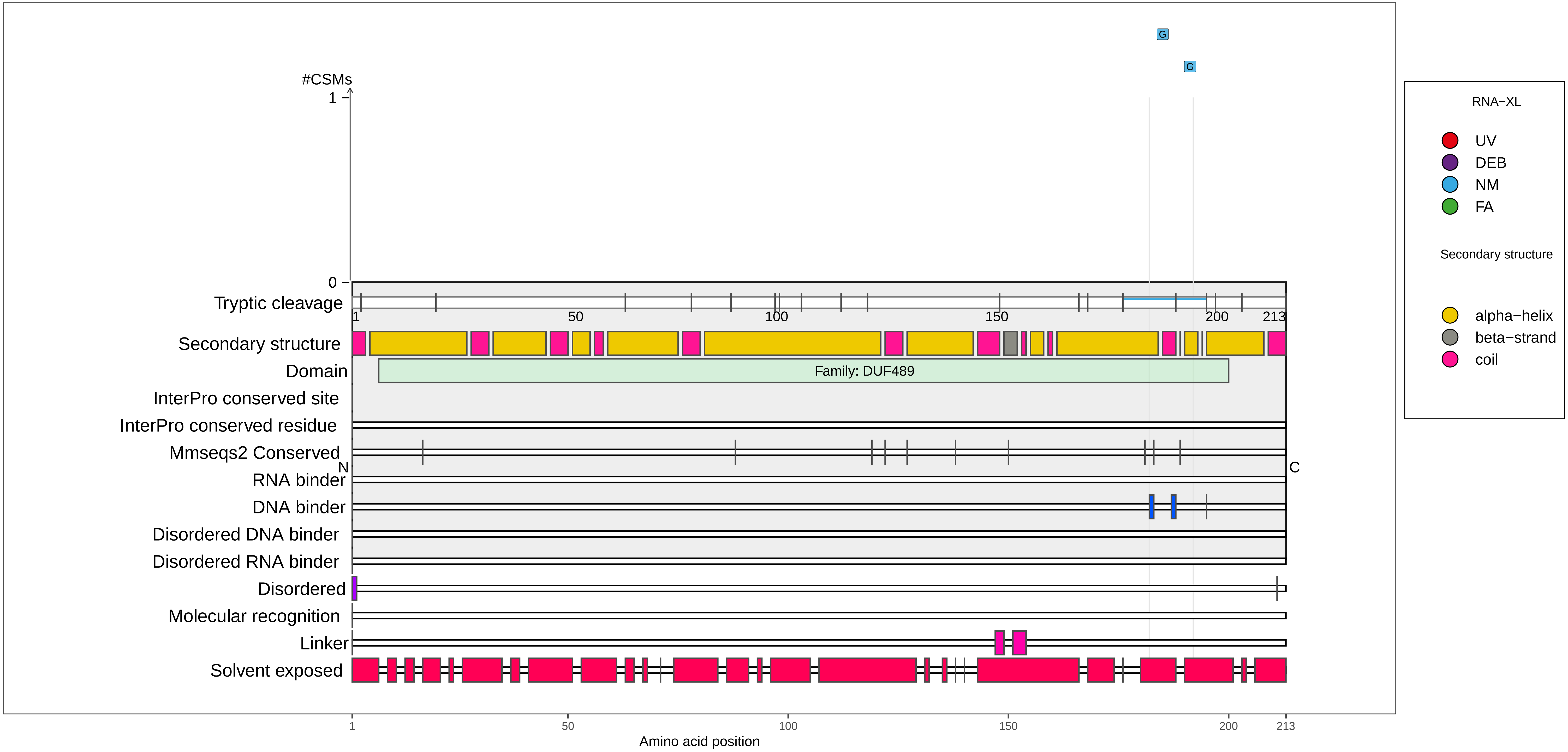

P25894
LOIP_ECOLI Metalloprotease LoiP

− Abundance:
tryptic [log10 Intensity]: 7.53 (Q 36)
PAXdb K12 strain [ppm]: 2.18 (Q 66)
PAXdb E.coli [ppm]: 1.75 (Q 66)

− RNA functions: not annotated

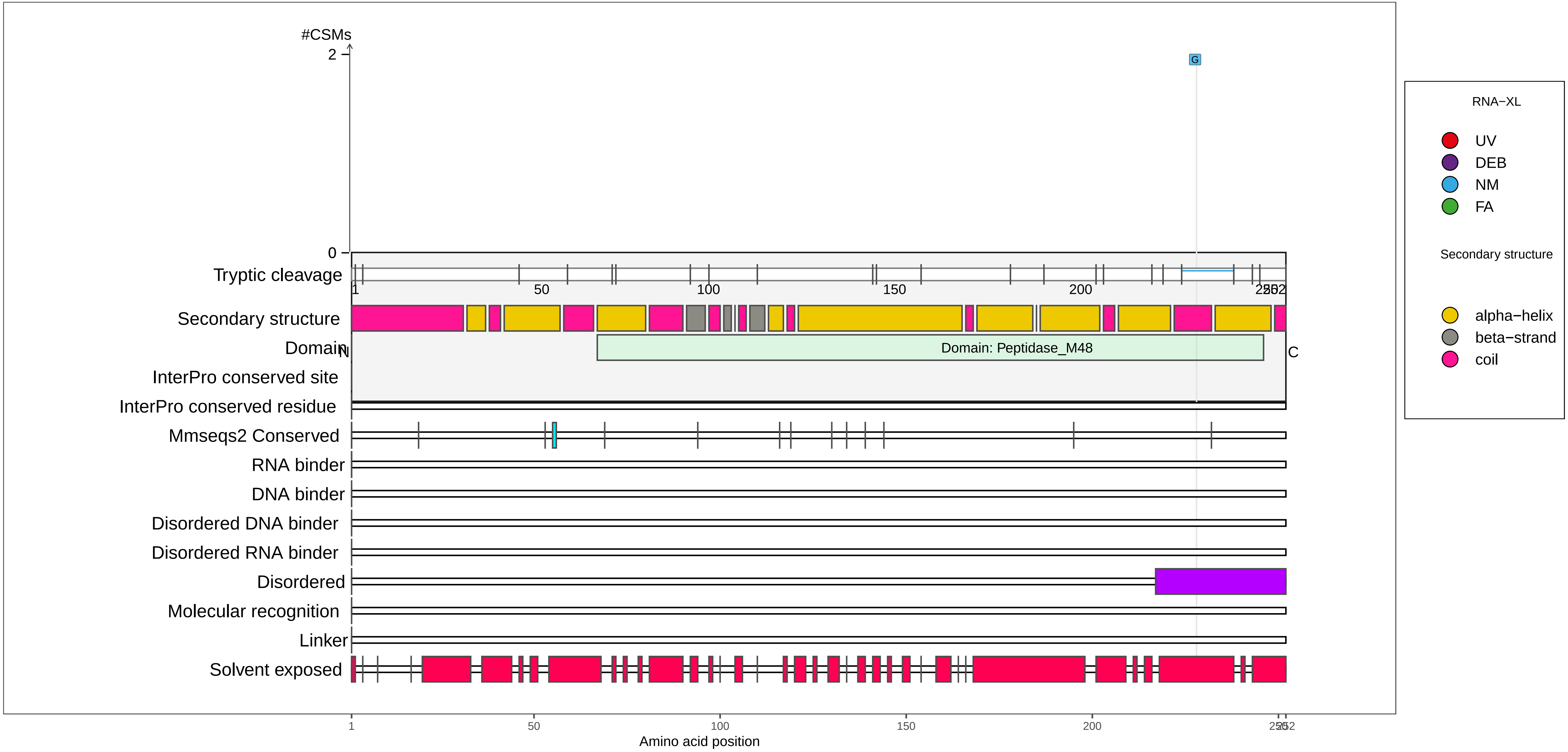

P27298
OPDA_ECOLI Oligopeptidase A

− Abundance:
tryptic [log10 Intensity]: 9.08 (Q 90)
PAXdb K12 strain [ppm]: 2.89 (Q 88)
PAXdb E.coli [ppm]: 2.52 (Q 86)

− RNA functions: not annotated

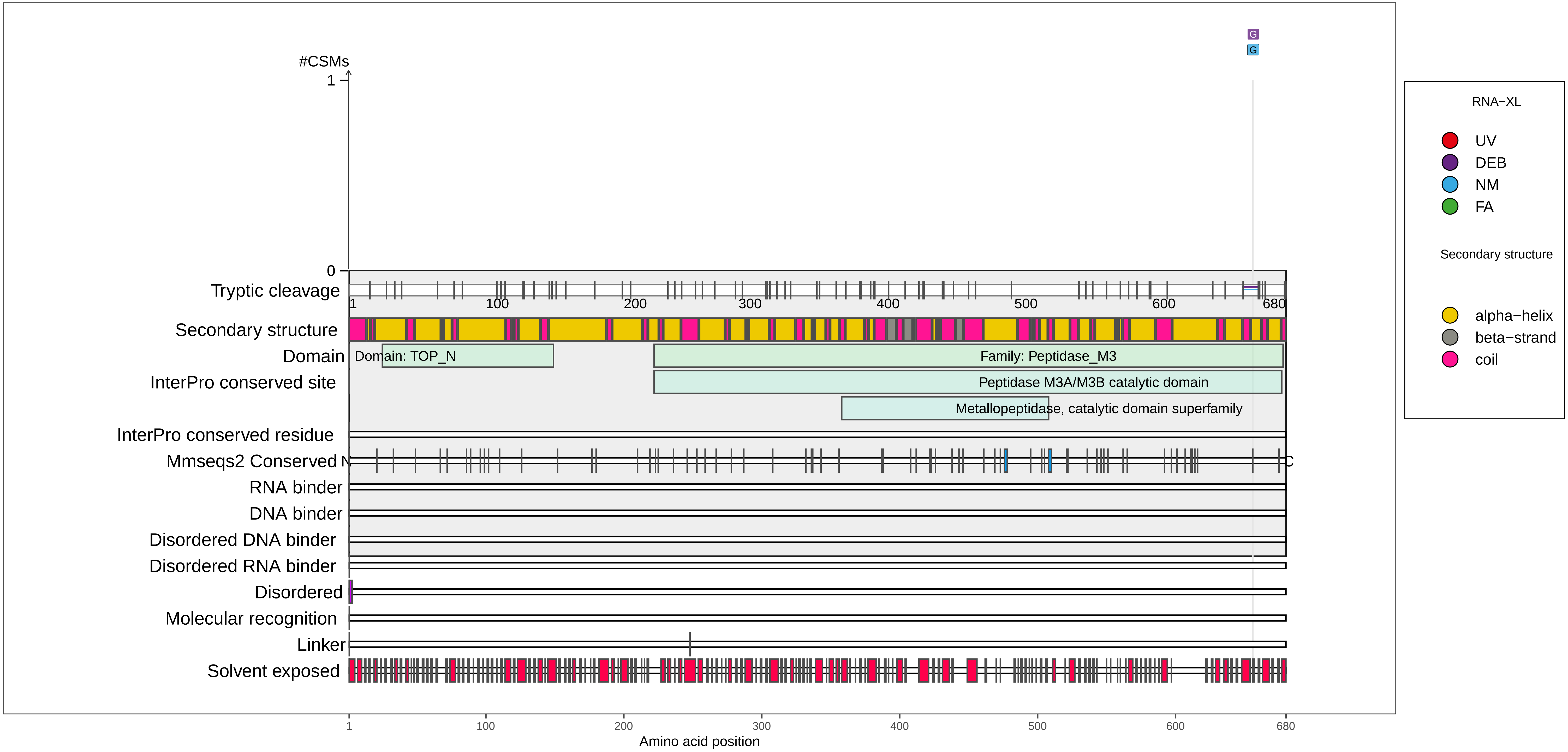

P27830
RMLB2_ECOLI dTDP−glucose 4,6−dehydratase 2

− Abundance:
tryptic [log10 Intensity]: 8.72 (Q 82)
PAXdb K12 strain [ppm]: 0.6 (Q 1)
PAXdb E.coli [ppm]: 0.73 (Q 41)

− RNA functions: not annotated

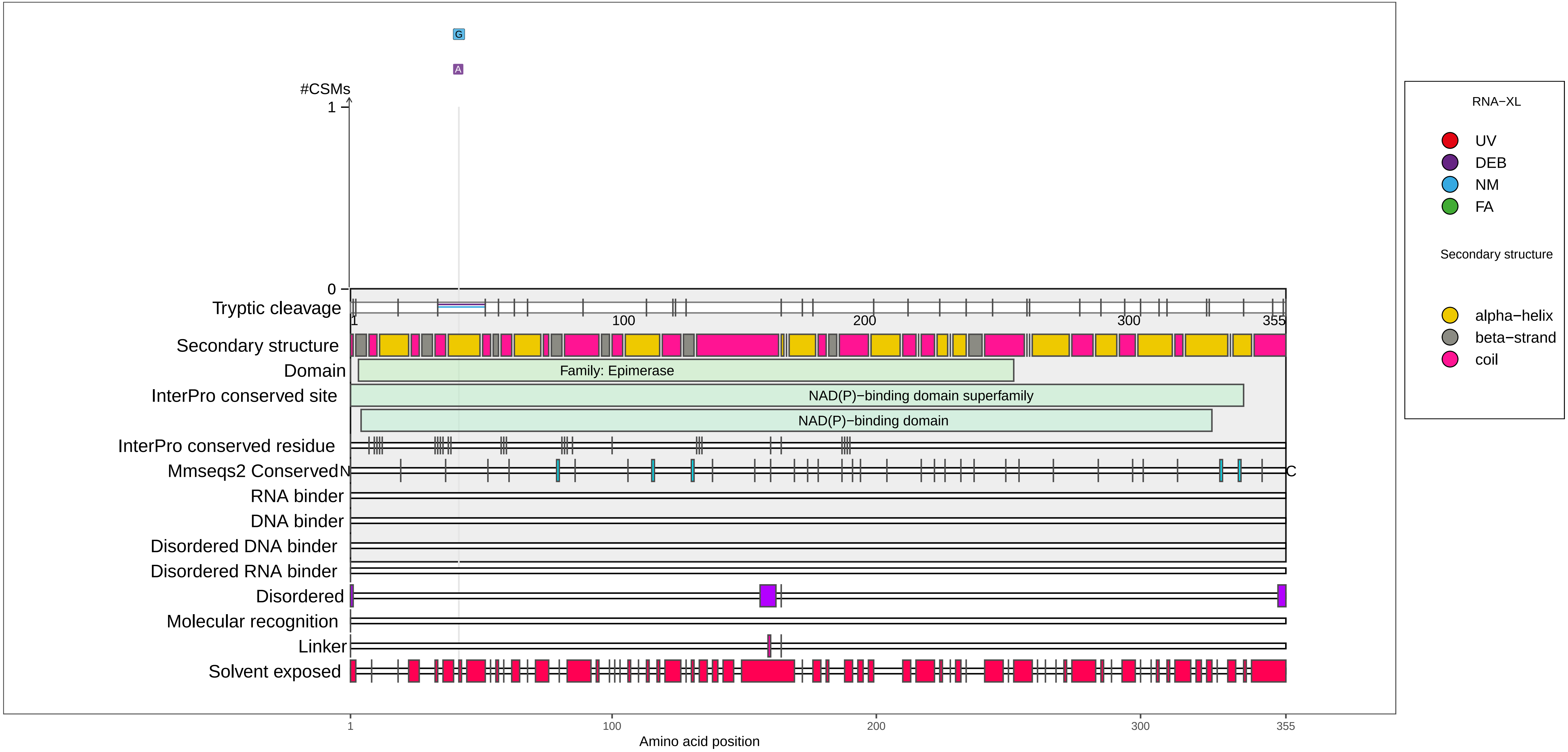

P30176
RIBX_ECOLI N−glycosidase YbiA

− Abundance:
tryptic [log10 Intensity]: NA (Q NA)
PAXdb K12 strain [ppm]: 1.27 (Q 24)
PAXdb E.coli [ppm]: −0.13 (Q 22)

− RNA functions: not annotated

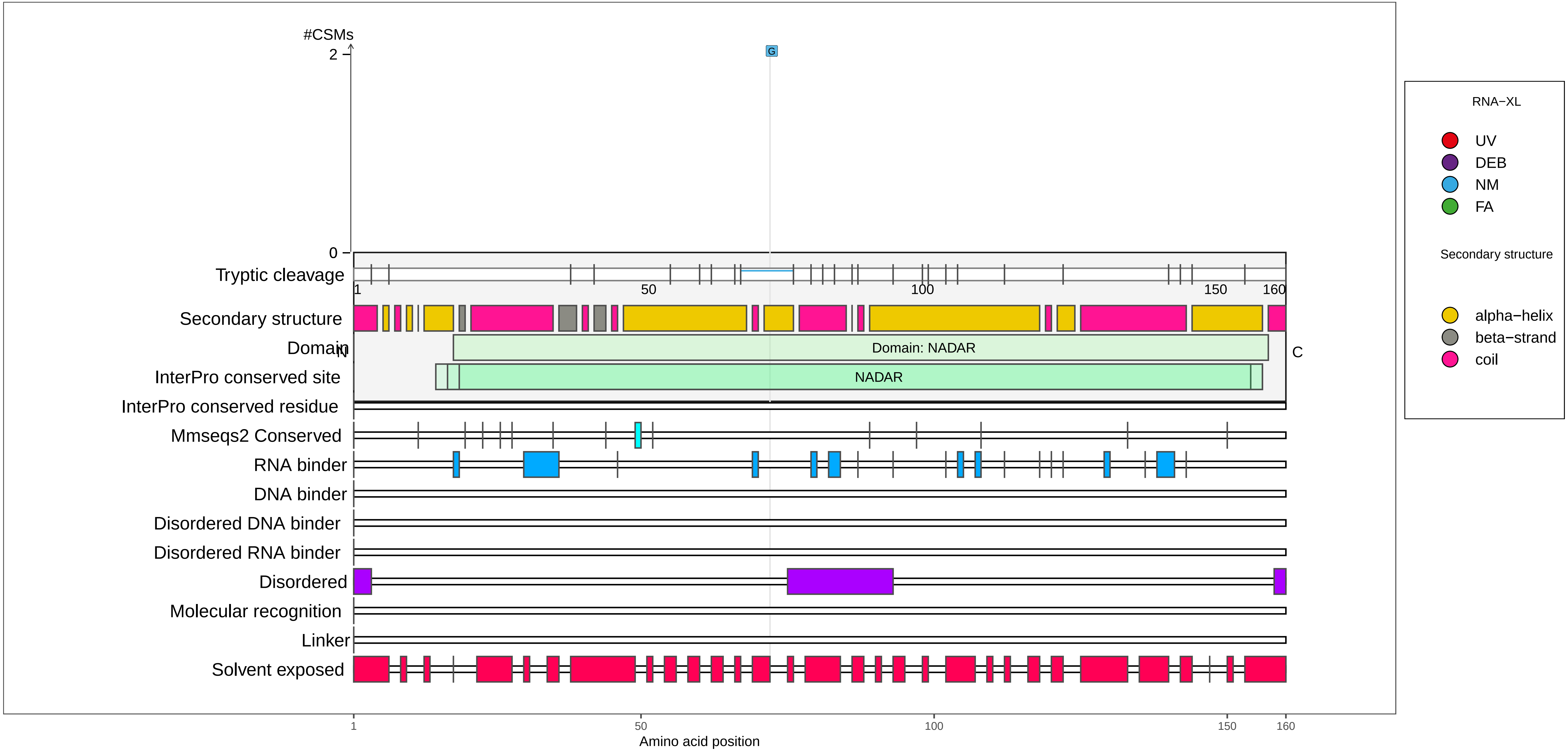

P30843
BASR_ECOLI Transcriptional regulatory protein BasR

− Abundance:
tryptic [log10 Intensity]: 8.74 (Q 82)
PAXdb K12 strain [ppm]: 2.18 (Q 66)
PAXdb E.coli [ppm]: 2.25 (Q 79)

− RNA functions: not annotated

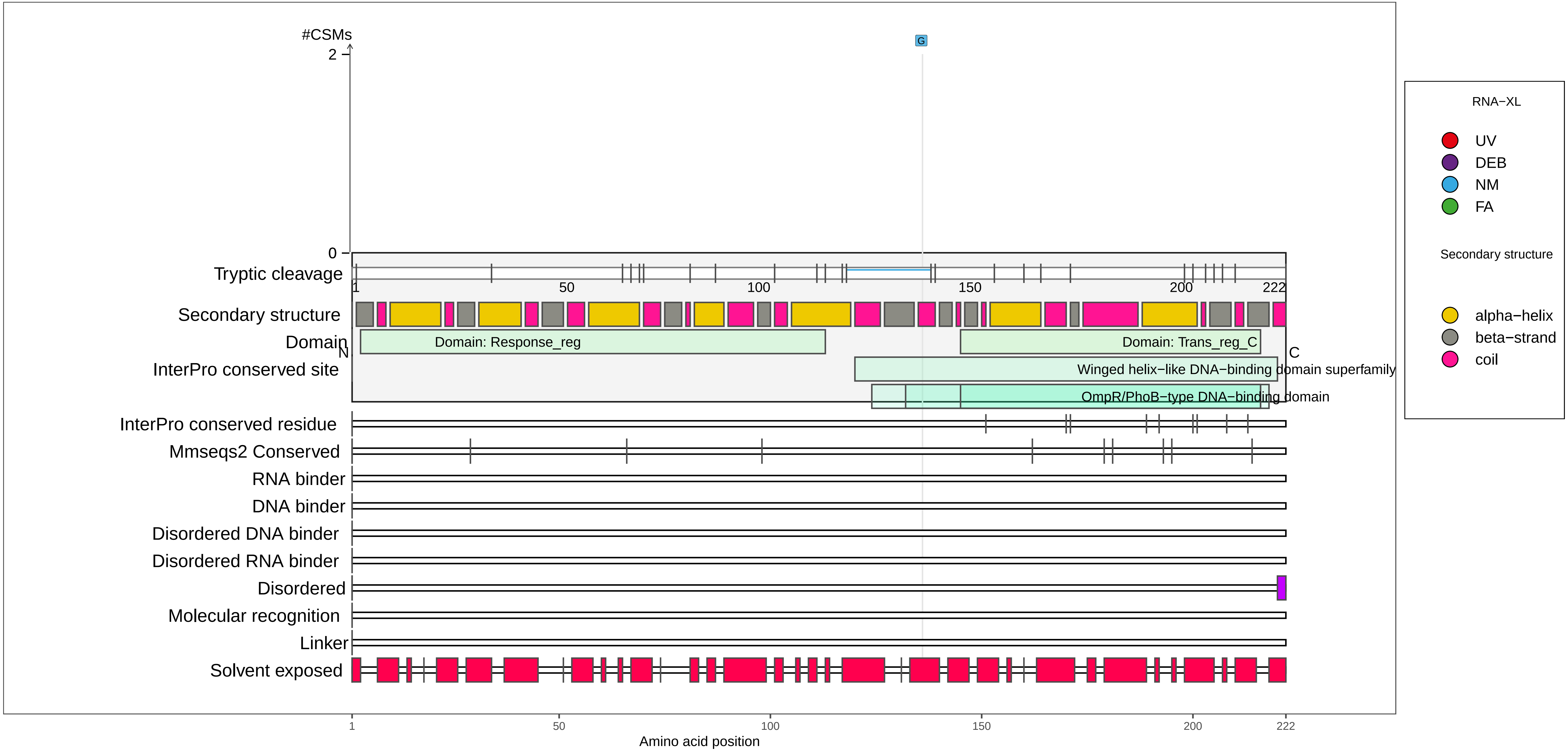

P30860
ARTJ_ECOLI ABC transporter arginine−binding protein 1

− Abundance:
tryptic [log10 Intensity]: 8.21 (Q 65)
PAXdb K12 strain [ppm]: 1.99 (Q 60)
PAXdb E.coli [ppm]: 2.67 (Q 89)

− RNA functions: not annotated

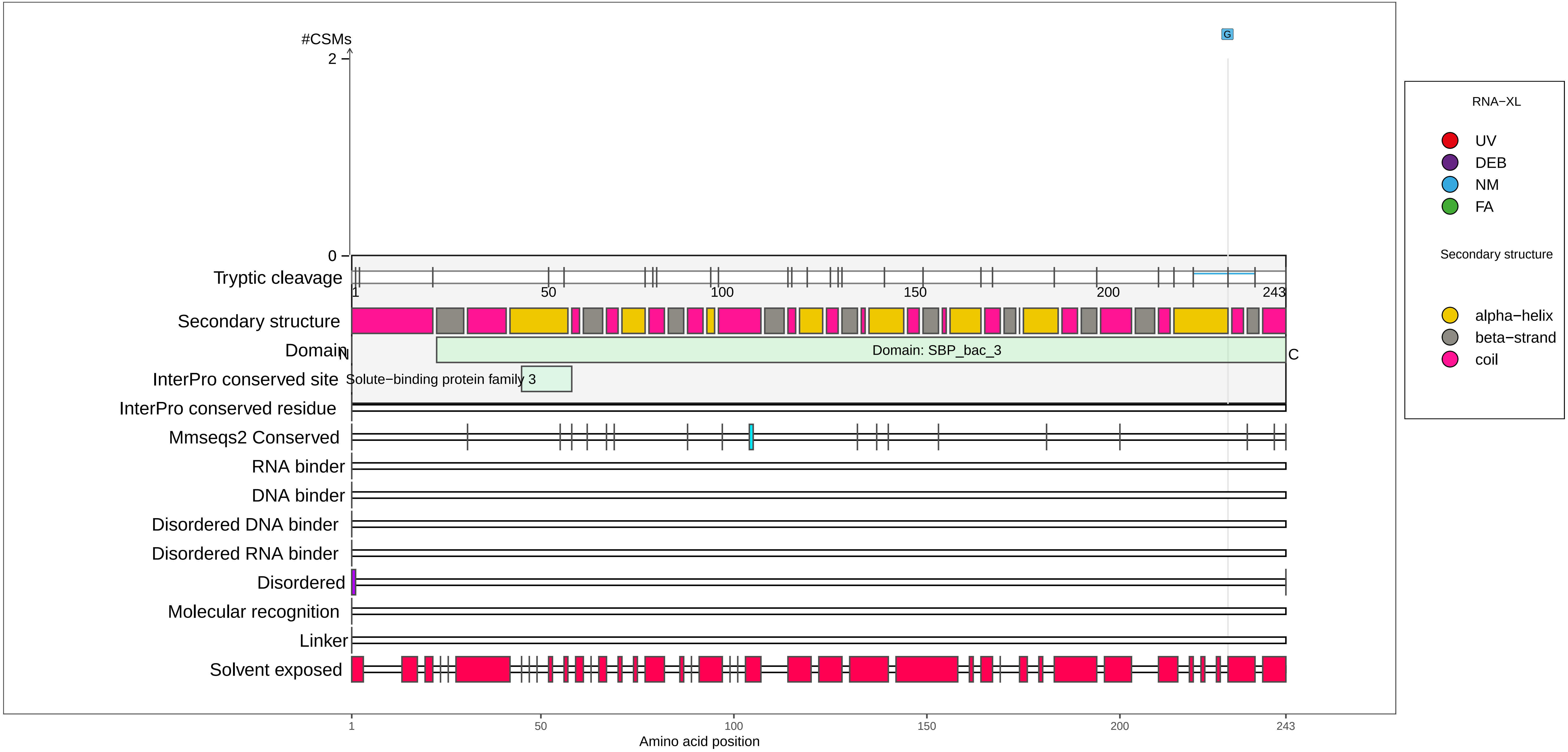

P31433
YICH_ECOLI AsmA family protein YicH

− Abundance:
tryptic [log10 Intensity]: 8.05 (Q 60)
PAXdb K12 strain [ppm]: NA (Q NA)
PAXdb E.coli [ppm]: 1.37 (Q 56)

− RNA functions: not annotated

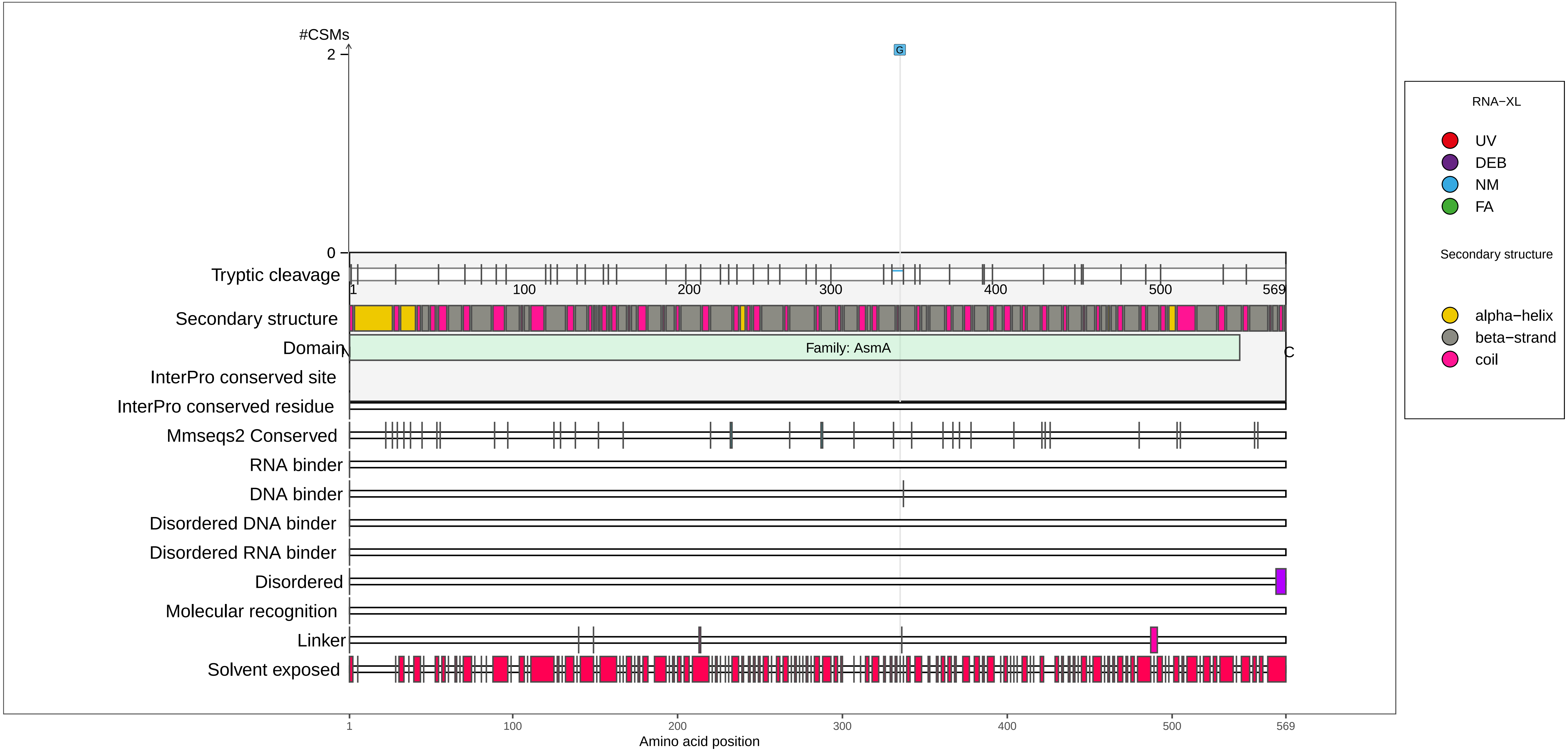

P31663
PANC_ECOLI Pantothenate synthetase

− Abundance:
tryptic [log10 Intensity]: 8.67 (Q 81)
PAXdb K12 strain [ppm]: 2.4 (Q 74)
PAXdb E.coli [ppm]: 2.78 (Q 91)

− RNA functions: not annotated

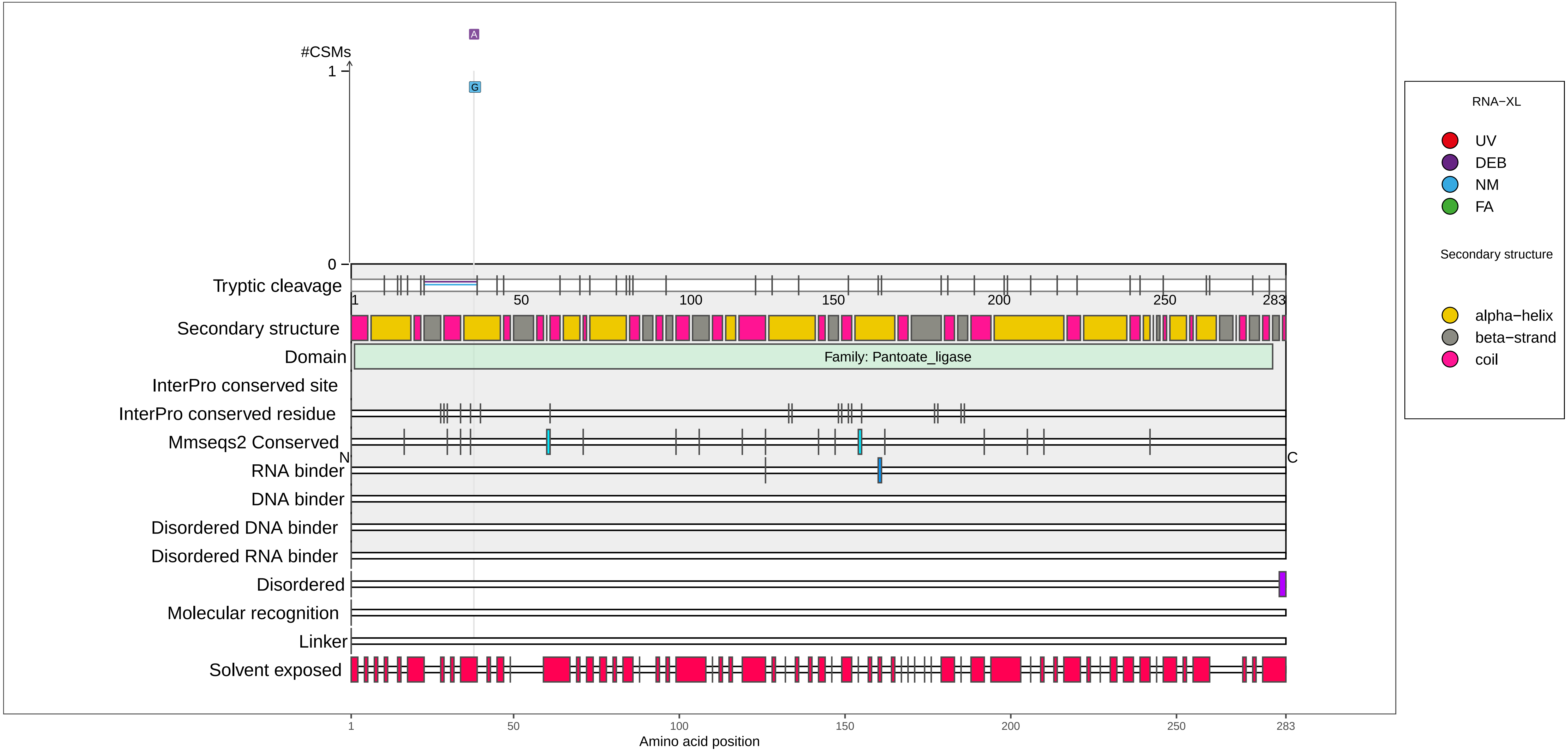

P31678
OTSB_ECOLI Trehalose−6−phosphate phosphatase
− Abundance:
tryptic [log10 Intensity]: NA (Q NA)
PAXdb K12 strain [ppm]: NA (Q NA)
PAXdb E.coli [ppm]: 1.5 (Q 60)

− RNA functions: not annotated

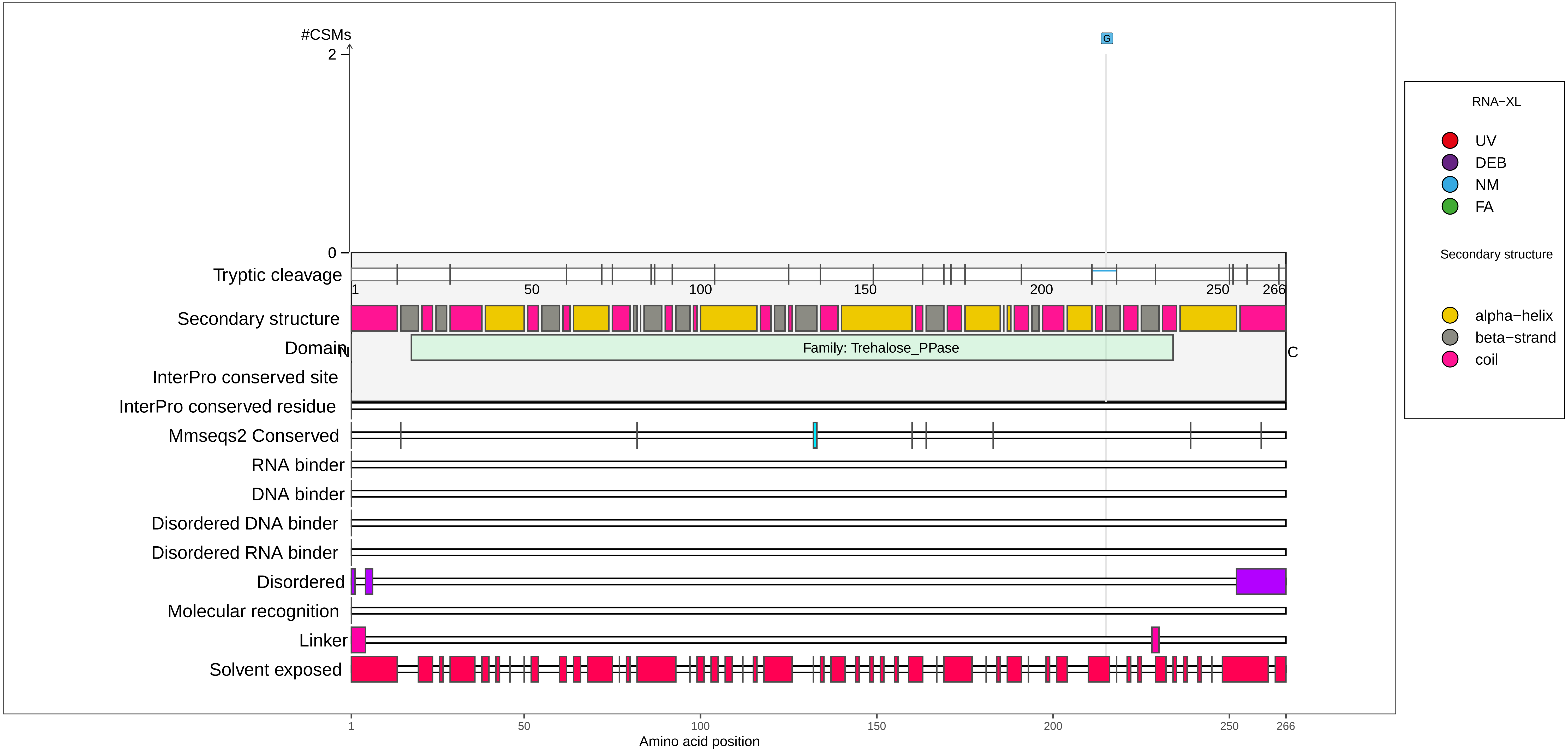

P33225
TORA_ECOLI Trimethylamine−N−oxide reductase 1

− Abundance:
tryptic [log10 Intensity]: NA (Q NA)
PAXdb K12 strain [ppm]: 0.88 (Q 7)
PAXdb E.coli [ppm]: 0.25 (Q 30)

− RNA functions: not annotated

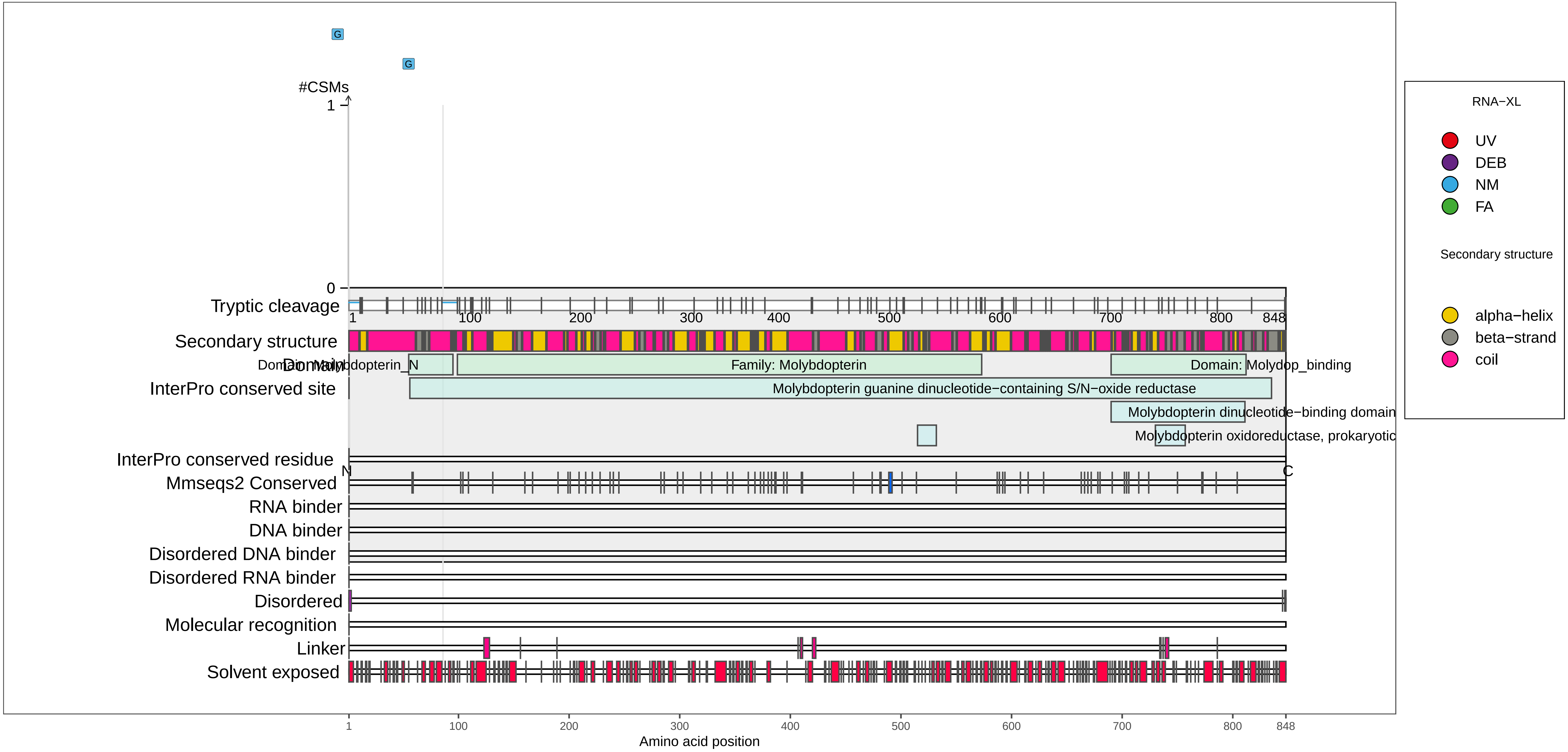

P33226
TORC_ECOLI Cytochrome c−type protein TorC
− Abundance:
tryptic [log10 Intensity]: NA (Q NA)
PAXdb K12 strain [ppm]: 1.74 (Q 51)
PAXdb E.coli [ppm]: −0.57 (Q 15)

− RNA functions: not annotated

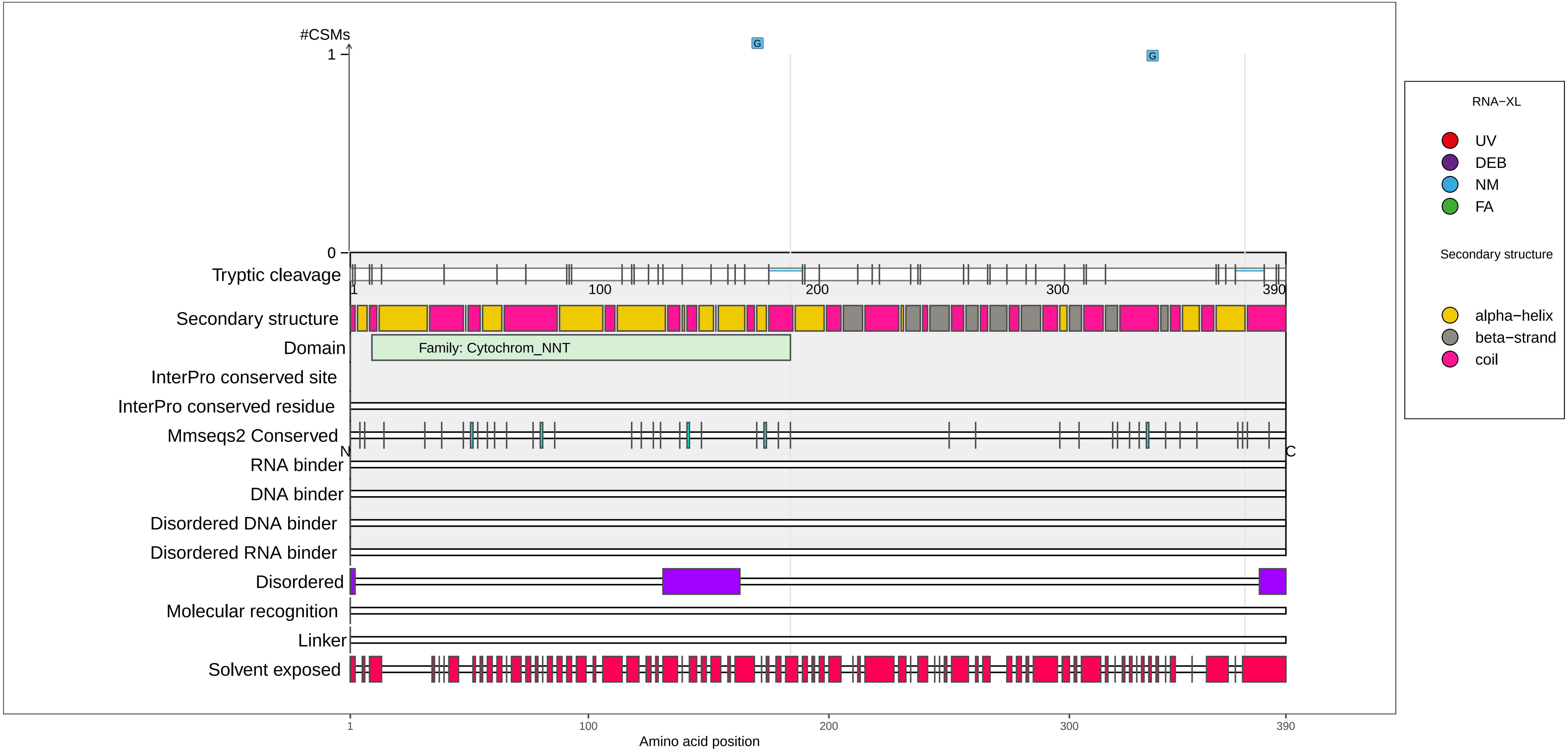

P33363
BGLX_ECOLI Periplasmic beta−glucosidase

− Abundance:
tryptic [log10 Intensity]: 8.21 (Q 66)
PAXdb K12 strain [ppm]: 1.73 (Q 50)
PAXdb E.coli [ppm]: 1.71 (Q 64)

− RNA functions: not annotated

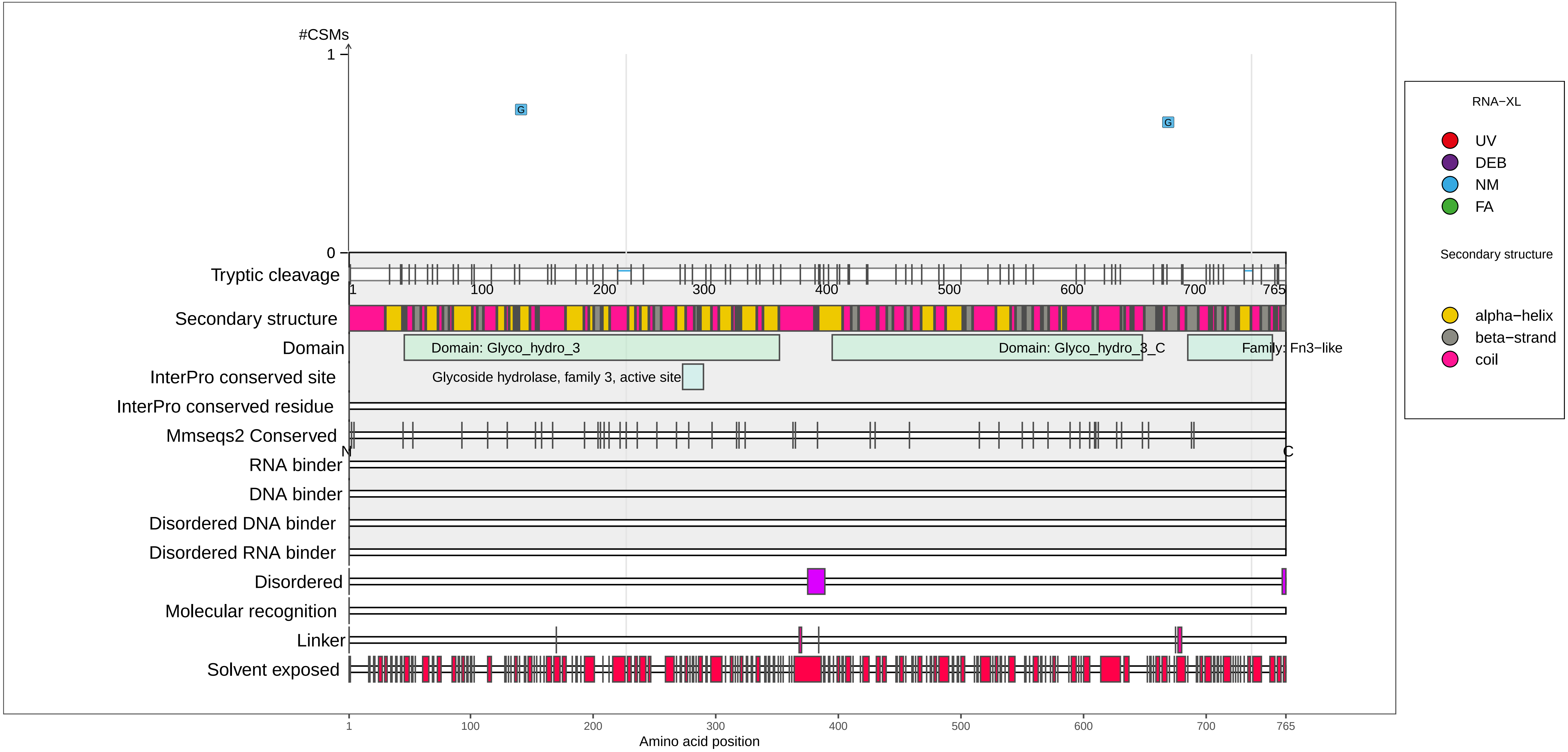

P37615
YHHM_ECOLI Uncharacterized protein YhhM

− Abundance:
tryptic [log10 Intensity]: 8.32 (Q 70)
PAXdb K12 strain [ppm]: NA (Q NA)
PAXdb E.coli [ppm]: 1.07 (Q 49)

− RNA functions: not annotated

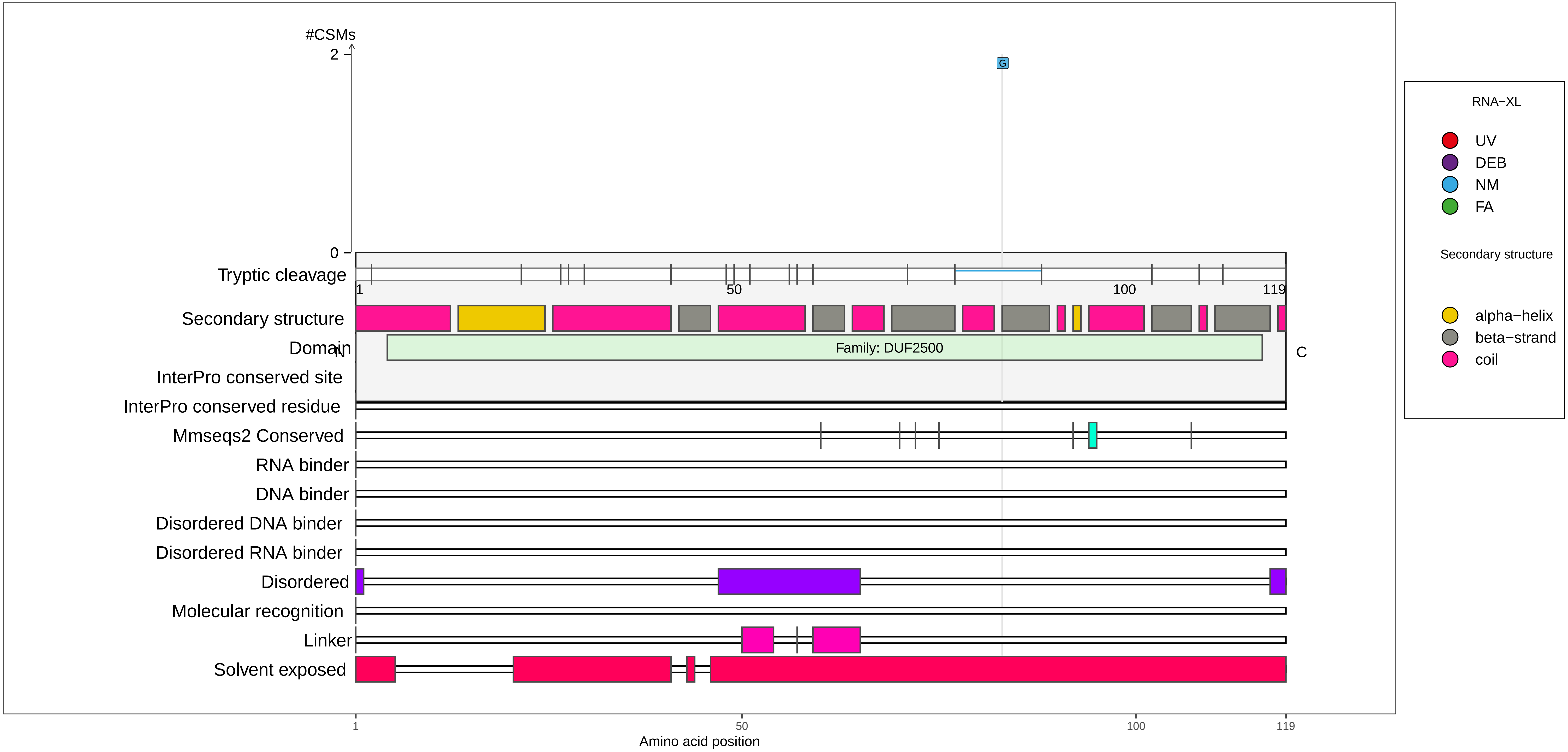

P37685
ALDB_ECOLI Aldehyde dehydrogenase B

− Abundance:
tryptic [log10 Intensity]: NA (Q NA)
PAXdb K12 strain [ppm]: 2.3 (Q 70)
PAXdb E.coli [ppm]: 2.95 (Q 94)

− RNA functions: not annotated

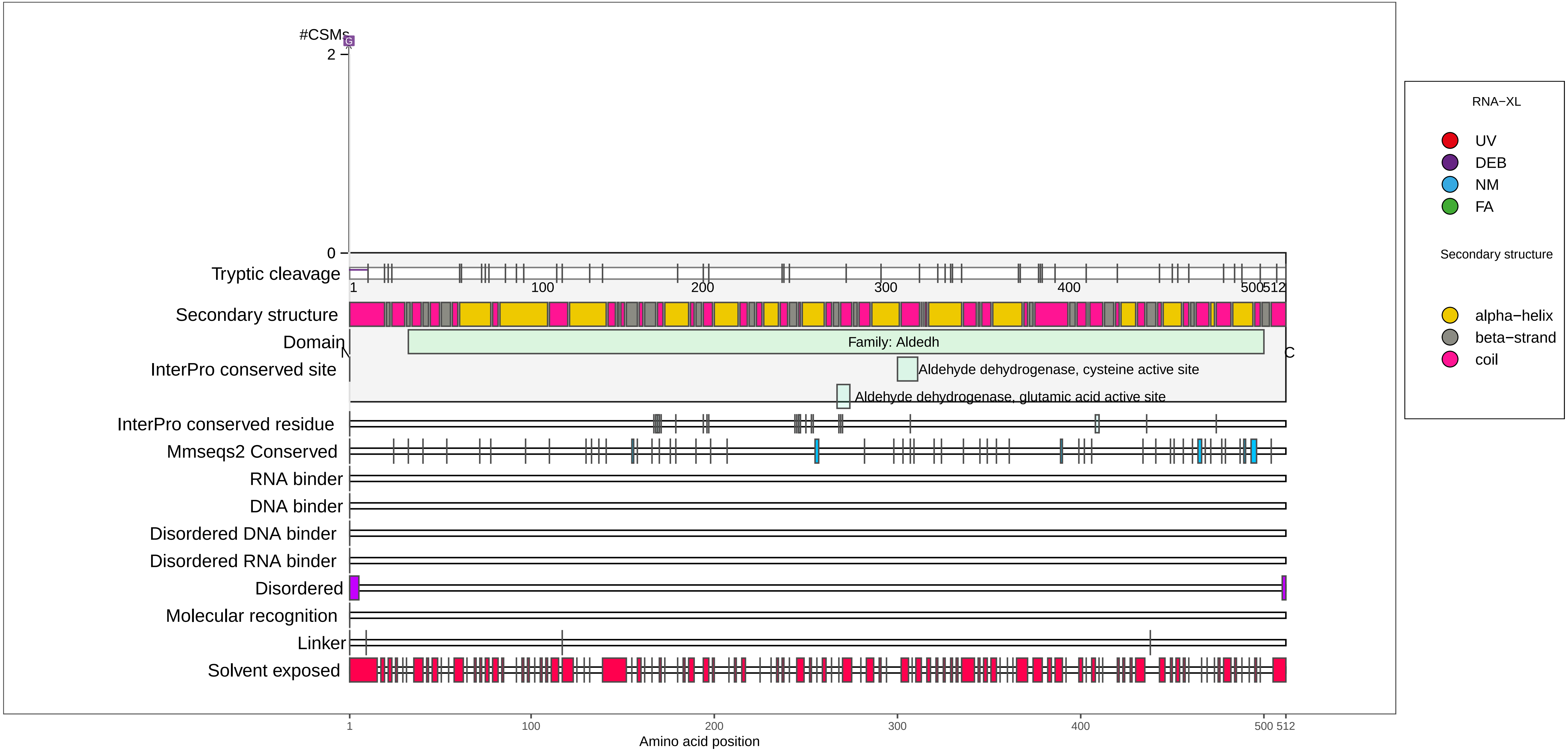

P39310
YTFB_ECOLI Cell division protein YtfB

− Abundance:
tryptic [log10 Intensity]: 7.51 (Q 35)
PAXdb K12 strain [ppm]: NA (Q NA)
PAXdb E.coli [ppm]: 1.91 (Q 70)

− RNA functions: not annotated

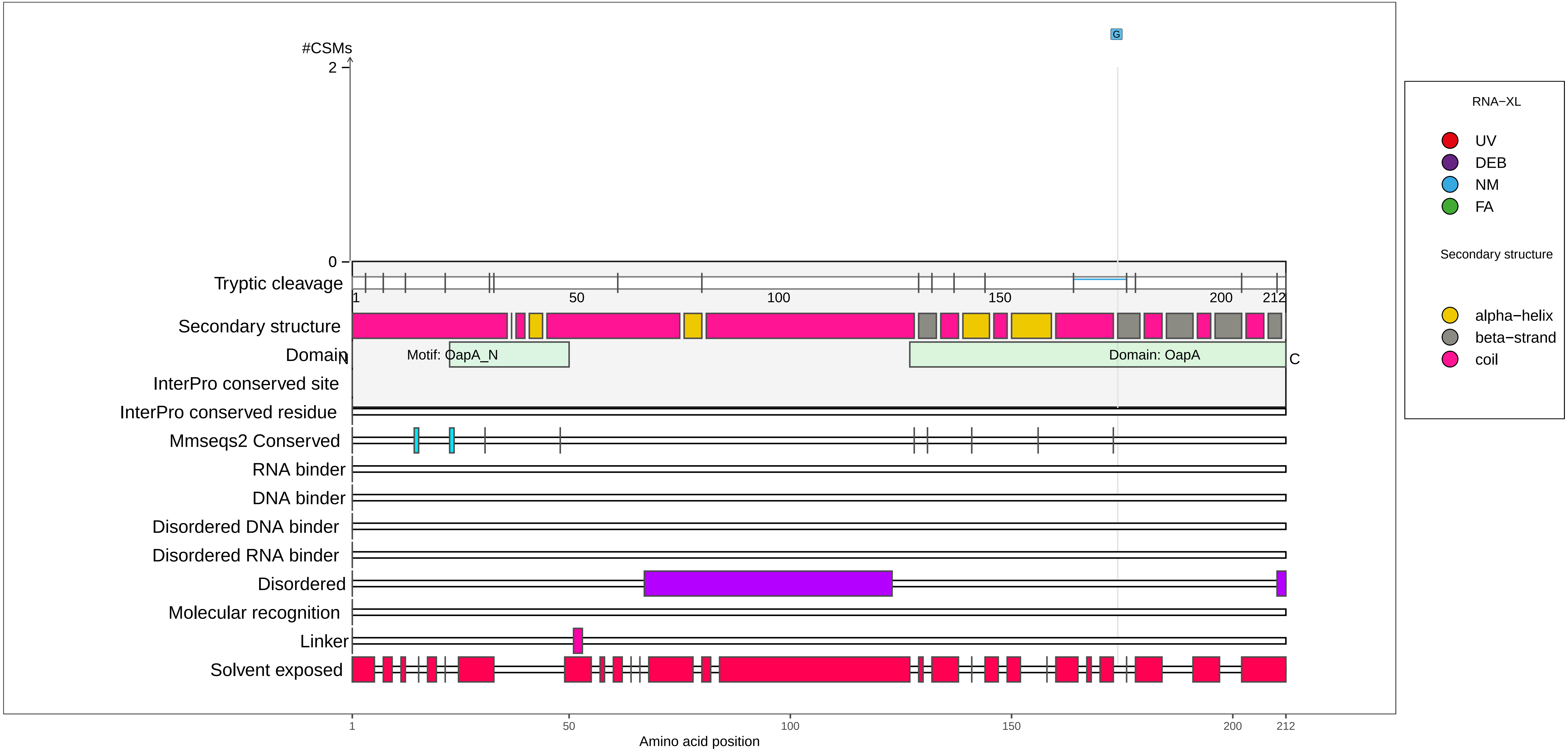

P39356
YJHU_ECOLI Uncharacterized transcriptional regulator YjhU
− Abundance:
tryptic [log10 Intensity]: NA (Q NA)
PAXdb K12 strain [ppm]: NA (Q NA)
PAXdb E.coli [ppm]: 1.76 (Q 66)

− RNA functions: not annotated

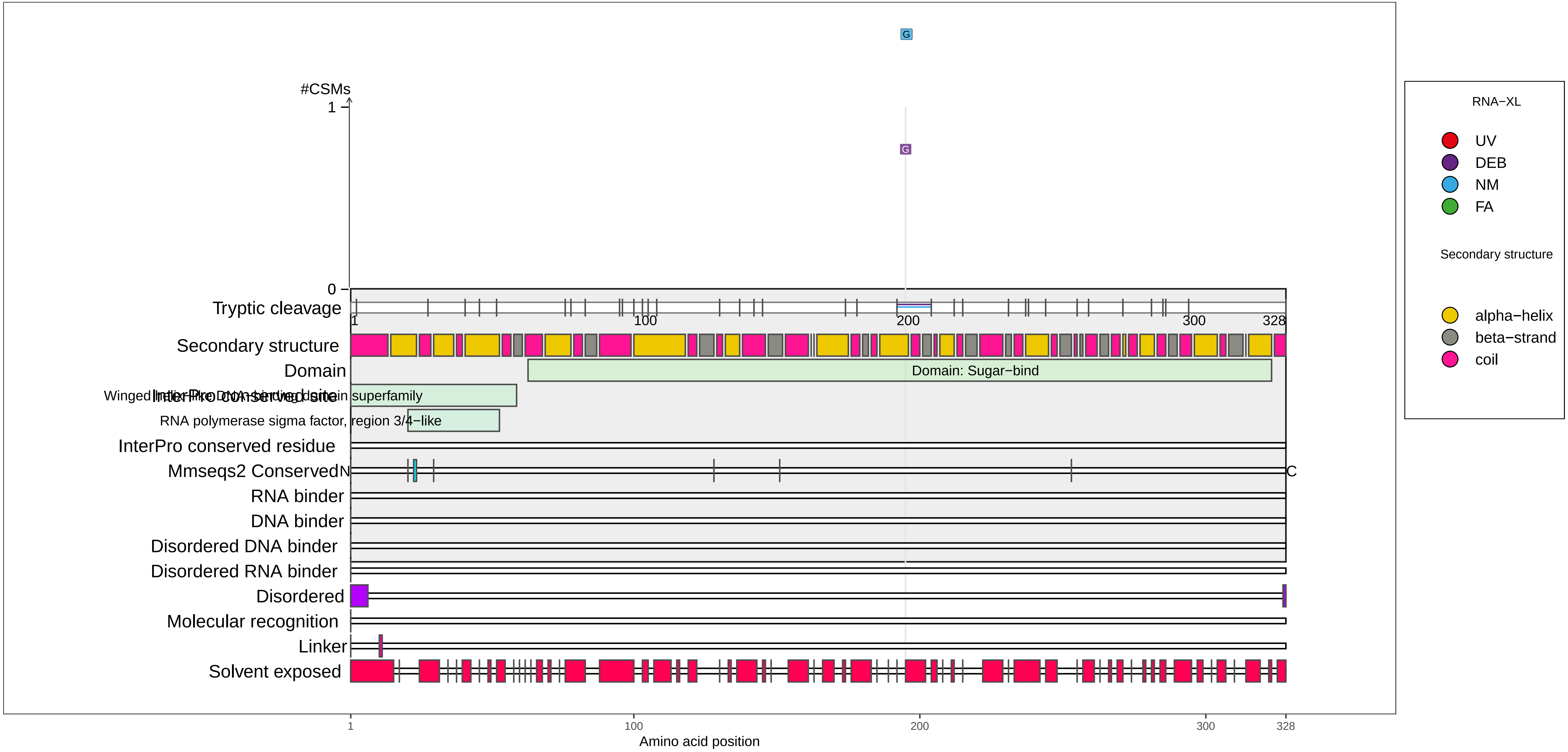

P45795
YRDB_ECOLI Uncharacterized protein YrdB

− Abundance:
tryptic [log10 Intensity]: 7.42 (Q 31)
PAXdb K12 strain [ppm]: 1.53 (Q 40)
PAXdb E.coli [ppm]: 0.77 (Q 42)

− RNA functions: not annotated

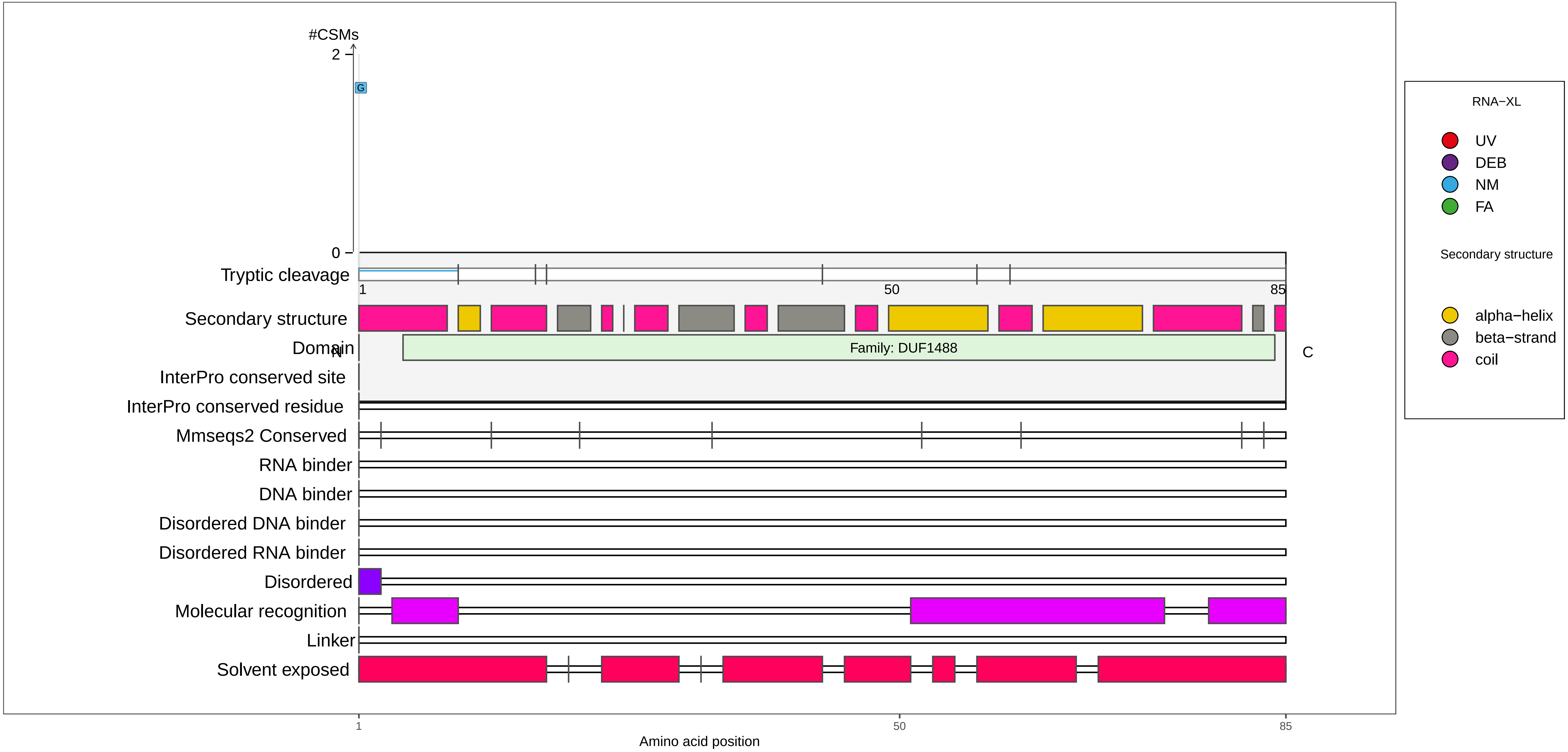

P45799
NUDE_ECOLI ADP compounds hydrolase NudE

− Abundance:
tryptic [log10 Intensity]: 8.23 (Q 67)
PAXdb K12 strain [ppm]: 2.36 (Q 72)
PAXdb E.coli [ppm]: 1.57 (Q 62)

− RNA functions: not annotated

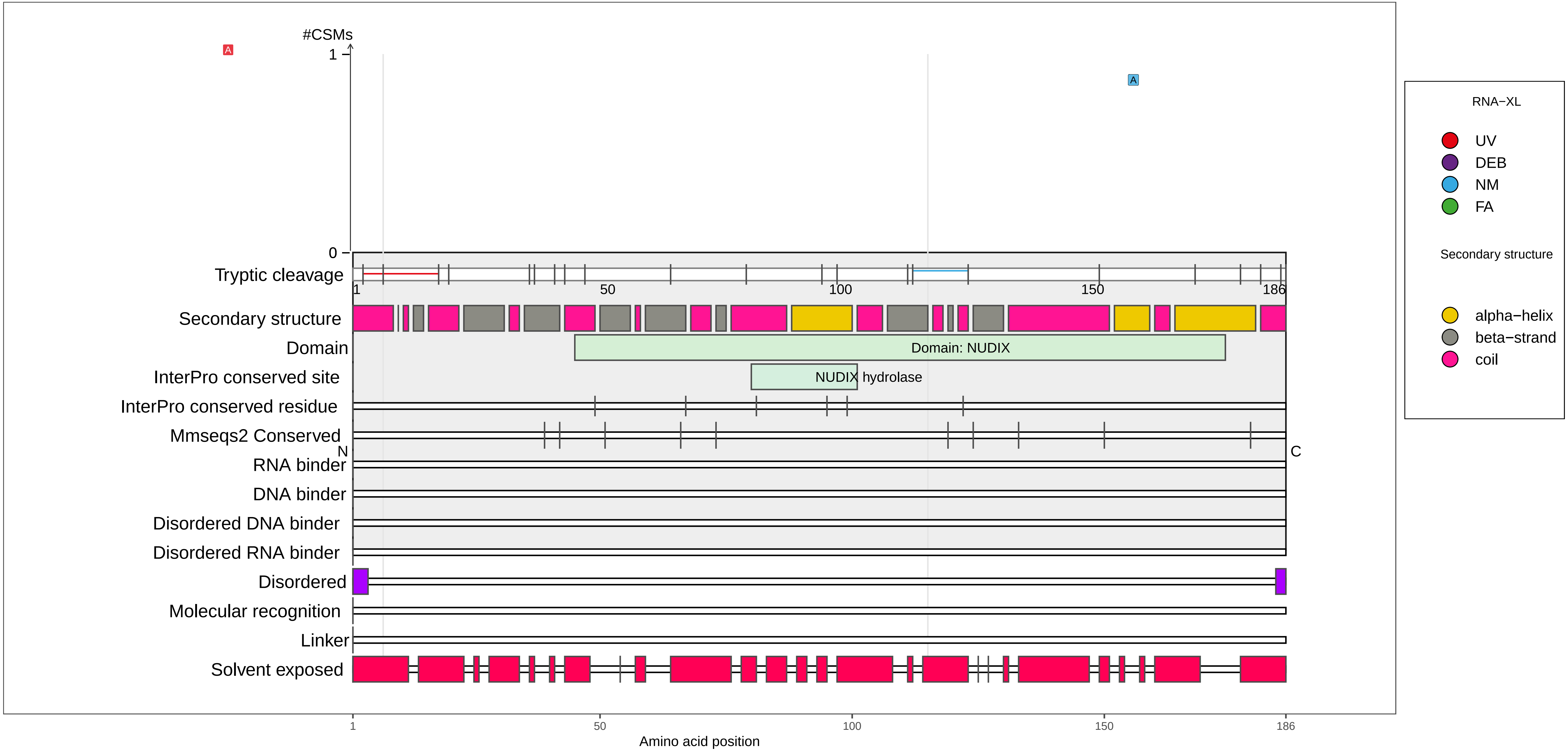

P51024
YAIL_ECOLI Uncharacterized protein YaiL

− Abundance:
tryptic [log10 Intensity]: 7.82 (Q 50)
PAXdb K12 strain [ppm]: 2.39 (Q 73)
PAXdb E.coli [ppm]: 1.7 (Q 64)

− RNA functions: not annotated

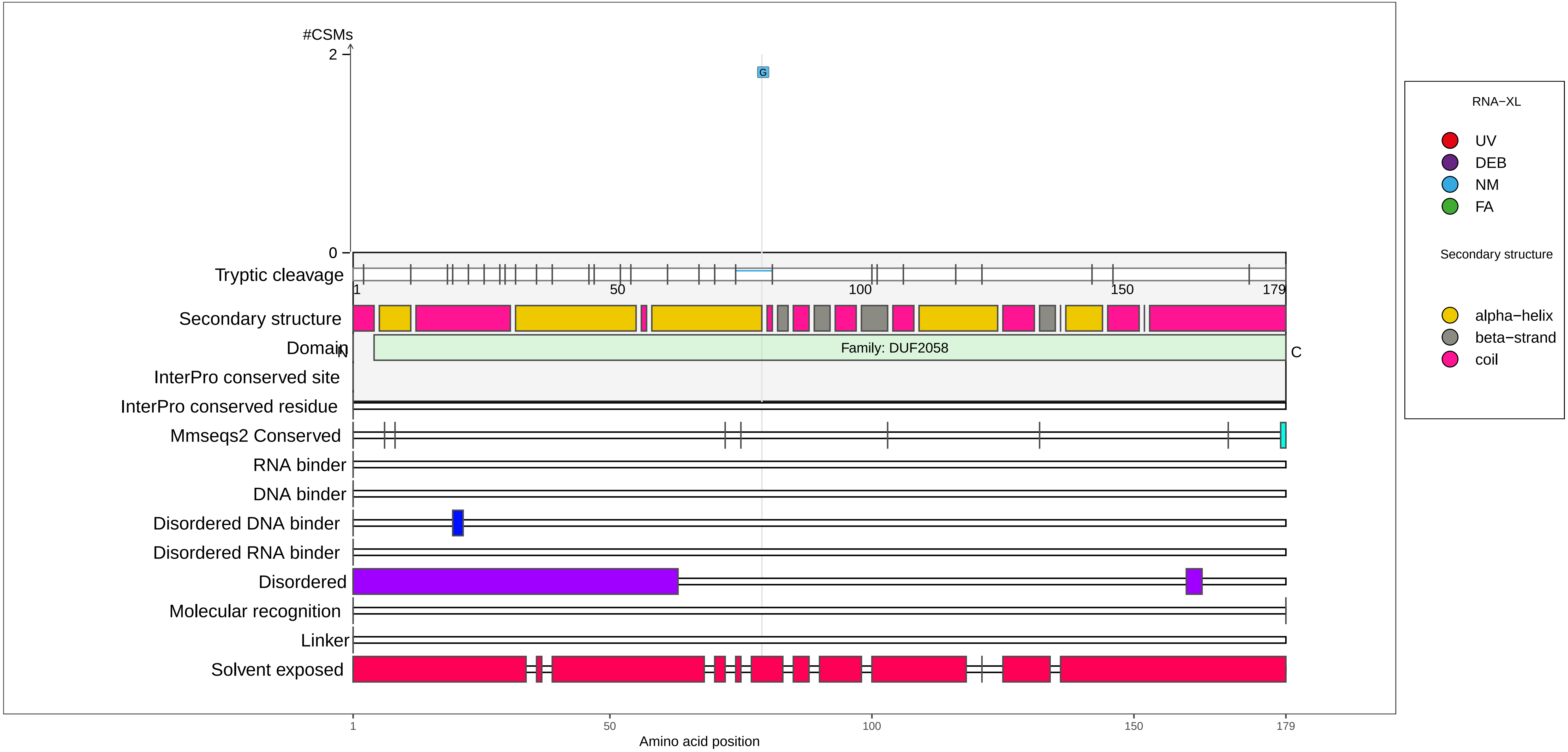

P52096
YAER_ECOLI Uncharacterized protein YaeR

− Abundance:
tryptic [log10 Intensity]: 7.95 (Q 56)
PAXdb K12 strain [ppm]: NA (Q NA)
PAXdb E.coli [ppm]: 2.05 (Q 73)

− RNA functions: not annotated

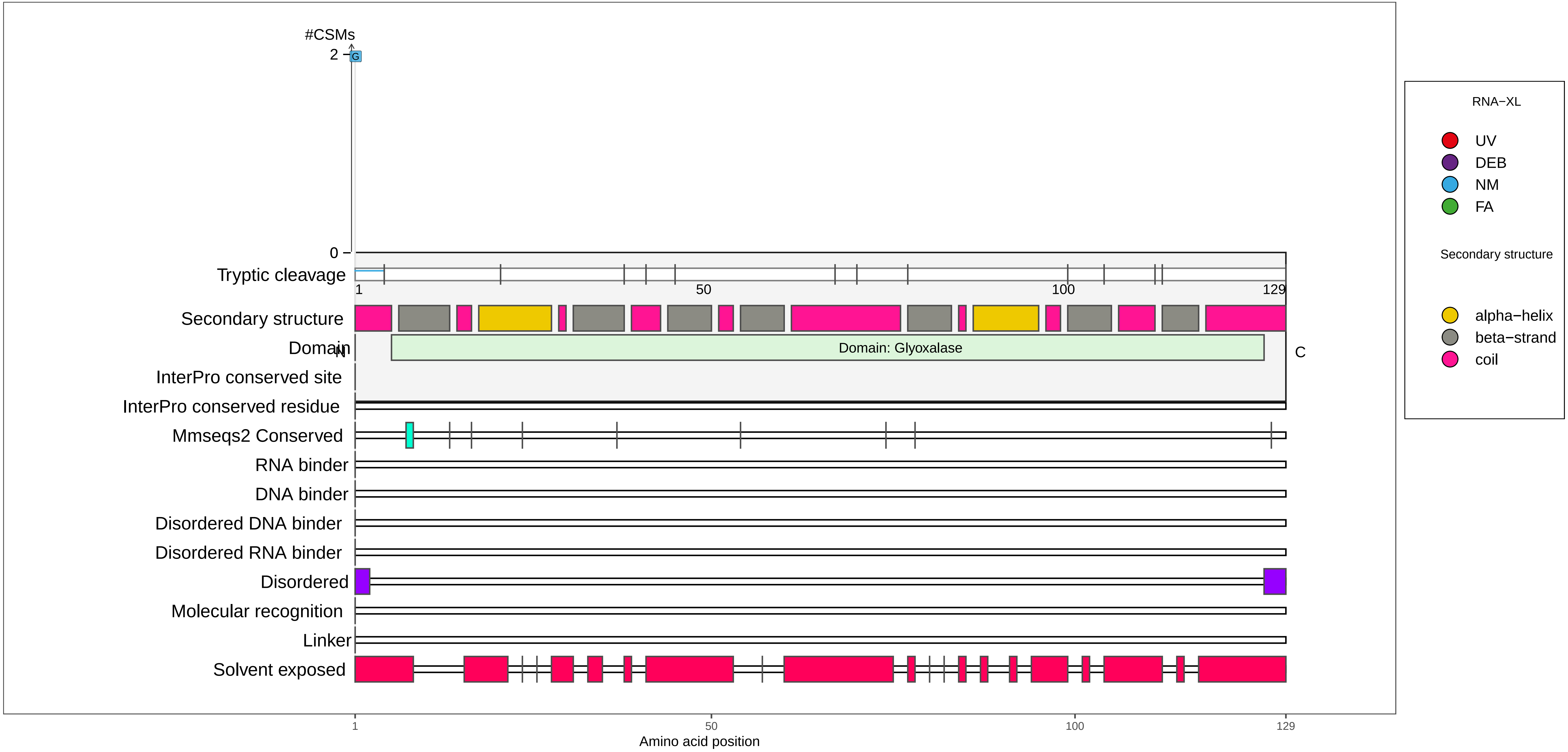

P52644
HSLJ_ECOLI Heat shock protein HslJ

− Abundance:
tryptic [log10 Intensity]: 8.11 (Q 62)
PAXdb K12 strain [ppm]: 1.48 (Q 37)
PAXdb E.coli [ppm]: 1.34 (Q 56)

− RNA functions: not annotated

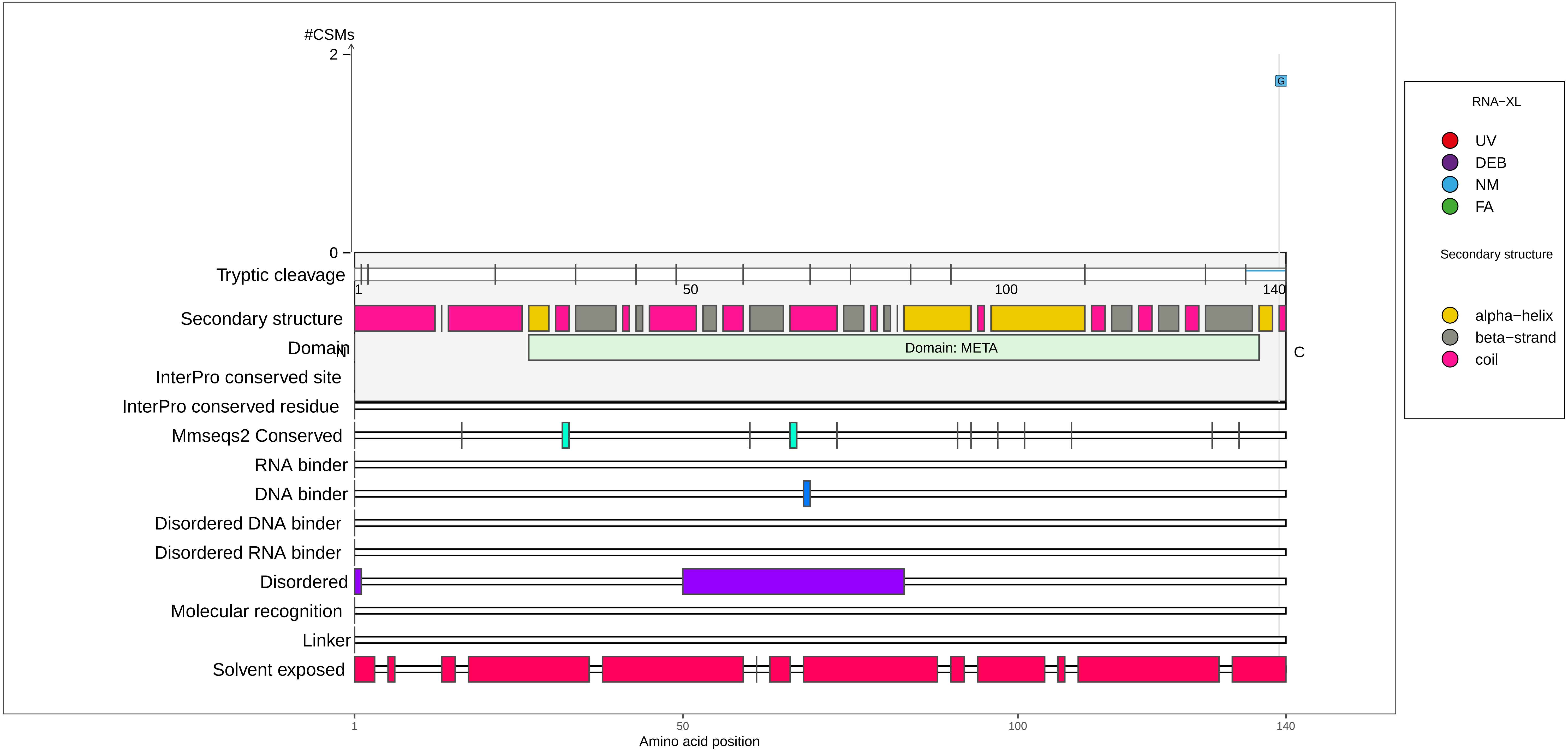

P56258
WECF_ECOLI TDP−N−acetylfucosamine:lipid II N−acetylfucosaminyltransferase

− Abundance:
tryptic [log10 Intensity]: 7.25 (Q 23)
PAXdb K12 strain [ppm]: 1.2 (Q 20)
PAXdb E.coli [ppm]: −0.15 (Q 21)

− RNA functions: not annotated

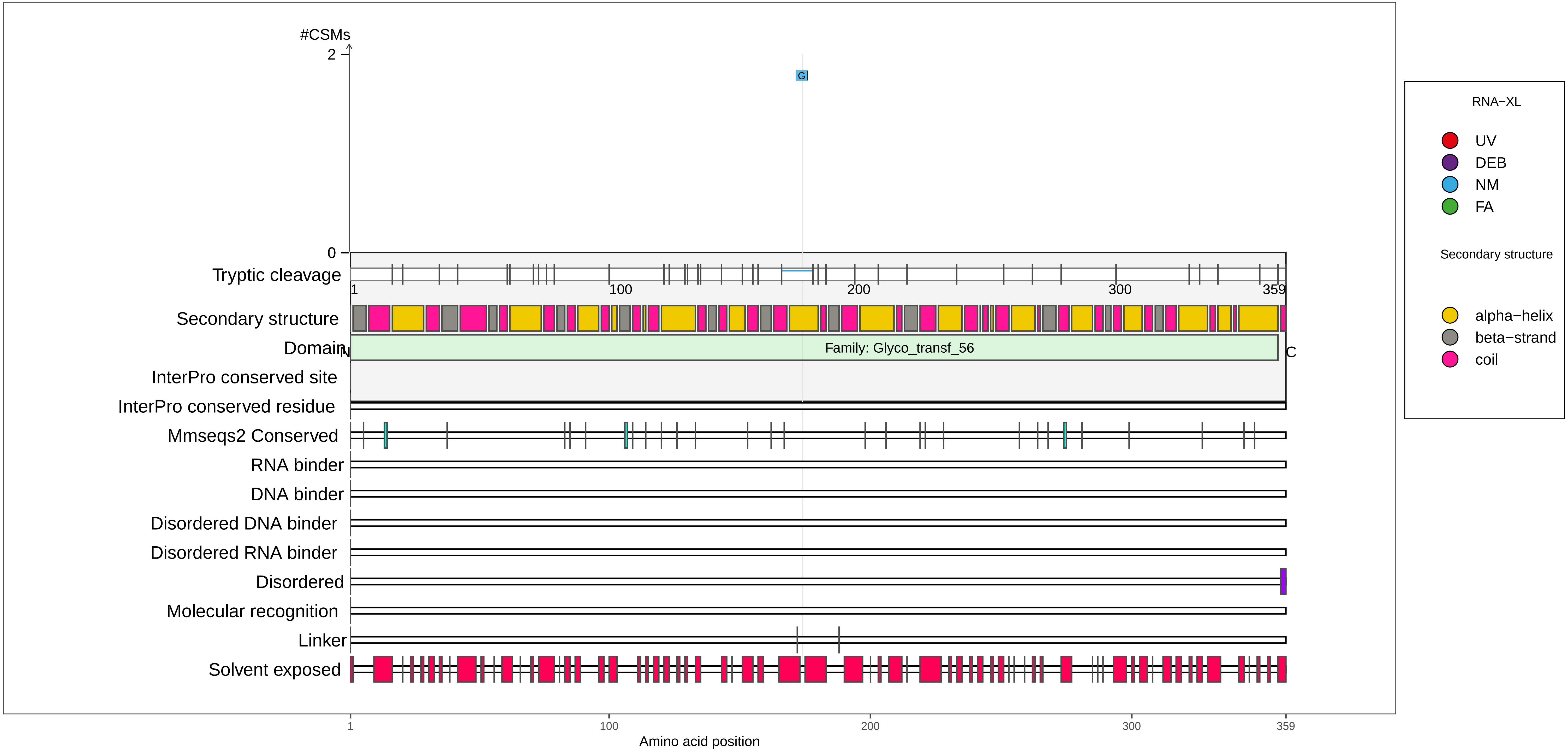

P60664
HIS6_ECOLI Imidazole glycerol phosphate synthase subunit HisF

− Abundance:
tryptic [log10 Intensity]: 8.54 (Q 77)
PAXdb K12 strain [ppm]: 1.5 (Q 38)
PAXdb E.coli [ppm]: 2.12 (Q 75)

− RNA functions: not annotated

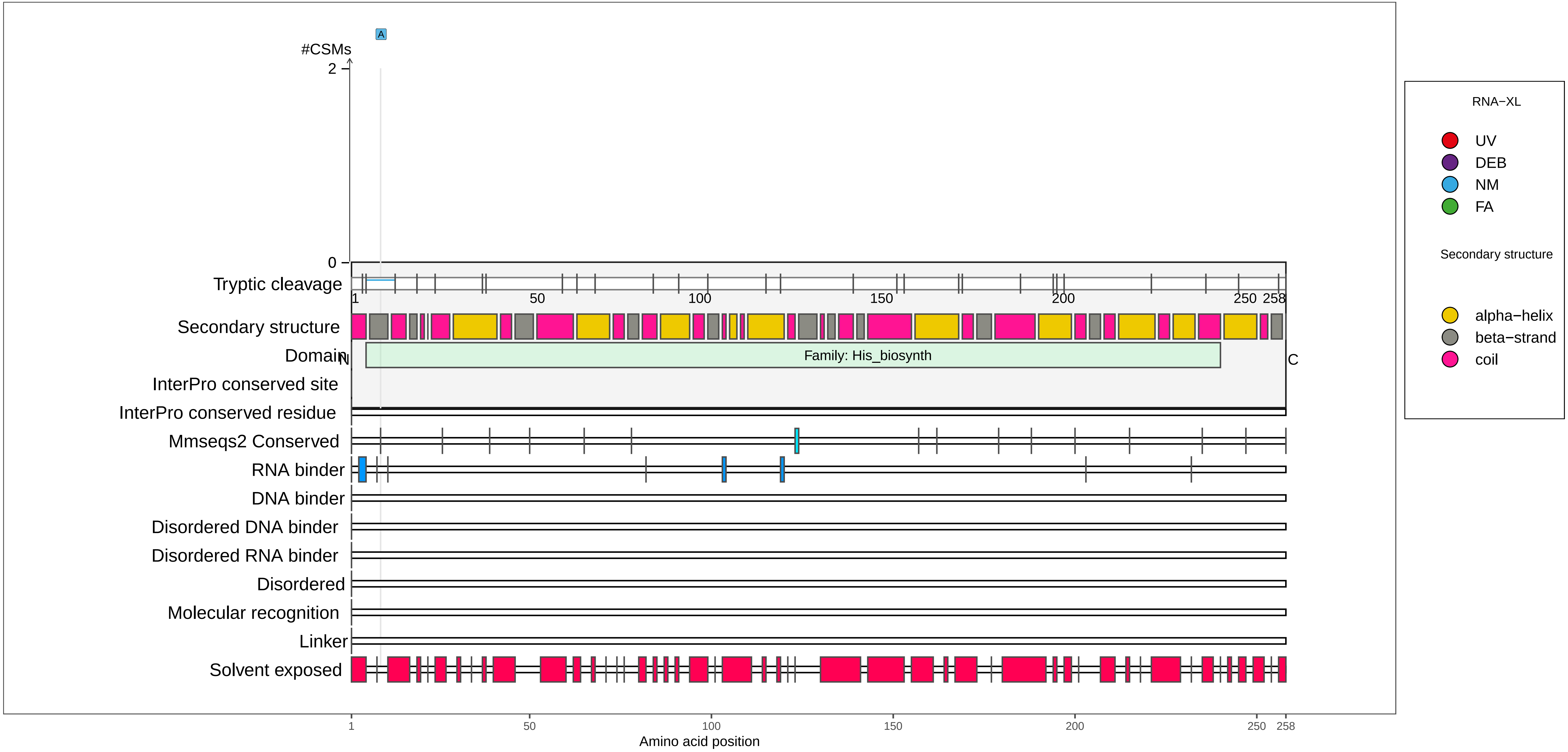

P64432
YPJD_ECOLI Inner membrane protein YpjD

− Abundance:
tryptic [log10 Intensity]: 7.19 (Q 20)
PAXdb K12 strain [ppm]: NA (Q NA)
PAXdb E.coli [ppm]: 0.48 (Q 36)

− RNA functions: not annotated

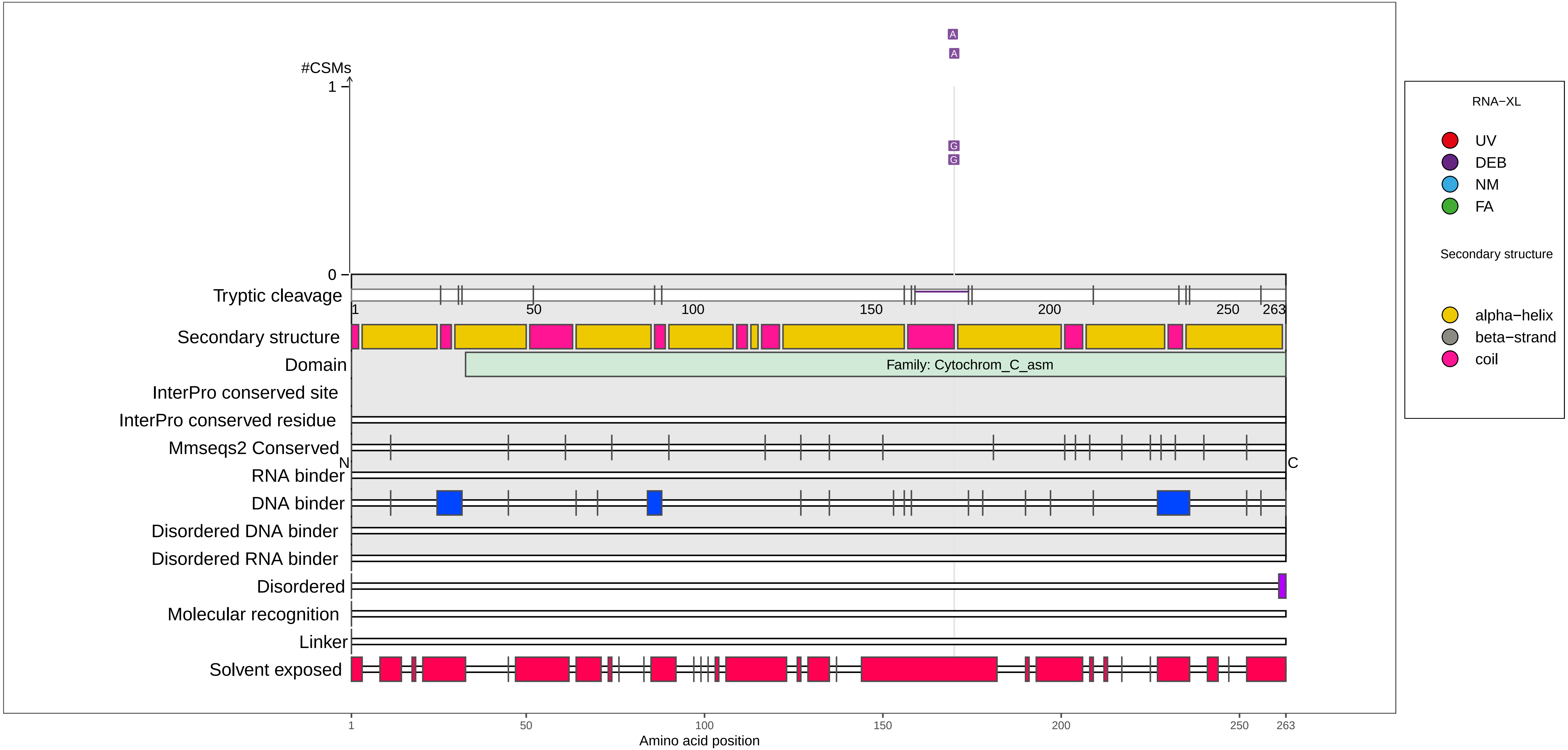

P64624
YHEO_ECOLI Uncharacterized protein YheO

− Abundance:
tryptic [log10 Intensity]: 7.61 (Q 40)
PAXdb K12 strain [ppm]: 2.6 (Q 80)
PAXdb E.coli [ppm]: 2.01 (Q 72)

− RNA functions: not annotated

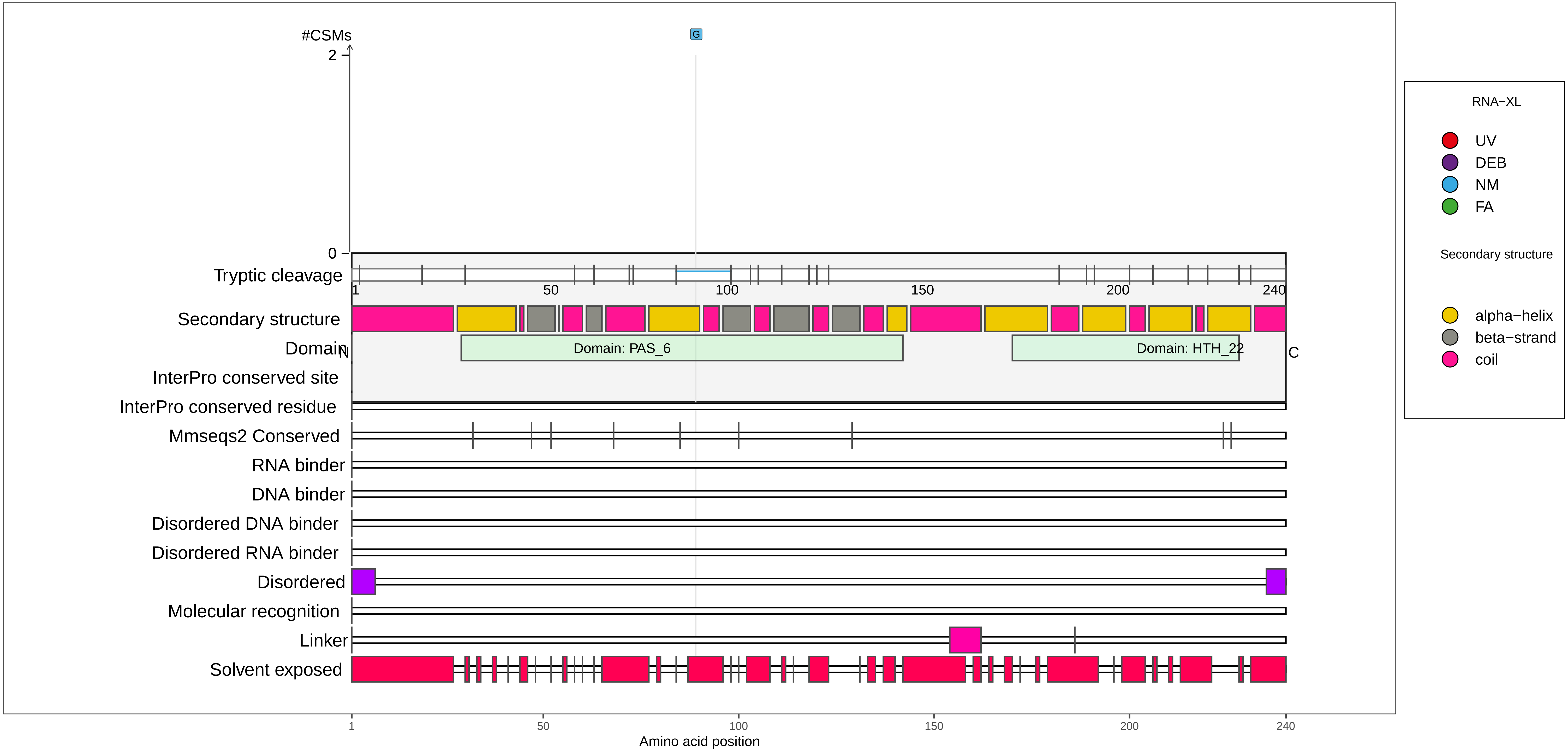

P67338
YOAH_ECOLI UPF0181 protein YoaH

− Abundance:
tryptic [log10 Intensity]: NA (Q NA)
PAXdb K12 strain [ppm]: NA (Q NA)
PAXdb E.coli [ppm]: −0.12 (Q 22)

− RNA functions: not annotated

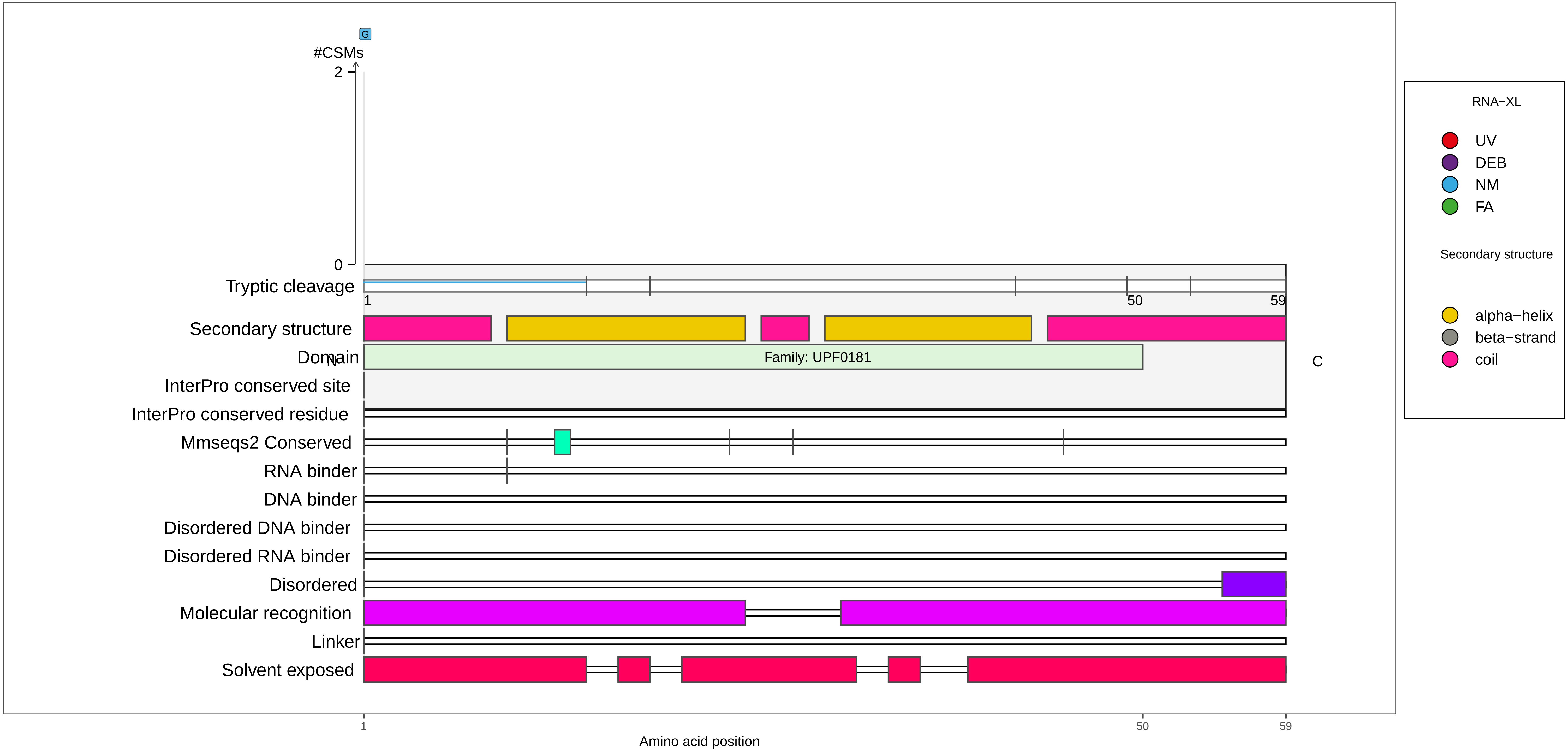

P67701
HIGA_ECOLI Antitoxin HigA

− Abundance:
tryptic [log10 Intensity]: 6.83 (Q 8)
PAXdb K12 strain [ppm]: NA (Q NA)
PAXdb E.coli [ppm]: 0.67 (Q 40)

− RNA functions: not annotated

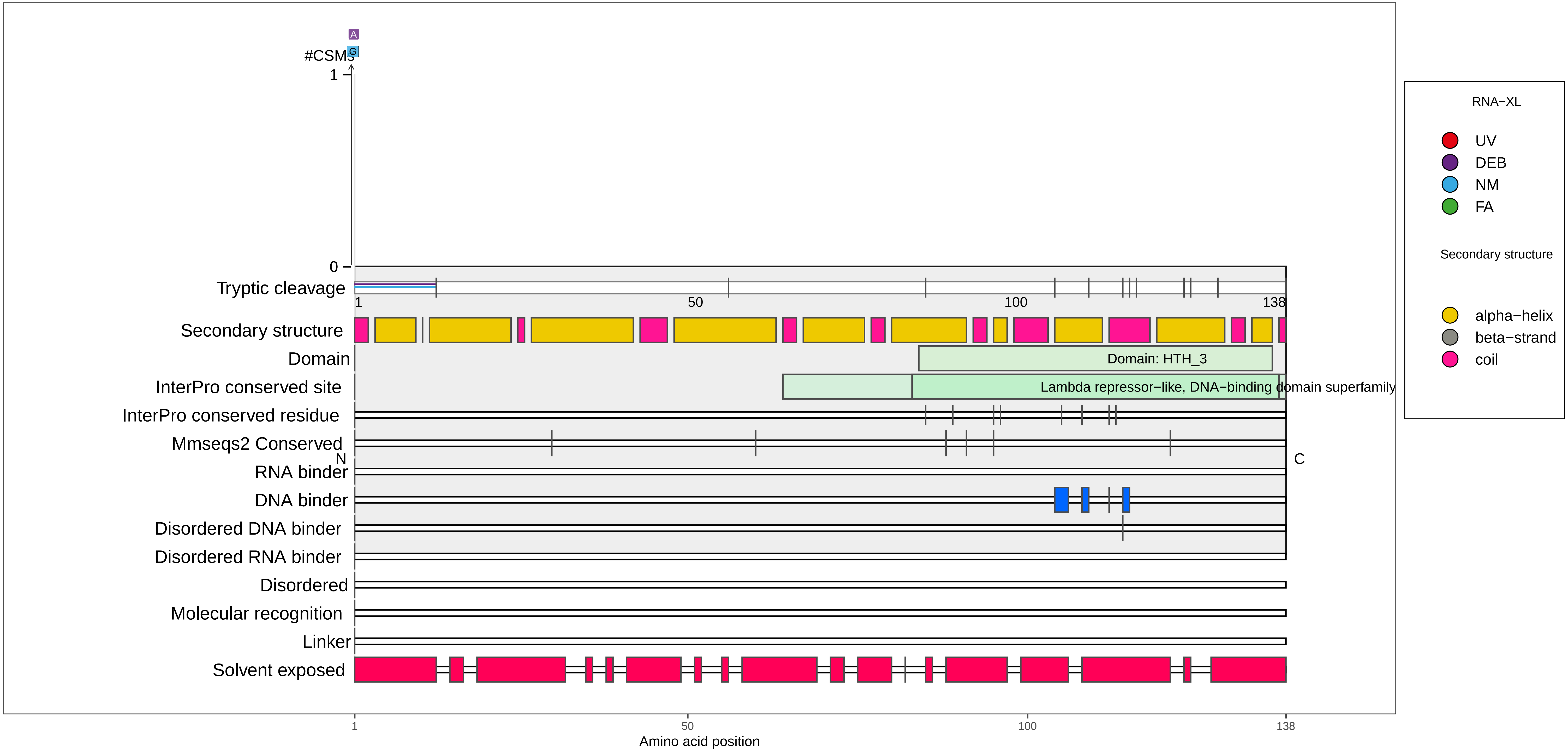

P69801
PTNC_ECOLI PTS system mannose−specific EIIC component

− Abundance:
tryptic [log10 Intensity]: 8.91 (Q 86)
PAXdb K12 strain [ppm]: NA (Q NA)
PAXdb E.coli [ppm]: 2.77 (Q 91)

− RNA functions: not annotated

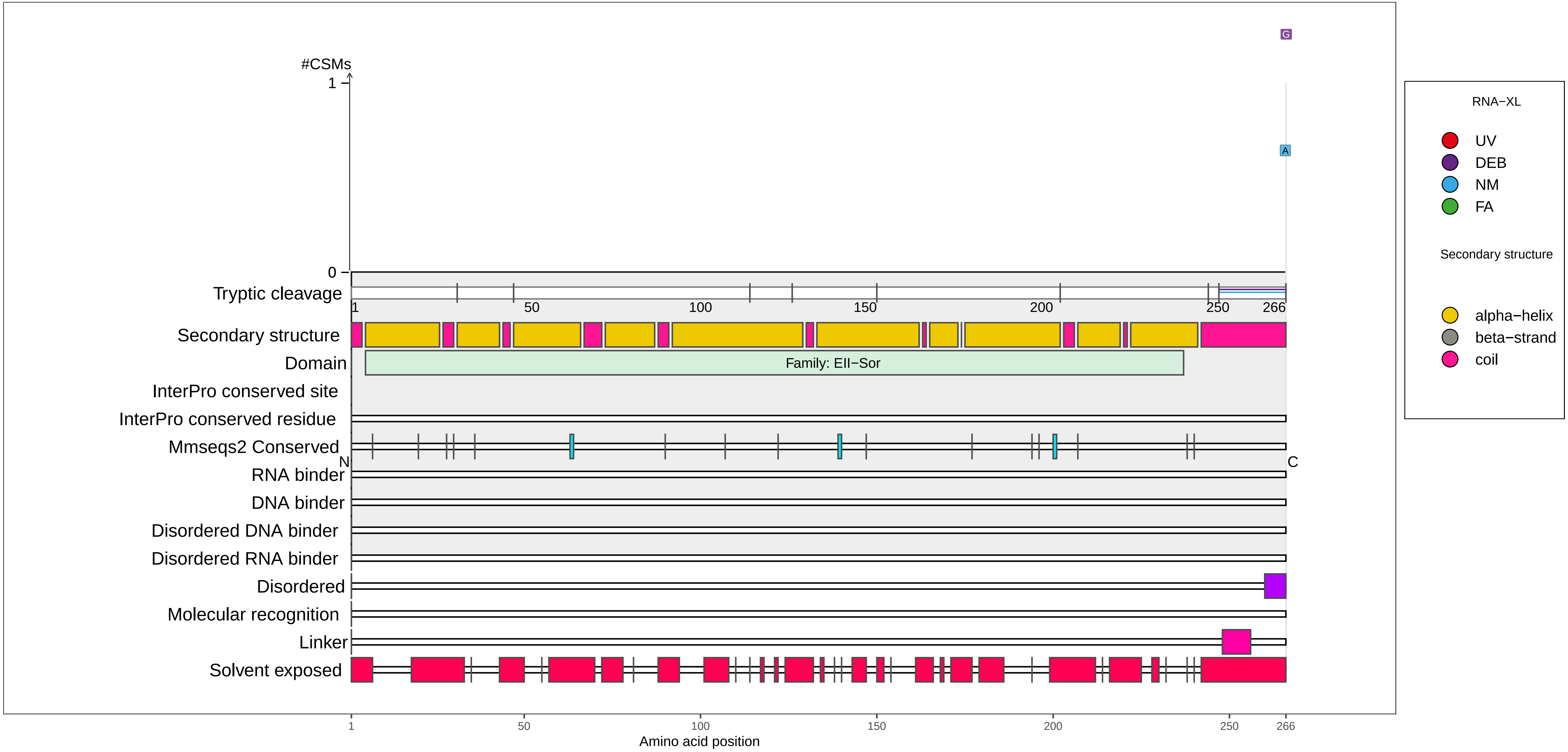

P75763
YBHI_ECOLI Inner membrane protein YbhI
− Abundance:
tryptic [log10 Intensity]: NA (Q NA)
PAXdb K12 strain [ppm]: NA (Q NA)
PAXdb E.coli [ppm]: −0.38 (Q 17)

− RNA functions: not annotated

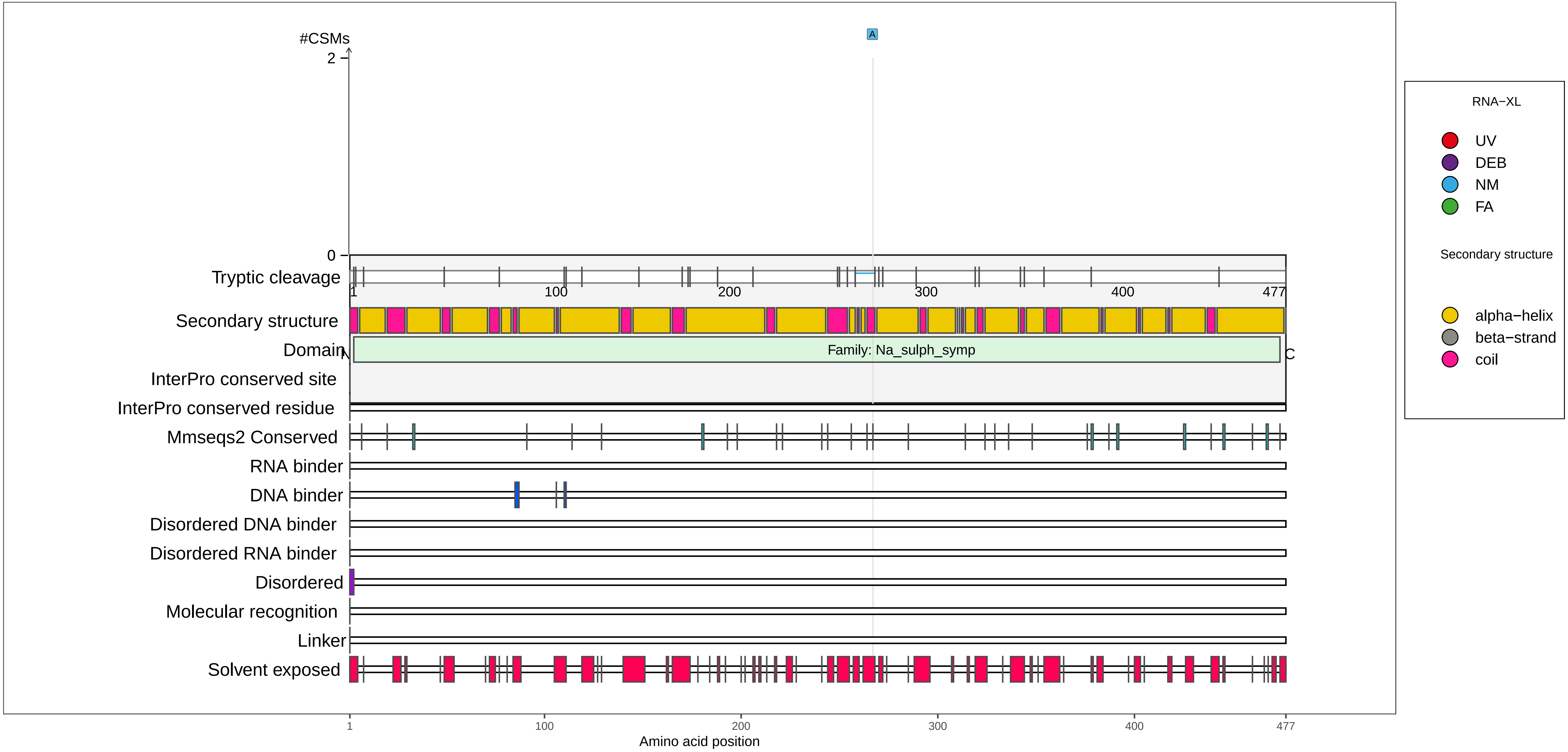

P75767
GNGF_ECOLI Putative gluconeogenesis factor

− Abundance:
tryptic [log10 Intensity]: 6.99 (Q 13)
PAXdb K12 strain [ppm]: NA (Q NA)
PAXdb E.coli [ppm]: 0.35 (Q 32)

− RNA functions: not annotated

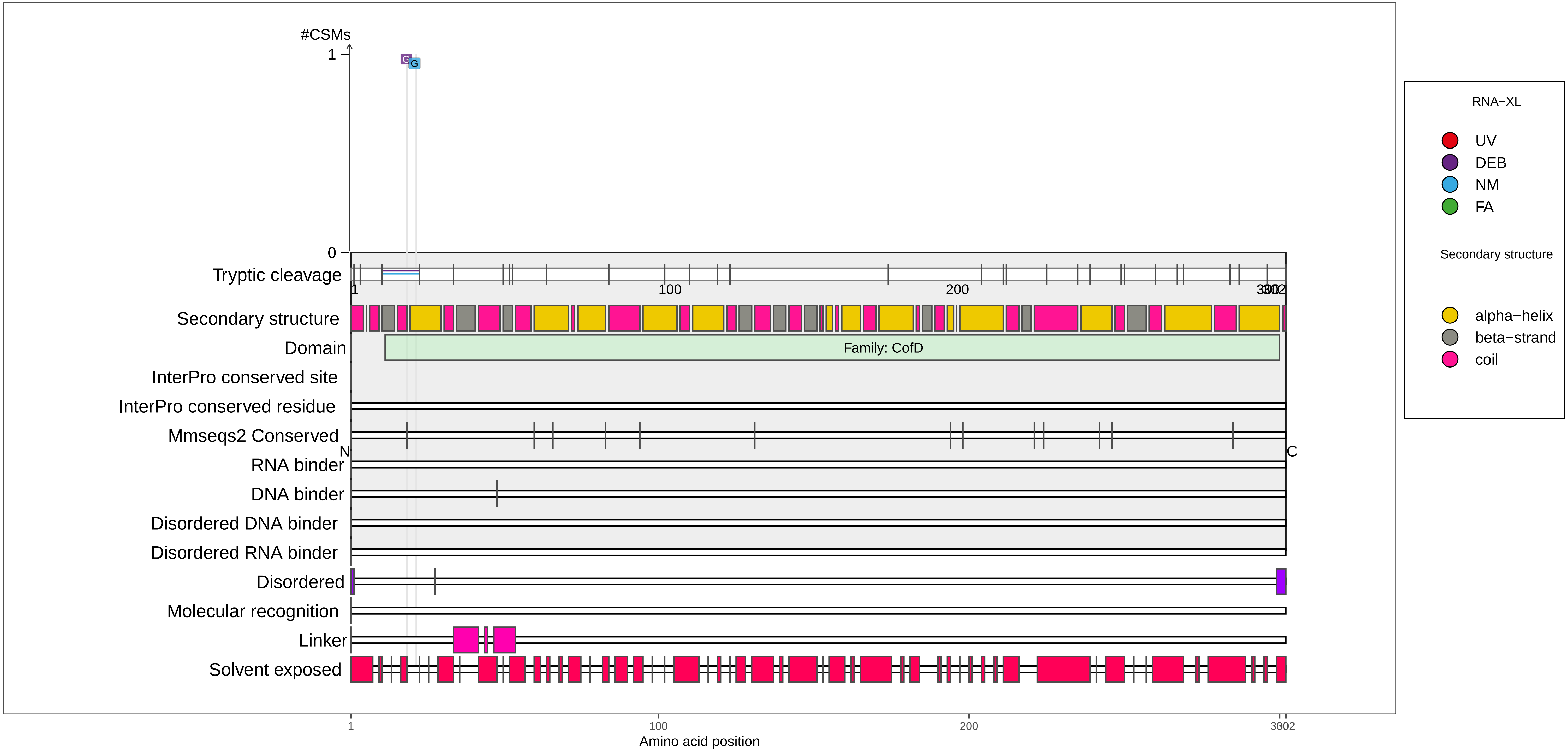

P75782
RLMF_ECOLI Ribosomal RNA large subunit methyltransferase F

− Abundance:
tryptic [log10 Intensity]: 6.58 (Q 3)
PAXdb K12 strain [ppm]: 1.49 (Q 38)
PAXdb E.coli [ppm]: 1.71 (Q 65)

− RNA functions:
23S rRNA (adenine(1618)−N(6))−methyltransferase activity; ncRNA metabolic process
ncRNA processing; RNA metabolic process; RNA methylation
RNA methyltransferase activity; RNA modification; RNA processing; rRNA (adenine−N6−)−methyltransferase activity
rRNA (adenine) methyltransferase activity; rRNA base methylation; rRNA metabolic process
rRNA methylation; rRNA methyltransferase activity; rRNA modification; rRNA processing

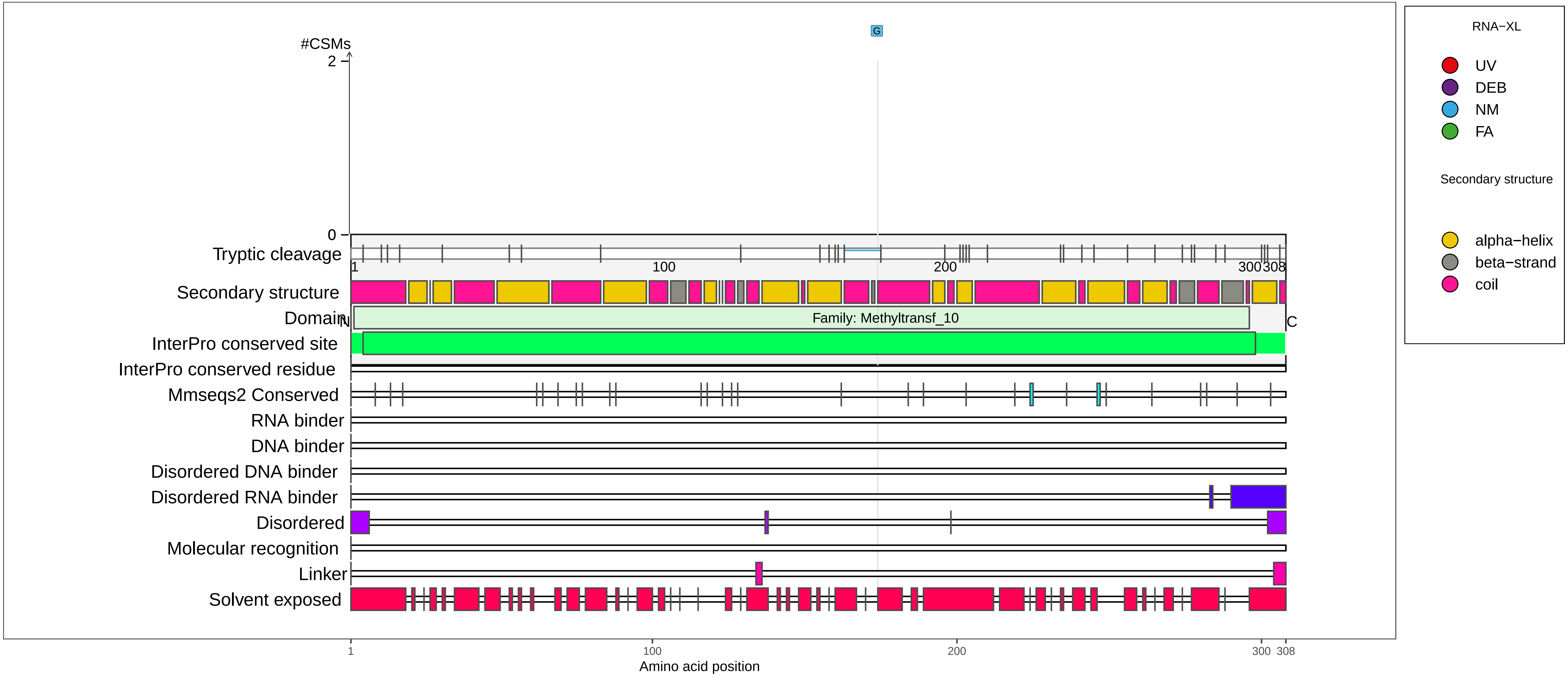

P76002
PLIG_ECOLI Inhibitor of g−type lysozyme

− Abundance:
tryptic [log10 Intensity]: 8.1 (Q 61)
PAXdb K12 strain [ppm]: 2.32 (Q 70)
PAXdb E.coli [ppm]: 2.05 (Q 73)

− RNA functions: not annotated

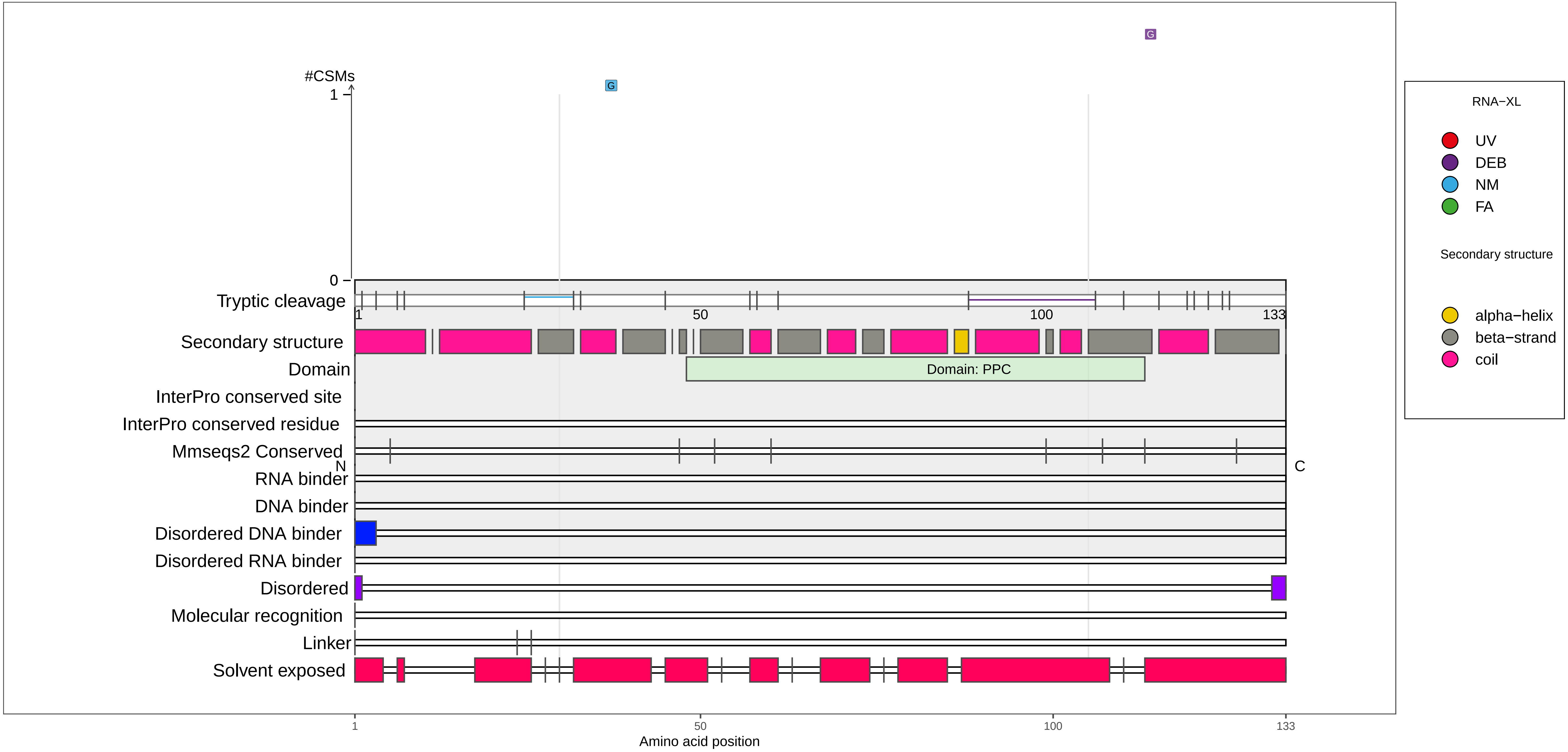

P76115
YNCD_ECOLI Probable TonB−dependent receptor YncD
− Abundance:
tryptic [log10 Intensity]: 8.97 (Q 87)
PAXdb K12 strain [ppm]: 0.88 (Q 7)
PAXdb E.coli [ppm]: 0.09 (Q 27)

− RNA functions: not annotated

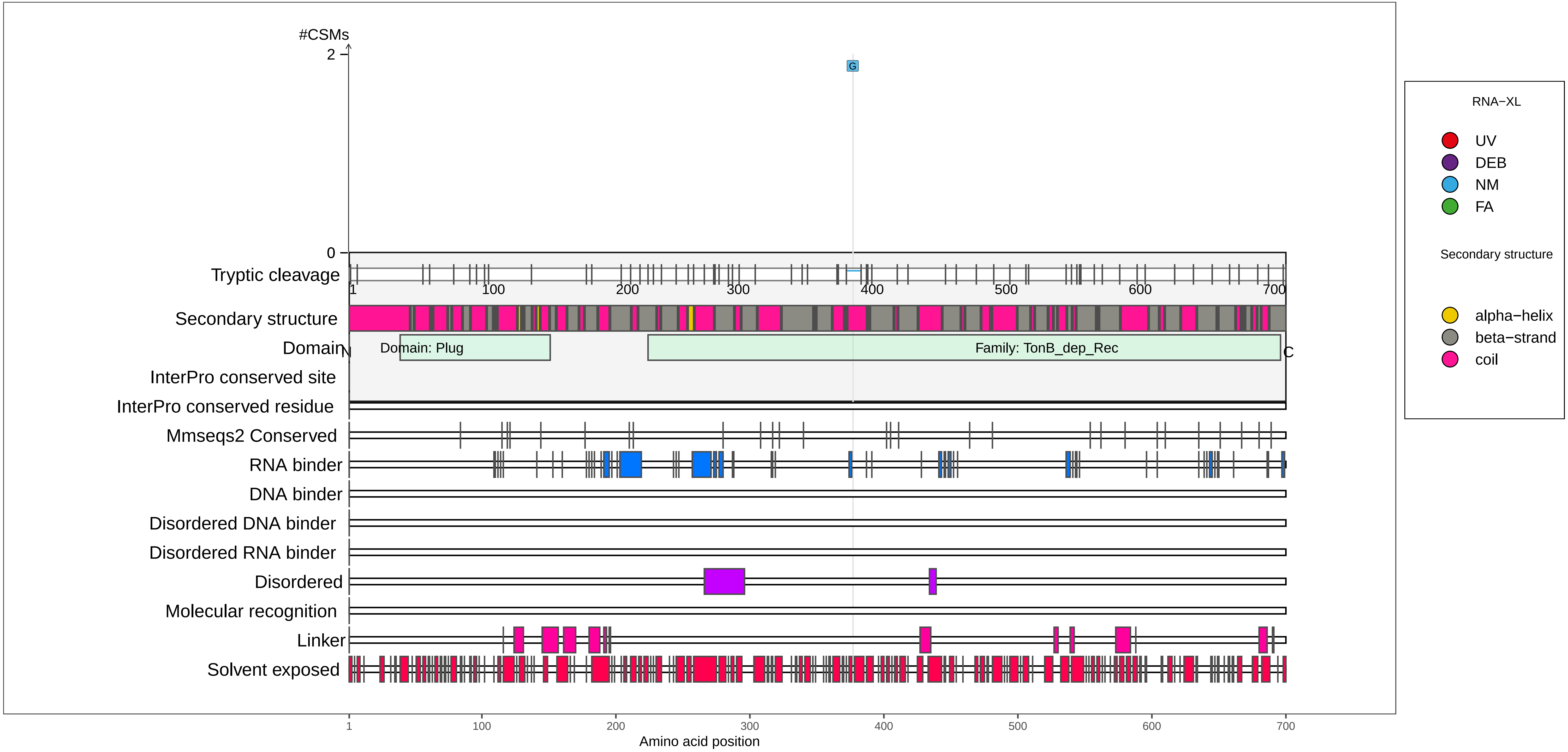

P76141
LSRR_ECOLI Transcriptional regulator LsrR

− Abundance:
tryptic [log10 Intensity]: NA (Q NA)
PAXdb K12 strain [ppm]: 1.2 (Q 20)
PAXdb E.coli [ppm]: 1.15 (Q 51)

− RNA functions:
RNA biosynthetic process; RNA metabolic process

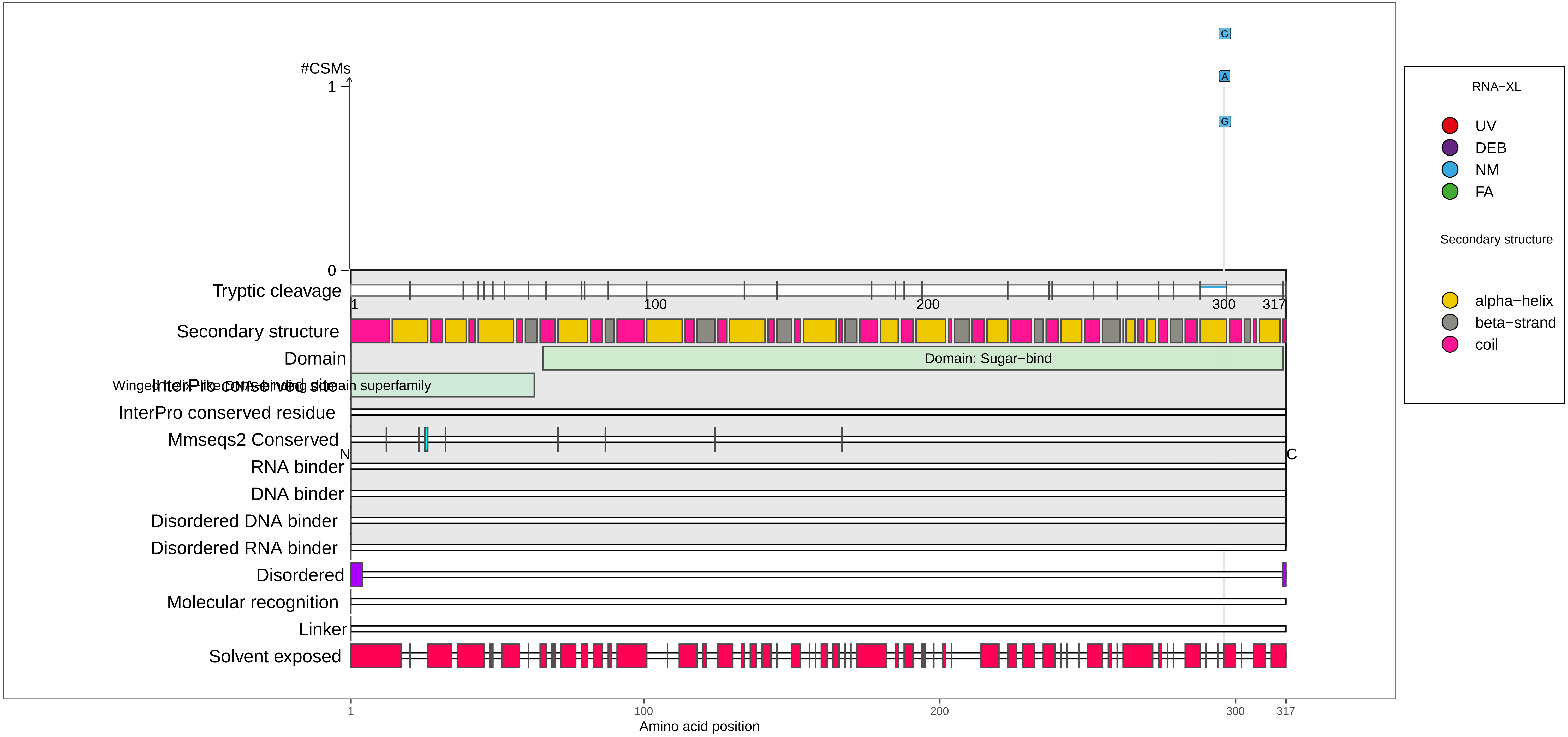

P76168
INTQ_ECOLI Putative defective protein IntQ
− RNA functions: not annotated

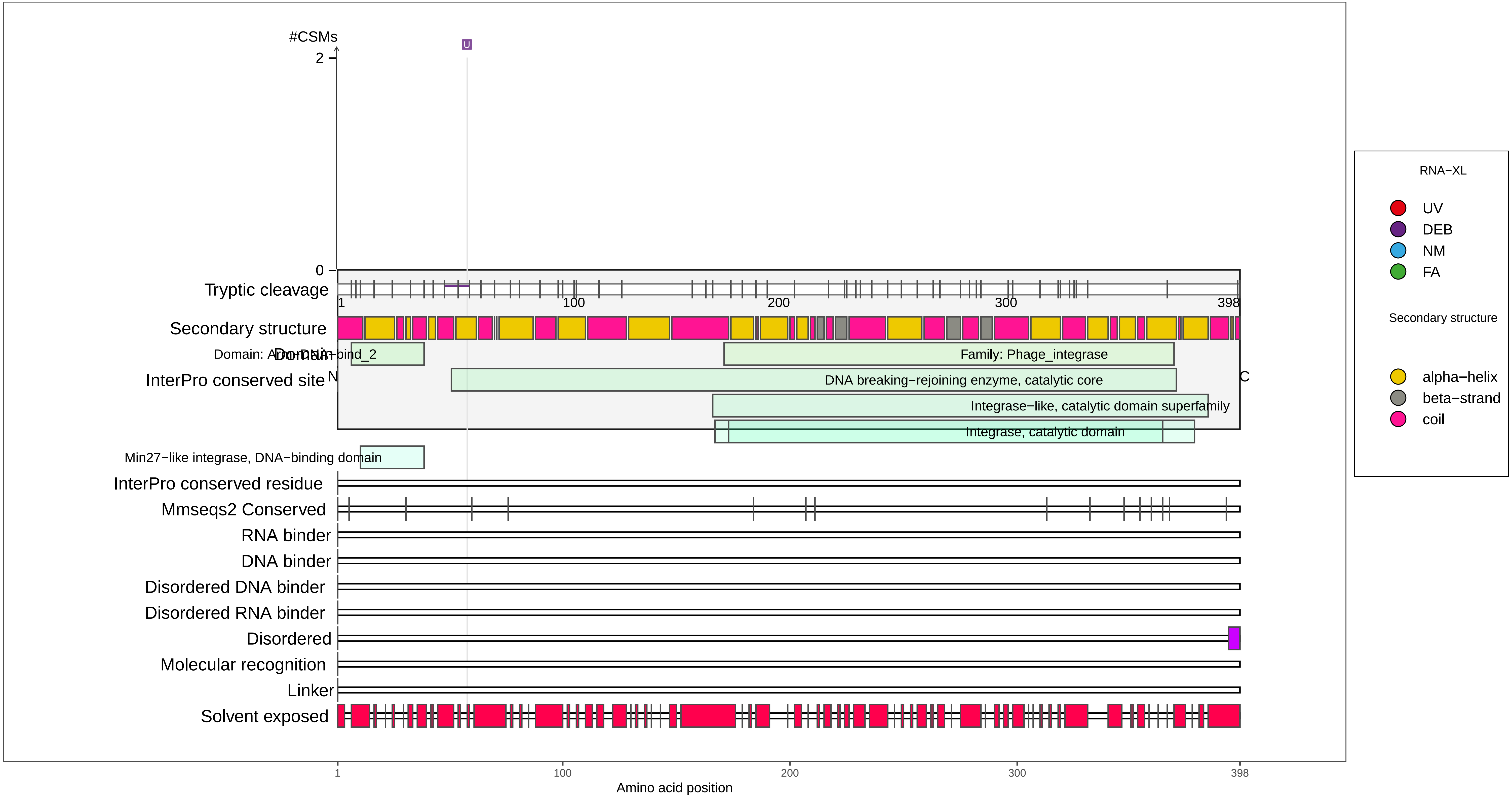

P76270
MSRC_ECOLI Free methionine−R−sulfoxide reductase

− Abundance:
tryptic [log10 Intensity]: 8.39 (Q 72)
PAXdb K12 strain [ppm]: 2.44 (Q 76)
PAXdb E.coli [ppm]: 2.31 (Q 81)

− RNA functions: not annotated

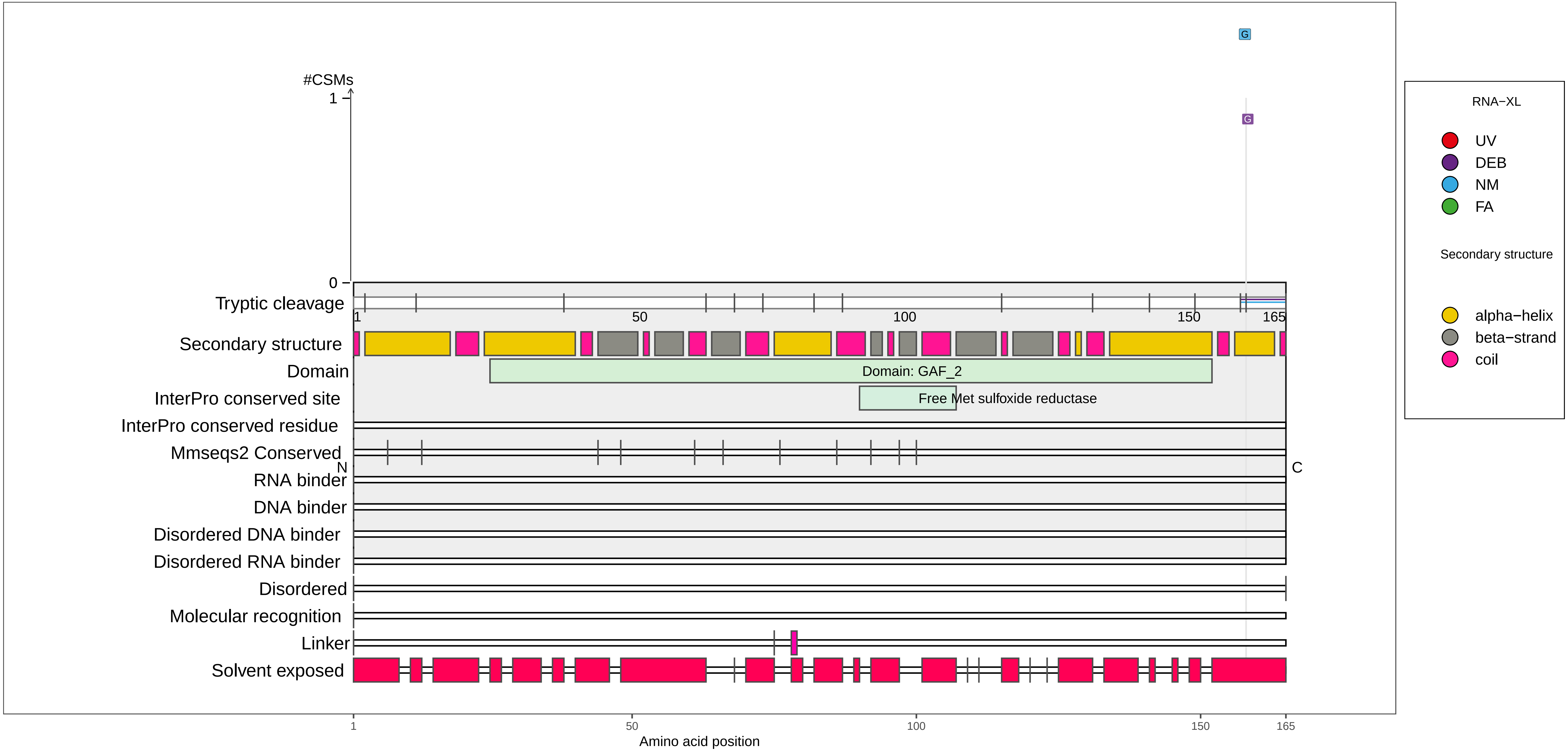

P76290
CMOA_ECOLI Carboxy−S−adenosyl−L−methionine synthase

− Abundance:
tryptic [log10 Intensity]: 7.55 (Q 37)
PAXdb K12 strain [ppm]: 1.7 (Q 49)
PAXdb E.coli [ppm]: 1.66 (Q 63)

− RNA functions:
ncRNA metabolic process; ncRNA processing; RNA metabolic process; RNA modification
RNA processing; tRNA metabolic process; tRNA modification; tRNA processing
tRNA wobble base modification; tRNA wobble uridine modification

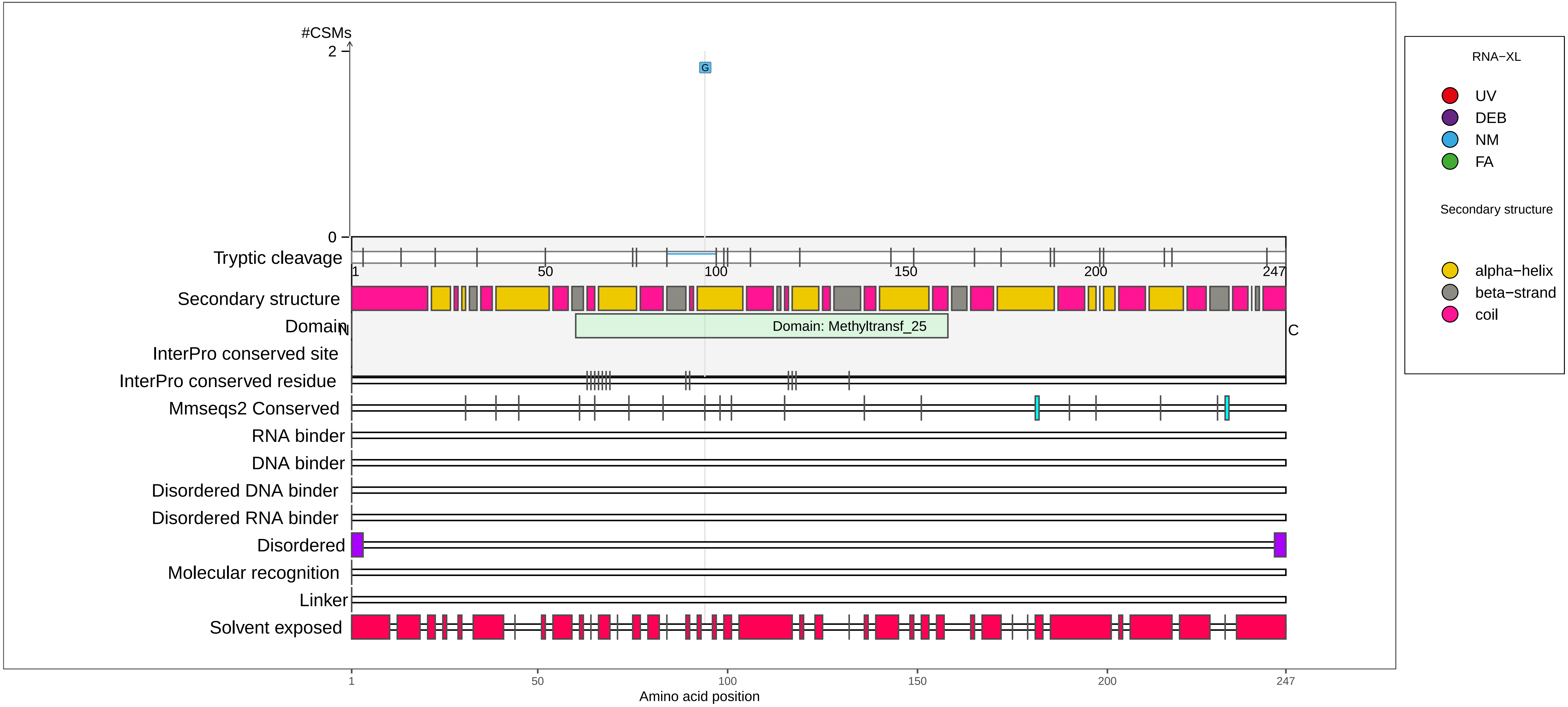

P76372
WZZB_ECOLI Chain length determinant protein

− Abundance:
tryptic [log10 Intensity]: 8.49 (Q 75)
PAXdb K12 strain [ppm]: 1.46 (Q 36)
PAXdb E.coli [ppm]: 2.42 (Q 84)

− RNA functions: not annotated

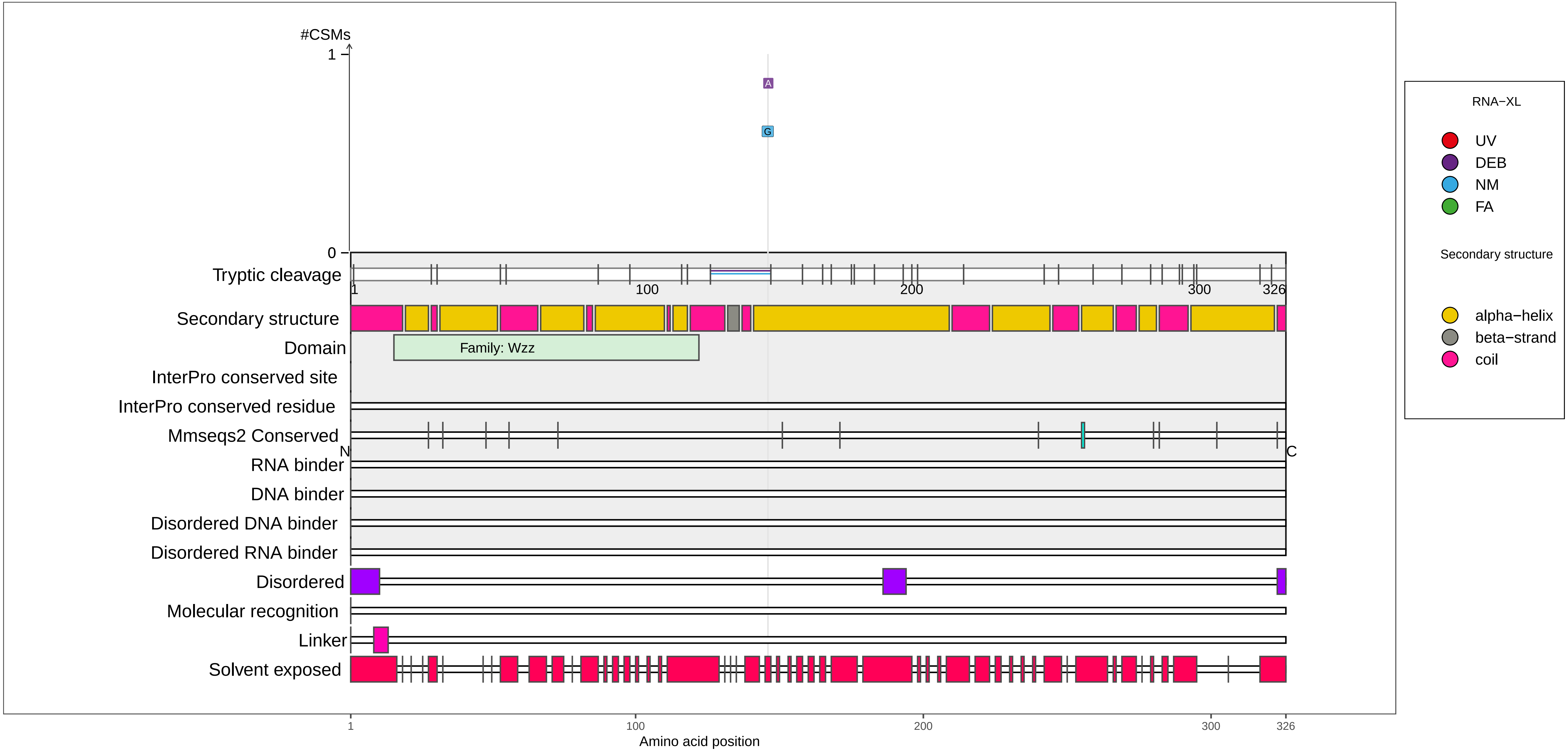

P76418
YEGU_ECOLI Uncharacterized protein YegU

− Abundance:
tryptic [log10 Intensity]: 8.16 (Q 63)
PAXdb K12 strain [ppm]: 0.77 (Q 4)
PAXdb E.coli [ppm]: 1.14 (Q 51)

− RNA functions: not annotated

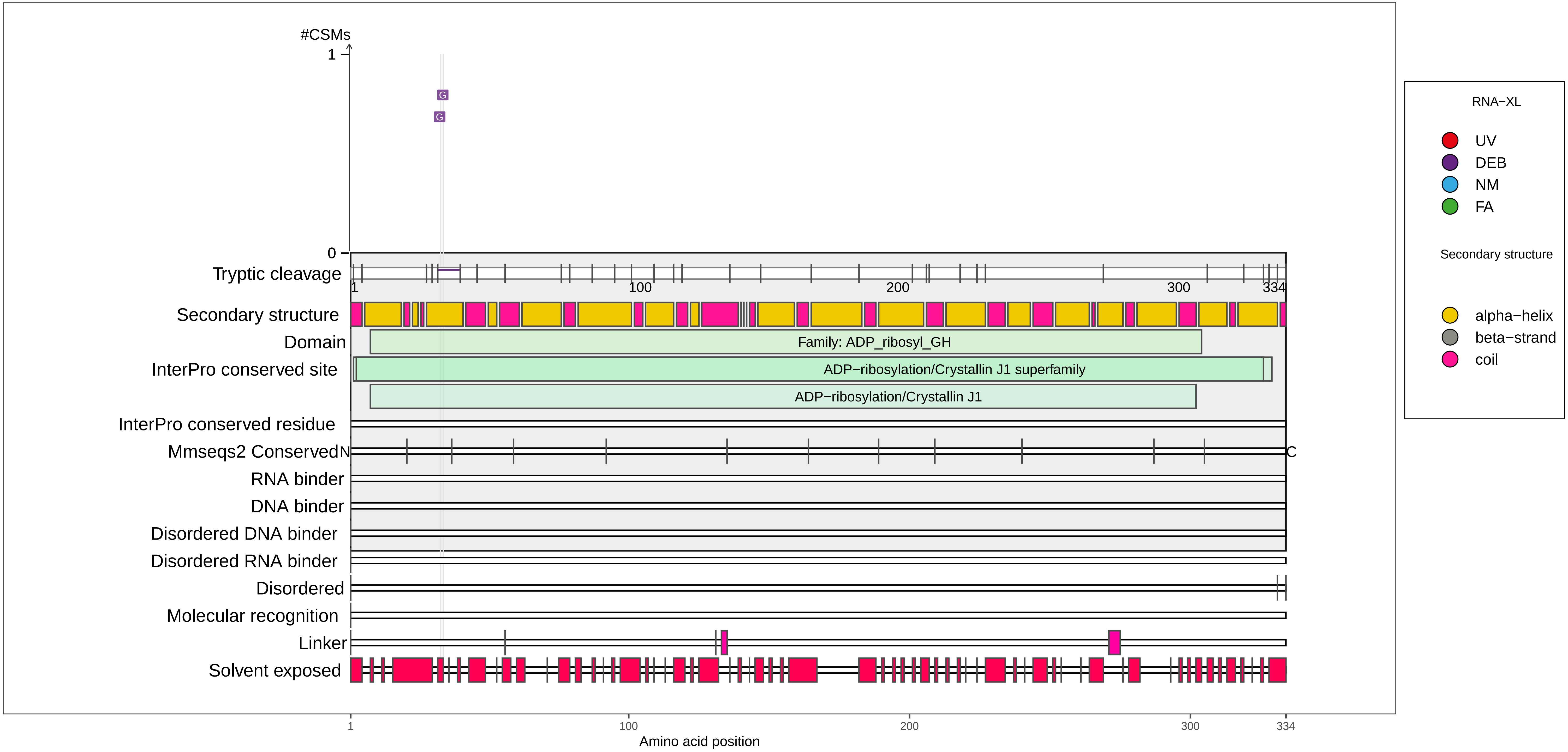

P76510
YFDN_ECOLI Uncharacterized protein YfdN

− Abundance:
tryptic [log10 Intensity]: NA (Q NA)
PAXdb K12 strain [ppm]: 1.99 (Q 60)
PAXdb E.coli [ppm]: −0.78 (Q 12)

− RNA functions: not annotated

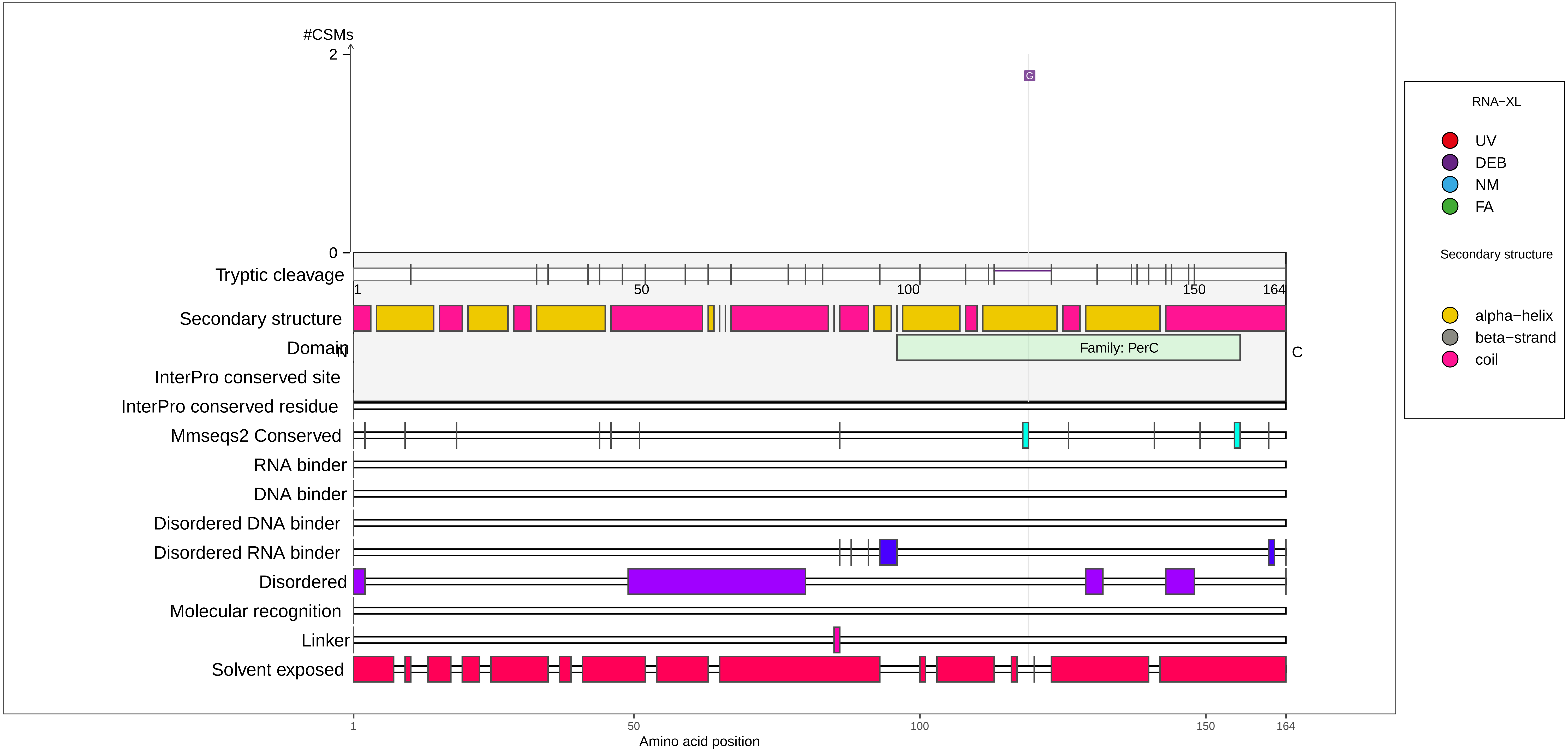

P76575
YFGJ_ECOLI Uncharacterized protein YfgJ

− Abundance:
tryptic [log10 Intensity]: NA (Q NA)
PAXdb K12 strain [ppm]: NA (Q NA)
PAXdb E.coli [ppm]: 0.7 (Q 41)

− RNA functions: not annotated

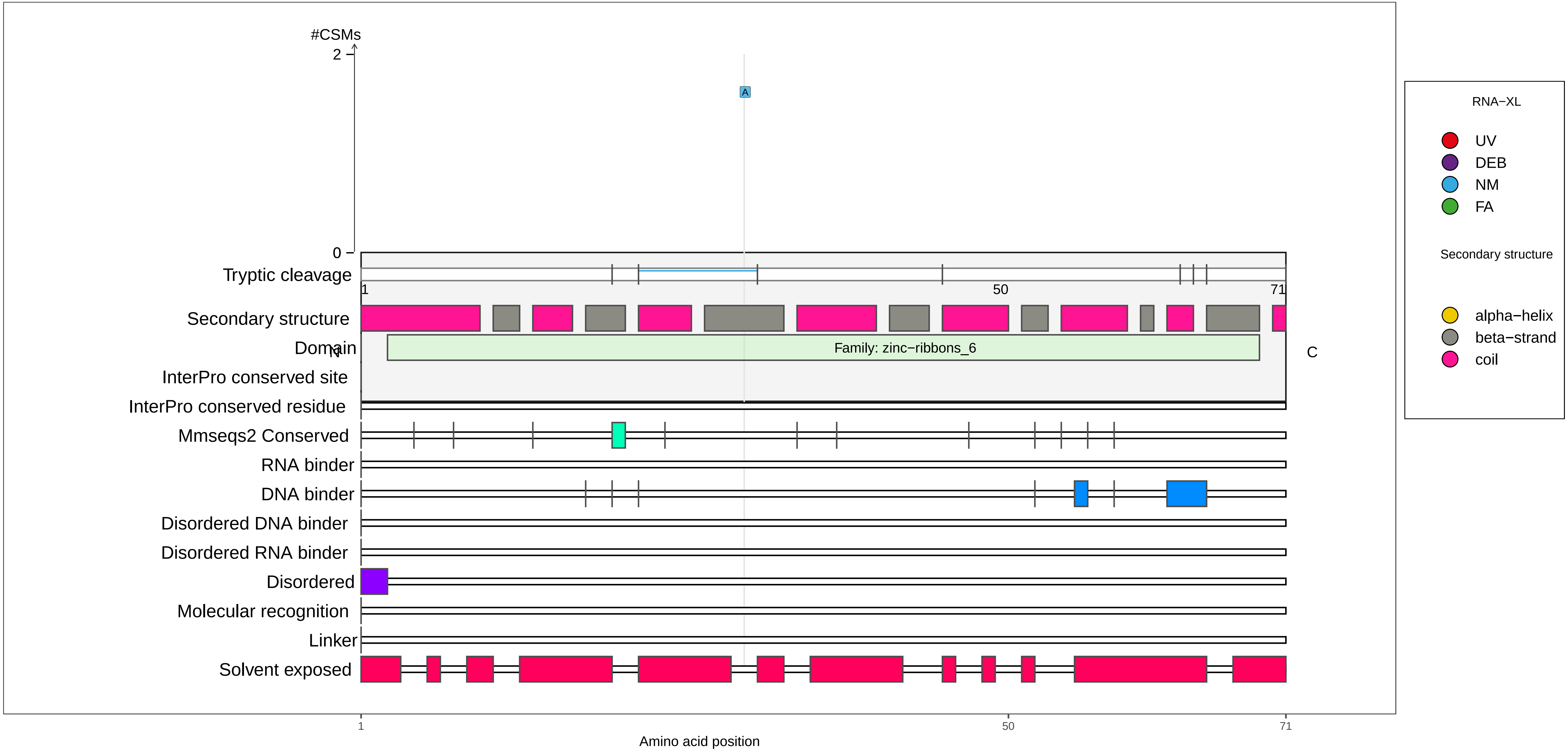

P76578
A2MG_ECOLI Alpha−2−macroglobulin

− Abundance:
tryptic [log10 Intensity]: 7.85 (Q 51)
PAXdb K12 strain [ppm]: 1.09 (Q 15)
PAXdb E.coli [ppm]: 1.21 (Q 53)

− RNA functions: not annotated

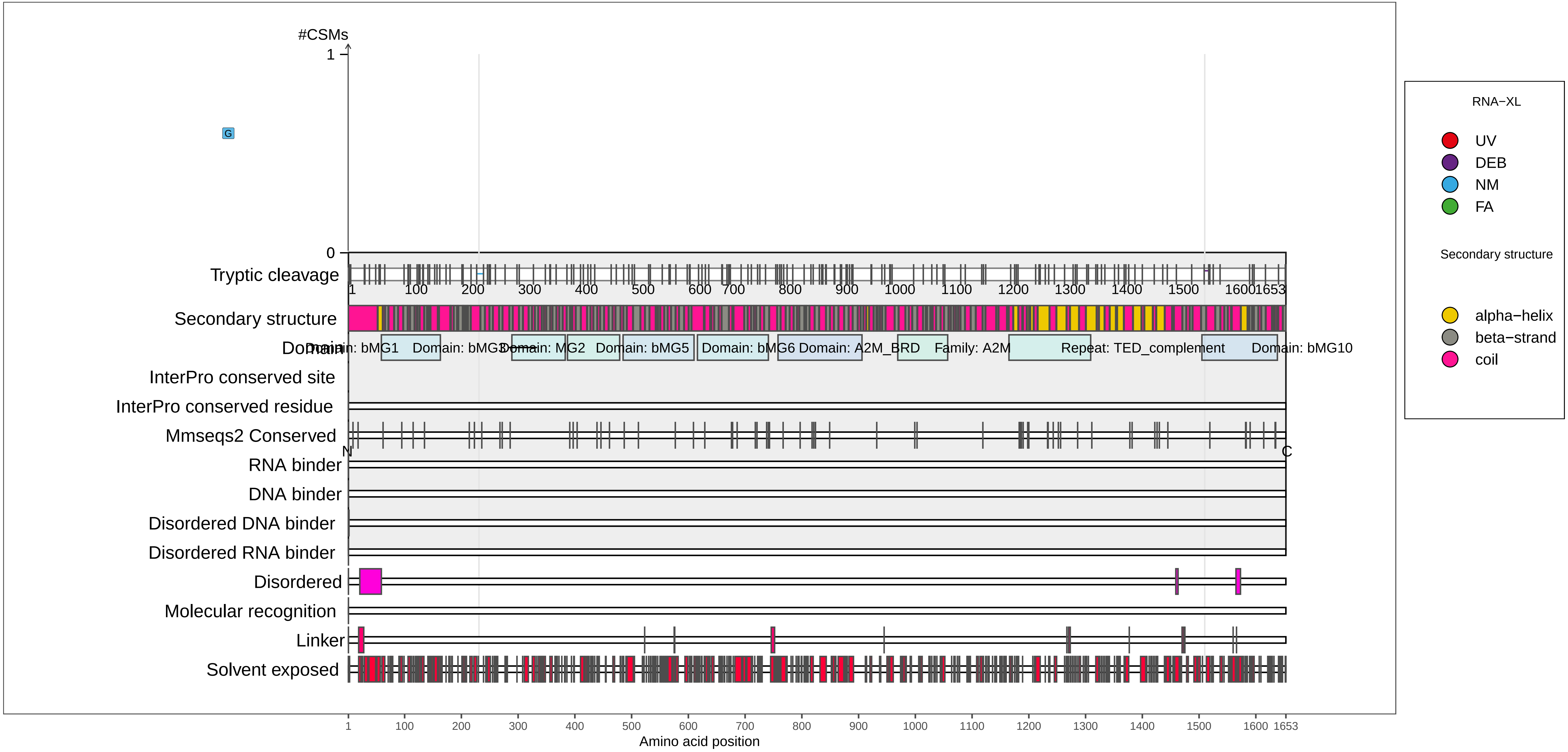

P76594
LYSAC_ECOLI Peptidyl−lysine N−acetyltransferase PatZ

− Abundance:
tryptic [log10 Intensity]: 7.44 (Q 32)
PAXdb K12 strain [ppm]: 1.5 (Q 38)
PAXdb E.coli [ppm]: 1.81 (Q 67)

− RNA functions: not annotated

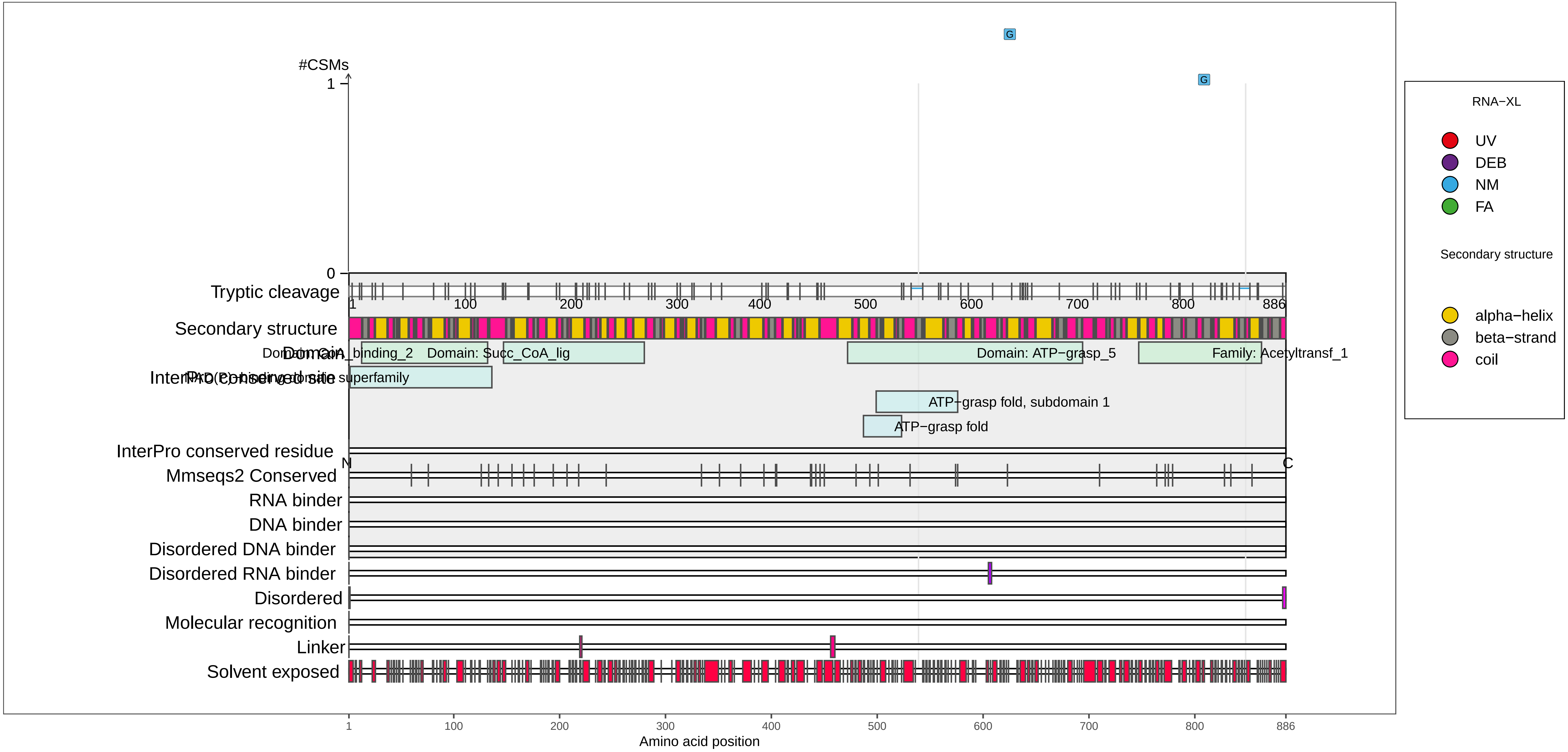

P77279
FETA_ECOLI Probable iron export ATP−binding protein FetA

− Abundance:
tryptic [log10 Intensity]: 7.01 (Q 13)
PAXdb K12 strain [ppm]: 1.62 (Q 46)
PAXdb E.coli [ppm]: 0.93 (Q 46)

− RNA functions: not annotated

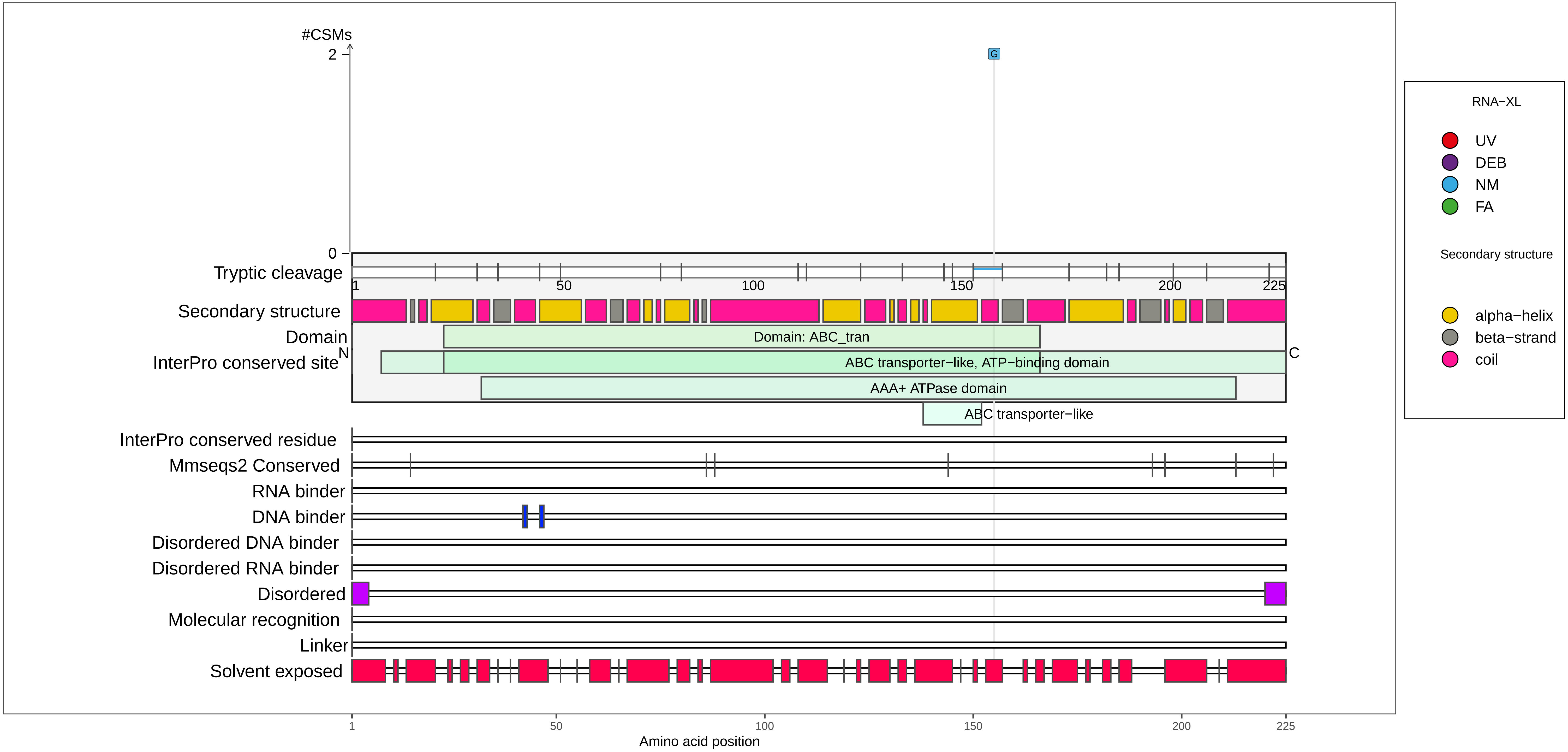

P77376
YDGJ_ECOLI Uncharacterized oxidoreductase YdgJ

− Abundance:
tryptic [log10 Intensity]: 7.89 (Q 53)
PAXdb K12 strain [ppm]: 1.38 (Q 31)
PAXdb E.coli [ppm]: 2.04 (Q 73)

− RNA functions: not annotated

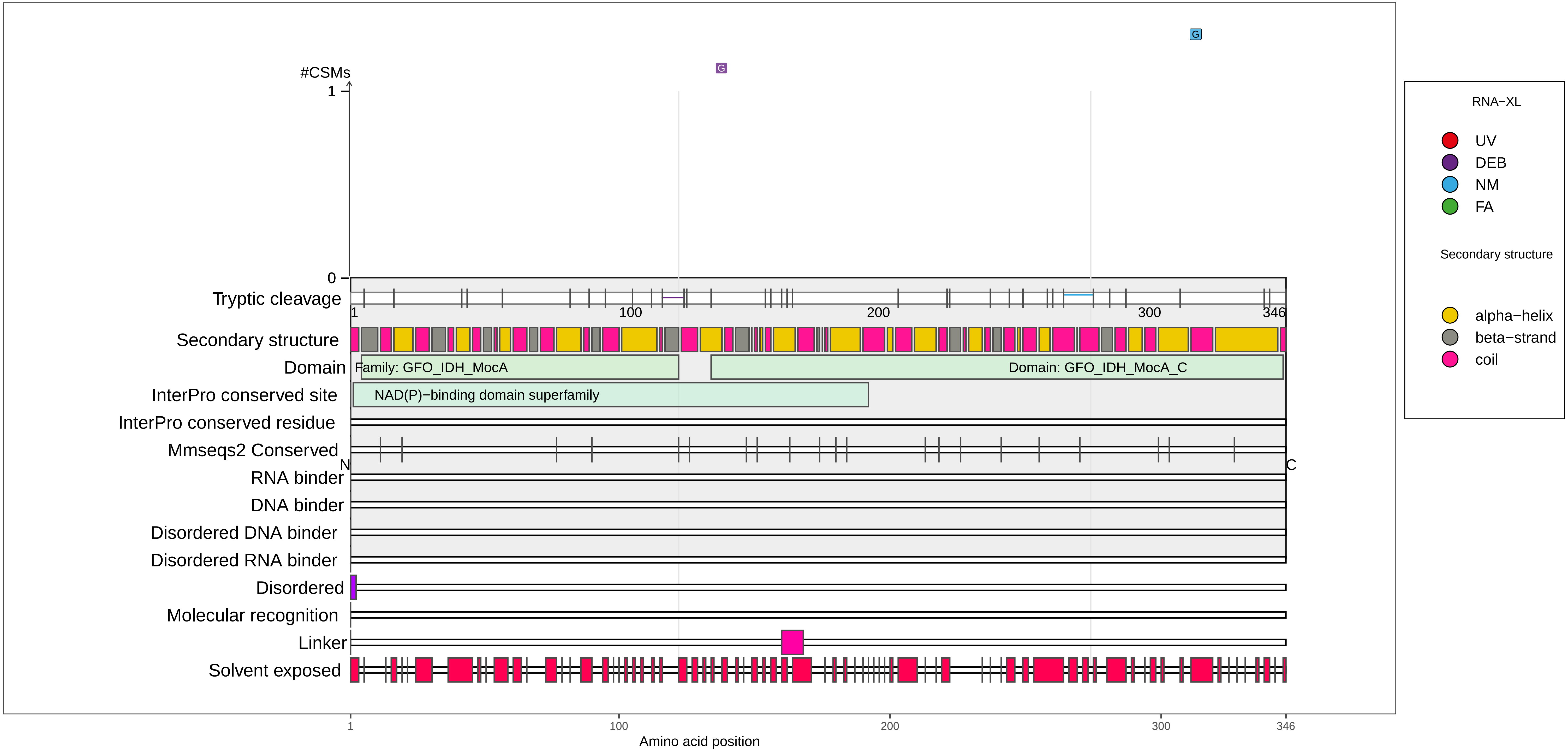

P77536
YKGF_ECOLI Uncharacterized electron transport protein YkgF
− Abundance:
tryptic [log10 Intensity]: 7.39 (Q 29)
PAXdb K12 strain [ppm]: 1.32 (Q 28)
PAXdb E.coli [ppm]: 0.67 (Q 40)

− RNA functions: not annotated

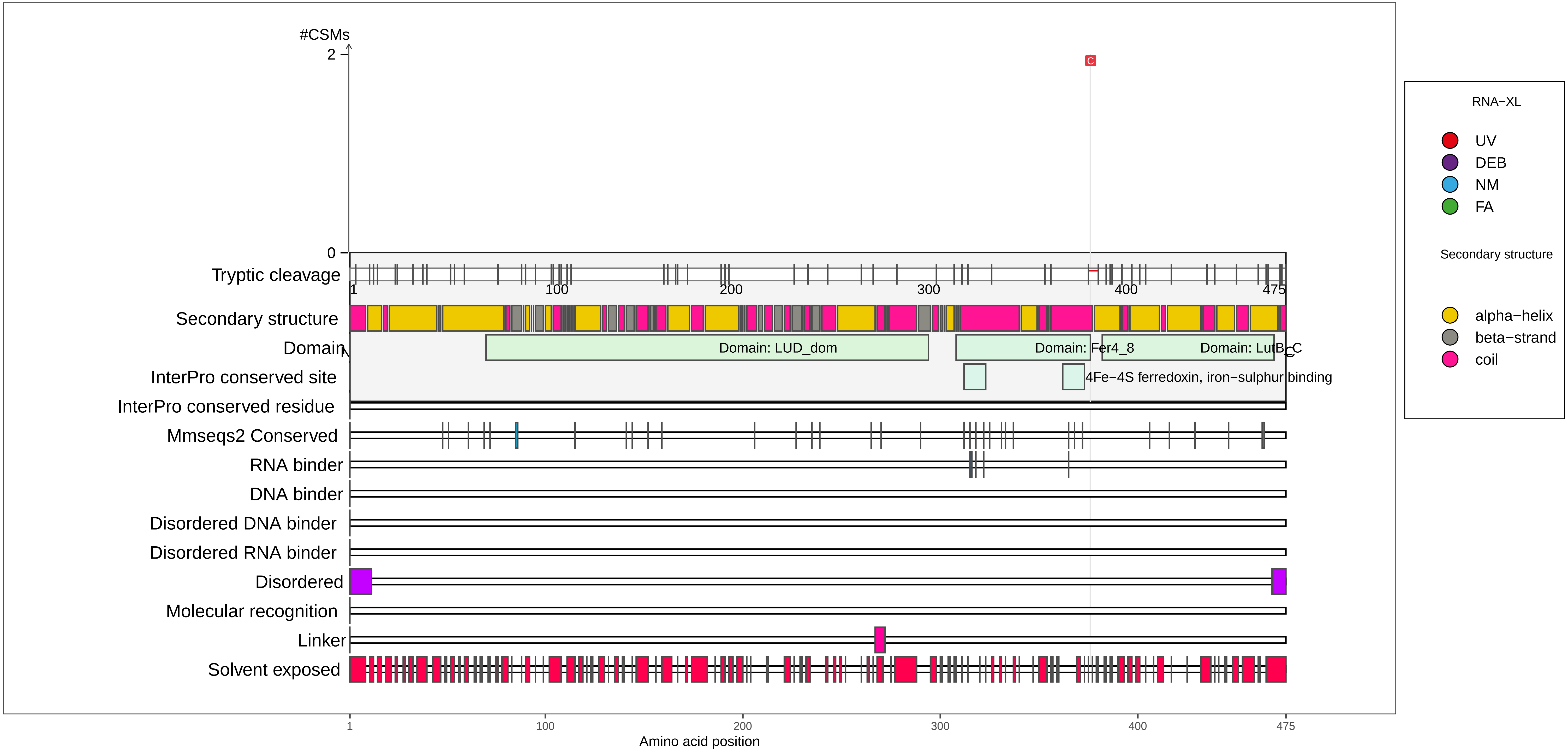

P77539
YDJL_ECOLI Uncharacterized zinc−type alcohol dehydrogenase−like protein YdjL

− Abundance:
tryptic [log10 Intensity]: 6.85 (Q 8)
PAXdb K12 strain [ppm]: NA (Q NA)
PAXdb E.coli [ppm]: 0.37 (Q 33)

− RNA functions: not annotated

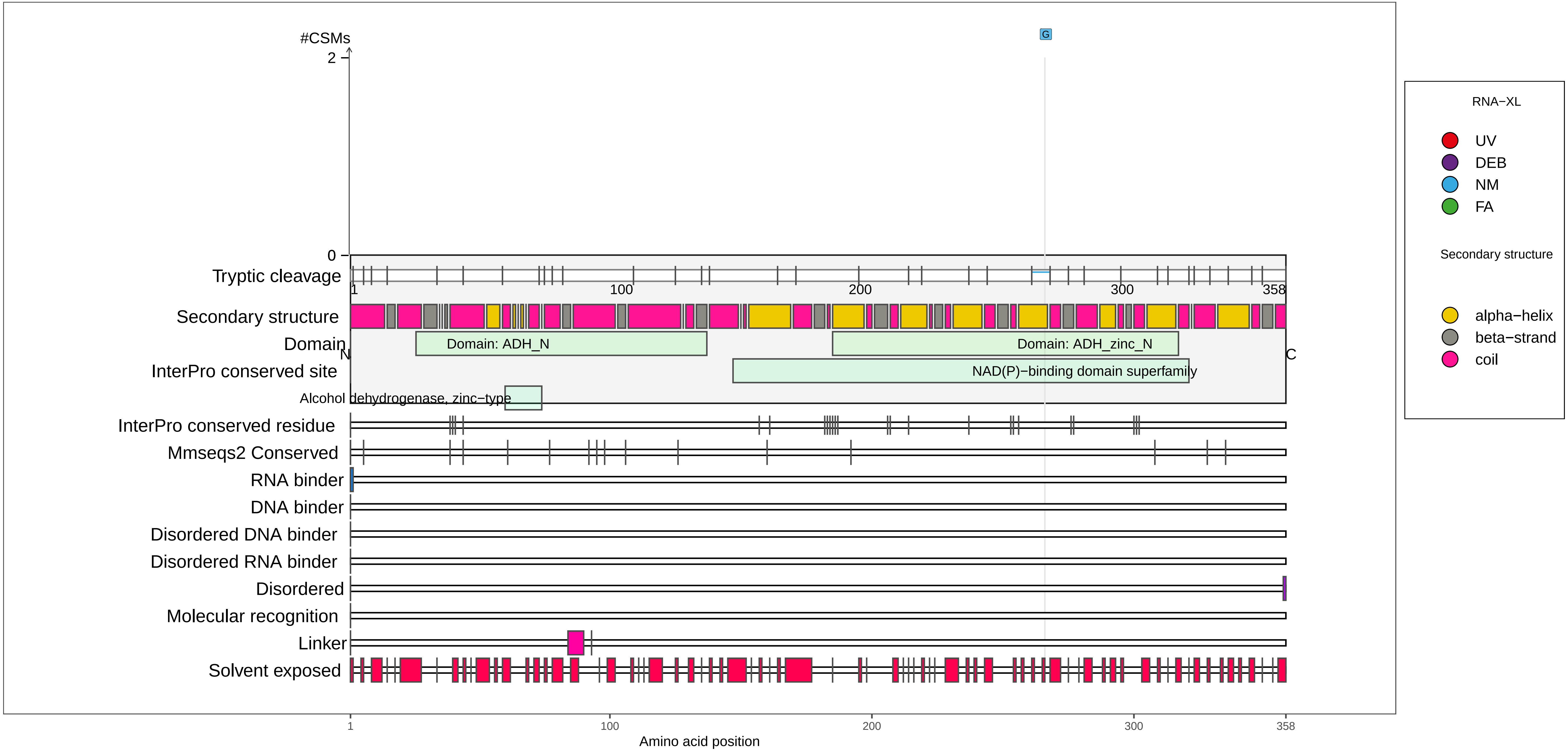

P77690
ARNB_ECOLI UDP−4−amino−4−deoxy−L−arabinose−−oxoglutarate aminotransferase

− Abundance:
tryptic [log10 Intensity]: 8.52 (Q 76)
PAXdb K12 strain [ppm]: 2.22 (Q 67)
PAXdb E.coli [ppm]: 2.07 (Q 74)

− RNA functions: not annotated

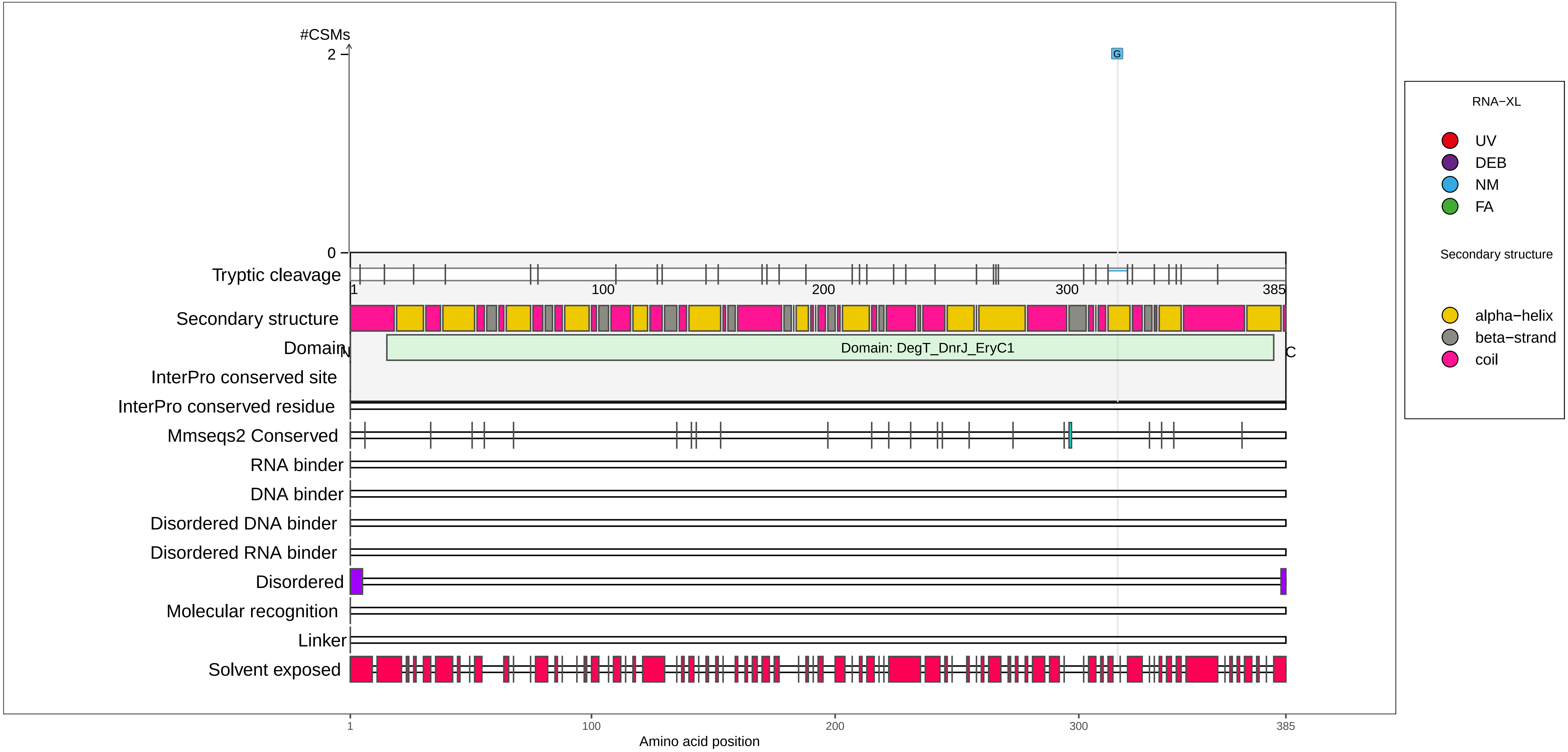

Q2M7R5
YIBT_ECOLI Uncharacterized protein YibT

− Abundance:
tryptic [log10 Intensity]: NA (Q NA)
PAXdb K12 strain [ppm]: NA (Q NA)
PAXdb E.coli [ppm]: 2.22 (Q 78)

− RNA functions: not annotated

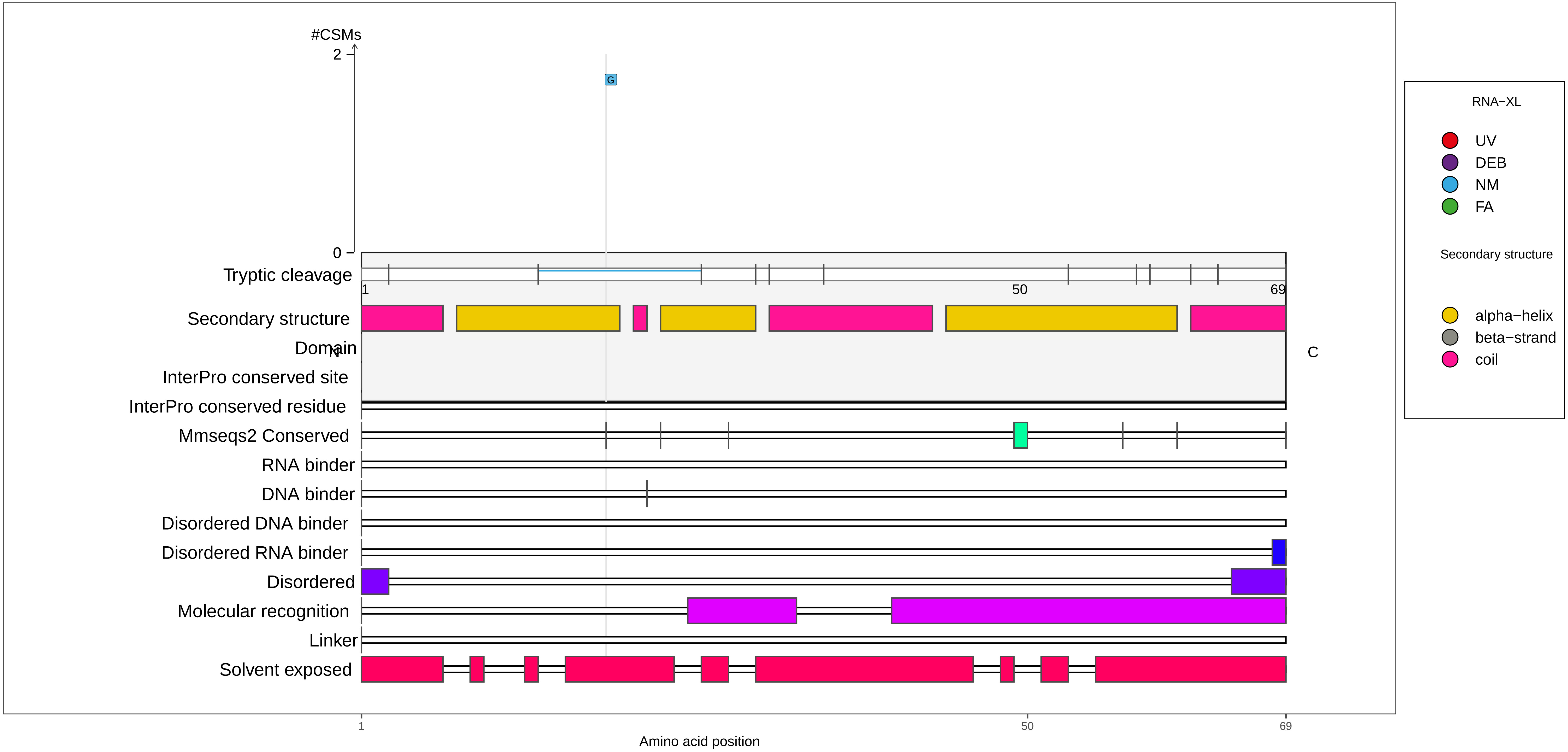

Q47622
SAPA_ECOLI Probable ABC transporter periplasmic−binding protein SapA

− Abundance:
tryptic [log10 Intensity]: 7.32 (Q 26)
PAXdb K12 strain [ppm]: 1.31 (Q 27)
PAXdb E.coli [ppm]: 1.05 (Q 49)

− RNA functions: not annotated

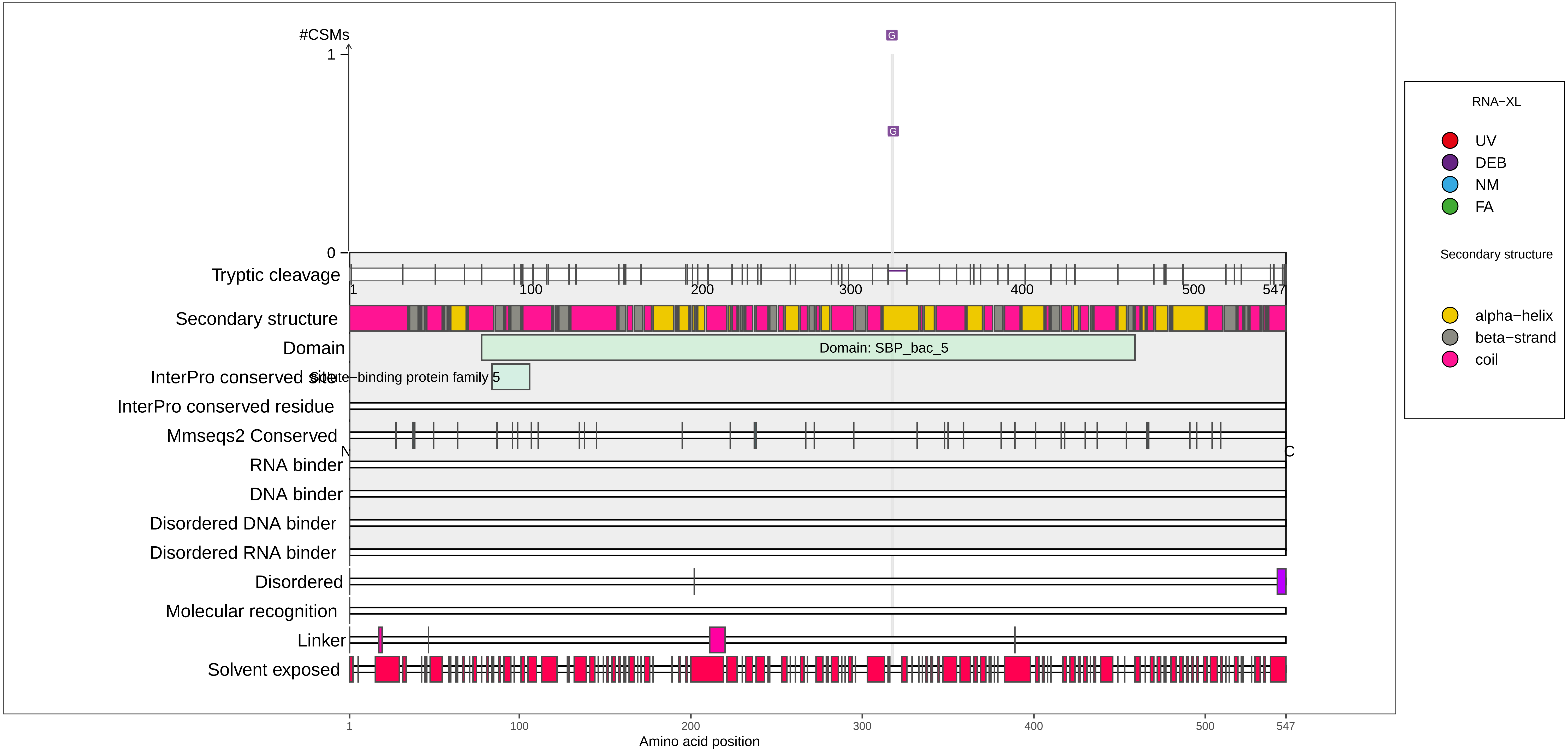

P00562
AK2H_ECOLI Bifunctional aspartokinase/homoserine dehydrogenase 2

− Abundance:
tryptic [log10 Intensity]: 7.5 (Q 35)
PAXdb K12 strain [ppm]: 2.06 (Q 62)
PAXdb E.coli [ppm]: 2.15 (Q 76)

− RNA functions: not annotated

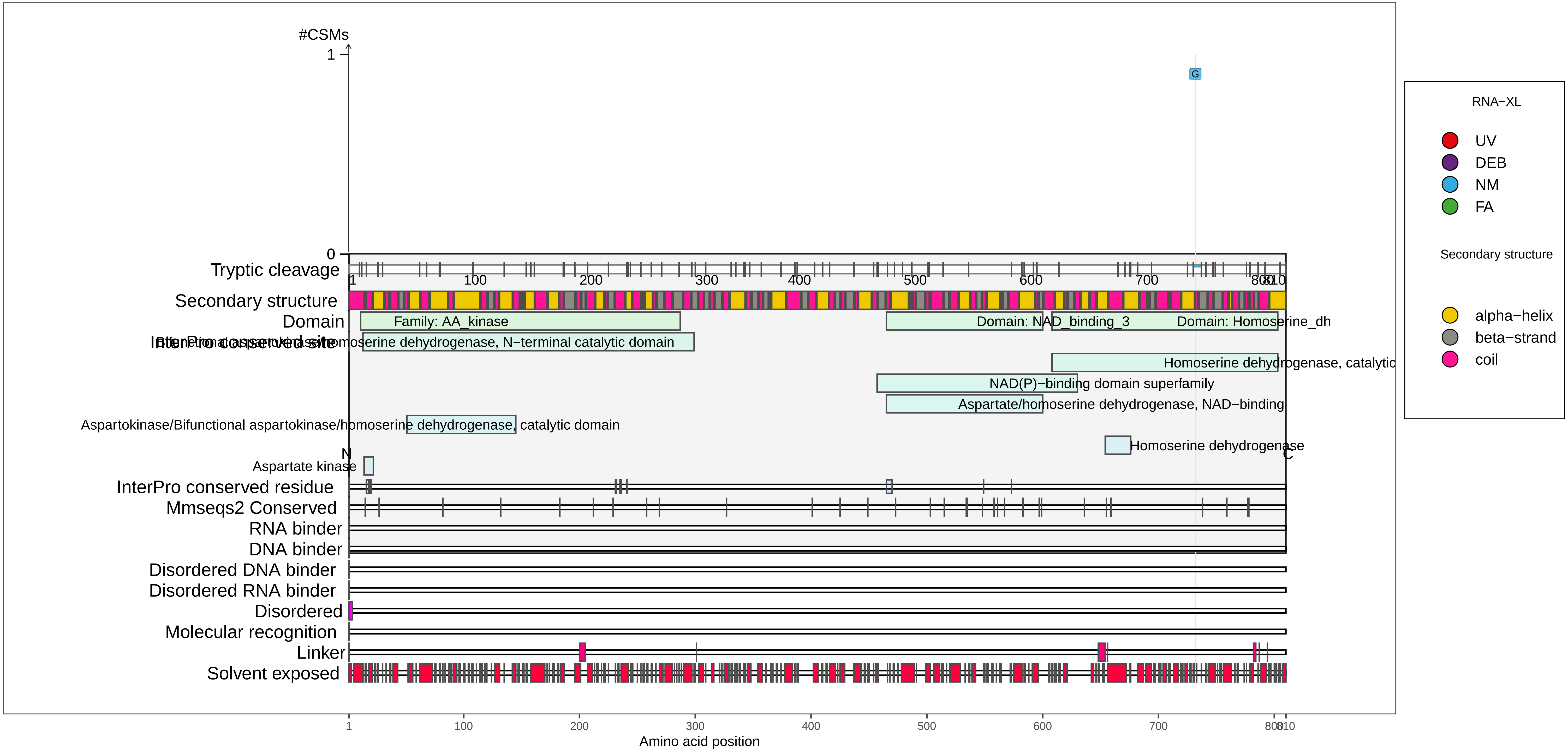

P03024
GALR_ECOLI HTH−type transcriptional regulator GalR
− Abundance:
tryptic [log10 Intensity]: 8.16 (Q 63)
PAXdb K12 strain [ppm]: 1.28 (Q 25)
PAXdb E.coli [ppm]: 0.28 (Q 31)

− RNA functions:
RNA biosynthetic process; RNA metabolic process

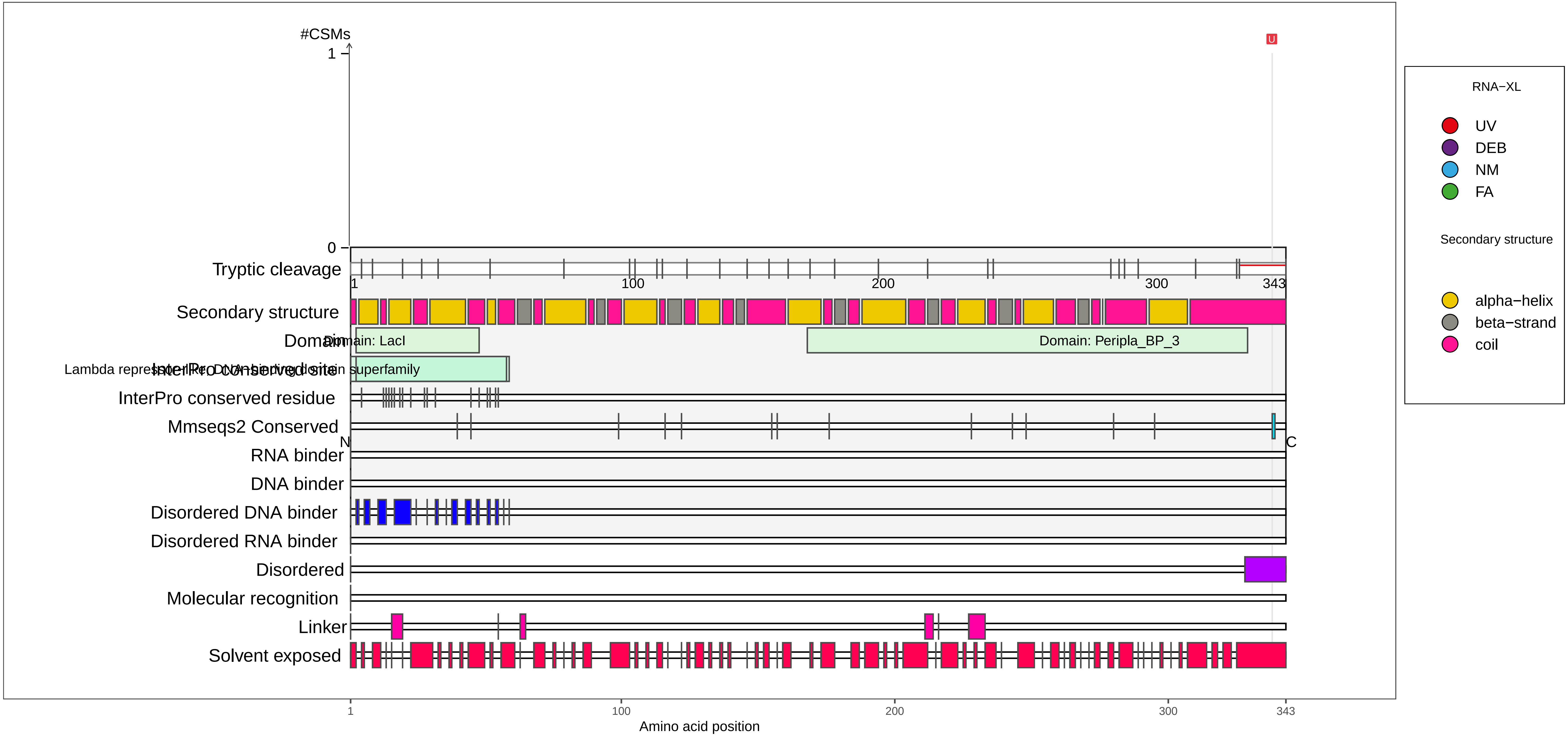

P03841
MALM_ECOLI Maltose operon periplasmic protein

− Abundance:
tryptic [log10 Intensity]: 7.64 (Q 42)
PAXdb K12 strain [ppm]: 3.06 (Q 92)
PAXdb E.coli [ppm]: 2.45 (Q 84)

− RNA functions: not annotated

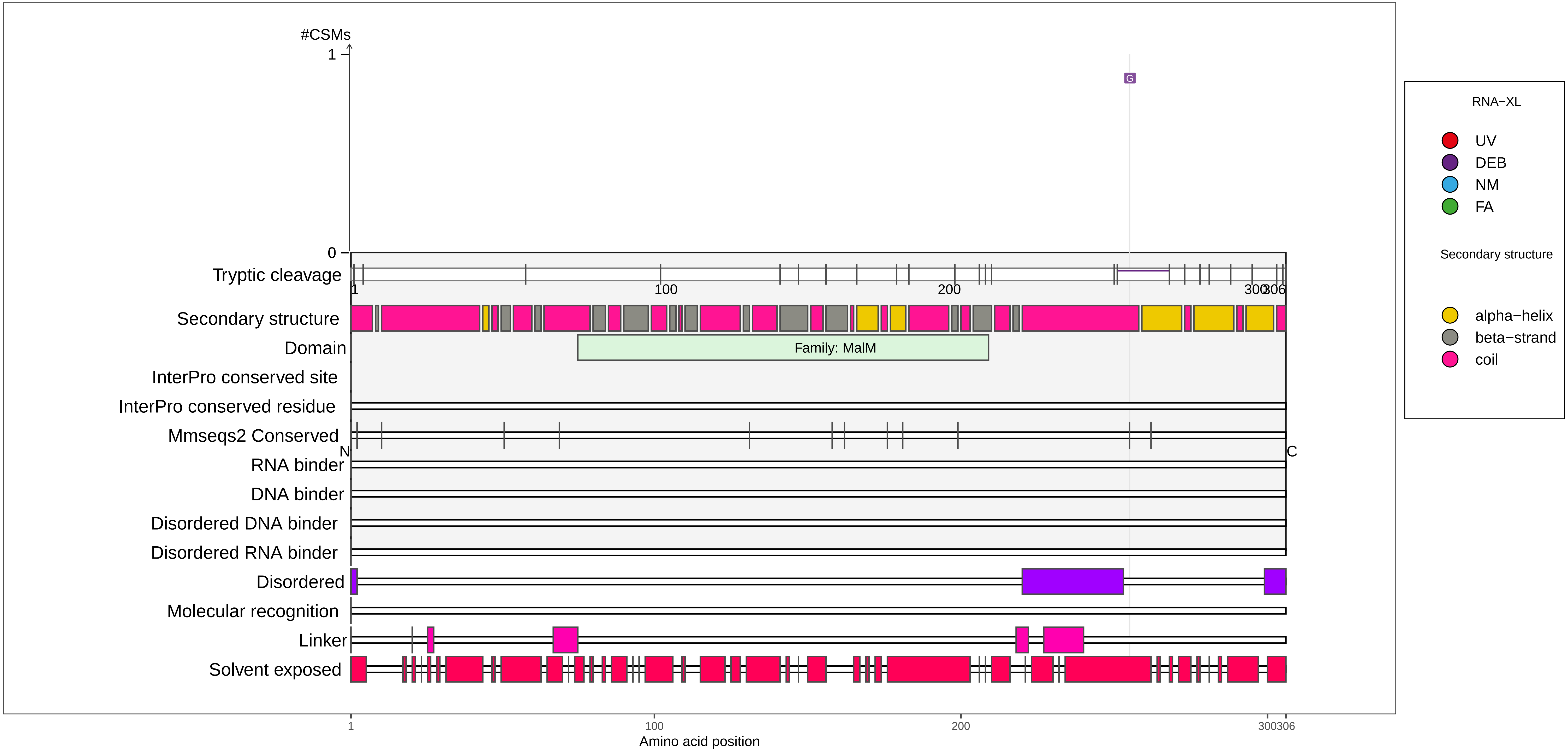

P04951
KDSB_ECOLI 3−deoxy−manno−octulosonate cytidylyltransferase

− Abundance:
tryptic [log10 Intensity]: 8.84 (Q 84)
PAXdb K12 strain [ppm]: 1.9 (Q 57)
PAXdb E.coli [ppm]: 2.45 (Q 84)

− RNA functions: not annotated

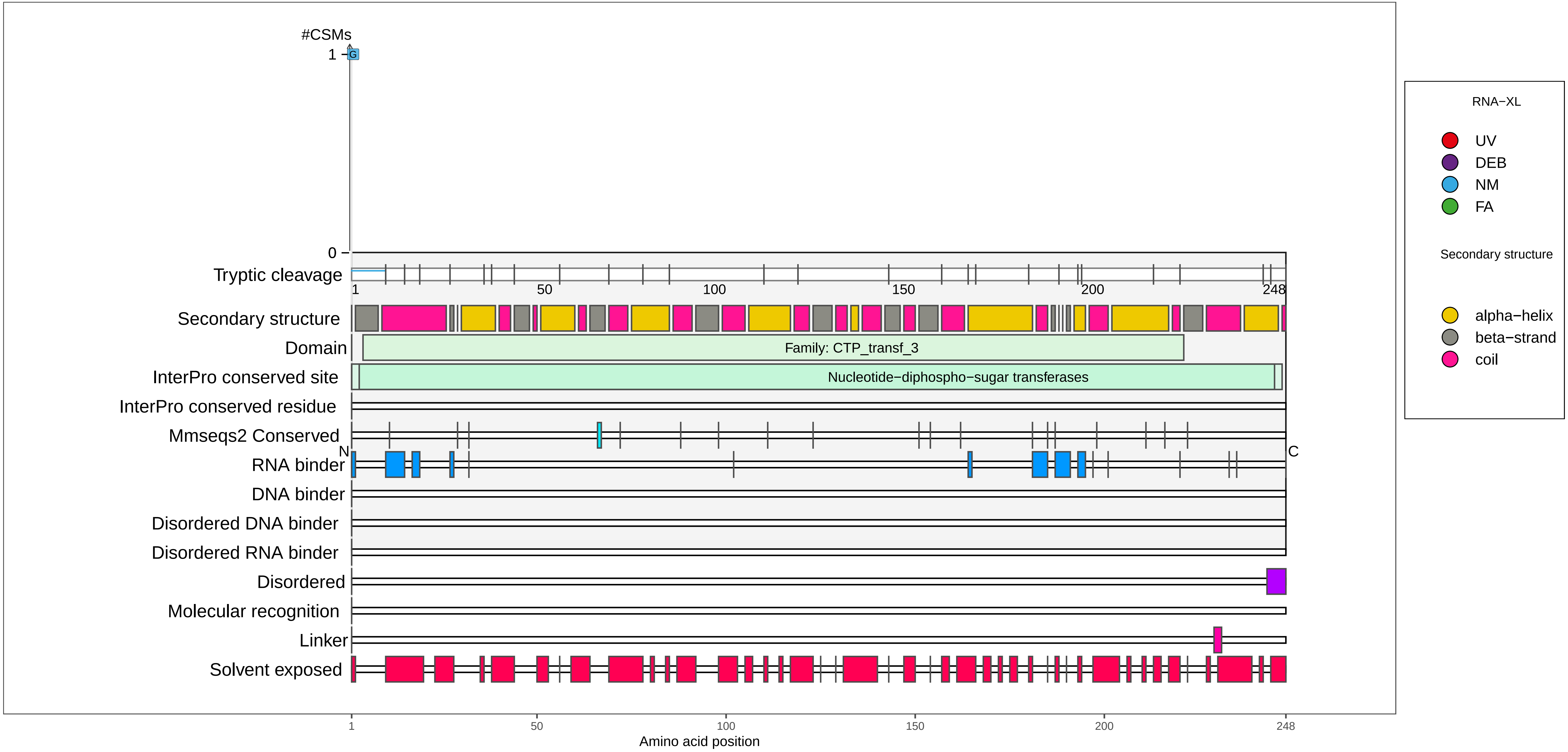

P05100
3MG1_ECOLI DNA−3−methyladenine glycosylase 1

− Abundance:
tryptic [log10 Intensity]: NA (Q NA)
PAXdb K12 strain [ppm]: NA (Q NA)
PAXdb E.coli [ppm]: 0.61 (Q 39)

− RNA functions: not annotated

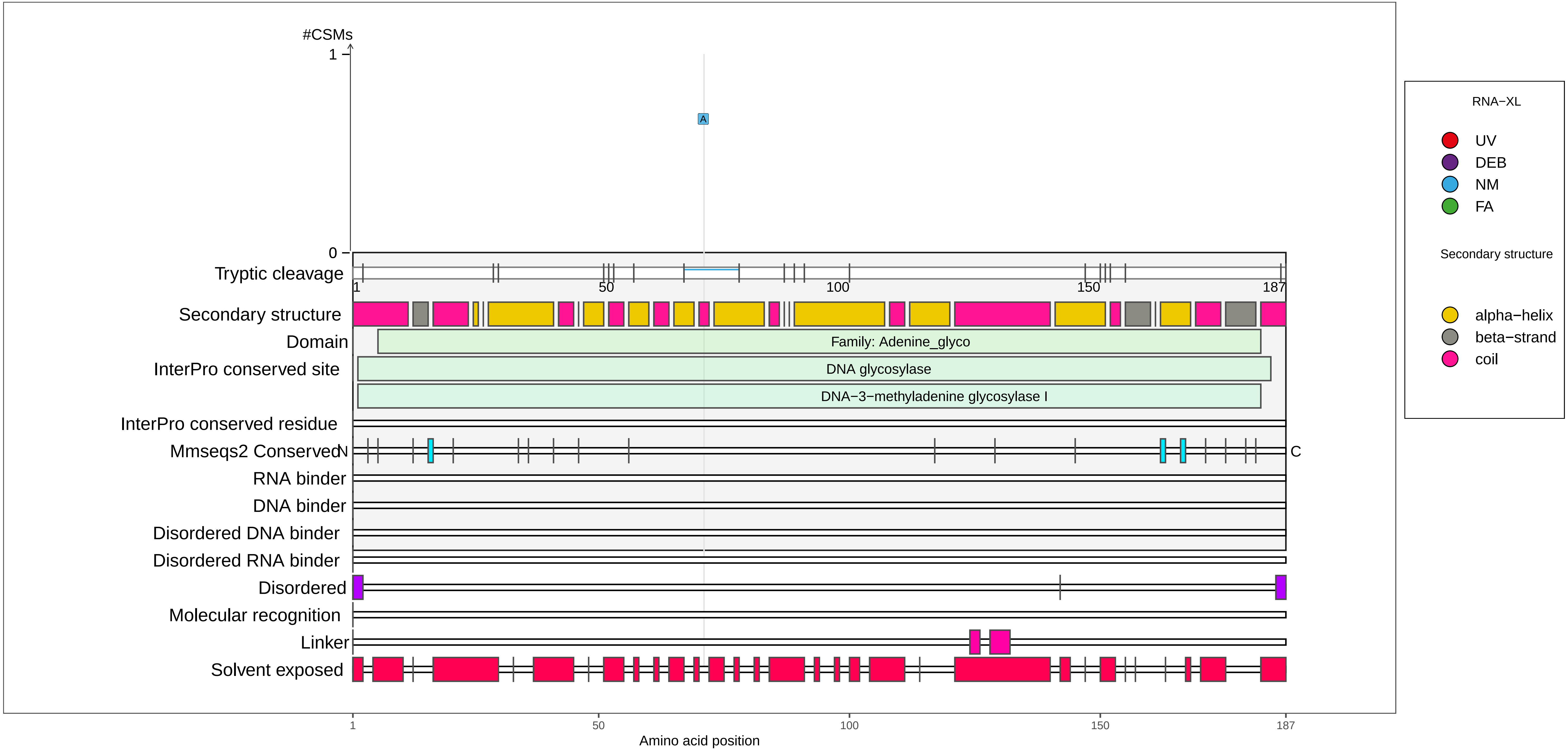

P05719
T1SK_ECOLI Type I restriction enzyme EcoKI specificity subunit

− Abundance:
tryptic [log10 Intensity]: NA (Q NA)
PAXdb K12 strain [ppm]: 1.18 (Q 19)
PAXdb E.coli [ppm]: 0.8 (Q 43)

− RNA functions: not annotated

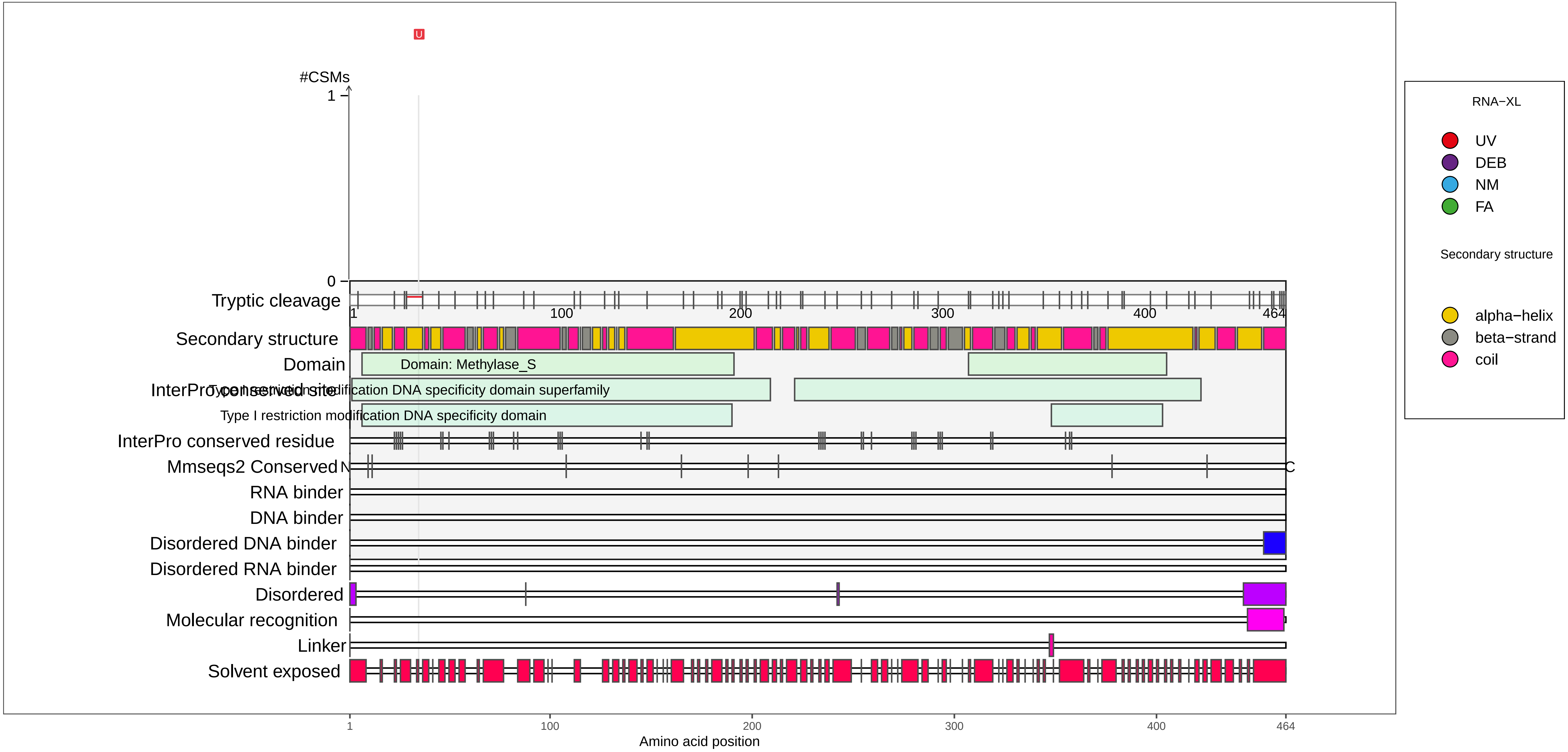

P07003
POXB_ECOLI Pyruvate dehydrogenase [ubiquinone]

− Abundance:
tryptic [log10 Intensity]: 7.16 (Q 19)
PAXdb K12 strain [ppm]: 2.91 (Q 89)
PAXdb E.coli [ppm]: 2.59 (Q 88)

− RNA functions: not annotated

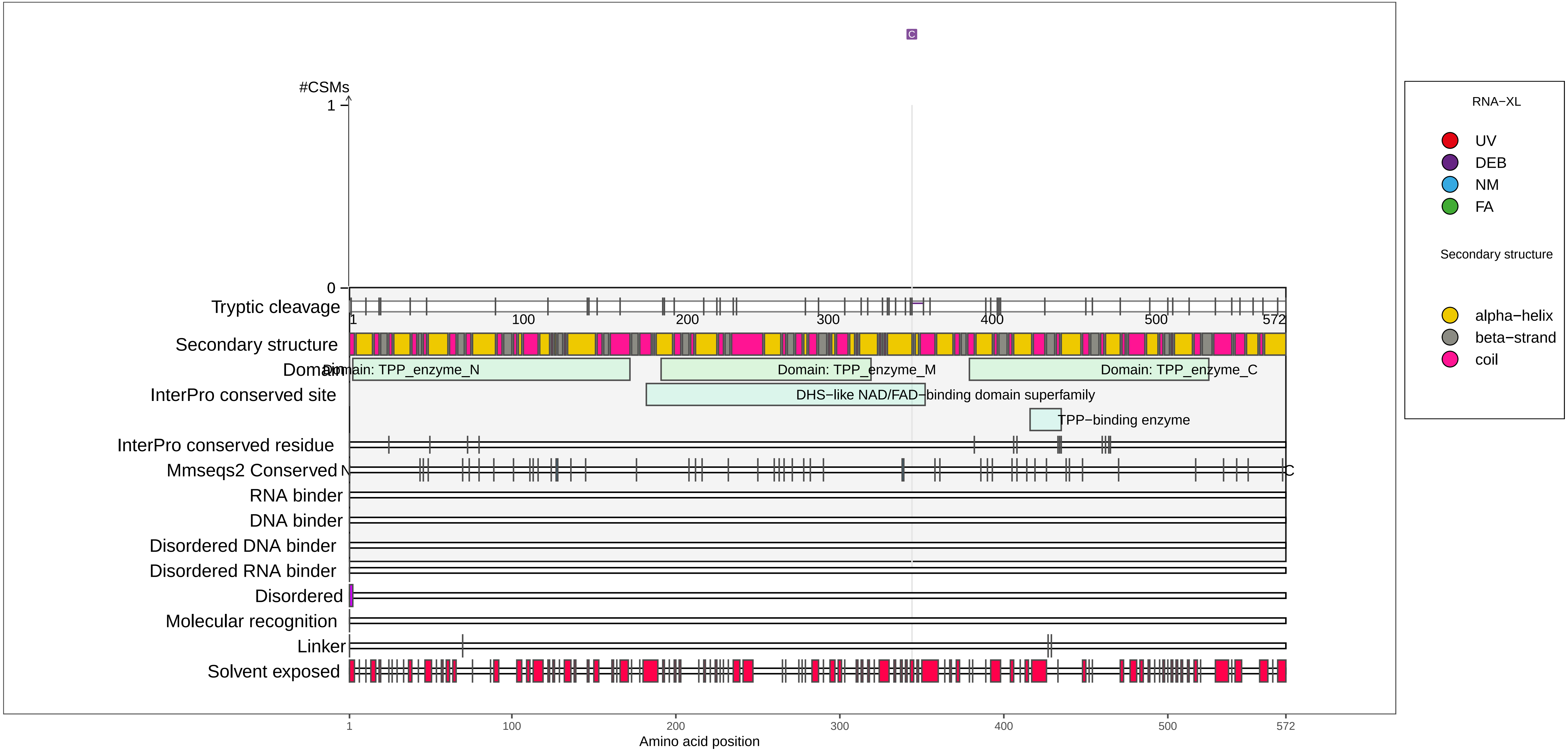

P07013
PRIB_ECOLI Primosomal replication protein N

− Abundance:
tryptic [log10 Intensity]: 7.68 (Q 44)
PAXdb K12 strain [ppm]: NA (Q NA)
PAXdb E.coli [ppm]: 0.31 (Q 32)

− RNA functions:
DNA replication, synthesis of RNA primer; RNA biosynthetic process
RNA metabolic process

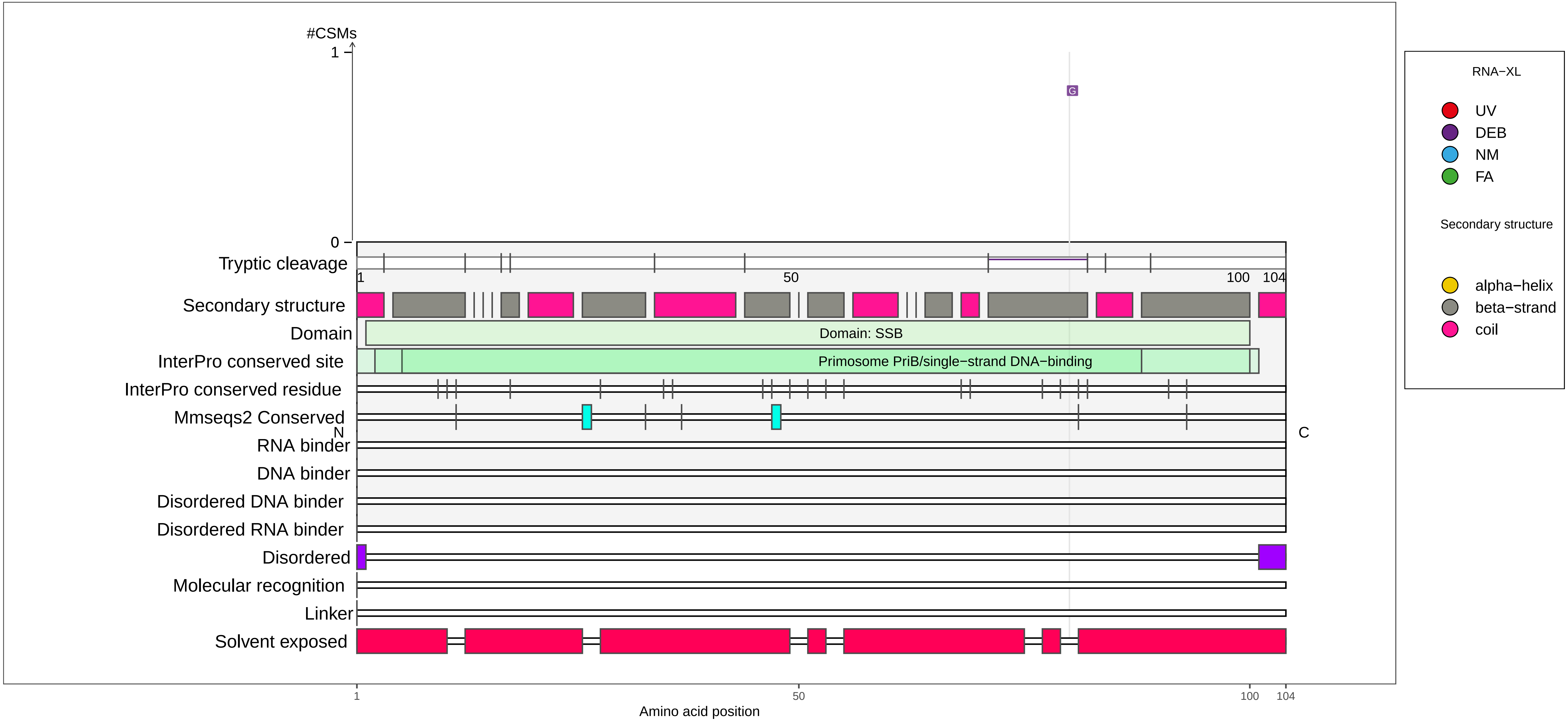

P07604
TYRR_ECOLI HTH−type transcriptional regulatory protein TyrR

− Abundance:
tryptic [log10 Intensity]: 6.43 (Q 2)
PAXdb K12 strain [ppm]: NA (Q NA)
PAXdb E.coli [ppm]: 1.78 (Q 66)

− RNA functions: not annotated

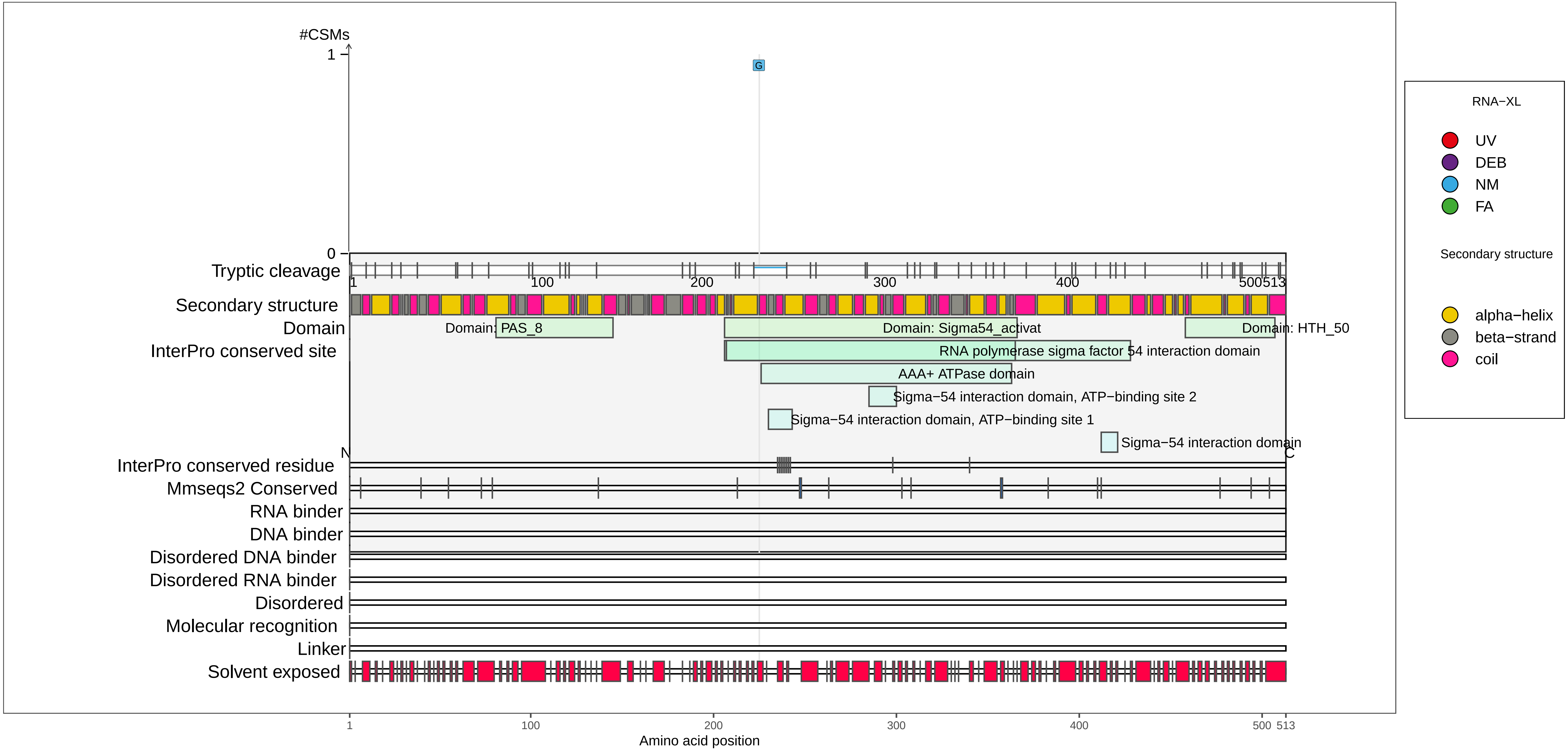

P07648
RECC_ECOLI RecBCD enzyme subunit RecC

− Abundance:
tryptic [log10 Intensity]: 7.06 (Q 15)
PAXdb K12 strain [ppm]: 1.16 (Q 18)
PAXdb E.coli [ppm]: 0.39 (Q 33)

− RNA functions: not annotated

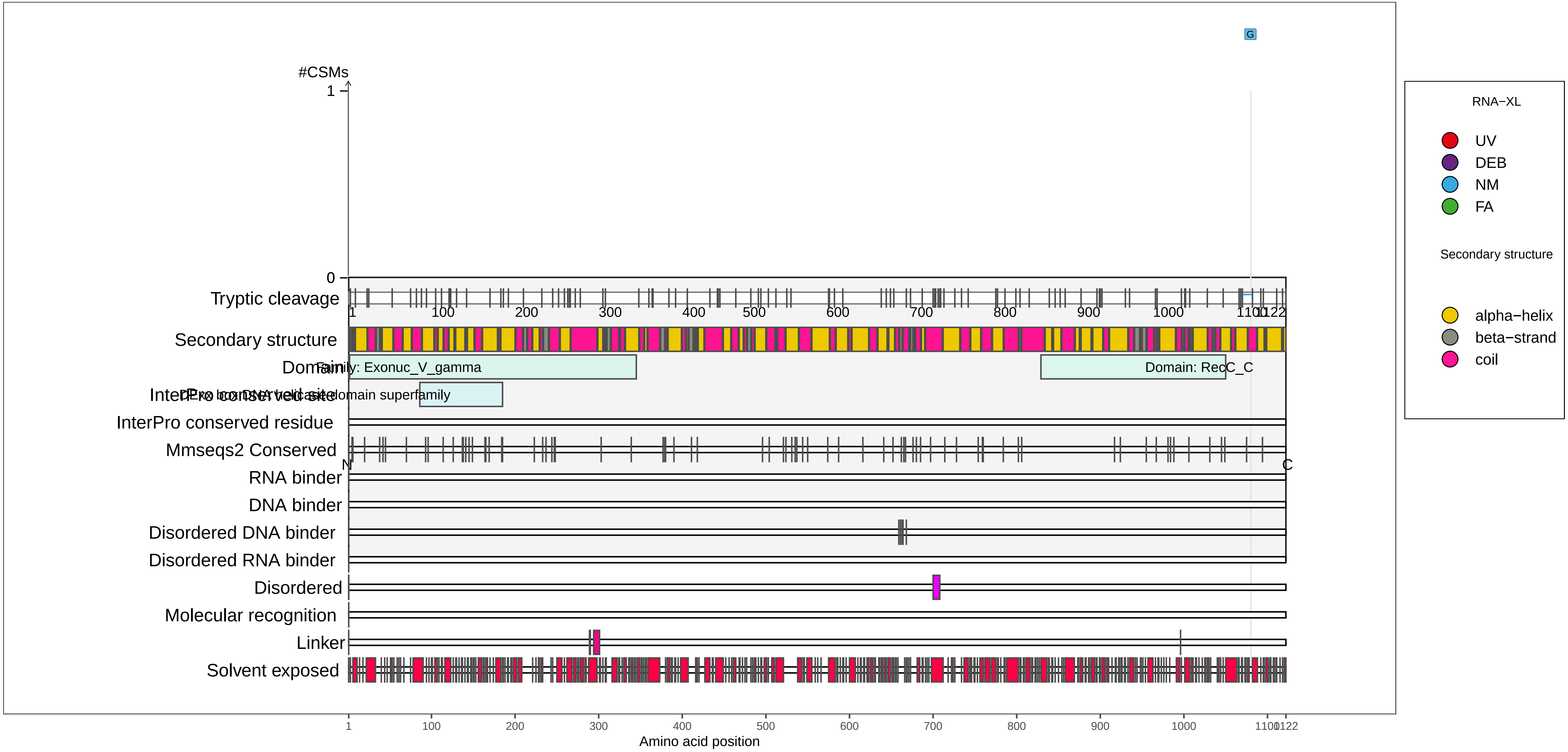

P08956
T1RK_ECOLI Type I restriction enzyme EcoKI endonuclease subunit

− Abundance:
tryptic [log10 Intensity]: 8.21 (Q 66)
PAXdb K12 strain [ppm]: 1.14 (Q 17)
PAXdb E.coli [ppm]: 1.16 (Q 51)

− RNA functions: not annotated

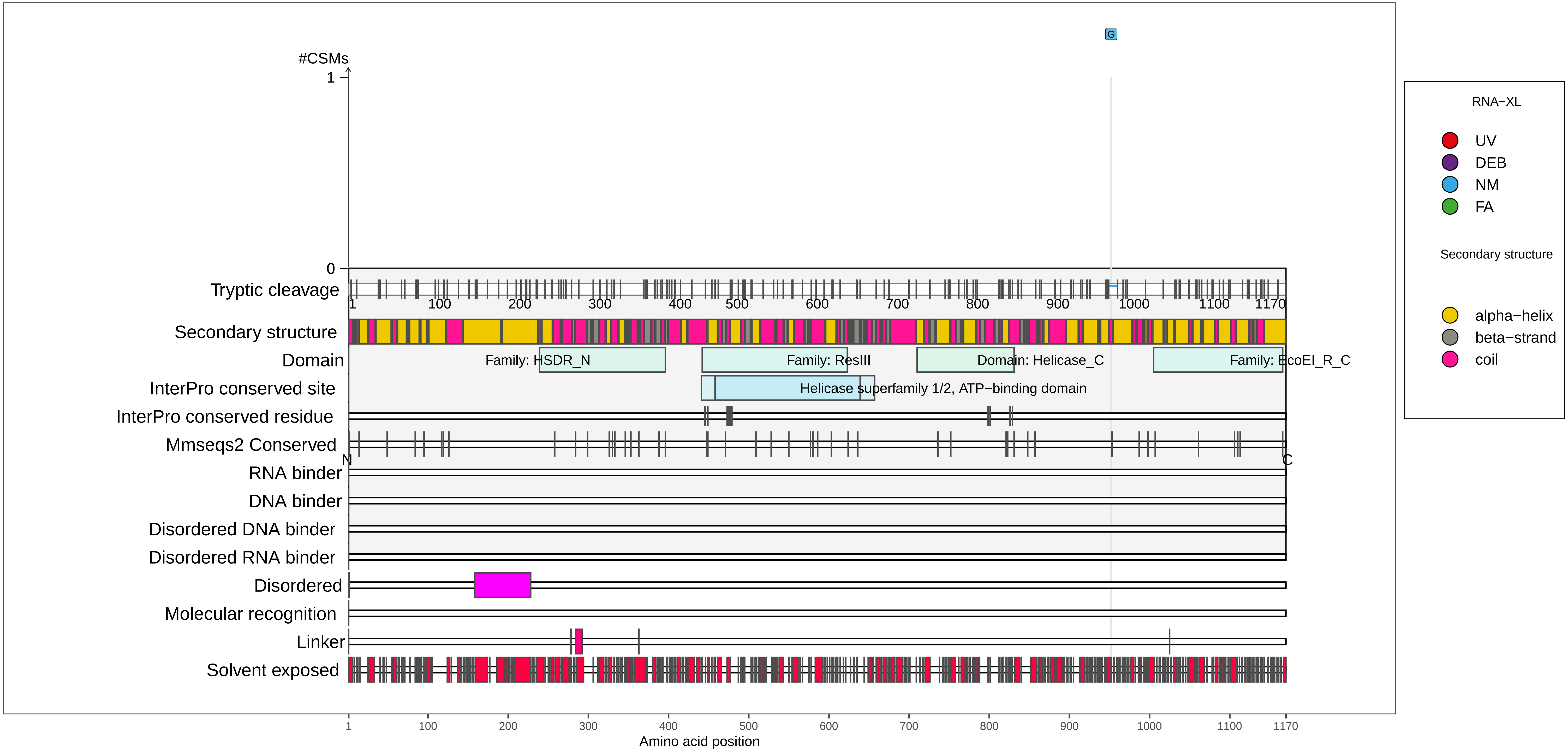

P09053
AVTA_ECOLI Valine−−pyruvate aminotransferase

− Abundance:
tryptic [log10 Intensity]: 8.28 (Q 69)
PAXdb K12 strain [ppm]: 1.93 (Q 58)
PAXdb E.coli [ppm]: 1.52 (Q 60)

− RNA functions: not annotated

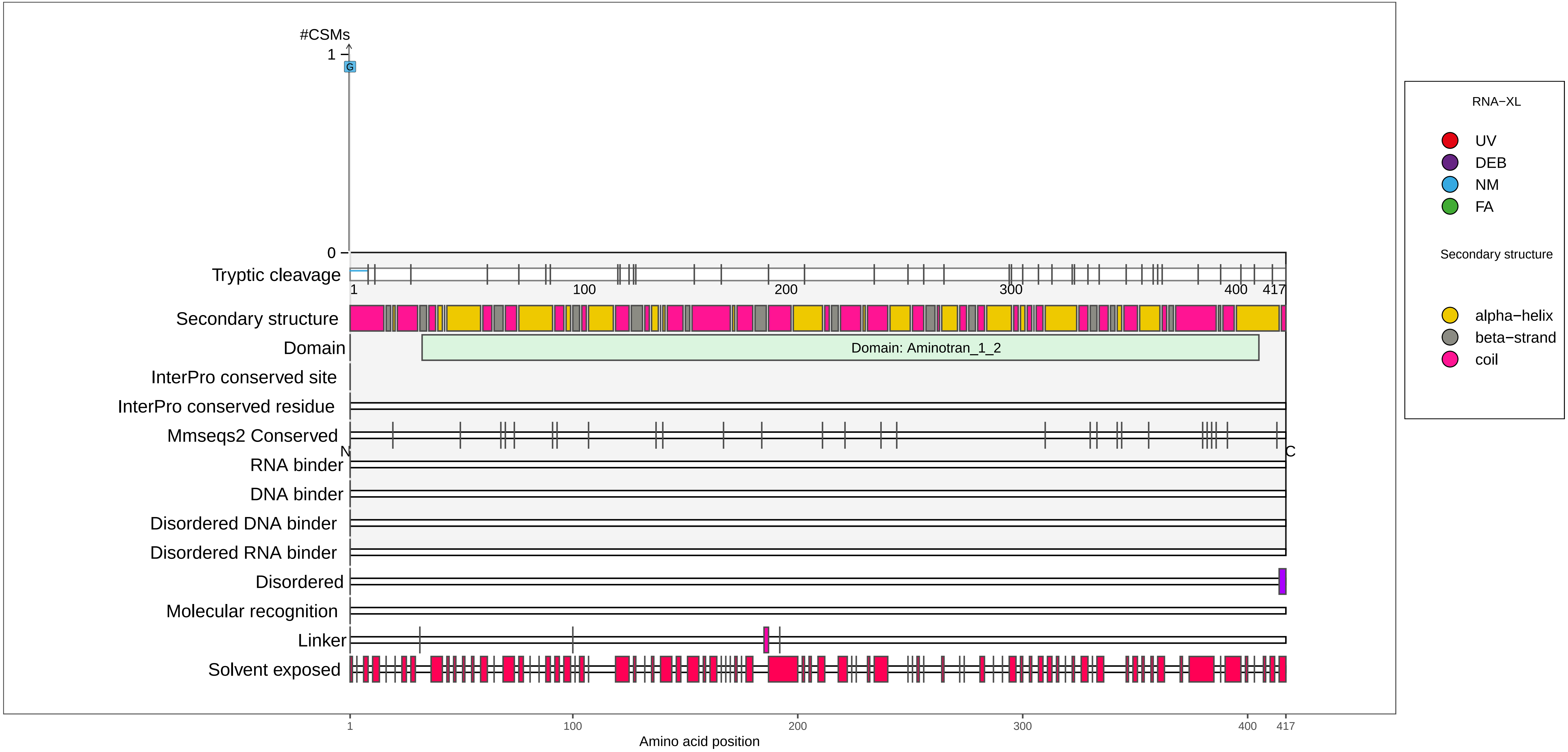

P09152
NARG_ECOLI Respiratory nitrate reductase 1 alpha chain

− Abundance:
tryptic [log10 Intensity]: NA (Q NA)
PAXdb K12 strain [ppm]: 0.96 (Q 9)
PAXdb E.coli [ppm]: 1.46 (Q 59)

− RNA functions: not annotated

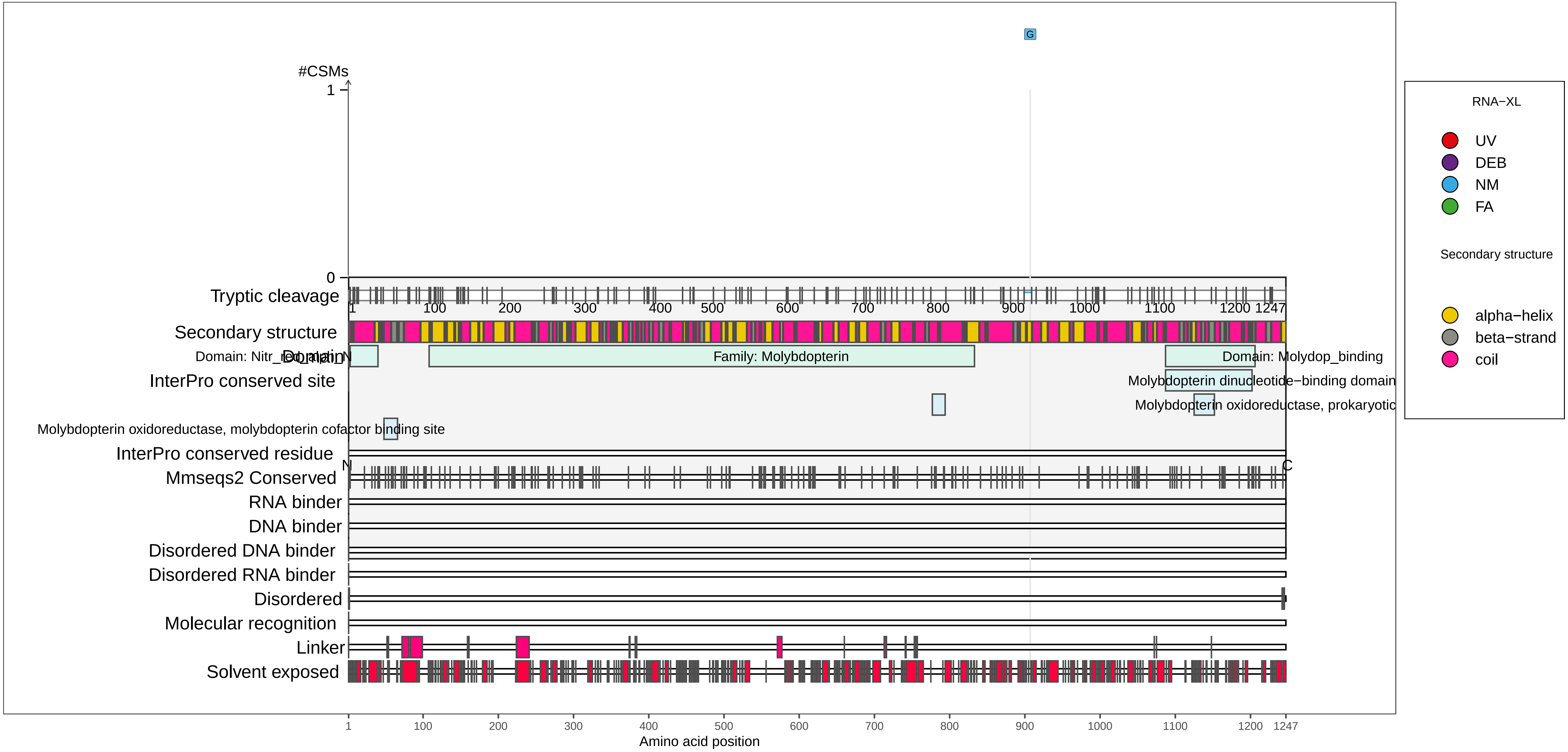

P0A6C8
ARGB_ECOLI Acetylglutamate kinase

− Abundance:
tryptic [log10 Intensity]: 8.78 (Q 83)
PAXdb K12 strain [ppm]: 2.83 (Q 87)
PAXdb E.coli [ppm]: 2.26 (Q 79)

− RNA functions: not annotated

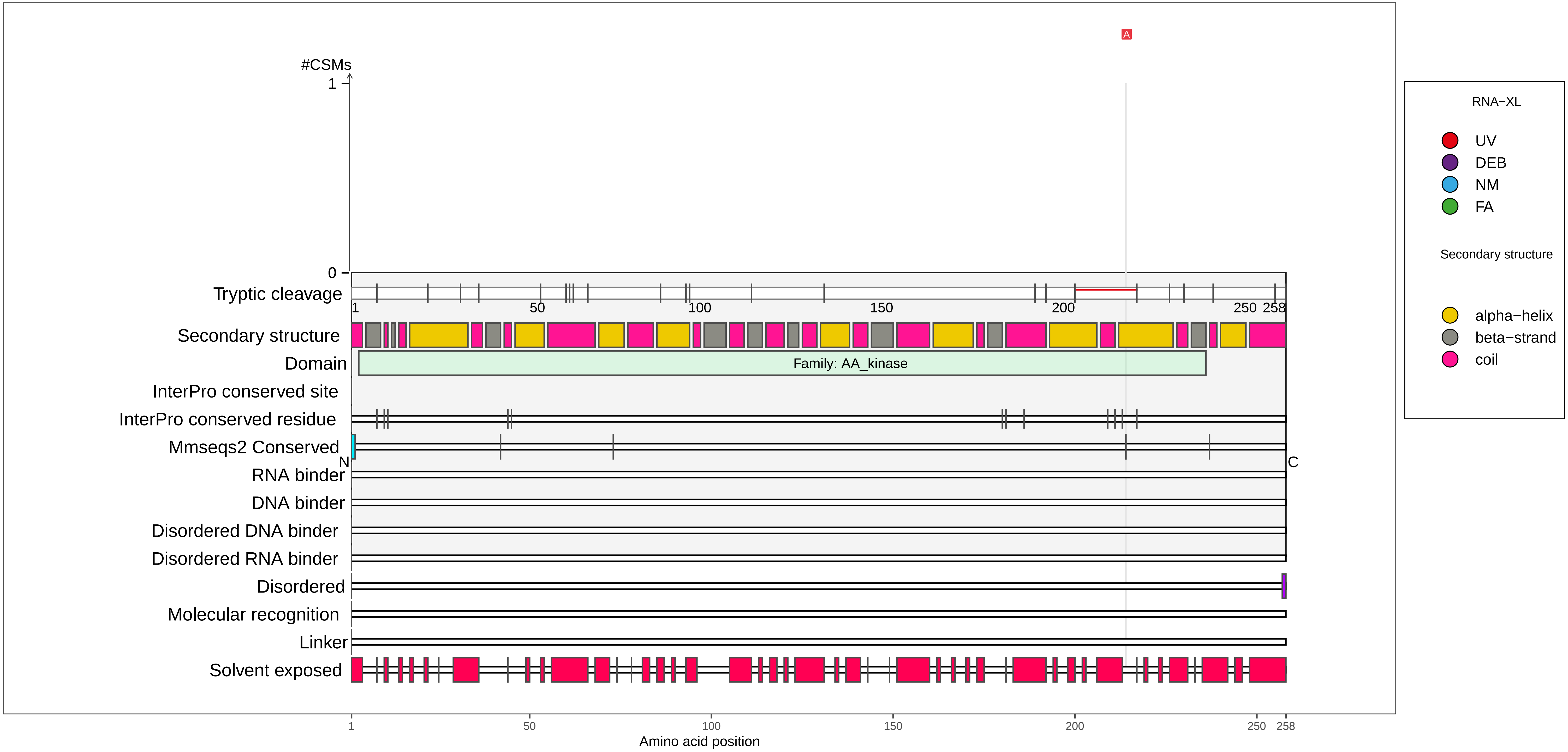

P0A6G7
CLPP_ECOLI ATP−dependent Clp protease proteolytic subunit

− Abundance:
tryptic [log10 Intensity]: 8.94 (Q 87)
PAXdb K12 strain [ppm]: 2.79 (Q 86)
PAXdb E.coli [ppm]: 3.17 (Q 96)

− RNA functions: not annotated

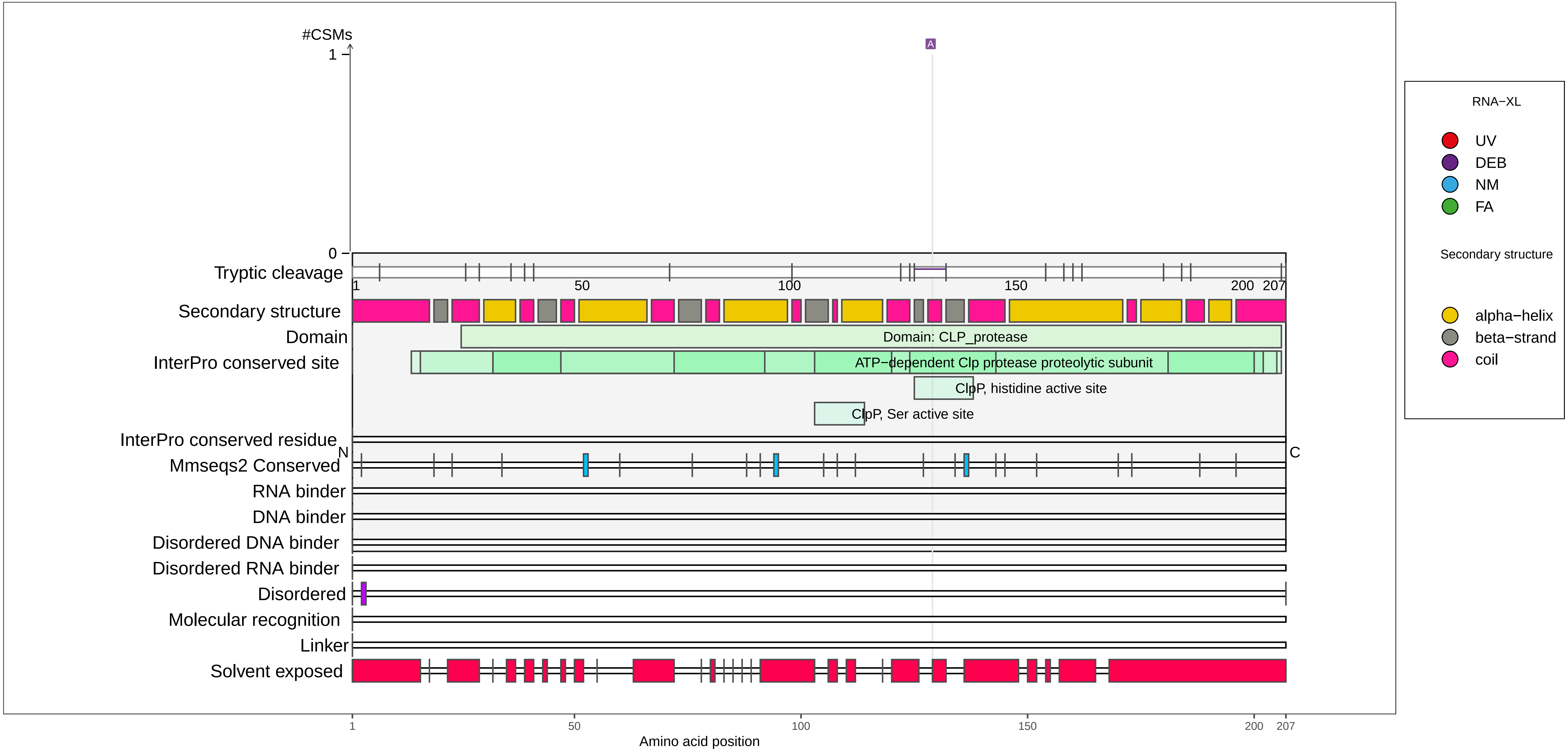

P0A6W0
GLSA2_ECOLI Glutaminase 2
− Abundance:
tryptic [log10 Intensity]: 7.47 (Q 33)
PAXdb K12 strain [ppm]: 1.69 (Q 49)
PAXdb E.coli [ppm]: 0.29 (Q 31)

− RNA functions: not annotated

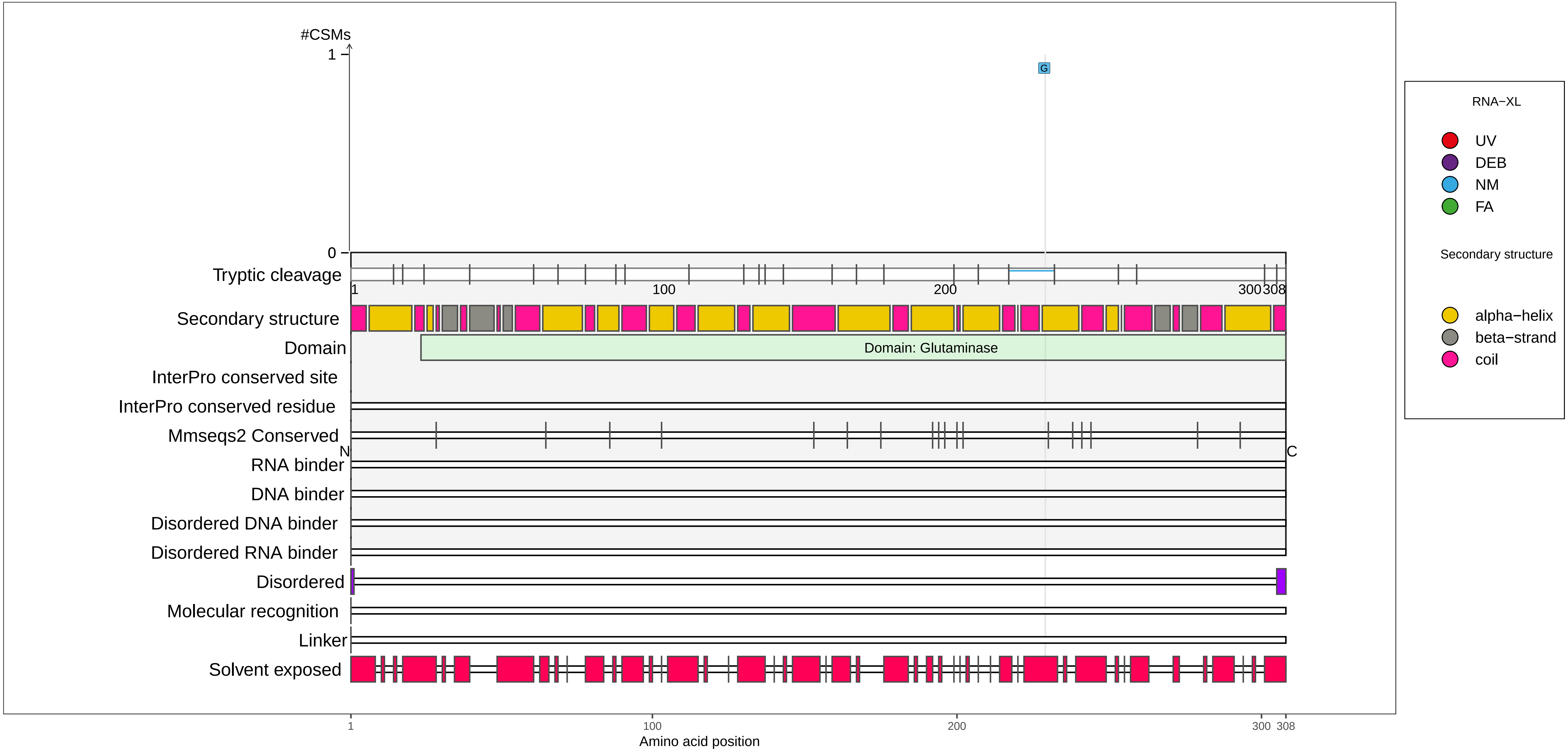

P0A6W3
MRAY_ECOLI Phospho−N−acetylmuramoyl−pentapeptide−transferase

− Abundance:
tryptic [log10 Intensity]: NA (Q NA)
PAXdb K12 strain [ppm]: NA (Q NA)
PAXdb E.coli [ppm]: 0.87 (Q 45)

− RNA functions: not annotated

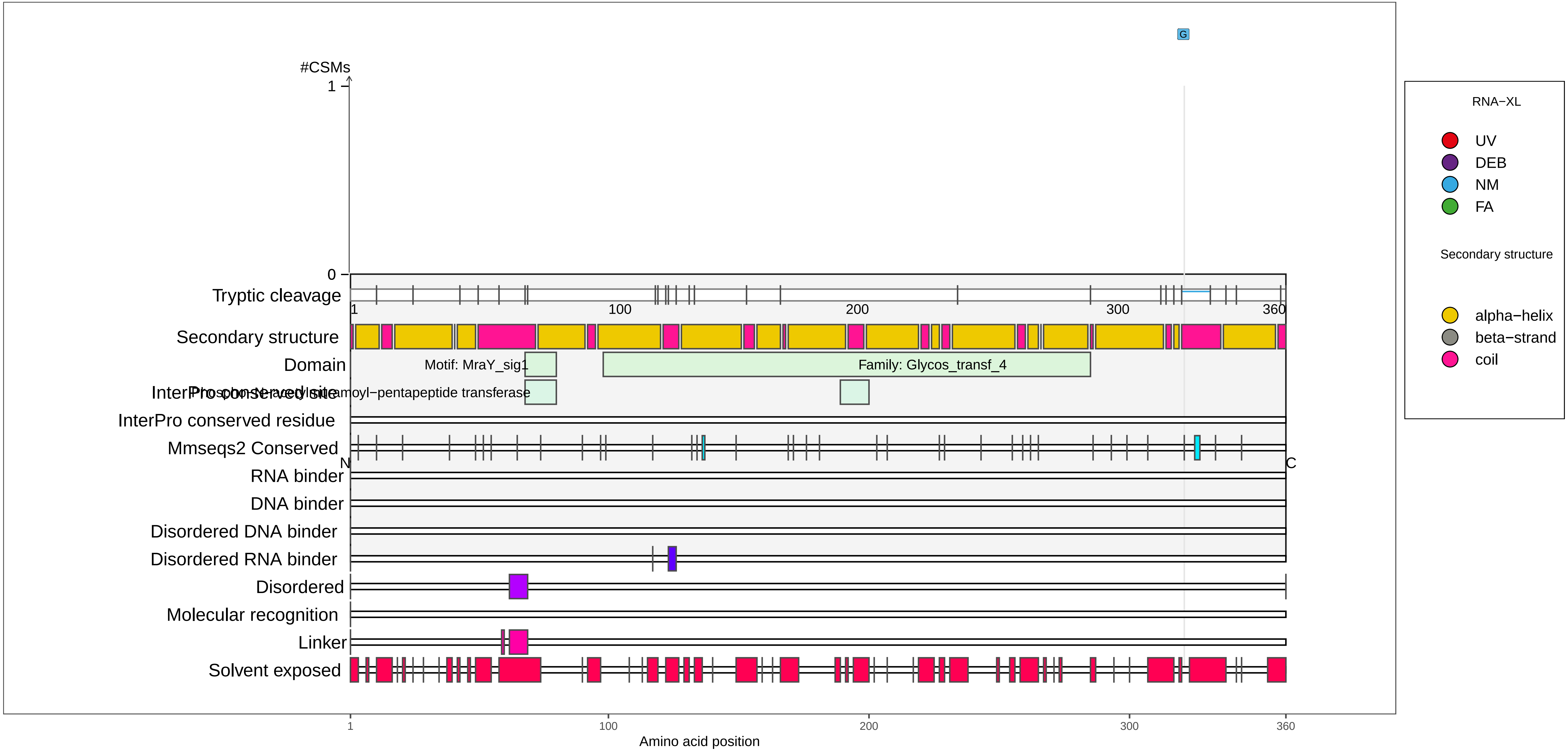

P0A725
LPXC_ECOLI UDP−3−O−acyl−N−acetylglucosamine deacetylase

− Abundance:
tryptic [log10 Intensity]: 7.4 (Q 30)
PAXdb K12 strain [ppm]: NA (Q NA)
PAXdb E.coli [ppm]: 1.55 (Q 61)

− RNA functions: not annotated

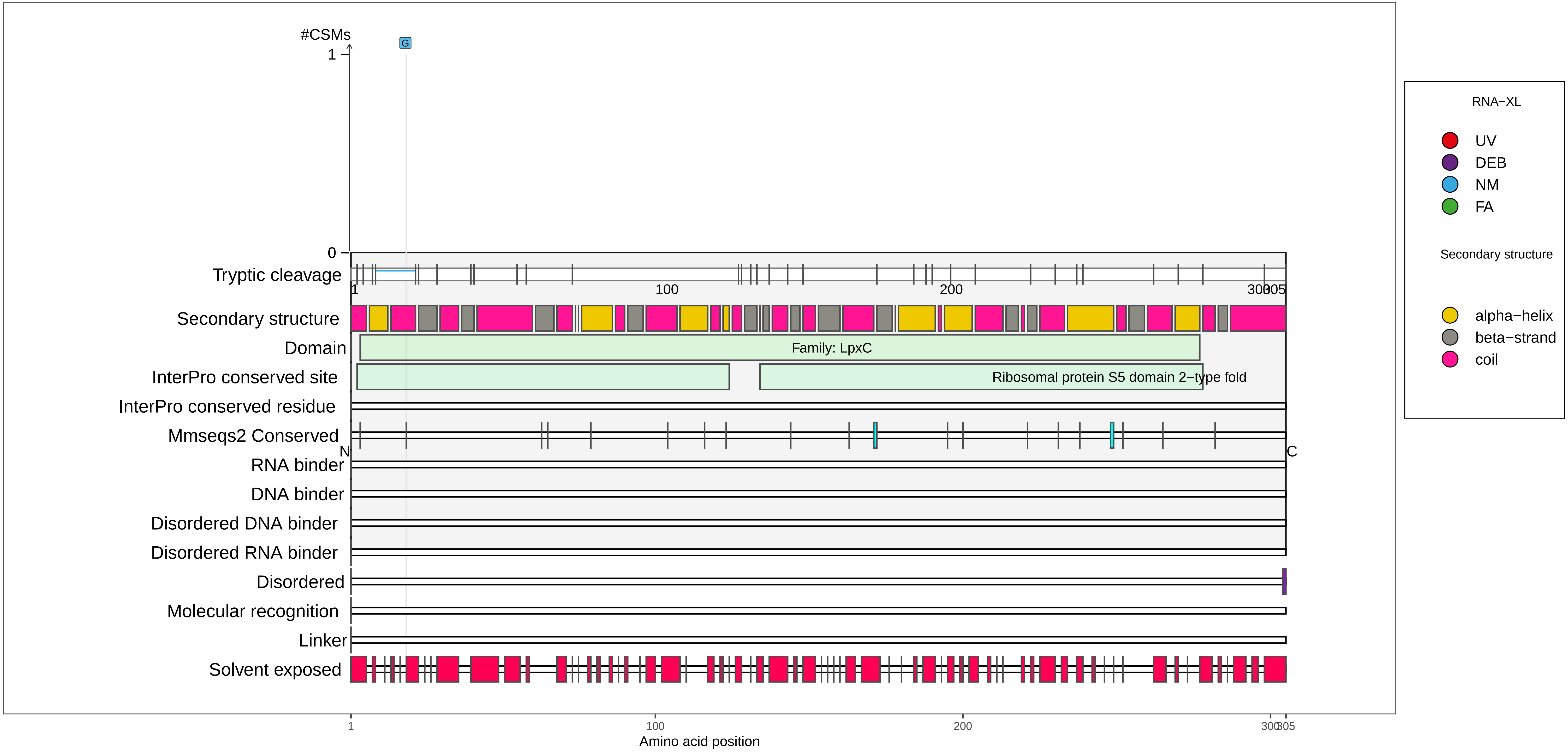

P0A742
MSCL_ECOLI Large−conductance mechanosensitive channel

− Abundance:
tryptic [log10 Intensity]: 8.08 (Q 61)
PAXdb K12 strain [ppm]: 1.87 (Q 56)
PAXdb E.coli [ppm]: 2.23 (Q 78)

− RNA functions: not annotated

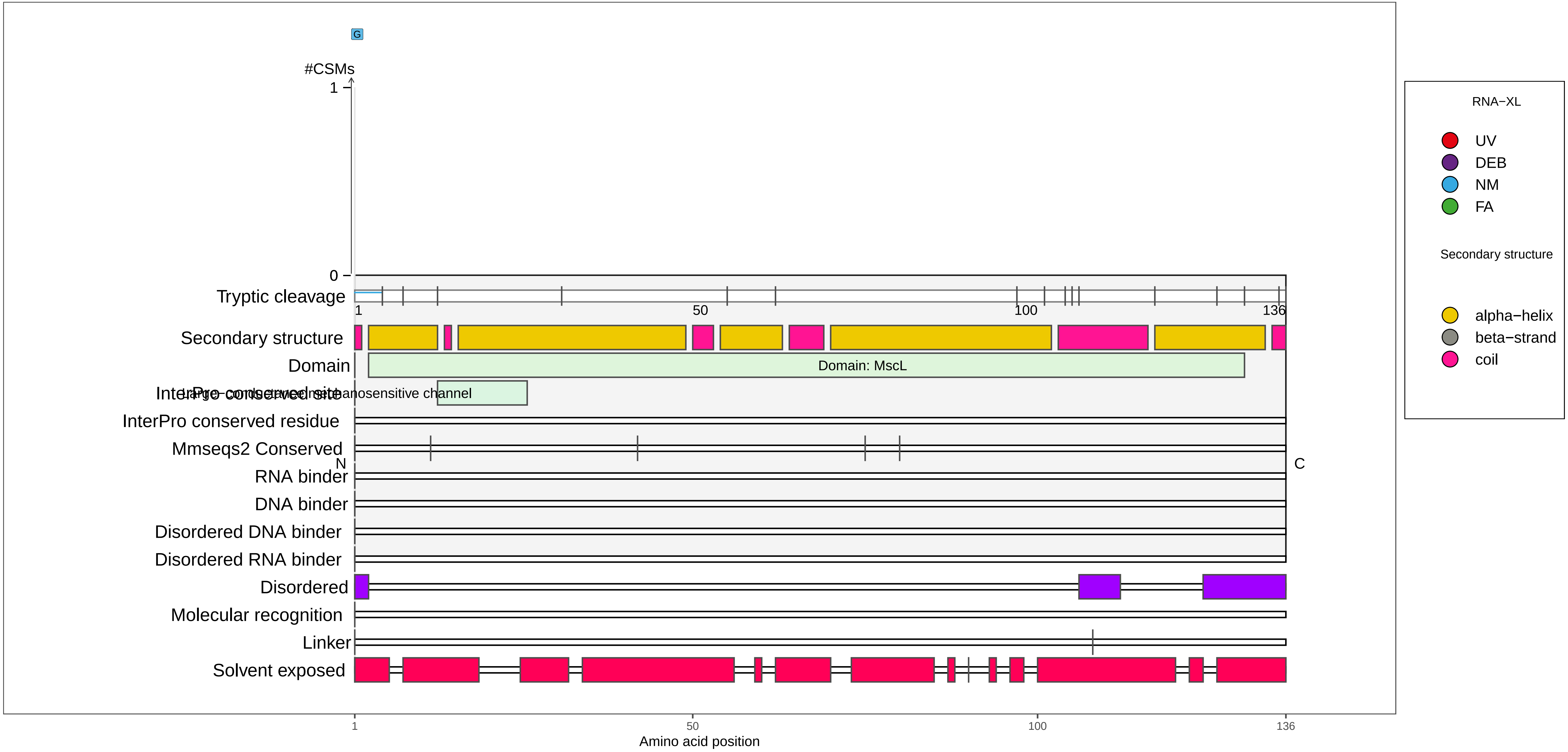

P0A7A7
PLSB_ECOLI Glycerol−3−phosphate acyltransferase

− Abundance:
tryptic [log10 Intensity]: 6.89 (Q 9)
PAXdb K12 strain [ppm]: 0.93 (Q 8)
PAXdb E.coli [ppm]: 2.3 (Q 80)

− RNA functions: not annotated

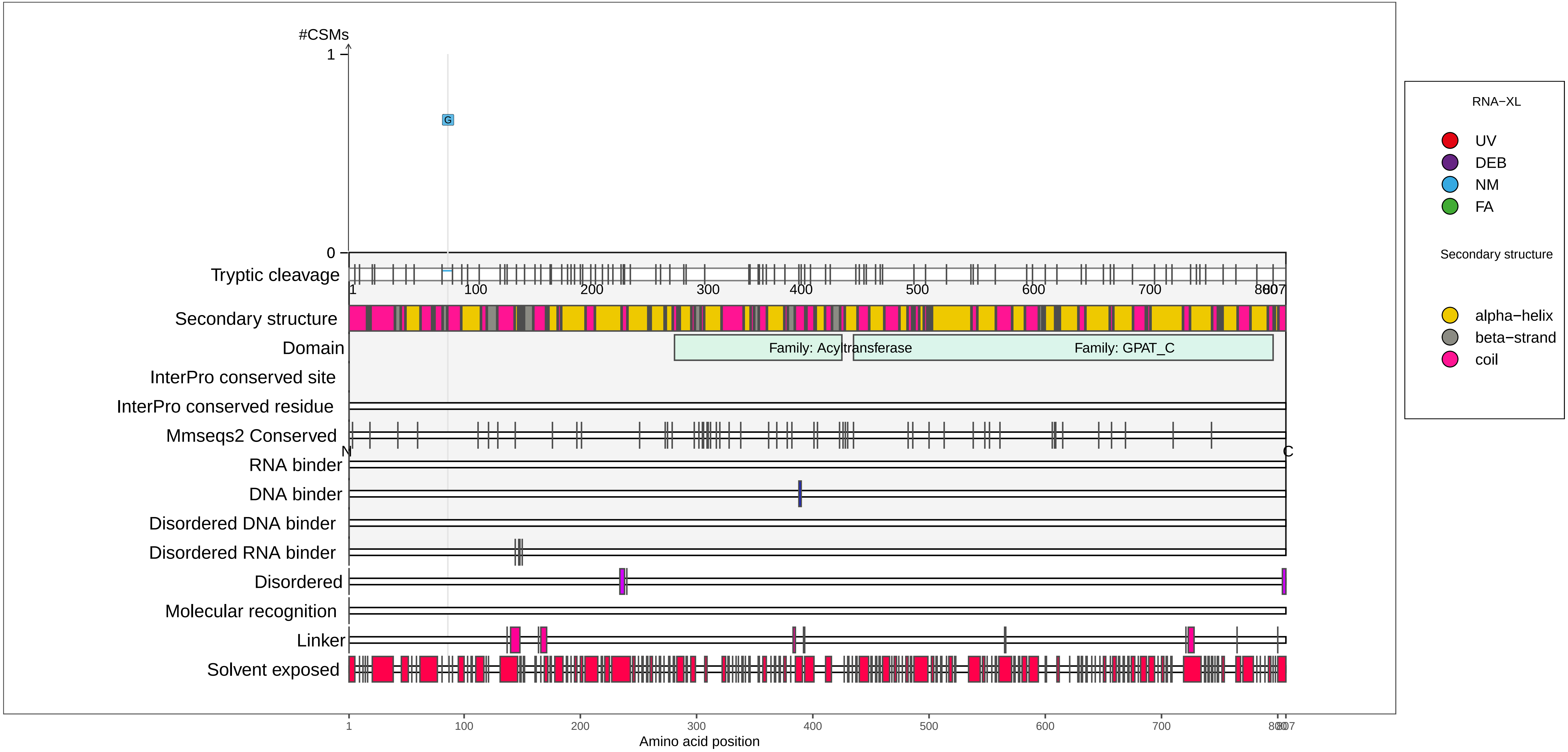

P0A7B1
PPK1_ECOLI Polyphosphate kinase

− Abundance:
tryptic [log10 Intensity]: 7.96 (Q 56)
PAXdb K12 strain [ppm]: 2.17 (Q 66)
PAXdb E.coli [ppm]: 1.98 (Q 72)

− RNA functions: not annotated

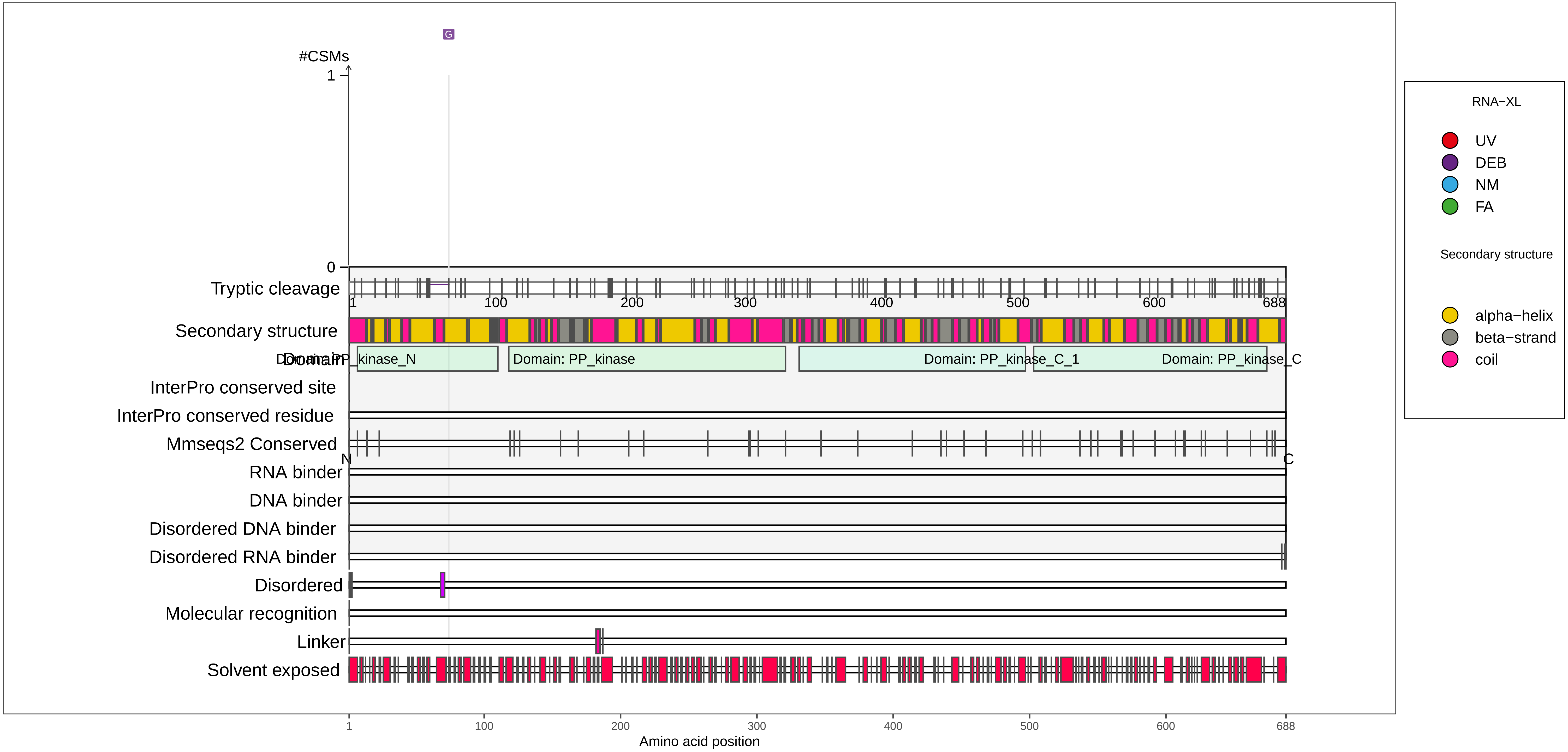

P0A7Y4
RNH_ECOLI Ribonuclease HI

− Abundance:
tryptic [log10 Intensity]: 7.73 (Q 46)
PAXdb K12 strain [ppm]: 1.33 (Q 28)
PAXdb E.coli [ppm]: 0.66 (Q 40)

− RNA functions:
DNA replication, removal of RNA primer; RNA catabolic process; RNA metabolic process
RNA phosphodiester bond hydrolysis
RNA phosphodiester bond hydrolysis, endonucleolytic; RNA−DNA hybrid ribonuclease activity

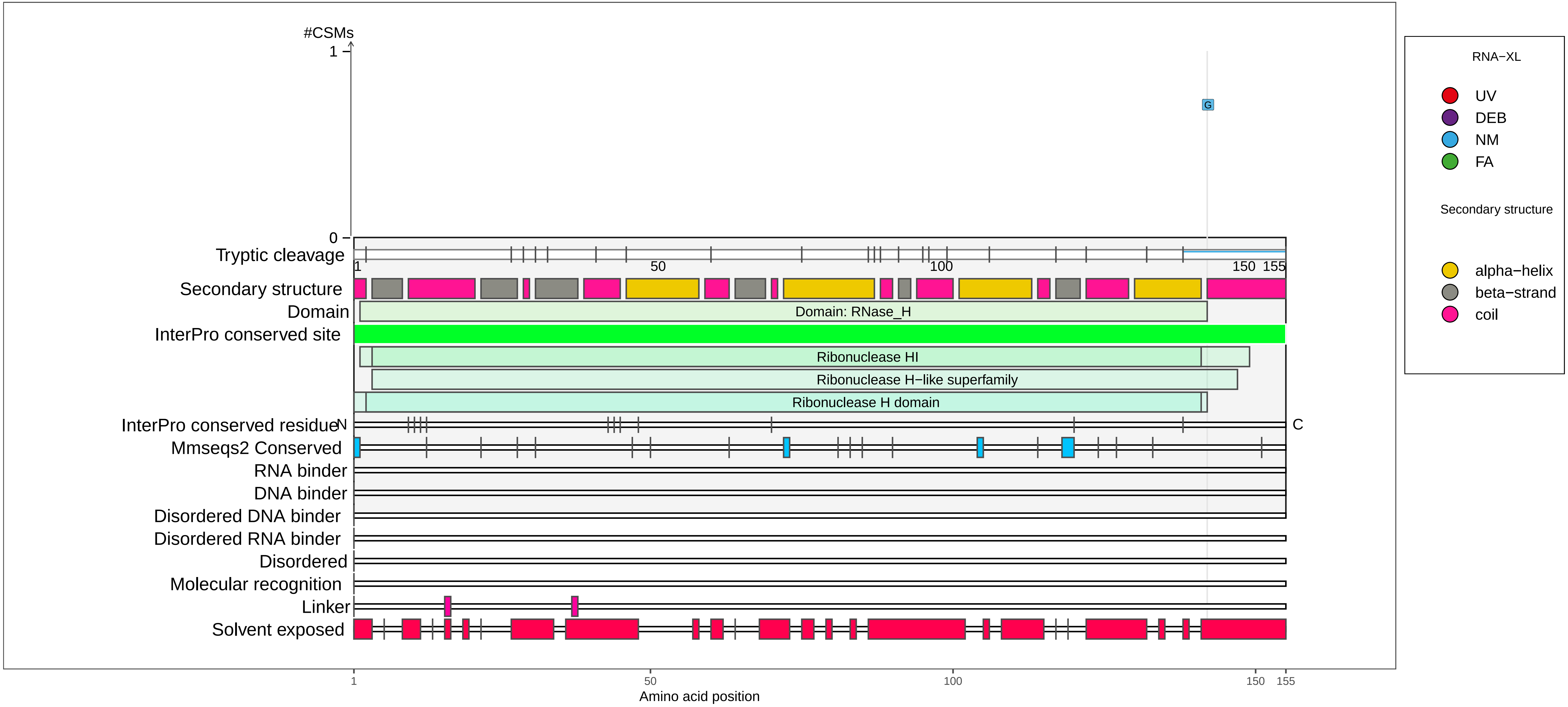

P0A7Y8
RNPA_ECOLI Ribonuclease P protein component

− Abundance:
tryptic [log10 Intensity]: 7.26 (Q 23)
PAXdb K12 strain [ppm]: NA (Q NA)
PAXdb E.coli [ppm]: 1.92 (Q 70)

− RNA functions:
3−tRNA processing endoribonuclease activity; ncRNA 3−end processing
ncRNA 5−end processing; ncRNA metabolic process; ncRNA processing; RNA 3−end processing
RNA 5−end processing; RNA binding; RNA metabolic process; RNA phosphodiester bond hydrolysis
RNA phosphodiester bond hydrolysis, endonucleolytic; RNA processing; tRNA 3−end processing
tRNA 3−trailer cleavage; tRNA 3−trailer cleavage, endonucleolytic
tRNA 5−end processing; tRNA 5−leader removal; tRNA binding; tRNA metabolic process; tRNA processing
tRNA−specific ribonuclease activity

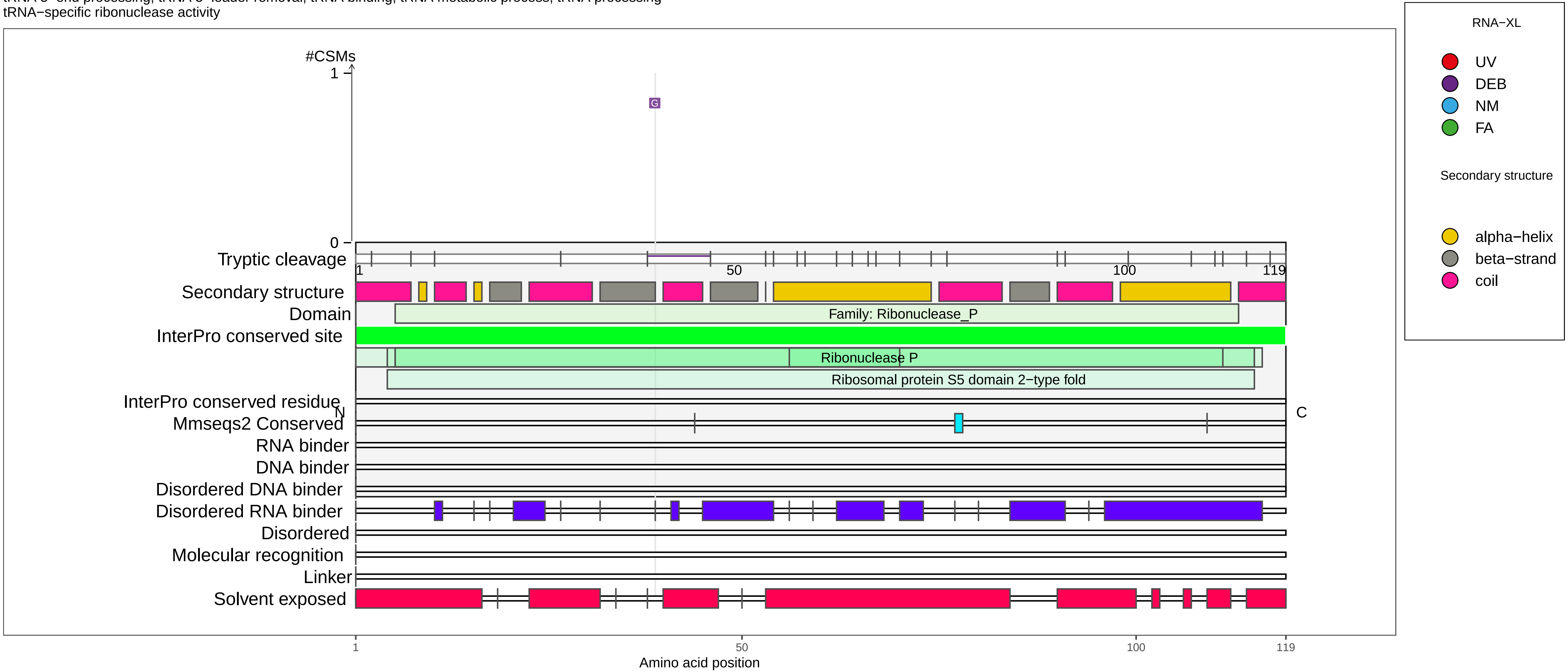

P0A823
SFSA_ECOLI Sugar fermentation stimulation protein A

− Abundance:
tryptic [log10 Intensity]: 7.54 (Q 37)
PAXdb K12 strain [ppm]: 1.41 (Q 33)
PAXdb E.coli [ppm]: 0.31 (Q 32)

− RNA functions: not annotated

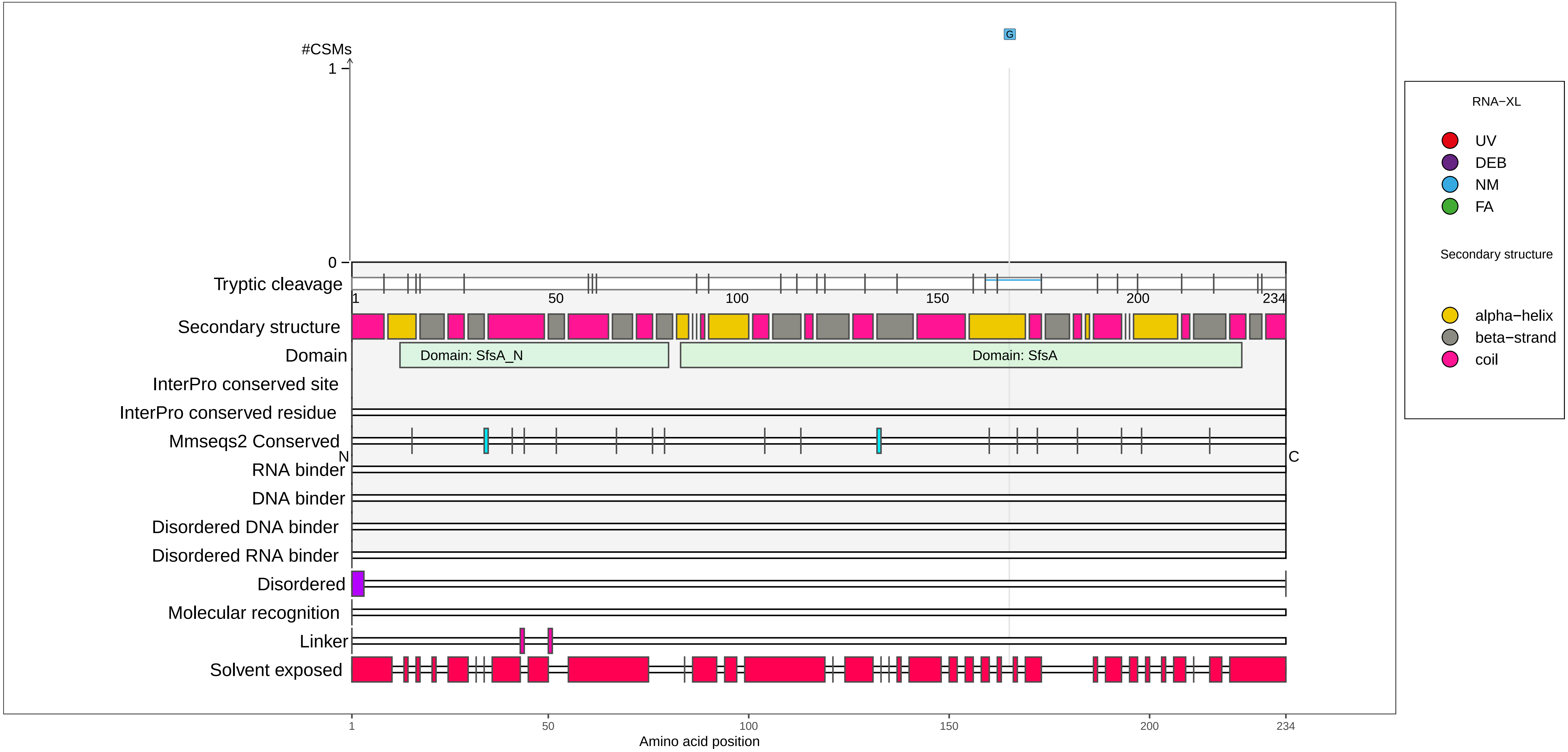

P0A873
TRMD_ECOLI tRNA (guanine−N(1)−)−methyltransferase

− Abundance:
tryptic [log10 Intensity]: 7.77 (Q 48)
PAXdb K12 strain [ppm]: 1.81 (Q 54)
PAXdb E.coli [ppm]: 1.21 (Q 52)

− RNA functions:
ncRNA metabolic process; ncRNA processing; RNA metabolic process; RNA methylation
RNA methyltransferase activity; RNA modification; RNA processing
tRNA (Guanine−1)−methyltransferase; tRNA (guanine−N1−)−methyltransferase activity
tRNA (guanine(37)−N(1))−methyltransferase activity; tRNA (guanine) methyltransferase activity
tRNA metabolic process; tRNA methylation; tRNA methyltransferase activity; tRNA modification
tRNA N1−guanine methylation; tRNA processing

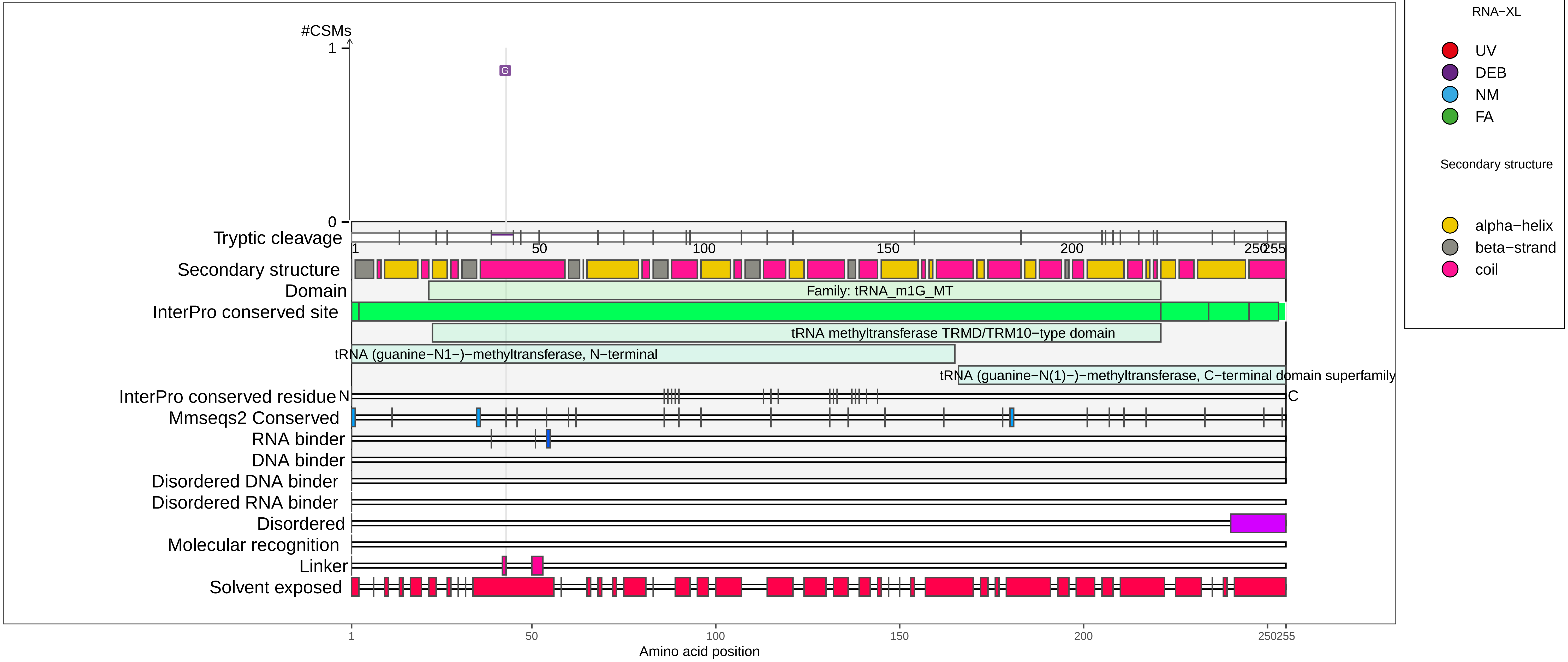

P0A877
TRPA_ECOLI Tryptophan synthase alpha chain
− Abundance:
tryptic [log10 Intensity]: 7.22 (Q 22)
PAXdb K12 strain [ppm]: 2.19 (Q 66)
PAXdb E.coli [ppm]: 2.82 (Q 92)

− RNA functions: not annotated

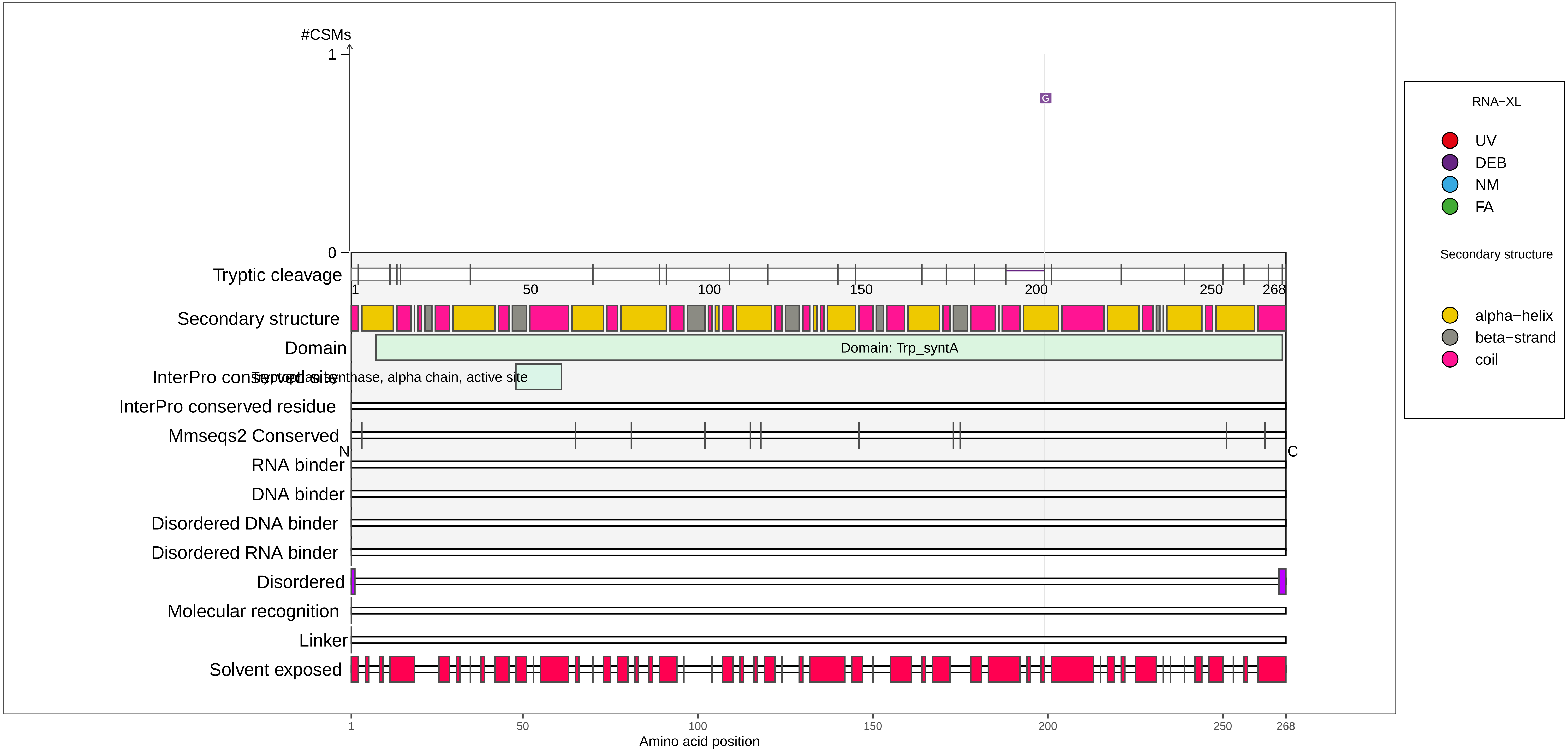

P0A8A4
PSRP_ECOLI Phosphoenolpyruvate synthase regulatory protein

− Abundance:
tryptic [log10 Intensity]: NA (Q NA)
PAXdb K12 strain [ppm]: NA (Q NA)
PAXdb E.coli [ppm]: 1.98 (Q 72)

− RNA functions: not annotated

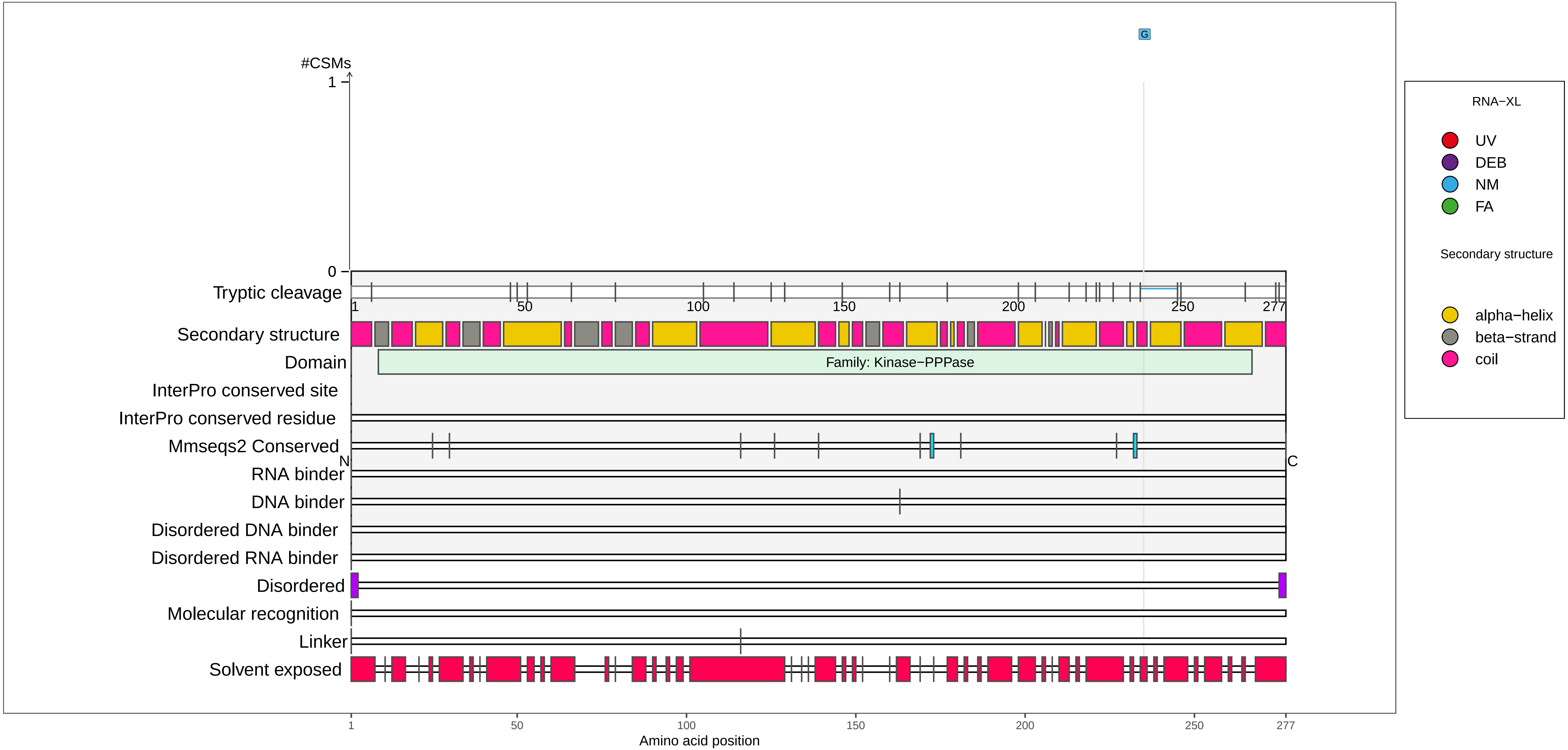

P0A8D0
NRDR_ECOLI Transcriptional repressor NrdR

− Abundance:
tryptic [log10 Intensity]: 7.3 (Q 25)
PAXdb K12 strain [ppm]: 2.5 (Q 77)
PAXdb E.coli [ppm]: 2.01 (Q 72)

− RNA functions: not annotated

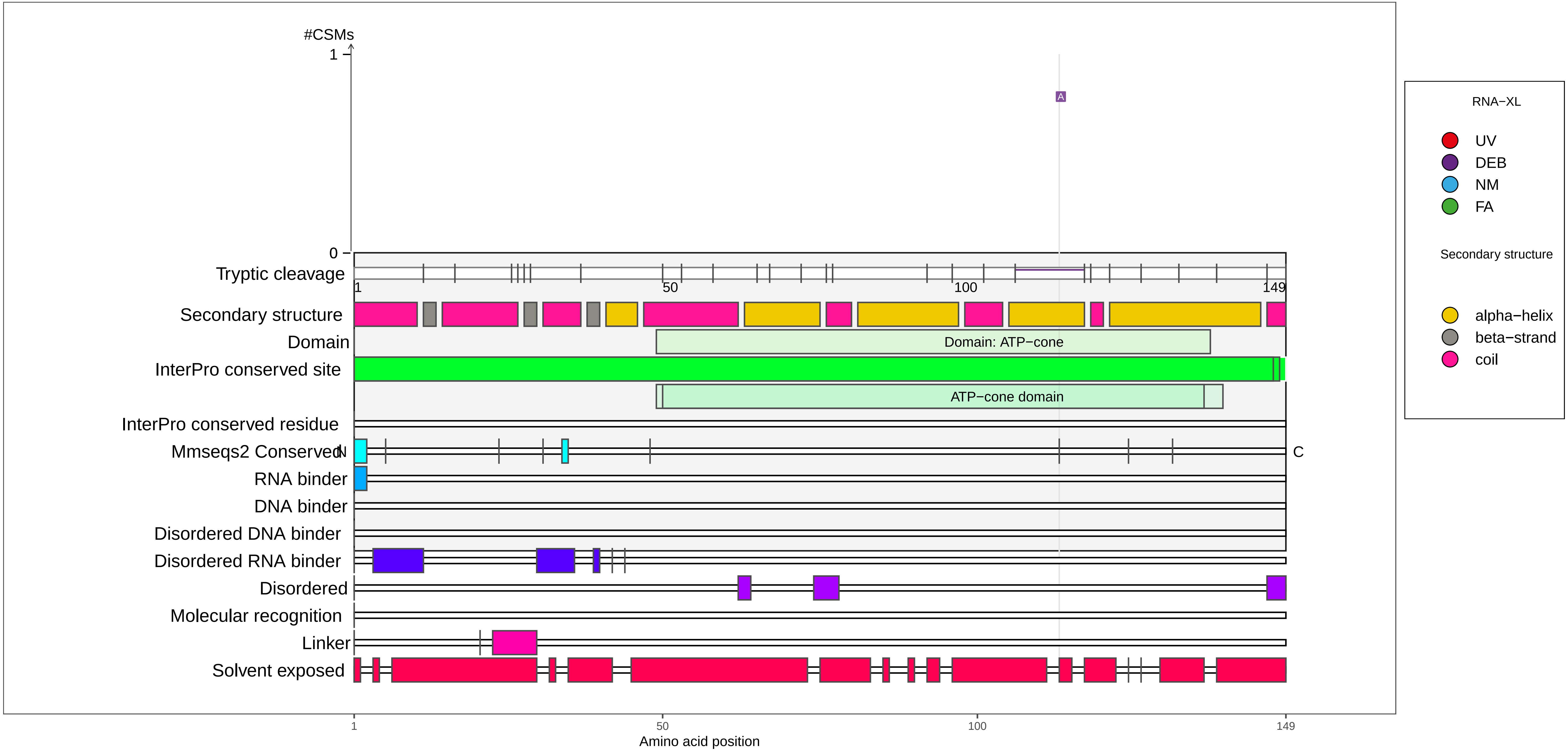

P0A8D6
YMDB_ECOLI O−acetyl−ADP−ribose deacetylase

− Abundance:
tryptic [log10 Intensity]: 6.87 (Q 9)
PAXdb K12 strain [ppm]: 1.85 (Q 55)
PAXdb E.coli [ppm]: 2.18 (Q 77)

− RNA functions: not annotated

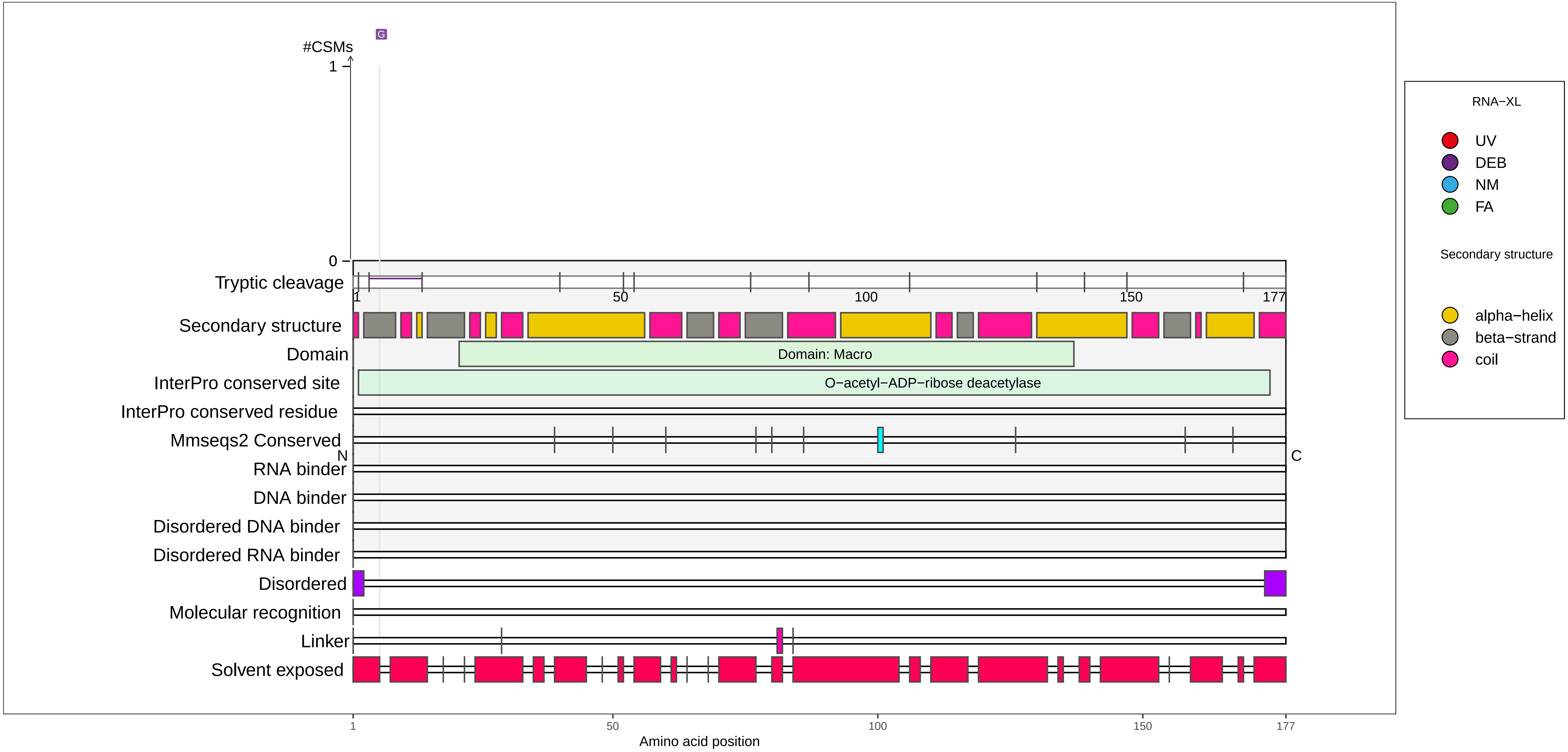

P0A8G3
UXAC_ECOLI Uronate isomerase

− Abundance:
tryptic [log10 Intensity]: 8.21 (Q 66)
PAXdb K12 strain [ppm]: 1.71 (Q 49)
PAXdb E.coli [ppm]: 2.11 (Q 75)

− RNA functions: not annotated

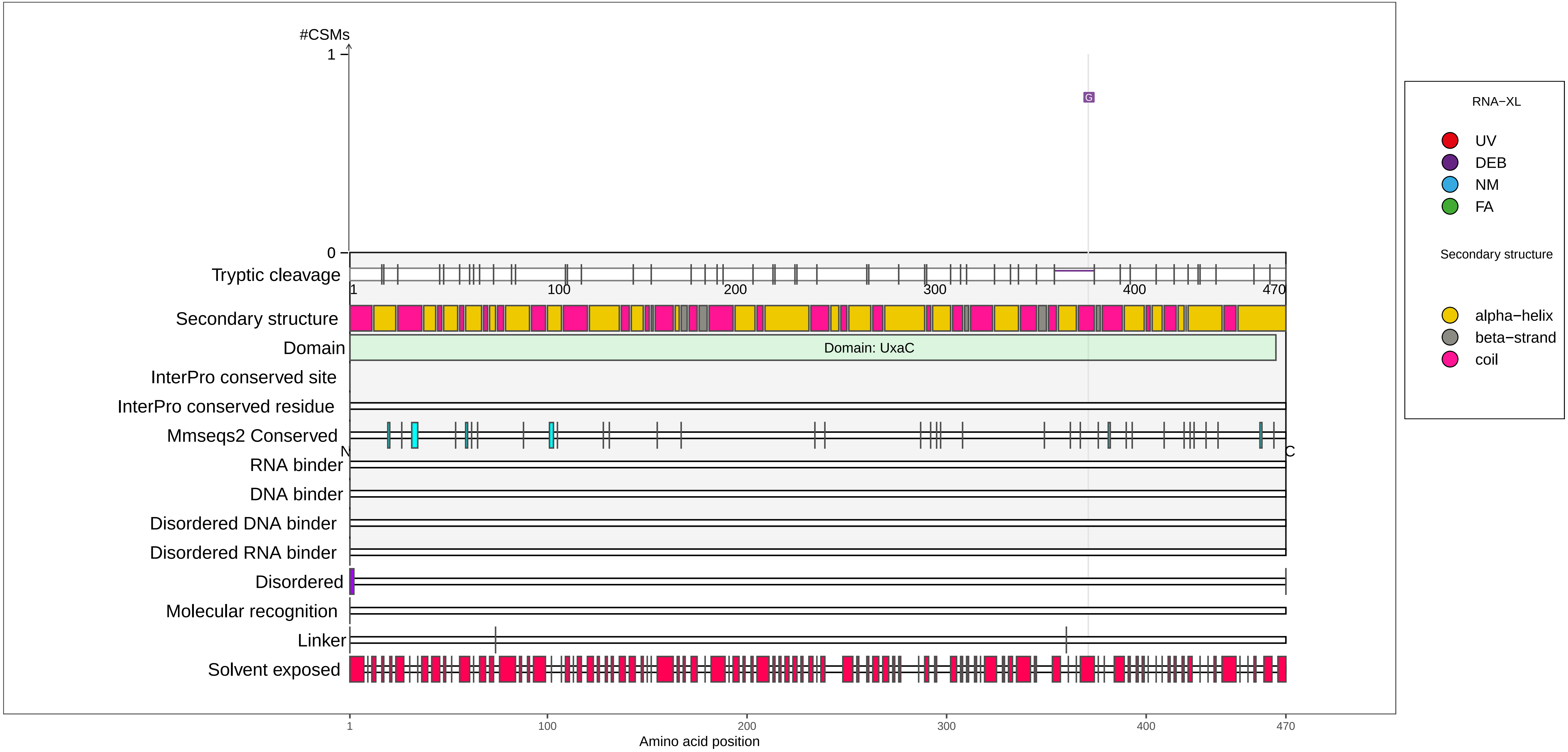

P0A8H6
YIHI_ECOLI Der GTPase−activating protein YihI
− Abundance:
tryptic [log10 Intensity]: 7.98 (Q 57)
PAXdb K12 strain [ppm]: 2.29 (Q 70)
PAXdb E.coli [ppm]: 2.58 (Q 88)

− RNA functions: not annotated

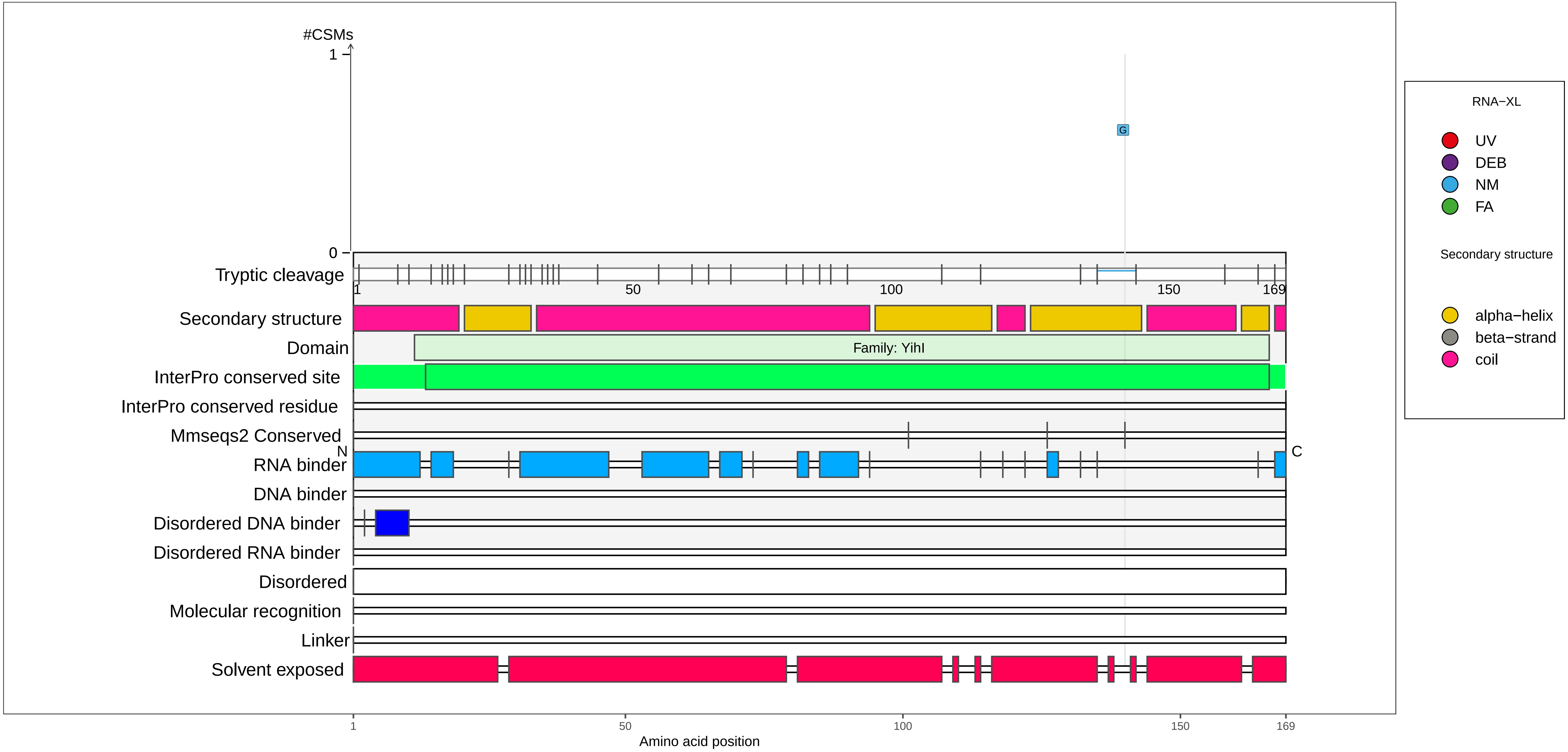

P0A8J2
DNAT_ECOLI Primosomal protein 1

− Abundance:
tryptic [log10 Intensity]: 7.47 (Q 33)
PAXdb K12 strain [ppm]: NA (Q NA)
PAXdb E.coli [ppm]: 1.56 (Q 61)

− RNA functions:
DNA replication, synthesis of RNA primer; RNA biosynthetic process
RNA metabolic process

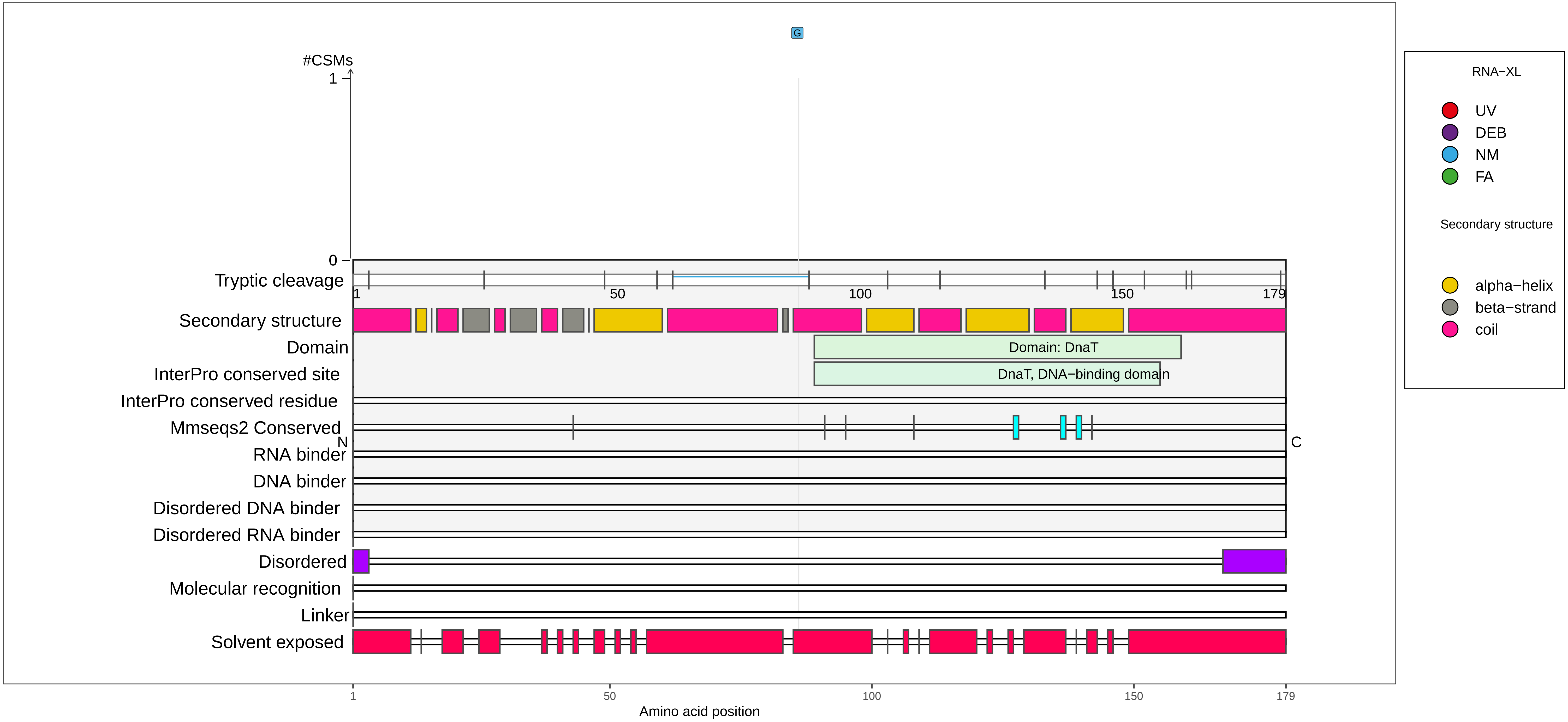

P0A8L7
YCIU_ECOLI UPF0263 protein YciU

− Abundance:
tryptic [log10 Intensity]: NA (Q NA)
PAXdb K12 strain [ppm]: NA (Q NA)
PAXdb E.coli [ppm]: 1.66 (Q 63)

− RNA functions: not annotated

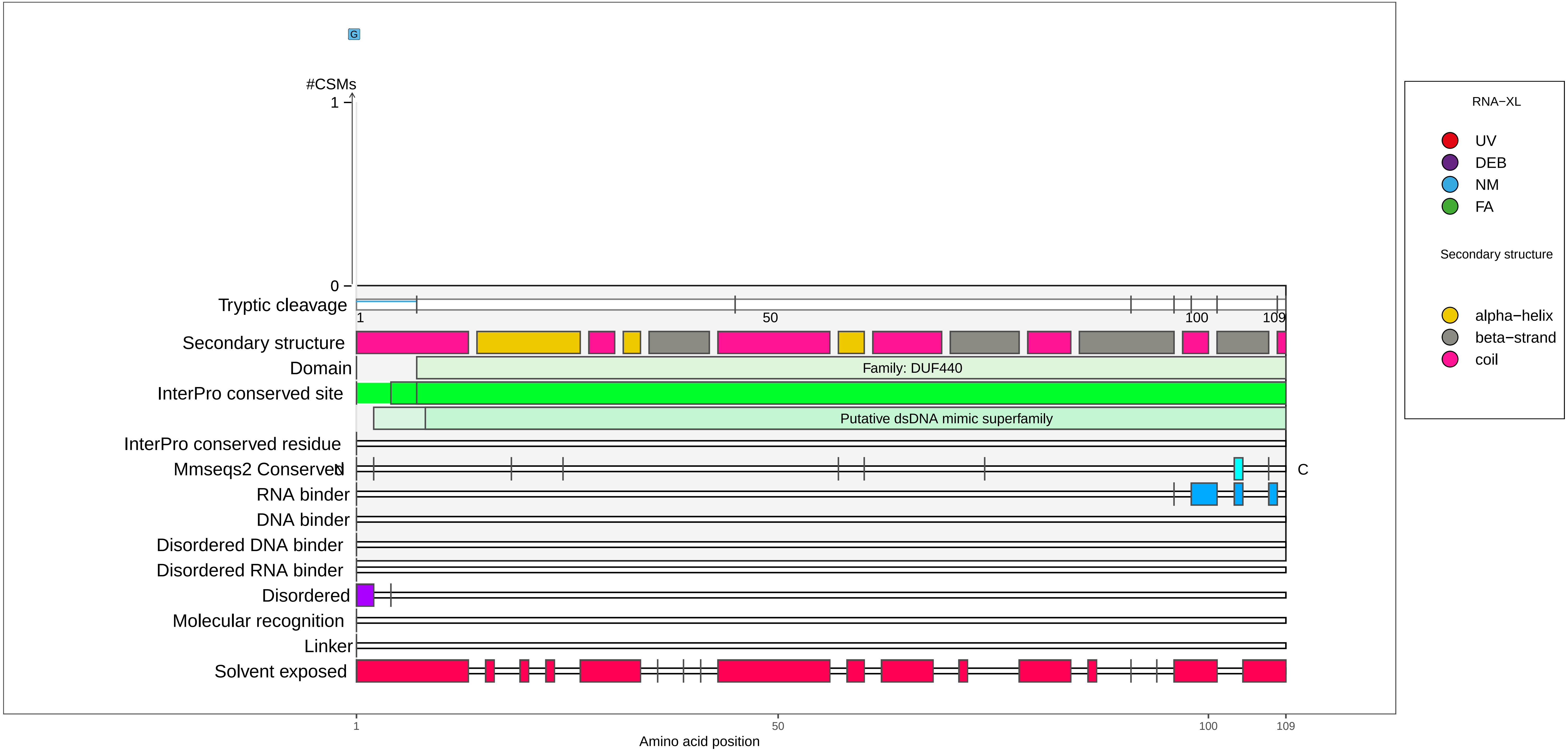

P0A8Q6
CLPS_ECOLI ATP−dependent Clp protease adapter protein ClpS

− Abundance:
tryptic [log10 Intensity]: 7.44 (Q 32)
PAXdb K12 strain [ppm]: 1.46 (Q 36)
PAXdb E.coli [ppm]: 0.9 (Q 45)

− RNA functions: not annotated

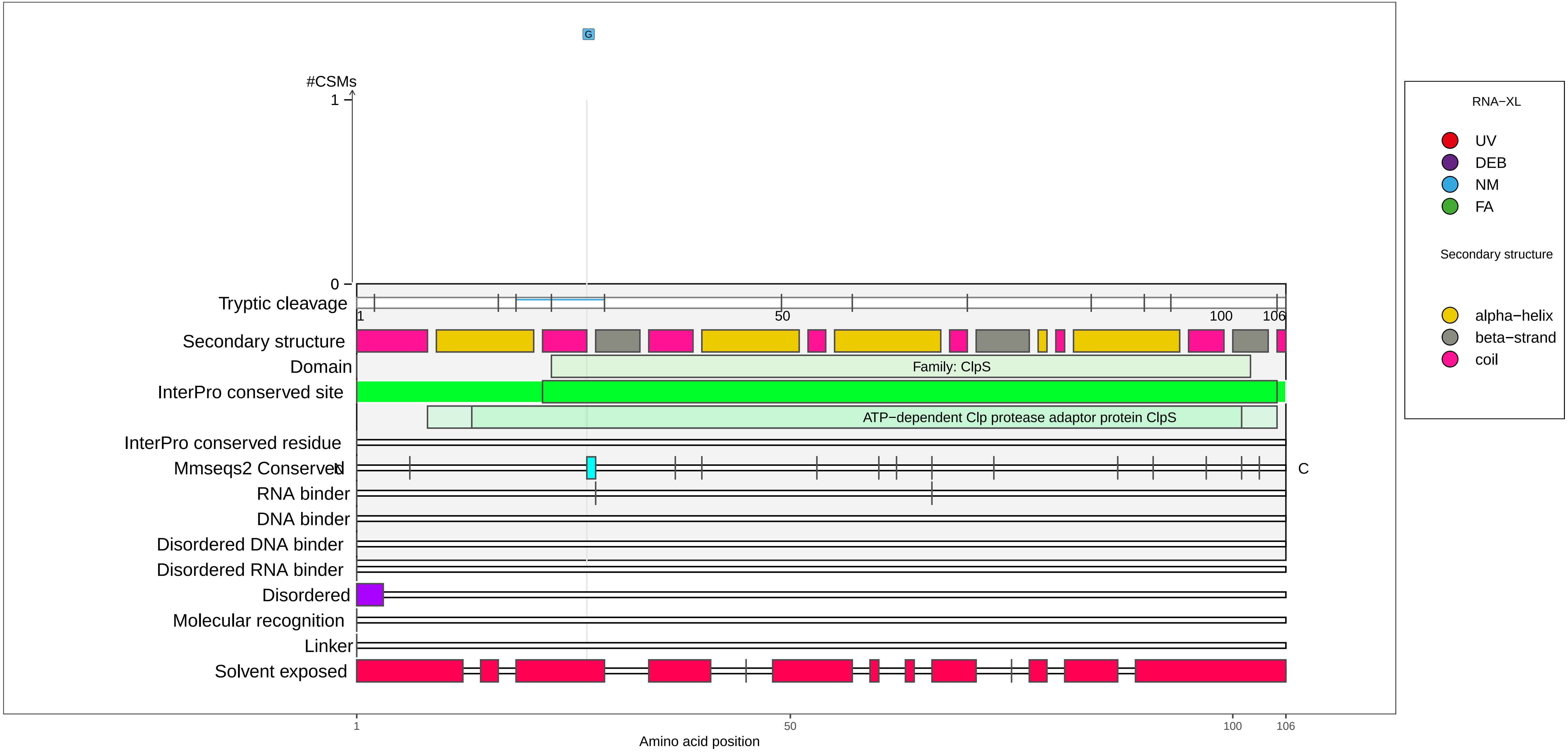

P0A8U6
METJ_ECOLI Met repressor

− Abundance:
tryptic [log10 Intensity]: 7.89 (Q 53)
PAXdb K12 strain [ppm]: 3.24 (Q 94)
PAXdb E.coli [ppm]: 2.86 (Q 93)

− RNA functions: not annotated

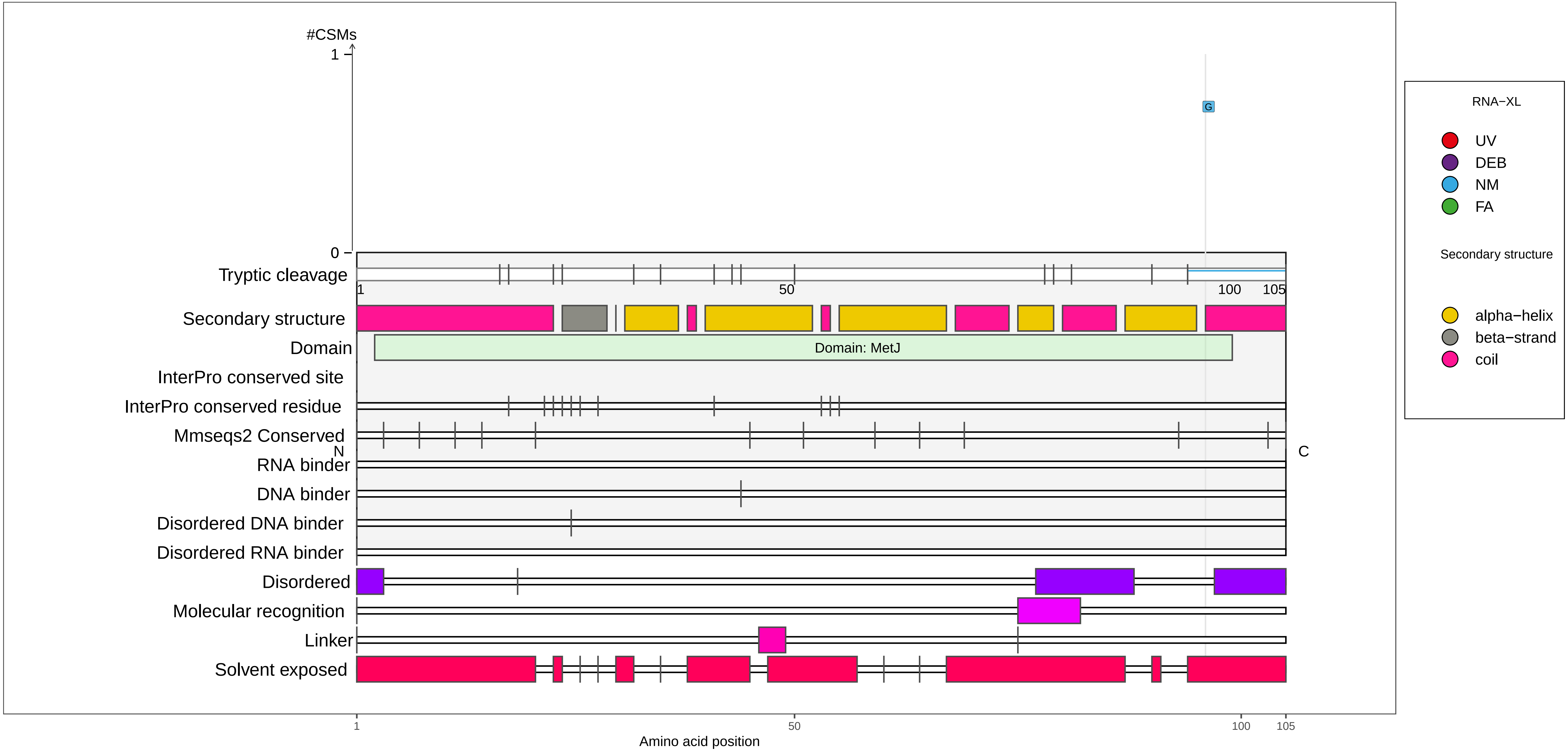

P0A8Z3
YBGC_ECOLI Acyl−CoA thioester hydrolase YbgC

− Abundance:
tryptic [log10 Intensity]: 7.56 (Q 38)
PAXdb K12 strain [ppm]: NA (Q NA)
PAXdb E.coli [ppm]: 1.93 (Q 70)

− RNA functions: not annotated

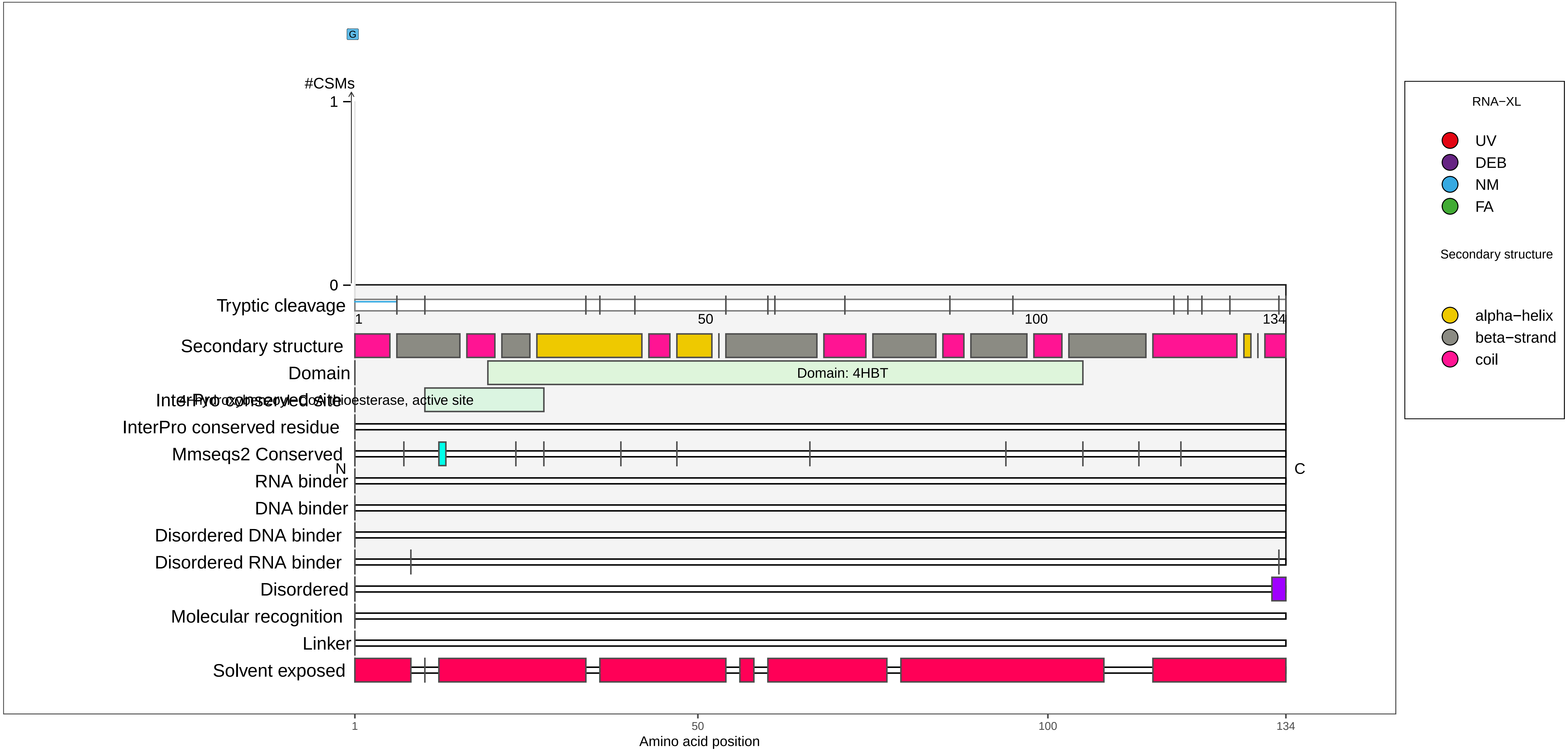

P0A903
BAMC_ECOLI Outer membrane protein assembly factor BamC

− Abundance:
tryptic [log10 Intensity]: 8.77 (Q 83)
PAXdb K12 strain [ppm]: 2.02 (Q 61)
PAXdb E.coli [ppm]: 2.45 (Q 84)

− RNA functions: not annotated

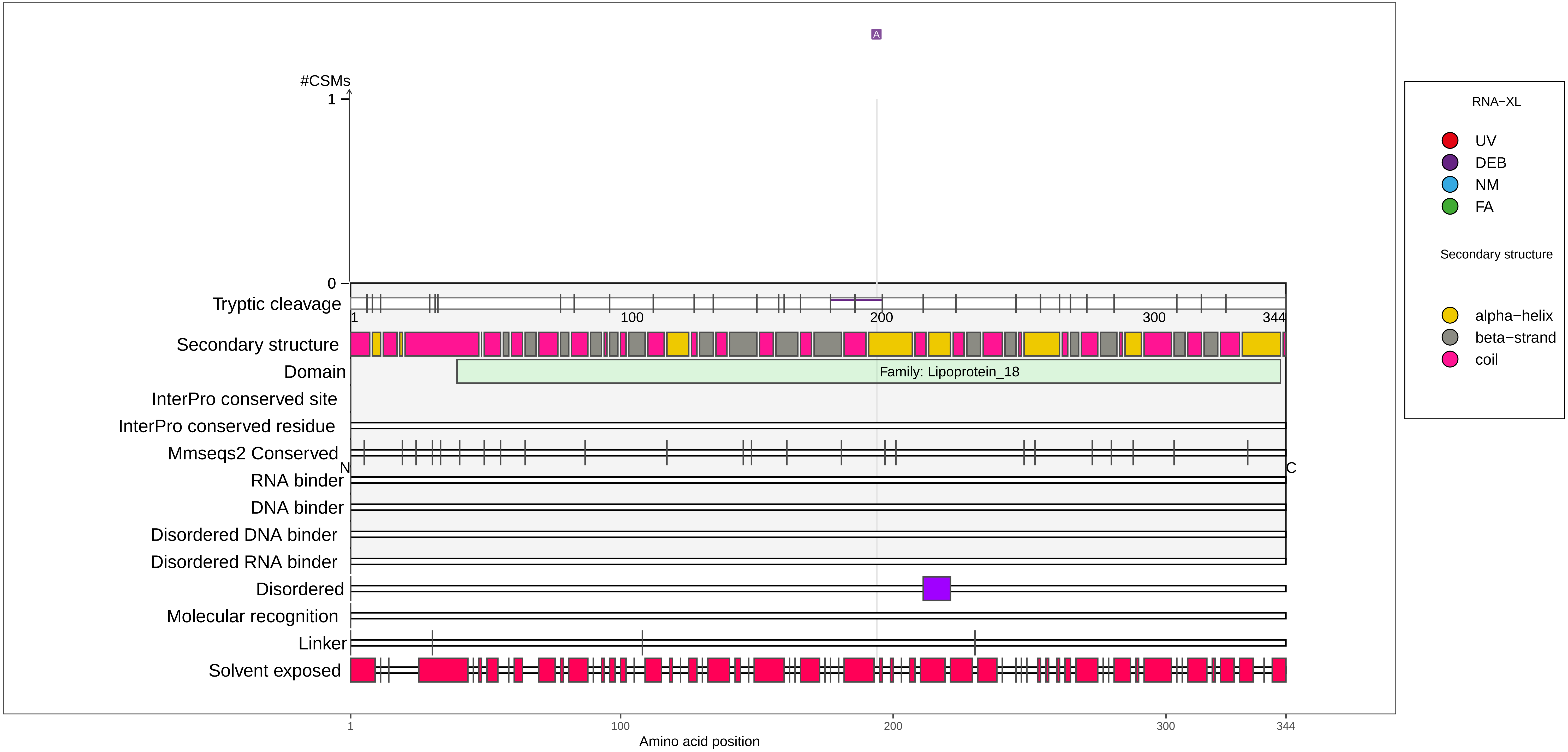

P0A917
OMPX_ECOLI Outer membrane protein X

− Abundance:
tryptic [log10 Intensity]: 9.6 (Q 97)
PAXdb K12 strain [ppm]: 1.54 (Q 40)
PAXdb E.coli [ppm]: 2.73 (Q 90)

− RNA functions: not annotated

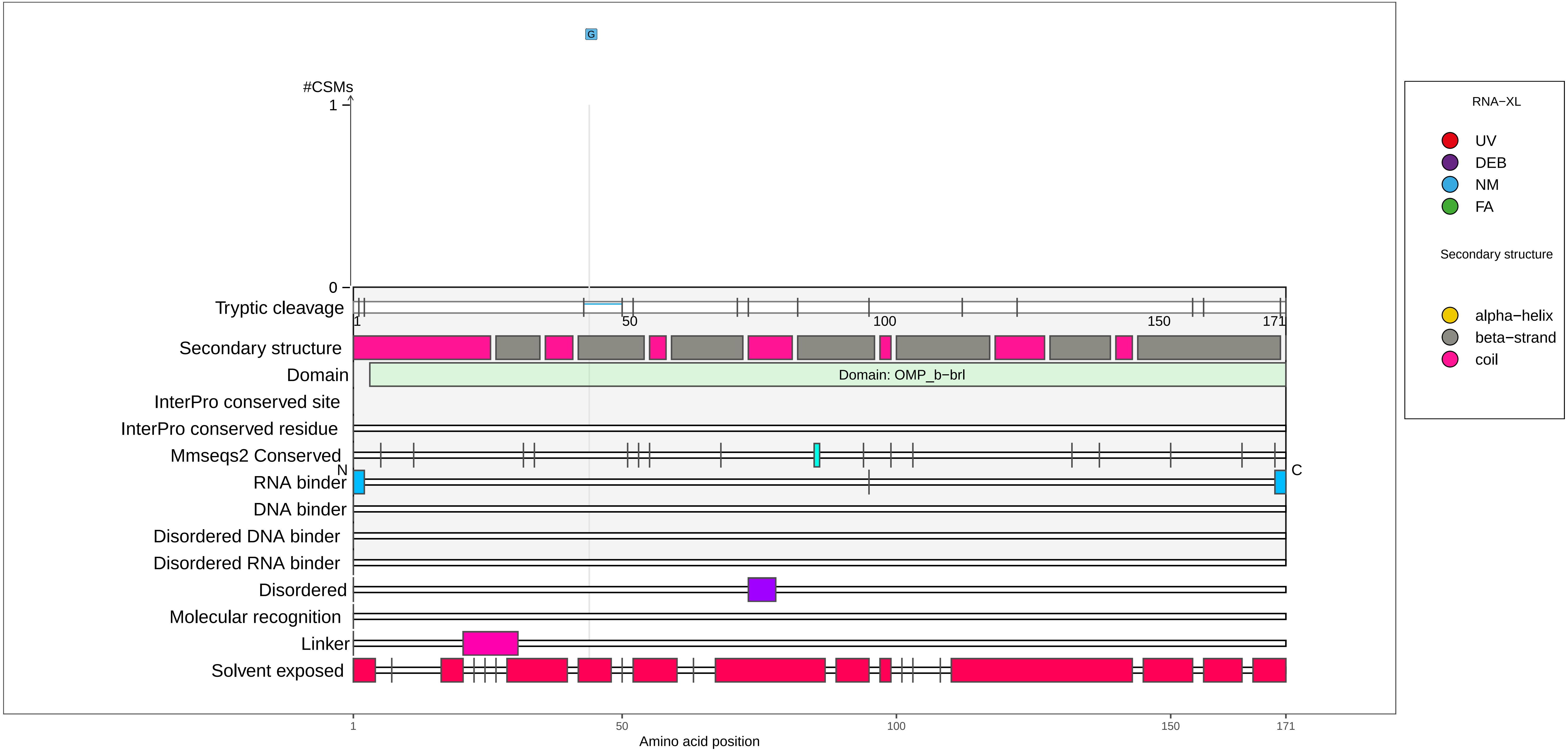

P0A935
MLTA_ECOLI Membrane−bound lytic murein transglycosylase A

− Abundance:
tryptic [log10 Intensity]: 8.17 (Q 64)
PAXdb K12 strain [ppm]: 0.85 (Q 6)
PAXdb E.coli [ppm]: 1.1 (Q 50)

− RNA functions: not annotated

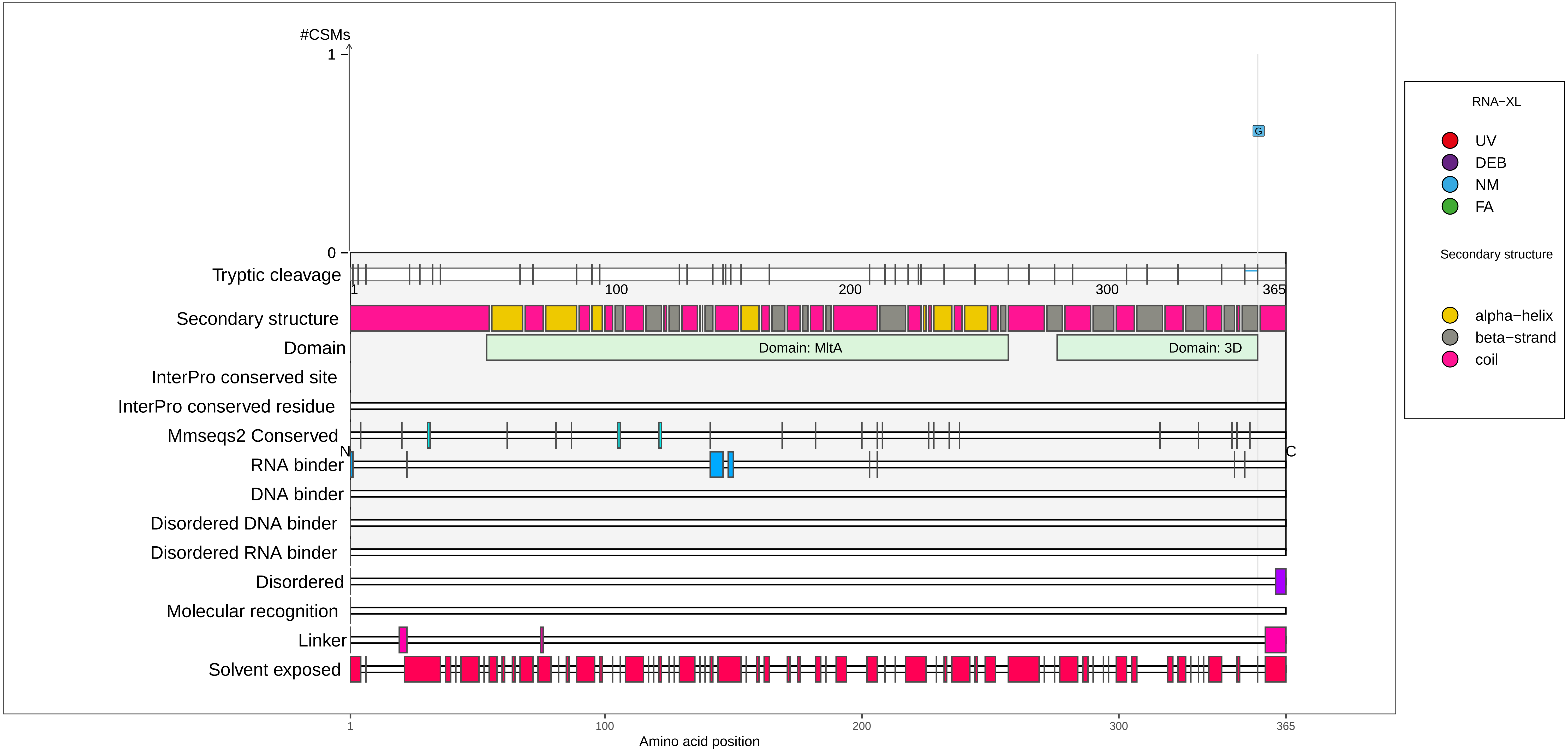

P0A951
ATDA_ECOLI Spermidine N(1)−acetyltransferase

− Abundance:
tryptic [log10 Intensity]: 7.93 (Q 55)
PAXdb K12 strain [ppm]: NA (Q NA)
PAXdb E.coli [ppm]: 1.6 (Q 62)

− RNA functions: not annotated

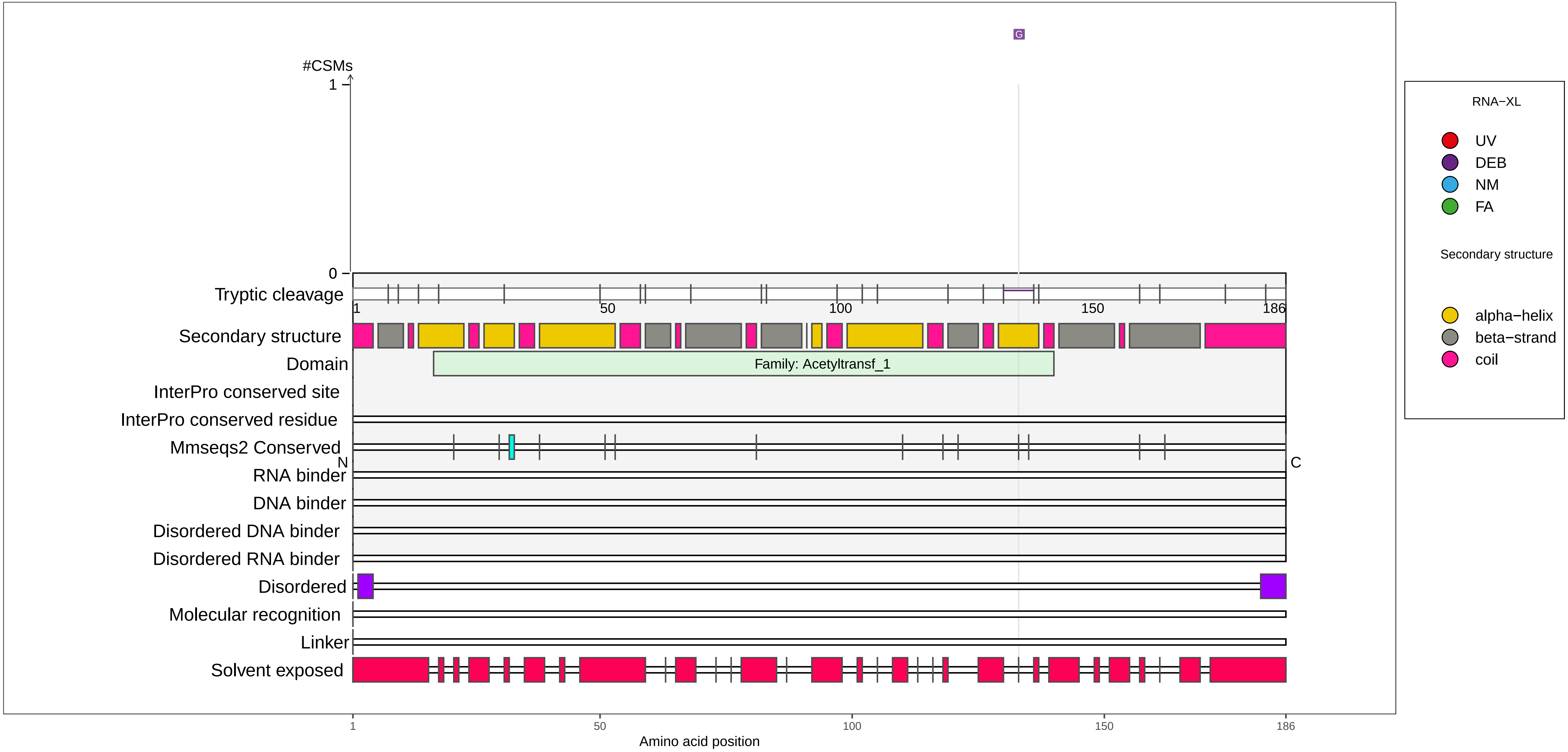

P0A991
ALF1_ECOLI Fructose−bisphosphate aldolase class 1

− Abundance:
tryptic [log10 Intensity]: 7.62 (Q 41)
PAXdb K12 strain [ppm]: 2.83 (Q 87)
PAXdb E.coli [ppm]: 2.92 (Q 94)

− RNA functions: not annotated

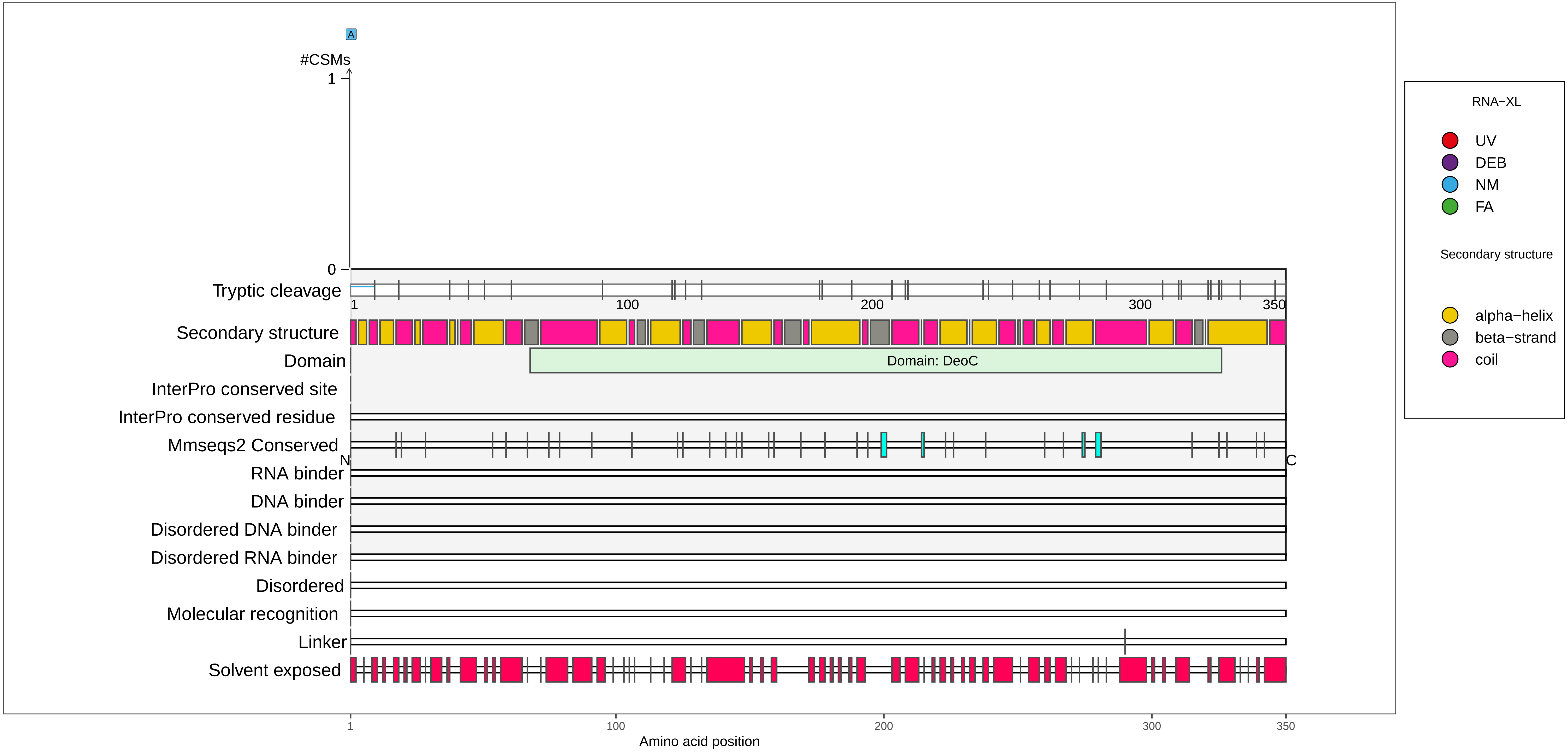

P0A9C0
GLPA_ECOLI Anaerobic glycerol−3−phosphate dehydrogenase subunit A

− Abundance:
tryptic [log10 Intensity]: 8.49 (Q 75)
PAXdb K12 strain [ppm]: 2.46 (Q 76)
PAXdb E.coli [ppm]: 2.04 (Q 73)

− RNA functions: not annotated

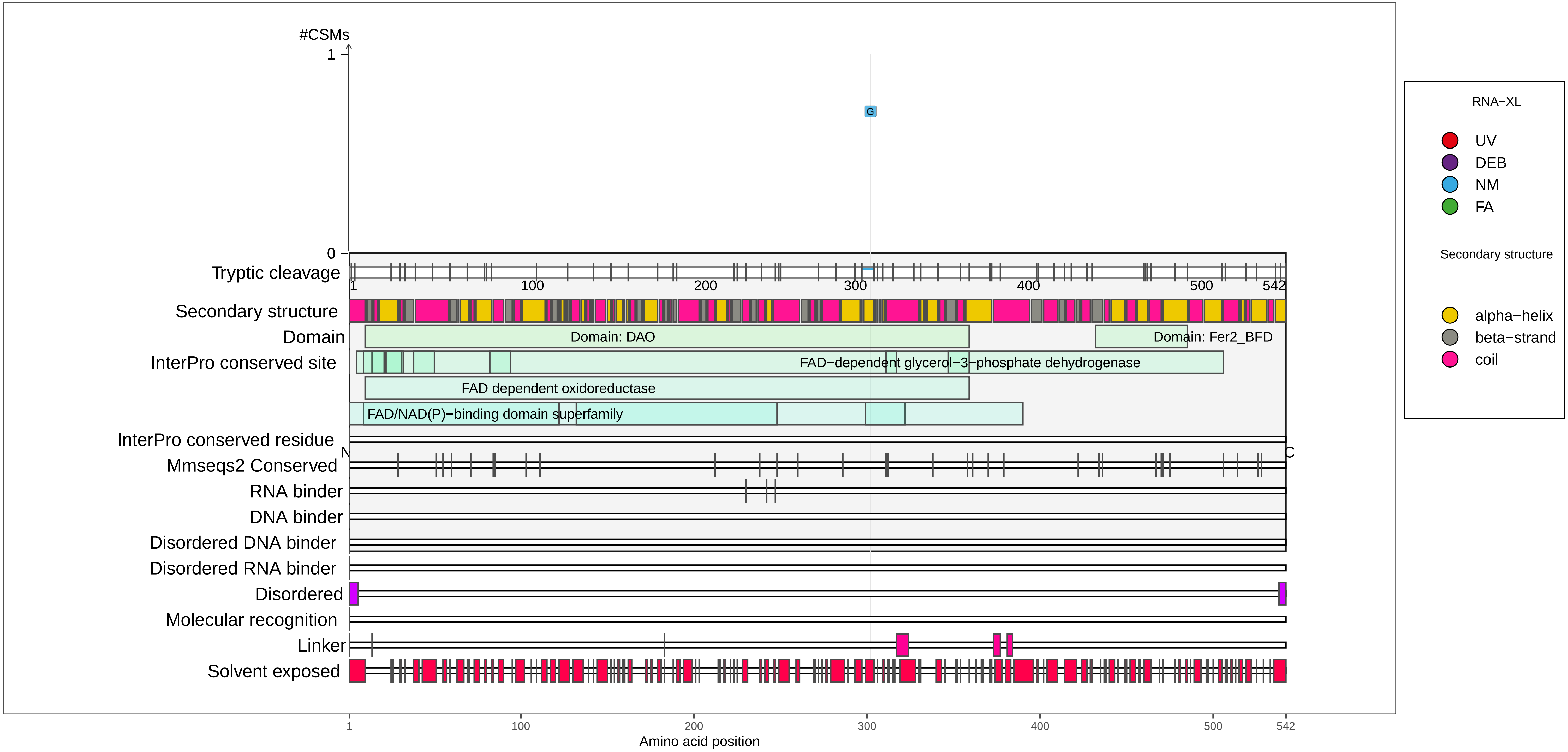

P0A9H1
MUG_ECOLI G/U mismatch−specific DNA glycosylase

− Abundance:
tryptic [log10 Intensity]: NA (Q NA)
PAXdb K12 strain [ppm]: 2.62 (Q 81)
PAXdb E.coli [ppm]: 2.03 (Q 73)

− RNA functions: not annotated

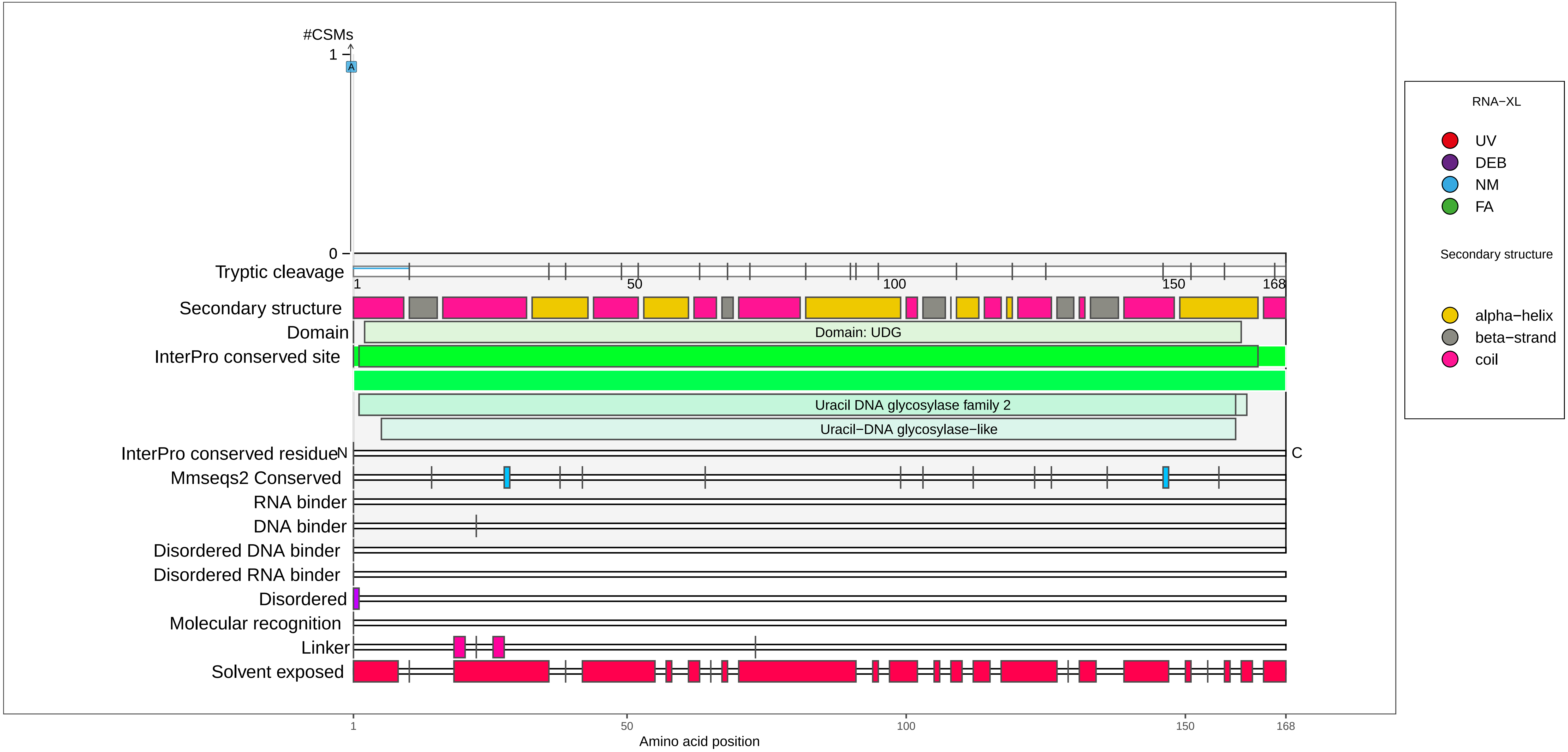

P0A9H7
CFA_ECOLI Cyclopropane−fatty−acyl−phospholipid synthase

− Abundance:
tryptic [log10 Intensity]: 8.66 (Q 80)
PAXdb K12 strain [ppm]: 2.12 (Q 65)
PAXdb E.coli [ppm]: 1.91 (Q 70)

− RNA functions: not annotated

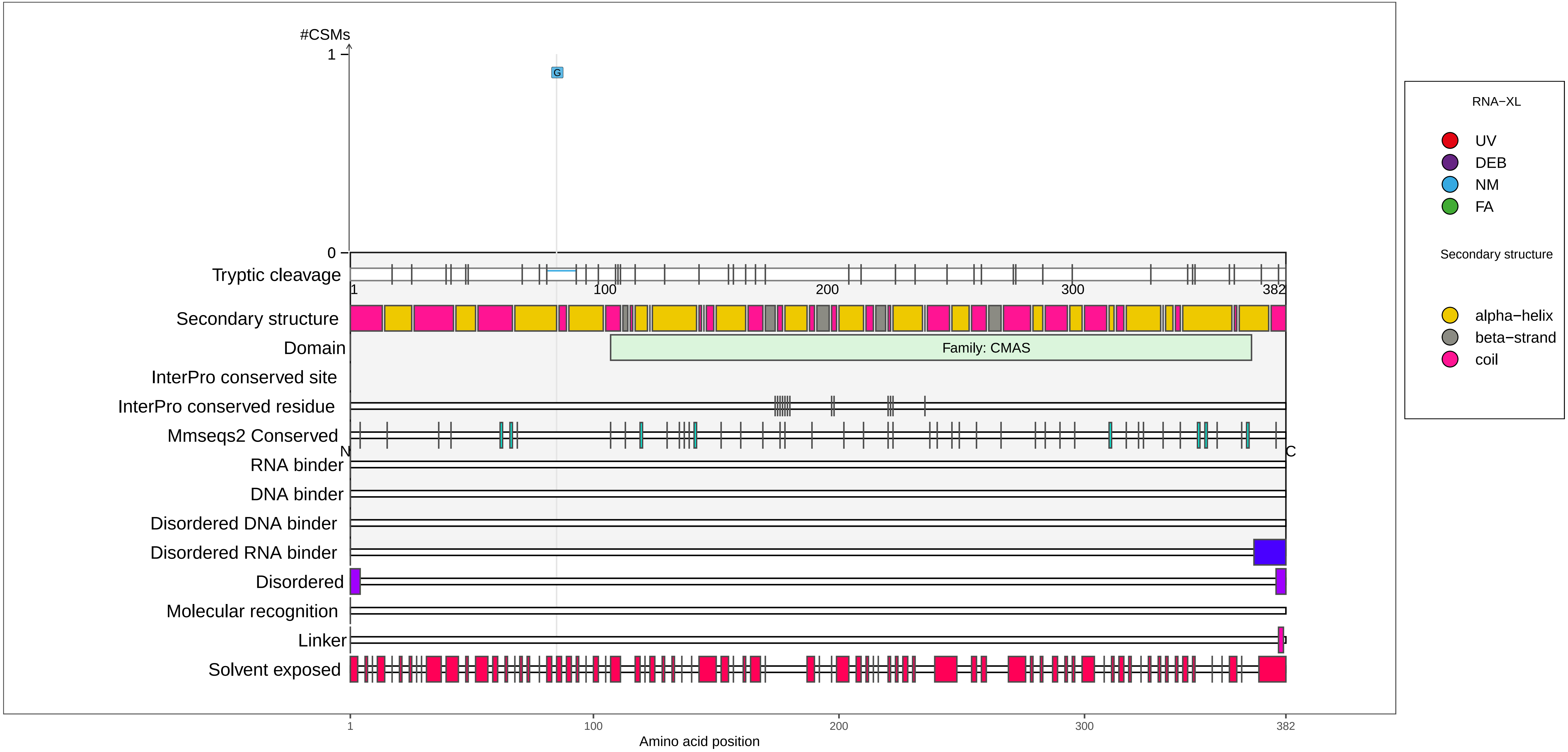

P0A9K1
PHOH_ECOLI Protein PhoH

− Abundance:
tryptic [log10 Intensity]: 6.72 (Q 6)
PAXdb K12 strain [ppm]: 0.81 (Q 5)
PAXdb E.coli [ppm]: 1.48 (Q 59)

− RNA functions: not annotated

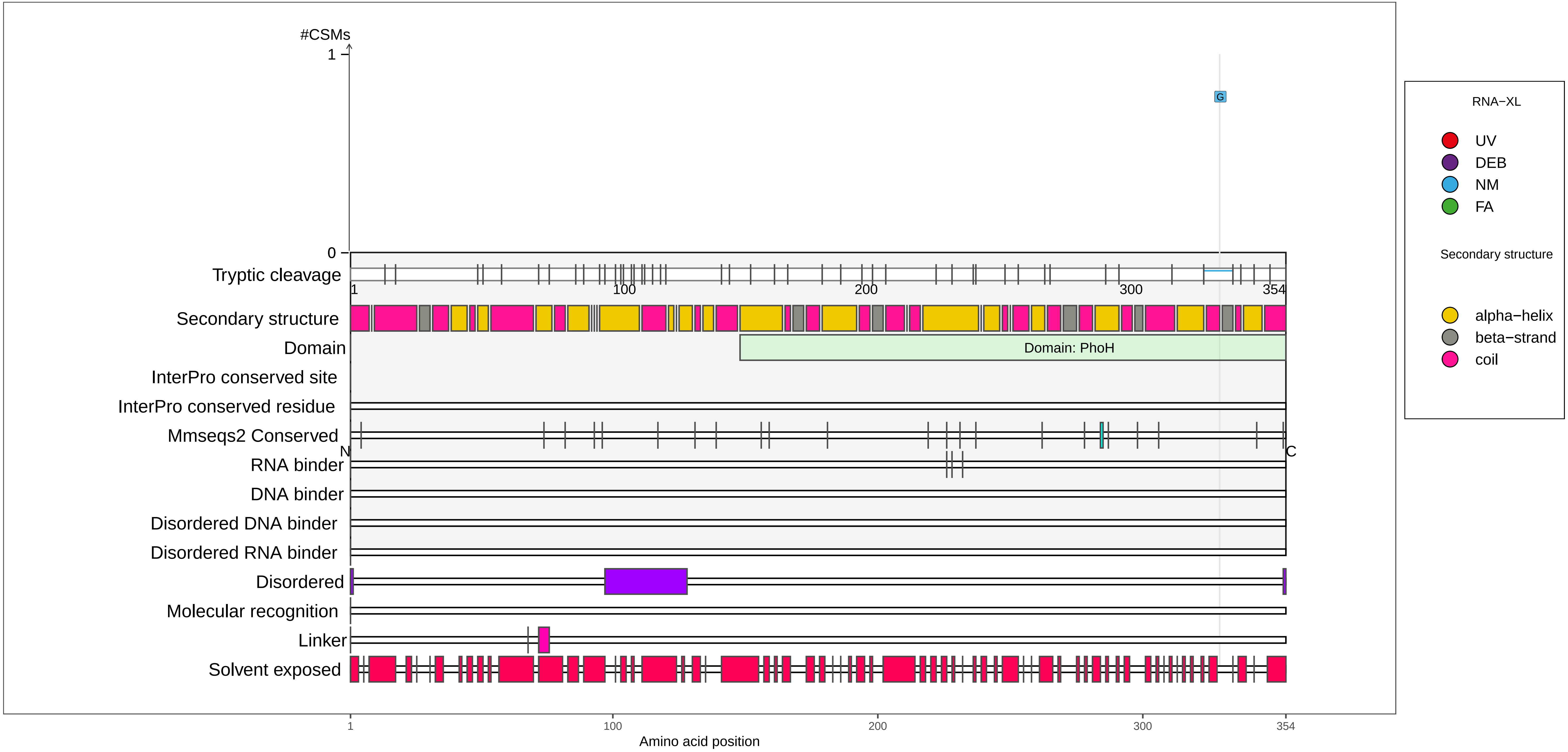

P0A9P9
IDNO_ECOLI 5−keto−D−gluconate 5−reductase

− Abundance:
tryptic [log10 Intensity]: 6.5 (Q 2)
PAXdb K12 strain [ppm]: NA (Q NA)
PAXdb E.coli [ppm]: 1.02 (Q 48)

− RNA functions: not annotated

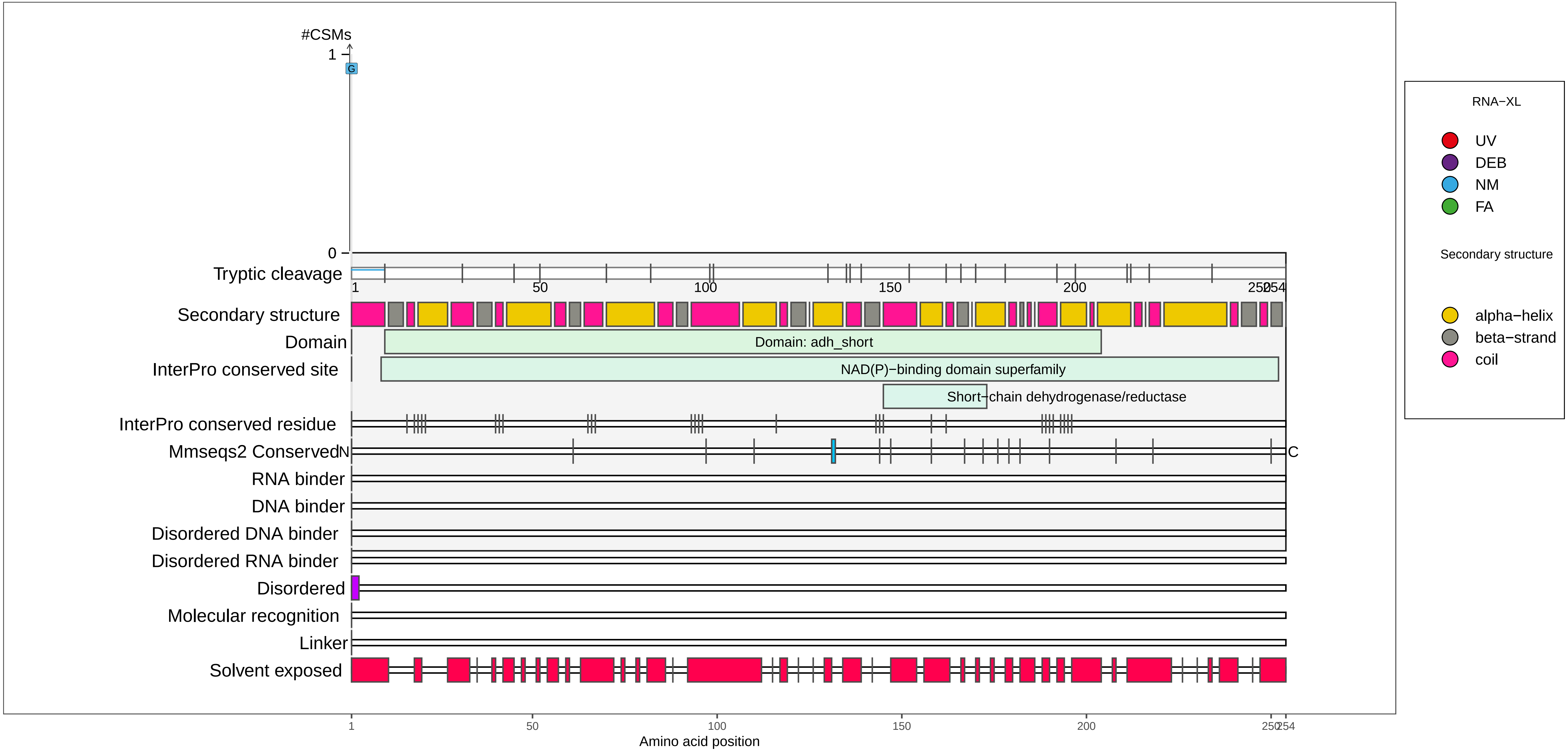

P0A9W3
ETTA_ECOLI Energy−dependent translational throttle protein EttA

− Abundance:
tryptic [log10 Intensity]: 8.58 (Q 78)
PAXdb K12 strain [ppm]: 2.79 (Q 86)
PAXdb E.coli [ppm]: 2.74 (Q 91)

− RNA functions:
RNA binding; rRNA binding; tRNA binding

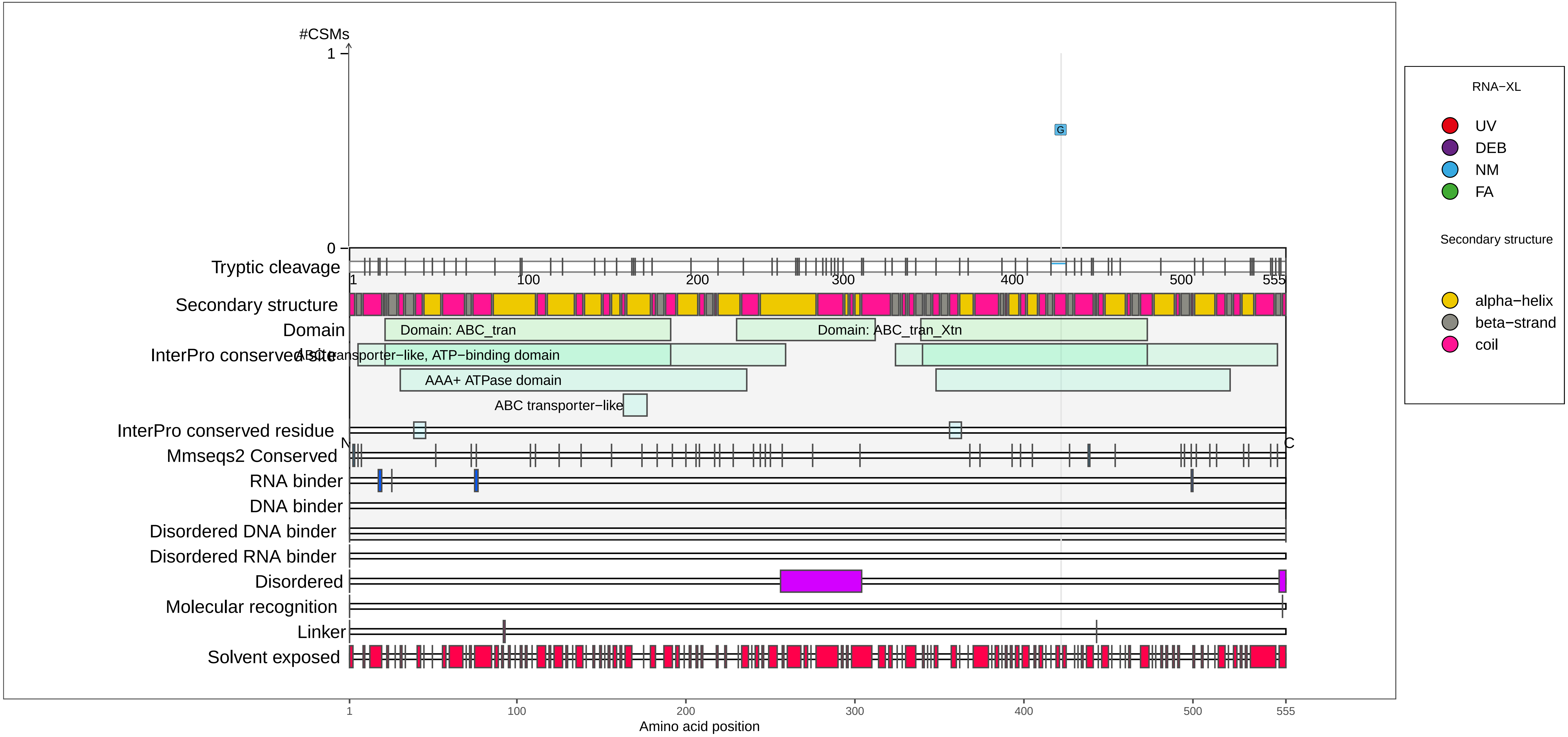

P0AA97
YAEQ_ECOLI Uncharacterized protein YaeQ

− Abundance:
tryptic [log10 Intensity]: NA (Q NA)
PAXdb K12 strain [ppm]: 1.36 (Q 30)
PAXdb E.coli [ppm]: 1.33 (Q 55)

− RNA functions: not annotated

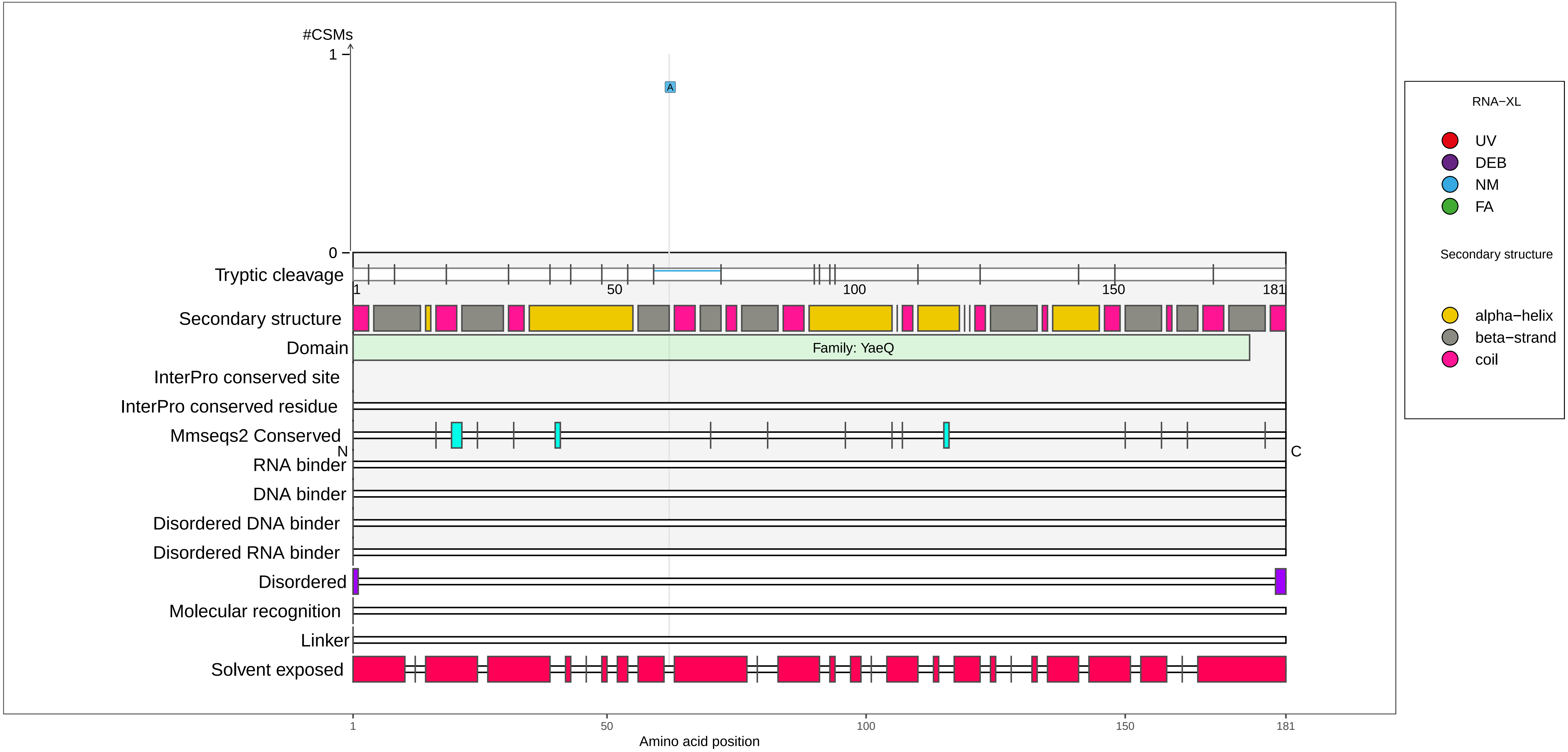

P0AAA1
YAGU_ECOLI Inner membrane protein YagU

− Abundance:
tryptic [log10 Intensity]: 7.9 (Q 54)
PAXdb K12 strain [ppm]: 1.18 (Q 19)
PAXdb E.coli [ppm]: 2.09 (Q 75)

− RNA functions: not annotated

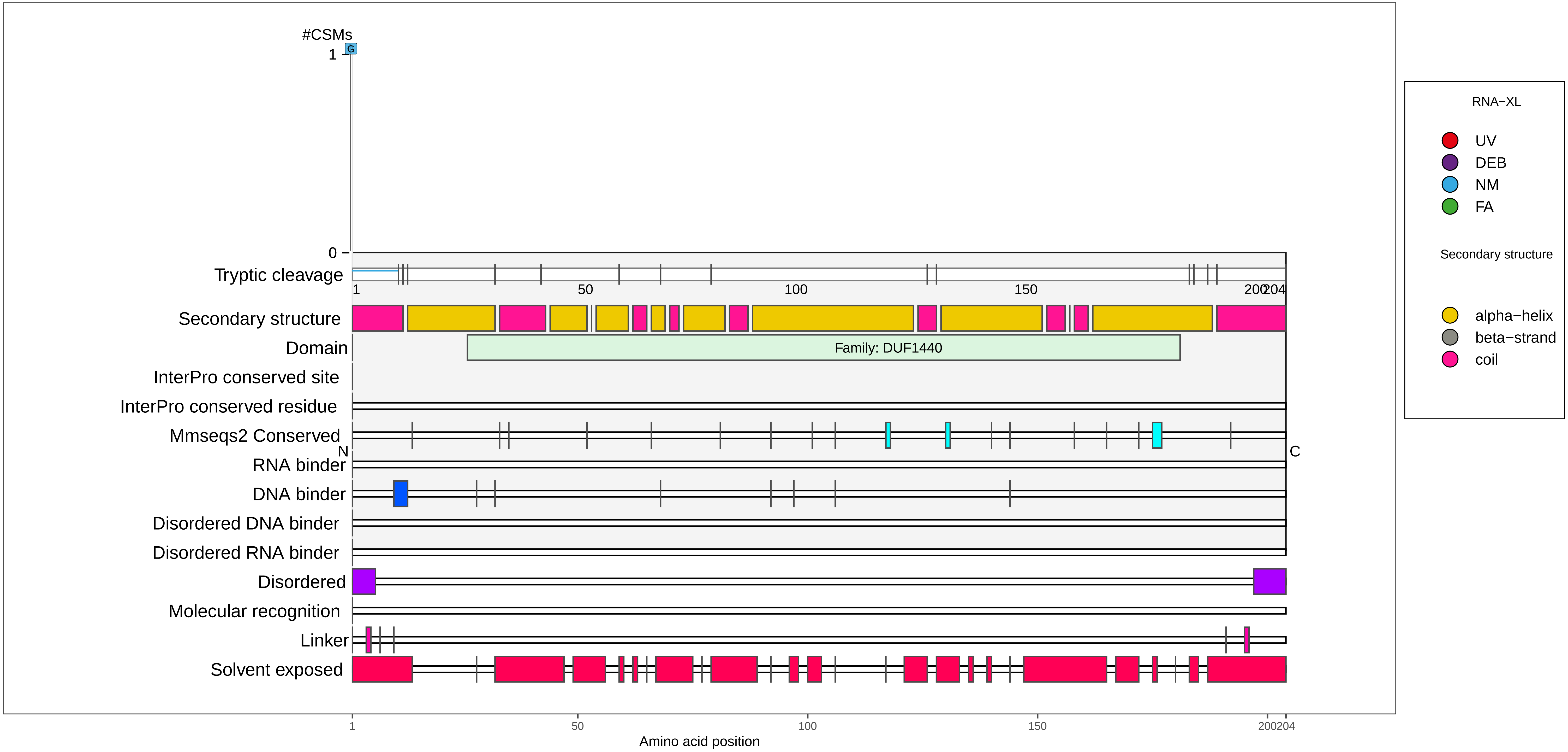

P0AAB4
UBID_ECOLI 3−octaprenyl−4−hydroxybenzoate carboxy−lyase

− Abundance:
tryptic [log10 Intensity]: 8.28 (Q 68)
PAXdb K12 strain [ppm]: 1.76 (Q 52)
PAXdb E.coli [ppm]: 1.7 (Q 64)

− RNA functions: not annotated

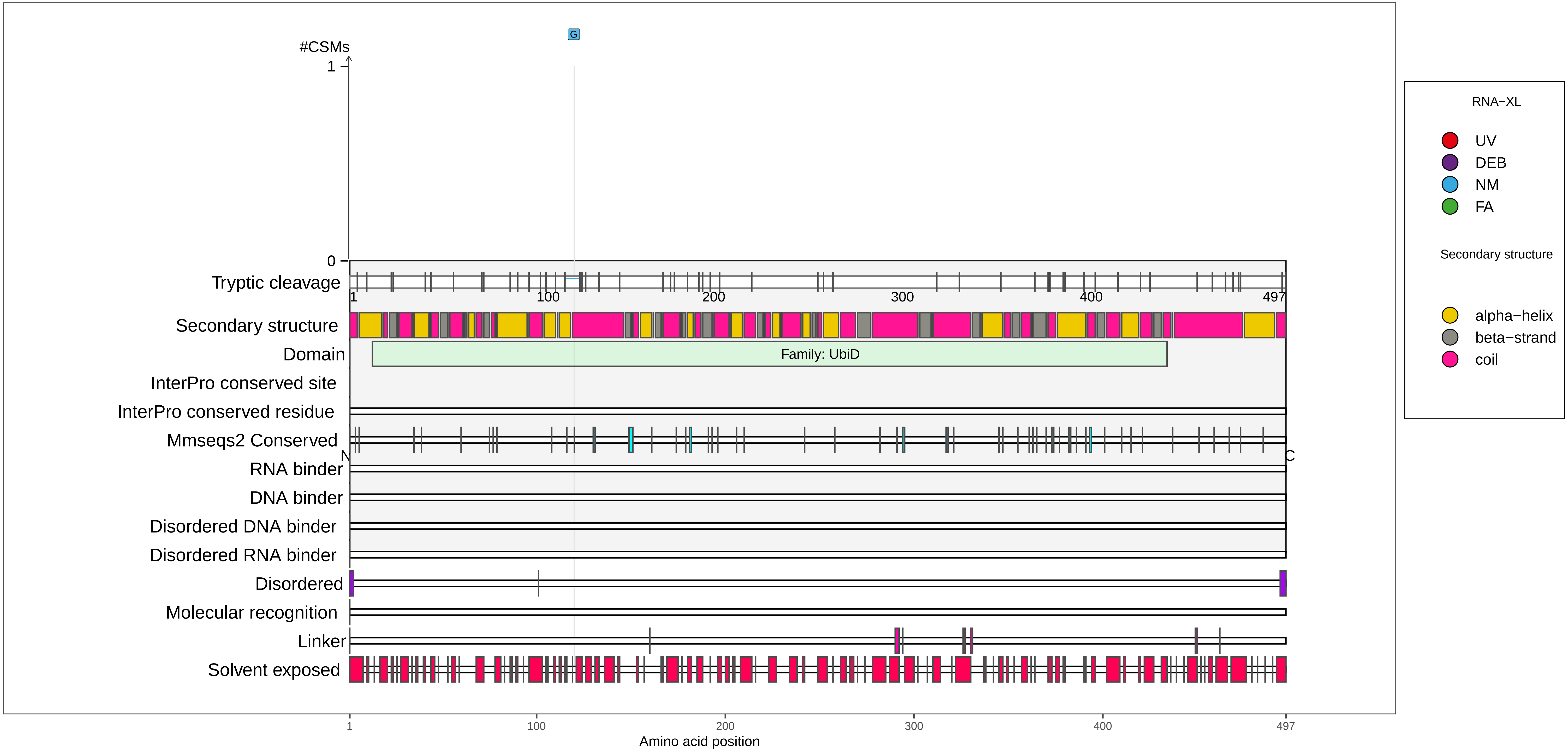

P0AAC4
YBHL_ECOLI Inner membrane protein YbhL

− Abundance:
tryptic [log10 Intensity]: 8.35 (Q 71)
PAXdb K12 strain [ppm]: 2.02 (Q 61)
PAXdb E.coli [ppm]: 0.57 (Q 38)

− RNA functions: not annotated

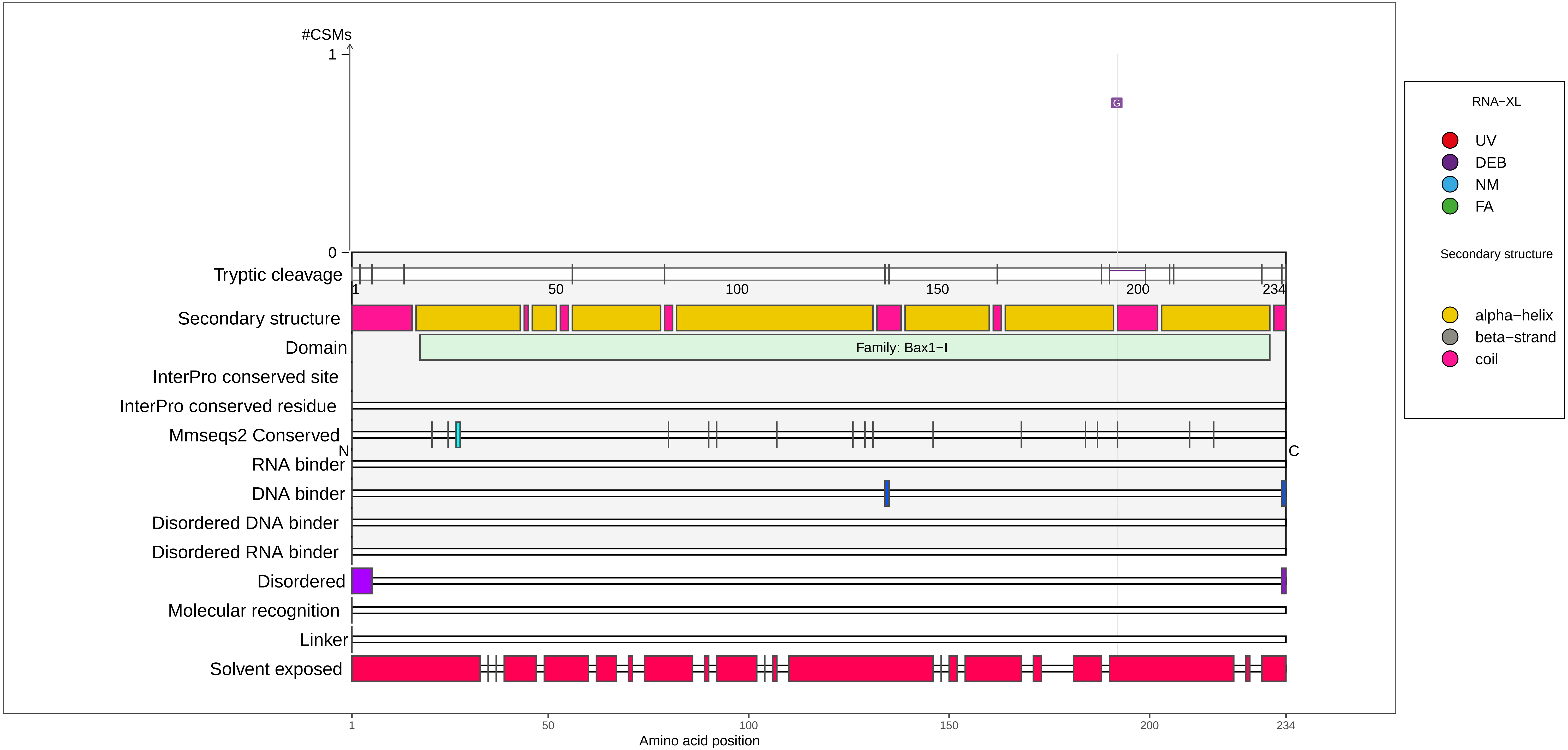

P0AAG8
MGLA_ECOLI Galactose/methyl galactoside import ATP−binding protein MglA
− Abundance:
tryptic [log10 Intensity]: 6.79 (Q 7)
PAXdb K12 strain [ppm]: 1.49 (Q 37)
PAXdb E.coli [ppm]: 2.17 (Q 77)

− RNA functions: not annotated

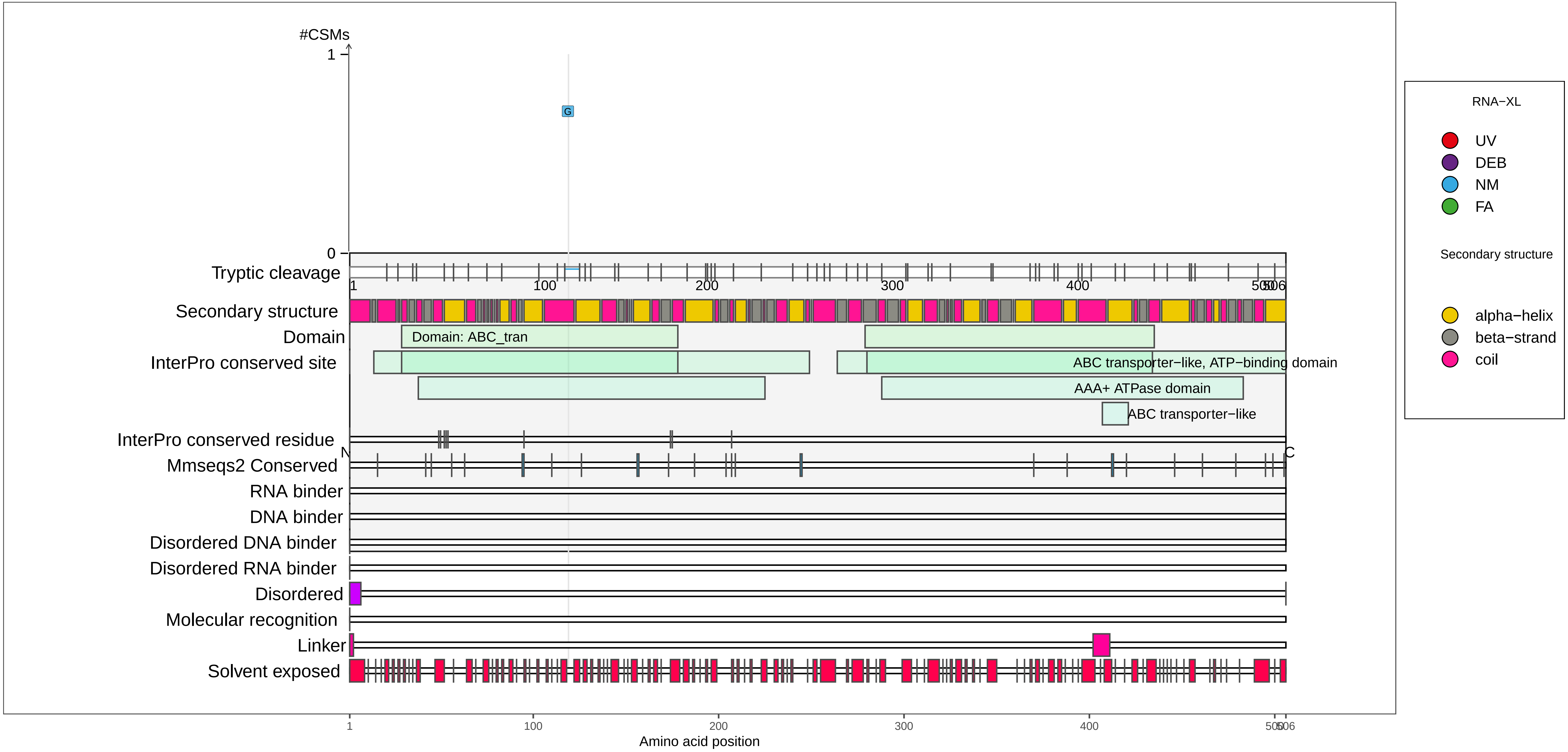

P0AAH4
SAPD_ECOLI Putrescine export system ATP−binding protein SapD

− Abundance:
tryptic [log10 Intensity]: NA (Q NA)
PAXdb K12 strain [ppm]: 1.68 (Q 48)
PAXdb E.coli [ppm]: −0.02 (Q 24)

− RNA functions: not annotated

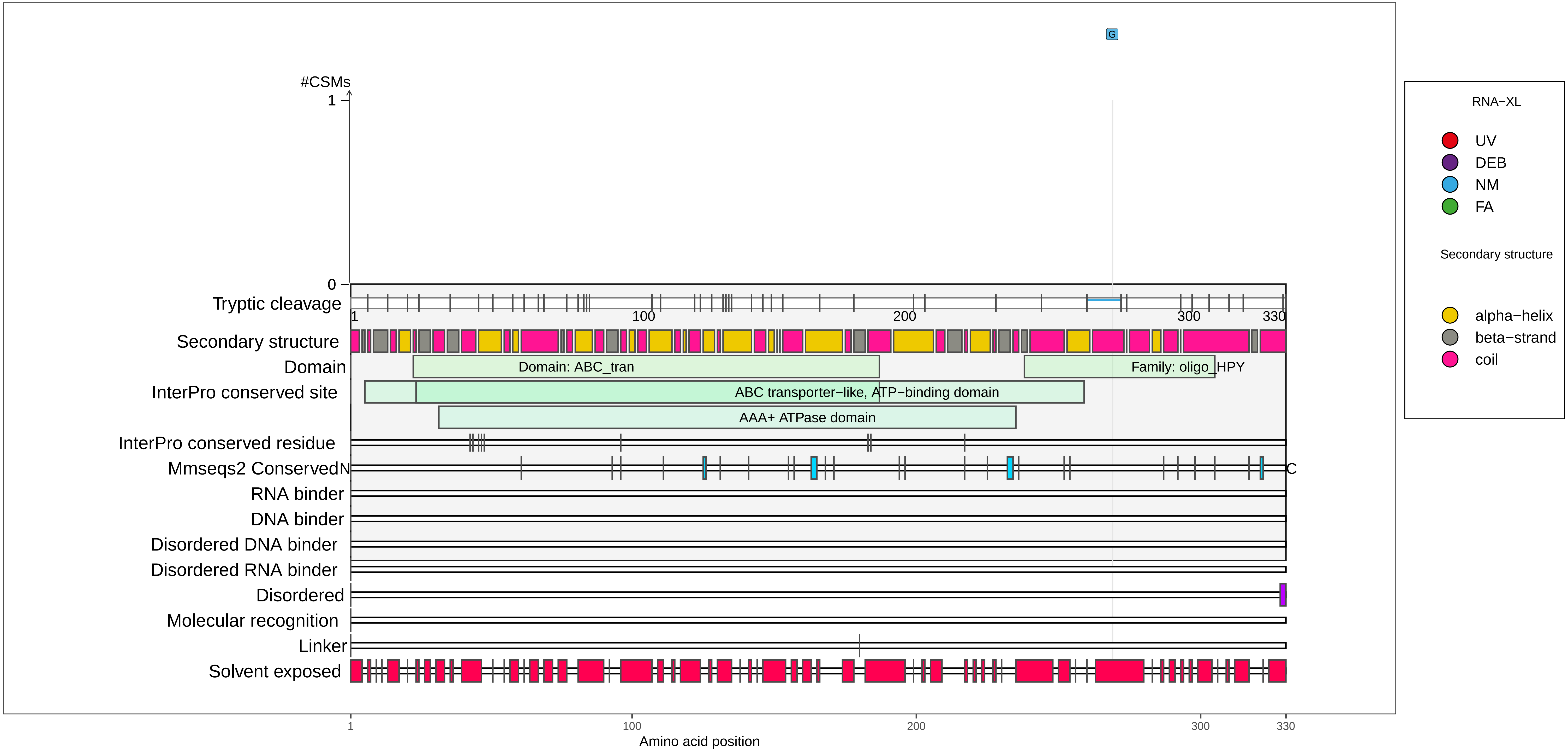

P0AAK4
HYDN_ECOLI Electron transport protein HydN

− Abundance:
tryptic [log10 Intensity]: NA (Q NA)
PAXdb K12 strain [ppm]: NA (Q NA)
PAXdb E.coli [ppm]: −2.7 (Q 1)

− RNA functions: not annotated

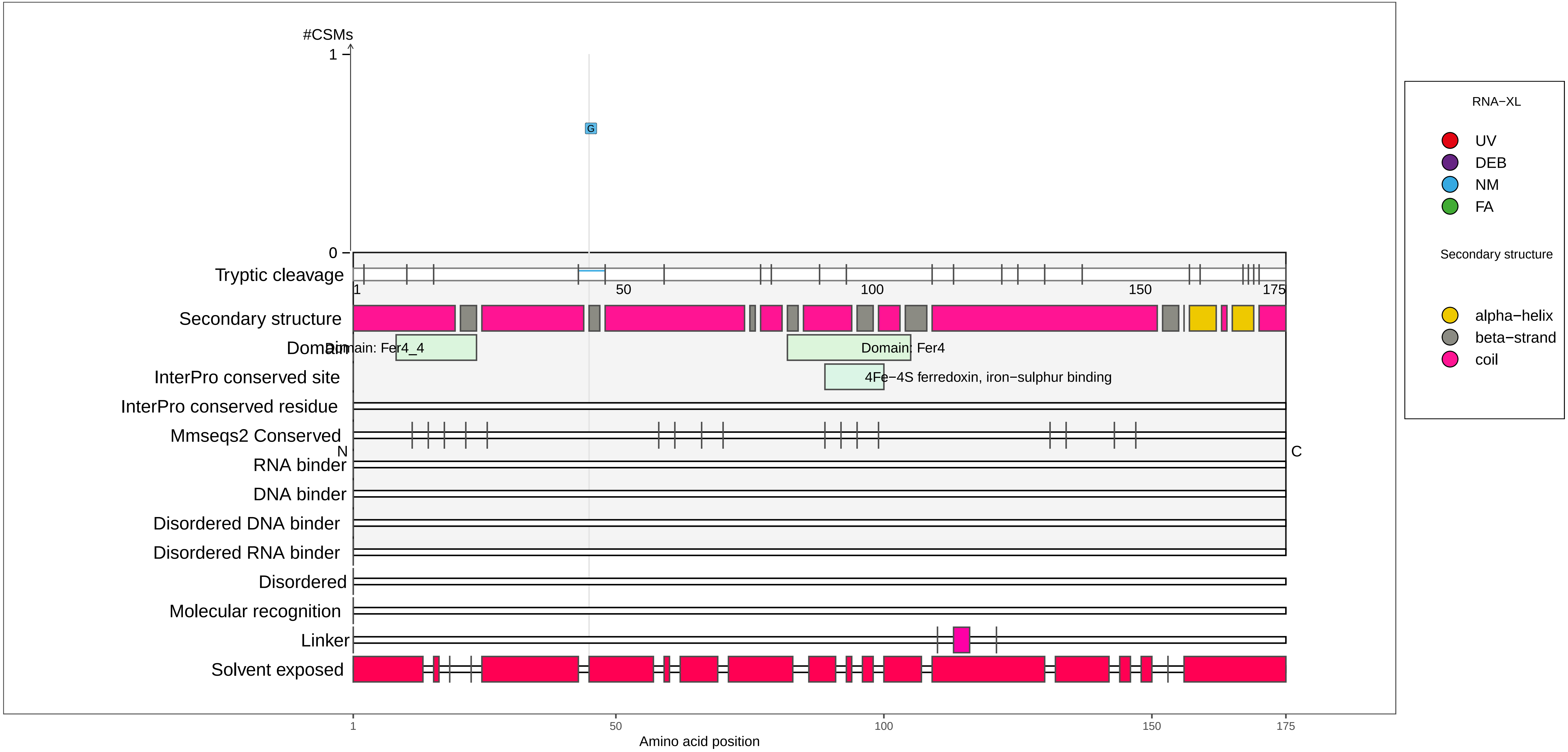

P0AAT4
YBDG_ECOLI Miniconductance mechanosensitive channel YbdG
− Abundance:
tryptic [log10 Intensity]: NA (Q NA)
PAXdb K12 strain [ppm]: NA (Q NA)
PAXdb E.coli [ppm]: 1.35 (Q 56)

− RNA functions: not annotated

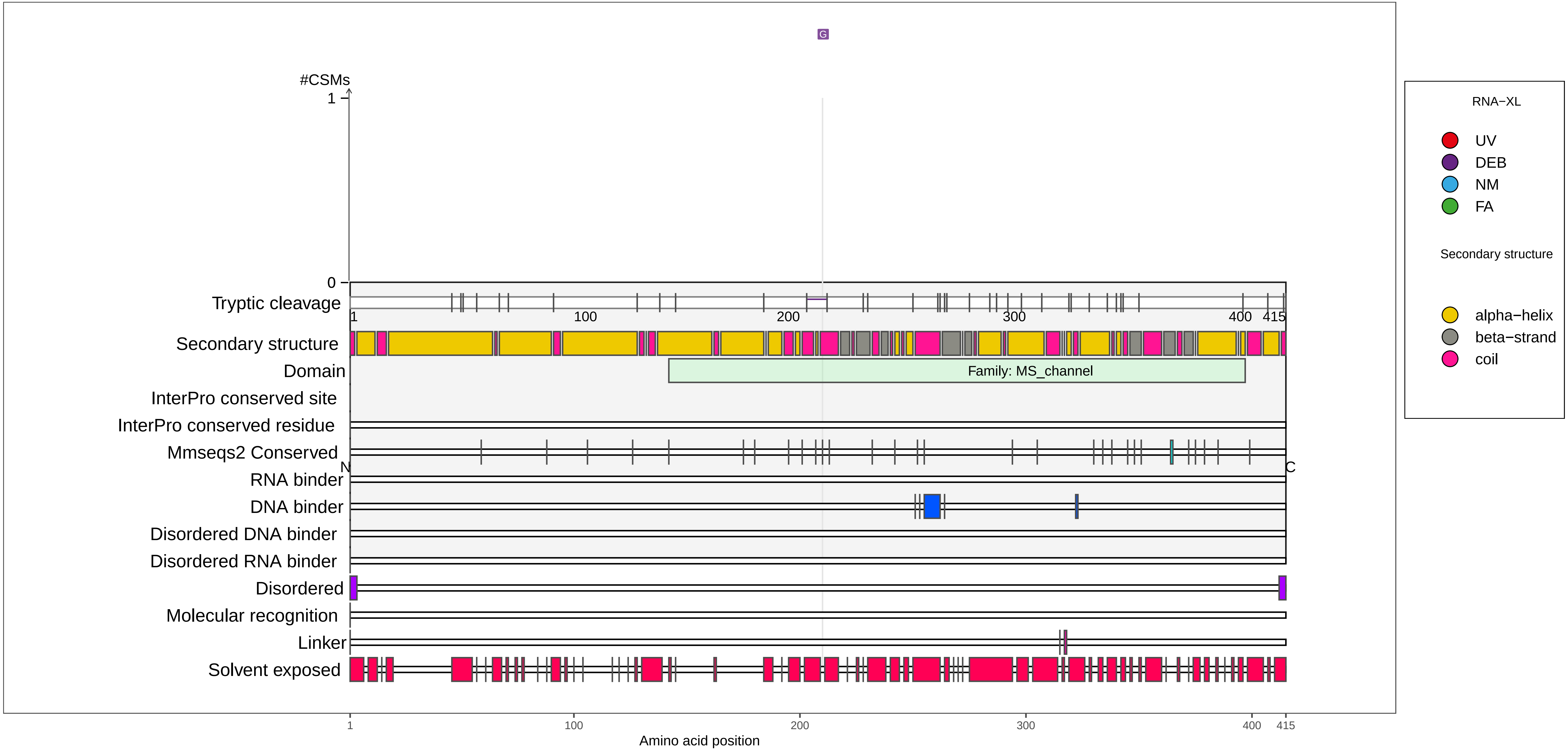

P0AAZ4
RARA_ECOLI Replication−associated recombination protein A

− Abundance:
tryptic [log10 Intensity]: NA (Q NA)
PAXdb K12 strain [ppm]: 1.79 (Q 53)
PAXdb E.coli [ppm]: 1.37 (Q 56)

− RNA functions: not annotated

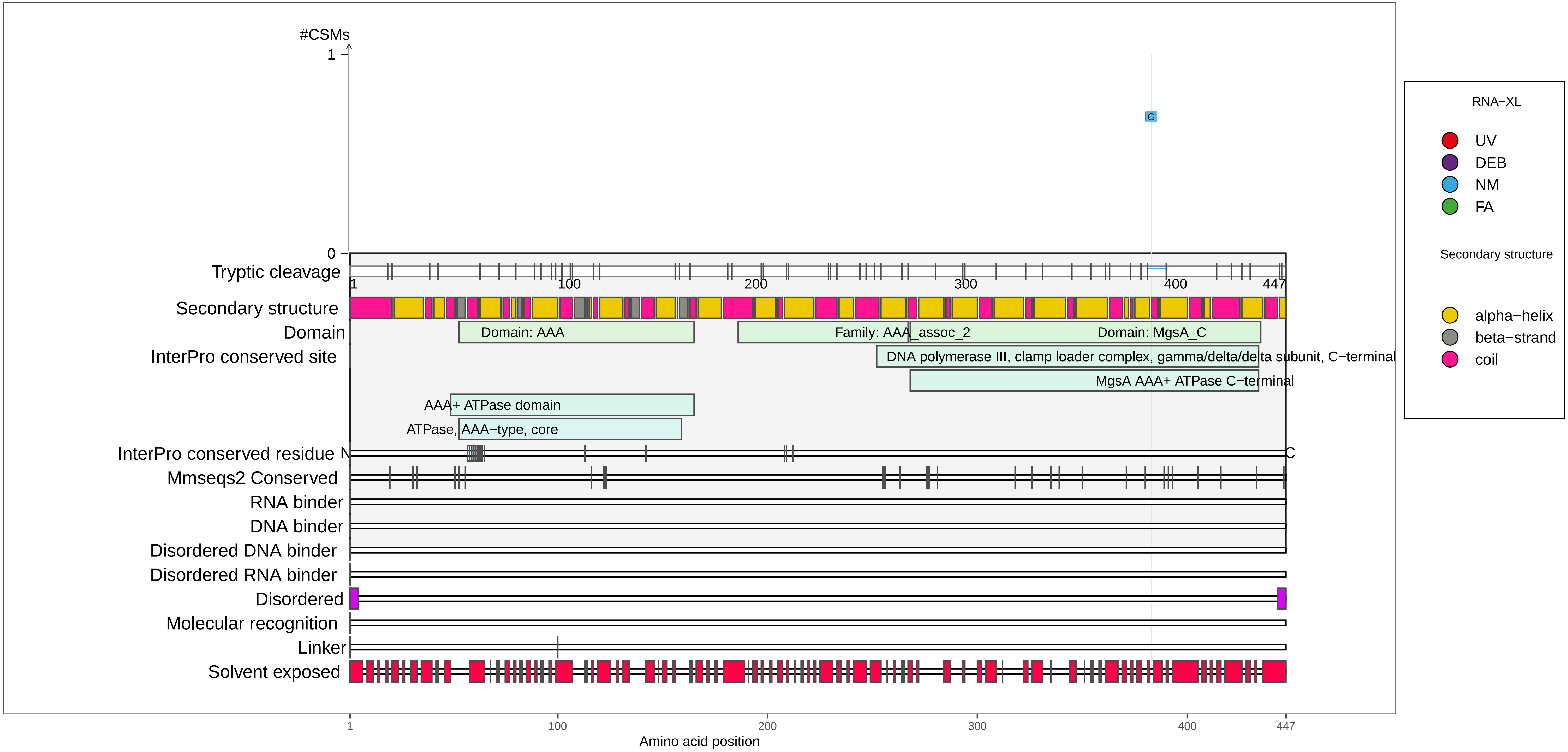

P0AB61
YCIN_ECOLI Protein YciN

− Abundance:
tryptic [log10 Intensity]: 8.44 (Q 73)
PAXdb K12 strain [ppm]: 3.41 (Q 96)
PAXdb E.coli [ppm]: 2.67 (Q 89)

− RNA functions: not annotated

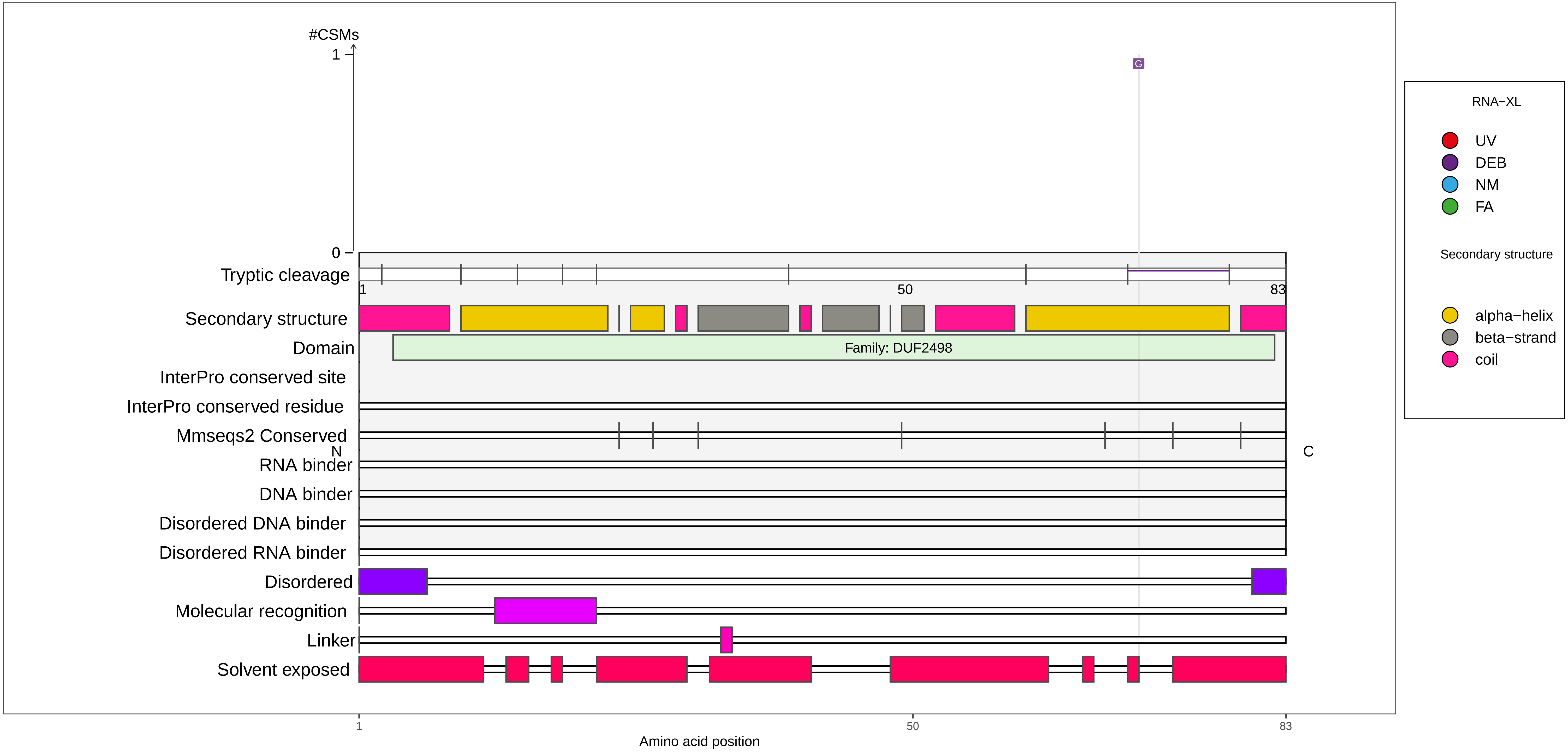

P0AB67
PNTB_ECOLI NAD(P) transhydrogenase subunit beta

− Abundance:
tryptic [log10 Intensity]: 7.34 (Q 27)
PAXdb K12 strain [ppm]: 1.97 (Q 59)
PAXdb E.coli [ppm]: 2.49 (Q 85)

− RNA functions: not annotated

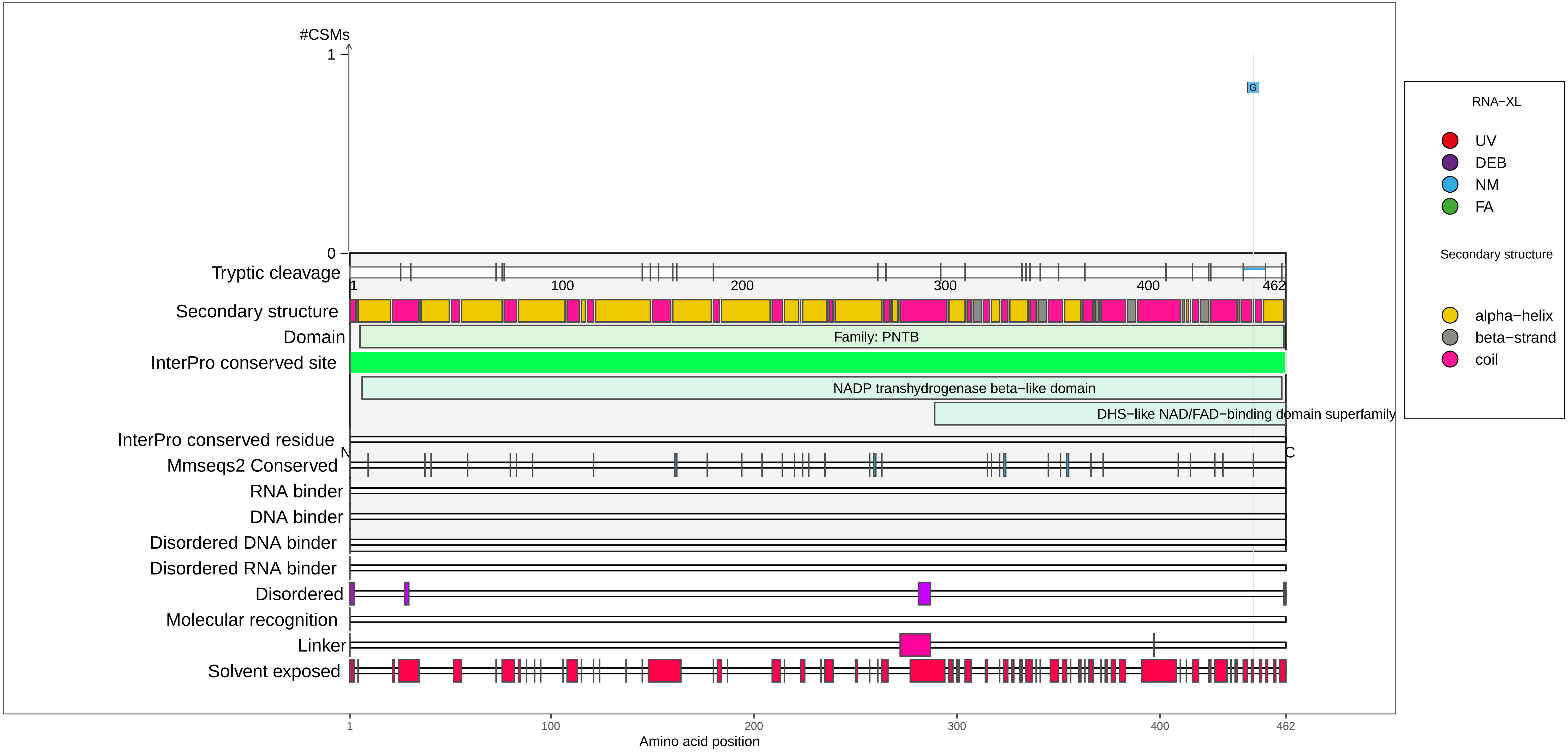

P0ABD8
BCCP_ECOLI Biotin carboxyl carrier protein of acetyl−CoA carboxylase

− Abundance:
tryptic [log10 Intensity]: 8.61 (Q 79)
PAXdb K12 strain [ppm]: 2.16 (Q 66)
PAXdb E.coli [ppm]: 2.47 (Q 85)

− RNA functions: not annotated

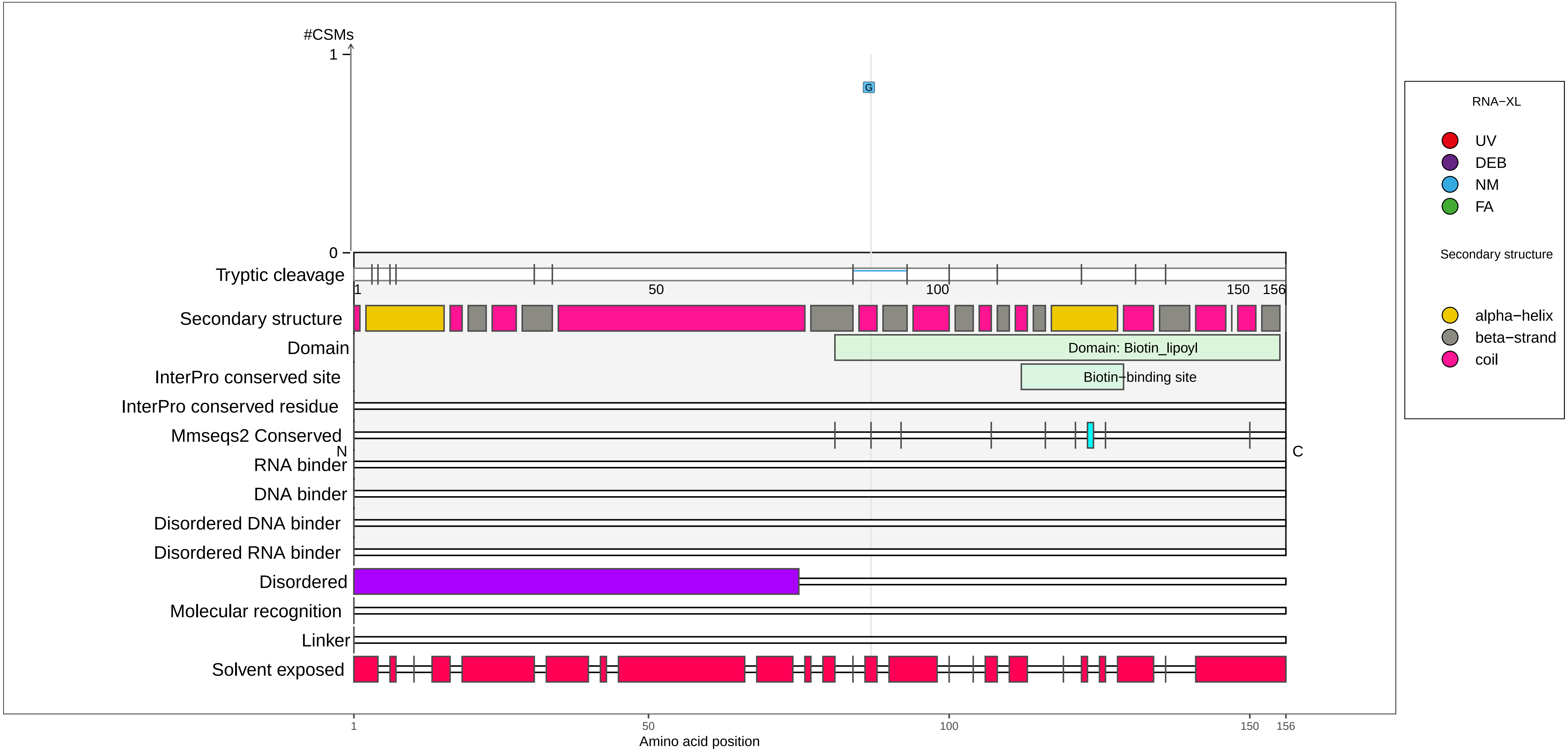

P0ABK2
CYDB_ECOLI Cytochrome bd−I ubiquinol oxidase subunit 2
− Abundance:
tryptic [log10 Intensity]: 9.89 (Q 99)
PAXdb K12 strain [ppm]: 2.65 (Q 82)
PAXdb E.coli [ppm]: 2.36 (Q 82)

− RNA functions: not annotated

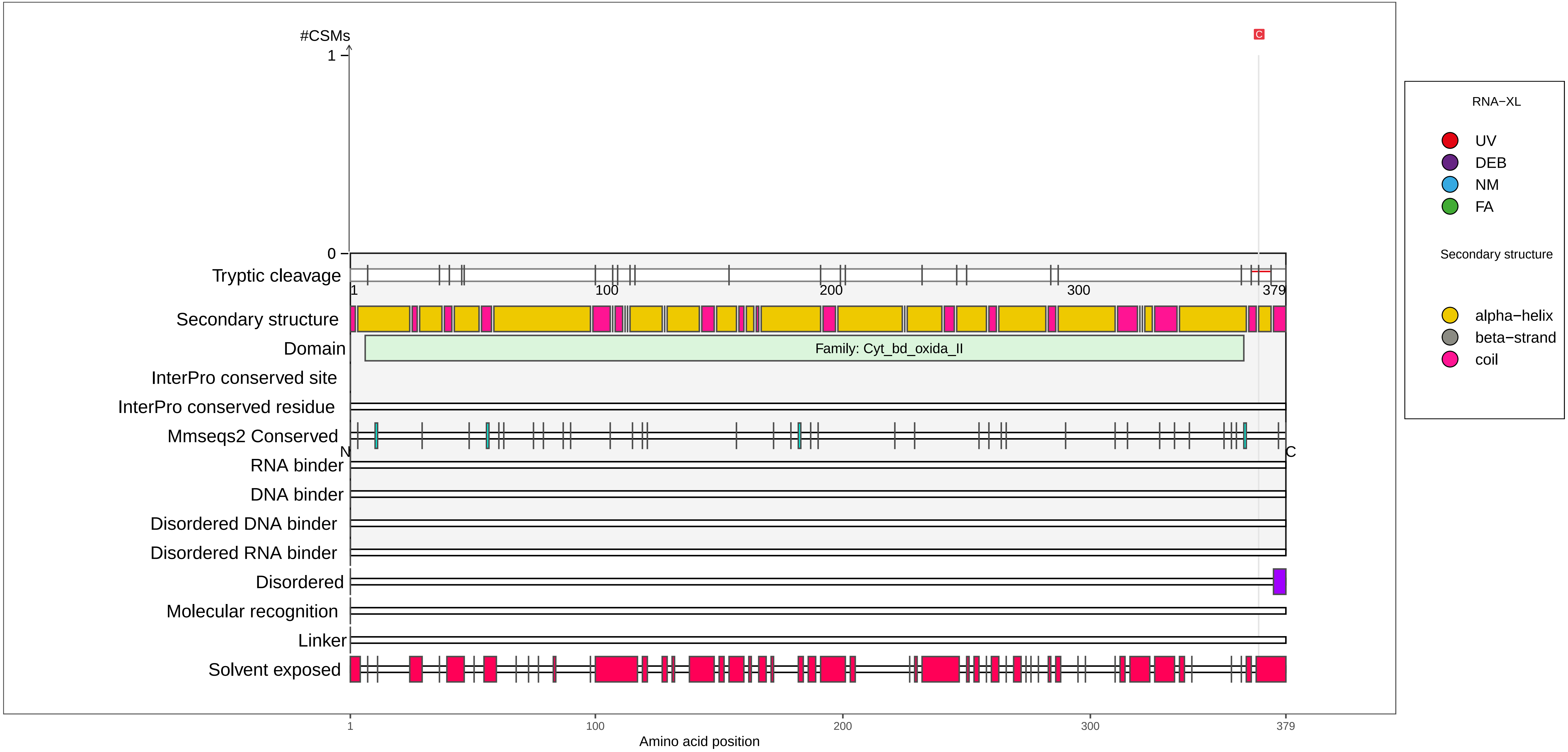

P0ABS8
HOLE_ECOLI DNA polymerase III subunit theta

− Abundance:
tryptic [log10 Intensity]: NA (Q NA)
PAXdb K12 strain [ppm]: 1.76 (Q 52)
PAXdb E.coli [ppm]: 1.04 (Q 49)

− RNA functions: not annotated

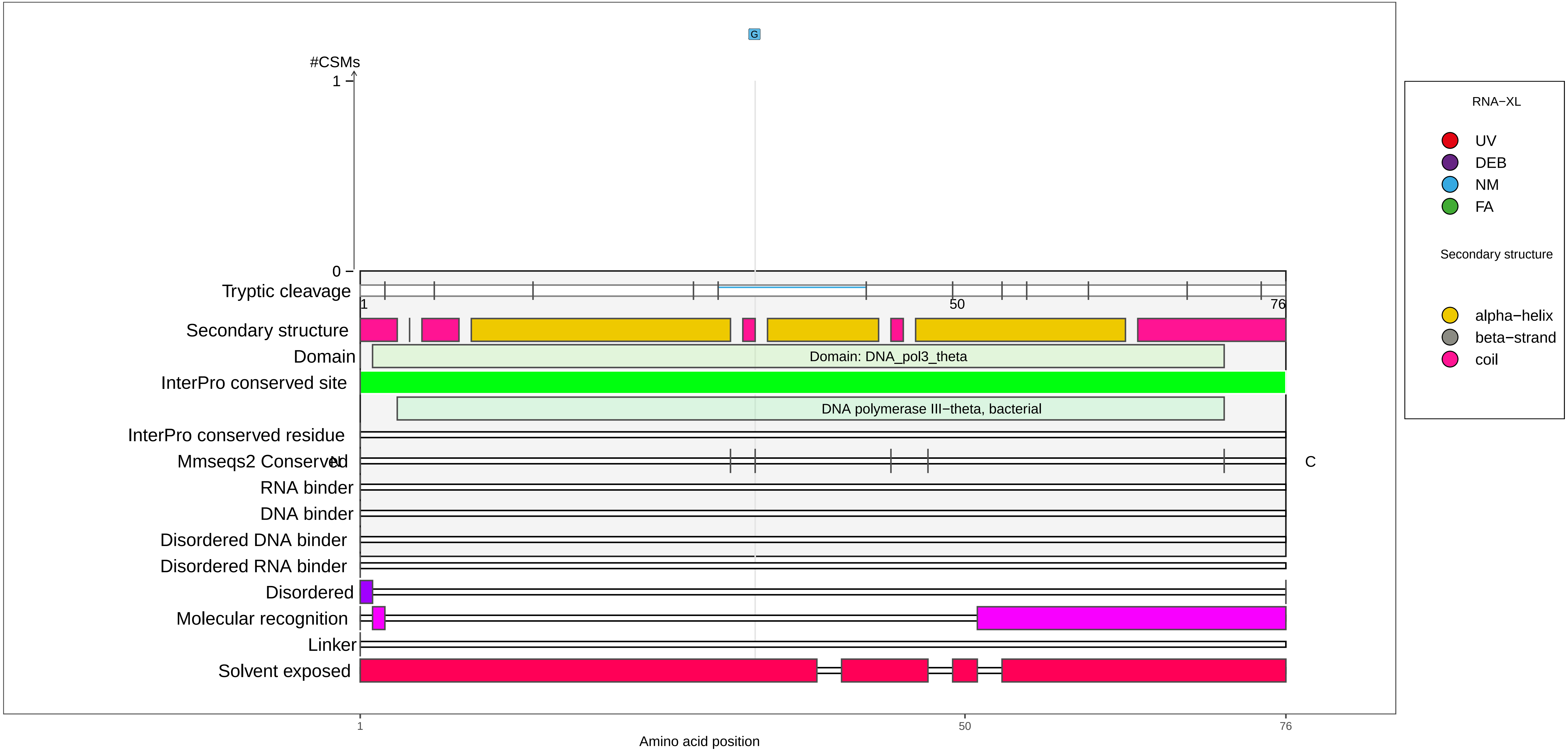

P0ABT2
DPS_ECOLI DNA protection during starvation protein

− Abundance:
tryptic [log10 Intensity]: 8.03 (Q 59)
PAXdb K12 strain [ppm]: 2.34 (Q 71)
PAXdb E.coli [ppm]: 3.49 (Q 99)

− RNA functions: not annotated

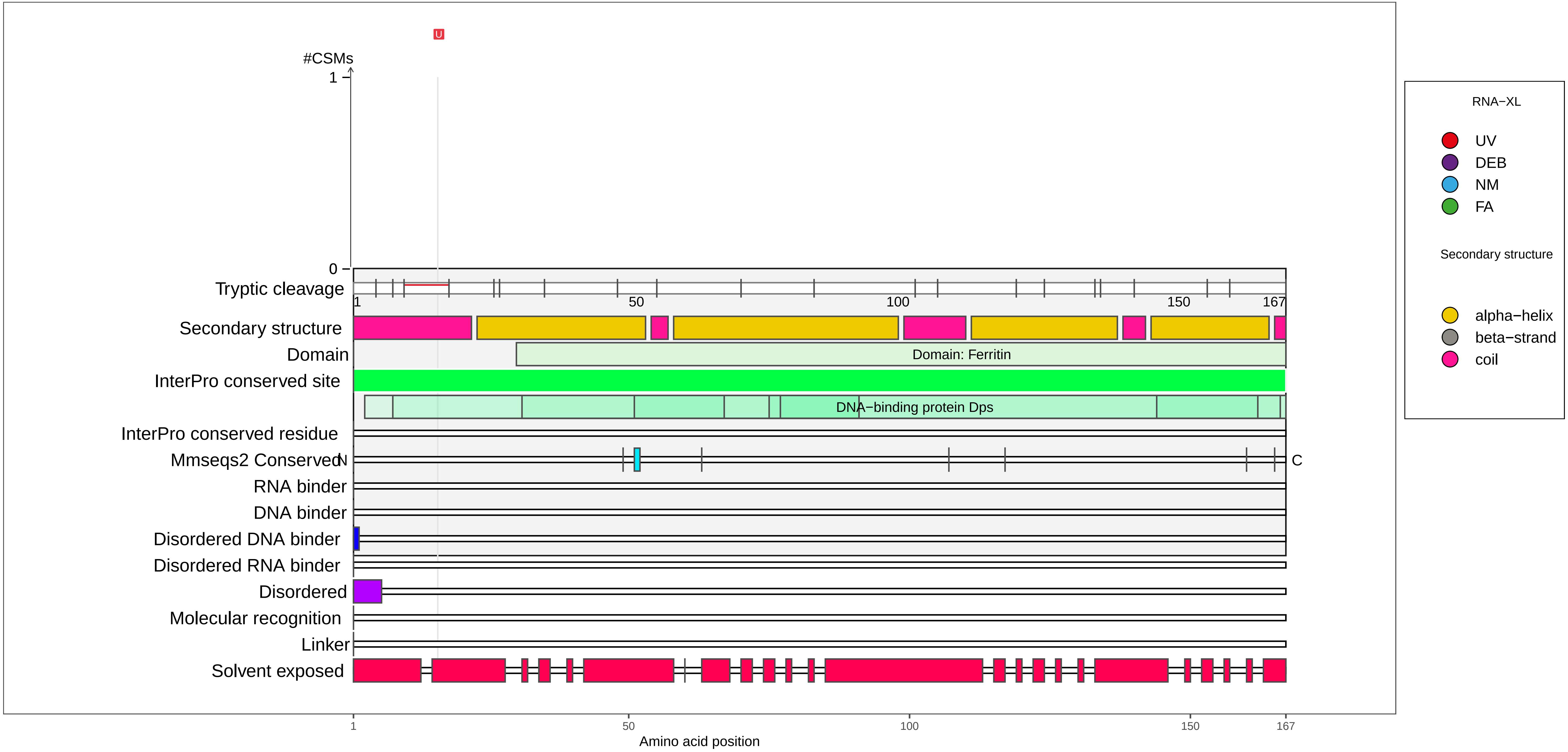

P0ABU5
ELBB_ECOLI Glyoxalase ElbB

− Abundance:
tryptic [log10 Intensity]: 8.08 (Q 60)
PAXdb K12 strain [ppm]: 1.72 (Q 50)
PAXdb E.coli [ppm]: 2.52 (Q 86)

− RNA functions: not annotated

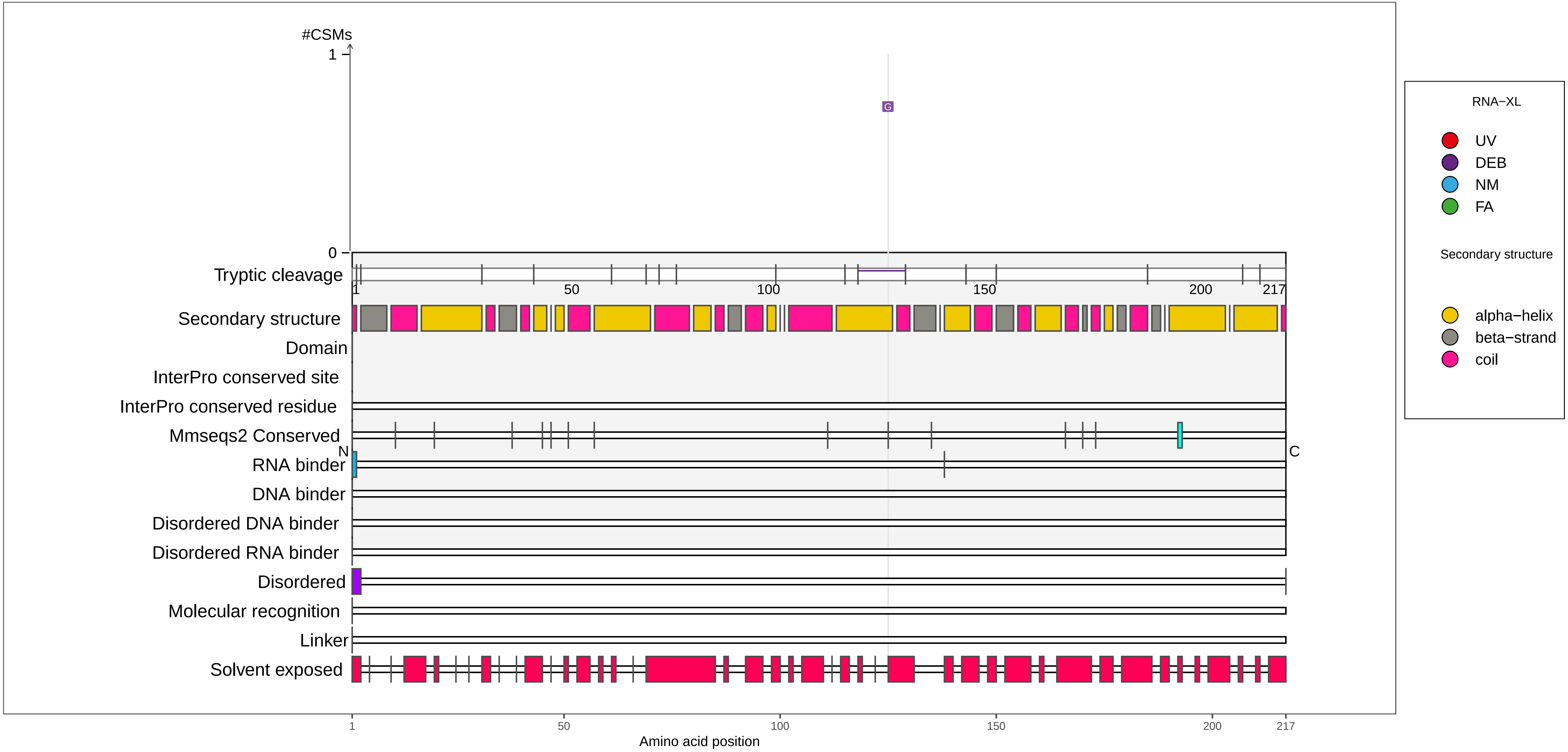

P0ABY2
FLIT_ECOLI Flagellar protein FliT
− Abundance:
tryptic [log10 Intensity]: NA (Q NA)
PAXdb K12 strain [ppm]: NA (Q NA)
PAXdb E.coli [ppm]: NA (Q NA)

− RNA functions: not annotated

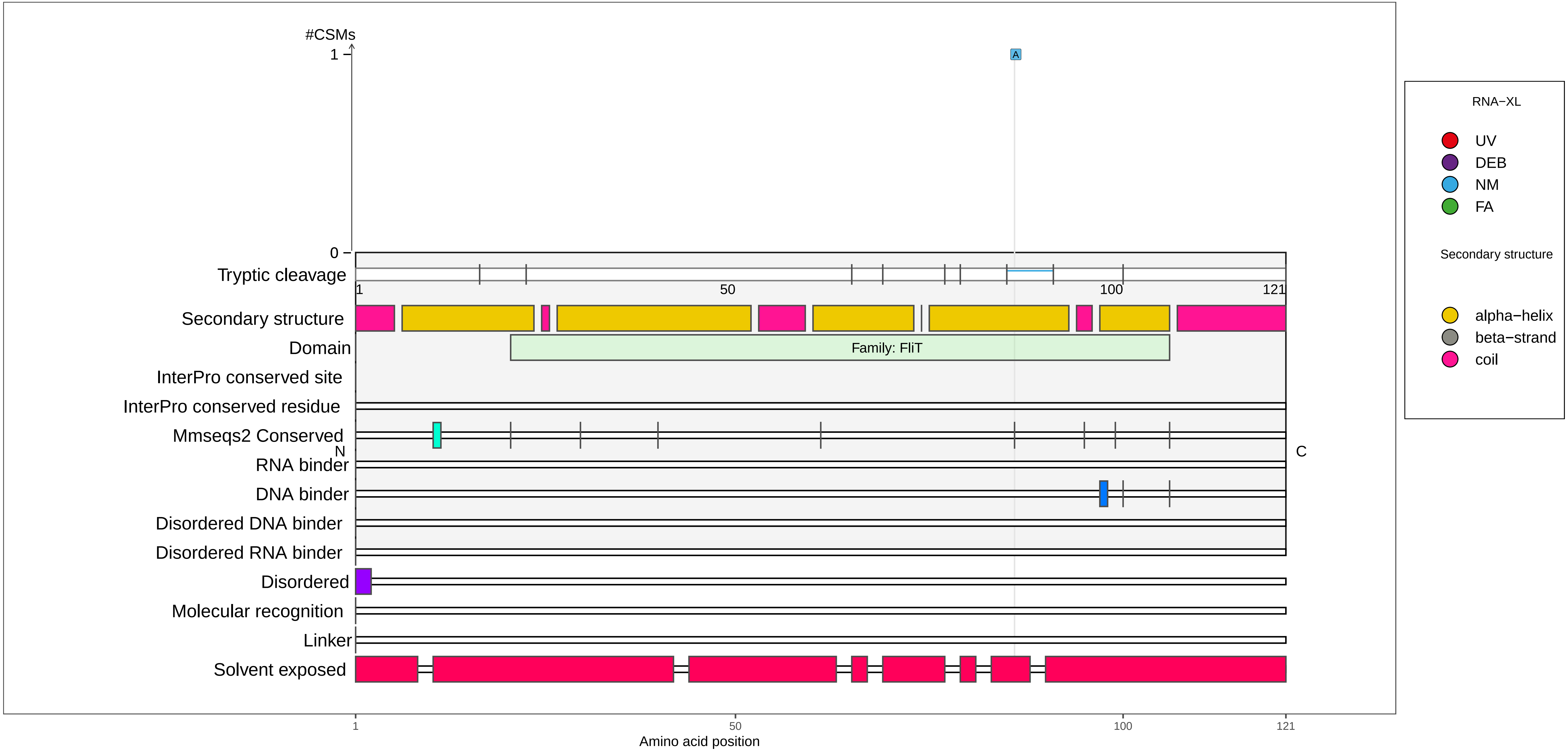

P0AC23
FOCA_ECOLI Probable formate transporter 1

− Abundance:
tryptic [log10 Intensity]: 7.78 (Q 48)
PAXdb K12 strain [ppm]: 2.06 (Q 62)
PAXdb E.coli [ppm]: 1.68 (Q 64)

− RNA functions: not annotated

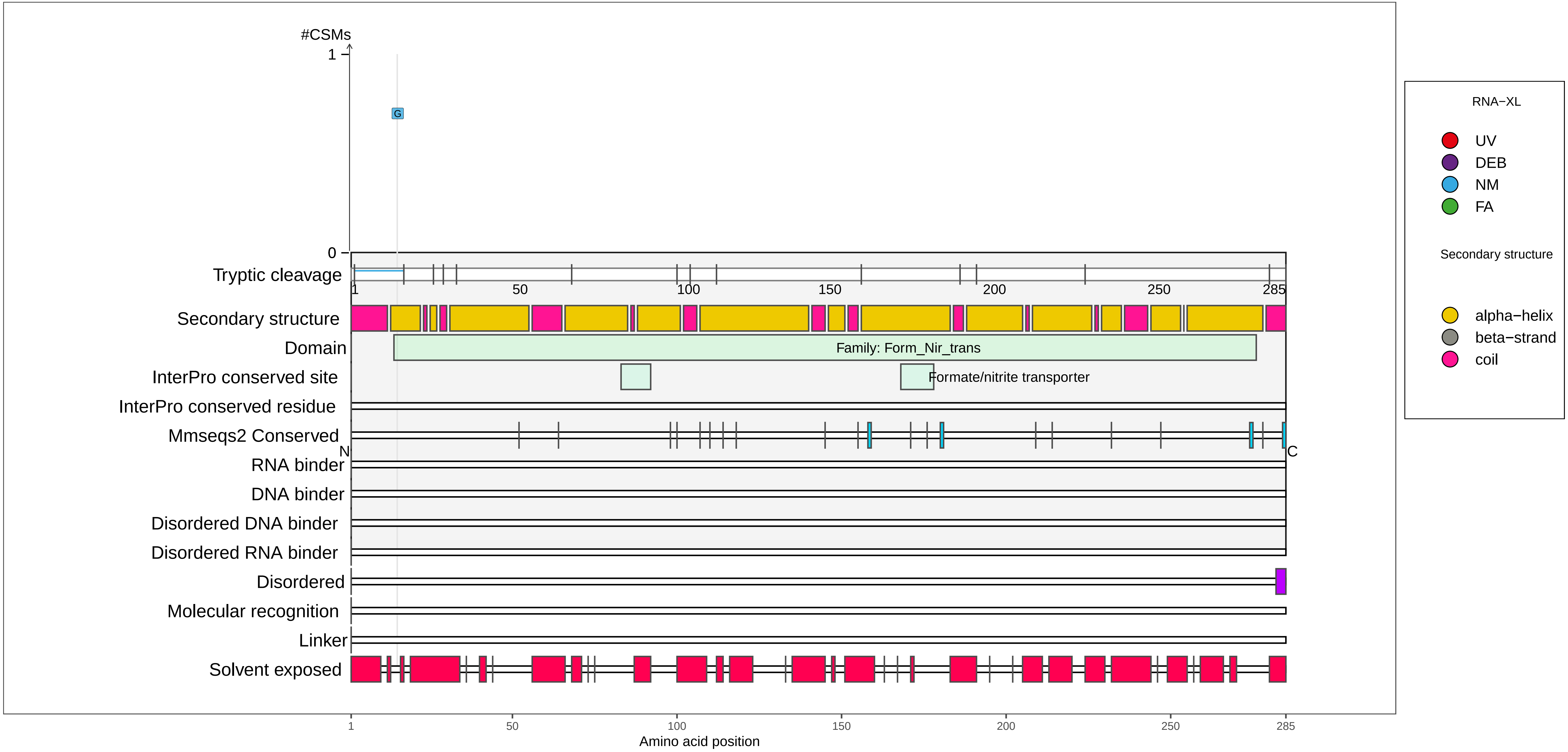

P0AC28
5FCL_ECOLI 5−formyltetrahydrofolate cyclo−ligase

− Abundance:
tryptic [log10 Intensity]: NA (Q NA)
PAXdb K12 strain [ppm]: NA (Q NA)
PAXdb E.coli [ppm]: 1.05 (Q 49)

− RNA functions: not annotated

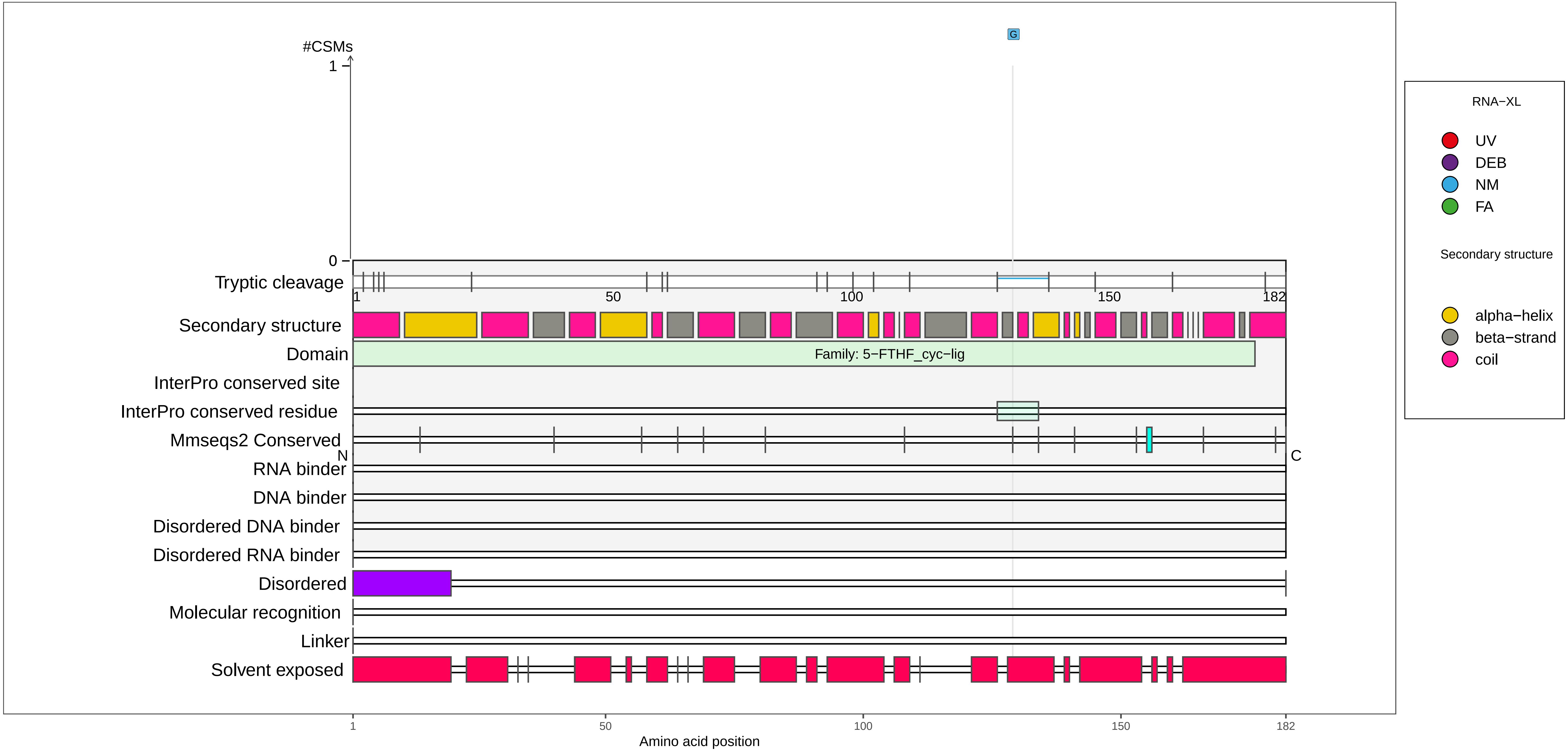

P0AC59
GLRX2_ECOLI Glutaredoxin 2

− Abundance:
tryptic [log10 Intensity]: 7.14 (Q 18)
PAXdb K12 strain [ppm]: 2.57 (Q 79)
PAXdb E.coli [ppm]: 2.69 (Q 90)

− RNA functions: not annotated

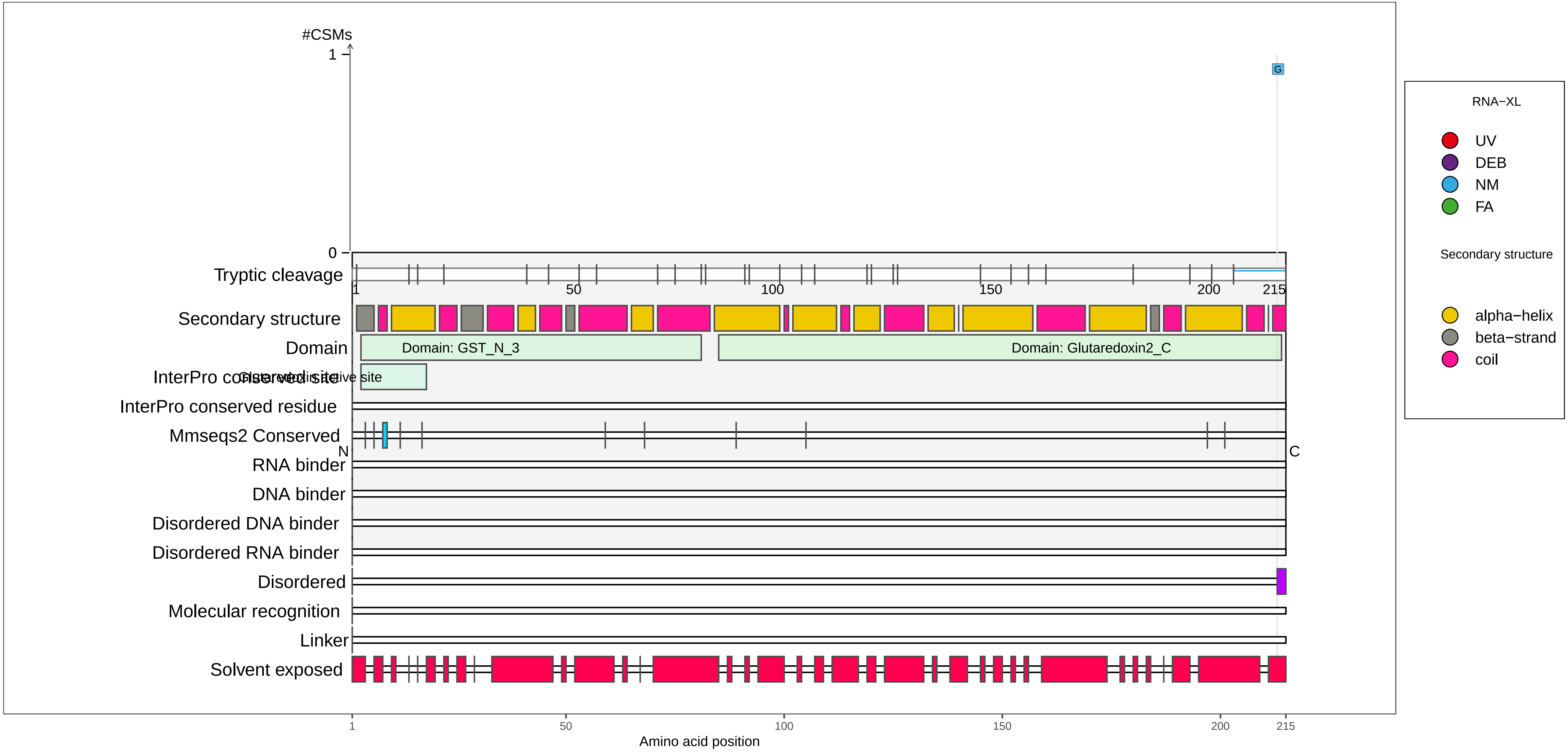

P0AC69
GLRX4_ECOLI Glutaredoxin 4

− Abundance:
tryptic [log10 Intensity]: 9.33 (Q 94)
PAXdb K12 strain [ppm]: 3.09 (Q 92)
PAXdb E.coli [ppm]: 3.21 (Q 97)

− RNA functions: not annotated

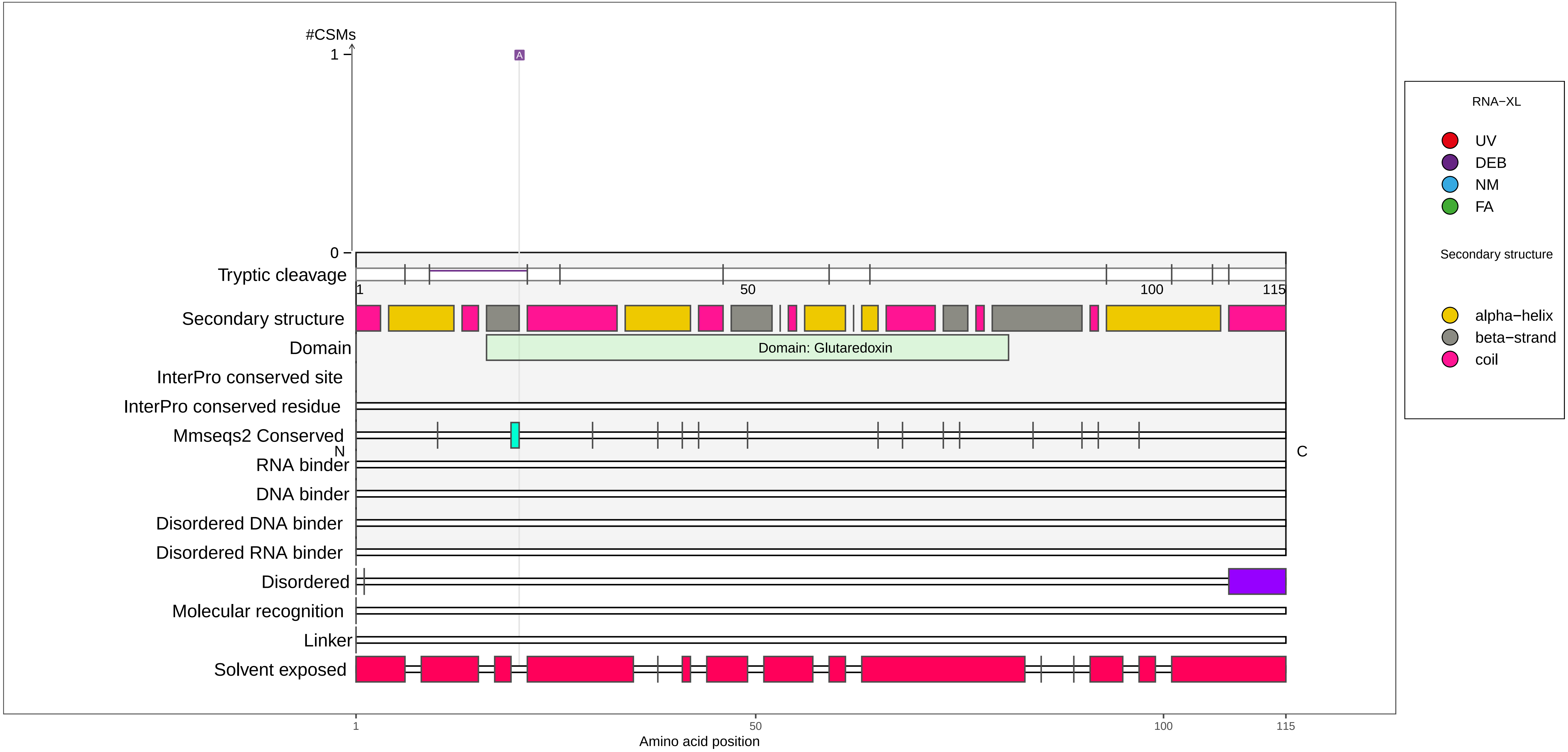

P0ACC3
ERPA_ECOLI Iron−sulfur cluster insertion protein ErpA

− Abundance:
tryptic [log10 Intensity]: 8.42 (Q 73)
PAXdb K12 strain [ppm]: 3.35 (Q 96)
PAXdb E.coli [ppm]: 2.79 (Q 91)

− RNA functions: not annotated

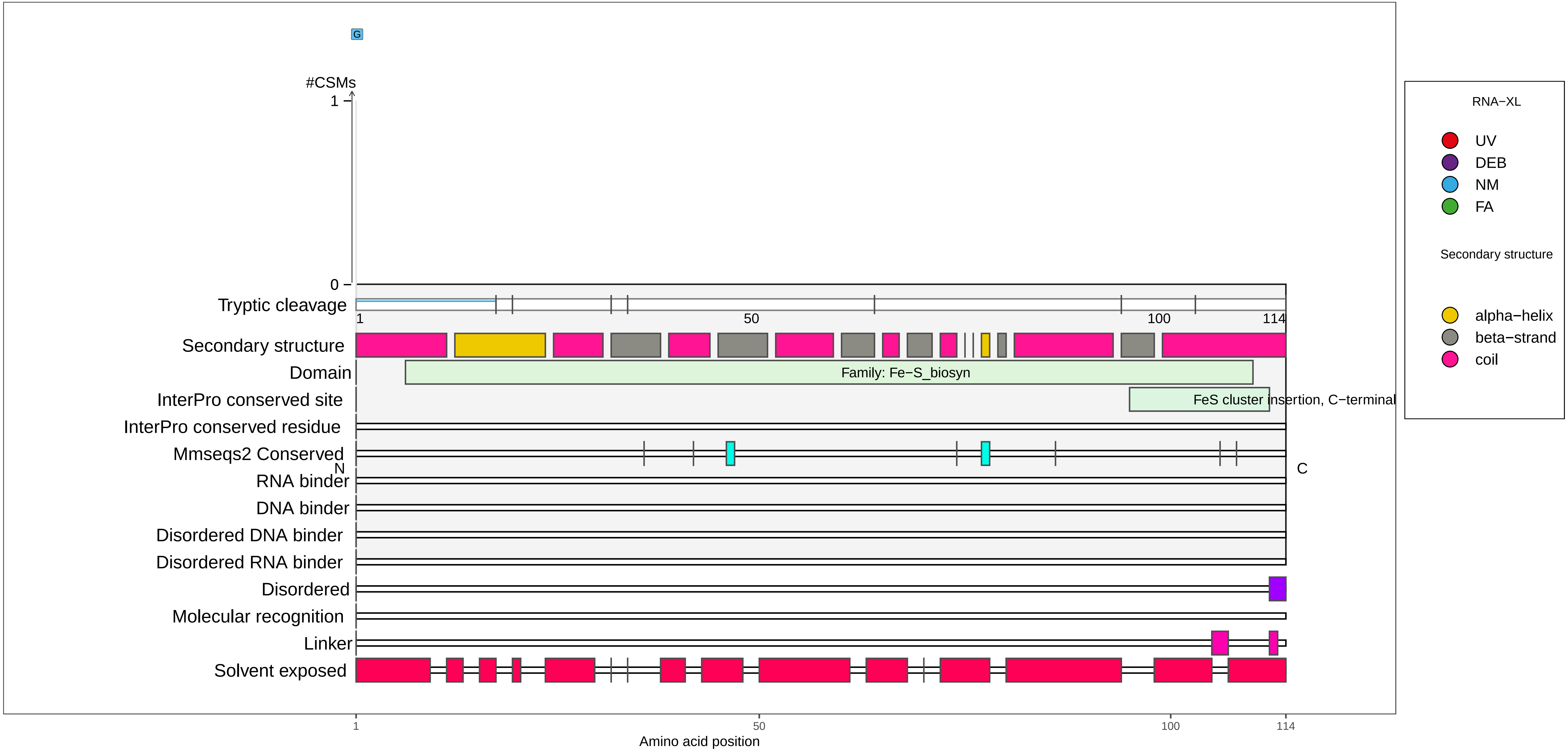

P0ACL2
EXUR_ECOLI Exu regulon transcriptional regulator

− Abundance:
tryptic [log10 Intensity]: 7.48 (Q 34)
PAXdb K12 strain [ppm]: 1.38 (Q 31)
PAXdb E.coli [ppm]: 1.43 (Q 58)

− RNA functions: not annotated

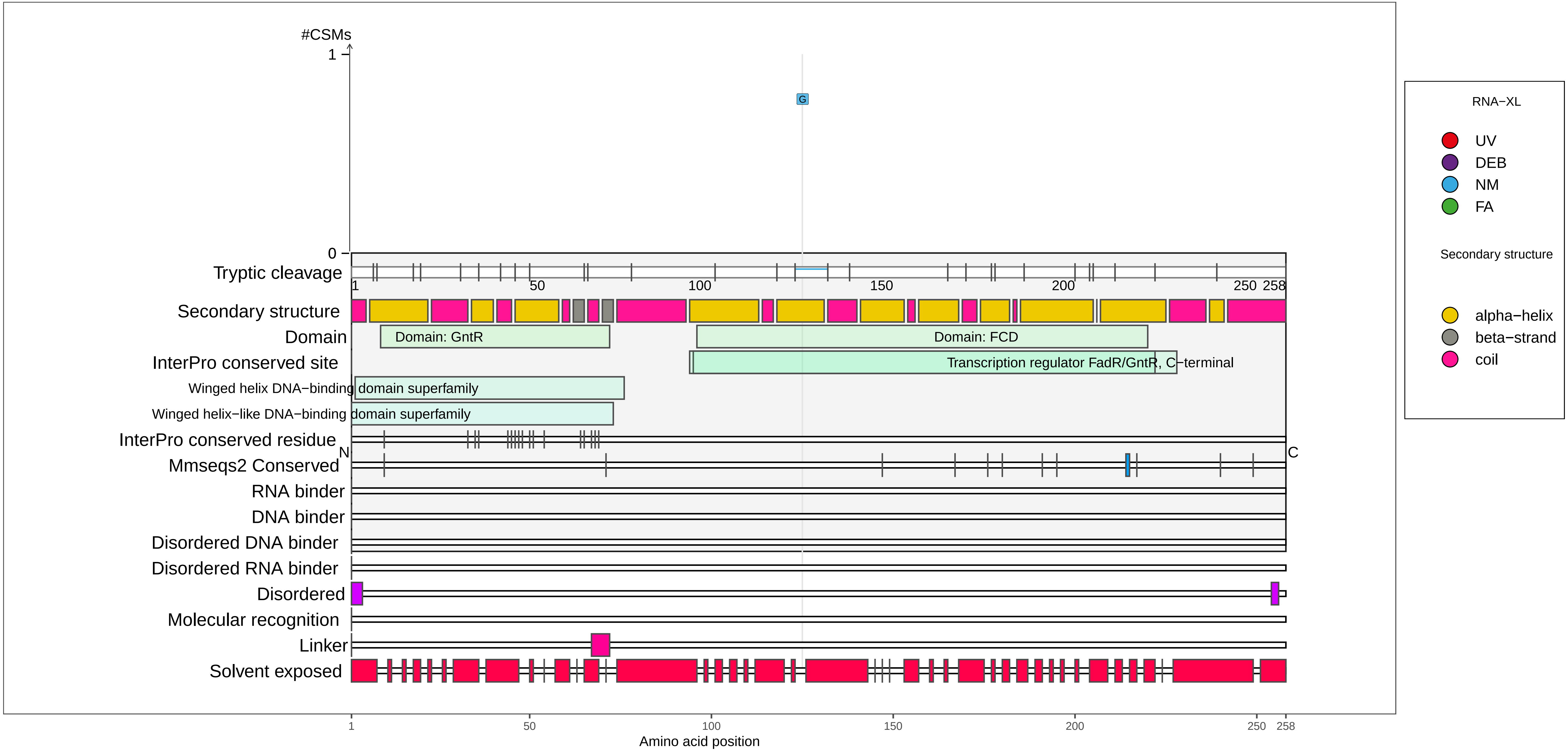

P0ACL9
PDHR_ECOLI Pyruvate dehydrogenase complex repressor

− Abundance:
tryptic [log10 Intensity]: NA (Q NA)
PAXdb K12 strain [ppm]: NA (Q NA)
PAXdb E.coli [ppm]: 1.19 (Q 52)

− RNA functions: not annotated

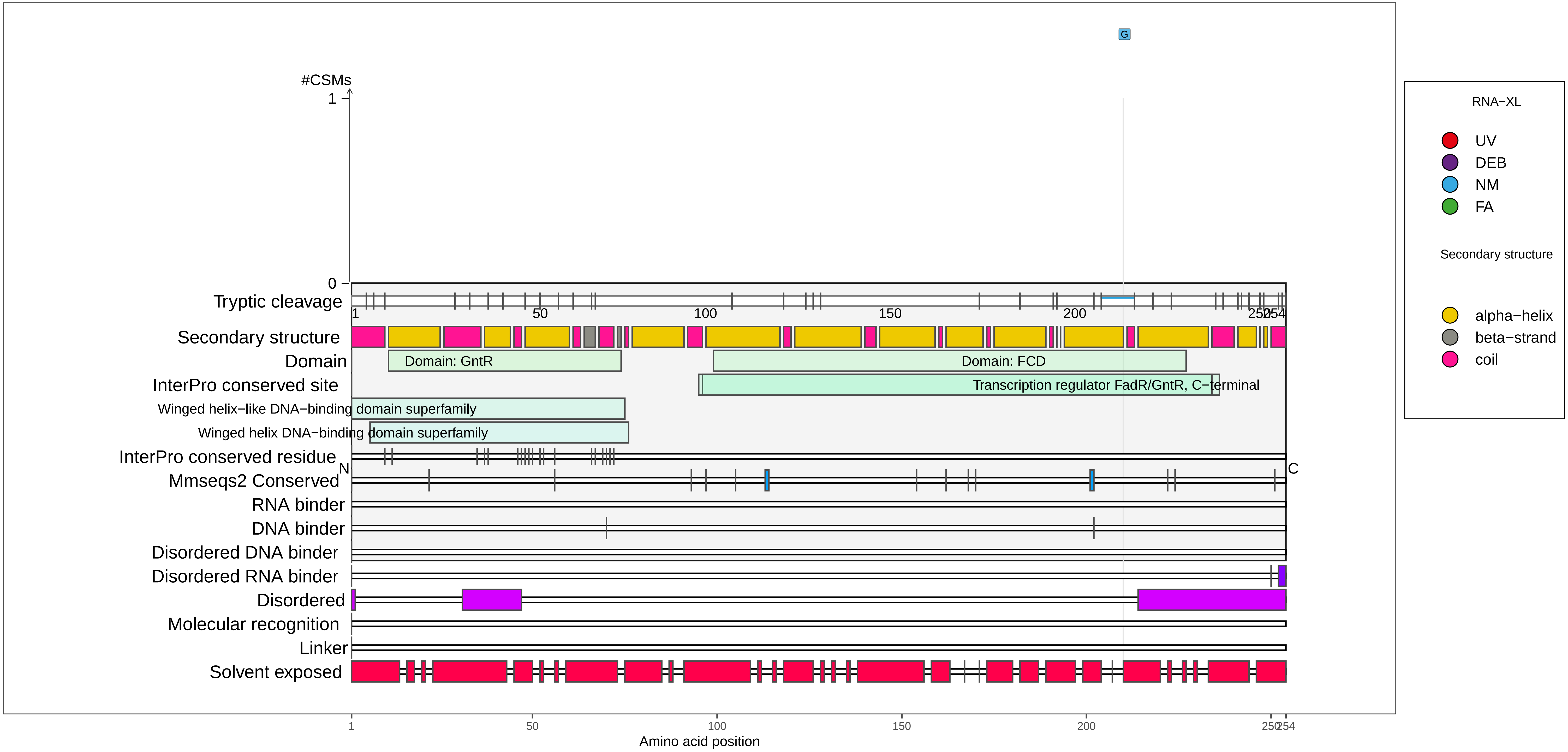

P0ACM9
YIHL_ECOLI Uncharacterized HTH−type transcriptional regulator YihL

− Abundance:
tryptic [log10 Intensity]: NA (Q NA)
PAXdb K12 strain [ppm]: 1.94 (Q 58)
PAXdb E.coli [ppm]: 0.17 (Q 28)

− RNA functions:
RNA biosynthetic process; RNA metabolic process

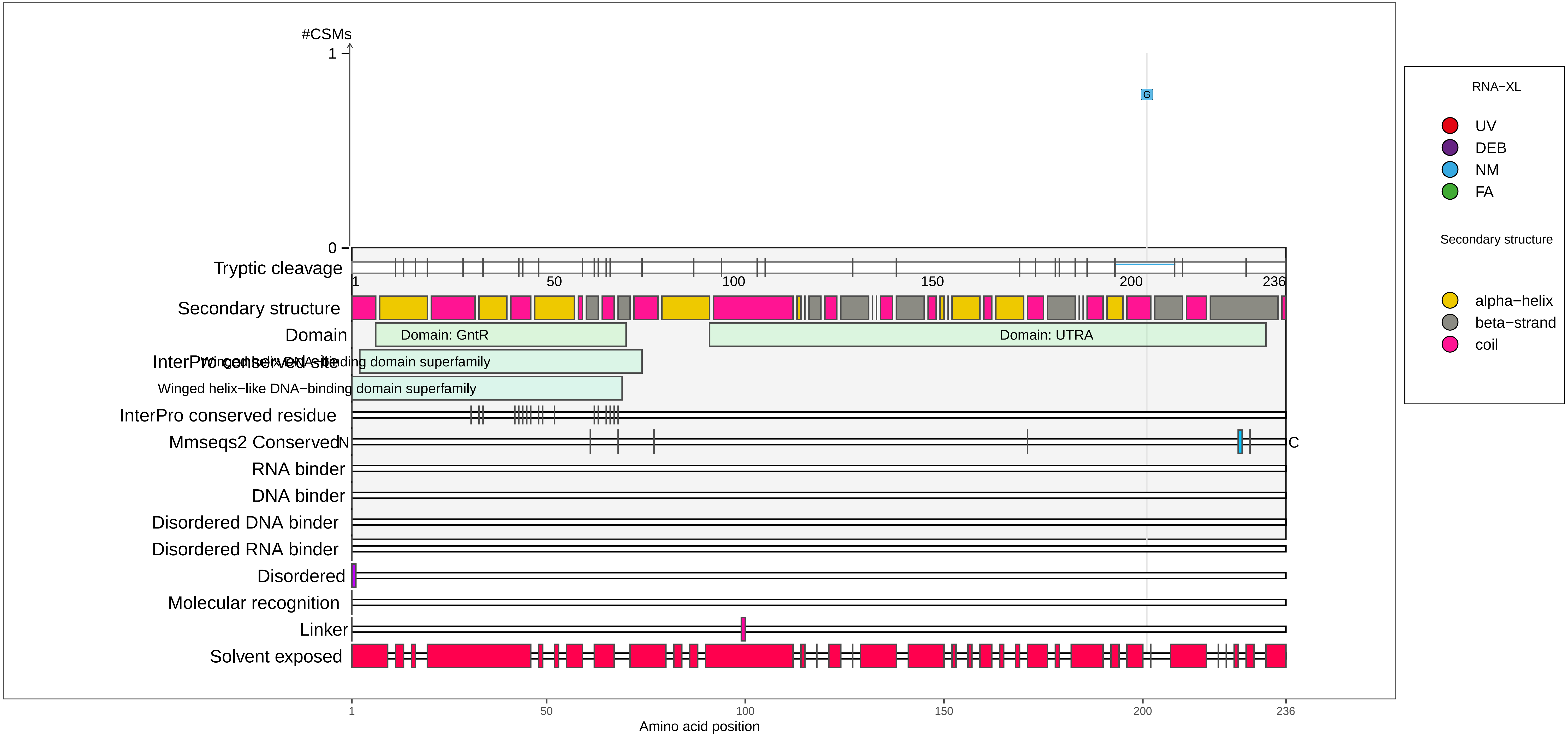

P0ACN4
ALLR_ECOLI HTH−type transcriptional repressor AllR

− Abundance:
tryptic [log10 Intensity]: 8.31 (Q 70)
PAXdb K12 strain [ppm]: 2.47 (Q 77)
PAXdb E.coli [ppm]: 1.78 (Q 66)

− RNA functions: not annotated

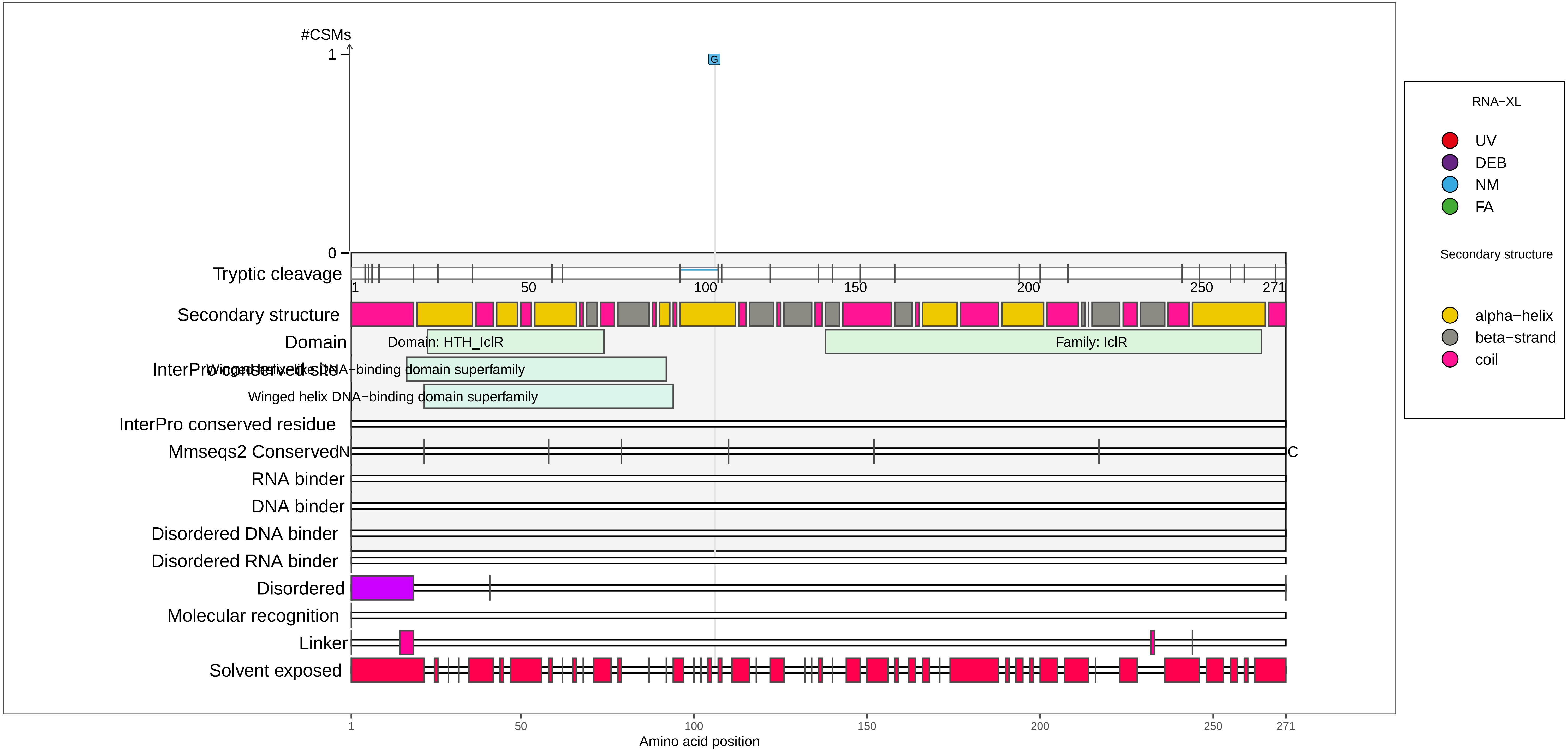

P0ACT2
ENVR_ECOLI Probable acrEF/envCD operon repressor

− Abundance:
tryptic [log10 Intensity]: NA (Q NA)
PAXdb K12 strain [ppm]: NA (Q NA)
PAXdb E.coli [ppm]: −1.8 (Q 2)

− RNA functions: not annotated

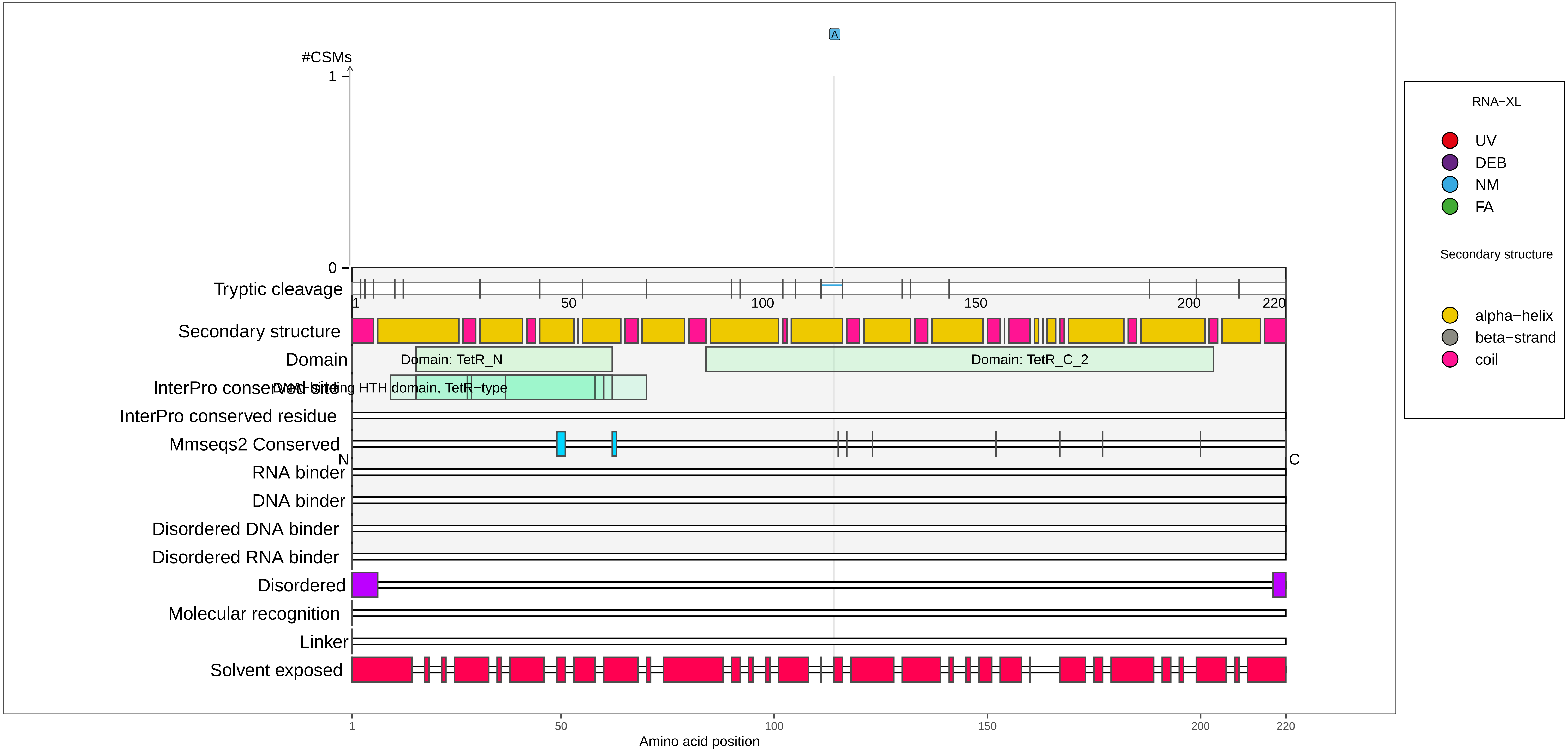

P0ACV0
LPXL_ECOLI Lipid A biosynthesis lauroyltransferase

− Abundance:
tryptic [log10 Intensity]: 7.69 (Q 44)
PAXdb K12 strain [ppm]: 0.99 (Q 10)
PAXdb E.coli [ppm]: 0.04 (Q 25)

− RNA functions: not annotated

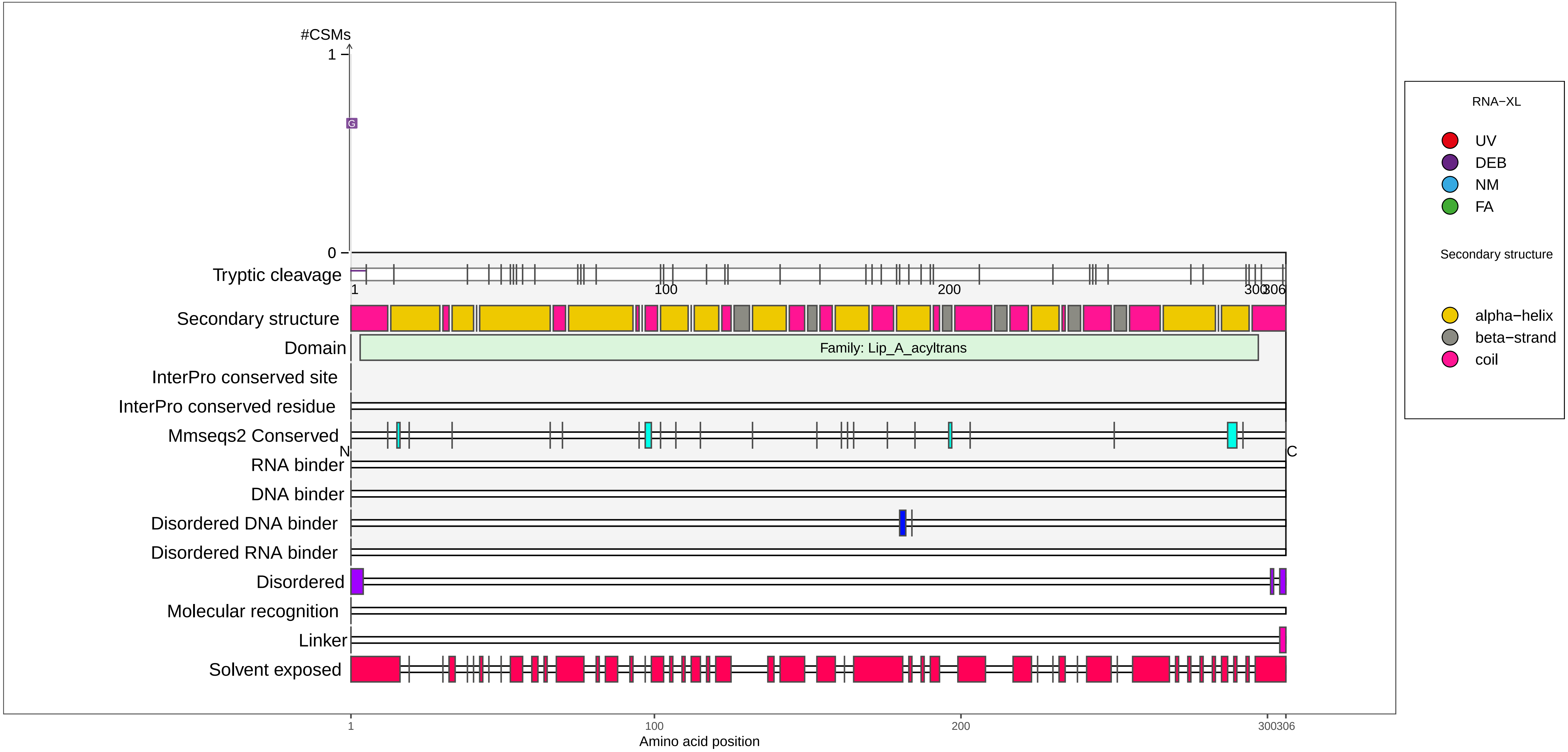

P0ACX3
YDHR_ECOLI Putative monooxygenase YdhR

− Abundance:
tryptic [log10 Intensity]: 8.45 (Q 73)
PAXdb K12 strain [ppm]: 2.44 (Q 76)
PAXdb E.coli [ppm]: 2.85 (Q 93)

− RNA functions: not annotated

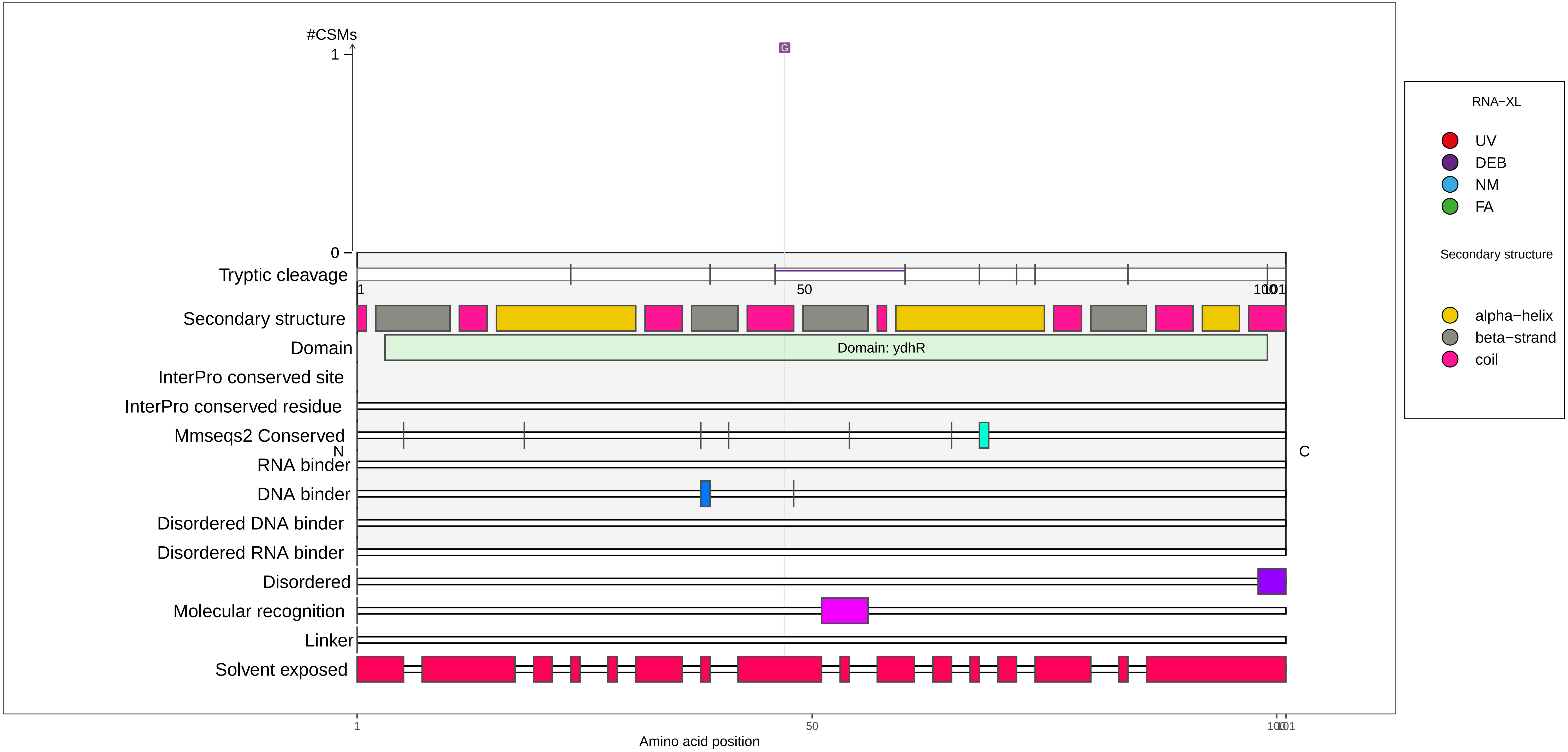

P0AD05
YECA_ECOLI Uncharacterized protein YecA

− Abundance:
tryptic [log10 Intensity]: 6.82 (Q 8)
PAXdb K12 strain [ppm]: 1.8 (Q 53)
PAXdb E.coli [ppm]: 2.45 (Q 84)

− RNA functions: not annotated

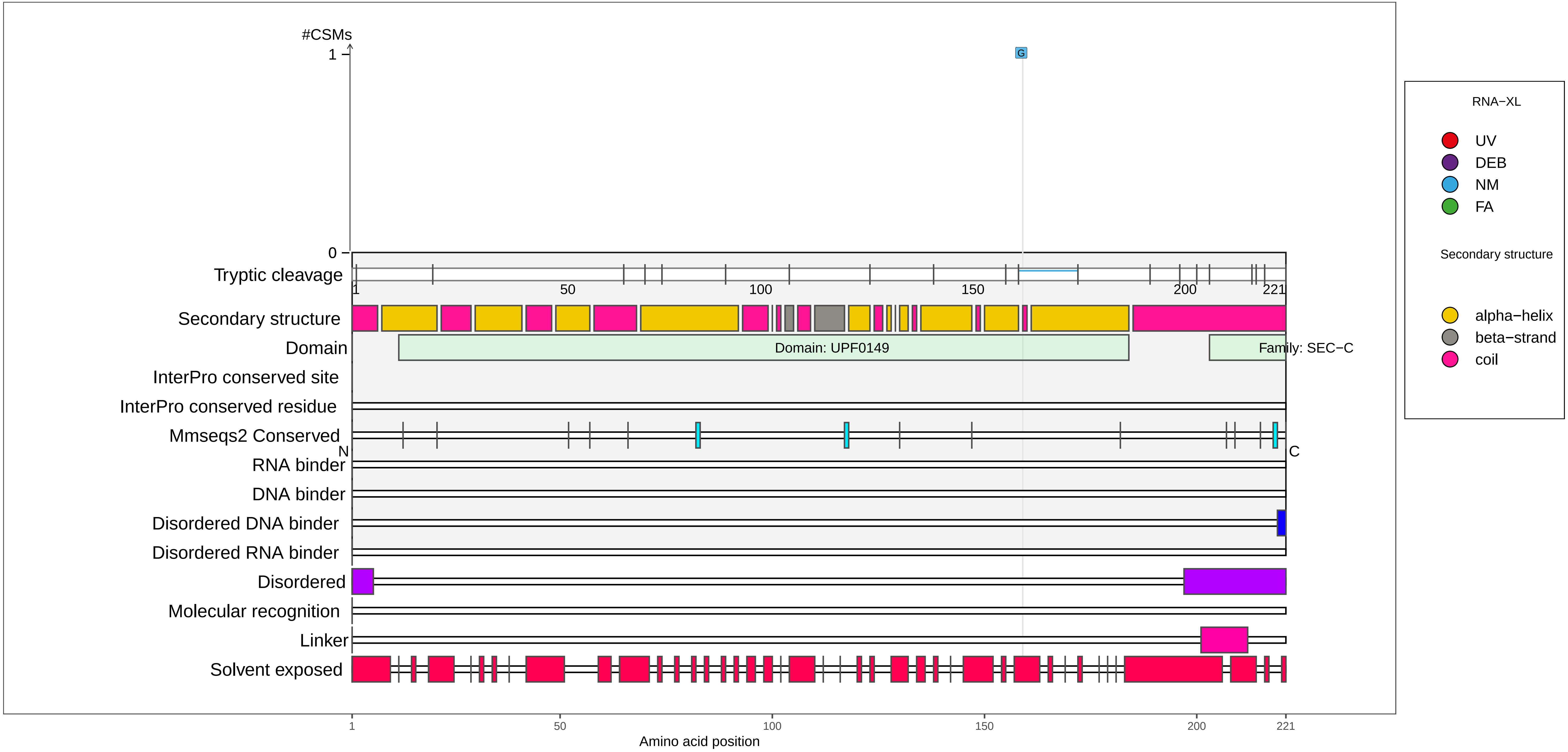

P0AD59
IVY_ECOLI Inhibitor of vertebrate lysozyme

− Abundance:
tryptic [log10 Intensity]: 7.75 (Q 47)
PAXdb K12 strain [ppm]: 2.44 (Q 76)
PAXdb E.coli [ppm]: 2.81 (Q 92)

− RNA functions: not annotated

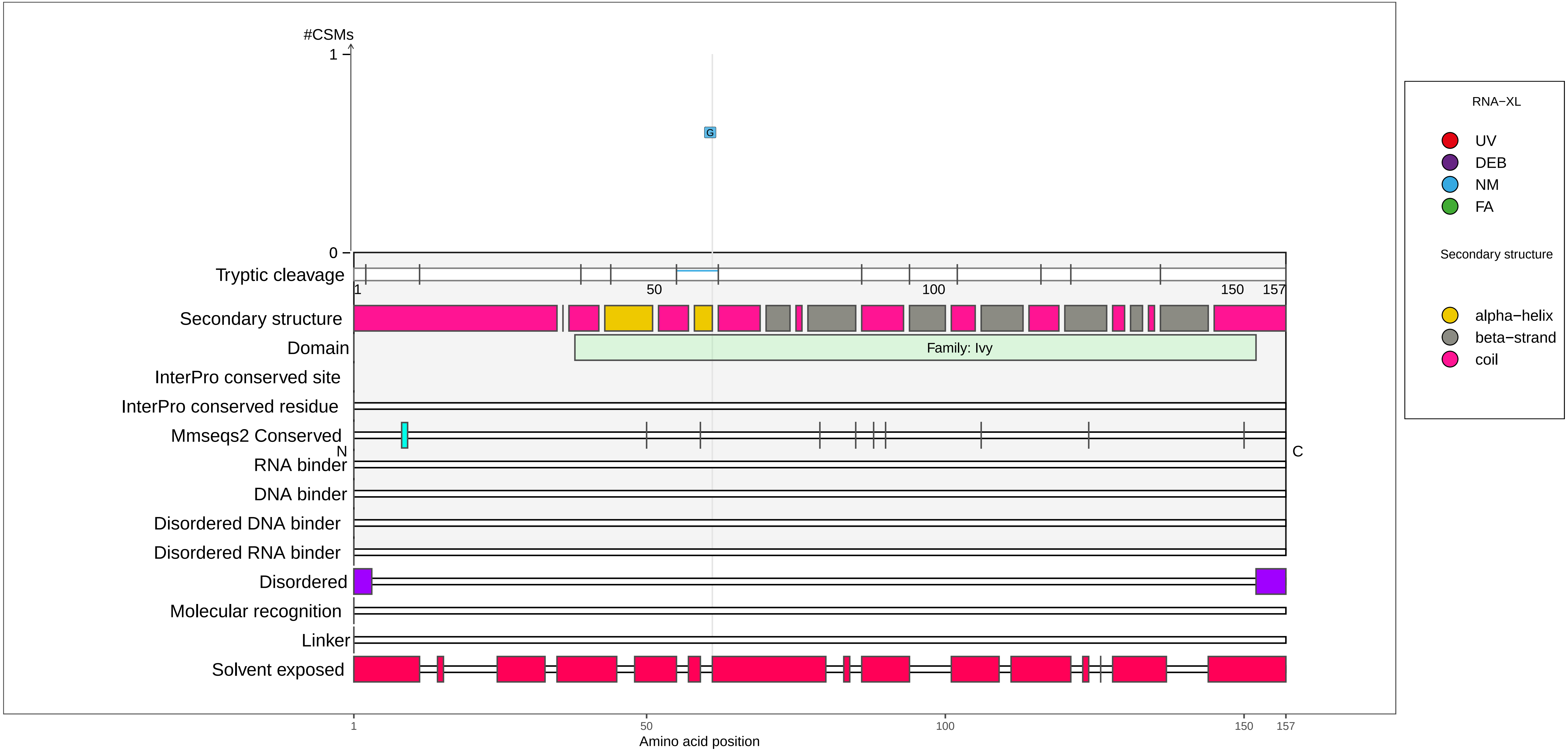

P0ADK0
YIAF_ECOLI Uncharacterized protein YiaF

− Abundance:
tryptic [log10 Intensity]: 8.33 (Q 70)
PAXdb K12 strain [ppm]: 1.37 (Q 31)
PAXdb E.coli [ppm]: 2.57 (Q 87)

− RNA functions: not annotated

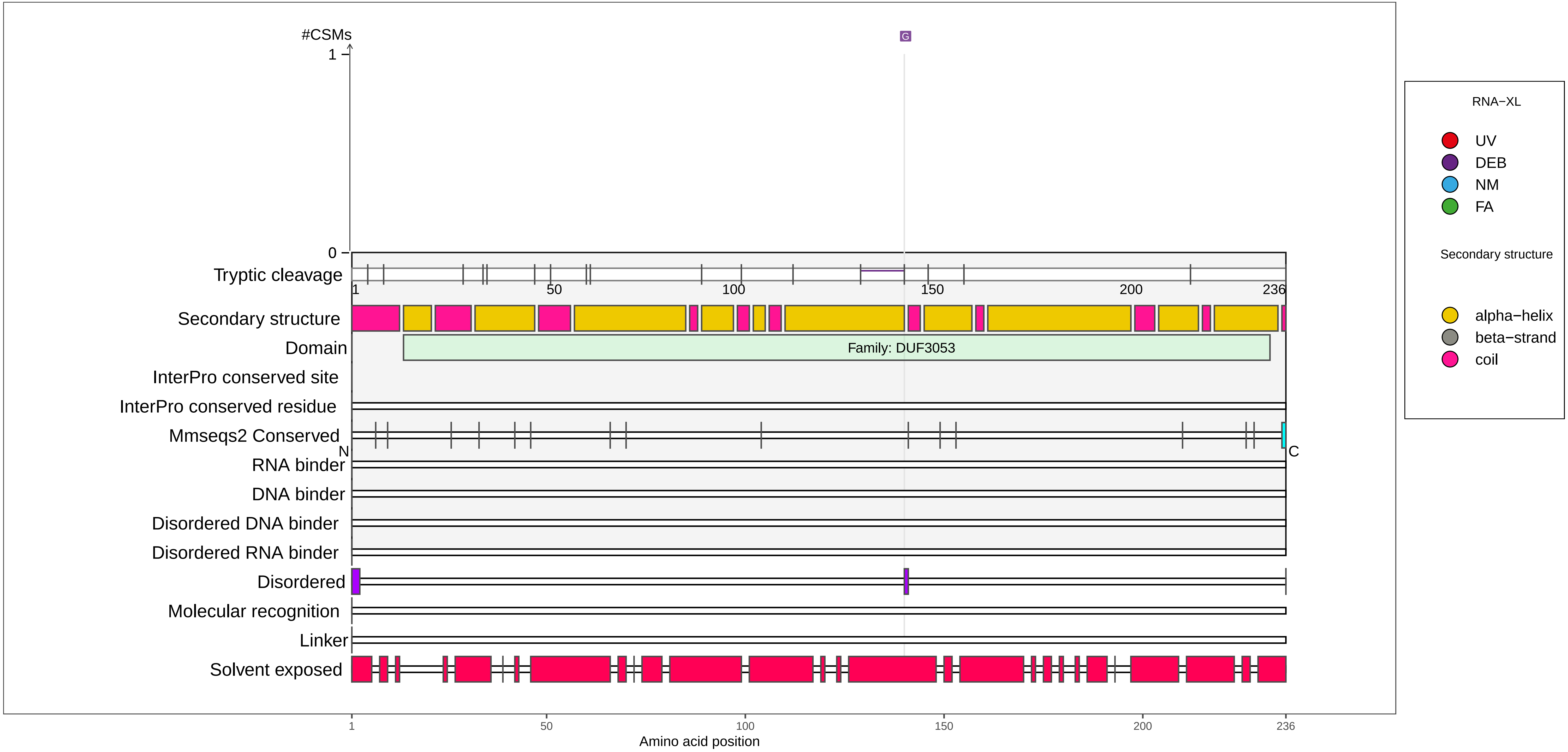

P0ADQ7
YGAM_ECOLI Uncharacterized protein YgaM

− Abundance:
tryptic [log10 Intensity]: NA (Q NA)
PAXdb K12 strain [ppm]: 2.52 (Q 78)
PAXdb E.coli [ppm]: 2.41 (Q 83)

− RNA functions: not annotated

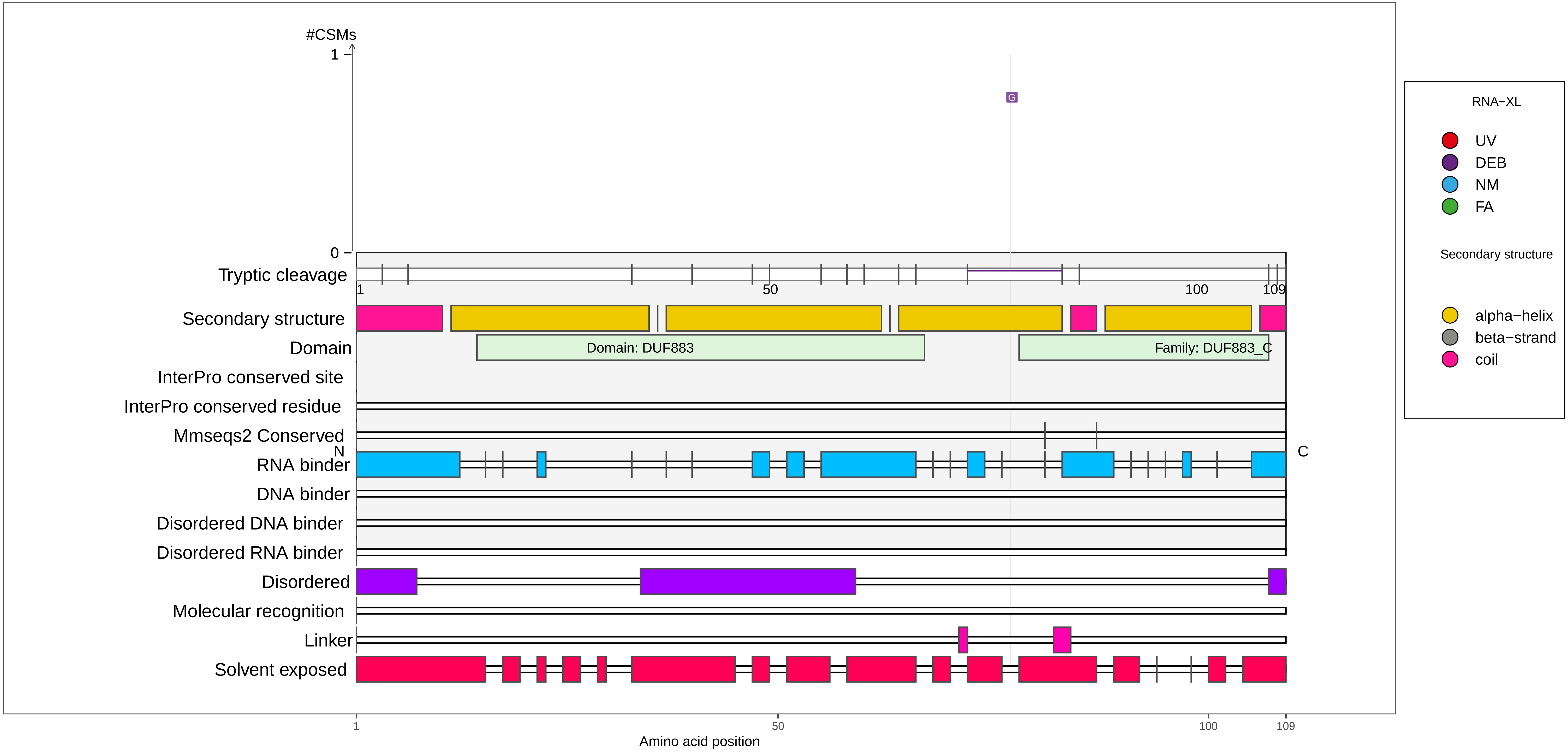

P0ADR6
RLMM_ECOLI Ribosomal RNA large subunit methyltransferase M

− Abundance:
tryptic [log10 Intensity]: 7.85 (Q 52)
PAXdb K12 strain [ppm]: 1.29 (Q 26)
PAXdb E.coli [ppm]: 1.18 (Q 52)

− RNA functions:
ncRNA metabolic process; ncRNA processing; RNA 2−O−methyltransferase activity
RNA metabolic process; RNA methylation; RNA methyltransferase activity; RNA modification
RNA processing; rRNA (cytosine−2−O−)−methyltransferase activity
rRNA (cytosine) methyltransferase activity; rRNA 2−O−methylation; rRNA metabolic process; rRNA methylation
rRNA methyltransferase activity; rRNA modification; rRNA processing

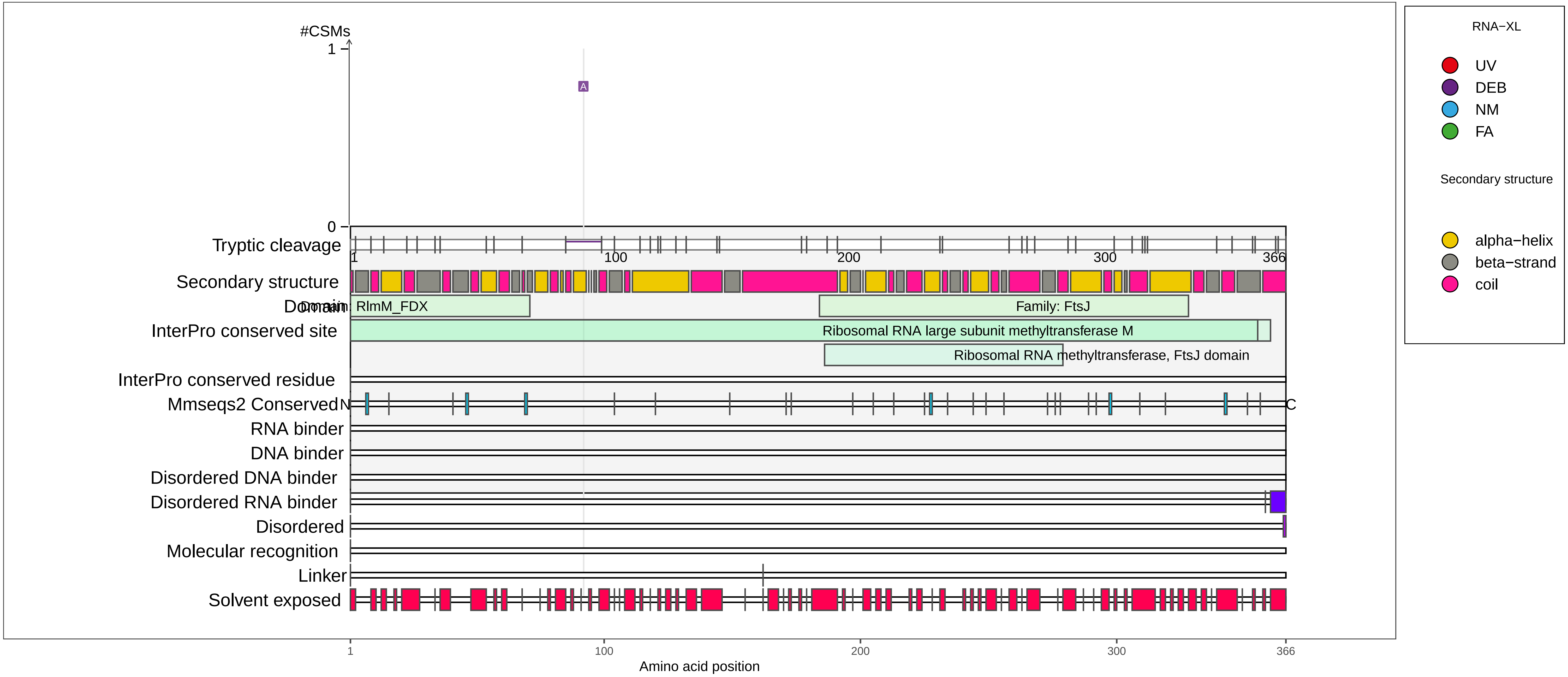

P0ADR8
PPNN_ECOLI Pyrimidine/purine nucleotide 5'−monophosphate nucleosidase

− Abundance:
tryptic [log10 Intensity]: 8.63 (Q 79)
PAXdb K12 strain [ppm]: 2.42 (Q 75)
PAXdb E.coli [ppm]: 1.7 (Q 64)

− RNA functions: not annotated

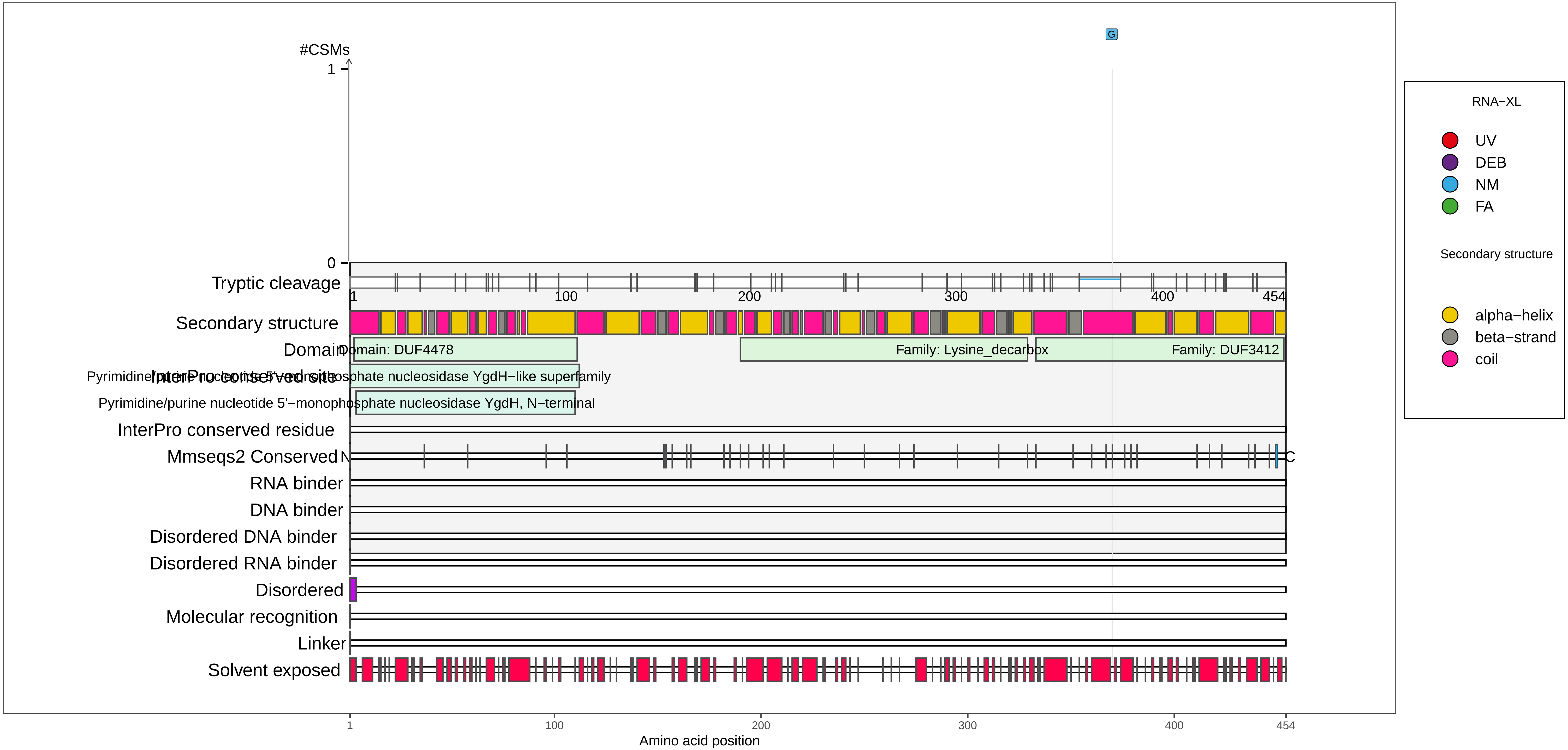

P0ADV1
LPTA_ECOLI Lipopolysaccharide export system protein LptA

− Abundance:
tryptic [log10 Intensity]: 8.17 (Q 64)
PAXdb K12 strain [ppm]: 2.58 (Q 80)
PAXdb E.coli [ppm]: 2.04 (Q 73)

− RNA functions: not annotated

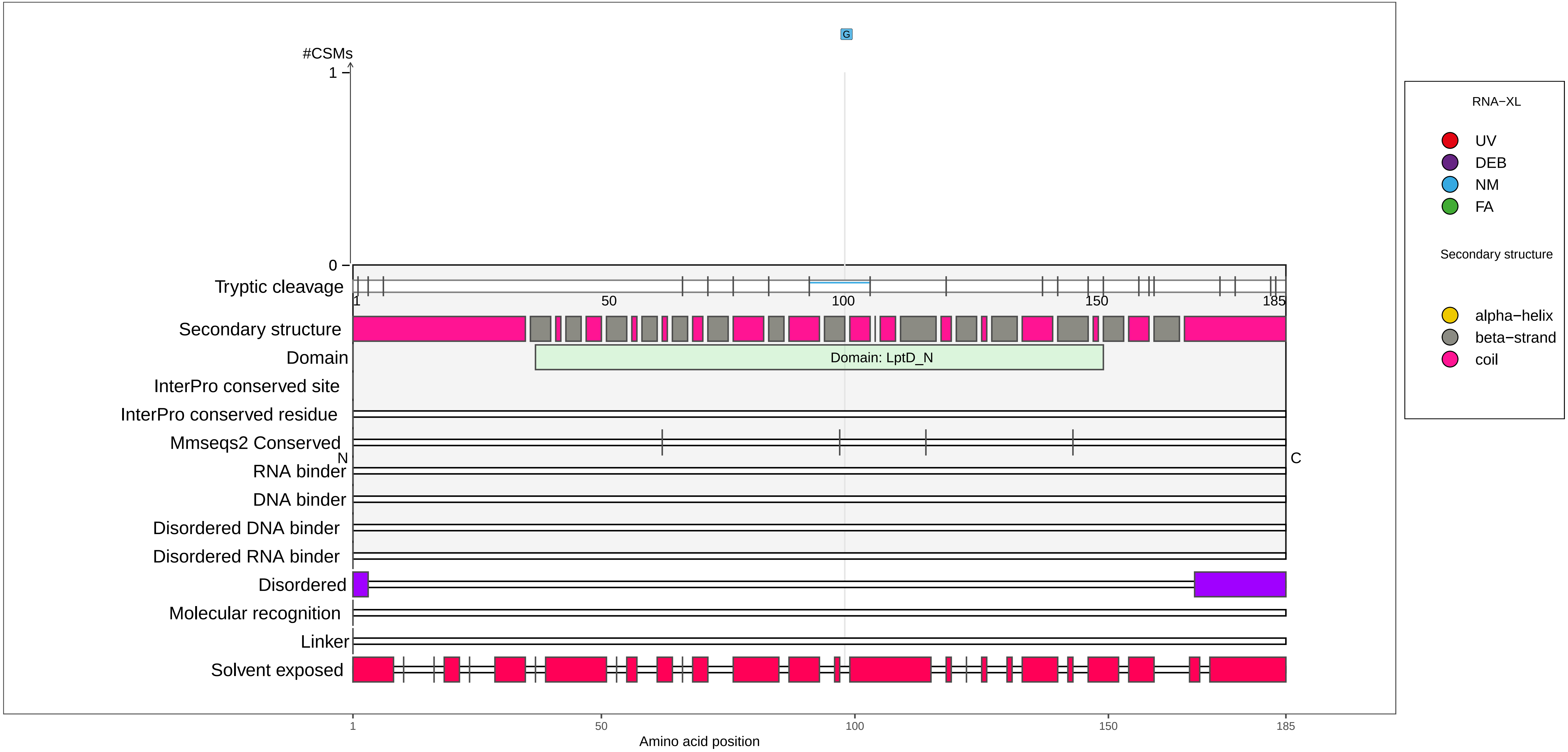

P0AE60
CEDA_ECOLI Cell division activator CedA

− Abundance:
tryptic [log10 Intensity]: 7.09 (Q 16)
PAXdb K12 strain [ppm]: 1.84 (Q 55)
PAXdb E.coli [ppm]: 1.4 (Q 57)

− RNA functions: not annotated

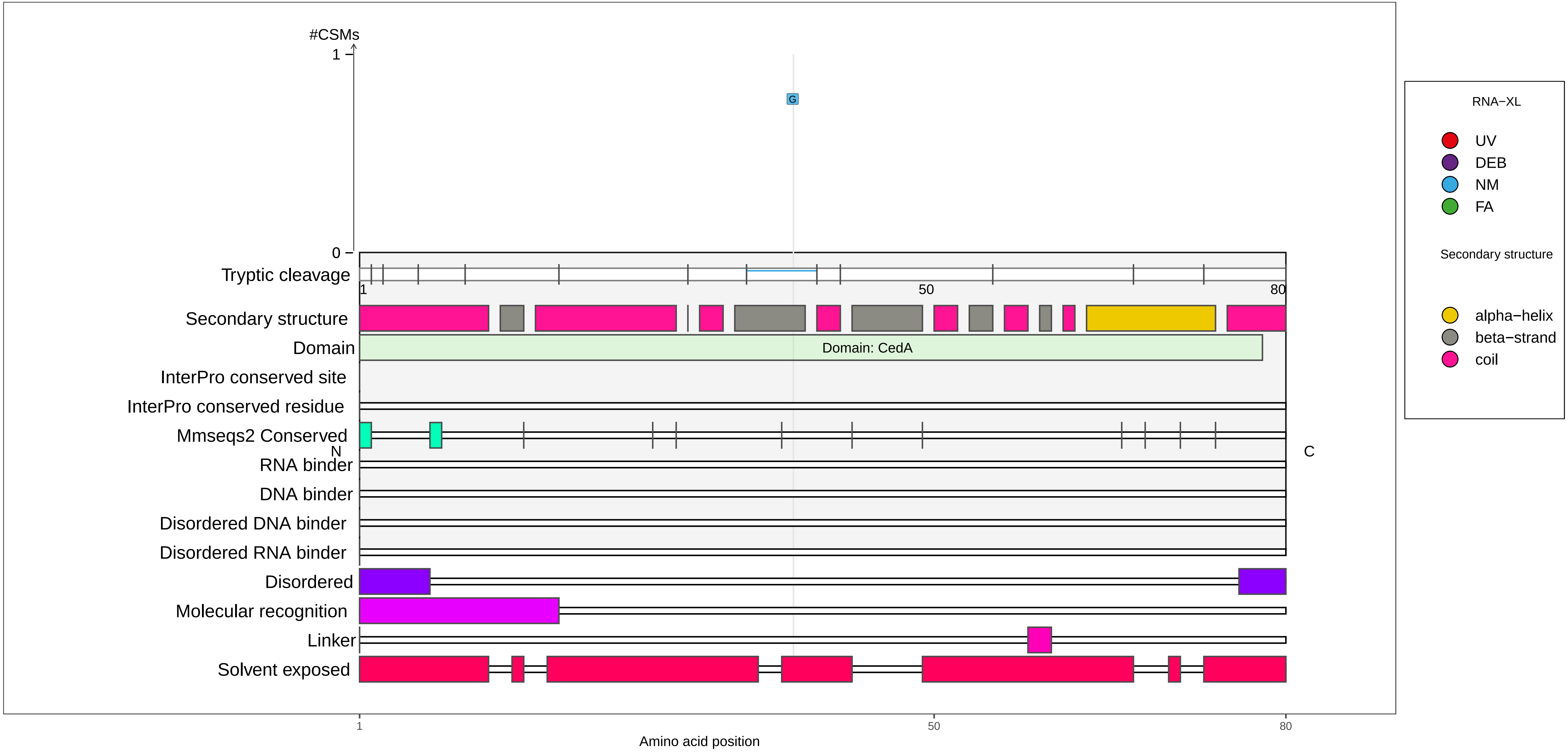

P0AEB7
YOAB_ECOLI RutC family protein YoaB

− Abundance:
tryptic [log10 Intensity]: 8.98 (Q 88)
PAXdb K12 strain [ppm]: NA (Q NA)
PAXdb E.coli [ppm]: 2.71 (Q 90)

− RNA functions: not annotated

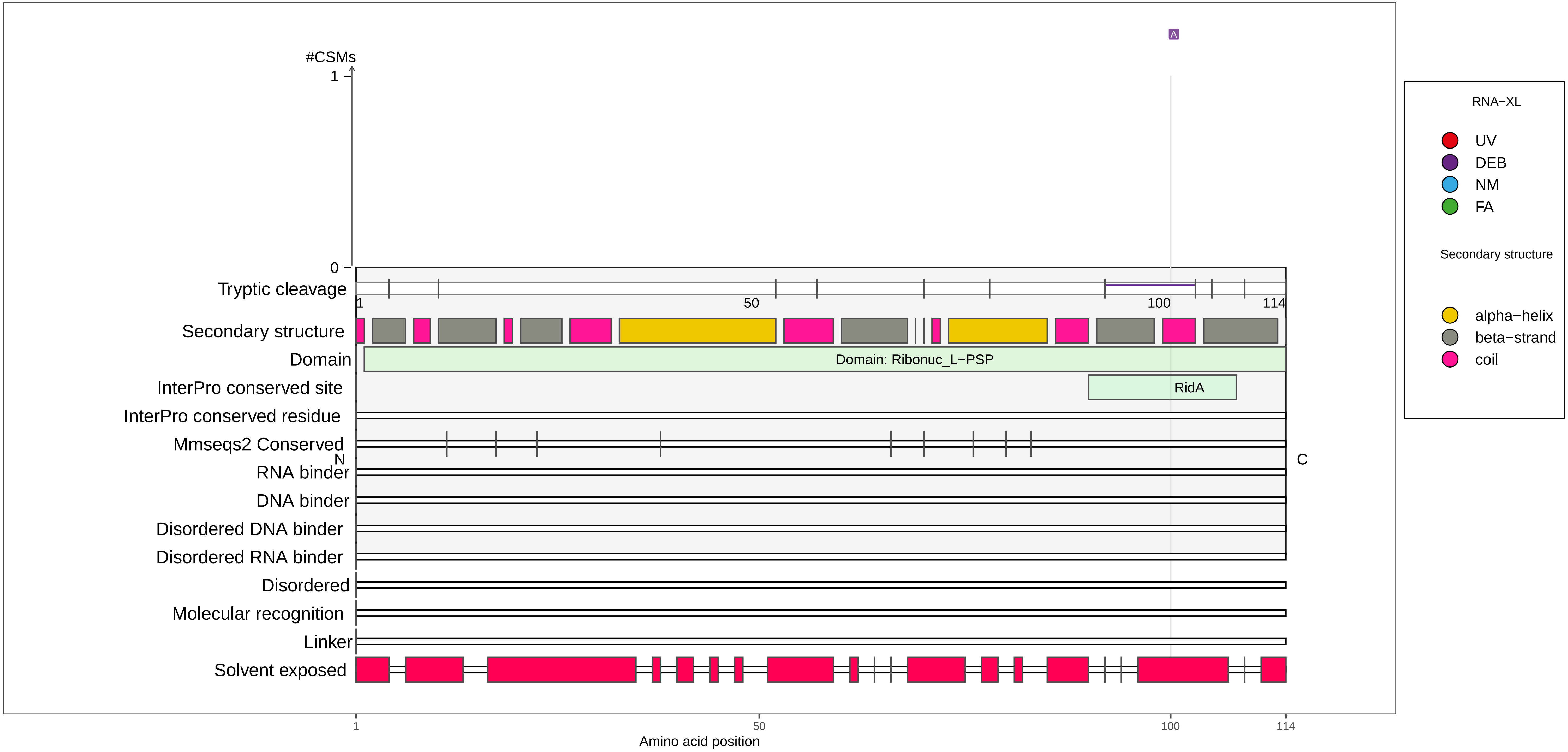

P0AEC3
ARCB_ECOLI Aerobic respiration control sensor protein ArcB

− Abundance:
tryptic [log10 Intensity]: 7.09 (Q 16)
PAXdb K12 strain [ppm]: 2.03 (Q 61)
PAXdb E.coli [ppm]: 2.18 (Q 77)

− RNA functions: not annotated

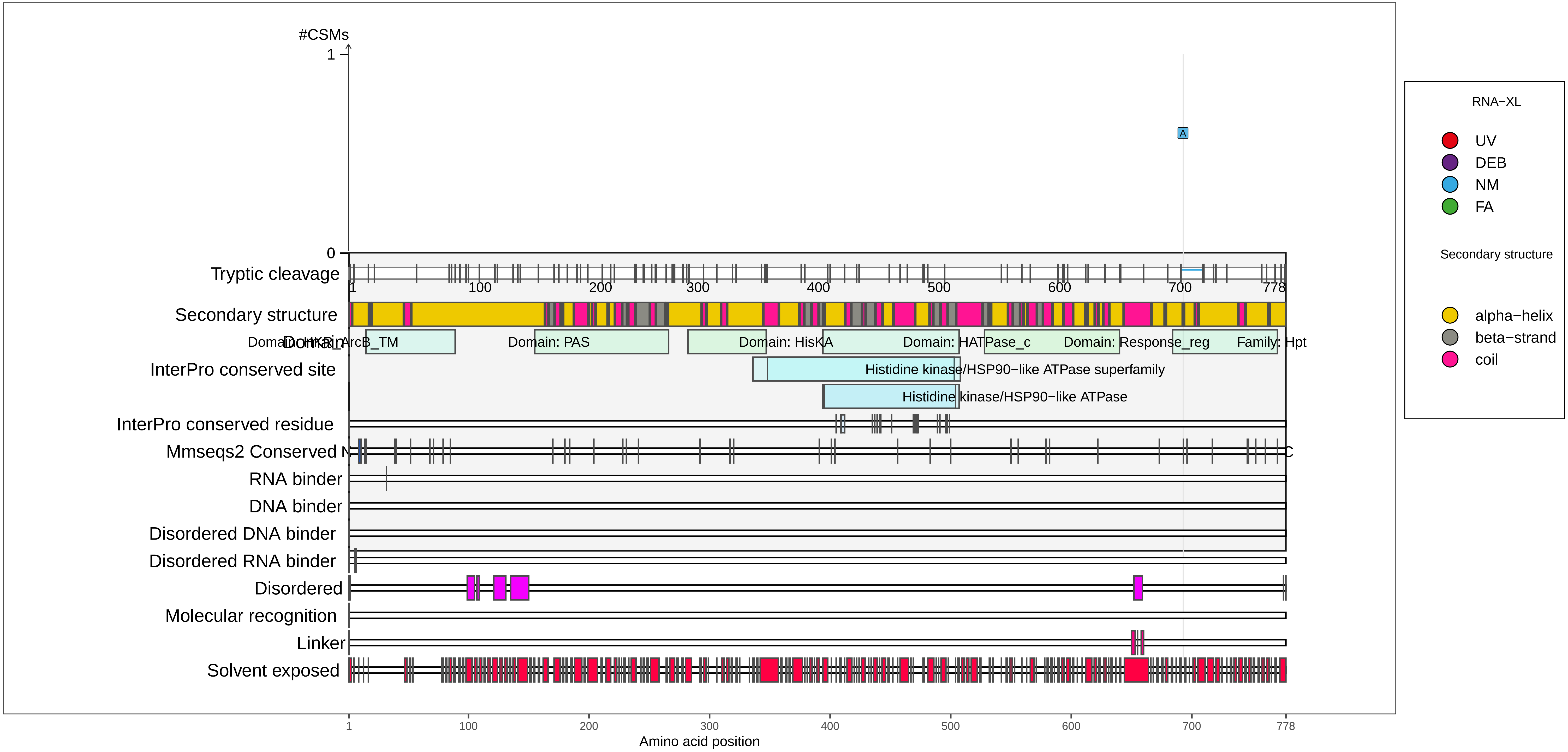

P0AEE5
DGAL_ECOLI D−galactose−binding periplasmic protein

− Abundance:
tryptic [log10 Intensity]: 7.18 (Q 20)
PAXdb K12 strain [ppm]: 3.02 (Q 91)
PAXdb E.coli [ppm]: 3.5 (Q 99)

− RNA functions: not annotated

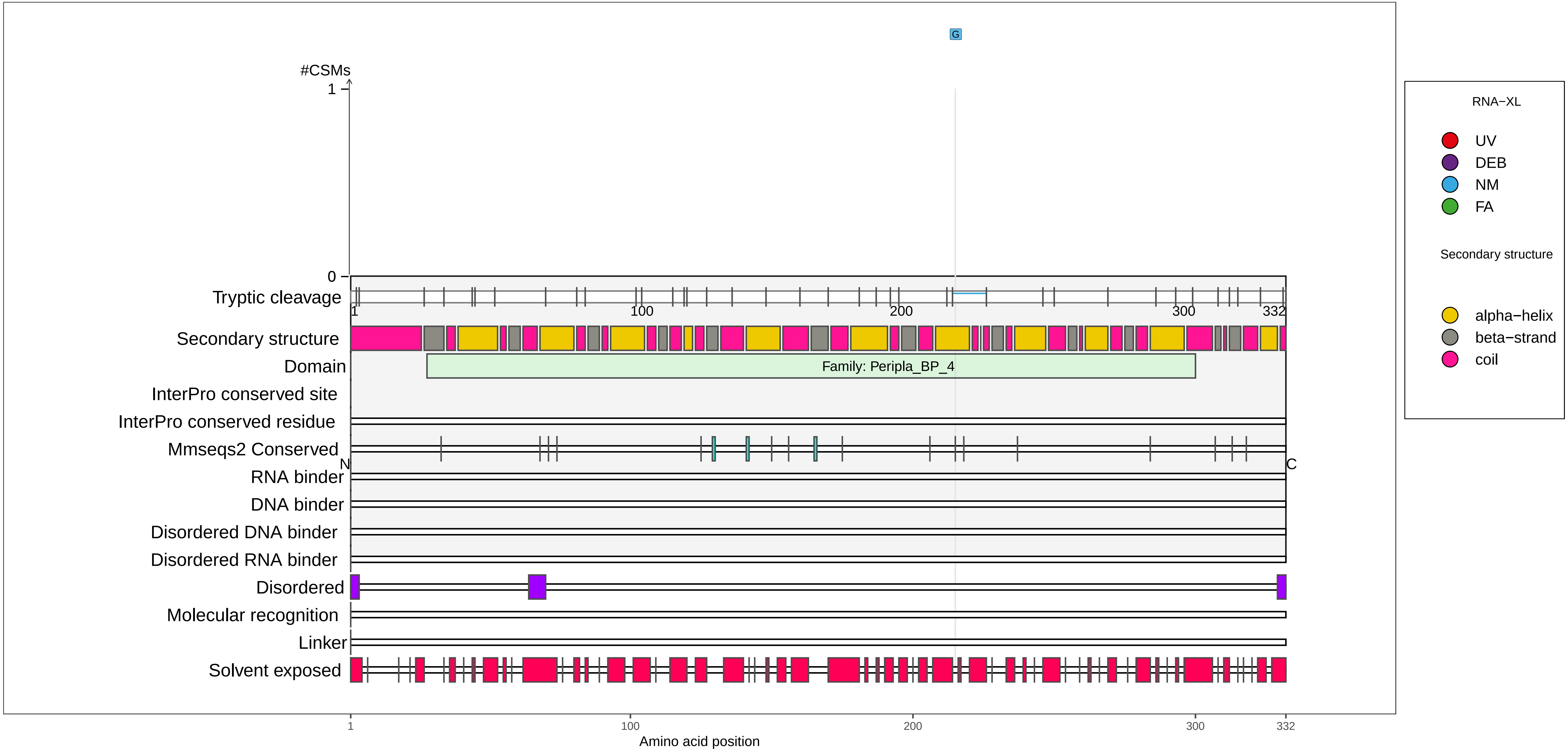

P0AEH1
RSEP_ECOLI Regulator of sigma−E protease RseP
− Abundance:
tryptic [log10 Intensity]: 7.52 (Q 36)
PAXdb K12 strain [ppm]: NA (Q NA)
PAXdb E.coli [ppm]: 0.73 (Q 41)

− RNA functions: not annotated

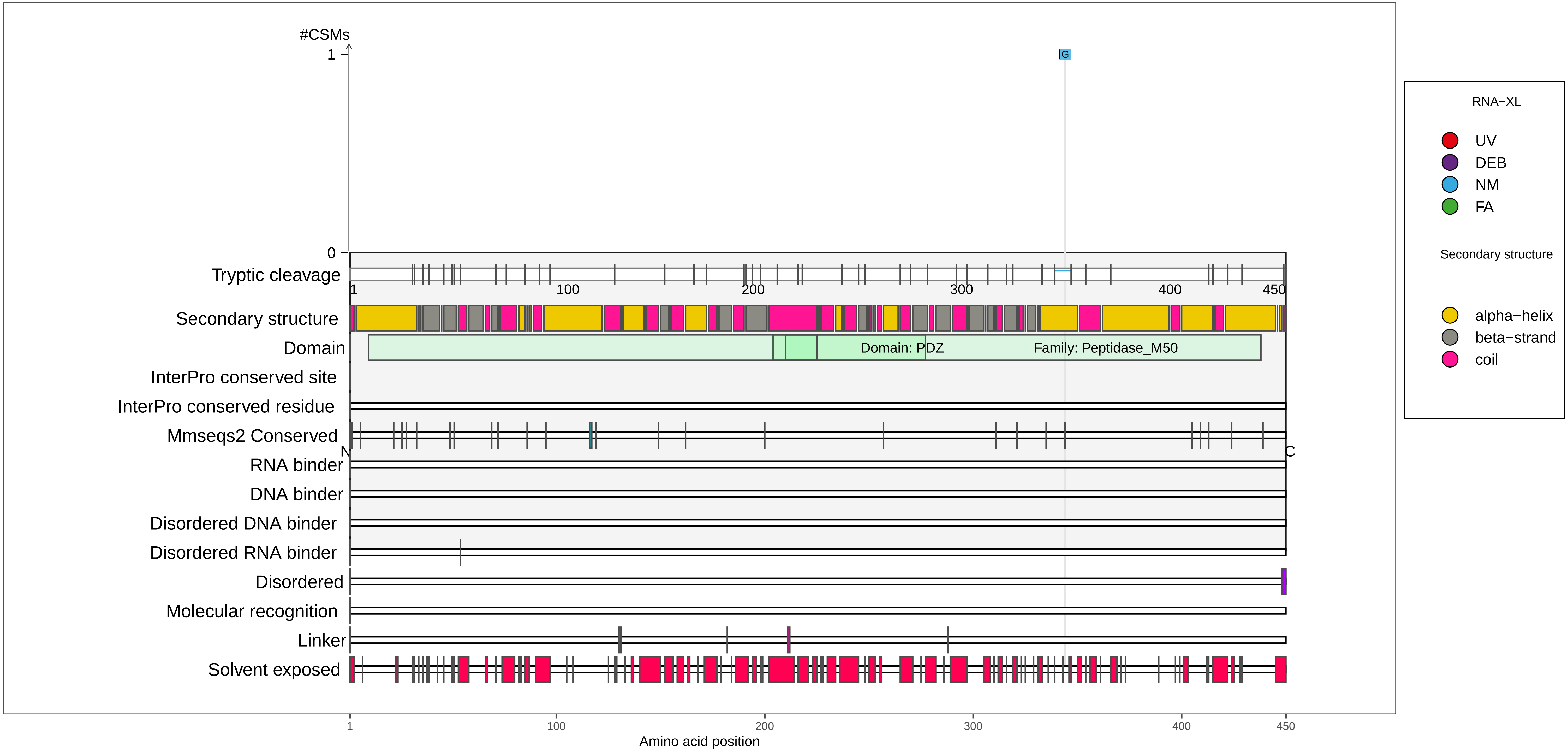

P0AEL3
FEOA_ECOLI Fe(2+) transport protein A

− Abundance:
tryptic [log10 Intensity]: NA (Q NA)
PAXdb K12 strain [ppm]: NA (Q NA)
PAXdb E.coli [ppm]: 2.56 (Q 87)

− RNA functions: not annotated

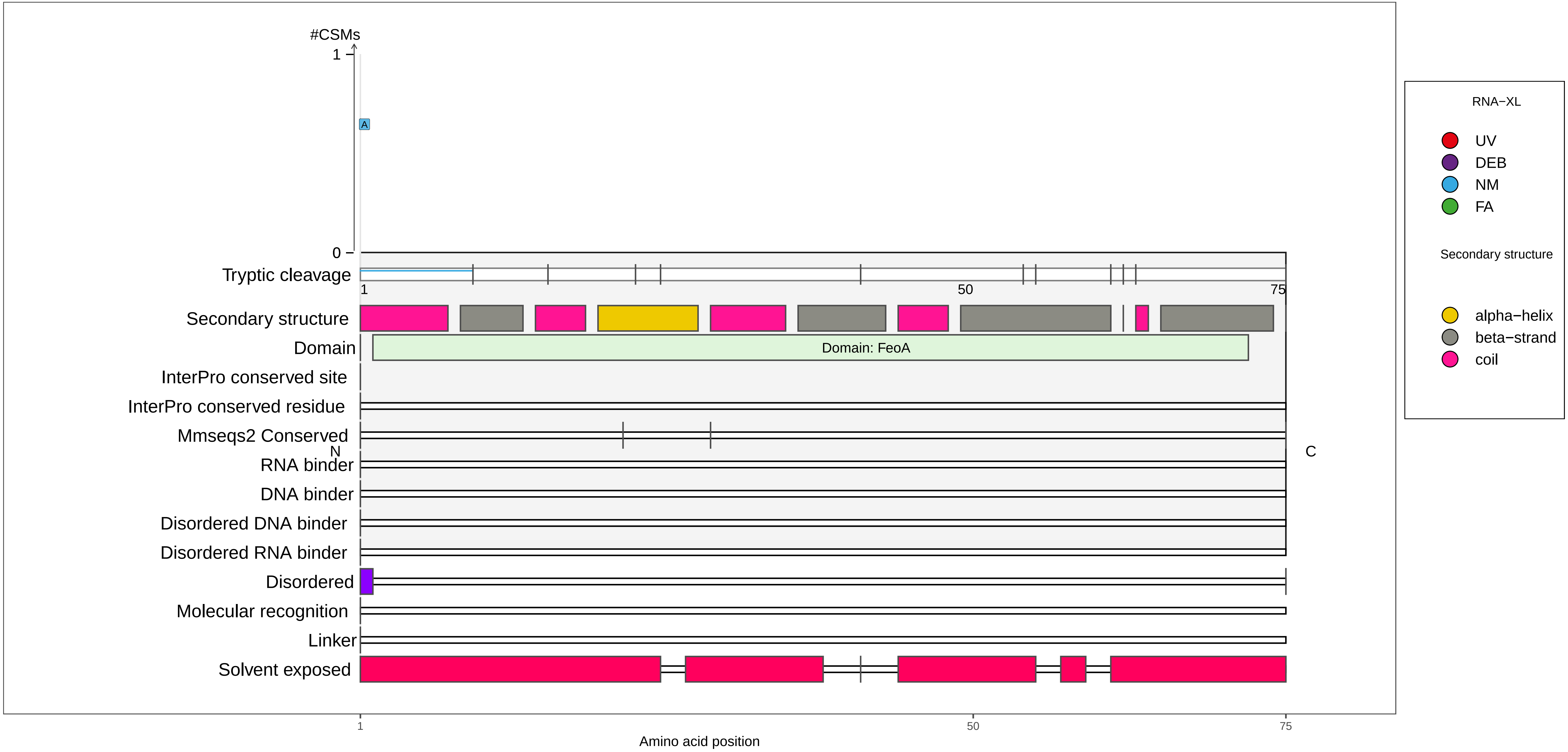

P0AEQ3
GLNH_ECOLI Glutamine−binding periplasmic protein

− Abundance:
tryptic [log10 Intensity]: 7.92 (Q 55)
PAXdb K12 strain [ppm]: 2.88 (Q 88)
PAXdb E.coli [ppm]: 3.47 (Q 99)

− RNA functions: not annotated

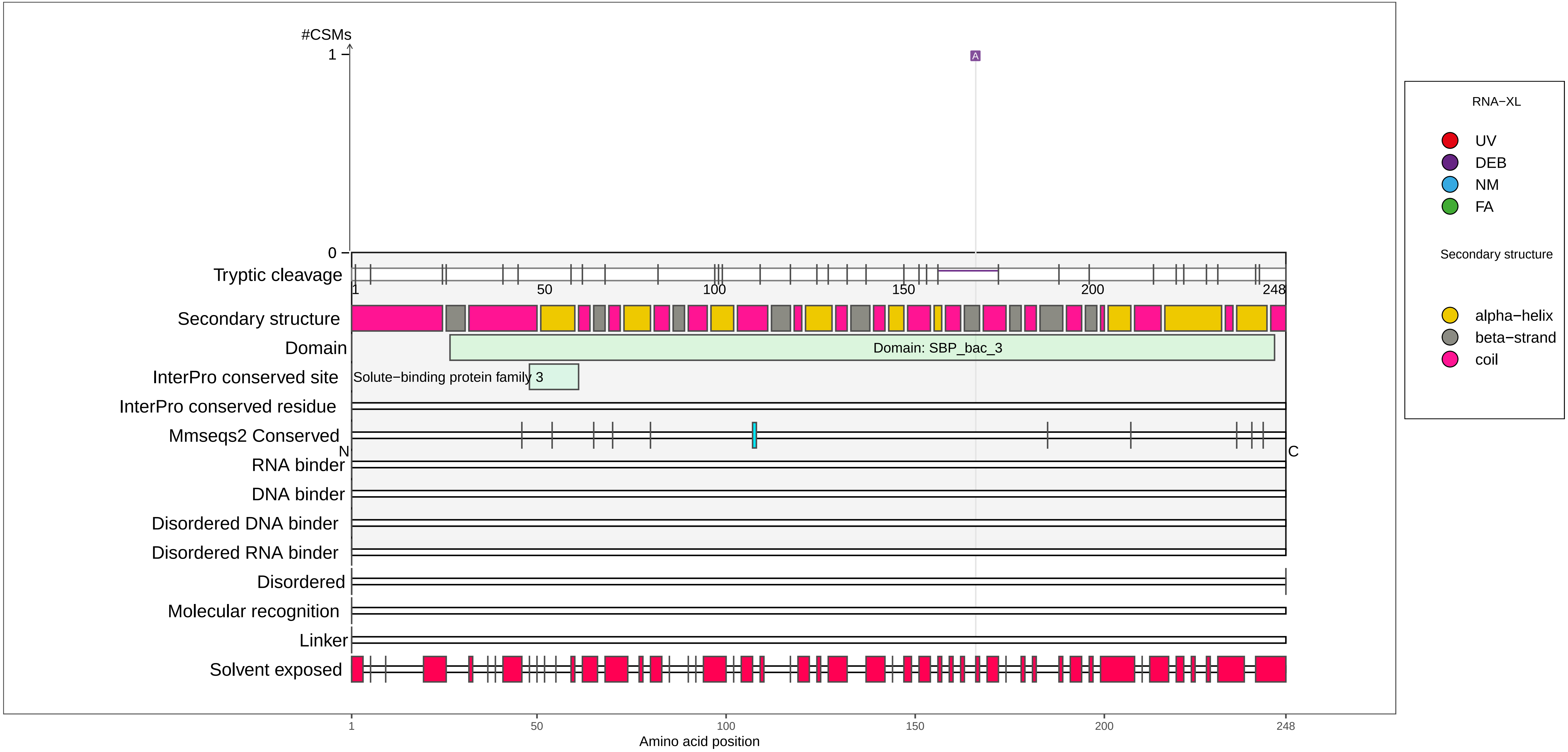

P0AEU0
HISJ_ECOLI Histidine−binding periplasmic protein

− Abundance:
tryptic [log10 Intensity]: 9.68 (Q 98)
PAXdb K12 strain [ppm]: 2.97 (Q 90)
PAXdb E.coli [ppm]: 2.97 (Q 94)

− RNA functions: not annotated

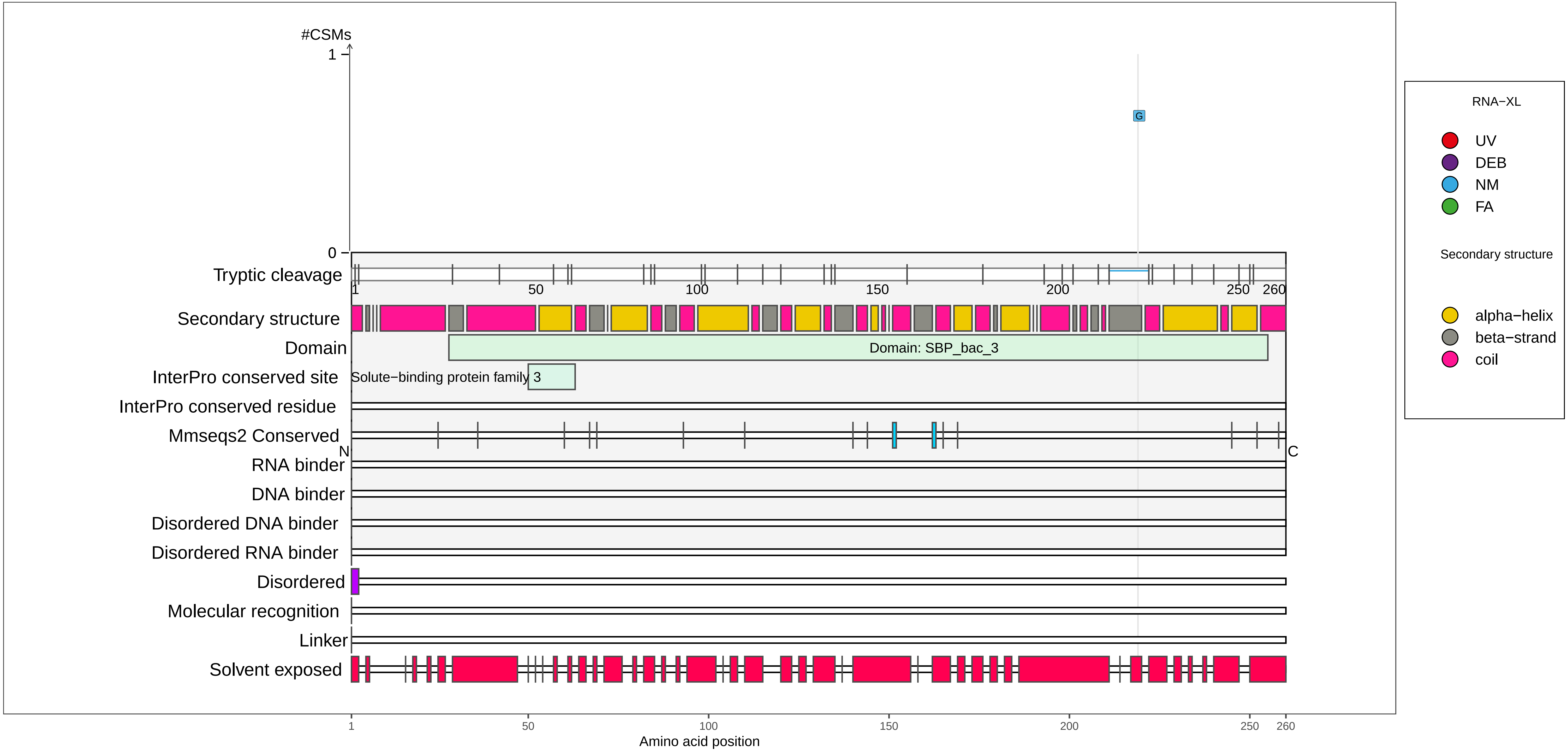

P0AEX5
KPPR_ECOLI Probable phosphoribulokinase

− Abundance:
tryptic [log10 Intensity]: 8.24 (Q 67)
PAXdb K12 strain [ppm]: 1.76 (Q 51)
PAXdb E.coli [ppm]: 1.85 (Q 68)

− RNA functions: not annotated

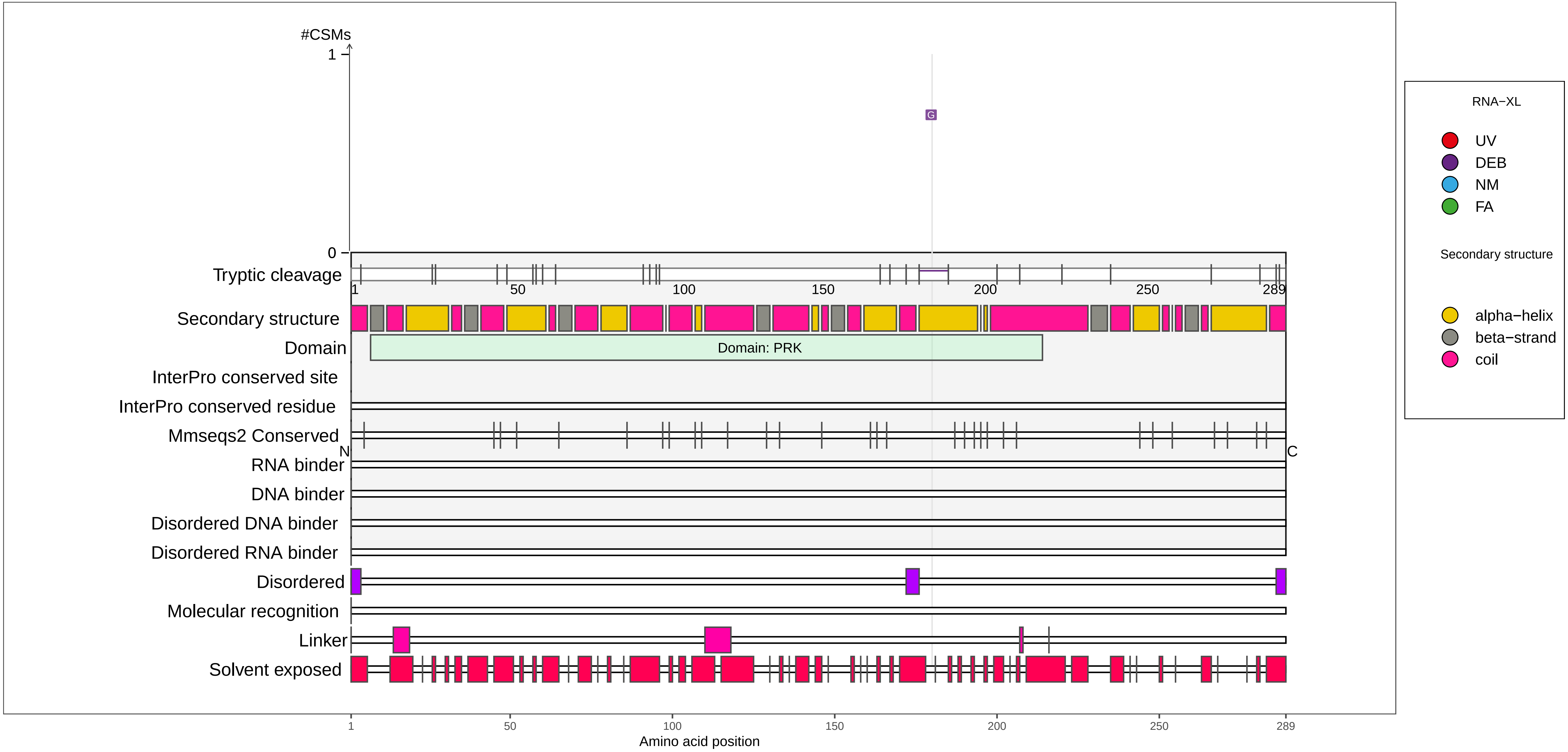

P0AEY1
MARC_ECOLI UPF0056 inner membrane protein MarC

− Abundance:
tryptic [log10 Intensity]: NA (Q NA)
PAXdb K12 strain [ppm]: NA (Q NA)
PAXdb E.coli [ppm]: −0.12 (Q 22)

− RNA functions: not annotated

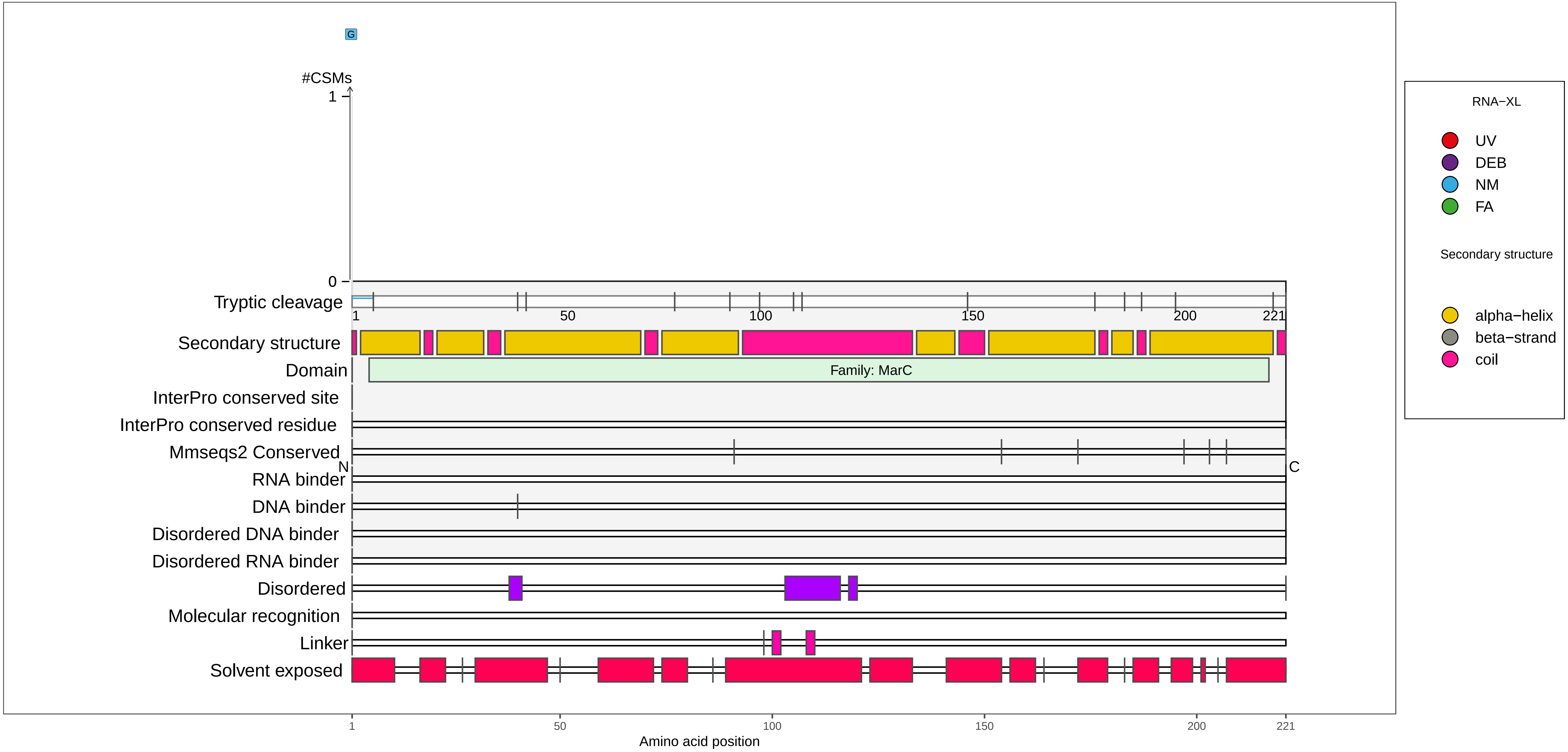

P0AEY3
MAZG_ECOLI Nucleoside triphosphate pyrophosphohydrolase

− Abundance:
tryptic [log10 Intensity]: 8.26 (Q 68)
PAXdb K12 strain [ppm]: NA (Q NA)
PAXdb E.coli [ppm]: 0.68 (Q 40)

− RNA functions: not annotated

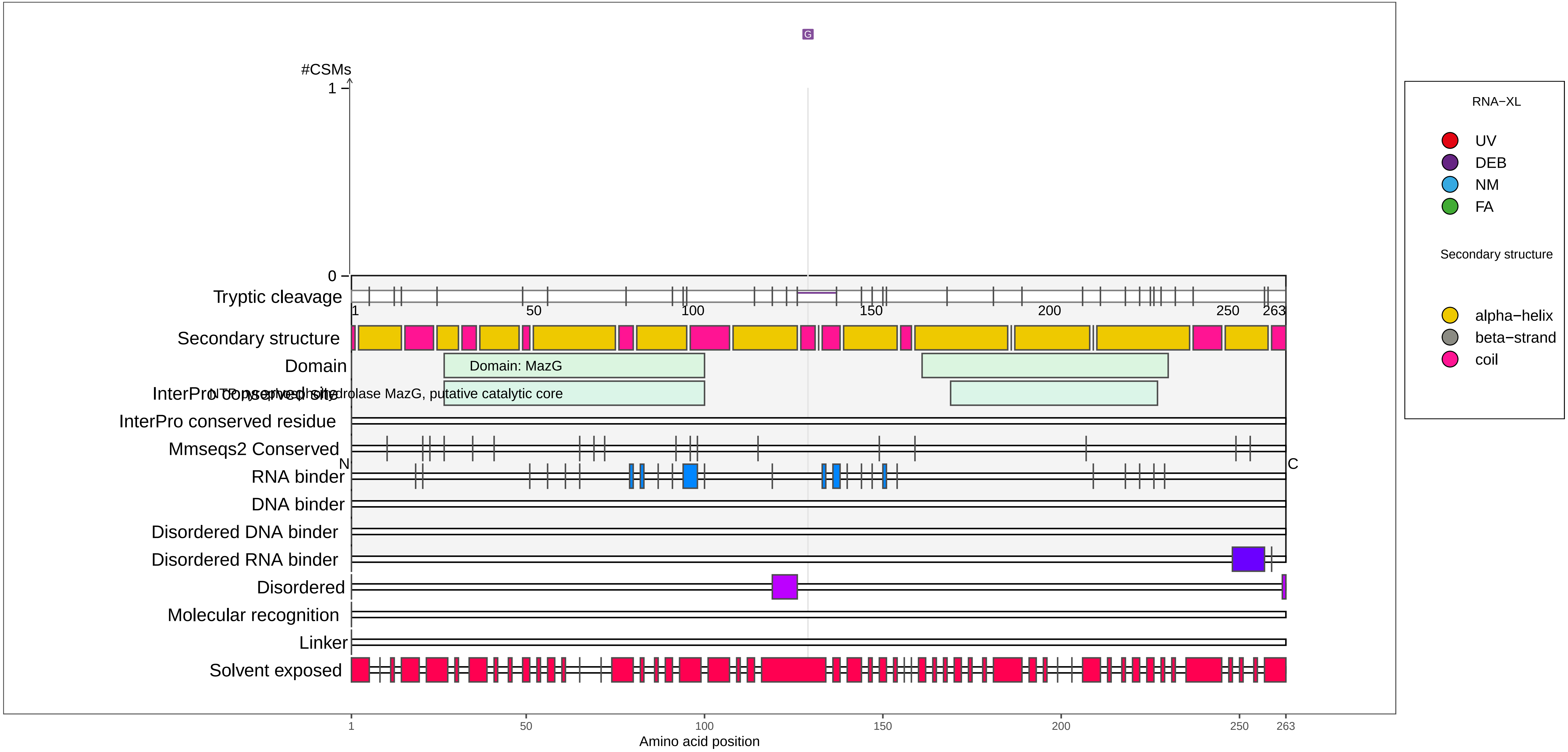

P0AF67
TSAE_ECOLI tRNA threonylcarbamoyladenosine biosynthesis protein TsaE

− Abundance:
tryptic [log10 Intensity]: 7.52 (Q 36)
PAXdb K12 strain [ppm]: 2.39 (Q 74)
PAXdb E.coli [ppm]: 2.07 (Q 74)

− RNA functions:
ncRNA metabolic process; ncRNA processing; RNA metabolic process; RNA modification
RNA processing; tRNA metabolic process; tRNA modification; tRNA processing
tRNA threonylcarbamoyladenosine metabolic process; tRNA threonylcarbamoyladenosine modification

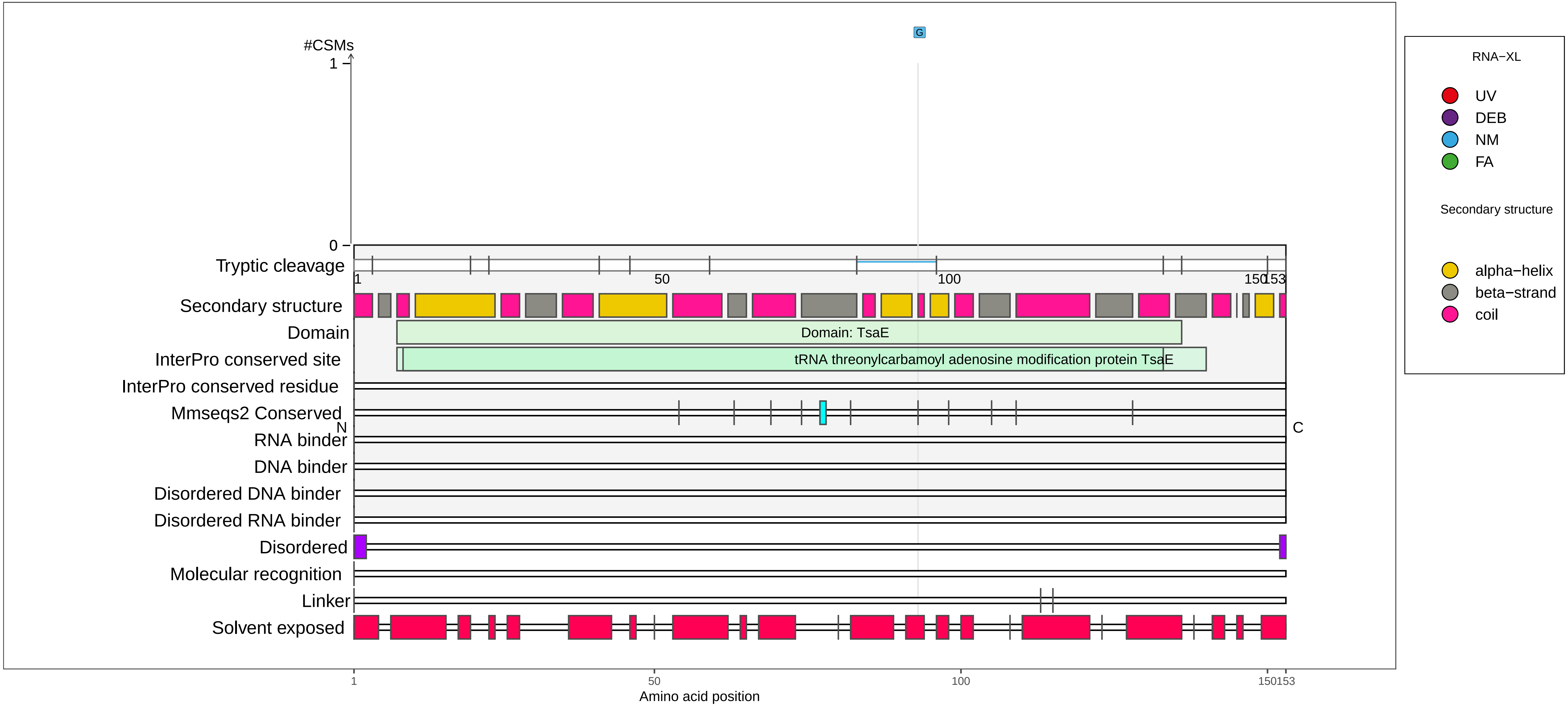

P0AF93
RIDA_ECOLI 2−iminobutanoate/2−iminopropanoate deaminase

− Abundance:
tryptic [log10 Intensity]: 8.31 (Q 70)
PAXdb K12 strain [ppm]: 2.92 (Q 89)
PAXdb E.coli [ppm]: 3.64 (Q 100)

− RNA functions: not annotated

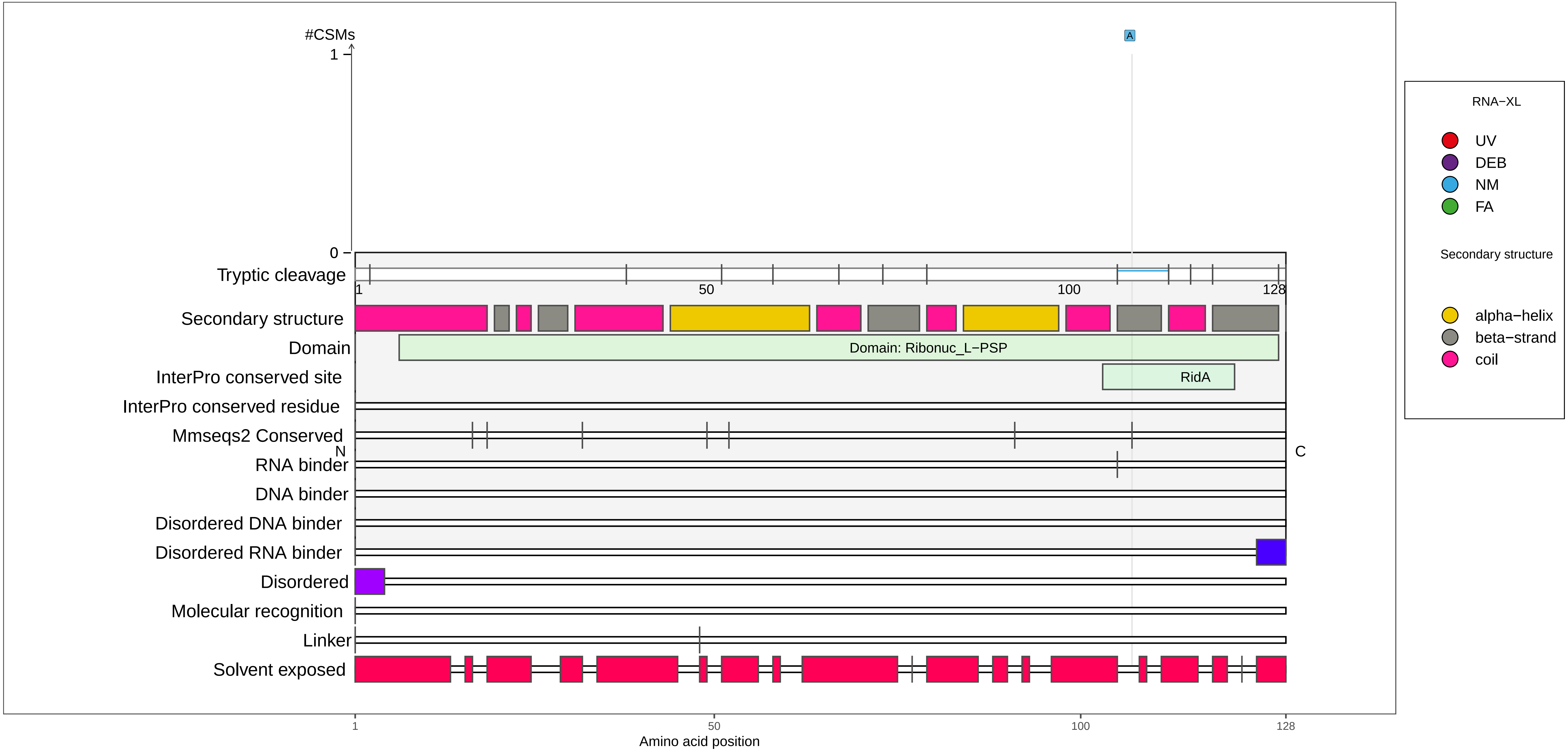

P0AFE4
NUOK_ECOLI NADH−quinone oxidoreductase subunit K

− Abundance:
tryptic [log10 Intensity]: 9.03 (Q 89)
PAXdb K12 strain [ppm]: NA (Q NA)
PAXdb E.coli [ppm]: 1.98 (Q 72)

− RNA functions: not annotated

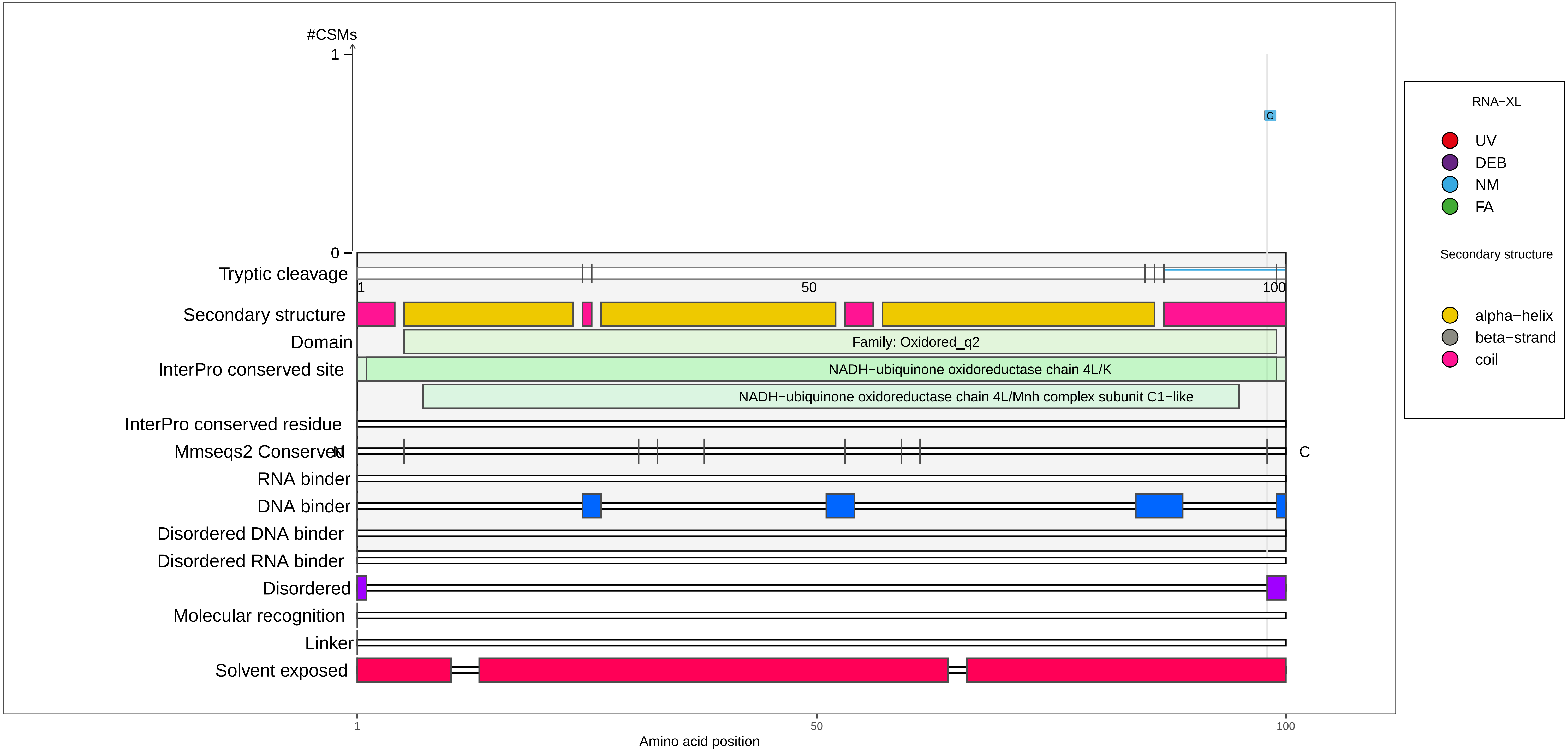

P0AFJ5
PHOB_ECOLI Phosphate regulon transcriptional regulatory protein PhoB

− Abundance:
tryptic [log10 Intensity]: 7.71 (Q 45)
PAXdb K12 strain [ppm]: NA (Q NA)
PAXdb E.coli [ppm]: 1.01 (Q 48)

− RNA functions: not annotated

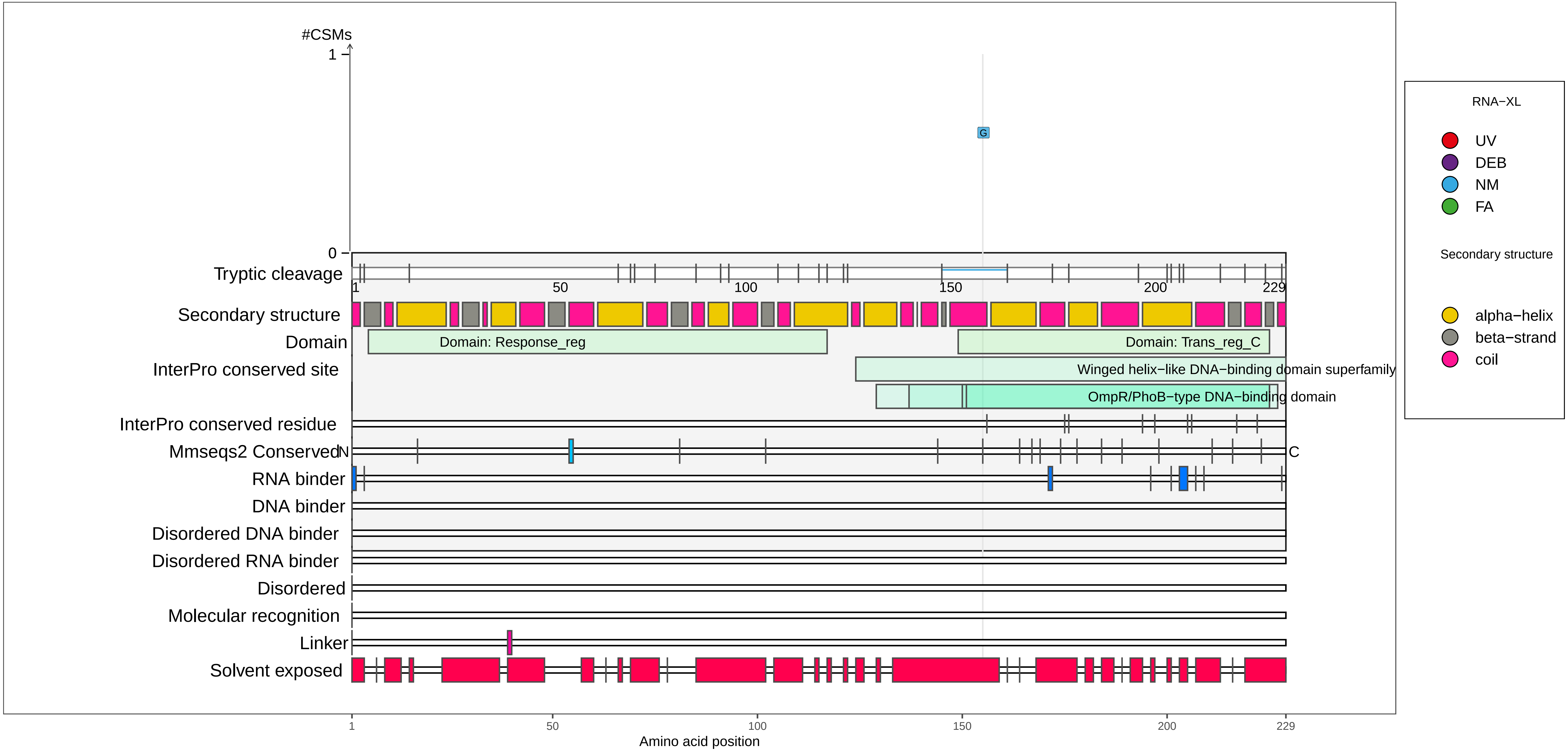

P0AFJ7
PITA_ECOLI Low−affinity inorganic phosphate transporter 1

− Abundance:
tryptic [log10 Intensity]: 6.78 (Q 7)
PAXdb K12 strain [ppm]: NA (Q NA)
PAXdb E.coli [ppm]: 1.46 (Q 59)

− RNA functions: not annotated

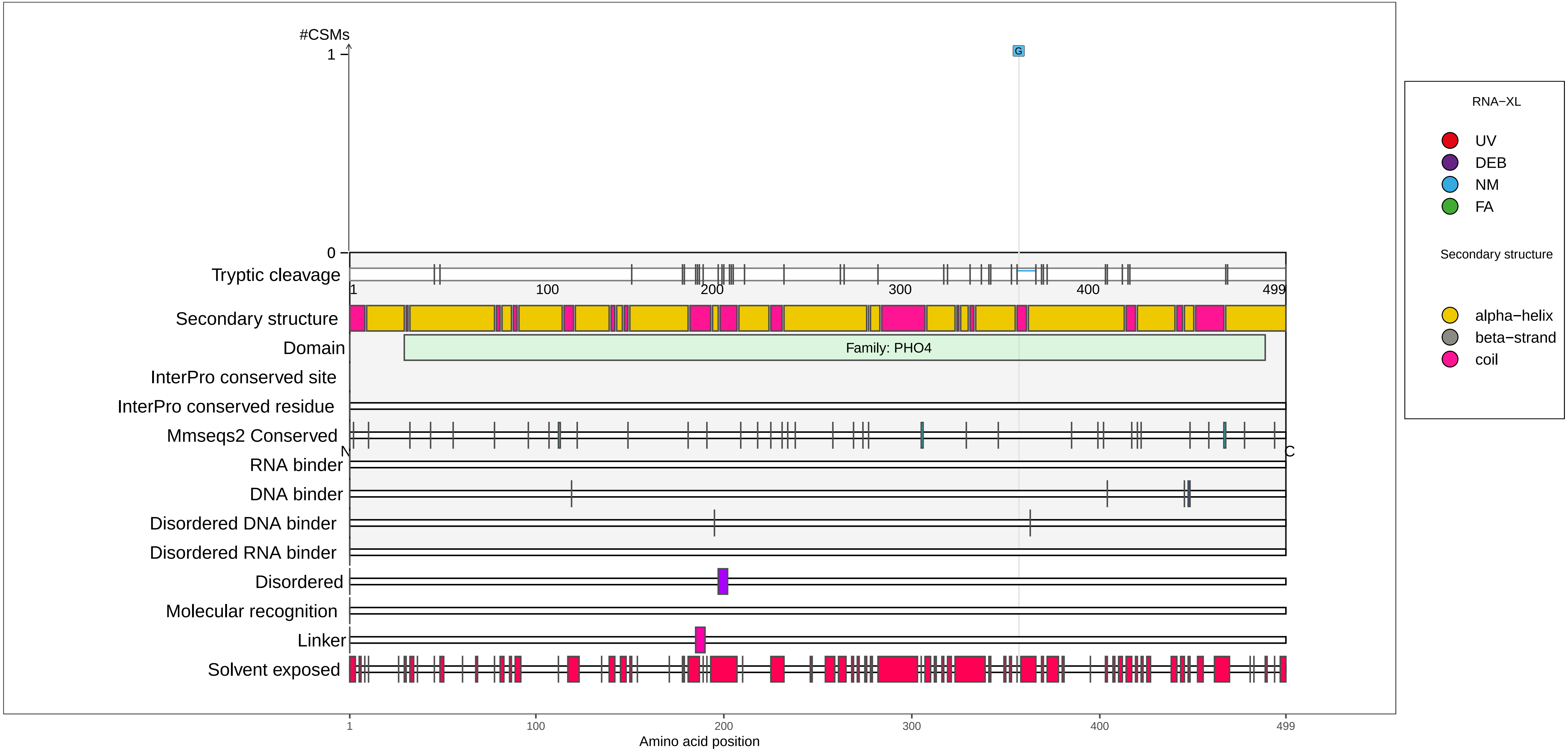

P0AFS3
FOLM_ECOLI Dihydromonapterin reductase

− Abundance:
tryptic [log10 Intensity]: 7.32 (Q 26)
PAXdb K12 strain [ppm]: NA (Q NA)
PAXdb E.coli [ppm]: −0.12 (Q 22)

− RNA functions: not annotated

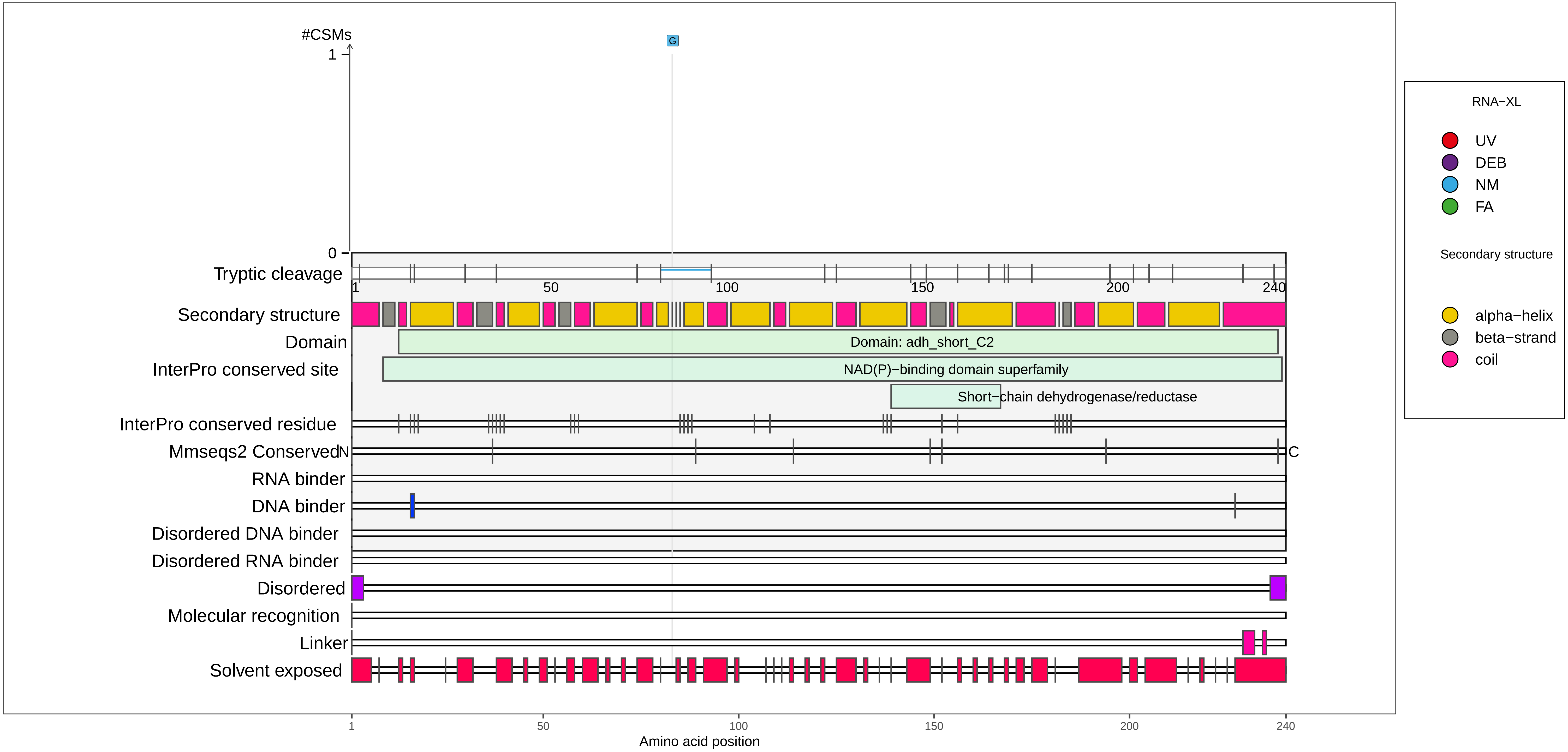

P0AFT5
BTSR_ECOLI Transcriptional regulatory protein BtsR

− Abundance:
tryptic [log10 Intensity]: 7.28 (Q 24)
PAXdb K12 strain [ppm]: 1.57 (Q 43)
PAXdb E.coli [ppm]: 0.82 (Q 43)

− RNA functions: not annotated

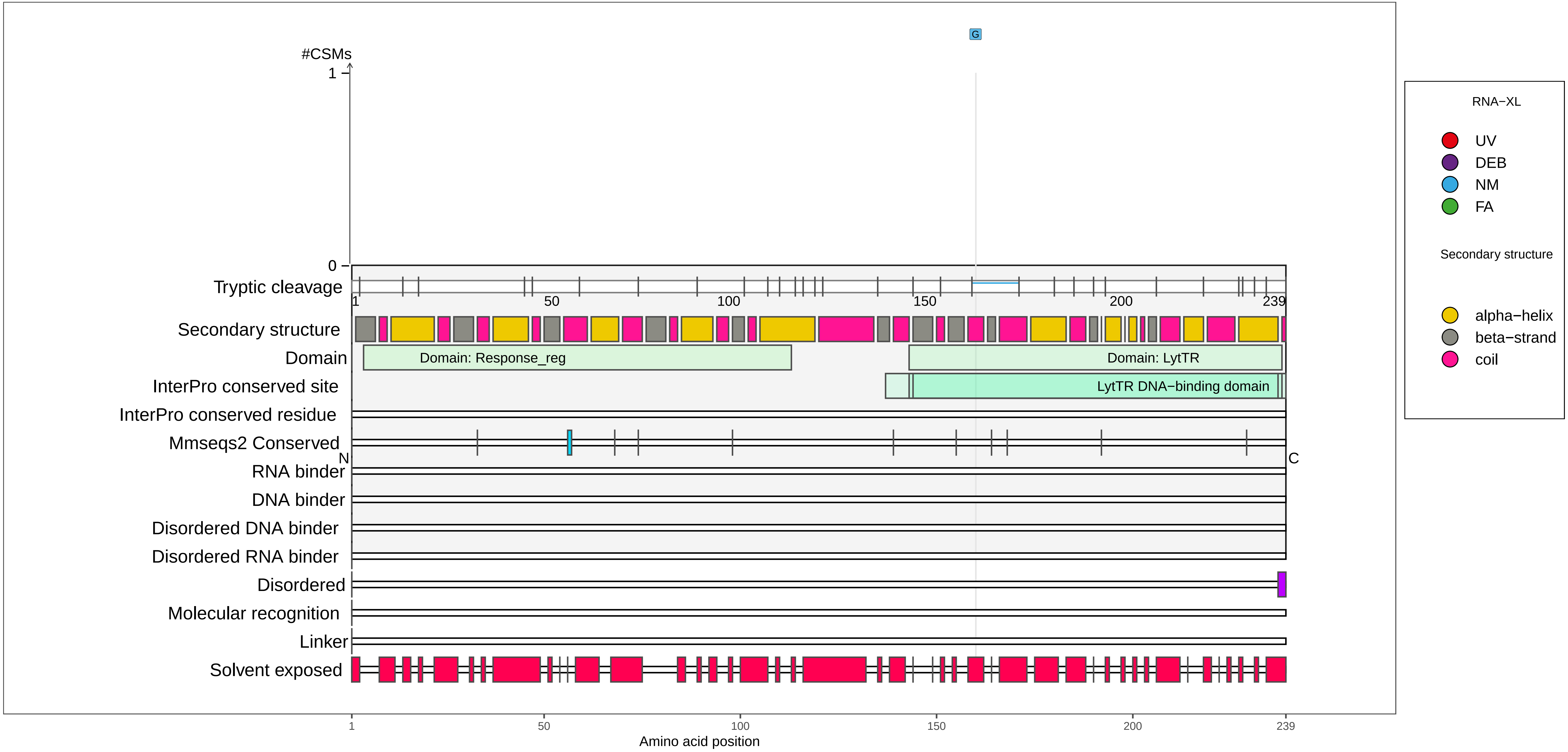

P0AFZ7
TRKH_ECOLI Trk system potassium uptake protein TrkH

− Abundance:
tryptic [log10 Intensity]: 6.36 (Q 1)
PAXdb K12 strain [ppm]: NA (Q NA)
PAXdb E.coli [ppm]: −1.59 (Q 3)

− RNA functions: not annotated

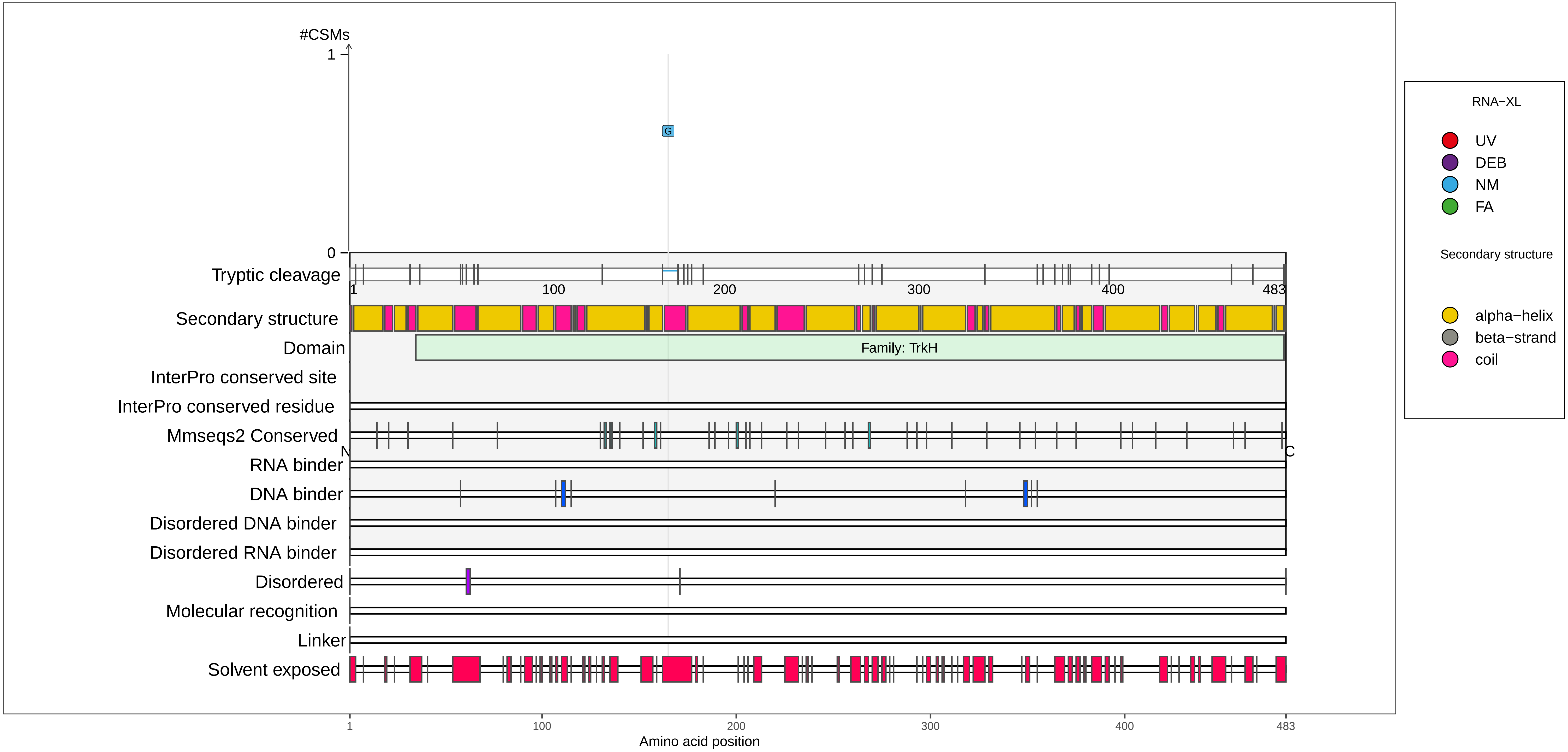

P0AG16
PUR1_ECOLI Amidophosphoribosyltransferase

− Abundance:
tryptic [log10 Intensity]: 7.3 (Q 25)
PAXdb K12 strain [ppm]: 2.22 (Q 68)
PAXdb E.coli [ppm]: 2.04 (Q 73)

− RNA functions: not annotated

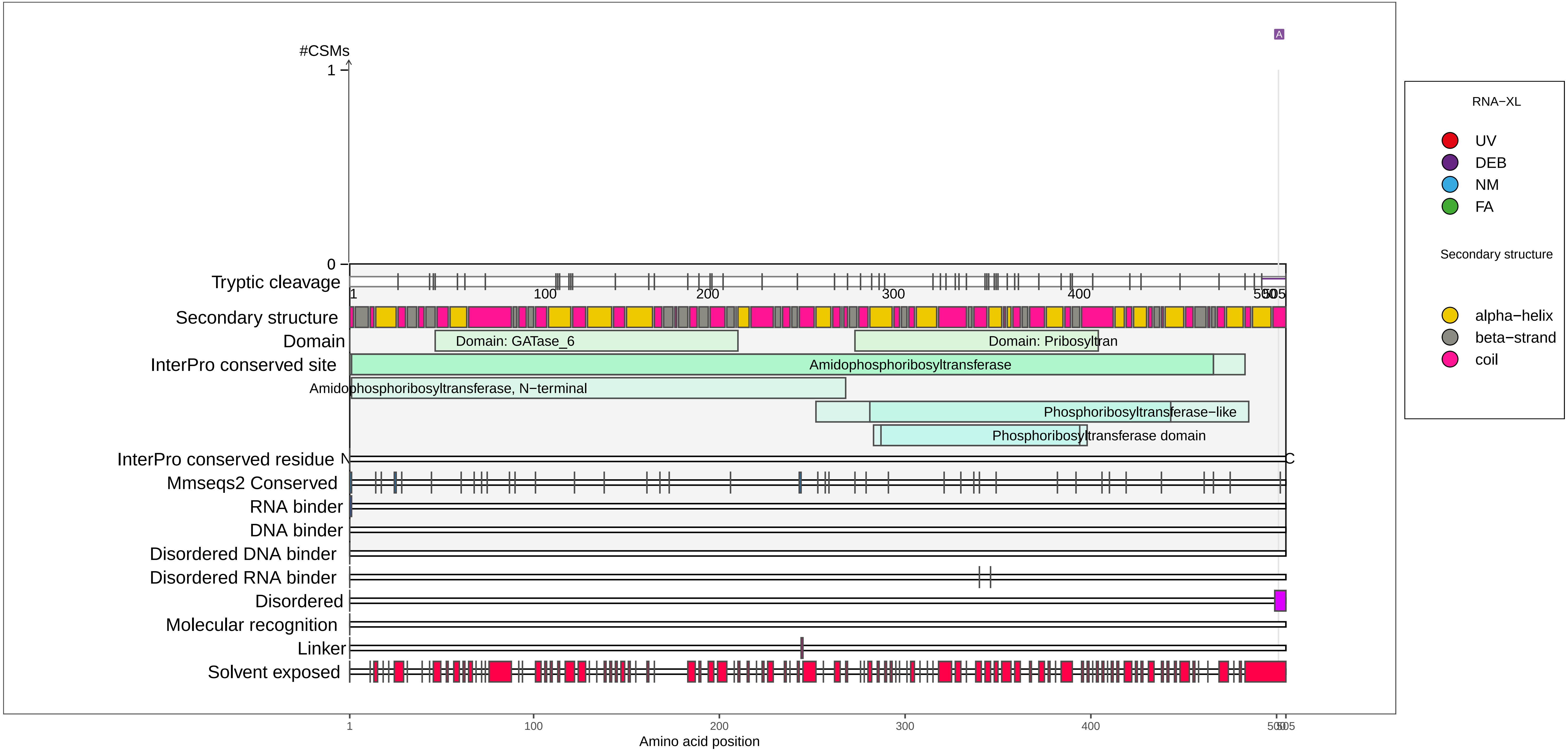

P0AG27
YIBN_ECOLI Uncharacterized protein YibN

− Abundance:
tryptic [log10 Intensity]: 8.42 (Q 73)
PAXdb K12 strain [ppm]: 2.35 (Q 72)
PAXdb E.coli [ppm]: 2.45 (Q 84)

− RNA functions: not annotated

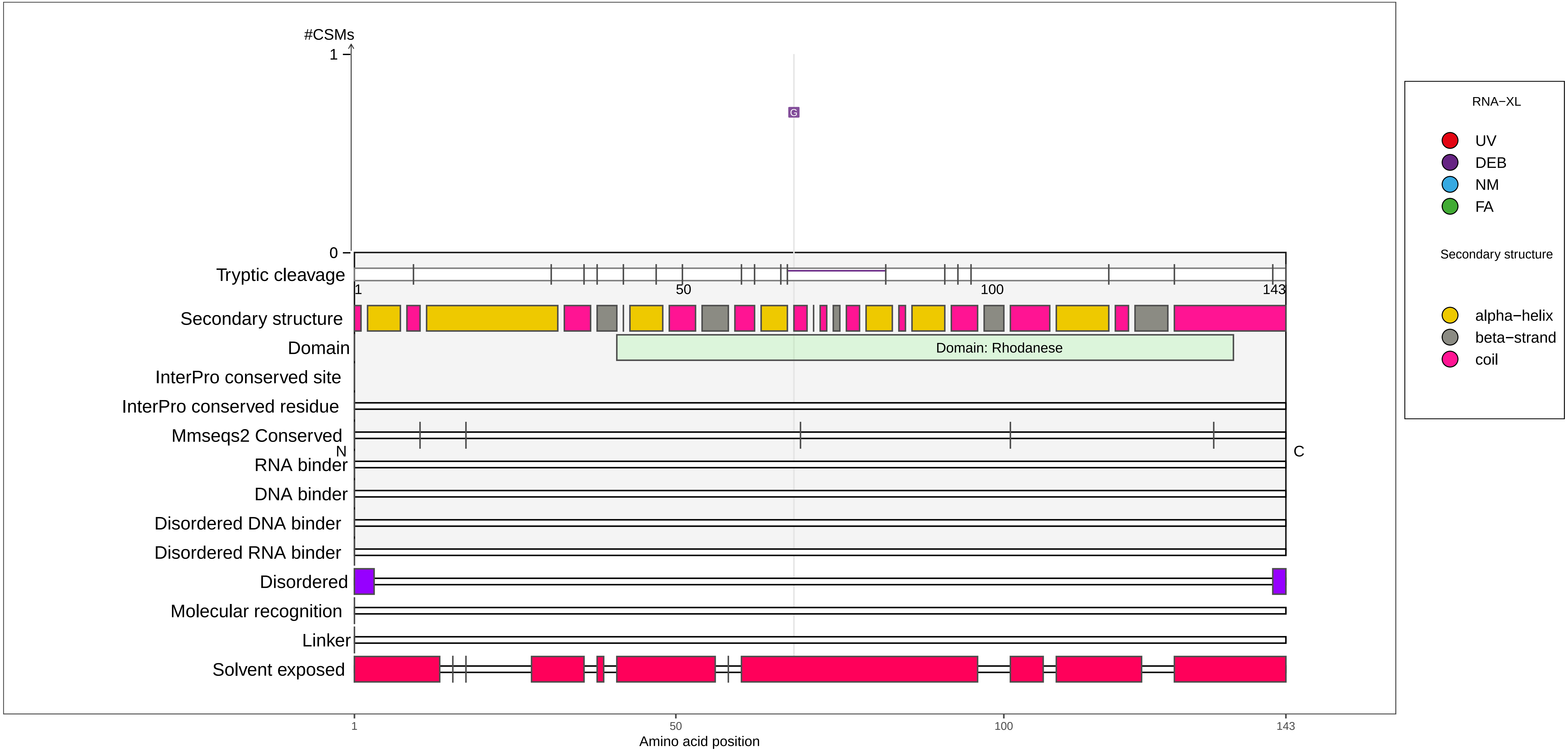

P0AGC3
SLT_ECOLI Soluble lytic murein transglycosylase

− Abundance:
tryptic [log10 Intensity]: 7.14 (Q 18)
PAXdb K12 strain [ppm]: 1.57 (Q 42)
PAXdb E.coli [ppm]: 1.89 (Q 69)

− RNA functions: not annotated

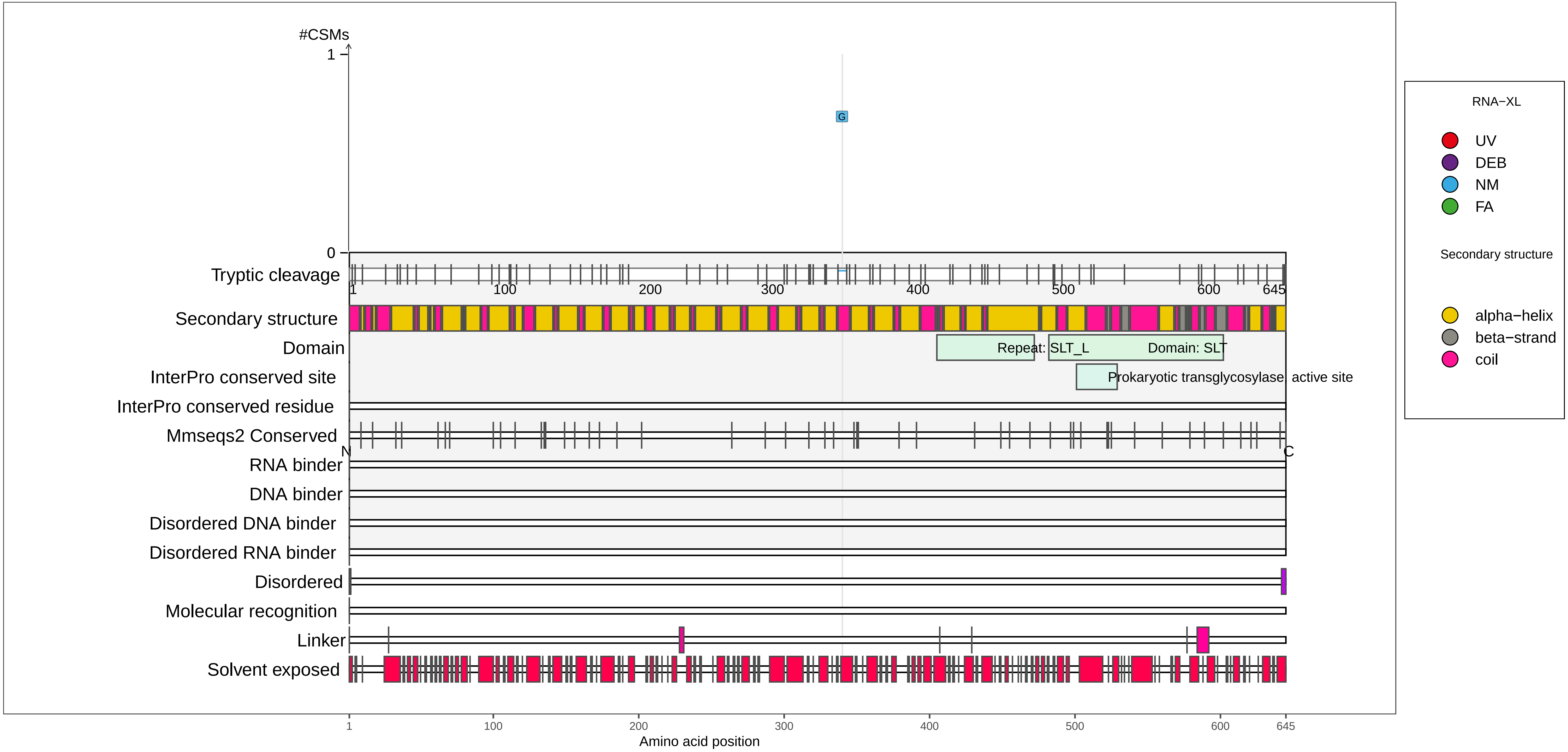

P0AGE6
CHRR_ECOLI Quinone reductase

− Abundance:
tryptic [log10 Intensity]: 8.05 (Q 59)
PAXdb K12 strain [ppm]: 1.37 (Q 31)
PAXdb E.coli [ppm]: 2.85 (Q 92)

− RNA functions: not annotated

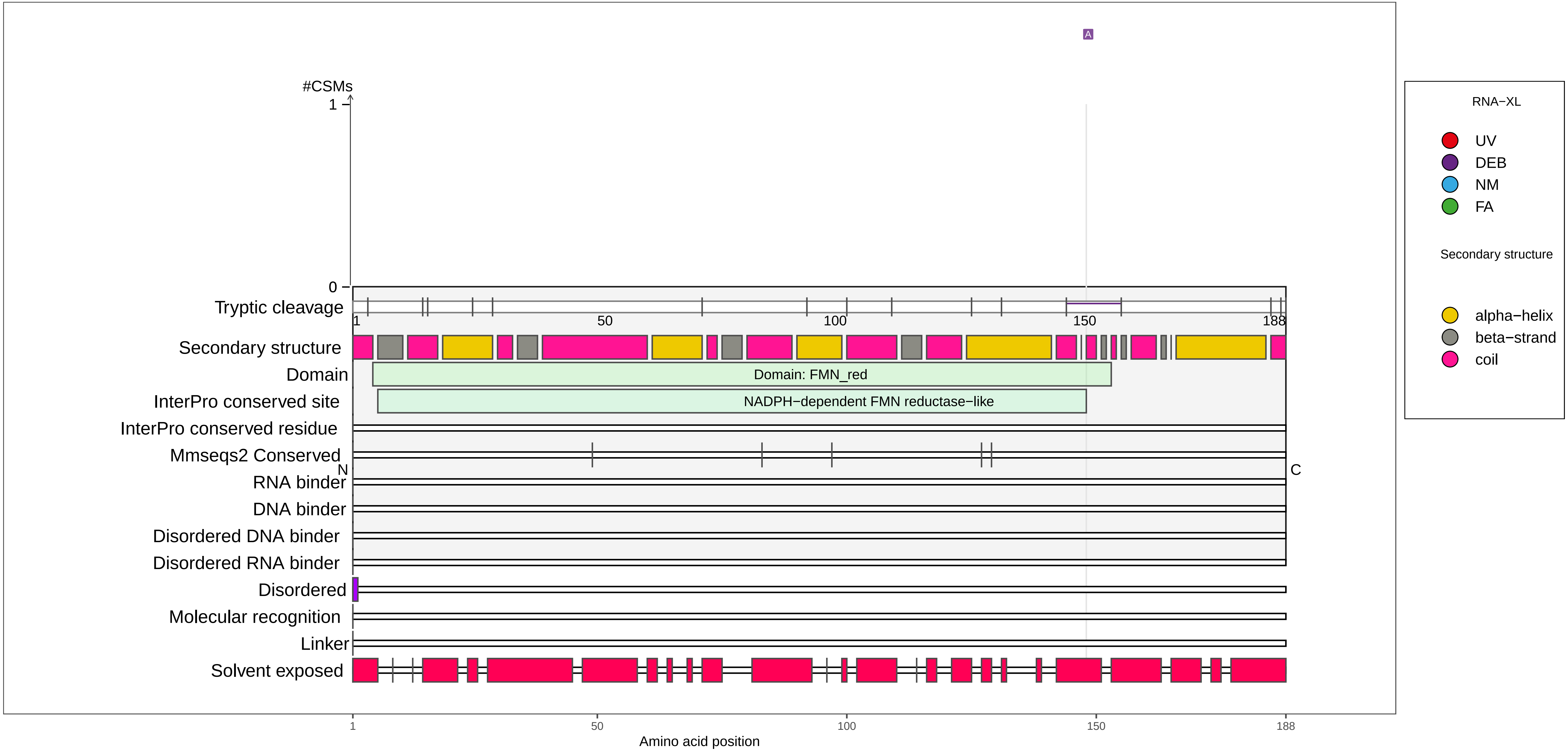

P0AGG4
THIO2_ECOLI Thioredoxin 2

− Abundance:
tryptic [log10 Intensity]: 7.92 (Q 54)
PAXdb K12 strain [ppm]: 2.8 (Q 86)
PAXdb E.coli [ppm]: 1.27 (Q 54)

− RNA functions: not annotated

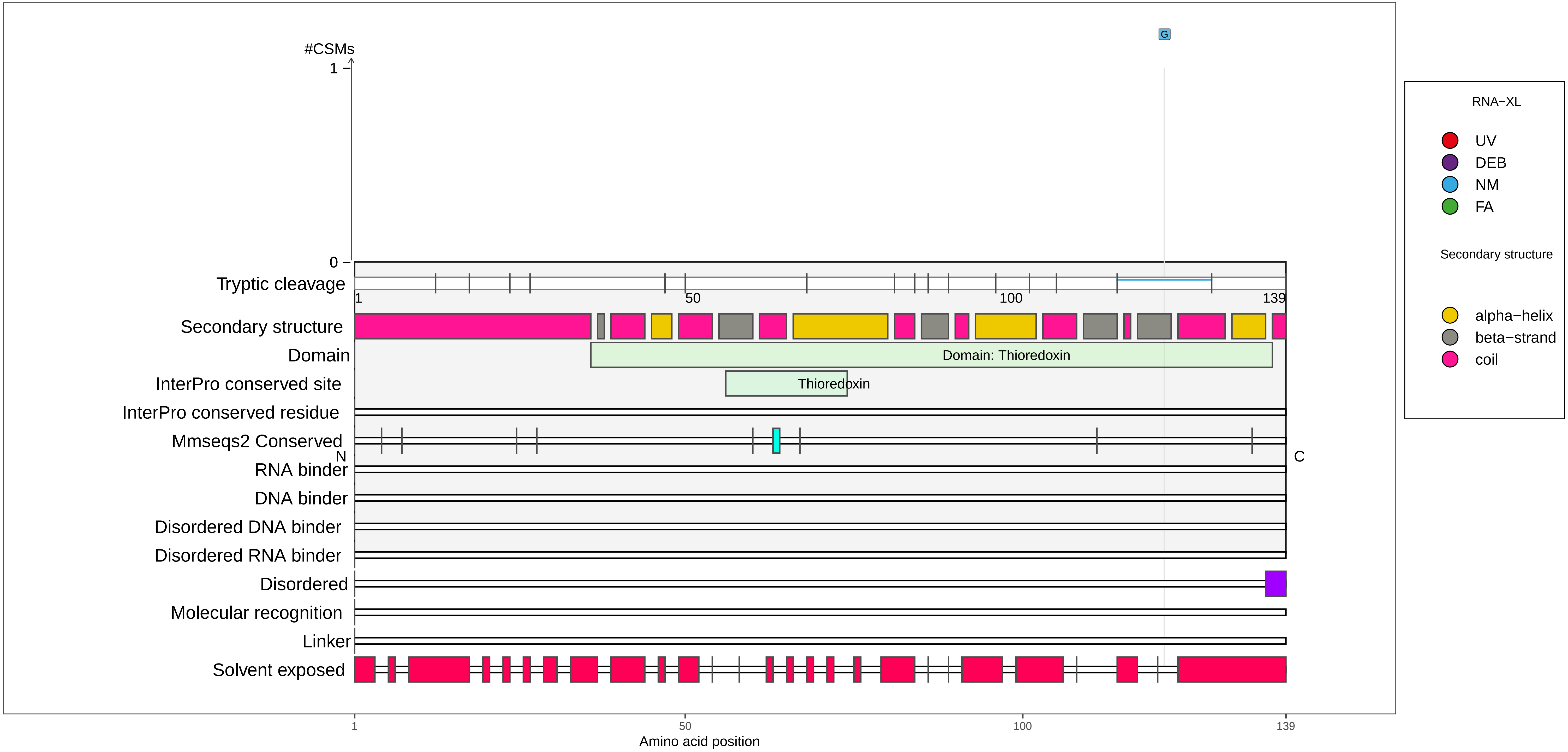

P0AGM0
YHHT_ECOLI Putative transport protein YhhT

− Abundance:
tryptic [log10 Intensity]: NA (Q NA)
PAXdb K12 strain [ppm]: NA (Q NA)
PAXdb E.coli [ppm]: −0.94 (Q 10)

− RNA functions: not annotated

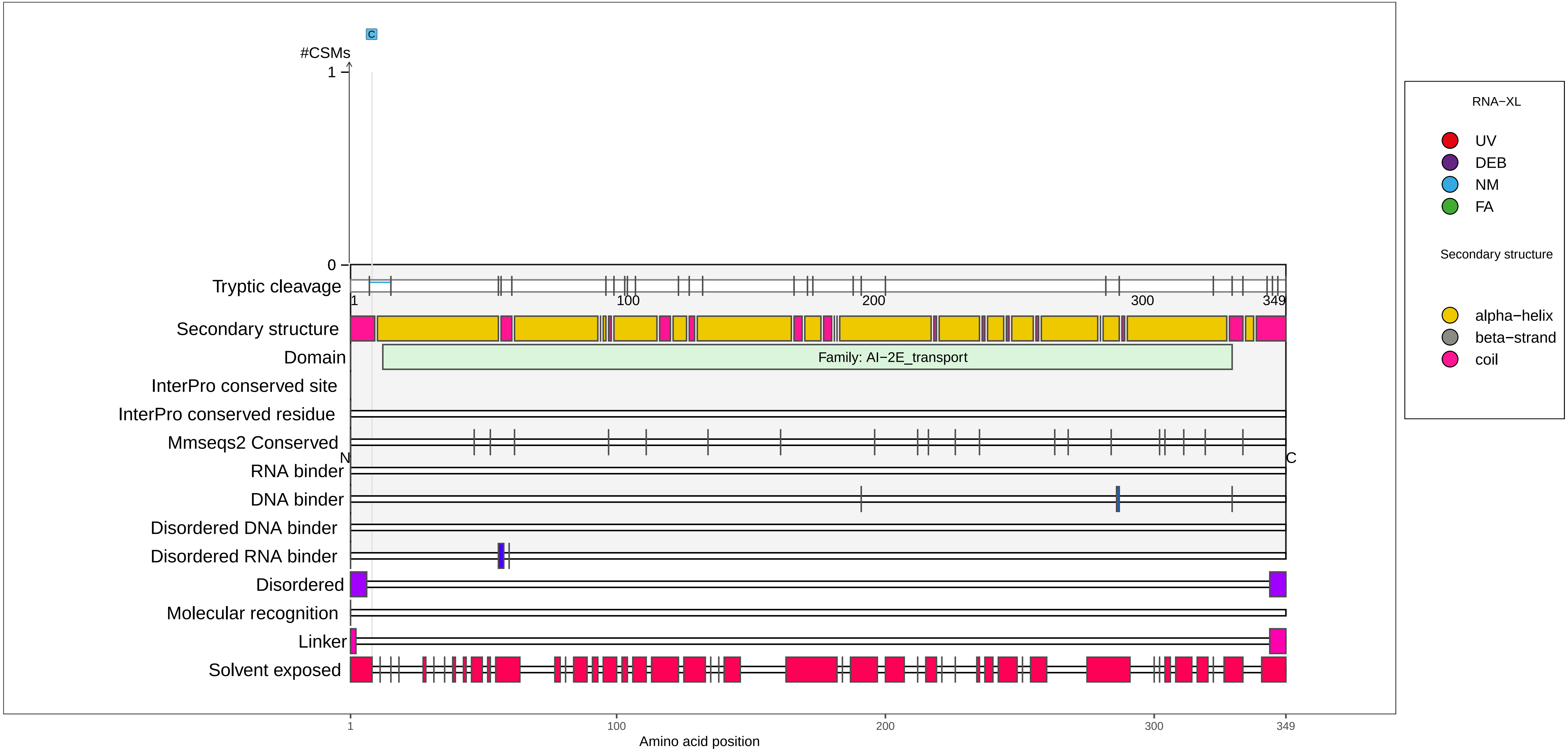

P0CB39
EPTC_ECOLI Phosphoethanolamine transferase EptC

− Abundance:
tryptic [log10 Intensity]: 7.82 (Q 50)
PAXdb K12 strain [ppm]: 2.16 (Q 66)
PAXdb E.coli [ppm]: 1.08 (Q 50)

− RNA functions: not annotated

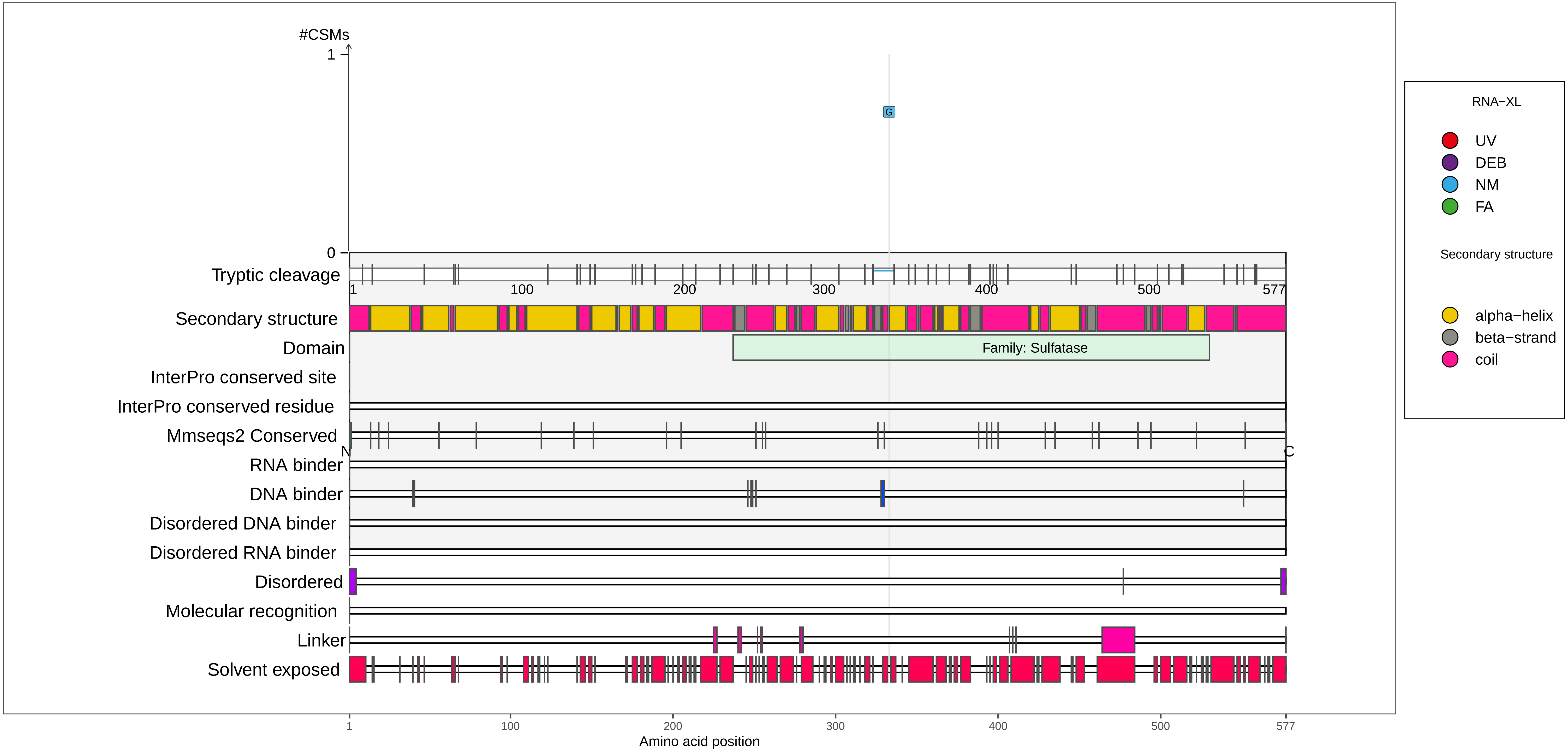

P0CF67
INSE2_ECOLI Transposase InsE for insertion sequence IS3B
− RNA functions: not annotated

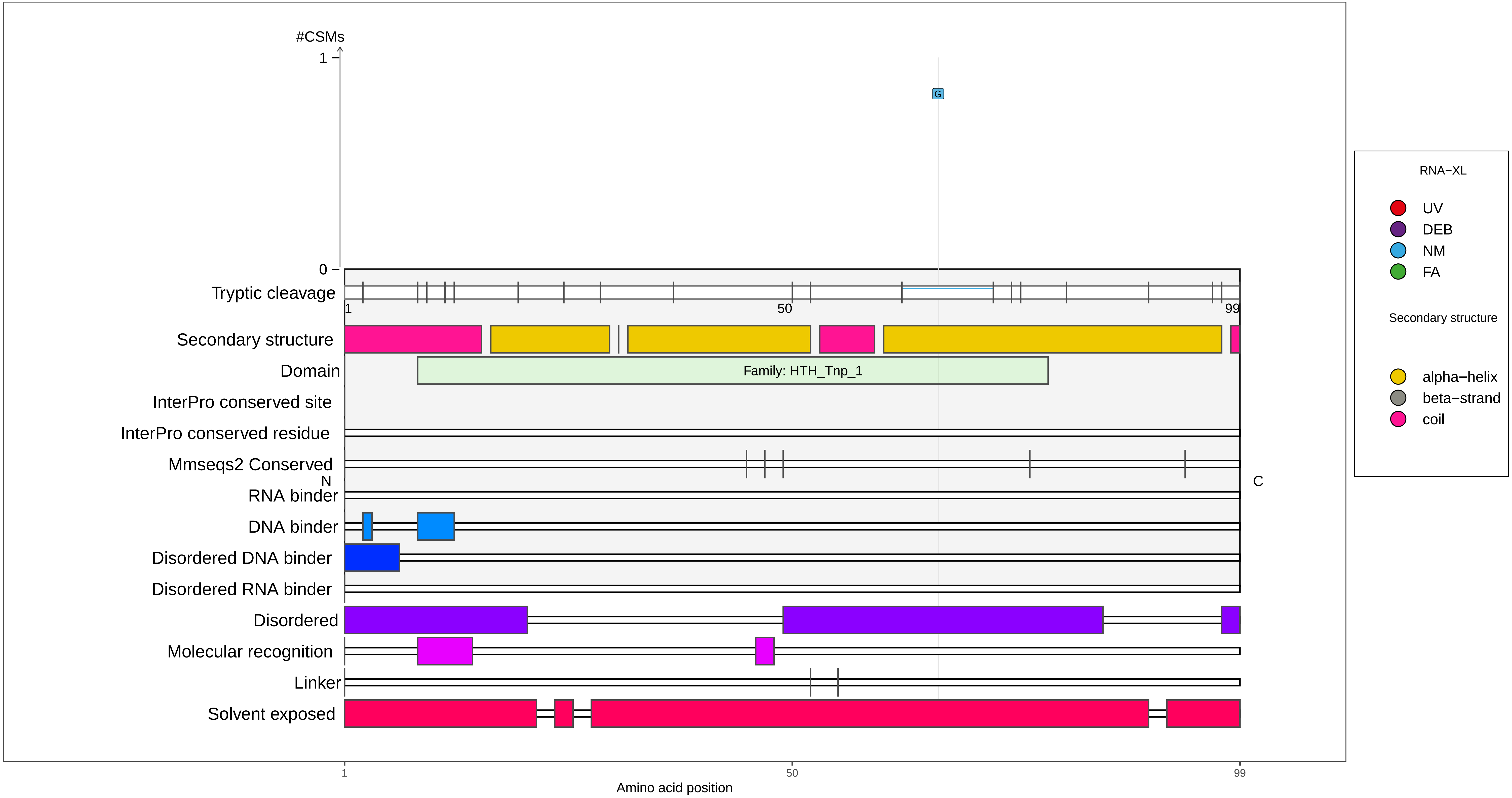

P0DP89
ILVGP_ECOLI Putative acetolactate synthase isozyme 2 large subunit
− RNA functions: not annotated

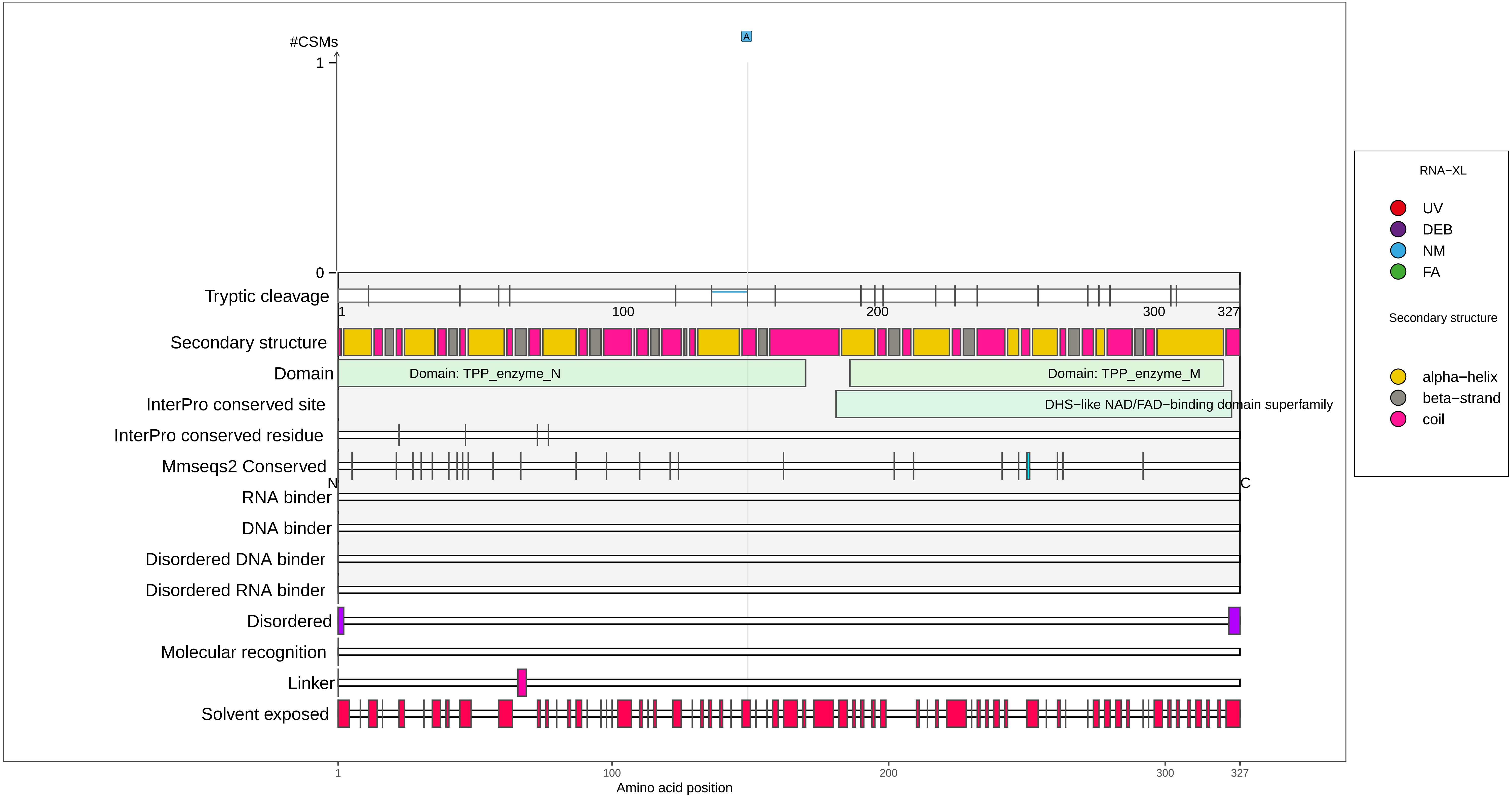

P10100
RLPA_ECOLI Endolytic peptidoglycan transglycosylase RlpA

− Abundance:
tryptic [log10 Intensity]: 7.7 (Q 44)
PAXdb K12 strain [ppm]: NA (Q NA)
PAXdb E.coli [ppm]: 1.58 (Q 62)

− RNA functions: not annotated

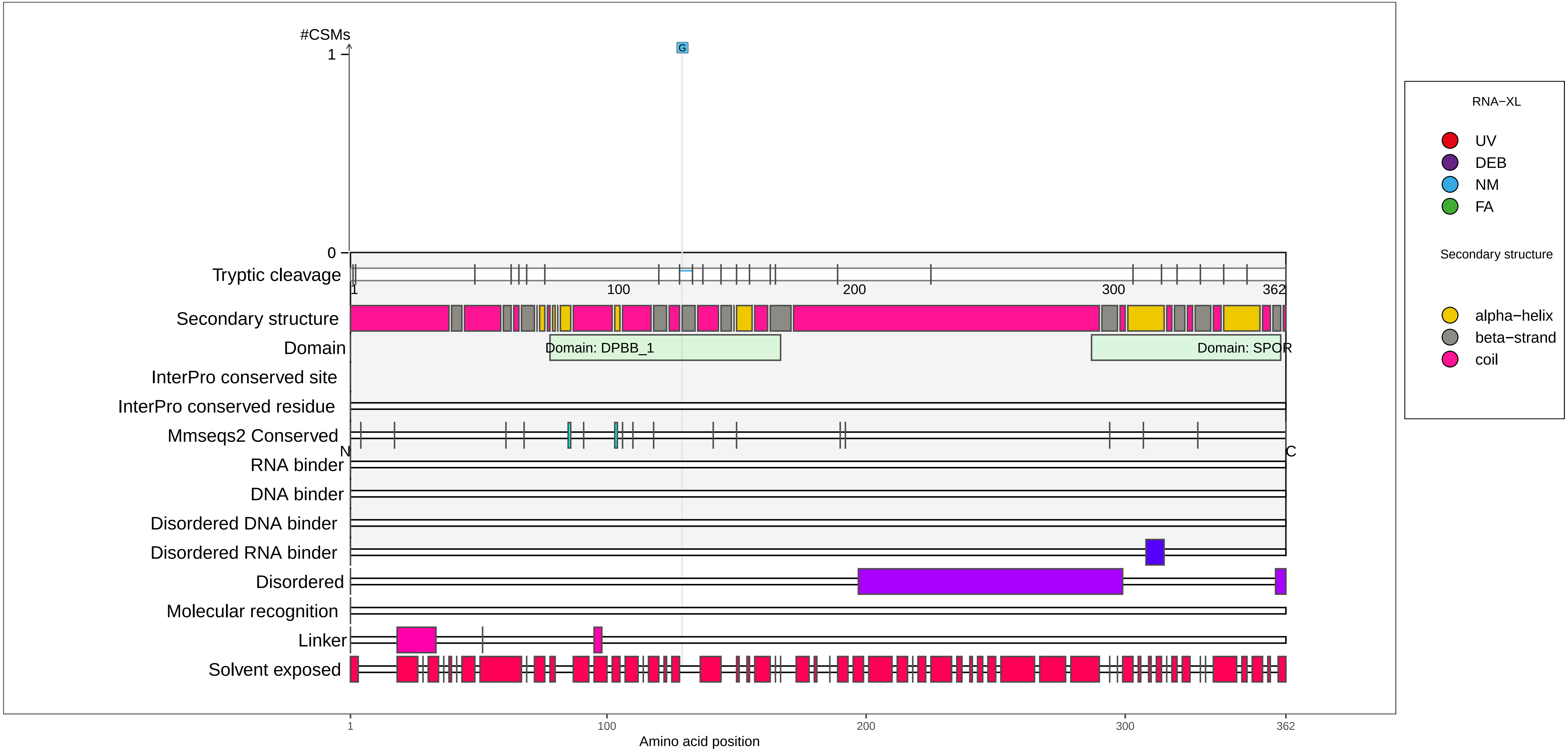

P10423
IAP_ECOLI Alkaline phosphatase isozyme conversion protein

− Abundance:
tryptic [log10 Intensity]: 8.83 (Q 84)
PAXdb K12 strain [ppm]: 1.39 (Q 32)
PAXdb E.coli [ppm]: 0.05 (Q 26)

− RNA functions: not annotated

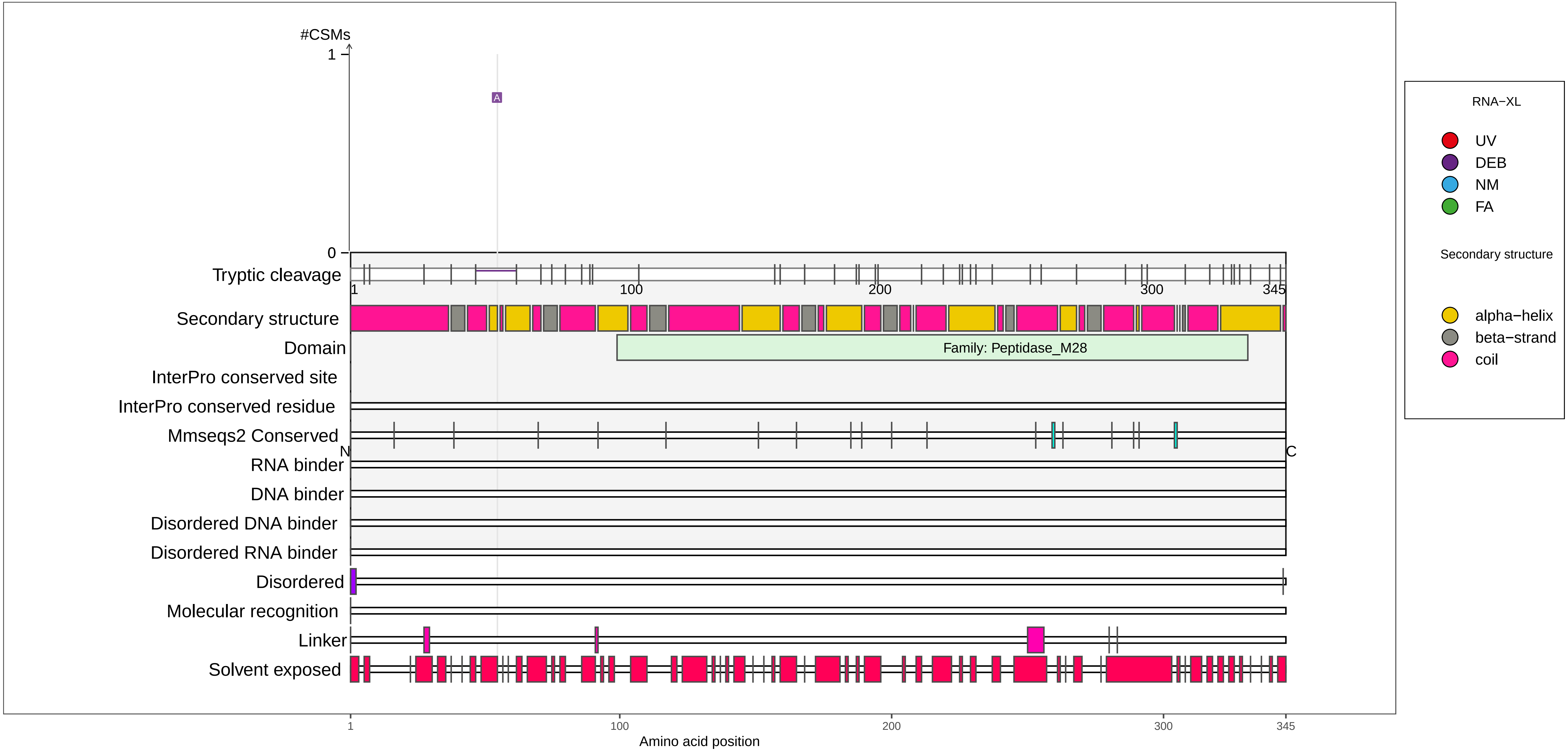

P10441
LPXB_ECOLI Lipid−A−disaccharide synthase

− Abundance:
tryptic [log10 Intensity]: NA (Q NA)
PAXdb K12 strain [ppm]: 0.79 (Q 5)
PAXdb E.coli [ppm]: 1.86 (Q 69)

− RNA functions: not annotated

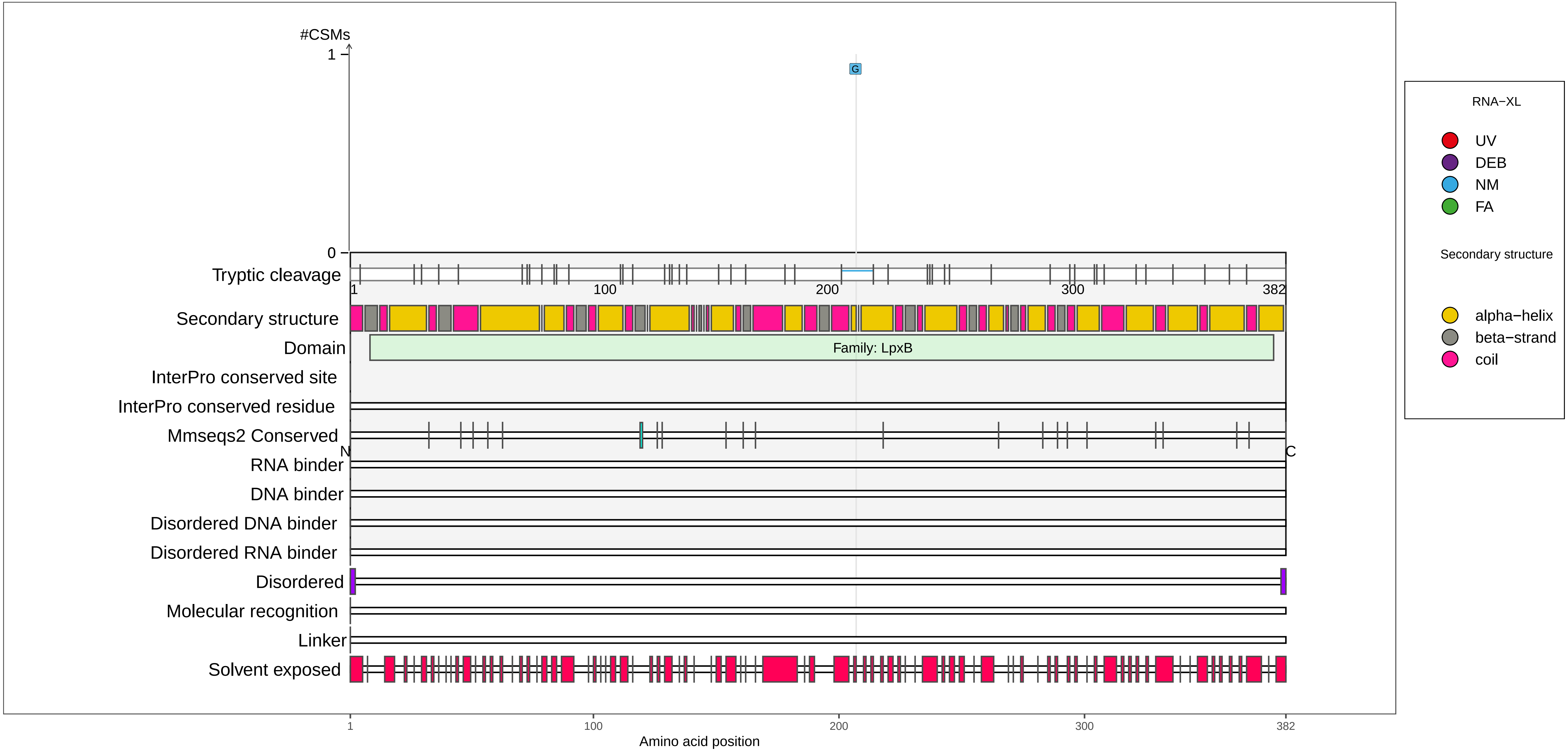

P10908
UGPQ_ECOLI Glycerophosphodiester phosphodiesterase, cytoplasmic

− Abundance:
tryptic [log10 Intensity]: 7.65 (Q 42)
PAXdb K12 strain [ppm]: 1.55 (Q 41)
PAXdb E.coli [ppm]: 1.53 (Q 60)

− RNA functions: not annotated

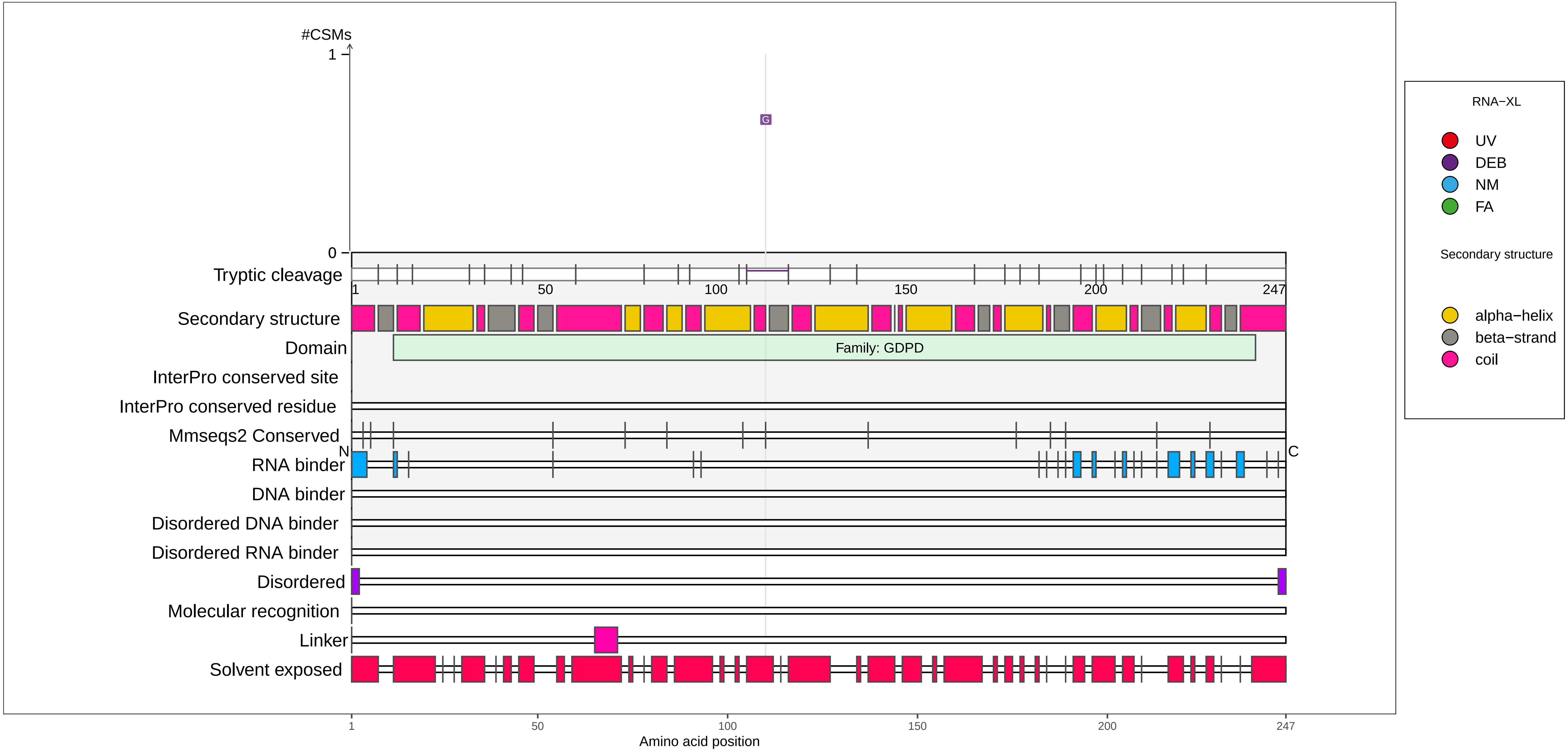

P11553
FUCK_ECOLI L−fuculokinase

− Abundance:
tryptic [log10 Intensity]: 7.71 (Q 45)
PAXdb K12 strain [ppm]: 1.39 (Q 32)
PAXdb E.coli [ppm]: 1.41 (Q 57)

− RNA functions: not annotated

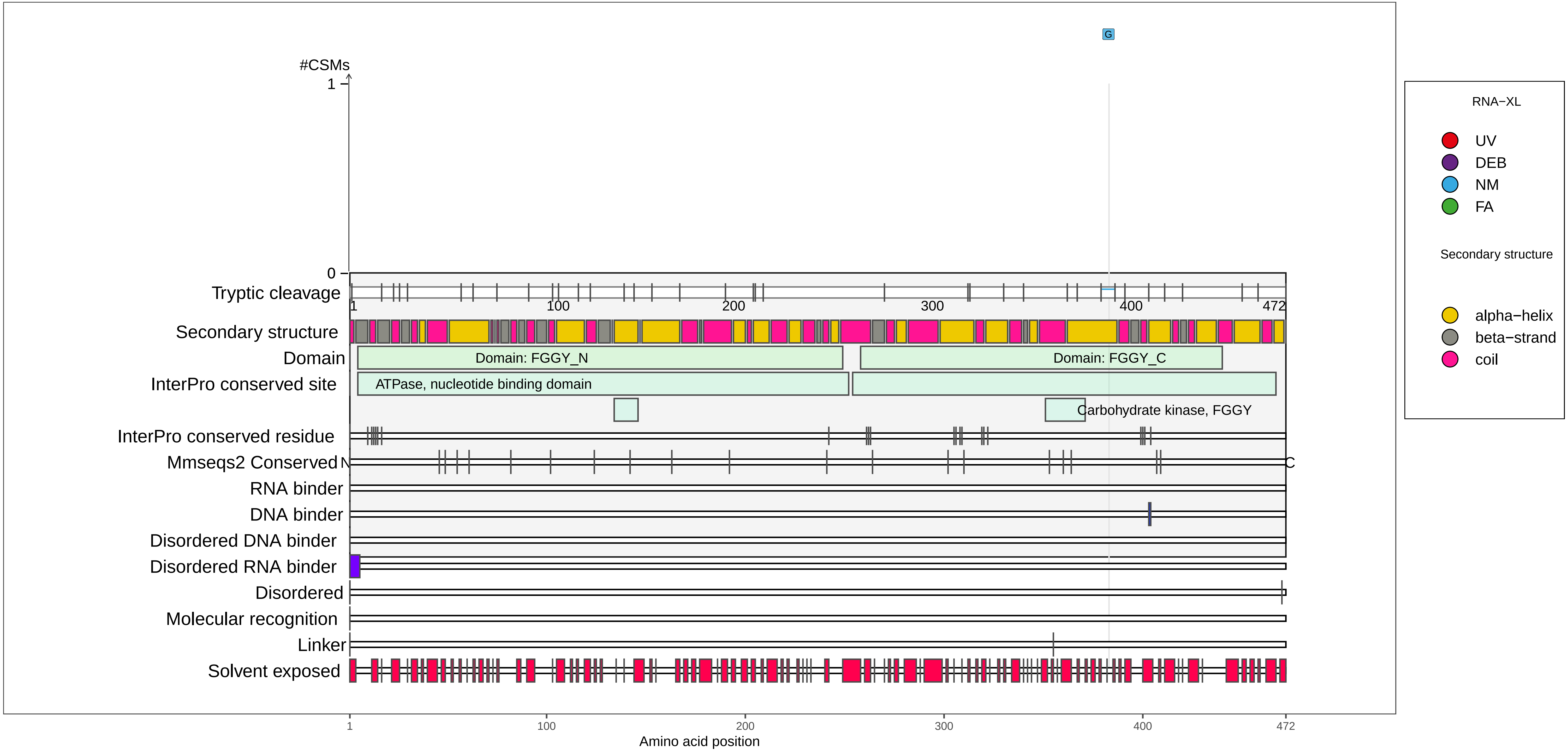

P13482
TREA_ECOLI Periplasmic trehalase

− Abundance:
tryptic [log10 Intensity]: NA (Q NA)
PAXdb K12 strain [ppm]: 1.62 (Q 46)
PAXdb E.coli [ppm]: 2.55 (Q 87)

− RNA functions: not annotated

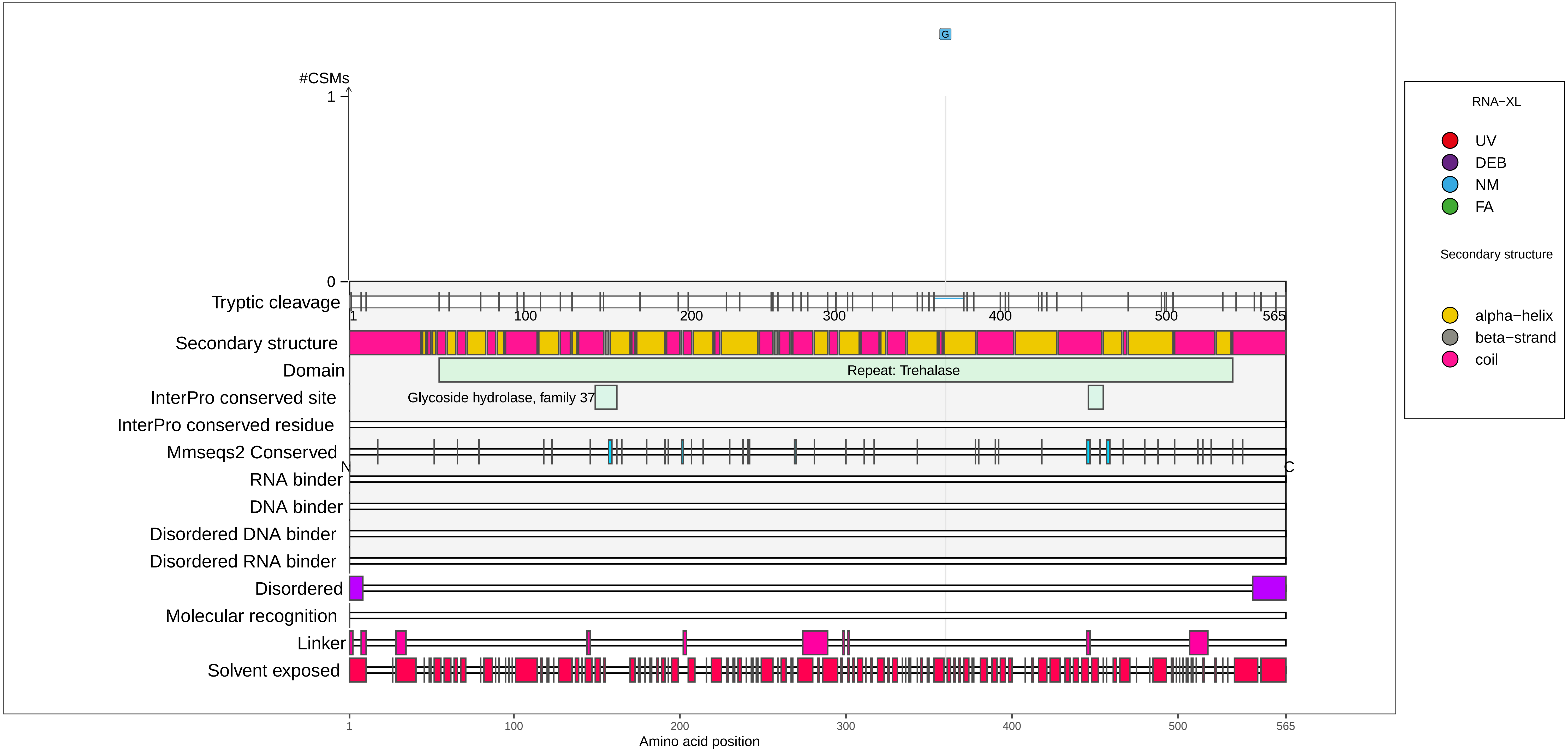

P13518
CSRD_ECOLI RNase E specificity factor CsrD

− Abundance:
tryptic [log10 Intensity]: NA (Q NA)
PAXdb K12 strain [ppm]: 0.73 (Q 3)
PAXdb E.coli [ppm]: 1.27 (Q 54)

− RNA functions:
ncRNA catabolic process; ncRNA metabolic process; RNA catabolic process
RNA metabolic process

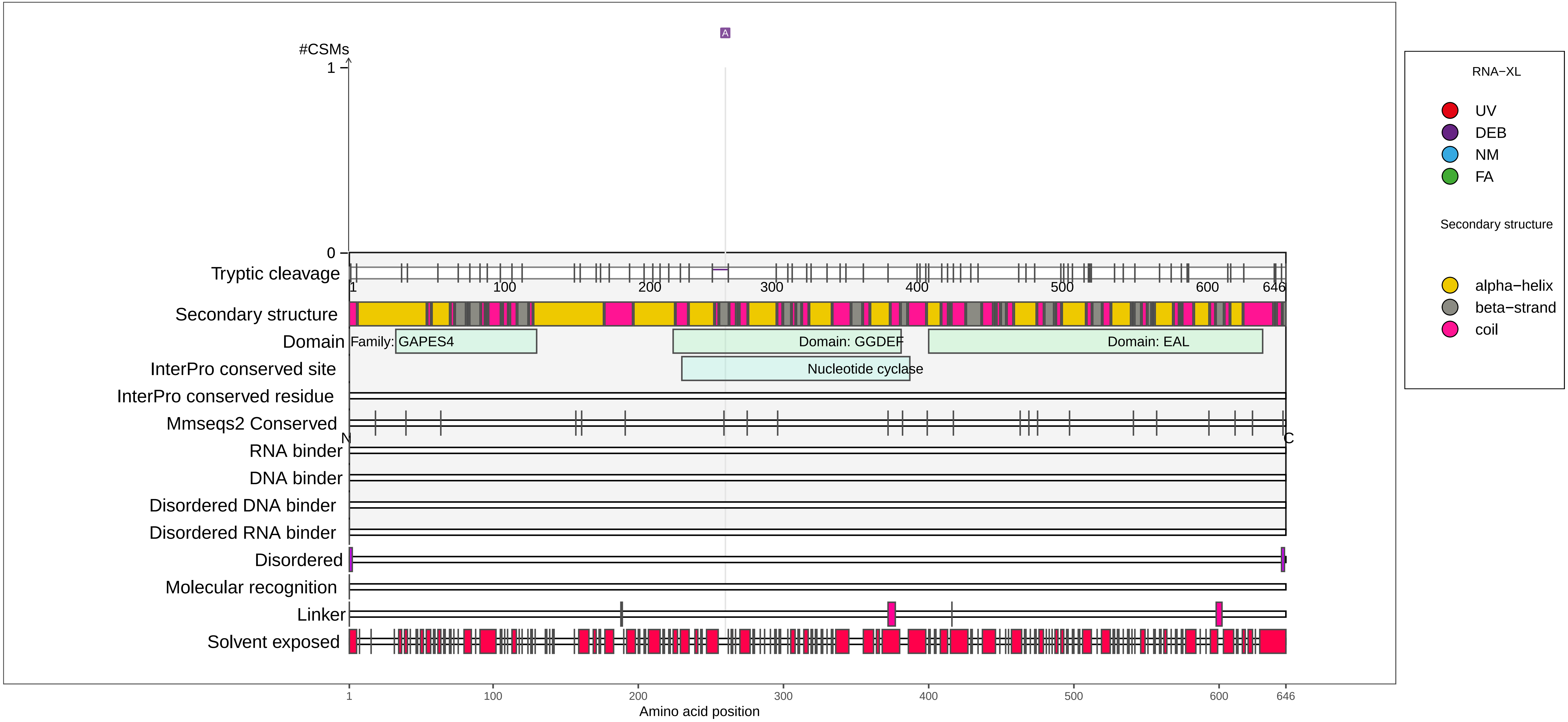

P13738
NHAA_ECOLI Na(+)/H(+) antiporter NhaA

− Abundance:
tryptic [log10 Intensity]: 7.27 (Q 24)
PAXdb K12 strain [ppm]: NA (Q NA)
PAXdb E.coli [ppm]: 0.72 (Q 41)

− RNA functions: not annotated

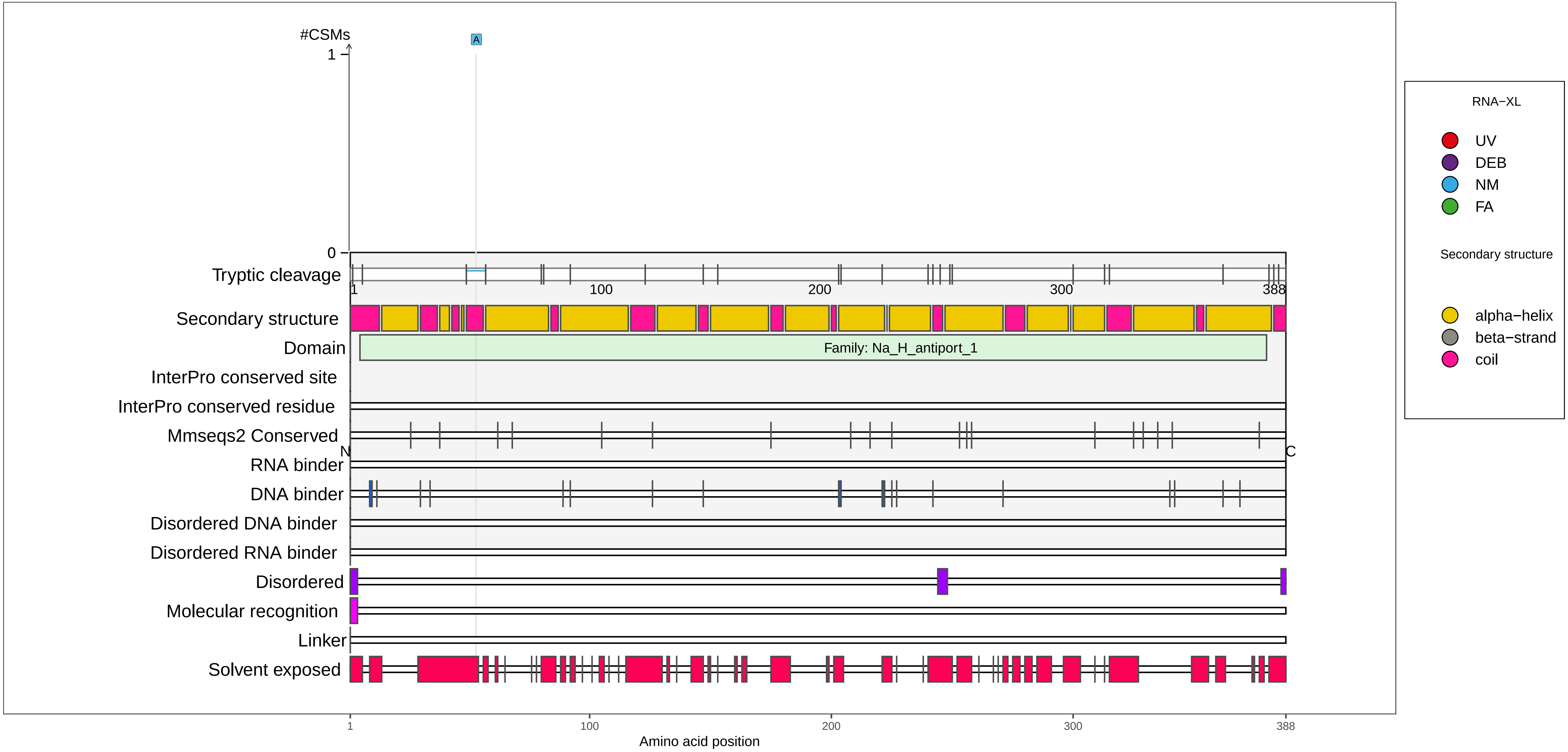

P15028
FECB_ECOLI Fe(3+) dicitrate−binding periplasmic protein

− Abundance:
tryptic [log10 Intensity]: 7.37 (Q 28)
PAXdb K12 strain [ppm]: NA (Q NA)
PAXdb E.coli [ppm]: −0.03 (Q 24)

− RNA functions: not annotated

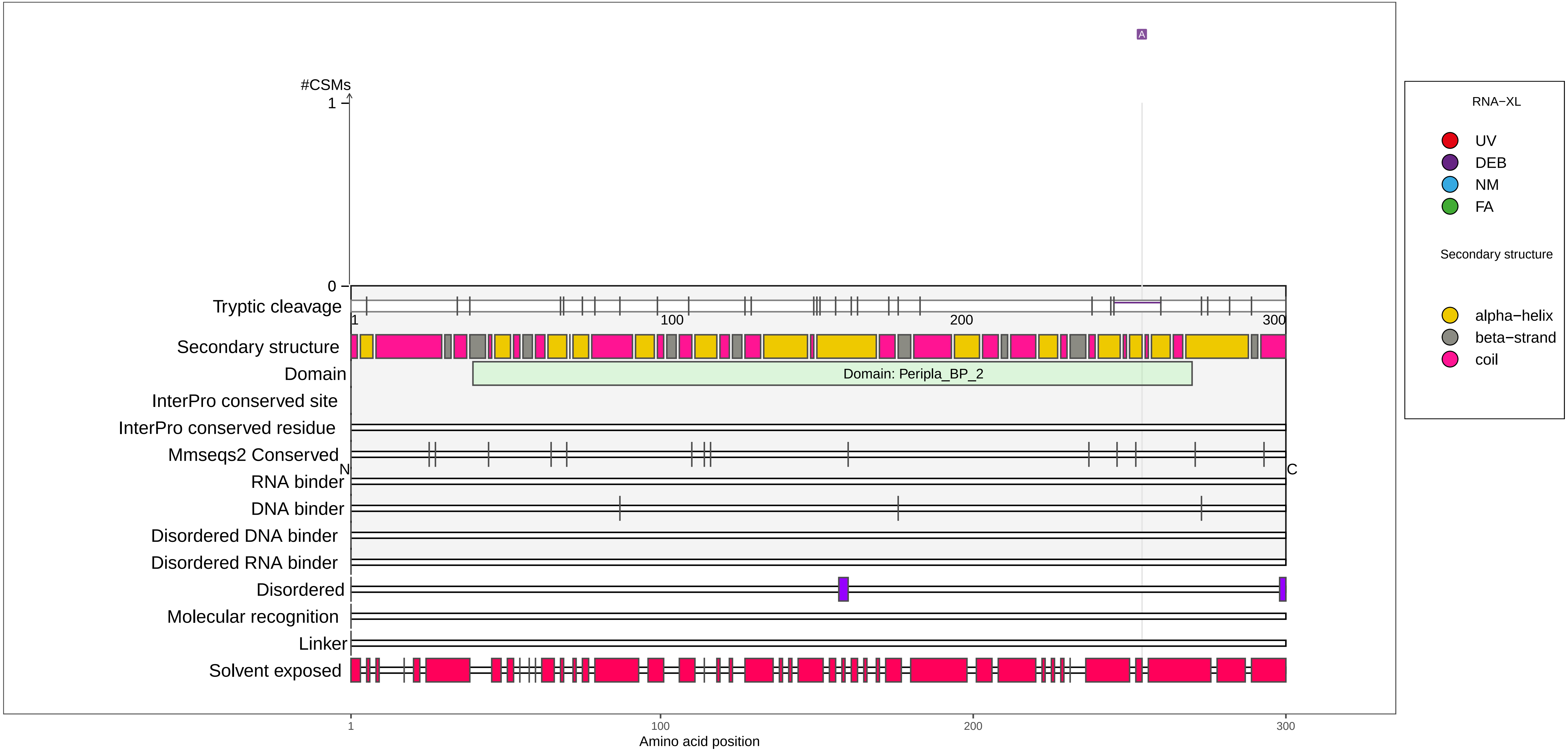

P16456
SELD_ECOLI Selenide, water dikinase

− Abundance:
tryptic [log10 Intensity]: 8.14 (Q 63)
PAXdb K12 strain [ppm]: 2.71 (Q 84)
PAXdb E.coli [ppm]: 2.37 (Q 82)

− RNA functions:
ncRNA metabolic process; ncRNA processing; RNA metabolic process; RNA modification
RNA processing; tRNA metabolic process; tRNA modification; tRNA processing
tRNA seleno−modification

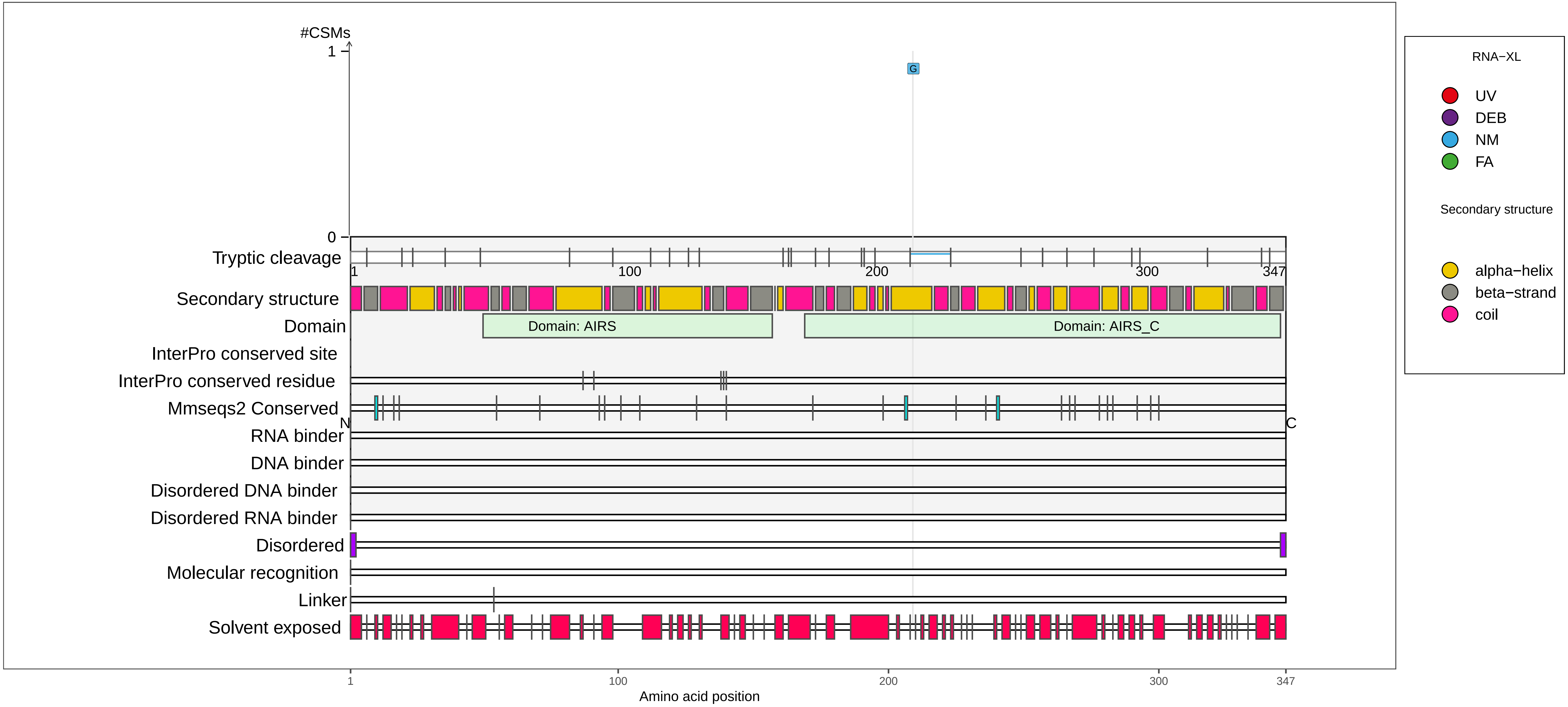

P16681
YJDN_ECOLI Protein YjdN

− Abundance:
tryptic [log10 Intensity]: 6.91 (Q 10)
PAXdb K12 strain [ppm]: 1.83 (Q 54)
PAXdb E.coli [ppm]: 2.53 (Q 86)

− RNA functions: not annotated

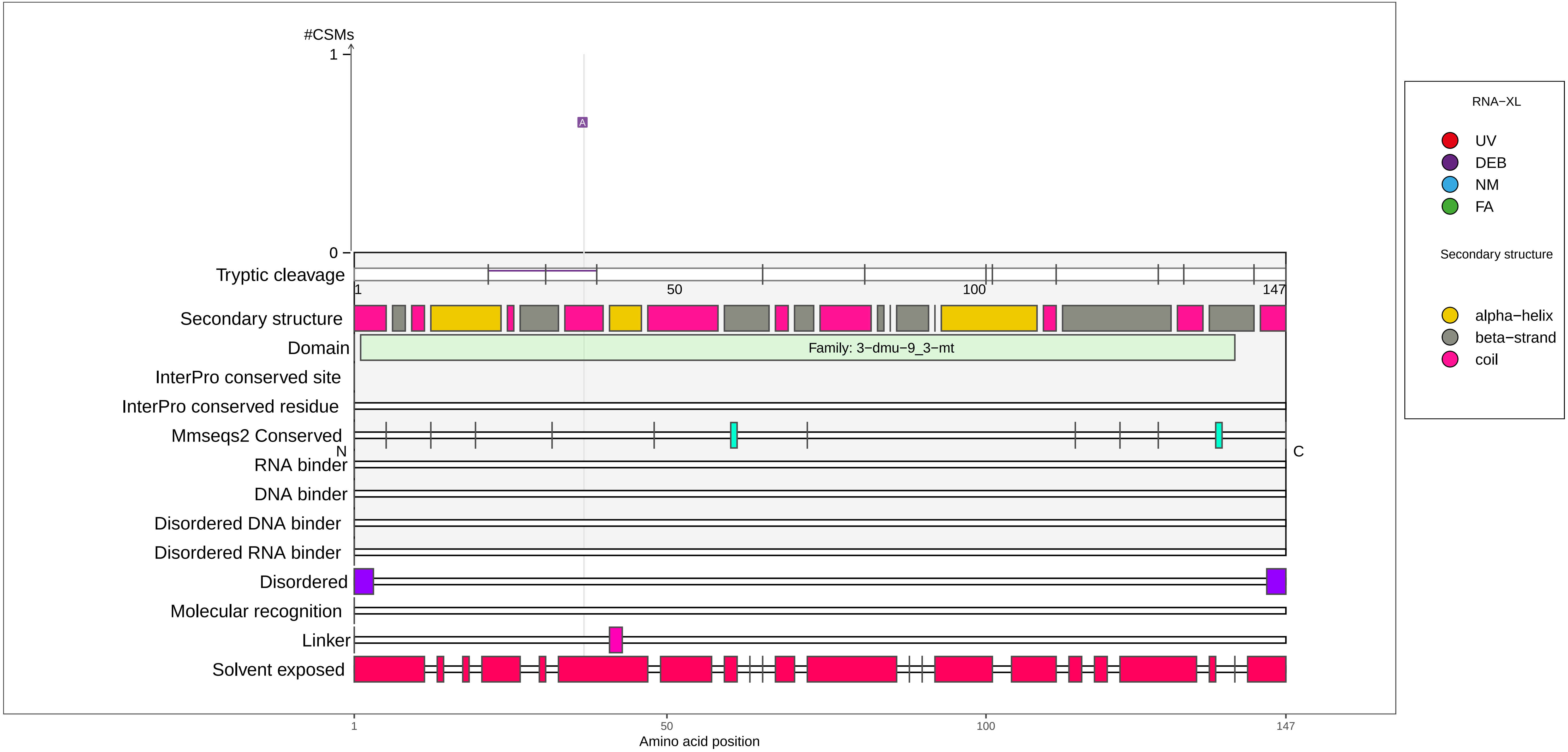

P16688
PHNJ_ECOLI Alpha−D−ribose 1−methylphosphonate 5−phosphate C−P lyase

− Abundance:
tryptic [log10 Intensity]: NA (Q NA)
PAXdb K12 strain [ppm]: NA (Q NA)
PAXdb E.coli [ppm]: 1.59 (Q 62)

− RNA functions: not annotated

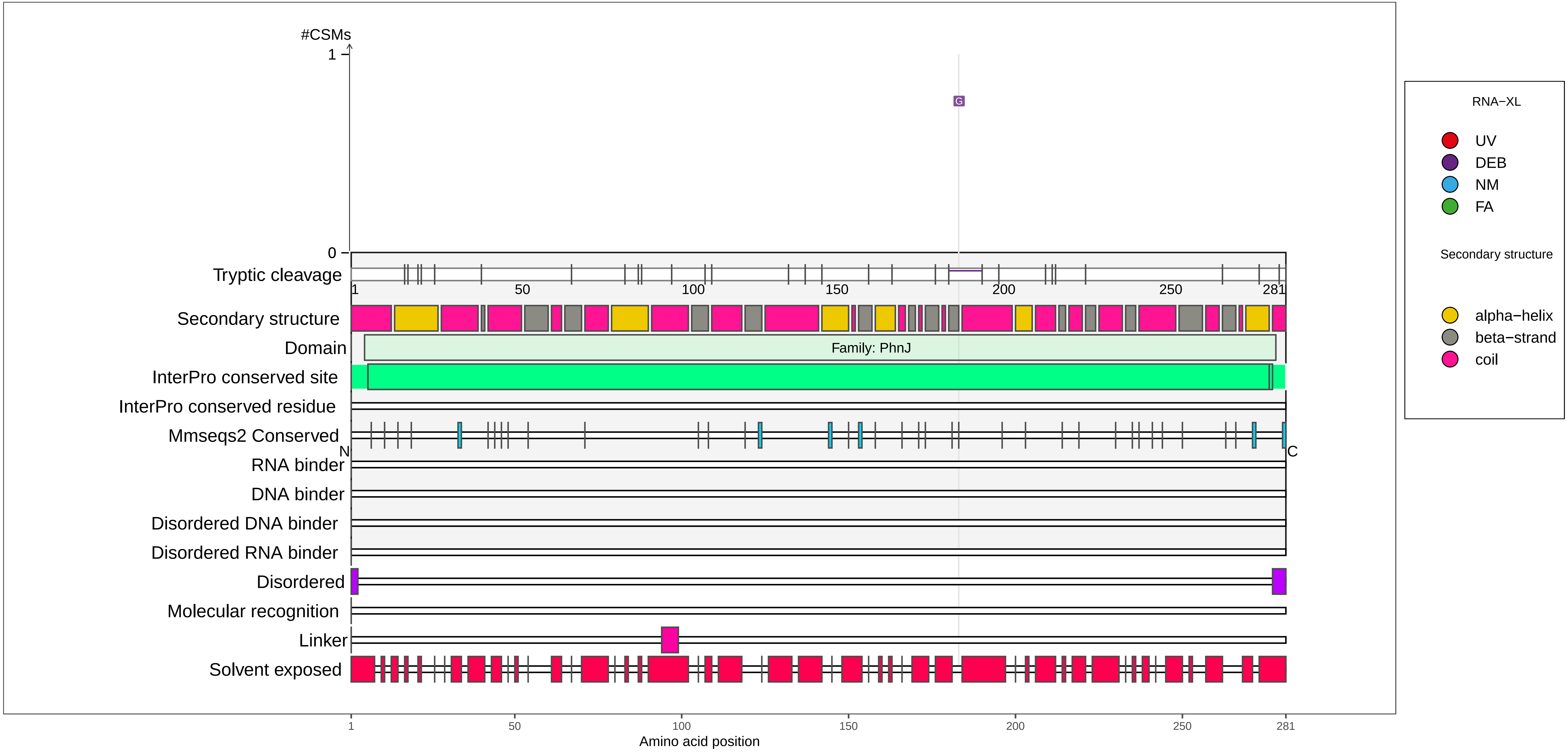

P17411
CHBF_ECOLI 6−phospho−beta−glucosidase

− Abundance:
tryptic [log10 Intensity]: NA (Q NA)
PAXdb K12 strain [ppm]: 1.25 (Q 23)
PAXdb E.coli [ppm]: −0.72 (Q 12)

− RNA functions: not annotated

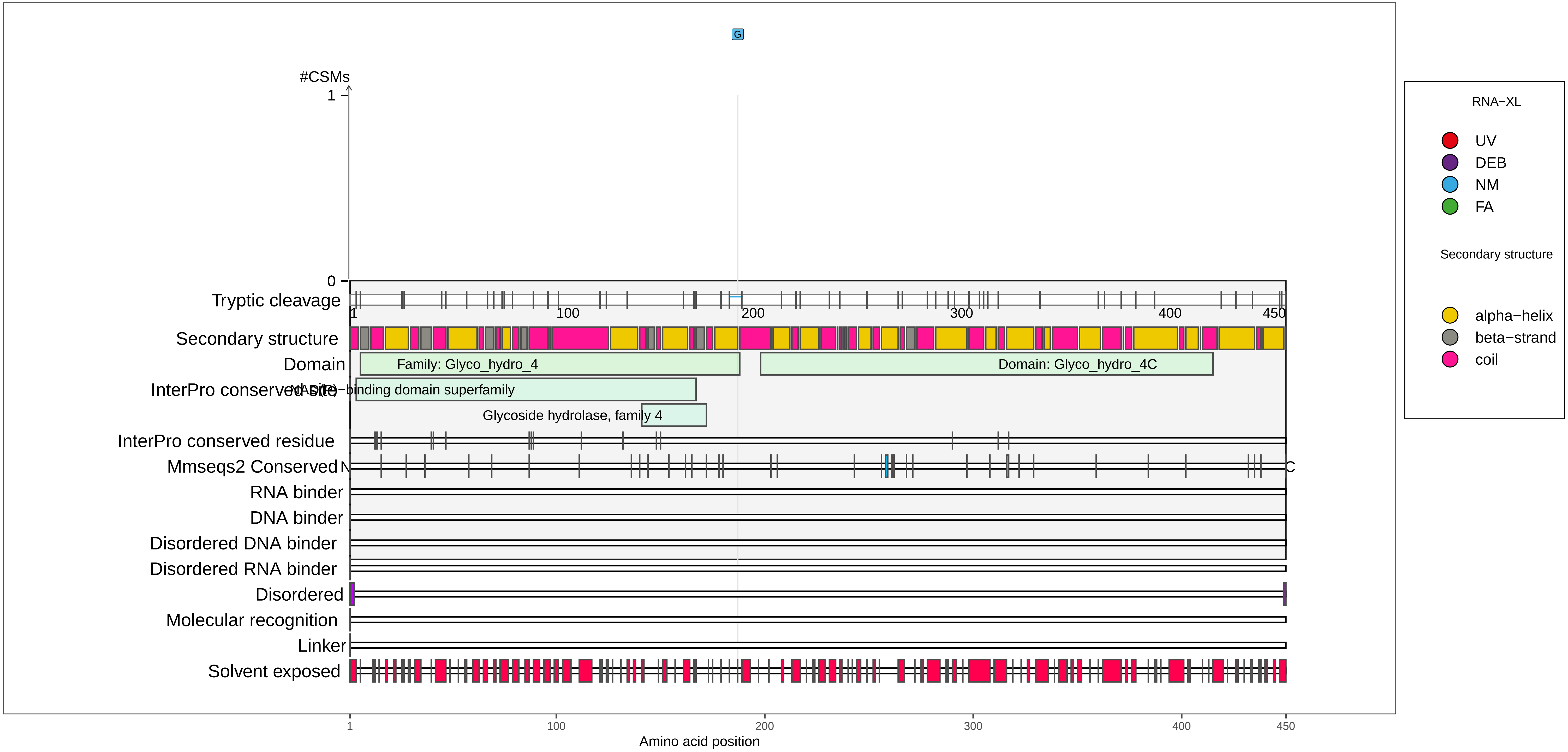

P17993
UBIG_ECOLI Ubiquinone biosynthesis O−methyltransferase

− Abundance:
tryptic [log10 Intensity]: 7.72 (Q 45)
PAXdb K12 strain [ppm]: 2.52 (Q 78)
PAXdb E.coli [ppm]: 2.06 (Q 74)

− RNA functions: not annotated

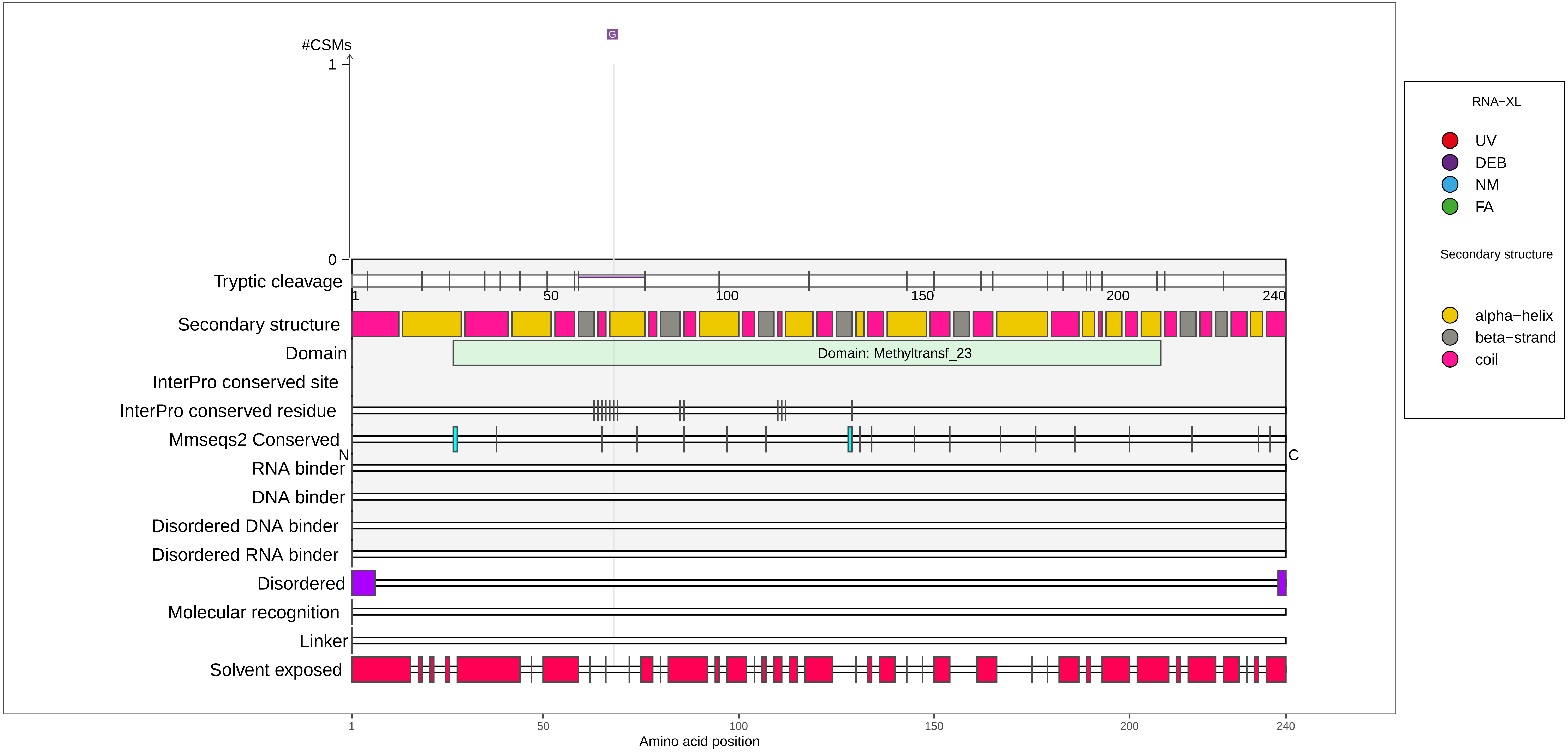

P18335
ARGD_ECOLI Acetylornithine/succinyldiaminopimelate aminotransferase

− Abundance:
tryptic [log10 Intensity]: 8.53 (Q 77)
PAXdb K12 strain [ppm]: 2.74 (Q 84)
PAXdb E.coli [ppm]: 2.51 (Q 86)

− RNA functions: not annotated

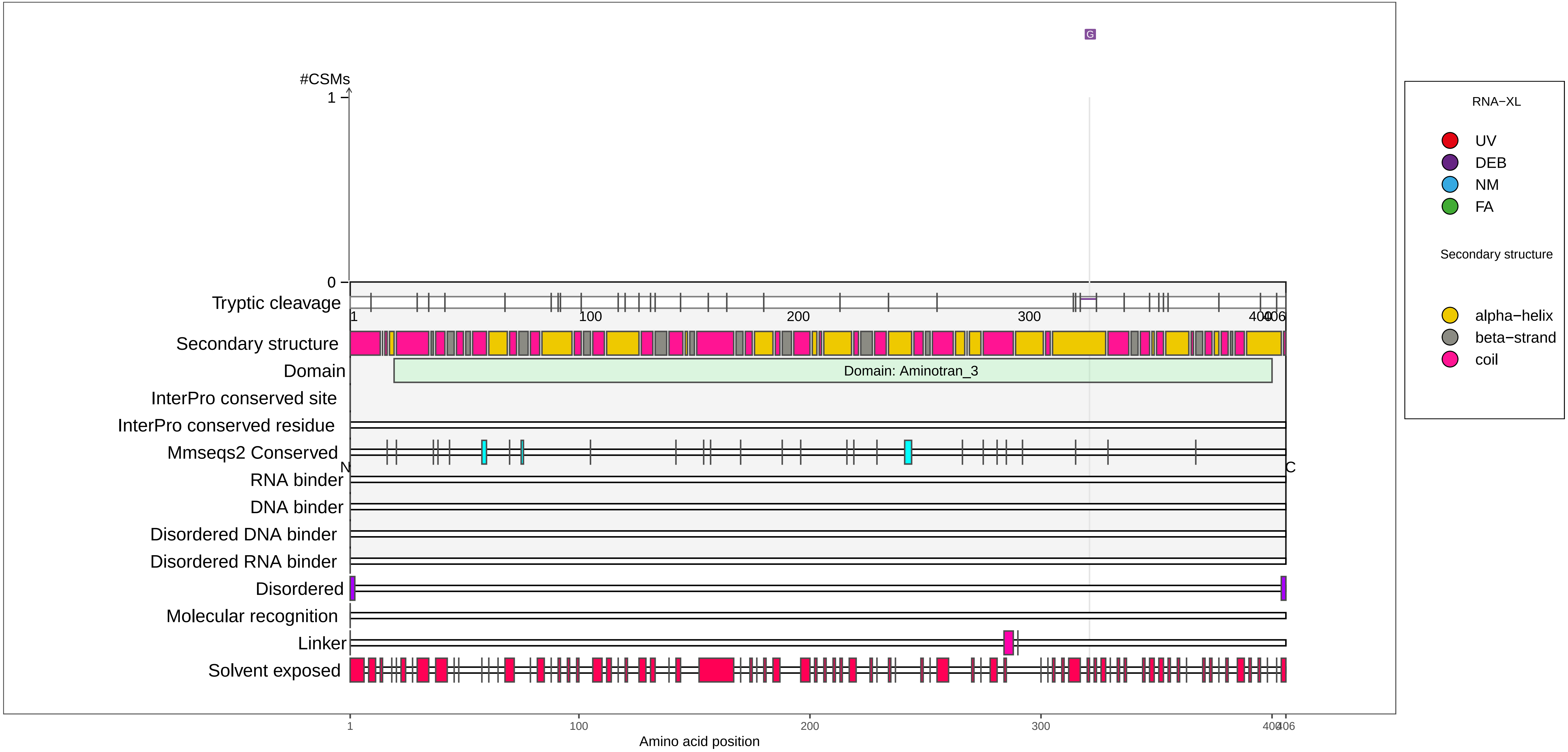

P21345
GLTP_ECOLI Proton/glutamate−aspartate symporter

− Abundance:
tryptic [log10 Intensity]: NA (Q NA)
PAXdb K12 strain [ppm]: NA (Q NA)
PAXdb E.coli [ppm]: 0.58 (Q 38)

− RNA functions: not annotated

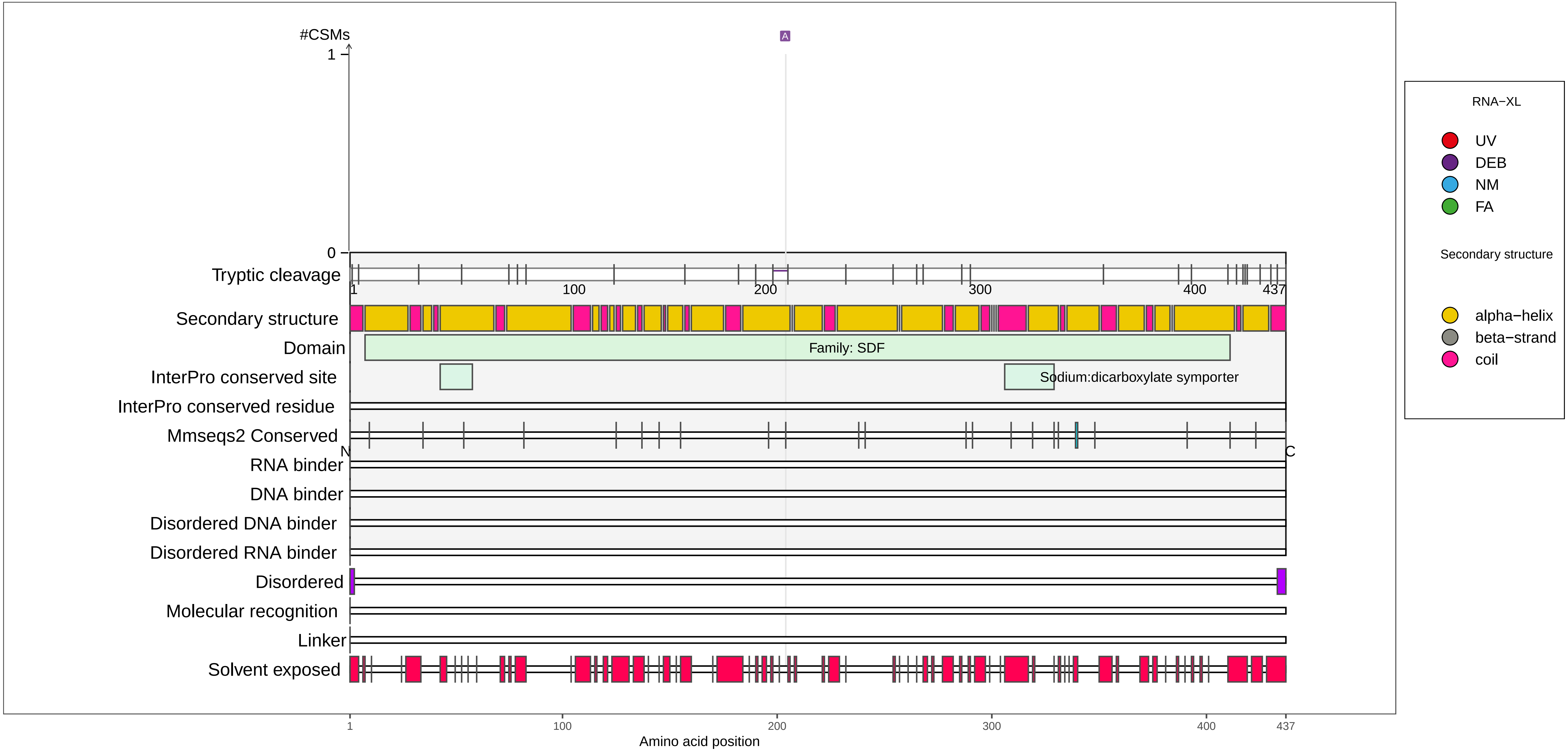

P22106
ASNB_ECOLI Asparagine synthetase B [glutamine−hydrolyzing]

− Abundance:
tryptic [log10 Intensity]: 8.19 (Q 65)
PAXdb K12 strain [ppm]: 2.44 (Q 76)
PAXdb E.coli [ppm]: 2.29 (Q 80)

− RNA functions: not annotated

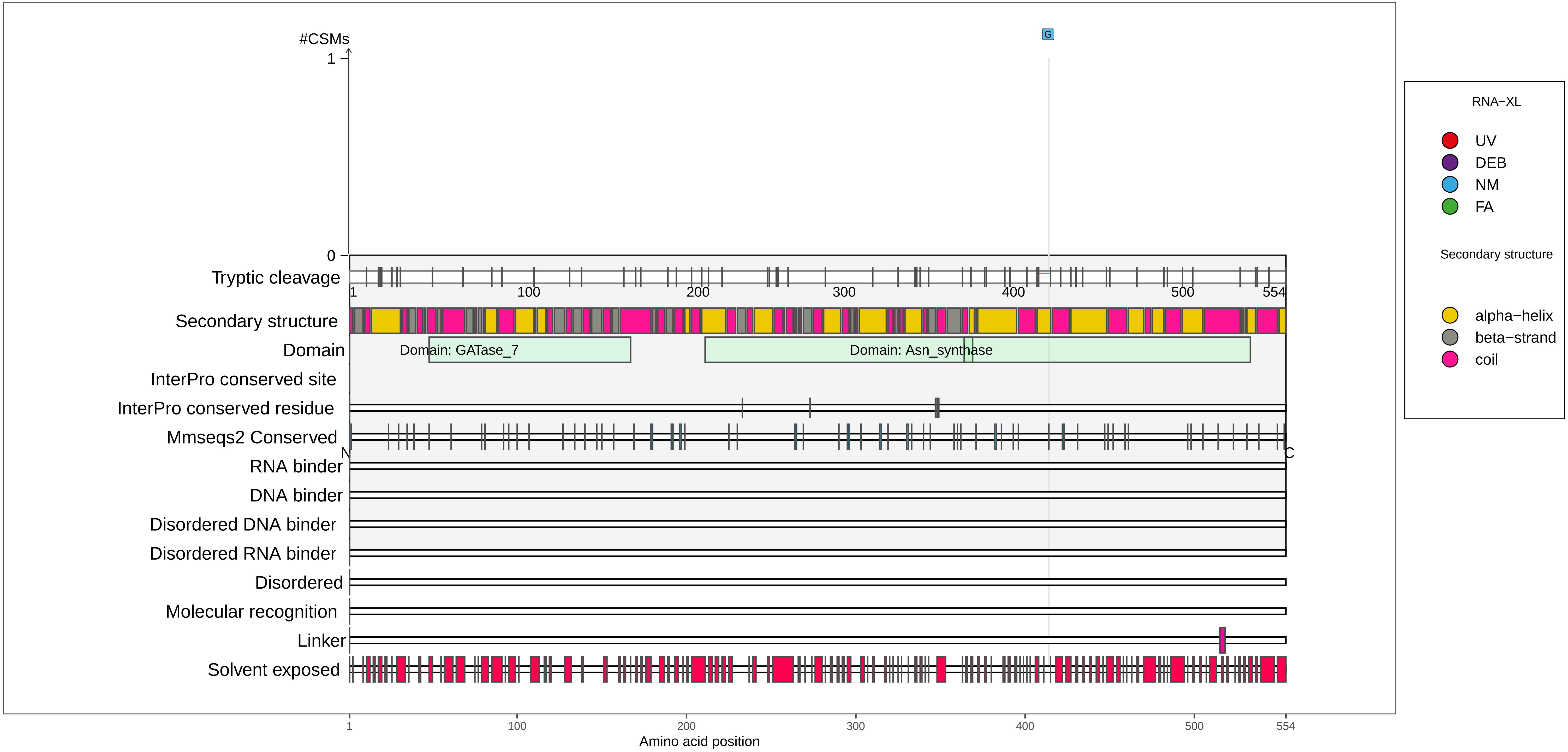

P22188
MURE_ECOLI UDP−N−acetylmuramoyl−L−alanyl−D−glutamate−−2,6−diaminopimelate ligase

− Abundance:
tryptic [log10 Intensity]: 8.53 (Q 76)
PAXdb K12 strain [ppm]: 1.94 (Q 58)
PAXdb E.coli [ppm]: 2.14 (Q 76)

− RNA functions: not annotated

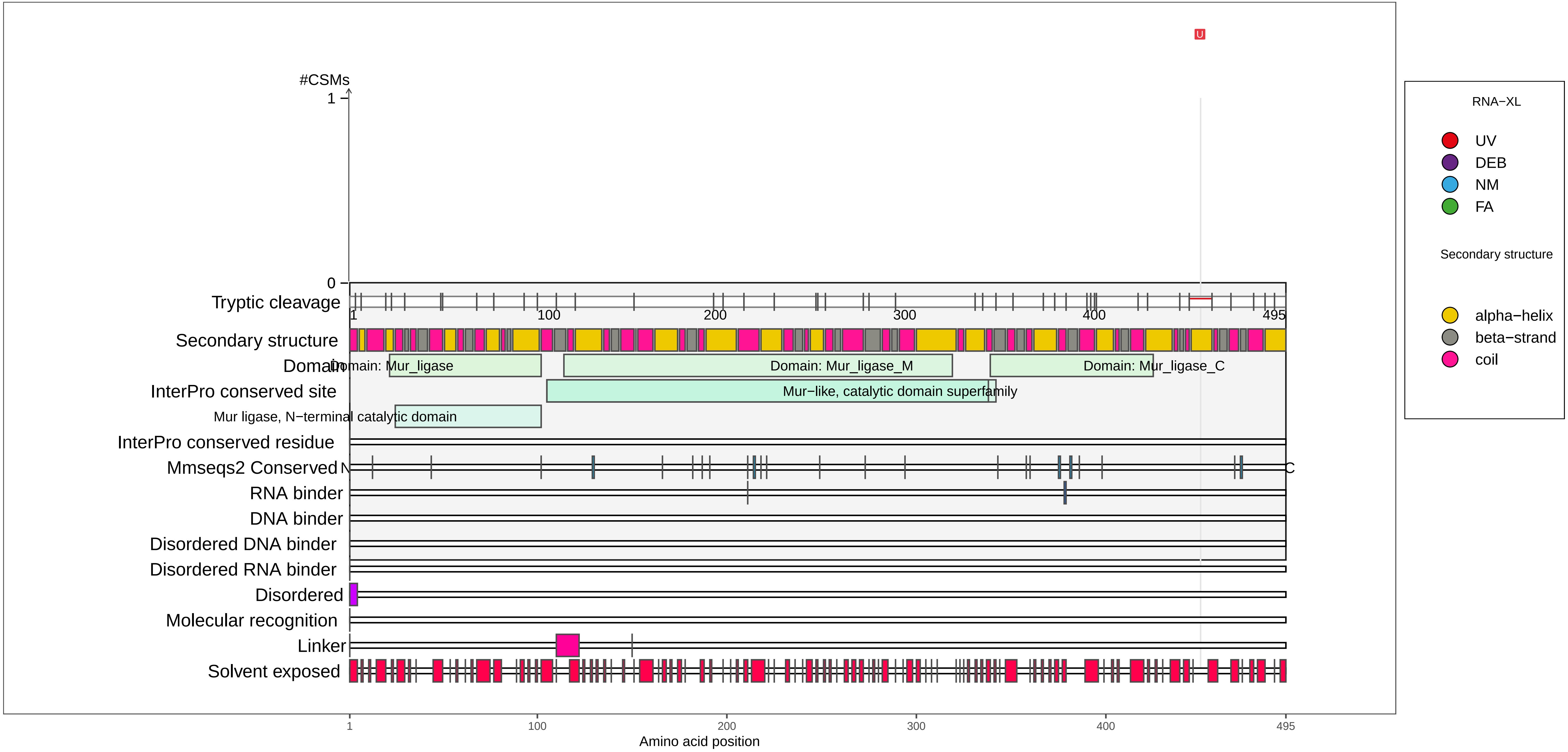

P23857
PSPE_ECOLI Thiosulfate sulfurtransferase PspE

− Abundance:
tryptic [log10 Intensity]: 9.21 (Q 93)
PAXdb K12 strain [ppm]: 2.94 (Q 89)
PAXdb E.coli [ppm]: 2.43 (Q 84)

− RNA functions: not annotated

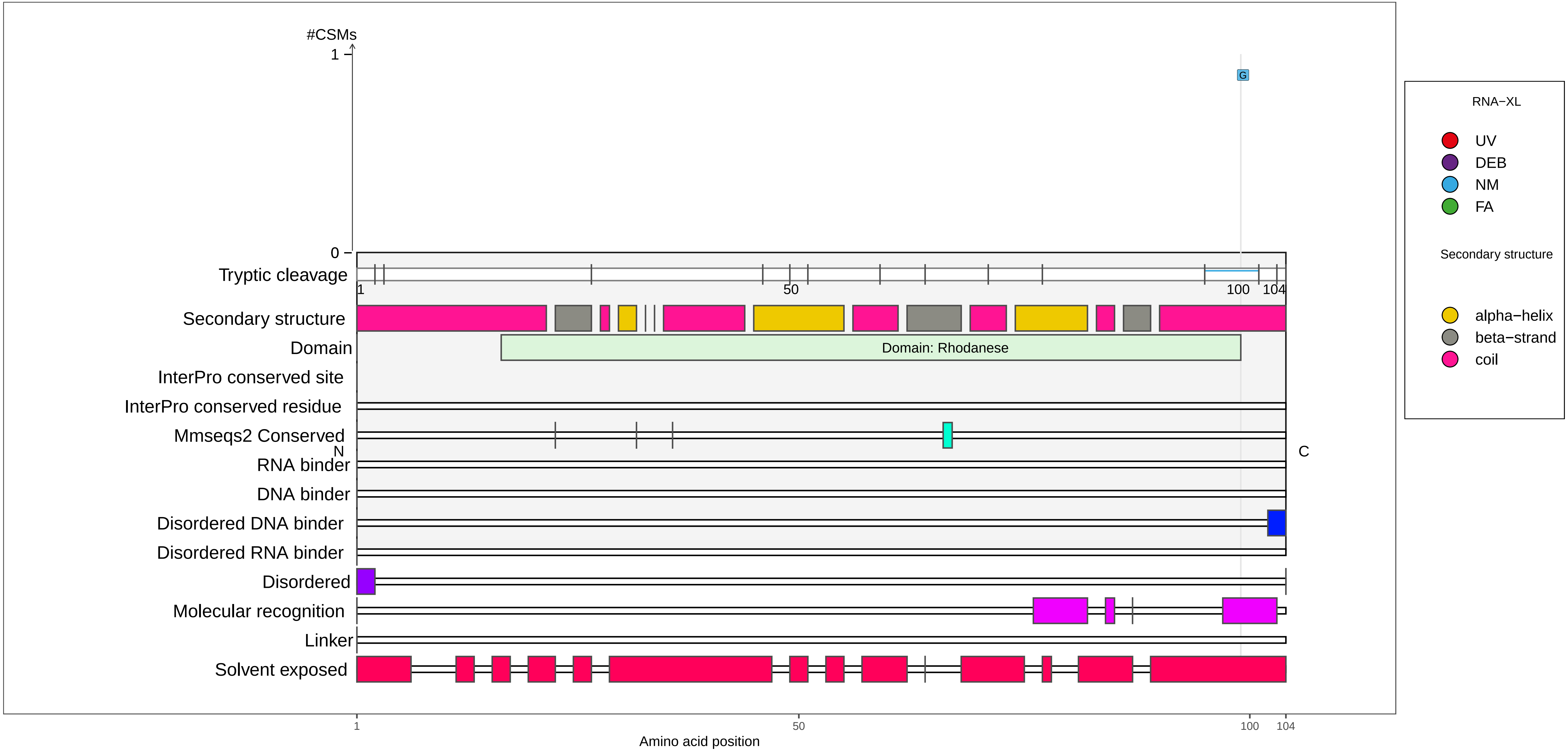

P23871
HEMH_ECOLI Ferrochelatase

− Abundance:
tryptic [log10 Intensity]: 7.02 (Q 14)
PAXdb K12 strain [ppm]: NA (Q NA)
PAXdb E.coli [ppm]: 0.71 (Q 41)

− RNA functions: not annotated

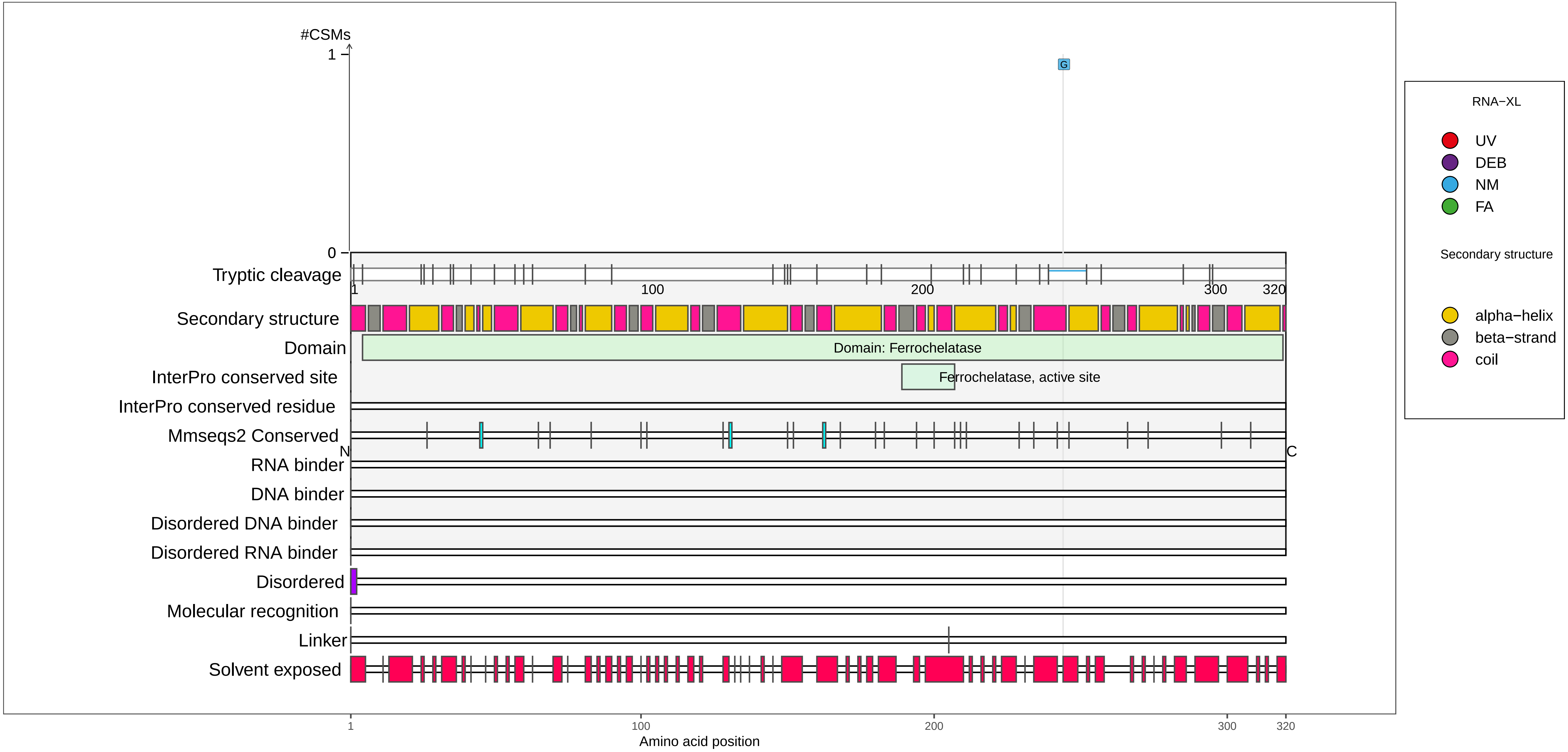

P24202
MRR_ECOLI Type IV methyl−directed restriction enzyme EcoKMrr

− Abundance:
tryptic [log10 Intensity]: 8.06 (Q 60)
PAXdb K12 strain [ppm]: 1.47 (Q 36)
PAXdb E.coli [ppm]: 1.24 (Q 53)

− RNA functions: not annotated

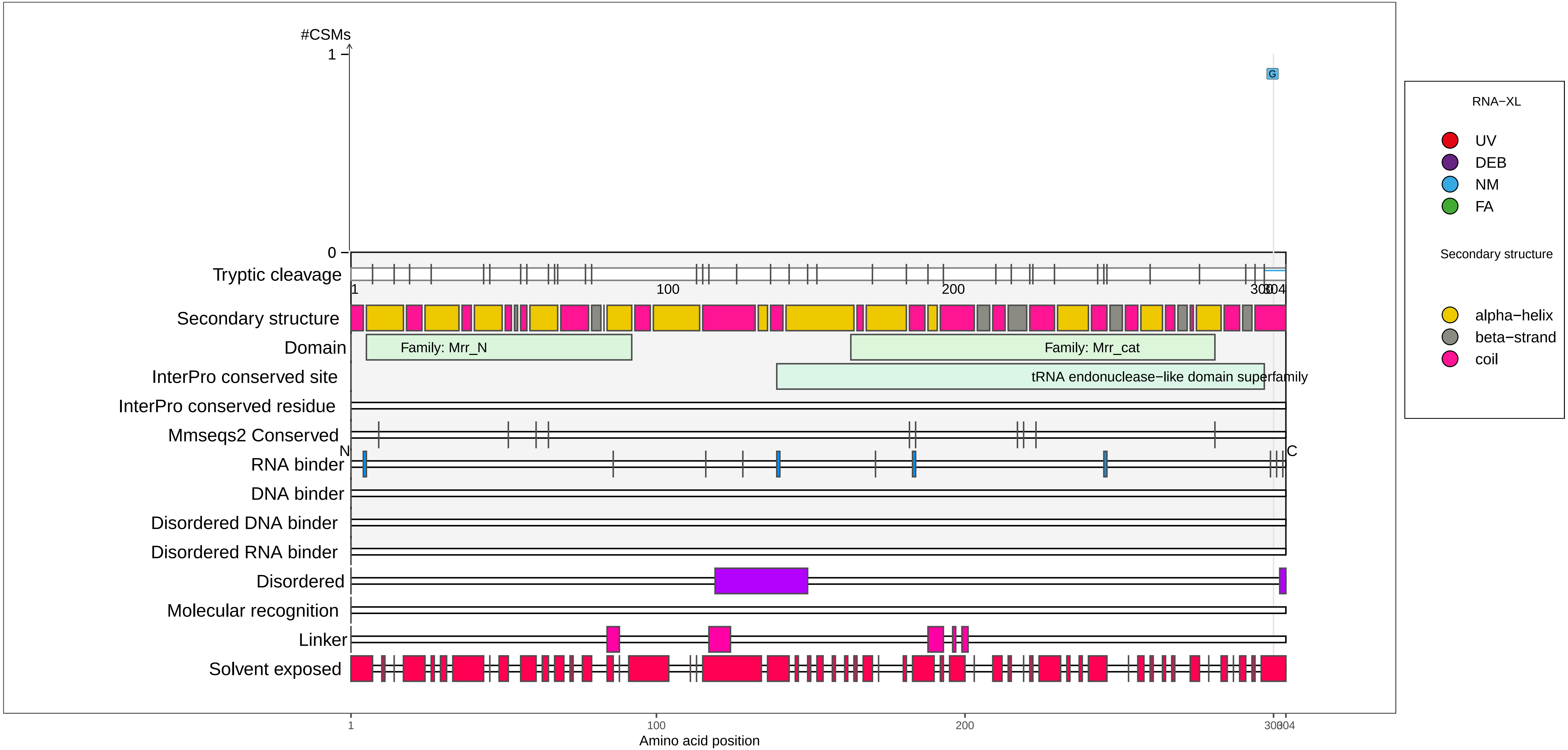

P24215
UXUA_ECOLI Mannonate dehydratase

− Abundance:
tryptic [log10 Intensity]: 7.69 (Q 44)
PAXdb K12 strain [ppm]: 2.21 (Q 67)
PAXdb E.coli [ppm]: 2.17 (Q 77)

− RNA functions: not annotated

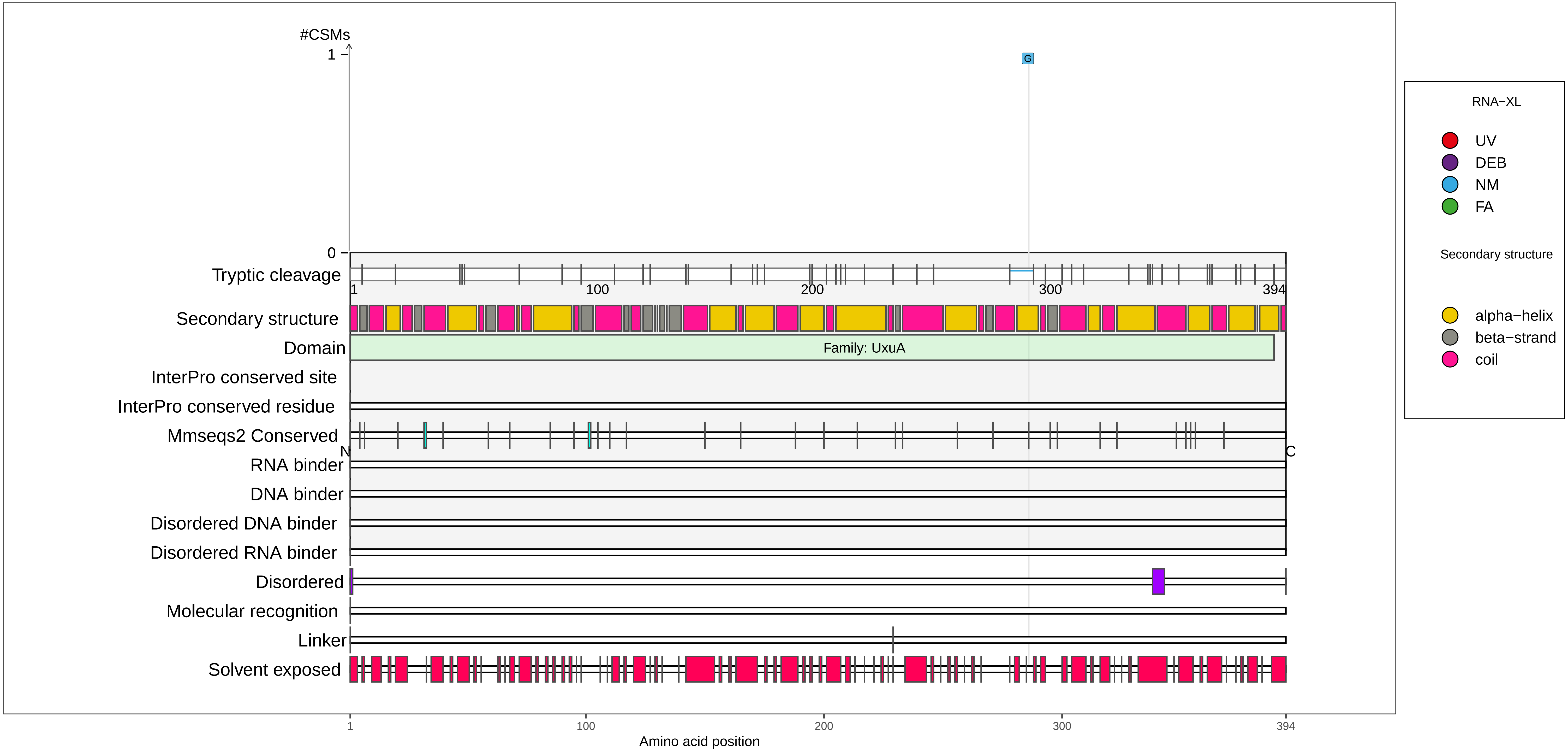

P24218
INTD_ECOLI Prophage integrase IntD

− Abundance:
tryptic [log10 Intensity]: NA (Q NA)
PAXdb K12 strain [ppm]: NA (Q NA)
PAXdb E.coli [ppm]: 0.75 (Q 42)

− RNA functions: not annotated

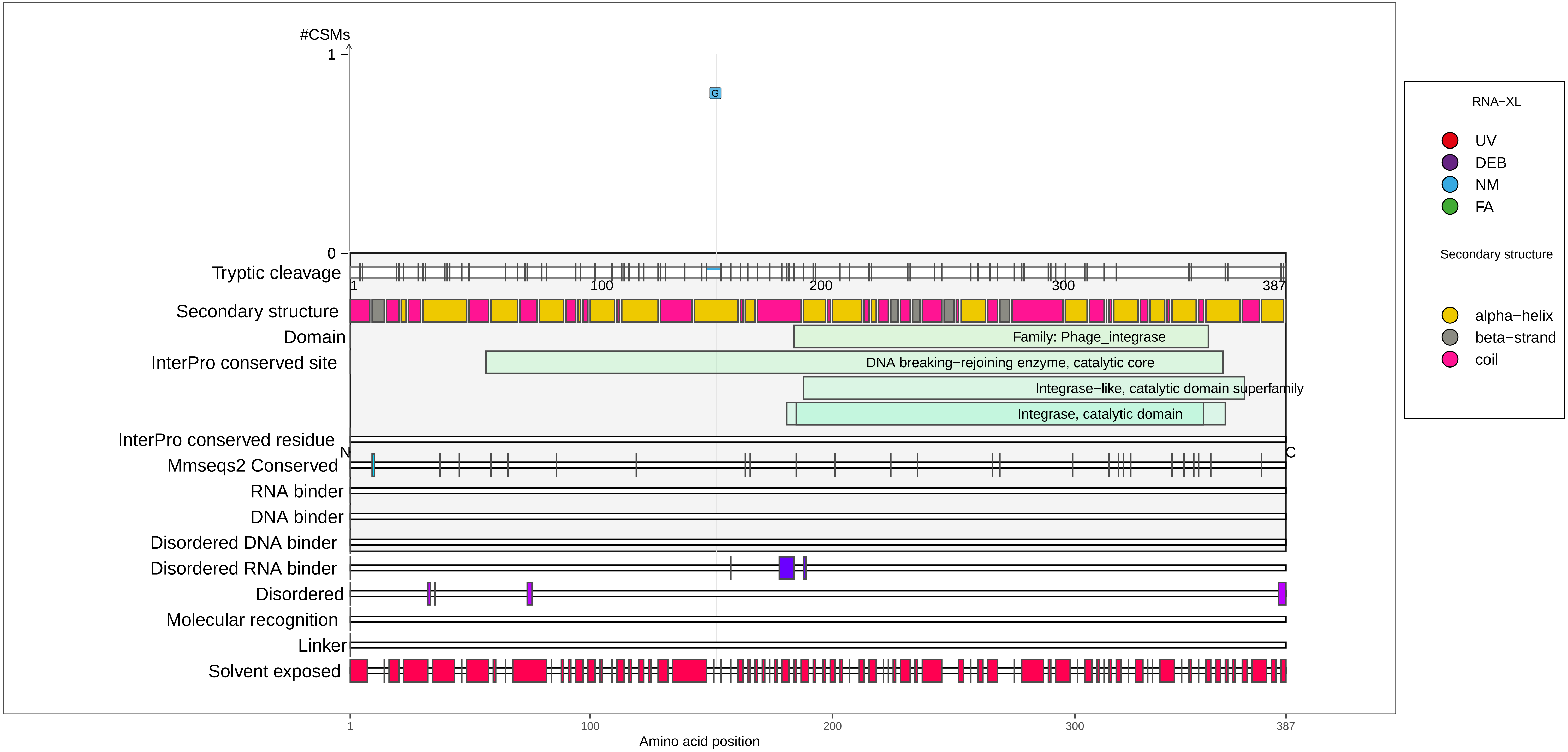

P24230
RECG_ECOLI ATP−dependent DNA helicase RecG

− Abundance:
tryptic [log10 Intensity]: 6.79 (Q 7)
PAXdb K12 strain [ppm]: 1.24 (Q 22)
PAXdb E.coli [ppm]: 1.23 (Q 53)

− RNA functions: not annotated

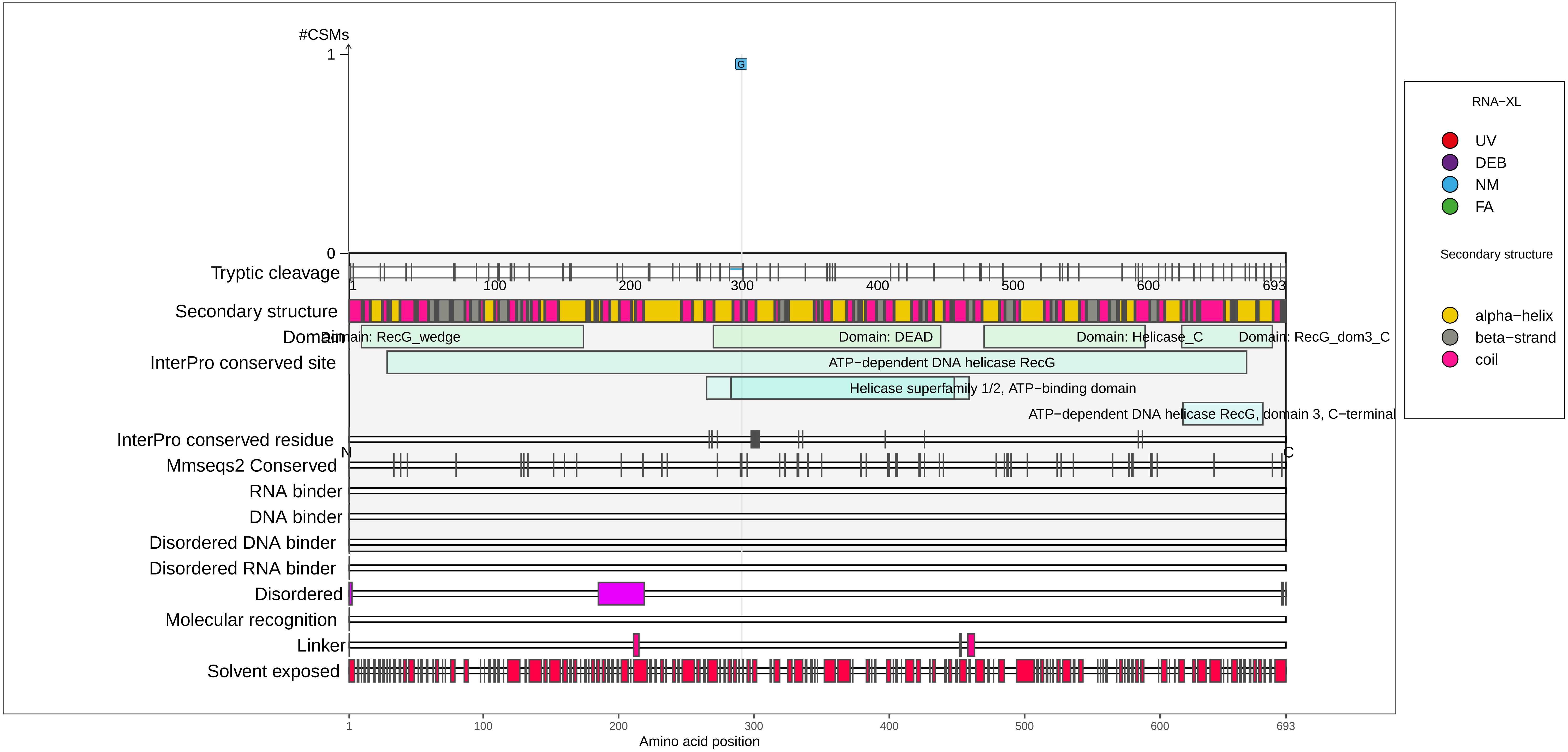

P24251
CRL_ECOLI Sigma factor−binding protein Crl
− RNA functions: not annotated

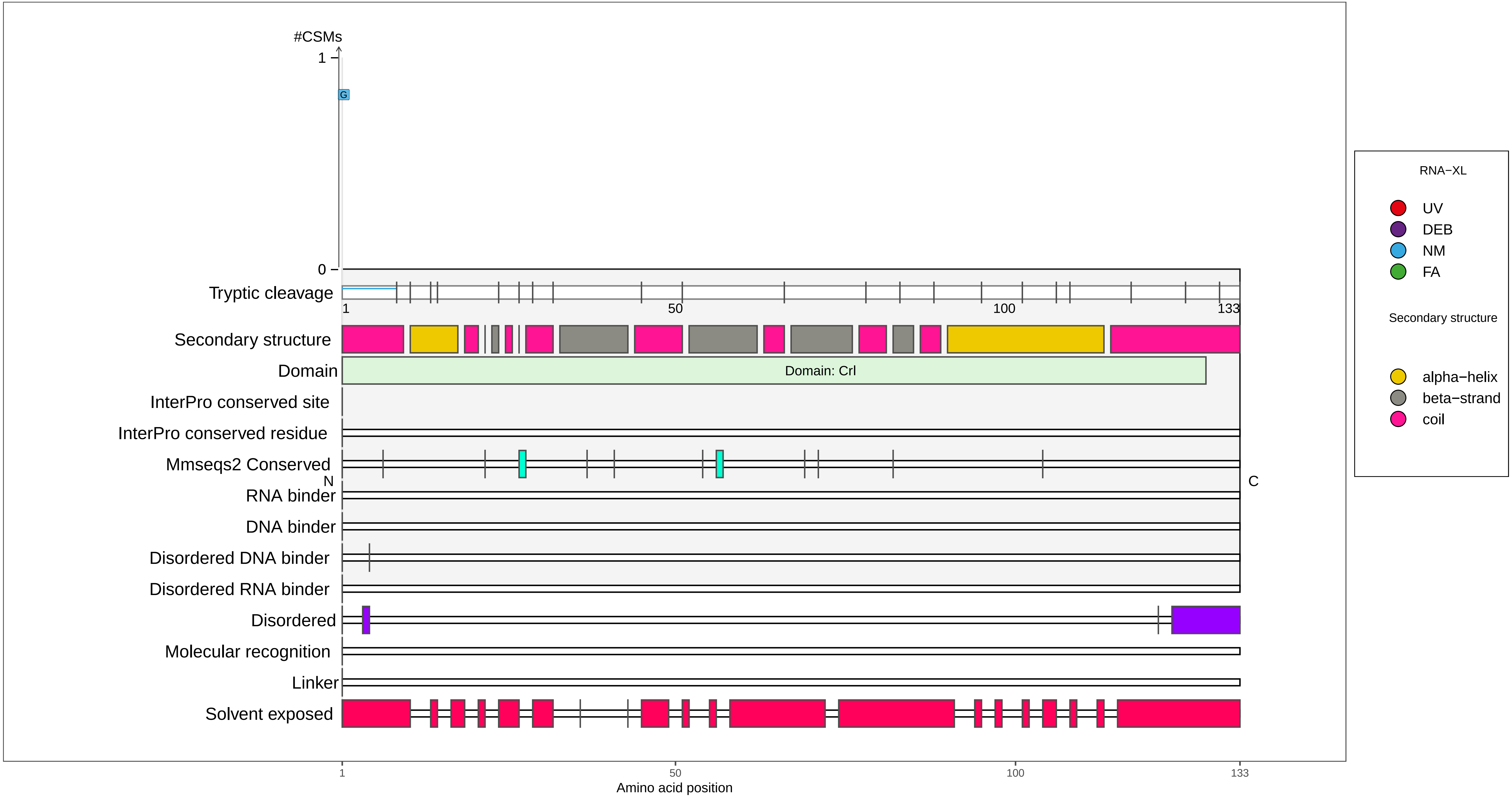

P24255
RP54_ECOLI RNA polymerase sigma−54 factor

− Abundance:
tryptic [log10 Intensity]: 7.58 (Q 39)
PAXdb K12 strain [ppm]: 1.61 (Q 45)
PAXdb E.coli [ppm]: 2.06 (Q 74)

− RNA functions:
5−3 RNA polymerase activity; DNA−directed 5−3 RNA polymerase activity
RNA biosynthetic process; RNA metabolic process; RNA polymerase activity

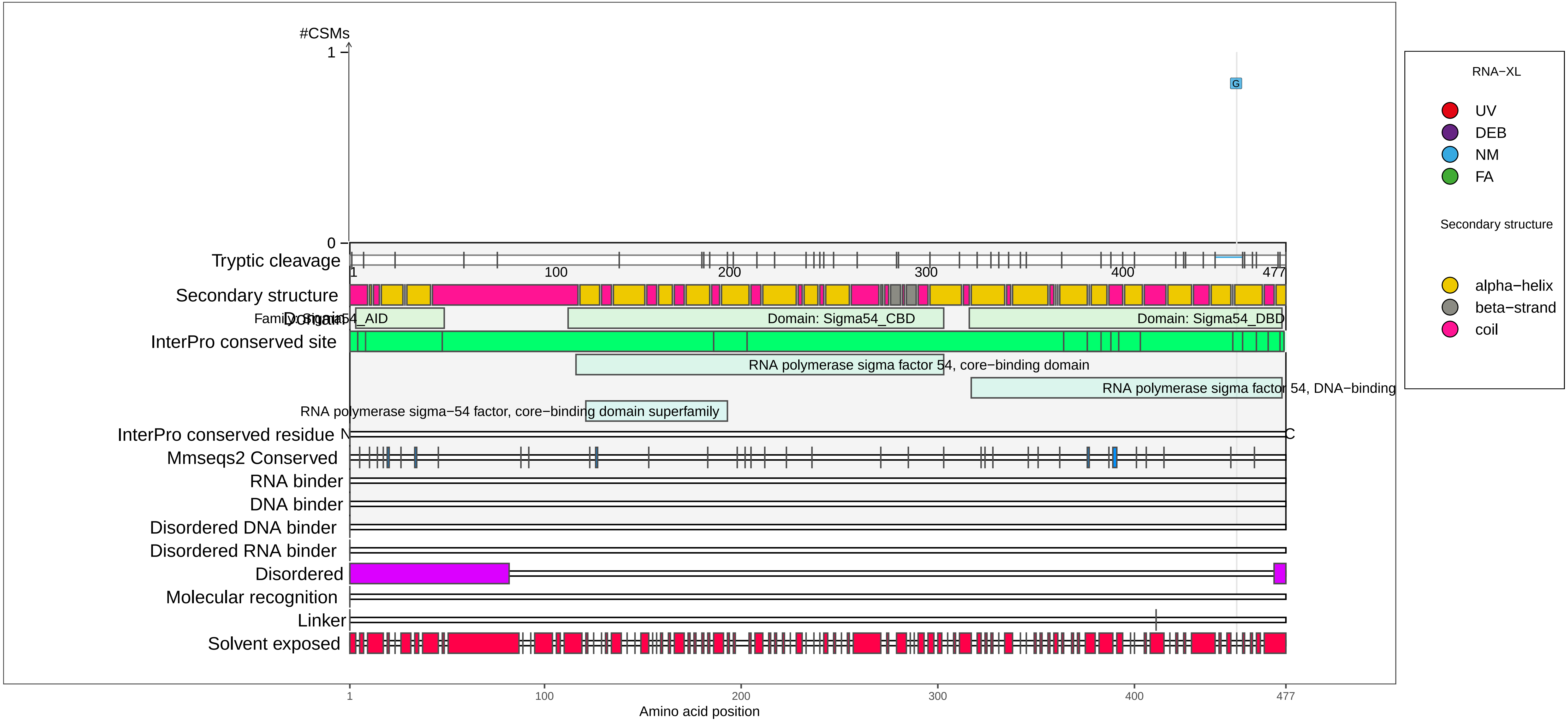

P25437
FRMA_ECOLI S−(hydroxymethyl)glutathione dehydrogenase

− Abundance:
tryptic [log10 Intensity]: 7.85 (Q 51)
PAXdb K12 strain [ppm]: 2.3 (Q 70)
PAXdb E.coli [ppm]: 2.25 (Q 79)

− RNA functions: not annotated

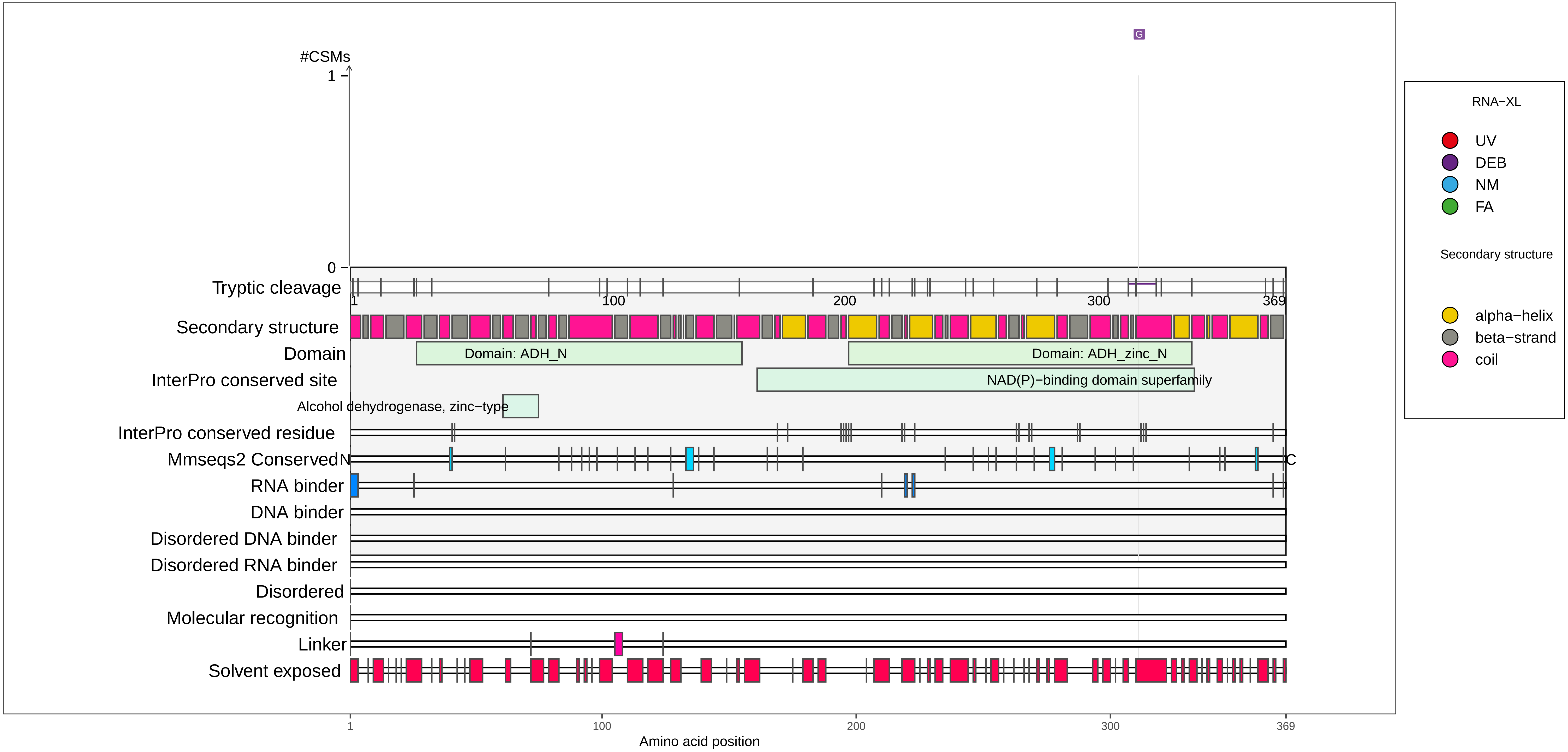

P25535
UBII_ECOLI 2−octaprenylphenol hydroxylase

− Abundance:
tryptic [log10 Intensity]: 7.67 (Q 43)
PAXdb K12 strain [ppm]: 1.67 (Q 48)
PAXdb E.coli [ppm]: 1.81 (Q 67)

− RNA functions: not annotated

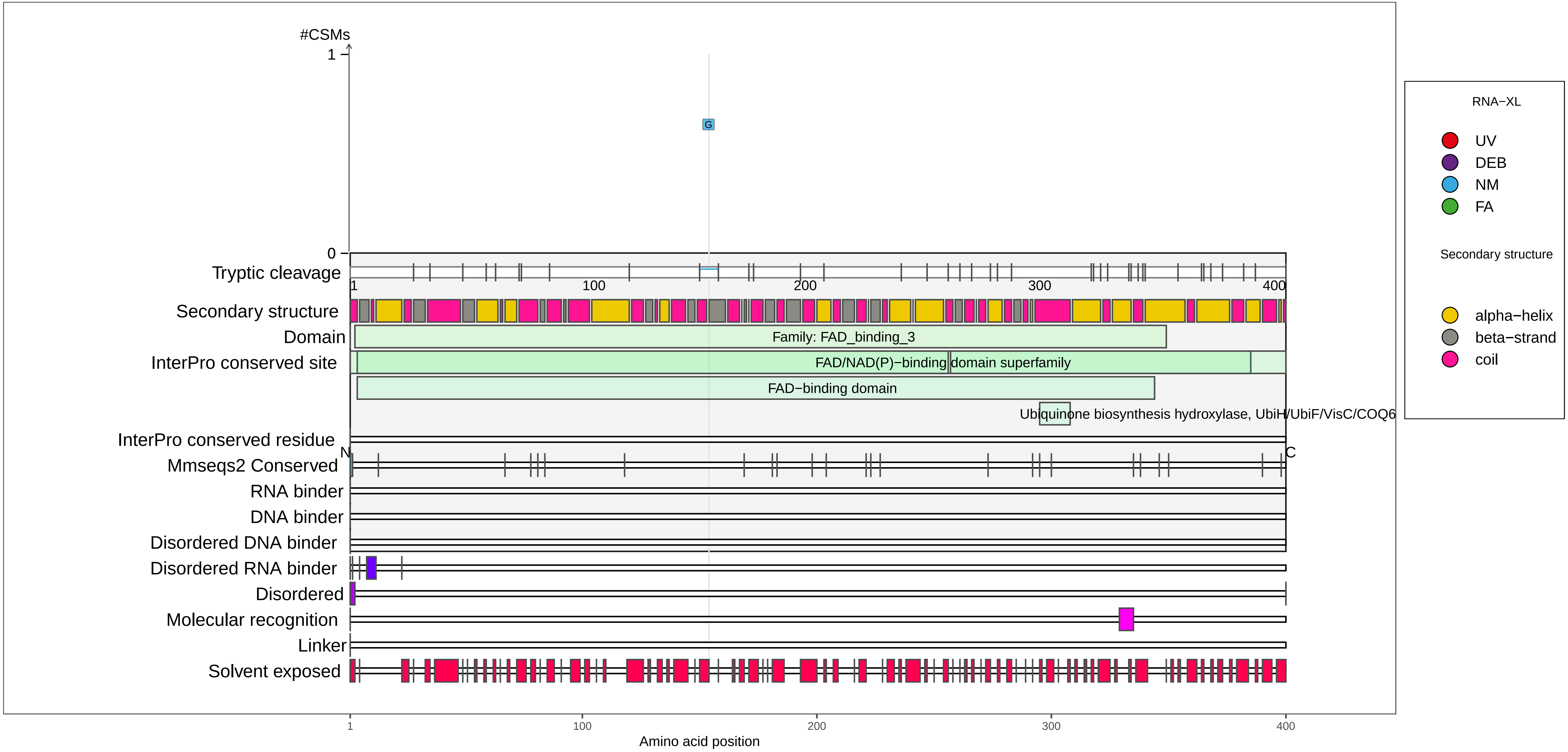

P25741
RFAP_ECOLI Lipopolysaccharide core heptose(I) kinase RfaP

− Abundance:
tryptic [log10 Intensity]: NA (Q NA)
PAXdb K12 strain [ppm]: 1.66 (Q 48)
PAXdb E.coli [ppm]: 0.8 (Q 43)

− RNA functions: not annotated

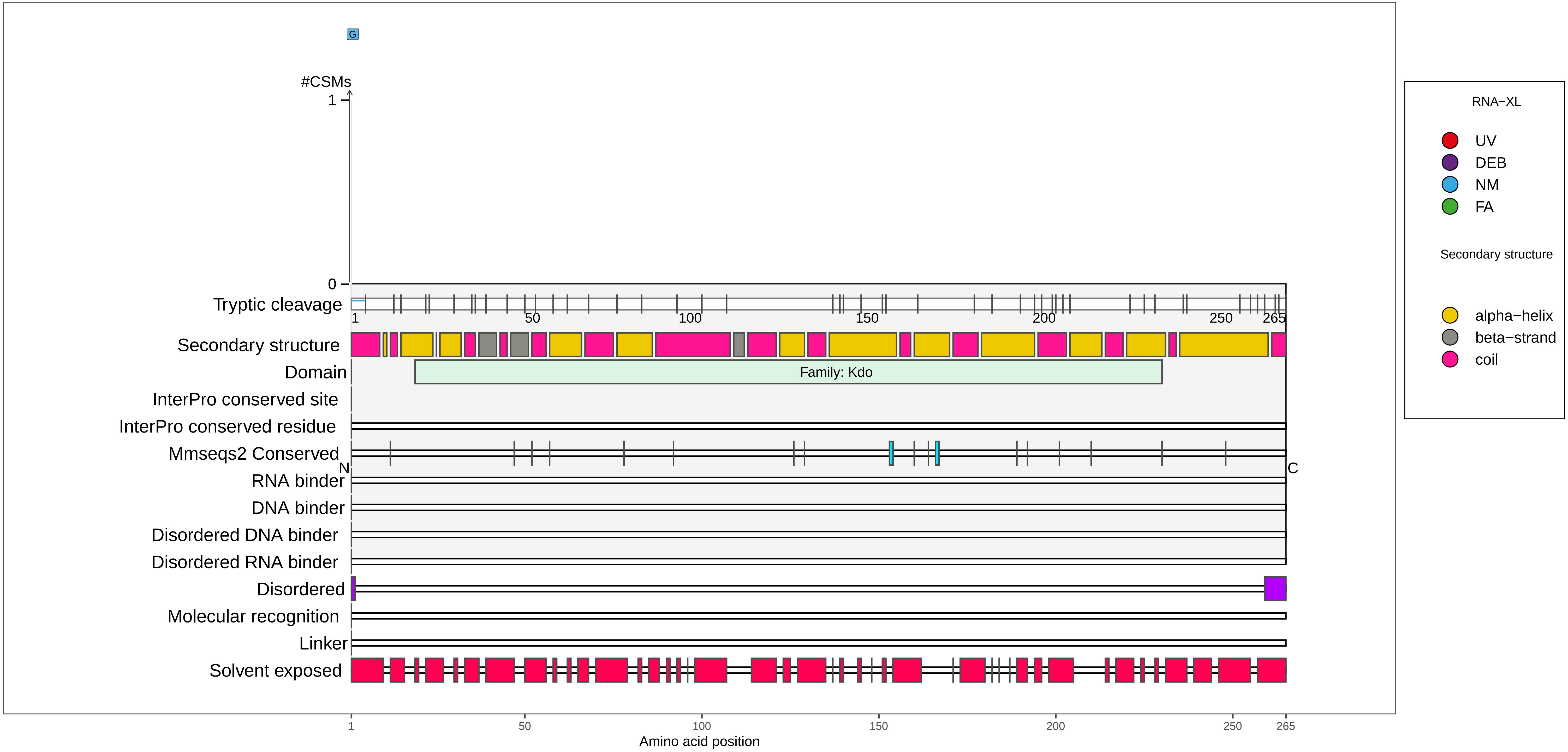

P25889

PREA_ECOLI NAD−dependent dihydropyrimidine dehydrogenase subunit PreA

− Abundance:
tryptic [log10 Intensity]: 7.86 (Q 52)
PAXdb K12 strain [ppm]: NA (Q NA)
PAXdb E.coli [ppm]: 1.57 (Q 62)

− RNA functions: not annotated

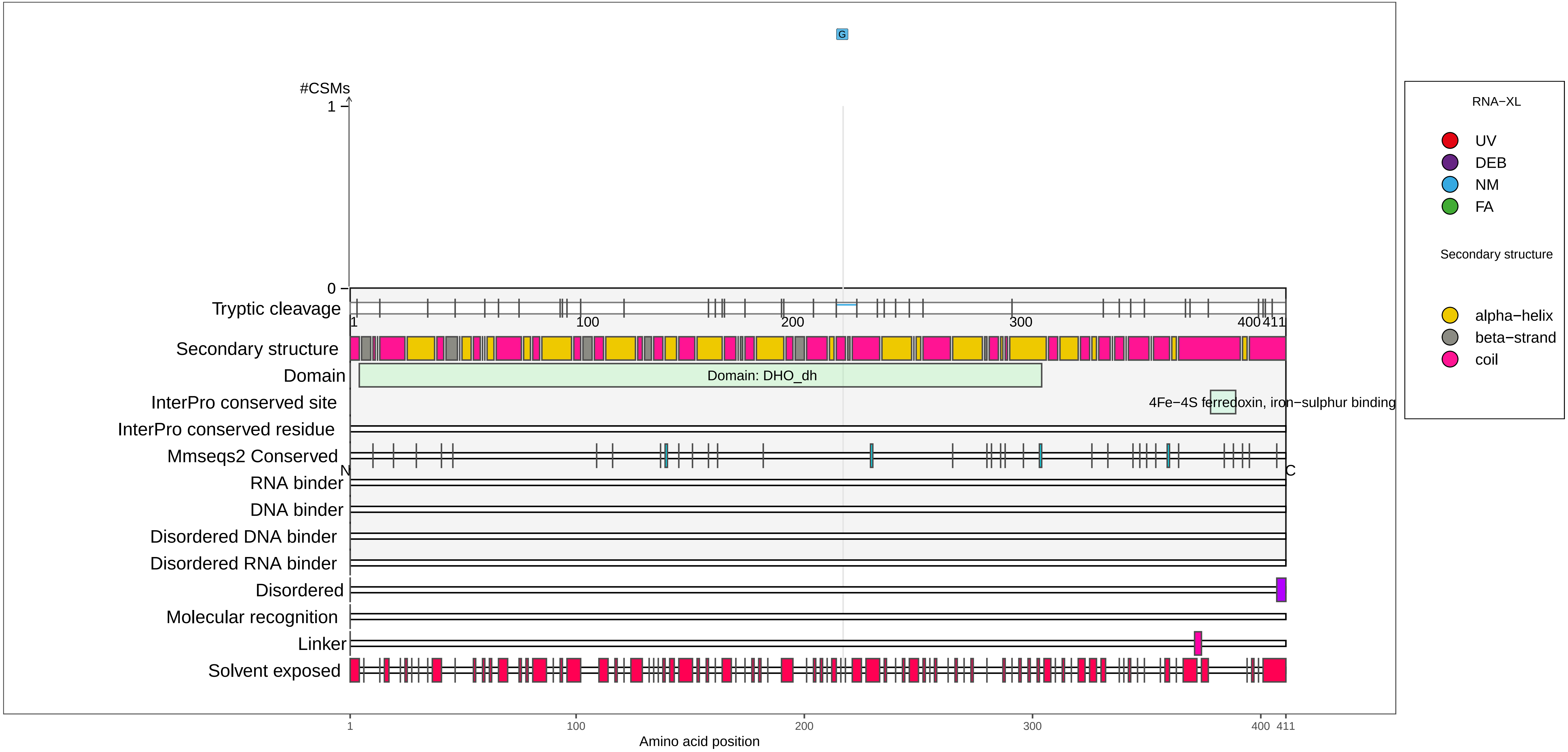

P26218
BGLH_ECOLI Cryptic outer membrane porin BglH

− Abundance:
tryptic [log10 Intensity]: NA (Q NA)
PAXdb K12 strain [ppm]: NA (Q NA)
PAXdb E.coli [ppm]: −2 (Q 1)

− RNA functions: not annotated

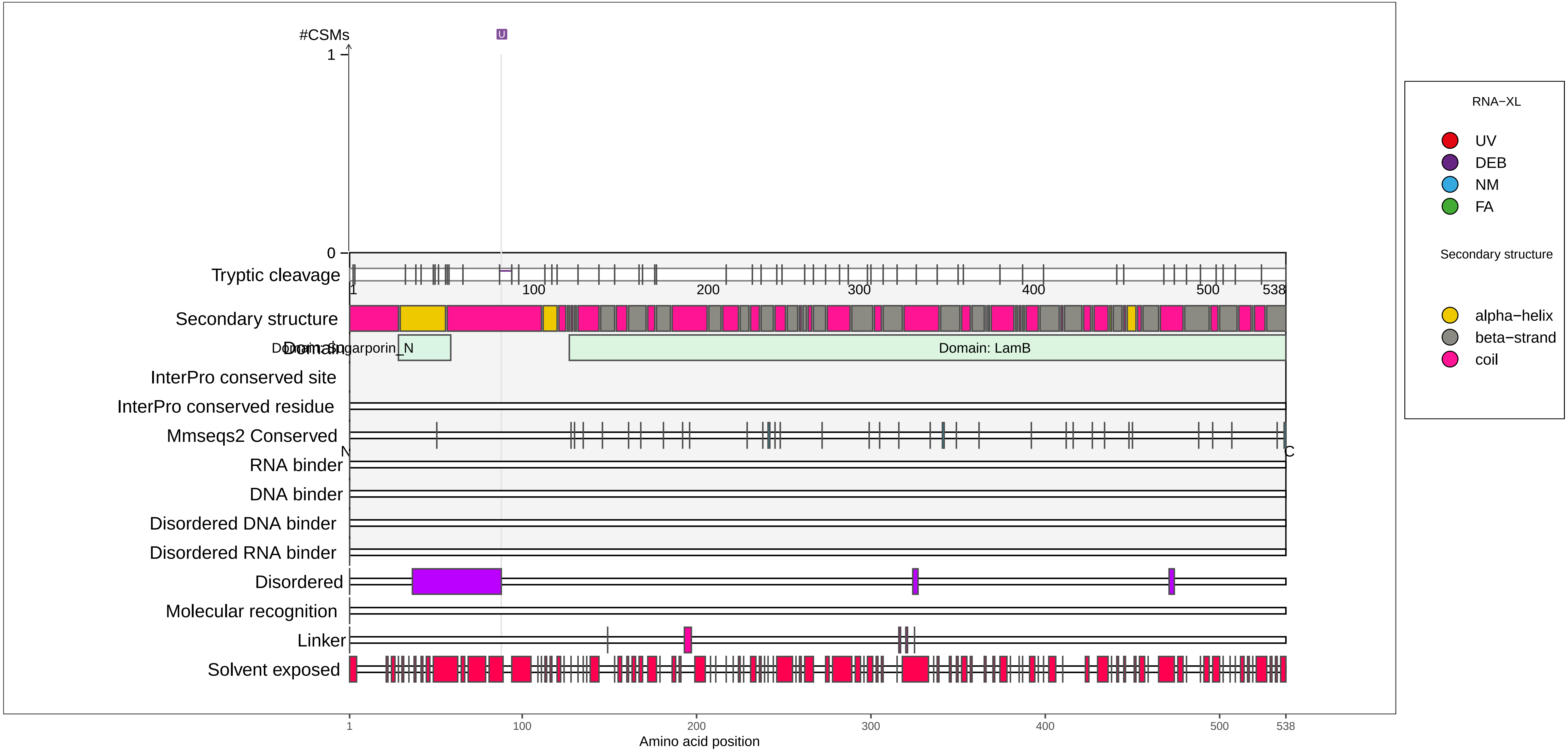

P28917
YDCC_ECOLI H repeat−associated putative transposase YdcC

− Abundance:
tryptic [log10 Intensity]: NA (Q NA)
PAXdb K12 strain [ppm]: NA (Q NA)
PAXdb E.coli [ppm]: −0.28 (Q 19)

− RNA functions: not annotated

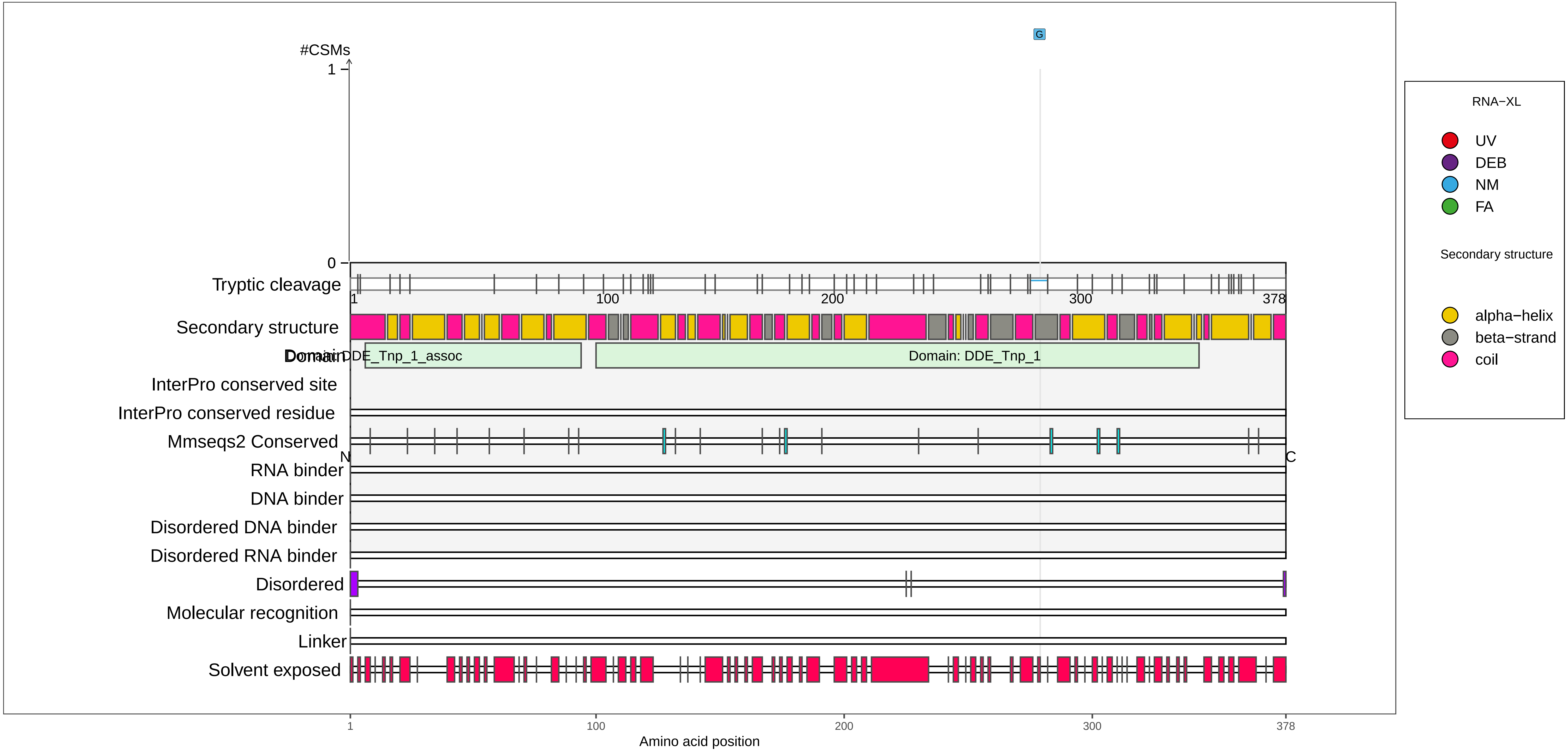

P29217
YCEH_ECOLI UPF0502 protein YceH

− Abundance:
tryptic [log10 Intensity]: 8 (Q 58)
PAXdb K12 strain [ppm]: 2.04 (Q 62)
PAXdb E.coli [ppm]: 2.46 (Q 85)

− RNA functions: not annotated

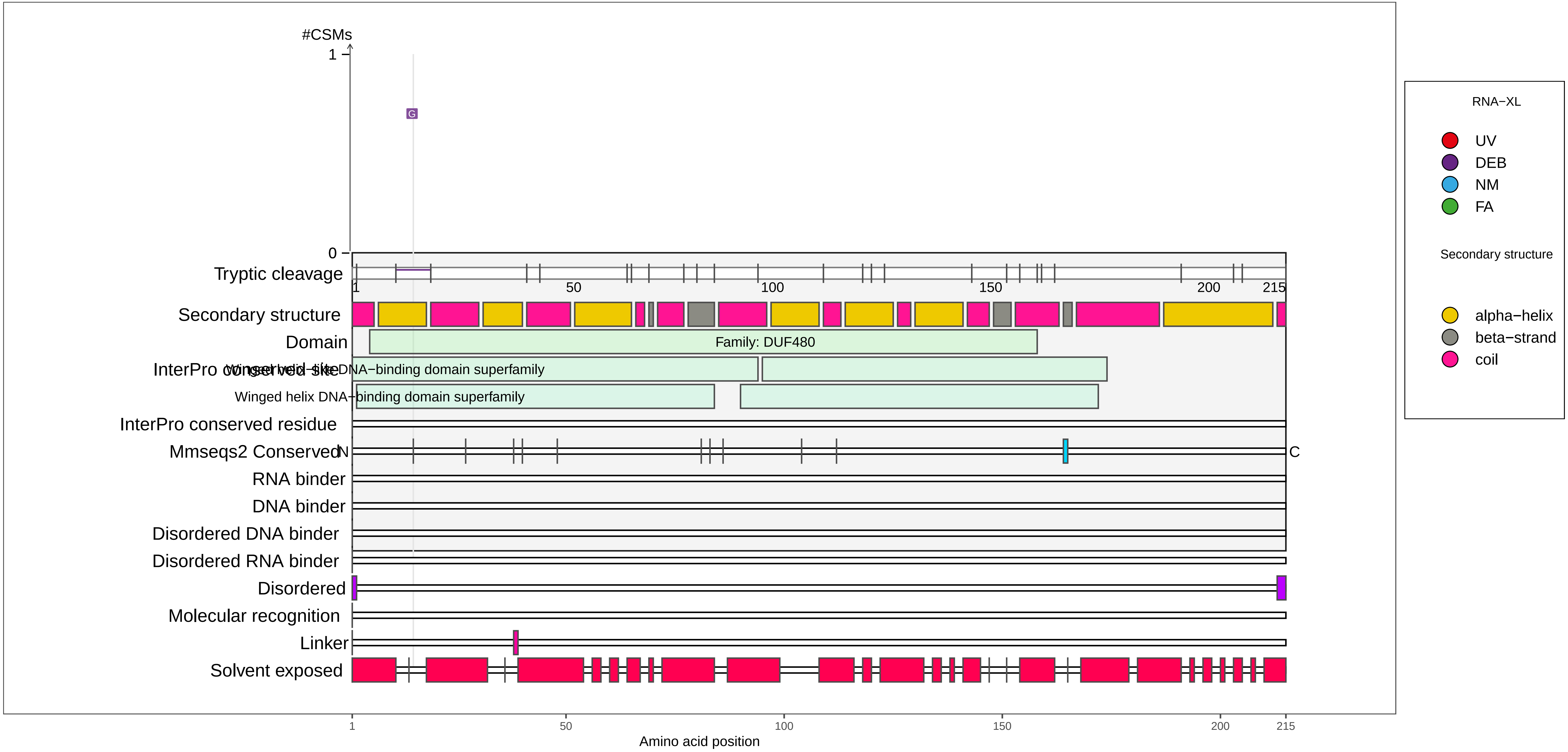

P30011
NADC_ECOLI Nicotinate−nucleotide pyrophosphorylase [carboxylating]

− Abundance:
tryptic [log10 Intensity]: 8.5 (Q 75)
PAXdb K12 strain [ppm]: 2 (Q 60)
PAXdb E.coli [ppm]: 2.37 (Q 82)

− RNA functions: not annotated

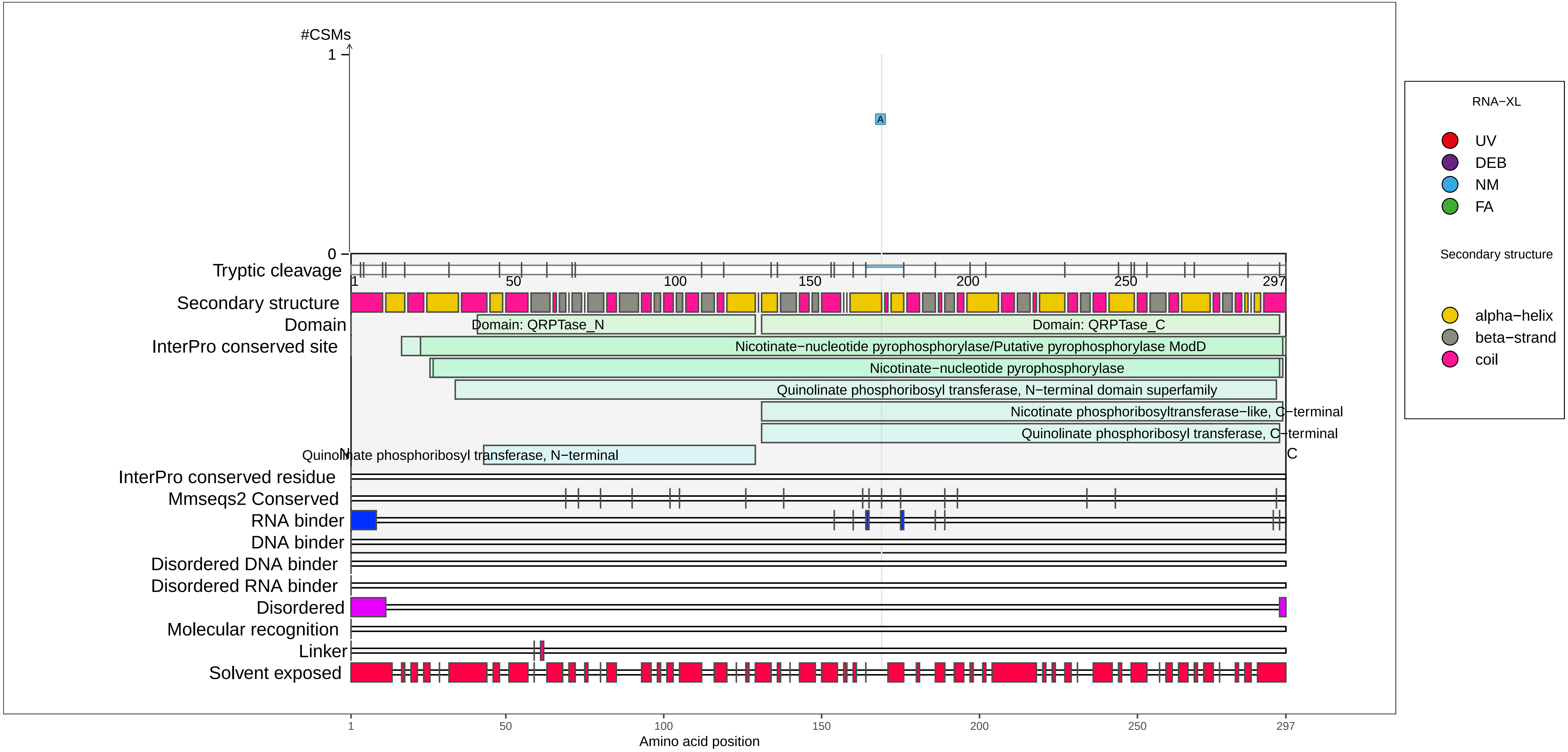

P30015
LHR_ECOLI Probable ATP−dependent helicase lhr

− Abundance:
tryptic [log10 Intensity]: NA (Q NA)
PAXdb K12 strain [ppm]: 1.51 (Q 39)
PAXdb E.coli [ppm]: 0.96 (Q 47)

− RNA functions:
RNA binding; RNA helicase activity

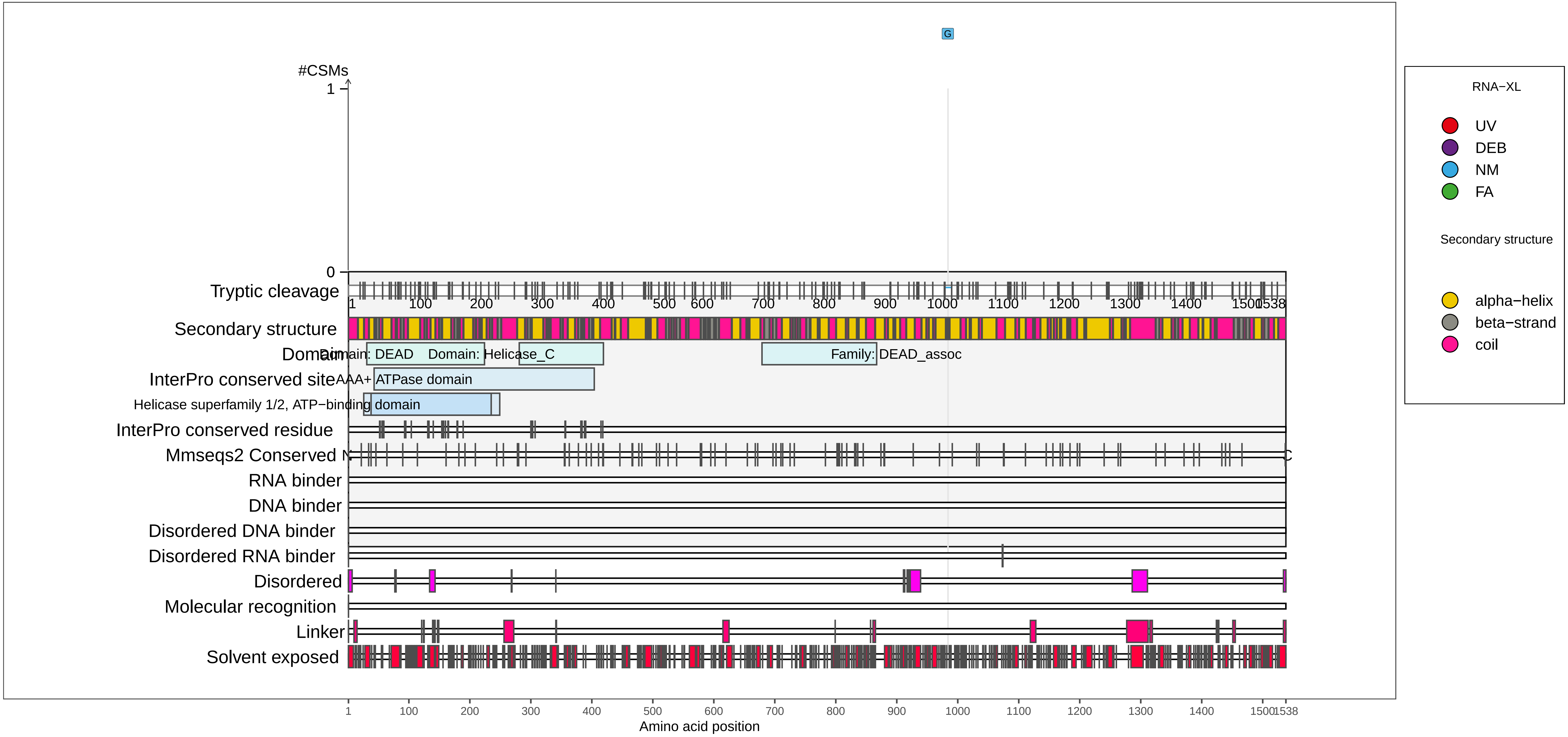

P30177
YBIB_ECOLI Uncharacterized protein YbiB

− Abundance:
tryptic [log10 Intensity]: 8.15 (Q 63)
PAXdb K12 strain [ppm]: 2.2 (Q 67)
PAXdb E.coli [ppm]: 2.48 (Q 85)

− RNA functions:
RNA binding

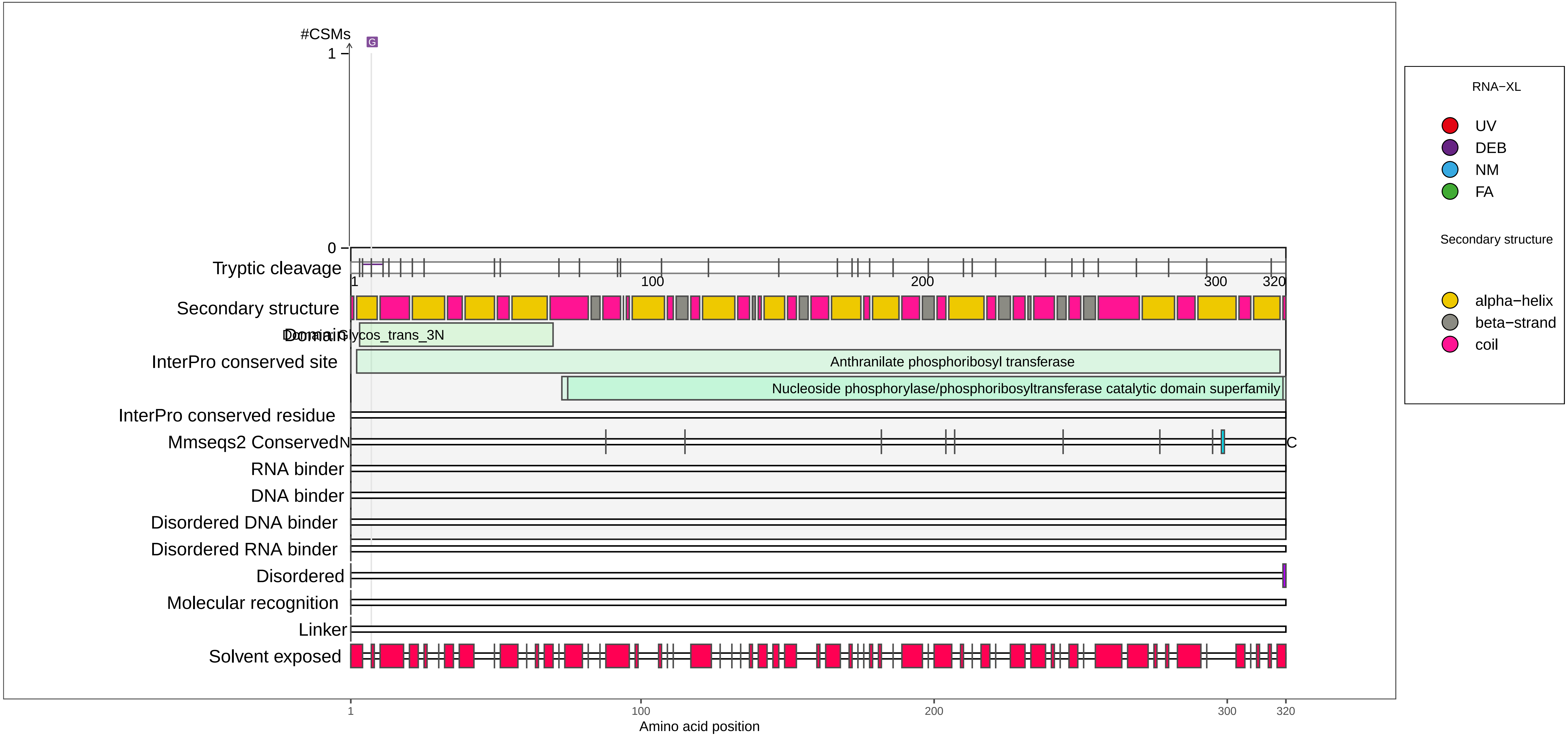

P30845
EPTA_ECOLI Phosphoethanolamine transferase EptA

− Abundance:
tryptic [log10 Intensity]: 8.2 (Q 65)
PAXdb K12 strain [ppm]: 1.91 (Q 57)
PAXdb E.coli [ppm]: −0.21 (Q 20)

− RNA functions: not annotated

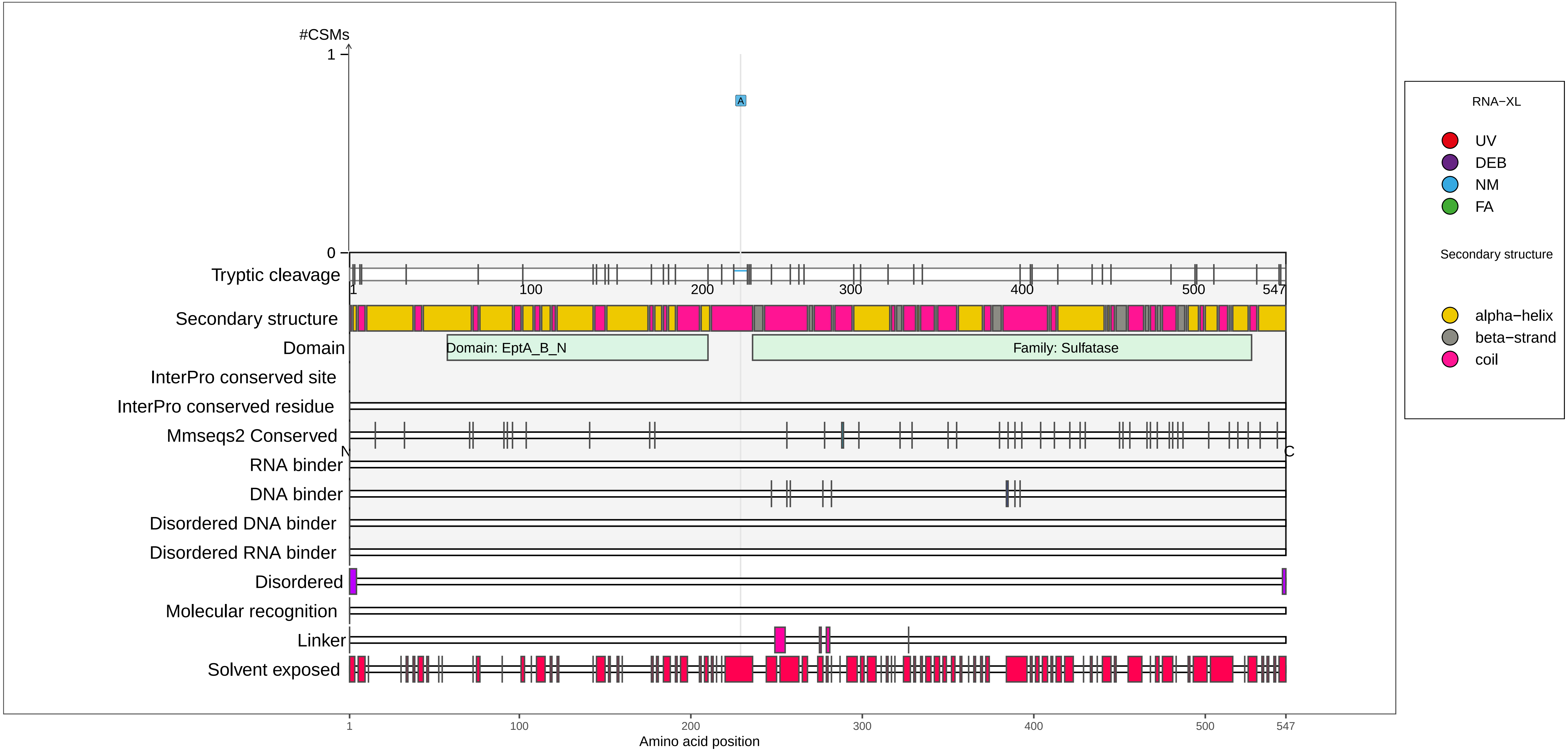

P30863
DKGB_ECOLI 2,5−diketo−D−gluconic acid reductase B

− Abundance:
tryptic [log10 Intensity]: 7.57 (Q 38)
PAXdb K12 strain [ppm]: 2.15 (Q 65)
PAXdb E.coli [ppm]: 1.88 (Q 69)

− RNA functions: not annotated

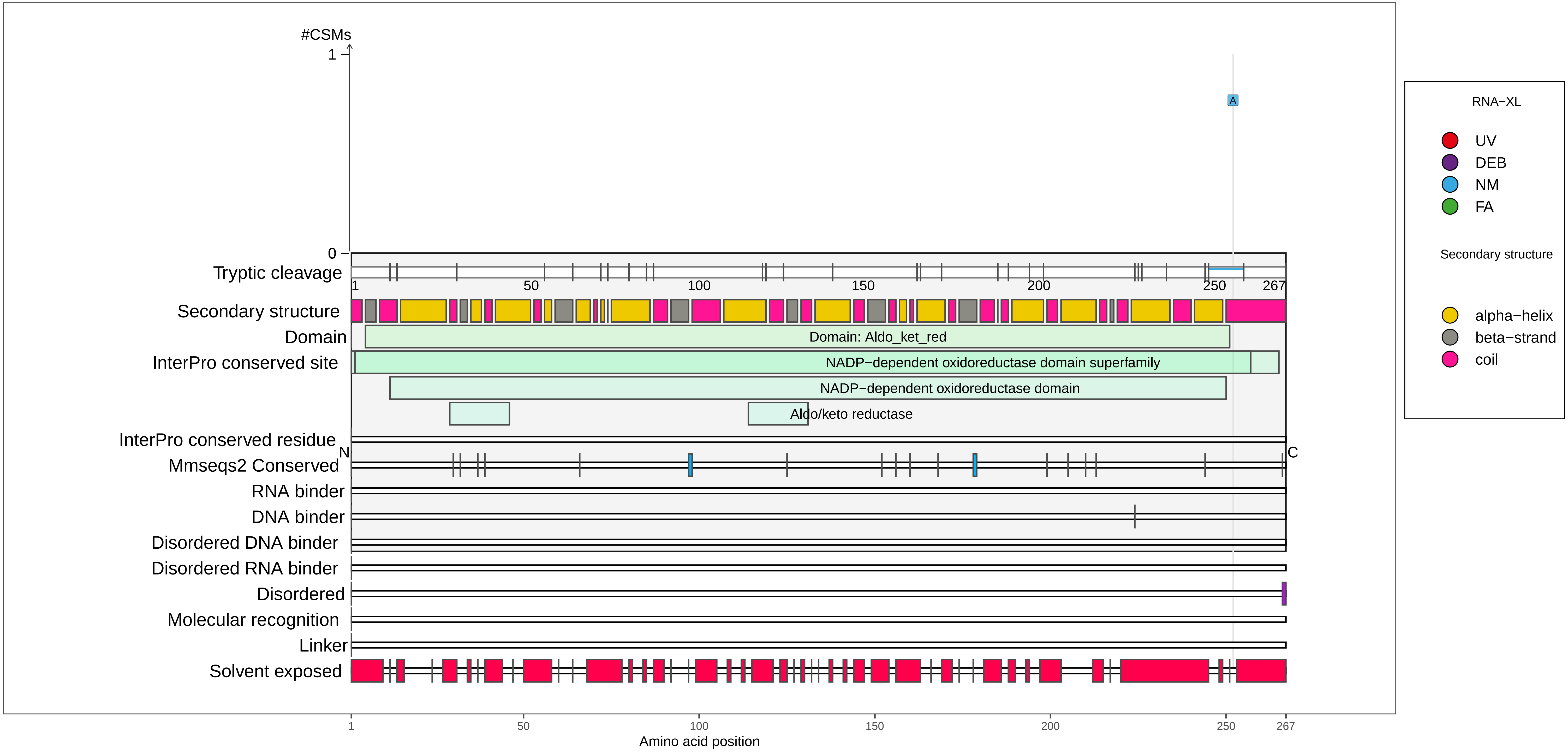

P30870
GLNE_ECOLI Bifunctional glutamine synthetase adenylyltransferase/adenylyl−removing enzyme

− Abundance:
tryptic [log10 Intensity]: 6.68 (Q 5)
PAXdb K12 strain [ppm]: 1.61 (Q 45)
PAXdb E.coli [ppm]: 1.66 (Q 63)

− RNA functions: not annotated

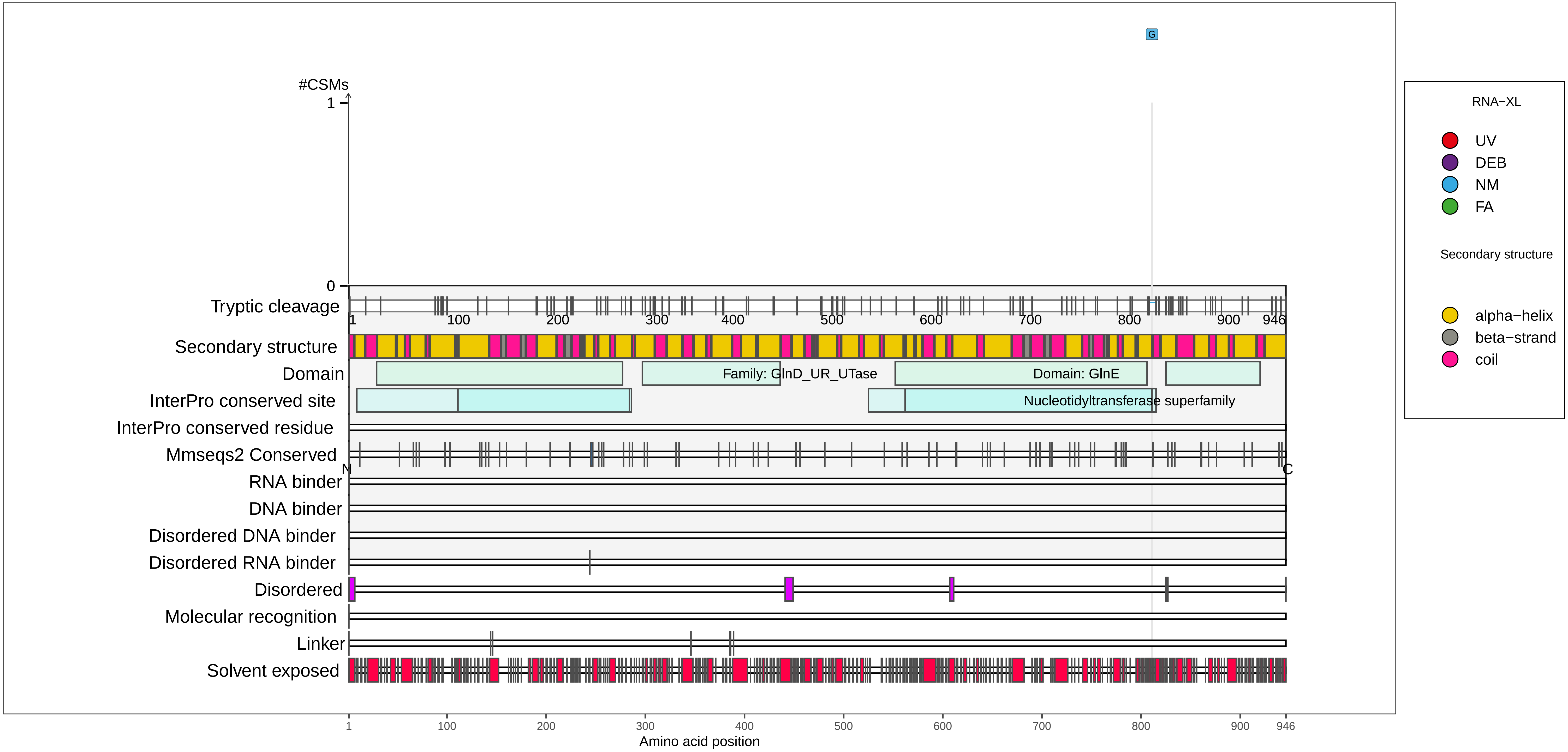

P31122
SOTB_ECOLI Sugar efflux transporter

− Abundance:
tryptic [log10 Intensity]: 7.81 (Q 49)
PAXdb K12 strain [ppm]: NA (Q NA)
PAXdb E.coli [ppm]: −1.39 (Q 5)

− RNA functions: not annotated

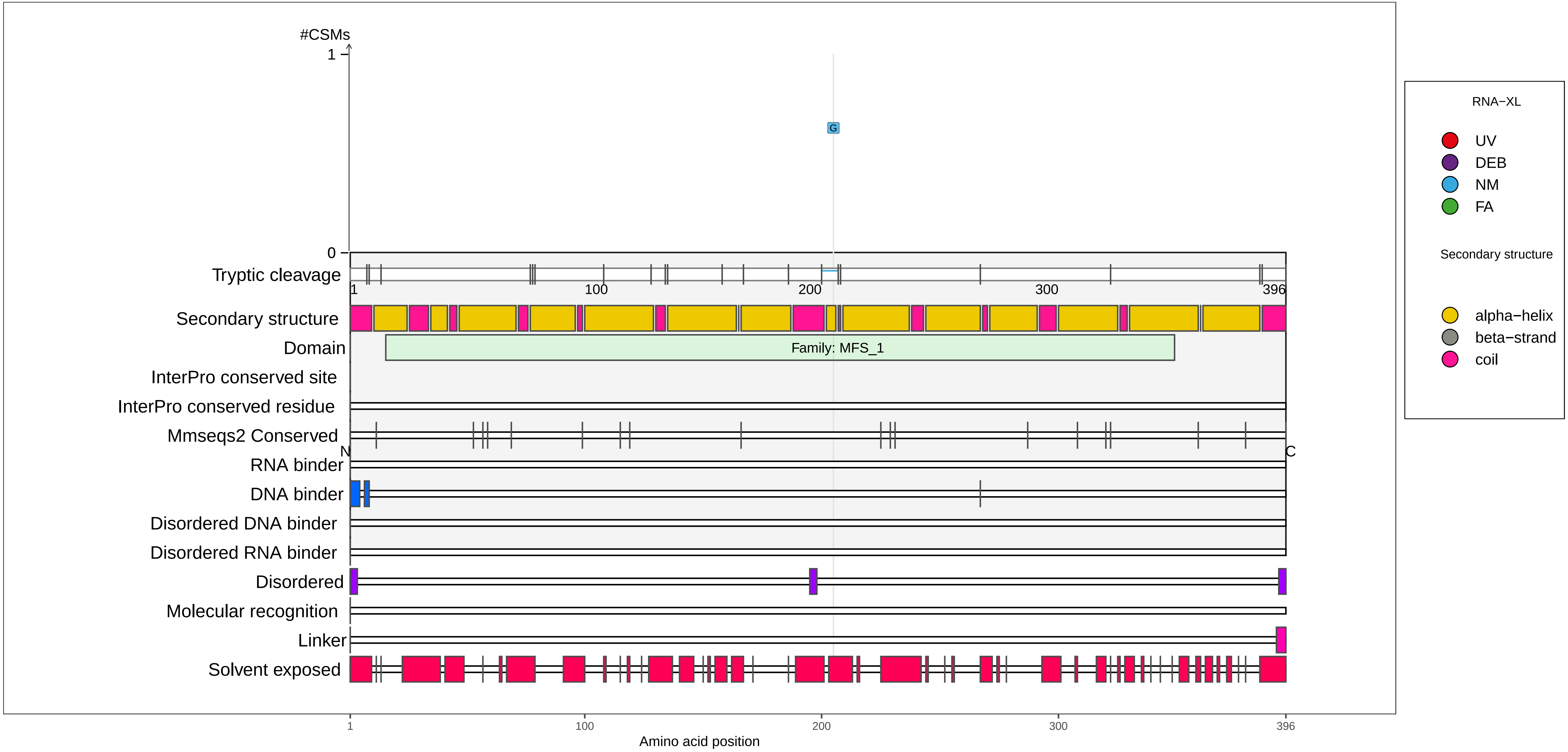

P31658
HCHA_ECOLI Protein/nucleic acid deglycase 1

− Abundance:
tryptic [log10 Intensity]: NA (Q NA)
PAXdb K12 strain [ppm]: 2.78 (Q 85)
PAXdb E.coli [ppm]: 2.75 (Q 91)

− RNA functions:
RNA metabolic process; RNA repair

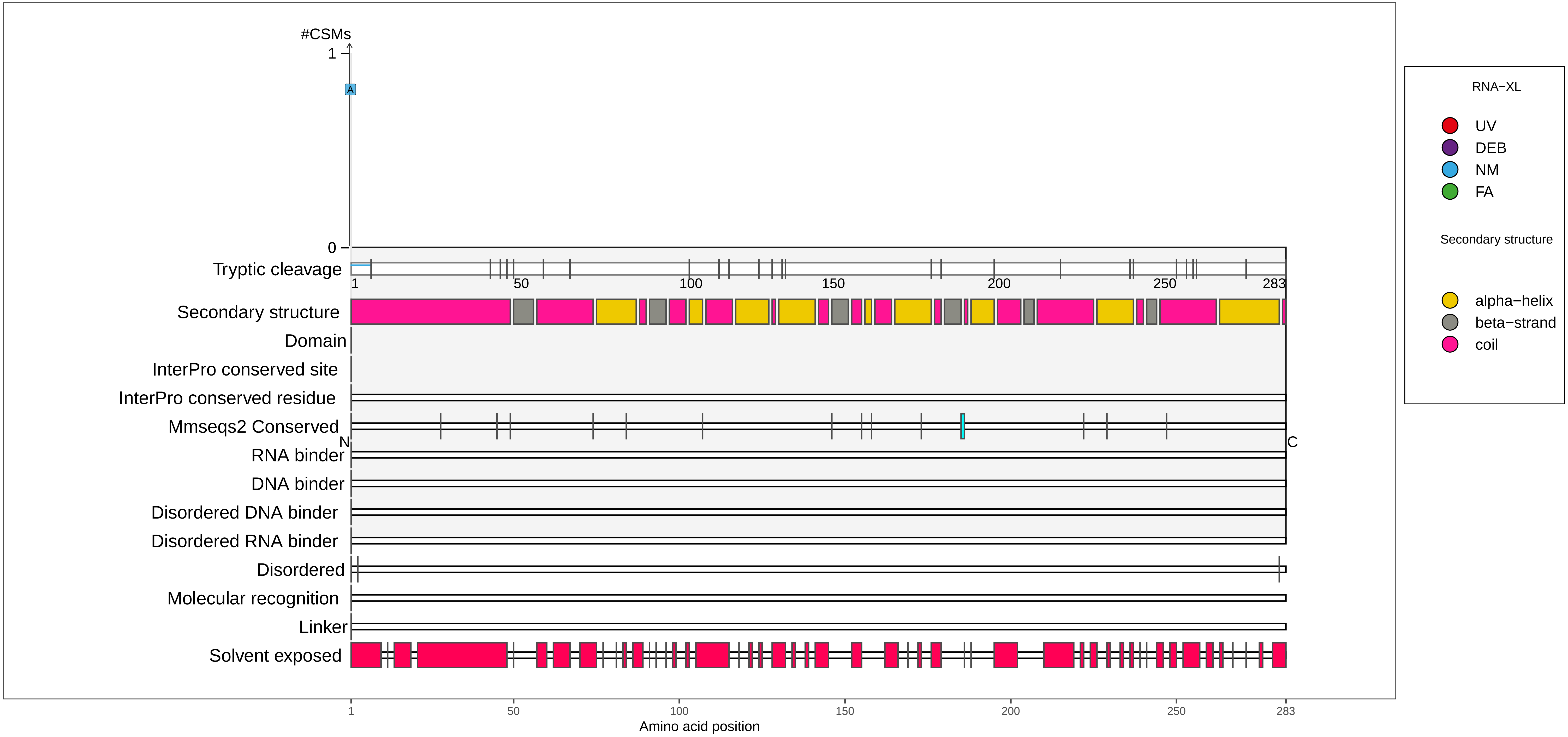

P31802
NARP_ECOLI Nitrate/nitrite response regulator protein NarP

− Abundance:
tryptic [log10 Intensity]: 7.84 (Q 51)
PAXdb K12 strain [ppm]: 1.35 (Q 30)
PAXdb E.coli [ppm]: 1.66 (Q 63)

− RNA functions: not annotated

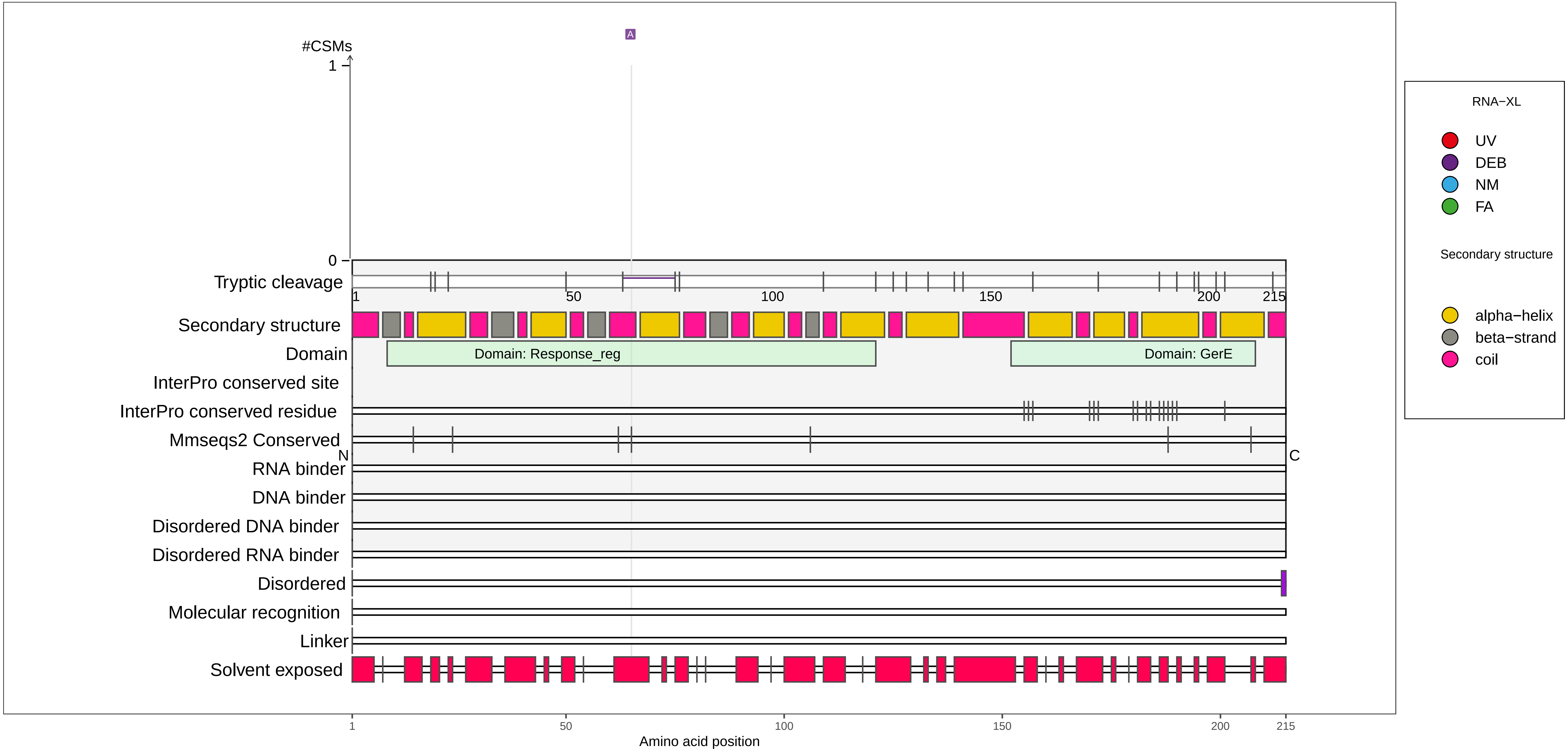

P32144
CSQR_ECOLI HTH−type transcriptional repressor CsqR

− Abundance:
tryptic [log10 Intensity]: 7.97 (Q 57)
PAXdb K12 strain [ppm]: 1.42 (Q 34)
PAXdb E.coli [ppm]: 1.69 (Q 64)

− RNA functions:
RNA biosynthetic process; RNA metabolic process

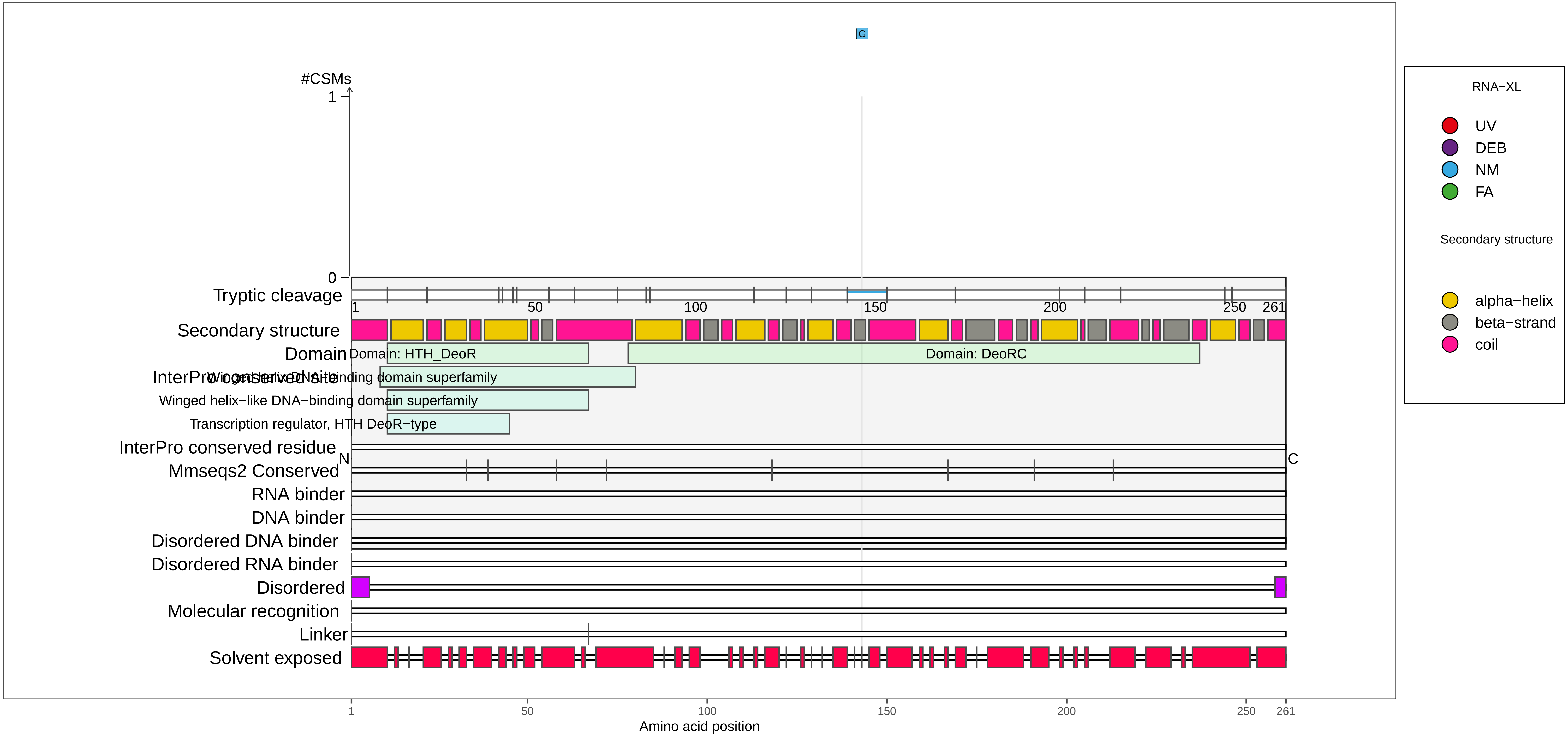

P32177
FDHD_ECOLI Sulfur carrier protein FdhD

− Abundance:
tryptic [log10 Intensity]: 7.29 (Q 25)
PAXdb K12 strain [ppm]: 1.59 (Q 44)
PAXdb E.coli [ppm]: 1.09 (Q 50)

− RNA functions: not annotated

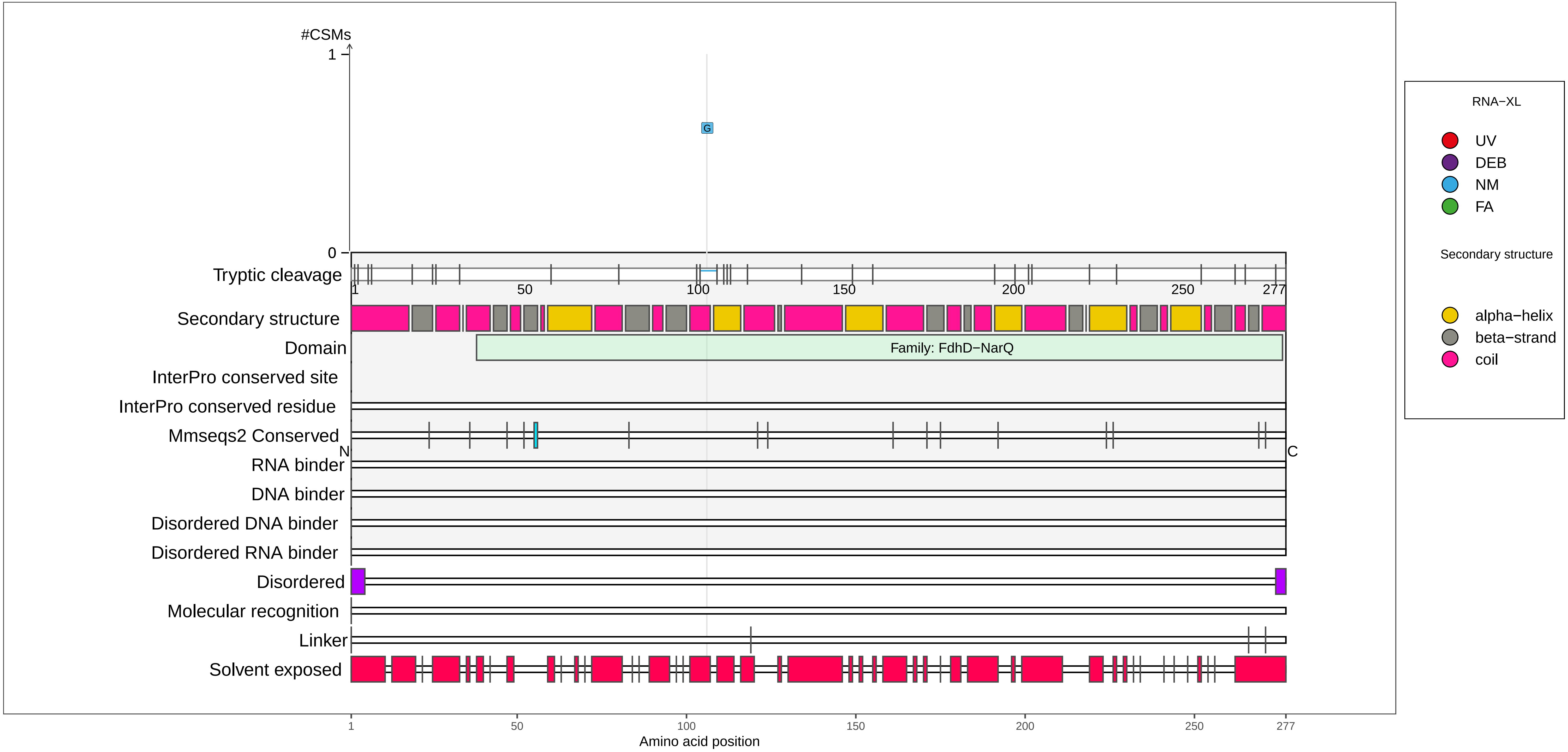

P32670
PTFX2_ECOLI Multiphosphoryl transfer protein 2

− Abundance:
tryptic [log10 Intensity]: 7.29 (Q 24)
PAXdb K12 strain [ppm]: 1.43 (Q 34)
PAXdb E.coli [ppm]: 0.11 (Q 27)

− RNA functions: not annotated

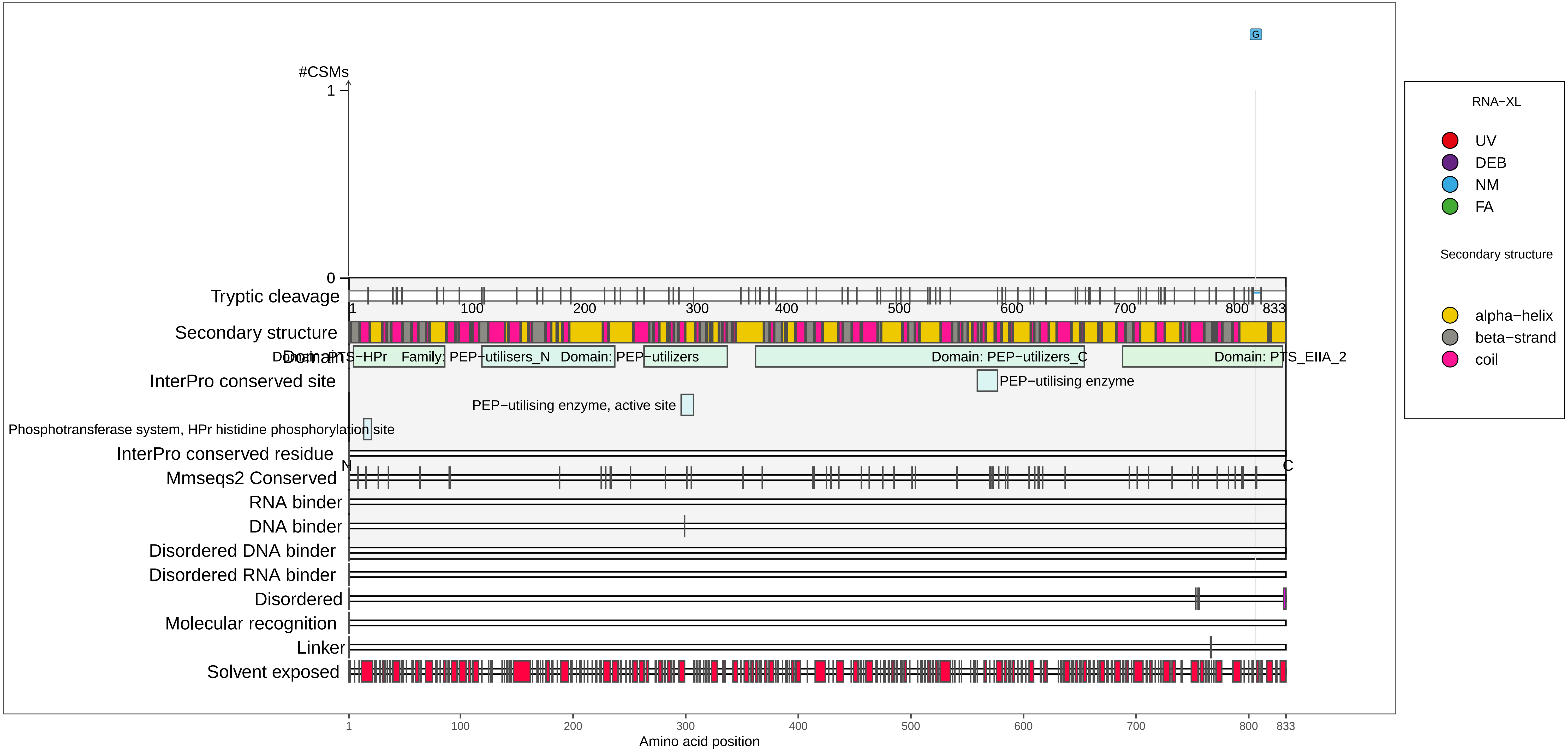

P32680
YJAG_ECOLI Uncharacterized protein YjaG

− Abundance:
tryptic [log10 Intensity]: 7.78 (Q 48)
PAXdb K12 strain [ppm]: NA (Q NA)
PAXdb E.coli [ppm]: 2.11 (Q 75)

− RNA functions: not annotated

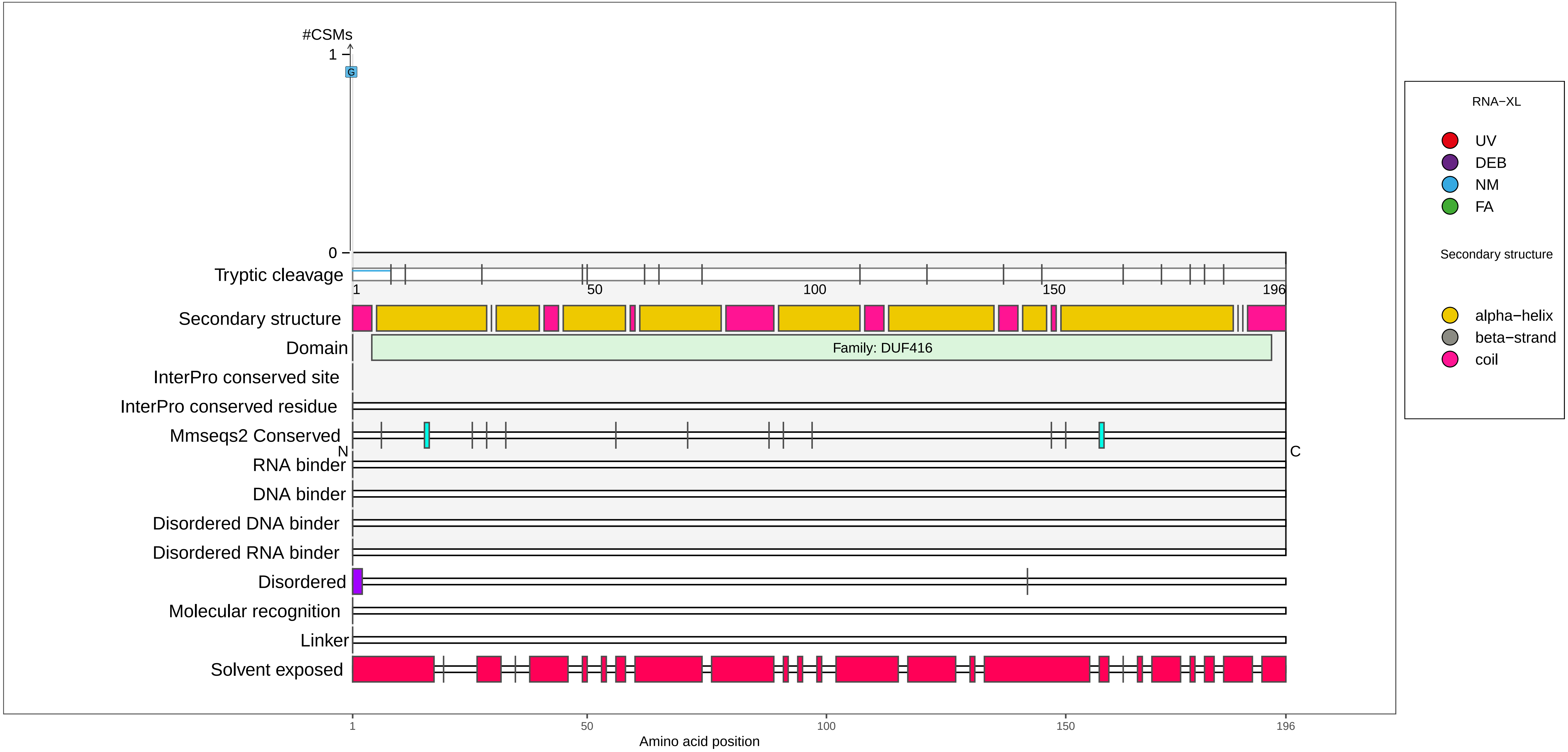

P32684
RLUF_ECOLI Dual−specificity RNA pseudouridine synthase RluF

− Abundance:
tryptic [log10 Intensity]: 7.76 (Q 47)
PAXdb K12 strain [ppm]: 0.83 (Q 6)
PAXdb E.coli [ppm]: 0.99 (Q 47)

− RNA functions:
ncRNA metabolic process; ncRNA processing; RNA binding; RNA metabolic process
RNA modification; RNA processing; RNA pseudouridylate synthase; rRNA metabolic process
rRNA modification; rRNA processing; rRNA pseudouridine synthase activity
rRNA pseudouridine synthesis; tRNA metabolic process; tRNA modification; tRNA processing
tRNA pseudouridine synthesis

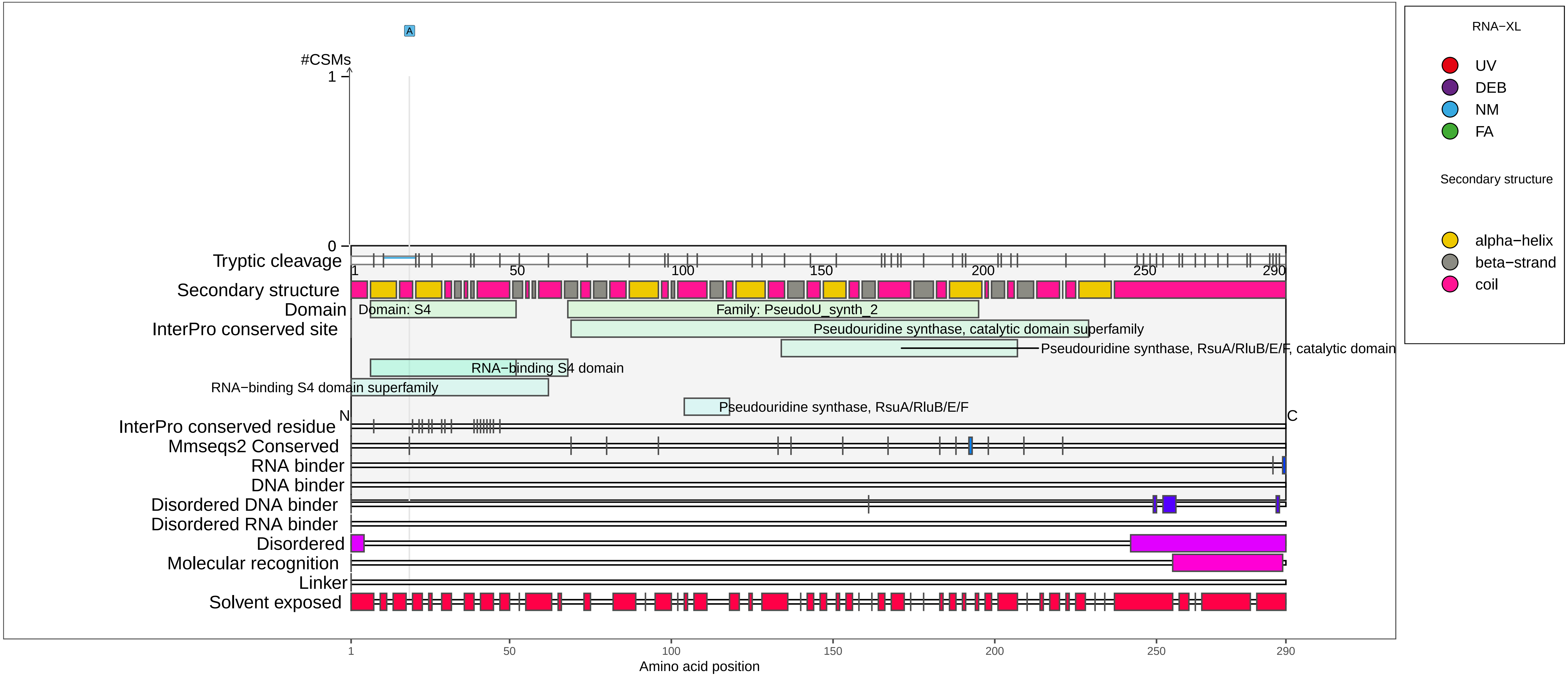

P32689
YJBH_ECOLI Uncharacterized lipoprotein YjbH
− Abundance:
tryptic [log10 Intensity]: NA (Q NA)
PAXdb K12 strain [ppm]: 0.83 (Q 6)
PAXdb E.coli [ppm]: −1.85 (Q 2)

− RNA functions: not annotated

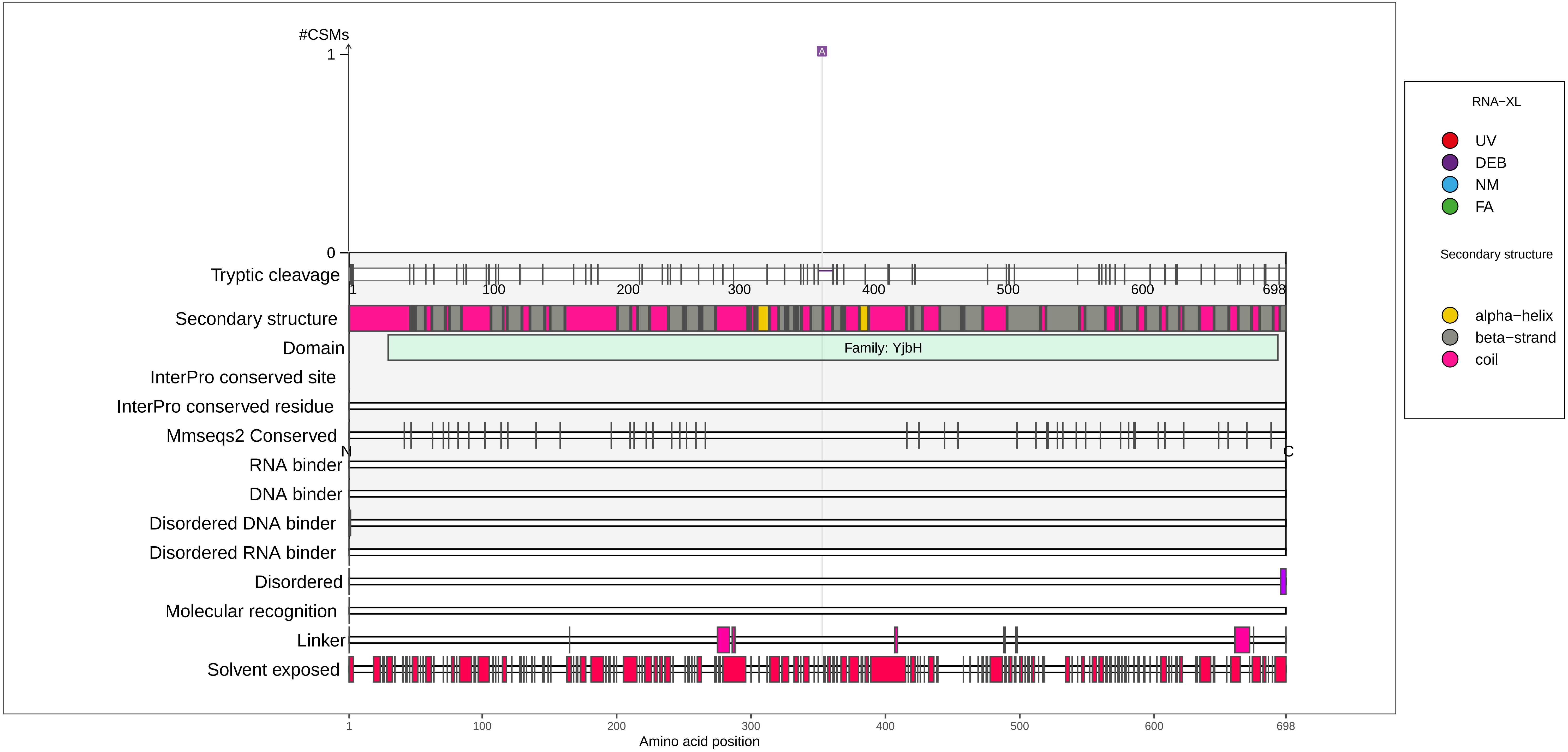

P32717
YJCS_ECOLI Putative alkyl/aryl−sulfatase YjcS
− Abundance:
tryptic [log10 Intensity]: NA (Q NA)
PAXdb K12 strain [ppm]: NA (Q NA)
PAXdb E.coli [ppm]: −0.74 (Q 12)

− RNA functions: not annotated

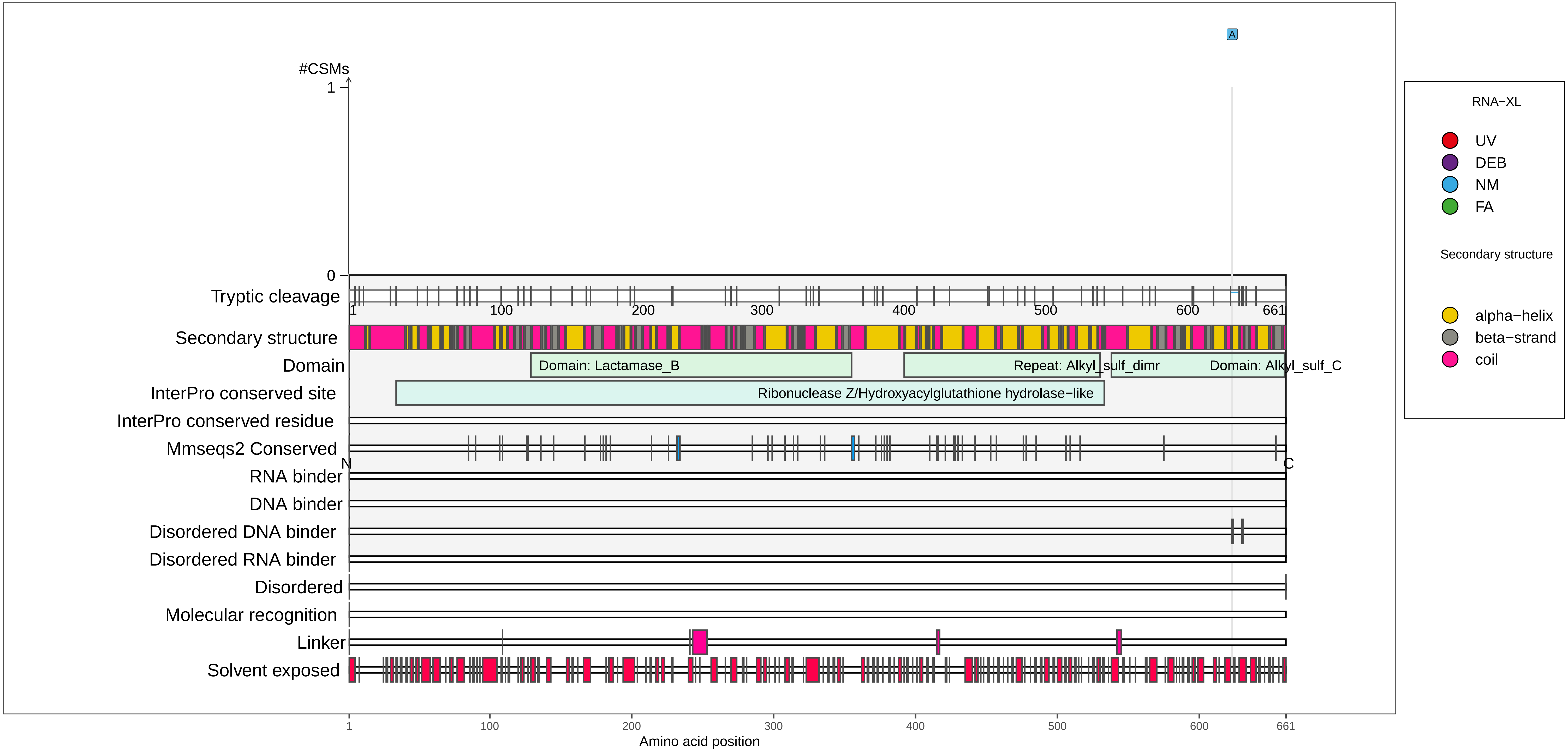

P32721
ALSA_ECOLI D−allose import ATP−binding protein AlsA

− Abundance:
tryptic [log10 Intensity]: NA (Q NA)
PAXdb K12 strain [ppm]: 1.64 (Q 46)
PAXdb E.coli [ppm]: 0.45 (Q 35)

− RNA functions: not annotated

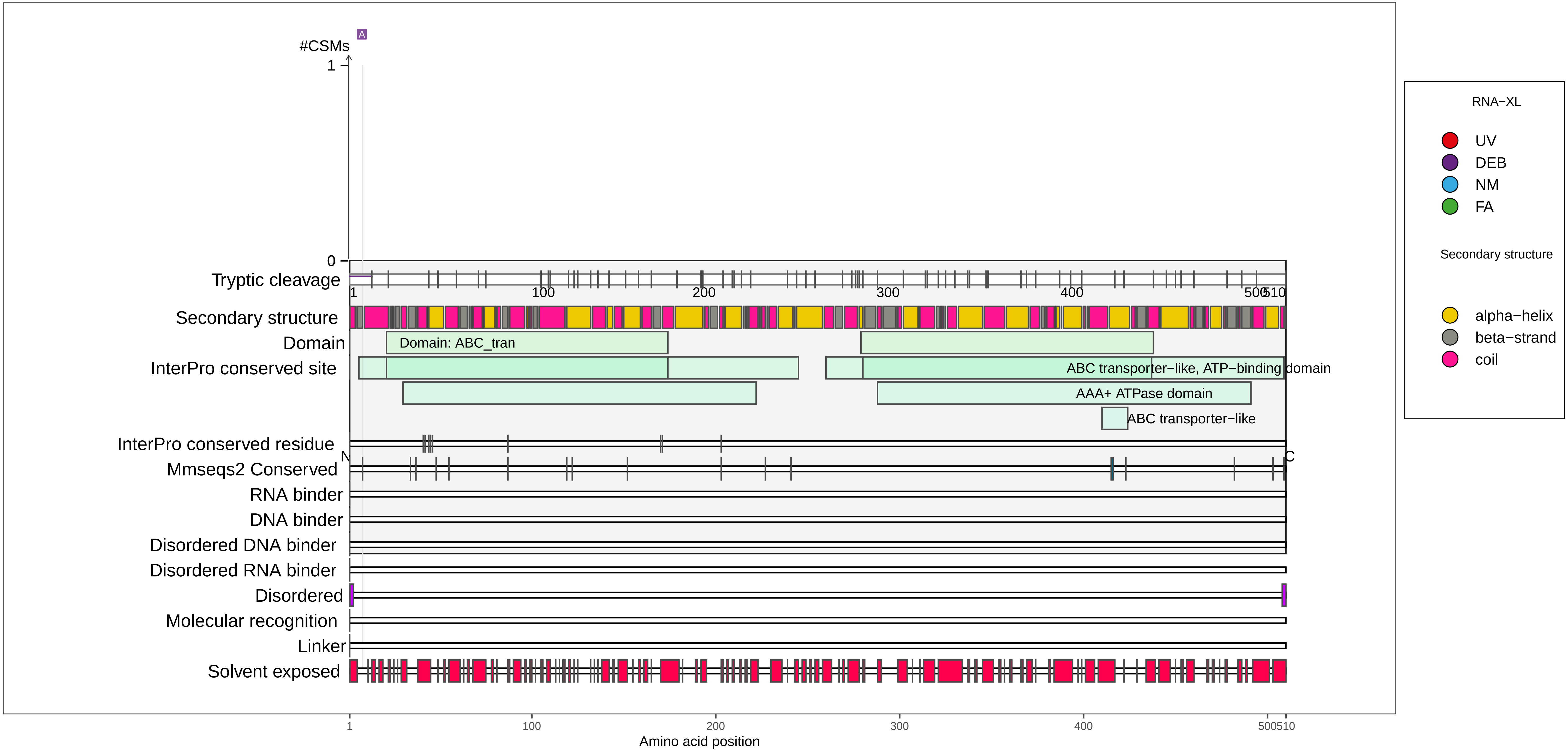

P33012
SBMC_ECOLI DNA gyrase inhibitor

− Abundance:
tryptic [log10 Intensity]: 8.12 (Q 62)
PAXdb K12 strain [ppm]: 1.37 (Q 31)
PAXdb E.coli [ppm]: 2.14 (Q 76)

− RNA functions: not annotated

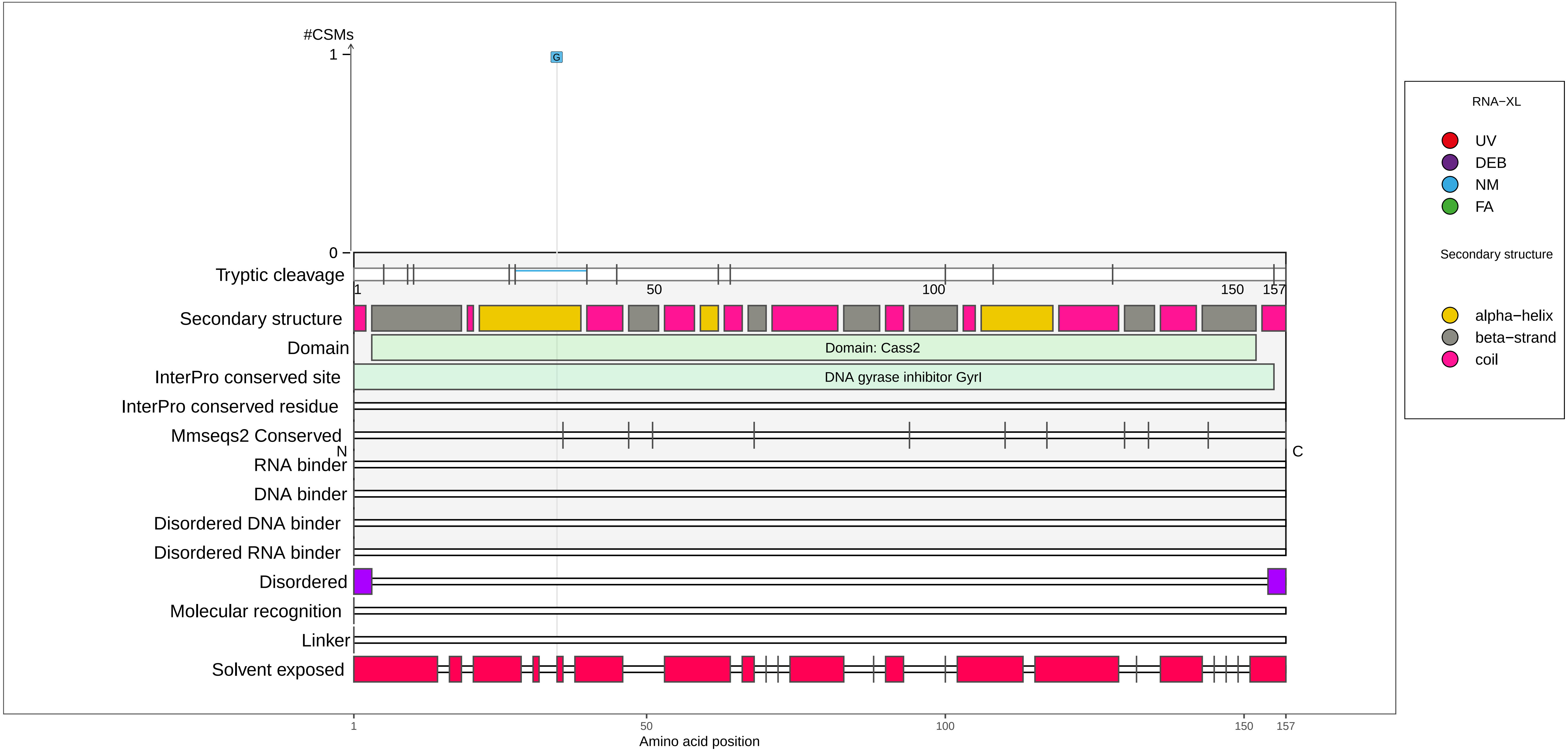

P33013
DACD_ECOLI D−alanyl−D−alanine carboxypeptidase DacD

− Abundance:
tryptic [log10 Intensity]: NA (Q NA)
PAXdb K12 strain [ppm]: NA (Q NA)
PAXdb E.coli [ppm]: −0.63 (Q 14)

− RNA functions: not annotated

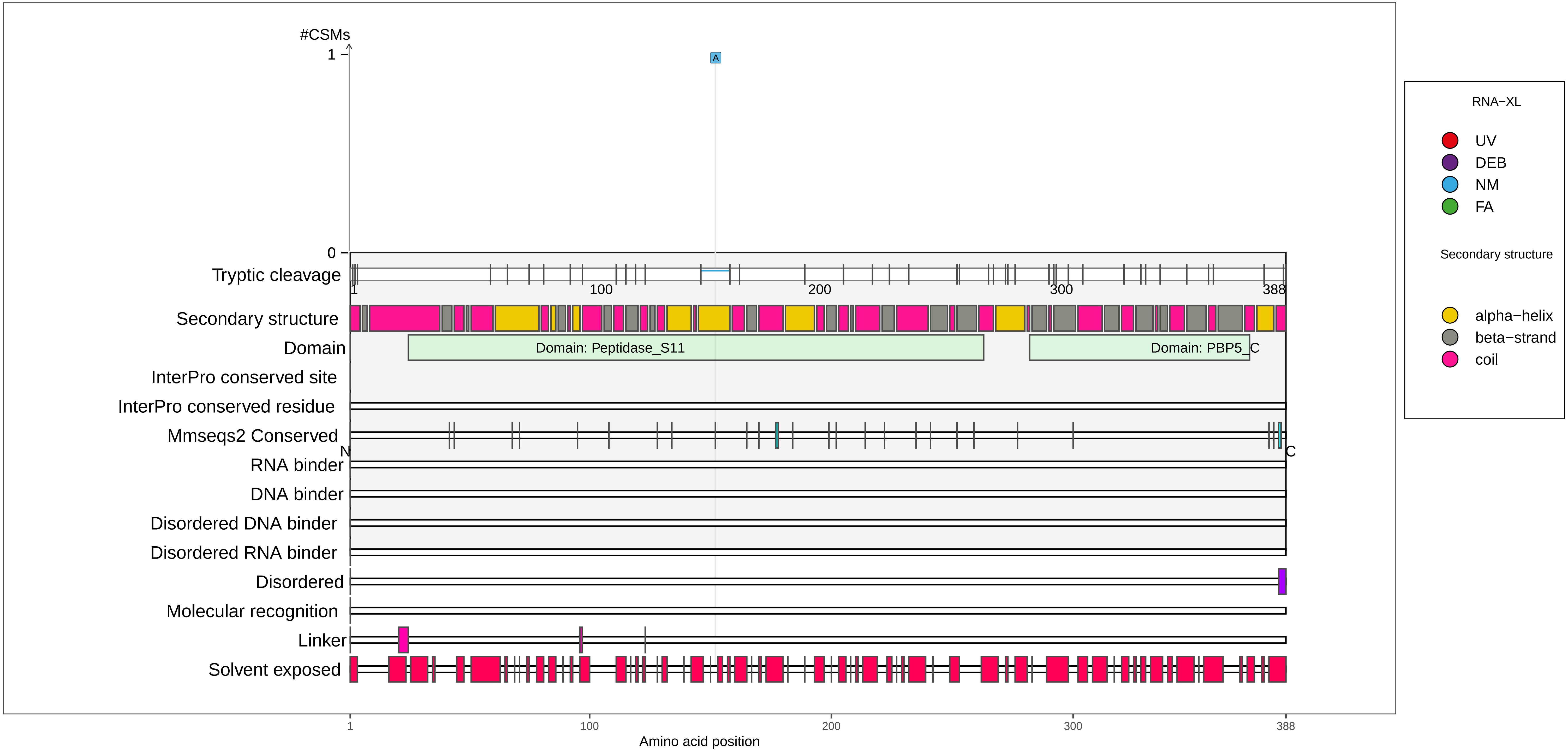

P33234
ADIY_ECOLI HTH−type transcriptional regulator AdiY

− Abundance:
tryptic [log10 Intensity]: NA (Q NA)
PAXdb K12 strain [ppm]: NA (Q NA)
PAXdb E.coli [ppm]: −1.7 (Q 2)

− RNA functions: not annotated

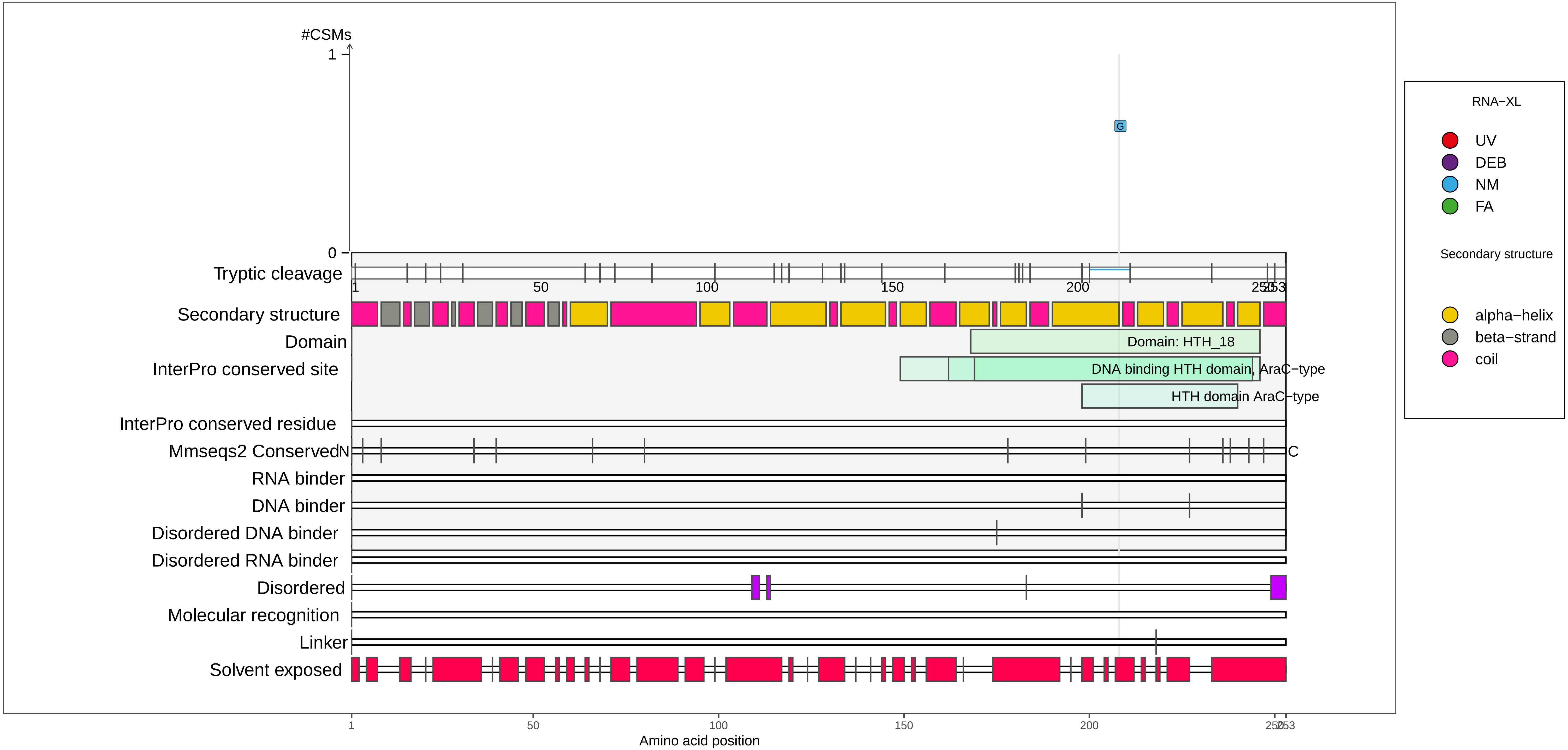

P33596
RECX_ECOLI Regulatory protein RecX

− Abundance:
tryptic [log10 Intensity]: 6.4 (Q 2)
PAXdb K12 strain [ppm]: NA (Q NA)
PAXdb E.coli [ppm]: 0.43 (Q 34)

− RNA functions: not annotated

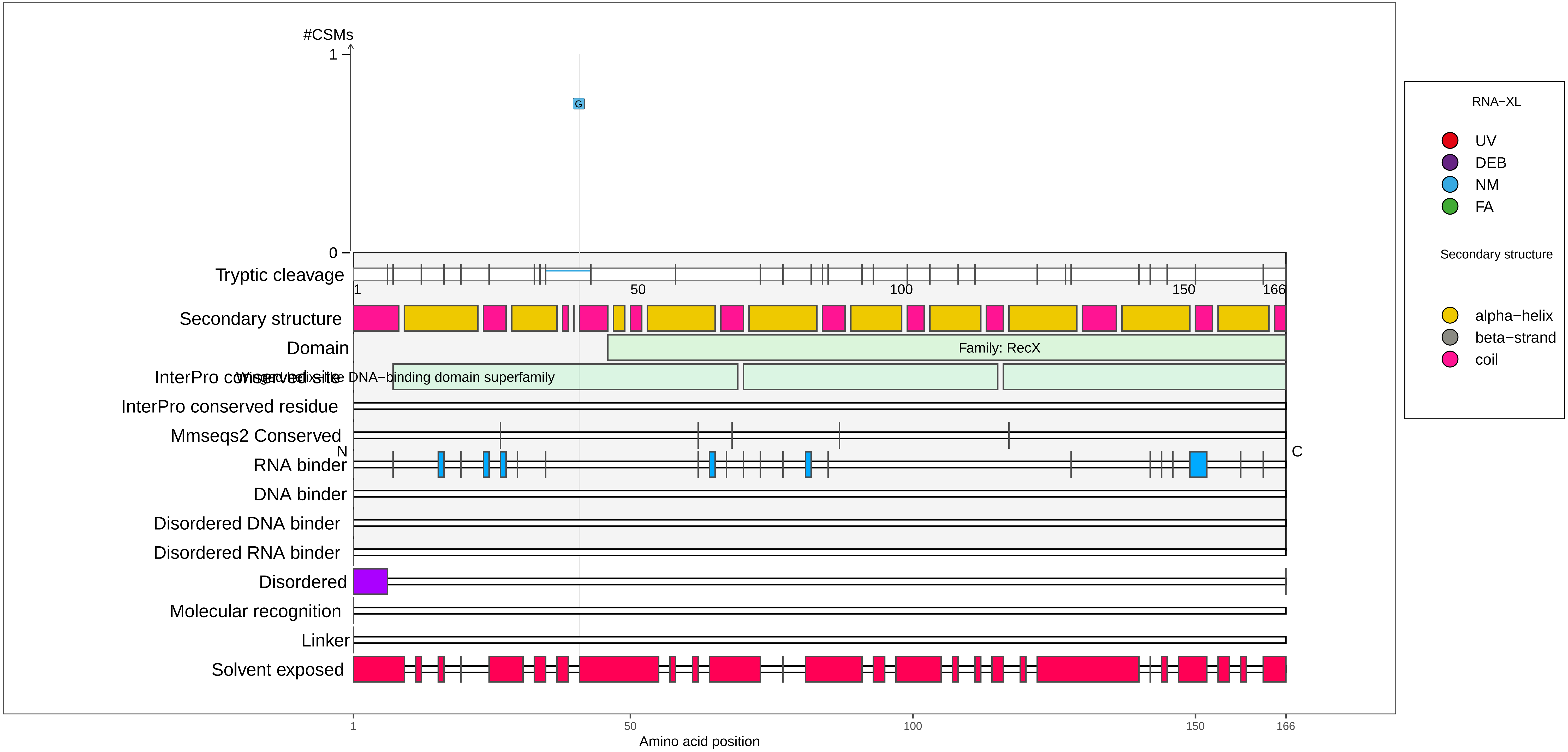

P36548
AMIA_ECOLI N−acetylmuramoyl−L−alanine amidase AmiA

− Abundance:
tryptic [log10 Intensity]: 7.17 (Q 20)
PAXdb K12 strain [ppm]: 1.15 (Q 18)
PAXdb E.coli [ppm]: 0.64 (Q 39)

− RNA functions: not annotated

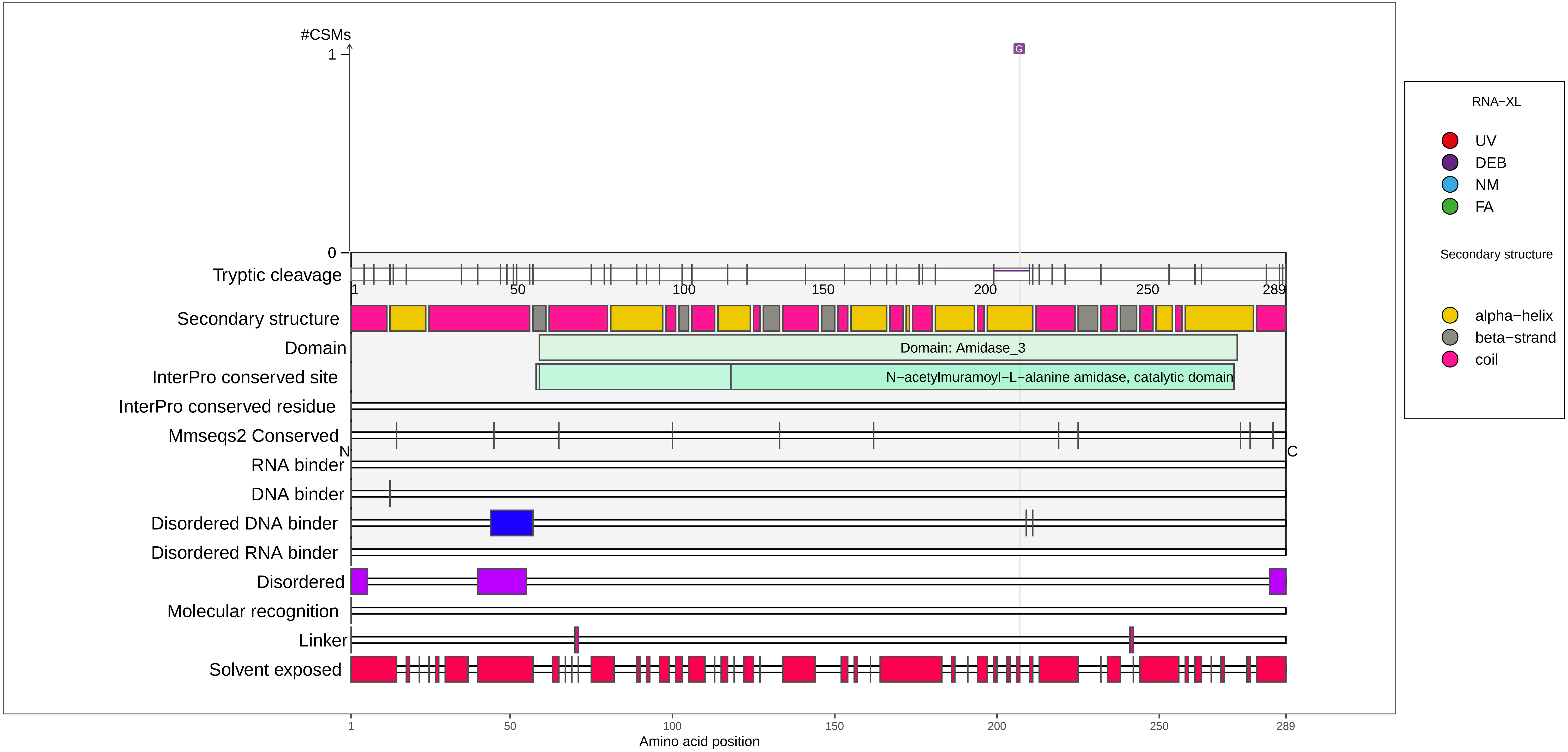

P36999
RLMA_ECOLI 23S rRNA (guanine(745)−N(1))−methyltransferase

− Abundance:
tryptic [log10 Intensity]: NA (Q NA)
PAXdb K12 strain [ppm]: NA (Q NA)
PAXdb E.coli [ppm]: 0.85 (Q 44)

− RNA functions:
23S rRNA (guanine(745)−N(1))−methyltransferase activity; ncRNA metabolic process
ncRNA processing; RNA metabolic process; RNA methylation; RNA methyltransferase activity
RNA modification; RNA processing; rRNA (guanine−N1−)−methyltransferase activity
rRNA (guanine) methyltransferase activity; rRNA base methylation; rRNA metabolic process
rRNA methylation; rRNA methyltransferase activity; rRNA modification; rRNA processing

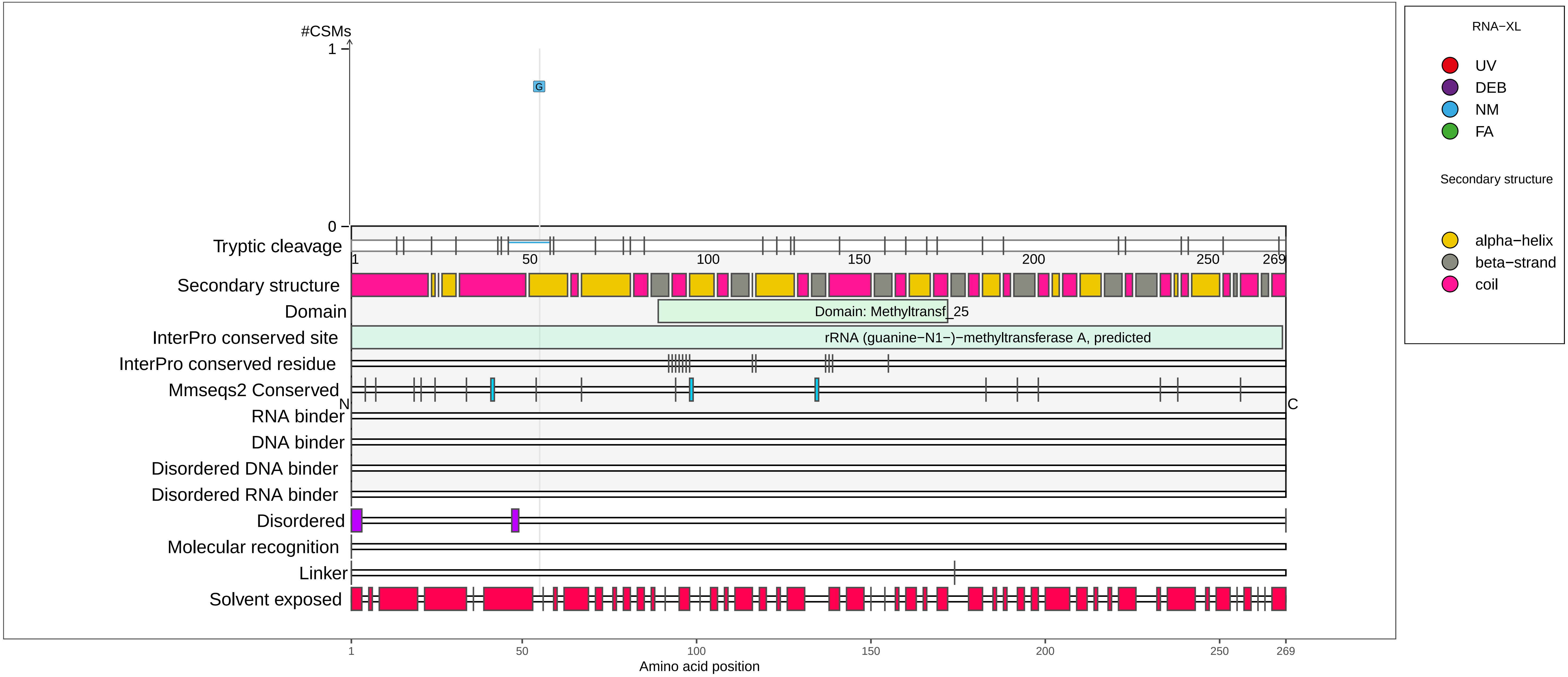

P37052
YCHJ_ECOLI UPF0225 protein YchJ

− Abundance:
tryptic [log10 Intensity]: 7.81 (Q 49)
PAXdb K12 strain [ppm]: NA (Q NA)
PAXdb E.coli [ppm]: 1.45 (Q 58)

− RNA functions: not annotated

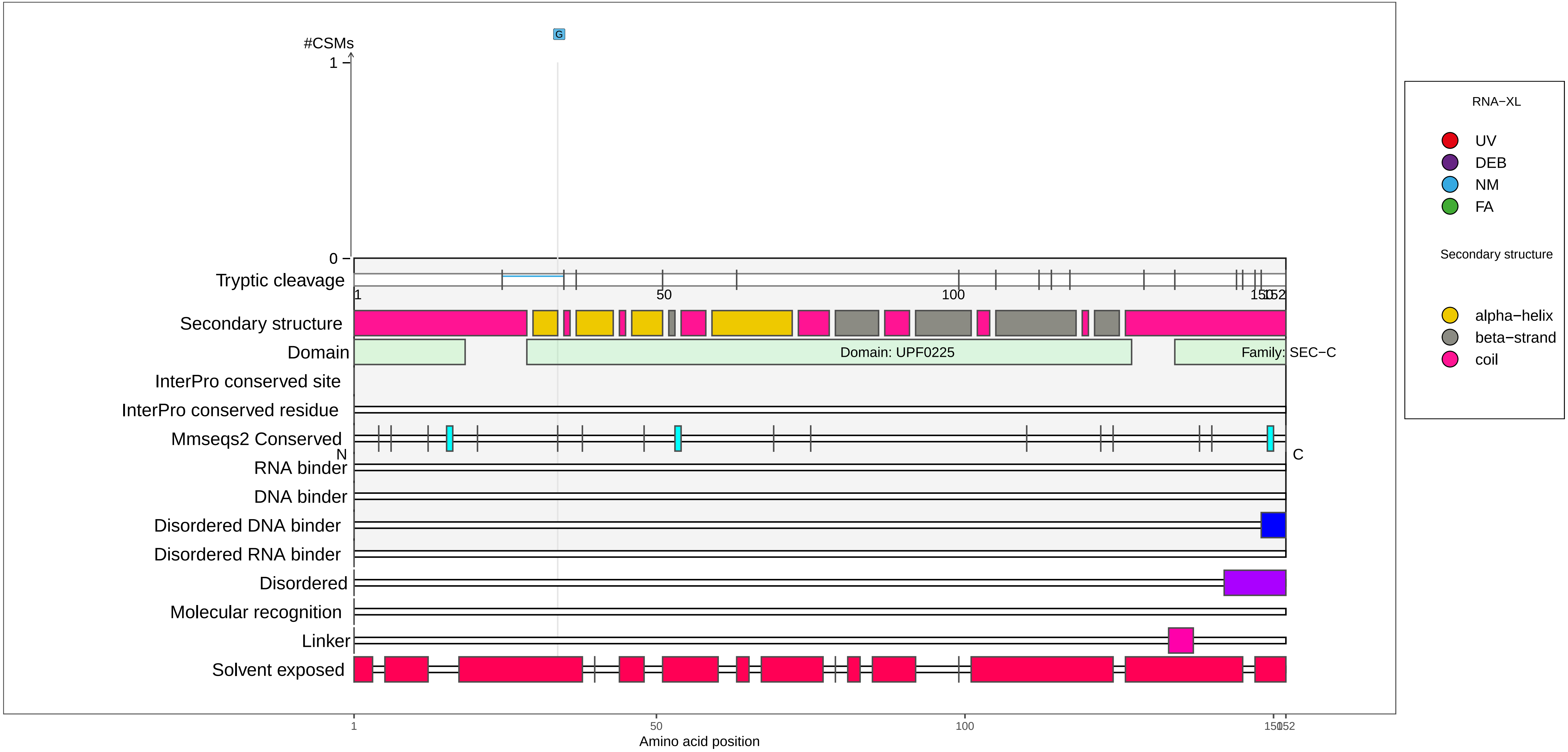

P37595
IAAA_ECOLI Isoaspartyl peptidase

− Abundance:
tryptic [log10 Intensity]: 7.74 (Q 46)
PAXdb K12 strain [ppm]: NA (Q NA)
PAXdb E.coli [ppm]: 0.97 (Q 47)

− RNA functions: not annotated

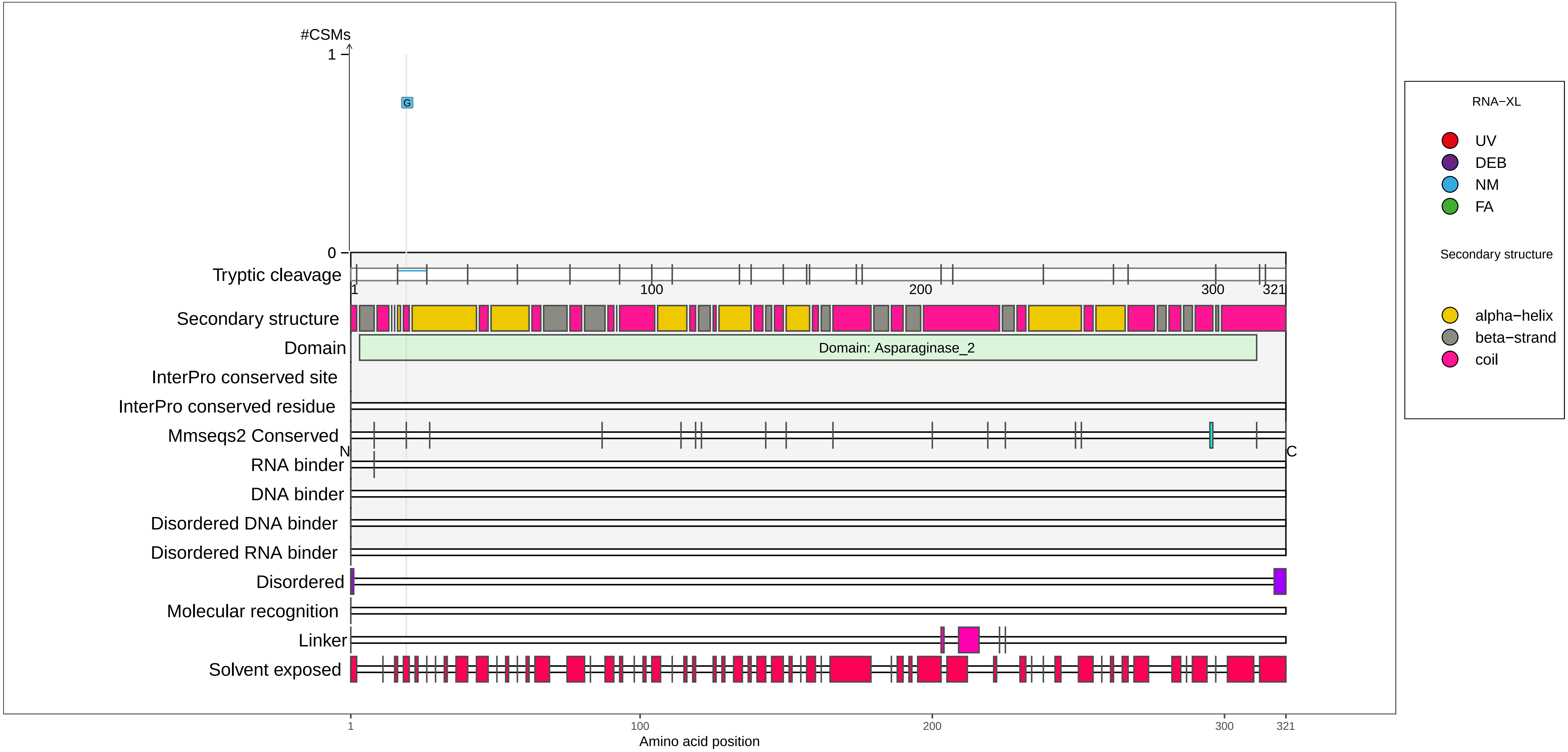

P37627
YHIJ_ECOLI Uncharacterized protein YhiJ

− Abundance:
tryptic [log10 Intensity]: NA (Q NA)
PAXdb K12 strain [ppm]: NA (Q NA)
PAXdb E.coli [ppm]: 0.24 (Q 30)

− RNA functions: not annotated

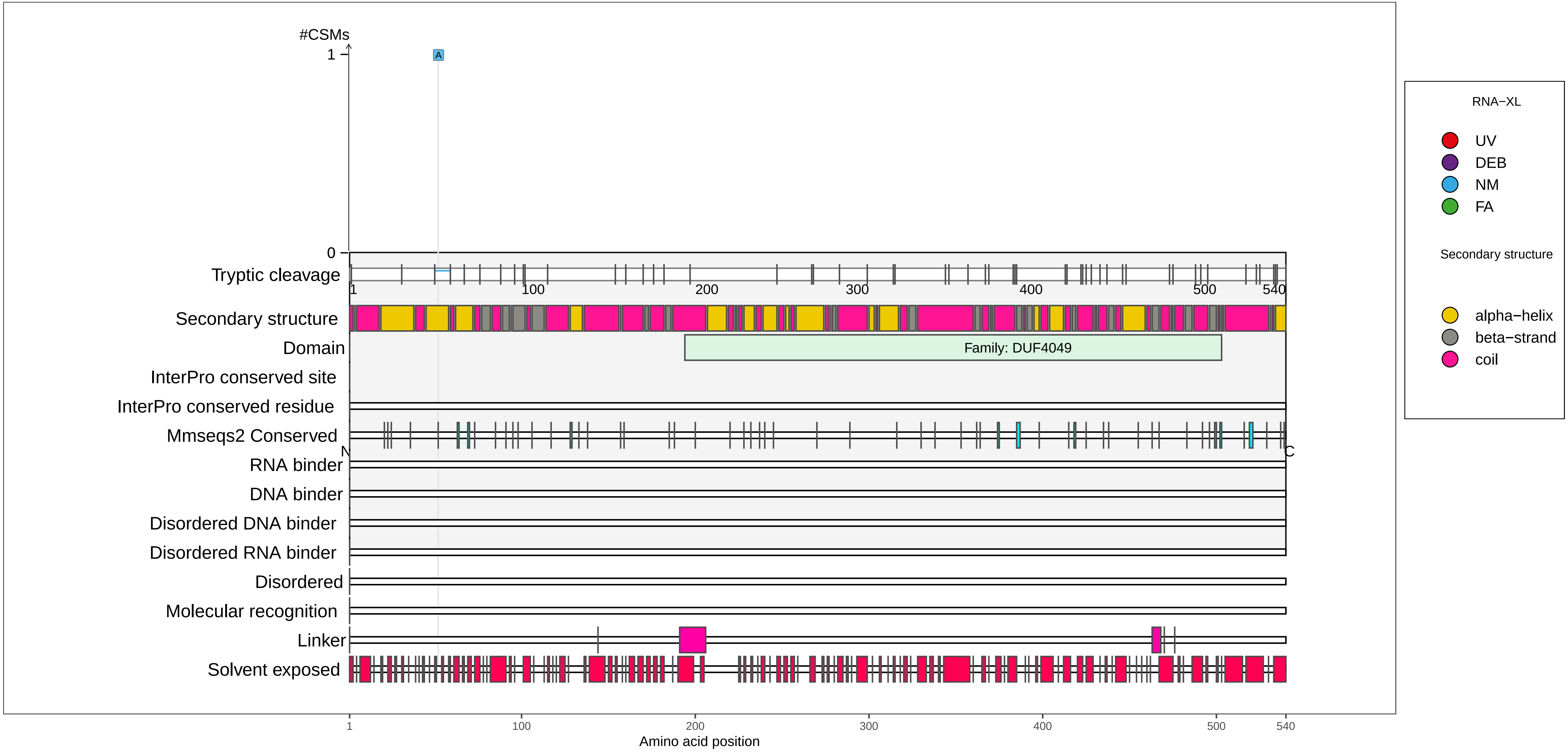

P37637
MDTF_ECOLI Multidrug resistance protein MdtF

− Abundance:
tryptic [log10 Intensity]: NA (Q NA)
PAXdb K12 strain [ppm]: 1.57 (Q 43)
PAXdb E.coli [ppm]: 0.65 (Q 40)

− RNA functions: not annotated

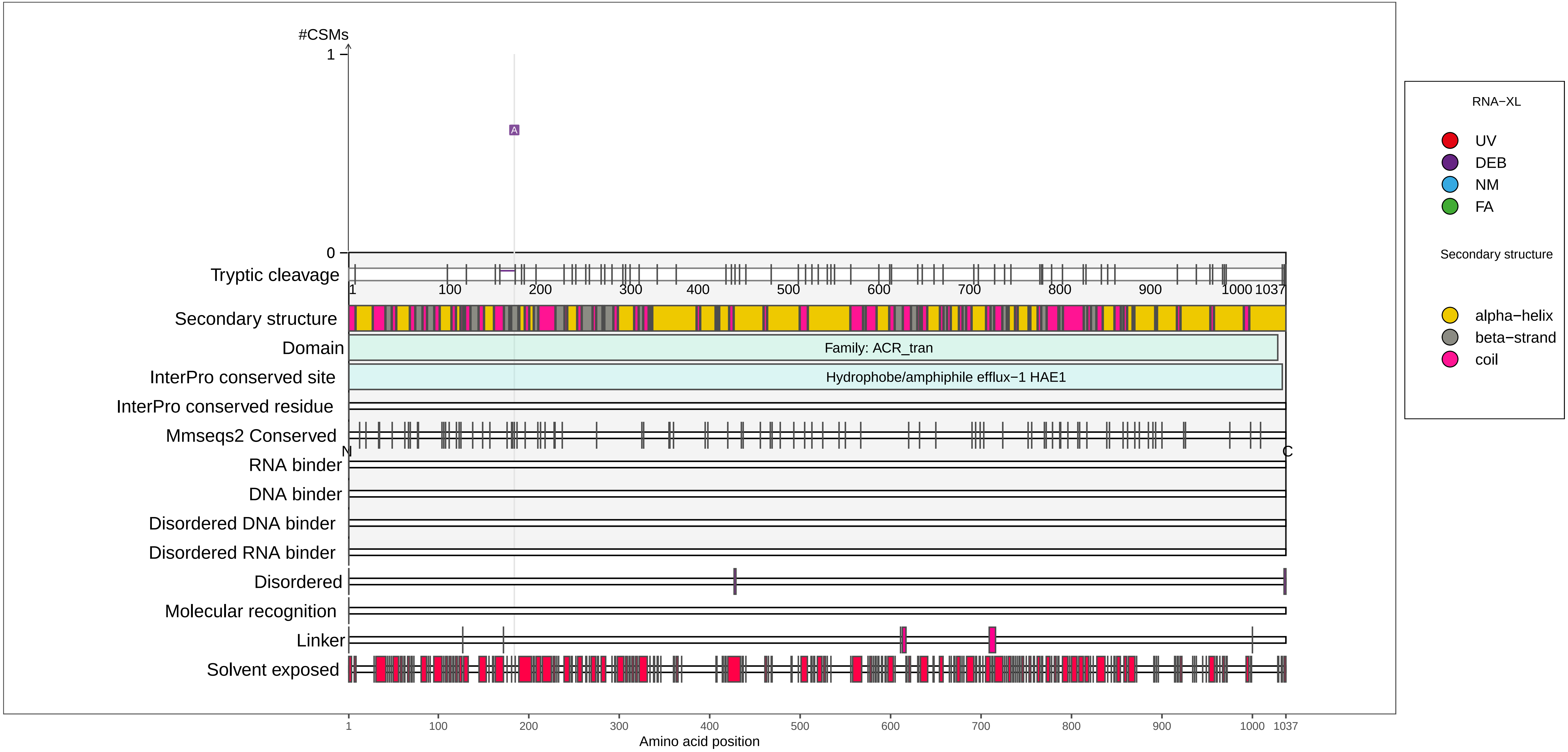

P37645
YHJG_ECOLI AsmA family protein YhjG

− Abundance:
tryptic [log10 Intensity]: NA (Q NA)
PAXdb K12 strain [ppm]: 1.68 (Q 48)
PAXdb E.coli [ppm]: 1.34 (Q 55)

− RNA functions: not annotated

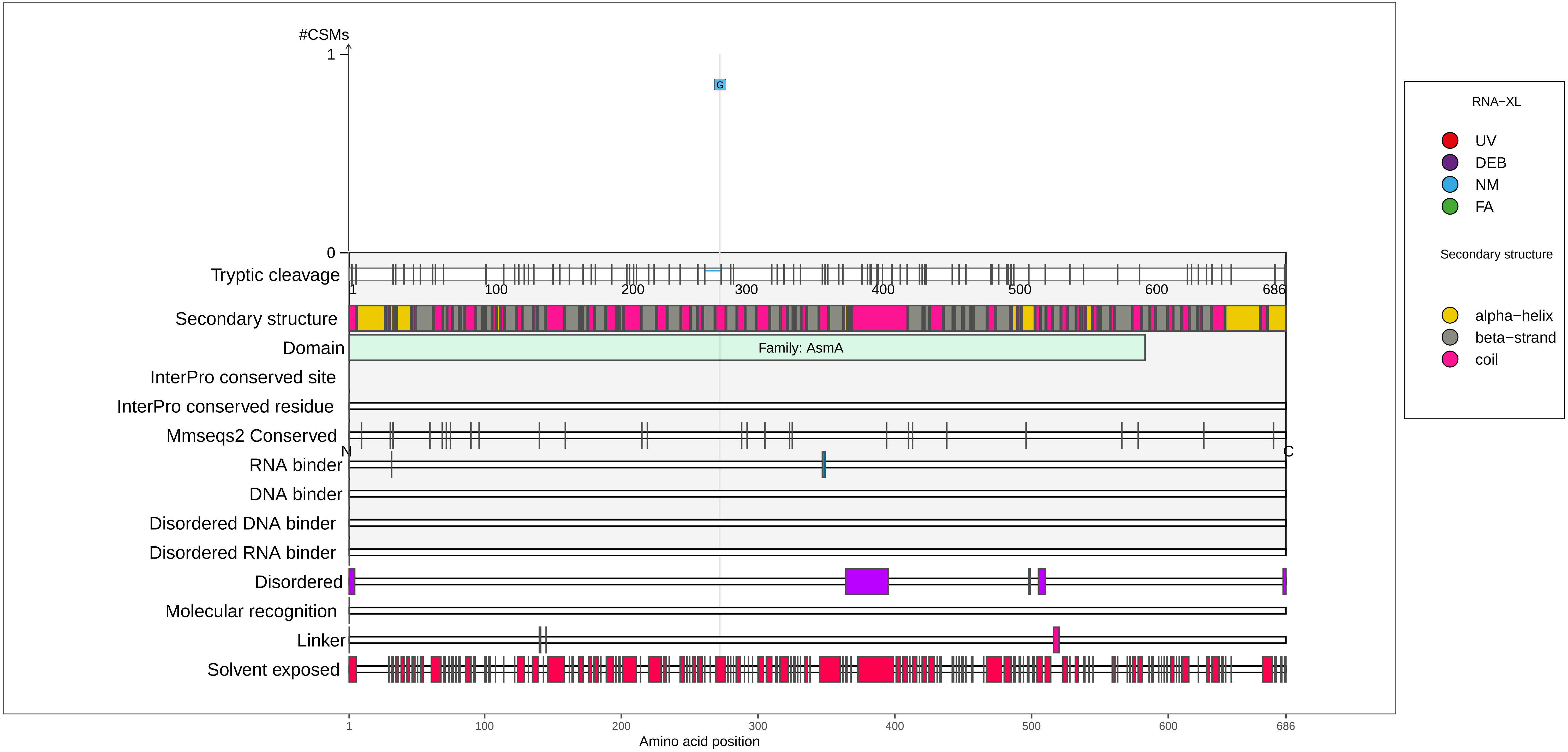

P37652
BCSB_ECOLI Cyclic di−GMP−binding protein

− Abundance:
tryptic [log10 Intensity]: 7.58 (Q 39)
PAXdb K12 strain [ppm]: NA (Q NA)
PAXdb E.coli [ppm]: −0.04 (Q 24)

− RNA functions: not annotated

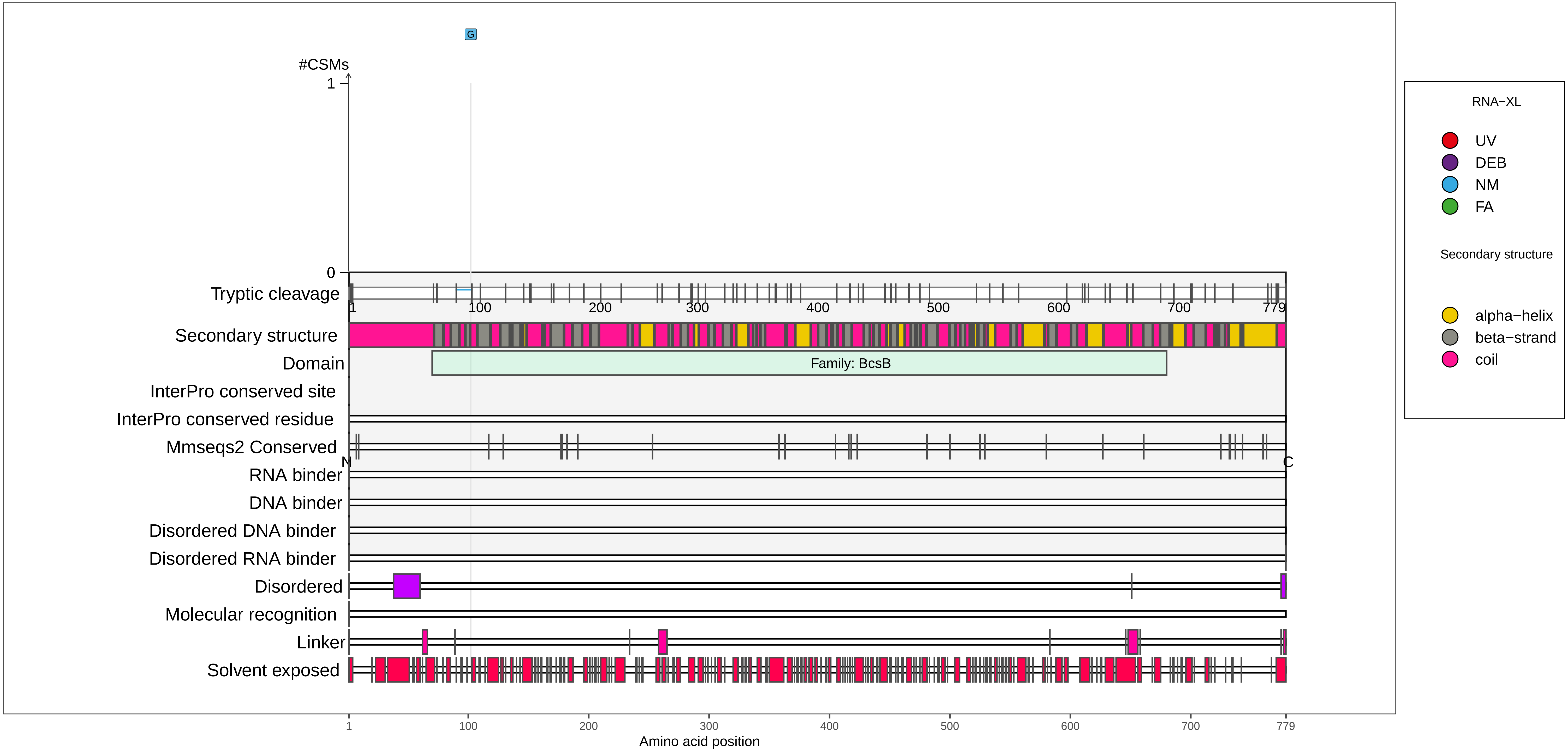

P37662
YHJX_ECOLI Uncharacterized MFS−type transporter YhjX

− Abundance:
tryptic [log10 Intensity]: NA (Q NA)
PAXdb K12 strain [ppm]: NA (Q NA)
PAXdb E.coli [ppm]: −1.64 (Q 3)

− RNA functions: not annotated

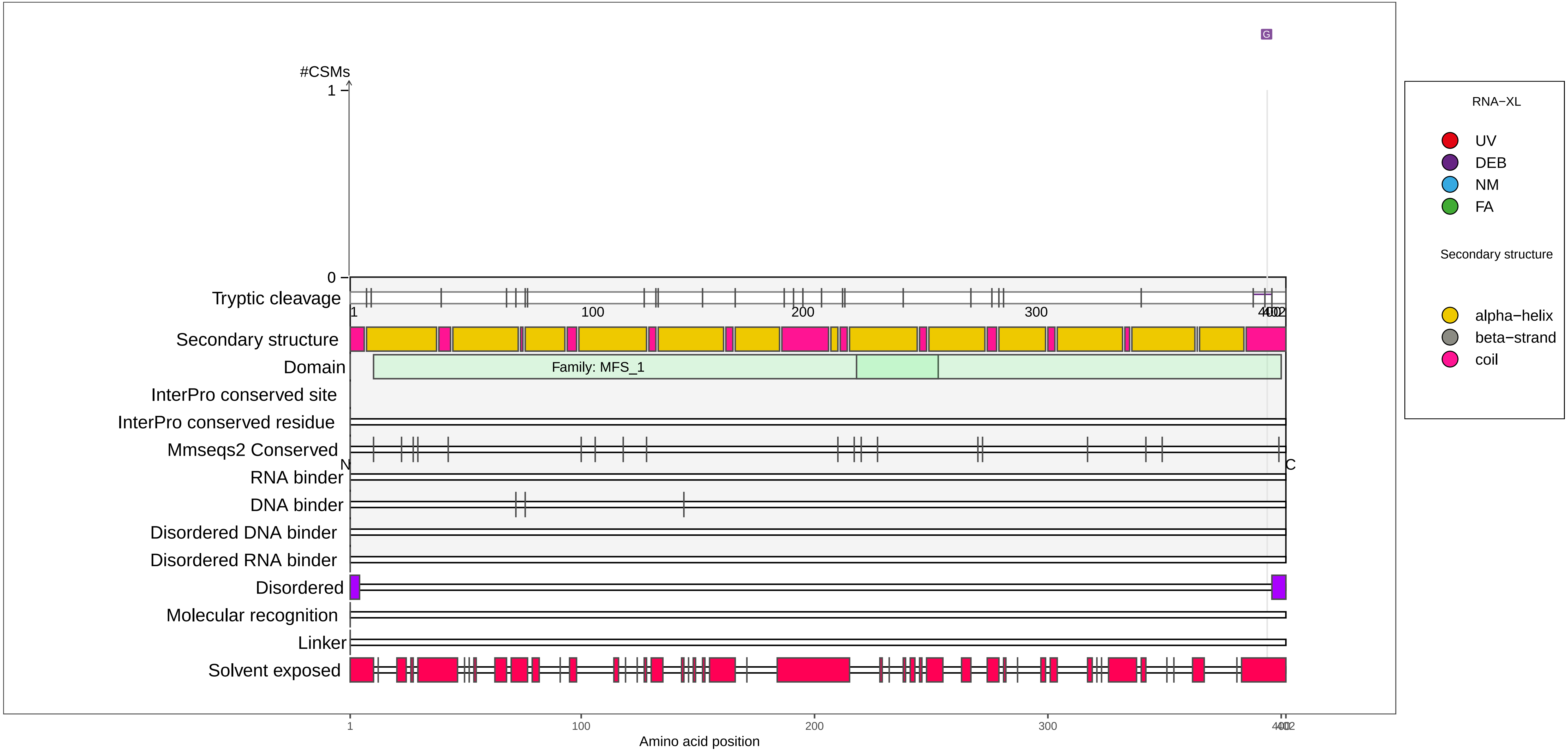

P37689
GPMI_ECOLI 2,3−bisphosphoglycerate−independent phosphoglycerate mutase
− Abundance:
tryptic [log10 Intensity]: 8.88 (Q 85)
PAXdb K12 strain [ppm]: 2.57 (Q 80)
PAXdb E.coli [ppm]: 2.67 (Q 89)

− RNA functions: not annotated

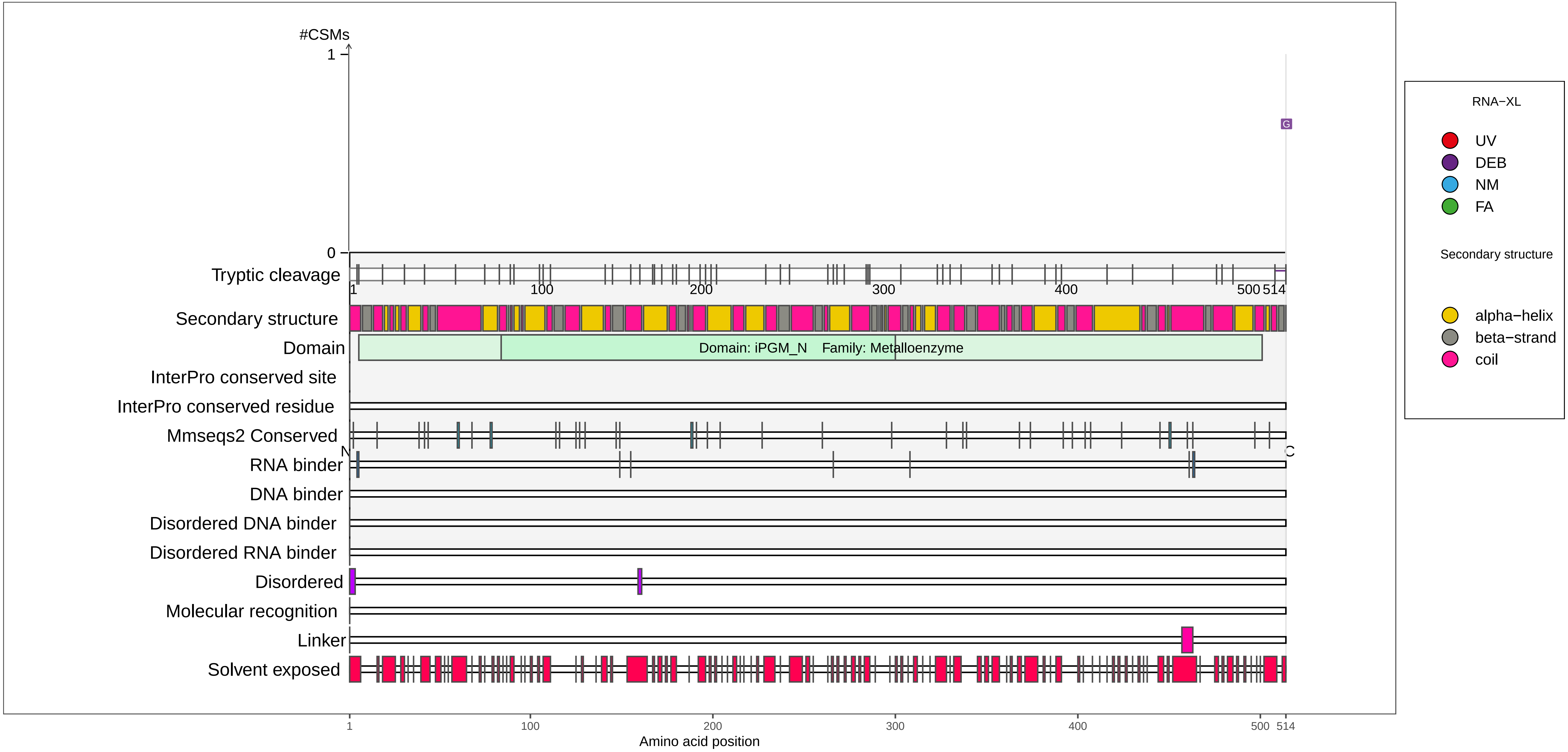

P38051
MENF_ECOLI Isochorismate synthase MenF
− Abundance:
tryptic [log10 Intensity]: 7.38 (Q 28)
PAXdb K12 strain [ppm]: 0.92 (Q 8)
PAXdb E.coli [ppm]: 0.12 (Q 27)

− RNA functions: not annotated

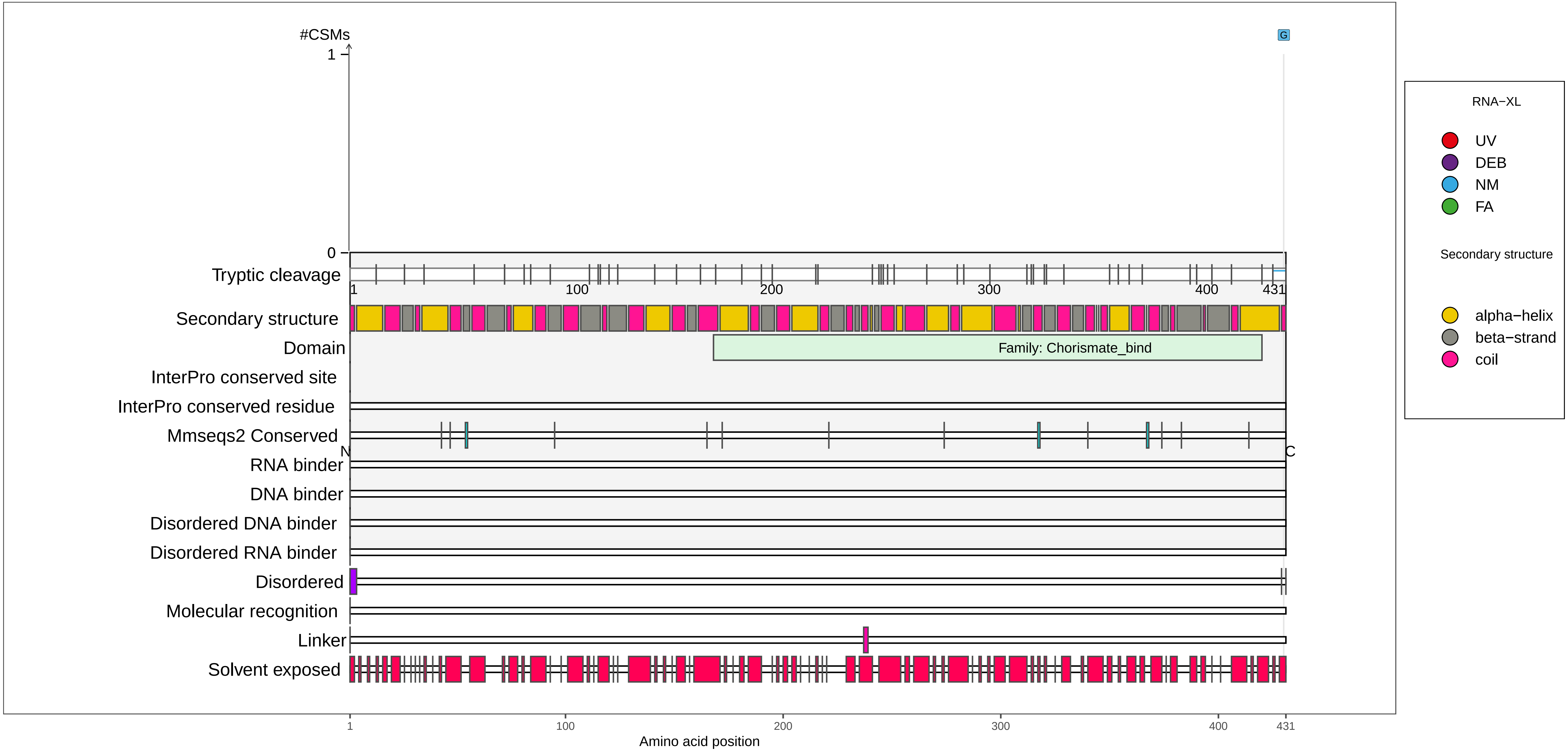

P38521
YGGL_ECOLI Uncharacterized protein YggL

− Abundance:
tryptic [log10 Intensity]: 7.95 (Q 56)
PAXdb K12 strain [ppm]: 2.32 (Q 71)
PAXdb E.coli [ppm]: 2.62 (Q 88)

− RNA functions: not annotated

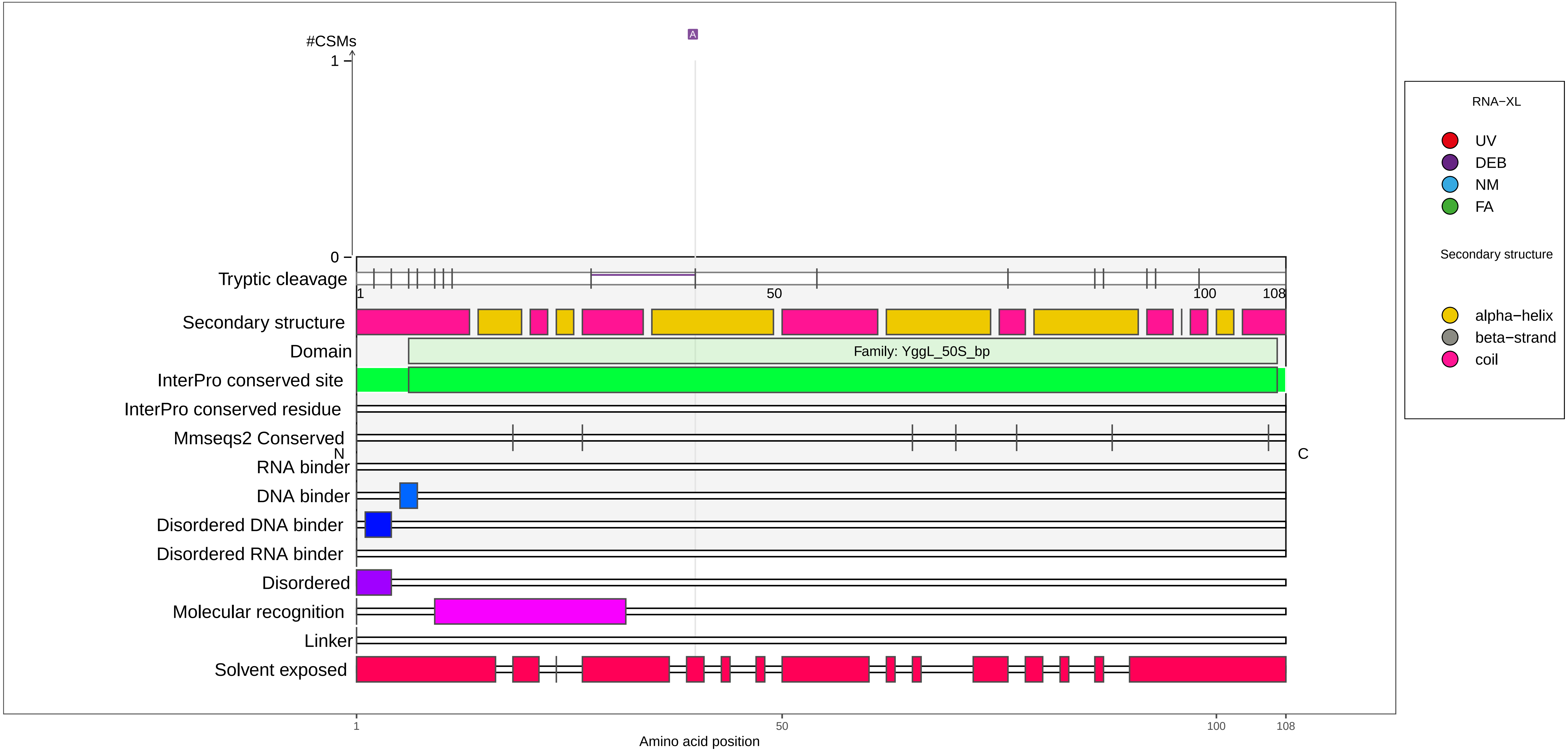

P39161
UXUR_ECOLI Uxu operon transcriptional regulator

− Abundance:
tryptic [log10 Intensity]: NA (Q NA)
PAXdb K12 strain [ppm]: NA (Q NA)
PAXdb E.coli [ppm]: 0.41 (Q 34)

− RNA functions:
RNA biosynthetic process; RNA metabolic process

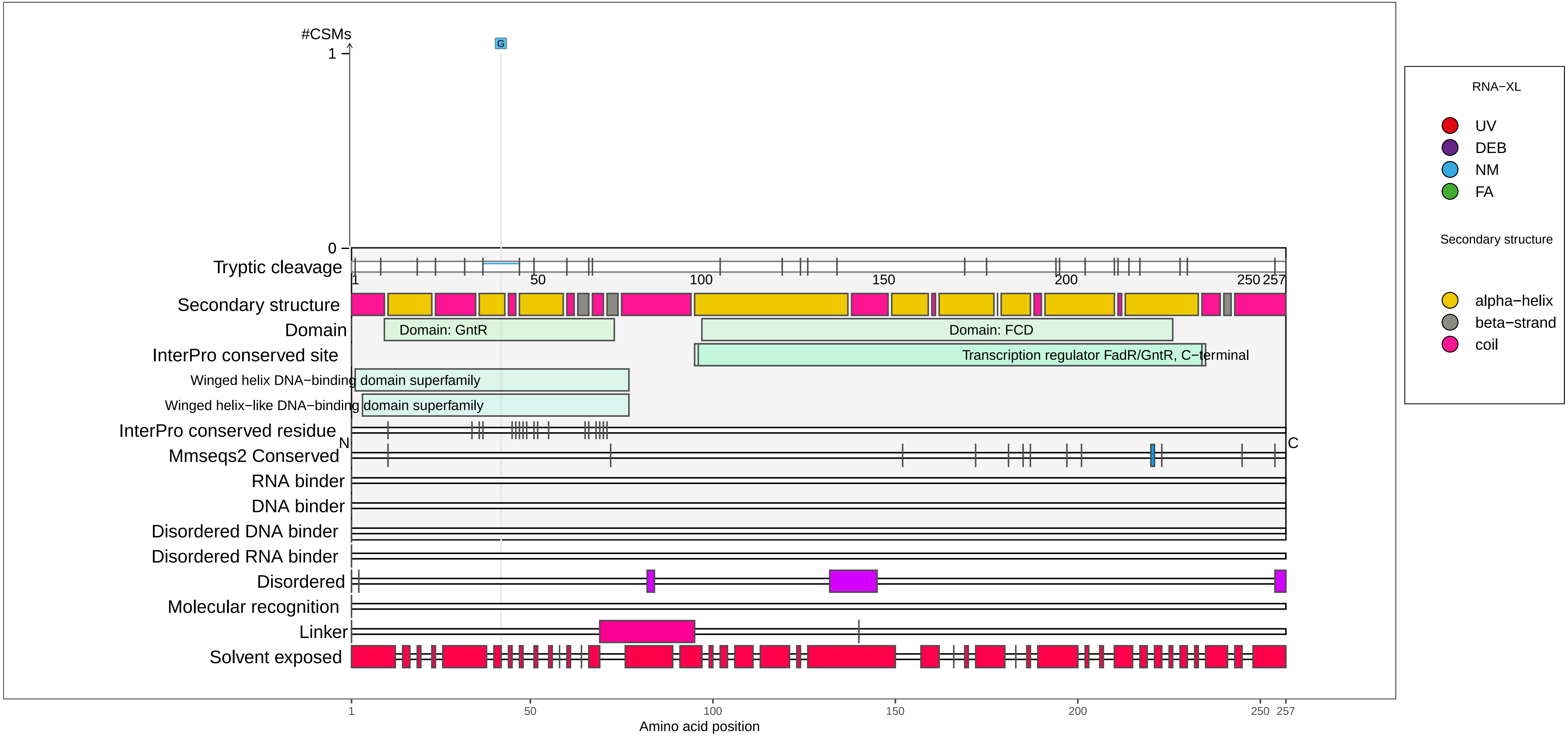

P39187
YTFJ_ECOLI Uncharacterized protein YtfJ

− Abundance:
tryptic [log10 Intensity]: 7.04 (Q 15)
PAXdb K12 strain [ppm]: 2.06 (Q 62)
PAXdb E.coli [ppm]: 2.52 (Q 86)

− RNA functions: not annotated

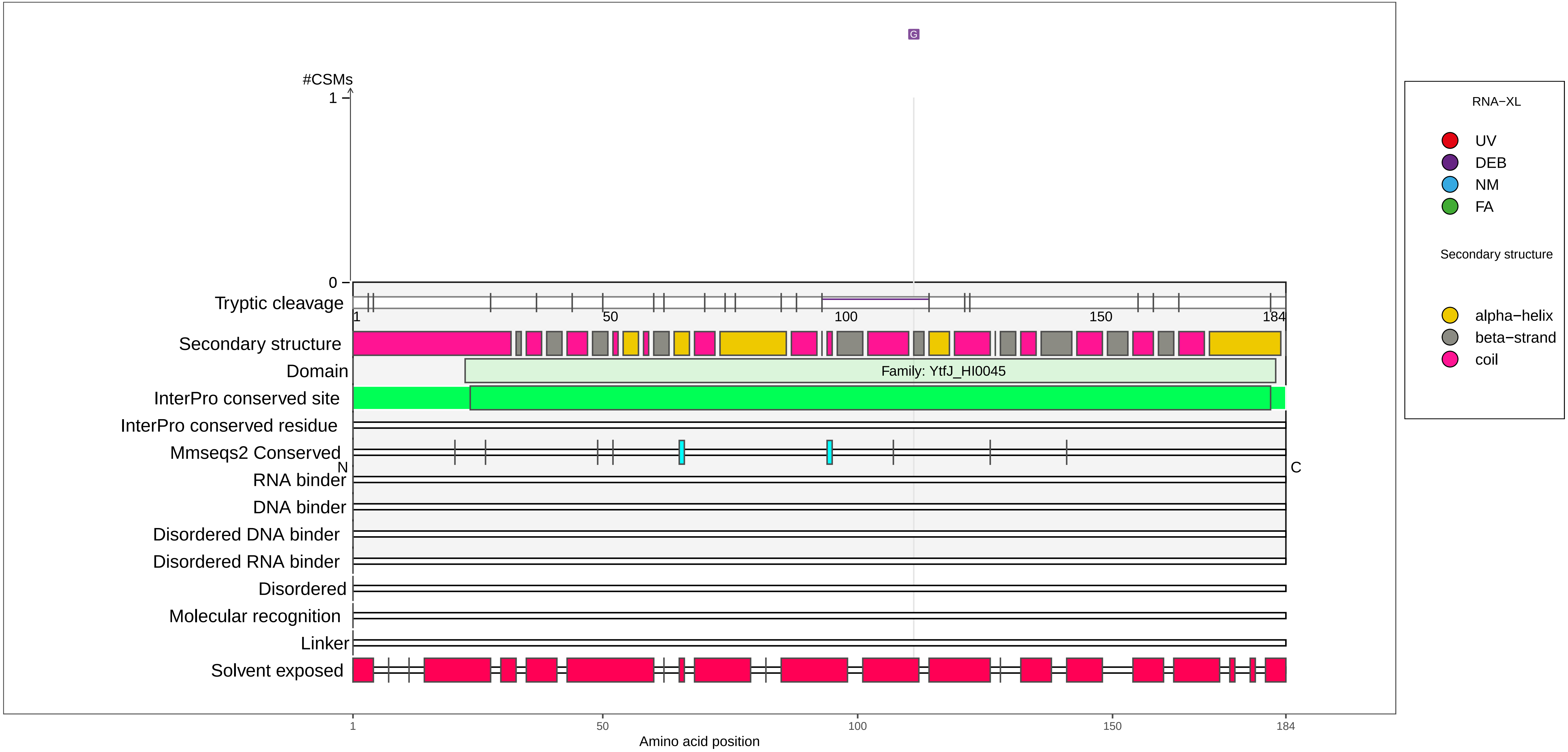

P39208
IDNK_ECOLI Thermosensitive gluconokinase

− Abundance:
tryptic [log10 Intensity]: NA (Q NA)
PAXdb K12 strain [ppm]: NA (Q NA)
PAXdb E.coli [ppm]: −0.61 (Q 14)

− RNA functions: not annotated

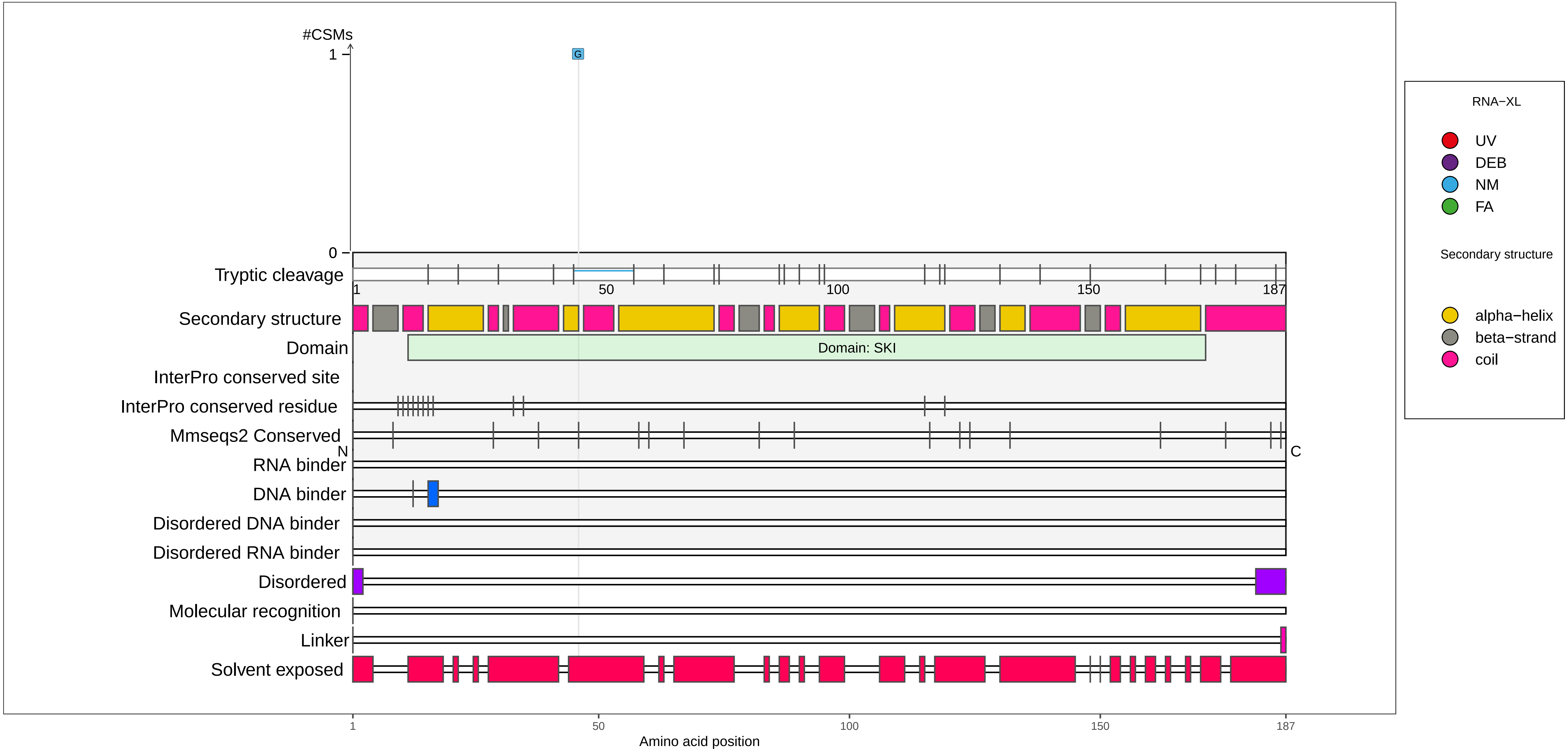

P39265
ALSB_ECOLI D−allose−binding periplasmic protein

− Abundance:
tryptic [log10 Intensity]: 7.41 (Q 30)
PAXdb K12 strain [ppm]: 2.43 (Q 75)
PAXdb E.coli [ppm]: 1.64 (Q 63)

− RNA functions: not annotated

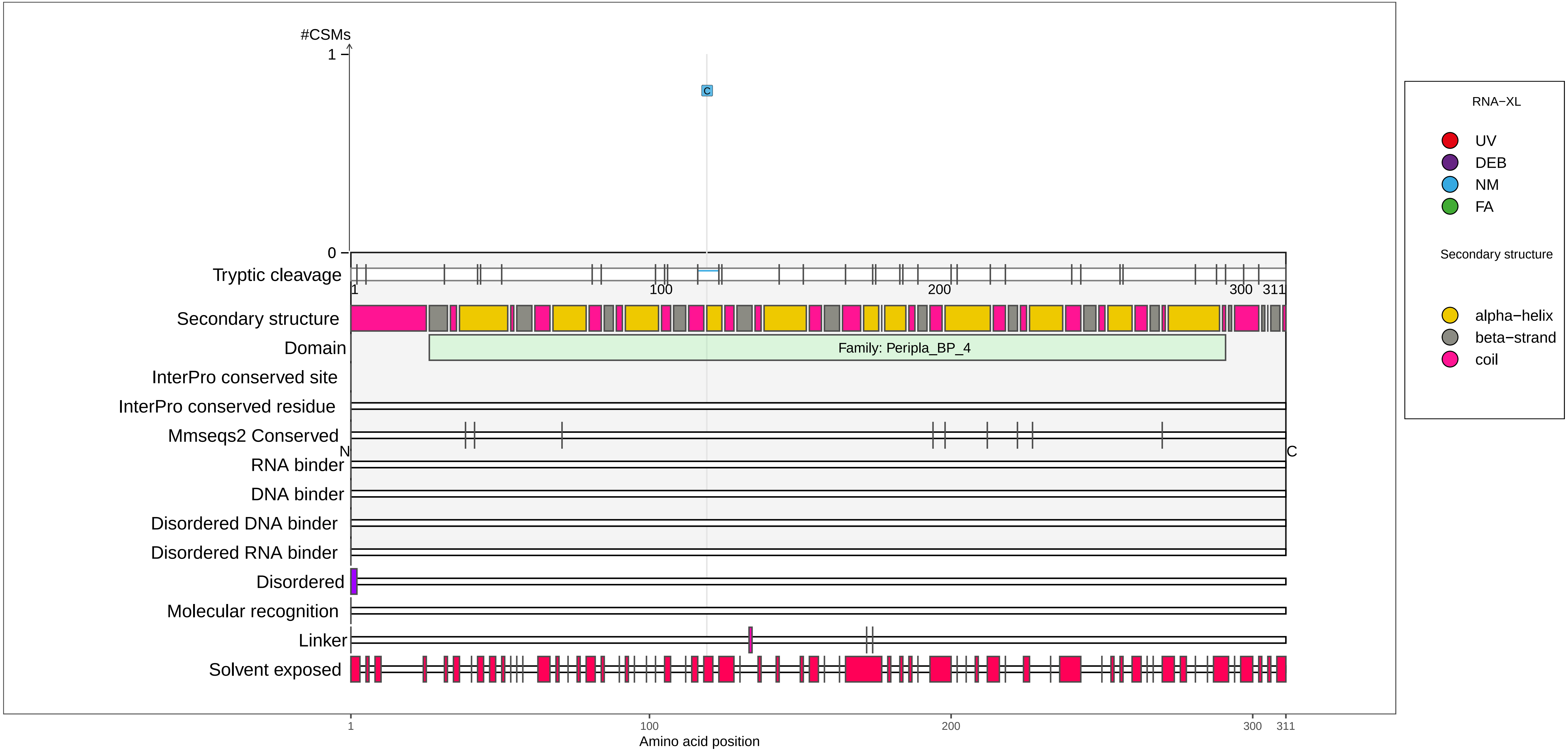

P39282
YJEM_ECOLI Inner membrane transporter YjeM

− Abundance:
tryptic [log10 Intensity]: 7.5 (Q 35)
PAXdb K12 strain [ppm]: NA (Q NA)
PAXdb E.coli [ppm]: NA (Q NA)

− RNA functions: not annotated

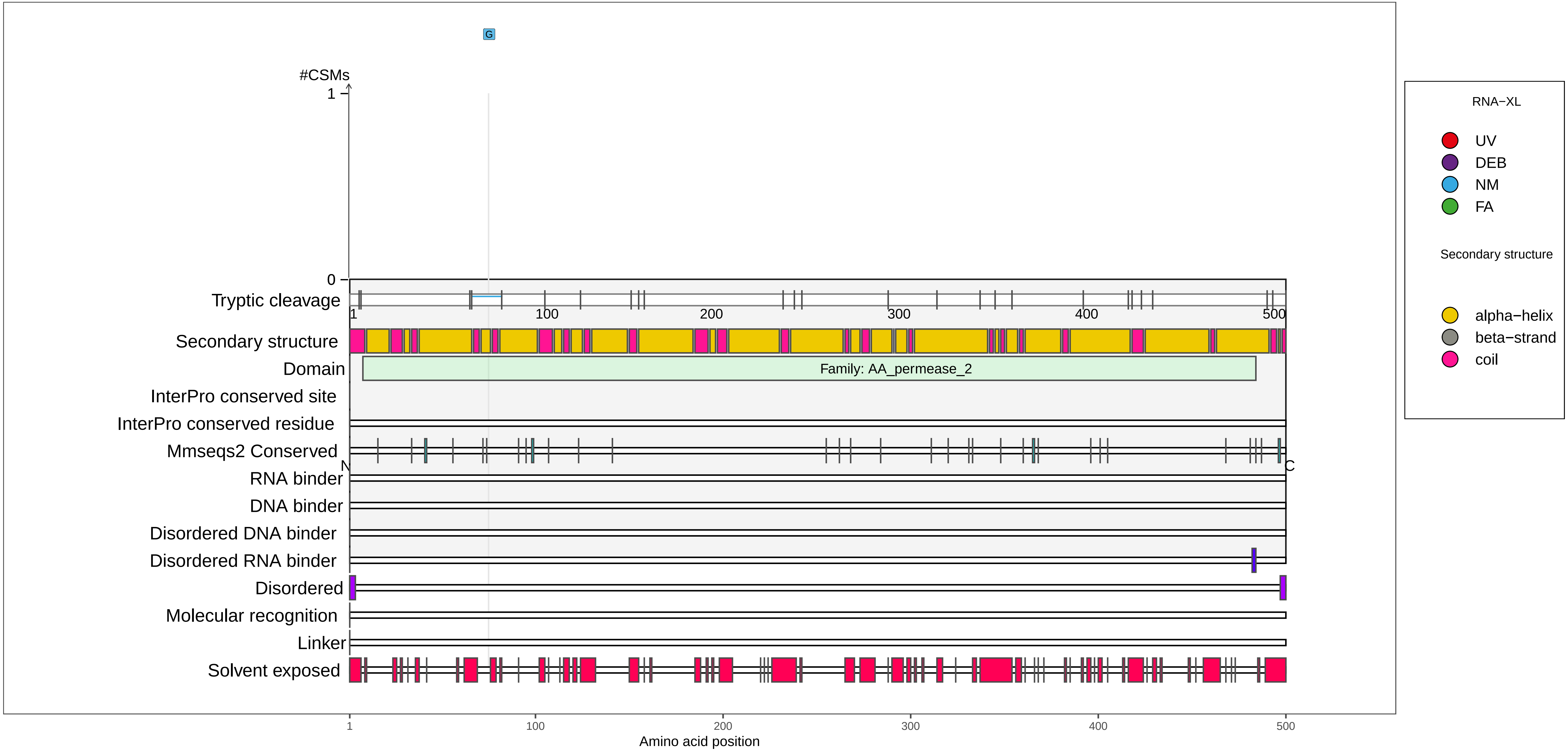

P39298
YJFP_ECOLI Esterase YjfP

− Abundance:
tryptic [log10 Intensity]: 7.54 (Q 37)
PAXdb K12 strain [ppm]: NA (Q NA)
PAXdb E.coli [ppm]: 1.15 (Q 51)

− RNA functions: not annotated

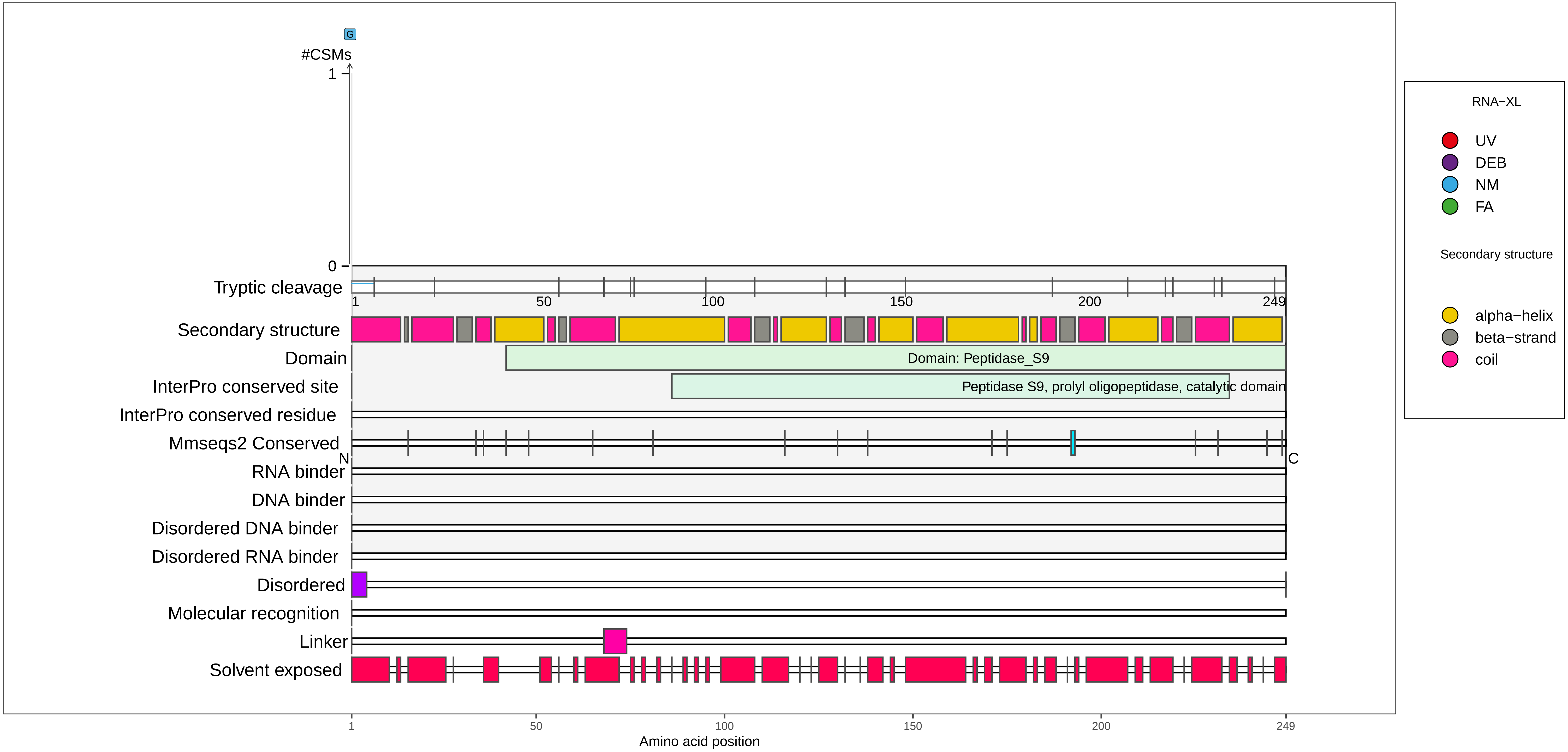

P39321
TAMB_ECOLI Translocation and assembly module subunit TamB

− Abundance:
tryptic [log10 Intensity]: 7.9 (Q 54)
PAXdb K12 strain [ppm]: 0.88 (Q 7)
PAXdb E.coli [ppm]: 0.53 (Q 37)

− RNA functions: not annotated

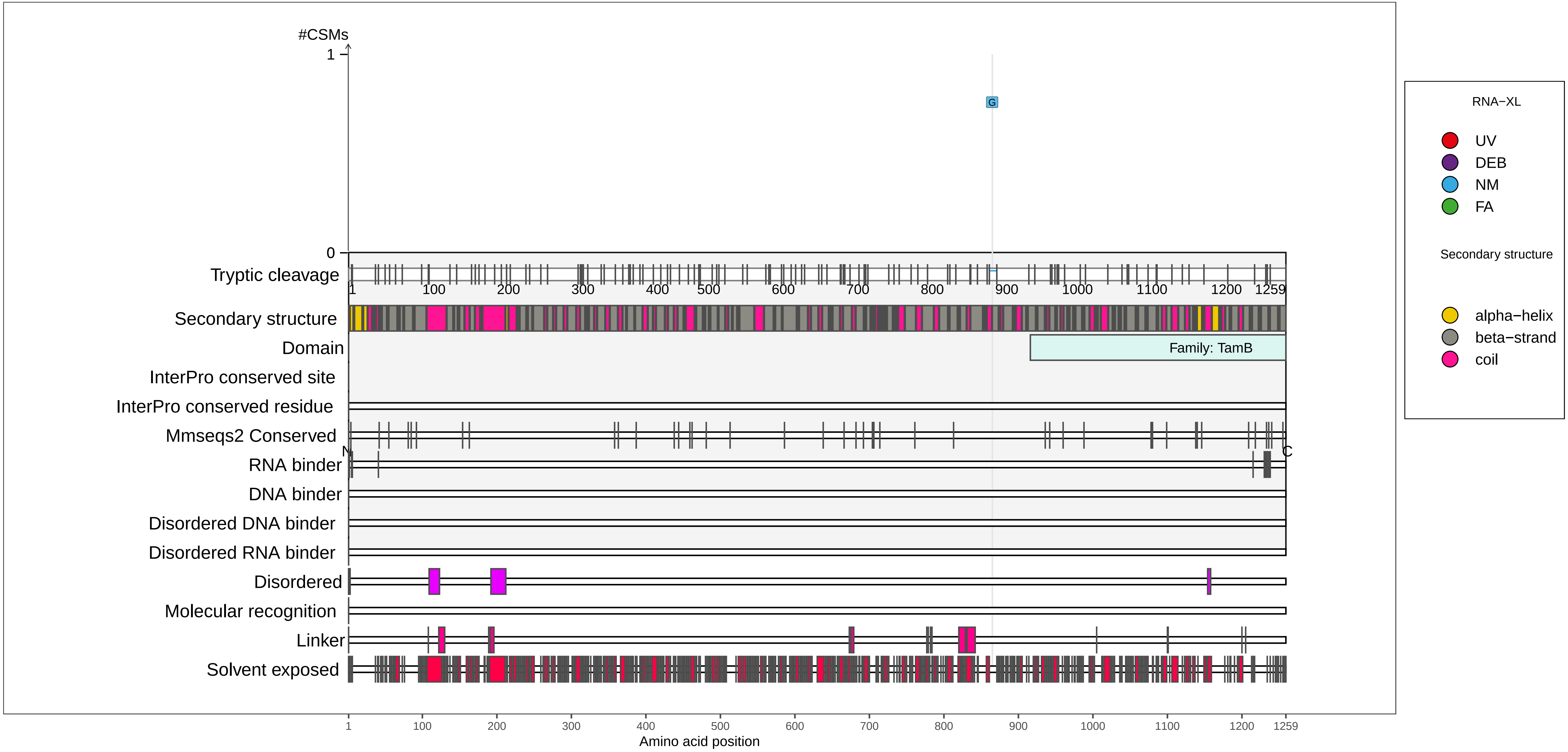

P39384
YJIM_ECOLI Putative dehydratase subunit YjiM
− Abundance:
tryptic [log10 Intensity]: NA (Q NA)
PAXdb K12 strain [ppm]: 0.88 (Q 7)
PAXdb E.coli [ppm]: −0.76 (Q 12)

− RNA functions: not annotated

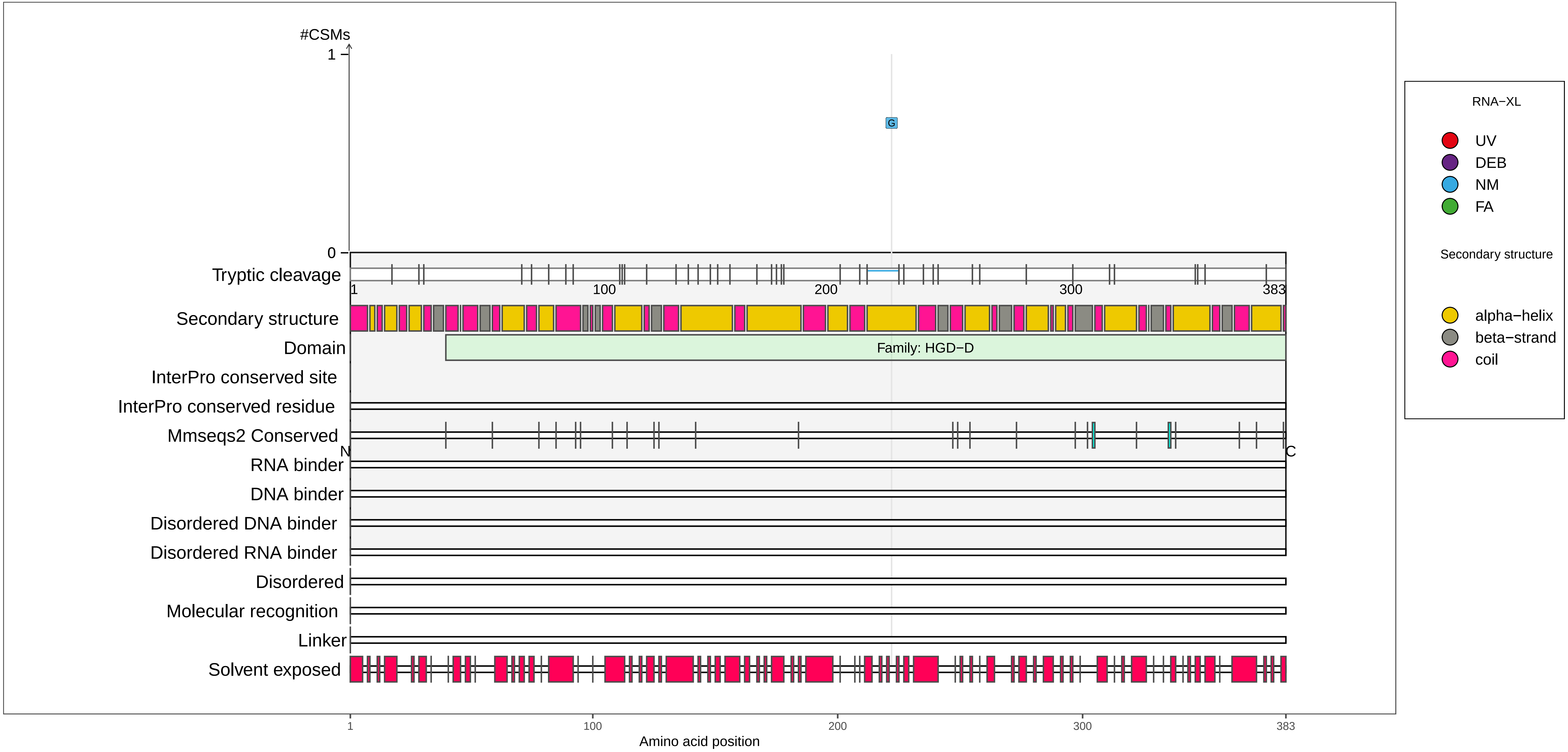

P39829
GARD_ECOLI Galactarate dehydratase (L−threo−forming)

− Abundance:
tryptic [log10 Intensity]: 7.64 (Q 42)
PAXdb K12 strain [ppm]: NA (Q NA)
PAXdb E.coli [ppm]: 0.43 (Q 34)

− RNA functions: not annotated

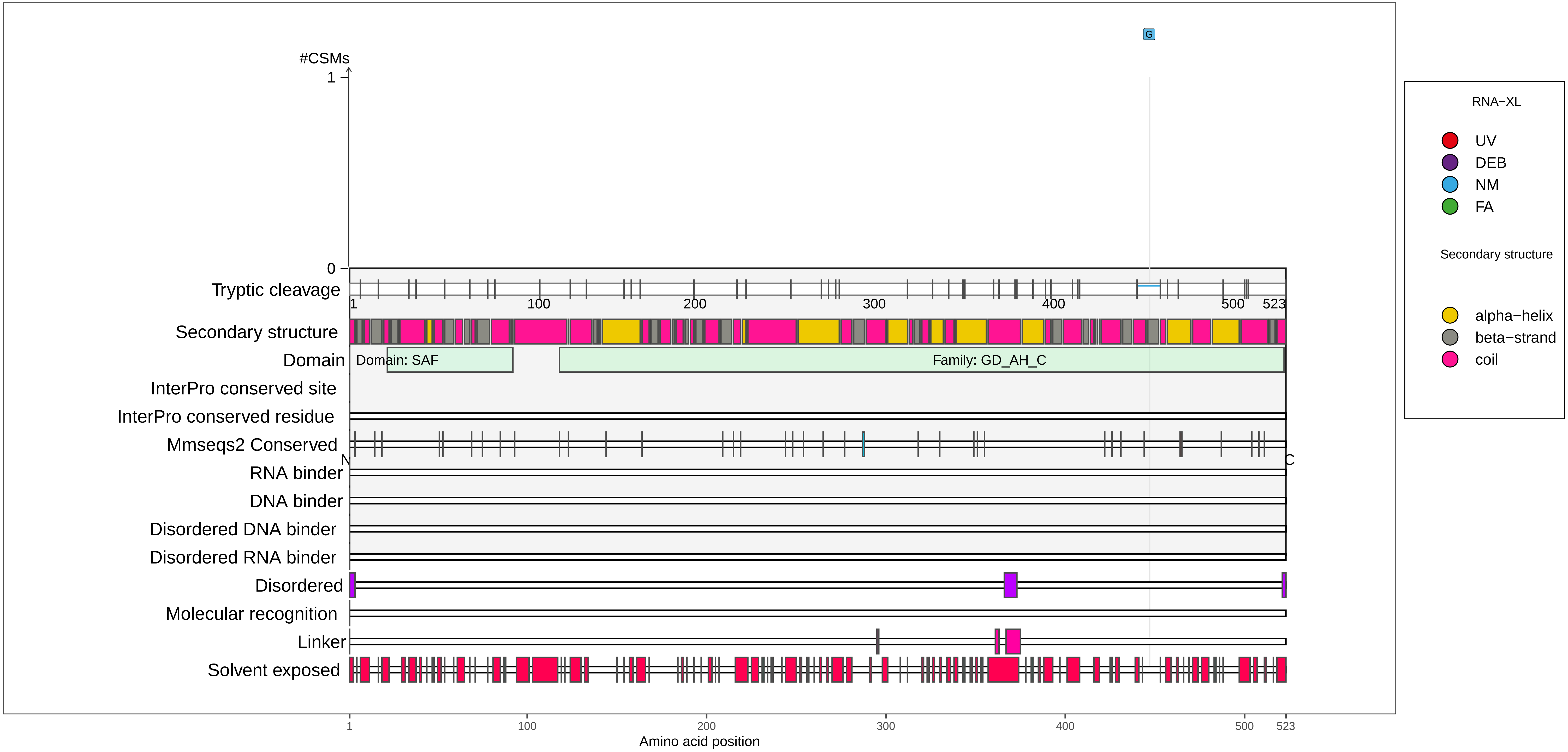

P40120
OPGD_ECOLI Glucans biosynthesis protein D

− Abundance:
tryptic [log10 Intensity]: 8.3 (Q 69)
PAXdb K12 strain [ppm]: 1.46 (Q 36)
PAXdb E.coli [ppm]: 1.37 (Q 56)

− RNA functions: not annotated

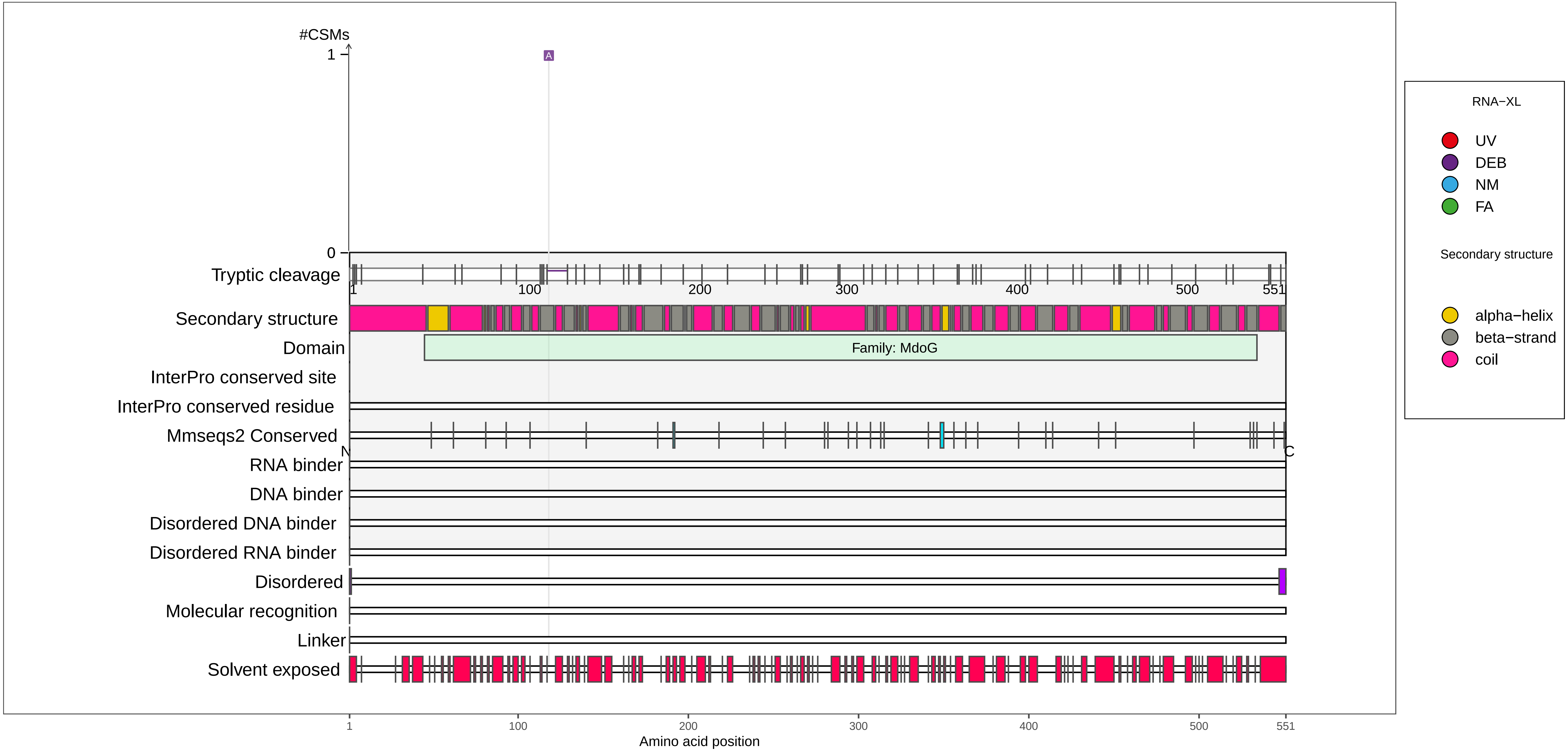

P41052
MLTB_ECOLI Membrane−bound lytic murein transglycosylase B

− Abundance:
tryptic [log10 Intensity]: 7.96 (Q 56)
PAXdb K12 strain [ppm]: 0.8 (Q 5)
PAXdb E.coli [ppm]: 0.98 (Q 47)

− RNA functions: not annotated

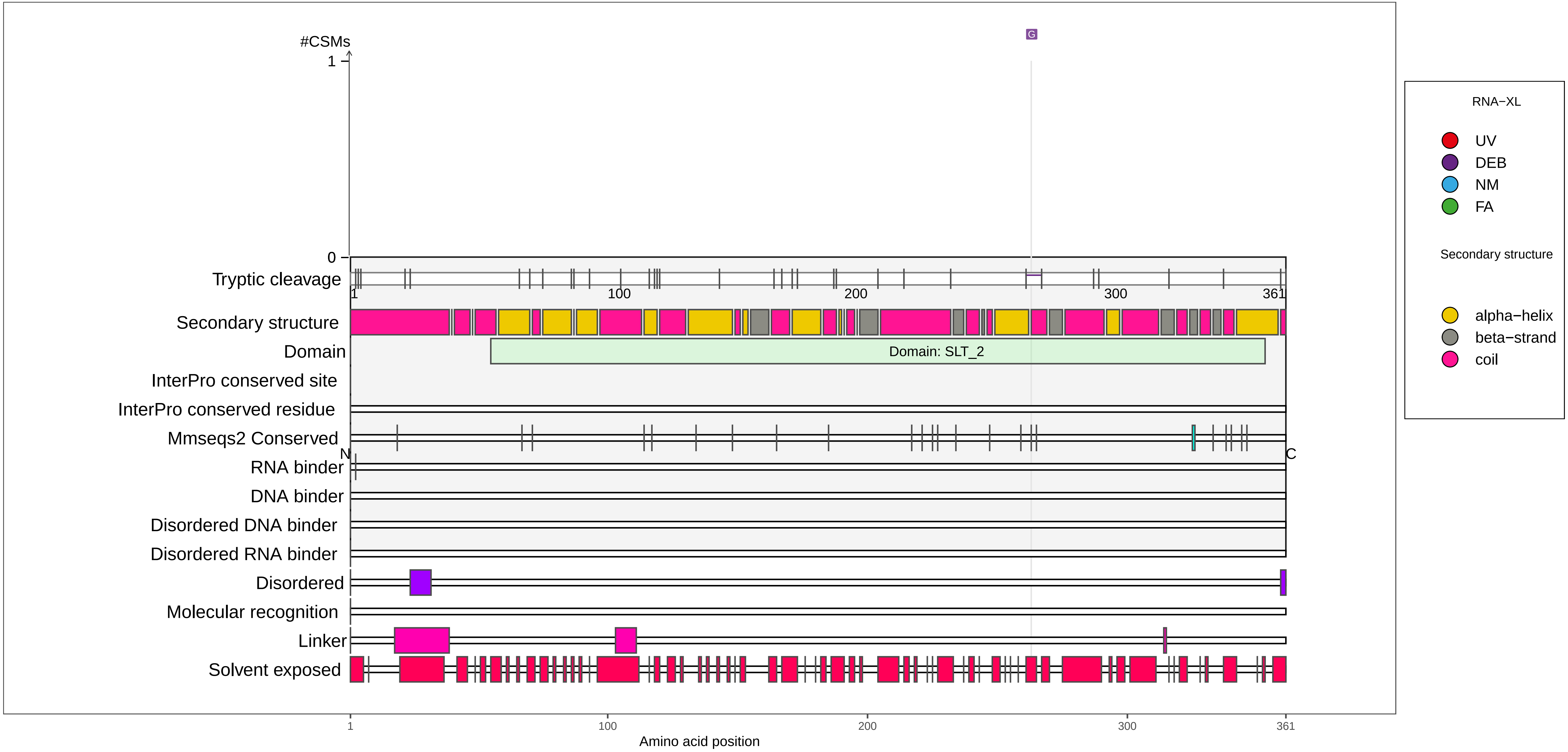

P42626
YHAM_ECOLI UPF0597 protein YhaM
− Abundance:
tryptic [log10 Intensity]: NA (Q NA)
PAXdb K12 strain [ppm]: NA (Q NA)
PAXdb E.coli [ppm]: −0.94 (Q 10)

− RNA functions: not annotated

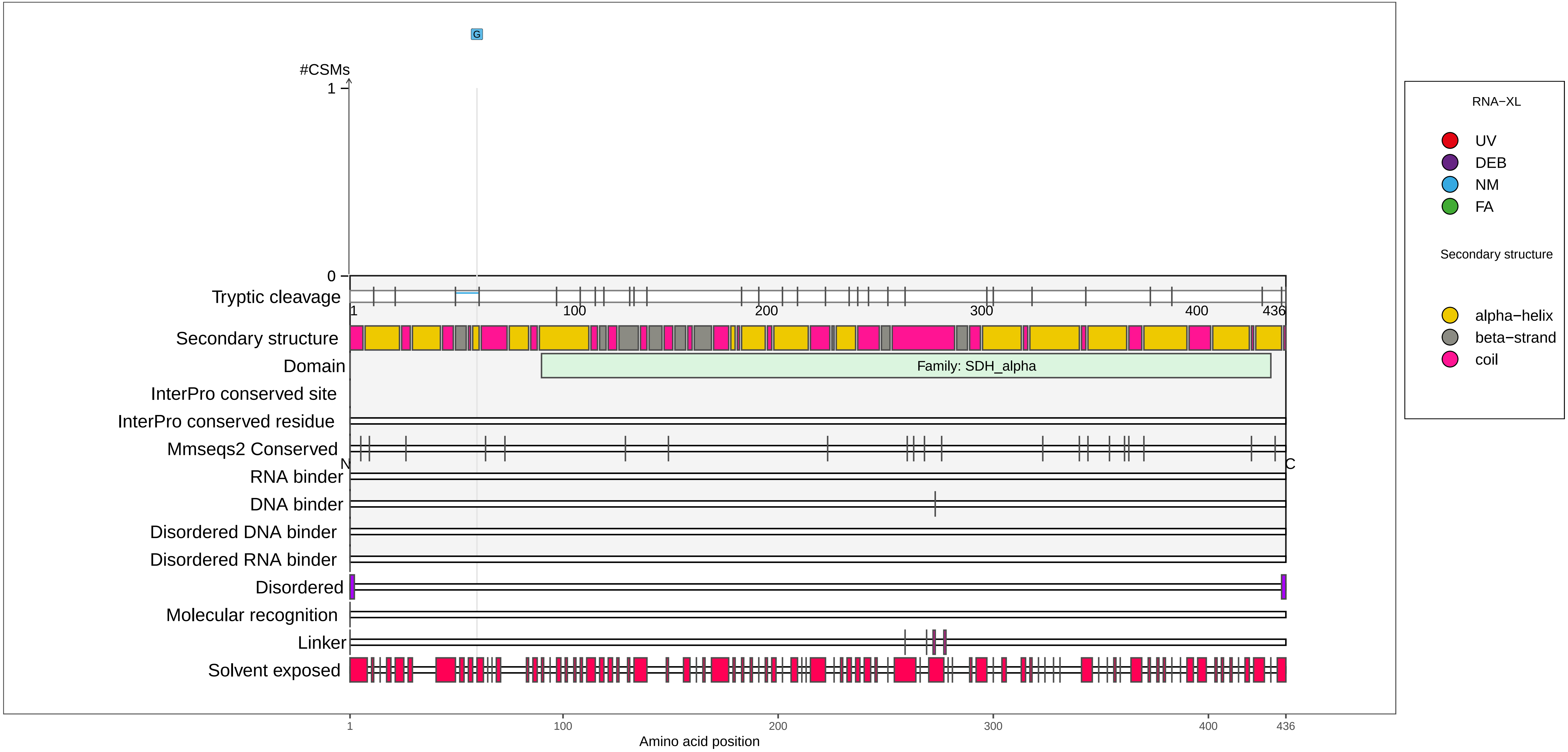

P43329
HRPA_ECOLI ATP−dependent RNA helicase HrpA

− Abundance:
tryptic [log10 Intensity]: 7.96 (Q 56)
PAXdb K12 strain [ppm]: 1.19 (Q 20)
PAXdb E.coli [ppm]: 1.43 (Q 58)

− RNA functions:
mRNA metabolic process; mRNA processing; RNA helicase activity; RNA metabolic process
RNA modification; RNA processing

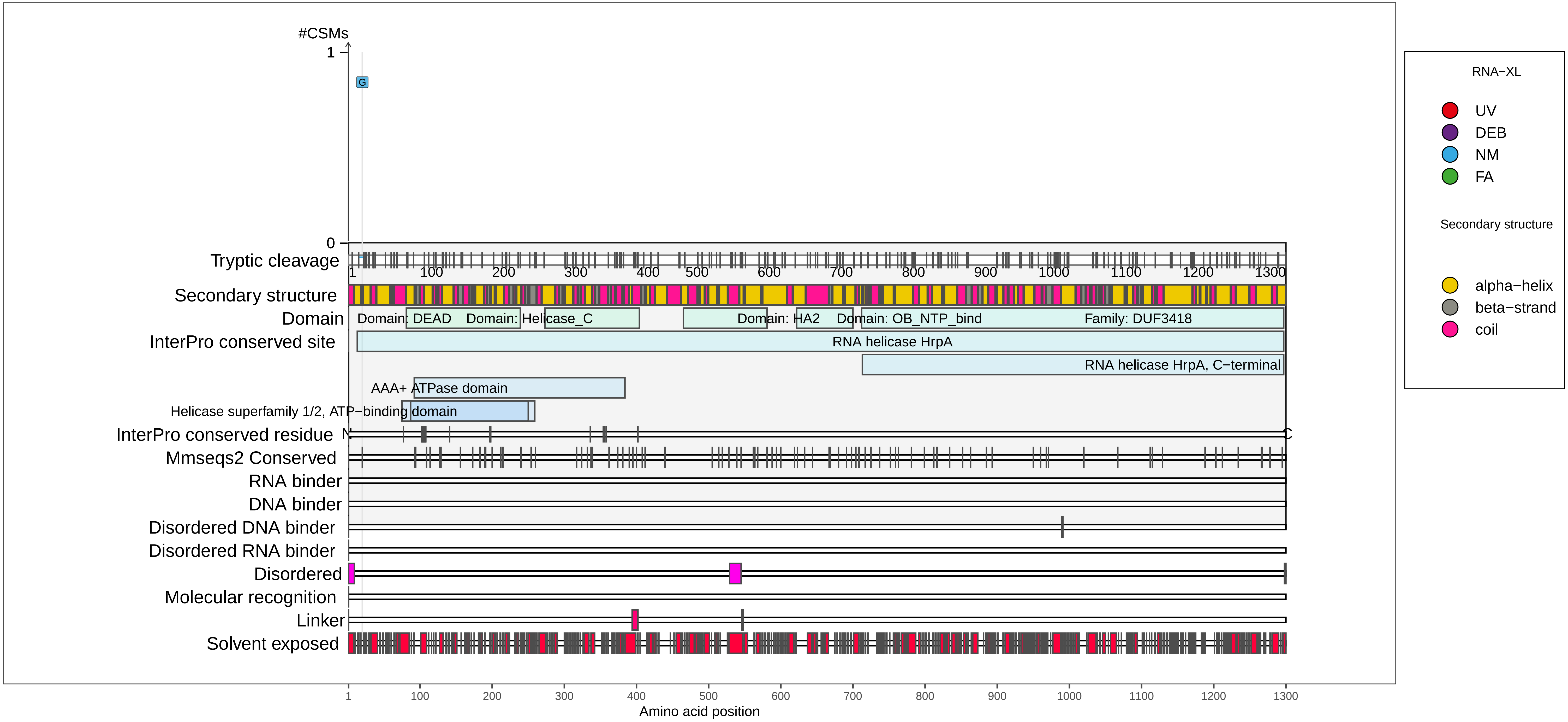

P45541
FRLC_ECOLI Fructoselysine 3−epimerase

− Abundance:
tryptic [log10 Intensity]: NA (Q NA)
PAXdb K12 strain [ppm]: NA (Q NA)
PAXdb E.coli [ppm]: −0.74 (Q 12)

− RNA functions: not annotated

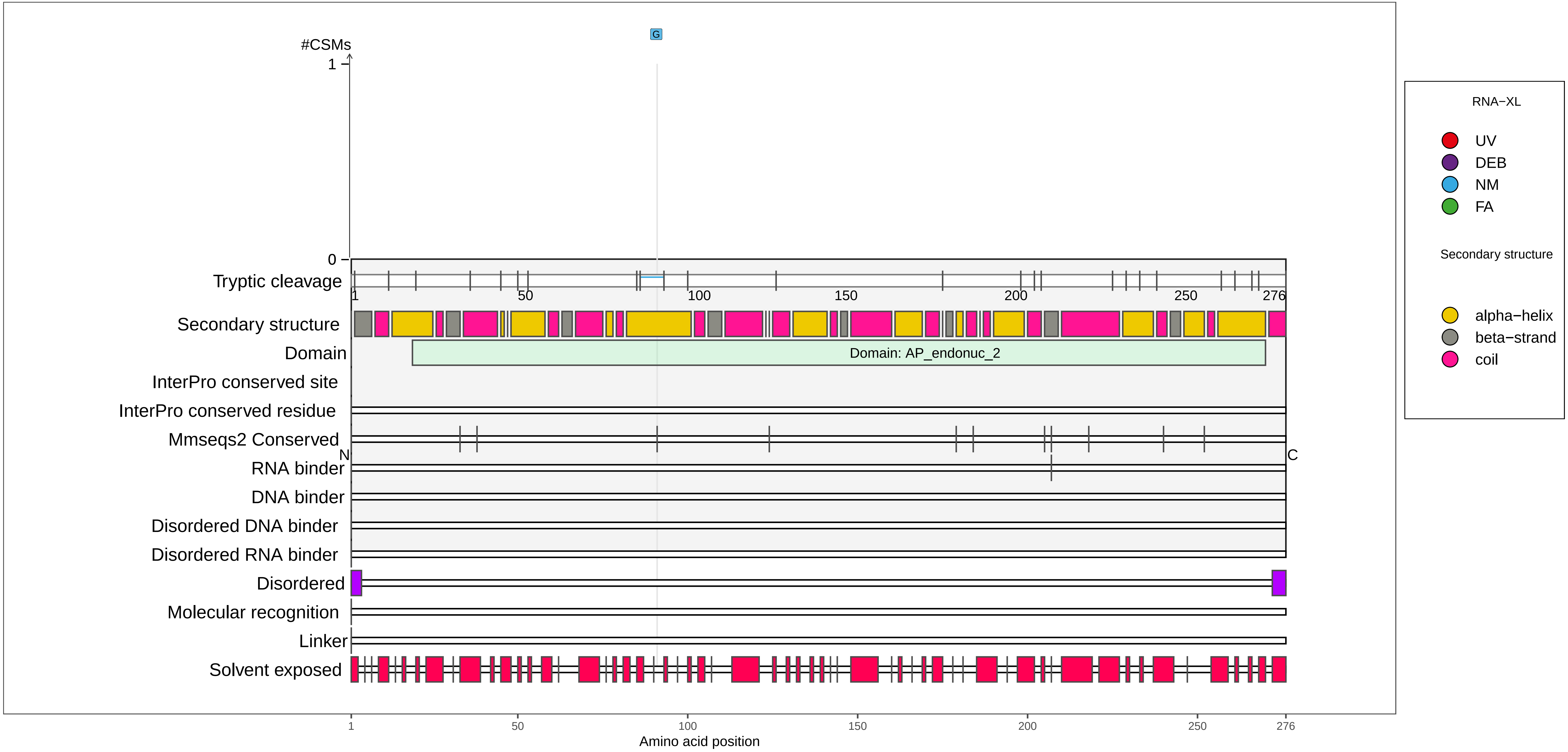

P45552
YHFZ_ECOLI Uncharacterized protein YhfZ

− Abundance:
tryptic [log10 Intensity]: 7.29 (Q 25)
PAXdb K12 strain [ppm]: NA (Q NA)
PAXdb E.coli [ppm]: −0.42 (Q 17)

− RNA functions: not annotated

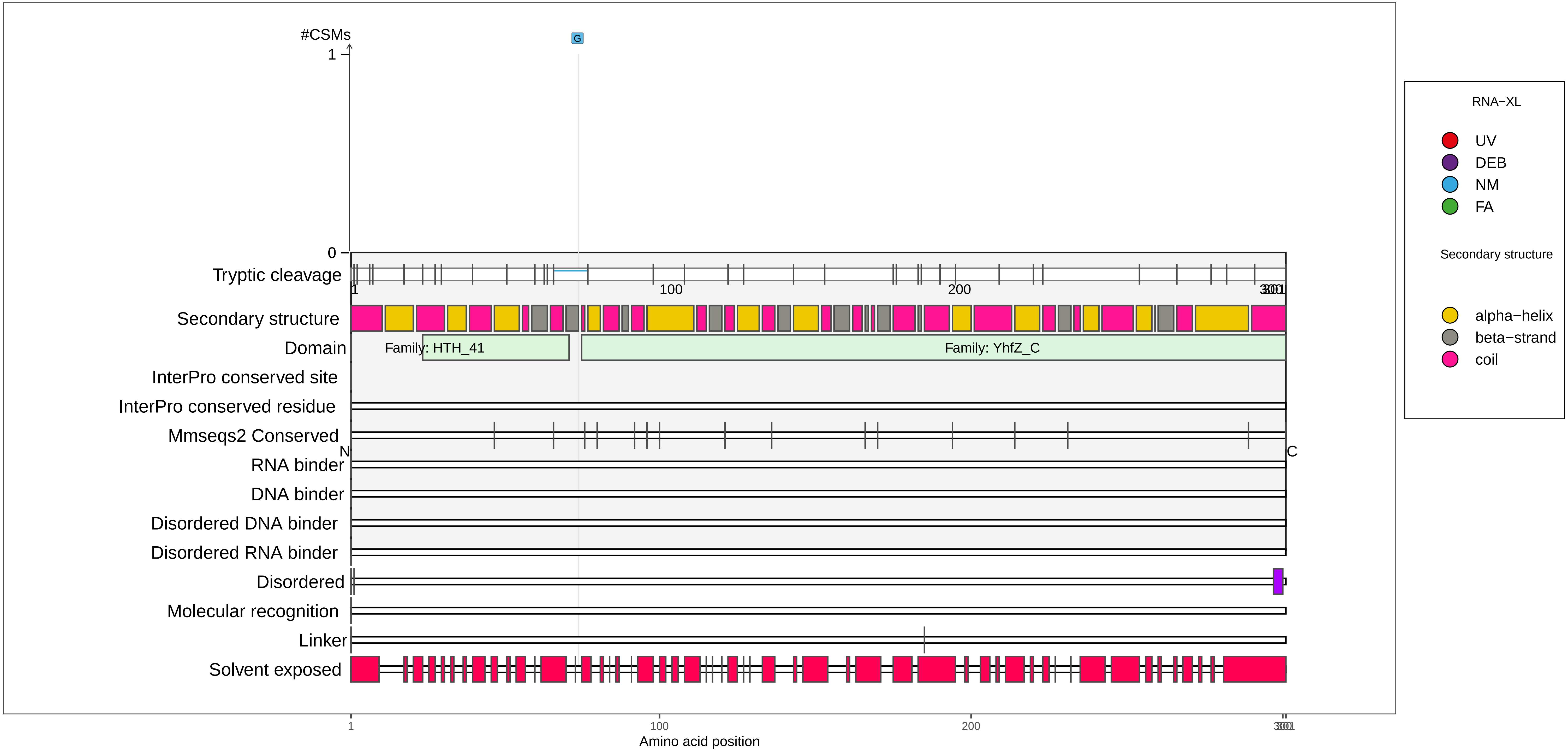

P46853
YHHX_ECOLI Uncharacterized oxidoreductase YhhX
− Abundance:
tryptic [log10 Intensity]: 8.92 (Q 86)
PAXdb K12 strain [ppm]: 1.56 (Q 42)
PAXdb E.coli [ppm]: 1.99 (Q 72)

− RNA functions: not annotated

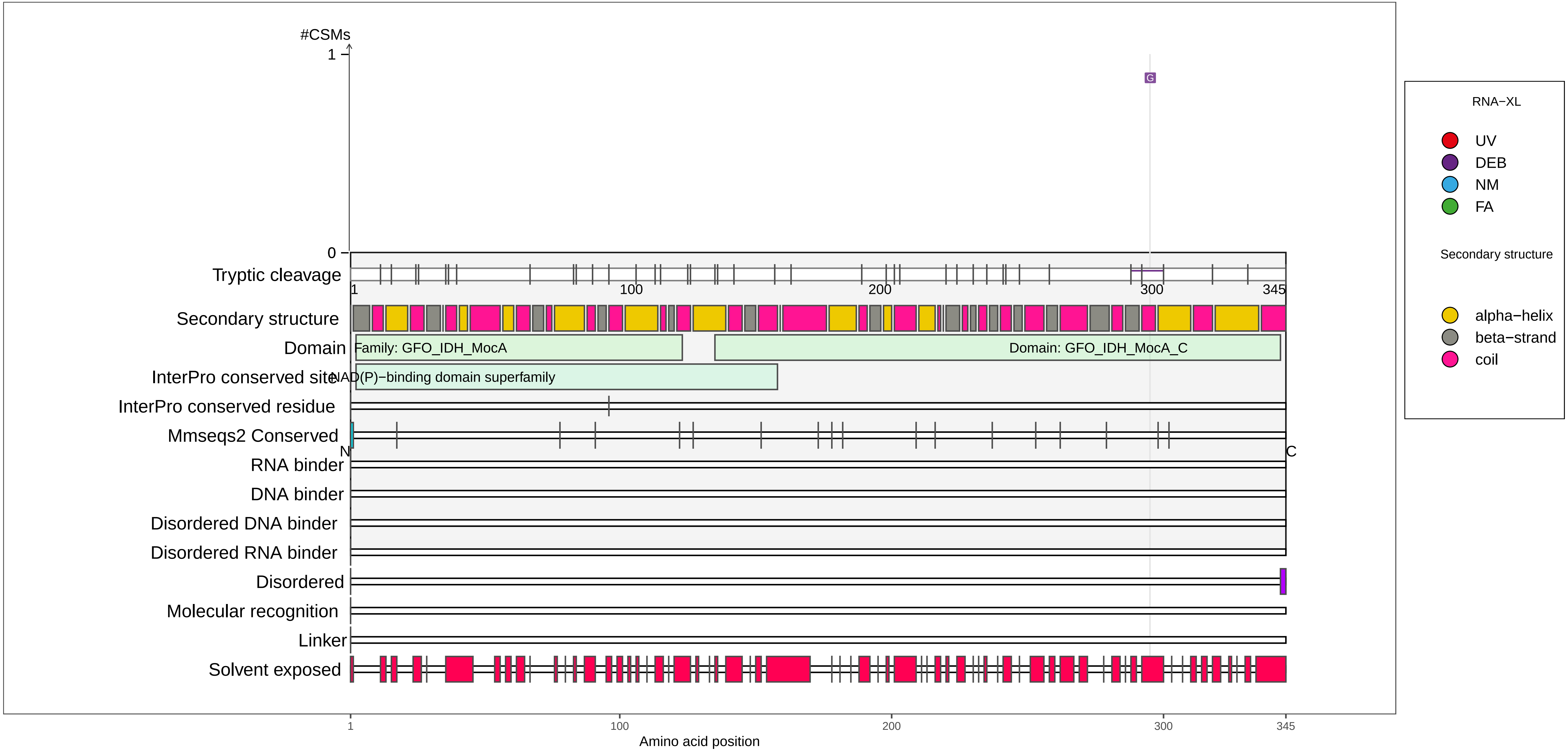

P52076
QSEB_ECOLI Transcriptional regulatory protein QseB

− Abundance:
tryptic [log10 Intensity]: 7.3 (Q 25)
PAXdb K12 strain [ppm]: NA (Q NA)
PAXdb E.coli [ppm]: 0.44 (Q 35)

− RNA functions:
RNA biosynthetic process; RNA metabolic process

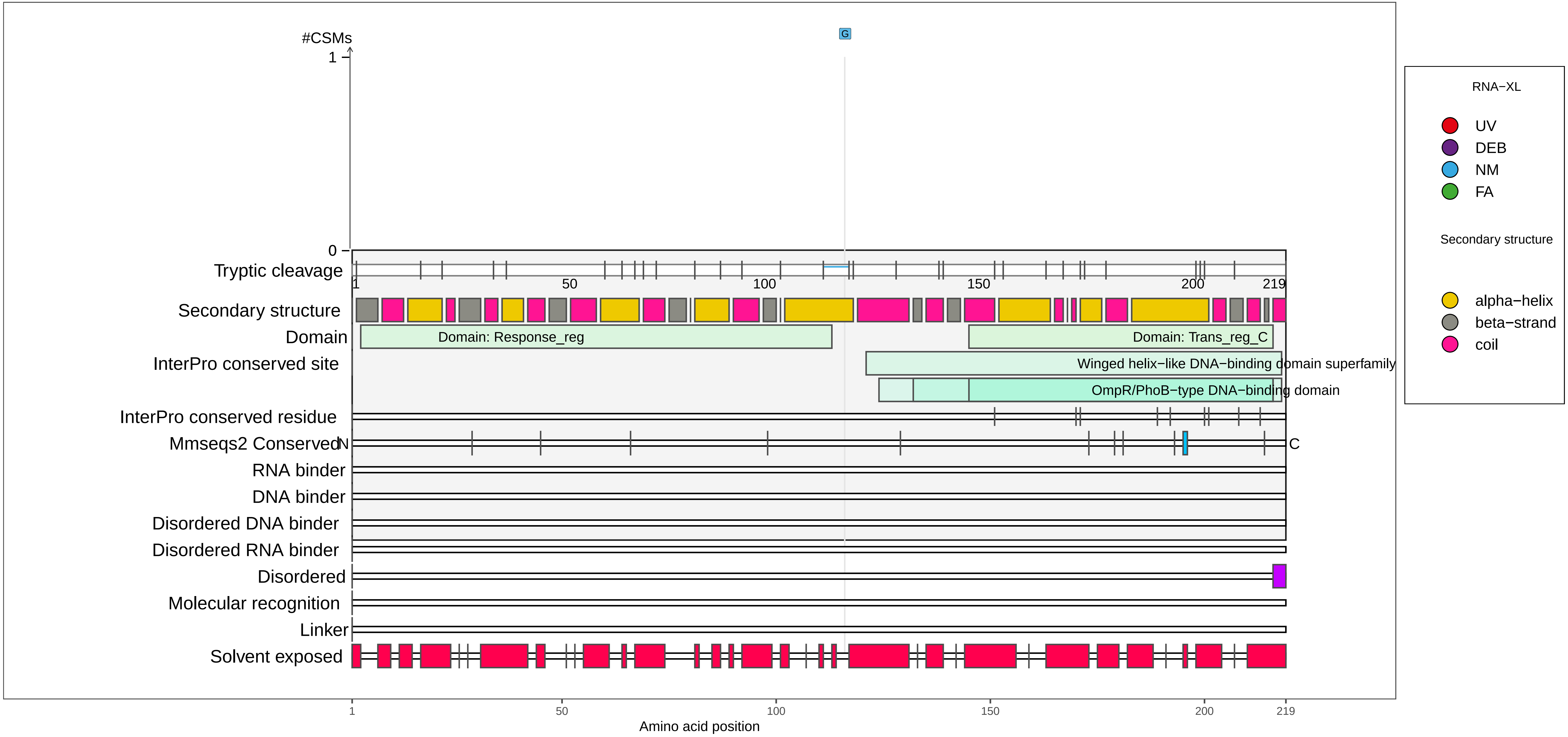

P52097
TILS_ECOLI tRNA(Ile)−lysidine synthase

− Abundance:
tryptic [log10 Intensity]: 7.64 (Q 42)
PAXdb K12 strain [ppm]: 1.83 (Q 54)
PAXdb E.coli [ppm]: 1.1 (Q 50)

− RNA functions:
ncRNA metabolic process; ncRNA processing; RNA metabolic process; RNA modification
RNA processing; tRNA metabolic process; tRNA modification; tRNA processing
tRNA wobble base modification; tRNA wobble cytosine modification

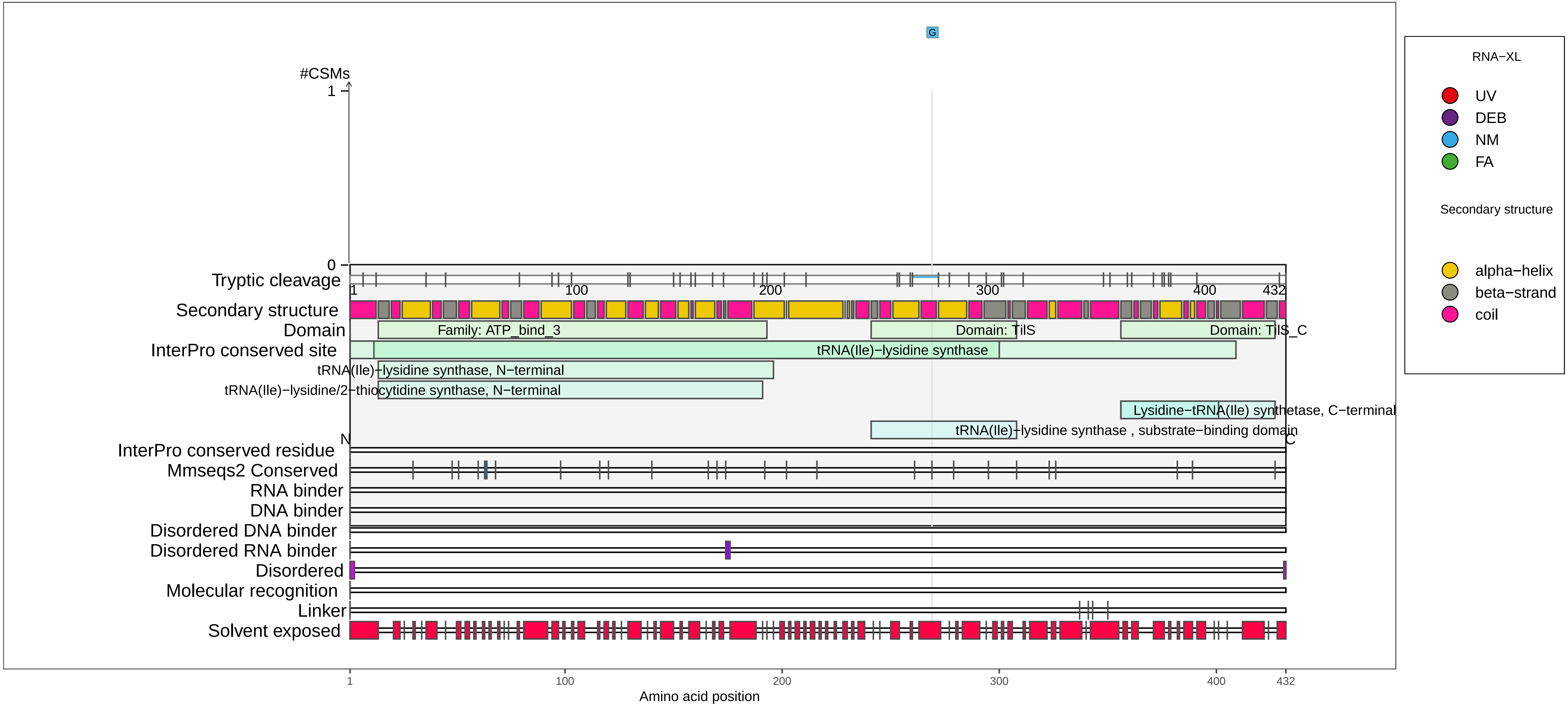

P52101
GLRK_ECOLI Sensor histidine kinase GlrK

− Abundance:
tryptic [log10 Intensity]: NA (Q NA)
PAXdb K12 strain [ppm]: 1.28 (Q 25)
PAXdb E.coli [ppm]: −0.63 (Q 14)

− RNA functions: not annotated

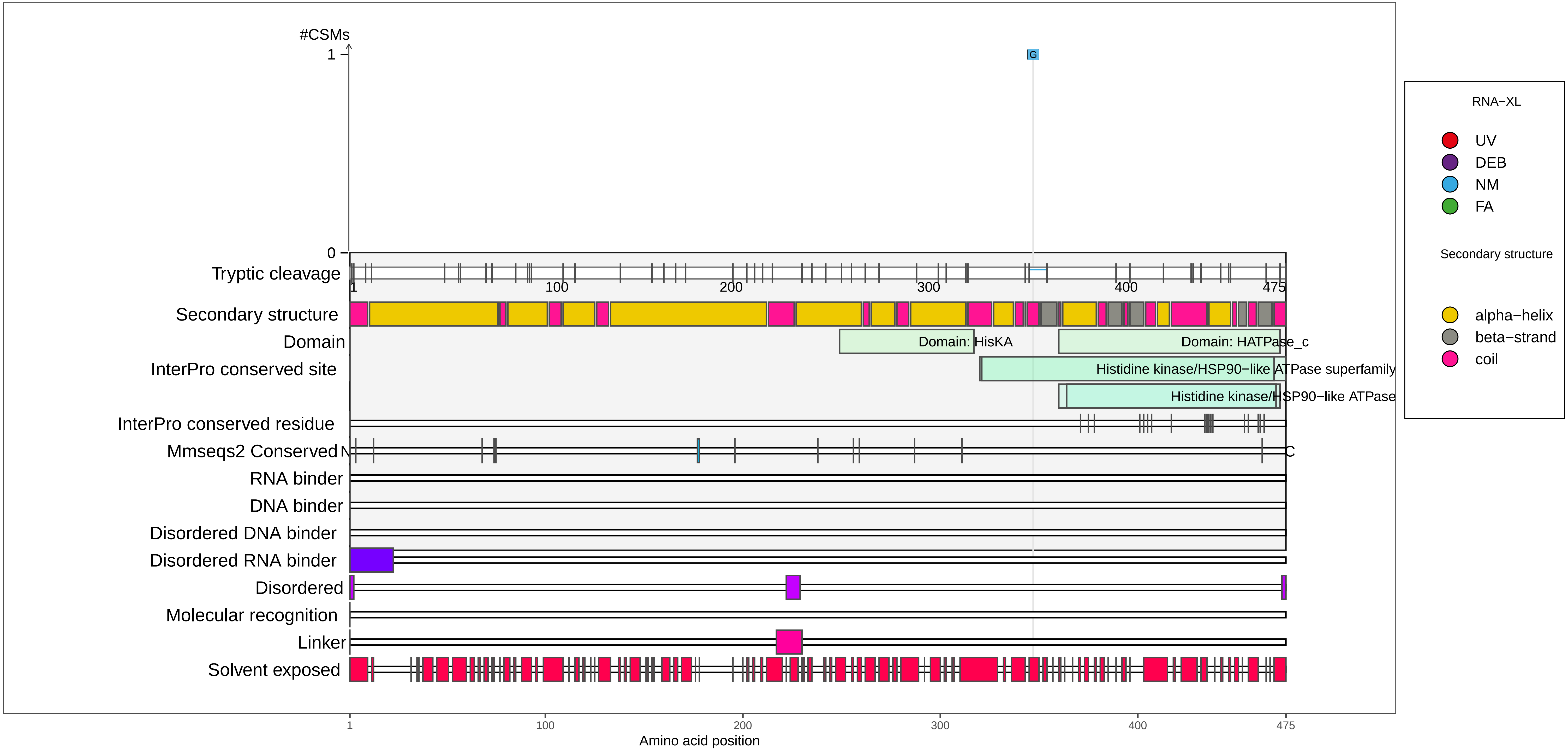

P52102
YFHL_ECOLI Ferredoxin YfhL

− Abundance:
tryptic [log10 Intensity]: 7.37 (Q 28)
PAXdb K12 strain [ppm]: NA (Q NA)
PAXdb E.coli [ppm]: 1.89 (Q 69)

− RNA functions: not annotated

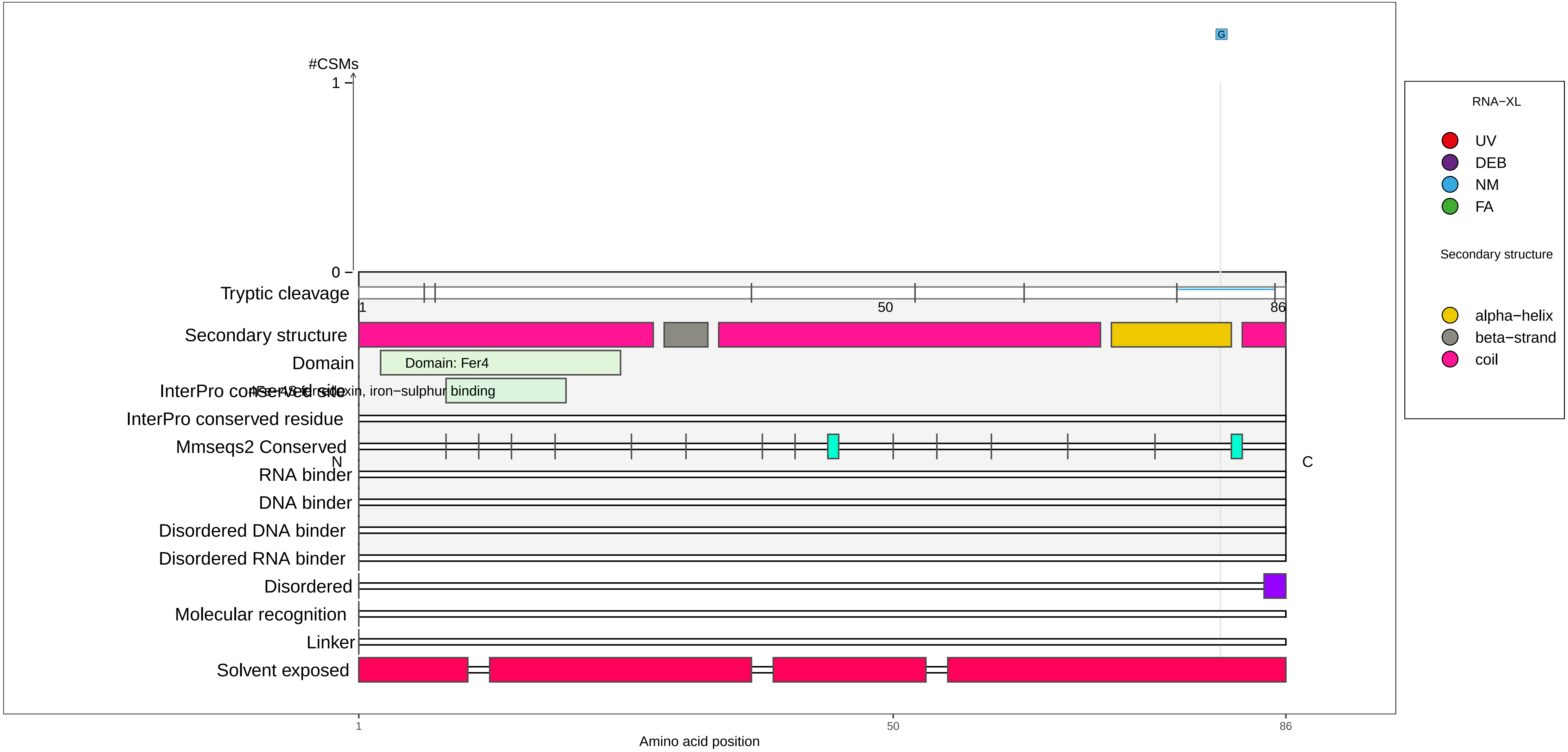

P54746
MNGB_ECOLI Mannosylglycerate hydrolase

− Abundance:
tryptic [log10 Intensity]: NA (Q NA)
PAXdb K12 strain [ppm]: 1.32 (Q 27)
PAXdb E.coli [ppm]: −0.22 (Q 20)

− RNA functions: not annotated

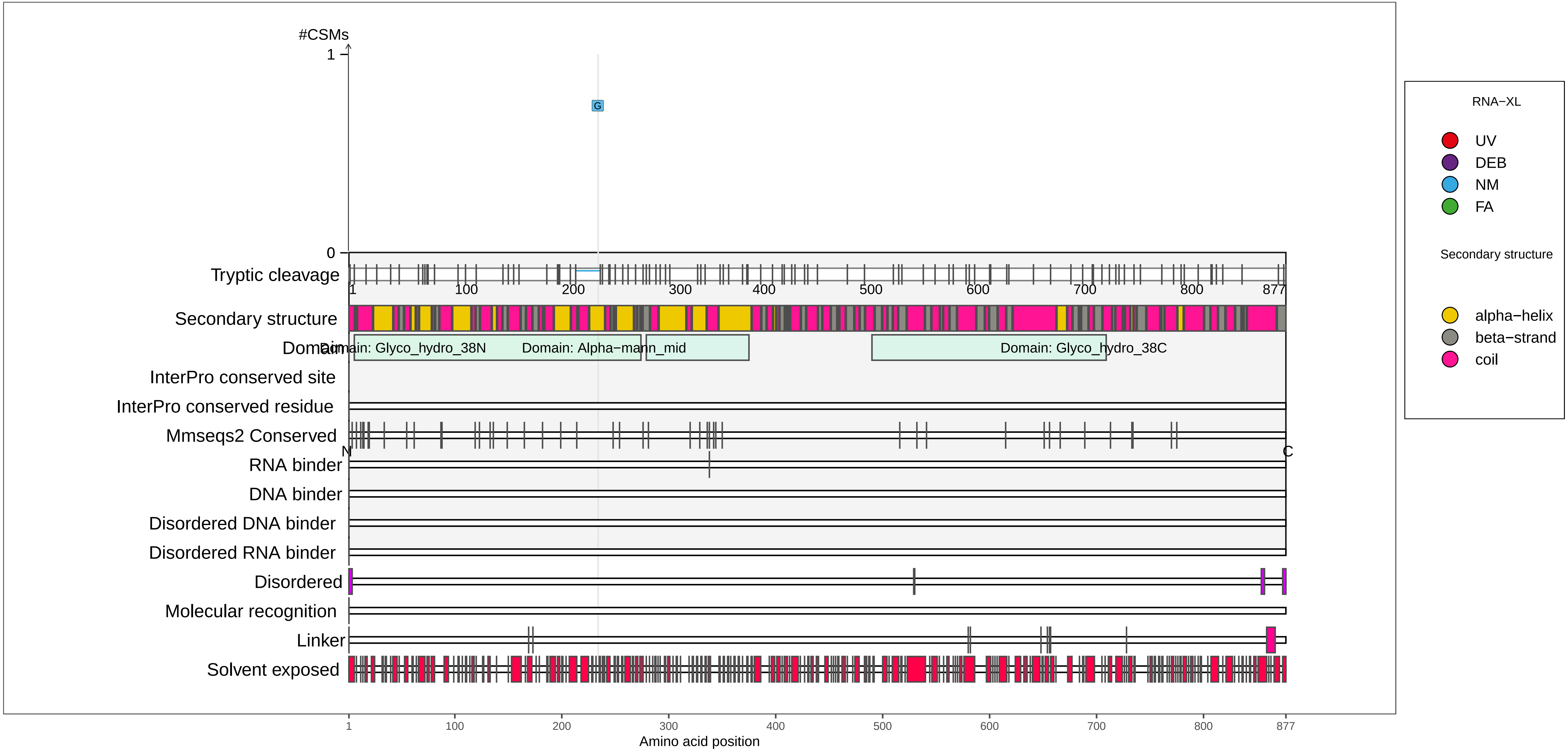

P60720
LIPB_ECOLI Octanoyltransferase

− Abundance:
tryptic [log10 Intensity]: 7.62 (Q 41)
PAXdb K12 strain [ppm]: 1.59 (Q 44)
PAXdb E.coli [ppm]: 1.19 (Q 52)

− RNA functions: not annotated

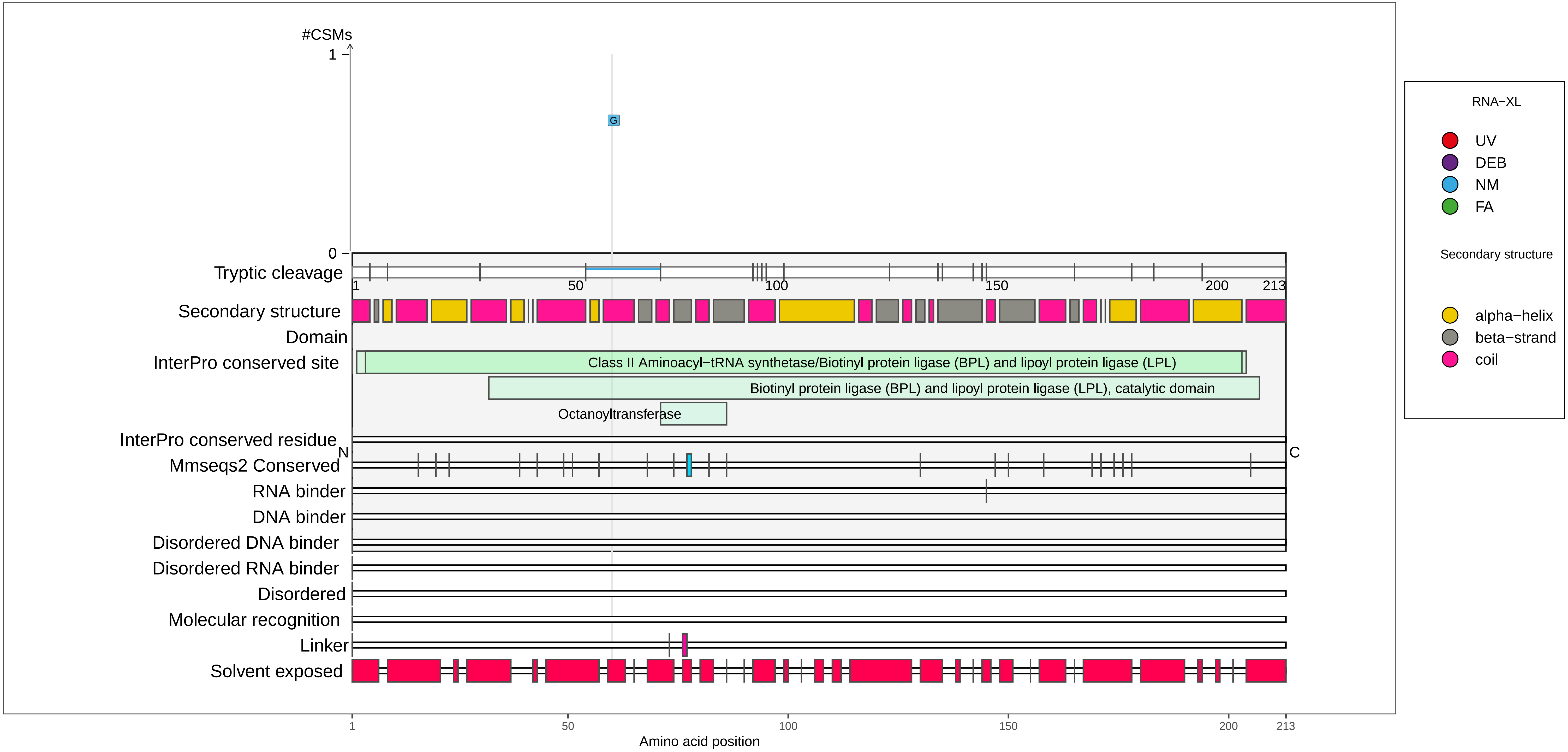

P61887
RMLA2_ECOLI Glucose−1−phosphate thymidylyltransferase 2

− Abundance:
tryptic [log10 Intensity]: 7.99 (Q 57)
PAXdb K12 strain [ppm]: NA (Q NA)
PAXdb E.coli [ppm]: 1.1 (Q 50)

− RNA functions: not annotated

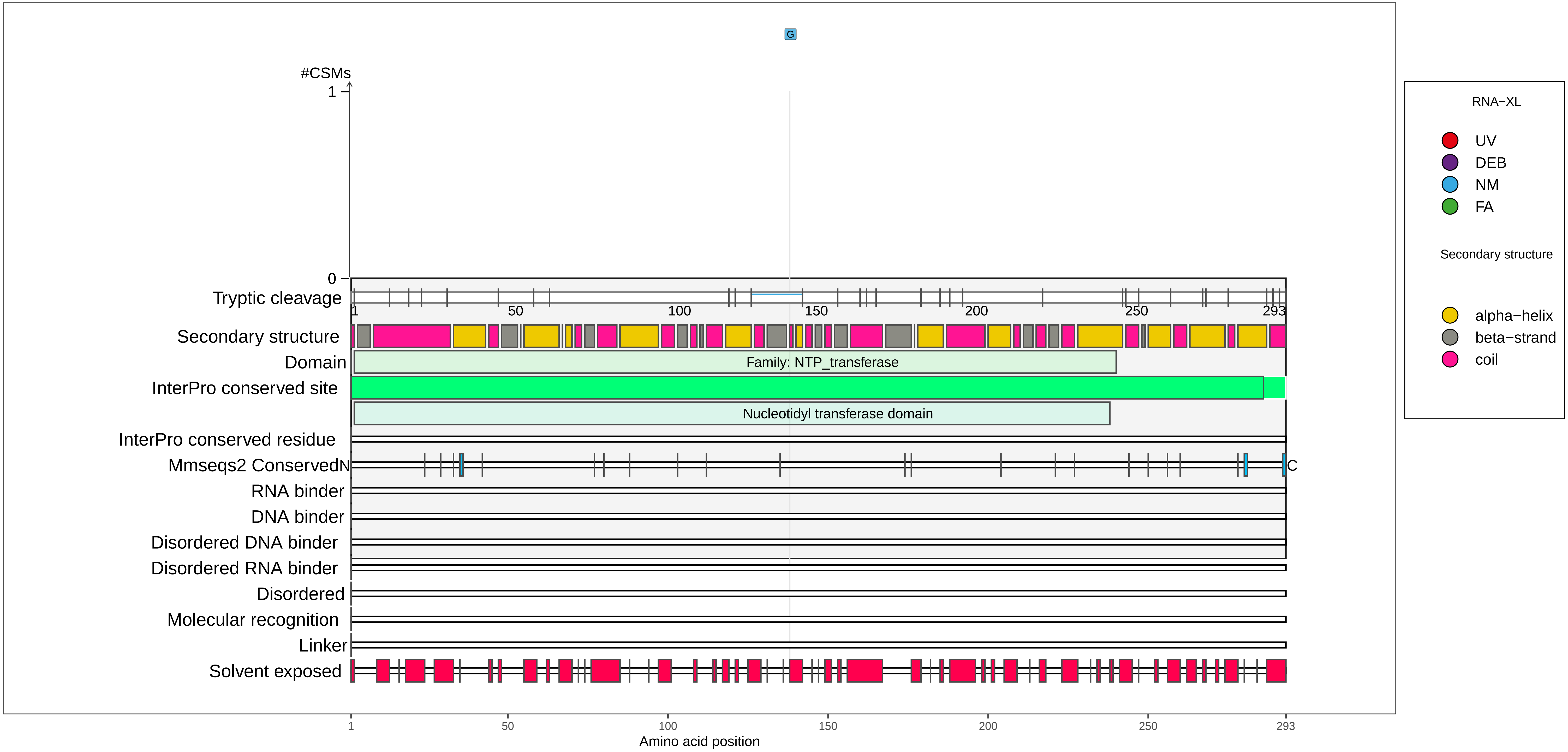

P64467
CNU_ECOLI OriC−binding nucleoid−associated protein

− Abundance:
tryptic [log10 Intensity]: 6.79 (Q 7)
PAXdb K12 strain [ppm]: NA (Q NA)
PAXdb E.coli [ppm]: 2.01 (Q 72)

− RNA functions: not annotated

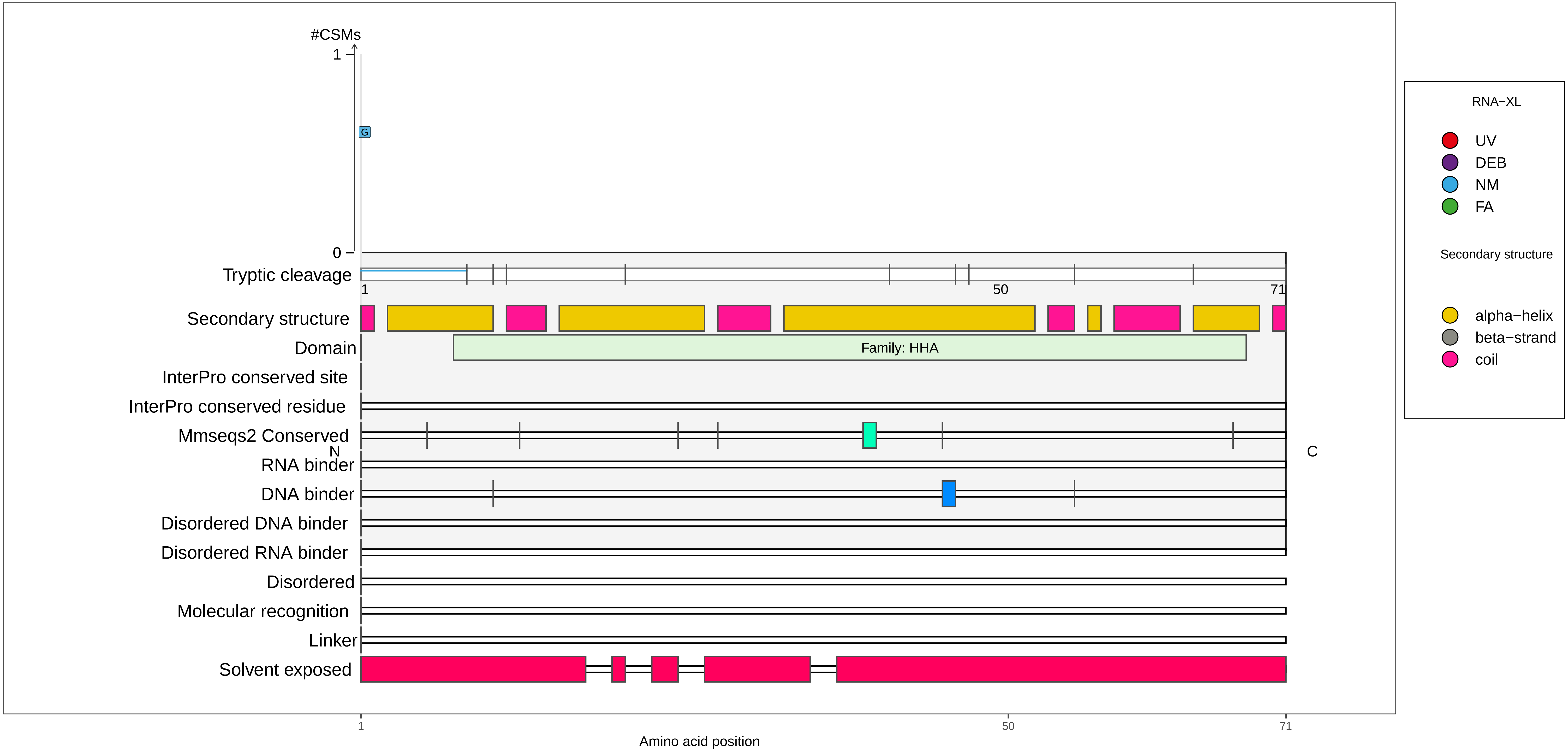

P67244
YQHA_ECOLI UPF0114 protein YqhA

− Abundance:
tryptic [log10 Intensity]: 8.07 (Q 60)
PAXdb K12 strain [ppm]: NA (Q NA)
PAXdb E.coli [ppm]: 1.26 (Q 54)

− RNA functions: not annotated

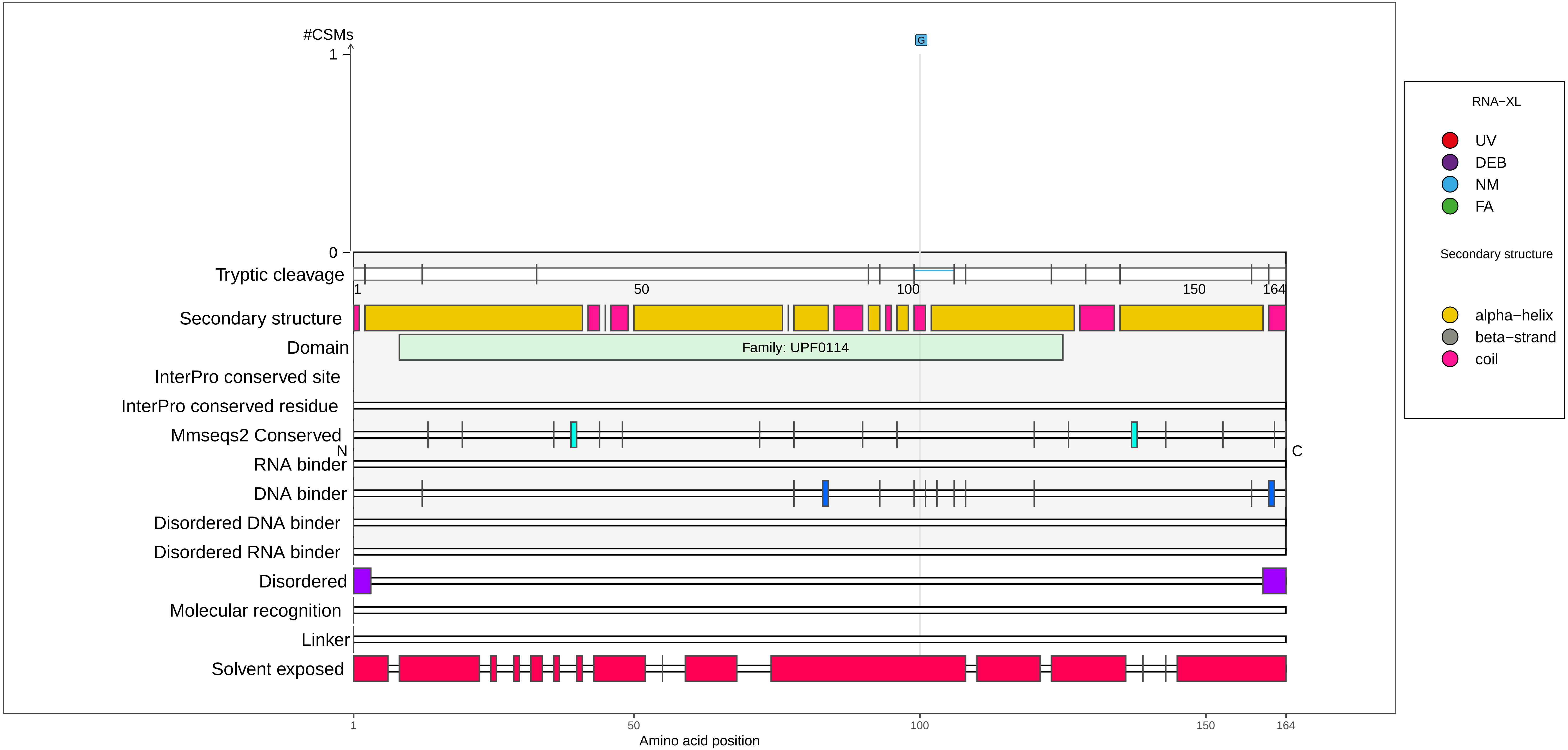

P67826
CUTC_ECOLI PF03932 family protein CutC

− Abundance:
tryptic [log10 Intensity]: 7.17 (Q 20)
PAXdb K12 strain [ppm]: 1.04 (Q 13)
PAXdb E.coli [ppm]: 1.68 (Q 64)

− RNA functions: not annotated

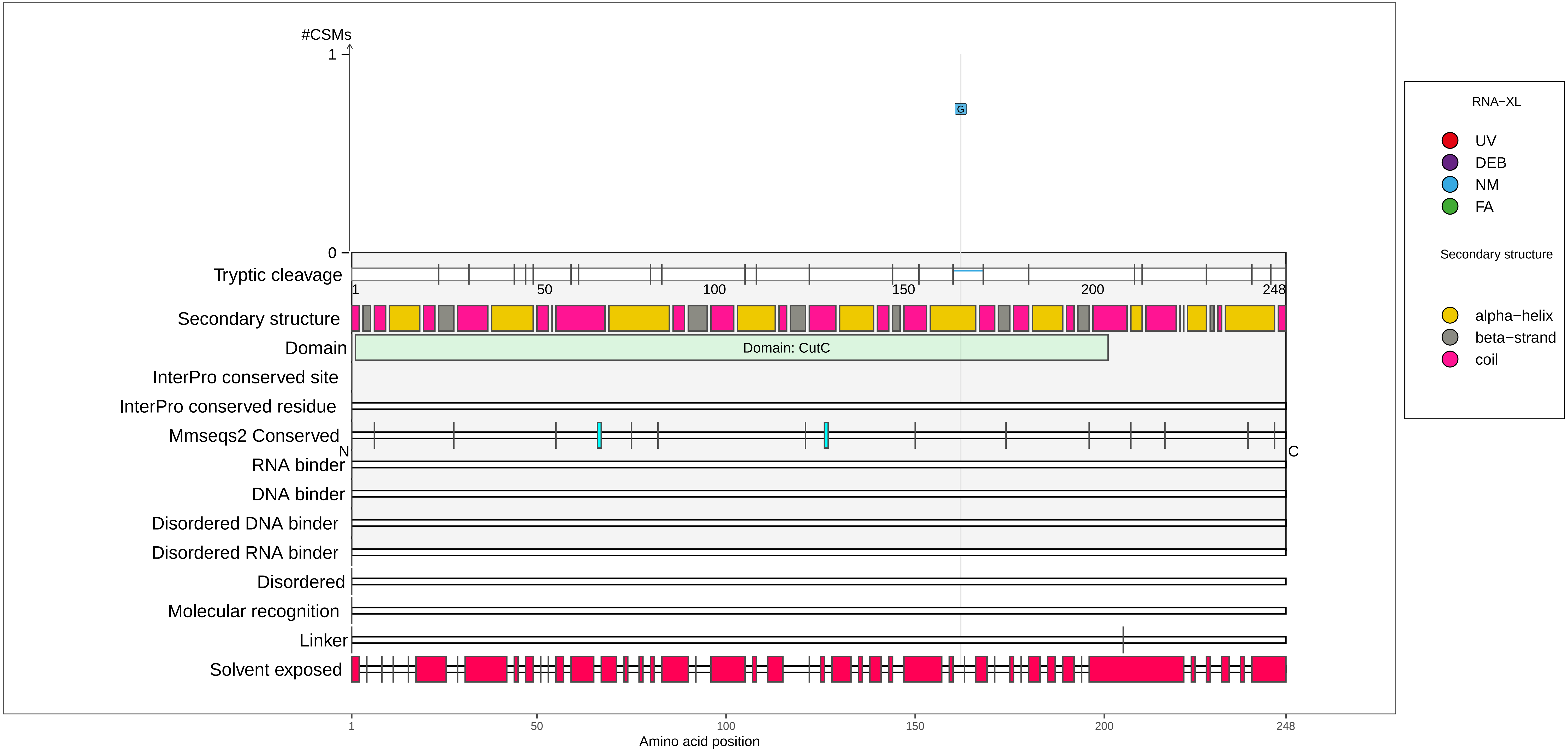

P67910
HLDD_ECOLI ADP−L−glycero−D−manno−heptose−6−epimerase

− Abundance:
tryptic [log10 Intensity]: 8.47 (Q 74)
PAXdb K12 strain [ppm]: 3.18 (Q 94)
PAXdb E.coli [ppm]: 2.98 (Q 94)

− RNA functions: not annotated

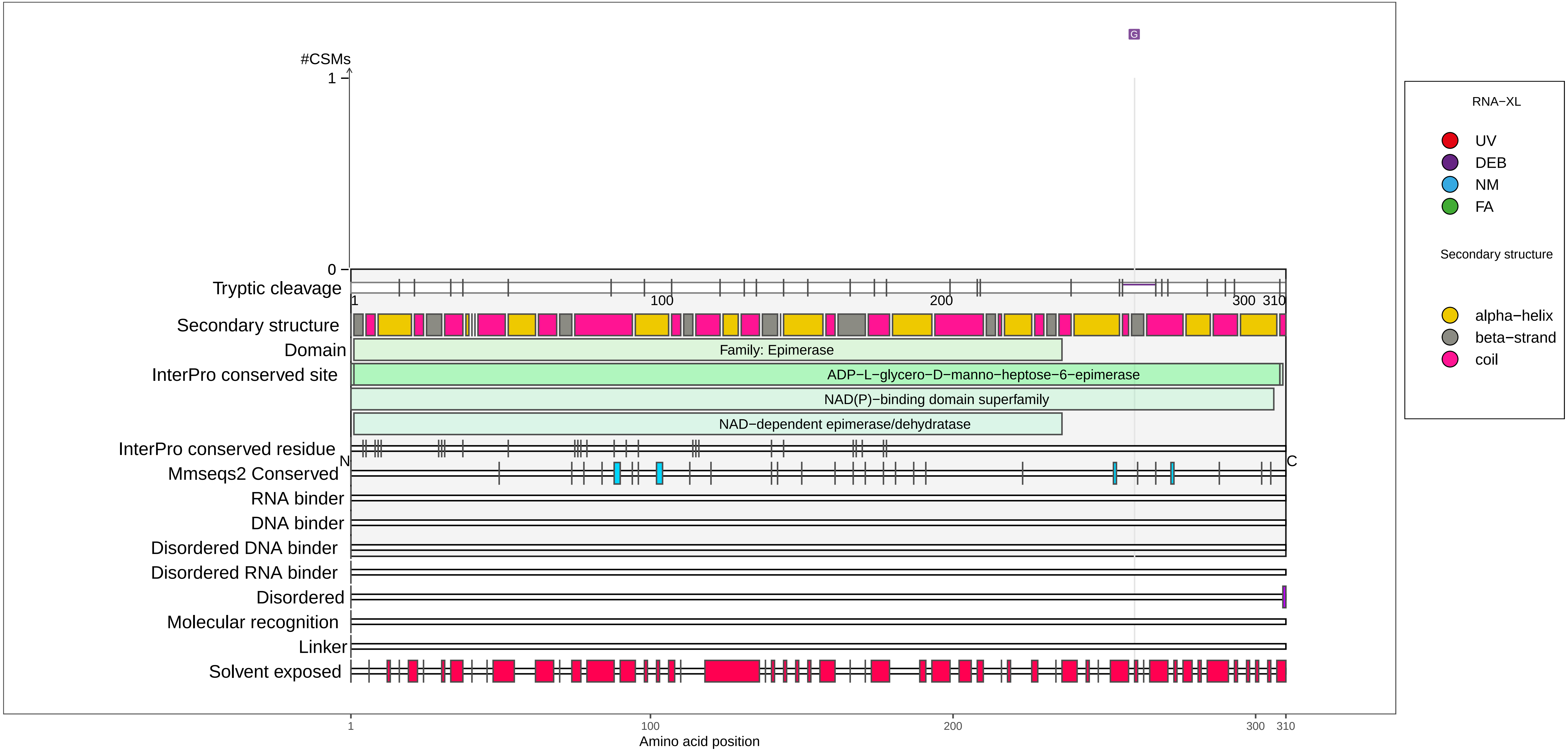

P68398
TADA_ECOLI tRNA−specific adenosine deaminase

− Abundance:
tryptic [log10 Intensity]: 7.38 (Q 29)
PAXdb K12 strain [ppm]: NA (Q NA)
PAXdb E.coli [ppm]: 1.33 (Q 55)

− RNA functions:
ncRNA metabolic process; ncRNA processing; RNA metabolic process; RNA modification
RNA processing; tRNA metabolic process; tRNA modification; tRNA processing
tRNA wobble adenosine to inosine editing; tRNA wobble base modification
tRNA−specific adenosine deaminase activity; tRNA−specific adenosine−34 deaminase activity
tRNA−specific adenosine−34 deaminase complex

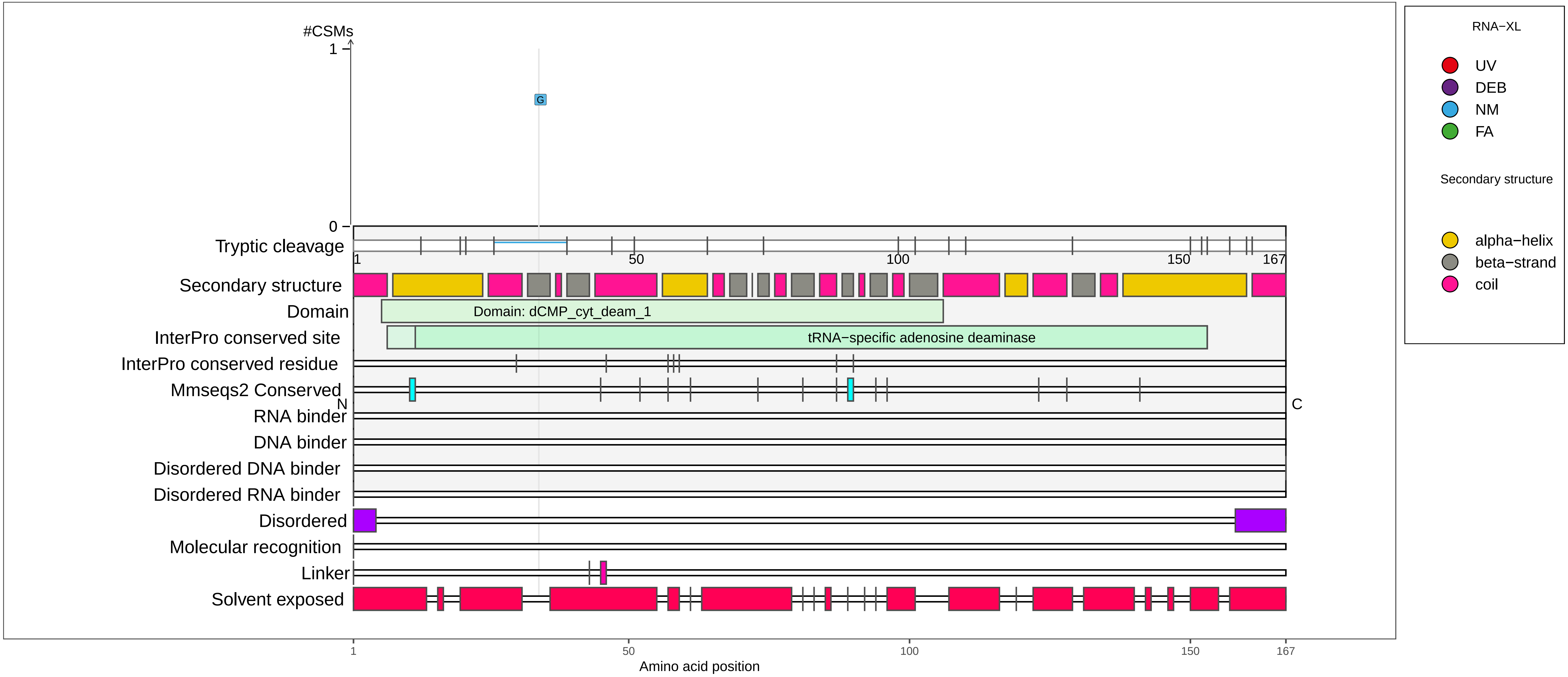

P69451
LCFA_ECOLI Long−chain−fatty−acid−−CoA ligase

− Abundance:
tryptic [log10 Intensity]: 8 (Q 57)
PAXdb K12 strain [ppm]: 1.87 (Q 56)
PAXdb E.coli [ppm]: 1.59 (Q 62)

− RNA functions: not annotated

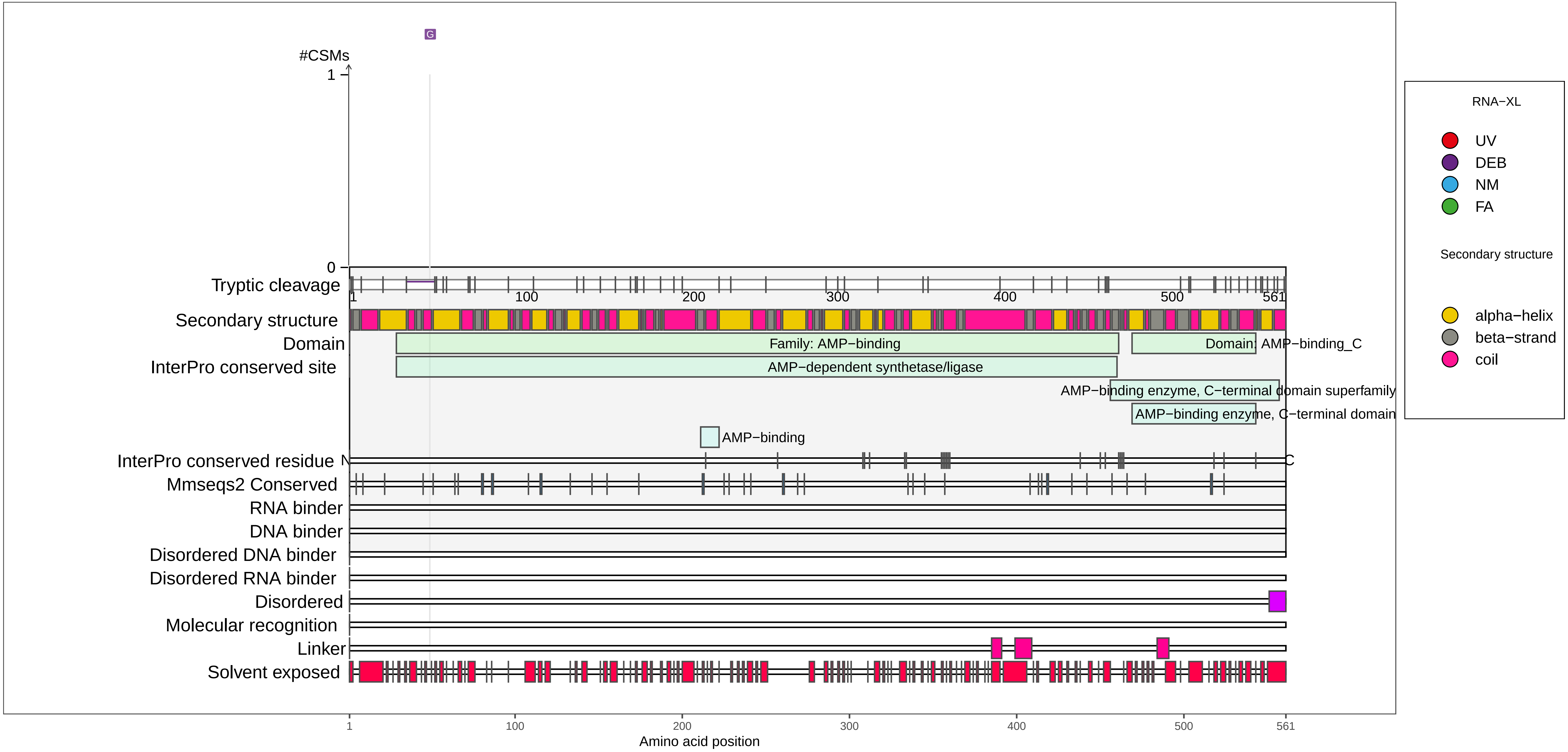

P69822
ULAB_ECOLI Ascorbate−specific PTS system EIIB component

− Abundance:
tryptic [log10 Intensity]: 7.92 (Q 54)
PAXdb K12 strain [ppm]: NA (Q NA)
PAXdb E.coli [ppm]: 1.43 (Q 58)

− RNA functions: not annotated

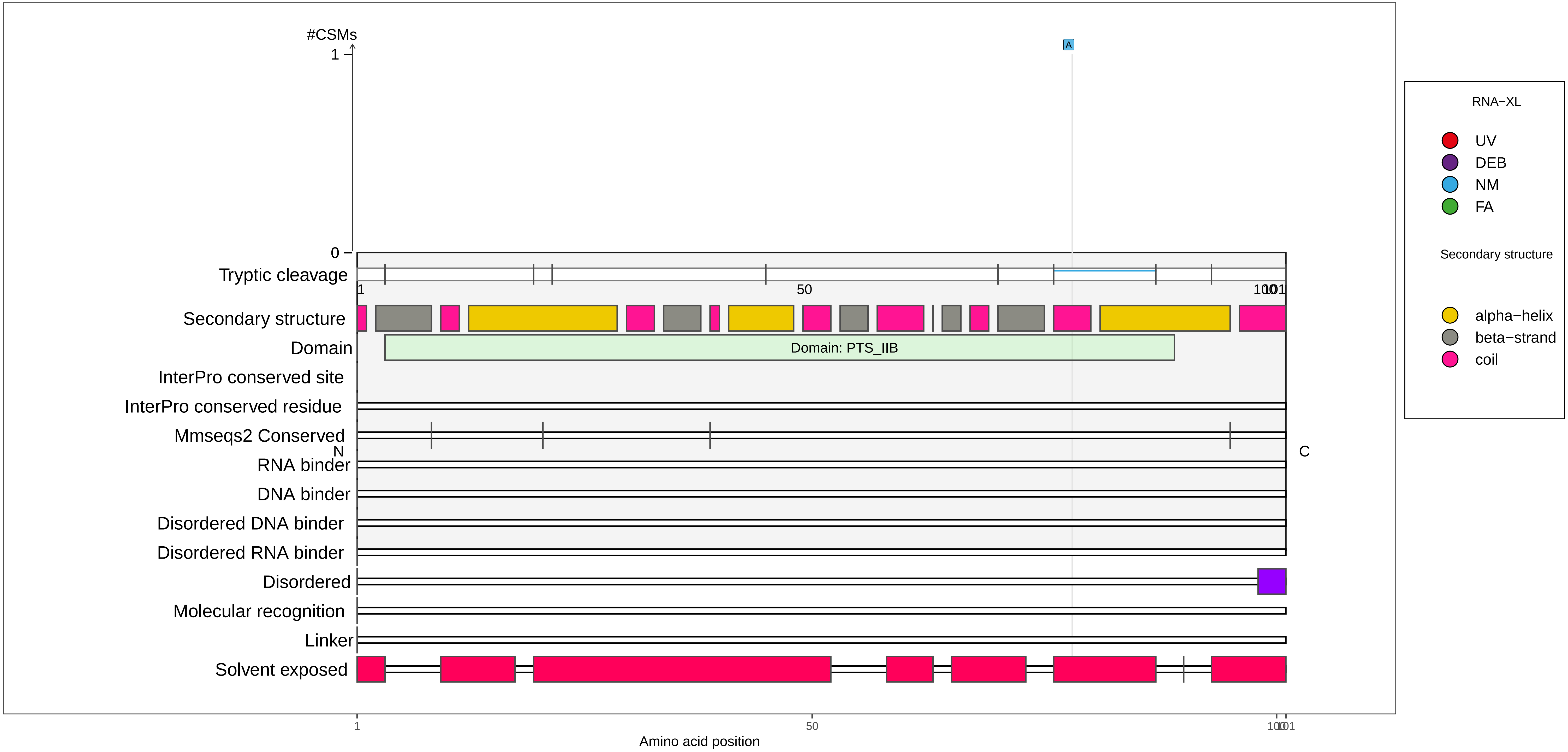

P69922
FUCI_ECOLI L−fucose isomerase

− Abundance:
tryptic [log10 Intensity]: 8.11 (Q 62)
PAXdb K12 strain [ppm]: 1.64 (Q 47)
PAXdb E.coli [ppm]: 0.49 (Q 36)

− RNA functions: not annotated

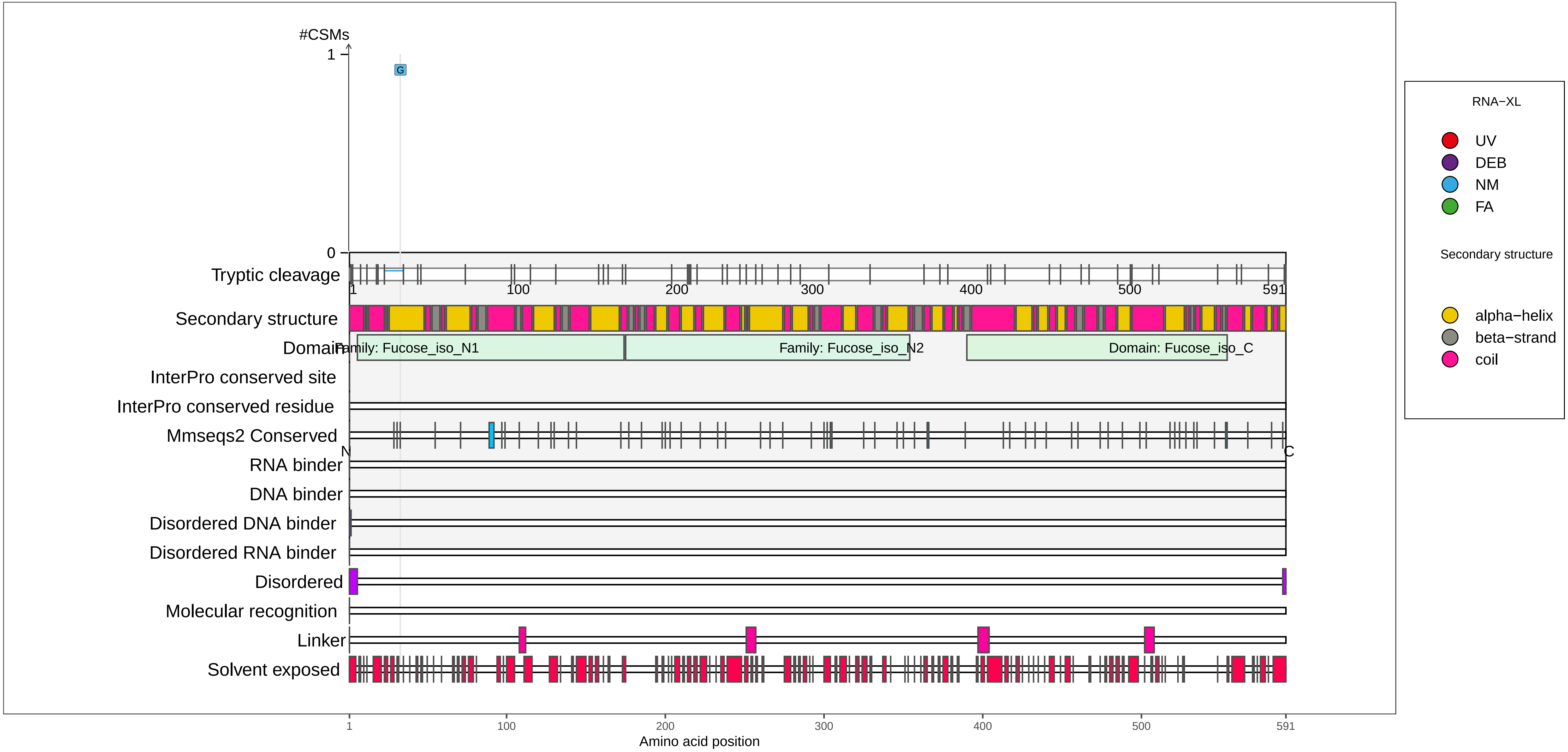

P71242
WCAK_ECOLI Colanic acid biosynthesis protein WcaK

− Abundance:
tryptic [log10 Intensity]: 7.33 (Q 26)
PAXdb K12 strain [ppm]: 0.93 (Q 9)
PAXdb E.coli [ppm]: −0.46 (Q 16)

− RNA functions: not annotated

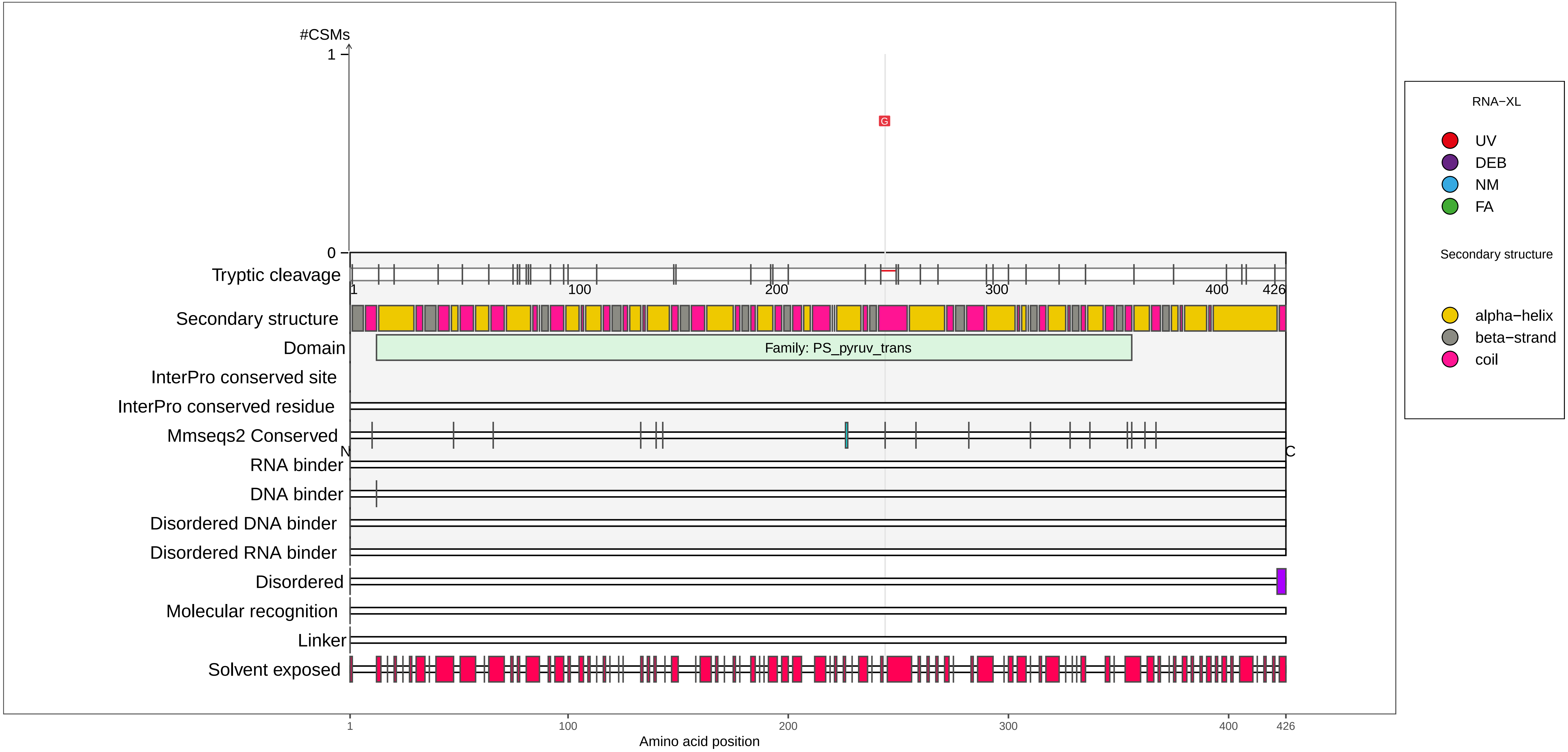

P75672
YAFS_ECOLI Uncharacterized protein YafS

− Abundance:
tryptic [log10 Intensity]: NA (Q NA)
PAXdb K12 strain [ppm]: NA (Q NA)
PAXdb E.coli [ppm]: −0.95 (Q 9)

− RNA functions: not annotated

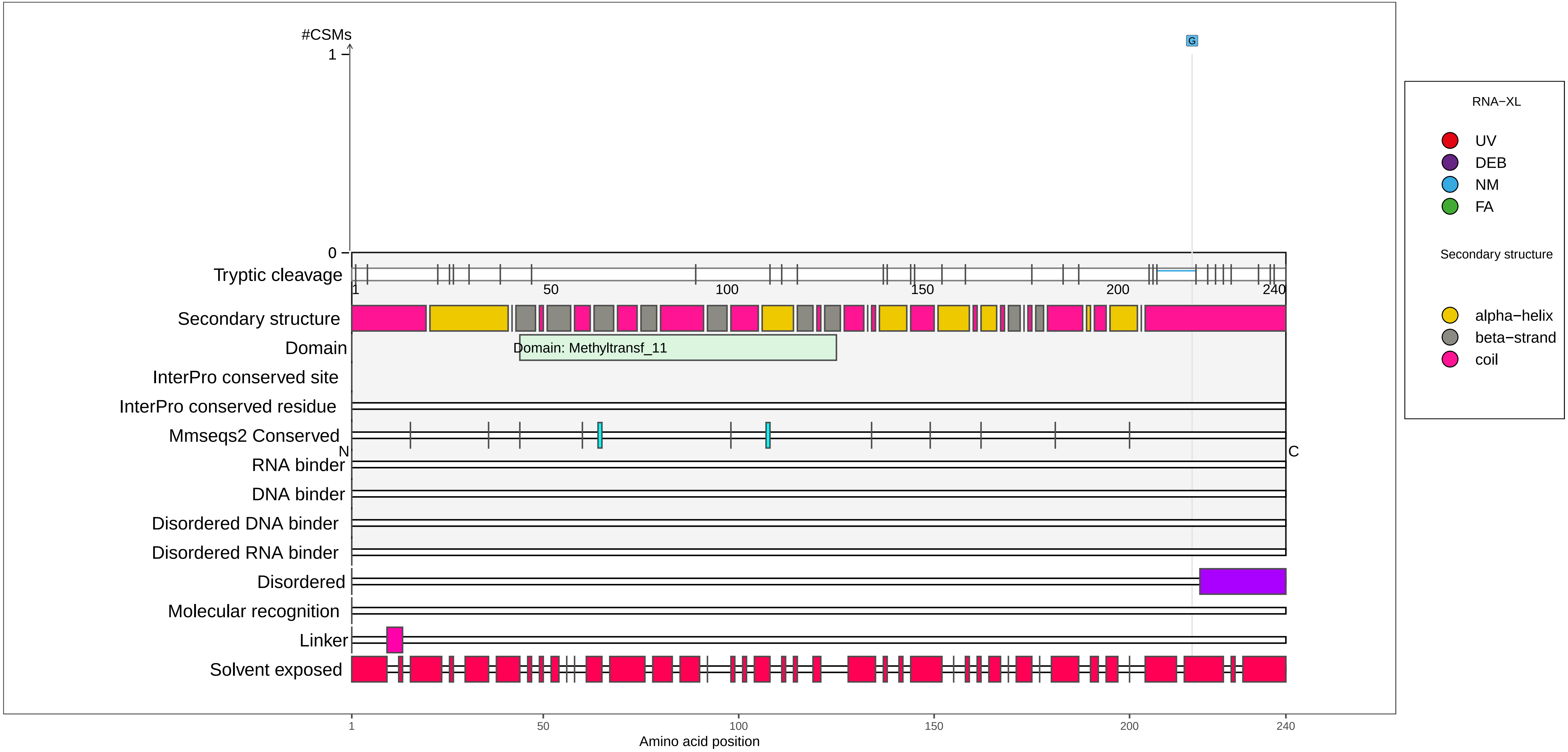

P75811
RCDA_ECOLI HTH−type transcriptional regulator RcdA

− Abundance:
tryptic [log10 Intensity]: NA (Q NA)
PAXdb K12 strain [ppm]: 1.31 (Q 27)
PAXdb E.coli [ppm]: 0.18 (Q 29)

− RNA functions:
RNA biosynthetic process; RNA metabolic process

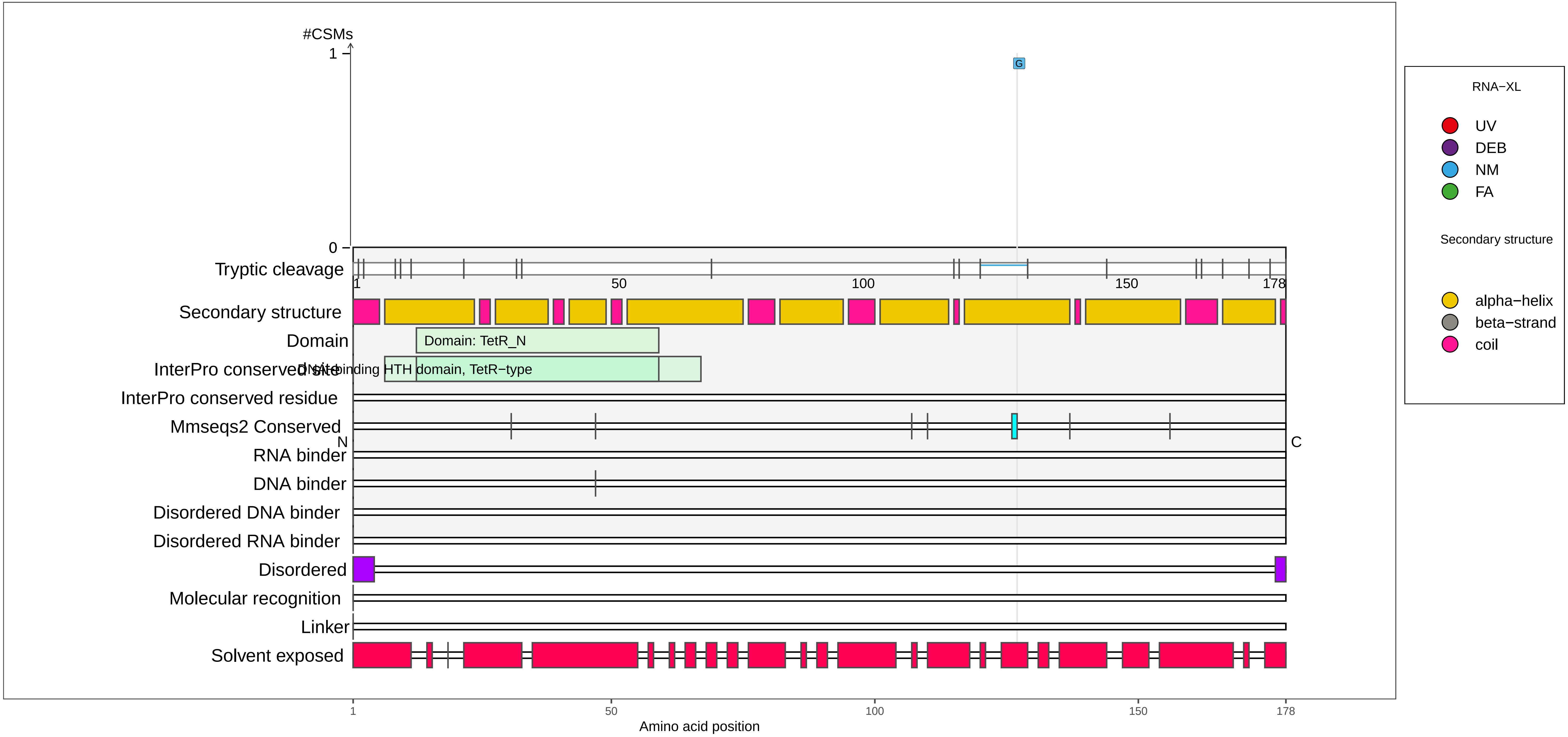

P75828
YBJD_ECOLI Uncharacterized protein YbjD

− Abundance:
tryptic [log10 Intensity]: 6.64 (Q 4)
PAXdb K12 strain [ppm]: 0.98 (Q 10)
PAXdb E.coli [ppm]: 0.76 (Q 42)

− RNA functions: not annotated

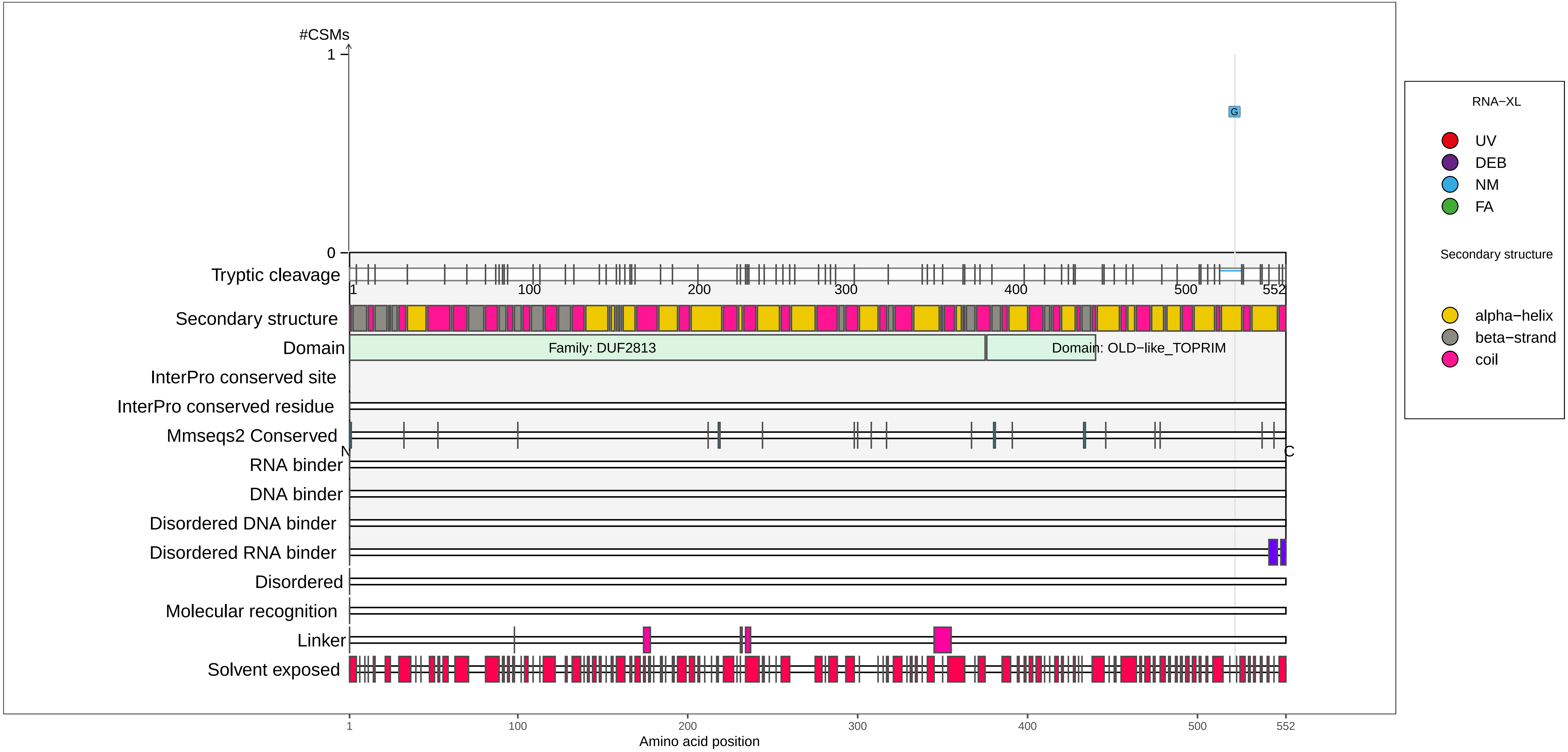

P75831
MACB_ECOLI Macrolide export ATP−binding/permease protein MacB

− Abundance:
tryptic [log10 Intensity]: 6.92 (Q 10)
PAXdb K12 strain [ppm]: 1.19 (Q 20)
PAXdb E.coli [ppm]: 0.35 (Q 32)

− RNA functions: not annotated

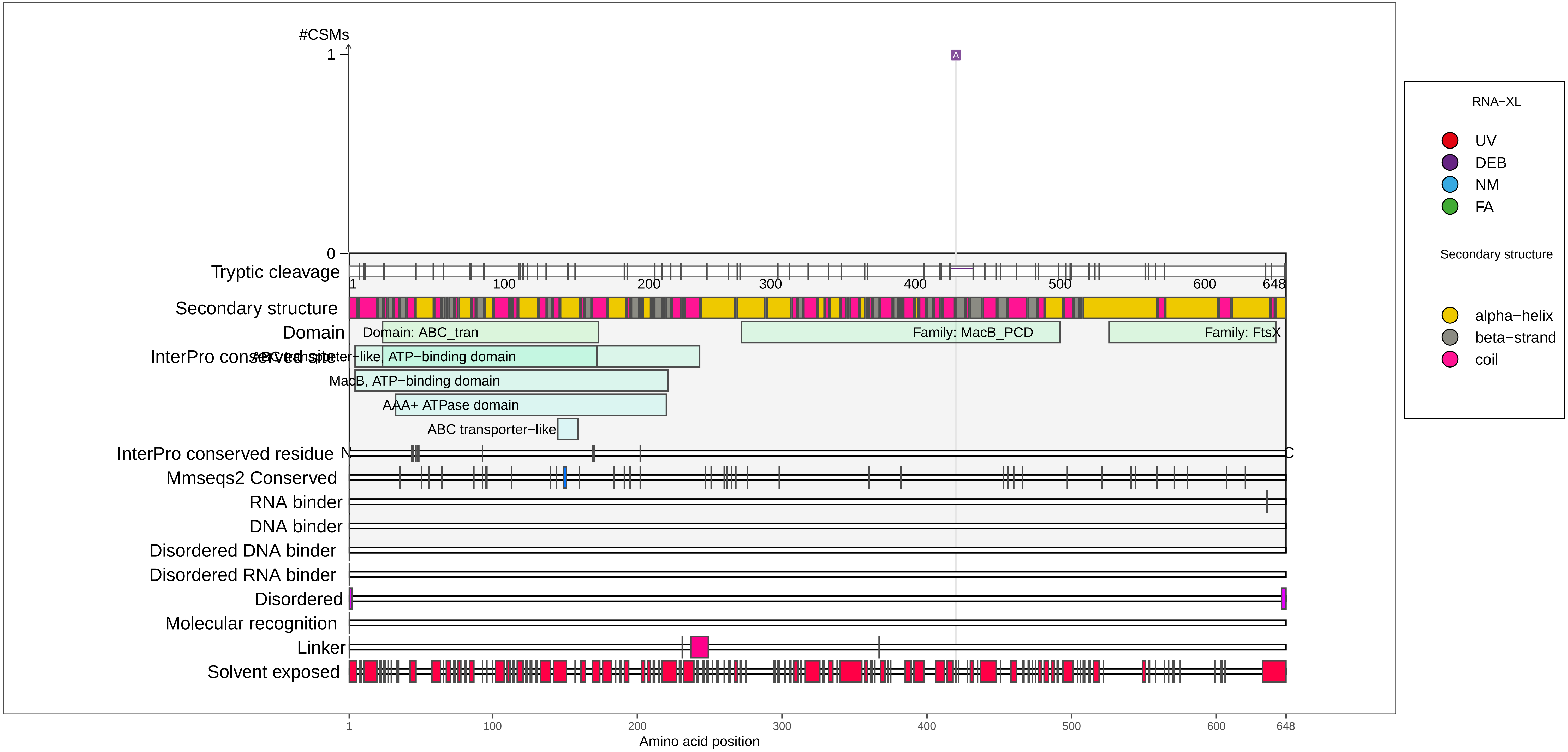

P75906
PGAB_ECOLI Poly−beta−1,6−N−acetyl−D−glucosamine N−deacetylase

− Abundance:
tryptic [log10 Intensity]: NA (Q NA)
PAXdb K12 strain [ppm]: NA (Q NA)
PAXdb E.coli [ppm]: −1.17 (Q 7)

− RNA functions: not annotated

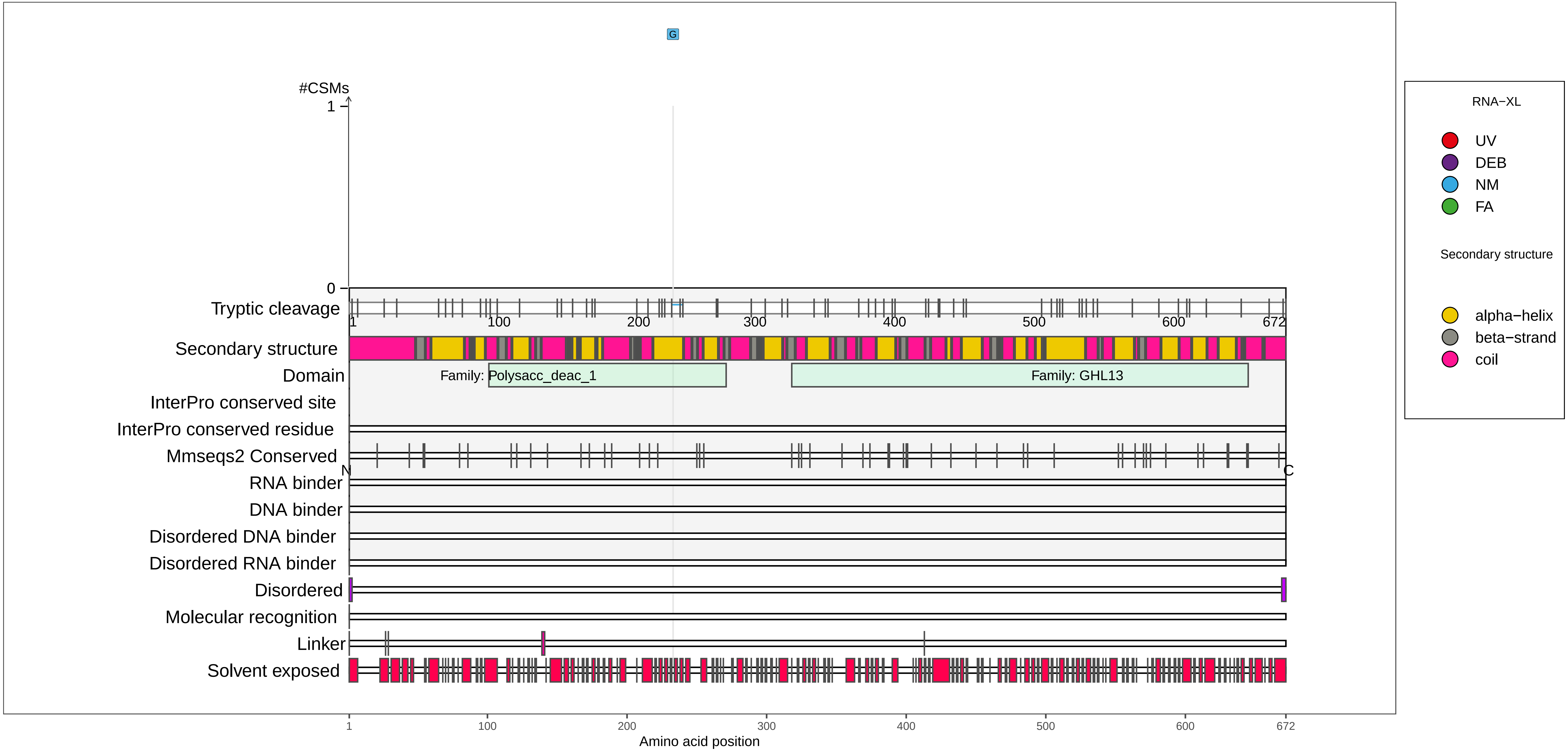

P75914
YCDX_ECOLI Probable phosphatase YcdX

− Abundance:
tryptic [log10 Intensity]: 8.26 (Q 68)
PAXdb K12 strain [ppm]: 2.14 (Q 65)
PAXdb E.coli [ppm]: 1.6 (Q 62)

− RNA functions: not annotated

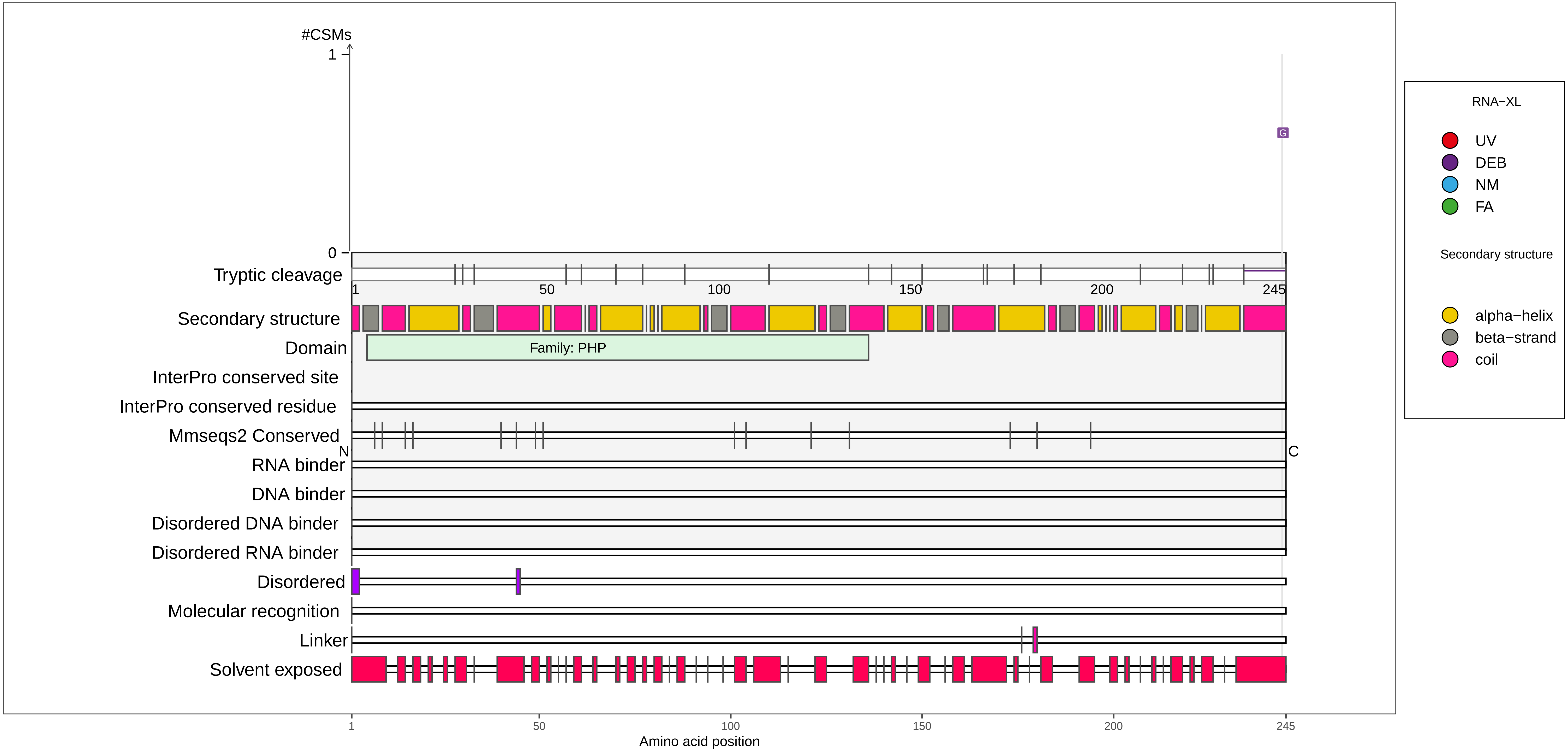

P75931
YCEM_ECOLI Putative oxidoreductase YceM

− Abundance:
tryptic [log10 Intensity]: 7.01 (Q 14)
PAXdb K12 strain [ppm]: NA (Q NA)
PAXdb E.coli [ppm]: 1.67 (Q 64)

− RNA functions: not annotated

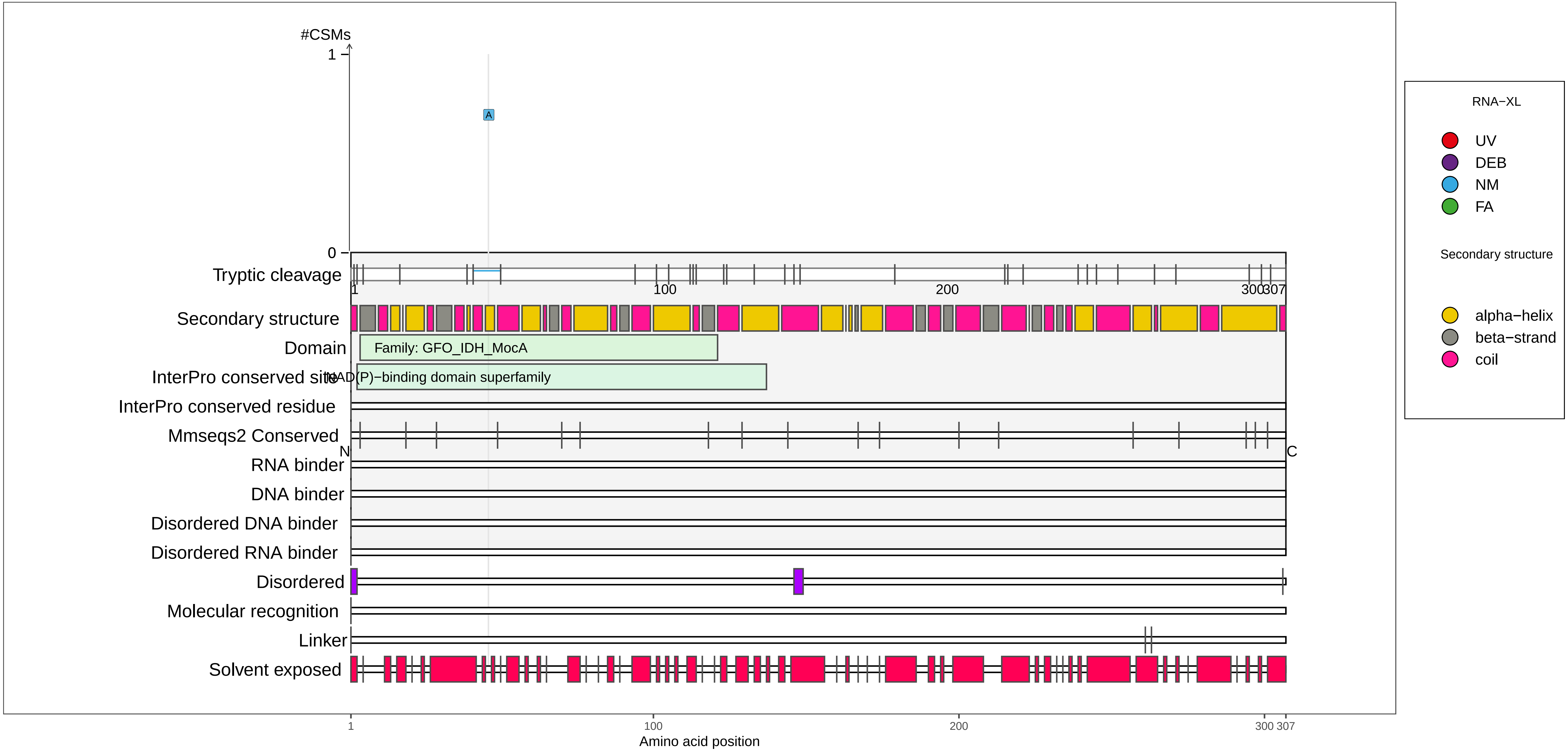

P75949
NAGZ_ECOLI Beta−hexosaminidase

− Abundance:
tryptic [log10 Intensity]: 7.59 (Q 39)
PAXdb K12 strain [ppm]: 2.49 (Q 77)
PAXdb E.coli [ppm]: 1.53 (Q 60)

− RNA functions: not annotated

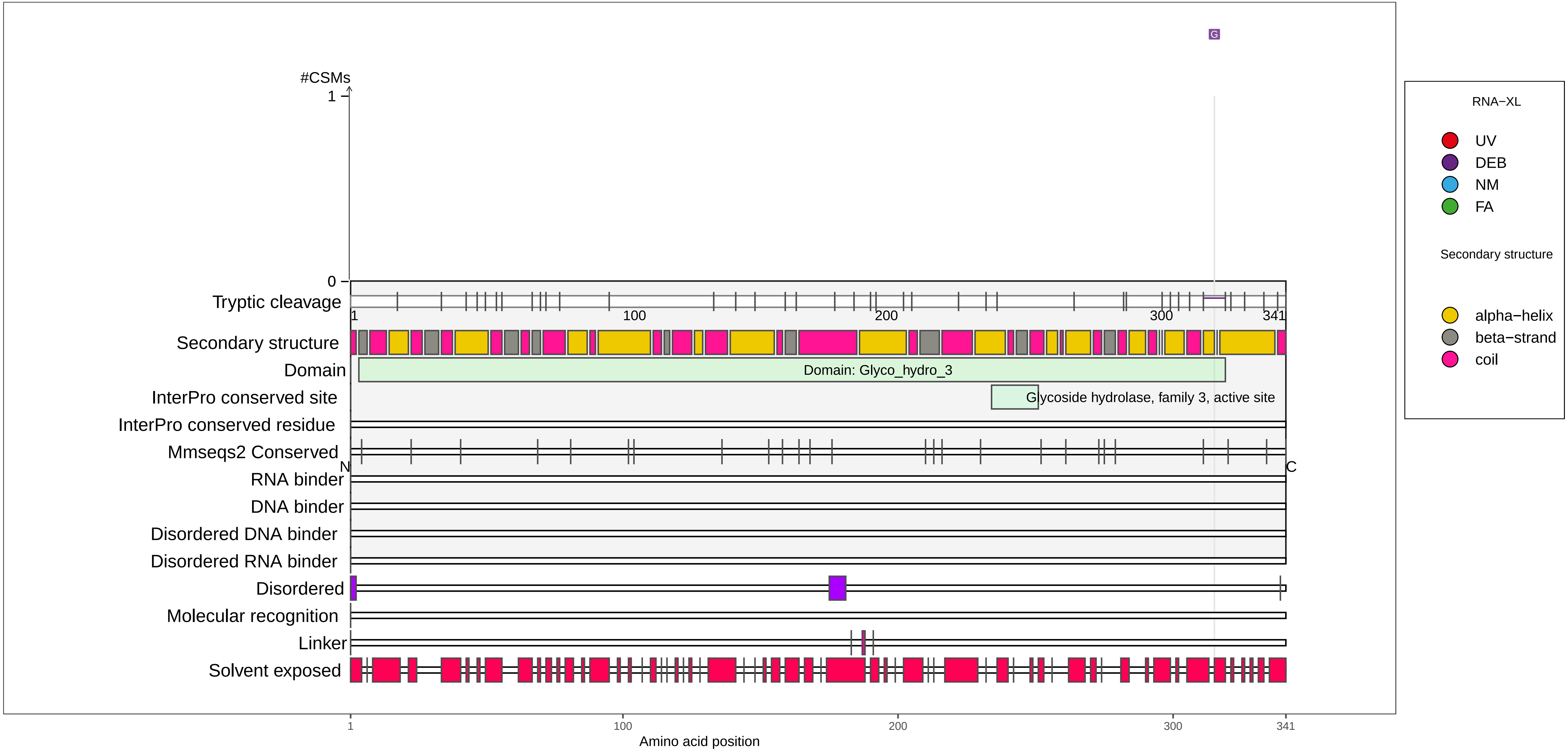

P75952
COMR_ECOLI HTH−type transcriptional repressor ComR

− Abundance:
tryptic [log10 Intensity]: 7.19 (Q 21)
PAXdb K12 strain [ppm]: NA (Q NA)
PAXdb E.coli [ppm]: 0.77 (Q 42)

− RNA functions: not annotated

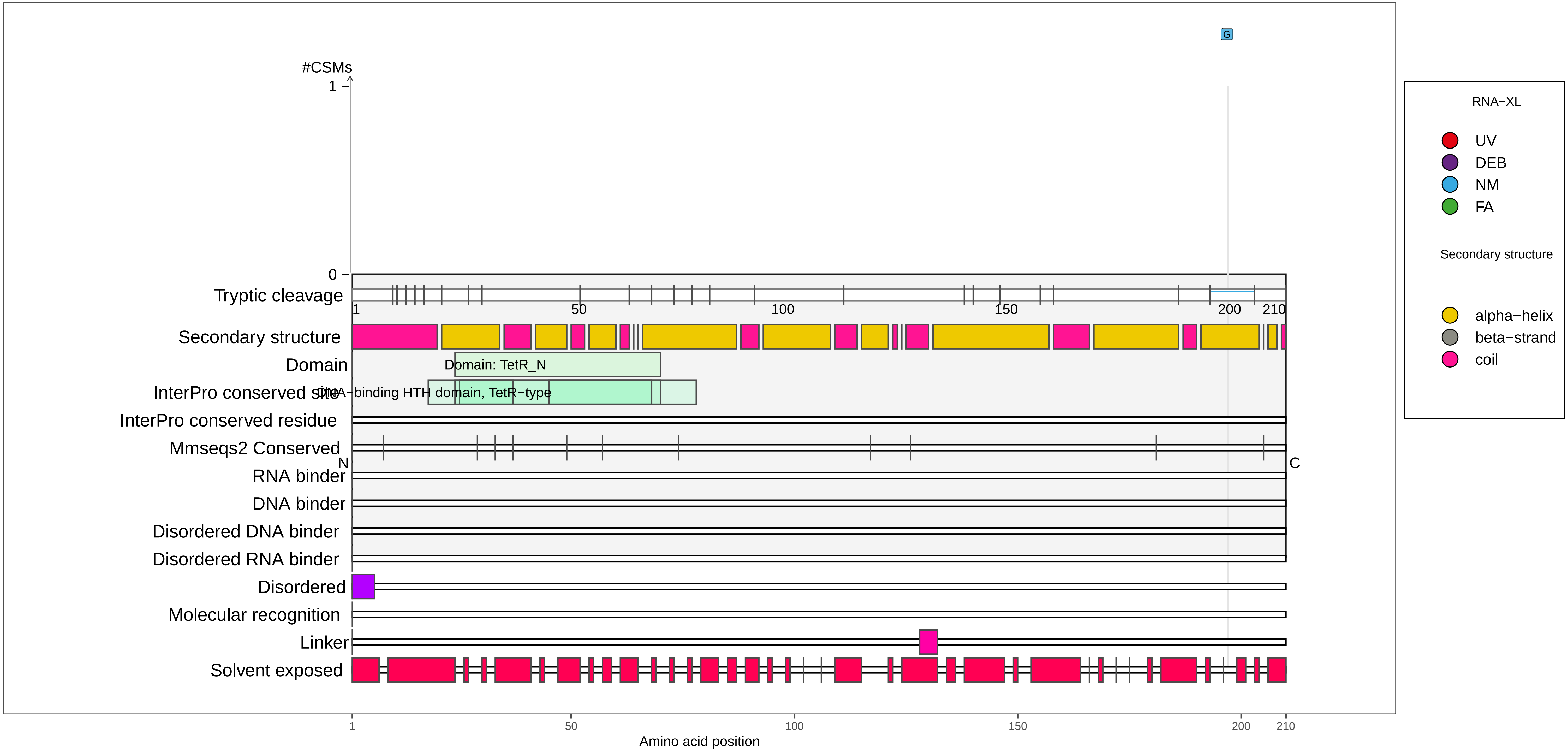

P75960
NPD_ECOLI NAD−dependent protein deacylase

− Abundance:
tryptic [log10 Intensity]: 7.44 (Q 32)
PAXdb K12 strain [ppm]: 1.06 (Q 13)
PAXdb E.coli [ppm]: 1.97 (Q 72)

− RNA functions: not annotated

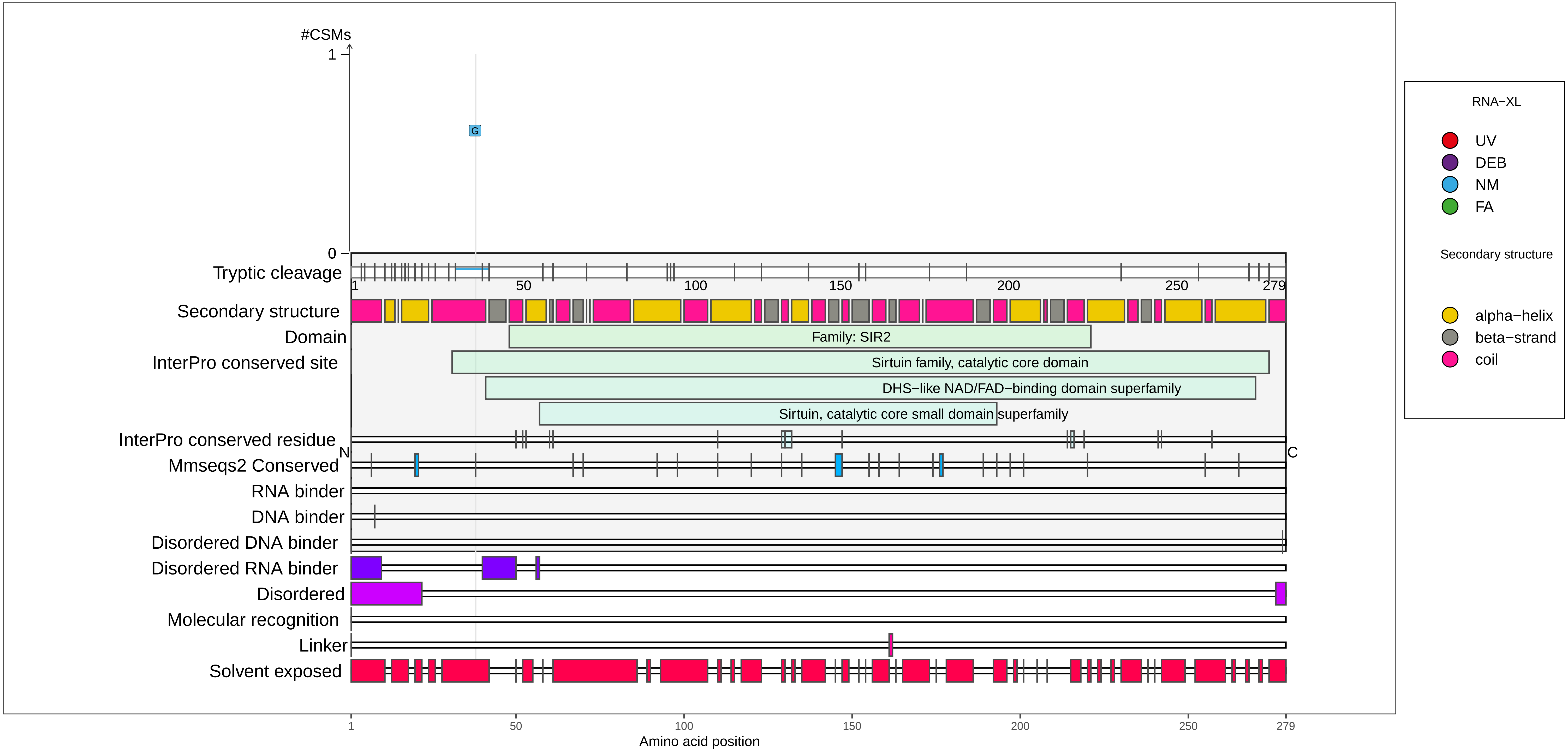

P75966
RLUE_ECOLI Ribosomal large subunit pseudouridine synthase E

− Abundance:
tryptic [log10 Intensity]: NA (Q NA)
PAXdb K12 strain [ppm]: 1.01 (Q 11)
PAXdb E.coli [ppm]: 0.68 (Q 40)

− RNA functions:
ncRNA metabolic process; ncRNA processing; RNA binding; RNA metabolic process
RNA modification; RNA processing; RNA pseudouridylate synthase; rRNA metabolic process
rRNA modification; rRNA processing; rRNA pseudouridine synthase activity
rRNA pseudouridine synthesis

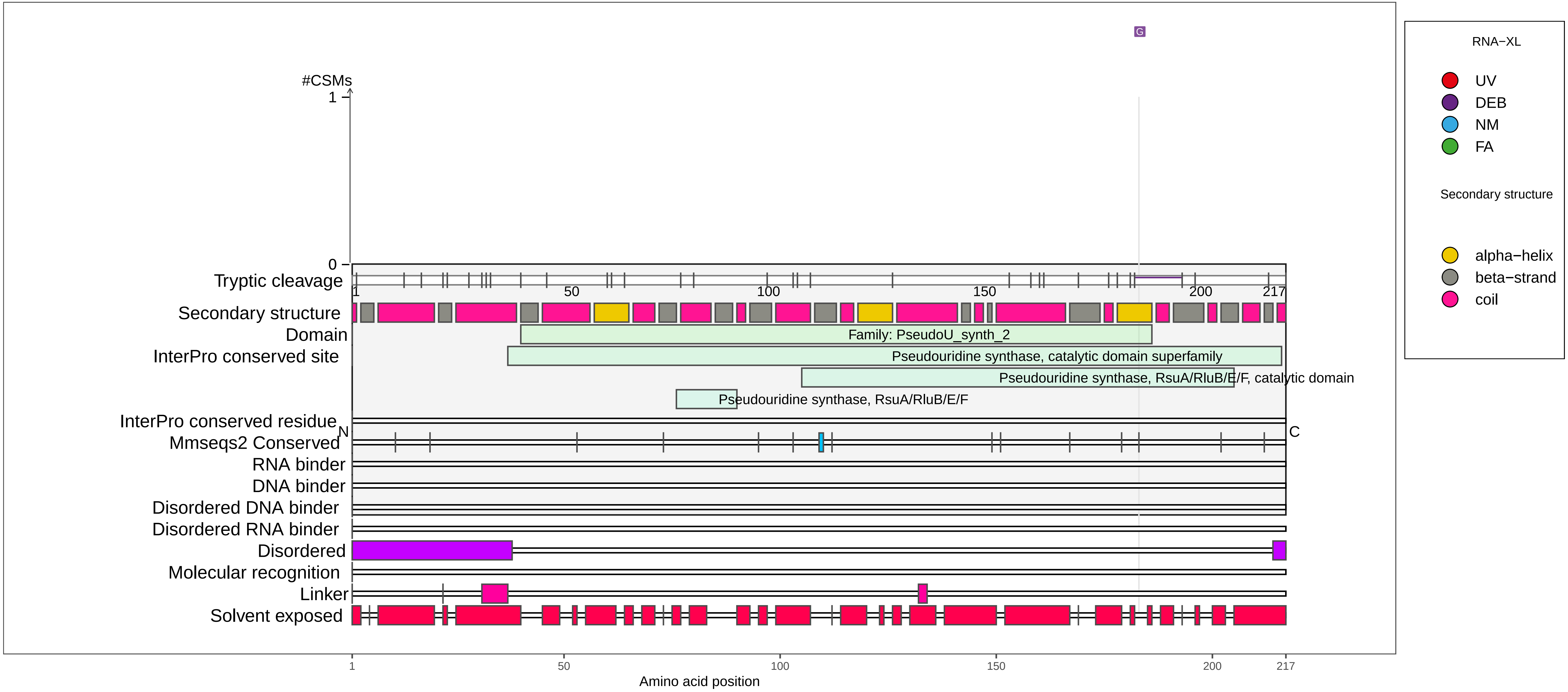

P75989
BLUR_ECOLI HTH−type transcriptional repressor BluR

− Abundance:
tryptic [log10 Intensity]: 7.16 (Q 19)
PAXdb K12 strain [ppm]: NA (Q NA)
PAXdb E.coli [ppm]: 1.42 (Q 57)

− RNA functions: not annotated

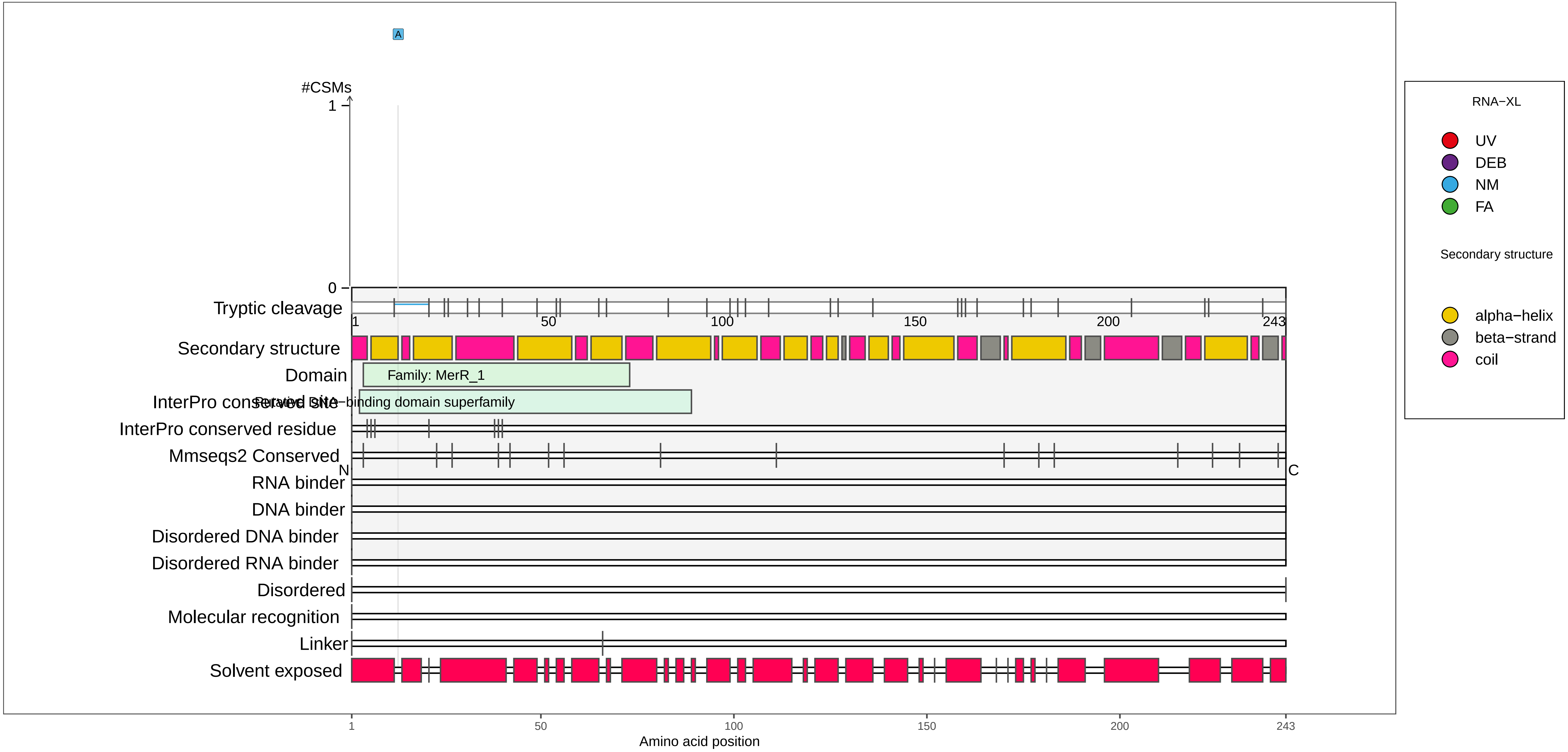

P76027
OPPD_ECOLI Oligopeptide transport ATP−binding protein OppD

− Abundance:
tryptic [log10 Intensity]: 6.25 (Q 1)
PAXdb K12 strain [ppm]: 1.31 (Q 27)
PAXdb E.coli [ppm]: 1.57 (Q 62)

− RNA functions: not annotated

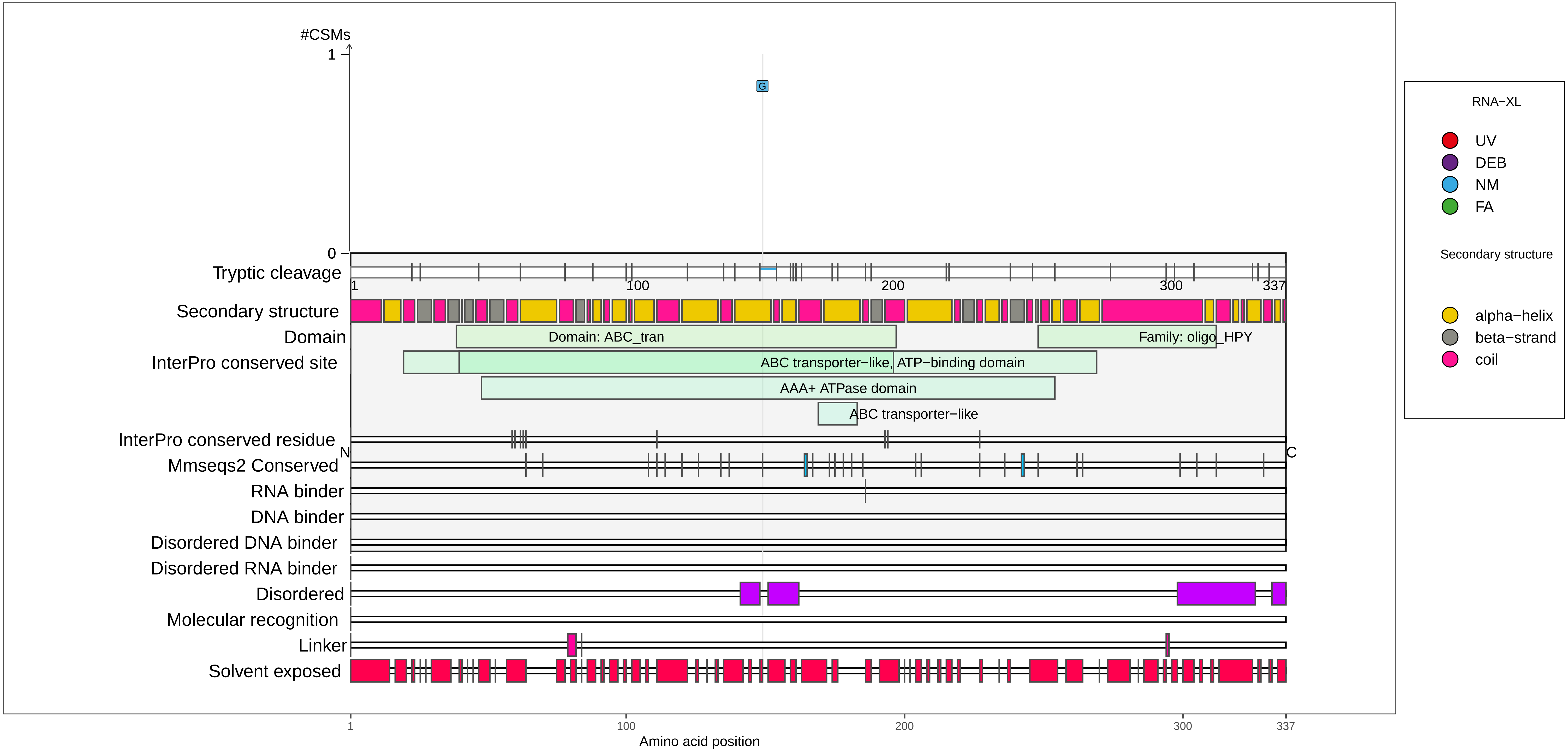

P76041
GGAP_ECOLI Glucosylglycerate phosphorylase

− Abundance:
tryptic [log10 Intensity]: NA (Q NA)
PAXdb K12 strain [ppm]: 0.73 (Q 3)
PAXdb E.coli [ppm]: −0.69 (Q 13)

− RNA functions: not annotated

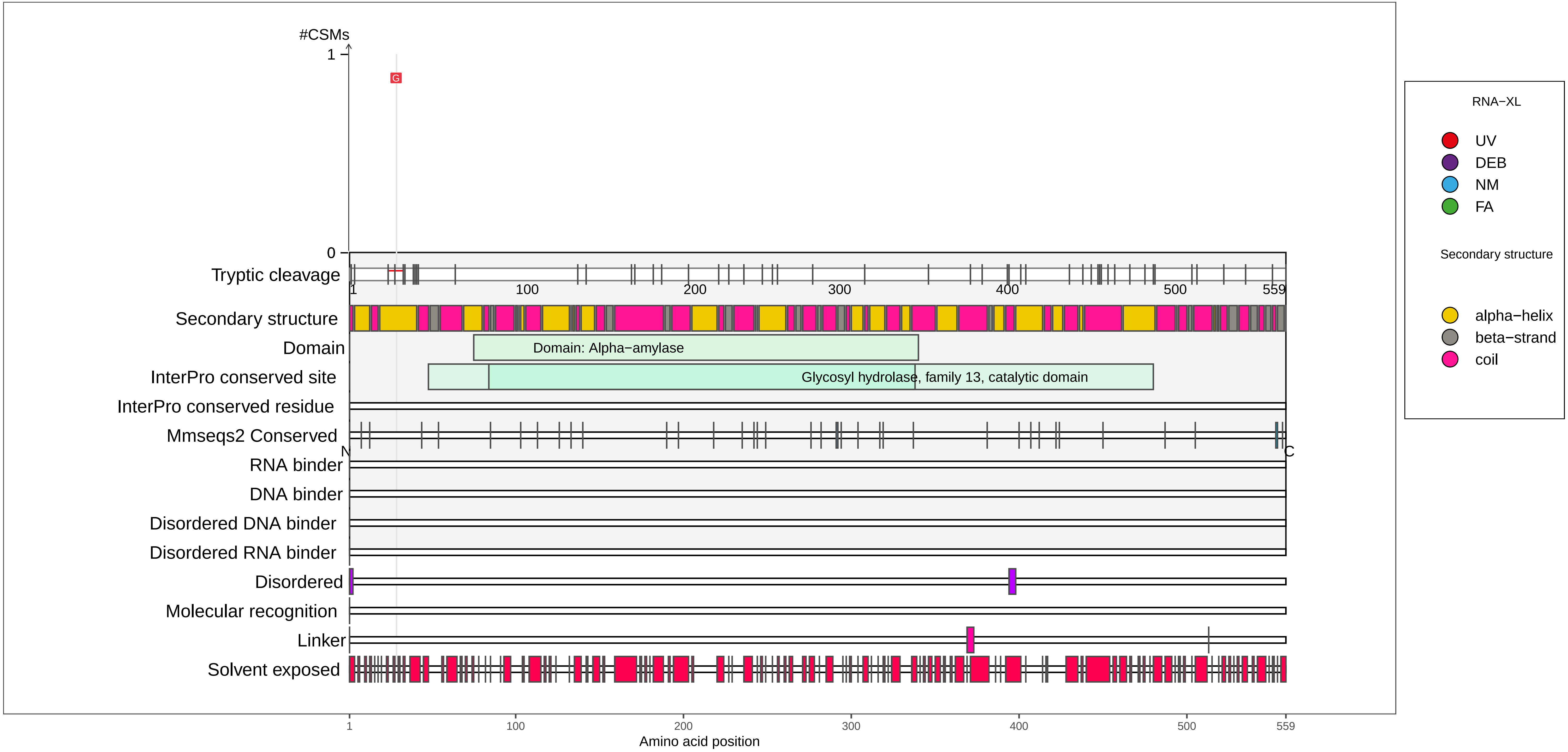

P76056
INTR_ECOLI Prophage integrase IntR

− Abundance:
tryptic [log10 Intensity]: NA (Q NA)
PAXdb K12 strain [ppm]: 0.77 (Q 4)
PAXdb E.coli [ppm]: −0.66 (Q 13)

− RNA functions: not annotated

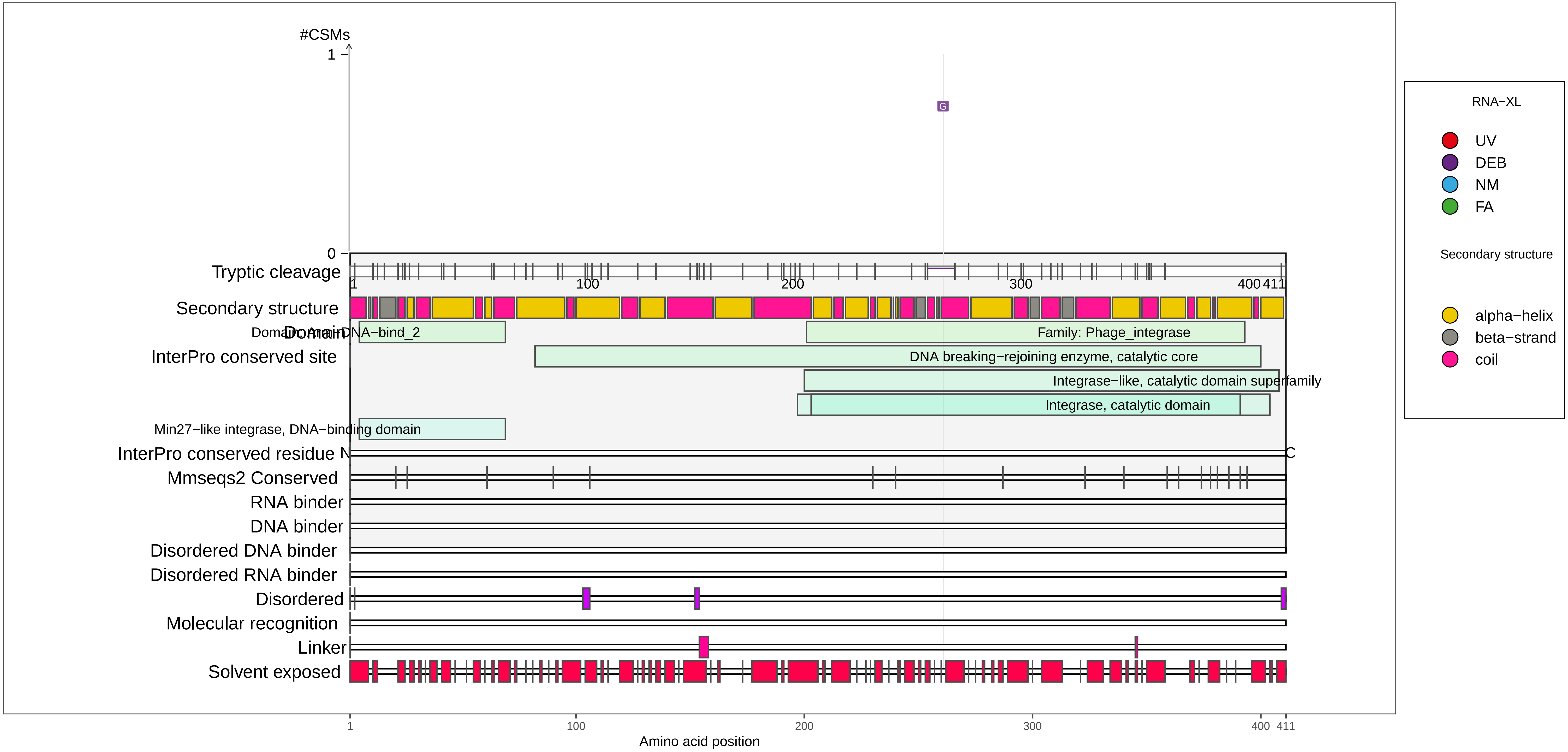

P76116
YNCE_ECOLI Uncharacterized protein YncE

− Abundance:
tryptic [log10 Intensity]: 8.85 (Q 85)
PAXdb K12 strain [ppm]: 1.46 (Q 36)
PAXdb E.coli [ppm]: 2.61 (Q 88)

− RNA functions: not annotated

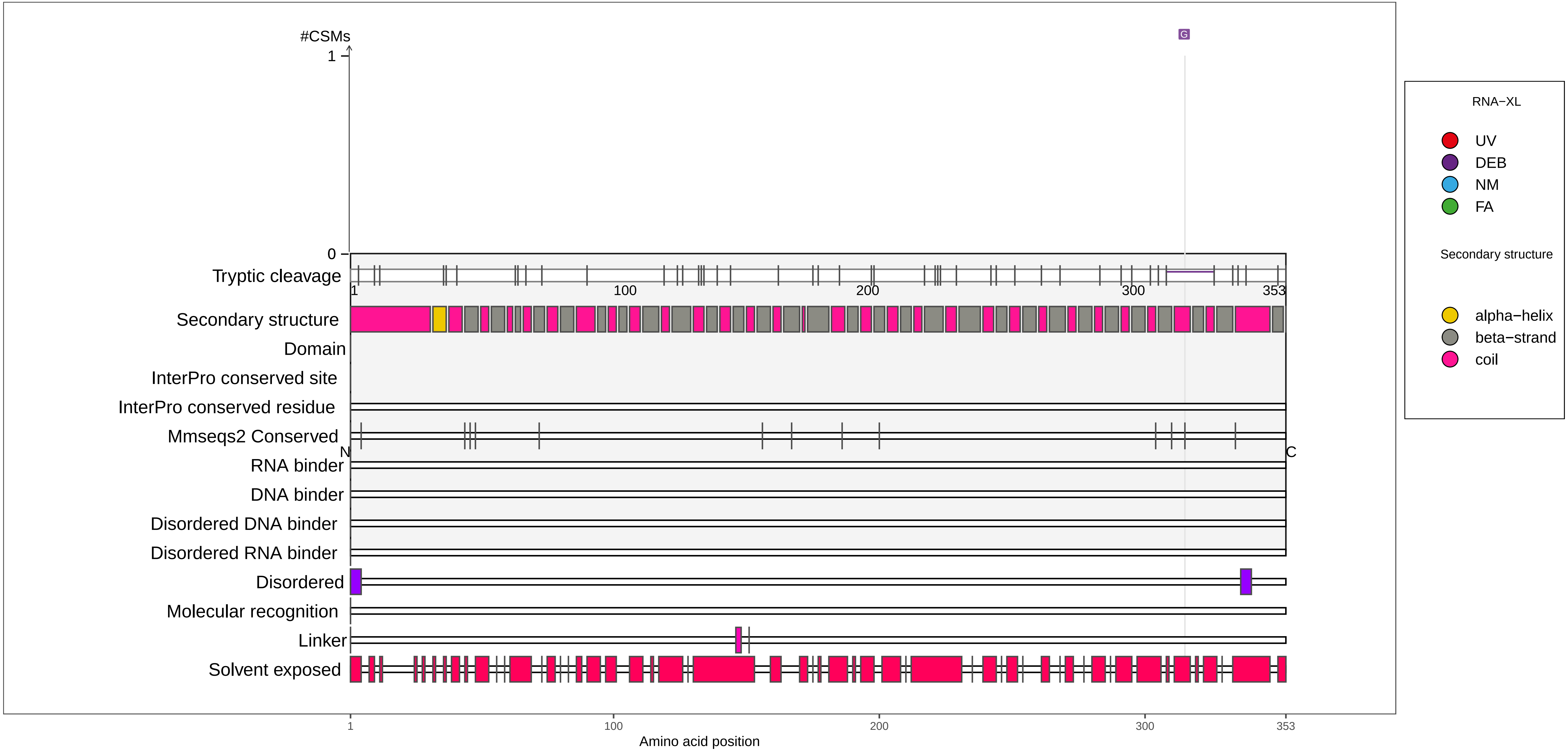

P76147
DGCF_ECOLI Probable diguanylate cyclase DgcF

− Abundance:
tryptic [log10 Intensity]: NA (Q NA)
PAXdb K12 strain [ppm]: NA (Q NA)
PAXdb E.coli [ppm]: −0.95 (Q 9)

− RNA functions: not annotated

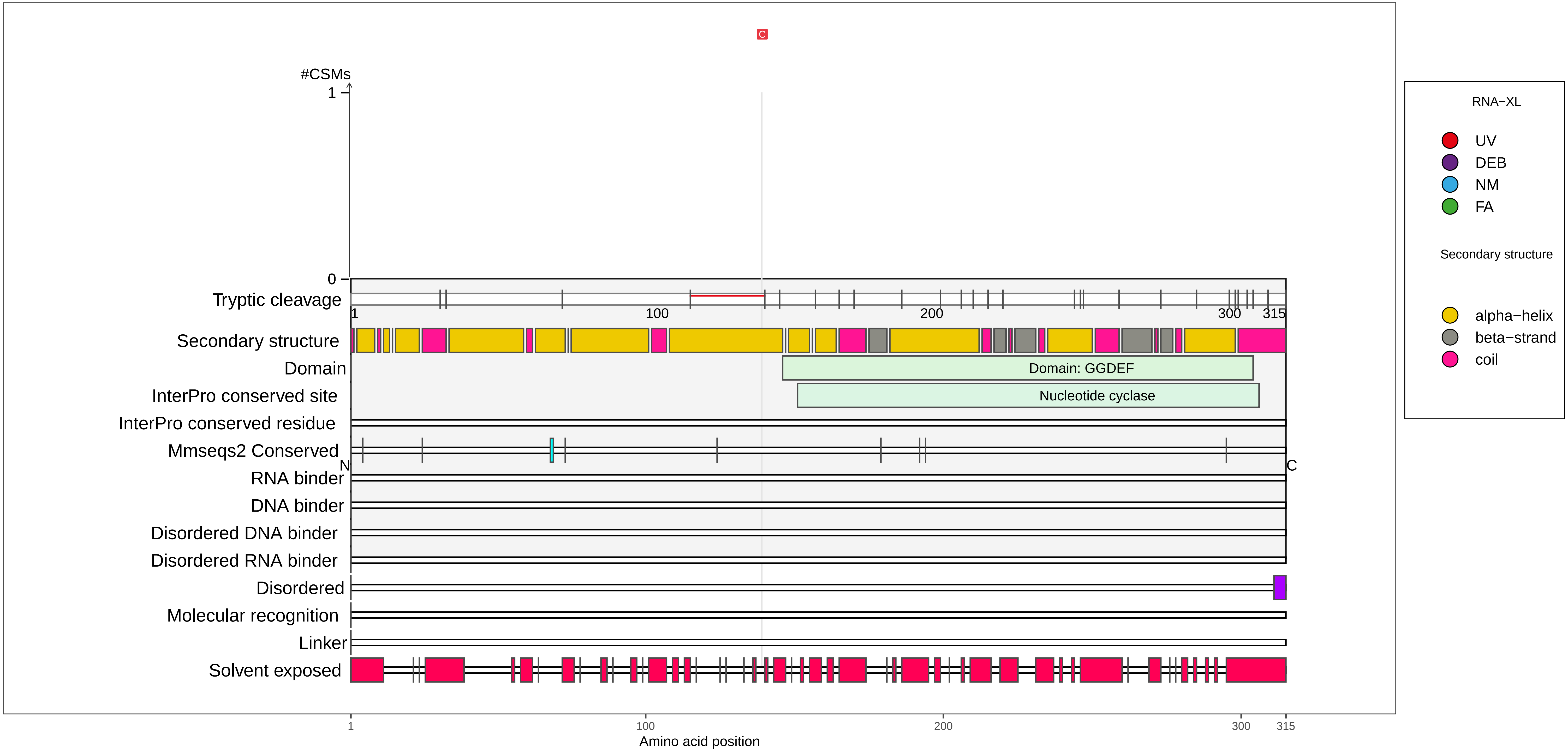

P76192
YDHV_ECOLI Uncharacterized oxidoreductase YdhV

− Abundance:
tryptic [log10 Intensity]: 8.97 (Q 88)
PAXdb K12 strain [ppm]: NA (Q NA)
PAXdb E.coli [ppm]: 0.46 (Q 35)

− RNA functions: not annotated

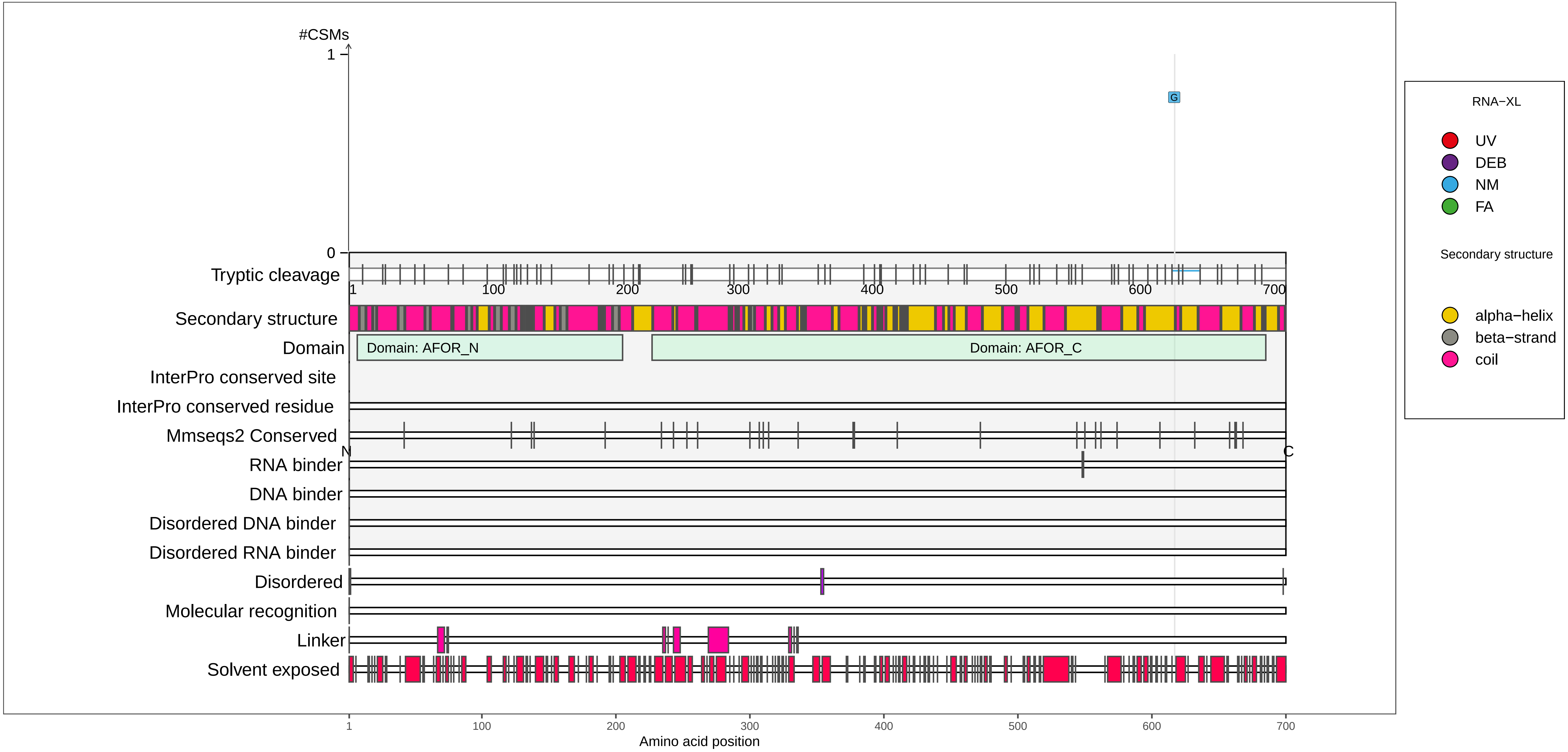

P76223
YNJB_ECOLI Protein YnjB

− Abundance:
tryptic [log10 Intensity]: 6.88 (Q 9)
PAXdb K12 strain [ppm]: NA (Q NA)
PAXdb E.coli [ppm]: 0.48 (Q 36)

− RNA functions: not annotated

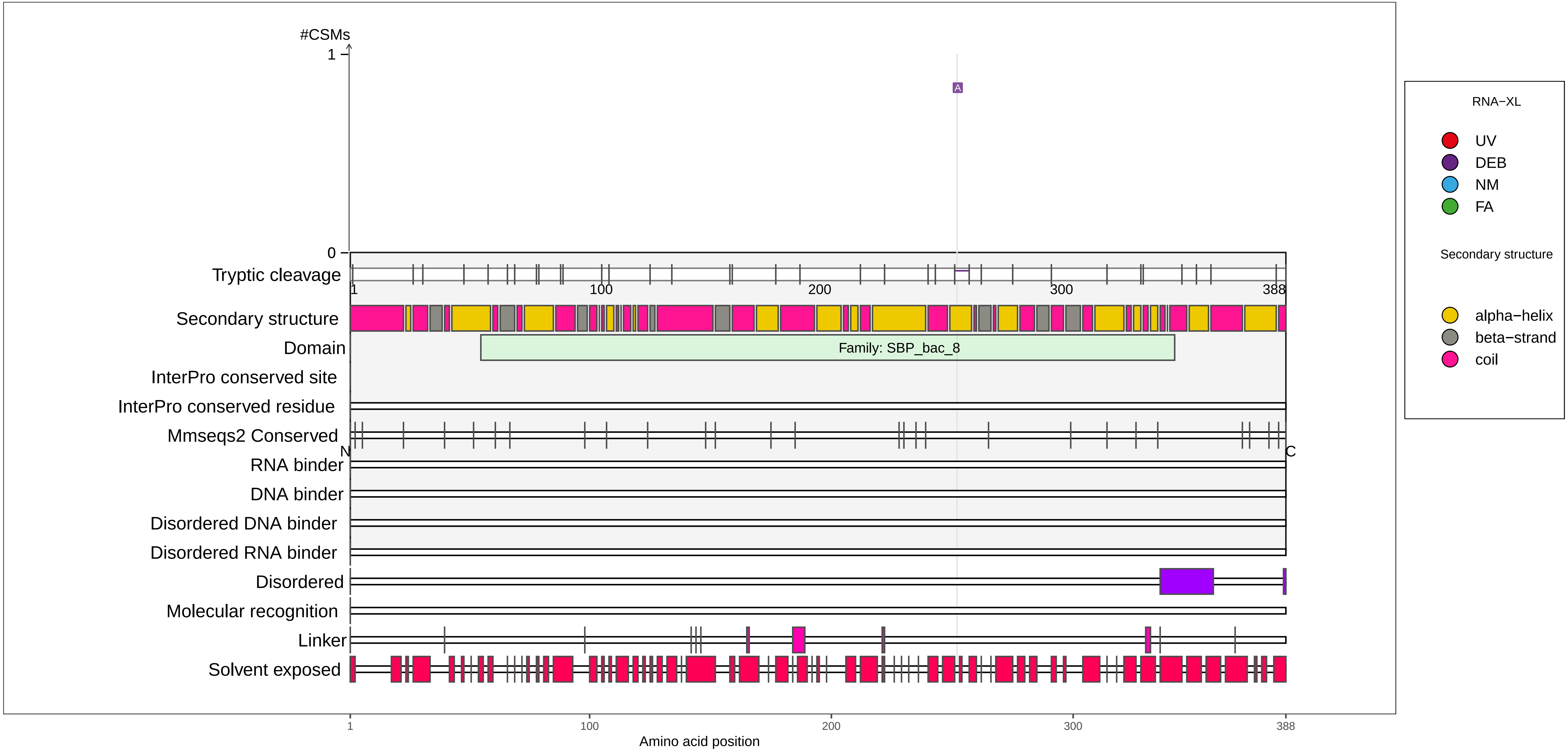

P76235
YEAH_ECOLI UPF0229 protein YeaH

− Abundance:
tryptic [log10 Intensity]: NA (Q NA)
PAXdb K12 strain [ppm]: NA (Q NA)
PAXdb E.coli [ppm]: 0.8 (Q 43)

− RNA functions: not annotated

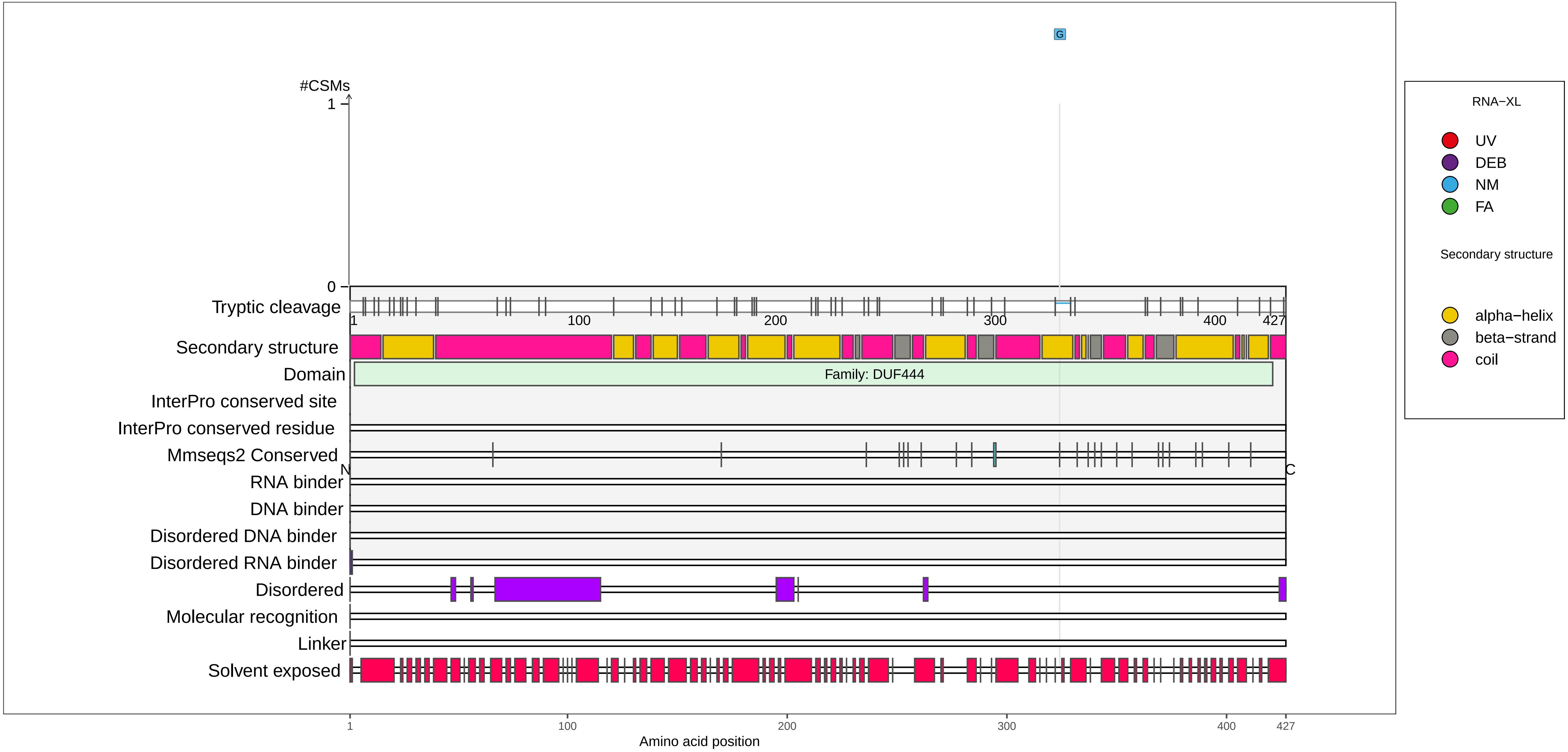

P76387
WZC_ECOLI Tyrosine−protein kinase wzc

− Abundance:
tryptic [log10 Intensity]: NA (Q NA)
PAXdb K12 strain [ppm]: 0.66 (Q 2)
PAXdb E.coli [ppm]: −1.21 (Q 6)

− RNA functions: not annotated

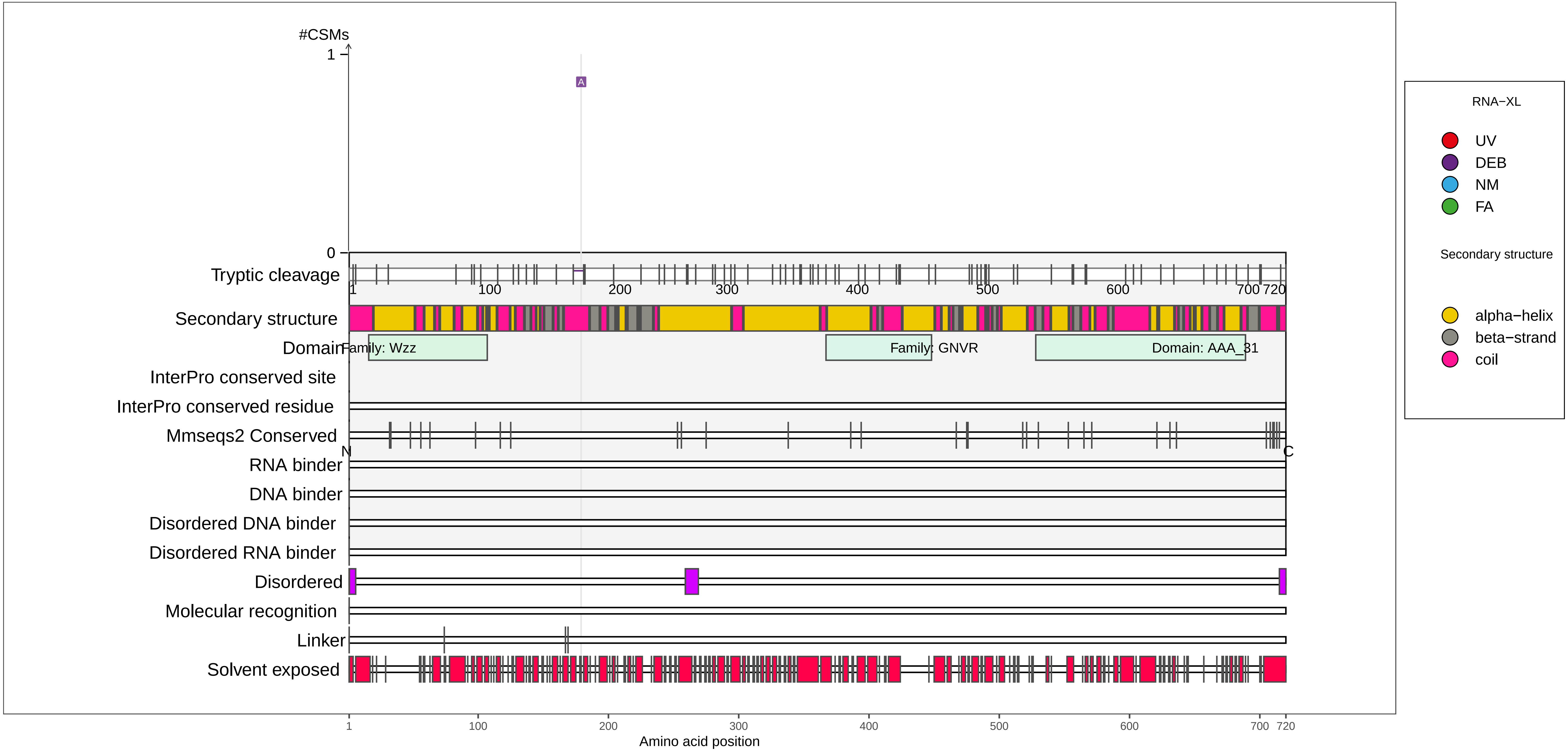

P76473
ARNT_ECOLI Undecaprenyl phosphate−alpha−4−amino−4−deoxy−L−arabinose arabinosyl transferase

− Abundance:
tryptic [log10 Intensity]: 8.1 (Q 61)
PAXdb K12 strain [ppm]: 1.09 (Q 15)
PAXdb E.coli [ppm]: −1.49 (Q 4)

− RNA functions: not annotated

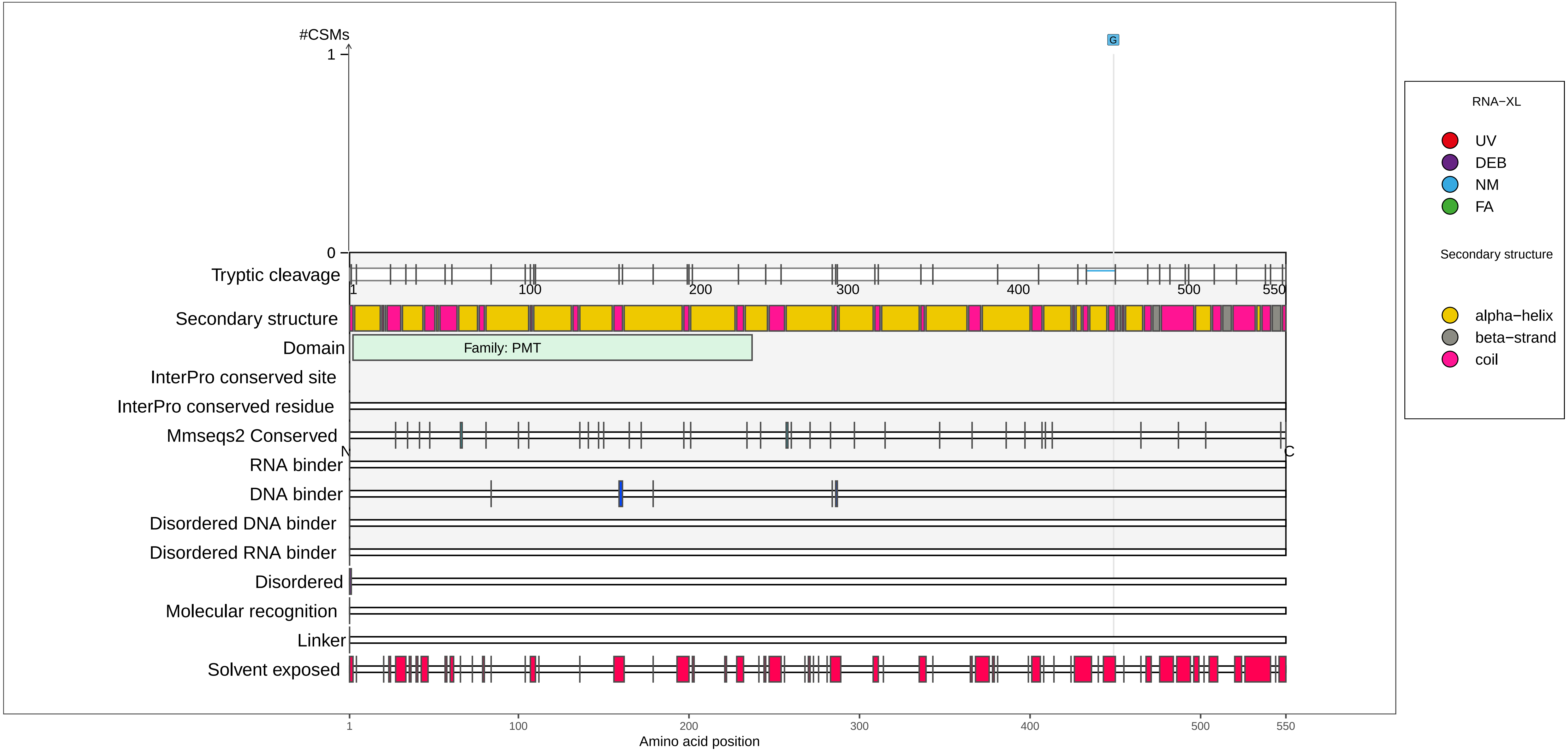

P76520
YFDX_ECOLI Protein YfdX

− Abundance:
tryptic [log10 Intensity]: NA (Q NA)
PAXdb K12 strain [ppm]: 1.28 (Q 25)
PAXdb E.coli [ppm]: −0.6 (Q 14)

− RNA functions: not annotated

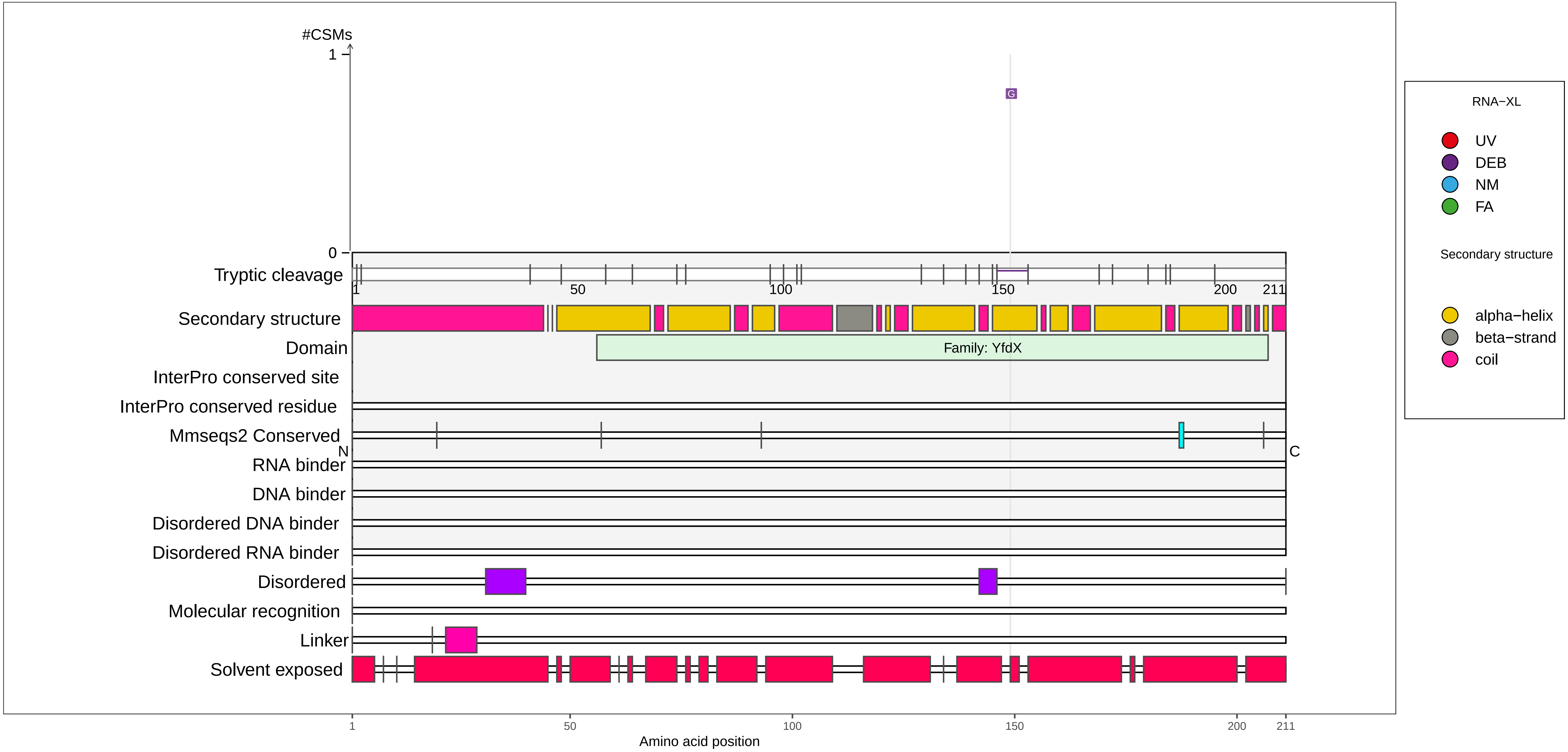

P76537
YFEY_ECOLI Uncharacterized protein YfeY

− Abundance:
tryptic [log10 Intensity]: 7.65 (Q 42)
PAXdb K12 strain [ppm]: 1.81 (Q 53)
PAXdb E.coli [ppm]: 1.22 (Q 53)

− RNA functions: not annotated

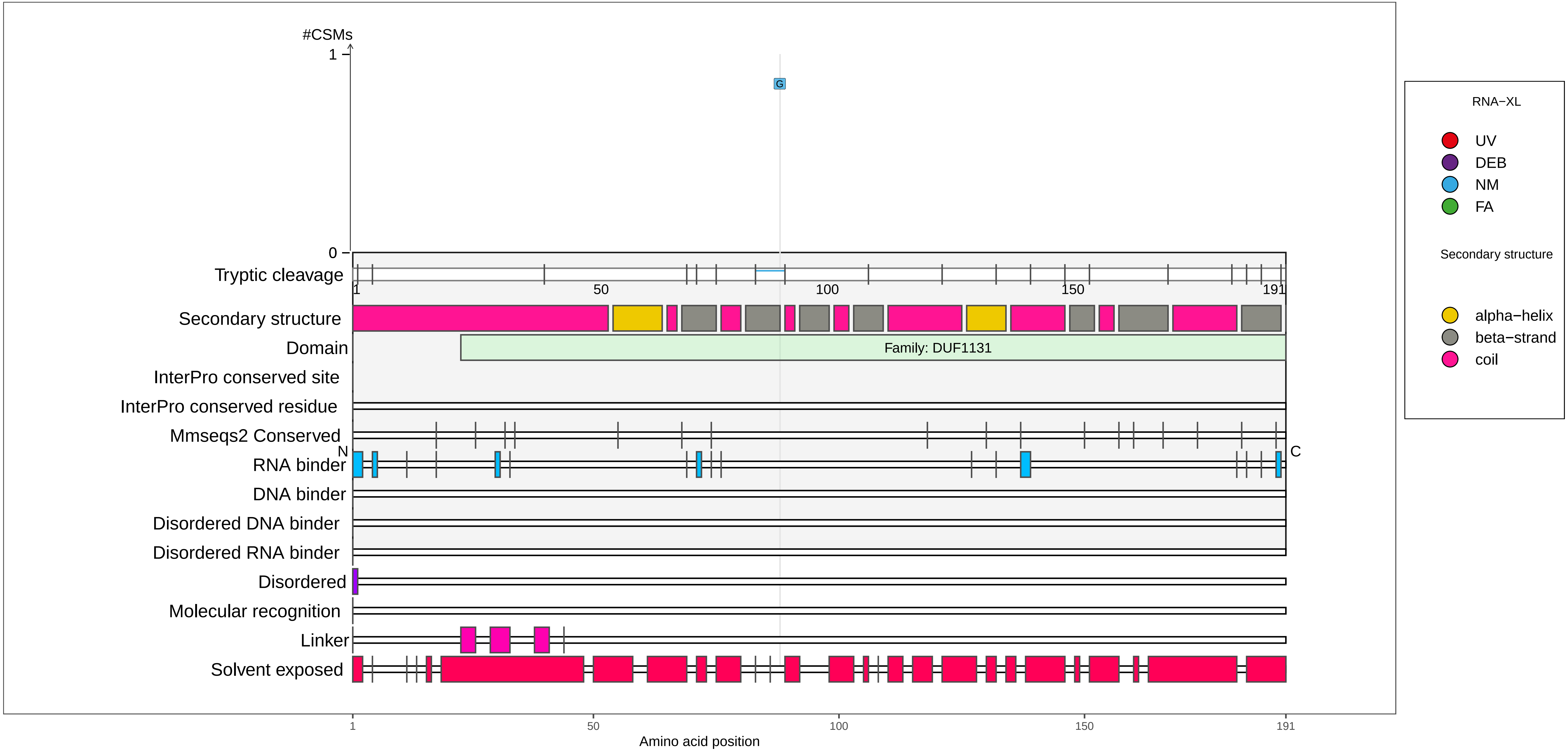

P76639
YGEH_ECOLI Uncharacterized protein YgeH

− Abundance:
tryptic [log10 Intensity]: NA (Q NA)
PAXdb K12 strain [ppm]: NA (Q NA)
PAXdb E.coli [ppm]: NA (Q NA)

− RNA functions: not annotated

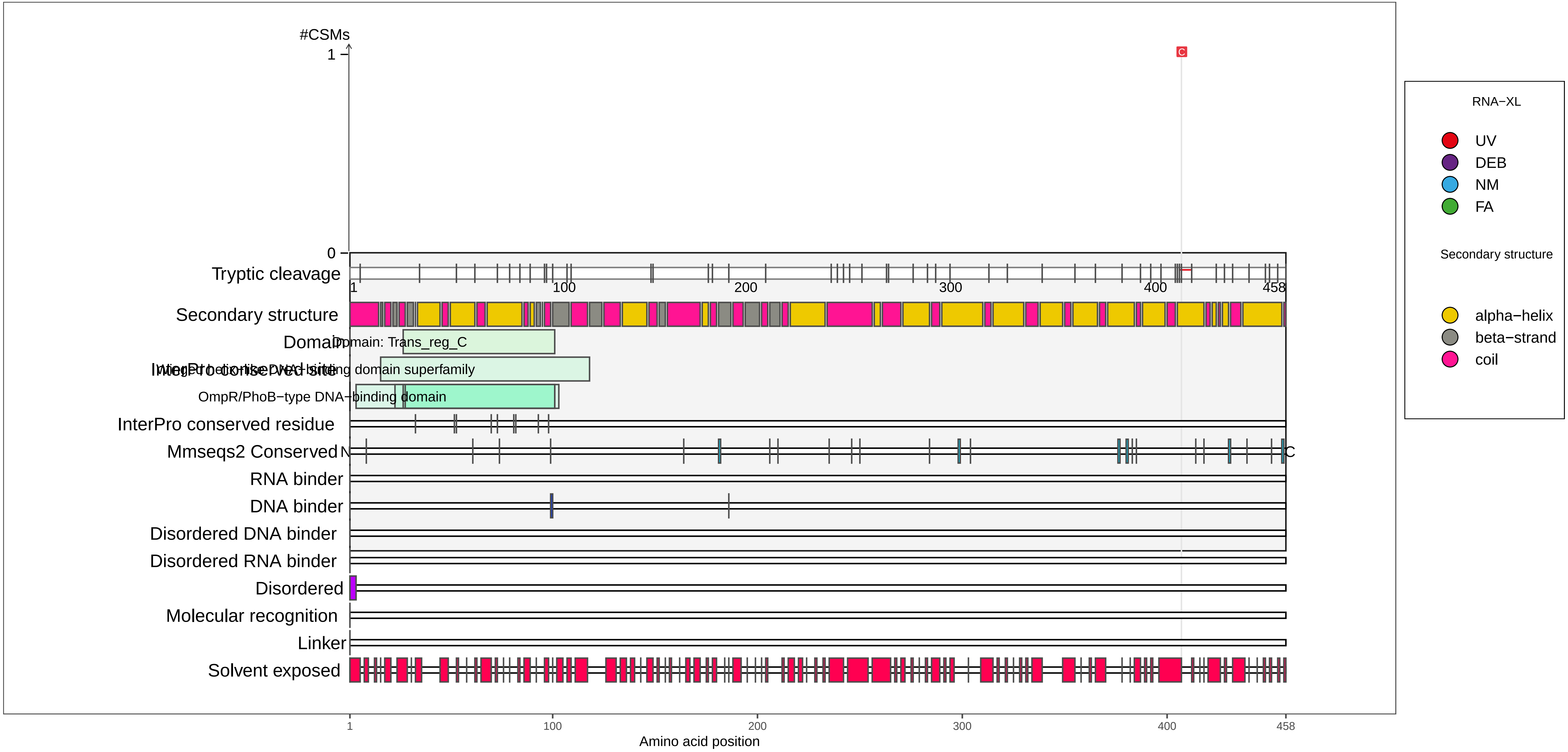

P76655
YQIG_ECOLI Putative outer membrane usher protein YqiG
− RNA functions: not annotated

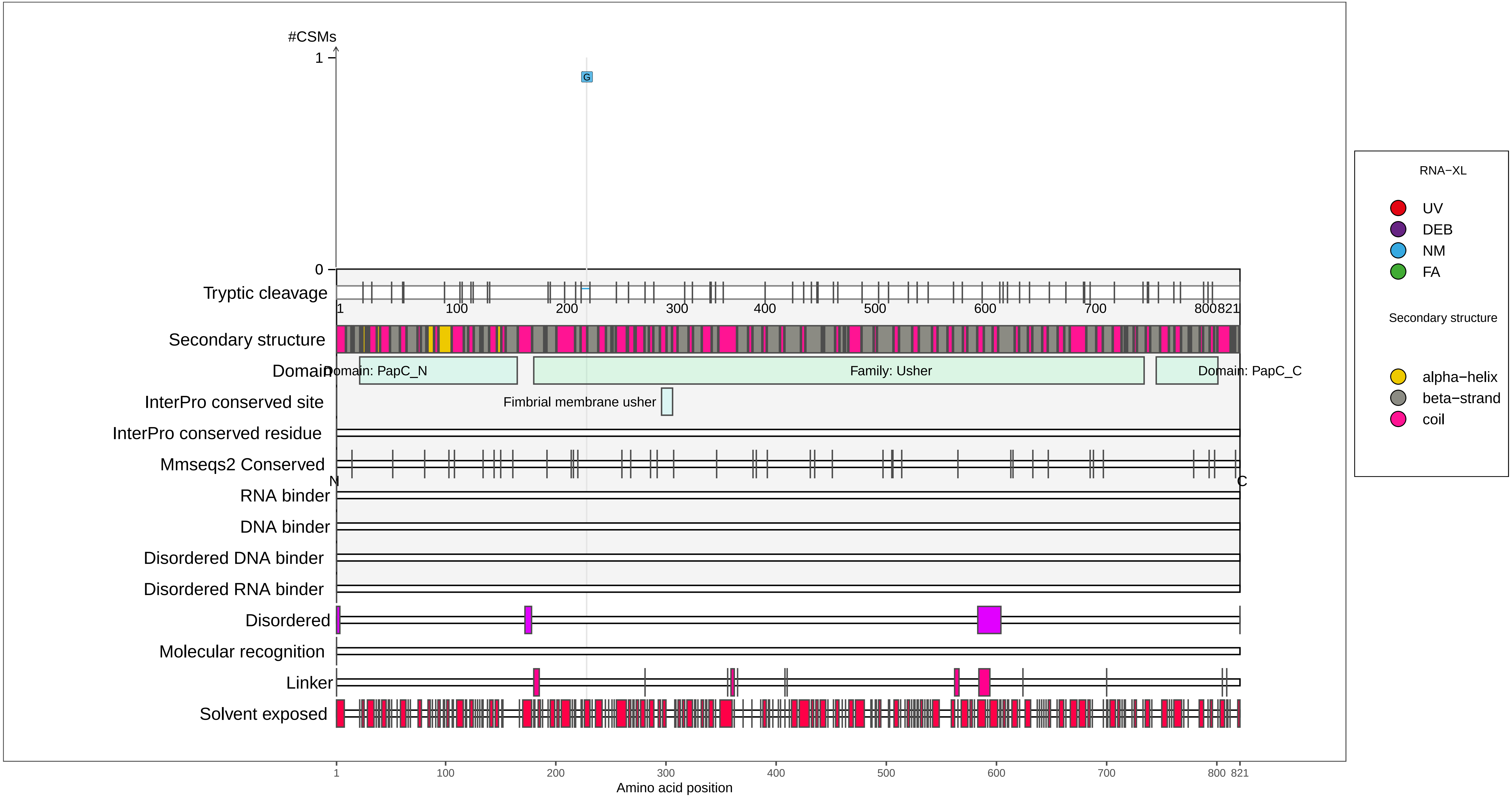

P77216
YBDN_ECOLI Uncharacterized protein YbdN

− Abundance:
tryptic [log10 Intensity]: NA (Q NA)
PAXdb K12 strain [ppm]: 0.91 (Q 8)
PAXdb E.coli [ppm]: 0.64 (Q 39)

− RNA functions: not annotated

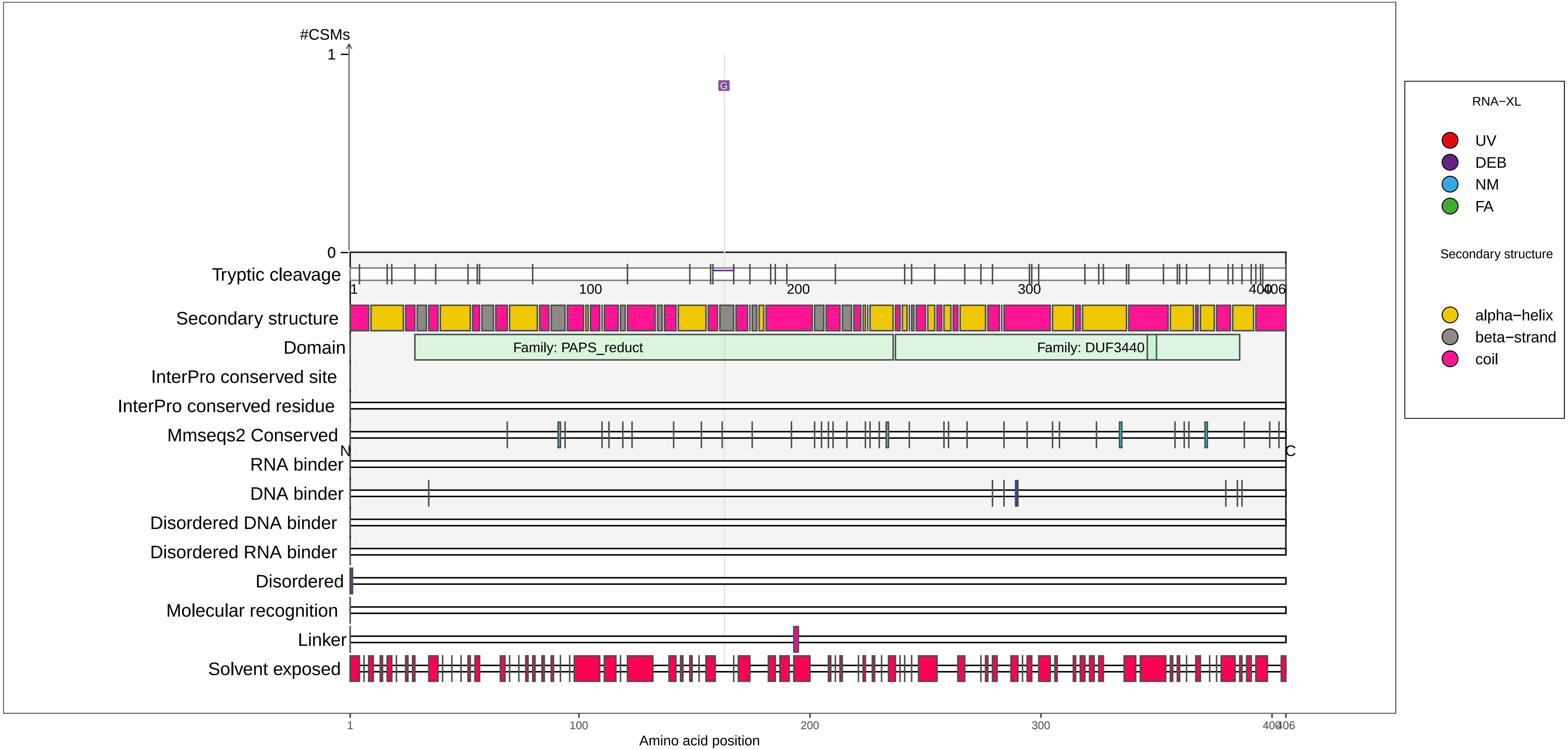

P77243
PRPD_ECOLI 2−methylcitrate dehydratase

− Abundance:
tryptic [log10 Intensity]: 8.87 (Q 85)
PAXdb K12 strain [ppm]: NA (Q NA)
PAXdb E.coli [ppm]: 2.19 (Q 77)

− RNA functions: not annotated

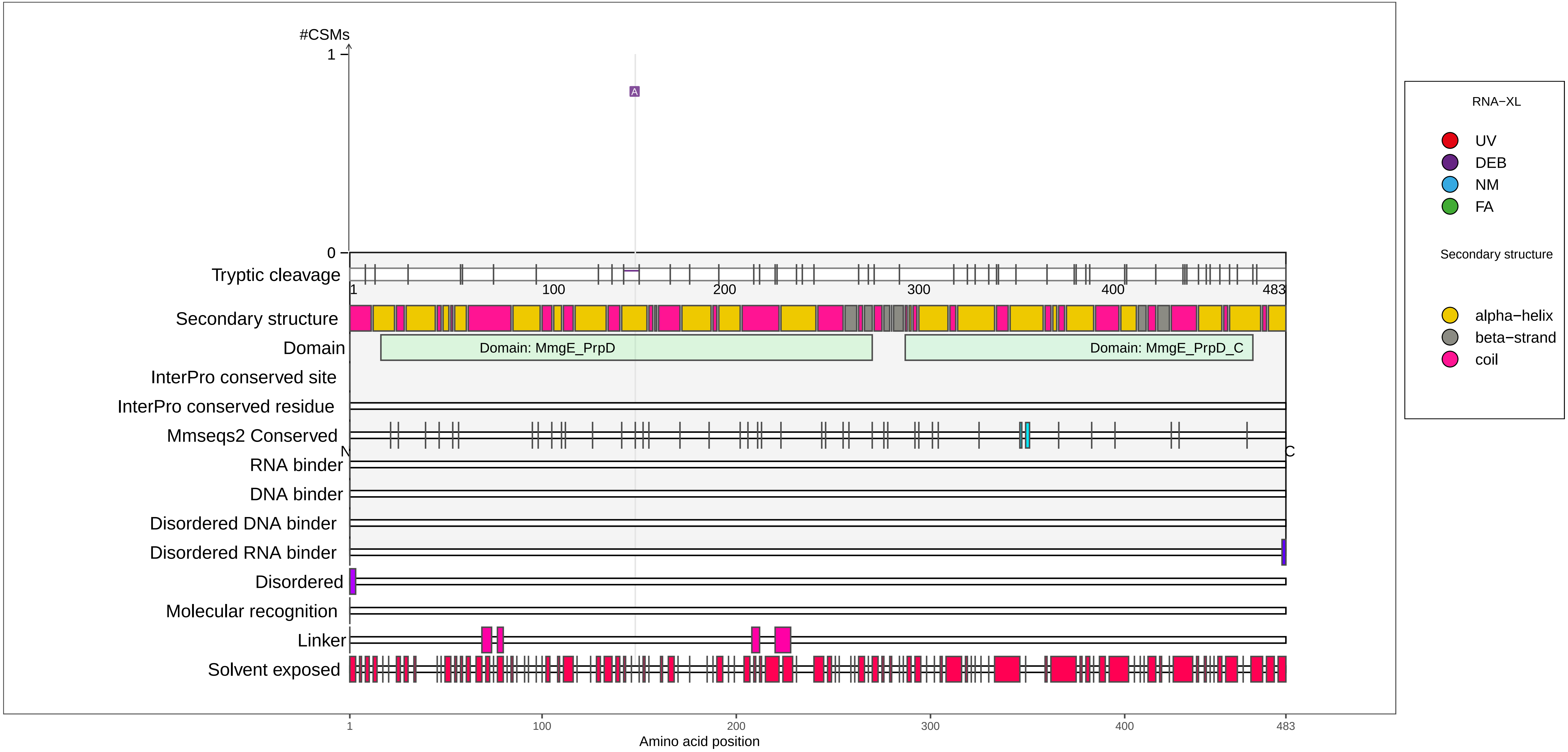

P77285
RSXG_ECOLI Ion−translocating oxidoreductase complex subunit G

− Abundance:
tryptic [log10 Intensity]: 7.05 (Q 15)
PAXdb K12 strain [ppm]: 1.95 (Q 59)
PAXdb E.coli [ppm]: 0.82 (Q 44)

− RNA functions: not annotated

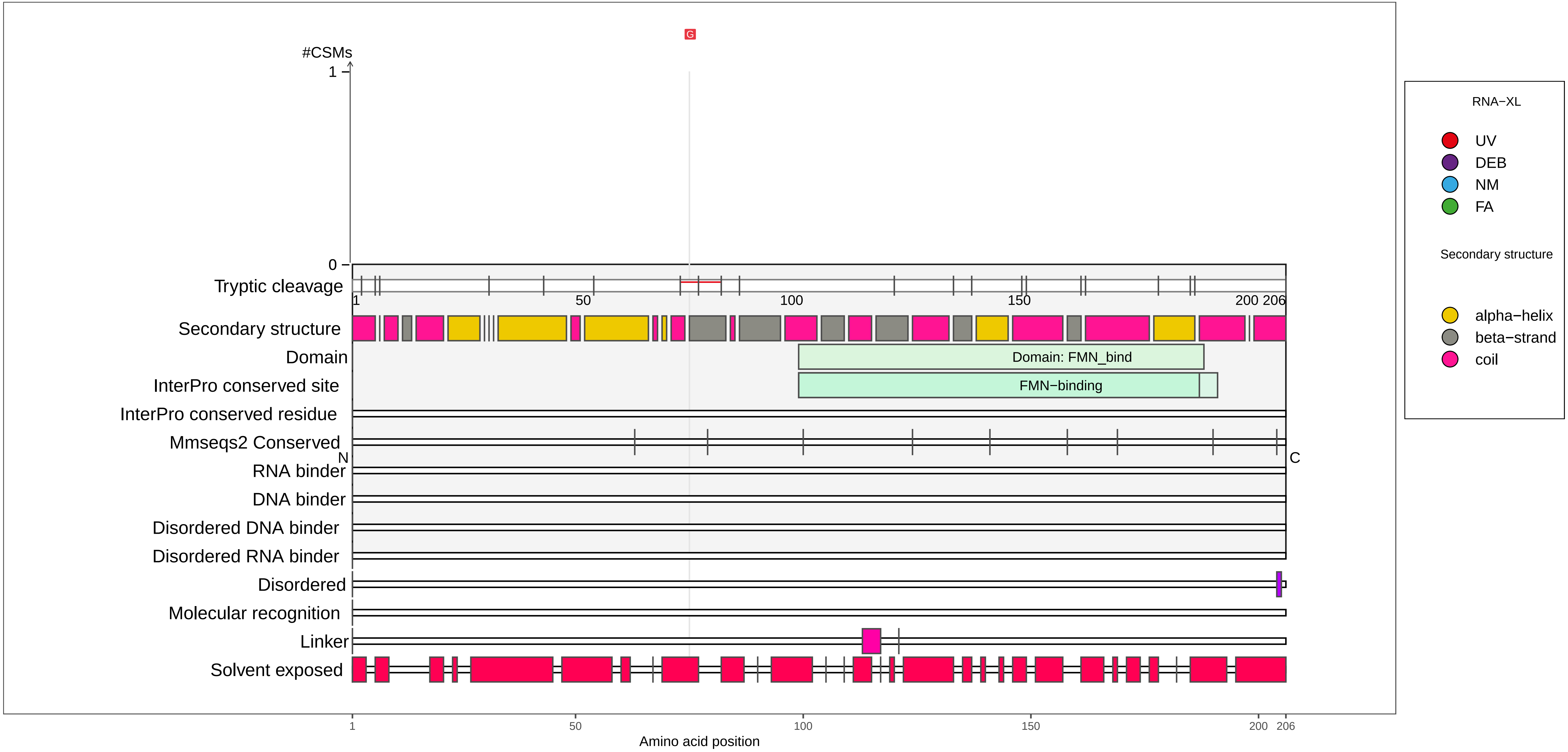

P77300
XYNR_ECOLI HTH−type transcriptional regulator XynR
− Abundance:
tryptic [log10 Intensity]: NA (Q NA)
PAXdb K12 strain [ppm]: NA (Q NA)
PAXdb E.coli [ppm]: 0.83 (Q 44)

− RNA functions:
RNA biosynthetic process; RNA metabolic process

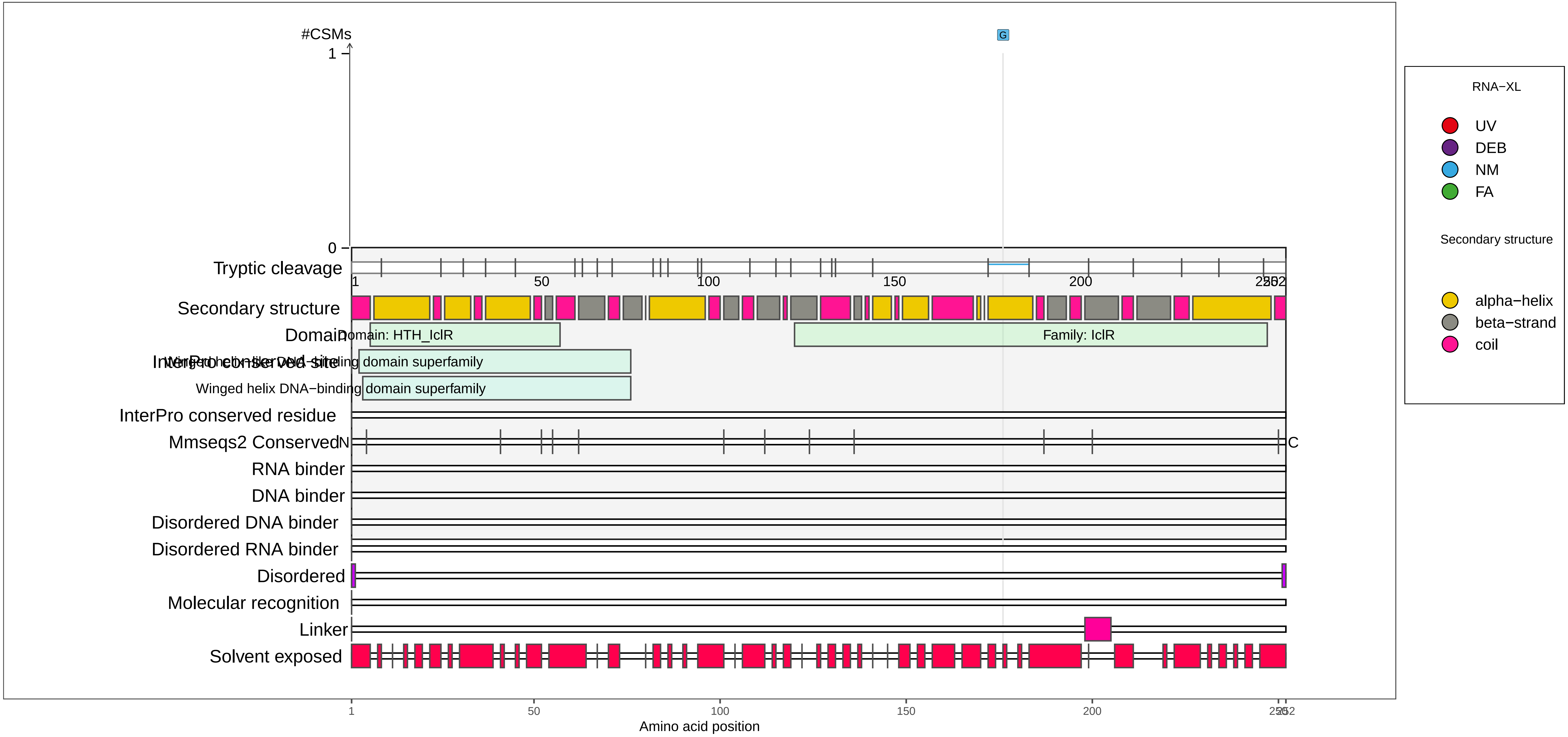

P77337
YDIS_ECOLI Probable electron transfer flavoprotein−quinone oxidoreductase YdiS

− Abundance:
tryptic [log10 Intensity]: NA (Q NA)
PAXdb K12 strain [ppm]: 0.9 (Q 8)
PAXdb E.coli [ppm]: −1.85 (Q 2)

− RNA functions: not annotated

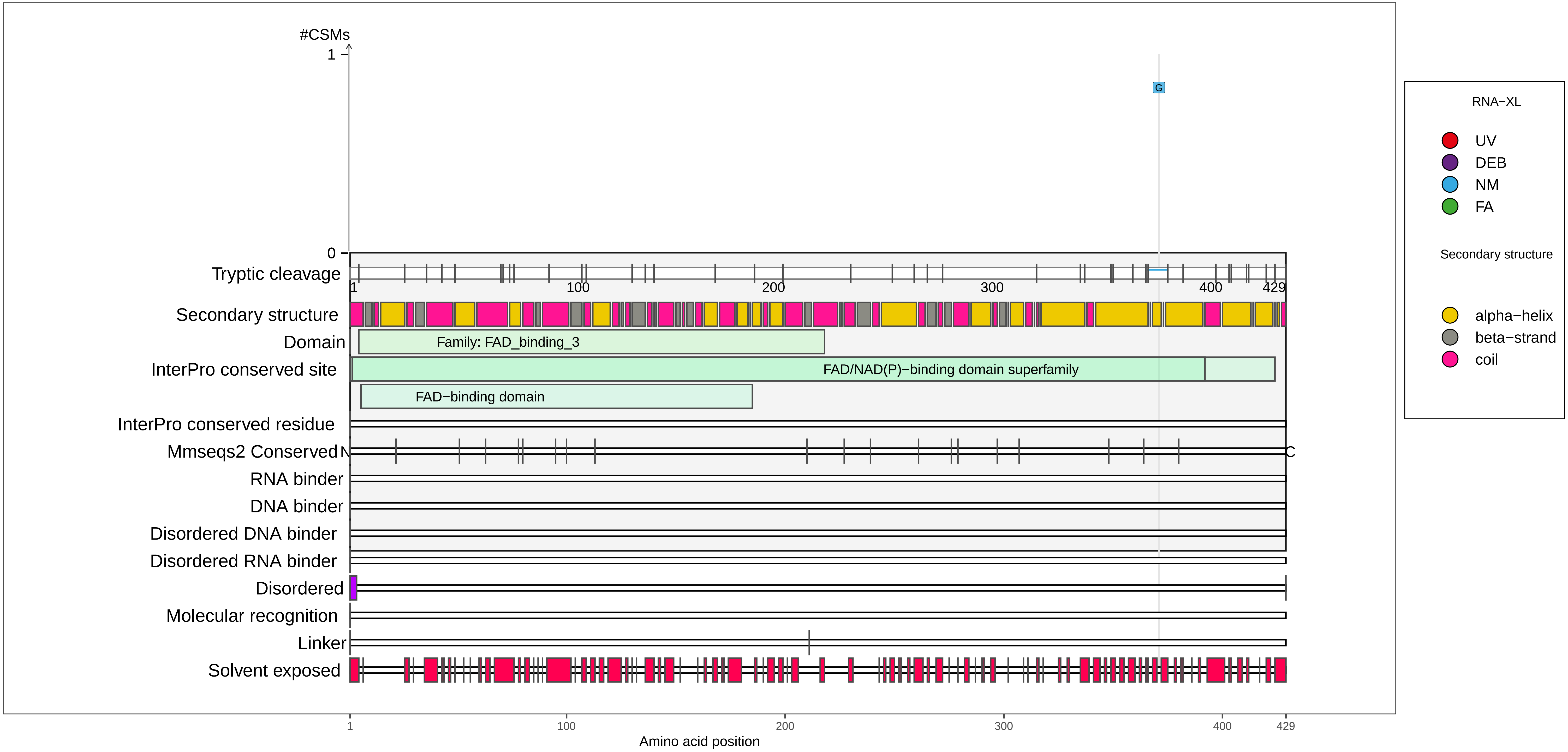

P77434
ALAC_ECOLI Glutamate−pyruvate aminotransferase AlaC

− Abundance:
tryptic [log10 Intensity]: 8.23 (Q 67)
PAXdb K12 strain [ppm]: 1.08 (Q 15)
PAXdb E.coli [ppm]: 1.55 (Q 61)

− RNA functions: not annotated

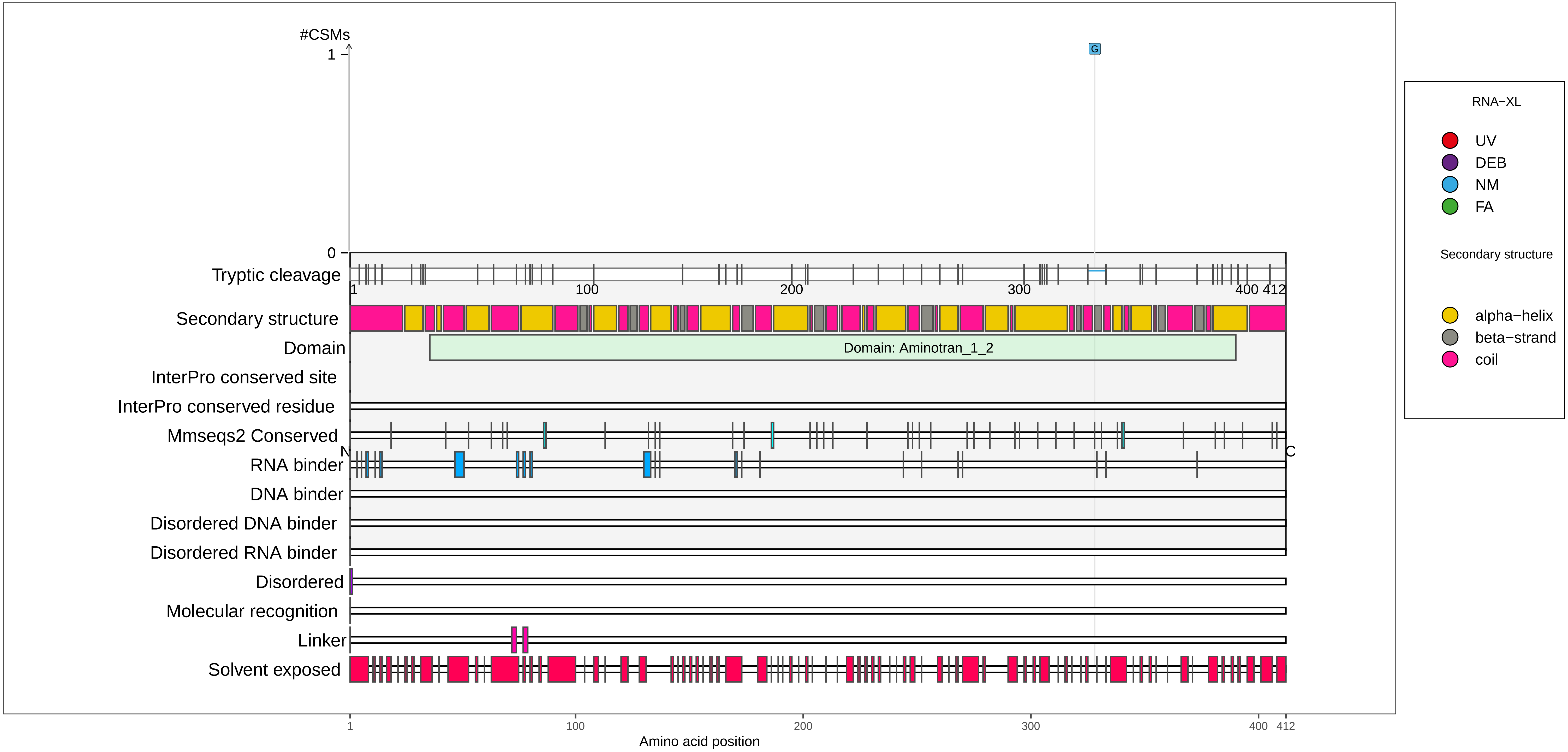

P77504
YBBP_ECOLI Uncharacterized ABC transporter permease YbbP
− Abundance:
tryptic [log10 Intensity]: NA (Q NA)
PAXdb K12 strain [ppm]: 0.72 (Q 3)
PAXdb E.coli [ppm]: −0.04 (Q 24)

− RNA functions: not annotated

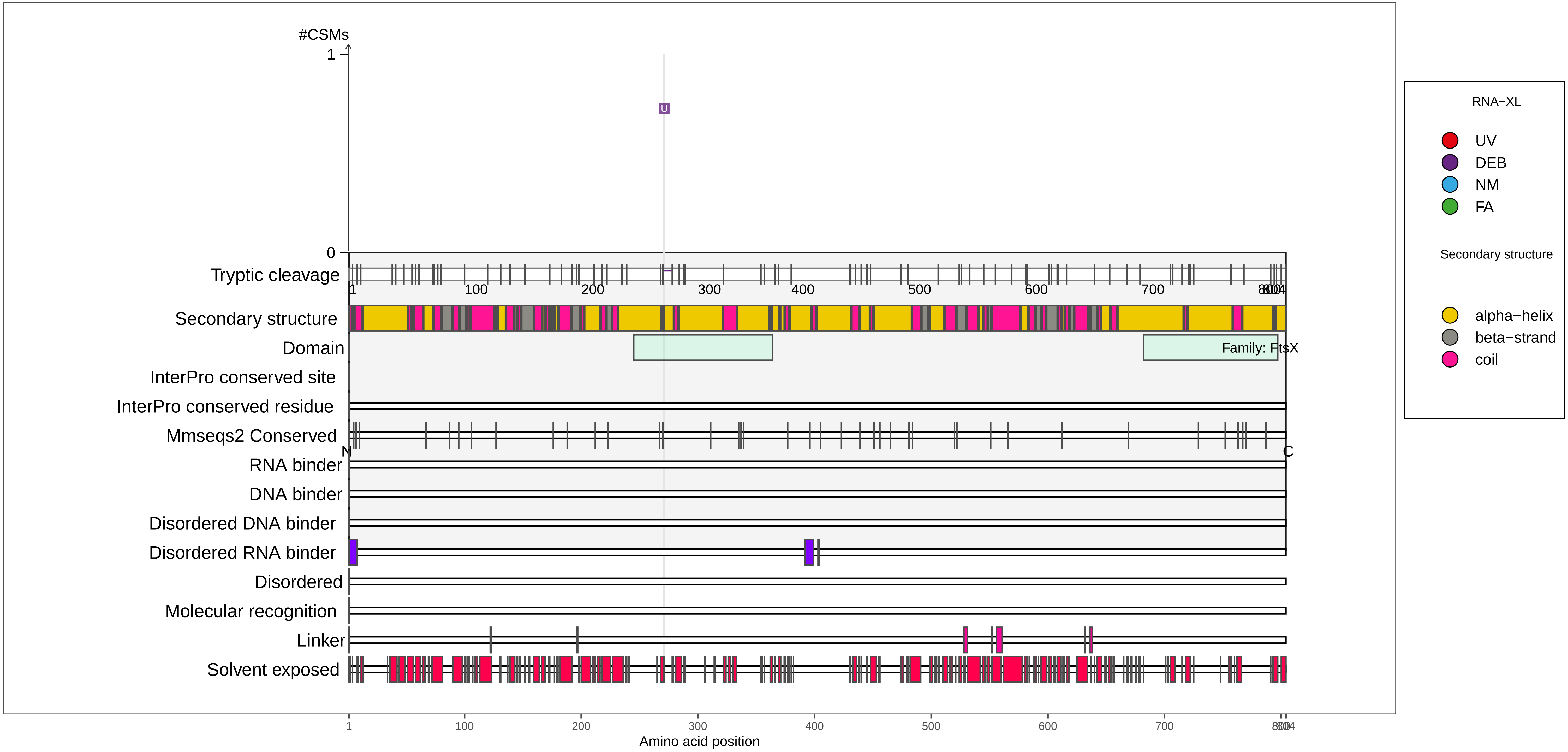

P77601
YKGA_ECOLI Putative HTH−type transcriptional regulator YkgA
− RNA functions: not annotated

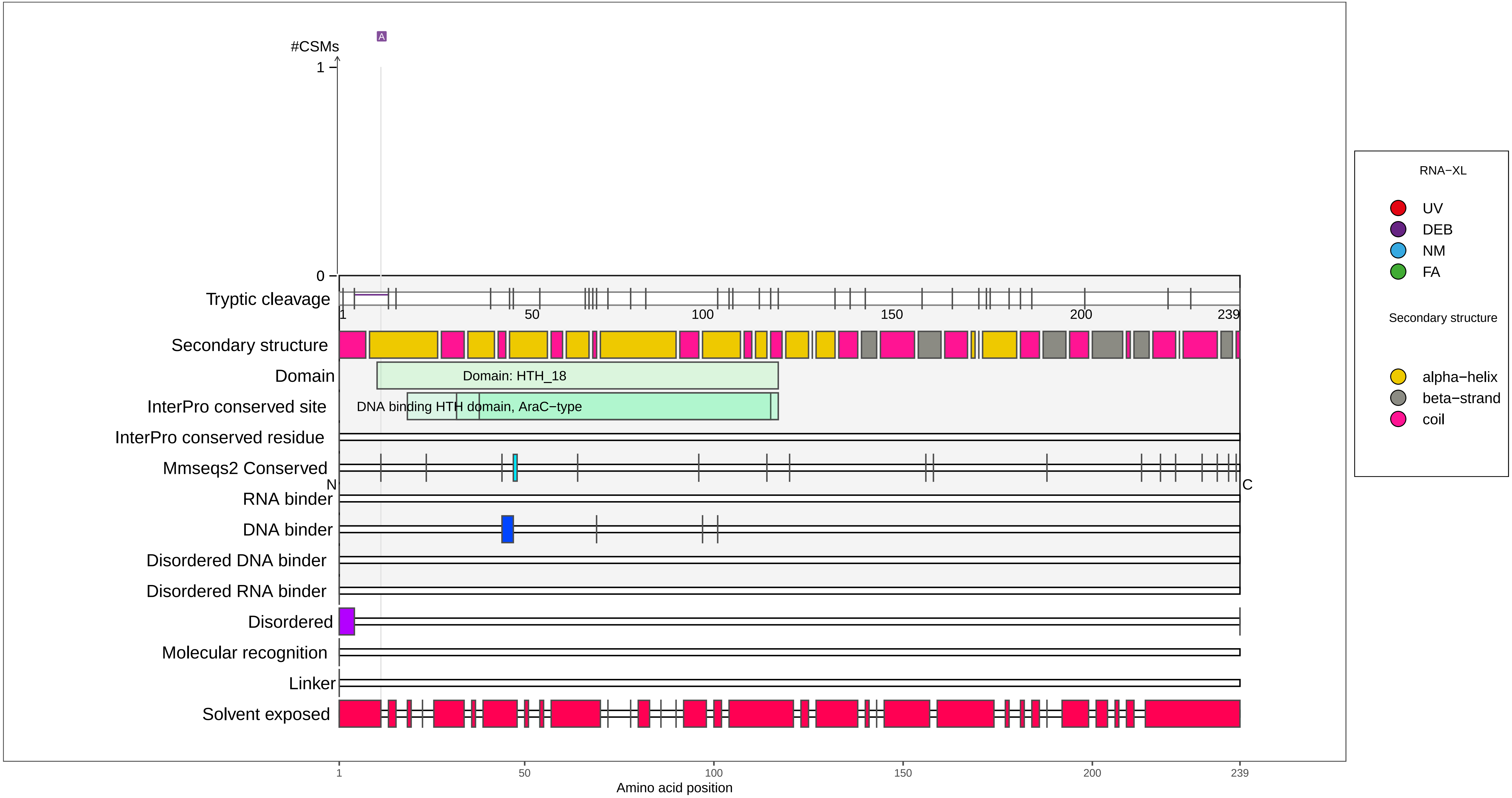

P77737
OPPF_ECOLI Oligopeptide transport ATP−binding protein OppF

− Abundance:
tryptic [log10 Intensity]: NA (Q NA)
PAXdb K12 strain [ppm]: 1.39 (Q 32)
PAXdb E.coli [ppm]: 1.51 (Q 60)

− RNA functions: not annotated

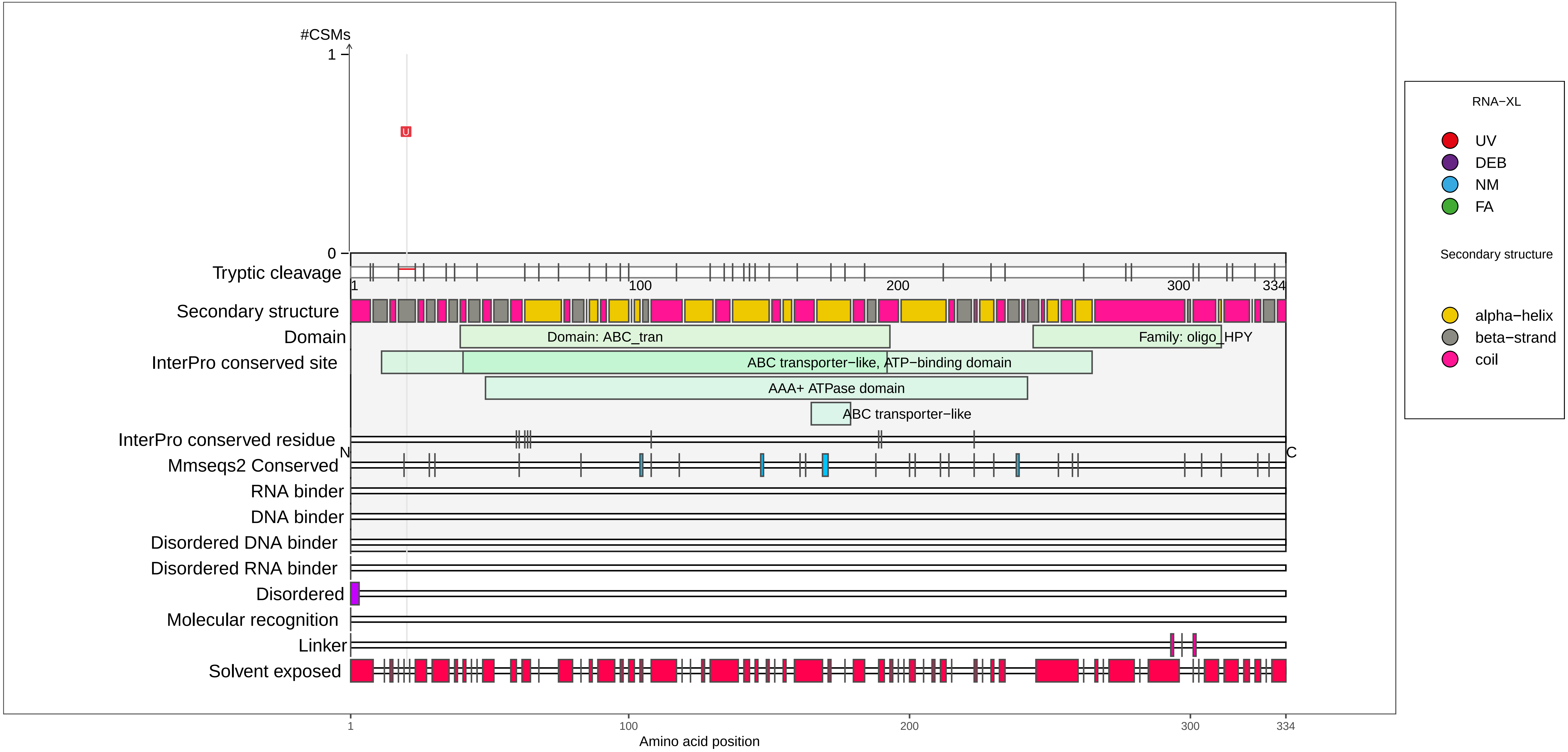

P77739
KT3K_ECOLI Probable ketoamine kinase YniA

− Abundance:
tryptic [log10 Intensity]: 8 (Q 58)
PAXdb K12 strain [ppm]: NA (Q NA)
PAXdb E.coli [ppm]: 2.26 (Q 79)

− RNA functions: not annotated

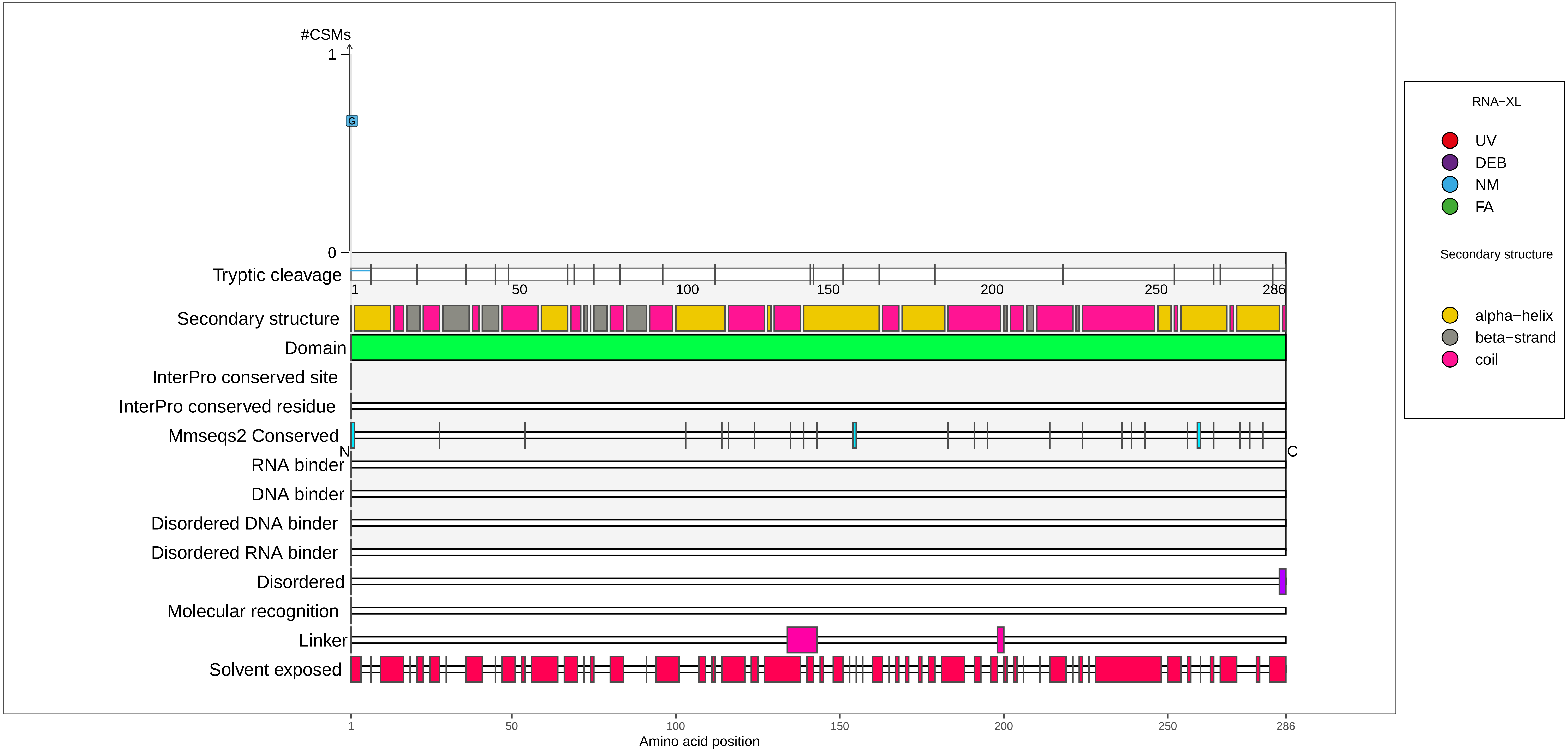

P77757
ARNC_ECOLI Undecaprenyl−phosphate 4−deoxy−4−formamido−L−arabinose transferase

− Abundance:
tryptic [log10 Intensity]: 7.01 (Q 14)
PAXdb K12 strain [ppm]: NA (Q NA)
PAXdb E.coli [ppm]: 1.31 (Q 55)

− RNA functions: not annotated

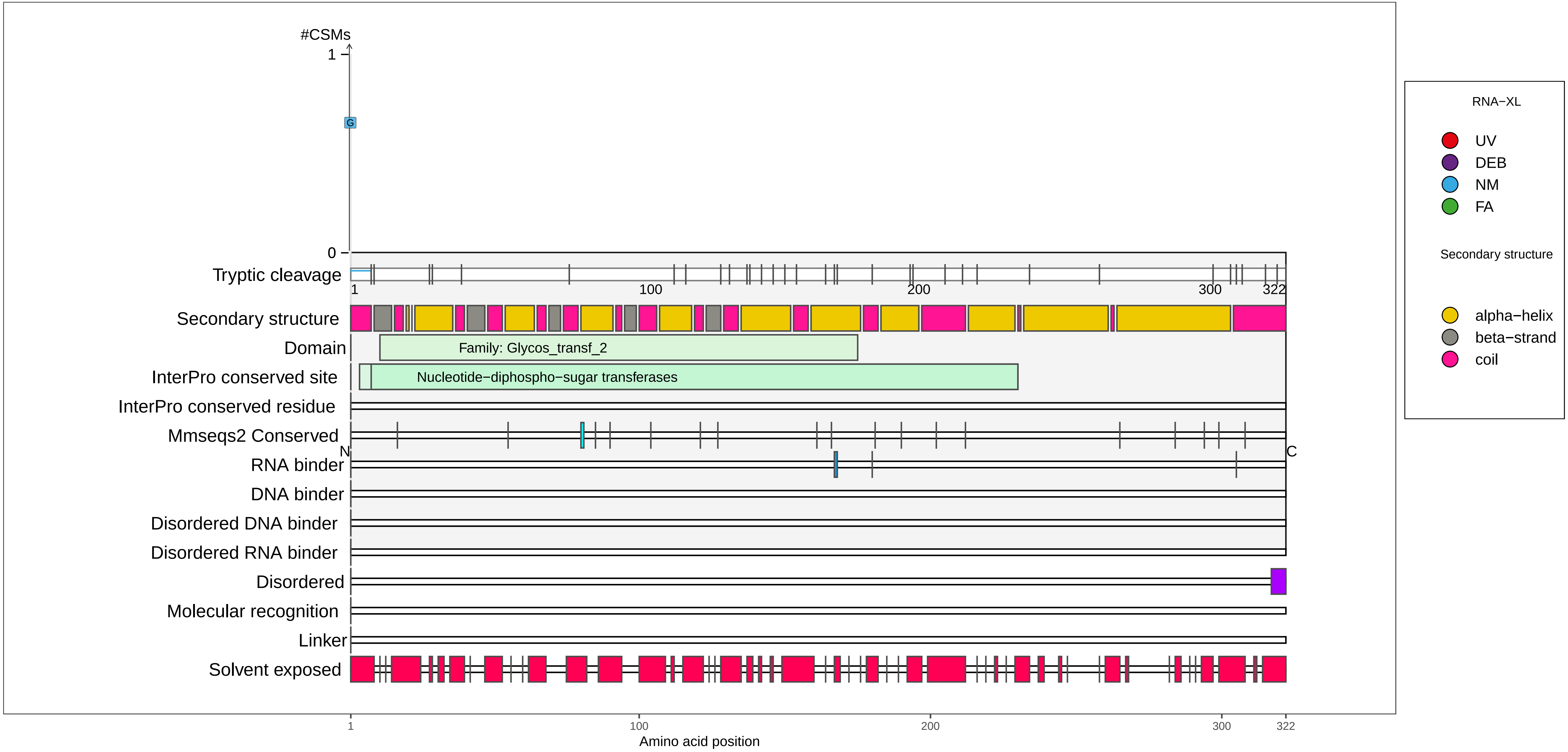

Q46796
YGEP_ECOLI Putative protein YgeP
− RNA functions: not annotated

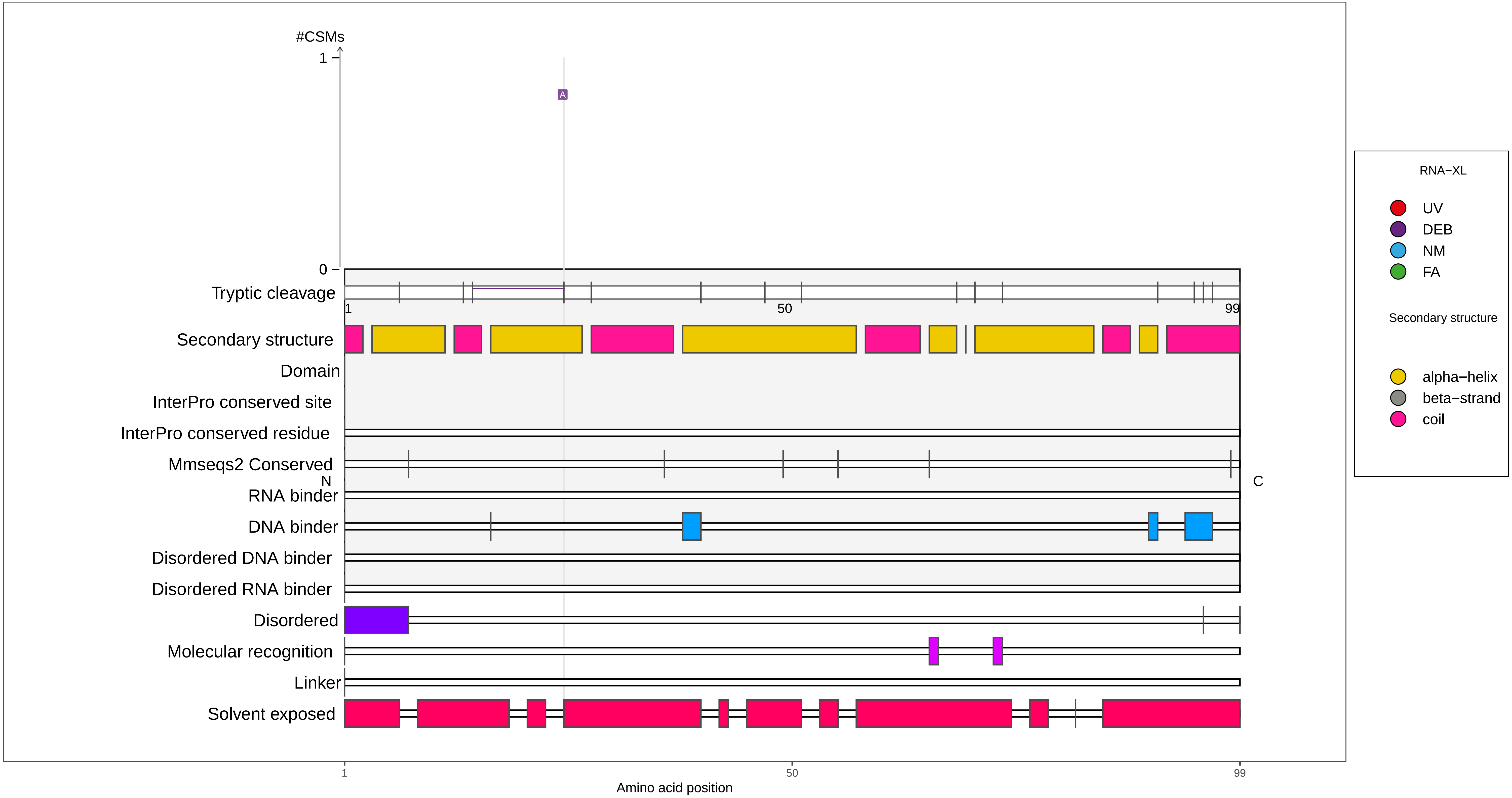

Q46807
ARCL_ECOLI Carbamate kinase−like protein YqeA
− Abundance:
tryptic [log10 Intensity]: NA (Q NA)
PAXdb K12 strain [ppm]: NA (Q NA)
PAXdb E.coli [ppm]: 1.16 (Q 51)

− RNA functions: not annotated

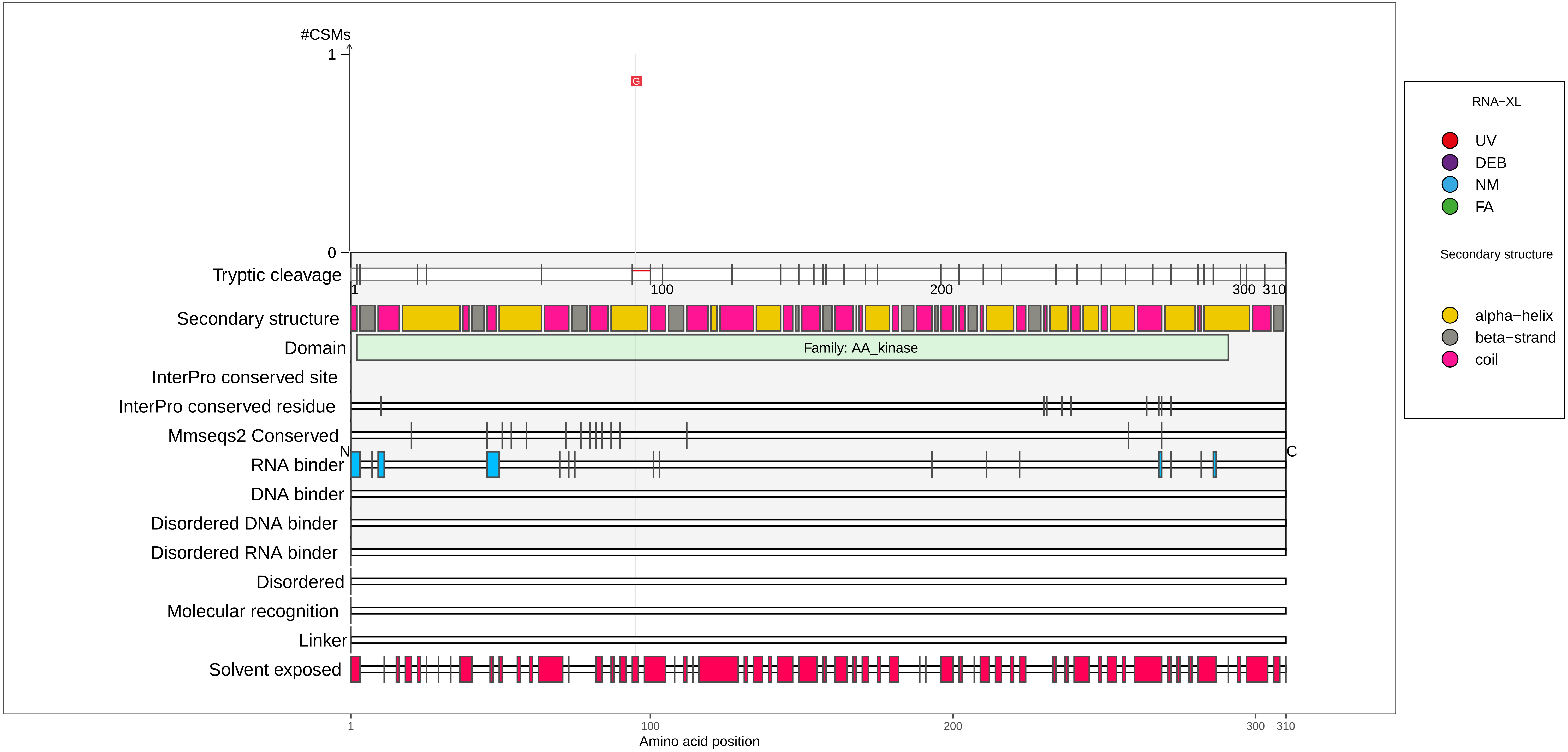

Q46822
IDI_ECOLI Isopentenyl−diphosphate Delta−isomerase

− Abundance:
tryptic [log10 Intensity]: 7.36 (Q 27)
PAXdb K12 strain [ppm]: NA (Q NA)
PAXdb E.coli [ppm]: 0.73 (Q 41)

− RNA functions: not annotated

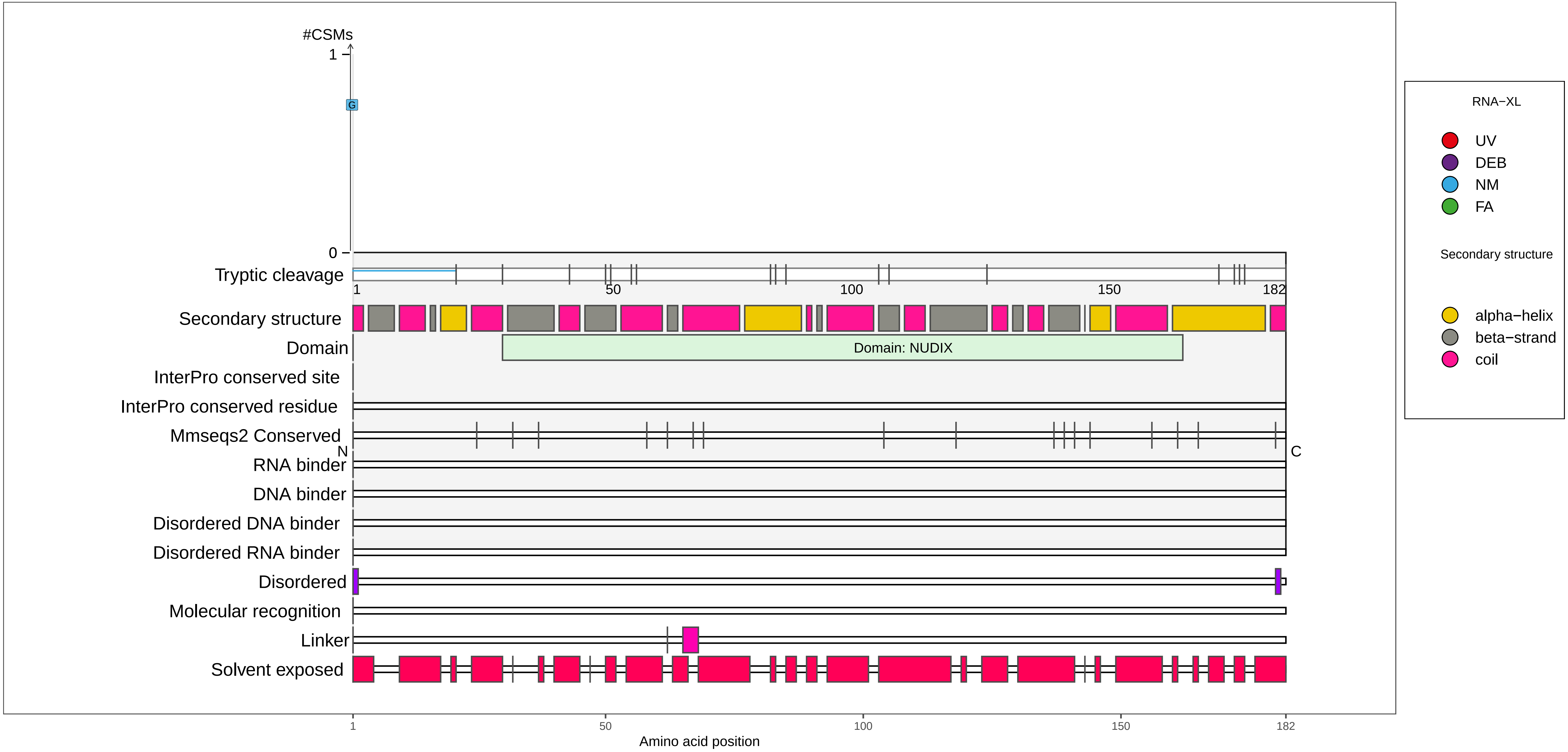

Q46845
YGHU_ECOLI Disulfide−bond oxidoreductase YghU

− Abundance:
tryptic [log10 Intensity]: 8.33 (Q 70)
PAXdb K12 strain [ppm]: 2.53 (Q 78)
PAXdb E.coli [ppm]: 2.57 (Q 87)

− RNA functions: not annotated

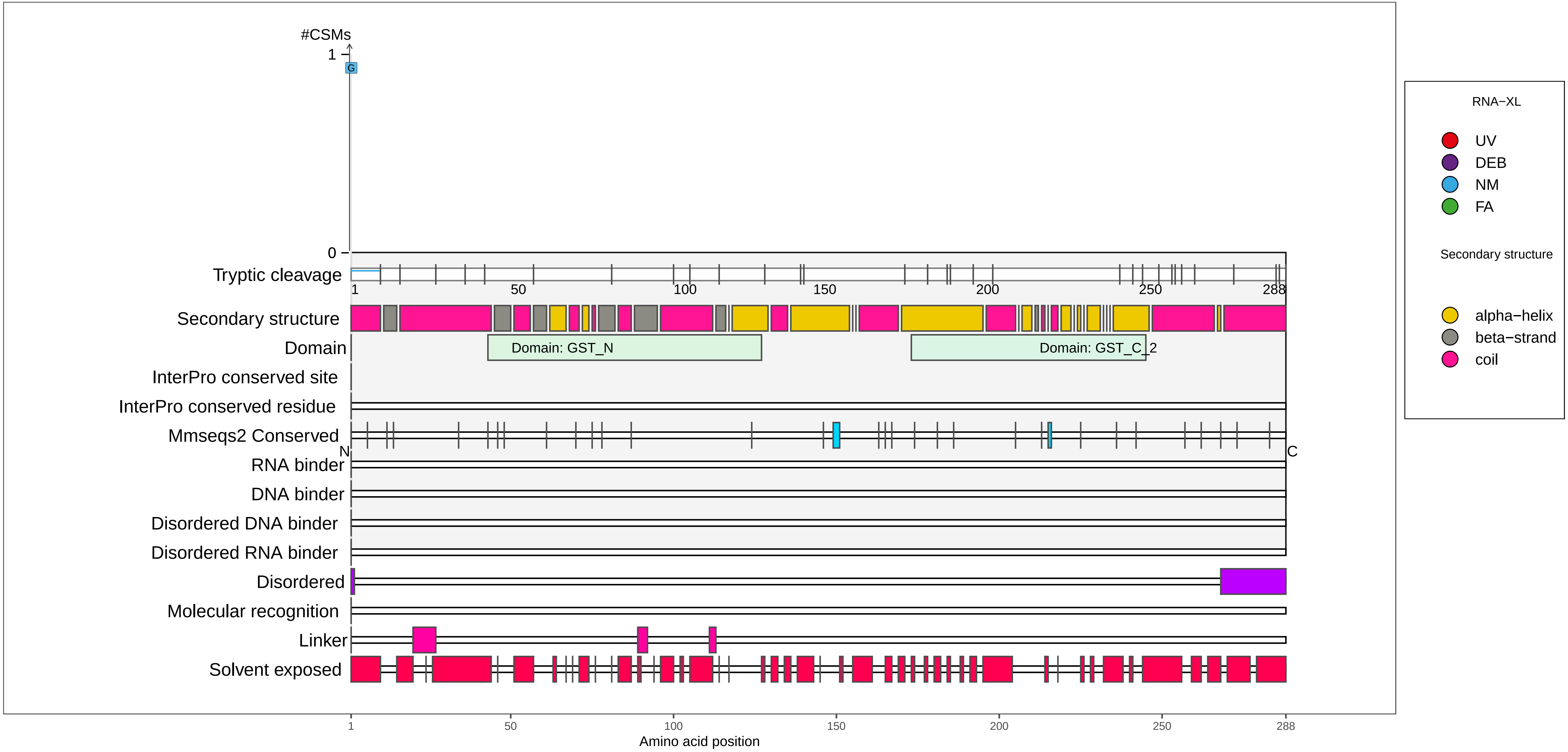

Q46925
CSDA_ECOLI Cysteine desulfurase CsdA

− Abundance:
tryptic [log10 Intensity]: 7.58 (Q 39)
PAXdb K12 strain [ppm]: 1.14 (Q 17)
PAXdb E.coli [ppm]: 1.43 (Q 58)

− RNA functions: not annotated

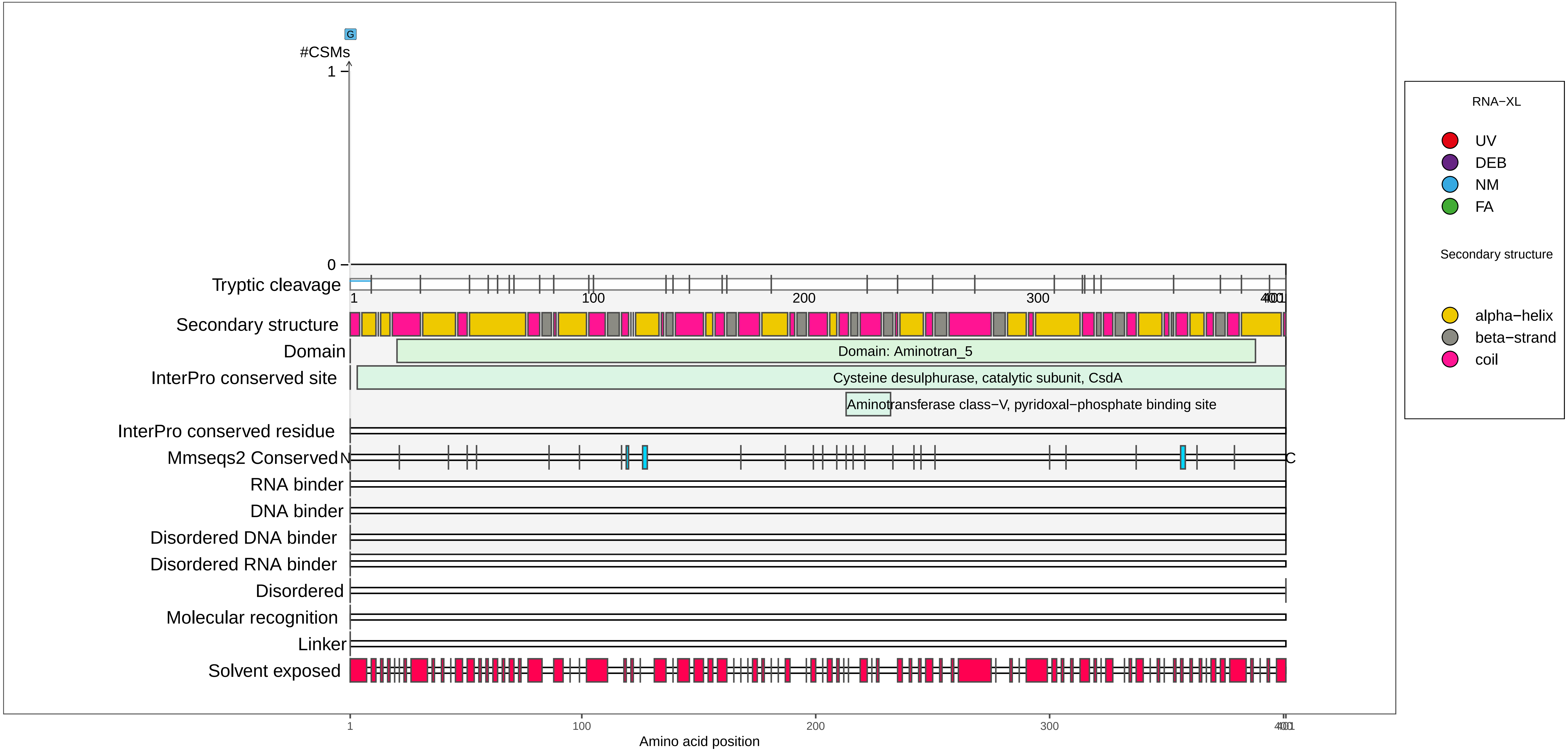

Q47706
UIDC_ECOLI Membrane−associated protein UidC

− Abundance:
tryptic [log10 Intensity]: 6.59 (Q 3)
PAXdb K12 strain [ppm]: 1.2 (Q 20)
PAXdb E.coli [ppm]: −0.99 (Q 9)

− RNA functions: not annotated

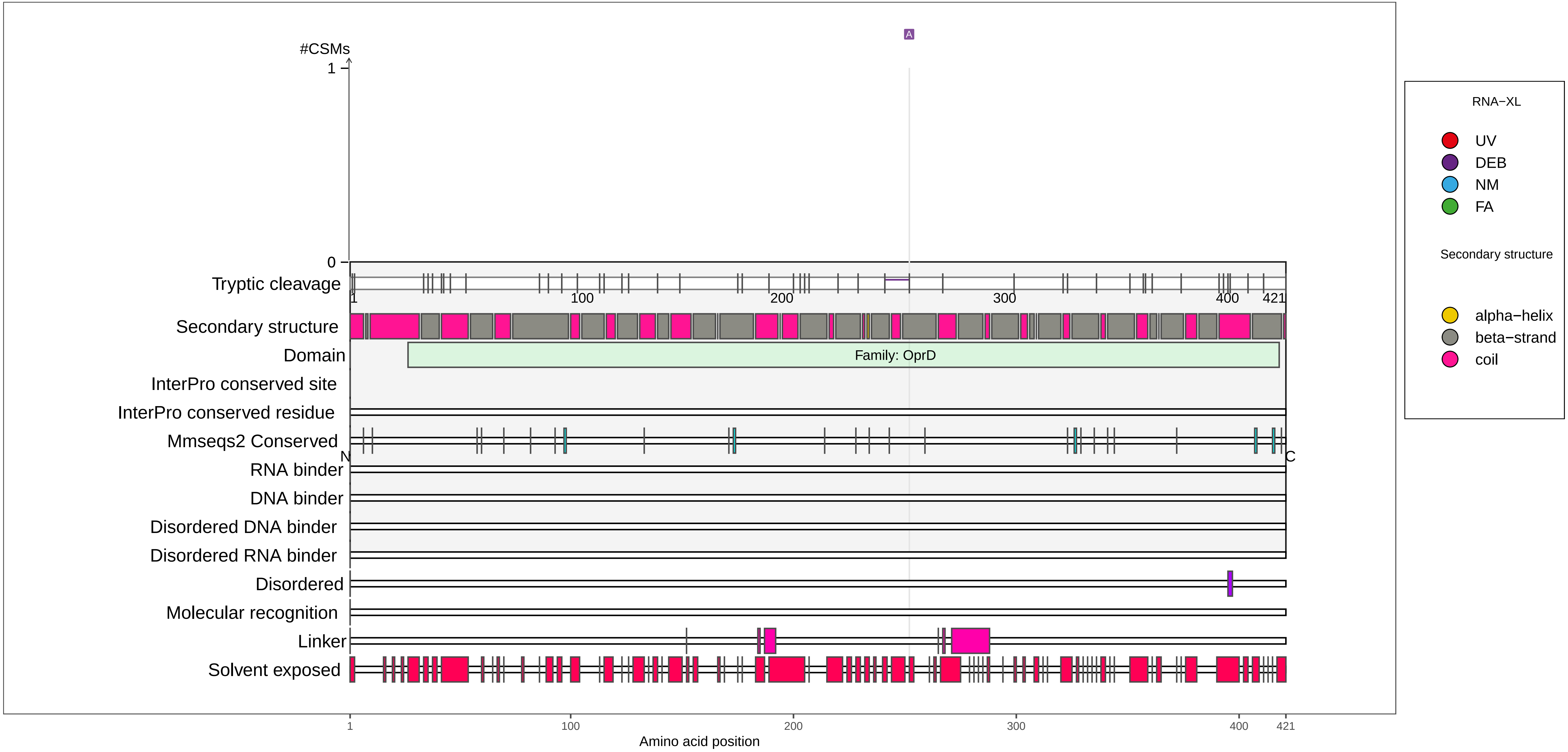

Q59385
COPA_ECOLI Copper−exporting P−type ATPase

− Abundance:
tryptic [log10 Intensity]: 7.45 (Q 33)
PAXdb K12 strain [ppm]: 2.17 (Q 66)
PAXdb E.coli [ppm]: 1.97 (Q 71)

− RNA functions: not annotated

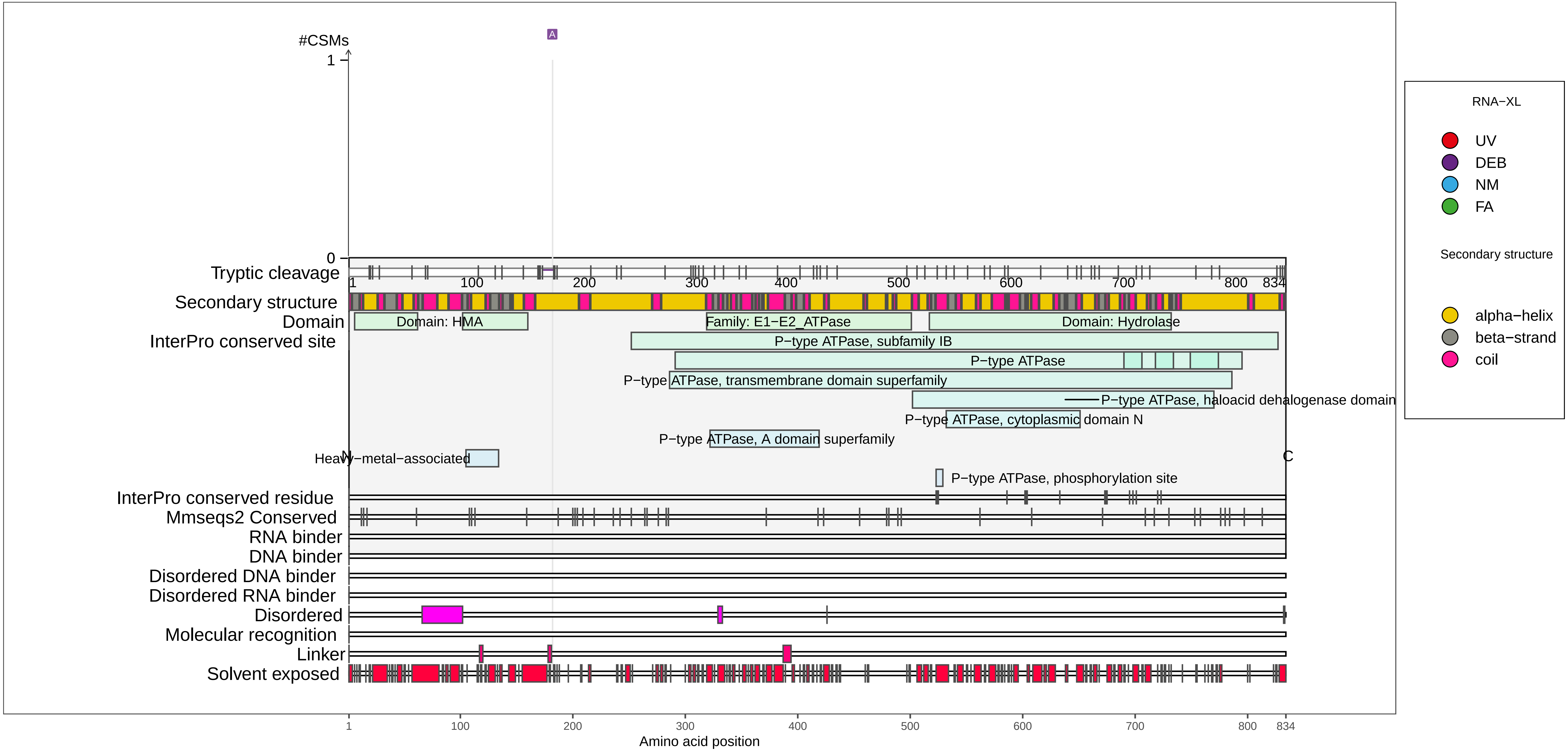
