## Supplementary File S1 for "Chemical crosslinking extends and complements UV crosslinking in analysis of RNA/DNA nucleic acid–protein interaction sites by mass spectrometry"

Supplementary Data S2. Exemplary mass spectra of UV-induced E. coli nucleotide-peptide crosslinks (S30 fraction, 2 replicate) identified by NuXL, but not by MSFragger-Labile workflow, visualized by OpenMS ToppView from NuXL output

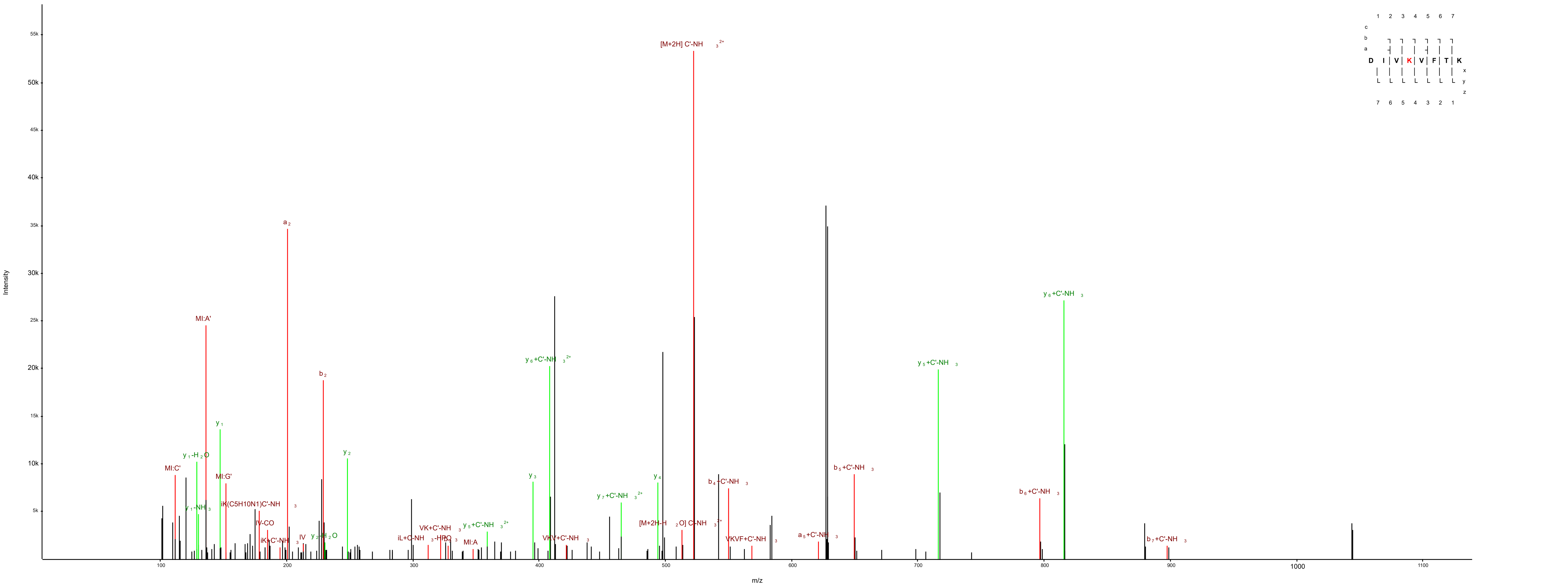


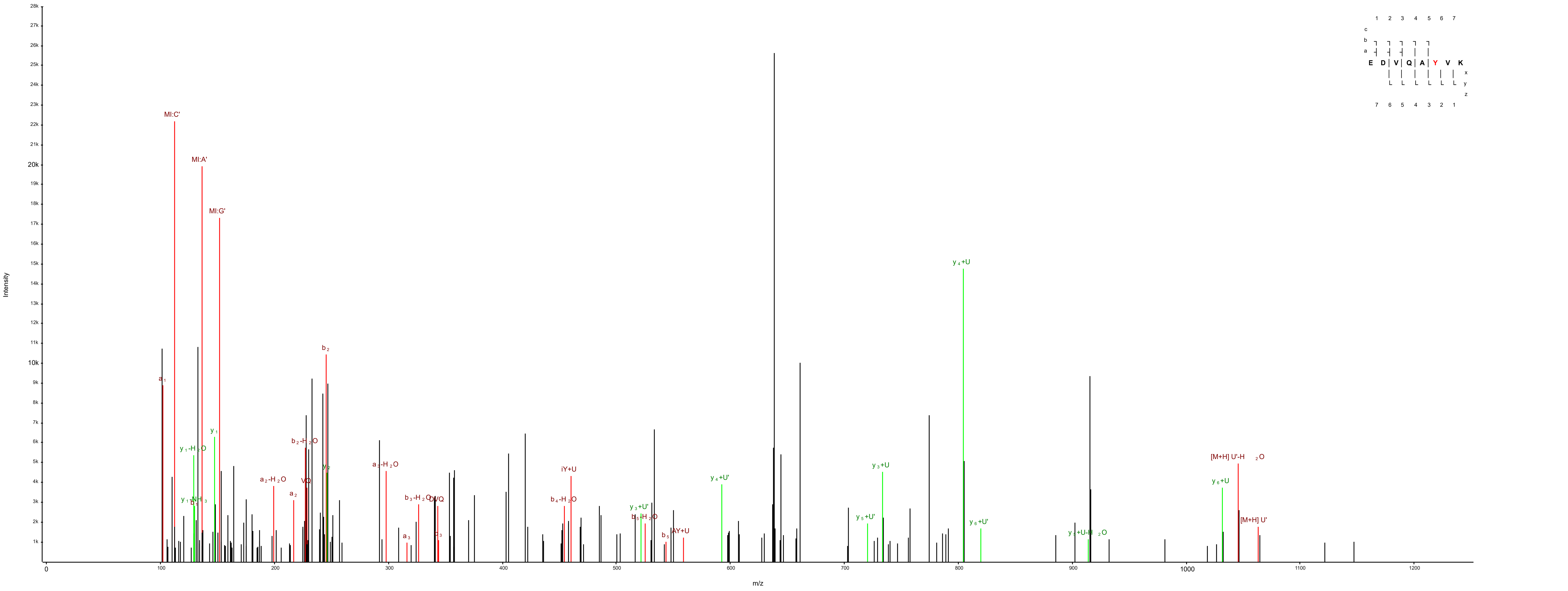


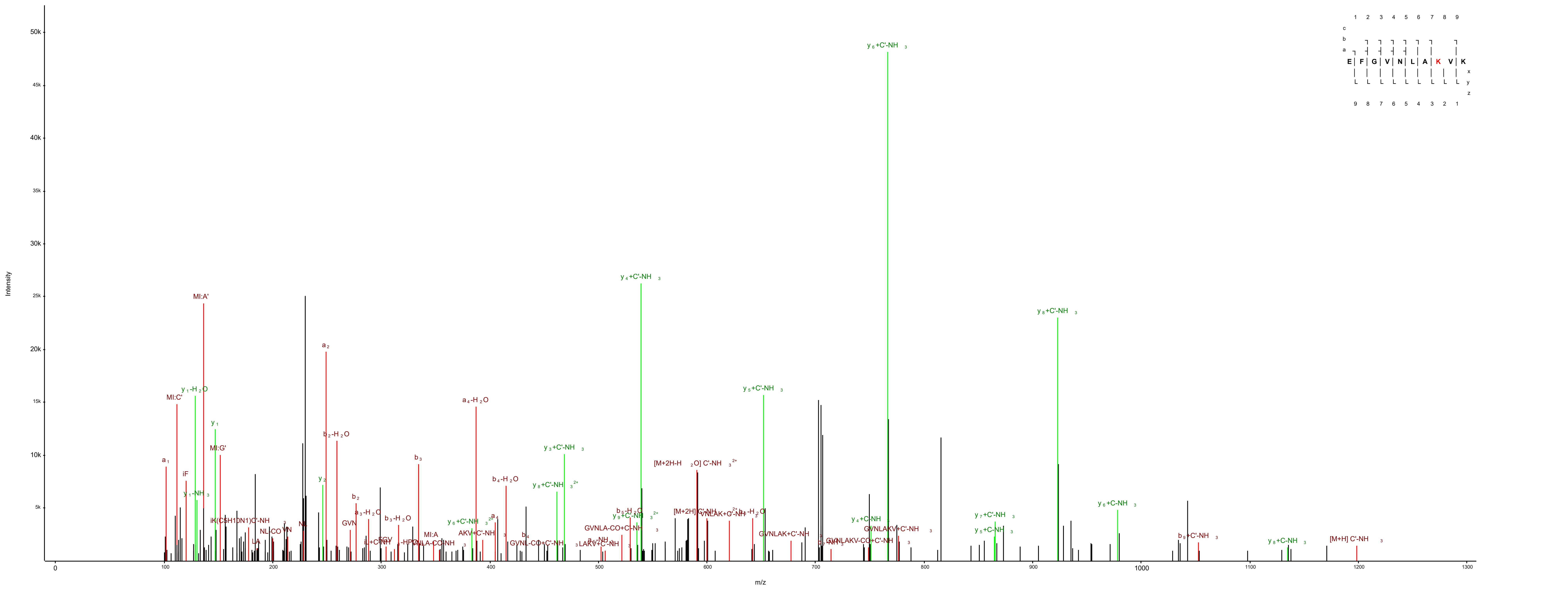
